# *Aspergillus fumigatus* versus Genus *Aspergillus*: Conservation, adaptive evolution and specific virulence genes

**DOI:** 10.1101/2020.12.01.406157

**Authors:** Shishir K Gupta, Mugdha Srivastava, Özge Osmanoglu, Zhuofei Xu, Axel A Brakhage, Thomas Dandekar

## Abstract

*Aspergillus* is an important fungal genus containing economically important species, as well as pathogenic species of animals and plants. Using eighteen fungal species of the genus *Aspergillus*, we conducted a comprehensive investigation of conserved genes and their evolution. This also allows to investigate the selection pressure driving the adaptive evolution in the pathogenic species *A. fumigatus.* Among single-copy orthologs (SCOs) for *A. fumigatus* and the closely related species *A. fischeri*, we identified 122 versus 50 positively selected genes (PSGs), respectively. Moreover, twenty conserved genes of unknown function were established to be clearly positively selected and thus important for adaption. *A. fumigatus* PSGs interacting with human host proteins show over-representation of adaptive, symbiosis-related, immunomodulatory, and virulence related pathways such as the TGF-β pathway, insulin receptor signaling, IL1 pathway and interfering with phagosomal of GTPase signaling. Additionally, among the virulence factor coding genes, secretory and membrane protein coding genes in multi-copy gene families, 212 genes underwent positive selection and also suggest increased adaptation such as fungal immune evasion mechanisms (*aspf2*), siderophore biosynthesis (*sidD*), fumarylalanine production (*sidE*), stress tolerance (*atfA*) and thermotolerance (*sodA*). These genes presumably contribute to host adaptation strategies. Genes for the biosynthesis of gliotoxin are shared among all the close relatives of *A. fumigatus* as ancient defense mechanism. Positive selection plays a crucial role in the adaptive evolution of *A. fumigatus*. The genome-wide profile of PSGs provides valuable targets for further research on the mechanisms of immune evasion, for antimycotic targeting and understanding fundamental processes of virulence.

## Introduction

Fungal diseases and specifically invasive fungal infections lead to an estimated 1.5 to 2 million deaths each year, which exceeds the global mortality estimates for either malaria or tuberculosis (Brown, et al. 2012). During 200 million years of evolution, aspergilli developed as a group of ubiquitous fungi (Galagan, et al. 2005). Among the known aspergilli, environmentally acquired pathogen *Aspergillus fumigatus* is the predominant, ubiquitous, opportunistic pathogenic species causing life-threatening invasive aspergillosis (IA) and chronic pulmonary aspergillosis (CPA) in immunodeficient patients as well as allergic disease in immunoreactive patients. *A. fumigatus* has superior ability to survive and grow in a wide range of environmental conditions. The conidia of *A. fumigatus* released from the conidiophores are dispersed in the environment and remain dormant until encountering the environmental conditions that allow metabolic activation. Once metabolically active, conidia swell and germinate into hyphal filaments that form mycelia and produce conidiophores (Sugui, et al. 2014). The inhalation of conidia, fungal growth and tissue invasion in immunocompromised patients can result in the establishment of invasive disease and represents a major cause of morbidity and mortality.

The availability of the sequenced genomes from several species of the genus *Aspergillus* opens up the possibility to examine the demarcation of fungal species at the whole-genome level. These include economically important fungi (*A. oryzae*, *A. niger*), plant pathogens (*A. zonatus*), animal pathogens (*A. fumigatus*, *A. clavatus*, *A. fischeri*, *A. brasiliensis*, *A. sydowii*, *A. tubingenesis*, *A. terreus*), plant and animal dual pathogens (*A. glaucus*, *A. versicolor*, *A. niger*, *A. flavus*, *A. wentii*) and non-pathogens (*A. nidulans*, *A. kawachii*, *A. oryzae*, *A. carbonarius*, *A. luchuensis*). Among these is *A. fischeri* that is a close homothallic sexual relative to *A. fumigatus*. This fungus can also cause keratitis and possibly pulmonary aspergillosis in transplant patients but is an extremely rare invasive pathogen (Chim, et al. 1998; Gerber, et al. 1973).

A recent comparative study emphasizes the high genomic and functional diversity within the genus *Aspergillus* (de Vries, et al. 2017). In the study, strong conservation was observed for central biological functions while high diversity was observed for many other physiological traits, such as secondary metabolism, carbon utilization, and stress response (de Vries, et al. 2017). However, a genome-wide scan indicating evolutionary forces operating on *Aspergillus* genes has not been performed yet. The analysis we performed here addresses this and provides a general resource to study gene evolution in the genus *Aspergillus*.

Homologous recombination and positive selection are major forces for the evolution of microorganisms that drive adaptation to new hosts, antimycotics and promote survival of pathogens in hostile environments including the human host (Gladieux, et al. 2014). Evolution experiments showed that recombination can accelerate species adaptation in stressful environments (Winkler and Kao 2012) by combining advantageous mutations and thereby assisting in their fixation (Shapiro, et al. 2009). Additionally, natural selection is the central force shaping the diversity of genotypes by acting on resulting phenotypes. Study of natural selection provides significant insights into the possible functional alterations during gene evolution and important nucleotide substitutions involved in adaptation. Adaptive changes of the protein-coding genes are ultimately responsible for evolutionary innovations that can have a significant impact on the adaptation of species to their environment and generate an overview of diversity (MacColl 2011). Fungal gene products which play a significant role in environmental adaptation including survival in the human body are more likely to be positively selected.

The evolutionary origin of virulence of *A. fumigatus* is not well known. Nevertheless, a well-accepted hypothesis suggests that interaction with free living predator amoeba during saprophytic existence may have posed a selection pressure on *A. fumigatus* to optimize survival that later endorsed accidental virulence in human host (Casadevall and Pirofski 2007; Hillmann, et al. 2015). Thus, we investigated whether evolutionary driving forces and positive selection is visible for virulence-related genes and how far in general *A. fumigatus* protein-coding genes are associated with a genome-wide signature of positive selection. We hence first compare the genomic features and then in detail the conserved gene families of aspergilli, species tree estimation followed by recombination and evolutionary analysis of single copy orthologs (SCOs). Next, we show the results indicating positive selected genes (PSGs) involved in host-pathogen interactions and environmental adaptations. Furthermore, the screen identified twenty conserved genes of unknown function which are clearly positively selected and hence conveying important uncharacterized adaptive functions. However, for a more complete picture on virulence evolution in multi-gene families have to be included and we point out positive selection in a number of virulence factors from these. Hence, we finally considered analysis of multi-gene families containing annotated virulence factor coding genes, secretory and membrane proteion coding genes in *A. fumigatus*. For each of these categories we establish here clear virulence, infection, or growth-mediating factors with positive selection. These PSGs illuminate virulence for *A. fumigatus* with genes for conidia germination, signal transduction, metabolism, mitochondrial activity, and transcriptional regulation. Note that the singletons and species-specific duplicated genes in *A. fumigatus* which could be associated with the increased virulence of the species (Table S1) were not analysed for species-level selection signature. Finally, we show that genes involve in biosynthesis of a very old and strong virulence factor gliotoxin are essential inventory, conserved among most Aspergilli since long time and hence do not share a species-resolved selection.

## Results

### Genomic features and quality assessment

The general and genomic features of aspergilli analysed in this study are listed in Table S2. The size and G+C content of the genomes ranged from 26.09 to 37.45 Mb and 48 to 52.8% respectively. *A. luchuensis* represented the largest *Aspergillus* genome among the analyzed genus. Since many of the fungal genomes are poorly annotated (Okagaki, et al. 2016), we assessed the quality of the genome annotation by testing for the presence of universally conserved fungal SCOs before performing the sequence comparisons (see Materials and methods). Almost all aspergilli genomes for the analysis had accepTable quality and completeness (Figure 1). This approach here is superior to large-scale genome analysis where the nonappearance of a gene in a particular genome might result from a poor quality of gene structure annotation rather than its true absence.

**Figure 1:**
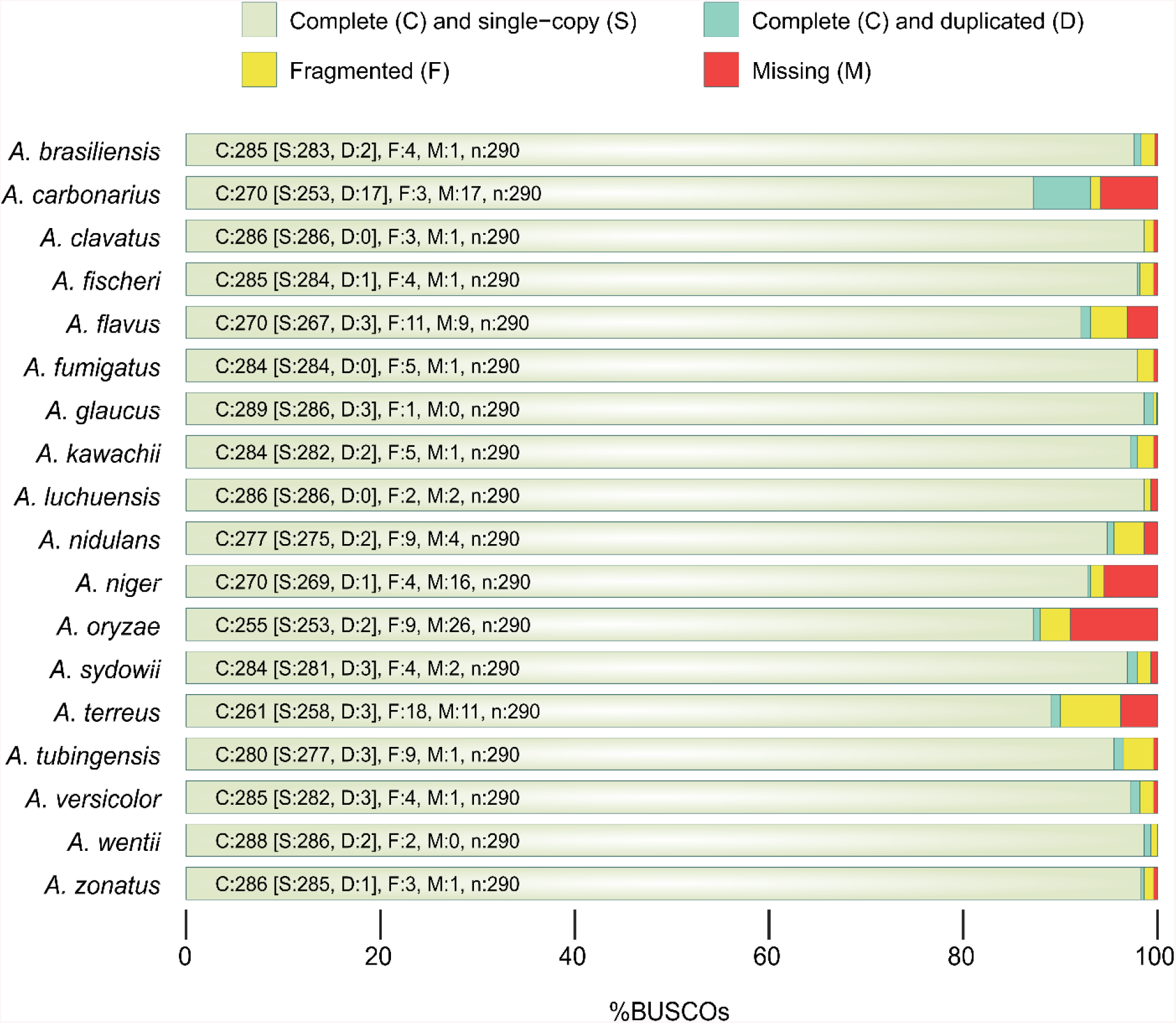
Genome completeness in the Aspergilli compared. Statistics and assessment of Aspergilli protein coding gene sets and their completeness in BUSCO notation (C:complete [S:single copy] and [D:duplicated], F:fragmented, M:missing, n: gene number).

### Orthologs and unique genes

Genome-wise comparisons revealed the similarity and differences among the different organisms (see Materials and Methods for details). The clustering of proteomes yield 18,474 groups from 189,010 protein sequences (Additional data file 1). The core genome spans ~23-35% of the total proteins encoded by each genome, revealing that core genome has a rather homogeneous gene content (Additional data file 2). We found that 3175 protein-coding genes are present as single copies in each genome, or 17% of the total ortholog clusters. These clusters comprise proteins involved in core biological processes, such as cell growth, cell division, protein folding and translation. Interestingly, sporocarp development involved in sexual reproduction was among the top five statistically over-represented (GO:0000909, FDR 4.79E-05) terms indicating aspergilli have the potential of undergoing sexual reproduction, which seems to be a conserved process. The full list of over-represented Gene Ontology (GO) terms in the three GO categories (i.e. ‘Biological Processes’, BP; ‘Molecular Function’, MF; ‘Cellular Component’, CC) is listed in Table S3.

Our comparative analysis also revealed that each genome also encodes species-specific proteins that have no homologues in other species. A total of ~2.5%-17% of the sequences in each *Aspergillus* genome were estimated to be so-called singletons at a conservative clustering threshold of OrthoMCL using a 1.5 inflation index. The largest number of singletons was present in *A. glaucus,* followed by *A. wentii*. Interestingly, out of a total of 517 species-specific sequences found in *A. fumigatus*, 29 contain secretion signals and 7 were annotated as cell rescue, defense, and virulence-related proteins by SignalP (Nielsen 2017) and FunCat (Ruepp, et al. 2004). It is likely that such proteins may also participate in some biological processes unique for *A. fumigatus* or play a role in the survival and propagation of this pathogen. However, in this regard more *in silico* and *in vitro* functional characterization is required. For instance, analysis of lineage-specific genes involved in adaptation of a species to a particular environment suggests that such genes evolved rapidly since their deduced proteins have a substrate to act on and therefore a number of such genes have been postulated to play a role and confer an adaptive advantage on a particular species (Nielsen, et al. 2005). In contrast to this hypothesis, studies on *Drosophila* suggested instead of being fast-evolving genes, singletons seem to have largely originated by *de novo* synthesis from non-coding regions like intergenic sequences (Zhou, et al. 2008). The further analysis of the species-specific proteins is beyond the scope of this manuscript but will be the subject of future studies. Figure 2 depicts the percentage of clustered and ‘species-specific proteins in each analysed proteome’. A quantitative summary of orthology assignments is given in Table S4.

**Figure 2:**
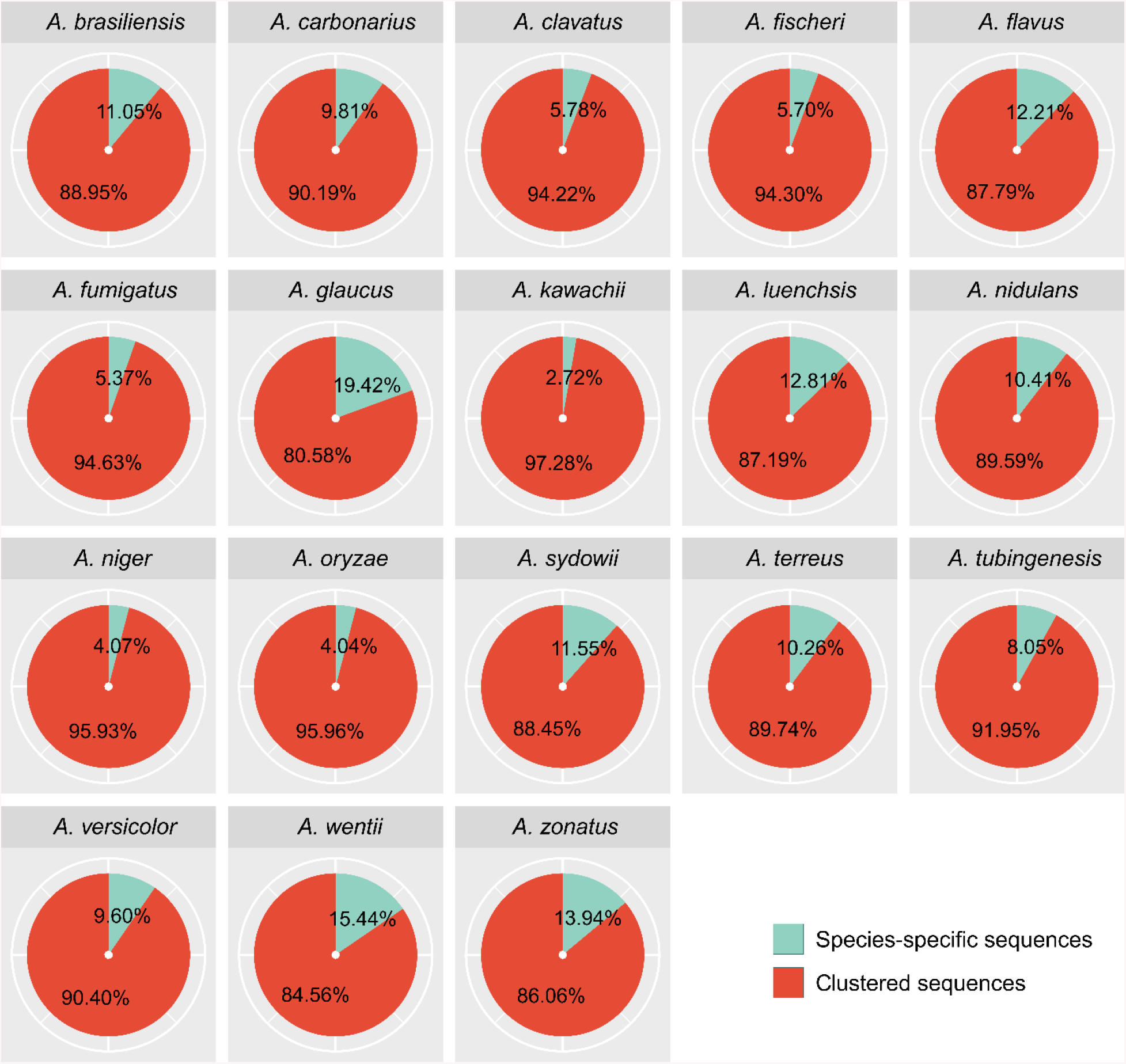
Distribution of Aspergilli proteins as unique or shared relative to the entire set of proteins. Partitioning Aspergilli gene sets by their traceable orthology reveals a spectrum of conservation from widespread orthologues found across the Aspergilli to species-specific genes with no recognizable homologues. The pie chart shows the percentage of unique and conserved proteins coded by among all eighteen sequenced Aspergilli identified by OrthoMCL analysis. Species-specific sequences include both the singletons and same species inparalog clusters.

### Reconciling of species tree and gene trees

Molecular phylogenies based on single genes often yield apparently conflicting tree structures and often produce incongruences (Jeffroy, et al. 2006). To overcome this problem, genome-scale methods for phylogenetic inference by combining multiple genes are well established to resolve incongruences (Gee 2003). Here, we reconstructed a species-tree using a supermatrix approach (Figure 3). We see strong bootstrap values that support the relationships among analyzed species. The proposed relationships agree with those recently reported by de Vries et al. (2017) (de Vries, et al. 2017). As expected, *A. fumigatus* and *A. fischeri* represented sister groups. However, in *A. fumigatus* comparatively more genetic change occurred. Moreover, *A. clavatus* was found as the closest ancestor of these two aspergilli but human infections due to *A. clavatus* are extremely rare case reports (Riddle, et al. 1968). Finally the three species *A. zonatus*, *A. glaucus* and *A. wentii* that are phylogenetically distant from *A. fumigatus*, also displayed the highest number of species-specific proteins (Figure 2).

**Figure 3:**
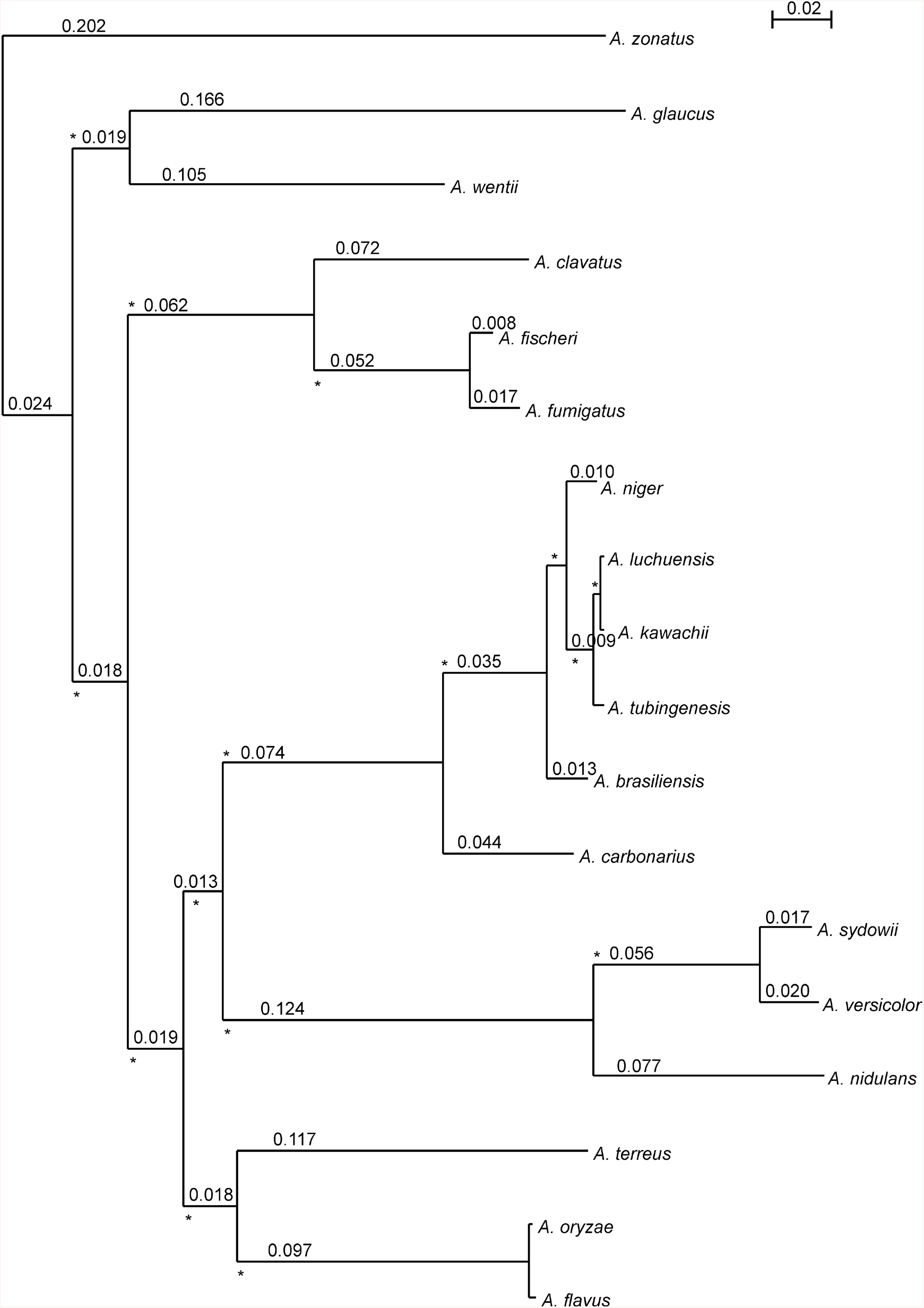
Aspergillus species tree. The maximum likelihood tree was obtained after concatenating the 3175 congruent orthologous proteins or protein fragments shared by all 18 genomes. On the branches of the tree are reported the branch length (black). Each node of the tree is supported by bootstrap value of 100. The scale bar (s/s) indicates number of substitutions per site.

### Positive selection in *A. fumigatus* single-copy ortholog protein-coding genes

In our analysis, recombination events were excluded to avoid spurious signals by these (see Materials and methods). Here, for *A. fumigatus* we identified 122 PSGs and for *A. fischeri* 50 PSGs (see below). The distribution of SCOs and *A. fumigatus* PSGs over the chromosomes is shown in Figure 4.

**Figure 4:**
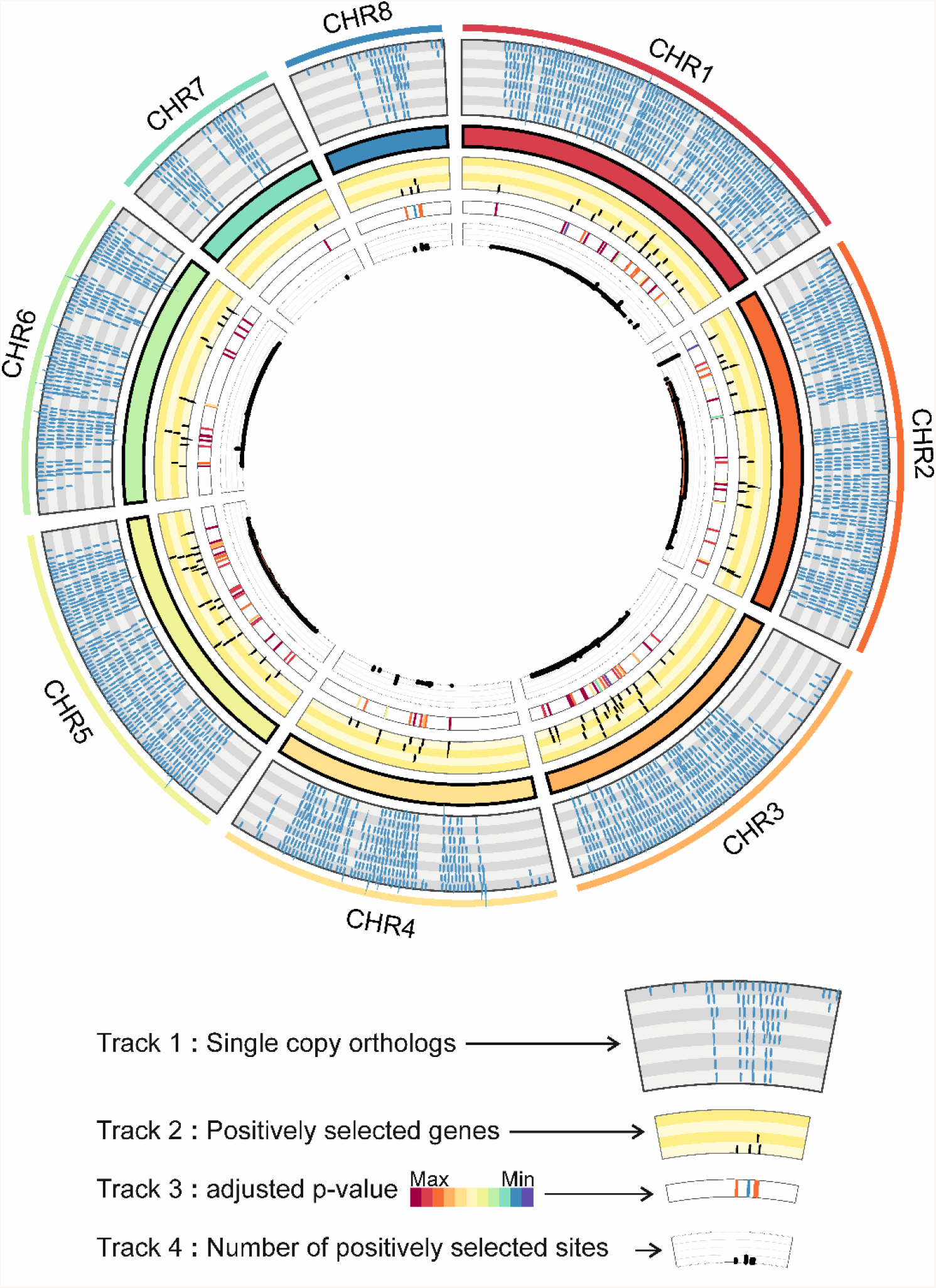
Chromosomal distribution of single-copy orthologs (SCOs) and positive selected genes (PSGs) in *A. fumigatus*. The circus plot shows the distribution of SCOs and PSGs over the different *A. fumigatus* chromosomes. Different color track shows the different information. Track1: SCOs in eight grey shaded subtracks, Track2: PSGs in four yellow shaded subtracks, Track3: heatmap of the q-values of PSGs and Track4: number of sites under selection in corresponding PSGs.

### Host interacting single-copy ortholog PSGs

The encounter between host and pathogen is mediated by host-pathogen protein-protein interactions (HP-PPI) which play crucial roles in infections, as they define the balance either in favour of the spread of the pathogen or their clearance (Nicod, et al. 2017). Genes involved in host-pathogen interactions are a frequent substrate of positive selection. We addressed the question whether the identified PSGs interact with host proteins and interfere with host defence processes. To answer this we verified the interacting partners of *A. fumigatus* protein products of PSGs in a previously established interspecies-interolog-based *A. fumigatus*-host interaction network (Remmele, et al. 2015) and the interspecies-interolog-based *A. fumigatus*-human interaction network reconstructed in this study. In total, 36% of SCO PSGs were identified to be involved in host-pathogen interactions (chi-squared = 306.18, p-value < 0.01). The resulting host-pathogen protein-protein interaction network consists of 44 *A. fumigatus* proteins coded by PSGs and 228 human proteins (Figure S1). The 40S ribosomal protein S9 Afua_3g06970 (Rps9A) was identified as top pathogen hub with 35 interactions followed by putative DEAD/DEAH box helicase Afua_5g02410 (Fal1) and putative proteasome component protein Afua_3g11300 (Prs2) (Table S5). Of note, proteins involved in pathogenicity with a higher degree of host interactions are involved in increasing the fitness of the pathogen in the host environment. We further performed a pathway and over-representation analysis (ORA) of targeted host proteins, which showed the over-representation of several immune-related pathways (Table S6). Signaling proteins are preferentially targeted by pathogens because they globally regulate many cellular processes. The pathway over-representation analysis of PSGs interacting human proteins resulted in significant over-representation of several signaling pathways (Table S6; Figure 5). Highly significant over-representation was observed for the transforming growth factor beta (TGF-β) pathway (pathway ORA corrected p-value 3.95E-12). Other top over-represented pathways include the insulin signaling pathway and the IL-1 signaling pathway. Notably, other potent immune pathways against *A. fumigatus* such as Toll-like receptor (TLR) pathway, T-cell and B-cell pathway (Hohl and Feldmesser 2007) were also significantly enriched. The adaptive evolution of several host interacting *A. fumigatus* genes might be due to environmental pressure, and these could contribute towards the various fungal strategies to adapt and evade recognition by amoeba as well as in the rare event of infection of the human host’s immune system.

**Figure 5:**
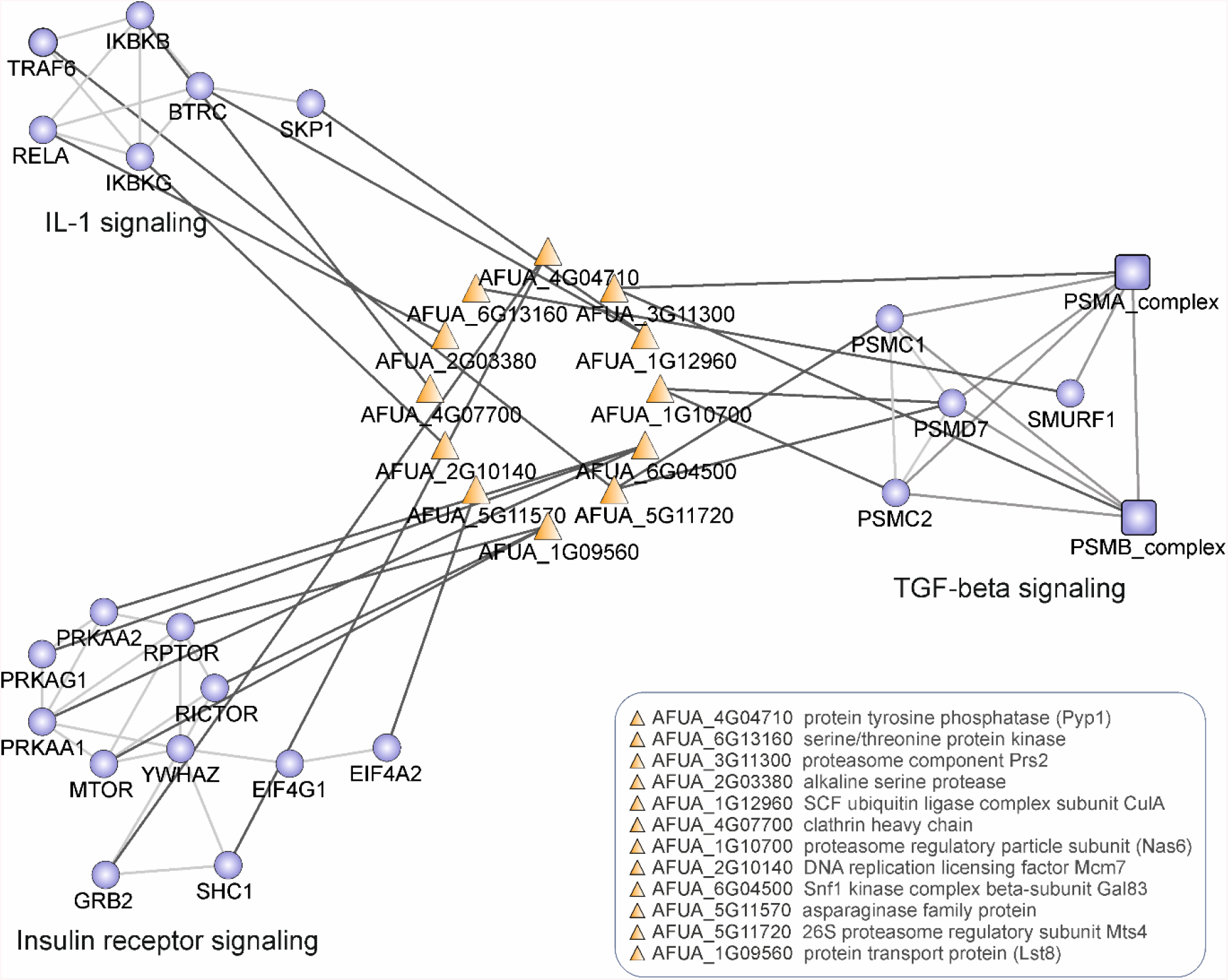
*A. fumigatus*-human host proteins-protein interactions (PPIs) map. PPIs between *A. fumigatus* proteins coded by positively selected genes (PSGs) and human proteins involved in three pathways are illustrated: IL1-signaling, Insulin receptor signaling and TGF-beta signaling. Orange -*A. fumigatus* proteins and blue - human proteins.

Table S7 summarizes the significantly enriched GO terms for PSGs interacting host proteins. Since the human host niche is in comparison to its global abundance in saprophytic niches rarely used by *A. fumigatus*, the selection pressure on *A. fumigatus* should be rather imposed by free-living amoebae and other microrganisms (e.g. (Stroe, et al. 2020)) during their co-survival in the soil niche. *A. fumigatus* possesses a molecular arsenal to survive in most environments including the human host (Marcos, et al. 2016). This requires powerful strategies for initial colonization (Paulussen, et al. 2017) and biofilm formation (Loussert, et al. 2010) which in the human host then permits to establish infection.

### Functional classification of single-copy ortholog PSGs

We further categorized the *A. fumigatus* PSGs in euKaryotic Orthologous Groups (KOG) categories based on a blast-based sequence similarity search. The most suiTable categories were identified, and multiple categories were used if genes were significantly similar to multiple query sequences with different KOG categories associated with them. If no annotation was retrieved for any of the genes, we scanned sequences against EGGNOGs (Huerta-Cepas, et al. 2016) HHM profiles for classification. In total, we were able to assign functional KOG categories to 107 PSGs (Figure 6). These genes are our prime candidates for having been shaped by positive selection during evolution and adaptation of *A. fumigatus* and were assigned to three large functional categories (as well as to the corresponding functional classes) according to the KOG database: (i) metabolism, (ii) cellular processes and signaling, and (iii) information storage and processing. In each category we studied the most interesting genes in greater detail to further investigate the probable causes of positive selection (see Additional file 1 Supplemental Note "Positive selection in genes classified under KOG functional categories"). Table S8 contains a full list of the *A. fumigatus* genes in single copy orthogroups evolved under positive selection.

**Figure 6:**
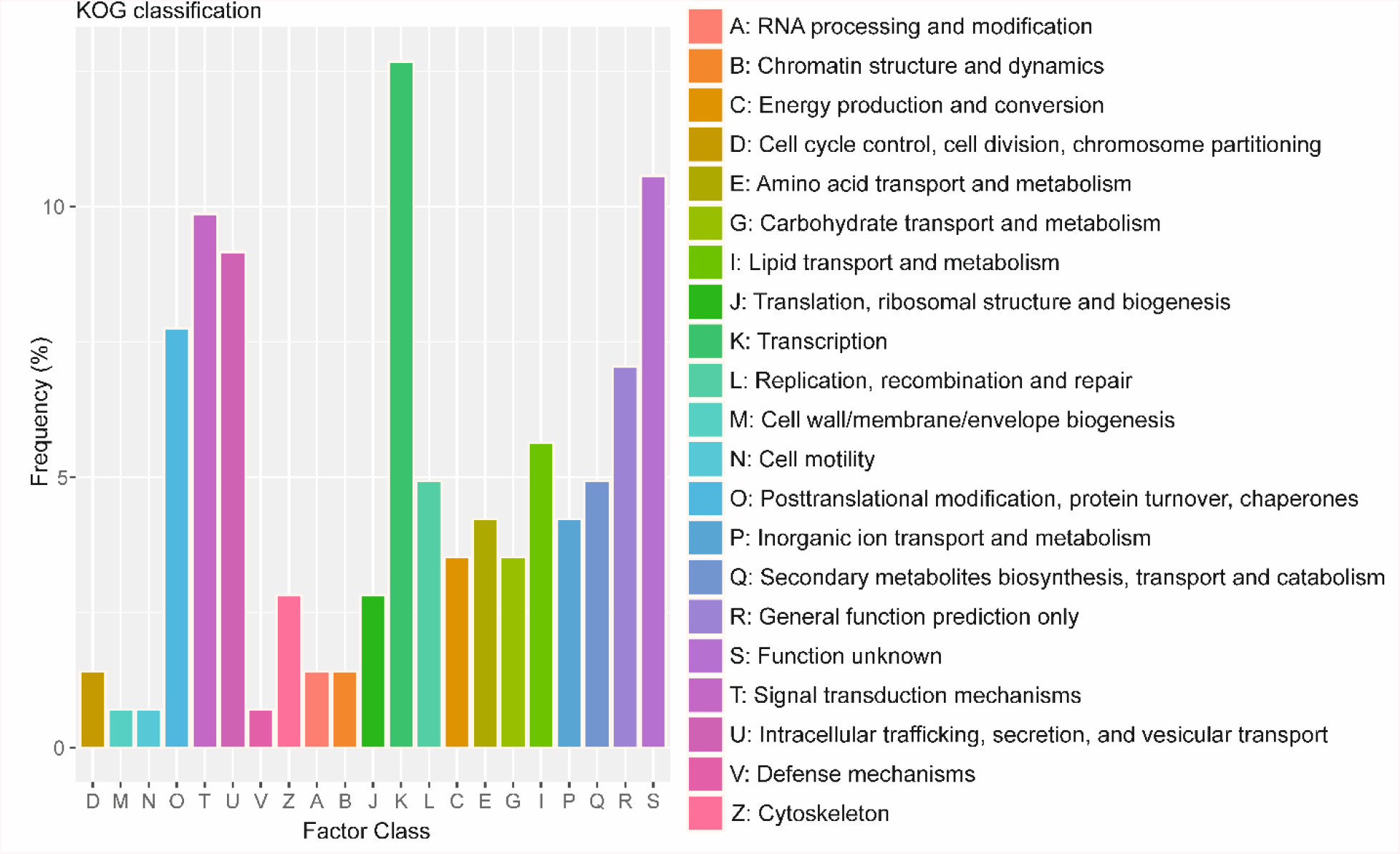
Functional categorization of positive selected genes (PSGs) in *A. fumigatus*. Distribution of the KOG function classification of PSGs. The categories of the KOG are shown on the horizontal axis and frequency are plotted on the vertical axis. The number of PSGs involved in transcription 18; in signal transduction mechanisms is 14; in intracellular trafficking, secretion, and vesicular transport is 13; in posttranslational modification, protein turnover, and chaperones is 11; in general function prediction is 10; in lipid transport and metabolism is 8; in replication, recombination, and repair is 7; in secondary metabolites biosynthesis, transport and catabolism is 7; in inorganic ion transport and metabolism is 6; in amino acid transport and metabolism is 6 and in unknown function is 15. The number of PSGs in other subgroups is ≤5.

### Positive selection in *A. fumigatus* in multi-gene families

Our analysis above focused only on SCOs; therefore, many secretory, membrane proteion coding genes and virulence associcated genes were involuntarily excluded by this criterion. To overcome this limitation, we collected multi-gene families from the literature (Puertolas-Balint, et al. 2019; Vivek-Ananth, et al. 2018) and analysed the positive selection for these genes. We focused on virulence associated genes in multi-gene families. i.e. non single-copy orthogroups, in which any of the eighteen Aspergilli we selected (Table S2) can have more than one paralog for a given gene. Out of 1498 non-SCOs genes analysed, we found 212 genes that underwent positive selection in *A. fumigatus* (Table S9). Major virulence-related genes under positive selection included genes involved in immunoreactivity, nutrient uptake, resistance to immune response, signaling and toxin and secondary metabolite biosynthesis and metabolism. Key examples are given in Table 1.

**Table 1:** Positive selection in *A. fumigatus* virulence genes^1^.

| Gene | Gene name | Annotation | Category |
| --- | --- | --- | --- |
| Afua_4g0958<br>0 | *aspf2* | Allergen Asp f2 | Allergens |
| Afua_5g0352<br>0 |  | Immunoreactive secreted protein |  |
| Afua_3g0342<br>0 | *sidD* | Nonribosomal peptide synthetase 4 | Nutrient uptake |
| Afua_1g1039<br>0 | *abcB* | Putative ABC multidrug transporter | Resistance to immune response |
| Afua_1g1744<br>0 | *abcA* | ABC drug exporter |  |
| Afua_5g0924<br>0 | *sodA* | Cu/Zn superoxide dismutase |  |
| Afua_7g0048<br>0 | *abcE* | Putative ABC transporter |  |
| Afua_2g0066<br>0 | *tcsB* | Putative sensor histidine kinase/response regulator | Signaling and regulation |
| Afua_2g1806<br>0 | *fgaM*<br>*T* | 4-dimethylallyltryptophan N-methyltransferase | Toxins and secondary metabolites |
| Afua_4g1449<br>0 | *tpcJ* | Putative dihydrogeodin oxidase |  |
| Afua_8g0037<br>0 | *fma-*<br>*PKS* | Fumagillin biosynthesis polyketide synthase |  |
| Afua_8g0044<br>0 |  | Dual-functional monooxygenase/methyltransferase |  |
| Afua_8g0046<br>0 |  | Methionine aminopeptidase type I, putative |  |

<sup>1</sup>See Table S9 for the complete list of PSGs in multi-gene family secretory, membrane protein coding genes and virulence associated genes (as defined by Puertolas-Balint, et al. 2019).

41 PSGs encoded proteins interact with human proteins. This was estabilsed by the analysis of an interolog and domain-domain interactions-based host pathogen interaction network. Enrichment analysis revealed the the significant over representation of pathogen interaction relevant GO terms such as GTPase mediated signaling, regulatory activity and response to stimuli (Figure S2; FDR ≤ 0.05):

Studies have suggested that pathogens can exploit membrane trafficking events and a variety of host signaling cascades to facilitate invasion, colonization, and proliferation by targeting host GTPases (Popoff 2014). Since during the saprophytic life-style *A. fumigatus* can interact with free living amoeba we speculate that selection in host GTPases interacting genes might be the result of conflict between *A. fumigatus* and amoeba. To investigate this, we have collected the unique genes belonging to the most enriched three categories (regulation of GTPases) and analyzed their orthology relations with two amoeba species: *Dictyostelium discoideum* and *Acanthamoeba castellanii*. Our analysis found that 45 of the GTPase-related genes had orthologs in one or both of the Amoeba species (Table S10) including homologs from direct BLAST (Altschul, et al. 1990) hits between human and *A. castellanii* and, *D. Discoideum*; and orthologs from Orthofinder (Emms and Kelly 2019) orthogroups of human, mouse, *A. castellanii* and *D. Discoideum*. The evidence for orthology relationship with amoeba of these genes that take a prominent role in host-pathogen interactions in human host may point to the similarity between mechanisms employed by *A. fumigatus* in human and amoeba host. These mechanisms may have evolved under the predation pressure from amoeba and can now be utilized to survive in the human host.

### Ancient defenses: Example gliotoxin biosynthesis cluster

Environmental virulence theories suggest that virulence traits of *A. fumigatus* have evolved as adaptation to natural predators in soil, such as free-living amoeba (Hillmann, et al. 2015). *A. fumigatus* is an organism with a wide range of niches, and it has natural predators and competitors. The selective pressure from predators is suggested to have given rise to several virulence mechanisms, especially to avoid the phagocytic killing. Indeed, *A. fumigatus* conidia were shown to follow the same phagocytic pathway in amoeba and macrophages (Bidochka, et al. 2010). The similarities between mechanisms used by *A. fumigatus*, especially at conidial stage, to counteract against phagocytes both in host and the environment points to the hypothesis that selective pressure exerted by amoeba predation caused evolution of virulence strategies in the mammalian host (Novohradska, et al. 2017). Furthermore, *A. fumigatus* was also shown to decrease viability in amoeba not only by germinating in the cell after its uptake but also via secretion of mainly gliotoxin (Hillmann, et al. 2015). Since *A. fumigatus* is an opportunistic pathogen and does not need a host to survive or replicate, it is suggested that these virulence mechanisms involving products of secondary metabolism such as gliotoxin, fumagillin, fumarylalanine, fumitremorgin, verruculogen, fumigaclavine, helvolic acid, sphingofungins and DHN-melanin evolved against their competitors and their predators in soil rather than host selection (Knowles, et al. 2020).

To investigate this example, we collected thirteen genes from gliotoxin gene cluster including biosynthesis genes, regulatory genes, transport genes etc. (Knowles, et al. 2020) and studied their conservation in sequenced *A. fumigatus* strains and eighteen Aspergilli. We found that all the thirteen gliotoxin gene cluster genes are conserved in all six sequenced *A. fumigatus* strains and are single copy orthologs except *gliA*, that had two paralogs in each strain (Table 2).

**Table 2:**
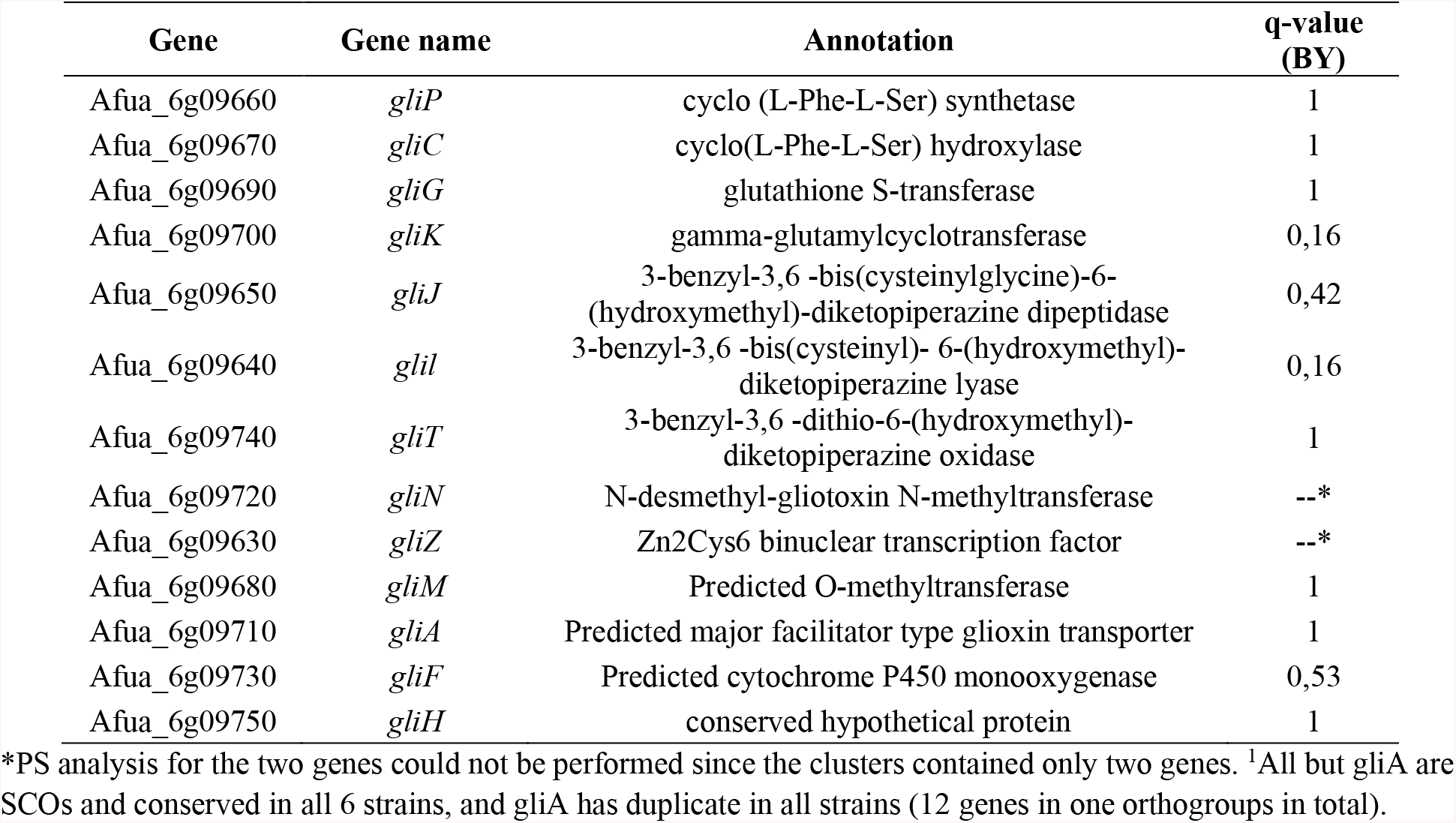
Gliotoxin biosynthesis genes^1^.

Comparison of eighteen Aspergillus species, on the other hand, showed high conservation of the gliotoxin gene cluster among *A. fumigatus* and its close relatives including *A. fischeri* and *A. clavatus* as well as the more distant species *A. flavus*, *A. oryzae* and *A. zonatus*. Indeed, *A. fischeri*, a ‘very close nonpathogenic relative of *A. fumigatus*’, is the only species that has orthologs of all the gliotoxin biosynthesis gene cluster (Figure 7) and was also shown to biosynthesize gliotoxin (Knowles, et al. 2020). Nevertheless, gliotoxin biosynthesis was not as important for the virulence of *A. fischeri* and loss of *laeA*, major regulator of secondary metabolism, did not affect its virulence (Knowles, et al. 2020).

**Figure 7:**
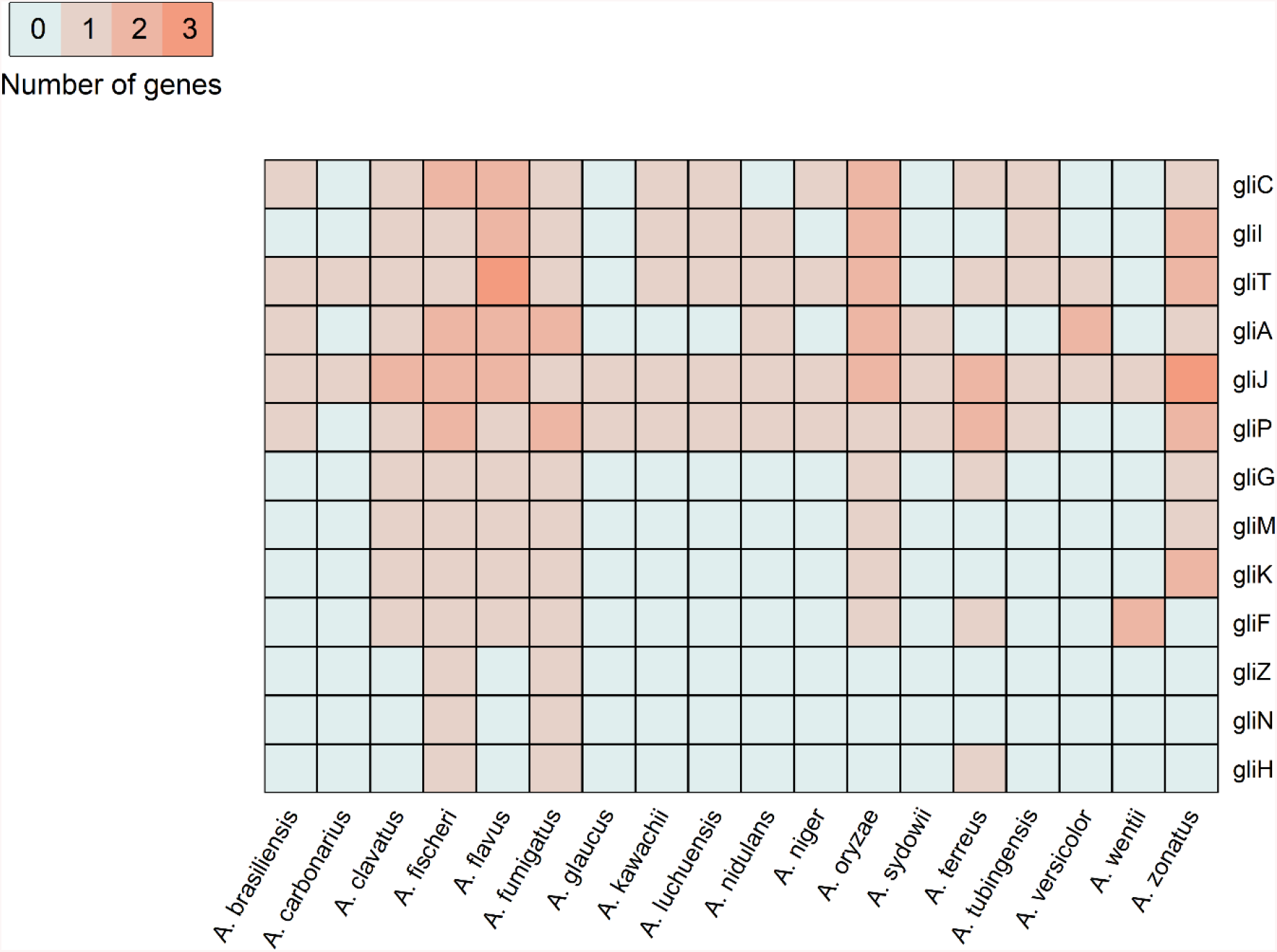
Conservation of gliotoxin biosynthesis gene cluster in Aspergillus species. Counts for each Aspergillus species in gliotoxin biosynthesis orthogroups are shown. Species that do not have an orthologs in a given gene orthogroup are represented gray.

We have also analyzed the traces of positive selection in gliotoxin biosynthesis genes of *A. fumigatus* and none of the genes showed signs of positive selection (Table 2). Gliotoxin was found to be the major amoebicidal factor in *A. fumigatus* (Hillmann, et al. 2015; Knowles, et al. 2020), which explains the lack of positive selection in gliotoxin biosynthesis pathway genes. Since gliotoxin evolved already long time ago against amoeba predation and now serves in “accidental virulence” against mammalian host, it’s plausible to find that there is no recent positive selection on those genes in an analysis compaling selection in current Aspergilli species.

## Discussion

The effectiveness of natural selection acting on an advantageous mutation and the likelihood of its long-term fixation is proportional to its fitness effect (Duret 2008). Fighting an infection in immunocompromised patients has to cope with versatile adaptation strategies of *A. fumigatus* (van de Veerdonk, et al. 2017) most likely resulting from its saprophytic life-style and its interaction with soil microorganisms including amoeba.

Previous work in the field includes the description of a toxin gene cluster evolved under positive selection in *A. parasiticus* (Carbone, et al. 2007). Yang et al. (2012) identified the duplicated gene pairs in *A. fumigatus* evolved under positive selection (Yang, et al. 2012). Species-wide comparisons of the genus Aspergillus provide a broad view of the biological diversity of the aspergilli and highlighted protein versatility and conservation (de Vries, et al. 2017). However, these studies did not offer the genome-wide signatures of recent selection in *A. fumigatus* investigated here. Indeed, positive selection is the evolutionary force to drive divergence of species from its close relatives (Wu and Ting 2004). To examine how positive selection may have operated during *A. fumigatus* evolution, we took advantage of the branch-site model’s ability to detect selection on the branch of interest. Maximimum likelihood (ML) codon-specific estimation of dN/dS ratio may highlight other fungal adaptation strategies. The data provided here further emphasize the functional importance of PSGs in multiple functional categories.

### Pathway and GO over-representation of *A. fumigatus* SCO proteins interacting with human host proteins

Our analysis indicates the over-representation of SCO PSGs interacting with host proteins down regulating the TGF-β (Letterio and Roberts 1998), insulin receptor and IL-1 signaling pathways (Karpac, et al. 2011). Pro-inflammatory cytokines are crucial for stimulating an effective immune response to *A. fumigatus* infection, which includes recruitment of neutrophils to the alveolar spaces, where they constitute more than 90% of the phagocytic cells (Balloy and Chignard 2009). The GO terms are given in Table S7.

The GO term ‘symbiosis, encompassing mutualism through parasitism’ was significantly enriched for PSGs targeted human proteins. This finding is consistent with earlier data showing the significant enrichment of this category in *A. fumigatus* targeted human proteins (Remmele, et al. 2015). The GO term ‘viral process’ and parent GO term ‘symbiont process’ were also over-represented. Together with other GO terms this indicates that in *A. fumigatus,* positive selection occurs mostly on genes involved in the initial stage of infection, where the host attempts to avoid pathogen establishment.

Among the multi-gene families PSGs interacting with the host, we identified many genes involved in regulation of GTPase cascade were targeted by *A. fumigatus*. Notably, the *D. discoideum* genome contains an unexpectedly large number or GTPases which involves complex signaling mechanisms in a wide range of biological processes (Wilkins and Insall 2001). Rho subfamily GTPases are also involved in controlling the reorganization of actin, which is essential for both phagocytosis (Swanson 2008) and subsequent maturation of phagosomes (Kruppa, et al. 2016). The kinetics and regulation of phagosomal maturation is very similar in *D. discoideum* and mammals (Duhon and Cardelli 2002). Once the pathogen is internalized in the host cell, fission and fusion events occur in the phagosomal membrane and the phagosomal membrane interacts with elements of the endolysosomal system to obtain proteases and lysosomal enzymes which is regulated by GTPases. It is likely that the natural selection in GTPase regulator interacting genes is the consequence of the conflict of fungi and amoeba.

We suggest that these host-pathogen interactions help *A. fumigatus* to survive also in the human host (see also (Hillmann, et al. 2015)). As a further example, host-pathogen interactions critical for survival of *Cryptococcus neoformans* in *A. castellanii* were shown to contribute to survival of the fungus inside macrophages (Steenbergen, et al. 2001). A recent review of such conserved virulence determinants focusses on this dual use of fungal adaptations beneficial to survive innate immune cells but also soil amoebae as the natural phagocytes (Novohradska, et al. 2017).

### Positive selection in other functionally annotated categories

All PSGs classified under KOG categories are discussed in the Additional file 1 Supplemental Note “Positive selection in genes classified under KOG functional categories”.

#### Conserved hypothetical protein-coding genes

Certain exceptions notwithstanding, conserved hypothetical proteins present in several sufficiently different genomes, are not truly ‘hypothetical’ anymore (Galperin and Koonin 2004). Twenty SCO PSGs code for conserved hypothetical proteins, expressed to survive hypoxic conditions (Afua_2g09800, Afua_3g13930, Afua_8g05450 and Afua_5g07480) or in early conidial development (Afua_6g13670, Afua_3g13930, Afua_5g06310, Afua_4g07030) in (Kroll, et al. 2014). Two are differentially expressed during dendritic cell infection (Afua_5g02110 and Afua_5g08770), and one is differentially expressed during neutrophil infection (Afua_4g07030). Moreover, two conserved hypothetical PSGs were annotated to be involved in signal transduction mechanisms (Afua_4g09890 and Afua_4g09910) by KOG categorization. These previously ignored conserved hypothetical genes are important as a general resource to understand the complex biology of Aspergillus better. Conserved hypothetical PSGs could be targets for novel and broad antimycotics as their positive selection and broad conservation suggests important functions for aspergilli.

#### Hypoxia-responsive genes

Hypoxia-tolerance is necessary for *A. fumigatus* to survive in host tissue but also during its saprophytic lifestyle. Oxygen levels drop from 21% in the atmosphere to 14% in the lung alveoli (Jain and Sznajder 2005), in surrounding tissue, oxygen availability is further reduced to 2 to 4% and in inflamed tissue to less than 1% (Lewis, et al. 1999). A broad range of metabolic pathways maintains energy levels under oxygen-limiting conditions. Compared with the gene expression profile of hypoxia-inducible target genes of *A. fumigatus* (Kroll, et al. 2014), 27% of the PSGs identified in our study significantly change their expression behaviour during low-oxygen conditions (Table S11).

#### Genes involved in early development of A. fumigatus

Conidia of *A. fumigatus* ingested or inhaled by immunocompromised patients can germinate in the lung and produce hyphae within 6-8 h to trigger infection (Lamarre, et al. 2008) and adaptation to aerobic metabolism and growth (Cagas, et al. 2011). Among the PSGs we identified, ~25% of the genes were significantly expressed at 8 h and 18% were significantly expressed at 16 h of growth. Highly downregulated PSGs at 8 h was the bZIP transcription factor Afua_3g11330 (*atfA*), followed by ammonium transporter Afua_1g10930 (*mep2*), while highly upregulated PSGs were 40S ribosomal protein S9-coding gene Afua_3g06970 (*rps9a*), followed by GDP-mannose pyrophosphorylase A (GMPP)-coding gene Afua_6g07620 (*srb1*). In *A. fumigatus* expression of 63% of the conidia-associated genes is controlled by *atfA* which is an essential gene for viability of this fungus in various environmental condition (Hagiwara, et al. 2014). It positively regulates conidial stress-related genes and negatively regulates the genes for germination, suggesting a role for *atfA* in conidial dormancy (Hagiwara, et al. 2016). *atfA* was also found to be pivotal for heat and oxidative stress tolerance in conidia by regulating the conidia-related genes responsible for stress protection (Hagiwara, et al. 2014). *mep2* coded proteins function as ammonium sensors in fungal development (Van Zeebroeck, et al. 2011) and induction of invasive filamentous growth (Joardar, et al. 2012). *srb1* gene encodes the GMPP in *A. fumigatus* and is essential for its viability. A highly downregulated PSG at 16 h was acyl-CoA thioesterase II-coding gene Afua_1g15170 (*acot2*), followed by *atfA*, while highly upregulated PSG was meiosis induction protein kinase Ime2, followed by an integral membrane protein 25D9-6 coding gene. *acot2* catalyzes the hydrolysis of acyl-CoAs to the free fatty acids and coenzyme A (CoASH), providing the potential to regulate intracellular levels of acyl-CoAs, free fatty acids, and CoASH and has important functions in lipid metabolism and other cellular processes (Hunt, et al. 2012). Overexpression of *acot2* can significantly modulate the mitochondrial fatty acid oxidation (Moffat, et al. 2014). Positive selection in *acot2* suggests respiratory activity was improved during the evolution of *A. fumigatus*. Other important genes in this category included serine/threonine protein kinase Afua_2g13140 (*ime2*) which is critical for proper initiation of meiotic progression and sporulation. In *A. nidulans*, *ime2* is required for the light-dependent control of mycotoxin production (Bayram, et al. 2009). Integral membrane protein 25D9-6 coding gene Afua_2g17080 which is a major component of peroxisomal membranes was also found to be evolved under positive selection.

#### Essential protein-coding genes

The essential genes of an organism constitute the minimal gene set required for the survival and growth of an organism, in particular defined as to sustain a functioning cellular life under most favourable culture conditions (Koonin 2003). Typically, such genes are more evolutionarily conserved than the non-essential genes (Jordan, et al. 2002), and rapid evolution is usually not expected among essential genes (Luo, et al. 2015) with few excepetions (Fang, et al. 2005). Several reports identified the essential genes in *A. fumigatus* using both the functional experimental methods and bioinformatics approaches (Carr, et al. 2010; Hu, et al. 2007; Lu, et al. 2014; Nierman, et al. 2005; Thykaer, et al. 2009). By comparing these with our results we noticed that among the experimentally identified essential genes only spliceosomal protein coding gene Afua_3g12290 (*dib1*) (Carr, et al. 2010) and mitochondrial import receptor subunit Afua_3g11860 (*tom22*) (Hu, et al. 2007) evolved under positive selection while among computationally identified essential genes by sequence similarity against essential eukaryotic genes (Nierman, et al. 2005), four genes (transcription initiation factor TFIID subunit 2 Afua_8g04950, DNA-directed RNA polymerase III subunit 22.9 kDa Afua_8g04350, DNA replication licensing factor Afua_2g10140 (*mcm7*) and Ccr4-Not transcription complex subunit Afua_3g10240 (*not1*)) evolved under positive selection.

Likewise, positive selection is often more prevalent in peripheral proteins while purifying selection is more stringent among central network proteins (hubs) in the interactome of organism (Fraser, et al. 2002; Kim, et al. 2007). To identify whether the PSGs follow the same criteria, we analysed the *A. fumigatus* interactome (Kaltdorf, et al. 2016) and compared the top 20% hubs with PSGs. In contrast to the general tendency, fifteen *A. fumigatus* hub proteins (≥5 interaction partners) were found to have evolved under adaptive evolution (Table S12). Notably, twelve of these proteins are also involved in host interactions (Table S13; e.g. asparagine synthase Afua_4g06900 (*asn2*), lon protease homolog Afua_2g11740 (*pim1*)), as detailed in result section "Host interacting PSGs" (see above). The importance of proteins during the infection is directly related to their number of interactions with the host (Crua Asensio, et al. 2017). This suggests that the observed positive selection at these hubs may be due first, to environmental imposed pressure to affect the pathogen fitness during amoeba attack and second, useful for survival in macrophages.

### Positive selection in *A. fischeri* protein-coding genes

*A. fischeri* is a sister-species to *A. fumigatus* but generally saprophytic and a pathogen only under rare circumstances (Chim, et al. 1998; Gerber, et al. 1973). The species tree confirms the similarity of the *A. fischeri* genome with *A. fumigatus* (Figure 3). Both species share 8536 one-to-one orthologs (Additional data file 3). To assess the extent of genome rearrangements more closely, the complete chromosome sequences of *A. fumigatus* and *A. fischeri* were aligned using the Mauve genome aligner (Darling, et al. 2004). The GC (guanine-cytosine) content was seen to be highly homogenous (49.2%) and genome sequences were relatively well-conserved. The alignment consisted of 318 Local Collinear Blocks (LCBs) with a minimum weight of 999. A genome synteny map is shown in Figure S3. We find several instances of genome rearrangements (reverse complement of the reference sequence) since *A. fumigatus* and *A. fischeri* separated from each other. Possibly the genome rearrangements occurred because of recombination that can be advantageous for fungal pathogens in stressful environments such as inside a human host (Goddard, et al. 2005). Furthermore, using the BS model (see methods), we identified 50 SCO PSGs in *A. fischeri* (Table S14). Among the orthologs of these genes in *A. fumigatus* only eight genes evolved under positive selection, although the sites where selection occurs were not the same for these genes. Taken together, this analysis shows clear differences regarding PSGs between both organisms and the adaptation of both aspergilli. In particular, *A. fischeri* has less than half as many PSGs and shares only eight PSGs with *A. fumigatus.* This difference suggests much less potential of *A. fischeri* to adapt even to a human host and is in line with only rarely observed clinical infections.

### *A. fumigatus* multi-copy virulence genes with positive selection

The analysis of only SCOs excluded many virulence related genes that have more than one copy in aspergilli genomes. Therefore, it is important and necessary to analyze positive selection in virulence genes that do not belong to SCOs. Our analysis of in these orthogroups identified 212 virulence genes that showed signatures of positive selection in *A. fumigatus* (Table S9). Some of these genes such as major allergen Afua_4g09580, genetic name *aspf2,* the nonribosomal peptide synthases and several ABC transporters were involved in processes such as immunoreactivity, nutrient uptake, resistance to immune response, signaling and toxin and secondary metabolite biosynthesis/metabolism (Table 1). In particular *aspf2* is found in eleven Aspergilli species from the 18 compared (not found in *A. brasiliensis, A. carbonarius, A. kawachii, A. luchuensis, A. niger, A. tubingensis*, and *A. zonatus*). Positive selection of *aspf2* in *A. fumigatus* and its conservation in so many *Aspergilli* may point to evolution of an anti-amoebal strategy employed by these fungal species based on inhibiting phagocytosis by protein binding. Thus, during human infection conidia of *A. fumigatus* start germinating and activate human immune responses from the complement system. However, *A. fumigatus* evades this immune response by regulating the complement system. For immune evasion, the allergen *aspf2* recruits immune regulators such as Factor-H, that is normally a factor to oppose complement activation. Furthermore, *aspf2* was shown to bind plasminogen and its activated version plasmin cleaves fibrinogen and this process aids damage of lung epithelial cells and helps with early infection. More importantly to confirm a more generally conserved evasion strategy against phagocytosis, conidia with mutant *aspf2* were shown to be more effectively phagocytosed and killed by neutrophils (Dasari, et al. 2018).

Despite the insufficient nutrient in host phagolysosome environment, *A. fumigatus* can swell up and initiate growth by utilizing their siderophore machinery to overcome the limited iron supply (Schrettl, et al. 2010). This process is also well documented when *A. fumigatus* interacts with *A. castellanii* and *D. discoideum* (Hillmann, et al. 2015; Van Waeyenberghe, et al. 2013). Nonribosomal peptide synthases Afua_3g03420 (*sidD*) involves in siderophore biosynthesis that facilitates survival in iron-scarce environments, complemented by other nutrient transporters (Table 1) was found to be positively selected. Decreasing iron content during infection is a strategy employed by the host during infections and pathogens evolve mechanisms to “obtain iron from the host”. Therefore, *A. fumigatus* uses two iron-uptake methods, one is via siderophores. Mutants of *sidD* among with other enzymes of the siderophore biosynthesis were shown decreased conidiation and attenuated virulence in iron-scarcity, indicating the induction of siderophore biosynthesis pathway during infection (McDonagh, et al. 2008; Slater, et al. 2011). Of note, we also observed positive selection in nonribosomal peptide synthetases Afua_3g03350 (*sidE*). Although, *sidE* do not contribute in siderophore biosynthesis it inolve in the production of fumarylalanine (Steinchen, et al. 2013). Fumarylalanine acts as immunomodulatory metabolite and *sidE* highly upregulates in murine lung infection (McDonagh, et al. 2008; Steinchen, et al. 2013).

Another result that came out from our study is that multifunctional Cu/Zn superoxide dismutase Afua_5g09240 *sodA* evolved under positive selection. During iron deplete conditions and host colonization, high expression of *sodA* in *A. fumigatus* provides self protection against own fungal oxidants and gliotoxin levels (Bruns, et al. 2010; Carberry, et al. 2012; Lambou, et al. 2010; Oberegger, et al. 2000). s*odA* is also recognized by sera from patients with confirmed *A. fumigatus* infections (Holdom, et al. 2000). Due to high expression of *sodA* during pathogenic growth it represent a valuable immunodiagnostic marker for *A. fumigatus* infections (Holdom, et al. 2000). In compare to wild-type, high susceptiblity to reactive oxygen species (ROS) and high temperature has also been observed in *sodA* mutants (Lambou, et al. 2010).

Moreover, three ABC transporters; Afua_1g17440 *abcA*, Afua_1g10390 *abcB*, and Afua_7g00480 *abcE* were found to be under positive selection in *A. fumigatus*. It was shown that all five ABC transporters (abcA-E) are induced by voriconazole and take part in azole resistance (Meneau, et al. 2016). Especially, *abcB* was found to be related to azole resistance and its loss contributed relatively more to increased susceptibility to azole indicating that *abcB* is essential for virulence and azole resistance while *abcA* while not being required may increase resistance in infection (Abad, et al. 2010; Paul, et al. 2013). The annotated virulence genes and their function constitute highly sought out data to understand pathogenic fungi.

### Evolutionary overview

Positive selection on *A. fumigatus* genes that are conserved among eighteen Aspergilli species can underline the *A. fumigatus*-specific functions of their protein products and consequently may be important in pathogenicity of *A. fumigatus*. The same applies for *A. fumigatus*-specific genes; however, it is not possible to run a positive selection analysis on these genes. Nevertheless, although not covered in the study, next step may involve analyzing positive selection in the genes in comparison with other sequenced *A. fumigatus* strains, since we have shown some of these genes are conserved in all strains and related to virulence (Table S1).

PSGs identified both in SCOs and multi-gene families can be related to recent virulence mechanisms evolved in *A. fumigatus* and show its unique virulence strategies against the host. Although it is assumed that *A. fumigatus* virulence mainly emerged accidentally upon selective pressure from natural predators such as amoeba, positive selection detected in these genes shows that host pressure may also have led to evolution of some strategies employed by *A. fumigatus*.

However, we show that one of the major virulence mechanisms of *A. fumigatus*, gliotoxin biosynthesis pathway, does not have signs of positive selection in our analysis set. This finding demonstrates that old and settled virulence mechanisms of *A. fumigatus* such as gliotoxin toxicity have in fact evolved under long term, global and continuous predation pressure as survival tricks and not as more recent and more species-specific virulence strategies.

Taken together, our work provides a detailed evolutionary perspective on *A. fumigatus* biology. For the first time the currently available extensive genome information (18 genomes) is not just analysed in terms of annotation or phylogenetic comparison but rather we show how positively selected genes in the pathogen *A. fumigatus* are identified, analyse their connection to virulence and add even detailed interactome analysis to reveal interactions with host. Blind spots in Aspergillus biology are illuminated, revealing highly conserved genes in the genus from our detailed positive selection analysis. Finally, we provide all data and scripts of the whole analysis as a resource to the community..

## Conclusion

We suggest selection plays a crucial role for aspergilli in general and specifically for the adaptive evolution of *A. fumigatus* from soil microorganism to human opportunistic pathogen. Eighteen fungal species of the genus *Aspergillus* were compared and analysed to identify PSGs. We identified 122 well annotated genes of *A. fumigatus* that evolved under positive selection pressure including signal transduction, metabolism, mitochondrial activity, regulation of transcription and conidia growth-related genes. This includes conserved genes of unknown function which we establish here to be clearly positively selected. Moreover, we detected positive selection signals in 212 secretory, membrane and virulence related multigene families. Genes analyzed here also point to immune evasion mechanism (*aspf2*, one of the central immune evasion gene was identified to evolve under positive selection). Futher positive selection was also identified in genes involved in siderophore biosynthesis (*sidD*), metabolite fumarylalanine production (*sidE*), stress tolerance controlling transcription factor (*atfA*) and multifunctional thermotolerance and self-protection regulating gene (*sodA,* immunodiagnostic marker for *A. fumigatus* infections). We found many PSGs interact with host GTPase regulation mechanisms, adaptive and immunomodulatory pathways. These are also medically relevant as they mitigate human host defences. Moreover, proteins coded by PSGs may provide targets for new antifungals. Taken together, our results identify *A. fumigatus* genes, which are strong candidates for having functional effects during adaptation to human host.

## Materials and methods

### Initial dataset, quality assessment and filtering

The complete set of protein-coding gene sequences and corresponding amino acid sequences of the eighteen *Aspergillus* genomes were were retrieved for GenBank database at NCBI (Benson, et al. 2005). BUSCO (Waterhouse, et al. 2018) was used to access the completeness of proteomes based on the presence of a benchmarking set of universal fungal single copy orthologs. Table S2 lists the version and details of the used fungal genomic assembly. BUSCO (Benchmarking Universal Single-Copy Orthologs) was used to assess genome quality. We hypothesize that if the genes are well demarcated in the sequenced genome, they should have an ortholog to the corresponding BUSCO set (Waterhouse, et al. 2018). Nearly all aspergilli genomes could be selected for the analysis with acceptable quality and completeness (Figure 1). Basic sequence quality features were first controlled as in (Hambuch and Parsch 2005). CDS (coding sequences) whose length was not a multiple of 3 or did not correspond to the length of the predicted protein, or that contained an internal stop codon, were eliminated; CDS shorter than 90 nt were eliminated; similarly, short peptides with unsuitable length (< 30 aa) for orthology analysis were eliminated.

### Orthologs and unique genes

The OrthoMCL algorithm, which uses multiple steps including BLASTP (Altschul, et al. 1990) and Markov clustering (Enright, et al. 2002) to group proteins into likely orthologous clusters, was used for identifying the orthologs between eighteen Aspergilli proteomes. Additionally, the proteins that are not shared with any other species but contain intra-species very close homologs, were extracted from the orthologous clusters. We also found single species clusters for the analysed aspergilli. Sequences in such clusters could be the consequence of gene duplications and together with singletons represent the species-specific proteins. This should be interpreted with caution, as an increase of the number of species in the analysis can change the number of proposed species-specific sequences. Although the evolutionary information seems to be lost for species-specific sequences, but these sequences can define the specific character of each species.

### Reconciling of gene trees and species trees

Sequences of single copy ortholog clusters were further used for estimating the species tree using super-matrix method (de Queiroz and Gatesy 2007). The filtered alignments were concatenated, and species tree was constructed using RaxML (Stamatakis 2014) under the probabilistic maximum-likelihood (ML) framework. We ran RaxML analyses to find the optimal ML estimate with 1000 bootstrap replicates. The resulting topologies of the species trees were applied to the selection analyses (Diekmann and Pereira-Leal 2015). For this purpose, we first produced the multiple sequence alignment (MSA) of 3175 SCOs clusters shared by all eighteen Aspergilli and concatenated them into a single alignment. The alignment quality was improved first by selecting conserved blocks, then by removing sequence parts with multiple substitutions. This allowed us to reduce their potentially misleading effects due to alignment error. We started with the 2,112,647 columns in the alignment and ended up with 1,246,185 columns to produce a giant phylogenetic matrix. Such extensive data removal is often impractical in single-gene analyses because too few positions remain available and produce a poorly resolved tree (Philippe, et al. 2000). Further, using the evidence from all characters in the derived phylogenetic matrix, the best probabilistic maximum likelihood (ML) species tree was estimated under the assumption that each character provides independent evidence of relationships.

### Single-Copy Orthologs Gene Families Data Set

Gene families were obtained from a custom run of the OrthoMCL pipeline (Li, et al. 2003) which implements Reverse Best Hit (RBH) Blast (Moreno-Hagelsieb and Latimer 2008) and MCL clustering algorithm (van Dongen and Abreu-Goodger 2012) to identify the ortholog clusters based on all-against-all Smith–Waterman protein sequence comparisons with an e-value cut off of 1e-5. Gene families including strictly one ortholog in each of the 18 species were selected (3175 gene families).

### Multiple sequence alignment (MSA)

Nucleotide and amino acid sequences of all orthologous gene groups were extracted from the initial sequence dataset. To reduce the effect of incorrect insertions/deletions (indels) on the codon alignments, MSA were initially performed for amino acid sequences of each ortholog group using T-Coffee (Notredame, et al. 2000), which combines the output of different aligners. The aligned amino acid sequences together with the corresponding nucleotide sequences of each ortholog group were converted into DNA alignments at the codon level using the program PAL2NAL (Suyama, et al. 2006). The misalignment errors can be an important source of false positives in genome-wide scans for positive selection in coding sequences (Jordan and Goldman 2012) and the removal of unreliable regions increases the power to detect positive selection (Privman, et al. 2012). Therefore, carefully filtered the potentially problematic sites in the alignments. To further improve alignment quality, we used a stringent Gblocks filtering (type = codons; minimum length of a block = 4; no gaps allowed) (Talavera and Castresana 2007) to remove gap-rich regions from the alignments, as these are problematic for positive selection inference (Fletcher and Yang 2010).

### Test for recombination (see Supplementary Methods)

#### Positive selection in *A. fumigatus* protein-coding genes

To explicitly test for positive selection in *A. fumigatus* among the orthologous clusters (alignment data available at https://funginet.hki-jena.de/data_files/76?version=1), we used a branch-site model implemented in PAML (Yang 2007), which allows four classes of codons: a strictly conserved class (ω < 1), a class that is conserved in the ‘background’ lineages but under positive selection in the ‘foreground’ lineage of interest, a class that is strictly neutral (ω = 1), and a class that is neutral on the ‘background’ lineages but under positive selection in the ‘foreground’ lineage of interest. When compared to the null model, which does not allow positive selection in the foreground lineage, this model provides a robust test for positive selection in a subset of codons on a particular lineage (Yang and dos Reis 2011). We applied this branch-site model to two sets of foreground lineages; the *A. fumigatus* terminal branch and *A. fischeri*. The significance of test was assessed using standard asymptotic assumptions (Self and Liang 1987). The calculations were performed using the codeml program from the PAML 4.2b package (Yang 2007). The Bayes empirical Bayes approach was employed to estimate the probabilities of positive selection for specific codons under the likelihood framework (Yang, et al. 2005).

To check whether the selection worked on the branch containing *A. fischeri* we similarly designated the corresponding branch as foreground and the rest as background. The log-likelihoods from each test were compared in a likelihood ratio test assuming a χ2 distribution of the test statistic. Bonferroni correction (Benjamini and Hochberg 1995) and FDR of 5% (Benjamini and Yekutieli 2001) were used to correct for multiple testing.

### Tests of functional category enrichment (see Supplementary methods)

#### Interolog network

The experimentally derived dataset of host-pathogen protein-protein interaction from the HPIDB (Ammari, et al. 2016) and PHISTO database (Durmus Tekir, et al. 2013) was used as a template to reconstruct interologs (Yu, et al. 2004) based PPIs network of PSGs coding proteins and human proteins. To determine the orthologs we used the Inparanoid algorithm (Sonnhammer and Ostlund 2015) which integrates all-versus-all Blast similarity results and Markov single linkage binding to construct putative orthologous groups in two proteomes. Blast e-value and was set to 1e-05 for clustering. Only the seed orthologs were considered to increase the prediction confidence. For, the pathogenic genes we also calculated interactions based on interacting domain profile pairs between host and pathogen as mentioned in (Gupta 2020).

### Inference of positive selection in *A. fumigatus* virulence genes (see Supplemental Methods)

#### Statistical analysis of PSGs

Multiple testing correction was performed to control for Type I errors according to the approach presented by Benjamini and Hochberg (Benjamini and Hochberg 1995). For the analyses of homologous recombination, recombination breakpoints were inferred by the Shimodaira-Hasegawa test (Shimodaira and Hasegawa 1999) with Bonferroni-corrected p-value and the threshold for significance was set at 0.05. For all genes tested for positive selection, the false discovery rate (FDR) was controlled by using the Benjamini-Yekutieli (BY) method (Benjamini and Yekutieli 2001) and the significance level was set to 10%. For all genes tested for recombination and positive selection, q-values were calculated from p-values using the R package q-value with the proportion of true null hypothesis set to 1 (π0 = 1) (Storey and Tibshirani 2003). Chi-square test and Fisher-exact test where appropriate were used to assess associations for the gene count data in the individual KOG categories; Bonferroni corrections for multiple testing were applied. All statistical analyses were carried out using R 3.4.4 (Turner 2011).

## Author contributions

S.K.G., M.S., Ö.O. and Z.X. performed the bioinformatics analyses, M.S. and T.D. conceived the project, S.K.G., M.S. and Ö.O. were involved in drafting the manuscript, TD supervised the work and discussed and finalized the manuscript with A.B. All authors analyzed the data, read and agreed to the final version of the manuscript.

## Acknowledgements

The authors gratefully acknowledge the support by the Deutsche Forschungsgemeinschaft (DFG) project number 210879364 (CRC/Transregio124 FungiNet; Projects A1– AB, project B1 - SKG and TD). MS would like to thank for financial support by Frauenbeauftragten Büro, University of Wuerzburg, Germany. We thank Dr. Alan Horowitz and Dr. Ulrike Rapp-Galmiche for stylistic suggestions and native speaker corrections. This publication was supported by the open access fund of the University of Würzburg.

## Availability of data and materials

All results are contained in the manuscript and its supplementary files. The tools used for the analysis of evolution on gene families in the genus *Aspergillus* with focus on *A. fumigatus* are made fully available by us including all codes and steps used. These analysis tools and methods can now be applied to any genus and phenotype of interest. The dataset and codes underlying our positive selection pipeline, as well as other additional data files are available at https://github.com/ShishirGupta-Wu/aspergillus_ps.

## Ethics approval and consent to participate

No ethical approval was required.

## Additional files

**Additional file 1:** Supplemental notes, Supplementary figures S1-S3, large supporting additional data files 1-3. (PDF 16,061 kb)

**Additional file 2:** Supplementary tables S1-S15. (XLSX 191 kb)

**Additional file 3:** Supplemental methods. (PDF 333 kb)

