## Additional file 1 for "*Aspergillus fumigatus* versus Genus *Aspergillus*: Conservation, adaptive evolution and specific virulence genes"

#### This PDF file includes:

**1. Supplemental note**

**(i) Recombination breakpoints**
**(ii) Positive selection in genes classified under KOG functional categories**
***- Metabolism***
***- Cellular processes and signaling***
***- Information storage and processing***
**References**

**2. Supplementary figures**
**- Figs. S1, S2 and S3**

**3. Additional data file 1**
**4. Additional data file 2**
**5. Additional data file 3**

#### Other Additional file for this manuscript includes the following:

**Additional file 2**
- **Captions for Tables S1 to S15 (Sheet in excel file)**
- **Tables S1 to S15 (Sheets in excel file)**

* Corresponding author

Note: We recommend to use bookmarks for navigation.

### Supplemental Note

#### Recombination breakpoints

Recombination is a phenomenon that occurs in the genomes of many organisms through the exchange or transfer of genomic fragments demarcated by recombination breakpoints. In the evolution of fungi, recombination is documented as one of the major forces driving diversity and adaptation [1, 2] subsequently resulting in the creation of mosaic sequences where the evolutionary history at each site may be different. With recombination, nucleotide sites in a sequence do not evolve along a single tree, but instead along a set of correlated trees [3]. Violating this tree-like assumption of evolution can lead to serious consequences when performing positive selection analysis [4]. Thus, the evolution of homologous recombinant sequences must be modelled by several phylogenies, one for each nonrecombinant fragment in the alignment [5]. The widely used likelihood methods for quantifying selection pressure on codon alignments [6] may suffer from high rates of false positives when the sequences being analyzed have undergone recombination [4, 7]. Therefore, we analyze the presence of recombination breakpoints in SCOs clusters. We used the Genetic Algorithm Recombination Detection (GARD) tool that implements the likelihood-based model selection procedure and uses the genetic algorithm to screen MSA for recombination breakpoints and quantifies the strength of support for their location [5, 8]. We implemented the Shimodaira-Hasegawa (SH) test to reject the potentially recombinant areas which might be due to substitution rate variation and not the recombination, since only a subset of recombination breakpoints convey any selective advantage, and the majority may be selectively neutral with limited biological significance [9]. We found that significant recombination events occurred in 879 single copy ortholog clusters (Additional file 2: Table S15).

We further conducted GO over-representation analysis of proteins that possibly experienced recombination. In the BP terms, the top two enriched GO terms were response to stress and purine ribonucleotide biosynthetic process, the MF terms were structural constituents of ribosome and phospholipase D (PLD) activity, and the CC terms were related to organelle and mitochondria. Here we have highlighted the role of proteins under BP and MF categories. Detailed alignments are listed in (Additional file 2: Table S5) including the low-level over-representation of recombination breakpoints. Studies have shown the positive correlation between stress and recombination activity [10, 11]. In *Drosophila melanogaster*, it has also been suggested that stress-induced recombination might be a mechanism of evolvability as certain types of stress, for instance heat shock stress, have the capacity to stimulate evolutionary changes [12]. Several stress-related proteins of *A. fumigatus* and *A. nidulans* have previously shown to be involved in recombination [13-16]. Highly significant over-representation of the GO category, cellular response to stress (GO:0033554, FDR 1.44E-03), suggest recombination might be a general stress response in aspergilli. The increased genetic flexibility by recombination might facilitate evolutionary adaptation of fungi to stressful environments.

Purines are essential nutrients required for functions as diverse as signal transduction, energy metabolism, and DNA synthesis. They are also required by the pathogenic fungi to enable growth and dissemination in the host. Due to varying concentrations of purines in their environmental niches and the host, most of the fungal pathogens can degrade, salvage, and synthesize de novo purines through several conserved enzymatic steps [17]. The purine nucleotide biosynthesis pathway is conserved between yeast and humans and is critically important for the normal functioning of cells. Additionally, dysregulation of this pathway can result in cytotoxicity [18]. High-frequency genetic variation in two purine biosynthesis genes influences survival of bacterial pathogen *Campylobacter jejuni* under stress and promotes its success as a pathogen [19]. We found that the over-representation of the GO category, purine ribonucleotide biosynthetic process (GO:0009152, FDR 1.82E-03), suggests the recombination in these genes could be the adaptive strategy of aspergilli.

Highly transcribed genes, such as the ribosomal genes, are more susceptible to recombination [20]. In agreement with this, we also found over-representation of several ribosome-related GO terms such as ribosome biogenesis (GO:0042254, FDR 2.54E-02), ribosome (GO:0005840, FDR 2.83E-02) and structural constituent of ribosome (GO:0003735, 2.77E-02) (Additional file 1: Table S5). Other than their primary functions in protein translation, ribosomal proteins (RPs) are found to have extra-ribosomal functions that are related to regulation of apoptosis, cell cycle arrest, cell proliferation, cell migration and invasion, and DNA damage repair [21, 22]. In response to DNA damage, RPs can help to maintain genomic stability by regulating the activity of the relevant key transcription factors [23]. To support the hyphal growth, the number of ribosomes in an ungerminated spore is insufficient, therefore, ribosome production is a major synthetic process during the early minutes of germination in filamentous fungi [24]. This process results in a 10-fold increase in the number of ribosomes actively engaged in translation within the first few minutes [25-27]. The degree of transcription involved enhances the opportunity for recombination. Thus, the over-representation of ribosome related GO terms in recombination breakpoints containing proteins is not surprising as there the transcription is strong.

Phospholipases are ubiquitous enzymes that participate in multiple cellular functions such as vesicular trafficking, receptor signaling and signal transduction. Phospholipase D (PLD) catalyzes the hydrolysis of phosphatidylcholine into phosphatidic acid (PA). The PLD encoding gene modulates the internalization of this pathogen into epithelial cells and represents a virulence factor of *A. fumigatus* in IA under immunosuppression by corticosteroids [28]. PLD activity and its presence in the nucleus are necessary for signaling and the completion of meiosis [29, 30]. Therefore, over-representation of phospholipase D activity is not surprising.

#### Positive selection in genes classified under KOG functional categories

##### *Metabolism*

Inhaled resting conidia in immunocompromised individuals become metabolically active within 30 min of ingestion. Cellular energy is essential for the multiplication and stress tolerance. Aspergillus species have an remarkable capacity for energy generation, and high efficiency to acquire and utilize a wide range of resources and can store considerable quantities of nutrients within the cell; all traits which contribute to their energy-generating capacity and competitive ability [31]. *A. fumigatus* and some other aspergilli possess and express multiple genes for the enzymes that regulate pathways which allow fungi to adapt their primary carbon metabolism requirement in the niche they inhabit and confer a selective advantage [32]. Also, the possibilities of positive selection signatures in other metabolic genes can be expected. Our scan revealed that several genes involved in metabolic processes evolve under positive selection pressure. This also includes eleven differentially expressed metabolic genes during conidial/hyphae growth. Positive selection pressure was observed in five genes (Afua_2g15730, Afua_2g17080, Afua_6g04500, Afua_1g07220 and Afua_3g08480) in KOG class carbohydrate transport and metabolism. Interestingly, a highly significant positive selection signal (Benjamini-Yekutieli, q-value 0) was observed in a key regulatory glycolytic enzyme 6-phosphofructo-2-kinase 1 coding gene (Afua_1g07220) which catalyzes the conversion of fructose 6-phosphate and ATP to fructose 1,6-bisphosphate and ADP. In response to the cells energy requirements, this enzyme can increase or decrease the rate of glycolysis through allosteric inhibition.

Active fatty acid and lipid metabolism ensure the capability of *A. fumigatus* for using alternative carbon sources in the stressful host environment in infected tissue during invasive aspergillosis [33, 34]. Additionally, also serve as the energy source for intracellular growth of *A. fumigatus* conidia [35]. Triacylglycerides (TAGs) are a class of neutral lipids that represent the most important primary storage form of energy for eukaryotic cells, which is often synthesized via multiple pathways. Phospholipid:DAG acyltransferase (PDAT) that transfer a fatty acyl moiety from a phospholipid (PL) to diacylglycerol (DAG) to form TAG represents one such acyl-CoA–independent TAG synthesis pathway. This membrane-localized enzyme coding gene Afua_3g09950 was identified as PSG by our analysis. The artificial microRNA silencing of PDAT suggests that membrane lipid turnover and TAG synthesis mediated by PDAT is essential for vigorous growth of alga *Chlamydomonas reinhardtii* under favorable culture conditions and for membrane lipid degradation with concomitant production of TAG for survival under stress [36]. BlastP based similarity search shows *A. fumigatus* PDAT shares 43% sequence similarity with *C. reinhardtii* PDAT. Further, having similar domains and similar GO annotations of both the proteins indicate their similar functions. Five genes involved in oxidative stress response (Afua_4g04710, Afua_3g11330, Afua_1g10930, Afua_3g09950 and Afua_2g11510) were found to be positively selected. We also found the signal of positive selection in acyl-CoA thioesterase II coding gene Afua_1g15170 which highly downregulates during 16 h of conidia growth. The enzyme is involved in lipid metabolism [37] as mentioned in later in the manuscript however, its role in *A. fumigatus* is not yet characterized. The Afua_5g08930 *ivdA* gene codes for the isovaleryl-CoA dehydrogenase enzyme, which participates in branched-chain amino acid leucine catabolic process and fatty acid beta-oxidation, (and) was found to be evolved under positive selection. The distant bacterial homolog *Mycobacterium tuberculosis* acyl-CoA dehydrogenase *mbtN*, which shows about 25% sequence identity to *A. fumigatus* Isovaleryl-CoA dehydrogenase and contains similar domain composition and GO annotation, is considered a virulence factor because of its involvement in the production of mycobactins and possibly vital for the iron acquisition within the macrophage [38, 39]. Another mitochondrial enzyme 3-methylcrotonyl-CoA carboxylase subunit alpha coding gene Afua_5g08910 *mccA*, involved in leucine degradation, was evolved under positive selection. This enzyme showed differential expression during conidia growth [40], oxygen-limitation [41], and host neutrophil interaction [42]. In oxygen-limiting conditions many pathogens are able to use leucine as the sole carbon or nitrogen source through the leucine degradation pathway leading to acetyl-CoA and acetoacetate, an important end-product of energy metabolism [43].

In fact, metabolism cannot proceed without energy, and thus adaptive evolution is expected in genes involved in the energy production and conversion subcategory, if the host encounters pathogen adaptation. *A. fumigatus* generate energy in vivo during fungal pathogenesis to drive the metabolic pathways required for fungal growth, host tissue invasion, and disease development using different mechanisms [44]. Mitochondrial respiration is active in hypoxia and critical for fungal pathogenesis [45, 46]. Interruption of mitochondrial respiration in *A. fumigatus* significantly impaired growth in normoxia and hypoxia, conidial germination, and displayed attenuated virulence [47]. Positive selection was observed for electron transfer flavoprotein-ubiquinone oxidoreductase (ETF:QO) coding gene Afua_3g10110, which is critical for linking central metabolism with the respiratory chain [48]. The Afua_3g08770 *grim-19* gene codes for the component of mitochondrial respiratory chain NADH:ubiquinone oxidoreductase complex (complex I), which catalyzes the first step of electron transfer in the oxidative phosphorylation system (OXPHOS), was positively selected. Elimination of *grim-19* destroys the assembly and electron transfer activity of complex I and influences the other complexes in the mitochondrial respiratory chain [49, 50]. It also influences the Warburg effect, favors cell protection and/or prevent aberrant cell growth, apoptosis and also appears to regulate innate and acquired immune responses in mouse models [50-52]. The subunit of mitochondrial matrix pyruvate dehydrogenase (PDH) complex, E1 component subunit alpha coding gene Afua_1g06960, was found to be rapidly evolved. The PDH complex catalyzes the overall conversion of pyruvate into acetyl-CoA and CO2, and thereby provides the primary link between the glycolytic pathway to the tricarboxylic acid (TCA) cycle. The product of PSG acetate kinase Afua_3g10750 *ack* gene converts acetyl phosphate to acetate along with the phosphorylation of ADP and is one of the most prominent ATP-generating reactions in anaerobic respiration. Therefore, *ack* is involved in glucose fermentation to acetate. The growth of a bacterial mutant lacking *ack* has been shown to be inhibited by glucose, suggesting that *ack* is involved in excretion of excess carbohydrate [53]. Positive selection of *A. fumigatus Ack* may therefore be the pathogens strategy to escape with the host interference during anaerobic respiration.

The amino acid asparagine is required for the synthesis of many important cellular proteins in normal cells. Differential expression of asparagine synthetase Afua_4g06900 *asn2* has been observed during early developmental stages of *A. fumigatus* conidia [40]. *asn2* and an another asparaginase family protein Afua_5g11570 showed the signature of positive selection. The positive selection was seen for the gene Afua_5g11450 which encodes for the regulatory protein of arginine metabolism. Arginine can be used as a nitrogen compound to repress the secretion of the alkaline protease that *A. fumigatus* uses to degrade and destroy the host complement system ability to make fungal hyphae susceptible to phagocytosis [54]. The biosynthesis of arginine is important for pathogenicity, hyphal growth, and siderophore biosynthesis, which is coordinated by the supply of ornithine that is produced either by mitochondrial glutamate or  in the cytosol by hydrolysis of arginine [55].

##### *Cellular processes and signaling*

A high number of PSGs annotated are involved in cellular processes and signaling (Additional file 2: Table S8). Majorly involved functional subcategories were signal transduction mechanisms; intracellular trafficking, secretion, and vesicular transport, posttranslational modification, protein turnover, chaperones, and others (Fig. 6). During the conidial growth, like other filamentous fungi, a germ tube apical extension of *A. fumigatus*, a mode of growth that involves the establishment of a stable axis of polarity, followed by the maintenance of growth in the same direction, is regulated by several polarity maintenance genes [56]. The ability to sustain polarization requires a constant stream of new cell wall and plasma membrane material to the hyphal apex [57]. The Centrin-binding protein coding gene Afua_1g13700 *sfi1*, essential for the initiation of spindle pole body duplication [58] and  critical  in the cell division cycle, was found to be positively selected. PSG Afua_3g14340 *mad2* which codes for the evolutionarily conserved central component of the mitotic checkpoint complex, is essential for the spindle assembly checkpoint mechanism [59]. PSG Afua_2g06110 *arp4* codes for a cytoskeletal protein involved in histone acetylation, histone exchange, and regulation of nucleosome density. In *C. albicans*, a mutant phenotype of *arp4* resulted in a decrease of cell size [60]. Several genes related to vesicles were positively selected (KOG class - 'U', Additional file 2: Table S8). Membrane-enclosed vesicles of the secretory system transport here components. The latter are required for membrane and cell wall biogenesis to the growing tip cell [61]. Besides involvement in growth, vesicles participate in importing nutrients from the host and exporting virulence factors to extracellular space. Differentially expressed PSGs during the conidia growth include the  clathrin heavy chain, mitochondrial import receptor subunit Afua_1g11860 *tom22*, and Rab small monomeric GTPase Afua_5g12130 *rab7*. The clathrin heavy chain is involved in the formation of coated vesicles and clathrin-dependent endocytosis. *tom22* mediates protein import into the mitochondrial matrix and mitochondrial outer membrane [62]. *rab7* plays a central role in the transport steps connecting late endosomes to upstream and downstream endocytic compartments and possibly to the phagocytic route [63]. The components of the cytoplasmic dynein complex, dynein family protein coding gene Afua_5g08510 *nudN*, showed the signature of positive selection. The cytoplasmic dynein complex play roles in mitotic regulation and intracellular transport of vesicles and organelles

[64]. Selection pressure was also observed on several genes involved in signal transduction mechanisms (KOG class - 'T', Additional file 2: Table S8). For instance, Afua_4g04710 *pyp1*, which has an ortholog in *Schizosaccharomyces pombe,* is involved in MAP kinase tyrosine phosphatase activity, negative regulation of G2/M transition of mitotic cell cycle, stress response, and osmo-sensory signaling pathway [65] and was found to be evolved under positive selection. Since *pyp1* is annotated as negative regulator of stress-activated MAPK cascade, it probably can control the functional consequence of MAPK activity in *A. fumigatus,* such as cell wall integrity, fungal morphology, and oxidative stress [66]. Protein phosphatase 2C (Afua_1g15800) mutants of *A. fumigatus* grow better than wild type in iron excess conditions [67]. Besides serine/threonine phosphatase activity, protein phosphatase 2C orthologs in fungi are involved in diverse functions (reviewed in [68]), including the virulence [69] in human fungal pathogen *Candida albicans*. Another important gene that evolved under adaptive evolution includes Afua_4g08050 *sac1*, which is required for virulence of *C. albicans* and its mutants. This gene also has induces attenuated filamentous growth and biofilm formation, hypersensitivity to cell wall stresses, and changed content and distribution of chitin [70]. Furthermore, several genes coding for proteins involved in posttranslational modification, protein turnover, and chaperone activity were positively selected (KOG class - 'O', Additional file 1: Table S8). This subcategory also includes Afua_3g12290 *dim1*, which is an essential gene for the viability of *A. fumigatus* [71]. The ortholog of *dim1* is involved in mRNA splicing, regulation of DNA methylation, mitotic spindle formation, and chromosome segregation [72].

##### *Information storage and processing*

In this category 33 PSGs were identified and among them most participate in transcription and related processes (Additional file 1: Table S8). PSG Afua_3g10240 *not1* codes a scaffold protein which is a critical component of Ccr4-Not complex regulating gene expression, mRNA turnover, translational repression, and transcription [73]. Not protein subunits have been found to repress transcription of promoters without a canonical TATA sequence and among the different components of Ccr4-Not complex, only *not1* is known to be essential for yeast viability [73] and growth [74]. The complex in general is regulatory and it regulates cellular processes upon sensing the nutrient levels and other signals and it is thought to be required to keep cells in the appropriate state and to respond to extracellular signals [75]. The ortholog of Afua_2g05020 *dopA* codes a regulatory protein which has been discovered to play a role in several processes, including transport from Golgi to endosome, hyphal growth, cell morphogenesis, spatiotemporal organization of multicellular structures, conidium formation and endoplasmic reticulum organization and viability [76]. PSG Afua_5g04200 *rpc82* codes for the RNA polymerase C subunit protein, which has been identified to be essential for yeast cell viability [77]. Additionally, this protein was also annotated with the GO term 'pathogenesis' (GO:0009405). The ortholog of C6 transcription factor (Afua_6g11110) regulates cell wall architecture, hyphal morphogenesis, virulence, and drug sensitivity in *C. albicans* [78]. In yeast, this protein participates in the regulation of glucose metabolism and fermentation [79]. Afua_5g12190 *taf30* codes a hub protein in the *A. fumigatus* interactome [80], which interacts with many complexes, such as general transcription factors that make up the RNA polymerase II preinitiation complex, chromatin remodeling complexes, and the histone modification enzymes. The highly conserved domain "Yeats" containing the *taf30* ortholog of yeast was found to affect the cell growth in a negative way, in which cells grow better than wild-type in stress conditions when this domain is absent [81]. Some genes involved in the initiation of transcription for instance, Afua_8g04950 *tsm1* and Afua_8g04350, were also found to be evolved under positive selection. Moreover, positive selected genes are involved in DNA replication, recombination, and repair. For instance, gene Afua_2g10140 *mcm7* that codes a part of MCM complex, which plays a role in initiation and regulation of DNA replication, was found to be positively selected. Higher amounts of evolutionarily conserved MCM proteins has been observed in proliferating cells [82]. Afua_5g03320 *ssl2* provides DNA helicase functions and participates in transcription and DNA repair. Furthermore, the ortholog of *ssl2* leads to growth defects when mutated in yeast [83] and is also essential for cell viability [84]. Under the subcategory RNA processing and modification, Afua_1g14310 *pan3*, the required subunit of the PAN enzyme the function of which is to shorten mRNA poly(A) tails [85] and regulate the activity of post replication repair complexes [86], was identified.

**Supplementary figures**

**Figure S1: Proteins-protein interactions (PPIs) map.** PPIs between *A. fumigatus* proteins coded by positively selected genes (PSGs) and human proteins. Orange - *A. fumigatus* proteins and blue - human proteins.

**Figure S2: Gene Ontology (GO) overrepresentation analysis.** Biological process (GO) significantly overrepresented in human proteins targeted by *A. fumigatus* virulence related proteins coded by multi-copy category positively selected genes (PSGs).

**Figure S3: *A. fumigatus* and *A. fischeri* genome synteny map**. Image was generated using the Mauve genome alignment tool. Upper and lower lines of the genomes correspond to *A. fumigatus* and *A. fischeri* respectively. Red vertical bars indicate concatenated chromosomal boundaries. Color-coded syntenic blocks indicate conserved segments (LCBs; Locally Collinear Blocks) identified by Mauve (minimum LCB weight = 999). Plots of sequence similarity are shown within each syntenic block. Regions with no color indicate no detectable homology between the two genomes with the settings used in Mauve. Unmatched regions (white area) within an LCBs indicate the presence of strain-specific sequence. The connecting lines between blocks indicate the location of each block in two genomes. Homologous regions with possible rearrangements are shown in first row.

### Figure S1

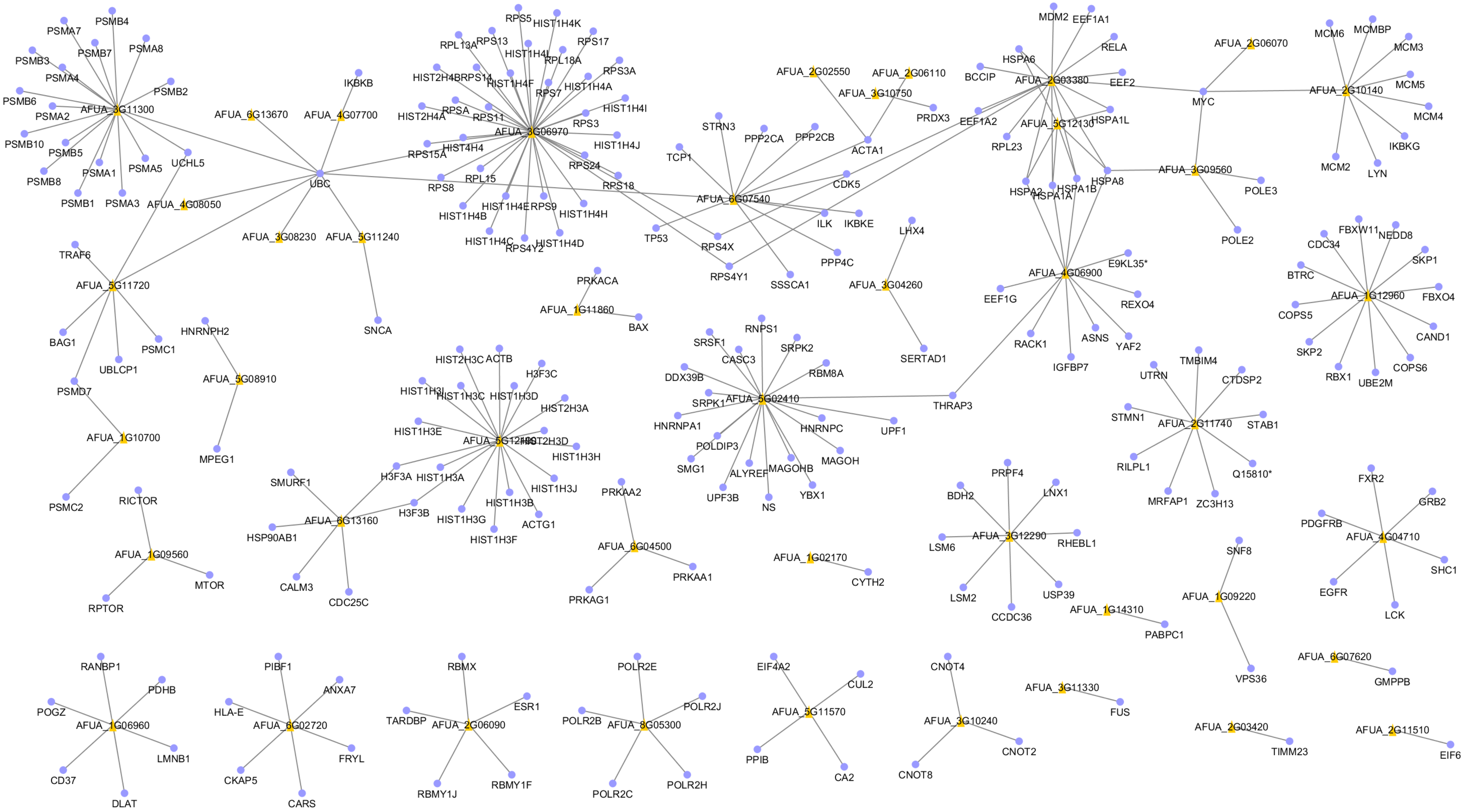

PSMB4
PSMA7
PSMB7
PSMA8
PSMB3
PSMA4
PSMB2
PSMB6
AFUA_3G11300
PSMA2
PSMB10
PSMB5
PSMA1
PSMA5
UCHL5
PSMB8
PSMB1
PSMA3
AFUA_4G08050
IKBKB
AFUA_6G13670
AFUA_4G07700
UBC
AFUA_3G08230
AFUA_5G11240
SNCA
TRAF6
AFUA_5G11720
BAG1
PSMC1
UBLCP1
PSMD7
AFUA_1G10700
PSMC2
HNRNPH2
AFUA_5G08910
MPEG1
RPS5
HIST1H4K
RPL13A
RPS13
RPS17
HIST1H4L
RPL18A
RPS3A
HIST2H4B
RPS14
HIST1H4F
RPS7
HIST1H4A
HIST2H4A
RPSA
RPS11
HIST1H4I
AFUA_3G06970
RPS3
RPS15A
HIST4H4
HIST1H4J
RPS24
RPS8
RPL15
RPS18
HIST1H4B
HIST1H4E
RPS9
HIST1H4H
HIST1H4C
RPS4Y2
HIST1H4D
AFUA_2G02550
AFUA_2G06110
AFUA_3G10750
PRDX3
STRN3
PPP2CA
PPP2CB
ACTA1
TCP1
CDK5
AFUA_6G07540
ILK
IKBKE
TP53
RPS4X
RPS4Y1
SSSCA1
PPP4C
MDM2
EEF1A1
HSPA6
RELA
BCCIP
AFUA_2G03380
EEF2
EEF1A2
AFUA_5G12130
HSPA1L
RPL23
HSPA2
HSPA1B
HSPA1A
HSPA8
AFUA_2G06070
MYC
AFUA_3G09560
POLE3
POLE2
MCM6
MCMBP
MCM3
AFUA_2G10140
MCM5
MCM4
IKBKG
MCM2
LYN
LHX4
AFUA_3G04260
SERTAD1
E9KL35*
AFUA_4G06900
EEF1G
REXO4
RACK1
ASNS
YAF2
IGFBP7
FBXW11
NEDD8
CDC34
SKP1
BTRC
AFUA_1G12960
FBXO4
COPS5
CAND1
SKP2
RBX1
UBE2M
COPS6
PRKACA
AFUA_1G11860
BAX
RNPS1
SRSF1
SRPK2
CASC3
RBM8A
DDX39B
SRPK1
AFUA_5G02410
THRAP3
HNRNPA1
HNRNPC
UPF1
POLDIP3
MAGOH
SMG1
ALYREF
MAGOHB
UPF3B
NS
YBX1
HIST2H3C
ACTB
H3F3C
HIST1H3I
HIST1H3C
HIST1H3D
HIST2H3A
HIST1H3E
AFUA_5G12180
HIST2H3D
HIST1H3H
H3F3A
HIST1H3A
HIST1H3J
H3F3B
HIST1H3B
HIST1H3G
ACTG1
HIST1H3F
SMURF1
HSP90AB1
AFUA_6G13160
CALM3
CDC25C
RICTOR
AFUA_1G09560
MTOR
RPTOR
PRKAA2
AFUA_6G04500
PRKAA1
PRKAG1
AFUA_1G02170
CYTH2
UTRN
TMBIM4
CTDSP2
STMN1
AFUA_2G11740
STAB1
RILPL1
Q15810*
MRFAP1
ZC3H13
PRPF4
BDH2
LNX1
AFUA_3G12290
LSM6
RHEBL1
LSM2
USP39
CCDC36
AFUA_1G14310
PABPC1
SNF8
AFUA_1G09220
VPS36
FXR2
GRB2
PDGFRB
AFUA_4G04710
SHC1
EGFR
LCK
AFUA_6G07620
GMPPB
RANBP1
PDHB
POGZ
AFUA_1G06960
LMNB1
CD37
DLAT
PIBF1
ANXA7
HLA-E
AFUA_6G02720
FRYL
CKAP5
CARS
RBMX
ESR1
TARDBP
AFUA_2G06090
RBMY1J
RBMY1F
POLR2E
POLR2J
POLR2B
AFUA_8G05300
POLR2C
POLR2H
EIF4A2
CUL2
AFUA_5G11570
PPIB
CA2
CNOT4
AFUA_3G10240
CNOT2
CNOT8
AFUA_3G11330
FUS
AFUA_2G03420
TIMM23
AFUA_2G11510
EIF6

**Figure S2**

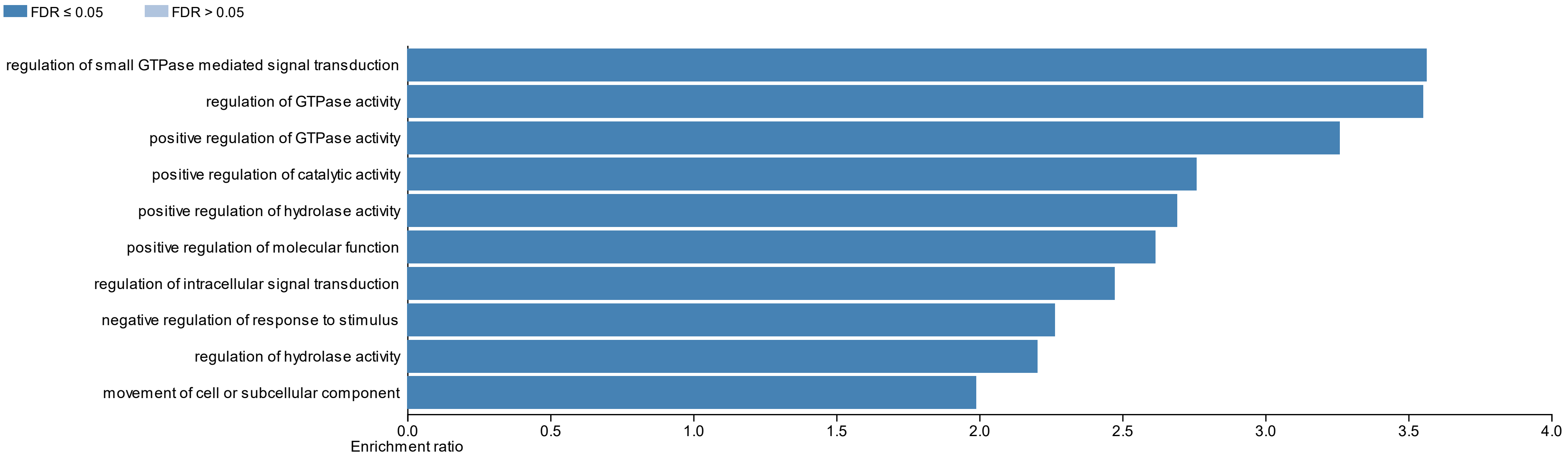
FDR ≤ 0.05
FDR > 0.05
regulation of small GTPase mediated signal transduction
regulation of GTPase activity
positive regulation of GTPase activity
positive regulation of catalytic activity
positive regulation of hydrolase activity
positive regulation of molecular function
regulation of intracellular signal transduction
negative regulation of response to stimulus
regulation of hydrolase activity
movement of cell or subcellular component
0.0
0.5
1.0
1.5
2.0
2.5
3.0
3.5
4.0
Enrichment ratio

### Figure S3

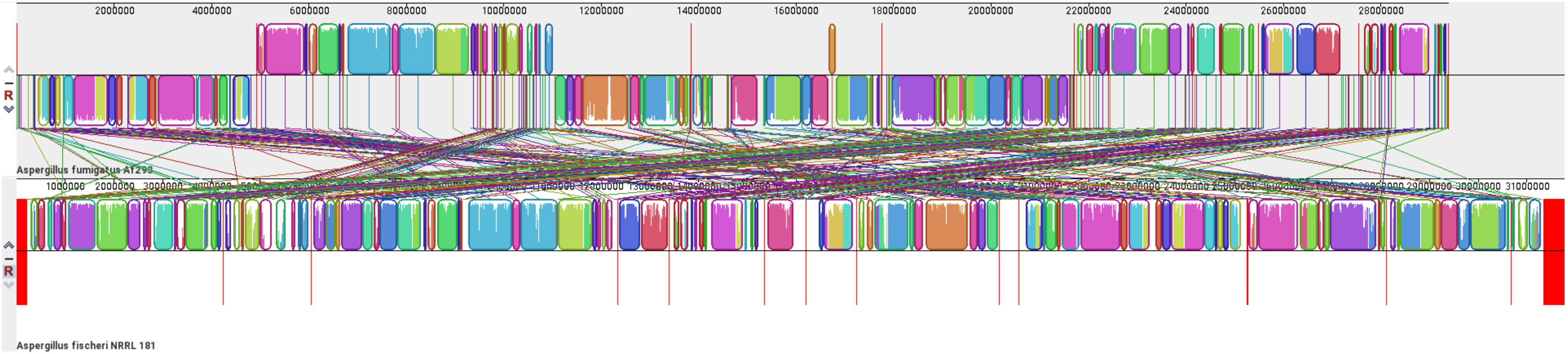

Aspergillus fumigatus Af293
Aspergillus fischeri NRRL 181

**Additional data file 1:** Clusters of orthologues identified by OrthoMCL (inflation index: 1.5). Abbreviations - abra : *Aspergillus brasiliensis*, acar : *A. carbonarius*, acla : *A. clavatus*, afis : *A. fischeri*, afla : *A. flavus*, afum : *A. fumigatus*, agla : *A. glaucus*, akaw : *A. kawachii*, alue : *A. luchuensis*, anid : *A. nidulans*, anig : *A. niger*, aora : *A. oryzae*, asyd : *A. sydowii*, ater : *A. terreus*, atub : *A. tubingenesis*, aver : *A. versicolor*, awen : *A. wentii*, azon : *A. zonatus*.

Group1: abra|OJJ66207.1 abra|OJJ67944.1 abra|OJJ69734.1 abra|OJJ73921.1 abra|OJJ75305.1 abra|OJJ75840.1 abra|OJJ73699.1 abra|OJJ69537.1 abra|OJJ78094.1 acar|OOF90653.1 acar|OOF94187.1 acar|OOG00776.1 acar|OOG01052.1 acar|OOF92566.1 acar|OOF93189.1 acar|OOF97512.1 acar|OOF98777.1 acla|EAW07798.1 acla|EAW08899.1 acla|EAW11681.1 akaw|GAA89528.1 akaw|GAA90187.1 akaw|GAA91252.1 aluc|OJZ80511.1 aluc|OJZ82816.1 aluc|OJZ88125.1 aluc|OJZ89282.1 aluc|OJZ83162.1 anid|CBF73441.1 anid|CBF90374.1 anid|CBF87872.1 anig|EHA17891.1 anig|EHA21256.1 anig|EHA21434.1 anig|EHA25711.1 anig|EHA28226.1 anig|EHA18088.1 anig|EHA19858.1 anig|EHA25622.1 anig|EHA25682.1 anig|EHA23283.1 aory|EIT74664.1 aory|EIT83097.1 aory|EIT80579.1 aory|EIT81467.1 aory|EIT83286.1 asyd|OJJ56148.1 asyd|OJJ63431.1 atub|OJI87775.1 atub|OJI89047.1 atub|OJI90732.1 atub|OJI88419.1 aver|OJI97506.1 aver|OJI99587.1 aver|OJJ03355.1 aver|OJJ05088.1 aver|OJJ08114.1 aver|OJJ03487.1 awen|OJJ29812.1 awen|OJJ35864.1 awen|OJJ38889.1 azon|OJJ42489.1 azon|OJJ44494.1 azon|OJJ46205.1 azon|OJJ47369.1 azon|OJJ45854.1 afis|EAW22835.1 afla|EED50110.1 afla|EED51745.1 afum|EAL90367.2 agla|OJJ78560.1 akaw|GAA89080.1 aory|EIT81736.1 ater|EAU35730.1 abra|OJJ67593.1 akaw|GAA89112.1 atub|OJI88453.1 acar|OOF91540.1 abra|OJJ70223.1 afla|EED52040.1 afum|EAL91343.1 akaw|GAA85904.1 aluc|OJZ91163.1 anid|CBF80212.1 atub|OJI83179.1 acar|OOF96620.1 akaw|GAA84550.1 atub|OJI85517.1 afla|EED52433.1 acla|EAW10758.1 afis|EAW16932.1 afla|EED48405.1 afla|EED51820.1 afla|EED51822.1 afla|EED52154.1 aory|EIT74440.1 anid|CBF73982.1 ater|EAU33387.1

Group2: abra|OJJ67588.1 abra|OJJ73922.1 abra|OJJ75304.1 acar|OOF99263.1 acar|OOG01053.1 acar|OOF99291.1 acar|OOF99559.1 acla|EAW07086.1 acla|EAW08665.1 acla|EAW08900.1 acla|EAW11679.1 acla|EAW15286.1 acla|EAW07799.1 afis|EAW16180.1 afis|EAW16446.1 afis|EAW20684.1 afla|EED44993.1 afla|EED47804.1 afla|EED49850.1 afla|EED50111.1 afla|EED52159.1 afum|EAL89049.1 afum|EAL91342.1 akaw|GAA82422.1 akaw|GAA89107.1 akaw|GAA89529.1 anid|CBF73453.1 anid|CBF87069.1 anid|CBF76036.1 anid|CBF87869.1 anid|CBF90372.1 anig|EHA17890.1 anig|EHA25707.1 anig|EHA20896.1 anig|EHA21433.1 anig|EHA24938.1 aory|EIT73739.1 aory|EIT74412.1 aory|EIT74615.1 aory|EIT80956.1 asyd|OJJ56147.1 asyd|OJJ63429.1 ater|EAU29302.1 ater|EAU30156.1 ater|EAU37759.1 ater|EAU30201.1 atub|OJI86150.1 atub|OJI88448.1 atub|OJI88518.1 atub|OJI90731.1 aver|OJI97508.1 aver|OJI99595.1 aver|OJJ03909.1 aver|OJJ05087.1 aver|OJI99581.1 aver|OJI99596.1 aver|OJJ07364.1 aver|OJJ08120.1 aver|OJJ08464.1 aver|OJI99625.1 aver|OJJ07311.1 aver|OJJ08781.1 awen|OJJ36076.1 awen|OJJ38362.1 awen|OJJ32929.1 awen|OJJ38888.1 azon|OJJ42030.1 azon|OJJ42483.1 azon|OJJ46579.1 azon|OJJ48580.1 afis|EAW22836.1 afum|EAL90366.1 agla|OJJ80563.1 aluc|OJZ85583.1 aluc|OJZ88124.1 asyd|OJJ61536.1 ater|EAU38874.1 awen|OJJ32292.1 abra|OJJ67652.1 afis|EAW16453.1 afis|EAW25605.1 agla|OJJ79232.1 akaw|GAA89168.1 aluc|OJZ81665.1 anig|EHA25778.1 ater|EAU33874.1 aver|OJJ07366.1 azon|OJJ50363.1 acla|EAW14636.1 ater|EAU36802.1 awen|OJJ34576.1 afum|EAL88812.1 aver|OJI99592.1 asyd|OJJ52456.1 aver|OJJ03957.1 awen|OJJ34650.1

Group3: abra|OJJ67584.1 abra|OJJ67975.1 abra|OJJ75778.1 abra|OJJ77754.1 acar|OOF92816.1 acar|OOF96723.1 acla|EAW08895.1 acla|EAW11760.1 acla|EAW12170.1 acla|EAW12336.1 acla|EAW14856.1 afla|EED47354.1 afla|EED52019.1 afla|EED47890.1 afla|EED48583.1 agla|OJJ79933.1 agla|OJJ80111.1 agla|OJJ83619.1 agla|OJJ86349.1 akaw|GAA84491.1 akaw|GAA85345.1 akaw|GAA87318.1 akaw|GAA91228.1 akaw|GAA92525.1 akaw|GAA90346.1 akaw|GAA85925.1 aluc|OJZ82842.1 aluc|OJZ85886.1 aluc|OJZ89214.1 aluc|OJZ90707.1 aluc|OJZ91539.1 aluc|OJZ91139.1 anid|CBF71467.1 anid|CBF83055.1 anid|CBF86052.1 anid|CBF87072.1 anid|CBF88289.1 anig|EHA19567.1 anig|EHA20193.1 anig|EHA21280.1 anig|EHA22576.1 anig|EHA24574.1 anig|EHA24700.1 anig|EHA28244.1 aory|EIT73118.1 aory|EIT76300.1 aory|EIT77964.1 aory|EIT81441.1 aory|EIT82867.1 ater|EAU29417.1 ater|EAU31921.1 ater|EAU32663.1 ater|EAU36720.1 atub|OJI79672.1 atub|OJI83150.1 atub|OJI85993.1 atub|OJI91114.1 atub|OJI82134.1 atub|OJI85578.1 atub|OJI89072.1 atub|OJI82802.1 aver|OJI96730.1 aver|OJJ05906.1 awen|OJJ29758.1 awen|OJJ34705.1 awen|OJJ40111.1 awen|OJJ29917.1 awen|OJJ30910.1 awen|OJJ38044.1 awen|OJJ39589.1 awen|OJJ29944.1 afis|EAW23793.1 afum|EAL87227.2 asyd|OJJ62861.1 aver|OJI96914.1 asyd|OJJ61528.1 abra|OJJ70374.1 acar|OOF91895.1 afis|EAW16886.1 afum|EAL86424.2 ater|EAU30225.1 azon|OJJ48577.1 acar|OOF93599.1 azon|OJJ49991.1 afla|EED46560.1 akaw|GAA87467.1 aluc|OJZ88529.1

Group4: abra|OJJ67662.1 abra|OJJ77815.1 abra|OJJ78181.1 abra|OJJ74373.1 abra|OJJ78098.1 abra|OJJ76619.1 acar|OOF91828.1 acar|OOF91847.1 acar|OOF92934.1 acar|OOF94993.1 acar|OOF97632.1 acar|OOF95055.1 acar|OOF95122.1 acla|EAW07064.1 afis|EAW19759.1 afis|EAW23638.1 akaw|GAA83602.1 akaw|GAA84851.1 akaw|GAA92094.1 akaw|GAA83831.1 akaw|GAA85928.1 akaw|GAA89177.1 akaw|GAA86655.1 akaw|GAA85575.1 akaw|GAA85859.1 aluc|OJZ81656.1 aluc|OJZ83631.1 aluc|OJZ90887.1 aluc|OJZ91221.1 aluc|OJZ81972.1 aluc|OJZ86820.1 aluc|OJZ89749.1 aluc|OJZ92392.1 aluc|OJZ83511.1 aluc|OJZ89332.1 aluc|OJZ91137.1 anid|CBF80211.1 anid|CBF82304.1 anig|EHA19289.1 anig|EHA24757.1 anig|EHA19606.1 anig|EHA25789.1 anig|EHA26583.1 aory|EIT74370.1 aory|EIT76972.1 ater|EAU35165.1 ater|EAU36856.1 atub|OJI79585.1 atub|OJI79696.1 atub|OJI83697.1 atub|OJI85053.1 atub|OJI86822.1 atub|OJI82881.1 atub|OJI88527.1 atub|OJI90000.1 atub|OJI83148.1 aver|OJJ04870.1 aver|OJJ06306.1

aver|OJJ07785.1 awen|OJJ29662.1 awen|OJJ29874.1 awen|OJJ33672.1 agla|OJJ83634.1 asyd|OJJ52727.1 aver|OJJ02001.1 azon|OJJ47400.1 afla|EED51023.1 abra|OJJ74811.1 acar|OOF95802.1 afla|EED52130.1 afum|EAL85129.1 atub|OJI85468.1 abra|OJJ77455.1 afla|EED49595.1 anig|EHA19288.1 asyd|OJJ56406.1 ater|EAU32666.1 ater|EAU38928.1 ater|EAU39559.1 anid|CBF79115.1 afis|EAW19771.1 akaw|GAA84588.1 akaw|GAA84589.1 aluc|OJZ89334.1 atub|OJI82404.1

Group5: abra|OJJ67238.1 abra|OJJ68732.1 acar|OOF92227.1 acar|OOF98313.1 acla|EAW12167.1 acla|EAW12516.1 afis|EAW15603.1 afis|EAW17072.1 afis|EAW19893.1 afis|EAW21005.1 afis|EAW24131.1 afis|EAW24651.1 afis|EAW22083.1 afis|EAW24734.1 afla|EED46089.1 afla|EED46801.1 afla|EED48098.1 afla|EED52234.1 afla|EED48355.1 afla|EED49615.1 afla|EED55083.1 afla|EED56403.1 afum|EAL85692.1 afum|EAL89381.1 afum|EAL91987.1 afum|EAL85730.1 afum|EAL86560.1 agla|OJJ79085.1 agla|OJJ82566.1 agla|OJJ86784.1 agla|OJJ87207.1 akaw|GAA89881.1 akaw|GAA91663.1 aluc|OJZ81798.1 aluc|OJZ87737.1 anid|CBF75787.1 anid|CBF86562.1 anid|CBF86663.1 anig|EHA20229.1 anig|EHA22418.1 aory|EIT72725.1 aory|EIT74466.1 aory|EIT76413.1 aory|EIT80974.1 aory|EIT81644.1 aory|EIT83015.1 asyd|OJJ59270.1 asyd|OJJ64851.1 asyd|OJJ60947.1 asyd|OJJ64782.1 asyd|OJJ65051.1 ater|EAU30561.1 ater|EAU31344.1 ater|EAU38103.1 atub|OJI80903.1 atub|OJI84369.1 aver|OJI98893.1 aver|OJI99174.1 aver|OJJ00274.1 aver|OJJ08481.1 aver|OJI98963.1 aver|OJJ01883.1 aver|OJJ07136.1 awen|OJJ30103.1 awen|OJJ40222.1 awen|OJJ41358.1 awen|OJJ31256.1 awen|OJJ37981.1 azon|OJJ46971.1 azon|OJJ48822.1 abra|OJJ73660.1 acar|OOG00838.1 akaw|GAA88498.1 aluc|OJZ88393.1 anig|EHA18121.1 aory|EIT72296.1 atub|OJI90988.1 awen|OJJ38335.1 anid|CBF74724.1 aluc|OJZ80164.1 aory|EIT76217.1 ater|EAU32520.1

Group6: abra|OJJ65900.1 abra|OJJ66009.1 abra|OJJ67667.1 abra|OJJ67833.1 abra|OJJ70432.1 abra|OJJ73134.1 abra|OJJ74399.1 abra|OJJ77760.1 abra|OJJ66336.1 abra|OJJ75222.1 abra|OJJ75882.1 acla|EAW13531.1 acla|EAW13625.1 afis|EAW16387.1 afis|EAW17052.1 afla|EED45665.1 afla|EED48448.1 afla|EED47332.1 afla|EED53171.1 afla|EED57354.1 akaw|GAA83600.1 akaw|GAA86003.1 akaw|GAA89183.1 akaw|GAA90148.1 akaw|GAA92662.1 akaw|GAA84585.1 akaw|GAA90211.1 akaw|GAA90881.1 aluc|OJZ81649.1 aluc|OJZ82246.1 aluc|OJZ82952.1 aluc|OJZ90228.1 aluc|OJZ92394.1 aluc|OJZ89329.1 aluc|OJZ83585.1 aluc|OJZ83788.1 aluc|OJZ90116.1 anig|EHA19545.1 anig|EHA19698.1 anig|EHA26581.1 anig|EHA21367.1 anig|EHA23246.1 anig|EHA25794.1 anig|EHA28496.1 aory|EIT80163.1 aory|EIT82886.1 asyd|OJJ53336.1 asyd|OJJ54299.1 ater|EAU30237.1 ater|EAU33224.1 atub|OJI80755.1 atub|OJI84681.1 atub|OJI86824.1 atub|OJI87738.1 atub|OJI88533.1 atub|OJI89152.1 atub|OJI89184.1 atub|OJI85471.1 atub|OJI88192.1 atub|OJI87802.1 aver|OJI96712.1 aver|OJJ08408.1 awen|OJJ30102.1 awen|OJJ41501.1 awen|OJJ41603.1 awen|OJJ31974.1 acar|OOF99302.1 afis|EAW21094.1 aory|EIT75368.1 ater|EAU31345.1 acar|OOF92089.1 aver|OJJ07905.1 aluc|OJZ83134.1 awen|OJJ35031.1 acar|OOF98127.1 afum|EAL87813.1 afum|EAL92117.1 anid|CBF75778.1 afis|EAW17483.1 anid|CBF84683.1 akaw|GAA83278.1

Group7: abra|OJJ65820.1 abra|OJJ66803.1 acar|OOF90643.1 acar|OOF94182.1 afis|EAW16457.1 afis|EAW21767.1 afla|EED55145.1 afla|EED55917.1 akaw|GAA92417.1 akaw|GAA93242.1 anid|CBF75176.1 anid|CBF84668.1 anid|CBF85153.1 anid|CBF88209.1 anid|CBF75912.1 anid|CBF80593.1 anid|CBF80303.1 anid|CBF80770.1 anig|EHA18419.1 anig|EHA24965.1 anig|EHA21352.1 aory|EIT76564.1 aory|EIT81636.1 aory|EIT81823.1 asyd|OJJ52880.1 asyd|OJJ54446.1 asyd|OJJ60235.1 asyd|OJJ61610.1 ater|EAU32131.1 ater|EAU35090.1 ater|EAU37273.1 aver|OJJ04030.1 aver|OJJ08401.1 aver|OJJ08851.1 aver|OJJ07396.1 azon|OJJ42892.1 azon|OJJ45918.1 azon|OJJ48221.1 azon|OJJ48680.1 azon|OJJ49289.1 agla|OJJ83550.1 anig|EHA26118.1 awen|OJJ41193.1 azon|OJJ42522.1 abra|OJJ67032.1 acla|EAW08788.1 afis|EAW19897.1 afis|EAW23754.1 agla|OJJ82646.1 akaw|GAA92151.1 aluc|OJZ85558.1 ater|EAU30865.1 atub|OJI79347.1 abra|OJJ76132.1 acar|OOF99791.1 acla|EAW09686.1 afis|EAW21453.1 afla|EED55883.1 afum|EAL92472.1 agla|OJJ85599.1 akaw|GAA88835.1 aluc|OJZ89593.1 anid|CBF76549.1 anig|EHA23018.1 aory|EIT76608.1 asyd|OJJ61156.1 atub|OJI85209.1 aver|OJJ00478.1 awen|OJJ39247.1 azon|OJJ51608.1 abra|OJJ77926.1 acar|OOF89902.1 afum|EAL87190.2 akaw|GAA83738.1 aluc|OJZ90994.1 atub|OJI82984.1 aver|OJJ07910.1 aver|OJJ00933.1

Group8: abra|OJJ66275.1 abra|OJJ77982.1 abra|OJJ70821.1 abra|OJJ71091.1 abra|OJJ75537.1 abra|OJJ78258.1 acar|OOF97571.1 acar|OOF98050.1 acar|OOG00735.1 acar|OOF98604.1 acla|EAW07624.1 acla|EAW08950.1 acla|EAW09019.1 acla|EAW09117.1 acla|EAW13520.1 acla|EAW15062.1 afla|EED45116.1 afla|EED47649.1 afla|EED49862.1 afla|EED51183.1 akaw|GAA82870.1 akaw|GAA91957.1 akaw|GAA85998.1 aluc|OJZ82661.1 aluc|OJZ84344.1 aluc|OJZ87437.1 aluc|OJZ91057.1 aluc|OJZ91332.1 anig|EHA18001.1 anig|EHA18935.1 anig|EHA20663.1 anig|EHA27435.1 anig|EHA27680.1 anig|EHA19760.1 anig|EHA23147.1 anig|EHA23656.1 anig|EHA20014.1 aory|EIT73707.1 aory|EIT80187.1 aory|EIT83384.1 asyd|OJJ57246.1 asyd|OJJ60063.1 asyd|OJJ63687.1 atub|OJI81211.1 atub|OJI83130.1 atub|OJI83325.1 atub|OJI88893.1 atub|OJI89724.1 atub|OJI83053.1 atub|OJI89947.1 aver|OJI99011.1 aver|OJJ08178.1 awen|OJJ29725.1 awen|OJJ32235.1 awen|OJJ34622.1 afla|EED49876.1 afum|EAL85113.2 akaw|GAA87094.1 aory|EIT83412.1 ater|EAU38971.1 afis|EAW16804.1 agla|OJJ85462.1 anid|CBF80487.1 ater|EAU29410.1 azon|OJJ42361.1 acla|EAW10019.1 afis|EAW20415.1 afum|EAL89230.2 anid|CBF85555.1 ater|EAU29808.1 azon|OJJ43495.1 aver|OJI97248.1 afum|EAL91103.2 akaw|GAA83290.1 akaw|GAA83291.1 afla|EED45117.1 aver|OJJ08105.1

Group9: abra|OJJ68001.1 abra|OJJ69377.1 abra|OJJ77432.1 abra|OJJ78055.1 acar|OOF92044.1 acar|OOF94347.1 afis|EAW19338.1 afis|EAW24682.1 afis|EAW20700.1 afis|EAW24697.1 afis|EAW25724.1 afla|EED45393.1 afla|EED47975.1 afla|EED51175.1 afla|EED57518.1 afum|EAL84397.1 afum|EAL84875.1 afum|EAL89339.1 afum|EAL94057.1 agla|OJJ83163.1 agla|OJJ85718.1 agla|OJJ88362.1 akaw|GAA85937.1 akaw|GAA88581.1 akaw|GAA88246.1 akaw|GAA90949.1 aluc|OJZ82213.1 aluc|OJZ82875.1 aluc|OJZ88485.1

aluc|OJZ91125.1 anid|CBF70387.1 anid|CBF79143.1 anid|CBF90097.1 anid|CBF74114.1 anid|CBF80184.1 anig|EHA21301.1 anig|EHA28527.1 aory|EIT75060.1 aory|EIT80443.1 aory|EIT79828.1 aory|EIT81356.1 aory|EIT81872.1 aory|EIT82404.1 asyd|OJJ53903.1 asyd|OJJ56275.1 asyd|OJJ57401.1 asyd|OJJ57276.1 asyd|OJJ58638.1 asyd|OJJ62031.1 ater|EAU31624.1 ater|EAU37396.1 ater|EAU31762.1 ater|EAU38791.1 atub|OJI79831.1 atub|OJI83136.1 atub|OJI88222.1 atub|OJI89099.1 aver|OJI96122.1 aver|OJJ01245.1 aver|OJI99530.1 aver|OJJ08000.1 aver|OJJ08490.1 awen|OJJ32995.1 awen|OJJ37839.1 awen|OJJ40791.1 azon|OJJ48191.1 azon|OJJ48520.1 acla|EAW14609.1 azon|OJJ43922.1 acla|EAW13612.1 afla|EED53479.1 anig|EHA20150.1 anig|EHA19826.1 afla|EED51738.1 afla|EED51739.1 ater|EAU33967.1

Group10: abra|OJJ73857.1 abra|OJJ76179.1 acar|OOF94216.1 acar|OOF99761.1 acar|OOG01005.1 acla|EAW07747.1 acla|EAW09647.1 afis|EAW17299.1 afis|EAW22784.1 afis|EAW20842.1 afis|EAW21493.1 afla|EED48870.1 afla|EED50046.1 afum|EAL84790.1 afum|EAL92512.1 afum|EAL90415.1 agla|OJJ83596.1 agla|OJJ85655.1 akaw|GAA82124.1 akaw|GAA89472.1 akaw|GAA92487.1 aluc|OJZ80154.1 aluc|OJZ88191.1 aluc|OJZ92051.1 aluc|OJZ91499.1 anid|CBF76107.1 anid|CBF87969.1 anid|CBF76478.1 anid|CBF80461.1 anid|CBF83927.1 anig|EHA17943.1 anig|EHA22980.1 anig|EHA22539.1 aory|EIT74611.1 aory|EIT76730.1 aory|EIT80155.1 aory|EIT81260.1 asyd|OJJ53335.1 asyd|OJJ61118.1 asyd|OJJ55439.1 asyd|OJJ60355.1 asyd|OJJ63488.1 ater|EAU29437.1 ater|EAU35045.1 ater|EAU38925.1 ater|EAU29951.1 atub|OJI80378.1 atub|OJI90790.1 atub|OJI85952.1 atub|OJI86482.1 aver|OJI97441.1 aver|OJJ06514.1 aver|OJI99693.1 aver|OJJ08409.1 aver|OJJ00436.1 awen|OJJ36113.1 awen|OJJ38951.1 awen|OJJ39202.1 awen|OJJ40168.1 azon|OJJ49724.1 azon|OJJ51661.1 abra|OJJ70289.1 acar|OOF91742.1 akaw|GAA92859.1 aluc|OJZ82554.1 anig|EHA22518.1 atub|OJI88795.1 awen|OJJ38795.1 afla|EED57355.1 afla|EED55928.1

Group11: abra|OJJ66566.1 abra|OJJ74913.1 acar|OOF91007.1 acar|OOF95904.1 acla|EAW08458.1 acla|EAW10929.1 afis|EAW20932.1 afis|EAW23503.1 afla|EED50286.1 afla|EED57404.1 afum|EAL84702.1 afum|EAL87958.1 agla|OJJ83956.1 agla|OJJ86749.1 akaw|GAA83509.1 akaw|GAA91151.1 aluc|OJZ81479.1 aluc|OJZ92486.1 anid|CBF82508.1 anid|CBF89725.1 anig|EHA18724.1 anig|EHA18871.1 anig|EHA25948.1 anig|EHA26487.1 aory|EIT77296.1 aory|EIT80109.1 asyd|OJJ54080.1 asyd|OJJ62235.1 ater|EAU35292.1 ater|EAU38577.1 atub|OJI86917.1 atub|OJI88701.1 aver|OJI96344.1 aver|OJJ08658.1 awen|OJJ34386.1 awen|OJJ39829.1 azon|OJJ47952.1 azon|OJJ48118.1 acar|OOF91006.1 acla|EAW10930.1 afis|EAW20931.1 afla|EED50285.1 afum|EAL84703.2 agla|OJJ83957.1 akaw|GAA91152.1 anid|CBF82510.1 anig|EHA25947.1 aory|EIT77323.1 asyd|OJJ54081.1 ater|EAU38576.1 aver|OJJ08657.1 awen|OJJ34387.1 azon|OJJ48119.1 abra|OJJ68224.1 acar|OOF95905.1 acla|EAW08457.1 afis|EAW23498.1 afla|EED57403.1 afum|EAL87965.1 akaw|GAA83219.1 aluc|OJZ83721.1 anig|EHA18870.1 aory|EIT80136.1 ater|EAU35291.1 atub|OJI90095.1 awen|OJJ39827.1 anig|EHA26486.1 atub|OJI86918.1 agla|OJJ83079.1

Group12: abra|OJJ67365.1 abra|OJJ70373.1 abra|OJJ72695.1 abra|OJJ77105.1 abra|OJJ78039.1 abra|OJJ77002.1 acar|OOF90082.1 acar|OOF91896.1 acar|OOF92990.1 acar|OOF91448.1 acla|EAW08775.1 acla|EAW12337.1 afis|EAW16885.1 afis|EAW20732.1 afis|EAW25626.1 afla|EED47350.1 afla|EED50231.1 afla|EED54427.1 afum|EAL84314.2 afum|EAL86423.2 agla|OJJ78903.1 agla|OJJ79932.1 agla|OJJ85297.1 akaw|GAA86951.1 akaw|GAA91977.1 aluc|OJZ80600.1 aluc|OJZ84401.1 aluc|OJZ82637.1 aluc|OJZ84514.1 anid|CBF80656.1 anid|CBF82934.1 anid|CBF83105.1 anig|EHA27442.1 anig|EHA27812.1 aory|EIT75862.1 aory|EIT77345.1 aory|EIT81172.1 aory|EIT81924.1 aory|EIT82860.1 asyd|OJJ53449.1 asyd|OJJ54612.1 asyd|OJJ61456.1 ater|EAU30228.1 ater|EAU38239.1 atub|OJI88868.1 atub|OJI89567.1 atub|OJI89671.1 aver|OJJ00792.1 aver|OJJ05324.1 aver|OJJ07546.1 aver|OJJ08303.1 awen|OJJ29598.1 awen|OJJ29759.1 awen|OJJ31482.1 awen|OJJ34518.1 awen|OJJ36047.1 azon|OJJ42524.1 afla|EED55182.1 akaw|GAA87046.1 anig|EHA27731.1 aory|EIT81588.1 asyd|OJJ55521.1 ater|EAU32705.1 aver|OJJ06599.1 azon|OJJ50398.1 afla|EED48862.1 ater|EAU32222.1

Group13: abra|OJJ74143.1 abra|OJJ77919.1 acar|OOF95048.1 acar|OOF95115.1 acar|OOF95586.1 acar|OOF98862.1 acar|OOF99364.1 afla|EED47372.1 afla|EED48594.1 akaw|GAA83744.1 akaw|GAA89741.1 akaw|GAA92822.1 akaw|GAA92977.1 aluc|OJZ80240.1 aluc|OJZ91576.1 aluc|OJZ80643.1 aluc|OJZ90987.1 anid|CBF75107.1 anid|CBF76688.1 anid|CBF85354.1 anid|CBF86773.1 anig|EHA19692.1 anig|EHA27189.1 aory|EIT73093.1 aory|EIT82880.1 asyd|OJJ59996.1 asyd|OJJ61226.1 asyd|OJJ64036.1 asyd|OJJ64686.1 ater|EAU32382.1 ater|EAU33556.1 atub|OJI82978.1 atub|OJI87498.1 aver|OJI98121.1 aver|OJJ00549.1 aver|OJJ02295.1 aver|OJI98789.1 abra|OJJ69652.1 acar|OOF98861.1 acla|EAW06691.1 afis|EAW16074.1 afla|EED48593.1 afum|EAL88706.2 agla|OJJ86094.1 akaw|GAA91631.1 aluc|OJZ82458.1 anid|CBF75109.1 anig|EHA28317.1 aory|EIT73047.1 ater|EAU33557.1 atub|OJI87980.1 awen|OJJ30359.1 azon|OJJ45380.1 acla|EAW06690.1 afis|EAW16075.1 afum|EAL88707.1 awen|OJJ30358.1 abra|OJJ76634.1 afla|EED53145.1 agla|OJJ82986.1 anig|EHA22603.1 aory|EIT75089.1 ater|EAU37809.1 atub|OJI86025.1 awen|OJJ31622.1 azon|OJJ47320.1

Group14: abra|OJJ65763.1 abra|OJJ65764.1 abra|OJJ68458.1 abra|OJJ70187.1 acar|OOF91603.1 acar|OOF93288.1 acar|OOF99308.1 acar|OOF99313.1 afis|EAW16734.1 afis|EAW19114.1 afla|EED45012.1 afla|EED57380.1 afla|EED57383.1 afum|EAL86261.1 afum|EAL86308.1 afum|EAL86264.1 afum|EAL93846.1 akaw|GAA83845.1 akaw|GAA86588.1 akaw|GAA92381.1 aluc|OJZ83240.1 aluc|OJZ85135.1 aluc|OJZ90870.1 anid|CBF74944.1 anid|CBF83824.1 anid|CBF84265.1 anig|EHA19590.1 anig|EHA25367.1 anig|EHA21095.1 anig|EHA22863.1 aory|EIT73582.1 aory|EIT75916.1 aory|EIT80067.1 aory|EIT80069.1 asyd|OJJ57214.1 asyd|OJJ58070.1 asyd|OJJ63941.1 asyd|OJJ60903.1 asyd|OJJ63609.1 asyd|OJJ63938.1 atub|OJI80457.1 atub|OJI82868.1 atub|OJI87697.1 aver|OJI97001.1 aver|OJJ00229.1 aver|OJJ03075.1 aver|OJJ04337.1 aver|OJJ07009.1 awen|OJJ30374.1 awen|OJJ30814.1 awen|OJJ32530.1 acla|EAW14406.1 agla|OJJ89198.1

azon|OJJ47391.1 aver|OJI97004.1 aver|OJI97328.1 awen|OJJ40814.1 azon|OJJ50090.1 ater|EAU37377.1 azon|OJJ42904.1 ater|EAU38814.1 asyd|OJJ55207.1

Group15: abra|OJJ69361.1 abra|OJJ71787.1 abra|OJJ75622.1 acar|OOF94364.1 acar|OOF96466.1 acar|OOG00218.1 acla|EAW08453.1 acla|EAW09357.1 acla|EAW14751.1 afis|EAW21780.1 afis|EAW23502.1 afis|EAW24282.1 afla|EED46347.1 afla|EED47994.1 afum|EAL85951.1 afum|EAL87959.1 afum|EAL92809.1 agla|OJJ80637.1 agla|OJJ84344.1 akaw|GAA84369.1 akaw|GAA85124.1 akaw|GAA88230.1 aluc|OJZ82194.1 aluc|OJZ86517.1 aluc|OJZ89073.1 anig|EHA22129.1 anig|EHA23464.1 anig|EHA28543.1 aory|EIT79847.1 aory|EIT81244.1 asyd|OJJ53917.1 asyd|OJJ56885.1 asyd|OJJ61185.1 asyd|OJJ62326.1 ater|EAU31230.1 ater|EAU38460.1 atub|OJI84007.1 atub|OJI85713.1 atub|OJI88241.1 aver|OJI96431.1 aver|OJJ00509.1 aver|OJJ08015.1 awen|OJJ33015.1 awen|OJJ34069.1 awen|OJJ39823.1 azon|OJJ47947.1 azon|OJJ48162.1 azon|OJJ48502.1 afla|EED57398.1 anid|CBF76607.1 agla|OJJ86745.1 abra|OJJ74918.1 acar|OOF95909.1 akaw|GAA83505.1 aluc|OJZ92489.1 anid|CBF89735.1 anig|EHA26483.1 aory|EIT80098.1 asyd|OJJ62232.1 ater|EAU35287.1 atub|OJI86922.1 aver|OJI96341.1

Group16: abra|OJJ69532.1 abra|OJJ73245.1 abra|OJJ75746.1 abra|OJJ71793.1 acar|OOF90431.1 acar|OOF92663.1 acar|OOG00484.1 acar|OOF96473.1 acla|EAW06919.1 acla|EAW08051.1 afis|EAW16536.1 afis|EAW23087.1 afla|EED48561.1 afla|EED48728.1 afla|EED56133.1 afum|EAL88392.1 afum|EAL89133.1 agla|OJJ82786.1 agla|OJJ89597.1 akaw|GAA84479.1 akaw|GAA85117.1 akaw|GAA87935.1 akaw|GAA88383.1 aluc|OJZ81182.1 aluc|OJZ82371.1 aluc|OJZ86523.1 aluc|OJZ89199.1 anid|CBF75152.1 anid|CBF81741.1 anid|CBF82143.1 anig|EHA18500.1 anig|EHA23358.1 anig|EHA28393.1 anig|EHA22123.1 aory|EIT73145.1 aory|EIT73234.1 aory|EIT76572.1 aory|EIT79614.1 asyd|OJJ54322.1 asyd|OJJ59597.1 asyd|OJJ59975.1 asyd|OJJ56163.1 atub|OJI83999.1 atub|OJI91379.1 atub|OJI85591.1 aver|OJJ02316.1 aver|OJJ05102.1 aver|OJJ05276.1 aver|OJJ02671.1 awen|OJJ30245.1 awen|OJJ39992.1 awen|OJJ40286.1 ater|EAU36211.1 ater|EAU36599.1 atub|OJI88071.1 awen|OJJ36697.1 azon|OJJ47500.1 azon|OJJ49147.1 afla|EED56590.1 agla|OJJ79942.1 anig|EHA23359.1

Group17: abra|OJJ67548.1 abra|OJJ67738.1 abra|OJJ69732.1 abra|OJJ75389.1 abra|OJJ67871.1 acar|OOF90253.1 acar|OOF90937.1 acar|OOF94189.1 acar|OOF94221.1 acar|OOF94457.1 acla|EAW11677.1 acla|EAW12103.1 afla|EED45500.1 afla|EED48836.1 afla|EED51171.1 afla|EED52156.1 akaw|GAA88112.1 akaw|GAA90185.1 akaw|GAA92429.1 akaw|GAA92752.1 akaw|GAA91880.1 aluc|OJZ81382.1 aluc|OJZ81593.1 aluc|OJZ83164.1 aluc|OJZ92960.1 aluc|OJZ82019.1 anid|CBF73431.1 anid|CBF87554.1 anig|EHA18677.1 anig|EHA25848.1 anig|EHA26015.1 anig|EHA28228.1 anig|EHA22237.1 aory|EIT74393.1 aory|EIT75554.1 aory|EIT81271.1 aory|EIT82420.1 aory|EIT81347.1 ater|EAU30799.1 ater|EAU34705.1 atub|OJI84570.1 atub|OJI87418.1 atub|OJI87773.1 atub|OJI88590.1 atub|OJI91588.1 aver|OJJ08108.1 aver|OJJ08968.1 azon|OJJ42487.1 azon|OJJ45532.1 afis|EAW17283.1 afum|EAL86701.1 agla|OJJ81086.1 anid|CBF82821.1 asyd|OJJ52999.1 asyd|OJJ61577.1 aver|OJJ00899.1 awen|OJJ35752.1 afla|EED45454.1 anid|CBF80162.1 aver|OJI99518.1

Group18: abra|OJJ67797.1 abra|OJJ73515.1 abra|OJJ68679.1 abra|OJJ70328.1 abra|OJJ76402.1 acar|OOF91783.1 acar|OOF92181.1 acar|OOG00703.1 acla|EAW07449.1 acla|EAW12568.1 afis|EAW18727.1 afis|EAW22482.1 afis|EAW24082.1 afla|EED45270.1 afla|EED51391.1 afum|EAL90712.1 afum|EAL91937.1 agla|OJJ79953.1 agla|OJJ82286.1 agla|OJJ87548.1 akaw|GAA82322.1 akaw|GAA89832.1 akaw|GAA92583.1 akaw|GAA87464.1 aluc|OJZ81855.1 aluc|OJZ82597.1 aluc|OJZ88532.1 aluc|OJZ91810.1 anig|EHA18233.1 anig|EHA22369.1 anig|EHA27480.1 anig|EHA22803.1 aory|EIT73742.1 aory|EIT78822.1 aory|EIT80707.1 asyd|OJJ53887.1 asyd|OJJ62829.1 ater|EAU30215.1 ater|EAU30505.1 ater|EAU39231.1 ater|EAU38177.1 atub|OJI84423.1 atub|OJI88833.1 atub|OJI91117.1 atub|OJI86266.1 aver|OJI96872.1 aver|OJJ07985.1 awen|OJJ29718.1 awen|OJJ32268.1 awen|OJJ35101.1 acla|EAW14021.1 aluc|OJZ90857.1 anid|CBF88889.1 atub|OJI82853.1 azon|OJJ50007.1 afla|EED50684.1 afla|EED54508.1 acar|OOF99644.1 afum|EAL93452.1 aory|EIT78049.1

Group19: abra|OJJ67556.1 abra|OJJ67739.1 abra|OJJ69721.1 abra|OJJ75388.1 abra|OJJ67870.1 acar|OOF90043.1 acar|OOF90938.1 acar|OOF94195.1 acar|OOF94224.1 acar|OOF94458.1 acla|EAW11676.1 acla|EAW12104.1 afla|EED45513.1 afla|EED48837.1 afla|EED51172.1 afla|EED52163.1 akaw|GAA88108.1 akaw|GAA90174.1 akaw|GAA92430.1 akaw|GAA92751.1 akaw|GAA91879.1 aluc|OJZ81378.1 aluc|OJZ81592.1 aluc|OJZ92959.1 aluc|OJZ82020.1 aluc|OJZ83176.1 anid|CBF73444.1 anid|CBF82822.1 anid|CBF87553.1 anig|EHA18673.1 anig|EHA25849.1 anig|EHA26016.1 anig|EHA28250.1 anig|EHA22236.1 aory|EIT74418.1 aory|EIT75385.1 aory|EIT81285.1 aory|EIT82421.1 aory|EIT81323.1 asyd|OJJ52998.1 asyd|OJJ61578.1 atub|OJI84572.1 atub|OJI87417.1 atub|OJI87761.1 atub|OJI91584.1 atub|OJI88591.1 aver|OJJ00900.1 aver|OJJ08115.1 aver|OJJ08967.1 azon|OJJ42478.1 azon|OJJ45531.1 afis|EAW17282.1 afum|EAL86700.1 agla|OJJ81085.1 ater|EAU30800.1 ater|EAU34706.1 awen|OJJ35753.1 anid|CBF80164.1 aver|OJI99519.1

Group20: abra|OJJ67608.1 abra|OJJ70431.1 acar|OOF89840.1 acar|OOF91955.1 acla|EAW11023.1 acla|EAW14999.1 afis|EAW18722.1 afis|EAW19401.1 afis|EAW20837.1 afis|EAW24523.1 afla|EED45470.1 afla|EED46489.1 afla|EED50679.1 afum|EAL84797.1 afum|EAL86188.1 afum|EAL93446.1 agla|OJJ79209.1 agla|OJJ83714.1 akaw|GAA89124.1 akaw|GAA90149.1 aluc|OJZ81714.1 aluc|OJZ83199.1 anid|CBF74338.1 anid|CBF82877.1 aory|EIT72354.1 aory|EIT75736.1 asyd|OJJ57954.1 asyd|OJJ59255.1 asyd|OJJ61569.1 ater|EAU30785.1 ater|EAU36144.1 ater|EAU36801.1 atub|OJI87739.1 atub|OJI88468.1 aver|OJJ00889.1 aver|OJJ01866.1 awen|OJJ32698.1 awen|OJJ34547.1 azon|OJJ42767.1 azon|OJJ43003.1 anig|EHA25725.1 agla|OJJ78774.1 anig|EHA21062.1 abra|OJJ76408.1 acar|OOF99650.1 acla|EAW14020.1 agla|OJJ80462.1

akaw|GAA82327.1 aluc|OJZ91804.1 anid|CBF70690.1 anig|EHA22798.1 asyd|OJJ58469.1 ater|EAU38182.1 atub|OJI86258.1 aver|OJJ01070.1 awen|OJJ32266.1 azon|OJJ46779.1 azon|OJJ51803.1 asyd|OJJ51967.1

Group21: abra|OJJ67735.1 abra|OJJ68593.1 abra|OJJ73309.1 abra|OJJ77765.1 acar|OOF90932.1 acar|OOF97505.1 acla|EAW12049.1 acla|EAW12177.1 afis|EAW17018.1 afis|EAW19983.1 afla|EED45460.1 afla|EED45667.1 afla|EED48486.1 afla|EED52023.1 afum|EAL84933.1 afum|EAL86540.1 akaw|GAA89817.1 akaw|GAA92425.1 aluc|OJZ81597.1 aluc|OJZ81873.1 anid|CBF69451.1 anid|CBF80428.1 anid|CBF82812.1 anid|CBF83139.1 anid|CBF88295.1 anid|CBF89312.1 anid|CBF86046.1 anig|EHA20911.1 anig|EHA25844.1 anig|EHA28237.1 aory|EIT73076.1 aory|EIT75517.1 aory|EIT81451.1 aory|EIT82410.1 aory|EIT79328.1 asyd|OJJ61573.1 asyd|OJJ62373.1 ater|EAU29529.1 ater|EAU30794.1 ater|EAU31923.1 ater|EAU35431.1 ater|EAU36706.1 atub|OJI80342.1 atub|OJI87755.1 aver|OJI96507.1 aver|OJJ00893.1 aver|OJJ07788.1 aver|OJJ06454.1 agla|OJJ79464.1 atub|OJI88585.1 awen|OJJ38079.1 azon|OJJ46787.1 azon|OJJ47411.1 akaw|GAA90162.1 aluc|OJZ83185.1 abra|OJJ70061.1 anig|EHA21035.1 acar|OOF91380.1 anig|EHA28257.1

Group22: abra|OJJ67988.1 abra|OJJ68663.1 abra|OJJ71084.1 abra|OJJ76673.1 acar|OOF92030.1 acar|OOF92164.1 acar|OOF98052.1 acar|OOF99336.1 akaw|GAA82877.1 akaw|GAA83886.1 akaw|GAA89737.1 akaw|GAA89804.1 akaw|GAA90960.1 akaw|GAA92519.1 akaw|GAA92933.1 aluc|OJZ80647.1 aluc|OJZ81887.1 aluc|OJZ82526.1 aluc|OJZ82862.1 aluc|OJZ87444.1 aluc|OJZ91142.1 aluc|OJZ91532.1 anig|EHA18444.1 anig|EHA19537.1 anig|EHA19580.1 anig|EHA20668.1 anig|EHA21292.1 anig|EHA22353.1 anig|EHA22571.1 anig|EHA27186.1 anig|EHA27490.1 anig|EHA22492.1 atub|OJI81204.1 atub|OJI82798.1 atub|OJI83155.1 atub|OJI84440.1 atub|OJI85986.1 atub|OJI87502.1 atub|OJI89087.1 atub|OJI87377.1 atub|OJI87376.1 awen|OJJ30975.1 awen|OJJ34464.1 azon|OJJ42197.1 azon|OJJ43917.1 azon|OJJ50236.1 azon|OJJ45149.1 azon|OJJ45207.1 afis|EAW19725.1 afla|EED53110.1 afum|EAL86801.1 agla|OJJ84005.1 aory|EIT74790.1 ater|EAU37613.1 abra|OJJ77748.1 aluc|OJZ90809.1 agla|OJJ88178.1 azon|OJJ46566.1 azon|OJJ42198.1

Group23: abra|OJJ70376.1 abra|OJJ76944.1 acar|OOF91394.1 acar|OOF91893.1 acar|OOF94281.1 acla|EAW09924.1 acla|EAW12333.1 acla|EAW12657.1 acla|EAW13468.1 afis|EAW16889.1 afis|EAW18023.1 afis|EAW20581.1 afis|EAW21172.1 afla|EED47357.1 afla|EED52279.1 afla|EED54755.1 afum|EAL84483.1 afum|EAL86427.2 afum|EAL92192.1 afum|EAL91835.1 agla|OJJ79937.1 agla|OJJ80007.1 agla|OJJ88693.1 akaw|GAA87096.1 akaw|GAA88352.1 akaw|GAA91976.1 aluc|OJZ82303.1 aluc|OJZ84342.1 aluc|OJZ82638.1 aory|EIT79353.1 aory|EIT82885.1 asyd|OJJ54585.1 asyd|OJJ55452.1 ater|EAU35839.1 ater|EAU37103.1 atub|OJI88134.1 atub|OJI89726.1 atub|OJI88871.1 aver|OJJ06525.1 aver|OJJ07569.1 awen|OJJ29755.1 awen|OJJ33285.1 awen|OJJ36510.1 azon|OJJ42518.1 azon|OJJ46073.1 azon|OJJ50660.1 abra|OJJ69460.1 anid|CBF83321.1 anig|EHA28452.1 asyd|OJJ60544.1 aver|OJI99870.1 azon|OJJ44385.1 anid|CBF76070.1 anid|CBF80669.1 anig|EHA27432.1 ater|EAU29931.1 ater|EAU30227.1 anig|EHA27678.1 aory|EIT74323.1

Group24: abra|OJJ69144.1 abra|OJJ70352.1 abra|OJJ77612.1 acar|OOF91158.1 acar|OOF99116.1 acla|EAW06570.1 acla|EAW08667.1 acla|EAW13589.1 afis|EAW15744.1 afis|EAW16026.1 afis|EAW20014.1 afum|EAL84904.1 afum|EAL85604.1 afum|EAL88655.1 agla|OJJ81906.1 agla|OJJ86054.1 agla|OJJ87984.1 akaw|GAA84019.1 akaw|GAA86313.1 aluc|OJZ80922.1 aluc|OJZ86130.1 anid|CBF71034.1 anid|CBF77968.1 anid|CBF84719.1 anig|EHA19425.1 anig|EHA24491.1 aory|EIT75778.1 aory|EIT81521.1 aory|EIT83013.1 asyd|OJJ52664.1 asyd|OJJ53328.1 asyd|OJJ55266.1 asyd|OJJ60058.1 ater|EAU29366.1 ater|EAU30124.1 ater|EAU32401.1 atub|OJI79926.1 atub|OJI82658.1 aver|OJJ02231.1 aver|OJJ04691.1 aver|OJJ06960.1 aver|OJJ08416.1 awen|OJJ30028.1 awen|OJJ38444.1 azon|OJJ43750.1 azon|OJJ45430.1 azon|OJJ48427.1 ater|EAU33615.1 awen|OJJ30431.1 awen|OJJ41731.1 afla|EED46021.1 afla|EED56407.1 anid|CBF75022.1 acla|EAW12683.1 afis|EAW17846.1 afum|EAL91655.1 ater|EAU33048.1 afla|EED51306.1

Group25: abra|OJJ70961.1 abra|OJJ76031.1 acar|OOF98169.1 acar|OOF99897.1 acla|EAW09101.1 acla|EAW09782.1 acla|EAW10522.1 afis|EAW21356.1 afis|EAW25044.1 afis|EAW25699.1 afis|EAW25729.1 afla|EED45847.1 afla|EED55407.1 afla|EED55771.1 afum|EAL84400.2 afum|EAL89712.1 afum|EAL92371.2 agla|OJJ78663.1 agla|OJJ87377.1 agla|OJJ89150.1 agla|OJJ88275.1 akaw|GAA82986.1 akaw|GAA91201.1 aluc|OJZ81536.1 aluc|OJZ87571.1 aluc|OJZ89483.1 anid|CBF76249.1 anid|CBF85912.1 anig|EHA20773.1 anig|EHA23111.1 anig|EHA25900.1 aory|EIT75461.1 aory|EIT80822.1 aory|EIT75500.1 aory|EIT76538.1 asyd|OJJ59042.1 asyd|OJJ60969.1 asyd|OJJ64398.1 ater|EAU35202.1 ater|EAU35794.1 ater|EAU36445.1 ater|EAU37742.1 atub|OJI81078.1 atub|OJI85318.1 aver|OJI98494.1 aver|OJJ00294.1 aver|OJJ01637.1 awen|OJJ37057.1 awen|OJJ39342.1 awen|OJJ40655.1 azon|OJJ51688.1 ater|EAU36998.1 afla|EED54858.1 anid|CBF69709.1 atub|OJI88650.1 aory|EIT79380.1

Group26: abra|OJJ71513.1 abra|OJJ73019.1 abra|OJJ77809.1 acar|OOF96188.1 acar|OOF96228.1 acar|OOF96266.1 acar|OOF98958.1 acla|EAW07995.1 acla|EAW11308.1 acla|EAW15070.1 afis|EAW18195.1 afis|EAW23030.1 afis|EAW20042.1 afla|EED45337.1 afla|EED54986.1 afla|EED56290.1 afum|EAL85162.1 afum|EAL88452.1 afum|EAL87556.1 agla|OJJ85896.1 agla|OJJ89448.1 agla|OJJ88920.1 akaw|GAA83840.1 akaw|GAA85596.1 akaw|GAA89403.1 aluc|OJZ83533.1 aluc|OJZ86251.1 aluc|OJZ90876.1 anid|CBF80914.1 anid|CBF86102.1 anig|EHA19596.1 anig|EHA24355.1 anig|EHA21614.1 aory|EIT73286.1 aory|EIT80288.1 asyd|OJJ56326.1 asyd|OJJ64481.1 ater|EAU29627.1 ater|EAU33801.1 atub|OJI82424.1 atub|OJI82873.1 atub|OJI84291.1 aver|OJI98582.1 aver|OJJ06387.1 awen|OJJ30557.1 awen|OJJ35716.1 azon|OJJ44174.1 azon|OJJ45510.1 azon|OJJ46319.1 anid|CBF84784.1 anid|CBF84786.1 aory|EIT73629.1 asyd|OJJ63232.1 ater|EAU38649.1 aver|OJI97724.1 awen|OJJ31798.1

Group27: abra|OJJ67095.1 abra|OJJ67686.1 abra|OJJ69631.1 abra|OJJ76432.1 acar|OOF92679.1 acar|OOF94074.1 acar|OOF94144.1 acar|OOF99676.1 acla|EAW10021.1 acla|EAW12377.1 acla|EAW13992.1 afis|EAW16823.1 afis|EAW18700.1 afis|EAW22316.1 afla|EED50648.1 afla|EED55075.1 afum|EAL86374.1 afum|EAL90874.1 afum|EAL93422.1 agla|OJJ79102.1 agla|OJJ82361.1 akaw|GAA82347.1 akaw|GAA89198.1 akaw|GAA91066.1 akaw|GAA91611.1 aluc|OJZ81632.1 aluc|OJZ82438.1 aluc|OJZ87880.1 aluc|OJZ91780.1 anid|CBF70637.1 anid|CBF82989.1 anig|EHA20183.1 anig|EHA25811.1 anig|EHA28335.1 aory|EIT78077.1 aory|EIT81953.1 aory|EIT82431.1 asyd|OJJ52074.1 asyd|OJJ54406.1 asyd|OJJ58493.1 asyd|OJJ64734.1 ater|EAU29368.1 ater|EAU35424.1 ater|EAU38206.1 atub|OJI80771.1 atub|OJI86236.1 atub|OJI88003.1 atub|OJI88548.1 aver|OJI98838.1 aver|OJJ01096.1 aver|OJJ03029.1 awen|OJJ32183.1 awen|OJJ40928.1 azon|OJJ46764.1 anig|EHA22777.1

Group28: abra|OJJ68117.1 abra|OJJ77078.1 abra|OJJ68815.1 acar|OOF90616.1 acar|OOF92960.1 acar|OOF93911.1 afis|EAW17741.1 afis|EAW17974.1 afla|EED45883.1 afla|EED55321.1 afum|EAL91555.1 afum|EAL91783.1 agla|OJJ81756.1 agla|OJJ85465.1 agla|OJJ89268.1 akaw|GAA86898.1 akaw|GAA86975.1 akaw|GAA90440.1 aluc|OJZ84488.1 aluc|OJZ85787.1 aluc|OJZ88708.1 aluc|OJZ90021.1 anid|CBF71442.1 anid|CBF84572.1 anig|EHA24771.1 anig|EHA27397.1 anig|EHA27792.1 aory|EIT75581.1 aory|EIT76204.1 asyd|OJJ53389.1 asyd|OJJ54954.1 ater|EAU30960.1 ater|EAU35114.1 ater|EAU36891.1 atub|OJI79571.1 atub|OJI84786.1 atub|OJI89592.1 atub|OJI91296.1 aver|OJI99661.1 aver|OJJ03907.1 aver|OJJ07242.1 aver|OJJ08354.1 awen|OJJ32944.1 awen|OJJ39601.1 azon|OJJ42107.1 azon|OJJ42757.1 azon|OJJ42786.1 azon|OJJ44596.1 azon|OJJ47560.1 acla|EAW11105.1 afla|EED52131.1 acla|EAW12809.1 ater|EAU36942.1 awen|OJJ32177.1 aory|EIT74465.1

Group29: anid|CBF71484.1 anid|CBF71852.1 anid|CBF76028.1 anid|CBF80836.1 anid|CBF81153.1 anid|CBF81850.1 anid|CBF84162.1 anid|CBF84238.1 anid|CBF86037.1 anid|CBF89606.1 anid|CBF80194.1 anid|CBF80847.1 anid|CBF84163.1 anid|CBF76085.1 anid|CBF71848.1 anid|CBF80228.1 anid|CBF81852.1 anid|CBF80226.1 anid|CBF82237.1 anid|CBF81848.1 anid|CBF86356.1 anid|CBF80843.1 anid|CBF80845.1 anid|CBF76165.1 anid|CBF80196.1 anid|CBF76161.1 anid|CBF80272.1 anid|CBF80253.1 anid|CBF80230.1 anid|CBF82238.1 anid|CBF80252.1 anid|CBF80219.1 anid|CBF89288.1 anid|CBF81155.1 anid|CBF76163.1 anid|CBF84359.1 anid|CBF89285.1 anid|CBF80218.1 anid|CBF82246.1 anid|CBF87221.1 anid|CBF81844.1 anid|CBF89286.1 anid|CBF81853.1 anid|CBF71850.1 anid|CBF74006.1 anid|CBF78409.1 anid|CBF87223.1 anid|CBF81157.1 anid|CBF79347.1 acar|OOF89759.1 afis|EAW17204.1 afis|EAW17205.1 afis|EAW22029.1 agla|OJJ82974.1 aory|EIT76232.1

Group30: abra|OJJ68096.1 abra|OJJ70248.1 abra|OJJ73110.1 abra|OJJ74007.1 acar|OOF92551.1 acar|OOF93899.1 acar|OOG00337.1 acar|OOG01129.1 afla|EED44907.1 afla|EED44932.1 afla|EED56562.1 agla|OJJ78703.1 agla|OJJ88418.1 akaw|GAA86542.1 akaw|GAA86880.1 akaw|GAA88056.1 akaw|GAA92925.1 akaw|GAA87488.1 aluc|OJZ81318.1 aluc|OJZ82516.1 aluc|OJZ85090.1 aluc|OJZ88037.1 aluc|OJZ90004.1 anid|CBF74975.1 anid|CBF78035.1 anid|CBF81473.1 anig|EHA17810.1 anig|EHA18624.1 anig|EHA25311.1 anig|EHA27379.1 anig|EHA20903.1 aory|EIT73500.1 aory|EIT73507.1 asyd|OJJ55224.1 asyd|OJJ59448.1 asyd|OJJ60089.1 asyd|OJJ61384.1 asyd|OJJ57439.1 ater|EAU39548.1 ater|EAU39587.1 atub|OJI80509.1 atub|OJI84804.1 atub|OJI91521.1 atub|OJI90644.1 aver|OJJ00711.1 aver|OJJ04176.1 aver|OJJ05739.1 aver|OJJ02824.1 awen|OJJ33639.1 awen|OJJ40047.1 acla|EAW07039.1 afis|EAW16665.1 afum|EAL89269.1 azon|OJJ50123.1

Group31: abra|OJJ69067.1 abra|OJJ70519.1 abra|OJJ77965.1 acar|OOF91229.1 acar|OOF97409.1 acar|OOF98850.1 acla|EAW08156.1 acla|EAW08763.1 afis|EAW15666.1 afis|EAW16089.1 afis|EAW23194.1 afla|EED46889.1 afla|EED47060.1 afla|EED48625.1 afum|EAL85682.1 afum|EAL88289.1 afum|EAL88721.1 agla|OJJ81804.1 agla|OJJ86130.1 agla|OJJ86345.1 akaw|GAA83366.1 akaw|GAA83698.1 akaw|GAA90597.1 aluc|OJZ83864.1 aluc|OJZ91035.1 aluc|OJZ86055.1 anid|CBF71178.1 anid|CBF74704.1 anid|CBF75129.1 anig|EHA19005.1 anig|EHA19734.1 anig|EHA24556.1 aory|EIT72496.1 aory|EIT73097.1 asyd|OJJ55863.1 asyd|OJJ60218.1 asyd|OJJ59982.1 ater|EAU32505.1 ater|EAU33161.1 ater|EAU36321.1 atub|OJI79852.1 atub|OJI83031.1 atub|OJI90232.1 aver|OJJ02309.1 aver|OJJ04050.1 aver|OJJ04804.1 awen|OJJ30341.1 awen|OJJ38341.1 awen|OJJ36585.1 azon|OJJ42517.1 azon|OJJ43041.1 aory|EIT72708.1 acla|EAW06676.1 azon|OJJ47622.1

Group32: abra|OJJ66128.1 abra|OJJ70165.1 abra|OJJ77944.1 acar|OOF90574.1 acar|OOF94003.1 afis|EAW24242.1 afis|EAW25195.1 afum|EAL85910.2 afum|EAL89210.1 anid|CBF71201.1 anid|CBF74964.1 asyd|OJJ52805.1 asyd|OJJ59802.1 ater|EAU29607.1 ater|EAU33975.1 ater|EAU34693.1 atub|OJI87381.1 atub|OJI87443.1 aver|OJI96126.1 aver|OJJ03002.1 aver|OJI98905.1 aver|OJI99605.1 aver|OJJ03852.1 afla|EED47392.1 akaw|GAA83719.1 aluc|OJZ91014.1 anid|CBF89341.1 aory|EIT73813.1 aory|EIT76002.1 atub|OJI83005.1 aver|OJJ05262.1 awen|OJJ36264.1 azon|OJJ48048.1 agla|OJJ85483.1 akaw|GAA86612.1 akaw|GAA92274.1 aluc|OJZ85163.1 aluc|OJZ90168.1 atub|OJI80439.1 atub|OJI81596.1 anig|EHA19716.1 ater|EAU36085.1 aver|OJJ08785.1 afum|EAL91911.1 acar|OOF93333.1 acar|OOF98718.1 asyd|OJJ55517.1 aver|OJJ06595.1 afla|EED45160.1 afla|EED54076.1 anid|CBF82991.1 azon|OJJ46519.1 asyd|OJJ52549.1

Group33: abra|OJJ66203.1 abra|OJJ68990.1 abra|OJJ74346.1 acar|OOF91561.1 acar|OOF91930.1 acar|OOF95383.1 acla|EAW07400.1 acla|EAW14202.1 afis|EAW19080.1 afis|EAW22426.1 afla|EED50168.1 afla|EED52504.1 afum|EAL91073.1 afum|EAL90765.1 afum|EAL93809.1 agla|OJJ84526.1 agla|OJJ87597.1 agla|OJJ84718.1 akaw|GAA86684.1 akaw|GAA89082.1 aluc|OJZ80508.1 aluc|OJZ85975.1 aluc|OJZ89784.1 anid|CBF76870.1 anid|CBF79672.1 anid|CBF88817.1 anig|EHA24631.1 anig|EHA26999.1 anig|EHA25684.1

aory|EIT76001.1 aory|EIT82781.1 asyd|OJJ52200.1 asyd|OJJ58034.1 asyd|OJJ61321.1 asyd|OJJ62887.1 ater|EAU32126.1 ater|EAU32863.1 ater|EAU32522.1 ater|EAU39281.1 atub|OJI85021.1 atub|OJI88421.1 aver|OJI96939.1 aver|OJJ00646.1 aver|OJJ04365.1 aver|OJJ05950.1 aver|OJJ07745.1 awen|OJJ32472.1 awen|OJJ35005.1 awen|OJJ34090.1 azon|OJJ45647.1 azon|OJJ46232.1 akaw|GAA90510.1 atub|OJI79765.1

Group34: abra|OJJ71614.1 abra|OJJ77069.1 acar|OOF92952.1 acar|OOF96330.1 acla|EAW13507.1 acla|EAW15156.1 afis|EAW20123.1 afis|EAW21377.1 afis|EAW20124.1 afis|EAW20605.1 afla|EED47726.1 afla|EED57252.1 afum|EAL84455.1 afum|EAL85242.1 agla|OJJ88636.1 agla|OJJ88823.1 akaw|GAA86984.1 akaw|GAA89313.1 aluc|OJZ84477.1 aluc|OJZ86355.1 anid|CBF84947.1 anid|CBF86944.1 anig|EHA21535.1 anig|EHA27785.1 aory|EIT73980.1 aory|EIT77148.1 asyd|OJJ55590.1 asyd|OJJ63148.1 ater|EAU35261.1 ater|EAU38740.1 atub|OJI84179.1 atub|OJI89602.1 aver|OJI97806.1 aver|OJJ06669.1 awen|OJJ31893.1 awen|OJJ33448.1 azon|OJJ46243.1 azon|OJJ50705.1 abra|OJJ71611.1 acar|OOF96327.1 acla|EAW15155.1 afla|EED57253.1 afum|EAL85241.2 agla|OJJ88828.1 akaw|GAA89316.1 anid|CBF84945.1 aory|EIT74161.1 asyd|OJJ63149.1 atub|OJI84182.1 aver|OJI97805.1 awen|OJJ31892.1 azon|OJJ42421.1 azon|OJJ46244.1

Group35: afis|EAW16487.1 afis|EAW19960.1 afla|EED48721.1 afla|EED49671.1 afum|EAL84955.1 afum|EAL89081.1 agla|OJJ79324.1 agla|OJJ82857.1 agla|OJJ88164.1 aory|EIT73140.1 aory|EIT80950.1 aory|EIT75645.1 aory|EIT79202.1 ater|EAU35270.1 ater|EAU36551.1 abra|OJJ68065.1 acla|EAW06870.1 afla|EED45908.1 akaw|GAA90894.1 aluc|OJZ82935.1 anid|CBF75512.1 anid|CBF84598.1 anig|EHA21355.1 anig|EHA22760.1 asyd|OJJ53379.1 atub|OJI89164.1 aver|OJJ02547.1 awen|OJJ29993.1 azon|OJJ49197.1 abra|OJJ71770.1 akaw|GAA85137.1 aluc|OJZ86500.1 anig|EHA22144.1 atub|OJI84026.1 azon|OJJ48613.1 abra|OJJ78207.1 acar|OOF98647.1 acla|EAW11241.1 afis|EAW18262.1 afum|EAL87626.1 akaw|GAA85819.1 aluc|OJZ91280.1 anid|CBF79531.1 anig|EHA19965.1 asyd|OJJ57895.1 ater|EAU33270.1 atub|OJI83270.1 aver|OJJ04500.1 awen|OJJ30636.1 afla|EED47227.1 asyd|OJJ59741.1 awen|OJJ40346.1 aver|OJJ08364.1

Group36: abra|OJJ72217.1 abra|OJJ77905.1 abra|OJJ77236.1 abra|OJJ77528.1 acar|OOF93116.1 acar|OOF98878.1 acar|OOF99197.1 acar|OOF96839.1 acla|EAW06700.1 acla|EAW13243.1 afis|EAW16233.1 afis|EAW16245.1 afis|EAW17393.1 afla|EED49461.1 afla|EED53306.1 afla|EED53348.1 afum|EAL88856.1 afum|EAL88873.1 afum|EAL91231.1 akaw|GAA83757.1 akaw|GAA84092.1 akaw|GAA90707.1 aluc|OJZ81007.1 aluc|OJZ84646.1 aluc|OJZ90974.1 anig|EHA19353.1 anig|EHA19676.1 anig|EHA27930.1 aory|EIT74908.1 aory|EIT75241.1 aory|EIT79470.1 atub|OJI82578.1 atub|OJI82965.1 atub|OJI89434.1 awen|OJJ30211.1 awen|OJJ30227.1 awen|OJJ37422.1 afla|EED48174.1 agla|OJJ80414.1 akaw|GAA84718.1 aluc|OJZ86969.1 anig|EHA21681.1 aory|EIT83042.1 atub|OJI83552.1 awen|OJJ31302.1 agla|OJJ86254.1 anid|CBF78900.1 asyd|OJJ64603.1 ater|EAU37455.1 aver|OJI98700.1 acla|EAW06515.1 ater|EAU33711.1

Group37: abra|OJJ72940.1 abra|OJJ77077.1 acar|OOF92763.1 acar|OOF92959.1 acar|OOF97146.1 afis|EAW18980.1 afis|EAW21099.1 afis|EAW24628.1 afla|EED53187.1 afla|EED54887.1 agla|OJJ80287.1 agla|OJJ82940.1 akaw|GAA86976.1 akaw|GAA91285.1 aluc|OJZ82784.1 aluc|OJZ84487.1 anig|EHA18922.1 anig|EHA21226.1 aory|EIT75171.1 aory|EIT79411.1 asyd|OJJ58358.1 asyd|OJJ60157.1 ater|EAU34862.1 ater|EAU37837.1 atub|OJI89016.1 atub|OJI89593.1 aver|OJJ02049.1 aver|OJJ04107.1 awen|OJJ31588.1 awen|OJJ37322.1 azon|OJJ47304.1 azon|OJJ47677.1 abra|OJJ70575.1 acla|EAW08374.1 afis|EAW23403.1 afla|EED47010.1 afum|EAL88067.1 agla|OJJ86391.1 akaw|GAA83419.1 aluc|OJZ83919.1 anid|CBF74791.1 anig|EHA19052.1 aory|EIT72601.1 ater|EAU36372.1 atub|OJI90290.1 awen|OJJ39474.1 acla|EAW14304.1 afum|EAL93716.2 anid|CBF69653.1 asyd|OJJ59075.1 aver|OJJ01668.1 anig|EHA27791.1

Group38: abra|OJJ65887.1 abra|OJJ73728.1 abra|OJJ73840.1 abra|OJJ75062.1 acar|OOF94948.1 acar|OOF97272.1 acar|OOG00894.1 acar|OOG00991.1 acla|EAW07735.1 acla|EAW09321.1 afis|EAW21824.1 afis|EAW24409.1 afla|EED45566.1 afla|EED50029.1 afum|EAL86078.1 afum|EAL92840.1 agla|OJJ83601.1 agla|OJJ85455.1 agla|OJJ89020.1 akaw|GAA86017.1 akaw|GAA88434.1 akaw|GAA88674.1 akaw|GAA89458.1 aluc|OJZ88209.1 aluc|OJZ88322.1 aluc|OJZ90245.1 aluc|OJZ92628.1 anid|CBF84638.1 anid|CBF87998.1 anig|EHA17956.1 anig|EHA18058.1 anig|EHA26362.1 aory|EIT74733.1 aory|EIT75750.1 asyd|OJJ53366.1 asyd|OJJ63506.1 ater|EAU38301.1 ater|EAU38950.1 atub|OJI81673.1 atub|OJI87058.1 atub|OJI90810.1 atub|OJI90917.1 aver|OJI97424.1 aver|OJJ08381.1 awen|OJJ32729.1 awen|OJJ37942.1 awen|OJJ38966.1 awen|OJJ41210.1 azon|OJJ46022.1 azon|OJJ48676.1 aory|EIT76918.1

Group39: abra|OJJ66342.1 abra|OJJ71162.1 abra|OJJ75572.1 acar|OOF89825.1 acar|OOF94094.1 acar|OOF94165.1 acar|OOF97998.1 acla|EAW09994.1 acla|EAW10337.1 afis|EAW21096.1 afis|EAW25252.1 afla|EED51678.1 afla|EED53719.1 afla|EED54679.1 afum|EAL86170.1 afum|EAL89914.1 afum|EAL92119.1 agla|OJJ80114.1 agla|OJJ89225.1 akaw|GAA82817.1 akaw|GAA84323.1 akaw|GAA90216.1 aluc|OJZ83127.1 aluc|OJZ87378.1 aluc|OJZ89020.1 anid|CBF80117.1 anid|CBF85345.1 anig|EHA20610.1 anig|EHA23510.1 anig|EHA28188.1 aory|EIT75350.1 aory|EIT78900.1 aory|EIT80684.1 asyd|OJJ54726.1 asyd|OJJ64027.1 atub|OJI81268.1 atub|OJI85769.1 atub|OJI87806.1 aver|OJI98113.1 aver|OJJ07441.1 awen|OJJ36352.1 awen|OJJ40966.1 awen|OJJ38784.1 ater|EAU34396.1 azon|OJJ44550.1 abra|OJJ67575.1 acar|OOF90092.1 akaw|GAA89104.1 anig|EHA25706.1 atub|OJI88443.1 afis|EAW24500.1

Group40: abra|OJJ73400.1 abra|OJJ73868.1 acar|OOG00598.1 acar|OOG01011.1 acla|EAW07007.1 acla|EAW07756.1 afis|EAW16623.1 afis|EAW19494.1 afis|EAW22793.1 afla|EED44972.1 afla|EED48978.1 afla|EED50061.1 afla|EED56714.1 afum|EAL87000.1 afum|EAL89214.1 afum|EAL90406.1 agla|OJJ83569.1 agla|OJJ85298.1 akaw|GAA83092.1 akaw|GAA89474.1 akaw|GAA89692.1 aluc|OJZ80692.1 aluc|OJZ88189.1 aluc|OJZ88639.1 anid|CBF81984.1 anid|CBF87954.1 anig|EHA17940.1 anig|EHA18328.1 anig|EHA27323.1

aory|EIT73515.1 aory|EIT73856.1 aory|EIT74647.1 aory|EIT79983.1 asyd|OJJ56154.1 asyd|OJJ59777.1 asyd|OJJ63479.1 ater|EAU34806.1 ater|EAU36732.1 ater|EAU38918.1 atub|OJI87552.1 atub|OJI90788.1 atub|OJI91225.1 aver|OJI97451.1 aver|OJJ02509.1 aver|OJJ05094.1 awen|OJJ36912.1 awen|OJJ38940.1 azon|OJJ45543.1 azon|OJJ49714.1 anid|CBF75442.1

Group41: abra|OJJ65606.1 abra|OJJ68131.1 abra|OJJ68372.1 abra|OJJ69834.1 acar|OOF90085.1 acar|OOF93362.1 acar|OOG00535.1 afla|EED45615.1 afla|EED45682.1 afla|EED54532.1 agla|OJJ80470.1 agla|OJJ80842.1 agla|OJJ83418.1 akaw|GAA86911.1 akaw|GAA87826.1 aluc|OJZ85462.1 aluc|OJZ90034.1 anig|EHA25004.1 anig|EHA25073.1 anig|EHA27410.1 aory|EIT75511.1 aory|EIT75540.1 aory|EIT78869.1 aory|EIT81830.1 asyd|OJJ52349.1 asyd|OJJ56735.1 asyd|OJJ60489.1 atub|OJI80101.1 atub|OJI84772.1 aver|OJI99815.1 aver|OJJ05882.1 aver|OJJ05988.1 awen|OJJ32535.1 awen|OJJ33535.1 awen|OJJ34624.1 awen|OJJ35129.1 awen|OJJ40106.1 acla|EAW07472.1 afis|EAW17054.1 afum|EAL86551.2 akaw|GAA83834.1 aluc|OJZ90883.1 anid|CBF86106.1 ater|EAU30279.1 atub|OJI82467.1 anig|EHA19603.1 atub|OJI82878.1 azon|OJJ47403.1 afla|EED57508.1

Group42: abra|OJJ65781.1 abra|OJJ69519.1 abra|OJJ69805.1 acar|OOF93570.1 acar|OOF94236.1 acla|EAW09965.1 acla|EAW15041.1 afis|EAW17149.1 afis|EAW21123.1 afis|EAW24564.1 afla|EED50957.1 afla|EED51796.1 afla|EED54117.1 afla|EED57909.1 afum|EAL86225.1 afum|EAL92146.1 agla|OJJ78698.1 agla|OJJ79809.1 agla|OJJ80088.1 akaw|GAA87854.1 akaw|GAA88307.1 akaw|GAA90882.1 aluc|OJZ82358.1 aluc|OJZ82950.1 aluc|OJZ85496.1 anid|CBF77670.1 anid|CBF87020.1 anig|EHA21430.1 anig|EHA25029.1 anig|EHA28405.1 aory|EIT76140.1 aory|EIT77042.1 aory|EIT81685.1 aory|EIT82334.1 asyd|OJJ53106.1 asyd|OJJ60460.1 ater|EAU32188.1 ater|EAU36777.1 atub|OJI80069.1 atub|OJI88083.1 atub|OJI89182.1 aver|OJI99780.1 aver|OJJ03746.1 awen|OJJ36453.1 awen|OJJ37008.1 awen|OJJ40041.1 azon|OJJ44424.1 azon|OJJ50127.1 aver|OJI99293.1

Group43: abra|OJJ67343.1 abra|OJJ74161.1 abra|OJJ74904.1 acar|OOF92471.1 acar|OOF95566.1 acar|OOF95902.1 afis|EAW18497.1 afis|EAW23515.1 afis|EAW24769.1 afum|EAL89410.1 afum|EAL93187.1 agla|OJJ83565.1 agla|OJJ85096.1 agla|OJJ86763.1 agla|OJJ88214.1 akaw|GAA83518.1 akaw|GAA86474.1 akaw|GAA89723.1 aluc|OJZ80660.1 aluc|OJZ85015.1 aluc|OJZ92477.1 anid|CBF70261.1 anid|CBF89705.1 anig|EHA25236.1 anig|EHA26493.1 anig|EHA27172.1 aory|EIT77595.1 aory|EIT78639.1 aory|EIT80122.1 asyd|OJJ58753.1 asyd|OJJ62252.1 asyd|OJJ62467.1 atub|OJI80582.1 atub|OJI86908.1 atub|OJI87516.1 aver|OJI96362.1 aver|OJJ01361.1 awen|OJJ31051.1 awen|OJJ37928.1 awen|OJJ39840.1 acla|EAW08468.1 afla|EED49731.1 afum|EAL87948.1 anid|CBF89382.1 ater|EAU35304.1 azon|OJJ43252.1 ater|EAU34034.1 aver|OJI96602.1 afla|EED57415.1

Group44: abra|OJJ68123.1 abra|OJJ69603.1 abra|OJJ72218.1 acar|OOF92701.1 acar|OOF93917.1 acar|OOF96841.1 acla|EAW12866.1 acla|EAW14975.1 afis|EAW17815.1 afis|EAW24499.1 afla|EED45599.1 afla|EED51849.1 afla|EED54703.1 afum|EAL86169.1 afum|EAL91624.1 agla|OJJ78971.1 agla|OJJ85550.1 agla|OJJ81686.1 akaw|GAA84717.1 akaw|GAA86904.1 akaw|GAA91591.1 aluc|OJZ82414.1 aluc|OJZ86970.1 aluc|OJZ90027.1 anig|EHA21680.1 anig|EHA27403.1 anig|EHA28353.1 aory|EIT75721.1 aory|EIT78771.1 aory|EIT81733.1 asyd|OJJ54895.1 asyd|OJJ58303.1 asyd|OJJ56516.1 asyd|OJJ56725.1 atub|OJI83551.1 atub|OJI84780.1 atub|OJI88027.1 aver|OJJ02109.1 aver|OJJ05998.1 aver|OJJ06206.1 awen|OJJ32626.1 awen|OJJ36932.1 anid|CBF73850.1 ater|EAU30287.1 azon|OJJ50345.1 anid|CBF80089.1 ater|EAU31562.1 awen|OJJ33657.1 ater|EAU29952.1

Group45: abra|OJJ68536.1 abra|OJJ72481.1 acar|OOF90150.1 acar|OOF94753.1 acla|EAW10385.1 acla|EAW12826.1 afis|EAW17874.1 afis|EAW17991.1 afis|EAW25211.1 afla|EED45114.1 afla|EED51912.1 afla|EED46241.1 afum|EAL89872.1 afum|EAL91801.1 afum|EAL91682.1 agla|OJJ80251.1 agla|OJJ83074.1 akaw|GAA86216.1 akaw|GAA91315.1 aluc|OJZ82750.1 aluc|OJZ90468.1 anid|CBF79986.1 anid|CBF89933.1 anig|EHA21199.1 anig|EHA23927.1 aory|EIT73554.1 aory|EIT76181.1 aory|EIT76362.1 ater|EAU31453.1 ater|EAU37217.1 ater|EAU31502.1 ater|EAU33093.1 atub|OJI81892.1 atub|OJI88986.1 aver|OJI96221.1 aver|OJJ02185.1 awen|OJJ41004.1 awen|OJJ41694.1 azon|OJJ45116.1 azon|OJJ46814.1 azon|OJJ48270.1 azon|OJJ48393.1 acla|EAW12711.1 asyd|OJJ58240.1 asyd|OJJ62114.1 awen|OJJ41555.1 anid|CBF69461.1 aver|OJJ01892.1 asyd|OJJ59277.1

Group46: abra|OJJ72575.1 abra|OJJ77969.1 acar|OOF94684.1 acar|OOF98846.1 akaw|GAA82423.1 akaw|GAA87621.1 akaw|GAA83693.1 akaw|GAA85201.1 aluc|OJZ90556.1 aluc|OJZ91039.1 anig|EHA19738.1 anig|EHA24003.1 asyd|OJJ54419.1 asyd|OJJ59096.1 atub|OJI81979.1 atub|OJI83035.1 acla|EAW06671.1 afis|EAW16094.1 afla|EED48630.1 afla|EED56583.1 afum|EAL88726.1 agla|OJJ88578.1 anid|CBF85458.1 aory|EIT73102.1 aory|EIT79674.1 asyd|OJJ58704.1 asyd|OJJ64100.1 ater|EAU30075.1 ater|EAU33170.1 aver|OJI98183.1 aver|OJJ01316.1 awen|OJJ30336.1 awen|OJJ40588.1 azon|OJJ42310.1 azon|OJJ45364.1 abra|OJJ76510.1 acar|OOF99461.1 afla|EED54610.1 agla|OJJ87529.1 aluc|OJZ91695.1 anig|EHA22708.1 aory|EIT78921.1 ater|EAU38073.1 atub|OJI86149.1 aver|OJJ01687.1 aver|OJJ06547.1 awen|OJJ36612.1 agla|OJJ86127.1 anid|CBF70342.1

Group47: acar|OOF91492.1 acar|OOF98333.1 acar|OOF92665.1 acar|OOF93877.1 afis|EAW15510.1 afis|EAW17137.1 afis|EAW16415.1 afis|EAW16568.1 afis|EAW17138.1 afis|EAW25618.1 afum|EAL89039.1 afum|EAL89165.1 anid|CBF80635.1 anid|CBF89254.1 ater|EAU31707.1 ater|EAU36836.1 abra|OJJ74339.1 acla|EAW07389.1 afis|EAW22415.1 afla|EED52493.1 afum|EAL90777.1 agla|OJJ87679.1 akaw|GAA86691.1 aluc|OJZ89792.1 anid|CBF88800.1 anig|EHA27005.1 aory|EIT82748.1 asyd|OJJ62896.1 ater|EAU39292.1

atub|OJI85014.1 aver|OJI96948.1 awen|OJJ34994.1 azon|OJJ45637.1 agla|OJJ82753.1 anig|EHA22499.1 atub|OJI84627.1 afis|EAW19287.1 acla|EAW06953.1 awen|OJJ40251.1 azon|OJJ49111.1 acar|OOG00127.1 akaw|GAA86019.1 aluc|OJZ90247.1 asyd|OJJ61228.1 aver|OJJ00551.1 anid|CBF76692.1 akaw|GAA91465.1 aluc|OJZ82070.1 afum|EAL84308.2

Group48: abra|OJJ66388.1 abra|OJJ69074.1 acla|EAW08755.1 acla|EAW11763.1 afis|EAW15677.1 afis|EAW23796.1 afis|EAW25170.1 afla|EED46854.1 afla|EED51603.1 afla|EED53628.1 afum|EAL85670.1 afum|EAL87230.1 afum|EAL89843.1 akaw|GAA90254.1 akaw|GAA90605.1 aluc|OJZ83086.1 aluc|OJZ86063.1 anid|CBF71157.1 anid|CBF79858.1 anid|CBF85251.1 aory|EIT72587.1 aory|EIT74962.1 aory|EIT80702.1 asyd|OJJ51987.1 asyd|OJJ54461.1 asyd|OJJ54551.1 asyd|OJJ55842.1 asyd|OJJ58159.1 asyd|OJJ59172.1 asyd|OJJ62925.1 asyd|OJJ64763.1 ater|EAU31334.1 ater|EAU32484.1 ater|EAU34334.1 atub|OJI79860.1 atub|OJI87858.1 aver|OJI98064.1 aver|OJJ04243.1 aver|OJJ04785.1 aver|OJJ07708.1 awen|OJJ30141.1 awen|OJJ33753.1 awen|OJJ38354.1 azon|OJJ42861.1 azon|OJJ43857.1 acar|OOF91216.1 agla|OJJ81826.1 anig|EHA24550.1

Group49: abra|OJJ66054.1 abra|OJJ67991.1 abra|OJJ68633.1 acar|OOF92033.1 acar|OOF92137.1 acla|EAW07055.1 acla|EAW12619.1 afis|EAW16409.1 afis|EAW16681.1 afis|EAW18065.1 afla|EED45379.1 afla|EED47440.1 afla|EED51440.1 afum|EAL84867.1 afum|EAL89033.1 afum|EAL89285.1 afum|EAL91877.1 akaw|GAA89772.1 akaw|GAA90957.1 aluc|OJZ81922.1 aluc|OJZ82865.1 anid|CBF73413.1 anid|CBF86422.1 anid|CBF87201.1 anig|EHA21295.1 anig|EHA22325.1 aory|EIT80529.1 aory|EIT81893.1 aory|EIT82380.1 asyd|OJJ59980.1 asyd|OJJ64938.1 ater|EAU30466.1 ater|EAU35871.1 atub|OJI84474.1 atub|OJI89090.1 aver|OJI99059.1 aver|OJJ02311.1 awen|OJJ36169.1 awen|OJJ41452.1 azon|OJJ47178.1 azon|OJJ48910.1 agla|OJJ87352.1 anid|CBF75131.1 aver|OJJ01785.1 asyd|OJJ59979.1 anid|CBF75132.1 aver|OJJ02312.1

Group50: abra|OJJ69511.1 abra|OJJ74314.1 acar|OOF94242.1 acar|OOF95413.1 acla|EAW07365.1 acla|EAW09962.1 afis|EAW21133.1 afis|EAW22392.1 afla|EED46140.1 afla|EED56055.1 afum|EAL90799.1 afum|EAL92151.1 agla|OJJ80083.1 agla|OJJ87654.1 akaw|GAA86711.1 akaw|GAA88313.1 aluc|OJZ82349.1 aluc|OJZ89812.1 anid|CBF87003.1 anid|CBF88741.1 anig|EHA27022.1 anig|EHA28411.1 aory|EIT76474.1 aory|EIT76698.1 asyd|OJJ60468.1 asyd|OJJ63953.1 ater|EAU36765.1 ater|EAU39326.1 atub|OJI84993.1 atub|OJI88091.1 aver|OJI96988.1 aver|OJI99788.1 awen|OJJ34969.1 awen|OJJ36463.1 acar|OOF94243.1 acla|EAW09961.1 afis|EAW21134.1 afum|EAL92153.1 agla|OJJ80077.1 anid|CBF87002.1 aory|EIT76417.1 ater|EAU36766.1 awen|OJJ36462.1 azon|OJJ44415.1 azon|OJJ44422.1 azon|OJJ45618.1 afla|EED46139.1

Group51: abra|OJJ66047.1 abra|OJJ76934.1 abra|OJJ77452.1 acar|OOF91381.1 acar|OOF99274.1 afis|EAW19956.1 afis|EAW22115.1 afla|EED52041.1 afla|EED54167.1 afla|EED55064.1 afum|EAL84958.1 afum|EAL91070.1 agla|OJJ80818.1 agla|OJJ80927.1 akaw|GAA84156.1 akaw|GAA87105.1 aluc|OJZ81077.1 aluc|OJZ84333.1 anig|EHA19285.1 anig|EHA27667.1 aory|EIT73694.1 aory|EIT75905.1 aory|EIT81463.1 aory|EIT81952.1 asyd|OJJ52197.1 asyd|OJJ52737.1 asyd|OJJ53817.1 asyd|OJJ54624.1 ater|EAU29271.1 ater|EAU29317.1 ater|EAU36080.1 atub|OJI82508.1 atub|OJI89736.1 aver|OJI95757.1 aver|OJI99556.1 aver|OJJ02993.1 aver|OJJ07236.1 aver|OJJ07532.1 aver|OJJ07918.1 awen|OJJ35197.1 awen|OJJ41937.1 anid|CBF78465.1 asyd|OJJ54956.1 azon|OJJ42548.1 afla|EED45216.1 agla|OJJ85358.1

Group52: abra|OJJ67587.1 abra|OJJ72232.1 abra|OJJ68022.1 abra|OJJ71087.1 acar|OOF96856.1 acar|OOF98832.1 afis|EAW19900.1 afis|EAW25740.1 afla|EED45923.1 afla|EED46506.1 afla|EED55031.1 afla|EED50270.1 afla|EED55600.1 akaw|GAA82874.1 akaw|GAA83260.1 akaw|GAA90929.1 aluc|OJZ80366.1 aluc|OJZ82896.1 aluc|OJZ83766.1 anig|EHA20401.1 anig|EHA20666.1 anig|EHA21320.1 aory|EIT72370.1 aory|EIT77313.1 aory|EIT81962.1 aory|EIT82672.1 ater|EAU30170.1 ater|EAU30810.1 ater|EAU34044.1 atub|OJI81207.1 atub|OJI89121.1 atub|OJI90137.1 afum|EAL85004.1 agla|OJJ80468.1 akaw|GAA84705.1 aluc|OJZ86982.1 anid|CBF73897.1 anig|EHA21665.1 aory|EIT75502.1 atub|OJI83537.1 acar|OOF92065.1 afis|EAW17166.1 awen|OJJ30813.1 aluc|OJZ87441.1 afla|EED45753.1 ater|EAU38566.1

Group53: acla|EAW06287.1 acla|EAW08342.1 acla|EAW12933.1 acla|EAW13032.1 afis|EAW17635.1 afis|EAW17710.1 afis|EAW18392.1 afis|EAW23377.1 afla|EED45712.1 afla|EED50529.1 afla|EED53015.1 afla|EED55580.1 afum|EAL88094.1 afum|EAL91525.1 afum|EAL93068.1 afum|EAL91450.1 akaw|GAA83332.1 akaw|GAA86352.1 akaw|GAA90807.1 akaw|GAA83394.1 anid|CBF70766.1 anid|CBF74845.1 anid|CBF76926.1 anid|CBF90134.1 aory|EIT75805.1 aory|EIT77402.1 aory|EIT82175.1 aory|EIT82691.1 ater|EAU31902.1 ater|EAU32280.1 ater|EAU35631.1 ater|EAU37071.1 awen|OJJ37126.1 awen|OJJ37209.1 acar|OOF97170.1 agla|OJJ86446.1 azon|OJJ50825.1 agla|OJJ81264.1 aluc|OJZ84881.1 anig|EHA27829.1 asyd|OJJ57069.1 atub|OJI90195.1 aver|OJJ03229.1 azon|OJJ50909.1 aluc|OJZ83828.1 asyd|OJJ62014.1

Group54: abra|OJJ65575.1 abra|OJJ65577.1 abra|OJJ66394.1 abra|OJJ68072.1 abra|OJJ78158.1 acar|OOF90635.1 acar|OOF94051.1 acar|OOF94121.1 agla|OJJ81248.1 agla|OJJ85122.1 agla|OJJ89317.1 akaw|GAA90259.1 akaw|GAA92419.1 aluc|OJZ80816.1 aluc|OJZ83081.1 aluc|OJZ87399.1 anid|CBF80614.1 anid|CBF80695.1 anig|EHA19835.1 anig|EHA28144.1 anig|EHA19917.1 anig|EHA20628.1 anig|EHA26671.1 ater|EAU29371.1 ater|EAU30114.1 atub|OJI81247.1 atub|OJI87864.1 aver|OJI97590.1 aver|OJJ07256.1 awen|OJJ29741.1 awen|OJJ32237.1 awen|OJJ32260.1 afis|EAW17155.1 afum|EAL86727.1 aory|EIT76930.1 azon|OJJ42341.1 afla|EED50167.1 asyd|OJJ52055.1 atub|OJI88902.1 abra|OJJ66294.1 abra|OJJ71136.1 akaw|GAA82838.1 azon|OJJ45212.1 awen|OJJ36390.1 azon|OJJ42821.1

Group55: abra|OJJ66953.1 abra|OJJ70025.1 acar|OOF93535.1 acar|OOF93655.1 agla|OJJ79210.1 agla|OJJ85149.1 akaw|GAA87635.1 akaw|GAA92208.1 aluc|OJZ85251.1 aluc|OJZ85625.1 anig|EHA24903.1 anig|EHA25451.1 aory|EIT80438.1 aory|EIT82112.1 asyd|OJJ55882.1 asyd|OJJ65094.1 ater|EAU32531.1 ater|EAU37243.1 atub|OJI79418.1 atub|OJI80318.1 aver|OJI99218.1 aver|OJJ04827.1 awen|OJJ30173.1 awen|OJJ38273.1 awen|OJJ41598.1 acla|EAW08800.1 afis|EAW15597.1 afla|EED50419.1 afum|EBA27239.1 anid|CBF78088.1 azon|OJJ43909.1 asyd|OJJ55868.1 acar|OOF93534.1 acla|EAW08799.1 afis|EAW15598.1 afum|EAL85734.1 agla|OJJ88612.1 akaw|GAA87636.1 anid|CBF78056.1 anig|EHA25452.1 aory|EIT82031.1 ater|EAU32530.1 atub|OJI80317.1 aver|OJJ04811.1 azon|OJJ43905.1

Group56: agla|OJJ81571.1 agla|OJJ85255.1 asyd|OJJ54206.1 asyd|OJJ59064.1 asyd|OJJ54427.1 ater|EAU31423.1 ater|EAU38101.1 aver|OJJ01658.1 aver|OJJ01906.1 awen|OJJ30145.1 awen|OJJ31520.1 abra|OJJ76240.1 acar|OOF99485.1 acla|EAW14198.1 afis|EAW18923.1 afla|EED50910.1 afum|EAL93655.1 agla|OJJ85250.1 akaw|GAA82174.1 aluc|OJZ91991.1 anid|CBF69674.1 anig|EHA22937.1 aory|EIT79045.1 asyd|OJJ59065.1 ater|EAU38098.1 atub|OJI86426.1 awen|OJJ31518.1 azon|OJJ47234.1 abra|OJJ76243.1 acar|OOF99488.1 acla|EAW06837.1 afis|EAW18926.1 afla|EED50916.1 afum|EAL93657.1 akaw|GAA82176.1 aluc|OJZ91988.1 anid|CBF69676.1 anig|EHA22935.1 aory|EIT78980.1 atub|OJI86423.1 azon|OJJ47236.1 afis|EAW16385.1 afla|EED48287.1 afum|EAL89013.2 aory|EIT72910.1

Group57: abra|OJJ69646.1 abra|OJJ77015.1 abra|OJJ77136.1 acar|OOF90285.1 acar|OOF91423.1 acar|OOF91458.1 acla|EAW09001.1 acla|EAW10860.1 acla|EAW12947.1 acla|EAW13503.1 afis|EAW17722.1 afis|EAW20602.1 afis|EAW20998.1 afum|EAL84459.1 afum|EAL91537.1 agla|OJJ78915.1 agla|OJJ79354.1 agla|OJJ88647.1 akaw|GAA87035.1 akaw|GAA90793.1 akaw|GAA91624.1 aluc|OJZ82450.1 aluc|OJZ84416.1 aluc|OJZ84546.1 anig|EHA27739.1 anig|EHA27843.1 anig|EHA28324.1 aory|EIT75439.1 aory|EIT77113.1 asyd|OJJ52539.1 asyd|OJJ55599.1 atub|OJI87989.1 atub|OJI89534.1 atub|OJI89658.1 aver|OJJ03865.1 aver|OJJ06679.1 awen|OJJ33394.1 awen|OJJ35790.1 afla|EED45724.1 afla|EED47731.1 anid|CBF86932.1 ater|EAU35247.1 azon|OJJ43330.1 agla|OJJ84075.1

Group58: acar|OOF92599.1 acar|OOF92620.1 abra|OJJ72332.1 acar|OOF94882.1 acla|EAW11990.1 afis|EAW19630.1 afla|EED56982.1 afum|EAL86892.1 agla|OJJ80775.1 akaw|GAA86084.1 aluc|OJZ90316.1 anid|CBF71323.1 anig|EHA23803.1 aory|EIT77586.1 asyd|OJJ55109.1 ater|EAU32896.1 atub|OJI81744.1 aver|OJJ07097.1 awen|OJJ41874.1 azon|OJJ47072.1 abra|OJJ72333.1 acar|OOF94880.1 acla|EAW11989.1 afis|EAW19629.1 afla|EED56981.1 afum|EAL86893.1 akaw|GAA86085.1 anid|CBF71321.1 aory|EIT77447.1 ater|EAU32897.1 azon|OJJ47071.1 abra|OJJ72335.1 acla|EAW11987.1 afis|EAW19627.1 afla|EED56979.1 afum|EAL86894.1 agla|OJJ80773.1 akaw|GAA86087.1 anid|CBF71319.1 aory|EIT77480.1 asyd|OJJ55111.1 ater|EAU32899.1 aver|OJJ07095.1 azon|OJJ47069.1

Group59: acar|OOF92600.1 acar|OOF92621.1 abra|OJJ76130.1 acar|OOF99793.1 acla|EAW09689.1 afis|EAW21450.1 afla|EED55879.1 afum|EAL92469.1 agla|OJJ85598.1 akaw|GAA88837.1 aluc|OJZ89591.1 anid|CBF76555.1 anig|EHA23020.1 aory|EIT76712.1 asyd|OJJ61157.1 ater|EAU35092.1 atub|OJI85211.1 aver|OJJ00479.1 awen|OJJ39248.1 azon|OJJ51607.1 abra|OJJ76131.1 acar|OOF99792.1 acla|EAW09687.1 afis|EAW21452.1 afla|EED55881.1 afum|EAL92471.1 akaw|GAA88836.1 aluc|OJZ89592.1 anid|CBF76551.1 anig|EHA23019.1 aory|EIT76739.1 ater|EAU35091.1 atub|OJI85210.1 abra|OJJ77110.1 acla|EAW09688.1 afis|EAW21451.1 afla|EED55880.1 afum|EAL92470.1 akaw|GAA90818.1 aluc|OJZ84519.1 anid|CBF76553.1 anig|EHA27817.1 aory|EIT76614.1 atub|OJI89561.1

Group60: afis|EAW15375.1 afis|EAW19388.1 afis|EAW19745.1 afis|EAW19746.1 afis|EAW20418.1 afis|EAW20688.1 afis|EAW21002.1 afis|EAW24636.1 afis|EAW25758.1 afis|EAW25761.1 afis|EAW17293.1 afis|EAW15376.1 afis|EAW17294.1 afis|EAW24637.1 afis|EAW15377.1 afis|EAW24638.1 afis|EAW15378.1 afis|EAW19381.1 afis|EAW22065.1 afis|EAW22066.1 afis|EAW24635.1 afis|EAW24639.1 afis|EAW24640.1 afis|EAW25759.1 afis|EAW22048.1 afis|EAW22062.1 afis|EAW23656.1 afis|EAW23657.1 afis|EAW25774.1 afis|EAW25760.1 anid|CBF76095.1 anid|CBF82248.1 ater|EAU29403.1 ater|EAU37740.1 acar|OOF89756.1 afum|EAL89306.1 asyd|OJJ55339.1 aver|OJJ07727.1 ater|EAU31660.1 aory|EIT81932.1 asyd|OJJ56289.1 afla|EED45889.1 aory|EIT75613.1 aluc|OJZ79928.1

Group61: abra|OJJ65992.1 abra|OJJ66994.1 abra|OJJ68819.1 acar|OOF90628.1 acar|OOF92880.1 afis|EAW20003.1 afis|EAW20740.1 afla|EED46008.1 afla|EED47482.1 afla|EED49530.1 agla|OJJ87844.1 agla|OJJ87962.1 akaw|GAA90432.1 akaw|GAA90866.1 akaw|GAA92170.1 aluc|OJZ82967.1 aluc|OJZ85578.1 aluc|OJZ85795.1 anig|EHA18425.1 anig|EHA21383.1 anig|EHA24761.1 anig|EHA24941.1 aory|EIT75392.1 aory|EIT79965.1 aory|EIT82595.1 asyd|OJJ52756.1 asyd|OJJ53312.1 asyd|OJJ54834.1 atub|OJI79374.1 atub|OJI79581.1 atub|OJI89200.1 atub|OJI91621.1 aver|OJJ07327.1 aver|OJJ08431.1 aver|OJJ08738.1 azon|OJJ43639.1 azon|OJJ50209.1 acla|EAW13602.1 afum|EAL84916.1 anid|CBF80344.1 anid|CBF84693.1 ater|EAU37349.1 awen|OJJ30010.1

Group62: abra|OJJ68708.1 abra|OJJ69824.1 acar|OOF92210.1 acar|OOF93339.1 acla|EAW08971.1 acla|EAW12537.1 afis|EAW15450.1 afis|EAW24112.1 afla|EED46821.1 afla|EED56823.1 akaw|GAA87836.1 akaw|GAA89862.1 aluc|OJZ81821.1 aluc|OJZ85472.1 anig|EHA22396.1 anig|EHA25013.1 atub|OJI80091.1 atub|OJI84392.1 awen|OJJ41377.1 awen|OJJ41767.1 abra|OJJ68707.1 acar|OOF92209.1 acla|EAW12538.1 afis|EAW24111.1 afum|EAL91965.1 agla|OJJ87188.1 akaw|GAA89861.1 aluc|OJZ81823.1 anid|CBF86525.1 anig|EHA22395.1 aory|EIT72582.1 asyd|OJJ64873.1 ater|EAU30539.1 atub|OJI84393.1 aver|OJI98986.1

azon|OJJ48844.1 afum|EAL91966.1 agla|OJJ87189.1 anid|CBF86527.1 aory|EIT72529.1 asyd|OJJ64872.1 ater|EAU30540.1 aver|OJI98984.1

Group63: agla|OJJ80048.1 agla|OJJ86306.1 agla|OJJ89670.1 agla|OJJ81524.1 agla|OJJ84920.1 agla|OJJ81876.1 agla|OJJ88808.1 akaw|GAA92616.1 akaw|GAA92655.1 akaw|GAA93101.1 akaw|GAA93218.1 akaw|GAA93326.1 akaw|GAA93341.1 akaw|GAA93349.1 akaw|GAA92654.1 anid|CBF73635.1 anid|CBF78260.1 anid|CBF84144.1 anid|CBF84167.1 anid|CBF74958.1 anid|CBF73637.1 anid|CBF75922.1 anid|CBF82905.1 anid|CBF83997.1 anid|CBF84165.1 anid|CBF89628.1 anid|CBF84258.1 anid|CBF83995.1 anid|CBF89115.1 anid|CBF79097.1 anid|CBF80200.1 anid|CBF79095.1 anid|CBF77664.1 anid|CBF80044.1 anid|CBF78884.1 anid|CBF78717.1 anid|CBF74110.1 anid|CBF74956.1 anid|CBF83687.1 acar|OOG00276.1 aluc|OJZ79659.1 afum|EAL90122.1 akaw|GAA93209.1

Group64: agla|OJJ84606.1 agla|OJJ86279.1 anid|CBF78809.1 anid|CBF85739.1 asyd|OJJ64288.1 asyd|OJJ65260.1 aver|OJI98381.1 aver|OJI99411.1 awen|OJJ30086.1 awen|OJJ30833.1 awen|OJJ38668.1 azon|OJJ44554.1 azon|OJJ44649.1 abra|OJJ67129.1 acar|OOF98424.1 afis|EAW19329.1 afla|EED47551.1 afum|EAL94046.1 akaw|GAA91039.1 aluc|OJZ87849.1 anig|EHA20152.1 aory|EIT74544.1 ater|EAU29453.1 atub|OJI80799.1 abra|OJJ67137.1 acar|OOF98421.1 acla|EAW14610.1 afis|EAW19339.1 afla|EED47549.1 afum|EAL94058.1 akaw|GAA91032.1 aluc|OJZ87840.1 anid|CBF78807.1 anig|EHA20314.1 aory|EIT74530.1 asyd|OJJ65259.1 ater|EAU29452.1 atub|OJI80806.1 aver|OJI99410.1 agla|OJJ83890.1 awen|OJJ41067.1 agla|OJJ89228.1 awen|OJJ40968.1

Group65: abra|OJJ65919.1 abra|OJJ71178.1 acar|OOF94968.1 acar|OOF97988.1 acla|EAW10328.1 acla|EAW13616.1 afis|EAW19973.1 afis|EAW25266.1 afla|EED50456.1 afla|EED53730.1 afum|EAL84937.1 afum|EAL89929.1 agla|OJJ80057.1 agla|OJJ89242.1 akaw|GAA82807.1 akaw|GAA92313.1 aluc|OJZ87365.1 aluc|OJZ90211.1 anid|CBF84421.1 anid|CBF85372.1 anig|EHA20600.1 anig|EHA23709.1 aory|EIT75304.1 aory|EIT82028.1 asyd|OJJ53545.1 asyd|OJJ57927.1 asyd|OJJ64047.1 ater|EAU31473.1 ater|EAU34407.1 atub|OJI81279.1 atub|OJI81642.1 aver|OJI97591.1 aver|OJI98132.1 aver|OJI98886.1 aver|OJJ08208.1 awen|OJJ36011.1 awen|OJJ40979.1 azon|OJJ42397.1 azon|OJJ44566.1 azon|OJJ45171.1 azon|OJJ50333.1 asyd|OJJ54388.1

Group66: abra|OJJ66757.1 abra|OJJ68421.1 abra|OJJ70242.1 acar|OOF92554.1 acar|OOF93838.1 afis|EAW19495.1 afis|EAW21208.1 afum|EAL86999.1 afum|EAL92219.1 agla|OJJ78585.1 agla|OJJ81224.1 agla|OJJ86343.1 agla|OJJ87022.1 akaw|GAA86545.1 akaw|GAA89691.1 akaw|GAA92527.1 aluc|OJZ80693.1 aluc|OJZ85093.1 aluc|OJZ90076.1 anig|EHA20997.1 anig|EHA27322.1 anig|EHA25314.1 aory|EIT73935.1 aory|EIT77370.1 asyd|OJJ59201.1 asyd|OJJ59780.1 ater|EAU34261.1 ater|EAU34807.1 atub|OJI80506.1 atub|OJI87553.1 atub|OJI84632.1 atub|OJI84730.1 aver|OJJ01814.1 aver|OJJ02507.1 awen|OJJ36122.1 awen|OJJ36911.1 azon|OJJ42905.1 azon|OJJ45981.1 acla|EAW11883.1 afla|EED50220.1 afla|EED56713.1 anid|CBF75444.1

Group67: abra|OJJ67357.1 abra|OJJ69620.1 abra|OJJ69925.1 acar|OOF92464.1 acar|OOF92686.1 acar|OOF93436.1 acla|EAW06385.1 acla|EAW08905.1 afis|EAW15491.1 afis|EAW18491.1 afla|EED49005.1 afla|EED52913.1 afum|EAL85832.1 afum|EAL93181.1 agla|OJJ80168.1 agla|OJJ85086.1 akaw|GAA86464.1 akaw|GAA87732.1 akaw|GAA91604.1 aluc|OJZ82429.1 aluc|OJZ85004.1 aluc|OJZ85363.1 anig|EHA25223.1 anig|EHA25546.1 anig|EHA28342.1 aory|EIT78593.1 aory|EIT82534.1 asyd|OJJ53723.1 asyd|OJJ56020.1 asyd|OJJ58801.1 ater|EAU29917.1 ater|EAU32628.1 atub|OJI80207.1 atub|OJI80594.1 atub|OJI88012.1 awen|OJJ31044.1 awen|OJJ38147.1 azon|OJJ43140.1 azon|OJJ50555.1 anid|CBF70161.1 aver|OJJ01405.1 aver|OJJ04967.1

Group68: abra|OJJ69659.1 abra|OJJ71032.1 acar|OOF92597.1 acar|OOF92618.1 acar|OOF92644.1 acar|OOF98104.1 aluc|OJZ82466.1 aluc|OJZ87497.1 anig|EHA20713.1 anig|EHA28309.1 atub|OJI81151.1 atub|OJI87972.1 acla|EAW08698.1 acla|EAW12766.1 afis|EAW15737.1 afis|EAW17928.1 afla|EED49112.1 afla|EED51315.1 afum|EAL85611.1 afum|EAL91739.1 agla|OJJ81894.1 agla|OJJ83005.1 akaw|GAA91638.1 akaw|GAA91639.1 aluc|OJZ82465.1 anid|CBF71049.1 anid|CBF80811.1 anig|EHA28310.1 aory|EIT78548.1 aory|EIT81553.1 asyd|OJJ52673.1 asyd|OJJ54450.1 ater|EAU32405.1 ater|EAU33543.1 atub|OJI87973.1 aver|OJJ04699.1 aver|OJJ07704.1 awen|OJJ38433.1 awen|OJJ41618.1 azon|OJJ43810.1 azon|OJJ48339.1 akaw|GAA82922.1

Group69: abra|OJJ70622.1 abra|OJJ71755.1 abra|OJJ72061.1 abra|OJJ76523.1 acar|OOF96715.1 acar|OOF96808.1 acar|OOF99444.1 acla|EAW13671.1 acla|EAW14172.1 afis|EAW15618.1 afis|EAW18886.1 afla|EED50879.1 afla|EED53904.1 afla|EED57853.1 agla|OJJ80215.1 agla|OJJ80629.1 akaw|GAA82436.1 akaw|GAA84874.1 aluc|OJZ84834.1 aluc|OJZ86797.1 aluc|OJZ91685.1 anig|EHA21886.1 anig|EHA22697.1 anig|EHA25744.1 aory|EIT78491.1 aory|EIT78970.1 aory|EIT81115.1 ater|EAU31982.1 ater|EAU32969.1 ater|EAU38058.1 atub|OJI83721.1 atub|OJI86137.1 aver|OJJ01614.1 aver|OJJ02222.1 awen|OJJ32444.1 awen|OJJ38463.1 azon|OJJ47202.1 azon|OJJ47982.1 afum|EAL93619.1 anid|CBF69896.1 asyd|OJJ59001.1

Group70: agla|OJJ81088.1 agla|OJJ81098.1 agla|OJJ84079.1 agla|OJJ81610.1 agla|OJJ85958.1 agla|OJJ84407.1 anid|CBF70684.1 anid|CBF83505.1 anid|CBF86544.1 anid|CBF70985.1 anid|CBF73878.1 anid|CBF76072.1 anid|CBF76571.1 anid|CBF77622.1 anid|CBF78018.1 anid|CBF78361.1 anid|CBF79546.1 anid|CBF82083.1 anid|CBF84126.1 anid|CBF88413.1 anid|CBF88708.1 anid|CBF89022.1 anid|CBF89295.1 ater|EAU29721.1 ater|EAU32284.1 ater|EAU32866.1 ater|EAU33186.1 ater|EAU30981.1 ater|EAU30143.1

ater|EAU35995.1 abra|OJJ69716.1 aluc|OJZ83181.1 anig|EHA28255.1 aory|EIT76332.1 atub|OJI87756.1 awen|OJJ29588.1 acar|OOF89784.1 aluc|OJZ82089.1 asyd|OJJ55193.1 agla|OJJ85314.1 agla|OJJ88024.1

Group71: abra|OJJ65673.1 abra|OJJ68238.1 abra|OJJ66038.1 abra|OJJ68515.1 abra|OJJ74212.1 abra|OJJ74240.1 acar|OOF92683.1 acar|OOF95490.1 acar|OOF95508.1 afla|EED47448.1 afla|EED57720.1 akaw|GAA86780.1 akaw|GAA86806.1 akaw|GAA91332.1 aluc|OJZ79540.1 aluc|OJZ89922.1 aluc|OJZ82729.1 aluc|OJZ89892.1 anig|EHA21182.1 anig|EHA27091.1 anig|EHA27122.1 aory|EIT79930.1 aory|EIT81920.1 atub|OJI84883.1 atub|OJI84915.1 atub|OJI88962.1 aver|OJI97104.1 aver|OJJ03419.1 acla|EAW07265.1 afum|EAL90895.1 ater|EAU34298.1 agla|OJJ87719.1 anid|CBF88605.1 awen|OJJ34900.1 azon|OJJ50067.1 afis|EAW22286.1 anid|CBF88560.1 asyd|OJJ63833.1 ater|EAU34323.1 asyd|OJJ63857.1

Group72: abra|OJJ66697.1 abra|OJJ68381.1 acar|OOF93705.1 acar|OOF93791.1 acar|OOF93987.1 acla|EAW11832.1 acla|EAW12357.1 afis|EAW16780.1 afis|EAW19443.1 afla|EED56190.1 afla|EED56651.1 afum|EAL86351.1 afum|EAL87052.1 agla|OJJ79770.1 agla|OJJ81157.1 akaw|GAA89641.1 akaw|GAA92564.1 aluc|OJZ80748.1 aluc|OJZ90112.1 anid|CBF80588.1 anid|CBF81398.1 anig|EHA20960.1 anig|EHA27274.1 aory|EIT73439.1 aory|EIT79687.1 asyd|OJJ54689.1 asyd|OJJ59403.1 ater|EAU30272.1 ater|EAU34745.1 atub|OJI84692.1 atub|OJI87197.1 atub|OJI87605.1 aver|OJI99850.1 aver|OJJ02870.1 aver|OJJ07480.1 aver|OJJ07987.1 awen|OJJ29847.1 awen|OJJ36844.1 azon|OJJ42956.1 azon|OJJ50350.1

Group73: abra|OJJ68502.1 abra|OJJ73653.1 acar|OOF98748.1 acar|OOG00831.1 acla|EAW07563.1 acla|EAW12139.1 acla|EAW15367.1 afis|EAW17197.1 afis|EAW17227.1 afis|EAW22597.1 afla|EED45179.1 afla|EED51197.1 afla|EED51520.1 afla|EED54722.1 afum|EAL86625.1 afum|EAL86649.1 afum|EAL90601.1 agla|OJJ80961.1 agla|OJJ86657.1 akaw|GAA88505.1 akaw|GAA92769.1 aluc|OJZ82715.1 aluc|OJZ88400.1 anid|CBF69884.1 anid|CBF89553.1 anig|EHA18128.1 anig|EHA21170.1 aory|EIT73539.1 aory|EIT78945.1 aory|EIT80690.1 asyd|OJJ59011.1 asyd|OJJ65377.1 ater|EAU31751.1 ater|EAU39106.1 atub|OJI88949.1 atub|OJI90995.1 aver|OJI99545.1 aver|OJJ01604.1 awen|OJJ35225.1 azon|OJJ51827.1

Group74: abra|OJJ66378.1 abra|OJJ68574.1 abra|OJJ75363.1 acar|OOF89872.1 acar|OOF92870.1 akaw|GAA90247.1 akaw|GAA91914.1 aluc|OJZ81966.1 aluc|OJZ83093.1 atub|OJI84517.1 atub|OJI87392.1 atub|OJI87843.1 acla|EAW08967.1 agla|OJJ89584.1 anid|CBF87061.1 anig|EHA22281.1 anig|EHA28155.1 asyd|OJJ60430.1 ater|EAU36814.1 aver|OJI99753.1 afla|EED48897.1 afum|EAL93812.1 akaw|GAA92059.1 aluc|OJZ92932.1 azon|OJJ46231.1 afis|EAW19083.1 agla|OJJ81069.1 anig|EHA26039.1 aory|EIT81242.1 ater|EAU36156.1 awen|OJJ35776.1 azon|OJJ45936.1 acla|EAW14375.1 afla|EED54315.1 aory|EIT76109.1 ater|EAU32130.1 awen|OJJ32478.1 afla|EED46053.1 aory|EIT83489.1

Group75: abra|OJJ66645.1 abra|OJJ77622.1 acar|OOF93665.1 acar|OOF93749.1 acar|OOF99113.1 afis|EAW19398.1 afis|EAW25441.1 afla|EED46395.1 afla|EED56341.1 afum|EAL87096.1 afum|EAL90099.1 agla|OJJ81109.1 agla|OJJ84218.1 akaw|GAA84009.1 akaw|GAA89598.1 aluc|OJZ80804.1 aluc|OJZ80911.1 anid|CBF74274.1 anid|CBF85786.1 anig|EHA19435.1 anig|EHA27230.1 aory|EIT72345.1 aory|EIT73378.1 aory|EIT83124.1 ater|EAU33678.1 ater|EAU36124.1 atub|OJI82668.1 atub|OJI87652.1 acla|EAW10153.1 acla|EAW11788.1 afla|EED50354.1 asyd|OJJ51908.1 asyd|OJJ64317.1 aver|OJJ05362.1 awen|OJJ36793.1 azon|OJJ43006.1 azon|OJJ43662.1 aver|OJI98410.1 awen|OJJ30031.1

Group76: abra|OJJ68339.1 abra|OJJ77792.1 acar|OOF89870.1 acar|OOF90688.1 acar|OOF90718.1 agla|OJJ81064.1 agla|OJJ85571.1 aluc|OJZ80535.1 aluc|OJZ83619.1 anig|EHA18777.1 anig|EHA25660.1 asyd|OJJ53057.1 asyd|OJJ61781.1 atub|OJI88396.1 atub|OJI89992.1 aver|OJI95881.1 aver|OJJ03787.1 awen|OJJ29921.1 awen|OJJ33519.1 awen|OJJ38011.1 abra|OJJ66230.1 afis|EAW16867.1 afla|EED45515.1 afla|EED46127.1 afum|EAL90432.1 akaw|GAA89059.1 akaw|GAA92106.1 anid|CBF80612.1 aory|EIT76508.1 asyd|OJJ64304.1 ater|EAU31983.1 aver|OJI98397.1 awen|OJJ35964.1 azon|OJJ42115.1 ater|EAU33486.1 aory|EIT73494.1 acar|OOF90354.1 afla|EED44960.1 awen|OJJ35329.1

Group77: abra|OJJ68971.1 abra|OJJ74920.1 acar|OOF91548.1 acar|OOF95911.1 acla|EAW08451.1 acla|EAW14216.1 afis|EAW19065.1 afis|EAW23496.1 afla|EED54431.1 afla|EED57396.1 afum|EAL87967.1 afum|EAL93794.1 agla|OJJ83427.1 agla|OJJ84542.1 agla|OJJ86743.1 akaw|GAA83503.1 akaw|GAA90498.1 aluc|OJZ85958.1 aluc|OJZ92491.1 anid|CBF79648.1 anid|CBF89739.1 anig|EHA24643.1 anig|EHA26481.1 aory|EIT75884.1 aory|EIT80127.1 asyd|OJJ52407.1 asyd|OJJ58022.1 asyd|OJJ62230.1 ater|EAU32108.1 ater|EAU35285.1 atub|OJI79746.1 atub|OJI86924.1 aver|OJI96339.1 aver|OJJ04377.1 awen|OJJ31497.1 awen|OJJ39821.1 azon|OJJ46209.1 azon|OJJ47945.1 anid|CBF76133.1

Group78: abra|OJJ72527.1 abra|OJJ73074.1 abra|OJJ74512.1 acar|OOF95231.1 acar|OOG00307.1 afis|EAW15886.1 afis|EAW22192.1 afla|EED49069.1 afla|EED49846.1 afla|EED56486.1 afum|EAL88520.1 afum|EAL90999.1 akaw|GAA82087.1 akaw|GAA86250.1 akaw|GAA88087.1 aluc|OJZ81351.1 aluc|OJZ90507.1 aluc|OJZ92093.1 anig|EHA18653.1 anig|EHA23964.1 anig|EHA26848.1 aory|EIT72810.1 aory|EIT82565.1 ater|EAU31708.1 ater|EAU34200.1 atub|OJI81931.1 atub|OJI86526.1 atub|OJI91555.1 aver|OJI97191.1 aver|OJJ02116.1 aver|OJJ08595.1 awen|OJJ34773.1 awen|OJJ36745.1 azon|OJJ45756.1 azon|OJJ45757.1 acla|EAW11155.1 anid|CBF88404.1 asyd|OJJ63749.1 aory|EIT83409.1

Group79: abra|OJJ69282.1 abra|OJJ71129.1 acar|OOF90163.1 acar|OOF98023.1 acar|OOF94440.1 acla|EAW10369.1 acla|EAW14671.1 afis|EAW24202.1 afis|EAW25223.1 afla|EED48070.1 afla|EED53683.1 afum|EAL85872.1 afum|EAL89884.1 agla|OJJ79190.1 agla|OJJ79675.1 akaw|GAA82845.1 akaw|GAA88154.1 aluc|OJZ82110.1 aluc|OJZ87406.1 anid|CBF74328.1 anid|CBF85271.1 anig|EHA20636.1 anig|EHA28620.1

aory|EIT75297.1 aory|EIT83329.1 asyd|OJJ53993.1 asyd|OJJ62910.1 ater|EAU31153.1 ater|EAU34382.1 atub|OJI81240.1 atub|OJI88326.1 aver|OJI98079.1 aver|OJJ08091.1 awen|OJJ33096.1 awen|OJJ40889.1 azon|OJJ42425.1 azon|OJJ44518.1 azon|OJJ44851.1

Group80: abra|OJJ69429.1 abra|OJJ76880.1 abra|OJJ74397.1 abra|OJJ75836.1 acar|OOF94306.1 acar|OOF95338.1 akaw|GAA84546.1 akaw|GAA87295.1 aluc|OJZ89279.1 aluc|OJZ89723.1 asyd|OJJ56900.1 asyd|OJJ58390.1 ater|EAU31031.1 ater|EAU36089.1 ater|EAU36364.1 ater|EAU36837.1 ater|EAU36966.1 ater|EAU39346.1 atub|OJI85077.1 atub|OJI85520.1 aver|OJJ02028.1 aver|OJJ03392.1 agla|OJJ83397.1 akaw|GAA88295.1 aluc|OJZ82266.1 anig|EHA28477.1 atub|OJI88172.1 anid|CBF82791.1 anig|EHA23290.1 anig|EHA26951.1 aory|EIT80997.1 aver|OJI99456.1 acla|EAW13699.1 afum|EAL86287.1 anid|CBF80711.1 awen|OJJ32049.1 afla|EED52718.1 afla|EED52719.1

Group81: abra|OJJ72988.1 abra|OJJ77279.1 acar|OOF93153.1 acar|OOF96164.1 acla|EAW08023.1 acla|EAW13278.1 afis|EAW17339.1 afis|EAW23057.1 afum|EAL88423.2 afum|EAL91183.1 akaw|GAA85562.1 akaw|GAA90670.1 aluc|OJZ83496.1 aluc|OJZ84689.1 anig|EHA24331.1 anig|EHA27964.1 asyd|OJJ54336.1 asyd|OJJ56391.1 ater|EAU33849.1 ater|EAU37607.1 atub|OJI82390.1 atub|OJI89393.1 aver|OJJ05288.1 aver|OJJ06322.1 awen|OJJ35679.1 awen|OJJ40816.1 afla|EED51042.1 afla|EED54960.1 agla|OJJ79345.1 agla|OJJ89418.1 anid|CBF81037.1 aory|EIT80299.1 azon|OJJ44147.1 azon|OJJ50224.1 anid|CBF84215.1 aory|EIT77041.1 asyd|OJJ57175.1 aver|OJJ03139.1

Group82: abra|OJJ73798.1 abra|OJJ76038.1 acar|OOF99891.1 acar|OOG00949.1 acla|EAW07105.1 acla|EAW07691.1 afla|EED45540.1 afla|EED49992.1 agla|OJJ86348.1 agla|OJJ87936.1 akaw|GAA88927.1 akaw|GAA89416.1 aluc|OJZ88257.1 aluc|OJZ89493.1 anid|CBF71497.1 anid|CBF88069.1 anig|EHA18002.1 anig|EHA23106.1 aory|EIT74740.1 aory|EIT75450.1 asyd|OJJ54934.1 asyd|OJJ60320.1 asyd|OJJ63552.1 ater|EAU32047.1 ater|EAU38995.1 atub|OJI85309.1 atub|OJI90856.1 aver|OJI97382.1 aver|OJI99669.1 aver|OJJ07264.1 awen|OJJ34497.1 awen|OJJ34697.1 awen|OJJ35343.1 azon|OJJ45277.1 azon|OJJ49485.1 azon|OJJ49775.1 afis|EAW22706.1 afum|EAL90486.1

Group83: abra|OJJ74861.1 abra|OJJ78108.1 aluc|OJZ91173.1 aluc|OJZ92439.1 anig|EHA19866.1 anig|EHA26528.1 atub|OJI83187.1 atub|OJI86868.1 acar|OOF95861.1 agla|OJJ86823.1 akaw|GAA83559.1 akaw|GAA85898.1 awen|OJJ39889.1 azon|OJJ43220.1 acar|OOF98757.1 acla|EAW06608.1 afis|EAW16215.1 afla|EED48709.1 afum|EAL88842.1 agla|OJJ86220.1 anid|CBF79542.1 aory|EIT73181.1 asyd|OJJ57900.1 ater|EAU33259.1 aver|OJJ04494.1 awen|OJJ30257.1 azon|OJJ43953.1 abra|OJJ78109.1 acla|EAW06607.1 afis|EAW16216.1 afla|EED48710.1 afum|EAL88843.1 akaw|GAA85897.1 anid|CBF79540.1 aory|EIT73231.1 ater|EAU33260.1 awen|OJJ30256.1 azon|OJJ43954.1

Group84: afla|EED45777.1 afla|EED56579.1 afla|EED46479.1 afla|EED47420.1 aory|EIT72361.1 aory|EIT81890.1 aory|EIT75763.1 aory|EIT79626.1 asyd|OJJ54274.1 asyd|OJJ58712.1 asyd|OJJ61516.1 asyd|OJJ61954.1 aver|OJI96045.1 aver|OJJ00852.1 aver|OJJ01325.1 aver|OJJ05230.1 abra|OJJ67908.1 afum|EAL86295.1 agla|OJJ85785.1 aluc|OJZ81978.1 anig|EHA22277.1 atub|OJI84530.1 awen|OJJ29991.1 azon|OJJ42128.1 abra|OJJ70236.1 akaw|GAA86549.1 aluc|OJZ85097.1 anid|CBF73730.1 anig|EHA25319.1 asyd|OJJ53339.1 ater|EAU37949.1 atub|OJI80501.1 aver|OJJ08406.1 awen|OJJ31368.1 awen|OJJ35920.1 asyd|OJJ65167.1 aver|OJI99319.1 awen|OJJ39978.1

Group85: abra|OJJ69585.1 acar|OOF91569.1 acla|EAW14381.1 acla|EAW14383.1 acla|EAW14384.1 afis|EAW19087.1 afis|EAW19089.1 afis|EAW19090.1 afla|EED54319.1 afla|EED54321.1 afla|EED54322.1 afum|EAL93817.1 afum|EAL93819.1 afum|EAL93820.1 agla|OJJ84514.1 agla|OJJ84515.1 agla|OJJ84517.1 akaw|GAA91580.1 akaw|GAA91581.1 akaw|GAA91583.1 aluc|OJZ82402.1 anid|CBF79686.1 anid|CBF79690.1 anid|CBF79692.1 anig|EHA28363.1 aory|EIT76048.1 aory|EIT76070.1 aory|EIT76088.1 asyd|OJJ58048.1 ater|EAU32135.1 ater|EAU32138.1 ater|EAU32139.1 atub|OJI88039.1 aver|OJJ04355.1 awen|OJJ32484.1 azon|OJJ46222.1 azon|OJJ46228.1 azon|OJJ46230.1

Group86: abra|OJJ65665.1 abra|OJJ65727.1 abra|OJJ70427.1 abra|OJJ73414.1 acar|OOG00335.1 acar|OOG00616.1 acla|EAW07048.1 acla|EAW10826.1 agla|OJJ78693.1 agla|OJJ85646.1 aluc|OJZ81154.1 aluc|OJZ88623.1 anid|CBF79793.1 anid|CBF82339.1 asyd|OJJ56350.1 asyd|OJJ56512.1 asyd|OJJ58126.1 aver|OJJ04280.1 aver|OJJ06363.1 awen|OJJ30203.1 awen|OJJ36125.1 azon|OJJ43588.1 azon|OJJ45284.1 afis|EAW21006.1 afum|EAL84365.1 agla|OJJ85438.1 akaw|GAA93193.1 atub|OJI88767.1 afla|EED56769.1 akaw|GAA83077.1 akaw|GAA93192.1 aory|EIT73951.1 atub|OJI91208.1 ater|EAU37240.1 afla|EED48948.1 aory|EIT79979.1 aory|EIT74379.1

Group87: abra|OJJ73172.1 abra|OJJ75139.1 acar|OOF97197.1 acar|OOG00411.1 acla|EAW06855.1 acla|EAW08318.1 afis|EAW16472.1 afis|EAW23353.1 afla|EED47207.1 afla|EED55602.1 afum|EAL88123.1 afum|EAL89066.1 agla|OJJ82874.1 agla|OJJ86473.1 akaw|GAA87999.1 akaw|GAA88746.1 aluc|OJZ81255.1 aluc|OJZ92706.1 anid|CBF74897.1 anid|CBF81437.1 anig|EHA18563.1 anig|EHA26294.1 aory|EIT79222.1 aory|EIT82665.1 asyd|OJJ59426.1 asyd|OJJ60103.1 ater|EAU35596.1 ater|EAU36570.1 atub|OJI87132.1 atub|OJI91451.1 aver|OJJ02845.1 aver|OJJ04159.1 awen|OJJ39536.1 awen|OJJ40364.1 azon|OJJ47731.1 azon|OJJ49213.1 anid|CBF74899.1

Group88: afis|EAW16767.1 afis|EAW20787.1 afis|EAW20816.1 aluc|OJZ89742.1 aluc|OJZ90793.1 atub|OJI82782.1 atub|OJI85059.1 aver|OJJ04748.1 aver|OJJ08594.1 awen|OJJ33411.1 awen|OJJ35564.1 afla|EED55613.1 anig|EHA22174.1 afum|EAL84837.1 afum|EAL86763.1 anid|CBF71491.1 asyd|OJJ52867.1

asyd|OJJ53645.1 asyd|OJJ54618.1 ater|EAU29406.1 awen|OJJ33444.1 azon|OJJ43208.1 afla|EED51750.1 agla|OJJ87003.1 akaw|GAA87278.1 anig|EHA19527.1 aory|EIT80503.1 asyd|OJJ56543.1 azon|OJJ47397.1 awen|OJJ33735.1 anid|CBF80305.1 aver|OJJ07540.1 awen|OJJ40176.1 agla|OJJ79599.1 akaw|GAA83902.1 anid|CBF86026.1 anid|CBF80765.1

Group89: aory|EIT73009.1 aory|EIT82639.1 abra|OJJ75430.1 acar|OOF90776.1 acla|EAW13800.1 acla|EAW13801.1 acla|EAW13802.1 afis|EAW18330.1 afis|EAW18331.1 afis|EAW18332.1 afla|EED48794.1 afla|EED48795.1 afla|EED49574.1 afum|EAL87695.1 afum|EAL87696.1 afum|EAL87697.1 agla|OJJ88054.1 akaw|GAA84206.1 akaw|GAA84207.1 aluc|OJZ88883.1 aluc|OJZ88884.1 anid|CBF79398.1 anid|CBF79400.1 anid|CBF79402.1 anig|EHA23623.1 aory|EIT72994.1 aory|EIT73029.1 asyd|OJJ57827.1 ater|EAU33343.1 ater|EAU33344.1 atub|OJI85907.1 atub|OJI85908.1 atub|OJI85909.1 aver|OJJ04565.1 awen|OJJ30712.1 azon|OJJ43305.1 afla|EED48796.1

Group90: aory|EIT73779.1 aory|EIT77467.1 awen|OJJ29827.1 awen|OJJ36939.1 abra|OJJ65853.1 acar|OOF95014.1 acla|EAW11610.1 afis|EAW23843.1 afla|EED56987.1 afum|EAL93778.1 agla|OJJ78690.1 akaw|GAA92291.1 aluc|OJZ90188.1 anig|EHA23687.1 ater|EAU31468.1 atub|OJI81615.1 aver|OJI99116.1 awen|OJJ29704.1 azon|OJJ46986.1 abra|OJJ77924.1 acar|OOF90880.1 afla|EED45127.1 akaw|GAA83740.1 aluc|OJZ90991.1 anid|CBF86319.1 anig|EHA19699.1 asyd|OJJ64996.1 atub|OJI82982.1 acar|OOG00327.1 asyd|OJJ55713.1 atub|OJI87196.1 aver|OJJ06799.1 afum|EAL87274.1 agla|OJJ86442.1 asyd|OJJ62873.1 ater|EAU31778.1 aver|OJI96926.1

Group91: abra|OJJ67337.1 acar|OOF92476.1 acla|EAW06397.1 afis|EAW18501.1 afla|EED52900.1 afum|EAL93192.1 agla|OJJ85126.1 akaw|GAA86479.1 aluc|OJZ85020.1 anid|CBF75182.1 anig|EHA25241.1 aory|EIT78666.1 asyd|OJJ59951.1 ater|EAU30070.1 atub|OJI80577.1 aver|OJJ02338.1 awen|OJJ31055.1 azon|OJJ50565.1 abra|OJJ69524.1 acar|OOF94230.1 acla|EAW11627.1 afis|EAW23829.1 afla|EED54410.1 afum|EAL87261.1 agla|OJJ82483.1 akaw|GAA88301.1 aluc|OJZ82364.1 anid|CBF75186.1 anig|EHA28401.1 aory|EIT75866.1 ater|EAU31384.1 atub|OJI88078.1 awen|OJJ38733.1 azon|OJJ50221.1 afla|EED51976.1 aory|EIT81396.1 agla|OJJ78561.1

Group92: abra|OJJ66053.1 abra|OJJ66142.1 abra|OJJ74823.1 abra|OJJ77498.1 acar|OOF98679.1 acar|OOF99228.1 afla|EED49691.1 afla|EED54455.1 afla|EED57013.1 agla|OJJ78526.1 agla|OJJ80805.1 agla|OJJ83730.1 akaw|GAA84123.1 akaw|GAA88361.1 aluc|OJZ81040.1 aluc|OJZ82294.1 anig|EHA18383.1 anig|EHA19327.1 anig|EHA25403.1 aory|EIT81151.1 aory|EIT82306.1 ater|EAU29270.1 ater|EAU38253.1 atub|OJI82544.1 atub|OJI88145.1 aver|OJJ01748.1 aver|OJJ01775.1 aver|OJJ08353.1 awen|OJJ34671.1 awen|OJJ41918.1 afis|EAW19512.1 afum|EAL86984.1 anid|CBF89390.1 asyd|OJJ59158.1 azon|OJJ42220.1 aory|EIT77599.1

Group93: abra|OJJ67791.1 abra|OJJ73324.1 abra|OJJ75373.1 acar|OOF94472.1 acar|OOF95395.1 afis|EAW17263.1 afis|EAW17790.1 afla|EED48868.1 afla|EED50759.1 afum|EAL86680.1 afum|EAL91599.1 akaw|GAA92735.1 akaw|GAA92794.1 aluc|OJZ88714.1 aluc|OJZ92943.1 aluc|OJZ88755.1 anig|EHA18417.1 anig|EHA26030.1 atub|OJI87402.1 atub|OJI91301.1 atub|OJI91325.1 awen|OJJ32920.1 awen|OJJ35770.1 acla|EAW12886.1 agla|OJJ84166.1 akaw|GAA92711.1 anid|CBF75220.1 aory|EIT78062.1 asyd|OJJ53461.1 ater|EAU29261.1 aver|OJJ08294.1 abra|OJJ67793.1 anid|CBF75222.1 aory|EIT78107.1 ater|EAU29259.1 awen|OJJ32922.1

Group94: abra|OJJ68844.1 abra|OJJ73357.1 acar|OOF90545.1 acar|OOG00565.1 acla|EAW06977.1 acla|EAW15277.1 afis|EAW16591.1 afis|EAW20248.1 afla|EED45005.1 afla|EED57097.1 afum|EAL85366.1 afum|EAL89187.1 akaw|GAA83127.1 akaw|GAA90404.1 aluc|OJZ85824.1 aluc|OJZ88683.1 anid|CBF78712.1 anid|CBF81894.1 anig|EHA18363.1 anig|EHA24737.1 aory|EIT73755.1 aory|EIT82298.1 asyd|OJJ59680.1 asyd|OJJ63029.1 ater|EAU32039.1 ater|EAU36671.1 atub|OJI79611.1 atub|OJI91263.1 aver|OJI97939.1 aver|OJJ02608.1 awen|OJJ31246.1 awen|OJJ40224.1 azon|OJJ48439.1 azon|OJJ49082.1 agla|OJJ82668.1 azon|OJJ45914.1

Group95: abra|OJJ69553.1 abra|OJJ70394.1 aory|EIT78629.1 aory|EIT82274.1 acar|OOF91880.1 afla|EED57090.1 agla|OJJ82606.1 akaw|GAA91965.1 aluc|OJZ82652.1 anid|CBF75146.1 anig|EHA27420.1 asyd|OJJ59972.1 ater|EAU30264.1 atub|OJI88884.1 aver|OJJ02319.1 awen|OJJ41493.1 abra|OJJ71088.1 acar|OOF98047.1 acla|EAW06408.1 afis|EAW18512.1 afla|EED52888.1 afum|EAL93207.1 agla|OJJ85018.1 akaw|GAA82873.1 aluc|OJZ87440.1 anid|CBF70237.1 anig|EHA20665.1 asyd|OJJ58768.1 ater|EAU29455.1 atub|OJI81208.1 aver|OJJ01377.1 awen|OJJ31066.1 azon|OJJ46021.1 acar|OOF98419.1 acar|OOF99514.1 azon|OJJ42344.1

Group96: abra|OJJ71749.1 abra|OJJ72735.1 acar|OOF90204.1 acar|OOF95966.1 afla|EED52022.1 afla|EED52611.1 afum|EAL85108.1 afum|EAL87310.1 agla|OJJ78437.1 agla|OJJ82556.1 akaw|GAA85163.1 akaw|GAA85341.1 aluc|OJZ86473.1 aluc|OJZ90703.1 anig|EHA22158.1 anig|EHA24129.1 aory|EIT73811.1 aory|EIT74178.1 aory|EIT81402.1 asyd|OJJ54106.1 asyd|OJJ56840.1 aver|OJJ03498.1 aver|OJJ08629.1 awen|OJJ31677.1 awen|OJJ33864.1 afis|EAW23882.1 agla|OJJ80999.1 anid|CBF77189.1 ater|EAU33464.1 atub|OJI82130.1 atub|OJI84056.1 azon|OJJ42521.1 azon|OJJ48689.1 acla|EAW11571.1 afis|EAW19784.1 afla|EED45206.1

Group97: abra|OJJ71850.1 abra|OJJ73479.1 acar|OOF96520.1 acar|OOG00674.1 acla|EAW06803.1 acla|EAW13882.1 afis|EAW16352.1 afis|EAW18594.1 afla|EED52753.1 afla|EED53519.1 afum|EAL88977.1

afum|EAL93302.1 agla|OJJ81646.1 agla|OJJ88248.1 akaw|GAA85067.1 akaw|GAA87429.1 aluc|OJZ86584.1 aluc|OJZ88568.1 anid|CBF70389.1 anid|CBF89223.1 anig|EHA18265.1 anig|EHA22076.1 aory|EIT75113.1 aory|EIT78157.1 asyd|OJJ58637.1 asyd|OJJ62519.1 ater|EAU34109.1 ater|EAU34888.1 atub|OJI83939.1 atub|OJI91154.1 aver|OJI96638.1 aver|OJJ01243.1 awen|OJJ31157.1 awen|OJJ37755.1 azon|OJJ45016.1 azon|OJJ46632.1

Group98: abra|OJJ72143.1 abra|OJJ77114.1 acar|OOF93003.1 acar|OOF96781.1 acla|EAW12957.1 acla|EAW14937.1 afis|EAW17702.1 afis|EAW24453.1 afum|EAL86122.1 afum|EAL91517.1 agla|OJJ79467.1 agla|OJJ80300.1 akaw|GAA84780.1 akaw|GAA90815.1 aluc|OJZ84523.1 aluc|OJZ86900.1 anid|CBF73820.1 anid|CBF89927.1 anig|EHA21742.1 anig|EHA27821.1 aory|EIT75637.1 aory|EIT83612.1 asyd|OJJ56569.1 asyd|OJJ62120.1 atub|OJI83619.1 atub|OJI89556.1 aver|OJI96228.1 aver|OJJ06153.1 awen|OJJ32744.1 awen|OJJ37133.1 azon|OJJ45294.1 azon|OJJ50831.1 afla|EED55221.1 ater|EAU29992.1 ater|EAU31909.1 afla|EED45698.1

Group99: abra|OJJ72622.1 abra|OJJ76815.1 acar|OOF91310.1 acar|OOF94641.1 acla|EAW11602.1 acla|EAW13375.1 afis|EAW20489.1 afis|EAW23856.1 afum|EAL84566.1 afum|EAL87284.1 agla|OJJ82510.1 agla|OJJ84827.1 akaw|GAA85243.1 akaw|GAA87206.1 aluc|OJZ84225.1 aluc|OJZ90600.1 anid|CBF77244.1 anid|CBF79220.1 anig|EHA24038.1 anig|EHA27574.1 aory|EIT74289.1 aory|EIT77786.1 asyd|OJJ55735.1 asyd|OJJ56812.1 ater|EAU29567.1 ater|EAU33499.1 atub|OJI82024.1 atub|OJI89859.1 aver|OJJ03523.1 aver|OJJ06821.1 awen|OJJ33189.1 awen|OJJ33817.1 azon|OJJ46391.1 azon|OJJ48719.1 afla|EED52576.1 afla|EED44774.1

Group100: abra|OJJ73827.1 abra|OJJ74268.1 acar|OOF95463.1 acar|OOG00979.1 acla|EAW07312.1 acla|EAW07722.1 afis|EAW22334.1 afis|EAW22754.1 afla|EED50015.1 afla|EED57748.1 afum|EAL90445.1 afum|EAL90858.1 agla|OJJ83671.1 agla|OJJ87714.1 akaw|GAA86755.1 akaw|GAA89445.1 aluc|OJZ88222.1 aluc|OJZ89865.1 anid|CBF88019.1 anid|CBF88645.1 anig|EHA17969.1 anig|EHA27065.1 aory|EIT74682.1 aory|EIT81059.1 asyd|OJJ63522.1 asyd|OJJ63881.1 ater|EAU38966.1 ater|EAU39471.1 atub|OJI84943.1 atub|OJI90822.1 aver|OJI97058.1 aver|OJI97408.1 awen|OJJ34912.1 awen|OJJ35426.1 azon|OJJ45572.1 azon|OJJ49750.1

Group101: abra|OJJ76503.1 abra|OJJ76504.1 acar|OOF99462.1 acar|OOF99465.1 afis|EAW18906.1 afis|EAW18907.1 afla|EED50840.1 afla|EED50841.1 afum|EAL93638.1 afum|EAL93639.1 agla|OJJ81511.1 agla|OJJ81512.1 akaw|GAA82419.1 akaw|GAA82420.1 aluc|OJZ91697.1 aluc|OJZ91699.1 anid|CBF69713.1 anid|CBF69715.1 anig|EHA22712.1 anig|EHA22713.1 aory|EIT78986.1 aory|EIT79030.1 asyd|OJJ59043.1 asyd|OJJ59044.1 ater|EAU38075.1 ater|EAU38076.1 atub|OJI86152.1 atub|OJI86154.1 aver|OJJ01638.1 aver|OJJ01639.1 awen|OJJ32463.1 awen|OJJ32464.1 azon|OJJ47220.1 azon|OJJ47221.1 acla|EAW14191.1 acla|EAW14192.1

Group102: afis|EAW17317.1 afis|EAW24567.1 afla|EED50210.1 afla|EED51099.1 afla|EED56609.1 afla|EED57630.1 anid|CBF82898.1 anid|CBF82901.1 aory|EIT75441.1 aory|EIT77262.1 aory|EIT79641.1 aory|EIT81362.1 aory|EIT82960.1 aory|EIT82255.1 aory|EIT81401.1 ater|EAU30305.1 ater|EAU39298.1 awen|OJJ35064.1 awen|OJJ38659.1 abra|OJJ69840.1 acar|OOF93363.1 acla|EAW13292.1 afla|EED57061.1 afum|EAL91153.1 akaw|GAA87820.1 aluc|OJZ85456.1 anig|EHA24998.1 asyd|OJJ62103.1 asyd|OJJ63346.1 atub|OJI80107.1 aver|OJI96209.1 aver|OJI97614.1 afla|EED52057.1 afum|EAL86230.1 acla|EAW15044.1 azon|OJJ48710.1

Group103: afla|EED46026.1 afla|EED49987.1 akaw|GAA89413.1 akaw|GAA90839.1 aluc|OJZ82994.1 aluc|OJZ88260.1 aory|EIT74653.1 aory|EIT75776.1 atub|OJI89228.1 atub|OJI90859.1 awen|OJJ30039.1 awen|OJJ35338.1 abra|OJJ65964.1 acar|OOF92910.1 acla|EAW13583.1 afis|EAW20018.1 afis|EAW22701.1 afum|EAL84899.2 afum|EAL90491.1 agla|OJJ87998.1 anid|CBF84731.1 anig|EHA21408.1 asyd|OJJ53303.1 ater|EAU29378.1 ater|EAU39001.1 aver|OJJ08438.1 azon|OJJ43685.1 abra|OJJ73794.1 acar|OOG00953.1 acla|EAW07686.1 agla|OJJ79008.1 anid|CBF88080.1 anig|EHA18005.1 asyd|OJJ63557.1 aver|OJI97375.1 azon|OJJ49780.1

Group104: agla|OJJ79992.1 agla|OJJ79999.1 azon|OJJ42536.1 azon|OJJ46701.1 abra|OJJ72154.1 acar|OOF95729.1 afis|EAW21487.1 afla|EED50612.1 agla|OJJ88432.1 anid|CBF86065.1 anig|EHA21734.1 aory|EIT78148.1 asyd|OJJ57414.1 ater|EAU29971.1 awen|OJJ31235.1 abra|OJJ77874.1 agla|OJJ85397.1 akaw|GAA83778.1 aluc|OJZ90949.1 anig|EHA19656.1 atub|OJI82940.1 azon|OJJ42437.1 afis|EAW18426.1 aluc|OJZ92434.1 awen|OJJ38296.1 akaw|GAA83564.1 ater|EAU38826.1 afla|EED48407.1 afla|EED49503.1 aory|EIT82647.1 awen|OJJ38298.1 agla|OJJ81249.1 aver|OJJ05768.1 asyd|OJJ52949.1 aver|OJI97602.1 azon|OJJ46505.1

Group105: abra|OJJ76497.1 acar|OOF99470.1 acla|EAW14363.1 afis|EAW18913.1 afla|EED50836.1 afum|EAL93644.1 agla|OJJ81517.1 akaw|GAA82414.1 aluc|OJZ91703.1 anid|CBF69695.1 anig|EHA22717.1 aory|EIT79010.1 asyd|OJJ59053.1 ater|EAU38080.1 atub|OJI86163.1 aver|OJJ01647.1 awen|OJJ31506.1 azon|OJJ41989.1 azon|OJJ47222.1 abra|OJJ76500.1 acar|OOF99467.1 acla|EAW14196.1 afis|EAW18912.1 afla|EED50837.1 afum|EAL93643.1 akaw|GAA82416.1 aluc|OJZ91702.1 anid|CBF69697.1 anig|EHA22716.1 aory|EIT79064.1 asyd|OJJ59051.1 ater|EAU38079.1 atub|OJI86161.1 awen|OJJ31505.1 azon|OJJ47223.1 azon|OJJ41991.1

Group106: abra|OJJ72950.1 abra|OJJ74416.1 acla|EAW06831.1 acla|EAW11196.1 afis|EAW15926.1 afis|EAW16381.1 afis|EAW19938.1 afum|EAL88562.1 afum|EAL89006.1 akaw|GAA85531.1 akaw|GAA87307.1 aluc|OJZ83460.1 aluc|OJZ89710.1 anid|CBF81021.1 anid|CBF89371.1 atub|OJI82352.1 atub|OJI85092.1 aver|OJI96612.1 aver|OJJ06329.1 awen|OJJ35654.1 awen|OJJ37914.1 acar|OOF96132.1 afla|EED54935.1 agla|OJJ89379.1 anig|EHA24304.1 aory|EIT80345.1 asyd|OJJ56383.1 ater|EAU35529.1 azon|OJJ48008.1 acar|OOF95327.1 afla|EED49933.1 anig|EHA26941.1 aory|EIT83444.1 asyd|OJJ62476.1 ater|EAU34071.1

Group107: abra|OJJ68266.1 acar|OOF97546.1 acla|EAW06826.1 afis|EAW16377.1 afla|EED53460.1 afum|EAL89002.1 agla|OJJ78595.1 akaw|GAA91930.1 aluc|OJZ82692.1 anid|CBF74547.1 anig|EHA27369.1 aory|EIT75115.1 asyd|OJJ57687.1 ater|EAU34075.1 atub|OJI88924.1 aver|OJJ05493.1 awen|OJJ37911.1 azon|OJJ45090.1 abra|OJJ68268.1 acar|OOF97548.1 acla|EAW07026.1 afis|EAW16652.1 afla|EED44948.1 afum|EAL89256.1 agla|OJJ78685.1 akaw|GAA91932.1 aluc|OJZ82690.1 anid|CBF74545.1 anig|EHA27367.1 aory|EIT73496.1 ater|EAU39576.1 atub|OJI88922.1 aver|OJJ05494.1 awen|OJJ40062.1 azon|OJJ45565.1

Group108: abra|OJJ69218.1 acar|OOF91100.1 acla|EAW08627.1 afis|EAW15804.1 afla|EED51241.1 afum|EAL85546.1 agla|OJJ81971.1 akaw|GAA92227.1 aluc|OJZ86200.1 anid|CBF70912.1 anig|EHA24423.1 aory|EIT83194.1 asyd|OJJ52602.1 ater|EAU33444.1 atub|OJI79993.1 aver|OJJ04628.1 awen|OJJ38514.1 azon|OJJ43738.1 abra|OJJ69219.1 acar|OOF91099.1 acla|EAW08626.1 afis|EAW15805.1 afla|EED51240.1 afum|EAL85545.1 akaw|GAA92228.1 aluc|OJZ86201.1 anid|CBF70910.1 anig|EHA24422.1 aory|EIT83177.1 asyd|OJJ52601.1 ater|EAU33443.1 atub|OJI79994.1 aver|OJJ04627.1 awen|OJJ38516.1 azon|OJJ43737.1

Group109: abra|OJJ73774.1 acar|OOG00931.1 acla|EAW07658.1 afis|EAW22676.1 afla|EED49957.1 afum|EAL90516.1 agla|OJJ79041.1 akaw|GAA88402.1 aluc|OJZ88280.1 anid|CBF88133.1 anig|EHA18025.1 aory|EIT83452.1 asyd|OJJ63590.1 ater|EAU39033.1 atub|OJI90880.1 aver|OJI97343.1 awen|OJJ35305.1 azon|OJJ49811.1 abra|OJJ73775.1 acar|OOG00932.1 acla|EAW07660.1 afis|EAW22677.1 afla|EED49958.1 afum|EAL90515.1 agla|OJJ79040.1 akaw|GAA88401.1 aluc|OJZ88279.1 anid|CBF88131.1 anig|EHA18024.1 aory|EIT83440.1 asyd|OJJ63589.1 ater|EAU39032.1 atub|OJI90879.1 aver|OJI97344.1 azon|OJJ49810.1

Group110: abra|OJJ74284.1 acar|OOF89817.1 acla|EAW07328.1 afis|EAW22360.1 afla|EED57768.1 afum|EAL90831.1 agla|OJJ87609.1 akaw|GAA86740.1 aluc|OJZ89849.1 anid|CBF88679.1 anig|EHA27050.1 aory|EIT81097.1 asyd|OJJ63900.1 ater|EAU39494.1 atub|OJI84957.1 aver|OJI97040.1 awen|OJJ34934.1 azon|OJJ45587.1 abra|OJJ74285.1 acar|OOF95446.1 acla|EAW07329.1 afis|EAW22361.1 afla|EED57769.1 afum|EAL90830.1 agla|OJJ87610.1 akaw|GAA86739.1 aluc|OJZ89848.1 anid|CBF88686.1 anig|EHA27049.1 aory|EIT81110.1 asyd|OJJ63902.1 ater|EAU39495.1 atub|OJI84958.1 aver|OJI97038.1 awen|OJJ34935.1

Group111: abra|OJJ74383.1 acar|OOF95354.1 acla|EAW07424.1 afis|EAW22450.1 afla|EED54465.1 afum|EAL90741.1 agla|OJJ87572.1 akaw|GAA87284.1 aluc|OJZ89735.1 anid|CBF88925.1 anig|EHA26961.1 aory|EIT81125.1 asyd|OJJ62853.1 ater|EAU39257.1 atub|OJI85066.1 aver|OJI96906.1 awen|OJJ35075.1 azon|OJJ50032.1 abra|OJJ74384.1 acar|OOF95353.1 acla|EAW07423.1 afis|EAW22449.1 afla|EED54467.1 afum|EAL90742.1 akaw|GAA87285.1 aluc|OJZ89734.1 anid|CBF88927.1 anig|EHA26960.1 aory|EIT81127.1 asyd|OJJ62852.1 ater|EAU39255.1 atub|OJI85067.1 aver|OJI96905.1 awen|OJJ35076.1 azon|OJJ50031.1

Group112: abra|OJJ75811.1 acar|OOG00066.1 acla|EAW14862.1 afis|EAW24390.1 afla|EED47820.1 afum|EAL86062.1 agla|OJJ78870.1 akaw|GAA84525.1 aluc|OJZ89258.1 anid|CBF73709.1 anig|EHA23310.1 aory|EIT77118.1 asyd|OJJ56644.1 ater|EAU30344.1 atub|OJI85541.1 aver|OJJ06082.1 awen|OJJ32814.1 azon|OJJ49018.1 abra|OJJ75812.1 acar|OOG00063.1 acla|EAW14861.1 afis|EAW24389.1 afum|EAL86061.1 agla|OJJ83285.1 akaw|GAA84526.1 aluc|OJZ89259.1 anid|CBF73707.1 anig|EHA23309.1 aory|EIT77172.1 asyd|OJJ56645.1 ater|EAU30345.1 atub|OJI85540.1 aver|OJJ06081.1 awen|OJJ32815.1 azon|OJJ49017.1

Group113: abra|OJJ77674.1 acar|OOF99062.1 acla|EAW06583.1 afis|EAW16009.1 afla|EED56386.1 afum|EAL88639.1 agla|OJJ86034.1 akaw|GAA83958.1 aluc|OJZ80845.1 anid|CBF74571.1 anig|EHA19478.1 aory|EIT83011.1 asyd|OJJ57651.1 ater|EAU33634.1 atub|OJI82722.1 aver|OJJ05511.1 awen|OJJ30456.1 azon|OJJ45447.1 abra|OJJ77675.1 acar|OOF99061.1 acla|EAW06587.1 afis|EAW16010.1 afla|EED56387.1 afum|EAL88640.1 agla|OJJ86035.1 akaw|GAA83957.1 aluc|OJZ80844.1 anid|CBF74569.1 anig|EHA19479.1 aory|EIT82998.1 ater|EAU33633.1 atub|OJI82723.1 aver|OJJ05510.1 awen|OJJ30455.1 azon|OJJ45446.1

Group114: abra|OJJ65981.1 abra|OJJ73719.1 acar|OOF92892.1 acar|OOG00881.1 acla|EAW07608.1 acla|EAW13600.1 afis|EAW20005.1 afis|EAW22638.1 afla|EED46007.1 afla|EED49889.1 afum|EAL84913.1 afum|EAL90554.2 akaw|GAA88444.1 akaw|GAA90854.1 aluc|OJZ82979.1 aluc|OJZ88332.1 anid|CBF84697.1 anid|CBF88205.1 anig|EHA18067.1 anig|EHA21394.1 asyd|OJJ53314.1 asyd|OJJ63639.1 atub|OJI89212.1 atub|OJI90927.1 aver|OJI97297.1 aver|OJJ08429.1 awen|OJJ30019.1 awen|OJJ35261.1 agla|OJJ80980.1 aory|EIT72460.1 aory|EIT75522.1 ater|EAU29347.1 ater|EAU39068.1 azon|OJJ44553.1

Group115: abra|OJJ66667.1 abra|OJJ72896.1 acar|OOF93682.1 acar|OOF93767.1 acar|OOF96087.1 acla|EAW11355.1 acla|EAW11808.1 afis|EAW18139.1 afis|EAW19418.1 afla|EED50375.1 afla|EED58065.1 afum|EAL87076.1 afum|EAL87496.1 agla|OJJ81131.1 agla|OJJ83822.1 akaw|GAA85483.1 akaw|GAA89616.1 aluc|OJZ80780.1 aluc|OJZ83405.1 anig|EHA24256.1 anig|EHA27249.1 aory|EIT79750.1 aory|EIT83148.1 atub|OJI82294.1 atub|OJI87632.1 awen|OJJ35581.1 awen|OJJ36816.1 azon|OJJ42986.1 azon|OJJ46941.1 anid|CBF74365.1 asyd|OJJ57784.1 ater|EAU34724.1 aver|OJJ05380.1 ater|EAU35462.1

Group116: abra|OJJ69228.1 abra|OJJ71754.1 asyd|OJJ52597.1 asyd|OJJ53400.1 aver|OJJ04201.1 aver|OJJ04624.1 abra|OJJ69224.1 acar|OOF91096.1 acla|EAW08622.1 afis|EAW15810.1 afla|EED51237.1 afum|EAL85541.1 agla|OJJ81976.1 akaw|GAA92232.1 aluc|OJZ86204.1 anid|CBF70905.1 anig|EHA24418.1 aory|EIT83188.1 ater|EAU33440.1 atub|OJI79998.1 awen|OJJ38521.1 azon|OJJ43734.1 acar|OOF91092.1 acla|EAW08620.1 afis|EAW15813.1 afla|EED51233.1 afum|EAL85538.1 akaw|GAA92236.1 aluc|OJZ86208.1 anid|CBF70903.1 aory|EIT83195.1 ater|EAU33435.1 atub|OJI80002.1 awen|OJJ38526.1

Group117: abra|OJJ69289.1 abra|OJJ70324.1 acar|OOF91780.1 acar|OOF94433.1 acla|EAW12318.1 acla|EAW14678.1 afis|EAW16903.1 afis|EAW24209.1 afla|EED45264.1 afla|EED48062.1 afum|EAL85879.1 afum|EAL86439.1 akaw|GAA88161.1 akaw|GAA92580.1 aluc|OJZ82118.1 aluc|OJZ82593.1 anig|EHA27482.1 anig|EHA28613.1 asyd|OJJ53986.1 asyd|OJJ54375.1 ater|EAU30213.1 ater|EAU31159.1 atub|OJI88319.1 atub|OJI88828.1 aver|OJJ05329.1 aver|OJJ08084.1 awen|OJJ29716.1 awen|OJJ33087.1 azon|OJJ42612.1 azon|OJJ44859.1 agla|OJJ79662.1 anid|CBF74267.1 aory|EIT83358.1 aory|EIT73698.1

Group118: abra|OJJ69846.1 abra|OJJ72321.1 acar|OOF94891.1 acar|OOF97566.1 afis|EAW19582.1 afis|EAW19642.1 afla|EED47805.1 afla|EED56993.1 agla|OJJ79220.1 agla|OJJ80784.1 akaw|GAA86072.1 akaw|GAA87813.1 aluc|OJZ85448.1 aluc|OJZ90307.1 anig|EHA23794.1 anig|EHA24991.1 aory|EIT77130.1 aory|EIT77504.1 atub|OJI80119.1 atub|OJI81731.1 aver|OJJ05597.1 aver|OJJ07110.1 acla|EAW12002.1 afum|EAL86880.1 anid|CBF71346.1 asyd|OJJ55096.1 ater|EAU32886.1 awen|OJJ41887.1 azon|OJJ42708.1 azon|OJJ46844.1 acla|EAW07815.1 anid|CBF74628.1 asyd|OJJ57585.1 ater|EAU30349.1

Group119: abra|OJJ70036.1 abra|OJJ72033.1 acla|EAW08504.1 acla|EAW09509.1 afis|EAW17096.1 afis|EAW21624.1 afla|EED48527.1 afla|EED57027.1 afla|EED57851.1 akaw|GAA84899.1 akaw|GAA88365.1 aluc|OJZ82290.1 aluc|OJZ86772.1 anig|EHA18733.1 anig|EHA21909.1 anig|EHA26569.1 aory|EIT72294.1 aory|EIT73160.1 aory|EIT81085.1 ater|EAU38312.1 ater|EAU38711.1 atub|OJI83756.1 atub|OJI88149.1 aver|OJJ03867.1 aver|OJJ08171.1 acar|OOF96691.1 afum|EAL92660.1 agla|OJJ79695.1 anid|CBF71556.1 asyd|OJJ52536.1 awen|OJJ33874.1 azon|OJJ48675.1 afla|EED46381.1 afla|EED46382.1

Group120: abra|OJJ72283.1 abra|OJJ78033.1 acar|OOF92932.1 acar|OOF94938.1 afla|EED49548.1 afla|EED50919.1 aluc|OJZ90262.1 aluc|OJZ91590.1 aory|EIT79028.1 aory|EIT82573.1 asyd|OJJ53419.1 asyd|OJJ54983.1 asyd|OJJ55080.1 asyd|OJJ65156.1 aver|OJJ07128.1 aver|OJJ08332.1 awen|OJJ32213.1 awen|OJJ32251.1 awen|OJJ41909.1 afis|EAW20797.1 afum|EAL84832.1 agla|OJJ85015.1 akaw|GAA86035.1 anid|CBF71371.1 anig|EHA23758.1 ater|EAU32868.1 atub|OJI81688.1 azon|OJJ47360.1 abra|OJJ76621.1 afla|EED51770.1 akaw|GAA92835.1 anig|EHA22614.1 aory|EIT81741.1 atub|OJI86039.1

Group121: abra|OJJ74623.1 abra|OJJ74837.1 acar|OOF95085.1 acar|OOF95151.1 acar|OOF95841.1 acla|EAW10761.1 acla|EAW14613.1 afis|EAW19343.1 afis|EAW24815.1 afla|EED47168.1 afla|EED50136.1 afla|EED56891.1 afum|EAL89459.1 afum|EAL94062.1 akaw|GAA81986.1 akaw|GAA83583.1 aluc|OJZ92204.1 aluc|OJZ92414.1 anig|EHA26551.1 anig|EHA26754.1 aory|EIT74736.1 aory|EIT79324.1 aory|EIT77583.1 ater|EAU36492.1 ater|EAU39073.1 atub|OJI86632.1 atub|OJI86844.1 azon|OJJ48665.1 azon|OJJ49253.1 agla|OJJ82911.1 anid|CBF85881.1 asyd|OJJ64378.1 aver|OJI98474.1 awen|OJJ40401.1

Group122: agla|OJJ84376.1 agla|OJJ85709.1 asyd|OJJ57959.1 asyd|OJJ61061.1 abra|OJJ71937.1 acar|OOF96594.1 acar|OOF96595.1 acla|EAW09592.1 acla|EAW09593.1 afis|EAW21545.1 afis|EAW21546.1 afla|EED55996.1 afla|EED55997.1 afum|EAL92565.1 afum|EAL92566.1 agla|OJJ85708.1 akaw|GAA84987.1 aluc|OJZ86674.1 anid|CBF76366.1 anid|CBF76368.1 anig|EHA21999.1 anig|EHA22000.1 aory|EIT76685.1 aory|EIT76776.1 asyd|OJJ61062.1 ater|EAU34993.1 ater|EAU34994.1 atub|OJI83854.1 aver|OJJ00378.1 aver|OJJ00379.1 awen|OJJ39144.1 awen|OJJ39146.1 azon|OJJ51476.1 azon|OJJ51477.1

Group123: abra|OJJ67331.1 acar|OOF92482.1 acla|EAW06403.1 afis|EAW18507.1 afla|EED52893.1 afum|EAL93201.1 agla|OJJ85114.1 akaw|GAA86485.1 aluc|OJZ85026.1 anid|CBF70247.1 anig|EHA25247.1 aory|EIT78591.1 asyd|OJJ58761.1 ater|EAU30064.1 atub|OJI80571.1 aver|OJJ01371.1 awen|OJJ31060.1 azon|OJJ50571.1 abra|OJJ67332.1 acar|OOF92481.1 acla|EAW06402.1 afis|EAW18506.1 afla|EED52895.1 afum|EAL93200.1 akaw|GAA86484.1 aluc|OJZ85025.1 anig|EHA25246.1 aory|EIT78664.1 asyd|OJJ58760.1 ater|EAU30065.1 atub|OJI80572.1 aver|OJJ01368.1 awen|OJJ31059.1 azon|OJJ50569.1

Group124: abra|OJJ68518.1 acar|OOF92724.1 acla|EAW12845.1 afis|EAW18008.1 afla|EED51887.1 afum|EAL91820.1 agla|OJJ80274.1 akaw|GAA91330.1 aluc|OJZ82732.1 anid|CBF80034.1 anig|EHA21184.1 aory|EIT81755.1 asyd|OJJ58272.1 ater|EAU31524.1 atub|OJI88965.1 aver|OJJ02152.1 awen|OJJ41532.1 azon|OJJ48252.1 atub|OJI79785.1 abra|OJJ69012.1 acar|OOF92723.1 acla|EAW12846.1 afis|EAW18009.1 afla|EED51886.1 afum|EAL91821.1 agla|OJJ80275.1 akaw|GAA90529.1 aluc|OJZ85998.1 anid|CBF80035.1 anig|EHA24612.1 aory|EIT81715.1 ater|EAU31525.1 awen|OJJ41531.1 azon|OJJ48251.1

Group125: abra|OJJ69958.1 acar|OOF93468.1 acla|EAW08829.1 afis|EAW15568.1 afla|EED49078.1 afum|EAL85765.1 agla|OJJ85182.1 akaw|GAA87702.1 aluc|OJZ85324.1 anid|CBF78110.1 anig|EHA25514.1 aory|EIT82568.1 ater|EAU32562.1 atub|OJI80248.1 abra|OJJ69959.1 acla|EAW08833.1 afis|EAW15567.1 afla|EED49076.1 afla|EED49077.1 afum|EAL85766.1 agla|OJJ85184.1 akaw|GAA87701.1 anid|CBF78112.1 anid|CBF78114.1 aory|EIT82498.1 aory|EIT82525.1 asyd|OJJ55899.1 ater|EAU32566.1 atub|OJI80249.1 aver|OJJ04844.1 awen|OJJ38227.1 azon|OJJ43068.1 acla|EAW08832.1 azon|OJJ47853.1

Group126: abra|OJJ71382.1 acar|OOF97787.1 acla|EAW10157.1 afis|EAW25437.1 afla|EED53925.1 afum|EAL90095.1 agla|OJJ87082.1 akaw|GAA82612.1 aluc|OJZ87158.1 anid|CBF77596.1 anig|EHA20417.1 aory|EIT78347.1 asyd|OJJ53142.1 ater|EAU34609.1 atub|OJI81482.1 aver|OJJ03710.1 awen|OJJ41165.1 azon|OJJ44743.1 abra|OJJ71383.1 acar|OOF97786.1 acla|EAW10158.1 afis|EAW25436.1 afla|EED53924.1 afum|EAL90094.1 agla|OJJ87081.1 akaw|GAA82611.1 aluc|OJZ87157.1 anid|CBF77594.1 anig|EHA20416.1 aory|EIT78262.1 asyd|OJJ53143.1 ater|EAU34608.1 atub|OJI81483.1 aver|OJJ03709.1

Group127: abra|OJJ72469.1 acar|OOF94768.1 acla|EAW12702.1 afis|EAW17865.1 afla|EED46230.1 afum|EAL91673.1 agla|OJJ83099.1 akaw|GAA86207.1 aluc|OJZ90456.1 anid|CBF76910.1 anig|EHA23918.1 aory|EIT76409.1 asyd|OJJ57089.1 ater|EAU33080.1 atub|OJI81883.1 aver|OJJ03211.1 awen|OJJ41706.1 azon|OJJ48402.1 abra|OJJ72470.1 acar|OOF94765.1 acla|EAW12703.1 afis|EAW17866.1 afla|EED46231.1 afum|EAL91674.1 agla|OJJ83086.1 akaw|GAA86208.1 aluc|OJZ90457.1 anid|CBF76912.1 anig|EHA23919.1 aory|EIT76527.1 ater|EAU33081.1 atub|OJI81884.1 awen|OJJ41705.1 azon|OJJ48401.1

Group128: abra|OJJ72932.1 acar|OOF96112.1 acla|EAW06458.1 afis|EAW15954.1 afla|EED58104.1 afum|EAL88587.1 agla|OJJ89359.1 akaw|GAA85514.1 aluc|OJZ83442.1 anid|CBF81063.1 anig|EHA24288.1 aory|EIT79164.1 asyd|OJJ56456.1 ater|EAU35503.1 atub|OJI82333.1 aver|OJJ06266.1 awen|OJJ35622.1 azon|OJJ46902.1 abra|OJJ72934.1 acar|OOF96115.1 acla|EAW06457.1 afis|EAW15953.1 afla|EED58105.1 afum|EAL88586.1 akaw|GAA85515.1 aluc|OJZ83443.1 anid|CBF81061.1 anig|EHA24289.1 aory|EIT79130.1 asyd|OJJ56455.1 ater|EAU35504.1 atub|OJI82334.1 aver|OJJ06267.1 awen|OJJ35623.1

Group129: abra|OJJ76465.1 acar|OOF99703.1 acla|EAW13966.1 afis|EAW18675.1 afla|EED50624.1 afum|EAL93395.2 agla|OJJ82401.1 aluc|OJZ91745.1 anid|CBF70588.1 anig|EHA22748.1 aory|EIT78145.1 asyd|OJJ58523.1 ater|EAU38235.1 atub|OJI86203.1 aver|OJJ01124.1 awen|OJJ32147.1 azon|OJJ46731.1 abra|OJJ76466.1 acar|OOF99704.1 acla|EAW13965.1 afis|EAW18674.1 afla|EED50625.1 afum|EAL93394.1 akaw|GAA82377.1 aluc|OJZ91744.1 anid|CBF70586.1 anig|EHA22747.1 aory|EIT77989.1 asyd|OJJ58524.1 ater|EAU38236.1 atub|OJI86202.1 aver|OJJ01125.1 awen|OJJ32146.1 azon|OJJ46730.1

Group130: abra|OJJ66702.1 abra|OJJ76322.1 acar|OOF93706.1 acar|OOF93792.1 afla|EED45227.1 afla|EED56652.1 anid|CBF78509.1 anid|CBF81402.1 anig|EHA22872.1 anig|EHA27277.1 aory|EIT73751.1 aory|EIT79654.1 asyd|OJJ59404.1 asyd|OJJ65313.1 ater|EAU29738.1 ater|EAU34746.1 atub|OJI86345.1 atub|OJI87602.1 acla|EAW11833.1 afis|EAW19444.1 afum|EAL87051.1 agla|OJJ81158.1 akaw|GAA89644.1 aluc|OJZ80745.1 aver|OJJ02869.1 awen|OJJ36845.1 azon|OJJ42319.1 azon|OJJ42955.1 acar|OOF97765.1 akaw|GAA82251.1 aluc|OJZ91893.1 aver|OJI99451.1 awen|OJJ40922.1

Group131: abra|OJJ68714.1 abra|OJJ76939.1 acar|OOF91389.1 acar|OOF92214.1 afum|EAL84486.2 afum|EAL91972.2 aluc|OJZ81814.1 aluc|OJZ84338.1 anig|EHA22401.1 anig|EHA27672.1 atub|OJI84386.1 atub|OJI89730.1 awen|OJJ33278.1 awen|OJJ41371.1 azon|OJJ48840.1 azon|OJJ50652.1 acla|EAW12532.1 acla|EAW13466.1 afis|EAW20578.1 afis|EAW24117.1 afla|EED46815.1 afla|EED52313.1 agla|OJJ88717.1 akaw|GAA87100.1 akaw|GAA89867.1 anid|CBF86537.1 aory|EIT74384.1 asyd|OJJ64866.1 ater|EAU37106.1 aver|OJI98979.1 anid|CBF78940.1 asyd|OJJ55646.1 aver|OJJ06731.1

Group132: abra|OJJ69799.1 abra|OJJ75720.1 acar|OOF90260.1 acar|OOF93578.1 afis|EAW21759.1 afis|EAW24741.1 afum|EAL89387.1 afum|EAL92789.1 akaw|GAA84456.1 akaw|GAA87860.1 aluc|OJZ85501.1 aluc|OJZ89172.1 anid|CBF80400.1 anid|CBF83678.1 anig|EHA23383.1 anig|EHA25034.1 aory|EIT75373.1 aory|EIT79769.1 asyd|OJJ54761.1 asyd|OJJ55920.1 asyd|OJJ60823.1 atub|OJI80064.1 atub|OJI85618.1 aver|OJJ00149.1 aver|OJJ04873.1 aver|OJJ07409.1 azon|OJJ45021.1 azon|OJJ48200.1 acla|EAW09376.1 afla|EED45538.1 agla|OJJ84391.1 ater|EAU38431.1 awen|OJJ34043.1

Group133: abra|OJJ71453.1 abra|OJJ72857.1 acar|OOF96054.1 acar|OOF97693.1 acar|OOF97744.1 acar|OOF98763.1 afis|EAW22749.1 afis|EAW24033.1 afla|EED45033.1 afla|EED49120.1 afum|EAL87464.1 afum|EAL90450.1 akaw|GAA82558.1 akaw|GAA85456.1 aluc|OJZ83364.1 aluc|OJZ87090.1 anig|EHA20361.1 anig|EHA24229.1 aory|EIT73568.1 aory|EIT78580.1 asyd|OJJ59869.1 asyd|OJJ65349.1 ater|EAU29414.1 ater|EAU29711.1 atub|OJI81548.1 atub|OJI82255.1 aver|OJI99497.1 aver|OJJ02418.1 acla|EAW11389.1 agla|OJJ86999.1 anid|CBF75320.1 awen|OJJ35505.1 azon|OJJ46979.1

Group134: abra|OJJ72895.1 abra|OJJ73247.1 acar|OOF96086.1 acar|OOG00487.1 acla|EAW06921.1 acla|EAW11356.1 afis|EAW16538.1 afis|EAW18138.1 afla|EED56131.1 afla|EED58064.1 afum|EAL87495.1 afum|EAL89135.1 agla|OJJ82784.1 agla|OJJ83821.1 akaw|GAA85482.1 akaw|GAA87933.1 aluc|OJZ81180.1 aluc|OJZ83404.1 anig|EHA18498.1 anig|EHA24255.1 aory|EIT76761.1 aory|EIT79757.1 ater|EAU35461.1 ater|EAU36601.1 atub|OJI82293.1 atub|OJI91377.1 awen|OJJ35580.1 awen|OJJ40283.1 azon|OJJ46943.1 azon|OJJ49145.1 anid|CBF81745.1 asyd|OJJ59599.1 aver|OJJ02669.1

Group135: anid|CBF74543.1 anid|CBF80800.1 asyd|OJJ54487.1 asyd|OJJ57688.1 aver|OJJ05495.1 aver|OJJ07692.1 awen|OJJ30771.1 awen|OJJ31710.1 abra|OJJ67440.1 acar|OOF92387.1 acar|OOF92388.1 acla|EAW06324.1 afis|EAW18438.1 afla|EED52977.1 afum|EAL93120.1 agla|OJJ85004.1 akaw|GAA86390.1 aluc|OJZ84924.1 anid|CBF70039.1 anig|EHA25142.1 anig|EHA25143.1 aory|EIT78691.1 aory|EIT78713.1 asyd|OJJ58866.1 ater|EAU29862.1 atub|OJI80675.1 aver|OJJ01467.1 azon|OJJ50475.1 acla|EAW06323.1 afis|EAW18437.1 afum|EAL93119.1 ater|EAU29854.1 afla|EED52975.1

Group136: aory|EIT76219.1 abra|OJJ66160.1 acla|EAW07091.1 ater|EAU30184.1 abra|OJJ76943.1 acar|OOF91392.1 acla|EAW14645.1 afis|EAW23759.1 afla|EED45469.1 afum|EAL87193.2 agla|OJJ78794.1 akaw|GAA87097.1 aluc|OJZ84341.1 anid|CBF86208.1 anig|EHA27675.1 anig|EHA27676.1 aory|EIT75611.1 asyd|OJJ64548.1 atub|OJI89727.1 aver|OJI98644.1 awen|OJJ40873.1 afis|EAW21192.1 anid|CBF78488.1 asyd|OJJ58352.1 ater|EAU36866.1 aver|OJJ02055.1 awen|OJJ31298.1 afis|EAW22111.1 afla|EED48116.1 aver|OJJ07112.1 azon|OJJ50585.1 awen|OJJ29841.1 awen|OJJ30060.1

Group137: abra|OJJ68073.1 acar|OOF92659.1 acla|EAW14013.1 afis|EAW18719.1 afla|EED50674.1 afum|EAL93443.2 agla|OJJ82339.1 akaw|GAA92398.1 aluc|OJZ83259.1 anid|CBF70677.1 anig|EHA21113.1 aory|EIT78139.1 asyd|OJJ58472.1 ater|EAU38185.1 atub|OJI87680.1 aver|OJJ01073.1 awen|OJJ32224.1 azon|OJJ46775.1 abra|OJJ68477.1 acar|OOF92676.1 acla|EAW07215.1 afis|EAW22228.1 afla|EED57619.1 afum|EAL90966.1 agla|OJJ87821.1 anid|CBF88467.1 anig|EHA21112.1 aory|EIT82940.1 asyd|OJJ63779.1 ater|EAU34233.1 aver|OJI97158.1 awen|OJJ34810.1 azon|OJJ45740.1

Group138: abra|OJJ71306.1 acar|OOF97864.1 acla|EAW10234.1 afis|EAW25358.1 afla|EED53840.1 afum|EAL90020.1 agla|OJJ83908.1 akaw|GAA82685.1 aluc|OJZ87238.1 anid|CBF77463.1 anig|EHA20484.1 aory|EIT78472.1 asyd|OJJ53218.1 ater|EAU34506.1 atub|OJI81404.1 aver|OJJ03634.1 awen|OJJ41084.1 azon|OJJ44664.1 abra|OJJ71307.1 acla|EAW10233.1 afis|EAW25359.1 afla|EED53841.1 afum|EAL90021.1 agla|OJJ83909.1 akaw|GAA82684.1 aluc|OJZ87237.1 anid|CBF77461.1 aory|EIT78435.1 asyd|OJJ53217.1 ater|EAU34507.1 atub|OJI81405.1 awen|OJJ41085.1 azon|OJJ44665.1

Group139: abra|OJJ71587.1 acar|OOF96317.1 acla|EAW15133.1 afis|EAW20105.1 afla|EED53369.1 afum|EAL85222.1 agla|OJJ88850.1 akaw|GAA89336.1 aluc|OJZ86327.1 anid|CBF84906.1 anig|EHA21551.1 aory|EIT75191.1 asyd|OJJ63169.1 ater|EAU38724.1 atub|OJI84207.1 aver|OJI97785.1 awen|OJJ31876.1 azon|OJJ46259.1 abra|OJJ71588.1 acla|EAW15132.1 afis|EAW20104.1 afum|EAL85221.1 agla|OJJ88849.1 akaw|GAA89335.1 aluc|OJZ86328.1 anid|CBF84908.1 anig|EHA21550.1 aory|EIT75224.1 asyd|OJJ63168.1 atub|OJI84206.1 aver|OJI97786.1 awen|OJJ31877.1 azon|OJJ46257.1

Group140: abra|OJJ72666.1 acar|OOF94609.1 acla|EAW11529.1 afis|EAW23925.1 afla|EED50596.1 afum|EAL87350.1 agla|OJJ79403.1 akaw|GAA85287.1 aluc|OJZ90645.1 anid|CBF77250.1 anig|EHA24080.1 aory|EIT82072.1 asyd|OJJ56809.1 ater|EAU32310.1 atub|OJI82075.1 aver|OJJ03525.1 awen|OJJ39029.1 azon|OJJ51358.1 abra|OJJ72668.1 acla|EAW11530.1 afis|EAW23924.1 afla|EED50597.1 afum|EAL87349.1 agla|OJJ79402.1 akaw|GAA85288.1 aluc|OJZ90646.1 anid|CBF77248.1 anig|EHA24081.1 aory|EIT78140.1 asyd|OJJ56810.1 ater|EAU32312.1 awen|OJJ39031.1 azon|OJJ51359.1

Group141: abra|OJJ74273.1 acar|OOF95457.1 acla|EAW07318.1 afis|EAW22350.1 afla|EED57754.1 afum|EAL90842.1 agla|OJJ87709.1 akaw|GAA86750.1 aluc|OJZ89860.1 anid|CBF88653.1 anig|EHA27060.1 aory|EIT81054.1 ater|EAU39489.1 aver|OJI97054.1 awen|OJJ34904.1 azon|OJJ45578.1 abra|OJJ74279.1 acar|OOF95452.1 acla|EAW07323.1 afis|EAW22355.1 afla|EED57761.1 afum|EAL90836.1 agla|OJJ87608.1 akaw|GAA86745.1 aluc|OJZ89854.1 anid|CBF88666.1 anig|EHA27055.1 aory|EIT81031.1 asyd|OJJ63892.1 atub|OJI84952.1 aver|OJI97047.1 awen|OJJ34929.1 azon|OJJ45582.1

Group142: abra|OJJ68218.1 abra|OJJ69789.1 acar|OOF93590.1 acar|OOF97507.1 agla|OJJ78902.1 agla|OJJ86249.1 akaw|GAA83228.1 akaw|GAA87869.1 aluc|OJZ83728.1 aluc|OJZ85510.1 anid|CBF84393.1 anid|CBF86693.1 atub|OJI80056.1 atub|OJI90104.1 acla|EAW09054.1 acla|EAW12123.1 afis|EAW25629.1 afla|EED55183.1 afum|EAL84317.1 anig|EHA18876.1 aory|EIT80665.1 aory|EIT81643.1 asyd|OJJ55507.1 ater|EAU32706.1 aver|OJJ06584.1 aver|OJJ08175.1 awen|OJJ36048.1 azon|OJJ43184.1 afla|EED51722.1 anig|EHA25042.1 asyd|OJJ53581.1 anid|CBF83169.1

Group143: abra|OJJ68305.1 abra|OJJ68976.1 afla|EED55146.1 afla|EED57528.1 agla|OJJ80112.1 agla|OJJ84537.1 akaw|GAA83169.1 akaw|GAA90501.1 aluc|OJZ83668.1 aluc|OJZ85964.1 aory|EIT81602.1 aory|EIT81847.1 asyd|OJJ59831.1 asyd|OJJ65307.1 atub|OJI79752.1 atub|OJI90039.1 awen|OJJ31501.1 awen|OJJ36340.1 azon|OJJ46215.1 azon|OJJ48729.1 acar|OOF91551.1 acar|OOF96683.1 acla|EAW14213.1 afis|EAW19068.1 afum|EAL93797.1 anid|CBF78527.1 anig|EHA18823.1 anig|EHA24640.1 aver|OJI99445.1 afla|EED54434.1 aory|EIT75875.1 ater|EAU32117.1

Group144: abra|OJJ69780.1 abra|OJJ73124.1 acar|OOF90902.1 acar|OOF91910.1 afis|EAW16809.1 afis|EAW20667.1 afis|EAW16810.1 atub|OJI88609.1 atub|OJI91509.1 awen|OJJ29822.1 awen|OJJ37778.1 azon|OJJ42117.1 azon|OJJ42607.1 afis|EAW17206.1 afis|EAW23422.1 afla|EED49098.1 afum|EAL85399.1 afum|EAL86742.1 afum|EBA27333.1 agla|OJJ84598.1 akaw|GAA88041.1 akaw|GAA92683.1 aluc|OJZ81300.1 aluc|OJZ85605.1 anig|EHA18609.1 ater|EAU32778.1 anig|EHA25864.1 afum|EAL86743.1 afis|EAW20625.1 afla|EED49487.1 atub|OJI83712.1 aluc|OJZ90138.1

Group145: abra|OJJ70543.1 abra|OJJ73773.1 acar|OOF94608.1 acar|OOG00929.1 afis|EAW15411.1 afis|EAW22675.1 afum|EAL84235.1 afum|EAL90517.1 akaw|GAA83389.1 akaw|GAA88403.1 aluc|OJZ83888.1 aluc|OJZ88281.1 anid|CBF74491.1 anid|CBF88137.1 anig|EHA18026.1 anig|EHA19025.1 aory|EIT72732.1 aory|EIT83438.1 asyd|OJJ57716.1 asyd|OJJ63598.1 ater|EAU36301.1 ater|EAU39034.1 atub|OJI90255.1 atub|OJI90881.1 azon|OJJ45068.1 azon|OJJ49812.1 acla|EAW07657.1 afla|EED49956.1 agla|OJJ79042.1 aver|OJI97338.1 aver|OJJ05451.1 awen|OJJ35304.1

Group146: abra|OJJ70875.1 abra|OJJ77724.1 acar|OOF98245.1 acar|OOF99016.1 acla|EAW10463.1 acla|EAW10819.1 acla|EAW11708.1 afis|EAW23731.1 afis|EAW25104.1 afum|EAL87170.1 afum|EAL89775.1 akaw|GAA83911.1 akaw|GAA91738.1 akaw|GAA93340.1 aluc|OJZ87647.1 aluc|OJZ90786.1 ater|EAU29552.1 ater|EAU31452.1 atub|OJI80998.1 atub|OJI82773.1 afla|EED55342.1 anid|CBF82779.1 anid|CBF86021.1 anig|EHA20840.1 asyd|OJJ55916.1 asyd|OJJ64453.1 ater|EAU39370.1 aver|OJI98556.1 aver|OJJ04856.1 awen|OJJ40718.1 anig|EHA19519.1 azon|OJJ48020.1

Group147: abra|OJJ73071.1 abra|OJJ77012.1 acla|EAW09005.1 acla|EAW15322.1 afis|EAW20323.1 afis|EAW25545.1 akaw|GAA87037.1 akaw|GAA88090.1 aluc|OJZ81354.1 aluc|OJZ84412.1 asyd|OJJ54270.1 asyd|OJJ55537.1 ater|EAU32181.1 ater|EAU39595.1 atub|OJI89661.1 atub|OJI91558.1 aver|OJJ05226.1 aver|OJJ06615.1 acar|OOG00304.1 afla|EED44875.1 afum|EAL85448.1 agla|OJJ78910.1 anid|CBF82101.1 anid|CBF83180.1 anig|EHA18655.1 aory|EIT83261.1 awen|OJJ36038.1 azon|OJJ51343.1 afla|EED54130.1 afum|EAL84263.1 anig|EHA27738.1 aory|EIT76050.1

Group148: abra|OJJ76652.1 abra|OJJ77533.1 acar|OOF99194.1 acar|OOF99350.1 afla|EED53116.1 afla|EED53300.1 akaw|GAA84088.1 akaw|GAA92992.1 aluc|OJZ81003.1 aluc|OJZ91553.1 anid|CBF82367.1 anid|CBF86914.1 anig|EHA19357.1 anig|EHA22588.1 asyd|OJJ54198.1 asyd|OJJ64607.1 ater|EAU30240.1 ater|EAU37451.1 ater|EAU37789.1 atub|OJI82582.1 atub|OJI86006.1 aver|OJI96893.1 aver|OJJ08526.1 aver|OJI98704.1 awen|OJJ31647.1 awen|OJJ37655.1 acla|EAW06705.1 agla|OJJ88350.1 aory|EIT74767.1 aory|EIT74899.1 azon|OJJ44918.1 asyd|OJJ53794.1

Group149: acar|OOF94510.1 acar|OOG00617.1 agla|OJJ78704.1 agla|OJJ82589.1 agla|OJJ88673.1 awen|OJJ33431.1 awen|OJJ40048.1 abra|OJJ73366.1 acar|OOG00573.1 acla|EAW12876.1 afis|EAW17800.1 afum|EAL91608.1 akaw|GAA83118.1 aluc|OJZ88674.1 anid|CBF87487.1 anig|EHA18356.1 asyd|OJJ52958.1 ater|EAU31398.1 atub|OJI91255.1 aver|OJJ08926.1 abra|OJJ73415.1 afla|EED44896.1 akaw|GAA83076.1 aluc|OJZ88622.1 anig|EHA18315.1 aory|EIT83248.1 asyd|OJJ62646.1 atub|OJI91207.1 aver|OJJ07782.1 azon|OJJ50122.1 afis|EAW20384.1 asyd|OJJ54467.1

Group150: acar|OOF95065.1 acar|OOF95131.1 acar|OOF95066.1 acar|OOF95132.1 abra|OJJ74648.1 acla|EAW10738.1 acla|EAW10739.1 afis|EAW24835.1 afis|EAW24836.1 afla|EED47139.1 afla|EED47140.1 afum|EAL89480.1 afum|EAL89481.2 agla|OJJ82028.1 akaw|GAA81962.1 akaw|GAA81963.1 aluc|OJZ92226.1 anid|CBF74584.1 anid|CBF74586.1 anig|EHA26730.1 aory|EIT79207.1 aory|EIT79226.1 asyd|OJJ57656.1 ater|EAU36469.1 ater|EAU36470.1 atub|OJI86656.1 aver|OJJ05516.1 aver|OJJ05517.1 awen|OJJ40422.1 awen|OJJ40423.1 azon|OJJ49280.1 azon|OJJ49281.1

Group151: acar|OOF95215.1 acar|OOG00549.1 afla|EED45224.1 afla|EED52255.1 afla|EED57542.1 afum|EAL86491.1 afum|EAL91022.1 aory|EIT73609.1 aory|EIT74331.1 aory|EIT81831.1 asyd|OJJ52204.1 asyd|OJJ54774.1 asyd|OJJ64753.1 ater|EAU30189.1 ater|EAU32767.1 ater|EAU34175.1 aver|OJI98855.1 aver|OJJ07392.1 aver|OJJ07305.1 abra|OJJ74541.1 acla|EAW07155.1 acla|EAW12276.1 afis|EAW16964.1 agla|OJJ87885.1 akaw|GAA82061.1 aluc|OJZ92121.1 anid|CBF86686.1 anig|EHA26824.1 atub|OJI86554.1 awen|OJJ34744.1 azon|OJJ42779.1 anid|CBF80465.1

Group152: anig|EHA18119.1 anig|EHA26153.1 awen|OJJ35539.1 awen|OJJ38334.1 acla|EAW13628.1 afis|EAW24672.1 acar|OOG00839.1 afla|EED48354.1 afum|EAL92114.1 akaw|GAA88497.1 aluc|OJZ88392.1 anid|CBF87369.1 anig|EHA18120.1 aory|EIT72297.1 asyd|OJJ53356.1 atub|OJI90987.1 aver|OJJ08385.1 acar|OOG00840.1 acla|EAW07713.1 afis|EAW20752.1 afla|EED48353.1 afum|EAL84865.1 agla|OJJ88184.1 akaw|GAA88496.1 anid|CBF82358.1 aory|EIT72300.1 asyd|OJJ54748.1 azon|OJJ43889.1 afum|EAL86291.2 afis|EAW21091.1 aver|OJJ08174.1 asyd|OJJ54203.1

Group153: asyd|OJJ60702.1 asyd|OJJ64821.1 aver|OJI98934.1 aver|OJJ00028.1 abra|OJJ72028.1 acar|OOF96677.1 afis|EAW21619.1 afla|EED57848.1 akaw|GAA84904.1 aluc|OJZ86765.1 anid|CBF83493.1 anig|EHA21914.1 aory|EIT81067.1 ater|EAU38309.1 atub|OJI83762.1 awen|OJJ37096.1 azon|OJJ48684.1 abra|OJJ77154.1 acar|OOF93040.1 acla|EAW12917.1 afis|EAW17740.1 afla|EED45741.1 afum|EAL91554.1 agla|OJJ81711.1 akaw|GAA90776.1 aluc|OJZ84568.1 anid|CBF86610.1 anig|EHA27862.1 aory|EIT75552.1 ater|EAU31867.1 atub|OJI89514.1 azon|OJJ50798.1

Group154: atub|OJI81651.1 atub|OJI86062.1 abra|OJJ65911.1 acar|OOF94961.1 acla|EAW12054.1 afis|EAW19697.1 afla|EED57032.1 afum|EAL86830.1 agla|OJJ80819.1 akaw|GAA92321.1 aluc|OJZ90219.1 anid|CBF71392.1 anig|EHA23717.1 aory|EIT82283.1 asyd|OJJ55041.1 ater|EAU33691.1 aver|OJI98387.1 awen|OJJ41939.1 azon|OJJ50749.1 abra|OJJ76597.1 akaw|GAA82501.1 aluc|OJZ91610.1 anig|EHA22639.1 acla|EAW11329.1 afis|EAW18166.1 afla|EED56329.1 afum|EAL87526.1 agla|OJJ85952.1 anid|CBF85749.1 aory|EIT73387.1 asyd|OJJ64295.1 aver|OJJ07167.1

Group155: abra|OJJ66622.1 acar|OOF91045.1 acla|EAW10875.1 afis|EAW20983.1 afla|EED50338.1 afum|EAL84647.2 agla|OJJ84060.1 akaw|GAA91103.1 aluc|OJZ81427.1 anid|CBF82618.1 anig|EHA25995.1 aory|EIT77260.1 asyd|OJJ54020.1 ater|EAU38633.1 atub|OJI88753.1 aver|OJJ08720.1 awen|OJJ34328.1 azon|OJJ48073.1 afla|EED50340.1 akaw|GAA91102.1 aluc|OJZ81426.1 acar|OOF91046.1 acla|EAW10874.1 afis|EAW20984.1 afla|EED50339.1 afum|EAL84646.1 agla|OJJ84063.1 anid|CBF82622.1 aory|EIT77350.1 atub|OJI88754.1 awen|OJJ34327.1 azon|OJJ48069.1

Group156: abra|OJJ68143.1 acar|OOF97481.1 acla|EAW13069.1 afis|EAW17595.1 afla|EED49225.1 afum|EAL91408.1 agla|OJJ81333.1 akaw|GAA92375.1 aluc|OJZ83234.1 anid|CBF90206.1 anig|EHA21091.1 aory|EIT83153.1 asyd|OJJ61974.1 ater|EAU37033.1 atub|OJI87702.1 aver|OJI96067.1 awen|OJJ37251.1 azon|OJJ50946.1 abra|OJJ68185.1 acla|EAW13066.1 afis|EAW17598.1 afla|EED49223.1 afum|EAL91411.1 agla|OJJ81335.1 akaw|GAA83261.1 aluc|OJZ83767.1 anid|CBF90203.1 asyd|OJJ61976.1 ater|EAU37035.1 atub|OJI90138.1 aver|OJI96069.1 awen|OJJ37248.1

Group157: abra|OJJ70602.1 acar|OOF97117.1 acla|EAW08180.1 afis|EAW23215.1 afla|EED47037.1 afum|EAL88269.1 akaw|GAA89942.1 aluc|OJZ83947.1 anig|EHA19076.1 aory|EIT72544.1 asyd|OJJ60180.1 ater|EAU36345.1 atub|OJI90318.1 awen|OJJ39449.1 abra|OJJ70604.1 acar|OOF97116.1 acla|EAW08178.1 afis|EAW23213.1 afum|EBA27340.1 agla|OJJ78731.1 akaw|GAA89944.1 aluc|OJZ83949.1 anid|CBF74742.1 anig|EHA19078.1 aory|EIT72666.1 asyd|OJJ60182.1 ater|EAU36343.1 atub|OJI90320.1 aver|OJJ04082.1 awen|OJJ39447.1 azon|OJJ47649.1 anid|CBF74745.1

Group158: abra|OJJ71268.1 acar|OOF97905.1 acar|OOF97907.1 acla|EAW10269.1 acla|EAW10270.1 afis|EAW25324.1 afis|EAW25325.1 afla|EED53799.1 afla|EED53800.1 afum|EAL89985.1 afum|EAL89986.1 agla|OJJ83862.1 agla|OJJ83863.1 akaw|GAA82722.1 aluc|OJZ87276.1 anid|CBF75258.1 anid|CBF75260.1 anig|EHA20517.1 anig|EHA20518.1 aory|EIT78367.1 aory|EIT78397.1 asyd|OJJ59900.1 asyd|OJJ59901.1 ater|EAU34470.1 atub|OJI81365.1 aver|OJJ02385.1 aver|OJJ02386.1 awen|OJJ41038.1 awen|OJJ41039.1 azon|OJJ44627.1 ater|EAU34472.1 ater|EAU34471.1

Group159: abra|OJJ72382.1 acar|OOF94837.1 acla|EAW11953.1 acla|EAW11954.1 afis|EAW19590.1 afis|EAW19591.1 afla|EED56945.1 afum|EAL86927.1 afum|EAL86928.1 agla|OJJ80730.1 agla|OJJ80732.1 akaw|GAA86129.1 akaw|GAA86131.1 aluc|OJZ90369.1 anid|CBF71264.1 anid|CBF71266.1 anig|EHA23843.1 anig|EHA23844.1 aory|EIT77519.1 aory|EIT77568.1 asyd|OJJ55144.1 asyd|OJJ55146.1 ater|EAU32934.1 ater|EAU32936.1 atub|OJI81795.1 aver|OJJ07061.1 aver|OJJ07063.1 awen|OJJ41805.1 awen|OJJ41808.1 azon|OJJ47024.1 azon|OJJ47026.1 acar|OOF94836.1

Group160: abra|OJJ72953.1 acar|OOF96134.1 acla|EAW11198.1 afis|EAW15928.1 afla|EED54933.1 afum|EAL88564.1 agla|OJJ89377.1 akaw|GAA85534.1 aluc|OJZ83463.1 anid|CBF81024.1 anig|EHA24306.1 aory|EIT80331.1 asyd|OJJ56385.1 ater|EAU35524.1 atub|OJI82355.1 aver|OJJ06327.1 awen|OJJ35652.1 azon|OJJ46886.1 abra|OJJ72954.1 acar|OOF96135.1 acla|EAW11199.1 afis|EAW15929.1 afla|EED54932.1 afum|EAL88565.1 akaw|GAA85535.1 anid|CBF81026.1 anig|EHA24307.1 aory|EIT80358.1 asyd|OJJ56386.1 ater|EAU35523.1 awen|OJJ35651.1 azon|OJJ46887.1

Group161: abra|OJJ74876.1 acar|OOF95878.1 acar|OOF95879.1 acla|EAW08494.1 acla|EAW08495.1 afis|EAW23540.1 afis|EAW23541.1 afla|EED57438.1 afla|EED57439.1 afum|EAL87925.1 afum|EAL87926.1 agla|OJJ86795.1 akaw|GAA83544.1 akaw|GAA83545.1 aluc|OJZ92453.1 anid|CBF89661.1 anid|CBF89663.1 anig|EHA26514.1 anig|EHA26515.1 aory|EIT80116.1 aory|EIT80129.1 asyd|OJJ62276.1 asyd|OJJ62277.1 ater|EAU35326.1 ater|EAU35327.1 atub|OJI86881.1 atub|OJI86882.1 aver|OJI96385.1 aver|OJI96386.1 awen|OJJ39871.1 awen|OJJ39872.1 azon|OJJ43230.1

Group162: abra|OJJ75341.1 acar|OOF89856.1 acla|EAW09853.1 afis|EAW21282.1 afla|EED54833.1 afum|EAL92299.1 akaw|GAA92034.1 aluc|OJZ92904.1 anid|CBF83359.1 anig|EHA26059.1 aory|EIT79370.1 ater|EAU35911.1 atub|OJI87363.1 awen|OJJ39417.1 abra|OJJ75940.1 acar|OOF99970.1 acla|EAW09854.1 afis|EAW21281.1 afla|EED54832.1 afum|EAL92298.1 agla|OJJ87478.1 akaw|GAA84642.1 aluc|OJZ89387.1 anid|CBF83358.1 anig|EHA23197.1 aory|EIT79445.1 asyd|OJJ60618.1 ater|EAU35910.1 atub|OJI85410.1 aver|OJI99942.1 awen|OJJ39418.1 azon|OJJ44300.1

Group163: abra|OJJ75791.1 acar|OOG00087.1 acar|OOG00097.1 acla|EAW14890.1 acla|EAW14891.1 afis|EAW24412.1 afis|EAW24413.1 afla|EED47763.1 afla|EED47764.1 afum|EAL86081.1 afum|EAL86082.1 agla|OJJ79215.1 agla|OJJ79216.1 akaw|GAA84506.1 akaw|GAA84510.1 aluc|OJZ89235.1 aluc|OJZ89239.1 anid|CBF73743.1 anid|CBF73745.1 anig|EHA23328.1 aory|EIT77117.1 aory|EIT77194.1 asyd|OJJ56623.1 ater|EAU30316.1 ater|EAU30317.1 atub|OJI85559.1 aver|OJJ06102.1 aver|OJJ06103.1 awen|OJJ32796.1 awen|OJJ32798.1 azon|OJJ49036.1 azon|OJJ49037.1

Group164: abra|OJJ76816.1 acar|OOF91311.1 acla|EAW13376.1 afis|EAW20490.1 afla|EED44773.1 afum|EAL84564.1 agla|OJJ84837.1 akaw|GAA87205.1 aluc|OJZ84226.1 anig|EHA27575.1 aory|EIT77777.1 asyd|OJJ55730.1 atub|OJI89858.1 aver|OJJ06816.1 awen|OJJ33190.1 azon|OJJ46392.1 abra|OJJ76822.1 acar|OOF91317.1 acla|EAW13382.1 afis|EAW20496.1 afla|EED44772.1 afum|EAL84558.1 agla|OJJ84806.1 akaw|GAA87200.1 aluc|OJZ84233.1 anid|CBF79209.1 anig|EHA27580.1 aory|EIT77769.1 ater|EAU37190.1 atub|OJI89852.1 awen|OJJ33196.1 azon|OJJ46400.1

Group165: abra|OJJ78214.1 acla|EAW11236.1 afis|EAW18269.1 afla|EED48735.1 afum|EAL87634.1 agla|OJJ88158.1 akaw|GAA85813.1 aluc|OJZ91286.1 anid|CBF79514.1 anig|EHA19971.1 aory|EIT73143.1 asyd|OJJ57887.1 ater|EAU33276.1 atub|OJI83276.1 aver|OJJ04507.1 awen|OJJ30643.1 azon|OJJ47972.1 abra|OJJ78215.1 acar|OOF98641.1 acla|EAW11235.1 afis|EAW18270.1 afla|EED48736.1 afum|EAL87635.1 agla|OJJ88157.1 akaw|GAA85812.1 aluc|OJZ91287.1 anid|CBF79512.1 anig|EHA19972.1 aory|EIT73147.1 asyd|OJJ57886.1 atub|OJI83277.1 aver|OJJ04508.1

Group166: abra|OJJ65897.1 abra|OJJ72196.1 acar|OOF94957.1 acar|OOF96818.1 agla|OJJ80319.1 agla|OJJ83741.1 agla|OJJ82619.1 akaw|GAA84738.1 akaw|GAA86008.1 aluc|OJZ86948.1 aluc|OJZ90234.1 anig|EHA21698.1 anig|EHA23734.1 asyd|OJJ53466.1 asyd|OJJ56558.1 ater|EAU29975.1 ater|EAU30764.1 atub|OJI81664.1 atub|OJI83575.1 aver|OJJ06162.1 aver|OJJ08289.1 azon|OJJ44535.1 azon|OJJ49495.1 acla|EAW07004.1 afis|EAW16621.1 afis|EAW23621.1 afla|EED55210.1 afum|EAL89216.2 anid|CBF73835.1 awen|OJJ32732.1 awen|OJJ40080.1

Group167: abra|OJJ66384.1 abra|OJJ76979.1 acar|OOF91425.1 acar|OOF94058.1 acar|OOF94128.1 acla|EAW11869.1 acla|EAW15042.1 afis|EAW19480.1 afis|EAW24565.1 afla|EED48343.1 afla|EED56690.1 afum|EAL86228.1 afum|EAL87015.1 akaw|GAA87066.1 akaw|GAA90250.1 aluc|OJZ83090.1 aluc|OJZ84377.1 anig|EHA27710.1 anig|EHA28152.1 aory|EIT73915.1 aory|EIT76297.1 atub|OJI87853.1 atub|OJI89692.1 azon|OJJ42424.1 azon|OJJ42921.1 asyd|OJJ56521.1 ater|EAU34790.1 aver|OJJ06201.1 awen|OJJ36886.1 anid|CBF84609.1 ater|EAU37362.1

Group168: abra|OJJ66675.1 abra|OJJ75504.1 acar|OOF90835.1 acar|OOF93687.1 acar|OOF93772.1 afis|EAW16470.1 afis|EAW19425.1 afum|EAL87069.1 afum|EAL89064.1 akaw|GAA84268.1 akaw|GAA89623.1 aluc|OJZ80773.1 aluc|OJZ88960.1 anig|EHA23560.1 anig|EHA27257.1 atub|OJI85829.1 atub|OJI87625.1 acla|EAW06851.1 afla|EED50383.1 agla|OJJ81137.1 anid|CBF74378.1 aory|EIT83131.1 asyd|OJJ57778.1 ater|EAU34729.1 aver|OJJ05387.1 awen|OJJ36823.1 azon|OJJ42980.1 afla|EED49104.1 asyd|OJJ57938.1 aver|OJJ04456.1 aory|EIT82490.1

Group169: abra|OJJ67685.1 abra|OJJ69630.1 acar|OOF92680.1 acar|OOF94075.1 acar|OOF94145.1 acar|OOF96015.1 afum|EAL86373.1 afum|EAL87420.1 akaw|GAA89197.1 akaw|GAA91610.1 aluc|OJZ81633.1 aluc|OJZ82435.1 anig|EHA25810.1 anig|EHA28336.1 atub|OJI88006.1 atub|OJI88547.1 afla|EED55077.1 aory|EIT81950.1 acla|EAW12376.1 afis|EAW16822.1 agla|OJJ86924.1 anid|CBF82987.1 asyd|OJJ64732.1 ater|EAU35425.1 aver|OJI98836.1 awen|OJJ35458.1 azon|OJJ51285.1 anid|CBF77369.1 asyd|OJJ53260.1 aver|OJJ03591.1 acla|EAW11432.1

Group170: acar|OOF91973.1 acar|OOF94079.1 acar|OOF94149.1 acar|OOF99292.1 afla|EED45949.1 afla|EED47803.1 aluc|OJZ79501.1 aluc|OJZ79636.1 aluc|OJZ81691.1 aluc|OJZ80076.1 aory|EIT75701.1 aory|EIT77131.1 asyd|OJJ52720.1 asyd|OJJ61396.1 asyd|OJJ61537.1 ater|EAU29987.1 ater|EAU30155.1 aver|OJJ00725.1 aver|OJJ04752.1 awen|OJJ32293.1 awen|OJJ35892.1 abra|OJJ67625.1 acla|EAW15287.1 afis|EAW20776.1 afum|EAL84844.1 agla|OJJ85083.1 akaw|GAA89145.1 anid|CBF87439.1 anig|EHA25753.1 atub|OJI88489.1 azon|OJJ42146.1

Group171: acar|OOF92319.1 acar|OOF92361.1 abra|OJJ67475.1 acla|EAW06297.1 afis|EAW18404.1 afla|EED53004.1 afum|EAL93084.1 agla|OJJ84961.1 akaw|GAA86362.1 aluc|OJZ84889.1 anid|CBF70745.1 anig|EHA25118.1 aory|EIT77437.1 asyd|OJJ58438.1 ater|EAU29821.1 atub|OJI80708.1 aver|OJJ01038.1 awen|OJJ31746.1 abra|OJJ67478.1 acla|EAW06294.1 afis|EAW18400.1 afla|EED53006.1 afum|EAL93078.1 agla|OJJ84971.1 akaw|GAA86360.1 anid|CBF70749.1 aory|EIT77424.1 ater|EAU29816.1 atub|OJI80711.1 awen|OJJ31749.1 azon|OJJ50474.1

Group172: acar|OOF98484.1 acar|OOF98497.1 abra|OJJ78390.1 acar|OOF98498.1 acla|EAW09142.1 acla|EAW09143.1 afis|EAW22004.1 afis|EAW22005.1 afla|EED46729.1 afum|EAL93019.1 afum|EAL93020.1 agla|OJJ84093.1 agla|OJJ84094.1 akaw|GAA85641.1 akaw|GAA85642.1 aluc|OJZ91469.1 anid|CBF82660.1 anid|CBF82662.1 anig|EHA20131.1 aory|EIT77865.1 aory|EIT77967.1 asyd|OJJ61666.1 asyd|OJJ61667.1 ater|EAU31123.1 ater|EAU31125.1 atub|OJI83462.1 aver|OJJ00987.1 awen|OJJ34299.1 awen|OJJ34300.1 azon|OJJ43338.1 azon|OJJ43339.1

Group173: afis|EAW21024.1 afis|EAW22408.1 afum|EAL90784.1 afum|EAL92063.1 awen|OJJ31391.1 abra|OJJ74333.1 acar|OOF95396.1 acla|EAW07383.1 afla|EED56072.1 akaw|GAA86696.1 aluc|OJZ89796.1 anid|CBF88792.1 anig|EHA27010.1 aory|EIT76782.1 asyd|OJJ62898.1 ater|EAU39300.1 atub|OJI85009.1 aver|OJI96952.1 awen|OJJ34988.1 acar|OOF95397.1 acla|EAW07385.1 afis|EAW22410.1 afla|EED52490.1 afum|EAL90782.1 akaw|GAA86694.1 anid|CBF88794.1 aory|EIT82779.1 ater|EAU39295.1 azon|OJJ50553.1 anig|EHA27008.1 aver|OJI96951.1

Group174: afla|EED46122.1 afla|EED53117.1 afla|EED48931.1 afla|EED51100.1 akaw|GAA86153.1 akaw|GAA92990.1 aory|EIT74792.1 aory|EIT79985.1 aory|EIT81307.1 atub|OJI81824.1 atub|OJI86011.1 abra|OJJ72410.1 acar|OOF94819.1 acla|EAW14266.1 afis|EAW19012.1 afum|EAL93746.1 agla|OJJ81770.1 aluc|OJZ90394.1 anig|EHA23863.1 aory|EIT76367.1 ater|EAU37790.1 awen|OJJ31644.1 abra|OJJ76649.1 acar|OOF99352.1 aluc|OJZ91559.1 anid|CBF86747.1 anig|EHA22591.1 asyd|OJJ64703.1 aver|OJI98807.1 azon|OJJ47337.1 afis|EAW16798.1

Group175: afum|EAL87729.1 afum|EAL88789.1 abra|OJJ65746.1 afis|EAW16109.1 afla|EED49345.1 agla|OJJ83935.1 akaw|GAA93207.1 aory|EIT79717.1 asyd|OJJ57625.1 ater|EAU33192.1 awen|OJJ41813.1 azon|OJJ42378.1 abra|OJJ77889.1 acar|OOF98892.1 afla|EED48269.1 akaw|GAA83769.1 aluc|OJZ90959.1 anig|EHA19664.1 asyd|OJJ64307.1 atub|OJI82950.1 aver|OJJ00097.1 aory|EIT72931.1 ater|EAU33625.1 acla|EAW06656.1 aluc|OJZ79714.1 asyd|OJJ60763.1 afla|EED57955.1 agla|OJJ86139.1 awen|OJJ38786.1 aory|EIT79719.1 aver|OJJ05561.1

Group176: ater|EAU33722.1 ater|EAU38738.1 abra|OJJ71609.1 acar|OOF96326.1 acla|EAW15150.1 acla|EAW15152.1 afis|EAW20118.1 afis|EAW20120.1 afla|EED53353.1 afla|EED57256.1 afum|EAL85235.1 afum|EBA27224.1 agla|OJJ88826.1 agla|OJJ88831.1 akaw|GAA89318.1 akaw|GAA89319.1 aluc|OJZ86351.1 aluc|OJZ86353.1 anid|CBF84937.1 anid|CBF84939.1 anig|EHA21536.1 aory|EIT74846.1 asyd|OJJ63156.1 ater|EAU38737.1 atub|OJI84184.1 aver|OJI97800.1 aver|OJI97801.1 awen|OJJ31888.1 azon|OJJ46246.1 aory|EIT74143.1 asyd|OJJ63154.1

Group177: awen|OJJ33301.1 awen|OJJ37090.1 abra|OJJ76956.1 acar|OOF91397.1 acla|EAW13475.1 afla|EED52244.1 agla|OJJ88687.1 akaw|GAA87087.1 aluc|OJZ84351.1 anid|CBF78932.1 anig|EHA27688.1 aory|EIT74404.1 asyd|OJJ62348.1 ater|EAU37096.1 atub|OJI89715.1 azon|OJJ50794.1 abra|OJJ77159.1 acar|OOF93045.1 acla|EAW12913.1 afis|EAW17747.1 afla|EED45747.1 afum|EAL91559.1 agla|OJJ81707.1 akaw|GAA90772.1 aluc|OJZ84572.1 anid|CBF86617.1 aory|EIT75770.1 asyd|OJJ64815.1 ater|EAU31860.1 atub|OJI89510.1 aver|OJI98928.1

Group178: abra|OJJ65971.1 acar|OOF92902.1 acla|EAW13592.1 afis|EAW20012.1 afla|EED46018.1 afum|EAL84906.1 akaw|GAA90846.1 aluc|OJZ82987.1 anid|CBF84721.1 anig|EHA21402.1 aory|EIT75680.1 asyd|OJJ53329.1 ater|EAU29354.1 atub|OJI89221.1 aver|OJJ08415.1 awen|OJJ30025.1 abra|OJJ65972.1 acar|OOF92900.1 acla|EAW13593.1 afis|EAW20011.1 afla|EED46017.1 afum|EAL84907.1 agla|OJJ87976.1 anid|CBF83963.1 anig|EHA21401.1 aory|EIT75560.1 asyd|OJJ61425.1 ater|EAU29353.1 aver|OJJ00759.1 awen|OJJ30024.1 azon|OJJ43649.1

Group179: abra|OJJ66528.1 acar|OOF90981.1 acla|EAW10962.1 afis|EAW20897.1 afla|EED50247.1 afum|EAL84735.1 agla|OJJ84007.1 akaw|GAA91185.1 aluc|OJZ81518.1 anid|CBF82460.1 anig|EHA25916.1 aory|EIT77290.1 ater|EAU38531.1 atub|OJI88667.1 awen|OJJ34450.1 azon|OJJ51738.1 abra|OJJ66529.1 acar|OOF90980.1 acla|EAW10961.1 afis|EAW20898.1 afla|EED50248.1 afum|EAL84734.2 agla|OJJ84008.1 akaw|GAA91184.1 aluc|OJZ81519.1 anid|CBF82458.1 aory|EIT77356.1 asyd|OJJ54115.1 atub|OJI88668.1 aver|OJJ08620.1 azon|OJJ51737.1

Group180: abra|OJJ66543.1 acar|OOF90995.1 afla|EED50933.1 agla|OJJ78717.1 akaw|GAA91168.1 aluc|OJZ81496.1 anid|CBF71517.1 anig|EHA25934.1 aory|EIT79048.1 asyd|OJJ54917.1 ater|EAU29874.1 atub|OJI88685.1 aver|OJJ07286.1 awen|OJJ30168.1 abra|OJJ67418.1 acar|OOF92403.1 acla|EAW06346.1 afis|EAW18457.1 afla|EED52951.1 afum|EAL93142.1 agla|OJJ85040.1 akaw|GAA86410.1 aluc|OJZ84950.1 anid|CBF70055.1 anig|EHA25165.1 aory|EIT78701.1 asyd|OJJ58858.1 atub|OJI80652.1 aver|OJJ01459.1 awen|OJJ31009.1 azon|OJJ50523.1

Group181: abra|OJJ67409.1 acar|OOF92406.1 acla|EAW06349.1 afis|EAW18461.1 afla|EED52947.1 afum|EAL93147.1 agla|OJJ85045.1 akaw|GAA86418.1 aluc|OJZ84959.1 anid|CBF70089.1 anig|EHA25173.1 aory|EIT78700.1 asyd|OJJ58841.1 ater|EAU29879.1 atub|OJI80641.1 aver|OJJ01443.1 awen|OJJ31013.1 azon|OJJ50526.1 abra|OJJ67410.1 acar|OOF92404.1 afis|EAW18460.1 afla|EED52948.1 afum|EAL93146.1 akaw|GAA86417.1 aluc|OJZ84958.1 anid|CBF70087.1 anig|EHA25172.1 aory|EIT78733.1 ater|EAU29878.1 atub|OJI80642.1 awen|OJJ31012.1

Group182: abra|OJJ67954.1 acar|OOF92799.1 acla|EAW11644.1 afis|EAW23813.1 afis|EAW23814.1 afla|EED51991.1 afla|EED52002.1 afum|EAL87248.1 agla|OJJ82453.1 akaw|GAA91242.1 akaw|GAA91243.1 aluc|OJZ82827.1 anid|CBF79881.1 anig|EHA21264.1 anig|EHA21265.1 aory|EIT81469.1 aory|EIT81487.1 asyd|OJJ58172.1 ater|EAU31366.1 atub|OJI89057.1 aver|OJJ04231.1 awen|OJJ33769.1 azon|OJJ46009.1 anid|CBF79883.1 asyd|OJJ58173.1 acla|EAW11643.1 afum|EAL87249.1 agla|OJJ82454.1 ater|EAU31369.1 awen|OJJ33770.1 azon|OJJ42873.1

Group183: abra|OJJ68614.1 acar|OOF92124.1 acar|OOF92125.1 acla|EAW12605.1 acla|EAW12606.1 afis|EAW18078.1 afis|EAW18079.1 afla|EED51472.1 afla|EED51473.1 afum|EAL91890.1 afum|EAL91891.1 agla|OJJ87372.1 akaw|GAA92907.1 akaw|GAA92908.1 aluc|OJZ81940.1 aluc|OJZ81941.1 anid|CBF86395.1 anid|CBF86397.1 anig|EHA22308.1 aory|EIT80638.1 aory|EIT80738.1 asyd|OJJ64952.1 asyd|OJJ64954.1 ater|EAU30443.1 ater|EAU30445.1 atub|OJI84493.1 atub|OJI84494.1 aver|OJI99074.1 aver|OJI99076.1 awen|OJJ41467.1 azon|OJJ48924.1

Group184: abra|OJJ71429.1 acar|OOF90402.1 acla|EAW10109.1 afis|EAW25478.1 afla|EED53973.1 afum|EAL90141.1 akaw|GAA82574.1 aluc|OJZ87109.1 anid|CBF77705.1 anig|EHA20376.1 asyd|OJJ53075.1 ater|EAU34650.1 atub|OJI81530.1 aver|OJJ03772.1 awen|OJJ41218.1 azon|OJJ44787.1 abra|OJJ71430.1 acla|EAW10108.1 afis|EAW25479.1 afla|EED53974.1 afum|EAL90144.1 akaw|GAA82573.1 aluc|OJZ87108.1 anid|CBF77703.1 anig|EHA20375.1 aory|EIT78327.1 asyd|OJJ53074.1 ater|EAU34651.1 atub|OJI81531.1 awen|OJJ41220.1 azon|OJJ44786.1

Group185: abra|OJJ73132.1 acar|OOF90742.1 acla|EAW13627.1 afis|EAW21092.1 afla|EED48284.1 afum|EAL92115.1 akaw|GAA92664.1 aluc|OJZ90118.1 anid|CBF83072.1 anig|EHA19695.1 anig|EHA19696.1 asyd|OJJ52355.1 ater|EAU32835.1 atub|OJI84679.1 aver|OJJ00828.1 awen|OJJ31224.1 azon|OJJ46149.1 acar|OOF90935.1 anid|CBF82817.1 aver|OJJ00896.1 acar|OOF90741.1 afis|EAW15628.1 afum|EAL85706.1 aory|EIT80551.1 asyd|OJJ61575.1 acar|OOF98130.1 acar|OOG00091.1 acla|EAW08502.1 afla|EED51590.1 asyd|OJJ61486.1 aory|EIT72957.1

Group186: abra|OJJ74487.1 acar|OOF95262.1 acla|EAW07208.1 afis|EAW22220.1 afla|EED57613.1 afum|EAL90974.1 agla|OJJ87824.1 akaw|GAA87381.1 aluc|OJZ89631.1 anig|EHA26874.1 ater|EAU34230.1 atub|OJI85171.1 aver|OJI97165.1 awen|OJJ34805.1 abra|OJJ74488.1 acar|OOF95261.1 acla|EAW07207.1 afis|EAW22219.1 afla|EED57612.1 afum|EAL90975.1 agla|OJJ87825.1 akaw|GAA87382.1 aluc|OJZ89630.1 anid|CBF88457.1 anig|EHA26873.1 aory|EIT82983.1 asyd|OJJ63772.1 ater|EAU34229.1 atub|OJI85172.1 aver|OJI97166.1 azon|OJJ45743.1

Group187: abra|OJJ74835.1 acar|OOF95828.1 acla|EAW09102.1 afis|EAW25706.1 afla|EED55155.1 afum|EAL84379.1 akaw|GAA83585.1 aluc|OJZ92412.1 anid|CBF83935.1 anig|EHA26553.1 aory|EIT81620.1 asyd|OJJ55434.1 ater|EAU32723.1 atub|OJI86842.1 aver|OJJ06511.1 awen|OJJ36213.1 azon|OJJ43614.1 abra|OJJ76623.1 acar|OOF99375.1 acla|EAW10020.1 afis|EAW22744.1 afla|EED50498.1 afum|EAL90452.1 akaw|GAA92834.1 aluc|OJZ91589.1 anig|EHA22613.1 aory|EIT82109.1 ater|EAU34860.1 atub|OJI86038.1 awen|OJJ40151.1 azon|OJJ48726.1

Group188: abra|OJJ74864.1 acar|OOF95864.1 acar|OOF95866.1 acla|EAW08515.1 acla|EAW08517.1 afis|EAW23553.1 afis|EAW23555.1 afla|EED57448.1 afla|EED57450.1 afum|EAL87911.1 afum|EAL87913.1 agla|OJJ86813.1 agla|OJJ86815.1 akaw|GAA83554.1 akaw|GAA83556.1 aluc|OJZ92442.1 anid|CBF89641.1 anid|CBF89643.1 anig|EHA26525.1 aory|EIT80074.1 aory|EIT80092.1 asyd|OJJ62288.1 asyd|OJJ62290.1 ater|EAU35336.1 ater|EAU35338.1 atub|OJI86871.1 aver|OJI96396.1 aver|OJI96398.1 awen|OJJ39884.1 awen|OJJ39886.1 azon|OJJ43221.1

Group189: abra|OJJ75158.1 acar|OOF97180.1 acar|OOF97181.1 acla|EAW08333.1 acla|EAW08334.1 afis|EAW23368.1 afis|EAW23369.1 afla|EED55588.1 afla|EED55589.1 afum|EAL88104.2 afum|EAL88105.1 agla|OJJ86455.1 agla|OJJ86456.1 akaw|GAA88764.1 akaw|GAA88769.1 aluc|OJZ92728.1 anid|CBF74819.1 anid|CBF74821.1 anig|EHA26273.1 aory|EIT82662.1 aory|EIT82702.1 asyd|OJJ60141.1 asyd|OJJ60142.1 ater|EAU35619.1 ater|EAU35620.1 atub|OJI87152.1 aver|OJJ04121.1 awen|OJJ39518.1 awen|OJJ39519.1 azon|OJJ47713.1 azon|OJJ47715.1

Group190: abra|OJJ75500.1 acar|OOF90833.1 acla|EAW09294.1 afis|EAW21853.1 afla|EED46457.1 afum|EAL92866.1 agla|OJJ84283.1 akaw|GAA84266.1 aluc|OJZ88959.1 anid|CBF76739.1 anig|EHA23561.1 aory|EIT72412.1 asyd|OJJ61254.1 ater|EAU30852.1 atub|OJI85830.1 aver|OJJ00577.1 awen|OJJ34137.1 azon|OJJ44207.1 abra|OJJ75502.1 acar|OOF90834.1 acla|EAW09295.1 afis|EAW21852.1 afla|EED46458.1 afum|EAL92865.1 akaw|GAA84267.1 anid|CBF76737.1 aory|EIT72344.1 asyd|OJJ61253.1 ater|EAU30851.1 awen|OJJ34135.1 azon|OJJ44209.1

Group191: abra|OJJ76457.1 acar|OOF99697.1 acla|EAW13972.1 acla|EAW13973.1 afis|EAW18681.1 afis|EAW18682.1 afla|EED50628.1 afum|EAL93402.1 afum|EAL93403.1 agla|OJJ82393.1 agla|OJJ82394.1 akaw|GAA82369.1 akaw|GAA82370.1 aluc|OJZ91752.1 aluc|OJZ91753.1 anid|CBF70601.1 anid|CBF70603.1 anig|EHA22755.1 anig|EHA22756.1 aory|EIT78067.1 asyd|OJJ58516.1 asyd|OJJ58517.1 ater|EAU38228.1 atub|OJI86210.1 atub|OJI86211.1 aver|OJJ01117.1 aver|OJJ01118.1 awen|OJJ32153.1 azon|OJJ46737.1 afla|EED50629.1 aory|EIT78047.1

Group192: abra|OJJ77384.1 acla|EAW09036.1 afla|EED56897.1 akaw|GAA91377.1 aluc|OJZ84792.1 anig|EHA28060.1 aory|EIT77450.1 asyd|OJJ57351.1 ater|EAU38783.1 atub|OJI89290.1 aver|OJJ05814.1 awen|OJJ35827.1 abra|OJJ78137.1 acar|OOF99382.1 acla|EAW14634.1 afis|EAW22732.1 afla|EED55765.1 afum|EAL90463.1 agla|OJJ78522.1 aluc|OJZ82078.1 anid|CBF78393.1 ater|EAU30739.1 awen|OJJ32040.1 azon|OJJ47350.1 acar|OOF96813.1 anig|EHA25049.1 aluc|OJZ82079.1 atub|OJI88447.1 akaw|GAA87875.1 aluc|OJZ85516.1 atub|OJI80050.1

Group193: abra|OJJ67081.1 abra|OJJ68837.1 abra|OJJ69761.1 agla|OJJ84620.1 agla|OJJ89301.1 aory|EIT74438.1 aory|EIT79603.1 asyd|OJJ52546.1 asyd|OJJ57148.1 asyd|OJJ63033.1 ater|EAU29247.1 ater|EAU29286.1 ater|EAU32050.1 ater|EAU35700.1 aver|OJI97935.1 aver|OJJ03858.1 awen|OJJ30061.1 awen|OJJ31262.1 acar|OOF90552.1 acla|EAW15294.1 afis|EAW20307.1 afla|EED52202.1 afla|EED56612.1 afum|EAL85435.2 akaw|GAA90413.1 aluc|OJZ85816.1 anid|CBF85212.1 atub|OJI79602.1 awen|OJJ29673.1 azon|OJJ45897.1

Group194: abra|OJJ67792.1 abra|OJJ73325.1 anig|EHA18415.1 anig|EHA18416.1 acla|EAW12887.1 afis|EAW17789.1 afla|EED50756.1 afum|EAL91598.1 agla|OJJ84165.1 akaw|GAA92793.1 aluc|OJZ88713.1 anid|CBF75218.1 aory|EIT78111.1 asyd|OJJ53462.1 ater|EAU29260.1 atub|OJI91300.1 aver|OJJ08293.1 awen|OJJ32921.1 acar|OOF95394.1 abra|OJJ75374.1 acar|OOF94473.1 afis|EAW17262.1 afla|EED48869.1 afum|EAL86679.1 akaw|GAA92734.1 aluc|OJZ92942.1 anig|EHA26031.1 aory|EIT81203.1 atub|OJI87401.1 awen|OJJ35771.1

Group195: abra|OJJ68217.1 abra|OJJ69796.1 acar|OOF92535.1 acar|OOF97501.1 akaw|GAA83229.1 akaw|GAA87863.1 aluc|OJZ83729.1 aluc|OJZ85504.1 anig|EHA18877.1 anig|EHA25036.1 aory|EIT80435.1 aory|EIT81864.1 atub|OJI80061.1 atub|OJI90106.1 awen|OJJ31685.1 awen|OJJ32019.1 azon|OJJ42231.1 azon|OJJ46071.1 acla|EAW15175.1 afis|EAW20145.1 afla|EED51644.1 afum|EAL85260.1 agla|OJJ89015.1 anid|CBF81813.1 asyd|OJJ59648.1 ater|EAU39446.1 aver|OJJ02634.1 asyd|OJJ58357.1 ater|EAU33077.1 aver|OJJ02050.1

Group196: abra|OJJ68439.1 abra|OJJ71007.1 acar|OOF93955.1 acar|OOF98135.1 afis|EAW19801.1 afis|EAW25026.1 agla|OJJ79158.1 agla|OJJ87950.1 akaw|GAA82946.1 akaw|GAA86933.1 aluc|OJZ87524.1 aluc|OJZ90058.1 anid|CBF81549.1 anid|CBF82379.1 anig|EHA20734.1 anig|EHA21013.1 asyd|OJJ54138.1 asyd|OJJ59494.1 ater|EAU30748.1 ater|EAU35761.1 atub|OJI81124.1 atub|OJI84749.1 aver|OJJ02780.1 aver|OJJ08598.1 acla|EAW10559.1 afla|EED55448.1 afum|EAL89669.1 aory|EIT80786.1 awen|OJJ40613.1 azon|OJJ49462.1

Group197: abra|OJJ69676.1 abra|OJJ72100.1 acar|OOF90056.1 acar|OOF96745.1 aluc|OJZ82484.1 aluc|OJZ86855.1 anig|EHA21784.1 anig|EHA28290.1 asyd|OJJ60799.1 asyd|OJJ62562.1 atub|OJI83664.1 atub|OJI87948.1 awen|OJJ31984.1 awen|OJJ34027.1 afis|EAW21744.1 afum|EAL92777.1 agla|OJJ81779.1 agla|OJJ84428.1 akaw|GAA84821.1 akaw|GAA88801.1 ater|EAU32512.1 aver|OJI96686.1 azon|OJJ42257.1 acla|EAW09389.1 afla|EED58033.1 anid|CBF83660.1 aory|EIT79754.1 ater|EAU38444.1 aver|OJJ00132.1 acar|OOF95939.1

Group198: abra|OJJ72836.1 abra|OJJ73712.1 agla|OJJ78798.1 agla|OJJ86951.1 akaw|GAA85433.1 akaw|GAA88451.1 aluc|OJZ83336.1 aluc|OJZ88338.1 anig|EHA18073.1 anig|EHA24212.1 aory|EIT75956.1 aory|EIT78575.1 asyd|OJJ53233.1 asyd|OJJ54785.1 ater|EAU29729.1 ater|EAU32826.1 atub|OJI82233.1 atub|OJI90934.1 aver|OJJ03618.1 aver|OJJ07380.1 awen|OJJ35489.1 awen|OJJ38098.1 acar|OOF96038.1 acla|EAW11404.1 afis|EAW24017.1 afla|EED49142.1 afum|EAL87447.1 anid|CBF77418.1 azon|OJJ47000.1 afla|EED54085.1

Group199: acar|OOF90457.1 acar|OOF90464.1 abra|OJJ71887.1 acla|EAW09633.1 afis|EAW21507.1 afla|EED55944.1 afum|EAL92527.1 agla|OJJ85669.1 akaw|GAA85029.1 aluc|OJZ86625.1 anid|CBF76445.1 anig|EHA22041.1 asyd|OJJ61103.1 ater|EAU35031.1 atub|OJI83900.1 aver|OJJ00420.1 awen|OJJ39188.1 azon|OJJ51505.1 acla|EAW09634.1 afis|EAW21506.1 afla|EED55943.1 afum|EAL92526.1 anid|CBF76447.1 asyd|OJJ61104.1 ater|EAU35032.1 aver|OJJ00421.1 awen|OJJ39189.1 acar|OOF96550.1 anig|EHA22040.1 afum|EAL92525.1

Group200: acar|OOF98095.1 acar|OOF99166.1 acla|EAW06529.1 acla|EAW10590.1 afis|EAW16066.1 afis|EAW24978.1 afla|EED48587.1 afla|EED55481.1 afum|EAL88698.1 afum|EAL89638.1 anid|CBF75098.1 anid|CBF81606.1 asyd|OJJ59522.1 asyd|OJJ60007.1 ater|EAU33563.1 ater|EAU35727.1 aver|OJJ02278.1 aver|OJJ02750.1 awen|OJJ30369.1 awen|OJJ40575.1 azon|OJJ45388.1 azon|OJJ49435.1 abra|OJJ71044.1 agla|OJJ79139.1 akaw|GAA82912.1 aluc|OJZ87482.1 anig|EHA20704.1 aory|EIT73215.1 aory|EIT80785.1 atub|OJI81166.1

Group201: afla|EED48129.1 afla|EED48474.1 anid|CBF75189.1 anid|CBF83027.1 aory|EIT73105.1 aory|EIT76193.1 aory|EIT79635.1 asyd|OJJ55485.1 asyd|OJJ56197.1 asyd|OJJ59940.1 ater|EAU31592.1 ater|EAU31786.1 aver|OJJ02347.1 aver|OJJ05124.1 aver|OJJ06564.1 aver|OJJ08128.1 azon|OJJ42131.1 azon|OJJ45782.1 azon|OJJ42807.1 abra|OJJ70953.1 acar|OOF98179.1 asyd|OJJ60083.1 asyd|OJJ64199.1 aver|OJI98288.1 aver|OJJ07880.1 afla|EED56568.1 aver|OJJ04169.1 anid|CBF87321.1 asyd|OJJ59110.1 aver|OJJ01702.1

Group202: agla|OJJ79264.1 agla|OJJ81588.1 asyd|OJJ56968.1 asyd|OJJ61442.1 aver|OJJ00778.1 aver|OJJ03319.1 azon|OJJ43207.1 azon|OJJ51818.1 abra|OJJ67615.1 acar|OOF91964.1 afla|EED44878.1 akaw|GAA89133.1 aluc|OJZ81703.1 anig|EHA25732.1 aory|EIT83259.1 atub|OJI88479.1 awen|OJJ36276.1 abra|OJJ68338.1 acar|OOF97638.1 akaw|GAA92100.1 aluc|OJZ83625.1 anid|CBF83952.1 anig|EHA18778.1 atub|OJI89994.1 awen|OJJ35809.1 azon|OJJ47565.1 akaw|GAA92300.1 aluc|OJZ90197.1 anig|EHA23695.1 atub|OJI81624.1

Group203: asyd|OJJ55488.1 asyd|OJJ61510.1 asyd|OJJ61953.1 asyd|OJJ60082.1 aver|OJI96044.1 aver|OJJ00847.1 aver|OJJ06567.1 awen|OJJ33588.1 awen|OJJ35927.1 abra|OJJ70268.1 afla|EED46478.1 agla|OJJ78786.1 akaw|GAA86529.1 aluc|OJZ85077.1 anid|CBF83020.1 anig|EHA25292.1 aory|EIT72376.1 aory|EIT82615.1 ater|EAU38779.1 atub|OJI80521.1 azon|OJJ42129.1 abra|OJJ70950.1 acar|OOF98180.1 akaw|GAA82994.1 aluc|OJZ87579.1 anig|EHA20781.1 atub|OJI81070.1 afla|EED49569.1 aver|OJJ04168.1 azon|OJJ42134.1

Group204: asyd|OJJ62594.1 awen|OJJ38006.1 awen|OJJ39405.1 abra|OJJ75729.1 acar|OOF90233.1 afis|EAW24067.1 afla|EED54453.1 afum|EAL91923.1 agla|OJJ87782.1 akaw|GAA84464.1 aluc|OJZ89181.1 anid|CBF83672.1 anig|EHA23374.1 aory|EIT81134.1 asyd|OJJ60812.1 ater|EAU32054.1 atub|OJI85608.1 aver|OJJ00139.1 abra|OJJ75956.1 acar|OOF99957.1 acla|EAW09841.1 akaw|GAA84658.1 aluc|OJZ89405.1 anid|CBF83380.1 anig|EHA23182.1 aory|EIT79378.1 ater|EAU35927.1 atub|OJI85392.1 afis|EAW21294.1 afum|EAL92311.1

Group205: awen|OJJ32368.1 awen|OJJ32676.1 azon|OJJ42336.1 azon|OJJ46795.1 abra|OJJ68299.1 acar|OOF95350.1 acar|OOF97574.1 acla|EAW14263.1 afla|EED54478.1 akaw|GAA83175.1 aluc|OJZ83673.1 anid|CBF85201.1 anig|EHA18829.1 aory|EIT80470.1 asyd|OJJ63024.1 ater|EAU32059.1 atub|OJI90044.1 aver|OJJ00341.1 awen|OJJ32193.1 abra|OJJ74389.1 akaw|GAA87289.1 aluc|OJZ89730.1 aory|EIT81129.1 atub|OJI85071.1 aory|EIT79607.1 asyd|OJJ55922.1 afla|EED56619.1 aver|OJI97944.1 afla|EED51613.1 agla|OJJ81483.1

Group206: abra|OJJ66942.1 acar|OOF91706.1 acar|OOF91708.1 acla|EAW14500.1 acla|EAW14501.1 afis|EAW19214.1 afis|EAW19215.1 afla|EED47685.1 afla|EED47686.1 afum|EAL93947.1 afum|EAL93948.1 agla|OJJ88482.1 akaw|GAA91478.1 akaw|GAA91479.1 aluc|OJZ85640.1 anid|CBF78591.1 anig|EHA24894.1 aory|EIT80213.1 aory|EIT80244.1 asyd|OJJ65083.1 ater|EAU37253.1 atub|OJI79431.1 aver|OJI99209.1 awen|OJJ37572.1 azon|OJJ51231.1 anid|CBF78589.1 ater|EAU37254.1 aver|OJI99208.1 awen|OJJ37571.1 asyd|OJJ65084.1

Group207: abra|OJJ67140.1 acar|OOF98417.1 acla|EAW14599.1 afis|EAW19328.1 afum|EAL94045.1 agla|OJJ86277.1 akaw|GAA91029.1 aluc|OJZ87832.1 anid|CBF83940.1 anig|EHA20311.1 aory|EIT74592.1 asyd|OJJ53609.1 ater|EAU29456.1 atub|OJI80809.1 aver|OJJ08139.1 awen|OJJ30097.1 azon|OJJ44450.1 abra|OJJ67993.1 acar|OOF92035.1 acla|EAW13634.1 afis|EAW23673.1 afla|EED47394.1 afum|EAL87116.1 akaw|GAA90955.1 aluc|OJZ82867.1 anig|EHA21297.1 aory|EIT81925.1 atub|OJI89092.1 azon|OJJ45993.1 afla|EED47553.1

Group208: abra|OJJ70660.1 acar|OOF97061.1 acla|EAW15046.1 afis|EAW23152.1 afla|EED48515.1 afum|EAL86237.1 agla|OJJ89650.1 akaw|GAA89996.1 aluc|OJZ84010.1 anid|CBF74423.1 anig|EHA19132.1 aory|EIT73250.1 asyd|OJJ57754.1 ater|EAU36259.1 atub|OJI90379.1 aver|OJJ05413.1 awen|OJJ36648.1 azon|OJJ47567.1 acla|EAW08098.1 afum|EAL88329.1 agla|OJJ89649.1 anig|EHA19133.1 azon|OJJ47566.1 acar|OOF97062.1 afis|EAW24607.1 anid|CBF87133.1 aory|EIT73278.1 asyd|OJJ60373.1 ater|EAU36260.1 aver|OJI99708.1

Group209: abra|OJJ71126.1 acar|OOF98025.1 acar|OOF98026.1 acla|EAW10372.1 acla|EAW10373.1 afis|EAW25219.1 afis|EAW25220.1 afla|EED53686.1 afla|EED53687.1 afum|EAL89880.1 afum|EAL89881.1 agla|OJJ79187.1 akaw|GAA82848.1 akaw|GAA82849.1 aluc|OJZ87409.1 anid|CBF85301.1 anig|EHA20639.1 aory|EIT75287.1 aory|EIT75345.1 asyd|OJJ62915.1 asyd|OJJ62916.1 ater|EAU34378.1 ater|EAU34379.1 atub|OJI81236.1 atub|OJI81237.1 aver|OJI98074.1 awen|OJJ40885.1 awen|OJJ40886.1 azon|OJJ44513.1 azon|OJJ44514.1

Group210: abra|OJJ73704.1 afis|EAW17806.1 afla|EED53530.1 afum|EBA27423.1 agla|OJJ78725.1 akaw|GAA88459.1 aluc|OJZ88348.1 ater|EAU30902.1 aver|OJI99268.1 awen|OJJ40823.1 afis|EAW16815.1 afis|EAW17324.1 afla|EED44724.1 afla|EED45247.1 akaw|GAA89045.1 aory|EIT72906.1 aory|EIT73594.1 asyd|OJJ53750.1 ater|EAU36770.1 azon|OJJ45102.1 aver|OJI98047.1 agla|OJJ86073.1 aory|EIT75237.1 asyd|OJJ57660.1 atub|OJI88380.1 aver|OJJ05522.1 asyd|OJJ62943.1 agla|OJJ80314.1 aluc|OJZ80551.1 anig|EHA27689.1

Group211: abra|OJJ74957.1 acar|OOF97356.1 acla|EAW08414.1 afis|EAW23455.1 afla|EED57346.1 afum|EAL88011.1 agla|OJJ86695.1 akaw|GAA83465.1 akaw|GAA83466.1 aluc|OJZ92529.1 anid|CBF89806.1 anig|EHA26445.1 anig|EHA26446.1 aory|EIT80091.1 asyd|OJJ62193.1 ater|EAU36921.1 atub|OJI86965.1 atub|OJI86966.1 aver|OJI96302.1 awen|OJJ39774.1 azon|OJJ47901.1 acla|EAW08415.1 afis|EAW23454.1 afla|EED57349.1 afum|EAL88012.1 agla|OJJ86696.1 anid|CBF89804.1 ater|EAU36920.1 aver|OJI96305.1 awen|OJJ39775.1

Group212: abra|OJJ75967.1 acar|OOF99952.1 acla|EAW09831.1 afis|EAW21302.1 afla|EED57262.1 afum|EAL92319.1 agla|OJJ87435.1 akaw|GAA88989.1 aluc|OJZ89414.1 anid|CBF83395.1 anig|EHA23173.1 aory|EIT74086.1 asyd|OJJ60638.1 ater|EAU35935.1 atub|OJI85381.1 aver|OJI99960.1 awen|OJJ39396.1 azon|OJJ44280.1 acla|EAW09826.1 afis|EAW21307.1 afum|EAL92323.1 anid|CBF83397.1 asyd|OJJ60645.1 ater|EAU35937.1 aver|OJI99963.1 afis|EAW21308.1 afum|EAL92324.1 anid|CBF83399.1 asyd|OJJ60644.1 ater|EAU35938.1

Group213: abra|OJJ76386.1 acar|OOF99627.1 acla|EAW14037.1 acla|EAW14038.1 afis|EAW18743.1 afis|EAW18744.1 afla|EED50699.1 afla|EED50700.1 afum|EAL93469.1 afum|EAL93470.1 agla|OJJ82309.1 akaw|GAA82305.1 akaw|GAA82306.1 aluc|OJZ91827.1 aluc|OJZ91828.1 anid|CBF69994.1 anid|CBF69996.1 anig|EHA22816.1 aory|EIT78103.1 aory|EIT78138.1 asyd|OJJ58888.1 asyd|OJJ58889.1 ater|EAU37870.1 ater|EAU37873.1 atub|OJI86282.1 atub|OJI86283.1 aver|OJJ01489.1 awen|OJJ32308.1 awen|OJJ32309.1 azon|OJJ42718.1

Group214: abra|OJJ65770.1 abra|OJJ69062.1 acar|OOF91221.1 acar|OOG00531.1 afla|EED46866.1 afla|EED56884.1 akaw|GAA90592.1 akaw|GAA90890.1 aluc|OJZ82939.1 aluc|OJZ86050.1 anig|EHA21359.1 anig|EHA24561.1 aory|EIT72522.1 aory|EIT77460.1 atub|OJI79847.1 atub|OJI89171.1 aver|OJJ03011.1 aver|OJJ07939.1 afis|EAW15670.1 afum|EAL85677.1 anid|CBF81578.1 asyd|OJJ53834.1 ater|EAU32494.1 awen|OJJ38346.1 acla|EAW11671.1 agla|OJJ81808.1 agla|OJJ85313.1 awen|OJJ38760.1 azon|OJJ42640.1

Group215: abra|OJJ66046.1 abra|OJJ74366.1 abra|OJJ77282.1 acar|OOF95364.1 acar|OOG00557.1 afis|EAW16747.1 afis|EAW22442.1 afla|EED54458.1 afla|EED54746.1 afum|EAL86322.1 afum|EAL90749.1 akaw|GAA86664.1 akaw|GAA90667.1 aluc|OJZ84692.1 aluc|OJZ89761.1 anig|EHA26981.1 anig|EHA27968.1 aory|EIT81130.1 aory|EIT81408.1 atub|OJI85041.1 atub|OJI89390.1 awen|OJJ35068.1 awen|OJJ37925.1 acla|EAW07417.1 agla|OJJ87580.1 anid|CBF88847.1 asyd|OJJ62866.1 ater|EAU39262.1 aver|OJI96919.1

Group216: abra|OJJ67846.1 abra|OJJ69869.1 acar|OOF90337.1 acar|OOF92827.1 acla|EAW11753.1 acla|EAW15317.1 akaw|GAA87787.1 akaw|GAA91854.1 aluc|OJZ82045.1 aluc|OJZ85422.1 anig|EHA22212.1 anig|EHA25601.1 asyd|OJJ52765.1 asyd|OJJ58155.1 ater|EAU31327.1 ater|EAU32679.1 aver|OJJ04248.1

aver|OJJ08748.1 awen|OJJ31347.1 awen|OJJ33731.1 afis|EAW23786.1 afla|EED51595.1 afum|EAL87220.2 agla|OJJ78939.1 anid|CBF79847.1 atub|OJI84604.1 azon|OJJ42847.1 atub|OJI80153.1 aory|EIT80419.1

Group217: abra|OJJ69627.1 abra|OJJ76818.1 acar|OOF91313.1 acar|OOF92684.1 acla|EAW08738.1 acla|EAW13378.1 agla|OJJ84834.1 agla|OJJ86329.1 akaw|GAA87204.1 akaw|GAA91607.1 anig|EHA27577.1 anig|EHA28339.1 atub|OJI88009.1 atub|OJI89856.1 awen|OJJ33192.1 awen|OJJ38390.1 azon|OJJ43848.1 azon|OJJ46477.1 afis|EAW15698.1 afla|EED51354.1 afum|EAL85650.1 aluc|OJZ82432.1 anid|CBF78282.1 aory|EIT80486.1 asyd|OJJ56072.1 ater|EAU39440.1 aver|OJJ05015.1 aluc|OJZ84228.1 asyd|OJJ52533.1

Group218: abra|OJJ69629.1 abra|OJJ74245.1 acar|OOF92681.1 acar|OOF95485.1 afla|EED55078.1 afla|EED57736.1 aluc|OJZ82434.1 aluc|OJZ89887.1 asyd|OJJ63867.1 asyd|OJJ64731.1 acla|EAW07299.1 agla|OJJ87688.1 akaw|GAA91609.1 anig|EHA28337.1 aory|EIT81976.1 atub|OJI84920.1 atub|OJI88007.1 aver|OJI98835.1 awen|OJJ34926.1 afis|EAW22346.1 afum|EAL90846.1 akaw|GAA86775.1 anid|CBF82985.1 anig|EHA27086.1 ater|EAU39484.1 aver|OJI97072.1 azon|OJJ50069.1 ater|EAU35426.1 aory|EIT79895.1

Group219: abra|OJJ69794.1 abra|OJJ76964.1 acar|OOF93584.1 acar|OOG00383.1 agla|OJJ81809.1 agla|OJJ84793.1 agla|OJJ85103.1 aluc|OJZ84359.1 aluc|OJZ85505.1 anig|EHA25037.1 anig|EHA27695.1 atub|OJI80060.1 atub|OJI89709.1 awen|OJJ30169.1 awen|OJJ37984.1 azon|OJJ46123.1 azon|OJJ46412.1 acla|EAW11252.1 afis|EAW18245.1 afla|EED44938.1 afum|EAL87610.1 akaw|GAA87864.1 aory|EIT73480.1 asyd|OJJ60485.1 ater|EAU30027.1 aver|OJI99811.1 acar|OOF91408.1 ater|EAU29360.1 akaw|GAA87080.1

Group220: abra|OJJ72131.1 abra|OJJ76584.1 acar|OOF96771.1 acar|OOF99402.1 agla|OJJ80546.1 agla|OJJ86323.1 agla|OJJ89016.1 akaw|GAA82487.1 akaw|GAA84791.1 aluc|OJZ86889.1 aluc|OJZ91622.1 anig|EHA21754.1 anig|EHA22647.1 asyd|OJJ57662.1 asyd|OJJ58966.1 atub|OJI83630.1 atub|OJI86075.1 aver|OJJ01560.1 aver|OJJ05524.1 awen|OJJ32389.1 awen|OJJ32572.1 awen|OJJ33544.1 acla|EAW14111.1 afis|EAW18813.1 afum|EAL93549.1 anid|CBF69799.1 aory|EIT77991.1 ater|EAU37980.1 azon|OJJ47150.1

Group221: acar|OOF92608.1 acar|OOF92629.1 abra|OJJ67199.1 acar|OOF98365.1 acla|EAW14525.1 acla|EAW14526.1 afis|EAW19243.1 afis|EAW19244.1 afla|EED47633.1 afla|EED47634.1 afum|EAL93972.1 afum|EAL93973.1 agla|OJJ88454.1 akaw|GAA90979.1 akaw|GAA90980.1 aluc|OJZ87775.1 anid|CBF78652.1 anig|EHA20259.1 aory|EIT80236.1 asyd|OJJ65195.1 ater|EAU29500.1 atub|OJI80870.1 aver|OJI99344.1 awen|OJJ37599.1 azon|OJJ51254.1 agla|OJJ88453.1 anid|CBF78654.1 azon|OJJ51255.1 ater|EAU29499.1

Group222: afis|EAW21841.1 afis|EAW25659.1 agla|OJJ84294.1 agla|OJJ87901.1 aory|EIT72356.1 aory|EIT79167.1 asyd|OJJ52337.1 asyd|OJJ61239.1 asyd|OJJ54547.1 aver|OJI96725.1 aver|OJI96959.1 aver|OJJ07630.1 aver|OJJ00562.1 abra|OJJ68267.1 acar|OOF97547.1 acla|EAW09304.1 afla|EED46529.1 afum|EAL92853.1 akaw|GAA91931.1 aluc|OJZ82691.1 anid|CBF76711.1 anig|EHA27368.1 ater|EAU38518.1 atub|OJI88923.1 awen|OJJ34124.1 azon|OJJ44222.1 aluc|OJZ80200.1 afla|EED47266.1 anid|CBF85164.1

Group223: afla|EED48230.1 afla|EED58028.1 anid|CBF83667.1 anid|CBF86363.1 aory|EIT72902.1 aory|EIT79778.1 asyd|OJJ60808.1 asyd|OJJ64969.1 aver|OJI99093.1 aver|OJJ00136.1 awen|OJJ30134.1 awen|OJJ35830.1 abra|OJJ72106.1 acar|OOF96749.1 acla|EAW09385.1 afis|EAW21750.1 afum|EAL92781.2 agla|OJJ84424.1 akaw|GAA84816.1 aluc|OJZ86861.1 anig|EHA21779.1 ater|EAU38440.1 atub|OJI83659.1 awen|OJJ34031.1 azon|OJJ48204.1 anid|CBF83471.1 ater|EAU34242.1 aver|OJJ00014.1 asyd|OJJ60689.1

Group224: azon|OJJ42769.1 azon|OJJ50372.1 abra|OJJ71617.1 acar|OOF96333.1 afla|EED57118.1 agla|OJJ80138.1 akaw|GAA89310.1 aluc|OJZ86358.1 anig|EHA21532.1 aory|EIT73985.1 atub|OJI84176.1 awen|OJJ40249.1 abra|OJJ77100.1 acar|OOF92984.1 acla|EAW12282.1 afis|EAW16959.1 afla|EED45239.1 afum|EAL86486.1 akaw|GAA86956.1 aluc|OJZ84508.1 anid|CBF80512.1 anig|EHA27807.1 aory|EIT73744.1 asyd|OJJ54736.1 ater|EAU30185.1 atub|OJI89572.1 aver|OJJ07435.1 agla|OJJ85349.1 awen|OJJ30386.1

Group225: azon|OJJ46793.1 azon|OJJ51796.1 abra|OJJ77127.1 acar|OOF93013.1 acla|EAW12902.1 afis|EAW16407.1 afla|EED51674.1 agla|OJJ81607.1 akaw|GAA90801.1 aluc|OJZ84537.1 anid|CBF82038.1 anig|EHA27835.1 aory|EIT79575.1 asyd|OJJ59200.1 ater|EAU36805.1 atub|OJI89543.1 aver|OJJ01812.1 awen|OJJ36343.1 acar|OOF93300.1 afla|EED49754.1 aory|EIT81671.1 asyd|OJJ54221.1 acar|OOF99299.1 aory|EIT77631.1 afla|EED48439.1 awen|OJJ38768.1 aory|EIT80498.1 afum|EAL91133.1 aver|OJJ05173.1

Group226: abra|OJJ67152.1 acar|OOF90071.1 acar|OOF98406.1 acla|EAW14572.1 acla|EAW14573.1 afis|EAW19295.1 afis|EAW19296.1 afla|EED47590.1 afla|EED47591.1 afum|EAL94014.1 afum|EAL94015.1 agla|OJJ88385.1 akaw|GAA91020.1 akaw|GAA91021.1 aluc|OJZ87823.1 aluc|OJZ87824.1 anid|CBF78768.1 anid|CBF78770.1 anig|EHA20303.1 aory|EIT74518.1 aory|EIT74526.1 asyd|OJJ65238.1 ater|EAU29476.1 ater|EAU29477.1 atub|OJI80820.1 atub|OJI80821.1 aver|OJI99388.1 awen|OJJ37646.1 azon|OJJ44911.1

Group227: abra|OJJ67842.1 acar|OOF92019.1 afis|EAW19539.1 afla|EED54092.1 afum|EAL86968.1 akaw|GAA91848.1 aluc|OJZ82050.1 anid|CBF75400.1 anig|EHA22208.1 aory|EIT75915.1 asyd|OJJ63641.1 ater|EAU35153.1 atub|OJI84608.1 aver|OJI97295.1 azon|OJJ48646.1 abra|OJJ76922.1 acar|OOF91369.1 afla|EED54074.1 akaw|GAA87116.1 aluc|OJZ84322.1 anid|CBF71754.1 anig|EHA27657.1 asyd|OJJ61735.1 atub|OJI89746.1 aver|OJI95828.1 awen|OJJ33258.1 anid|CBF71752.1 azon|OJJ44017.1 aory|EIT75930.1

Group228: abra|OJJ68668.1 acar|OOF92168.1 acla|EAW12580.1 afis|EAW24068.1 afla|EED51407.1 afum|EAL91924.1 agla|OJJ87316.1 aluc|OJZ81882.1 anid|CBF86457.1 anig|EHA22358.1 aory|EIT80674.1

asyd|OJJ64915.1 ater|EAU30490.1 atub|OJI84435.1 aver|OJI99035.1 awen|OJJ41422.1 azon|OJJ48887.1 acla|EAW12579.1 afis|EAW24069.1 afla|EED51406.1 afum|EAL91925.1 agla|OJJ87315.1 akaw|GAA89809.1 aluc|OJZ81881.1 anid|CBF86458.1 aory|EIT80695.1 asyd|OJJ64914.1 awen|OJJ41421.1 azon|OJJ48886.1

Group229: abra|OJJ68715.1 acar|OOF92215.1 acar|OOF92216.1 acla|EAW12530.1 acla|EAW12531.1 afis|EAW24118.1 afis|EAW24119.1 afla|EED46814.1 afum|EAL91973.1 afum|EAL91974.1 agla|OJJ87193.1 akaw|GAA89868.1 aluc|OJZ81813.1 anid|CBF86539.1 anid|CBF86540.1 anig|EHA22402.1 anig|EHA22403.1 aory|EIT72603.1 aory|EIT72683.1 asyd|OJJ64863.1 asyd|OJJ64865.1 ater|EAU30546.1 ater|EAU30547.1 atub|OJI84385.1 aver|OJI98974.1 aver|OJI98978.1 awen|OJJ41370.1 azon|OJJ48839.1 afla|EED46813.1

Group230: abra|OJJ68757.1 acar|OOF92245.1 acla|EAW12494.1 afis|EAW24151.1 afla|EED46780.1 afum|EAL92006.1 akaw|GAA89901.1 aluc|OJZ81770.1 anid|CBF77913.1 anig|EHA22446.1 aory|EIT72705.1 asyd|OJJ55298.1 ater|EAU30583.1 atub|OJI84342.1 aver|OJJ06928.1 awen|OJJ41327.1 abra|OJJ68758.1 acla|EAW12493.1 afis|EAW24152.1 afla|EED46779.1 afum|EAL92007.1 agla|OJJ89543.1 akaw|GAA89902.1 aluc|OJZ81769.1 anid|CBF77911.1 aory|EIT72741.1 atub|OJI84341.1 awen|OJJ41326.1 azon|OJJ48800.1

Group231: abra|OJJ70392.1 acar|OOF90061.1 acla|EAW12387.1 acla|EAW12388.1 afis|EAW16834.1 afis|EAW16835.1 afla|EED47328.1 afum|EAL86381.1 afum|EAL86382.2 agla|OJJ81545.1 agla|OJJ89426.1 akaw|GAA91967.1 aluc|OJZ82649.1 anid|CBF74687.1 anid|CBF74689.1 anig|EHA27422.1 anig|EHA27423.1 aory|EIT82824.1 asyd|OJJ60210.1 ater|EAU30239.1 atub|OJI88881.1 aver|OJJ04058.1 awen|OJJ35685.1 azon|OJJ44153.1 acar|OOF90060.1 afla|EED47327.1 aory|EIT82846.1 ater|EAU30242.1 acar|OOF91882.1

Group232: abra|OJJ70540.1 acar|OOF97393.1 acla|EAW08138.1 acla|EAW08139.1 afis|EAW23174.1 afis|EAW23175.1 afla|EED46944.1 afla|EED46945.1 afum|EAL88308.1 afum|EAL88309.1 agla|OJJ89686.1 agla|OJJ89687.1 akaw|GAA83385.1 akaw|GAA83386.1 aluc|OJZ83885.1 anid|CBF74496.1 anid|CBF74498.1 anig|EHA19022.1 aory|EIT72667.1 aory|EIT72762.1 asyd|OJJ57719.1 asyd|OJJ57720.1 ater|EAU36303.1 ater|EAU36304.1 atub|OJI90252.1 aver|OJJ05448.1 awen|OJJ36603.1 azon|OJJ47603.1 azon|OJJ47604.1

Group233: abra|OJJ73887.1 acar|OOG01025.1 acar|OOG01026.1 acla|EAW07769.1 acla|EAW07771.1 afis|EAW22805.1 afis|EAW22807.1 afla|EED50075.1 afum|EAL90394.1 afum|EBA27407.1 agla|OJJ83582.1 akaw|GAA89498.1 akaw|GAA89499.1 aluc|OJZ88162.1 anid|CBF87924.1 anig|EHA17916.1 anig|EHA17917.1 aory|EIT74704.1 asyd|OJJ63459.1 asyd|OJJ63460.1 ater|EAU38903.1 ater|EAU38904.1 atub|OJI90761.1 aver|OJI97472.1 aver|OJI97473.1 awen|OJJ38923.1 azon|OJJ49700.1 aory|EIT74626.1 afla|EED50076.1

Group234: abra|OJJ74769.1 acar|OOF95760.1 acla|EAW10642.1 afis|EAW24924.1 afla|EED55549.1 afum|EAL89579.1 agla|OJJ82153.1 akaw|GAA83643.1 aluc|OJZ92352.1 anid|CBF75699.1 anig|EHA26622.1 aory|EIT82671.1 asyd|OJJ57496.1 atub|OJI86780.1 aver|OJJ05684.1 awen|OJJ40521.1 azon|OJJ49380.1 abra|OJJ74770.1 acla|EAW10641.1 afis|EAW24925.1 afla|EED55548.1 afum|EAL89580.1 akaw|GAA83642.1 anid|CBF75701.1 aory|EIT82683.1 ater|EAU35655.1 atub|OJI86781.1 aver|OJJ05683.1 azon|OJJ49381.1

Group235: abra|OJJ74915.1 acar|OOF95906.1 acar|OOF95907.1 acla|EAW08455.1 acla|EAW08456.1 afis|EAW23499.1 afis|EAW23500.1 afla|EED57400.1 afla|EED57401.1 afum|EAL87962.2 afum|EAL87963.1 agla|OJJ86747.1 agla|OJJ86748.1 akaw|GAA83507.1 akaw|GAA83508.1 aluc|OJZ92487.1 anid|CBF89730.1 anid|CBF89731.1 anig|EHA26485.1 aory|EIT80151.1 aory|EIT80164.1 asyd|OJJ62234.1 ater|EAU35289.1 ater|EAU35290.1 atub|OJI86920.1 aver|OJI96343.1 awen|OJJ39825.1 azon|OJJ47949.1 azon|OJJ47950.1

Group236: abra|OJJ75075.1 acar|OOF97258.1 acar|OOF97259.1 acla|EAW08256.1 afis|EAW23290.1 afla|EED55682.1 afum|EAL88189.1 agla|OJJ86548.1 akaw|GAA88687.1 akaw|GAA88688.1 aluc|OJZ92641.1 anid|CBF89509.1 anig|EHA26351.1 aory|EIT77074.1 asyd|OJJ62397.1 ater|EAU37583.1 atub|OJI87072.1 aver|OJI96531.1 awen|OJJ39667.1 azon|OJJ47797.1 aory|EIT77075.1 acla|EAW08258.1 afis|EAW23291.1 afla|EED55680.1 afum|EAL88188.1 anid|CBF89508.1 asyd|OJJ62398.1 ater|EAU37582.1 azon|OJJ47796.1

Group237: abra|OJJ75137.1 acar|OOF97199.1 acar|OOF97200.1 afis|EAW23348.1 afis|EAW23350.1 afla|EED55617.1 afum|EAL88126.1 afum|EAL88128.1 agla|OJJ86478.1 akaw|GAA88743.1 aluc|OJZ92703.1 anid|CBF74910.1 anig|EHA26296.1 anig|EHA26297.1 aory|EIT81793.1 asyd|OJJ60100.1 ater|EAU35593.1 atub|OJI87130.1 aver|OJJ04162.1 awen|OJJ39539.1 awen|OJJ39543.1 azon|OJJ47741.1 agla|OJJ80588.1 aver|OJJ04161.1 anid|CBF74914.1 aory|EIT81802.1 ater|EAU35592.1 azon|OJJ47736.1 afla|EED55618.1

Group238: abra|OJJ75492.1 acar|OOF90824.1 acar|OOF90825.1 acla|EAW11224.1 acla|EAW11225.1 afis|EAW18278.1 afis|EAW18279.1 afla|EED48745.1 afla|EED48746.1 afum|EAL87644.1 afum|EAL87645.1 agla|OJJ88147.1 agla|OJJ88148.1 akaw|GAA84257.1 akaw|GAA84258.1 aluc|OJZ88951.1 anid|CBF79492.1 anid|CBF79494.1 anig|EHA23569.1 anig|EHA23570.1 aory|EIT73072.1 aory|EIT73086.1 asyd|OJJ57877.1 ater|EAU33284.1 ater|EAU33285.1 atub|OJI85838.1 aver|OJJ04517.1 awen|OJJ30651.1 azon|OJJ47964.1

Group239: abra|OJJ75717.1 acar|OOG00139.1 acar|OOG00141.1 acla|EAW09392.1 acla|EAW09394.1 afis|EAW21739.1 afis|EAW21741.1 afla|EED58014.1 afla|EED58016.1 afum|EAL92773.1 afum|EAL92774.1 agla|OJJ84432.1 agla|OJJ84449.1 akaw|GAA84452.1 akaw|GAA84453.1 aluc|OJZ89169.1 anid|CBF83696.1 anid|CBF83699.1 anig|EHA23386.1 aory|EIT79752.1 aory|EIT79775.1 asyd|OJJ60825.1 ater|EAU38420.1 ater|EAU38429.1 atub|OJI85621.1 aver|OJJ00151.1 awen|OJJ34021.1 awen|OJJ34023.1 azon|OJJ48535.1

Group240: abra|OJJ77333.1 acar|OOF93207.1 acla|EAW13139.1 afis|EAW17511.1 afla|EED49310.1 afum|EAL91331.1 agla|OJJ81447.1 akaw|GAA91425.1 aluc|OJZ84741.1 anid|CBF71568.1 anig|EHA28015.1 aory|EIT82226.1 asyd|OJJ61881.1 ater|EAU36952.1 atub|OJI89340.1 aver|OJI95979.1 awen|OJJ37333.1 azon|OJJ51021.1 afla|EED49307.1 afum|EAL91330.1 anid|CBF71531.1 asyd|OJJ61893.1 aver|OJI95989.1 azon|OJJ51020.1 acla|EAW13140.1 afis|EAW17512.1 anig|EHA28014.1 aory|EIT82236.1 awen|OJJ37334.1

Group241: abra|OJJ65980.1 abra|OJJ66239.1 acar|OOF90701.1 acar|OOF92893.1 acla|EAW11609.1 acla|EAW15289.1 afis|EAW23842.1 afis|EAW24576.1 akaw|GAA89050.1 akaw|GAA90853.1 aluc|OJZ80544.1 aluc|OJZ82980.1 anig|EHA21395.1 anig|EHA25651.1 asyd|OJJ54925.1 asyd|OJJ65267.1 ater|EAU32172.1 ater|EAU33781.1 atub|OJI88387.1 atub|OJI89213.1 aver|OJI99419.1 aver|OJJ07270.1 afla|EED45390.1 afum|EAL87273.1 agla|OJJ87625.1 anid|CBF78824.1 aory|EIT82439.1 azon|OJJ46551.1

Group242: abra|OJJ66056.1 abra|OJJ77098.1 afla|EED49909.1 afla|EED53379.1 akaw|GAA86958.1 akaw|GAA93126.1 aluc|OJZ81108.1 aluc|OJZ84506.1 anid|CBF80783.1 anid|CBF84621.1 anig|EHA19261.1 anig|EHA27805.1 aory|EIT72449.1 aory|EIT75126.1 asyd|OJJ54497.1 asyd|OJJ55641.1 ater|EAU30218.1 ater|EAU30988.1 atub|OJI82471.1 atub|OJI89574.1 aver|OJJ06724.1 aver|OJJ07683.1 awen|OJJ29737.1 awen|OJJ33568.1 acar|OOF92980.1 afis|EAW17111.1 afum|EAL86575.1 agla|OJJ79951.1

Group243: abra|OJJ66383.1 abra|OJJ71952.1 acar|OOF94059.1 acar|OOF94129.1 acar|OOF96603.1 afla|EED51643.1 afla|EED53089.1 akaw|GAA84976.1 akaw|GAA90249.1 aluc|OJZ83091.1 aluc|OJZ86686.1 aory|EIT74787.1 aory|EIT80741.1 atub|OJI83839.1 atub|OJI87845.1 azon|OJJ48234.1 azon|OJJ51069.1 acla|EAW12654.1 afis|EAW18031.1 afum|EAL91842.1 agla|OJJ85401.1 anid|CBF76050.1 anig|EHA21987.1 asyd|OJJ52485.1 ater|EAU36811.1 aver|OJJ03921.1 awen|OJJ41506.1 ater|EAU35537.1

Group244: abra|OJJ67133.1 abra|OJJ72705.1 akaw|GAA85309.1 akaw|GAA91036.1 aluc|OJZ87846.1 aluc|OJZ90669.1 atub|OJI80802.1 atub|OJI82099.1 aver|OJI96726.1 aver|OJJ07629.1 acar|OOF95940.1 anig|EHA24103.1 asyd|OJJ64079.1 azon|OJJ42255.1 afis|EAW22517.1 afum|EAL90676.1 aory|EIT73492.1 azon|OJJ49956.1 abra|OJJ75758.1 afla|EED44952.1 akaw|GAA84490.1 aluc|OJZ89211.1 anid|CBF85166.1 anig|EHA23347.1 asyd|OJJ54548.1 ater|EAU33498.1 atub|OJI85581.1 awen|OJJ40261.1

Group245: abra|OJJ68302.1 abra|OJJ74425.1 acar|OOF90436.1 acar|OOF90443.1 acar|OOF96685.1 afis|EAW24469.1 afis|EAW25516.1 aluc|OJZ80145.1 aluc|OJZ83671.1 aluc|OJZ89698.1 atub|OJI85105.1 atub|OJI90042.1 awen|OJJ34908.1 awen|OJJ41259.1 acla|EAW14952.1 afla|EED55056.1 afum|EAL86141.1 afum|EAL90181.1 agla|OJJ87287.1 akaw|GAA83172.1 anid|CBF77786.1 anig|EHA18826.1 aory|EIT81972.1 asyd|OJJ56269.1 ater|EAU33362.1 aver|OJJ08496.1 azon|OJJ42850.1 asyd|OJJ54439.1

Group246: abra|OJJ68357.1 abra|OJJ70141.1 afla|EED46535.1 afla|EED53487.1 akaw|GAA86624.1 akaw|GAA92129.1 aluc|OJZ83593.1 aluc|OJZ85179.1 anig|EHA18753.1 anig|EHA25391.1 aory|EIT72381.1 aory|EIT74971.1 asyd|OJJ56981.1 asyd|OJJ60301.1 ater|EAU38514.1 ater|EAU39003.1 atub|OJI80418.1 atub|OJI89965.1 awen|OJJ34116.1 awen|OJJ34635.1 acar|OOF93592.1 afis|EAW19844.1 afum|EAL85050.1 agla|OJJ84299.1 anid|CBF87147.1 aver|OJI99641.1 aver|OJJ03306.1 azon|OJJ50352.1

Group247: abra|OJJ71231.1 abra|OJJ72448.1 acar|OOF94788.1 acar|OOF97944.1 afla|EED46190.1 afla|EED56273.1 akaw|GAA82757.1 akaw|GAA86186.1 aluc|OJZ87316.1 aluc|OJZ90432.1 ater|EAU33047.1 ater|EAU33789.1 atub|OJI81329.1 atub|OJI81864.1 awen|OJJ31638.1 awen|OJJ41732.1 acla|EAW12682.1 afis|EAW17845.1 afum|EAL91654.1 agla|OJJ81772.1 agla|OJJ83123.1 anig|EHA20550.1 asyd|OJJ55268.1 aver|OJJ06959.1 azon|OJJ48428.1 anid|CBF77966.1 anig|EHA23901.1 aory|EIT76458.1

Group248: acar|OOF95098.1 acar|OOF95164.1 acar|OOF96577.1 afis|EAW21529.1 afis|EAW24614.1 agla|OJJ80793.1 agla|OJJ81846.1 agla|OJJ85693.1 agla|OJJ84478.1 aory|EIT76721.1 aory|EIT79947.1 asyd|OJJ52471.1 asyd|OJJ61079.1 ater|EAU31519.1 ater|EAU35012.1 abra|OJJ71920.1 acla|EAW09610.1 afla|EED47452.1 afla|EED55978.1 afum|EAL92550.1 akaw|GAA85002.1 aluc|OJZ86657.1 anid|CBF76400.1 anig|EHA22014.1 atub|OJI83870.1 aver|OJJ00395.1 awen|OJJ39164.1 azon|OJJ51493.1

Group249: afis|EAW16443.1 afis|EAW16700.1 afla|EED45811.1 afla|EED46054.1 anid|CBF78974.1 anid|CBF82132.1 anid|CBF88938.1 aory|EIT75703.1 aory|EIT80899.1 asyd|OJJ54312.1 asyd|OJJ55669.1 asyd|OJJ62858.1 aver|OJI95760.1 aver|OJI96910.1 aver|OJJ05266.1 aver|OJJ06750.1 abra|OJJ67169.1 acar|OOF97010.1 afla|EED55529.1 afum|EAL89297.1 akaw|GAA91005.1 aluc|OJZ87806.1 anig|EHA20288.1 ater|EAU36196.1 atub|OJI80839.1 azon|OJJ50602.1 afla|EED49535.1 aory|EIT82649.1

Group250: afis|EAW18309.1 afis|EAW25696.1 awen|OJJ30683.1 awen|OJJ32214.1 abra|OJJ75441.1 acar|OOF90786.1 acla|EAW13822.1 afla|EED53243.1 afum|EAL87675.1 agla|OJJ88078.1 akaw|GAA84216.1 aluc|OJZ88894.1 anid|CBF79436.1 anig|EHA23612.1 aory|EIT72989.1 asyd|OJJ54980.1 asyd|OJJ57846.1 ater|EAU33319.1 atub|OJI85895.1 aver|OJJ04318.1 aver|OJJ04546.1 awen|OJJ32538.1 azon|OJJ43103.1 afla|EED50918.1 aory|EIT74822.1 aory|EIT78997.1 asyd|OJJ52725.1 ater|EAU32817.1

Group251: afla|EED51654.1 afla|EED52464.1 aluc|OJZ84281.1 aluc|OJZ87033.1 anig|EHA25215.1 anig|EHA26913.1 anig|EHA27626.1 aory|EIT80434.1 aory|EIT82812.1 asyd|OJJ54529.1 asyd|OJJ62992.1 atub|OJI83482.1 atub|OJI89800.1 aver|OJI97976.1 aver|OJJ07658.1 abra|OJJ76870.1 acla|EAW10087.1 afis|EAW16918.1 akaw|GAA87153.1 anid|CBF80752.1 awen|OJJ30986.1 afis|EAW25500.1 afla|EED50854.1 afum|EAL90166.1 agla|OJJ87304.1 aory|EIT79059.1 awen|OJJ41242.1 azon|OJJ44808.1

Group252: agla|OJJ82640.1 agla|OJJ85276.1 asyd|OJJ57347.1 asyd|OJJ65114.1 aver|OJI99240.1 aver|OJJ05818.1 aver|OJJ00352.1 aver|OJJ05340.1 awen|OJJ30078.1 awen|OJJ32593.1 abra|OJJ78114.1 acar|OOF98754.1 acla|EAW07003.1 afis|EAW16620.1 afla|EED48645.1 afum|EAL89217.1 agla|OJJ84764.1 akaw|GAA85893.1 aluc|OJZ91179.1 anid|CBF81971.1 anig|EHA19870.1 aory|EIT73182.1 ater|EAU36705.1 atub|OJI83191.1 azon|OJJ42184.1 aory|EIT73798.1 asyd|OJJ61034.1 ater|EAU37620.1

Group253: asyd|OJJ62591.1 asyd|OJJ64803.1 abra|OJJ77288.1 acar|OOF93165.1 acla|EAW12880.1 afis|EAW17797.1 afla|EED45790.1 afum|EAL91605.2 agla|OJJ88361.1 akaw|GAA90661.1 aluc|OJZ84699.1 anid|CBF86643.1 anig|EHA27974.1 aory|EIT75720.1 ater|EAU31827.1 atub|OJI89383.1 aver|OJI98916.1 awen|OJJ37068.1 acar|OOF93164.1 afis|EAW17799.1 afla|EED45791.1 afum|EAL91607.1 ater|EAU31828.1 anid|CBF86641.1 aver|OJI98917.1 agla|OJJ78851.1 awen|OJJ37061.1 aory|EIT75595.1

Group254: abra|OJJ67087.1 afum|EAL86535.1 agla|OJJ83616.1 anid|CBF75171.1 anig|EHA20191.1 abra|OJJ67973.1 acar|OOF92814.1 acla|EAW11761.1 afis|EAW23794.1 afla|EED48581.1 akaw|GAA91229.1 aluc|OJZ82841.1 anig|EHA21278.1 aory|EIT73168.1 asyd|OJJ62859.1 atub|OJI89071.1 aver|OJI96912.1 awen|OJJ38042.1 azon|OJJ44002.1 akaw|GAA90147.1 aluc|OJZ91537.1 anig|EHA22574.1 atub|OJI87737.1 akaw|GAA92523.1 asyd|OJJ60253.1 atub|OJI85991.1 azon|OJJ46498.1 azon|OJJ45663.1

Group255: abra|OJJ67268.1 acar|OOF98298.1 acar|OOF98299.1 afis|EAW17109.1 afla|EED55276.1 afum|EAL86573.1 agla|OJJ88998.1 akaw|GAA91683.1 aluc|OJZ87711.1 anid|CBF85644.1 anig|EHA20205.1 anig|EHA20206.1 aory|EIT83585.1 asyd|OJJ64238.1 asyd|OJJ64239.1 ater|EAU29281.1 ater|EAU39434.1 atub|OJI80931.1 aver|OJI98331.1 aver|OJI98332.1 awen|OJJ40796.1 ater|EAU39435.1 afis|EAW17112.1 afum|EAL86576.1 anid|CBF85642.1 aory|EIT83532.1 afla|EED55275.1 aory|EIT83583.1

Group256: abra|OJJ67322.1 acar|OOF92491.1 acar|OOF92492.1 acla|EAW06413.1 acla|EAW06414.1 afis|EAW18517.1 afis|EAW18518.1 afla|EED52880.1 afla|EED52881.1 afum|EAL93212.1 afum|EAL93213.1 agla|OJJ85130.1 akaw|GAA86495.1 akaw|GAA86496.1 aluc|OJZ85037.1 aluc|OJZ85038.1 anid|CBF70228.1 anid|CBF70230.1 anig|EHA25256.1 aory|EIT78607.1 aory|EIT78620.1 asyd|OJJ58772.1 ater|EAU30052.1 ater|EAU30053.1 atub|OJI80559.1 aver|OJJ01381.1 awen|OJJ31070.1 azon|OJJ50580.1

Group257: abra|OJJ67358.1 acar|OOF92463.1 acla|EAW06384.1 afis|EAW18490.1 afla|EED52916.1 afum|EAL93180.1 agla|OJJ85085.1 akaw|GAA86463.1 aluc|OJZ85003.1 anid|CBF70157.1 anig|EHA25222.1 aory|EIT78645.1 asyd|OJJ58803.1 ater|EAU29914.1 atub|OJI80595.1 aver|OJJ01407.1 awen|OJJ31043.1 azon|OJJ50554.1 abra|OJJ67360.1 acla|EAW06383.1 afis|EAW18489.1 afla|EED52917.1 afum|EAL93179.1 akaw|GAA86461.1 anid|CBF70155.1 anig|EHA25220.1 aory|EIT78598.1 asyd|OJJ58804.1

Group258: abra|OJJ68582.1 atub|OJI91460.1 azon|OJJ47723.1 abra|OJJ75562.1 acar|OOF94716.1 acla|EAW13645.1 asyd|OJJ52397.1 ater|EAU35611.1 aver|OJJ01926.1 awen|OJJ38360.1 azon|OJJ47155.1 abra|OJJ76231.1 afis|EAW24533.1 afla|EED47408.1 afum|EAL86199.1 anid|CBF69447.1 anig|EHA18750.1 acla|EAW08325.1 afis|EAW23360.1 afla|EED55598.1 afum|EBA27335.1 agla|OJJ86466.1 aluc|OJZ92717.1 anig|EHA26282.1 aory|EIT82668.1 asyd|OJJ60112.1 ater|EAU35434.1 anig|EHA22166.1

Group259: abra|OJJ68687.1 acar|OOF92189.1 acar|OOF92190.1 acla|EAW12558.1 acla|EAW12560.1 afis|EAW24090.1 afis|EAW24092.1 afla|EED46847.1 afla|EED46849.1 afum|EAL91945.1 afum|EAL91947.2 agla|OJJ87160.1 agla|OJJ87166.1 akaw|GAA89840.1 akaw|GAA89842.1 aluc|OJZ81846.1 anid|CBF86494.1 anig|EHA22376.1 aory|EIT72742.1 asyd|OJJ64893.1 ater|EAU30514.1 ater|EAU30516.1 atub|OJI84415.1 aver|OJI99014.1 awen|OJJ41397.1 awen|OJJ41399.1 azon|OJJ48865.1 aory|EIT72593.1

Group260: abra|OJJ68761.1 acar|OOF92248.1 acla|EAW12490.1 afis|EAW24155.1 afla|EED46777.1 afum|EAL92010.1 akaw|GAA89905.1 aluc|OJZ81765.1 anid|CBF77904.1 anig|EHA22449.1 asyd|OJJ55301.1 atub|OJI84337.1 abra|OJJ68762.1 acla|EAW12489.1 afis|EAW24156.1 afum|EAL92011.1 agla|OJJ89525.1 akaw|GAA89906.1 aluc|OJZ81764.1 anid|CBF77902.1 anig|EHA22450.1 aory|EIT72652.1 asyd|OJJ55302.1 ater|EAU30587.1 atub|OJI84336.1 aver|OJJ06924.1 awen|OJJ41322.1 azon|OJJ48795.1

Group261: abra|OJJ69707.1 acar|OOF94204.1 acla|EAW14911.1 afla|EED55248.1 agla|OJJ79247.1 akaw|GAA90156.1 aluc|OJZ83191.1 anid|CBF73771.1 anig|EHA28261.1 aory|EIT77697.1 asyd|OJJ56592.1 atub|OJI87748.1 awen|OJJ35897.1 azon|OJJ50407.1 abra|OJJ72111.1 acar|OOF90025.1 afis|EAW24432.1 afum|EAL86100.1 akaw|GAA84811.1 aluc|OJZ86866.1 anig|EHA21775.1 ater|EAU30013.1 atub|OJI83654.1 aver|OJJ06133.1 awen|OJJ32770.1 aory|EIT83627.1 azon|OJJ44000.1 afla|EED49732.1

Group262: abra|OJJ69819.1 acar|OOF93344.1 afis|EAW15617.1 afla|EED52777.1 afum|EAL85718.1 agla|OJJ79351.1 akaw|GAA87841.1 aluc|OJZ85477.1 anid|CBF76275.1 anig|EHA25018.1 aory|EIT78176.1 asyd|OJJ60985.1 ater|EAU37197.1 atub|OJI80086.1 aver|OJJ00304.1 abra|OJJ74486.1 acar|OOF95263.1 acla|EAW08782.1 afla|EED57614.1 akaw|GAA87379.1 aluc|OJZ89633.1 anig|EHA26875.1 aory|EIT82993.1 atub|OJI85170.1 awen|OJJ40133.1 ater|EAU36083.1 asyd|OJJ52418.1 aver|OJJ03363.1

Group263: abra|OJJ70208.1 acar|OOF90365.1 acla|EAW13177.1 afis|EAW22620.1 afla|EED51827.1 afum|EAL87187.2 agla|OJJ85385.1 akaw|GAA86568.1 aluc|OJZ85119.1 anid|CBF84539.1 anig|EHA25339.1 aory|EIT81723.1 asyd|OJJ53483.1 ater|EAU32773.1 atub|OJI80474.1 aver|OJJ08276.1 awen|OJJ33335.1 azon|OJJ51798.1 acla|EAW11725.1 afis|EAW23750.1 asyd|OJJ64104.1 aver|OJI98187.1 azon|OJJ51793.1 afum|EAL90576.1 afla|EED53104.1 anid|CBF85467.1 agla|OJJ87544.1 aory|EIT74796.1

Group264: anig|EHA20613.1 atub|OJI81264.1 abra|OJJ71157.1 acar|OOF98002.1 acla|EAW10341.1 afis|EAW25250.1 afla|EED53714.1 afum|EAL89912.1 agla|OJJ89220.1 akaw|GAA82821.1 aluc|OJZ87382.1 anid|CBF85305.1 aory|EIT74933.1 asyd|OJJ63998.1 ater|EAU34356.1 aver|OJI98087.1 awen|OJJ40962.1 azon|OJJ44543.1 acla|EAW10342.1 afis|EAW25247.1 afla|EED53713.1 afum|EAL89909.1 agla|OJJ89214.1 akaw|GAA82822.1 aluc|OJZ87383.1 anid|CBF85306.1 awen|OJJ40961.1 azon|OJJ44529.1

Group265: abra|OJJ71667.1 acar|OOF96377.1 acar|OOF96378.1 acla|EAW15205.1 acla|EAW15206.1 afis|EAW20170.1 afis|EAW20171.1 afla|EED57177.1 afla|EED57179.1 afum|EAL85288.1 afum|EAL85289.1 agla|OJJ88759.1 akaw|GAA89264.1 akaw|GAA89265.1 aluc|OJZ86409.1 anid|CBF85026.1 anid|CBF85028.1 anig|EHA21490.1 aory|EIT73988.1 aory|EIT74058.1 asyd|OJJ63103.1 ater|EAU35205.1 ater|EAU35206.1 atub|OJI84122.1 aver|OJI97858.1 aver|OJI97859.1 awen|OJJ32067.1 azon|OJJ46099.1

Group266: abra|OJJ74363.1 acar|OOF95367.1 agla|OJJ88403.1 akaw|GAA86667.1 aluc|OJZ89764.1 anid|CBF83899.1 anig|EHA26984.1 asyd|OJJ55412.1 atub|OJI85038.1 aver|OJJ06490.1 awen|OJJ32658.1 abra|OJJ75265.1 acar|OOF94538.1 afis|EAW20807.1 afla|EED46108.1 afum|EAL84821.1 akaw|GAA91801.1 aluc|OJZ92823.1 anid|CBF87420.1 anig|EHA26186.1 aory|EIT76498.1 atub|OJI87279.1 atub|OJI84647.1 afla|EED56188.1 anig|EHA20917.1 aory|EIT73391.1 akaw|GAA92694.1 aluc|OJZ90152.1

Group267: abra|OJJ75148.1 acar|OOF97189.1 acar|OOF97190.1 acla|EAW08321.1 acla|EAW08322.1 afis|EAW23356.1 afis|EAW23357.1 afla|EED55606.1 afla|EED55607.1 afum|EAL88119.2 afum|EAL88120.2 agla|OJJ86469.1 akaw|GAA88753.1 akaw|GAA88754.1 aluc|OJZ92713.1 aluc|OJZ92714.1 anid|CBF74884.1 anig|EHA26286.1 aory|EIT81807.1 asyd|OJJ60109.1 ater|EAU35607.1 ater|EAU35608.1 atub|OJI87139.1 atub|OJI87140.1 aver|OJJ04153.1 awen|OJJ39530.1 azon|OJJ47726.1 azon|OJJ47728.1

Group268: abra|OJJ75712.1 acar|OOG00147.1 acla|EAW09400.1 afis|EAW21736.1 afla|EED58008.1 afum|EAL92769.1 agla|OJJ79847.1 akaw|GAA84446.1 aluc|OJZ89163.1 anig|EHA23391.1 aory|EIT79742.1 ater|EAU38417.1 atub|OJI85627.1 awen|OJJ34009.1 abra|OJJ75713.1 acar|OOG00146.1 acla|EAW09399.1 afla|EED58009.1 afum|EAL92770.1 agla|OJJ88777.1 akaw|GAA84447.1 aluc|OJZ89164.1 anig|EHA23390.1 aory|EIT79726.1 asyd|OJJ65106.1 atub|OJI85626.1 aver|OJI99233.1 azon|OJJ48541.1

Group269: abra|OJJ76491.1 acar|OOF99722.1 acar|OOF99723.1 acla|EAW13941.1 acla|EAW13944.1 afis|EAW18652.1 afis|EAW18653.1 afla|EED52680.1 afla|EED52681.1 afum|EAL93368.1 afum|EAL93369.1 agla|OJJ82422.1 akaw|GAA82406.1 akaw|GAA82407.1 aluc|OJZ91713.1 aluc|OJZ91714.1 anid|CBF70539.1 anid|CBF70540.1 anig|EHA22724.1 aory|EIT74312.1 aory|EIT74318.1 asyd|OJJ58549.1 ater|EAU38259.1 ater|EAU38260.1 atub|OJI86172.1 aver|OJJ01147.1 awen|OJJ32120.1 azon|OJJ46706.1

Group270: abra|OJJ76517.1 acar|OOF99456.1 acla|EAW14184.1 acla|EAW14185.1 afis|EAW18899.1 afis|EAW18900.1 afla|EED50848.1 afla|EED50853.1 afum|EAL93631.1 afum|EAL93632.1 agla|OJJ81505.1 akaw|GAA82429.1 akaw|GAA82430.1 aluc|OJZ91689.1 anid|CBF69917.1 anid|CBF69919.1 anig|EHA22702.1 aory|EIT78983.1 aory|EIT79046.1 asyd|OJJ58989.1 ater|EAU38067.1 ater|EAU38068.1 atub|OJI86142.1 aver|OJJ01626.1 aver|OJJ01627.1 awen|OJJ32457.1 azon|OJJ47214.1 azon|OJJ47215.1

Group271: abra|OJJ76807.1 acar|OOF91299.1 acla|EAW13366.1 afis|EAW20480.1 afla|EED44787.1 afum|EAL84574.1 agla|OJJ84832.1 akaw|GAA87216.1 aluc|OJZ84214.1 anid|CBF79227.1 anig|EHA27567.1 aory|EIT77778.1 asyd|OJJ55739.1 ater|EAU29559.1 atub|OJI89869.1 aver|OJJ06825.1 awen|OJJ33179.1 azon|OJJ46382.1 afla|EED44788.1 aory|EIT77780.1 awen|OJJ33175.1 azon|OJJ46828.1 acar|OOF91301.1 acla|EAW13367.1 afis|EAW20481.1 afum|EAL84573.1 akaw|GAA87215.1 aluc|OJZ84215.1

Group272: abra|OJJ77199.1 acar|OOF93082.1 acar|OOF93087.1 acla|EAW13188.1 acla|EAW13189.1 afis|EAW17452.1 afis|EAW17453.1 afla|EED49369.1 afla|EED49370.1 afum|EAL91280.1 afum|EAL91281.2 agla|OJJ80350.1 agla|OJJ80360.1 akaw|GAA90742.1 aluc|OJZ84606.1 anid|CBF71675.1 anid|CBF71677.1 anig|EHA27895.1 anig|EHA27896.1 aory|EIT79469.1 aory|EIT79540.1 asyd|OJJ61825.1 ater|EAU33932.1 ater|EAU33933.1 atub|OJI89473.1 aver|OJI95925.1 awen|OJJ37394.1 azon|OJJ51067.1

Group273: abra|OJJ77369.1 acar|OOF93239.1 acla|EAW13106.1 acla|EAW13107.1 afis|EAW17558.1 afla|EED49266.1 afla|EED49267.1 afum|EAL91372.1 agla|OJJ81410.1 akaw|GAA91392.1 akaw|GAA91393.1 aluc|OJZ84778.1 aluc|OJZ84779.1 anid|CBF90279.1 anig|EHA28046.1 aory|EIT82182.1 aory|EIT82245.1 asyd|OJJ61916.1 ater|EAU36989.1 atub|OJI89303.1 aver|OJI96009.1 awen|OJJ37296.1 azon|OJJ50989.1 afis|EAW17557.1 afum|EAL91371.1 agla|OJJ81411.1 anid|CBF90281.1 asyd|OJJ61917.1

Group274: abra|OJJ66221.1 abra|OJJ69595.1 aluc|OJZ80527.1 aluc|OJZ90304.1 anig|EHA19754.1 anig|EHA25667.1 aver|OJI98854.1 aver|OJJ06288.1 acar|OOF90660.1 acar|OOF90661.1 acla|EAW08925.1 afis|EAW17322.1 afum|EAL91158.1 agla|OJJ87386.1 akaw|GAA89065.1 anig|EHA25668.1 asyd|OJJ56432.1 ater|EAU32950.1 atub|OJI88403.1 awen|OJJ39606.1 azon|OJJ46461.1 akaw|GAA86069.1 awen|OJJ39353.1 acla|EAW09791.1 afis|EAW21345.1 afum|EAL92362.1 azon|OJJ51694.1

Group275: abra|OJJ68039.1 abra|OJJ73342.1 acar|OOF91804.1 acar|OOF94627.1 akaw|GAA83140.1 akaw|GAA90913.1 aluc|OJZ82913.1 aluc|OJZ88697.1 anig|EHA18377.1 anig|EHA21336.1 asyd|OJJ53721.1 asyd|OJJ60902.1 atub|OJI89139.1 atub|OJI91278.1 aver|OJJ00228.1 aver|OJJ07846.1 awen|OJJ29994.1 awen|OJJ33889.1 azon|OJJ42783.1 azon|OJJ50111.1 acla|EAW06969.1 afis|EAW24632.1 afla|EED55052.1 afum|EAL86254.1 anid|CBF83822.1 aory|EIT81944.1 ater|EAU36659.1

Group276: acar|OOF92615.1 acar|OOF92636.1 abra|OJJ69082.1 acar|OOF91209.1 acla|EAW08752.1 afis|EAW15684.1 afla|EED51370.1 afum|EAL85664.1 agla|OJJ81833.1 akaw|GAA90611.1 akaw|GAA90612.1 aluc|OJZ86070.1 anid|CBF71148.1 anig|EHA24544.1 aory|EIT80661.1 asyd|OJJ55837.1 ater|EAU32473.1 atub|OJI79866.1 atub|OJI79867.1 aver|OJJ04780.1 awen|OJJ38376.1 azon|OJJ43860.1 acla|EAW08751.1 afis|EAW15685.1 afla|EED51369.1 afum|EAL85663.1 anid|CBF71146.1

Group277: acar|OOF93141.1 acar|OOF96894.1 asyd|OJJ55471.1 asyd|OJJ61030.1 asyd|OJJ57185.1 abra|OJJ72261.1 acla|EAW13265.1 afis|EAW16849.1 afla|EED49789.1 afum|EAL86395.1 agla|OJJ88181.1 akaw|GAA84669.1 aluc|OJZ87020.1 anid|CBF84231.1 anig|EHA21635.1 aory|EIT77692.1 ater|EAU33865.1 atub|OJI83498.1 aver|OJJ00350.1 aver|OJJ03105.1 awen|OJJ36910.1 azon|OJJ50296.1 acla|EAW12401.1 afla|EED51180.1 aory|EIT81355.1 anid|CBF89260.1 azon|OJJ50917.1

Group278: acar|OOF94414.1 acar|OOF97591.1 afis|EAW24229.1 afis|EAW24933.1 afla|EED46551.1 afla|EED48042.1 afum|EAL85898.1 afum|EAL89589.1 aory|EIT77904.1 aory|EIT83324.1 asyd|OJJ53967.1 asyd|OJJ64678.1 ater|EAU29958.1 ater|EAU31178.1 aver|OJI98780.1 aver|OJJ08063.1 anig|EHA28592.1 abra|OJJ69311.1 acla|EAW14698.1 agla|OJJ79642.1 akaw|GAA88180.1 aluc|OJZ82140.1 anid|CBF74228.1 anig|EHA24084.1 atub|OJI88297.1 awen|OJJ33068.1 azon|OJJ48455.1

Group279: acla|EAW07586.1 acla|EAW11103.1 afis|EAW17139.1 afis|EAW25619.1 anid|CBF80637.1 anid|CBF89256.1 acar|OOF93375.1 abra|OJJ69171.1 acar|OOF91133.1 afla|EED55556.1 agla|OJJ80465.1 akaw|GAA86292.1 aluc|OJZ86158.1 anig|EHA24466.1 asyd|OJJ61355.1 ater|EAU32675.1 atub|OJI79950.1 aver|OJJ00682.1 awen|OJJ38095.1 azon|OJJ42539.1 aluc|OJZ82069.1 anid|CBF87524.1 ater|EAU31706.1 atub|OJI84626.1 akaw|GAA91464.1 anig|EHA22500.1 aory|EIT82677.1

Group280: afis|EAW18048.1 afis|EAW24488.1 afum|EAL86157.1 afum|EAL86730.1 afum|EAL91860.1 abra|OJJ68602.1 acar|OOF92112.1 acla|EAW12637.1 afla|EED51498.1 agla|OJJ85419.1 akaw|GAA91926.1 aluc|OJZ81954.1 anid|CBF79800.1 anig|EHA22293.1 aory|EIT80704.1 asyd|OJJ58129.1 ater|EAU31275.1 atub|OJI84505.1 aver|OJJ04276.1 awen|OJJ32709.1 awen|OJJ41486.1 azon|OJJ48216.1 afla|EED48171.1 agla|OJJ83081.1 aory|EIT83032.1 ater|EAU38954.1 aver|OJI98885.1

Group281: afla|EED45594.1 afla|EED53079.1 aory|EIT74816.1 aory|EIT75766.1 abra|OJJ66499.1 asyd|OJJ54572.1 aver|OJJ07582.1 azon|OJJ50366.1 abra|OJJ72643.1 acar|OOF94628.1 akaw|GAA85264.1 aluc|OJZ90622.1 anid|CBF69513.1 anig|EHA24059.1 asyd|OJJ56879.1 ater|EAU39542.1 atub|OJI82048.1 aver|OJJ08508.1 afla|EED47413.1 aory|EIT81918.1 aver|OJI99560.1 afla|EED52187.1 aory|EIT74424.1 ater|EAU31590.1 awen|OJJ31457.1 asyd|OJJ56253.1 ater|EAU31815.1

Group282: afla|EED51568.1 afla|EED51716.1 afla|EED56540.1 agla|OJJ78952.1 agla|OJJ80278.1 aory|EIT79700.1 aory|EIT80654.1 aory|EIT80755.1 asyd|OJJ57607.1 asyd|OJJ58114.1 aver|OJJ04291.1 aver|OJJ05577.1 awen|OJJ33707.1 awen|OJJ34440.1 abra|OJJ67863.1 acar|OOF92843.1 acla|EAW11735.1 afis|EAW23768.1 afum|EAL87202.1 agla|OJJ85386.1 akaw|GAA91870.1 aluc|OJZ82029.1 anid|CBF79783.1 anig|EHA22227.1 ater|EAU31312.1 atub|OJI84583.1 azon|OJJ42833.1

Group283: afum|EAL85403.1 afum|EAL87814.1 aory|EIT74045.1 aory|EIT79062.1 ater|EAU31644.1 ater|EAU36768.1 atub|OJI87665.1 atub|OJI90002.1 abra|OJJ65728.1 acar|OOF93412.1 afis|EAW19529.1 afla|EED50913.1 akaw|GAA93255.1 aluc|OJZ83633.1 asyd|OJJ62934.1 azon|OJJ46829.1 awen|OJJ30425.1 afis|EAW16772.1 akaw|GAA92091.1 anid|CBF73717.1 anig|EHA18786.1 asyd|OJJ64570.1 aver|OJI97354.1 aver|OJI98056.1 aver|OJI98667.1 afla|EED57223.1 asyd|OJJ63582.1

Group284: afum|EAL86514.1 afum|EAL91321.1 asyd|OJJ52759.1 asyd|OJJ54478.1 asyd|OJJ61874.1 azon|OJJ42558.1 azon|OJJ51475.1 azon|OJJ51028.1 abra|OJJ74834.1 acar|OOF95824.1 acla|EAW13149.1 afis|EAW17498.1 afla|EED49318.1 agla|OJJ81459.1 akaw|GAA83586.1 aluc|OJZ92411.1 anid|CBF71582.1 anig|EHA26554.1 aory|EIT82205.1 ater|EAU34444.1 ater|EAU36943.1 atub|OJI86841.1 aver|OJI95972.1 awen|OJJ37343.1 acla|EAW12237.1 afis|EAW16990.1 aver|OJJ08741.1

Group285: agla|OJJ79575.1 agla|OJJ86866.1 agla|OJJ88176.1 agla|OJJ89239.1 asyd|OJJ62473.1 asyd|OJJ64049.1 ater|EAU34049.1 ater|EAU34409.1 aver|OJI96608.1 aver|OJI98135.1 awen|OJJ38034.1 awen|OJJ40680.1 awen|OJJ40981.1 abra|OJJ74157.1 acar|OOF95570.1 acla|EAW12279.1 afis|EAW16961.1 afla|EED49740.1 afum|EAL86487.2 akaw|GAA89727.1 aluc|OJZ80656.1 anid|CBF89363.1 anig|EHA27176.1 aory|EIT75292.1 atub|OJI87511.1 azon|OJJ44568.1 anid|CBF85376.1

Group286: akaw|GAA84498.1 akaw|GAA89765.1 anig|EHA23338.1 anig|EHA27214.1 aver|OJJ00853.1 aver|OJJ07847.1 abra|OJJ75781.1 acar|OOF91480.1 acla|EAW10808.1 afis|EAW24770.1 afla|EED55185.1 afum|EAL89411.1 agla|OJJ83340.1 aluc|OJZ80618.1 aluc|OJZ89223.1 anid|CBF86704.1 aory|EIT83621.1 asyd|OJJ64740.1 ater|EAU37121.1 atub|OJI85569.1 atub|OJI87472.1 aver|OJI98844.1 awen|OJJ29862.1 anid|CBF83000.1 afla|EED53623.1 aory|EIT75056.1 asyd|OJJ61517.1

Group287: asyd|OJJ54631.1 asyd|OJJ65150.1 asyd|OJJ58326.1 abra|OJJ68110.1 afla|EED53398.1 akaw|GAA86893.1 aluc|OJZ90016.1 anid|CBF80628.1 anig|EHA27391.1 aory|EIT74923.1 ater|EAU36928.1 atub|OJI84791.1 aver|OJJ07525.1 awen|OJJ33176.1 abra|OJJ71207.1 acar|OOF97964.1 afla|EED54683.1 akaw|GAA82777.1 aluc|OJZ87336.1 anid|CBF80110.1 anig|EHA20573.1 aory|EIT78795.1 atub|OJI81308.1 aver|OJJ02085.1 afla|EED46389.1 aory|EIT72418.1 atub|OJI84522.1

Group288: awen|OJJ30753.1 awen|OJJ31728.1 awen|OJJ30755.1 awen|OJJ31726.1 abra|OJJ67455.1 acar|OOF92379.1 afis|EAW18419.1 afum|EAL93103.1 akaw|GAA86376.1 aluc|OJZ84910.1 anid|CBF70711.1 asyd|OJJ58453.1 ater|EAU29836.1 atub|OJI80688.1 aver|OJJ01052.1 azon|OJJ50489.1 acar|OOF92377.1 asyd|OJJ58451.1 acla|EAW06310.1 afla|EED52989.1 agla|OJJ84987.1 anid|CBF70709.1 aory|EIT78704.1 ater|EAU29837.1 aver|OJJ01053.1 ater|EAU29835.1 asyd|OJJ58452.1

Group289: azon|OJJ42747.1 azon|OJJ51180.1 abra|OJJ76401.1 acar|OOF99642.1 acla|EAW14023.1 afis|EAW18729.1 afla|EED50685.1 afum|EAL93454.1 agla|OJJ82287.1 akaw|GAA82321.1 aluc|OJZ91811.1 anid|CBF70016.1 anig|EHA22804.1 aory|EIT78004.1 asyd|OJJ58877.1 ater|EAU38175.1 atub|OJI86267.1 aver|OJJ01479.1 awen|OJJ32270.1 akaw|GAA82320.1 aluc|OJZ91812.1 acar|OOF99643.1 acla|EAW14022.1 afis|EAW18728.1 afum|EAL93453.1 anid|CBF70018.1 ater|EAU38176.1

Group290: abra|OJJ65930.1 acar|OOF94991.1 acla|EAW12067.1 afis|EAW19710.1 afla|EED57049.1 afum|EAL86817.2 agla|OJJ82841.1 akaw|GAA92305.1 akaw|GAA92306.1 aluc|OJZ90202.1 anid|CBF84278.1 anig|EHA23699.1 aory|EIT82284.1 asyd|OJJ57230.1 ater|EAU32834.1 atub|OJI81631.1 aver|OJJ03059.1 awen|OJJ32205.1 azon|OJJ42677.1 afla|EED57054.1 acar|OOF94992.1 acla|EAW12066.1 afis|EAW19709.1 afum|EAL86818.1 aory|EIT82258.1 ater|EAU32836.1 azon|OJJ42678.1

Group291: abra|OJJ66585.1 acar|OOF91022.1 acla|EAW10906.1 acla|EAW10907.1 afis|EAW20951.1 afis|EAW20952.1 afla|EED50307.1 afla|EED50308.1 afum|EAL84682.1 afum|EAL84683.2 agla|OJJ83929.1 akaw|GAA91132.1 akaw|GAA91134.1 aluc|OJZ81461.1 anid|CBF82552.1 anid|CBF82554.1 anig|EHA25968.1 aory|EIT77335.1 aory|EIT77337.1 asyd|OJJ54058.1 ater|EAU38602.1 ater|EAU38603.1 atub|OJI88718.1 aver|OJJ08683.1 awen|OJJ34364.1 awen|OJJ34365.1 azon|OJJ48100.1

Group292: abra|OJJ66932.1 acar|OOF91699.1 acla|EAW14495.1 afis|EAW19209.1 afla|EED52263.1 afum|EAL93939.2 agla|OJJ85720.1 akaw|GAA91486.1 aluc|OJZ85648.1 anid|CBF69569.1 anig|EHA24885.1 asyd|OJJ59150.1 ater|EAU34159.1 atub|OJI79439.1 aver|OJJ02018.1 awen|OJJ34677.1 azon|OJJ45456.1 acar|OOF91700.1 agla|OJJ88500.1 anid|CBF78572.1 anig|EHA24886.1 asyd|OJJ65074.1 ater|EAU37259.1 aver|OJI99199.1 awen|OJJ37563.1 azon|OJJ51226.1 aver|OJJ01740.1

Group293: abra|OJJ67205.1 acar|OOF98357.1 acar|OOF98358.1 acla|EAW14519.1 afis|EAW19237.1 afla|EED47640.1 afum|EAL93965.2 agla|OJJ88462.1 akaw|GAA90972.1 akaw|GAA90973.1 aluc|OJZ87769.1 anid|CBF78640.1 anig|EHA20253.1 aory|EIT80182.1 asyd|OJJ65189.1 ater|EAU37226.1 atub|OJI80876.1 aver|OJI99338.1 awen|OJJ37593.1 azon|OJJ51248.1 acla|EAW14518.1 afis|EAW19236.1 afla|EED47646.1 afum|EAL93964.1 anid|CBF78638.1 aory|EIT80263.1 ater|EAU37227.1

Group294: abra|OJJ67284.1 afla|EED54338.1 afum|EAL87110.1 agla|OJJ82024.1 akaw|GAA86525.1 aluc|OJZ85073.1 anid|CBF71655.1 anig|EHA25287.1 aory|EIT76146.1 asyd|OJJ61837.1 ater|EAU32860.1 atub|OJI80525.1 aver|OJI95936.1 awen|OJJ36119.1 abra|OJJ68987.1 acar|OOF91558.1 afla|EED48927.1 akaw|GAA90508.1 aluc|OJZ85973.1 anid|CBF87485.1 anig|EHA24632.1 aory|EIT79999.1 ater|EAU31715.1 atub|OJI79761.1 asyd|OJJ52961.1 aver|OJJ08927.1 awen|OJJ41925.1

Group295: abra|OJJ68390.1 afla|EED49558.1 agla|OJJ87147.1 aory|EIT82645.1 atub|OJI84701.1 abra|OJJ77648.1 acar|OOF99088.1 acla|EAW12091.1 afis|EAW19985.1 afum|EBA27209.1 akaw|GAA83987.1 aluc|OJZ80881.1 anid|CBF87267.1 anig|EHA19455.1 asyd|OJJ62371.1 ater|EAU33659.1 atub|OJI82696.1 aver|OJI98437.1 awen|OJJ38077.1 azon|OJJ42589.1 anid|CBF73965.1 aver|OJI96505.1 afla|EED52196.1 aory|EIT74391.1 ater|EAU29811.1 aluc|OJZ83787.1 anid|CBF85559.1

Group296: abra|OJJ69188.1 acar|OOF91124.1 acla|EAW08657.1 afis|EAW15774.1 afla|EED51274.1 afum|EAL85575.1 agla|OJJ81937.1 akaw|GAA86276.1 akaw|GAA86277.1 aluc|OJZ86174.1 anid|CBF70974.1 anig|EHA24450.1 anig|EHA24451.1 aory|EIT81499.1 asyd|OJJ52632.1 ater|EAU32367.1 atub|OJI79966.1 aver|OJJ04660.1 awen|OJJ38481.1 azon|OJJ43782.1 acla|EAW08658.1 afis|EAW15773.1 afla|EED49465.1 anid|CBF70972.1 ater|EAU32368.1 azon|OJJ43781.1 aory|EIT79571.1

Group297: abra|OJJ69236.1 acar|OOF91082.1 acar|OOF91084.1 acla|EAW08611.1 acla|EAW08613.1 afis|EAW15821.1 afis|EAW15822.1 afla|EED51224.1 afla|EED51226.1 afum|EAL85529.1 afum|EAL85530.1 agla|OJJ81985.1 akaw|GAA92244.1 akaw|GAA92246.1 aluc|OJZ86216.1 anid|CBF70886.1 anid|CBF70887.1 anig|EHA24410.1 aory|EIT83181.1 aory|EIT83189.1 asyd|OJJ52589.1 ater|EAU33426.1 ater|EAU33427.1 atub|OJI80011.1 aver|OJJ04616.1 awen|OJJ38535.1 azon|OJJ43727.1

Group298: abra|OJJ70462.1 acar|OOF97462.1 acar|OOF97465.1 acla|EAW13051.1 acla|EAW13053.1 afis|EAW17613.1 afis|EAW17616.1 afla|EED49204.1 afla|EED49207.1 afum|EAL91426.1 afum|EAL91429.1 agla|OJJ81324.1 akaw|GAA83308.1 akaw|GAA83311.1 aluc|OJZ83805.1 anid|CBF90169.1 anid|CBF90172.1 anig|EHA18952.1 aory|EIT78583.1 aory|EIT83172.1 asyd|OJJ61991.1 ater|EAU37048.1 ater|EAU37050.1 atub|OJI90172.1 aver|OJI96083.1 awen|OJJ37229.1 azon|OJJ50932.1

Group299: abra|OJJ71130.1 acar|OOF98022.1 acla|EAW10367.1 acla|EAW10368.1 afis|EAW25224.1 afis|EAW25225.1 afla|EED53669.1 afla|EED53682.1 afum|EAL89885.1 afum|EAL89886.1 akaw|GAA82844.1 aluc|OJZ87405.1 anid|CBF85264.1 anid|CBF85265.1 anig|EHA20634.1 anig|EHA20635.1 aory|EIT75307.1 aory|EIT75353.1 ater|EAU34383.1 ater|EAU34384.1 atub|OJI81241.1 aver|OJI98082.1 aver|OJI98083.1 agla|OJJ79183.1 asyd|OJJ62906.1 awen|OJJ40890.1 azon|OJJ44519.1

Group300: abra|OJJ72036.1 acar|OOF94174.1 acla|EAW09507.1 afla|EED57855.1 agla|OJJ82582.1 akaw|GAA84897.1 aluc|OJZ86774.1 anid|CBF83512.1 anig|EHA21907.1 aory|EIT81060.1 asyd|OJJ53748.1 asyd|OJJ60708.1 ater|EAU38317.1 atub|OJI83747.1 aver|OJJ00034.1 awen|OJJ33880.1 abra|OJJ72037.1 afla|EED56917.1 akaw|GAA84896.1 aluc|OJZ86775.1 anid|CBF73981.1 anig|EHA21906.1 aory|EIT77541.1 asyd|OJJ55360.1 atub|OJI83746.1 aver|OJJ06443.1 aver|OJJ07863.1

Group301: abra|OJJ72938.1 acar|OOF96120.1 acar|OOF96121.1 acla|EAW06452.1 acla|EAW06453.1 afis|EAW15949.1 afla|EED58109.1 afla|EED58110.1 afum|EAL88582.1 agla|OJJ89365.1 akaw|GAA85519.1 aluc|OJZ83447.1 anid|CBF81052.1 anig|EHA24293.1 anig|EHA24294.1 aory|EIT79139.1 aory|EIT79141.1 asyd|OJJ56450.1 ater|EAU35509.1 atub|OJI82338.1 atub|OJI82339.1 aver|OJJ06272.1 awen|OJJ35627.1 azon|OJJ46898.1 anid|CBF81054.1 ater|EAU35508.1 aver|OJJ06271.1

Group302: abra|OJJ73010.1 acar|OOF96182.1 acar|OOF96183.1 acla|EAW08002.1 acla|EAW08003.1 afis|EAW23037.1 afis|EAW23038.1 afla|EED54980.1 afum|EAL88443.1 afum|EAL88444.1 agla|OJJ89440.1 akaw|GAA85588.1 akaw|GAA85589.1 aluc|OJZ83525.1 anid|CBF80929.1 anid|CBF80931.1 anig|EHA24348.1 aory|EIT80318.1 aory|EIT80383.1 asyd|OJJ56316.1 ater|EAU29618.1 ater|EAU29621.1 atub|OJI82414.1 aver|OJJ06398.1 awen|OJJ35704.1 azon|OJJ44166.1 afla|EED54979.1

Group303: abra|OJJ73242.1 acar|OOG00479.1 acar|OOG00481.1 acla|EAW06915.1 acla|EAW06916.1 afis|EAW16532.1 afis|EAW16533.1 afla|EED56136.1 afla|EED56140.1 afum|EAL89129.1 afum|EAL89130.2 agla|OJJ82788.1 akaw|GAA87938.1 akaw|GAA87939.1 aluc|OJZ81186.1 anid|CBF81734.1 anid|CBF81736.1 anig|EHA18503.1 aory|EIT76580.1 aory|EIT76630.1 asyd|OJJ59594.1 ater|EAU36595.1 ater|EAU36596.1 atub|OJI91382.1 aver|OJJ02674.1 awen|OJJ40289.1 azon|OJJ49150.1

Group304: abra|OJJ73503.1 acar|OOG00688.1 acla|EAW07435.1 acla|EAW07436.1 afis|EAW22461.1 afis|EAW22462.1 afla|EED54482.1 afla|EED54483.1 afum|EAL90729.1 afum|EAL90730.1 agla|OJJ87561.1 akaw|GAA87450.1 akaw|GAA87451.1 aluc|OJZ88545.1 anid|CBF88911.1 anid|CBF88912.1 anig|EHA18245.1 aory|EIT81150.1 asyd|OJJ62841.1 ater|EAU39244.1 atub|OJI91130.1 aver|OJI96884.1 aver|OJI96885.1 awen|OJJ35087.1 awen|OJJ35088.1 azon|OJJ50020.1 aory|EIT81157.1

Group305: abra|OJJ74418.1 acar|OOF95322.1 acar|OOF95325.1 acla|EAW06834.1 afla|EED53447.1 afla|EED53451.1 agla|OJJ88208.1 akaw|GAA87309.1 akaw|GAA87312.1 aluc|OJZ89705.1 aluc|OJZ89708.1 anid|CBF89366.1 anid|CBF89368.1 anig|EHA26939.1 aory|EIT75073.1 asyd|OJJ62474.1 ater|EAU34066.1 ater|EAU34069.1 atub|OJI85094.1 aver|OJI96610.1 awen|OJJ37916.1 awen|OJJ37920.1 azon|OJJ45806.1 acar|OOF90541.1 acla|EAW11778.1 azon|OJJ50188.1 aory|EIT75099.1

Group306: abra|OJJ76286.1 acar|OOF99527.1 acla|EAW14330.1 afis|EAW18958.1 afum|EAL93691.1 akaw|GAA82215.1 aory|EIT76999.1 asyd|OJJ59136.1 awen|OJJ31557.1 azon|OJJ47274.1 abra|OJJ76287.1 acar|OOF99530.1 acla|EAW14329.1 afis|EAW18959.1 afla|EED51005.1 afum|EAL93692.1 agla|OJJ85336.1 akaw|GAA82216.1 aluc|OJZ91935.1 anid|CBF69598.1 anig|EHA22895.1 aory|EIT76941.1 ater|EAU38141.1 atub|OJI86377.1 aver|OJJ01725.1 awen|OJJ31558.1 azon|OJJ47275.1

Group307: abra|OJJ77576.1 acar|OOF99151.1 acla|EAW06541.1 acla|EAW06542.1 afis|EAW16053.1 afis|EAW16054.1 afla|EED56444.1 afla|EED56445.1 afum|EAL88685.1 afum|EAL88686.1 agla|OJJ86087.1 akaw|GAA84049.1 akaw|GAA84050.1 aluc|OJZ80961.1 anid|CBF75070.1 anid|CBF75072.1 anig|EHA19396.1 aory|EIT72818.1 asyd|OJJ60021.1 ater|EAU33580.1 ater|EAU33584.1 atub|OJI82625.1 aver|OJJ02264.1 awen|OJJ30385.1 awen|OJJ30389.1 azon|OJJ45401.1 aory|EIT72866.1

Group308: abra|OJJ77839.1 acar|OOF98932.1 acla|EAW11284.1 afis|EAW18216.1 afla|EED56254.1 afum|EAL87582.1 akaw|GAA83808.1 aory|EIT73366.1 ater|EAU33774.1 azon|OJJ50174.1 abra|OJJ77840.1 acar|OOF98931.1 acla|EAW11283.1 afis|EAW18217.1 afla|EED56253.1 afum|EAL87583.1 agla|OJJ85865.1 akaw|GAA83807.1 aluc|OJZ90915.1 anid|CBF86146.1 anig|EHA19627.1 aory|EIT73438.1 asyd|OJJ64511.1 atub|OJI82907.1 aver|OJI98609.1 awen|OJJ30595.1 azon|OJJ50175.1

Group309: abra|OJJ65742.1 abra|OJJ77048.1 acar|OOF91730.1 acar|OOF92937.1 aluc|OJZ84452.1 aluc|OJZ86322.1 atub|OJI84212.1 atub|OJI89624.1 acla|EAW13550.1 afis|EAW20627.1 afla|EED49777.1 afum|EAL84441.1 agla|OJJ86946.1 akaw|GAA87005.1 anid|CBF83228.1 anig|EHA27767.1 aory|EIT77606.1 asyd|OJJ55581.1 ater|EAU32164.1 aver|OJJ06660.1 awen|OJJ33462.1 azon|OJJ50718.1 afla|EED52520.1 anig|EHA19838.1 aory|EIT82787.1 awen|OJJ33741.1

Group310: abra|OJJ66443.1 abra|OJJ68021.1 abra|OJJ68923.1 acar|OOF91508.1 acar|OOF92064.1 agla|OJJ78513.1 agla|OJJ81294.1 agla|OJJ85809.1 akaw|GAA90325.1 akaw|GAA90930.1 aluc|OJZ82895.1 aluc|OJZ85908.1 anig|EHA21319.1 anig|EHA24684.1 aory|EIT75910.1 aory|EIT81247.1 asyd|OJJ59089.1 asyd|OJJ63774.1 atub|OJI79692.1 atub|OJI89120.1 aver|OJI97163.1 aver|OJJ01679.1 acla|EAW09049.1 afla|EED46360.1 ater|EAU34525.1 awen|OJJ36242.1

Group311: abra|OJJ68133.1 abra|OJJ69675.1 acar|OOF90055.1 acar|OOF90087.1 acla|EAW12871.1 afis|EAW17819.1 afla|EED45605.1 afum|EAL91629.1 agla|OJJ81780.1 akaw|GAA86913.1 akaw|GAA88800.1 aluc|OJZ90036.1 anid|CBF71190.1 anig|EHA27411.1 aory|EIT75710.1 asyd|OJJ55855.1 atub|OJI84770.1 atub|OJI87949.1 aver|OJJ04798.1 awen|OJJ38287.1 acla|EAW08770.1 afis|EAW15658.1 afum|EAL85688.2 aluc|OJZ82483.1 anig|EHA28291.1 ater|EAU32513.1

Group312: abra|OJJ68841.1 abra|OJJ73289.1 afis|EAW20222.1 afis|EAW20406.1 afla|EED48455.1 afla|EED50145.1 anig|EHA24185.1 anig|EHA24740.1 aory|EIT74688.1 aory|EIT83472.1 asyd|OJJ57669.1 asyd|OJJ58613.1 asyd|OJJ59804.1 asyd|OJJ64211.1 aver|OJI97339.1 aver|OJJ01220.1 aver|OJJ02487.1 acar|OOF90549.1 agla|OJJ82202.1 akaw|GAA90408.1 aluc|OJZ85820.1 anid|CBF70434.1 ater|EAU37675.1 atub|OJI79606.1 awen|OJJ30933.1 acar|OOG00326.1

Group313: abra|OJJ71059.1 abra|OJJ73085.1 akaw|GAA82900.1 akaw|GAA88075.1 anig|EHA18644.1 anig|EHA20691.1 ater|EAU29720.1 ater|EAU35713.1 atub|OJI81179.1 atub|OJI91543.1 aver|OJJ02730.1 aver|OJJ06544.1 awen|OJJ34513.1 awen|OJJ40561.1 acar|OOF98083.1 acla|EAW10600.1 afis|EAW24968.1 afum|EAL89627.1 agla|OJJ82234.1 aluc|OJZ87469.1 anid|CBF81643.1 aory|EIT80808.1 asyd|OJJ59538.1 azon|OJJ49422.1 afla|EED57716.1 aory|EIT79920.1

Group314: abra|OJJ77512.1 abra|OJJ78209.1 acar|OOF98645.1 acar|OOF99212.1 agla|OJJ84617.1 agla|OJJ88162.1 akaw|GAA84108.1 akaw|GAA85817.1 aluc|OJZ81024.1 aluc|OJZ91282.1 atub|OJI82559.1 atub|OJI83272.1 awen|OJJ30638.1 awen|OJJ31266.1 afis|EAW18264.1 anid|CBF79534.1 anig|EHA19340.1 aory|EIT75007.1 asyd|OJJ57897.1 aver|OJJ04498.1 azon|OJJ47978.1 afla|EED48723.1 afum|EAL87628.1 anig|EHA19967.1 ater|EAU33271.1 aory|EIT73106.1

Group315: acar|OOF95616.1 acar|OOF95668.1 afum|EAL86712.1 afum|EAL89365.2 aluc|OJZ83797.1 aluc|OJZ91020.1 atub|OJI83012.1 atub|OJI90166.1 abra|OJJ70451.1 afis|EAW19928.1 afis|EAW24742.1 agla|OJJ88966.1 akaw|GAA83713.1 anid|CBF78706.1 anig|EHA18945.1 asyd|OJJ64219.1 ater|EAU37496.1 aver|OJI98040.1 aver|OJI98302.1 awen|OJJ33410.1 azon|OJJ45929.1 afla|EED48357.1 afis|EAW16196.1 afum|EAL89388.1 asyd|OJJ61748.1 aver|OJI95845.1

Group316: acla|EAW06814.1 acla|EAW11026.1 afis|EAW16363.1 afis|EAW20840.1 afum|EAL84793.1 afum|EAL88988.1 agla|OJJ78575.1 agla|OJJ80753.1 aver|OJI96276.1 aver|OJI96627.1 awen|OJJ37768.1 awen|OJJ40172.1 abra|OJJ73491.1 acar|OOG00684.1 afla|EED53503.1 agla|OJJ79965.1 akaw|GAA87440.1 aluc|OJZ88557.1 anid|CBF89245.1 anig|EHA18255.1 aory|EIT74859.1 asyd|OJJ62507.1 ater|EAU34089.1 atub|OJI91142.1 azon|OJJ45066.1 asyd|OJJ62165.1

Group317: acla|EAW11040.1 acla|EAW11477.1 afla|EED47466.1 afla|EED49623.1 afla|EED50532.1 agla|OJJ82839.1 agla|OJJ86906.1 aory|EIT79957.1 aory|EIT81005.1 aory|EIT82111.1 awen|OJJ31954.1 awen|OJJ35441.1 azon|OJJ45954.1 azon|OJJ51301.1 abra|OJJ72795.1 acar|OOF89880.1 afis|EAW23976.1 afum|EAL87406.1 akaw|GAA85398.1 aluc|OJZ83292.1 anid|CBF77345.1 anig|EHA24176.1 asyd|OJJ53272.1 ater|EAU29776.1 atub|OJI82193.1 aver|OJJ03577.1

Group318: afla|EED44930.1 afla|EED51540.1 ater|EAU31301.1 ater|EAU38559.1 awen|OJJ33684.1 awen|OJJ36221.1 abra|OJJ67873.1 acar|OOF92853.1 acla|EAW11724.1 afis|EAW23749.1 afum|EAL87183.1 agla|OJJ78962.1 agla|OJJ89691.1 akaw|GAA91882.1 aluc|OJZ82015.1 anid|CBF79817.1 anig|EHA22241.1 aory|EIT80515.1 asyd|OJJ58137.1 atub|OJI84564.1 aver|OJJ04268.1 azon|OJJ42827.1 aver|OJJ03949.1 afis|EAW24762.1 afum|EAL89404.1 asyd|OJJ52466.1

Group319: afla|EED50195.1 afla|EED57633.1 anid|CBF78576.1 anid|CBF84581.1 anid|CBF88483.1 aory|EIT77388.1 aory|EIT82958.1 asyd|OJJ59798.1 asyd|OJJ63787.1 asyd|OJJ65070.1 aver|OJI97151.1 aver|OJI99195.1 aver|OJJ02491.1 abra|OJJ74472.1 acar|OOF95272.1 acla|EAW07225.1 afis|EAW22239.1 afum|EAL90953.1 agla|OJJ87812.1 akaw|GAA87366.1 aluc|OJZ89647.1 anig|EHA26885.1 ater|EAU34246.1 atub|OJI85157.1 awen|OJJ34817.1 azon|OJJ45732.1

Group320: aluc|OJZ79781.1 aluc|OJZ80029.1 aluc|OJZ80242.1 aluc|OJZ79859.1 aluc|OJZ80430.1 abra|OJJ65440.1 acla|EAW11311.1 afis|EAW16412.1 afla|EED56291.1 afum|EAL87548.1 akaw|GAA92640.1 anid|CBF73419.1 anig|EHA17691.1 asyd|OJJ52220.1 ater|EAU33807.1 aver|OJI98367.1 awen|OJJ29583.1 afis|EAW18185.1 akaw|GAA92960.1 anid|CBF85719.1 aory|EIT73430.1 asyd|OJJ64275.1 awen|OJJ30543.1 akaw|GAA89012.1 akaw|GAA93064.1 akaw|GAA93159.1

Group321: aluc|OJZ81129.1 aluc|OJZ82063.1 awen|OJJ30180.1 awen|OJJ35549.1 awen|OJJ30187.1 abra|OJJ65739.1 acar|OOF99878.1 acla|EAW14576.1 afla|EED53062.1 agla|OJJ86268.1 akaw|GAA91460.1 anid|CBF78777.1 aory|EIT74772.1 asyd|OJJ65241.1 atub|OJI84621.1 aver|OJI99392.1 azon|OJJ44977.1 abra|OJJ72613.1 afla|EED51141.1 anig|EHA24034.1 atub|OJI82018.1 awen|OJJ38779.1 aory|EIT81339.1 aory|EIT74785.1 akaw|GAA85238.1 aluc|OJZ90594.1

Group322: asyd|OJJ56859.1 abra|OJJ67208.1 acar|OOF98353.1 acla|EAW14728.1 afla|EED53515.1 akaw|GAA90969.1 aluc|OJZ87766.1 anid|CBF73997.1 anig|EHA20250.1 asyd|OJJ51948.1 ater|EAU33288.1 atub|OJI80879.1 aver|OJI96457.1 awen|OJJ36200.1 azon|OJJ50142.1 azon|OJJ46441.1 acar|OOF99565.1 afla|EED49927.1 agla|OJJ84266.1 akaw|GAA82243.1 aluc|OJZ91902.1 ater|EAU35687.1 acla|EAW09973.1 acla|EAW10724.1 agla|OJJ84410.1 agla|OJJ79135.1

Group323: abra|OJJ66489.1 acar|OOF90965.1 acar|OOF90966.1 acla|EAW10979.1 acla|EAW10980.1 afis|EAW20879.1 afis|EAW20880.1 afla|EED45565.1 afum|EAL84752.1 afum|EAL84753.1 agla|OJJ83989.1 akaw|GAA92472.1 akaw|GAA92473.1 aluc|OJZ81546.1 anid|CBF82862.1 anid|CBF82864.1 anig|EHA25889.1 aory|EIT75388.1 aory|EIT75432.1 asyd|OJJ61598.1 ater|EAU30811.1 atub|OJI88638.1 aver|OJJ00922.1 awen|OJJ34480.1 azon|OJJ51757.1 afla|EED45564.1

Group324: abra|OJJ67172.1 acar|OOF98391.1 acla|EAW14551.1 afis|EAW19268.1 afla|EED47606.1 afum|EAL93995.1 agla|OJJ88427.1 akaw|GAA91002.1 aluc|OJZ87803.1 anid|CBF78699.1 anig|EHA20284.1 aory|EIT80198.1 asyd|OJJ65220.1 ater|EAU29549.1 atub|OJI80844.1 aver|OJI99369.1 awen|OJJ37629.1 azon|OJJ47017.1 afla|EED47607.1 anid|CBF78697.1 aory|EIT80260.1 acar|OOF98392.1 acla|EAW14552.1 afis|EAW19269.1 afum|EAL93996.1 anig|EHA20285.1

Group325: abra|OJJ67303.1 acar|OOF92510.1 acla|EAW06432.1 afis|EAW18538.1 afla|EED52858.1 afum|EAL93236.1 agla|OJJ84448.1 akaw|GAA86513.1 aluc|OJZ85058.1 anig|EHA25273.1 aory|EIT78604.1 asyd|OJJ52110.1 atub|OJI80542.1 aver|OJJ05256.1 awen|OJJ36329.1 azon|OJJ42393.1 agla|OJJ82374.1 asyd|OJJ52384.1 aver|OJI96494.1 afis|EAW24603.1 afla|EED52462.1 aory|EIT82755.1 atub|OJI80421.1 aver|OJJ01939.1 asyd|OJJ53464.1 aver|OJJ08291.1

Group326: abra|OJJ68235.1 acar|OOF93581.1 acla|EAW10392.1 afis|EAW25199.1 afum|EBA27389.1 agla|OJJ85249.1 anid|CBF85213.1 asyd|OJJ54665.1 ater|EAU34341.1 atub|OJI83115.1 aver|OJI97964.1 awen|OJJ33408.1 abra|OJJ71100.1 afla|EED53646.1 akaw|GAA82868.1 aluc|OJZ87435.1 anig|EHA20657.1 aory|EIT75355.1 atub|OJI81213.1 azon|OJJ47678.1 aver|OJJ02024.1 awen|OJJ31517.1 asyd|OJJ63002.1 agla|OJJ86390.1 awen|OJJ39475.1 agla|OJJ89313.1

Group327: abra|OJJ68548.1 acar|OOF92744.1 acla|EAW12819.1 acla|EAW12820.1 afis|EAW17984.1 afis|EAW17985.1 afla|EED51920.1 afla|EED51921.1 afum|EAL91793.1 afum|EAL91794.1 agla|OJJ80241.1 akaw|GAA91305.1 akaw|GAA91306.1 aluc|OJZ82765.1 anid|CBF79988.1 anid|CBF79990.1 anig|EHA21208.1 aory|EIT81392.1 aory|EIT81447.1 asyd|OJJ58239.1 ater|EAU31494.1 ater|EAU31495.1 atub|OJI88997.1 aver|OJJ02186.1 awen|OJJ41561.1 azon|OJJ48276.1

Group328: abra|OJJ68713.1 acar|OOF92213.1 acla|EAW11027.1 afis|EAW20841.1 afla|EED46816.1 afum|EAL84792.1 agla|OJJ80754.1 akaw|GAA89866.1 aluc|OJZ81815.1 anid|CBF86535.1 anig|EHA22400.1 aory|EIT72698.1 asyd|OJJ62166.1 ater|EAU30544.1 atub|OJI84387.1 aver|OJI96277.1 awen|OJJ40171.1 azon|OJJ50392.1 afis|EAW24116.1 anid|CBF84526.1 acla|EAW12533.1 agla|OJJ88735.1 afum|EAL91971.1 awen|OJJ41372.1 aver|OJI98980.1 asyd|OJJ64867.1

Group329: abra|OJJ70297.1 acar|OOF91751.1 afis|EAW16949.1 afla|EED54756.1 afum|EAL86476.1 agla|OJJ79025.1 akaw|GAA93023.1 aluc|OJZ82562.1 anid|CBF73556.1 anig|EHA27157.1 aory|EIT79421.1 asyd|OJJ56717.1 atub|OJI88804.1 aver|OJJ06008.1 awen|OJJ32083.1 azon|OJJ45848.1 acar|OOF95544.1 agla|OJJ86358.1 akaw|GAA86838.1 aluc|OJZ89957.1 atub|OJI84849.1 agla|OJJ82216.1 ater|EAU31076.1 azon|OJJ46662.1 ater|EAU29923.1 awen|OJJ38719.1

Group330: abra|OJJ70396.1 acar|OOG01144.1 acla|EAW11244.1 afis|EAW19549.1 afla|EED49895.1 afum|EAL89219.1 agla|OJJ88525.1 akaw|GAA91962.1 aluc|OJZ82655.1 anig|EHA24571.1 asyd|OJJ65105.1 ater|EAU33680.1 atub|OJI88886.1 aver|OJI99232.1 awen|OJJ31981.1 azon|OJJ47790.1 afis|EAW18105.1 akaw|GAA92884.1 aory|EIT77372.1 afis|EAW16617.1 afum|EAL86960.1 anig|EHA27418.1 afla|EED50268.1 anid|CBF84502.1 asyd|OJJ53512.1 aver|OJI96187.1

Group331: abra|OJJ71980.1 acar|OOF96631.1 acla|EAW09559.1 acla|EAW09561.1 afis|EAW21576.1 afis|EAW21578.1 afla|EED57795.1 afum|EAL92595.1 afum|EAL92596.1 agla|OJJ85761.1 akaw|GAA84949.1 akaw|GAA84950.1 aluc|OJZ86717.1 anid|CBF76292.1 anid|CBF76294.1 anig|EHA21962.1 aory|EIT81026.1 aory|EIT81117.1 asyd|OJJ61004.1 ater|EAU34955.1 ater|EAU34956.1 atub|OJI83812.1 aver|OJJ00322.1 awen|OJJ39103.1 azon|OJJ51431.1 afla|EED57794.1

Group332: abra|OJJ75736.1 acar|OOF89930.1 acla|EAW14905.1 afis|EAW24426.1 afla|EED55252.1 afum|EAL86094.1 agla|OJJ79240.1 akaw|GAA84469.1 aluc|OJZ89188.1 anid|CBF73780.1 anig|EHA23369.1 aory|EIT83550.1 asyd|OJJ56598.1 ater|EAU30018.1 atub|OJI85602.1 aver|OJJ06127.1 awen|OJJ32778.1 azon|OJJ44006.1 acla|EAW14906.1 afis|EAW24427.1 afla|EED55251.1 afum|EAL86095.1 aluc|OJZ89187.1 anig|EHA23370.1 ater|EAU30019.1 awen|OJJ32779.1

Group333: abra|OJJ75938.1 acar|OOF99972.1 acar|OOF99973.1 acla|EAW09855.1 acla|EAW09856.1 afis|EAW21271.1 afis|EAW21280.1 afla|EED54831.1 afum|EAL92286.1 afum|EAL92296.1 agla|OJJ87477.1 akaw|GAA84639.1 akaw|GAA84640.1 aluc|OJZ89385.1 anid|CBF83352.1 anid|CBF83354.1 anig|EHA23199.1 aory|EIT79381.1 aory|EIT79404.1 asyd|OJJ60616.1 ater|EAU35907.1 ater|EAU35908.1 atub|OJI85413.1 aver|OJI99940.1 awen|OJJ39421.1 azon|OJJ44303.1

Group334: abra|OJJ76077.1 acar|OOF99848.1 acar|OOF99849.1 acla|EAW09743.1 acla|EAW09744.1 afis|EAW21398.1 afis|EAW21399.1 afla|EED55817.1 afla|EED55818.1 afum|EAL92412.1 afum|EAL92413.1 agla|OJJ85524.1 akaw|GAA88889.1 akaw|GAA88890.1 aluc|OJZ89533.1 anid|CBF83833.1 anig|EHA23070.1 aory|EIT76532.1 aory|EIT76875.1 asyd|OJJ60909.1 ater|EAU35159.1 atub|OJI85272.1 aver|OJJ00236.1 awen|OJJ39304.1 azon|OJJ51548.1 ater|EAU35160.1

Group335: abra|OJJ78321.1 acar|OOF98553.1 acla|EAW09204.1 acla|EAW09205.1 afis|EAW21942.1 afis|EAW21943.1 afla|EED46659.1 afla|EED46660.1 afum|EAL92956.1 afum|EAL92957.1 agla|OJJ84181.1 akaw|GAA85707.1 akaw|GAA85708.1 aluc|OJZ91396.1 anid|CBF84053.1 anid|CBF84055.1 anig|EHA20073.1 aory|EIT77914.1 aory|EIT77936.1 asyd|OJJ61344.1 ater|EAU30947.1 ater|EAU30991.1 atub|OJI83388.1 aver|OJJ00669.1 awen|OJJ34234.1 azon|OJJ43402.1

Group336: abra|OJJ65726.1 abra|OJJ73290.1 abra|OJJ77740.1 akaw|GAA83889.1 akaw|GAA92713.1 aluc|OJZ88758.1 aluc|OJZ90806.1 acar|OOF98988.1 acla|EAW12295.1 agla|OJJ85902.1 anig|EHA19533.1 aory|EIT79001.1 asyd|OJJ59393.1 ater|EAU30964.1 atub|OJI82795.1 atub|OJI91329.1 awen|OJJ36235.1 afis|EAW16926.1 afla|EED50932.1 anid|CBF79557.1 anig|EHA18455.1 aver|OJJ02879.1 afum|EAL87839.1 afis|EAW17484.1 agla|OJJ87319.1

Group337: abra|OJJ66243.1 abra|OJJ72838.1 afla|EED45789.1 afla|EED49140.1 aluc|OJZ80548.1 aluc|OJZ83338.1 anig|EHA24214.1 anig|EHA25648.1 aory|EIT75668.1 aory|EIT78573.1 atub|OJI82235.1 atub|OJI88383.1 acar|OOF96040.1 acla|EAW11403.1 afis|EAW24018.1 afum|EAL87448.1 akaw|GAA85435.1 akaw|GAA89048.1 anid|CBF87091.1 asyd|OJJ55451.1 ater|EAU29728.1 awen|OJJ35491.1 azon|OJJ46999.1 aver|OJJ06524.1 agla|OJJ78562.1

Group338: abra|OJJ66379.1 abra|OJJ67896.1 acar|OOF92868.1 acar|OOF94061.1 acar|OOF94131.1 akaw|GAA90248.1 akaw|GAA91900.1 aluc|OJZ81994.1 aluc|OJZ83092.1 anid|CBF82331.1 anid|CBF84191.1 anig|EHA22262.1 anig|EHA28154.1 atub|OJI84544.1 atub|OJI87844.1 afis|EAW20565.1 afla|EED45271.1 afum|EAL84496.1 agla|OJJ81768.1 aory|EIT73641.1 asyd|OJJ57136.1 ater|EAU32199.1 aver|OJJ03167.1 awen|OJJ36239.1 azon|OJJ45336.1

Group339: abra|OJJ66548.1 abra|OJJ77152.1 aluc|OJZ81501.1 aluc|OJZ84566.1 anig|EHA25929.1 anig|EHA27860.1 atub|OJI88680.1 atub|OJI89516.1 awen|OJJ34406.1 awen|OJJ37097.1 acar|OOF89954.1 afis|EAW17738.1 afla|EED50272.1 afum|EAL91553.1 agla|OJJ84037.1 akaw|GAA90778.1 akaw|GAA91173.1 anid|CBF82482.1 aory|EIT77312.1 asyd|OJJ54103.1 ater|EAU31869.1 aver|OJJ08632.1 azon|OJJ48148.1 acar|OOF93038.1 aory|EIT75734.1

Group340: abra|OJJ69049.1 abra|OJJ77527.1 akaw|GAA84093.1 akaw|GAA90580.1 aluc|OJZ81008.1 aluc|OJZ86038.1 anig|EHA19352.1 anig|EHA24573.1 asyd|OJJ51977.1 asyd|OJJ54328.1 atub|OJI79825.1 atub|OJI82576.1 acar|OOF99199.1 acla|EAW06702.1 afis|EAW16247.1 afum|EAL88875.1 agla|OJJ86256.1 anid|CBF82150.1 aory|EIT75005.1 ater|EAU37457.1 aver|OJJ05282.1 awen|OJJ30209.1 azon|OJJ43933.1 afla|EED53308.1 asyd|OJJ57961.1

Group341: abra|OJJ69066.1 abra|OJJ69440.1 acar|OOF91217.1 acar|OOF94300.1 afla|EED46891.1 afla|EED47923.1 akaw|GAA88378.1 akaw|GAA90596.1 aluc|OJZ82276.1 aluc|OJZ86054.1 anig|EHA24557.1 anig|EHA28470.1 aory|EIT72607.1 aory|EIT83373.1 atub|OJI79851.1 atub|OJI88163.1 awen|OJJ32949.1 awen|OJJ38342.1 acla|EAW08762.1 afis|EAW24340.1 afum|EAL86011.1 agla|OJJ83225.1 ater|EAU30407.1 aver|OJI96736.1 azon|OJJ48971.1

Group342: abra|OJJ70211.1 abra|OJJ76495.1 acar|OOF92571.1 acar|OOF99727.1 akaw|GAA82411.1 akaw|GAA86565.1 aluc|OJZ85115.1 aluc|OJZ91709.1 azon|OJJ46702.1 azon|OJJ50101.1 acla|EAW13937.1 afis|EAW18648.1 afla|EED52691.1 afum|EAL93364.1 agla|OJJ82426.1 anid|CBF70533.1 anig|EHA22720.1 anig|EHA25335.1 aory|EIT74236.1 asyd|OJJ58553.1 atub|OJI80479.1 atub|OJI86168.1 aver|OJJ01151.1 awen|OJJ32116.1 ater|EAU38263.1

Group343: abra|OJJ74685.1 abra|OJJ77443.1 acar|OOF95646.1 acar|OOF95697.1 asyd|OJJ52766.1 asyd|OJJ57547.1 asyd|OJJ56548.1 aver|OJJ05631.1 aver|OJJ06174.1 aver|OJJ08749.1 awen|OJJ30371.1 awen|OJJ40449.1 acla|EAW10711.1 afis|EAW24859.1 afla|EED47099.1 afum|EAL89507.1 agla|OJJ82077.1 akaw|GAA81929.1 aluc|OJZ92262.1 anid|CBF75821.1 anig|EHA26697.1 aory|EIT72503.1 ater|EAU36441.1 atub|OJI86696.1 azon|OJJ49305.1

Group344: abra|OJJ75156.1 abra|OJJ77714.1 acar|OOF97183.1 acar|OOF99026.1 akaw|GAA83918.1 akaw|GAA88762.1 aluc|OJZ90779.1 aluc|OJZ92725.1 anig|EHA19512.1 anig|EHA26276.1 ater|EAU35617.1 ater|EAU38821.1 atub|OJI82766.1 atub|OJI87150.1 acla|EAW08331.1 afis|EAW23366.1 afla|EED55592.1 afum|EAL88107.1 agla|OJJ86458.1 anid|CBF74825.1 aory|EIT82696.1 asyd|OJJ60139.1 aver|OJJ04123.1 awen|OJJ39521.1 azon|OJJ47717.1

Group345: acar|OOF90695.1 acar|OOF91261.1 afla|EED44834.1 afla|EED46120.1 agla|OJJ84890.1 agla|OJJ88358.1 anid|CBF79319.1 anid|CBF85127.1 asyd|OJJ55782.1 asyd|OJJ63353.1 aver|OJI97608.1 aver|OJJ06871.1 aver|OJJ07891.1 abra|OJJ76767.1 acla|EAW13321.1 afis|EAW20436.1 afum|EAL84616.1 akaw|GAA87259.1 aluc|OJZ84170.1 anig|EHA27531.1 aory|EIT77716.1 ater|EAU37722.1 atub|OJI89914.1 awen|OJJ33127.1 azon|OJJ46343.1

Group346: acar|OOG00252.1 acar|OOG00270.1 abra|OJJ66461.1 afis|EAW24599.1 agla|OJJ80217.1 aver|OJI96133.1 abra|OJJ67546.1 akaw|GAA88114.1 aluc|OJZ81386.1 anig|EHA18679.1 atub|OJI91591.1 abra|OJJ70572.1 acar|OOF97149.1 afla|EED47007.1 akaw|GAA83416.1 aluc|OJZ83916.1 anid|CBF74799.1 anig|EHA19049.1 aory|EIT72694.1 asyd|OJJ60153.1 ater|EAU36375.1 atub|OJI90286.1 aver|OJJ04111.1 awen|OJJ39478.1 awen|OJJ33480.1

Group347: acla|EAW06619.1 acla|EAW13553.1 awen|OJJ30278.1 awen|OJJ33599.1 azon|OJJ42199.1 azon|OJJ50720.1 abra|OJJ77044.1 acar|OOF90272.1 afis|EAW16146.1 afis|EAW20629.1 afla|EED49763.1 afum|EAL84438.1 afum|EAL88775.1 agla|OJJ80445.1 agla|OJJ86199.1 akaw|GAA87009.1 aluc|OJZ84447.1 anid|CBF84471.1 anig|EHA27764.1 aory|EIT77658.1 asyd|OJJ53537.1 ater|EAU32165.1 atub|OJI89628.1 aver|OJJ08214.1 azon|OJJ43971.1

Group348: acla|EAW07744.1 acla|EAW10138.1 afis|EAW20748.1 afis|EAW22781.1 afla|EED47130.1 afla|EED50040.1 afum|EAL84866.1 afum|EAL90418.1 anid|CBF71564.1 anid|CBF87981.1 ater|EAU33992.1 ater|EAU38933.1 awen|OJJ36890.1 awen|OJJ38956.1 abra|OJJ73849.1 acar|OOG00999.1 agla|OJJ83643.1 akaw|GAA89466.1 aluc|OJZ88200.1 anig|EHA17948.1 aory|EIT79245.1 asyd|OJJ63496.1 atub|OJI90798.1 aver|OJI97433.1 azon|OJJ49728.1

Group349: afis|EAW15424.1 afis|EAW16124.1 afum|EAL84220.1 afum|EAL88754.1 aory|EIT73054.1 aory|EIT80955.1 abra|OJJ69804.1 acar|OOF93572.1 acla|EAW06641.1 afla|EED48664.1 agla|OJJ86166.1 akaw|GAA87855.1 aluc|OJZ85497.1 anid|CBF85532.1 anig|EHA25030.1 asyd|OJJ64135.1 ater|EAU37174.1 atub|OJI80068.1 aver|OJI98218.1 awen|OJJ30306.1 azon|OJJ48681.1 acar|OOF91913.1 afla|EED47503.1 aory|EIT74578.1 ater|EAU33207.1

Group350: afis|EAW20695.1 afis|EAW25202.1 afum|EAL86271.1 afum|EAL89865.1 anid|CBF80363.1 anid|CBF85220.1 asyd|OJJ54819.1 asyd|OJJ60277.1 asyd|OJJ62998.1 ater|EAU34342.1 ater|EAU35428.1 aver|OJI97968.1 aver|OJJ07342.1 abra|OJJ71102.1 acar|OOF98041.1 acla|EAW10389.1 afla|EED53649.1 agla|OJJ88973.1 akaw|GAA82866.1 aluc|OJZ87433.1 anig|EHA20655.1 aory|EIT75301.1 atub|OJI81215.1 awen|OJJ40837.1 azon|OJJ46857.1

Group351: afla|EED45001.1 afla|EED45531.1 akaw|GAA83108.1 akaw|GAA90569.1 anig|EHA18348.1 anig|EHA22621.1 aory|EIT73636.1 aory|EIT75377.1 abra|OJJ68866.1 akaw|GAA90387.1 aluc|OJZ85844.1 anig|EHA24720.1 asyd|OJJ56361.1 ater|EAU36829.1 atub|OJI79632.1 aver|OJJ06353.1 abra|OJJ73381.1 acar|OOF90417.1 agla|OJJ82635.1 aluc|OJZ88657.1 anid|CBF90081.1 atub|OJI91243.1 awen|OJJ40195.1 azon|OJJ49062.1 awen|OJJ33525.1

Group352: afla|EED45010.1 afla|EED45410.1 anig|EHA21017.1 anig|EHA22867.1 aory|EIT73809.1 aory|EIT82470.1 aver|OJI96998.1 aver|OJJ03416.1 azon|OJJ43608.1 azon|OJJ47388.1 abra|OJJ68444.1 acar|OOF93950.1 afis|EAW19806.1 afla|EED50514.1 afum|EAL85087.1 akaw|GAA86929.1 aluc|OJZ90053.1 anid|CBF86126.1 asyd|OJJ63944.1 ater|EAU30743.1 atub|OJI84753.1 aver|OJJ08602.1 awen|OJJ34562.1 anid|CBF82389.1 asyd|OJJ54133.1

Group353: afla|EED45238.1 afla|EED49404.1 aory|EIT73665.1 aory|EIT79465.1 asyd|OJJ56263.1 asyd|OJJ61698.1 asyd|OJJ56732.1 aver|OJI95784.1 aver|OJJ05991.1 aver|OJJ08503.1 awen|OJJ33602.1 awen|OJJ36942.1 abra|OJJ77040.1 acar|OOF90076.1 acla|EAW13556.1 afis|EAW24047.1 agla|OJJ78446.1 akaw|GAA87012.1 aluc|OJZ84442.1 anid|CBF83192.1 anig|EHA27761.1 ater|EAU32167.1 atub|OJI89631.1 azon|OJJ50725.1 agla|OJJ78918.1

Group354: afla|EED45547.1 afla|EED48860.1 afum|EAL84770.1 afum|EAL86315.1 aory|EIT75539.1 aory|EIT81273.1 awen|OJJ29954.1 awen|OJJ34505.1 abra|OJJ67746.1 acar|OOF90945.1 acla|EAW12433.1 afis|EAW20865.1 agla|OJJ83914.1 akaw|GAA92437.1 aluc|OJZ81583.1 anid|CBF80387.1 anig|EHA25856.1 asyd|OJJ61074.1 ater|EAU30808.1 atub|OJI88598.1 aver|OJJ00391.1 azon|OJJ51781.1 acla|EAW10996.1 agla|OJJ79837.1 afis|EAW16740.1

Group355: afla|EED45554.1 afla|EED57298.1 aory|EIT74121.1 aory|EIT75629.1 asyd|OJJ56283.1 asyd|OJJ61592.1 asyd|OJJ61824.1 ater|EAU30081.1 ater|EAU30816.1 aver|OJI95924.1 aver|OJJ00914.1 aver|OJJ08478.1 abra|OJJ67752.1 acar|OOF90956.1 acla|EAW10988.1 afis|EAW20872.1 afum|EAL84762.1 agla|OJJ83979.1 akaw|GAA92442.1 aluc|OJZ81577.1 anid|CBF82847.1 anig|EHA25861.1 atub|OJI88603.1 awen|OJJ34489.1 azon|OJJ51766.1

Group356: afla|EED55154.1 afla|EED56856.1 aory|EIT77459.1 aory|EIT81607.1 asyd|OJJ52781.1 asyd|OJJ53407.1 asyd|OJJ55050.1 aver|OJI99562.1 aver|OJJ04194.1 aver|OJJ08759.1 abra|OJJ76674.1 acar|OOF99335.1 acla|EAW09104.1 afis|EAW25708.1 afum|EAL84381.1 agla|OJJ85490.1 akaw|GAA92518.1 aluc|OJZ91531.1 anid|CBF83922.1 anig|EHA22570.1 ater|EAU32722.1 atub|OJI85985.1 anig|EHA19936.1 asyd|OJJ55429.1 aver|OJJ06508.1

Group357: afum|EAL90938.1 afum|EAL91118.1 abra|OJJ70518.1 acar|OOF97412.1 acla|EAW08157.1 afis|EAW23195.1 afla|EED57873.1 afum|EAL88288.2 agla|OJJ86332.1 akaw|GAA83365.1 aluc|OJZ83863.1 anid|CBF74706.1 anig|EHA19003.1 aory|EIT83278.1 asyd|OJJ60219.1 ater|EAU36323.1 atub|OJI90231.1 aver|OJJ04049.1 awen|OJJ39425.1 azon|OJJ47626.1 afis|EAW20299.1 agla|OJJ84725.1 asyd|OJJ62722.1 aver|OJJ01792.1 azon|OJJ49994.1

Group358: agla|OJJ79814.1 agla|OJJ81857.1 agla|OJJ83394.1 asyd|OJJ52362.1 asyd|OJJ57936.1 asyd|OJJ60297.1 aver|OJI99638.1 aver|OJJ01984.1 aver|OJJ04458.1 awen|OJJ30856.1 awen|OJJ34435.1 awen|OJJ40033.1 acla|EAW09090.1 afis|EAW25682.1 afla|EED48118.1 afum|EAL84352.1 anid|CBF71525.1 aory|EIT76242.1 acla|EAW09112.1 afis|EAW25714.1 afum|EAL84386.2 anid|CBF74681.1 anid|CBF87142.1 ater|EAU32480.1 azon|OJJ46476.1

Group359: agla|OJJ83698.1 agla|OJJ89439.1 akaw|GAA85587.1 akaw|GAA85825.1 aluc|OJZ83524.1 aluc|OJZ91276.1 anig|EHA19962.1 anig|EHA24347.1 asyd|OJJ56315.1 asyd|OJJ58149.1 atub|OJI82413.1 atub|OJI83265.1 abra|OJJ73009.1 acar|OOF96181.1 acla|EAW08004.1 afis|EAW23039.1 afla|EED45399.1 afum|EAL88442.1 anid|CBF80933.1 aory|EIT82471.1 ater|EAU29619.1 aver|OJJ06399.1 awen|OJJ35703.1 azon|OJJ44165.1 aver|OJJ04255.1

Group360: aluc|OJZ83601.1 aluc|OJZ88595.1 anig|EHA18289.1 anig|EHA18760.1 asyd|OJJ56054.1 asyd|OJJ61370.1 atub|OJI89973.1 atub|OJI91180.1 aver|OJJ00697.1 aver|OJJ04995.1 awen|OJJ35022.1 awen|OJJ37732.1 abra|OJJ73445.1 acar|OOG00650.1 acla|EAW06781.1 afis|EAW16331.1 afla|EED53547.1 afum|EAL88956.1 agla|OJJ88261.1 akaw|GAA83049.1 anid|CBF84027.1 aory|EIT75029.1 ater|EAU34127.1 azon|OJJ45215.1 akaw|GAA92121.1

Group361: anid|CBF79016.1 anid|CBF89281.1 anid|CBF89401.1 anid|CBF82943.1 abra|OJJ69545.1 acla|EAW15297.1 afis|EAW17240.1 afla|EED45268.1 agla|OJJ88965.1 akaw|GAA91212.1 aluc|OJZ82859.1 aory|EIT72628.1 asyd|OJJ65384.1 ater|EAU37653.1 aver|OJI95878.1 awen|OJJ31342.1 atub|OJI81111.1 anig|EHA23888.1 aory|EIT73575.1 asyd|OJJ53722.1 anig|EHA19683.1 awen|OJJ36938.1 anid|CBF75970.1 ater|EAU39534.1 ater|EAU39535.1

Group362: anid|CBF80391.1 anid|CBF83286.1 asyd|OJJ54788.1 asyd|OJJ57255.1 aver|OJJ03039.1 aver|OJJ07376.1 abra|OJJ68496.1 acar|OOF98751.1 acla|EAW15364.1 afis|EAW17224.1 afla|EED51521.1 afum|EAL86646.1 agla|OJJ86656.1 akaw|GAA92774.1 aluc|OJZ82709.1 anid|CBF80759.1 anig|EHA21164.1 aory|EIT80558.1 ater|EAU37673.1 atub|OJI88942.1 azon|OJJ42623.1 azon|OJJ42665.1 asyd|OJJ57905.1 aver|OJJ04489.1 azon|OJJ45203.1

Group363: asyd|OJJ54205.1 asyd|OJJ55560.1 ater|EAU32170.1 ater|EAU32943.1 abra|OJJ75335.1 acar|OOF94500.1 acla|EAW13560.1 afis|EAW20637.1 afla|EED49750.1 afum|EAL84431.1 akaw|GAA92029.1 aluc|OJZ92899.1 anid|CBF83194.1 anig|EHA26064.1 aory|EIT77711.1 atub|OJI87357.1 awen|OJJ33608.1 abra|OJJ77031.1 acar|OOF91467.1 agla|OJJ78916.1 akaw|GAA87020.1 aluc|OJZ84434.1 anig|EHA27753.1 atub|OJI89639.1 aver|OJJ06641.1

Group364: atub|OJI82939.1 atub|OJI88437.1 abra|OJJ67568.1 afla|EED50521.1 akaw|GAA89098.1 anig|EHA25700.1 ater|EAU33843.1 abra|OJJ72071.1 acla|EAW14867.1 afis|EAW19913.1 afum|EAL84993.2 agla|OJJ81787.1 aory|EIT77284.1 ater|EAU32297.1 abra|OJJ77873.1 acar|OOF98898.1 aluc|OJZ90946.1 anig|EHA19655.1 aory|EIT82039.1 acar|OOG00619.1 afla|EED50223.1 anig|EHA19654.1 afla|EED50520.1 awen|OJJ34149.1 ater|EAU39457.1

Group365: awen|OJJ31642.1 awen|OJJ39457.1 abra|OJJ70593.1 acar|OOF97125.1 acar|OOF97948.1 acla|EAW08187.1 afis|EAW23221.1 afla|EED47029.1 afum|EAL88263.1 agla|OJJ86378.1 akaw|GAA83438.1 aluc|OJZ83939.1 anid|CBF76684.1 anig|EHA19069.1 ater|EAU36352.1 atub|OJI90312.1 abra|OJJ71237.1 afla|EED56272.1 akaw|GAA82751.1 aluc|OJZ87308.1 anig|EHA20544.1 aory|EIT73409.1 asyd|OJJ56499.1 atub|OJI81337.1 aver|OJJ06220.1

Group366: abra|OJJ66082.1 acar|OOF99287.1 acla|EAW06431.1 afis|EAW24735.1 afla|EED52785.1 afum|EAL89382.1 akaw|GAA84164.1 aluc|OJZ81089.1 anid|CBF79598.1 anig|EHA19281.1 aory|EIT78193.1 asyd|OJJ57988.1 ater|EAU31606.1 atub|OJI82495.1 aver|OJJ04414.1 awen|OJJ40223.1 afis|EAW18537.1 afla|EED56576.1 afum|EAL93235.1 anid|CBF82125.1 aory|EIT79676.1 awen|OJJ36227.1 ater|EAU32072.1 asyd|OJJ54295.1 aver|OJJ05250.1

Group367: abra|OJJ67359.1 acar|OOF93335.1 acla|EAW10444.1 afla|EED55318.1 agla|OJJ89047.1 akaw|GAA86462.1 anid|CBF87506.1 anig|EHA25221.1 aory|EIT76259.1 asyd|OJJ58362.1 ater|EAU39395.1 aver|OJJ02044.1 awen|OJJ40742.1 abra|OJJ70853.1 acar|OOF98263.1 afis|EAW25123.1 afum|EAL89794.1 akaw|GAA91720.1 aluc|OJZ87668.1 anid|CBF85688.1 anig|EHA20859.1 asyd|OJJ64259.1 atub|OJI80975.1 aver|OJI98351.1 azon|OJJ44113.1

Group368: abra|OJJ67850.1 acar|OOF92831.1 acla|EAW11749.1 afis|EAW23782.1 afum|EAL87216.1 agla|OJJ78934.1 akaw|GAA91858.1 aluc|OJZ82041.1 anig|EHA22215.1 atub|OJI84600.1 awen|OJJ33726.1 abra|OJJ75896.1 acar|OOG00008.1 afla|EED49453.1 agla|OJJ80043.1 akaw|GAA84602.1 aluc|OJZ89346.1 anig|EHA23235.1 aory|EIT79472.1 asyd|OJJ57259.1 ater|EAU39366.1 atub|OJI85452.1 aver|OJJ03037.1 awen|OJJ36056.1 azon|OJJ50429.1

Group369: abra|OJJ68472.1 acar|OOF94211.1 acla|EAW13768.1 afis|EAW19790.1 afla|EED45871.1 afum|EAL94073.1 akaw|GAA92393.1 aluc|OJZ83253.1 anid|CBF79916.1 anig|EHA21107.1 asyd|OJJ58193.1 ater|EAU36161.1 atub|OJI87684.1 aver|OJJ04212.1 awen|OJJ31450.1 acla|EAW14615.1 afis|EAW19353.1 afla|EED46572.1 anid|CBF76648.1 asyd|OJJ61206.1 aver|OJJ00530.1 acar|OOF94212.1 ater|EAU38489.1 aory|EIT77883.1 afum|EAL85101.1

Group370: abra|OJJ68801.1 acar|OOF91599.1 acla|EAW13559.1 afis|EAW20636.1 afum|EAL84432.1 akaw|GAA90452.1 aluc|OJZ85772.1 anig|EHA24781.1 aory|EIT77665.1 ater|EAU37652.1 atub|OJI79557.1 awen|OJJ33440.1 azon|OJJ50788.1 abra|OJJ70265.1 afla|EED54696.1 akaw|GAA86532.1 aluc|OJZ85080.1 anid|CBF85668.1 anig|EHA25295.1 aory|EIT78807.1 asyd|OJJ64247.1 aver|OJJ04749.1 afla|EED50779.1 aver|OJI98338.1 aory|EIT77987.1

Group371: abra|OJJ68927.1 acar|OOF90675.1 agla|OJJ86552.1 akaw|GAA90318.1 aluc|OJZ85915.1 anig|EHA24679.1 atub|OJI79705.1 aver|OJJ00751.1 abra|OJJ69600.1 acar|OOF93267.1 afis|EAW19224.1 afla|EED48204.1 anid|CBF83972.1 asyd|OJJ61415.1 awen|OJJ38318.1 abra|OJJ77028.1 afum|EAL93955.1 akaw|GAA87023.1 aluc|OJZ84430.1 anig|EHA27749.1 aory|EIT72916.1 atub|OJI89644.1 aver|OJJ08327.1 aver|OJJ02995.1 awen|OJJ29960.1

Group372: abra|OJJ70648.1 acar|OOF97073.1 acla|EAW08111.1 acla|EAW08112.1 afis|EAW23139.1 afis|EAW23140.1 afla|EED48502.1 afum|EAL88341.1 afum|EAL88342.1 agla|OJJ89661.1 akaw|GAA89984.1 aluc|OJZ83998.1 anid|CBF74444.1 anid|CBF74446.1 anig|EHA19121.1 aory|EIT73212.1 aory|EIT73270.1 asyd|OJJ57742.1 ater|EAU36277.1 atub|OJI90367.1 aver|OJJ05425.1 awen|OJJ36636.1 azon|OJJ47580.1 afla|EED48503.1 ater|EAU36278.1

Group373: abra|OJJ70697.1 acar|OOF90045.1 acla|EAW08067.1 afis|EAW23103.1 afla|EED48544.1 afum|EAL88376.1 agla|OJJ89639.1 akaw|GAA90028.1 akaw|GAA90029.1 aluc|OJZ84044.1 anid|CBF81363.1 anig|EHA19166.1 aory|EIT73061.1 asyd|OJJ59383.1 ater|EAU36232.1 atub|OJI90416.1 aver|OJJ02890.1 awen|OJJ36677.1 azon|OJJ47521.1 afla|EED48545.1 acla|EAW08068.1 afis|EAW23104.1 afum|EAL88375.1 anid|CBF81365.1 aory|EIT73082.1

Group374: abra|OJJ71208.1 acar|OOF97959.1 afla|EED54682.1 akaw|GAA82776.1 aluc|OJZ87335.1 anid|CBF80112.1 anig|EHA20571.1 anig|EHA20572.1 aory|EIT78810.1 asyd|OJJ65149.1 ater|EAU36930.1 atub|OJI81309.1 aver|OJJ02086.1 azon|OJJ44458.1 acar|OOF96688.1 afla|EED54681.1 asyd|OJJ65153.1 ater|EAU36931.1 acar|OOF97963.1 aory|EIT78922.1 afla|EED46387.1 afla|EED46388.1 aory|EIT72360.1 asyd|OJJ58325.1 atub|OJI84521.1

Group375: abra|OJJ72278.1 acar|OOF94942.1 afis|EAW20802.1 afla|EED56693.1 afum|EBA27208.1 agla|OJJ85014.1 akaw|GAA86030.1 aluc|OJZ90257.1 anid|CBF71378.1 anig|EHA23752.1 aory|EIT73887.1 asyd|OJJ55075.1 ater|EAU32872.1 atub|OJI81683.1 aver|OJJ07133.1 awen|OJJ41914.1 azon|OJJ47367.1 afis|EAW22078.1 afum|EAL91092.1 anid|CBF82802.1 asyd|OJJ57118.1 ater|EAU30146.1 afla|EED49551.1 aory|EIT82585.1 aory|EIT82590.1

Group376: abra|OJJ72657.1 acar|OOF94617.1 acla|EAW11521.1 afis|EAW23933.1 afla|EED50585.1 afum|EAL87357.1 agla|OJJ79413.1 akaw|GAA85279.1 akaw|GAA85280.1 aluc|OJZ90638.1 anid|CBF77273.1 anig|EHA24073.1 aory|EIT82169.1 asyd|OJJ56797.1 ater|EAU32304.1 atub|OJI82066.1 aver|OJJ03537.1 awen|OJJ39022.1 azon|OJJ51347.1 acla|EAW11520.1 afum|EAL87358.1 aory|EIT82139.1 afis|EAW23932.1 afla|EED50586.1 azon|OJJ51348.1

Group377: abra|OJJ72828.1 acar|OOF93602.1 acla|EAW14973.1 afis|EAW24498.1 afla|EED56620.1 afum|EAL86168.1 agla|OJJ79958.1 anid|CBF85575.1 aory|EIT79659.1 asyd|OJJ64180.1 ater|EAU31051.1 aver|OJJ07247.1 awen|OJJ34897.1 abra|OJJ75484.1 akaw|GAA84250.1 aluc|OJZ88942.1 anid|CBF80279.1 anig|EHA23578.1 atub|OJI85847.1 ater|EAU34059.1 anid|CBF71459.1 asyd|OJJ54948.1 afla|EED52220.1 aory|EIT74386.1 aver|OJI98257.1

Group378: abra|OJJ74406.1 acar|OOF95334.1 acla|EAW06822.1 afis|EAW16185.1 afla|EED53465.1 afum|EAL88816.2 agla|OJJ78591.1 akaw|GAA87300.1 aluc|OJZ89718.1 anid|CBF89935.1 anig|EHA26946.1 aory|EIT74868.1 asyd|OJJ62121.1 ater|EAU34079.1 atub|OJI85082.1 aver|OJI96229.1 awen|OJJ37907.1 azon|OJJ45085.1 afis|EAW16370.1 acla|EAW11051.1 afla|EED49634.1 aory|EIT80960.1 azon|OJJ45955.1 ater|EAU36747.1 azon|OJJ45949.1

Group379: abra|OJJ76313.1 acar|OOF99561.1 acar|OOF99562.1 acla|EAW14106.1 afis|EAW18806.1 afis|EAW18807.1 afla|EED50770.1 afum|EAL93542.1 afum|EAL93543.1 agla|OJJ80536.1 akaw|GAA82241.1 aluc|OJZ91904.1 anid|CBF69790.1 anig|EHA22880.1 aory|EIT78083.1 asyd|OJJ58961.1 ater|EAU37958.1 atub|OJI86354.1 aver|OJJ01555.1 awen|OJJ32382.1 azon|OJJ47143.1 agla|OJJ80538.1 acla|EAW14105.1 anid|CBF69788.1 awen|OJJ32381.1

Group380: abra|OJJ76772.1 acar|OOF91266.1 acla|EAW13327.1 afis|EAW20442.1 afla|EED44828.1 afum|EAL84610.2 agla|OJJ84883.1 akaw|GAA87253.1 akaw|GAA87254.1 aluc|OJZ84176.1 anid|CBF79308.1 anig|EHA27535.1 aory|EIT77723.1 asyd|OJJ55777.1 ater|EAU37717.1 atub|OJI89908.1 atub|OJI89909.1 aver|OJJ06866.1 awen|OJJ33132.1 azon|OJJ46347.1 acla|EAW13326.1 afis|EAW20441.1 afla|EED44829.1 afum|EAL84611.1 anid|CBF79310.1

Group381: abra|OJJ77345.1 acar|OOF93215.1 acla|EAW13129.1 acla|EAW13130.1 afis|EAW17533.1 afis|EAW17534.1 afla|EED49295.1 afla|EED49296.1 afum|EAL91347.1 afum|EAL91348.1 agla|OJJ81437.1 akaw|GAA91415.1 akaw|GAA91416.1 aluc|OJZ84756.1 anid|CBF90328.1 anid|CBF90329.1 anig|EHA28025.1 aory|EIT82179.1 asyd|OJJ61906.1 ater|EAU36962.1 ater|EAU36963.1 atub|OJI89328.1 aver|OJI95999.1 awen|OJJ37319.1 azon|OJJ51011.1

Group382: abra|OJJ66008.1 abra|OJJ66036.1 abra|OJJ72701.1 agla|OJJ79097.1 agla|OJJ82950.1 asyd|OJJ54673.1 asyd|OJJ57511.1 aver|OJJ05668.1 aver|OJJ07496.1 acar|OOF92090.1 akaw|GAA90880.1 aluc|OJZ82953.1 anig|EHA21368.1 aory|EIT75821.1 atub|OJI89185.1 awen|OJJ30435.1 atub|OJI87431.1 abra|OJJ67817.1 afum|EAL86776.1 anid|CBF79156.1 ater|EAU38014.1 afla|EED45927.1 afla|EED45926.1 anid|CBF75733.1

Group383: abra|OJJ66089.1 abra|OJJ76716.1 acar|OOF95538.1 afis|EAW21215.1 afla|EED53625.1 afum|EAL92226.1 aluc|OJZ91496.1 anig|EHA22536.1 aory|EIT75031.1 ater|EAU38549.1 atub|OJI85949.1 aver|OJI98307.1 aver|OJI99282.1 awen|OJJ30167.1 abra|OJJ66099.1 acla|EAW14986.1 afis|EAW24508.1 afum|EBA27258.1 agla|OJJ86285.1 anid|CBF78328.1 asyd|OJJ56085.1 aver|OJJ05028.1 akaw|GAA92484.1 awen|OJJ36979.1

Group384: abra|OJJ66370.1 abra|OJJ77693.1 acar|OOF98983.1 acar|OOF99046.1 akaw|GAA83937.1 akaw|GAA90239.1 atub|OJI82743.1 atub|OJI87834.1 acla|EAW06572.1 afis|EAW16024.1 afla|EED56405.1 afum|EAL88653.1 agla|OJJ81572.1 aluc|OJZ83101.1 anid|CBF75012.1 anig|EHA19493.1 anig|EHA28166.1 aory|EIT83000.1 asyd|OJJ60056.1 ater|EAU33617.1 aver|OJJ02233.1 awen|OJJ30433.1 azon|OJJ45432.1 aluc|OJZ80819.1

Group385: abra|OJJ67097.1 abra|OJJ77658.1 aluc|OJZ80870.1 aluc|OJZ87876.1 anig|EHA19465.1 anig|EHA20181.1 atub|OJI80773.1 atub|OJI82707.1 acar|OOF98457.1 acar|OOF99078.1 acla|EAW06500.1 afis|EAW15996.1 afla|EED56370.1 afum|EAL88628.2 agla|OJJ86011.1 akaw|GAA83977.1 akaw|GAA91064.1 anid|CBF81691.1 aory|EIT73327.1 asyd|OJJ59562.1 ater|EAU33651.1 aver|OJJ02706.1 awen|OJJ30473.1 azon|OJJ45465.1

Group386: abra|OJJ67844.1 abra|OJJ76158.1 acar|OOF92021.1 acar|OOF99774.1 akaw|GAA82106.1 akaw|GAA91850.1 aluc|OJZ82048.1 aluc|OJZ92070.1 anig|EHA22210.1 anig|EHA22999.1 atub|OJI84606.1 atub|OJI86502.1 awen|OJJ37808.1 awen|OJJ39219.1 acla|EAW09660.1 afis|EAW21479.1 afla|EED55911.1 afum|EAL92499.1 agla|OJJ85632.1 anid|CBF76506.1 aory|EIT76810.1 asyd|OJJ61132.1 aver|OJJ00453.1 azon|OJJ51636.1

Group387: abra|OJJ68864.1 abra|OJJ75043.1 acar|OOF90525.1 acar|OOF97290.1 akaw|GAA88660.1 akaw|GAA90389.1 aluc|OJZ85842.1 aluc|OJZ92609.1 atub|OJI79630.1 atub|OJI87043.1 awen|OJJ31662.1 awen|OJJ39696.1 acla|EAW08232.1 afis|EAW23266.1 afla|EED55747.1 afum|EAL88213.1 agla|OJJ86586.1 anid|CBF84130.1 anig|EHA24722.1 aory|EIT83228.1 asyd|OJJ57114.1 ater|EAU37532.1 aver|OJJ03186.1 azon|OJJ47828.1

Group388: abra|OJJ70275.1 abra|OJJ75558.1 acar|OOF90876.1 acar|OOF98673.1 agla|OJJ82823.1 agla|OJJ88521.1 akaw|GAA84313.1 akaw|GAA92874.1 aluc|OJZ82537.1 aluc|OJZ89008.1 anig|EHA22505.1 anig|EHA23520.1 asyd|OJJ55019.1 asyd|OJJ59700.1 atub|OJI85781.1 atub|OJI88781.1 aver|OJJ02587.1 aver|OJJ07186.1 awen|OJJ32269.1 awen|OJJ32616.1 anid|CBF78793.1 afla|EED45595.1 aory|EIT75779.1 aory|EIT75483.1

Group389: abra|OJJ70579.1 abra|OJJ75835.1 akaw|GAA83423.1 akaw|GAA84545.1 aluc|OJZ83923.1 aluc|OJZ89278.1 anig|EHA19056.1 anig|EHA23291.1 atub|OJI85521.1 atub|OJI90295.1 awen|OJJ32048.1 awen|OJJ39470.1 acar|OOF97142.1 acla|EAW08201.1 afis|EAW23235.1 afla|EED47014.1 afum|EAL88247.1 agla|OJJ86395.1 anid|CBF75150.1 aory|EIT72630.1 asyd|OJJ59974.1 ater|EAU36368.1 aver|OJJ02317.1 azon|OJJ47672.1

Group390: abra|OJJ70867.1 abra|OJJ72108.1 acar|OOF90121.1 acar|OOF96751.1 aluc|OJZ86863.1 aluc|OJZ87652.1 anig|EHA20846.1 anig|EHA21778.1 aory|EIT75800.1 aory|EIT76249.1 atub|OJI80991.1 atub|OJI83657.1 acla|EAW10459.1 afis|EAW25108.1 afla|EED55334.1 afum|EAL89779.1 anid|CBF86068.1 asyd|OJJ64458.1 ater|EAU39378.1 aver|OJI98561.1 awen|OJJ37055.1 azon|OJJ44547.1 akaw|GAA84814.1 afla|EED45959.1

Group391: acar|OOF90694.1 acar|OOF95969.1 afla|EED46110.1 afla|EED52607.1 agla|OJJ82549.1 agla|OJJ84416.1 awen|OJJ31372.1 awen|OJJ33859.1 abra|OJJ72741.1 acla|EAW11574.1 afis|EAW23880.1 afum|EAL87308.1 akaw|GAA85352.1 aluc|OJZ90714.1 anid|CBF77193.1 anig|EHA24133.1 aory|EIT74176.1 asyd|OJJ56836.1 ater|EAU33467.1 atub|OJI82140.1 aver|OJJ03501.1 azon|OJJ48693.1 asyd|OJJ58816.1 aory|EIT76373.1

Group392: acar|OOF92653.1 acar|OOG00348.1 abra|OJJ69030.1 afla|EED56357.1 akaw|GAA90546.1 aluc|OJZ86016.1 anig|EHA24595.1 aory|EIT73420.1 ater|EAU31400.1 atub|OJI79805.1 awen|OJJ34779.1 abra|OJJ73125.1 acla|EAW07183.1 afis|EAW22196.1 afum|EAL90995.1 agla|OJJ87849.1 akaw|GAA88040.1 aluc|OJZ81299.1 anid|CBF88408.1 anig|EHA18608.1 asyd|OJJ63753.1 atub|OJI91508.1 aver|OJI97188.1 azon|OJJ45760.1

Group393: acar|OOF94038.1 acar|OOF94108.1 abra|OJJ77702.1 acar|OOF99038.1 acla|EAW12405.1 afis|EAW16864.1 afla|EED51093.1 afum|EAL86403.2 agla|OJJ78684.1 akaw|GAA83928.1 aluc|OJZ90767.1 anig|EHA19502.1 aory|EIT81997.1 asyd|OJJ62647.1 ater|EAU37649.1 atub|OJI82755.1 awen|OJJ36987.1 azon|OJJ45261.1 acla|EAW15311.1 akaw|GAA90280.1 aluc|OJZ83060.1 anig|EHA28126.1 atub|OJI87886.1 awen|OJJ38066.1

Group394: acar|OOF95062.1 acar|OOF95129.1 afla|EED47135.1 afla|EED49651.1 anid|CBF74606.1 anid|CBF84541.1 asyd|OJJ53476.1 asyd|OJJ57659.1 aver|OJJ05520.1 aver|OJJ08280.1 abra|OJJ74650.1 acla|EAW10735.1 afis|EAW24839.1 afum|EAL89485.1 agla|OJJ82030.1 akaw|GAA81960.1 aluc|OJZ92229.1 anig|EHA26728.1 aory|EIT79283.1 ater|EAU36464.1 atub|OJI86659.1 awen|OJJ40425.1 azon|OJJ49283.1 ater|EAU29279.1

Group395: afis|EAW18417.1 afis|EAW22052.1 afum|EAL86752.2 afum|EAL93101.1 aory|EIT78709.1 aory|EIT83477.1 asyd|OJJ51851.1 asyd|OJJ58449.1 awen|OJJ30750.1 awen|OJJ31731.1 abra|OJJ65787.1 acar|OOF92376.1 acla|EAW06308.1 agla|OJJ84984.1 akaw|GAA86375.1 anid|CBF70714.1 anig|EHA25131.1 ater|EAU29833.1 atub|OJI80690.1 aver|OJJ01050.1 azon|OJJ50491.1 abra|OJJ67456.1 aluc|OJZ84908.1 akaw|GAA92636.1

Group396: afis|EAW19022.1 afis|EAW24624.1 afis|EAW25668.1 afum|EAL84338.1 afum|EAL93753.1 ater|EAU32064.1 ater|EAU33941.1 ater|EAU38525.1 abra|OJJ68917.1 acar|OOF91505.1 acla|EAW14258.1 afla|EED56624.1 agla|OJJ82736.1 akaw|GAA90332.1 aluc|OJZ85900.1 anid|CBF79090.1 anid|CBF85580.1 anig|EHA24692.1 aory|EIT79597.1 asyd|OJJ64176.1 atub|OJI79685.1 aver|OJI98251.1 awen|OJJ31400.1 aver|OJJ08543.1

Group397: afla|EED46864.1 afla|EED51110.1 aory|EIT72646.1 aory|EIT81348.1 ater|EAU32491.1 ater|EAU37681.1 abra|OJJ65858.1 acar|OOF95019.1 acla|EAW14585.1 agla|OJJ81814.1 akaw|GAA92287.1 aluc|OJZ90184.1 anig|EHA23683.1 atub|OJI81611.1 awen|OJJ38347.1 abra|OJJ76688.1 acar|OOF99324.1 akaw|GAA92506.1 aluc|OJZ91519.1 anid|CBF73586.1 anig|EHA22557.1 asyd|OJJ56216.1 atub|OJI85974.1 aver|OJJ05148.1

Group398: afla|EED49844.1 afla|EED50421.1 anid|CBF71215.1 anid|CBF78131.1 aory|EIT82058.1 aory|EIT83396.1 asyd|OJJ55867.1 asyd|OJJ57404.1 azon|OJJ43885.1 azon|OJJ46240.1 abra|OJJ70022.1 acar|OOF93528.1 acla|EAW08798.1 afis|EAW15601.1 afum|EAL85732.1 akaw|GAA87639.1 aluc|OJZ85254.1 anig|EHA25455.1 ater|EAU32528.1 atub|OJI80314.1 aver|OJJ02118.1 aver|OJJ04809.1 awen|OJJ38276.1 asyd|OJJ57345.1

Group399: agla|OJJ82200.1 agla|OJJ84366.1 abra|OJJ67531.1 afis|EAW24553.1 afla|EED51537.1 afum|EAL86218.1 aluc|OJZ87028.1 anid|CBF78469.1 anig|EHA21629.1 asyd|OJJ57285.1 ater|EAU37648.1 atub|OJI83486.1 aver|OJJ05871.1 awen|OJJ38297.1 abra|OJJ68399.1 acar|OOF93974.1 akaw|GAA92548.1 aluc|OJZ90097.1 anig|EHA20979.1 aory|EIT74555.1 atub|OJI84709.1 awen|OJJ37870.1 awen|OJJ40901.1 ater|EAU36047.1

Group400: agla|OJJ84117.1 agla|OJJ84488.1 ater|EAU31647.1 ater|EAU32102.1 ater|EAU32790.1 awen|OJJ34277.1 awen|OJJ34655.1 awen|OJJ38014.1 awen|OJJ38039.1 abra|OJJ74152.1 acar|OOF95573.1 afla|EED54552.1 akaw|GAA89731.1 aluc|OJZ80652.1 anig|EHA27180.1 aory|EIT78872.1 asyd|OJJ61410.1 atub|OJI87507.1 aver|OJJ00740.1 akaw|GAA89130.1 anig|EHA25730.1 atub|OJI88476.1 anid|CBF73465.1 aluc|OJZ81708.1

Group401: akaw|GAA85083.1 akaw|GAA91235.1 aluc|OJZ82835.1 aluc|OJZ86559.1 atub|OJI83963.1 atub|OJI89065.1 abra|OJJ67962.1 acar|OOF92808.1 acla|EAW11652.1 afis|EAW23801.1 afla|EED51619.1 afum|EAL87235.2 agla|OJJ78848.1 anig|EHA21273.1 aory|EIT80604.1 asyd|OJJ61019.1 ater|EAU31355.1 awen|OJJ33760.1 azon|OJJ42864.1 abra|OJJ71827.1 acar|OOF96504.1 anid|CBF85398.1 anig|EHA22090.1 ater|EAU39385.1

Group402: akaw|GAA91381.1 akaw|GAA92466.1 aluc|OJZ81553.1 aluc|OJZ84788.1 anig|EHA25881.1 anig|EHA28056.1 atub|OJI88632.1 atub|OJI89294.1 abra|OJJ66481.1 acar|OOF93248.1 afla|EED46099.1 anid|CBF82158.1 aory|EIT76444.1 asyd|OJJ54333.1 ater|EAU37661.1 aver|OJJ05287.1 awen|OJJ41952.1 abra|OJJ77379.1 afla|EED46096.1 anid|CBF82155.1 afla|EED46097.1 akaw|GAA92464.1 anid|CBF82156.1 aory|EIT76468.1

Group403: aory|EIT74183.1 aory|EIT82897.1 ater|EAU31229.1 ater|EAU37935.1 aver|OJJ03325.1 aver|OJJ08019.1 abra|OJJ69359.1 acar|OOF94365.1 acla|EAW14750.1 afis|EAW24281.1 afla|EED47995.1 afum|EAL85950.1 agla|OJJ80635.1 akaw|GAA88229.1 aluc|OJZ82191.1 anid|CBF74148.1 anig|EHA28544.1 asyd|OJJ53921.1 asyd|OJJ56960.1 atub|OJI88244.1 awen|OJJ33016.1 azon|OJJ48501.1 acar|OOF90185.1 afla|EED52650.1

Group404: awen|OJJ30773.1 awen|OJJ31708.1 abra|OJJ68030.1 acar|OOF92075.1 afla|EED45369.1 agla|OJJ84153.1 akaw|GAA90923.1 aluc|OJZ82903.1 aory|EIT82429.1 ater|EAU36888.1 atub|OJI89129.1 abra|OJJ70336.1 acla|EAW11117.1 afla|EED51703.1 aluc|OJZ82606.1 anig|EHA27474.1 aory|EIT81577.1 asyd|OJJ64590.1 ater|EAU34064.1 atub|OJI88841.1 aver|OJI98688.1 azon|OJJ45907.1 aver|OJI98018.1 afla|EED55092.1

Group405: abra|OJJ66091.1 acla|EAW07082.1 afla|EED53627.1 aory|EIT75030.1 asyd|OJJ64735.1 ater|EAU38556.1 aver|OJJ06182.1 awen|OJJ30165.1 abra|OJJ77781.1 afis|EAW19895.1 afum|EAL92224.2 agla|OJJ86287.1 akaw|GAA83866.1 aluc|OJZ90834.1 anid|CBF78324.1 anig|EHA19557.1 atub|OJI82820.1 aver|OJI98311.1 awen|OJJ36980.1 afis|EAW21213.1 asyd|OJJ56540.1 asyd|OJJ56087.1 aver|OJJ05030.1 awen|OJJ33354.1

Group406: abra|OJJ66593.1 acar|OOF91028.1 acla|EAW10898.1 afis|EAW20960.1 afum|EAL84673.1 agla|OJJ83918.1 akaw|GAA91124.1 aluc|OJZ81454.1 anid|CBF82567.1 anig|EHA25974.1 aory|EIT77369.1 asyd|OJJ54050.1 ater|EAU38610.1 atub|OJI88727.1 aver|OJJ08692.1 awen|OJJ34357.1 azon|OJJ48093.1 acla|EAW10899.1 afis|EAW20959.1 afla|EED50315.1 afum|EAL84674.1 akaw|GAA91125.1 aluc|OJZ81455.1 aory|EIT77298.1

Group407: anid|CBF79149.1 abra|OJJ68851.1 acar|OOF93604.1 afis|EAW23435.1 afla|EED49617.1 akaw|GAA87401.1 aluc|OJZ87639.1 anid|CBF79147.1 anig|EHA25624.1 aory|EIT80987.1 ater|EAU36062.1 atub|OJI79619.1 abra|OJJ76884.1 afla|EED53582.1 anid|CBF85241.1 asyd|OJJ56896.1 afis|EAW23434.1 aory|EIT80692.1 ater|EAU31633.1 atub|OJI88357.1 afla|EED51736.1 aory|EIT80667.1 aory|EIT75082.1 atub|OJI81007.1

Group408: abra|OJJ66957.1 acar|OOF93651.1 acla|EAW14508.1 afis|EAW19221.1 afla|EED47668.1 agla|OJJ88465.1 akaw|GAA92204.1 aluc|OJZ85619.1 anid|CBF78606.1 anig|EHA24907.1 azon|OJJ51238.1 abra|OJJ69809.1 acar|OOF93568.1 akaw|GAA87851.1 aluc|OJZ85491.1 aory|EIT82845.1 atub|OJI80074.1 awen|OJJ37584.1 azon|OJJ46584.1 afla|EED52824.1 asyd|OJJ65099.1 aver|OJI99225.1 afla|EED47373.1 atub|OJI79412.1

Group409: abra|OJJ66987.1 acar|OOF93625.1 acla|EAW08983.1 afis|EAW24689.1 afla|EED48347.1 afum|EAL89330.1 agla|OJJ83266.1 akaw|GAA92176.1 aluc|OJZ85586.1 anid|CBF73671.1 anig|EHA24935.1 aory|EIT76285.1 asyd|OJJ56664.1 ater|EAU31416.1 atub|OJI79381.1 aver|OJJ06059.1 awen|OJJ32833.1 azon|OJJ49002.1 acla|EAW14852.1 afis|EAW24379.1 afla|EED47832.1 afum|EAL86051.1 aory|EIT77109.1 ater|EAU30357.1

Group410: abra|OJJ67104.1 acar|OOF95836.1 afla|EED47889.1 aluc|OJZ82016.1 anig|EHA22240.1 aory|EIT77218.1 atub|OJI84565.1 abra|OJJ75997.1 acar|OOF99928.1 acla|EAW09808.1 afis|EAW21328.1 afla|EED57286.1 afum|EAL92346.2 agla|OJJ87411.1 akaw|GAA88962.1 aluc|OJZ89443.1 anid|CBF83430.1 anig|EHA23143.1 aory|EIT74039.1 asyd|OJJ60660.1 ater|EAU35953.1 aver|OJI99983.1 awen|OJJ39370.1 azon|OJJ51710.1

Group411: abra|OJJ67606.1 acar|OOF93337.1 acla|EAW11030.1 afis|EAW20824.1 afla|EED45471.1 afum|EAL84805.1 akaw|GAA89123.1 aluc|OJZ81715.1 anid|CBF77690.1 anig|EHA25724.1 aory|EIT75480.1 asyd|OJJ60009.1 ater|EAU37676.1 atub|OJI88467.1 aver|OJJ04442.1 awen|OJJ36217.1 afla|EED52784.1 aory|EIT78156.1 atub|OJI87922.1 acar|OOF98999.1 akaw|GAA83906.1 anig|EHA19524.1 aver|OJJ02276.1 azon|OJJ50341.1

Group412: afis|EAW15422.1 afum|EAL84222.1 abra|OJJ74863.1 acar|OOF95863.1 acla|EAW08764.1 afla|EED57451.1 agla|OJJ81802.1 akaw|GAA83557.1 aluc|OJZ92441.1 anid|CBF89639.1 anig|EHA26526.1 aory|EIT80148.1 asyd|OJJ62291.1 ater|EAU32509.1 atub|OJI86870.1 aver|OJI96399.1 awen|OJJ39887.1 azon|OJJ50235.1 afum|EAL85683.1 afis|EAW15665.1 afla|EED46890.1 awen|OJJ38340.1 acla|EAW08989.1 aory|EIT72560.1

Group413: abra|OJJ67909.1 acar|OOF98177.1 afla|EED46476.1 agla|OJJ79399.1 anig|EHA22276.1 aory|EIT72342.1 asyd|OJJ61955.1 ater|EAU37942.1 aver|OJI96047.1 awen|OJJ33590.1 azon|OJJ42127.1 abra|OJJ70267.1 akaw|GAA86530.1 aluc|OJZ85078.1 anid|CBF83033.1 anig|EHA25293.1 asyd|OJJ61489.1 atub|OJI80520.1 aver|OJJ00830.1 aory|EIT73700.1 asyd|OJJ55487.1 aver|OJJ06566.1 afla|EED45145.1 aory|EIT82578.1

Group414: abra|OJJ68074.1 acar|OOF96032.1 acla|EAW11411.1 afis|EAW24010.1 afla|EED49151.1 afum|EAL87440.1 agla|OJJ86942.1 akaw|GAA92399.1 aluc|OJZ83260.1 anid|CBF77404.1 anig|EHA21114.1 aory|EIT79051.1 asyd|OJJ53240.1 ater|EAU29741.1 atub|OJI87679.1 aver|OJJ03610.1 awen|OJJ35482.1 azon|OJJ51266.1 aory|EIT78571.1 afla|EED50884.1 ater|EAU38050.1 asyd|OJJ59006.1 aver|OJJ01610.1 awen|OJJ37601.1

Group415: abra|OJJ68223.1 acla|EAW07100.1 afis|EAW20575.1 afum|EAL84490.1 akaw|GAA83221.1 aluc|OJZ83722.1 anig|EHA18872.1 atub|OJI90097.1 azon|OJJ50650.1 abra|OJJ76992.1 acar|OOF91437.1 afla|EED52310.1 agla|OJJ88718.1 akaw|GAA87054.1 aluc|OJZ84392.1 anid|CBF78937.1 anig|EHA27721.1 aory|EIT74487.1 asyd|OJJ55645.1 ater|EAU37109.1 atub|OJI89679.1 aver|OJJ06730.1 awen|OJJ33275.1 acla|EAW13463.1

Group416: abra|OJJ68471.1 acar|OOF94213.1 acla|EAW13767.1 afis|EAW19791.1 afis|EAW22422.1 afla|EED45872.1 afum|EAL85100.2 afum|EAL90770.1 agla|OJJ87604.1 akaw|GAA92392.1 aluc|OJZ83252.1 anid|CBF88815.1 anig|EHA21105.1 aory|EIT75744.1 asyd|OJJ62890.1 ater|EAU36158.1 atub|OJI87685.1 awen|OJJ35001.1 azon|OJJ50741.1 acla|EAW07396.1 ater|EAU39285.1 anig|EHA21106.1 afla|EED52500.1 aory|EIT82777.1

Group417: abra|OJJ69765.1 afis|EAW19682.1 afla|EED54789.1 afum|EAL86840.1 agla|OJJ80273.1 akaw|GAA87888.1 aluc|OJZ85533.1 anid|CBF74487.1 anig|EHA25064.1 aory|EIT79365.1 asyd|OJJ57713.1 awen|OJJ37833.1 abra|OJJ73953.1 acar|OOG01081.1 akaw|GAA89558.1 aluc|OJZ88091.1 anig|EHA17861.1 asyd|OJJ58685.1 ater|EAU36023.1 atub|OJI90699.1 anid|CBF87757.1 aver|OJI97605.1 aory|EIT73667.1 asyd|OJJ63356.1

Group418: abra|OJJ70549.1 afla|EED53198.1 agla|OJJ89090.1 akaw|GAA83393.1 aluc|OJZ83893.1 asyd|OJJ55203.1 atub|OJI90260.1 aver|OJJ07013.1 abra|OJJ70550.1 acar|OOF97171.1 acla|EAW08341.1 afis|EAW23376.1 afla|EED55581.1 afum|EAL88095.1 agla|OJJ86447.1 anid|CBF74843.1 anig|EHA19029.1 asyd|OJJ60130.1 ater|EAU35630.1 atub|OJI90261.1 aver|OJJ04131.1 awen|OJJ39507.1 azon|OJJ47706.1 aory|EIT74938.1

Group419: agla|OJJ85379.1 asyd|OJJ59944.1 aver|OJJ02344.1 abra|OJJ77167.1 acar|OOF93057.1 acla|EAW12893.1 afis|EAW17769.1 afla|EED45780.1 afum|EAL91577.1 agla|OJJ81693.1 akaw|GAA90765.1 aluc|OJZ84580.1 anid|CBF86634.1 anig|EHA27873.1 asyd|OJJ64807.1 ater|EAU31848.1 atub|OJI89500.1 aver|OJI98920.1 awen|OJJ37072.1 acar|OOF91588.1 afla|EED45781.1 aory|EIT75674.1 afla|EED45782.1 aory|EIT75774.1

Group420: acar|OOF98107.1 akaw|GAA82924.1 anig|EHA20714.1 atub|OJI81150.1 abra|OJJ71031.1 acar|OOF98105.1 afis|EAW24984.1 afla|EED55473.1 afum|EBA27385.1 agla|OJJ82277.1 akaw|GAA82923.1 anid|CBF81625.1 aory|EIT80888.1 asyd|OJJ59528.1 ater|EAU35736.1 aver|OJJ02742.1 awen|OJJ40581.1 azon|OJJ49444.1 acla|EAW10584.1 afis|EAW24985.1 afum|EBA27386.1 anid|CBF81605.1 aory|EIT80797.1 aver|OJJ02751.1

Group421: abra|OJJ71639.1 acar|OOF96356.1 acla|EAW15180.1 afis|EAW20150.1 afla|EED57200.1 afum|EAL85264.1 agla|OJJ88784.1 akaw|GAA89286.1 akaw|GAA89287.1 aluc|OJZ86382.1 anid|CBF84986.1 anig|EHA21512.1 aory|EIT73993.1 asyd|OJJ63125.1 ater|EAU38765.1 atub|OJI84151.1 aver|OJI97830.1 awen|OJJ32023.1 azon|OJJ46067.1 acla|EAW15179.1 afis|EAW20149.1 anid|CBF84984.1 aory|EIT74054.1 afum|EAL85265.1

Group422: abra|OJJ71666.1 acar|OOF96376.1 acla|EAW15203.1 acla|EAW15204.1 afis|EAW20168.1 afis|EAW20169.1 afla|EED57180.1 afum|EAL85286.1 afum|EAL85287.1 agla|OJJ88761.1 akaw|GAA89266.1 akaw|GAA89267.1 aluc|OJZ86408.1 anid|CBF85015.1 anid|CBF85017.1 anig|EHA21491.1 aory|EIT74089.1 asyd|OJJ63104.1 ater|EAU38800.1 atub|OJI84123.1 aver|OJI97857.1 awen|OJJ32065.1 awen|OJJ32066.1 azon|OJJ46098.1

Group423: abra|OJJ72573.1 acar|OOF94686.1 akaw|GAA85199.1 aluc|OJZ90554.1 anig|EHA24001.1 atub|OJI81977.1 abra|OJJ77405.1 acar|OOF93271.1 acla|EAW13764.1 afis|EAW19795.1 afla|EED52177.1 afum|EAL85095.2 agla|OJJ83624.1 akaw|GAA91360.1 aluc|OJZ84812.1 anig|EHA28076.1 aory|EIT74472.1 asyd|OJJ54440.1 asyd|OJJ56546.1 ater|EAU36883.1 atub|OJI89273.1 aver|OJJ06176.1 awen|OJJ36480.1 azon|OJJ42299.1

Group424: abra|OJJ73313.1 acar|OOG00380.1 afis|EAW20348.1 afla|EED51166.1 agla|OJJ89330.1 akaw|GAA92810.1 aluc|OJZ88733.1 anid|CBF85228.1 anig|EHA18436.1 aory|EIT82315.1 asyd|OJJ62986.1 atub|OJI91309.1 aver|OJI97982.1 awen|OJJ29882.1 afis|EAW17164.1 anid|CBF76678.1 asyd|OJJ61222.1 aver|OJI96139.1 anid|CBF80144.1 aory|EIT81349.1 aver|OJI99535.1 afla|EED57011.1 asyd|OJJ61869.1 aver|OJI95967.1

Group425: abra|OJJ73412.1 acar|OOG00609.1 akaw|GAA83079.1 aluc|OJZ88625.1 anid|CBF80318.1 anig|EHA18317.1 asyd|OJJ57587.1 atub|OJI91210.1 aver|OJJ05595.1 awen|OJJ38061.1 abra|OJJ76670.1 acar|OOF99340.1 acla|EAW09094.1 afis|EAW25688.1 afla|EED55156.1 afum|EAL84357.1 agla|OJJ78885.1 akaw|GAA92522.1 aluc|OJZ91536.1 anig|EHA22573.1 aory|EIT81660.1 ater|EAU32946.1 atub|OJI85990.1 awen|OJJ41728.1

Group426: abra|OJJ74271.1 acar|OOF95459.1 acla|EAW07315.1 afis|EAW22331.1 afla|EED57752.1 afum|EAL90861.1 agla|OJJ87711.1 akaw|GAA86752.1 aluc|OJZ89862.1 anid|CBF88649.1 anig|EHA27062.1 aory|EIT81123.1 asyd|OJJ63884.1 ater|EAU39460.1 atub|OJI84946.1 aver|OJI97056.1 awen|OJJ34909.1 azon|OJJ45576.1 abra|OJJ74564.1 acar|OOF95196.1 akaw|GAA82037.1 aluc|OJZ92145.1 anig|EHA26804.1 atub|OJI86574.1

Group427: abra|OJJ74337.1 acar|OOF95390.1 acla|EAW07388.1 afis|EAW22412.1 afla|EED52492.1 afum|EAL90780.1 agla|OJJ80127.1 akaw|GAA86692.1 aluc|OJZ89793.1 anid|CBF88798.1 anig|EHA27006.1 aory|EIT82746.1 asyd|OJJ64791.1 ater|EAU39293.1 atub|OJI85013.1 aver|OJI98903.1 awen|OJJ34993.1 azon|OJJ45636.1 afis|EAW15417.1 afla|EED50613.1 afum|EAL84228.1 aory|EIT78052.1 ater|EAU37467.1 awen|OJJ32510.1

Group428: abra|OJJ76931.1 acar|OOF90090.1 acla|EAW13434.1 afis|EAW20550.1 afla|EED45551.1 afum|EAL84510.1 akaw|GAA87109.1 aluc|OJZ84329.1 anid|CBF73888.1 anig|EHA27663.1 anig|EHA27664.1 aory|EIT75407.1 asyd|OJJ53735.1 ater|EAU31070.1 atub|OJI89739.1 aver|OJJ07850.1 awen|OJJ33265.1 azon|OJJ50631.1 acar|OOF99064.1 acla|EAW11193.1 afis|EAW15923.1 afum|EAL88560.2 ater|EAU32036.1 awen|OJJ35657.1

Group429: abra|OJJ77163.1 acar|OOF93049.1 acla|EAW11721.1 afis|EAW23742.1 afla|EED45436.1 afum|EAL87179.1 agla|OJJ81702.1 akaw|GAA90769.1 aluc|OJZ84576.1 anid|CBF86622.1 anig|EHA27869.1 aory|EIT82382.1 asyd|OJJ65088.1 ater|EAU37973.1 atub|OJI89506.1 aver|OJI99214.1 awen|OJJ37083.1 azon|OJJ42895.1 anid|CBF87116.1 awen|OJJ35821.1 ater|EAU31855.1 asyd|OJJ64812.1 aver|OJI98925.1 anid|CBF71823.1

Group430: abra|OJJ65741.1 abra|OJJ71669.1 asyd|OJJ61538.1 asyd|OJJ63100.1 aver|OJI97862.1 aver|OJI98822.1 aver|OJJ00865.1 acar|OOF96381.1 acla|EAW15209.1 afis|EAW20174.1 afla|EED57174.1 afum|EAL85292.1 agla|OJJ88756.1 akaw|GAA89262.1 aluc|OJZ86412.1 anid|CBF85033.1 anig|EHA21487.1 aory|EIT74095.1 ater|EAU35210.1 atub|OJI84119.1 awen|OJJ41764.1 azon|OJJ46103.1 afla|EED54878.1

Group431: abra|OJJ65836.1 abra|OJJ67758.1 abra|OJJ72366.1 afum|EAL87737.1 afum|EAL88791.1 agla|OJJ79294.1 agla|OJJ80455.1 akaw|GAA87903.1 akaw|GAA92853.1 akaw|GAA92883.1 akaw|GAA93148.1 anig|EHA21873.1 anig|EHA26120.1 azon|OJJ46212.1 azon|OJJ47661.1 acar|OOF97545.1 aluc|OJZ80010.1 aluc|OJZ90353.1 anid|CBF79035.1 asyd|OJJ52047.1 atub|OJI81780.1 agla|OJJ84748.1 akaw|GAA86116.1

Group432: abra|OJJ65984.1 abra|OJJ67112.1 abra|OJJ73081.1 acar|OOF92889.1 acar|OOF98440.1 akaw|GAA88080.1 akaw|GAA90857.1 akaw|GAA91052.1 aluc|OJZ81343.1 aluc|OJZ82976.1 aluc|OJZ87863.1 anig|EHA18648.1 anig|EHA20169.1 anig|EHA21392.1 atub|OJI80785.1 atub|OJI89209.1 atub|OJI91548.1 azon|OJJ42157.1 azon|OJJ42333.1 azon|OJJ42440.1 azon|OJJ42744.1 anid|CBF71426.1 awen|OJJ31434.1

Group433: abra|OJJ66045.1 abra|OJJ73392.1 abra|OJJ77643.1 afla|EED46392.1 afla|EED49513.1 atub|OJI82691.1 atub|OJI87256.1 atub|OJI87448.1 atub|OJI91231.1 acar|OOF99092.1 agla|OJJ85351.1 akaw|GAA83097.1 akaw|GAA83991.1 aluc|OJZ80886.1 aluc|OJZ88644.1 anig|EHA18335.1 anig|EHA19451.1 aory|EIT72357.1 asyd|OJJ54995.1 aver|OJJ07206.1 awen|OJJ40082.1 afis|EAW17049.1 aver|OJI95758.1

Group434: abra|OJJ67714.1 abra|OJJ68753.1 akaw|GAA89213.1 akaw|GAA89897.1 aluc|OJZ81616.1 aluc|OJZ81774.1 anig|EHA22442.1 anig|EHA25827.1 atub|OJI84347.1 atub|OJI88567.1 acar|OOF92242.1 acla|EAW12498.1 afis|EAW24147.1 afla|EED46784.1 afum|EAL92002.1 agla|OJJ87227.1 anid|CBF77921.1 aory|EIT72760.1 asyd|OJJ55295.1 ater|EAU30579.1 aver|OJJ06931.1 awen|OJJ41332.1 azon|OJJ48802.1

Group435: abra|OJJ70999.1 abra|OJJ75834.1 acar|OOF98141.1 acla|EAW10556.1 afis|EAW25023.1 afla|EED55445.1 afum|EAL89672.1 agla|OJJ89191.1 akaw|GAA82949.1 aluc|OJZ87528.1 anid|CBF81543.1 anig|EHA20738.1 aory|EIT80817.1 asyd|OJJ59491.1 ater|EAU35764.1 atub|OJI81120.1 aver|OJJ02783.1 awen|OJJ40617.1 azon|OJJ49465.1 akaw|GAA84544.1 aluc|OJZ89277.1 anig|EHA23292.1 atub|OJI85522.1

Group436: abra|OJJ73227.1 abra|OJJ77180.1 acar|OOF91798.1 acar|OOG00465.1 agla|OJJ82802.1 agla|OJJ84567.1 asyd|OJJ52864.1 asyd|OJJ59702.1 acla|EAW06901.1 afis|EAW16518.1 afla|EED56153.1 afum|EAL89113.1 akaw|GAA87953.1 aluc|OJZ81201.1 anid|CBF75551.1 anig|EHA18516.1 aory|EIT76710.1 ater|EAU36584.1 atub|OJI91396.1 aver|OJJ02585.1 aver|OJJ08843.1 awen|OJJ40305.1 azon|OJJ49166.1

Group437: abra|OJJ75244.1 abra|OJJ75269.1 acar|OOF94534.1 acar|OOF94549.1 akaw|GAA91787.1 akaw|GAA91807.1 aluc|OJZ92806.1 aluc|OJZ92829.1 anig|EHA26180.1 anig|EHA26203.1 atub|OJI87249.1 atub|OJI87286.1 acla|EAW12239.1 afis|EAW16988.1 afla|EED48988.1 afum|EAL86512.1 agla|OJJ79828.1 anid|CBF87433.1 aory|EIT80045.1 ater|EAU31739.1 aver|OJJ08882.1 awen|OJJ35990.1 asyd|OJJ52913.1

Group438: abra|OJJ75609.1 abra|OJJ76068.1 akaw|GAA84359.1 akaw|GAA88899.1 aluc|OJZ89061.1 aluc|OJZ89522.1 anig|EHA23079.1 anig|EHA23477.1 atub|OJI85281.1 atub|OJI85725.1 acar|OOG00229.1 acla|EAW09371.1 afis|EAW21764.1 afla|EED46335.1 afum|EAL92794.1 agla|OJJ84387.1 anid|CBF83753.1 aory|EIT81195.1 asyd|OJJ60858.1 ater|EAU38451.1 aver|OJJ00182.1 awen|OJJ34047.1 azon|OJJ48184.1

Group439: acar|OOF90104.1 acar|OOF90211.1 acar|OOF90646.1 akaw|GAA86640.1 akaw|GAA87321.1 akaw|GAA88463.1 akaw|GAA90279.1 aluc|OJZ83061.1 aluc|OJZ88352.1 aluc|OJZ85197.1 anig|EHA18084.1 anig|EHA28127.1 atub|OJI80404.1 atub|OJI87885.1 atub|OJI90947.1 azon|OJJ42159.1 azon|OJJ48921.1 azon|OJJ43897.1 azon|OJJ46414.1 abra|OJJ73701.1 afla|EED57243.1 aory|EIT74140.1 awen|OJJ30907.1

Group440: acar|OOF90258.1 acar|OOF90411.1 acar|OOF94537.1 akaw|GAA90238.1 akaw|GAA91802.1 aluc|OJZ83102.1 aluc|OJZ92824.1 atub|OJI87281.1 atub|OJI87833.1 awen|OJJ33606.1 awen|OJJ39914.1 abra|OJJ75266.1 acla|EAW08995.1 afla|EED48390.1 afum|EAL91138.1 anig|EHA26183.1 aory|EIT83109.1 ater|EAU30261.1 ater|EAU37025.1 azon|OJJ45138.1 anig|EHA27462.1 ater|EAU34045.1 acar|OOF95569.1

Group441: acar|OOF94062.1 acar|OOF94132.1 abra|OJJ66364.1 acla|EAW08975.1 afis|EAW15454.1 afla|EED54752.1 afum|EAL84191.1 agla|OJJ80556.1 akaw|GAA90237.1 aluc|OJZ83103.1 anid|CBF80456.1 anig|EHA28167.1 aory|EIT78886.1 asyd|OJJ54768.1 ater|EAU32265.1 atub|OJI87832.1 aver|OJJ07399.1 awen|OJJ36431.1 azon|OJJ43598.1 atub|OJI80330.1 afla|EED48876.1 aory|EIT81202.1 awen|OJJ31672.1

Group442: acla|EAW10867.1 acla|EAW11076.1 afla|EED50345.1 afla|EED55539.1 afum|EAL84638.1 afum|EAL87765.1 aory|EIT77302.1 aory|EIT80884.1 asyd|OJJ54018.1 asyd|OJJ56874.1 abra|OJJ66629.1 acar|OOF91053.1 afis|EAW20991.1 agla|OJJ84069.1 akaw|GAA91096.1 aluc|OJZ81419.1 anid|CBF82616.1 anig|EHA26002.1 ater|EAU38640.1 atub|OJI88762.1 aver|OJJ08722.1 awen|OJJ34319.1 azon|OJJ48065.1

Group443: afis|EAW17232.1 afis|EAW21390.1 afum|EAL86652.1 afum|EAL92404.2 aory|EIT72927.1 aory|EIT76529.1 azon|OJJ48879.1 azon|OJJ50259.1 azon|OJJ51536.1 abra|OJJ65877.1 abra|OJJ76067.1 acar|OOF99863.1 acla|EAW09752.1 afla|EED48188.1 akaw|GAA88900.1 aluc|OJZ89521.1 anid|CBF83840.1 anig|EHA23080.1 asyd|OJJ60914.1 ater|EAU35174.1 atub|OJI85282.1 aver|OJJ00241.1 awen|OJJ30796.1

Group444: afis|EAW19359.1 afis|EAW20492.1 aver|OJJ07259.1 aver|OJJ07872.1 awen|OJJ33502.1 awen|OJJ34946.1 abra|OJJ66132.1 acar|OOF94018.1 acla|EAW14627.1 afla|EED56212.1 afum|EAL84562.1 agla|OJJ80511.1 akaw|GAA92689.1 aluc|OJZ90147.1 anid|CBF73880.1 anig|EHA20923.1 aory|EIT73321.1 asyd|OJJ64747.1 ater|EAU37645.1 atub|OJI84652.1 awen|OJJ38031.1 asyd|OJJ53765.1 asyd|OJJ54937.1

Group445: afis|EAW20796.1 afis|EAW22075.1 afum|EAL84833.1 afum|EAL91095.1 anid|CBF71369.1 anid|CBF82800.1 ater|EAU30149.1 ater|EAU32867.1 abra|OJJ72284.1 acar|OOF94937.1 afla|EED49547.1 agla|OJJ85016.1 akaw|GAA86036.1 aluc|OJZ90263.1 anig|EHA23759.1 aory|EIT73956.1 asyd|OJJ55081.1 asyd|OJJ57119.1 atub|OJI81689.1 aver|OJJ07127.1 awen|OJJ41908.1 azon|OJJ47364.1 afla|EED56697.1

Group446: afla|EED44969.1 afla|EED46569.1 afla|EED51147.1 aory|EIT73475.1 aory|EIT77864.1 aory|EIT81352.1 asyd|OJJ56870.1 asyd|OJJ61208.1 abra|OJJ75570.1 acar|OOF90888.1 acla|EAW09352.1

afis|EAW21817.1 afum|EAL92833.1 agla|OJJ84325.1 akaw|GAA84321.1 aluc|OJZ89018.1 anid|CBF76652.1 anig|EHA23512.1 ater|EAU38486.1 atub|OJI85771.1 aver|OJJ00532.1 awen|OJJ34091.1 azon|OJJ44255.1

Group447: afla|EED47292.1 afla|EED54844.1 agla|OJJ87461.1 agla|OJJ89110.1 aory|EIT79178.1 aory|EIT79336.1 asyd|OJJ60317.1 asyd|OJJ60627.1 asyd|OJJ62712.1 abra|OJJ75953.1 acar|OOF99959.1 acla|EAW09843.1 afis|EAW21292.1 afum|EAL92309.1 akaw|GAA84656.1 aluc|OJZ89402.1 anid|CBF83376.1 anig|EHA23185.1 ater|EAU35925.1 atub|OJI85395.1 aver|OJI99951.1 awen|OJJ39407.1 azon|OJJ44290.1

Group448: afla|EED48892.1 afla|EED55381.1 aory|EIT80798.1 aory|EIT81174.1 asyd|OJJ56534.1 asyd|OJJ64421.1 aver|OJI98520.1 aver|OJJ06188.1 abra|OJJ70928.1 acar|OOF98198.1 acla|EAW10494.1 afis|EAW25073.1 afum|EAL89742.1 agla|OJJ89119.1 akaw|GAA83014.1 aluc|OJZ87599.1 anid|CBF85962.1 anig|EHA20799.1 ater|EAU39552.1 atub|OJI81048.1 awen|OJJ40684.1 azon|OJJ44061.1 awen|OJJ34526.1

Group449: afla|EED54572.1 afla|EED56566.1 asyd|OJJ53770.1 asyd|OJJ62761.1 ater|EAU30266.1 ater|EAU39167.1 aver|OJI96804.1 aver|OJJ07878.1 abra|OJJ73591.1 acar|OOG00775.1 acla|EAW07510.1 afis|EAW22541.1 afum|EAL90653.1 agla|OJJ80900.1 akaw|GAA88561.1 aluc|OJZ88459.1 anid|CBF89038.1 anig|EHA18179.1 aory|EIT78798.1 aory|EIT79646.1 atub|OJI91057.1 awen|OJJ35170.1 azon|OJJ49937.1

Group450: agla|OJJ78857.1 agla|OJJ87323.1 abra|OJJ66001.1 acar|OOF92098.1 acla|EAW11077.1 afla|EED45932.1 akaw|GAA90873.1 aluc|OJZ82959.1 anid|CBF84691.1 anig|EHA21375.1 aory|EIT75815.1 asyd|OJJ53330.1 ater|EAU32160.1 atub|OJI89191.1 aver|OJJ08414.1 azon|OJJ46594.1 afla|EED53485.1 aory|EIT75146.1 asyd|OJJ61788.1 aver|OJI95887.1 ater|EAU32697.1 asyd|OJJ55442.1 ater|EAU35122.1

Group451: agla|OJJ81417.1 agla|OJJ84332.1 abra|OJJ77363.1 acar|OOF93232.1 acla|EAW13113.1 afis|EAW17551.1 afla|EED49274.1 afum|EAL91365.1 akaw|GAA91399.1 aluc|OJZ84772.1 anid|CBF82159.1 anig|EHA28040.1 aory|EIT82213.1 asyd|OJJ54334.1 ater|EAU36983.1 atub|OJI89309.1 awen|OJJ32551.1 azon|OJJ50995.1 afis|EAW17400.1 afla|EED46104.1 afum|EAL91238.1 anid|CBF79798.1 aory|EIT76423.1

Group452: akaw|GAA83965.1 akaw|GAA90739.1 aluc|OJZ80854.1 aluc|OJZ84609.1 abra|OJJ77200.1 acar|OOF93088.1 acla|EAW13191.1 afis|EAW17450.1 afla|EED49372.1 afum|EAL91278.2 agla|OJJ80362.1 akaw|GAA90738.1 aluc|OJZ84610.1 anid|CBF71683.1 anig|EHA27899.1 aory|EIT79587.1 asyd|OJJ61822.1 ater|EAU33930.1 atub|OJI89471.1 aver|OJI95922.1 awen|OJJ37396.1 azon|OJJ51068.1 anig|EHA27898.1

Group453: anid|CBF69431.1 anid|CBF73986.1 abra|OJJ66331.1 acar|OOG00530.1 acla|EAW09349.1 afis|EAW21804.1 afla|EED56853.1 afum|EAL92830.1 agla|OJJ79164.1 akaw|GAA90208.1 aluc|OJZ83137.1 anig|EHA28196.1 aory|EIT77503.1 asyd|OJJ65096.1 ater|EAU38484.1 atub|OJI87797.1 aver|OJI99221.1 azon|OJJ47555.1 aver|OJJ01938.1 afla|EED46583.1 aory|EIT77946.1 asyd|OJJ52385.1 agla|OJJ83529.1

Group454: anid|CBF73961.1 anid|CBF84419.1 asyd|OJJ53557.1 asyd|OJJ53743.1 asyd|OJJ57304.1 ater|EAU32191.1 ater|EAU37274.1 aver|OJJ05855.1 aver|OJJ07856.1 aver|OJJ08197.1 abra|OJJ66428.1 acla|EAW13742.1 afis|EAW19838.1 afla|EED45944.1 afum|EAL85055.2 agla|OJJ84458.1 akaw|GAA90293.1 aluc|OJZ83045.1 anig|EHA28114.1 aory|EIT75577.1 atub|OJI87903.1 awen|OJJ29654.1 azon|OJJ42315.1

Group455: aory|EIT75378.1 aory|EIT78539.1 awen|OJJ34056.1 awen|OJJ35494.1 azon|OJJ46995.1 azon|OJJ50589.1 abra|OJJ72842.1 acar|OOF96043.1 acla|EAW11400.1 afis|EAW24021.1 afla|EED45752.1 afla|EED49134.1 afla|EED49135.1 afum|EAL87451.1 agla|OJJ86954.1 akaw|GAA85444.1 aluc|OJZ83350.1 anid|CBF77424.1 anig|EHA24217.1 asyd|OJJ53230.1 ater|EAU29725.1 atub|OJI82243.1 aver|OJJ03622.1

Group456: asyd|OJJ51978.1 asyd|OJJ52250.1 asyd|OJJ56074.1 abra|OJJ76969.1 acar|OOF91414.1 acla|EAW13273.1 acla|EAW13494.1 afis|EAW20594.1 afla|EED47642.1 afum|EAL84467.1 agla|OJJ88662.1 akaw|GAA87075.1 aluc|OJZ84365.1 anid|CBF78279.1 anig|EHA27700.1 aory|EIT80239.1 ater|EAU35240.1 atub|OJI89703.1 aver|OJJ05017.1 awen|OJJ33318.1 azon|OJJ50690.1 afis|EAW17345.1 afum|EAL91189.2

Group457: asyd|OJJ52031.1 asyd|OJJ52140.1 abra|OJJ68385.1 acar|OOF93982.1 acla|EAW12147.1 afla|EED51420.1 afum|EAL86589.1 anid|CBF81959.1 aory|EIT80646.1 aver|OJJ05804.1 abra|OJJ68386.1 afis|EAW17129.1 akaw|GAA92560.1 aluc|OJZ90106.1 anig|EHA20964.1 ater|EAU31612.1 atub|OJI84696.1 awen|OJJ32473.1 azon|OJJ46830.1 afis|EAW25756.1 akaw|GAA93234.1 aluc|OJZ79642.1 asyd|OJJ57363.1

Group458: asyd|OJJ57402.1 asyd|OJJ58377.1 abra|OJJ75419.1 acar|OOF90764.1 acla|EAW13791.1 afis|EAW18341.1 afla|EED54057.1 afum|EAL87704.1 agla|OJJ88043.1 akaw|GAA84194.1 aluc|OJZ88871.1 anid|CBF79375.1 anig|EHA23633.1 asyd|OJJ57817.1 ater|EAU34676.1 atub|OJI85920.1 aver|OJJ04178.1 awen|OJJ35725.1 azon|OJJ42531.1 acar|OOF90590.1 ater|EAU35690.1 awen|OJJ36256.1 asyd|OJJ60091.1

Group459: asyd|OJJ59828.1 asyd|OJJ60302.1 asyd|OJJ60331.1 aver|OJI99642.1 aver|OJJ02459.1 aver|OJI99677.1 awen|OJJ29685.1 awen|OJJ30621.1 awen|OJJ36172.1 abra|OJJ77900.1 acar|OOF98884.1 acla|EAW11251.1 afis|EAW18246.1 afla|EED56217.1 afum|EAL87611.1 agla|OJJ85824.1 akaw|GAA83762.1 aluc|OJZ90969.1 anid|CBF87149.1 anig|EHA19671.1 aory|EIT73372.1 ater|EAU33739.1 atub|OJI82960.1

Group460: asyd|OJJ60480.1 asyd|OJJ61028.1 awen|OJJ32562.1 awen|OJJ36305.1 azon|OJJ43188.1 azon|OJJ47398.1 afum|EAL86366.1 abra|OJJ75145.1 akaw|GAA88750.1 aluc|OJZ92710.1 anig|EHA26289.1 acar|OOF98762.1 ater|EAU36669.1 acla|EAW09016.1 aver|OJI99800.1 afla|EED54373.1 aluc|OJZ81261.1 aory|EIT75853.1 ater|EAU36009.1 aver|OJJ07827.1 atub|OJI87136.1 ater|EAU33600.1 asyd|OJJ53703.1

Group461: ater|EAU29737.1 ater|EAU36828.1 abra|OJJ68505.1 acar|OOF90180.1 acar|OOF91789.1 acla|EAW11100.1 afis|EAW20763.1 afla|EED53200.1 afum|EAL84856.1 agla|OJJ84006.1 akaw|GAA92768.1 aluc|OJZ82717.1 anid|CBF80354.1 anig|EHA21171.1 aory|EIT75122.1 asyd|OJJ54828.1 atub|OJI88951.1 aver|OJJ07333.1 awen|OJJ29787.1 azon|OJJ42797.1 asyd|OJJ58563.1 aver|OJJ01162.1 azon|OJJ42773.1

Group462: atub|OJI83219.1 atub|OJI83220.1 abra|OJJ73330.1 afla|EED57225.1 agla|OJJ88240.1 akaw|GAA92790.1 anid|CBF74610.1 anig|EHA18401.1 aory|EIT73990.1 ater|EAU30210.1 atub|OJI91293.1 aver|OJJ05551.1 awen|OJJ36971.1 abra|OJJ78150.1 acar|OOF91819.1 agla|OJJ79110.1 akaw|GAA85875.1 aluc|OJZ91202.1 anig|EHA19907.1 asyd|OJJ55391.1 aver|OJJ08266.1 agla|OJJ78827.1 azon|OJJ42621.1

Group463: azon|OJJ45795.1 azon|OJJ48029.1 abra|OJJ67774.1 acla|EAW12350.1 afis|EAW16775.1 afla|EED45004.1 afum|EAL86344.1 agla|OJJ83713.1 aory|EIT73669.1 asyd|OJJ60273.1 ater|EAU32446.1 atub|OJI80491.1 aver|OJJ07654.1 awen|OJJ34537.1 acla|EAW13294.1 afla|EED47565.1 afum|EAL86272.1 aory|EIT74524.1 awen|OJJ35954.1 ater|EAU33716.1 aver|OJJ06720.1 afis|EAW20694.1 ater|EAU36512.1

Group464: abra|OJJ65849.1 acla|EAW08786.1 afis|EAW15938.1 aluc|OJZ80301.1 ater|EAU31058.1 atub|OJI87454.1 aver|OJI96479.1 awen|OJJ39979.1 abra|OJJ70264.1 agla|OJJ88746.1 akaw|GAA86533.1 aluc|OJZ85081.1 anig|EHA25296.1 aory|EIT73096.1 asyd|OJJ55484.1 atub|OJI80518.1 aver|OJJ06563.1 abra|OJJ70920.1 acar|OOF98204.1 akaw|GAA83020.1 aluc|OJZ87606.1 atub|OJI81042.1 asyd|OJJ52755.1

Group465: abra|OJJ66209.1 acar|OOF90651.1 acla|EAW06843.1 afis|EAW20271.1 afum|EAL85405.2 akaw|GAA89078.1 aluc|OJZ80513.1 anig|EHA25680.1 aory|EIT83312.1 asyd|OJJ60801.1 ater|EAU35068.1 atub|OJI88417.1 abra|OJJ67048.1 afla|EED52068.1 anig|EHA18595.1 azon|OJJ42510.1 afla|EED52437.1 aluc|OJZ90229.1 anig|EHA23731.1 ater|EAU37218.1 afla|EED52417.1 akaw|GAA86004.1 azon|OJJ50736.1

Group466: anid|CBF73652.1 abra|OJJ78129.1 acar|OOF90670.1 acla|EAW14637.1 afla|EED45059.1 akaw|GAA85878.1 aluc|OJZ91191.1 anid|CBF78296.1 anig|EHA19887.1 asyd|OJJ56106.1 atub|OJI83210.1 aver|OJJ05048.1 azon|OJJ45964.1 acla|EAW11632.1 asyd|OJJ52073.1 ater|EAU31025.1 atub|OJI91230.1 aver|OJJ03030.1 afla|EED45199.1 aory|EIT73607.1 asyd|OJJ59264.1 awen|OJJ35370.1 aver|OJI99658.1

Group467: abra|OJJ66772.1 acar|OOF93853.1 acla|EAW08992.1 afis|EAW15416.1 afla|EED48724.1 afum|EAL84229.2 agla|OJJ79941.1 anid|CBF81992.1 anig|EHA27337.1 aory|EIT73206.1 asyd|OJJ56162.1 ater|EAU32712.1 aver|OJJ05101.1 awen|OJJ39993.1 azon|OJJ42917.1 abra|OJJ75745.1 akaw|GAA84478.1 aluc|OJZ89198.1 anig|EHA23360.1 atub|OJI85592.1 afla|EED55174.1 aory|EIT81585.1 awen|OJJ36897.1

Group468: abra|OJJ67101.1 acar|OOF98451.1 afis|EAW20274.1 afla|EED57102.1 afum|EAL85408.1 agla|OJJ87503.1 akaw|GAA91060.1 aluc|OJZ87872.1 anid|CBF78242.1 anig|EHA20178.1 aory|EIT82822.1 asyd|OJJ56196.1 ater|EAU31785.1 atub|OJI80777.1 aver|OJJ05123.1 awen|OJJ35879.1 afla|EED48128.1 aory|EIT76261.1 agla|OJJ85549.1 asyd|OJJ61684.1 ater|EAU38792.1 aver|OJJ03092.1 awen|OJJ29992.1

Group469: abra|OJJ67813.1 acar|OOF89857.1 afis|EAW19368.1 afla|EED48200.1 afum|EAL94087.1 akaw|GAA87912.1 aluc|OJZ85623.1 anid|CBF87299.1 anig|EHA18476.1 aory|EIT72963.1 asyd|OJJ57229.1 ater|EAU30775.1 atub|OJI79416.1 aver|OJJ03060.1 awen|OJJ32619.1 agla|OJJ83617.1 afis|EAW24515.1 afla|EED47664.1 afum|EAL86181.1 aluc|OJZ88782.1 anid|CBF84276.1 aory|EIT80178.1 atub|OJI91354.1

Group470: abra|OJJ67869.1 acar|OOF92849.1 afla|EED51544.1 agla|OJJ78960.1 akaw|GAA91876.1 aluc|OJZ82023.1 anid|CBF79791.1 anig|EHA22232.1 aory|EIT80600.1 asyd|OJJ58125.1 ater|EAU31305.1 atub|OJI84576.1 aver|OJJ04281.1 awen|OJJ33687.1 abra|OJJ68892.1 acla|EAW11730.1 afis|EAW23762.1 afum|EAL87195.1 agla|OJJ82460.1 akaw|GAA90357.1 aluc|OJZ85875.1 anig|EHA24715.1 atub|OJI79661.1

Group471: abra|OJJ67925.1 acar|OOF92776.1 afis|EAW17948.1 afum|EAL91758.1 agla|OJJ89340.1 akaw|GAA91272.1 aluc|OJZ82797.1 anid|CBF80065.1 anig|EHA21238.1 ater|EAU31422.1 atub|OJI89028.1 aver|OJJ02091.1 awen|OJJ41595.1 azon|OJJ48310.1 abra|OJJ74125.1 acla|EAW14963.1 afla|EED48217.1 akaw|GAA89760.1 aluc|OJZ80623.1 anig|EHA27208.1 aory|EIT72959.1 asyd|OJJ58320.1 awen|OJJ31633.1

Group472: abra|OJJ67981.1 acla|EAW10293.1 afis|EAW16666.1 afla|EED50863.1 afum|EAL89270.1 agla|OJJ79452.1 akaw|GAA91221.1 aluc|OJZ82849.1 anid|CBF83064.1 anig|EHA21286.1 aory|EIT78981.1 asyd|OJJ53493.1 ater|EAU38052.1 atub|OJI89079.1 aver|OJJ08255.1 awen|OJJ41013.1 abra|OJJ74173.1 acar|OOF95553.1 akaw|GAA86843.1 aluc|OJZ89962.1 anig|EHA27162.1 atub|OJI84843.1 agla|OJJ89278.1

Group473: awen|OJJ41504.1 abra|OJJ71797.1 acar|OOF96477.1 acla|EAW13728.1 afis|EAW18033.1 afum|EAL91844.1 agla|OJJ85408.1 akaw|GAA85113.1 aluc|OJZ86527.1 anid|CBF81681.1 anig|EHA22119.1 aory|EIT81720.1 asyd|OJJ53625.1 asyd|OJJ58292.1 atub|OJI83995.1 aver|OJJ05942.1 awen|OJJ31297.1 azon|OJJ50337.1 acar|OOF92709.1 afla|EED51862.1 anid|CBF80076.1 ater|EAU31550.1 aver|OJJ02132.1

Group474: abra|OJJ68397.1 acar|OOF94205.1 acla|EAW12189.1 afis|EAW23755.1 afla|EED52086.1 afum|EAL87191.1 akaw|GAA92551.1 aluc|OJZ90099.1 anid|CBF84318.1 anig|EHA20976.1 aory|EIT76905.1 asyd|OJJ55649.1 ater|EAU29520.1 atub|OJI84707.1 aver|OJJ06734.1 awen|OJJ34639.1 acla|EAW14123.1 afla|EED45259.1 aory|EIT73569.1 ater|EAU38832.1 afis|EAW18834.1 afum|EAL93570.1 afis|EAW20404.1

Group475: abra|OJJ68432.1 acar|OOF93958.1 afis|EAW25721.1 afla|EED52101.1 afum|EAL84395.2 agla|OJJ87948.1 akaw|GAA86939.1 akaw|GAA86940.1 aluc|OJZ90065.1 anid|CBF82387.1 anig|EHA21009.1

aory|EIT76909.1 asyd|OJJ54134.1 ater|EAU30751.1 atub|OJI84742.1 aver|OJJ08601.1 awen|OJJ29682.1 afla|EED52100.1 anid|CBF82386.1 aory|EIT76914.1 afla|EED45119.1 aory|EIT73766.1 ater|EAU30752.1

Group476: abra|OJJ68512.1 akaw|GAA91334.1 aluc|OJZ82727.1 anig|EHA21180.1 atub|OJI88960.1 abra|OJJ69632.1 acar|OOF92677.1 acla|EAW10562.1 afis|EAW25029.1 afla|EED55452.1 afum|EAL89666.1 agla|OJJ79155.1 akaw|GAA91612.1 aluc|OJZ82439.1 anid|CBF81553.1 anig|EHA28334.1 aory|EIT80876.1 asyd|OJJ59496.1 ater|EAU35758.1 atub|OJI88002.1 aver|OJJ02778.1 awen|OJJ40610.1 azon|OJJ49459.1

Group477: afum|EAL94100.1 abra|OJJ73872.1 afla|EED45842.1 akaw|GAA89484.1 aluc|OJZ88178.1 anid|CBF80737.1 anig|EHA17930.1 aory|EIT75520.1 asyd|OJJ55544.1 ater|EAU35671.1 atub|OJI90777.1 aver|OJJ06622.1 awen|OJJ38075.1 abra|OJJ78163.1 akaw|GAA85862.1 aluc|OJZ91217.1 anig|EHA19923.1 asyd|OJJ61397.1 atub|OJI83235.1 afis|EAW20414.1 afum|EBA27371.1 ater|EAU29532.1 aver|OJJ00726.1

Group478: abra|OJJ72302.1 acar|OOF94922.1 acla|EAW10039.1 afis|EAW24476.1 afla|EED56937.1 afum|EAL86149.1 agla|OJJ86412.1 akaw|GAA86053.1 aluc|OJZ90283.1 anid|CBF78363.1 anig|EHA23773.1 aory|EIT77524.1 asyd|OJJ57323.1 ater|EAU29927.1 atub|OJI81707.1 aver|OJJ05837.1 awen|OJJ40788.1 acla|EAW12030.1 afis|EAW24671.1 afum|EAL89324.1 anig|EHA28157.1 azon|OJJ48190.1 awen|OJJ34571.1

Group479: abra|OJJ72891.1 acar|OOF96081.1 acla|EAW11359.1 afis|EAW18135.1 afla|EED58060.1 afum|EAL87492.1 akaw|GAA85478.1 aluc|OJZ83399.1 anid|CBF75378.1 anid|CBF84404.1 aory|EIT72894.1 aory|EIT79796.1 asyd|OJJ53570.1 asyd|OJJ59841.1 ater|EAU35458.1 atub|OJI82285.1 aver|OJJ02448.1 aver|OJJ08185.1 awen|OJJ35577.1 afla|EED56220.1 aory|EIT73370.1 afla|EED48228.1 ater|EAU33738.1

Group480: abra|OJJ73797.1 acar|OOG00952.1 acla|EAW07688.1 afis|EAW22703.1 afla|EED49989.1 afum|EAL90489.1 agla|OJJ79006.1 akaw|GAA89415.1 aluc|OJZ88258.1 anid|CBF88074.1 anig|EHA18003.1 aory|EIT73485.1 asyd|OJJ63555.1 ater|EAU38999.1 atub|OJI90857.1 aver|OJI97379.1 awen|OJJ35340.1 azon|OJJ49778.1 aory|EIT74604.1 afis|EAW21081.1 afum|EAL92102.1 agla|OJJ79284.1 afla|EED44920.1

Group481: abra|OJJ74536.1 acar|OOF96722.1 afla|EED46724.1 akaw|GAA82065.1 aluc|OJZ92117.1 anig|EHA26827.1 aory|EIT77932.1 asyd|OJJ61662.1 ater|EAU31119.1 atub|OJI86550.1 abra|OJJ78387.1 acar|OOF98501.1 acla|EAW09146.1 afis|EAW22000.1 afum|EAL93015.1 agla|OJJ84097.1 akaw|GAA85645.1 aluc|OJZ91466.1 anid|CBF82670.1 atub|OJI83459.1 aver|OJJ00983.1 awen|OJJ34296.1 azon|OJJ43342.1

Group482: abra|OJJ76072.1 acar|OOF99854.1 afis|EAW16839.1 afla|EED55812.1 afum|EAL86386.1 agla|OJJ80806.1 akaw|GAA88895.1 aluc|OJZ89526.1 anid|CBF73895.1 anig|EHA23075.1 aory|EIT76740.1 asyd|OJJ53659.1 ater|EAU36679.1 atub|OJI85277.1 aver|OJJ07767.1 abra|OJJ76505.1 akaw|GAA92900.1 anig|EHA22711.1 acar|OOF99003.1 agla|OJJ79991.1 awen|OJJ36307.1 acla|EAW12392.1 anig|EHA25400.1

Group483: abra|OJJ76831.1 acar|OOF91325.1 acla|EAW13393.1 afis|EAW16462.1 afla|EED45207.1 afum|EAL89058.1 agla|OJJ84801.1 akaw|GAA87191.1 aluc|OJZ84243.1 anid|CBF75808.1 anig|EHA27588.1 aory|EIT73549.1 asyd|OJJ55721.1 atub|OJI89840.1 aver|OJJ06807.1 awen|OJJ33206.1 azon|OJJ46480.1 awen|OJJ33434.1 acla|EAW06841.1 asyd|OJJ57540.1 aver|OJJ05638.1 afis|EAW20505.1 afum|EAL84551.1

Group484: abra|OJJ76940.1 acar|OOF91390.1 acla|EAW09287.1 afis|EAW21860.1 afla|EED45974.1 afum|EAL92873.1 agla|OJJ79533.1 akaw|GAA87099.1 aluc|OJZ84339.1 anid|CBF89674.1 anig|EHA27673.1 aory|EIT75604.1 asyd|OJJ62270.1 ater|EAU29774.1 atub|OJI89729.1 aver|OJI96379.1 awen|OJJ34144.1 azon|OJJ50245.1 afla|EED46450.1 agla|OJJ84275.1 aory|EIT72411.1 ater|EAU30859.1 azon|OJJ43482.1

Group485: abra|OJJ77386.1 acar|OOF93255.1 acla|EAW11088.1 afis|EAW20767.1 afla|EED50191.1 afum|EAL84852.1 akaw|GAA91375.1 aluc|OJZ84794.1 anid|CBF73862.1 anig|EHA28062.1 aory|EIT77384.1 asyd|OJJ56520.1 ater|EAU31812.1 atub|OJI89288.1 aver|OJJ06202.1 awen|OJJ33710.1 azon|OJJ50068.1 awen|OJJ34924.1 acla|EAW07300.1 aory|EIT79889.1 ater|EAU39485.1 asyd|OJJ57568.1 aver|OJJ05611.1

Group486: abra|OJJ78343.1 acar|OOF98537.1 acla|EAW13688.1 afis|EAW19912.1 afla|EED47242.1 afum|EAL84995.1 agla|OJJ84607.1 akaw|GAA85690.1 aluc|OJZ91415.1 anid|CBF87843.1 anig|EHA20088.1 aory|EIT79169.1 asyd|OJJ61361.1 ater|EAU31064.1 atub|OJI83410.1 aver|OJJ00688.1 awen|OJJ33622.1 azon|OJJ43388.1 afla|EED46677.1 anid|CBF84039.1 aory|EIT77968.1 awen|OJJ34250.1 agla|OJJ88201.1

Group487: abra|OJJ65482.1 abra|OJJ70272.1 awen|OJJ34689.1 awen|OJJ34690.1 acar|OOF93269.1 afum|EAL92101.1 agla|OJJ82948.1 aluc|OJZ82533.1 abra|OJJ68088.1 acla|EAW15262.1 afla|EED50507.1 akaw|GAA86873.1 aluc|OJZ89997.1 anig|EHA27372.1 ater|EAU33813.1 atub|OJI84811.1 afis|EAW21079.1 aory|EIT82101.1 acar|OOF90259.1 agla|OJJ86366.1 awen|OJJ32169.1 anig|EHA17942.1

Group488: abra|OJJ67142.1 abra|OJJ74374.1 acar|OOF95362.1 acar|OOF98414.1 akaw|GAA87274.1 akaw|GAA91027.1 aluc|OJZ87830.1 aluc|OJZ89747.1 atub|OJI80811.1 atub|OJI85055.1 acla|EAW13670.1 afis|EAW19949.1 afla|EED45542.1 afum|EAL84966.1 anid|CBF80394.1 anig|EHA20309.1 aory|EIT75592.1 asyd|OJJ54728.1 ater|EAU31211.1 aver|OJJ07440.1 azon|OJJ45120.1 anig|EHA26970.1

Group489: abra|OJJ68479.1 abra|OJJ69702.1 akaw|GAA88821.1 akaw|GAA92402.1 aluc|OJZ82505.1 aluc|OJZ83263.1 atub|OJI87674.1 atub|OJI87926.1 acar|OOF92670.1 acla|EAW15184.1 afis|EAW20154.1 afla|EED57195.1 afum|EAL85269.1 agla|OJJ88775.1 anid|CBF84996.1 anig|EHA28265.1 aory|EIT74166.1 asyd|OJJ63119.1 ater|EAU38769.1 aver|OJI97839.1 awen|OJJ32030.1 azon|OJJ46082.1

Group490: abra|OJJ69764.1 abra|OJJ76049.1 acla|EAW08772.1 acla|EAW09780.1 aluc|OJZ85534.1 aluc|OJZ89505.1 ater|EAU29246.1 ater|EAU35201.1 acar|OOF99880.1 afis|EAW21358.1 afla|EED55773.1 afum|EAL92373.1 agla|OJJ85477.1 akaw|GAA88916.1 anid|CBF76263.1 anig|EHA23095.1 aory|EIT76750.1 asyd|OJJ60971.1 atub|OJI85298.1 aver|OJJ00296.1 awen|OJJ39340.1 azon|OJJ51686.1

Group491: abra|OJJ74560.1 abra|OJJ78116.1 asyd|OJJ52502.1 asyd|OJJ54384.1 atub|OJI83194.1 atub|OJI86158.1 acla|EAW12888.1 afis|EAW17788.1 afum|EAL91597.1 acar|OOF98752.1 afla|EED48713.1 agla|OJJ82644.1 akaw|GAA85891.1 aluc|OJZ91181.1 aluc|OJZ92144.1 anid|CBF84655.1 anig|EHA26805.1 aory|EIT73117.1 ater|EAU33264.1 awen|OJJ36279.1 anig|EHA19872.1 anig|EHA19902.1

Group492: acar|OOF93735.1 acar|OOF93823.1 abra|OJJ66740.1 acla|EAW07107.1 afis|EAW19478.1 afla|EED56686.1 afum|EAL87018.1 agla|OJJ78581.1 akaw|GAA89678.1 aluc|OJZ80708.1 anid|CBF69805.1 anig|EHA27309.1 aory|EIT73937.1 asyd|OJJ58969.1 ater|EAU30765.1 atub|OJI87566.1 aver|OJJ01564.1 awen|OJJ40135.1 azon|OJJ45831.1 afis|EAW22119.1 agla|OJJ81206.1 afum|EAL91067.1

Group493: acar|OOF95045.1 acar|OOF95112.1 abra|OJJ73294.1 acla|EAW14608.1 afis|EAW19337.1 afum|EAL94055.1 agla|OJJ79423.1 akaw|GAA92709.1 aluc|OJZ88753.1 anid|CBF90107.1 anig|EHA18454.1 asyd|OJJ59230.1 ater|EAU30249.1 atub|OJI91323.1 aver|OJJ07884.1 awen|OJJ40776.1 afis|EAW24675.1 afla|EED54362.1 asyd|OJJ57396.1 ater|EAU31824.1 aver|OJJ01840.1 asyd|OJJ53780.1

Group494: acar|OOF95079.1 acar|OOF95144.1 afla|EED48018.1 afla|EED48886.1 anig|EHA24784.1 anig|EHA26744.1 aory|EIT81173.1 aory|EIT82914.1 awen|OJJ31657.1 awen|OJJ37608.1 abra|OJJ74633.1 acla|EAW14723.1 afis|EAW24259.1 afum|EAL85927.2 akaw|GAA81976.1 aluc|OJZ92213.1 asyd|OJJ62240.1 ater|EAU31205.1 atub|OJI86644.1 aver|OJI96349.1 awen|OJJ34660.1 azon|OJJ51810.1

Group495: acar|OOG00243.1 acar|OOG00261.1 acar|OOG00244.1 acar|OOG00262.1 abra|OJJ67538.1 acla|EAW06974.1 afis|EAW16588.1 afla|EED56077.1 afum|EAL89183.1 agla|OJJ78537.1 akaw|GAA88121.1 aluc|OJZ81394.1 anid|CBF78530.1 anig|EHA18687.1 aory|EIT76791.1 asyd|OJJ65303.1 atub|OJI91599.1 aver|OJI99441.1 awen|OJJ37035.1 azon|OJJ49092.1 awen|OJJ37036.1 awen|OJJ40227.1

Group496: acla|EAW12269.1 acla|EAW15163.1 agla|OJJ78609.1 agla|OJJ88803.1 ater|EAU31506.1 ater|EAU38748.1 aver|OJI97813.1 aver|OJI98043.1 abra|OJJ71622.1 acar|OOF96339.1 afis|EAW20132.1 afla|EED57220.1 afum|EAL85248.1 akaw|GAA89303.1 aluc|OJZ86365.1 anid|CBF84960.1 anig|EHA21527.1 aory|EIT74164.1 asyd|OJJ63141.1 atub|OJI84169.1 awen|OJJ31998.1 azon|OJJ50773.1

Group497: afis|EAW15552.1 afis|EAW16853.1 anid|CBF75924.1 anid|CBF80262.1 anid|CBF83068.1 abra|OJJ72655.1 akaw|GAA83700.1 aluc|OJZ91032.1 aory|EIT81911.1 ater|EAU36759.1 atub|OJI83029.1 aver|OJJ04021.1 awen|OJJ30091.1 ater|EAU37833.1 aluc|OJZ90302.1 agla|OJJ89297.1 afla|EED47474.1 awen|OJJ38684.1 agla|OJJ80033.1 asyd|OJJ61018.1 ater|EAU29857.1 atub|OJI81728.1

Group498: afis|EAW16688.1 afis|EAW25102.1 ater|EAU31451.1 ater|EAU31645.1 ater|EAU39369.1 awen|OJJ29684.1 awen|OJJ39500.1 awen|OJJ40716.1 abra|OJJ70876.1 acar|OOF98244.1 acla|EAW10465.1 afla|EED55343.1 afum|EAL89773.1 agla|OJJ85448.1 akaw|GAA91739.1 aluc|OJZ87646.1 anid|CBF86019.1 anig|EHA20839.1 asyd|OJJ64452.1 atub|OJI80999.1 aver|OJI98555.1 azon|OJJ42298.1

Group499: afis|EAW17401.1 afis|EAW23311.1 agla|OJJ82761.1 agla|OJJ86536.1 asyd|OJJ52975.1 asyd|OJJ54697.1 asyd|OJJ62410.1 aver|OJI96543.1 aver|OJJ08941.1 abra|OJJ75086.1 acar|OOF97247.1 acla|EAW08278.1 afla|EED55666.1 afum|EAL88166.1 akaw|GAA88698.1 aluc|OJZ92652.1 anid|CBF89498.1 aory|EIT81808.1 ater|EAU35550.1 atub|OJI87083.1 awen|OJJ39656.1 azon|OJJ47784.1

Group500: afis|EAW18163.1 afis|EAW21088.1 afum|EAL87523.1 afum|EAL92111.1 abra|OJJ77694.1 acar|OOF99045.1 acla|EAW11332.1 afla|EED56330.1 agla|OJJ85962.1 akaw|GAA83936.1 aluc|OJZ90757.1 anid|CBF85753.1 anig|EHA19494.1 aory|EIT73367.1 asyd|OJJ64297.1 ater|EAU33690.1 atub|OJI82744.1 aver|OJI98390.1 awen|OJJ30508.1 azon|OJJ50317.1 acla|EAW10845.1 acla|EAW13632.1

Group501: afla|EED45701.1 afla|EED55177.1 aory|EIT75733.1 aory|EIT81570.1 ater|EAU31908.1 ater|EAU32716.1 azon|OJJ50388.1 azon|OJJ50758.1 abra|OJJ68251.1 acar|OOF97530.1 acla|EAW09063.1 afis|EAW25635.1 afum|EAL84323.1 agla|OJJ78897.1 akaw|GAA83210.1 aluc|OJZ83712.1 anid|CBF83153.1 anig|EHA18863.1 asyd|OJJ55515.1 atub|OJI90087.1 aver|OJJ06592.1 awen|OJJ36092.1

Group502: afla|EED49422.1 afla|EED56716.1 afla|EED53576.1 aory|EIT73869.1 aory|EIT74843.1 aory|EIT79474.1 abra|OJJ66759.1 acar|OOF93842.1 acla|EAW11885.1 afis|EAW19497.1 afum|EAL86996.1 agla|OJJ81661.1 akaw|GAA89696.1 aluc|OJZ80689.1 anid|CBF75434.1 anig|EHA27326.1 asyd|OJJ59785.1 ater|EAU34809.1 atub|OJI87548.1 aver|OJJ02503.1 awen|OJJ36914.1 azon|OJJ42897.1

Group503: afla|EED49854.1 afla|EED57953.1 asyd|OJJ60093.1 asyd|OJJ60346.1 ater|EAU29423.1 ater|EAU34802.1 aver|OJI97568.1 aver|OJI99687.1 aver|OJJ03906.1 aver|OJJ04180.1 aver|OJI98275.1 aver|OJJ03018.1 acla|EAW11879.1 afis|EAW19490.1 afla|EED45383.1 afum|EAL87004.1 anid|CBF87414.1 aory|EIT82446.1 aory|EIT83413.1 asyd|OJJ65387.1 awen|OJJ36906.1 azon|OJJ47353.1

Group504: afla|EED52027.1 afla|EED55465.1 aory|EIT80878.1 aory|EIT81484.1 awen|OJJ35115.1 awen|OJJ40599.1 abra|OJJ77085.1 acar|OOF92969.1 acla|EAW10573.1 afis|EAW24994.1 afum|EAL89655.1

agla|OJJ82261.1 akaw|GAA86970.1 aluc|OJZ84494.1 anid|CBF81588.1 anig|EHA27797.1 asyd|OJJ59513.1 ater|EAU35745.1 atub|OJI89586.1 aver|OJJ02760.1 azon|OJJ49449.1 azon|OJJ49978.1

Group505: afla|EED54894.1 afla|EED55162.1 aory|EIT79416.1 aory|EIT81659.1 abra|OJJ69688.1 acar|OOF93996.1 afis|EAW25764.1 akaw|GAA88812.1 aluc|OJZ82495.1 anig|EHA28277.1 aory|EIT79352.1 asyd|OJJ53577.1 atub|OJI87936.1 aver|OJJ08180.1 awen|OJJ29839.1 azon|OJJ45270.1 acla|EAW12187.1 afis|EAW17124.1 afum|EAL86585.1 anid|CBF83965.1 asyd|OJJ61426.1 aver|OJJ00760.1

Group506: afum|EAL86746.1 afum|EAL86767.1 afum|EAL90935.1 afum|EAL91121.1 awen|OJJ29553.1 awen|OJJ29590.1 awen|OJJ29640.1 abra|OJJ65612.1 afis|EAW15508.1 agla|OJJ81552.1 aver|OJJ03119.1 abra|OJJ66326.1 afis|EAW20713.1 afla|EED52357.1 agla|OJJ80478.1 akaw|GAA87890.1 anid|CBF84445.1 anig|EHA21856.1 asyd|OJJ52037.1 azon|OJJ42065.1 asyd|OJJ62703.1 asyd|OJJ51872.1

Group507: agla|OJJ78579.1 agla|OJJ86328.1 agla|OJJ87482.1 agla|OJJ87483.1 azon|OJJ42717.1 azon|OJJ45989.1 abra|OJJ77701.1 acar|OOF99039.1 acla|EAW09904.1 afis|EAW17120.1 afla|EED48004.1 akaw|GAA83929.1 aluc|OJZ90766.1 anig|EHA19500.1 aory|EIT82898.1 asyd|OJJ57236.1 ater|EAU33755.1 atub|OJI82754.1 aver|OJJ00941.1 awen|OJJ35399.1 anid|CBF87717.1 aver|OJJ03049.1

Group508: agla|OJJ79099.1 agla|OJJ80685.1 abra|OJJ65682.1 afis|EAW18187.1 afum|EAL87550.1 abra|OJJ67677.1 acar|OOF92161.1 afla|EED45129.1 agla|OJJ82590.1 akaw|GAA83877.1 aluc|OJZ81891.1 anig|EHA22351.1 ater|EAU32846.1 atub|OJI84444.1 afis|EAW17780.1 akaw|GAA89800.1 aver|OJJ08350.1 awen|OJJ31361.1 acla|EAW13858.1 anid|CBF83961.1 aory|EIT73733.1 ater|EAU34039.1

Group509: agla|OJJ79128.1 agla|OJJ84540.1 asyd|OJJ53810.1 asyd|OJJ58024.1 aver|OJJ04376.1 aver|OJJ07911.1 awen|OJJ31498.1 awen|OJJ36278.1 abra|OJJ68972.1 acar|OOF91549.1 acla|EAW14215.1 afis|EAW19066.1 afla|EED54432.1 afum|EAL93795.1 akaw|GAA90499.1 aluc|OJZ85961.1 anid|CBF79650.1 anig|EHA24642.1 aory|EIT75890.1 ater|EAU32114.1 atub|OJI79748.1 azon|OJJ46211.1

Group510: agla|OJJ81570.1 agla|OJJ84260.1 asyd|OJJ54207.1 asyd|OJJ61270.1 awen|OJJ30146.1 awen|OJJ34161.1 abra|OJJ78235.1 acar|OOF98625.1 acla|EAW09275.1 afis|EAW21872.1 afla|EED46436.1 afum|EAL92886.1 akaw|GAA85790.1 aluc|OJZ91309.1 anid|CBF76772.1 anig|EHA19993.1 aory|EIT72312.1 ater|EAU30870.1 atub|OJI83302.1 aver|OJJ00599.1 aver|OJJ01907.1 azon|OJJ43468.1

Group511: agla|OJJ82328.1 agla|OJJ89180.1 ater|EAU35772.1 ater|EAU39458.1 atub|OJI81101.1 atub|OJI87185.1 awen|OJJ30919.1 awen|OJJ40626.1 abra|OJJ70986.1 acar|OOF98145.1 acla|EAW10546.1 afis|EAW25014.1 afla|EED55435.1 afum|EAL89682.1 akaw|GAA82963.1 aluc|OJZ87544.1 anid|CBF81525.1 anig|EHA20748.1 aory|EIT80926.1 asyd|OJJ59480.1 aver|OJJ02794.1 azon|OJJ49057.1

Group512: agla|OJJ83608.1 agla|OJJ87442.1 anig|EHA21812.1 anig|EHA23178.1 ater|EAU35930.1 ater|EAU39415.1 awen|OJJ36034.1 awen|OJJ39401.1 abra|OJJ75961.1 acar|OOF99954.1 acla|EAW09837.1 afis|EAW21297.1 afla|EED54848.1 afum|EAL92314.1 akaw|GAA84661.1 aluc|OJZ89409.1 anid|CBF83385.1 aory|EIT79420.1 asyd|OJJ60632.1 atub|OJI85387.1 aver|OJI99955.1 azon|OJJ44285.1

Group513: akaw|GAA83114.1 akaw|GAA92362.1 aluc|OJZ79905.1 aluc|OJZ81127.1 aluc|OJZ88669.1 aluc|OJZ88850.1 abra|OJJ73371.1 acar|OOG00577.1 acla|EAW06992.1 afis|EAW16606.1 afla|EED45025.1 afum|EAL89203.1 agla|OJJ82647.1 anid|CBF81913.1 anig|EHA18352.1 aory|EIT73821.1 asyd|OJJ59694.1 ater|EAU36688.1 atub|OJI91251.1 aver|OJJ02593.1 awen|OJJ40200.1 azon|OJJ49067.1

Group514: akaw|GAA89013.1 akaw|GAA93063.1 akaw|GAA93173.1 aluc|OJZ79783.1 aluc|OJZ79855.1 aluc|OJZ80033.1 aluc|OJZ80379.1 acar|OOF93307.1 acla|EAW11112.1 afis|EAW25575.1 afla|EED45095.1 afum|EAL84285.2 agla|OJJ78554.1 anid|CBF80123.1 anig|EHA17692.1 aory|EIT73824.1 asyd|OJJ58342.1 ater|EAU31353.1 aver|OJJ02069.1 awen|OJJ38810.1 agla|OJJ85585.1 awen|OJJ40900.1

Group515: asyd|OJJ54189.1 asyd|OJJ55024.1 ater|EAU32854.1 ater|EAU36887.1 aver|OJJ07182.1 aver|OJJ08550.1 abra|OJJ69557.1 acar|OOF91593.1 acla|EAW12881.1 afis|EAW17796.1 afla|EED57033.1 afum|EAL91604.2 agla|OJJ89112.1 akaw|GAA91566.1 aluc|OJZ82384.1 anid|CBF71395.1 anig|EHA28379.1 aory|EIT82249.1 atub|OJI88056.1 awen|OJJ41608.1 azon|OJJ48327.1 agla|OJJ84736.1

Group516: asyd|OJJ54603.1 asyd|OJJ56125.1 asyd|OJJ60498.1 aver|OJI99822.1 aver|OJJ05067.1 aver|OJJ07552.1 abra|OJJ67246.1 acar|OOF98311.1 acla|EAW08524.1 afis|EAW23563.1 afum|EAL87905.1 agla|OJJ80849.1 akaw|GAA91666.1 aluc|OJZ87731.1 anid|CBF69379.1 anig|EHA20223.1 aory|EIT74378.1 ater|EAU37656.1 atub|OJI80909.1 awen|OJJ30069.1 awen|OJJ37809.1 anid|CBF78333.1

Group517: asyd|OJJ55069.1 asyd|OJJ55250.1 ater|EAU31464.1 ater|EAU32149.1 aver|OJJ06973.1 aver|OJJ07142.1 abra|OJJ77233.1 acar|OOF93115.1 acla|EAW12666.1 afis|EAW17830.1 afla|EED46172.1 afum|EAL91640.1 agla|OJJ80651.1 akaw|GAA90709.1 aluc|OJZ84643.1 anid|CBF77996.1 anig|EHA27929.1 aory|EIT74935.1 atub|OJI89438.1 awen|OJJ41759.1 azon|OJJ48451.1 afla|EED53219.1

Group518: aver|OJJ00309.1 aver|OJJ08547.1 abra|OJJ66043.1 afis|EAW24627.1 afum|EAL86253.1 abra|OJJ69961.1 acar|OOF99004.1 acla|EAW11755.1 afla|EED51597.1 agla|OJJ78815.1 akaw|GAA87698.1 aluc|OJZ85322.1 anid|CBF76281.1 anig|EHA25512.1 aory|EIT80452.1 asyd|OJJ60989.1 ater|EAU31328.1 atub|OJI80251.1 awen|OJJ31328.1 afis|EAW23788.1 afum|EAL87222.1 awen|OJJ33732.1

Group519: awen|OJJ30937.1 awen|OJJ35649.1 acar|OOF98703.1 afla|EED50782.1 aory|EIT78014.1 abra|OJJ75482.1 afis|EAW24629.1 agla|OJJ82330.1 anid|CBF87140.1 asyd|OJJ53426.1 ater|EAU36583.1 aver|OJJ08319.1 acar|OOF93299.1 anid|CBF82039.1 asyd|OJJ54222.1 aver|OJJ05174.1 anig|EHA20951.1 atub|OJI84677.1 acar|OOF99300.1 ater|EAU34863.1 ater|EAU32944.1 awen|OJJ32536.1

Group520: azon|OJJ42806.1 azon|OJJ43585.1 asyd|OJJ55440.1 abra|OJJ73119.1 acla|EAW13712.1 afum|EAL91905.2 aluc|OJZ80612.1 asyd|OJJ57580.1 aver|OJJ05600.1 abra|OJJ75285.1 acar|OOF91874.1 afla|EED47775.1 akaw|GAA91823.1 aluc|OJZ92846.1 anig|EHA26164.1 atub|OJI87302.1 awen|OJJ38175.1 akaw|GAA85218.1 aver|OJJ06515.1 afis|EAW18097.1 aory|EIT77197.1 aory|EIT77132.1

Group521: abra|OJJ65490.1 anig|EHA18426.1 atub|OJI91620.1 anid|CBF78136.1 abra|OJJ70409.1 acla|EAW12074.1 afis|EAW19715.1 afla|EED47481.1 afum|EAL86812.1 agla|OJJ83008.1 akaw|GAA91946.1 aluc|OJZ82672.1 anid|CBF82263.1 anig|EHA21039.1 aory|EIT79956.1 asyd|OJJ52475.1 atub|OJI88908.1 aver|OJJ03940.1 awen|OJJ33572.1 acar|OOF90658.1 afla|EED47480.1 aory|EIT79950.1

Group522: abra|OJJ65676.1 acar|OOF97626.1 afla|EED58063.1 agla|OJJ89300.1 aluc|OJZ79543.1 aory|EIT79786.1 ater|EAU33067.1 aver|OJJ03420.1 awen|OJJ35949.1 azon|OJJ46841.1 abra|OJJ72223.1 acar|OOF96846.1 afum|EAL86781.1 agla|OJJ85961.1 akaw|GAA84714.1 aluc|OJZ86973.1 anid|CBF77158.1 anig|EHA21675.1 asyd|OJJ56903.1 atub|OJI83548.1 aver|OJJ03390.1 awen|OJJ36080.1

Group523: abra|OJJ65756.1 acla|EAW07338.1 afis|EAW19764.1 afla|EED46307.1 afum|EAL84477.1 agla|OJJ89164.1 anid|CBF85553.1 aory|EIT76353.1 asyd|OJJ64152.1 ater|EAU36860.1 aver|OJI99267.1 abra|OJJ66793.1 acar|OOF93870.1 akaw|GAA86858.1 aluc|OJZ89981.1 anig|EHA27353.1 aory|EIT73825.1 atub|OJI84829.1 azon|OJJ50413.1 afum|EAL85138.1 aver|OJI98232.1 afla|EED44981.1

Group524: abra|OJJ65922.1 acar|OOF94983.1 afla|EED52093.1 aluc|OJZ90209.1 anig|EHA23706.1 atub|OJI81639.1 aver|OJJ07181.1 awen|OJJ29695.1 abra|OJJ68440.1 acar|OOF93953.1 afis|EAW19803.1 afla|EED52094.1 agla|OJJ87952.1 akaw|GAA86932.1 aluc|OJZ90057.1 anid|CBF82416.1 anig|EHA21014.1 aory|EIT76923.1 asyd|OJJ54150.1 ater|EAU30746.1 atub|OJI84750.1 aver|OJJ08591.1

Group525: asyd|OJJ65354.1 ater|EAU38218.1 awen|OJJ32091.1 azon|OJJ46668.1 abra|OJJ67699.1 afis|EAW16912.1 afla|EED51126.1 agla|OJJ81314.1 akaw|GAA89208.1 aluc|OJZ81621.1 anid|CBF84302.1 anig|EHA25822.1 aory|EIT81344.1 asyd|OJJ55578.1 atub|OJI88559.1 aver|OJJ06657.1 akaw|GAA90566.1 anid|CBF90346.1 ater|EAU37575.1 asyd|OJJ62616.1 aver|OJJ03451.1 aver|OJI99503.1

Group526: acla|EAW15302.1 anid|CBF84723.1 abra|OJJ76196.1 acar|OOF99748.1 afla|EED52713.1 agla|OJJ81243.1 akaw|GAA82138.1 aluc|OJZ92033.1 anig|EHA22966.1 aory|EIT78196.1 ater|EAU38283.1 atub|OJI86466.1 awen|OJJ32095.1 azon|OJJ46673.1 afis|EAW20227.1 afum|EAL85344.1 aver|OJJ08433.1 afis|EAW25742.1 afla|EED46022.1 aory|EIT75659.1 asyd|OJJ53309.1 awen|OJJ30032.1

Group527: abra|OJJ66248.1 acar|OOF90703.1 afis|EAW15427.1 afla|EED56771.1 afum|EAL84216.1 agla|OJJ83506.1 akaw|GAA89041.1 aluc|OJZ80554.1 anid|CBF78298.1 anig|EHA25643.1 aory|EIT73857.1 asyd|OJJ61726.1 ater|EAU33504.1 atub|OJI88377.1 aver|OJJ05045.1 awen|OJJ38047.1 atub|OJI82848.1 aver|OJJ05231.1 afla|EED52547.1 aory|EIT74224.1 asyd|OJJ56102.1 aluc|OJZ90847.1

Group528: abra|OJJ67067.1 acar|OOF90566.1 afis|EAW22299.1 afla|EED53134.1 afum|EAL91113.1 agla|OJJ82720.1 anid|CBF85181.1 anig|EHA18939.1 aory|EIT75127.1 ater|EAU32823.1 atub|OJI88353.1 aver|OJJ05787.1 awen|OJJ31414.1 abra|OJJ77976.1 akaw|GAA83682.1 aluc|OJZ91051.1 anig|EHA19748.1 atub|OJI83047.1 azon|OJJ42193.1 anid|CBF87385.1 akaw|GAA89027.1 aluc|OJZ80577.1

Group529: abra|OJJ67286.1 afla|EED54791.1 anid|CBF87758.1 anig|EHA25285.1 aory|EIT79425.1 asyd|OJJ63355.1 atub|OJI80526.1 aver|OJI97606.1 abra|OJJ70023.1 acar|OOF93529.1 afis|EAW15602.1 afla|EED50420.1 afum|EAL85731.1 agla|OJJ81554.1 akaw|GAA87638.1 aluc|OJZ85253.1 anid|CBF78955.1 anig|EHA25454.1 aory|EIT82173.1 ater|EAU32529.1 atub|OJI80315.1 awen|OJJ38274.1

Group530: abra|OJJ67324.1 acar|OOF92489.1 acla|EAW06411.1 afis|EAW18515.1 afla|EED52883.1 afum|EAL93210.1 agla|OJJ85106.1 akaw|GAA86492.1 aluc|OJZ85034.1 anid|CBF84636.1 anig|EHA25254.1 aory|EIT78633.1 atub|OJI80563.1 aver|OJJ08377.1 awen|OJJ31069.1 azon|OJJ50578.1 acar|OOF91935.1 afla|EED47190.1 akaw|GAA89092.1 anig|EHA25694.1 aory|EIT79183.1 atub|OJI88431.1

Group531: abra|OJJ67569.1 akaw|GAA89099.1 anig|EHA25701.1 aver|OJJ07705.1 abra|OJJ70824.1 acar|OOF91940.1 acla|EAW09022.1 afis|EAW17020.1 afla|EED57105.1 agla|OJJ89322.1 akaw|GAA89419.1 aluc|OJZ88254.1 anid|CBF86050.1 anig|EHA17999.1 aory|EIT74056.1 atub|OJI90852.1 awen|OJJ32256.1 azon|OJJ49710.1 acla|EAW10946.1 akaw|GAA89819.1 aluc|OJZ81871.1 aver|OJJ07786.1

Group532: abra|OJJ67571.1 acar|OOF91945.1 acla|EAW07104.1 afis|EAW17308.1 afla|EED57039.1 afum|EAL91143.1 agla|OJJ83702.1 akaw|GAA84673.1 aluc|OJZ87016.1 anid|CBF79774.1 aory|EIT77654.1 asyd|OJJ58117.1 atub|OJI88439.1 aver|OJJ04288.1 awen|OJJ30697.1 acar|OOG00400.1 afla|EED49713.1 agla|OJJ88113.1 asyd|OJJ58093.1 atub|OJI83503.1 asyd|OJJ53589.1 aver|OJJ08159.1

Group533: abra|OJJ67572.1 acar|OOF91946.1 afla|EED57712.1 agla|OJJ83703.1 akaw|GAA89101.1 anid|CBF79773.1 anig|EHA25703.1 aory|EIT79942.1 asyd|OJJ58116.1 ater|EAU33168.1 atub|OJI88440.1

aver|OJJ04289.1 awen|OJJ30698.1 abra|OJJ77771.1 acar|OOF93071.1 agla|OJJ86555.1 akaw|GAA83875.1 aluc|OJZ90821.1 anig|EHA19547.1 atub|OJI82809.1 azon|OJJ50365.1 azon|OJJ42498.1

Group534: abra|OJJ67769.1 acla|EAW13437.1 afis|EAW17522.1 agla|OJJ79313.1 aory|EIT83023.1 ater|EAU39522.1 awen|OJJ32751.1 azon|OJJ42601.1 abra|OJJ74180.1 acar|OOF95540.1 akaw|GAA86835.1 aluc|OJZ89954.1 anig|EHA27155.1 atub|OJI84852.1 abra|OJJ78121.1 akaw|GAA89008.1 aluc|OJZ80000.1 anig|EHA19877.1 atub|OJI83202.1 aver|OJI97603.1 azon|OJJ43572.1 awen|OJJ34683.1

Group535: abra|OJJ67872.1 acla|EAW11678.1 akaw|GAA91881.1 aluc|OJZ82018.1 anid|CBF73446.1 anig|EHA22238.1 atub|OJI84569.1 aver|OJJ08116.1 abra|OJJ69722.1 acar|OOF94194.1 afla|EED52162.1 akaw|GAA90175.1 aluc|OJZ83175.1 anig|EHA28249.1 aory|EIT74416.1 asyd|OJJ54298.1 atub|OJI87762.1 awen|OJJ40352.1 azon|OJJ42479.1 acar|OOF99262.1 acar|OOF99303.1 atub|OJI80754.1

Group536: abra|OJJ68322.1 acar|OOF97579.1 acla|EAW08933.1 afis|EAW15466.1 afla|EED51782.1 afum|EAL85856.1 akaw|GAA92073.1 aluc|OJZ83651.1 anid|CBF83126.1 anig|EHA18802.1 aory|EIT81734.1 asyd|OJJ55850.1 ater|EAU34052.1 atub|OJI90020.1 awen|OJJ29732.1 azon|OJJ48706.1 anig|EHA24765.1 afla|EED45870.1 aory|EIT75363.1 aver|OJJ04793.1 awen|OJJ38111.1 akaw|GAA83679.1

Group537: abra|OJJ68433.1 acar|OOF93957.1 afla|EED52099.1 agla|OJJ87949.1 akaw|GAA86938.1 aluc|OJZ90064.1 anid|CBF82384.1 anig|EHA21010.1 aory|EIT76917.1 asyd|OJJ54135.1 ater|EAU30750.1 atub|OJI84743.1 aver|OJJ08600.1 abra|OJJ68434.1 afla|EED52098.1 akaw|GAA86937.1 aluc|OJZ90063.1 anid|CBF82382.1 asyd|OJJ54136.1 atub|OJI84744.1 awen|OJJ29691.1 azon|OJJ48597.1

Group538: abra|OJJ68774.1 acar|OOF92259.1 acla|EAW12478.1 afla|EED46766.1 afum|EAL92024.1 agla|OJJ89513.1 akaw|GAA89917.1 aluc|OJZ81752.1 anid|CBF77878.1 anig|EHA22460.1 asyd|OJJ55313.1 ater|EAU30598.1 atub|OJI84323.1 aver|OJJ06913.1 awen|OJJ41310.1 azon|OJJ48783.1 azon|OJJ48784.1 acar|OOF92261.1 acla|EAW12477.1 agla|OJJ89512.1 akaw|GAA89918.1 ater|EAU30599.1

Group539: abra|OJJ69330.1 acar|OOF94397.1 acla|EAW14710.1 afis|EAW24245.1 afla|EED48029.1 afum|EAL85913.2 agla|OJJ79629.1 akaw|GAA88201.1 aluc|OJZ82162.1 anid|CBF86924.1 anig|EHA28570.1 aory|EIT83325.1 asyd|OJJ55608.1 ater|EAU32738.1 atub|OJI88275.1 aver|OJJ06683.1 awen|OJJ38017.1 azon|OJJ51787.1 asyd|OJJ53731.1 afis|EAW23670.1 afum|EAL87115.1 agla|OJJ81340.1

Group540: abra|OJJ70397.1 acar|OOF91879.1 acla|EAW07052.1 afis|EAW25614.1 afla|EED45236.1 agla|OJJ79504.1 akaw|GAA91961.1 aluc|OJZ82656.1 anig|EHA27417.1 aory|EIT73635.1 ater|EAU31421.1 atub|OJI88887.1 awen|OJJ37774.1 azon|OJJ44467.1 acar|OOF98993.1 agla|OJJ88672.1 asyd|OJJ54458.1 ater|EAU30186.1 aver|OJJ07698.1 awen|OJJ33156.1 afla|EED53590.1 aory|EIT75094.1

Group541: abra|OJJ70452.1 acar|OOF95031.1 acla|EAW08562.1 afis|EAW23604.1 afla|EED52317.1 afum|EAL86541.1 agla|OJJ83593.1 akaw|GAA90131.1 aluc|OJZ83218.1 anid|CBF81471.1 anig|EHA21076.1 aory|EIT74491.1 asyd|OJJ52470.1 atub|OJI87719.1 aver|OJJ03017.1 awen|OJJ36161.1 azon|OJJ45785.1 aver|OJJ03945.1 agla|OJJ79566.1 awen|OJJ39928.1 asyd|OJJ59447.1 aver|OJJ02825.1

Group542: abra|OJJ70674.1 acar|OOF97053.1 acla|EAW08090.1 afis|EAW23125.1 afla|EED48525.1 afum|EAL88355.1 agla|OJJ87945.1 akaw|GAA90006.1 aluc|OJZ84022.1 anid|CBF74410.1 anig|EHA19143.1 aory|EIT73109.1 asyd|OJJ57761.1 ater|EAU36253.1 atub|OJI90393.1 aver|OJJ05405.1 awen|OJJ36655.1 azon|OJJ47551.1 awen|OJJ33435.1 afla|EED47672.1 aory|EIT80214.1 ater|EAU37239.1

Group543: abra|OJJ71147.1 acar|OOF98008.1 acla|EAW10357.1 afis|EAW25237.1 afla|EED53704.1 afum|EAL89898.1 agla|OJJ89205.1 akaw|GAA82830.1 aluc|OJZ87391.1 anid|CBF85312.1 anig|EHA20620.1 aory|EIT75340.1 asyd|OJJ64005.1 ater|EAU34361.1 atub|OJI81256.1 aver|OJI98093.1 awen|OJJ40948.1 azon|OJJ44541.1 afis|EAW25236.1 afum|EAL89897.1 awen|OJJ40953.1 azon|OJJ44540.1

Group544: abra|OJJ72457.1 acar|OOF94781.1 acla|EAW12689.1 afis|EAW17853.1 afla|EED46207.1 afum|EAL91661.1 agla|OJJ83117.1 akaw|GAA86193.1 aluc|OJZ90443.1 anid|CBF77956.1 anig|EHA23907.1 aory|EIT76431.1 asyd|OJJ61702.1 ater|EAU33055.1 atub|OJI81872.1 aver|OJI95791.1 awen|OJJ41720.1 azon|OJJ48418.1 afla|EED45133.1 aory|EIT73615.1 awen|OJJ30412.1 agla|OJJ86835.1

Group545: abra|OJJ73099.1 acar|OOG00332.1 acla|EAW07049.1 afis|EAW16675.1 afla|EED49659.1 afum|EBA27373.1 agla|OJJ78688.1 akaw|GAA88064.1 aluc|OJZ81327.1 anid|CBF89880.1 anig|EHA18631.1 aory|EIT80967.1 asyd|OJJ54564.1 ater|EAU33734.1 atub|OJI91530.1 aver|OJJ07590.1 awen|OJJ40031.1 azon|OJJ50640.1 acla|EAW08376.1 afis|EAW23405.1 afum|EAL88065.2 ater|EAU37518.1

Group546: abra|OJJ74582.1 acar|OOF90133.1 acla|EAW13424.1 afis|EAW16727.1 afla|EED45977.1 afum|EAL86301.1 agla|OJJ78669.1 akaw|GAA82023.1 aluc|OJZ92160.1 anid|CBF85294.1 anig|EHA26789.1 aory|EIT75394.1 asyd|OJJ58681.1 ater|EAU38428.1 atub|OJI86588.1 aver|OJJ01293.1 azon|OJJ50620.1 acar|OOF98979.1 afla|EED51702.1 aory|EIT80478.1 aver|OJI99084.1 anid|CBF71192.1

Group547: abra|OJJ76094.1 acar|OOF99830.1 acar|OOF99831.1 acla|EAW09725.1 afis|EAW21416.1 afla|EED55842.1 afum|EAL92431.1 agla|OJJ85546.1 akaw|GAA88872.1 akaw|GAA88873.1 aluc|OJZ89550.1 anid|CBF83794.1 anig|EHA23056.1 aory|EIT76615.1 asyd|OJJ60885.1 ater|EAU35132.1 atub|OJI85255.1 atub|OJI85256.1 aver|OJJ00209.1 awen|OJJ39286.1 azon|OJJ51568.1 aory|EIT76662.1

Group548: abra|OJJ76170.1 acar|OOF99769.1 acla|EAW13698.1 afis|EAW21484.1 afla|EED55916.1 afum|EAL92504.1 agla|OJJ85638.1 akaw|GAA82116.1 aluc|OJZ92060.1 anid|CBF76494.1 anig|EHA22989.1 aory|EIT76674.1 asyd|OJJ61126.1 ater|EAU35054.1 atub|OJI86492.1 aver|OJJ00447.1 awen|OJJ39213.1 azon|OJJ51641.1 acla|EAW09655.1 afla|EED49764.1 aory|EIT77609.1 awen|OJJ29776.1

Group549: abra|OJJ66058.1 abra|OJJ74755.1 akaw|GAA83656.1 akaw|GAA93128.1 aluc|OJZ81105.1 aluc|OJZ92336.1 atub|OJI82473.1 atub|OJI86765.1 awen|OJJ33564.1 awen|OJJ36363.1 afla|EED47092.1 agla|OJJ80118.1 anig|EHA19264.1 ater|EAU30986.1 acar|OOF95748.1 anid|CBF84619.1 anig|EHA26635.1 aory|EIT72514.1 asyd|OJJ55640.1 aver|OJJ06723.1 aory|EIT72452.1

Group550: abra|OJJ68047.1 abra|OJJ70802.1 acar|OOF96908.1 acar|OOF96943.1 aory|EIT72863.1 aory|EIT81579.1 acla|EAW11122.1 afis|EAW15851.1 afla|EED56522.1 afum|EAL88487.1 agla|OJJ87257.1 akaw|GAA90118.1 aluc|OJZ84148.1 anid|CBF81176.1 anig|EHA21341.1 asyd|OJJ59292.1 ater|EAU31666.1 atub|OJI90520.1 aver|OJJ02983.1 awen|OJJ36776.1 azon|OJJ47422.1

Group551: abra|OJJ69621.1 abra|OJJ74815.1 abra|OJJ78091.1 afis|EAW21065.1 afis|EAW22323.1 afla|EED44992.1 afla|EED49898.1 akaw|GAA85057.1 akaw|GAA85907.1 aluc|OJZ86595.1 aluc|OJZ91160.1 anig|EHA19855.1 anig|EHA26580.1 aory|EIT72455.1 aory|EIT73686.1 atub|OJI83175.1 atub|OJI83928.1 acar|OOF98779.1 afum|EAL86268.1 aver|OJJ04473.1 awen|OJJ40160.1

Group552: abra|OJJ72113.1 abra|OJJ72712.1 anig|EHA21773.1 anig|EHA24109.1 atub|OJI82107.1 atub|OJI83652.1 acar|OOF96753.1 acla|EAW14914.1 afis|EAW24435.1 afla|EED55246.1 afum|EAL86103.1 agla|OJJ79250.1 akaw|GAA84809.1 aluc|OJZ86868.1 anid|CBF73765.1 aory|EIT83588.1 asyd|OJJ56590.1 ater|EAU30011.1 aver|OJJ06135.1 awen|OJJ32764.1 azon|OJJ43998.1

Group553: abra|OJJ73065.1 abra|OJJ73716.1 acar|OOG00294.1 acar|OOG00876.1 akaw|GAA88095.1 akaw|GAA88447.1 aluc|OJZ81361.1 aluc|OJZ88334.1 anig|EHA18070.1 anig|EHA18661.1 atub|OJI90930.1 atub|OJI91565.1 azon|OJJ45231.1 azon|OJJ46835.1 azon|OJJ51792.1 acla|EAW10825.1 afis|EAW24747.1 afla|EED47445.1 afum|EAL89394.1 aory|EIT81906.1 awen|OJJ29808.1

Group554: acar|OOF91686.1 acar|OOF91797.1 ater|EAU30834.1 ater|EAU37279.1 awen|OJJ37554.1 awen|OJJ37864.1 abra|OJJ66921.1 acla|EAW14487.1 afis|EAW19196.1 afla|EED47712.1 afum|EAL93928.1 agla|OJJ88511.1 akaw|GAA91498.1 aluc|OJZ85659.1 anid|CBF78550.1 anig|EHA24874.1 aory|EIT80197.1 asyd|OJJ65057.1 atub|OJI79452.1 aver|OJI99182.1 azon|OJJ51215.1

Group555: acar|OOF93691.1 acar|OOF93775.1 abra|OJJ66681.1 acla|EAW11818.1 afis|EAW19429.1 afla|EED56633.1 afum|EAL87065.1 agla|OJJ81141.1 akaw|GAA89626.1 aluc|OJZ80768.1 anid|CBF74384.1 anig|EHA27260.1 aory|EIT79639.1 asyd|OJJ57774.1 ater|EAU34732.1 atub|OJI87621.1 awen|OJJ36828.1 azon|OJJ42976.1 aver|OJJ05392.1 aver|OJJ05391.1 awen|OJJ36827.1

Group556: acar|OOF94034.1 acar|OOF94104.1 afis|EAW20233.1 afis|EAW24546.1 afum|EAL85350.1 afum|EAL86210.1 ater|EAU29965.1 ater|EAU37764.1 abra|OJJ66417.1 afla|EED55131.1 akaw|GAA90283.1 aluc|OJZ83057.1 anig|EHA28123.1 aory|EIT81603.1 asyd|OJJ56039.1 atub|OJI87890.1 aver|OJJ04985.1 awen|OJJ41447.1 azon|OJJ47374.1 abra|OJJ67779.1 ater|EAU37374.1

Group557: acar|OOG00254.1 acar|OOG01226.1 acar|OOG01227.1 abra|OJJ70128.1 acla|EAW13482.1 afis|EAW20590.1 afla|EED52238.1 afum|EAL84471.1 agla|OJJ88668.1 akaw|GAA86642.1 aluc|OJZ85199.1 anid|CBF85436.1 anig|EHA25406.1 aory|EIT74451.1 asyd|OJJ64088.1 ater|EAU35229.1 atub|OJI80402.1 aver|OJI98172.1 awen|OJJ36225.1 acar|OOF91412.1 awen|OJJ33308.1

Group558: acla|EAW09303.1 acla|EAW15329.1 afum|EAL85377.2 afum|EAL92855.1 asyd|OJJ53789.1 asyd|OJJ61241.1 abra|OJJ75519.1 acar|OOF90846.1 afis|EAW21843.1 afla|EED46527.1 agla|OJJ84291.1 akaw|GAA84283.1 aluc|OJZ88976.1 anid|CBF76715.1 anig|EHA23549.1 aory|EIT72431.1 ater|EAU38520.1 atub|OJI85813.1 awen|OJJ34127.1 azon|OJJ44216.1 azon|OJJ51717.1

Group559: acla|EAW09341.1 acla|EAW11685.1 afis|EAW21796.1 afis|EAW23700.1 afum|EAL87141.1 afum|EAL92821.1 abra|OJJ75576.1 acar|OOF90894.1 afla|EED46599.1 agla|OJJ84330.1 akaw|GAA84328.1 aluc|OJZ89025.1 anid|CBF76627.1 anig|EHA23507.1 aory|EIT77837.1 asyd|OJJ61195.1 ater|EAU38478.1 atub|OJI85765.1 aver|OJJ00519.1 awen|OJJ34081.1 azon|OJJ44259.1

Group560: afis|EAW16155.1 afis|EAW23200.1 afum|EAL88027.1 afum|EAL88283.1 ater|EAU36328.1 ater|EAU36668.1 abra|OJJ70513.1 acar|OOF97416.1 acla|EAW08162.1 afla|EED47054.1 agla|OJJ78748.1 akaw|GAA83360.1 aluc|OJZ83858.1 anid|CBF74719.1 anig|EHA18999.1 aory|EIT72491.1 asyd|OJJ60202.1 atub|OJI90225.1 aver|OJJ04064.1 awen|OJJ39430.1 azon|OJJ47634.1

Group561: afis|EAW17104.1 afis|EAW25308.1 agla|OJJ88961.1 agla|OJJ89287.1 abra|OJJ71251.1 acar|OOF97919.1 acla|EAW10284.1 afum|EAL89971.1 akaw|GAA82736.1 aluc|OJZ87291.1 anid|CBF75291.1 anig|EHA20532.1 aory|EIT78474.1 asyd|OJJ59887.1 ater|EAU34456.1 atub|OJI81350.1 aver|OJJ02399.1 awen|OJJ41022.1 azon|OJJ44609.1 asyd|OJJ65171.1 afla|EED53785.1

Group562: afis|EAW19712.1 afis|EAW25702.1 afum|EAL84375.1 afum|EAL86815.1 abra|OJJ77041.1 acar|OOF90274.1 afla|EED50904.1 akaw|GAA87011.1 aluc|OJZ84444.1 anig|EHA27762.1 aory|EIT79058.1

asyd|OJJ63378.1 ater|EAU32932.1 atub|OJI89630.1 aver|OJJ05795.1 awen|OJJ39868.1 anid|CBF81946.1 agla|OJJ84461.1 aver|OJI97557.1 ater|EAU34785.1 asyd|OJJ57373.1

Group563: afis|EAW22109.1 afis|EAW22582.1 afum|EAL90617.1 afum|EAL91075.1 abra|OJJ73636.1 acar|OOG00816.1 acla|EAW07548.1 afla|EED54627.1 agla|OJJ80942.1 akaw|GAA88519.1 aluc|OJZ88414.1 anid|CBF88345.1 anig|EHA18141.1 aory|EIT78908.1 asyd|OJJ63718.1 ater|EAU39122.1 atub|OJI91010.1 aver|OJI97222.1 awen|OJJ35210.1 azon|OJJ49899.1 anid|CBF89563.1

Group564: afla|EED44994.1 afla|EED49089.1 asyd|OJJ52855.1 asyd|OJJ55885.1 aver|OJJ04830.1 aver|OJJ08832.1 awen|OJJ36406.1 awen|OJJ39156.1 abra|OJJ70006.1 acar|OOF93512.1 acla|EAW15361.1 agla|OJJ83218.1 akaw|GAA87655.1 aluc|OJZ85273.1 anid|CBF78094.1 anig|EHA25471.1 aory|EIT82516.1 ater|EAU34442.1 atub|OJI80297.1 aory|EIT73678.1 aver|OJJ01999.1

Group565: afla|EED45878.1 afla|EED45984.1 afla|EED47653.1 afum|EAL84420.1 afum|EAL86318.1 aory|EIT75536.1 aory|EIT75748.1 aory|EIT80201.1 acar|OOF99716.1 acla|EAW12434.1 afis|EAW16743.1 akaw|GAA82399.1 aluc|OJZ91723.1 anid|CBF89574.1 anig|EHA22730.1 asyd|OJJ62337.1 ater|EAU30784.1 atub|OJI86180.1 aver|OJI96442.1 awen|OJJ29697.1 anid|CBF83502.1

Group566: afla|EED45981.1 afla|EED55382.1 aory|EIT75435.1 aory|EIT80896.1 ater|EAU33990.1 ater|EAU39553.1 abra|OJJ70929.1 acar|OOF98196.1 acla|EAW10495.1 afis|EAW25072.1 afum|EAL89741.1 agla|OJJ89122.1 akaw|GAA83013.1 aluc|OJZ87598.1 anid|CBF85960.1 anig|EHA20798.1 asyd|OJJ64420.1 atub|OJI81049.1 aver|OJI98519.1 awen|OJJ40683.1 azon|OJJ44060.1

Group567: afla|EED48911.1 afla|EED51260.1 aory|EIT81176.1 aory|EIT81506.1 awen|OJJ29671.1 awen|OJJ38491.1 abra|OJJ69199.1 acar|OOF91116.1 acla|EAW08647.1 afis|EAW15784.1 afum|EAL85566.1 agla|OJJ81947.1 akaw|GAA92212.1 aluc|OJZ86184.1 anid|CBF70950.1 anig|EHA24440.1 asyd|OJJ52622.1 ater|EAU32357.1 atub|OJI79976.1 aver|OJJ04649.1 azon|OJJ43792.1

Group568: afla|EED51201.1 afla|EED53566.1 asyd|OJJ62648.1 asyd|OJJ65376.1 aver|OJI96748.1 aver|OJI99544.1 abra|OJJ73167.1 acar|OOG00407.1 acla|EAW06285.1 afis|EAW22140.1 afum|EAL91050.1 agla|OJJ87935.1 akaw|GAA88004.1 aluc|OJZ81260.1 anid|CBF89140.1 anig|EHA18567.1 aory|EIT75205.1 ater|EAU34142.1 atub|OJI91456.1 awen|OJJ34714.1 azon|OJJ50465.1

Group569: afum|EAL87769.1 afum|EAL87803.1 akaw|GAA92345.1 akaw|GAA93055.1 akaw|GAA92650.1 akaw|GAA93090.1 awen|OJJ29538.1 awen|OJJ29541.1 abra|OJJ65531.1 acla|EAW13472.1 aluc|OJZ79518.1 anid|CBF73631.1 anig|EHA21825.1 abra|OJJ65574.1 agla|OJJ85769.1 abra|OJJ65598.1 ater|EAU30115.1 acar|OOF90498.1 ater|EAU30117.1 acla|EAW11437.1 anig|EHA21861.1

Group570: agla|OJJ78645.1 agla|OJJ81233.1 agla|OJJ83403.1 abra|OJJ71870.1 acar|OOF96539.1 acla|EAW13900.1 afis|EAW18612.1 afla|EED52733.1 afum|EAL93322.1 akaw|GAA85045.1 aluc|OJZ86609.1 anid|CBF70422.1 anig|EHA22057.1 aory|EIT78158.1 asyd|OJJ58619.1 ater|EAU34909.1 atub|OJI83917.1 aver|OJJ01226.1 awen|OJJ32073.1 azon|OJJ46654.1 agla|OJJ81232.1

Group571: agla|OJJ82854.1 agla|OJJ88960.1 abra|OJJ71795.1 acar|OOF96475.1 acla|EAW12077.1 afis|EAW19718.1 afla|EED52815.1 afum|EAL86808.1 akaw|GAA85115.1 aluc|OJZ86525.1 anid|CBF78801.1 anig|EHA22121.1 aory|EIT78250.1 asyd|OJJ55181.1 ater|EAU34844.1 atub|OJI83997.1 aver|OJJ07026.1 acar|OOG00120.1 anid|CBF84151.1 asyd|OJJ65073.1 aver|OJI99198.1

Group572: agla|OJJ83728.1 agla|OJJ88980.1 abra|OJJ65910.1 acar|OOF90095.1 afla|EED53605.1 afum|EAL88809.2 akaw|GAA92322.1 aluc|OJZ90220.1 anid|CBF70180.1 anig|EHA23718.1 aory|EIT75102.1 atub|OJI81652.1 aver|OJJ03066.1 abra|OJJ66145.1 afis|EAW16403.1 anid|CBF86674.1 aver|OJJ05884.1 awen|OJJ29890.1 afis|EAW16176.1 aver|OJJ01772.1 asyd|OJJ52559.1

Group573: akaw|GAA82012.1 akaw|GAA83300.1 anig|EHA23471.1 anig|EHA26778.1 azon|OJJ46821.1 azon|OJJ49225.1 abra|OJJ74596.1 acar|OOF95173.1 acla|EAW10785.1 afis|EAW24793.1 afla|EED47193.1 afum|EAL89435.1 agla|OJJ82887.1 aluc|OJZ92171.1 anid|CBF81462.1 aory|EIT79250.1 asyd|OJJ59441.1 ater|EAU36515.1 atub|OJI86603.1 aver|OJJ02832.1 awen|OJJ40374.1

Group574: aluc|OJZ79634.1 aluc|OJZ79637.1 aluc|OJZ79681.1 aluc|OJZ90921.1 abra|OJJ77846.1 acar|OOF98926.1 acla|EAW11277.1 afis|EAW18223.1 afla|EED56246.1 afum|EAL87589.2 agla|OJJ85859.1 akaw|GAA83801.1 anid|CBF86151.1 anig|EHA19633.1 aory|EIT73368.1 asyd|OJJ64517.1 ater|EAU33768.1 atub|OJI82913.1 aver|OJI98615.1 awen|OJJ30601.1 azon|OJJ50167.1

Group575: aluc|OJZ79645.1 aluc|OJZ80467.1 anid|CBF81159.1 anid|CBF87224.1 anid|CBF82244.1 agla|OJJ86031.1 akaw|GAA88990.1 awen|OJJ36065.1 abra|OJJ70212.1 afla|EED52528.1 akaw|GAA86564.1 anig|EHA25334.1 aory|EIT82764.1 atub|OJI80480.1 acar|OOG00877.1 asyd|OJJ57261.1 aver|OJJ03035.1 azon|OJJ47412.1 awen|OJJ30459.1 azon|OJJ45449.1 aluc|OJZ85114.1

Group576: aluc|OJZ80315.1 aluc|OJZ86416.1 anid|CBF76119.1 anid|CBF85040.1 atub|OJI80382.1 atub|OJI84115.1 abra|OJJ71674.1 acar|OOF96385.1 acla|EAW15213.1 afis|EAW20178.1 afla|EED57170.1 afum|EAL85296.1 agla|OJJ84688.1 akaw|GAA89258.1 anig|EHA21483.1 aory|EIT74051.1 asyd|OJJ63096.1 ater|EAU35214.1 aver|OJI97866.1 awen|OJJ31177.1 azon|OJJ46108.1

Group577: aluc|OJZ87121.1 aluc|OJZ90552.1 anig|EHA20385.1 anig|EHA23999.1 atub|OJI81520.1 atub|OJI81975.1 awen|OJJ29935.1 awen|OJJ40118.1 awen|OJJ31429.1 awen|OJJ33906.1 abra|OJJ70173.1 acar|OOF90667.1 afla|EED52274.1 akaw|GAA82584.1 asyd|OJJ53431.1 aver|OJJ04439.1 azon|OJJ42631.1 abra|OJJ72571.1 aory|EIT80573.1 afla|EED51481.1 aory|EIT83049.1

Group578: aluc|OJZ89485.1 aluc|OJZ91533.1 asyd|OJJ52882.1 asyd|OJJ54759.1 atub|OJI85315.1 atub|OJI85987.1 abra|OJJ76672.1 acar|OOF99337.1 afla|EED47468.1 akaw|GAA92520.1 anid|CBF87401.1 anig|EHA18623.1 aory|EIT79967.1 ater|EAU31454.1 aver|OJJ08854.1 awen|OJJ38083.1 agla|OJJ86847.1 ater|EAU36924.1 awen|OJJ41649.1 aver|OJJ00789.1 aver|OJJ07411.1

Group579: anid|CBF70118.1 anid|CBF76209.1 awen|OJJ38751.1 awen|OJJ40905.1 abra|OJJ66448.1 acla|EAW11664.1 afis|EAW15397.1 agla|OJJ79144.1 asyd|OJJ56278.1 ater|EAU32821.1 aver|OJJ08485.1 awen|OJJ37492.1 abra|OJJ67824.1 acar|OOF93612.1 akaw|GAA91839.1 anig|EHA22193.1 atub|OJI84618.1 acla|EAW11665.1 awen|OJJ38753.1 aver|OJJ05914.1 aluc|OJZ82059.1

Group580: anid|CBF71344.1 anid|CBF71770.1 asyd|OJJ55097.1 asyd|OJJ55529.1 aver|OJJ07109.1 aver|OJJ07313.1 abra|OJJ72319.1 acar|OOF94892.1 acla|EAW12001.1 afis|EAW19641.1 afla|EED56992.1 afum|EAL86881.1 agla|OJJ80783.1 akaw|GAA86071.1 aluc|OJZ90306.1 anig|EHA23792.1 aory|EIT77578.1 ater|EAU32888.1 atub|OJI81730.1 awen|OJJ41886.1 azon|OJJ50727.1

Group581: anid|CBF80560.1 anid|CBF82261.1 abra|OJJ66944.1 akaw|GAA91476.1 aluc|OJZ85637.1 atub|OJI79427.1 abra|OJJ71064.1 acar|OOF90585.1 acla|EAW07015.1 afis|EAW25726.1 afla|EED53906.1 aory|EIT78492.1 asyd|OJJ52818.1 ater|EAU32822.1 aver|OJJ07278.1 awen|OJJ38756.1 azon|OJJ42176.1 aver|OJJ08797.1 awen|OJJ32537.1 afum|EAL87822.1 acla|EAW11668.1

Group582: asyd|OJJ52091.1 asyd|OJJ52556.1 asyd|OJJ55512.1 asyd|OJJ65388.1 abra|OJJ68248.1 acar|OOF97527.1 acla|EAW09059.1 afis|EAW25632.1 afla|EED55180.1 afum|EAL84320.1 agla|OJJ78899.1 akaw|GAA83213.1 aluc|OJZ83715.1 anid|CBF83158.1 anig|EHA18866.1 aory|EIT81642.1 ater|EAU32718.1 atub|OJI90091.1 aver|OJJ06589.1 awen|OJJ36052.1 aver|OJJ03845.1

Group583: asyd|OJJ52420.1 asyd|OJJ54955.1 abra|OJJ76861.1 acar|OOF91348.1 acla|EAW13422.1 afis|EAW20534.1 afla|EED52483.1 afum|EAL84524.1 agla|OJJ84698.1 akaw|GAA87161.1 aluc|OJZ84273.1 anid|CBF71440.1 anig|EHA27615.1 aory|EIT82754.1 asyd|OJJ55687.1 ater|EAU37151.1 atub|OJI89810.1 aver|OJJ06771.1 aver|OJJ07241.1 awen|OJJ33240.1 anid|CBF78998.1

Group584: asyd|OJJ53054.1 asyd|OJJ62331.1 aver|OJI96437.1 aver|OJJ03790.1 abra|OJJ67212.1 acar|OOF98347.1 acla|EAW08554.1 afis|EAW23596.1 afla|EED55272.1 afum|EAL87871.1 agla|OJJ86856.1 akaw|GAA90965.1 aluc|OJZ87761.1 anid|CBF89583.1 anig|EHA20246.1 aory|EIT83575.1 ater|EAU35356.1 atub|OJI80883.1 awen|OJJ39918.1 azon|OJJ45892.1 anid|CBF77738.1

Group585: asyd|OJJ53773.1 asyd|OJJ64196.1 aver|OJI98285.1 aver|OJJ01986.1 abra|OJJ71111.1 acar|OOF98036.1 acla|EAW10381.1 afis|EAW25212.1 afla|EED53666.1 afum|EAL89873.1 agla|OJJ88949.1 akaw|GAA82858.1 aluc|OJZ87423.1 anid|CBF85604.1 anig|EHA19933.1 anig|EHA20646.1 aory|EIT75278.1 ater|EAU34369.1 atub|OJI81225.1 awen|OJJ40874.1 azon|OJJ46813.1

Group586: asyd|OJJ54182.1 asyd|OJJ55740.1 aver|OJJ06826.1 aver|OJJ08558.1 abra|OJJ76808.1 acar|OOF91302.1 acla|EAW13368.1 afis|EAW20482.1 afla|EED44786.1 afum|EAL84572.1 akaw|GAA87214.1 aluc|OJZ84216.1 anid|CBF79229.1 anig|EHA27568.1 aory|EIT73548.1 aory|EIT77772.1 ater|EAU29560.1 atub|OJI89868.1 awen|OJJ33180.1 azon|OJJ46383.1 afla|EED45102.1

Group587: asyd|OJJ54581.1 asyd|OJJ54583.1 asyd|OJJ59186.1 awen|OJJ29889.1 awen|OJJ35053.1 awen|OJJ38450.1 azon|OJJ44478.1 azon|OJJ45681.1 abra|OJJ67703.1 acar|OOF97596.1 agla|OJJ81404.1 akaw|GAA89211.1 aluc|OJZ81618.1 anig|EHA25825.1 ater|EAU38935.1 atub|OJI88562.1 acar|OOG00088.1 afla|EED51583.1 aory|EIT80517.1 afis|EAW20205.1 afla|EED51443.1

Group588: asyd|OJJ60296.1 asyd|OJJ62116.1 aver|OJI96223.1 aver|OJI99634.1 aver|OJJ02653.1 aver|OJJ06318.1 abra|OJJ68285.1 acar|OOF90110.1 afis|EAW22106.1 afla|EED55144.1 afum|EAL91085.1 agla|OJJ82696.1 akaw|GAA83187.1 aluc|OJZ83686.1 anid|CBF87172.1 anig|EHA18840.1 aory|EIT81583.1 asyd|OJJ56395.1 ater|EAU32730.1 atub|OJI90057.1 awen|OJJ40235.1

Group589: ater|EAU35203.1 ater|EAU36444.1 abra|OJJ76030.1 acar|OOF99899.1 acla|EAW09783.1 afis|EAW21355.1 afla|EED55770.1 afum|EAL92370.2 agla|OJJ87376.1 aluc|OJZ89481.1 anid|CBF76251.1 anig|EHA23112.1 aory|EIT76609.1 asyd|OJJ60968.1 atub|OJI85319.1 aver|OJJ00292.1 awen|OJJ39343.1 azon|OJJ51689.1 afla|EED53075.1 aory|EIT74793.1 asyd|OJJ59168.1

Group590: azon|OJJ46530.1 azon|OJJ50393.1 acla|EAW13391.1 afla|EED49406.1 agla|OJJ78968.1 aory|EIT79482.1 asyd|OJJ52710.1 aver|OJJ04739.1 awen|OJJ35362.1 acar|OOF92857.1 afis|EAW24526.1 agla|OJJ81446.1 akaw|GAA82748.1 afum|EAL86190.1 ater|EAU37241.1 atub|OJI83744.1 afis|EAW22564.1 afum|EAL94067.1 agla|OJJ81434.1 aluc|OJZ87304.1 ater|EAU36467.1

Group591: abra|OJJ65586.1 acla|EAW06849.1 afum|EAL86639.1 anid|CBF87353.1 asyd|OJJ53531.1 ater|EAU31407.1 aver|OJJ08219.1 abra|OJJ68254.1 acar|OOF97533.1 afla|EED51066.1 akaw|GAA83207.1

aluc|OJZ83709.1 anig|EHA18860.1 aory|EIT77028.1 ater|EAU32153.1 atub|OJI90081.1 aver|OJJ00862.1 awen|OJJ38280.1 azon|OJJ43291.1 asyd|OJJ61534.1 azon|OJJ43503.1

Group592: abra|OJJ65854.1 acar|OOF95015.1 acla|EAW13528.1 afis|EAW20729.1 afla|EED48906.1 afum|EAL84879.1 agla|OJJ84616.1 akaw|GAA92290.1 aluc|OJZ90187.1 anid|CBF85584.1 anig|EHA23686.1 aory|EIT81194.1 asyd|OJJ64174.1 atub|OJI81614.1 aver|OJJ06463.1 awen|OJJ29911.1 azon|OJJ42620.1 afla|EED55134.1 aory|EIT81587.1 ater|EAU32788.1 aver|OJI98249.1

Group593: abra|OJJ66409.1 acar|OOF97558.1 acla|EAW15053.1 afla|EED47219.1 aluc|OJZ81110.1 anid|CBF76141.1 aory|EIT79310.1 ater|EAU32457.1 aver|OJJ03023.1 awen|OJJ37014.1 abra|OJJ67968.1 afis|EAW17012.1 afla|EED55205.1 agla|OJJ86640.1 aory|EIT83525.1 azon|OJJ42140.1 aory|EIT79570.1 akaw|GAA86290.1 awen|OJJ29897.1 aver|OJJ06536.1 asyd|OJJ54884.1

Group594: abra|OJJ66784.1 afla|EED47234.1 anid|CBF87107.1 aory|EIT79229.1 abra|OJJ75747.1 acar|OOG00125.1 acla|EAW09092.1 afla|EED52418.1 agla|OJJ82218.1 akaw|GAA84480.1 aluc|OJZ89200.1 anig|EHA23357.1 aory|EIT83316.1 asyd|OJJ56738.1 ater|EAU31807.1 atub|OJI85590.1 aver|OJJ05987.1 awen|OJJ41471.1 azon|OJJ44445.1 afum|EAL84355.1 afis|EAW25686.1

Group595: abra|OJJ67354.1 akaw|GAA86466.1 aluc|OJZ85007.1 asyd|OJJ58136.1 ater|EAU35299.1 atub|OJI80590.1 aver|OJJ04269.1 awen|OJJ33479.1 abra|OJJ69839.1 akaw|GAA87821.1 aluc|OJZ85457.1 anid|CBF84465.1 anig|EHA24999.1 atub|OJI80106.1 azon|OJJ47408.1 afla|EED57088.1 aory|EIT82252.1 awen|OJJ29832.1 asyd|OJJ53544.1 aver|OJJ08209.1 awen|OJJ37797.1

Group596: abra|OJJ67576.1 acla|EAW07712.1 afis|EAW19707.1 afla|EED52185.1 afum|EAL86292.2 agla|OJJ80363.1 akaw|GAA89105.1 anid|CBF82328.1 aory|EIT74319.1 asyd|OJJ59047.1 ater|EAU32212.1 atub|OJI88444.1 aver|OJJ01643.1 awen|OJJ35443.1 azon|OJJ49488.1 asyd|OJJ52892.1 aver|OJJ08863.1 afum|EBA27291.1 afis|EAW25591.1 awen|OJJ40142.1 agla|OJJ81186.1

Group597: abra|OJJ67718.1 afis|EAW16742.1 afla|EED49725.1 afum|EAL86317.2 aory|EIT77616.1 abra|OJJ76618.1 akaw|GAA86494.1 aluc|OJZ85036.1 anig|EHA22616.1 aory|EIT76392.1 asyd|OJJ54900.1 ater|EAU30402.1 atub|OJI86041.1 awen|OJJ32259.1 asyd|OJJ60225.1 atub|OJI80561.1 aver|OJJ04042.1 afla|EED46080.1 afla|EED46081.1 agla|OJJ80285.1 awen|OJJ32907.1

Group598: abra|OJJ67990.1 acar|OOF92032.1 afis|EAW22103.1 afla|EED45380.1 akaw|GAA90958.1 aluc|OJZ82864.1 anig|EHA21294.1 aory|EIT82399.1 atub|OJI89089.1 awen|OJJ36170.1 abra|OJJ71233.1 akaw|GAA82755.1 aluc|OJZ87312.1 anid|CBF84724.1 anig|EHA20548.1 asyd|OJJ53307.1 atub|OJI81333.1 aver|OJJ08434.1 azon|OJJ43637.1 abra|OJJ76888.1 afis|EAW16705.1

Group599: abra|OJJ68494.1 acar|OOF98749.1 acla|EAW12142.1 afis|EAW17194.1 afla|EED45176.1 afum|EAL86622.1 agla|OJJ86660.1 akaw|GAA92776.1 aluc|OJZ82707.1 anid|CBF69890.1 anig|EHA21162.1 aory|EIT73731.1 asyd|OJJ59007.1 ater|EAU34144.1 atub|OJI88940.1 aver|OJJ01608.1 awen|OJJ34716.1 agla|OJJ87933.1 anid|CBF89139.1 asyd|OJJ62650.1 aver|OJI96750.1

Group600: abra|OJJ69337.1 acar|OOF94389.1 acla|EAW06601.1 afis|EAW16003.1 afla|EED48032.1 afum|EAL88635.2 agla|OJJ79631.1 akaw|GAA88207.1 aluc|OJZ82169.1 anid|CBF85846.1 anig|EHA28563.1 aory|EIT83338.1 asyd|OJJ53958.1 ater|EAU31185.1 atub|OJI88268.1 aver|OJJ08054.1 awen|OJJ33053.1 azon|OJJ48464.1 asyd|OJJ64357.1 ater|EAU33640.1 aver|OJI98453.1

Group601: abra|OJJ69426.1 acar|OOF94310.1 acla|EAW08550.1 afis|EAW23591.1 afum|EAL87875.1 akaw|GAA88292.1 aluc|OJZ82263.1 anid|CBF73805.1 anig|EHA28480.1 asyd|OJJ56617.1 aver|OJJ06107.1 awen|OJJ35870.1 abra|OJJ69775.1 acar|OOF93598.1 afla|EED45633.1 akaw|GAA87878.1 aluc|OJZ85519.1 anig|EHA25052.1 aory|EIT75817.1 atub|OJI80047.1 azon|OJJ47676.1

Group602: abra|OJJ69887.1 acar|OOF93848.1 acla|EAW11893.1 afis|EAW19504.1 afla|EED46651.1 agla|OJJ81680.1 akaw|GAA87765.1 aluc|OJZ85400.1 anid|CBF87939.1 anig|EHA25582.1 aory|EIT77814.1 asyd|OJJ63469.1 ater|EAU38715.1 atub|OJI80171.1 aver|OJI97463.1 awen|OJJ36926.1 azon|OJJ45056.1 akaw|GAA89704.1 aluc|OJZ80680.1 anig|EHA27332.1 atub|OJI87539.1

Group603: abra|OJJ70214.1 acar|OOF92565.1 afla|EED46373.1 akaw|GAA86562.1 aluc|OJZ85112.1 anid|CBF78840.1 anig|EHA25332.1 aory|EIT77546.1 ater|EAU29773.1 atub|OJI80482.1 azon|OJJ45196.1 afis|EAW23745.1 anid|CBF78842.1 afla|EED52200.1 asyd|OJJ65270.1 aver|OJI99422.1 awen|OJJ33680.1 akaw|GAA93170.1 aluc|OJZ83029.1 aory|EIT74347.1 afla|EED56929.1

Group604: abra|OJJ70243.1 anig|EHA25313.1 abra|OJJ71272.1 acar|OOF92553.1 afis|EAW17415.1 afla|EED48947.1 agla|OJJ88526.1 akaw|GAA82718.1 aluc|OJZ87272.1 anid|CBF75478.1 anig|EHA20513.1 asyd|OJJ63978.1 ater|EAU39296.1 atub|OJI81370.1 aver|OJI96975.1 awen|OJJ31635.1 azon|OJJ46586.1 afla|EED48946.1 agla|OJJ87753.1 aory|EIT79982.1 aory|EIT80005.1

Group605: abra|OJJ70312.1 acar|OOF93014.1 acla|EAW13439.1 afis|EAW20553.1 afla|EED45906.1 afum|EAL84508.1 agla|OJJ80823.1 akaw|GAA93011.1 aluc|OJZ82574.1 anid|CBF78981.1 anig|EHA27499.1 aory|EIT75600.1 asyd|OJJ55678.1 ater|EAU37124.1 atub|OJI88818.1 aver|OJJ06761.1 awen|OJJ41921.1 azon|OJJ42685.1 anig|EHA23882.1 afla|EED52345.1 aory|EIT74410.1

Group606: abra|OJJ70650.1 acar|OOF97072.1 acla|EAW08109.1 afis|EAW23142.1 afla|EED47570.1 afum|EAL88339.1 agla|OJJ89659.1 akaw|GAA89986.1 aluc|OJZ84000.1 anid|CBF74440.1 anig|EHA19122.1 aory|EIT73130.1 asyd|OJJ57744.1 ater|EAU36275.1 atub|OJI90369.1 aver|OJJ05423.1 awen|OJJ36638.1 azon|OJJ47578.1 afla|EED44702.1 aluc|OJZ80351.1 aory|EIT79586.1

Group607: abra|OJJ70731.1 acar|OOF97007.1 acla|EAW11182.1 afis|EAW15913.1 afla|EED57767.1 afum|EAL88548.2 agla|OJJ89577.1 akaw|GAA90059.1 aluc|OJZ84076.1 anid|CBF81305.1 anig|EHA19195.1 aory|EIT81027.1 asyd|OJJ59352.1 ater|EAU36192.1 atub|OJI90448.1 aver|OJJ02922.1 awen|OJJ36717.1 azon|OJJ47476.1 acla|EAW10824.1 afum|EAL89393.1 awen|OJJ36142.1

Group608: abra|OJJ70975.1 acar|OOF98156.1 acla|EAW10534.1 afis|EAW25001.1 afla|EED55421.1 afum|EAL89697.1 agla|OJJ89163.1 akaw|GAA82974.1 aluc|OJZ87557.1 anid|CBF81488.1 anig|EHA20760.1 atub|OJI81090.1 awen|OJJ40638.1 abra|OJJ71753.1 aluc|OJZ84833.1 anig|EHA25743.1 asyd|OJJ53401.1 ater|EAU35784.1 aver|OJJ04200.1 asyd|OJJ61749.1 aver|OJI95846.1

Group609: abra|OJJ71034.1 acar|OOF98103.1 acla|EAW10585.1 afla|EED55474.1 afum|EAL89644.1 agla|OJJ82278.1 akaw|GAA93319.1 aluc|OJZ87495.1 anid|CBF81619.1 anig|EHA20712.1 asyd|OJJ59527.1 ater|EAU35735.1 atub|OJI81154.1 aver|OJJ02744.1 awen|OJJ40580.1 azon|OJJ49440.1 afis|EAW24983.1 akaw|GAA82920.1 acla|EAW12399.1 afis|EAW16847.1 afum|EAL86393.1

Group610: abra|OJJ71163.1 acar|OOF97997.1 acla|EAW14151.1 afis|EAW18864.1 afla|EED53721.1 afum|EAL93596.1 agla|OJJ85239.1 akaw|GAA82816.1 aluc|OJZ87377.1 anid|CBF85347.1 anig|EHA20609.1 aory|EIT75273.1 asyd|OJJ64028.1 ater|EAU34397.1 atub|OJI81269.1 aver|OJI98114.1 awen|OJJ32721.1 azon|OJJ50355.1 afla|EED45395.1 aory|EIT82387.1 awen|OJJ39952.1

Group611: abra|OJJ71567.1 acar|OOF96301.1 acla|EAW12384.1 afis|EAW16831.1 afla|EED47160.1 afum|EAL86379.1 akaw|GAA89357.1 aluc|OJZ86304.1 anid|CBF69502.1 anig|EHA21568.1 aory|EIT79185.1 asyd|OJJ63349.1 ater|EAU36074.1 atub|OJI84232.1 aver|OJI97611.1 awen|OJJ31351.1 azon|OJJ50260.1 asyd|OJJ59227.1 aver|OJJ01837.1 azon|OJJ48154.1 awen|OJJ35570.1

Group612: abra|OJJ71741.1 acar|OOF96432.1 acla|EAW06445.1 afis|EAW18550.1 afla|EED52825.1 afum|EAL93250.1 agla|OJJ81594.1 akaw|GAA85155.1 aluc|OJZ86482.1 anid|CBF70273.1 anig|EHA22165.1 aory|EIT78166.1 asyd|OJJ58746.1 ater|EAU33222.1 atub|OJI84048.1 aver|OJJ01356.1 awen|OJJ31108.1 azon|OJJ46554.1 ater|EAU31802.1 afla|EED56761.1 aory|EIT73941.1

Group613: abra|OJJ72042.1 acar|OOF96701.1 acla|EAW09503.1 afis|EAW21633.1 afla|EED57861.1 afum|EAL92666.2 agla|OJJ79697.1 akaw|GAA84891.1 aluc|OJZ86780.1 anid|CBF83510.1 anig|EHA21902.1 aory|EIT81075.1 asyd|OJJ60709.1 ater|EAU38319.1 atub|OJI83740.1 aver|OJJ00035.1 awen|OJJ33882.1 azon|OJJ48672.1 afum|EAL91062.1 acla|EAW07114.1 afis|EAW22126.1

Group614: abra|OJJ72229.1 acar|OOF96852.1 acla|EAW12118.1 afis|EAW17264.1 afla|EED48866.1 afum|EAL86681.1 agla|OJJ84958.1 akaw|GAA84708.1 aluc|OJZ86979.1 anig|EHA21670.1 aory|EIT81253.1 atub|OJI83541.1 azon|OJJ50249.1 abra|OJJ75321.1 acar|OOF94506.1 akaw|GAA92016.1 aluc|OJZ92882.1 anig|EHA26078.1 atub|OJI87341.1 acar|OOF94471.1 ater|EAU36154.1

Group615: abra|OJJ72267.1 acar|OOG00128.1 acla|EAW10037.1 afis|EAW23420.1 afla|EED49100.1 afum|EAL84260.1 akaw|GAA86022.1 anig|EHA23745.1 ater|EAU37850.1 atub|OJI81675.1 abra|OJJ73122.1 acar|OOF92500.1 afis|EAW22027.1 akaw|GAA88043.1 aluc|OJZ81302.1 anig|EHA18611.1 aory|EIT82499.1 atub|OJI91511.1 afum|EAL88049.1 acla|EAW12426.1 afis|EAW15611.1

Group616: abra|OJJ72269.1 acar|OOF90351.1 afis|EAW19947.1 afla|EED46130.1 afum|EAL84968.2 agla|OJJ85569.1 akaw|GAA86024.1 aluc|OJZ90251.1 anid|CBF84249.1 anig|EHA23747.1 aory|EIT76460.1 asyd|OJJ57200.1 ater|EAU30029.1 atub|OJI81677.1 aver|OJJ03090.1 awen|OJJ32569.1 afum|EAL90430.1 akaw|GAA84794.1 aluc|OJZ86886.1 atub|OJI83633.1 awen|OJJ31250.1

Group617: abra|OJJ72859.1 acar|OOF96056.1 acla|EAW11387.1 afis|EAW18109.1 afla|EED49118.1 afum|EAL87466.1 agla|OJJ86997.1 akaw|GAA85458.1 aluc|OJZ83366.1 anid|CBF75326.1 anig|EHA24231.1 aory|EIT78544.1 asyd|OJJ59867.1 ater|EAU29709.1 atub|OJI82257.1 aver|OJJ02421.1 awen|OJJ35507.1 azon|OJJ46977.1 afla|EED55499.1 agla|OJJ87955.1 aory|EIT80816.1

Group618: abra|OJJ72915.1 acar|OOF96097.1 acla|EAW11688.1 afis|EAW15970.1 afla|EED58087.1 afum|EAL87149.1 agla|OJJ83846.1 aluc|OJZ83425.1 anig|EHA24277.1 aory|EIT79158.1 asyd|OJJ56475.1 ater|EAU35478.1 atub|OJI82311.1 aver|OJJ06243.1 awen|OJJ35603.1 azon|OJJ46922.1 anid|CBF69504.1 asyd|OJJ59223.1 aver|OJJ01833.1 acla|EAW11689.1 afis|EAW23705.1

Group619: abra|OJJ73104.1 acar|OOG00342.1 acla|EAW12431.1 afis|EAW16738.1 afla|EED44914.1 afum|EAL86313.2 agla|OJJ82713.1 akaw|GAA88061.1 aluc|OJZ81323.1 anid|CBF82088.1 anig|EHA18629.1 aory|EIT73472.1 asyd|OJJ54256.1 ater|EAU39589.1 atub|OJI91526.1 aver|OJJ05209.1 awen|OJJ39338.1 anid|CBF75878.1 asyd|OJJ60230.1 aver|OJJ04036.1 azon|OJJ51650.1

Group620: abra|OJJ73308.1 acar|OOG00533.1 acla|EAW12048.1 afis|EAW17017.1 afla|EED45666.1 akaw|GAA89816.1 aluc|OJZ81874.1 anig|EHA21034.1 aory|EIT81376.1 asyd|OJJ54465.1 ater|EAU36711.1

aver|OJJ07789.1 awen|OJJ31272.1 azon|OJJ47410.1 afla|EED48480.1 anid|CBF86045.1 anig|EHA28230.1 afla|EED52021.1 anid|CBF73486.1 aver|OJJ06468.1 anid|CBF89301.1

Group621: abra|OJJ74126.1 acar|OOF95610.1 afis|EAW24721.1 afla|EED53234.1 afum|EAL89370.1 agla|OJJ87996.1 akaw|GAA89758.1 aluc|OJZ80625.1 anid|CBF82175.1 anig|EHA27207.1 aory|EIT74852.1 asyd|OJJ54960.1 atub|OJI87480.1 aver|OJJ08805.1 awen|OJJ30037.1 azon|OJJ43678.1 asyd|OJJ64485.1 atub|OJI88492.1 aver|OJI98586.1 aver|OJJ07231.1 asyd|OJJ52829.1

Group622: abra|OJJ74136.1 acar|OOF95594.1 acla|EAW07094.1 afis|EAW22104.1 afla|EED54116.1 afum|EAL91087.1 akaw|GAA89750.1 aluc|OJZ80634.1 anid|CBF79913.1 anig|EHA27198.1 aory|EIT76054.1 asyd|OJJ58191.1 ater|EAU32259.1 atub|OJI87489.1 aver|OJJ04214.1 awen|OJJ34666.1 azon|OJJ50218.1 acla|EAW11902.1 afis|EAW19515.1 afum|EAL86981.2 anid|CBF86632.1

Group623: abra|OJJ74191.1 acar|OOF95530.1 acla|EAW07243.1 afis|EAW22264.1 afla|EED57656.1 afum|EAL90916.1 agla|OJJ87791.1 akaw|GAA86826.1 aluc|OJZ89944.1 anid|CBF88515.1 anig|EHA27143.1 aory|EIT80760.1 asyd|OJJ63805.1 ater|EAU34274.1 atub|OJI84862.1 aver|OJI97133.1 awen|OJJ34837.1 azon|OJJ45710.1 afla|EED51484.1 asyd|OJJ58154.1 aver|OJJ04249.1

Group624: abra|OJJ74556.1 acar|OOF95199.1 acla|EAW12292.1 afis|EAW16945.1 afla|EED46082.1 afum|EAL86472.2 agla|OJJ89555.1 akaw|GAA82040.1 aluc|OJZ92139.1 anid|CBF88435.1 anig|EHA26810.1 aory|EIT76401.1 asyd|OJJ63762.1 ater|EAU35227.1 atub|OJI86571.1 aver|OJI97178.1 awen|OJJ36332.1 azon|OJJ46487.1 acla|EAW11680.1 afis|EAW23697.1 anid|CBF76117.1

Group625: abra|OJJ74819.1 acar|OOF95807.1 acla|EAW08501.1 afis|EAW16872.1 afum|EAL86410.2 akaw|GAA83596.1 aluc|OJZ92398.1 anid|CBF86626.1 anig|EHA26567.1 asyd|OJJ64810.1 ater|EAU32516.1 atub|OJI86828.1 aver|OJI98923.1 awen|OJJ32502.1 azon|OJJ43488.1 acar|OOF90627.1 afis|EAW17135.1 afla|EED45618.1 anig|EHA24760.1 aory|EIT75444.1 ater|EAU38152.1

Group626: abra|OJJ76271.1 acar|OOF99519.1 afis|EAW24673.1 akaw|GAA82204.1 aluc|OJZ91949.1 anid|CBF90103.1 anig|EHA22908.1 atub|OJI86394.1 aver|OJI96119.1 abra|OJJ77743.1 afis|EAW19876.1 afla|EED56872.1 agla|OJJ82021.1 anid|CBF74093.1 asyd|OJJ65119.1 ater|EAU36111.1 aver|OJI99248.1 awen|OJJ34656.1 awen|OJJ38125.1 aory|EIT77510.1 asyd|OJJ62028.1

Group627: abra|OJJ76451.1 acar|OOF99694.1 acla|EAW13977.1 afis|EAW18686.1 afla|EED53481.1 afum|EAL89344.2 akaw|GAA82363.1 aluc|OJZ91759.1 anig|EHA22761.1 aory|EIT75172.1 asyd|OJJ58512.1 ater|EAU38222.1 atub|OJI86217.1 aver|OJJ01113.1 awen|OJJ32157.1 azon|OJJ46741.1 acla|EAW13614.1 afis|EAW16454.1 anig|EHA20148.1 afum|EAL93407.1 afis|EAW24702.1

Group628: abra|OJJ76531.1 acar|OOF99442.1 acla|EAW12929.1 afis|EAW17727.1 afla|EED48895.1 afum|EAL91542.1 agla|OJJ87872.1 akaw|GAA82443.1 aluc|OJZ91673.1 anid|CBF90286.1 anig|EHA22691.1 aory|EIT81184.1 asyd|OJJ61920.1 ater|EAU36831.1 atub|OJI86128.1 aver|OJI96012.1 awen|OJJ39163.1 azon|OJJ44435.1 acar|OOF93188.1 acla|EAW12526.1 awen|OJJ40114.1

Group629: abra|OJJ76817.1 acar|OOF91312.1 acla|EAW13377.1 afis|EAW19711.1 afla|EED47757.1 afum|EAL86816.2 agla|OJJ84836.1 aluc|OJZ84227.1 anid|CBF79211.1 anig|EHA27576.1 aory|EIT77795.1 asyd|OJJ55731.1 ater|EAU30448.1 atub|OJI89857.1 aver|OJJ06817.1 awen|OJJ33191.1 azon|OJJ46393.1 aory|EIT77115.1 afla|EED44738.1 asyd|OJJ54596.1 aver|OJJ07556.1

Group630: abra|OJJ76842.1 acar|OOF89970.1 acla|EAW13405.1 afis|EAW20516.1 afla|EED52466.1 afum|EAL84541.1 agla|OJJ84774.1 akaw|GAA87180.1 aluc|OJZ84253.1 anid|CBF79059.1 anig|EHA27598.1 aory|EIT82819.1 asyd|OJJ55708.1 ater|EAU37185.1 atub|OJI89830.1 aver|OJJ06794.1 awen|OJJ33222.1 azon|OJJ45121.1 aluc|OJZ87023.1 anig|EHA21634.1 atub|OJI83495.1

Group631: abra|OJJ77007.1 acar|OOF91453.1 acla|EAW15047.1 afis|EAW24611.1 afla|EED47377.1 afum|EAL86241.1 agla|OJJ78906.1 akaw|GAA87041.1 aluc|OJZ84406.1 anid|CBF83177.1 anig|EHA27736.1 aory|EIT75955.1 asyd|OJJ55523.1 ater|EAU32183.1 atub|OJI89666.1 aver|OJJ06601.1 awen|OJJ36042.1 azon|OJJ45129.1 acla|EAW15048.1 aory|EIT82833.1 afla|EED54123.1

Group632: abra|OJJ77033.1 acar|OOF91472.1 acla|EAW07271.1 afis|EAW22292.1 afla|EED57692.1 afum|EAL90889.1 agla|OJJ87756.1 akaw|GAA87018.1 aluc|OJZ84436.1 anid|CBF88573.1 anig|EHA27755.1 aory|EIT79900.1 asyd|OJJ63839.1 ater|EAU34305.1 atub|OJI89637.1 aver|OJI97098.1 awen|OJJ34866.1 azon|OJJ45662.1 afla|EED51864.1 aory|EIT81771.1 awen|OJJ30800.1

Group633: abra|OJJ77037.1 acar|OOF90079.1 acla|EAW13558.1 afis|EAW20634.1 afla|EED49758.1 afum|EAL84434.1 agla|OJJ86283.1 akaw|GAA87014.1 aluc|OJZ84440.1 anid|CBF83213.1 anig|EHA27759.1 aory|EIT77698.1 asyd|OJJ65203.1 ater|EAU32169.1 atub|OJI89633.1 aver|OJJ06646.1 awen|OJJ33605.1 azon|OJJ50726.1 asyd|OJJ55565.1 azon|OJJ42610.1 aver|OJI99352.1

Group634: abra|OJJ77109.1 acar|OOF92994.1 afla|EED51851.1 agla|OJJ86977.1 akaw|GAA90819.1 aluc|OJZ84518.1 anid|CBF89943.1 anig|EHA27816.1 aory|EIT81728.1 asyd|OJJ62111.1 ater|EAU31034.1 atub|OJI89562.1 aver|OJJ03877.1 awen|OJJ33658.1 acla|EAW14977.1 aver|OJI96219.1 asyd|OJJ57090.1 ater|EAU34328.1 awen|OJJ33488.1 aver|OJJ03210.1 asyd|OJJ62627.1

Group635: abra|OJJ77191.1 acar|OOF93077.1 acla|EAW13182.1 afis|EAW17458.1 afla|EED49364.1 afum|EAL91286.1 agla|OJJ80049.1 akaw|GAA90749.1 aluc|OJZ84599.1 anid|CBF71666.1 anig|EHA27889.1 aory|EIT79494.1 asyd|OJJ59263.1 ater|EAU33943.1 atub|OJI89480.1 aver|OJJ01874.1 awen|OJJ37389.1 azon|OJJ51062.1 agla|OJJ80359.1 asyd|OJJ61831.1 aver|OJI95930.1

Group636: abra|OJJ77325.1 acar|OOF93201.1 acla|EAW13147.1 afis|EAW17500.1 afla|EED45479.1 afum|EAL91323.1 agla|OJJ81453.1 akaw|GAA91431.1 aluc|OJZ84735.1 anid|CBF71578.1 anig|EHA28008.1 aory|EIT82181.1 asyd|OJJ61876.1 ater|EAU36946.1 atub|OJI89346.1 aver|OJI95974.1 awen|OJJ37341.1 azon|OJJ51026.1 agla|OJJ78829.1 aory|EIT75365.1 asyd|OJJ62947.1

Group637: abra|OJJ77475.1 acar|OOF99250.1 acla|EAW06757.1 afis|EAW16307.1 afla|EED53242.1 afum|EAL88929.1 agla|OJJ88269.1 akaw|GAA84143.1 aluc|OJZ81063.1 anid|CBF69567.1 anig|EHA19305.1 aory|EIT75142.1 asyd|OJJ59151.1 ater|EAU29335.1 atub|OJI82522.1 aver|OJJ08866.1 awen|OJJ34880.1 azon|OJJ44967.1 aver|OJJ01741.1 awen|OJJ36323.1 asyd|OJJ52895.1

Group638: abra|OJJ78236.1 acar|OOF98624.1 acla|EAW09274.1 afis|EAW21874.1 afla|EED46435.1 afum|EAL92888.1 agla|OJJ84259.1 akaw|GAA85789.1 aluc|OJZ91310.1 anid|CBF76774.1 anig|EHA19994.1 aory|EIT72385.1 asyd|OJJ61272.1 ater|EAU30873.1 atub|OJI83303.1 aver|OJJ00600.1 awen|OJJ34162.1 azon|OJJ43467.1 afla|EED46434.1 awen|OJJ34163.1 asyd|OJJ61271.1

Group639: abra|OJJ65541.1 abra|OJJ65833.1 abra|OJJ67278.1 akaw|GAA92069.1 akaw|GAA92846.1 asyd|OJJ52003.1 asyd|OJJ62485.1 acar|OOF95207.1 afla|EED55152.1 agla|OJJ86434.1 anid|CBF76638.1 anig|EHA18761.1 aory|EIT81617.1 aver|OJJ07599.1 awen|OJJ31646.1 aluc|OJZ88789.1 acar|OOG00873.1 agla|OJJ79830.1 asyd|OJJ54555.1 awen|OJJ30844.1

Group640: abra|OJJ65699.1 abra|OJJ77440.1 aluc|OJZ81081.1 aluc|OJZ88716.1 asyd|OJJ59917.1 asyd|OJJ61765.1 aver|OJI96472.1 aver|OJJ02369.1 afla|EED56833.1 afum|EAL84269.2 agla|OJJ80658.1 akaw|GAA84158.1 anid|CBF85281.1 anig|EHA18818.1 aory|EIT73911.1 ater|EAU34168.1 atub|OJI82505.1 awen|OJJ30259.1 akaw|GAA92795.1 anig|EHA28199.1

Group641: abra|OJJ66191.1 abra|OJJ68084.1 acar|OOF91939.1 acar|OOF93882.1 akaw|GAA86869.1 akaw|GAA89096.1 anig|EHA25698.1 anig|EHA27364.1 atub|OJI84816.1 atub|OJI88435.1 acla|EAW09010.1 afis|EAW25589.1 afum|EAL84297.2 aluc|OJZ89993.1 asyd|OJJ61550.1 aver|OJJ00875.1 awen|OJJ41959.1 afla|EED48865.1 anid|CBF82995.1 aory|EIT81254.1

Group642: abra|OJJ68152.1 abra|OJJ75347.1 aluc|OJZ83228.1 aluc|OJZ92909.1 atub|OJI87369.1 atub|OJI87708.1 acar|OOF94489.1 acla|EAW08546.1 afis|EAW23583.1 afla|EED48907.1 afum|EAL87881.1 akaw|GAA92040.1 anid|CBF87513.1 anig|EHA21084.1 anig|EHA26055.1 aory|EIT81206.1 ater|EAU36167.1 awen|OJJ30073.1 asyd|OJJ52977.1 aver|OJJ08943.1

Group643: abra|OJJ70063.1 abra|OJJ75532.1 akaw|GAA84171.1 akaw|GAA87613.1 akaw|GAA91952.1 aluc|OJZ81096.1 aluc|OJZ82666.1 aluc|OJZ85234.1 anig|EHA19275.1 anig|EHA25438.1 anig|EHA27440.1 atub|OJI80344.1 atub|OJI82486.1 atub|OJI88899.1 acar|OOF93551.1 afis|EAW25548.1 afla|EED45174.1 afla|EED46370.1 anid|CBF82283.1 aory|EIT81213.1

Group644: abra|OJJ74604.1 abra|OJJ77441.1 acar|OOF95100.1 acar|OOF95166.1 acla|EAW10775.1 afis|EAW24801.1 afla|EED47186.1 afum|EAL89442.1 agla|OJJ82895.1 akaw|GAA82004.1 aluc|OJZ92183.1 anid|CBF81510.1 anig|EHA26770.1 aory|EIT79225.1 asyd|OJJ59469.1 ater|EAU36506.1 atub|OJI86611.1 aver|OJJ02805.1 awen|OJJ40384.1 azon|OJJ49233.1

Group645: acar|OOF90070.1 acar|OOF90633.1 akaw|GAA85210.1 akaw|GAA92948.1 abra|OJJ65532.1 anig|EHA21123.1 abra|OJJ67562.1 afla|EED57361.1 akaw|GAA84330.1 aluc|OJZ89029.1 aory|EIT78652.1 ater|EAU36945.1 atub|OJI85762.1 awen|OJJ30886.1 abra|OJJ72584.1 afla|EED44973.1 aluc|OJZ90565.1 anig|EHA24011.1 atub|OJI81988.1 afla|EED52856.1

Group646: acar|OOF90425.1 acar|OOF90429.1 acar|OOF93386.1 abra|OJJ69872.1 acla|EAW08934.1 afis|EAW15465.1 afla|EED46918.1 afum|EAL85857.1 agla|OJJ78859.1 akaw|GAA87784.1 aluc|OJZ85419.1 anid|CBF71773.1 anig|EHA25598.1 aory|EIT72659.1 asyd|OJJ61719.1 ater|EAU32668.1 atub|OJI80156.1 aver|OJI95813.1 awen|OJJ38110.1 azon|OJJ43167.1

Group647: acar|OOF90437.1 acar|OOF90444.1 abra|OJJ68818.1 acla|EAW13660.1 afis|EAW19540.1 agla|OJJ84857.1 akaw|GAA90433.1 aluc|OJZ85794.1 anid|CBF73461.1 anig|EHA24767.1 aory|EIT74867.1 asyd|OJJ55924.1 ater|EAU34189.1 atub|OJI79580.1 azon|OJJ42666.1 afum|EAL84970.1 azon|OJJ45282.1 afis|EAW19943.1 agla|OJJ79424.1 agla|OJJ89038.1

Group648: acar|OOF90442.1 acar|OOF90448.1 abra|OJJ78364.1 acar|OOF98523.1 acla|EAW09171.1 afis|EAW21975.1 afla|EED46698.1 afum|EAL92990.1 agla|OJJ84127.1 akaw|GAA85669.1 aluc|OJZ91440.1 anid|CBF82718.1 anig|EHA20104.1 aory|EIT77954.1 asyd|OJJ61636.1 ater|EAU31093.1 atub|OJI83434.1 aver|OJJ00959.1 awen|OJJ34268.1 azon|OJJ43367.1

Group649: acar|OOF90479.1 acar|OOF90486.1 acar|OOF96185.1 abra|OJJ73013.1 acla|EAW07999.1 afis|EAW23034.1 afla|EED54982.1 afum|EAL88447.1 agla|OJJ89443.1 akaw|GAA85592.1 aluc|OJZ83528.1

anid|CBF80924.1 anig|EHA24351.1 aory|EIT80284.1 asyd|OJJ56320.1 ater|EAU29623.1 atub|OJI82417.1 aver|OJJ06394.1 awen|OJJ35708.1 azon|OJJ44168.1

Group650: acar|OOF90594.1 acar|OOG00347.1 abra|OJJ73121.1 afis|EAW19817.1 afla|EED53226.1 afum|EAL85074.1 akaw|GAA88044.1 aluc|OJZ81304.1 anid|CBF82022.1 anig|EHA18612.1 aory|EIT75068.1 asyd|OJJ56203.1 ater|EAU29416.1 atub|OJI91512.1 aver|OJJ05130.1 awen|OJJ36147.1 azon|OJJ42172.1 azon|OJJ50764.1 aver|OJJ06466.1 ater|EAU34240.1

Group651: acar|OOF90625.1 acar|OOF90863.1 acar|OOF92572.1 acar|OOF94857.1 akaw|GAA86114.1 akaw|GAA86569.1 akaw|GAA90436.1 aluc|OJZ85120.1 aluc|OJZ85791.1 aluc|OJZ90350.1 atub|OJI79576.1 atub|OJI80473.1 atub|OJI81777.1 azon|OJJ43546.1 abra|OJJ70235.1 afla|EED57683.1 akaw|GAA86570.1 anig|EHA25341.1 aory|EIT79899.1 anig|EHA23836.1

Group652: acar|OOF92603.1 acar|OOF92624.1 asyd|OJJ51990.1 asyd|OJJ56630.1 abra|OJJ68460.1 acla|EAW14885.1 afis|EAW24404.1 afla|EED47809.1 afum|EAL86075.1 agla|OJJ89341.1 akaw|GAA92382.1 aluc|OJZ83241.1 anid|CBF73735.1 anig|EHA21096.1 aory|EIT77126.1 ater|EAU30327.1 atub|OJI87696.1 aver|OJJ06096.1 awen|OJJ32804.1 azon|OJJ49024.1

Group653: acar|OOF92614.1 acar|OOF92635.1 awen|OJJ30756.1 awen|OJJ31725.1 abra|OJJ67454.1 acla|EAW06311.1 afis|EAW18420.1 afla|EED52988.1 afum|EAL93104.1 agla|OJJ84988.1 akaw|GAA86377.1 aluc|OJZ84911.1 anid|CBF70700.1 anig|EHA25132.1 aory|EIT78732.1 asyd|OJJ58461.1 ater|EAU29838.1 atub|OJI80687.1 aver|OJJ01061.1 azon|OJJ50488.1

Group654: acar|OOF93661.1 acar|OOF93745.1 abra|OJJ66641.1 acla|EAW11784.1 afis|EAW19394.1 afum|EAL87100.1 agla|OJJ81105.1 akaw|GAA89594.1 aluc|OJZ80808.1 anid|CBF74281.1 anig|EHA27226.1 aory|EIT83121.1 asyd|OJJ56700.1 ater|EAU36057.1 atub|OJI87656.1 aver|OJJ05358.1 awen|OJJ36788.1 azon|OJJ43011.1 asyd|OJJ51861.1 aver|OJJ06025.1

Group655: acar|OOF93670.1 acar|OOF93754.1 abra|OJJ66651.1 acla|EAW11794.1 afis|EAW19404.1 afla|EED50359.1 afum|EAL87091.1 agla|OJJ81118.1 akaw|GAA89603.1 aluc|OJZ80795.1 anid|CBF74343.1 anig|EHA27236.1 aory|EIT83145.1 asyd|OJJ51900.1 asyd|OJJ57797.1 ater|EAU36141.1 atub|OJI87646.1 aver|OJJ05367.1 awen|OJJ36800.1 azon|OJJ42998.1

Group656: acar|OOF93714.1 acar|OOF93801.1 agla|OJJ81170.1 agla|OJJ82654.1 abra|OJJ66712.1 acla|EAW11843.1 afis|EAW19454.1 afla|EED56662.1 afum|EAL87042.1 akaw|GAA89654.1 aluc|OJZ80735.1 anid|CBF81419.1 anig|EHA27286.1 aory|EIT79629.1 asyd|OJJ59414.1 ater|EAU34757.1 atub|OJI87592.1 aver|OJJ02859.1 awen|OJJ36855.1 azon|OJJ42945.1

Group657: acar|OOF94063.1 acar|OOF94133.1 abra|OJJ66361.1 acla|EAW10010.1 afis|EAW21046.1 afla|EED51624.1 afum|EAL92082.1 agla|OJJ82364.1 akaw|GAA90234.1 aluc|OJZ83107.1 anid|CBF87132.1 anig|EHA28170.1 aory|EIT80496.1 asyd|OJJ60365.1 atub|OJI87827.1 aver|OJI99702.1 awen|OJJ32176.1 afla|EED48902.1 aory|EIT81167.1 awen|OJJ36966.1

Group658: acar|OOF94066.1 acar|OOF94136.1 abra|OJJ66356.1 acla|EAW10006.1 afis|EAW21050.1 afla|EED51620.1 afum|EAL92086.1 akaw|GAA90230.1 aluc|OJZ83111.1 anid|CBF87104.1 anig|EHA28174.1 aory|EIT76079.1 asyd|OJJ60388.1 ater|EAU36822.1 atub|OJI87823.1 aver|OJI99717.1 awen|OJJ32173.1 afla|EED54348.1 asyd|OJJ53360.1 aver|OJJ07646.1

Group659: acar|OOF95099.1 acar|OOF95165.1 asyd|OJJ59458.1 asyd|OJJ62484.1 abra|OJJ74605.1 acla|EAW10774.1 afis|EAW24802.1 afla|EED47184.1 afum|EAL89443.1 agla|OJJ82896.1 akaw|GAA82003.1 aluc|OJZ92184.1 anid|CBF81490.1 anig|EHA26769.1 aory|EIT79286.1 ater|EAU36505.1 atub|OJI86612.1 aver|OJJ02815.1 awen|OJJ40385.1 azon|OJJ49235.1

Group660: acar|OOF95626.1 acar|OOF95677.1 abra|OJJ74662.1 acla|EAW10965.1 afis|EAW20769.1 afla|EED50244.1 afum|EAL84738.2 agla|OJJ84002.1 akaw|GAA81949.1 aluc|OJZ92239.1 anig|EHA26715.1 asyd|OJJ54118.1 ater|EAU32725.1 atub|OJI86673.1 aver|OJJ08618.1 awen|OJJ34453.1 acar|OOF90978.1 afis|EAW20895.1 aory|EIT77360.1 ater|EAU38529.1

Group661: acar|OOF95637.1 acar|OOF95688.1 aver|OJJ05622.1 aver|OJJ07803.1 abra|OJJ74674.1 acla|EAW10723.1 afis|EAW24850.1 afla|EED47121.1 afum|EAL89496.1 agla|OJJ82064.1 akaw|GAA81939.1 aluc|OJZ92251.1 anid|CBF75837.1 anig|EHA26705.1 aory|EIT79282.1 asyd|OJJ57556.1 ater|EAU36453.1 atub|OJI86684.1 awen|OJJ40435.1 azon|OJJ49296.1

Group662: acar|OOF96241.1 acar|OOF96278.1 afum|EAL85174.2 afum|EAL87772.2 abra|OJJ71525.1 acla|EAW15082.1 afis|EAW20054.1 afla|EED45326.1 agla|OJJ88906.1 akaw|GAA89392.1 aluc|OJZ86265.1 anid|CBF84810.1 anig|EHA21603.1 aory|EIT73801.1 asyd|OJJ63220.1 ater|EAU38663.1 atub|OJI84278.1 aver|OJI97735.1 awen|OJJ31812.1 azon|OJJ46310.1

Group663: acar|OOF96244.1 acar|OOF96281.1 afum|EAL85177.1 afum|EAL87761.1 abra|OJJ71529.1 acla|EAW15085.1 afis|EAW20057.1 afla|EED45323.1 agla|OJJ88903.1 akaw|GAA89389.1 aluc|OJZ86269.1 anid|CBF84842.1 anig|EHA21601.1 aory|EIT73791.1 asyd|OJJ63216.1 ater|EAU38666.1 atub|OJI84273.1 aver|OJI97739.1 awen|OJJ31816.1 azon|OJJ46307.1

Group664: acar|OOF96246.1 acar|OOF96283.1 afum|EAL85179.1 afum|EAL87759.1 abra|OJJ71533.1 acla|EAW15087.1 afis|EAW20059.1 afla|EED45321.1 agla|OJJ88901.1 akaw|GAA89387.1 aluc|OJZ86271.1 anid|CBF84838.1 anig|EHA21598.1 aory|EIT73620.1 asyd|OJJ63214.1 ater|EAU38668.1 atub|OJI84270.1 aver|OJI97741.1 awen|OJJ31819.1 azon|OJJ46305.1

Group665: acar|OOF96247.1 acar|OOF96284.1 afum|EAL85180.1 afum|EAL87758.1 abra|OJJ71534.1 acla|EAW15088.1 afis|EAW20060.1 afla|EED45319.1 agla|OJJ88899.1 akaw|GAA89386.1 aluc|OJZ86272.1 anid|CBF84836.1 anig|EHA21597.1 aory|EIT73579.1 asyd|OJJ63213.1 ater|EAU38669.1 atub|OJI84268.1 aver|OJI97742.1 awen|OJJ31820.1 azon|OJJ46304.1

Group666: acar|OOF97654.1 acar|OOF97701.1 abra|OJJ71503.1 acla|EAW10045.1 afis|EAW25539.1 afla|EED44716.1 afum|EAL90202.1 agla|OJJ87266.1 akaw|GAA82517.1 aluc|OJZ87044.1 anid|CBF77824.1 anig|EHA20317.1 aory|EIT83461.1 asyd|OJJ53006.1 ater|EAU33380.1 atub|OJI81591.1 aver|OJJ03837.1 awen|OJJ41281.1 azon|OJJ49499.1 afla|EED44715.1

Group667: acar|OOF97687.1 acar|OOF97737.1 abra|OJJ71461.1 acla|EAW10090.1 afis|EAW25497.1 afla|EED53997.1 afum|EAL90163.1 agla|OJJ87307.1 akaw|GAA82552.1 aluc|OJZ87083.1 anid|CBF77751.1 anig|EHA20354.1 aory|EIT78321.1 asyd|OJJ53046.1 asyd|OJJ54800.1 ater|EAU34668.1 atub|OJI81554.1 aver|OJI95782.1 aver|OJJ03799.1 awen|OJJ41240.1

Group668: acar|OOF99309.1 acar|OOF99314.1 abra|OJJ74376.1 akaw|GAA87275.1 aluc|OJZ89746.1 anig|EHA26968.1 atub|OJI85056.1 abra|OJJ78032.1 akaw|GAA85950.1 aluc|OJZ91112.1 anig|EHA19809.1 aory|EIT82873.1 asyd|OJJ64719.1 ater|EAU30104.1 atub|OJI83109.1 aver|OJJ05112.1 awen|OJJ33401.1 azon|OJJ45152.1 acla|EAW11016.1 awen|OJJ38250.1

Group669: acla|EAW06566.1 acla|EAW13723.1 afis|EAW16030.1 afis|EAW19868.1 afum|EAL85029.1 afum|EAL88661.1 agla|OJJ86058.1 agla|OJJ87705.1 awen|OJJ30418.1 awen|OJJ39976.1 abra|OJJ67634.1 acar|OOF91979.1 afla|EED48196.1 akaw|GAA89153.1 aluc|OJZ81683.1 anig|EHA25759.1 aory|EIT72945.1 asyd|OJJ52747.1 atub|OJI88498.1 azon|OJJ45425.1

Group670: acla|EAW07085.1 acla|EAW08664.1 abra|OJJ67836.1 afum|EAL87810.1 agla|OJJ83639.1 akaw|GAA84845.1 aluc|OJZ86826.1 anig|EHA28243.1 asyd|OJJ62960.1 ater|EAU36707.1 atub|OJI89149.1 aver|OJJ06460.1 aver|OJJ03911.1 anig|EHA28234.1 asyd|OJJ64572.1 atub|OJI83692.1 aver|OJI98007.1 anid|CBF78723.1 aver|OJI98669.1 aver|OJJ06464.1

Group671: acla|EAW10425.1 acla|EAW11189.1 afum|EAL88556.2 afum|EAL89817.2 abra|OJJ70818.1 acar|OOF98284.1 afis|EAW25144.1 afla|EED55301.1 agla|OJJ89017.1 akaw|GAA91697.1 aluc|OJZ87695.1 anid|CBF85681.1 anig|EHA20883.1 aory|EIT83608.1 asyd|OJJ64255.1 ater|EAU39416.1 atub|OJI80947.1 aver|OJI98346.1 awen|OJJ40769.1 azon|OJJ46879.1

Group672: afis|EAW16434.1 afis|EAW20510.1 agla|OJJ84787.1 agla|OJJ85600.1 abra|OJJ76837.1 acar|OOF91329.1 acla|EAW13399.1 afla|EED52451.1 afum|EAL84546.1 akaw|GAA87186.1 aluc|OJZ84248.1 anid|CBF79187.1 anig|EHA27593.1 aory|EIT82791.1 asyd|OJJ55716.1 ater|EAU37204.1 atub|OJI89835.1 aver|OJJ06802.1 awen|OJJ33211.1 azon|OJJ46484.1

Group673: afis|EAW17153.1 afis|EAW21838.1 afum|EAL87745.2 afum|EBA27475.1 abra|OJJ75520.1 acar|OOF90847.1 acla|EAW09307.1 afla|EED46537.1 agla|OJJ84300.1 akaw|GAA84284.1 aluc|OJZ88977.1 anid|CBF76703.1 anig|EHA23548.1 aory|EIT72350.1 asyd|OJJ61234.1 ater|EAU38513.1 atub|OJI85812.1 aver|OJJ00557.1 awen|OJJ34114.1 azon|OJJ44226.1

Group674: afis|EAW17990.1 afis|EAW20711.1 afum|EAL91127.1 afum|EAL91800.1 abra|OJJ68535.1 acar|OOF92737.1 acla|EAW12825.1 afla|EED51913.1 agla|OJJ80250.1 akaw|GAA91316.1 aluc|OJZ82748.1 anid|CBF79983.1 anig|EHA21198.1 aory|EIT81369.1 asyd|OJJ58245.1 ater|EAU31501.1 atub|OJI88983.1 aver|OJJ02181.1 awen|OJJ41556.1 azon|OJJ48271.1

Group675: afis|EAW19531.1 afis|EAW22425.1 afum|EAL86975.1 afum|EAL90766.1 abra|OJJ74345.1 acar|OOF95384.1 acla|EAW07399.1 afla|EED52503.1 agla|OJJ87607.1 akaw|GAA86685.1 aluc|OJZ89785.1 anid|CBF88811.1 anig|EHA27000.1 asyd|OJJ62888.1 ater|EAU39282.1 atub|OJI85020.1 aver|OJI96940.1 awen|OJJ35004.1 azon|OJJ42263.1 azon|OJJ45640.1

Group676: afis|EAW23730.1 afis|EAW25103.1 awen|OJJ30012.1 awen|OJJ40717.1 abra|OJJ65991.1 acar|OOF92881.1 acla|EAW10464.1 acla|EAW11709.1 afla|EED46010.1 afum|EAL87168.1 akaw|GAA90865.1 aluc|OJZ82968.1 anid|CBF78451.1 anid|CBF84695.1 anig|EHA21384.1 aory|EIT75669.1 ater|EAU34822.1 atub|OJI89201.1 aver|OJJ08430.1 azon|OJJ48019.1

Group677: afla|EED45834.1 afla|EED51243.1 aory|EIT75702.1 aory|EIT83187.1 abra|OJJ69216.1 acar|OOF91102.1 acla|EAW08629.1 afis|EAW15802.1 afum|EAL85548.1 agla|OJJ81969.1 akaw|GAA92225.1 aluc|OJZ86198.1 anid|CBF70916.1 anig|EHA24425.1 asyd|OJJ52604.1 ater|EAU33446.1 atub|OJI79990.1 aver|OJJ04630.1 awen|OJJ38512.1 azon|OJJ43740.1

Group678: afla|EED46513.1 afla|EED55226.1 aory|EIT72395.1 aory|EIT83617.1 abra|OJJ72126.1 acar|OOF96766.1 acla|EAW14932.1 afis|EAW24447.1 afum|EAL86116.1 agla|OJJ80293.1 akaw|GAA84797.1

aluc|OJZ86883.1 anid|CBF73809.1 anig|EHA21761.1 asyd|OJJ56577.1 ater|EAU29999.1 atub|OJI83636.1 aver|OJJ06146.1 awen|OJJ32750.1 azon|OJJ43988.1

Group679: afla|EED46574.1 afla|EED49749.1 afla|EED52838.1 akaw|GAA87766.1 akaw|GAA91614.1 aluc|OJZ82440.1 aluc|OJZ85401.1 aluc|OJZ82904.1 anig|EHA21327.1 anig|EHA28333.1 aory|EIT77940.1 aory|EIT78630.1 abra|OJJ69634.1 agla|OJJ80224.1 asyd|OJJ52421.1 ater|EAU37501.1 atub|OJI88000.1 awen|OJJ31941.1 atub|OJI89130.1 akaw|GAA90922.1

Group680: afla|EED47735.1 afla|EED54121.1 abra|OJJ66976.1 acar|OOF93637.1 acla|EAW08563.1 afis|EAW23605.1 afum|EAL87861.1 agla|OJJ86863.1 akaw|GAA92189.1 aluc|OJZ85601.1 anid|CBF89577.1 anig|EHA24923.1 aory|EIT75914.1 aory|EIT77123.1 asyd|OJJ62333.1 ater|EAU35372.1 atub|OJI79396.1 aver|OJI96439.1 awen|OJJ39929.1 ater|EAU39302.1

Group681: agla|OJJ78415.1 agla|OJJ84293.1 abra|OJJ69363.1 acar|OOF94362.1 acla|EAW14753.1 afis|EAW24284.1 afla|EED47992.1 afum|EAL85953.1 akaw|GAA88232.1 aluc|OJZ82196.1 anid|CBF74135.1 anig|EHA28541.1 aory|EIT79812.1 asyd|OJJ53915.1 ater|EAU31232.1 atub|OJI88239.1 aver|OJJ08013.1 awen|OJJ33012.1 azon|OJJ48503.1 agla|OJJ83146.1

Group682: agla|OJJ78543.1 agla|OJJ79518.1 anid|CBF74064.1 anid|CBF77736.1 anid|CBF83503.1 anid|CBF87508.1 anid|CBF88991.1 anid|CBF89857.1 abra|OJJ69705.1 acla|EAW09195.1 afis|EAW18410.1 afla|EED46570.1 afum|EAL93090.1 aluc|OJZ80203.1 aory|EIT77911.1 asyd|OJJ53853.1 awen|OJJ41393.1 asyd|OJJ62070.1 anid|CBF73616.1 asyd|OJJ56787.1

Group683: agla|OJJ79096.1 agla|OJJ82949.1 abra|OJJ65777.1 acar|OOF92088.1 afla|EED45916.1 akaw|GAA90885.1 aluc|OJZ82946.1 anid|CBF84425.1 anig|EHA21364.1 aory|EIT75550.1 atub|OJI89178.1 abra|OJJ72700.1 afis|EAW19995.1 afum|EAL84921.1 asyd|OJJ54671.1 aver|OJJ07499.1 awen|OJJ30436.1 afla|EED49831.1 asyd|OJJ53641.1 aver|OJJ01165.1

Group684: agla|OJJ79930.1 agla|OJJ82748.1 awen|OJJ29762.1 awen|OJJ33576.1 abra|OJJ70370.1 acar|OOF91900.1 acla|EAW12338.1 afis|EAW16884.1 afla|EED47358.1 afum|EAL86420.1 akaw|GAA91980.1 aluc|OJZ82634.1 anid|CBF80682.1 anig|EHA27445.1 aory|EIT82887.1 asyd|OJJ54592.1 ater|EAU30230.1 atub|OJI88865.1 aver|OJJ07560.1 azon|OJJ50386.1

Group685: akaw|GAA85714.1 akaw|GAA87141.1 aluc|OJZ84296.1 aluc|OJZ91389.1 aory|EIT81041.1 aory|EIT81749.1 atub|OJI83382.1 atub|OJI89773.1 abra|OJJ78316.1 acar|OOF90169.1 acla|EAW09510.1 afis|EAW21622.1 afla|EED51839.1 afum|EAL92658.1 anig|EHA20067.1 asyd|OJJ65401.1 ater|EAU38311.1 aver|OJI95769.1 awen|OJJ38680.1 azon|OJJ47988.1

Group686: aluc|OJZ79488.1 aluc|OJZ79629.1 abra|OJJ68492.1 acar|OOF90053.1 acla|EAW12140.1 afis|EAW17196.1 afla|EED45178.1 afum|EAL86624.1 agla|OJJ86658.1 akaw|GAA92778.1 aluc|OJZ82705.1 anid|CBF69887.1 anig|EHA21160.1 aory|EIT73771.1 asyd|OJJ59009.1 ater|EAU31750.1 atub|OJI88938.1 aver|OJJ01606.1 azon|OJJ51828.1 asyd|OJJ53646.1

Group687: anid|CBF84338.1 anid|CBF85236.1 abra|OJJ66254.1 acar|OOF90725.1 acla|EAW13654.1 afis|EAW19951.1 afum|EAL84964.1 aluc|OJZ80562.1 atub|OJI88369.1 aver|OJI97993.1 abra|OJJ68406.1 afla|EED50983.1 akaw|GAA92542.1 aluc|OJZ90090.1 anig|EHA20984.1 aory|EIT75403.1 asyd|OJJ62976.1 atub|OJI84715.1 afla|EED45866.1 aory|EIT77017.1

Group688: aory|EIT72414.1 aory|EIT75970.1 abra|OJJ66829.1 acar|OOF91607.1 acla|EAW14412.1 afis|EAW19120.1 afla|EED46512.1 afla|EED54294.1 afum|EAL93852.1 agla|OJJ80435.1 akaw|GAA90466.1 aluc|OJZ85758.1 anid|CBF75403.1 anig|EHA24798.1 asyd|OJJ59823.1 ater|EAU37383.1 atub|OJI79543.1 aver|OJJ02464.1 awen|OJJ37451.1 azon|OJJ51128.1

Group689: aory|EIT73542.1 aory|EIT78881.1 awen|OJJ31263.1 awen|OJJ35154.1 awen|OJJ38678.1 abra|OJJ73576.1 acar|OOG00757.1 acla|EAW07494.1 afis|EAW22527.1 afla|EED54555.1 afum|EAL90667.1 agla|OJJ88671.1 akaw|GAA88574.1 aluc|OJZ88476.1 anig|EHA18193.1 asyd|OJJ54459.1 ater|EAU39184.1 atub|OJI91071.1 aver|OJJ07697.1 azon|OJJ49947.1

Group690: asyd|OJJ52010.1 asyd|OJJ52511.1 asyd|OJJ58584.1 abra|OJJ76189.1 acar|OOF99753.1 acla|EAW13913.1 afis|EAW18624.1 afum|EAL93335.1 agla|OJJ81236.1 akaw|GAA82133.1 aluc|OJZ92039.1 anid|CBF70490.1 anig|EHA22972.1 aory|EIT78164.1 ater|EAU34924.1 atub|OJI86473.1 aver|OJJ01188.1 awen|OJJ32087.1 azon|OJJ46666.1 afla|EED52723.1

Group691: asyd|OJJ52365.1 asyd|OJJ62740.1 aver|OJI96783.1 aver|OJJ01982.1 abra|OJJ74531.1 acar|OOF95221.1 acla|EAW07161.1 afis|EAW22176.1 afla|EED57555.1 afum|EAL91013.1 agla|OJJ87878.1 akaw|GAA82069.1 aluc|OJZ92113.1 anid|CBF89075.1 anig|EHA26832.1 aory|EIT81844.1 ater|EAU34181.1 atub|OJI86546.1 awen|OJJ34753.1 azon|OJJ45807.1

Group692: asyd|OJJ52442.1 asyd|OJJ59129.1 ater|EAU33481.1 ater|EAU38150.1 abra|OJJ76293.1 acar|OOF99538.1 acla|EAW14323.1 afis|EAW18965.1 afla|EED51014.1 afum|EAL93697.2 agla|OJJ85347.1 akaw|GAA82221.1 aluc|OJZ91929.1 anid|CBF69609.1 anig|EHA22891.1 aory|EIT76953.1 atub|OJI86372.1 aver|OJJ01717.1 awen|OJJ31565.1 azon|OJJ46798.1

Group693: asyd|OJJ52683.1 asyd|OJJ58527.1 aver|OJJ01128.1 aver|OJJ08669.1 abra|OJJ76471.1 acar|OOF99708.1 acla|EAW13962.1 afis|EAW18671.1 afla|EED50620.1 afum|EAL93390.1 agla|OJJ82404.1 akaw|GAA82383.1 aluc|OJZ91738.1 anid|CBF70581.1 anig|EHA22744.1 aory|EIT78086.1 ater|EAU38240.1 atub|OJI86196.1 awen|OJJ32143.1 azon|OJJ46724.1

Group694: asyd|OJJ52787.1 asyd|OJJ58807.1 aver|OJJ01410.1 aver|OJJ08762.1 abra|OJJ67374.1 acar|OOF92437.1 acla|EAW06381.1 afis|EAW18487.1 afla|EED52921.1 afum|EAL93176.1 agla|OJJ85082.1 akaw|GAA86448.1 aluc|OJZ84991.1 anid|CBF70149.1 anig|EHA25203.1 aory|EIT78597.1 ater|EAU29912.1 atub|OJI80609.1 awen|OJJ31041.1 azon|OJJ48423.1

Group695: asyd|OJJ54188.1 asyd|OJJ58628.1 aver|OJJ01234.1 aver|OJJ08551.1 abra|OJJ71860.1 acar|OOF96532.1 acla|EAW13891.1 afis|EAW18603.1 afla|EED52744.1 afum|EAL93311.1 agla|OJJ81655.1 akaw|GAA85055.1 aluc|OJZ86597.1 anid|CBF70407.1 anig|EHA22065.1 aory|EIT78216.1 ater|EAU34898.1 atub|OJI83926.1 awen|OJJ31169.1 azon|OJJ46646.1

Group696: asyd|OJJ54443.1 asyd|OJJ57339.1 awen|OJJ33856.1 awen|OJJ40919.1 abra|OJJ66256.1 acar|OOF90727.1 aluc|OJZ80564.1 anig|EHA25634.1 atub|OJI88367.1 awen|OJJ38260.1 abra|OJJ77747.1 akaw|GAA83887.1 aluc|OJZ90808.1 anid|CBF81976.1 anig|EHA19536.1 aory|EIT82874.1 atub|OJI82797.1 aver|OJJ05823.1 azon|OJJ50277.1 afla|EED47338.1

Group697: asyd|OJJ55918.1 asyd|OJJ60822.1 aver|OJJ00148.1 aver|OJJ04871.1 abra|OJJ75721.1 acar|OOF90261.1 acla|EAW09377.1 afis|EAW21758.1 afla|EED58019.1 afum|EAL92788.2 agla|OJJ84392.1 akaw|GAA84457.1 aluc|OJZ89173.1 anid|CBF83680.1 anig|EHA23382.1 aory|EIT79759.1 ater|EAU38432.1 atub|OJI85617.1 awen|OJJ34042.1 azon|OJJ48199.1

Group698: asyd|OJJ57403.1 asyd|OJJ62790.1 aver|OJI96837.1 aver|OJJ07752.1 abra|OJJ73561.1 acar|OOG00743.1 acla|EAW07478.1 afis|EAW22510.1 afla|EED54538.1 afum|EAL90683.1 agla|OJJ80868.1 akaw|GAA88593.1 aluc|OJZ88500.1 anid|CBF88979.1 anig|EHA18209.1 aory|EIT78737.1 ater|EAU39204.1 atub|OJI91087.1 awen|OJJ35136.1 azon|OJJ49963.1

Group699: asyd|OJJ61696.1 asyd|OJJ63657.1 aver|OJI95780.1 aver|OJI97277.1 abra|OJJ73686.1 acar|OOG00858.1 acla|EAW07601.1 afis|EAW22632.1 afla|EED49882.1 afum|EAL90561.1 agla|OJJ80987.1 akaw|GAA88473.1 aluc|OJZ88363.1 anid|CBF88232.1 anig|EHA18095.1 aory|EIT72462.1 ater|EAU39076.1 atub|OJI90959.1 awen|OJJ35254.1 azon|OJJ49852.1

Group700: asyd|OJJ63591.1 asyd|OJJ65132.1 abra|OJJ65990.1 acar|OOF92883.1 acla|EAW13601.1 afis|EAW20004.1 afla|EED46009.1 afum|EAL84915.1 agla|OJJ87963.1 akaw|GAA90864.1 aluc|OJZ82969.1 anig|EHA21385.1 aory|EIT75787.1 ater|EAU29342.1 atub|OJI89202.1 aver|OJJ07612.1 awen|OJJ30013.1 azon|OJJ43642.1 aver|OJI97342.1 ater|EAU30135.1

Group701: ater|EAU29422.1 ater|EAU29976.1 awen|OJJ32730.1 awen|OJJ40859.1 abra|OJJ72200.1 acar|OOF96820.1 acla|EAW14947.1 afis|EAW24462.1 afla|EED55204.1 afum|EAL86133.1 agla|OJJ80320.1 akaw|GAA84736.1 aluc|OJZ86950.1 anid|CBF73838.1 anig|EHA21697.1 aory|EIT83625.1 asyd|OJJ56557.1 atub|OJI83573.1 aver|OJJ06163.1 azon|OJJ45283.1

Group702: aver|OJI99622.1 aver|OJJ08782.1 abra|OJJ69941.1 acar|OOF93451.1 acla|EAW08881.1 afis|EAW15514.1 afla|EED49021.1 agla|OJJ80185.1 akaw|GAA87718.1 aluc|OJZ85345.1 anid|CBF87265.1 anig|EHA25533.1 asyd|OJJ60386.1 ater|EAU32610.1 atub|OJI80225.1 aver|OJI99715.1 awen|OJJ38168.1 azon|OJJ43128.1 afum|EAL85817.1 aory|EIT82522.1

Group703: aver|OJJ00302.1 aver|OJJ08608.1 abra|OJJ68423.1 acar|OOF93930.1 acla|EAW11595.1 afis|EAW23861.1 afla|EED51802.1 afum|EAL87289.1 agla|OJJ87920.1 akaw|GAA86949.1 aluc|OJZ90074.1 anid|CBF82425.1 anig|EHA20999.1 aory|EIT81707.1 asyd|OJJ54128.1 asyd|OJJ60984.1 ater|EAU30753.1 atub|OJI84732.1 awen|OJJ36149.1 azon|OJJ43896.1

Group704: aver|OJJ03177.1 aver|OJJ07034.1 abra|OJJ72236.1 acar|OOF96858.1 acla|EAW14993.1 afis|EAW24519.1 afla|EED47314.1 afum|EAL86185.1 akaw|GAA84703.1 aluc|OJZ86984.1 anid|CBF71222.1 anig|EHA21663.1 aory|EIT82888.1 asyd|OJJ55173.1 ater|EAU29959.1 atub|OJI83535.1 awen|OJJ32602.1 afla|EED57070.1 aory|EIT82265.1 azon|OJJ48731.1

Group705: azon|OJJ42149.1 azon|OJJ43551.1 abra|OJJ67976.1 acar|OOF92817.1 acla|EAW11759.1 afis|EAW23792.1 afla|EED51600.1 afum|EAL87226.1 agla|OJJ78821.1 akaw|GAA91227.1 aluc|OJZ82843.1 anid|CBF79854.1 anig|EHA21281.1 aory|EIT80489.1 asyd|OJJ58158.1 ater|EAU31332.1 atub|OJI89073.1 aver|OJJ04245.1 awen|OJJ33749.1 awen|OJJ34607.1

Group706: azon|OJJ42457.1 azon|OJJ44531.1 abra|OJJ77870.1 acar|OOF98903.1 afla|EED56222.1 akaw|GAA83780.1 aluc|OJZ90944.1 anid|CBF86194.1 anig|EHA19652.1 aory|EIT73290.1 asyd|OJJ64539.1 ater|EAU33742.1 atub|OJI82936.1 aver|OJI98635.1 awen|OJJ38717.1 acla|EAW11254.1 afis|EAW19361.1 afum|EAL94081.1 afla|EED49734.1 aory|EIT77649.1

Group707: azon|OJJ43566.1 azon|OJJ47404.1 azon|OJJ43567.1 abra|OJJ73130.1 acla|EAW12351.1 afis|EAW16776.1 afla|EED50851.1 afum|EAL86345.1 agla|OJJ79835.1 anig|EHA18704.1 aory|EIT78959.1

ater|EAU34771.1 azon|OJJ42286.1 akaw|GAA89139.1 aluc|OJZ81697.1 aory|EIT81792.1 atub|OJI88483.1 awen|OJJ29815.1 afla|EED55657.1 awen|OJJ34470.1

Group708: abra|OJJ65712.1 acla|EAW11438.1 afis|EAW24094.1 awen|OJJ32718.1 azon|OJJ51154.1 abra|OJJ68624.1 acla|EAW12144.1 aluc|OJZ81931.1 aory|EIT78678.1 asyd|OJJ58603.1 atub|OJI84484.1 abra|OJJ70046.1 acar|OOF98851.1 acla|EAW07758.1 akaw|GAA93052.1 anig|EHA25445.1 atub|OJI80326.1 ater|EAU38181.1 azon|OJJ51651.1 awen|OJJ36405.1

Group709: abra|OJJ65771.1 acar|OOG00532.1 afis|EAW15669.1 afum|EAL85679.1 agla|OJJ81807.1 akaw|GAA90889.1 aluc|OJZ82940.1 anig|EHA21360.1 asyd|OJJ53833.1 ater|EAU32495.1 atub|OJI89172.1 aver|OJJ07938.1 awen|OJJ38345.1 abra|OJJ69063.1 acar|OOF91220.1 akaw|GAA90593.1 aluc|OJZ86051.1 anig|EHA24560.1 atub|OJI79848.1 aver|OJJ03010.1

Group710: abra|OJJ65912.1 acar|OOF94962.1 afla|EED52047.1 agla|OJJ80820.1 akaw|GAA92320.1 aluc|OJZ90218.1 anig|EHA23716.1 aory|EIT81383.1 atub|OJI81650.1 abra|OJJ78044.1 agla|OJJ88747.1 aluc|OJZ80304.1 anid|CBF82048.1 asyd|OJJ54231.1 ater|EAU30133.1 atub|OJI87457.1 aver|OJJ05184.1 awen|OJJ39981.1 awen|OJJ41941.1 ater|EAU32853.1

Group711: abra|OJJ66027.1 acar|OOF98054.1 akaw|GAA84509.1 aluc|OJZ89238.1 atub|OJI87439.1 awen|OJJ39904.1 azon|OJJ45880.1 abra|OJJ68242.1 acar|OOG00392.1 acla|EAW08538.1 afis|EAW23576.1 afla|EED48210.1 afum|EAL87889.1 anid|CBF86753.1 aory|EIT72922.1 asyd|OJJ64708.1 ater|EAU32766.1 aver|OJI98810.1 awen|OJJ29877.1 acar|OOG00089.1

Group712: abra|OJJ66125.1 acar|OOF94014.1 acla|EAW12396.1 afis|EAW16844.1 afla|EED54593.1 afum|EAL86390.1 akaw|GAA92685.1 aluc|OJZ90143.1 anid|CBF82122.1 anig|EHA20926.1 aory|EIT78860.1 asyd|OJJ54293.1 ater|EAU29537.1 ater|EAU37948.1 atub|OJI84656.1 aver|OJJ08125.1 awen|OJJ41474.1 anid|CBF87727.1 awen|OJJ36145.1 aver|OJJ05248.1

Group713: abra|OJJ66251.1 acar|OOF90722.1 acla|EAW13684.1 afis|EAW19922.1 afla|EED50973.1 afum|EAL84987.2 agla|OJJ79334.1 akaw|GAA89036.1 aluc|OJZ80559.1 anid|CBF78449.1 anig|EHA25638.1 aory|EIT77050.1 asyd|OJJ52716.1 ater|EAU39414.1 atub|OJI88372.1 aver|OJJ04745.1 awen|OJJ29958.1 ater|EAU38132.1 awen|OJJ36275.1 anid|CBF71116.1

Group714: abra|OJJ66260.1 acar|OOF90729.1 afla|EED53425.1 agla|OJJ88623.1 aluc|OJZ80567.1 anid|CBF87285.1 anig|EHA25631.1 aory|EIT75214.1 asyd|OJJ56365.1 atub|OJI88364.1 aver|OJJ06349.1 awen|OJJ40804.1 azon|OJJ42356.1 abra|OJJ68264.1 acar|OOF97543.1 akaw|GAA83201.1 aluc|OJZ83702.1 anig|EHA18853.1 atub|OJI90074.1 acla|EAW12046.1

Group715: abra|OJJ66444.1 acar|OOG00548.1 acla|EAW08961.1 afis|EAW15440.1 afla|EED49520.1 afum|EAL84206.1 agla|OJJ89304.1 akaw|GAA86649.1 aluc|OJZ85207.1 anid|CBF80104.1 anig|EHA26091.1 aory|EIT82592.1 asyd|OJJ58321.1 aver|OJI98310.1 awen|OJJ32245.1 azon|OJJ50232.1 asyd|OJJ56181.1 aver|OJJ02088.1 asyd|OJJ52211.1 awen|OJJ36078.1

Group716: abra|OJJ66628.1 acar|OOF91051.1 acla|EAW10868.1 afis|EAW20990.1 afla|EED46119.1 afum|EAL84639.1 agla|OJJ84068.1 akaw|GAA91097.1 aluc|OJZ81420.1 anid|CBF82620.1 anig|EHA26001.1 aory|EIT77263.1 asyd|OJJ54019.1 ater|EAU38639.1 atub|OJI88761.1 aver|OJJ08721.1 awen|OJJ34320.1 azon|OJJ48067.1 aory|EIT76479.1 afla|EED50344.1

Group717: abra|OJJ66789.1 acar|OOF93863.1 afla|EED49543.1 akaw|GAA86855.1 aluc|OJZ89976.1 anig|EHA27350.1 aory|EIT82644.1 atub|OJI84832.1 acla|EAW08552.1 afis|EAW23594.1 afum|EAL87873.1 agla|OJJ86851.1 aver|OJJ05126.1 awen|OJJ39916.1 azon|OJJ45890.1 afla|EED49567.1 aory|EIT82571.1 aver|OJI96425.1 anid|CBF82020.1 asyd|OJJ56199.1

Group718: abra|OJJ66926.1 acar|OOF91694.1 akaw|GAA91491.1 anig|EHA24880.1 atub|OJI79444.1 abra|OJJ67524.1 acar|OOG00111.1 agla|OJJ87236.1 awen|OJJ29656.1 aluc|OJZ85159.1 anid|CBF84480.1 aver|OJI96719.1 awen|OJJ39541.1 afis|EAW23743.1 asyd|OJJ53652.1 aver|OJI98697.1 asyd|OJJ64600.1 agla|OJJ88829.1 aver|OJJ04862.1 ater|EAU31016.1

Group719: abra|OJJ66927.1 acar|OOF91695.1 acla|EAW14492.1 afis|EAW19206.1 afla|EED47706.1 afum|EAL93936.1 agla|OJJ88505.1 akaw|GAA91490.1 aluc|OJZ85652.1 anid|CBF78561.1 anig|EHA24881.1 aory|EIT80267.1 asyd|OJJ65063.1 ater|EAU37266.1 atub|OJI79443.1 aver|OJI99187.1 awen|OJJ37559.1 azon|OJJ51221.1 ater|EAU31748.1 awen|OJJ37930.1

Group720: abra|OJJ66958.1 acar|OOF93650.1 acla|EAW14962.1 afis|EAW24485.1 afla|EED45449.1 afum|EAL86154.1 agla|OJJ88956.1 akaw|GAA92203.1 aluc|OJZ85618.1 anid|CBF82826.1 anig|EHA24908.1 aory|EIT82465.1 asyd|OJJ55929.1 ater|EAU37956.1 atub|OJI79411.1 aver|OJJ04879.1 awen|OJJ34538.1 azon|OJJ51794.1 agla|OJJ83743.1 aver|OJJ00902.1

Group721: abra|OJJ67036.1 acla|EAW07070.1 afis|EAW20317.1 afum|EAL85445.1 agla|OJJ78448.1 akaw|GAA93114.1 abra|OJJ72609.1 acar|OOF94646.1 akaw|GAA85235.1 aluc|OJZ90591.1 anig|EHA24030.1 atub|OJI82014.1 azon|OJJ44436.1 aver|OJJ06054.1 acla|EAW08310.1 asyd|OJJ57393.1 azon|OJJ45053.1 acla|EAW15313.1 afla|EED55819.1 aory|EIT76576.1

Group722: abra|OJJ67530.1 afis|EAW20793.1 anig|EHA18697.1 ater|EAU31981.1 atub|OJI91610.1 aver|OJJ05285.1 abra|OJJ74028.1 acar|OOG01145.1 afla|EED54908.1 agla|OJJ83374.1 akaw|GAA87511.1 aluc|OJZ88009.1 anid|CBF87701.1 anig|EHA17789.1 aory|EIT80304.1 asyd|OJJ63330.1 atub|OJI90618.1 awen|OJJ38650.1 aver|OJI97632.1 anid|CBF84596.1

Group723: abra|OJJ67629.1 acar|OOF91975.1 acla|EAW08578.1 afis|EAW19829.1 afla|EED55476.1 afum|EAL85063.1 akaw|GAA89149.1 aluc|OJZ81687.1 anid|CBF81789.1 anig|EHA25756.1 aory|EIT80791.1 asyd|OJJ59633.1 ater|EAU38817.1 atub|OJI88494.1 aver|OJJ08870.1 azon|OJJ44622.1 acar|OOF94209.1 anid|CBF87415.1 asyd|OJJ52901.1 acla|EAW12655.1

Group724: abra|OJJ67755.1 agla|OJJ81288.1 aver|OJJ06529.1 awen|OJJ36303.1 abra|OJJ68240.1 acar|OOF91925.1 afis|EAW22251.1 afla|EED45387.1 anig|EHA27220.1 aory|EIT82443.1 asyd|OJJ64210.1 atub|OJI82856.1 awen|OJJ29879.1 afla|EED49833.1 agla|OJJ89342.1 aory|EIT83402.1 awen|OJJ29870.1 anid|CBF88940.1 aver|OJI99604.1 asyd|OJJ56544.1

Group725: abra|OJJ68054.1 acla|EAW07812.1 afla|EED48275.1 agla|OJJ87900.1 akaw|GAA90900.1 aluc|OJZ82926.1 anid|CBF87851.1 anig|EHA21348.1 aory|EIT72960.1 asyd|OJJ56525.1 ater|EAU29450.1 atub|OJI89156.1 aver|OJJ06198.1 awen|OJJ31590.1 aluc|OJZ80199.1 asyd|OJJ63418.1 ater|EAU38861.1 aver|OJI97520.1 awen|OJJ38874.1 awen|OJJ39633.1

Group726: abra|OJJ68111.1 acla|EAW15307.1 afla|EED55277.1 agla|OJJ83148.1 akaw|GAA92450.1 aluc|OJZ81568.1 anig|EHA27392.1 aory|EIT75017.1 asyd|OJJ62945.1 ater|EAU32219.1 atub|OJI88617.1 aver|OJI98045.1 awen|OJJ35781.1 afla|EED53156.1 awen|OJJ29938.1 ater|EAU30130.1 asyd|OJJ62946.1 aver|OJI98044.1 aver|OJJ07921.1 aver|OJJ03358.1

Group727: abra|OJJ68833.1 acar|OOF90556.1 acla|EAW11325.1 afis|EAW18170.1 afla|EED56323.1 afum|EAL87532.1 agla|OJJ85940.1 akaw|GAA90418.1 aluc|OJZ85812.1 anid|CBF82352.1 anig|EHA24747.1 aory|EIT73358.1 asyd|OJJ54213.1 ater|EAU33984.1 atub|OJI79598.1 aver|OJJ08518.1 awen|OJJ31290.1 ater|EAU33834.1 awen|OJJ30517.1 aluc|OJZ80361.1

Group728: abra|OJJ68942.1 acar|OOF91523.1 acla|EAW14240.1 afis|EAW19040.1 afla|EED50395.1 afum|EAL93769.1 agla|OJJ84572.1 akaw|GAA90311.1 aluc|OJZ85928.1 anig|EHA24668.1 aory|EIT82123.1 asyd|OJJ65324.1 ater|EAU32079.1 atub|OJI79718.1 aver|OJI99469.1 azon|OJJ46176.1 afla|EED56613.1 agla|OJJ84621.1 aory|EIT79679.1 awen|OJJ31261.1

Group729: abra|OJJ69052.1 acar|OOF91238.1 afis|EAW20344.1 afla|EED45064.1 afum|EAL85464.1 agla|OJJ85556.1 akaw|GAA90584.1 aluc|OJZ86042.1 anid|CBF85628.1 anig|EHA24569.1 aory|EIT73572.1 asyd|OJJ57132.1 ater|EAU37144.1 atub|OJI79829.1 aver|OJJ03172.1 awen|OJJ38366.1 azon|OJJ45225.1 ater|EAU37679.1 afla|EED56599.1 aory|EIT79596.1

Group730: abra|OJJ69229.1 acar|OOF91091.1 acla|EAW12717.1 afis|EAW15814.1 afla|EED52103.1 afum|EAL85537.1 agla|OJJ83067.1 akaw|GAA92237.1 aluc|OJZ86209.1 anid|CBF70901.1 anig|EHA24417.1 asyd|OJJ52596.1 ater|EAU33434.1 atub|OJI80003.1 aver|OJJ04623.1 awen|OJJ38527.1 afis|EAW17879.1 afum|EAL91687.1 aver|OJJ05547.1 awen|OJJ41685.1

Group731: abra|OJJ69242.1 acar|OOF91077.1 acla|EAW08606.1 afis|EAW15827.1 afla|EED51218.1 afum|EAL85524.1 agla|OJJ81990.1 akaw|GAA92251.1 aluc|OJZ86222.1 anid|CBF70872.1 anig|EHA24405.1 aory|EIT80412.1 asyd|OJJ52582.1 ater|EAU33421.1 atub|OJI80016.1 aver|OJJ04610.1 awen|OJJ38541.1 azon|OJJ43722.1 asyd|OJJ54750.1 aver|OJJ07417.1

Group732: abra|OJJ69380.1 acar|OOF94345.1 acla|EAW08419.1 afis|EAW23450.1 afla|EED57353.1 afum|EBA27331.1 agla|OJJ86147.1 akaw|GAA88248.1 aluc|OJZ82215.1 anid|CBF89797.1 anig|EHA28525.1 aory|EIT80130.1 asyd|OJJ53729.1 atub|OJI88220.1 aver|OJI98284.1 awen|OJJ31384.1 azon|OJJ47909.1 asyd|OJJ62197.1 aver|OJI96309.1 awen|OJJ39779.1

Group733: abra|OJJ69833.1 acar|OOF93358.1 afla|EED54260.1 agla|OJJ79197.1 akaw|GAA87827.1 aluc|OJZ85463.1 anid|CBF88684.1 anig|EHA25005.1 aory|EIT76164.1 asyd|OJJ56863.1 atub|OJI80100.1 aver|OJJ03462.1 awen|OJJ32851.1 abra|OJJ70031.1 acla|EAW15278.1 afis|EAW16787.1 afum|EAL86357.1 anig|EHA18716.1 ater|EAU36812.1 atub|OJI79320.1

Group734: abra|OJJ69864.1 acar|OOF90332.1 afis|EAW22301.1 afla|EED45628.1 agla|OJJ82723.1 akaw|GAA87792.1 aluc|OJZ85427.1 anid|CBF85184.1 anig|EHA25606.1 aory|EIT75771.1 ater|EAU32684.1 atub|OJI80148.1 aver|OJI96605.1 awen|OJJ31411.1 azon|OJJ42404.1 acar|OOF91868.1 afla|EED49499.1 aory|EIT82625.1 asyd|OJJ62470.1 ater|EAU38702.1

Group735: abra|OJJ70266.1 afla|EED54695.1 akaw|GAA86531.1 aluc|OJZ85079.1 anig|EHA25294.1 aory|EIT78911.1 atub|OJI80519.1 abra|OJJ78041.1 acla|EAW08785.1 afis|EAW15939.1 afla|EED54424.1 afum|EAL88574.1 agla|OJJ80367.1 aluc|OJZ80302.1 asyd|OJJ52753.1 ater|EAU31057.1 atub|OJI87455.1 aver|OJI96478.1 awen|OJJ39980.1 ater|EAU37127.1

Group736: abra|OJJ70395.1 acar|OOF90083.1 afla|EED52271.1 agla|OJJ85325.1 akaw|GAA91963.1 aluc|OJZ82654.1 anig|EHA27419.1 aory|EIT74349.1 asyd|OJJ61023.1 atub|OJI88885.1 aver|OJJ00344.1

awen|OJJ33584.1 afla|EED48143.1 aory|EIT74445.1 asyd|OJJ58659.1 afla|EED52287.1 afla|EED55148.1 aory|EIT81619.1 awen|OJJ34668.1 awen|OJJ37795.1

Group737: abra|OJJ70402.1 acar|OOF93249.1 acla|EAW14348.1 afis|EAW20782.1 afla|EED52329.1 afum|EAL84841.1 agla|OJJ85851.1 aory|EIT74419.1 asyd|OJJ62498.1 ater|EAU32955.1 aver|OJI96618.1 azon|OJJ44872.1 abra|OJJ77381.1 akaw|GAA91380.1 aluc|OJZ84789.1 anig|EHA28057.1 atub|OJI89293.1 aory|EIT74407.1 ater|EAU31843.1 azon|OJJ50673.1

Group738: acla|EAW07697.1 afis|EAW22714.1 afum|EAL90480.1 agla|OJJ78998.1 azon|OJJ49768.1 abra|OJJ76709.1 acar|OOF98067.1 akaw|GAA92489.1 aluc|OJZ91501.1 anid|CBF75580.1 anig|EHA22541.1 atub|OJI85954.1 awen|OJJ37863.1 afla|EED45147.1 aory|EIT73529.1 asyd|OJJ55486.1 aver|OJJ06565.1 awen|OJJ35349.1 awen|OJJ33591.1 azon|OJJ42132.1

Group739: abra|OJJ70566.1 acla|EAW08365.1 afis|EAW23395.1 afla|EED55562.1 afum|EAL88075.1 agla|OJJ86422.1 akaw|GAA83410.1 aluc|OJZ83909.1 anid|CBF74809.1 anig|EHA19043.1 aory|EIT82686.1 asyd|OJJ60147.1 ater|EAU35648.1 atub|OJI90280.1 aver|OJJ04116.1 awen|OJJ39484.1 azon|OJJ47686.1 acla|EAW12884.1 afis|EAW17793.1 afum|EAL91602.1

Group740: abra|OJJ70960.1 acar|OOF98170.1 acla|EAW10521.1 afis|EAW25045.1 afla|EED48351.1 afum|EAL89713.1 agla|OJJ89149.1 akaw|GAA82987.1 aluc|OJZ87572.1 anid|CBF85914.1 anig|EHA20774.1 aory|EIT80893.1 asyd|OJJ64399.1 ater|EAU35795.1 atub|OJI81077.1 aver|OJI98495.1 awen|OJJ40656.1 azon|OJJ44034.1 aory|EIT72298.1 afla|EED55406.1

Group741: abra|OJJ71146.1 acar|OOF98009.1 acar|OOF98010.1 acla|EAW10360.1 afis|EAW25233.1 afla|EED53679.1 afum|EAL89894.1 agla|OJJ79174.1 akaw|GAA82831.1 aluc|OJZ87392.1 anid|CBF85327.1 anig|EHA20621.1 anig|EHA20622.1 aory|EIT75303.1 asyd|OJJ64017.1 ater|EAU34392.1 atub|OJI81254.1 aver|OJI98103.1 awen|OJJ40899.1 azon|OJJ44526.1

Group742: abra|OJJ71723.1 acar|OOF96427.1 acla|EAW10027.1 afis|EAW15606.1 afum|EAL85727.1 akaw|GAA85167.1 aluc|OJZ86469.1 anig|EHA21441.1 atub|OJI84060.1 azon|OJJ45251.1 afla|EED55033.1 agla|OJJ82925.1 aory|EIT76156.1 asyd|OJJ53836.1 afla|EED48194.1 aory|EIT72936.1 azon|OJJ45977.1 afla|EED54138.1 aory|EIT81964.1 asyd|OJJ56435.1

Group743: abra|OJJ71853.1 acar|OOF96523.1 afla|EED52750.1 afum|EAL93305.1 agla|OJJ81650.1 akaw|GAA85064.1 aluc|OJZ86587.1 anid|CBF70395.1 anig|EHA22073.1 aory|EIT78183.1 asyd|OJJ58634.1 atub|OJI83936.1 aver|OJJ01240.1 awen|OJJ31160.1 acla|EAW13885.1 afis|EAW18597.1 ater|EAU34891.1 azon|OJJ46636.1 asyd|OJJ54226.1 aver|OJJ05179.1

Group744: abra|OJJ72081.1 acar|OOF90349.1 afla|EED45358.1 agla|OJJ82915.1 aluc|OJZ86835.1 aory|EIT82374.1 asyd|OJJ56205.1 atub|OJI83683.1 aver|OJJ05132.1 awen|OJJ35919.1 azon|OJJ42174.1 afla|EED45415.1 akaw|GAA92568.1 aluc|OJZ82578.1 anig|EHA22490.1 aory|EIT73709.1 asyd|OJJ65181.1 aver|OJI99335.1 aory|EIT82376.1 awen|OJJ39611.1

Group745: abra|OJJ72207.1 acar|OOF96828.1 acla|EAW14969.1 afis|EAW24492.1 afla|EED54697.1 afum|EAL86161.1 agla|OJJ80329.1 akaw|GAA84727.1 aluc|OJZ86959.1 anid|CBF71478.1 anig|EHA21689.1 aory|EIT78783.1 asyd|OJJ54943.1 ater|EAU29944.1 atub|OJI83564.1 aver|OJJ07252.1 awen|OJJ33962.1 azon|OJJ42613.1 azon|OJJ48606.1 awen|OJJ32703.1

Group746: abra|OJJ72535.1 acar|OOF94717.1 acla|EAW12742.1 afis|EAW17905.1 afla|EED46297.1 afum|EAL91714.1 agla|OJJ83037.1 akaw|GAA86257.1 aluc|OJZ90515.1 anid|CBF76985.1 anig|EHA23970.1 aory|EIT76354.1 asyd|OJJ57035.1 ater|EAU33123.1 atub|OJI81939.1 aver|OJJ03257.1 awen|OJJ41645.1 azon|OJJ48367.1 aluc|OJZ90514.1 aory|EIT76342.1

Group747: abra|OJJ72848.1 acar|OOF96049.1 acla|EAW11395.1 afis|EAW24026.1 afla|EED49128.1 afum|EAL87457.1 agla|OJJ86959.1 akaw|GAA85449.1 aluc|OJZ83357.1 anid|CBF84482.1 anig|EHA24222.1 aory|EIT78568.1 asyd|OJJ53530.1 ater|EAU29718.1 atub|OJI82248.1 aver|OJJ08220.1 awen|OJJ35499.1 azon|OJJ43175.1 acla|EAW12378.1 awen|OJJ40008.1

Group748: abra|OJJ73145.1 acar|OOG00357.1 acla|EAW07021.1 afis|EAW16647.1 afla|EED45442.1 afum|EAL89251.1 agla|OJJ82595.1 akaw|GAA88027.1 aluc|OJZ81286.1 anid|CBF82058.1 anig|EHA18585.1 aory|EIT82414.1 asyd|OJJ54238.1 ater|EAU39572.1 atub|OJI91493.1 aver|OJJ05190.1 awen|OJJ40067.1 azon|OJJ45557.1 aver|OJJ01178.1 afla|EED50972.1

Group749: abra|OJJ73181.1 acar|OOG00417.1 afla|EED47213.1 afum|EAL89070.1 akaw|GAA87994.1 aluc|OJZ81249.1 anig|EHA18557.1 ater|EAU36561.1 awen|OJJ40360.1 azon|OJJ49208.1 acla|EAW06859.1 afis|EAW16476.1 agla|OJJ82869.1 anid|CBF75495.1 aory|EIT79277.1 asyd|OJJ59750.1 atub|OJI91445.1 aver|OJJ02537.1 acar|OOG00418.1 atub|OJI91444.1

Group750: abra|OJJ73280.1 acar|OOG00516.1 acla|EAW06955.1 afis|EAW16570.1 afla|EED56094.1 afum|EAL89167.1 agla|OJJ82745.1 akaw|GAA92725.1 aluc|OJZ88769.1 anid|CBF81824.1 anig|EHA18465.1 aory|EIT76687.1 asyd|OJJ59656.1 ater|EAU36638.1 atub|OJI91342.1 aver|OJJ02626.1 awen|OJJ40248.1 azon|OJJ49110.1 afum|EAL87777.1 aory|EIT72969.1

Group751: abra|OJJ73388.1 acar|OOG00590.1 acla|EAW07000.1 afis|EAW16615.1 afla|EED44990.1 afum|EAL89221.1 agla|OJJ82629.1 akaw|GAA83101.1 aluc|OJZ88649.1 anid|CBF82980.1 anig|EHA18340.1 aory|EIT73600.1 asyd|OJJ53668.1 ater|EAU36700.1 atub|OJI91236.1 aver|OJJ07774.1 awen|OJJ40187.1 azon|OJJ49474.1 anig|EHA18404.1 ater|EAU37639.1

Group752: abra|OJJ73407.1 acar|OOG00604.1 acla|EAW07012.1 afis|EAW16632.1 afla|EED48324.1 afum|EAL89228.1 agla|OJJ82617.1 akaw|GAA83084.1 aluc|OJZ88631.1 anid|CBF82011.1 anig|EHA18321.1 aory|EIT73498.1 asyd|OJJ56190.1 ater|EAU36752.1 atub|OJI91215.1 aver|OJJ05118.1 awen|OJJ40077.1 azon|OJJ43490.1 aory|EIT76284.1 afla|EED48295.1

Group753: abra|OJJ73423.1 acar|OOG00627.1 acla|EAW07036.1 afis|EAW16662.1 afla|EED44895.1 afum|EAL89266.1 agla|OJJ78707.1 akaw|GAA83069.1 aluc|OJZ88620.1 anid|CBF84550.1 anig|EHA18311.1 aory|EIT83238.1 asyd|OJJ53473.1 ater|EAU31782.1 atub|OJI91205.1 aver|OJJ08283.1 awen|OJJ40051.1 azon|OJJ47485.1 afla|EED56569.1 aory|EIT79633.1

Group754: abra|OJJ73685.1 acar|OOG00855.1 acar|OOG00856.1 acla|EAW07600.1 afis|EAW22631.1 afla|EED49880.1 afla|EED49881.1 afum|EBA27410.1 agla|OJJ80989.1 akaw|GAA88474.1 aluc|OJZ88364.1 anid|CBF88234.1 anig|EHA18096.1 aory|EIT72461.1 ater|EAU39077.1 atub|OJI90961.1 aver|OJI97276.1 awen|OJJ35252.1 azon|OJJ49854.1 asyd|OJJ63658.1

Group755: abra|OJJ73793.1 acar|OOG00954.1 acla|EAW07685.1 afis|EAW22700.1 afla|EED49986.1 afum|EAL90492.1 agla|OJJ79009.1 akaw|GAA89412.1 aluc|OJZ88261.1 anid|CBF88081.1 anig|EHA18006.1 aory|EIT74617.1 asyd|OJJ63558.1 ater|EAU39002.1 atub|OJI90860.1 aver|OJI97374.1 awen|OJJ35337.1 azon|OJJ49781.1 ater|EAU30260.1 awen|OJJ38024.1

Group756: abra|OJJ73803.1 acar|OOG00958.1 acla|EAW07675.1 afis|EAW22690.1 afla|EED49983.1 afum|EAL90502.1 agla|OJJ79015.1 akaw|GAA89422.1 aluc|OJZ88250.1 anid|CBF88089.1 anig|EHA17993.1 aory|EIT74622.1 asyd|OJJ63564.1 ater|EAU39009.1 atub|OJI90849.1 aver|OJI97369.1 awen|OJJ33510.1 azon|OJJ49784.1 acar|OOG00352.1 afla|EED46078.1

Group757: abra|OJJ73866.1 acar|OOF97590.1 acla|EAW06976.1 afis|EAW16590.1 afla|EED46107.1 afum|EAL89185.1 agla|OJJ83704.1 akaw|GAA89476.1 aluc|OJZ88187.1 anid|CBF76004.1 anig|EHA17938.1 aory|EIT76398.1 asyd|OJJ57426.1 ater|EAU29326.1 atub|OJI90786.1 aver|OJJ03990.1 awen|OJJ40225.1 azon|OJJ49083.1 anid|CBF75976.1 asyd|OJJ60263.1

Group758: abra|OJJ74117.1 acar|OOG01222.1 acla|EAW07959.1 afis|EAW22994.1 afla|EED58187.1 afum|EAL90205.1 agla|OJJ83290.1 akaw|GAA87590.1 aluc|OJZ87915.1 anid|CBF87568.1 anig|EHA17710.1 aory|EIT79090.1 asyd|OJJ63243.1 ater|EAU30624.1 atub|OJI90527.1 aver|OJI97713.1 awen|OJJ38567.1 azon|OJJ50434.1 anig|EHA17709.1 afum|EAL90206.1

Group759: abra|OJJ74269.1 acar|OOF95462.1 acla|EAW07313.1 afis|EAW22333.1 afla|EED57749.1 afum|EAL90859.1 agla|OJJ87713.1 akaw|GAA86754.1 aluc|OJZ89864.1 anid|CBF78344.1 anig|EHA27064.1 aory|EIT81050.1 asyd|OJJ56131.1 ater|EAU39462.1 atub|OJI84944.1 aver|OJJ05072.1 awen|OJJ34911.1 asyd|OJJ61756.1 aver|OJI95855.1 agla|OJJ78853.1

Group760: abra|OJJ74757.1 acar|OOF95750.1 acla|EAW10654.1 afis|EAW24913.1 afla|EED46998.1 afum|EAL89567.1 agla|OJJ82924.1 akaw|GAA83654.1 aluc|OJZ92341.1 anid|CBF75675.1 anig|EHA26633.1 aory|EIT72680.1 asyd|OJJ57485.1 ater|EAU36385.1 atub|OJI86767.1 aver|OJJ05695.1 awen|OJJ40509.1 azon|OJJ49368.1 afla|EED48195.1 aory|EIT72956.1

Group761: abra|OJJ74761.1 acar|OOF95754.1 acla|EAW10650.1 afis|EAW24917.1 afla|EED47002.1 afum|EAL89572.2 agla|OJJ82145.1 akaw|GAA83650.1 aluc|OJZ92345.1 anid|CBF75683.1 anig|EHA26629.1 aory|EIT72569.1 asyd|OJJ57489.1 ater|EAU36381.1 atub|OJI86771.1 aver|OJJ05691.1 awen|OJJ40513.1 azon|OJJ49372.1 anid|CBF75685.1 atub|OJI86772.1

Group762: abra|OJJ74833.1 acar|OOF95823.1 acla|EAW11247.1 afis|EAW18253.1 afla|EED56199.1 afum|EAL87616.1 agla|OJJ88191.1 akaw|GAA83587.1 aluc|OJZ92410.1 anid|CBF86203.1 anig|EHA26555.1 aory|EIT73440.1 asyd|OJJ58331.1 ater|EAU33725.1 atub|OJI86840.1 aver|OJJ02080.1 awen|OJJ30627.1 azon|OJJ47986.1 aver|OJI98640.1 asyd|OJJ64544.1

Group763: abra|OJJ74891.1 acar|OOF95894.1 acla|EAW12408.1 afis|EAW17228.1 afla|EED45013.1 agla|OJJ86664.1 akaw|GAA83528.1 aluc|OJZ92468.1 anig|EHA26500.1 aory|EIT73647.1 asyd|OJJ62261.1 ater|EAU35311.1 atub|OJI86897.1 aver|OJI96370.1 awen|OJJ39852.1 afla|EED54064.1 aory|EIT76126.1 asyd|OJJ56095.1 aver|OJJ05039.1 afla|EED46550.1

Group764: abra|OJJ74964.1 acar|OOF97350.1 acla|EAW08410.1 afis|EAW23459.1 afla|EED57342.1 afum|EAL88007.2 agla|OJJ86690.1 akaw|GAA83462.1 aluc|OJZ92534.1 anid|CBF89818.1 anig|EHA26441.1 aory|EIT80102.1 asyd|OJJ62188.1 ater|EAU37472.1 atub|OJI86970.1 aver|OJI96297.1 awen|OJJ39771.1 azon|OJJ47905.1 afla|EED57341.1 aory|EIT80055.1

Group765: abra|OJJ75326.1 acar|OOF90139.1 acla|EAW12017.1 afis|EAW19657.1 afla|EED57008.1 afum|EAL86866.1 agla|OJJ80802.1 akaw|GAA92021.1 aluc|OJZ92889.1 anid|CBF82008.1 anig|EHA26074.1

aory|EIT82264.1 asyd|OJJ55074.1 asyd|OJJ65337.1 ater|EAU32873.1 atub|OJI87347.1 aver|OJI99483.1 aver|OJJ07134.1 awen|OJJ41915.1 agla|OJJ85377.1

Group766: abra|OJJ75618.1 acar|OOG00217.1 acla|EAW09356.1 afis|EAW21782.1 afla|EED46350.1 afum|EAL92810.1 agla|OJJ84341.1 akaw|GAA84367.1 aluc|OJZ89071.1 anid|CBF76611.1 anig|EHA23465.1 aory|EIT81207.1 asyd|OJJ61187.1 ater|EAU38459.1 atub|OJI85717.1 aver|OJJ00510.1 awen|OJJ34070.1 azon|OJJ48159.1 afum|EAL85431.1 anid|CBF76609.1

Group767: abra|OJJ75770.1 acla|EAW09729.1 afis|EAW21415.1 afla|EED54885.1 afum|EAL92430.1 agla|OJJ78973.1 akaw|GAA89736.1 aluc|OJZ82585.1 anid|CBF82747.1 aory|EIT79345.1 asyd|OJJ54779.1 ater|EAU34053.1 atub|OJI87188.1 awen|OJJ30924.1 azon|OJJ44473.1 afis|EAW17192.1 afum|EAL86620.1 akaw|GAA92574.1 anid|CBF80434.1 agla|OJJ79605.1

Group768: abra|OJJ75774.1 acla|EAW12650.1 afis|EAW18035.1 afla|EED45683.1 afum|EAL91846.1 agla|OJJ82507.1 anid|CBF81936.1 anig|EHA18442.1 aory|EIT76073.1 asyd|OJJ58678.1 ater|EAU31610.1 atub|OJI87184.1 aver|OJJ01290.1 awen|OJJ30920.1 acar|OOG00708.1 afla|EED54070.1 asyd|OJJ57410.1 aver|OJJ05771.1 awen|OJJ35374.1 aory|EIT75690.1

Group769: abra|OJJ76070.1 acar|OOF99859.1 acla|EAW09749.1 afis|EAW21393.1 afla|EED55810.1 afum|EAL92407.1 agla|OJJ85517.1 akaw|GAA88897.1 aluc|OJZ89524.1 anid|CBF71432.1 anig|EHA23077.1 aory|EIT76869.1 asyd|OJJ61713.1 ater|EAU35164.1 atub|OJI85279.1 aver|OJJ07221.1 awen|OJJ39311.1 azon|OJJ51541.1 asyd|OJJ54970.1 ater|EAU32747.1

Group770: abra|OJJ76073.1 acar|OOF99853.1 afis|EAW16840.1 afla|EED47451.1 afum|EAL86387.1 akaw|GAA88894.1 aluc|OJZ89527.1 anig|EHA23074.1 aory|EIT81896.1 atub|OJI85276.1 aver|OJJ06456.1 awen|OJJ36309.1 abra|OJJ76507.1 agla|OJJ80808.1 akaw|GAA92901.1 anid|CBF73893.1 anig|EHA22710.1 asyd|OJJ53660.1 ater|EAU36678.1 aver|OJJ07768.1

Group771: abra|OJJ76117.1 acar|OOF99808.1 acla|EAW09703.1 afis|EAW21437.1 afla|EED45636.1 afum|EAL92452.2 agla|OJJ85581.1 akaw|GAA88850.1 aluc|OJZ89574.1 anid|CBF76585.1 anig|EHA23033.1 aory|EIT76558.1 asyd|OJJ61172.1 ater|EAU35108.1 atub|OJI85230.1 aver|OJJ00497.1 awen|OJJ39261.1 azon|OJJ51593.1 aory|EIT75602.1 afla|EED55865.1

Group772: abra|OJJ76529.1 acar|OOF99449.1 acla|EAW14176.1 afis|EAW18891.1 afla|EED50873.1 afum|EBA27495.1 agla|OJJ81497.1 akaw|GAA82440.1 akaw|GAA82441.1 aluc|OJZ91675.1 aluc|OJZ91676.1 anid|CBF69904.1 anig|EHA22693.1 aory|EIT79015.1 asyd|OJJ58997.1 ater|EAU38063.1 atub|OJI86130.1 aver|OJJ01619.1 awen|OJJ32450.1 azon|OJJ47208.1

Group773: abra|OJJ77055.1 acar|OOF92945.1 acla|EAW13514.1 afis|EAW19907.1 afla|EED54205.1 afum|EAL84999.1 agla|OJJ88625.1 akaw|GAA86997.1 aluc|OJZ84460.1 anid|CBF86958.1 anig|EHA27775.1 aory|EIT76071.1 asyd|OJJ55597.1 ater|EAU35255.1 atub|OJI89615.1 aver|OJJ06677.1 awen|OJJ33456.1 azon|OJJ50712.1 afis|EAW20612.1 acla|EAW13697.1

Group774: abra|OJJ77217.1 acar|OOF93104.1 acla|EAW09073.1 afis|EAW16463.1 afla|EED45621.1 afum|EAL89059.1 akaw|GAA90721.1 aluc|OJZ84628.1 anid|CBF82099.1 anig|EHA27915.1 aory|EIT75489.1 ater|EAU35691.1 atub|OJI89451.1 azon|OJJ51362.1 awen|OJJ34525.1 acla|EAW06842.1 afis|EAW25646.1 asyd|OJJ54269.1 agla|OJJ79388.1 afum|EAL84332.1

Group775: abra|OJJ77859.1 acar|OOF98914.1 acla|EAW11267.1 afis|EAW18231.1 afla|EED52136.1 afum|EAL87599.1 agla|OJJ85846.1 akaw|GAA83790.1 aluc|OJZ90933.1 anid|CBF86176.1 anig|EHA19643.1 aory|EIT73335.1 asyd|OJJ64527.1 ater|EAU33760.1 atub|OJI82925.1 aver|OJI98625.1 awen|OJJ30611.1 azon|OJJ48005.1 azon|OJJ42776.1 awen|OJJ36209.1

Group776: abra|OJJ77974.1 acar|OOF98840.1 acla|EAW06664.1 afis|EAW16101.1 afla|EED48636.1 afum|EAL88733.1 agla|OJJ86121.1 akaw|GAA83686.1 aluc|OJZ91047.1 anid|CBF85446.1 anig|EHA19744.1 aory|EIT73060.1 asyd|OJJ64094.1 ater|EAU33175.1 aver|OJI98176.1 awen|OJJ30331.1 azon|OJJ45359.1 anid|CBF85444.1 asyd|OJJ64092.1 atub|OJI83043.1

Group777: abra|OJJ78228.1 acar|OOF98631.1 afis|EAW21866.1 afla|EED46443.1 afum|EAL92880.2 agla|OJJ84269.1 akaw|GAA85796.1 aluc|OJZ91303.1 anid|CBF76759.1 anig|EHA19985.1 aory|EIT72387.1 asyd|OJJ61264.1 ater|EAU30863.1 atub|OJI83294.1 aver|OJJ00590.1 awen|OJJ34155.1 azon|OJJ43474.1 afla|EED54738.1 anid|CBF76761.1 aver|OJJ00591.1

Group778: abra|OJJ78259.1 acar|OOF98601.1 acla|EAW09251.1 afis|EAW21896.1 afla|EED46412.1 afum|EBA27479.1 agla|OJJ84236.1 akaw|GAA85766.1 aluc|OJZ91333.1 anig|EHA20015.1 aory|EIT72351.1 asyd|OJJ61292.1 ater|EAU30891.1 atub|OJI83326.1 aver|OJJ00618.1 awen|OJJ34185.1 azon|OJJ43448.1 aluc|OJZ91334.1 akaw|GAA85767.1 anid|CBF76812.1

Group779: abra|OJJ66219.1 abra|OJJ77447.1 awen|OJJ34603.1 awen|OJJ39635.1 acar|OOF90664.1 afla|EED48277.1 afla|EED48278.1 akaw|GAA89067.1 aluc|OJZ80525.1 anig|EHA25670.1 anig|EHA25671.1 aory|EIT72908.1 aory|EIT72950.1 ater|EAU37355.1 atub|OJI88405.1 anid|CBF74667.1 acar|OOF90665.1 anid|CBF74669.1 awen|OJJ34600.1

Group780: abra|OJJ67019.1 abra|OJJ74812.1 akaw|GAA83601.1 akaw|GAA93299.1 anig|EHA25612.1 anig|EHA26582.1 afis|EAW20313.1 afla|EED56766.1 afum|EAL85436.1 agla|OJJ83095.1 aluc|OJZ92393.1 ater|EAU31040.1 atub|OJI86823.1 awen|OJJ33515.1 acla|EAW07069.1 aory|EIT73952.1 asyd|OJJ56123.1 aver|OJJ05065.1 azon|OJJ42468.1

Group781: abra|OJJ67077.1 abra|OJJ70209.1 akaw|GAA86567.1 akaw|GAA91080.1 aluc|OJZ85118.1 aluc|OJZ87895.1 anig|EHA20194.1 anig|EHA25338.1 aory|EIT75167.1 aory|EIT80564.1 atub|OJI80475.1 atub|OJI80757.1 acar|OOF90366.1 afis|EAW16753.1 afla|EED51374.1 afum|EAL86325.1 agla|OJJ78769.1 aver|OJJ05553.1 afla|EED53531.1

Group782: abra|OJJ68020.1 abra|OJJ77410.1 acar|OOF92063.1 acar|OOF93277.1 akaw|GAA90931.1 akaw|GAA91354.1 anig|EHA21318.1 anig|EHA28081.1 atub|OJI89119.1 atub|OJI89267.1 acla|EAW09048.1 afla|EED46361.1 agla|OJJ81295.1 aluc|OJZ82894.1 aory|EIT81187.1 asyd|OJJ63773.1 aver|OJI97164.1 awen|OJJ30865.1 aluc|OJZ84818.1

Group783: abra|OJJ68371.1 abra|OJJ72091.1 acar|OOF96736.1 acla|EAW09472.1 afis|EAW21661.1 afla|EED57928.1 afum|EAL92696.1 agla|OJJ79735.1 akaw|GAA84829.1 aluc|OJZ86846.1 anid|CBF83566.1 anig|EHA21792.1 aory|EIT82318.1 asyd|OJJ60745.1 ater|EAU38349.1 atub|OJI83673.1 aver|OJJ00077.1 awen|OJJ33927.1 azon|OJJ48635.1

Group784: acar|OOF90452.1 acar|OOF90459.1 abra|OJJ71882.1 acla|EAW13911.1 afis|EAW18623.1 afla|EED52721.1 afum|EAL93333.1 agla|OJJ81237.1 akaw|GAA85034.1 aluc|OJZ86620.1 anid|CBF70493.1 anig|EHA22046.1 aory|EIT78225.1 asyd|OJJ58583.1 ater|EAU34923.1 atub|OJI83905.1 aver|OJJ01187.1 awen|OJJ32086.1 azon|OJJ46665.1

Group785: acar|OOF90454.1 acar|OOF90462.1 abra|OJJ71884.1 acla|EAW09637.1 afis|EAW21503.1 afla|EED55939.1 afum|EAL92522.1 agla|OJJ85666.1 akaw|GAA85032.1 aluc|OJZ86622.1 anid|CBF76453.1 anig|EHA22044.1 aory|EIT76619.1 asyd|OJJ61107.1 ater|EAU35035.1 atub|OJI83903.1 aver|OJJ00424.1 awen|OJJ39192.1 azon|OJJ51508.1

Group786: acar|OOF90455.1 acar|OOF90461.1 abra|OJJ71885.1 acla|EAW09636.1 afis|EAW21504.1 afla|EED55940.1 afum|EAL92523.1 agla|OJJ85667.1 akaw|GAA85031.1 aluc|OJZ86623.1 anid|CBF76451.1 anig|EHA22043.1 aory|EIT76886.1 asyd|OJJ61106.1 ater|EAU35034.1 atub|OJI83902.1 aver|OJJ00423.1 awen|OJJ39191.1 azon|OJJ51507.1

Group787: acar|OOF90456.1 acar|OOF90463.1 abra|OJJ71886.1 acla|EAW09635.1 afis|EAW21505.1 afla|EED55941.1 afum|EAL92524.1 agla|OJJ85668.1 akaw|GAA85030.1 aluc|OJZ86624.1 anid|CBF76449.1 anig|EHA22042.1 aory|EIT76597.1 asyd|OJJ61105.1 ater|EAU35033.1 atub|OJI83901.1 aver|OJJ00422.1 awen|OJJ39190.1 azon|OJJ51506.1

Group788: acar|OOF90909.1 acar|OOF90910.1 abra|OJJ67526.1 afla|EED49685.1 anig|EHA18706.1 aory|EIT81018.1 asyd|OJJ57249.1 ater|EAU38522.1 aver|OJJ03045.1 awen|OJJ31267.1 abra|OJJ75516.1 acar|OOF90843.1 afla|EED46523.1 akaw|GAA84280.1 aluc|OJZ88973.1 anig|EHA23552.1 atub|OJI85817.1 aory|EIT72379.1 agla|OJJ87127.1

Group789: acar|OOF92001.1 acar|OOF92003.1 abra|OJJ67845.1 acla|EAW11054.1 afis|EAW20822.1 afla|EED52866.1 afum|EAL84807.1 agla|OJJ88409.1 akaw|GAA91851.1 aluc|OJZ82047.1 anid|CBF78348.1 anig|EHA22211.1 aory|EIT78637.1 asyd|OJJ56141.1 atub|OJI84605.1 aver|OJJ05080.1 awen|OJJ29709.1 azon|OJJ45742.1 awen|OJJ34808.1

Group790: acar|OOF92286.1 acar|OOF92327.1 abra|OJJ67516.1 acla|EAW06255.1 afis|EAW18362.1 afla|EED53043.1 afum|EAL93033.1 agla|OJJ84926.1 akaw|GAA86324.1 aluc|OJZ84848.1 anid|CBF70825.1 anig|EHA25082.1 aory|EIT77408.1 asyd|OJJ58399.1 ater|EAU32230.1 atub|OJI80748.1 aver|OJJ01000.1 awen|OJJ31788.1 azon|OJJ50438.1

Group791: acar|OOF92287.1 acar|OOF92328.1 abra|OJJ67515.1 acla|EAW06254.1 afis|EAW18361.1 afla|EED53044.1 afum|EAL93032.2 agla|OJJ84925.1 akaw|GAA86325.1 aluc|OJZ84849.1 anid|CBF70827.1 anig|EHA25083.1 aory|EIT77438.1 asyd|OJJ58398.1 ater|EAU32229.1 atub|OJI80747.1 aver|OJJ00999.1 awen|OJJ31789.1 azon|OJJ42001.1

Group792: acar|OOF92288.1 acar|OOF92329.1 abra|OJJ67514.1 acla|EAW06260.1 afis|EAW18367.1 afla|EED53038.1 afum|EAL93041.1 agla|OJJ84931.1 akaw|GAA86326.1 aluc|OJZ84850.1 anid|CBF70815.1 anig|EHA25084.1 aory|EIT77399.1 asyd|OJJ58404.1 ater|EAU32236.1 atub|OJI80746.1 aver|OJJ01005.1 awen|OJJ31783.1 azon|OJJ50443.1

Group793: acar|OOF92289.1 acar|OOF92330.1 abra|OJJ67513.1 acla|EAW06259.1 afis|EAW18366.1 afla|EED53039.1 afum|EAL93040.1 agla|OJJ84930.1 akaw|GAA86327.1 aluc|OJZ84851.1 anid|CBF70817.1 anig|EHA25085.1 aory|EIT77396.1 asyd|OJJ58403.1 ater|EAU32235.1 atub|OJI80745.1 aver|OJJ01004.1 awen|OJJ31784.1 azon|OJJ50442.1

Group794: acar|OOF92291.1 acar|OOF92332.1 abra|OJJ67510.1 afis|EAW18368.1 afla|EED53037.1 afum|EAL93043.1 agla|OJJ84933.1 akaw|GAA86329.1 aluc|OJZ84854.1 anid|CBF70814.1 anig|EHA25087.1

aory|EIT77401.1 asyd|OJJ58405.1 ater|EAU32238.1 atub|OJI80742.1 aver|OJJ01006.1 awen|OJJ31781.1 azon|OJJ43320.1 acla|EAW06262.1

Group795: acar|OOF92293.1 acar|OOF92334.1 abra|OJJ67507.1 acla|EAW06264.1 afis|EAW18370.1 afla|EED53036.1 afum|EAL93045.1 agla|OJJ84935.1 akaw|GAA86331.1 aluc|OJZ84856.1 anid|CBF70808.1 anig|EHA25089.1 aory|EIT77405.1 asyd|OJJ58408.1 ater|EAU32240.1 atub|OJI80740.1 aver|OJJ01010.1 awen|OJJ31779.1 azon|OJJ50445.1

Group796: acar|OOF92297.1 acar|OOF92338.1 abra|OJJ67503.1 acla|EAW06267.1 afis|EAW18373.1 afla|EED53033.1 afum|EAL93048.1 agla|OJJ84937.1 akaw|GAA86335.1 aluc|OJZ84860.1 anid|CBF70800.1 anig|EHA25093.1 aory|EIT77403.1 asyd|OJJ58411.1 ater|EAU32243.1 atub|OJI80736.1 aver|OJJ01013.1 awen|OJJ31777.1 azon|OJJ50448.1

Group797: acar|OOF92298.1 acar|OOF92339.1 abra|OJJ67502.1 acla|EAW06268.1 afis|EAW18374.1 afla|EED53032.1 afum|EAL93049.1 agla|OJJ84938.1 akaw|GAA86336.1 aluc|OJZ84861.1 anid|CBF70798.1 anig|EHA25094.1 aory|EIT77421.1 asyd|OJJ58412.1 ater|EAU32244.1 atub|OJI80735.1 aver|OJJ01014.1 awen|OJJ31766.1 azon|OJJ50449.1

Group798: acar|OOF92299.1 acar|OOF92340.1 abra|OJJ67501.1 acla|EAW06269.1 afis|EAW18375.1 afla|EED53031.1 afum|EAL93050.1 agla|OJJ84939.1 akaw|GAA86337.1 aluc|OJZ84862.1 anid|CBF70797.1 anig|EHA25095.1 aory|EIT77415.1 asyd|OJJ58413.1 ater|EAU32245.1 atub|OJI80734.1 aver|OJJ01015.1 awen|OJJ31767.1 azon|OJJ50450.1

Group799: acar|OOF92301.1 acar|OOF92342.1 abra|OJJ67499.1 acla|EAW06271.1 afis|EAW18377.1 afla|EED53029.1 afum|EAL93052.1 agla|OJJ84941.1 akaw|GAA86339.1 aluc|OJZ84864.1 anid|CBF70793.1 anig|EHA25096.1 aory|EIT77426.1 asyd|OJJ58415.1 ater|EAU32247.1 atub|OJI80732.1 aver|OJJ01017.1 awen|OJJ31769.1 azon|OJJ50452.1

Group800: acar|OOF92303.1 acar|OOF92343.1 abra|OJJ67498.1 acla|EAW06272.1 afis|EAW18378.1 afla|EED53028.1 afum|EAL93053.1 agla|OJJ84942.1 akaw|GAA86340.1 aluc|OJZ84865.1 anid|CBF70791.1 anig|EHA25097.1 aory|EIT77397.1 asyd|OJJ58416.1 ater|EAU32248.1 atub|OJI80731.1 aver|OJJ01018.1 awen|OJJ31770.1 azon|OJJ50453.1

Group801: acar|OOF92304.1 acar|OOF92345.1 abra|OJJ67496.1 acla|EAW06274.1 afis|EAW18380.1 afla|EED53026.1 afum|EAL93055.1 agla|OJJ84944.1 akaw|GAA86342.1 aluc|OJZ84867.1 anid|CBF70787.1 anig|EHA25099.1 aory|EIT77406.1 asyd|OJJ58418.1 ater|EAU32249.1 atub|OJI80729.1 aver|OJJ01020.1 awen|OJJ31772.1 azon|OJJ50455.1

Group802: acar|OOF92305.1 acar|OOF92346.1 abra|OJJ67495.1 acla|EAW06275.1 afis|EAW18381.1 afla|EED53025.1 afum|EAL93056.1 agla|OJJ84945.1 akaw|GAA86343.1 aluc|OJZ84869.1 anid|CBF70785.1 anig|EHA25100.1 aory|EIT77407.1 asyd|OJJ58419.1 ater|EAU32250.1 atub|OJI80728.1 aver|OJJ01021.1 awen|OJJ31773.1 azon|OJJ50456.1

Group803: acar|OOF92306.1 acar|OOF92347.1 abra|OJJ67494.1 acla|EAW06276.1 afis|EAW18382.1 afla|EED53024.1 afum|EAL93057.1 agla|OJJ84946.1 akaw|GAA86344.1 aluc|OJZ84870.1 anid|CBF70783.1 anig|EHA25101.1 aory|EIT77418.1 asyd|OJJ58420.1 ater|EAU32251.1 atub|OJI80727.1 aver|OJJ01022.1 awen|OJJ31774.1 azon|OJJ50457.1

Group804: acar|OOF92307.1 acar|OOF92348.1 abra|OJJ67492.1 acla|EAW06278.1 afis|EAW18384.1 afla|EED53022.1 afum|EAL93059.1 agla|OJJ84948.1 akaw|GAA86346.1 aluc|OJZ84872.1 anid|CBF70781.1 anig|EHA25103.1 aory|EIT77395.1 asyd|OJJ58422.1 ater|EAU32253.1 atub|OJI80726.1 aver|OJJ01023.1 awen|OJJ31775.1 azon|OJJ50458.1

Group805: acar|OOF92308.1 acar|OOF92349.1 abra|OJJ67491.1 acla|EAW06279.1 afis|EAW18385.1 afla|EED53021.1 afum|EAL93060.1 agla|OJJ84949.1 akaw|GAA86347.1 aluc|OJZ84874.1 anid|CBF70779.1 anig|EHA25104.1 aory|EIT77414.1 asyd|OJJ58423.1 ater|EAU32254.1 atub|OJI80725.1 aver|OJJ01024.1 awen|OJJ31776.1 azon|OJJ45926.1

Group806: acar|OOF92310.1 acar|OOF92351.1 abra|OJJ67490.1 acla|EAW06282.1 afis|EAW18388.1 afla|EED53019.1 afum|EAL93063.1 agla|OJJ84951.1 akaw|GAA86348.1 aluc|OJZ84875.1 anid|CBF70772.1 anig|EHA25106.1 aory|EIT77433.1 asyd|OJJ58427.1 ater|EAU32261.1 atub|OJI80723.1 aver|OJJ01028.1 awen|OJJ31763.1 azon|OJJ50461.1

Group807: acar|OOF92311.1 acar|OOF92352.1 abra|OJJ67489.1 acla|EAW06284.1 afis|EAW18390.1 afla|EED53017.1 afum|EAL93065.1 agla|OJJ84953.1 akaw|GAA86350.1 aluc|OJZ84878.1 anid|CBF70776.1 anig|EHA25107.1 aory|EIT77439.1 asyd|OJJ58425.1 ater|EAU32263.1 atub|OJI80721.1 aver|OJJ01026.1 awen|OJJ31761.1 azon|OJJ50464.1

Group808: acar|OOF92312.1 acar|OOF92353.1 abra|OJJ67488.1 acla|EAW06286.1 afis|EAW18391.1 afla|EED53016.1 afum|EAL93066.1 agla|OJJ84955.1 akaw|GAA86351.1 aluc|OJZ84880.1 anid|CBF70777.1 anig|EHA25110.1 aory|EIT77398.1 asyd|OJJ58424.1 ater|EAU32278.1 atub|OJI80719.1 aver|OJJ01025.1 awen|OJJ31758.1 azon|OJJ50466.1

Group809: acar|OOF92314.1 acar|OOF92355.1 abra|OJJ67485.1 acla|EAW06289.1 afis|EAW18394.1 afla|EED53013.1 afum|EAL93070.1 agla|OJJ84966.1 akaw|GAA86354.1 aluc|OJZ84883.1 anid|CBF70762.1 anig|EHA25112.1 aory|EIT77419.1 asyd|OJJ58430.1 ater|EAU32282.1 atub|OJI80717.1 aver|OJJ01032.1 awen|OJJ31756.1 azon|OJJ50472.1

Group810: acar|OOF92317.1 acar|OOF92359.1 abra|OJJ67482.1 acla|EAW06291.1 afis|EAW18397.1 afla|EED53009.1 afum|EAL93074.1 agla|OJJ84968.1 akaw|GAA86357.1 aluc|OJZ84886.1 anid|CBF70756.1 anig|EHA25115.1 aory|EIT77428.1 asyd|OJJ58434.1 ater|EAU32285.1 atub|OJI80714.1 aver|OJJ01035.1 awen|OJJ31754.1 azon|OJJ50470.1

Group811: acar|OOF92318.1 acar|OOF92360.1 afum|EAL85117.1 afum|EAL93075.1 abra|OJJ67480.1 acla|EAW06292.1 afis|EAW18398.1 afla|EED53008.1 akaw|GAA86358.1 aluc|OJZ84887.1 anid|CBF70755.1 anig|EHA25116.1 aory|EIT77431.1 asyd|OJJ58436.1 ater|EAU29814.1 atub|OJI80713.1 aver|OJJ01036.1 awen|OJJ31752.1 azon|OJJ50468.1

Group812: acar|OOF92320.1 acar|OOF92362.1 abra|OJJ67474.1 acla|EAW06298.1 afis|EAW18405.1 afla|EED53003.1 afum|EAL93085.1 agla|OJJ84956.1 akaw|GAA86363.1 aluc|OJZ84891.1 anid|CBF69754.1 anig|EHA25120.1 aory|EIT77409.1 asyd|OJJ58942.1 ater|EAU29822.1 atub|OJI80707.1 aver|OJJ01538.1 awen|OJJ31745.1 azon|OJJ50505.1

Group813: acar|OOF92596.1 acar|OOF92617.1 abra|OJJ68969.1 acla|EAW14218.1 afis|EAW19063.1 afla|EED54429.1 afum|EAL93792.1 agla|OJJ84544.1 akaw|GAA90496.1 aluc|OJZ85956.1 anid|CBF79645.1 anig|EHA24645.1 aory|EIT75848.1 asyd|OJJ58020.1 ater|EAU32106.1 atub|OJI79744.1 aver|OJJ04379.1 awen|OJJ31494.1 azon|OJJ46207.1

Group814: acar|OOF92598.1 acar|OOF92619.1 abra|OJJ68487.1 acla|EAW06807.1 afis|EAW16356.1 afla|EED53514.1 afum|EAL88981.1 agla|OJJ88237.1 akaw|GAA92784.1 aluc|OJZ82699.1 anid|CBF89230.1 anig|EHA21155.1 aory|EIT74940.1 asyd|OJJ62515.1 ater|EAU34103.1 atub|OJI88931.1 aver|OJI96634.1 awen|OJJ37760.1 azon|OJJ45058.1

Group815: acar|OOF92601.1 acar|OOF92622.1 abra|OJJ77122.1 acla|EAW12935.1 afis|EAW17711.1 afla|EED45713.1 afum|EAL91526.1 agla|OJJ81737.1 akaw|GAA90806.1 aluc|OJZ84532.1 anid|CBF76922.1 anig|EHA27830.1 aory|EIT75682.1 asyd|OJJ57071.1 ater|EAU31890.1 atub|OJI89548.1 aver|OJJ03227.1 awen|OJJ37125.1 azon|OJJ50824.1

Group816: acar|OOF92602.1 acar|OOF92623.1 abra|OJJ75110.1 acla|EAW08316.1 afis|EAW23324.1 afla|EED55642.1 afum|EAL88155.1 agla|OJJ86519.1 akaw|GAA88719.1 aluc|OJZ92676.1 anid|CBF89451.1 anig|EHA26322.1 aory|EIT81779.1 asyd|OJJ62432.1 ater|EAU35572.1 atub|OJI87104.1 aver|OJI96568.1 awen|OJJ39570.1 azon|OJJ47763.1

Group817: acar|OOF92604.1 acar|OOF92625.1 abra|OJJ78212.1 acla|EAW11239.1 afis|EAW18266.1 afla|EED48732.1 afum|EAL87631.1 agla|OJJ88160.1 akaw|GAA85815.1 aluc|OJZ91284.1 anid|CBF79520.1 anig|EHA19969.1 aory|EIT73261.1 asyd|OJJ57890.1 ater|EAU33273.1 atub|OJI83274.1 aver|OJJ04504.1 awen|OJJ30640.1 azon|OJJ47975.1

Group818: acar|OOF92606.1 acar|OOF92627.1 abra|OJJ72009.1 acla|EAW09531.1 afis|EAW21601.1 afla|EED57828.1 afum|EAL92620.1 agla|OJJ85796.1 akaw|GAA84924.1 aluc|OJZ86745.1 anid|CBF83460.1 anig|EHA21934.1 aory|EIT81025.1 asyd|OJJ60683.1 ater|EAU38291.1 atub|OJI83782.1 aver|OJJ00008.1 awen|OJJ39081.1 azon|OJJ51413.1

Group819: acar|OOF92607.1 acar|OOF92628.1 abra|OJJ71278.1 acla|EAW10260.1 afis|EAW25333.1 afla|EED53813.1 afum|EAL89996.1 agla|OJJ83874.1 akaw|GAA82710.1 aluc|OJZ87265.1 anid|CBF75238.1 anig|EHA20507.1 aory|EIT78421.1 asyd|OJJ59912.1 ater|EAU34479.1 atub|OJI81377.1 aver|OJJ02374.1 awen|OJJ41027.1 azon|OJJ44637.1

Group820: acar|OOF92609.1 acar|OOF92630.1 abra|OJJ74612.1 acla|EAW10770.1 afis|EAW24806.1 afla|EED47177.1 afum|EAL89448.1 agla|OJJ82901.1 akaw|GAA81996.1 aluc|OJZ92192.1 anid|CBF85894.1 anig|EHA26762.1 aory|EIT79234.1 asyd|OJJ64389.1 ater|EAU36500.1 atub|OJI86621.1 aver|OJI98483.1 awen|OJJ40390.1 azon|OJJ49240.1

Group821: acar|OOF92610.1 acar|OOF92631.1 abra|OJJ71181.1 acla|EAW10323.1 afis|EAW25269.1 afla|EED53734.1 afum|EAL89932.1 agla|OJJ89237.1 akaw|GAA82804.1 aluc|OJZ87362.1 anid|CBF85380.1 anig|EHA20597.1 aory|EIT75356.1 asyd|OJJ64050.1 ater|EAU34412.1 atub|OJI81283.1 aver|OJI98136.1 awen|OJJ40983.1 azon|OJJ44569.1

Group822: acar|OOF92611.1 acar|OOF92632.1 abra|OJJ72794.1 acla|EAW11478.1 afis|EAW23975.1 afla|EED50533.1 afum|EAL87405.1 agla|OJJ86905.1 akaw|GAA85397.1 aluc|OJZ83290.1 anid|CBF77343.1 anig|EHA24175.1 aory|EIT82080.1 asyd|OJJ53273.1 ater|EAU29777.1 atub|OJI82192.1 aver|OJJ03576.1 awen|OJJ35440.1 azon|OJJ51303.1

Group823: acar|OOF92612.1 acar|OOF92633.1 abra|OJJ66525.1 acla|EAW10966.1 afis|EAW20894.1 afla|EED50243.1 afum|EAL84739.1 agla|OJJ84001.1 akaw|GAA91188.1 aluc|OJZ81515.1 anid|CBF82444.1

anig|EHA25913.1 aory|EIT77371.1 asyd|OJJ54119.1 ater|EAU38528.1 atub|OJI88664.1 aver|OJJ08617.1 awen|OJJ34454.1 azon|OJJ51741.1

Group824: acar|OOF92616.1 acar|OOF92637.1 abra|OJJ76860.1 acla|EAW13421.1 afis|EAW20532.1 afla|EED52482.1 afum|EAL84525.1 agla|OJJ84699.1 akaw|GAA87162.1 aluc|OJZ84272.1 anid|CBF79000.1 anig|EHA27614.1 aory|EIT82763.1 asyd|OJJ55688.1 ater|EAU37152.1 atub|OJI89811.1 aver|OJJ06772.1 awen|OJJ33239.1 azon|OJJ50618.1

Group825: acar|OOF93657.1 acar|OOF93741.1 abra|OJJ66636.1 acla|EAW11779.1 afis|EAW19389.1 afla|EED50347.1 afum|EAL87106.1 agla|OJJ81100.1 akaw|GAA89589.1 aluc|OJZ80813.1 anid|CBF74291.1 anig|EHA27221.1 aory|EIT81936.1 asyd|OJJ51912.1 ater|EAU36052.1 atub|OJI87662.1 aver|OJJ05353.1 awen|OJJ36782.1 azon|OJJ43016.1

Group826: acar|OOF93659.1 acar|OOF93743.1 abra|OJJ66639.1 acla|EAW11782.1 afis|EAW19392.1 afla|EED50350.1 afum|EAL87102.1 agla|OJJ81103.1 akaw|GAA89592.1 aluc|OJZ80810.1 anid|CBF74285.1 anig|EHA27224.1 aory|EIT81937.1 asyd|OJJ51914.1 ater|EAU36055.1 atub|OJI87658.1 aver|OJJ05356.1 awen|OJJ36785.1 azon|OJJ43013.1

Group827: acar|OOF93660.1 acar|OOF93744.1 abra|OJJ66640.1 acla|EAW11783.1 afis|EAW19393.1 afla|EED50351.1 agla|OJJ81104.1 akaw|GAA89593.1 aluc|OJZ80809.1 anig|EHA27225.1 aory|EIT81941.1 asyd|OJJ51863.1 ater|EAU36056.1 atub|OJI87657.1 aver|OJJ05357.1 awen|OJJ36787.1 azon|OJJ43012.1 afum|EAL87101.1 anid|CBF74283.1

Group828: acar|OOF93663.1 acar|OOF93747.1 abra|OJJ66643.1 acla|EAW11786.1 afis|EAW19396.1 afla|EED50352.1 afum|EAL87098.1 agla|OJJ81107.1 akaw|GAA89596.1 aluc|OJZ80806.1 anid|CBF74278.1 anig|EHA27228.1 aory|EIT83119.1 asyd|OJJ51859.1 ater|EAU36122.1 atub|OJI87654.1 aver|OJJ05360.1 awen|OJJ36791.1 azon|OJJ43008.1

Group829: acar|OOF93667.1 acar|OOF93751.1 abra|OJJ66647.1 acla|EAW11790.1 afis|EAW19400.1 afla|EED50355.1 afum|EAL87094.1 agla|OJJ81111.1 akaw|GAA89600.1 aluc|OJZ80802.1 anid|CBF74336.1 anig|EHA27232.1 aory|EIT83130.1 asyd|OJJ51906.1 ater|EAU36126.1 atub|OJI87650.1 aver|OJJ05363.1 awen|OJJ36795.1 azon|OJJ43004.1

Group830: acar|OOF93669.1 acar|OOF93753.1 abra|OJJ66649.1 acla|EAW11792.1 afis|EAW19403.1 afla|EED50358.1 afum|EAL87092.1 agla|OJJ81117.1 akaw|GAA89602.1 aluc|OJZ80800.1 anid|CBF74298.1 anig|EHA27234.1 aory|EIT83146.1 asyd|OJJ51902.1 ater|EAU36143.1 atub|OJI87648.1 aver|OJJ05365.1 awen|OJJ36798.1 azon|OJJ43001.1

Group831: acar|OOF93671.1 acar|OOF93755.1 abra|OJJ66653.1 acla|EAW11795.1 afis|EAW19405.1 afla|EED50361.1 afum|EAL87090.1 agla|OJJ81119.1 akaw|GAA89604.1 aluc|OJZ80794.1 anid|CBF74345.1 anig|EHA27237.1 aory|EIT83126.1 asyd|OJJ57796.1 ater|EAU36140.1 atub|OJI87645.1 aver|OJJ05368.1 awen|OJJ36801.1 azon|OJJ42997.1

Group832: acar|OOF93672.1 acar|OOF93756.1 abra|OJJ66655.1 acla|EAW11796.1 afis|EAW19406.1 afla|EED50362.1 afum|EAL87088.1 agla|OJJ81120.1 akaw|GAA89605.1 aluc|OJZ80793.1 anid|CBF74347.1 anig|EHA27238.1 aory|EIT83144.1 asyd|OJJ57795.1 ater|EAU36139.1 atub|OJI87644.1 aver|OJJ05369.1 awen|OJJ36802.1 azon|OJJ42996.1

Group833: acar|OOF93673.1 acar|OOF93757.1 abra|OJJ66656.1 acla|EAW11797.1 afis|EAW19407.1 afla|EED50363.1 afum|EAL87087.1 agla|OJJ81121.1 akaw|GAA89606.1 aluc|OJZ80792.1 anid|CBF74349.1 anig|EHA27239.1 aory|EIT83134.1 asyd|OJJ57794.1 ater|EAU36138.1 atub|OJI87643.1 aver|OJJ05370.1 awen|OJJ36803.1 azon|OJJ42995.1

Group834: acar|OOF93674.1 acar|OOF93758.1 abra|OJJ66657.1 acla|EAW11798.1 afis|EAW19408.1 afla|EED50364.1 afum|EAL87086.1 agla|OJJ81122.1 akaw|GAA89607.1 aluc|OJZ80791.1 anid|CBF74356.1 anig|EHA27240.1 aory|EIT83115.1 asyd|OJJ57793.1 ater|EAU34715.1 atub|OJI87642.1 aver|OJJ05371.1 awen|OJJ36804.1 azon|OJJ42994.1

Group835: acar|OOF93676.1 acar|OOF93761.1 abra|OJJ66660.1 acla|EAW11802.1 afis|EAW19412.1 afla|EED50367.1 afum|EAL87082.1 agla|OJJ81126.1 akaw|GAA89610.1 aluc|OJZ80788.1 anid|CBF74358.1 anig|EHA27242.1 aory|EIT83138.1 asyd|OJJ57789.1 ater|EAU34718.1 atub|OJI87639.1 aver|OJJ05375.1 awen|OJJ36808.1 azon|OJJ42992.1

Group836: acar|OOF93677.1 acar|OOF93762.1 abra|OJJ66661.1 acla|EAW11803.1 afis|EAW19413.1 afla|EED50368.1 afum|EAL87081.1 agla|OJJ81127.1 akaw|GAA89611.1 aluc|OJZ80787.1 anid|CBF74360.1 anig|EHA27243.1 aory|EIT83132.1 asyd|OJJ57788.1 ater|EAU34719.1 atub|OJI87638.1 aver|OJJ05376.1 awen|OJJ36809.1 azon|OJJ42991.1

Group837: acar|OOF93678.1 acar|OOF93763.1 abra|OJJ66662.1 acla|EAW11804.1 afis|EAW19414.1 afla|EED50369.1 afum|EAL87080.2 agla|OJJ81128.1 akaw|GAA89612.1 aluc|OJZ80786.1 anid|CBF74362.1 anig|EHA27244.1 aory|EIT83123.1 asyd|OJJ57787.1 ater|EAU34720.1 atub|OJI87636.1 aver|OJJ05377.1 awen|OJJ36810.1 azon|OJJ42990.1

Group838: acar|OOF93679.1 acar|OOF93765.1 abra|OJJ66664.1 acla|EAW11805.1 afis|EAW19415.1 afla|EED50371.1 afum|EAL87079.1 agla|OJJ81129.1 akaw|GAA89613.1 aluc|OJZ80783.1 anid|CBF74364.1 anig|EHA27246.1 aory|EIT83116.1 asyd|OJJ57785.1 ater|EAU34721.1 atub|OJI87635.1 aver|OJJ05379.1 awen|OJJ36812.1 azon|OJJ42989.1

Group839: acar|OOF93681.1 acar|OOF93764.1 abra|OJJ66666.1 acla|EAW11807.1 afis|EAW19417.1 afla|EED50373.1 afum|EAL87077.1 agla|OJJ81130.1 akaw|GAA89615.1 aluc|OJZ80781.1 anid|CBF86348.1 anig|EHA27248.1 aory|EIT83140.1 asyd|OJJ64978.1 ater|EAU34723.1 atub|OJI87633.1 aver|OJI99100.1 awen|OJJ36815.1 azon|OJJ42987.1

Group840: acar|OOF93685.1 acar|OOF93770.1 abra|OJJ66671.1 acla|EAW11812.1 afis|EAW19422.1 afla|EED50379.1 afum|EAL87072.1 agla|OJJ81134.1 akaw|GAA89620.1 aluc|OJZ80776.1 anid|CBF74373.1 anig|EHA27253.1 aory|EIT83120.1 asyd|OJJ57781.1 ater|EAU34726.1 atub|OJI87628.1 aver|OJJ05384.1 awen|OJJ36820.1 azon|OJJ42982.1

Group841: acar|OOF93686.1 acar|OOF93771.1 abra|OJJ66672.1 acla|EAW11813.1 afis|EAW19423.1 afla|EED50380.1 afum|EAL87071.1 agla|OJJ81135.1 akaw|GAA89621.1 aluc|OJZ80775.1 anid|CBF74375.1 anig|EHA27254.1 aory|EIT83122.1 asyd|OJJ57780.1 ater|EAU34727.1 atub|OJI87627.1 aver|OJJ05385.1 awen|OJJ36821.1 azon|OJJ42981.1

Group842: acar|OOF93690.1 acar|OOF93776.1 abra|OJJ66680.1 acla|EAW11817.1 afis|EAW19428.1 afla|EED56632.1 afum|EAL87066.1 agla|OJJ81140.1 akaw|GAA89625.1 aluc|OJZ80769.1 anid|CBF74382.1 anig|EHA27259.1 aory|EIT79645.1 asyd|OJJ57775.1 ater|EAU34731.1 atub|OJI87622.1 aver|OJJ05390.1 awen|OJJ36826.1 azon|OJJ42977.1

Group843: acar|OOF93692.1 acar|OOF93778.1 abra|OJJ66683.1 acla|EAW11820.1 afis|EAW19431.1 afla|EED56635.1 afum|EAL87063.1 agla|OJJ81143.1 akaw|GAA89628.1 aluc|OJZ80766.1 anid|CBF81374.1 anig|EHA27262.1 aory|EIT79669.1 asyd|OJJ59388.1 ater|EAU34734.1 atub|OJI87619.1 aver|OJJ02885.1 awen|OJJ36832.1 azon|OJJ42974.1

Group844: acar|OOF93693.1 acar|OOF93777.1 abra|OJJ66682.1 acla|EAW11819.1 afis|EAW19430.1 afla|EED56634.1 afum|EAL87064.1 agla|OJJ81142.1 akaw|GAA89627.1 aluc|OJZ80767.1 anid|CBF74386.1 anig|EHA27261.1 aory|EIT79632.1 asyd|OJJ57773.1 ater|EAU34733.1 atub|OJI87620.1 aver|OJJ05393.1 awen|OJJ36829.1 azon|OJJ42975.1

Group845: acar|OOF93696.1 acar|OOF93781.1 abra|OJJ66687.1 acla|EAW11822.1 afis|EAW19433.1 afla|EED56638.1 afum|EAL87062.1 agla|OJJ81145.1 akaw|GAA89631.1 aluc|OJZ80762.1 anid|CBF81380.1 anig|EHA27265.1 aory|EIT79671.1 asyd|OJJ59391.1 ater|EAU34735.1 atub|OJI87616.1 aver|OJJ02881.1 awen|OJJ36833.1 azon|OJJ42972.1

Group846: acar|OOF93699.1 acar|OOF93784.1 abra|OJJ66690.1 acla|EAW11825.1 afis|EAW19436.1 afla|EED56641.1 afum|EAL87059.1 agla|OJJ81148.1 akaw|GAA89634.1 aluc|OJZ80759.1 anid|CBF81383.1 anig|EHA27268.1 aory|EIT79678.1 asyd|OJJ59395.1 ater|EAU34738.1 atub|OJI87613.1 aver|OJJ02877.1 awen|OJJ36834.1 azon|OJJ42969.1

Group847: acar|OOF93701.1 acar|OOF93787.1 abra|OJJ66693.1 acla|EAW11828.1 afis|EAW19440.1 afla|EED56647.1 afum|EAL87056.1 agla|OJJ81155.1 akaw|GAA89637.1 aluc|OJZ80752.1 anid|CBF79760.1 anig|EHA27271.1 aory|EIT79660.1 asyd|OJJ58101.1 ater|EAU34741.1 atub|OJI87609.1 aver|OJJ04303.1 awen|OJJ36840.1 azon|OJJ42961.1

Group848: acar|OOF93703.1 acar|OOF93790.1 abra|OJJ66695.1 acla|EAW11830.1 afis|EAW19442.1 afla|EED56649.1 afum|EAL87054.1 agla|OJJ81153.1 akaw|GAA89639.1 aluc|OJZ80750.1 anid|CBF81395.1 anig|EHA27272.1 aory|EIT79630.1 asyd|OJJ59401.1 ater|EAU34743.1 atub|OJI87607.1 aver|OJJ02872.1 awen|OJJ36842.1 azon|OJJ42958.1

Group849: acar|OOF93707.1 acar|OOF93793.1 abra|OJJ66703.1 acla|EAW11834.1 afis|EAW19445.1 afla|EED56653.1 afum|EAL87050.1 agla|OJJ81160.1 akaw|GAA89645.1 aluc|OJZ80744.1 anid|CBF81404.1 anig|EHA27278.1 aory|EIT79624.1 asyd|OJJ59405.1 ater|EAU34747.1 atub|OJI87601.1 aver|OJJ02868.1 awen|OJJ36846.1 azon|OJJ42954.1

Group850: acar|OOF93708.1 acar|OOF93794.1 abra|OJJ66704.1 acla|EAW11835.1 afis|EAW19446.1 afla|EED56654.1 afum|EAL87049.1 agla|OJJ81161.1 akaw|GAA89646.1 aluc|OJZ80743.1 anid|CBF81405.1 anig|EHA27279.1 aory|EIT79662.1 asyd|OJJ59406.1 ater|EAU34748.1 atub|OJI87600.1 aver|OJJ02867.1 awen|OJJ36847.1 azon|OJJ42953.1

Group851: acar|OOF93709.1 acar|OOF93795.1 abra|OJJ66706.1 acla|EAW11837.1 afis|EAW19448.1 afla|EED56656.1 afum|EBA27299.1 agla|OJJ81163.1 akaw|GAA89648.1 aluc|OJZ80741.1 anid|CBF81409.1 anig|EHA27281.1 aory|EIT79680.1 asyd|OJJ59408.1 ater|EAU34750.1 atub|OJI87598.1 aver|OJJ02865.1 awen|OJJ36849.1 azon|OJJ42951.1

Group852: acar|OOF93710.1 acar|OOF93796.1 abra|OJJ66708.1 acla|EAW11839.1 afis|EAW19450.1 afla|EED56658.1 afum|EAL87046.1 agla|OJJ81165.1 akaw|GAA89650.1 aluc|OJZ80739.1 anid|CBF81412.1

anig|EHA27283.1 aory|EIT79655.1 asyd|OJJ59410.1 ater|EAU34752.1 atub|OJI87596.1 aver|OJJ02863.1 awen|OJJ36851.1 azon|OJJ42949.1

Group853: acar|OOF93711.1 acar|OOF93797.1 abra|OJJ66709.1 acla|EAW11840.1 afis|EAW19451.1 afla|EED56659.1 afum|EAL87045.1 agla|OJJ81166.1 akaw|GAA89651.1 aluc|OJZ80738.1 anid|CBF81414.1 anig|EHA27284.1 aory|EIT79675.1 asyd|OJJ59411.1 ater|EAU34753.1 atub|OJI87595.1 aver|OJJ02862.1 awen|OJJ36852.1 azon|OJJ42948.1

Group854: acar|OOF93715.1 acar|OOF93802.1 abra|OJJ66713.1 acla|EAW11844.1 afis|EAW19455.1 afla|EED56663.1 afum|EAL87041.1 agla|OJJ81171.1 akaw|GAA89655.1 aluc|OJZ80734.1 anid|CBF81421.1 anig|EHA27287.1 aory|EIT79656.1 asyd|OJJ59415.1 ater|EAU34758.1 atub|OJI87591.1 aver|OJJ02858.1 awen|OJJ36856.1 azon|OJJ42944.1

Group855: acar|OOF93716.1 acar|OOF93803.1 abra|OJJ66714.1 acla|EAW11845.1 afis|EAW19456.1 afla|EED56664.1 afum|EAL87040.1 agla|OJJ81173.1 akaw|GAA89656.1 aluc|OJZ80733.1 anid|CBF81423.1 anig|EHA27288.1 aory|EIT79663.1 asyd|OJJ59416.1 ater|EAU34759.1 atub|OJI87590.1 aver|OJJ02857.1 awen|OJJ36858.1 azon|OJJ42943.1

Group856: acar|OOF93717.1 acar|OOF93804.1 abra|OJJ66715.1 acla|EAW11846.1 afis|EAW19457.1 afla|EED56665.1 afum|EAL87039.1 agla|OJJ81174.1 akaw|GAA89657.1 aluc|OJZ80732.1 anid|CBF81425.1 anig|EHA27289.1 aory|EIT79668.1 asyd|OJJ59417.1 ater|EAU34760.1 atub|OJI87589.1 aver|OJJ02856.1 awen|OJJ36859.1 azon|OJJ42942.1

Group857: acar|OOF93718.1 acar|OOF93805.1 abra|OJJ66716.1 acla|EAW11847.1 afis|EAW19458.1 afla|EED56669.1 afum|EAL87038.1 agla|OJJ81175.1 akaw|GAA89658.1 aluc|OJZ80731.1 anid|CBF81427.1 anig|EHA27290.1 aory|EIT79631.1 asyd|OJJ59418.1 ater|EAU34761.1 atub|OJI87588.1 aver|OJJ02851.1 awen|OJJ36860.1 azon|OJJ42941.1

Group858: acar|OOF93721.1 acar|OOF93808.1 abra|OJJ66719.1 acla|EAW11850.1 afis|EAW19461.1 afla|EED56672.1 afum|EAL87035.1 agla|OJJ81178.1 akaw|GAA89661.1 aluc|OJZ80728.1 anid|CBF74529.1 anig|EHA27293.1 aory|EIT82995.1 asyd|OJJ57737.1 ater|EAU34764.1 atub|OJI87584.1 aver|OJJ05430.1 awen|OJJ36863.1 azon|OJJ42938.1

Group859: acar|OOF93723.1 acar|OOF93810.1 abra|OJJ66721.1 acla|EAW11852.1 afis|EAW19463.1 afla|EED56674.1 afum|EAL87033.1 agla|OJJ81180.1 akaw|GAA89663.1 aluc|OJZ80726.1 anid|CBF75491.1 anig|EHA27294.1 aory|EIT73836.1 asyd|OJJ59753.1 ater|EAU34766.1 atub|OJI87582.1 aver|OJJ02535.1 awen|OJJ36865.1 azon|OJJ42936.1

Group860: acar|OOF93724.1 acar|OOF93811.1 abra|OJJ66722.1 acla|EAW11853.1 afis|EAW19464.1 afla|EED56675.1 afum|EAL87032.1 agla|OJJ81181.1 akaw|GAA89664.1 aluc|OJZ80725.1 anid|CBF75490.1 anig|EHA27295.1 aory|EIT73882.1 asyd|OJJ59754.1 ater|EAU34768.1 atub|OJI87581.1 aver|OJJ02534.1 awen|OJJ36866.1 azon|OJJ42935.1

Group861: acar|OOF93726.1 acar|OOF93814.1 abra|OJJ66726.1 acla|EAW11856.1 afis|EAW19467.1 afla|EED56677.1 afum|EAL87029.1 agla|OJJ81187.1 akaw|GAA89667.1 aluc|OJZ80722.1 anid|CBF75486.1 anig|EHA27298.1 aory|EIT73931.1 asyd|OJJ59756.1 ater|EAU34777.1 atub|OJI87578.1 aver|OJJ02532.1 awen|OJJ36872.1 azon|OJJ42932.1

Group862: acar|OOF93727.1 acar|OOF93815.1 abra|OJJ66727.1 acla|EAW11857.1 afis|EAW19468.1 afla|EED56678.1 afum|EAL87028.1 agla|OJJ81188.1 akaw|GAA89668.1 aluc|OJZ80721.1 anid|CBF75484.1 anig|EHA27299.1 aory|EIT73958.1 asyd|OJJ59757.1 ater|EAU34778.1 atub|OJI87577.1 aver|OJJ02531.1 awen|OJJ36873.1 azon|OJJ42931.1

Group863: acar|OOF93728.1 acar|OOF93816.1 abra|OJJ66728.1 acla|EAW11858.1 afis|EAW19469.1 afla|EED56679.1 afum|EAL87027.1 agla|OJJ81190.1 akaw|GAA89669.1 aluc|OJZ80720.1 anid|CBF75482.1 anig|EHA27300.1 aory|EIT73843.1 asyd|OJJ59758.1 ater|EAU34779.1 atub|OJI87576.1 aver|OJJ02530.1 awen|OJJ36874.1 azon|OJJ42930.1

Group864: acar|OOF93729.1 acar|OOF93817.1 abra|OJJ66731.1 acla|EAW11859.1 afis|EAW19470.1 afla|EED56680.1 afum|EAL87026.1 agla|OJJ81191.1 akaw|GAA89670.1 aluc|OJZ80718.1 anid|CBF75480.1 anig|EHA27301.1 aory|EIT73854.1 asyd|OJJ59759.1 ater|EAU34780.1 atub|OJI87574.1 aver|OJJ02529.1 awen|OJJ36875.1 azon|OJJ42929.1

Group865: acar|OOF93731.1 acar|OOF93819.1 abra|OJJ66733.1 acla|EAW11861.1 afis|EAW19472.1 afla|EED56682.1 afum|EAL87024.1 agla|OJJ81193.1 akaw|GAA89672.1 aluc|OJZ80716.1 anid|CBF75474.1 anig|EHA27303.1 aory|EIT73961.1 asyd|OJJ59761.1 ater|EAU34782.1 atub|OJI87572.1 aver|OJJ02527.1 awen|OJJ36877.1 azon|OJJ42927.1

Group866: acar|OOF93734.1 acar|OOF93822.1 abra|OJJ66736.1 acla|EAW11864.1 afis|EAW19475.1 afla|EED56683.1 afum|EAL87021.1 agla|OJJ81198.1 akaw|GAA89675.1 aluc|OJZ80711.1 anid|CBF75470.1 anig|EHA27306.1 aory|EIT73904.1 asyd|OJJ59764.1 ater|EAU34787.1 atub|OJI87569.1 aver|OJJ02524.1 awen|OJJ36880.1 azon|OJJ42925.1

Group867: acar|OOF93736.1 acar|OOF93825.1 abra|OJJ66739.1 acla|EAW11866.1 afis|EAW19477.1 afla|EED56685.1 afum|EAL87019.1 agla|OJJ81207.1 akaw|GAA89677.1 aluc|OJZ80709.1 anid|CBF71519.1 anig|EHA27308.1 aory|EIT73831.1 asyd|OJJ54916.1 ater|EAU34789.1 atub|OJI87567.1 aver|OJJ07287.1 awen|OJJ36882.1 azon|OJJ42923.1

Group868: acar|OOF93739.1 acar|OOF93829.1 abra|OJJ66743.1 acla|EAW11872.1 afis|EAW19483.1 afla|EED56701.1 afum|EAL87012.1 agla|OJJ81208.1 akaw|GAA89681.1 aluc|OJZ80704.1 anid|CBF75464.1 anig|EHA27313.1 aory|EIT73864.1 asyd|OJJ59767.1 ater|EAU34796.1 atub|OJI87562.1 aver|OJJ02520.1 awen|OJJ36895.1 azon|OJJ42918.1

Group869: acar|OOF94046.1 acar|OOF94116.1 agla|OJJ78942.1 agla|OJJ84364.1 agla|OJJ88800.1 asyd|OJJ55478.1 asyd|OJJ56071.1 aver|OJJ05014.1 aver|OJJ06558.1 akaw|GAA90264.1 aluc|OJZ83077.1 anig|EHA28141.1 ater|EAU36656.1 atub|OJI87869.1 awen|OJJ30395.1 aory|EIT78619.1 azon|OJJ48036.1 anig|EHA28514.1 awen|OJJ34834.1

Group870: acar|OOF94064.1 acar|OOF94134.1 abra|OJJ66360.1 acla|EAW10009.1 afis|EAW21047.1 afla|EED51623.1 afum|EAL92083.1 agla|OJJ82365.1 akaw|GAA90233.1 aluc|OJZ83108.1 anid|CBF87099.1 anig|EHA28171.1 aory|EIT80605.1 asyd|OJJ60391.1 ater|EAU36819.1 atub|OJI87826.1 aver|OJI99720.1 awen|OJJ32175.1 azon|OJJ46754.1

Group871: acar|OOF94087.1 acar|OOF94157.1 abra|OJJ66351.1 acla|EAW06467.1 afis|EAW15962.1 afla|EED58095.1 afum|EAL88595.1 agla|OJJ89350.1 akaw|GAA90224.1 aluc|OJZ83117.1 anid|CBF81081.1 anig|EHA28180.1 aory|EIT79147.1 asyd|OJJ56464.1 ater|EAU35492.1 atub|OJI87817.1 aver|OJJ06256.1 awen|OJJ35612.1 azon|OJJ46912.1

Group872: acar|OOF94090.1 acar|OOF94160.1 abra|OJJ66347.1 acla|EAW09988.1 afis|EAW21105.1 afla|EED51687.1 afum|EAL92126.1 agla|OJJ80110.1 akaw|GAA90221.1 aluc|OJZ83120.1 anid|CBF87045.1 anig|EHA28184.1 aory|EIT80607.1 asyd|OJJ60445.1 ater|EAU36794.1 atub|OJI87813.1 aver|OJI99763.1 awen|OJJ36366.1 azon|OJJ43506.1

Group873: acar|OOF94095.1 acar|OOF94164.1 abra|OJJ66343.1 acla|EAW09991.1 afis|EAW21101.1 afla|EED48394.1 afum|EAL92122.1 agla|OJJ80116.1 akaw|GAA90217.1 aluc|OJZ83126.1 anid|CBF87046.1 atub|OJI87807.1 aver|OJI99762.1 awen|OJJ37901.1 azon|OJJ44446.1 acla|EAW09992.1 awen|OJJ36357.1 afla|EED51683.1 asyd|OJJ60444.1

Group874: acar|OOF95064.1 acar|OOF95130.1 abra|OJJ74649.1 acla|EAW10737.1 afis|EAW24838.1 afla|EED47138.1 afum|EAL89484.1 agla|OJJ82029.1 akaw|GAA81961.1 aluc|OJZ92227.1 anid|CBF74603.1 anig|EHA26729.1 aory|EIT79201.1 asyd|OJJ57657.1 ater|EAU36468.1 atub|OJI86657.1 aver|OJJ05518.1 awen|OJJ40424.1 azon|OJJ49282.1

Group875: acar|OOF95067.1 acar|OOF95133.1 abra|OJJ74647.1 acla|EAW10740.1 afis|EAW24834.1 afla|EED47141.1 afum|EAL89479.1 agla|OJJ82027.1 akaw|GAA81964.1 aluc|OJZ92225.1 anid|CBF74582.1 anig|EHA26731.1 aory|EIT79244.1 asyd|OJJ57655.1 ater|EAU36471.1 atub|OJI86655.1 aver|OJJ05515.1 awen|OJJ40421.1 azon|OJJ49279.1

Group876: acar|OOF95068.1 acar|OOF95134.1 abra|OJJ74646.1 acla|EAW10741.1 afis|EAW24833.1 afla|EED47142.1 afum|EAL89478.1 agla|OJJ82026.1 akaw|GAA81965.1 aluc|OJZ92224.1 anid|CBF74580.1 anig|EHA26732.1 aory|EIT79261.1 asyd|OJJ57654.1 ater|EAU36472.1 atub|OJI86654.1 aver|OJJ05514.1 awen|OJJ40420.1 azon|OJJ49278.1

Group877: acar|OOF95069.1 acar|OOF95135.1 abra|OJJ68490.1 acla|EAW10742.1 afis|EAW24832.1 afla|EED47143.1 afum|EAL89477.1 agla|OJJ82025.1 akaw|GAA92780.1 aluc|OJZ82703.1 anid|CBF74578.1 anig|EHA21158.1 aory|EIT79304.1 asyd|OJJ57653.1 ater|EAU36473.1 atub|OJI88936.1 aver|OJJ05513.1 awen|OJJ40419.1 azon|OJJ49277.1

Group878: acar|OOF95070.1 acar|OOF95136.1 abra|OJJ74645.1 acla|EAW10743.1 afis|EAW24831.1 afla|EED47144.1 afum|EAL89476.1 agla|OJJ82019.1 akaw|GAA81966.1 aluc|OJZ92223.1 anid|CBF74576.1 anig|EHA26733.1 aory|EIT79280.1 asyd|OJJ57652.1 ater|EAU36474.1 atub|OJI86653.1 aver|OJJ05512.1 awen|OJJ40418.1 azon|OJJ49276.1

Group879: acar|OOF95076.1 acar|OOF95140.1 abra|OJJ74640.1 acla|EAW10748.1 afis|EAW24826.1 afla|EED47149.1 afum|EAL89470.1 agla|OJJ82015.1 akaw|GAA81971.1 aluc|OJZ92219.1 anid|CBF85858.1 anig|EHA26739.1 aory|EIT79249.1 asyd|OJJ64367.1 ater|EAU36479.1 atub|OJI86650.1 aver|OJI98462.1 awen|OJJ40412.1 azon|OJJ49262.1

Group880: acar|OOF95080.1 acar|OOF95145.1 abra|OJJ74631.1 acla|EAW10753.1 afis|EAW24822.1 afla|EED47153.1 afum|EAL89466.1 agla|OJJ82920.1 akaw|GAA81978.1 aluc|OJZ92212.1 anid|CBF85851.1 anig|EHA26746.1 aory|EIT79200.1 asyd|OJJ64363.1 ater|EAU36483.1 atub|OJI86641.1 aver|OJI98458.1 awen|OJJ40408.1 azon|OJJ49257.1

Group881: acar|OOF95083.1 acar|OOF95148.1 abra|OJJ74626.1 acla|EAW10756.1 afis|EAW24818.1 afla|EED47162.1 afum|EAL91338.1 akaw|GAA81982.1 aluc|OJZ92208.1 anid|CBF85876.1 anig|EHA26750.1

aory|EIT79206.1 asyd|OJJ64375.1 ater|EAU36488.1 atub|OJI86637.1 aver|OJI98471.1 awen|OJJ40404.1 afum|EAL89462.1 afis|EAW17523.1

Group882: acar|OOF95084.1 acar|OOF95149.1 abra|OJJ74624.1 acla|EAW10760.1 afis|EAW24816.1 afla|EED47167.1 afum|EAL89460.2 agla|OJJ82910.1 akaw|GAA81984.1 aluc|OJZ92206.1 anid|CBF85879.1 anig|EHA26752.1 aory|EIT79187.1 asyd|OJJ64377.1 ater|EAU39452.1 atub|OJI86635.1 aver|OJI98473.1 awen|OJJ40402.1 azon|OJJ43613.1

Group883: acar|OOF95086.1 acar|OOF95152.1 abra|OJJ74618.1 acla|EAW10764.1 afis|EAW24812.1 afla|EED47171.1 afum|EAL89455.1 agla|OJJ82907.1 akaw|GAA81988.1 aluc|OJZ92201.1 anid|CBF85885.1 anig|EHA26756.1 aory|EIT79186.1 asyd|OJJ64381.1 ater|EAU36495.1 atub|OJI86629.1 aver|OJI98477.1 awen|OJJ40396.1 azon|OJJ49249.1

Group884: acar|OOF95087.1 acar|OOF95153.1 abra|OJJ74617.1 acla|EAW10765.1 afis|EAW24811.1 afla|EED47172.1 afum|EAL89454.1 agla|OJJ82906.1 akaw|GAA81989.1 aluc|OJZ92200.1 anid|CBF85886.1 anig|EHA26757.1 aory|EIT79291.1 asyd|OJJ64382.1 ater|EAU36496.1 atub|OJI86627.1 aver|OJI98478.1 awen|OJJ40395.1 azon|OJJ49248.1

Group885: acar|OOF95089.1 acar|OOF95155.1 abra|OJJ74615.1 acla|EAW10767.1 afis|EAW24809.1 afla|EED47174.1 afum|EAL89452.1 agla|OJJ82904.1 akaw|GAA81991.1 aluc|OJZ92198.1 anid|CBF85888.1 anig|EHA26759.1 aory|EIT79300.1 asyd|OJJ64384.1 ater|EAU36497.1 atub|OJI86625.1 aver|OJI98480.1 awen|OJJ40393.1 azon|OJJ49246.1

Group886: acar|OOF95090.1 acar|OOF95156.1 abra|OJJ74614.1 acla|EAW10768.1 afis|EAW24808.1 afla|EED47175.1 afum|EAL89450.2 agla|OJJ82903.1 akaw|GAA81992.1 aluc|OJZ92197.1 anid|CBF85890.1 anig|EHA26760.1 aory|EIT79254.1 asyd|OJJ64385.1 ater|EAU36498.1 atub|OJI86624.1 aver|OJI98481.1 awen|OJJ40392.1 azon|OJJ49245.1

Group887: acar|OOF95091.1 acar|OOF95157.1 abra|OJJ74613.1 acla|EAW10769.1 afis|EAW24807.1 afla|EED47176.1 afum|EAL89449.1 agla|OJJ82902.1 akaw|GAA81995.1 aluc|OJZ92193.1 anid|CBF85892.1 anig|EHA26761.1 aory|EIT79197.1 asyd|OJJ64387.1 ater|EAU36499.1 atub|OJI86622.1 aver|OJI98482.1 awen|OJJ40391.1 azon|OJJ49241.1

Group888: acar|OOF95092.1 acar|OOF95158.1 abra|OJJ74611.1 acla|EAW10771.1 afis|EAW24805.1 afla|EED47180.1 afum|EAL89447.1 agla|OJJ82900.1 akaw|GAA81997.1 aluc|OJZ92191.1 anid|CBF81492.1 anig|EHA26763.1 aory|EIT79217.1 asyd|OJJ59459.1 ater|EAU36502.1 atub|OJI86620.1 aver|OJJ02814.1 awen|OJJ40389.1 azon|OJJ49239.1

Group889: acar|OOF95093.1 acar|OOF95159.1 abra|OJJ74610.1 acla|EAW10772.1 afis|EAW24804.1 afla|EED47181.1 afum|EAL89446.1 agla|OJJ82899.1 akaw|GAA81998.1 aluc|OJZ92190.1 anid|CBF81493.1 anig|EHA26764.1 aory|EIT79264.1 asyd|OJJ59460.1 ater|EAU36503.1 atub|OJI86619.1 aver|OJJ02813.1 awen|OJJ40388.1 azon|OJJ49238.1

Group890: acar|OOF95095.1 acar|OOF95162.1 abra|OJJ74608.1 acla|EAW10773.1 afis|EAW24803.1 afla|EED47183.1 afum|EAL89444.1 agla|OJJ82897.1 akaw|GAA82000.1 aluc|OJZ92188.1 anid|CBF81495.1 anig|EHA26765.1 aory|EIT79220.1 asyd|OJJ59463.1 ater|EAU36504.1 atub|OJI86617.1 aver|OJJ02811.1 azon|OJJ49236.1 awen|OJJ40386.1

Group891: acar|OOF95101.1 acar|OOF95167.1 abra|OJJ74603.1 acla|EAW10776.1 afis|EAW24798.1 afla|EED47189.1 afum|EAL89440.1 agla|OJJ82894.1 akaw|GAA82005.1 aluc|OJZ92182.1 anid|CBF81508.1 anig|EHA26771.1 aory|EIT79258.1 asyd|OJJ59468.1 ater|EAU36507.1 atub|OJI86610.1 aver|OJJ02806.1 awen|OJJ40383.1 azon|OJJ49232.1

Group892: acar|OOF95102.1 acar|OOF95168.1 afis|EAW19312.1 afis|EAW24797.1 awen|OJJ34628.1 awen|OJJ40382.1 abra|OJJ74602.1 acla|EAW10777.1 afla|EED47188.1 afum|EAL89439.1 akaw|GAA82006.1 aluc|OJZ92181.1 anid|CBF81506.1 anig|EHA26772.1 asyd|OJJ60343.1 ater|EAU36508.1 atub|OJI86609.1 aver|OJI99684.1 aory|EIT79270.1

Group893: acar|OOF95104.1 acar|OOF95170.1 abra|OJJ74600.1 acla|EAW10779.1 afis|EAW24795.1 afla|EED47187.1 afum|EAL89437.1 agla|OJJ82891.1 akaw|GAA82008.1 aluc|OJZ92179.1 anid|CBF81503.1 anig|EHA26774.1 aory|EIT79230.1 asyd|OJJ59466.1 ater|EAU36510.1 atub|OJI86607.1 aver|OJJ02808.1 awen|OJJ40380.1 azon|OJJ49230.1

Group894: acar|OOF95106.1 acar|OOF95172.1 abra|OJJ74598.1 acla|EAW10781.1 afis|EAW24788.1 afla|EED47191.1 afum|EAL89427.1 agla|OJJ82890.1 akaw|GAA82010.1 aluc|OJZ92176.1 anid|CBF81466.1 anig|EHA26776.1 aory|EIT79298.1 asyd|OJJ59443.1 ater|EAU36513.1 atub|OJI86605.1 aver|OJJ02830.1 awen|OJJ40378.1 azon|OJJ49229.1

Group895: acar|OOF95629.1 acar|OOF95680.1 abra|OJJ74664.1 acla|EAW10731.1 afis|EAW24844.1 afla|EED47127.1 afum|EAL89490.1 agla|OJJ82044.1 akaw|GAA81947.1 aluc|OJZ92241.1 anid|CBF75849.1 anig|EHA26713.1 aory|EIT79274.1 asyd|OJJ57571.1 ater|EAU36460.1 atub|OJI86675.1 aver|OJJ05608.1 awen|OJJ40429.1 azon|OJJ49290.1

Group896: acar|OOF95630.1 acar|OOF95681.1 abra|OJJ74665.1 acla|EAW10730.1 afis|EAW24845.1 afla|EED47126.1 afum|EAL89491.1 agla|OJJ82045.1 akaw|GAA81946.1 aluc|OJZ92242.1 anid|CBF75847.1 anig|EHA26712.1 aory|EIT79168.1 asyd|OJJ57562.1 ater|EAU36459.1 atub|OJI86676.1 aver|OJJ05616.1 awen|OJJ40430.1 azon|OJJ49291.1

Group897: acar|OOF95631.1 acar|OOF95683.1 abra|OJJ74666.1 acla|EAW10729.1 afis|EAW24846.1 afla|EED47125.1 afum|EAL89492.1 agla|OJJ82058.1 akaw|GAA81945.1 aluc|OJZ92244.1 anid|CBF75845.1 anig|EHA26711.1 aory|EIT79177.1 asyd|OJJ57561.1 ater|EAU36457.1 atub|OJI86678.1 aver|OJJ05617.1 awen|OJJ40431.1 azon|OJJ49292.1

Group898: acar|OOF95632.1 acar|OOF95684.1 abra|OJJ74671.1 acla|EAW10726.1 afis|EAW24849.1 afla|EED47122.1 afum|EAL89495.1 agla|OJJ82061.1 akaw|GAA81942.1 aluc|OJZ92248.1 anid|CBF75839.1 anig|EHA26708.1 aory|EIT79246.1 asyd|OJJ57557.1 ater|EAU36454.1 atub|OJI86681.1 aver|OJJ05621.1 awen|OJJ40434.1 azon|OJJ49295.1

Group899: acar|OOF95633.1 acar|OOF95685.1 abra|OJJ74668.1 acla|EAW10728.1 afis|EAW24847.1 afla|EED47124.1 afum|EAL89493.1 agla|OJJ82059.1 akaw|GAA81944.1 aluc|OJZ92246.1 anid|CBF75843.1 anig|EHA26710.1 aory|EIT79296.1 asyd|OJJ57560.1 ater|EAU36456.1 atub|OJI86679.1 aver|OJJ05618.1 awen|OJJ40432.1 azon|OJJ49293.1

Group900: acar|OOF95638.1 acar|OOF95689.1 abra|OJJ74675.1 acla|EAW10722.1 afis|EAW24851.1 afla|EED47120.1 afum|EAL89498.1 agla|OJJ82065.1 akaw|GAA81938.1 aluc|OJZ92252.1 anid|CBF75835.1 anig|EHA26704.1 aory|EIT79290.1 asyd|OJJ57555.1 ater|EAU36452.1 atub|OJI86685.1 aver|OJJ05623.1 awen|OJJ40436.1 azon|OJJ49297.1

Group901: acar|OOF95639.1 acar|OOF95690.1 abra|OJJ74677.1 acla|EAW10720.1 afis|EAW24853.1 afla|EED47118.1 afum|EAL89500.1 agla|OJJ82067.1 akaw|GAA81937.1 aluc|OJZ92254.1 anid|CBF75831.1 anig|EHA26703.1 aory|EIT79297.1 asyd|OJJ57553.1 ater|EAU36449.1 atub|OJI86687.1 aver|OJJ05625.1 awen|OJJ40438.1 azon|OJJ49299.1

Group902: acar|OOF95640.1 acar|OOF95691.1 abra|OJJ74678.1 acla|EAW10719.1 afis|EAW24854.1 afla|EED47117.1 afum|EAL89501.1 agla|OJJ82068.1 akaw|GAA81936.1 aluc|OJZ92255.1 anid|CBF90310.1 anig|EHA26702.1 aory|EIT79215.1 asyd|OJJ61915.1 ater|EAU36448.1 atub|OJI86688.1 aver|OJI96008.1 awen|OJJ40439.1 azon|OJJ46990.1

Group903: acar|OOF95641.1 acar|OOF95692.1 abra|OJJ74679.1 acla|EAW10716.1 afis|EAW24855.1 afla|EED47094.1 afum|EAL89503.2 agla|OJJ82069.1 akaw|GAA81935.1 aluc|OJZ92256.1 anid|CBF75829.1 anig|EHA26701.1 aory|EIT72488.1 asyd|OJJ57552.1 ater|EAU36447.1 atub|OJI86690.1 aver|OJJ05626.1 awen|OJJ40443.1 azon|OJJ49301.1

Group904: acar|OOF95643.1 acar|OOF95694.1 abra|OJJ74681.1 acla|EAW10714.1 afis|EAW24856.1 afla|EED47096.1 afum|EAL89504.1 agla|OJJ82071.1 akaw|GAA81933.1 aluc|OJZ92258.1 anid|CBF75827.1 anig|EHA26700.1 aory|EIT72489.1 asyd|OJJ57550.1 ater|EAU36446.1 atub|OJI86692.1 aver|OJJ05628.1 awen|OJJ40445.1 azon|OJJ49302.1

Group905: acar|OOF95644.1 acar|OOF95695.1 abra|OJJ74683.1 acla|EAW10713.1 afis|EAW24857.1 afla|EED47097.1 afum|EAL89505.2 agla|OJJ82072.1 akaw|GAA81931.1 aluc|OJZ92260.1 anid|CBF75825.1 anig|EHA26699.1 aory|EIT72494.1 asyd|OJJ57549.1 ater|EAU36443.1 atub|OJI86694.1 aver|OJJ05629.1 awen|OJJ40446.1 azon|OJJ49303.1

Group906: acar|OOF95645.1 acar|OOF95696.1 abra|OJJ74684.1 acla|EAW10712.1 afis|EAW24858.1 afla|EED47098.1 afum|EAL89506.2 agla|OJJ82073.1 akaw|GAA81930.1 aluc|OJZ92261.1 anid|CBF75823.1 anig|EHA26698.1 aory|EIT72492.1 asyd|OJJ57548.1 ater|EAU36442.1 atub|OJI86695.1 aver|OJJ05630.1 awen|OJJ40447.1 azon|OJJ49304.1

Group907: acar|OOF95647.1 acar|OOF95698.1 abra|OJJ74686.1 acla|EAW10710.1 afis|EAW24860.1 afla|EED47100.1 afum|EAL89509.1 agla|OJJ82078.1 akaw|GAA81928.1 aluc|OJZ92263.1 anid|CBF75819.1 anig|EHA26696.1 aory|EIT72779.1 asyd|OJJ57546.1 ater|EAU36440.1 atub|OJI86697.1 aver|OJJ05632.1 awen|OJJ40450.1 azon|OJJ49306.1

Group908: acar|OOF95648.1 acar|OOF95699.1 abra|OJJ74687.1 acla|EAW10709.1 afis|EAW24861.1 afla|EED47101.1 afum|EAL89510.1 agla|OJJ82079.1 akaw|GAA81927.1 aluc|OJZ92264.1 anid|CBF75817.1 anig|EHA26695.1 aory|EIT72784.1 asyd|OJJ57545.1 ater|EAU36439.1 atub|OJI86698.1 aver|OJJ05633.1 awen|OJJ40451.1 azon|OJJ49307.1

Group909: acar|OOF95649.1 acar|OOF95700.1 abra|OJJ74688.1 acla|EAW10708.1 afis|EAW24862.1 afla|EED47102.1 afum|EAL89511.1 agla|OJJ82080.1 akaw|GAA81926.1 aluc|OJZ92265.1 anid|CBF75815.1 anig|EHA26694.1 aory|EIT72778.1 asyd|OJJ57544.1 ater|EAU36438.1 atub|OJI86699.1 aver|OJJ05634.1 awen|OJJ40452.1 azon|OJJ49308.1

Group910: acar|OOF95650.1 acar|OOF95701.1 abra|OJJ74689.1 acla|EAW10707.1 afis|EAW24863.1 afla|EED47103.1 afum|EAL89513.1 agla|OJJ82081.1 akaw|GAA81925.1 aluc|OJZ92266.1 anid|CBF75813.1

anig|EHA26693.1 aory|EIT72782.1 asyd|OJJ57543.1 ater|EAU36437.1 atub|OJI86700.1 aver|OJJ05635.1 awen|OJJ40453.1 azon|OJJ49309.1

Group911: acar|OOF95651.1 acar|OOF95702.1 abra|OJJ74690.1 acla|EAW10706.1 afis|EAW24864.1 afla|EED47104.1 afum|EAL89514.1 agla|OJJ82082.1 akaw|GAA81924.1 aluc|OJZ92267.1 anid|CBF75811.1 anig|EHA26692.1 aory|EIT72780.1 asyd|OJJ57542.1 ater|EAU36436.1 atub|OJI86701.1 aver|OJJ05636.1 awen|OJJ40454.1 azon|OJJ49310.1

Group912: acar|OOF95653.1 acar|OOF95704.1 abra|OJJ74693.1 acla|EAW10705.1 afis|EAW24865.1 afla|EED47106.1 afum|EAL89515.1 agla|OJJ82083.1 akaw|GAA81923.1 aluc|OJZ92270.1 anid|CBF75809.1 anig|EHA26691.1 aory|EIT72781.1 asyd|OJJ57541.1 ater|EAU36435.1 atub|OJI86704.1 aver|OJJ05637.1 awen|OJJ40455.1 azon|OJJ49311.1

Group913: acar|OOF95660.1 acar|OOF95711.1 abra|OJJ68465.1 acla|EAW10698.1 afis|EAW24872.1 afla|EED47112.1 afum|EAL89522.1 agla|OJJ82093.1 akaw|GAA92385.1 aluc|OJZ83244.1 anid|CBF75794.1 anig|EHA21099.1 aory|EIT79170.1 asyd|OJJ57530.1 ater|EAU36429.1 atub|OJI87692.1 aver|OJJ05648.1 awen|OJJ40463.1 azon|OJJ49323.1

Group914: acar|OOF95661.1 acar|OOF95712.1 abra|OJJ68466.1 acla|EAW10697.1 afis|EAW24873.1 afla|EED47113.1 afum|EAL89523.1 agla|OJJ82094.1 akaw|GAA92386.1 aluc|OJZ83245.1 anid|CBF75792.1 anig|EHA21100.1 aory|EIT79294.1 asyd|OJJ57531.1 ater|EAU36428.1 atub|OJI87691.1 aver|OJJ05647.1 awen|OJJ40464.1 azon|OJJ49324.1

Group915: acar|OOF95662.1 acar|OOF95713.1 abra|OJJ68467.1 acla|EAW10696.1 afis|EAW24874.1 afla|EED47114.1 afum|EAL89524.1 agla|OJJ82095.1 akaw|GAA92388.1 aluc|OJZ83248.1 anid|CBF75790.1 anig|EHA21101.1 aory|EIT79175.1 asyd|OJJ57532.1 ater|EAU36427.1 atub|OJI87689.1 aver|OJJ05646.1 awen|OJJ40465.1 azon|OJJ49325.1

Group916: acar|OOF95663.1 acar|OOF95714.1 abra|OJJ74702.1 acla|EAW10695.1 afis|EAW24875.1 afla|EED47115.1 afum|EAL89525.1 agla|OJJ82096.1 akaw|GAA81917.1 aluc|OJZ92279.1 anid|CBF75789.1 anig|EHA26685.1 aory|EIT79273.1 asyd|OJJ57533.1 ater|EAU36426.1 atub|OJI86712.1 aver|OJJ05645.1 awen|OJJ40466.1 azon|OJJ49328.1

Group917: acar|OOF95665.1 acar|OOF95716.1 abra|OJJ74704.1 acla|EAW10693.1 afis|EAW24877.1 afla|EED47089.1 afum|EAL89527.1 agla|OJJ82098.1 akaw|GAA81914.1 aluc|OJZ92284.1 anid|CBF75758.1 anig|EHA26684.1 aory|EIT72484.1 asyd|OJJ57527.1 ater|EAU36424.1 atub|OJI86715.1 aver|OJJ05651.1 awen|OJJ40467.1 azon|OJJ49330.1

Group918: acar|OOF96224.1 acar|OOF96262.1 abra|OJJ71508.1 acla|EAW15066.1 afis|EAW20038.1 afla|EED45342.1 afum|EAL85159.1 agla|OJJ88925.1 akaw|GAA89405.1 aluc|OJZ86247.1 anid|CBF84777.1 anig|EHA21617.1 aory|EIT73643.1 asyd|OJJ63237.1 ater|EAU38653.1 atub|OJI84295.1 aver|OJI97719.1 awen|OJJ31793.1 azon|OJJ43703.1

Group919: acar|OOF96229.1 acar|OOF96267.1 abra|OJJ71515.1 acla|EAW15072.1 afis|EAW20044.1 afla|EED45336.1 afum|EAL85164.2 agla|OJJ88918.1 akaw|GAA89401.1 aluc|OJZ86253.1 anid|CBF84789.1 anig|EHA21612.1 aory|EIT73570.1 asyd|OJJ63230.1 ater|EAU38648.1 atub|OJI84290.1 aver|OJI97726.1 awen|OJJ31800.1 azon|OJJ46321.1

Group920: acar|OOF96233.1 acar|OOF96270.1 abra|OJJ71517.1 acla|EAW15075.1 afis|EAW20046.1 afla|EED45334.1 afum|EAL85167.1 agla|OJJ88914.1 akaw|GAA89400.1 aluc|OJZ86256.1 anid|CBF84795.1 anig|EHA21610.1 aory|EIT73786.1 asyd|OJJ63227.1 ater|EAU38656.1 atub|OJI84287.1 aver|OJI97728.1 awen|OJJ31804.1 azon|OJJ46318.1

Group921: acar|OOF96234.1 acar|OOF96271.1 abra|OJJ71518.1 acla|EAW15076.1 afis|EAW20047.1 afla|EED45333.1 afum|EAL85168.2 agla|OJJ88913.1 akaw|GAA89399.1 aluc|OJZ86257.1 anid|CBF84797.1 anig|EHA21609.1 aory|EIT73741.1 asyd|OJJ63226.1 ater|EAU38657.1 atub|OJI84286.1 aver|OJI97729.1 awen|OJJ31805.1 azon|OJJ46317.1

Group922: acar|OOF96235.1 acar|OOF96272.1 abra|OJJ71519.1 acla|EAW15077.1 afis|EAW20048.1 afla|EED45332.1 afum|EAL85169.1 agla|OJJ88912.1 akaw|GAA89398.1 aluc|OJZ86258.1 anid|CBF84799.1 anig|EHA21608.1 aory|EIT73661.1 asyd|OJJ63225.1 ater|EAU38658.1 atub|OJI84285.1 aver|OJI97730.1 awen|OJJ31806.1 azon|OJJ46316.1

Group923: acar|OOF96236.1 acar|OOF96273.1 abra|OJJ71520.1 acla|EAW15078.1 afis|EAW20049.1 afum|EAL85170.2 agla|OJJ88911.1 akaw|GAA89397.1 aluc|OJZ86259.1 anid|CBF84801.1 anig|EHA21607.1 aory|EIT73827.1 asyd|OJJ63224.1 ater|EAU38659.1 atub|OJI84284.1 aver|OJI97731.1 awen|OJJ31807.1 azon|OJJ46315.1 afla|EED45331.1

Group924: acar|OOF96239.1 acar|OOF96276.1 abra|OJJ71523.1 acla|EAW15080.1 afis|EAW20052.1 afla|EED45328.1 afum|EAL85172.1 agla|OJJ88908.1 akaw|GAA89394.1 aluc|OJZ86262.1 anid|CBF84806.1 anig|EHA21605.1 aory|EIT73685.1 asyd|OJJ63222.1 ater|EAU38661.1 atub|OJI84281.1 aver|OJI97733.1 awen|OJJ31810.1 azon|OJJ46312.1

Group925: acar|OOF96240.1 acar|OOF96277.1 abra|OJJ71524.1 acla|EAW15081.1 afis|EAW20053.1 afla|EED45327.1 afum|EAL85173.2 agla|OJJ88907.1 akaw|GAA89393.1 aluc|OJZ86263.1 anid|CBF84808.1 anig|EHA21604.1 aory|EIT73613.1 asyd|OJJ63221.1 ater|EAU38662.1 atub|OJI84280.1 aver|OJI97734.1 awen|OJJ31811.1 azon|OJJ46311.1

Group926: acar|OOF96242.1 acar|OOF96279.1 abra|OJJ71527.1 acla|EAW15083.1 afis|EAW20055.1 afla|EED45325.1 afum|EAL85175.2 agla|OJJ88905.1 akaw|GAA89391.1 aluc|OJZ86266.1 anid|CBF84813.1 anig|EHA21602.1 aory|EIT73716.1 asyd|OJJ63218.1 ater|EAU38664.1 atub|OJI84275.1 aver|OJI97737.1 awen|OJJ31814.1 azon|OJJ46309.1

Group927: acar|OOF96243.1 acar|OOF96280.1 afum|EAL85176.2 afum|EAL87762.1 abra|OJJ71528.1 acla|EAW15084.1 afis|EAW20056.1 afla|EED45324.1 agla|OJJ88904.1 akaw|GAA89390.1 aluc|OJZ86268.1 anid|CBF84817.1 aory|EIT73595.1 asyd|OJJ63217.1 ater|EAU38665.1 atub|OJI84274.1 aver|OJI97738.1 awen|OJJ31815.1 azon|OJJ46308.1

Group928: acar|OOF96245.1 acar|OOF96282.1 abra|OJJ71531.1 acla|EAW15086.1 afis|EAW20058.1 afla|EED45322.1 afum|EAL85178.1 agla|OJJ88902.1 akaw|GAA89388.1 aluc|OJZ86270.1 anid|CBF84840.1 anig|EHA21599.1 aory|EIT73696.1 asyd|OJJ63215.1 ater|EAU38667.1 atub|OJI84271.1 aver|OJI97740.1 awen|OJJ31817.1 azon|OJJ46306.1

Group929: acar|OOF96248.1 acar|OOF96285.1 afum|EAL85181.2 afum|EAL87756.2 abra|OJJ71535.1 acla|EAW15089.1 afis|EAW20061.1 afla|EED45318.1 agla|OJJ88898.1 akaw|GAA89385.1 aluc|OJZ86273.1 anid|CBF84834.1 anig|EHA21596.1 aory|EIT73796.1 asyd|OJJ63212.1 ater|EAU38670.1 atub|OJI84267.1 aver|OJI97743.1 awen|OJJ31821.1

Group930: acar|OOF96249.1 acar|OOF96286.1 abra|OJJ71537.1 acla|EAW15090.1 afis|EAW20062.1 afla|EED45317.1 afum|EAL85182.2 agla|OJJ88897.1 akaw|GAA89384.1 aluc|OJZ86274.1 anid|CBF84832.1 anig|EHA21595.1 aory|EIT73715.1 asyd|OJJ63211.1 ater|EAU38671.1 atub|OJI84266.1 aver|OJI97744.1 awen|OJJ31822.1 azon|OJJ46303.1

Group931: acar|OOF96250.1 acar|OOF96287.1 abra|OJJ71540.1 acla|EAW15095.1 afis|EAW20064.1 afla|EED45313.1 afum|EAL85184.1 agla|OJJ88893.1 akaw|GAA89381.1 aluc|OJZ86278.1 anid|CBF84828.1 anig|EHA21593.1 aory|EIT73604.1 asyd|OJJ63209.1 ater|EAU38673.1 atub|OJI84261.1 aver|OJI97746.1 awen|OJJ31825.1 azon|OJJ46300.1

Group932: acar|OOF96251.1 acar|OOF96290.1 abra|OJJ71543.1 acla|EAW15097.1 afis|EAW20069.1 afla|EED45310.1 afum|EAL85187.1 agla|OJJ88891.1 akaw|GAA89379.1 aluc|OJZ86281.1 anid|CBF84823.1 anig|EHA21590.1 aory|EIT73765.1 asyd|OJJ63206.1 ater|EAU38675.1 atub|OJI84258.1 aver|OJI97749.1 awen|OJJ31828.1 azon|OJJ46296.1

Group933: acar|OOF96252.1 acar|OOF96291.1 abra|OJJ71545.1 acla|EAW15099.1 afis|EAW20071.1 afla|EED45308.1 afum|EAL85189.1 agla|OJJ88889.1 akaw|GAA89377.1 aluc|OJZ86283.1 anid|CBF84845.1 anig|EHA21588.1 aory|EIT73815.1 asyd|OJJ63204.1 ater|EAU38676.1 atub|OJI84255.1 aver|OJI97751.1 awen|OJJ31830.1 azon|OJJ46294.1

Group934: acar|OOF96254.1 acar|OOF96293.1 abra|OJJ71549.1 acla|EAW15103.1 afis|EAW20075.1 afla|EED45304.1 afum|EAL85193.2 agla|OJJ88885.1 akaw|GAA89373.1 aluc|OJZ86287.1 anid|CBF84853.1 anig|EHA21584.1 aory|EIT73551.1 asyd|OJJ63200.1 ater|EAU38683.1 atub|OJI84251.1 aver|OJI97755.1 awen|OJJ31834.1 azon|OJJ46290.1

Group935: acar|OOF96255.1 acar|OOF96294.1 abra|OJJ71550.1 acla|EAW15104.1 afis|EAW20076.1 afla|EED45302.1 afum|EAL85194.1 agla|OJJ88884.1 akaw|GAA89372.1 aluc|OJZ86288.1 anid|CBF84855.1 anig|EHA21583.1 aory|EIT73645.1 asyd|OJJ63199.1 ater|EAU38684.1 atub|OJI84250.1 aver|OJI97756.1 awen|OJJ31835.1 azon|OJJ46289.1

Group936: acar|OOF96256.1 acar|OOF96295.1 abra|OJJ71551.1 acla|EAW15105.1 afis|EAW20077.1 afla|EED45298.1 afum|EAL85195.1 agla|OJJ88883.1 akaw|GAA89371.1 aluc|OJZ86289.1 anid|CBF84867.1 anig|EHA21582.1 aory|EIT73681.1 asyd|OJJ63198.1 ater|EAU38685.1 atub|OJI84249.1 aver|OJI97757.1 awen|OJJ31836.1 azon|OJJ46288.1

Group937: acar|OOF96257.1 acar|OOF96296.1 abra|OJJ71554.1 acla|EAW15108.1 afis|EAW20079.1 afla|EED45296.1 afum|EAL85198.1 agla|OJJ88880.1 akaw|GAA89368.1 aluc|OJZ86292.1 anid|CBF84858.1 anig|EHA21579.1 aory|EIT73808.1 asyd|OJJ63195.1 ater|EAU38687.1 atub|OJI84246.1 aver|OJI97760.1 awen|OJJ31839.1 azon|OJJ46284.1

Group938: acar|OOF96906.1 acar|OOF96942.1 abra|OJJ70803.1 acla|EAW11121.1 afis|EAW15850.1 afla|EED56523.1 afum|EAL88486.1 agla|OJJ87258.1 akaw|GAA90119.1 aluc|OJZ84149.1 anid|CBF81174.1 anig|EHA19253.1 aory|EIT72791.1 asyd|OJJ59291.1 ater|EAU31665.1 atub|OJI90521.1 aver|OJJ02984.1 awen|OJJ36777.1 azon|OJJ47421.1

Group939: acar|OOF96913.1 acar|OOF96948.1 abra|OJJ70797.1 acla|EAW11126.1 afis|EAW15855.1 afla|EED56517.1 afum|EAL88491.1 agla|OJJ87252.1 akaw|GAA90113.1 aluc|OJZ84142.1 anid|CBF81185.1

anig|EHA19250.1 aory|EIT72794.1 asyd|OJJ59297.1 ater|EAU31670.1 atub|OJI90516.1 aver|OJJ02978.1 awen|OJJ36772.1 azon|OJJ47426.1

Group940: acar|OOF96914.1 acar|OOF96949.1 abra|OJJ70796.1 acla|EAW11127.1 afis|EAW15856.1 afla|EED56516.1 afum|EAL88492.1 agla|OJJ87251.1 akaw|GAA90112.1 aluc|OJZ84141.1 anid|CBF81187.1 anig|EHA19249.1 aory|EIT72847.1 asyd|OJJ59298.1 ater|EAU31671.1 atub|OJI90515.1 aver|OJJ02977.1 awen|OJJ36771.1 azon|OJJ47427.1

Group941: acar|OOF96916.1 acar|OOF96951.1 abra|OJJ70794.1 acla|EAW11129.1 afis|EAW15858.1 afla|EED56514.1 afum|EAL88494.1 agla|OJJ87249.1 akaw|GAA90110.1 aluc|OJZ84139.1 anid|CBF81191.1 anig|EHA19247.1 aory|EIT72852.1 asyd|OJJ59300.1 ater|EAU31673.1 atub|OJI90513.1 aver|OJJ02975.1 awen|OJJ36769.1 azon|OJJ47429.1

Group942: acar|OOF96917.1 acar|OOF96952.1 abra|OJJ70793.1 acla|EAW11130.1 afis|EAW15861.1 afla|EED56513.1 afum|EAL88496.1 agla|OJJ87248.1 akaw|GAA90109.1 aluc|OJZ84138.1 anid|CBF81193.1 anig|EHA19246.1 aory|EIT72867.1 asyd|OJJ59301.1 ater|EAU31674.1 atub|OJI90512.1 aver|OJJ02974.1 awen|OJJ36768.1 azon|OJJ47430.1

Group943: acar|OOF96918.1 acar|OOF96953.1 abra|OJJ70792.1 acla|EAW11131.1 afis|EAW15862.1 afla|EED56512.1 afum|EAL88497.1 agla|OJJ87247.1 akaw|GAA90108.1 aluc|OJZ84137.1 anid|CBF81194.1 anig|EHA19245.1 aory|EIT72817.1 asyd|OJJ59302.1 ater|EAU31675.1 atub|OJI90511.1 aver|OJJ02973.1 awen|OJJ36767.1 azon|OJJ48009.1

Group944: acar|OOF96919.1 acar|OOF96954.1 abra|OJJ70791.1 acla|EAW11132.1 afis|EAW15863.1 afla|EED56511.1 afum|EAL88498.1 agla|OJJ87246.1 akaw|GAA90107.1 aluc|OJZ84136.1 anid|CBF81196.1 anig|EHA19244.1 aory|EIT72814.1 asyd|OJJ59303.1 ater|EAU31676.1 atub|OJI90510.1 aver|OJJ02972.1 awen|OJJ36766.1 azon|OJJ47431.1

Group945: acar|OOF96921.1 acar|OOF96956.1 abra|OJJ70789.1 acla|EAW11133.1 afis|EAW15865.1 afla|EED56510.1 afum|EAL88499.1 agla|OJJ87244.1 akaw|GAA90105.1 aluc|OJZ84134.1 anid|CBF81200.1 anig|EHA19242.1 aory|EIT72849.1 asyd|OJJ59304.1 ater|EAU31678.1 atub|OJI90508.1 aver|OJJ02971.1 awen|OJJ36765.1 azon|OJJ47433.1

Group946: acar|OOF96924.1 acar|OOF96959.1 abra|OJJ70786.1 acla|EAW11135.1 afis|EAW15867.1 afla|EED56508.1 afum|EAL88501.1 agla|OJJ87241.1 akaw|GAA90103.1 aluc|OJZ84131.1 anid|CBF81205.1 anig|EHA19239.1 aory|EIT72789.1 asyd|OJJ59307.1 ater|EAU31680.1 atub|OJI90505.1 aver|OJJ02968.1 awen|OJJ36762.1 azon|OJJ47436.1

Group947: acar|OOF96927.1 acar|OOF96962.1 abra|OJJ70781.1 acla|EAW11138.1 afis|EAW15869.1 afla|EED56505.1 afum|EAL88503.1 agla|OJJ87238.1 akaw|GAA90100.1 aluc|OJZ84128.1 anid|CBF81211.1 anig|EHA19236.1 aory|EIT72793.1 asyd|OJJ59310.1 ater|EAU31686.1 atub|OJI90502.1 aver|OJJ02965.1 awen|OJJ36760.1 azon|OJJ47439.1

Group948: acar|OOF96928.1 acar|OOF96963.1 abra|OJJ70780.1 acla|EAW11139.1 afis|EAW15870.1 afla|EED56504.1 afum|EAL88504.1 agla|OJJ87237.1 akaw|GAA90099.1 aluc|OJZ84127.1 anid|CBF81213.1 anig|EHA19235.1 aory|EIT72861.1 asyd|OJJ59311.1 ater|EAU31687.1 atub|OJI90501.1 aver|OJJ02964.1 awen|OJJ36759.1 azon|OJJ47440.1

Group949: acar|OOF96930.1 acar|OOF96965.1 abra|OJJ70778.1 acla|EAW11141.1 afis|EAW15872.1 afla|EED56503.1 afum|EAL88506.1 agla|OJJ87235.1 akaw|GAA90097.1 aluc|OJZ84125.1 anid|CBF81219.1 anig|EHA19233.1 aory|EIT72819.1 asyd|OJJ59312.1 ater|EAU31689.1 atub|OJI90499.1 aver|OJJ02963.1 awen|OJJ36758.1 azon|OJJ47442.1

Group950: acar|OOF96931.1 acar|OOF96966.1 abra|OJJ70777.1 acla|EAW11142.1 afis|EAW15873.1 afla|EED56502.1 afum|EBA27350.1 akaw|GAA90096.1 aluc|OJZ84124.1 anid|CBF81221.1 anig|EHA19232.1 aory|EIT72844.1 asyd|OJJ59313.1 ater|EAU31690.1 atub|OJI90497.1 aver|OJJ02962.1 awen|OJJ36757.1 azon|OJJ47443.1 afum|EBA27351.1

Group951: acar|OOF96932.1 acar|OOF96967.1 abra|OJJ70776.1 acla|EAW11143.1 afis|EAW15874.1 afla|EED56501.1 afum|EAL88508.1 agla|OJJ87234.1 akaw|GAA90095.1 aluc|OJZ84123.1 anid|CBF81222.1 anig|EHA19231.1 aory|EIT72827.1 asyd|OJJ59314.1 ater|EAU31691.1 atub|OJI90495.1 aver|OJJ02961.1 awen|OJJ36756.1 azon|OJJ47444.1

Group952: acar|OOF97657.1 acar|OOF97704.1 abra|OJJ71500.1 acla|EAW10048.1 afis|EAW25536.1 afla|EED44712.1 afum|EAL90200.1 agla|OJJ87267.1 akaw|GAA82520.1 aluc|OJZ87047.1 anid|CBF77819.1 anig|EHA20320.1 aory|EIT83462.1 asyd|OJJ53009.1 ater|EAU33377.1 atub|OJI81588.1 aver|OJJ03834.1 awen|OJJ41278.1 azon|OJJ49501.1

Group953: acar|OOF97658.1 acar|OOF97705.1 abra|OJJ71499.1 acla|EAW10050.1 afis|EAW25534.1 afla|EED44709.1 afum|EAL90198.1 agla|OJJ87269.1 akaw|GAA82521.1 aluc|OJZ87048.1 anid|CBF77817.1 anig|EHA20321.1 aory|EIT83458.1 asyd|OJJ53010.1 ater|EAU33376.1 atub|OJI81587.1 aver|OJJ03833.1 awen|OJJ41276.1 azon|OJJ49508.1

Group954: acar|OOF97660.1 acar|OOF97707.1 abra|OJJ71497.1 acla|EAW10052.1 afis|EAW25532.1 afla|EED54033.1 afum|EAL90196.2 agla|OJJ87270.1 akaw|GAA82523.1 aluc|OJZ87050.1 anid|CBF77813.1 anig|EHA20323.1 aory|EIT78299.1 asyd|OJJ53012.1 ater|EAU33374.1 atub|OJI81585.1 aver|OJJ03831.1 awen|OJJ41274.1 azon|OJJ49506.1

Group955: acar|OOF97661.1 acar|OOF97708.1 abra|OJJ71496.1 acla|EAW10053.1 afis|EAW25531.1 afla|EED54032.1 afum|EBA27398.1 agla|OJJ87271.1 akaw|GAA82524.1 aluc|OJZ87051.1 anid|CBF77811.1 anig|EHA20324.1 aory|EIT78333.1 asyd|OJJ53013.1 ater|EAU33373.1 atub|OJI81584.1 aver|OJJ03830.1 awen|OJJ41273.1 azon|OJJ49505.1

Group956: acar|OOF97663.1 acar|OOF97710.1 abra|OJJ71494.1 acla|EAW10055.1 afis|EAW25529.1 afla|EED54030.1 afum|EAL90194.1 agla|OJJ87273.1 akaw|GAA82526.1 aluc|OJZ87053.1 anid|CBF77821.1 anig|EHA20326.1 aory|EIT78259.1 asyd|OJJ53015.1 ater|EAU33372.1 atub|OJI81582.1 aver|OJJ03828.1 awen|OJJ41271.1 azon|OJJ49509.1

Group957: acar|OOF97664.1 acar|OOF97711.1 abra|OJJ71493.1 acla|EAW10057.1 afis|EAW25527.1 afla|EED54028.1 afum|EAL90192.1 agla|OJJ87275.1 akaw|GAA82527.1 aluc|OJZ87054.1 anid|CBF77805.1 anig|EHA20327.1 aory|EIT78292.1 asyd|OJJ53017.1 ater|EAU33370.1 atub|OJI81581.1 aver|OJJ03826.1 awen|OJJ41269.1 azon|OJJ49511.1

Group958: acar|OOF97666.1 acar|OOF97713.1 abra|OJJ71491.1 acla|EAW10059.1 afis|EAW25525.1 afla|EED55229.1 afum|EAL90190.2 agla|OJJ81067.1 akaw|GAA82529.1 aluc|OJZ87056.1 anid|CBF77802.1 anig|EHA20329.1 aory|EIT83576.1 asyd|OJJ53019.1 ater|EAU32314.1 atub|OJI81579.1 aver|OJJ03824.1 awen|OJJ40768.1 azon|OJJ44041.1

Group959: acar|OOF97669.1 acar|OOF97715.1 abra|OJJ71489.1 acla|EAW10061.1 afis|EAW25523.1 afla|EED54026.1 afum|EAL90188.2 agla|OJJ87278.1 akaw|GAA82531.1 aluc|OJZ87058.1 anid|CBF77800.1 anig|EHA20331.1 aory|EIT78257.1 asyd|OJJ53021.1 ater|EAU33368.1 atub|OJI81577.1 aver|OJJ03822.1 awen|OJJ41267.1 azon|OJJ44832.1

Group960: acar|OOF97671.1 acar|OOF97718.1 abra|OJJ71486.1 acla|EAW10064.1 afis|EAW25521.1 afla|EED54023.1 afum|EAL90186.2 agla|OJJ87281.1 akaw|GAA82534.1 aluc|OJZ87061.1 anid|CBF77796.1 anig|EHA20334.1 aory|EIT78303.1 asyd|OJJ53024.1 ater|EAU33367.1 atub|OJI81574.1 aver|OJJ03819.1 awen|OJJ41264.1 azon|OJJ44829.1

Group961: acar|OOF97672.1 acar|OOF97719.1 abra|OJJ71485.1 acla|EAW10065.1 afis|EAW25520.1 afla|EED54022.1 afum|EAL90185.1 agla|OJJ87282.1 akaw|GAA82535.1 aluc|OJZ87062.1 anid|CBF77794.1 anig|EHA20335.1 aory|EIT78334.1 asyd|OJJ53025.1 ater|EAU33366.1 atub|OJI81573.1 aver|OJJ03818.1 awen|OJJ41263.1 azon|OJJ44828.1

Group962: acar|OOF97673.1 acar|OOF97720.1 abra|OJJ71481.1 acla|EAW10068.1 afis|EAW25517.1 afla|EED54017.1 afum|EAL90182.1 agla|OJJ87286.1 akaw|GAA82538.1 aluc|OJZ87066.1 anid|CBF77788.1 anig|EHA20338.1 aory|EIT78273.1 asyd|OJJ53028.1 ater|EAU33363.1 atub|OJI81570.1 aver|OJJ03816.1 awen|OJJ41260.1 azon|OJJ44825.1

Group963: acar|OOF97674.1 acar|OOF97721.1 abra|OJJ71478.1 acla|EAW10071.1 afis|EAW25515.1 afla|EED54016.1 afum|EAL90180.1 agla|OJJ87288.1 akaw|GAA82539.1 aluc|OJZ87067.1 anid|CBF77785.1 anig|EHA20339.1 aory|EIT78267.1 asyd|OJJ53029.1 ater|EAU33361.1 atub|OJI81569.1 aver|OJJ03815.1 awen|OJJ41258.1 azon|OJJ44824.1

Group964: acar|OOF97675.1 acar|OOF97723.1 abra|OJJ71476.1 acla|EAW10073.1 afis|EAW25513.1 afla|EED54014.1 afum|EAL90178.1 agla|OJJ87291.1 akaw|GAA82540.1 aluc|OJZ87070.1 anid|CBF77781.1 anig|EHA20340.1 aory|EIT78254.1 asyd|OJJ53031.1 ater|EAU33359.1 atub|OJI81567.1 aver|OJJ03813.1 awen|OJJ41256.1 azon|OJJ44821.1

Group965: acar|OOF97678.1 acar|OOF97726.1 abra|OJJ71474.1 acla|EAW10076.1 afis|EAW25510.1 afla|EED54011.1 afum|EAL90176.2 agla|OJJ87295.1 akaw|GAA82542.1 aluc|OJZ87072.1 anid|CBF77775.1 anig|EHA20342.1 aory|EIT78253.1 asyd|OJJ53034.1 ater|EAU33357.1 atub|OJI81565.1 aver|OJJ03810.1 awen|OJJ41252.1 azon|OJJ44819.1

Group966: acar|OOF97679.1 acar|OOF97727.1 abra|OJJ71472.1 acla|EAW10078.1 afis|EAW25508.1 afla|EED54008.1 afum|EAL90174.1 agla|OJJ87297.1 akaw|GAA82544.1 aluc|OJZ87074.1 anid|CBF77772.1 anig|EHA20343.1 aory|EIT78280.1 asyd|OJJ53036.1 ater|EAU33355.1 atub|OJI81563.1 aver|OJJ03808.1 awen|OJJ41250.1 azon|OJJ44817.1

Group967: acar|OOF97680.1 acar|OOF97729.1 abra|OJJ71470.1 acla|EAW10081.1 afis|EAW25506.1 afla|EED54006.1 afum|EAL90172.1 agla|OJJ87298.1 akaw|GAA82545.1 aluc|OJZ87075.1 anid|CBF77768.1 anig|EHA20345.1 aory|EIT78338.1 asyd|OJJ53037.1 ater|EAU33354.1 atub|OJI81561.1 aver|OJJ03807.1 awen|OJJ41249.1 azon|OJJ44816.1

Group968: acar|OOF97681.1 acar|OOF97730.1 abra|OJJ71469.1 acla|EAW10082.1 afis|EAW25505.1 afla|EED54005.1 afum|EAL90171.1 agla|OJJ87299.1 akaw|GAA82546.1 aluc|OJZ87076.1 anid|CBF77766.1

anig|EHA20346.1 aory|EIT78265.1 asyd|OJJ53038.1 ater|EAU33353.1 atub|OJI81560.1 aver|OJJ03806.1 awen|OJJ41248.1 azon|OJJ44813.1

Group969: acar|OOF97682.1 acar|OOF97731.1 abra|OJJ71468.1 acla|EAW10083.1 afis|EAW25504.1 afla|EED54003.1 afum|EAL90170.1 agla|OJJ87300.1 akaw|GAA82547.1 aluc|OJZ87077.1 anid|CBF77764.1 anig|EHA20347.1 aory|EIT78261.1 asyd|OJJ53039.1 ater|EAU33352.1 atub|OJI81559.1 aver|OJJ03805.1 awen|OJJ41247.1 azon|OJJ44812.1

Group970: acar|OOF97684.1 acar|OOF97734.1 abra|OJJ71464.1 acla|EAW10086.1 afis|EAW25501.1 afla|EED54000.1 afum|EAL90167.1 agla|OJJ87303.1 akaw|GAA82550.1 aluc|OJZ87080.1 anid|CBF77756.1 anig|EHA20351.1 aory|EIT78290.1 asyd|OJJ53043.1 ater|EAU34671.1 atub|OJI81557.1 aver|OJJ03801.1 awen|OJJ41243.1 azon|OJJ44809.1

Group971: acar|OOF97685.1 acar|OOF97735.1 abra|OJJ71465.1 acla|EAW10085.1 afis|EAW25502.1 afla|EED54001.1 afum|EAL90168.1 agla|OJJ87302.1 akaw|GAA82549.1 aluc|OJZ87079.1 anid|CBF77758.1 anig|EHA20350.1 aory|EIT78293.1 asyd|OJJ53042.1 ater|EAU34672.1 atub|OJI81558.1 aver|OJJ03802.1 awen|OJJ41244.1 azon|OJJ44810.1

Group972: acar|OOF97686.1 acar|OOF97736.1 abra|OJJ71463.1 acla|EAW10088.1 afis|EAW25499.1 afla|EED53999.1 afum|EAL90165.1 agla|OJJ87305.1 akaw|GAA82551.1 aluc|OJZ87081.1 anid|CBF77754.1 anig|EHA20352.1 aory|EIT78297.1 asyd|OJJ53044.1 ater|EAU34670.1 atub|OJI81556.1 aver|OJJ03800.1 awen|OJJ41241.1 azon|OJJ44807.1

Group973: acar|OOF97690.1 acar|OOF97740.1 abra|OJJ71458.1 acla|EAW10091.1 afis|EAW25496.1 afla|EED53996.1 afum|EAL90162.1 agla|OJJ87309.1 akaw|GAA82553.1 aluc|OJZ87085.1 anid|CBF77749.1 anig|EHA20355.1 aory|EIT78288.1 asyd|OJJ53047.1 ater|EAU34667.1 atub|OJI81553.1 aver|OJJ03798.1 awen|OJJ41238.1 azon|OJJ44804.1

Group974: acar|OOF97691.1 acar|OOF97741.1 abra|OJJ71456.1 acla|EAW10093.1 afis|EAW25494.1 afla|EED53994.1 afum|EAL90160.1 agla|OJJ87311.1 akaw|GAA82555.1 aluc|OJZ87087.1 anid|CBF77745.1 anig|EHA20357.1 aory|EIT78318.1 asyd|OJJ53049.1 ater|EAU34665.1 atub|OJI81551.1 aver|OJJ03796.1 awen|OJJ41236.1 azon|OJJ44802.1

Group975: acar|OOF97692.1 acar|OOF97742.1 abra|OJJ71455.1 acla|EAW10094.1 afis|EAW25493.1 afla|EED53993.1 afum|EAL90158.1 akaw|GAA82556.1 aluc|OJZ87088.1 anig|EHA20359.1 aory|EIT78309.1 asyd|OJJ53050.1 ater|EAU34664.1 atub|OJI81550.1 aver|OJJ03795.1 awen|OJJ41235.1 azon|OJJ44801.1 afum|EAL90159.1 anig|EHA20358.1

Group976: acar|OOF97694.1 acar|OOF97743.1 abra|OJJ71454.1 acla|EAW10095.1 afis|EAW25492.1 afla|EED53992.1 afum|EAL90157.1 agla|OJJ87140.1 akaw|GAA82557.1 aluc|OJZ87089.1 anid|CBF77743.1 anig|EHA20360.1 aory|EIT78276.1 asyd|OJJ53051.1 ater|EAU34663.1 atub|OJI81549.1 aver|OJJ03794.1 awen|OJJ41234.1 azon|OJJ44799.1

Group977: acar|OOF97695.1 acar|OOF97745.1 abra|OJJ71451.1 acla|EAW10096.1 afis|EAW25491.1 afla|EED53991.1 afum|EAL90156.1 agla|OJJ87139.1 akaw|GAA82559.1 aluc|OJZ87091.1 anid|CBF77741.1 anig|EHA20362.1 aory|EIT78277.1 asyd|OJJ53052.1 ater|EAU34662.1 atub|OJI81547.1 aver|OJJ03792.1 awen|OJJ41233.1 azon|OJJ44798.1

Group978: acar|OOF97697.1 acar|OOF97747.1 abra|OJJ71447.1 acla|EAW10099.1 afis|EAW25488.1 afla|EED53986.1 afum|EAL90153.1 agla|OJJ87136.1 akaw|GAA82562.1 aluc|OJZ87094.1 anid|CBF77724.1 anig|EHA20365.1 aory|EIT78270.1 asyd|OJJ53061.1 ater|EAU34659.1 atub|OJI81544.1 aver|OJJ03783.1 awen|OJJ41229.1 azon|OJJ44796.1

Group979: acar|OOF97698.1 acar|OOF97748.1 abra|OJJ71446.1 acla|EAW10100.1 afis|EAW25487.1 afla|EED53985.1 afum|EAL90152.1 agla|OJJ87135.1 akaw|GAA82563.1 aluc|OJZ87095.1 anid|CBF77722.1 anig|EHA20366.1 aory|EIT78345.1 asyd|OJJ53062.1 ater|EAU34658.1 atub|OJI81543.1 aver|OJJ03782.1 awen|OJJ41228.1 azon|OJJ44795.1

Group980: acar|OOF97699.1 acar|OOF97749.1 abra|OJJ71444.1 acla|EAW10102.1 afis|EAW25485.1 afla|EED53983.1 afum|EAL90150.1 agla|OJJ87133.1 akaw|GAA82565.1 aluc|OJZ87097.1 anid|CBF77713.1 anig|EHA20368.1 aory|EIT78331.1 asyd|OJJ53067.1 ater|EAU34656.1 atub|OJI81541.1 aver|OJJ03778.1 awen|OJJ41226.1 azon|OJJ44781.1

Group981: acar|OOF98471.1 acar|OOF98473.1 abra|OJJ78401.1 acla|EAW09131.1 afis|EAW22015.1 afla|EED46740.1 afum|EAL93030.1 agla|OJJ84082.1 akaw|GAA85630.1 aluc|OJZ91480.1 anid|CBF82625.1 anig|EHA20142.1 aory|EIT77878.1 asyd|OJJ54012.1 ater|EAU31136.1 atub|OJI83473.1 aver|OJJ08730.1 awen|OJJ34311.1 azon|OJJ43327.1

Group982: acar|OOF98472.1 acar|OOF98486.1 abra|OJJ78402.1 acla|EAW09130.1 afis|EAW22016.1 afla|EED46741.1 afum|EAL93031.1 agla|OJJ84081.1 akaw|GAA85629.1 aluc|OJZ91481.1 anid|CBF82627.1 anig|EHA20143.1 aory|EIT77969.1 asyd|OJJ54011.1 ater|EAU31137.1 atub|OJI83474.1 aver|OJJ08731.1 awen|OJJ34312.1 azon|OJJ43328.1

Group983: acar|OOF98475.1 acar|OOF98487.1 abra|OJJ78399.1 acla|EAW09133.1 afis|EAW22013.1 afla|EED46738.1 afum|EAL93028.1 agla|OJJ84084.1 akaw|GAA85632.1 aluc|OJZ91478.1 anid|CBF82644.1 anig|EHA20140.1 aory|EIT77900.1 asyd|OJJ61676.1 ater|EAU31135.1 atub|OJI83471.1 aver|OJJ00997.1 awen|OJJ34309.1 azon|OJJ48062.1

Group984: acar|OOF98476.1 acar|OOF98489.1 abra|OJJ78398.1 acla|EAW09134.1 afis|EAW22012.1 afla|EED46737.1 afum|EAL93027.1 agla|OJJ84085.1 akaw|GAA85633.1 aluc|OJZ91477.1 anid|CBF82646.1 anig|EHA20139.1 aory|EIT77930.1 asyd|OJJ61675.1 ater|EAU31134.1 atub|OJI83470.1 aver|OJJ00996.1 awen|OJJ34308.1 azon|OJJ48061.1

Group985: acar|OOF98478.1 acar|OOF98491.1 abra|OJJ78396.1 acla|EAW09136.1 afis|EAW22011.1 afla|EED46735.1 afum|EAL93026.1 agla|OJJ84087.1 akaw|GAA85635.1 aluc|OJZ91475.1 anid|CBF82649.1 anig|EHA20137.1 aory|EIT77821.1 asyd|OJJ61673.1 ater|EAU31132.1 atub|OJI83468.1 aver|OJJ00994.1 awen|OJJ34306.1 azon|OJJ43333.1

Group986: acar|OOF98481.1 acar|OOF98494.1 abra|OJJ78393.1 acla|EAW09139.1 afis|EAW22008.1 afla|EED46732.1 afum|EAL93023.1 agla|OJJ84090.1 akaw|GAA85638.1 aluc|OJZ91472.1 anid|CBF82655.1 anig|EHA20134.1 aory|EIT77839.1 asyd|OJJ61670.1 ater|EAU31130.1 atub|OJI83465.1 aver|OJJ00991.1 awen|OJJ34303.1 azon|OJJ43335.1

Group987: acar|OOF98482.1 acar|OOF98495.1 abra|OJJ78391.1 acla|EAW09141.1 afis|EAW22006.1 afla|EED46730.1 afum|EAL93021.2 agla|OJJ84092.1 akaw|GAA85640.1 aluc|OJZ91470.1 anid|CBF82659.1 anig|EHA20132.1 aory|EIT77908.1 asyd|OJJ61668.1 ater|EAU31128.1 atub|OJI83463.1 aver|OJJ00989.1 awen|OJJ34301.1 azon|OJJ43337.1

Group988: acar|OOF98483.1 acar|OOF98496.1 abra|OJJ78392.1 acla|EAW09140.1 afis|EAW22007.1 afla|EED46731.1 afum|EAL93022.1 agla|OJJ84091.1 akaw|GAA85639.1 aluc|OJZ91471.1 anid|CBF82657.1 anig|EHA20133.1 aory|EIT77818.1 asyd|OJJ61669.1 ater|EAU31129.1 atub|OJI83464.1 aver|OJJ00990.1 awen|OJJ34302.1 azon|OJJ43336.1

Group989: acar|OOF98531.1 acar|OOF98532.1 abra|OJJ78355.1 acla|EAW09180.1 afis|EAW21966.1 afla|EED46687.1 afum|EAL92980.1 agla|OJJ84137.1 akaw|GAA85679.1 aluc|OJZ91430.1 anid|CBF82734.1 anig|EHA20099.1 aory|EIT77938.1 asyd|OJJ61626.1 ater|EAU31086.1 atub|OJI83423.1 aver|OJJ00951.1 awen|OJJ34260.1 azon|OJJ43377.1

Group990: acla|EAW12052.1 acla|EAW15326.1 afis|EAW19692.1 afis|EAW20347.1 afla|EED45214.1 afla|EED48467.1 afum|EAL85467.1 afum|EAL86834.1 aory|EIT73589.1 aory|EIT76272.1 asyd|OJJ53694.1 asyd|OJJ62962.1 aver|OJI98005.1 aver|OJJ07810.1 acar|OOF91909.1 anid|CBF75992.1 ater|EAU37786.1 awen|OJJ38808.1 azon|OJJ45237.1

Group991: afis|EAW15984.1 afis|EAW22037.1 abra|OJJ77639.1 acar|OOF99107.1 acla|EAW06489.1 afla|EED56354.1 afum|EAL88616.1 agla|OJJ85987.1 akaw|GAA83995.1 aluc|OJZ80891.1 anid|CBF85810.1 anig|EHA19447.1 aory|EIT73316.1 asyd|OJJ64340.1 ater|EAU33668.1 atub|OJI82687.1 aver|OJI98433.1 awen|OJJ30486.1 azon|OJJ45477.1

Group992: afis|EAW16169.1 afis|EAW20696.1 abra|OJJ78079.1 acar|OOF98785.1 acla|EAW06612.1 afla|EED48696.1 afum|EAL88803.1 agla|OJJ86214.1 akaw|GAA85915.1 aluc|OJZ91151.1 anid|CBF79561.1 anig|EHA19849.1 aory|EIT73253.1 asyd|OJJ57924.1 ater|EAU33245.1 atub|OJI83165.1 aver|OJJ04468.1 awen|OJJ30269.1 azon|OJJ43965.1

Group993: afis|EAW16186.1 afis|EAW21276.1 afum|EAL88817.1 afum|EAL92291.2 abra|OJJ66405.1 acla|EAW11035.1 afla|EED49635.1 agla|OJJ80815.1 akaw|GAA90273.1 aluc|OJZ83068.1 anig|EHA28132.1 aory|EIT80977.1 asyd|OJJ54301.1 ater|EAU36744.1 atub|OJI87878.1 azon|OJJ42729.1 ater|EAU31600.1 azon|OJJ45943.1 anid|CBF80686.1

Group994: afis|EAW16703.1 afis|EAW22511.1 abra|OJJ73562.1 acar|OOG00744.1 acla|EAW07479.1 afla|EED54539.1 afum|EAL90682.1 agla|OJJ80876.1 akaw|GAA88592.1 aluc|OJZ88499.1 anid|CBF88981.1 anig|EHA18208.1 aory|EIT78923.1 asyd|OJJ62789.1 ater|EAU39198.1 atub|OJI91086.1 aver|OJI96836.1 awen|OJJ35137.1 azon|OJJ49962.1

Group995: afis|EAW17302.1 afis|EAW21312.1 abra|OJJ75975.1 acar|OOF99944.1 acla|EAW09822.1 afla|EED57269.1 afum|EAL92328.1 agla|OJJ87430.1 akaw|GAA88981.1 aluc|OJZ89422.1 anid|CBF83405.1 anig|EHA23165.1 aory|EIT74144.1 asyd|OJJ60646.1 ater|EAU35941.1 atub|OJI85371.1 aver|OJI99964.1 awen|OJJ39388.1 azon|OJJ44276.1

Group996: afis|EAW20681.1 afis|EAW24898.1 abra|OJJ74730.1 acar|OOF89795.1 acla|EAW10669.1 afla|EED46977.1 afum|EAL89552.2 agla|OJJ82121.1 akaw|GAA81887.1 aluc|OJZ92314.1 anid|CBF75653.1 anig|EHA26655.1 aory|EIT72604.1 asyd|OJJ57466.1 ater|EAU36401.1 atub|OJI86743.1 aver|OJJ05715.1 awen|OJJ40490.1 azon|OJJ49353.1

Group997: afis|EAW21461.1 afis|EAW22039.1 abra|OJJ76153.1 acar|OOF89976.1 acla|EAW09678.1 afla|EED55893.1 afum|EAL92480.1 agla|OJJ85609.1 akaw|GAA82102.1 aluc|OJZ92076.1 anid|CBF76534.1

anig|EHA23002.1 aory|EIT76550.1 asyd|OJJ61148.1 ater|EAU35074.1 atub|OJI86506.1 aver|OJJ00470.1 awen|OJJ39238.1 azon|OJJ51616.1

Group998: afis|EAW24479.1 afis|EAW24670.1 abra|OJJ67630.1 acar|OOF91977.1 afis|EAW16900.1 afla|EED52360.1 afla|EED53180.1 afum|EAL86435.1 akaw|GAA89150.1 aluc|OJZ81686.1 anig|EHA25757.1 aory|EIT74904.1 atub|OJI88495.1 azon|OJJ45915.1 abra|OJJ70817.1 acla|EAW12031.1 afum|EAL86150.1 anig|EHA28156.1 awen|OJJ40789.1

Group999: afla|EED44958.1 afla|EED54307.1 aory|EIT73482.1 aory|EIT76036.1 abra|OJJ68802.1 acar|OOF91598.1 acla|EAW12204.1 afis|EAW17076.1 afum|EAL86562.1 agla|OJJ84498.1 akaw|GAA90451.1 aluc|OJZ85773.1 anid|CBF78312.1 anig|EHA24780.1 asyd|OJJ52532.1 ater|EAU37366.1 atub|OJI79558.1 aver|OJJ03869.1 awen|OJJ32505.1

Group1000: afla|EED47569.1 afla|EED49784.1 abra|OJJ77270.1 acar|OOF93140.1 acla|EAW13268.1 afis|EAW17349.1 afum|EAL91192.2 agla|OJJ88227.1 akaw|GAA90678.1 aluc|OJZ84677.1 anid|CBF84233.1 anig|EHA27955.1 aory|EIT77705.1 asyd|OJJ57184.1 ater|EAU33864.1 atub|OJI89403.1 aver|OJJ03106.1 awen|OJJ34563.1 azon|OJJ50311.1

Group1001: afla|EED51012.1 afla|EED56994.1 aory|EIT76959.1 aory|EIT77587.1 abra|OJJ72322.1 acar|OOF94890.1 acla|EAW12003.1 afis|EAW19643.1 afum|EAL86879.1 agla|OJJ80785.1 akaw|GAA86073.1 aluc|OJZ90308.1 anig|EHA23795.1 asyd|OJJ58365.1 ater|EAU32885.1 atub|OJI81732.1 aver|OJJ02041.1 awen|OJJ41894.1 azon|OJJ42706.1

Group1002: afla|EED51028.1 afla|EED52463.1 aver|OJJ01603.1 aver|OJJ05985.1 abra|OJJ72678.1 acar|OOF97379.1 acla|EAW12138.1 afis|EAW17198.1 afum|EAL86626.2 akaw|GAA85296.1 aluc|OJZ90655.1 anid|CBF69883.1 anig|EHA24091.1 aory|EIT82808.1 asyd|OJJ56742.1 asyd|OJJ59012.1 atub|OJI82085.1 afla|EED45180.1 aory|EIT73605.1

Group1003: agla|OJJ78416.1 agla|OJJ80642.1 abra|OJJ72427.1 acar|OOF94806.1 acla|EAW12660.1 afis|EAW17824.1 afla|EED46164.1 afum|EAL91634.1 akaw|GAA86168.1 aluc|OJZ90411.1 anid|CBF78007.1 anig|EHA23877.1 aory|EIT76471.1 asyd|OJJ55244.1 ater|EAU33025.1 atub|OJI81842.1 aver|OJJ06980.1 awen|OJJ41754.1 azon|OJJ48444.1

Group1004: agla|OJJ78484.1 agla|OJJ87510.1 abra|OJJ75909.1 acar|OOF99994.1 acla|EAW09880.1 afis|EAW21246.1 afla|EED54806.1 afum|EAL92259.1 akaw|GAA84615.1 aluc|OJZ89359.1 anid|CBF83238.1 anig|EHA23222.1 aory|EIT79446.1 asyd|OJJ60599.1 ater|EAU35886.1 atub|OJI85439.1 aver|OJI99922.1 awen|OJJ36559.1 azon|OJJ44332.1

Group1005: agla|OJJ78485.1 agla|OJJ87511.1 abra|OJJ75908.1 acar|OOF99995.1 acla|EAW09881.1 afis|EAW21245.1 afla|EED54805.1 afum|EAL92258.1 akaw|GAA84614.1 aluc|OJZ89358.1 anid|CBF83236.1 anig|EHA23223.1 aory|EIT79346.1 asyd|OJJ60600.1 ater|EAU35885.1 atub|OJI85440.1 aver|OJI99923.1 awen|OJJ36558.1 azon|OJJ44333.1

Group1006: agla|OJJ78486.1 agla|OJJ87512.1 abra|OJJ75907.1 acar|OOF99996.1 acla|EAW09882.1 afis|EAW21244.1 afla|EED54804.1 afum|EAL92257.1 akaw|GAA84613.1 aluc|OJZ89357.1 anid|CBF83253.1 anig|EHA23224.1 aory|EIT79338.1 asyd|OJJ60588.1 ater|EAU35884.1 atub|OJI85441.1 aver|OJI99911.1 awen|OJJ36555.1 azon|OJJ44334.1

Group1007: agla|OJJ78634.1 agla|OJJ84721.1 asyd|OJJ56854.1 asyd|OJJ56884.1 aver|OJJ03469.1 aver|OJJ07611.1 afla|EED52628.1 aory|EIT74179.1 afis|EAW23894.1 afum|EAL87322.1 acar|OOF92970.1 acla|EAW11559.1 akaw|GAA86969.1 aluc|OJZ84495.1 anid|CBF77167.1 ater|EAU33448.1 atub|OJI89585.1 awen|OJJ39055.1 azon|OJJ51386.1

Group1008: agla|OJJ80738.1 agla|OJJ86280.1 abra|OJJ77802.1 acar|OOF90102.1 agla|OJJ86304.1 akaw|GAA83847.1 aluc|OJZ90868.1 anig|EHA19588.1 aory|EIT75382.1 asyd|OJJ55038.1 atub|OJI82866.1 aver|OJJ07235.1 awen|OJJ36240.1 awen|OJJ37877.1 afla|EED45523.1 asyd|OJJ54957.1 agla|OJJ79081.1 akaw|GAA91814.1 anig|EHA23349.1

Group1009: agla|OJJ82043.1 agla|OJJ82886.1 abra|OJJ74586.1 acar|OOF95180.1 acla|EAW10792.1 afis|EAW24784.1 afla|EED47201.1 afum|EAL89423.1 akaw|GAA82019.1 aluc|OJZ92164.1 anid|CBF81451.1 anig|EHA26785.1 aory|EIT79216.1 asyd|OJJ59433.1 ater|EAU36525.1 atub|OJI86593.1 aver|OJJ02838.1 awen|OJJ40369.1 azon|OJJ49218.1

Group1010: agla|OJJ82106.1 agla|OJJ85287.1 abra|OJJ74712.1 acar|OOF95721.1 acla|EAW10686.1 afis|EAW24884.1 afla|EED47082.1 afum|EAL89534.1 akaw|GAA81905.1 aluc|OJZ92293.1 anid|CBF75745.1 anig|EHA26676.1 aory|EIT72490.1 asyd|OJJ57520.1 ater|EAU36417.1 atub|OJI86723.1 aver|OJJ05658.1 awen|OJJ40474.1 azon|OJJ49338.1

Group1011: agla|OJJ82204.1 agla|OJJ88613.1 abra|OJJ66850.1 acar|OOF91626.1 acla|EAW14428.1 afis|EAW19135.1 afla|EED54279.1 afum|EAL93867.1 akaw|GAA91556.1 aluc|OJZ85734.1 anid|CBF86332.1 anig|EHA24816.1 aory|EIT76149.1 asyd|OJJ64990.1 ater|EAU37406.1 atub|OJI79519.1 aver|OJI99110.1 awen|OJJ37471.1 azon|OJJ51151.1

Group1012: agla|OJJ87407.1 agla|OJJ87526.1 abra|OJJ76001.1 acar|OOF99924.1 acla|EAW09804.1 afis|EAW21332.1 afla|EED57290.1 afum|EAL92350.1 akaw|GAA88959.1 aluc|OJZ89446.1 anid|CBF83435.1 anig|EHA23140.1 aory|EIT74092.1 asyd|OJJ60664.1 ater|EAU35958.1 atub|OJI85347.1 aver|OJI99987.1 awen|OJJ39366.1 azon|OJJ51705.1

Group1013: akaw|GAA87623.1 akaw|GAA91524.1 abra|OJJ66888.1 acar|OOF91659.1 acla|EAW14460.1 afis|EAW19168.1 afla|EED54229.1 afum|EAL93898.1 agla|OJJ88556.1 aluc|OJZ85694.1 anid|CBF86275.1 anig|EHA24847.1 aory|EIT75898.1 asyd|OJJ65023.1 ater|EAU37305.1 atub|OJI79484.1 aver|OJI99142.1 awen|OJJ37521.1 azon|OJJ51187.1

Group1014: akaw|GAA90564.1 akaw|GAA91190.1 abra|OJJ66522.1 acar|OOF90976.1 acla|EAW10968.1 afis|EAW20892.1 afla|EED45587.1 afum|EAL84741.1 agla|OJJ84000.1 aluc|OJZ81513.1 anid|CBF82751.1 anig|EHA25911.1 aory|EIT75801.1 asyd|OJJ61613.1 ater|EAU30831.1 atub|OJI88662.1 aver|OJJ00936.1 awen|OJJ34455.1 azon|OJJ51742.1

Group1015: akaw|GAA91161.1 akaw|GAA92850.1 atub|OJI87463.1 atub|OJI88691.1 abra|OJJ66554.1 acar|OOF90999.1 acla|EAW10940.1 afla|EED50275.1 afum|EAL84713.1 agla|OJJ83968.1 aluc|OJZ81489.1 anid|CBF82446.1 ater|EAU38568.1 aver|OJJ08627.1 awen|OJJ34399.1 azon|OJJ48135.1 afis|EAW20921.1 aory|EIT77308.1 aluc|OJZ79652.1

Group1016: aluc|OJZ80175.1 aluc|OJZ91283.1 abra|OJJ78210.1 acla|EAW11240.1 afis|EAW18265.1 afum|EAL87630.1 agla|OJJ88161.1 akaw|GAA85816.1 anid|CBF79522.1 anig|EHA19968.1 aory|EIT73099.1 asyd|OJJ57891.1 ater|EAU33272.1 atub|OJI83273.1 aver|OJJ04503.1 awen|OJJ30639.1 azon|OJJ47977.1 afla|EED48730.1 afla|EED48731.1

Group1017: anid|CBF75896.1 anid|CBF87037.1 abra|OJJ68334.1 acar|OOF97603.1 acla|EAW09978.1 afis|EAW21111.1 afla|EED51763.1 afum|EAL92134.1 agla|OJJ80102.1 akaw|GAA92087.1 aluc|OJZ83637.1 anig|EHA18789.1 aory|EIT79989.1 asyd|OJJ60451.1 ater|EAU36787.1 atub|OJI90006.1 aver|OJI99769.1 awen|OJJ36440.1 azon|OJJ44433.1

Group1018: anid|CBF76123.1 anid|CBF81569.1 abra|OJJ71017.1 acar|OOF98123.1 acla|EAW10567.1 afis|EAW25033.1 afla|EED55458.1 afum|EAL89661.1 agla|OJJ82245.1 akaw|GAA82938.1 aluc|OJZ87515.1 anig|EHA20726.1 aory|EIT80849.1 asyd|OJJ59506.1 ater|EAU35752.1 atub|OJI81135.1 aver|OJJ02768.1 awen|OJJ40606.1 azon|OJJ49455.1

Group1019: anid|CBF84679.1 anid|CBF86911.1 asyd|OJJ53342.1 asyd|OJJ64609.1 awen|OJJ34698.1 awen|OJJ40005.1 abra|OJJ69889.1 acar|OOF93409.1 afla|EED48936.1 agla|OJJ87937.1 akaw|GAA85843.1 akaw|GAA87761.1 aluc|OJZ85395.1 anig|EHA25579.1 aory|EIT80037.1 ater|EAU30714.1 atub|OJI80176.1 aver|OJJ08403.1 aver|OJI98705.1

Group1020: anid|CBF85624.1 anid|CBF85626.1 abra|OJJ69051.1 acar|OOF91239.1 afis|EAW20343.1 afla|EED45065.1 afum|EAL85460.1 akaw|GAA90583.1 aluc|OJZ86041.1 anig|EHA24570.1 aory|EIT73625.1 asyd|OJJ57131.1 ater|EAU37680.1 atub|OJI79828.1 aver|OJJ03173.1 awen|OJJ38367.1 azon|OJJ45226.1 awen|OJJ38122.1 agla|OJJ87449.1

Group1021: aory|EIT72521.1 aory|EIT77023.1 abra|OJJ69072.1 acar|OOF91232.1 acla|EAW08768.1 afis|EAW15662.1 afla|EED46885.1 afum|EAL85686.1 agla|OJJ81784.1 akaw|GAA90603.1 aluc|OJZ86061.1 anid|CBF71182.1 anig|EHA24552.1 asyd|OJJ55861.1 atub|OJI79858.1 aver|OJJ04802.1 awen|OJJ38338.1 azon|OJJ43025.1 afla|EED51086.1

Group1022: aory|EIT72608.1 aory|EIT74906.1 asyd|OJJ54447.1 asyd|OJJ56044.1 aver|OJJ04989.1 aver|OJJ07712.1 abra|OJJ68724.1 acar|OOF92223.1 afla|EED46807.1 akaw|GAA89875.1 aluc|OJZ81804.1 anig|EHA22412.1 asyd|OJJ56930.1 atub|OJI84376.1 awen|OJJ34986.1 azon|OJJ48830.1 acla|EAW12522.1 afla|EED53315.1 ater|EAU31747.1

Group1023: aory|EIT76145.1 aory|EIT78526.1 abra|OJJ72823.1 acar|OOF96028.1 acla|EAW11415.1 afis|EAW24006.1 afla|EED49155.1 afum|EAL87436.1 agla|OJJ86937.1 akaw|GAA85424.1 aluc|OJZ83325.1 anid|CBF77396.1 anig|EHA24203.1 asyd|OJJ53244.1 ater|EAU29745.1 atub|OJI82222.1 aver|OJJ03606.1 awen|OJJ35478.1 azon|OJJ51270.1

Group1024: aory|EIT76394.1 aory|EIT77963.1 abra|OJJ78294.1 acar|OOF98573.1 acla|EAW09222.1 afis|EAW21926.1 afla|EED46618.1 afum|EAL92939.1 agla|OJJ84196.1 akaw|GAA85733.1 aluc|OJZ91368.1 anid|CBF76880.1 anig|EHA20046.1 asyd|OJJ61326.1 ater|EAU30932.1 atub|OJI83361.1 aver|OJJ00651.1 awen|OJJ34216.1 azon|OJJ43418.1

Group1025: aory|EIT81453.1 aory|EIT82110.1 abra|OJJ76662.1 acar|OOF99341.1 afis|EAW20834.1 afla|EED52049.1 afum|EAL84798.1 agla|OJJ80143.1 akaw|GAA93003.1 aluc|OJZ91540.1 anid|CBF88060.1 anig|EHA22577.1 asyd|OJJ54521.1 atub|OJI85994.1 aver|OJJ07664.1 awen|OJJ36032.1 afla|EED50429.1 aver|OJI97388.1 asyd|OJJ63546.1

Group1026: asyd|OJJ51877.1 asyd|OJJ54109.1 abra|OJJ66556.1 acar|OOF91001.1 acla|EAW10938.1 afis|EAW20923.1 afla|EED50277.1 afum|EAL84711.1 agla|OJJ83966.1 akaw|GAA91159.1 aluc|OJZ81487.1

anid|CBF82451.1 anig|EHA25940.1 aory|EIT77347.1 ater|EAU38570.1 atub|OJI88693.1 aver|OJJ08625.1 awen|OJJ34397.1 azon|OJJ48133.1

Group1027: asyd|OJJ51975.1 asyd|OJJ60994.1 abra|OJJ71988.1 acar|OOF96641.1 acla|EAW09550.1 afis|EAW21586.1 afla|EED57807.1 afum|EAL92604.1 agla|OJJ85779.1 akaw|GAA84941.1 aluc|OJZ86725.1 anid|CBF76287.1 anig|EHA21954.1 aory|EIT81038.1 ater|EAU34945.1 atub|OJI83803.1 aver|OJJ00314.1 awen|OJJ39095.1 azon|OJJ51428.1

Group1028: asyd|OJJ52256.1 asyd|OJJ61135.1 abra|OJJ76154.1 acar|OOF99776.1 acla|EAW09663.1 afis|EAW21474.1 afla|EED46116.1 afum|EAL92493.1 agla|OJJ85624.1 akaw|GAA82103.1 aluc|OJZ92074.1 anid|CBF76511.1 anig|EHA23001.1 aory|EIT76380.1 ater|EAU35059.1 atub|OJI86505.1 aver|OJJ00457.1 awen|OJJ39223.1 azon|OJJ51633.1

Group1029: asyd|OJJ52749.1 asyd|OJJ64102.1 abra|OJJ77967.1 acar|OOF98848.1 acla|EAW06673.1 afis|EAW16092.1 afla|EED48628.1 afum|EAL88724.1 agla|OJJ86129.1 akaw|GAA83695.1 aluc|OJZ91037.1 anid|CBF85462.1 anig|EHA19736.1 aory|EIT73247.1 ater|EAU33164.1 atub|OJI83033.1 aver|OJI98185.1 awen|OJJ30338.1 azon|OJJ45366.1

Group1030: asyd|OJJ53223.1 asyd|OJJ57663.1 aver|OJJ03629.1 aver|OJJ05525.1 abra|OJJ71299.1 acar|OOF97871.1 afis|EAW25353.1 afla|EED53834.1 afum|EAL90015.1 agla|OJJ83901.1 akaw|GAA82690.1 aluc|OJZ87244.1 anid|CBF77438.1 anig|EHA20489.1 aory|EIT78444.1 ater|EAU34501.1 atub|OJI81398.1 awen|OJJ41076.1 azon|OJJ44658.1

Group1031: asyd|OJJ53278.1 asyd|OJJ56762.1 abra|OJJ72786.1 acar|OOF96003.1 acla|EAW11483.1 afis|EAW23970.1 afla|EED50538.1 afum|EAL87400.1 agla|OJJ86899.1 akaw|GAA85393.1 aluc|OJZ83284.1 anid|CBF77331.1 anig|EHA24170.1 aory|EIT82142.1 ater|EAU29782.1 atub|OJI82187.1 aver|OJJ03570.1 awen|OJJ38973.1 azon|OJJ51308.1

Group1032: asyd|OJJ53282.1 asyd|OJJ56766.1 abra|OJJ72783.1 acar|OOF96001.1 acla|EAW11484.1 afis|EAW23969.1 afla|EED50540.1 afum|EAL87399.1 agla|OJJ86898.1 akaw|GAA85391.1 aluc|OJZ83282.1 anid|CBF77330.1 anig|EHA24169.1 aory|EIT82164.1 ater|EAU29783.1 atub|OJI82186.1 aver|OJJ03568.1 awen|OJJ38974.1 azon|OJJ51309.1

Group1033: asyd|OJJ53391.1 asyd|OJJ55131.1 abra|OJJ72369.1 acar|OOF94851.1 acla|EAW11966.1 afis|EAW19602.1 afla|EED56958.1 afum|EAL86916.1 agla|OJJ80744.1 akaw|GAA86119.1 aluc|OJZ90357.1 anid|CBF71283.1 anig|EHA23833.1 aory|EIT77537.1 ater|EAU32923.1 atub|OJI81785.1 aver|OJJ07072.1 awen|OJJ41838.1 azon|OJJ47049.1

Group1034: asyd|OJJ53445.1 asyd|OJJ58376.1 abra|OJJ68456.1 acar|OOF93941.1 afla|EED51144.1 akaw|GAA86918.1 aluc|OJZ90041.1 anid|CBF87350.1 anig|EHA21029.1 aory|EIT81354.1 asyd|OJJ61782.1 ater|EAU30292.1 atub|OJI84765.1 aver|OJI95882.1 awen|OJJ41192.1 azon|OJJ51544.1 asyd|OJJ60364.1 aver|OJI99700.1 aver|OJJ08306.1

Group1035: asyd|OJJ53802.1 asyd|OJJ61125.1 abra|OJJ76171.1 acar|OOF99768.1 acla|EAW09654.1 afis|EAW21485.1 afla|EED55918.1 afum|EAL92505.1 agla|OJJ85644.1 akaw|GAA82117.1 aluc|OJZ92059.1 anid|CBF76492.1 anig|EHA22988.1 aory|EIT76579.1 ater|EAU35053.1 atub|OJI86491.1 aver|OJJ00446.1 awen|OJJ39212.1 azon|OJJ51642.1

Group1036: asyd|OJJ55564.1 asyd|OJJ65204.1 aver|OJI99353.1 aver|OJJ06645.1 abra|OJJ77036.1 acar|OOF92935.1 acla|EAW13557.1 afis|EAW20633.1 afla|EED49759.1 afum|EAL84435.1 agla|OJJ86284.1 akaw|GAA87015.1 aluc|OJZ84439.1 anid|CBF83215.1 anig|EHA27758.1 aory|EIT77675.1 ater|EAU32168.1 atub|OJI89634.1 awen|OJJ33604.1

Group1037: asyd|OJJ55692.1 asyd|OJJ57944.1 aver|OJJ04452.1 aver|OJJ06777.1 abra|OJJ76856.1 acar|OOF91342.1 acla|EAW13417.1 afis|EAW20528.1 afla|EED52477.1 afum|EAL84529.1 agla|OJJ84707.1 akaw|GAA87166.1 aluc|OJZ84268.1 anid|CBF79013.1 anig|EHA27610.1 aory|EIT82744.1 ater|EAU37160.1 atub|OJI89815.1 awen|OJJ33235.1

Group1038: asyd|OJJ57870.1 asyd|OJJ61020.1 abra|OJJ75464.1 acar|OOF90276.1 acla|EAW13831.1 afis|EAW18300.1 afla|EED48769.1 afum|EAL87666.1 agla|OJJ88123.1 akaw|GAA84237.1 aluc|OJZ88920.1 anid|CBF79478.1 anig|EHA23592.1 aory|EIT73032.1 ater|EAU33308.1 atub|OJI85872.1 aver|OJJ04524.1 awen|OJJ30675.1 azon|OJJ43275.1

Group1039: ater|EAU29462.1 ater|EAU34080.1 ater|EAU38788.1 abra|OJJ74396.1 acar|OOF95340.1 acla|EAW06820.1 afis|EAW16368.1 afla|EED53468.1 afum|EAL88993.1 agla|OJJ78589.1 akaw|GAA87294.1 aluc|OJZ89724.1 anid|CBF89352.1 anig|EHA26952.1 aory|EIT74981.1 asyd|OJJ62501.1 atub|OJI85076.1 aver|OJI96622.1 awen|OJJ37773.1

Group1040: ater|EAU30138.1 ater|EAU38860.1 abra|OJJ73936.1 acar|OOG01067.1 acla|EAW07813.1 afis|EAW22849.1 afla|EED50124.1 afum|EAL90351.1 agla|OJJ83475.1 akaw|GAA89542.1 aluc|OJZ88108.1 anid|CBF87853.1 anig|EHA17877.1 aory|EIT74737.1 asyd|OJJ63417.1 atub|OJI90715.1 aver|OJI97521.1 awen|OJJ38872.1 azon|OJJ49662.1

Group1041: ater|EAU30572.1 ater|EAU36084.1 abra|OJJ68744.1 acar|OOF90309.1 acla|EAW14290.1 afis|EAW18993.1 afla|EED46791.1 afum|EAL93728.1 agla|OJJ82981.1 akaw|GAA89891.1 aluc|OJZ81782.1 anid|CBF77932.1 anig|EHA22433.1 aory|EIT72766.1 asyd|OJJ52417.1 atub|OJI84356.1 aver|OJJ03362.1 awen|OJJ31615.1 azon|OJJ47315.1

Group1042: ater|EAU30835.1 ater|EAU31368.1 abra|OJJ68417.1 acar|OOF91799.1 acla|EAW08827.1 afis|EAW15570.1 afla|EED51627.1 afum|EAL85763.1 agla|OJJ85441.1 akaw|GAA92531.1 aluc|OJZ90080.1 anig|EHA20993.1 aory|EIT80506.1 asyd|OJJ57633.1 atub|OJI84726.1 aver|OJJ04438.1 awen|OJJ38058.1 awen|OJJ38229.1 aver|OJI95759.1

Group1043: atub|OJI80331.1 atub|OJI90561.1 abra|OJJ74085.1 acar|OOG01195.1 acla|EAW07929.1 afis|EAW22967.1 afla|EED54404.1 afum|EAL90230.1 agla|OJJ83320.1 akaw|GAA87565.1 aluc|OJZ87950.1 anid|CBF87610.1 anig|EHA17736.1 aory|EIT75873.1 asyd|OJJ63271.1 ater|EAU30655.1 aver|OJI97686.1 awen|OJJ38596.1 azon|OJJ47331.1

Group1044: aver|OJI96628.1 aver|OJJ08269.1 abra|OJJ73489.1 acar|OOG00683.1 acla|EAW06813.1 afis|EAW16362.1 afla|EED53506.1 afum|EAL88987.1 agla|OJJ79111.1 akaw|GAA87438.1 aluc|OJZ88559.1 anid|CBF89241.1 anig|EHA18256.1 aory|EIT75032.1 asyd|OJJ62508.1 ater|EAU34090.1 atub|OJI91144.1 awen|OJJ37766.1 asyd|OJJ55388.1

Group1045: aver|OJI96684.1 aver|OJJ05918.1 abra|OJJ73439.1 acar|OOG00645.1 acla|EAW06773.1 afis|EAW16324.1 afla|EED53558.1 afum|EAL88947.1 agla|OJJ88286.1 akaw|GAA83054.1 aluc|OJZ88601.1 anid|CBF89160.1 anig|EHA18295.1 aory|EIT74902.1 asyd|OJJ62559.1 ater|EAU34133.1 atub|OJI91186.1 awen|OJJ37724.1 azon|OJJ44983.1

Group1046: aver|OJI98887.1 aver|OJJ06757.1 abra|OJJ77001.1 acar|OOF91446.1 acla|EAW13450.1 afis|EAW20561.1 afla|EED52343.1 afum|EAL84500.1 agla|OJJ88729.1 akaw|GAA87047.1 aluc|OJZ84400.1 anid|CBF78975.1 anig|EHA27730.1 aory|EIT74333.1 asyd|OJJ55674.1 ater|EAU37119.1 atub|OJI89672.1 awen|OJJ33268.1 azon|OJJ50642.1

Group1047: aver|OJI98940.1 aver|OJI99461.1 abra|OJJ69903.1 acar|OOF93421.1 acla|EAW08920.1 afis|EAW15474.1 afla|EED46925.1 afum|EAL85847.1 agla|OJJ89052.1 akaw|GAA87749.1 aluc|OJZ85382.1 anid|CBF86601.1 anig|EHA25565.1 aory|EIT72600.1 asyd|OJJ64827.1 ater|EAU32654.1 atub|OJI80188.1 awen|OJJ36395.1 azon|OJJ44109.1

Group1048: aver|OJJ00758.1 aver|OJJ05227.1 abra|OJJ75869.1 acar|OOG00025.1 acla|EAW09931.1 afis|EAW21178.1 afla|EED54765.1 afum|EAL92198.1 agla|OJJ79995.1 akaw|GAA84575.1 aluc|OJZ89312.1 anid|CBF82104.1 anig|EHA23255.1 aory|EIT79387.1 asyd|OJJ54271.1 ater|EAU35844.1 atub|OJI85487.1 awen|OJJ36517.1 afla|EED54766.1

Group1049: aver|OJJ01895.1 aver|OJJ04283.1 abra|OJJ68598.1 acar|OOF92108.1 acla|EAW12641.1 afis|EAW18044.1 afla|EED51502.1 afum|EAL91855.2 agla|OJJ85415.1 akaw|GAA91922.1 aluc|OJZ81958.1 anid|CBF79781.1 anig|EHA22289.1 aory|EIT80736.1 asyd|OJJ58122.1 ater|EAU31279.1 atub|OJI84509.1 awen|OJJ41490.1 azon|OJJ48220.1

Group1050: aver|OJJ05206.1 aver|OJJ06610.1 aluc|OJZ88621.1 anig|EHA18312.1 atub|OJI91206.1 abra|OJJ73422.1 acar|OOG00625.1 acla|EAW07037.1 afis|EAW16663.1 afla|EED44906.1 afum|EAL89267.1 agla|OJJ78706.1 akaw|GAA83070.1 anid|CBF82078.1 aory|EIT73489.1 asyd|OJJ54252.1 ater|EAU39585.1 awen|OJJ40050.1 azon|OJJ50117.1

Group1051: aver|OJJ07706.1 aver|OJJ08297.1 abra|OJJ76161.1 acar|OOF95039.1 afis|EAW19929.1 afum|EAL84979.1 agla|OJJ80644.1 akaw|GAA82108.1 aluc|OJZ92068.1 anid|CBF84561.1 anig|EHA22995.1 aory|EIT79289.1 asyd|OJJ53459.1 ater|EAU31756.1 atub|OJI86499.1 awen|OJJ33359.1 awen|OJJ33575.1 acla|EAW13677.1 afla|EED47286.1

Group1052: awen|OJJ29714.1 awen|OJJ31500.1 abra|OJJ72682.1 acar|OOF95931.1 acla|EAW12316.1 afis|EAW16904.1 afla|EED45262.1 afum|EAL86440.1 agla|OJJ84538.1 akaw|GAA85300.1 aluc|OJZ90660.1 anid|CBF80774.1 anig|EHA24095.1 aory|EIT73651.1 asyd|OJJ54502.1 ater|EAU30211.1 atub|OJI82089.1 aver|OJJ07679.1 azon|OJJ46214.1

Group1053: awen|OJJ30059.1 awen|OJJ32960.1 abra|OJJ69424.1 acar|OOF94312.1 acla|EAW14803.1 afis|EAW24334.1 afla|EED47930.1 afum|EAL86004.1 agla|OJJ83203.1 akaw|GAA88290.1 aluc|OJZ82260.1 anig|EHA28482.1 aory|EIT79819.1 ater|EAU30414.1 atub|OJI88178.1 anid|CBF74037.1 aver|OJJ07954.1 azon|OJJ48958.1 asyd|OJJ53852.1

Group1054: awen|OJJ30747.1 awen|OJJ31734.1 abra|OJJ67459.1 acar|OOF92371.1 acla|EAW06307.1 afis|EAW18416.1 afla|EED52993.1 afum|EAL93098.1 agla|OJJ84982.1 akaw|GAA86374.1 aluc|OJZ84905.1 anid|CBF70716.1 anig|EHA25129.1 aory|EIT77436.1 asyd|OJJ58446.1 ater|EAU29831.1 atub|OJI80693.1 aver|OJJ01046.1 azon|OJJ50492.1

Group1055: awen|OJJ30760.1 awen|OJJ31721.1 abra|OJJ67449.1 acar|OOF90341.1 acla|EAW06315.1 afis|EAW18429.1 afla|EED52984.1 afum|EAL93109.1 agla|OJJ84994.1 akaw|GAA86381.1 aluc|OJZ84915.1

anid|CBF70683.1 anig|EHA25136.1 aory|EIT78705.1 asyd|OJJ58466.1 ater|EAU29845.1 atub|OJI80683.1 aver|OJJ01065.1 azon|OJJ50482.1

Group1056: awen|OJJ30764.1 awen|OJJ31717.1 abra|OJJ67446.1 acar|OOF90343.1 acla|EAW06317.1 afis|EAW18431.1 afla|EED52982.1 afum|EAL93111.2 agla|OJJ84997.1 akaw|GAA86383.1 aluc|OJZ84917.1 anid|CBF70025.1 anig|EHA25137.1 aory|EIT78725.1 asyd|OJJ58872.1 ater|EAU29847.1 atub|OJI80681.1 aver|OJJ01475.1 azon|OJJ50481.1

Group1057: awen|OJJ30768.1 awen|OJJ31713.1 abra|OJJ67442.1 acar|OOF92386.1 acla|EAW06321.1 afis|EAW18435.1 afla|EED52980.1 afum|EAL93116.1 agla|OJJ85002.1 akaw|GAA86387.1 aluc|OJZ84922.1 anid|CBF70033.1 anig|EHA25140.1 aory|EIT78731.1 asyd|OJJ58868.1 ater|EAU29851.1 atub|OJI80677.1 aver|OJJ01470.1 azon|OJJ50478.1

Group1058: awen|OJJ30787.1 awen|OJJ31694.1 abra|OJJ75327.1 acar|OOF90141.1 acla|EAW15368.1 afla|EED45606.1 akaw|GAA92022.1 aluc|OJZ92890.1 anig|EHA26073.1 aory|EIT75814.1 ater|EAU36093.1 atub|OJI87348.1 aver|OJI99803.1 akaw|GAA86921.1 aluc|OJZ90045.1 anig|EHA21025.1 ater|EAU38089.1 atub|OJI84761.1 asyd|OJJ52000.1

Group1059: awen|OJJ31367.1 awen|OJJ39791.1 awen|OJJ33381.1 awen|OJJ35034.1 azon|OJJ43925.1 azon|OJJ44485.1 abra|OJJ70024.1 acar|OOF93532.1 afla|EED45807.1 agla|OJJ82719.1 akaw|GAA87637.1 aluc|OJZ85252.1 anig|EHA25453.1 atub|OJI80316.1 aver|OJJ07005.1 acar|OOF93530.1 agla|OJJ78997.1 aver|OJJ04358.1 asyd|OJJ58045.1

Group1060: awen|OJJ32955.1 awen|OJJ34548.1 abra|OJJ69452.1 acar|OOF94289.1 acla|EAW14816.1 afis|EAW24346.1 afla|EED47917.1 afum|EAL86016.1 agla|OJJ83207.1 akaw|GAA88367.1 aluc|OJZ82288.1 anid|CBF74019.1 anig|EHA28460.1 aory|EIT83370.1 asyd|OJJ53841.1 ater|EAU30400.1 atub|OJI88151.1 aver|OJJ07944.1 azon|OJJ48966.1

Group1061: awen|OJJ35632.1 awen|OJJ41410.1 abra|OJJ68673.1 acar|OOF92174.1 acla|EAW12575.1 afis|EAW24073.1 afla|EED51400.1 afum|EAL91929.1 agla|OJJ87142.1 akaw|GAA89826.1 aluc|OJZ81862.1 anid|CBF86467.1 anig|EHA22364.1 aory|EIT80717.1 asyd|OJJ64908.1 ater|EAU30495.1 atub|OJI84429.1 aver|OJI99030.1 azon|OJJ48882.1

Group1062: awen|OJJ35956.1 awen|OJJ39245.1 abra|OJJ76134.1 acar|OOF99789.1 acla|EAW09684.1 afis|EAW21455.1 afla|EED55885.1 afum|EAL92474.1 agla|OJJ85603.1 akaw|GAA88833.1 aluc|OJZ89596.1 anid|CBF76546.1 anig|EHA23016.1 aory|EIT76856.1 asyd|OJJ61154.1 ater|EAU35080.1 atub|OJI85207.1 aver|OJJ00476.1 azon|OJJ51610.1

Group1063: awen|OJJ35989.1 awen|OJJ38023.1 abra|OJJ75270.1 acar|OOF94533.1 acla|EAW12238.1 afis|EAW16989.1 afla|EED48968.1 afum|EAL86513.1 agla|OJJ79827.1 akaw|GAA91808.1 aluc|OJZ92830.1 anid|CBF87435.1 anig|EHA26179.1 aory|EIT79996.1 asyd|OJJ52914.1 ater|EAU31738.1 atub|OJI87287.1 aver|OJJ08884.1 azon|OJJ50274.1

Group1064: awen|OJJ36336.1 awen|OJJ40727.1 abra|OJJ70865.1 acar|OOF98251.1 acla|EAW10457.1 afis|EAW25110.1 afla|EED55332.1 afum|EAL89781.1 agla|OJJ89071.1 akaw|GAA91732.1 aluc|OJZ87654.1 anid|CBF86071.1 anig|EHA20848.1 aory|EIT76213.1 asyd|OJJ64460.1 ater|EAU39380.1 atub|OJI80988.1 aver|OJI98563.1 azon|OJJ44099.1

Group1065: awen|OJJ38099.1 awen|OJJ40555.1 abra|OJJ74809.1 acar|OOF95801.1 acla|EAW10605.1 afis|EAW24962.1 afla|EED55500.1 afum|EAL89620.1 agla|OJJ82197.1 akaw|GAA83603.1 aluc|OJZ92391.1 anid|CBF81656.1 anig|EHA26584.1 aory|EIT80890.1 asyd|OJJ59547.1 ater|EAU35693.1 atub|OJI86821.1 aver|OJJ02721.1 azon|OJJ49417.1

Group1066: awen|OJJ40158.1 awen|OJJ41199.1 abra|OJJ71411.1 acar|OOF97761.1 acla|EAW10126.1 afis|EAW25462.1 afla|EED53951.1 afum|EAL90125.1 agla|OJJ87114.1 akaw|GAA82589.1 aluc|OJZ87129.1 anid|CBF77649.1 anig|EHA20391.1 aory|EIT78320.1 asyd|OJJ53118.1 ater|EAU34631.1 atub|OJI81511.1 aver|OJJ03736.1 azon|OJJ44770.1

Group1067: azon|OJJ42233.1 azon|OJJ45236.1 acla|EAW12053.1 afis|EAW19693.1 afla|EED48466.1 afum|EAL86833.1 aory|EIT76269.1 asyd|OJJ62963.1 aver|OJI98004.1 acla|EAW15332.1 afis|EAW20372.1 afla|EED45213.1 afum|EAL85486.1 anid|CBF75994.1 aory|EIT73663.1 asyd|OJJ53695.1 ater|EAU37785.1 aver|OJJ07811.1 awen|OJJ38807.1

Group1068: azon|OJJ43179.1 azon|OJJ47539.1 abra|OJJ70678.1 acar|OOF97049.1 acla|EAW08086.1 afis|EAW23122.1 afla|EED48531.1 afum|EAL88358.1 agla|OJJ89617.1 akaw|GAA90010.1 aluc|OJZ84026.1 anid|CBF74402.1 anig|EHA19147.1 aory|EIT73068.1 asyd|OJJ57764.1 ater|EAU36250.1 atub|OJI90398.1 aver|OJJ05402.1 awen|OJJ36659.1

Group1069: azon|OJJ44484.1 azon|OJJ48556.1 abra|OJJ75695.1 acar|OOG00157.1 acla|EAW09413.1 afis|EAW21723.1 afla|EED57992.1 afum|EAL92756.1 agla|OJJ79861.1 akaw|GAA84431.1 aluc|OJZ89146.1 anid|CBF83716.1 anig|EHA23402.1 aory|EIT79772.1 asyd|OJJ60837.1 ater|EAU38407.1 atub|OJI85642.1 aver|OJJ00161.1 awen|OJJ33996.1

Group1070: azon|OJJ45306.1 azon|OJJ46721.1 abra|OJJ76474.1 acar|OOF99711.1 acla|EAW13959.1 afis|EAW18668.1 afla|EED52660.1 afum|EAL93386.1 agla|OJJ82407.1 akaw|GAA82386.1 aluc|OJZ91735.1 anid|CBF70575.1 anig|EHA22741.1 aory|EIT74271.1 asyd|OJJ58530.1 ater|EAU38243.1 atub|OJI86193.1 aver|OJJ01132.1 awen|OJJ32140.1

Group1071: azon|OJJ45770.1 azon|OJJ47370.1 abra|OJJ72219.1 agla|OJJ78976.1 anig|EHA21679.1 awen|OJJ32400.1 abra|OJJ74055.1 acar|OOG01168.1 acla|EAW13644.1 afla|EED58134.1 afum|EAL84922.1 akaw|GAA87537.1 aluc|OJZ87980.1 anig|EHA17763.1 aory|EIT79161.1 asyd|OJJ57210.1 ater|EAU30685.1 atub|OJI90590.1 aver|OJJ03079.1

Group1072: azon|OJJ45841.1 azon|OJJ45942.1 abra|OJJ69362.1 acar|OOF94363.1 acla|EAW14752.1 afis|EAW24283.1 afla|EED47993.1 afum|EAL85952.1 agla|OJJ83145.1 akaw|GAA88231.1 aluc|OJZ82195.1 anid|CBF74136.1 anig|EHA28542.1 aory|EIT79842.1 asyd|OJJ53916.1 ater|EAU31231.1 atub|OJI88240.1 aver|OJJ08014.1 awen|OJJ33013.1

Group1073: azon|OJJ50179.1 azon|OJJ50395.1 abra|OJJ68426.1 acar|OOF93927.1 acla|EAW11928.1 afis|EAW19562.1 afla|EED50583.1 afum|EAL86950.1 agla|OJJ81713.1 akaw|GAA86946.1 aluc|OJZ90071.1 anid|CBF89380.1 anig|EHA21002.1 aory|EIT82053.1 asyd|OJJ62464.1 ater|EAU31050.1 atub|OJI84735.1 aver|OJI96599.1 awen|OJJ35963.1

Group1074: abra|OJJ65489.1 acar|OOF90657.1 afis|EAW25622.1 afla|EED47479.1 afum|EAL85425.1 agla|OJJ86171.1 akaw|GAA88141.1 aluc|OJZ85402.1 anig|EHA18427.1 aory|EIT79955.1 asyd|OJJ55499.1 ater|EAU36065.1 atub|OJI91619.1 aver|OJJ06577.1 afla|EED49747.1 afum|EAL84310.1 akaw|GAA87767.1 aory|EIT77702.1 aluc|OJZ79947.1

Group1075: abra|OJJ65876.1 aory|EIT76450.1 abra|OJJ68863.1 acar|OOF90526.1 afis|EAW20215.1 akaw|GAA90390.1 aluc|OJZ85841.1 anig|EHA24723.1 atub|OJI79629.1 awen|OJJ31221.1 abra|OJJ77712.1 acar|OOF99028.1 agla|OJJ85976.1 aluc|OJZ90777.1 asyd|OJJ59565.1 atub|OJI82764.1 awen|OJJ41677.1 aver|OJJ02704.1 azon|OJJ46135.1

Group1076: afla|EED47310.1 aory|EIT82826.1 abra|OJJ77715.1 acar|OOF99024.1 acla|EAW06603.1 afis|EAW16220.1 afum|EAL88848.1 agla|OJJ86232.1 akaw|GAA83917.1 aluc|OJZ90780.1 anig|EHA19513.1 asyd|OJJ59566.1 atub|OJI82767.1 aver|OJJ02703.1 awen|OJJ29818.1 azon|OJJ42618.1 acar|OOF98224.1 ater|EAU33263.1 awen|OJJ30252.1

Group1077: abra|OJJ65944.1 acar|OOF92929.1 acla|EAW13567.1 afis|EAW20033.1 afla|EED46044.1 afum|EAL84883.1 agla|OJJ88021.1 akaw|GAA90822.1 aluc|OJZ83013.1 anid|CBF84765.1 anig|EHA21428.1 aory|EIT75479.1 asyd|OJJ53284.1 ater|EAU29399.1 atub|OJI89248.1 aver|OJJ08457.1 awen|OJJ30056.1 azon|OJJ46327.1 afla|EED46043.1

Group1078: abra|OJJ65999.1 acar|OOF92101.1 afla|EED56561.1 agla|OJJ88417.1 akaw|GAA90871.1 aluc|OJZ82962.1 anig|EHA21378.1 asyd|OJJ57315.1 atub|OJI89194.1 aver|OJJ05846.1 awen|OJJ33380.1 azon|OJJ44468.1 anid|CBF79162.1 akaw|GAA92926.1 aluc|OJZ82517.1 anig|EHA20902.1 ater|EAU38011.1 asyd|OJJ62803.1 aver|OJI96849.1

Group1079: abra|OJJ66158.1 acar|OOG00538.1 acla|EAW08523.1 afis|EAW23562.1 afum|EAL87906.1 agla|OJJ80848.1 asyd|OJJ55543.1 aver|OJJ06621.1 abra|OJJ67247.1 acar|OOF89887.1 afla|EED52167.1 akaw|GAA91667.1 aluc|OJZ87730.1 anig|EHA20222.1 aory|EIT74456.1 ater|EAU35672.1 atub|OJI80910.1 afla|EED47300.1 aory|EIT82893.1

Group1080: abra|OJJ66244.1 acla|EAW14177.1 afis|EAW16160.1 afla|EED50871.1 afum|EBA27496.1 agla|OJJ81498.1 akaw|GAA89047.1 aluc|OJZ80549.1 anid|CBF69906.1 anig|EHA25647.1 aory|EIT79021.1 asyd|OJJ58996.1 ater|EAU33887.1 atub|OJI88382.1 aver|OJJ01620.1 azon|OJJ47209.1 acar|OOF99450.1 afis|EAW18892.1 afla|EED48908.1

Group1081: abra|OJJ66266.1 acar|OOF97511.1 akaw|GAA89028.1 aluc|OJZ80576.1 anig|EHA25621.1 atub|OJI88355.1 abra|OJJ69860.1 afla|EED45622.1 agla|OJJ84525.1 akaw|GAA87796.1 aluc|OJZ85431.1 anid|CBF86345.1 anig|EHA25610.1 aory|EIT75614.1 asyd|OJJ64976.1 ater|EAU32692.1 atub|OJI80144.1 aver|OJI99098.1 awen|OJJ32475.1

Group1082: abra|OJJ66514.1 acar|OOF90929.1 acla|EAW15358.1 afis|EAW20353.1 afla|EED45813.1 afum|EAL85471.1 agla|OJJ82840.1 akaw|GAA91197.1 aluc|OJZ81532.1 anid|CBF78206.1 anig|EHA25904.1 aory|EIT75838.1 atub|OJI88655.1 aver|OJJ04934.1 awen|OJJ30867.1 azon|OJJ42291.1 afla|EED50237.1 aluc|OJZ81531.1 anig|EHA25905.1

Group1083: abra|OJJ66594.1 acar|OOF91029.1 acla|EAW10897.1 afis|EAW20961.1 afla|EED50316.1 afum|EAL84672.1 agla|OJJ83917.1 akaw|GAA91123.1 aluc|OJZ81453.1 anid|CBF82569.1 anig|EHA25975.1 aory|EIT77366.1 asyd|OJJ54049.1 ater|EAU38611.1 atub|OJI88728.1 aver|OJJ08694.1 awen|OJJ34356.1 azon|OJJ48092.1 aory|EIT77330.1

Group1084: abra|OJJ66648.1 acar|OOF93752.1 acla|EAW11791.1 afis|EAW19402.1 afla|EED50356.1 afum|EAL87093.1 agla|OJJ81115.1 akaw|GAA89601.1 aluc|OJZ80801.1 anid|CBF74340.1 anig|EHA27233.1

aory|EIT83133.1 asyd|OJJ51904.1 ater|EAU36127.1 atub|OJI87649.1 aver|OJJ05364.1 awen|OJJ36797.1 azon|OJJ43002.1 acar|OOF93668.1

Group1085: abra|OJJ66851.1 acar|OOF91627.1 acla|EAW14429.1 afis|EAW19136.1 afla|EED54277.1 afum|EAL93868.1 agla|OJJ88615.1 akaw|GAA91555.1 aluc|OJZ85733.1 anid|CBF86327.1 anig|EHA24817.1 aory|EIT75982.1 asyd|OJJ64992.1 ater|EAU37407.1 atub|OJI79518.1 aver|OJI99111.1 awen|OJJ37473.1 azon|OJJ51152.1 anid|CBF86329.1

Group1086: abra|OJJ66857.1 acar|OOF91634.1 acla|EAW14436.1 afis|EAW19144.1 afla|EED54269.1 afum|EAL93875.1 agla|OJJ88587.1 akaw|GAA91548.1 aluc|OJZ85726.1 anid|CBF86314.1 anig|EHA24822.1 aory|EIT75975.1 asyd|OJJ64999.1 ater|EAU37416.1 atub|OJI79512.1 aver|OJI99119.1 awen|OJJ37483.1 azon|OJJ51160.1 aory|EIT76087.1

Group1087: abra|OJJ66865.1 acar|OOF91641.1 acla|EAW14441.1 afis|EAW19149.1 afla|EED54264.1 afum|EAL93880.1 agla|OJJ88575.1 akaw|GAA91543.1 aluc|OJZ85717.1 anid|CBF86306.1 anig|EHA24827.1 aory|EIT76024.1 asyd|OJJ65005.1 ater|EAU37422.1 atub|OJI79507.1 aver|OJI99124.1 awen|OJJ37502.1 azon|OJJ51167.1 afla|EED54263.1

Group1088: abra|OJJ67116.1 acar|OOF98436.1 acla|EAW13746.1 afis|EAW19825.1 afla|EED56208.1 afum|EAL85067.1 agla|OJJ83144.1 akaw|GAA91049.1 aluc|OJZ87859.1 anid|CBF83120.1 anig|EHA20165.1 aory|EIT73304.1 asyd|OJJ61473.1 ater|EAU31280.1 atub|OJI80788.1 aver|OJJ00814.1 awen|OJJ36528.1 azon|OJJ47562.1 agla|OJJ79980.1

Group1089: abra|OJJ67209.1 acar|OOF98351.1 afis|EAW19743.1 afum|EAL86783.1 akaw|GAA90968.1 aluc|OJZ87765.1 anig|EHA20249.1 ater|EAU29472.1 atub|OJI80880.1 abra|OJJ75764.1 afla|EED56526.1 aory|EIT82926.1 asyd|OJJ58672.1 atub|OJI87194.1 aver|OJJ01283.1 awen|OJJ30928.1 azon|OJJ42672.1 awen|OJJ34642.1 aver|OJJ07429.1

Group1090: abra|OJJ67296.1 acar|OOF92515.1 afla|EED45654.1 agla|OJJ79150.1 akaw|GAA86517.1 aluc|OJZ85063.1 anid|CBF81698.1 anig|EHA25277.1 aory|EIT75443.1 asyd|OJJ59570.1 atub|OJI80535.1 aver|OJJ02697.1 acar|OOF90954.1 afla|EED50408.1 aory|EIT82054.1 asyd|OJJ59578.1 ater|EAU32270.1 aver|OJJ02691.1 awen|OJJ36407.1

Group1091: abra|OJJ67369.1 acla|EAW10405.1 afis|EAW25166.1 afla|EED44928.1 afum|EAL89839.1 anid|CBF85620.1 aory|EIT77682.1 asyd|OJJ64202.1 ater|EAU31038.1 aver|OJI98291.1 abra|OJJ68396.1 akaw|GAA92552.1 aluc|OJZ90100.1 anig|EHA20975.1 aory|EIT83236.1 atub|OJI84706.1 awen|OJJ34638.1 aory|EIT73481.1 afla|EED49793.1

Group1092: abra|OJJ67422.1 acar|OOF91907.1 acla|EAW06342.1 afis|EAW18454.1 afla|EED52954.1 afum|EAL93139.1 agla|OJJ84146.1 akaw|GAA86407.1 anig|EHA25162.1 aory|EIT78718.1 asyd|OJJ58854.1 ater|EAU29871.1 atub|OJI80655.1 aver|OJJ01455.1 azon|OJJ50520.1 anid|CBF71560.1 asyd|OJJ54214.1 aver|OJJ08517.1 agla|OJJ85037.1

Group1093: abra|OJJ67523.1 awen|OJJ30978.1 abra|OJJ69776.1 acar|OOF93595.1 acla|EAW11058.1 afis|EAW20815.1 afla|EED53102.1 afum|EBA27207.1 agla|OJJ81282.1 akaw|GAA87877.1 aluc|OJZ85518.1 anid|CBF86687.1 anig|EHA25051.1 aory|EIT74763.1 asyd|OJJ64752.1 ater|EAU38934.1 atub|OJI80048.1 aver|OJI98853.1 azon|OJJ45137.1

Group1094: abra|OJJ67596.1 acla|EAW12346.1 afis|EAW20358.1 afla|EED57868.1 afum|EAL90941.1 agla|OJJ88440.1 akaw|GAA89115.1 aluc|OJZ81724.1 anid|CBF77124.1 anig|EHA25715.1 aory|EIT81103.1 asyd|OJJ62720.1 ater|EAU34084.1 atub|OJI88457.1 aver|OJI99735.1 awen|OJJ36944.1 afum|EAL85475.1 afum|EAL91116.1 asyd|OJJ60405.1

Group1095: abra|OJJ67622.1 acar|OOF91970.1 acla|EAW12407.1 afis|EAW16866.1 afla|EED45897.1 afum|EAL86405.2 akaw|GAA89142.1 aluc|OJZ81694.1 anig|EHA25737.1 aory|EIT75474.1 asyd|OJJ56695.1 atub|OJI88486.1 aver|OJJ06029.1 awen|OJJ33543.1 azon|OJJ42803.1 acar|OOF91765.1 awen|OJJ34441.1 azon|OJJ50737.1 afla|EED52164.1

Group1096: abra|OJJ67694.1 acar|OOF90173.1 acla|EAW12072.1 afis|EAW19713.1 afla|EED56920.1 afum|EAL86814.1 akaw|GAA89203.1 aluc|OJZ81627.1 anid|CBF82745.1 anig|EHA25816.1 aory|EIT72578.1 asyd|OJJ61617.1 atub|OJI88553.1 aver|OJJ00940.1 awen|OJJ31364.1 azon|OJJ47340.1 acar|OOF91834.1 afla|EED46824.1 aory|EIT77570.1

Group1097: abra|OJJ67728.1 acar|OOF89960.1 afis|EAW20574.1 afla|EED52326.1 afum|EAL84491.1 agla|OJJ89291.1 akaw|GAA89222.1 aluc|OJZ81606.1 anid|CBF87431.1 anig|EHA25839.1 aory|EIT74400.1 asyd|OJJ52907.1 atub|OJI88579.1 aver|OJJ08876.1 awen|OJJ30548.1 awen|OJJ29983.1 akaw|GAA85320.1 aluc|OJZ90681.1 anig|EHA27494.1

Group1098: abra|OJJ67796.1 agla|OJJ84477.1 abra|OJJ75071.1 acar|OOF97263.1 acla|EAW08252.1 afis|EAW23286.1 afla|EED55688.1 afum|EAL88193.1 akaw|GAA88683.1 aluc|OJZ92637.1 anid|CBF89516.1 anig|EHA26354.1 aory|EIT77084.1 asyd|OJJ62394.1 ater|EAU37586.1 atub|OJI87068.1 aver|OJI96527.1 awen|OJJ39670.1 azon|OJJ47800.1

Group1099: aory|EIT77379.1 abra|OJJ71274.1 acar|OOF97900.1 acla|EAW10254.1 afis|EAW25337.1 afla|EED53807.1 afum|EAL90000.1 agla|OJJ83868.1 akaw|GAA82716.1 aluc|OJZ87270.1 anid|CBF75246.1 anig|EHA20511.1 asyd|OJJ59907.1 ater|EAU34477.1 atub|OJI81372.1 aver|OJJ02378.1 awen|OJJ41034.1 azon|OJJ44631.1 aory|EIT78377.1

Group1100: abra|OJJ67904.1 acar|OOF92874.1 acla|EAW11719.1 afis|EAW23740.1 afla|EED51530.1 afum|EAL87177.1 agla|OJJ78966.1 akaw|GAA91909.1 aluc|OJZ81983.1 anid|CBF73900.1 anig|EHA22271.1 aory|EIT80572.1 asyd|OJJ65360.1 atub|OJI84535.1 aver|OJI99509.1 awen|OJJ33676.1 azon|OJJ44443.1 afla|EED57578.1 aory|EIT82975.1

Group1101: ater|EAU31288.1 awen|OJJ33664.1 abra|OJJ75236.1 acar|OOF94556.1 acla|EAW11702.1 afis|EAW23723.1 afla|EED53191.1 afum|EAL87163.1 akaw|GAA91779.1 aluc|OJZ92797.1 anid|CBF86826.1 anig|EHA26211.1 aory|EIT74820.1 asyd|OJJ56249.1 atub|OJI87241.1 aver|OJJ08511.1 azon|OJJ44225.1 afla|EED51505.1 aory|EIT80632.1

Group1102: abra|OJJ67986.1 acar|OOF92028.1 acla|EAW13238.1 afis|EAW21052.1 afla|EED48858.1 akaw|GAA91216.1 aluc|OJZ82854.1 anid|CBF73882.1 anig|EHA21290.1 aory|EIT79551.1 asyd|OJJ57026.1 ater|EAU38426.1 atub|OJI89084.1 aver|OJJ06477.1 afum|EAL92088.2 asyd|OJJ55377.1 aory|EIT81277.1 anid|CBF77000.1 aver|OJJ03264.1

Group1103: abra|OJJ68002.1 akaw|GAA90948.1 aluc|OJZ82876.1 anig|EHA21302.1 atub|OJI89100.1 abra|OJJ77430.1 afis|EAW24679.1 afum|EAL89347.1 anid|CBF90095.1 asyd|OJJ62032.1 awen|OJJ40790.1 azon|OJJ48192.1 afla|EED49910.1 aory|EIT72471.1 asyd|OJJ57385.1 ater|EAU31630.1 anid|CBF88170.1 aver|OJI97314.1 awen|OJJ32480.1

Group1104: abra|OJJ68219.1 acar|OOF97509.1 acla|EAW09056.1 afis|EAW25630.1 afla|EED55187.1 afum|EAL84318.1 agla|OJJ78900.1 akaw|GAA83227.1 aluc|OJZ83727.1 anid|CBF83164.1 anig|EHA18875.1 aory|EIT83560.1 asyd|OJJ55506.1 ater|EAU32707.1 atub|OJI90103.1 aver|OJJ06581.1 awen|OJJ36050.1 azon|OJJ50396.1 acla|EAW09057.1

Group1105: abra|OJJ68227.1 afla|EED48923.1 anid|CBF73570.1 aory|EIT80015.1 asyd|OJJ56694.1 aver|OJJ06030.1 abra|OJJ72608.1 acar|OOF94647.1 afis|EAW20314.1 afla|EED52548.1 afum|EAL85437.1 akaw|GAA85234.1 aluc|OJZ90590.1 anig|EHA24029.1 ater|EAU33505.1 atub|OJI82013.1 awen|OJJ33813.1 azon|OJJ48723.1 aory|EIT74186.1

Group1106: abra|OJJ68422.1 acar|OOF97621.1 acla|EAW10999.1 afis|EAW20862.1 afla|EED45444.1 afum|EAL84773.1 agla|OJJ83749.1 akaw|GAA92526.1 aluc|OJZ90075.1 anid|CBF71444.1 anig|EHA20998.1 aory|EIT82447.1 asyd|OJJ60710.1 ater|EAU30805.1 atub|OJI84731.1 aver|OJJ00036.1 awen|OJJ34508.1 azon|OJJ51786.1 agla|OJJ89303.1

Group1107: abra|OJJ68428.1 acar|OOF94918.1 afis|EAW23747.1 afla|EED51539.1 afum|EAL87182.1 agla|OJJ79306.1 akaw|GAA86943.1 aluc|OJZ90068.1 anid|CBF81672.1 anig|EHA21006.1 aory|EIT80703.1 asyd|OJJ59555.1 ater|EAU31300.1 atub|OJI84739.1 aver|OJI97564.1 awen|OJJ33683.1 azon|OJJ44480.1 anig|EHA25691.1 aver|OJJ02713.1

Group1108: abra|OJJ68521.1 acar|OOF92726.1 acla|EAW12843.1 afis|EAW18006.1 afla|EED51889.1 afum|EAL91818.1 agla|OJJ80269.1 akaw|GAA91328.1 aluc|OJZ82735.1 anid|CBF80030.1 anig|EHA21186.1 aory|EIT82919.1 asyd|OJJ58269.1 ater|EAU31522.1 atub|OJI88969.1 aver|OJJ02154.1 awen|OJJ41536.1 azon|OJJ48254.1 agla|OJJ80268.1

Group1109: abra|OJJ68529.1 acar|OOF92733.1 acla|EAW12835.1 afis|EAW18000.1 afla|EED51896.1 afum|EAL91811.1 agla|OJJ80263.1 akaw|GAA91321.1 aluc|OJZ82742.1 anid|CBF80021.1 anig|EHA21193.1 aory|EIT76177.1 asyd|OJJ58263.1 ater|EAU31512.1 atub|OJI88977.1 aver|OJJ02161.1 awen|OJJ41542.1 azon|OJJ48260.1 aory|EIT76174.1

Group1110: abra|OJJ68552.1 acar|OOF92747.1 acla|EAW12815.1 afis|EAW17980.1 afla|EED51925.1 afum|EAL91789.1 agla|OJJ80237.1 akaw|GAA91301.1 aluc|OJZ82769.1 anid|CBF79979.1 anig|EHA21211.1 aory|EIT81371.1 asyd|OJJ58235.1 ater|EAU31490.1 atub|OJI89000.1 aver|OJJ02190.1 awen|OJJ41565.1 azon|OJJ48280.1 akaw|GAA92005.1

Group1111: abra|OJJ68565.1 acar|OOF92758.1 acla|EAW12801.1 afis|EAW17967.1 afla|EED51939.1 afum|EAL91776.1 agla|OJJ80213.1 akaw|GAA91289.1 aluc|OJZ82780.1 anid|CBF79959.1 anig|EHA21223.1 aory|EIT81465.1 asyd|OJJ58221.1 ater|EAU31477.1 atub|OJI89013.1 aver|OJJ02201.1 awen|OJJ41577.1 azon|OJJ48290.1 akaw|GAA91288.1

Group1112: abra|OJJ68634.1 acar|OOF92138.1 acla|EAW12618.1 afis|EAW18066.1 afla|EED51450.1 afum|EBA27443.1 agla|OJJ87353.1 akaw|GAA89773.1 aluc|OJZ81921.1 anid|CBF86420.1 anig|EHA22326.1 anig|EHA22327.1 aory|EIT80631.1 asyd|OJJ64939.1 ater|EAU30462.1 atub|OJI84473.1 aver|OJI99061.1 awen|OJJ41453.1 azon|OJJ48911.1

Group1113: abra|OJJ68759.1 acar|OOF92246.1 acla|EAW12492.1 afis|EAW24153.1 afla|EED46778.1 afum|EAL92008.1 agla|OJJ89544.1 akaw|GAA89903.1 aluc|OJZ81767.1 anid|CBF77909.1 anig|EHA22447.1

aory|EIT72606.1 asyd|OJJ55299.1 ater|EAU30585.1 atub|OJI84340.1 aver|OJJ06926.1 awen|OJJ41324.1 azon|OJJ48798.1 ater|EAU30584.1

Group1114: abra|OJJ68992.1 acar|OOF92704.1 acla|EAW12644.1 afis|EAW18041.1 afla|EED51855.1 afla|EED51856.1 afum|EAL91852.1 agla|OJJ85413.1 akaw|GAA90512.1 aluc|OJZ85977.1 anid|CBF80101.1 anig|EHA24629.1 aory|EIT81757.1 asyd|OJJ58297.1 ater|EAU31556.1 atub|OJI79767.1 aver|OJJ02127.1 awen|OJJ41496.1 azon|OJJ48224.1

Group1115: abra|OJJ69111.1 acar|OOF91183.1 acla|EAW08721.1 afis|EAW15726.1 afla|EED51338.1 afum|EAL85621.1 agla|OJJ81865.1 akaw|GAA90636.1 aluc|OJZ86100.1 anid|CBF71089.1 anig|EHA24519.1 aory|EIT81534.1 asyd|OJJ52700.1 ater|EAU32431.1 atub|OJI79895.1 aver|OJJ04728.1 awen|OJJ38410.1 azon|OJJ43832.1 afla|EED51337.1

Group1116: abra|OJJ69231.1 acar|OOF91089.1 acla|EAW08618.1 afis|EAW15816.1 afla|EED51231.1 afum|EAL85535.1 agla|OJJ81980.1 akaw|GAA92239.1 aluc|OJZ86211.1 anid|CBF70897.1 anig|EHA24415.1 aory|EIT83178.1 asyd|OJJ52594.1 ater|EAU33432.1 atub|OJI80005.1 aver|OJJ04621.1 awen|OJJ38530.1 azon|OJJ43732.1 atub|OJI80006.1

Group1117: abra|OJJ69301.1 acar|OOF94424.1 acla|EAW14687.1 afis|EAW24218.1 afla|EED48053.1 afum|EAL85887.1 agla|OJJ79652.1 akaw|GAA88170.1 aluc|OJZ82129.1 anid|CBF74250.1 anig|EHA28602.1 aory|EIT83328.1 asyd|OJJ53976.1 ater|EAU31168.1 atub|OJI88307.1 aver|OJJ08075.1 awen|OJJ33078.1 azon|OJJ44868.1 aory|EIT83336.1

Group1118: abra|OJJ69328.1 acar|OOF94399.1 acla|EAW14712.1 afis|EAW24247.1 afla|EED48026.1 afum|EAL85915.1 agla|OJJ79626.1 akaw|GAA88199.1 aluc|OJZ82160.1 anid|CBF74202.1 anig|EHA28572.1 aory|EIT83332.1 asyd|OJJ53955.1 ater|EAU31188.1 atub|OJI88277.1 aver|OJJ08051.1 awen|OJJ33059.1 azon|OJJ48467.1 afla|EED48027.1

Group1119: abra|OJJ69488.1 acar|OOF94260.1 acla|EAW09905.1 afis|EAW21154.1 afla|EED54658.1 afum|EAL92172.1 agla|OJJ80039.1 akaw|GAA88332.1 aluc|OJZ82327.1 anid|CBF86989.1 anig|EHA28431.1 aory|EIT78787.1 asyd|OJJ60510.1 ater|EAU39556.1 atub|OJI88111.1 aver|OJI99836.1 awen|OJJ36482.1 azon|OJJ44363.1 anig|EHA21818.1

Group1120: abra|OJJ69514.1 acar|OOF94239.1 acla|EAW09954.1 afis|EAW21142.1 afla|EED46141.1 afum|EAL92161.1 agla|OJJ80069.1 akaw|GAA88310.1 aluc|OJZ82352.1 anid|CBF87008.1 anig|EHA28408.1 aory|EIT76511.1 asyd|OJJ60471.1 ater|EAU36763.1 atub|OJI88088.1 aver|OJI99791.1 awen|OJJ36466.1 azon|OJJ44412.1 azon|OJJ42893.1

Group1121: abra|OJJ69523.1 acar|OOF94231.1 acla|EAW09971.1 afis|EAW21117.1 afla|EED51790.1 afum|EAL92141.1 agla|OJJ80094.1 akaw|GAA88302.1 aluc|OJZ82363.1 anid|CBF87028.1 anig|EHA28402.1 aory|EIT81753.1 asyd|OJJ60456.1 ater|EAU36782.1 atub|OJI88079.1 aver|OJJ06754.1 awen|OJJ36447.1 azon|OJJ44428.1 aver|OJI99774.1

Group1122: abra|OJJ69615.1 acar|OOF92691.1 acla|EAW12075.1 afis|EAW19716.1 afla|EED53109.1 afum|EAL86811.1 agla|OJJ80862.1 akaw|GAA91599.1 aluc|OJZ82423.1 anid|CBF80080.1 anig|EHA28346.1 aory|EIT74768.1 asyd|OJJ58309.1 ater|EAU37777.1 atub|OJI88017.1 aver|OJJ02103.1 awen|OJJ35106.1 azon|OJJ45858.1 awen|OJJ34696.1

Group1123: abra|OJJ69668.1 acar|OOF95768.1 acla|EAW10632.1 afis|EAW24935.1 afla|EED55537.1 afum|EAL87763.1 agla|OJJ82162.1 akaw|GAA91646.1 aluc|OJZ82473.1 anid|CBF75718.1 anig|EHA28300.1 aory|EIT80819.1 asyd|OJJ57503.1 ater|EAU35664.1 atub|OJI87964.1 aver|OJJ05675.1 awen|OJJ40530.1 azon|OJJ49390.1 afum|EAL89592.1

Group1124: abra|OJJ69841.1 acar|OOF93364.1 acla|EAW13293.1 afis|EAW24568.1 afla|EED56610.1 afum|EAL91152.1 akaw|GAA87819.1 aluc|OJZ85455.1 anid|CBF82896.1 anig|EHA24997.1 aory|EIT81324.1 asyd|OJJ62102.1 ater|EAU39297.1 atub|OJI80109.1 aver|OJI96208.1 afis|EAW17316.1 aory|EIT82251.1 ater|EAU30304.1 aory|EIT79627.1

Group1125: abra|OJJ69857.1 acar|OOF93382.1 acla|EAW08537.1 afis|EAW23575.1 afla|EED57541.1 afum|EAL87890.1 agla|OJJ85306.1 akaw|GAA87799.1 aluc|OJZ85434.1 anig|EHA24980.1 aory|EIT81841.1 asyd|OJJ56731.1 atub|OJI80134.1 awen|OJJ38171.1 azon|OJJ45131.1 agla|OJJ86836.1 awen|OJJ39903.1 azon|OJJ45879.1 afla|EED54164.1

Group1126: abra|OJJ70048.1 acar|OOF93540.1 afla|EED47673.1 agla|OJJ79802.1 akaw|GAA87627.1 aluc|OJZ85243.1 anid|CBF84432.1 anig|EHA25443.1 aory|EIT80242.1 asyd|OJJ61842.1 ater|EAU34191.1 atub|OJI80328.1 aver|OJJ05342.1 awen|OJJ36937.1 aver|OJI95942.1 agla|OJJ80121.1 asyd|OJJ53636.1 aver|OJJ01170.1 ater|EAU29232.1

Group1127: abra|OJJ70093.1 acla|EAW12322.1 afis|EAW17167.1 afla|EED51057.1 afum|EAL89052.1 agla|OJJ78763.1 akaw|GAA87602.1 aluc|OJZ85222.1 anig|EHA25426.1 aory|EIT76987.1 ater|EAU34026.1 atub|OJI80361.1 azon|OJJ42125.1 akaw|GAA87480.1 aluc|OJZ88046.1 anig|EHA17818.1 ater|EAU29611.1 atub|OJI90654.1 awen|OJJ36983.1

Group1128: abra|OJJ70121.1 acar|OOF92449.1 acla|EAW11044.1 afis|EAW16194.1 afla|EED48259.1 afum|EAL88825.1 akaw|GAA86648.1 aluc|OJZ85206.1 anid|CBF74996.1 anig|EHA25410.1 aory|EIT72896.1 ater|EAU34204.1 atub|OJI80396.1 aver|OJJ05082.1 azon|OJJ46585.1 afla|EED53310.1 aory|EIT74931.1 azon|OJJ45951.1 afla|EED49627.1

Group1129: abra|OJJ70144.1 acar|OOF96811.1 afla|EED52394.1 akaw|GAA90567.1 akaw|GAA90568.1 aluc|OJZ86936.1 anig|EHA25649.1 aory|EIT83282.1 asyd|OJJ53406.1 atub|OJI88384.1 aver|OJJ04195.1 awen|OJJ35567.1 azon|OJJ42608.1 akaw|GAA84747.1 anig|EHA21708.1 atub|OJI83585.1 afla|EED52395.1 anig|EHA25751.1 akaw|GAA91340.1

Group1130: abra|OJJ70231.1 acar|OOF91736.1 acla|EAW11658.1 afis|EAW20417.1 afla|EED53976.1 agla|OJJ82616.1 aluc|OJZ90859.1 aver|OJJ06455.1 awen|OJJ40076.1 azon|OJJ45549.1 abra|OJJ73408.1 acar|OOG00605.1 akaw|GAA83083.1 aluc|OJZ88630.1 anig|EHA18320.1 aory|EIT74373.1 atub|OJI91214.1 afla|EED52225.1 agla|OJJ88203.1

Group1131: abra|OJJ70466.1 acar|OOF97458.1 acla|EAW13048.1 afis|EAW17619.1 afla|EED49199.1 afla|EED49200.1 afum|EAL91432.1 agla|OJJ81306.1 akaw|GAA83315.1 aluc|OJZ83810.1 anid|CBF90163.1 anig|EHA18955.1 aory|EIT78513.1 asyd|OJJ61994.1 ater|EAU37053.1 atub|OJI90177.1 aver|OJI96086.1 awen|OJJ37225.1 azon|OJJ50928.1

Group1132: abra|OJJ70467.1 acar|OOF97457.1 acla|EAW13047.1 afis|EAW17620.1 afla|EED49197.1 afla|EED49198.1 afum|EAL91433.2 agla|OJJ81305.1 akaw|GAA83316.1 aluc|OJZ83811.1 anid|CBF90161.1 anig|EHA18956.1 aory|EIT78563.1 asyd|OJJ61995.1 ater|EAU37054.1 atub|OJI90178.1 aver|OJI96087.1 awen|OJJ37224.1 azon|OJJ50927.1

Group1133: abra|OJJ70612.1 acar|OOF97110.1 acla|EAW13015.1 afis|EAW17652.1 afla|EED50506.1 afum|EAL91467.1 agla|OJJ79552.1 akaw|GAA89951.1 aluc|OJZ83958.1 anid|CBF90075.1 anig|EHA19086.1 aory|EIT82132.1 asyd|OJJ62041.1 ater|EAU37088.1 atub|OJI90330.1 aver|OJI96147.1 awen|OJJ37193.1 azon|OJJ50892.1 aver|OJJ07605.1

Group1134: abra|OJJ70647.1 acar|OOF97074.1 acla|EAW08113.1 afis|EAW23138.1 afla|EED48501.1 afum|EAL88343.1 agla|OJJ89662.1 akaw|GAA89983.1 aluc|OJZ83997.1 anid|CBF74448.1 anig|EHA19120.1 aory|EIT73103.1 asyd|OJJ57741.1 ater|EAU36279.1 atub|OJI90366.1 aver|OJJ05426.1 awen|OJJ36635.1 azon|OJJ47581.1 awen|OJJ36634.1

Group1135: abra|OJJ70656.1 acar|OOF97066.1 acla|EAW08103.1 afis|EAW23148.1 afla|EED48511.1 afum|EAL88333.1 agla|OJJ89654.1 akaw|GAA89992.1 aluc|OJZ84006.1 anid|CBF74430.1 anig|EHA19128.1 aory|EIT73155.1 asyd|OJJ57750.1 ater|EAU32975.1 atub|OJI90375.1 aver|OJJ05417.1 awen|OJJ36643.1 azon|OJJ47571.1 acar|OOG00309.1

Group1136: abra|OJJ70745.1 acar|OOF96999.1 acla|EAW11175.1 afis|EAW15906.1 afla|EED56463.1 afum|EAL88541.1 agla|OJJ89566.1 akaw|GAA90066.1 aluc|OJZ84090.1 anid|CBF81270.1 anig|EHA19203.1 aory|EIT72857.1 asyd|OJJ59333.1 ater|EAU36183.1 atub|OJI90461.1 aver|OJJ02941.1 awen|OJJ36725.1 azon|OJJ47484.1 ater|EAU30153.1

Group1137: abra|OJJ70747.1 acar|OOF96998.1 acla|EAW11174.1 afis|EAW15905.1 afla|EED56464.1 afum|EAL88540.1 agla|OJJ89565.1 akaw|GAA90067.1 aluc|OJZ84091.1 anid|CBF81274.1 anig|EHA19204.1 aory|EIT72825.1 asyd|OJJ59334.1 ater|EAU36182.1 atub|OJI90462.1 aver|OJJ02940.1 awen|OJJ36726.1 azon|OJJ47471.1 aory|EIT72838.1

Group1138: abra|OJJ70748.1 acar|OOF96997.1 acla|EAW11173.1 afis|EAW15904.1 afla|EED56466.1 afum|EAL88539.1 agla|OJJ89564.1 akaw|GAA90068.1 aluc|OJZ84092.1 anid|CBF81276.1 anig|EHA19205.1 aory|EIT72390.1 asyd|OJJ59335.1 ater|EAU36181.1 atub|OJI90463.1 aver|OJJ02939.1 awen|OJJ36727.1 azon|OJJ47470.1 afla|EED46475.1

Group1139: abra|OJJ70830.1 afis|EAW19787.1 afla|EED45955.1 afum|EAL85107.2 agla|OJJ88990.1 anid|CBF74550.1 anig|EHA20974.1 aory|EIT75768.1 asyd|OJJ57685.1 ater|EAU34824.1 aver|OJJ04314.1 awen|OJJ33838.1 abra|OJJ76587.1 acar|OOF99400.1 akaw|GAA82491.1 aluc|OJZ91619.1 anig|EHA22646.1 atub|OJI86072.1 asyd|OJJ52721.1

Group1140: abra|OJJ70840.1 acar|OOF98274.1 acla|EAW13281.1 afis|EAW17336.1 afla|EED46105.1 afum|EAL91180.1 agla|OJJ82057.1 akaw|GAA91708.1 aluc|OJZ87681.1 anid|CBF85633.1 anig|EHA20871.1 aory|EIT76521.1 asyd|OJJ56056.1 ater|EAU33888.1 atub|OJI80962.1 aver|OJJ04998.1 awen|OJJ33506.1 azon|OJJ50300.1 azon|OJJ51817.1

Group1141: abra|OJJ70845.1 acar|OOF98270.1 acla|EAW10436.1 afis|EAW25131.1 afla|EED55314.1 afum|EAL89802.1 agla|OJJ89035.1 akaw|GAA91713.1 aluc|OJZ87676.1 anid|CBF86816.1 anig|EHA20866.1 aory|EIT83519.1 asyd|OJJ64657.1 ater|EAU39403.1 atub|OJI80967.1 aver|OJI98760.1 awen|OJJ40756.1 azon|OJJ44121.1 aver|OJJ02113.1

Group1142: abra|OJJ70882.1 acar|OOF98239.1 acla|EAW10470.1 afis|EAW25097.1 afla|EED55351.1 afum|EAL89768.1 agla|OJJ85476.1 akaw|GAA91744.1 aluc|OJZ87640.1 anid|CBF86010.1 anig|EHA20834.1

aory|EIT80916.1 asyd|OJJ64447.1 ater|EAU39355.1 atub|OJI81005.1 aver|OJI98550.1 awen|OJJ40711.1 azon|OJJ44087.1 ater|EAU29372.1

Group1143: abra|OJJ70985.1 acar|OOF98146.1 acla|EAW10545.1 afis|EAW25013.1 afla|EED55432.1 afum|EAL89683.1 agla|OJJ89177.1 akaw|GAA82964.1 aluc|OJZ87545.1 aluc|OJZ87546.1 anid|CBF81523.1 anig|EHA20749.1 aory|EIT80770.1 asyd|OJJ59478.1 ater|EAU35773.1 atub|OJI81100.1 aver|OJJ02795.1 awen|OJJ40627.1 azon|OJJ49056.1

Group1144: abra|OJJ71009.1 acar|OOF98133.1 acla|EAW10561.1 afis|EAW25028.1 afla|EED55451.1 afum|EAL89667.1 agla|OJJ79156.1 agla|OJJ82560.1 akaw|GAA82944.1 aluc|OJZ87522.1 anid|CBF81555.1 anig|EHA20732.1 aory|EIT80765.1 asyd|OJJ59497.1 ater|EAU35759.1 atub|OJI81127.1 aver|OJJ02777.1 awen|OJJ40611.1 azon|OJJ49460.1

Group1145: abra|OJJ71029.1 acar|OOF98110.1 acla|EAW10582.1 afis|EAW24987.1 afla|EED55471.1 afum|EAL89647.1 agla|OJJ82275.1 akaw|GAA82926.1 aluc|OJZ87501.1 anid|CBF81601.1 anig|EHA20716.1 aory|EIT80792.1 asyd|OJJ59520.1 ater|EAU35738.1 atub|OJI81148.1 aver|OJJ02753.1 awen|OJJ40583.1 azon|OJJ41984.1 azon|OJJ49442.1

Group1146: abra|OJJ71043.1 acar|OOF98096.1 acla|EAW10591.1 afis|EAW24977.1 afla|EED55482.1 afum|EAL89637.1 agla|OJJ79140.1 akaw|GAA82913.1 aluc|OJZ87485.1 anid|CBF81627.1 anig|EHA20706.1 aory|EIT80881.1 asyd|OJJ59529.1 ater|EAU35726.1 atub|OJI81163.1 aver|OJJ02741.1 awen|OJJ40574.1 azon|OJJ49436.1 afla|EED55483.1

Group1147: abra|OJJ71134.1 acar|OOF98018.1 acla|EAW10363.1 afis|EAW25229.1 afla|EED53673.1 afum|EAL89890.2 agla|OJJ79179.1 akaw|GAA82840.1 aluc|OJZ87401.1 anid|CBF85338.1 anig|EHA20630.1 aory|EIT75324.1 asyd|OJJ64023.1 ater|EAU34387.1 atub|OJI81245.1 aver|OJI98109.1 awen|OJJ40894.1 azon|OJJ44523.1 aory|EIT75321.1

Group1148: abra|OJJ71194.1 acar|OOF97972.1 acla|EAW10310.1 afis|EAW25283.1 afla|EED53749.1 afum|EAL89946.1 agla|OJJ89257.1 akaw|GAA82789.1 aluc|OJZ87348.1 anid|CBF85422.1 anig|EHA20584.1 aory|EIT78394.1 asyd|OJJ64075.1 ater|EAU34425.1 atub|OJI81295.1 aver|OJI98162.1 awen|OJJ40996.1 azon|OJJ44586.1 asyd|OJJ60350.1

Group1149: abra|OJJ71328.1 acar|OOF97846.1 acla|EAW10211.1 afis|EAW25383.1 afla|EED53860.1 afum|EAL90041.1 agla|OJJ83775.1 akaw|GAA82664.1 aluc|OJZ87214.1 anid|CBF77488.1 anig|EHA20465.1 aory|EIT78406.1 asyd|OJJ53195.1 ater|EAU34529.1 atub|OJI81425.1 aver|OJJ03655.1 awen|OJJ41109.1 azon|OJJ44687.1 aory|EIT78415.1

Group1150: abra|OJJ71401.1 acar|OOF97769.1 acla|EAW10133.1 afis|EAW25454.1 afla|EED53942.1 afla|EED53943.1 afum|EAL90113.1 agla|OJJ87107.1 akaw|GAA82596.1 aluc|OJZ87141.1 anid|CBF77635.1 anig|EHA20399.1 aory|EIT78263.1 asyd|OJJ53125.1 ater|EAU34624.1 atub|OJI81501.1 aver|OJJ03729.1 awen|OJJ41186.1 azon|OJJ44762.1

Group1151: abra|OJJ71618.1 acar|OOF96335.1 acla|EAW13285.1 afis|EAW17333.1 afla|EED57246.1 afum|EAL91170.2 akaw|GAA89309.1 aluc|OJZ86359.1 anid|CBF84952.1 anig|EHA21531.1 aory|EIT74025.1 asyd|OJJ63145.1 ater|EAU38742.1 atub|OJI84175.1 aver|OJI97809.1 azon|OJJ50869.1 acla|EAW10844.1 afla|EED47443.1 aory|EIT81909.1

Group1152: abra|OJJ71994.1 acar|OOF96648.1 acla|EAW09538.1 afis|EAW21594.1 afla|EED57816.1 afum|EAL92612.1 agla|OJJ85787.1 akaw|GAA84934.1 aluc|OJZ86732.1 anid|CBF83446.1 anig|EHA21947.1 aory|EIT81030.1 asyd|OJJ60671.1 ater|EAU34930.1 atub|OJI83796.1 aver|OJI99994.1 awen|OJJ39087.1 azon|OJJ51418.1 ater|EAU32744.1

Group1153: abra|OJJ72045.1 acar|OOF96703.1 acla|EAW09501.1 afis|EAW21635.1 afla|EED48487.1 afum|EAL92669.1 agla|OJJ79700.1 akaw|GAA84889.1 aluc|OJZ86782.1 anid|CBF83498.1 anig|EHA21900.1 aory|EIT83266.1 asyd|OJJ60712.1 ater|EAU38321.1 atub|OJI83738.1 aver|OJJ00045.1 awen|OJJ33884.1 azon|OJJ48669.1 aory|EIT73220.1

Group1154: abra|OJJ72230.1 acar|OOF96853.1 acla|EAW14990.1 afis|EAW25547.1 afla|EED47847.1 afum|EAL86183.1 agla|OJJ80332.1 akaw|GAA84707.1 aluc|OJZ86980.1 anid|CBF74013.1 anig|EHA21669.1 aory|EIT77206.1 asyd|OJJ56425.1 atub|OJI83540.1 aver|OJJ06292.1 awen|OJJ32605.1 afis|EAW24517.1 asyd|OJJ53838.1 aver|OJJ07941.1

Group1155: abra|OJJ72280.1 acar|OOF94941.1 afis|EAW20800.1 afla|EED49545.1 afum|EAL84829.1 agla|OJJ85012.1 akaw|GAA86032.1 aluc|OJZ90259.1 anid|CBF71374.1 anig|EHA23754.1 aory|EIT82577.1 asyd|OJJ55077.1 ater|EAU32870.1 atub|OJI81685.1 aver|OJJ07131.1 awen|OJJ41912.1 azon|OJJ47358.1 anig|EHA23755.1 afum|EAL84830.1

Group1156: abra|OJJ72423.1 acar|OOF90372.1 acla|EAW12860.1 afis|EAW17809.1 afla|EED46159.1 afum|EAL91617.1 agla|OJJ80653.1 akaw|GAA86165.1 aluc|OJZ90406.1 anid|CBF78020.1 anig|EHA23874.1 aory|EIT76396.1 asyd|OJJ55235.1 ater|EAU33015.1 atub|OJI81837.1 aver|OJJ06988.1 awen|OJJ41763.1 azon|OJJ44903.1 afla|EED46160.1

Group1157: abra|OJJ72434.1 acar|OOF94801.1 acla|EAW12667.1 afis|EAW17831.1 afla|EED46173.1 afum|EAL91641.1 agla|OJJ80640.1 akaw|GAA86174.1 aluc|OJZ90417.1 anid|CBF77994.1 anig|EHA23885.1 aory|EIT76438.1 asyd|OJJ55252.1 ater|EAU33034.1 atub|OJI81850.1 aver|OJI99643.1 awen|OJJ41750.1 azon|OJJ48442.1 aver|OJJ06972.1

Group1158: abra|OJJ72484.1 acar|OOF94751.1 acla|EAW12715.1 afis|EAW17877.1 afla|EED46293.1 afum|EAL91685.1 agla|OJJ83069.1 akaw|GAA86219.1 aluc|OJZ90471.1 anig|EHA23930.1 aory|EIT76328.1 asyd|OJJ61521.1 ater|EAU33097.1 atub|OJI81895.1 aver|OJJ05549.1 awen|OJJ41687.1 asyd|OJJ57661.1 aver|OJJ05523.1 aver|OJJ01273.1

Group1159: abra|OJJ72549.1 acar|OOF94703.1 acla|EAW12754.1 afis|EAW17917.1 afla|EED46320.1 afum|EBA27439.1 agla|OJJ83026.1 akaw|GAA85181.1 aluc|OJZ90537.1 anid|CBF76959.1 anig|EHA23985.1 aory|EIT81197.1 atub|OJI81958.1 awen|OJJ41628.1 asyd|OJJ57052.1 aver|OJJ03244.1 azon|OJJ48353.1 ater|EAU33139.1 asyd|OJJ57051.1

Group1160: acla|EAW12414.1 acar|OOF98070.1 afla|EED52231.1 agla|OJJ84011.1 anid|CBF69489.1 aory|EIT74442.1 asyd|OJJ59240.1 ater|EAU37441.1 awen|OJJ35818.1 azon|OJJ49326.1 acar|OOG00738.1 asyd|OJJ60713.1 ater|EAU30842.1 aver|OJJ00046.1 afla|EED47470.1 aory|EIT79973.1 ater|EAU31019.1 azon|OJJ48614.1 aver|OJJ01850.1

Group1161: abra|OJJ72629.1 acar|OOF94635.1 acla|EAW11593.1 afis|EAW23864.1 afla|EED52589.1 afum|EAL87292.1 agla|OJJ82532.1 akaw|GAA85249.1 aluc|OJZ90606.1 anid|CBF77222.1 anig|EHA24044.1 aory|EIT74309.1 asyd|OJJ56820.1 ater|EAU33487.1 atub|OJI82030.1 aver|OJJ03515.1 awen|OJJ33839.1 azon|OJJ48708.1 awen|OJJ33432.1

Group1162: abra|OJJ72716.1 acar|OOF95949.1 acla|EAW11552.1 afis|EAW23901.1 afla|EED52636.1 afum|EAL87328.1 agla|OJJ79367.1 akaw|GAA85324.1 aluc|OJZ90685.1 anid|CBF77143.1 anig|EHA24113.1 aory|EIT74168.1 asyd|OJJ56912.1 ater|EAU32337.1 atub|OJI82111.1 aver|OJJ03380.1 awen|OJJ39048.1 azon|OJJ51378.1 afis|EAW25771.1

Group1163: abra|OJJ72751.1 acar|OOF95974.1 acla|EAW11510.1 afis|EAW23943.1 afla|EED50573.1 afum|EAL87368.1 agla|OJJ79431.1 akaw|GAA85363.1 aluc|OJZ90725.1 anid|CBF77256.1 anig|EHA24143.1 aory|EIT82138.1 asyd|OJJ56806.1 ater|EAU36036.1 atub|OJI82153.1 aver|OJJ03528.1 awen|OJJ39012.1 azon|OJJ51337.1 awen|OJJ40567.1

Group1164: abra|OJJ72900.1 acar|OOF89981.1 acla|EAW11351.1 afis|EAW18143.1 afla|EED58069.1 afum|EAL87500.1 agla|OJJ80189.1 akaw|GAA85487.1 aluc|OJZ83409.1 anid|CBF81125.1 anig|EHA24260.1 aory|EIT79709.1 asyd|OJJ56489.1 atub|OJI82297.1 aver|OJJ06229.1 awen|OJJ35585.1 awen|OJJ38003.1 aver|OJJ03876.1 ater|EAU30131.1

Group1165: abra|OJJ72905.1 acar|OOF89821.1 acla|EAW11349.1 afis|EAW18146.1 afla|EED58074.1 afum|EAL87503.1 agla|OJJ83830.1 akaw|GAA85490.1 aluc|OJZ83415.1 anid|CBF81118.1 anig|EHA24264.1 asyd|OJJ56487.1 ater|EAU35469.1 atub|OJI82302.1 aver|OJJ06232.1 awen|OJJ35589.1 azon|OJJ46936.1 afla|EED58073.1 afum|EAL87504.1

Group1166: abra|OJJ72907.1 acar|OOF96090.1 acla|EAW11346.1 afis|EAW18148.1 afla|EED58076.1 afum|EAL87506.1 agla|OJJ83833.1 akaw|GAA85492.1 aluc|OJZ83417.1 anid|CBF81114.1 anig|EHA24266.1 aory|EIT79127.1 ater|EAU35470.1 atub|OJI82304.1 aver|OJJ06234.1 awen|OJJ35591.1 azon|OJJ46913.1 asyd|OJJ56485.1 asyd|OJJ57077.1

Group1167: abra|OJJ73268.1 acar|OOG00508.1 acla|EAW06945.1 afis|EAW16559.1 afla|EED56102.1 afum|EAL89157.1 agla|OJJ82763.1 akaw|GAA87914.1 aluc|OJZ81161.1 anid|CBF81806.1 anig|EHA18478.1 aory|EIT76895.1 asyd|OJJ59644.1 ater|EAU36627.1 atub|OJI91356.1 aver|OJJ02638.1 awen|OJJ40257.1 azon|OJJ49121.1 acar|OOF91712.1

Group1168: abra|OJJ73284.1 acar|OOG00520.1 acla|EAW06959.1 afis|EAW16574.1 afla|EED56090.1 afum|EAL89171.2 agla|OJJ82706.1 akaw|GAA92721.1 aluc|OJZ88765.1 anid|CBF81830.1 anig|EHA18461.1 aory|EIT76622.1 asyd|OJJ59660.1 ater|EAU36642.1 atub|OJI91338.1 aver|OJJ02622.1 awen|OJJ40244.1 azon|OJJ49105.1 asyd|OJJ52085.1

Group1169: abra|OJJ73346.1 acar|OOG00556.1 acla|EAW06973.1 afis|EAW16587.1 afla|EED56078.1 afum|EAL89182.1 agla|OJJ82679.1 akaw|GAA83136.1 aluc|OJZ88693.1 anid|CBF81890.1 anig|EHA18373.1 aory|EIT76830.1 asyd|OJJ59676.1 ater|EAU36664.1 ater|EAU39117.1 atub|OJI91273.1 aver|OJJ02610.1 awen|OJJ40228.1 azon|OJJ49093.1

Group1170: abra|OJJ73352.1 acar|OOG00563.1 acla|EAW12784.1 afis|EAW19851.1 afla|EED52646.1 afum|EAL85042.1 agla|OJJ81301.1 akaw|GAA83131.1 aluc|OJZ88687.1 anid|CBF73979.1 anig|EHA18367.1 aory|EIT74299.1 asyd|OJJ54702.1 ater|EAU32479.1 atub|OJI91267.1 aver|OJJ07471.1 awen|OJJ32014.1 azon|OJJ46075.1 afis|EAW17946.1

Group1171: abra|OJJ73409.1 acar|OOG00606.1 acla|EAW07013.1 afis|EAW16640.1 afla|EED44962.1 afum|EAL89242.1 agla|OJJ82610.1 akaw|GAA83082.1 aluc|OJZ88629.1 anid|CBF82018.1 anig|EHA18319.1

aory|EIT73512.1 asyd|OJJ56191.1 ater|EAU36751.1 atub|OJI91213.1 aver|OJJ05120.1 awen|OJJ40074.1 azon|OJJ45550.1 anid|CBF76269.1

Group1172: abra|OJJ73438.1 acar|OOG00644.1 acla|EAW06774.1 afis|EAW16325.1 afla|EED53556.1 afum|EAL88949.1 agla|OJJ88288.1 akaw|GAA83055.1 aluc|OJZ88602.1 anid|CBF89162.1 anig|EHA18296.1 aory|EIT75182.1 asyd|OJJ62547.1 ater|EAU34132.1 atub|OJI91187.1 aver|OJI96673.1 awen|OJJ37725.1 azon|OJJ44985.1 afla|EED53557.1

Group1173: abra|OJJ73449.1 acar|OOG00652.1 acla|EAW11065.1 afis|EAW20809.1 afla|EED50135.1 afum|EAL84819.1 akaw|GAA83047.1 aluc|OJZ88592.1 anid|CBF69930.1 anig|EHA18288.1 aory|EIT74700.1 asyd|OJJ58924.1 ater|EAU34826.1 atub|OJI91178.1 aver|OJJ01520.1 awen|OJJ29971.1 azon|OJJ46931.1 asyd|OJJ52486.1 aver|OJJ03920.1

Group1174: abra|OJJ73545.1 acar|OOG00719.1 acla|EAW07464.1 afis|EAW22495.1 afla|EED54524.1 afum|EAL90698.1 agla|OJJ82639.1 akaw|GAA88607.1 aluc|OJZ88515.1 anid|CBF78415.1 anig|EHA18221.1 aory|EIT78926.1 asyd|OJJ58988.1 atub|OJI91100.1 aver|OJJ01628.1 afis|EAW23820.1 afla|EED45632.1 afum|EAL87255.1 aory|EIT75782.1

Group1175: abra|OJJ73649.1 acar|OOG00830.1 acla|EAW07562.1 afis|EAW22596.1 afla|EED54646.1 afum|EAL90602.1 agla|OJJ80960.1 akaw|GAA88506.1 aluc|OJZ88401.1 anid|CBF88373.1 anig|EHA18129.1 aory|EIT78830.1 asyd|OJJ63733.1 ater|EAU39107.1 atub|OJI90997.1 aver|OJI97208.1 awen|OJJ35224.1 azon|OJJ49885.1 afla|EED54647.1

Group1176: abra|OJJ73717.1 acla|EAW14401.1 afis|EAW16390.1 afla|EED52126.1 afum|EAL89016.2 agla|OJJ84496.1 akaw|GAA88446.1 aluc|OJZ88333.1 anid|CBF79713.1 anig|EHA18069.1 aory|EIT74447.1 asyd|OJJ58060.1 ater|EAU37368.1 atub|OJI90929.1 aver|OJJ04343.1 awen|OJJ32507.1 azon|OJJ45230.1 afis|EAW19108.1 afum|EAL93839.1

Group1177: abra|OJJ73743.1 acar|OOG00906.1 acla|EAW07637.1 afis|EAW22657.1 afla|EED49925.1 afum|EAL90535.1 agla|OJJ81040.1 akaw|GAA88422.1 aluc|OJZ88309.1 anid|CBF80225.1 anig|EHA18046.1 aory|EIT72274.1 asyd|OJJ54399.1 ater|EAU39050.1 atub|OJI90904.1 aver|OJJ05335.1 awen|OJJ35266.1 azon|OJJ49829.1 ater|EAU30036.1

Group1178: abra|OJJ73792.1 acar|OOG00955.1 afis|EAW22699.1 afla|EED49985.1 afum|EAL90493.1 agla|OJJ79011.1 akaw|GAA89411.1 aluc|OJZ88262.1 anid|CBF88085.1 anig|EHA18007.1 aory|EIT74716.1 asyd|OJJ63561.1 ater|EAU39007.1 atub|OJI90861.1 aver|OJI97372.1 awen|OJJ35335.1 acar|OOF98064.1 anid|CBF78395.1 acla|EAW07684.1

Group1179: abra|OJJ73822.1 acar|OOG00974.1 acla|EAW07711.1 afis|EAW22727.1 afla|EED50013.1 afum|EAL90468.2 agla|OJJ83675.1 akaw|GAA89441.1 aluc|OJZ88226.1 anid|CBF88026.1 anig|EHA17974.1 aory|EIT74607.1 asyd|OJJ63528.1 ater|EAU38970.1 atub|OJI90827.1 aver|OJI97403.1 awen|OJJ35422.1 azon|OJJ49754.1 aory|EIT74669.1

Group1180: abra|OJJ73906.1 acar|OOG01039.1 acla|EAW07784.1 afis|EAW22820.1 afla|EED50095.1 afum|EBA27405.1 agla|OJJ83518.1 akaw|GAA89513.1 aluc|OJZ88141.1 anid|CBF87897.1 anig|EHA17904.1 aory|EIT74756.1 asyd|OJJ63443.1 ater|EAU38890.1 atub|OJI90744.1 aver|OJI97493.1 awen|OJJ38907.1 azon|OJJ49688.1 afla|EED50094.1

Group1181: abra|OJJ73972.1 acar|OOG01099.1 acla|EAW07843.1 afis|EAW22879.1 afla|EED50182.1 afum|EAL90320.1 akaw|GAA89575.1 aluc|OJZ88072.1 anid|CBF87810.1 anig|EHA17845.1 aory|EIT83503.1 asyd|OJJ63389.1 ater|EAU38843.1 atub|OJI90681.1 aver|OJI97546.1 awen|OJJ38843.1 azon|OJJ49640.1 afla|EED46055.1 azon|OJJ45935.1

Group1182: abra|OJJ73974.1 acar|OOG01101.1 acla|EAW07845.1 afis|EAW22881.1 afla|EED50179.1 afum|EAL90318.1 agla|OJJ83449.1 akaw|GAA89577.1 aluc|OJZ88070.1 anid|CBF87807.1 anig|EHA17843.1 aory|EIT77377.1 asyd|OJJ63387.1 ater|EAU38845.1 atub|OJI90679.1 aver|OJI97548.1 awen|OJJ38841.1 azon|OJJ49638.1 agla|OJJ83448.1

Group1183: abra|OJJ73990.1 acar|OOG01116.1 acla|EAW07861.1 afis|EAW22896.1 afla|EED54867.1 afum|EAL90302.1 agla|OJJ83416.1 akaw|GAA87473.1 aluc|OJZ88055.1 anid|CBF87777.1 anig|EHA17827.1 aory|EIT79357.1 asyd|OJJ63370.1 atub|OJI90663.1 aver|OJI97576.1 awen|OJJ38822.1 azon|OJJ49621.1 ater|EAU35980.1 awen|OJJ38821.1

Group1184: abra|OJJ74134.1 acar|OOF95597.1 acla|EAW08432.1 afis|EAW23475.1 afla|EED57371.1 afum|EAL87990.1 agla|OJJ86714.1 akaw|GAA89752.1 aluc|OJZ80631.1 anid|CBF89773.1 anig|EHA27200.1 aory|EIT80150.1 asyd|OJJ62211.1 ater|EAU36905.1 atub|OJI87487.1 aver|OJI96322.1 awen|OJJ37876.1 asyd|OJJ52090.1 aver|OJJ03003.1

Group1185: abra|OJJ74469.1 acar|OOF95278.1 acla|EAW07228.1 afis|EAW22242.1 afla|EED57635.1 afum|EAL90951.1 agla|OJJ87809.1 akaw|GAA87362.1 aluc|OJZ89651.1 anid|CBF88489.1 anig|EHA26889.1 aory|EIT82991.1 asyd|OJJ63790.1 ater|EAU34249.1 atub|OJI85153.1 aver|OJI97147.1 awen|OJJ34820.1 azon|OJJ45730.1 aver|OJJ06299.1

Group1186: acar|OOF99264.1 awen|OJJ30905.1 agla|OJJ80074.1 akaw|GAA89762.1 aluc|OJZ80621.1 anid|CBF86745.1 anig|EHA27210.1 aory|EIT75558.1 asyd|OJJ64702.1 ater|EAU30235.1 atub|OJI87475.1 aver|OJI98806.1 acar|OOF99265.1 afla|EED49606.1 aory|EIT80966.1 azon|OJJ42634.1 asyd|OJJ53572.1 aory|EIT80991.1 aver|OJI99007.1

Group1187: abra|OJJ74764.1 acar|OOF95757.1 acla|EAW10646.1 afis|EAW24920.1 afla|EED55553.1 afum|EAL89575.1 agla|OJJ82149.1 akaw|GAA83647.1 aluc|OJZ92348.1 anig|EHA26626.1 aory|EIT82708.1 asyd|OJJ57492.1 asyd|OJJ62481.1 ater|EAU35652.1 atub|OJI86775.1 aver|OJJ05688.1 awen|OJJ40517.1 azon|OJJ49376.1 anid|CBF75691.1

Group1188: abra|OJJ74777.1 acar|OOF95766.1 acla|EAW10634.1 afis|EAW24932.1 afla|EED55541.1 afum|EAL89588.1 agla|OJJ82160.1 akaw|GAA83635.1 aluc|OJZ92358.1 anid|CBF75714.1 anig|EHA26615.1 aory|EIT82707.1 asyd|OJJ57501.1 ater|EAU35662.1 atub|OJI86788.1 aver|OJJ05677.1 awen|OJJ40528.1 azon|OJJ49389.1 afum|EAL87767.1

Group1189: abra|OJJ74785.1 acar|OOF95775.1 acla|EAW10626.1 afis|EAW24941.1 afla|EED55524.1 afum|EAL89598.1 agla|OJJ82168.1 akaw|GAA83628.1 aluc|OJZ92365.1 anig|EHA26608.1 aory|EIT80862.1 asyd|OJJ57516.1 ater|EAU35668.1 atub|OJI86795.1 aver|OJJ05663.1 awen|OJJ40535.1 azon|OJJ46849.1 azon|OJJ49396.1 afla|EED47694.1

Group1190: abra|OJJ74880.1 acar|OOF95883.1 acla|EAW08490.1 afis|EAW23536.1 afla|EED57434.1 afum|EAL87930.1 agla|OJJ86791.1 akaw|GAA83540.1 aluc|OJZ92457.1 anid|CBF89670.1 anig|EHA26510.1 aory|EIT80126.1 asyd|OJJ62272.1 ater|EAU35322.1 atub|OJI86886.1 aver|OJI96381.1 awen|OJJ39866.1 azon|OJJ43234.1 afla|EED57433.1

Group1191: abra|OJJ74903.1 acar|OOF95901.1 acla|EAW08469.1 afis|EAW23516.1 afla|EED57417.1 afum|EAL87947.2 agla|OJJ86764.1 akaw|GAA83519.1 aluc|OJZ92476.1 anid|CBF89704.1 anig|EHA26494.1 aory|EIT80064.1 asyd|OJJ62253.1 ater|EAU35305.1 atub|OJI86907.1 aver|OJI96363.1 awen|OJJ39841.1 azon|OJJ43251.1 aory|EIT80158.1

Group1192: abra|OJJ74942.1 acar|OOF97373.1 acla|EAW08428.1 afis|EAW23472.1 afla|EED57368.1 afum|EAL87994.1 agla|OJJ86711.1 akaw|GAA83479.1 aluc|OJZ92516.1 anid|CBF89779.1 anig|EHA26459.1 aory|EIT80072.1 asyd|OJJ62207.1 ater|EAU36908.1 atub|OJI86949.1 aver|OJI96319.1 awen|OJJ39798.1 azon|OJJ47919.1 awen|OJJ40002.1

Group1193: abra|OJJ75019.1 acar|OOF89964.1 akaw|GAA88640.1 aluc|OJZ92586.1 anid|CBF84112.1 anig|EHA26399.1 atub|OJI87018.1 abra|OJJ75020.1 acla|EAW08212.1 afis|EAW23246.1 afla|EED55727.1 afum|EAL88235.1 agla|OJJ86614.1 aory|EIT83220.1 asyd|OJJ57103.1 ater|EAU37528.1 aver|OJJ03196.1 awen|OJJ39724.1 azon|OJJ47850.1

Group1194: abra|OJJ75054.1 acar|OOF97283.1 acla|EAW08239.1 afis|EAW23273.1 afla|EED51546.1 afum|EAL88206.1 agla|OJJ86575.1 akaw|GAA88667.1 aluc|OJZ92619.1 anid|CBF89542.1 anig|EHA26369.1 aory|EIT77094.1 asyd|OJJ62380.1 ater|EAU37558.1 atub|OJI87053.1 aver|OJI96514.1 awen|OJJ39684.1 azon|OJJ47821.1 aory|EIT80554.1

Group1195: abra|OJJ75063.1 acar|OOF97271.1 acla|EAW08246.1 afis|EAW23280.1 afla|EED55696.1 afum|EAL88199.1 agla|OJJ86567.1 akaw|GAA88675.1 aluc|OJZ92629.1 anid|CBF89529.1 anig|EHA26361.1 aory|EIT77085.1 asyd|OJJ62387.1 ater|EAU37578.1 atub|OJI87059.1 aver|OJI96520.1 awen|OJJ39677.1 azon|OJJ43147.1 awen|OJJ31335.1

Group1196: abra|OJJ75119.1 acar|OOF97214.1 acla|EAW09724.1 afis|EAW21417.1 afum|EAL92432.2 aluc|OJZ92686.1 anig|EHA26314.1 atub|OJI87114.1 azon|OJJ50750.1 abra|OJJ75768.1 afla|EED52344.1 agla|OJJ78972.1 aory|EIT74507.1 asyd|OJJ58674.1 atub|OJI87190.1 aver|OJJ01285.1 awen|OJJ30922.1 asyd|OJJ52504.1 aver|OJJ03875.1

Group1197: aver|OJJ04160.1 afis|EAW19670.1 afla|EED46493.1 afum|EAL92627.1 agla|OJJ86969.1 anid|CBF74912.1 anig|EHA22175.1 aory|EIT72425.1 asyd|OJJ60375.1 ater|EAU35595.1 atub|OJI91472.1 aver|OJJ05939.1 awen|OJJ39537.1 azon|OJJ51156.1 agla|OJJ86474.1 asyd|OJJ53618.1 awen|OJJ30584.1 asyd|OJJ60102.1 asyd|OJJ53622.1

Group1198: abra|OJJ75142.1 acar|OOF97194.1 anid|CBF74890.1 anig|EHA26291.1 asyd|OJJ60105.1 ater|EAU35601.1 aver|OJJ04157.1 awen|OJJ39534.1 abra|OJJ76897.1 afla|EED44885.1 agla|OJJ86440.1 akaw|GAA87147.1 aluc|OJZ84290.1 anig|EHA27630.1 aory|EIT83262.1 asyd|OJJ57332.1 atub|OJI89779.1 aver|OJJ05829.1 awen|OJJ40829.1

Group1199: abra|OJJ75173.1 acar|OOF94605.1 acla|EAW13003.1 afis|EAW17665.1 afla|EED50492.1 afum|EAL91479.1 agla|OJJ79538.1 akaw|GAA88780.1 aluc|OJZ92741.1 anid|CBF90049.1 anig|EHA26262.1 aory|EIT82090.1 asyd|OJJ62054.1 ater|EAU31988.1 atub|OJI87166.1 aver|OJI96160.1 awen|OJJ37176.1 azon|OJJ50876.1 asyd|OJJ61465.1

Group1200: abra|OJJ75218.1 acar|OOF94566.1 acla|EAW12260.1 afis|EAW16969.1 afla|EED48292.1 afum|EAL86497.1 agla|OJJ79785.1 akaw|GAA91768.1 aluc|OJZ92782.1 anig|EHA26228.1 aory|EIT76281.1

asyd|OJJ64772.1 atub|OJI87224.1 aver|OJI98872.1 awen|OJJ35998.1 azon|OJJ50271.1 abra|OJJ76878.1 afum|EAL87809.1 ater|EAU38553.1

Group1201: abra|OJJ75219.1 acar|OOF94565.1 acla|EAW12259.1 afis|EAW16978.1 afla|EED48290.1 afum|EAL86504.1 agla|OJJ79786.1 akaw|GAA91769.1 aluc|OJZ92783.1 anig|EHA26227.1 aory|EIT76317.1 asyd|OJJ64771.1 atub|OJI87225.1 aver|OJI98871.1 awen|OJJ35997.1 azon|OJJ50269.1 anig|EHA26226.1 afum|EAL86503.1 afla|EED48291.1

Group1202: abra|OJJ75302.1 acar|OOF92541.1 acla|EAW09084.1 afis|EAW25675.1 afla|EED47643.1 afum|EAL84345.1 agla|OJJ86330.1 akaw|GAA91835.1 aluc|OJZ92857.1 anid|CBF78280.1 anig|EHA26149.1 aory|EIT76311.1 asyd|OJJ52225.1 ater|EAU36081.1 atub|OJI87314.1 aver|OJJ05016.1 awen|OJJ35856.1 azon|OJJ42600.1 asyd|OJJ56073.1

Group1203: abra|OJJ75308.1 acar|OOF94521.1 acla|EAW12232.1 afis|EAW16996.1 afla|EED48959.1 afum|EAL86520.1 agla|OJJ79823.1 akaw|GAA91992.1 aluc|OJZ92862.1 anid|CBF87466.1 anig|EHA26103.1 aory|EIT80011.1 asyd|OJJ52936.1 ater|EAU31732.1 atub|OJI87320.1 aver|OJJ08903.1 awen|OJJ35985.1 azon|OJJ47407.1 azon|OJJ50280.1

Group1204: abra|OJJ75752.1 acar|OOG00116.1 acla|EAW09398.1 afis|EAW16786.1 afla|EED47412.1 afum|EAL86356.1 agla|OJJ78549.1 akaw|GAA84485.1 aluc|OJZ89206.1 anid|CBF84608.1 anig|EHA23353.1 aory|EIT81898.1 asyd|OJJ60434.1 ater|EAU37659.1 atub|OJI85586.1 aver|OJI99756.1 awen|OJJ40856.1 azon|OJJ42408.1 ater|EAU30953.1

Group1205: abra|OJJ75970.1 acar|OOF99950.1 acla|EAW09828.1 afis|EAW21305.1 afla|EED57265.1 afum|EAL92321.1 akaw|GAA88986.1 aluc|OJZ89417.1 anid|CBF71434.1 anig|EHA23171.1 aory|EIT73994.1 asyd|OJJ65347.1 ater|EAU35936.1 atub|OJI85377.1 aver|OJJ07224.1 awen|OJJ39393.1 azon|OJJ50301.1 asyd|OJJ54967.1 ater|EAU32173.1

Group1206: abra|OJJ75996.1 acar|OOF99929.1 acla|EAW09809.1 afis|EAW21327.1 afla|EED57285.1 afum|EAL92345.1 agla|OJJ87412.1 akaw|GAA88963.1 aluc|OJZ89442.1 anid|CBF83428.1 anig|EHA23144.1 aory|EIT74010.1 asyd|OJJ60659.1 ater|EAU35952.1 atub|OJI85351.1 aver|OJI99982.1 awen|OJJ39371.1 azon|OJJ51711.1 afla|EED57284.1

Group1207: abra|OJJ76229.1 acar|OOF99477.1 acla|EAW07053.1 afis|EAW16679.1 afla|EED45240.1 afum|EAL89283.1 akaw|GAA82167.1 aluc|OJZ92000.1 anid|CBF82050.1 aory|EIT73588.1 asyd|OJJ54232.1 ater|EAU29467.1 atub|OJI86434.1 aver|OJJ05185.1 awen|OJJ36977.1 azon|OJJ42100.1 akaw|GAA92942.1 atub|OJI86436.1 akaw|GAA82166.1

Group1208: abra|OJJ76593.1 acar|OOF99394.1 acla|EAW14308.1 afis|EAW18977.1 afla|EED53193.1 afum|EAL93713.2 agla|OJJ82935.1 akaw|GAA82497.1 aluc|OJZ91613.1 anid|CBF69648.1 anig|EHA22642.1 aory|EIT74974.1 asyd|OJJ59078.1 ater|EAU38168.1 atub|OJI86066.1 aver|OJJ01671.1 awen|OJJ31583.1 azon|OJJ47298.1 afla|EED53194.1

Group1209: abra|OJJ76704.1 acar|OOF89909.1 acla|EAW07574.1 afis|EAW22608.1 afla|EED49806.1 afum|EAL90590.1 agla|OJJ87452.1 akaw|GAA92495.1 aluc|OJZ91507.1 anid|CBF80167.1 aory|EIT77615.1 asyd|OJJ63679.1 atub|OJI85959.1 aver|OJI97260.1 awen|OJJ35236.1 azon|OJJ49875.1 agla|OJJ80978.1 anid|CBF88267.1 aver|OJI99521.1

Group1210: abra|OJJ76824.1 acar|OOF91318.1 acla|EAW13384.1 afis|EAW20498.1 afla|EED44740.1 afum|EAL84556.1 akaw|GAA87198.1 aluc|OJZ84235.1 anid|CBF79206.1 anig|EHA27582.1 aory|EIT77799.1 asyd|OJJ55728.1 ater|EAU37192.1 atub|OJI89850.1 aver|OJJ06814.1 awen|OJJ33198.1 azon|OJJ46398.1 aory|EIT77801.1 afla|EED44741.1

Group1211: abra|OJJ76859.1 acar|OOF91347.1 acla|EAW13420.1 afis|EAW20531.1 afla|EED52481.1 afum|EAL84526.1 agla|OJJ84700.1 akaw|GAA87163.1 aluc|OJZ84271.1 anid|CBF79002.1 anig|EHA27613.1 aory|EIT82813.1 asyd|OJJ55689.1 ater|EAU32506.1 atub|OJI89812.1 aver|OJJ06773.1 awen|OJJ33238.1 azon|OJJ50610.1 afis|EAW15383.1

Group1212: abra|OJJ76952.1 acar|OOF91399.1 acla|EAW13478.1 afis|EAW20585.1 afla|EED52259.1 afum|EAL84480.1 agla|OJJ80829.1 akaw|GAA87090.1 aluc|OJZ84348.1 anid|CBF78926.1 anig|EHA27685.1 aory|EIT76185.1 asyd|OJJ55629.1 ater|EAU37094.1 atub|OJI89720.1 aver|OJJ06706.1 awen|OJJ33298.1 azon|OJJ50675.1 afis|EAW16394.1

Group1213: abra|OJJ76962.1 acla|EAW13481.1 afis|EAW20589.1 afla|EED52239.1 afum|EAL84472.1 akaw|GAA87081.1 aluc|OJZ84357.1 anid|CBF80645.1 anig|EHA27694.1 aory|EIT74320.1 asyd|OJJ54615.1 ater|EAU35228.1 atub|OJI89710.1 aver|OJJ07544.1 azon|OJJ44240.1 acla|EAW11256.1 afis|EAW18243.1 afum|EAL87607.1 ater|EAU38946.1

Group1214: abra|OJJ77019.1 acla|EAW12022.1 afis|EAW20396.1 afla|EED51074.1 afum|EAL86857.1 akaw|GAA87033.1 aluc|OJZ84419.1 anig|EHA27741.1 aory|EIT83235.1 asyd|OJJ61467.1 ater|EAU34588.1 atub|OJI89655.1 aver|OJJ07304.1 awen|OJJ32287.1 afis|EAW19666.1 aory|EIT77057.1 ater|EAU30085.1 ater|EAU31682.1 aver|OJJ00806.1

Group1215: abra|OJJ77072.1 acar|OOF96452.1 acla|EAW11107.1 afis|EAW23663.1 afla|EED48431.1 afum|EAL85154.2 agla|OJJ81757.1 aluc|OJZ84480.1 anid|CBF75932.1 aory|EIT81669.1 atub|OJI89599.1 aver|OJI99663.1 awen|OJJ32900.1 afla|EED51081.1 aory|EIT77059.1 agla|OJJ80484.1 awen|OJJ40921.1 awen|OJJ40116.1 awen|OJJ32943.1

Group1216: akaw|GAA86981.1 anig|EHA27470.1 azon|OJJ46863.1 acla|EAW08585.1 afis|EAW16399.1 afum|EAL89027.1 azon|OJJ47065.1 acla|EAW11106.1 afla|EED48430.1 agla|OJJ80486.1 aory|EIT76978.1 atub|OJI88846.1 awen|OJJ32901.1 afla|EED51080.1 agla|OJJ88498.1 akaw|GAA92594.1 aluc|OJZ82611.1 afla|EED54189.1 awen|OJJ34609.1

Group1217: abra|OJJ77092.1 acar|OOG00596.1 afis|EAW17058.1 afla|EED52120.1 afum|EAL86554.1 akaw|GAA86963.1 aluc|OJZ84501.1 anid|CBF88306.1 anig|EHA27801.1 asyd|OJJ62781.1 ater|EAU31929.1 atub|OJI89579.1 aver|OJI96826.1 azon|OJJ45055.1 acar|OOF90672.1 aory|EIT74489.1 azon|OJJ43507.1 afla|EED52121.1 anid|CBF88998.1

Group1218: abra|OJJ77232.1 acar|OOF93114.1 afla|EED48701.1 agla|OJJ83280.1 akaw|GAA90710.1 aluc|OJZ84642.1 anid|CBF78734.1 anig|EHA27928.1 aory|EIT73111.1 asyd|OJJ55609.1 atub|OJI89439.1 aver|OJJ06684.1 awen|OJJ30084.1 abra|OJJ77423.1 acla|EAW13221.1 afis|EAW20821.1 afum|EAL84808.1 ater|EAU31194.1 awen|OJJ31931.1

Group1219: abra|OJJ77364.1 acar|OOF93233.1 acla|EAW13112.1 afis|EAW17552.1 afla|EED49273.1 afum|EAL91366.1 agla|OJJ81416.1 akaw|GAA91398.1 aluc|OJZ84773.1 anid|CBF90298.1 anig|EHA28041.1 aory|EIT82235.1 asyd|OJJ61925.1 ater|EAU36982.1 atub|OJI89308.1 aver|OJI96017.1 awen|OJJ37301.1 azon|OJJ50994.1 anid|CBF87295.1

Group1220: abra|OJJ77453.1 acar|OOF99273.1 acla|EAW12649.1 afis|EAW18036.1 afla|EED54071.1 afum|EAL91847.1 agla|OJJ78571.1 akaw|GAA84155.1 aluc|OJZ81076.1 anid|CBF81938.1 anig|EHA19286.1 aory|EIT76067.1 asyd|OJJ57409.1 ater|EAU29318.1 ater|EAU31609.1 atub|OJI82509.1 aver|OJJ05772.1 awen|OJJ34693.1 azon|OJJ48230.1

Group1221: abra|OJJ77467.1 acar|OOF99267.1 acla|EAW07127.1 afla|EED54176.1 agla|OJJ87943.1 akaw|GAA84150.1 aluc|OJZ81071.1 anig|EHA19298.1 aory|EIT75990.1 asyd|OJJ52794.1 ater|EAU29321.1 atub|OJI82515.1 aver|OJJ08772.1 awen|OJJ34711.1 akaw|GAA91805.1 atub|OJI87284.1 asyd|OJJ64953.1 aver|OJI99075.1 aluc|OJZ92827.1

Group1222: abra|OJJ77471.1 acar|OOF99253.1 acla|EAW06761.1 afis|EAW16311.1 afla|EED54184.1 afum|EAL88933.1 agla|OJJ88270.1 akaw|GAA84146.1 aluc|OJZ81067.1 anid|CBF69560.1 anig|EHA19302.1 asyd|OJJ59156.1 ater|EAU29331.1 atub|OJI88508.1 aver|OJJ01746.1 awen|OJJ37711.1 azon|OJJ44970.1 aluc|OJZ81673.1 anig|EHA25767.1

Group1223: abra|OJJ77555.1 acar|OOF99172.1 acla|EAW06522.1 afis|EAW16238.1 afla|EED53339.1 afum|EAL88859.1 agla|OJJ86247.1 akaw|GAA84068.1 aluc|OJZ80982.1 anid|CBF78861.1 anig|EHA19378.1 aory|EIT78064.1 asyd|OJJ64575.1 atub|OJI82604.1 aver|OJI98672.1 awen|OJJ30219.1 azon|OJJ43941.1 afla|EED50785.1 agla|OJJ79233.1

Group1224: abra|OJJ77564.1 acar|OOF99168.1 acla|EAW06526.1 afis|EAW16070.1 afla|EED48589.1 afum|EAL88702.1 agla|OJJ86098.1 akaw|GAA84062.1 aluc|OJZ80974.1 anid|CBF75103.1 anig|EHA19385.1 aory|EIT83270.1 asyd|OJJ60005.1 ater|EAU33561.1 atub|OJI82611.1 aver|OJJ02281.1 awen|OJJ30365.1 azon|OJJ45384.1 afla|EED57884.1

Group1225: abra|OJJ77597.1 acar|OOF99129.1 acla|EAW06561.1 afis|EAW16035.1 afla|EED56421.1 afum|EAL88666.2 agla|OJJ86065.1 akaw|GAA84031.1 aluc|OJZ80937.1 anig|EHA19414.1 aory|EIT72879.1 ater|EAU33605.1 atub|OJI82644.1 awen|OJJ34579.1 azon|OJJ45420.1 aluc|OJZ83622.1 akaw|GAA92103.1 anig|EHA27514.1 atub|OJI87849.1

Group1226: abra|OJJ77599.1 acar|OOF99127.1 acla|EAW06563.1 afis|EAW16033.1 afla|EED56415.1 afla|EED56416.1 afum|EAL88664.1 agla|OJJ86061.1 akaw|GAA84029.1 aluc|OJZ80934.1 anid|CBF75034.1 anig|EHA19416.1 aory|EIT72886.1 asyd|OJJ60050.1 ater|EAU33608.1 atub|OJI82646.1 aver|OJJ02239.1 awen|OJJ30414.1 azon|OJJ45422.1

Group1227: abra|OJJ77664.1 acar|OOF99071.1 acla|EAW06508.1 afis|EAW24531.1 afla|EED56377.1 agla|OJJ85724.1 akaw|GAA83971.1 aluc|OJZ80864.1 anig|EHA19471.1 aory|EIT73456.1 ater|EAU33642.1 atub|OJI82713.1 aver|OJJ02022.1 awen|OJJ30465.1 azon|OJJ45455.1 afum|EAL86197.1 asyd|OJJ54667.1 afum|EAL86196.1 agla|OJJ86406.1

Group1228: abra|OJJ77796.1 acar|OOF90711.1 akaw|GAA89038.1 aluc|OJZ80557.1 anig|EHA25640.1 atub|OJI88374.1 aver|OJJ02063.1 awen|OJJ38007.1 abra|OJJ77916.1 afla|EED51608.1 agla|OJJ82626.1 anid|CBF73906.1 anig|EHA18406.1 aory|EIT80458.1 asyd|OJJ53666.1 ater|EAU37641.1 aver|OJJ07773.1 awen|OJJ40190.1 agla|OJJ83009.1

Group1229: abra|OJJ77806.1 acar|OOF98961.1 afis|EAW18189.1 afla|EED56286.1 afum|EAL87553.1 agla|OJJ86044.1 akaw|GAA83843.1 aluc|OJZ90873.1 anid|CBF86100.1 anig|EHA19593.1 aory|EIT73443.1

asyd|OJJ61783.1 ater|EAU33804.1 atub|OJI82872.1 aver|OJI95883.1 awen|OJJ30553.1 azon|OJJ45513.1 aver|OJI98581.1 asyd|OJJ64479.1

Group1230: abra|OJJ77818.1 acar|OOF98950.1 acla|EAW11305.1 afis|EAW18192.1 afla|EED56282.1 afum|EAL87560.1 agla|OJJ85885.1 akaw|GAA83828.1 aluc|OJZ90890.1 anid|CBF86109.1 anig|EHA19609.1 aory|EIT73431.1 asyd|OJJ64488.1 ater|EAU33797.1 atub|OJI82884.1 aver|OJI98589.1 awen|OJJ30564.1 azon|OJJ45507.1 aory|EIT73332.1

Group1231: abra|OJJ77831.1 acar|OOF98939.1 acla|EAW11295.1 afis|EAW18207.1 afla|EED56264.1 afum|EAL87572.1 agla|OJJ85875.1 akaw|GAA83816.1 aluc|OJZ90905.1 anid|CBF86133.1 anig|EHA19620.1 aory|EIT73347.1 asyd|OJJ64503.1 ater|EAU33782.1 atub|OJI82898.1 aver|OJI98602.1 awen|OJJ30579.1 azon|OJJ45497.1 afla|EED56263.1

Group1232: abra|OJJ77887.1 acar|OOF90364.1 afis|EAW17355.1 afla|EED49462.1 agla|OJJ82474.1 akaw|GAA82230.1 aluc|OJZ91918.1 anig|EHA19662.1 aory|EIT79498.1 asyd|OJJ56938.1 ater|EAU29950.1 aver|OJJ03347.1 awen|OJJ32673.1 azon|OJJ46453.1 asyd|OJJ57438.1 aver|OJJ05740.1 awen|OJJ31966.1 asyd|OJJ57599.1 anig|EHA23994.1

Group1233: abra|OJJ77901.1 acar|OOF98882.1 acla|EAW06511.1 afis|EAW16229.1 afla|EED53352.1 afum|EAL88852.1 agla|OJJ86237.1 akaw|GAA83761.1 aluc|OJZ90970.1 anid|CBF86221.1 anig|EHA19672.1 aory|EIT75062.1 asyd|OJJ64562.1 ater|EAU33715.1 atub|OJI82961.1 aver|OJI98659.1 awen|OJJ30233.1 azon|OJJ43950.1 anid|CBF86220.1

Group1234: abra|OJJ78153.1 acar|OOF98713.1 acla|EAW12149.1 afis|EAW17172.1 afla|EED45420.1 afum|EAL86603.1 akaw|GAA85873.1 aluc|OJZ91205.1 anig|EHA19910.1 aory|EIT82401.1 ater|EAU31964.1 atub|OJI83222.1 aver|OJJ03361.1 awen|OJJ31986.1 azon|OJJ49155.1 akaw|GAA92449.1 atub|OJI88616.1 aluc|OJZ81569.1 anig|EHA25894.1

Group1235: abra|OJJ78155.1 acar|OOF90381.1 acla|EAW11243.1 afis|EAW18260.1 afla|EED48719.1 afum|EAL87623.1 agla|OJJ88167.1 akaw|GAA85871.1 aluc|OJZ91207.1 anid|CBF79523.1 anig|EHA19911.1 aory|EIT73142.1 asyd|OJJ57893.1 ater|EAU33268.1 atub|OJI83223.1 aver|OJJ04502.1 awen|OJJ30632.1 awen|OJJ30633.1 azon|OJJ47984.1

Group1236: abra|OJJ78254.1 acar|OOF98608.1 acla|EAW09255.1 afis|EAW21892.1 afla|EED46416.1 afum|EAL92906.1 agla|OJJ84240.1 akaw|GAA85772.1 aluc|OJZ91328.1 anid|CBF76804.1 anig|EHA20010.1 aory|EIT72374.1 asyd|OJJ61288.1 atub|OJI83321.1 aver|OJJ00614.1 awen|OJJ34181.1 azon|OJJ43451.1 acar|OOF99280.1 ater|EAU36657.1

Group1237: abra|OJJ78362.1 acar|OOF98525.1 acla|EAW09173.1 afis|EAW21973.1 afla|EED46696.1 afum|EAL92988.1 agla|OJJ84129.1 akaw|GAA85671.1 aluc|OJZ91436.1 anid|CBF82714.1 anig|EHA20102.1 aory|EIT77832.1 asyd|OJJ61638.1 ater|EAU31091.1 atub|OJI83430.1 aver|OJJ00961.1 awen|OJJ34266.1 azon|OJJ43370.1 aory|EIT77819.1

Group1238: abra|OJJ68129.1 abra|OJJ70102.1 acla|EAW12416.1 akaw|GAA87598.1 akaw|GAA86909.1 aluc|OJZ85218.1 aluc|OJZ90032.1 anig|EHA18784.1 anig|EHA27408.1 anig|EHA25422.1 atub|OJI80367.1 atub|OJI84774.1 awen|OJJ39118.1 anid|CBF75781.1 ater|EAU32451.1 azon|OJJ42520.1 afla|EED47323.1 aory|EIT82878.1

Group1239: abra|OJJ73597.1 abra|OJJ74454.1 acar|OOF95298.1 acar|OOG00783.1 akaw|GAA87348.1 akaw|GAA88555.1 aluc|OJZ88453.1 aluc|OJZ89666.1 anig|EHA18174.1 anig|EHA26903.1 atub|OJI85134.1 atub|OJI91051.1 afla|EED49261.1 aory|EIT82199.1 asyd|OJJ62434.1 ater|EAU29287.1 aver|OJI96570.1 aver|OJI98026.1

Group1240: acar|OOF90451.1 acar|OOF90458.1 abra|OJJ71881.1 acla|EAW13910.1 afis|EAW18622.1 afla|EED52720.1 afum|EAL93332.1 agla|OJJ81238.1 akaw|GAA85035.1 aluc|OJZ86619.1 anid|CBF70495.1 anig|EHA22047.1 asyd|OJJ58582.1 ater|EAU34922.1 atub|OJI83907.1 aver|OJJ01186.1 awen|OJJ32085.1 azon|OJJ46664.1

Group1241: acar|OOF90710.1 acar|OOF98737.1 atub|OJI84596.1 atub|OJI89205.1 abra|OJJ65987.1 afis|EAW19527.1 afla|EED49678.1 agla|OJJ87966.1 akaw|GAA90861.1 aluc|OJZ82972.1 anid|CBF78948.1 anig|EHA21389.1 aory|EIT80959.1 asyd|OJJ55653.1 ater|EAU29344.1 aver|OJJ06737.1 awen|OJJ30016.1 azon|OJJ43646.1

Group1242: acar|OOF91721.1 acar|OOF91725.1 abra|OJJ73107.1 afis|EAW19523.1 afla|EED54039.1 afum|EAL84403.1 akaw|GAA88059.1 aluc|OJZ81321.1 anid|CBF87177.1 anig|EHA18627.1 aory|EIT74598.1 asyd|OJJ63343.1 ater|EAU32279.1 atub|OJI91524.1 aver|OJI97618.1 azon|OJJ45982.1 afis|EAW25732.1 ater|EAU37760.1

Group1243: acar|OOF92284.1 acar|OOF92325.1 abra|OJJ67518.1 acla|EAW06257.1 afis|EAW18364.1 afla|EED53041.1 afum|EAL93037.1 agla|OJJ84927.1 aluc|OJZ84846.1 anid|CBF70821.1 anig|EHA25081.1 aory|EIT77411.1 asyd|OJJ58401.1 ater|EAU32232.1 atub|OJI80749.1 aver|OJJ01002.1 awen|OJJ31787.1 azon|OJJ42024.1

Group1244: acar|OOF92296.1 acar|OOF92337.1 abra|OJJ67504.1 acla|EAW06266.1 afis|EAW18372.1 afla|EED53034.1 afum|EAL93047.1 agla|OJJ84936.1 akaw|GAA86334.1 aluc|OJZ84859.1 anid|CBF70802.1 anig|EHA25092.1 asyd|OJJ58410.1 ater|EAU32242.1 atub|OJI80737.1 aver|OJJ01012.1 awen|OJJ31778.1 azon|OJJ50447.1

Group1245: acar|OOF92300.1 acar|OOF92341.1 abra|OJJ67500.1 acla|EAW06270.1 afis|EAW18376.1 afla|EED53030.1 afum|EAL93051.1 agla|OJJ84940.1 akaw|GAA86338.1 aluc|OJZ84863.1 anid|CBF70795.1 aory|EIT77422.1 asyd|OJJ58414.1 ater|EAU32246.1 atub|OJI80733.1 aver|OJJ01016.1 awen|OJJ31768.1 azon|OJJ50451.1

Group1246: acar|OOF92302.1 acar|OOF92344.1 abra|OJJ67497.1 acla|EAW06273.1 afis|EAW18379.1 afla|EED53027.1 afum|EAL93054.1 agla|OJJ84943.1 akaw|GAA86341.1 aluc|OJZ84866.1 anid|CBF70789.1 anig|EHA25098.1 aory|EIT77417.1 asyd|OJJ58417.1 atub|OJI80730.1 aver|OJJ01019.1 awen|OJJ31771.1 azon|OJJ50454.1

Group1247: acar|OOF92313.1 acar|OOF92354.1 abra|OJJ67486.1 acla|EAW06288.1 afis|EAW18393.1 afla|EED53014.1 afum|EAL93069.1 agla|OJJ84960.1 akaw|GAA86353.1 aluc|OJZ84882.1 anid|CBF70764.1 anig|EHA25111.1 asyd|OJJ58429.1 ater|EAU32281.1 atub|OJI80718.1 aver|OJJ01031.1 azon|OJJ50467.1 awen|OJJ31757.1

Group1248: acar|OOF92613.1 acar|OOF92634.1 abra|OJJ68141.1 acla|EAW07634.1 afis|EAW22654.1 afla|EED49922.1 afum|EAL90538.1 agla|OJJ81037.1 akaw|GAA92377.1 aluc|OJZ83236.1 anid|CBF88199.1 anig|EHA21093.1 asyd|OJJ63635.1 ater|EAU39052.1 atub|OJI87700.1 aver|OJI97301.1 awen|OJJ35269.1 azon|OJJ49838.1

Group1249: acar|OOF93658.1 acar|OOF93742.1 abra|OJJ66637.1 acla|EAW11780.1 afis|EAW19390.1 afla|EED50348.1 afum|EAL87105.1 agla|OJJ81101.1 akaw|GAA89590.1 aluc|OJZ80812.1 anig|EHA27222.1 aory|EIT81938.1 asyd|OJJ51911.1 ater|EAU36053.1 atub|OJI87661.1 aver|OJJ05352.1 awen|OJJ36783.1 azon|OJJ43015.1

Group1250: acar|OOF93662.1 acar|OOF93746.1 abra|OJJ66642.1 acla|EAW11785.1 afis|EAW19395.1 afum|EAL87099.1 agla|OJJ81106.1 akaw|GAA89595.1 aluc|OJZ80807.1 anid|CBF74279.1 anig|EHA27227.1 aory|EIT83136.1 asyd|OJJ51860.1 ater|EAU36121.1 atub|OJI87655.1 aver|OJJ05359.1 awen|OJJ36789.1 azon|OJJ43009.1

Group1251: acar|OOF93664.1 acar|OOF93748.1 abra|OJJ66644.1 acla|EAW11787.1 afis|EAW19397.1 afla|EED50353.1 afum|EAL87097.1 agla|OJJ81108.1 akaw|GAA89597.1 aluc|OJZ80805.1 anid|CBF74276.1 anig|EHA27229.1 aory|EIT83127.1 ater|EAU36123.1 atub|OJI87653.1 aver|OJJ05361.1 awen|OJJ36792.1 azon|OJJ43007.1

Group1252: acar|OOF93684.1 acar|OOF93769.1 abra|OJJ66669.1 acla|EAW11810.1 afis|EAW19420.1 afla|EED50377.1 afum|EAL87074.1 agla|OJJ81132.1 akaw|GAA89618.1 aluc|OJZ80778.1 anid|CBF74369.1 anig|EHA27251.1 aory|EIT83135.1 asyd|OJJ57783.1 atub|OJI87630.1 aver|OJJ05382.1 awen|OJJ36818.1 azon|OJJ42984.1

Group1253: acar|OOF93697.1 acar|OOF93782.1 abra|OJJ66688.1 acla|EAW11823.1 afis|EAW19434.1 afla|EED56639.1 afum|EAL87061.1 agla|OJJ81146.1 akaw|GAA89632.1 aluc|OJZ80761.1 anid|CBF81381.1 anig|EHA27266.1 aory|EIT79620.1 asyd|OJJ59392.1 ater|EAU34736.1 atub|OJI87615.1 aver|OJJ02880.1 azon|OJJ42971.1

Group1254: acar|OOF93702.1 acar|OOF93788.1 abra|OJJ66694.1 acla|EAW11829.1 afis|EAW19441.1 afla|EED56648.1 afum|EAL87055.1 agla|OJJ81154.1 akaw|GAA89638.1 aluc|OJZ80751.1 anid|CBF81393.1 aory|EIT79649.1 asyd|OJJ59399.1 ater|EAU34742.1 atub|OJI87608.1 aver|OJJ02873.1 awen|OJJ36841.1 azon|OJJ42959.1

Group1255: acar|OOF93712.1 acar|OOF93798.1 abra|OJJ66710.1 acla|EAW11841.1 afis|EAW19452.1 afla|EED56660.1 afum|EAL87044.1 agla|OJJ81167.1 akaw|GAA89652.1 aluc|OJZ80737.1 anid|CBF81416.1 anig|EHA27285.1 asyd|OJJ59412.1 ater|EAU34754.1 atub|OJI87594.1 aver|OJJ02861.1 azon|OJJ42947.1 awen|OJJ36853.1

Group1256: acar|OOF93719.1 acar|OOF93806.1 abra|OJJ66717.1 acla|EAW11848.1 afis|EAW19459.1 afla|EED56670.1 afum|EAL87037.1 agla|OJJ81176.1 akaw|GAA89659.1 aluc|OJZ80730.1 anig|EHA27291.1 aory|EIT79651.1 asyd|OJJ59419.1 ater|EAU34762.1 atub|OJI87586.1 aver|OJJ02850.1 awen|OJJ36861.1 azon|OJJ42940.1

Group1257: acar|OOF93720.1 acar|OOF93807.1 abra|OJJ66718.1 acla|EAW11849.1 afis|EAW19460.1 afla|EED56671.1 afum|EAL87036.1 agla|OJJ81177.1 akaw|GAA89660.1 aluc|OJZ80729.1 anid|CBF81429.1 anig|EHA27292.1 asyd|OJJ59420.1 ater|EAU34763.1 atub|OJI87585.1 aver|OJJ02849.1 awen|OJJ36862.1 azon|OJJ42939.1

Group1258: acar|OOF93730.1 acar|OOF93818.1 abra|OJJ66732.1 acla|EAW11860.1 afis|EAW19471.1 afla|EED56681.1 afum|EAL87025.1 agla|OJJ81192.1 akaw|GAA89671.1 aluc|OJZ80717.1 anid|CBF75472.1

anig|EHA27302.1 asyd|OJJ59760.1 ater|EAU34781.1 atub|OJI87573.1 aver|OJJ02528.1 awen|OJJ36876.1 azon|OJJ42928.1

Group1259: acar|OOF93732.1 acar|OOF93820.1 abra|OJJ66734.1 acla|EAW11862.1 afis|EAW19473.1 afla|EED57650.1 afum|EAL87023.1 agla|OJJ81195.1 akaw|GAA89673.1 aluc|OJZ80715.1 anid|CBF71137.1 anig|EHA27304.1 asyd|OJJ55833.1 ater|EAU34783.1 atub|OJI87571.1 aver|OJJ04775.1 awen|OJJ36878.1 azon|OJJ48920.1

Group1260: acar|OOF93733.1 acar|OOF93821.1 abra|OJJ66735.1 acla|EAW11863.1 afis|EAW19474.1 afum|EAL87022.1 agla|OJJ81194.1 akaw|GAA89674.1 aluc|OJZ80713.1 anid|CBF75476.1 anig|EHA27305.1 aory|EIT73892.1 asyd|OJJ59763.1 ater|EAU34784.1 atub|OJI87570.1 aver|OJJ02525.1 awen|OJJ36879.1 azon|OJJ42926.1

Group1261: acar|OOF93738.1 acar|OOF93827.1 abra|OJJ66742.1 acla|EAW11870.1 afis|EAW19481.1 afla|EED56691.1 afum|EAL87014.1 akaw|GAA89679.1 aluc|OJZ80707.1 anid|CBF75468.1 anig|EHA27311.1 aory|EIT73840.1 asyd|OJJ59765.1 ater|EAU34793.1 atub|OJI87564.1 aver|OJJ02523.1 awen|OJJ36887.1 azon|OJJ42920.1

Group1262: acar|OOF94028.1 acar|OOF94098.1 abra|OJJ66425.1 acla|EAW06805.1 afis|EAW16354.1 afla|EED53518.1 afum|EAL88979.1 akaw|GAA90289.1 aluc|OJZ83051.1 anid|CBF89224.1 anig|EHA28115.1 aory|EIT74854.1 asyd|OJJ62518.1 ater|EAU34107.1 atub|OJI87897.1 aver|OJI96637.1 awen|OJJ37758.1 azon|OJJ47345.1

Group1263: acar|OOF94037.1 acar|OOF94107.1 abra|OJJ71135.1 afla|EED48894.1 akaw|GAA82839.1 aluc|OJZ87400.1 anig|EHA20629.1 aory|EIT81248.1 atub|OJI81246.1 abra|OJJ74706.1 acar|OOF89904.1 akaw|GAA81911.1 aluc|OJZ92286.1 anig|EHA26682.1 atub|OJI86717.1 awen|OJJ33632.1 abra|OJJ76740.1 asyd|OJJ65386.1

Group1264: acar|OOF94049.1 acar|OOF94119.1 akaw|GAA85929.1 akaw|GAA90360.1 aluc|OJZ85872.1 aluc|OJZ91136.1 atub|OJI79658.1 atub|OJI83147.1 abra|OJJ78065.1 acla|EAW14269.1 afis|EAW19008.1 afla|EED49855.1 afum|EAL93742.1 anid|CBF71642.1 anig|EHA24713.1 asyd|OJJ61653.1 anid|CBF82888.1 awen|OJJ32641.1

Group1265: acar|OOF94065.1 acar|OOF94135.1 abra|OJJ66358.1 acla|EAW10008.1 afis|EAW21048.1 afla|EED51622.1 afum|EAL92084.1 akaw|GAA90232.1 aluc|OJZ83109.1 anid|CBF87100.1 anig|EHA28172.1 aory|EIT80748.1 asyd|OJJ60390.1 ater|EAU36820.1 atub|OJI87825.1 aver|OJI99719.1 awen|OJJ32174.1 azon|OJJ46753.1

Group1266: acar|OOF94088.1 acar|OOF94158.1 abra|OJJ66350.1 acla|EAW09997.1 afis|EAW21076.1 afla|EED51677.1 afum|EAL92099.1 akaw|GAA90223.1 aluc|OJZ83118.1 anid|CBF87050.1 anig|EHA28181.1 aory|EIT80671.1 asyd|OJJ60442.1 ater|EAU36800.1 atub|OJI87816.1 aver|OJI99760.1 awen|OJJ36350.1 azon|OJJ45249.1

Group1267: acar|OOF95043.1 acar|OOF95110.1 abra|OJJ70456.1 awen|OJJ40126.1 abra|OJJ75241.1 acla|EAW14126.1 afis|EAW18829.1 afla|EED52811.1 afum|EAL93563.1 akaw|GAA91784.1 aluc|OJZ92803.1 anig|EHA26206.1 aory|EIT77227.1 ater|EAU33497.1 atub|OJI87246.1 awen|OJJ39969.1 aory|EIT73400.1 awen|OJJ40159.1

Group1268: acar|OOF95074.1 acar|OOF95139.1 abra|OJJ74641.1 acla|EAW10747.1 afis|EAW24827.1 afla|EED47148.1 afum|EAL89472.1 agla|OJJ82016.1 akaw|GAA81970.1 aluc|OJZ92220.1 anid|CBF85861.1 anig|EHA26738.1 asyd|OJJ64368.1 ater|EAU36478.1 atub|OJI86651.1 aver|OJI98463.1 awen|OJJ40413.1 azon|OJJ49263.1

Group1269: acar|OOF95075.1 acar|OOF95141.1 abra|OJJ74639.1 acla|EAW10749.1 afis|EAW24825.1 afla|EED47150.1 afum|EAL89469.1 agla|OJJ82013.1 akaw|GAA81972.1 aluc|OJZ92218.1 anid|CBF85857.1 anig|EHA26740.1 asyd|OJJ64366.1 ater|EAU36480.1 atub|OJI86649.1 aver|OJI98461.1 awen|OJJ40411.1 azon|OJJ49260.1

Group1270: acar|OOF95077.1 acar|OOF95142.1 abra|OJJ74637.1 acla|EAW10754.1 afis|EAW24823.1 afla|EED47152.1 afum|EAL89467.1 agla|OJJ82011.1 akaw|GAA81974.1 aluc|OJZ92216.1 anid|CBF85853.1 anig|EHA26742.1 aory|EIT79248.1 asyd|OJJ64364.1 atub|OJI86647.1 aver|OJI98459.1 awen|OJJ40409.1 azon|OJJ49258.1

Group1271: acar|OOF95081.1 acar|OOF95146.1 abra|OJJ74628.1 acla|EAW10751.1 afis|EAW24820.1 afla|EED47156.1 afum|EAL89464.1 agla|OJJ82918.1 akaw|GAA81980.1 aluc|OJZ92210.1 anid|CBF85872.1 anig|EHA26748.1 asyd|OJJ64373.1 ater|EAU36485.1 atub|OJI86639.1 aver|OJI98469.1 awen|OJJ40406.1 azon|OJJ49255.1

Group1272: acar|OOF95103.1 acar|OOF95169.1 abra|OJJ74601.1 acla|EAW10778.1 afis|EAW24796.1 afum|EAL89438.1 agla|OJJ82892.1 akaw|GAA82007.1 aluc|OJZ92180.1 anid|CBF81504.1 anig|EHA26773.1 aory|EIT79228.1 asyd|OJJ59467.1 ater|EAU36509.1 atub|OJI86608.1 aver|OJJ02807.1 awen|OJJ40381.1 azon|OJJ49231.1

Group1273: acar|OOF95628.1 acar|OOF95679.1 abra|OJJ74663.1 acla|EAW10732.1 afis|EAW24843.1 afla|EED47128.1 afum|EAL89489.1 agla|OJJ82033.1 akaw|GAA81948.1 aluc|OJZ92240.1 anid|CBF75851.1 anig|EHA26714.1 aory|EIT79238.1 asyd|OJJ57572.1 ater|EAU36461.1 atub|OJI86674.1 aver|OJJ05607.1 awen|OJJ40428.1

Group1274: acar|OOF95634.1 acar|OOF95682.1 abra|OJJ74669.1 acla|EAW10727.1 afis|EAW24848.1 afla|EED47123.1 afum|EAL89494.1 agla|OJJ82060.1 akaw|GAA81943.1 aluc|OJZ92247.1 anid|CBF75841.1 anig|EHA26709.1 aory|EIT79325.1 asyd|OJJ57559.1 ater|EAU36455.1 atub|OJI86680.1 aver|OJJ05619.1 awen|OJJ40433.1

Group1275: acar|OOF95654.1 acar|OOF95705.1 abra|OJJ74694.1 acla|EAW10704.1 afis|EAW24866.1 afla|EED47107.1 afum|EAL89516.1 agla|OJJ82084.1 akaw|GAA81922.1 aluc|OJZ92271.1 anid|CBF75806.1 anig|EHA26690.1 asyd|OJJ57539.1 ater|EAU36434.1 atub|OJI86706.1 aver|OJJ05639.1 awen|OJJ40457.1 azon|OJJ49316.1

Group1276: acar|OOF95656.1 acar|OOF95707.1 abra|OJJ74696.1 acla|EAW10702.1 afis|EAW24868.1 afla|EED47108.1 afum|EAL89518.1 agla|OJJ82086.1 akaw|GAA81921.1 aluc|OJZ92273.1 anid|CBF75804.1 anig|EHA26689.1 asyd|OJJ57537.1 ater|EAU36432.1 atub|OJI86708.1 aver|OJJ05641.1 awen|OJJ40459.1 azon|OJJ49318.1

Group1277: acar|OOF95657.1 acar|OOF95708.1 abra|OJJ74697.1 acla|EAW10701.1 afis|EAW24869.1 afla|EED47109.1 afum|EAL89519.1 agla|OJJ82088.1 akaw|GAA81920.1 aluc|OJZ92276.1 anid|CBF75800.1 anig|EHA26688.1 asyd|OJJ57528.1 ater|EAU36431.1 atub|OJI86709.1 aver|OJJ05650.1 awen|OJJ40460.1 azon|OJJ49320.1

Group1278: acar|OOF95658.1 acar|OOF95709.1 abra|OJJ74699.1 acla|EAW10700.1 afis|EAW24870.1 afla|EED47110.1 afum|EAL89520.1 agla|OJJ82091.1 akaw|GAA81919.1 aluc|OJZ92277.1 anid|CBF75798.1 anig|EHA26687.1 asyd|OJJ57529.1 ater|EAU36430.1 atub|OJI86710.1 aver|OJJ05649.1 awen|OJJ40461.1 azon|OJJ49322.1

Group1279: acar|OOF96222.1 acar|OOF96260.1 abra|OJJ71506.1 acla|EAW15064.1 afis|EAW20036.1 afum|EAL85157.1 agla|OJJ88927.1 akaw|GAA89407.1 aluc|OJZ86245.1 anid|CBF84773.1 anig|EHA21619.1 aory|EIT73713.1 asyd|OJJ63239.1 ater|EAU38654.1 atub|OJI84297.1 aver|OJI97717.1 awen|OJJ31791.1 azon|OJJ46325.1

Group1280: acar|OOF96232.1 acar|OOF96269.1 abra|OJJ71516.1 acla|EAW15074.1 afis|EAW20045.1 afla|EED45335.1 afum|EAL85166.1 agla|OJJ88916.1 aluc|OJZ86255.1 anid|CBF84793.1 anig|EHA21611.1 aory|EIT73562.1 asyd|OJJ63228.1 ater|EAU38655.1 atub|OJI84288.1 aver|OJI97727.1 awen|OJJ31802.1 azon|OJJ46323.1

Group1281: acar|OOF96253.1 acar|OOF96292.1 abra|OJJ71548.1 acla|EAW15102.1 afis|EAW20074.1 afla|EED45305.1 afum|EAL85192.1 agla|OJJ88886.1 akaw|GAA89374.1 aluc|OJZ86286.1 anig|EHA21585.1 aory|EIT73597.1 asyd|OJJ63201.1 ater|EAU38682.1 atub|OJI84252.1 aver|OJI97754.1 awen|OJJ31833.1 azon|OJJ46291.1

Group1282: acar|OOF96903.1 acar|OOF96938.1 abra|OJJ70807.1 acla|EAW10044.1 afis|EAW25540.1 afum|EAL90203.1 agla|OJJ87262.1 akaw|GAA90121.1 aluc|OJZ84153.1 anid|CBF77829.1 anig|EHA19257.1 aory|EIT82996.1 asyd|OJJ53004.1 ater|EAU31663.1 atub|OJI90525.1 aver|OJJ03839.1 awen|OJJ36781.1 azon|OJJ43018.1

Group1283: acar|OOF96912.1 acar|OOF96947.1 abra|OJJ70798.1 acla|EAW11125.1 afis|EAW15854.1 afla|EED56518.1 afum|EAL88490.1 agla|OJJ87253.1 akaw|GAA90114.1 aluc|OJZ84143.1 anid|CBF81183.1 anig|EHA19251.1 asyd|OJJ59296.1 ater|EAU31669.1 atub|OJI90517.1 aver|OJJ02979.1 awen|OJJ36773.1 azon|OJJ47425.1

Group1284: acar|OOF96922.1 acar|OOF96957.1 abra|OJJ70788.1 acla|EAW11134.1 afis|EAW15866.1 afla|EED56509.1 afum|EAL88500.1 agla|OJJ87243.1 akaw|GAA90104.1 aluc|OJZ84133.1 anid|CBF81202.1 anig|EHA19241.1 asyd|OJJ59305.1 ater|EAU31679.1 atub|OJI90507.1 aver|OJJ02970.1 awen|OJJ36764.1 azon|OJJ47434.1

Group1285: acar|OOF96933.1 acar|OOF96968.1 asyd|OJJ59315.1 abra|OJJ70775.1 acla|EAW11144.1 afis|EAW15875.1 afla|EED56500.1 afum|EAL88509.2 agla|OJJ87233.1 aluc|OJZ84122.1 anid|CBF81224.1 anig|EHA19230.1 aory|EIT72804.1 ater|EAU31692.1 atub|OJI90494.1 aver|OJJ02960.1 awen|OJJ36755.1 azon|OJJ47445.1

Group1286: acar|OOF96934.1 acar|OOF96969.1 azon|OJJ42010.1 azon|OJJ45917.1 abra|OJJ68484.1 acla|EAW11145.1 afis|EAW15876.1 afla|EED56499.1 afum|EAL88510.1 agla|OJJ87232.1 akaw|GAA91928.1 aluc|OJZ82694.1 anid|CBF81226.1 anig|EHA21141.1 aory|EIT72869.1 atub|OJI88926.1 aver|OJJ02959.1 awen|OJJ36754.1

Group1287: acar|OOF96935.1 acar|OOF96970.1 abra|OJJ68483.1 acla|EAW11146.1 afis|EAW15877.1 afla|EED56498.1 afum|EAL88511.1 agla|OJJ87231.1 akaw|GAA91929.1 aluc|OJZ82693.1 anid|CBF81228.1

anig|EHA21140.1 aory|EIT72870.1 ater|EAU31697.1 atub|OJI88925.1 aver|OJJ02958.1 awen|OJJ36753.1 azon|OJJ47446.1

Group1288: acar|OOF96937.1 acar|OOF96973.1 abra|OJJ70773.1 acla|EAW11149.1 afis|EAW15880.1 afla|EED56492.1 afum|EAL88514.1 agla|OJJ89541.1 akaw|GAA90092.1 aluc|OJZ84119.1 anid|CBF81234.1 anig|EHA19228.1 aory|EIT72835.1 asyd|OJJ59317.1 atub|OJI90492.1 aver|OJJ02956.1 awen|OJJ36751.1 azon|OJJ47448.1

Group1289: acar|OOF97655.1 acar|OOF97702.1 abra|OJJ71502.1 acla|EAW10046.1 afis|EAW25538.1 afla|EED44714.1 afum|EAL90201.1 akaw|GAA82518.1 aluc|OJZ87045.1 anid|CBF77822.1 anig|EHA20318.1 aory|EIT83463.1 asyd|OJJ53007.1 ater|EAU33379.1 atub|OJI81590.1 aver|OJJ03836.1 awen|OJJ41280.1 azon|OJJ49503.1

Group1290: acar|OOF97659.1 acar|OOF97706.1 abra|OJJ71498.1 acla|EAW10051.1 afis|EAW25533.1 afla|EED44708.1 afum|EAL90197.1 akaw|GAA82522.1 aluc|OJZ87049.1 anid|CBF77815.1 anig|EHA20322.1 aory|EIT83464.1 asyd|OJJ53011.1 ater|EAU33375.1 atub|OJI81586.1 aver|OJJ03832.1 awen|OJJ41275.1 azon|OJJ49507.1

Group1291: acar|OOF97662.1 acar|OOF97709.1 abra|OJJ71495.1 acla|EAW10054.1 afis|EAW25530.1 afla|EED54031.1 afum|EBA27397.1 agla|OJJ87272.1 akaw|GAA82525.1 aluc|OJZ87052.1 anid|CBF77809.1 anig|EHA20325.1 aory|EIT78310.1 asyd|OJJ53014.1 atub|OJI81583.1 aver|OJJ03829.1 awen|OJJ41272.1 azon|OJJ49504.1

Group1292: acar|OOF97667.1 acar|OOF97714.1 abra|OJJ71490.1 acla|EAW10060.1 afis|EAW25524.1 afla|EED54027.1 afum|EAL90189.1 agla|OJJ87277.1 akaw|GAA82530.1 aluc|OJZ87057.1 anig|EHA20330.1 aory|EIT78322.1 asyd|OJJ53020.1 ater|EAU33369.1 atub|OJI81578.1 aver|OJJ03823.1 awen|OJJ41268.1 azon|OJJ49513.1

Group1293: acar|OOF97670.1 acar|OOF97717.1 abra|OJJ71487.1 acla|EAW10063.1 afis|EAW25522.1 afla|EED54024.1 afum|EAL90187.1 agla|OJJ87280.1 akaw|GAA82533.1 aluc|OJZ87060.1 anid|CBF77798.1 anig|EHA20333.1 aory|EIT78336.1 asyd|OJJ53023.1 atub|OJI81575.1 aver|OJJ03820.1 awen|OJJ41265.1 azon|OJJ44830.1

Group1294: acar|OOF97677.1 acar|OOF97725.1 abra|OJJ71475.1 acla|EAW10075.1 afis|EAW25511.1 afla|EED54012.1 afum|EAL90177.1 agla|OJJ87294.1 akaw|GAA82541.1 aluc|OJZ87071.1 anid|CBF77777.1 aory|EIT78286.1 asyd|OJJ53033.1 ater|EAU33358.1 atub|OJI81566.1 aver|OJJ03811.1 awen|OJJ41253.1 azon|OJJ44820.1

Group1295: acar|OOF97683.1 acar|OOF97732.1 abra|OJJ71466.1 acla|EAW10084.1 afis|EAW25503.1 afla|EED54002.1 afum|EAL90169.2 agla|OJJ87301.1 akaw|GAA82548.1 aluc|OJZ87078.1 anid|CBF77762.1 anig|EHA20348.1 aory|EIT78296.1 asyd|OJJ53041.1 ater|EAU33351.1 aver|OJJ03803.1 awen|OJJ41245.1 azon|OJJ44811.1

Group1296: acar|OOF97696.1 acar|OOF97746.1 ater|EAU30101.1 ater|EAU30103.1 abra|OJJ71449.1 afla|EED53988.1 agla|OJJ87138.1 akaw|GAA82561.1 aluc|OJZ87092.1 anid|CBF77728.1 anig|EHA20364.1 aory|EIT78348.1 asyd|OJJ53059.1 atub|OJI81546.1 aver|OJJ03785.1 awen|OJJ41230.1 acla|EAW14376.1 aver|OJI99265.1

Group1297: acar|OOF98470.1 acar|OOF98485.1 abra|OJJ78403.1 acla|EAW14650.1 afis|EAW17292.1 afla|EED46742.1 agla|OJJ84080.1 akaw|GAA85628.1 aluc|OJZ91482.1 anid|CBF82629.1 anig|EHA20144.1 aory|EIT77955.1 asyd|OJJ54010.1 ater|EAU31138.1 atub|OJI83475.1 aver|OJJ08732.1 awen|OJJ34313.1 azon|OJJ43329.1

Group1298: acar|OOF98474.1 acar|OOF98488.1 abra|OJJ78400.1 acla|EAW09132.1 afis|EAW22014.1 afla|EED46739.1 afum|EAL93029.1 agla|OJJ84083.1 akaw|GAA85631.1 aluc|OJZ91479.1 anid|CBF82642.1 anig|EHA20141.1 aory|EIT77943.1 asyd|OJJ61677.1 atub|OJI83472.1 aver|OJJ00998.1 awen|OJJ34310.1 azon|OJJ48063.1

Group1299: acar|OOF98480.1 acar|OOF98493.1 abra|OJJ78394.1 acla|EAW09138.1 afis|EAW22009.1 afla|EED46733.1 afum|EAL93024.1 agla|OJJ84089.1 akaw|GAA85637.1 aluc|OJZ91473.1 anid|CBF82653.1 anig|EHA20135.1 aory|EIT77921.1 asyd|OJJ61671.1 atub|OJI83466.1 aver|OJJ00992.1 awen|OJJ34304.1 azon|OJJ48064.1

Group1300: acar|OOG00033.1 acar|OOG00123.1 abra|OJJ75863.1 acla|EAW14825.1 afis|EAW24354.1 afla|EED47909.1 afum|EAL86025.2 agla|OJJ83234.1 akaw|GAA84569.1 aluc|OJZ89305.1 anig|EHA23261.1 aory|EIT83374.1 asyd|OJJ53696.1 ater|EAU30389.1 atub|OJI85494.1 aver|OJJ07812.1 awen|OJJ32871.1 azon|OJJ48979.1

Group1301: acla|EAW14649.1 afis|EAW22112.1 afis|EAW24562.1 afum|EAL86223.1 asyd|OJJ54831.1 asyd|OJJ65331.1 aver|OJJ07476.1 afla|EED54620.1 agla|OJJ78532.1 anid|CBF80350.1 aory|EIT78818.1 ater|EAU31010.1 ater|EAU33401.1 ater|EAU36517.1 aver|OJJ07330.1 acla|EAW15031.1 aver|OJI99476.1 afis|EAW21218.1

Group1302: acla|EAW08761.1 acla|EAW12263.1 abra|OJJ69061.1 acar|OOF91222.1 afis|EAW15671.1 afla|EED46862.1 afum|EAL85676.2 akaw|GAA90591.1 aluc|OJZ86049.1 anid|CBF71172.1 anig|EHA24562.1 aory|EIT72618.1 asyd|OJJ55852.1 ater|EAU32490.1 atub|OJI79846.1 aver|OJJ04795.1 awen|OJJ38348.1 azon|OJJ43869.1

Group1303: acla|EAW09069.1 acla|EAW11073.1 afla|EED55166.1 afla|EED57249.1 anid|CBF83136.1 anid|CBF84254.1 aory|EIT74130.1 aory|EIT81645.1 asyd|OJJ55425.1 asyd|OJJ57203.1 ater|EAU32710.1 ater|EAU33927.1 aver|OJJ03086.1 aver|OJJ06504.1 afis|EAW25639.1 afum|EAL84326.1 agla|OJJ78892.1 awen|OJJ36107.1

Group1304: acla|EAW09129.1 acla|EAW11451.1 aory|EIT74374.1 abra|OJJ77018.1 acar|OOF95335.1 afla|EED56545.1 akaw|GAA90906.1 aluc|OJZ82920.1 atub|OJI89146.1 awen|OJJ36105.1 acar|OOF91778.1 afla|EED52330.1 agla|OJJ85629.1 ater|EAU39199.1 awen|OJJ32501.1 azon|OJJ46534.1 aory|EIT79688.1 anid|CBF87732.1

Group1305: afis|EAW16696.1 afis|EAW24556.1 afum|EAL86220.1 afum|EAL89293.1 abra|OJJ70132.1 acar|OOF96170.1 afla|EED46901.1 agla|OJJ88675.1 akaw|GAA86632.1 aluc|OJZ85189.1 anid|CBF81791.1 anig|EHA25402.1 aory|EIT72585.1 asyd|OJJ60097.1 ater|EAU32525.1 atub|OJI80408.1 aver|OJJ04183.1 awen|OJJ36331.1

Group1306: afis|EAW17245.1 afis|EAW25661.1 abra|OJJ68041.1 acar|OOF90403.1 acla|EAW12126.1 afla|EED56768.1 afum|EAL86664.2 akaw|GAA90912.1 aluc|OJZ82914.1 anid|CBF84467.1 anig|EHA21337.1 aory|EIT73930.1 asyd|OJJ53543.1 atub|OJI89140.1 aver|OJJ08210.1 awen|OJJ40087.1 ater|EAU38594.1 aver|OJJ03052.1

Group1307: afla|EED44734.1 afla|EED49486.1 agla|OJJ78680.1 agla|OJJ85924.1 abra|OJJ66288.1 aory|EIT79948.1 aver|OJJ07505.1 abra|OJJ77218.1 akaw|GAA90720.1 aluc|OJZ84629.1 atub|OJI89450.1 afis|EAW23523.1 afla|EED47473.1 azon|OJJ42569.1 aory|EIT77798.1 ater|EAU36203.1 aver|OJJ07255.1 anid|CBF78331.1

Group1308: afla|EED47371.1 afla|EED50262.1 afla|EED50950.1 ater|EAU30176.1 ater|EAU32383.1 ater|EAU38563.1 afis|EAW17332.1 agla|OJJ84032.1 anid|CBF85356.1 afum|EAL91169.1 aory|EIT76964.1 asyd|OJJ62637.1 asyd|OJJ62661.1 aver|OJI96759.1 aver|OJI99292.1 awen|OJJ34422.1 aory|EIT82881.1 anid|CBF89127.1

Group1309: afla|EED50259.1 afla|EED52529.1 aory|EIT74308.1 aory|EIT77339.1 abra|OJJ66540.1 acar|OOF90993.1 acla|EAW10950.1 afis|EAW20911.1 afum|EAL84723.1 akaw|GAA91175.1 aluc|OJZ81504.1 anid|CBF82490.1 anig|EHA25926.1 asyd|OJJ54107.1 ater|EAU38544.1 atub|OJI88677.1 aver|OJJ08628.1 awen|OJJ34431.1

Group1310: agla|OJJ78539.1 agla|OJJ78727.1 abra|OJJ72757.1 acar|OOF95978.1 acla|EAW11507.1 afla|EED52511.1 afum|EAL91104.2 akaw|GAA85367.1 aluc|OJZ90731.1 anid|CBF77292.1 anig|EHA24147.1 aory|EIT82146.1 asyd|OJJ56788.1 ater|EAU31605.1 atub|OJI82157.1 aver|OJJ03548.1 awen|OJJ39008.1 aory|EIT82761.1

Group1311: agla|OJJ78573.1 agla|OJJ78791.1 asyd|OJJ53623.1 asyd|OJJ55815.1 awen|OJJ34401.1 awen|OJJ38082.1 afla|EED55668.1 aory|EIT77088.1 asyd|OJJ64785.1 aver|OJI98897.1 anid|CBF79150.1 akaw|GAA92796.1 aluc|OJZ88717.1 anig|EHA28200.1 anid|CBF85279.1 asyd|OJJ62636.1 aver|OJJ05940.1 asyd|OJJ61766.1

Group1312: agla|OJJ78989.1 agla|OJJ83687.1 abra|OJJ73813.1 acar|OOG00968.1 acla|EAW07700.1 afis|EAW22717.1 afla|EED50003.1 afum|EAL90477.1 akaw|GAA89432.1 aluc|OJZ88237.1 anid|CBF88050.1 anig|EHA17984.1 asyd|OJJ63539.1 ater|EAU38986.1 atub|OJI90839.1 aver|OJI97394.1 awen|OJJ35411.1 azon|OJJ49765.1

Group1313: agla|OJJ80769.1 agla|OJJ80770.1 abra|OJJ72336.1 acar|OOF94879.1 acla|EAW11986.1 afis|EAW19626.1 afla|EED56978.1 afum|EAL86895.1 akaw|GAA86088.1 aluc|OJZ90317.1 anid|CBF71317.1 asyd|OJJ55112.1 ater|EAU32900.1 atub|OJI81745.1 aver|OJJ07094.1 awen|OJJ41873.1 azon|OJJ47068.1 aory|EIT77552.1

Group1314: agla|OJJ80830.1 agla|OJJ84729.1 awen|OJJ32928.1 awen|OJJ33337.1 abra|OJJ74236.1 afla|EED48423.1 aluc|OJZ89898.1 anig|EHA27098.1 aory|EIT80749.1 atub|OJI84909.1 acla|EAW13274.1 afis|EAW17343.1 afum|EBA27425.1 anid|CBF71489.1 asyd|OJJ51916.1 afla|EED51567.1 aory|EIT83078.1 ater|EAU30102.1

Group1315: agla|OJJ81338.1 agla|OJJ84780.1 awen|OJJ36016.1 awen|OJJ38019.1 abra|OJJ73446.1 acar|OOF98836.1 afis|EAW24692.1 afla|EED45076.1 afum|EAL89334.1 akaw|GAA89024.1 aluc|OJZ80580.1 anid|CBF76032.1 anig|EHA25618.1 aory|EIT73772.1 asyd|OJJ52453.1 ater|EAU33313.1 atub|OJI88350.1 aver|OJJ03963.1

Group1316: agla|OJJ83725.1 agla|OJJ88209.1 awen|OJJ30544.1 awen|OJJ37831.1 azon|OJJ42347.1 azon|OJJ45910.1 ater|EAU34086.1 acar|OOF92105.1 aver|OJJ07715.1 acar|OOF99481.1 afis|EAW19566.1

afis|EAW19382.1 aluc|OJZ85537.1 ater|EAU36043.1 afis|EAW19567.1 anig|EHA22173.1 aver|OJJ07008.1 asyd|OJJ55209.1

Group1317: agla|OJJ84608.1 agla|OJJ88488.1 abra|OJJ68915.1 acar|OOF91503.1 acla|EAW14256.1 afis|EAW19024.1 afla|EED56627.1 afum|EAL93755.1 akaw|GAA90334.1 aluc|OJZ85898.1 anid|CBF73887.1 anig|EHA24694.1 aory|EIT79613.1 asyd|OJJ53658.1 ater|EAU32066.1 atub|OJI79683.1 aver|OJJ07766.1 awen|OJJ31397.1

Group1318: akaw|GAA84492.1 akaw|GAA92643.1 abra|OJJ68086.1 acar|OOF93885.1 afis|EAW23619.1 afum|EAL87843.1 akaw|GAA86871.1 aluc|OJZ89995.1 anig|EHA27365.1 asyd|OJJ55804.1 atub|OJI84813.1 aver|OJJ03908.1 awen|OJJ33610.1 abra|OJJ75773.1 afla|EED51751.1 aory|EIT80721.1 ater|EAU34085.1 azon|OJJ50739.1

Group1319: akaw|GAA88994.1 akaw|GAA93237.1 aluc|OJZ79544.1 aluc|OJZ80344.1 abra|OJJ70263.1 acar|OOF91460.1 afla|EED57905.1 agla|OJJ83807.1 akaw|GAA86534.1 aluc|OJZ85082.1 anid|CBF78690.1 aory|EIT82332.1 asyd|OJJ53892.1 ater|EAU33679.1 atub|OJI80517.1 aver|OJJ07990.1 afla|EED57907.1 aory|EIT82357.1

Group1320: akaw|GAA89097.1 akaw|GAA93124.1 abra|OJJ70832.1 acla|EAW13725.1 afla|EED45954.1 agla|OJJ89628.1 aluc|OJZ83078.1 anig|EHA20972.1 aory|EIT73606.1 asyd|OJJ52723.1 atub|OJI87867.1 aver|OJJ04315.1 awen|OJJ32035.1 azon|OJJ46438.1 awen|OJJ34430.1 akaw|GAA90262.1 aory|EIT75383.1 awen|OJJ34591.1

Group1321: aluc|OJZ85535.1 aluc|OJZ86787.1 anig|EHA19890.1 anig|EHA21895.1 atub|OJI83732.1 atub|OJI89790.1 abra|OJJ69763.1 ater|EAU29245.1 abra|OJJ72050.1 acar|OOF96707.1 afla|EED57904.1 agla|OJJ78987.1 akaw|GAA84884.1 aory|EIT82360.1 asyd|OJJ61024.1 aver|OJJ01910.1 awen|OJJ33896.1 azon|OJJ45975.1

Group1322: anid|CBF69554.1 anid|CBF74620.1 asyd|OJJ57601.1 asyd|OJJ59165.1 aver|OJJ01755.1 aver|OJJ05583.1 abra|OJJ77468.1 acar|OOF99266.1 acla|EAW07128.1 afis|EAW22137.1 afla|EED54177.1 afum|EAL91053.1 akaw|GAA84149.1 aluc|OJZ81070.1 anig|EHA19299.1 ater|EAU29324.1 atub|OJI82516.1 awen|OJJ34710.1

Group1323: anid|CBF71205.1 anid|CBF74966.1 anid|CBF89272.1 abra|OJJ69173.1 acar|OOF91131.1 afla|EED55558.1 agla|OJJ84719.1 akaw|GAA86289.1 aluc|OJZ86160.1 anig|EHA24464.1 asyd|OJJ61621.1 ater|EAU32677.1 aver|OJJ00945.1 awen|OJJ38097.1 azon|OJJ42541.1 aory|EIT82704.1 atub|OJI79952.1 ater|EAU34906.1

Group1324: anid|CBF77312.1 anid|CBF84491.1 abra|OJJ72772.1 acar|OOF95991.1 afis|EAW23959.1 afla|EED50553.1 afum|EAL87388.1 agla|OJJ86884.1 akaw|GAA85381.1 aluc|OJZ90749.1 anig|EHA24160.1 aory|EIT82046.1 asyd|OJJ56774.1 ater|EAU29793.1 atub|OJI82175.1 aver|OJJ03559.1 awen|OJJ38988.1 azon|OJJ51320.1

Group1325: anid|CBF87056.1 anid|CBF87058.1 abra|OJJ66247.1 acar|OOF90705.1 acla|EAW07050.1 afis|EAW17102.1 afla|EED51639.1 afum|EAL89282.1 akaw|GAA89043.1 aluc|OJZ80553.1 anig|EHA25644.1 aory|EIT80540.1 atub|OJI88378.1 aver|OJI99751.1 awen|OJJ37983.1 afis|EAW16677.1 afla|EED47132.1 aory|EIT79173.1

Group1326: anig|EHA19581.1 anig|EHA25318.1 awen|OJJ33562.1 awen|OJJ34654.1 abra|OJJ70237.1 acar|OOF92557.1 afis|EAW21623.1 afla|EED53598.1 afum|EAL92659.1 agla|OJJ88653.1 akaw|GAA86548.1 aluc|OJZ85096.1 anid|CBF79555.1 aory|EIT74955.1 asyd|OJJ57931.1 atub|OJI80502.1 aver|OJJ04466.1 azon|OJJ45043.1

Group1327: anig|EHA20767.1 anig|EHA21813.1 atub|OJI81082.1 abra|OJJ70967.1 acar|OOF98164.1 acla|EAW10527.1 afis|EAW25039.1 afla|EED55413.1 afum|EAL89707.1 agla|OJJ89154.1 akaw|GAA82981.1 aluc|OJZ87565.1 anid|CBF85903.1 aory|EIT80777.1 asyd|OJJ64393.1 ater|EAU35789.1 aver|OJI98488.1 awen|OJJ40645.1

Group1328: anig|EHA20833.1 afum|EAL89767.1 abra|OJJ70883.1 acar|OOF98234.1 acla|EAW10471.1 afis|EAW25096.1 afla|EED55352.1 agla|OJJ89081.1 akaw|GAA87402.1 aluc|OJZ87638.1 anid|CBF86008.1 aory|EIT80853.1 asyd|OJJ64446.1 ater|EAU39354.1 atub|OJI81009.1 aver|OJI98549.1 awen|OJJ40710.1 azon|OJJ44086.1

Group1329: anig|EHA21819.1 anig|EHA24599.1 abra|OJJ69025.1 acar|OOF92661.1 acla|EAW07121.1 afis|EAW22131.1 afla|EED51689.1 afum|EAL91059.1 akaw|GAA90542.1 aluc|OJZ86012.1 aory|EIT80445.1 ater|EAU33620.1 atub|OJI79801.1 awen|OJJ32638.1 azon|OJJ42095.1 afla|EED56398.1 aory|EIT83004.1 aver|OJJ06436.1

Group1330: anig|EHA25931.1 anig|EHA28231.1 abra|OJJ66546.1 acla|EAW07078.1 afla|EED45458.1 akaw|GAA91171.1 aluc|OJZ81499.1 aory|EIT82415.1 ater|EAU30796.1 atub|OJI88682.1 aver|OJJ06470.1 akaw|GAA92427.1 anig|EHA25846.1 atub|OJI88588.1 acar|OOF90934.1 anid|CBF82815.1 asyd|OJJ61574.1 aver|OJJ00895.1

Group1331: aory|EIT72308.1 aory|EIT72311.1 abra|OJJ69636.1 acar|OOF92673.1 acla|EAW10864.1 afis|EAW20994.1 afum|EAL84634.1 agla|OJJ84072.1 akaw|GAA91616.1 aluc|OJZ82442.1 anid|CBF82637.1 anig|EHA28331.1 asyd|OJJ54015.1 ater|EAU38643.1 atub|OJI87998.1 aver|OJJ08726.1 awen|OJJ34316.1 azon|OJJ48058.1

Group1332: aory|EIT72309.1 aory|EIT72310.1 abra|OJJ66632.1 acar|OOF91054.1 acla|EAW10863.1 afis|EAW20995.1 afum|EAL84633.1 agla|OJJ84073.1 akaw|GAA91095.1 aluc|OJZ81416.1 anid|CBF82633.1 anig|EHA26004.1 asyd|OJJ54013.1 ater|EAU38644.1 atub|OJI88764.1 aver|OJJ08728.1 awen|OJJ34315.1 azon|OJJ48060.1

Group1333: asyd|OJJ52021.1 asyd|OJJ52275.1 abra|OJJ73341.1 acar|OOG00528.1 acla|EAW13716.1 afis|EAW19870.1 afla|EED53094.1 afum|EAL85027.1 akaw|GAA83141.1 aluc|OJZ88698.1 anid|CBF81882.1 anig|EHA18378.1 aory|EIT74789.1 asyd|OJJ59668.1 ater|EAU30732.1 atub|OJI91279.1 aver|OJJ02615.1 ater|EAU36648.1

Group1334: asyd|OJJ52762.1 asyd|OJJ62749.1 abra|OJJ73605.1 acar|OOG00789.1 acla|EAW07522.1 afis|EAW22552.1 afla|EED54584.1 afum|EAL90643.1 agla|OJJ80914.1 akaw|GAA88548.1 aluc|OJZ88444.1 anid|CBF89060.1 anig|EHA18168.1 aory|EIT78947.1 ater|EAU39152.1 atub|OJI91042.1 aver|OJI96789.1 awen|OJJ35183.1

Group1335: asyd|OJJ54165.1 asyd|OJJ59819.1 aver|OJJ02469.1 aver|OJJ08576.1 abra|OJJ76687.1 acar|OOF99325.1 acla|EAW12040.1 afis|EAW19689.1 afla|EED56767.1 afum|EAL86837.2 akaw|GAA92507.1 aluc|OJZ91520.1 anid|CBF75398.1 anig|EHA22558.1 aory|EIT73885.1 ater|EAU37774.1 atub|OJI85975.1 awen|OJJ31435.1

Group1336: asyd|OJJ54481.1 asyd|OJJ64266.1 abra|OJJ70844.1 acar|OOF98271.1 acla|EAW10435.1 afis|EAW25133.1 afla|EED55313.1 afum|EAL89804.1 agla|OJJ89034.1 akaw|GAA91712.1 aluc|OJZ87677.1 anid|CBF85703.1 anig|EHA20867.1 aory|EIT83632.1 ater|EAU39404.1 atub|OJI80966.1 aver|OJI98358.1 awen|OJJ40757.1

Group1337: asyd|OJJ55684.1 asyd|OJJ56117.1 aver|OJJ05058.1 aver|OJJ06768.1 abra|OJJ76907.1 acar|OOF91355.1 afis|EAW20540.1 afla|EED52439.1 afum|EAL84518.2 akaw|GAA87131.1 aluc|OJZ84306.1 anid|CBF78991.1 anig|EHA27642.1 aory|EIT82783.1 ater|EAU37140.1 atub|OJI89762.1 awen|OJJ33246.1 azon|OJJ50626.1

Group1338: asyd|OJJ56868.1 asyd|OJJ60506.1 aver|OJI99831.1 aver|OJJ03458.1 aver|OJJ05527.1 abra|OJJ65536.1 afis|EAW19680.1 afum|EAL86843.1 aluc|OJZ79939.1 anig|EHA19938.1 abra|OJJ65772.1 atub|OJI89174.1 awen|OJJ39572.1 awen|OJJ39573.1 ater|EAU37595.1 anid|CBF80809.1 aory|EIT76448.1 ater|EAU39331.1

Group1339: asyd|OJJ58286.1 asyd|OJJ62494.1 abra|OJJ69000.1 acar|OOF92712.1 afis|EAW18028.1 afla|EED51869.1 afum|EAL91839.1 agla|OJJ85399.1 akaw|GAA90519.1 aluc|OJZ85984.1 anid|CBF80060.1 anig|EHA24622.1 aory|EIT81740.1 ater|EAU31540.1 atub|OJI79774.1 aver|OJJ02138.1 awen|OJJ41513.1 azon|OJJ48929.1

Group1340: asyd|OJJ60628.1 abra|OJJ75954.1 acar|OOF99958.1 acla|EAW09842.1 afis|EAW21293.1 afla|EED54845.1 afum|EAL92310.1 agla|OJJ87460.1 akaw|GAA84657.1 aluc|OJZ89403.1 anid|CBF83378.1 anig|EHA23184.1 aory|EIT79413.1 ater|EAU35926.1 atub|OJI85394.1 aver|OJI99952.1 awen|OJJ39406.1 azon|OJJ44289.1

Group1341: ater|EAU29424.1 ater|EAU33846.1 abra|OJJ77281.1 acar|OOF93155.1 acla|EAW13280.1 afis|EAW17337.1 afla|EED51045.1 afum|EAL91181.1 agla|OJJ79349.1 akaw|GAA90668.1 aluc|OJZ84691.1 anid|CBF87334.1 anig|EHA27966.1 aory|EIT77013.1 asyd|OJJ56362.1 atub|OJI89391.1 aver|OJJ06352.1 azon|OJJ44714.1

Group1342: ater|EAU29807.1 ater|EAU31871.1 abra|OJJ77150.1 acar|OOF93036.1 acla|EAW12919.1 afis|EAW17736.1 afla|EED45737.1 afum|EAL91551.1 agla|OJJ81714.1 akaw|GAA90780.1 aluc|OJZ84564.1 anid|CBF86592.1 anig|EHA27858.1 asyd|OJJ64830.1 atub|OJI89518.1 aver|OJI98943.1 awen|OJJ37099.1 azon|OJJ50800.1

Group1343: ater|EAU31940.1 ater|EAU38199.1 abra|OJJ76424.1 acar|OOF99668.1 acla|EAW14001.1 afis|EAW18709.1 afla|EED50658.1 afum|EAL93432.1 akaw|GAA82339.1 aluc|OJZ91788.1 anid|CBF70654.1 anig|EHA22785.1 aory|EIT78023.1 asyd|OJJ58483.1 atub|OJI86244.1 aver|OJJ01086.1 awen|OJJ32192.1 azon|OJJ46757.1

Group1344: ater|EAU35144.1 ater|EAU37867.1 abra|OJJ77032.1 acar|OOF91470.1 acla|EAW13563.1 afis|EAW20685.1 afla|EED49742.1 afum|EAL84413.1 agla|OJJ80325.1 akaw|GAA87019.1 aluc|OJZ84435.1 anid|CBF83195.1 anig|EHA27754.1 aory|EIT77691.1 asyd|OJJ55559.1 atub|OJI89638.1 aver|OJJ06640.1 awen|OJJ34542.1

Group1345: ater|EAU35442.1 ater|EAU37095.1 abra|OJJ76968.1 acar|OOF91413.1 acla|EAW13484.1 afis|EAW20592.1 afla|EED52233.1 afum|EAL84469.1 agla|OJJ79954.1 akaw|GAA87076.1 aluc|OJZ84364.1

anid|CBF78915.1 anig|EHA27699.1 asyd|OJJ55617.1 ater|EAU35238.1 atub|OJI89704.1 aver|OJJ06692.1 awen|OJJ33552.1

Group1346: ater|EAU35971.1 ater|EAU38852.1 abra|OJJ76019.1 acar|OOF99906.1 acla|EAW09787.1 afis|EAW21351.1 afla|EED55763.1 afum|EAL92366.1 agla|OJJ80812.1 akaw|GAA88941.1 aluc|OJZ89466.1 anid|CBF76199.1 anig|EHA23122.1 aory|EIT76697.1 asyd|OJJ60381.1 atub|OJI85329.1 aver|OJI99710.1 awen|OJJ39349.1

Group1347: aver|OJI97203.1 aver|OJJ08497.1 abra|OJJ74529.1 acar|OOF95222.1 acla|EAW07163.1 afis|EAW22178.1 afla|EED57557.1 afum|EAL91011.1 agla|OJJ87877.1 akaw|GAA82071.1 aluc|OJZ92111.1 anid|CBF88382.1 aory|EIT81848.1 asyd|OJJ63737.1 ater|EAU34183.1 atub|OJI86544.1 awen|OJJ34758.1 azon|OJJ45803.1

Group1348: awen|OJJ29894.1 awen|OJJ38081.1 abra|OJJ76942.1 akaw|GAA87098.1 aluc|OJZ84340.1 anig|EHA27674.1 atub|OJI89728.1 acla|EAW12080.1 afis|EAW19721.1 agla|OJJ86006.1 aver|OJI96480.1 acar|OOF91391.1 afla|EED57523.1 afum|EAL86805.2 anig|EHA19572.1 ater|EAU30973.1 atub|OJI83020.1 ater|EAU34067.1

Group1349: awen|OJJ30757.1 awen|OJJ31724.1 abra|OJJ67453.1 acar|OOF92380.1 acla|EAW06312.1 afis|EAW18421.1 afla|EED52987.1 afum|EAL93105.1 akaw|GAA86378.1 aluc|OJZ84912.1 anid|CBF70698.1 anig|EHA25133.1 aory|EIT78706.1 asyd|OJJ58462.1 ater|EAU29839.1 atub|OJI80686.1 aver|OJJ01062.1 azon|OJJ50486.1

Group1350: awen|OJJ30769.1 awen|OJJ31712.1 abra|OJJ67441.1 acar|OOF92385.1 acla|EAW06322.1 afis|EAW18436.1 afla|EED52979.1 afum|EAL93117.1 agla|OJJ85003.1 akaw|GAA86388.1 aluc|OJZ84923.1 anid|CBF70035.1 anig|EHA25141.1 asyd|OJJ58867.1 ater|EAU29852.1 atub|OJI80676.1 aver|OJJ01469.1 azon|OJJ50477.1

Group1351: awen|OJJ30784.1 awen|OJJ31697.1 abra|OJJ77412.1 acar|OOF93279.1 acla|EAW08581.1 afis|EAW17309.1 afla|EED48984.1 afum|EAL91144.1 agla|OJJ85394.1 akaw|GAA91352.1 aluc|OJZ84821.1 anid|CBF89993.1 anig|EHA28083.1 asyd|OJJ62080.1 ater|EAU35447.1 atub|OJI89265.1 aver|OJI96188.1 azon|OJJ44474.1

Group1352: awen|OJJ30785.1 awen|OJJ31696.1 abra|OJJ67989.1 acar|OOF92031.1 afis|EAW24766.1 afla|EED47575.1 afum|EAL89408.2 agla|OJJ88067.1 akaw|GAA90959.1 aluc|OJZ82863.1 anid|CBF90079.1 anig|EHA21293.1 aory|EIT74521.1 asyd|OJJ62039.1 ater|EAU29463.1 atub|OJI89088.1 aver|OJI96145.1 azon|OJJ46302.1

Group1353: awen|OJJ32597.1 awen|OJJ38114.1 abra|OJJ77103.1 acar|OOF92988.1 afla|EED45185.1 akaw|GAA86953.1 aluc|OJZ84512.1 anid|CBF81557.1 anig|EHA27810.1 aory|EIT73710.1 asyd|OJJ59498.1 ater|EAU38093.1 atub|OJI89569.1 aver|OJJ02776.1 azon|OJJ42741.1 afla|EED47336.1 aory|EIT82859.1 aver|OJJ03900.1

Group1354: awen|OJJ32803.1 awen|OJJ35367.1 abra|OJJ75800.1 acar|OOG00076.1 acla|EAW14886.1 afis|EAW24405.1 afla|EED47807.1 afum|EAL86076.1 agla|OJJ78882.1 akaw|GAA84516.1 aluc|OJZ89248.1 anid|CBF75173.1 anig|EHA23319.1 aory|EIT77184.1 asyd|OJJ59956.1 ater|EAU30320.1 atub|OJI85550.1 aver|OJJ02333.1

Group1355: awen|OJJ34458.1 awen|OJJ34686.1 abra|OJJ66071.1 acar|OOG00583.1 acla|EAW08522.1 afis|EAW16392.1 afla|EED52127.1 afum|EAL89031.1 akaw|GAA84173.1 aluc|OJZ81098.1 anid|CBF81685.1 anig|EHA19273.1 aory|EIT74371.1 asyd|OJJ62625.1 ater|EAU29596.1 atub|OJI82482.1 aver|OJI96704.1 azon|OJJ43880.1

Group1356: azon|OJJ42761.1 azon|OJJ42890.1 azon|OJJ42564.1 azon|OJJ50002.1 azon|OJJ44488.1 azon|OJJ42513.1 azon|OJJ43493.1 azon|OJJ42351.1 azon|OJJ42537.1 azon|OJJ48473.1 azon|OJJ43882.1 azon|OJJ44466.1 azon|OJJ51813.1 azon|OJJ42495.1 azon|OJJ46515.1 afla|EED48610.1 afla|EED57034.1 aory|EIT82256.1

Group1357: azon|OJJ42597.1 azon|OJJ49266.1 abra|OJJ70118.1 acla|EAW12241.1 afis|EAW21131.1 agla|OJJ80450.1 asyd|OJJ55221.1 atub|OJI80395.1 aver|OJI97990.1 awen|OJJ37684.1 asyd|OJJ61887.1 afum|EAL84504.1 agla|OJJ80675.1 afis|EAW20557.1 awen|OJJ35176.1 azon|OJJ42354.1 asyd|OJJ59262.1 anig|EHA25411.1

Group1358: azon|OJJ43518.1 azon|OJJ49486.1 abra|OJJ66404.1 acla|EAW12367.1 afis|EAW20845.1 agla|OJJ86411.1 akaw|GAA90272.1 aluc|OJZ83069.1 anid|CBF78174.1 anig|EHA28133.1 asyd|OJJ55965.1 atub|OJI87877.1 aver|OJJ04911.1 awen|OJJ36259.1 abra|OJJ68228.1 afum|EAL84788.1 acar|OOG00612.1 azon|OJJ45925.1

Group1359: azon|OJJ44012.1 azon|OJJ45468.1 abra|OJJ69814.1 acar|OOF93561.1 acla|EAW11673.1 afis|EAW23690.1 afla|EED51830.1 afum|EAL87135.1 agla|OJJ79265.1 akaw|GAA87843.1 aluc|OJZ85481.1 anid|CBF80383.1 anig|EHA25021.1 aory|EIT81761.1 asyd|OJJ54786.1 atub|OJI80083.1 aver|OJJ07379.1 awen|OJJ38091.1

Group1360: abra|OJJ65585.1 afum|EBA27422.1 abra|OJJ65589.1 asyd|OJJ55957.1 abra|OJJ68807.1 afla|EED46563.1 aluc|OJZ85779.1 anid|CBF70722.1 aory|EIT77899.1 atub|OJI79564.1 awen|OJJ35847.1 acla|EAW13291.1 afis|EAW17318.1 afum|EAL91154.1 azon|OJJ50647.1 aver|OJJ04903.1 afum|EAL89333.1 azon|OJJ50595.1

Group1361: abra|OJJ65623.1 acar|OOF97131.1 acla|EAW12840.1 afum|EAL87789.1 agla|OJJ86050.1 akaw|GAA92893.1 anid|CBF78703.1 asyd|OJJ62680.1 ater|EAU36754.1 awen|OJJ36388.1 azon|OJJ43355.1 afla|EED54881.1 aory|EIT79373.1 acar|OOF98798.1 acla|EAW11601.1 aluc|OJZ85600.1 atub|OJI79395.1 anig|EHA24924.1

Group1362: abra|OJJ65672.1 afis|EAW16875.1 afum|EAL86412.1 aluc|OJZ79539.1 ater|EAU30978.1 aver|OJJ03435.1 awen|OJJ36144.1 azon|OJJ46815.1 abra|OJJ67798.1 aluc|OJZ90858.1 atub|OJI82852.1 abra|OJJ68250.1 acar|OOF97529.1 akaw|GAA83211.1 aluc|OJZ83713.1 anig|EHA18864.1 atub|OJI90088.1 ater|EAU37972.1

Group1363: abra|OJJ65773.1 acar|OOF92085.1 agla|OJJ81015.1 akaw|GAA90888.1 aluc|OJZ82942.1 anid|CBF86755.1 anig|EHA21361.1 asyd|OJJ61469.1 ater|EAU35421.1 atub|OJI89175.1 aver|OJJ00810.1 awen|OJJ34702.1 afla|EED45830.1 akaw|GAA85168.1 aluc|OJZ86468.1 anig|EHA21442.1 aory|EIT75507.1 azon|OJJ48930.1

Group1364: abra|OJJ65864.1 acar|OOF95036.1 akaw|GAA92281.1 aluc|OJZ90178.1 anid|CBF80753.1 anig|EHA23677.1 asyd|OJJ54523.1 atub|OJI81605.1 aver|OJJ07660.1 abra|OJJ69850.1 acar|OOF93373.1 afla|EED55058.1 akaw|GAA87808.1 aluc|OJZ85443.1 anig|EHA24987.1 aory|EIT81967.1 atub|OJI80124.1 azon|OJJ46475.1

Group1365: abra|OJJ65896.1 acar|OOF94956.1 acla|EAW11696.1 afis|EAW23713.1 afla|EED44871.1 afum|EAL87157.1 agla|OJJ81717.1 akaw|GAA86009.1 aluc|OJZ90235.1 anid|CBF83004.1 anig|EHA23735.1 aory|EIT82717.1 asyd|OJJ61502.1 ater|EAU31796.1 atub|OJI81665.1 aver|OJJ00840.1 awen|OJJ37102.1 azon|OJJ50803.1

Group1366: abra|OJJ65913.1 acar|OOF94964.1 acla|EAW12019.1 afis|EAW19660.1 afla|EED57030.1 afum|EAL86863.1 agla|OJJ80822.1 akaw|GAA92319.1 aluc|OJZ90217.1 anid|CBF71390.1 anig|EHA23715.1 aory|EIT82277.1 asyd|OJJ55042.1 ater|EAU32852.1 atub|OJI81649.1 aver|OJJ07166.1 awen|OJJ41922.1 azon|OJJ42684.1

Group1367: abra|OJJ65945.1 acar|OOF92928.1 acla|EAW13569.1 afis|EAW20032.1 afla|EED46042.1 afum|EAL84884.1 agla|OJJ88019.1 akaw|GAA90823.1 aluc|OJZ83012.1 anid|CBF84764.1 anig|EHA21427.1 aory|EIT75566.1 asyd|OJJ53285.1 ater|EAU29398.1 atub|OJI89247.1 aver|OJJ08456.1 awen|OJJ30055.1 azon|OJJ43701.1

Group1368: abra|OJJ65946.1 acar|OOF92925.1 acla|EAW13570.1 afis|EAW20031.1 afla|EED46041.1 afum|EAL84885.1 agla|OJJ88018.1 akaw|GAA90824.1 aluc|OJZ83011.1 anid|CBF84760.1 anig|EHA21426.1 aory|EIT75583.1 asyd|OJJ53286.1 ater|EAU29395.1 atub|OJI89246.1 aver|OJJ08455.1 awen|OJJ30054.1 azon|OJJ43702.1

Group1369: abra|OJJ65948.1 acar|OOF92924.1 acla|EAW13571.1 afis|EAW20030.1 afla|EED46039.1 afum|EAL84887.1 agla|OJJ88016.1 akaw|GAA90826.1 aluc|OJZ83009.1 anid|CBF84756.1 anig|EHA21424.1 aory|EIT75781.1 asyd|OJJ53288.1 ater|EAU29393.1 atub|OJI89244.1 aver|OJJ08453.1 awen|OJJ30052.1 azon|OJJ43700.1

Group1370: abra|OJJ65949.1 acar|OOF92923.1 acla|EAW13572.1 afis|EAW20029.1 afla|EED46038.1 afum|EAL84888.1 agla|OJJ88015.1 akaw|GAA90827.1 aluc|OJZ83008.1 anid|CBF84754.1 anig|EHA21423.1 aory|EIT75542.1 asyd|OJJ53289.1 ater|EAU29392.1 atub|OJI89243.1 aver|OJJ08452.1 awen|OJJ30051.1 azon|OJJ43698.1

Group1371: abra|OJJ65951.1 acar|OOF92921.1 acla|EAW13573.1 afis|EAW20028.1 afla|EED46037.1 afum|EAL84889.1 agla|OJJ88000.1 akaw|GAA90828.1 aluc|OJZ83006.1 anid|CBF84753.1 anig|EHA21422.1 aory|EIT75829.1 asyd|OJJ53290.1 ater|EAU29391.1 atub|OJI89241.1 aver|OJJ08450.1 awen|OJJ30042.1 azon|OJJ43688.1

Group1372: abra|OJJ65952.1 acar|OOF92920.1 acla|EAW13574.1 afis|EAW20027.1 afla|EED46036.1 afum|EAL84890.1 agla|OJJ88001.1 akaw|GAA90829.1 aluc|OJZ83005.1 anid|CBF84751.1 anig|EHA21421.1 aory|EIT75375.1 asyd|OJJ53291.1 ater|EAU29390.1 atub|OJI89240.1 aver|OJJ08449.1 awen|OJJ30043.1 azon|OJJ43689.1

Group1373: abra|OJJ65954.1 acar|OOF92918.1 acla|EAW13575.1 afis|EAW20026.1 afla|EED46034.1 afum|EAL84891.1 agla|OJJ88004.1 akaw|GAA90831.1 aluc|OJZ83003.1 anid|CBF84749.1 anig|EHA21419.1 aory|EIT75468.1 asyd|OJJ53293.1 ater|EAU29388.1 atub|OJI89237.1 aver|OJJ08447.1 awen|OJJ30045.1 azon|OJJ43691.1

Group1374: abra|OJJ65956.1 acar|OOF92916.1 acla|EAW13577.1 afis|EAW20024.1 afla|EED46032.1 afum|EAL84893.1 agla|OJJ88006.1 akaw|GAA90833.1 aluc|OJZ83001.1 anid|CBF84745.1 anig|EHA21417.1

aory|EIT75584.1 asyd|OJJ53295.1 ater|EAU29386.1 atub|OJI89235.1 aver|OJJ08445.1 awen|OJJ30046.1 azon|OJJ43693.1

Group1375: abra|OJJ65957.1 acar|OOF92915.1 acla|EAW13578.1 afis|EAW20023.1 afla|EED46031.1 afum|EAL84894.1 agla|OJJ88007.1 akaw|GAA90834.1 aluc|OJZ83000.1 anid|CBF84743.1 anig|EHA21416.1 aory|EIT75718.1 asyd|OJJ53296.1 ater|EAU29385.1 atub|OJI89234.1 aver|OJJ08444.1 awen|OJJ30047.1 azon|OJJ43694.1

Group1376: abra|OJJ65958.1 acar|OOF92914.1 acla|EAW13579.1 afis|EAW20022.1 afla|EED46030.1 afum|EAL84895.1 agla|OJJ88011.1 akaw|GAA90835.1 aluc|OJZ82999.1 anid|CBF84741.1 anig|EHA21415.1 aory|EIT75399.1 asyd|OJJ53297.1 ater|EAU29384.1 atub|OJI89233.1 aver|OJJ08443.1 awen|OJJ30048.1 azon|OJJ43695.1

Group1377: abra|OJJ65959.1 acar|OOF92913.1 acla|EAW13580.1 afis|EAW20021.1 afla|EED46029.1 afum|EAL84896.2 agla|OJJ88012.1 akaw|GAA90836.1 aluc|OJZ82998.1 anid|CBF84739.1 anig|EHA21414.1 aory|EIT75790.1 asyd|OJJ53298.1 ater|EAU29383.1 atub|OJI89232.1 aver|OJJ08442.1 awen|OJJ30049.1 azon|OJJ43696.1

Group1378: abra|OJJ65963.1 acar|OOF92911.1 acla|EAW13582.1 afis|EAW20019.1 afla|EED46027.1 afum|EAL84898.1 agla|OJJ87997.1 akaw|GAA90838.1 aluc|OJZ82995.1 anid|CBF84733.1 anig|EHA21410.1 aory|EIT75391.1 asyd|OJJ53302.1 ater|EAU29380.1 atub|OJI89229.1 aver|OJJ08439.1 awen|OJJ30040.1 azon|OJJ43684.1

Group1379: abra|OJJ65969.1 acar|OOF92904.1 acla|EAW13590.1 afis|EAW20013.1 afla|EED46020.1 afum|EAL84905.1 agla|OJJ87978.1 akaw|GAA90844.1 aluc|OJZ82989.1 anid|CBF84715.1 anig|EHA21404.1 aory|EIT75372.1 asyd|OJJ53326.1 ater|EAU29355.1 atub|OJI89223.1 aver|OJJ08417.1 awen|OJJ30027.1 azon|OJJ43661.1

Group1380: abra|OJJ65976.1 acar|OOF92896.1 acla|EAW13597.1 afis|EAW20007.1 afla|EED46013.1 afum|EAL84911.1 agla|OJJ87972.1 akaw|GAA90850.1 aluc|OJZ82983.1 anid|CBF84706.1 anig|EHA21397.1 aory|EIT75456.1 asyd|OJJ53321.1 ater|EAU29349.1 atub|OJI89217.1 aver|OJJ08422.1 awen|OJJ30020.1 azon|OJJ49472.1

Group1381: abra|OJJ65993.1 acar|OOF92879.1 acla|EAW13603.1 afis|EAW15644.1 afla|EED45983.1 agla|OJJ87961.1 akaw|GAA90867.1 aluc|OJZ82966.1 anid|CBF80346.1 anig|EHA21382.1 aory|EIT75657.1 asyd|OJJ54833.1 atub|OJI89199.1 aver|OJJ07328.1 awen|OJJ30009.1 azon|OJJ46985.1 asyd|OJJ52075.1 aver|OJJ03028.1

Group1382: abra|OJJ65998.1 acar|OOF92102.1 akaw|GAA90870.1 aluc|OJZ82963.1 anig|EHA21379.1 atub|OJI89195.1 abra|OJJ66040.1 acla|EAW10004.1 afis|EAW21054.1 afum|EAL92090.1 agla|OJJ81308.1 anid|CBF83224.1 aory|EIT77099.1 asyd|OJJ55572.1 ater|EAU37572.1 aver|OJJ06651.1 awen|OJJ33474.1 afla|EED55697.1

Group1383: abra|OJJ66217.1 acar|OOF89906.1 acla|EAW11713.1 afis|EAW19322.1 afla|EED48416.1 afum|EAL94040.1 agla|OJJ88754.1 akaw|GAA89069.1 aluc|OJZ80523.1 anid|CBF74663.1 anig|EHA25672.1 aory|EIT83055.1 asyd|OJJ54973.1 atub|OJI88407.1 aver|OJJ07218.1 awen|OJJ33251.1 azon|OJJ50629.1 awen|OJJ34601.1

Group1384: abra|OJJ66228.1 acar|OOF90678.1 acla|EAW07009.1 afis|EAW16625.1 afla|EED44976.1 afum|EAL89212.2 akaw|GAA89061.1 aluc|OJZ80533.1 anig|EHA25662.1 aory|EIT73723.1 asyd|OJJ56749.1 ater|EAU36726.1 atub|OJI88398.1 aver|OJJ05982.1 awen|OJJ40148.1 azon|OJJ42882.1 acar|OOG00595.1 azon|OJJ42590.1

Group1385: abra|OJJ66483.1 acar|OOF90960.1 acla|EAW10986.1 afis|EAW20874.1 afla|EED45556.1 afum|EAL84760.1 agla|OJJ83981.1 akaw|GAA92468.1 aluc|OJZ81551.1 anid|CBF82851.1 anig|EHA25883.1 aory|EIT75688.1 asyd|OJJ61594.1 ater|EAU30815.1 atub|OJI88634.1 aver|OJJ00916.1 awen|OJJ34486.1 azon|OJJ51763.1

Group1386: akaw|GAA92469.1 aluc|OJZ81550.1 atub|OJI88635.1 abra|OJJ66485.1 acar|OOF90962.1 acla|EAW10984.1 afis|EAW20876.1 afla|EED45558.1 afum|EAL84757.1 agla|OJJ83983.1 anid|CBF82855.1 anig|EHA25885.1 aory|EIT75434.1 asyd|OJJ61595.1 ater|EAU30813.1 aver|OJJ00918.1 awen|OJJ34484.1 azon|OJJ51761.1

Group1387: abra|OJJ66488.1 acar|OOF90964.1 acla|EAW10981.1 afis|EAW20878.1 afla|EED45563.1 afum|EAL84754.1 agla|OJJ83988.1 akaw|GAA92471.1 aluc|OJZ81547.1 anid|CBF82860.1 anig|EHA25888.1 aory|EIT75724.1 asyd|OJJ61597.1 ater|EAU38706.1 atub|OJI88637.1 aver|OJJ00921.1 awen|OJJ34481.1 azon|OJJ51758.1

Group1388: abra|OJJ66491.1 acar|OOF90967.1 acla|EAW10978.1 afis|EAW20881.1 afla|EED45568.1 afum|EAL84751.1 agla|OJJ83990.1 akaw|GAA91209.1 aluc|OJZ81544.1 anid|CBF82866.1 anig|EHA25891.1 aory|EIT75376.1 asyd|OJJ61599.1 ater|EAU30823.1 atub|OJI88640.1 aver|OJJ00923.1 awen|OJJ34479.1 azon|OJJ51756.1

Group1389: abra|OJJ66504.1 acar|OOF90968.1 acla|EAW10977.1 afis|EAW20882.1 afla|EED45567.1 afum|EAL84750.1 agla|OJJ83991.1 akaw|GAA91206.1 aluc|OJZ81541.1 anid|CBF82764.1 anig|EHA25895.1 aory|EIT75654.1 asyd|OJJ61604.1 ater|EAU30824.1 atub|OJI88645.1 aver|OJJ00927.1 awen|OJJ34478.1 azon|OJJ51755.1

Group1390: abra|OJJ66506.1 acar|OOF90970.1 acla|EAW10975.1 afis|EAW20885.1 afla|EED45570.1 afum|EAL84748.1 agla|OJJ83992.1 akaw|GAA91204.1 aluc|OJZ81539.1 anid|CBF82758.1 anig|EHA25897.1 aory|EIT75723.1 asyd|OJJ61607.1 ater|EAU30825.1 atub|OJI88647.1 aver|OJJ00930.1 awen|OJJ34462.1 azon|OJJ51752.1

Group1391: abra|OJJ66507.1 acar|OOF90971.1 acla|EAW10974.1 afis|EAW20886.1 afla|EED45571.1 afum|EAL84747.1 agla|OJJ83993.1 akaw|GAA91203.1 aluc|OJZ81538.1 anid|CBF82760.1 anig|EHA25898.1 aory|EIT75837.1 asyd|OJJ61606.1 ater|EAU30826.1 atub|OJI88648.1 aver|OJJ00929.1 awen|OJJ34461.1 azon|OJJ51750.1

Group1392: abra|OJJ66508.1 acar|OOF90972.1 acla|EAW10973.1 afis|EAW20887.1 afla|EED45572.1 afum|EAL84746.1 agla|OJJ83994.1 akaw|GAA91202.1 aluc|OJZ81537.1 anid|CBF82762.1 anig|EHA25899.1 aory|EIT75742.1 asyd|OJJ61605.1 ater|EAU30827.1 atub|OJI88649.1 aver|OJJ00928.1 awen|OJJ34460.1 azon|OJJ51749.1

Group1393: abra|OJJ66518.1 acar|OOF90973.1 acla|EAW10971.1 afis|EAW20888.1 afla|EED45576.1 afum|EAL84745.1 agla|OJJ83996.1 akaw|GAA91193.1 aluc|OJZ81526.1 anid|CBF82768.1 anig|EHA25908.1 aory|EIT75514.1 asyd|OJJ61601.1 ater|EAU30828.1 atub|OJI88659.1 aver|OJJ00925.1 awen|OJJ34459.1 azon|OJJ51748.1

Group1394: abra|OJJ66532.1 acar|OOF90983.1 acla|EAW10960.1 afis|EAW20899.1 afla|EED50249.1 afum|EAL84733.1 agla|OJJ84012.1 akaw|GAA91183.1 aluc|OJZ81520.1 anid|CBF82455.1 anig|EHA25917.1 aory|EIT77292.1 asyd|OJJ54114.1 ater|EAU38532.1 atub|OJI88669.1 aver|OJJ08621.1 awen|OJJ34449.1 azon|OJJ51735.1

Group1395: abra|OJJ66534.1 acar|OOF90986.1 acla|EAW10958.1 afis|EAW20901.1 afla|EED50251.1 afum|EAL84730.1 agla|OJJ84015.1 akaw|GAA91181.1 aluc|OJZ81522.1 anid|CBF82453.1 anig|EHA25920.1 aory|EIT77336.1 asyd|OJJ54112.1 ater|EAU38534.1 atub|OJI88671.1 aver|OJJ08623.1 awen|OJJ34447.1 azon|OJJ51733.1

Group1396: abra|OJJ66536.1 acar|OOF90987.1 acla|EAW10955.1 afis|EAW20906.1 afla|EED50254.1 afum|EAL84727.1 agla|OJJ84022.1 akaw|GAA91179.1 aluc|OJZ81524.1 anid|CBF82467.1 anig|EHA25922.1 aory|EIT77306.1 asyd|OJJ54098.1 ater|EAU38536.1 atub|OJI88673.1 aver|OJJ08640.1 awen|OJJ34445.1 azon|OJJ51728.1

Group1397: abra|OJJ66538.1 acar|OOF90989.1 acla|EAW10953.1 afis|EAW20908.1 afla|EED50256.1 afum|EAL84725.1 agla|OJJ84024.1 akaw|GAA91177.1 aluc|OJZ81509.1 anid|CBF82471.1 anig|EHA25924.1 aory|EIT77316.1 asyd|OJJ54100.1 ater|EAU38538.1 atub|OJI88675.1 aver|OJJ08638.1 awen|OJJ34443.1 azon|OJJ51720.1

Group1398: abra|OJJ66542.1 acar|OOF90994.1 acla|EAW09002.1 afis|EAW25543.1 afla|EED50261.1 afum|EAL84261.1 agla|OJJ84031.1 akaw|GAA91167.1 aluc|OJZ81495.1 anid|CBF80805.1 anig|EHA25935.1 aory|EIT77304.1 asyd|OJJ54485.1 ater|EAU38562.1 atub|OJI88686.1 aver|OJJ07694.1 awen|OJJ34423.1 azon|OJJ49761.1

Group1399: abra|OJJ66544.1 acar|OOF90996.1 acla|EAW10945.1 afis|EAW20913.1 afla|EED50263.1 afum|EAL84721.1 agla|OJJ79574.1 akaw|GAA91169.1 aluc|OJZ81497.1 anid|CBF84410.1 anig|EHA25933.1 aory|EIT77352.1 asyd|OJJ53562.1 ater|EAU38564.1 atub|OJI88684.1 aver|OJJ08190.1 awen|OJJ34421.1 azon|OJJ48150.1

Group1400: abra|OJJ66552.1 acar|OOF89951.1 acla|EAW10941.1 afis|EAW20919.1 afla|EED50273.1 afum|EAL84715.1 agla|OJJ80887.1 akaw|GAA91165.1 aluc|OJZ81492.1 anid|CBF82484.1 anig|EHA25937.1 aory|EIT77325.1 asyd|OJJ54104.1 atub|OJI88688.1 aver|OJJ08631.1 awen|OJJ34402.1 azon|OJJ49948.1 awen|OJJ35153.1

Group1401: abra|OJJ66558.1 acar|OOF90013.1 acla|EAW10937.1 afis|EAW20924.1 afla|EED50278.1 afum|EAL84710.1 agla|OJJ83965.1 akaw|GAA91158.1 aluc|OJZ81486.1 anid|CBF82449.1 anig|EHA25941.1 aory|EIT77317.1 asyd|OJJ54111.1 ater|EAU38571.1 atub|OJI88694.1 aver|OJJ08624.1 awen|OJJ34396.1 azon|OJJ48132.1

Group1402: abra|OJJ66562.1 acar|OOF91003.1 acla|EAW10934.1 afis|EAW20927.1 afla|EED50281.1 afum|EAL84707.1 agla|OJJ83961.1 akaw|GAA91155.1 aluc|OJZ81483.1 anid|CBF82497.1 anig|EHA25944.1 aory|EIT77267.1 asyd|OJJ54088.1 ater|EAU38573.1 atub|OJI88698.1 aver|OJJ08650.1 awen|OJJ34392.1 azon|OJJ48123.1

Group1403: abra|OJJ66563.1 acar|OOF91004.1 acla|EAW10933.1 afis|EAW20928.1 afla|EED50282.1 afum|EAL84706.1 agla|OJJ83960.1 akaw|GAA91154.1 aluc|OJZ81482.1 anid|CBF82515.1 anig|EHA25945.1

aory|EIT77363.1 asyd|OJJ54084.1 ater|EAU38574.1 atub|OJI88699.1 aver|OJJ08654.1 awen|OJJ34390.1 azon|OJJ48122.1

Group1404: abra|OJJ66564.1 acar|OOF91005.1 acla|EAW10932.1 afis|EAW20929.1 afla|EED50283.1 afum|EAL84705.1 agla|OJJ83959.1 akaw|GAA91153.1 aluc|OJZ81481.1 anid|CBF82513.1 anig|EHA25946.1 aory|EIT77279.1 asyd|OJJ54083.1 ater|EAU38575.1 atub|OJI88700.1 aver|OJJ08655.1 awen|OJJ34389.1 azon|OJJ48121.1

Group1405: abra|OJJ66568.1 acar|OOF91009.1 acla|EAW10927.1 afis|EAW20402.1 afla|EED50288.1 afum|EAL85501.1 agla|OJJ83952.1 akaw|GAA91149.1 aluc|OJZ81476.1 anid|CBF82504.1 anig|EHA25950.1 aory|EIT77254.1 asyd|OJJ54078.1 ater|EAU38579.1 atub|OJI88703.1 aver|OJJ08660.1 awen|OJJ34384.1 azon|OJJ48115.1

Group1406: abra|OJJ66570.1 acar|OOF91010.1 acla|EAW10924.1 afis|EAW20934.1 afla|EED50291.1 afum|EAL84700.1 agla|OJJ83951.1 akaw|GAA91148.1 aluc|OJZ81474.1 anid|CBF82532.1 anig|EHA25951.1 aory|EIT77301.1 asyd|OJJ54076.1 ater|EAU38580.1 atub|OJI88704.1 aver|OJJ08663.1 awen|OJJ34382.1 azon|OJJ48114.1

Group1407: abra|OJJ66573.1 acar|OOF91014.1 acla|EAW10917.1 afis|EAW20941.1 afla|EED50296.1 afum|EAL84693.1 agla|OJJ83946.1 akaw|GAA91145.1 aluc|OJZ81471.1 anid|CBF82534.1 anig|EHA25956.1 aory|EIT77257.1 asyd|OJJ54069.1 ater|EAU38586.1 atub|OJI88707.1 aver|OJJ08673.1 awen|OJJ34376.1 azon|OJJ50768.1

Group1408: abra|OJJ66576.1 acar|OOF91017.1 acla|EAW10914.1 afis|EAW20944.1 afla|EED50298.1 afum|EAL84690.2 agla|OJJ83943.1 akaw|GAA91142.1 aluc|OJZ81468.1 anid|CBF82539.1 anig|EHA25959.1 aory|EIT77376.1 asyd|OJJ54066.1 ater|EAU38588.1 atub|OJI88710.1 aver|OJJ08676.1 awen|OJJ34372.1 azon|OJJ48108.1

Group1409: abra|OJJ66577.1 acar|OOF91018.1 acla|EAW10913.1 afis|EAW20945.1 afla|EED50299.1 afum|EAL84689.2 agla|OJJ83941.1 akaw|GAA91141.1 aluc|OJZ81467.1 anid|CBF82541.1 anig|EHA25960.1 aory|EIT77303.1 asyd|OJJ54065.1 ater|EAU38596.1 atub|OJI88711.1 aver|OJJ08677.1 awen|OJJ34371.1 azon|OJJ48107.1

Group1410: abra|OJJ66579.1 acar|OOF91019.1 acla|EAW10911.1 afis|EAW20947.1 afla|EED50301.1 afum|EAL84687.1 agla|OJJ83939.1 akaw|GAA91139.1 aluc|OJZ81465.1 anid|CBF82545.1 anig|EHA25962.1 aory|EIT77341.1 asyd|OJJ54063.1 ater|EAU38598.1 atub|OJI88713.1 aver|OJJ08679.1 awen|OJJ34369.1 azon|OJJ48105.1

Group1411: abra|OJJ66580.1 acar|OOF91020.1 acla|EAW10910.1 afis|EAW20948.1 afla|EED50302.1 afum|EAL84686.1 agla|OJJ83938.1 akaw|GAA91138.1 aluc|OJZ81464.1 anid|CBF82547.1 anig|EHA25963.1 aory|EIT77278.1 asyd|OJJ54062.1 ater|EAU38599.1 atub|OJI88714.1 aver|OJJ08680.1 awen|OJJ34368.1 azon|OJJ48104.1

Group1412: abra|OJJ66581.1 acar|OOF89819.1 acla|EAW10909.1 afis|EAW20949.1 afla|EED50303.1 afum|EAL84685.1 agla|OJJ83936.1 akaw|GAA91137.1 aluc|OJZ81463.1 anid|CBF82549.1 anig|EHA25964.1 aory|EIT77276.1 asyd|OJJ54061.1 ater|EAU38600.1 atub|OJI88715.1 aver|OJJ08681.1 awen|OJJ34367.1 azon|OJJ48103.1

Group1413: abra|OJJ66583.1 acar|OOF91021.1 acla|EAW10908.1 afis|EAW20950.1 afla|EED50305.1 afum|EAL84684.1 agla|OJJ83930.1 akaw|GAA91135.1 aluc|OJZ81462.1 anid|CBF82550.1 anig|EHA25966.1 aory|EIT77342.1 asyd|OJJ54060.1 ater|EAU38601.1 atub|OJI88716.1 aver|OJJ08682.1 awen|OJJ34366.1 azon|OJJ48101.1

Group1414: abra|OJJ66587.1 acar|OOF91024.1 acla|EAW10904.1 afis|EAW20954.1 afla|EED50310.1 afum|EAL84680.2 agla|OJJ83925.1 akaw|GAA91130.1 aluc|OJZ81459.1 anid|CBF82558.1 anig|EHA25970.1 aory|EIT77289.1 asyd|OJJ54056.1 ater|EAU38605.1 atub|OJI88720.1 aver|OJJ08685.1 awen|OJJ34362.1 azon|OJJ48098.1

Group1415: abra|OJJ66588.1 acar|OOF91025.1 acla|EAW10903.1 afis|EAW20955.1 afla|EED50311.1 afum|EAL84679.1 agla|OJJ83924.1 akaw|GAA91129.1 aluc|OJZ81458.1 anid|CBF82560.1 anig|EHA25971.1 aory|EIT77266.1 asyd|OJJ54054.1 ater|EAU38606.1 atub|OJI88721.1 aver|OJJ08687.1 awen|OJJ34361.1 azon|OJJ48097.1

Group1416: abra|OJJ66589.1 acar|OOF91026.1 acla|EAW10902.1 afis|EAW20956.1 afla|EED50312.1 afum|EAL84677.1 agla|OJJ83922.1 akaw|GAA91128.1 aluc|OJZ81457.1 anid|CBF82562.1 anig|EHA25972.1 aory|EIT77295.1 asyd|OJJ54052.1 ater|EAU38607.1 atub|OJI88722.1 aver|OJJ08688.1 awen|OJJ34360.1 azon|OJJ48096.1

Group1417: abra|OJJ66590.1 acar|OOF91027.1 acla|EAW10901.1 afis|EAW20957.1 afla|EED50313.1 afum|EAL84676.1 agla|OJJ83920.1 akaw|GAA91127.1 aluc|OJZ81456.1 anid|CBF82564.1 anig|EHA25973.1 aory|EIT77362.1 asyd|OJJ54051.1 ater|EAU38608.1 atub|OJI88723.1 aver|OJJ08689.1 awen|OJJ34359.1 azon|OJJ48095.1

Group1418: abra|OJJ66596.1 acar|OOF91030.1 acla|EAW10896.1 afis|EAW20962.1 afla|EED50317.1 afum|EAL84671.1 agla|OJJ83916.1 akaw|GAA91122.1 aluc|OJZ81450.1 anid|CBF82571.1 anig|EHA25976.1 aory|EIT77307.1 asyd|OJJ54048.1 ater|EAU38612.1 atub|OJI88729.1 aver|OJJ08695.1 awen|OJJ34354.1 azon|OJJ48091.1

Group1419: abra|OJJ66598.1 acar|OOF91031.1 acla|EAW10895.1 afis|EAW20963.1 afla|EED50318.1 afum|EAL84670.1 agla|OJJ83915.1 akaw|GAA91121.1 aluc|OJZ81449.1 anid|CBF82572.1 anig|EHA25977.1 aory|EIT77348.1 asyd|OJJ54047.1 ater|EAU38613.1 atub|OJI88730.1 aver|OJJ08696.1 awen|OJJ34352.1 azon|OJJ48089.1

Group1420: abra|OJJ66600.1 acar|OOF91033.1 acla|EAW10892.1 afis|EAW20966.1 afla|EED50320.1 afum|EAL84667.1 agla|OJJ84043.1 akaw|GAA91119.1 aluc|OJZ81446.1 anid|CBF82578.1 anig|EHA25979.1 aory|EIT77255.1 asyd|OJJ54042.1 ater|EAU38616.1 atub|OJI88732.1 aver|OJJ08699.1 awen|OJJ34349.1 azon|OJJ48086.1

Group1421: abra|OJJ66601.1 acar|OOF91034.1 acla|EAW10891.1 afis|EAW20967.1 afla|EED50321.1 afum|EAL84666.1 agla|OJJ84044.1 akaw|GAA91118.1 aluc|OJZ81445.1 anid|CBF82580.1 anig|EHA25980.1 aory|EIT77332.1 asyd|OJJ54041.1 ater|EAU38617.1 atub|OJI88733.1 aver|OJJ08700.1 awen|OJJ34348.1 azon|OJJ48085.1

Group1422: abra|OJJ66604.1 acar|OOF90114.1 acla|EAW10889.1 afis|EAW20969.1 afla|EED50323.1 afum|EAL84665.1 agla|OJJ84046.1 akaw|GAA91116.1 aluc|OJZ81442.1 anid|CBF82584.1 anig|EHA25982.1 aory|EIT77349.1 asyd|OJJ54039.1 ater|EAU38619.1 atub|OJI88736.1 aver|OJJ08701.1 awen|OJJ34346.1 azon|OJJ48084.1

Group1423: abra|OJJ66606.1 acar|OOF90112.1 acla|EAW10887.1 afis|EAW20971.1 afla|EED50325.1 afum|EAL84663.1 agla|OJJ84048.1 akaw|GAA91114.1 aluc|OJZ81440.1 anid|CBF82585.1 anig|EHA25984.1 aory|EIT77291.1 asyd|OJJ54037.1 ater|EAU38620.1 atub|OJI88738.1 aver|OJJ08703.1 awen|OJJ34344.1 azon|OJJ48083.1

Group1424: abra|OJJ66610.1 acar|OOF91035.1 acla|EAW10886.1 afis|EAW20972.1 afla|EED50326.1 afum|EAL84661.1 agla|OJJ84050.1 akaw|GAA91113.1 aluc|OJZ81439.1 anid|CBF82587.1 anig|EHA25985.1 aory|EIT77361.1 asyd|OJJ54036.1 ater|EAU38622.1 atub|OJI88742.1 aver|OJJ08704.1 awen|OJJ34341.1 azon|OJJ48081.1

Group1425: abra|OJJ66612.1 acar|OOF91036.1 acla|EAW10884.1 afis|EAW20974.1 afla|EED50329.1 afum|EAL84659.1 agla|OJJ84052.1 akaw|GAA91111.1 aluc|OJZ81437.1 anid|CBF82593.1 anig|EHA25987.1 aory|EIT77331.1 asyd|OJJ54033.1 ater|EAU38624.1 atub|OJI88744.1 aver|OJJ08707.1 awen|OJJ34339.1 azon|OJJ48079.1

Group1426: abra|OJJ66613.1 acar|OOF91038.1 acla|EAW10883.1 afis|EAW20975.1 afla|EED50330.1 afum|EAL84658.1 agla|OJJ84053.1 akaw|GAA91110.1 aluc|OJZ81436.1 anid|CBF82595.1 anig|EHA25988.1 aory|EIT77270.1 asyd|OJJ54032.1 ater|EAU38625.1 atub|OJI88745.1 aver|OJJ08708.1 awen|OJJ34337.1 azon|OJJ48078.1

Group1427: abra|OJJ66617.1 acar|OOF91040.1 acla|EAW10880.1 afis|EAW20978.1 afla|EED50333.1 afum|EAL84655.1 agla|OJJ84055.1 akaw|GAA91108.1 aluc|OJZ81432.1 anid|CBF82598.1 anig|EHA25990.1 aory|EIT77374.1 asyd|OJJ54030.1 ater|EAU38627.1 atub|OJI88748.1 aver|OJJ08710.1 awen|OJJ34334.1 azon|OJJ48076.1

Group1428: abra|OJJ66619.1 acar|OOF91042.1 acla|EAW10878.1 afis|EAW20980.1 afla|EED50335.1 afum|EAL84651.2 agla|OJJ84057.1 akaw|GAA91106.1 aluc|OJZ81430.1 anid|CBF71772.1 anig|EHA25992.1 aory|EIT77286.1 asyd|OJJ61720.1 ater|EAU38628.1 atub|OJI88750.1 aver|OJI95814.1 awen|OJJ34332.1 azon|OJJ48074.1

Group1429: abra|OJJ66624.1 acar|OOF91048.1 acla|EAW10872.1 afis|EAW20986.1 afla|EED50341.1 afum|EAL84644.1 agla|OJJ84065.1 akaw|GAA91101.1 aluc|OJZ81425.1 anid|CBF82611.1 anig|EHA25997.1 aory|EIT77327.1 asyd|OJJ54022.1 ater|EAU38635.1 atub|OJI88756.1 aver|OJJ08718.1 awen|OJJ34325.1 azon|OJJ48071.1

Group1430: abra|OJJ66627.1 acar|OOF91052.1 acla|EAW10869.1 afis|EAW20989.1 afla|EED50343.1 afum|EAL84640.2 agla|OJJ84067.1 akaw|GAA91098.1 aluc|OJZ81421.1 anid|CBF82605.1 anig|EHA26000.1 aory|EIT77281.1 asyd|OJJ54026.1 ater|EAU38638.1 atub|OJI88760.1 aver|OJJ08714.1 awen|OJJ34321.1 azon|OJJ48066.1

Group1431: abra|OJJ66749.1 acar|OOF93832.1 acla|EAW11875.1 afis|EAW19486.1 afla|EED56704.1 afum|EAL87008.1 agla|OJJ81211.1 akaw|GAA89685.1 aluc|OJZ80700.1 anid|CBF75460.1 anig|EHA27316.1 aory|EIT73946.1 asyd|OJJ59769.1 ater|EAU34799.1 atub|OJI87559.1 aver|OJJ02516.1 awen|OJJ36902.1 azon|OJJ42915.1

Group1432: abra|OJJ66751.1 acar|OOF93833.1 acla|EAW11876.1 afis|EAW19487.1 afla|EED56705.1 afum|EAL87007.1 agla|OJJ81213.1 akaw|GAA89686.1 aluc|OJZ80699.1 anid|CBF75458.1 anig|EHA27317.1

aory|EIT73903.1 asyd|OJJ59771.1 ater|EAU34800.1 atub|OJI87558.1 aver|OJJ02515.1 awen|OJJ36903.1 azon|OJJ42913.1

Group1433: abra|OJJ66752.1 acar|OOF93834.1 acla|EAW11877.1 afis|EAW19488.1 afla|EED56706.1 afum|EAL87006.1 agla|OJJ81214.1 akaw|GAA89687.1 aluc|OJZ80698.1 anid|CBF75456.1 anig|EHA27318.1 aory|EIT73834.1 asyd|OJJ59772.1 ater|EAU34801.1 atub|OJI87557.1 aver|OJJ02514.1 awen|OJJ36904.1 azon|OJJ42912.1

Group1434: abra|OJJ66753.1 acar|OOF93835.1 acla|EAW11880.1 afis|EAW19491.1 afla|EED56710.1 afum|EAL87003.1 agla|OJJ81220.1 akaw|GAA89688.1 aluc|OJZ80697.1 anid|CBF75452.1 anig|EHA27319.1 aory|EIT73847.1 asyd|OJJ59774.1 ater|EAU34803.1 atub|OJI87556.1 aver|OJJ02512.1 awen|OJJ36907.1 azon|OJJ42909.1

Group1435: abra|OJJ66755.1 acar|OOF93836.1 acla|EAW11881.1 afis|EAW19492.1 afla|EED56711.1 afum|EAL87002.1 agla|OJJ81221.1 akaw|GAA89689.1 aluc|OJZ80696.1 anid|CBF75450.1 anig|EHA27320.1 aory|EIT73907.1 asyd|OJJ59775.1 ater|EAU34804.1 atub|OJI87555.1 aver|OJJ02511.1 awen|OJJ36908.1 azon|OJJ42908.1

Group1436: abra|OJJ66773.1 acar|OOF91583.1 acla|EAW15030.1 afis|EAW20305.1 agla|OJJ85746.1 akaw|GAA87178.1 aluc|OJZ84255.1 asyd|OJJ60493.1 atub|OJI89828.1 awen|OJJ36138.1 acla|EAW10581.1 afla|EED52812.1 aory|EIT78206.1 awen|OJJ37047.1 afis|EAW17210.1 azon|OJJ49313.1 afum|EAL85433.1 awen|OJJ32513.1

Group1437: abra|OJJ66819.1 acar|OOF91602.1 acla|EAW14405.1 afis|EAW19113.1 afla|EED54301.1 afum|EAL93845.1 agla|OJJ84489.1 akaw|GAA90457.1 aluc|OJZ85767.1 anid|CBF79734.1 anig|EHA24786.1 aory|EIT76106.1 asyd|OJJ58069.1 ater|EAU37372.1 atub|OJI79552.1 aver|OJJ04338.1 awen|OJJ32523.1 azon|OJJ50791.1

Group1438: abra|OJJ66828.1 acar|OOF91606.1 acla|EAW14411.1 afis|EAW19119.1 afla|EED54295.1 afum|EAL93851.1 agla|OJJ80434.1 akaw|GAA90464.1 aluc|OJZ85759.1 anid|CBF79724.1 anig|EHA24797.1 aory|EIT76143.1 asyd|OJJ58075.1 ater|EAU37382.1 atub|OJI79544.1 aver|OJJ04332.1 awen|OJJ37450.1 azon|OJJ51127.1

Group1439: abra|OJJ66830.1 acar|OOF91608.1 acla|EAW14413.1 afis|EAW19121.1 afla|EED54293.1 afum|EAL93853.1 agla|OJJ80436.1 akaw|GAA90467.1 aluc|OJZ85757.1 anid|CBF79722.1 anig|EHA24799.1 aory|EIT76028.1 asyd|OJJ58077.1 ater|EAU37384.1 atub|OJI79542.1 aver|OJJ04330.1 awen|OJJ37452.1 azon|OJJ51129.1

Group1440: abra|OJJ66832.1 acar|OOF91609.1 acla|EAW14414.1 afis|EAW19122.1 afla|EED54292.1 afum|EAL93854.1 agla|OJJ80437.1 akaw|GAA90468.1 aluc|OJZ85756.1 anid|CBF79720.1 anig|EHA24800.1 aory|EIT76096.1 asyd|OJJ58078.1 ater|EAU37385.1 atub|OJI79541.1 aver|OJJ04329.1 awen|OJJ37453.1 azon|OJJ51130.1

Group1441: abra|OJJ66833.1 acar|OOF91610.1 acla|EAW14415.1 afis|EAW19123.1 afla|EED54291.1 afum|EAL93855.1 agla|OJJ80438.1 akaw|GAA90469.1 aluc|OJZ85755.1 anid|CBF79718.1 anig|EHA24801.1 aory|EIT76111.1 asyd|OJJ58079.1 ater|EAU37386.1 atub|OJI79540.1 aver|OJJ04328.1 awen|OJJ37454.1 azon|OJJ51131.1

Group1442: abra|OJJ66834.1 acar|OOF91611.1 acla|EAW14416.1 afis|EAW19124.1 afla|EED54290.1 afum|EAL93856.1 agla|OJJ80439.1 akaw|GAA90470.1 aluc|OJZ85754.1 anid|CBF79717.1 anig|EHA24802.1 aory|EIT76047.1 asyd|OJJ58080.1 ater|EAU37387.1 atub|OJI79538.1 aver|OJJ04327.1 awen|OJJ37455.1 azon|OJJ51134.1

Group1443: abra|OJJ66835.1 acar|OOF91613.1 acla|EAW14419.1 afis|EAW19126.1 afla|EED54288.1 afum|EAL93858.1 agla|OJJ80441.1 akaw|GAA90472.1 aluc|OJZ85752.1 anid|CBF79737.1 anig|EHA24804.1 aory|EIT75920.1 asyd|OJJ58083.1 ater|EAU37393.1 atub|OJI79535.1 aver|OJJ04325.1 awen|OJJ37457.1 azon|OJJ51137.1

Group1444: abra|OJJ66836.1 acar|OOF91614.1 acla|EAW14420.1 afis|EAW19127.1 afla|EED54287.1 afum|EAL93859.1 agla|OJJ88596.1 akaw|GAA90473.1 aluc|OJZ85751.1 anid|CBF79739.1 anig|EHA24805.1 aory|EIT75974.1 asyd|OJJ58084.1 ater|EAU37394.1 atub|OJI79534.1 aver|OJJ04324.1 awen|OJJ37459.1 azon|OJJ51138.1

Group1445: abra|OJJ66840.1 acar|OOF91617.1 agla|OJJ85152.1 akaw|GAA90476.1 aluc|OJZ85747.1 anig|EHA24808.1 atub|OJI79532.1 aver|OJI98647.1 awen|OJJ33714.1 asyd|OJJ59197.1 azon|OJJ44469.1 acla|EAW12945.1 afla|EED50142.1 aluc|OJZ85744.1 aory|EIT74655.1 asyd|OJJ64551.1 atub|OJI79529.1 azon|OJJ49271.1

Group1446: abra|OJJ66843.1 acar|OOF91620.1 acla|EAW14417.1 afis|EAW19130.1 afla|EED54284.1 afum|EAL93862.1 agla|OJJ88602.1 akaw|GAA90479.1 aluc|OJZ85740.1 anid|CBF79758.1 anig|EHA24811.1 aory|EIT76041.1 asyd|OJJ58100.1 ater|EAU37398.1 atub|OJI79525.1 aver|OJJ04304.1 awen|OJJ37465.1 azon|OJJ51143.1

Group1447: abra|OJJ66846.1 acar|OOF91624.1 acla|EAW14425.1 afis|EAW19132.1 afla|EED54281.1 afum|EAL93864.1 agla|OJJ88607.1 akaw|GAA91559.1 aluc|OJZ85737.1 anid|CBF86338.1 anig|EHA24813.1 aory|EIT76060.1 asyd|OJJ64988.1 ater|EAU37405.1 atub|OJI79522.1 aver|OJI99108.1 awen|OJJ37468.1 azon|OJJ51146.1

Group1448: abra|OJJ66847.1 acar|OOF91625.1 acla|EAW14426.1 afis|EAW19133.1 afla|EED54280.1 afum|EAL93865.1 agla|OJJ88608.1 akaw|GAA91558.1 aluc|OJZ85736.1 anid|CBF86336.1 anig|EHA24814.1 aory|EIT76085.1 asyd|OJJ64989.1 ater|EAU37401.1 atub|OJI79521.1 aver|OJI99109.1 awen|OJJ37469.1 azon|OJJ51147.1

Group1449: abra|OJJ66853.1 acar|OOF91629.1 acla|EAW14431.1 afis|EAW19138.1 afla|EED54275.1 afum|EAL93870.1 agla|OJJ88617.1 akaw|GAA91553.1 aluc|OJZ85731.1 anid|CBF86324.1 anig|EHA24818.1 aory|EIT75921.1 asyd|OJJ64993.1 ater|EAU37408.1 atub|OJI79516.1 aver|OJI99112.1 awen|OJJ37475.1 azon|OJJ51155.1

Group1450: abra|OJJ66854.1 acar|OOF91630.1 acla|EAW14432.1 afis|EAW19139.1 afla|EED54274.1 afum|EAL93871.2 agla|OJJ88591.1 akaw|GAA91552.1 aluc|OJZ85730.1 anid|CBF86322.1 anig|EHA24819.1 aory|EIT75977.1 asyd|OJJ64994.1 ater|EAU37410.1 atub|OJI79515.1 aver|OJI99113.1 awen|OJJ37476.1 azon|OJJ51157.1

Group1451: abra|OJJ66855.1 acar|OOF91632.1 acla|EAW14433.1 afis|EAW19141.1 afla|EED54272.1 afum|EAL93872.1 agla|OJJ88589.1 akaw|GAA91551.1 aluc|OJZ85728.1 anid|CBF86321.1 anig|EHA24820.1 aory|EIT76058.1 asyd|OJJ64995.1 ater|EAU37412.1 atub|OJI79514.1 aver|OJI99115.1 awen|OJJ37481.1 azon|OJJ51158.1

Group1452: abra|OJJ66856.1 acar|OOF91633.1 acla|EAW14434.1 afis|EAW19142.1 afla|EED54271.1 afum|EAL93873.1 agla|OJJ88588.1 akaw|GAA91550.1 aluc|OJZ85727.1 anid|CBF86317.1 anig|EHA24821.1 aory|EIT76153.1 asyd|OJJ64997.1 ater|EAU37413.1 atub|OJI79513.1 aver|OJI99117.1 awen|OJJ37482.1 azon|OJJ51159.1

Group1453: abra|OJJ66858.1 acar|OOF91636.1 acla|EAW14437.1 afis|EAW19145.1 afla|EED54268.1 afum|EAL93876.1 agla|OJJ88586.1 akaw|GAA91547.1 aluc|OJZ85725.1 anid|CBF86313.1 anig|EHA24823.1 aory|EIT76154.1 asyd|OJJ65000.1 ater|EAU37417.1 atub|OJI79511.1 aver|OJI99120.1 awen|OJJ37484.1 azon|OJJ51161.1

Group1454: abra|OJJ66861.1 acar|OOF91638.1 acla|EAW14438.1 afis|EAW19146.1 afla|EED54267.1 afum|EAL93877.1 agla|OJJ88584.1 akaw|GAA91546.1 aluc|OJZ85721.1 anid|CBF86311.1 anig|EHA24824.1 aory|EIT76119.1 asyd|OJJ65001.1 ater|EAU37420.1 atub|OJI79510.1 aver|OJI99121.1 awen|OJJ37485.1 azon|OJJ51163.1

Group1455: abra|OJJ66864.1 acar|OOF91640.1 acla|EAW14440.1 afis|EAW19148.1 afla|EED54265.1 afum|EAL93879.1 agla|OJJ88576.1 akaw|GAA91544.1 aluc|OJZ85719.1 anid|CBF86308.1 anig|EHA24826.1 aory|EIT75979.1 asyd|OJJ65004.1 ater|EAU37421.1 atub|OJI79508.1 aver|OJI99123.1 awen|OJJ37501.1 azon|OJJ51166.1

Group1456: abra|OJJ66866.1 acar|OOF91643.1 acla|EAW14442.1 afis|EAW19150.1 afla|EED54257.1 afum|EAL93881.1 agla|OJJ88572.1 akaw|GAA91542.1 aluc|OJZ85716.1 anid|CBF86305.1 anig|EHA24828.1 aory|EIT76051.1 asyd|OJJ65006.1 ater|EAU37424.1 atub|OJI79506.1 aver|OJI99125.1 awen|OJJ37503.1 azon|OJJ51168.1

Group1457: abra|OJJ66869.1 acar|OOF91646.1 acla|EAW14445.1 afis|EAW19153.1 afla|EED54252.1 afum|EAL93884.1 agla|OJJ88569.1 akaw|GAA91539.1 aluc|OJZ85712.1 anid|CBF86299.1 anig|EHA24831.1 aory|EIT76023.1 asyd|OJJ65009.1 ater|EAU37426.1 atub|OJI79503.1 aver|OJI99128.1 awen|OJJ37506.1 azon|OJJ51172.1

Group1458: abra|OJJ66871.1 acar|OOF91647.1 acla|EAW14446.1 afis|EAW19154.1 afla|EED54251.1 afum|EAL93885.1 agla|OJJ88568.1 akaw|GAA91538.1 aluc|OJZ85711.1 anid|CBF86297.1 anig|EHA24832.1 aory|EIT75939.1 asyd|OJJ65010.1 ater|EAU37427.1 atub|OJI79502.1 aver|OJI99129.1 awen|OJJ37507.1 azon|OJJ51173.1

Group1459: abra|OJJ66873.1 acar|OOF91649.1 acla|EAW14448.1 afis|EAW19156.1 afla|EED54246.1 afum|EAL93887.1 agla|OJJ88566.1 akaw|GAA91536.1 aluc|OJZ85708.1 anid|CBF86294.1 anig|EHA24834.1 aory|EIT75988.1 asyd|OJJ65012.1 ater|EAU37429.1 atub|OJI79499.1 aver|OJI99131.1 awen|OJJ37509.1 azon|OJJ51174.1

Group1460: abra|OJJ66874.1 acar|OOF91650.1 acla|EAW14449.1 afis|EAW19157.1 afla|EED54245.1 afum|EBA27505.1 agla|OJJ88565.1 akaw|GAA91535.1 aluc|OJZ85707.1 anid|CBF86292.1 anig|EHA24835.1 aory|EIT75937.1 asyd|OJJ65013.1 ater|EAU37430.1 atub|OJI79498.1 aver|OJI99132.1 awen|OJJ37510.1 azon|OJJ51175.1

Group1461: abra|OJJ66875.1 acar|OOF91651.1 acla|EAW14450.1 afis|EAW19158.1 afla|EED54244.1 afum|EBA27506.1 agla|OJJ88564.1 akaw|GAA91534.1 aluc|OJZ85706.1 anid|CBF86291.1 anig|EHA24836.1

aory|EIT76133.1 asyd|OJJ65014.1 ater|EAU37431.1 atub|OJI79497.1 aver|OJI99133.1 awen|OJJ37511.1 azon|OJJ51176.1

Group1462: abra|OJJ66879.1 acar|OOF91654.1 acla|EAW14454.1 afis|EAW19162.1 afla|EED54240.1 afum|EAL93892.2 agla|OJJ88562.1 akaw|GAA91530.1 aluc|OJZ85703.1 anid|CBF86285.1 anig|EHA24840.1 aory|EIT76157.1 asyd|OJJ65017.1 ater|EAU37311.1 atub|OJI79493.1 aver|OJI99135.1 awen|OJJ37515.1 azon|OJJ51178.1

Group1463: abra|OJJ66880.1 acar|OOF91655.1 acla|EAW14455.1 afis|EAW19163.1 afla|EED54239.1 afum|EAL93893.1 agla|OJJ88561.1 akaw|GAA91529.1 aluc|OJZ85702.1 anid|CBF86282.1 anig|EHA24841.1 aory|EIT76014.1 asyd|OJJ65018.1 ater|EAU37310.1 atub|OJI79492.1 aver|OJI99136.1 awen|OJJ37516.1 azon|OJJ51181.1

Group1464: abra|OJJ66883.1 acar|OOF91657.1 acla|EAW14457.1 afis|EAW19165.1 afla|EED54232.1 afum|EAL93895.1 agla|OJJ88559.1 akaw|GAA91527.1 aluc|OJZ85699.1 anid|CBF86279.1 anig|EHA24843.1 aory|EIT75967.1 asyd|OJJ65020.1 ater|EAU37308.1 atub|OJI79489.1 aver|OJI99138.1 awen|OJJ37518.1 azon|OJJ51185.1

Group1465: abra|OJJ66887.1 acar|OOF91660.1 acla|EAW14459.1 afis|EAW19167.1 afla|EED54230.1 afum|EAL93897.1 agla|OJJ88557.1 akaw|GAA91525.1 aluc|OJZ85696.1 anid|CBF86276.1 anig|EHA24846.1 aory|EIT75971.1 asyd|OJJ65022.1 ater|EAU37306.1 atub|OJI79485.1 aver|OJI99141.1 awen|OJJ37520.1 azon|OJJ51186.1

Group1466: abra|OJJ66889.1 acar|OOF91661.1 acla|EAW14461.1 afis|EAW19169.1 afla|EED54228.1 afum|EAL93899.1 agla|OJJ88555.1 akaw|GAA91523.1 aluc|OJZ85693.1 anid|CBF86273.1 anig|EHA24848.1 aory|EIT76026.1 asyd|OJJ65024.1 ater|EAU37304.1 atub|OJI79483.1 aver|OJI99143.1 awen|OJJ37522.1 azon|OJJ51188.1

Group1467: abra|OJJ66890.1 acar|OOF91662.1 acla|EAW14462.1 afis|EAW19170.1 afla|EED54226.1 afum|EAL93900.1 agla|OJJ88554.1 akaw|GAA91522.1 aluc|OJZ85692.1 anid|CBF86271.1 anig|EHA24849.1 aory|EIT75952.1 asyd|OJJ65025.1 ater|EAU37303.1 atub|OJI79482.1 aver|OJI99144.1 awen|OJJ37523.1 azon|OJJ51189.1

Group1468: abra|OJJ66891.1 acar|OOF91663.1 acla|EAW14463.1 afis|EAW19171.1 afla|EED54224.1 afum|EAL93901.1 agla|OJJ88553.1 akaw|GAA91521.1 aluc|OJZ85691.1 anid|CBF86270.1 anig|EHA24850.1 aory|EIT75965.1 asyd|OJJ65026.1 ater|EAU37302.1 atub|OJI79481.1 aver|OJI99145.1 awen|OJJ37524.1 azon|OJJ51190.1

Group1469: abra|OJJ66893.1 acar|OOF91664.1 acla|EAW14465.1 afis|EAW19173.1 afla|EED54222.1 afum|EAL93903.1 agla|OJJ88551.1 akaw|GAA91519.1 aluc|OJZ85689.1 anid|CBF86266.1 anig|EHA24852.1 aory|EIT76009.1 asyd|OJJ65029.1 ater|EAU37300.1 atub|OJI79479.1 aver|OJI99148.1 awen|OJJ37526.1 azon|OJJ51193.1

Group1470: abra|OJJ66894.1 acar|OOF91665.1 acla|EAW14466.1 afis|EAW19174.1 afla|EED54221.1 afum|EAL93904.1 agla|OJJ88550.1 akaw|GAA91518.1 aluc|OJZ85688.1 anid|CBF86265.1 anig|EHA24853.1 aory|EIT76027.1 asyd|OJJ65030.1 ater|EAU37299.1 atub|OJI79478.1 aver|OJI99149.1 awen|OJJ37527.1 azon|OJJ51194.1

Group1471: abra|OJJ66895.1 acar|OOF91666.1 acla|EAW14467.1 afis|EAW19175.1 afla|EED54220.1 afum|EAL93905.1 agla|OJJ88548.1 akaw|GAA91517.1 aluc|OJZ85687.1 anid|CBF86263.1 anig|EHA24854.1 aory|EIT75931.1 asyd|OJJ65031.1 ater|EAU37298.1 atub|OJI79477.1 aver|OJI99150.1 awen|OJJ37528.1 azon|OJJ51195.1

Group1472: abra|OJJ66896.1 acar|OOF91667.1 acla|EAW14468.1 afis|EAW19176.1 afla|EED54219.1 afum|EAL93906.1 agla|OJJ88547.1 akaw|GAA91516.1 aluc|OJZ85686.1 anid|CBF86262.1 anig|EHA24855.1 aory|EIT76150.1 asyd|OJJ65032.1 ater|EAU37297.1 atub|OJI79476.1 aver|OJI99151.1 awen|OJJ37530.1 azon|OJJ51196.1

Group1473: abra|OJJ66897.1 acar|OOF91668.1 acla|EAW14469.1 afis|EAW19177.1 afla|EED54218.1 afum|EAL93907.1 agla|OJJ88546.1 akaw|GAA91515.1 aluc|OJZ85685.1 anid|CBF86260.1 anig|EHA24856.1 aory|EIT75968.1 asyd|OJJ65033.1 ater|EAU37296.1 atub|OJI79475.1 aver|OJI99152.1 awen|OJJ37531.1 azon|OJJ51197.1

Group1474: abra|OJJ66899.1 acar|OOF91671.1 acla|EAW14470.1 afis|EAW19179.1 afla|EED54217.1 afum|EAL93909.1 agla|OJJ88544.1 akaw|GAA91513.1 aluc|OJZ85683.1 anid|CBF86259.1 anig|EHA24858.1 aory|EIT76147.1 asyd|OJJ65035.1 ater|EAU37294.1 atub|OJI79473.1 aver|OJI99154.1 awen|OJJ37533.1 azon|OJJ51199.1

Group1475: abra|OJJ66900.1 acar|OOF91672.1 acla|EAW14471.1 afis|EAW19180.1 afla|EED54216.1 afum|EAL93910.1 agla|OJJ88543.1 akaw|GAA91512.1 aluc|OJZ85682.1 anid|CBF86257.1 anig|EHA24859.1 aory|EIT76034.1 asyd|OJJ65036.1 ater|EAU37293.1 atub|OJI79472.1 aver|OJI99155.1 awen|OJJ37534.1 azon|OJJ51200.1

Group1476: abra|OJJ66903.1 acar|OOF91673.1 acla|EAW14473.1 afis|EAW19182.1 afla|EED54213.1 afum|EAL93912.1 agla|OJJ88540.1 akaw|GAA91510.1 aluc|OJZ85680.1 anid|CBF86253.1 anig|EHA24861.1 aory|EIT76125.1 asyd|OJJ65038.1 ater|EAU37291.1 atub|OJI79470.1 aver|OJI99157.1 awen|OJJ37536.1 azon|OJJ51202.1

Group1477: abra|OJJ66905.1 acar|OOF91675.1 acla|EAW14474.1 afis|EAW19183.1 afla|EED54212.1 afum|EAL93913.1 agla|OJJ88539.1 akaw|GAA91509.1 aluc|OJZ85676.1 anid|CBF86252.1 anig|EHA24864.1 aory|EIT75985.1 asyd|OJJ65041.1 ater|EAU37290.1 atub|OJI79467.1 aver|OJI99159.1 awen|OJJ37538.1 azon|OJJ51203.1

Group1478: abra|OJJ66906.1 acar|OOF91676.1 acla|EAW14475.1 afis|EAW19184.1 afla|EED54211.1 afum|EAL93915.1 akaw|GAA91508.1 aluc|OJZ85675.1 anid|CBF86250.1 anig|EHA24865.1 aory|EIT76099.1 asyd|OJJ65042.1 ater|EAU37289.1 atub|OJI79466.1 aver|OJI99160.1 awen|OJJ37539.1 azon|OJJ51204.1 agla|OJJ88538.1

Group1479: abra|OJJ66907.1 acar|OOF91677.1 acla|EAW14476.1 afla|EED54210.1 afum|EAL93916.1 agla|OJJ88537.1 akaw|GAA91507.1 aluc|OJZ85673.1 anid|CBF86248.1 anig|EHA24866.1 aory|EIT76017.1 asyd|OJJ65043.1 ater|EAU37287.1 atub|OJI79465.1 aver|OJI99162.1 awen|OJJ37542.1 afis|EAW19185.1 azon|OJJ51206.1

Group1480: abra|OJJ66909.1 acar|OOF91679.1 acla|EAW14478.1 afis|EAW19187.1 afla|EED47724.1 afum|EAL93918.1 agla|OJJ88534.1 akaw|GAA91505.1 aluc|OJZ85671.1 anid|CBF86246.1 anig|EHA24868.1 aory|EIT77160.1 asyd|OJJ65044.1 ater|EAU37286.1 atub|OJI79463.1 aver|OJI99165.1 awen|OJJ37545.1 azon|OJJ51210.1

Group1481: abra|OJJ66911.1 acar|OOF91680.1 acla|EAW14480.1 afis|EAW19189.1 afla|EED47722.1 afum|EAL93921.1 agla|OJJ88532.1 akaw|GAA91504.1 aluc|OJZ85670.1 anid|CBF86243.1 anig|EHA24869.1 aory|EIT77116.1 asyd|OJJ65046.1 ater|EAU37285.1 atub|OJI79462.1 aver|OJI99167.1 awen|OJJ37549.1 azon|OJJ51211.1

Group1482: abra|OJJ66915.1 acar|OOF91682.1 acla|EAW14482.1 afis|EAW19191.1 afla|EED47718.1 afum|EAL93922.1 agla|OJJ88518.1 akaw|GAA91503.1 aluc|OJZ85665.1 anid|CBF86239.1 anig|EHA24870.1 aory|EIT80246.1 asyd|OJJ65049.1 ater|EAU37284.1 atub|OJI79458.1 aver|OJI99171.1 awen|OJJ37551.1 azon|OJJ51212.1

Group1483: abra|OJJ66918.1 acar|OOF91685.1 acla|EAW14485.1 afis|EAW19194.1 afla|EED47714.1 afum|EAL93925.2 agla|OJJ88515.1 akaw|GAA91500.1 aluc|OJZ85661.1 anid|CBF78544.1 anig|EHA24872.1 aory|EIT80273.1 asyd|OJJ65054.1 ater|EAU37281.1 atub|OJI79455.1 aver|OJI99177.1 awen|OJJ37552.1 azon|OJJ51213.1

Group1484: abra|OJJ66919.1 acar|OOF91687.1 acla|EAW14486.1 afis|EAW19195.1 afla|EED47713.1 afum|EAL93927.2 agla|OJJ88514.1 akaw|GAA91499.1 aluc|OJZ85660.1 anid|CBF78546.1 anig|EHA24873.1 aory|EIT80279.1 asyd|OJJ65055.1 ater|EAU37280.1 atub|OJI79454.1 aver|OJI99178.1 awen|OJJ37553.1 azon|OJJ51214.1

Group1485: abra|OJJ66923.1 acar|OOF91690.1 acla|EAW14489.1 afis|EAW19203.1 afla|EED47709.1 afum|EAL93933.1 agla|OJJ88509.1 akaw|GAA91496.1 aluc|OJZ85657.1 anid|CBF78554.1 anig|EHA24876.1 aory|EIT80207.1 asyd|OJJ65059.1 ater|EAU37277.1 atub|OJI79449.1 aver|OJI99184.1 awen|OJJ37556.1 azon|OJJ51219.1

Group1486: abra|OJJ66924.1 acar|OOF91692.1 acla|EAW14490.1 afis|EAW19204.1 afla|EED47708.1 afum|EAL93934.1 agla|OJJ88508.1 akaw|GAA91493.1 aluc|OJZ85654.1 anid|CBF78558.1 anig|EHA24878.1 aory|EIT80227.1 asyd|OJJ65061.1 ater|EAU37268.1 atub|OJI79446.1 aver|OJI99185.1 awen|OJJ37557.1 azon|OJJ51220.1

Group1487: abra|OJJ66928.1 acar|OOF91696.1 acla|EAW14493.1 afis|EAW19207.1 afla|EED47705.1 afum|EAL93937.1 agla|OJJ88504.1 akaw|GAA91489.1 aluc|OJZ85651.1 anid|CBF78563.1 anig|EHA24882.1 aory|EIT80245.1 asyd|OJJ65064.1 ater|EAU37265.1 atub|OJI79442.1 aver|OJI99188.1 awen|OJJ37560.1 azon|OJJ51222.1

Group1488: abra|OJJ66931.1 acar|OOF91698.1 acla|EAW14494.1 afis|EAW19208.1 afla|EED47703.1 afum|EAL93938.2 agla|OJJ88501.1 akaw|GAA91487.1 aluc|OJZ85649.1 anid|CBF78566.1 anig|EHA24884.1 aory|EIT80217.1 asyd|OJJ65067.1 ater|EAU37260.1 atub|OJI79440.1 aver|OJI99191.1 awen|OJJ37562.1 azon|OJJ51225.1

Group1489: abra|OJJ66934.1 acar|OOF91701.1 acla|EAW14496.1 afis|EAW19210.1 afla|EED47702.1 afum|EAL93940.1 agla|OJJ88490.1 akaw|GAA91485.1 aluc|OJZ85647.1 anid|CBF78570.1 anig|EHA24887.1 aory|EIT80269.1 asyd|OJJ65075.1 ater|EAU37258.1 atub|OJI79438.1 aver|OJI99200.1 awen|OJJ37564.1 azon|OJJ51227.1

Group1490: afum|EAL92928.1 abra|OJJ78283.1 acar|OOF98585.1 acla|EAW09232.1 afis|EAW21916.1 afla|EED46633.1 agla|OJJ84207.1 akaw|GAA85744.1 aluc|OJZ91357.1 anig|EHA20035.1 aory|EIT77912.1

asyd|OJJ61312.1 ater|EAU30918.1 atub|OJI83350.1 aver|OJJ00637.1 awen|OJJ34206.1 azon|OJJ43428.1 anid|CBF76852.1

Group1491: abra|OJJ66937.1 acar|OOF91703.1 acla|EAW15005.1 afis|EAW24537.1 afla|EED47700.1 afum|EAL86201.1 agla|OJJ86874.1 akaw|GAA91483.1 aluc|OJZ85645.1 anig|EHA24890.1 aory|EIT80258.1 asyd|OJJ63536.1 ater|EAU39600.1 atub|OJI79436.1 aver|OJI96723.1 awen|OJJ38055.1 azon|OJJ42331.1 agla|OJJ81013.1

Group1492: abra|OJJ66938.1 acar|OOF91704.1 acla|EAW14497.1 afis|EAW19211.1 afla|EED47699.1 afum|EAL93941.1 agla|OJJ88486.1 akaw|GAA91482.1 aluc|OJZ85644.1 anid|CBF78583.1 anig|EHA24891.1 aory|EIT80265.1 asyd|OJJ65078.1 ater|EAU37256.1 atub|OJI79435.1 aver|OJI99203.1 awen|OJJ37565.1 azon|OJJ51228.1

Group1493: abra|OJJ66940.1 acar|OOF91705.1 acla|EAW14498.1 afis|EAW19212.1 afla|EED47698.1 afum|EBA27508.1 agla|OJJ88485.1 akaw|GAA91481.1 aluc|OJZ85642.1 anid|CBF78581.1 anig|EHA24892.1 aory|EIT80243.1 asyd|OJJ65076.1 ater|EAU37255.1 atub|OJI79433.1 aver|OJI99201.1 awen|OJJ37568.1 azon|OJJ51229.1

Group1494: abra|OJJ66948.1 acar|OOF91714.1 acla|EAW14503.1 afis|EAW19217.1 afla|EED47682.1 afum|EAL93950.1 agla|OJJ88478.1 akaw|GAA91472.1 aluc|OJZ85632.1 anid|CBF78594.1 anig|EHA24899.1 aory|EIT80231.1 asyd|OJJ65091.1 ater|EAU37251.1 atub|OJI79422.1 aver|OJI99216.1 awen|OJJ37574.1 azon|OJJ51234.1

Group1495: abra|OJJ66952.1 acar|OOF93656.1 acla|EAW14505.1 afis|EAW19218.1 afla|EED47681.1 afum|EAL93951.2 akaw|GAA91468.1 aluc|OJZ85626.1 anid|CBF78597.1 anig|EHA24902.1 aory|EIT80228.1 asyd|OJJ65093.1 ater|EAU37250.1 atub|OJI79419.1 aver|OJI99217.1 azon|OJJ51235.1 agla|OJJ88477.1 awen|OJJ37576.1

Group1496: abra|OJJ66955.1 acar|OOF93653.1 acla|EAW14506.1 afis|EAW19219.1 afla|EED47670.1 afum|EAL93952.1 agla|OJJ88476.1 akaw|GAA92206.1 aluc|OJZ85621.1 anid|CBF78603.1 anig|EHA24905.1 aory|EIT80271.1 asyd|OJJ65097.1 ater|EAU37238.1 atub|OJI79414.1 aver|OJI99223.1 awen|OJJ37577.1 azon|OJJ51236.1

Group1497: abra|OJJ66956.1 acar|OOF93652.1 acla|EAW14507.1 afis|EAW19220.1 afla|EED47669.1 afum|EAL93953.1 agla|OJJ88464.1 akaw|GAA92205.1 aluc|OJZ85620.1 anid|CBF78605.1 anig|EHA24906.1 aory|EIT80172.1 asyd|OJJ65098.1 ater|EAU37237.1 atub|OJI79413.1 aver|OJI99224.1 awen|OJJ37583.1 azon|OJJ51237.1

Group1498: abra|OJJ66962.1 acar|OOF93645.1 acla|EAW14512.1 afis|EAW19230.1 afla|EED47658.1 afum|EAL93959.1 agla|OJJ88473.1 akaw|GAA92199.1 aluc|OJZ85614.1 anid|CBF78614.1 anig|EHA24912.1 aory|EIT80215.1 asyd|OJJ65178.1 ater|EAU37233.1 atub|OJI79407.1 aver|OJI99331.1 awen|OJJ37588.1 azon|OJJ51242.1

Group1499: abra|OJJ66980.1 acar|OOF93631.1 acla|EAW11770.1 afis|EAW23678.1 afla|EED52197.1 afum|EAL87121.1 agla|OJJ78538.1 akaw|GAA92184.1 aluc|OJZ85594.1 anid|CBF89572.1 anig|EHA24928.1 aory|EIT74397.1 asyd|OJJ62335.1 ater|EAU35375.1 atub|OJI79390.1 aver|OJI96441.1 awen|OJJ35575.1 azon|OJJ43197.1

Group1500: abra|OJJ66988.1 acar|OOF93624.1 acla|EAW09074.1 afis|EAW25663.1 afla|EED46073.1 afum|EAL84335.1 akaw|GAA92175.1 aluc|OJZ85585.1 anid|CBF84674.1 anig|EHA24936.1 aory|EIT83504.1 asyd|OJJ53347.1 atub|OJI79380.1 aver|OJJ08397.1 awen|OJJ35373.1 azon|OJJ45069.1 afla|EED55027.1 aory|EIT81986.1

Group1501: abra|OJJ66992.1 acar|OOF93622.1 acla|EAW15343.1 afis|EAW20364.1 afla|EED51949.1 afum|EAL85480.2 agla|OJJ81790.1 akaw|GAA92172.1 aluc|OJZ85580.1 anid|CBF69463.1 anig|EHA24939.1 aory|EIT81480.1 asyd|OJJ59276.1 ater|EAU36758.1 atub|OJI79376.1 aver|OJJ01891.1 awen|OJJ32524.1 azon|OJJ50243.1

Group1502: abra|OJJ67094.1 acla|EAW11007.1 akaw|GAA91067.1 aluc|OJZ87881.1 anig|EHA20184.1 atub|OJI80770.1 abra|OJJ69053.1 acar|OOF91237.1 afum|EAL86660.1 abra|OJJ72679.1 acar|OOF95928.1 afis|EAW19541.1 agla|OJJ88408.1 anig|EHA24092.1 ater|EAU39521.1 atub|OJI82086.1 acar|OOF91465.1 afum|EAL86967.1

Group1503: abra|OJJ67118.1 acar|OOF98434.1 acla|EAW11710.1 afis|EAW23729.1 afla|EED47523.1 afum|EAL87167.1 agla|OJJ88177.1 akaw|GAA91047.1 aluc|OJZ87857.1 anid|CBF69394.1 anig|EHA20163.1 aory|EIT74567.1 asyd|OJJ52918.1 ater|EAU30755.1 atub|OJI80790.1 aver|OJJ08888.1 awen|OJJ32546.1 azon|OJJ51795.1

Group1504: abra|OJJ67150.1 acar|OOF98408.1 acla|EAW14575.1 afis|EAW19299.1 afla|EED47588.1 afum|EAL94018.1 agla|OJJ88383.1 akaw|GAA91023.1 aluc|OJZ87826.1 anid|CBF78774.1 anig|EHA20305.1 aory|EIT74538.1 asyd|OJJ65240.1 ater|EAU29474.1 atub|OJI80817.1 aver|OJI99390.1 awen|OJJ37648.1 azon|OJJ44913.1

Group1505: abra|OJJ67151.1 acar|OOF98407.1 acla|EAW14574.1 afis|EAW19297.1 afla|EED47589.1 afum|EAL94016.1 agla|OJJ88384.1 akaw|GAA91022.1 aluc|OJZ87825.1 anid|CBF78772.1 anig|EHA20304.1 aory|EIT74535.1 asyd|OJJ65239.1 ater|EAU29475.1 atub|OJI80819.1 aver|OJI99389.1 awen|OJJ37647.1 azon|OJJ44912.1

Group1506: abra|OJJ67153.1 acar|OOF98405.1 acla|EAW14571.1 afis|EAW19294.1 afla|EED47592.1 afum|EAL94013.1 agla|OJJ88386.1 akaw|GAA91019.1 aluc|OJZ87821.1 anid|CBF78767.1 anig|EHA20302.1 aory|EIT74534.1 asyd|OJJ65237.1 ater|EAU29478.1 atub|OJI80823.1 aver|OJI99386.1 awen|OJJ37645.1 azon|OJJ44910.1

Group1507: abra|OJJ67154.1 acar|OOF98404.1 acla|EAW14570.1 afis|EAW19283.1 afla|EED47593.1 afum|EAL94009.1 agla|OJJ88395.1 akaw|GAA91018.1 aluc|OJZ87820.1 anid|CBF78763.1 anig|EHA20301.1 aory|EIT80189.1 asyd|OJJ65235.1 ater|EAU29479.1 atub|OJI80824.1 aver|OJI99384.1 awen|OJJ37644.1 azon|OJJ44909.1

Group1508: abra|OJJ67157.1 acar|OOF98402.1 acla|EAW14568.1 afis|EAW19281.1 afla|EED47596.1 afum|EAL94007.1 agla|OJJ88393.1 akaw|GAA91015.1 aluc|OJZ87816.1 anid|CBF78760.1 anig|EHA20297.1 aory|EIT80256.1 asyd|OJJ65233.1 ater|EAU29482.1 atub|OJI80828.1 aver|OJI99382.1 awen|OJJ37642.1 azon|OJJ44906.1

Group1509: abra|OJJ67159.1 acar|OOF98400.1 acla|EAW14567.1 afis|EAW19280.1 afla|EED47598.1 afum|EAL94006.1 agla|OJJ88391.1 akaw|GAA91013.1 aluc|OJZ87814.1 anid|CBF78756.1 anig|EHA20295.1 aory|EIT80259.1 asyd|OJJ65231.1 ater|EAU29483.1 atub|OJI80830.1 aver|OJI99380.1 awen|OJJ37640.1 azon|OJJ44904.1

Group1510: abra|OJJ67163.1 acar|OOF98397.1 acla|EAW14565.1 afis|EAW19278.1 afla|EED47600.1 afum|EAL94004.1 agla|OJJ88389.1 akaw|GAA91011.1 aluc|OJZ87812.1 anid|CBF78752.1 anig|EHA20294.1 aory|EIT80253.1 asyd|OJJ65229.1 ater|EAU29485.1 atub|OJI80832.1 aver|OJI99378.1 awen|OJJ37638.1 azon|OJJ44882.1

Group1511: abra|OJJ67164.1 acar|OOF98398.1 acla|EAW14564.1 afis|EAW19277.1 afla|EED47601.1 afum|EAL94003.1 agla|OJJ88396.1 akaw|GAA91010.1 aluc|OJZ87811.1 anid|CBF78750.1 anig|EHA20293.1 aory|EIT80205.1 asyd|OJJ65228.1 ater|EAU29486.1 atub|OJI80833.1 aver|OJI99377.1 awen|OJJ37637.1 azon|OJJ44883.1

Group1512: abra|OJJ67173.1 acar|OOF98390.1 acla|EAW14550.1 afis|EAW19267.1 afla|EED47608.1 afum|EAL93994.1 agla|OJJ88426.1 akaw|GAA91001.1 aluc|OJZ87802.1 anid|CBF78695.1 anig|EHA20283.1 aory|EIT80277.1 asyd|OJJ65218.1 ater|EAU29548.1 atub|OJI80845.1 aver|OJI99368.1 awen|OJJ37628.1 azon|OJJ47016.1

Group1513: abra|OJJ67175.1 acar|OOF98388.1 acla|EAW14548.1 afis|EAW19265.1 afla|EED47609.1 afum|EAL93992.1 agla|OJJ88424.1 akaw|GAA90999.1 aluc|OJZ87800.1 anid|CBF78691.1 anig|EHA20281.1 aory|EIT80181.1 asyd|OJJ65216.1 ater|EAU29546.1 atub|OJI80847.1 aver|OJI99366.1 awen|OJJ37626.1 azon|OJJ47015.1

Group1514: abra|OJJ67176.1 acar|OOF98386.1 acla|EAW14547.1 afis|EAW19264.1 afla|EED47610.1 afum|EAL93991.1 agla|OJJ88423.1 akaw|GAA90998.1 aluc|OJZ87799.1 anid|CBF78688.1 anig|EHA20280.1 aory|EIT80241.1 asyd|OJJ65214.1 ater|EAU29545.1 atub|OJI80848.1 aver|OJI99364.1 awen|OJJ37625.1 azon|OJJ47014.1

Group1515: abra|OJJ67178.1 acar|OOF98385.1 acla|EAW14545.1 afis|EAW19263.1 afla|EED47612.1 afum|EAL93990.1 agla|OJJ88421.1 aluc|OJZ87797.1 anid|CBF78684.1 anig|EHA20278.1 aory|EIT80200.1 asyd|OJJ65212.1 ater|EAU29543.1 atub|OJI80850.1 aver|OJI99362.1 awen|OJJ37623.1 azon|OJJ47012.1 aory|EIT80261.1

Group1516: abra|OJJ67179.1 acar|OOF98384.1 acla|EAW14544.1 afis|EAW19262.1 afla|EED47613.1 afum|EAL93989.1 agla|OJJ88420.1 akaw|GAA90997.1 aluc|OJZ87796.1 anid|CBF78682.1 anig|EHA20277.1 aory|EIT80281.1 asyd|OJJ65211.1 ater|EAU29542.1 atub|OJI80851.1 aver|OJI99361.1 awen|OJJ37622.1 azon|OJJ47011.1

Group1517: abra|OJJ67180.1 acar|OOF98382.1 acla|EAW14543.1 afis|EAW19261.1 afla|EED47614.1 afum|EAL93988.1 agla|OJJ88433.1 akaw|GAA90996.1 aluc|OJZ87795.1 anid|CBF78680.1 anig|EHA20276.1 aory|EIT80274.1 asyd|OJJ65210.1 ater|EAU29514.1 atub|OJI80852.1 aver|OJI99360.1 awen|OJJ37621.1 azon|OJJ47010.1

Group1518: abra|OJJ67182.1 acar|OOF98380.1 acla|EAW14541.1 afis|EAW19259.1 afla|EED47616.1 afum|EAL93986.1 agla|OJJ88436.1 akaw|GAA90994.1 aluc|OJZ87793.1 anid|CBF78678.1 anig|EHA20274.1 aory|EIT80183.1 asyd|OJJ65209.1 ater|EAU29516.1 atub|OJI80854.1 aver|OJI99358.1 awen|OJJ37620.1 azon|OJJ47009.1

Group1519: abra|OJJ67183.1 acar|OOF98379.1 acla|EAW14540.1 afis|EAW19258.1 afla|EED47618.1 afum|EAL93985.1 agla|OJJ88437.1 akaw|GAA90993.1 aluc|OJZ87792.1 anid|CBF78676.1 anig|EHA20273.1

aory|EIT80276.1 asyd|OJJ65208.1 ater|EAU29517.1 atub|OJI80855.1 aver|OJI99357.1 awen|OJJ37619.1 azon|OJJ47008.1

Group1520: abra|OJJ67187.1 acar|OOF98377.1 acla|EAW14537.1 afis|EAW19253.1 afla|EED47621.1 afum|EAL93982.1 agla|OJJ88444.1 akaw|GAA90990.1 aluc|OJZ87788.1 anid|CBF78671.1 anig|EHA20270.1 aory|EIT80226.1 asyd|OJJ65205.1 ater|EAU29510.1 atub|OJI80859.1 aver|OJI99354.1 awen|OJJ37616.1 azon|OJJ47005.1

Group1521: abra|OJJ67189.1 acar|OOF98374.1 acla|EAW14534.1 afis|EAW19252.1 afla|EED47626.1 afum|EAL93981.1 agla|OJJ88445.1 akaw|GAA90988.1 aluc|OJZ87786.1 anid|CBF78669.1 anig|EHA20266.1 aory|EIT80202.1 asyd|OJJ65202.1 ater|EAU29506.1 atub|OJI80861.1 aver|OJI99351.1 awen|OJJ37615.1 azon|OJJ47004.1

Group1522: abra|OJJ67190.1 acar|OOF98373.1 acla|EAW14533.1 afis|EAW19251.1 afla|EED47627.1 afum|EAL93980.1 agla|OJJ88446.1 akaw|GAA90987.1 aluc|OJZ87785.1 anid|CBF78667.1 anig|EHA20265.1 aory|EIT80224.1 asyd|OJJ65201.1 ater|EAU29505.1 atub|OJI80862.1 aver|OJI99350.1 awen|OJJ37614.1 azon|OJJ51262.1

Group1523: abra|OJJ67192.1 acar|OOF98372.1 acla|EAW14532.1 afis|EAW19250.1 afla|EED47628.1 afum|EAL93979.1 agla|OJJ88447.1 akaw|GAA90986.1 aluc|OJZ87784.1 anid|CBF78663.1 anig|EHA20264.1 aory|EIT80188.1 asyd|OJJ65199.1 ater|EAU29504.1 atub|OJI80863.1 aver|OJI99348.1 awen|OJJ37612.1 azon|OJJ51261.1

Group1524: abra|OJJ67193.1 acar|OOF98371.1 acla|EAW14531.1 afis|EAW19249.1 afla|EED47629.1 afum|EAL93978.1 agla|OJJ88448.1 akaw|GAA90985.1 aluc|OJZ87782.1 anid|CBF78665.1 anig|EHA20263.1 aory|EIT80199.1 asyd|OJJ65200.1 ater|EAU29503.1 atub|OJI80865.1 aver|OJI99349.1 awen|OJJ37611.1 azon|OJJ51260.1

Group1525: abra|OJJ67195.1 acar|OOF98369.1 acla|EAW14529.1 afis|EAW19247.1 afla|EED47631.1 afum|EAL93976.1 agla|OJJ88450.1 akaw|GAA90983.1 aluc|OJZ87780.1 anid|CBF78658.1 anig|EHA20262.1 aory|EIT80177.1 asyd|OJJ65197.1 ater|EAU29501.1 atub|OJI80867.1 aver|OJI99346.1 awen|OJJ37604.1 azon|OJJ51257.1

Group1526: abra|OJJ67200.1 acar|OOF98364.1 acla|EAW14524.1 afis|EAW19242.1 afla|EED47635.1 afum|EAL93971.2 agla|OJJ88455.1 akaw|GAA90978.1 aluc|OJZ87774.1 anid|CBF78650.1 anig|EHA20258.1 aory|EIT80223.1 asyd|OJJ65194.1 ater|EAU29498.1 atub|OJI80871.1 aver|OJI99343.1 awen|OJJ37598.1 azon|OJJ51253.1

Group1527: abra|OJJ67201.1 acar|OOF98362.1 acla|EAW14523.1 afis|EAW19241.1 afla|EED47636.1 afum|EAL93970.2 agla|OJJ88456.1 akaw|GAA90977.1 aluc|OJZ87773.1 anid|CBF78648.1 anig|EHA20257.1 aory|EIT80218.1 asyd|OJJ65193.1 ater|EAU29497.1 atub|OJI80872.1 aver|OJI99342.1 awen|OJJ37597.1 azon|OJJ51252.1

Group1528: abra|OJJ67202.1 acar|OOF98361.1 acla|EAW14522.1 afis|EAW19240.1 afla|EED47637.1 afum|EAL93969.1 agla|OJJ88458.1 akaw|GAA90976.1 aluc|OJZ87772.1 anid|CBF78646.1 anig|EHA20256.1 aory|EIT80185.1 asyd|OJJ65192.1 ater|EAU37223.1 atub|OJI80873.1 aver|OJI99341.1 awen|OJJ37596.1 azon|OJJ51251.1

Group1529: abra|OJJ67203.1 acar|OOF98360.1 acla|EAW14521.1 afis|EAW19239.1 afla|EED47638.1 afum|EBA27509.1 agla|OJJ88459.1 akaw|GAA90975.1 aluc|OJZ87771.1 anid|CBF78644.1 anig|EHA20255.1 aory|EIT80222.1 asyd|OJJ65191.1 ater|EAU37224.1 atub|OJI80874.1 aver|OJI99340.1 awen|OJJ37595.1 azon|OJJ51250.1

Group1530: abra|OJJ67204.1 acar|OOF98359.1 acla|EAW14520.1 afis|EAW19238.1 afla|EED47639.1 afum|EAL93966.1 agla|OJJ88461.1 akaw|GAA90974.1 aluc|OJZ87770.1 anid|CBF78642.1 anig|EHA20254.1 aory|EIT80175.1 asyd|OJJ65190.1 ater|EAU37225.1 atub|OJI80875.1 aver|OJI99339.1 awen|OJJ37594.1 azon|OJJ51249.1

Group1531: abra|OJJ67213.1 acar|OOF98346.1 acla|EAW08553.1 afis|EAW23595.1 afla|EED47505.1 afum|EAL87872.1 agla|OJJ86855.1 akaw|GAA90964.1 aluc|OJZ87760.1 anid|CBF89585.1 anig|EHA20245.1 aory|EIT74581.1 asyd|OJJ62318.1 ater|EAU35355.1 atub|OJI80884.1 aver|OJI96426.1 awen|OJJ39917.1 azon|OJJ45891.1

Group1532: abra|OJJ67215.1 acar|OOF98343.1 acla|EAW10167.1 afis|EAW25427.1 afla|EED53915.1 afum|EAL90086.1 agla|OJJ87071.1 akaw|GAA90962.1 aluc|OJZ87757.1 anid|CBF77551.1 anig|EHA20243.1 aory|EIT78350.1 asyd|OJJ53152.1 ater|EAU34597.1 atub|OJI80886.1 aver|OJJ03699.1 awen|OJJ41156.1 azon|OJJ44734.1

Group1533: abra|OJJ67225.1 acar|OOF98326.1 acla|EAW10408.1 afis|EAW25160.1 afla|EED53622.1 afum|EAL89835.1 agla|OJJ88637.1 akaw|GAA91652.1 aluc|OJZ87749.1 anid|CBF80654.1 anig|EHA20237.1 aory|EIT75132.1 asyd|OJJ54613.1 ater|EAU34333.1 atub|OJI80894.1 aver|OJJ07545.1 awen|OJJ33447.1 azon|OJJ50704.1

Group1534: abra|OJJ67239.1 acar|OOF89889.1 acla|EAW13993.1 afis|EAW18701.1 afla|EED50650.1 afum|EAL93424.1 agla|OJJ82360.1 akaw|GAA91664.1 aluc|OJZ87736.1 anid|CBF70640.1 anig|EHA20228.1 aory|EIT78150.1 asyd|OJJ58491.1 ater|EAU38205.1 atub|OJI80904.1 aver|OJJ01094.1 awen|OJJ32184.1 azon|OJJ48705.1

Group1535: abra|OJJ67298.1 acar|OOF92514.1 acla|EAW15241.1 afis|EAW20208.1 afla|EED57135.1 afum|EAL85325.1 agla|OJJ84656.1 akaw|GAA86516.1 aluc|OJZ85061.1 anid|CBF85098.1 anig|EHA25276.1 aory|EIT74134.1 asyd|OJJ63059.1 ater|EAU32018.1 atub|OJI80536.1 aver|OJI97904.1 awen|OJJ31212.1 azon|OJJ46148.1

Group1536: abra|OJJ67306.1 acar|OOF92507.1 acla|EAW06427.1 afis|EAW18531.1 afla|EED52870.1 afum|EAL93228.1 agla|OJJ85144.1 akaw|GAA86509.1 aluc|OJZ85052.1 anid|CBF70204.1 anig|EHA25269.1 aory|EIT78588.1 asyd|OJJ58784.1 ater|EAU30039.1 atub|OJI80546.1 aver|OJJ01392.1 awen|OJJ31089.1 azon|OJJ50594.1

Group1537: abra|OJJ67308.1 acar|OOF92504.1 acla|EAW06424.1 afis|EAW18528.1 afla|EED52873.1 afum|EAL93225.1 agla|OJJ85141.1 akaw|GAA86507.1 aluc|OJZ85050.1 anid|CBF70208.1 anig|EHA25267.1 aory|EIT78611.1 asyd|OJJ58781.1 ater|EAU30041.1 atub|OJI80548.1 aver|OJJ01389.1 awen|OJJ31086.1 azon|OJJ50591.1

Group1538: abra|OJJ67310.1 acar|OOF92502.1 acla|EAW06422.1 afis|EAW18526.1 afla|EED52875.1 afum|EAL93221.1 agla|OJJ85140.1 akaw|GAA86505.1 aluc|OJZ85048.1 anid|CBF70211.1 anig|EHA25265.1 aory|EIT78615.1 asyd|OJJ58779.1 ater|EAU30043.1 atub|OJI80550.1 aver|OJJ01388.1 awen|OJJ31080.1 azon|OJJ50588.1

Group1539: abra|OJJ67311.1 acar|OOF92501.1 acla|EAW06421.1 afis|EAW18525.1 afla|EED52876.1 afum|EAL93220.1 agla|OJJ85139.1 akaw|GAA86504.1 aluc|OJZ85047.1 anid|CBF70213.1 anig|EHA25264.1 aory|EIT78600.1 asyd|OJJ58778.1 ater|EAU30044.1 atub|OJI80551.1 aver|OJJ01387.1 awen|OJJ31079.1 azon|OJJ50587.1

Group1540: abra|OJJ67319.1 acar|OOF92496.1 acla|EAW06417.1 afis|EAW18521.1 afla|EED52877.1 afum|EAL93216.1 agla|OJJ85133.1 akaw|GAA86499.1 aluc|OJZ85041.1 anid|CBF70223.1 anig|EHA25259.1 aory|EIT78623.1 asyd|OJJ58774.1 ater|EAU30049.1 atub|OJI80556.1 aver|OJJ01383.1 awen|OJJ31073.1 azon|OJJ49993.1

Group1541: abra|OJJ67321.1 acar|OOF92493.1 acla|EAW06415.1 afis|EAW18519.1 afla|EED52879.1 afum|EAL93214.1 agla|OJJ85131.1 akaw|GAA86497.1 aluc|OJZ85039.1 anid|CBF70226.1 anig|EHA25257.1 aory|EIT78673.1 asyd|OJJ58773.1 ater|EAU30051.1 atub|OJI80558.1 aver|OJJ01382.1 awen|OJJ31071.1 azon|OJJ50581.1

Group1542: abra|OJJ67323.1 acar|OOF92490.1 acla|EAW06412.1 afis|EAW18516.1 afla|EED52882.1 afum|EAL93211.1 agla|OJJ85129.1 akaw|GAA86493.1 aluc|OJZ85035.1 anid|CBF70232.1 anig|EHA25255.1 aory|EIT78656.1 asyd|OJJ58771.1 ater|EAU30054.1 atub|OJI80562.1 aver|OJJ01380.1 awen|OJJ31592.1 azon|OJJ50579.1

Group1543: abra|OJJ67325.1 acar|OOF92488.1 acla|EAW06410.1 afis|EAW18514.1 afla|EED52884.1 afum|EAL93209.1 agla|OJJ85107.1 akaw|GAA86491.1 aluc|OJZ85033.1 anid|CBF70234.1 anig|EHA25253.1 aory|EIT78594.1 asyd|OJJ58770.1 ater|EAU30055.1 atub|OJI80564.1 aver|OJJ01379.1 awen|OJJ31068.1 azon|OJJ50577.1

Group1544: abra|OJJ67326.1 acar|OOF92487.1 acla|EAW06409.1 afis|EAW18513.1 afla|EED52887.1 afum|EAL93208.1 agla|OJJ85108.1 akaw|GAA86490.1 aluc|OJZ85032.1 anid|CBF70236.1 anig|EHA25252.1 aory|EIT78631.1 asyd|OJJ58769.1 ater|EAU30057.1 atub|OJI80565.1 aver|OJJ01378.1 awen|OJJ31067.1 azon|OJJ50576.1

Group1545: abra|OJJ67327.1 acar|OOF92486.1 acla|EAW06407.1 afis|EAW18511.1 afla|EED52889.1 afum|EAL93206.1 agla|OJJ85109.1 akaw|GAA86489.1 aluc|OJZ85030.1 anid|CBF70239.1 anig|EHA25251.1 aory|EIT78676.1 asyd|OJJ58767.1 ater|EAU30058.1 atub|OJI80567.1 aver|OJJ01376.1 awen|OJJ31065.1 azon|OJJ50575.1

Group1546: abra|OJJ67328.1 acar|OOF92485.1 acla|EAW06406.1 afis|EAW18510.1 afla|EED52890.1 afum|EAL93205.1 agla|OJJ85110.1 akaw|GAA86488.1 aluc|OJZ85029.1 anid|CBF70241.1 anig|EHA25250.1 aory|EIT78606.1 asyd|OJJ58766.1 ater|EAU30059.1 atub|OJI80568.1 aver|OJJ01375.1 awen|OJJ31064.1 azon|OJJ50574.1

Group1547: abra|OJJ67329.1 acar|OOF92483.1 acla|EAW06405.1 afis|EAW18509.1 afla|EED52891.1 afum|EAL93204.1 agla|OJJ85111.1 akaw|GAA86487.1 aluc|OJZ85028.1 anid|CBF70243.1 anig|EHA25249.1 aory|EIT78599.1 asyd|OJJ58765.1 ater|EAU30060.1 atub|OJI80569.1 aver|OJJ01374.1 awen|OJJ31063.1 azon|OJJ50573.1

Group1548: abra|OJJ67330.1 acar|OOF92484.1 acla|EAW06404.1 afis|EAW18508.1 afla|EED52894.1 afum|EAL93203.1 agla|OJJ85112.1 akaw|GAA86486.1 aluc|OJZ85027.1 anid|CBF70245.1 anig|EHA25248.1

aory|EIT78590.1 asyd|OJJ58762.1 ater|EAU30063.1 atub|OJI80570.1 aver|OJJ01372.1 awen|OJJ31061.1 azon|OJJ50572.1

Group1549: abra|OJJ67334.1 acar|OOF92479.1 acla|EAW06400.1 afis|EAW18504.1 afla|EED52897.1 afum|EAL93197.1 agla|OJJ85119.1 akaw|GAA86482.1 aluc|OJZ85023.1 anid|CBF70249.1 anig|EHA25244.1 aory|EIT78613.1 asyd|OJJ58759.1 ater|EAU30067.1 atub|OJI80574.1 aver|OJJ01367.1 awen|OJJ31058.1 azon|OJJ50567.1

Group1550: abra|OJJ67335.1 acar|OOF92478.1 acla|EAW06399.1 afis|EAW18503.1 afla|EED52898.1 afum|EAL93196.2 agla|OJJ85118.1 akaw|GAA86481.1 aluc|OJZ85022.1 anid|CBF70251.1 anig|EHA25243.1 aory|EIT78654.1 asyd|OJJ58758.1 ater|EAU30068.1 atub|OJI80575.1 aver|OJJ01366.1 awen|OJJ31057.1 azon|OJJ50568.1

Group1551: abra|OJJ67336.1 acar|OOF92477.1 acla|EAW06398.1 afis|EAW18502.1 afla|EED52899.1 afum|EAL93195.1 agla|OJJ85123.1 akaw|GAA86480.1 aluc|OJZ85021.1 anid|CBF70253.1 anig|EHA25242.1 aory|EIT78660.1 asyd|OJJ58757.1 ater|EAU30069.1 atub|OJI80576.1 aver|OJJ01365.1 awen|OJJ31056.1 azon|OJJ50566.1

Group1552: abra|OJJ67339.1 acar|OOF92474.1 acla|EAW06395.1 afis|EAW18499.1 afla|EED52902.1 afum|EAL93190.1 agla|OJJ85128.1 akaw|GAA86477.1 aluc|OJZ85018.1 anid|CBF70257.1 anig|EHA25239.1 aory|EIT78634.1 asyd|OJJ58755.1 ater|EAU30071.1 atub|OJI80579.1 aver|OJJ01363.1 awen|OJJ31053.1 azon|OJJ50563.1

Group1553: abra|OJJ67340.1 acar|OOF92473.1 acla|EAW06394.1 afis|EAW18498.1 afla|EED52903.1 afum|EAL93188.1 agla|OJJ85105.1 akaw|GAA86476.1 aluc|OJZ85017.1 anid|CBF70259.1 anig|EHA25238.1 aory|EIT78622.1 asyd|OJJ58754.1 ater|EAU30072.1 atub|OJI80580.1 aver|OJJ01362.1 awen|OJJ31052.1 azon|OJJ50564.1

Group1554: abra|OJJ67346.1 acar|OOF92469.1 acla|EAW06390.1 afis|EAW18496.1 afla|EED52907.1 afum|EAL93186.1 agla|OJJ85091.1 akaw|GAA86472.1 aluc|OJZ85013.1 anid|CBF70170.1 anig|EHA25234.1 aory|EIT78612.1 asyd|OJJ58795.1 ater|EAU30073.1 atub|OJI80584.1 aver|OJJ01400.1 awen|OJJ31049.1 azon|OJJ50560.1

Group1555: abra|OJJ67347.1 acar|OOF92468.1 acla|EAW06389.1 afis|EAW18495.1 afla|EED52908.1 afum|EAL93185.1 agla|OJJ85090.1 akaw|GAA86471.1 aluc|OJZ85012.1 anid|CBF70168.1 anig|EHA25233.1 aory|EIT78663.1 asyd|OJJ58797.1 ater|EAU29921.1 atub|OJI80585.1 aver|OJJ01401.1 awen|OJJ31048.1 azon|OJJ50559.1

Group1556: abra|OJJ67350.1 acar|OOF92466.1 acla|EAW06386.1 afis|EAW18492.1 afla|EED52912.1 afum|EAL93182.1 agla|OJJ85087.1 akaw|GAA86468.1 aluc|OJZ85009.1 anid|CBF70162.1 anig|EHA25230.1 aory|EIT78650.1 asyd|OJJ58800.1 ater|EAU29918.1 atub|OJI80588.1 aver|OJJ01404.1 awen|OJJ31045.1 azon|OJJ50556.1

Group1557: abra|OJJ67375.1 acar|OOF92436.1 acla|EAW06380.1 afis|EAW18486.1 afla|EED52922.1 afum|EAL93175.1 agla|OJJ85081.1 akaw|GAA86447.1 aluc|OJZ84990.1 anid|CBF70147.1 anig|EHA25202.1 aory|EIT78638.1 asyd|OJJ58808.1 ater|EAU29911.1 atub|OJI80610.1 aver|OJJ01411.1 awen|OJJ31033.1 azon|OJJ50552.1

Group1558: abra|OJJ67377.1 acar|OOF92435.1 acla|EAW06379.1 afis|EAW18485.1 afla|EED52923.1 afum|EBA27486.1 agla|OJJ85074.1 akaw|GAA86446.1 aluc|OJZ84988.1 anid|CBF70145.1 anig|EHA25201.1 aory|EIT78668.1 asyd|OJJ58809.1 ater|EAU29910.1 atub|OJI80611.1 aver|OJJ01412.1 awen|OJJ31035.1 azon|OJJ50550.1

Group1559: abra|OJJ67380.1 acar|OOF92432.1 acla|EAW06377.1 afis|EAW18483.1 afla|EED52925.1 afum|EAL93172.1 agla|OJJ85072.1 akaw|GAA86444.1 aluc|OJZ84986.1 anid|CBF86341.1 anig|EHA25199.1 aory|EIT78658.1 asyd|OJJ64985.1 ater|EAU29909.1 atub|OJI80614.1 aver|OJI99105.1 awen|OJJ31037.1 azon|OJJ50548.1

Group1560: abra|OJJ67381.1 acar|OOF92433.1 acla|EAW06376.1 afis|EAW18482.1 afla|EED52926.1 afum|EAL93171.1 agla|OJJ85063.1 akaw|GAA86443.1 aluc|OJZ84985.1 anid|CBF86343.1 anig|EHA25198.1 aory|EIT78671.1 asyd|OJJ64984.1 ater|EAU29908.1 atub|OJI80615.1 aver|OJI99104.1 awen|OJJ31038.1 azon|OJJ50547.1

Group1561: abra|OJJ67385.1 acar|OOF92428.1 acla|EAW06374.1 afis|EAW18480.1 afla|EED52929.1 afum|EAL93169.1 agla|OJJ85066.1 akaw|GAA86439.1 aluc|OJZ84981.1 anid|CBF70139.1 anig|EHA25194.1 aory|EIT78702.1 asyd|OJJ58812.1 ater|EAU29896.1 atub|OJI80620.1 aver|OJJ01415.1 awen|OJJ31040.1 azon|OJJ50545.1

Group1562: abra|OJJ67387.1 acar|OOF92427.1 acla|EAW06372.1 afis|EAW18478.1 afla|EED52932.1 afum|EAL93166.1 agla|OJJ85068.1 akaw|GAA86437.1 aluc|OJZ84980.1 anid|CBF70136.1 anig|EHA25192.1 aory|EIT78730.1 asyd|OJJ58814.1 ater|EAU29895.1 atub|OJI80622.1 aver|OJJ01417.1 awen|OJJ31031.1 azon|OJJ50544.1

Group1563: abra|OJJ67393.1 acar|OOF92423.1 acla|EAW06370.1 afis|EAW18476.1 afla|EED52934.1 afum|EAL93163.1 agla|OJJ85071.1 akaw|GAA86434.1 aluc|OJZ84976.1 anid|CBF70132.1 anig|EHA25188.1 aory|EIT78693.1 asyd|OJJ58817.1 ater|EAU29893.1 atub|OJI80625.1 aver|OJJ01421.1 awen|OJJ31029.1 azon|OJJ50542.1

Group1564: abra|OJJ67394.1 acar|OOF92422.1 acla|EAW06369.1 afis|EAW18475.1 afla|EED52935.1 afum|EAL93162.1 agla|OJJ85062.1 akaw|GAA86433.1 aluc|OJZ84975.1 anid|CBF70116.1 anig|EHA25187.1 aory|EIT78723.1 asyd|OJJ58819.1 ater|EAU29892.1 atub|OJI80626.1 aver|OJJ01423.1 awen|OJJ31028.1 azon|OJJ50541.1

Group1565: abra|OJJ67395.1 acar|OOF92421.1 acla|EAW06368.1 afis|EAW18474.1 afla|EED52936.1 afum|EAL93161.1 agla|OJJ85061.1 akaw|GAA86432.1 aluc|OJZ84974.1 anid|CBF70108.1 anig|EHA25186.1 aory|EIT78688.1 asyd|OJJ58823.1 ater|EAU29891.1 atub|OJI80627.1 aver|OJJ01427.1 awen|OJJ31026.1 azon|OJJ50540.1

Group1566: abra|OJJ67396.1 acar|OOF92419.1 acla|EAW06364.1 afis|EAW18473.1 afla|EED52937.1 afum|EAL93160.2 agla|OJJ85060.1 akaw|GAA86431.1 aluc|OJZ84973.1 anid|CBF70107.1 anig|EHA25185.1 aory|EIT78722.1 asyd|OJJ58827.1 ater|EAU29890.1 atub|OJI80628.1 aver|OJJ01430.1 awen|OJJ31025.1 azon|OJJ50539.1

Group1567: abra|OJJ67398.1 acar|OOF92418.1 acla|EAW06362.1 afis|EAW18471.1 afla|EED52939.1 afum|EAL93158.1 agla|OJJ85056.1 akaw|GAA86429.1 aluc|OJZ84971.1 anid|CBF70105.1 anig|EHA25183.1 aory|EIT78698.1 asyd|OJJ58829.1 ater|EAU29888.1 atub|OJI80630.1 aver|OJJ01432.1 awen|OJJ31023.1 azon|OJJ50537.1

Group1568: abra|OJJ67399.1 acar|OOF92417.1 acla|EAW06359.1 afis|EAW18470.1 afla|EED52940.1 afum|EAL93156.1 agla|OJJ85055.1 akaw|GAA86428.1 aluc|OJZ84969.1 anid|CBF70103.1 anig|EHA25182.1 aory|EIT78679.1 asyd|OJJ58830.1 ater|EAU29887.1 atub|OJI80631.1 aver|OJJ01433.1 awen|OJJ31022.1 azon|OJJ50536.1

Group1569: abra|OJJ67400.1 acar|OOF92416.1 acla|EAW06358.1 afis|EAW18469.1 afla|EED52941.1 afum|EAL93155.1 agla|OJJ85054.1 akaw|GAA86427.1 aluc|OJZ84968.1 anid|CBF70101.1 anig|EHA25181.1 aory|EIT78712.1 asyd|OJJ58831.1 ater|EAU29886.1 atub|OJI80632.1 aver|OJJ01434.1 awen|OJJ31021.1 azon|OJJ50535.1

Group1570: abra|OJJ67401.1 acar|OOF92415.1 acla|EAW06357.1 afis|EAW18468.1 afla|EED52942.1 afum|EAL93154.1 agla|OJJ85053.1 akaw|GAA86426.1 aluc|OJZ84967.1 anid|CBF70100.1 anig|EHA25180.1 aory|EIT78694.1 asyd|OJJ58832.1 ater|EAU29885.1 atub|OJI80633.1 aver|OJJ01435.1 awen|OJJ31020.1 azon|OJJ50534.1

Group1571: abra|OJJ67403.1 acar|OOF92413.1 acla|EAW06355.1 afis|EAW18466.1 afla|EED52944.1 afum|EAL93152.1 agla|OJJ85051.1 akaw|GAA86424.1 aluc|OJZ84965.1 anid|CBF70096.1 anig|EHA25178.1 aory|EIT78707.1 asyd|OJJ58834.1 ater|EAU29883.1 atub|OJI80635.1 aver|OJJ01437.1 awen|OJJ31018.1 azon|OJJ50532.1

Group1572: abra|OJJ67411.1 acar|OOF89994.1 acla|EAW06348.1 afis|EAW18459.1 afla|EED52949.1 afum|EAL93145.1 agla|OJJ85042.1 akaw|GAA86416.1 aluc|OJZ84957.1 anid|CBF70085.1 anig|EHA25171.1 aory|EIT78721.1 asyd|OJJ58842.1 ater|EAU29877.1 atub|OJI80643.1 aver|OJJ01444.1 awen|OJJ31011.1 azon|OJJ50525.1

Group1573: abra|OJJ67412.1 acar|OOF89992.1 acla|EAW06347.1 afis|EAW18458.1 afla|EED52950.1 afum|EAL93143.1 agla|OJJ85041.1 akaw|GAA86415.1 aluc|OJZ84956.1 anid|CBF70084.1 anig|EHA25170.1 aory|EIT78692.1 asyd|OJJ58843.1 ater|EAU29876.1 atub|OJI80644.1 aver|OJJ01445.1 awen|OJJ31010.1 azon|OJJ50524.1

Group1574: abra|OJJ67419.1 acar|OOF92402.1 acla|EAW06345.1 afis|EAW18456.1 afla|EED52952.1 afum|EAL93141.1 agla|OJJ85039.1 akaw|GAA86409.1 aluc|OJZ84949.1 anid|CBF70053.1 anig|EHA25164.1 aory|EIT78736.1 asyd|OJJ58857.1 ater|EAU29873.1 atub|OJI80653.1 aver|OJJ01458.1 awen|OJJ31008.1 azon|OJJ50522.1

Group1575: abra|OJJ67421.1 acar|OOF92401.1 acla|EAW06344.1 afis|EAW18455.1 afla|EED52953.1 afum|EAL93140.1 agla|OJJ85038.1 akaw|GAA86408.1 aluc|OJZ84947.1 anid|CBF70057.1 anig|EHA25163.1 aory|EIT78696.1 asyd|OJJ58855.1 ater|EAU29872.1 atub|OJI80654.1 aver|OJJ01456.1 awen|OJJ31007.1 azon|OJJ50521.1

Group1576: abra|OJJ67423.1 acar|OOF89873.1 acla|EAW06341.1 afis|EAW18453.1 afla|EED52955.1 afum|EAL93138.1 agla|OJJ85032.1 akaw|GAA86406.1 aluc|OJZ84944.1 anid|CBF70061.1 anig|EHA25161.1 aory|EIT78714.1 asyd|OJJ58853.1 ater|EAU29870.1 atub|OJI80656.1 aver|OJJ01454.1 awen|OJJ31005.1 azon|OJJ50519.1

Group1577: abra|OJJ67424.1 acar|OOF89874.1 acla|EAW06340.1 afis|EAW18452.1 afla|EED52956.1 afum|EAL93137.1 agla|OJJ85030.1 akaw|GAA86405.1 aluc|OJZ84943.1 anid|CBF70062.1 anig|EHA25160.1

aory|EIT78680.1 asyd|OJJ58852.1 ater|EAU29869.1 atub|OJI80657.1 aver|OJJ01453.1 awen|OJJ31004.1 azon|OJJ50517.1

Group1578: abra|OJJ67426.1 acar|OOF92399.1 acla|EAW06338.1 afis|EAW18450.1 afla|EED52958.1 afum|EAL93135.1 agla|OJJ85028.1 akaw|GAA86404.1 aluc|OJZ84941.1 anid|CBF70064.1 anig|EHA25158.1 aory|EIT78699.1 asyd|OJJ58850.1 ater|EAU29868.1 atub|OJI80659.1 aver|OJJ01451.1 awen|OJJ31002.1 azon|OJJ50516.1

Group1579: abra|OJJ67429.1 acar|OOF92397.1 acla|EAW06335.1 afis|EAW18447.1 afla|EED52963.1 afum|EAL93132.1 agla|OJJ85024.1 akaw|GAA86401.1 aluc|OJZ84937.1 anid|CBF70070.1 anig|EHA25155.1 aory|EIT78728.1 asyd|OJJ58847.1 ater|EAU29866.1 atub|OJI80662.1 aver|OJJ01448.1 awen|OJJ30999.1 azon|OJJ50513.1

Group1580: abra|OJJ67431.1 anig|EHA25151.1 atub|OJI80666.1 abra|OJJ69856.1 acar|OOF97498.1 akaw|GAA87800.1 aluc|OJZ85435.1 anig|EHA24981.1 atub|OJI80133.1 abra|OJJ75776.1 acar|OOF95028.1 acla|EAW07821.1 afis|EAW22857.1 afla|EED47892.1 afum|EAL90343.1 akaw|GAA86398.1 aory|EIT77247.1 aluc|OJZ84933.1

Group1581: abra|OJJ67434.1 acar|OOF92394.1 acla|EAW06332.1 afis|EAW18445.1 afla|EED52967.1 afum|EAL93130.1 agla|OJJ85021.1 akaw|GAA86396.1 aluc|OJZ84930.1 anid|CBF70041.1 anig|EHA25149.1 aory|EIT78729.1 asyd|OJJ58865.1 ater|EAU29864.1 atub|OJI80669.1 aver|OJJ01466.1 awen|OJJ30993.1 azon|OJJ50510.1

Group1582: abra|OJJ67435.1 acar|OOF92393.1 acla|EAW06331.1 afis|EAW18444.1 afla|EED52968.1 afum|EAL93128.1 agla|OJJ85020.1 akaw|GAA86395.1 aluc|OJZ84929.1 anid|CBF70043.1 anig|EHA25148.1 aory|EIT78686.1 asyd|OJJ58864.1 ater|EAU29863.1 atub|OJI80670.1 aver|OJJ01465.1 awen|OJJ30992.1 azon|OJJ50511.1

Group1583: abra|OJJ67437.1 acar|OOF92391.1 acla|EAW06327.1 afis|EAW18441.1 afla|EED52972.1 afum|EAL93123.1 agla|OJJ85006.1 akaw|GAA86393.1 aluc|OJZ84927.1 anid|CBF70049.1 anig|EHA25146.1 aory|EIT78690.1 asyd|OJJ58861.1 ater|EAU29860.1 atub|OJI80672.1 aver|OJJ01462.1 awen|OJJ30989.1 azon|OJJ50507.1

Group1584: abra|OJJ67438.1 acar|OOF92389.1 acla|EAW06328.1 afis|EAW18442.1 afla|EED52971.1 afum|EAL93125.1 agla|OJJ85008.1 akaw|GAA86392.1 aluc|OJZ84926.1 anid|CBF70047.1 anig|EHA25145.1 aory|EIT78682.1 asyd|OJJ58862.1 ater|EAU29859.1 atub|OJI80673.1 aver|OJJ01463.1 awen|OJJ30990.1 azon|OJJ50508.1

Group1585: abra|OJJ67439.1 acar|OOF92390.1 acla|EAW06329.1 afis|EAW18443.1 afla|EED52970.1 afum|EAL93126.1 agla|OJJ85009.1 akaw|GAA86391.1 aluc|OJZ84925.1 anid|CBF70045.1 anig|EHA25144.1 aory|EIT78727.1 asyd|OJJ58863.1 ater|EAU29858.1 atub|OJI80674.1 aver|OJJ01464.1 awen|OJJ30991.1 azon|OJJ50509.1

Group1586: abra|OJJ67452.1 acla|EAW06313.1 afla|EED52986.1 akaw|GAA86379.1 aluc|OJZ84913.1 anid|CBF70694.1 anig|EHA25134.1 aory|EIT78726.1 ater|EAU29843.1 atub|OJI80685.1 azon|OJJ50483.1 afis|EAW18422.1 afum|EAL93106.1 asyd|OJJ58464.1 aver|OJJ01063.1 awen|OJJ31723.1 agla|OJJ84990.1 awen|OJJ30758.1

Group1587: abra|OJJ67471.1 acar|OOF89932.1 acla|EAW06300.1 afis|EAW18407.1 afla|EED52997.1 afum|EAL93087.1 agla|OJJ84959.1 akaw|GAA86365.1 aluc|OJZ84895.1 anid|CBF70741.1 anig|EHA25122.1 aory|EIT77423.1 asyd|OJJ58439.1 ater|EAU29825.1 atub|OJI80705.1 aver|OJJ01039.1 awen|OJJ31743.1 azon|OJJ50501.1

Group1588: abra|OJJ67609.1 acar|OOF91956.1 acla|EAW11006.1 afis|EAW20847.1 afla|EED45466.1 afum|EAL84786.1 akaw|GAA89125.1 aluc|OJZ81713.1 anid|CBF82875.1 anig|EHA25726.1 aory|EIT75524.1 asyd|OJJ61570.1 ater|EAU30786.1 atub|OJI88471.1 aver|OJJ00890.1 awen|OJJ34544.1 azon|OJJ42460.1 ater|EAU37631.1

Group1589: abra|OJJ67742.1 acar|OOF90949.1 acla|EAW10991.1 afis|EAW20869.1 afla|EED45539.1 afum|EAL84765.1 agla|OJJ83977.1 akaw|GAA92433.1 aluc|OJZ81588.1 anid|CBF82842.1 anig|EHA25852.1 aory|EIT75440.1 asyd|OJJ61589.1 ater|EAU30819.1 atub|OJI88594.1 aver|OJJ00911.1 awen|OJJ34495.1 azon|OJJ51769.1

Group1590: abra|OJJ67743.1 acar|OOF90948.1 acla|EAW10992.1 afis|EAW20868.1 afla|EED45543.1 afum|EAL84766.2 agla|OJJ83976.1 akaw|GAA92434.1 aluc|OJZ81586.1 anid|CBF82840.1 anig|EHA25853.1 aory|EIT75711.1 asyd|OJJ61588.1 ater|EAU30820.1 atub|OJI88595.1 aver|OJJ00910.1 awen|OJJ34498.1 azon|OJJ51773.1

Group1591: abra|OJJ67744.1 acar|OOF90947.1 acla|EAW10993.1 afis|EAW20867.1 afla|EED45544.1 afum|EAL84767.1 agla|OJJ83975.1 akaw|GAA92435.1 aluc|OJZ81585.1 anid|CBF82838.1 anig|EHA25854.1 aory|EIT75422.1 asyd|OJJ61587.1 ater|EAU30821.1 atub|OJI88596.1 aver|OJJ00909.1 awen|OJJ34499.1 azon|OJJ51774.1

Group1592: abra|OJJ67747.1 acar|OOF90944.1 acla|EAW10997.1 afis|EAW20864.1 afla|EED45548.1 afum|EAL84771.1 agla|OJJ83751.1 akaw|GAA92438.1 aluc|OJZ81582.1 anid|CBF82835.1 anig|EHA25857.1 aory|EIT75692.1 asyd|OJJ61584.1 ater|EAU30807.1 atub|OJI88599.1 aver|OJJ00906.1 awen|OJJ34506.1 azon|OJJ51784.1

Group1593: abra|OJJ67748.1 acar|OOF90943.1 acla|EAW10998.1 afis|EAW20863.1 afla|EED45549.1 afum|EAL84772.1 agla|OJJ83750.1 akaw|GAA92439.1 aluc|OJZ81581.1 anid|CBF82833.1 anig|EHA25858.1 aory|EIT75610.1 asyd|OJJ61583.1 ater|EAU30806.1 atub|OJI88600.1 aver|OJJ00905.1 awen|OJJ34507.1 azon|OJJ51785.1

Group1594: abra|OJJ67749.1 acar|OOF90950.1 acla|EAW10990.1 afis|EAW20870.1 afla|EED45552.1 afum|EAL84764.1 agla|OJJ83978.1 akaw|GAA92440.1 aluc|OJZ81580.1 anid|CBF82844.1 anig|EHA25859.1 aory|EIT75541.1 asyd|OJJ61590.1 ater|EAU30818.1 atub|OJI88601.1 aver|OJJ00912.1 awen|OJJ34494.1 azon|OJJ51768.1

Group1595: abra|OJJ67799.1 acar|OOF94206.1 agla|OJJ79105.1 anid|CBF81915.1 asyd|OJJ59697.1 ater|EAU35300.1 aver|OJJ04270.1 awen|OJJ41729.1 azon|OJJ43893.1 asyd|OJJ65109.1 aver|OJI99236.1 asyd|OJJ55389.1 aver|OJJ08268.1 aver|OJJ02590.1 awen|OJJ34589.1 asyd|OJJ54350.1 aver|OJJ05302.1 asyd|OJJ58135.1

Group1596: abra|OJJ67847.1 acar|OOF92828.1 acla|EAW11752.1 afis|EAW23785.1 afla|EED51594.1 afum|EAL87219.2 agla|OJJ78938.1 akaw|GAA91855.1 aluc|OJZ82044.1 anid|CBF79845.1 anig|EHA22213.1 aory|EIT80570.1 asyd|OJJ58153.1 ater|EAU31326.1 atub|OJI84603.1 aver|OJJ04251.1 awen|OJJ33730.1 azon|OJJ42852.1

Group1597: abra|OJJ67848.1 acar|OOF92829.1 acla|EAW11751.1 afis|EAW23784.1 afla|EED51593.1 afum|EAL87218.2 agla|OJJ78937.1 akaw|GAA91856.1 aluc|OJZ82043.1 anid|CBF79841.1 anig|EHA22214.1 aory|EIT80650.1 asyd|OJJ58152.1 ater|EAU31325.1 atub|OJI84602.1 aver|OJJ04252.1 awen|OJJ33729.1 azon|OJJ42853.1

Group1598: abra|OJJ67851.1 acar|OOF92832.1 acla|EAW11748.1 afis|EAW23781.1 afla|EED51585.1 afum|EAL87215.1 agla|OJJ78933.1 akaw|GAA91859.1 aluc|OJZ82040.1 anid|CBF79839.1 anig|EHA22216.1 aory|EIT80522.1 asyd|OJJ58148.1 ater|EAU31322.1 atub|OJI84599.1 aver|OJJ04256.1 awen|OJJ33724.1 azon|OJJ42856.1

Group1599: abra|OJJ67859.1 acar|OOF92840.1 acla|EAW11739.1 afis|EAW23772.1 afla|EED51573.1 afum|EAL87206.1 agla|OJJ78947.1 akaw|GAA91866.1 aluc|OJZ82033.1 anid|CBF79823.1 anig|EHA22223.1 aory|EIT80715.1 asyd|OJJ58141.1 ater|EAU31316.1 atub|OJI84588.1 aver|OJJ04264.1 awen|OJJ33703.1 azon|OJJ42837.1

Group1600: abra|OJJ67861.1 acar|OOF92839.1 acla|EAW11737.1 afis|EAW23770.1 afla|EED51571.1 afum|EAL87204.1 agla|OJJ78949.1 anid|CBF79821.1 anig|EHA22225.1 aory|EIT80633.1 asyd|OJJ58138.1 ater|EAU31314.1 aver|OJJ04267.1 awen|OJJ33705.1 azon|OJJ42836.1 akaw|GAA91868.1 aluc|OJZ82031.1 atub|OJI84586.1

Group1601: abra|OJJ67865.1 acar|OOF92845.1 acla|EAW11734.1 afis|EAW23766.1 afla|EED51552.1 afum|EAL87199.1 agla|OJJ78956.1 akaw|GAA91872.1 aluc|OJZ82027.1 anid|CBF79770.1 anig|EHA22229.1 aory|EIT80651.1 asyd|OJJ58112.1 ater|EAU31310.1 atub|OJI84581.1 aver|OJJ04293.1 awen|OJJ33692.1 azon|OJJ42832.1

Group1602: abra|OJJ67866.1 acar|OOF92846.1 acla|EAW11733.1 afis|EAW23765.1 afla|EED51550.1 afum|EAL87198.1 agla|OJJ78957.1 akaw|GAA91873.1 aluc|OJZ82026.1 anid|CBF79772.1 anig|EHA22230.1 aory|EIT80718.1 asyd|OJJ58110.1 ater|EAU31311.1 atub|OJI84579.1 aver|OJJ04294.1 awen|OJJ33689.1 azon|OJJ42830.1

Group1603: abra|OJJ67881.1 acar|OOF92865.1 acla|EAW11722.1 afis|EAW23744.1 afla|EED51536.1 afum|EAL87180.1 agla|OJJ78963.1 akaw|GAA91888.1 aluc|OJZ82009.1 anid|CBF81714.1 anig|EHA22248.1 aory|EIT80594.1 asyd|OJJ59582.1 ater|EAU31299.1 atub|OJI84558.1 aver|OJJ02687.1 awen|OJJ33679.1 azon|OJJ42820.1

Group1604: abra|OJJ67913.1 acar|OOF92766.1 acla|EAW12796.1 afis|EAW17959.1 afla|EED51948.1 afum|EAL91770.1 agla|OJJ80207.1 akaw|GAA91283.1 aluc|OJZ82786.1 anid|CBF79950.1 anig|EHA21227.1 aory|EIT81390.1 asyd|OJJ58216.1 ater|EAU31448.1 atub|OJI89018.1 aver|OJJ02206.1 awen|OJJ41583.1 azon|OJJ48295.1

Group1605: abra|OJJ67915.1 acar|OOF92767.1 acla|EAW12794.1 afis|EAW17957.1 afla|EED51950.1 afum|EAL91768.1 agla|OJJ80204.1 akaw|GAA91281.1 aluc|OJZ82788.1 anid|CBF79947.1 anig|EHA21229.1 aory|EIT81406.1 asyd|OJJ58214.1 ater|EAU31445.1 atub|OJI89020.1 aver|OJJ02208.1 awen|OJJ41585.1 azon|OJJ48297.1

Group1606: abra|OJJ67917.1 acar|OOF92769.1 acla|EAW12792.1 afis|EAW17955.1 afla|EED51952.1 afum|EAL91766.1 agla|OJJ80202.1 akaw|GAA91279.1 aluc|OJZ82790.1 anid|CBF79943.1 anig|EHA21231.1

aory|EIT81414.1 asyd|OJJ58212.1 ater|EAU31443.1 atub|OJI89021.1 aver|OJJ02210.1 awen|OJJ41587.1 azon|OJJ48302.1

Group1607: abra|OJJ67918.1 acar|OOF92770.1 acla|EAW12791.1 afis|EAW17951.1 afla|EED51953.1 afum|EAL91761.1 agla|OJJ80201.1 akaw|GAA91278.1 aluc|OJZ82791.1 anid|CBF79941.1 anig|EHA21232.1 aory|EIT81400.1 asyd|OJJ58211.1 ater|EAU31442.1 atub|OJI89022.1 aver|OJJ02211.1 awen|OJJ41588.1 azon|OJJ48303.1

Group1608: abra|OJJ67919.1 acar|OOF92771.1 acla|EAW12790.1 afis|EAW17952.1 afla|EED51954.1 afum|EAL91762.1 agla|OJJ80200.1 akaw|GAA91277.1 aluc|OJZ82792.1 anid|CBF79939.1 anig|EHA21233.1 aory|EIT81379.1 asyd|OJJ58210.1 ater|EAU31441.1 atub|OJI89023.1 aver|OJJ02212.1 awen|OJJ41589.1 azon|OJJ48304.1

Group1609: abra|OJJ67920.1 acar|OOF92772.1 acla|EAW12789.1 afis|EAW17953.1 afla|EED51955.1 afum|EAL91763.1 agla|OJJ80199.1 akaw|GAA91276.1 aluc|OJZ82793.1 anid|CBF79934.1 anig|EHA21234.1 aory|EIT81461.1 asyd|OJJ58207.1 ater|EAU31440.1 atub|OJI89024.1 aver|OJJ02215.1 awen|OJJ41590.1 azon|OJJ48305.1

Group1610: abra|OJJ67922.1 acar|OOF92773.1 acla|EAW12788.1 afis|EAW17954.1 afla|EED51956.1 afum|EAL91765.1 agla|OJJ80198.1 akaw|GAA91275.1 aluc|OJZ82794.1 anid|CBF79932.1 anig|EHA21235.1 aory|EIT81403.1 asyd|OJJ58206.1 ater|EAU31439.1 atub|OJI89025.1 aver|OJJ02216.1 awen|OJJ41591.1 azon|OJJ48306.1

Group1611: abra|OJJ67923.1 acar|OOF92774.1 acla|EAW12787.1 afis|EAW17950.1 afla|EED51957.1 afum|EAL91760.1 agla|OJJ80197.1 akaw|GAA91274.1 aluc|OJZ82795.1 anid|CBF79931.1 anig|EHA21236.1 aory|EIT81374.1 asyd|OJJ58205.1 ater|EAU31438.1 atub|OJI89026.1 aver|OJJ02217.1 awen|OJJ41592.1 azon|OJJ48308.1

Group1612: abra|OJJ67926.1 acar|OOF92775.1 acla|EAW12785.1 afis|EAW17947.1 afla|EED51959.1 afum|EAL91757.1 agla|OJJ80195.1 akaw|GAA91271.1 aluc|OJZ82798.1 anid|CBF79929.1 anig|EHA21239.1 aory|EIT81382.1 asyd|OJJ58204.1 ater|EAU31431.1 atub|OJI89029.1 aver|OJJ04202.1 awen|OJJ41596.1 azon|OJJ48312.1

Group1613: abra|OJJ67928.1 acar|OOF92777.1 acla|EAW12781.1 afis|EAW17943.1 afla|EED51961.1 afum|EAL91754.1 agla|OJJ82989.1 akaw|GAA91269.1 aluc|OJZ82800.1 anid|CBF79925.1 anig|EHA21240.1 aory|EIT81385.1 asyd|OJJ58202.1 ater|EAU31396.1 atub|OJI89031.1 aver|OJJ04207.1 awen|OJJ41600.1 azon|OJJ48323.1

Group1614: abra|OJJ67929.1 acar|OOF92779.1 acla|EAW11622.1 afis|EAW23836.1 afla|EED51962.1 afum|EAL87267.1 agla|OJJ82498.1 akaw|GAA91268.1 aluc|OJZ82801.1 anid|CBF79923.1 anig|EHA21242.1 aory|EIT81370.1 asyd|OJJ58200.1 ater|EAU31395.1 atub|OJI89032.1 aver|OJJ04208.1 awen|OJJ33798.1 azon|OJJ48745.1

Group1615: abra|OJJ67936.1 acar|OOF92785.1 acla|EAW11625.1 afis|EAW23831.1 afla|EED51974.1 afum|EAL87263.1 agla|OJJ82481.1 akaw|GAA91260.1 aluc|OJZ82808.1 anid|CBF79904.1 anig|EHA21248.1 aory|EIT81407.1 asyd|OJJ58187.1 ater|EAU31385.1 atub|OJI89039.1 aver|OJJ04218.1 awen|OJJ33789.1 azon|OJJ48757.1

Group1616: abra|OJJ67938.1 acar|OOF92787.1 acla|EAW11628.1 afis|EAW23828.1 afla|EED51977.1 afum|EAL87260.1 agla|OJJ82484.1 akaw|GAA91258.1 aluc|OJZ82810.1 anid|CBF79908.1 anig|EHA21249.1 aory|EIT81387.1 asyd|OJJ58189.1 ater|EAU31383.1 atub|OJI89040.1 aver|OJJ04216.1 awen|OJJ33787.1 azon|OJJ48759.1

Group1617: abra|OJJ67940.1 acar|OOF92789.1 acla|EAW11630.1 afis|EAW23826.1 afla|EED51980.1 afum|EAL87258.1 agla|OJJ82486.1 akaw|GAA91256.1 aluc|OJZ82812.1 anid|CBF79911.1 anig|EHA21252.1 aory|EIT81452.1 asyd|OJJ58190.1 ater|EAU31381.1 atub|OJI89043.1 aver|OJJ04215.1 awen|OJJ33784.1 azon|OJJ42886.1

Group1618: abra|OJJ67942.1 acar|OOF92791.1 acla|EAW11636.1 afis|EAW23821.1 afla|EED51983.1 afum|EAL87256.1 agla|OJJ82468.1 akaw|GAA91254.1 aluc|OJZ82814.1 anid|CBF79894.1 anig|EHA21254.1 aory|EIT81391.1 asyd|OJJ58182.1 ater|EAU31373.1 atub|OJI89045.1 aver|OJJ04224.1 awen|OJJ33777.1 azon|OJJ42883.1

Group1619: abra|OJJ67951.1 acar|OOF92798.1 acla|EAW11641.1 afis|EAW23816.1 afla|EED51989.1 afum|EAL87251.1 agla|OJJ82462.1 akaw|GAA91246.1 aluc|OJZ82824.1 anid|CBF79887.1 anig|EHA21262.1 aory|EIT81439.1 asyd|OJJ58175.1 ater|EAU31371.1 atub|OJI89053.1 aver|OJJ04229.1 awen|OJJ33773.1 azon|OJJ42877.1

Group1620: abra|OJJ67957.1 acar|OOF92802.1 acla|EAW11646.1 afis|EAW23810.1 afla|EED52005.1 afum|EAL87244.1 agla|OJJ82450.1 akaw|GAA91239.1 aluc|OJZ82830.1 anid|CBF79876.1 anig|EHA21268.1 aory|EIT81434.1 asyd|OJJ58169.1 ater|EAU31363.1 atub|OJI89060.1 aver|OJJ04234.1 awen|OJJ33766.1 azon|OJJ42870.1

Group1621: abra|OJJ67958.1 acar|OOF92803.1 acla|EAW11647.1 afis|EAW23809.1 afla|EED52006.1 afum|EAL87243.1 agla|OJJ82449.1 akaw|GAA91238.1 aluc|OJZ82831.1 anid|CBF79874.1 anig|EHA21269.1 aory|EIT81404.1 asyd|OJJ58168.1 ater|EAU31362.1 atub|OJI89061.1 aver|OJJ04235.1 awen|OJJ33765.1 azon|OJJ42868.1

Group1622: abra|OJJ67977.1 acar|OOF92819.1 acla|EAW11758.1 afis|EAW23791.1 afla|EED51599.1 afum|EAL87225.1 agla|OJJ78819.1 akaw|GAA91226.1 aluc|OJZ82844.1 anid|CBF79853.1 anig|EHA21282.1 aory|EIT80507.1 asyd|OJJ58157.1 ater|EAU31330.1 atub|OJI89074.1 aver|OJJ04246.1 awen|OJJ33746.1 azon|OJJ42843.1

Group1623: abra|OJJ67978.1 acar|OOF92818.1 acla|EAW11757.1 afis|EAW23790.1 afla|EED51598.1 afum|EAL87224.1 agla|OJJ78817.1 akaw|GAA91225.1 aluc|OJZ82845.1 anid|CBF79851.1 anig|EHA21283.1 aory|EIT80526.1 asyd|OJJ58156.1 ater|EAU31329.1 atub|OJI89075.1 aver|OJJ04247.1 awen|OJJ33745.1 azon|OJJ42844.1

Group1624: abra|OJJ68015.1 acar|OOF92057.1 acla|EAW08795.1 afis|EAW15605.1 afla|EED55861.1 afum|EAL85728.1 agla|OJJ79064.1 akaw|GAA90936.1 aluc|OJZ82888.1 anid|CBF69625.1 anig|EHA21314.1 aory|EIT76549.1 asyd|OJJ59120.1 ater|EAU37248.1 atub|OJI89113.1 aver|OJJ01709.1 awen|OJJ32398.1 azon|OJJ43874.1

Group1625: abra|OJJ68016.1 acar|OOF92058.1 afla|EED54104.1 agla|OJJ80413.1 akaw|GAA90935.1 aluc|OJZ82889.1 anid|CBF83706.1 anig|EHA21315.1 aory|EIT77159.1 asyd|OJJ53740.1 atub|OJI89114.1 aver|OJJ07853.1 awen|OJJ35049.1 azon|OJJ42272.1 acla|EAW14887.1 awen|OJJ34541.1 aory|EIT75976.1 awen|OJJ30142.1

Group1626: abra|OJJ68060.1 acar|OOF92081.1 acla|EAW13655.1 afis|EAW19935.1 afla|EED45904.1 afum|EAL84975.2 agla|OJJ79327.1 akaw|GAA90896.1 aluc|OJZ82932.1 anid|CBF84602.1 anig|EHA21353.1 aory|EIT75822.1 asyd|OJJ53375.1 ater|EAU36898.1 atub|OJI89161.1 aver|OJJ08369.1 awen|OJJ29978.1 azon|OJJ50215.1

Group1627: abra|OJJ68075.1 acar|OOF94470.1 acla|EAW12117.1 afis|EAW17265.1 afla|EED48857.1 afum|EAL86682.1 agla|OJJ85378.1 akaw|GAA92736.1 aluc|OJZ92944.1 anid|CBF87528.1 anig|EHA26028.1 aory|EIT81286.1 asyd|OJJ52984.1 ater|EAU36128.1 atub|OJI87403.1 aver|OJJ08953.1 awen|OJJ35769.1 azon|OJJ50248.1

Group1628: abra|OJJ68089.1 acar|OOF93893.1 acla|EAW15283.1 afis|EAW17127.1 afla|EED48286.1 afum|EAL86587.1 agla|OJJ78597.1 akaw|GAA86874.1 aluc|OJZ89998.1 anid|CBF80813.1 anig|EHA27373.1 aory|EIT72919.1 asyd|OJJ54451.1 ater|EAU34073.1 atub|OJI84810.1 aver|OJJ07703.1 awen|OJJ29885.1 azon|OJJ50148.1

Group1629: abra|OJJ68125.1 acar|OOF93919.1 acla|EAW12869.1 afis|EAW17817.1 afla|EED45601.1 afum|EAL91627.1 agla|OJJ81689.1 akaw|GAA86906.1 aluc|OJZ90029.1 anid|CBF80748.1 anig|EHA27405.1 aory|EIT75765.1 asyd|OJJ54539.1 ater|EAU30285.1 atub|OJI84778.1 aver|OJJ07642.1 awen|OJJ36934.1 azon|OJJ42822.1

Group1630: abra|OJJ68126.1 acar|OOF93920.1 acla|EAW12870.1 afis|EAW17818.1 afla|EED45602.1 afum|EAL91628.1 agla|OJJ85762.1 akaw|GAA86907.1 aluc|OJZ90030.1 anid|CBF80750.1 anig|EHA27406.1 asyd|OJJ62953.1 atub|OJI84777.1 aver|OJJ07643.1 awen|OJJ31294.1 azon|OJJ50609.1 aver|OJI98031.1 asyd|OJJ54538.1

Group1631: abra|OJJ68139.1 acar|OOF98356.1 acla|EAW14517.1 afis|EAW19235.1 afla|EED47647.1 afum|EAL93963.1 agla|OJJ88463.1 akaw|GAA92378.1 aluc|OJZ83237.1 anid|CBF78634.1 anig|EHA21094.1 aory|EIT80221.1 asyd|OJJ65123.1 ater|EAU37228.1 atub|OJI87699.1 aver|OJI99251.1 awen|OJJ37592.1 azon|OJJ51245.1

Group1632: abra|OJJ68145.1 acar|OOF89988.1 acla|EAW07390.1 afis|EAW22416.1 afla|EED52494.1 afum|EAL90776.1 agla|OJJ87680.1 akaw|GAA92373.1 aluc|OJZ83231.1 anid|CBF88053.1 anig|EHA21089.1 aory|EIT82805.1 asyd|OJJ63540.1 ater|EAU39291.1 atub|OJI87705.1 aver|OJI97393.1 awen|OJJ34995.1 azon|OJJ45638.1

Group1633: agla|OJJ86892.1 akaw|GAA88060.1 aluc|OJZ81322.1 abra|OJJ68375.1 agla|OJJ79451.1 aluc|OJZ90115.1 anig|EHA18628.1 ater|EAU37955.1 atub|OJI84684.1 awen|OJJ35546.1 azon|OJJ50297.1 acar|OOF90164.1 atub|OJI82870.1 awen|OJJ29729.1 acar|OOF90659.1 afla|EED57502.1 ater|EAU37868.1 atub|OJI91525.1

Group1634: abra|OJJ68168.1 acar|OOF97466.1 acla|EAW13054.1 afis|EAW17612.1 afla|EED49203.1 afum|EAL91425.1 agla|OJJ81325.1 akaw|GAA83275.1 aluc|OJZ83781.1 anid|CBF90174.1 anig|EHA18921.1 aory|EIT78579.1 asyd|OJJ61990.1 ater|EAU37047.1 atub|OJI90151.1 aver|OJI96082.1 awen|OJJ37230.1 azon|OJJ50933.1

Group1635: abra|OJJ68169.1 acar|OOF97467.1 acla|EAW13055.1 afis|EAW17611.1 afla|EED49202.1 afum|EAL91424.1 agla|OJJ81326.1 akaw|GAA83274.1 aluc|OJZ83780.1 anid|CBF90175.1 anig|EHA18920.1

aory|EIT78562.1 asyd|OJJ61989.1 ater|EAU37046.1 atub|OJI90150.1 aver|OJI96081.1 awen|OJJ37231.1 azon|OJJ50934.1

Group1636: abra|OJJ68171.1 acar|OOF97468.1 acla|EAW13056.1 afis|EAW17610.1 afla|EED49201.1 afum|EAL91423.1 agla|OJJ81331.1 akaw|GAA83273.1 aluc|OJZ83779.1 anid|CBF90177.1 anig|EHA18919.1 aory|EIT78514.1 asyd|OJJ61988.1 ater|EAU37045.1 atub|OJI90149.1 aver|OJI96080.1 awen|OJJ37232.1 azon|OJJ50935.1

Group1637: abra|OJJ68172.1 acar|OOF97470.1 acla|EAW13057.1 afis|EAW17609.1 afla|EED49213.1 afum|EAL91422.1 agla|OJJ81356.1 akaw|GAA83272.1 aluc|OJZ83778.1 anid|CBF90179.1 anig|EHA18918.1 aory|EIT83168.1 asyd|OJJ61987.1 ater|EAU37044.1 atub|OJI90148.1 aver|OJI96079.1 awen|OJJ37233.1 azon|OJJ50936.1

Group1638: abra|OJJ68173.1 acar|OOF97471.1 acla|EAW13058.1 afis|EAW17608.1 afla|EED49214.1 afum|EAL91421.1 agla|OJJ81355.1 akaw|GAA83271.1 aluc|OJZ83777.1 anid|CBF90181.1 anig|EHA18917.1 aory|EIT83162.1 asyd|OJJ61986.1 ater|EAU37043.1 atub|OJI90147.1 aver|OJI96078.1 awen|OJJ37234.1 azon|OJJ50937.1

Group1639: abra|OJJ68176.1 acar|OOF97473.1 acla|EAW13060.1 afis|EAW17604.1 afla|EED49217.1 afum|EAL91417.1 agla|OJJ81352.1 akaw|GAA83268.1 aluc|OJZ83774.1 anid|CBF90186.1 anig|EHA18914.1 aory|EIT83159.1 asyd|OJJ61983.1 ater|EAU37041.1 atub|OJI90145.1 aver|OJI96075.1 awen|OJJ37237.1 azon|OJJ50939.1

Group1640: abra|OJJ68177.1 acar|OOF97474.1 acla|EAW13061.1 afis|EAW17603.1 afla|EED49218.1 afum|EAL91416.1 agla|OJJ81351.1 akaw|GAA83267.1 aluc|OJZ83773.1 anid|CBF90188.1 anig|EHA18913.1 aory|EIT83167.1 asyd|OJJ61982.1 ater|EAU37040.1 atub|OJI90144.1 aver|OJI96074.1 awen|OJJ37238.1 azon|OJJ50940.1

Group1641: abra|OJJ68181.1 acar|OOF97477.1 acla|EAW13063.1 afis|EAW17601.1 afla|EED49220.1 afum|EBA27434.1 agla|OJJ81348.1 akaw|GAA83265.1 aluc|OJZ83771.1 anid|CBF90190.1 anig|EHA18911.1 aory|EIT83161.1 asyd|OJJ61981.1 ater|EAU37038.1 atub|OJI90142.1 aver|OJI96073.1 awen|OJJ37244.1 azon|OJJ50942.1

Group1642: abra|OJJ68182.1 acar|OOF97476.1 acla|EAW13064.1 afis|EAW17600.1 afla|EED49221.1 afum|EAL91414.1 agla|OJJ81347.1 akaw|GAA83264.1 aluc|OJZ83770.1 anid|CBF90192.1 anig|EHA18910.1 aory|EIT83165.1 asyd|OJJ61980.1 ater|EAU37037.1 atub|OJI90141.1 aver|OJI96072.1 awen|OJJ37245.1 azon|OJJ50943.1

Group1643: abra|OJJ68184.1 acar|OOF97479.1 acla|EAW13065.1 afis|EAW17599.1 afla|EED49222.1 afum|EAL91412.2 agla|OJJ81345.1 akaw|GAA83262.1 aluc|OJZ83768.1 anig|EHA18908.1 aory|EIT83170.1 asyd|OJJ61978.1 ater|EAU37036.1 atub|OJI90139.1 aver|OJI96070.1 awen|OJJ37247.1 azon|OJJ50945.1 anid|CBF90194.1

Group1644: abra|OJJ68190.1 acar|OOF97486.1 acla|EAW13072.1 afis|EAW17592.1 afla|EED49230.1 afum|EAL91405.1 agla|OJJ81357.1 akaw|GAA83255.1 aluc|OJZ83756.1 anid|CBF90212.1 anig|EHA18903.1 aory|EIT83166.1 asyd|OJJ61969.1 ater|EAU37029.1 atub|OJI90131.1 aver|OJI96062.1 awen|OJJ37254.1 azon|OJJ50949.1

Group1645: abra|OJJ68193.1 acar|OOF97489.1 acla|EAW13076.1 afis|EAW17588.1 afla|EED49234.1 afum|EAL91401.1 agla|OJJ81364.1 akaw|GAA83252.1 aluc|OJZ83753.1 anid|CBF90220.1 anig|EHA18900.1 aory|EIT83171.1 asyd|OJJ61966.1 ater|EAU37023.1 atub|OJI90128.1 aver|OJI96058.1 awen|OJJ37258.1 azon|OJJ50953.1

Group1646: abra|OJJ68194.1 acar|OOF97490.1 acla|EAW13077.1 afis|EAW17587.1 afla|EED49235.1 afum|EAL91400.1 agla|OJJ81365.1 akaw|GAA83251.1 aluc|OJZ83752.1 anid|CBF90222.1 anig|EHA18899.1 aory|EIT83154.1 asyd|OJJ61965.1 ater|EAU37022.1 atub|OJI90127.1 aver|OJI96057.1 awen|OJJ37259.1 azon|OJJ50954.1

Group1647: abra|OJJ68196.1 acar|OOF97491.1 acla|EAW13079.1 afis|EAW17585.1 afla|EED49237.1 afum|EAL91398.2 agla|OJJ81368.1 akaw|GAA83249.1 aluc|OJZ83750.1 anid|CBF90225.1 anig|EHA18897.1 aory|EIT83169.1 asyd|OJJ61963.1 ater|EAU37020.1 atub|OJI90125.1 aver|OJI96055.1 awen|OJJ37261.1 azon|OJJ50956.1

Group1648: abra|OJJ68197.1 acar|OOF89886.1 acla|EAW13080.1 afis|EAW17584.1 afla|EED49239.1 afum|EAL91397.1 agla|OJJ81369.1 akaw|GAA83248.1 aluc|OJZ83749.1 anid|CBF90227.1 anig|EHA18896.1 aory|EIT82222.1 asyd|OJJ61962.1 ater|EAU37019.1 atub|OJI90124.1 aver|OJI96054.1 awen|OJJ37262.1 azon|OJJ50957.1

Group1649: abra|OJJ68199.1 acar|OOF90119.1 acla|EAW13081.1 afis|EAW17583.1 afla|EED49240.1 afum|EAL91396.1 agla|OJJ81370.1 akaw|GAA83247.1 aluc|OJZ83748.1 anid|CBF90229.1 anig|EHA18895.1 aory|EIT82239.1 asyd|OJJ61961.1 ater|EAU37018.1 atub|OJI90123.1 aver|OJI96053.1 awen|OJJ37265.1 azon|OJJ50959.1

Group1650: abra|OJJ68200.1 acar|OOF90117.1 acla|EAW13083.1 afis|EAW17581.1 afla|EED49241.1 agla|OJJ81372.1 akaw|GAA83245.1 aluc|OJZ83747.1 anid|CBF90232.1 anig|EHA18893.1 aory|EIT82216.1 asyd|OJJ61959.1 ater|EAU37016.1 atub|OJI90122.1 aver|OJI96051.1 awen|OJJ37267.1 azon|OJJ50960.1 afum|EAL91394.1

Group1651: abra|OJJ68201.1 acar|OOF90116.1 acla|EAW13084.1 afis|EAW17580.1 afla|EED49242.1 afum|EAL91393.1 agla|OJJ81379.1 akaw|GAA83244.1 aluc|OJZ83746.1 anid|CBF90234.1 anig|EHA18892.1 aory|EIT82232.1 asyd|OJJ61958.1 ater|EAU37015.1 atub|OJI90121.1 aver|OJI96050.1 awen|OJJ37268.1 azon|OJJ50961.1

Group1652: abra|OJJ68203.1 acar|OOF90234.1 acla|EAW13086.1 afis|EAW17579.1 afla|EED49243.1 afum|EAL91392.1 agla|OJJ81378.1 akaw|GAA83243.1 aluc|OJZ83744.1 anid|CBF90238.1 anig|EHA18891.1 aory|EIT82211.1 asyd|OJJ61957.1 ater|EAU37014.1 atub|OJI90120.1 aver|OJI96049.1 awen|OJJ37269.1 azon|OJJ50962.1

Group1653: abra|OJJ68204.1 acar|OOF90236.1 acla|EAW13087.1 afis|EAW17577.1 afla|EED49244.1 afum|EAL91390.1 agla|OJJ81377.1 akaw|GAA83242.1 aluc|OJZ83741.1 anid|CBF90245.1 anig|EHA18890.1 aory|EIT82198.1 asyd|OJJ61951.1 ater|EAU37013.1 atub|OJI90119.1 aver|OJI96042.1 awen|OJJ37270.1 azon|OJJ50969.1

Group1654: abra|OJJ68205.1 acar|OOF90235.1 acla|EAW13088.1 afis|EAW17576.1 afla|EED49245.1 afum|EAL91389.1 agla|OJJ81381.1 akaw|GAA83241.1 aluc|OJZ83740.1 anid|CBF90247.1 anig|EHA18889.1 aory|EIT82237.1 asyd|OJJ61950.1 ater|EAU37012.1 atub|OJI90118.1 aver|OJI96041.1 awen|OJJ37271.1 azon|OJJ50971.1

Group1655: abra|OJJ68207.1 acar|OOF97493.1 acla|EAW13090.1 afis|EAW17574.1 afla|EED49247.1 afum|EAL91387.1 agla|OJJ81384.1 akaw|GAA83239.1 aluc|OJZ83738.1 anid|CBF90250.1 anig|EHA18887.1 aory|EIT82189.1 asyd|OJJ61948.1 ater|EAU37010.1 atub|OJI90116.1 aver|OJI96040.1 awen|OJJ37273.1 azon|OJJ50973.1

Group1656: abra|OJJ68211.1 acar|OOF97495.1 acla|EAW12973.1 afis|EAW17691.1 afla|EED50453.1 afum|EAL91505.1 agla|OJJ79488.1 akaw|GAA83235.1 aluc|OJZ83734.1 anid|CBF89980.1 anig|EHA18883.1 aory|EIT82051.1 asyd|OJJ62090.1 ater|EAU31953.1 atub|OJI90112.1 aver|OJI96196.1 awen|OJJ37145.1 azon|OJJ50845.1

Group1657: abra|OJJ68212.1 acar|OOF97496.1 acla|EAW12972.1 afis|EAW17692.1 afla|EED50452.1 afum|EAL91506.1 agla|OJJ79487.1 akaw|GAA83234.1 aluc|OJZ83733.1 anid|CBF89978.1 anig|EHA18882.1 aory|EIT82145.1 asyd|OJJ62091.1 ater|EAU31958.1 atub|OJI90111.1 aver|OJI96197.1 awen|OJJ37144.1 azon|OJJ50844.1

Group1658: abra|OJJ68249.1 acar|OOF97528.1 acla|EAW09061.1 afis|EAW25634.1 afla|EED55178.1 afum|EAL84322.1 agla|OJJ78898.1 akaw|GAA83212.1 aluc|OJZ83714.1 anid|CBF83155.1 anig|EHA18865.1 aory|EIT81648.1 asyd|OJJ55514.1 ater|EAU32715.1 atub|OJI90089.1 aver|OJJ06591.1 awen|OJJ36053.1 azon|OJJ50759.1

Group1659: abra|OJJ68257.1 acar|OOF97534.1 acla|EAW09064.1 afis|EAW25636.1 afla|EED55176.1 afum|EAL84324.1 agla|OJJ78896.1 akaw|GAA83205.1 aluc|OJZ83707.1 anid|CBF83151.1 anig|EHA18858.1 aory|EIT81641.1 asyd|OJJ55516.1 ater|EAU32717.1 atub|OJI90079.1 aver|OJJ06593.1 awen|OJJ36093.1 azon|OJJ50756.1

Group1660: abra|OJJ68259.1 acar|OOF97536.1 acla|EAW14109.1 afis|EAW18811.1 afla|EED50761.1 afum|EAL93547.1 agla|OJJ80544.1 akaw|GAA92415.1 aluc|OJZ83273.1 anid|CBF69795.1 anig|EHA18856.1 aory|EIT78026.1 asyd|OJJ58964.1 ater|EAU37966.1 atub|OJI87668.1 aver|OJJ01558.1 awen|OJJ32387.1 azon|OJJ47147.1

Group1661: abra|OJJ68274.1 acar|OOF97552.1 acla|EAW09071.1 agla|OJJ78841.1 akaw|GAA83195.1 aluc|OJZ83697.1 anid|CBF78459.1 anig|EHA18848.1 asyd|OJJ64190.1 ater|EAU32708.1 atub|OJI90068.1 aver|OJI98267.1 awen|OJJ35541.1 azon|OJJ44995.1 acar|OOF92523.1 azon|OJJ42182.1 afla|EED55164.1 azon|OJJ45941.1

Group1662: abra|OJJ68281.1 acar|OOF97555.1 acla|EAW09093.1 afis|EAW25687.1 afla|EED55157.1 afum|EAL84356.1 agla|OJJ78886.1 akaw|GAA83191.1 aluc|OJZ83690.1 anid|CBF83933.1 anig|EHA18844.1 aory|EIT81633.1 asyd|OJJ55435.1 ater|EAU32721.1 atub|OJI90062.1 aver|OJJ06512.1 awen|OJJ36120.1 azon|OJJ50742.1

Group1663: abra|OJJ68301.1 acar|OOF96686.1 acla|EAW12631.1 afis|EAW18053.1 afla|EED56195.1 afum|EAL91864.1 agla|OJJ87336.1 akaw|GAA83173.1 aluc|OJZ83672.1 anid|CBF83350.1 anig|EHA18827.1 aory|EIT73338.1 asyd|OJJ60615.1 ater|EAU34830.1 atub|OJI90043.1 aver|OJI99939.1 awen|OJJ41435.1 azon|OJJ48899.1

Group1664: abra|OJJ68398.1 acar|OOF93975.1 acla|EAW08574.1 afis|EAW23634.1 afla|EED47506.1 afum|EAL87826.1 akaw|GAA92549.1 aluc|OJZ90098.1 anig|EHA20978.1 aory|EIT74517.1 asyd|OJJ54836.1

atub|OJI84708.1 aver|OJJ07325.1 azon|OJJ50231.1 aory|EIT74543.1 asyd|OJJ53612.1 aver|OJJ07720.1 awen|OJJ29655.1

Group1665: abra|OJJ68415.1 acar|OOG00093.1 acla|EAW11055.1 afis|EAW20382.1 afum|EAL85495.1 agla|OJJ83841.1 akaw|GAA92533.1 aluc|OJZ90082.1 anid|CBF70730.1 anig|EHA20991.1 asyd|OJJ56111.1 ater|EAU30781.1 atub|OJI84724.1 aver|OJJ05054.1 awen|OJJ30077.1 azon|OJJ48238.1 anid|CBF78288.1 acla|EAW15020.1

Group1666: abra|OJJ68449.1 acar|OOF93946.1 acla|EAW11772.1 afis|EAW23677.1 afla|EED54146.1 afum|EAL87120.1 agla|OJJ78443.1 akaw|GAA86924.1 aluc|OJZ90048.1 anid|CBF75094.1 anig|EHA21022.1 aory|EIT75945.1 asyd|OJJ60010.1 ater|EAU29439.1 atub|OJI84758.1 aver|OJJ02275.1 awen|OJJ38063.1 azon|OJJ50775.1

Group1667: abra|OJJ68459.1 acar|OOF92643.1 acla|EAW08639.1 afis|EAW15792.1 afla|EED51253.1 afum|EAL85558.1 agla|OJJ81955.1 akaw|GAA92421.1 aluc|OJZ80815.1 anid|CBF70968.1 anig|EHA21125.1 aory|EIT81555.1 asyd|OJJ52614.1 ater|EAU32350.1 atub|OJI87664.1 aver|OJJ04641.1 awen|OJJ38500.1 azon|OJJ43801.1

Group1668: abra|OJJ68464.1 acar|OOF94169.1 acla|EAW10056.1 afis|EAW25528.1 afla|EED54029.1 afum|EAL90193.2 agla|OJJ87274.1 akaw|GAA92384.1 aluc|OJZ83243.1 anid|CBF77807.1 anig|EHA21098.1 aory|EIT78344.1 asyd|OJJ53016.1 ater|EAU33371.1 atub|OJI87693.1 aver|OJJ03827.1 awen|OJJ41270.1 azon|OJJ49510.1

Group1669: abra|OJJ68468.1 acar|OOF89893.1 acla|EAW10429.1 afis|EAW25140.1 afla|EED55305.1 afum|EAL89812.1 agla|OJJ89027.1 akaw|GAA92389.1 aluc|OJZ83249.1 anid|CBF86095.1 anig|EHA21102.1 aory|EIT83530.1 asyd|OJJ64474.1 ater|EAU39409.1 atub|OJI87688.1 aver|OJI98578.1 awen|OJJ40764.1 azon|OJJ44129.1

Group1670: abra|OJJ68475.1 acar|OOF92975.1 acla|EAW06477.1 afis|EAW15972.1 afla|EED58084.1 afum|EAL88604.1 agla|OJJ83843.1 akaw|GAA92396.1 aluc|OJZ83256.1 anid|CBF81100.1 anig|EHA21110.1 aory|EIT79134.1 asyd|OJJ56477.1 ater|EAU35476.1 atub|OJI87682.1 aver|OJJ06241.1 awen|OJJ35600.1 azon|OJJ46920.1

Group1671: abra|OJJ68476.1 acar|OOF99927.1 acla|EAW09807.1 afis|EAW21329.1 afla|EED57287.1 afum|EAL92347.1 agla|OJJ87410.1 akaw|GAA92397.1 aluc|OJZ83258.1 anid|CBF83431.1 anig|EHA21111.1 aory|EIT74125.1 asyd|OJJ60661.1 ater|EAU35955.1 atub|OJI87681.1 aver|OJI99984.1 awen|OJJ39369.1 azon|OJJ51709.1

Group1672: abra|OJJ68478.1 acar|OOF96519.1 acla|EAW13881.1 afis|EAW18593.1 afla|EED52754.1 afum|EAL93301.1 agla|OJJ81645.1 akaw|GAA92401.1 aluc|OJZ83262.1 anid|CBF70381.1 anig|EHA21115.1 aory|EIT78198.1 asyd|OJJ58642.1 ater|EAU34887.1 atub|OJI87675.1 aver|OJJ01248.1 awen|OJJ31156.1 azon|OJJ46631.1

Group1673: abra|OJJ68485.1 acar|OOG00353.1 acla|EAW07017.1 afis|EAW16643.1 afla|EED44966.1 afum|EAL89246.1 agla|OJJ82604.1 akaw|GAA92786.1 aluc|OJZ82697.1 anid|CBF82045.1 anig|EHA21146.1 aory|EIT73490.1 asyd|OJJ54225.1 ater|EAU39568.1 atub|OJI88929.1 aver|OJJ05178.1 awen|OJJ40071.1 azon|OJJ45553.1

Group1674: abra|OJJ68486.1 acar|OOF95442.1 acla|EAW07333.1 afis|EAW22365.1 afla|EED57773.1 afum|EBA27416.1 agla|OJJ87616.1 akaw|GAA92785.1 aluc|OJZ82698.1 anid|CBF88690.1 anig|EHA21153.1 aory|EIT81113.1 asyd|OJJ63908.1 ater|EAU39499.1 atub|OJI88930.1 aver|OJI97033.1 awen|OJJ34939.1 azon|OJJ45591.1

Group1675: abra|OJJ68497.1 acar|OOF99362.1 acla|EAW14277.1 afis|EAW18997.1 afla|EED53139.1 afum|EAL93732.1 agla|OJJ80191.1 akaw|GAA92773.1 aluc|OJZ82710.1 anid|CBF86779.1 anig|EHA21165.1 aory|EIT74918.1 asyd|OJJ64682.1 ater|EAU37806.1 atub|OJI88943.1 aver|OJI98784.1 awen|OJJ31625.1 azon|OJJ47325.1

Group1676: abra|OJJ68499.1 acar|OOF93648.1 acla|EAW14509.1 afis|EAW19222.1 afla|EED47663.1 afum|EAL93954.1 agla|OJJ88466.1 akaw|GAA92772.1 aluc|OJZ82712.1 anid|CBF78608.1 anig|EHA21167.1 aory|EIT80278.1 asyd|OJJ65103.1 ater|EAU37236.1 atub|OJI88945.1 aver|OJI99230.1 awen|OJJ37585.1 azon|OJJ51239.1

Group1677: abra|OJJ68500.1 acar|OOF92674.1 acla|EAW08059.1 afis|EAW23095.1 afla|EED48553.1 afum|EAL88383.1 agla|OJJ89607.1 akaw|GAA92771.1 aluc|OJZ82713.1 anid|CBF75180.1 anig|EHA21168.1 aory|EIT73191.1 asyd|OJJ59952.1 ater|EAU36223.1 atub|OJI88946.1 aver|OJJ02337.1 awen|OJJ36686.1 azon|OJJ47510.1

Group1678: abra|OJJ68501.1 acar|OOF89894.1 acla|EAW11963.1 afis|EAW19600.1 afla|EED56956.1 afum|EAL86918.1 agla|OJJ80742.1 akaw|GAA92770.1 aluc|OJZ82714.1 anid|CBF71291.1 anig|EHA21169.1 aory|EIT77492.1 asyd|OJJ55128.1 ater|EAU32925.1 atub|OJI88947.1 aver|OJJ07076.1 awen|OJJ41841.1 azon|OJJ47038.1

Group1679: abra|OJJ68507.1 acar|OOF92954.1 acla|EAW07233.1 afis|EAW22254.1 afla|EED57640.1 afum|EAL90924.1 agla|OJJ87802.1 akaw|GAA92766.1 aluc|OJZ82719.1 anid|CBF88500.1 anig|EHA21173.1 aory|EIT82931.1 asyd|OJJ63795.1 ater|EAU34255.1 atub|OJI88953.1 aver|OJI97142.1 awen|OJJ34826.1 azon|OJJ45723.1

Group1680: abra|OJJ68510.1 acar|OOF92672.1 acla|EAW07690.1 afis|EAW22705.1 afla|EED49991.1 afum|EAL90487.1 agla|OJJ79004.1 akaw|GAA92761.1 aluc|OJZ82724.1 anid|CBF88071.1 anig|EHA21178.1 aory|EIT74613.1 asyd|OJJ63553.1 ater|EAU38996.1 atub|OJI88958.1 aver|OJI97381.1 awen|OJJ35342.1 azon|OJJ49776.1

Group1681: abra|OJJ68511.1 acar|OOF92678.1 acla|EAW14940.1 afis|EAW24455.1 afla|EED55219.1 afum|EAL86124.1 agla|OJJ80302.1 akaw|GAA91335.1 aluc|OJZ82725.1 anid|CBF75392.1 anig|EHA21179.1 aory|EIT83615.1 asyd|OJJ59834.1 ater|EAU29438.1 atub|OJI88959.1 aver|OJJ02454.1 awen|OJJ32742.1 azon|OJJ45292.1

Group1682: abra|OJJ68517.1 acar|OOF99154.1 acla|EAW06536.1 afis|EAW16059.1 afla|EED48573.1 afum|EAL88692.1 agla|OJJ86106.1 akaw|GAA91331.1 aluc|OJZ82731.1 anid|CBF75082.1 anig|EHA21183.1 aory|EIT73064.1 asyd|OJJ60019.1 ater|EAU33575.1 atub|OJI88964.1 aver|OJJ02266.1 awen|OJJ30380.1 azon|OJJ45396.1

Group1683: abra|OJJ68524.1 acar|OOF92729.1 acla|EAW12839.1 afis|EAW18004.1 afla|EED51892.1 afum|EAL91815.2 agla|OJJ80272.1 akaw|GAA91325.1 aluc|OJZ82738.1 anid|CBF80032.1 anig|EHA21189.1 aory|EIT76182.1 asyd|OJJ58271.1 ater|EAU31516.1 atub|OJI88973.1 aver|OJJ02153.1 awen|OJJ41538.1 azon|OJJ48256.1

Group1684: abra|OJJ68526.1 acar|OOF92730.1 acla|EAW12838.1 afis|EAW18003.1 afla|EED51893.1 afum|EAL91814.2 agla|OJJ80266.1 akaw|GAA91324.1 aluc|OJZ82739.1 anid|CBF80026.1 anig|EHA21190.1 aory|EIT76172.1 asyd|OJJ58266.1 ater|EAU31515.1 atub|OJI88974.1 aver|OJJ02157.1 awen|OJJ41539.1 azon|OJJ48257.1

Group1685: abra|OJJ68527.1 acar|OOF92731.1 acla|EAW12837.1 afis|EAW18002.1 afla|EED51894.1 afum|EAL91813.1 agla|OJJ80265.1 akaw|GAA91323.1 aluc|OJZ82740.1 anid|CBF80025.1 anig|EHA21191.1 aory|EIT76169.1 asyd|OJJ58265.1 ater|EAU31514.1 atub|OJI88975.1 aver|OJJ02158.1 awen|OJJ41540.1 azon|OJJ48258.1

Group1686: abra|OJJ68528.1 acar|OOF92732.1 acla|EAW12836.1 afis|EAW18001.1 afla|EED51895.1 afum|EAL91812.1 agla|OJJ80264.1 akaw|GAA91322.1 aluc|OJZ82741.1 anid|CBF80023.1 anig|EHA21192.1 aory|EIT76176.1 asyd|OJJ58264.1 ater|EAU31513.1 atub|OJI88976.1 aver|OJJ02160.1 awen|OJJ41541.1 azon|OJJ48259.1

Group1687: abra|OJJ68531.1 acar|OOF92734.1 acla|EAW12834.1 afis|EAW17999.1 afla|EED51899.1 afum|EAL91810.1 agla|OJJ80262.1 akaw|GAA91320.1 aluc|OJZ82743.1 anid|CBF80019.1 anig|EHA21194.1 aory|EIT76173.1 asyd|OJJ58262.1 ater|EAU31511.1 atub|OJI88978.1 aver|OJJ02162.1 awen|OJJ41543.1 azon|OJJ48261.1

Group1688: abra|OJJ68532.1 acar|OOF92735.1 acla|EAW12833.1 afis|EAW17998.1 afla|EED51900.1 afum|EAL91809.2 agla|OJJ80261.1 akaw|GAA91319.1 aluc|OJZ82744.1 anid|CBF80017.1 anig|EHA21195.1 aory|EIT76171.1 asyd|OJJ58261.1 ater|EAU31510.1 atub|OJI88979.1 aver|OJJ02163.1 awen|OJJ41544.1 azon|OJJ48262.1

Group1689: abra|OJJ68533.1 acar|OOF92736.1 afis|EAW17997.1 afla|EED51903.1 afum|EAL91807.2 akaw|GAA91318.1 aluc|OJZ82746.1 anig|EHA21196.1 aory|EIT76183.1 asyd|OJJ58260.1 atub|OJI88980.1 aver|OJJ02164.1 awen|OJJ41545.1 acla|EAW12832.1 agla|OJJ80257.1 ater|EAU31509.1 azon|OJJ48263.1 anid|CBF80015.1

Group1690: abra|OJJ68537.1 acar|OOF90151.1 acla|EAW12827.1 afis|EAW17992.1 afla|EED51911.1 afum|EAL91802.1 agla|OJJ80253.1 akaw|GAA91314.1 aluc|OJZ82751.1 anid|CBF80004.1 anig|EHA21200.1 aory|EIT76179.1 asyd|OJJ58250.1 ater|EAU31503.1 atub|OJI88987.1 aver|OJJ02174.1 awen|OJJ41554.1 azon|OJJ48269.1

Group1691: abra|OJJ68538.1 acar|OOF90152.1 acla|EAW12828.1 afis|EAW17993.1 afla|EED51908.1 afum|EAL91803.1 agla|OJJ80254.1 akaw|GAA91313.1 aluc|OJZ82752.1 anid|CBF80005.1 anig|EHA21201.1 aory|EIT76170.1 asyd|OJJ58251.1 ater|EAU31504.1 atub|OJI88988.1 aver|OJJ02173.1 awen|OJJ41553.1 azon|OJJ48268.1

Group1692: abra|OJJ68541.1 acar|OOF92739.1 acla|EAW12829.1 afis|EAW17994.1 afla|EED51907.1 afum|EAL91804.1 agla|OJJ80255.1 akaw|GAA91312.1 aluc|OJZ82753.1 anid|CBF80009.1 anig|EHA21202.1 aory|EIT76184.1 asyd|OJJ58256.1 ater|EAU31505.1 atub|OJI88990.1 aver|OJJ02167.1 awen|OJJ41551.1 azon|OJJ48267.1

Group1693: abra|OJJ68542.1 acar|OOF92740.1 acla|EAW12830.1 afis|EAW17995.1 afla|EED51906.1 afum|EAL91805.1 agla|OJJ80259.1 akaw|GAA91311.1 aluc|OJZ82755.1 anid|CBF80011.1 anig|EHA21203.1

aory|EIT76180.1 asyd|OJJ58257.1 ater|EAU31507.1 atub|OJI88991.1 aver|OJJ02166.1 awen|OJJ41550.1 azon|OJJ48266.1

Group1694: abra|OJJ68544.1 acar|OOF92742.1 acla|EAW12823.1 afis|EAW17988.1 afla|EED51915.1 afum|EAL91797.1 agla|OJJ80248.1 akaw|GAA91309.1 aluc|OJZ82759.1 anid|CBF79996.1 anig|EHA21205.1 aory|EIT81365.1 asyd|OJJ58247.1 ater|EAU31498.1 atub|OJI88993.1 aver|OJJ02179.1 awen|OJJ41558.1 azon|OJJ48273.1

Group1695: abra|OJJ68546.1 acar|OOF92743.1 acla|EAW12822.1 afis|EAW17987.1 afla|EED51916.1 afum|EAL91796.1 agla|OJJ80242.1 akaw|GAA91308.1 aluc|OJZ82762.1 anid|CBF80000.1 anig|EHA21206.1 aory|EIT81367.1 asyd|OJJ58248.1 ater|EAU31497.1 atub|OJI88995.1 aver|OJJ02176.1 awen|OJJ41559.1 azon|OJJ48274.1

Group1696: abra|OJJ68549.1 acar|OOF92745.1 acla|EAW12818.1 afis|EAW17983.1 afla|EED51922.1 afum|EAL91792.1 agla|OJJ80240.1 akaw|GAA91304.1 aluc|OJZ82766.1 anid|CBF79992.1 anig|EHA21209.1 aory|EIT81388.1 asyd|OJJ58238.1 ater|EAU31493.1 atub|OJI88998.1 aver|OJJ02187.1 awen|OJJ41562.1 azon|OJJ48277.1

Group1697: abra|OJJ68554.1 acar|OOF92749.1 acla|EAW12813.1 afis|EAW17978.1 afla|EED51927.1 afum|EAL91787.1 agla|OJJ80235.1 akaw|GAA91299.1 aluc|OJZ82771.1 anid|CBF79976.1 anig|EHA21213.1 aory|EIT81454.1 asyd|OJJ58232.1 ater|EAU31489.1 atub|OJI89002.1 aver|OJJ02192.1 awen|OJJ41567.1 azon|OJJ44836.1

Group1698: abra|OJJ68555.1 acar|OOF92750.1 acla|EAW12812.1 afis|EAW17977.1 afla|EED51928.1 afum|EAL91786.1 agla|OJJ80234.1 akaw|GAA91298.1 aluc|OJZ82772.1 anid|CBF79975.1 anig|EHA21214.1 aory|EIT81416.1 asyd|OJJ58231.1 ater|EAU31488.1 atub|OJI89003.1 aver|OJJ02193.1 awen|OJJ41568.1 azon|OJJ44837.1

Group1699: abra|OJJ68556.1 acar|OOF92751.1 acla|EAW12811.1 afis|EAW17976.1 afla|EED51929.1 afum|EAL91785.1 agla|OJJ80233.1 akaw|GAA91297.1 aluc|OJZ82773.1 anid|CBF79973.1 anig|EHA21215.1 aory|EIT81423.1 asyd|OJJ58230.1 ater|EAU31487.1 atub|OJI89004.1 aver|OJJ02194.1 awen|OJJ41569.1 azon|OJJ44838.1

Group1700: abra|OJJ68557.1 acar|OOF92752.1 acla|EAW12810.1 afis|EAW17975.1 afla|EED51930.1 afum|EAL91784.1 agla|OJJ80232.1 akaw|GAA91296.1 aluc|OJZ82774.1 anid|CBF79972.1 anig|EHA21216.1 aory|EIT81446.1 asyd|OJJ58229.1 ater|EAU31486.1 atub|OJI89005.1 aver|OJJ02195.1 awen|OJJ41570.1 azon|OJJ48283.1

Group1701: abra|OJJ68559.1 acar|OOF92754.1 acla|EAW12806.1 afis|EAW17972.1 afla|EED51932.1 afum|EAL91781.1 agla|OJJ80230.1 akaw|GAA91294.1 aluc|OJZ82775.1 anid|CBF79968.1 anig|EHA21218.1 aory|EIT81462.1 asyd|OJJ58227.1 ater|EAU31482.1 atub|OJI89007.1 aver|OJJ02197.1 awen|OJJ41572.1 azon|OJJ48285.1

Group1702: abra|OJJ68561.1 acar|OOF92755.1 acla|EAW12804.1 afis|EAW17970.1 afla|EED51934.1 afum|EAL91779.1 agla|OJJ80228.1 akaw|GAA91292.1 aluc|OJZ82777.1 anid|CBF79964.1 anig|EHA21220.1 aory|EIT81427.1 asyd|OJJ58224.1 ater|EAU31480.1 atub|OJI89009.1 aver|OJJ02198.1 awen|OJJ41574.1 azon|OJJ48287.1

Group1703: abra|OJJ68564.1 acar|OOF92757.1 acla|EAW12802.1 afis|EAW17968.1 afla|EED51937.1 afum|EAL91777.1 agla|OJJ80226.1 akaw|GAA91290.1 aluc|OJZ82779.1 anid|CBF79961.1 anig|EHA21222.1 aory|EIT81398.1 asyd|OJJ58222.1 ater|EAU31478.1 atub|OJI89012.1 aver|OJJ02200.1 awen|OJJ41576.1 azon|OJJ48289.1

Group1704: abra|OJJ68567.1 acar|OOF92761.1 acla|EAW12798.1 afis|EAW17964.1 afla|EED51946.1 afum|EAL91774.1 agla|OJJ80209.1 akaw|GAA91286.1 aluc|OJZ82782.1 anid|CBF79954.1 anig|EHA21225.1 aory|EIT81431.1 asyd|OJJ58218.1 ater|EAU31450.1 atub|OJI89015.1 aver|OJJ02204.1 awen|OJJ41580.1 azon|OJJ48293.1

Group1705: abra|OJJ68599.1 acar|OOF92109.1 acla|EAW12640.1 afis|EAW18045.1 afla|EED51501.1 afum|EAL91857.1 agla|OJJ85416.1 akaw|GAA91923.1 aluc|OJZ81957.1 anid|CBF79779.1 anig|EHA22290.1 aory|EIT80534.1 asyd|OJJ58121.1 ater|EAU31278.1 atub|OJI84508.1 aver|OJJ04284.1 awen|OJJ41489.1 azon|OJJ48219.1

Group1706: abra|OJJ68601.1 acar|OOF92111.1 acla|EAW12638.1 afis|EAW18047.1 afla|EED51499.1 afum|EAL91859.1 agla|OJJ85418.1 akaw|GAA91925.1 aluc|OJZ81955.1 anid|CBF79776.1 anig|EHA22292.1 aory|EIT80455.1 asyd|OJJ58119.1 ater|EAU31276.1 atub|OJI84506.1 aver|OJJ04286.1 awen|OJJ41487.1 azon|OJJ48217.1

Group1707: abra|OJJ68605.1 acar|OOF92114.1 acla|EAW12588.1 afis|EAW18087.1 afla|EED51483.1 afum|EAL91899.1 agla|OJJ85421.1 akaw|GAA92916.1 aluc|OJZ81952.1 anid|CBF79764.1 anig|EHA22301.1 aory|EIT80442.1 asyd|OJJ58103.1 ater|EAU30435.1 atub|OJI84502.1 aver|OJJ04301.1 awen|OJJ41480.1 azon|OJJ48212.1

Group1708: abra|OJJ68606.1 acar|OOF92117.1 acla|EAW12587.1 afis|EAW18086.1 afla|EED51480.1 afum|EAL91898.1 agla|OJJ85422.1 akaw|GAA92915.1 aluc|OJZ81948.1 anid|CBF79762.1 anig|EHA22302.1 aory|EIT80466.1 asyd|OJJ58102.1 ater|EAU30436.1 atub|OJI84501.1 aver|OJJ04302.1 awen|OJJ41479.1 azon|OJJ48211.1

Group1709: abra|OJJ68607.1 acar|OOF92118.1 acla|EAW12593.1 afis|EAW18085.1 afla|EED51479.1 afum|EAL91897.1 agla|OJJ85423.1 akaw|GAA92914.1 aluc|OJZ81947.1 anid|CBF86384.1 aory|EIT80735.1 asyd|OJJ64960.1 ater|EAU30437.1 atub|OJI84500.1 aver|OJI99082.1 awen|OJJ41478.1 azon|OJJ48210.1 aory|EIT81994.1

Group1710: abra|OJJ68608.1 acar|OOF92119.1 acla|EAW12594.1 afis|EAW18084.1 afla|EED51478.1 afum|EAL91896.1 agla|OJJ85424.1 akaw|GAA92913.1 aluc|OJZ81946.1 anid|CBF86386.1 anig|EHA22303.1 aory|EIT80678.1 asyd|OJJ64959.1 ater|EAU30438.1 atub|OJI84499.1 aver|OJI99081.1 awen|OJJ41477.1 azon|OJJ48209.1

Group1711: abra|OJJ68610.1 acar|OOF92120.1 acla|EAW12601.1 afis|EAW18083.1 afla|EED51477.1 afum|EAL91895.1 agla|OJJ85446.1 akaw|GAA92912.1 aluc|OJZ81945.1 anid|CBF86388.1 anig|EHA22304.1 aory|EIT80541.1 asyd|OJJ64958.1 ater|EAU30439.1 atub|OJI84498.1 aver|OJI99080.1 awen|OJJ41472.1 azon|OJJ48931.1

Group1712: abra|OJJ68611.1 acar|OOF92121.1 acla|EAW12602.1 afis|EAW18082.1 afla|EED51476.1 afum|EAL91894.1 agla|OJJ85447.1 akaw|GAA92911.1 aluc|OJZ81944.1 anid|CBF86390.1 anig|EHA22305.1 aory|EIT80460.1 asyd|OJJ64957.1 ater|EAU30440.1 atub|OJI84497.1 aver|OJI99079.1 awen|OJJ41470.1 azon|OJJ48927.1

Group1713: abra|OJJ68612.1 acar|OOF92122.1 acla|EAW12603.1 afis|EAW18081.1 afla|EED51475.1 afum|EAL91893.1 agla|OJJ87374.1 akaw|GAA92910.1 aluc|OJZ81943.1 anid|CBF86391.1 anig|EHA22306.1 aory|EIT80723.1 asyd|OJJ64956.1 ater|EAU30441.1 atub|OJI84496.1 aver|OJI99078.1 awen|OJJ41469.1 azon|OJJ48926.1

Group1714: abra|OJJ68613.1 acar|OOF92123.1 acla|EAW12604.1 afis|EAW18080.1 afla|EED51474.1 afum|EAL91892.1 agla|OJJ87373.1 akaw|GAA92909.1 aluc|OJZ81942.1 anid|CBF86393.1 anig|EHA22307.1 aory|EIT80697.1 asyd|OJJ64955.1 ater|EAU30442.1 atub|OJI84495.1 aver|OJI99077.1 awen|OJJ41468.1 azon|OJJ48925.1

Group1715: abra|OJJ68615.1 acar|OOF92126.1 acla|EAW12607.1 afis|EAW18077.1 afla|EED51471.1 afum|EAL91889.1 agla|OJJ87370.1 akaw|GAA92906.1 aluc|OJZ81939.1 anid|CBF86404.1 anig|EHA22309.1 aory|EIT80562.1 asyd|OJJ64951.1 ater|EAU30446.1 atub|OJI84492.1 aver|OJI99073.1 awen|OJJ41466.1 azon|OJJ48923.1

Group1716: abra|OJJ68617.1 acar|OOF92128.1 acla|EAW12609.1 afis|EAW18075.1 afla|EED51467.1 afum|EAL91887.1 agla|OJJ87367.1 akaw|GAA92904.1 aluc|OJZ81937.1 anid|CBF86400.1 anig|EHA22311.1 aory|EIT80468.1 asyd|OJJ64949.1 ater|EAU30452.1 atub|OJI84490.1 aver|OJI99071.1 awen|OJJ41464.1 azon|OJJ48919.1

Group1717: abra|OJJ68618.1 acar|OOF92129.1 acla|EAW12610.1 afis|EAW18074.1 afla|EED51464.1 afum|EAL91886.1 agla|OJJ87366.1 akaw|GAA92903.1 aluc|OJZ81936.1 anid|CBF86398.1 anig|EHA22312.1 aory|EIT80711.1 asyd|OJJ64948.1 ater|EAU30454.1 atub|OJI84489.1 aver|OJI99070.1 awen|OJJ41463.1 azon|OJJ48918.1

Group1718: abra|OJJ68619.1 acar|OOF92130.1 acla|EAW12611.1 afis|EAW18073.1 afla|EED51463.1 afum|EAL91885.1 agla|OJJ87365.1 akaw|GAA92899.1 aluc|OJZ81935.1 anid|CBF86406.1 anig|EHA22313.1 aory|EIT80619.1 asyd|OJJ64947.1 ater|EAU30455.1 atub|OJI84488.1 aver|OJI99069.1 awen|OJJ41462.1 azon|OJJ48917.1

Group1719: abra|OJJ68620.1 acar|OOF92131.1 acla|EAW13703.1 afis|EAW19885.1 afla|EED57031.1 afum|EAL85012.1 agla|OJJ89079.1 akaw|GAA92898.1 aluc|OJZ81934.1 anid|CBF82756.1 anig|EHA22314.1 aory|EIT82313.1 asyd|OJJ61608.1 ater|EAU33697.1 atub|OJI84487.1 aver|OJJ00931.1 awen|OJJ36996.1 azon|OJJ42189.1

Group1720: abra|OJJ68623.1 acar|OOF92133.1 acla|EAW12612.1 afis|EAW18072.1 afla|EED51461.1 afum|EAL91884.1 agla|OJJ87364.1 akaw|GAA92897.1 aluc|OJZ81932.1 anid|CBF86411.1 anig|EHA22316.1 aory|EIT80603.1 asyd|OJJ64944.1 ater|EAU30457.1 atub|OJI84485.1 aver|OJI99066.1 awen|OJJ41459.1 azon|OJJ48916.1

Group1721: abra|OJJ68626.1 acar|OOF92136.1 acla|EAW12613.1 afis|EAW18071.1 afla|EED51460.1 afum|EAL91883.1 agla|OJJ87363.1 akaw|GAA92895.1 aluc|OJZ81929.1 anid|CBF86413.1 anig|EHA22318.1 aory|EIT80647.1 asyd|OJJ64943.1 ater|EAU30458.1 atub|OJI84481.1 aver|OJI99065.1 awen|OJJ41458.1 azon|OJJ48915.1

Group1722: abra|OJJ68636.1 acar|OOF92140.1 acla|EAW12615.1 afis|EAW18069.1 afla|EED51458.1 afum|EAL91881.2 agla|OJJ87356.1 akaw|GAA89775.1 aluc|OJZ81919.1 anid|CBF86416.1 anig|EHA22328.1

aory|EIT80491.1 asyd|OJJ64941.1 ater|EAU30460.1 atub|OJI84471.1 aver|OJI99063.1 awen|OJJ41456.1 azon|OJJ48913.1

Group1723: abra|OJJ68637.1 acar|OOF92141.1 acla|EAW12614.1 afis|EAW18070.1 afla|EED51459.1 afum|EAL91882.1 agla|OJJ87357.1 akaw|GAA89776.1 aluc|OJZ81918.1 anid|CBF86415.1 anig|EHA22329.1 aory|EIT80742.1 asyd|OJJ64942.1 ater|EAU30459.1 atub|OJI84470.1 aver|OJI99064.1 awen|OJJ41457.1 azon|OJJ48914.1

Group1724: abra|OJJ68642.1 acar|OOF92146.1 acla|EAW12620.1 afis|EAW18064.1 afla|EED51433.1 afum|EAL91876.1 akaw|GAA89782.1 aluc|OJZ81910.1 anid|CBF86424.1 anig|EHA22334.1 aory|EIT80728.1 asyd|OJJ64937.1 ater|EAU30467.1 atub|OJI84463.1 aver|OJI99058.1 awen|OJJ41451.1 azon|OJJ48909.1 agla|OJJ87350.1

Group1725: abra|OJJ68643.1 acar|OOF92145.1 acla|EAW12621.1 afis|EAW18063.1 afla|EED51432.1 afum|EAL91875.1 agla|OJJ87349.1 akaw|GAA89783.1 aluc|OJZ81908.1 anid|CBF86426.1 anig|EHA22335.1 aory|EIT80621.1 asyd|OJJ64936.1 ater|EAU30468.1 atub|OJI84461.1 aver|OJI99057.1 awen|OJJ41449.1 azon|OJJ48908.1

Group1726: abra|OJJ68647.1 acar|OOF92149.1 acla|EAW12623.1 afis|EAW18061.1 afla|EED51429.1 afum|EAL91872.1 agla|OJJ87325.1 akaw|GAA89787.1 aluc|OJZ81904.1 anid|CBF86430.1 anig|EHA22339.1 aory|EIT80637.1 asyd|OJJ64934.1 ater|EAU30473.1 atub|OJI84457.1 aver|OJI99054.1 awen|OJJ41445.1 azon|OJJ48907.1

Group1727: abra|OJJ68648.1 acar|OOF92150.1 acla|EAW12624.1 afis|EAW18060.1 afla|EED51428.1 afum|EAL91871.1 agla|OJJ87326.1 akaw|GAA89788.1 aluc|OJZ81903.1 anid|CBF86432.1 anig|EHA22340.1 aory|EIT80552.1 asyd|OJJ64933.1 ater|EAU30474.1 atub|OJI84456.1 aver|OJI99053.1 awen|OJJ41444.1 azon|OJJ48905.1

Group1728: abra|OJJ68649.1 acar|OOF92151.1 acla|EAW12625.1 afis|EAW18059.1 afla|EED51427.1 afum|EAL91870.1 agla|OJJ87327.1 akaw|GAA89789.1 aluc|OJZ81902.1 anid|CBF86434.1 anig|EHA22341.1 aory|EIT80495.1 asyd|OJJ64932.1 ater|EAU30475.1 atub|OJI84455.1 aver|OJI99051.1 awen|OJJ41442.1 azon|OJJ48904.1

Group1729: abra|OJJ68650.1 acar|OOF92152.1 acla|EAW12626.1 afis|EAW18058.1 afla|EED51426.1 afum|EBA27442.1 agla|OJJ87328.1 akaw|GAA89790.1 aluc|OJZ81901.1 anid|CBF86435.1 anig|EHA22342.1 aory|EIT80761.1 asyd|OJJ64931.1 ater|EAU30476.1 atub|OJI84454.1 aver|OJI99050.1 awen|OJJ41441.1 azon|OJJ48903.1

Group1730: abra|OJJ68652.1 acar|OOF92153.1 acla|EAW12628.1 afis|EAW18056.1 afla|EED51423.1 afum|EAL91867.1 agla|OJJ87330.1 akaw|GAA89792.1 aluc|OJZ81899.1 anid|CBF86443.1 anig|EHA22344.1 aory|EIT80509.1 asyd|OJJ64928.1 ater|EAU30478.1 atub|OJI84452.1 aver|OJI99047.1 awen|OJJ41439.1 azon|OJJ48901.1

Group1731: abra|OJJ68655.1 acar|OOF92155.1 acla|EAW12630.1 afis|EAW18054.1 afla|EED51421.1 afum|EAL91865.1 agla|OJJ87335.1 akaw|GAA89794.1 aluc|OJZ81897.1 anid|CBF86438.1 anig|EHA22345.1 aory|EIT80746.1 asyd|OJJ64927.1 ater|EAU30479.1 atub|OJI84450.1 aver|OJI99045.1 awen|OJJ41436.1 azon|OJJ48900.1

Group1732: abra|OJJ68656.1 acar|OOF92156.1 acla|EAW12632.1 afis|EAW18052.1 afla|EED51418.1 afum|EAL91863.1 agla|OJJ87339.1 akaw|GAA89795.1 aluc|OJZ81896.1 anid|CBF86439.1 anig|EHA22346.1 aory|EIT80553.1 asyd|OJJ64925.1 ater|EAU30480.1 atub|OJI84449.1 aver|OJI99044.1 awen|OJJ41433.1 azon|OJJ48898.1

Group1733: abra|OJJ68657.1 acar|OOF92157.1 acla|EAW12633.1 afis|EAW18051.1 afla|EED51417.1 afum|EAL91862.1 agla|OJJ87340.1 akaw|GAA89796.1 aluc|OJZ81895.1 anid|CBF86440.1 anig|EHA22347.1 aory|EIT80630.1 asyd|OJJ64924.1 ater|EAU30481.1 atub|OJI84448.1 aver|OJI99043.1 awen|OJJ41432.1 azon|OJJ48897.1

Group1734: abra|OJJ68659.1 acar|OOF92159.1 acla|EAW12590.1 afis|EAW24059.1 afla|EED51416.1 afum|EAL91914.1 agla|OJJ87341.1 akaw|GAA89797.1 aluc|OJZ81894.1 anid|CBF86444.1 anig|EHA22348.1 aory|EIT80652.1 asyd|OJJ64923.1 ater|EAU30482.1 atub|OJI84447.1 aver|OJI99042.1 awen|OJJ41431.1 azon|OJJ48896.1

Group1735: abra|OJJ68660.1 acar|OOF92158.1 acla|EAW12591.1 afis|EAW24060.1 afla|EED51415.1 afum|EAL91915.1 agla|OJJ87342.1 akaw|GAA89798.1 aluc|OJZ81893.1 anid|CBF86446.1 anig|EHA22349.1 aory|EIT80731.1 asyd|OJJ64922.1 ater|EAU30483.1 atub|OJI84446.1 aver|OJI99041.1 awen|OJJ41430.1 azon|OJJ48895.1

Group1736: abra|OJJ68662.1 acar|OOF92162.1 acla|EAW12586.1 afis|EAW24062.1 afla|EED51412.1 afum|EAL91917.1 agla|OJJ87344.1 akaw|GAA89802.1 aluc|OJZ81889.1 anid|CBF86449.1 anig|EHA22352.1 aory|EIT80533.1 asyd|OJJ64920.1 ater|EAU30484.1 atub|OJI84442.1 aver|OJI99040.1 awen|OJJ41428.1 azon|OJJ48893.1

Group1737: abra|OJJ68664.1 acar|OOF92165.1 acla|EAW12585.1 afis|EAW24063.1 afla|EED51411.1 afum|EAL91918.1 agla|OJJ87345.1 akaw|GAA89805.1 aluc|OJZ81886.1 anid|CBF86451.1 anig|EHA22354.1 aory|EIT80561.1 asyd|OJJ64919.1 ater|EAU30485.1 atub|OJI84439.1 aver|OJI99039.1 awen|OJJ41426.1 azon|OJJ48891.1

Group1738: abra|OJJ68665.1 acar|OOF92167.1 acla|EAW12584.1 afis|EAW24064.1 afla|EED51410.1 afum|EAL91919.1 agla|OJJ87346.1 akaw|GAA89806.1 aluc|OJZ81885.1 anid|CBF86452.1 anig|EHA22355.1 aory|EIT80617.1 asyd|OJJ64918.1 ater|EAU30486.1 atub|OJI84438.1 aver|OJI99038.1 awen|OJJ41425.1 azon|OJJ48890.1

Group1739: abra|OJJ68666.1 acar|OOF92166.1 acla|EAW12583.1 afis|EAW24065.1 afla|EED51409.1 afum|EAL91920.1 agla|OJJ87347.1 akaw|GAA89807.1 aluc|OJZ81884.1 anid|CBF86454.1 anig|EHA22356.1 aory|EIT80471.1 asyd|OJJ64917.1 ater|EAU30487.1 atub|OJI84437.1 aver|OJI99037.1 awen|OJJ41424.1 azon|OJJ48889.1

Group1740: abra|OJJ68670.1 acar|OOF92170.1 acla|EAW12577.1 afis|EAW24071.1 afla|EED51404.1 afum|EAL91927.1 agla|OJJ87312.1 akaw|GAA89811.1 aluc|OJZ81879.1 anid|CBF86462.1 anig|EHA22359.1 aory|EIT80762.1 asyd|OJJ64912.1 ater|EAU30492.1 atub|OJI84433.1 aver|OJI99033.1 awen|OJJ41419.1 azon|OJJ48884.1

Group1741: abra|OJJ68677.1 acar|OOF92179.1 acla|EAW12570.1 afis|EAW24080.1 afla|EED51393.1 afum|EAL91935.1 agla|OJJ87151.1 akaw|GAA89830.1 aluc|OJZ81857.1 anid|CBF86478.1 anig|EHA22367.1 aory|EIT80640.1 asyd|OJJ64902.1 ater|EAU30501.1 atub|OJI84425.1 aver|OJI99024.1 awen|OJJ41418.1 azon|OJJ48875.1

Group1742: abra|OJJ68678.1 acar|OOF92180.1 acla|EAW12569.1 afis|EAW24081.1 afla|EED51392.1 afum|EAL91936.1 agla|OJJ87152.1 akaw|GAA89831.1 aluc|OJZ81856.1 anid|CBF86480.1 anig|EHA22368.1 aory|EIT80524.1 asyd|OJJ64901.1 ater|EAU30504.1 atub|OJI84424.1 aver|OJI99022.1 awen|OJJ41407.1 azon|OJJ48873.1

Group1743: abra|OJJ68681.1 acar|OOF92183.1 acla|EAW12566.1 afis|EAW24084.1 afla|EED51389.1 afum|EAL91939.1 agla|OJJ87154.1 akaw|GAA89834.1 aluc|OJZ81852.1 anid|CBF86484.1 anig|EHA22370.1 aory|EIT80477.1 asyd|OJJ64899.1 ater|EAU30507.1 atub|OJI84421.1 aver|OJI99020.1 awen|OJJ41405.1 azon|OJJ48871.1

Group1744: abra|OJJ68682.1 acar|OOF92184.1 acla|EAW12565.1 afis|EAW24085.1 afla|EED51388.1 afum|EAL91940.1 agla|OJJ87155.1 akaw|GAA89835.1 aluc|OJZ81851.1 anid|CBF86486.1 anig|EHA22371.1 aory|EIT80535.1 asyd|OJJ64898.1 ater|EAU30508.1 atub|OJI84420.1 aver|OJI99019.1 awen|OJJ41404.1 azon|OJJ48870.1

Group1745: abra|OJJ68684.1 acar|OOF92186.1 acla|EAW12563.1 afis|EAW24087.1 afla|EED51386.1 afum|EAL91942.1 agla|OJJ87157.1 akaw|GAA89837.1 aluc|OJZ81849.1 anid|CBF86489.1 anig|EHA22373.1 aory|EIT80476.1 asyd|OJJ64896.1 ater|EAU30511.1 atub|OJI84418.1 aver|OJI99017.1 awen|OJJ41402.1 azon|OJJ48868.1

Group1746: abra|OJJ68685.1 acar|OOF92187.1 acla|EAW12562.1 afis|EAW24088.1 afla|EED46851.1 afum|EAL91943.1 agla|OJJ87158.1 akaw|GAA89838.1 aluc|OJZ81848.1 anid|CBF86491.1 anig|EHA22374.1 aory|EIT72584.1 asyd|OJJ64895.1 ater|EAU30512.1 atub|OJI84417.1 aver|OJI99016.1 awen|OJJ41401.1 azon|OJJ48867.1

Group1747: abra|OJJ68686.1 acar|OOF92188.1 acla|EAW12561.1 afis|EAW24089.1 afla|EED46850.1 afum|EAL91944.1 agla|OJJ87159.1 akaw|GAA89839.1 aluc|OJZ81847.1 anid|CBF86493.1 anig|EHA22375.1 aory|EIT72520.1 asyd|OJJ64894.1 ater|EAU30513.1 atub|OJI84416.1 aver|OJI99015.1 awen|OJJ41400.1 azon|OJJ48866.1

Group1748: abra|OJJ68689.1 acar|OOF92193.1 acla|EAW12555.1 afis|EAW24096.1 afla|EED46838.1 afum|EAL91950.1 agla|OJJ87169.1 akaw|GAA89844.1 aluc|OJZ81843.1 anid|CBF86498.1 anig|EHA22379.1 aory|EIT72590.1 asyd|OJJ64889.1 ater|EAU30523.1 atub|OJI84411.1 aver|OJI99003.1 awen|OJJ41394.1 azon|OJJ48862.1

Group1749: abra|OJJ68690.1 acar|OOF92194.1 acla|EAW12554.1 afis|EAW24097.1 afla|EED46837.1 afum|EAL91951.2 agla|OJJ87171.1 akaw|GAA89845.1 aluc|OJZ81842.1 anid|CBF86500.1 anig|EHA22380.1 aory|EIT72693.1 asyd|OJJ64888.1 ater|EAU30524.1 atub|OJI84410.1 aver|OJI99002.1 awen|OJJ41392.1 azon|OJJ48861.1

Group1750: abra|OJJ68691.1 acar|OOF92195.1 acla|EAW12553.1 afis|EAW24098.1 afla|EED46836.1 afum|EAL91952.1 agla|OJJ87172.1 akaw|GAA89846.1 aluc|OJZ81841.1 anid|CBF86501.1 anig|EHA22381.1 aory|EIT72579.1 asyd|OJJ64887.1 ater|EAU30525.1 atub|OJI84409.1 aver|OJI99001.1 awen|OJJ41391.1 azon|OJJ48860.1

Group1751: abra|OJJ68692.1 acar|OOF92196.1 acla|EAW12552.1 afis|EAW24099.1 afla|EED46835.1 afum|EAL91953.2 agla|OJJ87173.1 akaw|GAA89847.1 aluc|OJZ81840.1 anid|CBF86503.1 anig|EHA22382.1

aory|EIT72533.1 asyd|OJJ64886.1 ater|EAU30526.1 atub|OJI84408.1 aver|OJI99000.1 awen|OJJ41390.1 azon|OJJ48859.1

Group1752: abra|OJJ68693.1 acar|OOF92197.1 acla|EAW12551.1 afis|EAW24100.1 afla|EED46834.1 afum|EAL91954.1 agla|OJJ87174.1 akaw|GAA89848.1 aluc|OJZ81838.1 anid|CBF86505.1 anig|EHA22383.1 aory|EIT72706.1 asyd|OJJ64885.1 ater|EAU30527.1 atub|OJI84407.1 aver|OJI98999.1 awen|OJJ41389.1 azon|OJJ48858.1

Group1753: abra|OJJ68695.1 acar|OOF92199.1 acla|EAW12549.1 afis|EAW24101.1 afla|EED46832.1 afum|EAL91955.1 agla|OJJ87176.1 akaw|GAA89850.1 aluc|OJZ81836.1 anid|CBF86507.1 anig|EHA22385.1 aory|EIT72575.1 asyd|OJJ64883.1 ater|EAU30529.1 atub|OJI84405.1 aver|OJI98997.1 awen|OJJ41387.1 azon|OJJ48856.1

Group1754: abra|OJJ68696.1 acar|OOF92200.1 acla|EAW12548.1 afis|EAW24102.1 afla|EED46831.1 afum|EAL91956.1 agla|OJJ87177.1 akaw|GAA89851.1 aluc|OJZ81835.1 anid|CBF86509.1 anig|EHA22386.1 aory|EIT72546.1 asyd|OJJ64882.1 ater|EAU30530.1 atub|OJI84404.1 aver|OJI98996.1 awen|OJJ41386.1 azon|OJJ48855.1

Group1755: abra|OJJ68697.1 acar|OOF92201.1 acla|EAW12547.1 afis|EAW24103.1 afla|EED46830.1 afum|EAL91957.1 agla|OJJ87178.1 akaw|GAA89852.1 aluc|OJZ81834.1 anid|CBF86511.1 anig|EHA22387.1 aory|EIT72771.1 asyd|OJJ64881.1 ater|EAU30531.1 atub|OJI84403.1 aver|OJI98995.1 awen|OJJ41385.1 azon|OJJ48854.1

Group1756: abra|OJJ68698.1 acar|OOF92202.1 acla|EAW12546.1 afis|EAW24104.1 afla|EED46829.1 afum|EAL91958.1 agla|OJJ87179.1 akaw|GAA89853.1 aluc|OJZ81833.1 anid|CBF86513.1 anig|EHA22388.1 aory|EIT72709.1 asyd|OJJ64880.1 ater|EAU30532.1 atub|OJI84402.1 aver|OJI98994.1 awen|OJJ41384.1 azon|OJJ48853.1

Group1757: abra|OJJ68703.1 acar|OOF92205.1 acla|EAW12542.1 afis|EAW24107.1 afla|EED46826.1 afum|EAL91961.1 agla|OJJ87182.1 akaw|GAA89857.1 aluc|OJZ81828.1 anid|CBF86518.1 anig|EHA22391.1 aory|EIT72757.1 asyd|OJJ64877.1 ater|EAU30535.1 atub|OJI84398.1 aver|OJI98990.1 awen|OJJ41381.1 azon|OJJ48850.1

Group1758: abra|OJJ68704.1 acar|OOF92206.1 acla|EAW12541.1 afis|EAW24108.1 afla|EED46825.1 afum|EAL91962.1 agla|OJJ87183.1 akaw|GAA89858.1 aluc|OJZ81826.1 anid|CBF86520.1 anig|EHA22392.1 aory|EIT72547.1 asyd|OJJ64876.1 ater|EAU30536.1 atub|OJI84396.1 aver|OJI98989.1 awen|OJJ41380.1 azon|OJJ48848.1

Group1759: abra|OJJ68706.1 acar|OOF92208.1 acla|EAW12539.1 afis|EAW24110.1 afla|EED46822.1 afum|EAL91964.1 agla|OJJ87185.1 akaw|GAA89860.1 aluc|OJZ81824.1 anid|CBF86523.1 anig|EHA22394.1 aory|EIT72674.1 asyd|OJJ64874.1 ater|EAU30538.1 atub|OJI84394.1 aver|OJI98987.1 awen|OJJ41378.1 azon|OJJ48846.1

Group1760: abra|OJJ68709.1 acar|OOF92211.1 acla|EAW12536.1 afis|EAW24113.1 afla|EED46820.1 afum|EAL91967.1 agla|OJJ87190.1 akaw|GAA89863.1 aluc|OJZ81818.1 anid|CBF86529.1 anig|EHA22397.1 aory|EIT72599.1 asyd|OJJ64871.1 ater|EAU30541.1 atub|OJI84391.1 aver|OJI98983.1 awen|OJJ41375.1 azon|OJJ48843.1

Group1761: abra|OJJ68710.1 acar|OOF92212.1 acla|EAW12535.1 afis|EAW24114.1 afla|EED46819.1 afum|EAL91968.1 agla|OJJ87191.1 akaw|GAA89864.1 aluc|OJZ81817.1 anid|CBF86531.1 anig|EHA22398.1 aory|EIT72684.1 asyd|OJJ64870.1 ater|EAU30542.1 atub|OJI84390.1 aver|OJI98982.1 awen|OJJ41374.1 azon|OJJ48842.1

Group1762: abra|OJJ68717.1 acar|OOF92217.1 acla|EAW12529.1 afis|EAW24120.1 afla|EED46812.1 afum|EAL91975.1 agla|OJJ87194.1 akaw|GAA89869.1 aluc|OJZ81812.1 anid|CBF86542.1 anig|EHA22405.1 aory|EIT72532.1 asyd|OJJ64862.1 ater|EAU30548.1 atub|OJI84384.1 aver|OJI98973.1 awen|OJJ41369.1 azon|OJJ48836.1

Group1763: abra|OJJ68719.1 acar|OOF92218.1 acla|EAW12528.1 afis|EAW24121.1 afla|EED46810.1 afum|EAL91977.1 agla|OJJ87196.1 akaw|GAA89870.1 aluc|OJZ81810.1 anid|CBF86546.1 anig|EHA22407.1 aory|EIT72617.1 asyd|OJJ64860.1 ater|EAU30549.1 atub|OJI84382.1 aver|OJI98972.1 awen|OJJ41368.1 azon|OJJ48835.1

Group1764: abra|OJJ68722.1 acar|OOF92221.1 acla|EAW12524.1 afis|EAW24124.1 afla|EED46809.1 afum|EAL91980.1 agla|OJJ87199.1 akaw|GAA89872.1 aluc|OJZ81807.1 anid|CBF86551.1 anig|EHA22410.1 aory|EIT72759.1 asyd|OJJ64857.1 ater|EAU30551.1 atub|OJI84379.1 aver|OJI98969.1 awen|OJJ41365.1 azon|OJJ48832.1

Group1765: abra|OJJ68723.1 acar|OOF92222.1 acla|EAW12523.1 afis|EAW24125.1 afla|EED46808.1 afum|EAL91981.1 agla|OJJ87200.1 akaw|GAA89873.1 aluc|OJZ81806.1 anid|CBF86553.1 anig|EHA22411.1 aory|EIT72530.1 asyd|OJJ64856.1 ater|EAU30552.1 atub|OJI84378.1 aver|OJI98968.1 awen|OJJ41364.1 azon|OJJ48831.1

Group1766: abra|OJJ68725.1 acar|OOF92224.1 acla|EAW12521.1 afis|EAW24126.1 afla|EED46806.1 afum|EAL91982.1 agla|OJJ87202.1 akaw|GAA89876.1 aluc|OJZ81803.1 anid|CBF86555.1 anig|EHA22413.1 aory|EIT72636.1 asyd|OJJ64855.1 ater|EAU30553.1 atub|OJI84375.1 aver|OJI98967.1 awen|OJJ41363.1 azon|OJJ48829.1

Group1767: abra|OJJ68728.1 acar|OOF92225.1 acla|EAW12520.1 afis|EAW24127.1 afla|EED46805.1 afum|EAL91983.1 agla|OJJ87203.1 akaw|GAA89877.1 aluc|OJZ81802.1 anid|CBF86557.1 anig|EHA22414.1 aory|EIT72714.1 asyd|OJJ64854.1 ater|EAU30557.1 atub|OJI84374.1 aver|OJI98966.1 awen|OJJ41362.1 azon|OJJ48827.1

Group1768: abra|OJJ68729.1 acar|OOF92226.1 acla|EAW12519.1 afis|EAW24128.1 afla|EED46804.1 afum|EAL91984.1 agla|OJJ87204.1 akaw|GAA89878.1 aluc|OJZ81801.1 anid|CBF86559.1 anig|EHA22415.1 aory|EIT72675.1 asyd|OJJ64853.1 ater|EAU30558.1 atub|OJI84372.1 aver|OJI98965.1 awen|OJJ41361.1 azon|OJJ48825.1

Group1769: abra|OJJ68730.1 acar|OOF92228.1 acla|EAW12518.1 afis|EAW24129.1 afla|EED46803.1 afum|EAL91985.1 agla|OJJ87205.1 akaw|GAA89879.1 aluc|OJZ81800.1 anid|CBF86560.1 anig|EHA22416.1 aory|EIT72669.1 asyd|OJJ64852.1 ater|EAU30559.1 atub|OJI84371.1 aver|OJI98964.1 awen|OJJ41360.1 azon|OJJ48824.1

Group1770: abra|OJJ68734.1 acar|OOF92230.1 acla|EAW12513.1 afis|EAW24133.1 afla|EED46800.1 afum|EAL91988.1 agla|OJJ87210.1 akaw|GAA89882.1 aluc|OJZ81796.1 anid|CBF86566.1 anig|EHA22420.1 aory|EIT72682.1 asyd|OJJ64849.1 ater|EAU30562.1 atub|OJI84367.1 aver|OJI98961.1 awen|OJJ41356.1 azon|OJJ48819.1

Group1771: abra|OJJ68735.1 acar|OOF92231.1 acla|EAW12512.1 afis|EAW24134.1 afla|EED46799.1 afum|EAL91989.2 agla|OJJ87211.1 akaw|GAA89883.1 aluc|OJZ81794.1 anid|CBF86568.1 anig|EHA22421.1 aory|EIT72755.1 asyd|OJJ64848.1 ater|EAU30563.1 atub|OJI84366.1 aver|OJI98960.1 awen|OJJ41355.1 azon|OJJ48818.1

Group1772: abra|OJJ68736.1 acar|OOF92232.1 acla|EAW12511.1 afis|EAW24135.1 afla|EED46798.1 afum|EAL91990.1 agla|OJJ87212.1 akaw|GAA89884.1 aluc|OJZ81793.1 anid|CBF86570.1 anig|EHA22422.1 aory|EIT72542.1 asyd|OJJ64847.1 ater|EAU30564.1 atub|OJI84365.1 aver|OJI98959.1 awen|OJJ41354.1 azon|OJJ48817.1

Group1773: abra|OJJ68738.1 acar|OOF92234.1 afis|EAW24137.1 afla|EED46797.1 afum|EAL91992.1 agla|OJJ87215.1 akaw|GAA89886.1 aluc|OJZ81789.1 anid|CBF77943.1 anig|EHA22426.1 aory|EIT72631.1 asyd|OJJ55282.1 ater|EAU30566.1 atub|OJI84363.1 aver|OJJ06946.1 awen|OJJ41352.1 azon|OJJ48815.1 acla|EAW12509.1

Group1774: abra|OJJ68739.1 acar|OOF92235.1 acla|EAW12508.1 afis|EAW24138.1 afla|EED46796.1 afum|EAL91993.1 agla|OJJ87216.1 akaw|GAA89887.1 aluc|OJZ81788.1 anid|CBF77941.1 anig|EHA22427.1 aory|EIT72772.1 asyd|OJJ55283.1 ater|EAU30567.1 atub|OJI84362.1 aver|OJJ06945.1 awen|OJJ41351.1 azon|OJJ48814.1

Group1775: abra|OJJ68742.1 acar|OOF90307.1 acla|EAW12506.1 afis|EAW24141.1 afla|EED46793.1 afum|EAL91996.1 agla|OJJ87220.1 akaw|GAA89889.1 aluc|OJZ81784.1 anid|CBF77936.1 anig|EHA22431.1 aory|EIT72574.1 asyd|OJJ55286.1 ater|EAU30570.1 atub|OJI84358.1 aver|OJJ06942.1 awen|OJJ41347.1 azon|OJJ48811.1

Group1776: abra|OJJ68743.1 acar|OOF90308.1 acla|EAW12505.1 afis|EAW24142.1 afla|EED46792.1 afum|EAL91997.1 agla|OJJ87221.1 akaw|GAA89890.1 aluc|OJZ81783.1 anid|CBF77934.1 anig|EHA22432.1 aory|EIT72571.1 asyd|OJJ55287.1 ater|EAU30571.1 atub|OJI84357.1 aver|OJJ06941.1 awen|OJJ41346.1 azon|OJJ48810.1

Group1777: abra|OJJ68748.1 acar|OOF92236.1 acla|EAW12503.1 afis|EAW24143.1 afla|EED46789.1 afum|EAL91998.1 agla|OJJ87222.1 akaw|GAA89892.1 aluc|OJZ81779.1 anid|CBF77930.1 anig|EHA22436.1 aory|EIT72633.1 asyd|OJJ55290.1 ater|EAU30574.1 atub|OJI84353.1 aver|OJJ06939.1 awen|OJJ41345.1 azon|OJJ48809.1

Group1778: abra|OJJ68749.1 acar|OOF92237.1 acla|EAW12502.1 afis|EAW24144.1 afla|EED46788.1 afum|EAL91999.1 agla|OJJ87223.1 akaw|GAA89893.1 aluc|OJZ81778.1 anid|CBF77928.1 anig|EHA22438.1 aory|EIT72562.1 asyd|OJJ55291.1 ater|EAU30575.1 atub|OJI84351.1 aver|OJJ06936.1 awen|OJJ41336.1 azon|OJJ48806.1

Group1779: abra|OJJ68750.1 acar|OOF92238.1 acla|EAW12501.1 afis|EAW24145.1 afla|EED46787.1 afum|EAL92000.1 agla|OJJ87224.1 akaw|GAA89894.1 aluc|OJZ81777.1 anid|CBF77926.1 anig|EHA22439.1 aory|EIT72718.1 asyd|OJJ55292.1 ater|EAU30576.1 atub|OJI84350.1 aver|OJJ06935.1 awen|OJJ41335.1 azon|OJJ48804.1

Group1780: abra|OJJ68752.1 acar|OOF92239.1 acla|EAW12499.1 afis|EAW24146.1 afla|EED46783.1 afum|EAL92001.1 agla|OJJ87226.1 akaw|GAA89896.1 aluc|OJZ81775.1 anid|CBF77923.1 anig|EHA22441.1

aory|EIT72696.1 asyd|OJJ55294.1 ater|EAU30578.1 atub|OJI84348.1 aver|OJJ06932.1 awen|OJJ41333.1 azon|OJJ48803.1

Group1781: abra|OJJ68756.1 acar|OOF92244.1 acla|EAW12495.1 afis|EAW24150.1 afla|EED46781.1 afum|EAL92005.1 agla|OJJ89542.1 akaw|GAA89900.1 aluc|OJZ81771.1 anid|CBF77915.1 anig|EHA22445.1 aory|EIT72518.1 asyd|OJJ55297.1 ater|EAU30582.1 atub|OJI84344.1 aver|OJJ06929.1 awen|OJJ41328.1 azon|OJJ48801.1

Group1782: abra|OJJ68763.1 acar|OOF92249.1 acla|EAW12488.1 afis|EAW24157.1 afla|EED46776.1 afum|EAL92012.2 agla|OJJ89524.1 akaw|GAA89907.1 aluc|OJZ81762.1 anid|CBF77900.1 anig|EHA22451.1 aory|EIT72774.1 asyd|OJJ55303.1 ater|EAU30588.1 atub|OJI84334.1 aver|OJJ06923.1 awen|OJJ41321.1 azon|OJJ48794.1

Group1783: abra|OJJ68766.1 acar|OOF92252.1 acla|EAW12485.1 afis|EAW24160.1 afla|EED46773.1 afum|EAL92016.1 agla|OJJ89521.1 akaw|GAA89910.1 aluc|OJZ81759.1 anid|CBF77894.1 anig|EHA22453.1 aory|EIT72644.1 asyd|OJJ55306.1 ater|EAU30591.1 atub|OJI84331.1 aver|OJJ06920.1 awen|OJJ41318.1 azon|OJJ48791.1

Group1784: abra|OJJ68768.1 acar|OOF92253.1 acla|EAW12484.1 afis|EAW24161.1 afla|EED46772.1 afum|EAL92017.1 agla|OJJ89519.1 akaw|GAA89911.1 aluc|OJZ81758.1 anid|CBF77892.1 anig|EHA22454.1 aory|EIT72707.1 asyd|OJJ55307.1 ater|EAU30592.1 atub|OJI84329.1 aver|OJJ06919.1 awen|OJJ41316.1 azon|OJJ48790.1

Group1785: abra|OJJ68769.1 acar|OOF92254.1 acla|EAW12483.1 afis|EAW24162.1 afla|EED46771.1 afum|EAL92018.1 agla|OJJ89518.1 akaw|GAA89912.1 aluc|OJZ81757.1 anid|CBF77890.1 anig|EHA22455.1 aory|EIT72701.1 asyd|OJJ55308.1 ater|EAU30593.1 atub|OJI84328.1 aver|OJJ06918.1 awen|OJJ41315.1 azon|OJJ48789.1

Group1786: abra|OJJ68770.1 acar|OOF92255.1 acla|EAW12482.1 afis|EAW24163.1 afla|EED46770.1 afum|EAL92020.1 agla|OJJ89517.1 akaw|GAA89913.1 aluc|OJZ81756.1 anid|CBF77888.1 anig|EHA22456.1 aory|EIT72763.1 asyd|OJJ55309.1 ater|EAU30594.1 atub|OJI84327.1 aver|OJJ06917.1 awen|OJJ41314.1 azon|OJJ48788.1

Group1787: abra|OJJ68785.1 acar|OOF92269.1 acla|EAW12469.1 afis|EAW24172.1 afla|EED46757.1 afum|EAL92034.1 agla|OJJ89503.1 akaw|GAA89927.1 aluc|OJZ81743.1 anid|CBF77861.1 anig|EHA22469.1 aory|EIT72255.1 asyd|OJJ55323.1 ater|EAU30607.1 atub|OJI84314.1 aver|OJJ06905.1 awen|OJJ41300.1 azon|OJJ48774.1

Group1788: abra|OJJ68788.1 acar|OOF92272.1 acla|EAW12466.1 afis|EAW24175.1 afla|EED46754.1 afum|EAL92037.1 agla|OJJ89499.1 akaw|GAA89929.1 aluc|OJZ81740.1 anid|CBF77855.1 anig|EHA22472.1 aory|EIT72259.1 asyd|OJJ55326.1 ater|EAU30610.1 atub|OJI84311.1 aver|OJJ06902.1 awen|OJJ41295.1 azon|OJJ48771.1

Group1789: abra|OJJ68790.1 acar|OOF92274.1 acla|EAW12463.1 afis|EAW24178.1 afla|EED46751.1 afum|EAL92040.1 agla|OJJ89496.1 akaw|GAA89932.1 aluc|OJZ81736.1 anid|CBF77849.1 anig|EHA22475.1 aory|EIT72252.1 asyd|OJJ55329.1 ater|EAU30613.1 atub|OJI84308.1 aver|OJJ06899.1 awen|OJJ41291.1 azon|OJJ48765.1

Group1790: abra|OJJ68793.1 acar|OOF92278.1 acla|EAW12460.1 afis|EAW24182.1 afla|EED46749.1 afum|EAL92042.1 agla|OJJ89493.1 akaw|GAA89933.1 aluc|OJZ81734.1 anid|CBF77844.1 anig|EHA22478.1 aory|EIT72258.1 asyd|OJJ55332.1 ater|EAU30615.1 atub|OJI84305.1 aver|OJJ06896.1 awen|OJJ41288.1 azon|OJJ48762.1

Group1791: abra|OJJ68794.1 acar|OOF92277.1 acla|EAW12459.1 afis|EAW24183.1 afla|EED46748.1 afum|EAL92043.1 agla|OJJ89492.1 akaw|GAA89934.1 aluc|OJZ81733.1 anid|CBF77842.1 anig|EHA22479.1 aory|EIT72260.1 asyd|OJJ55333.1 ater|EAU30616.1 atub|OJI84304.1 aver|OJJ06895.1 awen|OJJ41287.1 azon|OJJ48769.1

Group1792: abra|OJJ68797.1 acar|OOF92281.1 acla|EAW14657.1 afis|EAW24186.1 afla|EED46746.1 afum|EAL92046.1 agla|OJJ89489.1 akaw|GAA89936.1 aluc|OJZ81730.1 anid|CBF77836.1 anig|EHA22482.1 aory|EIT72257.1 asyd|OJJ55336.1 ater|EAU30619.1 atub|OJI84301.1 aver|OJJ06891.1 awen|OJJ41284.1 azon|OJJ43704.1

Group1793: abra|OJJ68827.1 acla|EAW12865.1 afis|EAW17814.1 afla|EED45598.1 afum|EAL91623.1 agla|OJJ81685.1 akaw|GAA90426.1 aluc|OJZ85803.1 anid|CBF79113.1 anig|EHA24752.1 aory|EIT75707.1 asyd|OJJ54395.1 atub|OJI79588.1 aver|OJJ07596.1 awen|OJJ31341.1 anid|CBF80733.1 awen|OJJ36931.1 asyd|OJJ54560.1

Group1794: abra|OJJ68855.1 acar|OOF90533.1 acla|EAW15256.1 afis|EAW20230.1 afla|EED57124.1 afum|EAL85347.2 agla|OJJ84632.1 akaw|GAA90398.1 aluc|OJZ85833.1 anid|CBF85122.1 anig|EHA24730.1 aory|EIT74034.1 asyd|OJJ63044.1 ater|EAU32032.1 atub|OJI79621.1 aver|OJI97919.1 awen|OJJ31233.1 azon|OJJ46156.1

Group1795: abra|OJJ68856.1 acar|OOF90532.1 afla|EED57125.1 afum|EAL85346.1 agla|OJJ84633.1 akaw|GAA90397.1 aluc|OJZ85834.1 anid|CBF85120.1 anig|EHA24729.1 aory|EIT74077.1 ater|EAU32031.1 atub|OJI79622.1 aver|OJI97918.1 awen|OJJ31232.1 azon|OJJ46155.1 acla|EAW15255.1 afis|EAW20229.1 asyd|OJJ63045.1

Group1796: abra|OJJ68859.1 acar|OOF90530.1 acla|EAW15250.1 afis|EAW20218.1 afla|EED57129.1 afum|EAL85333.1 agla|OJJ84638.1 akaw|GAA90394.1 aluc|OJZ85837.1 anid|CBF85115.1 anig|EHA24726.1 aory|EIT74090.1 asyd|OJJ63049.1 ater|EAU32027.1 atub|OJI79625.1 aver|OJI97914.1 awen|OJJ31228.1 azon|OJJ46151.1

Group1797: abra|OJJ68871.1 acar|OOF89965.1 acla|EAW15248.1 afis|EAW20214.1 afla|EED57130.1 afum|EAL85331.1 agla|OJJ84640.1 akaw|GAA90383.1 aluc|OJZ85848.1 anid|CBF85111.1 anig|EHA28098.1 aory|EIT74018.1 asyd|OJJ63051.1 ater|EAU32024.1 atub|OJI79636.1 aver|OJI97912.1 awen|OJJ31220.1 azon|OJJ46137.1

Group1798: abra|OJJ68913.1 acar|OOF91500.1 acla|EAW15323.1 afis|EAW20327.1 afla|EED57093.1 afum|EAL85453.1 agla|OJJ86344.1 akaw|GAA90336.1 aluc|OJZ85896.1 anid|CBF87781.1 anig|EHA24696.1 aory|EIT82282.1 asyd|OJJ63373.1 ater|EAU32055.1 atub|OJI79681.1 aver|OJI97573.1 awen|OJJ39915.1 azon|OJJ45889.1

Group1799: abra|OJJ68918.1 acar|OOF91506.1 acla|EAW14259.1 afis|EAW19021.1 afla|EED56623.1 afum|EAL93752.1 agla|OJJ84614.1 akaw|GAA90331.1 aluc|OJZ85901.1 anid|CBF85579.1 anig|EHA24691.1 aory|EIT79672.1 asyd|OJJ64177.1 ater|EAU32063.1 atub|OJI79686.1 aver|OJI98253.1 awen|OJJ31311.1 azon|OJJ46160.1

Group1800: abra|OJJ68919.1 acar|OOF91507.1 acla|EAW14260.1 afis|EAW19020.1 afla|EED56622.1 afum|EAL93751.1 agla|OJJ84615.1 akaw|GAA90330.1 aluc|OJZ85902.1 anid|CBF85577.1 anig|EHA24690.1 aory|EIT79600.1 asyd|OJJ64178.1 ater|EAU32062.1 atub|OJI79687.1 aver|OJI98254.1 awen|OJJ31310.1 azon|OJJ46161.1

Group1801: abra|OJJ68926.1 acar|OOF91512.1 acla|EAW14248.1 afis|EAW19032.1 afla|EED50385.1 afum|EAL93761.1 agla|OJJ84590.1 akaw|GAA90322.1 aluc|OJZ85911.1 anid|CBF79596.1 anig|EHA24681.1 aory|EIT83141.1 asyd|OJJ57985.1 ater|EAU32071.1 atub|OJI79701.1 aver|OJJ04415.1 awen|OJJ31461.1 azon|OJJ46163.1

Group1802: abra|OJJ68929.1 acar|OOF90676.1 afis|EAW19225.1 afum|EAL93956.1 agla|OJJ89343.1 akaw|GAA90320.1 aluc|OJZ85913.1 anid|CBF90358.1 anig|EHA24677.1 aory|EIT72912.1 asyd|OJJ55570.1 ater|EAU37576.1 atub|OJI79703.1 aver|OJJ06649.1 awen|OJJ29830.1 afla|EED51132.1 aory|EIT81305.1 agla|OJJ81307.1

Group1803: abra|OJJ68932.1 acar|OOF91515.1 acla|EAW14247.1 afis|EAW19033.1 afla|EED50387.1 afum|EAL93762.2 agla|OJJ84588.1 akaw|GAA90317.1 aluc|OJZ85916.1 anid|CBF79600.1 anig|EHA24675.1 aory|EIT83129.1 asyd|OJJ57990.1 ater|EAU32073.1 atub|OJI79707.1 aver|OJJ04412.1 awen|OJJ31463.1 azon|OJJ46164.1

Group1804: abra|OJJ68933.1 acar|OOF91516.1 acla|EAW14246.1 afla|EED50389.1 agla|OJJ84587.1 akaw|GAA90316.1 aluc|OJZ85917.1 anig|EHA24674.1 aory|EIT83117.1 asyd|OJJ57991.1 ater|EAU32074.1 atub|OJI79710.1 aver|OJJ04411.1 awen|OJJ31464.1 azon|OJJ46166.1 afum|EAL93763.1 anid|CBF79601.1 afis|EAW19034.1

Group1805: abra|OJJ68937.1 acar|OOF91518.1 acla|EAW14243.1 afis|EAW19037.1 afla|EED50391.1 afum|EAL93766.1 agla|OJJ84584.1 akaw|GAA90314.1 aluc|OJZ85922.1 anid|CBF79605.1 anig|EHA24672.1 aory|EIT82128.1 asyd|OJJ57993.1 ater|EAU32076.1 atub|OJI79712.1 aver|OJJ04409.1 awen|OJJ31466.1 azon|OJJ46169.1

Group1806: abra|OJJ68941.1 acar|OOF91521.1 acla|EAW14242.1 afis|EAW19038.1 afla|EED50393.1 afum|EAL93767.1 agla|OJJ84582.1 akaw|GAA90312.1 aluc|OJZ85927.1 anid|CBF79609.1 anig|EHA24670.1 aory|EIT82042.1 asyd|OJJ57994.1 ater|EAU32077.1 atub|OJI79716.1 aver|OJJ04408.1 awen|OJJ31468.1 azon|OJJ46172.1

Group1807: abra|OJJ68943.1 acar|OOF91522.1 acla|EAW14241.1 afis|EAW19039.1 afla|EED50394.1 afum|EAL93768.1 agla|OJJ84573.1 akaw|GAA90310.1 aluc|OJZ85929.1 anid|CBF79610.1 anig|EHA24669.1 aory|EIT82167.1 asyd|OJJ57995.1 ater|EAU32078.1 atub|OJI79717.1 aver|OJJ04407.1 awen|OJJ31469.1 azon|OJJ46173.1

Group1808: abra|OJJ68944.1 acar|OOF91524.1 acla|EAW14239.1 afis|EAW19041.1 afla|EED50396.1 afum|EAL93770.1 agla|OJJ84564.1 akaw|GAA90309.1 aluc|OJZ85930.1 anid|CBF79612.1 anig|EHA24667.1 aory|EIT82062.1 asyd|OJJ57999.1 ater|EAU32080.1 atub|OJI79719.1 aver|OJJ04403.1 awen|OJJ31470.1 azon|OJJ46179.1

Group1809: abra|OJJ68945.1 acar|OOF91525.1 acla|EAW14238.1 afis|EAW19042.1 afla|EED50397.1 afum|EAL93771.1 agla|OJJ84563.1 akaw|GAA90308.1 aluc|OJZ85931.1 anid|CBF79613.1 anig|EHA24666.1

aory|EIT82121.1 asyd|OJJ57998.1 ater|EAU32081.1 atub|OJI79720.1 aver|OJJ04404.1 awen|OJJ31471.1 azon|OJJ46181.1

Group1810: abra|OJJ68948.1 acar|OOF91526.1 acla|EAW14236.1 afis|EAW19044.1 afla|EED50401.1 afum|EAL93773.1 agla|OJJ84561.1 akaw|GAA90306.1 aluc|OJZ85934.1 anig|EHA24664.1 aory|EIT82075.1 ater|EAU32083.1 atub|OJI79723.1 azon|OJJ46184.1 anid|CBF79617.1 asyd|OJJ58000.1 aver|OJJ04402.1 awen|OJJ31473.1

Group1811: abra|OJJ68949.1 acar|OOF91527.1 acla|EAW14235.1 afis|EAW19045.1 afla|EED50402.1 afum|EAL93774.1 agla|OJJ84560.1 akaw|GAA90305.1 aluc|OJZ85935.1 anid|CBF79619.1 anig|EHA24663.1 aory|EIT82000.1 asyd|OJJ58001.1 ater|EAU32084.1 atub|OJI79724.1 aver|OJJ04400.1 awen|OJJ31474.1 azon|OJJ46188.1

Group1812: abra|OJJ68952.1 acar|OOF91530.1 acla|EAW14234.1 afis|EAW19046.1 afla|EED50405.1 afum|EAL93775.1 agla|OJJ84559.1 akaw|GAA90481.1 aluc|OJZ85938.1 anid|CBF79621.1 anig|EHA24661.1 aory|EIT82119.1 asyd|OJJ58003.1 ater|EAU32085.1 atub|OJI79726.1 aver|OJJ04398.1 awen|OJJ31475.1 azon|OJJ46189.1

Group1813: abra|OJJ68953.1 acar|OOF91531.1 acla|EAW14233.1 afis|EAW19047.1 afla|EED50406.1 afum|EAL93776.1 agla|OJJ84558.1 akaw|GAA90482.1 aluc|OJZ85939.1 anid|CBF79622.1 anig|EHA24660.1 aory|EIT82158.1 asyd|OJJ58004.1 ater|EAU32086.1 atub|OJI79727.1 aver|OJJ04397.1 awen|OJJ31477.1 azon|OJJ46190.1

Group1814: abra|OJJ68956.1 acar|OOF91535.1 acla|EAW14229.1 afis|EAW19052.1 afla|EED54409.1 afum|EAL93781.1 agla|OJJ84555.1 akaw|GAA90484.1 aluc|OJZ85942.1 anid|CBF79627.1 anig|EHA24657.1 aory|EIT75857.1 asyd|OJJ58009.1 ater|EAU32089.1 atub|OJI79730.1 aver|OJJ04391.1 awen|OJJ31483.1 azon|OJJ46193.1

Group1815: abra|OJJ68957.1 acar|OOF91534.1 acla|EAW14228.1 afis|EAW19053.1 afla|EED54411.1 afum|EAL93782.1 agla|OJJ84554.1 akaw|GAA90485.1 aluc|OJZ85943.1 anid|CBF79628.1 anig|EHA24656.1 aory|EIT75871.1 asyd|OJJ58010.1 ater|EAU32090.1 atub|OJI79731.1 aver|OJJ04390.1 awen|OJJ31484.1 azon|OJJ46194.1

Group1816: abra|OJJ68959.1 acar|OOF91537.1 acla|EAW14226.1 afis|EAW19055.1 afla|EED54413.1 afum|EAL93784.1 agla|OJJ84552.1 akaw|GAA90487.1 aluc|OJZ85945.1 anid|CBF79630.1 anig|EHA24654.1 aory|EIT75877.1 asyd|OJJ58012.1 ater|EAU32091.1 atub|OJI79733.1 aver|OJJ04388.1 awen|OJJ31486.1 azon|OJJ46196.1

Group1817: abra|OJJ68960.1 acar|OOF91538.1 acla|EAW14225.1 afis|EAW19056.1 afla|EED54414.1 afum|EAL93785.1 agla|OJJ84551.1 akaw|GAA90488.1 aluc|OJZ85946.1 anid|CBF79632.1 anig|EHA24653.1 aory|EIT75852.1 asyd|OJJ58013.1 ater|EAU32092.1 atub|OJI79734.1 aver|OJJ04387.1 awen|OJJ31487.1 azon|OJJ46197.1

Group1818: abra|OJJ68961.1 acar|OOF91539.1 acla|EAW14224.1 afis|EAW19057.1 afla|EED54415.1 afum|EAL93786.1 agla|OJJ84550.1 akaw|GAA90489.1 aluc|OJZ85947.1 anid|CBF79634.1 anig|EHA24652.1 aory|EIT75893.1 asyd|OJJ58014.1 ater|EAU32093.1 atub|OJI79735.1 aver|OJJ04386.1 awen|OJJ31488.1 azon|OJJ46199.1

Group1819: abra|OJJ68963.1 acar|OOF91541.1 acla|EAW14223.1 afis|EAW19058.1 afla|EED54418.1 afum|EAL93787.1 agla|OJJ84549.1 akaw|GAA90491.1 aluc|OJZ85950.1 anid|CBF79638.1 anig|EHA24650.1 aory|EIT75892.1 asyd|OJJ58015.1 ater|EAU32094.1 atub|OJI79738.1 aver|OJJ04385.1 awen|OJJ31489.1 azon|OJJ46201.1

Group1820: abra|OJJ68965.1 acar|OOF91542.1 acla|EAW14222.1 afis|EAW19059.1 afla|EED54421.1 afum|EAL93788.1 agla|OJJ84548.1 akaw|GAA90492.1 aluc|OJZ85952.1 anid|CBF79639.1 anig|EHA24649.1 aory|EIT75888.1 asyd|OJJ58016.1 ater|EAU32095.1 atub|OJI79740.1 aver|OJJ04384.1 awen|OJJ31490.1 azon|OJJ46202.1

Group1821: abra|OJJ68966.1 acar|OOF91544.1 acla|EAW14221.1 afis|EAW19060.1 afla|EED54422.1 afum|EAL93789.1 agla|OJJ84547.1 akaw|GAA90493.1 aluc|OJZ85953.1 anid|CBF79640.1 anig|EHA24648.1 aory|EIT75846.1 asyd|OJJ58017.1 ater|EAU32098.1 atub|OJI79741.1 aver|OJJ04382.1 awen|OJJ31491.1 azon|OJJ46203.1

Group1822: abra|OJJ68967.1 acar|OOF91545.1 acla|EAW14220.1 afis|EAW19061.1 afla|EED54423.1 afum|EAL93790.1 agla|OJJ84546.1 akaw|GAA90494.1 aluc|OJZ85954.1 anid|CBF79641.1 anig|EHA24647.1 aory|EIT75876.1 asyd|OJJ58018.1 ater|EAU32099.1 atub|OJI79742.1 aver|OJJ04381.1 awen|OJJ31492.1 azon|OJJ46204.1

Group1823: abra|OJJ68968.1 acar|OOF91546.1 acla|EAW14219.1 afis|EAW19062.1 afla|EED54428.1 afum|EAL93791.1 agla|OJJ84545.1 akaw|GAA90495.1 aluc|OJZ85955.1 anid|CBF79643.1 anig|EHA24646.1 aory|EIT75878.1 asyd|OJJ58019.1 ater|EAU32105.1 atub|OJI79743.1 aver|OJJ04380.1 awen|OJJ31493.1 azon|OJJ46206.1

Group1824: abra|OJJ68970.1 acar|OOF91547.1 acla|EAW14217.1 afis|EAW19064.1 afla|EED54430.1 afum|EAL93793.2 agla|OJJ84543.1 akaw|GAA90497.1 aluc|OJZ85957.1 anid|CBF79646.1 anig|EHA24644.1 aory|EIT75886.1 asyd|OJJ58021.1 ater|EAU32107.1 atub|OJI79745.1 aver|OJJ04378.1 awen|OJJ31496.1 azon|OJJ46208.1

Group1825: abra|OJJ68973.1 acar|OOF91550.1 acla|EAW14214.1 afis|EAW19067.1 afla|EED54433.1 afum|EAL93796.1 agla|OJJ84539.1 akaw|GAA90500.1 aluc|OJZ85962.1 anid|CBF79652.1 anig|EHA24641.1 aory|EIT75842.1 asyd|OJJ58025.1 ater|EAU32115.1 atub|OJI79749.1 aver|OJJ04374.1 awen|OJJ31499.1 azon|OJJ46213.1

Group1826: abra|OJJ68978.1 acar|OOF91553.1 acla|EAW14210.1 afis|EAW19071.1 afla|EED54436.1 afum|EAL93799.1 agla|OJJ84534.1 akaw|GAA90503.1 aluc|OJZ85966.1 anid|CBF79663.1 anig|EHA24638.1 aory|EIT75872.1 asyd|OJJ58028.1 ater|EAU32119.1 atub|OJI79754.1 aver|OJJ04371.1 awen|OJJ31503.1 azon|OJJ46218.1

Group1827: abra|OJJ68982.1 acar|OOF91554.1 acla|EAW14209.1 afis|EAW19072.1 afla|EED54437.1 afum|EAL93800.1 agla|OJJ84532.1 akaw|GAA90504.1 aluc|OJZ85968.1 anid|CBF79661.1 anig|EHA24637.1 aory|EIT75860.1 asyd|OJJ58029.1 ater|EAU32120.1 atub|OJI79756.1 aver|OJJ04370.1 awen|OJJ32466.1 azon|OJJ46219.1

Group1828: abra|OJJ68984.1 acar|OOF91555.1 acla|EAW14207.1 afis|EAW19074.1 afla|EED54439.1 afum|EAL93802.1 agla|OJJ84529.1 akaw|GAA90506.1 aluc|OJZ85970.1 anid|CBF79667.1 anig|EHA24635.1 aory|EIT75870.1 asyd|OJJ58031.1 ater|EAU32122.1 atub|OJI79758.1 aver|OJJ04368.1 awen|OJJ32468.1 azon|OJJ46238.1

Group1829: abra|OJJ68991.1 acar|OOF92703.1 acla|EAW12643.1 afis|EAW18042.1 afla|EED51854.1 afum|EAL91853.1 agla|OJJ85414.1 akaw|GAA90511.1 aluc|OJZ85976.1 anid|CBF80097.1 anig|EHA24630.1 aory|EIT81766.1 asyd|OJJ58299.1 ater|EAU31557.1 atub|OJI79766.1 aver|OJJ02125.1 awen|OJJ41492.1 azon|OJJ48223.1

Group1830: abra|OJJ68994.1 acar|OOF92705.1 acla|EAW12645.1 afis|EAW18040.1 afla|EED51857.1 afum|EAL91851.1 agla|OJJ85412.1 akaw|GAA90513.1 aluc|OJZ85978.1 anid|CBF80103.1 anig|EHA24628.1 aory|EIT81744.1 asyd|OJJ58296.1 ater|EAU31555.1 atub|OJI79768.1 aver|OJJ02128.1 awen|OJJ41497.1 azon|OJJ48225.1

Group1831: abra|OJJ68996.1 acar|OOF92707.1 acla|EAW12647.1 afis|EAW18038.1 afla|EED51860.1 afum|EAL91849.1 agla|OJJ85410.1 akaw|GAA90515.1 aluc|OJZ85980.1 anid|CBF80073.1 anig|EHA24626.1 aory|EIT81689.1 asyd|OJJ58294.1 ater|EAU31552.1 atub|OJI79770.1 aver|OJJ02130.1 awen|OJJ41499.1 azon|OJJ48227.1

Group1832: abra|OJJ68997.1 acar|OOF92708.1 acla|EAW12648.1 afis|EAW18037.1 afla|EED51861.1 afum|EAL91848.1 agla|OJJ85409.1 akaw|GAA90516.1 aluc|OJZ85981.1 anid|CBF80074.1 anig|EHA24625.1 aory|EIT81730.1 asyd|OJJ58293.1 ater|EAU31551.1 atub|OJI79771.1 aver|OJJ02131.1 awen|OJJ41500.1 azon|OJJ48228.1

Group1833: abra|OJJ69004.1 acar|OOF92717.1 acla|EAW12855.1 afis|EAW18018.1 afla|EED51873.1 afum|EAL91830.1 agla|OJJ85372.1 akaw|GAA90522.1 aluc|OJZ85990.1 anid|CBF80052.1 anig|EHA24620.1 aory|EIT81725.1 asyd|OJJ58280.1 ater|EAU31534.1 atub|OJI79779.1 aver|OJJ02143.1 awen|OJJ41521.1 azon|OJJ48243.1

Group1834: abra|OJJ69005.1 acar|OOF92718.1 acla|EAW12854.1 afla|EED51874.1 afum|EAL91829.1 akaw|GAA90523.1 aluc|OJZ85991.1 anig|EHA24619.1 aory|EIT81686.1 ater|EAU31533.1 atub|OJI79780.1 afis|EAW18017.1 asyd|OJJ58279.1 aver|OJJ02144.1 awen|OJJ41522.1 agla|OJJ85371.1 anid|CBF80050.1 azon|OJJ48244.1

Group1835: abra|OJJ69008.1 acar|OOF92719.1 acla|EAW12850.1 afis|EAW18013.1 afla|EED51883.1 afum|EAL91825.1 agla|OJJ80286.1 akaw|GAA90525.1 aluc|OJZ85994.1 anid|CBF80040.1 anig|EHA24616.1 aory|EIT81726.1 asyd|OJJ58275.1 ater|EAU31529.1 atub|OJI79782.1 aver|OJJ02149.1 awen|OJJ41528.1 azon|OJJ48248.1

Group1836: abra|OJJ69010.1 acar|OOF92721.1 acla|EAW12848.1 afis|EAW18011.1 afla|EED51885.1 afum|EAL91823.1 agla|OJJ80283.1 akaw|GAA90527.1 aluc|OJZ85996.1 anid|CBF80037.1 anig|EHA24614.1 aory|EIT81760.1 asyd|OJJ58273.1 ater|EAU31527.1 atub|OJI79784.1 aver|OJJ02151.1 awen|OJJ41530.1 azon|OJJ48250.1

Group1837: azon|OJJ51098.1 abra|OJJ74349.1 acar|OOF95381.1 acla|EAW07403.1 afis|EAW22429.1 afla|EED52507.1 afum|EAL90762.1 agla|OJJ87594.1 akaw|GAA86681.1 aluc|OJZ89781.1 anid|CBF88824.1 anig|EHA26996.1 aory|EIT82792.1 asyd|OJJ62884.1 ater|EAU39278.1 atub|OJI85025.1 aver|OJI96936.1 awen|OJJ35008.1

Group1838: abra|OJJ69024.1 acar|OOF92985.1 acla|EAW06466.1 afis|EAW15961.1 afla|EED58096.1 afum|EAL88594.1 agla|OJJ89351.1 akaw|GAA90541.1 aluc|OJZ86011.1 anid|CBF81079.1 anig|EHA24600.1

aory|EIT79116.1 asyd|OJJ56463.1 ater|EAU35493.1 atub|OJI79800.1 aver|OJJ06258.1 awen|OJJ35613.1 azon|OJJ46911.1

Group1839: abra|OJJ69026.1 acar|OOF92660.1 acla|EAW12556.1 afis|EAW24095.1 afla|EED46839.1 afum|EAL91949.1 agla|OJJ87168.1 akaw|GAA92787.1 aluc|OJZ82696.1 anid|CBF75138.1 anig|EHA21145.1 aory|EIT72775.1 asyd|OJJ59978.1 ater|EAU30520.1 atub|OJI88928.1 aver|OJJ02313.1 awen|OJJ41395.1 azon|OJJ48863.1

Group1840: abra|OJJ69029.1 acar|OOF92658.1 acla|EAW10072.1 afis|EAW25514.1 afla|EED54015.1 afum|EAL90179.1 agla|OJJ87289.1 akaw|GAA90545.1 aluc|OJZ86015.1 anid|CBF77783.1 anig|EHA24596.1 aory|EIT78264.1 asyd|OJJ53030.1 ater|EAU33360.1 atub|OJI79804.1 aver|OJJ03814.1 awen|OJJ41257.1 azon|OJJ44822.1

Group1841: abra|OJJ69034.1 acar|OOF89828.1 acla|EAW08636.1 afis|EAW15795.1 afla|EED51250.1 afum|EAL85555.1 agla|OJJ81958.1 akaw|GAA90549.1 aluc|OJZ86019.1 anid|CBF70929.1 anig|EHA24592.1 aory|EIT81525.1 asyd|OJJ52611.1 ater|EAU32347.1 atub|OJI79808.1 aver|OJJ04638.1 awen|OJJ38503.1 azon|OJJ43748.1

Group1842: abra|OJJ69036.1 acar|OOF92649.1 acla|EAW12687.1 afis|EAW17850.1 afla|EED46204.1 afum|EAL91659.1 agla|OJJ83119.1 akaw|GAA90551.1 aluc|OJZ86021.1 anid|CBF77960.1 anig|EHA24590.1 aory|EIT76404.1 asyd|OJJ55272.1 ater|EAU33052.1 atub|OJI79810.1 aver|OJJ06955.1 awen|OJJ41723.1 azon|OJJ48421.1

Group1843: abra|OJJ69037.1 acar|OOF92647.1 acla|EAW14104.1 afis|EAW18805.1 afla|EED50771.1 afum|EAL93541.1 agla|OJJ80535.1 akaw|GAA90552.1 aluc|OJZ86022.1 anid|CBF88044.1 anig|EHA24589.1 aory|EIT78003.1 asyd|OJJ63545.1 ater|EAU37943.1 atub|OJI79811.1 aver|OJI97389.1 awen|OJJ32380.1 azon|OJJ47142.1

Group1844: abra|OJJ69038.1 acar|OOF92646.1 acla|EAW06768.1 afis|EAW16318.1 afla|EED53563.1 afum|EAL88942.1 agla|OJJ88279.1 akaw|GAA90553.1 aluc|OJZ86023.1 anid|CBF89147.1 anig|EHA24588.1 aory|EIT74896.1 asyd|OJJ62568.1 ater|EAU34139.1 atub|OJI79812.1 aver|OJI96692.1 awen|OJJ37718.1 azon|OJJ44975.1

Group1845: abra|OJJ69039.1 acar|OOF92645.1 acla|EAW11330.1 afis|EAW18165.1 afla|EED53435.1 afum|EAL87525.2 agla|OJJ85953.1 akaw|GAA90554.1 aluc|OJZ86024.1 anid|CBF85751.1 anig|EHA24587.1 aory|EIT74836.1 asyd|OJJ64296.1 ater|EAU36013.1 atub|OJI79813.1 aver|OJI98389.1 awen|OJJ30512.1 azon|OJJ50291.1

Group1846: abra|OJJ69057.1 acar|OOF91226.1 acla|EAW08757.1 afis|EAW15674.1 afla|EED46856.1 afum|EAL85673.1 agla|OJJ81821.1 akaw|GAA90587.1 aluc|OJZ86045.1 anid|CBF71161.1 anig|EHA24566.1 aory|EIT72754.1 asyd|OJJ55844.1 ater|EAU32486.1 atub|OJI79842.1 aver|OJJ04787.1 awen|OJJ38352.1 azon|OJJ42790.1

Group1847: abra|OJJ69058.1 acar|OOF91225.1 acla|EAW08756.1 afis|EAW15673.1 afla|EED46857.1 afum|EAL85675.1 agla|OJJ81820.1 akaw|GAA90588.1 aluc|OJZ86046.1 anid|CBF71163.1 anig|EHA24565.1 aory|EIT72609.1 asyd|OJJ55845.1 ater|EAU32487.1 atub|OJI79843.1 aver|OJJ04788.1 awen|OJJ38351.1 azon|OJJ42791.1

Group1848: abra|OJJ69064.1 acar|OOF91219.1 acla|EAW08760.1 afis|EAW15668.1 afla|EED46876.1 afum|EAL85680.1 agla|OJJ81806.1 akaw|GAA90594.1 aluc|OJZ86052.1 anid|CBF71174.1 anig|EHA24559.1 aory|EIT72756.1 asyd|OJJ55853.1 ater|EAU32503.1 atub|OJI79849.1 aver|OJJ04796.1 awen|OJJ38344.1 azon|OJJ43043.1

Group1849: abra|OJJ69065.1 acar|OOF91218.1 acla|EAW08759.1 afis|EAW15667.1 afla|EED46877.1 afum|EAL85681.1 agla|OJJ81805.1 akaw|GAA90595.1 aluc|OJZ86053.1 anid|CBF71176.1 anig|EHA24558.1 aory|EIT72761.1 asyd|OJJ55854.1 ater|EAU32504.1 atub|OJI79850.1 aver|OJJ04797.1 awen|OJJ38343.1 azon|OJJ43042.1

Group1850: abra|OJJ69071.1 acar|OOF91231.1 acla|EAW08767.1 afis|EAW15663.1 afla|EED46886.1 afum|EAL85685.1 agla|OJJ81785.1 akaw|GAA90602.1 aluc|OJZ86060.1 anid|CBF71180.1 anig|EHA24553.1 aory|EIT72594.1 asyd|OJJ55862.1 ater|EAU32510.1 atub|OJI79857.1 aver|OJJ04803.1 awen|OJJ38339.1 azon|OJJ43024.1

Group1851: abra|OJJ69078.1 acar|OOF91213.1 afis|EAW15680.1 afla|EED46852.1 afum|EAL85668.1 akaw|GAA90607.1 aluc|OJZ86066.1 anid|CBF71154.1 anig|EHA24548.1 aory|EIT72663.1 asyd|OJJ59218.1 ater|EAU32482.1 atub|OJI79862.1 aver|OJJ01828.1 awen|OJJ38357.1 azon|OJJ48601.1 asyd|OJJ55840.1 aver|OJJ04783.1

Group1852: abra|OJJ69080.1 acar|OOF91211.1 acla|EAW08754.1 afis|EAW15682.1 afla|EED51372.1 afum|EAL85666.1 agla|OJJ81831.1 akaw|GAA90609.1 aluc|OJZ86068.1 anid|CBF71152.1 anig|EHA24546.1 aory|EIT80462.1 asyd|OJJ55839.1 ater|EAU32475.1 atub|OJI79864.1 aver|OJJ04782.1 awen|OJJ38374.1 azon|OJJ43858.1

Group1853: abra|OJJ69081.1 acar|OOF91210.1 acla|EAW08753.1 afis|EAW15683.1 afla|EED51371.1 afum|EAL85665.1 agla|OJJ81832.1 akaw|GAA90610.1 aluc|OJZ86069.1 anid|CBF71150.1 anig|EHA24545.1 aory|EIT80441.1 asyd|OJJ55838.1 ater|EAU32474.1 atub|OJI79865.1 aver|OJJ04781.1 awen|OJJ38375.1 azon|OJJ43859.1

Group1854: abra|OJJ69084.1 acar|OOF91208.1 acla|EAW08750.1 afis|EAW15686.1 afla|EED51368.1 afum|EAL85662.1 agla|OJJ81834.1 akaw|GAA90613.1 aluc|OJZ86071.1 anid|CBF71144.1 anig|EHA24543.1 aory|EIT80616.1 asyd|OJJ55836.1 ater|EAU32467.1 atub|OJI79868.1 aver|OJJ04779.1 awen|OJJ38377.1 azon|OJJ43861.1

Group1855: abra|OJJ69089.1 acar|OOF91204.1 acla|EAW08743.1 afis|EAW15693.1 afla|EED51360.1 afum|EAL85655.1 agla|OJJ81840.1 akaw|GAA90618.1 aluc|OJZ86078.1 anid|CBF71129.1 anig|EHA24538.1 aory|EIT80527.1 asyd|OJJ55829.1 ater|EAU32462.1 atub|OJI79875.1 aver|OJJ04771.1 awen|OJJ38384.1 azon|OJJ43853.1

Group1856: abra|OJJ69090.1 acar|OOF91203.1 acla|EAW08742.1 afis|EAW15694.1 afla|EED51359.1 afum|EAL85654.1 agla|OJJ81841.1 akaw|GAA90619.1 aluc|OJZ86079.1 anid|CBF71127.1 anig|EHA24537.1 aory|EIT80431.1 asyd|OJJ55828.1 ater|EAU32461.1 atub|OJI79876.1 aver|OJJ04770.1 awen|OJJ38385.1 azon|OJJ43852.1

Group1857: abra|OJJ69091.1 acar|OOF91202.1 acla|EAW08741.1 afis|EAW15695.1 afla|EED51358.1 afum|EAL85653.1 agla|OJJ81842.1 akaw|GAA90620.1 aluc|OJZ86080.1 anid|CBF71119.1 anig|EHA24535.1 aory|EIT80729.1 asyd|OJJ55827.1 ater|EAU32460.1 atub|OJI79877.1 aver|OJJ04769.1 awen|OJJ38386.1 azon|OJJ43850.1

Group1858: abra|OJJ69093.1 acar|OOF91201.1 acla|EAW08740.1 afis|EAW15696.1 afla|EED51357.1 afum|EAL85652.2 agla|OJJ81850.1 akaw|GAA90621.1 aluc|OJZ86081.1 anid|CBF71121.1 anig|EHA24534.1 aory|EIT80641.1 asyd|OJJ55826.1 ater|EAU32455.1 atub|OJI79878.1 aver|OJJ04768.1 awen|OJJ38389.1 azon|OJJ43849.1

Group1859: abra|OJJ69094.1 acar|OOF91200.1 acla|EAW08737.1 afis|EAW15699.1 afla|EED51353.1 afum|EAL85649.1 agla|OJJ81849.1 akaw|GAA90622.1 aluc|OJZ86083.1 anid|CBF71123.1 anig|EHA24532.1 aory|EIT80448.1 asyd|OJJ55825.1 ater|EAU32452.1 atub|OJI79879.1 aver|OJJ04767.1 awen|OJJ38391.1 azon|OJJ43847.1

Group1860: abra|OJJ69099.1 acar|OOF91197.1 acla|EAW08734.1 afis|EAW15702.1 afla|EED51349.1 afum|EAL85645.1 agla|OJJ81844.1 akaw|GAA90625.1 aluc|OJZ86088.1 anid|CBF71108.1 anig|EHA24529.1 aory|EIT80559.1 asyd|OJJ52712.1 ater|EAU32442.1 atub|OJI79884.1 aver|OJJ04741.1 awen|OJJ38396.1 azon|OJJ43845.1

Group1861: abra|OJJ69100.1 acar|OOF91196.1 acla|EAW08733.1 afis|EAW15707.1 afla|EED51348.1 afum|EAL85640.1 agla|OJJ81851.1 akaw|GAA90626.1 aluc|OJZ86089.1 anid|CBF71106.1 anig|EHA24528.1 aory|EIT80499.1 asyd|OJJ52711.1 atub|OJI79885.1 aver|OJJ04740.1 awen|OJJ38397.1 ater|EAU32441.1 azon|OJJ43844.1

Group1862: abra|OJJ69104.1 acar|OOF91194.1 acla|EAW08728.1 afis|EAW15703.1 afla|EED51345.1 afum|EAL85644.1 agla|OJJ81855.1 akaw|GAA90629.1 aluc|OJZ86092.1 anid|CBF71100.1 anig|EHA24525.1 aory|EIT80418.1 asyd|OJJ52707.1 ater|EAU32438.1 atub|OJI79888.1 aver|OJJ04736.1 awen|OJJ38402.1 azon|OJJ43841.1

Group1863: abra|OJJ69108.1 acar|OOF91181.1 acla|EAW08724.1 afis|EAW15732.1 afla|EED51341.1 afum|EAL85615.2 agla|OJJ81861.1 akaw|GAA90633.1 aluc|OJZ86096.1 anid|CBF71094.1 anig|EHA24522.1 aory|EIT81523.1 asyd|OJJ52704.1 ater|EAU32434.1 atub|OJI79892.1 aver|OJJ04733.1 awen|OJJ38406.1 azon|OJJ43836.1

Group1864: abra|OJJ69110.1 acar|OOF91184.1 acla|EAW08722.1 afis|EAW15730.1 afla|EED51339.1 afum|EAL85617.1 agla|OJJ81864.1 akaw|GAA90635.1 aluc|OJZ86098.1 anid|CBF71090.1 anig|EHA24520.1 aory|EIT81546.1 asyd|OJJ52701.1 ater|EAU32432.1 atub|OJI79894.1 aver|OJJ04729.1 awen|OJJ38409.1 azon|OJJ43833.1

Group1865: abra|OJJ69113.1 acar|OOF91185.1 acla|EAW08719.1 afis|EAW15728.1 afla|EED51335.1 afum|EAL85619.1 agla|OJJ81867.1 akaw|GAA90638.1 aluc|OJZ86102.1 anid|CBF71085.1 anig|EHA24517.1 aory|EIT81524.1 asyd|OJJ52698.1 ater|EAU32430.1 atub|OJI79897.1 aver|OJJ04726.1 awen|OJJ38412.1 azon|OJJ43830.1

Group1866: abra|OJJ69116.1 acar|OOF91188.1 acla|EAW08716.1 afis|EAW15724.1 afla|EED51332.1 afum|EAL85623.1 agla|OJJ81870.1 akaw|GAA90641.1 aluc|OJZ86105.1 anid|CBF71079.1 anig|EHA24515.1 aory|EIT81503.1 asyd|OJJ52695.1 ater|EAU32427.1 atub|OJI79900.1 aver|OJJ04723.1 awen|OJJ38415.1 azon|OJJ43827.1

Group1867: abra|OJJ69119.1 acar|OOF91191.1 acla|EAW08713.1 afis|EAW15721.1 afla|EED51329.1 afum|EAL85626.1 agla|OJJ81873.1 akaw|GAA90644.1 aluc|OJZ86108.1 anid|CBF71075.1 anig|EHA24513.1

aory|EIT81515.1 asyd|OJJ52692.1 ater|EAU32424.1 atub|OJI79903.1 aver|OJJ04720.1 awen|OJJ38418.1 azon|OJJ43824.1

Group1868: abra|OJJ69120.1 acar|OOF91192.1 acla|EAW08712.1 afis|EAW15720.1 afla|EED51328.1 afum|EAL85628.1 agla|OJJ81874.1 akaw|GAA90645.1 aluc|OJZ86109.1 anid|CBF71073.1 anig|EHA24512.1 aory|EIT81554.1 asyd|OJJ52691.1 ater|EAU32423.1 atub|OJI79904.1 aver|OJJ04719.1 awen|OJJ38419.1 azon|OJJ43823.1

Group1869: abra|OJJ69121.1 acar|OOF91193.1 acla|EAW08711.1 afis|EAW15719.1 afla|EED51327.1 afum|EAL85629.2 agla|OJJ81875.1 akaw|GAA90646.1 aluc|OJZ86110.1 anid|CBF71071.1 anig|EHA24511.1 aory|EIT81517.1 asyd|OJJ52690.1 ater|EAU32420.1 atub|OJI79905.1 aver|OJJ04718.1 awen|OJJ38420.1 azon|OJJ43822.1

Group1870: abra|OJJ69122.1 acar|OOF91178.1 acla|EAW08710.1 afis|EAW15718.1 afla|EED51326.1 afum|EAL85630.2 agla|OJJ81877.1 akaw|GAA90647.1 aluc|OJZ86111.1 anid|CBF71070.1 anig|EHA24510.1 aory|EIT81526.1 asyd|OJJ52689.1 ater|EAU32421.1 atub|OJI79906.1 aver|OJJ04717.1 awen|OJJ38421.1 azon|OJJ43821.1

Group1871: abra|OJJ69123.1 acar|OOF91177.1 acla|EAW08709.1 afis|EAW15717.1 afla|EED51325.1 afum|EAL85631.1 agla|OJJ81878.1 akaw|GAA90648.1 aluc|OJZ86112.1 anid|CBF71068.1 anig|EHA24509.1 aory|EIT81505.1 asyd|OJJ52688.1 ater|EAU32422.1 atub|OJI79907.1 aver|OJJ04716.1 awen|OJJ38422.1 azon|OJJ43820.1

Group1872: abra|OJJ69125.1 acar|OOF91175.1 acla|EAW08707.1 afis|EAW15715.1 afla|EED51323.1 afum|EAL85633.1 agla|OJJ81886.1 akaw|GAA90650.1 aluc|OJZ86114.1 anid|CBF71064.1 anig|EHA24507.1 aory|EIT81539.1 asyd|OJJ52685.1 ater|EAU32419.1 atub|OJI79909.1 aver|OJJ04713.1 awen|OJJ38425.1 azon|OJJ43818.1

Group1873: abra|OJJ69126.1 acar|OOF91174.1 acla|EAW08706.1 afis|EAW15714.1 afla|EED51322.1 afum|EAL85634.1 agla|OJJ81887.1 akaw|GAA90651.1 aluc|OJZ86115.1 anid|CBF71058.1 anig|EHA24506.1 aory|EIT81529.1 asyd|OJJ52680.1 ater|EAU32418.1 atub|OJI79910.1 aver|OJJ04706.1 awen|OJJ38426.1 azon|OJJ43817.1

Group1874: abra|OJJ69127.1 acar|OOF91173.1 acla|EAW08705.1 afis|EAW15713.1 afla|EED51321.1 afum|EAL85635.1 agla|OJJ81888.1 akaw|GAA90652.1 aluc|OJZ86116.1 anid|CBF71060.1 anig|EHA24505.1 aory|EIT81530.1 asyd|OJJ52679.1 ater|EAU32417.1 atub|OJI79911.1 aver|OJJ04705.1 awen|OJJ38427.1 azon|OJJ43816.1

Group1875: abra|OJJ69128.1 acar|OOF91172.1 acla|EAW08704.1 afis|EAW15712.1 afla|EED51320.1 afum|EAL85636.1 agla|OJJ81889.1 akaw|GAA90653.1 aluc|OJZ86117.1 anid|CBF71057.1 anig|EHA24504.1 aory|EIT81550.1 asyd|OJJ52678.1 ater|EAU32416.1 atub|OJI79912.1 aver|OJJ04704.1 awen|OJJ38428.1 azon|OJJ43815.1

Group1876: abra|OJJ69131.1 acar|OOF91169.1 acla|EAW08701.1 afis|EAW15709.1 afla|EED51317.1 afum|EAL85638.1 agla|OJJ81892.1 akaw|GAA90656.1 aluc|OJZ86120.1 anid|CBF71053.1 anig|EHA24501.1 aory|EIT81516.1 asyd|OJJ52675.1 ater|EAU32414.1 atub|OJI79916.1 aver|OJJ04701.1 awen|OJJ38431.1 azon|OJJ43812.1

Group1877: abra|OJJ69132.1 acar|OOF91168.1 acla|EAW08700.1 afis|EAW15708.1 afla|EED51316.1 afum|EAL85639.1 agla|OJJ81893.1 akaw|GAA90657.1 aluc|OJZ86121.1 anid|CBF71051.1 anig|EHA24500.1 aory|EIT81557.1 asyd|OJJ52674.1 ater|EAU32413.1 atub|OJI79917.1 aver|OJJ04700.1 awen|OJJ38432.1 azon|OJJ43811.1

Group1878: abra|OJJ69136.1 acar|OOF91164.1 acla|EAW08697.1 afis|EAW15738.1 afla|EED51314.1 afum|EAL85610.1 agla|OJJ81895.1 akaw|GAA86320.1 aluc|OJZ86124.1 anid|CBF71047.1 anig|EHA24497.1 aory|EIT81566.1 asyd|OJJ52672.1 ater|EAU32406.1 atub|OJI79920.1 aver|OJJ04698.1 awen|OJJ38434.1 azon|OJJ43808.1

Group1879: abra|OJJ69138.1 acar|OOF91163.1 acla|EAW08696.1 afis|EAW15739.1 afla|EED51313.1 afum|EAL85609.1 agla|OJJ81900.1 akaw|GAA86319.1 aluc|OJZ86125.1 anid|CBF71045.1 anig|EHA24496.1 aory|EIT81532.1 asyd|OJJ52671.1 ater|EAU32408.1 atub|OJI79921.1 aver|OJJ04697.1 awen|OJJ38438.1 azon|OJJ43807.1

Group1880: abra|OJJ69143.1 acar|OOF91159.1 acla|EAW08666.1 afis|EAW15743.1 afla|EED51309.1 afum|EAL85605.1 agla|OJJ81904.1 akaw|GAA86314.1 aluc|OJZ86129.1 anid|CBF71038.1 anig|EHA24492.1 aory|EIT81535.1 asyd|OJJ52666.1 ater|EAU32402.1 atub|OJI79925.1 aver|OJJ04693.1 awen|OJJ38443.1 azon|OJJ43749.1

Group1881: abra|OJJ69147.1 acar|OOF91156.1 acla|EAW08669.1 afis|EAW15757.1 afla|EED51302.1 afum|EAL85591.1 agla|OJJ81905.1 akaw|GAA86311.1 aluc|OJZ86133.1 anid|CBF71025.1 anig|EHA24488.1 aory|EIT81545.1 asyd|OJJ52660.1 ater|EAU32400.1 atub|OJI79929.1 aver|OJJ04688.1 awen|OJJ38445.1 azon|OJJ43751.1

Group1882: abra|OJJ69154.1 acar|OOF91149.1 acla|EAW08671.1 afis|EAW15753.1 afla|EED51298.1 afum|EAL85596.1 agla|OJJ81909.1 akaw|GAA86308.1 aluc|OJZ86141.1 anid|CBF71023.1 anig|EHA24483.1 aory|EIT81538.1 asyd|OJJ52659.1 ater|EAU32397.1 atub|OJI79933.1 aver|OJJ04687.1 awen|OJJ38453.1 azon|OJJ43753.1

Group1883: abra|OJJ69155.1 acar|OOF91148.1 acla|EAW08672.1 afis|EAW15752.1 afla|EED51297.1 afum|EAL85597.1 agla|OJJ81910.1 akaw|GAA86307.1 aluc|OJZ86142.1 anid|CBF71021.1 anig|EHA24482.1 aory|EIT81540.1 asyd|OJJ52658.1 ater|EAU32396.1 atub|OJI79934.1 aver|OJJ04686.1 awen|OJJ38454.1 azon|OJJ43754.1

Group1884: abra|OJJ69156.1 acar|OOF91147.1 acla|EAW08673.1 afis|EAW15751.1 afla|EED51296.1 afum|EAL85598.1 agla|OJJ81911.1 akaw|GAA86306.1 aluc|OJZ86143.1 anid|CBF71019.1 anig|EHA24481.1 aory|EIT81520.1 asyd|OJJ52657.1 ater|EAU32395.1 atub|OJI79935.1 aver|OJJ04685.1 awen|OJJ38455.1 azon|OJJ43755.1

Group1885: abra|OJJ69157.1 acar|OOF91146.1 acla|EAW08674.1 afis|EAW15750.1 afum|EAL85599.1 agla|OJJ81913.1 akaw|GAA86305.1 aluc|OJZ86144.1 anid|CBF71017.1 anig|EHA24480.1 aory|EIT81492.1 asyd|OJJ52656.1 ater|EAU32394.1 atub|OJI79936.1 aver|OJJ04684.1 awen|OJJ38456.1 azon|OJJ43756.1 afla|EED51295.1

Group1886: abra|OJJ69158.1 acar|OOF91145.1 acla|EAW08675.1 afis|EAW15749.1 afla|EED51294.1 afum|EAL85600.1 agla|OJJ81914.1 akaw|GAA86304.1 aluc|OJZ86145.1 anid|CBF71015.1 anig|EHA24479.1 aory|EIT81541.1 asyd|OJJ52655.1 ater|EAU32393.1 atub|OJI79937.1 aver|OJJ04683.1 awen|OJJ38457.1 azon|OJJ43757.1

Group1887: abra|OJJ69159.1 acar|OOF91144.1 acla|EAW08676.1 afis|EAW15748.1 afla|EED51293.1 afum|EAL85601.1 agla|OJJ81915.1 akaw|GAA86303.1 aluc|OJZ86146.1 anid|CBF71013.1 anig|EHA24478.1 aory|EIT81547.1 asyd|OJJ52654.1 ater|EAU32392.1 atub|OJI79938.1 aver|OJJ04682.1 awen|OJJ38458.1 azon|OJJ43759.1

Group1888: abra|OJJ69161.1 acar|OOF91141.1 acla|EAW08678.1 afis|EAW15746.1 afla|EED51291.1 afum|EAL85602.1 agla|OJJ81917.1 akaw|GAA86301.1 aluc|OJZ86148.1 anid|CBF71009.1 anig|EHA24476.1 aory|EIT81488.1 asyd|OJJ52651.1 ater|EAU32390.1 atub|OJI79940.1 aver|OJJ04680.1 awen|OJJ38460.1 azon|OJJ43761.1

Group1889: abra|OJJ69162.1 acar|OOF91140.1 acla|EAW08679.1 afis|EAW15754.1 afla|EED51290.1 afum|EAL85595.1 agla|OJJ81918.1 akaw|GAA86300.1 aluc|OJZ86149.1 anid|CBF71007.1 anig|EHA24475.1 aory|EIT81493.1 asyd|OJJ52650.1 ater|EAU32389.1 atub|OJI79941.1 aver|OJJ04679.1 awen|OJJ38462.1 azon|OJJ43762.1

Group1890: abra|OJJ69164.1 acar|OOF91139.1 acla|EAW08680.1 afis|EAW15758.1 afla|EED51289.1 afum|EAL85590.1 agla|OJJ81920.1 akaw|GAA86299.1 aluc|OJZ86151.1 anid|CBF71006.1 anig|EHA24474.1 aory|EIT81496.1 asyd|OJJ52648.1 ater|EAU32388.1 atub|OJI79943.1 aver|OJJ04677.1 awen|OJJ38466.1 azon|OJJ43764.1

Group1891: abra|OJJ69165.1 acar|OOF91138.1 acla|EAW08681.1 afis|EAW15759.1 afla|EED51288.1 afum|EAL85589.1 agla|OJJ81921.1 akaw|GAA86298.1 aluc|OJZ86152.1 anid|CBF71004.1 anig|EHA24473.1 aory|EIT81511.1 asyd|OJJ52647.1 ater|EAU32387.1 atub|OJI79944.1 aver|OJJ04676.1 awen|OJJ38467.1 azon|OJJ43765.1

Group1892: abra|OJJ69166.1 acar|OOF91137.1 acla|EAW08682.1 afis|EAW15760.1 afla|EED51287.1 afum|EAL85588.1 agla|OJJ81922.1 akaw|GAA86297.1 aluc|OJZ86153.1 anid|CBF71002.1 anig|EHA24472.1 aory|EIT81510.1 asyd|OJJ52646.1 ater|EAU32386.1 atub|OJI79945.1 aver|OJJ04675.1 awen|OJJ38468.1 azon|OJJ43766.1

Group1893: abra|OJJ69167.1 acar|OOF91136.1 acla|EAW08683.1 afis|EAW15761.1 afla|EED51286.1 afum|EAL85587.1 agla|OJJ81923.1 akaw|GAA86296.1 aluc|OJZ86154.1 anid|CBF71000.1 anig|EHA24471.1 aory|EIT81497.1 asyd|OJJ52645.1 ater|EAU32385.1 atub|OJI79946.1 aver|OJJ04674.1 awen|OJJ38469.1 azon|OJJ43767.1

Group1894: abra|OJJ69168.1 acar|OOF91135.1 acla|EAW08684.1 afis|EAW15762.1 afla|EED51285.1 afum|EAL85586.1 agla|OJJ81924.1 akaw|GAA86295.1 aluc|OJZ86155.1 anid|CBF70998.1 anig|EHA24470.1 aory|EIT81522.1 asyd|OJJ52644.1 ater|EAU32384.1 atub|OJI79947.1 aver|OJJ04673.1 awen|OJJ38470.1 azon|OJJ43769.1

Group1895: abra|OJJ69177.1 acar|OOF91129.1 acla|EAW08686.1 afis|EAW15763.1 afla|EED51284.1 afum|EAL85585.1 agla|OJJ81925.1 akaw|GAA86287.1 aluc|OJZ86163.1 anid|CBF70996.1 anig|EHA24461.1 aory|EIT81489.1 asyd|OJJ52643.1 ater|EAU32378.1 atub|OJI79955.1 aver|OJJ04672.1 awen|OJJ38471.1 azon|OJJ43770.1

Group1896: abra|OJJ69178.1 acar|OOF90317.1 acla|EAW08687.1 afis|EAW15764.1 afla|EED51283.1 afum|EAL85584.1 agla|OJJ81927.1 akaw|GAA86286.1 aluc|OJZ86164.1 anid|CBF70994.1 anig|EHA24460.1

aory|EIT81513.1 asyd|OJJ52642.1 ater|EAU32377.1 atub|OJI79956.1 aver|OJJ04670.1 awen|OJJ38472.1 azon|OJJ43772.1

Group1897: abra|OJJ69179.1 acar|OOF90318.1 acla|EAW08688.1 afis|EAW15765.1 afla|EED51282.1 afum|EAL85583.1 agla|OJJ81928.1 akaw|GAA86285.1 aluc|OJZ86165.1 anid|CBF70993.1 anig|EHA24459.1 aory|EIT81558.1 asyd|OJJ52641.1 ater|EAU32376.1 atub|OJI79957.1 aver|OJJ04669.1 awen|OJJ38473.1 azon|OJJ43773.1

Group1898: abra|OJJ69180.1 acar|OOF90319.1 acla|EAW08689.1 afis|EAW15766.1 afla|EED51281.1 afum|EAL85582.1 agla|OJJ81930.1 akaw|GAA86284.1 aluc|OJZ86166.1 anig|EHA24458.1 aory|EIT81544.1 asyd|OJJ52640.1 ater|EAU32375.1 atub|OJI79958.1 awen|OJJ38474.1 anid|CBF70991.1 aver|OJJ04668.1 azon|OJJ43774.1

Group1899: abra|OJJ69181.1 acar|OOF90321.1 acla|EAW08690.1 afis|EAW15767.1 afla|EED51280.1 afum|EAL85581.1 agla|OJJ81931.1 akaw|GAA86283.1 aluc|OJZ86167.1 anid|CBF70989.1 anig|EHA24457.1 aory|EIT81536.1 asyd|OJJ52639.1 ater|EAU32374.1 atub|OJI79959.1 aver|OJJ04667.1 awen|OJJ38475.1 azon|OJJ43775.1

Group1900: abra|OJJ69183.1 acar|OOF90320.1 acla|EAW08691.1 afis|EAW15768.1 afla|EED51279.1 afum|EAL85580.1 agla|OJJ81932.1 akaw|GAA86282.1 aluc|OJZ86168.1 anid|CBF70987.1 anig|EHA24456.1 aory|EIT81531.1 asyd|OJJ52638.1 ater|EAU32373.1 atub|OJI79960.1 aver|OJJ04666.1 awen|OJJ38476.1 azon|OJJ43776.1

Group1901: abra|OJJ69184.1 acar|OOF91128.1 acla|EAW08662.1 afis|EAW15769.1 afla|EED51278.1 afum|EAL85579.1 agla|OJJ81933.1 akaw|GAA86281.1 aluc|OJZ86170.1 anid|CBF70983.1 anig|EHA24455.1 aory|EIT81490.1 asyd|OJJ52636.1 ater|EAU32372.1 atub|OJI79962.1 aver|OJJ04664.1 awen|OJJ38477.1 azon|OJJ43777.1

Group1902: abra|OJJ69186.1 acar|OOF91126.1 acla|EAW08660.1 afis|EAW15771.1 afla|EED51276.1 afum|EAL85577.1 agla|OJJ81935.1 akaw|GAA86279.1 aluc|OJZ86172.1 anid|CBF70979.1 anig|EHA24453.1 aory|EIT81507.1 asyd|OJJ52634.1 ater|EAU32370.1 atub|OJI79964.1 aver|OJJ04662.1 awen|OJJ38479.1 azon|OJJ43779.1

Group1903: abra|OJJ69187.1 acar|OOF91125.1 acla|EAW08659.1 afis|EAW15772.1 afla|EED51275.1 afum|EAL85576.1 agla|OJJ81936.1 akaw|GAA86278.1 aluc|OJZ86173.1 anid|CBF70977.1 anig|EHA24452.1 aory|EIT81560.1 asyd|OJJ52633.1 ater|EAU32369.1 atub|OJI79965.1 aver|OJJ04661.1 awen|OJJ38480.1 azon|OJJ43780.1

Group1904: abra|OJJ69189.1 acar|OOF91122.1 acla|EAW08656.1 afis|EAW15775.1 afla|EED51271.1 afum|EAL85574.1 agla|OJJ81938.1 akaw|GAA86275.1 aluc|OJZ86175.1 anid|CBF70976.1 anig|EHA24449.1 aory|EIT81533.1 asyd|OJJ52631.1 ater|EAU32366.1 atub|OJI79967.1 aver|OJJ04659.1 awen|OJJ38482.1 azon|OJJ43783.1

Group1905: abra|OJJ69191.1 acar|OOF91120.1 acla|EAW08654.1 afis|EAW15777.1 afla|EED51268.1 afum|EAL85572.1 agla|OJJ81940.1 akaw|GAA86273.1 aluc|OJZ86177.1 anid|CBF70933.1 anig|EHA24447.1 aory|EIT81563.1 asyd|OJJ52629.1 ater|EAU32364.1 atub|OJI79969.1 aver|OJJ04657.1 awen|OJJ38484.1 azon|OJJ43785.1

Group1906: abra|OJJ69192.1 acar|OOF91121.1 acla|EAW08653.1 afis|EAW15778.1 afla|EED51266.1 afum|EAL85571.1 agla|OJJ81941.1 akaw|GAA86272.1 aluc|OJZ86178.1 anid|CBF70935.1 anig|EHA24446.1 aory|EIT81565.1 asyd|OJJ52628.1 ater|EAU32363.1 atub|OJI79970.1 aver|OJJ04656.1 awen|OJJ38485.1 azon|OJJ43786.1

Group1907: abra|OJJ69193.1 acar|OOF91119.1 acla|EAW08652.1 afis|EAW15779.1 afla|EED51265.1 afum|EAL85570.1 agla|OJJ81942.1 akaw|GAA86271.1 aluc|OJZ86179.1 anid|CBF70937.1 anig|EHA24445.1 aory|EIT81542.1 asyd|OJJ52627.1 ater|EAU32362.1 atub|OJI79971.1 aver|OJJ04655.1 awen|OJJ38486.1 azon|OJJ43787.1

Group1908: abra|OJJ69194.1 acar|OOF91118.1 acla|EAW08651.1 afis|EAW15780.1 afla|EED51264.1 afum|EBA27235.1 agla|OJJ81943.1 akaw|GAA86270.1 aluc|OJZ86180.1 anid|CBF70939.1 anig|EHA24444.1 aory|EIT81528.1 asyd|OJJ52626.1 ater|EAU32361.1 atub|OJI79972.1 aver|OJJ04654.1 awen|OJJ38487.1 azon|OJJ43788.1

Group1909: abra|OJJ69196.1 acar|OOF90027.1 acla|EAW08650.1 afis|EAW15781.1 afla|EED51263.1 afum|EAL85569.1 agla|OJJ81944.1 akaw|GAA92209.1 aluc|OJZ86181.1 anid|CBF70941.1 anig|EHA24443.1 aory|EIT81567.1 asyd|OJJ52625.1 ater|EAU32360.1 atub|OJI79973.1 aver|OJJ04652.1 awen|OJJ38488.1 azon|OJJ43789.1

Group1910: abra|OJJ69197.1 acar|OOF90026.1 acla|EAW08649.1 afis|EAW15782.1 afla|EED51262.1 afum|EAL85568.1 agla|OJJ81945.1 akaw|GAA92210.1 aluc|OJZ86182.1 anid|CBF70945.1 anig|EHA24442.1 aory|EIT81562.1 asyd|OJJ52624.1 ater|EAU32359.1 atub|OJI79974.1 aver|OJJ04651.1 awen|OJJ38489.1 azon|OJJ43790.1

Group1911: abra|OJJ69198.1 acar|OOF91117.1 acla|EAW08648.1 afis|EAW15783.1 afla|EED51261.1 afum|EAL85567.1 agla|OJJ81946.1 akaw|GAA92211.1 aluc|OJZ86183.1 anid|CBF70946.1 anig|EHA24441.1 aory|EIT81552.1 asyd|OJJ52623.1 ater|EAU32358.1 atub|OJI79975.1 aver|OJJ04650.1 awen|OJJ38490.1 azon|OJJ43791.1

Group1912: abra|OJJ69200.1 acar|OOF91115.1 acla|EAW08646.1 afis|EAW15785.1 afla|EED51259.1 afum|EAL85565.1 agla|OJJ81948.1 akaw|GAA92213.1 aluc|OJZ86185.1 anid|CBF70954.1 anig|EHA24439.1 aory|EIT81568.1 asyd|OJJ52621.1 ater|EAU32356.1 atub|OJI79977.1 aver|OJJ04648.1 awen|OJJ38492.1 azon|OJJ43794.1

Group1913: abra|OJJ69204.1 acar|OOF91112.1 acla|EAW08643.1 afis|EAW15788.1 afla|EED51257.1 afum|EAL85562.1 agla|OJJ81951.1 akaw|GAA92216.1 aluc|OJZ86188.1 anid|CBF70960.1 anig|EHA24435.1 aory|EIT81501.1 asyd|OJJ52618.1 ater|EAU32353.1 atub|OJI79980.1 aver|OJJ04645.1 awen|OJJ38496.1 azon|OJJ43797.1

Group1914: abra|OJJ69205.1 acar|OOF91109.1 acla|EAW08642.1 afis|EAW15789.1 afla|EED51256.1 afum|EAL85561.1 agla|OJJ81952.1 akaw|GAA92217.1 aluc|OJZ86189.1 anid|CBF70962.1 anig|EHA24434.1 aory|EIT81537.1 asyd|OJJ52617.1 ater|EAU32352.1 atub|OJI79981.1 aver|OJJ04644.1 awen|OJJ38497.1 azon|OJJ43798.1

Group1915: abra|OJJ69206.1 acar|OOF91111.1 acla|EAW08641.1 afis|EAW15790.1 afla|EED51255.1 afum|EAL85560.1 agla|OJJ81953.1 akaw|GAA92218.1 aluc|OJZ86190.1 anid|CBF70964.1 anig|EHA24433.1 aory|EIT81514.1 asyd|OJJ52616.1 ater|EAU32351.1 atub|OJI79982.1 aver|OJJ04643.1 awen|OJJ38498.1 azon|OJJ43799.1

Group1916: abra|OJJ69208.1 acar|OOF91108.1 acla|EAW08638.1 afis|EAW15793.1 afla|EED51252.1 afum|EAL85557.1 agla|OJJ81956.1 akaw|GAA92219.1 aluc|OJZ86191.1 anid|CBF70970.1 anig|EHA24431.1 aory|EIT81504.1 asyd|OJJ52613.1 ater|EAU32349.1 atub|OJI79983.1 aver|OJJ04640.1 awen|OJJ38501.1 azon|OJJ43802.1

Group1917: abra|OJJ69209.1 acar|OOF91107.1 acla|EAW08637.1 afis|EAW15794.1 afla|EED51251.1 afum|EAL85556.1 agla|OJJ81957.1 akaw|GAA92220.1 aluc|OJZ86192.1 anid|CBF70931.1 anig|EHA24430.1 aory|EIT81500.1 asyd|OJJ52612.1 ater|EAU32348.1 atub|OJI79984.1 aver|OJJ04639.1 awen|OJJ38502.1 azon|OJJ43803.1

Group1918: abra|OJJ69210.1 acar|OOF91106.1 acla|EAW08635.1 afis|EAW15796.1 afla|EED51249.1 afum|EAL85554.1 agla|OJJ81960.1 akaw|GAA92221.1 aluc|OJZ86193.1 anid|CBF70927.1 anig|EHA24429.1 aory|EIT81508.1 asyd|OJJ52610.1 ater|EAU32346.1 atub|OJI79985.1 aver|OJJ04637.1 awen|OJJ38504.1 azon|OJJ43747.1

Group1919: abra|OJJ69211.1 acar|OOF91105.1 acla|EAW08634.1 afis|EAW15797.1 afla|EED51248.1 afum|EAL85553.2 agla|OJJ81961.1 akaw|GAA92222.1 aluc|OJZ86194.1 anid|CBF70925.1 anig|EHA24428.1 aory|EIT81561.1 asyd|OJJ52609.1 ater|EAU32345.1 atub|OJI79986.1 aver|OJJ04636.1 awen|OJJ38505.1 azon|OJJ43746.1

Group1920: abra|OJJ69212.1 acar|OOF91104.1 acla|EAW08633.1 afis|EAW15798.1 afla|EED51247.1 afum|EAL85552.1 agla|OJJ81962.1 akaw|GAA92223.1 aluc|OJZ86195.1 anid|CBF70923.1 anig|EHA24427.1 aory|EIT81502.1 asyd|OJJ52608.1 ater|EAU32344.1 atub|OJI79988.1 aver|OJJ04635.1 awen|OJJ38506.1 azon|OJJ43745.1

Group1921: abra|OJJ69215.1 acar|OOF91103.1 acla|EAW08630.1 afis|EAW15801.1 afla|EED51244.1 afum|EAL85549.2 agla|OJJ81968.1 akaw|GAA92224.1 aluc|OJZ86196.1 anid|CBF70918.1 anig|EHA24426.1 aory|EIT83186.1 asyd|OJJ52605.1 ater|EAU33447.1 atub|OJI79989.1 aver|OJJ04632.1 awen|OJJ38510.1 azon|OJJ43742.1

Group1922: abra|OJJ69217.1 acar|OOF91101.1 acla|EAW08628.1 afis|EAW15803.1 afla|EED51242.1 afum|EAL85547.1 agla|OJJ81970.1 akaw|GAA92226.1 aluc|OJZ86199.1 anid|CBF70914.1 anig|EHA24424.1 aory|EIT83176.1 asyd|OJJ52603.1 ater|EAU33445.1 atub|OJI79991.1 aver|OJJ04629.1 awen|OJJ38513.1 azon|OJJ43739.1

Group1923: abra|OJJ69222.1 acar|OOF91098.1 acla|EAW12739.1 afis|EAW17902.1 afla|EED51238.1 afum|EAL91710.1 agla|OJJ81973.1 akaw|GAA92230.1 aluc|OJZ86202.1 anid|CBF70908.1 anig|EHA24420.1 aory|EIT83180.1 asyd|OJJ52599.1 ater|EAU33438.1 atub|OJI79996.1 aver|OJJ04626.1 awen|OJJ38518.1 azon|OJJ43736.1

Group1924: abra|OJJ69230.1 acar|OOF91090.1 acla|EAW08619.1 afis|EAW15815.1 afla|EED51232.1 afum|EAL85536.1 agla|OJJ81979.1 akaw|GAA92238.1 aluc|OJZ86210.1 anid|CBF70899.1 anig|EHA24416.1 aory|EIT83190.1 asyd|OJJ52595.1 ater|EAU33433.1 atub|OJI80004.1 aver|OJJ04622.1 awen|OJJ38528.1 azon|OJJ43733.1

Group1925: abra|OJJ69232.1 acar|OOF91088.1 acla|EAW08617.1 afis|EAW15817.1 afla|EED51230.1 afum|EAL85534.1 agla|OJJ81981.1 akaw|GAA92240.1 aluc|OJZ86212.1 anid|CBF70895.1 anig|EHA24414.1

aory|EIT83179.1 asyd|OJJ52593.1 ater|EAU33431.1 atub|OJI80007.1 aver|OJJ04620.1 awen|OJJ38531.1 azon|OJJ43731.1

Group1926: abra|OJJ69234.1 acar|OOF91086.1 acla|EAW08615.1 afis|EAW15819.1 afla|EED51228.1 afum|EAL85532.1 agla|OJJ81983.1 akaw|GAA92242.1 aluc|OJZ86214.1 anid|CBF70891.1 anig|EHA24412.1 aory|EIT83174.1 asyd|OJJ52591.1 ater|EAU33429.1 atub|OJI80009.1 aver|OJJ04618.1 awen|OJJ38533.1 azon|OJJ43729.1

Group1927: abra|OJJ69235.1 acar|OOF91085.1 acla|EAW08614.1 afis|EAW15820.1 afla|EED51227.1 afum|EAL85531.1 agla|OJJ81984.1 akaw|GAA92243.1 aluc|OJZ86215.1 anid|CBF70889.1 anig|EHA24411.1 aory|EIT83196.1 asyd|OJJ52590.1 ater|EAU33428.1 atub|OJI80010.1 aver|OJJ04617.1 awen|OJJ38534.1 azon|OJJ43728.1

Group1928: abra|OJJ69237.1 acar|OOF91081.1 acla|EAW08610.1 afis|EAW15823.1 afla|EED51223.1 afum|EAL85528.1 agla|OJJ81986.1 akaw|GAA92247.1 aluc|OJZ86217.1 anid|CBF70884.1 anig|EHA24409.1 aory|EIT83191.1 asyd|OJJ52588.1 ater|EAU33425.1 atub|OJI80012.1 aver|OJJ04615.1 awen|OJJ38536.1 azon|OJJ43726.1

Group1929: abra|OJJ69238.1 acar|OOF91080.1 acla|EAW08609.1 afis|EAW15824.1 afla|EED51221.1 afum|EAL85527.1 agla|OJJ81987.1 akaw|GAA92248.1 aluc|OJZ86218.1 anid|CBF70878.1 anig|EHA24408.1 aory|EIT80400.1 asyd|OJJ52586.1 ater|EAU33424.1 atub|OJI80013.1 aver|OJJ04613.1 awen|OJJ38538.1 azon|OJJ43725.1

Group1930: abra|OJJ69239.1 acar|OOF91079.1 acla|EAW08608.1 afis|EAW15825.1 afla|EED51220.1 afum|EAL85526.1 agla|OJJ81988.1 akaw|GAA92249.1 aluc|OJZ86220.1 anid|CBF70876.1 anig|EHA24407.1 aory|EIT80394.1 asyd|OJJ52584.1 ater|EAU33423.1 atub|OJI80014.1 aver|OJJ04612.1 awen|OJJ38539.1 azon|OJJ43724.1

Group1931: abra|OJJ69240.1 acar|OOF91078.1 acla|EAW08607.1 afis|EAW15826.1 afla|EED51219.1 afum|EAL85525.1 agla|OJJ81989.1 akaw|GAA92250.1 aluc|OJZ86221.1 anid|CBF70874.1 anig|EHA24406.1 aory|EIT80397.1 asyd|OJJ52583.1 ater|EAU33422.1 atub|OJI80015.1 aver|OJJ04611.1 awen|OJJ38540.1 azon|OJJ43723.1

Group1932: abra|OJJ69244.1 acar|OOF91076.1 acla|EAW08605.1 afis|EAW15828.1 afla|EED51217.1 afum|EAL85523.1 agla|OJJ81992.1 akaw|GAA92252.1 aluc|OJZ86225.1 anid|CBF70869.1 anig|EHA24403.1 aory|EIT80398.1 asyd|OJJ52580.1 ater|EAU33419.1 atub|OJI80018.1 aver|OJJ04608.1 awen|OJJ38542.1 azon|OJJ43721.1

Group1933: abra|OJJ69247.1 acla|EAW08602.1 afis|EAW15831.1 afla|EED51215.1 afum|EAL85521.1 agla|OJJ81995.1 akaw|GAA92254.1 aluc|OJZ86228.1 anid|CBF70863.1 anig|EHA24401.1 aory|EIT80403.1 asyd|OJJ52577.1 ater|EAU33417.1 atub|OJI80021.1 aver|OJJ04606.1 awen|OJJ38544.1 azon|OJJ43718.1 aory|EIT80399.1

Group1934: abra|OJJ69248.1 acar|OOF91073.1 acla|EAW08601.1 afis|EAW15832.1 afla|EED51214.1 afum|EAL85520.1 agla|OJJ81996.1 akaw|GAA92255.1 aluc|OJZ86229.1 anid|CBF70861.1 anig|EHA24400.1 aory|EIT80409.1 asyd|OJJ52576.1 ater|EAU33416.1 atub|OJI80022.1 aver|OJJ04605.1 awen|OJJ38545.1 azon|OJJ43717.1

Group1935: abra|OJJ69249.1 acar|OOF91072.1 acla|EAW08600.1 afis|EAW15833.1 afla|EED51213.1 afum|EAL85519.1 agla|OJJ81997.1 akaw|GAA92256.1 aluc|OJZ86230.1 anid|CBF70859.1 anig|EHA24399.1 aory|EIT80405.1 asyd|OJJ52575.1 ater|EAU33415.1 atub|OJI80023.1 aver|OJJ04604.1 awen|OJJ38546.1 azon|OJJ43716.1

Group1936: abra|OJJ69250.1 acar|OOF91071.1 acla|EAW08599.1 afis|EAW15834.1 afla|EED51212.1 afum|EAL85518.1 agla|OJJ81998.1 akaw|GAA92257.1 aluc|OJZ86231.1 anid|CBF70858.1 anig|EHA24398.1 aory|EIT80395.1 asyd|OJJ52574.1 ater|EAU33414.1 atub|OJI80024.1 aver|OJJ04603.1 awen|OJJ38547.1 azon|OJJ43715.1

Group1937: abra|OJJ69252.1 acar|OOF91069.1 acla|EAW08597.1 afis|EAW15836.1 afla|EED51210.1 afum|EAL85515.1 agla|OJJ82000.1 akaw|GAA92259.1 aluc|OJZ86232.1 anid|CBF70854.1 anig|EHA24397.1 aory|EIT80401.1 asyd|OJJ52573.1 ater|EAU33412.1 atub|OJI80026.1 aver|OJJ04602.1 awen|OJJ38550.1 azon|OJJ43713.1

Group1938: atub|OJI80028.1 abra|OJJ69256.1 acar|OOF91065.1 acla|EAW08593.1 afis|EAW15841.1 afla|EED51208.1 afum|EAL85510.1 agla|OJJ84917.1 akaw|GAA92262.1 aluc|OJZ86236.1 anid|CBF70845.1 anig|EHA24392.1 aory|EIT80413.1 asyd|OJJ52568.1 ater|EAU33394.1 aver|OJJ04597.1 awen|OJJ38555.1 azon|OJJ43711.1

Group1939: abra|OJJ69257.1 acar|OOF91064.1 acla|EAW08592.1 afis|EAW15842.1 afla|EED51207.1 afum|EAL85509.1 agla|OJJ84916.1 akaw|GAA92263.1 aluc|OJZ86237.1 anid|CBF70843.1 anig|EHA24391.1 aory|EIT80396.1 asyd|OJJ52567.1 ater|EAU33393.1 atub|OJI80029.1 aver|OJJ04596.1 awen|OJJ38556.1 azon|OJJ43710.1

Group1940: abra|OJJ69259.1 acar|OOF91063.1 acla|EAW08590.1 afis|EAW15844.1 afla|EED51205.1 afum|EAL85507.1 agla|OJJ84915.1 akaw|GAA92265.1 aluc|OJZ86239.1 anid|CBF70839.1 anig|EHA24390.1 asyd|OJJ52565.1 ater|EAU33391.1 atub|OJI80031.1 aver|OJJ04594.1 awen|OJJ38559.1 azon|OJJ43708.1 awen|OJJ38558.1

Group1941: abra|OJJ69262.1 acar|OOF91060.1 acla|EAW08589.1 afis|EAW15846.1 afla|EED51204.1 afum|EAL85505.1 agla|OJJ84913.1 akaw|GAA92267.1 aluc|OJZ86241.1 anid|CBF70835.1 anig|EHA24388.1 aory|EIT80414.1 asyd|OJJ52564.1 ater|EAU33390.1 atub|OJI80033.1 aver|OJJ04592.1 awen|OJJ38561.1 azon|OJJ48761.1

Group1942: abra|OJJ69263.1 acar|OOF91059.1 acla|EAW08588.1 afis|EAW15847.1 afla|EED51203.1 afum|EAL85504.1 agla|OJJ84911.1 akaw|GAA92268.1 aluc|OJZ86242.1 anid|CBF70833.1 anig|EHA24387.1 aory|EIT80410.1 asyd|OJJ52563.1 ater|EAU33389.1 atub|OJI80034.1 aver|OJJ04591.1 awen|OJJ38562.1 azon|OJJ43615.1

Group1943: abra|OJJ69268.1 acar|OOF89861.1 acla|EAW14653.1 afis|EAW24188.1 afla|EED48081.1 afum|EAL85861.1 agla|OJJ79691.1 akaw|GAA88143.1 aluc|OJZ82097.1 anid|CBF74306.1 anig|EHA28632.1 aory|EIT82476.1 asyd|OJJ54008.1 ater|EAU31140.1 atub|OJI88340.1 aver|OJJ08103.1 awen|OJJ33111.1 azon|OJJ41999.1

Group1944: abra|OJJ69275.1 acar|OOF94447.1 acla|EAW14663.1 afis|EAW24194.1 afla|EED48077.1 afum|EAL85866.1 agla|OJJ79684.1 akaw|GAA88148.1 aluc|OJZ82102.1 anid|CBF74313.1 anig|EHA28627.1 aory|EIT82481.1 asyd|OJJ54002.1 ater|EAU31145.1 atub|OJI88333.1 aver|OJJ08099.1 awen|OJJ33105.1 azon|OJJ44844.1

Group1945: abra|OJJ69276.1 acar|OOF94446.1 acla|EAW14664.1 afis|EAW24195.1 afla|EED48076.1 afum|EAL85867.1 agla|OJJ79683.1 akaw|GAA88149.1 aluc|OJZ82103.1 anid|CBF74315.1 anig|EHA28626.1 aory|EIT82479.1 asyd|OJJ54001.1 ater|EAU31146.1 atub|OJI88332.1 aver|OJJ08098.1 awen|OJJ33104.1 azon|OJJ44845.1

Group1946: abra|OJJ69277.1 acar|OOF94445.1 acla|EAW14665.1 afis|EAW24196.1 afla|EED48075.1 afum|EAL85868.1 agla|OJJ79682.1 akaw|GAA88150.1 aluc|OJZ82104.1 anid|CBF74317.1 anig|EHA28625.1 aory|EIT82482.1 asyd|OJJ54000.1 ater|EAU31147.1 atub|OJI88331.1 aver|OJJ08097.1 awen|OJJ33103.1 azon|OJJ44846.1

Group1947: abra|OJJ69278.1 acar|OOF94444.1 acla|EAW14666.1 afis|EAW24197.1 afla|EED48074.1 afum|EAL85869.1 agla|OJJ79681.1 akaw|GAA88151.1 aluc|OJZ82105.1 anid|CBF74319.1 anig|EHA28624.1 aory|EIT82477.1 asyd|OJJ53999.1 ater|EAU31148.1 atub|OJI88330.1 aver|OJJ08096.1 awen|OJJ33102.1 azon|OJJ44847.1

Group1948: abra|OJJ69279.1 acar|OOF94443.1 acla|EAW14668.1 afis|EAW24199.1 afla|EED48073.1 afum|EAL85871.1 agla|OJJ79678.1 akaw|GAA88152.1 aluc|OJZ82106.1 anid|CBF74323.1 anig|EHA28623.1 aory|EIT83321.1 asyd|OJJ53996.1 ater|EAU31150.1 atub|OJI88329.1 aver|OJJ08094.1 awen|OJJ33099.1 azon|OJJ44848.1

Group1949: abra|OJJ69281.1 acar|OOF94441.1 acla|EAW14670.1 afis|EAW24201.1 afla|EED48071.1 afum|EBA27248.1 agla|OJJ79676.1 akaw|GAA88153.1 aluc|OJZ82109.1 anid|CBF74326.1 anig|EHA28621.1 aory|EIT83320.1 asyd|OJJ53994.1 ater|EAU31152.1 atub|OJI88327.1 aver|OJJ08092.1 awen|OJJ33097.1 azon|OJJ44850.1

Group1950: abra|OJJ69283.1 acar|OOF94439.1 acla|EAW14672.1 afis|EAW24203.1 afla|EED48069.1 afum|EAL85873.2 agla|OJJ79674.1 akaw|GAA88155.1 aluc|OJZ82112.1 anid|CBF74330.1 anig|EHA28619.1 aory|EIT83340.1 asyd|OJJ53992.1 ater|EAU31154.1 atub|OJI88325.1 aver|OJJ08090.1 awen|OJJ33095.1 azon|OJJ44852.1

Group1951: abra|OJJ69284.1 acar|OOF94438.1 acla|EAW14673.1 afis|EAW24204.1 afla|EED48067.1 afum|EAL85874.2 agla|OJJ79673.1 akaw|GAA88156.1 aluc|OJZ82113.1 anid|CBF74332.1 anig|EHA28618.1 aory|EIT83335.1 asyd|OJJ53991.1 ater|EAU31155.1 atub|OJI88324.1 aver|OJJ08089.1 awen|OJJ33094.1 azon|OJJ44854.1

Group1952: abra|OJJ69285.1 acar|OOF94436.1 acla|EAW14674.1 afis|EAW24205.1 afla|EED48066.1 afum|EAL85875.1 agla|OJJ79672.1 akaw|GAA88157.1 aluc|OJZ82114.1 anid|CBF74272.1 anig|EHA28617.1 aory|EIT83356.1 asyd|OJJ53990.1 ater|EAU31156.1 atub|OJI88323.1 aver|OJJ08088.1 awen|OJJ33093.1 azon|OJJ44855.1

Group1953: abra|OJJ69286.1 acar|OOF94437.1 acla|EAW14675.1 afis|EAW24206.1 afla|EED48065.1 afum|EAL85876.1 agla|OJJ79671.1 akaw|GAA88158.1 aluc|OJZ82115.1 anid|CBF74271.1 anig|EHA28616.1 aory|EIT83359.1 asyd|OJJ53989.1 ater|EAU31157.1 atub|OJI88322.1 aver|OJJ08087.1 awen|OJJ33092.1 azon|OJJ44856.1

Group1954: abra|OJJ69288.1 acar|OOF94434.1 acla|EAW14677.1 afis|EAW24208.1 afla|EED48063.1 afum|EAL85878.1 agla|OJJ79669.1 akaw|GAA88160.1 aluc|OJZ82117.1 anid|CBF74269.1 anig|EHA28614.1

aory|EIT83351.1 asyd|OJJ53987.1 ater|EAU31158.1 atub|OJI88320.1 aver|OJJ08085.1 awen|OJJ33089.1 azon|OJJ44858.1

Group1955: abra|OJJ69292.1 acar|OOF94431.1 acla|EAW14680.1 afis|EAW24211.1 afla|EED48060.1 afum|EAL85880.1 agla|OJJ79654.1 akaw|GAA88163.1 aluc|OJZ82121.1 anid|CBF74263.1 anig|EHA28611.1 aory|EIT83350.1 asyd|OJJ53983.1 ater|EAU31161.1 atub|OJI88317.1 aver|OJJ08082.1 awen|OJJ33085.1 azon|OJJ44861.1

Group1956: abra|OJJ69293.1 acar|OOF94430.1 acla|EAW14681.1 afis|EAW24212.1 afla|EED48059.1 afum|EAL85881.1 agla|OJJ79663.1 akaw|GAA88164.1 aluc|OJZ82122.1 anid|CBF74261.1 anig|EHA28610.1 aory|EIT83323.1 asyd|OJJ53982.1 ater|EAU31162.1 atub|OJI88316.1 aver|OJJ08081.1 awen|OJJ33084.1 azon|OJJ44862.1

Group1957: abra|OJJ69294.1 acar|OOF94429.1 acla|EAW14682.1 afis|EAW24213.1 afla|EED48058.1 afum|EAL85882.1 agla|OJJ79665.1 akaw|GAA88165.1 aluc|OJZ82123.1 anid|CBF74260.1 anig|EHA28609.1 aory|EIT83345.1 asyd|OJJ53981.1 ater|EAU31163.1 atub|OJI88315.1 aver|OJJ08080.1 awen|OJJ33083.1 azon|OJJ44863.1

Group1958: abra|OJJ69296.1 acar|OOF94428.1 acla|EAW14683.1 afis|EAW24214.1 afla|EED48057.1 afum|EAL85883.1 agla|OJJ79666.1 akaw|GAA88166.1 aluc|OJZ82125.1 anid|CBF74258.1 anig|EHA28607.1 aory|EIT83357.1 asyd|OJJ53980.1 ater|EAU31164.1 atub|OJI88313.1 aver|OJJ08079.1 awen|OJJ33082.1 azon|OJJ44864.1

Group1959: abra|OJJ69297.1 acar|OOF94427.1 acla|EAW14684.1 afis|EAW24215.1 afla|EED48056.1 afum|EAL85884.1 agla|OJJ79667.1 akaw|GAA88167.1 aluc|OJZ82126.1 anid|CBF74256.1 anig|EHA28606.1 aory|EIT83352.1 asyd|OJJ53979.1 ater|EAU31165.1 atub|OJI88312.1 aver|OJJ08078.1 awen|OJJ33081.1 azon|OJJ44865.1

Group1960: abra|OJJ69298.1 acar|OOF94425.1 acla|EAW14685.1 afis|EAW24216.1 afla|EED48055.1 afum|EAL85885.1 agla|OJJ79668.1 akaw|GAA88168.1 aluc|OJZ82127.1 anid|CBF74254.1 anig|EHA28605.1 aory|EIT83339.1 asyd|OJJ53978.1 ater|EAU31166.1 atub|OJI88311.1 aver|OJJ08077.1 awen|OJJ33080.1 azon|OJJ44866.1

Group1961: abra|OJJ69299.1 acar|OOF94426.1 acla|EAW14686.1 afis|EAW24217.1 afla|EED48054.1 afum|EAL85886.1 agla|OJJ79653.1 akaw|GAA88169.1 aluc|OJZ82128.1 anid|CBF74252.1 anig|EHA28603.1 aory|EIT83360.1 asyd|OJJ53977.1 ater|EAU31167.1 atub|OJI88310.1 aver|OJJ08076.1 awen|OJJ33079.1 azon|OJJ44867.1

Group1962: abra|OJJ69302.1 acar|OOF94423.1 acla|EAW14688.1 afis|EAW24219.1 afla|EED48052.1 afum|EAL85888.1 agla|OJJ79651.1 akaw|GAA88171.1 aluc|OJZ82130.1 anid|CBF74248.1 anig|EHA28601.1 aory|EIT83354.1 asyd|OJJ53975.1 ater|EAU31169.1 atub|OJI88306.1 aver|OJJ08074.1 awen|OJJ33077.1 azon|OJJ44869.1

Group1963: abra|OJJ69305.1 acar|OOF94420.1 acla|EAW14691.1 afis|EAW24222.1 afla|EED48050.1 afum|EAL85891.2 agla|OJJ79648.1 akaw|GAA88174.1 aluc|OJZ82134.1 anid|CBF74239.1 anig|EHA28597.1 aory|EIT83349.1 asyd|OJJ53971.1 ater|EAU31172.1 atub|OJI88303.1 aver|OJJ08070.1 awen|OJJ33074.1 azon|OJJ44873.1

Group1964: abra|OJJ69307.1 acar|OOF94419.1 acla|EAW14694.1 afis|EAW24225.1 afla|EED48046.1 afum|EAL85893.1 agla|OJJ79646.1 akaw|GAA88176.1 aluc|OJZ82136.1 anid|CBF74234.1 anig|EHA28595.1 aory|EIT83343.1 asyd|OJJ53969.1 ater|EAU31174.1 atub|OJI88301.1 aver|OJJ08067.1 awen|OJJ33071.1 azon|OJJ44875.1

Group1965: abra|OJJ69308.1 acar|OOF94418.1 acla|EAW14695.1 afis|EAW24226.1 afla|EED48045.1 afum|EAL85894.1 agla|OJJ79645.1 akaw|GAA88177.1 aluc|OJZ82137.1 anid|CBF74232.1 anig|EHA28594.1 aory|EIT83341.1 asyd|OJJ53968.1 ater|EAU31175.1 atub|OJI88300.1 aver|OJJ08066.1 awen|OJJ33070.1 azon|OJJ44877.1

Group1966: abra|OJJ69312.1 acar|OOF94415.1 acla|EAW14738.1 afis|EAW24269.1 afla|EED48008.1 afum|EAL85937.1 agla|OJJ79596.1 akaw|GAA88181.1 aluc|OJZ82141.1 anid|CBF74167.1 anig|EHA28591.1 aory|EIT82915.1 asyd|OJJ53936.1 ater|EAU31214.1 atub|OJI88296.1 aver|OJJ08034.1 awen|OJJ33031.1 azon|OJJ48488.1

Group1967: abra|OJJ69313.1 acar|OOF94413.1 acla|EAW14737.1 afis|EAW24268.1 afla|EED48009.1 afum|EAL85936.1 agla|OJJ79597.1 akaw|GAA88182.1 aluc|OJZ82142.1 anid|CBF74169.1 anig|EHA28589.1 aory|EIT82903.1 asyd|OJJ53937.1 ater|EAU31213.1 atub|OJI88295.1 aver|OJJ08035.1 awen|OJJ33032.1 azon|OJJ48487.1

Group1968: abra|OJJ69314.1 acar|OOF94412.1 acla|EAW14736.1 afis|EAW24267.1 afla|EED48010.1 afum|EAL85935.1 agla|OJJ79598.1 akaw|GAA88183.1 aluc|OJZ82143.1 anid|CBF74171.1 anig|EHA28588.1 aory|EIT82909.1 asyd|OJJ53938.1 ater|EAU31212.1 atub|OJI88294.1 aver|OJJ08036.1 awen|OJJ33033.1 azon|OJJ48486.1

Group1969: abra|OJJ69315.1 acar|OOF90302.1 acla|EAW14734.1 afis|EAW24266.1 afla|EED48011.1 afum|EAL85934.1 agla|OJJ79606.1 akaw|GAA88184.1 aluc|OJZ82144.1 anid|CBF74173.1 anig|EHA28586.1 aory|EIT82916.1 asyd|OJJ53940.1 ater|EAU31210.1 atub|OJI88292.1 aver|OJJ08037.1 awen|OJJ33034.1 azon|OJJ48485.1

Group1970: abra|OJJ69316.1 acar|OOF90303.1 acla|EAW14732.1 afis|EAW24265.1 afla|EED48012.1 afum|EAL85933.1 agla|OJJ79607.1 akaw|GAA88185.1 aluc|OJZ82145.1 anid|CBF74175.1 anig|EHA28585.1 aory|EIT82905.1 asyd|OJJ53941.1 ater|EAU31209.1 atub|OJI88291.1 aver|OJJ08038.1 awen|OJJ33035.1 azon|OJJ48484.1

Group1971: abra|OJJ69319.1 acar|OOF94411.1 acla|EAW14726.1 afis|EAW24262.1 akaw|GAA88188.1 aluc|OJZ82148.1 ater|EAU31207.1 atub|OJI88287.1 aver|OJJ08043.1 awen|OJJ33039.1 azon|OJJ48479.1 afum|EAL85930.1 agla|OJJ79610.1 anid|CBF74186.1 anig|EHA28582.1 aory|EIT82904.1 asyd|OJJ53946.1 afla|EED48015.1

Group1972: abra|OJJ69329.1 acar|OOF94398.1 acla|EAW14711.1 afis|EAW24246.1 afla|EED48028.1 afum|EAL85914.1 agla|OJJ79627.1 akaw|GAA88200.1 aluc|OJZ82161.1 anid|CBF74210.1 anig|EHA28571.1 aory|EIT83355.1 asyd|OJJ53956.1 ater|EAU31187.1 atub|OJI88276.1 aver|OJJ08052.1 awen|OJJ33058.1 azon|OJJ48466.1

Group1973: abra|OJJ69332.1 acar|OOF94395.1 acla|EAW14708.1 afis|EAW24239.1 afla|EED48031.1 afum|EAL85907.1 agla|OJJ79630.1 akaw|GAA88203.1 aluc|OJZ82164.1 anid|CBF74213.1 anig|EHA28568.1 aory|EIT83322.1 asyd|OJJ53957.1 ater|EAU31186.1 atub|OJI88273.1 aver|OJJ08053.1 awen|OJJ33054.1 azon|OJJ48465.1

Group1974: anig|EHA28560.1 abra|OJJ69342.1 acar|OOF94385.1 acla|EAW14704.1 afis|EAW24235.1 afla|EED48036.1 afum|EAL85904.1 agla|OJJ79636.1 akaw|GAA88211.1 aluc|OJZ82174.1 anid|CBF74219.1 aory|EIT83346.1 asyd|OJJ53961.1 ater|EAU31183.1 atub|OJI88263.1 aver|OJJ08057.1 awen|OJJ33063.1 azon|OJJ48461.1

Group1975: abra|OJJ69343.1 acar|OOF94386.1 acla|EAW14703.1 afis|EAW24234.1 afla|EED48037.1 afum|EAL85903.1 agla|OJJ79637.1 akaw|GAA88212.1 aluc|OJZ82175.1 anid|CBF74221.1 anig|EHA28559.1 aory|EIT83326.1 asyd|OJJ53962.1 ater|EAU31182.1 atub|OJI88262.1 aver|OJJ08058.1 awen|OJJ33064.1 azon|OJJ48460.1

Group1976: abra|OJJ69345.1 acar|OOF94379.1 acla|EAW14701.1 afis|EAW24232.1 afla|EED48039.1 afum|EAL85901.2 agla|OJJ79639.1 akaw|GAA88214.1 aluc|OJZ82177.1 anid|CBF74223.1 anig|EHA28557.1 aory|EIT83353.1 asyd|OJJ53964.1 ater|EAU31180.1 atub|OJI88260.1 aver|OJJ08060.1 awen|OJJ33065.1 azon|OJJ48458.1

Group1977: abra|OJJ69346.1 acar|OOF94378.1 acla|EAW14700.1 afis|EAW24231.1 afla|EED48040.1 afum|EAL85900.1 agla|OJJ79640.1 akaw|GAA88215.1 aluc|OJZ82178.1 anid|CBF74224.1 anig|EHA28556.1 aory|EIT83342.1 asyd|OJJ53965.1 ater|EAU31179.1 atub|OJI88259.1 aver|OJJ08061.1 awen|OJJ33066.1 azon|OJJ48457.1

Group1978: abra|OJJ69349.1 acar|OOF94376.1 acla|EAW14739.1 afis|EAW24270.1 afla|EED48007.1 afum|EAL85938.1 agla|OJJ79595.1 akaw|GAA88219.1 aluc|OJZ82181.1 anid|CBF74165.1 anig|EHA28553.1 aory|EIT82896.1 asyd|OJJ53935.1 ater|EAU31215.1 atub|OJI88254.1 aver|OJJ08033.1 awen|OJJ33030.1 azon|OJJ48489.1

Group1979: abra|OJJ69351.1 acar|OOF94375.1 acla|EAW14740.1 afis|EAW24271.1 afla|EED48006.1 afum|EAL85939.1 agla|OJJ79594.1 akaw|GAA88220.1 aluc|OJZ82183.1 anid|CBF74163.1 anig|EHA28552.1 aory|EIT82918.1 asyd|OJJ53933.1 ater|EAU31216.1 atub|OJI88253.1 aver|OJJ08031.1 awen|OJJ33028.1 azon|OJJ48490.1

Group1980: abra|OJJ69353.1 acar|OOF94372.1 acla|EAW14743.1 afis|EAW24274.1 afla|EED48002.1 afum|EAL85942.1 agla|OJJ79587.1 akaw|GAA88222.1 aluc|OJZ82186.1 anid|CBF74158.1 anig|EHA28550.1 aory|EIT82912.1 asyd|OJJ53927.1 ater|EAU31221.1 atub|OJI88251.1 aver|OJJ08026.1 awen|OJJ33023.1 azon|OJJ48494.1

Group1981: abra|OJJ69355.1 acar|OOF94370.1 acla|EAW14745.1 afis|EAW24276.1 afla|EED48000.1 afum|EAL85945.1 agla|OJJ79585.1 akaw|GAA88224.1 aluc|OJZ82187.1 anid|CBF74156.1 anig|EHA28548.1 aory|EIT82907.1 asyd|OJJ53925.1 ater|EAU31219.1 atub|OJI88249.1 aver|OJJ08024.1 awen|OJJ33021.1 azon|OJJ48496.1

Group1982: abra|OJJ69357.1 acar|OOF94367.1 acla|EAW14748.1 afis|EAW24279.1 afla|EED47997.1 afum|EAL85948.1 agla|OJJ80633.1 akaw|GAA88227.1 aluc|OJZ82189.1 anid|CBF74151.1 anig|EHA28546.1 aory|EIT82902.1 asyd|OJJ53923.1 ater|EAU31228.1 atub|OJI88246.1 aver|OJJ08021.1 awen|OJJ33018.1 azon|OJJ48499.1

Group1983: abra|OJJ69366.1 acar|OOF94359.1 acla|EAW14756.1 afis|EAW24287.1 afla|EED47989.1 afum|EAL85956.1 agla|OJJ83152.1 akaw|GAA88235.1 aluc|OJZ82200.1 anid|CBF74129.1 anig|EHA28539.1

aory|EIT79810.1 asyd|OJJ53912.1 ater|EAU31238.1 atub|OJI88236.1 aver|OJJ08010.1 awen|OJJ33009.1 azon|OJJ48506.1

Group1984: abra|OJJ69367.1 acar|OOF94358.1 acla|EAW14757.1 afis|EAW24288.1 afla|EED47988.1 afum|EAL85957.1 agla|OJJ83153.1 akaw|GAA88236.1 aluc|OJZ82201.1 anid|CBF74127.1 anig|EHA28538.1 aory|EIT79844.1 asyd|OJJ53911.1 ater|EAU31239.1 atub|OJI88235.1 aver|OJJ08009.1 awen|OJJ33008.1 azon|OJJ48507.1

Group1985: abra|OJJ69369.1 acar|OOF89837.1 acla|EAW14759.1 afis|EAW24290.1 afla|EED47986.1 afum|EAL85958.1 agla|OJJ83155.1 akaw|GAA88238.1 aluc|OJZ82203.1 anid|CBF74125.1 anig|EHA28536.1 aory|EIT79849.1 asyd|OJJ53909.1 ater|EAU31240.1 atub|OJI88233.1 aver|OJJ08007.1 awen|OJJ33006.1 azon|OJJ48509.1

Group1986: abra|OJJ69370.1 acar|OOF94357.1 acla|EAW14760.1 afis|EAW24291.1 afla|EED47985.1 afum|EAL85959.1 agla|OJJ83156.1 akaw|GAA88239.1 aluc|OJZ82204.1 anid|CBF74123.1 anig|EHA28535.1 aory|EIT79854.1 asyd|OJJ53908.1 ater|EAU31241.1 atub|OJI88232.1 aver|OJJ08006.1 awen|OJJ33005.1 azon|OJJ48510.1

Group1987: abra|OJJ69371.1 acar|OOF94355.1 acla|EAW14761.1 afis|EAW24292.1 afla|EED47984.1 afum|EAL85960.1 agla|OJJ83157.1 akaw|GAA88240.1 aluc|OJZ82205.1 anid|CBF74121.1 anig|EHA28534.1 aory|EIT79808.1 asyd|OJJ53907.1 ater|EAU31242.1 atub|OJI88231.1 aver|OJJ08005.1 awen|OJJ33004.1 azon|OJJ48511.1

Group1988: abra|OJJ69373.1 acar|OOF94354.1 acla|EAW14763.1 afis|EAW24294.1 afla|EED47982.1 afum|EAL85962.1 agla|OJJ83159.1 akaw|GAA88242.1 aluc|OJZ82208.1 anid|CBF74118.1 anig|EHA28532.1 aory|EIT79845.1 asyd|OJJ53905.1 ater|EAU31244.1 atub|OJI88228.1 aver|OJJ08003.1 awen|OJJ33001.1 azon|OJJ48513.1

Group1989: abra|OJJ69375.1 acar|OOF94351.1 acla|EAW14765.1 afis|EAW24296.1 afla|EED47980.1 afum|EAL85964.1 agla|OJJ83161.1 akaw|GAA88244.1 aluc|OJZ82210.1 anid|CBF75402.1 anig|EHA28530.1 aory|EIT79827.1 asyd|OJJ59822.1 ater|EAU31246.1 atub|OJI88226.1 aver|OJJ02465.1 awen|OJJ32998.1 azon|OJJ48514.1

Group1990: abra|OJJ69376.1 acar|OOF94349.1 acla|EAW14766.1 afis|EAW24297.1 afla|EED47979.1 afum|EAL85965.1 agla|OJJ83162.1 akaw|GAA88245.1 aluc|OJZ82211.1 anid|CBF74116.1 anig|EHA28529.1 aory|EIT79833.1 asyd|OJJ53904.1 ater|EAU31248.1 atub|OJI88225.1 aver|OJJ08001.1 awen|OJJ32997.1 azon|OJJ48515.1

Group1991: abra|OJJ69381.1 acar|OOF94344.1 acla|EAW14768.1 afis|EAW24299.1 afla|EED47973.1 afum|EAL85967.1 agla|OJJ83167.1 akaw|GAA88249.1 aluc|OJZ82216.1 anid|CBF74108.1 anig|EHA28524.1 aory|EIT79831.1 asyd|OJJ53901.1 ater|EAU31250.1 atub|OJI88219.1 aver|OJJ07998.1 awen|OJJ32993.1 azon|OJJ48523.1

Group1992: abra|OJJ69383.1 acar|OOF94343.1 acla|EAW14770.1 afis|EAW24301.1 afla|EED47971.1 afum|EAL85969.1 agla|OJJ83169.1 akaw|GAA88251.1 aluc|OJZ82217.1 anid|CBF74104.1 anig|EHA28523.1 aory|EIT79840.1 asyd|OJJ53899.1 ater|EAU31252.1 atub|OJI88218.1 aver|OJJ07996.1 awen|OJJ32991.1 azon|OJJ48525.1

Group1993: abra|OJJ69384.1 acar|OOF94342.1 acla|EAW14771.1 afis|EAW24302.1 afla|EED47970.1 afum|EAL85970.1 agla|OJJ83170.1 akaw|GAA88252.1 aluc|OJZ82218.1 anid|CBF74102.1 anig|EHA28522.1 aory|EIT79832.1 asyd|OJJ53898.1 ater|EAU31253.1 atub|OJI88217.1 aver|OJJ07995.1 awen|OJJ32990.1 azon|OJJ45840.1

Group1994: abra|OJJ69385.1 acar|OOF94341.1 acla|EAW14772.1 afis|EAW24303.1 afla|EED47969.1 afum|EAL85971.1 agla|OJJ83171.1 akaw|GAA88253.1 aluc|OJZ82219.1 anid|CBF74100.1 anig|EHA28521.1 aory|EIT79817.1 asyd|OJJ53897.1 ater|EAU31256.1 atub|OJI88216.1 aver|OJJ07994.1 awen|OJJ32989.1 azon|OJJ48526.1

Group1995: abra|OJJ69386.1 acar|OOF94340.1 acla|EAW14773.1 afis|EAW24304.1 afla|EED47968.1 afum|EAL85972.1 agla|OJJ83172.1 akaw|GAA88254.1 aluc|OJZ82220.1 anid|CBF79548.1 anig|EHA28520.1 aory|EIT79824.1 asyd|OJJ62162.1 ater|EAU31257.1 atub|OJI88215.1 aver|OJI96272.1 awen|OJJ32988.1 azon|OJJ48527.1

Group1996: abra|OJJ69387.1 acar|OOF94339.1 acla|EAW14774.1 afis|EAW24305.1 afla|EED47965.1 afum|EAL85973.1 agla|OJJ83173.1 akaw|GAA88255.1 aluc|OJZ82221.1 anid|CBF74098.1 anig|EHA28519.1 aory|EIT79856.1 asyd|OJJ53896.1 ater|EAU31258.1 atub|OJI88214.1 aver|OJJ07993.1 awen|OJJ32987.1 azon|OJJ48932.1

Group1997: abra|OJJ69398.1 acar|OOF94332.1 acla|EAW14780.1 afis|EAW24311.1 afla|EED47956.1 afum|EAL85979.1 agla|OJJ83179.1 akaw|GAA88264.1 aluc|OJZ82234.1 anid|CBF74081.1 anig|EHA28507.1 aory|EIT79851.1 asyd|OJJ53882.1 ater|EAU31264.1 atub|OJI88204.1 aver|OJJ07980.1 awen|OJJ32981.1 azon|OJJ48936.1

Group1998: abra|OJJ69400.1 acar|OOF94330.1 acla|EAW14782.1 afis|EAW24313.1 afla|EED47954.1 afum|EAL85981.1 agla|OJJ83181.1 akaw|GAA88266.1 aluc|OJZ82236.1 anid|CBF74079.1 anig|EHA28505.1 aory|EIT79862.1 asyd|OJJ53880.1 ater|EAU31266.1 atub|OJI88202.1 aver|OJJ07978.1 awen|OJJ32979.1 azon|OJJ48938.1

Group1999: abra|OJJ69402.1 acar|OOF94329.1 acla|EAW14784.1 afis|EAW24315.1 afla|EED47952.1 afum|EAL85983.1 agla|OJJ83183.1 akaw|GAA88267.1 aluc|OJZ82238.1 anid|CBF74078.1 anig|EHA28503.1 aory|EIT79841.1 asyd|OJJ53878.1 ater|EAU31267.1 atub|OJI88200.1 aver|OJJ07976.1 awen|OJJ32977.1 azon|OJJ48940.1

Group2000: abra|OJJ69404.1 acar|OOF94327.1 acla|EAW14786.1 afis|EAW24317.1 afla|EED47950.1 afum|EAL85985.1 agla|OJJ83185.1 akaw|GAA88269.1 aluc|OJZ82240.1 anid|CBF74074.1 anig|EHA28501.1 aory|EIT79859.1 asyd|OJJ53876.1 ater|EAU31270.1 atub|OJI88198.1 aver|OJJ07974.1 awen|OJJ32975.1 azon|OJJ48942.1

Group2001: abra|OJJ69405.1 acar|OOF94326.1 acla|EAW14787.1 afis|EAW24318.1 afla|EED47949.1 afum|EAL85987.1 agla|OJJ83186.1 akaw|GAA88270.1 aluc|OJZ82242.1 anid|CBF74073.1 anig|EHA28500.1 aory|EIT79816.1 asyd|OJJ53875.1 ater|EAU31271.1 atub|OJI88196.1 aver|OJJ07973.1 awen|OJJ32974.1 azon|OJJ48943.1

Group2002: abra|OJJ69410.1 acar|OOF94323.1 afla|EED47946.1 afum|EAL85990.1 akaw|GAA88277.1 aluc|OJZ82247.1 anid|CBF74069.1 anig|EHA28494.1 aory|EIT79850.1 asyd|OJJ53873.1 ater|EAU31273.1 atub|OJI88191.1 aver|OJJ07971.1 azon|OJJ48945.1 acla|EAW14789.1 afis|EAW24320.1 agla|OJJ83189.1 awen|OJJ32972.1

Group2003: abra|OJJ69411.1 acar|OOF94322.1 acla|EAW14790.1 afis|EAW24321.1 afla|EED47945.1 afum|EAL85991.1 agla|OJJ83190.1 akaw|GAA88278.1 aluc|OJZ82248.1 anid|CBF74068.1 anig|EHA28493.1 aory|EIT79836.1 asyd|OJJ53872.1 ater|EAU31274.1 atub|OJI88190.1 aver|OJJ07970.1 awen|OJJ32971.1 azon|OJJ48946.1

Group2004: abra|OJJ69412.1 acar|OOF94321.1 acla|EAW14791.1 afis|EAW24322.1 afla|EED47944.1 afum|EAL85992.1 agla|OJJ83191.1 akaw|GAA88279.1 aluc|OJZ82249.1 anid|CBF74066.1 anig|EHA28492.1 aory|EIT79820.1 asyd|OJJ53871.1 ater|EAU30430.1 atub|OJI88189.1 aver|OJJ07969.1 awen|OJJ32970.1 azon|OJJ48947.1

Group2005: abra|OJJ69413.1 acar|OOF94320.1 acla|EAW14793.1 afis|EAW24324.1 afla|EED47942.1 afum|EAL85994.1 akaw|GAA88281.1 aluc|OJZ82250.1 anid|CBF74060.1 anig|EHA28491.1 aory|EIT79823.1 asyd|OJJ53870.1 atub|OJI88187.1 aver|OJJ07968.1 awen|OJJ33577.1 azon|OJJ42851.1 afla|EED51658.1 aory|EIT80622.1

Group2006: abra|OJJ69414.1 acar|OOF94318.1 acla|EAW14795.1 afis|EAW24326.1 afla|EED47940.1 afum|EAL85996.1 agla|OJJ83193.1 akaw|GAA88282.1 aluc|OJZ82251.1 anid|CBF74056.1 anig|EHA28490.1 aory|EIT79834.1 asyd|OJJ53868.1 ater|EAU30427.1 atub|OJI88186.1 aver|OJJ07966.1 awen|OJJ32968.1 azon|OJJ48949.1

Group2007: abra|OJJ69415.1 acar|OOF94317.1 acla|EAW14796.1 afis|EAW24327.1 afla|EED47939.1 afum|EAL85997.1 agla|OJJ83194.1 akaw|GAA88283.1 aluc|OJZ82252.1 anid|CBF74054.1 anig|EHA28489.1 aory|EIT79852.1 asyd|OJJ53867.1 ater|EAU30426.1 atub|OJI88185.1 aver|OJJ07965.1 awen|OJJ32967.1 azon|OJJ48950.1

Group2008: abra|OJJ69417.1 acar|OOF94315.1 acla|EAW14798.1 afis|EAW24329.1 afla|EED47937.1 afum|EAL85999.1 agla|OJJ83196.1 akaw|GAA88285.1 aluc|OJZ82254.1 anid|CBF74052.1 anig|EHA28487.1 aory|EIT79861.1 asyd|OJJ53865.1 atub|OJI88183.1 aver|OJJ07963.1 awen|OJJ32965.1 azon|OJJ48953.1 ater|EAU30424.1

Group2009: abra|OJJ69419.1 acar|OOF90227.1 acla|EAW14799.1 afis|EAW24330.1 afla|EED47935.1 afum|EAL86000.1 agla|OJJ83197.1 akaw|GAA88286.1 aluc|OJZ82255.1 anid|CBF74050.1 anig|EHA28486.1 aory|EIT79863.1 asyd|OJJ53863.1 ater|EAU30422.1 atub|OJI88182.1 aver|OJJ07962.1 awen|OJJ32964.1 azon|OJJ48954.1

Group2010: abra|OJJ69420.1 acar|OOF90228.1 acla|EAW14800.1 afis|EAW24331.1 afla|EED47933.1 afum|EAL86001.1 agla|OJJ83198.1 akaw|GAA88287.1 aluc|OJZ82256.1 anid|CBF74048.1 anig|EHA28485.1 aory|EIT79848.1 asyd|OJJ53862.1 ater|EAU30418.1 atub|OJI88181.1 aver|OJJ07961.1 awen|OJJ32963.1 azon|OJJ48955.1

Group2011: abra|OJJ69421.1 acar|OOF90226.1 acla|EAW14801.1 afis|EAW24332.1 afla|EED47932.1 afum|EAL86002.1 agla|OJJ83199.1 akaw|GAA88288.1 aluc|OJZ82258.1 anid|CBF74045.1 anig|EHA28484.1 aory|EIT79839.1 asyd|OJJ53857.1 ater|EAU30417.1 atub|OJI88180.1 aver|OJJ07958.1 awen|OJJ32962.1 azon|OJJ48960.1

Group2012: abra|OJJ69423.1 acar|OOF94313.1 acla|EAW14802.1 afis|EAW24333.1 afla|EED47931.1 afum|EAL86003.1 agla|OJJ83202.1 akaw|GAA88289.1 aluc|OJZ82259.1 anid|CBF74039.1 anig|EHA28483.1

aory|EIT79843.1 asyd|OJJ53859.1 ater|EAU30416.1 atub|OJI88179.1 aver|OJJ07959.1 awen|OJJ32961.1 azon|OJJ48959.1

Group2013: abra|OJJ69446.1 acar|OOF94294.1 acla|EAW14811.1 afis|EAW24341.1 afla|EED47922.1 afum|EAL86012.1 agla|OJJ83224.1 akaw|GAA88372.1 aluc|OJZ82282.1 anid|CBF74027.1 anig|EHA28465.1 aory|EIT83372.1 asyd|OJJ53846.1 ater|EAU30406.1 atub|OJI88157.1 aver|OJJ07948.1 awen|OJJ32950.1 azon|OJJ48970.1

Group2014: abra|OJJ69448.1 acar|OOF94293.1 acla|EAW14812.1 afis|EAW24342.1 afla|EED47920.1 afum|EAL86013.1 agla|OJJ83223.1 akaw|GAA88371.1 aluc|OJZ82283.1 anid|CBF74025.1 anig|EHA28464.1 aory|EIT83362.1 asyd|OJJ53845.1 ater|EAU30404.1 atub|OJI88155.1 aver|OJJ07947.1 awen|OJJ32951.1 azon|OJJ48969.1

Group2015: abra|OJJ69449.1 acar|OOF94292.1 acla|EAW14813.1 afis|EAW24343.1 afla|EED47919.1 afum|EAL86014.1 agla|OJJ83210.1 akaw|GAA88370.1 aluc|OJZ82285.1 anid|CBF74028.1 anig|EHA28463.1 aory|EIT83364.1 asyd|OJJ53847.1 ater|EAU30403.1 atub|OJI88154.1 aver|OJJ07949.1 awen|OJJ32952.1 azon|OJJ48968.1

Group2016: abra|OJJ69453.1 acar|OOF94288.1 acla|EAW14817.1 afis|EAW24347.1 afla|EED47916.1 afum|EAL86017.1 agla|OJJ83206.1 akaw|GAA88359.1 aluc|OJZ82296.1 anid|CBF74017.1 anig|EHA28459.1 aory|EIT83378.1 asyd|OJJ53840.1 ater|EAU30399.1 atub|OJI88143.1 aver|OJJ07943.1 awen|OJJ32956.1 azon|OJJ48964.1

Group2017: abra|OJJ69456.1 acar|OOF94285.1 acla|EAW14821.1 afis|EAW24349.1 afla|EED47914.1 afum|EAL86019.2 agla|OJJ81762.1 akaw|GAA88356.1 aluc|OJZ82299.1 anid|CBF76155.1 anig|EHA28456.1 aory|EIT83376.1 asyd|OJJ52360.1 ater|EAU30395.1 atub|OJI88140.1 aver|OJJ01988.1 awen|OJJ32875.1 azon|OJJ48976.1

Group2018: abra|OJJ69457.1 acar|OOF94284.1 acla|EAW14820.1 afis|EAW24350.1 afla|EED47913.1 afum|EAL86020.2 agla|OJJ81761.1 akaw|GAA88355.1 aluc|OJZ82300.1 anid|CBF76157.1 anig|EHA28455.1 aory|EIT83375.1 asyd|OJJ52359.1 ater|EAU30394.1 atub|OJI88139.1 aver|OJJ01990.1 awen|OJJ32874.1 azon|OJJ48975.1

Group2019: abra|OJJ69458.1 acar|OOF94283.1 acla|EAW14819.1 afis|EAW24351.1 afla|EED47912.1 afum|EAL86021.1 agla|OJJ81760.1 akaw|GAA88354.1 aluc|OJZ82301.1 anid|CBF76159.1 anig|EHA28454.1 aory|EIT83371.1 asyd|OJJ52358.1 ater|EAU30393.1 atub|OJI88137.1 aver|OJJ01991.1 awen|OJJ32873.1 azon|OJJ48974.1

Group2020: abra|OJJ69459.1 acar|OOF94282.1 acla|EAW09925.1 afis|EAW21173.1 afla|EED54758.1 afum|EAL92193.1 agla|OJJ80005.1 akaw|GAA88353.1 aluc|OJZ82302.1 anid|CBF83317.1 anig|EHA28453.1 aory|EIT79382.1 asyd|OJJ60559.1 ater|EAU35840.1 atub|OJI88135.1 aver|OJI99885.1 awen|OJJ36511.1 azon|OJJ44387.1

Group2021: abra|OJJ69463.1 acar|OOF94278.1 acla|EAW09922.1 afis|EAW21170.1 afla|EED54735.1 afum|EAL92189.2 agla|OJJ80010.1 akaw|GAA88349.1 aluc|OJZ82306.1 anid|CBF83232.1 anig|EHA28450.1 aory|EIT78745.1 asyd|OJJ60541.1 ater|EAU35837.1 atub|OJI88132.1 aver|OJI99868.1 awen|OJJ36507.1 azon|OJJ44379.1

Group2022: abra|OJJ69467.1 acar|OOF94275.1 acla|EAW09920.1 afis|EAW21168.1 afla|EED54732.1 afum|EAL92187.1 agla|OJJ80012.1 akaw|GAA88346.1 aluc|OJZ82311.1 anid|CBF86960.1 anig|EHA28447.1 aory|EIT78891.1 asyd|OJJ60539.1 ater|EAU35835.1 atub|OJI88129.1 aver|OJI99866.1 awen|OJJ36505.1 azon|OJJ44377.1

Group2023: abra|OJJ69468.1 acar|OOF94274.1 acla|EAW09919.1 afis|EAW21167.1 afla|EED54731.1 afum|EAL92186.1 agla|OJJ80013.1 akaw|GAA88345.1 aluc|OJZ82312.1 anid|CBF86961.1 anig|EHA28446.1 aory|EIT78775.1 asyd|OJJ60537.1 ater|EAU35834.1 atub|OJI88128.1 aver|OJI99864.1 awen|OJJ36504.1 azon|OJJ44376.1

Group2024: abra|OJJ69469.1 acar|OOF94273.1 acla|EAW09918.1 afis|EAW21166.1 afla|EED54730.1 afum|EAL92185.1 agla|OJJ80015.1 akaw|GAA88344.1 aluc|OJZ82313.1 anid|CBF86963.1 anig|EHA28445.1 aory|EIT78843.1 asyd|OJJ60536.1 ater|EAU35833.1 atub|OJI88127.1 aver|OJI99863.1 awen|OJJ36503.1 azon|OJJ44375.1

Group2025: abra|OJJ69471.1 acar|OOF94272.1 acla|EAW09917.1 afis|EAW21165.1 afla|EED54729.1 afum|EAL92184.1 agla|OJJ80016.1 akaw|GAA88343.1 aluc|OJZ82314.1 anid|CBF86964.1 anig|EHA28444.1 aory|EIT78934.1 asyd|OJJ60535.1 ater|EAU35832.1 atub|OJI88126.1 aver|OJI99862.1 awen|OJJ36500.1 azon|OJJ44374.1

Group2026: abra|OJJ69473.1 acar|OOF94270.1 acla|EAW09915.1 afis|EAW21163.1 afla|EED54726.1 afum|EAL92182.1 agla|OJJ80020.1 akaw|GAA88342.1 aluc|OJZ82316.1 anid|CBF86968.1 anig|EHA28442.1 aory|EIT78844.1 asyd|OJJ60532.1 ater|EAU35829.1 atub|OJI88125.1 aver|OJI99859.1 awen|OJJ36497.1 azon|OJJ44372.1

Group2027: abra|OJJ69474.1 acar|OOF94268.1 acla|EAW09914.1 afis|EAW21162.1 afla|EED54725.1 afum|EAL92181.1 agla|OJJ80021.1 akaw|GAA88341.1 aluc|OJZ82317.1 anid|CBF86970.1 anig|EHA28441.1 aory|EIT78833.1 asyd|OJJ60531.1 ater|EAU35828.1 atub|OJI88124.1 aver|OJI99858.1 awen|OJJ36496.1 azon|OJJ44371.1

Group2028: abra|OJJ69475.1 acar|OOF94269.1 acla|EAW09913.1 afis|EAW21161.1 afla|EED54724.1 afum|EAL92180.1 agla|OJJ80022.1 akaw|GAA88340.1 aluc|OJZ82318.1 anid|CBF86972.1 anig|EHA28440.1 aory|EIT78951.1 asyd|OJJ60530.1 ater|EAU35827.1 atub|OJI88123.1 aver|OJI99857.1 awen|OJJ36495.1 azon|OJJ44370.1

Group2029: abra|OJJ69477.1 acar|OOF94265.1 acla|EAW09911.1 afis|EAW21160.1 afla|EED54649.1 afum|EAL92178.1 agla|OJJ80028.1 akaw|GAA88338.1 aluc|OJZ82320.1 anid|CBF86977.1 anig|EHA28438.1 aory|EIT78896.1 asyd|OJJ60527.1 ater|EAU35822.1 atub|OJI88120.1 aver|OJI99855.1 awen|OJJ36492.1 azon|OJJ44365.1

Group2030: abra|OJJ69481.1 acar|OOF94263.1 acla|EAW09908.1 akaw|GAA88335.1 aluc|OJZ82323.1 anig|EHA28435.1 atub|OJI88117.1 awen|OJJ36488.1 agla|OJJ80036.1 afis|EAW21157.1 afum|EAL92175.1 afla|EED54653.1 aory|EIT78820.1 ater|EAU35819.1 anid|CBF86982.1 asyd|OJJ60523.1 aver|OJI99847.1 azon|OJJ44359.1

Group2031: abra|OJJ69483.1 acar|OOF94262.1 acla|EAW09907.1 afis|EAW21156.1 afla|EED54652.1 afum|EAL92174.1 agla|OJJ80037.1 akaw|GAA88334.1 aluc|OJZ82324.1 anid|CBF86984.1 anig|EHA28434.1 aory|EIT78871.1 asyd|OJJ60518.1 ater|EAU35818.1 atub|OJI88116.1 aver|OJI99845.1 awen|OJJ36487.1 azon|OJJ44360.1

Group2032: abra|OJJ69485.1 acar|OOF94261.1 acla|EAW09906.1 afis|EAW21155.1 afla|EED54661.1 afum|EAL92173.1 agla|OJJ80038.1 akaw|GAA88333.1 aluc|OJZ82326.1 anid|CBF86986.1 anig|EHA28432.1 aory|EIT78925.1 asyd|OJJ60520.1 ater|EAU35817.1 atub|OJI88114.1 aver|OJI99846.1 awen|OJJ36486.1 azon|OJJ44362.1

Group2033: abra|OJJ69492.1 acar|OOF94251.1 acla|EAW09902.1 afis|EAW21150.1 afla|EED54662.1 afum|EAL92169.1 agla|OJJ78417.1 akaw|GAA88328.1 aluc|OJZ82332.1 anid|CBF75144.1 anig|EHA28428.1 aory|EIT78748.1 asyd|OJJ59971.1 ater|EAU36738.1 atub|OJI88108.1 aver|OJJ02320.1 awen|OJJ36477.1 asyd|OJJ54978.1

Group2034: abra|OJJ69496.1 acar|OOF94254.1 acla|EAW09950.1 afis|EAW21146.1 afla|EED46146.1 afum|EAL92165.1 agla|OJJ80070.1 akaw|GAA88325.1 aluc|OJZ82336.1 anid|CBF87010.1 anig|EHA28424.1 aory|EIT76496.1 asyd|OJJ60472.1 ater|EAU36761.1 atub|OJI88104.1 aver|OJI99792.1 awen|OJJ36469.1 azon|OJJ44354.1

Group2035: abra|OJJ69497.1 acar|OOF94256.1 acla|EAW09951.1 afis|EAW21145.1 afla|EED46147.1 afum|EAL92164.1 agla|OJJ80071.1 akaw|GAA88324.1 aluc|OJZ82337.1 anid|CBF86995.1 anig|EHA28423.1 aory|EIT76462.1 asyd|OJJ60473.1 ater|EAU36757.1 atub|OJI88103.1 aver|OJI99793.1 awen|OJJ36470.1 azon|OJJ44355.1

Group2036: abra|OJJ69498.1 acar|OOF94257.1 acla|EAW09952.1 afis|EAW21144.1 afla|EED46148.1 afum|EAL92163.1 agla|OJJ80072.1 akaw|GAA88323.1 aluc|OJZ82338.1 anid|CBF86997.1 anig|EHA28422.1 aory|EIT76514.1 asyd|OJJ60474.1 ater|EAU36756.1 atub|OJI88102.1 aver|OJI99794.1 awen|OJJ36471.1 azon|OJJ44356.1

Group2037: abra|OJJ69505.1 acar|OOF94247.1 acla|EAW09957.1 afis|EAW21136.1 afla|EED51811.1 afum|EAL92155.1 agla|OJJ80076.1 akaw|GAA88318.1 aluc|OJZ82344.1 anid|CBF87016.1 anig|EHA28416.1 aory|EIT81758.1 asyd|OJJ60463.1 ater|EAU36774.1 atub|OJI88096.1 aver|OJI99783.1 awen|OJJ36456.1 azon|OJJ44420.1

Group2038: abra|OJJ69506.1 acar|OOF94246.1 acla|EAW09958.1 afis|EAW21137.1 afla|EED51812.1 afum|EAL92156.1 agla|OJJ80081.1 akaw|GAA88317.1 aluc|OJZ82345.1 anid|CBF87014.1 anig|EHA28415.1 aory|EIT81705.1 asyd|OJJ60464.1 ater|EAU36773.1 atub|OJI88095.1 aver|OJI99784.1 awen|OJJ36457.1 azon|OJJ44418.1

Group2039: abra|OJJ69508.1 acar|OOF94245.1 acla|EAW09959.1 afis|EAW21138.1 afla|EED51813.1 afum|EAL92157.2 agla|OJJ80080.1 akaw|GAA88316.1 aluc|OJZ82346.1 anid|CBF87012.1 anig|EHA28414.1 aory|EIT81694.1 asyd|OJJ60465.1 ater|EAU36772.1 atub|OJI88094.1 aver|OJI99785.1 awen|OJJ36458.1 azon|OJJ44419.1

Group2040: abra|OJJ69510.1 acar|OOF94244.1 acla|EAW09960.1 afis|EAW21135.1 afla|EED46138.1 afum|EAL92154.1 agla|OJJ80078.1 akaw|GAA88314.1 aluc|OJZ82348.1 anid|CBF87000.1 anig|EHA28412.1 aory|EIT76494.1 asyd|OJJ60467.1 ater|EAU36767.1 atub|OJI88092.1 aver|OJI99787.1 awen|OJJ36461.1 azon|OJJ44416.1

Group2041: abra|OJJ69513.1 acar|OOF94240.1 acla|EAW09955.1 afis|EAW21141.1 afla|EED46142.1 afum|EAL92160.1 agla|OJJ80068.1 akaw|GAA88311.1 aluc|OJZ82351.1 anid|CBF87007.1 anig|EHA28409.1

aory|EIT76415.1 asyd|OJJ60470.1 ater|EAU36762.1 atub|OJI88089.1 aver|OJI99790.1 awen|OJJ36465.1 azon|OJJ44413.1

Group2042: abra|OJJ69516.1 acar|OOF94238.1 acla|EAW09964.1 afis|EAW21124.1 afla|EED51799.1 afum|EAL92147.1 agla|OJJ80086.1 akaw|GAA88308.1 aluc|OJZ82354.1 anid|CBF87018.1 anig|EHA28406.1 aory|EIT81724.1 asyd|OJJ60461.1 ater|EAU36776.1 atub|OJI88086.1 aver|OJI99781.1 awen|OJJ36454.1 azon|OJJ44423.1

Group2043: abra|OJJ69520.1 acar|OOF94237.1 acla|EAW09966.1 afis|EAW21122.1 afla|EED51795.1 afum|EAL92145.1 agla|OJJ80089.1 akaw|GAA88306.1 aluc|OJZ82359.1 anid|CBF87021.1 anig|EHA28404.1 aory|EIT81713.1 asyd|OJJ60459.1 ater|EAU36778.1 atub|OJI88082.1 aver|OJI99779.1 awen|OJJ36452.1 azon|OJJ44425.1

Group2044: abra|OJJ69522.1 acar|OOF94233.1 acla|EAW09968.1 afis|EAW21120.1 afla|EED51793.1 afum|EBA27448.1 agla|OJJ80091.1 akaw|GAA88305.1 aluc|OJZ82362.1 anid|CBF87025.1 anig|EHA28403.1 aory|EIT81754.1 asyd|OJJ60457.1 ater|EAU36779.1 atub|OJI88080.1 aver|OJI99777.1 awen|OJJ36450.1 azon|OJJ44427.1

Group2045: abra|OJJ69525.1 acar|OOF94229.1 acla|EAW09972.1 afis|EAW21116.1 afla|EED51767.1 afum|EAL92139.1 agla|OJJ80095.1 akaw|GAA88300.1 aluc|OJZ82365.1 anid|CBF87030.1 anig|EHA28400.1 aory|EIT79993.1 asyd|OJJ60455.1 ater|EAU36783.1 atub|OJI88077.1 aver|OJI99773.1 awen|OJJ36446.1 azon|OJJ44429.1

Group2046: abra|OJJ69529.1 afis|EAW22089.1 afla|EED53382.1 aory|EIT74986.1 asyd|OJJ58665.1 ater|EAU39308.1 azon|OJJ42020.1 abra|OJJ77940.1 acar|OOF93336.1 acla|EAW10026.1 akaw|GAA83724.1 aluc|OJZ91007.1 asyd|OJJ53344.1 atub|OJI82999.1 aver|OJJ08400.1 afla|EED53383.1 azon|OJJ45239.1 anid|CBF70508.1

Group2047: abra|OJJ69535.1 acar|OOF95965.1 acla|EAW11570.1 afis|EAW23883.1 afla|EED52612.1 afum|EAL87311.1 agla|OJJ82570.1 akaw|GAA88385.1 aluc|OJZ82373.1 anid|CBF77187.1 anig|EHA28391.1 aory|EIT74250.1 asyd|OJJ56841.1 ater|EAU33463.1 atub|OJI88069.1 aver|OJJ03497.1 awen|OJJ33865.1 azon|OJJ48688.1

Group2048: abra|OJJ69536.1 acar|OOF91604.1 acla|EAW14409.1 afis|EAW19117.1 afla|EED54297.1 afum|EAL93849.1 agla|OJJ80432.1 akaw|GAA91927.1 aluc|OJZ82695.1 anid|CBF79728.1 anig|EHA21143.1 aory|EIT75966.1 asyd|OJJ58073.1 ater|EAU37379.1 atub|OJI88927.1 aver|OJJ04334.1 awen|OJJ37447.1 azon|OJJ51125.1

Group2049: abra|OJJ69575.1 acar|OOF91580.1 acla|EAW14394.1 afis|EAW19102.1 afla|EED54308.1 afum|EAL93834.1 agla|OJJ84500.1 akaw|GAA91572.1 aluc|OJZ82392.1 anid|CBF79696.1 anig|EHA28372.1 aory|EIT75942.1 asyd|OJJ58050.1 ater|EAU32151.1 atub|OJI88049.1 aver|OJJ04353.1 awen|OJJ32495.1 azon|OJJ46034.1

Group2050: abra|OJJ69580.1 acar|OOF91573.1 acla|EAW14388.1 afis|EAW19095.1 afla|EED54328.1 afum|EAL93826.1 agla|OJJ84506.1 akaw|GAA91576.1 aluc|OJZ82398.1 anid|CBF79705.1 anig|EHA28367.1 aory|EIT75940.1 asyd|OJJ58055.1 ater|EAU32143.1 atub|OJI88044.1 aver|OJJ04348.1 awen|OJJ32488.1 azon|OJJ46043.1

Group2051: abra|OJJ69583.1 acar|OOF91571.1 acla|EAW14386.1 afis|EAW19092.1 afla|EED54327.1 afum|EAL93823.1 agla|OJJ84509.1 akaw|GAA91578.1 aluc|OJZ82400.1 anid|CBF79709.1 anig|EHA28365.1 asyd|OJJ58057.1 ater|EAU32141.1 atub|OJI88041.1 aver|OJJ04346.1 awen|OJJ32486.1 azon|OJJ46045.1 afum|EAL93822.1

Group2052: abra|OJJ69584.1 acar|OOF91570.1 acla|EAW14385.1 afis|EAW19091.1 afla|EED54326.1 afum|EAL93821.1 agla|OJJ84512.1 akaw|GAA91579.1 aluc|OJZ82401.1 anid|CBF79711.1 anig|EHA28364.1 aory|EIT76092.1 asyd|OJJ58058.1 ater|EAU32140.1 atub|OJI88040.1 aver|OJJ04345.1 awen|OJJ32485.1 azon|OJJ46046.1

Group2053: abra|OJJ69587.1 acar|OOF91568.1 acla|EAW14380.1 afis|EAW19086.1 afla|EED54318.1 afum|EAL93816.1 agla|OJJ84518.1 akaw|GAA91584.1 aluc|OJZ82404.1 anid|CBF79684.1 anig|EHA28362.1 aory|EIT76159.1 asyd|OJJ58047.1 ater|EAU32134.1 atub|OJI88037.1 aver|OJJ04356.1 awen|OJJ32483.1 azon|OJJ46227.1

Group2054: abra|OJJ69588.1 acar|OOF91567.1 acla|EAW14379.1 afis|EAW19085.1 afla|EED54317.1 afum|EAL93815.1 agla|OJJ84521.1 akaw|GAA91585.1 aluc|OJZ82406.1 anid|CBF79683.1 anig|EHA28361.1 aory|EIT75897.1 asyd|OJJ58046.1 ater|EAU32133.1 atub|OJI88036.1 aver|OJJ04357.1 awen|OJJ32482.1 azon|OJJ46226.1

Group2055: abra|OJJ69590.1 acar|OOF91565.1 acla|EAW14378.1 afis|EAW19084.1 afla|EED54316.1 afum|EAL93814.1 agla|OJJ84522.1 akaw|GAA91586.1 aluc|OJZ82408.1 anid|CBF79676.1 anig|EHA28360.1 aory|EIT75902.1 asyd|OJJ58038.1 ater|EAU32132.1 atub|OJI88034.1 aver|OJJ04362.1 awen|OJJ32481.1 azon|OJJ46224.1

Group2056: abra|OJJ69592.1 anid|CBF82180.1 anig|EHA28357.1 ater|EAU30193.1 abra|OJJ73493.1 acar|OOG00685.1 afis|EAW20408.1 afum|EAL89236.1 agla|OJJ88380.1 akaw|GAA87442.1 aluc|OJZ88555.1 anig|EHA18253.1 atub|OJI91140.1 awen|OJJ31333.1 azon|OJJ49482.1 afis|EAW19984.1 afum|EAL84932.1 anid|CBF84172.1

Group2057: abra|OJJ69602.1 acar|OOF92702.1 acla|EAW10013.1 afis|EAW21041.1 afla|EED51373.1 afum|EAL92080.1 agla|OJJ84213.1 akaw|GAA91590.1 aluc|OJZ82413.1 anid|CBF80091.1 anig|EHA28354.1 aory|EIT80745.1 asyd|OJJ58302.1 ater|EAU31725.1 atub|OJI88028.1 aver|OJJ02110.1 awen|OJJ32696.1 azon|OJJ50700.1

Group2058: abra|OJJ69616.1 acar|OOF92690.1 acla|EAW12076.1 afis|EAW19717.1 afla|EED53108.1 afum|EAL86810.1 agla|OJJ80861.1 akaw|GAA91600.1 aluc|OJZ82424.1 anid|CBF80078.1 anig|EHA28345.1 aory|EIT74760.1 asyd|OJJ58310.1 ater|EAU37776.1 atub|OJI88016.1 aver|OJJ02101.1 awen|OJJ35107.1 azon|OJJ49987.1

Group2059: abra|OJJ69628.1 afla|EED55079.1 agla|OJJ86032.1 aluc|OJZ82433.1 anid|CBF82983.1 anig|EHA28338.1 asyd|OJJ64730.1 ater|EAU30171.1 atub|OJI88008.1 aver|OJI98834.1 awen|OJJ30457.1 azon|OJJ45448.1 acar|OOF92682.1 afis|EAW16821.1 afum|EAL86371.1 acla|EAW12375.1 aory|EIT81573.1 akaw|GAA91608.1

Group2060: abra|OJJ69635.1 acar|OOF93616.1 acla|EAW15185.1 afis|EAW20155.1 afla|EED55122.1 afum|EAL85270.1 agla|OJJ89111.1 akaw|GAA91615.1 aluc|OJZ82441.1 anid|CBF69484.1 anig|EHA28332.1 aory|EIT79219.1 asyd|OJJ59246.1 atub|OJI87999.1 aver|OJJ01857.1 awen|OJJ36185.1 aory|EIT81639.1 afla|EED47294.1

Group2061: abra|OJJ69637.1 acar|OOF92675.1 acla|EAW12571.1 afis|EAW24079.1 afla|EED51394.1 afum|EAL91934.1 agla|OJJ87150.1 akaw|GAA91617.1 aluc|OJZ82443.1 anid|CBF86476.1 anig|EHA28330.1 aory|EIT80502.1 asyd|OJJ64903.1 ater|EAU30500.1 atub|OJI87997.1 aver|OJI99025.1 awen|OJJ41417.1 azon|OJJ48876.1

Group2062: azon|OJJ44974.1 abra|OJJ73161.1 acar|OOG00404.1 acla|EAW06767.1 afis|EAW16317.1 afla|EED53564.1 afum|EAL88940.1 agla|OJJ88278.1 akaw|GAA88011.1 aluc|OJZ81269.1 anid|CBF89145.1 anig|EHA18569.1 aory|EIT75175.1 asyd|OJJ62569.1 ater|EAU34140.1 atub|OJI91476.1 aver|OJI96693.1 awen|OJJ37717.1

Group2063: abra|OJJ69655.1 acar|OOG00750.1 acla|EAW07485.1 afis|EAW22518.1 afla|EED54547.1 afum|EAL90675.1 agla|OJJ80878.1 akaw|GAA91634.1 aluc|OJZ82461.1 anid|CBF88995.1 anig|EHA28314.1 aory|EIT78864.1 asyd|OJJ62783.1 ater|EAU39191.1 atub|OJI87977.1 aver|OJI96828.1 awen|OJJ35145.1 azon|OJJ49955.1

Group2064: abra|OJJ69657.1 acar|OOF95821.1 acla|EAW11286.1 afis|EAW18213.1 afla|EED56258.1 afum|EAL87579.1 agla|OJJ85869.1 akaw|GAA91636.1 aluc|OJZ82463.1 anid|CBF75160.1 anig|EHA28312.1 aory|EIT73422.1 asyd|OJJ59962.1 ater|EAU33776.1 atub|OJI87975.1 aver|OJJ02328.1 awen|OJJ30591.1 azon|OJJ50185.1

Group2065: abra|OJJ69663.1 acar|OOF98113.1 acla|EAW10577.1 afis|EAW24990.1 afla|EED55469.1 afum|EAL89650.1 agla|OJJ82264.1 akaw|GAA91641.1 aluc|OJZ82468.1 anid|CBF81596.1 anig|EHA28307.1 aory|EIT80882.1 asyd|OJJ59517.1 ater|EAU35741.1 atub|OJI87970.1 aver|OJJ02756.1 awen|OJJ40595.1 azon|OJJ49446.1

Group2066: abra|OJJ69664.1 acar|OOG00819.1 acla|EAW07551.1 afis|EAW22585.1 afla|EED54630.1 afum|EAL90614.1 agla|OJJ80945.1 akaw|GAA91642.1 aluc|OJZ82469.1 anid|CBF88352.1 anig|EHA28306.1 aory|EIT78765.1 asyd|OJJ63721.1 ater|EAU39114.1 atub|OJI87968.1 aver|OJI97219.1 awen|OJJ35213.1 azon|OJJ49896.1

Group2067: abra|OJJ69665.1 acar|OOG00018.1 acla|EAW09937.1 afis|EAW21182.1 afla|EED54777.1 afum|EAL92203.1 agla|OJJ79988.1 akaw|GAA91643.1 aluc|OJZ82470.1 anid|CBF83310.1 anig|EHA28303.1 aory|EIT79439.1 asyd|OJJ60549.1 ater|EAU35848.1 atub|OJI87967.1 aver|OJI99875.1 awen|OJJ36521.1 azon|OJJ44397.1

Group2068: abra|OJJ69666.1 acar|OOG00019.1 acla|EAW09936.1 afis|EAW21181.1 afla|EED54778.1 afum|EAL92202.1 agla|OJJ79998.1 akaw|GAA91644.1 aluc|OJZ82471.1 anid|CBF83308.1 anig|EHA28302.1 aory|EIT79454.1 asyd|OJJ60550.1 ater|EAU35847.1 atub|OJI87966.1 aver|OJI99876.1 awen|OJJ36520.1 azon|OJJ44396.1

Group2069: abra|OJJ69667.1 acar|OOF97488.1 acla|EAW13075.1 afis|EAW17589.1 afla|EED49233.1 afum|EAL91402.1 agla|OJJ81360.1 akaw|GAA91645.1 aluc|OJZ82472.1 anid|CBF90218.1 anig|EHA28301.1 aory|EIT83160.1 asyd|OJJ61967.1 ater|EAU37024.1 atub|OJI87965.1 aver|OJI96059.1 awen|OJJ37257.1 azon|OJJ50952.1

Group2070: abra|OJJ69669.1 acar|OOF95822.1 acla|EAW06663.1 afis|EAW16102.1 afla|EED48637.1 afum|EAL88734.1 agla|OJJ86131.1 akaw|GAA91647.1 aluc|OJZ82474.1 anid|CBF85442.1 anig|EHA28299.1

aory|EIT73069.1 asyd|OJJ64091.1 ater|EAU33179.1 atub|OJI87963.1 aver|OJI98175.1 awen|OJJ30330.1 azon|OJJ45358.1

Group2071: abra|OJJ69677.1 acar|OOF92983.1 acla|EAW14808.1 afis|EAW24337.1 afla|EED47927.1 afum|EAL86008.1 agla|OJJ83205.1 akaw|GAA88802.1 aluc|OJZ82485.1 anid|CBF74030.1 anig|EHA28289.1 aory|EIT79853.1 asyd|OJJ53849.1 ater|EAU30411.1 atub|OJI87947.1 aver|OJJ07950.1 awen|OJJ32957.1 azon|OJJ48963.1

Group2072: abra|OJJ69682.1 acar|OOF93221.1 acla|EAW13124.1 afis|EAW17540.1 afla|EED49287.1 afum|EAL91354.1 agla|OJJ81429.1 akaw|GAA92416.1 aluc|OJZ83274.1 anid|CBF90315.1 anig|EHA28284.1 aory|EIT82215.1 asyd|OJJ61912.1 ater|EAU36970.1 atub|OJI87666.1 aver|OJI96005.1 awen|OJJ37313.1 azon|OJJ51005.1

Group2073: abra|OJJ69687.1 acar|OOF92648.1 acla|EAW10319.1 afis|EAW25273.1 afla|EED53738.1 afum|EAL89936.1 agla|OJJ89247.1 akaw|GAA88811.1 aluc|OJZ82494.1 anid|CBF85387.1 anig|EHA28278.1 aory|EIT75306.1 asyd|OJJ64055.1 ater|EAU34416.1 atub|OJI87938.1 aver|OJI98140.1 awen|OJJ40987.1 azon|OJJ44574.1

Group2074: abra|OJJ69698.1 acar|OOF90521.1 acla|EAW15242.1 afis|EAW20209.1 afla|EED57134.1 afum|EAL85326.1 agla|OJJ84655.1 akaw|GAA88816.1 aluc|OJZ82500.1 anid|CBF85100.1 anig|EHA28270.1 aory|EIT74032.1 asyd|OJJ63057.1 ater|EAU32019.1 atub|OJI87931.1 aver|OJI97905.1 awen|OJJ31213.1 azon|OJJ46146.1

Group2075: abra|OJJ69699.1 acar|OOF95410.1 acla|EAW07370.1 afis|EAW22395.1 afla|EED56059.1 afum|EAL90796.1 agla|OJJ87658.1 akaw|GAA88818.1 aluc|OJZ82502.1 anid|CBF88747.1 anig|EHA28269.1 aory|EIT76693.1 asyd|OJJ63956.1 ater|EAU39322.1 atub|OJI87929.1 aver|OJI96985.1 awen|OJJ34973.1 azon|OJJ45622.1

Group2076: abra|OJJ69701.1 acar|OOF93222.1 acla|EAW13123.1 afis|EAW17541.1 afla|EED49286.1 afum|EAL91355.1 agla|OJJ81428.1 akaw|GAA88820.1 aluc|OJZ82504.1 anid|CBF90313.1 anig|EHA28266.1 aory|EIT82202.1 asyd|OJJ61913.1 ater|EAU36971.1 atub|OJI87927.1 aver|OJI96006.1 awen|OJJ37312.1 azon|OJJ51004.1

Group2077: abra|OJJ69703.1 acar|OOF98863.1 acla|EAW06689.1 afis|EAW16076.1 afla|EED48596.1 afum|EAL88708.1 agla|OJJ86093.1 akaw|GAA88822.1 aluc|OJZ82506.1 anid|CBF75115.1 anig|EHA28263.1 aory|EIT73056.1 asyd|OJJ59995.1 ater|EAU33555.1 atub|OJI87925.1 aver|OJJ02296.1 awen|OJJ30357.1 azon|OJJ45379.1

Group2078: abra|OJJ69718.1 acar|OOF94196.1 acla|EAW09976.1 afis|EAW21113.1 afla|EED51764.1 afum|EAL92136.1 agla|OJJ80098.1 akaw|GAA90171.1 aluc|OJZ83179.1 anid|CBF87036.1 anig|EHA28253.1 aory|EIT80019.1 asyd|OJJ60452.1 ater|EAU36786.1 atub|OJI87758.1 aver|OJI99770.1 awen|OJJ36441.1 azon|OJJ44432.1

Group2079: abra|OJJ69746.1 acar|OOF94176.1 acla|EAW09985.1 afis|EAW21108.1 afla|EED51696.1 afum|EAL92129.1 agla|OJJ80107.1 akaw|GAA90198.1 aluc|OJZ83150.1 anid|CBF87041.1 anig|EHA28216.1 aory|EIT80588.1 asyd|OJJ60448.1 ater|EAU36792.1 atub|OJI87787.1 aver|OJI99766.1 awen|OJJ36382.1 azon|OJJ51534.1

Group2080: aver|OJJ05453.1 abra|OJJ70544.1 acar|OOF94607.1 acla|EAW08135.1 afis|EAW23171.1 afla|EED46949.1 afum|EAL88312.2 agla|OJJ89683.1 akaw|GAA83390.1 aluc|OJZ83889.1 anid|CBF74485.1 anig|EHA19026.1 aory|EIT72627.1 asyd|OJJ57712.1 ater|EAU36300.1 atub|OJI90256.1 awen|OJJ36606.1 azon|OJJ47601.1

Group2081: abra|OJJ69784.1 acar|OOF95612.1 afum|EAL89018.1 anid|CBF84864.1 anig|EHA25046.1 asyd|OJJ56179.1 atub|OJI80053.1 aver|OJJ05815.1 abra|OJJ77383.1 acar|OOF93253.1 agla|OJJ82715.1 akaw|GAA91378.1 aluc|OJZ84791.1 anig|EHA28059.1 atub|OJI89291.1 awen|OJJ36484.1 aver|OJI99823.1 asyd|OJJ57350.1

Group2082: abra|OJJ69822.1 acar|OOF93341.1 acla|EAW08969.1 afis|EAW15448.1 afla|EED56825.1 afum|EAL84198.2 agla|OJJ86271.1 akaw|GAA87838.1 aluc|OJZ85474.1 anid|CBF78305.1 anig|EHA25015.1 aory|EIT73942.1 asyd|OJJ56090.1 ater|EAU30838.1 atub|OJI80089.1 aver|OJJ05033.1 awen|OJJ31223.1 azon|OJJ45071.1

Group2083: abra|OJJ69827.1 acar|OOF93352.1 afla|EED55192.1 akaw|GAA87833.1 aluc|OJZ85469.1 anid|CBF78301.1 anig|EHA25010.1 aory|EIT83545.1 asyd|OJJ56100.1 atub|OJI80094.1 aver|OJJ05043.1 acla|EAW08979.1 afis|EAW15430.1 afum|EAL84213.1 ater|EAU32701.1 awen|OJJ33446.1 afla|EED47528.1 aory|EIT74571.1

Group2084: abra|OJJ69888.1 acar|OOF93408.1 acla|EAW10806.1 afis|EAW24768.1 afla|EED49741.1 afum|EAL89409.1 agla|OJJ82608.1 akaw|GAA87764.1 aluc|OJZ85399.1 anid|CBF69441.1 anig|EHA25581.1 aory|EIT77620.1 ater|EAU36289.1 atub|OJI80172.1 awen|OJJ39939.1 acar|OOF92530.1 asyd|OJJ60681.1 aver|OJJ00006.1

Group2085: abra|OJJ69898.1 acar|OOF93417.1 acla|EAW08922.1 afis|EAW15471.1 afla|EED46922.1 afum|EAL85850.1 agla|OJJ80151.1 akaw|GAA87752.1 aluc|OJZ85385.1 anid|CBF78234.1 anig|EHA25569.1 aory|EIT72668.1 asyd|OJJ56036.1 ater|EAU32657.1 atub|OJI80185.1 aver|OJJ04982.1 awen|OJJ38121.1 azon|OJJ48022.1

Group2086: abra|OJJ69901.1 acar|OOF93419.1 acla|EAW08921.1 afis|EAW15473.1 afla|EED46923.1 afum|EAL85848.1 akaw|GAA87750.1 aluc|OJZ85383.1 anid|CBF78228.1 anig|EHA25567.1 aory|EIT72551.1 asyd|OJJ56034.1 ater|EAU32656.1 atub|OJI80187.1 aver|OJJ04980.1 awen|OJJ38126.1 azon|OJJ43160.1 agla|OJJ80154.1

Group2087: abra|OJJ69906.1 acar|OOF93423.1 acla|EAW08918.1 afis|EAW15476.1 afla|EED46939.1 afum|EAL85845.1 agla|OJJ80155.1 akaw|GAA87747.1 aluc|OJZ85380.1 anid|CBF78226.1 anig|EHA25563.1 aory|EIT72564.1 asyd|OJJ56033.1 ater|EAU32651.1 atub|OJI80190.1 aver|OJJ04979.1 awen|OJJ38133.1 azon|OJJ43159.1

Group2088: abra|OJJ69914.1 acar|OOF93426.1 acla|EAW08915.1 afis|EAW15479.1 afla|EED46934.1 afum|EAL85842.1 agla|OJJ80158.1 akaw|GAA87742.1 aluc|OJZ85374.1 anid|CBF78221.1 anig|EHA25558.1 aory|EIT72728.1 asyd|OJJ56028.1 ater|EAU32648.1 atub|OJI80195.1 aver|OJJ04975.1 awen|OJJ38138.1 azon|OJJ43152.1

Group2089: abra|OJJ69916.1 acar|OOF93428.1 acla|EAW08912.1 afis|EAW15483.1 afla|EED46930.1 afla|EED48092.1 afum|EAL85839.1 agla|OJJ80162.1 akaw|GAA87740.1 aluc|OJZ85372.1 anig|EHA25556.1 aory|EIT72736.1 asyd|OJJ56026.1 ater|EAU32647.1 atub|OJI80198.1 aver|OJJ04973.1 awen|OJJ38142.1 azon|OJJ43148.1

Group2090: abra|OJJ69940.1 acar|OOF93449.1 acla|EAW08883.1 afis|EAW15506.1 afla|EED49019.1 afum|EAL85818.1 agla|OJJ80183.1 akaw|GAA87719.1 aluc|OJZ85347.1 anid|CBF84473.1 anig|EHA25534.1 aory|EIT82494.1 asyd|OJJ53540.1 ater|EAU32612.1 atub|OJI80223.1 aver|OJJ08213.1 awen|OJJ38166.1 azon|OJJ45800.1

Group2091: abra|OJJ69952.1 acar|OOF93462.1 acla|EAW08870.1 afis|EAW15524.1 afla|EED49034.1 afum|EAL85804.1 agla|OJJ85222.1 akaw|GAA87707.1 aluc|OJZ85330.1 anig|EHA25519.1 aory|EIT82486.1 asyd|OJJ55992.1 ater|EAU32599.1 atub|OJI80242.1 aver|OJJ04940.1 awen|OJJ38188.1 azon|OJJ43107.1 afla|EED49033.1

Group2092: abra|OJJ69953.1 acar|OOF93463.1 acla|EAW08869.1 afis|EAW15525.1 afla|EED49036.1 afum|EAL85803.1 agla|OJJ85221.1 akaw|GAA87706.1 aluc|OJZ85329.1 anid|CBF78217.1 anig|EHA25518.1 aory|EIT82537.1 asyd|OJJ55991.1 ater|EAU32598.1 atub|OJI80243.1 aver|OJJ04939.1 awen|OJJ38189.1 azon|OJJ43106.1

Group2093: abra|OJJ69955.1 acar|OOF93465.1 acla|EAW08867.1 afis|EAW15527.1 afla|EED49038.1 afum|EAL85801.1 agla|OJJ85219.1 akaw|GAA87704.1 aluc|OJZ85327.1 anid|CBF78212.1 anig|EHA25517.1 aory|EIT82564.1 asyd|OJJ55989.1 ater|EAU32597.1 atub|OJI80245.1 aver|OJJ04937.1 awen|OJJ38191.1 azon|OJJ43104.1

Group2094: abra|OJJ69962.1 acar|OOF93470.1 acla|EAW08834.1 afis|EAW15566.1 afla|EED49075.1 afum|EAL85767.2 agla|OJJ85185.1 akaw|GAA87697.1 aluc|OJZ85321.1 anid|CBF78116.1 anig|EHA25511.1 aory|EIT82536.1 asyd|OJJ55900.1 ater|EAU32567.1 atub|OJI80252.1 aver|OJJ04845.1 awen|OJJ38226.1 azon|OJJ43069.1

Group2095: abra|OJJ69963.1 acar|OOF93471.1 acla|EAW08835.1 afis|EAW15565.1 afla|EED49074.1 agla|OJJ85186.1 akaw|GAA87696.1 aluc|OJZ85320.1 anid|CBF78118.1 anig|EHA25510.1 aory|EIT82524.1 asyd|OJJ55901.1 ater|EAU32568.1 atub|OJI80253.1 aver|OJJ04846.1 awen|OJJ38225.1 azon|OJJ43070.1 afum|EAL85768.1

Group2096: abra|OJJ69971.1 acar|OOF93475.1 acla|EAW08842.1 afis|EAW15559.1 afla|EED49063.1 afum|EAL85773.1 agla|OJJ85195.1 akaw|GAA87690.1 aluc|OJZ85311.1 anid|CBF78159.1 anig|EHA25506.1 aory|EIT82560.1 asyd|OJJ55960.1 ater|EAU32575.1 atub|OJI80260.1 aver|OJJ04906.1 awen|OJJ38217.1 azon|OJJ43077.1

Group2097: abra|OJJ69974.1 acar|OOF93479.1 acla|EAW08844.1 afis|EAW15556.1 afla|EED49061.1 afum|EAL85776.1 agla|OJJ85198.1 akaw|GAA87687.1 aluc|OJZ85307.1 anid|CBF78163.1 anig|EHA25503.1 aory|EIT82517.1 asyd|OJJ55964.1 ater|EAU32577.1 atub|OJI80264.1 aver|OJJ04910.1 awen|OJJ38214.1 azon|OJJ43081.1

Group2098: abra|OJJ69975.1 acar|OOF93481.1 acla|EAW08846.1 afis|EAW15554.1 afla|EED49057.1 afum|EAL85779.1 agla|OJJ85201.1 akaw|GAA87686.1 aluc|OJZ85306.1 anid|CBF78167.1 anig|EHA25502.1 aory|EIT82497.1 asyd|OJJ55968.1 ater|EAU32580.1 atub|OJI80265.1 aver|OJJ04915.1 awen|OJJ38210.1 azon|OJJ43084.1

Group2099: abra|OJJ69978.1 acar|OOF93484.1 acla|EAW08847.1 afis|EAW15549.1 afla|EED49055.1 afum|EAL85783.1 agla|OJJ85202.1 akaw|GAA87683.1 aluc|OJZ85302.1 anid|CBF78165.1 anig|EHA25499.1

aory|EIT82558.1 asyd|OJJ55969.1 ater|EAU32583.1 atub|OJI80268.1 aver|OJJ04916.1 awen|OJJ38209.1 azon|OJJ43086.1

Group2100: abra|OJJ69979.1 acar|OOF93483.1 acla|EAW08849.1 afis|EAW15547.1 afla|EED49054.1 afum|EAL85785.1 agla|OJJ85205.1 akaw|GAA87682.1 aluc|OJZ85301.1 anid|CBF78176.1 anig|EHA25498.1 aory|EIT82504.1 asyd|OJJ55970.1 ater|EAU32582.1 atub|OJI80269.1 aver|OJJ04917.1 awen|OJJ38208.1 azon|OJJ43087.1

Group2101: abra|OJJ69982.1 acar|OOF93486.1 acla|EAW08852.1 afis|EAW15544.1 afla|EED49051.1 afum|EAL85787.1 agla|OJJ85207.1 akaw|GAA87680.1 aluc|OJZ85298.1 anid|CBF78182.1 anig|EHA25496.1 aory|EIT82527.1 asyd|OJJ55973.1 ater|EAU32584.1 atub|OJI80272.1 aver|OJJ04921.1 awen|OJJ38205.1 azon|OJJ43090.1

Group2102: abra|OJJ69983.1 acar|OOF93487.1 acla|EAW08853.1 afis|EAW15543.1 afla|EED49050.1 afum|EAL85788.1 agla|OJJ85208.1 akaw|GAA87679.1 aluc|OJZ85297.1 anid|CBF78183.1 anig|EHA25495.1 aory|EIT82528.1 asyd|OJJ55974.1 ater|EAU32585.1 atub|OJI80273.1 aver|OJJ04922.1 awen|OJJ38204.1 azon|OJJ43091.1

Group2103: abra|OJJ69984.1 acar|OOF93489.1 acla|EAW08855.1 afis|EAW15541.1 afla|EED49049.1 afum|EAL85789.1 agla|OJJ85209.1 akaw|GAA87677.1 aluc|OJZ85295.1 anid|CBF78187.1 anig|EHA25493.1 aory|EIT82548.1 asyd|OJJ55976.1 ater|EAU32586.1 atub|OJI80275.1 aver|OJJ04924.1 awen|OJJ38202.1 azon|OJJ43093.1

Group2104: abra|OJJ69985.1 acar|OOF93490.1 acla|EAW08856.1 afis|EAW15540.1 afla|EED49048.1 afum|EAL85790.1 agla|OJJ85210.1 akaw|GAA87676.1 aluc|OJZ85294.1 anid|CBF78189.1 anig|EHA25492.1 aory|EIT82530.1 asyd|OJJ55977.1 ater|EAU32587.1 atub|OJI80276.1 aver|OJJ04925.1 awen|OJJ38201.1 azon|OJJ43094.1

Group2105: abra|OJJ69987.1 acar|OOF93491.1 acla|EAW08858.1 afis|EAW15538.1 afla|EED49046.1 afum|EAL85792.1 agla|OJJ85211.1 akaw|GAA87674.1 aluc|OJZ85292.1 anid|CBF78193.1 anig|EHA25490.1 aory|EIT82570.1 asyd|OJJ55979.1 ater|EAU32589.1 atub|OJI80278.1 aver|OJJ04927.1 awen|OJJ38200.1 azon|OJJ43095.1

Group2106: abra|OJJ69989.1 acar|OOF93494.1 acla|EAW08861.1 afis|EAW15533.1 afla|EED49044.1 afum|EAL85795.1 agla|OJJ85214.1 akaw|GAA87672.1 aluc|OJZ85290.1 anid|CBF78199.1 anig|EHA25488.1 aory|EIT82552.1 asyd|OJJ55983.1 ater|EAU32592.1 atub|OJI80280.1 aver|OJJ04930.1 awen|OJJ38197.1 azon|OJJ43096.1

Group2107: abra|OJJ69990.1 acar|OOF93496.1 acla|EAW08863.1 afis|EAW15531.1 afla|EED49042.1 afum|EAL85797.1 agla|OJJ85216.1 akaw|GAA87671.1 aluc|OJZ85289.1 anid|CBF78202.1 anig|EHA25486.1 aory|EIT82550.1 asyd|OJJ55985.1 ater|EAU32594.1 atub|OJI80281.1 aver|OJJ04932.1 awen|OJJ38195.1 azon|OJJ43099.1

Group2108: abra|OJJ69991.1 acar|OOF93497.1 acla|EAW08864.1 afis|EAW15530.1 afla|EED49041.1 afum|EAL85798.1 agla|OJJ85217.1 akaw|GAA87670.1 aluc|OJZ85288.1 anid|CBF78204.1 anig|EHA25485.1 aory|EIT82567.1 asyd|OJJ55986.1 ater|EAU32595.1 atub|OJI80282.1 aver|OJJ04933.1 awen|OJJ38194.1 azon|OJJ43100.1

Group2109: abra|OJJ69995.1 acar|OOF93502.1 acla|EAW08824.1 afis|EAW15573.1 afla|EED49082.1 afum|EAL85761.1 agla|OJJ85179.1 akaw|GAA87666.1 aluc|OJZ85284.1 anid|CBF78105.1 anig|EHA25481.1 aory|EIT82549.1 asyd|OJJ55895.1 ater|EAU32558.1 atub|OJI80286.1 aver|OJJ04840.1 awen|OJJ38232.1 azon|OJJ43064.1

Group2110: abra|OJJ70001.1 acar|OOF93507.1 acla|EAW08818.1 afis|EAW15579.1 afla|EED49093.1 afum|EAL85755.1 agla|OJJ85173.1 akaw|GAA87660.1 aluc|OJZ85278.1 anid|CBF78100.1 anig|EHA25475.1 aory|EIT82531.1 asyd|OJJ55889.1 ater|EAU32551.1 atub|OJI80292.1 aver|OJJ04834.1 awen|OJJ38241.1 azon|OJJ43058.1

Group2111: abra|OJJ70003.1 acar|OOF93509.1 acla|EAW08816.1 afis|EAW15581.1 afla|EED49090.1 afum|EAL85753.2 agla|OJJ85170.1 akaw|GAA87657.1 aluc|OJZ85276.1 anid|CBF78096.1 anig|EHA25473.1 aory|EIT82539.1 asyd|OJJ55886.1 ater|EAU32549.1 atub|OJI80294.1 aver|OJJ04831.1 awen|OJJ38243.1 azon|OJJ43055.1

Group2112: abra|OJJ70008.1 acar|OOF93514.1 acla|EAW08815.1 afis|EAW15582.1 afla|EED49088.1 afum|EAL85752.1 agla|OJJ85169.1 akaw|GAA87654.1 aluc|OJZ85271.1 anid|CBF78090.1 anig|EHA25470.1 aory|EIT82507.1 asyd|OJJ55883.1 ater|EAU32547.1 atub|OJI80300.1 aver|OJJ04828.1 awen|OJJ38246.1 azon|OJJ43052.1

Group2113: abra|OJJ70011.1 acar|OOF93517.1 acla|EAW08813.1 afis|EAW15584.1 afla|EED54445.1 afum|EAL85748.2 agla|OJJ85162.1 akaw|GAA87651.1 aluc|OJZ85267.1 anid|CBF78060.1 anig|EHA25466.1 aory|EIT81147.1 ater|EAU32545.1 atub|OJI80303.1 aver|OJJ04813.1 awen|OJJ38249.1 azon|OJJ43030.1 anig|EHA25467.1

Group2114: abra|OJJ70013.1 acar|OOF93519.1 acla|EAW08811.1 afis|EAW15586.1 afla|EED54443.1 afum|EAL85746.1 agla|OJJ85166.1 akaw|GAA87649.1 aluc|OJZ85264.1 anid|CBF78063.1 anig|EHA25464.1 aory|EIT81154.1 asyd|OJJ55870.1 ater|EAU32543.1 atub|OJI80305.1 aver|OJJ04815.1 awen|OJJ38252.1 azon|OJJ43032.1

Group2115: abra|OJJ70014.1 acar|OOF93520.1 acla|EAW08809.1 afis|EAW15587.1 afla|EED50410.1 afum|EAL85745.1 agla|OJJ85161.1 akaw|GAA87648.1 aluc|OJZ85263.1 anid|CBF78069.1 anig|EHA25463.1 aory|EIT82131.1 asyd|OJJ55872.1 ater|EAU32541.1 atub|OJI80306.1 aver|OJJ04817.1 awen|OJJ38254.1 azon|OJJ43033.1

Group2116: abra|OJJ70015.1 acar|OOF93521.1 acla|EAW08808.1 afis|EAW15588.1 afla|EED50411.1 afum|EAL85744.1 agla|OJJ85160.1 akaw|GAA87646.1 aluc|OJZ85261.1 anid|CBF78071.1 anig|EHA25462.1 aory|EIT82015.1 asyd|OJJ55873.1 ater|EAU32539.1 atub|OJI80307.1 aver|OJJ04818.1 awen|OJJ38255.1 azon|OJJ43038.1

Group2117: abra|OJJ70016.1 acar|OOF93522.1 acla|EAW08807.1 afis|EAW15589.1 afla|EED50412.1 afum|EAL85743.1 agla|OJJ85159.1 akaw|GAA87645.1 aluc|OJZ85260.1 anid|CBF78072.1 anig|EHA25461.1 aory|EIT82097.1 asyd|OJJ55874.1 ater|EAU32538.1 atub|OJI80308.1 aver|OJJ04819.1 awen|OJJ38262.1 azon|OJJ43023.1

Group2118: abra|OJJ70018.1 acar|OOF93524.1 acla|EAW08805.1 afis|EAW15591.1 afla|EED50414.1 afum|EAL85741.1 agla|OJJ85157.1 akaw|GAA87643.1 aluc|OJZ85258.1 anid|CBF78076.1 anig|EHA25459.1 aory|EIT82095.1 asyd|OJJ55876.1 ater|EAU32537.1 atub|OJI80310.1 aver|OJJ04821.1 awen|OJJ38264.1 azon|OJJ43021.1

Group2119: abra|OJJ70026.1 acar|OOF93536.1 acla|EAW08802.1 afis|EAW15595.1 afla|EED50418.1 afum|EAL85736.2 agla|OJJ85151.1 akaw|GAA87634.1 aluc|OJZ85250.1 anid|CBF78087.1 anig|EHA25450.1 aory|EIT82070.1 asyd|OJJ55881.1 ater|EAU32532.1 atub|OJI80319.1 aver|OJJ04826.1 awen|OJJ38271.1 azon|OJJ43910.1

Group2120: abra|OJJ70042.1 acar|OOF93537.1 acla|EAW08803.1 afis|EAW15594.1 afla|EED50417.1 afum|EAL85737.2 agla|OJJ85153.1 akaw|GAA87633.1 aluc|OJZ85249.1 anid|CBF78085.1 anig|EHA25449.1 aory|EIT82162.1 asyd|OJJ55880.1 ater|EAU32533.1 atub|OJI80320.1 aver|OJJ04825.1 awen|OJJ38270.1 azon|OJJ43912.1

Group2121: abra|OJJ70083.1 acla|EAW11426.1 afis|EAW20385.1 afla|EED45578.1 afum|EAL85496.1 agla|OJJ83253.1 akaw|GAA87604.1 aluc|OJZ85224.1 anid|CBF86057.1 anig|EHA25428.1 aory|EIT75627.1 asyd|OJJ54466.1 ater|EAU32965.1 atub|OJI80357.1 aver|OJJ07783.1 awen|OJJ30093.1 afis|EAW16167.1 aory|EIT75578.1

Group2122: abra|OJJ70153.1 acar|OOF93582.1 afis|EAW17230.1 agla|OJJ86975.1 akaw|GAA84496.1 aluc|OJZ89221.1 anig|EHA23341.1 asyd|OJJ53738.1 ater|EAU34915.1 atub|OJI85571.1 aver|OJI98051.1 afis|EAW19779.1 afla|EED45371.1 afum|EAL85119.2 anid|CBF71521.1 aory|EIT82393.1 asyd|OJJ62940.1 afis|EAW20309.1

Group2123: abra|OJJ70159.1 acar|OOF93314.1 acla|EAW06525.1 afis|EAW16241.1 afla|EED57104.1 afum|EAL88863.1 agla|OJJ88977.1 akaw|GAA86606.1 aluc|OJZ85156.1 anid|CBF78253.1 anig|EHA25372.1 aory|EIT73977.1 asyd|OJJ56050.1 ater|EAU34336.1 atub|OJI80433.1 aver|OJJ04993.1 awen|OJJ36019.1 azon|OJJ42535.1

Group2124: abra|OJJ70295.1 acar|OOF91749.1 acla|EAW10279.1 afis|EAW25315.1 afla|EED49708.1 afum|EAL89976.1 agla|OJJ88748.1 aluc|OJZ82560.1 anig|EHA22521.1 aory|EIT77651.1 asyd|OJJ62816.1 ater|EAU32274.1 atub|OJI88802.1 aver|OJI96860.1 awen|OJJ40751.1 azon|OJJ44610.1 awen|OJJ41048.1 agla|OJJ80540.1

Group2125: abra|OJJ70321.1 acar|OOF91776.1 acla|EAW12313.1 afis|EAW16939.1 afla|EED45249.1 afum|EAL86466.1 agla|OJJ79973.1 akaw|GAA92577.1 aluc|OJZ82590.1 anid|CBF75409.1 anig|EHA27485.1 aory|EIT73687.1 asyd|OJJ59815.1 ater|EAU30208.1 atub|OJI88825.1 aver|OJJ02473.1 awen|OJJ29713.1 azon|OJJ45106.1

Group2126: abra|OJJ70342.1 acar|OOF91814.1 abra|OJJ74452.1 acla|EAW12256.1 afla|EED48966.1 afum|EAL86656.1 agla|OJJ87924.1 akaw|GAA87346.1 aluc|OJZ89668.1 anig|EHA26906.1 aory|EIT79988.1 asyd|OJJ52239.1 atub|OJI85132.1 awen|OJJ31495.1 afla|EED48965.1 anig|EHA26905.1 aver|OJJ02177.1 aory|EIT83308.1

Group2127: abra|OJJ70367.1 acar|OOF91904.1 acla|EAW12332.1 afis|EAW16890.1 afla|EED53384.1 afum|EAL86428.2 agla|OJJ79944.1 akaw|GAA91984.1 aluc|OJZ82629.1 anid|CBF80798.1 anig|EHA27448.1 aory|EIT75200.1 asyd|OJJ54488.1 ater|EAU30224.1 atub|OJI88862.1 aver|OJJ07691.1 awen|OJJ29748.1 azon|OJJ50203.1

Group2128: abra|OJJ70389.1 acar|OOF90063.1 acla|EAW12390.1 afis|EAW16837.1 afla|EED47319.1 afum|EAL86385.1 agla|OJJ83707.1 akaw|GAA91969.1 aluc|OJZ82647.1 anid|CBF80639.1 anig|EHA27425.1

aory|EIT82834.1 asyd|OJJ54621.1 ater|EAU30244.1 atub|OJI88879.1 aver|OJJ07537.1 awen|OJJ33871.1 azon|OJJ43960.1

Group2129: abra|OJJ70449.1 acar|OOF91831.1 acla|EAW09124.1 afis|EAW24708.1 afla|EED52336.1 afum|EAL89350.1 agla|OJJ82814.1 akaw|GAA90132.1 aluc|OJZ83215.1 anid|CBF83955.1 anig|EHA21075.1 aory|EIT74337.1 asyd|OJJ61431.1 ater|EAU37947.1 atub|OJI87721.1 aver|OJJ00766.1 awen|OJJ39949.1 azon|OJJ51122.1

Group2130: abra|OJJ70463.1 acar|OOF97461.1 acla|EAW13050.1 afis|EAW17617.1 afla|EED49208.1 afum|EAL91430.1 agla|OJJ81323.1 akaw|GAA83312.1 aluc|OJZ83806.1 anid|CBF90167.1 anig|EHA18953.1 aory|EIT83163.1 asyd|OJJ61992.1 ater|EAU37051.1 atub|OJI90173.1 aver|OJI96084.1 awen|OJJ37228.1 azon|OJJ50931.1

Group2131: abra|OJJ70464.1 acar|OOF97459.1 acla|EAW13049.1 afis|EAW17618.1 afla|EED49209.1 afum|EAL91431.1 agla|OJJ81322.1 akaw|GAA83313.1 aluc|OJZ83807.1 anid|CBF90165.1 anig|EHA18954.1 aory|EIT83149.1 asyd|OJJ61993.1 ater|EAU37052.1 atub|OJI90174.1 aver|OJI96085.1 awen|OJJ37227.1 azon|OJJ50930.1

Group2132: abra|OJJ70468.1 acar|OOF97456.1 acla|EAW13046.1 afis|EAW17621.1 afla|EED49196.1 afum|EAL91434.1 agla|OJJ81304.1 akaw|GAA83317.1 aluc|OJZ83812.1 anid|CBF90159.1 anig|EHA18957.1 aory|EIT78509.1 asyd|OJJ61996.1 ater|EAU37055.1 atub|OJI90179.1 aver|OJI96088.1 awen|OJJ37223.1 azon|OJJ50926.1

Group2133: abra|OJJ70469.1 acar|OOF97455.1 acla|EAW13045.1 afis|EAW17622.1 afla|EED49195.1 afum|EAL91435.1 agla|OJJ81303.1 akaw|GAA83318.1 aluc|OJZ83813.1 anid|CBF90157.1 anig|EHA18958.1 aory|EIT78576.1 asyd|OJJ61997.1 ater|EAU37056.1 atub|OJI90180.1 aver|OJI96089.1 awen|OJJ37222.1 azon|OJJ50925.1

Group2134: abra|OJJ70470.1 acar|OOF97453.1 acla|EAW13044.1 afis|EAW17623.1 afla|EED49194.1 afum|EAL91437.1 agla|OJJ81302.1 akaw|GAA83319.1 aluc|OJZ83814.1 anid|CBF90156.1 anig|EHA18959.1 aory|EIT78552.1 asyd|OJJ61998.1 ater|EAU37057.1 atub|OJI90181.1 aver|OJI96090.1 awen|OJJ37221.1 azon|OJJ50924.1

Group2135: abra|OJJ70473.1 acar|OOF97452.1 acla|EAW13041.1 afis|EAW17626.1 afla|EED49192.1 afum|EAL91440.1 agla|OJJ81276.1 akaw|GAA83322.1 aluc|OJZ83817.1 anid|CBF90150.1 anig|EHA18962.1 aory|EIT78505.1 asyd|OJJ62001.1 ater|EAU37060.1 atub|OJI90184.1 aver|OJI96093.1 awen|OJJ37218.1 azon|OJJ50921.1

Group2136: abra|OJJ70475.1 acar|OOF97450.1 acla|EAW13039.1 afis|EAW17628.1 afla|EED49190.1 afum|EAL91442.1 agla|OJJ81275.1 akaw|GAA83324.1 aluc|OJZ83819.1 anid|CBF90147.1 anig|EHA18964.1 aory|EIT78550.1 asyd|OJJ62002.1 ater|EAU37062.1 atub|OJI90186.1 aver|OJI96094.1 awen|OJJ37216.1 azon|OJJ50919.1

Group2137: abra|OJJ70476.1 acar|OOF97449.1 acla|EAW13038.1 afis|EAW17629.1 afla|EED49189.1 afum|EAL91443.1 agla|OJJ81272.1 akaw|GAA83325.1 aluc|OJZ83820.1 anid|CBF90141.1 anig|EHA18965.1 aory|EIT78529.1 asyd|OJJ62006.1 ater|EAU37063.1 atub|OJI90187.1 aver|OJI96098.1 awen|OJJ37215.1 azon|OJJ50916.1

Group2138: abra|OJJ70477.1 acar|OOF97448.1 acla|EAW13037.1 afis|EAW17630.1 afla|EED49188.1 afum|EAL91445.1 agla|OJJ81271.1 akaw|GAA83326.1 aluc|OJZ83822.1 anid|CBF90143.1 anig|EHA18966.1 aory|EIT78536.1 asyd|OJJ62005.1 ater|EAU37065.1 atub|OJI90189.1 aver|OJI96096.1 awen|OJJ37214.1 azon|OJJ50914.1

Group2139: abra|OJJ70478.1 acar|OOF97447.1 acla|EAW13036.1 afis|EAW17631.1 afla|EED49187.1 afum|EAL91446.1 agla|OJJ81270.1 akaw|GAA83327.1 aluc|OJZ83823.1 anid|CBF90145.1 anig|EHA18967.1 aory|EIT78534.1 asyd|OJJ62004.1 ater|EAU37066.1 atub|OJI90190.1 aver|OJI96095.1 awen|OJJ37213.1 azon|OJJ50913.1

Group2140: abra|OJJ70481.1 acar|OOF97444.1 acla|EAW13035.1 afis|EAW17632.1 afla|EED49186.1 afum|EAL91447.1 agla|OJJ81269.1 akaw|GAA83329.1 aluc|OJZ83825.1 anid|CBF90139.1 anig|EHA18969.1 aory|EIT78523.1 asyd|OJJ62011.1 ater|EAU37067.1 atub|OJI90192.1 aver|OJI96103.1 awen|OJJ37212.1 azon|OJJ50912.1

Group2141: abra|OJJ70482.1 acar|OOF97443.1 acla|EAW13034.1 afis|EAW17633.1 afla|EED50531.1 afum|EAL91448.1 agla|OJJ81266.1 akaw|GAA83330.1 aluc|OJZ83826.1 anid|CBF90137.1 anig|EHA18970.1 aory|EIT82093.1 asyd|OJJ62012.1 ater|EAU37069.1 atub|OJI90193.1 aver|OJI96104.1 awen|OJJ37211.1 azon|OJJ50911.1

Group2142: abra|OJJ70483.1 acar|OOF97442.1 acla|EAW13033.1 afis|EAW17634.1 afla|EED50530.1 afum|EAL91449.1 agla|OJJ81265.1 akaw|GAA83331.1 aluc|OJZ83827.1 anid|CBF90135.1 anig|EHA18971.1 aory|EIT82030.1 asyd|OJJ62013.1 ater|EAU37070.1 atub|OJI90194.1 aver|OJI96105.1 awen|OJJ37210.1 azon|OJJ50910.1

Group2143: abra|OJJ70484.1 acar|OOF97441.1 acla|EAW13031.1 afis|EAW17636.1 afla|EED50528.1 afum|EAL91451.2 agla|OJJ81263.1 akaw|GAA83333.1 aluc|OJZ83829.1 anid|CBF90132.1 anig|EHA18972.1 aory|EIT82038.1 asyd|OJJ62015.1 ater|EAU37072.1 atub|OJI90196.1 aver|OJI96106.1 awen|OJJ37208.1 azon|OJJ50908.1

Group2144: abra|OJJ70488.1 acar|OOF97438.1 acla|EAW13028.1 afis|EAW17639.1 afla|EED50524.1 afum|EAL91454.1 agla|OJJ79567.1 akaw|GAA83336.1 aluc|OJZ83832.1 anid|CBF90125.1 anig|EHA18975.1 aory|EIT82049.1 asyd|OJJ62018.1 ater|EAU37075.1 atub|OJI90199.1 aver|OJI96109.1 awen|OJJ37206.1 azon|OJJ50905.1

Group2145: abra|OJJ70489.1 acar|OOF97437.1 acla|EAW13027.1 afis|EAW17640.1 afla|EED50523.1 afum|EAL91455.1 agla|OJJ79568.1 akaw|GAA83337.1 aluc|OJZ83833.1 anig|EHA18976.1 aory|EIT82102.1 asyd|OJJ62019.1 ater|EAU37076.1 atub|OJI90200.1 aver|OJI96110.1 awen|OJJ37205.1 azon|OJJ50904.1 anid|CBF90123.1

Group2146: abra|OJJ70490.1 acar|OOF97436.1 acla|EAW13026.1 afis|EAW17641.1 afla|EED50522.1 afum|EAL91456.1 agla|OJJ79569.1 akaw|GAA83338.1 aluc|OJZ83834.1 anid|CBF90121.1 anig|EHA18977.1 aory|EIT82016.1 asyd|OJJ62020.1 ater|EAU37077.1 atub|OJI90201.1 aver|OJI96111.1 awen|OJJ37204.1 azon|OJJ50903.1

Group2147: abra|OJJ70492.1 acar|OOF97435.1 acla|EAW13025.1 afis|EAW17642.1 afla|EED50518.1 afum|EAL91457.1 agla|OJJ79562.1 akaw|GAA83340.1 aluc|OJZ83836.1 anid|CBF90119.1 anig|EHA18979.1 aory|EIT82059.1 asyd|OJJ62021.1 ater|EAU37078.1 atub|OJI90203.1 aver|OJI96112.1 awen|OJJ37203.1 azon|OJJ50902.1

Group2148: abra|OJJ70493.1 acar|OOF97432.1 acla|EAW13024.1 afis|EAW17643.1 afla|EED50517.1 afum|EAL91458.1 agla|OJJ79561.1 akaw|GAA83341.1 aluc|OJZ83837.1 anid|CBF90118.1 anig|EHA18980.1 aory|EIT82149.1 asyd|OJJ62022.1 ater|EAU37079.1 atub|OJI90204.1 aver|OJI96113.1 awen|OJJ37202.1 azon|OJJ50901.1

Group2149: abra|OJJ70495.1 acar|OOF97431.1 acla|EAW13022.1 afis|EAW17645.1 afla|EED50512.1 afum|EAL91460.1 agla|OJJ79559.1 akaw|GAA83343.1 aluc|OJZ83840.1 anid|CBF90114.1 anig|EHA18982.1 aory|EIT82064.1 asyd|OJJ62024.1 ater|EAU37081.1 atub|OJI90206.1 aver|OJI96115.1 awen|OJJ37200.1 azon|OJJ50899.1

Group2150: abra|OJJ70498.1 acar|OOF97428.1 acla|EAW08176.1 afis|EAW23212.1 afla|EED47039.1 afum|EAL88272.1 agla|OJJ78733.1 akaw|GAA83346.1 aluc|OJZ83843.1 anid|CBF74740.1 anig|EHA18985.1 aory|EIT72595.1 asyd|OJJ60184.1 ater|EAU36340.1 atub|OJI90209.1 aver|OJJ04080.1 awen|OJJ39445.1 azon|OJJ47647.1

Group2151: abra|OJJ70499.1 acar|OOF97427.1 acla|EAW08175.1 afis|EAW23211.1 afla|EED47040.1 afum|EAL88273.1 agla|OJJ78734.1 akaw|GAA83347.1 aluc|OJZ83844.1 anid|CBF74738.1 anig|EHA18986.1 aory|EIT72653.1 asyd|OJJ60185.1 ater|EAU36339.1 atub|OJI90210.1 aver|OJJ04079.1 awen|OJJ39444.1 azon|OJJ47646.1

Group2152: abra|OJJ70500.1 acar|OOF97425.1 acla|EAW08174.1 afis|EAW23210.1 afla|EED47041.1 afum|EAL88274.1 agla|OJJ78735.1 akaw|GAA83348.1 aluc|OJZ83845.1 anid|CBF74736.1 anig|EHA18987.1 aory|EIT72592.1 asyd|OJJ60186.1 ater|EAU36338.1 atub|OJI90211.1 aver|OJJ04078.1 awen|OJJ39443.1 azon|OJJ47645.1

Group2153: abra|OJJ70504.1 acar|OOF97421.1 acla|EAW08170.1 afis|EAW23206.1 afla|EED47047.1 afum|EAL88277.1 agla|OJJ78739.1 akaw|GAA83353.1 aluc|OJZ83849.1 anid|CBF74734.1 anig|EHA18991.1 aory|EIT72747.1 asyd|OJJ60189.1 ater|EAU36334.1 atub|OJI90215.1 aver|OJJ04075.1 awen|OJJ39438.1 azon|OJJ47641.1

Group2154: abra|OJJ70509.1 acar|OOF97420.1 acla|EAW08166.1 afis|EAW23204.1 afla|EED47050.1 afum|EAL88279.1 agla|OJJ78743.1 akaw|GAA83356.1 aluc|OJZ83854.1 anid|CBF74730.1 anig|EHA18995.1 aory|EIT72509.1 asyd|OJJ60194.1 ater|EAU36332.1 atub|OJI90221.1 aver|OJJ04071.1 awen|OJJ39434.1 azon|OJJ47638.1

Group2155: abra|OJJ70510.1 acar|OOF97419.1 acla|EAW08165.1 afis|EAW23203.1 afla|EED47051.1 afum|EAL88280.1 agla|OJJ78745.1 akaw|GAA83357.1 aluc|OJZ83855.1 anid|CBF74728.1 anig|EHA18996.1 aory|EIT72511.1 asyd|OJJ60195.1 ater|EAU36331.1 atub|OJI90222.1 aver|OJJ04070.1 awen|OJJ39433.1 azon|OJJ47637.1

Group2156: abra|OJJ70512.1 acar|OOF97417.1 acla|EAW08163.1 afis|EAW23201.1 afla|EED47053.1 afum|EAL88282.1 agla|OJJ78747.1 akaw|GAA83359.1 aluc|OJZ83857.1 anid|CBF74721.1 anig|EHA18998.1 aory|EIT72498.1 asyd|OJJ60201.1 ater|EAU36329.1 atub|OJI90224.1 aver|OJJ04065.1 awen|OJJ39431.1 azon|OJJ47635.1

Group2157: abra|OJJ70514.1 acar|OOF97415.1 acla|EAW08161.1 afis|EAW23199.1 afla|EED47055.1 afum|EAL88284.1 agla|OJJ78749.1 akaw|GAA83361.1 aluc|OJZ83859.1 anid|CBF74717.1 anig|EHA19000.1

aory|EIT72508.1 asyd|OJJ60203.1 ater|EAU36327.1 atub|OJI90226.1 aver|OJJ04063.1 awen|OJJ39429.1 azon|OJJ47633.1

Group2158: abra|OJJ70515.1 acar|OOF97414.1 acla|EAW08160.1 afis|EAW23198.1 afla|EED47056.1 afum|EAL88285.1 agla|OJJ86331.1 akaw|GAA83362.1 aluc|OJZ83860.1 anid|CBF74715.1 anig|EHA19001.1 aory|EIT72476.1 asyd|OJJ60205.1 ater|EAU36325.1 atub|OJI90228.1 aver|OJJ04062.1 awen|OJJ39428.1 azon|OJJ47632.1

Group2159: abra|OJJ70521.1 acar|OOF97407.1 acla|EAW08154.1 afis|EAW23185.1 afla|EED47062.1 afum|EAL88298.1 agla|OJJ86360.1 akaw|GAA83368.1 aluc|OJZ83866.1 anid|CBF74698.1 anig|EHA19007.1 aory|EIT72505.1 asyd|OJJ60215.1 ater|EAU36320.1 atub|OJI90234.1 aver|OJJ04053.1 awen|OJJ36587.1 azon|OJJ47619.1

Group2160: abra|OJJ70522.1 acar|OOF97406.1 acla|EAW08153.1 afis|EAW23186.1 afla|EED47064.1 afum|EAL88297.1 agla|OJJ86361.1 akaw|GAA83369.1 aluc|OJZ83867.1 anid|CBF74696.1 anig|EHA19008.1 aory|EIT72495.1 asyd|OJJ60214.1 ater|EAU36319.1 atub|OJI90235.1 aver|OJJ04054.1 awen|OJJ36588.1 azon|OJJ47618.1

Group2161: abra|OJJ70529.1 acar|OOF97402.1 acla|EAW08142.1 afis|EAW23178.1 afla|EED49000.1 afum|EAL88305.1 agla|OJJ89690.1 akaw|GAA83374.1 aluc|OJZ83872.1 anid|CBF74512.1 anig|EHA19013.1 aory|EIT80028.1 asyd|OJJ57730.1 ater|EAU36311.1 atub|OJI90240.1 aver|OJJ05439.1 awen|OJJ36601.1 azon|OJJ47607.1

Group2162: abra|OJJ70532.1 acar|OOF97400.1 acla|EAW08145.1 afis|EAW23181.1 afla|EED47073.1 afum|EAL88302.1 akaw|GAA83377.1 aluc|OJZ83875.1 anid|CBF74516.1 anig|EHA19015.1 aory|EIT72487.1 asyd|OJJ57732.1 ater|EAU36314.1 atub|OJI90243.1 aver|OJJ05437.1 awen|OJJ36597.1 azon|OJJ47611.1 agla|OJJ86370.1

Group2163: abra|OJJ70536.1 acar|OOF97397.1 acla|EAW08148.1 afis|EAW23191.1 afla|EED47070.1 afum|EAL88292.1 agla|OJJ86367.1 akaw|GAA83381.1 aluc|OJZ83880.1 anid|CBF74521.1 anig|EHA19018.1 aory|EIT72473.1 asyd|OJJ57733.1 ater|EAU36315.1 atub|OJI90247.1 aver|OJJ05434.1 awen|OJJ36593.1 azon|OJJ47612.1

Group2164: abra|OJJ70537.1 acar|OOF97396.1 acla|EAW08149.1 afis|EAW23190.1 afla|EED47069.1 afum|EAL88293.1 agla|OJJ86365.1 akaw|GAA83382.1 aluc|OJZ83881.1 anid|CBF74523.1 anig|EHA19019.1 aory|EIT72486.1 asyd|OJJ57734.1 ater|EAU36309.1 atub|OJI90249.1 aver|OJJ05433.1 awen|OJJ36592.1 azon|OJJ47614.1

Group2165: abra|OJJ70539.1 acar|OOF97394.1 acla|EAW08151.1 afis|EAW23188.1 afla|EED47067.1 afum|EAL88295.1 agla|OJJ86363.1 akaw|GAA83384.1 aluc|OJZ83884.1 anid|CBF74527.1 anig|EHA19021.1 aory|EIT72500.1 asyd|OJJ57736.1 ater|EAU36310.1 atub|OJI90251.1 aver|OJJ05431.1 awen|OJJ36590.1 azon|OJJ47616.1

Group2166: abra|OJJ70541.1 acar|OOF97392.1 acla|EAW08137.1 afis|EAW23173.1 afla|EED46946.1 afum|EAL88310.1 agla|OJJ89685.1 akaw|GAA83387.1 aluc|OJZ83886.1 anid|CBF74495.1 anig|EHA19023.1 aory|EIT72576.1 asyd|OJJ57718.1 ater|EAU36302.1 atub|OJI90253.1 aver|OJJ05449.1 awen|OJJ36604.1 azon|OJJ47602.1

Group2167: abra|OJJ70552.1 acar|OOF97168.1 acla|EAW08344.1 afis|EAW23379.1 afla|EED55578.1 afum|EAL88092.1 agla|OJJ86444.1 akaw|GAA83396.1 aluc|OJZ83895.1 anid|CBF74849.1 anig|EHA19031.1 aory|EIT82685.1 asyd|OJJ60128.1 ater|EAU35633.1 atub|OJI90263.1 aver|OJJ04133.1 awen|OJJ39503.1 azon|OJJ47704.1

Group2168: abra|OJJ70554.1 acar|OOF97165.1 acla|EAW08350.1 afis|EAW23383.1 afla|EED55571.1 afum|EAL88088.1 agla|OJJ86437.1 akaw|GAA83399.1 aluc|OJZ83897.1 anid|CBF74854.1 anig|EHA19034.1 aory|EIT82660.1 asyd|OJJ60126.1 ater|EAU35636.1 atub|OJI90267.1 aver|OJJ04136.1 awen|OJJ39498.1 azon|OJJ47700.1

Group2169: abra|OJJ70555.1 acar|OOF97164.1 acla|EAW08353.1 afis|EAW23384.1 afla|EED55572.1 afum|EAL88087.1 agla|OJJ86436.1 akaw|GAA83400.1 aluc|OJZ83898.1 anid|CBF74856.1 anig|EHA19035.1 aory|EIT82663.1 asyd|OJJ60125.1 ater|EAU35637.1 atub|OJI90268.1 aver|OJJ04137.1 awen|OJJ39497.1 azon|OJJ47699.1

Group2170: abra|OJJ70556.1 acar|OOF97163.1 acla|EAW08354.1 afis|EAW23385.1 afla|EED55573.1 afum|EAL88086.2 agla|OJJ86435.1 akaw|GAA83401.1 aluc|OJZ83899.1 anid|CBF74875.1 anig|EHA19036.1 aory|EIT82690.1 asyd|OJJ60114.1 ater|EAU35638.1 atub|OJI90269.1 aver|OJJ04148.1 awen|OJJ39496.1 azon|OJJ47698.1

Group2171: abra|OJJ70557.1 acar|OOF97162.1 acla|EAW08355.1 afis|EAW23386.1 afla|EED55574.1 afum|EAL88085.1 agla|OJJ86432.1 akaw|GAA83402.1 aluc|OJZ83900.1 anid|CBF74873.1 anig|EHA19037.1 aory|EIT82673.1 asyd|OJJ60115.1 ater|EAU35639.1 atub|OJI90270.1 aver|OJJ04147.1 awen|OJJ39495.1 azon|OJJ47696.1

Group2172: abra|OJJ70559.1 acar|OOF97161.1 acla|EAW08357.1 afis|EAW23388.1 afla|EED55576.1 afum|EAL88083.1 agla|OJJ86430.1 akaw|GAA83404.1 aluc|OJZ83903.1 anid|CBF74870.1 anig|EHA19038.1 aory|EIT82706.1 asyd|OJJ60117.1 ater|EAU35640.1 atub|OJI90272.1 aver|OJJ04145.1 awen|OJJ39493.1 azon|OJJ47694.1

Group2173: abra|OJJ70560.1 acar|OOF97160.1 acla|EAW08358.1 afis|EAW23389.1 afla|EED55568.1 afum|EAL88082.1 agla|OJJ86428.1 akaw|GAA83405.1 aluc|OJZ83904.1 anid|CBF74868.1 anig|EHA19039.1 aory|EIT82664.1 asyd|OJJ60118.1 ater|EAU35642.1 atub|OJI90273.1 aver|OJJ04144.1 awen|OJJ39492.1 azon|OJJ47693.1

Group2174: abra|OJJ70561.1 acar|OOF97158.1 acla|EAW08360.1 afis|EAW23391.1 afla|EED55566.1 afum|EAL88080.1 agla|OJJ86426.1 akaw|GAA83406.1 aluc|OJZ83905.1 anid|CBF74864.1 anig|EHA19040.1 aory|EIT82703.1 asyd|OJJ60121.1 ater|EAU35644.1 atub|OJI90276.1 aver|OJJ04141.1 awen|OJJ39488.1 azon|OJJ47691.1

Group2175: abra|OJJ70564.1 acar|OOF97156.1 acla|EAW08362.1 afis|EAW23393.1 afla|EED55564.1 afum|EAL88077.1 agla|OJJ86420.1 akaw|GAA83408.1 aluc|OJZ83907.1 anid|CBF74860.1 anig|EHA19042.1 aory|EIT82692.1 asyd|OJJ60123.1 ater|EAU35646.1 atub|OJI90278.1 aver|OJJ04139.1 awen|OJJ39486.1 azon|OJJ47689.1

Group2176: abra|OJJ70567.1 acar|OOF97155.1 acla|EAW08366.1 afis|EAW23396.1 afla|EED55560.1 afum|EAL88074.1 agla|OJJ86410.1 akaw|GAA83411.1 aluc|OJZ83910.1 anid|CBF74807.1 anig|EHA19044.1 aory|EIT82680.1 asyd|OJJ60148.1 ater|EAU35649.1 atub|OJI90281.1 aver|OJJ04115.1 awen|OJJ39483.1 azon|OJJ48236.1

Group2177: abra|OJJ70568.1 acar|OOF97154.1 acla|EAW08367.1 afis|EAW23397.1 afla|EED55559.1 afum|EAL88073.1 agla|OJJ86409.1 akaw|GAA83412.1 aluc|OJZ83911.1 anid|CBF74805.1 anig|EHA19045.1 aory|EIT82705.1 asyd|OJJ60149.1 ater|EAU35650.1 atub|OJI90282.1 aver|OJJ04114.1 awen|OJJ39482.1 azon|OJJ47684.1

Group2178: abra|OJJ70569.1 acar|OOF97152.1 acla|EAW08371.1 afis|EAW23398.1 afla|EED55554.1 afum|EAL88072.1 agla|OJJ86403.1 akaw|GAA83413.1 aluc|OJZ83913.1 anid|CBF74803.1 anig|EHA19046.1 aory|EIT82712.1 asyd|OJJ60150.1 ater|EAU35651.1 atub|OJI90283.1 aver|OJJ04113.1 awen|OJJ39481.1 azon|OJJ47683.1

Group2179: abra|OJJ70570.1 acar|OOF97151.1 acla|EAW08370.1 afis|EAW23399.1 afla|EED47005.1 afum|EAL88071.1 agla|OJJ86402.1 akaw|GAA83414.1 aluc|OJZ83914.1 anid|CBF74801.1 anig|EHA19047.1 aory|EIT72537.1 asyd|OJJ60152.1 ater|EAU36378.1 atub|OJI90284.1 aver|OJJ04112.1 awen|OJJ39480.1 azon|OJJ47682.1

Group2180: abra|OJJ70571.1 acar|OOF97150.1 acla|EAW08369.1 afis|EAW23400.1 afla|EED47006.1 afum|EAL88070.1 agla|OJJ86401.1 akaw|GAA83415.1 aluc|OJZ83915.1 anid|CBF73601.1 anig|EHA19048.1 aory|EIT72638.1 asyd|OJJ56680.1 ater|EAU36377.1 atub|OJI90285.1 aver|OJJ06043.1 awen|OJJ39479.1 azon|OJJ47681.1

Group2181: abra|OJJ70574.1 acar|OOF97147.1 acla|EAW08373.1 afis|EAW23402.1 afla|EED47009.1 afum|EAL88068.1 agla|OJJ86389.1 akaw|GAA83418.1 aluc|OJZ83918.1 anid|CBF74795.1 anig|EHA19051.1 aory|EIT72635.1 asyd|OJJ60155.1 ater|EAU36373.1 atub|OJI90289.1 aver|OJJ04109.1 awen|OJJ39476.1 azon|OJJ47679.1

Group2182: abra|OJJ70576.1 acar|OOF97145.1 acla|EAW08375.1 afis|EAW23404.1 afla|EED47011.1 afum|EAL88066.1 agla|OJJ86392.1 akaw|GAA83420.1 aluc|OJZ83920.1 anid|CBF74793.1 anig|EHA19053.1 aory|EIT72673.1 asyd|OJJ60156.1 ater|EAU36371.1 atub|OJI90292.1 aver|OJJ04108.1 awen|OJJ39473.1 azon|OJJ47675.1

Group2183: abra|OJJ70577.1 acar|OOF97144.1 acla|EAW08203.1 afis|EAW23237.1 afla|EED47012.1 afum|EAL88245.1 agla|OJJ86393.1 akaw|GAA83421.1 aluc|OJZ83921.1 anid|CBF74787.1 anig|EHA19054.1 aory|EIT72690.1 asyd|OJJ60159.1 ater|EAU36370.1 atub|OJI90293.1 aver|OJJ04105.1 awen|OJJ39472.1 azon|OJJ47674.1

Group2184: abra|OJJ70578.1 acar|OOF97143.1 acla|EAW08202.1 afis|EAW23236.1 afla|EED47013.1 afum|EAL88246.1 agla|OJJ86394.1 akaw|GAA83422.1 aluc|OJZ83922.1 anid|CBF74785.1 anig|EHA19055.1 aory|EIT72768.1 asyd|OJJ60160.1 ater|EAU36369.1 atub|OJI90294.1 aver|OJJ04104.1 awen|OJJ39471.1 azon|OJJ47673.1

Group2185: abra|OJJ70580.1 acar|OOF97138.1 acla|EAW08200.1 afis|EAW23234.1 afla|EED47015.1 afum|EAL88248.1 agla|OJJ86396.1 akaw|GAA83424.1 aluc|OJZ83925.1 anid|CBF74783.1 anig|EHA19057.1 aory|EIT72553.1 asyd|OJJ60161.1 ater|EAU36367.1 atub|OJI90297.1 aver|OJJ04103.1 awen|OJJ39469.1 azon|OJJ47669.1

Group2186: abra|OJJ70582.1 acar|OOF97137.1 acla|EAW08198.1 afis|EAW23232.1 afla|EED47016.1 afum|EAL88250.1 agla|OJJ86398.1 akaw|GAA83426.1 aluc|OJZ83927.1 anid|CBF74779.1 anig|EHA19059.1

aory|EIT72651.1 asyd|OJJ60163.1 ater|EAU36361.1 atub|OJI90299.1 aver|OJJ04101.1 awen|OJJ39467.1 azon|OJJ47667.1

Group2187: abra|OJJ70584.1 acar|OOF97136.1 acla|EAW08197.1 afis|EAW23231.1 afla|EED47019.1 afum|EAL88252.1 agla|OJJ86399.1 akaw|GAA83427.1 aluc|OJZ83928.1 anid|CBF74777.1 anig|EHA19060.1 aory|EIT72752.1 asyd|OJJ60164.1 ater|EAU36360.1 atub|OJI90300.1 aver|OJJ04100.1 awen|OJJ39466.1 azon|OJJ47666.1

Group2188: abra|OJJ70585.1 acar|OOF97135.1 acla|EAW08196.1 afis|EAW23230.1 afla|EED47020.1 afum|EAL88253.1 agla|OJJ86400.1 akaw|GAA83428.1 aluc|OJZ83929.1 anid|CBF74775.1 anig|EHA19061.1 aory|EIT72543.1 asyd|OJJ60165.1 ater|EAU36359.1 atub|OJI90301.1 aver|OJJ04099.1 awen|OJJ39465.1 azon|OJJ47665.1

Group2189: abra|OJJ70589.1 acar|OOF97128.1 acla|EAW08192.1 afis|EAW23226.1 afla|EED47024.1 afum|EAL88257.1 agla|OJJ86383.1 akaw|GAA83433.1 aluc|OJZ83934.1 anid|CBF74767.1 anig|EHA19065.1 aory|EIT72765.1 asyd|OJJ60169.1 ater|EAU36356.1 atub|OJI90306.1 aver|OJJ04095.1 awen|OJJ39461.1 azon|OJJ47660.1

Group2190: abra|OJJ70592.1 acar|OOF97126.1 acla|EAW08188.1 afis|EAW23222.1 afla|EED47028.1 afum|EAL88262.1 agla|OJJ86379.1 akaw|GAA83437.1 aluc|OJZ83938.1 anid|CBF74759.1 anig|EHA19068.1 aory|EIT72692.1 asyd|OJJ60173.1 ater|EAU36353.1 atub|OJI90311.1 aver|OJJ04091.1 awen|OJJ39458.1 azon|OJJ47656.1

Group2191: abra|OJJ70594.1 acar|OOF97124.1 acla|EAW08186.1 afis|EAW23220.1 afla|EED47030.1 afum|EAL88264.1 agla|OJJ86377.1 akaw|GAA83439.1 aluc|OJZ83940.1 anid|CBF74757.1 anig|EHA19070.1 aory|EIT72550.1 asyd|OJJ60174.1 ater|EAU36351.1 atub|OJI90313.1 aver|OJJ04089.1 awen|OJJ39456.1 azon|OJJ47654.1

Group2192: abra|OJJ70595.1 acar|OOF97123.1 acla|EAW08185.1 afis|EAW23219.1 afla|EED47031.1 afum|EAL88265.1 agla|OJJ86376.1 akaw|GAA83440.1 aluc|OJZ83941.1 anid|CBF74755.1 anig|EHA19071.1 aory|EIT72573.1 asyd|OJJ60175.1 ater|EAU36350.1 atub|OJI90314.1 aver|OJJ04088.1 awen|OJJ39455.1 azon|OJJ47653.1

Group2193: abra|OJJ70600.1 acar|OOF97120.1 acla|EAW08182.1 afis|EAW23217.1 afla|EED47035.1 afum|EAL88267.1 agla|OJJ86373.1 akaw|GAA89940.1 aluc|OJZ83945.1 anid|CBF74749.1 anig|EHA19074.1 aory|EIT72655.1 asyd|OJJ60178.1 ater|EAU36347.1 atub|OJI90316.1 aver|OJJ04084.1 awen|OJJ39451.1 azon|OJJ47651.1

Group2194: abra|OJJ70606.1 acar|OOF97115.1 acla|EAW13020.1 afis|EAW17647.1 afla|EED50511.1 afum|EAL91462.1 agla|OJJ79557.1 akaw|GAA89946.1 aluc|OJZ83952.1 anid|CBF90111.1 anig|EHA19080.1 aory|EIT82166.1 asyd|OJJ62026.1 ater|EAU37083.1 atub|OJI90323.1 aver|OJI96117.1 awen|OJJ37198.1 azon|OJJ50897.1

Group2195: abra|OJJ70607.1 acar|OOF97114.1 acla|EAW13019.1 afis|EAW17648.1 afla|EED50510.1 afum|EAL91464.1 agla|OJJ79556.1 akaw|GAA89947.1 aluc|OJZ83954.1 anid|CBF90087.1 anig|EHA19081.1 aory|EIT82125.1 asyd|OJJ62036.1 ater|EAU37084.1 atub|OJI90324.1 aver|OJI96142.1 awen|OJJ37197.1 azon|OJJ50896.1

Group2196: abra|OJJ70608.1 acar|OOF97113.1 acla|EAW13018.1 afis|EAW17649.1 afla|EED50509.1 afum|EAL91465.1 agla|OJJ79555.1 akaw|GAA89948.1 aluc|OJZ83955.1 anid|CBF90085.1 anig|EHA19082.1 aory|EIT82043.1 asyd|OJJ62037.1 ater|EAU37085.1 atub|OJI90325.1 aver|OJI96143.1 awen|OJJ37196.1 azon|OJJ50895.1

Group2197: abra|OJJ70611.1 acar|OOF97111.1 acla|EAW13016.1 afis|EAW17651.1 afla|EED50508.1 afum|EAL91466.1 agla|OJJ79553.1 akaw|GAA89950.1 aluc|OJZ83957.1 anid|CBF90077.1 anig|EHA19084.1 aory|EIT82060.1 asyd|OJJ62040.1 ater|EAU37087.1 atub|OJI90327.1 aver|OJI96146.1 awen|OJJ37194.1 azon|OJJ50893.1

Group2198: abra|OJJ70613.1 acar|OOF97109.1 acla|EAW13014.1 afis|EAW17653.1 afla|EED50505.1 afum|EAL91468.1 agla|OJJ79551.1 akaw|GAA89952.1 aluc|OJZ83959.1 anid|CBF90074.1 anig|EHA19087.1 aory|EIT82010.1 asyd|OJJ62042.1 ater|EAU37089.1 atub|OJI90331.1 aver|OJI96148.1 awen|OJJ37192.1 azon|OJJ50891.1

Group2199: abra|OJJ70616.1 acar|OOF97105.1 acla|EAW13011.1 afis|EAW17656.1 afla|EED50502.1 afum|EAL91471.1 agla|OJJ79549.1 akaw|GAA89955.1 aluc|OJZ83962.1 anid|CBF90066.1 anig|EHA19090.1 aory|EIT82103.1 asyd|OJJ62045.1 ater|EAU31997.1 atub|OJI90334.1 aver|OJI96151.1 awen|OJJ37187.1 azon|OJJ50889.1

Group2200: abra|OJJ70617.1 acar|OOF97104.1 acla|EAW13010.1 afis|EAW17657.1 afla|EED50501.1 afum|EAL91472.1 agla|OJJ79548.1 akaw|GAA89956.1 aluc|OJZ83963.1 anid|CBF90065.1 anig|EHA19091.1 aory|EIT82023.1 asyd|OJJ62046.1 ater|EAU31996.1 atub|OJI90335.1 aver|OJI96152.1 awen|OJJ37186.1 azon|OJJ50888.1

Group2201: abra|OJJ70618.1 acar|OOF97103.1 acla|EAW13009.1 afis|EAW17658.1 afla|EED50500.1 afum|EAL91473.1 agla|OJJ79547.1 akaw|GAA89957.1 aluc|OJZ83964.1 anid|CBF90063.1 anig|EHA19092.1 aory|EIT82147.1 asyd|OJJ62048.1 ater|EAU31994.1 atub|OJI90337.1 aver|OJI96153.1 awen|OJJ37185.1 azon|OJJ50886.1

Group2202: abra|OJJ70619.1 acar|OOF97102.1 acla|EAW13008.1 afis|EAW17659.1 afla|EED50499.1 afum|EAL91474.1 agla|OJJ79546.1 akaw|GAA89958.1 aluc|OJZ83965.1 anid|CBF90061.1 anig|EHA19093.1 aory|EIT82055.1 asyd|OJJ62049.1 ater|EAU31993.1 atub|OJI90338.1 aver|OJI96154.1 awen|OJJ37184.1 azon|OJJ50885.1

Group2203: abra|OJJ70620.1 acar|OOF97101.1 acla|EAW13007.1 afis|EAW17660.1 afla|EED50497.1 afum|EAL91475.1 agla|OJJ79529.1 akaw|GAA89959.1 aluc|OJZ83966.1 anid|CBF90059.1 anig|EHA19094.1 aory|EIT82172.1 asyd|OJJ62050.1 ater|EAU31992.1 atub|OJI90339.1 aver|OJI96155.1 awen|OJJ37182.1 azon|OJJ50884.1

Group2204: abra|OJJ70625.1 acar|OOF97099.1 acla|EAW13005.1 afis|EAW17663.1 afla|EED50495.1 afum|EAL91477.2 agla|OJJ79532.1 akaw|GAA89963.1 aluc|OJZ83970.1 anid|CBF90056.1 anig|EHA19099.1 aory|EIT82160.1 asyd|OJJ62052.1 ater|EAU31990.1 atub|OJI90343.1 aver|OJI96158.1 awen|OJJ37179.1 azon|OJJ50881.1

Group2205: abra|OJJ70626.1 acar|OOF97098.1 acla|EAW08131.1 afis|EAW23168.1 afla|EED46953.1 afum|EAL88316.1 agla|OJJ89679.1 akaw|GAA89964.1 aluc|OJZ83971.1 anid|CBF74478.1 anig|EHA19100.1 aory|EIT72624.1 asyd|OJJ57707.1 ater|EAU36297.1 atub|OJI90344.1 aver|OJJ05457.1 awen|OJJ36615.1 azon|OJJ47598.1

Group2206: abra|OJJ70627.1 acar|OOF97097.1 acla|EAW08130.1 afis|EAW23167.1 afla|EED46954.1 afum|EAL88317.2 agla|OJJ89678.1 akaw|GAA89965.1 aluc|OJZ83972.1 anid|CBF74476.1 anig|EHA19101.1 aory|EIT72656.1 asyd|OJJ57706.1 ater|EAU36296.1 atub|OJI90345.1 aver|OJJ05458.1 awen|OJJ36616.1 azon|OJJ41979.1

Group2207: abra|OJJ70628.1 acar|OOF97096.1 acla|EAW08129.1 afis|EAW23166.1 afla|EED46955.1 afum|EAL88318.1 agla|OJJ89677.1 akaw|GAA89966.1 aluc|OJZ83974.1 anid|CBF74475.1 anig|EHA19102.1 aory|EIT72623.1 asyd|OJJ57705.1 ater|EAU36295.1 atub|OJI90347.1 aver|OJJ05459.1 awen|OJJ36617.1 azon|OJJ47597.1

Group2208: abra|OJJ70629.1 acar|OOF97095.1 acla|EAW08128.1 afis|EAW23165.1 afla|EED46957.1 afum|EBA27343.1 agla|OJJ89676.1 akaw|GAA89967.1 aluc|OJZ83976.1 anid|CBF74473.1 anig|EHA19103.1 aory|EIT72540.1 asyd|OJJ57704.1 ater|EAU36294.1 atub|OJI90349.1 aver|OJJ05460.1 awen|OJJ36618.1 azon|OJJ47595.1

Group2209: abra|OJJ70630.1 acar|OOF97093.1 acla|EAW08126.1 afis|EAW23163.1 afla|EED46958.1 afum|EAL88320.1 agla|OJJ89674.1 akaw|GAA89969.1 aluc|OJZ83978.1 anid|CBF74467.1 anig|EHA19105.1 aory|EIT72637.1 asyd|OJJ57700.1 ater|EAU36293.1 atub|OJI90351.1 aver|OJJ05464.1 awen|OJJ36620.1 azon|OJJ47593.1

Group2210: abra|OJJ70639.1 acar|OOF97080.1 acla|EAW08118.1 afis|EAW23133.1 afla|EED48995.1 afum|EAL88348.1 agla|OJJ89666.1 akaw|GAA89975.1 aluc|OJZ83986.1 anid|CBF74456.1 anig|EHA19112.1 aory|EIT80032.1 asyd|OJJ57677.1 ater|EAU36286.1 atub|OJI90358.1 aver|OJJ05483.1 awen|OJJ36627.1 azon|OJJ47586.1

Group2211: abra|OJJ70640.1 acar|OOF97081.1 acla|EAW08117.1 afis|EAW23134.1 afla|EED48497.1 afum|EAL88347.1 agla|OJJ89665.1 akaw|GAA89976.1 aluc|OJZ83987.1 anid|CBF74455.1 anig|EHA19113.1 aory|EIT73203.1 asyd|OJJ57676.1 ater|EAU36282.1 atub|OJI90359.1 aver|OJJ05482.1 awen|OJJ36630.1 azon|OJJ47584.1

Group2212: abra|OJJ70646.1 acar|OOF97075.1 acla|EAW08114.1 afis|EAW23137.1 afla|EED48500.1 afum|EAL88344.1 agla|OJJ89663.1 akaw|GAA89982.1 aluc|OJZ83996.1 anid|CBF74450.1 anig|EHA19119.1 aory|EIT73193.1 asyd|OJJ57740.1 ater|EAU36280.1 atub|OJI90365.1 aver|OJJ05427.1 awen|OJJ36633.1 azon|OJJ47582.1

Group2213: abra|OJJ70653.1 acar|OOF97070.1 acla|EAW08106.1 afis|EAW23145.1 afla|EED48508.1 afum|EAL88336.1 agla|OJJ89657.1 akaw|GAA89989.1 aluc|OJZ84003.1 anid|CBF74436.1 anig|EHA19125.1 aory|EIT73083.1 asyd|OJJ57747.1 ater|EAU36266.1 atub|OJI90372.1 aver|OJJ05420.1 awen|OJJ36640.1 azon|OJJ47574.1

Group2214: abra|OJJ70654.1 acar|OOF97068.1 acla|EAW08105.1 afis|EAW23146.1 afla|EED48509.1 afum|EAL88335.1 agla|OJJ89656.1 akaw|GAA89990.1 aluc|OJZ84004.1 anid|CBF74434.1 anig|EHA19126.1 aory|EIT73165.1 asyd|OJJ57748.1 ater|EAU36265.1 atub|OJI90373.1 aver|OJJ05419.1 awen|OJJ36641.1 azon|OJJ47573.1

Group2215: abra|OJJ70657.1 acar|OOF97065.1 acla|EAW08102.1 afis|EAW23149.1 afla|EED48512.1 afum|EAL88332.1 agla|OJJ89653.1 akaw|GAA89993.1 aluc|OJZ84007.1 anid|CBF74428.1 anig|EHA19129.1

aory|EIT73157.1 asyd|OJJ57751.1 ater|EAU36263.1 atub|OJI90376.1 aver|OJJ05416.1 awen|OJJ36645.1 azon|OJJ47570.1

Group2216: abra|OJJ70659.1 acar|OOF97063.1 acla|EAW08100.1 afis|EAW23151.1 afla|EED48514.1 afum|EAL88330.2 agla|OJJ89651.1 akaw|GAA89995.1 aluc|OJZ84009.1 anid|CBF74424.1 anig|EHA19131.1 aory|EIT73110.1 asyd|OJJ57753.1 ater|EAU36261.1 atub|OJI90378.1 aver|OJJ05414.1 awen|OJJ36647.1 azon|OJJ47568.1

Group2217: abra|OJJ70661.1 acar|OOF97060.1 acla|EAW08097.1 afis|EAW23153.1 afla|EED48516.1 afum|EAL88328.1 agla|OJJ89648.1 akaw|GAA89997.1 aluc|OJZ84011.1 anid|CBF74421.1 anig|EHA19134.1 aory|EIT73108.1 asyd|OJJ57755.1 ater|EAU36258.1 atub|OJI90380.1 aver|OJJ05412.1 awen|OJJ36649.1 azon|OJJ47553.1

Group2218: abra|OJJ70666.1 acar|OOF97058.1 acla|EAW08096.1 afis|EAW23154.1 afla|EED48519.1 afum|EAL88327.2 agla|OJJ89646.1 akaw|GAA90001.1 aluc|OJZ84016.1 anid|CBF74419.1 anig|EHA19138.1 aory|EIT73174.1 asyd|OJJ57756.1 ater|EAU36257.1 atub|OJI90387.1 aver|OJJ05411.1 awen|OJJ36650.1 azon|OJJ47545.1

Group2219: abra|OJJ70669.1 acar|OOF97057.1 acla|EAW08094.1 afis|EAW23129.1 afla|EED48521.1 afum|EAL88351.1 agla|OJJ89645.1 akaw|GAA90002.1 aluc|OJZ84017.1 anid|CBF74417.1 anig|EHA19139.1 aory|EIT73127.1 asyd|OJJ57757.1 ater|EAU36256.1 atub|OJI90389.1 aver|OJJ05410.1 awen|OJJ36651.1 azon|OJJ47547.1

Group2220: abra|OJJ70671.1 acar|OOF97055.1 acla|EAW08093.1 afis|EAW23128.1 afla|EED48522.1 afum|EAL88352.1 agla|OJJ89644.1 akaw|GAA90003.1 aluc|OJZ84019.1 anid|CBF74413.1 anig|EHA19140.1 aory|EIT73195.1 asyd|OJJ57758.1 ater|EAU36255.1 atub|OJI90390.1 aver|OJJ05408.1 awen|OJJ36652.1 azon|OJJ47548.1

Group2221: abra|OJJ70672.1 acar|OOF97054.1 acla|EAW08092.1 afis|EAW23127.1 afla|EED48523.1 afum|EAL88353.1 agla|OJJ89643.1 akaw|GAA90004.1 aluc|OJZ84020.1 anid|CBF74412.1 anig|EHA19141.1 aory|EIT73139.1 asyd|OJJ57759.1 ater|EAU36254.1 atub|OJI90391.1 aver|OJJ05407.1 awen|OJJ36653.1 azon|OJJ47549.1

Group2222: abra|OJJ70675.1 acar|OOF97052.1 acla|EAW08089.1 afis|EAW23124.1 afla|EED48528.1 afum|EAL88356.1 agla|OJJ89641.1 akaw|GAA90007.1 aluc|OJZ84023.1 anid|CBF74406.1 anig|EHA19144.1 aory|EIT73244.1 asyd|OJJ57762.1 ater|EAU36252.1 atub|OJI90395.1 aver|OJJ05404.1 awen|OJJ36656.1 azon|OJJ47552.1

Group2223: abra|OJJ70677.1 acar|OOF97050.1 acla|EAW08087.1 afis|EAW23123.1 afla|EED48530.1 afum|EAL88357.1 agla|OJJ89616.1 akaw|GAA90009.1 aluc|OJZ84025.1 anid|CBF74404.1 anig|EHA19146.1 aory|EIT73116.1 asyd|OJJ57763.1 ater|EAU36251.1 atub|OJI90397.1 aver|OJJ05403.1 awen|OJJ36658.1 azon|OJJ47540.1

Group2224: abra|OJJ70680.1 acar|OOF97047.1 acla|EAW08084.1 afis|EAW23120.1 afla|EED48533.1 afum|EAL88360.1 agla|OJJ89619.1 akaw|GAA90012.1 aluc|OJZ84028.1 anid|CBF74398.1 anig|EHA19149.1 aory|EIT73210.1 asyd|OJJ57766.1 ater|EAU36249.1 atub|OJI90400.1 aver|OJJ05400.1 awen|OJJ36661.1 azon|OJJ47537.1

Group2225: abra|OJJ70681.1 acar|OOF97046.1 acla|EAW08083.1 afis|EAW23119.1 afla|EED48534.1 afum|EAL88361.1 agla|OJJ89621.1 akaw|GAA90013.1 aluc|OJZ84029.1 anid|CBF74396.1 anig|EHA19150.1 aory|EIT73258.1 asyd|OJJ57767.1 ater|EAU36248.1 atub|OJI90401.1 aver|OJJ05399.1 awen|OJJ36662.1 azon|OJJ47536.1

Group2226: abra|OJJ70684.1 acar|OOF97043.1 acla|EAW08078.1 afis|EAW23117.1 afla|EED48535.1 afum|EAL88363.1 agla|OJJ89624.1 akaw|GAA90016.1 aluc|OJZ84032.1 anid|CBF74393.1 anig|EHA19153.1 aory|EIT73171.1 asyd|OJJ57769.1 ater|EAU36245.1 atub|OJI90404.1 aver|OJJ05397.1 awen|OJJ36665.1 azon|OJJ47534.1

Group2227: abra|OJJ70685.1 acar|OOF97042.1 acla|EAW08077.1 afis|EAW23116.1 afla|EED48536.1 afum|EAL88364.1 agla|OJJ89625.1 akaw|GAA90017.1 aluc|OJZ84033.1 anid|CBF74391.1 anig|EHA19154.1 aory|EIT73186.1 asyd|OJJ57770.1 ater|EAU36244.1 atub|OJI90405.1 aver|OJJ05396.1 awen|OJJ36666.1 azon|OJJ47533.1

Group2228: abra|OJJ70686.1 acar|OOF97041.1 afis|EAW23115.1 afla|EED53328.1 afum|EAL88365.1 agla|OJJ89626.1 akaw|GAA90018.1 aluc|OJZ84034.1 anid|CBF76024.1 anig|EHA19155.1 aory|EIT74989.1 asyd|OJJ52450.1 ater|EAU36243.1 atub|OJI90406.1 aver|OJJ03966.1 awen|OJJ36667.1 azon|OJJ47532.1 aory|EIT74930.1

Group2229: abra|OJJ70689.1 acar|OOF97038.1 acla|EAW08076.1 afis|EAW23111.1 afla|EED48537.1 afum|EAL88368.1 agla|OJJ89630.1 akaw|GAA90021.1 aluc|OJZ84037.1 anid|CBF74389.1 anig|EHA19158.1 aory|EIT73070.1 asyd|OJJ57771.1 ater|EAU36239.1 atub|OJI90409.1 aver|OJJ05395.1 awen|OJJ36670.1 azon|OJJ47529.1

Group2230: abra|OJJ70694.1 acar|OOF97035.1 acla|EAW08071.1 afis|EAW23107.1 afla|EED48541.1 afum|EAL88372.1 agla|OJJ89636.1 akaw|GAA90025.1 aluc|OJZ84041.1 anid|CBF81370.1 anig|EHA19163.1 aory|EIT73129.1 asyd|OJJ59386.1 ater|EAU36235.1 atub|OJI90413.1 aver|OJJ02887.1 awen|OJJ36674.1 azon|OJJ47524.1

Group2231: abra|OJJ70696.1 acar|OOF97033.1 acla|EAW08069.1 afis|EAW23105.1 afla|EED48543.1 afum|EAL88374.1 agla|OJJ89638.1 akaw|GAA90027.1 aluc|OJZ84043.1 anid|CBF81367.1 anig|EHA19165.1 aory|EIT73169.1 asyd|OJJ59384.1 ater|EAU36233.1 atub|OJI90415.1 aver|OJJ02889.1 awen|OJJ36676.1 azon|OJJ47522.1

Group2232: abra|OJJ70700.1 acar|OOF90048.1 acla|EAW08065.1 afis|EAW23101.1 afla|EED48549.1 afum|EAL88377.1 agla|OJJ89613.1 akaw|GAA90031.1 aluc|OJZ84046.1 anid|CBF81359.1 anig|EHA19167.1 aory|EIT73156.1 asyd|OJJ59381.1 ater|EAU36227.1 atub|OJI90418.1 aver|OJJ02892.1 awen|OJJ36680.1 azon|OJJ47517.1

Group2233: abra|OJJ70701.1 acar|OOF89978.1 acla|EAW08064.1 afis|EAW23100.1 afla|EED48548.1 afum|EAL88378.2 agla|OJJ89612.1 akaw|GAA90032.1 aluc|OJZ84047.1 anig|EHA19168.1 aory|EIT73049.1 asyd|OJJ59380.1 ater|EAU36228.1 atub|OJI90419.1 aver|OJJ02893.1 awen|OJJ36681.1 azon|OJJ47516.1 anid|CBF81358.1

Group2234: abra|OJJ70702.1 acar|OOF89980.1 acla|EAW08063.1 afis|EAW23099.1 afla|EED48547.1 afum|EAL88379.1 agla|OJJ89611.1 akaw|GAA90033.1 aluc|OJZ84049.1 anid|CBF81356.1 anig|EHA19169.1 aory|EIT73063.1 asyd|OJJ59379.1 ater|EAU36230.1 atub|OJI90420.1 aver|OJJ02894.1 awen|OJJ36682.1 azon|OJJ47515.1

Group2235: abra|OJJ70704.1 acar|OOF97032.1 acla|EAW08061.1 afis|EAW23097.1 afla|EED48551.1 afum|EAL88381.2 agla|OJJ89609.1 akaw|GAA90035.1 aluc|OJZ84051.1 anid|CBF81352.1 anig|EHA19171.1 aory|EIT73218.1 asyd|OJJ59377.1 ater|EAU36225.1 atub|OJI90422.1 aver|OJJ02896.1 awen|OJJ36684.1 azon|OJJ47512.1

Group2236: abra|OJJ70705.1 acar|OOF97031.1 acla|EAW08060.1 afis|EAW23096.1 afla|EED48552.1 afum|EAL88382.2 agla|OJJ89608.1 akaw|GAA90036.1 aluc|OJZ84052.1 anid|CBF81350.1 anig|EHA19172.1 aory|EIT73272.1 asyd|OJJ59376.1 ater|EAU36224.1 atub|OJI90423.1 aver|OJJ02897.1 awen|OJJ36685.1 azon|OJJ47511.1

Group2237: abra|OJJ70706.1 acar|OOF97029.1 acla|EAW08058.1 afis|EAW23094.1 afla|EED48554.1 afum|EAL88384.2 agla|OJJ89606.1 akaw|GAA90037.1 aluc|OJZ84053.1 anid|CBF87089.1 anig|EHA19173.1 aory|EIT73275.1 asyd|OJJ60399.1 atub|OJI90424.1 aver|OJI99728.1 awen|OJJ36688.1 azon|OJJ47509.1 aory|EIT73221.1

Group2238: abra|OJJ70708.1 acar|OOF97028.1 acla|EAW08056.1 afis|EAW23092.1 afla|EED48556.1 afum|EBA27345.1 agla|OJJ89604.1 akaw|GAA90039.1 aluc|OJZ84055.1 anid|CBF81348.1 anig|EHA19176.1 aory|EIT73227.1 asyd|OJJ59374.1 ater|EAU36215.1 atub|OJI90426.1 aver|OJJ02900.1 awen|OJJ36690.1 azon|OJJ47506.1

Group2239: abra|OJJ70710.1 acar|OOF97026.1 acla|EAW08054.1 afis|EAW23090.1 afla|EED48558.1 afum|EAL88389.1 agla|OJJ89602.1 akaw|GAA90041.1 aluc|OJZ84057.1 anid|CBF81345.1 anig|EHA19177.1 aory|EIT73059.1 asyd|OJJ59372.1 ater|EAU36213.1 atub|OJI90428.1 aver|OJJ02902.1 awen|OJJ36692.1 azon|OJJ47504.1

Group2240: abra|OJJ70711.1 acar|OOF97025.1 acla|EAW08053.1 afis|EAW23089.1 afla|EED48559.1 afum|EAL88390.1 agla|OJJ89601.1 akaw|GAA90042.1 aluc|OJZ84058.1 anid|CBF81343.1 anig|EHA19178.1 aory|EIT73173.1 asyd|OJJ59371.1 ater|EAU36212.1 atub|OJI90429.1 aver|OJJ02903.1 awen|OJJ36694.1 azon|OJJ47503.1

Group2241: abra|OJJ70714.1 acar|OOF97022.1 acla|EAW08049.1 afis|EAW23085.1 afla|EED48563.1 afum|EAL88394.1 agla|OJJ89595.1 akaw|GAA90045.1 aluc|OJZ84061.1 anid|CBF81339.1 anig|EHA19181.1 aory|EIT73232.1 asyd|OJJ59369.1 ater|EAU36209.1 atub|OJI90432.1 aver|OJJ02905.1 awen|OJJ36699.1 azon|OJJ47498.1

Group2242: abra|OJJ70715.1 acar|OOF97021.1 acla|EAW08048.1 afis|EAW23084.1 afla|EED48564.1 afum|EAL88395.1 agla|OJJ89594.1 akaw|GAA90046.1 aluc|OJZ84062.1 anid|CBF81337.1 anig|EHA19182.1 aory|EIT73208.1 asyd|OJJ59368.1 ater|EAU36208.1 atub|OJI90433.1 aver|OJJ02906.1 awen|OJJ36700.1 azon|OJJ47497.1

Group2243: abra|OJJ70716.1 acar|OOF97020.1 acla|EAW08047.1 afis|EAW23083.1 afla|EED48565.1 afum|EAL88396.1 agla|OJJ89592.1 akaw|GAA90047.1 aluc|OJZ84063.1 anid|CBF81335.1 anig|EHA19183.1 aory|EIT73176.1 asyd|OJJ59367.1 ater|EAU36207.1 atub|OJI90434.1 aver|OJJ02907.1 awen|OJJ36701.1 azon|OJJ47496.1

Group2244: abra|OJJ70717.1 acar|OOF97019.1 acla|EAW08046.1 afis|EAW23082.1 afla|EED48566.1 afum|EAL88397.1 agla|OJJ89591.1 akaw|GAA90048.1 aluc|OJZ84064.1 anid|CBF81333.1 anig|EHA19184.1

aory|EIT73236.1 asyd|OJJ59366.1 ater|EAU36206.1 atub|OJI90435.1 aver|OJJ02908.1 awen|OJJ36702.1 azon|OJJ47495.1

Group2245: abra|OJJ70718.1 acar|OOF97018.1 acla|EAW08044.1 afis|EAW23080.1 afla|EED48567.1 afum|EAL88399.1 agla|OJJ89588.1 akaw|GAA90049.1 aluc|OJZ84065.1 anid|CBF81330.1 anig|EHA19185.1 aory|EIT73202.1 asyd|OJJ59364.1 ater|EAU36205.1 atub|OJI90436.1 aver|OJJ02910.1 awen|OJJ36704.1 azon|OJJ47494.1

Group2246: abra|OJJ70719.1 acar|OOF97017.1 acla|EAW08043.1 afis|EAW23079.1 afla|EED48568.1 afum|EAL88400.1 agla|OJJ89587.1 akaw|GAA90050.1 aluc|OJZ84066.1 anid|CBF81328.1 anig|EHA19186.1 aory|EIT73095.1 asyd|OJJ59363.1 ater|EAU36204.1 atub|OJI90437.1 aver|OJJ02911.1 awen|OJJ36705.1 azon|OJJ47493.1

Group2247: abra|OJJ70720.1 acar|OOF97016.1 acla|EAW08042.1 afis|EAW23078.1 afla|EED48569.1 afum|EAL88401.1 agla|OJJ89586.1 akaw|GAA90051.1 aluc|OJZ84067.1 anid|CBF81326.1 anig|EHA19187.1 aory|EIT73149.1 asyd|OJJ59362.1 ater|EAU36202.1 atub|OJI90438.1 aver|OJJ02912.1 awen|OJJ36706.1 azon|OJJ47489.1

Group2248: abra|OJJ70722.1 acar|OOF97015.1 afis|EAW23077.1 afla|EED48570.1 afum|EAL88402.1 agla|OJJ89585.1 akaw|GAA90052.1 aluc|OJZ84068.1 anid|CBF81324.1 anig|EHA19188.1 aory|EIT73161.1 asyd|OJJ59361.1 ater|EAU36201.1 atub|OJI90440.1 aver|OJJ02913.1 awen|OJJ36707.1 azon|OJJ47490.1 aver|OJJ06633.1

Group2249: abra|OJJ70726.1 acar|OOF97013.1 acla|EAW08040.1 afis|EAW23075.1 afla|EED56447.1 afum|EAL88404.1 agla|OJJ89582.1 akaw|GAA90054.1 aluc|OJZ84071.1 anid|CBF81318.1 anig|EHA19190.1 aory|EIT72820.1 asyd|OJJ59359.1 ater|EAU36199.1 atub|OJI90443.1 aver|OJJ02915.1 awen|OJJ36709.1 azon|OJJ47492.1

Group2250: abra|OJJ70727.1 acar|OOF97012.1 acla|EAW08039.1 afis|EAW23074.1 afla|EED56449.1 afum|EAL88405.1 agla|OJJ89581.1 akaw|GAA90055.1 aluc|OJZ84072.1 anid|CBF81316.1 anig|EHA19191.1 aory|EIT72790.1 asyd|OJJ59358.1 ater|EAU36198.1 atub|OJI90444.1 aver|OJJ02916.1 awen|OJJ36710.1 azon|OJJ47487.1

Group2251: abra|OJJ70728.1 acar|OOF97011.1 acla|EAW08038.1 afis|EAW23073.1 afla|EED56450.1 afum|EAL88406.1 agla|OJJ89580.1 akaw|GAA90056.1 aluc|OJZ84073.1 anid|CBF81314.1 anig|EHA19192.1 aory|EIT72843.1 asyd|OJJ59357.1 ater|EAU36197.1 atub|OJI90445.1 aver|OJJ02917.1 awen|OJJ36711.1 azon|OJJ47488.1

Group2252: abra|OJJ70729.1 acar|OOF97009.1 acla|EAW11184.1 afis|EAW15915.1 afla|EED56455.1 afum|EAL88550.2 agla|OJJ89579.1 akaw|GAA90057.1 aluc|OJZ84074.1 anid|CBF81309.1 anig|EHA19193.1 aory|EIT72851.1 asyd|OJJ59354.1 ater|EAU36193.1 atub|OJI90446.1 aver|OJJ02920.1 awen|OJJ36715.1 azon|OJJ47474.1

Group2253: abra|OJJ70733.1 acar|OOF97005.1 acla|EAW11180.1 afis|EAW15911.1 afla|EED56457.1 afum|EAL88546.2 agla|OJJ89575.1 akaw|GAA90061.1 aluc|OJZ84078.1 anid|CBF81301.1 anig|EHA19197.1 aory|EIT72816.1 asyd|OJJ59350.1 ater|EAU36191.1 atub|OJI90450.1 aver|OJJ02924.1 awen|OJJ36719.1 azon|OJJ47478.1

Group2254: abra|OJJ70735.1 acar|OOF97004.1 acla|EAW11179.1 afis|EAW15910.1 afla|EED56458.1 afum|EAL88545.1 agla|OJJ89574.1 akaw|GAA90062.1 aluc|OJZ84080.1 anid|CBF81299.1 anig|EHA19198.1 aory|EIT72821.1 asyd|OJJ59349.1 ater|EAU36189.1 atub|OJI90451.1 aver|OJJ02925.1 awen|OJJ36720.1 azon|OJJ47479.1

Group2255: abra|OJJ70738.1 acar|OOF97003.1 acla|EAW11178.1 afis|EAW15909.1 afla|EED56459.1 afum|EAL88544.1 agla|OJJ89573.1 akaw|GAA90063.1 aluc|OJZ84083.1 anid|CBF81298.1 anig|EHA19199.1 aory|EIT72803.1 asyd|OJJ59348.1 ater|EAU36188.1 atub|OJI90453.1 aver|OJJ02926.1 awen|OJJ36721.1 azon|OJJ47480.1

Group2256: abra|OJJ70741.1 acar|OOF97002.1 acla|EAW11176.1 afis|EAW15907.1 afla|EED56461.1 afum|EAL88542.1 agla|OJJ78421.1 akaw|GAA90065.1 aluc|OJZ84086.1 anid|CBF81263.1 anig|EHA19202.1 aory|EIT72874.1 asyd|OJJ59332.1 ater|EAU36186.1 atub|OJI90457.1 aver|OJJ02942.1 awen|OJJ36723.1 azon|OJJ47482.1

Group2257: abra|OJJ70751.1 acar|OOF96994.1 acla|EAW11170.1 afis|EAW15901.1 afla|EED56470.1 afum|EAL88536.1 agla|OJJ89561.1 akaw|GAA90071.1 aluc|OJZ84095.1 anid|CBF81281.1 anig|EHA19208.1 aory|EIT72864.1 asyd|OJJ59338.1 ater|EAU36180.1 atub|OJI90468.1 aver|OJJ02936.1 awen|OJJ36730.1 azon|OJJ47467.1

Group2258: abra|OJJ70752.1 acar|OOF96992.1 acla|EAW11169.1 afis|EAW15900.1 afla|EED56471.1 afum|EAL88535.1 agla|OJJ89560.1 akaw|GAA90072.1 aluc|OJZ84097.1 anid|CBF81283.1 anig|EHA19209.1 aory|EIT72802.1 asyd|OJJ59340.1 ater|EAU36179.1 atub|OJI90470.1 aver|OJJ02935.1 awen|OJJ36731.1 azon|OJJ47466.1

Group2259: abra|OJJ70753.1 acar|OOF96991.1 acla|EAW11168.1 afis|EAW15899.1 afla|EED56472.1 afum|EAL88534.1 agla|OJJ89553.1 akaw|GAA90073.1 aluc|OJZ84098.1 anid|CBF81285.1 anig|EHA19210.1 aory|EIT72807.1 asyd|OJJ59341.1 ater|EAU36178.1 atub|OJI90471.1 aver|OJJ02934.1 awen|OJJ36732.1 azon|OJJ47465.1

Group2260: abra|OJJ70754.1 acar|OOF96990.1 acla|EAW11167.1 afis|EAW15898.1 afla|EED56473.1 afum|EAL88533.1 agla|OJJ89552.1 akaw|GAA90074.1 aluc|OJZ84099.1 anid|CBF81287.1 anig|EHA19211.1 aory|EIT72796.1 asyd|OJJ59342.1 ater|EAU36177.1 atub|OJI90472.1 aver|OJJ02933.1 awen|OJJ36733.1 azon|OJJ47464.1

Group2261: abra|OJJ70755.1 acar|OOF96989.1 acla|EAW11166.1 afis|EAW15897.1 afla|EED56474.1 afum|EAL88532.1 agla|OJJ89551.1 akaw|GAA90075.1 aluc|OJZ84100.1 anid|CBF81288.1 anig|EHA19212.1 aory|EIT72834.1 asyd|OJJ59343.1 ater|EAU36176.1 atub|OJI90473.1 aver|OJJ02931.1 awen|OJJ36734.1 azon|OJJ47463.1

Group2262: abra|OJJ70756.1 acar|OOF96988.1 acla|EAW11165.1 afis|EAW15896.1 afla|EED56475.1 afum|EAL88531.1 agla|OJJ89550.1 akaw|GAA90076.1 aluc|OJZ84101.1 anid|CBF81290.1 anig|EHA19213.1 aory|EIT72841.1 asyd|OJJ59344.1 ater|EAU36175.1 atub|OJI90474.1 aver|OJJ02930.1 awen|OJJ36735.1 azon|OJJ47462.1

Group2263: abra|OJJ70758.1 acar|OOF96986.1 acla|EAW11163.1 afis|EAW15894.1 afla|EED56477.1 afum|EAL88529.1 agla|OJJ89548.1 akaw|GAA90078.1 aluc|OJZ84103.1 anid|CBF81294.1 anig|EHA19215.1 aory|EIT72822.1 asyd|OJJ59346.1 ater|EAU36173.1 atub|OJI90476.1 aver|OJJ02928.1 awen|OJJ36737.1 azon|OJJ47460.1

Group2264: abra|OJJ70760.1 acar|OOF96984.1 acla|EAW11161.1 afis|EAW15892.1 afla|EED56479.1 afum|EAL88527.1 agla|OJJ89530.1 akaw|GAA90080.1 aluc|OJZ84105.1 anid|CBF81259.1 anig|EHA19217.1 aory|EIT72795.1 asyd|OJJ59330.1 ater|EAU31714.1 atub|OJI90479.1 aver|OJJ02944.1 awen|OJJ36739.1 azon|OJJ47459.1

Group2265: abra|OJJ70762.1 acar|OOF96982.1 acla|EAW11159.1 afis|EAW15890.1 afla|EED56482.1 afum|EAL88525.2 agla|OJJ89532.1 akaw|GAA90082.1 aluc|OJZ84107.1 anid|CBF81256.1 anig|EHA19219.1 aory|EIT72811.1 asyd|OJJ59328.1 ater|EAU31712.1 atub|OJI90481.1 aver|OJJ02946.1 awen|OJJ36740.1 azon|OJJ47457.1

Group2266: abra|OJJ70766.1 acar|OOF96979.1 acla|EAW11156.1 afis|EAW15887.1 afla|EED56485.1 afum|EAL88521.2 agla|OJJ89534.1 akaw|GAA90086.1 aluc|OJZ84112.1 anid|CBF81250.1 anig|EHA19222.1 aory|EIT72797.1 asyd|OJJ59325.1 ater|EAU31709.1 atub|OJI90485.1 aver|OJJ02949.1 awen|OJJ36743.1 azon|OJJ47454.1

Group2267: abra|OJJ70767.1 acar|OOF96977.1 acla|EAW11154.1 afis|EAW15885.1 afla|EED56487.1 afum|EAL88519.1 agla|OJJ89536.1 akaw|GAA90087.1 aluc|OJZ84113.1 anid|CBF81243.1 anig|EHA19223.1 aory|EIT72824.1 asyd|OJJ59321.1 ater|EAU31705.1 atub|OJI90486.1 aver|OJJ02952.1 awen|OJJ36746.1 azon|OJJ47453.1

Group2268: abra|OJJ70768.1 acar|OOF96978.1 acla|EAW11153.1 afis|EAW15884.1 afla|EED56488.1 afum|EAL88518.1 agla|OJJ89537.1 akaw|GAA90088.1 aluc|OJZ84114.1 anid|CBF81241.1 anig|EHA19224.1 aory|EIT72873.1 asyd|OJJ59320.1 ater|EAU31704.1 atub|OJI90487.1 aver|OJJ02953.1 awen|OJJ36747.1 azon|OJJ47452.1

Group2269: abra|OJJ70770.1 acar|OOF96976.1 acla|EAW11151.1 afis|EAW15882.1 afla|EED56489.1 afum|EAL88516.1 agla|OJJ89539.1 akaw|GAA90090.1 aluc|OJZ84116.1 anid|CBF81237.1 anig|EHA19226.1 aory|EIT72859.1 asyd|OJJ59319.1 ater|EAU31703.1 atub|OJI90489.1 aver|OJJ02954.1 awen|OJJ36749.1 azon|OJJ47450.1

Group2270: abra|OJJ70774.1 acar|OOF96936.1 acla|EAW11147.1 afis|EAW15878.1 afla|EED56497.1 afum|EAL88512.1 agla|OJJ87230.1 akaw|GAA90094.1 aluc|OJZ84121.1 anid|CBF81230.1 anig|EHA19229.1 aory|EIT72868.1 asyd|OJJ59316.1 ater|EAU31699.1 atub|OJI90493.1 aver|OJJ02957.1 awen|OJJ36752.1 azon|OJJ47447.1

Group2271: abra|OJJ70795.1 acar|OOF96950.1 acla|EAW11128.1 afis|EAW15857.1 afla|EED56515.1 afum|EAL88493.1 agla|OJJ87250.1 akaw|GAA90111.1 aluc|OJZ84140.1 anid|CBF81189.1 anig|EHA19248.1 aory|EIT72805.1 asyd|OJJ59299.1 ater|EAU31672.1 atub|OJI90514.1 aver|OJJ02976.1 awen|OJJ36770.1 azon|OJJ47428.1

Group2272: abra|OJJ70809.1 acar|OOF98293.1 acla|EAW10418.1 afis|EAW25151.1 afla|EED55282.1 afum|EAL89826.2 agla|OJJ89004.1 akaw|GAA91689.1 aluc|OJZ87705.1 anid|CBF81138.1 anig|EHA20888.1 aory|EIT83553.1 asyd|OJJ56507.1 ater|EAU39430.1 atub|OJI80938.1 aver|OJJ06212.1 awen|OJJ40780.1 azon|OJJ46872.1

Group2273: abra|OJJ70815.1 acar|OOF98285.1 acla|EAW10424.1 afis|EAW25145.1 afla|EED55294.1 afum|EAL89818.1 agla|OJJ89008.1 akaw|GAA91695.1 aluc|OJZ87698.1 anid|CBF85679.1 anig|EHA20885.1

aory|EIT83533.1 asyd|OJJ64254.1 ater|EAU39423.1 atub|OJI80945.1 aver|OJI98344.1 awen|OJJ40770.1 azon|OJJ46878.1

Group2274: abra|OJJ70833.1 acar|OOF98281.1 acla|EAW10427.1 afis|EAW25142.1 afla|EED55303.1 afum|EAL89814.1 agla|OJJ89025.1 akaw|GAA91701.1 aluc|OJZ87689.1 anid|CBF85686.1 anig|EHA20878.1 aory|EIT83517.1 asyd|OJJ64258.1 ater|EAU39411.1 atub|OJI80951.1 aver|OJI98350.1 awen|OJJ40766.1 azon|OJJ44131.1

Group2275: abra|OJJ70834.1 acar|OOF98280.1 acla|EAW10428.1 afis|EAW25141.1 afla|EED55304.1 afum|EAL89813.1 agla|OJJ89026.1 akaw|GAA91702.1 aluc|OJZ87688.1 anid|CBF86093.1 anig|EHA20877.1 aory|EIT83571.1 asyd|OJJ64473.1 ater|EAU39410.1 atub|OJI80952.1 aver|OJI98577.1 awen|OJJ40765.1 azon|OJJ44130.1

Group2276: abra|OJJ70837.1 acar|OOF98277.1 acla|EAW10432.1 afis|EAW25137.1 afla|EED55309.1 afum|EAL89808.2 agla|OJJ89030.1 akaw|GAA91705.1 aluc|OJZ87684.1 anid|CBF85710.1 anig|EHA20874.1 aory|EIT83613.1 asyd|OJJ64270.1 ater|EAU39407.1 atub|OJI80959.1 aver|OJI98362.1 awen|OJJ40761.1 azon|OJJ44125.1

Group2277: abra|OJJ70839.1 acar|OOF98275.1 acla|EAW10433.1 afis|EAW25135.1 afla|EED55311.1 afum|EAL89806.2 agla|OJJ89032.1 akaw|GAA91707.1 aluc|OJZ87682.1 anid|CBF85706.1 anig|EHA20872.1 aory|EIT83546.1 asyd|OJJ64268.1 ater|EAU39406.1 atub|OJI80961.1 aver|OJI98360.1 awen|OJJ40759.1 azon|OJJ44123.1

Group2278: abra|OJJ70843.1 acar|OOF98272.1 acla|EAW10434.1 afis|EAW25134.1 afla|EED55312.1 afum|EAL89805.1 agla|OJJ89033.1 akaw|GAA91711.1 aluc|OJZ87678.1 anid|CBF85705.1 anig|EHA20868.1 aory|EIT83614.1 asyd|OJJ64267.1 ater|EAU39405.1 atub|OJI80965.1 aver|OJI98359.1 awen|OJJ40758.1 azon|OJJ44122.1

Group2279: abra|OJJ70848.1 acar|OOF98266.1 acla|EAW10439.1 afis|EAW25128.1 afla|EED55316.1 afum|EAL89799.1 agla|OJJ89042.1 akaw|GAA91716.1 aluc|OJZ87672.1 anid|CBF85697.1 anig|EHA20863.1 aory|EIT83568.1 asyd|OJJ64263.1 ater|EAU39401.1 atub|OJI80970.1 aver|OJI98355.1 awen|OJJ40753.1 azon|OJJ44117.1

Group2280: abra|OJJ70851.1 acar|OOF98265.1 acla|EAW10441.1 afis|EAW25126.1 afla|EED55317.1 afum|EAL89797.1 agla|OJJ89044.1 akaw|GAA91718.1 aluc|OJZ87670.1 anid|CBF85694.1 anig|EHA20861.1 aory|EIT83542.1 asyd|OJJ64261.1 ater|EAU39399.1 atub|OJI80972.1 aver|OJI98353.1 awen|OJJ40750.1 azon|OJJ44115.1

Group2281: abra|OJJ70854.1 acar|OOF98262.1 acla|EAW10447.1 afis|EAW25120.1 afla|EED55322.1 afum|EAL89791.1 agla|OJJ89053.1 akaw|GAA91721.1 aluc|OJZ87667.1 anid|CBF86087.1 anig|EHA20858.1 aory|EIT76263.1 asyd|OJJ64469.1 ater|EAU39392.1 atub|OJI80976.1 aver|OJI98573.1 awen|OJJ40739.1 azon|OJJ44108.1

Group2282: abra|OJJ70855.1 acar|OOF98261.1 acla|EAW10446.1 afis|EAW25121.1 afla|EED55320.1 afum|EAL89792.1 agla|OJJ89049.1 akaw|GAA91722.1 aluc|OJZ87665.1 anid|CBF86089.1 anig|EHA20857.1 aory|EIT76239.1 asyd|OJJ64471.1 ater|EAU39393.1 atub|OJI80978.1 aver|OJI98575.1 awen|OJJ40740.1 azon|OJJ44110.1

Group2283: abra|OJJ70856.1 acar|OOF98260.1 acla|EAW10445.1 afis|EAW25122.1 afla|EED55319.1 afum|EAL89793.1 agla|OJJ89048.1 akaw|GAA91723.1 aluc|OJZ87664.1 anid|CBF86091.1 anig|EHA20856.1 aory|EIT76220.1 asyd|OJJ64472.1 ater|EAU39394.1 atub|OJI80979.1 aver|OJI98576.1 awen|OJJ40741.1 azon|OJJ44112.1

Group2284: abra|OJJ70857.1 acar|OOF98259.1 acla|EAW10449.1 afis|EAW25118.1 afla|EED55324.1 afum|EAL89789.1 agla|OJJ89059.1 akaw|GAA91724.1 aluc|OJZ87663.1 anid|CBF86086.1 anig|EHA20855.1 aory|EIT76226.1 asyd|OJJ64468.1 ater|EAU39386.1 atub|OJI80980.1 aver|OJI98572.1 awen|OJJ40737.1 azon|OJJ44107.1

Group2285: abra|OJJ70858.1 acar|OOF98258.1 acla|EAW10450.1 afis|EAW25117.1 afla|EED55325.1 afum|EAL89788.1 agla|OJJ89063.1 akaw|GAA91725.1 aluc|OJZ87662.1 anid|CBF86084.1 anig|EHA20854.1 aory|EIT76188.1 asyd|OJJ64467.1 ater|EAU39387.1 atub|OJI80981.1 aver|OJI98571.1 awen|OJJ40734.1 azon|OJJ44106.1

Group2286: abra|OJJ70859.1 acar|OOF98257.1 acla|EAW10451.1 afis|EAW25116.1 afla|EED55326.1 afum|EAL89787.1 agla|OJJ89064.1 akaw|GAA91726.1 aluc|OJZ87661.1 anid|CBF86082.1 anig|EHA20853.1 aory|EIT76238.1 asyd|OJJ64466.1 ater|EAU39388.1 atub|OJI80982.1 aver|OJI98570.1 awen|OJJ40733.1 azon|OJJ44105.1

Group2287: abra|OJJ70860.1 acar|OOF98256.1 acla|EAW10452.1 afis|EAW25115.1 afla|EED55327.1 afum|EAL89786.1 agla|OJJ89065.1 akaw|GAA91727.1 aluc|OJZ87660.1 anid|CBF86080.1 anig|EHA20852.1 aory|EIT76212.1 asyd|OJJ64465.1 ater|EAU39389.1 atub|OJI80983.1 aver|OJI98569.1 awen|OJJ40732.1 azon|OJJ44104.1

Group2288: abra|OJJ70861.1 acar|OOF98255.1 acla|EAW10453.1 afis|EAW25114.1 afla|EED55328.1 afum|EAL89785.1 agla|OJJ89066.1 akaw|GAA91728.1 aluc|OJZ87659.1 anid|CBF86079.1 anig|EHA20851.1 aory|EIT76255.1 asyd|OJJ64464.1 ater|EAU39390.1 atub|OJI80984.1 aver|OJI98568.1 awen|OJJ40731.1 azon|OJJ44103.1

Group2289: abra|OJJ70862.1 acar|OOF98254.1 acla|EAW10454.1 afis|EAW25113.1 afla|EED55329.1 afum|EAL89784.1 agla|OJJ89067.1 akaw|GAA91729.1 aluc|OJZ87658.1 anid|CBF86077.1 anig|EHA20850.1 aory|EIT76257.1 asyd|OJJ64463.1 ater|EAU39383.1 atub|OJI80985.1 aver|OJI98567.1 awen|OJJ40730.1 azon|OJJ44102.1

Group2290: abra|OJJ70864.1 acar|OOF98252.1 acla|EAW10456.1 afis|EAW25111.1 afla|EED55331.1 afum|EAL89782.2 agla|OJJ89070.1 akaw|GAA91731.1 aluc|OJZ87656.1 anid|CBF86073.1 anig|EHA20849.1 aory|EIT76253.1 asyd|OJJ64461.1 ater|EAU39381.1 atub|OJI80987.1 aver|OJI98564.1 awen|OJJ40728.1 azon|OJJ44100.1

Group2291: abra|OJJ70878.1 acar|OOF98242.1 acla|EAW10467.1 afis|EAW25100.1 afla|EED55345.1 afum|EAL89771.1 agla|OJJ85468.1 akaw|GAA91741.1 aluc|OJZ87644.1 anid|CBF86015.1 anig|EHA20837.1 aory|EIT80879.1 asyd|OJJ64450.1 ater|EAU39362.1 atub|OJI81001.1 aver|OJI98553.1 awen|OJJ40714.1 azon|OJJ44091.1

Group2292: abra|OJJ70880.1 acar|OOF98241.1 acla|EAW10468.1 afis|EAW25099.1 afla|EED55347.1 afum|EAL89770.1 agla|OJJ85472.1 akaw|GAA91742.1 aluc|OJZ87643.1 anid|CBF86013.1 anig|EHA20836.1 aory|EIT80895.1 asyd|OJJ64449.1 ater|EAU39361.1 atub|OJI81002.1 aver|OJI98552.1 awen|OJJ40713.1 azon|OJJ44090.1

Group2293: abra|OJJ70890.1 acar|OOF98227.1 acla|EAW10472.1 afis|EAW25095.1 afla|EED55353.1 afum|EAL89766.2 agla|OJJ89082.1 akaw|GAA87406.1 aluc|OJZ87634.1 anid|CBF86004.1 anig|EHA20829.1 aory|EIT80918.1 asyd|OJJ64445.1 ater|EAU39353.1 atub|OJI81015.1 aver|OJI98548.1 awen|OJJ40709.1 azon|OJJ44085.1

Group2294: abra|OJJ70892.1 acar|OOF90217.1 acla|EAW10474.1 afis|EAW25093.1 afla|EED55354.1 afum|EAL89764.1 agla|OJJ89084.1 akaw|GAA87408.1 aluc|OJZ87631.1 anid|CBF86001.1 anig|EHA20827.1 aory|EIT80886.1 asyd|OJJ64443.1 ater|EAU39351.1 atub|OJI81018.1 aver|OJI98545.1 awen|OJJ40707.1 azon|OJJ44083.1

Group2295: abra|OJJ70895.1 acar|OOF90219.1 acla|EAW10482.1 afis|EAW25085.1 afla|EED55362.1 afum|EAL89754.1 agla|OJJ89097.1 akaw|GAA87411.1 aluc|OJZ87628.1 anid|CBF85988.1 anig|EHA20824.1 aory|EIT80873.1 asyd|OJJ64434.1 ater|EAU39341.1 atub|OJI81022.1 aver|OJI98535.1 awen|OJJ40697.1 azon|OJJ44075.1

Group2296: abra|OJJ70897.1 acar|OOF90220.1 acla|EAW10481.1 afis|EAW25086.1 afla|EED55361.1 afum|EAL89755.1 agla|OJJ89096.1 akaw|GAA87412.1 aluc|OJZ87626.1 anid|CBF85990.1 anig|EHA20823.1 aory|EIT80880.1 asyd|OJJ64435.1 ater|EAU39342.1 atub|OJI81023.1 aver|OJI98536.1 awen|OJJ40698.1 azon|OJJ44076.1

Group2297: abra|OJJ70898.1 acar|OOF98228.1 acla|EAW10480.1 afis|EAW25087.1 afla|EED55360.1 afum|EAL89756.1 agla|OJJ89095.1 akaw|GAA87413.1 aluc|OJZ87625.1 anid|CBF85992.1 anig|EHA20822.1 aory|EIT80799.1 asyd|OJJ64436.1 ater|EAU39343.1 atub|OJI81024.1 aver|OJI98537.1 awen|OJJ40700.1 azon|OJJ44077.1

Group2298: abra|OJJ70899.1 acar|OOF98221.1 acla|EAW10479.1 afis|EAW25088.1 afla|EED55359.1 afum|EAL89757.1 agla|OJJ89094.1 akaw|GAA87414.1 aluc|OJZ87624.1 anid|CBF85994.1 anig|EHA20821.1 aory|EIT80835.1 asyd|OJJ64437.1 ater|EAU39344.1 atub|OJI81025.1 aver|OJI98538.1 awen|OJJ40701.1 azon|OJJ44078.1

Group2299: abra|OJJ70901.1 acar|OOF98218.1 acla|EAW10477.1 afis|EAW25090.1 afla|EED55357.1 afum|EAL89760.2 agla|OJJ89087.1 akaw|GAA87416.1 aluc|OJZ87622.1 anid|CBF85997.1 anig|EHA20819.1 aory|EIT80906.1 asyd|OJJ64440.1 ater|EAU39345.1 atub|OJI81027.1 aver|OJI98542.1 awen|OJJ40704.1 azon|OJJ44080.1

Group2300: abra|OJJ70906.1 acar|OOF98214.1 acla|EAW10484.1 afis|EAW25083.1 afla|EED55367.1 afum|EAL89752.1 agla|OJJ89102.1 akaw|GAA87420.1 aluc|OJZ87618.1 anid|CBF85985.1 anig|EHA20816.1 aory|EIT80861.1 asyd|OJJ64432.1 ater|EAU39339.1 atub|OJI81031.1 aver|OJI98533.1 awen|OJJ40695.1 azon|OJJ44072.1

Group2301: abra|OJJ70911.1 acar|OOF98211.1 acla|EAW10486.1 afis|EAW25081.1 afla|EED55369.1 afum|EAL89750.1 agla|OJJ89105.1 akaw|GAA87422.1 aluc|OJZ87614.1 anid|CBF85977.1 anig|EHA20811.1 aory|EIT80845.1 asyd|OJJ64429.1 ater|EAU39519.1 atub|OJI81035.1 aver|OJI98529.1 awen|OJJ40693.1 azon|OJJ44069.1

Group2302: abra|OJJ70912.1 acar|OOF98210.1 acla|EAW10487.1 afis|EAW25080.1 afla|EED55370.1 afum|EAL89749.1 agla|OJJ89106.1 akaw|GAA87423.1 aluc|OJZ87613.1 anid|CBF85975.1 anig|EHA20810.1

aory|EIT80866.1 asyd|OJJ64428.1 ater|EAU39539.1 atub|OJI81036.1 aver|OJI98528.1 awen|OJJ40692.1 azon|OJJ44068.1

Group2303: abra|OJJ70913.1 acar|OOF89971.1 acla|EAW10488.1 afis|EAW25079.1 afla|EED55371.1 afum|EAL89748.1 agla|OJJ89107.1 akaw|GAA87424.1 aluc|OJZ87612.1 anid|CBF85973.1 anig|EHA20809.1 aory|EIT80872.1 asyd|OJJ64427.1 ater|EAU39540.1 atub|OJI81037.1 aver|OJI98527.1 awen|OJJ40691.1 azon|OJJ44067.1

Group2304: abra|OJJ70915.1 acar|OOF98209.1 acla|EAW10489.1 afis|EAW25078.1 afla|EED55375.1 afum|EAL89747.1 agla|OJJ89115.1 akaw|GAA87425.1 aluc|OJZ87610.1 anid|CBF85970.1 anig|EHA20808.1 aory|EIT80925.1 asyd|OJJ64426.1 ater|EAU39545.1 atub|OJI81038.1 aver|OJI98526.1 awen|OJJ40689.1 azon|OJJ44066.1

Group2305: abra|OJJ70916.1 acar|OOF98208.1 acla|EAW10490.1 afis|EAW25077.1 afla|EED55376.1 afum|EAL89746.1 agla|OJJ89116.1 akaw|GAA87426.1 aluc|OJZ87609.1 anid|CBF85968.1 anig|EHA20807.1 aory|EIT80812.1 asyd|OJJ64425.1 ater|EAU39546.1 atub|OJI81039.1 aver|OJI98524.1 awen|OJJ40688.1 azon|OJJ44065.1

Group2306: abra|OJJ70918.1 acar|OOF98206.1 acla|EAW10492.1 afis|EAW25075.1 afla|EED55378.1 afum|EAL89744.1 agla|OJJ89117.1 akaw|GAA87428.1 aluc|OJZ87608.1 anid|CBF85964.1 anig|EHA20806.1 aory|EIT80830.1 asyd|OJJ64423.1 ater|EAU39550.1 atub|OJI81040.1 aver|OJI98522.1 awen|OJJ40686.1 azon|OJJ44063.1

Group2307: abra|OJJ70931.1 acar|OOF98195.1 acla|EAW10497.1 afis|EAW25070.1 afla|EED55384.1 afum|EAL89739.1 agla|OJJ89124.1 akaw|GAA83011.1 aluc|OJZ87596.1 anid|CBF85958.1 anig|EHA20796.1 aory|EIT80805.1 asyd|OJJ64418.1 ater|EAU39555.1 atub|OJI81051.1 aver|OJI98517.1 awen|OJJ40681.1 azon|OJJ44058.1

Group2308: abra|OJJ70933.1 acar|OOF89985.1 acla|EAW10499.1 afis|EAW25068.1 afla|EED55386.1 afum|EAL89737.2 agla|OJJ89126.1 akaw|GAA83009.1 aluc|OJZ87594.1 anid|CBF85954.1 anig|EHA20794.1 aory|EIT80870.1 asyd|OJJ64416.1 ater|EAU35815.1 atub|OJI81053.1 aver|OJI98515.1 awen|OJJ40675.1 azon|OJJ44056.1

Group2309: abra|OJJ70940.1 acar|OOF98192.1 acla|EAW10505.1 afis|EAW25062.1 afla|EED55391.1 afum|EAL89731.2 agla|OJJ89133.1 akaw|GAA83004.1 aluc|OJZ87588.1 anid|CBF85943.1 anig|EHA20789.1 aory|EIT80778.1 asyd|OJJ64412.1 ater|EAU35810.1 atub|OJI81060.1 aver|OJI98511.1 awen|OJJ40669.1 azon|OJJ44050.1

Group2310: abra|OJJ70941.1 acar|OOF98191.1 acla|EAW10506.1 afis|EAW25061.1 afla|EED55392.1 afum|EAL89729.1 agla|OJJ89134.1 akaw|GAA83003.1 aluc|OJZ87587.1 anid|CBF85941.1 anig|EHA20788.1 aory|EIT80784.1 asyd|OJJ64411.1 ater|EAU35809.1 atub|OJI81061.1 aver|OJI98510.1 awen|OJJ40668.1 azon|OJJ44049.1

Group2311: abra|OJJ70942.1 acar|OOF98190.1 acla|EAW10507.1 afis|EAW25060.1 afla|EED55393.1 afum|EAL89728.1 agla|OJJ89135.1 akaw|GAA83002.1 aluc|OJZ87586.1 anid|CBF85940.1 anig|EHA20787.1 aory|EIT80892.1 asyd|OJJ64410.1 ater|EAU35808.1 atub|OJI81062.1 aver|OJI98509.1 awen|OJJ40667.1 azon|OJJ44048.1

Group2312: abra|OJJ70944.1 acar|OOF98187.1 acla|EAW10512.1 afis|EAW25055.1 afla|EED55396.1 afum|EAL89723.2 agla|OJJ89137.1 akaw|GAA82999.1 aluc|OJZ87584.1 anid|CBF85933.1 anig|EHA20785.1 aory|EIT80802.1 asyd|OJJ64408.1 ater|EAU35805.1 atub|OJI81064.1 aver|OJI98506.1 awen|OJJ40665.1 azon|OJJ44044.1

Group2313: abra|OJJ70945.1 acar|OOF98186.1 acla|EAW10513.1 afis|EAW25054.1 afla|EED55397.1 afum|EAL89722.1 agla|OJJ89138.1 akaw|GAA82998.1 aluc|OJZ87583.1 anid|CBF85931.1 anig|EHA20784.1 aory|EIT80901.1 asyd|OJJ64407.1 ater|EAU35804.1 atub|OJI81065.1 aver|OJI98505.1 awen|OJJ40664.1 azon|OJJ44045.1

Group2314: abra|OJJ70946.1 acar|OOF98185.1 acla|EAW10514.1 afis|EAW25053.1 afla|EED55398.1 afum|EAL89721.1 agla|OJJ89139.1 akaw|GAA82997.1 aluc|OJZ87582.1 anid|CBF85929.1 anig|EHA20783.1 aory|EIT80773.1 asyd|OJJ64406.1 ater|EAU35803.1 atub|OJI81066.1 aver|OJI98504.1 awen|OJJ40663.1 azon|OJJ44043.1

Group2315: abra|OJJ70954.1 acar|OOF98176.1 acla|EAW10515.1 afis|EAW25051.1 afla|EED55400.1 afum|EAL89719.1 agla|OJJ89141.1 akaw|GAA82993.1 aluc|OJZ87578.1 anid|CBF85925.1 anig|EHA20780.1 aory|EIT80833.1 asyd|OJJ64405.1 ater|EAU35801.1 atub|OJI81071.1 aver|OJI98501.1 awen|OJJ40661.1 azon|OJJ44040.1

Group2316: abra|OJJ70955.1 acar|OOF98175.1 acla|EAW10516.1 afis|EAW25050.1 afla|EED55401.1 afum|EAL89718.1 agla|OJJ89142.1 akaw|GAA82992.1 aluc|OJZ87577.1 anid|CBF85923.1 anig|EHA20779.1 aory|EIT80841.1 asyd|OJJ64404.1 ater|EAU35800.1 atub|OJI81072.1 aver|OJI98500.1 awen|OJJ40660.1 azon|OJJ44039.1

Group2317: abra|OJJ70957.1 acar|OOF98173.1 acla|EAW10518.1 afis|EAW25048.1 afla|EED55403.1 afum|EAL89716.1 agla|OJJ89144.1 akaw|GAA82990.1 aluc|OJZ87575.1 anid|CBF85920.1 anig|EHA20777.1 aory|EIT80783.1 asyd|OJJ64402.1 ater|EAU35798.1 atub|OJI81074.1 aver|OJI98498.1 awen|OJJ40659.1 azon|OJJ44037.1

Group2318: abra|OJJ70958.1 acar|OOF98172.1 acla|EAW10519.1 afis|EAW25047.1 afla|EED55404.1 afum|EAL89715.1 agla|OJJ89145.1 akaw|GAA82989.1 aluc|OJZ87574.1 anid|CBF85918.1 anig|EHA20776.1 aory|EIT80839.1 asyd|OJJ64401.1 ater|EAU35797.1 atub|OJI81075.1 aver|OJI98497.1 awen|OJJ40658.1 azon|OJJ44036.1

Group2319: abra|OJJ70959.1 acar|OOF98171.1 acla|EAW10520.1 afis|EAW25046.1 afla|EED55405.1 afum|EAL89714.1 agla|OJJ89148.1 akaw|GAA82988.1 aluc|OJZ87573.1 anid|CBF85916.1 anig|EHA20775.1 aory|EIT80804.1 asyd|OJJ64400.1 ater|EAU35796.1 atub|OJI81076.1 aver|OJI98496.1 awen|OJJ40657.1 azon|OJJ44035.1

Group2320: abra|OJJ70963.1 acar|OOF98167.1 acla|EAW10524.1 afis|EAW25042.1 afla|EED55410.1 afum|EAL89710.1 agla|OJJ89152.1 akaw|GAA82984.1 aluc|OJZ87569.1 anid|CBF85908.1 anig|EHA20771.1 aory|EIT80900.1 asyd|OJJ64396.1 ater|EAU35792.1 atub|OJI81080.1 aver|OJI98492.1 awen|OJJ40647.1 azon|OJJ44032.1

Group2321: abra|OJJ70964.1 acar|OOF98166.1 acla|EAW10525.1 afis|EAW25041.1 afla|EED55411.1 afum|EAL89709.1 agla|OJJ89153.1 akaw|GAA82983.1 aluc|OJZ87568.1 anid|CBF85907.1 anig|EHA20770.1 aory|EIT80795.1 asyd|OJJ64395.1 ater|EAU35791.1 atub|OJI81081.1 aver|OJI98490.1 awen|OJJ40646.1 azon|OJJ44031.1

Group2322: abra|OJJ70968.1 acar|OOF98162.1 acla|EAW10528.1 afis|EAW25038.1 afla|EED55415.1 afum|EAL89706.1 agla|OJJ89155.1 akaw|GAA82980.1 aluc|OJZ87564.1 anid|CBF85901.1 anig|EHA20766.1 aory|EIT80889.1 asyd|OJJ64392.1 ater|EAU35788.1 atub|OJI81083.1 aver|OJI98487.1 awen|OJJ40644.1 azon|OJJ44028.1

Group2323: abra|OJJ70971.1 acar|OOF98159.1 acla|EAW10531.1 afis|EAW24998.1 afla|EED55418.1 afum|EAL89703.1 agla|OJJ89158.1 akaw|GAA82977.1 aluc|OJZ87561.1 anid|CBF85898.1 anig|EHA20763.1 aory|EIT80823.1 asyd|OJJ64390.1 ater|EAU35786.1 atub|OJI81086.1 aver|OJI98485.1 awen|OJJ40641.1 azon|OJJ44026.1

Group2324: abra|OJJ70973.1 acar|OOF98158.1 acla|EAW10532.1 afis|EAW24999.1 afla|EED55419.1 afum|EAL89699.1 agla|OJJ89161.1 akaw|GAA82976.1 aluc|OJZ87559.1 anid|CBF81499.1 anig|EHA20762.1 aory|EIT80834.1 asyd|OJJ59465.1 ater|EAU35785.1 atub|OJI81088.1 aver|OJJ02809.1 awen|OJJ40640.1 azon|OJJ44024.1

Group2325: abra|OJJ70976.1 acar|OOF98154.1 acla|EAW10535.1 afis|EAW25002.1 afla|EED55422.1 afum|EAL89696.1 agla|OJJ89165.1 akaw|GAA82973.1 aluc|OJZ87556.1 anid|CBF81486.1 anig|EHA20759.1 aory|EIT80883.1 ater|EAU35783.1 atub|OJI81091.1 awen|OJJ40637.1 azon|OJJ49046.1 asyd|OJJ59457.1 aver|OJJ02816.1

Group2326: abra|OJJ70977.1 acar|OOF98155.1 acla|EAW10536.1 afis|EAW25003.1 afla|EED55423.1 afum|EAL89695.1 agla|OJJ89166.1 akaw|GAA82972.1 aluc|OJZ87555.1 anid|CBF81484.1 anig|EHA20758.1 aory|EIT80840.1 asyd|OJJ59456.1 ater|EAU35782.1 atub|OJI81092.1 aver|OJJ02817.1 awen|OJJ40636.1 azon|OJJ49047.1

Group2327: abra|OJJ70979.1 acar|OOF98153.1 acla|EAW10538.1 afis|EAW25005.1 afla|EED55425.1 afum|EAL89693.1 agla|OJJ89168.1 akaw|GAA82970.1 aluc|OJZ87553.1 anid|CBF81481.1 anig|EHA20756.1 aory|EIT80871.1 asyd|OJJ59454.1 ater|EAU35780.1 atub|OJI81093.1 aver|OJJ02819.1 awen|OJJ40634.1 azon|OJJ49049.1

Group2328: abra|OJJ70980.1 acar|OOF98151.1 acla|EAW10539.1 afis|EAW25006.1 afla|EED55426.1 afum|EAL89691.2 agla|OJJ89169.1 akaw|GAA82969.1 aluc|OJZ87551.1 anid|CBF81512.1 anig|EHA20754.1 aory|EIT80772.1 asyd|OJJ59470.1 ater|EAU35779.1 atub|OJI81094.1 aver|OJJ02804.1 awen|OJJ40633.1 azon|OJJ49050.1

Group2329: abra|OJJ70981.1 acar|OOF98150.1 acla|EAW10540.1 afis|EAW25007.1 afla|EED55427.1 afum|EAL89690.1 agla|OJJ89170.1 akaw|GAA82968.1 aluc|OJZ87550.1 anid|CBF81514.1 anig|EHA20753.1 aory|EIT80826.1 asyd|OJJ59472.1 ater|EAU35778.1 atub|OJI81095.1 aver|OJJ02802.1 awen|OJJ40632.1 azon|OJJ49051.1

Group2330: abra|OJJ70982.1 acar|OOF98148.1 acla|EAW10541.1 afis|EAW25008.1 afla|EED55428.1 afum|EBA27387.1 agla|OJJ89173.1 akaw|GAA82967.1 aluc|OJZ87549.1 anid|CBF81516.1 anig|EHA20752.1 aory|EIT80794.1 asyd|OJJ59473.1 ater|EAU35777.1 atub|OJI81097.1 aver|OJJ02801.1 awen|OJJ40631.1 azon|OJJ49052.1

Group2331: abra|OJJ70983.1 acar|OOF98149.1 acla|EAW10542.1 afis|EAW25010.1 afla|EED55430.1 afum|EAL89686.1 agla|OJJ89174.1 akaw|GAA82966.1 aluc|OJZ87548.1 anid|CBF81518.1 anig|EHA20751.1

aory|EIT80803.1 asyd|OJJ59474.1 ater|EAU35776.1 atub|OJI81098.1 aver|OJJ02799.1 awen|OJJ40630.1 azon|OJJ49053.1

Group2332: abra|OJJ70984.1 acar|OOF98147.1 acla|EAW10543.1 afis|EAW25011.1 afla|EED55431.1 afum|EAL89685.1 agla|OJJ89175.1 akaw|GAA82965.1 aluc|OJZ87547.1 anid|CBF81519.1 anig|EHA20750.1 aory|EIT80824.1 asyd|OJJ59475.1 ater|EAU35775.1 atub|OJI81099.1 aver|OJJ02798.1 awen|OJJ40629.1 azon|OJJ49054.1

Group2333: abra|OJJ70989.1 acar|OOF98144.1 acla|EAW10547.1 afis|EAW25015.1 afla|EED55437.1 afum|EAL89681.1 agla|OJJ89181.1 akaw|GAA82962.1 aluc|OJZ87543.1 anid|CBF81527.1 anig|EHA20747.1 aory|EIT80927.1 asyd|OJJ59481.1 ater|EAU35771.1 atub|OJI81104.1 aver|OJJ02793.1 awen|OJJ40625.1 azon|OJJ49058.1

Group2334: abra|OJJ70991.1 acar|OOF98143.1 acla|EAW10548.1 afis|EAW25016.1 afla|EED55439.1 afum|EAL89680.1 agla|OJJ89182.1 akaw|GAA82960.1 aluc|OJZ87541.1 anid|CBF81529.1 anig|EHA20746.1 aory|EIT80844.1 asyd|OJJ59483.1 ater|EAU35770.1 atub|OJI81106.1 aver|OJJ02791.1 awen|OJJ40624.1 azon|OJJ49059.1

Group2335: abra|OJJ70996.1 acar|OOF90034.1 acla|EAW10552.1 afis|EAW25019.1 afla|EED55442.1 afum|EAL89676.1 agla|OJJ89187.1 akaw|GAA82952.1 aluc|OJZ87532.1 anid|CBF81538.1 anig|EHA20741.1 aory|EIT80858.1 asyd|OJJ59488.1 ater|EAU35767.1 atub|OJI81117.1 aver|OJJ02786.1 awen|OJJ40621.1 azon|OJJ49469.1

Group2336: abra|OJJ70998.1 acar|OOF98142.1 acla|EAW10555.1 afis|EAW25022.1 afla|EED55444.1 afum|EAL89673.1 agla|OJJ89190.1 akaw|GAA82950.1 aluc|OJZ87529.1 anid|CBF81541.1 anig|EHA20739.1 aory|EIT80864.1 asyd|OJJ59490.1 ater|EAU35765.1 atub|OJI81119.1 aver|OJJ02784.1 awen|OJJ40618.1 azon|OJJ49466.1

Group2337: abra|OJJ71001.1 acar|OOF98139.1 acla|EAW10558.1 afis|EAW25025.1 afla|EED55447.1 afum|EAL89670.1 agla|OJJ79159.1 akaw|GAA82947.1 aluc|OJZ87525.1 anid|CBF81547.1 anig|EHA20736.1 aory|EIT80809.1 asyd|OJJ59493.1 ater|EAU35762.1 atub|OJI81123.1 aver|OJJ02781.1 awen|OJJ40614.1 azon|OJJ49463.1

Group2338: abra|OJJ71008.1 acar|OOF98134.1 acla|EAW10560.1 afis|EAW25027.1 afla|EED55449.1 afum|EAL89668.1 agla|OJJ79157.1 akaw|GAA82945.1 aluc|OJZ87523.1 anid|CBF81551.1 anig|EHA20733.1 aory|EIT80929.1 asyd|OJJ59495.1 ater|EAU35760.1 atub|OJI81125.1 aver|OJJ02779.1 awen|OJJ40612.1 azon|OJJ49461.1

Group2339: abra|OJJ71013.1 acar|OOF98126.1 acla|EAW10564.1 afis|EAW25031.1 afla|EED55454.1 afum|EAL89664.1 agla|OJJ82242.1 akaw|GAA82941.1 aluc|OJZ87518.1 anid|CBF81564.1 anig|EHA20729.1 aory|EIT80774.1 asyd|OJJ59502.1 ater|EAU35756.1 atub|OJI81131.1 aver|OJJ02772.1 awen|OJJ40608.1 azon|OJJ49458.1

Group2340: abra|OJJ71016.1 acar|OOF98124.1 acla|EAW10566.1 afis|EAW25032.1 afla|EED55457.1 afum|EAL89662.1 agla|OJJ82244.1 akaw|GAA82939.1 aluc|OJZ87516.1 anid|CBF81567.1 anig|EHA20727.1 aory|EIT80779.1 asyd|OJJ59504.1 ater|EAU35755.1 atub|OJI81134.1 aver|OJJ02770.1 awen|OJJ40607.1 azon|OJJ49456.1

Group2341: abra|OJJ71019.1 acar|OOF98122.1 acla|EAW10568.1 afis|EAW25034.1 afla|EED55459.1 afum|EAL89660.1 agla|OJJ82255.1 akaw|GAA82936.1 aluc|OJZ87513.1 anid|CBF81574.1 anig|EHA20725.1 aory|EIT80788.1 asyd|OJJ59507.1 ater|EAU35751.1 atub|OJI81138.1 aver|OJJ02767.1 awen|OJJ40604.1 azon|OJJ49454.1

Group2342: abra|OJJ71021.1 acar|OOF98119.1 acla|EAW10570.1 afis|EAW25036.1 afla|EED55462.1 afum|EAL89658.1 agla|OJJ82257.1 akaw|GAA82934.1 aluc|OJZ87511.1 anid|CBF81583.1 anig|EHA20723.1 aory|EIT80855.1 asyd|OJJ59510.1 ater|EAU35747.1 atub|OJI81140.1 aver|OJJ02764.1 awen|OJJ40602.1 azon|OJJ49452.1

Group2343: abra|OJJ71022.1 acar|OOF98117.1 acla|EAW10571.1 afis|EAW25037.1 afla|EED55463.1 afum|EAL89657.1 agla|OJJ82258.1 akaw|GAA82933.1 aluc|OJZ87510.1 anid|CBF81585.1 anig|EHA20722.1 aory|EIT80922.1 asyd|OJJ59511.1 ater|EAU38720.1 atub|OJI81141.1 aver|OJJ02762.1 awen|OJJ40601.1 azon|OJJ49451.1

Group2344: abra|OJJ71023.1 acar|OOF98118.1 acla|EAW10572.1 afis|EAW24995.1 afla|EED55464.1 afum|EAL89656.1 agla|OJJ82259.1 akaw|GAA82932.1 aluc|OJZ87509.1 anid|CBF81586.1 anig|EHA20721.1 aory|EIT80875.1 asyd|OJJ59512.1 ater|EAU35746.1 atub|OJI81142.1 aver|OJJ02761.1 awen|OJJ40600.1 azon|OJJ49450.1

Group2345: abra|OJJ71028.1 acar|OOF98111.1 acla|EAW10579.1 afis|EAW24988.1 afla|EED55470.1 afum|EAL89648.1 agla|OJJ82274.1 akaw|GAA82927.1 aluc|OJZ87502.1 anid|CBF81599.1 anig|EHA20717.1 aory|EIT80811.1 asyd|OJJ59519.1 ater|EAU35739.1 atub|OJI81147.1 aver|OJJ02754.1 awen|OJJ40593.1 azon|OJJ49443.1

Group2346: abra|OJJ71030.1 acar|OOF98109.1 acla|EAW10583.1 afis|EAW24986.1 afla|EED55472.1 afum|EAL89646.1 agla|OJJ82276.1 akaw|GAA82925.1 aluc|OJZ87498.1 anid|CBF81603.1 anig|EHA20715.1 aory|EIT80838.1 asyd|OJJ59521.1 ater|EAU35737.1 atub|OJI81149.1 aver|OJJ02752.1 awen|OJJ40582.1 azon|OJJ49441.1

Group2347: abra|OJJ71036.1 acar|OOF98101.1 acla|EAW10586.1 afis|EAW24982.1 afla|EED55477.1 afum|EAL89642.1 agla|OJJ82279.1 akaw|GAA82919.1 aluc|OJZ87491.1 anid|CBF81614.1 anig|EHA20711.1 aory|EIT80831.1 asyd|OJJ59526.1 ater|EAU35734.1 atub|OJI81157.1 aver|OJJ02746.1 awen|OJJ40579.1 azon|OJJ49430.1

Group2348: abra|OJJ71038.1 acar|OOF98100.1 acla|EAW10587.1 afis|EAW24981.1 afla|EED55478.1 afum|EAL89641.1 agla|OJJ79136.1 akaw|GAA82918.1 aluc|OJZ87490.1 anid|CBF81612.1 anig|EHA20710.1 aory|EIT80854.1 asyd|OJJ59525.1 ater|EAU35733.1 atub|OJI81158.1 aver|OJJ02747.1 awen|OJJ40578.1 azon|OJJ49431.1

Group2349: abra|OJJ71040.1 acar|OOF98098.1 acla|EAW10589.1 afis|EAW24979.1 afla|EED55480.1 afum|EAL89639.1 agla|OJJ79138.1 akaw|GAA82916.1 aluc|OJZ87488.1 anid|CBF81608.1 anig|EHA20708.1 aory|EIT80915.1 asyd|OJJ59523.1 ater|EAU35729.1 atub|OJI81160.1 aver|OJJ02749.1 awen|OJJ40576.1 azon|OJJ49434.1

Group2350: abra|OJJ71042.1 acar|OOF98097.1 acla|EAW10592.1 afis|EAW24976.1 afla|EED55484.1 afum|EAL89636.1 agla|OJJ79141.1 akaw|GAA82914.1 aluc|OJZ87486.1 anid|CBF81629.1 anig|EHA20707.1 aory|EIT80868.1 asyd|OJJ59530.1 ater|EAU35725.1 atub|OJI81162.1 aver|OJJ02740.1 awen|OJJ40573.1 azon|OJJ49438.1

Group2351: abra|OJJ71045.1 acar|OOF98094.1 acla|EAW10593.1 afis|EAW24975.1 afla|EED55485.1 afum|EAL89635.1 agla|OJJ82241.1 akaw|GAA82911.1 aluc|OJZ87481.1 anid|CBF81630.1 anig|EHA20703.1 aory|EIT80859.1 asyd|OJJ59531.1 ater|EAU35723.1 atub|OJI81167.1 aver|OJJ02739.1 awen|OJJ40572.1 azon|OJJ49429.1

Group2352: abra|OJJ71047.1 acar|OOF98092.1 acla|EAW10595.1 afis|EAW24973.1 afla|EED55486.1 afum|EAL89633.1 agla|OJJ82239.1 akaw|GAA82909.1 aluc|OJZ87479.1 anid|CBF81634.1 anig|EHA20701.1 aory|EIT80885.1 asyd|OJJ59533.1 ater|EAU35721.1 atub|OJI81169.1 aver|OJJ02737.1 awen|OJJ40570.1 azon|OJJ49427.1

Group2353: abra|OJJ71048.1 acar|OOF98091.1 acla|EAW10596.1 afis|EAW24972.1 afla|EED55487.1 afum|EBA27384.1 agla|OJJ82238.1 akaw|GAA82908.1 aluc|OJZ87478.1 anid|CBF81636.1 anig|EHA20700.1 aory|EIT80911.1 asyd|OJJ59534.1 ater|EAU35720.1 atub|OJI81170.1 aver|OJJ02736.1 awen|OJJ40569.1 azon|OJJ49426.1

Group2354: abra|OJJ71049.1 acar|OOF98090.1 acla|EAW10597.1 afis|EAW24971.1 afla|EED55490.1 afum|EAL89630.2 agla|OJJ82237.1 akaw|GAA82907.1 aluc|OJZ87477.1 anid|CBF81638.1 anig|EHA20699.1 aory|EIT80796.1 asyd|OJJ59535.1 ater|EAU35719.1 atub|OJI81171.1 aver|OJJ02735.1 awen|OJJ40568.1 azon|OJJ49425.1

Group2355: abra|OJJ71052.1 acar|OOF98087.1 acla|EAW10603.1 afis|EAW24965.1 afla|EED55495.1 afum|EAL89623.1 agla|OJJ82224.1 akaw|GAA82905.1 aluc|OJZ87475.1 anid|CBF81647.1 anig|EHA20695.1 aory|EIT80847.1 asyd|OJJ59541.1 ater|EAU35709.1 atub|OJI81173.1 aver|OJJ02727.1 awen|OJJ40558.1 azon|OJJ49419.1

Group2356: abra|OJJ71054.1 acar|OOF98085.1 acla|EAW14735.1 afis|EAW19536.1 afla|EED54353.1 afum|EAL86970.2 agla|OJJ85406.1 akaw|GAA82902.1 aluc|OJZ87473.1 anid|CBF76891.1 anig|EHA20693.1 aory|EIT78008.1 asyd|OJJ61335.1 ater|EAU31542.1 atub|OJI81175.1 aver|OJJ00659.1 awen|OJJ34757.1 aory|EIT75941.1

Group2357: abra|OJJ71061.1 acar|OOF98082.1 acla|EAW10598.1 afis|EAW24970.1 afla|EED55491.1 afum|EAL89629.1 agla|OJJ82236.1 akaw|GAA82898.1 aluc|OJZ87467.1 anid|CBF81640.1 anig|EHA20689.1 aory|EIT80891.1 asyd|OJJ59536.1 ater|EAU35715.1 atub|OJI81181.1 aver|OJJ02733.1 awen|OJJ40564.1 azon|OJJ49424.1

Group2358: abra|OJJ71075.1 acar|OOF98069.1 acla|EAW10604.1 afis|EAW24964.1 afla|EED55497.1 afum|EAL89622.1 agla|OJJ82226.1 akaw|GAA82887.1 aluc|OJZ87455.1 anid|CBF81649.1 anig|EHA20676.1 aory|EIT80869.1 asyd|OJJ59542.1 ater|EAU35707.1 atub|OJI81193.1 aver|OJJ02726.1 awen|OJJ40556.1 azon|OJJ49418.1

Group2359: abra|OJJ71103.1 acar|OOF90221.1 acla|EAW10388.1 afis|EAW25205.1 afla|EED53664.1 afum|EAL89866.2 agla|OJJ88951.1 akaw|GAA82865.1 aluc|OJZ87432.1 anid|CBF85221.1 anig|EHA20653.1 aory|EIT75288.1 asyd|OJJ62997.1 ater|EAU34349.1 atub|OJI81216.1 aver|OJI97970.1 awen|OJJ40840.1 azon|OJJ46822.1

Group2360: abra|OJJ71104.1 acar|OOF90222.1 acla|EAW10387.1 afis|EAW25206.1 afla|EED53662.1 afum|EAL89867.1 agla|OJJ88952.1 akaw|GAA82864.1 aluc|OJZ87431.1 anid|CBF85223.1 anig|EHA20652.1

aory|EIT75326.1 asyd|OJJ62996.1 ater|EAU34348.1 atub|OJI81217.1 aver|OJI97971.1 awen|OJJ40841.1 azon|OJJ46823.1

Group2361: abra|OJJ71105.1 acar|OOF90223.1 acla|EAW10386.1 afis|EAW25207.1 afla|EED53660.1 afum|EAL89868.1 agla|OJJ79791.1 akaw|GAA82863.1 aluc|OJZ87430.1 anid|CBF85225.1 anig|EHA20651.1 aory|EIT75316.1 asyd|OJJ62995.1 ater|EAU34347.1 atub|OJI81218.1 aver|OJI97972.1 awen|OJJ35846.1 azon|OJJ45132.1

Group2362: abra|OJJ71112.1 acar|OOF98038.1 acla|EAW10380.1 afis|EAW25213.1 afla|EED53667.1 afum|EAL89874.1 agla|OJJ88948.1 akaw|GAA82857.1 aluc|OJZ87422.1 anid|CBF85606.1 anig|EHA20645.1 aory|EIT75347.1 asyd|OJJ64197.1 ater|EAU34370.1 atub|OJI81226.1 aver|OJI98286.1 awen|OJJ40875.1 azon|OJJ46812.1

Group2363: abra|OJJ71117.1 acar|OOF98033.1 acla|EAW10378.1 afis|EAW25215.1 afla|EED53692.1 afum|EAL89876.1 agla|OJJ88944.1 akaw|GAA82853.1 aluc|OJZ87418.1 anid|CBF85254.1 anig|EHA20642.1 aory|EIT75323.1 asyd|OJJ62921.1 ater|EAU34372.1 atub|OJI81230.1 aver|OJI98068.1 awen|OJJ40878.1 azon|OJJ44506.1

Group2364: abra|OJJ71119.1 acar|OOF98031.1 acla|EAW10376.1 afis|EAW25217.1 afla|EED53690.1 afum|EAL89878.1 agla|OJJ79184.1 akaw|GAA82851.1 aluc|OJZ87417.1 anid|CBF85273.1 anig|EHA20641.1 aory|EIT75328.1 asyd|OJJ62920.1 ater|EAU34374.1 atub|OJI81231.1 aver|OJI98070.1 awen|OJJ40880.1 azon|OJJ44508.1

Group2365: abra|OJJ71125.1 acar|OOF98028.1 acla|EAW10374.1 afis|EAW25218.1 afla|EED53688.1 afum|EAL89879.1 agla|OJJ79186.1 akaw|GAA82850.1 aluc|OJZ87410.1 anid|CBF85299.1 anig|EHA20640.1 aory|EIT75354.1 asyd|OJJ62917.1 ater|EAU34377.1 atub|OJI81235.1 aver|OJI98073.1 awen|OJJ40883.1 azon|OJJ44512.1

Group2366: abra|OJJ71128.1 acar|OOF98024.1 acla|EAW10370.1 afis|EAW25222.1 afla|EED53684.1 afum|EAL89883.1 agla|OJJ79189.1 akaw|GAA82846.1 aluc|OJZ87407.1 anid|CBF85258.1 anig|EHA20637.1 aory|EIT75281.1 asyd|OJJ62911.1 ater|EAU34381.1 atub|OJI81239.1 aver|OJI98078.1 awen|OJJ40888.1 azon|OJJ44517.1

Group2367: abra|OJJ71132.1 acar|OOF98020.1 acla|EAW10365.1 afis|EAW25227.1 afla|EED53671.1 afum|EAL89888.1 agla|OJJ79181.1 akaw|GAA82842.1 aluc|OJZ87403.1 anid|CBF85269.1 anig|EHA20632.1 aory|EIT75290.1 asyd|OJJ62908.1 ater|EAU34385.1 atub|OJI81243.1 aver|OJI98080.1 awen|OJJ40892.1 azon|OJJ44521.1

Group2368: abra|OJJ71133.1 acar|OOF98019.1 acla|EAW10364.1 afis|EAW25228.1 afla|EED53672.1 afum|EAL89889.1 agla|OJJ79180.1 akaw|GAA82841.1 aluc|OJZ87402.1 anid|CBF85262.1 anig|EHA20631.1 aory|EIT75302.1 asyd|OJJ62905.1 ater|EAU34386.1 atub|OJI81244.1 aver|OJI98084.1 awen|OJJ40893.1 azon|OJJ44522.1

Group2369: abra|OJJ71142.1 acar|OOF98014.1 acla|EAW10359.1 afis|EAW25234.1 afla|EED53675.1 afum|EAL89895.1 agla|OJJ89203.1 akaw|GAA82835.1 aluc|OJZ87396.1 anid|CBF85334.1 anig|EHA20626.1 aory|EIT75320.1 asyd|OJJ64021.1 ater|EAU34389.1 atub|OJI81250.1 aver|OJI98107.1 awen|OJJ40946.1 azon|OJJ44527.1

Group2370: abra|OJJ71143.1 acar|OOF98013.1 acla|EAW10358.1 afis|EAW25235.1 afla|EED53676.1 afum|EAL89896.2 agla|OJJ89204.1 akaw|GAA82834.1 aluc|OJZ87395.1 anid|CBF85332.1 anig|EHA20625.1 aory|EIT75309.1 asyd|OJJ64020.1 ater|EAU34390.1 atub|OJI81251.1 aver|OJI98106.1 awen|OJJ40947.1 azon|OJJ44528.1

Group2371: abra|OJJ71145.1 acar|OOF98011.1 acla|EAW10361.1 afis|EAW25232.1 afla|EED53678.1 afum|EAL89893.1 agla|OJJ79175.1 akaw|GAA82832.1 aluc|OJZ87393.1 anid|CBF85329.1 anig|EHA20623.1 aory|EIT75284.1 asyd|OJJ64018.1 ater|EAU34391.1 atub|OJI81253.1 aver|OJI98104.1 awen|OJJ40898.1 azon|OJJ44525.1

Group2372: abra|OJJ71150.1 acar|OOF98007.1 acla|EAW10352.1 afis|EAW25242.1 afla|EED53702.1 afum|EAL89904.1 agla|OJJ89208.1 akaw|GAA82827.1 aluc|OJZ87388.1 anid|CBF85319.1 anig|EHA20617.1 aory|EIT75296.1 asyd|OJJ64011.1 ater|EAU34367.1 atub|OJI81259.1 aver|OJI98099.1 awen|OJJ40955.1 azon|OJJ44538.1

Group2373: abra|OJJ71151.1 acar|OOF98006.1 acla|EAW10346.1 afis|EAW25243.1 afla|EED53700.1 afum|EAL89905.1 agla|OJJ89209.1 akaw|GAA82826.1 aluc|OJZ87387.1 anid|CBF85321.1 anig|EHA20616.1 aory|EIT75325.1 asyd|OJJ64012.1 ater|EAU34360.1 atub|OJI81260.1 aver|OJI98100.1 awen|OJJ40956.1 azon|OJJ44537.1

Group2374: abra|OJJ71158.1 acar|OOF98000.1 acla|EAW10339.1 afis|EAW25248.1 afla|EED53716.1 afum|EAL89910.1 agla|OJJ89222.1 akaw|GAA82820.1 aluc|OJZ87381.1 anid|CBF85341.1 anig|EHA20612.1 aory|EIT75359.1 asyd|OJJ64025.1 ater|EAU34355.1 atub|OJI81265.1 aver|OJI98111.1 awen|OJJ40964.1 azon|OJJ44545.1

Group2375: abra|OJJ71167.1 acar|OOF97996.1 acla|EAW10336.1 afis|EAW25253.1 afla|EED53722.1 afum|EAL89916.1 agla|OJJ89227.1 akaw|GAA82814.1 aluc|OJZ87374.1 anid|CBF85349.1 anig|EHA20607.1 aory|EIT75275.1 asyd|OJJ64033.1 ater|EAU34398.1 atub|OJI81272.1 aver|OJI98117.1 awen|OJJ40967.1 azon|OJJ44551.1

Group2376: abra|OJJ71169.1 acar|OOF97994.1 acla|EAW10333.1 afis|EAW25259.1 afla|EED53724.1 afum|EAL89921.1 agla|OJJ89230.1 akaw|GAA82812.1 aluc|OJZ87372.1 anid|CBF85360.1 anig|EHA20605.1 aory|EIT75308.1 asyd|OJJ64039.1 ater|EAU34400.1 atub|OJI81274.1 aver|OJI98123.1 awen|OJJ40972.1 azon|OJJ44558.1

Group2377: abra|OJJ71170.1 acar|OOF97993.1 acla|EAW10332.1 afla|EED53725.1 agla|OJJ89231.1 akaw|GAA82811.1 aluc|OJZ87371.1 anid|CBF85362.1 anig|EHA20604.1 aory|EIT75319.1 asyd|OJJ64040.1 ater|EAU34401.1 atub|OJI81275.1 aver|OJI98124.1 awen|OJJ40973.1 afis|EAW25260.1 afum|EAL89922.1 azon|OJJ44559.1

Group2378: abra|OJJ71173.1 acar|OOF97992.1 acla|EAW10331.1 afis|EAW25263.1 afla|EED53727.1 afum|EAL89926.2 agla|OJJ89233.1 akaw|GAA82810.1 aluc|OJZ87369.1 anid|CBF85365.1 anig|EHA20603.1 aory|EIT75277.1 asyd|OJJ64043.1 ater|EAU34402.1 atub|OJI81276.1 aver|OJI98127.1 awen|OJJ40975.1 azon|OJJ47355.1

Group2379: abra|OJJ71174.1 acar|OOF97991.1 acla|EAW10330.1 afis|EAW25264.1 afla|EED53728.1 afum|EAL89927.1 agla|OJJ89234.1 akaw|GAA82809.1 aluc|OJZ87368.1 anid|CBF85367.1 anig|EHA20602.1 aory|EIT75295.1 asyd|OJJ64045.1 ater|EAU34403.1 atub|OJI81277.1 aver|OJI98128.1 awen|OJJ40976.1 azon|OJJ44563.1

Group2380: abra|OJJ71179.1 acar|OOF97987.1 acla|EAW10327.1 afis|EAW25267.1 afla|EED53731.1 afum|EAL89930.1 agla|OJJ89241.1 akaw|GAA82806.1 aluc|OJZ87364.1 anid|CBF85374.1 anig|EHA20599.1 aory|EIT75331.1 asyd|OJJ64048.1 ater|EAU34408.1 atub|OJI81280.1 aver|OJI98133.1 awen|OJJ40980.1 azon|OJJ44567.1

Group2381: abra|OJJ71182.1 acar|OOF97984.1 acla|EAW10322.1 afis|EAW25270.1 afla|EED53735.1 afum|EAL89933.1 agla|OJJ89243.1 akaw|GAA82803.1 aluc|OJZ87361.1 anid|CBF85382.1 anig|EHA20596.1 aory|EIT75338.1 asyd|OJJ64051.1 ater|EAU34413.1 atub|OJI81284.1 aver|OJI98137.1 awen|OJJ40984.1 azon|OJJ44570.1

Group2382: abra|OJJ71183.1 acar|OOF97983.1 acla|EAW10321.1 afis|EAW25271.1 afla|EED53736.1 afum|EAL89934.1 agla|OJJ89244.1 akaw|GAA82802.1 aluc|OJZ87360.1 anid|CBF85384.1 anig|EHA20595.1 aory|EIT75335.1 asyd|OJJ64053.1 ater|EAU34414.1 atub|OJI81285.1 aver|OJI98138.1 awen|OJJ40985.1 azon|OJJ44572.1

Group2383: abra|OJJ71184.1 acar|OOF97982.1 acla|EAW10320.1 afis|EAW25272.1 afla|EED53737.1 afum|EAL89935.1 agla|OJJ89245.1 akaw|GAA82801.1 aluc|OJZ87359.1 anid|CBF85385.1 anig|EHA20594.1 aory|EIT75311.1 asyd|OJJ64054.1 ater|EAU34415.1 atub|OJI81286.1 aver|OJI98139.1 awen|OJJ40986.1 azon|OJJ44573.1

Group2384: abra|OJJ71185.1 acar|OOF97981.1 acla|EAW10318.1 afis|EAW25274.1 afla|EED53739.1 afum|EAL89937.1 agla|OJJ89248.1 akaw|GAA82800.1 aluc|OJZ87358.1 anid|CBF85389.1 anig|EHA20593.1 aory|EIT75291.1 asyd|OJJ64056.1 ater|EAU34417.1 atub|OJI81287.1 aver|OJI98141.1 awen|OJJ40988.1 azon|OJJ44575.1

Group2385: abra|OJJ71187.1 acar|OOF97980.1 acla|EAW10316.1 afis|EAW25276.1 afla|EED53742.1 afum|EAL89939.1 agla|OJJ89250.1 akaw|GAA82798.1 aluc|OJZ87357.1 anid|CBF85392.1 anig|EHA20591.1 aory|EIT75317.1 asyd|OJJ64058.1 ater|EAU34419.1 atub|OJI81288.1 aver|OJI98143.1 awen|OJJ40990.1 azon|OJJ44578.1

Group2386: abra|OJJ71188.1 acar|OOF97979.1 acla|EAW10315.1 afis|EAW25277.1 afla|EED53744.1 afum|EAL89940.1 agla|OJJ89251.1 akaw|GAA82797.1 aluc|OJZ87356.1 anid|CBF85394.1 anig|EHA20590.1 aory|EIT75351.1 asyd|OJJ64059.1 ater|EAU34420.1 atub|OJI81289.1 aver|OJI98145.1 awen|OJJ40991.1 azon|OJJ44580.1

Group2387: abra|OJJ71189.1 acar|OOF97978.1 acla|EAW10314.1 afis|EAW25278.1 afla|EED53745.1 afum|EAL89941.1 agla|OJJ89252.1 akaw|GAA82796.1 aluc|OJZ87355.1 anid|CBF85396.1 anig|EHA20589.1 aory|EIT75313.1 asyd|OJJ64060.1 ater|EAU34421.1 atub|OJI81290.1 aver|OJI98146.1 awen|OJJ40992.1 azon|OJJ44581.1

Group2388: abra|OJJ71192.1 acar|OOF97975.1 acla|EAW10312.1 afis|EAW25281.1 afla|EED53747.1 afum|EAL89944.1 agla|OJJ89255.1 akaw|GAA82793.1 aluc|OJZ87352.1 anid|CBF85418.1 anig|EHA20586.1 aory|EIT75318.1 asyd|OJJ64073.1 ater|EAU34422.1 atub|OJI81293.1 aver|OJI98160.1 awen|OJJ40994.1 azon|OJJ44583.1

Group2389: abra|OJJ71198.1 acar|OOF97970.1 acla|EAW10305.1 afis|EAW25287.1 afla|EED53752.1 afum|EAL89950.1 agla|OJJ89260.1 akaw|GAA82785.1 aluc|OJZ87344.1 anid|CBF85429.1 anig|EHA20581.1

asyd|OJJ64083.1 ater|EAU34428.1 atub|OJI81300.1 aver|OJI98168.1 awen|OJJ40999.1 azon|OJJ44591.1 aory|EIT78420.1

Group2390: abra|OJJ71203.1 acar|OOF97968.1 acla|EAW10304.1 afis|EAW25288.1 afla|EED53753.1 afum|EAL89951.1 agla|OJJ89261.1 akaw|GAA82781.1 aluc|OJZ87340.1 anid|CBF85431.1 anig|EHA20577.1 aory|EIT78400.1 asyd|OJJ64084.1 ater|EAU34429.1 atub|OJI81304.1 aver|OJI98169.1 awen|OJJ41000.1 azon|OJJ44594.1

Group2391: abra|OJJ71204.1 acar|OOF97967.1 acla|EAW10303.1 afis|EAW25289.1 afla|EED53754.1 afum|EAL89952.1 agla|OJJ89262.1 akaw|GAA82780.1 aluc|OJZ87339.1 anid|CBF85433.1 anig|EHA20576.1 aory|EIT78451.1 asyd|OJJ64085.1 ater|EAU34430.1 atub|OJI81305.1 aver|OJI98170.1 awen|OJJ41001.1 azon|OJJ44593.1

Group2392: abra|OJJ71205.1 acar|OOF97965.1 acla|EAW10302.1 afis|EAW25290.1 afla|EED53755.1 afum|EAL89953.1 agla|OJJ89263.1 akaw|GAA82779.1 aluc|OJZ87338.1 anid|CBF85435.1 anig|EHA20575.1 aory|EIT78386.1 asyd|OJJ64086.1 ater|EAU34431.1 atub|OJI81306.1 aver|OJI98171.1 awen|OJJ41002.1 azon|OJJ44592.1

Group2393: abra|OJJ71206.1 acar|OOF97966.1 acla|EAW10301.1 afis|EAW25291.1 afla|EED53756.1 afum|EAL89954.1 agla|OJJ89265.1 akaw|GAA82778.1 aluc|OJZ87337.1 anid|CBF85413.1 anig|EHA20574.1 aory|EIT78379.1 asyd|OJJ64069.1 ater|EAU34432.1 atub|OJI81307.1 aver|OJI98156.1 awen|OJJ41003.1 azon|OJJ44595.1

Group2394: abra|OJJ71210.1 acar|OOF97956.1 acla|EAW10300.1 afis|EAW25292.1 afla|EED53757.1 afum|EAL89955.1 agla|OJJ89269.1 akaw|GAA82774.1 aluc|OJZ87333.1 anid|CBF85411.1 anig|EHA20569.1 aory|EIT78389.1 asyd|OJJ64068.1 ater|EAU34433.1 atub|OJI81311.1 aver|OJI98155.1 awen|OJJ41006.1 azon|OJJ44602.1

Group2395: abra|OJJ71211.1 acar|OOF97955.1 acla|EAW10298.1 afis|EAW25294.1 afla|EED53759.1 afum|EAL89957.1 agla|OJJ89270.1 akaw|GAA82773.1 aluc|OJZ87332.1 anid|CBF85409.1 anig|EHA20568.1 aory|EIT78363.1 asyd|OJJ64067.1 ater|EAU34434.1 atub|OJI81312.1 aver|OJI98154.1 awen|OJJ41008.1 azon|OJJ44601.1

Group2396: abra|OJJ71227.1 acar|OOF97952.1 acla|EAW10295.1 afis|EAW25297.1 afla|EED53763.1 afum|EAL89960.1 agla|OJJ89273.1 akaw|GAA82762.1 aluc|OJZ87320.1 anid|CBF85405.1 anig|EHA20554.1 aory|EIT78455.1 asyd|OJJ64064.1 ater|EAU34438.1 atub|OJI81325.1 aver|OJI98151.1 awen|OJJ41011.1 azon|OJJ44598.1

Group2397: abra|OJJ71241.1 acar|OOF97926.1 acla|EAW09108.1 afis|EAW25717.1 afla|EED53769.1 afum|EAL84392.2 agla|OJJ85442.1 akaw|GAA82746.1 aluc|OJZ87301.1 anid|CBF77088.1 anig|EHA20541.1 aory|EIT78470.1 asyd|OJJ56975.1 ater|EAU34440.1 atub|OJI81339.1 aver|OJJ03313.1 awen|OJJ33601.1 azon|OJJ44472.1

Group2398: abra|OJJ71244.1 acar|OOF97924.1 acla|EAW10291.1 afis|EAW25300.1 afla|EED53774.1 afum|EAL89963.2 agla|OJJ89280.1 akaw|GAA82743.1 aluc|OJZ87298.1 anid|CBF75193.1 anig|EHA20539.1 aory|EIT78388.1 asyd|OJJ59938.1 ater|EAU34448.1 atub|OJI81342.1 aver|OJJ02349.1 awen|OJJ41015.1 azon|OJJ44603.1

Group2399: abra|OJJ71245.1 acar|OOF97923.1 acla|EAW10290.1 afis|EAW25301.1 afla|EED53776.1 afum|EAL89964.1 agla|OJJ89281.1 akaw|GAA82742.1 aluc|OJZ87297.1 anid|CBF75195.1 anig|EHA20538.1 aory|EIT78454.1 asyd|OJJ59937.1 ater|EAU34449.1 atub|OJI81343.1 aver|OJJ02350.1 awen|OJJ41016.1 azon|OJJ42625.1

Group2400: abra|OJJ71248.1 acar|OOF97920.1 acla|EAW10287.1 afis|EAW25305.1 afla|EED53783.1 afum|EAL89968.2 agla|OJJ89284.1 akaw|GAA82739.1 aluc|OJZ87294.1 anid|CBF75296.1 anig|EHA20535.1 aory|EIT78359.1 asyd|OJJ59884.1 ater|EAU34453.1 atub|OJI81346.1 aver|OJJ02402.1 awen|OJJ41019.1 azon|OJJ44606.1

Group2401: abra|OJJ71255.1 acar|OOF97916.1 acla|EAW10281.1 afis|EAW25313.1 afla|EED53788.1 afum|EAL89974.1 agla|OJJ83848.1 akaw|GAA82733.1 aluc|OJZ87288.1 anid|CBF75287.1 anig|EHA20529.1 aory|EIT78401.1 asyd|OJJ59889.1 ater|EAU34458.1 atub|OJI81353.1 aver|OJJ02397.1 awen|OJJ41053.1 azon|OJJ44614.1

Group2402: abra|OJJ71256.1 acar|OOF97915.1 acla|EAW10280.1 afis|EAW25314.1 afla|EED53789.1 afum|EAL89975.1 agla|OJJ83849.1 akaw|GAA82732.1 aluc|OJZ87287.1 anid|CBF75285.1 anig|EHA20528.1 aory|EIT78413.1 asyd|OJJ59890.1 ater|EAU34459.1 atub|OJI81354.1 aver|OJJ02396.1 awen|OJJ41050.1 azon|OJJ44612.1

Group2403: abra|OJJ71258.1 acar|OOF97914.1 acla|EAW10278.1 afis|EAW25316.1 afla|EED53791.1 afum|EAL89977.2 agla|OJJ83854.1 akaw|GAA82731.1 aluc|OJZ87285.1 anid|CBF75283.1 anig|EHA20527.1 aory|EIT78447.1 asyd|OJJ59892.1 ater|EAU34460.1 atub|OJI81356.1 aver|OJJ02394.1 awen|OJJ41047.1 azon|OJJ44616.1

Group2404: abra|OJJ71259.1 acar|OOF97913.1 acla|EAW10277.1 afis|EAW25317.1 afla|EED53792.1 afum|EAL89978.2 agla|OJJ83855.1 akaw|GAA82730.1 aluc|OJZ87284.1 anid|CBF75281.1 anig|EHA20526.1 aory|EIT78371.1 asyd|OJJ59893.1 ater|EAU34464.1 atub|OJI81357.1 aver|OJJ02393.1 awen|OJJ41046.1 azon|OJJ44617.1

Group2405: abra|OJJ71262.1 acar|OOF97912.1 acla|EAW10275.1 afis|EAW25319.1 afla|EED53794.1 afum|EAL89980.2 agla|OJJ83857.1 akaw|GAA82727.1 aluc|OJZ87281.1 anid|CBF75269.1 anig|EHA20523.1 aory|EIT78456.1 asyd|OJJ59895.1 ater|EAU34465.1 atub|OJI81360.1 aver|OJJ02391.1 awen|OJJ41044.1 azon|OJJ44619.1

Group2406: abra|OJJ71263.1 acar|OOF97911.1 acla|EAW10274.1 afis|EAW25320.1 afla|EED53795.1 afum|EAL89981.1 agla|OJJ83858.1 akaw|GAA82726.1 aluc|OJZ87280.1 anid|CBF75267.1 anig|EHA20522.1 aory|EIT78395.1 asyd|OJJ59896.1 ater|EAU34466.1 atub|OJI81361.1 aver|OJJ02390.1 awen|OJJ41043.1 azon|OJJ44620.1

Group2407: abra|OJJ71264.1 acar|OOF97910.1 acla|EAW10273.1 afis|EAW25321.1 afla|EED53796.1 afum|EAL89982.1 agla|OJJ83859.1 akaw|GAA82725.1 aluc|OJZ87279.1 anid|CBF75265.1 anig|EHA20521.1 aory|EIT78365.1 asyd|OJJ59897.1 ater|EAU34467.1 atub|OJI81362.1 aver|OJJ02389.1 awen|OJJ41042.1 azon|OJJ44621.1

Group2408: abra|OJJ71265.1 acar|OOF97909.1 acla|EAW10272.1 afis|EAW25322.1 afla|EED53797.1 afum|EAL89983.1 agla|OJJ83860.1 akaw|GAA82724.1 aluc|OJZ87278.1 anid|CBF75263.1 anig|EHA20520.1 aory|EIT78429.1 asyd|OJJ59898.1 ater|EAU34468.1 atub|OJI81363.1 aver|OJJ02388.1 awen|OJJ41041.1 azon|OJJ44624.1

Group2409: abra|OJJ71267.1 acar|OOF97908.1 acla|EAW10271.1 afis|EAW25323.1 afla|EED53798.1 afum|EAL89984.1 agla|OJJ83861.1 akaw|GAA82723.1 aluc|OJZ87277.1 anid|CBF75261.1 anig|EHA20519.1 aory|EIT78438.1 asyd|OJJ59899.1 ater|EAU34469.1 atub|OJI81364.1 aver|OJJ02387.1 awen|OJJ41040.1 azon|OJJ44626.1

Group2410: abra|OJJ71270.1 acar|OOF97902.1 acla|EAW10268.1 afis|EAW25326.1 afla|EED53803.1 afum|EAL89987.1 agla|OJJ83870.1 akaw|GAA82720.1 aluc|OJZ87274.1 anid|CBF75254.1 anig|EHA20515.1 aory|EIT78467.1 asyd|OJJ59904.1 ater|EAU34473.1 atub|OJI81368.1 aver|OJJ02381.1 awen|OJJ41031.1 azon|OJJ44634.1

Group2411: abra|OJJ71271.1 acar|OOF97903.1 acla|EAW10267.1 afis|EAW25327.1 afla|EED53804.1 afum|EAL89988.1 agla|OJJ83869.1 akaw|GAA82719.1 aluc|OJZ87273.1 anid|CBF75252.1 anig|EHA20514.1 aory|EIT78360.1 asyd|OJJ59905.1 ater|EAU34474.1 atub|OJI81369.1 aver|OJJ02380.1 awen|OJJ41032.1 azon|OJJ44633.1

Group2412: abra|OJJ71276.1 acar|OOF97897.1 acla|EAW10257.1 afis|EAW25336.1 afla|EED53810.1 afum|EAL89999.1 agla|OJJ83871.1 akaw|GAA82713.1 aluc|OJZ87267.1 anid|CBF75242.1 anig|EHA20509.1 aory|EIT78463.1 asyd|OJJ59910.1 atub|OJI81375.1 aver|OJJ02376.1 awen|OJJ41030.1 azon|OJJ44635.1 afla|EED53809.1

Group2413: abra|OJJ71277.1 acar|OOF97896.1 acla|EAW10258.1 afis|EAW25335.1 afla|EED53811.1 afum|EAL89998.1 agla|OJJ83872.1 akaw|GAA82712.1 aluc|OJZ87266.1 anid|CBF75240.1 anig|EHA20508.1 aory|EIT78414.1 asyd|OJJ59911.1 ater|EAU34478.1 atub|OJI81376.1 aver|OJJ02375.1 awen|OJJ41029.1 azon|OJJ44636.1

Group2414: abra|OJJ71279.1 acar|OOF97892.1 acla|EAW10261.1 afis|EAW25332.1 afla|EED53814.1 afum|EAL89994.1 agla|OJJ83875.1 akaw|GAA82709.1 aluc|OJZ87264.1 anid|CBF75236.1 anig|EHA20506.1 aory|EIT78424.1 asyd|OJJ59913.1 ater|EAU34480.1 atub|OJI81378.1 aver|OJJ02373.1 awen|OJJ41026.1 azon|OJJ44638.1

Group2415: abra|OJJ71280.1 acar|OOF97893.1 acla|EAW10262.1 afis|EAW25331.1 afla|EED53815.1 afum|EAL89993.1 agla|OJJ83877.1 akaw|GAA82708.1 aluc|OJZ87263.1 anid|CBF75234.1 anig|EHA20505.1 aory|EIT78448.1 asyd|OJJ59914.1 ater|EAU34481.1 atub|OJI81379.1 aver|OJJ02372.1 awen|OJJ41025.1 azon|OJJ44639.1

Group2416: abra|OJJ71286.1 acar|OOF97886.1 acla|EAW10251.1 afis|EAW25341.1 afla|EED53822.1 afum|EBA27392.1 agla|OJJ83884.1 akaw|GAA82703.1 aluc|OJZ87257.1 anid|CBF75213.1 anig|EHA20502.1 aory|EIT78427.1 asyd|OJJ59928.1 ater|EAU34487.1 atub|OJI81385.1 aver|OJJ02358.1 awen|OJJ41062.1 azon|OJJ44645.1

Group2417: abra|OJJ71287.1 acar|OOF97884.1 acla|EAW10250.1 afis|EAW25342.1 afla|EED53823.1 afum|EAL90005.2 agla|OJJ83885.1 akaw|GAA82702.1 aluc|OJZ87256.1 anid|CBF75211.1 anig|EHA20501.1 aory|EIT78440.1 asyd|OJJ59927.1 ater|EAU34488.1 atub|OJI81386.1 aver|OJJ02359.1 awen|OJJ41063.1 azon|OJJ44646.1

Group2418: abra|OJJ71288.1 acar|OOF97885.1 acla|EAW10249.1 afis|EAW25343.1 afla|EED53824.1 afum|EAL90006.1 agla|OJJ83886.1 akaw|GAA82701.1 aluc|OJZ87255.1 anid|CBF75209.1 anig|EHA20500.1

aory|EIT78410.1 asyd|OJJ59926.1 ater|EAU34489.1 atub|OJI81387.1 aver|OJJ02360.1 awen|OJJ41065.1 azon|OJJ44647.1

Group2419: abra|OJJ71289.1 acar|OOF97883.1 acla|EAW10248.1 afis|EAW25344.1 afla|EED53825.1 afum|EAL90007.1 agla|OJJ83887.1 akaw|GAA82700.1 aluc|OJZ87254.1 anid|CBF75207.1 anig|EHA20499.1 aory|EIT78419.1 asyd|OJJ59925.1 ater|EAU34490.1 atub|OJI81388.1 aver|OJJ02361.1 awen|OJJ41066.1 azon|OJJ44648.1

Group2420: abra|OJJ71290.1 acar|OOF97882.1 acla|EAW10247.1 afis|EAW25345.1 afla|EED53826.1 afum|EAL90008.2 agla|OJJ83891.1 akaw|GAA82699.1 aluc|OJZ87253.1 anid|CBF75205.1 anig|EHA20498.1 aory|EIT78441.1 asyd|OJJ59924.1 ater|EAU34491.1 atub|OJI81389.1 aver|OJJ02362.1 awen|OJJ41068.1 azon|OJJ44650.1

Group2421: abra|OJJ71291.1 acar|OOF97881.1 acla|EAW10246.1 afis|EAW25346.1 afla|EED53827.1 afum|EAL90009.1 agla|OJJ83892.1 akaw|GAA82698.1 aluc|OJZ87252.1 anid|CBF75203.1 anig|EHA20497.1 aory|EIT78374.1 asyd|OJJ59923.1 ater|EAU34492.1 atub|OJI81390.1 aver|OJJ02363.1 awen|OJJ41069.1 azon|OJJ44651.1

Group2422: abra|OJJ71293.1 acar|OOF97877.1 acla|EAW10245.1 afis|EAW25347.1 afla|EED53828.1 afum|EAL90010.2 agla|OJJ83893.1 akaw|GAA82696.1 aluc|OJZ87250.1 anid|CBF77426.1 anig|EHA20495.1 aory|EIT78392.1 asyd|OJJ53229.1 ater|EAU34495.1 atub|OJI81392.1 aver|OJJ03623.1 awen|OJJ41070.1 azon|OJJ44652.1

Group2423: abra|OJJ71294.1 acar|OOF97876.1 acla|EAW10244.1 afis|EAW25348.1 afla|EED53829.1 afum|EAL90011.1 agla|OJJ83894.1 akaw|GAA82695.1 aluc|OJZ87249.1 anid|CBF77428.1 anig|EHA20494.1 aory|EIT78450.1 asyd|OJJ53228.1 ater|EAU34496.1 atub|OJI81393.1 aver|OJJ03624.1 awen|OJJ41071.1 azon|OJJ44653.1

Group2424: abra|OJJ71295.1 acar|OOF97875.1 acla|EAW10243.1 afis|EAW25349.1 afla|EED53830.1 afum|EAL90012.1 agla|OJJ83895.1 akaw|GAA82694.1 aluc|OJZ87248.1 anid|CBF77430.1 anig|EHA20493.1 asyd|OJJ53227.1 ater|EAU34497.1 atub|OJI81394.1 aver|OJJ03625.1 awen|OJJ41072.1 azon|OJJ44654.1 aory|EIT78356.1

Group2425: abra|OJJ71296.1 acar|OOF97874.1 acla|EAW10242.1 afis|EAW25350.1 afla|EED53831.1 afum|EAL90013.1 agla|OJJ83896.1 akaw|GAA82693.1 aluc|OJZ87247.1 anid|CBF77432.1 anig|EHA20492.1 aory|EIT78425.1 asyd|OJJ53226.1 ater|EAU34498.1 atub|OJI81395.1 aver|OJJ03626.1 awen|OJJ41073.1 azon|OJJ44655.1

Group2426: abra|OJJ71297.1 acar|OOF97873.1 acla|EAW10241.1 afis|EAW25351.1 afla|EED53832.1 afum|EAL90014.1 agla|OJJ83897.1 akaw|GAA82692.1 aluc|OJZ87246.1 anid|CBF77434.1 anig|EHA20491.1 aory|EIT78366.1 asyd|OJJ53225.1 ater|EAU34499.1 atub|OJI81396.1 aver|OJJ03627.1 awen|OJJ41074.1 azon|OJJ44656.1

Group2427: abra|OJJ71300.1 acar|OOF97870.1 acla|EAW10238.1 afis|EAW25354.1 afla|EED53835.1 afum|EAL90016.1 agla|OJJ83902.1 akaw|GAA82689.1 aluc|OJZ87243.1 anid|CBF77471.1 anig|EHA20488.1 aory|EIT78370.1 asyd|OJJ53222.1 ater|EAU34502.1 atub|OJI81399.1 aver|OJJ03630.1 awen|OJJ41077.1 azon|OJJ44659.1

Group2428: abra|OJJ71302.1 acar|OOF97869.1 acla|EAW10237.1 afis|EAW25355.1 afla|EED53836.1 afum|EAL90017.1 agla|OJJ83903.1 akaw|GAA82688.1 aluc|OJZ87241.1 anid|CBF77469.1 anig|EHA20487.1 aory|EIT78369.1 asyd|OJJ53221.1 ater|EAU34503.1 atub|OJI81401.1 aver|OJJ03631.1 awen|OJJ41078.1 azon|OJJ44660.1

Group2429: abra|OJJ71303.1 acar|OOF97868.1 acla|EAW10236.1 afis|EAW25356.1 afla|EED53838.1 afum|EAL90018.1 agla|OJJ83905.1 akaw|GAA82687.1 aluc|OJZ87240.1 anid|CBF77467.1 anig|EHA20486.1 aory|EIT78449.1 asyd|OJJ53220.1 ater|EAU34504.1 atub|OJI81402.1 aver|OJJ03632.1 awen|OJJ41081.1 azon|OJJ44662.1

Group2430: abra|OJJ71304.1 acar|OOF97867.1 acla|EAW10235.1 afis|EAW25357.1 afla|EED53839.1 afum|EAL90019.1 agla|OJJ83906.1 akaw|GAA82686.1 aluc|OJZ87239.1 anid|CBF77465.1 anig|EHA20485.1 aory|EIT78442.1 asyd|OJJ53219.1 ater|EAU34505.1 atub|OJI81403.1 aver|OJJ03633.1 awen|OJJ41082.1 azon|OJJ44663.1

Group2431: abra|OJJ71308.1 acar|OOF97863.1 acla|EAW10232.1 afis|EAW25360.1 afum|EAL90022.1 agla|OJJ83910.1 akaw|GAA82683.1 aluc|OJZ87236.1 anid|CBF77459.1 anig|EHA20483.1 aory|EIT78368.1 asyd|OJJ53216.1 ater|EAU34508.1 atub|OJI81406.1 aver|OJJ03635.1 awen|OJJ41086.1 azon|OJJ44666.1 afla|EED53842.1

Group2432: abra|OJJ71310.1 acar|OOF97862.1 acla|EAW10230.1 afis|EAW25362.1 afla|EED53844.1 afum|EAL90024.1 agla|OJJ83912.1 akaw|GAA82681.1 aluc|OJZ87234.1 anid|CBF77455.1 anig|EHA20481.1 aory|EIT78471.1 asyd|OJJ53214.1 ater|EAU34509.1 atub|OJI81408.1 aver|OJJ03637.1 awen|OJJ41088.1 azon|OJJ44668.1

Group2433: abra|OJJ71311.1 acar|OOF97861.1 acla|EAW10229.1 afis|EAW25363.1 afla|EED53845.1 afum|EAL90025.1 agla|OJJ83913.1 akaw|GAA82680.1 aluc|OJZ87233.1 anid|CBF77453.1 anig|EHA20480.1 aory|EIT78383.1 asyd|OJJ53213.1 ater|EAU34510.1 atub|OJI81409.1 aver|OJJ03638.1 awen|OJJ41089.1 azon|OJJ44669.1

Group2434: abra|OJJ71312.1 acar|OOF97860.1 acla|EAW10228.1 afis|EAW25364.1 afla|EED53846.1 afum|EAL90026.1 agla|OJJ83752.1 akaw|GAA82679.1 aluc|OJZ87232.1 anid|CBF77452.1 anig|EHA20479.1 aory|EIT78459.1 asyd|OJJ53211.1 ater|EAU34511.1 atub|OJI81410.1 aver|OJJ03639.1 awen|OJJ41091.1 azon|OJJ44670.1

Group2435: abra|OJJ71315.1 acar|OOF97857.1 acla|EAW10224.1 afis|EAW25368.1 afla|EED53849.1 afum|EAL90028.2 agla|OJJ83755.1 akaw|GAA82676.1 aluc|OJZ87229.1 anid|CBF77446.1 anig|EHA20476.1 aory|EIT78461.1 asyd|OJJ53208.1 ater|EAU34513.1 atub|OJI81413.1 aver|OJJ03642.1 awen|OJJ41094.1 azon|OJJ44673.1

Group2436: abra|OJJ71316.1 acar|OOF97856.1 acla|EAW10223.1 afis|EAW25369.1 afla|EED53850.1 afum|EAL90029.2 agla|OJJ83756.1 akaw|GAA82675.1 aluc|OJZ87228.1 anid|CBF77444.1 anig|EHA20475.1 aory|EIT78384.1 asyd|OJJ53207.1 ater|EAU34514.1 atub|OJI81414.1 aver|OJJ03643.1 awen|OJJ41095.1 azon|OJJ44674.1

Group2437: abra|OJJ71317.1 acar|OOF97855.1 acla|EAW10222.1 afis|EAW25370.1 afla|EED53851.1 afum|EAL90030.1 agla|OJJ83757.1 akaw|GAA82674.1 aluc|OJZ87227.1 anid|CBF77442.1 anig|EHA20474.1 aory|EIT78393.1 asyd|OJJ53206.1 ater|EAU34515.1 atub|OJI81415.1 aver|OJJ03644.1 awen|OJJ41096.1 azon|OJJ44675.1

Group2438: abra|OJJ71320.1 acar|OOF97854.1 acla|EAW10219.1 afis|EAW25373.1 afla|EED53854.1 afum|EAL90033.1 agla|OJJ83760.1 akaw|GAA82671.1 aluc|OJZ87224.1 anid|CBF77475.1 anig|EHA20471.1 aory|EIT78376.1 asyd|OJJ53203.1 ater|EAU34518.1 atub|OJI81418.1 aver|OJJ03647.1 awen|OJJ41099.1 azon|OJJ44678.1

Group2439: abra|OJJ71321.1 acar|OOF97852.1 acla|EAW10218.1 afis|EAW25375.1 afla|EED53855.1 afum|EAL90034.1 agla|OJJ83761.1 akaw|GAA82670.1 aluc|OJZ87223.1 anid|CBF77477.1 anig|EHA20470.1 aory|EIT78390.1 asyd|OJJ53202.1 ater|EAU34519.1 atub|OJI81419.1 aver|OJJ03648.1 awen|OJJ41100.1 azon|OJJ44679.1

Group2440: abra|OJJ71322.1 acar|OOF97853.1 acla|EAW10217.1 afis|EAW25376.1 afla|EED53856.1 afum|EAL90035.1 agla|OJJ83765.1 akaw|GAA82669.1 aluc|OJZ87222.1 anid|CBF77479.1 anig|EHA20469.1 aory|EIT78453.1 asyd|OJJ53201.1 ater|EAU34520.1 atub|OJI81420.1 aver|OJJ03649.1 awen|OJJ41103.1 azon|OJJ44680.1

Group2441: abra|OJJ71324.1 acar|OOF97850.1 acla|EAW10215.1 afis|EAW25378.1 afla|EED53858.1 afum|EAL90037.1 agla|OJJ83767.1 akaw|GAA82667.1 aluc|OJZ87220.1 anid|CBF77483.1 anig|EHA20468.1 aory|EIT78426.1 asyd|OJJ53199.1 ater|EAU34522.1 atub|OJI81421.1 aver|OJJ03651.1 awen|OJJ41105.1 azon|OJJ44682.1

Group2442: abra|OJJ71325.1 acar|OOF97849.1 acla|EAW10214.1 afis|EAW25379.1 afla|EED53859.1 afum|EAL90038.1 agla|OJJ83772.1 akaw|GAA82666.1 aluc|OJZ87219.1 anid|CBF77485.1 anig|EHA20467.1 aory|EIT78378.1 asyd|OJJ53197.1 ater|EAU34527.1 atub|OJI81422.1 aver|OJJ03653.1 awen|OJJ41106.1 azon|OJJ44683.1

Group2443: abra|OJJ71329.1 acla|EAW10210.1 afis|EAW25384.1 afla|EED54692.1 afum|EAL90043.1 agla|OJJ83776.1 akaw|GAA82663.1 aluc|OJZ87213.1 anid|CBF77490.1 anig|EHA20464.1 asyd|OJJ53194.1 ater|EAU34530.1 atub|OJI81426.1 aver|OJJ03656.1 awen|OJJ41111.1 azon|OJJ44688.1 acar|OOF97845.1 afla|EED53861.1

Group2444: abra|OJJ71333.1 acar|OOF97842.1 acla|EAW10206.1 afis|EAW25389.1 afla|EED53866.1 afum|EAL90048.2 agla|OJJ83792.1 akaw|GAA82659.1 aluc|OJZ87209.1 anid|CBF77500.1 anig|EHA20461.1 aory|EIT78380.1 asyd|OJJ53190.1 ater|EAU34537.1 atub|OJI81430.1 aver|OJJ03660.1 awen|OJJ41116.1 azon|OJJ44692.1

Group2445: abra|OJJ71334.1 acar|OOF97838.1 acla|EAW10205.1 afis|EAW25390.1 afla|EED53867.1 afum|EAL90049.1 agla|OJJ87028.1 akaw|GAA82658.1 aluc|OJZ87208.1 anid|CBF77502.1 anig|EHA20459.1 aory|EIT78439.1 asyd|OJJ53189.1 ater|EAU34538.1 atub|OJI81431.1 aver|OJJ03661.1 awen|OJJ41117.1 azon|OJJ44693.1

Group2446: abra|OJJ71337.1 acar|OOF97835.1 acla|EAW10202.1 afis|EAW25393.1 afla|EED53870.1 afum|EAL90052.1 agla|OJJ87031.1 akaw|GAA82655.1 aluc|OJZ87205.1 anid|CBF77508.1 anig|EHA20458.1 aory|EIT78391.1 asyd|OJJ53186.1 ater|EAU34541.1 atub|OJI81435.1 aver|OJJ03665.1 awen|OJJ41119.1 azon|OJJ43212.1

Group2447: abra|OJJ71339.1 acar|OOF97833.1 acla|EAW10200.1 afis|EAW25395.1 afla|EED53872.1 afum|EAL90054.1 agla|OJJ87033.1 akaw|GAA82653.1 aluc|OJZ87203.1 anid|CBF77512.1 anig|EHA20456.1

aory|EIT78468.1 asyd|OJJ53185.1 ater|EAU34543.1 atub|OJI81437.1 aver|OJJ03666.1 awen|OJJ41121.1 azon|OJJ44696.1

Group2448: abra|OJJ71340.1 acar|OOF97832.1 acla|EAW10199.1 afis|EAW25396.1 afla|EED53873.1 afum|EAL90055.1 agla|OJJ87034.1 akaw|GAA82652.1 aluc|OJZ87202.1 anid|CBF77514.1 anig|EHA20455.1 aory|EIT78362.1 asyd|OJJ53184.1 ater|EAU34544.1 atub|OJI81438.1 aver|OJJ03667.1 awen|OJJ41122.1 azon|OJJ44697.1

Group2449: abra|OJJ71341.1 acar|OOF97831.1 acla|EAW10198.1 afis|EAW25397.1 afla|EED53874.1 afum|EAL90056.1 agla|OJJ87035.1 akaw|GAA82651.1 aluc|OJZ87201.1 anid|CBF77516.1 anig|EHA20454.1 aory|EIT78399.1 asyd|OJJ53183.1 ater|EAU34545.1 atub|OJI81439.1 aver|OJJ03668.1 awen|OJJ41123.1 azon|OJJ44698.1

Group2450: abra|OJJ71342.1 acar|OOF97830.1 acla|EAW10197.1 afis|EAW25398.1 afla|EED53875.1 afum|EAL90057.1 agla|OJJ87036.1 akaw|GAA82650.1 aluc|OJZ87200.1 anid|CBF77518.1 anig|EHA20453.1 aory|EIT78445.1 asyd|OJJ53182.1 ater|EAU34549.1 atub|OJI81440.1 aver|OJJ03669.1 awen|OJJ41124.1 azon|OJJ44699.1

Group2451: abra|OJJ71345.1 acar|OOF97827.1 acla|EAW10194.1 afis|EAW25401.1 afla|EED53879.1 afum|EAL90060.1 agla|OJJ87042.1 akaw|GAA82647.1 aluc|OJZ87196.1 anid|CBF77524.1 anig|EHA20451.1 aory|EIT78402.1 asyd|OJJ53180.1 ater|EAU34552.1 atub|OJI81443.1 aver|OJJ03671.1 awen|OJJ41128.1 azon|OJJ44702.1

Group2452: abra|OJJ71346.1 acar|OOF97824.1 acla|EAW10192.1 afis|EAW25403.1 afla|EED53882.1 afum|EAL90062.1 agla|OJJ87044.1 akaw|GAA82646.1 aluc|OJZ87193.1 anid|CBF77530.1 anig|EHA20450.1 aory|EIT78396.1 asyd|OJJ53177.1 ater|EAU34554.1 atub|OJI81445.1 aver|OJJ03675.1 awen|OJJ41130.1 azon|OJJ44704.1

Group2453: abra|OJJ71347.1 acar|OOF97823.1 acla|EAW10191.1 afis|EAW25404.1 afla|EED53883.1 afum|EAL90063.1 agla|OJJ87045.1 akaw|GAA82645.1 aluc|OJZ87192.1 anid|CBF77532.1 anig|EHA20449.1 aory|EIT78372.1 asyd|OJJ53176.1 ater|EAU34555.1 atub|OJI81446.1 aver|OJJ03676.1 awen|OJJ41131.1 azon|OJJ44705.1

Group2454: abra|OJJ71349.1 acar|OOF97821.1 acla|EAW10189.1 afis|EAW25406.1 afla|EED53885.1 afum|EAL90065.1 agla|OJJ87047.1 akaw|GAA82643.1 aluc|OJZ87190.1 anid|CBF77536.1 anig|EHA20447.1 aory|EIT78483.1 asyd|OJJ53174.1 ater|EAU34557.1 atub|OJI81448.1 aver|OJJ03678.1 awen|OJJ41133.1 azon|OJJ44707.1

Group2455: abra|OJJ71351.1 acar|OOF97818.1 acla|EAW10187.1 afis|EAW25408.1 afla|EED53887.1 afum|EAL90067.1 agla|OJJ87049.1 akaw|GAA82641.1 aluc|OJZ87188.1 anid|CBF77540.1 anig|EHA20445.1 aory|EIT78477.1 asyd|OJJ53172.1 ater|EAU34559.1 atub|OJI81450.1 aver|OJJ03680.1 awen|OJJ41135.1 azon|OJJ44709.1

Group2456: abra|OJJ71352.1 acar|OOF97819.1 acla|EAW10186.1 afis|EAW25409.1 afla|EED53888.1 afum|EAL90068.1 agla|OJJ87050.1 akaw|GAA82640.1 aluc|OJZ87187.1 anid|CBF77588.1 anig|EHA20444.1 aory|EIT78490.1 asyd|OJJ53171.1 ater|EAU34560.1 atub|OJI81451.1 aver|OJJ03681.1 awen|OJJ41136.1 azon|OJJ44715.1

Group2457: abra|OJJ71353.1 acar|OOF97817.1 acla|EAW10183.1 afis|EAW25411.1 afla|EED53890.1 afum|EAL90070.1 agla|OJJ87052.1 akaw|GAA82639.1 aluc|OJZ87186.1 anid|CBF77584.1 anig|EHA20443.1 aory|EIT78494.1 asyd|OJJ53169.1 ater|EAU34562.1 atub|OJI81452.1 aver|OJJ03683.1 awen|OJJ41138.1 azon|OJJ44717.1

Group2458: abra|OJJ71354.1 acar|OOF97816.1 acla|EAW10182.1 afis|EAW25412.1 afla|EED53891.1 afum|EAL90071.1 agla|OJJ87054.1 akaw|GAA82638.1 aluc|OJZ87185.1 anid|CBF77582.1 anig|EHA20442.1 aory|EIT78487.1 asyd|OJJ53168.1 ater|EAU34563.1 atub|OJI81453.1 aver|OJJ03684.1 awen|OJJ41139.1 azon|OJJ44719.1

Group2459: abra|OJJ71355.1 acar|OOF97815.1 acla|EAW10181.1 afis|EAW25413.1 afla|EED53892.1 afum|EAL90072.1 agla|OJJ87055.1 akaw|GAA82637.1 aluc|OJZ87184.1 anid|CBF77580.1 anig|EHA20441.1 aory|EIT78502.1 asyd|OJJ53167.1 ater|EAU34564.1 atub|OJI81455.1 aver|OJJ03685.1 awen|OJJ41140.1 azon|OJJ44720.1

Group2460: abra|OJJ71357.1 acar|OOF97813.1 acla|EAW10179.1 afis|EAW25415.1 afla|EED53894.1 afum|EAL90074.1 agla|OJJ87057.1 akaw|GAA82635.1 aluc|OJZ87182.1 anid|CBF77576.1 anig|EHA20440.1 aory|EIT78485.1 asyd|OJJ53165.1 ater|EAU34566.1 atub|OJI81457.1 aver|OJJ03687.1 awen|OJJ41142.1 azon|OJJ44722.1

Group2461: abra|OJJ71358.1 acar|OOF97812.1 acla|EAW10178.1 afis|EAW25416.1 afla|EED53895.1 afum|EAL90075.1 agla|OJJ87058.1 akaw|GAA82634.1 aluc|OJZ87181.1 anid|CBF77574.1 anig|EHA20439.1 aory|EIT78501.1 asyd|OJJ53164.1 ater|EAU34567.1 atub|OJI81458.1 aver|OJJ03688.1 awen|OJJ41144.1 azon|OJJ44723.1

Group2462: abra|OJJ71359.1 acar|OOF97811.1 acla|EAW10177.1 afis|EAW25417.1 afla|EED53896.1 afum|EAL90076.1 agla|OJJ87059.1 akaw|GAA82633.1 aluc|OJZ87180.1 anid|CBF77573.1 anig|EHA20438.1 aory|EIT78493.1 asyd|OJJ53163.1 ater|EAU34568.1 atub|OJI81459.1 aver|OJJ03689.1 awen|OJJ41145.1 azon|OJJ44724.1

Group2463: abra|OJJ71360.1 acar|OOF97810.1 acla|EAW10176.1 afis|EAW25418.1 afla|EED53897.1 afum|EAL90077.1 agla|OJJ87060.1 akaw|GAA82632.1 aluc|OJZ87179.1 anid|CBF77571.1 anig|EHA20437.1 aory|EIT78486.1 asyd|OJJ53162.1 ater|EAU34569.1 atub|OJI81460.1 aver|OJJ03690.1 awen|OJJ41146.1 azon|OJJ44725.1

Group2464: abra|OJJ71361.1 acar|OOF97809.1 acla|EAW10175.1 afis|EAW25419.1 afla|EED53898.1 afum|EAL90078.1 agla|OJJ87061.1 akaw|GAA82631.1 aluc|OJZ87178.1 anid|CBF77569.1 anig|EHA20436.1 aory|EIT78503.1 asyd|OJJ53161.1 ater|EAU34570.1 atub|OJI81461.1 aver|OJJ03691.1 awen|OJJ41147.1 azon|OJJ44726.1

Group2465: abra|OJJ71363.1 acar|OOF97808.1 acla|EAW10174.1 afis|EAW25420.1 afla|EED53899.1 afum|EAL90079.1 agla|OJJ87064.1 akaw|GAA82630.1 aluc|OJZ87177.1 anid|CBF77567.1 anig|EHA20435.1 aory|EIT78497.1 asyd|OJJ53160.1 ater|EAU34571.1 atub|OJI81462.1 aver|OJJ03692.1 awen|OJJ41148.1 azon|OJJ44727.1

Group2466: abra|OJJ71365.1 acar|OOF97807.1 acla|EAW10173.1 afis|EAW25421.1 afla|EED53900.1 afum|EAL90080.2 agla|OJJ87063.1 akaw|GAA82629.1 aluc|OJZ87175.1 anid|CBF77563.1 anig|EHA20433.1 aory|EIT78488.1 asyd|OJJ53158.1 ater|EAU34572.1 atub|OJI81463.1 aver|OJJ03693.1 awen|OJJ41149.1 azon|OJJ44728.1

Group2467: abra|OJJ71366.1 acar|OOF97806.1 acla|EAW10172.1 afis|EAW25422.1 afla|EED53901.1 afum|EAL90081.1 agla|OJJ87062.1 akaw|GAA82628.1 aluc|OJZ87174.1 anid|CBF77561.1 anig|EHA20432.1 aory|EIT78499.1 asyd|OJJ53157.1 ater|EAU34573.1 atub|OJI81464.1 aver|OJJ03694.1 awen|OJJ41150.1 azon|OJJ44729.1

Group2468: abra|OJJ71367.1 acar|OOF97805.1 acla|EAW10171.1 afis|EAW25423.1 afla|EED53902.1 afum|EAL90082.1 agla|OJJ87065.1 akaw|GAA82627.1 aluc|OJZ87173.1 anid|CBF77559.1 anig|EHA20431.1 aory|EIT78495.1 asyd|OJJ53156.1 ater|EAU34577.1 atub|OJI81465.1 aver|OJJ03695.1 awen|OJJ41151.1 azon|OJJ44730.1

Group2469: abra|OJJ71368.1 acar|OOF97804.1 acla|EAW10170.1 afis|EAW25424.1 afla|EED53903.1 afum|EAL90083.1 agla|OJJ87066.1 akaw|GAA82626.1 aluc|OJZ87172.1 anid|CBF77557.1 anig|EHA20430.1 aory|EIT78480.1 asyd|OJJ53155.1 ater|EAU34578.1 atub|OJI81466.1 aver|OJJ03696.1 awen|OJJ41152.1 azon|OJJ44731.1

Group2470: abra|OJJ71370.1 acar|OOF97798.1 acla|EAW10168.1 afis|EAW25426.1 afla|EED53914.1 afum|EAL90085.1 agla|OJJ87070.1 akaw|GAA82624.1 aluc|OJZ87170.1 anid|CBF77553.1 anig|EHA20428.1 aory|EIT78484.1 asyd|OJJ53153.1 ater|EAU34591.1 atub|OJI81469.1 aver|OJJ03698.1 awen|OJJ41155.1 azon|OJJ44733.1

Group2471: abra|OJJ71371.1 acar|OOF97797.1 acla|EAW10166.1 afis|EAW25428.1 afla|EED53916.1 afum|EAL90087.1 agla|OJJ87073.1 akaw|GAA82623.1 aluc|OJZ87169.1 anid|CBF77549.1 anig|EHA20427.1 aory|EIT78314.1 asyd|OJJ53151.1 ater|EAU34598.1 atub|OJI81470.1 aver|OJJ03700.1 awen|OJJ41157.1 azon|OJJ44735.1

Group2472: abra|OJJ71372.1 acar|OOF97796.1 acla|EAW10165.1 afis|EAW25429.1 afla|EED53917.1 afum|EAL90088.1 agla|OJJ87074.1 akaw|GAA82622.1 aluc|OJZ87168.1 anid|CBF77547.1 anig|EHA20426.1 aory|EIT78324.1 asyd|OJJ53150.1 ater|EAU34600.1 atub|OJI81471.1 aver|OJJ03701.1 awen|OJJ41158.1 azon|OJJ44736.1

Group2473: abra|OJJ71374.1 acar|OOF97794.1 acla|EAW10163.1 afis|EAW25431.1 afla|EED53919.1 afum|EAL90089.1 agla|OJJ87076.1 akaw|GAA82620.1 aluc|OJZ87166.1 anid|CBF77543.1 anig|EHA20425.1 aory|EIT78269.1 asyd|OJJ53148.1 ater|EAU34603.1 atub|OJI81473.1 aver|OJJ03703.1 awen|OJJ41160.1 azon|OJJ44738.1

Group2474: abra|OJJ71376.1 acar|OOF97792.1 acla|EAW10148.1 afis|EAW25445.1 afla|EED53931.1 afum|EAL90103.1 agla|OJJ87089.1 akaw|GAA82618.1 aluc|OJZ87164.1 anid|CBF77610.1 anig|EHA20423.1 aory|EIT78328.1 asyd|OJJ53135.1 ater|EAU34615.1 atub|OJI81475.1 aver|OJJ03717.1 awen|OJJ41174.1 azon|OJJ44750.1

Group2475: abra|OJJ71378.1 acar|OOF97791.1 acla|EAW10146.1 afis|EAW25447.1 afla|EED53929.1 afum|EAL90105.1 agla|OJJ87086.1 akaw|GAA82616.1 aluc|OJZ87162.1 anid|CBF77606.1 anig|EHA20421.1 aory|EIT78352.1 asyd|OJJ53138.1 ater|EAU34613.1 atub|OJI81477.1 aver|OJJ03714.1 awen|OJJ41169.1 azon|OJJ44747.1

Group2476: abra|OJJ71379.1 acar|OOF97790.1 acla|EAW10154.1 afis|EAW25440.1 afla|EED53928.1 afum|EAL90098.1 agla|OJJ87085.1 akaw|GAA82615.1 aluc|OJZ87161.1 anid|CBF77602.1 anig|EHA20420.1

aory|EIT78271.1 asyd|OJJ53139.1 ater|EAU34612.1 atub|OJI81478.1 aver|OJJ03713.1 awen|OJJ41168.1 azon|OJJ44746.1

Group2477: abra|OJJ71380.1 acar|OOF97789.1 acla|EAW10155.1 afis|EAW25439.1 afla|EED53927.1 afum|EAL90097.1 agla|OJJ87084.1 akaw|GAA82614.1 aluc|OJZ87160.1 anid|CBF77600.1 anig|EHA20419.1 aory|EIT78298.1 asyd|OJJ53140.1 ater|EAU34611.1 atub|OJI81480.1 aver|OJJ03712.1 awen|OJJ41167.1 azon|OJJ44745.1

Group2478: abra|OJJ71384.1 acar|OOF97785.1 acla|EAW10159.1 afis|EAW25435.1 afla|EED53923.1 afum|EAL90093.1 agla|OJJ87080.1 akaw|GAA82610.1 aluc|OJZ87156.1 anid|CBF77592.1 anig|EHA20415.1 aory|EIT78272.1 asyd|OJJ53144.1 ater|EAU34607.1 atub|OJI81484.1 aver|OJJ03708.1 awen|OJJ41164.1 azon|OJJ44742.1

Group2479: abra|OJJ71386.1 acar|OOF97783.1 acla|EAW10161.1 afis|EAW25433.1 afla|EED53921.1 afum|EAL90091.1 agla|OJJ87078.1 akaw|GAA82608.1 aluc|OJZ87154.1 anid|CBF77590.1 anig|EHA20413.1 aory|EIT78355.1 asyd|OJJ53146.1 ater|EAU34605.1 atub|OJI81486.1 aver|OJJ03706.1 awen|OJJ41162.1 azon|OJJ44740.1

Group2480: abra|OJJ71389.1 acar|OOF97779.1 acla|EAW10150.1 afis|EAW25443.1 afla|EED53933.1 afum|EAL90101.1 agla|OJJ87092.1 akaw|GAA82605.1 aluc|OJZ87151.1 anid|CBF77614.1 anig|EHA20410.1 aory|EIT78302.1 asyd|OJJ53133.1 ater|EAU34616.1 atub|OJI81489.1 aver|OJJ03719.1 awen|OJJ41176.1 azon|OJJ44752.1

Group2481: abra|OJJ71390.1 acar|OOF97778.1 acla|EAW10151.1 afis|EAW25442.1 afla|EED53934.1 afum|EAL90100.1 agla|OJJ87094.1 akaw|GAA82604.1 aluc|OJZ87150.1 anid|CBF77616.1 anig|EHA20409.1 aory|EIT78339.1 asyd|OJJ53132.1 ater|EAU34617.1 atub|OJI81490.1 aver|OJJ03720.1 awen|OJJ41177.1 azon|OJJ44753.1

Group2482: abra|OJJ71393.1 acar|OOF97775.1 acla|EAW10145.1 afis|EAW25448.1 afla|EED53935.1 afum|EAL90106.1 agla|OJJ87095.1 akaw|GAA82603.1 aluc|OJZ87148.1 anid|CBF77624.1 anig|EHA20407.1 aory|EIT78256.1 asyd|OJJ53131.1 ater|EAU34618.1 atub|OJI81492.1 aver|OJJ03723.1 awen|OJJ41179.1 azon|OJJ44755.1

Group2483: abra|OJJ71395.1 acar|OOF97774.1 acla|EAW10144.1 afis|EAW25449.1 afla|EED53936.1 afum|EAL90108.1 agla|OJJ87101.1 akaw|GAA82602.1 aluc|OJZ87147.1 anid|CBF77626.1 anig|EHA20406.1 aory|EIT78279.1 asyd|OJJ53130.1 ater|EAU34619.1 atub|OJI81494.1 aver|OJJ03724.1 awen|OJJ41180.1 azon|OJJ44756.1

Group2484: abra|OJJ71397.1 acar|OOF97772.1 acla|EAW10142.1 afis|EAW25451.1 afla|EED53938.1 afum|EAL90110.1 agla|OJJ87103.1 akaw|GAA82600.1 aluc|OJZ87145.1 anid|CBF77629.1 anig|EHA20404.1 aory|EIT78274.1 asyd|OJJ53128.1 ater|EAU34621.1 atub|OJI81496.1 aver|OJJ03726.1 awen|OJJ41182.1 azon|OJJ44758.1

Group2485: abra|OJJ71399.1 acar|OOF97771.1 acla|EAW10135.1 afis|EAW25452.1 afla|EED53940.1 afum|EAL90111.1 agla|OJJ87105.1 akaw|GAA82598.1 aluc|OJZ87143.1 anid|CBF77056.1 anig|EHA20403.1 aory|EIT78282.1 asyd|OJJ56993.1 ater|EAU34622.1 atub|OJI81498.1 aver|OJJ03294.1 awen|OJJ41184.1 azon|OJJ44760.1

Group2486: abra|OJJ71400.1 acar|OOF97770.1 acla|EAW10134.1 afis|EAW25453.1 afla|EED53941.1 afum|EAL90112.1 agla|OJJ87106.1 akaw|GAA82597.1 aluc|OJZ87142.1 anid|CBF77633.1 anig|EHA20402.1 aory|EIT78319.1 asyd|OJJ53126.1 ater|EAU34623.1 atub|OJI81500.1 aver|OJJ03728.1 awen|OJJ41185.1 azon|OJJ44761.1

Group2487: abra|OJJ71402.1 acar|OOF97768.1 acla|EAW10132.1 afis|EAW25455.1 afla|EED53944.1 afum|EAL90114.1 agla|OJJ87108.1 akaw|GAA82595.1 aluc|OJZ87139.1 anid|CBF77637.1 anig|EHA20398.1 aory|EIT78284.1 asyd|OJJ53124.1 ater|EAU34625.1 atub|OJI81502.1 aver|OJJ03730.1 awen|OJJ41187.1 azon|OJJ44763.1

Group2488: abra|OJJ71404.1 acar|OOF97767.1 acla|EAW10131.1 afis|EAW25456.1 afla|EED53945.1 afum|EAL90115.2 agla|OJJ87109.1 akaw|GAA82594.1 aluc|OJZ87138.1 anid|CBF77639.1 anig|EHA20397.1 aory|EIT78325.1 asyd|OJJ53123.1 ater|EAU34626.1 atub|OJI81503.1 aver|OJJ03731.1 awen|OJJ41188.1 azon|OJJ44764.1

Group2489: abra|OJJ71405.1 acar|OOF97766.1 acla|EAW10130.1 afis|EAW25457.1 afla|EED53946.1 afum|EAL90117.1 agla|OJJ87110.1 akaw|GAA82593.1 aluc|OJZ87137.1 anid|CBF77641.1 anig|EHA20396.1 aory|EIT78306.1 asyd|OJJ53122.1 ater|EAU34627.1 atub|OJI81504.1 aver|OJJ03732.1 awen|OJJ41189.1 azon|OJJ44767.1

Group2490: abra|OJJ71407.1 acar|OOF97764.1 acla|EAW10128.1 afis|EAW25459.1 afla|EED53948.1 afum|EAL90119.1 agla|OJJ87112.1 akaw|GAA82592.1 aluc|OJZ87135.1 anid|CBF77645.1 anig|EHA20394.1 aory|EIT78349.1 asyd|OJJ53120.1 ater|EAU34629.1 atub|OJI81507.1 aver|OJJ03734.1 awen|OJJ41191.1 azon|OJJ44768.1

Group2491: abra|OJJ71409.1 acla|EAW10127.1 afis|EAW25460.1 afla|EED53949.1 afum|EAL90121.1 agla|OJJ87113.1 akaw|GAA82591.1 aluc|OJZ87131.1 anid|CBF77647.1 anig|EHA20393.1 aory|EIT78304.1 asyd|OJJ53119.1 ater|EAU34630.1 atub|OJI81509.1 aver|OJJ03735.1 awen|OJJ41195.1 azon|OJJ44769.1 afum|EAL87795.1

Group2492: abra|OJJ71413.1 acar|OOF90159.1 acla|EAW10125.1 afis|EAW25463.1 afla|EED53955.1 afum|EAL90126.1 agla|OJJ87115.1 akaw|GAA82588.1 aluc|OJZ87128.1 anid|CBF77653.1 anig|EHA20389.1 aory|EIT78291.1 asyd|OJJ53114.1 ater|EAU34632.1 atub|OJI81512.1 aver|OJJ03741.1 awen|OJJ41200.1 azon|OJJ44771.1

Group2493: abra|OJJ71416.1 acar|OOF90160.1 acla|EAW10123.1 afis|EAW25464.1 afla|EED53958.1 afum|EAL90127.1 agla|OJJ87116.1 akaw|GAA82586.1 aluc|OJZ87124.1 anid|CBF77655.1 anig|EHA20387.1 aory|EIT78307.1 asyd|OJJ53115.1 ater|EAU34634.1 atub|OJI81516.1 aver|OJJ03740.1 awen|OJJ41201.1 azon|OJJ44774.1

Group2494: abra|OJJ71418.1 acar|OOF97758.1 acla|EAW10122.1 afis|EAW25465.1 afla|EED53960.1 afum|EAL90128.2 agla|OJJ87117.1 akaw|GAA82585.1 aluc|OJZ87122.1 anid|CBF77651.1 anig|EHA20386.1 aory|EIT78354.1 asyd|OJJ53117.1 ater|EAU34635.1 atub|OJI81519.1 aver|OJJ03737.1 awen|OJJ41205.1 azon|OJJ44775.1

Group2495: abra|OJJ71420.1 acar|OOF97757.1 acla|EAW10120.1 afis|EAW25467.1 afla|EED53963.1 afum|EAL90131.1 agla|OJJ87118.1 akaw|GAA82582.1 aluc|OJZ87118.1 anid|CBF77666.1 anig|EHA20384.1 aory|EIT78330.1 asyd|OJJ53102.1 ater|EAU34636.1 atub|OJI81522.1 aver|OJJ03751.1 awen|OJJ41206.1 azon|OJJ44777.1

Group2496: abra|OJJ71421.1 acar|OOF97756.1 acla|EAW10111.1 afis|EAW25476.1 afla|EED53964.1 afum|EAL90139.1 agla|OJJ87120.1 akaw|GAA82581.1 aluc|OJZ87117.1 anid|CBF77674.1 anig|EHA20383.1 aory|EIT78326.1 asyd|OJJ53091.1 ater|EAU34640.1 atub|OJI81523.1 aver|OJJ03755.1 awen|OJJ41207.1 azon|OJJ44779.1

Group2497: abra|OJJ71423.1 acar|OOF97755.1 acla|EAW10116.1 afis|EAW25471.1 afla|EED53968.1 afum|EAL90135.1 agla|OJJ87122.1 akaw|GAA82579.1 aluc|OJZ87115.1 anid|CBF77682.1 anig|EHA20381.1 aory|EIT78300.1 asyd|OJJ53088.1 ater|EAU34644.1 atub|OJI81525.1 aver|OJJ03758.1 awen|OJJ41212.1 azon|OJJ44793.1

Group2498: abra|OJJ71424.1 acar|OOF97754.1 acla|EAW10119.1 afis|EAW25470.1 afla|EED53969.1 afum|EAL90134.1 agla|OJJ87123.1 akaw|GAA82578.1 aluc|OJZ87114.1 anid|CBF77684.1 anig|EHA20380.1 aory|EIT78285.1 asyd|OJJ53087.1 ater|EAU34646.1 atub|OJI81526.1 aver|OJJ03759.1 awen|OJJ41213.1 azon|OJJ44792.1

Group2499: abra|OJJ71425.1 acar|OOF97753.1 acla|EAW10118.1 afis|EAW25469.1 afla|EED53970.1 afum|EAL90133.1 agla|OJJ87124.1 akaw|GAA82577.1 aluc|OJZ87113.1 anid|CBF77686.1 anig|EHA20379.1 aory|EIT78275.1 asyd|OJJ53086.1 ater|EAU34647.1 atub|OJI81527.1 aver|OJJ03760.1 awen|OJJ41214.1 azon|OJJ44791.1

Group2500: abra|OJJ71426.1 acar|OOF97752.1 acla|EAW10117.1 afis|EAW25468.1 afla|EED53971.1 afum|EAL90132.1 agla|OJJ87125.1 akaw|GAA82576.1 aluc|OJZ87112.1 anid|CBF77688.1 anig|EHA20378.1 aory|EIT78308.1 asyd|OJJ53085.1 ater|EAU34648.1 atub|OJI81528.1 aver|OJJ03761.1 awen|OJJ41215.1 azon|OJJ44790.1

Group2501: abra|OJJ71428.1 acar|OOF97750.1 acla|EAW10110.1 afis|EAW25477.1 afla|EED53972.1 afum|EAL90140.1 agla|OJJ87126.1 akaw|GAA82575.1 aluc|OJZ87110.1 anid|CBF77709.1 anig|EHA20377.1 aory|EIT78305.1 asyd|OJJ53077.1 ater|EAU34649.1 atub|OJI81529.1 aver|OJJ03771.1 awen|OJJ41216.1 azon|OJJ44789.1

Group2502: abra|OJJ71432.1 acar|OOF90401.1 acla|EAW10106.1 afis|EAW25481.1 afla|EED53979.1 afum|EAL90146.1 agla|OJJ87129.1 akaw|GAA82571.1 aluc|OJZ87106.1 anid|CBF77701.1 anig|EHA20374.1 aory|EIT78311.1 asyd|OJJ53072.1 ater|EAU34652.1 atub|OJI81532.1 aver|OJJ03773.1 awen|OJJ41222.1 azon|OJJ44785.1

Group2503: abra|OJJ71435.1 acar|OOF90399.1 acla|EAW10104.1 afis|EAW25483.1 afla|EED53981.1 afum|EAL90148.1 agla|OJJ87131.1 akaw|GAA82569.1 aluc|OJZ87103.1 anid|CBF77698.1 anig|EHA20372.1 aory|EIT78335.1 asyd|OJJ53070.1 ater|EAU34654.1 atub|OJI81535.1 aver|OJJ03775.1 awen|OJJ41224.1 azon|OJJ44783.1

Group2504: abra|OJJ71507.1 acar|OOF96261.1 acla|EAW15065.1 afis|EAW20037.1 afla|EED45343.1 afum|EAL85158.1 agla|OJJ88926.1 akaw|GAA89406.1 aluc|OJZ86246.1 anid|CBF84775.1 anig|EHA21618.1 aory|EIT73818.1 asyd|OJJ63238.1 ater|EAU38647.1 atub|OJI84296.1 aver|OJI97718.1 awen|OJJ31792.1 azon|OJJ46326.1

Group2505: abra|OJJ71542.1 acar|OOF96289.1 acla|EAW15096.1 afis|EAW20068.1 afla|EED45311.1 afum|EAL85186.1 agla|OJJ88892.1 akaw|GAA89380.1 aluc|OJZ86280.1 anid|CBF84825.1 anig|EHA21591.1

aory|EIT73564.1 asyd|OJJ63207.1 ater|EAU38674.1 atub|OJI84259.1 aver|OJI97748.1 awen|OJJ31827.1 azon|OJJ46297.1

Group2506: abra|OJJ71556.1 acar|OOF90418.1 acla|EAW15109.1 afis|EAW20080.1 afla|EED45295.1 afum|EAL85199.1 agla|OJJ88879.1 akaw|GAA89367.1 aluc|OJZ86293.1 anid|CBF84857.1 anig|EHA21578.1 aory|EIT73533.1 asyd|OJJ63194.1 ater|EAU38688.1 atub|OJI84243.1 aver|OJI97761.1 awen|OJJ31840.1 azon|OJJ46283.1

Group2507: abra|OJJ71560.1 acar|OOF90420.1 acla|EAW15112.1 afis|EAW20083.1 afla|EED45291.1 afum|EAL85202.1 agla|OJJ88875.1 akaw|GAA89364.1 aluc|OJZ86297.1 anid|CBF84871.1 anig|EHA21575.1 aory|EIT73626.1 asyd|OJJ63191.1 ater|EAU38691.1 atub|OJI84240.1 aver|OJI97765.1 awen|OJJ31843.1 azon|OJJ46280.1

Group2508: abra|OJJ71561.1 acar|OOF90421.1 acla|EAW15113.1 afis|EAW20084.1 afla|EED45290.1 afum|EAL85203.1 agla|OJJ88874.1 akaw|GAA89363.1 aluc|OJZ86298.1 anid|CBF84873.1 anig|EHA21574.1 aory|EIT73721.1 asyd|OJJ63190.1 ater|EAU38692.1 atub|OJI84239.1 aver|OJI97766.1 awen|OJJ31844.1 azon|OJJ46279.1

Group2509: abra|OJJ71562.1 acar|OOF96298.1 acla|EAW15114.1 afis|EAW20085.1 afla|EED45289.1 afum|EAL85204.1 agla|OJJ88873.1 akaw|GAA89362.1 aluc|OJZ86299.1 anid|CBF84875.1 anig|EHA21573.1 aory|EIT73546.1 asyd|OJJ63189.1 ater|EAU38693.1 atub|OJI84237.1 aver|OJI97767.1 awen|OJJ31845.1 azon|OJJ46278.1

Group2510: abra|OJJ71563.1 acar|OOF96299.1 acla|EAW15115.1 afis|EAW20086.1 afla|EED45288.1 afum|EAL85205.1 agla|OJJ88872.1 akaw|GAA89361.1 aluc|OJZ86300.1 anid|CBF84877.1 anig|EHA21572.1 aory|EIT73633.1 asyd|OJJ63188.1 ater|EAU38694.1 atub|OJI84236.1 aver|OJI97768.1 awen|OJJ31846.1 azon|OJJ46277.1

Group2511: abra|OJJ71568.1 acar|OOF96302.1 acla|EAW15118.1 afis|EAW20088.1 afla|EED45282.1 afum|EAL85207.1 agla|OJJ88868.1 akaw|GAA89356.1 aluc|OJZ86305.1 anid|CBF84880.1 anig|EHA21567.1 aory|EIT73561.1 asyd|OJJ63185.1 ater|EAU38697.1 atub|OJI84231.1 aver|OJI97770.1 awen|OJJ31849.1 azon|OJJ46274.1

Group2512: abra|OJJ71570.1 acar|OOF96303.1 acla|EAW15119.1 afis|EAW20089.1 afla|EED45281.1 afum|EAL85208.1 agla|OJJ88867.1 akaw|GAA89355.1 aluc|OJZ86307.1 anid|CBF84882.1 anig|EHA21566.1 aory|EIT73628.1 asyd|OJJ63184.1 ater|EAU38698.1 atub|OJI84229.1 aver|OJI97771.1 awen|OJJ31851.1 azon|OJJ46273.1

Group2513: abra|OJJ71573.1 acar|OOF96305.1 acla|EAW15122.1 afis|EAW20092.1 afla|EED45278.1 afum|EAL85211.1 agla|OJJ88862.1 akaw|GAA89351.1 aluc|OJZ86310.1 anid|CBF84897.1 anig|EHA21563.1 aory|EIT73632.1 asyd|OJJ63175.1 ater|EAU38701.1 atub|OJI84225.1 aver|OJI97779.1 awen|OJJ31855.1 azon|OJJ46270.1

Group2514: abra|OJJ71577.1 acar|OOF96306.1 acla|EAW15124.1 afis|EAW20094.1 afla|EED45276.1 afum|EAL85212.1 agla|OJJ88860.1 akaw|GAA89347.1 aluc|OJZ86314.1 anid|CBF84893.1 anig|EHA21561.1 aory|EIT73817.1 asyd|OJJ63177.1 ater|EAU38704.1 atub|OJI84221.1 aver|OJI97777.1 awen|OJJ31857.1 azon|OJJ46268.1

Group2515: abra|OJJ71580.1 acar|OOF96310.1 acla|EAW15127.1 afis|EAW20097.1 afla|EED45273.1 afum|EAL85215.1 agla|OJJ88857.1 akaw|GAA89344.1 aluc|OJZ86317.1 anid|CBF84888.1 anig|EHA21558.1 aory|EIT73668.1 asyd|OJJ63180.1 ater|EAU38705.1 atub|OJI84218.1 aver|OJI97774.1 awen|OJJ31860.1 azon|OJJ46265.1

Group2516: abra|OJJ71584.1 acar|OOF96314.1 acla|EAW15129.1 afis|EAW20101.1 afla|EED53372.1 afum|EAL85218.1 agla|OJJ88854.1 akaw|GAA89339.1 aluc|OJZ86324.1 anid|CBF84901.1 anig|EHA21554.1 aory|EIT74990.1 asyd|OJJ63172.1 ater|EAU38721.1 atub|OJI84210.1 aver|OJI97782.1 awen|OJJ31871.1 azon|OJJ46263.1

Group2517: abra|OJJ71585.1 acar|OOF96315.1 acla|EAW15130.1 afis|EAW20102.1 afla|EED53371.1 afum|EAL85219.1 agla|OJJ88853.1 akaw|GAA89338.1 aluc|OJZ86325.1 anid|CBF84903.1 anig|EHA21553.1 aory|EIT75070.1 asyd|OJJ63171.1 ater|EAU38722.1 atub|OJI84209.1 aver|OJI97783.1 awen|OJJ31872.1 azon|OJJ46262.1

Group2518: abra|OJJ71586.1 acar|OOF96316.1 acla|EAW15131.1 afis|EAW20103.1 afla|EED53370.1 afum|EAL85220.1 agla|OJJ88851.1 akaw|GAA89337.1 aluc|OJZ86326.1 anid|CBF84905.1 anig|EHA21552.1 aory|EIT75110.1 asyd|OJJ63170.1 ater|EAU38723.1 atub|OJI84208.1 aver|OJI97784.1 awen|OJJ31875.1 azon|OJJ46260.1

Group2519: abra|OJJ71593.1 acar|OOF96321.1 acla|EAW15138.1 afis|EAW20110.1 afla|EED53362.1 afum|EAL85227.1 agla|OJJ88842.1 akaw|GAA89330.1 aluc|OJZ86335.1 anid|CBF84919.1 anig|EHA21546.1 aory|EIT74851.1 asyd|OJJ63164.1 ater|EAU38728.1 atub|OJI84200.1 aver|OJI97791.1 awen|OJJ31881.1 azon|OJJ46253.1

Group2520: abra|OJJ71594.1 acar|OOF96322.1 acla|EAW15139.1 afis|EAW20111.1 afla|EED53361.1 afum|EAL85228.2 agla|OJJ88841.1 akaw|GAA89329.1 aluc|OJZ86336.1 anid|CBF84921.1 anig|EHA21545.1 aory|EIT75176.1 asyd|OJJ63163.1 ater|EAU38729.1 atub|OJI84199.1 aver|OJI97792.1 awen|OJJ31882.1 azon|OJJ46252.1

Group2521: abra|OJJ71595.1 acar|OOF96323.1 acla|EAW15140.1 afis|EAW20112.1 afla|EED53360.1 afum|EAL85229.1 agla|OJJ88840.1 akaw|GAA89328.1 aluc|OJZ86337.1 anid|CBF84923.1 anig|EHA21544.1 aory|EIT75201.1 asyd|OJJ63162.1 ater|EAU38730.1 atub|OJI84198.1 aver|OJI97793.1 awen|OJJ31883.1 azon|OJJ46251.1

Group2522: abra|OJJ71596.1 acar|OOF96324.1 acla|EAW15141.1 afis|EAW20113.1 afla|EED53357.1 afum|EAL85230.1 agla|OJJ82514.1 akaw|GAA89327.1 aluc|OJZ86338.1 anid|CBF84924.1 anig|EHA21543.1 aory|EIT75098.1 asyd|OJJ63161.1 ater|EAU38732.1 atub|OJI84197.1 aver|OJI97794.1 azon|OJJ46250.1 awen|OJJ31884.1

Group2523: abra|OJJ71615.1 acar|OOF96331.1 acla|EAW15157.1 afis|EAW20125.1 afla|EED57251.1 afum|EAL85243.1 agla|OJJ88822.1 akaw|GAA89312.1 aluc|OJZ86356.1 anid|CBF84949.1 anig|EHA21534.1 aory|EIT74081.1 asyd|OJJ63147.1 ater|EAU38741.1 atub|OJI84178.1 aver|OJI97807.1 awen|OJJ31894.1 azon|OJJ46242.1

Group2524: abra|OJJ71621.1 acar|OOF96338.1 acla|EAW15161.1 afis|EAW20130.1 afla|EED57222.1 afum|EAL85246.1 agla|OJJ88805.1 akaw|GAA89305.1 aluc|OJZ86364.1 anid|CBF84956.1 anig|EHA21528.1 aory|EIT74137.1 asyd|OJJ63143.1 ater|EAU38747.1 atub|OJI84170.1 aver|OJI97811.1 awen|OJJ31996.1 azon|OJJ46051.1

Group2525: abra|OJJ71623.1 acar|OOF96340.1 acla|EAW15164.1 afis|EAW20133.1 afla|EED57219.1 afum|EAL85249.1 agla|OJJ88802.1 akaw|GAA89302.1 aluc|OJZ86366.1 anid|CBF84962.1 anig|EHA21526.1 aory|EIT74020.1 asyd|OJJ63140.1 ater|EAU38749.1 atub|OJI84168.1 aver|OJI97814.1 awen|OJJ32001.1 azon|OJJ46054.1

Group2526: abra|OJJ71624.1 acar|OOF96341.1 acla|EAW15165.1 afis|EAW20134.1 afla|EED57218.1 afum|EAL85250.1 agla|OJJ88801.1 akaw|GAA89301.1 aluc|OJZ86367.1 anid|CBF84964.1 anig|EHA21525.1 aory|EIT74013.1 asyd|OJJ63139.1 ater|EAU38750.1 atub|OJI84167.1 aver|OJI97815.1 awen|OJJ32002.1 azon|OJJ46055.1

Group2527: abra|OJJ71626.1 acar|OOF96343.1 acla|EAW15166.1 afis|EAW20135.1 afla|EED57217.1 afum|EBA27225.1 agla|OJJ88799.1 akaw|GAA89300.1 aluc|OJZ86368.1 anid|CBF84965.1 anig|EHA21524.1 aory|EIT73997.1 asyd|OJJ63138.1 ater|EAU38758.1 atub|OJI84166.1 aver|OJI97816.1 awen|OJJ32003.1 azon|OJJ46056.1

Group2528: abra|OJJ71627.1 acar|OOF96345.1 acla|EAW15168.1 afis|EAW20137.1 afla|EED57212.1 afum|EAL85252.1 agla|OJJ88798.1 akaw|GAA89298.1 aluc|OJZ86369.1 anid|CBF84968.1 anig|EHA21523.1 aory|EIT74094.1 asyd|OJJ63136.1 ater|EAU38756.1 atub|OJI84165.1 aver|OJI97818.1 awen|OJJ32006.1 azon|OJJ46058.1

Group2529: abra|OJJ71628.1 acar|OOF96346.1 acla|EAW15169.1 afis|EAW20138.1 afla|EED57210.1 afum|EAL85253.1 agla|OJJ88797.1 akaw|GAA89297.1 aluc|OJZ86370.1 anid|CBF84970.1 anig|EHA21522.1 aory|EIT74150.1 asyd|OJJ63135.1 ater|EAU38755.1 atub|OJI84164.1 aver|OJI97819.1 awen|OJJ32007.1 azon|OJJ46063.1

Group2530: abra|OJJ71630.1 acar|OOF96347.1 acla|EAW15172.1 afis|EAW20141.1 afla|EED57209.1 afum|EAL85256.1 agla|OJJ88791.1 akaw|GAA89295.1 aluc|OJZ86372.1 anid|CBF84973.1 anig|EHA21520.1 aory|EIT74038.1 asyd|OJJ63133.1 ater|EAU38753.1 atub|OJI84162.1 aver|OJI97821.1 awen|OJJ32010.1 azon|OJJ46079.1

Group2531: abra|OJJ71633.1 acar|OOF96350.1 acla|EAW15174.1 afis|EAW20143.1 afla|EED57205.1 afum|EAL85258.1 agla|OJJ88789.1 akaw|GAA89293.1 aluc|OJZ86376.1 anid|CBF84978.1 anig|EHA21518.1 aory|EIT74107.1 asyd|OJJ63130.1 atub|OJI84158.1 aver|OJI97824.1 awen|OJJ32013.1 azon|OJJ46076.1 ater|EAU38760.1

Group2532: abra|OJJ71637.1 acar|OOF96354.1 acla|EAW15177.1 afis|EAW20147.1 afla|EED57202.1 afum|EAL85262.1 agla|OJJ88786.1 akaw|GAA89289.1 aluc|OJZ86380.1 anid|CBF84980.1 anig|EHA21513.1 aory|EIT73995.1 asyd|OJJ63127.1 ater|EAU38763.1 atub|OJI84153.1 aver|OJI97828.1 awen|OJJ32021.1 azon|OJJ46069.1

Group2533: abra|OJJ71643.1 acar|OOF96359.1 acla|EAW15183.1 afis|EAW20153.1 afla|EED57197.1 afum|EAL85268.1 agla|OJJ88781.1 akaw|GAA89283.1 aluc|OJZ86386.1 anid|CBF84993.1 anig|EHA21509.1 aory|EIT74014.1 asyd|OJJ63121.1 ater|EAU38767.1 atub|OJI84147.1 aver|OJI97837.1 awen|OJJ32026.1 azon|OJJ46080.1

Group2534: abra|OJJ71649.1 acar|OOF96363.1 acla|EAW15186.1 afis|EAW20156.1 afla|EED57194.1 afum|EAL85271.1 agla|OJJ88774.1 akaw|GAA89280.1 aluc|OJZ86393.1 anid|CBF84998.1 anig|EHA21505.1

aory|EIT74147.1 asyd|OJJ63118.1 ater|EAU38770.1 atub|OJI84140.1 aver|OJI97841.1 awen|OJJ32031.1 azon|OJJ46083.1

Group2535: abra|OJJ71655.1 acar|OOF96369.1 acla|EAW15192.1 afis|EAW20161.1 afla|EED57187.1 afum|EAL85279.1 agla|OJJ88769.1 akaw|GAA89274.1 aluc|OJZ86401.1 anid|CBF85009.1 anig|EHA21499.1 aory|EIT74084.1 asyd|OJJ63112.1 ater|EAU38778.1 atub|OJI84134.1 aver|OJI97847.1 awen|OJJ32056.1 azon|OJJ46090.1

Group2536: abra|OJJ71656.1 acar|OOF96370.1 acla|EAW15194.1 afis|EAW20163.1 afla|EED57186.1 afum|EAL85281.1 agla|OJJ88767.1 akaw|GAA89273.1 aluc|OJZ86402.1 anid|CBF85011.1 anig|EHA21498.1 aory|EIT74088.1 asyd|OJJ63111.1 ater|EAU38794.1 atub|OJI84133.1 aver|OJI97849.1 awen|OJJ32058.1 azon|OJJ46092.1

Group2537: abra|OJJ71663.1 acar|OOF96373.1 acla|EAW15199.1 afis|EAW20165.1 afla|EED57183.1 afum|EAL85283.1 agla|OJJ88764.1 akaw|GAA89270.1 aluc|OJZ86405.1 anid|CBF85022.1 anig|EHA21494.1 aory|EIT74158.1 asyd|OJJ63107.1 ater|EAU38797.1 atub|OJI84126.1 aver|OJI97854.1 awen|OJJ32062.1 azon|OJJ46095.1

Group2538: abra|OJJ71665.1 acar|OOF96375.1 acla|EAW15202.1 afis|EAW20167.1 afla|EED57181.1 afum|EAL85285.1 agla|OJJ88762.1 akaw|GAA89268.1 aluc|OJZ86407.1 anid|CBF85019.1 anig|EHA21492.1 aory|EIT74007.1 asyd|OJJ63105.1 ater|EAU38799.1 atub|OJI84124.1 aver|OJI97856.1 awen|OJJ32064.1 azon|OJJ46097.1

Group2539: abra|OJJ71668.1 acar|OOF96380.1 acla|EAW15208.1 afis|EAW20173.1 afla|EED57175.1 afum|EAL85291.1 agla|OJJ88757.1 akaw|GAA89263.1 aluc|OJZ86411.1 anid|CBF85031.1 anig|EHA21488.1 aory|EIT74021.1 asyd|OJJ63101.1 ater|EAU35209.1 atub|OJI84120.1 aver|OJI97861.1 awen|OJJ32068.1 azon|OJJ46101.1

Group2540: abra|OJJ71670.1 acar|OOF96382.1 acla|EAW15210.1 afis|EAW20175.1 afla|EED57173.1 afum|EAL85293.1 agla|OJJ84691.1 akaw|GAA89261.1 aluc|OJZ86413.1 anid|CBF85035.1 anig|EHA21486.1 aory|EIT73987.1 asyd|OJJ63099.1 ater|EAU35211.1 atub|OJI84118.1 aver|OJI97863.1 awen|OJJ31174.1 azon|OJJ46104.1

Group2541: abra|OJJ71671.1 acar|OOF96383.1 acla|EAW15211.1 afis|EAW20176.1 afla|EED57172.1 afum|EAL85294.1 agla|OJJ84690.1 akaw|GAA89260.1 aluc|OJZ86414.1 anid|CBF85037.1 anig|EHA21485.1 aory|EIT73964.1 asyd|OJJ63098.1 ater|EAU35212.1 atub|OJI84117.1 aver|OJI97864.1 awen|OJJ31175.1 azon|OJJ46105.1

Group2542: abra|OJJ71672.1 acar|OOF96384.1 acla|EAW15212.1 afis|EAW20177.1 afla|EED57171.1 afum|EAL85295.1 agla|OJJ84689.1 akaw|GAA89259.1 aluc|OJZ86415.1 anid|CBF85039.1 anig|EHA21484.1 aory|EIT74066.1 asyd|OJJ63097.1 ater|EAU35213.1 atub|OJI84116.1 aver|OJI97865.1 awen|OJJ31176.1 azon|OJJ46106.1

Group2543: abra|OJJ71675.1 acar|OOF96386.1 acla|EAW15214.1 afis|EAW20179.1 afla|EED57169.1 afum|EAL85297.1 agla|OJJ84687.1 akaw|GAA89257.1 aluc|OJZ86417.1 anid|CBF85042.1 anig|EHA21482.1 aory|EIT74151.1 asyd|OJJ63095.1 ater|EAU35215.1 atub|OJI84114.1 aver|OJI97867.1 awen|OJJ31178.1 azon|OJJ46109.1

Group2544: abra|OJJ71676.1 acar|OOF96387.1 acla|EAW15215.1 afis|EAW20180.1 afla|EED57168.1 afum|EAL85298.1 agla|OJJ84686.1 akaw|GAA89256.1 aluc|OJZ86418.1 anid|CBF85044.1 anig|EHA21481.1 aory|EIT73991.1 asyd|OJJ63094.1 ater|EAU35216.1 atub|OJI84113.1 aver|OJI97868.1 awen|OJJ31179.1 azon|OJJ46110.1

Group2545: abra|OJJ71677.1 acar|OOF96388.1 acla|EAW15216.1 afis|EAW20181.1 afla|EED57167.1 afum|EAL85299.1 agla|OJJ84685.1 akaw|GAA89255.1 aluc|OJZ86419.1 anid|CBF85046.1 anig|EHA21480.1 aory|EIT74127.1 asyd|OJJ63093.1 ater|EAU35217.1 atub|OJI84112.1 aver|OJI97869.1 awen|OJJ31180.1 azon|OJJ46111.1

Group2546: abra|OJJ71678.1 acar|OOF96389.1 acla|EAW15217.1 afis|EAW20182.1 afla|EED57166.1 afum|EAL85300.1 agla|OJJ84684.1 akaw|GAA89254.1 aluc|OJZ86420.1 anid|CBF85048.1 anig|EHA21479.1 aory|EIT74003.1 asyd|OJJ63092.1 ater|EAU35218.1 atub|OJI84111.1 aver|OJI97870.1 awen|OJJ31181.1 azon|OJJ46112.1

Group2547: abra|OJJ71679.1 acar|OOF96390.1 acla|EAW15218.1 afis|EAW20183.1 afla|EED57165.1 afum|EAL85301.1 agla|OJJ84683.1 akaw|GAA89253.1 aluc|OJZ86421.1 anid|CBF85050.1 anig|EHA21478.1 aory|EIT74002.1 asyd|OJJ63091.1 ater|EAU35219.1 atub|OJI84110.1 aver|OJI97871.1 awen|OJJ31182.1 azon|OJJ46113.1

Group2548: abra|OJJ71682.1 acar|OOF96393.1 acla|EAW15221.1 afis|EAW20186.1 afla|EED57164.1 afum|EAL85305.1 agla|OJJ84680.1 akaw|GAA89250.1 aluc|OJZ86424.1 anid|CBF85054.1 anig|EHA21474.1 aory|EIT74048.1 asyd|OJJ63088.1 ater|EAU35222.1 atub|OJI84107.1 aver|OJI97874.1 awen|OJJ31186.1 azon|OJJ46116.1

Group2549: abra|OJJ71685.1 acar|OOF96395.1 acla|EAW15223.1 afis|EAW20188.1 afla|EED57158.1 afum|EAL85307.1 agla|OJJ84678.1 akaw|GAA89248.1 aluc|OJZ86427.1 anid|CBF85058.1 anig|EHA21472.1 aory|EIT73974.1 asyd|OJJ63086.1 ater|EAU35223.1 atub|OJI84104.1 aver|OJI97876.1 awen|OJJ31188.1 azon|OJJ46118.1

Group2550: abra|OJJ71686.1 acar|OOF96396.1 acla|EAW15224.1 afis|EAW20189.1 afla|EED57157.1 afum|EAL85308.1 agla|OJJ84677.1 akaw|GAA89247.1 aluc|OJZ86428.1 anid|CBF85060.1 anig|EHA21471.1 aory|EIT74135.1 asyd|OJJ63085.1 ater|EAU35224.1 atub|OJI84103.1 aver|OJI97877.1 awen|OJJ31189.1 azon|OJJ46119.1

Group2551: abra|OJJ71687.1 acar|OOF96397.1 acla|EAW15225.1 afis|EAW20190.1 afla|EED57156.1 afum|EAL85309.1 agla|OJJ84676.1 akaw|GAA89246.1 aluc|OJZ86429.1 anid|CBF85062.1 anig|EHA21470.1 aory|EIT74033.1 asyd|OJJ63084.1 ater|EAU35225.1 atub|OJI84102.1 aver|OJI97879.1 awen|OJJ31190.1 azon|OJJ46124.1

Group2552: abra|OJJ71689.1 acar|OOF96398.1 acla|EAW15226.1 afis|EAW20191.1 afla|EED57155.1 afum|EAL85310.1 agla|OJJ84675.1 akaw|GAA89245.1 aluc|OJZ86431.1 anid|CBF85064.1 anig|EHA21469.1 aory|EIT74046.1 asyd|OJJ63083.1 ater|EAU35226.1 atub|OJI84100.1 aver|OJI97881.1 awen|OJJ31196.1 azon|OJJ46120.1

Group2553: abra|OJJ71691.1 acar|OOF96399.1 acla|EAW15228.1 afis|EAW20192.1 afla|EED57152.1 afum|EAL85312.2 agla|OJJ84673.1 akaw|GAA89244.1 aluc|OJZ86435.1 anid|CBF85065.1 anig|EHA21468.1 aory|EIT74165.1 asyd|OJJ63082.1 ater|EAU32000.1 atub|OJI84095.1 aver|OJI97882.1 awen|OJJ31197.1 azon|OJJ46125.1

Group2554: abra|OJJ71692.1 acar|OOF96400.1 acla|EAW15229.1 afis|EAW20193.1 afla|EED57151.1 afum|EAL85313.1 agla|OJJ84672.1 akaw|GAA89243.1 aluc|OJZ86436.1 anid|CBF85067.1 anig|EHA21467.1 aory|EIT74001.1 asyd|OJJ63081.1 ater|EAU32001.1 atub|OJI84094.1 aver|OJI97883.1 awen|OJJ31198.1 azon|OJJ46126.1

Group2555: abra|OJJ71693.1 acar|OOF96402.1 acla|EAW15230.1 afis|EAW20194.1 afla|EED57150.1 afum|EAL85314.1 agla|OJJ84671.1 akaw|GAA89242.1 aluc|OJZ86437.1 anid|CBF85069.1 anig|EHA21466.1 aory|EIT74019.1 asyd|OJJ63080.1 ater|EAU32002.1 atub|OJI84093.1 aver|OJI97884.1 awen|OJJ31199.1 azon|OJJ46127.1

Group2556: abra|OJJ71697.1 acar|OOF96405.1 acla|EAW15233.1 afis|EAW20197.1 afla|EED57147.1 afum|EAL85317.1 agla|OJJ84668.1 akaw|GAA89238.1 aluc|OJZ86440.1 anid|CBF85078.1 anig|EHA21461.1 aory|EIT74146.1 asyd|OJJ63076.1 ater|EAU32005.1 atub|OJI84088.1 aver|OJI97888.1 awen|OJJ31203.1 azon|OJJ46128.1

Group2557: abra|OJJ71699.1 acar|OOF96407.1 acla|EAW15235.1 afis|EAW20199.1 afla|EED57145.1 afum|EAL85319.1 agla|OJJ84666.1 akaw|GAA89236.1 aluc|OJZ86442.1 anid|CBF85082.1 anig|EHA21459.1 aory|EIT74005.1 asyd|OJJ63073.1 ater|EAU32007.1 atub|OJI84086.1 aver|OJI97890.1 awen|OJJ31205.1 azon|OJJ41971.1

Group2558: abra|OJJ71700.1 acar|OOF96408.1 acla|EAW15236.1 afis|EAW20200.1 afla|EED57144.1 afum|EAL85320.1 agla|OJJ84665.1 akaw|GAA89235.1 aluc|OJZ86443.1 anid|CBF85084.1 anig|EHA21458.1 aory|EIT74097.1 asyd|OJJ63072.1 ater|EAU32008.1 atub|OJI84085.1 aver|OJI97891.1 awen|OJJ31206.1 azon|OJJ46130.1

Group2559: abra|OJJ71701.1 acar|OOF96409.1 acla|EAW15237.1 afis|EAW20201.1 afla|EED57143.1 afum|EAL85321.1 agla|OJJ84663.1 akaw|GAA89234.1 aluc|OJZ86444.1 anid|CBF85085.1 anig|EHA21457.1 aory|EIT73998.1 asyd|OJJ63070.1 ater|EAU32009.1 atub|OJI84084.1 aver|OJI97892.1 awen|OJJ31207.1 azon|OJJ46131.1

Group2560: abra|OJJ71705.1 acar|OOF96415.1 afis|EAW20206.1 afla|EED57137.1 afum|EAL85324.1 agla|OJJ84661.1 akaw|GAA89232.1 aluc|OJZ86447.1 anig|EHA21454.1 aory|EIT74126.1 asyd|OJJ63064.1 ater|EAU32017.1 atub|OJI84081.1 aver|OJI97897.1 awen|OJJ31210.1 acla|EAW15239.1 anid|CBF85093.1 azon|OJJ46134.1

Group2561: abra|OJJ71714.1 acar|OOF96419.1 acla|EAW06442.1 afis|EAW18547.1 afla|EED52833.1 afum|EAL93247.1 agla|OJJ81584.1 akaw|GAA85175.1 aluc|OJZ86459.1 anid|CBF70177.1 anig|EHA21449.1 aory|EIT78161.1 asyd|OJJ58793.1 ater|EAU30022.1 atub|OJI84068.1 aver|OJJ01397.1 awen|OJJ31099.1 azon|OJJ46565.1

Group2562: abra|OJJ71715.1 acar|OOF96420.1 acla|EAW06441.1 afis|EAW18546.1 afla|EED52832.1 afum|EAL93246.1 agla|OJJ81587.1 akaw|GAA85174.1 aluc|OJZ86461.1 anid|CBF70265.1 anig|EHA21448.1 aory|EIT78207.1 asyd|OJJ58750.1 ater|EAU34832.1 atub|OJI84067.1 aver|OJJ01360.1 awen|OJJ31100.1 azon|OJJ46562.1

Group2563: abra|OJJ71716.1 acar|OOF96421.1 acla|EAW06440.1 afis|EAW18545.1 afla|EED52831.1 afum|EAL93245.1 agla|OJJ81589.1 akaw|GAA85173.1 aluc|OJZ86462.1 anid|CBF70267.1 anig|EHA21447.1

aory|EIT78194.1 asyd|OJJ58749.1 ater|EAU34833.1 atub|OJI84066.1 aver|OJJ01359.1 awen|OJJ31101.1 azon|OJJ46561.1

Group2564: abra|OJJ71719.1 acar|OOF96423.1 acla|EAW06439.1 afis|EAW18543.1 afla|EED52829.1 afum|EAL93243.1 agla|OJJ81591.1 akaw|GAA85171.1 aluc|OJZ86465.1 anid|CBF70269.1 anig|EHA21445.1 aory|EIT78205.1 asyd|OJJ58748.1 ater|EAU34834.1 atub|OJI84064.1 aver|OJJ01358.1 awen|OJJ31103.1 azon|OJJ46558.1

Group2565: abra|OJJ71720.1 acar|OOF96425.1 acla|EAW06444.1 afis|EAW18549.1 afla|EED52828.1 afum|EAL93249.1 agla|OJJ81592.1 akaw|GAA85170.1 aluc|OJZ86466.1 anid|CBF70271.1 anig|EHA21444.1 aory|EIT78197.1 asyd|OJJ58747.1 ater|EAU34836.1 atub|OJI84063.1 aver|OJJ01357.1 awen|OJJ31105.1 azon|OJJ46557.1

Group2566: abra|OJJ71742.1 acar|OOF96433.1 acla|EAW06446.1 afis|EAW18551.1 afla|EED52823.1 afum|EAL93251.1 agla|OJJ81605.1 akaw|GAA85156.1 aluc|OJZ86481.1 anid|CBF70277.1 anig|EHA22164.1 aory|EIT78237.1 asyd|OJJ58744.1 ater|EAU34839.1 atub|OJI84049.1 aver|OJJ01354.1 awen|OJJ31111.1 azon|OJJ46587.1

Group2567: abra|OJJ71745.1 acar|OOF96436.1 acla|EAW06449.1 afis|EAW18553.1 afla|EED52822.1 afum|EAL93253.1 agla|OJJ81603.1 akaw|GAA85159.1 aluc|OJZ86478.1 anid|CBF70280.1 anig|EHA22162.1 aory|EIT78230.1 asyd|OJJ58741.1 ater|EAU34841.1 atub|OJI84052.1 aver|OJJ01351.1 awen|OJJ31114.1 azon|OJJ46590.1

Group2568: abra|OJJ71803.1 acar|OOF96483.1 acla|EAW13844.1 afis|EAW18558.1 afla|EED52808.1 afum|EAL93258.1 agla|OJJ81601.1 akaw|GAA85107.1 aluc|OJZ86533.1 anid|CBF70286.1 anig|EHA22113.1 aory|EIT78195.1 asyd|OJJ58737.1 ater|EAU34847.1 atub|OJI83988.1 aver|OJJ01348.1 awen|OJJ31116.1 azon|OJJ46596.1

Group2569: abra|OJJ71806.1 acar|OOF96484.1 acla|EAW13845.1 afis|EAW18559.1 afla|EED52807.1 afum|EAL93259.1 agla|OJJ81600.1 akaw|GAA85106.1 aluc|OJZ86536.1 anid|CBF70288.1 anig|EHA22112.1 aory|EIT78169.1 asyd|OJJ58735.1 ater|EAU34848.1 atub|OJI83987.1 aver|OJJ01347.1 awen|OJJ31117.1 azon|OJJ46595.1

Group2570: abra|OJJ71809.1 acar|OOF96487.1 acla|EAW13848.1 afis|EAW18562.1 afla|EED52792.1 afum|EAL93263.1 agla|OJJ81609.1 akaw|GAA85103.1 aluc|OJZ86539.1 anid|CBF70294.1 anig|EHA22110.1 aory|EIT78177.1 asyd|OJJ58732.1 ater|EAU34850.1 atub|OJI83984.1 aver|OJJ01344.1 awen|OJJ31120.1 azon|OJJ46597.1

Group2571: abra|OJJ71810.1 acar|OOF96488.1 acla|EAW13849.1 afis|EAW18563.1 afla|EED52791.1 afum|EAL93265.2 agla|OJJ81612.1 akaw|GAA85102.1 aluc|OJZ86541.1 anid|CBF70298.1 anig|EHA22109.1 aory|EIT78239.1 asyd|OJJ58730.1 ater|EAU34851.1 atub|OJI83982.1 aver|OJJ01343.1 awen|OJJ31121.1 azon|OJJ46602.1

Group2572: abra|OJJ71811.1 acar|OOF96489.1 acla|EAW13850.1 afis|EAW18564.1 afla|EED52790.1 afum|EAL93266.1 agla|OJJ81613.1 akaw|GAA85101.1 aluc|OJZ86542.1 anid|CBF70300.1 anig|EHA22108.1 aory|EIT78209.1 asyd|OJJ58729.1 ater|EAU34852.1 atub|OJI83981.1 aver|OJJ01342.1 awen|OJJ31122.1 azon|OJJ46603.1

Group2573: abra|OJJ71813.1 acar|OOF96491.1 acla|EAW13853.1 afis|EAW18567.1 afla|EED52786.1 afum|EAL93269.2 agla|OJJ81616.1 akaw|GAA85097.1 aluc|OJZ86546.1 anid|CBF70305.1 anig|EHA22104.1 aory|EIT78200.1 asyd|OJJ58726.1 ater|EAU34854.1 atub|OJI83977.1 aver|OJJ01339.1 awen|OJJ31124.1 azon|OJJ46605.1

Group2574: abra|OJJ71816.1 acar|OOF96493.1 acla|EAW13856.1 afis|EAW18570.1 afla|EED52781.1 afum|EAL93273.1 agla|OJJ81621.1 akaw|GAA85094.1 aluc|OJZ86548.1 anid|CBF70311.1 anig|EHA22101.1 aory|EIT78224.1 asyd|OJJ58723.1 ater|EAU34855.1 atub|OJI83974.1 aver|OJJ01336.1 awen|OJJ31127.1 azon|OJJ46608.1

Group2575: abra|OJJ71817.1 acar|OOF96494.1 acla|EAW13857.1 afis|EAW18571.1 afla|EED52780.1 afum|EAL93274.1 agla|OJJ81622.1 akaw|GAA85093.1 aluc|OJZ86549.1 anid|CBF70313.1 anig|EHA22100.1 aory|EIT78222.1 asyd|OJJ58722.1 ater|EAU34856.1 atub|OJI83973.1 aver|OJJ01335.1 awen|OJJ31128.1 azon|OJJ46609.1

Group2576: abra|OJJ71818.1 acar|OOF96495.1 acla|EAW13860.1 afis|EAW18572.1 afla|EED52776.1 afum|EAL93275.1 agla|OJJ81623.1 akaw|GAA85092.1 aluc|OJZ86550.1 anid|CBF70315.1 anig|EHA22099.1 aory|EIT78170.1 asyd|OJJ58721.1 ater|EAU34857.1 atub|OJI83972.1 aver|OJJ01334.1 awen|OJJ31129.1 azon|OJJ46610.1

Group2577: abra|OJJ71819.1 acar|OOF96496.1 acla|EAW13861.1 afis|EAW18573.1 afla|EED52775.1 afum|EAL93276.1 agla|OJJ81624.1 akaw|GAA85091.1 aluc|OJZ86551.1 anid|CBF70317.1 anig|EHA22098.1 aory|EIT78219.1 asyd|OJJ58720.1 ater|EAU34858.1 atub|OJI83971.1 aver|OJJ01333.1 awen|OJJ31130.1 azon|OJJ46611.1

Group2578: abra|OJJ71820.1 acar|OOF96497.1 acla|EAW13862.1 afis|EAW18574.1 afla|EED52774.1 afum|EAL93277.1 agla|OJJ81625.1 akaw|GAA85090.1 aluc|OJZ86552.1 anid|CBF70319.1 anig|EHA22097.1 aory|EIT78236.1 asyd|OJJ58719.1 ater|EAU34869.1 atub|OJI83970.1 aver|OJJ01332.1 awen|OJJ31131.1 azon|OJJ46612.1

Group2579: abra|OJJ71821.1 acar|OOF96498.1 acla|EAW13863.1 afis|EAW18575.1 afla|EED52773.1 afum|EAL93278.1 agla|OJJ81626.1 akaw|GAA85089.1 aluc|OJZ86553.1 anid|CBF70320.1 anig|EHA22096.1 aory|EIT78159.1 asyd|OJJ58718.1 ater|EAU34868.1 atub|OJI83969.1 aver|OJJ01331.1 awen|OJJ31132.1 azon|OJJ46613.1

Group2580: abra|OJJ71823.1 acar|OOF96499.1 acla|EAW13864.1 afis|EAW18576.1 afla|EED52772.1 afum|EAL93279.1 agla|OJJ81627.1 akaw|GAA85087.1 aluc|OJZ86555.1 anid|CBF70322.1 anig|EHA22094.1 aory|EIT78160.1 asyd|OJJ58717.1 ater|EAU34867.1 atub|OJI83967.1 aver|OJJ01330.1 awen|OJJ31133.1 azon|OJJ46614.1

Group2581: abra|OJJ71824.1 acar|OOF96500.1 acla|EAW13865.1 afis|EAW18577.1 afla|EED52771.1 afum|EAL93280.1 agla|OJJ81628.1 akaw|GAA85086.1 aluc|OJZ86556.1 anid|CBF70324.1 anig|EHA22093.1 aory|EIT78235.1 asyd|OJJ58716.1 ater|EAU34866.1 atub|OJI83966.1 aver|OJJ01329.1 awen|OJJ31134.1 azon|OJJ46615.1

Group2582: abra|OJJ71829.1 acar|OOF96505.1 acla|EAW13866.1 afis|EAW18578.1 afla|EED52770.1 afum|EAL93281.1 agla|OJJ81629.1 akaw|GAA85082.1 aluc|OJZ86561.1 anid|CBF70354.1 anig|EHA22089.1 aory|EIT78184.1 asyd|OJJ58657.1 ater|EAU34861.1 atub|OJI83960.1 aver|OJJ01265.1 awen|OJJ31135.1 azon|OJJ46616.1

Group2583: abra|OJJ71837.1 acar|OOF90146.1 acla|EAW13868.1 afis|EAW18580.1 afla|EED52766.1 afum|EAL93284.1 agla|OJJ81631.1 akaw|GAA85078.1 aluc|OJZ86568.1 anid|CBF70360.1 anig|EHA22086.1 aory|EIT78154.1 asyd|OJJ58653.1 ater|EAU34873.1 atub|OJI83953.1 aver|OJJ01262.1 awen|OJJ31143.1 azon|OJJ46619.1

Group2584: abra|OJJ71840.1 acar|OOF90149.1 acla|EAW13871.1 afis|EAW18582.1 afla|EED52762.1 afum|EAL93286.1 agla|OJJ81634.1 akaw|GAA85077.1 aluc|OJZ86573.1 anid|CBF70362.1 anig|EHA22084.1 aory|EIT78249.1 asyd|OJJ58652.1 ater|EAU34876.1 atub|OJI83950.1 aver|OJJ01258.1 awen|OJJ31146.1 azon|OJJ46621.1

Group2585: abra|OJJ71842.1 acar|OOF96511.1 acla|EAW13873.1 afis|EAW18585.1 afla|EED52761.1 afum|EAL93289.1 agla|OJJ81636.1 akaw|GAA85075.1 aluc|OJZ86575.1 anid|CBF70366.1 anig|EHA22082.1 aory|EIT78178.1 asyd|OJJ58650.1 ater|EAU34878.1 atub|OJI83948.1 aver|OJJ01256.1 awen|OJJ31148.1 azon|OJJ46623.1

Group2586: abra|OJJ71843.1 acar|OOF96512.1 acla|EAW13874.1 afis|EAW18586.1 afla|EED52760.1 afum|EAL93290.1 agla|OJJ81637.1 akaw|GAA85074.1 aluc|OJZ86576.1 anid|CBF70370.1 anig|EHA22081.1 aory|EIT78214.1 asyd|OJJ58649.1 ater|EAU34879.1 atub|OJI83946.1 aver|OJJ01255.1 awen|OJJ31149.1 azon|OJJ46624.1

Group2587: abra|OJJ71844.1 acar|OOF96513.1 acla|EAW13875.1 afis|EAW18587.1 afla|EED52759.1 afum|EAL93294.2 agla|OJJ81638.1 akaw|GAA85073.1 aluc|OJZ86577.1 anid|CBF70372.1 anig|EHA22080.1 aory|EIT78199.1 asyd|OJJ58648.1 ater|EAU34880.1 atub|OJI83945.1 aver|OJJ01254.1 awen|OJJ31150.1 azon|OJJ46625.1

Group2588: abra|OJJ71845.1 acar|OOF96514.1 acla|EAW13876.1 afis|EAW18588.1 afla|EED52758.1 afum|EAL93295.2 agla|OJJ81640.1 akaw|GAA85072.1 aluc|OJZ86578.1 anid|CBF70374.1 anig|EHA22079.1 aory|EIT78218.1 asyd|OJJ58647.1 ater|EAU34881.1 atub|OJI83944.1 aver|OJJ01253.1 awen|OJJ31151.1 azon|OJJ46626.1

Group2589: abra|OJJ71846.1 acar|OOF96515.1 acla|EAW13877.1 afis|EAW18589.1 afla|EED52757.1 afum|EAL93296.2 agla|OJJ81641.1 akaw|GAA85071.1 aluc|OJZ86579.1 anid|CBF70375.1 anig|EHA22078.1 aory|EIT78231.1 asyd|OJJ58646.1 ater|EAU34882.1 atub|OJI83943.1 aver|OJJ01252.1 awen|OJJ31152.1 azon|OJJ46627.1

Group2590: abra|OJJ71851.1 acar|OOF96521.1 acla|EAW13883.1 afis|EAW18595.1 afla|EED52752.1 afum|EAL93303.1 agla|OJJ81647.1 akaw|GAA85066.1 aluc|OJZ86585.1 anid|CBF70391.1 anig|EHA22075.1 aory|EIT78172.1 asyd|OJJ58636.1 ater|EAU34889.1 atub|OJI83938.1 aver|OJJ01242.1 awen|OJJ31158.1 azon|OJJ46633.1

Group2591: abra|OJJ71854.1 acar|OOF96524.1 acla|EAW13886.1 afis|EAW18598.1 afla|EED52749.1 afum|EAL93306.1 agla|OJJ81651.1 akaw|GAA85063.1 aluc|OJZ86588.1 anid|CBF70397.1 anig|EHA22072.1 aory|EIT78201.1 asyd|OJJ58633.1 ater|EAU34892.1 atub|OJI83935.1 aver|OJJ01239.1 awen|OJJ31161.1 azon|OJJ46637.1

Group2592: abra|OJJ71855.1 acar|OOF96525.1 acla|EAW13887.1 afis|EAW18599.1 afla|EED52748.1 afum|EAL93307.1 agla|OJJ81652.1 akaw|GAA85062.1 aluc|OJZ86589.1 anid|CBF70399.1 anig|EHA22071.1

aory|EIT78171.1 asyd|OJJ58632.1 ater|EAU34893.1 atub|OJI83934.1 aver|OJJ01238.1 awen|OJJ31162.1 azon|OJJ46638.1

Group2593: abra|OJJ71857.1 acar|OOF96527.1 acla|EAW13889.1 afis|EAW18601.1 afla|EED52746.1 afum|EAL93309.1 agla|OJJ81653.1 akaw|GAA85060.1 aluc|OJZ86591.1 anid|CBF70403.1 anig|EHA22069.1 aory|EIT78188.1 asyd|OJJ58630.1 ater|EAU34895.1 atub|OJI83932.1 aver|OJJ01236.1 awen|OJJ31166.1 azon|OJJ46640.1

Group2594: abra|OJJ71858.1 acar|OOF96528.1 acla|EAW13890.1 afis|EAW18602.1 afla|EED52745.1 afum|EAL93310.1 agla|OJJ81654.1 akaw|GAA85059.1 aluc|OJZ86592.1 anid|CBF70405.1 anig|EHA22068.1 aory|EIT78217.1 asyd|OJJ58629.1 ater|EAU34896.1 atub|OJI83931.1 aver|OJJ01235.1 awen|OJJ31167.1 azon|OJJ46641.1

Group2595: abra|OJJ71861.1 acar|OOF96533.1 acla|EAW13892.1 afis|EAW18604.1 afla|EED52743.1 afum|EAL93312.1 agla|OJJ81656.1 akaw|GAA85054.1 aluc|OJZ86598.1 anid|CBF70409.1 anig|EHA22064.1 aory|EIT78243.1 asyd|OJJ58627.1 ater|EAU34899.1 atub|OJI83925.1 aver|OJJ01233.1 awen|OJJ31170.1 azon|OJJ46647.1

Group2596: abra|OJJ71864.1 acar|OOF96534.1 acla|EAW13894.1 afis|EAW18606.1 afla|EED52740.1 afum|EBA27488.1 agla|OJJ81660.1 akaw|GAA85052.1 aluc|OJZ86601.1 anig|EHA22062.1 aory|EIT78163.1 ater|EAU34901.1 atub|OJI83922.1 awen|OJJ31172.1 azon|OJJ46649.1 anid|CBF70412.1 asyd|OJJ58625.1 aver|OJJ01231.1

Group2597: abra|OJJ71865.1 acar|OOF96535.1 acla|EAW13895.1 afis|EAW18607.1 afla|EED52739.1 afum|EAL93317.1 agla|OJJ81666.1 akaw|GAA85051.1 aluc|OJZ86602.1 anid|CBF70414.1 anig|EHA22061.1 aory|EIT78220.1 asyd|OJJ58624.1 ater|EAU34902.1 atub|OJI83921.1 aver|OJJ01230.1 awen|OJJ31173.1 azon|OJJ46650.1

Group2598: abra|OJJ71866.1 acar|OOF96536.1 acla|EAW13896.1 afis|EAW18608.1 afla|EED52738.1 afum|EAL93318.1 agla|OJJ81665.1 akaw|GAA85049.1 aluc|OJZ86603.1 anid|CBF70416.1 anig|EHA22060.1 aory|EIT78223.1 asyd|OJJ58623.1 ater|EAU34903.1 atub|OJI83920.1 aver|OJJ01229.1 awen|OJJ32069.1 azon|OJJ46651.1

Group2599: abra|OJJ71869.1 acar|OOF96540.1 acla|EAW13899.1 afis|EAW18611.1 afla|EED52736.1 afum|EAL93321.1 agla|OJJ81234.1 akaw|GAA85046.1 aluc|OJZ86608.1 anid|CBF70420.1 anig|EHA22058.1 aory|EIT78190.1 asyd|OJJ58620.1 ater|EAU34908.1 atub|OJI83918.1 aver|OJJ01227.1 awen|OJJ32072.1 azon|OJJ46652.1

Group2600: abra|OJJ71873.1 acar|OOF96544.1 acla|EAW13903.1 afis|EAW18615.1 afla|EED48366.1 afum|EAL93325.1 agla|OJJ81229.1 akaw|GAA85042.1 aluc|OJZ86612.1 anid|CBF70428.1 anig|EHA22054.1 aory|EIT83110.1 asyd|OJJ58616.1 ater|EAU34912.1 atub|OJI83914.1 aver|OJJ01223.1 awen|OJJ32076.1 azon|OJJ46656.1

Group2601: abra|OJJ71874.1 acar|OOF96545.1 acla|EAW13904.1 afis|EAW18616.1 afla|EED52730.1 afum|EAL93326.1 agla|OJJ81228.1 akaw|GAA85041.1 aluc|OJZ86613.1 anid|CBF70426.1 anig|EHA22053.1 aory|EIT78192.1 asyd|OJJ58617.1 ater|EAU34913.1 atub|OJI83913.1 aver|OJJ01224.1 awen|OJJ32077.1 azon|OJJ46657.1

Group2602: abra|OJJ71876.1 acla|EAW13905.1 afis|EAW18617.1 afla|EED52728.1 afum|EAL93327.1 agla|OJJ81227.1 akaw|GAA85040.1 aluc|OJZ86614.1 anid|CBF70424.1 anig|EHA22052.1 aory|EIT78248.1 asyd|OJJ58618.1 ater|EAU34917.1 atub|OJI83912.1 aver|OJJ01225.1 awen|OJJ32078.1 azon|OJJ46658.1 afla|EED52729.1

Group2603: abra|OJJ71878.1 acar|OOF96546.1 acla|EAW13907.1 afis|EAW18619.1 afla|EED52726.1 afum|EAL93329.1 agla|OJJ81241.1 akaw|GAA85038.1 aluc|OJZ86616.1 anid|CBF70501.1 anig|EHA22050.1 aory|EIT78153.1 asyd|OJJ58579.1 ater|EAU34919.1 atub|OJI83910.1 aver|OJJ01183.1 awen|OJJ32081.1 azon|OJJ46660.1

Group2604: abra|OJJ71879.1 acar|OOF96547.1 acla|EAW13908.1 afis|EAW18620.1 afla|EED52725.1 afum|EAL93330.1 agla|OJJ81240.1 akaw|GAA85037.1 aluc|OJZ86617.1 anid|CBF70499.1 anig|EHA22049.1 aory|EIT78189.1 asyd|OJJ58580.1 ater|EAU34920.1 atub|OJI83909.1 aver|OJJ01184.1 awen|OJJ32082.1 azon|OJJ46661.1

Group2605: abra|OJJ71880.1 acar|OOF96548.1 acla|EAW13909.1 afis|EAW18621.1 afla|EED52724.1 afum|EAL93331.1 agla|OJJ81239.1 akaw|GAA85036.1 aluc|OJZ86618.1 anid|CBF70497.1 anig|EHA22048.1 aory|EIT78181.1 asyd|OJJ58581.1 ater|EAU34921.1 atub|OJI83908.1 aver|OJJ01185.1 awen|OJJ32084.1 azon|OJJ46663.1

Group2606: abra|OJJ71889.1 acar|OOF96551.1 acla|EAW09632.1 afis|EAW21508.1 afla|EED55945.1 afum|EAL92528.1 agla|OJJ85671.1 akaw|GAA85028.1 aluc|OJZ86626.1 anid|CBF76443.1 anig|EHA22039.1 aory|EIT76729.1 asyd|OJJ61102.1 ater|EAU35030.1 atub|OJI83899.1 aver|OJJ00419.1 awen|OJJ39187.1 azon|OJJ51504.1

Group2607: abra|OJJ71891.1 acar|OOF96552.1 acla|EAW09631.1 afis|EAW21509.1 afla|EED55947.1 afum|EAL92529.1 agla|OJJ85673.1 akaw|GAA85027.1 aluc|OJZ86630.1 anid|CBF76441.1 anig|EHA22038.1 aory|EIT76718.1 asyd|OJJ61101.1 ater|EAU35029.1 atub|OJI83898.1 aver|OJJ00416.1 awen|OJJ39185.1 azon|OJJ51503.1

Group2608: abra|OJJ71895.1 acar|OOF96556.1 acla|EAW09627.1 afis|EAW21513.1 afla|EED55951.1 afum|EAL92533.1 agla|OJJ85677.1 akaw|GAA85023.1 aluc|OJZ86634.1 anid|CBF76433.1 anig|EHA22035.1 aory|EIT76705.1 asyd|OJJ61096.1 ater|EAU35027.1 atub|OJI83894.1 aver|OJJ00411.1 awen|OJJ39181.1 azon|OJJ51512.1

Group2609: abra|OJJ71896.1 acar|OOF96557.1 acla|EAW09626.1 afis|EAW21514.1 afla|EED55952.1 afum|EAL92534.1 agla|OJJ85678.1 akaw|GAA85022.1 aluc|OJZ86635.1 anid|CBF76431.1 anig|EHA22034.1 aory|EIT76554.1 asyd|OJJ61095.1 ater|EAU35026.1 atub|OJI83893.1 aver|OJJ00410.1 awen|OJJ39180.1 azon|OJJ51513.1

Group2610: abra|OJJ71898.1 acar|OOF96559.1 acla|EAW09624.1 afis|EAW21516.1 afla|EED55954.1 afum|EAL92536.1 agla|OJJ85680.1 akaw|GAA85020.1 aluc|OJZ86637.1 anid|CBF76427.1 anig|EHA22032.1 aory|EIT76837.1 asyd|OJJ61093.1 ater|EAU35024.1 atub|OJI83891.1 aver|OJJ00408.1 awen|OJJ39178.1 azon|OJJ51516.1

Group2611: abra|OJJ71902.1 acar|OOF96560.1 acla|EAW09621.1 afis|EAW21518.1 afla|EED55956.1 afum|EAL92539.1 agla|OJJ85681.1 akaw|GAA85018.1 aluc|OJZ86639.1 anid|CBF76423.1 anig|EHA22030.1 aory|EIT76726.1 asyd|OJJ61091.1 ater|EAU35022.1 atub|OJI83888.1 aver|OJJ00406.1 awen|OJJ39175.1 azon|OJJ51515.1

Group2612: abra|OJJ71903.1 acar|OOF96561.1 acla|EAW09620.1 afis|EAW21519.1 afla|EED55957.1 afum|EAL92540.1 agla|OJJ85682.1 akaw|GAA85017.1 aluc|OJZ86640.1 anid|CBF76419.1 anig|EHA22029.1 aory|EIT76756.1 asyd|OJJ61090.1 ater|EAU35021.1 atub|OJI83887.1 aver|OJJ00405.1 awen|OJJ39174.1 azon|OJJ51514.1

Group2613: abra|OJJ71905.1 acar|OOF96562.1 acla|EAW09618.1 afis|EAW21521.1 afla|EED55959.1 afum|EAL92542.1 agla|OJJ85684.1 akaw|GAA85015.1 aluc|OJZ86642.1 anid|CBF76416.1 anig|EHA22027.1 aory|EIT76533.1 asyd|OJJ61088.1 ater|EAU35020.1 atub|OJI83885.1 aver|OJJ00403.1 awen|OJJ39172.1 azon|OJJ51501.1

Group2614: abra|OJJ71908.1 acar|OOF96568.1 acla|EAW09615.1 afis|EAW21524.1 afla|EED55962.1 afum|EAL92545.1 agla|OJJ85687.1 akaw|GAA85011.1 aluc|OJZ86646.1 anid|CBF76410.1 anig|EHA22024.1 aory|EIT76688.1 asyd|OJJ61085.1 ater|EAU35017.1 atub|OJI83881.1 aver|OJJ00400.1 awen|OJJ39169.1 azon|OJJ51498.1

Group2615: abra|OJJ71917.1 acar|OOF96574.1 acla|EAW09613.1 afis|EAW21526.1 afla|EED55965.1 afum|EAL92547.1 agla|OJJ85689.1 akaw|GAA85005.1 aluc|OJZ86654.1 anid|CBF76406.1 anig|EHA22017.1 aory|EIT76778.1 asyd|OJJ61082.1 ater|EAU35015.1 atub|OJI83873.1 aver|OJJ00398.1 awen|OJJ39167.1 azon|OJJ51496.1

Group2616: abra|OJJ71918.1 acar|OOF96575.1 acla|EAW09612.1 afis|EAW21527.1 afla|EED55966.1 afum|EAL92548.1 agla|OJJ85690.1 akaw|GAA85004.1 aluc|OJZ86655.1 anid|CBF76404.1 anig|EHA22016.1 aory|EIT76686.1 asyd|OJJ61081.1 ater|EAU35014.1 atub|OJI83872.1 aver|OJJ00397.1 awen|OJJ39166.1 azon|OJJ51495.1

Group2617: abra|OJJ71921.1 acar|OOF96578.1 acla|EAW09607.1 afis|EAW21530.1 afla|EED55979.1 afum|EAL92551.1 agla|OJJ85694.1 akaw|GAA85001.1 aluc|OJZ86658.1 anid|CBF76398.1 anig|EHA22013.1 aory|EIT76728.1 asyd|OJJ61078.1 ater|EAU35009.1 atub|OJI83869.1 aver|OJJ00394.1 awen|OJJ39162.1 azon|OJJ51492.1

Group2618: abra|OJJ71922.1 acar|OOF96579.1 acla|EAW09606.1 afis|EAW21531.1 afla|EED55980.1 afum|EAL92552.1 agla|OJJ85695.1 akaw|GAA85000.1 aluc|OJZ86659.1 anid|CBF76396.1 anig|EHA22012.1 aory|EIT76765.1 asyd|OJJ61077.1 ater|EAU35008.1 atub|OJI83868.1 aver|OJJ00393.1 awen|OJJ39161.1 azon|OJJ51491.1

Group2619: abra|OJJ71923.1 acar|OOF96580.1 acla|EAW09605.1 afis|EAW21532.1 afla|EED55981.1 afum|EAL92553.1 agla|OJJ85696.1 akaw|GAA84999.1 aluc|OJZ86660.1 anid|CBF76394.1 anig|EHA22011.1 aory|EIT76691.1 asyd|OJJ61076.1 ater|EAU35007.1 atub|OJI83867.1 aver|OJJ00392.1 awen|OJJ39160.1 azon|OJJ51490.1

Group2620: abra|OJJ71924.1 acar|OOF96581.1 acla|EAW09604.1 afis|EAW21533.1 afla|EED55982.1 afum|EAL92554.1 agla|OJJ85697.1 akaw|GAA84998.1 aluc|OJZ86661.1 anid|CBF76392.1 anig|EHA22010.1 aory|EIT76892.1 asyd|OJJ61073.1 ater|EAU35006.1 atub|OJI83864.1 aver|OJJ00390.1 awen|OJJ39159.1 azon|OJJ51489.1

Group2621: abra|OJJ71928.1 acar|OOF96585.1 afis|EAW21536.1 afla|EED55986.1 afum|EAL92556.2 agla|OJJ85700.1 anid|CBF76386.1 aory|EIT76707.1 aver|OJJ00386.1 acla|EAW09601.1 akaw|GAA84995.1

aluc|OJZ86665.1 anig|EHA22008.1 asyd|OJJ61070.1 ater|EAU35003.1 atub|OJI83861.1 awen|OJJ39155.1 azon|OJJ51486.1

Group2622: abra|OJJ71929.1 acar|OOF96586.1 acla|EAW09600.1 afis|EAW21537.1 afla|EED55989.1 afum|EAL92557.1 agla|OJJ85701.1 akaw|GAA84994.1 aluc|OJZ86666.1 anid|CBF76382.1 anig|EHA22007.1 aory|EIT76583.1 asyd|OJJ61068.1 ater|EAU35002.1 atub|OJI83860.1 aver|OJJ00384.1 awen|OJJ39154.1 azon|OJJ51485.1

Group2623: abra|OJJ71931.1 acar|OOF96588.1 acla|EAW09598.1 afis|EAW21539.1 afla|EED55991.1 afum|EAL92559.2 agla|OJJ85703.1 akaw|GAA84992.1 aluc|OJZ86668.1 anid|CBF76378.1 anig|EHA22005.1 aory|EIT76546.1 asyd|OJJ61066.1 ater|EAU35001.1 atub|OJI83859.1 aver|OJJ00382.1 awen|OJJ39152.1 azon|OJJ51478.1

Group2624: abra|OJJ71935.1 acar|OOF96591.1 acla|EAW09595.1 afis|EAW21542.1 afla|EED55993.1 afum|EAL92562.1 agla|OJJ85705.1 akaw|GAA84989.1 aluc|OJZ86672.1 anid|CBF76374.1 anig|EHA22002.1 aory|EIT76774.1 asyd|OJJ61064.1 ater|EAU34997.1 atub|OJI83856.1 aver|OJJ00381.1 awen|OJJ39148.1 azon|OJJ51482.1

Group2625: abra|OJJ71938.1 acar|OOF96596.1 acla|EAW09591.1 afis|EAW21547.1 afla|EED55999.1 afum|EAL92567.1 agla|OJJ85710.1 akaw|GAA84986.1 aluc|OJZ86675.1 anid|CBF76364.1 anig|EHA21998.1 aory|EIT76832.1 asyd|OJJ61060.1 ater|EAU34992.1 atub|OJI83853.1 aver|OJJ00377.1 awen|OJJ39143.1 azon|OJJ51464.1

Group2626: abra|OJJ71940.1 acar|OOF96598.1 acla|EAW09590.1 afis|EAW21548.1 afla|EED56001.1 afum|EAL92568.1 agla|OJJ85726.1 akaw|GAA84985.1 aluc|OJZ86676.1 anid|CBF76362.1 anig|EHA21997.1 aory|EIT76581.1 asyd|OJJ61059.1 ater|EAU34991.1 atub|OJI83852.1 aver|OJJ00376.1 awen|OJJ39142.1 azon|OJJ51465.1

Group2627: abra|OJJ71942.1 acar|OOF96599.1 acla|EAW09588.1 afis|EAW21550.1 afla|EED56002.1 afum|EAL92570.1 agla|OJJ85728.1 akaw|GAA84983.1 aluc|OJZ86678.1 anid|CBF76358.1 anig|EHA21995.1 aory|EIT76540.1 asyd|OJJ61057.1 ater|EAU34989.1 atub|OJI83849.1 aver|OJJ00374.1 awen|OJJ39140.1 azon|OJJ51467.1

Group2628: abra|OJJ71946.1 acar|OOF96600.1 acla|EAW09587.1 afis|EAW21551.1 afla|EED56003.1 afum|EAL92571.1 agla|OJJ85729.1 akaw|GAA84980.1 aluc|OJZ86682.1 anid|CBF76356.1 anig|EHA21992.1 aory|EIT76839.1 asyd|OJJ61056.1 ater|EAU34988.1 atub|OJI83845.1 aver|OJJ00373.1 awen|OJJ39139.1 azon|OJJ51470.1

Group2629: abra|OJJ71951.1 acar|OOF96602.1 acla|EAW09583.1 afis|EAW21554.1 afla|EED56006.1 afum|EAL92574.1 agla|OJJ85731.1 akaw|GAA84977.1 aluc|OJZ86685.1 anid|CBF76350.1 anig|EHA21988.1 aory|EIT76802.1 asyd|OJJ61053.1 ater|EAU34980.1 atub|OJI83841.1 aver|OJJ00370.1 awen|OJJ39136.1 azon|OJJ51474.1

Group2630: abra|OJJ71956.1 acar|OOF96615.1 acla|EAW09579.1 afis|EAW21558.1 afla|EED56010.1 afum|EAL92577.1 agla|OJJ85736.1 akaw|GAA84971.1 aluc|OJZ86691.1 anid|CBF76344.1 anig|EHA21982.1 aory|EIT76530.1 asyd|OJJ61049.1 ater|EAU34976.1 atub|OJI83833.1 aver|OJJ00365.1 awen|OJJ39132.1 azon|OJJ51460.1

Group2631: abra|OJJ71957.1 acar|OOF96613.1 acla|EAW09578.1 afis|EAW21559.1 afla|EED56011.1 afum|EAL92578.1 agla|OJJ85737.1 akaw|GAA84970.1 aluc|OJZ86692.1 anid|CBF76342.1 anig|EHA21981.1 aory|EIT76831.1 asyd|OJJ61048.1 ater|EAU34974.1 atub|OJI83832.1 aver|OJJ00364.1 awen|OJJ39131.1 azon|OJJ51459.1

Group2632: abra|OJJ71959.1 acar|OOF96611.1 acla|EAW09577.1 afis|EAW21560.1 afla|EED56012.1 afum|EAL92579.1 agla|OJJ85739.1 akaw|GAA84969.1 aluc|OJZ86693.1 anid|CBF76340.1 anig|EHA21980.1 aory|EIT76670.1 asyd|OJJ61046.1 atub|OJI83831.1 aver|OJJ00363.1 awen|OJJ39129.1 azon|OJJ51458.1 ater|EAU34973.1

Group2633: abra|OJJ71960.1 acar|OOF96610.1 acla|EAW09576.1 afis|EAW21561.1 afla|EED56014.1 afum|EAL92580.1 agla|OJJ85740.1 akaw|GAA84968.1 aluc|OJZ86694.1 anid|CBF76338.1 anig|EHA21979.1 aory|EIT76536.1 asyd|OJJ61045.1 ater|EAU34972.1 atub|OJI83830.1 aver|OJJ00362.1 awen|OJJ39128.1 azon|OJJ51456.1

Group2634: abra|OJJ71961.1 acar|OOF96609.1 acla|EAW09575.1 afis|EAW21562.1 afla|EED56015.1 afum|EAL92581.1 agla|OJJ85741.1 akaw|GAA84967.1 aluc|OJZ86696.1 anid|CBF76336.1 anig|EHA21978.1 aory|EIT76848.1 asyd|OJJ61044.1 ater|EAU34971.1 atub|OJI83829.1 aver|OJJ00361.1 awen|OJJ39127.1 azon|OJJ51455.1

Group2635: abra|OJJ71966.1 acar|OOF89890.1 acla|EAW09571.1 afis|EAW21565.1 afum|EAL92584.1 agla|OJJ85750.1 akaw|GAA84962.1 aluc|OJZ86702.1 anid|CBF76330.1 anig|EHA21974.1 aory|EIT76870.1 asyd|OJJ61038.1 ater|EAU34967.1 atub|OJI83824.1 aver|OJJ00356.1 awen|OJJ39120.1 azon|OJJ51450.1 afla|EED56022.1

Group2636: abra|OJJ71969.1 acar|OOF96604.1 acla|EAW09570.1 afis|EAW21566.1 afla|EED56031.1 afum|EAL92585.1 agla|OJJ85751.1 akaw|GAA84961.1 aluc|OJZ86704.1 anid|CBF76328.1 anig|EHA21972.1 aory|EIT76642.1 asyd|OJJ61016.1 ater|EAU34965.1 atub|OJI83823.1 aver|OJJ00336.1 awen|OJJ39115.1 azon|OJJ51449.1

Group2637: abra|OJJ71970.1 acar|OOF96621.1 acla|EAW09569.1 afis|EAW21567.1 afla|EED56034.1 afum|EAL92586.1 agla|OJJ85752.1 akaw|GAA84960.1 aluc|OJZ86705.1 anid|CBF76326.1 anig|EHA21971.1 aory|EIT76629.1 asyd|OJJ61015.1 ater|EAU34964.1 atub|OJI83822.1 aver|OJJ00334.1 awen|OJJ39113.1 azon|OJJ51448.1

Group2638: abra|OJJ71971.1 acar|OOF96622.1 acla|EAW09568.1 afis|EAW21568.1 afla|EED56035.1 afum|EAL92587.1 agla|OJJ85753.1 akaw|GAA84959.1 aluc|OJZ86706.1 anid|CBF76324.1 anig|EHA21970.1 aory|EIT76766.1 asyd|OJJ61014.1 ater|EAU34963.1 atub|OJI83821.1 aver|OJJ00333.1 awen|OJJ39112.1 azon|OJJ51447.1

Group2639: abra|OJJ71972.1 acar|OOF96623.1 acla|EAW09567.1 afis|EAW21569.1 afla|EED56036.1 afum|EBA27467.1 agla|OJJ85754.1 akaw|GAA84958.1 aluc|OJZ86707.1 anid|CBF76322.1 anig|EHA21969.1 aory|EIT76617.1 asyd|OJJ61013.1 ater|EAU34962.1 atub|OJI83820.1 aver|OJJ00332.1 awen|OJJ39111.1 azon|OJJ51446.1

Group2640: abra|OJJ71973.1 acar|OOF96625.1 acla|EAW09565.1 afis|EAW21571.1 afla|EED56037.1 afum|EAL92589.1 agla|OJJ85771.1 akaw|GAA84957.1 aluc|OJZ86708.1 anid|CBF76318.1 anig|EHA21968.1 aory|EIT76745.1 asyd|OJJ61011.1 ater|EAU34961.1 atub|OJI83819.1 aver|OJJ00330.1 awen|OJJ39110.1 azon|OJJ51445.1

Group2641: abra|OJJ71974.1 acar|OOF96624.1 acla|EAW09564.1 afis|EAW21572.1 afla|EED56038.1 afum|EAL92590.1 agla|OJJ85768.1 akaw|GAA84956.1 aluc|OJZ86709.1 anid|CBF76316.1 anig|EHA21967.1 aory|EIT76889.1 asyd|OJJ61010.1 ater|EAU34960.1 atub|OJI83818.1 aver|OJJ00329.1 awen|OJJ39109.1 azon|OJJ51444.1

Group2642: abra|OJJ71975.1 acar|OOF96626.1 acla|EAW09563.1 afis|EAW21573.1 afla|EED57792.1 afum|EAL92591.2 agla|OJJ85766.1 akaw|GAA84955.1 aluc|OJZ86710.1 anid|CBF76314.1 anig|EHA21966.1 aory|EIT81122.1 asyd|OJJ61009.1 ater|EAU34959.1 atub|OJI83817.1 aver|OJJ00328.1 awen|OJJ39108.1 azon|OJJ51443.1

Group2643: abra|OJJ71978.1 acar|OOF96628.1 acla|EAW09562.1 afis|EAW21575.1 afla|EED57793.1 afum|EAL92593.1 agla|OJJ85764.1 akaw|GAA84952.1 aluc|OJZ86715.1 anid|CBF76306.1 anig|EHA21963.1 aory|EIT81108.1 asyd|OJJ61006.1 ater|EAU34957.1 atub|OJI83814.1 aver|OJJ00324.1 awen|OJJ39105.1 azon|OJJ51438.1

Group2644: abra|OJJ71983.1 acar|OOF96634.1 acla|EAW09556.1 afis|EAW21581.1 afla|EED57799.1 afum|EAL92599.2 agla|OJJ85757.1 akaw|GAA84946.1 aluc|OJZ86720.1 anid|CBF76300.1 anig|EHA21959.1 aory|EIT81064.1 asyd|OJJ61001.1 ater|EAU34953.1 atub|OJI83809.1 aver|OJJ00319.1 awen|OJJ39100.1 azon|OJJ51434.1

Group2645: abra|OJJ71987.1 acar|OOF96642.1 acla|EAW09551.1 afis|EAW21585.1 afla|EED57806.1 afum|EAL92603.2 agla|OJJ85777.1 akaw|GAA84942.1 aluc|OJZ86724.1 anid|CBF76289.1 anig|EHA21955.1 aory|EIT81058.1 asyd|OJJ60995.1 ater|EAU34946.1 atub|OJI83804.1 aver|OJJ00315.1 awen|OJJ39096.1 azon|OJJ51429.1

Group2646: abra|OJJ71990.1 acar|OOF96644.1 acla|EAW09545.1 afis|EAW21588.1 afla|EED57809.1 afum|EAL92606.1 agla|OJJ85781.1 akaw|GAA84939.1 aluc|OJZ86727.1 anid|CBF76285.1 anig|EHA21952.1 aory|EIT81040.1 asyd|OJJ60992.1 ater|EAU34943.1 atub|OJI83801.1 aver|OJJ00312.1 awen|OJJ39093.1 azon|OJJ51426.1

Group2647: abra|OJJ71991.1 acar|OOF96645.1 acla|EAW09544.1 afis|EAW21589.1 afla|EED57811.1 afum|EAL92607.1 agla|OJJ85782.1 akaw|GAA84938.1 aluc|OJZ86728.1 anid|CBF76283.1 anig|EHA21951.1 aory|EIT81035.1 asyd|OJJ60991.1 ater|EAU34942.1 atub|OJI83800.1 aver|OJJ00311.1 awen|OJJ39092.1 azon|OJJ51425.1

Group2648: abra|OJJ71993.1 acar|OOF96647.1 acla|EAW09542.1 afis|EAW21591.1 afla|EED57813.1 afum|EAL92609.1 agla|OJJ85784.1 akaw|GAA84935.1 aluc|OJZ86731.1 anid|CBF83440.1 anig|EHA21948.1 aory|EIT81099.1 asyd|OJJ60668.1 ater|EAU34940.1 atub|OJI83797.1 aver|OJI99991.1 awen|OJJ39090.1 azon|OJJ51421.1

Group2649: abra|OJJ71995.1 acar|OOF96649.1 acla|EAW09540.1 afis|EAW21593.1 afla|EED57815.1 afum|EAL92611.1 agla|OJJ85788.1 akaw|GAA84933.1 aluc|OJZ86733.1 anid|CBF83444.1 anig|EHA21946.1 aory|EIT81101.1 asyd|OJJ60670.1 ater|EAU34931.1 atub|OJI83795.1 aver|OJI99993.1 awen|OJJ39088.1 azon|OJJ51419.1

Group2650: abra|OJJ71997.1 acar|OOF96651.1 acla|EAW09541.1 afis|EAW21592.1 afla|EED57814.1 afum|EAL92610.1 agla|OJJ85789.1 akaw|GAA84932.1 aluc|OJZ86735.1 anid|CBF83442.1 anig|EHA21944.1

aory|EIT81111.1 asyd|OJJ60669.1 ater|EAU34932.1 atub|OJI83793.1 aver|OJI99992.1 awen|OJJ39089.1 azon|OJJ51420.1

Group2651: abra|OJJ71998.1 acar|OOF96653.1 acla|EAW09536.1 afis|EAW21596.1 afla|EED57819.1 afum|EAL92614.1 agla|OJJ85790.1 akaw|GAA84931.1 aluc|OJZ86736.1 anid|CBF83453.1 anig|EHA21943.1 aory|EIT81091.1 asyd|OJJ60672.1 ater|EAU34928.1 atub|OJI83792.1 aver|OJI99995.1 awen|OJJ39086.1 azon|OJJ51417.1

Group2652: abra|OJJ72007.1 acar|OOF96657.1 acla|EAW09534.1 afis|EAW21599.1 afla|EED57824.1 afum|EAL92618.1 agla|OJJ85794.1 akaw|GAA84926.1 aluc|OJZ86743.1 anid|CBF83457.1 anig|EHA21936.1 aory|EIT81107.1 asyd|OJJ60679.1 ater|EAU38289.1 atub|OJI83784.1 aver|OJJ00004.1 awen|OJJ39083.1 azon|OJJ51415.1

Group2653: abra|OJJ72008.1 acla|EAW09533.1 anid|CBF83458.1 asyd|OJJ60682.1 ater|EAU38290.1 aver|OJJ00007.1 azon|OJJ51414.1 acla|EAW09532.1 afis|EAW21600.1 afla|EED57827.1 afum|EAL92619.1 agla|OJJ85795.1 akaw|GAA84925.1 aluc|OJZ86744.1 anig|EHA21935.1 aory|EIT81065.1 atub|OJI83783.1 awen|OJJ39082.1

Group2654: abra|OJJ72010.1 acar|OOF96658.1 acla|EAW09529.1 afis|EAW21603.1 afla|EED57829.1 afum|EAL92622.1 agla|OJJ85797.1 akaw|GAA84923.1 aluc|OJZ86746.1 anid|CBF83462.1 anig|EHA21933.1 aory|EIT81061.1 asyd|OJJ60684.1 ater|EAU38292.1 atub|OJI83781.1 aver|OJJ00009.1 awen|OJJ39080.1 azon|OJJ51412.1

Group2655: abra|OJJ72014.1 acar|OOF96663.1 acla|EAW09526.1 afis|EAW21606.1 afla|EED57833.1 afum|EAL92625.1 agla|OJJ85801.1 akaw|GAA84919.1 aluc|OJZ86750.1 anid|CBF83469.1 anig|EHA21929.1 aory|EIT81024.1 asyd|OJJ60688.1 ater|EAU38295.1 atub|OJI83777.1 aver|OJJ00013.1 awen|OJJ39076.1 azon|OJJ51408.1

Group2656: abra|OJJ72015.1 acar|OOF96664.1 acla|EAW09525.1 afis|EAW21607.1 afla|EED57834.1 afum|EAL92626.1 agla|OJJ85802.1 akaw|GAA84918.1 aluc|OJZ86751.1 anid|CBF83473.1 anig|EHA21928.1 aory|EIT81105.1 asyd|OJJ60690.1 ater|EAU38296.1 atub|OJI83775.1 aver|OJJ00015.1 awen|OJJ39075.1 azon|OJJ51407.1

Group2657: abra|OJJ72016.1 acar|OOF96666.1 acla|EAW09524.1 afis|EAW21608.1 afla|EED57837.1 afum|EAL92642.1 agla|OJJ85803.1 akaw|GAA84917.1 aluc|OJZ86752.1 anid|CBF83475.1 anig|EHA21927.1 aory|EIT81080.1 asyd|OJJ60691.1 ater|EAU38297.1 atub|OJI83774.1 aver|OJJ00016.1 awen|OJJ39074.1 azon|OJJ51406.1

Group2658: abra|OJJ72017.1 acar|OOF96665.1 acla|EAW09523.1 afis|EAW21609.1 afla|EED57838.1 afum|EAL92643.1 agla|OJJ85804.1 akaw|GAA84916.1 aluc|OJZ86753.1 anid|CBF83477.1 anig|EHA21926.1 aory|EIT81100.1 asyd|OJJ60692.1 ater|EAU38298.1 atub|OJI83773.1 aver|OJJ00017.1 awen|OJJ39073.1 azon|OJJ51405.1

Group2659: abra|OJJ72018.1 acar|OOF96667.1 acla|EAW09521.1 afis|EAW21611.1 afla|EED57840.1 afum|EAL92645.1 agla|OJJ78616.1 akaw|GAA84914.1 aluc|OJZ86755.1 anid|CBF83480.1 anig|EHA21924.1 aory|EIT81096.1 asyd|OJJ60694.1 ater|EAU38302.1 atub|OJI83772.1 aver|OJJ00020.1 awen|OJJ39071.1 azon|OJJ51402.1

Group2660: abra|OJJ72019.1 acar|OOF96669.1 acla|EAW09520.1 afis|EAW21612.1 afla|EED57841.1 afum|EAL92646.1 agla|OJJ78617.1 akaw|GAA84913.1 aluc|OJZ86756.1 anid|CBF83482.1 anig|EHA21923.1 aory|EIT81062.1 asyd|OJJ60695.1 ater|EAU38303.1 atub|OJI83771.1 aver|OJJ00021.1 awen|OJJ39070.1 azon|OJJ51401.1

Group2661: abra|OJJ72020.1 acar|OOF96668.1 acla|EAW09515.1 afis|EAW21617.1 afla|EED57842.1 afum|EAL92652.1 agla|OJJ78622.1 akaw|GAA84912.1 aluc|OJZ86757.1 anid|CBF83484.1 anig|EHA21922.1 aory|EIT81032.1 asyd|OJJ60696.1 ater|EAU38304.1 atub|OJI83770.1 aver|OJJ00022.1 awen|OJJ39069.1 azon|OJJ51400.1

Group2662: abra|OJJ72022.1 acar|OOF96671.1 acla|EAW09517.1 afis|EAW21615.1 afla|EED57844.1 afum|EAL92649.1 agla|OJJ78624.1 akaw|GAA84910.1 aluc|OJZ86759.1 anid|CBF83485.1 anig|EHA21920.1 aory|EIT81023.1 asyd|OJJ60698.1 ater|EAU38305.1 atub|OJI83768.1 aver|OJJ00024.1 awen|OJJ39067.1 azon|OJJ51398.1

Group2663: abra|OJJ72024.1 acar|OOF96673.1 acla|EAW09519.1 afis|EAW21613.1 afla|EED57846.1 afum|EAL92647.1 agla|OJJ78626.1 akaw|GAA84908.1 aluc|OJZ86761.1 anid|CBF83489.1 anig|EHA21918.1 aory|EIT81029.1 asyd|OJJ60700.1 ater|EAU38307.1 atub|OJI83766.1 aver|OJJ00026.1 awen|OJJ39065.1 azon|OJJ51396.1

Group2664: abra|OJJ72032.1 acar|OOF96682.1 acla|EAW09511.1 afis|EAW21621.1 afla|EED57850.1 afum|EAL92657.2 agla|OJJ82578.1 akaw|GAA84900.1 aluc|OJZ86770.1 anid|CBF83495.1 anig|EHA21910.1 aory|EIT81046.1 asyd|OJJ60704.1 ater|EAU38310.1 atub|OJI83757.1 aver|OJJ00029.1 awen|OJJ33873.1 azon|OJJ48677.1

Group2665: abra|OJJ72044.1 acar|OOF96702.1 acla|EAW09502.1 afis|EAW21634.1 afla|EED57876.1 afum|EAL92667.1 agla|OJJ79699.1 akaw|GAA84890.1 aluc|OJZ86781.1 anid|CBF83500.1 anig|EHA21901.1 aory|EIT83264.1 asyd|OJJ60711.1 ater|EAU38320.1 atub|OJI83739.1 aver|OJJ00038.1 awen|OJJ33883.1 azon|OJJ48670.1

Group2666: abra|OJJ72046.1 acar|OOF96704.1 acla|EAW09500.1 afis|EAW21636.1 afla|EED57879.1 afum|EAL92670.1 agla|OJJ79701.1 akaw|GAA84888.1 aluc|OJZ86783.1 anid|CBF83517.1 anig|EHA21899.1 aory|EIT83275.1 asyd|OJJ60714.1 ater|EAU38322.1 atub|OJI83737.1 aver|OJJ00047.1 awen|OJJ33885.1 azon|OJJ48668.1

Group2667: abra|OJJ72048.1 acar|OOF96706.1 acla|EAW09498.1 afis|EAW21638.1 afla|EED57881.1 afum|EAL92672.2 agla|OJJ79703.1 akaw|GAA84886.1 aluc|OJZ86785.1 anid|CBF83521.1 anig|EHA21897.1 aory|EIT83269.1 asyd|OJJ60716.1 ater|EAU38323.1 atub|OJI83735.1 aver|OJJ00049.1 awen|OJJ33887.1 azon|OJJ48666.1

Group2668: abra|OJJ72052.1 acar|OOF96709.1 acla|EAW09495.1 afis|EAW21641.1 afla|EED57902.1 afum|EAL92675.1 agla|OJJ79708.1 akaw|GAA84882.1 aluc|OJZ86789.1 anid|CBF83526.1 anig|EHA21894.1 aory|EIT82320.1 asyd|OJJ60718.1 ater|EAU38326.1 atub|OJI83730.1 aver|OJJ00051.1 awen|OJJ33898.1 azon|OJJ48662.1

Group2669: abra|OJJ72054.1 acar|OOF96711.1 acla|EAW09493.1 afis|EAW21643.1 afla|EED57900.1 afum|EAL92677.1 agla|OJJ79710.1 akaw|GAA84880.1 aluc|OJZ86791.1 anid|CBF83528.1 anig|EHA21892.1 aory|EIT82343.1 asyd|OJJ60720.1 ater|EAU38328.1 atub|OJI83728.1 aver|OJJ00053.1 awen|OJJ33900.1 azon|OJJ48660.1

Group2670: abra|OJJ72056.1 acar|OOF96713.1 acla|EAW09492.1 afis|EAW21644.1 afla|EED57899.1 afum|EAL92678.1 agla|OJJ79711.1 akaw|GAA84878.1 aluc|OJZ86793.1 anid|CBF83530.1 anig|EHA21890.1 aory|EIT82331.1 asyd|OJJ60721.1 ater|EAU38329.1 atub|OJI83726.1 aver|OJJ00054.1 awen|OJJ33901.1 azon|OJJ48658.1

Group2671: abra|OJJ72062.1 acar|OOF96716.1 acla|EAW09489.1 afis|EAW21647.1 afla|EED57896.1 afum|EAL92681.1 agla|OJJ79718.1 akaw|GAA84873.1 aluc|OJZ86798.1 anid|CBF83537.1 anig|EHA21885.1 aory|EIT82735.1 asyd|OJJ60725.1 ater|EAU38332.1 atub|OJI83720.1 aver|OJJ00058.1 awen|OJJ33907.1 azon|OJJ48655.1

Group2672: abra|OJJ72063.1 acar|OOF96717.1 acla|EAW09479.1 afis|EAW21657.1 afla|EED57895.1 afum|EAL92692.1 agla|OJJ79719.1 akaw|GAA84872.1 aluc|OJZ86799.1 anid|CBF83539.1 anig|EHA21884.1 aory|EIT82741.1 asyd|OJJ60727.1 ater|EAU38333.1 atub|OJI83719.1 aver|OJJ00059.1 awen|OJJ33908.1 azon|OJJ48653.1

Group2673: abra|OJJ72064.1 acar|OOF96718.1 acla|EAW09478.1 afis|EAW21658.1 afla|EED57894.1 afum|EAL92693.1 agla|OJJ79720.1 akaw|GAA84871.1 aluc|OJZ86800.1 anid|CBF83541.1 anig|EHA21883.1 aory|EIT82736.1 asyd|OJJ60728.1 ater|EAU38334.1 atub|OJI83718.1 aver|OJJ00060.1 awen|OJJ33909.1 azon|OJJ48652.1

Group2674: abra|OJJ72065.1 acar|OOF96719.1 acla|EAW09477.1 afis|EAW21659.1 afla|EED57893.1 afum|EAL92694.1 agla|OJJ79721.1 akaw|GAA84870.1 aluc|OJZ86801.1 anid|CBF83542.1 anig|EHA21882.1 aory|EIT82740.1 asyd|OJJ60729.1 ater|EAU38335.1 atub|OJI83717.1 aver|OJJ00061.1 awen|OJJ33910.1 azon|OJJ48651.1

Group2675: abra|OJJ72068.1 acar|OOF96724.1 acla|EAW09483.1 afis|EAW21653.1 afla|EED57886.1 afum|EAL92688.1 agla|OJJ79724.1 akaw|GAA84867.1 aluc|OJZ86803.1 anid|CBF83550.1 anig|EHA21879.1 aory|EIT83265.1 asyd|OJJ60733.1 ater|EAU38339.1 atub|OJI83715.1 aver|OJJ00065.1 awen|OJJ33914.1 azon|OJJ48647.1

Group2676: abra|OJJ72075.1 acar|OOF96727.1 acla|EAW09485.1 afis|EAW21651.1 afla|EED57918.1 afum|EAL92685.2 agla|OJJ79727.1 akaw|GAA84842.1 aluc|OJZ86829.1 anid|CBF83553.1 anig|EHA21806.1 aory|EIT82364.1 asyd|OJJ60735.1 ater|EAU38343.1 atub|OJI83689.1 aver|OJJ00067.1 awen|OJJ33917.1 azon|OJJ48643.1

Group2677: abra|OJJ72085.1 acar|OOF96730.1 acla|EAW09488.1 afis|EAW21648.1 afla|EED57923.1 afum|EAL92682.2 agla|OJJ79730.1 akaw|GAA84835.1 aluc|OJZ86839.1 anid|CBF83559.1 anig|EHA21798.1 aory|EIT82355.1 asyd|OJJ60738.1 ater|EAU38345.1 atub|OJI83679.1 aver|OJJ00070.1 awen|OJJ33919.1 azon|OJJ48645.1

Group2678: abra|OJJ72086.1 acar|OOF96732.1 acla|EAW09467.1 afis|EAW21669.1 afla|EED57933.1 afum|EAL92704.1 agla|OJJ79740.1 akaw|GAA84834.1 aluc|OJZ86840.1 anid|CBF83582.1 anig|EHA21797.1 aory|EIT82344.1 asyd|OJJ60781.1 ater|EAU38353.1 atub|OJI83678.1 aver|OJJ00114.1 awen|OJJ33932.1 azon|OJJ48640.1

Group2679: abra|OJJ72087.1 acar|OOF96731.1 acla|EAW09468.1 afis|EAW21668.1 afla|EED57932.1 afum|EAL92703.1 agla|OJJ79739.1 akaw|GAA84833.1 aluc|OJZ86841.1 anid|CBF83580.1 anig|EHA21796.1

aory|EIT82317.1 asyd|OJJ60749.1 ater|EAU38352.1 atub|OJI83677.1 aver|OJJ00081.1 awen|OJJ33931.1 azon|OJJ48639.1

Group2680: abra|OJJ72089.1 acar|OOF96735.1 acla|EAW09470.1 afis|EAW21666.1 afla|EED57930.1 afum|EAL92701.1 agla|OJJ79737.1 akaw|GAA84831.1 aluc|OJZ86843.1 anid|CBF83569.1 anig|EHA21794.1 aory|EIT82323.1 asyd|OJJ60747.1 ater|EAU38351.1 atub|OJI83675.1 aver|OJJ00079.1 awen|OJJ33930.1 azon|OJJ48637.1

Group2681: abra|OJJ72090.1 acar|OOF96734.1 acla|EAW09471.1 afis|EAW21665.1 afla|EED57929.1 afum|EAL92700.1 agla|OJJ79736.1 akaw|GAA84830.1 aluc|OJZ86844.1 anid|CBF83568.1 anig|EHA21793.1 aory|EIT82319.1 asyd|OJJ60746.1 ater|EAU38350.1 atub|OJI83674.1 aver|OJJ00078.1 awen|OJJ33929.1 azon|OJJ48636.1

Group2682: abra|OJJ72092.1 acar|OOF96737.1 acla|EAW09473.1 afis|EAW21662.1 afla|EED57927.1 afum|EAL92697.1 agla|OJJ79734.1 akaw|GAA84828.1 aluc|OJZ86847.1 anid|CBF83564.1 anig|EHA21791.1 aory|EIT82336.1 asyd|OJJ60744.1 ater|EAU38348.1 atub|OJI83672.1 aver|OJJ00076.1 awen|OJJ33926.1 azon|OJJ48634.1

Group2683: abra|OJJ72094.1 acar|OOF96739.1 acla|EAW09475.1 afis|EAW21664.1 afla|EED57925.1 afum|EAL92699.1 agla|OJJ79732.1 akaw|GAA84826.1 aluc|OJZ86849.1 anid|CBF83562.1 anig|EHA21789.1 aory|EIT82328.1 asyd|OJJ60742.1 ater|EAU38347.1 atub|OJI83670.1 aver|OJJ00074.1 awen|OJJ33924.1 azon|OJJ48632.1

Group2684: abra|OJJ72095.1 acar|OOF96740.1 acla|EAW09476.1 afis|EAW21660.1 afla|EED57924.1 afum|EAL92695.1 agla|OJJ79731.1 akaw|GAA84825.1 aluc|OJZ86850.1 anid|CBF83560.1 anig|EHA21788.1 aory|EIT82356.1 asyd|OJJ60741.1 ater|EAU38346.1 atub|OJI83669.1 aver|OJJ00073.1 awen|OJJ33923.1 azon|OJJ48631.1

Group2685: abra|OJJ72096.1 acar|OOF96741.1 acla|EAW09391.1 afis|EAW21742.1 afla|EED58036.1 afum|EAL92775.1 agla|OJJ84431.1 akaw|GAA84824.1 aluc|OJZ86851.1 anid|CBF83748.1 anig|EHA21787.1 aory|EIT79774.1 asyd|OJJ60855.1 ater|EAU38446.1 atub|OJI83668.1 aver|OJJ00179.1 awen|OJJ34024.1 azon|OJJ48534.1

Group2686: abra|OJJ72099.1 acar|OOF96744.1 acla|EAW09390.1 afis|EAW21743.1 afla|EED58034.1 afum|EAL92776.1 agla|OJJ84427.1 akaw|GAA84822.1 aluc|OJZ86854.1 anid|CBF83738.1 anig|EHA21785.1 aory|EIT79762.1 asyd|OJJ60853.1 ater|EAU38445.1 atub|OJI83665.1 aver|OJJ00177.1 awen|OJJ34026.1 azon|OJJ48530.1

Group2687: abra|OJJ72101.1 acar|OOF96746.1 acla|EAW09388.1 afis|EAW21746.1 afla|EED58032.1 afum|EAL92778.1 agla|OJJ84430.1 akaw|GAA84820.1 aluc|OJZ86856.1 anid|CBF83662.1 anig|EHA21783.1 aory|EIT79789.1 asyd|OJJ60804.1 ater|EAU38443.1 atub|OJI83663.1 aver|OJJ00133.1 awen|OJJ34028.1 azon|OJJ48201.1

Group2688: abra|OJJ72105.1 acar|OOF96748.1 acla|EAW09386.1 afis|EAW21749.1 afla|EED58029.1 afum|EAL92780.2 agla|OJJ84425.1 akaw|GAA84817.1 aluc|OJZ86860.1 anid|CBF83665.1 anig|EHA21780.1 aory|EIT79704.1 asyd|OJJ60806.1 ater|EAU38441.1 atub|OJI83660.1 aver|OJJ00135.1 awen|OJJ34030.1 azon|OJJ48203.1

Group2689: abra|OJJ72110.1 acar|OOF90022.1 acla|EAW14910.1 afis|EAW24431.1 afla|EED55244.1 afum|EAL86099.1 agla|OJJ79245.1 akaw|GAA84812.1 aluc|OJZ86865.1 anid|CBF73773.1 anig|EHA21776.1 aory|EIT83529.1 asyd|OJJ56593.1 ater|EAU30014.1 atub|OJI83655.1 aver|OJJ06132.1 awen|OJJ32771.1 azon|OJJ44001.1

Group2690: abra|OJJ72112.1 acar|OOF96752.1 acla|EAW14912.1 afis|EAW24433.1 afla|EED55247.1 afum|EAL86101.1 agla|OJJ79248.1 akaw|GAA84810.1 aluc|OJZ86867.1 anid|CBF73767.1 anig|EHA21774.1 aory|EIT83512.1 asyd|OJJ56591.1 ater|EAU30012.1 atub|OJI83653.1 aver|OJJ06134.1 awen|OJJ32768.1 azon|OJJ43999.1

Group2691: abra|OJJ72115.1 acar|OOF96755.1 acla|EAW14916.1 afis|EAW24438.1 afla|EED55241.1 afum|EAL86106.1 agla|OJJ78777.1 akaw|GAA84807.1 aluc|OJZ86870.1 anid|CBF73762.1 anig|EHA21771.1 aory|EIT83515.1 asyd|OJJ56588.1 ater|EAU30009.1 atub|OJI83649.1 aver|OJJ06137.1 awen|OJJ32761.1 azon|OJJ43996.1

Group2692: abra|OJJ72116.1 acar|OOF96756.1 acla|EAW14917.1 afis|EAW24439.1 afla|EED55239.1 afum|EAL86107.1 agla|OJJ78778.1 akaw|GAA84806.1 aluc|OJZ86871.1 anid|CBF73760.1 anig|EHA21770.1 aory|EIT83598.1 asyd|OJJ56587.1 ater|EAU30008.1 atub|OJI83648.1 aver|OJJ06138.1 awen|OJJ32760.1 azon|OJJ43995.1

Group2693: abra|OJJ72117.1 acar|OOF96758.1 acla|EAW14918.1 afis|EAW24440.1 afla|EED55238.1 afum|EAL86108.1 agla|OJJ78779.1 akaw|GAA84805.1 aluc|OJZ86872.1 anid|CBF73758.1 anig|EHA21769.1 aory|EIT83628.1 asyd|OJJ56586.1 ater|EAU30007.1 atub|OJI83647.1 aver|OJJ06139.1 awen|OJJ32759.1 azon|OJJ43994.1

Group2694: abra|OJJ72118.1 acar|OOF96759.1 acla|EAW14919.1 afis|EAW24441.1 afla|EED55237.1 afum|EAL86109.1 agla|OJJ78780.1 akaw|GAA84804.1 aluc|OJZ86873.1 anid|CBF73756.1 anig|EHA21768.1 aory|EIT83564.1 asyd|OJJ56585.1 ater|EAU30006.1 atub|OJI83645.1 aver|OJJ06140.1 awen|OJJ32758.1 azon|OJJ43993.1

Group2695: abra|OJJ72119.1 acar|OOF96760.1 acla|EAW14920.1 afis|EAW24442.1 afla|EED55236.1 afum|EAL86110.1 agla|OJJ78783.1 akaw|GAA84803.1 aluc|OJZ86875.1 anid|CBF73754.1 anig|EHA21767.1 aory|EIT83587.1 asyd|OJJ56584.1 ater|EAU30005.1 atub|OJI83643.1 aver|OJJ06141.1 awen|OJJ32757.1 azon|OJJ43991.1

Group2696: abra|OJJ72123.1 acar|OOF96763.1 acla|EAW14923.1 afis|EAW24445.1 afla|EED55232.1 afum|EAL86114.1 agla|OJJ80290.1 akaw|GAA84800.1 aluc|OJZ86880.1 anid|CBF73749.1 anig|EHA21764.1 aory|EIT83556.1 asyd|OJJ56580.1 ater|EAU30002.1 atub|OJI83639.1 aver|OJJ06144.1 awen|OJJ32754.1 azon|OJJ43989.1

Group2697: abra|OJJ72124.1 acar|OOF96764.1 acla|EAW14930.1 afis|EAW24446.1 afla|EED55231.1 afum|EAL86115.1 agla|OJJ80291.1 akaw|GAA84799.1 aluc|OJZ86881.1 anid|CBF73806.1 anig|EHA21763.1 aory|EIT83582.1 asyd|OJJ56579.1 ater|EAU30001.1 atub|OJI83638.1 aver|OJJ06145.1 awen|OJJ32753.1 azon|OJJ43992.1

Group2698: abra|OJJ72129.1 acar|OOF93633.1 acla|EAW13225.1 afis|EAW17419.1 afla|EED53977.1 agla|OJJ86132.1 akaw|GAA84793.1 aluc|OJZ86887.1 anid|CBF71783.1 aory|EIT78268.1 atub|OJI83632.1 aver|OJI95842.1 awen|OJJ40006.1 azon|OJJ48326.1 asyd|OJJ53805.1 ater|EAU32275.1 afis|EAW19998.1 ater|EAU38678.1

Group2699: abra|OJJ72139.1 acar|OOF96777.1 acla|EAW14933.1 afis|EAW24448.1 afla|EED55225.1 afum|EAL86117.1 agla|OJJ80294.1 akaw|GAA84783.1 aluc|OJZ86897.1 anid|CBF73811.1 anig|EHA21746.1 aory|EIT83551.1 asyd|OJJ56576.1 ater|EAU29998.1 atub|OJI83622.1 aver|OJJ06147.1 awen|OJJ32749.1 azon|OJJ45300.1

Group2700: abra|OJJ72140.1 acar|OOF96778.1 acla|EAW14934.1 afis|EAW24449.1 afla|EED55224.1 afum|EAL86118.1 agla|OJJ80295.1 akaw|GAA84784.1 aluc|OJZ86896.1 anid|CBF73813.1 anig|EHA21745.1 aory|EIT83509.1 asyd|OJJ56575.1 ater|EAU29997.1 atub|OJI83623.1 aver|OJJ06148.1 awen|OJJ32748.1 azon|OJJ45301.1

Group2701: abra|OJJ72141.1 acar|OOF96779.1 acla|EAW14935.1 afis|EAW24450.1 afla|EED55223.1 afum|EAL86119.1 agla|OJJ80296.1 akaw|GAA84782.1 aluc|OJZ86898.1 anid|CBF73815.1 anig|EHA21744.1 aory|EIT83569.1 asyd|OJJ56574.1 ater|EAU29996.1 atub|OJI83621.1 aver|OJJ06149.1 awen|OJJ32747.1 azon|OJJ45299.1

Group2702: abra|OJJ72145.1 acar|OOF96782.1 acla|EAW14939.1 afis|EAW24454.1 afla|EED55220.1 afum|EAL86123.1 agla|OJJ80301.1 akaw|GAA84779.1 aluc|OJZ86902.1 anid|CBF73822.1 anig|EHA21741.1 aory|EIT83594.1 asyd|OJJ56567.1 ater|EAU29991.1 atub|OJI83618.1 aver|OJJ06154.1 awen|OJJ32743.1 azon|OJJ45293.1

Group2703: abra|OJJ72146.1 acar|OOF96783.1 acla|EAW14941.1 afis|EAW24456.1 afla|EED55218.1 afum|EAL86126.1 agla|OJJ80303.1 akaw|GAA84778.1 aluc|OJZ86903.1 anid|CBF73824.1 anig|EHA21740.1 aory|EIT83584.1 asyd|OJJ56566.1 ater|EAU29988.1 atub|OJI83617.1 aver|OJJ06155.1 awen|OJJ32741.1 azon|OJJ45291.1

Group2704: abra|OJJ72147.1 acar|OOF96784.1 acla|EAW14942.1 afis|EAW24457.1 afla|EED55217.1 afum|EAL86127.1 agla|OJJ80304.1 akaw|GAA84777.1 aluc|OJZ86904.1 anid|CBF73826.1 anig|EHA21739.1 aory|EIT83603.1 asyd|OJJ56564.1 ater|EAU29985.1 atub|OJI83616.1 aver|OJJ06157.1 awen|OJJ32740.1 azon|OJJ45290.1

Group2705: abra|OJJ72148.1 acar|OOF96785.1 acla|EAW14943.1 afis|EAW24458.1 afla|EED55216.1 afum|EAL86128.1 agla|OJJ80305.1 akaw|GAA84776.1 aluc|OJZ86905.1 anid|CBF73828.1 anig|EHA21738.1 aory|EIT83601.1 asyd|OJJ56563.1 ater|EAU29984.1 atub|OJI83615.1 aver|OJJ06158.1 awen|OJJ32739.1 azon|OJJ45289.1

Group2706: abra|OJJ72149.1 acar|OOF96786.1 acla|EAW14944.1 afis|EAW24459.1 afla|EED55215.1 afum|EAL86129.1 agla|OJJ80306.1 akaw|GAA84775.1 aluc|OJZ86906.1 anid|CBF73830.1 anig|EHA21737.1 aory|EIT83574.1 asyd|OJJ56562.1 ater|EAU29983.1 atub|OJI83614.1 aver|OJJ06159.1 awen|OJJ32738.1 azon|OJJ45288.1

Group2707: abra|OJJ72150.1 acar|OOF96787.1 acla|EAW14945.1 afis|EAW24460.1 afla|EED55214.1 afum|EAL86130.1 agla|OJJ80308.1 akaw|GAA84774.1 aluc|OJZ86909.1 anid|CBF73832.1 anig|EHA21736.1 aory|EIT83521.1 asyd|OJJ56561.1 ater|EAU29982.1 atub|OJI83613.1 aver|OJJ06160.1 awen|OJJ32736.1 azon|OJJ45287.1

Group2708: abra|OJJ72209.1 acar|OOF96830.1 acla|EAW14967.1 afis|EAW24490.1 afla|EED54675.1 afum|EAL86159.1 agla|OJJ81567.1 akaw|GAA84725.1 aluc|OJZ86961.1 anid|CBF89709.1 anig|EHA21687.1

aory|EIT78852.1 asyd|OJJ62250.1 ater|EAU29966.1 atub|OJI83562.1 aver|OJI96360.1 awen|OJJ32706.1 azon|OJJ45248.1

Group2709: abra|OJJ72241.1 acar|OOF96866.1 acla|EAW07579.1 afis|EAW22615.1 afla|EED47379.1 afum|EAL90583.1 akaw|GAA84698.1 aluc|OJZ86989.1 anid|CBF84548.1 anig|EHA21657.1 aory|EIT82884.1 asyd|OJJ53472.1 ater|EAU39426.1 atub|OJI83530.1 aver|OJJ08284.1 awen|OJJ36695.1 azon|OJJ47502.1 aver|OJJ03406.1

Group2710: abra|OJJ72282.1 acar|OOF94939.1 afis|EAW20798.1 afla|EED49549.1 afum|EAL84831.1 akaw|GAA86034.1 aluc|OJZ90261.1 anig|EHA23757.1 asyd|OJJ55079.1 ater|EAU30148.1 atub|OJI81687.1 aver|OJJ07129.1 awen|OJJ41910.1 azon|OJJ47366.1 acar|OOF97630.1 afla|EED56695.1 aory|EIT73901.1 asyd|OJJ57120.1

Group2711: abra|OJJ72290.1 acar|OOF94933.1 acla|EAW12012.1 afis|EAW19652.1 afla|EED57003.1 afum|EAL86870.1 agla|OJJ80798.1 akaw|GAA86042.1 aluc|OJZ90271.1 anid|CBF71361.1 anig|EHA23763.1 aory|EIT77490.1 asyd|OJJ55084.1 ater|EAU32876.1 atub|OJI81695.1 aver|OJJ07123.1 awen|OJJ41903.1 azon|OJJ42697.1

Group2712: abra|OJJ72296.1 acar|OOF94928.1 acla|EAW12007.1 afis|EAW19647.1 afla|EED56998.1 afum|EAL86875.1 agla|OJJ80789.1 akaw|GAA86047.1 aluc|OJZ90276.1 anid|CBF71354.1 anig|EHA23768.1 aory|EIT77515.1 asyd|OJJ55089.1 ater|EAU32881.1 atub|OJI81700.1 aver|OJJ07118.1 awen|OJJ41898.1 azon|OJJ42702.1

Group2713: abra|OJJ72298.1 acar|OOF94926.1 acla|EAW12006.1 afis|EAW19646.1 afla|EED56997.1 afum|EAL86876.1 agla|OJJ80788.1 akaw|GAA86049.1 aluc|OJZ90278.1 anid|CBF71352.1 anig|EHA23769.1 aory|EIT77535.1 asyd|OJJ55091.1 ater|EAU32883.1 atub|OJI81703.1 aver|OJJ07117.1 awen|OJJ41897.1 azon|OJJ42703.1

Group2714: abra|OJJ72299.1 acar|OOF94924.1 acla|EAW12005.1 afis|EAW19645.1 afla|EED56996.1 afum|EAL86877.1 agla|OJJ80787.1 akaw|GAA86050.1 aluc|OJZ90280.1 anid|CBF71350.1 anig|EHA23770.1 aory|EIT77477.1 asyd|OJJ55092.1 ater|EAU32884.1 atub|OJI81704.1 aver|OJJ07116.1 awen|OJJ41896.1 azon|OJJ42704.1

Group2715: abra|OJJ72318.1 acar|OOF94893.1 acla|EAW12000.1 afis|EAW19640.1 afla|EED56991.1 afum|EAL86882.1 agla|OJJ80782.1 akaw|GAA86067.1 aluc|OJZ90301.1 anid|CBF71343.1 anig|EHA23789.1 aory|EIT77491.1 asyd|OJJ55098.1 ater|EAU32889.1 atub|OJI81727.1 aver|OJJ07108.1 awen|OJJ41885.1 azon|OJJ42710.1

Group2716: abra|OJJ72324.1 acar|OOF89799.1 acla|EAW11999.1 afis|EAW19639.1 afla|EED56990.1 afum|EAL86883.1 agla|OJJ80781.1 akaw|GAA86074.1 aluc|OJZ90309.1 anid|CBF71341.1 anig|EHA23796.1 aory|EIT77478.1 asyd|OJJ55099.1 ater|EAU32890.1 atub|OJI81734.1 aver|OJJ07107.1 awen|OJJ41884.1 azon|OJJ42711.1

Group2717: abra|OJJ72327.1 acar|OOF94887.1 acla|EAW11996.1 afis|EAW19636.1 afla|EED56988.1 afum|EAL86886.1 agla|OJJ80778.1 akaw|GAA86077.1 aluc|OJZ90312.1 anid|CBF71335.1 anig|EHA23799.1 aory|EIT77538.1 asyd|OJJ55103.1 ater|EAU32892.1 atub|OJI81737.1 aver|OJJ07104.1 awen|OJJ41881.1 azon|OJJ47079.1

Group2718: abra|OJJ72328.1 acar|OOF94886.1 acla|EAW11995.1 afis|EAW19635.1 afla|EED56986.1 afum|EAL86887.1 agla|OJJ80777.1 akaw|GAA86078.1 aluc|OJZ90313.1 anid|CBF71333.1 anig|EHA23800.1 aory|EIT77532.1 asyd|OJJ55104.1 atub|OJI81738.1 aver|OJJ07103.1 awen|OJJ41880.1 azon|OJJ47078.1 ater|EAU32893.1

Group2719: abra|OJJ72338.1 acar|OOF94878.1 acla|EAW11985.1 afis|EAW19625.1 afla|EED56977.1 afum|EBA27293.1 agla|OJJ80768.1 akaw|GAA86089.1 aluc|OJZ90318.1 anid|CBF71315.1 anig|EHA23804.1 aory|EIT77446.1 asyd|OJJ55113.1 ater|EAU32903.1 atub|OJI81747.1 aver|OJJ07093.1 awen|OJJ41872.1 azon|OJJ47067.1

Group2720: abra|OJJ72339.1 acar|OOF94877.1 acla|EAW11984.1 afis|EAW19624.1 afla|EED56976.1 afum|EAL86898.1 agla|OJJ80767.1 akaw|GAA86090.1 aluc|OJZ90319.1 anid|CBF71313.1 anig|EHA23805.1 aory|EIT77523.1 asyd|OJJ55114.1 ater|EAU32902.1 atub|OJI81748.1 aver|OJJ07092.1 awen|OJJ41871.1 azon|OJJ47066.1

Group2721: abra|OJJ72343.1 acar|OOF94874.1 acla|EAW11983.1 afis|EAW19621.1 afla|EED56973.1 afum|EAL86900.1 agla|OJJ80766.1 akaw|GAA86092.1 aluc|OJZ90322.1 anid|CBF71293.1 anig|EHA23808.1 aory|EIT77575.1 asyd|OJJ55117.1 ater|EAU32905.1 atub|OJI81749.1 aver|OJJ07089.1 awen|OJJ41868.1 azon|OJJ47061.1

Group2722: abra|OJJ72344.1 acar|OOF94873.1 acla|EAW11982.1 afis|EAW19620.1 afla|EED56972.1 afum|EAL86902.1 agla|OJJ80765.1 akaw|GAA86093.1 aluc|OJZ90323.1 anid|CBF71296.1 anig|EHA23809.1 aory|EIT77533.1 asyd|OJJ55118.1 ater|EAU32906.1 atub|OJI81750.1 aver|OJJ07088.1 awen|OJJ41867.1 azon|OJJ47060.1

Group2723: abra|OJJ72345.1 acar|OOF94872.1 acla|EAW11981.1 afis|EAW19619.1 afla|EED56967.1 afum|EAL86903.1 agla|OJJ80764.1 akaw|GAA86094.1 aluc|OJZ90324.1 anid|CBF71298.1 anig|EHA23810.1 aory|EIT77589.1 asyd|OJJ55119.1 ater|EAU32907.1 atub|OJI81751.1 aver|OJJ07087.1 awen|OJJ41866.1 azon|OJJ47059.1

Group2724: abra|OJJ72346.1 acar|OOF94871.1 acla|EAW11980.1 afis|EAW19618.1 afla|EED56968.1 afum|EAL86904.1 agla|OJJ80763.1 akaw|GAA86095.1 aluc|OJZ90325.1 anid|CBF71300.1 anig|EHA23811.1 aory|EIT77479.1 asyd|OJJ55120.1 ater|EAU32908.1 atub|OJI81752.1 aver|OJJ07086.1 awen|OJJ41865.1 azon|OJJ47058.1

Group2725: abra|OJJ72349.1 acar|OOF94869.1 acla|EAW11977.1 afis|EAW19609.1 afla|EED56969.1 afum|EAL86913.1 agla|OJJ80761.1 akaw|GAA86096.1 aluc|OJZ90328.1 anid|CBF71302.1 anig|EHA23812.1 aory|EIT77540.1 asyd|OJJ55121.1 ater|EAU32910.1 atub|OJI81755.1 aver|OJJ07083.1 awen|OJJ41862.1 azon|OJJ47055.1

Group2726: abra|OJJ72351.1 acar|OOF94867.1 acla|EAW11975.1 afis|EAW19611.1 afla|EED56966.1 afum|EAL86911.1 agla|OJJ80759.1 akaw|GAA86098.1 aluc|OJZ90330.1 anid|CBF71306.1 anig|EHA23814.1 aory|EIT77494.1 asyd|OJJ55123.1 ater|EAU32921.1 atub|OJI81757.1 aver|OJJ07080.1 awen|OJJ41859.1 azon|OJJ47053.1

Group2727: abra|OJJ72352.1 acar|OOF94866.1 acla|EAW11973.1 afis|EAW19613.1 afla|EED56961.1 afum|EAL86909.1 agla|OJJ80758.1 akaw|GAA86099.1 aluc|OJZ90331.1 anid|CBF71308.1 anig|EHA23815.1 aory|EIT77512.1 asyd|OJJ55126.1 ater|EAU32914.1 atub|OJI81758.1 aver|OJJ07078.1 awen|OJJ41857.1 azon|OJJ47051.1

Group2728: abra|OJJ72353.1 acar|OOF94865.1 acla|EAW11972.1 afis|EAW19614.1 afla|EED56962.1 afum|EAL86908.1 agla|OJJ80757.1 akaw|GAA86100.1 aluc|OJZ90332.1 anid|CBF71310.1 anig|EHA23816.1 aory|EIT77559.1 asyd|OJJ55127.1 ater|EAU32915.1 atub|OJI81759.1 aver|OJJ07077.1 awen|OJJ41856.1 azon|OJJ47050.1

Group2729: abra|OJJ72355.1 acar|OOF94863.1 acla|EAW11961.1 afis|EAW19598.1 afla|EED56954.1 afum|EAL86920.1 agla|OJJ80740.1 akaw|GAA86102.1 aluc|OJZ90334.1 anid|CBF71287.1 anig|EHA23818.1 aory|EIT77544.1 asyd|OJJ55129.1 ater|EAU32927.1 atub|OJI81761.1 aver|OJJ07074.1 awen|OJJ41844.1 azon|OJJ44888.1

Group2730: abra|OJJ72368.1 acar|OOF94849.1 acla|EAW11964.1 afis|EAW19601.1 afla|EED56957.1 afum|EAL86917.1 agla|OJJ80743.1 akaw|GAA86118.1 aluc|OJZ90356.1 anid|CBF71281.1 anig|EHA23834.1 aory|EIT77555.1 asyd|OJJ55133.1 ater|EAU32924.1 atub|OJI81783.1 aver|OJJ07071.1 awen|OJJ41839.1 azon|OJJ47044.1

Group2731: abra|OJJ72370.1 acar|OOF94852.1 afis|EAW25546.1 afla|EED51182.1 agla|OJJ80811.1 akaw|GAA86120.1 aluc|OJZ90359.1 anid|CBF80641.1 anig|EHA23832.1 aory|EIT81311.1 asyd|OJJ65320.1 ater|EAU36020.1 atub|OJI81786.1 aver|OJI99465.1 anid|CBF82274.1 asyd|OJJ59966.1 anid|CBF86654.1 aver|OJJ02324.1

Group2732: abra|OJJ72373.1 acar|OOF94855.1 acla|EAW11967.1 afis|EAW19617.1 afla|EED56965.1 afum|EAL86905.1 agla|OJJ80746.1 akaw|GAA86123.1 aluc|OJZ90362.1 anid|CBF71277.1 anig|EHA23829.1 aory|EIT77547.1 asyd|OJJ55139.1 ater|EAU32917.1 atub|OJI81789.1 aver|OJJ07066.1 awen|OJJ41850.1 azon|OJJ47046.1

Group2733: abra|OJJ72377.1 acar|OOF94841.1 acla|EAW11958.1 afis|EAW19595.1 afla|EED56951.1 afum|EAL86923.1 agla|OJJ80736.1 akaw|GAA86125.1 aluc|OJZ90364.1 anid|CBF71262.1 anig|EHA23839.1 aory|EIT77501.1 asyd|OJJ55150.1 ater|EAU32929.1 atub|OJI81791.1 aver|OJJ07057.1 awen|OJJ41849.1 azon|OJJ47020.1

Group2734: abra|OJJ72379.1 acar|OOF94840.1 acla|EAW11957.1 afis|EAW19594.1 afla|EED56950.1 afum|EAL86924.1 agla|OJJ80735.1 akaw|GAA86127.1 aluc|OJZ90366.1 anid|CBF71275.1 anig|EHA23841.1 aory|EIT77449.1 asyd|OJJ55149.1 ater|EAU32930.1 atub|OJI81793.1 aver|OJJ07058.1 awen|OJJ41811.1 azon|OJJ47021.1

Group2735: abra|OJJ72381.1 acar|OOF94838.1 acla|EAW11955.1 afis|EAW19592.1 afla|EED56948.1 afum|EAL86926.1 agla|OJJ80733.1 akaw|GAA86128.1 aluc|OJZ90368.1 anid|CBF71268.1 anig|EHA23842.1 aory|EIT77465.1 asyd|OJJ55147.1 ater|EAU32933.1 atub|OJI81794.1 aver|OJJ07060.1 awen|OJJ41809.1 azon|OJJ47023.1

Group2736: abra|OJJ72388.1 acar|OOF94835.1 acla|EAW11949.1 afis|EAW19586.1 afla|EED56914.1 afum|EAL86932.1 agla|OJJ80723.1 akaw|GAA86136.1 aluc|OJZ90374.1 anid|CBF71254.1 anig|EHA23848.1 aory|EIT77498.1 asyd|OJJ55153.1 ater|EAU32962.1 atub|OJI81800.1 aver|OJJ07054.1 awen|OJJ41800.1 azon|OJJ47031.1

Group2737: abra|OJJ72389.1 acar|OOF94834.1 acla|EAW11948.1 afis|EAW19585.1 afla|EED56913.1 afum|EAL86933.1 agla|OJJ80722.1 akaw|GAA86137.1 aluc|OJZ90376.1 anid|CBF71253.1 anig|EHA23849.1

aory|EIT77525.1 asyd|OJJ55155.1 ater|EAU32963.1 atub|OJI81801.1 aver|OJJ07053.1 awen|OJJ41799.1 azon|OJJ47032.1

Group2738: abra|OJJ72391.1 acar|OOF94833.1 acla|EAW11947.1 afis|EAW19584.1 afla|EED56912.1 afum|EAL86934.1 agla|OJJ80712.1 akaw|GAA86138.1 aluc|OJZ90378.1 anid|CBF71251.1 anig|EHA23850.1 aory|EIT77463.1 asyd|OJJ55156.1 ater|EAU32967.1 atub|OJI81803.1 aver|OJJ07052.1 awen|OJJ41798.1 azon|OJJ47033.1

Group2739: abra|OJJ72394.1 acar|OOF94832.1 acla|EAW11946.1 afis|EAW19583.1 afla|EED56911.1 afum|EAL86935.2 agla|OJJ80711.1 akaw|GAA86139.1 aluc|OJZ90379.1 anid|CBF71249.1 anig|EHA23851.1 aory|EIT77567.1 asyd|OJJ55157.1 ater|EAU32968.1 atub|OJI81804.1 aver|OJJ07051.1 awen|OJJ41797.1 azon|OJJ47034.1

Group2740: abra|OJJ72395.1 acar|OOF94831.1 acla|EAW11945.1 afis|EAW19581.1 afla|EED56910.1 afum|EAL86937.1 agla|OJJ80710.1 akaw|GAA86140.1 aluc|OJZ90380.1 anid|CBF71245.1 anig|EHA23852.1 aory|EIT77560.1 asyd|OJJ55159.1 ater|EAU32972.1 atub|OJI81806.1 aver|OJJ07049.1 awen|OJJ41795.1 azon|OJJ47036.1

Group2741: abra|OJJ72396.1 acar|OOF94830.1 acla|EAW11944.1 afis|EAW19580.1 afla|EED56909.1 afum|EAL86938.1 agla|OJJ80709.1 akaw|GAA86141.1 aluc|OJZ90381.1 anid|CBF71243.1 anig|EHA23853.1 aory|EIT77483.1 asyd|OJJ55160.1 ater|EAU32973.1 atub|OJI81807.1 aver|OJJ07047.1 awen|OJJ41794.1 azon|OJJ47037.1

Group2742: abra|OJJ72397.1 acar|OOF94829.1 acla|EAW11943.1 afis|EAW19579.1 afla|EED56804.1 afum|EAL86939.1 agla|OJJ80708.1 akaw|GAA86142.1 aluc|OJZ90382.1 anid|CBF71239.1 anig|EHA23854.1 aory|EIT73874.1 asyd|OJJ55163.1 ater|EAU32976.1 atub|OJI81808.1 aver|OJJ07044.1 awen|OJJ41792.1 azon|OJJ47041.1

Group2743: abra|OJJ72399.1 acar|OOF94828.1 acla|EAW11941.1 afis|EAW19577.1 afla|EED56805.1 afum|EAL86941.2 agla|OJJ80706.1 akaw|GAA86144.1 aluc|OJZ90384.1 anid|CBF71236.1 anig|EHA23856.1 aory|EIT73858.1 asyd|OJJ55165.1 ater|EAU33003.1 atub|OJI81811.1 aver|OJJ07042.1 awen|OJJ41790.1 azon|OJJ47039.1

Group2744: abra|OJJ72401.1 acar|OOF94827.1 acla|EAW11940.1 afis|EAW19575.1 afla|EED56808.1 afum|EAL86942.1 agla|OJJ80704.1 akaw|GAA86145.1 aluc|OJZ90386.1 anid|CBF71234.1 anig|EHA23857.1 aory|EIT73879.1 asyd|OJJ55166.1 ater|EAU33002.1 atub|OJI81814.1 aver|OJJ07040.1 awen|OJJ41788.1 azon|OJJ44890.1

Group2745: abra|OJJ72402.1 acar|OOF94826.1 acla|EAW11939.1 afis|EAW19574.1 afla|EED56809.1 afum|EAL86943.1 agla|OJJ80702.1 akaw|GAA86146.1 aluc|OJZ90387.1 anid|CBF71232.1 anig|EHA23858.1 aory|EIT73889.1 asyd|OJJ55167.1 ater|EAU33001.1 atub|OJI81815.1 aver|OJJ07039.1 awen|OJJ41787.1 azon|OJJ44891.1

Group2746: abra|OJJ72404.1 acar|OOF94824.1 acla|EAW11937.1 afis|EAW19572.1 afla|EED56811.1 afum|EAL86945.1 agla|OJJ80700.1 akaw|GAA86148.1 aluc|OJZ90389.1 anid|CBF71228.1 anig|EHA23859.1 aory|EIT73949.1 asyd|OJJ55170.1 ater|EAU32995.1 atub|OJI81818.1 aver|OJJ07037.1 awen|OJJ41785.1 azon|OJJ44893.1

Group2747: abra|OJJ72405.1 acar|OOF94823.1 acla|EAW11936.1 afis|EAW19571.1 afla|EED56812.1 afum|EAL86946.1 agla|OJJ80699.1 akaw|GAA86149.1 aluc|OJZ90390.1 anid|CBF71226.1 anig|EHA23860.1 aory|EIT73876.1 asyd|OJJ55171.1 ater|EAU32994.1 atub|OJI81819.1 aver|OJJ07036.1 awen|OJJ41784.1 azon|OJJ44894.1

Group2748: abra|OJJ72408.1 acar|OOF94821.1 acla|EAW11933.1 afis|EAW19569.1 afla|EED56814.1 afum|EAL86947.1 agla|OJJ80697.1 akaw|GAA86151.1 aluc|OJZ90392.1 anid|CBF78033.1 anig|EHA23862.1 aory|EIT73916.1 asyd|OJJ55227.1 ater|EAU32992.1 atub|OJI81821.1 aver|OJJ06994.1 awen|OJJ41782.1 azon|OJJ44896.1

Group2749: abra|OJJ72412.1 acar|OOF94817.1 acla|EAW11929.1 afis|EAW19563.1 afla|EED56817.1 afum|EAL86949.1 agla|OJJ80692.1 akaw|GAA86155.1 aluc|OJZ90396.1 anid|CBF78024.1 anig|EHA23866.1 aory|EIT73850.1 asyd|OJJ55233.1 ater|EAU32986.1 atub|OJI81827.1 aver|OJJ06990.1 awen|OJJ41778.1 azon|OJJ44899.1

Group2750: abra|OJJ72413.1 acar|OOF94816.1 acla|EAW12857.1 afis|EAW17821.1 afla|EED56818.1 afum|EAL91631.1 agla|OJJ80690.1 akaw|GAA86156.1 aluc|OJZ90397.1 anid|CBF78026.1 anig|EHA23867.1 aory|EIT73888.1 asyd|OJJ55232.1 ater|EAU32985.1 atub|OJI81828.1 aver|OJJ06991.1 awen|OJJ41777.1 azon|OJJ44900.1

Group2751: abra|OJJ72414.1 acar|OOF94815.1 acla|EAW12858.1 afis|EAW17820.1 afla|EED56819.1 afum|EAL91630.1 agla|OJJ80689.1 akaw|GAA86157.1 aluc|OJZ90398.1 anid|CBF78027.1 anig|EHA23868.1 aory|EIT73947.1 asyd|OJJ55231.1 ater|EAU32984.1 atub|OJI81829.1 aver|OJJ06992.1 awen|OJJ41776.1 azon|OJJ44901.1

Group2752: abra|OJJ72415.1 acar|OOF94814.1 acla|EAW12859.1 afis|EAW17808.1 afla|EED56820.1 afum|EAL91616.1 agla|OJJ80688.1 akaw|GAA86158.1 aluc|OJZ90399.1 anid|CBF78029.1 anig|EHA23869.1 aory|EIT73877.1 asyd|OJJ55230.1 ater|EAU32982.1 atub|OJI81830.1 aver|OJJ06993.1 awen|OJJ41775.1 azon|OJJ44902.1

Group2753: abra|OJJ72417.1 acar|OOF90199.1 acla|EAW13737.1 afis|EAW17067.1 afla|EED52009.1 afum|EAL87736.1 agla|OJJ83275.1 akaw|GAA86160.1 aluc|OJZ90401.1 aory|EIT81373.1 awen|OJJ36556.1 azon|OJJ46436.1 abra|OJJ73401.1 acar|OOF98722.1 anid|CBF84645.1 anig|EHA18326.1 ater|EAU36994.1 aver|OJJ04312.1

Group2754: abra|OJJ72421.1 acar|OOF90370.1 acla|EAW12862.1 afis|EAW17811.1 afla|EED46162.1 afum|EAL91619.1 agla|OJJ80655.1 akaw|GAA86163.1 aluc|OJZ90404.1 anid|CBF78013.1 anig|EHA23872.1 aory|EIT76383.1 asyd|OJJ55240.1 ater|EAU33017.1 atub|OJI81835.1 aver|OJJ06983.1 awen|OJJ41761.1 azon|OJJ44878.1

Group2755: abra|OJJ72422.1 acar|OOF90371.1 acla|EAW12861.1 afis|EAW17810.1 afla|EED46161.1 afum|EAL91618.1 agla|OJJ80654.1 akaw|GAA86164.1 aluc|OJZ90405.1 anid|CBF78014.1 anig|EHA23873.1 aory|EIT76437.1 asyd|OJJ55239.1 ater|EAU33016.1 atub|OJI81836.1 aver|OJJ06984.1 awen|OJJ41762.1 azon|OJJ44879.1

Group2756: abra|OJJ72424.1 acar|OOF90373.1 afis|EAW17822.1 afla|EED46167.1 agla|OJJ80652.1 akaw|GAA86166.1 aluc|OJZ90407.1 anid|CBF78011.1 anig|EHA23875.1 aory|EIT76519.1 asyd|OJJ63016.1 ater|EAU33023.1 atub|OJI81838.1 aver|OJI97948.1 awen|OJJ41760.1 azon|OJJ48452.1 acar|OOF98065.1 anid|CBF85197.1

Group2757: abra|OJJ72425.1 acar|OOF94810.1 acla|EAW12659.1 afis|EAW17823.1 afla|EED46166.1 afum|EAL91633.1 agla|OJJ80641.1 akaw|GAA86167.1 aluc|OJZ90408.1 anid|CBF78009.1 anig|EHA23876.1 aory|EIT76477.1 asyd|OJJ55242.1 ater|EAU33024.1 atub|OJI81840.1 aver|OJJ06982.1 awen|OJJ41751.1 azon|OJJ48443.1

Group2758: abra|OJJ72428.1 acar|OOF94804.1 acla|EAW12661.1 afis|EAW17825.1 afla|EED46163.1 afum|EAL91635.1 agla|OJJ80646.1 akaw|GAA86169.1 aluc|OJZ90412.1 anid|CBF78005.1 anig|EHA23878.1 aory|EIT76524.1 asyd|OJJ55245.1 ater|EAU33026.1 atub|OJI81843.1 aver|OJJ06979.1 awen|OJJ41755.1 azon|OJJ48445.1

Group2759: abra|OJJ72429.1 acar|OOF94805.1 acla|EAW12662.1 afis|EAW17826.1 afla|EED46168.1 afum|EAL91636.1 agla|OJJ80647.1 akaw|GAA86170.1 aluc|OJZ90413.1 anid|CBF78003.1 anig|EHA23879.1 aory|EIT76408.1 asyd|OJJ55246.1 ater|EAU33027.1 atub|OJI81845.1 aver|OJJ06978.1 awen|OJJ41756.1 azon|OJJ48446.1

Group2760: abra|OJJ72430.1 acar|OOF94803.1 acla|EAW12663.1 afis|EAW17827.1 afla|EED46169.1 afum|EAL91637.1 agla|OJJ80648.1 akaw|GAA86171.1 aluc|OJZ90414.1 anid|CBF78001.1 anig|EHA23880.1 aory|EIT76516.1 asyd|OJJ55247.1 ater|EAU33031.1 atub|OJI81846.1 aver|OJJ06977.1 awen|OJJ41757.1 azon|OJJ48447.1

Group2761: abra|OJJ72439.1 acar|OOF94798.1 acla|EAW12672.1 afis|EAW17835.1 afla|EED46180.1 afum|EAL91645.1 agla|OJJ83131.1 akaw|GAA86178.1 aluc|OJZ90423.1 anid|CBF77987.1 anig|EHA23892.1 aory|EIT76377.1 asyd|OJJ55256.1 ater|EAU33039.1 atub|OJI81855.1 aver|OJJ06968.1 awen|OJJ41741.1 azon|OJJ48437.1

Group2762: abra|OJJ72440.1 acar|OOF94797.1 acla|EAW12673.1 afis|EAW17836.1 afla|EED46181.1 afum|EAL91646.1 agla|OJJ83130.1 akaw|GAA86179.1 aluc|OJZ90424.1 anid|CBF77985.1 anig|EHA23893.1 aory|EIT76515.1 asyd|OJJ55257.1 ater|EAU33040.1 atub|OJI81856.1 aver|OJJ06967.1 awen|OJJ41740.1 azon|OJJ48436.1

Group2763: abra|OJJ72443.1 acar|OOF94793.1 acla|EAW12676.1 afis|EAW17839.1 afla|EED46184.1 afum|EAL91649.1 agla|OJJ83128.1 akaw|GAA86182.1 aluc|OJZ90427.1 anid|CBF77979.1 anig|EHA23897.1 aory|EIT76493.1 asyd|OJJ55260.1 ater|EAU33042.1 atub|OJI81859.1 aver|OJJ06965.1 awen|OJJ41737.1 azon|OJJ48434.1

Group2764: abra|OJJ72445.1 acar|OOF94791.1 acla|EAW12678.1 afis|EAW17841.1 afla|EED46186.1 afum|EAL91650.1 agla|OJJ83126.1 akaw|GAA86183.1 aluc|OJZ90429.1 anid|CBF77975.1 anig|EHA23898.1 aory|EIT76513.1 asyd|OJJ55262.1 ater|EAU33043.1 atub|OJI81861.1 aver|OJJ06963.1 awen|OJJ41735.1 azon|OJJ48432.1

Group2765: abra|OJJ72446.1 acar|OOF94790.1 acla|EAW12679.1 afis|EAW17842.1 afla|EED46187.1 afum|EAL91651.1 agla|OJJ83125.1 akaw|GAA86184.1 aluc|OJZ90430.1 anid|CBF77973.1 anig|EHA23899.1 aory|EIT76407.1 asyd|OJJ55263.1 ater|EAU33044.1 atub|OJI81862.1 aver|OJJ06962.1 awen|OJJ41734.1 azon|OJJ48431.1

Group2766: abra|OJJ72450.1 acar|OOF94786.1 acla|EAW12684.1 afis|EAW17847.1 afla|EED46201.1 afum|EAL91656.1 agla|OJJ83122.1 akaw|GAA86187.1 aluc|OJZ90435.1 anid|CBF77964.1 anig|EHA23902.1

aory|EIT76452.1 asyd|OJJ55269.1 ater|EAU33049.1 atub|OJI81866.1 aver|OJJ06958.1 awen|OJJ41726.1 azon|OJJ48425.1

Group2767: abra|OJJ72458.1 acar|OOF94779.1 acla|EAW12690.1 afis|EAW17854.1 afla|EED46208.1 afum|EAL91662.1 agla|OJJ83116.1 akaw|GAA86194.1 aluc|OJZ90445.1 anid|CBF77955.1 anig|EHA23908.1 aory|EIT76378.1 asyd|OJJ55275.1 ater|EAU33056.1 atub|OJI81873.1 aver|OJJ06952.1 awen|OJJ41719.1 azon|OJJ48416.1

Group2768: abra|OJJ72459.1 acar|OOF94780.1 acla|EAW12691.1 afis|EAW17855.1 afla|EED46209.1 afum|EAL91663.1 agla|OJJ83115.1 akaw|GAA86195.1 aluc|OJZ90446.1 anid|CBF77953.1 anig|EHA23909.1 aory|EIT76395.1 asyd|OJJ55276.1 ater|EAU33057.1 atub|OJI81874.1 aver|OJJ06951.1 awen|OJJ41718.1 azon|OJJ48415.1

Group2769: abra|OJJ72460.1 acar|OOF94778.1 acla|EAW12692.1 afis|EAW17856.1 afla|EED46210.1 afum|EAL91664.1 agla|OJJ83113.1 akaw|GAA86196.1 aluc|OJZ90447.1 anid|CBF77951.1 anig|EHA23910.1 aory|EIT76442.1 asyd|OJJ55277.1 ater|EAU33058.1 atub|OJI81875.1 aver|OJJ06950.1 awen|OJJ41716.1 azon|OJJ48411.1

Group2770: abra|OJJ72461.1 acar|OOF94777.1 acla|EAW12693.1 afis|EAW17857.1 afla|EED46212.1 afum|EAL91665.1 agla|OJJ83112.1 akaw|GAA86197.1 aluc|OJZ90448.1 anid|CBF77949.1 anig|EHA23911.1 aory|EIT76364.1 asyd|OJJ55278.1 ater|EAU33059.1 atub|OJI81876.1 aver|OJJ06949.1 awen|OJJ41715.1 azon|OJJ48410.1

Group2771: abra|OJJ72462.1 acar|OOF94775.1 acla|EAW12694.1 afis|EAW17858.1 afla|EED46213.1 afum|EAL91666.1 agla|OJJ83111.1 akaw|GAA86198.1 aluc|OJZ90449.1 anid|CBF77947.1 anig|EHA23912.1 aory|EIT76487.1 asyd|OJJ55279.1 ater|EAU33064.1 atub|OJI81877.1 aver|OJJ06948.1 awen|OJJ41714.1 azon|OJJ48409.1

Group2772: abra|OJJ72467.1 acar|OOF94773.1 acla|EAW12699.1 afis|EAW17862.1 afla|EED46225.1 afum|EAL91670.2 agla|OJJ83101.1 akaw|GAA86203.1 aluc|OJZ90454.1 anid|CBF88078.1 anig|EHA23916.1 aory|EIT76473.1 asyd|OJJ57087.1 ater|EAU33070.1 atub|OJI81881.1 aver|OJJ03214.1 awen|OJJ41709.1 azon|OJJ48404.1

Group2773: abra|OJJ72468.1 acar|OOF94772.1 acla|EAW12700.1 afis|EAW17863.1 afla|EED46226.1 afum|EAL91671.1 agla|OJJ83100.1 akaw|GAA86204.1 aluc|OJZ90455.1 anid|CBF84085.1 anig|EHA23917.1 aory|EIT76453.1 asyd|OJJ57088.1 ater|EAU33078.1 atub|OJI81882.1 aver|OJJ03213.1 awen|OJJ41707.1 azon|OJJ48403.1

Group2774: abra|OJJ72471.1 acar|OOF94764.1 acla|EAW12704.1 afis|EAW17867.1 afla|EED46236.1 afum|EAL91675.1 agla|OJJ83087.1 akaw|GAA86209.1 aluc|OJZ90458.1 anid|CBF89893.1 anig|EHA23920.1 aory|EIT76466.1 asyd|OJJ62139.1 ater|EAU33082.1 atub|OJI81885.1 aver|OJI96249.1 awen|OJJ41704.1 azon|OJJ48400.1

Group2775: abra|OJJ72476.1 acar|OOF94760.1 acla|EAW12707.1 afis|EAW17870.1 afla|EED46238.1 afum|EAL91678.1 agla|OJJ83090.1 akaw|GAA86212.1 aluc|OJZ90462.1 anid|CBF89899.1 anig|EHA23923.1 aory|EIT76429.1 asyd|OJJ62136.1 ater|EAU33088.1 atub|OJI81888.1 aver|OJI96246.1 awen|OJJ41700.1 azon|OJJ48397.1

Group2776: abra|OJJ72478.1 acar|OOF94756.1 acla|EAW12709.1 afis|EAW17872.1 afla|EED46239.1 afum|EAL91680.1 agla|OJJ83092.1 akaw|GAA86214.1 aluc|OJZ90465.1 anid|CBF89903.1 anig|EHA23925.1 aory|EIT76379.1 asyd|OJJ62134.1 ater|EAU33090.1 atub|OJI81890.1 aver|OJI96244.1 awen|OJJ41698.1 azon|OJJ48395.1

Group2777: abra|OJJ72482.1 acar|OOF94754.1 acla|EAW12712.1 afis|EAW17875.1 afla|EED46242.1 afum|EAL91683.1 agla|OJJ83071.1 akaw|GAA86217.1 aluc|OJZ90469.1 anid|CBF89904.1 anig|EHA23928.1 aory|EIT76495.1 asyd|OJJ59281.1 ater|EAU33095.1 atub|OJI81893.1 aver|OJJ01894.1 awen|OJJ41689.1 aver|OJI96243.1

Group2778: abra|OJJ72488.1 acar|OOF94747.1 acla|EAW12719.1 afis|EAW17882.1 afla|EED46269.1 afum|EAL91690.1 agla|OJJ83065.1 akaw|GAA86223.1 aluc|OJZ90475.1 anig|EHA23934.1 aory|EIT76427.1 asyd|OJJ57008.1 ater|EAU33102.1 atub|OJI81900.1 aver|OJJ03279.1 awen|OJJ41684.1 azon|OJJ48392.1 anid|CBF77035.1

Group2779: abra|OJJ72489.1 acar|OOF92642.1 acla|EAW12720.1 afis|EAW17883.1 afla|EED46270.1 afum|EAL91691.1 agla|OJJ83064.1 akaw|GAA86224.1 aluc|OJZ90476.1 anid|CBF77033.1 anig|EHA23935.1 aory|EIT76482.1 asyd|OJJ57009.1 ater|EAU33103.1 atub|OJI81901.1 aver|OJJ03278.1 awen|OJJ41682.1 azon|OJJ48391.1

Group2780: abra|OJJ72493.1 acar|OOF94745.1 acla|EAW12722.1 afis|EAW17885.1 afla|EED46272.1 afum|EAL91693.1 agla|OJJ83062.1 akaw|GAA86227.1 aluc|OJZ90479.1 anid|CBF77029.1 anig|EHA23938.1 aory|EIT76321.1 asyd|OJJ57011.1 ater|EAU33105.1 atub|OJI81904.1 aver|OJJ03276.1 awen|OJJ41674.1 azon|OJJ48389.1

Group2781: abra|OJJ72494.1 acar|OOF94744.1 acla|EAW12723.1 afis|EAW17886.1 afla|EED46273.1 afum|EAL91694.1 agla|OJJ83061.1 akaw|GAA86228.1 aluc|OJZ90480.1 anid|CBF77027.1 anig|EHA23939.1 aory|EIT76334.1 asyd|OJJ57012.1 ater|EAU33106.1 atub|OJI81905.1 aver|OJJ03275.1 awen|OJJ41673.1 azon|OJJ48388.1

Group2782: abra|OJJ72497.1 acar|OOF94742.1 acla|EAW12725.1 afis|EAW17888.1 afla|EED46275.1 afum|EAL91697.1 agla|OJJ83059.1 akaw|GAA86229.1 aluc|OJZ90481.1 anid|CBF77026.1 anig|EHA23940.1 aory|EIT76333.1 asyd|OJJ57013.1 ater|EAU33107.1 atub|OJI81906.1 aver|OJJ03274.1 awen|OJJ41671.1 azon|OJJ48386.1

Group2783: abra|OJJ72503.1 acar|OOF94738.1 acla|EAW12727.1 afis|EAW17890.1 afla|EED46282.1 afum|EAL91699.1 agla|OJJ83055.1 akaw|GAA86231.1 aluc|OJZ90485.1 anid|CBF77016.1 anig|EHA23943.1 aory|EIT76352.1 asyd|OJJ57017.1 ater|EAU33110.1 atub|OJI81910.1 aver|OJJ03272.1 awen|OJJ41667.1 azon|OJJ48384.1

Group2784: abra|OJJ72507.1 acar|OOF94735.1 acla|EAW12730.1 afis|EAW17893.1 afla|EED46286.1 afum|EAL91702.1 agla|OJJ83049.1 akaw|GAA86235.1 aluc|OJZ90490.1 anid|CBF77006.1 anig|EHA23948.1 aory|EIT76341.1 asyd|OJJ57021.1 ater|EAU33114.1 atub|OJI81914.1 aver|OJJ03268.1 awen|OJJ41659.1 azon|OJJ48379.1

Group2785: abra|OJJ72509.1 acar|OOF94733.1 acla|EAW12732.1 afis|EAW17895.1 afla|EED46267.1 afum|EAL91703.1 agla|OJJ83047.1 akaw|GAA86237.1 aluc|OJZ90492.1 anid|CBF77004.1 anig|EHA23950.1 aory|EIT76414.1 asyd|OJJ57023.1 ater|EAU33115.1 atub|OJI81916.1 aver|OJJ03266.1 awen|OJJ41657.1 azon|OJJ48377.1

Group2786: abra|OJJ72511.1 acar|OOF94732.1 acla|EAW12733.1 afis|EAW17896.1 afla|EED46266.1 afum|EAL91704.1 agla|OJJ83046.1 akaw|GAA86238.1 aluc|OJZ90493.1 anid|CBF77002.1 anig|EHA23951.1 aory|EIT76411.1 asyd|OJJ57025.1 ater|EAU33116.1 atub|OJI81917.1 aver|OJJ03265.1 awen|OJJ41655.1 azon|OJJ48375.1

Group2787: abra|OJJ72513.1 acar|OOF94730.1 acla|EAW12734.1 afis|EAW17897.1 afla|EED46265.1 afum|EAL91705.1 agla|OJJ83045.1 akaw|GAA86240.1 aluc|OJZ90495.1 anid|CBF76998.1 anig|EHA23954.1 aory|EIT76381.1 asyd|OJJ57027.1 ater|EAU33117.1 atub|OJI81920.1 aver|OJJ03263.1 awen|OJJ41654.1 azon|OJJ48374.1

Group2788: abra|OJJ72514.1 acar|OOF94729.1 acla|EAW12735.1 afis|EAW17898.1 afla|EED46264.1 afum|EAL91706.1 agla|OJJ83044.1 akaw|GAA86241.1 aluc|OJZ90496.1 anid|CBF76996.1 anig|EHA23955.1 aory|EIT76461.1 asyd|OJJ57028.1 ater|EAU33118.1 atub|OJI81921.1 aver|OJJ03262.1 awen|OJJ41653.1 azon|OJJ48373.1

Group2789: abra|OJJ72515.1 acar|OOF94728.1 afis|EAW17899.1 afla|EED46263.1 agla|OJJ83043.1 akaw|GAA86242.1 aluc|OJZ90497.1 anid|CBF76994.1 anig|EHA23956.1 aory|EIT76422.1 asyd|OJJ57029.1 atub|OJI81922.1 aver|OJJ03261.1 awen|OJJ41652.1 azon|OJJ48372.1 acla|EAW12736.1 afum|EAL91707.1 ater|EAU33119.1

Group2790: abra|OJJ72516.1 acar|OOF94727.1 acla|EAW12737.1 afis|EAW17900.1 afla|EED46262.1 afum|EAL91708.1 agla|OJJ83042.1 akaw|GAA86243.1 aluc|OJZ90498.1 anid|CBF76992.1 anig|EHA23957.1 aory|EIT76428.1 asyd|OJJ57030.1 ater|EAU33120.1 atub|OJI81923.1 aver|OJJ03260.1 awen|OJJ41651.1 azon|OJJ48371.1

Group2791: abra|OJJ72518.1 acar|OOF94726.1 acla|EAW12738.1 afis|EAW17901.1 afla|EED46261.1 afum|EAL91709.1 agla|OJJ83041.1 akaw|GAA86244.1 aluc|OJZ90499.1 anid|CBF76990.1 anig|EHA23958.1 aory|EIT76470.1 asyd|OJJ57032.1 ater|EAU33121.1 atub|OJI81924.1 aver|OJJ03259.1 awen|OJJ41648.1 azon|OJJ48370.1

Group2792: abra|OJJ72538.1 acar|OOF89947.1 acla|EAW12744.1 afis|EAW17907.1 afla|EED46301.1 afum|EAL91716.2 agla|OJJ83035.1 akaw|GAA86262.1 aluc|OJZ90520.1 anid|CBF76981.1 anig|EHA23974.1 aory|EIT76350.1 asyd|OJJ57037.1 ater|EAU33125.1 atub|OJI81944.1 aver|OJJ03255.1 awen|OJJ41643.1 azon|OJJ48364.1

Group2793: abra|OJJ72539.1 acar|OOF89942.1 acla|EAW12745.1 afis|EAW17908.1 afla|EED46302.1 afum|EAL91717.2 agla|OJJ83034.1 akaw|GAA86263.1 aluc|OJZ90522.1 anid|CBF76979.1 anig|EHA23975.1 aory|EIT76327.1 asyd|OJJ57038.1 ater|EAU33126.1 atub|OJI81945.1 aver|OJJ03253.1 awen|OJJ41641.1 azon|OJJ48363.1

Group2794: abra|OJJ72543.1 acar|OOF94712.1 acla|EAW12747.1 afis|EAW17911.1 afla|EED46313.1 afum|EAL91720.1 agla|OJJ83032.1 akaw|GAA86267.1 aluc|OJZ90527.1 anid|CBF76967.1 anig|EHA23979.1 aory|EIT76323.1 asyd|OJJ57041.1 ater|EAU33131.1 atub|OJI81949.1 aver|OJJ03252.1 awen|OJJ41638.1 azon|OJJ48360.1

Group2795: abra|OJJ72544.1 acar|OOF94711.1 acla|EAW12748.1 afis|EAW17912.1 afla|EED46314.1 afum|EAL91721.1 agla|OJJ83031.1 akaw|GAA86268.1 aluc|OJZ90528.1 anid|CBF76969.1 anig|EHA23980.1

aory|EIT76335.1 asyd|OJJ57042.1 ater|EAU33132.1 atub|OJI81950.1 aver|OJJ03251.1 awen|OJJ41637.1 azon|OJJ48359.1

Group2796: abra|OJJ72545.1 acar|OOF94710.1 acla|EAW12749.1 afis|EAW17913.1 afla|EED46315.1 afum|EAL91722.1 agla|OJJ83030.1 akaw|GAA86269.1 aluc|OJZ90529.1 anid|CBF76971.1 anig|EHA23981.1 aory|EIT76319.1 asyd|OJJ57043.1 ater|EAU33133.1 atub|OJI81951.1 aver|OJJ03250.1 awen|OJJ41636.1 azon|OJJ48356.1

Group2797: abra|OJJ72551.1 acar|OOF94702.1 acla|EAW12756.1 afis|EAW17919.1 afla|EED46322.1 afum|EAL91730.1 agla|OJJ83023.1 akaw|GAA85183.1 aluc|OJZ90538.1 anid|CBF76955.1 anig|EHA23986.1 aory|EIT81188.1 asyd|OJJ57054.1 ater|EAU33554.1 atub|OJI81959.1 aver|OJJ03243.1 awen|OJJ41627.1 azon|OJJ48350.1

Group2798: abra|OJJ72554.1 acar|OOF94700.1 acla|EAW12759.1 afis|EAW17921.1 afla|EED46324.1 afum|EAL91732.1 agla|OJJ83020.1 akaw|GAA85186.1 aluc|OJZ90541.1 anid|CBF76951.1 anig|EHA23989.1 aory|EIT81201.1 asyd|OJJ57057.1 ater|EAU33550.1 atub|OJI81963.1 aver|OJJ03241.1 awen|OJJ41624.1 azon|OJJ48347.1

Group2799: abra|OJJ72555.1 acar|OOF94698.1 acla|EAW12760.1 afis|EAW17922.1 afla|EED46325.1 afum|EAL91733.1 agla|OJJ83019.1 akaw|GAA85187.1 aluc|OJZ90542.1 anid|CBF76949.1 anig|EHA23990.1 aory|EIT81261.1 asyd|OJJ57058.1 ater|EAU33549.1 atub|OJI81964.1 aver|OJJ03240.1 awen|OJJ41623.1 azon|OJJ48346.1

Group2800: abra|OJJ72562.1 acar|OOF94694.1 acla|EAW11409.1 afis|EAW24012.1 afla|EED49148.1 afum|EAL87442.1 agla|OJJ86944.1 akaw|GAA85191.1 aluc|OJZ90546.1 anid|CBF77408.1 aory|EIT78554.1 asyd|OJJ53238.1 ater|EAU29735.1 atub|OJI81968.1 aver|OJJ03612.1 awen|OJJ35484.1 azon|OJJ51264.1 awen|OJJ37387.1

Group2801: abra|OJJ72564.1 acar|OOF94690.1 acla|EAW12764.1 afis|EAW17926.1 afla|EED46328.1 afum|EAL91737.1 agla|OJJ83013.1 akaw|GAA92409.1 aluc|OJZ83268.1 anid|CBF76942.1 anig|EHA21121.1 aory|EIT81217.1 asyd|OJJ57062.1 ater|EAU33545.1 atub|OJI87669.1 aver|OJJ03236.1 awen|OJJ41620.1 azon|OJJ48343.1

Group2802: abra|OJJ72565.1 acar|OOF94689.1 acla|EAW12765.1 afis|EAW17927.1 afla|EED58037.1 afum|EAL91738.1 agla|OJJ83007.1 akaw|GAA92406.1 aluc|OJZ83267.1 anid|CBF76940.1 anig|EHA21120.1 aory|EIT79801.1 asyd|OJJ57063.1 ater|EAU33544.1 atub|OJI87670.1 aver|OJJ03235.1 awen|OJJ41619.1 azon|OJJ48342.1

Group2803: abra|OJJ72567.1 acar|OOF94687.1 acla|EAW12767.1 afis|EAW17930.1 afla|EED49111.1 afum|EAL91741.1 agla|OJJ83003.1 akaw|GAA85194.1 aluc|OJZ90549.1 anid|CBF76936.1 anig|EHA23997.1 aory|EIT78530.1 asyd|OJJ57064.1 ater|EAU33540.1 atub|OJI81971.1 aver|OJJ03234.1 awen|OJJ41616.1 azon|OJJ48337.1

Group2804: abra|OJJ72568.1 acar|OOF97957.1 acla|EAW14929.1 afla|EED48920.1 afum|EAL85693.1 akaw|GAA85196.1 aluc|OJZ90551.1 anig|EHA23998.1 aory|EIT81168.1 ater|EAU36518.1 atub|OJI81973.1 aver|OJJ07138.1 afis|EAW22084.1 acla|EAW14284.1 afla|EED49819.1 aory|EIT77604.1 afla|EED55283.1 aory|EIT83580.1

Group2805: abra|OJJ72581.1 acar|OOF94671.1 acla|EAW12768.1 afis|EAW17931.1 afla|EED49110.1 afum|EAL91742.1 agla|OJJ83002.1 akaw|GAA85207.1 aluc|OJZ90562.1 anid|CBF76934.1 anig|EHA24008.1 aory|EIT78507.1 asyd|OJJ57065.1 ater|EAU33539.1 atub|OJI81985.1 aver|OJJ03233.1 awen|OJJ41615.1 azon|OJJ48336.1

Group2806: abra|OJJ72582.1 acar|OOF94670.1 acla|EAW12769.1 afis|EAW17932.1 afla|EED49109.1 afum|EAL91743.1 agla|OJJ83001.1 akaw|GAA85208.1 aluc|OJZ90563.1 anid|CBF76932.1 anig|EHA24009.1 aory|EIT78546.1 asyd|OJJ57066.1 ater|EAU33538.1 atub|OJI81986.1 aver|OJJ03232.1 awen|OJJ41614.1 azon|OJJ48335.1

Group2807: abra|OJJ72583.1 acar|OOF94669.1 acla|EAW12770.1 afis|EAW17933.1 afla|EED49108.1 afum|EAL91744.1 agla|OJJ83000.1 akaw|GAA85209.1 aluc|OJZ90564.1 anid|CBF76930.1 anig|EHA24010.1 aory|EIT78564.1 asyd|OJJ57067.1 ater|EAU33537.1 atub|OJI81987.1 aver|OJJ03231.1 awen|OJJ41613.1 azon|OJJ48333.1

Group2808: abra|OJJ72585.1 acar|OOF94667.1 acla|EAW12771.1 afis|EAW17934.1 afla|EED49107.1 afum|EAL91745.1 agla|OJJ82999.1 akaw|GAA85211.1 aluc|OJZ90566.1 anid|CBF77039.1 anig|EHA24012.1 aory|EIT78519.1 asyd|OJJ57002.1 ater|EAU33536.1 atub|OJI81989.1 aver|OJJ03285.1 awen|OJJ41612.1 azon|OJJ48331.1

Group2809: abra|OJJ72588.1 acar|OOF94662.1 acla|EAW12774.1 afis|EAW17937.1 afla|EED49105.1 afum|EAL91748.1 agla|OJJ82995.1 akaw|GAA85214.1 aluc|OJZ90569.1 anid|CBF77044.1 anig|EHA24014.1 aory|EIT78518.1 asyd|OJJ56999.1 ater|EAU33533.1 atub|OJI81992.1 aver|OJJ03288.1 awen|OJJ41609.1 azon|OJJ48328.1

Group2810: abra|OJJ72589.1 acar|OOF94661.1 acla|EAW12775.1 afis|EAW17938.1 afla|EED49103.1 afum|EAL91749.1 agla|OJJ82994.1 akaw|GAA85215.1 aluc|OJZ90570.1 anid|CBF77046.1 anig|EHA24015.1 aory|EIT82547.1 asyd|OJJ56998.1 ater|EAU33530.1 atub|OJI81993.1 aver|OJJ03289.1 awen|OJJ41607.1 azon|OJJ48325.1

Group2811: abra|OJJ72592.1 acar|OOF94658.1 acla|EAW12778.1 afis|EAW17941.1 afla|EED52574.1 afum|EAL91752.1 agla|OJJ82991.1 akaw|GAA85219.1 aluc|OJZ90573.1 anid|CBF77049.1 anig|EHA24018.1 aory|EIT74256.1 asyd|OJJ56996.1 ater|EAU33524.1 atub|OJI81997.1 aver|OJJ03291.1 awen|OJJ41602.1 azon|OJJ48320.1

Group2812: abra|OJJ72596.1 acar|OOF94657.1 acla|EAW11620.1 afla|EED52571.1 akaw|GAA85224.1 aluc|OJZ90577.1 anig|EHA24021.1 aory|EIT74280.1 ater|EAU33521.1 atub|OJI82002.1 azon|OJJ48744.1 afis|EAW23838.1 afum|EAL87269.1 asyd|OJJ56992.1 aver|OJJ03295.1 anid|CBF77058.1 agla|OJJ82497.1 awen|OJJ33801.1

Group2813: abra|OJJ72598.1 acar|OOF94656.1 acla|EAW11619.1 afis|EAW23853.1 afla|EED52570.1 afum|EAL87281.1 agla|OJJ82496.1 akaw|GAA85225.1 aluc|OJZ90579.1 anid|CBF77060.1 anig|EHA24022.1 aory|EIT74300.1 asyd|OJJ56991.1 ater|EAU33520.1 atub|OJI82004.1 aver|OJJ03296.1 awen|OJJ33802.1 azon|OJJ48743.1

Group2814: abra|OJJ72599.1 acar|OOF94655.1 acla|EAW11618.1 afis|EAW23852.1 afla|EED52569.1 afum|EAL87280.1 agla|OJJ82495.1 akaw|GAA85226.1 aluc|OJZ90580.1 anid|CBF77062.1 anig|EHA24023.1 aory|EIT74211.1 asyd|OJJ56990.1 ater|EAU33519.1 atub|OJI82005.1 aver|OJJ03297.1 awen|OJJ33803.1 azon|OJJ48742.1

Group2815: abra|OJJ72600.1 acar|OOF94653.1 acla|EAW11617.1 afis|EAW23851.1 afla|EED52568.1 afum|EAL87279.1 agla|OJJ82494.1 akaw|GAA85228.1 aluc|OJZ90583.1 anid|CBF77063.1 anig|EHA24024.1 aory|EIT74296.1 asyd|OJJ56989.1 ater|EAU33518.1 atub|OJI82006.1 aver|OJJ03298.1 awen|OJJ33804.1 azon|OJJ48740.1

Group2816: abra|OJJ72603.1 acar|OOF94650.1 acla|EAW11611.1 afis|EAW23844.1 afla|EED52553.1 afum|EAL87275.1 agla|OJJ82490.1 akaw|GAA85231.1 aluc|OJZ90586.1 anid|CBF77071.1 anig|EHA24026.1 aory|EIT74313.1 asyd|OJJ56986.1 ater|EAU33511.1 atub|OJI82009.1 aver|OJJ03301.1 awen|OJJ33807.1 azon|OJJ48737.1

Group2817: abra|OJJ72605.1 acar|OOF94649.1 afis|EAW23848.1 afla|EED52565.1 agla|OJJ82491.1 akaw|GAA85232.1 aluc|OJZ90588.1 anig|EHA24027.1 aory|EIT74169.1 atub|OJI82011.1 aver|OJJ03300.1 acla|EAW11614.1 afum|EAL87276.2 anid|CBF77067.1 asyd|OJJ56987.1 ater|EAU33515.1 awen|OJJ33806.1 azon|OJJ48727.1

Group2818: abra|OJJ72625.1 acar|OOF94639.1 acla|EAW11599.1 afis|EAW23858.1 afla|EED52583.1 afum|EAL87286.1 agla|OJJ82525.1 akaw|GAA85245.1 aluc|OJZ90602.1 anid|CBF77236.1 anig|EHA24039.1 aory|EIT74247.1 asyd|OJJ56814.1 ater|EAU33493.1 atub|OJI82026.1 aver|OJJ03521.1 awen|OJJ33819.1 azon|OJJ48716.1

Group2819: abra|OJJ72626.1 acar|OOF94638.1 acla|EAW11598.1 afis|EAW23859.1 afla|EED52584.1 afum|EAL87287.1 agla|OJJ82526.1 akaw|GAA85246.1 aluc|OJZ90603.1 anid|CBF77234.1 anig|EHA24040.1 aory|EIT74284.1 asyd|OJJ56815.1 ater|EAU33492.1 atub|OJI82027.1 aver|OJJ03520.1 awen|OJJ33820.1 azon|OJJ48715.1

Group2820: abra|OJJ72627.1 acar|OOF94637.1 acla|EAW11596.1 afis|EAW23860.1 afla|EED52585.1 afum|EAL87288.1 agla|OJJ82527.1 akaw|GAA85247.1 aluc|OJZ90604.1 anid|CBF77232.1 anig|EHA24041.1 aory|EIT74206.1 asyd|OJJ56816.1 ater|EAU33490.1 atub|OJI82028.1 awen|OJJ33821.1 azon|OJJ48714.1 aver|OJJ03519.1

Group2821: abra|OJJ72635.1 acar|OOF94633.1 acla|EAW11594.1 afis|EAW23862.1 afla|EED52586.1 agla|OJJ82528.1 akaw|GAA85256.1 aluc|OJZ90614.1 anid|CBF77230.1 anig|EHA24052.1 aory|EIT74306.1 asyd|OJJ56817.1 ater|EAU33489.1 atub|OJI82040.1 aver|OJJ03518.1 awen|OJJ33822.1 azon|OJJ48713.1 aluc|OJZ91238.1

Group2822: abra|OJJ72640.1 acar|OOF94631.1 acla|EAW11584.1 afis|EAW23869.1 afla|EED52597.1 afum|EAL87297.1 agla|OJJ82538.1 akaw|GAA85261.1 aluc|OJZ90619.1 anid|CBF77217.1 anig|EHA24056.1 aory|EIT74235.1 asyd|OJJ56823.1 ater|EAU33478.1 atub|OJI82045.1 aver|OJJ03512.1 awen|OJJ33845.1 azon|OJJ48702.1

Group2823: abra|OJJ72641.1 acar|OOF94630.1 acla|EAW11583.1 afis|EAW23870.1 afla|EED52598.1 afum|EAL87298.1 agla|OJJ82539.1 akaw|GAA85262.1 aluc|OJZ90620.1 anid|CBF77219.1 anig|EHA24057.1 aory|EIT74294.1 asyd|OJJ56822.1 ater|EAU33479.1 atub|OJI82046.1 aver|OJJ03513.1 awen|OJJ33846.1 azon|OJJ48701.1

Group2824: abra|OJJ72642.1 acar|OOF94629.1 acla|EAW11582.1 afis|EAW23871.1 afla|EED52599.1 afum|EAL87299.1 agla|OJJ82540.1 akaw|GAA85263.1 aluc|OJZ90621.1 anid|CBF77220.1 anig|EHA24058.1

aory|EIT74246.1 asyd|OJJ56821.1 ater|EAU33475.1 atub|OJI82047.1 aver|OJJ03514.1 awen|OJJ33847.1 azon|OJJ48700.1

Group2825: abra|OJJ72647.1 acar|OOF94623.1 acla|EAW11581.1 afis|EAW23872.1 afla|EED52600.1 afum|EAL87300.1 agla|OJJ82542.1 akaw|GAA85267.1 aluc|OJZ90625.1 anid|CBF84521.1 anig|EHA24062.1 aory|EIT74293.1 asyd|OJJ60278.1 ater|EAU33474.1 atub|OJI82052.1 aver|OJJ03978.1 awen|OJJ33848.1 azon|OJJ48699.1

Group2826: abra|OJJ72648.1 acar|OOF94622.1 acla|EAW11580.1 afis|EAW23874.1 afla|EED52602.1 afum|EAL87302.1 agla|OJJ82544.1 akaw|GAA85269.1 aluc|OJZ90627.1 anid|CBF77209.1 anig|EHA24063.1 aory|EIT74307.1 asyd|OJJ56829.1 ater|EAU33472.1 atub|OJI82053.1 aver|OJJ03507.1 awen|OJJ33850.1 azon|OJJ48698.1

Group2827: abra|OJJ72653.1 acar|OOF94620.1 afis|EAW23877.1 afla|EED52605.1 afum|EAL87305.1 agla|OJJ82547.1 akaw|GAA85272.1 aluc|OJZ90631.1 anid|CBF77201.1 anig|EHA24068.1 aory|EIT74181.1 asyd|OJJ56833.1 ater|EAU33469.1 atub|OJI82057.1 aver|OJJ03504.1 awen|OJJ33854.1 azon|OJJ48695.1 acla|EAW11577.1

Group2828: abra|OJJ72654.1 acar|OOF94619.1 acla|EAW11576.1 afis|EAW23878.1 afla|EED52606.1 afum|EAL87306.1 agla|OJJ82548.1 akaw|GAA85273.1 aluc|OJZ90632.1 anid|CBF77199.1 anig|EHA24069.1 aory|EIT74260.1 asyd|OJJ56834.1 ater|EAU33468.1 atub|OJI82059.1 aver|OJJ03503.1 awen|OJJ33858.1 azon|OJJ48694.1

Group2829: abra|OJJ72658.1 acar|OOF94615.1 acla|EAW11522.1 afis|EAW23931.1 afla|EED50587.1 afum|EAL87356.1 agla|OJJ79411.1 akaw|GAA85281.1 aluc|OJZ90639.1 anid|CBF77275.1 anig|EHA24074.1 aory|EIT82143.1 asyd|OJJ56796.1 ater|EAU32296.1 atub|OJI82067.1 aver|OJJ03538.1 awen|OJJ39023.1 azon|OJJ51349.1

Group2830: abra|OJJ72659.1 acar|OOF94614.1 acla|EAW11523.1 afis|EAW23930.1 afla|EED50588.1 afum|EAL87355.1 agla|OJJ79407.1 akaw|GAA85282.1 aluc|OJZ90640.1 anid|CBF77277.1 anig|EHA24075.1 aory|EIT82106.1 asyd|OJJ56795.1 ater|EAU32306.1 atub|OJI82068.1 aver|OJJ03539.1 awen|OJJ39024.1 azon|OJJ51350.1

Group2831: abra|OJJ72661.1 acar|OOF94612.1 acla|EAW11526.1 afis|EAW23928.1 afla|EED50590.1 afum|EAL87353.1 agla|OJJ79406.1 akaw|GAA85284.1 aluc|OJZ90642.1 anid|CBF77281.1 anig|EHA24077.1 aory|EIT82136.1 asyd|OJJ56793.1 ater|EAU32307.1 atub|OJI82070.1 aver|OJJ03541.1 awen|OJJ39026.1 azon|OJJ51352.1

Group2832: abra|OJJ72662.1 acar|OOF94611.1 acla|EAW11527.1 afis|EAW23927.1 afla|EED50591.1 afum|EAL87352.1 agla|OJJ79405.1 akaw|GAA85285.1 aluc|OJZ90643.1 anid|CBF77254.1 anig|EHA24078.1 aory|EIT82117.1 asyd|OJJ56807.1 ater|EAU32308.1 atub|OJI82071.1 aver|OJJ03527.1 awen|OJJ39027.1 azon|OJJ51353.1

Group2833: abra|OJJ72663.1 acar|OOF94610.1 acla|EAW11528.1 afis|EAW23926.1 afla|EED50594.1 afum|EAL87351.1 agla|OJJ79404.1 akaw|GAA85286.1 aluc|OJZ90644.1 anid|CBF77252.1 anig|EHA24079.1 aory|EIT82151.1 asyd|OJJ56808.1 ater|EAU32309.1 atub|OJI82072.1 aver|OJJ03526.1 awen|OJJ39028.1 azon|OJJ51354.1

Group2834: abra|OJJ72669.1 acar|OOF97389.1 acla|EAW11531.1 afis|EAW23923.1 afla|EED50599.1 afum|EAL87348.1 agla|OJJ79397.1 akaw|GAA85289.1 aluc|OJZ90647.1 anid|CBF77090.1 anig|EHA24082.1 aory|EIT78106.1 asyd|OJJ56959.1 ater|EAU32313.1 atub|OJI82077.1 aver|OJJ03326.1 awen|OJJ39032.1 azon|OJJ51360.1

Group2835: abra|OJJ72670.1 acar|OOF97388.1 acla|EAW11537.1 afis|EAW23919.1 afla|EED50608.1 afum|EAL87345.1 agla|OJJ79386.1 akaw|GAA85290.1 aluc|OJZ90648.1 anid|CBF77096.1 anig|EHA24083.1 aory|EIT78069.1 asyd|OJJ56955.1 ater|EAU32318.1 atub|OJI82078.1 aver|OJJ03330.1 awen|OJJ39036.1 azon|OJJ51364.1

Group2836: abra|OJJ72675.1 acar|OOF97380.1 acla|EAW11538.1 afis|EAW23913.1 afla|EED50618.1 afum|EAL87342.1 agla|OJJ79382.1 akaw|GAA85294.1 aluc|OJZ90653.1 anid|CBF77098.1 anig|EHA24088.1 aory|EIT78036.1 asyd|OJJ56954.1 ater|EAU32319.1 atub|OJI82082.1 aver|OJJ03331.1 awen|OJJ39038.1 azon|OJJ51365.1

Group2837: abra|OJJ72683.1 acar|OOF95932.1 acla|EAW13159.1 afis|EAW17487.1 afla|EED49329.1 afum|EAL91310.1 agla|OJJ81475.1 akaw|GAA85301.1 aluc|OJZ90661.1 anid|CBF71613.1 anig|EHA24096.1 aory|EIT79552.1 asyd|OJJ61862.1 ater|EAU36932.1 atub|OJI82090.1 aver|OJI95960.1 awen|OJJ37355.1 azon|OJJ51040.1

Group2838: abra|OJJ72684.1 acar|OOF95933.1 acla|EAW11540.1 afis|EAW23911.1 afla|EED52657.1 afum|EAL87340.1 agla|OJJ79380.1 akaw|GAA85302.1 aluc|OJZ90662.1 anid|CBF77104.1 anig|EHA24097.1 aory|EIT74175.1 asyd|OJJ56950.1 ater|EAU32321.1 atub|OJI82091.1 aver|OJJ03335.1 awen|OJJ39040.1 azon|OJJ51367.1

Group2839: abra|OJJ72685.1 acar|OOF95934.1 acla|EAW11541.1 afis|EAW23910.1 afla|EED52655.1 afum|EAL87339.1 agla|OJJ79379.1 akaw|GAA85303.1 aluc|OJZ90663.1 anid|CBF77106.1 anig|EHA24098.1 aory|EIT74315.1 asyd|OJJ56949.1 ater|EAU32322.1 atub|OJI82092.1 aver|OJJ03336.1 awen|OJJ39041.1 azon|OJJ51368.1

Group2840: abra|OJJ72687.1 acar|OOF95936.1 acla|EAW11542.1 afis|EAW23909.1 afla|EED52654.1 afum|EAL87337.1 agla|OJJ79378.1 akaw|GAA85305.1 aluc|OJZ90665.1 anid|CBF77108.1 anig|EHA24100.1 aory|EIT74231.1 asyd|OJJ56948.1 ater|EAU32324.1 atub|OJI82094.1 aver|OJJ03337.1 awen|OJJ39042.1 azon|OJJ51370.1

Group2841: abra|OJJ72706.1 acar|OOF95943.1 acla|EAW11546.1 afis|EAW23907.1 afla|EED52642.1 afum|EAL87335.1 agla|OJJ79369.1 akaw|GAA85310.1 aluc|OJZ90671.1 anid|CBF77127.1 anig|EHA24104.1 aory|EIT74245.1 asyd|OJJ56941.1 ater|EAU32331.1 atub|OJI82101.1 aver|OJJ03344.1 awen|OJJ39043.1 azon|OJJ51372.1

Group2842: abra|OJJ72707.1 acar|OOF95944.1 acla|EAW11548.1 afis|EAW23905.1 afla|EED52640.1 afum|EAL87333.1 agla|OJJ79370.1 akaw|GAA85311.1 aluc|OJZ90672.1 anid|CBF77125.1 anig|EHA24105.1 aory|EIT74243.1 asyd|OJJ56940.1 ater|EAU32332.1 atub|OJI82102.1 aver|OJJ03345.1 awen|OJJ39044.1 azon|OJJ51373.1

Group2843: abra|OJJ72714.1 acar|OOF95946.1 acla|EAW11550.1 afis|EAW23903.1 afla|EED52638.1 afum|EAL87331.1 agla|OJJ79372.1 akaw|GAA85322.1 aluc|OJZ90683.1 anid|CBF77139.1 anig|EHA24111.1 aory|EIT74295.1 asyd|OJJ56914.1 ater|EAU32335.1 atub|OJI82109.1 aver|OJJ03378.1 awen|OJJ39046.1 azon|OJJ51375.1

Group2844: abra|OJJ72715.1 acar|OOF95947.1 acla|EAW11551.1 afis|EAW23902.1 afla|EED52637.1 afum|EAL87329.1 agla|OJJ79373.1 akaw|GAA85323.1 aluc|OJZ90684.1 anid|CBF77141.1 anig|EHA24112.1 aory|EIT74272.1 asyd|OJJ56913.1 ater|EAU32336.1 atub|OJI82110.1 aver|OJJ03379.1 awen|OJJ39047.1 azon|OJJ51376.1

Group2845: abra|OJJ72719.1 acar|OOF95952.1 acla|EAW11555.1 afis|EAW23898.1 afla|EED52632.1 afum|EAL87325.1 agla|OJJ78640.1 akaw|GAA85326.1 aluc|OJZ90688.1 anid|CBF77148.1 anig|EHA24116.1 aory|EIT74173.1 asyd|OJJ56910.1 ater|EAU32339.1 atub|OJI82113.1 aver|OJJ03383.1 awen|OJJ39050.1 azon|OJJ51382.1

Group2846: abra|OJJ72721.1 acar|OOF95954.1 acla|EAW11557.1 afis|EAW23896.1 afla|EED52630.1 afum|EAL87324.1 agla|OJJ78637.1 akaw|GAA85327.1 aluc|OJZ90690.1 anid|CBF77152.1 anig|EHA24117.1 aory|EIT74255.1 asyd|OJJ56908.1 ater|EAU32340.1 atub|OJI82115.1 aver|OJJ03385.1 awen|OJJ39052.1 azon|OJJ51384.1

Group2847: abra|OJJ72722.1 acar|OOF95955.1 acla|EAW11558.1 afis|EAW23895.1 afla|EED52629.1 afum|EAL87323.1 agla|OJJ78636.1 akaw|GAA85328.1 aluc|OJZ90691.1 anid|CBF77154.1 anig|EHA24118.1 aory|EIT74200.1 asyd|OJJ56907.1 ater|EAU32341.1 atub|OJI82116.1 aver|OJJ03386.1 awen|OJJ39053.1 azon|OJJ51385.1

Group2848: abra|OJJ72725.1 acar|OOF95958.1 acla|EAW11562.1 afis|EAW23891.1 afla|EED52621.1 afum|EAL87319.1 agla|OJJ78631.1 akaw|GAA85331.1 aluc|OJZ90694.1 anid|CBF77171.1 anig|EHA24119.1 aory|EIT74302.1 asyd|OJJ56851.1 ater|EAU33451.1 atub|OJI82119.1 aver|OJJ03466.1 awen|OJJ39058.1 azon|OJJ51389.1

Group2849: abra|OJJ72727.1 acar|OOF95960.1 acla|EAW11564.1 afis|EAW23889.1 afla|EED52619.1 afum|EBA27309.1 agla|OJJ78629.1 akaw|GAA85333.1 aluc|OJZ90696.1 anid|CBF77175.1 anig|EHA24121.1 aory|EIT74266.1 asyd|OJJ56848.1 ater|EAU33454.1 atub|OJI82122.1 aver|OJJ03491.1 awen|OJJ39061.1 azon|OJJ51391.1

Group2850: abra|OJJ72729.1 acar|OOF95961.1 acla|EAW11565.1 afis|EAW23888.1 afla|EED52617.1 afum|EAL87315.1 agla|OJJ78628.1 akaw|GAA85334.1 aluc|OJZ90697.1 anid|CBF77177.1 anig|EHA24123.1 aory|EIT74304.1 asyd|OJJ56847.1 ater|EAU33456.1 atub|OJI82123.1 aver|OJJ03492.1 awen|OJJ39063.1 azon|OJJ51392.1

Group2851: abra|OJJ72733.1 acar|OOF95963.1 acla|EAW11568.1 afis|EAW23885.1 afla|EED52614.1 afum|EBA27307.1 agla|OJJ82572.1 akaw|GAA85339.1 aluc|OJZ90701.1 anid|CBF77183.1 anig|EHA24127.1 aory|EIT74261.1 asyd|OJJ56843.1 ater|EAU33461.1 atub|OJI82128.1 aver|OJJ03495.1 awen|OJJ33867.1 azon|OJJ48687.1

Group2852: abra|OJJ72736.1 acar|OOF95967.1 acla|EAW11572.1 afis|EAW23881.1 afla|EED52610.1 afum|EAL87309.1 agla|OJJ82558.1 akaw|GAA85342.1 aluc|OJZ90704.1 anid|CBF77191.1 anig|EHA24130.1 aory|EIT74269.1 asyd|OJJ56839.1 ater|EAU33465.1 atub|OJI82131.1 aver|OJJ03499.1 awen|OJJ33863.1 azon|OJJ48690.1

Group2853: abra|OJJ72748.1 acar|OOF95971.1 acla|EAW11515.1 afis|EAW23938.1 afla|EED50579.1 afum|EAL87363.1 agla|OJJ79426.1 akaw|GAA85360.1 aluc|OJZ90722.1 anid|CBF77265.1 anig|EHA24140.1

aory|EIT82037.1 asyd|OJJ56801.1 ater|EAU32295.1 atub|OJI82150.1 aver|OJJ03533.1 awen|OJJ39017.1 azon|OJJ51342.1

Group2854: abra|OJJ72749.1 acar|OOF95972.1 acla|EAW11514.1 afis|EAW23939.1 afla|EED50578.1 afum|EAL87364.1 agla|OJJ79427.1 akaw|GAA85361.1 aluc|OJZ90723.1 anid|CBF77263.1 anig|EHA24141.1 aory|EIT82108.1 asyd|OJJ56802.1 ater|EAU32294.1 atub|OJI82151.1 aver|OJJ03532.1 awen|OJJ39016.1 azon|OJJ51341.1

Group2855: abra|OJJ72750.1 acar|OOF95973.1 acla|EAW11513.1 afis|EAW23940.1 afla|EED50577.1 afum|EAL87365.1 agla|OJJ79428.1 akaw|GAA85362.1 aluc|OJZ90724.1 anid|CBF77262.1 anig|EHA24142.1 aory|EIT82017.1 asyd|OJJ56803.1 ater|EAU32293.1 atub|OJI82152.1 aver|OJJ03531.1 awen|OJJ39015.1 azon|OJJ51340.1

Group2856: abra|OJJ72755.1 acar|OOF95976.1 acla|EAW11512.1 afis|EAW23941.1 afla|EED50576.1 afum|EAL87366.1 agla|OJJ79429.1 akaw|GAA85365.1 aluc|OJZ90728.1 anid|CBF77260.1 anig|EHA24145.1 aory|EIT82066.1 asyd|OJJ56804.1 ater|EAU32292.1 atub|OJI82155.1 aver|OJJ03530.1 awen|OJJ39014.1 azon|OJJ51339.1

Group2857: abra|OJJ72756.1 acar|OOF95977.1 acla|EAW11508.1 afis|EAW23945.1 afla|EED50571.1 afum|EAL87370.1 agla|OJJ79436.1 akaw|GAA85366.1 aluc|OJZ90729.1 anid|CBF77290.1 anig|EHA24146.1 aory|EIT82165.1 asyd|OJJ56784.1 ater|EAU32289.1 atub|OJI82156.1 aver|OJJ03549.1 awen|OJJ39009.1 azon|OJJ51335.1

Group2858: abra|OJJ72761.1 acar|OOF95980.1 acla|EAW11503.1 afis|EAW23949.1 afla|EED50567.1 afum|EAL87375.1 agla|OJJ79442.1 akaw|GAA85371.1 aluc|OJZ90735.1 anid|CBF77283.1 anig|EHA24151.1 aory|EIT82002.1 asyd|OJJ56792.1 ater|EAU29803.1 atub|OJI82163.1 aver|OJJ03542.1 awen|OJJ39000.1 azon|OJJ51331.1

Group2859: abra|OJJ72763.1 acar|OOF95983.1 acla|EAW11501.1 afis|EAW23951.1 afla|EED50565.1 afum|EAL87379.1 agla|OJJ79444.1 akaw|GAA85373.1 aluc|OJZ90738.1 anid|CBF77296.1 anig|EHA24153.1 aory|EIT82176.1 asyd|OJJ56782.1 ater|EAU29801.1 atub|OJI82165.1 aver|OJJ03551.1 awen|OJJ38997.1 azon|OJJ51329.1

Group2860: abra|OJJ72764.1 acar|OOF95984.1 acla|EAW11500.1 afis|EAW23952.1 afla|EED50564.1 afum|EAL87380.1 agla|OJJ79445.1 akaw|GAA85374.1 aluc|OJZ90739.1 anid|CBF77298.1 anig|EHA24154.1 aory|EIT82174.1 asyd|OJJ56781.1 ater|EAU29800.1 atub|OJI82166.1 aver|OJJ03552.1 awen|OJJ38996.1 azon|OJJ51328.1

Group2861: abra|OJJ72765.1 acar|OOF95985.1 acla|EAW11499.1 afis|EAW23953.1 afla|EED50562.1 afum|EAL87381.1 agla|OJJ79446.1 akaw|GAA85375.1 aluc|OJZ90740.1 anid|CBF77300.1 anig|EHA24155.1 aory|EIT82008.1 asyd|OJJ56780.1 ater|EAU29799.1 atub|OJI82167.1 aver|OJJ03553.1 awen|OJJ38995.1 azon|OJJ51327.1

Group2862: abra|OJJ72767.1 acar|OOF95986.1 acla|EAW11498.1 afis|EAW23954.1 afla|EED50561.1 afum|EAL87382.1 agla|OJJ79447.1 akaw|GAA85376.1 aluc|OJZ90743.1 anid|CBF77302.1 anig|EHA24156.1 aory|EIT82091.1 asyd|OJJ56779.1 ater|EAU29798.1 atub|OJI82170.1 aver|OJJ03554.1 awen|OJJ38994.1 azon|OJJ51326.1

Group2863: abra|OJJ72768.1 acar|OOF95987.1 acla|EAW11497.1 afis|EAW23955.1 afla|EED50557.1 afum|EAL87383.1 agla|OJJ86878.1 akaw|GAA85377.1 aluc|OJZ90745.1 anid|CBF77304.1 anig|EHA24157.1 aory|EIT82133.1 asyd|OJJ56778.1 ater|EAU29797.1 atub|OJI82171.1 aver|OJJ03555.1 awen|OJJ38993.1 azon|OJJ51324.1

Group2864: abra|OJJ72769.1 acar|OOF95988.1 acla|EAW11496.1 afis|EAW23956.1 afla|EED50556.1 afum|EAL87384.1 agla|OJJ86879.1 akaw|GAA85378.1 aluc|OJZ90746.1 anid|CBF77306.1 anig|EHA24158.1 aory|EIT82152.1 asyd|OJJ56777.1 ater|EAU29796.1 atub|OJI82172.1 aver|OJJ03556.1 awen|OJJ38991.1 azon|OJJ51323.1

Group2865: abra|OJJ72770.1 acar|OOF95989.1 acla|EAW11495.1 afis|EAW23957.1 afla|EED50555.1 afum|EAL87385.1 agla|OJJ86880.1 akaw|GAA85379.1 aluc|OJZ90747.1 anid|CBF77308.1 anig|EHA24159.1 aory|EIT82065.1 asyd|OJJ56776.1 ater|EAU29795.1 atub|OJI82173.1 aver|OJJ03557.1 awen|OJJ38990.1 azon|OJJ51322.1

Group2866: abra|OJJ72773.1 acar|OOF95992.1 acla|EAW11493.1 afis|EAW23960.1 afla|EED50552.1 afum|EAL87389.1 agla|OJJ86885.1 akaw|GAA85382.1 aluc|OJZ90750.1 anid|CBF77314.1 anig|EHA24161.1 aory|EIT82096.1 asyd|OJJ56773.1 ater|EAU29792.1 atub|OJI82176.1 aver|OJJ03560.1 awen|OJJ38987.1 azon|OJJ51319.1

Group2867: abra|OJJ72775.1 acar|OOF95993.1 acla|EAW11491.1 afis|EAW23962.1 afla|EED50548.1 afum|EAL87391.2 agla|OJJ86887.1 akaw|GAA85384.1 anig|EHA24162.1 aory|EIT82083.1 ater|EAU29790.1 atub|OJI82177.1 awen|OJJ38985.1 azon|OJJ51317.1 anid|CBF83986.1 asyd|OJJ61403.1 aver|OJJ00731.1 aluc|OJZ90752.1

Group2868: abra|OJJ72776.1 acar|OOF95994.1 acla|EAW11490.1 afis|EAW23963.1 afla|EED50547.1 afum|EAL87393.1 agla|OJJ86888.1 akaw|GAA85385.1 aluc|OJZ90754.1 anid|CBF77316.1 anig|EHA24163.1 aory|EIT82035.1 asyd|OJJ56772.1 ater|EAU29789.1 atub|OJI82178.1 aver|OJJ03561.1 awen|OJJ38984.1 azon|OJJ51316.1

Group2869: abra|OJJ72779.1 acar|OOF95996.1 acla|EAW11488.1 afis|EAW23965.1 afla|EED50545.1 afum|EAL87395.1 agla|OJJ86894.1 akaw|GAA85387.1 aluc|OJZ83277.1 anid|CBF77322.1 anig|EHA24165.1 aory|EIT82007.1 asyd|OJJ56770.1 ater|EAU29787.1 atub|OJI82181.1 aver|OJJ03563.1 awen|OJJ38978.1 azon|OJJ51314.1

Group2870: abra|OJJ72780.1 acar|OOF95999.1 acla|EAW11487.1 afis|EAW23966.1 afla|EED50544.1 afum|EAL87396.1 agla|OJJ86895.1 akaw|GAA85388.1 aluc|OJZ83279.1 anid|CBF77324.1 anig|EHA24166.1 aory|EIT82088.1 asyd|OJJ56769.1 ater|EAU29786.1 atub|OJI82183.1 aver|OJJ03565.1 awen|OJJ38977.1 azon|OJJ51313.1

Group2871: abra|OJJ72781.1 acar|OOF96000.1 acla|EAW11486.1 afis|EAW23967.1 afla|EED50542.1 afum|EAL87397.1 agla|OJJ86896.1 akaw|GAA85389.1 aluc|OJZ83280.1 anid|CBF77326.1 anig|EHA24167.1 aory|EIT82086.1 asyd|OJJ56768.1 ater|EAU29785.1 atub|OJI82184.1 aver|OJJ03566.1 awen|OJJ38976.1 azon|OJJ51312.1

Group2872: abra|OJJ72782.1 acar|OOF96002.1 acla|EAW11485.1 afis|EAW23968.1 afla|EED50541.1 afum|EAL87398.2 agla|OJJ86897.1 akaw|GAA85390.1 aluc|OJZ83281.1 anid|CBF77328.1 anig|EHA24168.1 aory|EIT82024.1 asyd|OJJ56767.1 ater|EAU29784.1 atub|OJI82185.1 aver|OJJ03567.1 awen|OJJ38975.1 azon|OJJ51310.1

Group2873: abra|OJJ72792.1 acar|OOF90123.1 acla|EAW11480.1 afis|EAW23973.1 afla|EED50535.1 afum|EAL87403.1 agla|OJJ86902.1 akaw|GAA85395.1 aluc|OJZ83288.1 anid|CBF77339.1 anig|EHA24173.1 aory|EIT82084.1 asyd|OJJ53275.1 ater|EAU29779.1 atub|OJI82190.1 aver|OJJ03574.1 awen|OJJ35438.1 azon|OJJ51305.1

Group2874: abra|OJJ72797.1 acar|OOF96005.1 acla|EAW11475.1 afis|EAW23978.1 afla|EED49183.1 afum|EAL87408.1 agla|OJJ86908.1 akaw|GAA85400.1 aluc|OJZ83294.1 anid|CBF77349.1 anig|EHA24178.1 aory|EIT78581.1 asyd|OJJ53271.1 ater|EAU29772.1 atub|OJI82195.1 aver|OJJ03579.1 awen|OJJ35446.1 azon|OJJ51300.1

Group2875: abra|OJJ72798.1 acar|OOF96006.1 acla|EAW11474.1 afis|EAW23979.1 afla|EED49182.1 afum|EAL87409.1 agla|OJJ86910.1 akaw|GAA85401.1 aluc|OJZ83295.1 anid|CBF77351.1 anig|EHA24179.1 aory|EIT78531.1 asyd|OJJ53270.1 ater|EAU29771.1 atub|OJI82196.1 aver|OJJ03580.1 awen|OJJ35447.1 azon|OJJ51299.1

Group2876: abra|OJJ72800.1 acar|OOF96008.1 acla|EAW11444.1 afis|EAW23981.1 afla|EED49178.1 afum|EAL87411.1 agla|OJJ86914.1 akaw|GAA85403.1 aluc|OJZ83298.1 anid|CBF77356.1 anig|EHA24181.1 aory|EIT78527.1 asyd|OJJ53268.1 ater|EAU29770.1 atub|OJI82198.1 aver|OJJ03583.1 awen|OJJ35449.1 azon|OJJ51297.1

Group2877: abra|OJJ72801.1 acar|OOF96009.1 acla|EAW11443.1 afis|EAW23982.1 afla|EED49177.1 afum|EAL87412.1 agla|OJJ86916.1 akaw|GAA85404.1 aluc|OJZ83299.1 anid|CBF77358.1 anig|EHA24182.1 aory|EIT78537.1 asyd|OJJ53267.1 ater|EAU29769.1 atub|OJI82199.1 aver|OJJ03584.1 awen|OJJ35450.1 azon|OJJ51293.1

Group2878: abra|OJJ72803.1 acar|OOF96011.1 acla|EAW11441.1 afis|EAW23984.1 afla|EED49175.1 afum|EAL87413.1 agla|OJJ86918.1 akaw|GAA85406.1 aluc|OJZ83301.1 anid|CBF77362.1 anig|EHA24183.1 aory|EIT78558.1 asyd|OJJ53265.1 ater|EAU29767.1 atub|OJI82201.1 aver|OJJ03586.1 awen|OJJ35452.1 azon|OJJ51291.1

Group2879: abra|OJJ72804.1 acar|OOF96012.1 acla|EAW11440.1 afis|EAW23985.1 afla|EED49173.1 afum|EAL87414.1 agla|OJJ86919.1 akaw|GAA85407.1 aluc|OJZ83302.1 anid|CBF77364.1 anig|EHA24184.1 aory|EIT78557.1 asyd|OJJ53264.1 ater|EAU29766.1 atub|OJI82202.1 aver|OJJ03587.1 awen|OJJ35453.1 azon|OJJ51290.1

Group2880: abra|OJJ72808.1 acar|OOF96014.1 acla|EAW11433.1 afis|EAW23989.1 afla|EED49169.1 afum|EAL87419.1 agla|OJJ86923.1 akaw|GAA85411.1 aluc|OJZ83306.1 anid|CBF77367.1 anig|EHA24189.1 aory|EIT78567.1 asyd|OJJ53261.1 ater|EAU29761.1 atub|OJI82206.1 aver|OJJ03590.1 awen|OJJ35457.1 azon|OJJ51286.1

Group2881: abra|OJJ72810.1 acar|OOF96017.1 acla|EAW11430.1 afis|EAW23993.1 afla|EED49167.1 afum|EAL87423.1 agla|OJJ86926.1 akaw|GAA85413.1 aluc|OJZ83308.1 anid|CBF77373.1 anig|EHA24191.1 aory|EIT78512.1 asyd|OJJ53258.1 ater|EAU29758.1 atub|OJI82208.1 aver|OJJ03593.1 awen|OJJ35462.1 azon|OJJ51281.1

Group2882: abra|OJJ72811.1 acar|OOF96018.1 acla|EAW11429.1 afis|EAW23994.1 afla|EED49166.1 afum|EAL87424.1 agla|OJJ86927.1 akaw|GAA85414.1 aluc|OJZ83309.1 anid|CBF77375.1 anig|EHA24192.1

aory|EIT78551.1 asyd|OJJ53257.1 ater|EAU29757.1 atub|OJI82209.1 aver|OJJ03594.1 awen|OJJ35463.1 azon|OJJ51280.1

Group2883: abra|OJJ72814.1 acar|OOF96020.1 acla|EAW11428.1 afis|EAW23995.1 afla|EED49165.1 afum|EAL87425.1 agla|OJJ86928.1 akaw|GAA85415.1 aluc|OJZ83312.1 anid|CBF77378.1 anig|EHA24194.1 aory|EIT78566.1 asyd|OJJ53255.1 ater|EAU29756.1 atub|OJI82212.1 aver|OJJ03597.1 awen|OJJ35465.1 azon|OJJ51278.1

Group2884: abra|OJJ72815.1 acar|OOF96021.1 acla|EAW11427.1 afis|EAW23996.1 afla|EED49164.1 afum|EAL87426.1 agla|OJJ86929.1 akaw|GAA85416.1 aluc|OJZ83314.1 anid|CBF77379.1 anig|EHA24195.1 aory|EIT78572.1 asyd|OJJ53254.1 ater|EAU29755.1 atub|OJI82213.1 aver|OJJ03599.1 awen|OJJ35467.1 azon|OJJ51277.1

Group2885: abra|OJJ72818.1 acar|OOF96024.1 acla|EAW11422.1 afis|EAW23999.1 afla|EED49162.1 afum|EAL87429.1 agla|OJJ86933.1 akaw|GAA85419.1 aluc|OJZ83320.1 anid|CBF77385.1 anig|EHA24198.1 aory|EIT78542.1 asyd|OJJ53250.1 ater|EAU29750.1 atub|OJI82217.1 awen|OJJ35470.1 azon|OJJ51274.1 aver|OJJ03602.1

Group2886: abra|OJJ72820.1 acar|OOF96025.1 acla|EAW11419.1 afis|EAW24003.1 afla|EED49158.1 afum|EAL87433.1 agla|OJJ86935.1 akaw|GAA85421.1 aluc|OJZ83322.1 anid|CBF77390.1 anig|EHA24200.1 aory|EIT78547.1 asyd|OJJ53247.1 ater|EAU29748.1 atub|OJI82219.1 aver|OJJ03604.1 awen|OJJ35474.1 azon|OJJ51272.1

Group2887: abra|OJJ72822.1 acar|OOF96027.1 acla|EAW11417.1 afis|EAW24005.1 afla|EED49156.1 afum|EAL87435.1 agla|OJJ86936.1 akaw|GAA85423.1 aluc|OJZ83324.1 anid|CBF77394.1 anig|EHA24202.1 aory|EIT78532.1 asyd|OJJ53245.1 ater|EAU29746.1 atub|OJI82221.1 aver|OJJ03605.1 awen|OJJ35477.1 azon|OJJ51271.1

Group2888: abra|OJJ72824.1 acar|OOF96029.1 acla|EAW11414.1 afis|EAW24007.1 afla|EED49154.1 afum|EAL87437.1 agla|OJJ86938.1 akaw|GAA85425.1 aluc|OJZ83326.1 anid|CBF77398.1 anig|EHA24204.1 aory|EIT78570.1 asyd|OJJ53243.1 ater|EAU29744.1 atub|OJI82223.1 aver|OJJ03607.1 awen|OJJ35479.1 azon|OJJ51269.1

Group2889: abra|OJJ72825.1 acar|OOF96030.1 acla|EAW11413.1 afis|EAW24008.1 afla|EED49153.1 afum|EAL87438.1 agla|OJJ86939.1 akaw|GAA85426.1 aluc|OJZ83327.1 anid|CBF77400.1 anig|EHA24205.1 aory|EIT78506.1 asyd|OJJ53242.1 ater|EAU29743.1 atub|OJI82224.1 aver|OJJ03608.1 awen|OJJ35480.1 azon|OJJ51268.1

Group2890: abra|OJJ72826.1 acar|OOF96031.1 acla|EAW11412.1 afis|EAW24009.1 afla|EED49152.1 afum|EAL87439.1 agla|OJJ86940.1 akaw|GAA85427.1 aluc|OJZ83328.1 anid|CBF77402.1 anig|EHA24206.1 aory|EIT78555.1 asyd|OJJ53241.1 ater|EAU29742.1 atub|OJI82225.1 aver|OJJ03609.1 awen|OJJ35481.1 azon|OJJ51267.1

Group2891: abra|OJJ72827.1 acar|OOF96033.1 acla|EAW11410.1 afis|EAW24011.1 afla|EED49150.1 afum|EAL87441.1 agla|OJJ86943.1 akaw|GAA85428.1 aluc|OJZ83330.1 anid|CBF77406.1 anig|EHA24207.1 aory|EIT78559.1 asyd|OJJ53239.1 ater|EAU29740.1 atub|OJI82226.1 aver|OJJ03611.1 awen|OJJ35483.1 azon|OJJ51265.1

Group2892: abra|OJJ72831.1 acar|OOF96035.1 acla|EAW11408.1 afis|EAW24013.1 afla|EED49147.1 afum|EAL87443.1 agla|OJJ86945.1 akaw|GAA85430.1 aluc|OJZ83332.1 anid|CBF77410.1 anig|EHA24209.1 aory|EIT78522.1 asyd|OJJ53237.1 ater|EAU29734.1 atub|OJI82228.1 aver|OJJ03614.1 awen|OJJ35485.1 azon|OJJ51263.1

Group2893: abra|OJJ72833.1 acar|OOF96036.1 acla|EAW11406.1 afis|EAW24015.1 afla|EED49144.1 afum|EAL87445.1 agla|OJJ86949.1 akaw|GAA85431.1 aluc|OJZ83334.1 anid|CBF77414.1 anig|EHA24210.1 aory|EIT78508.1 asyd|OJJ53235.1 ater|EAU29731.1 atub|OJI82230.1 aver|OJJ03616.1 awen|OJJ35487.1 azon|OJJ47003.1

Group2894: abra|OJJ72835.1 acar|OOF96037.1 acla|EAW11405.1 afis|EAW24016.1 afla|EED49143.1 afum|EAL87446.1 agla|OJJ86950.1 akaw|GAA85432.1 aluc|OJZ83335.1 anid|CBF77416.1 anig|EHA24211.1 aory|EIT78565.1 asyd|OJJ53234.1 ater|EAU29730.1 atub|OJI82232.1 aver|OJJ03617.1 awen|OJJ35488.1 azon|OJJ47001.1

Group2895: abra|OJJ72837.1 acar|OOF96039.1 acla|EAW06923.1 afis|EAW16540.1 afla|EED53743.1 afum|EAL89137.1 agla|OJJ85576.1 akaw|GAA85434.1 aluc|OJZ83337.1 anid|CBF69989.1 anig|EHA24213.1 aory|EIT75293.1 asyd|OJJ58892.1 ater|EAU39054.1 atub|OJI82234.1 aver|OJJ01493.1 awen|OJJ40280.1 azon|OJJ49143.1

Group2896: abra|OJJ72840.1 acar|OOF96042.1 acla|EAW11402.1 afis|EAW24019.1 afla|EED49137.1 afum|EAL87449.1 agla|OJJ86952.1 akaw|GAA85442.1 aluc|OJZ83347.1 anid|CBF77420.1 anig|EHA24215.1 aory|EIT78560.1 asyd|OJJ53232.1 ater|EAU29727.1 atub|OJI82241.1 aver|OJJ03619.1 awen|OJJ35492.1 azon|OJJ46998.1

Group2897: abra|OJJ72844.1 acar|OOF96045.1 acla|EAW11399.1 afis|EAW24022.1 afla|EED49132.1 afum|EAL87453.1 agla|OJJ86955.1 akaw|GAA85445.1 aluc|OJZ83352.1 anid|CBF75300.1 anig|EHA24218.1 aory|EIT78561.1 asyd|OJJ59881.1 ater|EAU29724.1 atub|OJI82244.1 aver|OJJ02405.1 awen|OJJ35495.1 azon|OJJ46994.1

Group2898: abra|OJJ72847.1 acar|OOF96048.1 acla|EAW11396.1 afis|EAW24025.1 afla|EED49129.1 afum|EAL87456.1 agla|OJJ86958.1 akaw|GAA85448.1 aluc|OJZ83355.1 anid|CBF75305.1 anig|EHA24221.1 aory|EIT78574.1 asyd|OJJ59876.1 ater|EAU29719.1 atub|OJI82247.1 aver|OJJ02410.1 awen|OJJ35498.1 azon|OJJ46991.1

Group2899: abra|OJJ72850.1 acar|OOF96051.1 acla|EAW11393.1 afis|EAW24028.1 afla|EED49126.1 afum|EAL87459.1 agla|OJJ86961.1 akaw|GAA85451.1 aluc|OJZ83359.1 anid|CBF75310.1 anig|EHA24224.1 aory|EIT78543.1 asyd|OJJ59874.1 ater|EAU29716.1 atub|OJI82250.1 aver|OJJ02413.1 awen|OJJ35501.1 azon|OJJ46988.1

Group2900: abra|OJJ72853.1 acar|OOF96052.1 acla|EAW11391.1 afis|EAW24031.1 afla|EED49123.1 afum|EAL87462.1 agla|OJJ87001.1 akaw|GAA85454.1 aluc|OJZ83362.1 anid|CBF75316.1 anig|EHA24227.1 aory|EIT78584.1 asyd|OJJ59871.1 ater|EAU29713.1 atub|OJI82253.1 aver|OJJ02416.1 awen|OJJ35503.1 azon|OJJ46983.1

Group2901: abra|OJJ72854.1 acar|OOF96053.1 acla|EAW11390.1 afis|EAW24032.1 afla|EED49121.1 afum|EAL87463.1 agla|OJJ87000.1 akaw|GAA85455.1 aluc|OJZ83363.1 anid|CBF75318.1 anig|EHA24228.1 aory|EIT78515.1 asyd|OJJ59870.1 ater|EAU29712.1 atub|OJI82254.1 aver|OJJ02417.1 awen|OJJ35504.1 azon|OJJ46981.1

Group2902: abra|OJJ72858.1 acar|OOF96055.1 acla|EAW11388.1 afis|EAW24034.1 afla|EED49119.1 afum|EAL87465.1 agla|OJJ86998.1 akaw|GAA85457.1 aluc|OJZ83365.1 anid|CBF75324.1 anig|EHA24230.1 aory|EIT78541.1 asyd|OJJ59868.1 ater|EAU29710.1 atub|OJI82256.1 aver|OJJ02419.1 awen|OJJ35506.1 azon|OJJ46978.1

Group2903: abra|OJJ72860.1 acar|OOF96057.1 acla|EAW11386.1 afis|EAW18110.1 afla|EED49117.1 afum|EAL87467.1 agla|OJJ86996.1 akaw|GAA85459.1 aluc|OJZ83367.1 anid|CBF75328.1 anig|EHA24232.1 aory|EIT78582.1 asyd|OJJ59865.1 ater|EAU29708.1 atub|OJI82259.1 aver|OJJ02422.1 awen|OJJ35508.1 azon|OJJ47630.1

Group2904: abra|OJJ72861.1 acar|OOF96058.1 acla|EAW11385.1 afis|EAW18111.1 afla|EED49114.1 afum|EAL87468.1 agla|OJJ86995.1 akaw|GAA85460.1 aluc|OJZ83368.1 anid|CBF75330.1 anig|EHA24233.1 aory|EIT78525.1 asyd|OJJ59864.1 ater|EAU29707.1 atub|OJI82260.1 aver|OJJ02423.1 awen|OJJ35509.1 azon|OJJ46976.1

Group2905: abra|OJJ72862.1 acar|OOF96060.1 acla|EAW11384.1 afis|EAW18112.1 afla|EED49113.1 afum|EAL87469.1 agla|OJJ86994.1 akaw|GAA85461.1 aluc|OJZ83369.1 anid|CBF75332.1 anig|EHA24234.1 aory|EIT78586.1 asyd|OJJ59863.1 ater|EAU29706.1 atub|OJI82261.1 aver|OJJ02424.1 awen|OJJ35510.1 azon|OJJ46975.1

Group2906: abra|OJJ72863.1 acar|OOF96062.1 acla|EAW11379.1 afis|EAW18116.1 afla|EED58041.1 afum|EAL87473.1 agla|OJJ86980.1 akaw|GAA85462.1 aluc|OJZ83376.1 anid|CBF75340.1 anig|EHA24235.1 aory|EIT79770.1 asyd|OJJ59859.1 ater|EAU29701.1 atub|OJI82262.1 aver|OJJ02428.1 awen|OJJ35515.1 azon|OJJ46970.1

Group2907: abra|OJJ72865.1 acar|OOF96064.1 acla|EAW11382.1 afis|EAW18114.1 afla|EED58039.1 afum|EAL87471.1 agla|OJJ86982.1 akaw|GAA85464.1 aluc|OJZ83378.1 anid|CBF75336.1 anig|EHA24237.1 aory|EIT79795.1 asyd|OJJ59861.1 ater|EAU29703.1 atub|OJI82264.1 aver|OJJ02426.1 awen|OJJ35512.1 azon|OJJ46973.1

Group2908: abra|OJJ72866.1 acar|OOF96065.1 acla|EAW11383.1 afis|EAW18113.1 afla|EED58038.1 afum|EAL87470.1 agla|OJJ86991.1 akaw|GAA85465.1 aluc|OJZ83379.1 anid|CBF75334.1 anig|EHA24238.1 aory|EIT79794.1 asyd|OJJ59862.1 ater|EAU29705.1 atub|OJI82265.1 aver|OJJ02425.1 awen|OJJ35511.1 azon|OJJ46974.1

Group2909: abra|OJJ72869.1 acar|OOF96068.1 acla|EAW11376.1 afis|EAW18119.1 afla|EED58044.1 afum|EAL87477.1 agla|OJJ87011.1 akaw|GAA85468.1 aluc|OJZ83382.1 anid|CBF75346.1 anig|EHA24241.1 aory|EIT79722.1 asyd|OJJ59854.1 ater|EAU29698.1 atub|OJI82268.1 aver|OJJ02431.1 awen|OJJ35518.1 azon|OJJ46966.1

Group2910: abra|OJJ72871.1 acar|OOF96069.1 acla|EAW11375.1 afis|EAW18120.1 afla|EED58045.1 afum|EAL87478.1 agla|OJJ87014.1 akaw|GAA85469.1 aluc|OJZ83385.1 anid|CBF75348.1 anig|EHA24243.1 aory|EIT79798.1 asyd|OJJ59853.1 ater|EAU29697.1 atub|OJI82271.1 aver|OJJ02432.1 awen|OJJ35521.1 azon|OJJ46965.1

Group2911: abra|OJJ72874.1 acar|OOF96071.1 acla|EAW11374.1 afis|EAW18121.1 afla|EED58046.1 afum|EAL87479.1 agla|OJJ87015.1 akaw|GAA85470.1 aluc|OJZ83386.1 anid|CBF75352.1 anig|EHA24244.1

aory|EIT79741.1 asyd|OJJ59852.1 ater|EAU29695.1 atub|OJI82272.1 aver|OJJ02435.1 awen|OJJ35523.1 azon|OJJ46963.1

Group2912: abra|OJJ72880.1 acar|OOF96073.1 acla|EAW11366.1 afis|EAW18129.1 afla|EED58053.1 afum|EAL87486.1 agla|OJJ83795.1 akaw|GAA85472.1 aluc|OJZ83391.1 anid|CBF75366.1 anig|EHA24246.1 aory|EIT79707.1 asyd|OJJ59848.1 ater|EAU35448.1 atub|OJI82277.1 aver|OJJ02440.1 awen|OJJ35557.1 azon|OJJ46953.1

Group2913: abra|OJJ72886.1 acar|OOF96076.1 acla|EAW11364.1 afis|EAW18130.1 afla|EED58054.1 afum|EAL87487.1 agla|OJJ83802.1 akaw|GAA85473.1 aluc|OJZ83394.1 anid|CBF75370.1 anig|EHA24247.1 aory|EIT79721.1 asyd|OJJ59847.1 ater|EAU35452.1 atub|OJI82280.1 aver|OJJ02441.1 awen|OJJ35561.1 azon|OJJ46951.1

Group2914: abra|OJJ72887.1 acar|OOF96077.1 acla|EAW11363.1 afis|EAW18132.1 afla|EED58056.1 afum|EAL87489.1 agla|OJJ83814.1 akaw|GAA85474.1 aluc|OJZ83395.1 anid|CBF75372.1 anig|EHA24248.1 aory|EIT79771.1 asyd|OJJ59846.1 ater|EAU35454.1 atub|OJI82281.1 aver|OJJ02442.1 awen|OJJ35571.1 azon|OJJ46949.1

Group2915: abra|OJJ72890.1 acar|OOF96080.1 acla|EAW11360.1 afis|EAW18134.1 afla|EED58059.1 afum|EAL87491.1 agla|OJJ83818.1 akaw|GAA85477.1 aluc|OJZ83398.1 anid|CBF75380.1 anig|EHA24251.1 aory|EIT79745.1 asyd|OJJ59843.1 ater|EAU35457.1 atub|OJI82284.1 aver|OJJ02446.1 awen|OJJ35576.1 azon|OJJ46946.1

Group2916: abra|OJJ72892.1 acar|OOF96083.1 acla|EAW11358.1 afis|EAW18136.1 afla|EED58061.1 afum|EAL87493.1 agla|OJJ83819.1 akaw|GAA85479.1 aluc|OJZ83400.1 anid|CBF75376.1 anig|EHA24252.1 aory|EIT79756.1 asyd|OJJ59840.1 ater|EAU35459.1 atub|OJI82288.1 aver|OJJ02449.1 awen|OJJ35578.1 azon|OJJ46945.1

Group2917: abra|OJJ72894.1 acar|OOF96085.1 acla|EAW11357.1 afis|EAW18137.1 afla|EED58062.1 afum|EAL87494.1 agla|OJJ83820.1 akaw|GAA85481.1 aluc|OJZ83403.1 anid|CBF75384.1 anig|EHA24254.1 aory|EIT79803.1 asyd|OJJ59838.1 ater|EAU35460.1 atub|OJI82291.1 aver|OJJ02450.1 awen|OJJ35579.1 azon|OJJ46944.1

Group2918: abra|OJJ72898.1 acar|OOF96088.1 acla|EAW11353.1 afis|EAW18141.1 afla|EED58067.1 afum|EAL87498.1 agla|OJJ83826.1 akaw|GAA85485.1 aluc|OJZ83407.1 anid|CBF81129.1 anig|EHA24258.1 aory|EIT79748.1 asyd|OJJ56491.1 ater|EAU35464.1 atub|OJI82296.1 aver|OJJ06227.1 awen|OJJ35583.1 azon|OJJ46939.1

Group2919: abra|OJJ72904.1 acar|OOF89820.1 acla|EAW11350.1 afis|EAW18145.1 afla|EED58072.1 afum|EAL87502.1 agla|OJJ83828.1 akaw|GAA85489.1 aluc|OJZ83414.1 anid|CBF81120.1 anig|EHA24263.1 aory|EIT79155.1 asyd|OJJ56488.1 ater|EAU35468.1 atub|OJI82301.1 aver|OJJ06231.1 awen|OJJ35588.1 azon|OJJ46937.1

Group2920: abra|OJJ72906.1 acar|OOF96089.1 acla|EAW11348.1 afis|EAW18147.1 afla|EED58075.1 afum|EAL87505.1 agla|OJJ83832.1 akaw|GAA85491.1 aluc|OJZ83416.1 anid|CBF81116.1 anig|EHA24265.1 aory|EIT79152.1 asyd|OJJ56486.1 ater|EAU35446.1 atub|OJI82303.1 aver|OJJ06233.1 awen|OJJ35590.1 azon|OJJ46935.1

Group2921: abra|OJJ72909.1 acar|OOF96092.1 acla|EAW11344.1 afis|EAW18150.1 afla|EED58078.1 afum|EAL87508.1 agla|OJJ83835.1 akaw|GAA85494.1 aluc|OJZ83419.1 anid|CBF81110.1 anig|EHA24268.1 aory|EIT79111.1 asyd|OJJ56483.1 ater|EAU35472.1 atub|OJI82306.1 aver|OJJ06236.1 awen|OJJ35593.1 azon|OJJ46915.1

Group2922: abra|OJJ72910.1 acar|OOF96093.1 acla|EAW11343.1 afis|EAW18151.1 afla|EED58080.1 afum|EAL87509.1 agla|OJJ83836.1 akaw|GAA85495.1 aluc|OJZ83421.1 anid|CBF81108.1 anig|EHA24269.1 aory|EIT79142.1 asyd|OJJ56481.1 ater|EAU35473.1 atub|OJI82307.1 aver|OJJ06238.1 awen|OJJ35594.1 azon|OJJ46916.1

Group2923: abra|OJJ72912.1 acar|OOF96094.1 acla|EAW06479.1 afis|EAW15974.1 afla|EED58082.1 afum|EAL88606.1 agla|OJJ83838.1 akaw|GAA85496.1 aluc|OJZ83422.1 anid|CBF81105.1 anig|EHA24270.1 aory|EIT79133.1 asyd|OJJ56479.1 ater|EAU35474.1 atub|OJI82308.1 aver|OJJ06239.1 awen|OJJ35597.1 azon|OJJ46918.1

Group2924: abra|OJJ72913.1 acar|OOF96095.1 acla|EAW06478.1 afis|EAW15973.1 afla|EED58083.1 afum|EAL88605.1 agla|OJJ83842.1 akaw|GAA85497.1 aluc|OJZ83423.1 anid|CBF81101.1 anig|EHA24271.1 aory|EIT79117.1 asyd|OJJ56478.1 ater|EAU35475.1 atub|OJI82309.1 aver|OJJ06240.1 awen|OJJ35599.1 azon|OJJ46919.1

Group2925: abra|OJJ72914.1 acar|OOF96096.1 acla|EAW06476.1 afis|EAW15971.1 afla|EED58085.1 afum|EAL88603.1 agla|OJJ83844.1 akaw|GAA85498.1 aluc|OJZ83424.1 anid|CBF81098.1 anig|EHA24272.1 aory|EIT79112.1 asyd|OJJ56476.1 ater|EAU35477.1 atub|OJI82310.1 aver|OJJ06242.1 awen|OJJ35601.1 azon|OJJ46921.1

Group2926: abra|OJJ72917.1 acar|OOF96099.1 acla|EAW06474.1 afis|EAW15968.1 afla|EED58089.1 afum|EAL88601.1 agla|OJJ89290.1 akaw|GAA85500.1 aluc|OJZ83427.1 anid|CBF81094.1 anig|EHA24279.1 aory|EIT79146.1 asyd|OJJ56470.1 ater|EAU35480.1 atub|OJI82315.1 aver|OJJ06249.1 awen|OJJ35605.1 azon|OJJ46925.1

Group2927: abra|OJJ72920.1 acar|OOF96102.1 acla|EAW06471.1 afis|EAW15965.1 afla|EED58091.1 afum|EAL88598.1 agla|OJJ89346.1 akaw|GAA85503.1 aluc|OJZ83430.1 anid|CBF81089.1 anig|EHA24281.1 aory|EIT79110.1 asyd|OJJ56467.1 ater|EAU35487.1 atub|OJI82318.1 aver|OJJ06252.1 awen|OJJ35608.1 azon|OJJ46928.1

Group2928: abra|OJJ72922.1 acar|OOF96103.1 acla|EAW06469.1 afis|EAW15964.1 afla|EED58093.1 afum|EAL88597.1 agla|OJJ89348.1 akaw|GAA85504.1 aluc|OJZ83432.1 anid|CBF81085.1 anig|EHA24282.1 aory|EIT79125.1 asyd|OJJ56465.1 ater|EAU35488.1 atub|OJI82321.1 aver|OJJ06254.1 awen|OJJ35610.1 azon|OJJ46930.1

Group2929: abra|OJJ72930.1 acar|OOF96110.1 acla|EAW06460.1 afis|EAW15956.1 afla|EED58102.1 afum|EAL88589.1 agla|OJJ89357.1 akaw|GAA85512.1 aluc|OJZ83440.1 anid|CBF81067.1 anig|EHA24286.1 aory|EIT79135.1 asyd|OJJ56458.1 ater|EAU35501.1 atub|OJI82331.1 aver|OJJ06264.1 awen|OJJ35618.1 azon|OJJ46904.1

Group2930: abra|OJJ72931.1 acar|OOF96111.1 acla|EAW06459.1 afis|EAW15955.1 afla|EED58103.1 afum|EAL88588.1 agla|OJJ89358.1 akaw|GAA85513.1 aluc|OJZ83441.1 anid|CBF81065.1 anig|EHA24287.1 aory|EIT79109.1 asyd|OJJ56457.1 ater|EAU35502.1 atub|OJI82332.1 aver|OJJ06265.1 awen|OJJ35619.1 azon|OJJ46903.1

Group2931: abra|OJJ72935.1 acar|OOF96116.1 acla|EAW06456.1 afis|EAW15952.1 afla|EED58106.1 afum|EAL88585.1 agla|OJJ89362.1 akaw|GAA85516.1 aluc|OJZ83444.1 anid|CBF81059.1 anig|EHA24290.1 aory|EIT79114.1 asyd|OJJ56454.1 ater|EAU35505.1 atub|OJI82335.1 aver|OJJ06268.1 awen|OJJ35624.1 azon|OJJ46901.1

Group2932: abra|OJJ72936.1 acar|OOF96117.1 acla|EAW06455.1 afis|EAW15951.1 afla|EED58107.1 afum|EAL88584.1 agla|OJJ89363.1 akaw|GAA85517.1 aluc|OJZ83445.1 anid|CBF81057.1 anig|EHA24291.1 aory|EIT79166.1 asyd|OJJ56453.1 ater|EAU35506.1 atub|OJI82336.1 aver|OJJ06269.1 awen|OJJ35625.1 azon|OJJ46900.1

Group2933: abra|OJJ72937.1 acar|OOF96119.1 acla|EAW06454.1 afis|EAW15950.1 afla|EED58108.1 afum|EAL88583.1 agla|OJJ89364.1 akaw|GAA85518.1 aluc|OJZ83446.1 anid|CBF81056.1 anig|EHA24292.1 aory|EIT79143.1 asyd|OJJ56451.1 ater|EAU35507.1 atub|OJI82337.1 aver|OJJ06270.1 awen|OJJ35626.1 azon|OJJ46899.1

Group2934: abra|OJJ72942.1 acar|OOF96123.1 acla|EAW11209.1 afis|EAW15942.1 afla|EED58116.1 afum|EAL88576.1 agla|OJJ89368.1 akaw|GAA85524.1 aluc|OJZ83452.1 anid|CBF80959.1 anig|EHA24298.1 aory|EIT79144.1 asyd|OJJ56346.1 ater|EAU35514.1 atub|OJI82344.1 aver|OJJ06367.1 awen|OJJ35640.1 azon|OJJ46895.1

Group2935: abra|OJJ72945.1 acar|OOF96126.1 acla|EAW11205.1 afis|EAW15935.1 afla|EED58120.1 afum|EAL88571.1 agla|OJJ89371.1 akaw|GAA85527.1 aluc|OJZ83455.1 anid|CBF80966.1 anig|EHA24301.1 aory|EIT79156.1 asyd|OJJ56349.1 ater|EAU35517.1 atub|OJI82347.1 aver|OJJ06364.1 awen|OJJ35644.1 azon|OJJ46892.1

Group2936: abra|OJJ72949.1 acar|OOF96130.1 acla|EAW11195.1 afis|EAW15925.1 afla|EED54936.1 afum|EAL88561.1 agla|OJJ89380.1 akaw|GAA85530.1 aluc|OJZ83459.1 anid|CBF81019.1 anig|EHA24303.1 aory|EIT80380.1 asyd|OJJ56382.1 ater|EAU35530.1 atub|OJI82351.1 aver|OJJ06331.1 awen|OJJ35655.1 azon|OJJ46884.1

Group2937: abra|OJJ72956.1 acar|OOF96137.1 acla|EAW11201.1 afis|EAW15931.1 afla|EED54930.1 afum|EAL88567.1 agla|OJJ89375.1 akaw|GAA85537.1 aluc|OJZ83465.1 anid|CBF81029.1 anig|EHA24309.1 aory|EIT80323.1 asyd|OJJ56388.1 ater|EAU35521.1 atub|OJI82358.1 aver|OJJ06325.1 awen|OJJ35648.1 azon|OJJ46889.1

Group2938: abra|OJJ72960.1 acar|OOF96142.1 acla|EAW11192.1 afis|EAW15922.1 afla|EED54938.1 afum|EAL88559.1 agla|OJJ89386.1 akaw|GAA85540.1 aluc|OJZ83468.1 anid|CBF81013.1 anig|EHA24312.1 aory|EIT80338.1 asyd|OJJ56379.1 ater|EAU35533.1 atub|OJI82362.1 aver|OJJ06334.1 awen|OJJ35658.1 azon|OJJ46881.1

Group2939: abra|OJJ72967.1 acar|OOF96149.1 acla|EAW11187.1 afis|EAW15918.1 afla|EED54943.1 afum|EAL88554.1 agla|OJJ89393.1 akaw|GAA85546.1 aluc|OJZ83478.1 anid|CBF81002.1 anig|EHA24315.1 aory|EIT80355.1 asyd|OJJ56373.1 ater|EAU35538.1 atub|OJI82368.1 aver|OJJ06340.1 awen|OJJ35661.1 azon|OJJ44132.1

Group2940: abra|OJJ72971.1 acar|OOF96150.1 acla|EAW08037.1 afis|EAW23072.1 afla|EED54944.1 afum|EAL88407.1 agla|OJJ89394.1 akaw|GAA85547.1 aluc|OJZ83482.1 anid|CBF80992.1 anig|EHA24317.1

aory|EIT80389.1 asyd|OJJ56368.1 ater|EAU35539.1 atub|OJI82372.1 aver|OJJ06346.1 awen|OJJ35663.1 azon|OJJ44133.1

Group2941: abra|OJJ72972.1 acar|OOF96151.1 acla|EAW08036.1 afis|EAW23071.1 afla|EED54945.1 afum|EAL88408.1 agla|OJJ89395.1 akaw|GAA85548.1 aluc|OJZ83483.1 anid|CBF80994.1 anig|EHA24318.1 aory|EIT80359.1 asyd|OJJ56369.1 ater|EAU35540.1 atub|OJI82373.1 aver|OJJ06345.1 awen|OJJ35664.1 azon|OJJ44134.1

Group2942: abra|OJJ72973.1 acar|OOF96152.1 acla|EAW08035.1 afis|EAW23070.1 afla|EED54946.1 afum|EAL88409.1 agla|OJJ89396.1 akaw|GAA85549.1 aluc|OJZ83484.1 anid|CBF80996.1 anig|EHA24319.1 aory|EIT80382.1 asyd|OJJ56370.1 ater|EAU37593.1 atub|OJI82374.1 aver|OJJ06344.1 awen|OJJ35665.1 azon|OJJ44135.1

Group2943: abra|OJJ72975.1 acar|OOF96153.1 acla|EAW08034.1 afis|EAW23069.1 afla|EED54947.1 afum|EAL88410.1 agla|OJJ89397.1 akaw|GAA85550.1 aluc|OJZ83485.1 anid|CBF80998.1 anig|EHA24320.1 aory|EIT80368.1 asyd|OJJ56371.1 ater|EAU37596.1 atub|OJI82376.1 aver|OJJ06343.1 awen|OJJ35666.1 azon|OJJ44137.1

Group2944: abra|OJJ72976.1 acar|OOF96154.1 acla|EAW08033.1 afis|EAW23068.1 afla|EED54948.1 afum|EAL88411.1 agla|OJJ89398.1 akaw|GAA85551.1 aluc|OJZ83486.1 anid|CBF81000.1 anig|EHA24321.1 aory|EIT80295.1 asyd|OJJ56372.1 ater|EAU37597.1 atub|OJI82377.1 aver|OJJ06342.1 awen|OJJ35667.1 azon|OJJ44138.1

Group2945: abra|OJJ72979.1 acar|OOF96158.1 acla|EAW08031.1 afis|EAW23065.1 afla|EED54952.1 afum|EAL88414.1 agla|OJJ89409.1 akaw|GAA85554.1 aluc|OJZ83489.1 anid|CBF80981.1 anig|EHA24324.1 aory|EIT80321.1 asyd|OJJ56358.1 ater|EAU37601.1 atub|OJI82381.1 aver|OJJ06356.1 awen|OJJ35670.1 azon|OJJ44140.1

Group2946: abra|OJJ72980.1 acar|OOF96156.1 acla|EAW08030.1 afis|EAW23064.1 afla|EED54953.1 afum|EAL88415.1 agla|OJJ89410.1 akaw|GAA85555.1 aluc|OJZ83490.1 anid|CBF80979.1 anig|EHA24325.1 aory|EIT80361.1 asyd|OJJ56357.1 ater|EAU37602.1 atub|OJI82382.1 aver|OJJ06357.1 awen|OJJ35671.1 azon|OJJ44141.1

Group2947: abra|OJJ72982.1 acar|OOF96159.1 acla|EAW08028.1 afis|EAW23062.1 afla|EED54955.1 afum|EAL88417.1 agla|OJJ89412.1 akaw|GAA85557.1 aluc|OJZ83492.1 anid|CBF80975.1 anig|EHA24327.1 aory|EIT80363.1 asyd|OJJ56355.1 ater|EAU37603.1 atub|OJI82386.1 aver|OJJ06359.1 awen|OJJ35673.1 azon|OJJ44142.1

Group2948: abra|OJJ72983.1 acar|OOF96160.1 acla|EAW08027.1 afis|EAW23061.1 afla|EED54957.1 afum|EAL88418.1 agla|OJJ89414.1 akaw|GAA85558.1 aluc|OJZ83493.1 anid|CBF80973.1 anig|EHA24328.1 aory|EIT80353.1 asyd|OJJ56354.1 ater|EAU37604.1 atub|OJI82387.1 aver|OJJ06360.1 awen|OJJ35676.1 azon|OJJ44143.1

Group2949: abra|OJJ72984.1 acar|OOF96161.1 acla|EAW08026.1 afis|EAW23060.1 afla|EED54958.1 afum|EAL88419.1 agla|OJJ89415.1 akaw|GAA85559.1 aluc|OJZ83494.1 anid|CBF80972.1 anig|EHA24329.1 aory|EIT80387.1 asyd|OJJ56353.1 ater|EAU37605.1 atub|OJI82388.1 aver|OJJ06361.1 awen|OJJ35677.1 azon|OJJ44144.1

Group2950: abra|OJJ72985.1 acar|OOF96162.1 acla|EAW08025.1 afis|EAW23059.1 afla|EED54959.1 afum|EAL88420.1 agla|OJJ89417.1 akaw|GAA85560.1 aluc|OJZ83495.1 anid|CBF80970.1 anig|EHA24330.1 aory|EIT80388.1 asyd|OJJ56352.1 ater|EAU37606.1 atub|OJI82389.1 aver|OJJ06362.1 awen|OJJ35678.1 azon|OJJ44146.1

Group2951: abra|OJJ72989.1 acar|OOF96165.1 acla|EAW08022.1 afis|EAW23056.1 afla|EED54961.1 afum|EAL88424.1 agla|OJJ89419.1 akaw|GAA85563.1 aluc|OJZ83497.1 anid|CBF81039.1 anig|EHA24332.1 aory|EIT80362.1 asyd|OJJ56392.1 ater|EAU37608.1 atub|OJI82391.1 aver|OJJ06321.1 awen|OJJ35680.1 azon|OJJ44148.1

Group2952: abra|OJJ72990.1 acar|OOF96166.1 acla|EAW08021.1 afis|EAW23055.1 afla|EED54962.1 afum|EAL88425.1 agla|OJJ89420.1 akaw|GAA85564.1 aluc|OJZ83498.1 anid|CBF81041.1 anig|EHA24333.1 aory|EIT80303.1 asyd|OJJ56393.1 ater|EAU37609.1 atub|OJI82392.1 aver|OJJ06320.1 awen|OJJ35681.1 azon|OJJ44149.1

Group2953: abra|OJJ72991.1 acar|OOF96167.1 acla|EAW08020.1 afis|EAW23054.1 afla|EED54964.1 afum|EAL88426.1 agla|OJJ89422.1 akaw|GAA85565.1 aluc|OJZ83499.1 anid|CBF81043.1 anig|EHA24334.1 aory|EIT80378.1 asyd|OJJ56441.1 ater|EAU37610.1 atub|OJI82393.1 aver|OJJ06280.1 awen|OJJ35682.1 azon|OJJ44150.1

Group2954: abra|OJJ72997.1 acar|OOF96173.1 acla|EAW08016.1 afis|EAW23050.1 afla|EED54968.1 afum|EAL88430.1 agla|OJJ89429.1 akaw|GAA85571.1 aluc|OJZ83505.1 anid|CBF80953.1 anig|EHA24338.1 aory|EIT80340.1 asyd|OJJ56343.1 ater|EAU29602.1 atub|OJI82399.1 aver|OJJ06370.1 awen|OJJ35689.1 azon|OJJ44154.1

Group2955: abra|OJJ72998.1 acar|OOF89938.1 acla|EAW08015.1 afis|EAW23049.1 afla|EED54969.1 afum|EAL88431.1 agla|OJJ89430.1 akaw|GAA85572.1 aluc|OJZ83506.1 anid|CBF80951.1 anig|EHA24339.1 aory|EIT80346.1 asyd|OJJ56342.1 ater|EAU29601.1 atub|OJI82400.1 aver|OJJ06371.1 awen|OJJ35690.1 azon|OJJ44155.1

Group2956: abra|OJJ72999.1 acar|OOF89939.1 acla|EAW08014.1 afis|EAW23048.1 afla|EED54970.1 afum|EAL88432.1 agla|OJJ89431.1 akaw|GAA85573.1 aluc|OJZ83509.1 anid|CBF80949.1 anig|EHA24340.1 aory|EIT80379.1 asyd|OJJ56341.1 ater|EAU29600.1 atub|OJI82402.1 aver|OJJ06372.1 awen|OJJ35691.1 azon|OJJ44157.1

Group2957: abra|OJJ73002.1 acar|OOF96175.1 acla|EAW08011.1 afis|EAW23046.1 afla|EED54973.1 afum|EAL88435.1 agla|OJJ89432.1 akaw|GAA85580.1 aluc|OJZ83516.1 anid|CBF80946.1 anig|EHA24341.1 aory|EIT80343.1 asyd|OJJ56339.1 ater|EAU29614.1 atub|OJI82407.1 aver|OJJ06374.1 awen|OJJ35694.1 azon|OJJ44158.1

Group2958: abra|OJJ73003.1 acar|OOF96176.1 acla|EAW08010.1 afis|EAW23045.1 afla|EED54974.1 afum|EAL88436.1 agla|OJJ89433.1 akaw|GAA85581.1 aluc|OJZ83517.1 anid|CBF80944.1 anig|EHA24342.1 aory|EIT80339.1 asyd|OJJ56338.1 ater|EAU29613.1 atub|OJI82408.1 aver|OJJ06375.1 awen|OJJ35695.1 azon|OJJ44159.1

Group2959: abra|OJJ73006.1 acar|OOF96178.1 acla|EAW08007.1 afis|EAW23042.1 afla|EED54976.1 afum|EAL88439.1 agla|OJJ89436.1 akaw|GAA85584.1 aluc|OJZ83521.1 anid|CBF80938.1 anig|EHA24344.1 aory|EIT80316.1 asyd|OJJ56312.1 ater|EAU29617.1 atub|OJI82410.1 aver|OJJ06402.1 awen|OJJ35700.1 azon|OJJ44162.1

Group2960: abra|OJJ73008.1 acar|OOF96180.1 acla|EAW08005.1 afis|EAW23040.1 afla|EED54978.1 afum|EAL88441.1 agla|OJJ89438.1 akaw|GAA85586.1 aluc|OJZ83523.1 anid|CBF80935.1 anig|EHA24346.1 aory|EIT80392.1 asyd|OJJ56314.1 ater|EAU29603.1 atub|OJI82412.1 aver|OJJ06400.1 awen|OJJ35702.1 azon|OJJ44164.1

Group2961: abra|OJJ73016.1 acar|OOF96186.1 acla|EAW07997.1 afis|EAW23032.1 afla|EED54984.1 afum|EAL88450.1 agla|OJJ89445.1 akaw|GAA85594.1 aluc|OJZ83531.1 anid|CBF80910.1 anig|EHA24353.1 aory|EIT80327.1 asyd|OJJ56328.1 ater|EAU29625.1 atub|OJI82421.1 aver|OJJ06385.1 awen|OJJ35713.1 azon|OJJ44172.1

Group2962: abra|OJJ73022.1 acar|OOF96190.1 acla|EAW07993.1 afis|EAW23028.1 afla|EED54987.1 afum|EAL88453.2 agla|OJJ89451.1 akaw|GAA85597.1 aluc|OJZ83535.1 anid|CBF80920.1 anig|EHA24356.1 aory|EIT80371.1 asyd|OJJ56322.1 ater|EAU29630.1 atub|OJI82426.1 aver|OJJ06391.1 awen|OJJ35717.1 azon|OJJ44176.1

Group2963: abra|OJJ73023.1 acar|OOF96191.1 acla|EAW07992.1 afis|EAW23027.1 afla|EED54988.1 afum|EAL88455.1 agla|OJJ89452.1 akaw|GAA85598.1 aluc|OJZ83536.1 anid|CBF80918.1 anig|EHA24357.1 aory|EIT80332.1 asyd|OJJ56323.1 ater|EAU29629.1 atub|OJI82427.1 aver|OJJ06390.1 awen|OJJ35718.1 azon|OJJ44177.1

Group2964: abra|OJJ73025.1 acar|OOF96192.1 acla|EAW07991.1 afis|EAW23026.1 afla|EED54989.1 afum|EAL88456.1 agla|OJJ89453.1 akaw|GAA85599.1 aluc|OJZ83537.1 anid|CBF80916.1 anig|EHA24358.1 aory|EIT80337.1 asyd|OJJ56324.1 ater|EAU29628.1 atub|OJI82428.1 aver|OJJ06389.1 awen|OJJ35719.1 azon|OJJ44178.1

Group2965: abra|OJJ73029.1 acar|OOF96195.1 acla|EAW07988.1 afis|EAW23023.1 afla|EED54993.1 afum|EAL88459.1 agla|OJJ89456.1 akaw|GAA85602.1 aluc|OJZ83540.1 anid|CBF80905.1 anig|EHA24360.1 aory|EIT80347.1 asyd|OJJ56332.1 ater|EAU29651.1 atub|OJI82432.1 aver|OJJ06382.1 awen|OJJ30720.1 azon|OJJ44181.1

Group2966: abra|OJJ73030.1 acar|OOF96196.1 acla|EAW07987.1 afis|EAW23022.1 afla|EED54994.1 afum|EAL88460.1 agla|OJJ89457.1 akaw|GAA85603.1 aluc|OJZ83541.1 anid|CBF80903.1 anig|EHA24361.1 aory|EIT80374.1 asyd|OJJ56333.1 ater|EAU29253.1 atub|OJI82433.1 aver|OJJ06381.1 awen|OJJ30721.1 azon|OJJ44182.1

Group2967: abra|OJJ73031.1 acar|OOF96197.1 acla|EAW07986.1 afis|EAW23021.1 afla|EED54995.1 afum|EAL88461.1 agla|OJJ89458.1 akaw|GAA85604.1 aluc|OJZ83542.1 anid|CBF80901.1 anig|EHA24362.1 aory|EIT80385.1 asyd|OJJ56334.1 ater|EAU29652.1 atub|OJI82434.1 aver|OJJ06380.1 awen|OJJ30722.1 azon|OJJ44183.1

Group2968: abra|OJJ73032.1 acar|OOF96198.1 acla|EAW07985.1 afis|EAW23020.1 afla|EED54996.1 afum|EAL88462.1 agla|OJJ89459.1 akaw|GAA85605.1 aluc|OJZ83543.1 anid|CBF80900.1 anig|EHA24363.1 aory|EIT80298.1 asyd|OJJ56335.1 ater|EAU29653.1 atub|OJI82435.1 aver|OJJ06379.1 awen|OJJ30723.1 azon|OJJ44184.1

Group2969: abra|OJJ73037.1 acar|OOF96200.1 acla|EAW07982.1 afis|EAW23017.1 afla|EED54999.1 afum|EAL88464.1 agla|OJJ89463.1 akaw|GAA85608.1 aluc|OJZ83548.1 anig|EHA24366.1 aory|EIT80350.1

asyd|OJJ56309.1 ater|EAU29657.1 atub|OJI82439.1 aver|OJJ06405.1 awen|OJJ29535.1 azon|OJJ44187.1 anid|CBF80894.1

Group2970: abra|OJJ73038.1 acar|OOF96201.1 acla|EAW07981.1 afis|EAW23016.1 afla|EED55000.1 afum|EAL88465.1 agla|OJJ89464.1 akaw|GAA85609.1 aluc|OJZ83549.1 anid|CBF80896.1 anig|EHA24367.1 aory|EIT80294.1 asyd|OJJ56310.1 ater|EAU29658.1 atub|OJI82440.1 aver|OJJ06404.1 awen|OJJ29534.1 azon|OJJ44188.1

Group2971: abra|OJJ73039.1 acar|OOF96204.1 acla|EAW07980.1 afis|EAW23015.1 afla|EED55001.1 afum|EAL88466.1 agla|OJJ89465.1 akaw|GAA85610.1 aluc|OJZ83550.1 anid|CBF80890.1 anig|EHA24368.1 aory|EIT80296.1 asyd|OJJ56307.1 ater|EAU29659.1 atub|OJI82441.1 aver|OJJ06408.1 awen|OJJ30728.1 azon|OJJ44189.1

Group2972: abra|OJJ73040.1 acar|OOF96203.1 acla|EAW07979.1 afis|EAW23014.1 afla|EED55002.1 afum|EAL88467.2 agla|OJJ89466.1 akaw|GAA85611.1 aluc|OJZ83551.1 anid|CBF80888.1 anig|EHA24369.1 aory|EIT80286.1 asyd|OJJ56306.1 ater|EAU29660.1 atub|OJI82442.1 aver|OJJ06409.1 awen|OJJ30729.1 azon|OJJ44190.1

Group2973: abra|OJJ73041.1 acar|OOF96202.1 acla|EAW07978.1 afis|EAW23013.1 afla|EED55003.1 afum|EAL88468.1 agla|OJJ89467.1 akaw|GAA85612.1 aluc|OJZ83552.1 anid|CBF80886.1 anig|EHA24370.1 aory|EIT80391.1 asyd|OJJ56305.1 ater|EAU29661.1 atub|OJI82443.1 aver|OJJ06410.1 awen|OJJ30730.1 azon|OJJ44191.1

Group2974: abra|OJJ73042.1 acar|OOF96205.1 acla|EAW07977.1 afis|EAW23012.1 afla|EED55004.1 afum|EAL88469.1 agla|OJJ89468.1 akaw|GAA85613.1 aluc|OJZ83553.1 anid|CBF80885.1 anig|EHA24371.1 aory|EIT80335.1 asyd|OJJ56304.1 ater|EAU29662.1 atub|OJI82444.1 aver|OJJ06411.1 awen|OJJ30731.1 azon|OJJ44192.1

Group2975: abra|OJJ73046.1 acar|OOF96208.1 acla|EAW07973.1 afis|EAW23008.1 afla|EED55007.1 afum|EAL88472.1 agla|OJJ89472.1 akaw|GAA85617.1 aluc|OJZ83557.1 anid|CBF80879.1 anig|EHA24375.1 aory|EIT80310.1 asyd|OJJ56302.1 ater|EAU29665.1 atub|OJI82447.1 aver|OJJ06415.1 awen|OJJ30733.1 azon|OJJ44196.1

Group2976: abra|OJJ73048.1 acar|OOF96211.1 acla|EAW07971.1 afis|EAW23006.1 afla|EED55009.1 afum|EAL88474.1 agla|OJJ89473.1 akaw|GAA85619.1 aluc|OJZ83559.1 anid|CBF80875.1 anig|EHA24377.1 aory|EIT80349.1 asyd|OJJ56300.1 ater|EAU29667.1 atub|OJI82449.1 aver|OJJ06417.1 awen|OJJ30735.1 azon|OJJ44197.1

Group2977: abra|OJJ73049.1 acar|OOF96212.1 acla|EAW07970.1 afis|EAW23005.1 afla|EED55010.1 afum|EAL88475.1 agla|OJJ89474.1 akaw|GAA85620.1 aluc|OJZ83560.1 anid|CBF80873.1 anig|EHA24378.1 aory|EIT80326.1 asyd|OJJ56299.1 ater|EAU29668.1 atub|OJI82450.1 aver|OJJ06418.1 awen|OJJ30736.1 azon|OJJ44198.1

Group2978: abra|OJJ73050.1 acar|OOF96213.1 acla|EAW07969.1 afis|EAW23004.1 afla|EED55011.1 afum|EAL88476.2 agla|OJJ89475.1 akaw|GAA85621.1 aluc|OJZ83561.1 anid|CBF80871.1 anig|EHA24379.1 aory|EIT80312.1 asyd|OJJ56298.1 ater|EAU29669.1 atub|OJI82451.1 aver|OJJ06419.1 awen|OJJ30737.1 azon|OJJ44199.1

Group2979: abra|OJJ73051.1 acar|OOF96214.1 acla|EAW07968.1 afis|EAW23003.1 afla|EED55012.1 afum|EAL88477.1 agla|OJJ89476.1 akaw|GAA85622.1 aluc|OJZ83562.1 anid|CBF80869.1 anig|EHA24380.1 aory|EIT80369.1 asyd|OJJ56297.1 ater|EAU29670.1 atub|OJI82452.1 aver|OJJ06420.1 awen|OJJ30738.1 azon|OJJ44200.1

Group2980: abra|OJJ73052.1 acar|OOF96215.1 acla|EAW07967.1 afis|EAW23002.1 afla|EED55013.1 afum|EAL88478.1 agla|OJJ89477.1 akaw|GAA85623.1 aluc|OJZ83564.1 anid|CBF80868.1 anig|EHA24381.1 aory|EIT80329.1 asyd|OJJ56296.1 ater|EAU29671.1 atub|OJI82454.1 aver|OJJ06421.1 awen|OJJ30739.1 azon|OJJ44201.1

Group2981: abra|OJJ73056.1 acar|OOF96217.1 acla|EAW07965.1 afis|EAW23000.1 afla|EED55015.1 afum|EAL88480.1 agla|OJJ89479.1 akaw|GAA85624.1 aluc|OJZ83567.1 anid|CBF80864.1 anig|EHA24383.1 aory|EIT80314.1 asyd|OJJ56294.1 ater|EAU29673.1 atub|OJI82456.1 aver|OJJ06423.1 awen|OJJ30741.1 azon|OJJ44204.1

Group2982: abra|OJJ73058.1 acar|OOF96218.1 acla|EAW07963.1 afis|EAW22998.1 afla|EED55017.1 afum|EAL88482.1 agla|OJJ89481.1 akaw|GAA85626.1 aluc|OJZ83569.1 anid|CBF80860.1 anig|EHA24385.1 aory|EIT80381.1 asyd|OJJ56292.1 ater|EAU29674.1 atub|OJI82458.1 aver|OJJ06425.1 awen|OJJ30743.1 azon|OJJ44202.1

Group2983: abra|OJJ73059.1 acar|OOF96220.1 acla|EAW07962.1 afis|EAW22997.1 afla|EED55018.1 afum|EAL88483.1 agla|OJJ89482.1 akaw|GAA85627.1 aluc|OJZ83570.1 anid|CBF80856.1 anig|EHA24386.1 aory|EIT80291.1 asyd|OJJ56291.1 ater|EAU29675.1 atub|OJI82459.1 aver|OJJ06426.1 awen|OJJ30744.1 azon|OJJ44205.1

Group2984: abra|OJJ73062.1 acar|OOG00286.1 afla|EED50225.1 agla|OJJ79957.1 akaw|GAA88097.1 aluc|OJZ81364.1 anid|CBF85609.1 anig|EHA18664.1 aory|EIT76192.1 asyd|OJJ64207.1 ater|EAU30207.1 atub|OJI91568.1 aver|OJI98295.1 awen|OJJ39951.1 afla|EED45252.1 aory|EIT73530.1 agla|OJJ80099.1 aory|EIT77256.1

Group2985: abra|OJJ73142.1 acar|OOG00354.1 acla|EAW07018.1 afis|EAW16644.1 afla|EED44956.1 afum|EAL89248.1 agla|OJJ82603.1 akaw|GAA88030.1 aluc|OJZ81289.1 anid|CBF82053.1 anig|EHA18588.1 aory|EIT73503.1 asyd|OJJ54235.1 ater|EAU39569.1 atub|OJI91497.1 aver|OJJ05187.1 awen|OJJ40070.1 azon|OJJ45554.1

Group2986: abra|OJJ73143.1 acar|OOG00355.1 acla|EAW07019.1 afis|EAW16645.1 afla|EED44955.1 afum|EAL89249.1 agla|OJJ82602.1 akaw|GAA88029.1 aluc|OJZ81288.1 anid|CBF82055.1 anig|EHA18587.1 aory|EIT73491.1 asyd|OJJ54236.1 ater|EAU39570.1 atub|OJI91496.1 aver|OJJ05188.1 awen|OJJ40069.1 azon|OJJ45555.1

Group2987: abra|OJJ73147.1 acar|OOG00358.1 acla|EAW07023.1 afis|EAW16649.1 afla|EED44951.1 afum|EAL89253.1 agla|OJJ78458.1 akaw|GAA88025.1 aluc|OJZ81284.1 anid|CBF82061.1 anig|EHA18583.1 aory|EIT73464.1 asyd|OJJ54240.1 ater|EAU39573.1 atub|OJI91491.1 aver|OJJ05192.1 awen|OJJ40065.1 azon|OJJ45561.1

Group2988: abra|OJJ73148.1 acar|OOG00360.1 acla|EAW07024.1 afis|EAW16650.1 afla|EED44950.1 afum|EAL89254.1 agla|OJJ78459.1 akaw|GAA88024.1 aluc|OJZ81282.1 anid|CBF82063.1 anig|EHA18582.1 aory|EIT73478.1 asyd|OJJ54241.1 ater|EAU39574.1 atub|OJI91490.1 aver|OJJ05193.1 awen|OJJ40064.1 azon|OJJ45562.1

Group2989: abra|OJJ73149.1 acar|OOG00361.1 acla|EAW07025.1 afis|EAW16651.1 afla|EED44949.1 afum|EAL89255.1 agla|OJJ78460.1 akaw|GAA88023.1 aluc|OJZ81281.1 anid|CBF82064.1 anig|EHA18581.1 aory|EIT73505.1 asyd|OJJ54242.1 ater|EAU39575.1 atub|OJI91489.1 aver|OJJ05194.1 awen|OJJ40063.1 azon|OJJ45563.1

Group2990: abra|OJJ73151.1 acar|OOG00363.1 acla|EAW07028.1 afis|EAW16654.1 afla|EED44946.1 afum|EAL89258.1 agla|OJJ78687.1 akaw|GAA88021.1 aluc|OJZ81279.1 anid|CBF82067.1 anig|EHA18579.1 aory|EIT73521.1 asyd|OJJ54244.1 ater|EAU39578.1 atub|OJI91487.1 aver|OJJ05196.1 awen|OJJ40060.1 azon|OJJ45567.1

Group2991: abra|OJJ73156.1 acar|OOG00371.1 acla|EAW07030.1 afis|EAW16656.1 afla|EED44941.1 afum|EAL89260.1 agla|OJJ78716.1 akaw|GAA88016.1 aluc|OJZ81274.1 anid|CBF82070.1 anig|EHA18574.1 aory|EIT73470.1 asyd|OJJ54246.1 ater|EAU39579.1 atub|OJI91481.1 aver|OJJ05199.1 awen|OJJ40058.1 azon|OJJ50081.1

Group2992: abra|OJJ73158.1 acar|OOG00369.1 acla|EAW07032.1 afis|EAW16658.1 afla|EED44944.1 afum|EAL89262.1 agla|OJJ78714.1 akaw|GAA88014.1 aluc|OJZ81272.1 anid|CBF82072.1 anig|EHA18572.1 aory|EIT73479.1 asyd|OJJ54248.1 ater|EAU39581.1 atub|OJI91479.1 aver|OJJ05202.1 awen|OJJ40056.1 azon|OJJ50105.1

Group2993: abra|OJJ73159.1 acar|OOG00368.1 acla|EAW07033.1 afis|EAW16659.1 afla|EED44936.1 afum|EAL89263.1 agla|OJJ78713.1 akaw|GAA88013.1 aluc|OJZ81271.1 anid|CBF82073.1 anig|EHA18571.1 aory|EIT73463.1 asyd|OJJ54249.1 ater|EAU39582.1 atub|OJI91478.1 aver|OJJ05203.1 awen|OJJ40055.1 azon|OJJ50106.1

Group2994: abra|OJJ73160.1 acar|OOG00405.1 acla|EAW06769.1 afis|EAW16319.1 afla|EED53562.1 afum|EAL88943.1 agla|OJJ88280.1 akaw|GAA88012.1 aluc|OJZ81270.1 anid|CBF89151.1 anig|EHA18570.1 aory|EIT74869.1 asyd|OJJ62566.1 ater|EAU34138.1 atub|OJI91477.1 aver|OJI96690.1 awen|OJJ37720.1 azon|OJJ44978.1

Group2995: abra|OJJ73168.1 acar|OOG00409.1 acla|EAW07130.1 afis|EAW22141.1 afla|EED53567.1 afum|EAL91049.1 agla|OJJ87934.1 akaw|GAA88003.1 aluc|OJZ81259.1 anid|CBF89137.1 anig|EHA18566.1 aory|EIT75049.1 asyd|OJJ62649.1 ater|EAU34143.1 atub|OJI91455.1 aver|OJI96749.1 awen|OJJ34715.1 azon|OJJ45847.1

Group2996: abra|OJJ73170.1 acar|OOG00410.1 acla|EAW10796.1 afis|EAW24780.1 afla|EED47205.1 afum|EAL89419.1 agla|OJJ82876.1 akaw|GAA88001.1 aluc|OJZ81257.1 anid|CBF81439.1 anig|EHA18564.1 aory|EIT79182.1 asyd|OJJ59428.1 ater|EAU36528.1 atub|OJI91453.1 aver|OJJ02843.1 awen|OJJ40366.1 azon|OJJ49215.1

Group2997: abra|OJJ73173.1 acar|OOG00413.1 acla|EAW06856.1 afis|EAW16473.1 afla|EED47208.1 afum|EAL89067.1 agla|OJJ82873.1 akaw|GAA87998.1 aluc|OJZ81254.1 anid|CBF81435.1 anig|EHA18562.1 aory|EIT79309.1 asyd|OJJ59424.1 ater|EAU36569.1 atub|OJI91450.1 aver|OJJ02846.1 awen|OJJ40363.1 azon|OJJ49211.1

Group2998: abra|OJJ73176.1 acar|OOG00416.1 acla|EAW06858.1 afis|EAW16475.1 afla|EED47210.1 afum|EAL89069.1 agla|OJJ82871.1 akaw|GAA87996.1 aluc|OJZ81251.1 anid|CBF81431.1 anig|EHA18560.1

aory|EIT79330.1 asyd|OJJ59422.1 ater|EAU36562.1 atub|OJI91448.1 aver|OJJ02848.1 awen|OJJ40361.1 azon|OJJ49209.1

Group2999: abra|OJJ73183.1 acar|OOG00420.1 acla|EAW06861.1 afis|EAW16478.1 afla|EED47215.1 afum|EAL89072.1 agla|OJJ82867.1 akaw|GAA87992.1 aluc|OJZ81246.1 anid|CBF75501.1 anig|EHA18555.1 aory|EIT79241.1 asyd|OJJ59747.1 ater|EAU36560.1 atub|OJI91442.1 aver|OJJ02541.1 awen|OJJ40357.1 azon|OJJ49206.1

Group3000: abra|OJJ73184.1 acar|OOG00421.1 acla|EAW06862.1 afis|EAW16479.1 afla|EED47216.1 afum|EAL89073.1 agla|OJJ82866.1 akaw|GAA87991.1 aluc|OJZ81245.1 anid|CBF75499.1 anig|EHA18554.1 aory|EIT79319.1 asyd|OJJ59748.1 ater|EAU36559.1 atub|OJI91441.1 aver|OJJ02540.1 awen|OJJ40356.1 azon|OJJ49205.1

Group3001: abra|OJJ73185.1 acar|OOG00422.1 acla|EAW06863.1 afis|EAW16480.1 afla|EED47217.1 afum|EAL89074.1 agla|OJJ82865.1 akaw|GAA87990.1 aluc|OJZ81244.1 anid|CBF75503.1 anig|EHA18553.1 aory|EIT79172.1 asyd|OJJ59746.1 ater|EAU36558.1 atub|OJI91440.1 aver|OJJ02542.1 awen|OJJ40354.1 azon|OJJ49204.1

Group3002: abra|OJJ73189.1 acar|OOG00425.1 acla|EAW06865.1 afis|EAW16482.1 afla|EED47222.1 afum|EAL89076.1 agla|OJJ82862.1 akaw|GAA87988.1 aluc|OJZ81241.1 anid|CBF75506.1 anig|EHA18551.1 aory|EIT79303.1 asyd|OJJ59744.1 ater|EAU36556.1 atub|OJI91438.1 aver|OJJ02544.1 awen|OJJ40350.1 azon|OJJ49202.1

Group3003: abra|OJJ73190.1 acar|OOG00426.1 acla|EAW06866.1 afis|EAW16483.1 afla|EED47223.1 afum|EAL89077.1 agla|OJJ82861.1 akaw|GAA87987.1 aluc|OJZ81240.1 anid|CBF75508.1 anig|EHA18550.1 aory|EIT79190.1 asyd|OJJ59743.1 ater|EAU36555.1 atub|OJI91437.1 aver|OJJ02545.1 awen|OJJ40348.1 azon|OJJ49201.1

Group3004: abra|OJJ73191.1 acar|OOG00428.1 acla|EAW06867.1 afis|EAW16484.1 afla|EED47224.1 afum|EAL89078.1 agla|OJJ82860.1 akaw|GAA87986.1 aluc|OJZ81239.1 anid|CBF75520.1 anig|EHA18549.1 aory|EIT79323.1 asyd|OJJ59736.1 ater|EAU36554.1 atub|OJI91436.1 aver|OJJ02551.1 awen|OJJ40343.1 azon|OJJ49200.1

Group3005: abra|OJJ73193.1 acar|OOG00429.1 acla|EAW06869.1 afis|EAW16486.1 afla|EED47226.1 afum|EAL89080.1 agla|OJJ82858.1 akaw|GAA87984.1 aluc|OJZ81236.1 anid|CBF75516.1 anig|EHA18547.1 aory|EIT79231.1 asyd|OJJ59738.1 ater|EAU36552.1 atub|OJI91434.1 aver|OJJ02549.1 awen|OJJ40345.1 azon|OJJ49198.1

Group3006: abra|OJJ73194.1 acar|OOG00431.1 acla|EAW06871.1 afis|EAW16488.1 afla|EED47229.1 afum|EAL89082.1 agla|OJJ82852.1 akaw|GAA87983.1 aluc|OJZ81235.1 anid|CBF75510.1 anig|EHA18546.1 aory|EIT79266.1 asyd|OJJ59742.1 ater|EAU36548.1 atub|OJI91432.1 aver|OJJ02546.1 awen|OJJ40342.1 azon|OJJ49196.1

Group3007: abra|OJJ73196.1 acar|OOG00434.1 acla|EAW06873.1 afis|EAW16490.1 afla|EED47231.1 afum|EAL89084.1 agla|OJJ82829.1 akaw|GAA87981.1 aluc|OJZ81233.1 anid|CBF75522.1 anig|EHA18544.1 aory|EIT79209.1 asyd|OJJ59735.1 ater|EAU36547.1 atub|OJI91430.1 aver|OJJ02552.1 awen|OJJ40339.1 azon|OJJ49193.1

Group3008: abra|OJJ73199.1 acar|OOG00438.1 acla|EAW06876.1 afis|EAW16493.1 afla|EED47237.1 afum|EAL89087.1 agla|OJJ82832.1 akaw|GAA87978.1 aluc|OJZ81230.1 anid|CBF75527.1 anig|EHA18541.1 aory|EIT79313.1 asyd|OJJ59732.1 ater|EAU36545.1 atub|OJI91427.1 aver|OJJ02555.1 awen|OJJ40336.1 azon|OJJ49190.1

Group3009: abra|OJJ73202.1 acar|OOG00441.1 acla|EAW06879.1 afis|EAW16496.1 afla|EED47251.1 afum|EAL89090.1 agla|OJJ82835.1 akaw|GAA87975.1 aluc|OJZ81227.1 anid|CBF75606.1 anig|EHA18538.1 aory|EIT79275.1 asyd|OJJ59727.1 ater|EAU36543.1 atub|OJI91424.1 aver|OJJ02559.1 awen|OJJ40333.1 azon|OJJ49187.1

Group3010: abra|OJJ73203.1 acar|OOG00442.1 acla|EAW06880.1 afis|EAW16497.1 afla|EED56182.1 afum|EBA27367.1 agla|OJJ82836.1 akaw|GAA87974.1 aluc|OJZ81224.1 anid|CBF75605.1 anig|EHA18537.1 aory|EIT73365.1 asyd|OJJ59726.1 ater|EAU36542.1 atub|OJI91422.1 aver|OJJ02560.1 awen|OJJ40332.1 azon|OJJ49186.1

Group3011: abra|OJJ73204.1 acar|OOG00443.1 acla|EAW06881.1 afis|EAW16498.1 afla|EED56181.1 afum|EAL89093.1 agla|OJJ82838.1 akaw|GAA87973.1 aluc|OJZ81222.1 anid|CBF75604.1 anig|EHA18536.1 aory|EIT73379.1 asyd|OJJ59725.1 ater|EAU36541.1 atub|OJI91420.1 aver|OJJ02561.1 awen|OJJ40331.1 azon|OJJ49185.1

Group3012: abra|OJJ73205.1 acar|OOG00444.1 acla|EAW06882.1 afis|EAW16499.1 afla|EED56180.1 afum|EAL89094.1 agla|OJJ82826.1 akaw|GAA87972.1 aluc|OJZ81221.1 anid|CBF75602.1 anig|EHA18535.1 aory|EIT76650.1 asyd|OJJ59724.1 ater|EAU36540.1 atub|OJI91419.1 aver|OJJ02562.1 awen|OJJ40330.1 azon|OJJ49184.1

Group3013: abra|OJJ73206.1 acar|OOG00445.1 acla|EAW06883.1 afis|EAW16500.1 afla|EED56179.1 afum|EAL89095.1 agla|OJJ82827.1 akaw|GAA87971.1 aluc|OJZ81220.1 anid|CBF75600.1 anig|EHA18534.1 aory|EIT76763.1 asyd|OJJ59723.1 ater|EAU36539.1 atub|OJI91418.1 aver|OJJ02563.1 awen|OJJ40328.1 azon|OJJ49183.1

Group3014: abra|OJJ73207.1 acar|OOG00446.1 acla|EAW06884.1 afis|EAW16501.1 afla|EED56178.1 afum|EAL89096.1 agla|OJJ82828.1 akaw|GAA87970.1 aluc|OJZ81219.1 anid|CBF75598.1 anig|EHA18533.1 aory|EIT76626.1 asyd|OJJ59722.1 ater|EAU36538.1 atub|OJI91417.1 aver|OJJ02564.1 awen|OJJ40327.1 azon|OJJ49182.1

Group3015: abra|OJJ73209.1 acar|OOG00447.1 acla|EAW06885.1 afis|EAW16502.1 afla|EED56177.1 afum|EAL89097.1 agla|OJJ82815.1 akaw|GAA87969.1 aluc|OJZ81218.1 anid|CBF75589.1 anig|EHA18532.1 aory|EIT76868.1 asyd|OJJ59721.1 ater|EAU36537.1 atub|OJI91416.1 aver|OJJ02565.1 awen|OJJ40326.1 azon|OJJ49181.1

Group3016: abra|OJJ73210.1 acar|OOG00449.1 acla|EAW06886.1 afis|EAW16503.1 afla|EED56175.1 afum|EAL89098.1 agla|OJJ82816.1 akaw|GAA87968.1 aluc|OJZ81217.1 anid|CBF75591.1 anig|EHA18531.1 aory|EIT76792.1 asyd|OJJ59720.1 ater|EAU36536.1 atub|OJI91415.1 aver|OJJ02566.1 awen|OJJ40324.1 azon|OJJ49180.1

Group3017: abra|OJJ73212.1 acar|OOG00448.1 acla|EAW06888.1 afis|EAW16505.1 afla|EED56173.1 afum|EAL89100.1 agla|OJJ82818.1 akaw|GAA87966.1 aluc|OJZ81215.1 anid|CBF75594.1 anig|EHA18529.1 aory|EIT76612.1 asyd|OJJ59718.1 ater|EAU36535.1 atub|OJI91412.1 aver|OJJ02568.1 awen|OJJ40321.1 azon|OJJ49178.1

Group3018: abra|OJJ73216.1 acar|OOG00453.1 acla|EAW06891.1 afis|EAW16508.1 afla|EED56170.1 afum|EAL89103.1 agla|OJJ82822.1 akaw|GAA87963.1 aluc|OJZ81212.1 anid|CBF75578.1 anig|EHA18526.1 aory|EIT76719.1 asyd|OJJ59714.1 ater|EAU36572.1 atub|OJI91407.1 aver|OJJ02572.1 awen|OJJ40317.1 azon|OJJ47186.1

Group3019: abra|OJJ73218.1 acar|OOG00455.1 acla|EAW06892.1 afis|EAW16509.1 afla|EED56164.1 afum|EAL89104.2 agla|OJJ82812.1 akaw|GAA87961.1 aluc|OJZ81210.1 anid|CBF75572.1 anig|EHA18524.1 aory|EIT76824.1 asyd|OJJ59711.1 ater|EAU36575.1 atub|OJI91405.1 aver|OJJ02575.1 awen|OJJ40315.1 azon|OJJ49175.1

Group3020: abra|OJJ73219.1 acar|OOG00457.1 acla|EAW06894.1 afis|EAW16511.1 afla|EED56162.1 afum|EAL89106.1 agla|OJJ82811.1 akaw|GAA87959.1 aluc|OJZ81208.1 anid|CBF75568.1 anig|EHA18523.1 aory|EIT76692.1 asyd|OJJ59709.1 ater|EAU36577.1 atub|OJI91404.1 aver|OJJ02576.1 awen|OJJ40313.1 azon|OJJ49173.1

Group3021: abra|OJJ73220.1 acar|OOG00458.1 acla|EAW06895.1 afis|EAW16512.1 afla|EED56161.1 afum|EAL89107.2 agla|OJJ82810.1 akaw|GAA87958.1 aluc|OJZ81207.1 anid|CBF75566.1 anig|EHA18522.1 aory|EIT76654.1 asyd|OJJ59708.1 ater|EAU36578.1 atub|OJI91403.1 aver|OJJ02577.1 awen|OJJ40312.1 azon|OJJ49172.1

Group3022: abra|OJJ73222.1 acar|OOG00460.1 acla|EAW06897.1 afis|EAW16514.1 afla|EED56159.1 afum|EAL89109.1 agla|OJJ82808.1 akaw|GAA87956.1 aluc|OJZ81205.1 anid|CBF75562.1 anig|EHA18520.1 aory|EIT76696.1 asyd|OJJ59706.1 ater|EAU36580.1 atub|OJI91401.1 aver|OJJ02579.1 awen|OJJ40309.1 azon|OJJ49170.1

Group3023: abra|OJJ73223.1 acar|OOG00461.1 acla|EAW06898.1 afis|EAW16515.1 afla|EED56158.1 afum|EAL89110.2 agla|OJJ82807.1 akaw|GAA87955.1 aluc|OJZ81204.1 anid|CBF75560.1 anig|EHA18519.1 aory|EIT76553.1 asyd|OJJ59705.1 ater|EAU36581.1 atub|OJI91400.1 aver|OJJ02580.1 awen|OJJ40308.1 azon|OJJ49169.1

Group3024: abra|OJJ73224.1 acar|OOG00462.1 acla|EAW06899.1 afis|EAW16516.1 afla|EED56157.1 afum|EAL89111.2 agla|OJJ82806.1 akaw|GAA87954.1 aluc|OJZ81203.1 anid|CBF75558.1 anig|EHA18518.1 aory|EIT76604.1 asyd|OJJ59704.1 ater|EAU36582.1 atub|OJI91398.1 aver|OJJ02582.1 awen|OJJ40307.1 azon|OJJ49168.1

Group3025: abra|OJJ73228.1 acar|OOG00466.1 acla|EAW06902.1 afis|EAW16519.1 afla|EED56152.1 afum|EAL89114.1 agla|OJJ82801.1 akaw|GAA87952.1 aluc|OJZ81200.1 anid|CBF75555.1 anig|EHA18515.1 aory|EIT76840.1 asyd|OJJ57431.1 ater|EAU36585.1 atub|OJI91395.1 aver|OJJ05749.1 awen|OJJ40303.1 azon|OJJ49165.1

Group3026: abra|OJJ73229.1 acar|OOG00468.1 acla|EAW06904.1 afis|EAW16521.1 afla|EED56150.1 afum|EAL89116.1 agla|OJJ82800.1 akaw|GAA87951.1 aluc|OJZ81199.1 anid|CBF75547.1 anig|EHA18514.1 aory|EIT76779.1 asyd|OJJ57441.1 ater|EAU36586.1 atub|OJI91394.1 aver|OJJ05737.1 awen|OJJ40302.1 azon|OJJ49164.1

Group3027: abra|OJJ73230.1 acar|OOG00469.1 acla|EAW06905.1 afis|EAW16522.1 afla|EED56149.1 afum|EAL89117.1 agla|OJJ82799.1 akaw|GAA87950.1 aluc|OJZ81198.1 anid|CBF75545.1 anig|EHA18513.1

aory|EIT76528.1 asyd|OJJ57442.1 ater|EAU36587.1 atub|OJI91393.1 aver|OJJ05736.1 awen|OJJ40301.1 azon|OJJ49163.1

Group3028: abra|OJJ73231.1 acar|OOG00470.1 acla|EAW06906.1 afis|EAW16523.1 afla|EED56148.1 afum|EAL89118.1 agla|OJJ82798.1 akaw|GAA87949.1 aluc|OJZ81197.1 anid|CBF75543.1 anig|EHA18512.1 aory|EIT76854.1 asyd|OJJ57443.1 ater|EAU36588.1 atub|OJI91392.1 aver|OJJ05735.1 awen|OJJ40300.1 azon|OJJ49162.1

Group3029: abra|OJJ73232.1 acar|OOG00471.1 acla|EAW06907.1 afis|EAW16524.1 afla|EED56147.1 afum|EAL89119.1 agla|OJJ82797.1 akaw|GAA87948.1 aluc|OJZ81196.1 anid|CBF75541.1 anig|EHA18511.1 aory|EIT76769.1 asyd|OJJ57446.1 ater|EAU36589.1 atub|OJI91391.1 aver|OJJ05734.1 awen|OJJ40299.1 azon|OJJ49161.1

Group3030: abra|OJJ73234.1 acar|OOG00472.1 acla|EAW06909.1 afis|EAW16526.1 afla|EED56145.1 afum|EAL89122.1 agla|OJJ82796.1 akaw|GAA87946.1 aluc|OJZ81194.1 anid|CBF75537.1 anig|EHA18509.1 aory|EIT76537.1 asyd|OJJ57447.1 ater|EAU36590.1 atub|OJI91389.1 aver|OJJ05732.1 awen|OJJ40297.1 azon|OJJ49159.1

Group3031: abra|OJJ73236.1 acar|OOG00473.1 acla|EAW06910.1 afis|EAW16527.1 afla|EED56144.1 afum|EAL89123.2 agla|OJJ82795.1 akaw|GAA87945.1 aluc|OJZ81193.1 anid|CBF75535.1 anig|EHA18508.1 aory|EIT76806.1 asyd|OJJ57448.1 ater|EAU36591.1 atub|OJI91388.1 aver|OJJ05731.1 awen|OJJ40296.1 azon|OJJ49158.1

Group3032: abra|OJJ73237.1 acar|OOG00474.1 acla|EAW06911.1 afis|EAW16528.1 afla|EED56143.1 afum|EAL89124.1 agla|OJJ82793.1 akaw|GAA87944.1 aluc|OJZ81191.1 anid|CBF75533.1 anig|EHA18507.1 aory|EIT76880.1 asyd|OJJ57449.1 ater|EAU36592.1 atub|OJI91387.1 aver|OJJ05730.1 awen|OJJ40295.1 azon|OJJ49157.1

Group3033: abra|OJJ73239.1 acar|OOG00476.1 acla|EAW06912.1 afis|EAW16529.1 afla|EED56141.1 afum|EAL89125.1 agla|OJJ82791.1 akaw|GAA87942.1 aluc|OJZ81189.1 anid|CBF75529.1 anig|EHA18505.1 aory|EIT76699.1 asyd|OJJ57450.1 ater|EAU36593.1 atub|OJI91385.1 aver|OJJ05728.1 awen|OJJ40293.1 azon|OJJ49154.1

Group3034: abra|OJJ73240.1 acar|OOG00477.1 acla|EAW06913.1 afis|EAW16530.1 afla|EED56138.1 afum|EAL89127.1 agla|OJJ82790.1 akaw|GAA87941.1 aluc|OJZ81188.1 anid|CBF81731.1 anig|EHA18504.1 aory|EIT76798.1 asyd|OJJ59592.1 ater|EAU36594.1 atub|OJI91384.1 aver|OJJ02677.1 awen|OJJ40291.1 azon|OJJ49153.1

Group3035: abra|OJJ73246.1 acar|OOG00485.1 acla|EAW06920.1 afis|EAW16537.1 afla|EED56132.1 afum|EAL89134.1 agla|OJJ82785.1 akaw|GAA87934.1 aluc|OJZ81181.1 anid|CBF81743.1 anig|EHA18499.1 aory|EIT76834.1 asyd|OJJ59598.1 ater|EAU36600.1 atub|OJI91378.1 aver|OJJ02670.1 awen|OJJ40285.1 azon|OJJ49146.1

Group3036: abra|OJJ73248.1 acar|OOG00488.1 acla|EAW06922.1 afis|EAW16539.1 afla|EED56130.1 afum|EAL89136.1 agla|OJJ82783.1 akaw|GAA87932.1 aluc|OJZ81179.1 anid|CBF81746.1 anig|EHA18497.1 aory|EIT76822.1 asyd|OJJ59600.1 ater|EAU36602.1 atub|OJI91375.1 aver|OJJ02668.1 awen|OJJ40282.1 azon|OJJ49144.1

Group3037: abra|OJJ73249.1 acar|OOG00491.1 acla|EAW06924.1 afis|EAW16541.1 afla|EED56127.1 afum|EAL89138.1 agla|OJJ82782.1 akaw|GAA87931.1 aluc|OJZ81178.1 anid|CBF81748.1 anig|EHA18496.1 aory|EIT76785.1 asyd|OJJ59601.1 ater|EAU36603.1 atub|OJI91374.1 aver|OJJ02667.1 awen|OJJ40279.1 azon|OJJ49142.1

Group3038: abra|OJJ73251.1 acar|OOG00495.1 acla|EAW06929.1 afis|EAW16544.1 afla|EED56121.1 afum|EAL89142.2 agla|OJJ82779.1 akaw|GAA87929.1 aluc|OJZ81176.1 anid|CBF81754.1 anig|EHA18494.1 aory|EIT76628.1 asyd|OJJ59604.1 ater|EAU36607.1 atub|OJI91372.1 aver|OJJ02665.1 awen|OJJ40276.1 azon|OJJ49139.1

Group3039: abra|OJJ73252.1 acar|OOG00494.1 acla|EAW06930.1 afis|EAW16545.1 afla|EED56120.1 afum|EAL89143.1 agla|OJJ82778.1 akaw|GAA87928.1 aluc|OJZ81175.1 anid|CBF81755.1 anig|EHA18493.1 aory|EIT76556.1 asyd|OJJ59605.1 ater|EAU36608.1 atub|OJI91371.1 aver|OJJ02664.1 awen|OJJ40275.1 azon|OJJ49138.1

Group3040: abra|OJJ73253.1 acar|OOG00496.1 acla|EAW06931.1 afis|EAW16546.1 afum|EAL89144.1 agla|OJJ82777.1 akaw|GAA87927.1 aluc|OJZ81174.1 anid|CBF81757.1 anig|EHA18492.1 aory|EIT76862.1 asyd|OJJ59606.1 ater|EAU36609.1 atub|OJI91370.1 aver|OJJ02663.1 awen|OJJ40271.1 azon|OJJ49137.1 afla|EED56118.1

Group3041: abra|OJJ73256.1 acar|OOG00499.1 acla|EAW06933.1 afis|EAW16548.1 afla|EED56116.1 afum|EAL89146.1 agla|OJJ82775.1 akaw|GAA87925.1 aluc|OJZ81172.1 anid|CBF81761.1 anig|EHA18490.1 aory|EIT76624.1 asyd|OJJ59607.1 ater|EAU36611.1 atub|OJI91367.1 aver|OJJ02662.1 awen|OJJ40269.1 azon|OJJ49133.1

Group3042: abra|OJJ73257.1 acar|OOG00498.1 acla|EAW06934.1 afis|EAW16549.1 afla|EED56115.1 afum|EAL89147.1 agla|OJJ82774.1 akaw|GAA87924.1 aluc|OJZ81171.1 anid|CBF81763.1 anig|EHA18489.1 aory|EIT76873.1 asyd|OJJ59608.1 ater|EAU36612.1 atub|OJI91366.1 aver|OJJ02661.1 awen|OJJ40268.1 azon|OJJ49132.1

Group3043: abra|OJJ73258.1 acar|OOG00500.1 acla|EAW06935.1 afis|EAW16550.1 afla|EED56112.1 afum|EAL89148.1 agla|OJJ82773.1 akaw|GAA87923.1 aluc|OJZ81170.1 anid|CBF81765.1 anig|EHA18488.1 aory|EIT76657.1 ater|EAU36613.1 atub|OJI91365.1 awen|OJJ40267.1 azon|OJJ49131.1 asyd|OJJ59609.1 aver|OJJ02660.1

Group3044: abra|OJJ73259.1 acar|OOG00501.1 acla|EAW06936.1 afis|EAW16551.1 afla|EED56111.1 afum|EAL89149.1 agla|OJJ82772.1 akaw|GAA87922.1 aluc|OJZ81169.1 anid|CBF81766.1 anig|EHA18487.1 aory|EIT76585.1 asyd|OJJ59610.1 ater|EAU36617.1 atub|OJI91364.1 aver|OJJ02659.1 awen|OJJ40266.1 azon|OJJ49130.1

Group3045: abra|OJJ73262.1 acar|OOG00502.1 acla|EAW06939.1 afis|EAW16554.1 afla|EED56108.1 afum|EAL89152.1 agla|OJJ82769.1 akaw|GAA87920.1 aluc|OJZ81167.1 anid|CBF81795.1 anig|EHA18484.1 aory|EIT76711.1 asyd|OJJ59637.1 ater|EAU36619.1 atub|OJI91362.1 aver|OJJ02644.1 awen|OJJ40264.1 azon|OJJ49127.1

Group3046: abra|OJJ73263.1 acar|OOG00503.1 acla|EAW06940.1 afis|EAW16555.1 afla|EED56107.1 afum|EAL89153.1 agla|OJJ82768.1 akaw|GAA87919.1 aluc|OJZ81166.1 anid|CBF81796.1 anig|EHA18483.1 aory|EIT76566.1 asyd|OJJ59638.1 ater|EAU36620.1 atub|OJI91361.1 aver|OJJ02643.1 awen|OJJ40263.1 azon|OJJ49126.1

Group3047: abra|OJJ73264.1 acar|OOG00504.1 acla|EAW06941.1 afis|EAW16556.1 afla|EED56106.1 afum|EAL89154.1 agla|OJJ82767.1 akaw|GAA87918.1 aluc|OJZ81165.1 anid|CBF81798.1 anig|EHA18482.1 aory|EIT76736.1 asyd|OJJ59639.1 ater|EAU36621.1 atub|OJI91360.1 aver|OJJ02642.1 awen|OJJ40262.1 azon|OJJ49125.1

Group3048: abra|OJJ73265.1 acar|OOG00505.1 acla|EAW06942.1 afis|EAW16557.1 afla|EED56105.1 afum|EAL89155.1 agla|OJJ82766.1 akaw|GAA87917.1 aluc|OJZ81164.1 anid|CBF81802.1 anig|EHA18481.1 aory|EIT76677.1 asyd|OJJ59642.1 ater|EAU36624.1 atub|OJI91359.1 aver|OJJ02640.1 awen|OJJ40260.1 azon|OJJ49122.1

Group3049: abra|OJJ73266.1 acar|OOG00506.1 acla|EAW06943.1 afis|EAW16558.1 afla|EED56104.1 afum|EAL89156.1 agla|OJJ82765.1 akaw|GAA87916.1 aluc|OJZ81163.1 anid|CBF81800.1 anig|EHA18480.1 aory|EIT76573.1 asyd|OJJ59640.1 ater|EAU36625.1 atub|OJI91358.1 aver|OJJ02641.1 awen|OJJ40259.1 azon|OJJ49123.1

Group3050: abra|OJJ73269.1 acar|OOG00509.1 acla|EAW06946.1 afis|EAW16561.1 afla|EED56101.1 afum|EAL89159.1 agla|OJJ82762.1 akaw|GAA87913.1 aluc|OJZ81160.1 anid|CBF81808.1 anig|EHA18477.1 aory|EIT76636.1 asyd|OJJ59645.1 ater|EAU36628.1 atub|OJI91355.1 aver|OJJ02637.1 awen|OJJ40256.1 azon|OJJ49120.1

Group3051: abra|OJJ73275.1 acar|OOG00514.1 acla|EAW06950.1 afis|EAW16565.1 afla|EED56098.1 afum|EAL89162.1 agla|OJJ82756.1 akaw|GAA92730.1 aluc|OJZ88774.1 anid|CBF81817.1 anig|EHA18468.1 aory|EIT76722.1 asyd|OJJ59651.1 ater|EAU36634.1 atub|OJI91345.1 aver|OJJ02631.1 awen|OJJ40253.1 azon|OJJ49115.1

Group3052: abra|OJJ73278.1 acar|OOG00515.1 acla|EAW06954.1 afis|EAW16569.1 afla|EED56095.1 afum|EAL89166.1 agla|OJJ82751.1 akaw|GAA92727.1 aluc|OJZ88771.1 anid|CBF81822.1 anig|EHA18467.1 aory|EIT82359.1 asyd|OJJ59654.1 ater|EAU36637.1 atub|OJI91344.1 aver|OJJ02628.1 awen|OJJ40250.1 azon|OJJ41980.1

Group3053: abra|OJJ73281.1 acar|OOG00517.1 acla|EAW06956.1 afis|EAW16571.1 afla|EED56093.1 afum|EAL89168.1 agla|OJJ82703.1 akaw|GAA92724.1 aluc|OJZ88768.1 anid|CBF81826.1 anig|EHA18464.1 aory|EIT76754.1 asyd|OJJ59657.1 ater|EAU36639.1 atub|OJI91341.1 aver|OJJ02625.1 awen|OJJ40247.1 azon|OJJ49108.1

Group3054: abra|OJJ73283.1 acar|OOG00519.1 acla|EAW06958.1 afis|EAW16573.1 afla|EED56091.1 afum|EAL89170.2 agla|OJJ82705.1 akaw|GAA92722.1 aluc|OJZ88766.1 anid|CBF81828.1 anig|EHA18462.1 aory|EIT76851.1 asyd|OJJ59659.1 ater|EAU36641.1 atub|OJI91339.1 aver|OJJ02623.1 awen|OJJ40245.1 azon|OJJ49106.1

Group3055: abra|OJJ73287.1 acar|OOG00522.1 acla|EAW06962.1 afis|EAW16577.1 afla|EED56087.1 afum|EAL89173.1 agla|OJJ82700.1 akaw|GAA92718.1 aluc|OJZ88763.1 anid|CBF81876.1 anig|EHA18459.1 aory|EIT76884.1 asyd|OJJ59665.1 ater|EAU36645.1 atub|OJI91335.1 aver|OJJ02618.1 awen|OJJ40240.1 azon|OJJ49103.1

Group3056: abra|OJJ73331.1 acar|OOF91495.1 agla|OJJ78824.1 anig|EHA18400.1 abra|OJJ78190.1 acar|OOF98667.1 acla|EAW12173.1 akaw|GAA85839.1 aluc|OJZ91261.1 anig|EHA19949.1 asyd|OJJ52476.1

ater|EAU36116.1 atub|OJI83250.1 aver|OJJ03936.1 awen|OJJ39598.1 azon|OJJ42251.1 afla|EED54616.1 aory|EIT78824.1

Group3057: abra|OJJ73338.1 acar|OOG00523.1 acla|EAW06963.1 afis|EAW16578.1 afla|EED56086.1 afum|EAL89174.1 agla|OJJ82699.1 akaw|GAA83146.1 aluc|OJZ88703.1 anid|CBF81878.1 anig|EHA18387.1 aory|EIT76746.1 asyd|OJJ59666.1 ater|EAU36646.1 atub|OJI91284.1 aver|OJJ02617.1 awen|OJJ40239.1 azon|OJJ49102.1

Group3058: abra|OJJ73339.1 acar|OOG00524.1 acla|EAW06964.1 afis|EAW16579.1 afla|EED56085.1 afum|EAL89175.1 agla|OJJ82698.1 akaw|GAA83145.1 aluc|OJZ88702.1 anid|CBF81880.1 anig|EHA18386.1 aory|EIT76894.1 asyd|OJJ59667.1 ater|EAU36647.1 atub|OJI91283.1 aver|OJJ02616.1 awen|OJJ40238.1 azon|OJJ49101.1

Group3059: abra|OJJ73340.1 acar|OOG00527.1 acla|EAW06965.1 afis|EAW16580.1 afla|EED56082.1 afum|EAL89176.1 agla|OJJ82695.1 akaw|GAA83142.1 aluc|OJZ88699.1 anid|CBF81883.1 anig|EHA18379.1 aory|EIT76744.1 asyd|OJJ59669.1 ater|EAU36649.1 atub|OJI91280.1 aver|OJJ02614.1 awen|OJJ40234.1 azon|OJJ49100.1

Group3060: abra|OJJ73343.1 acar|OOG00553.1 acla|EAW06970.1 afis|EAW16584.1 afla|EED56081.1 afum|EAL89179.1 agla|OJJ82694.1 akaw|GAA83139.1 aluc|OJZ88696.1 anid|CBF81885.1 anig|EHA18376.1 aory|EIT76749.1 asyd|OJJ59671.1 ater|EAU36661.1 atub|OJI91277.1 aver|OJJ02613.1 awen|OJJ40232.1 azon|OJJ49096.1

Group3061: abra|OJJ73344.1 acar|OOG00554.1 acla|EAW06971.1 afis|EAW16585.1 afla|EED56080.1 afum|EAL89180.1 agla|OJJ82693.1 akaw|GAA83138.1 aluc|OJZ88695.1 anid|CBF81887.1 anig|EHA18375.1 aory|EIT76713.1 asyd|OJJ59672.1 ater|EAU36662.1 atub|OJI91275.1 aver|OJJ02612.1 awen|OJJ40231.1 azon|OJJ49095.1

Group3062: abra|OJJ73345.1 acar|OOG00555.1 acla|EAW06972.1 afis|EAW16586.1 afla|EED56079.1 afum|EAL89181.2 agla|OJJ82692.1 akaw|GAA83137.1 aluc|OJZ88694.1 anid|CBF81889.1 anig|EHA18374.1 aory|EIT76590.1 asyd|OJJ59673.1 ater|EAU36663.1 atub|OJI91274.1 aver|OJJ02611.1 awen|OJJ40230.1 azon|OJJ49094.1

Group3063: abra|OJJ73359.1 acar|OOG00567.1 acla|EAW06979.1 afis|EAW16593.1 afla|EED45008.1 afum|EAL89189.1 agla|OJJ82660.1 akaw|GAA83125.1 aluc|OJZ88681.1 anid|CBF81897.1 anig|EHA18362.1 aory|EIT73639.1 asyd|OJJ59682.1 ater|EAU36673.1 atub|OJI91262.1 aver|OJJ02606.1 awen|OJJ40214.1 azon|OJJ49080.1

Group3064: abra|OJJ73361.1 acar|OOG00568.1 acla|EAW06981.1 afis|EAW16595.1 afla|EED45014.1 afum|EAL89191.1 agla|OJJ82659.1 akaw|GAA83123.1 aluc|OJZ88679.1 anid|CBF81901.1 anig|EHA18360.1 aory|EIT73637.1 asyd|OJJ59685.1 ater|EAU36674.1 atub|OJI91260.1 aver|OJJ02602.1 awen|OJJ40213.1 azon|OJJ49079.1

Group3065: abra|OJJ73362.1 acar|OOG00569.1 acla|EAW06982.1 afis|EAW16596.1 afla|EED45015.1 afum|EAL89192.1 agla|OJJ82658.1 akaw|GAA83122.1 aluc|OJZ88678.1 anid|CBF81902.1 anig|EHA18359.1 aory|EIT73689.1 asyd|OJJ59686.1 ater|EAU36675.1 atub|OJI91259.1 aver|OJJ02601.1 awen|OJJ40212.1 azon|OJJ49078.1

Group3066: abra|OJJ73365.1 acar|OOG00572.1 acla|EAW06985.1 afis|EAW16599.1 afla|EED45018.1 afum|EAL89195.1 agla|OJJ82655.1 akaw|GAA83119.1 aluc|OJZ88675.1 anid|CBF81905.1 anig|EHA18357.1 aory|EIT73617.1 asyd|OJJ59688.1 ater|EAU36680.1 atub|OJI91256.1 aver|OJJ02599.1 awen|OJJ40209.1 azon|OJJ49075.1

Group3067: abra|OJJ73368.1 acar|OOG00574.1 acla|EAW06988.1 afis|EAW16602.1 afla|EED45021.1 afum|EAL89198.1 agla|OJJ82650.1 akaw|GAA83116.1 aluc|OJZ88672.1 anid|CBF81909.1 anig|EHA18354.1 aory|EIT73783.1 asyd|OJJ59690.1 ater|EAU36683.1 atub|OJI91253.1 aver|OJJ02597.1 awen|OJJ40204.1 azon|OJJ49071.1

Group3068: abra|OJJ73378.1 acar|OOF90413.1 acla|EAW06994.1 afis|EAW16609.1 afla|EED45027.1 afum|EAL89205.1 agla|OJJ82638.1 akaw|GAA83111.1 aluc|OJZ88665.1 anid|CBF81965.1 anig|EHA18350.1 aory|EIT73642.1 asyd|OJJ57355.1 ater|EAU36693.1 atub|OJI91247.1 aver|OJJ05810.1 awen|OJJ40198.1 azon|OJJ49069.1

Group3069: abra|OJJ73382.1 acar|OOG00581.1 acla|EAW06997.1 afis|EAW16611.1 afla|EED45000.1 afum|EAL89207.1 agla|OJJ82634.1 akaw|GAA83107.1 aluc|OJZ88656.1 anid|CBF81963.1 anig|EHA18347.1 aory|EIT73611.1 asyd|OJJ57358.1 ater|EAU36696.1 atub|OJI91242.1 aver|OJJ05808.1 awen|OJJ40193.1 azon|OJJ49061.1

Group3070: abra|OJJ73383.1 acar|OOG00582.1 acla|EAW06998.1 afis|EAW16613.1 afla|EED44999.1 afum|EAL89223.1 agla|OJJ82633.1 akaw|GAA83106.1 aluc|OJZ88654.1 anid|CBF81961.1 anig|EHA18346.1 aory|EIT73720.1 asyd|OJJ57359.1 ater|EAU36697.1 atub|OJI91241.1 aver|OJJ05807.1 awen|OJJ40192.1 azon|OJJ49060.1

Group3071: abra|OJJ73402.1 acar|OOG00600.1 acla|EAW07005.1 afis|EAW16622.1 afla|EED44970.1 afum|EAL89215.1 agla|OJJ82622.1 akaw|GAA83088.1 aluc|OJZ88635.1 anid|CBF81987.1 anig|EHA18325.1 aory|EIT73499.1 asyd|OJJ56156.1 ater|EAU36736.1 atub|OJI91219.1 aver|OJJ05096.1 awen|OJJ40086.1 azon|OJJ49496.1

Group3072: abra|OJJ73424.1 acar|OOG00628.1 acla|EAW07034.1 afis|EAW16660.1 afla|EED44893.1 afum|EAL89264.1 agla|OJJ78709.1 akaw|GAA83068.1 aluc|OJZ88619.1 anid|CBF82075.1 anig|EHA18310.1 aory|EIT83240.1 asyd|OJJ54250.1 ater|EAU39583.1 atub|OJI91204.1 aver|OJJ05204.1 awen|OJJ40053.1 azon|OJJ50109.1

Group3073: abra|OJJ73425.1 acar|OOG00629.1 acla|EAW07035.1 afis|EAW16661.1 afla|EED44894.1 afum|EAL89265.1 agla|OJJ78708.1 akaw|GAA83067.1 aluc|OJZ88618.1 anid|CBF82076.1 anig|EHA18309.1 aory|EIT83251.1 asyd|OJJ54251.1 ater|EAU39584.1 atub|OJI91203.1 aver|OJJ05205.1 awen|OJJ40052.1 azon|OJJ50116.1

Group3074: abra|OJJ73432.1 acar|OOG00639.1 acla|EAW06771.1 afis|EAW16322.1 afla|EED53560.1 afum|EAL88945.1 agla|OJJ88284.1 akaw|GAA83060.1 aluc|OJZ88609.1 anid|CBF89157.1 anig|EHA18302.1 aory|EIT75216.1 asyd|OJJ62563.1 ater|EAU34135.1 atub|OJI91194.1 aver|OJI96687.1 awen|OJJ37722.1 azon|OJJ44981.1

Group3075: abra|OJJ73437.1 acar|OOG00643.1 acla|EAW06775.1 afis|EAW16326.1 afla|EED53555.1 afum|EAL88950.1 agla|OJJ88266.1 akaw|GAA83056.1 aluc|OJZ88604.1 anid|CBF89164.1 anig|EHA18297.1 aory|EIT75038.1 asyd|OJJ62546.1 ater|EAU34131.1 atub|OJI91188.1 aver|OJI96672.1 awen|OJJ37726.1 azon|OJJ44986.1

Group3076: abra|OJJ73440.1 acar|OOG00646.1 acla|EAW06772.1 afis|EAW16323.1 afla|EED53559.1 afum|EAL88946.1 agla|OJJ88285.1 akaw|GAA83053.1 aluc|OJZ88600.1 anid|CBF89158.1 anig|EHA18294.1 aory|EIT74917.1 asyd|OJJ62560.1 ater|EAU34134.1 atub|OJI91185.1 aver|OJI96685.1 awen|OJJ37723.1 azon|OJJ44982.1

Group3077: abra|OJJ73442.1 acar|OOG00648.1 acla|EAW06777.1 afis|EAW16328.1 afla|EED53553.1 afum|EAL88952.1 agla|OJJ88264.1 akaw|GAA83052.1 aluc|OJZ88598.1 anid|CBF89168.1 anig|EHA18292.1 aory|EIT74949.1 asyd|OJJ62543.1 ater|EAU34129.1 atub|OJI91183.1 aver|OJI96670.1 awen|OJJ37729.1 azon|OJJ44988.1

Group3078: abra|OJJ73451.1 acar|OOG00654.1 acla|EAW06782.1 afis|EAW16332.1 afla|EED53545.1 afum|EAL88957.1 agla|OJJ88260.1 akaw|GAA83045.1 aluc|OJZ88590.1 anid|CBF89183.1 anig|EHA18285.1 aory|EIT75091.1 asyd|OJJ62537.1 ater|EAU34126.1 atub|OJI91176.1 aver|OJI96663.1 awen|OJJ37733.1 azon|OJJ44990.1

Group3079: abra|OJJ73456.1 acar|OOG00656.1 acla|EAW06783.1 afis|EAW16333.1 afla|EED53543.1 afum|EAL88958.1 agla|OJJ88259.1 akaw|GAA83042.1 aluc|OJZ88587.1 anid|CBF89188.1 anig|EHA18281.1 aory|EIT75083.1 asyd|OJJ62536.1 ater|EAU34125.1 atub|OJI91173.1 aver|OJI96661.1 awen|OJJ37734.1 azon|OJJ44993.1

Group3080: abra|OJJ73458.1 acar|OOG00659.1 acla|EAW06785.1 afis|EAW16335.1 afla|EED53541.1 afum|EAL88960.1 agla|OJJ88251.1 akaw|GAA83040.1 aluc|OJZ88585.1 anid|CBF89192.1 anig|EHA18280.1 aory|EIT74882.1 asyd|OJJ62534.1 ater|EAU34123.1 atub|OJI91171.1 aver|OJI96659.1 awen|OJJ37737.1 azon|OJJ44997.1

Group3081: abra|OJJ73460.1 acar|OOG00661.1 acla|EAW06787.1 afis|EAW16336.1 afla|EED53539.1 afum|EAL88962.1 agla|OJJ88254.1 akaw|GAA83038.1 aluc|OJZ88583.1 anid|CBF89196.1 anig|EHA18278.1 aory|EIT75013.1 asyd|OJJ62532.1 ater|EAU34121.1 atub|OJI91169.1 aver|OJI96657.1 awen|OJJ37739.1 azon|OJJ45000.1

Group3082: abra|OJJ73463.1 acar|OOG00664.1 acla|EAW06789.1 afis|EAW16339.1 afla|EED53536.1 afum|EAL88964.1 agla|OJJ88256.1 akaw|GAA83035.1 aluc|OJZ88581.1 anid|CBF89199.1 anig|EHA18277.1 aory|EIT74845.1 asyd|OJJ62530.1 ater|EAU34119.1 atub|OJI91167.1 aver|OJI96655.1 awen|OJJ37741.1 azon|OJJ45002.1

Group3083: abra|OJJ73464.1 acar|OOG00665.1 acla|EAW06790.1 afis|EAW16340.1 afla|EED53535.1 afum|EAL88965.1 agla|OJJ88257.1 akaw|GAA83034.1 aluc|OJZ88580.1 anid|CBF89201.1 anig|EHA18276.1 aory|EIT75162.1 asyd|OJJ62529.1 ater|EAU34117.1 atub|OJI91166.1 aver|OJI96654.1 awen|OJJ37742.1 azon|OJJ45003.1

Group3084: abra|OJJ73465.1 acar|OOG00666.1 acla|EAW06792.1 afis|EAW16341.1 afla|EED53534.1 afum|EAL88966.1 agla|OJJ88247.1 akaw|GAA83033.1 aluc|OJZ88579.1 anid|CBF89203.1 anig|EHA18275.1 aory|EIT75067.1 asyd|OJJ62528.1 ater|EAU34116.1 atub|OJI91165.1 aver|OJI96651.1 awen|OJJ37743.1 azon|OJJ45004.1

Group3085: abra|OJJ73466.1 acar|OOG00667.1 acla|EAW06793.1 afis|EAW16342.1 afla|EED53533.1 afum|EAL88967.1 agla|OJJ88246.1 akaw|GAA83032.1 aluc|OJZ88578.1 anid|CBF89205.1 anig|EHA18274.1

aory|EIT75133.1 asyd|OJJ62527.1 ater|EAU34115.1 atub|OJI91164.1 aver|OJI96650.1 awen|OJJ37744.1 azon|OJJ45005.1

Group3086: abra|OJJ73467.1 acar|OOG00668.1 acla|EAW06794.1 afis|EAW16343.1 afla|EED53532.1 afum|EAL88968.1 agla|OJJ88245.1 akaw|GAA83031.1 aluc|OJZ88577.1 anid|CBF89206.1 anig|EHA18273.1 aory|EIT74961.1 asyd|OJJ62526.1 ater|EAU34114.1 atub|OJI91163.1 aver|OJI96649.1 awen|OJJ37745.1 azon|OJJ45006.1

Group3087: abra|OJJ73470.1 acar|OOG00669.1 acla|EAW06795.1 afis|EAW16344.1 afla|EED53526.1 afum|EAL88969.1 agla|OJJ88244.1 akaw|GAA83030.1 aluc|OJZ88576.1 anid|CBF89217.1 anig|EHA18272.1 aory|EIT74890.1 asyd|OJJ62524.1 ater|EAU34113.1 atub|OJI91162.1 aver|OJI96645.1 awen|OJJ37747.1 azon|OJJ45008.1

Group3088: abra|OJJ73474.1 acar|OOG00670.1 acla|EAW06799.1 afis|EAW16348.1 afla|EED53523.1 afum|EAL88973.1 agla|OJJ88239.1 akaw|GAA83027.1 aluc|OJZ88573.1 anid|CBF89210.1 anig|EHA18271.1 aory|EIT75044.1 asyd|OJJ62522.1 ater|EAU34111.1 atub|OJI91159.1 aver|OJI96641.1 awen|OJJ37751.1 azon|OJJ45010.1

Group3089: abra|OJJ73476.1 acar|OOG00673.1 acla|EAW06802.1 afis|EAW16351.1 afla|EED53521.1 afum|EAL88976.1 agla|OJJ88249.1 akaw|GAA83025.1 aluc|OJZ88571.1 anid|CBF89221.1 anig|EHA18269.1 aory|EIT75100.1 asyd|OJJ62520.1 ater|EAU34110.1 atub|OJI91157.1 aver|OJI96639.1 awen|OJJ37754.1 azon|OJJ45013.1

Group3090: abra|OJJ73484.1 acar|OOG00678.1 acla|EAW06808.1 afis|EAW16357.1 afla|EED53513.1 afum|EAL88982.1 agla|OJJ88236.1 akaw|GAA87433.1 aluc|OJZ88564.1 anid|CBF89232.1 anig|EHA18261.1 aory|EIT75027.1 asyd|OJJ62514.1 ater|EAU34102.1 atub|OJI91150.1 aver|OJI96633.1 awen|OJJ37761.1 azon|OJJ45061.1

Group3091: abra|OJJ73485.1 acar|OOG00680.1 acla|EAW06809.1 afis|EAW16358.1 afla|EED53512.1 afum|EAL88983.1 agla|OJJ88235.1 akaw|GAA87434.1 aluc|OJZ88563.1 anid|CBF89234.1 anig|EHA18260.1 aory|EIT75048.1 asyd|OJJ62513.1 ater|EAU34101.1 atub|OJI91149.1 aver|OJI96632.1 awen|OJJ37762.1 azon|OJJ45062.1

Group3092: abra|OJJ73486.1 acar|OOG00679.1 acla|EAW06810.1 afis|EAW16359.1 afla|EED53510.1 afum|EAL88984.1 agla|OJJ88233.1 akaw|GAA87435.1 aluc|OJZ88562.1 anid|CBF89236.1 anig|EHA18259.1 aory|EIT75112.1 asyd|OJJ62512.1 ater|EAU34100.1 atub|OJI91147.1 aver|OJI96631.1 awen|OJJ37763.1 azon|OJJ45063.1

Group3093: abra|OJJ73487.1 acar|OOG00681.1 acla|EAW06811.1 afis|EAW16360.1 afla|EED53508.1 afum|EAL88985.1 agla|OJJ88232.1 akaw|GAA87436.1 aluc|OJZ88561.1 anid|CBF89237.1 anig|EHA18258.1 aory|EIT75230.1 asyd|OJJ62510.1 ater|EAU34092.1 atub|OJI91146.1 aver|OJI96630.1 awen|OJJ37764.1 azon|OJJ45064.1

Group3094: abra|OJJ73488.1 acar|OOG00682.1 acla|EAW06812.1 afis|EAW16361.1 afla|EED53507.1 afum|EAL88986.1 agla|OJJ88231.1 akaw|GAA87437.1 aluc|OJZ88560.1 anid|CBF89239.1 anig|EHA18257.1 aory|EIT75024.1 asyd|OJJ62509.1 ater|EAU34091.1 atub|OJI91145.1 aver|OJI96629.1 awen|OJJ37765.1 azon|OJJ45065.1

Group3095: abra|OJJ73496.1 acar|OOG00694.1 acla|EAW07430.1 afis|EAW22456.1 afla|EED54488.1 afum|EAL90735.1 agla|OJJ87568.1 akaw|GAA87445.1 aluc|OJZ88552.1 anid|CBF88921.1 anig|EHA18250.1 aory|EIT78863.1 asyd|OJJ62846.1 ater|EAU39249.1 atub|OJI91137.1 aver|OJI96890.1 awen|OJJ35081.1 azon|OJJ50026.1

Group3096: abra|OJJ73497.1 acar|OOG00693.1 acla|EAW07431.1 afis|EAW22457.1 afla|EED54487.1 afum|EAL90734.1 agla|OJJ87567.1 akaw|GAA87446.1 aluc|OJZ88551.1 anid|CBF88920.1 anig|EHA18249.1 aory|EIT78747.1 asyd|OJJ62845.1 ater|EAU39248.1 atub|OJI91136.1 aver|OJI96889.1 awen|OJJ35082.1 azon|OJJ50025.1

Group3097: abra|OJJ73507.1 acar|OOG00697.1 acla|EAW07440.1 afis|EAW22467.1 afla|EED54499.1 afum|EAL90724.1 agla|OJJ87557.1 akaw|GAA87455.1 aluc|OJZ88540.1 anid|CBF88903.1 anig|EHA18241.1 aory|EIT78913.1 asyd|OJJ62837.1 ater|EAU39241.1 atub|OJI91125.1 aver|OJI96880.1 awen|OJJ35093.1 azon|OJJ50016.1

Group3098: abra|OJJ73508.1 acar|OOG00698.1 acla|EAW07441.1 afis|EAW22468.1 afla|EED54500.1 afum|EAL90723.1 agla|OJJ87556.1 akaw|GAA87456.1 aluc|OJZ88539.1 anid|CBF88902.1 anig|EHA18240.1 aory|EIT78813.1 asyd|OJJ62836.1 ater|EAU39240.1 atub|OJI91124.1 aver|OJI96879.1 awen|OJJ35094.1 azon|OJJ50015.1

Group3099: abra|OJJ73510.1 acar|OOG00699.1 acla|EAW07443.1 afis|EAW22470.1 afla|EED54502.1 afum|EAL90721.1 agla|OJJ87554.1 akaw|GAA87458.1 aluc|OJZ88537.1 anid|CBF88900.1 anig|EHA18238.1 aory|EIT78910.1 asyd|OJJ62834.1 ater|EAU39238.1 atub|OJI91122.1 aver|OJI96877.1 awen|OJJ35096.1 azon|OJJ50013.1

Group3100: abra|OJJ73512.1 acar|OOG00701.1 acla|EAW07445.1 afis|EAW22472.1 afla|EED54504.1 afum|EAL90719.1 agla|OJJ87552.1 akaw|GAA87460.1 aluc|OJZ88535.1 anid|CBF88896.1 anig|EHA18236.1 aory|EIT78949.1 asyd|OJJ62832.1 ater|EAU39236.1 atub|OJI91120.1 aver|OJI96875.1 awen|OJJ35098.1 azon|OJJ50011.1

Group3101: abra|OJJ73513.1 acar|OOG00700.1 acla|EAW07447.1 afis|EAW22474.1 afla|EED54506.1 afum|EAL90717.1 agla|OJJ87550.1 akaw|GAA87462.1 aluc|OJZ88534.1 anid|CBF88892.1 anig|EHA18235.1 aory|EIT78850.1 asyd|OJJ62831.1 ater|EAU39235.1 atub|OJI91119.1 aver|OJI96874.1 awen|OJJ35099.1 azon|OJJ50010.1

Group3102: abra|OJJ73516.1 acar|OOG00706.1 acla|EAW07451.1 afis|EAW22483.1 afla|EED54509.1 afum|EAL90711.1 agla|OJJ87543.1 akaw|GAA87465.1 aluc|OJZ88531.1 anid|CBF88887.1 anig|EHA18232.1 aory|EIT78939.1 asyd|OJJ62826.1 ater|EAU39232.1 atub|OJI91116.1 aver|OJI96871.1 awen|OJJ35102.1 azon|OJJ50006.1

Group3103: abra|OJJ73529.1 acar|OOG00710.1 acla|EAW07454.1 afis|EAW22486.1 afla|EED54512.1 afum|EAL90707.1 agla|OJJ80863.1 akaw|GAA88617.1 aluc|OJZ88526.1 anid|CBF88942.1 anig|EHA18228.1 aory|EIT78750.1 asyd|OJJ62815.1 ater|EAU39228.1 atub|OJI91111.1 aver|OJI96859.1 awen|OJJ35105.1 azon|OJJ49989.1

Group3104: abra|OJJ73534.1 acar|OOG00712.1 acla|EAW07456.1 afis|EAW22487.1 afla|EED54516.1 afum|EAL90706.1 agla|OJJ80860.1 akaw|GAA88616.1 aluc|OJZ88525.1 anid|CBF88944.1 anig|EHA18227.1 aory|EIT78867.1 asyd|OJJ62814.1 ater|EAU39227.1 atub|OJI91110.1 aver|OJI96858.1 awen|OJJ35108.1 azon|OJJ49986.1

Group3105: abra|OJJ73537.1 acar|OOG00713.1 acla|EAW07459.1 afis|EAW22490.1 afla|EED54519.1 afum|EAL90703.1 agla|OJJ80856.1 akaw|GAA88613.1 aluc|OJZ88522.1 anid|CBF88950.1 anig|EHA18225.1 aory|EIT78823.1 asyd|OJJ62811.1 ater|EAU39225.1 atub|OJI91107.1 aver|OJI96857.1 awen|OJJ35110.1 azon|OJJ49983.1

Group3106: abra|OJJ73539.1 acar|OOG00715.1 acla|EAW07460.1 afis|EAW22491.1 afla|EED54520.1 afum|EAL90702.1 agla|OJJ80854.1 akaw|GAA88612.1 aluc|OJZ88520.1 anid|CBF88951.1 anig|EHA18224.1 aory|EIT78935.1 asyd|OJJ62809.1 ater|EAU39224.1 atub|OJI91105.1 aver|OJI96855.1 awen|OJJ35112.1 azon|OJJ49981.1

Group3107: abra|OJJ73541.1 acar|OOG00716.1 acla|EAW07461.1 afis|EAW22492.1 afla|EED54521.1 afum|EAL90701.1 agla|OJJ80853.1 akaw|GAA88610.1 aluc|OJZ88518.1 anid|CBF88953.1 anig|EHA18223.1 aory|EIT78920.1 asyd|OJJ62808.1 ater|EAU39223.1 atub|OJI91103.1 aver|OJI96854.1 awen|OJJ35113.1 azon|OJJ49980.1

Group3108: abra|OJJ73547.1 acar|OOG00721.1 acla|EAW07466.1 afis|EAW22499.1 afla|EED54526.1 afum|EAL90694.1 agla|OJJ80847.1 akaw|GAA88605.1 aluc|OJZ88513.1 anid|CBF88960.1 anig|EHA18219.1 aory|EIT78800.1 asyd|OJJ62804.1 ater|EAU39219.1 atub|OJI91098.1 aver|OJI96850.1 awen|OJJ35122.1 azon|OJJ49976.1

Group3109: abra|OJJ73549.1 acar|OOG00723.1 acla|EAW07468.1 afis|EAW22501.1 afla|EED54529.1 afum|EAL90692.1 agla|OJJ80851.1 akaw|GAA88603.1 aluc|OJZ88511.1 anid|CBF88962.1 anig|EHA18217.1 aory|EIT78773.1 asyd|OJJ62799.1 ater|EAU39217.1 atub|OJI91096.1 aver|OJI96846.1 awen|OJJ35124.1 azon|OJJ49974.1

Group3110: abra|OJJ73550.1 acar|OOG00724.1 acla|EAW07469.1 afis|EAW22502.1 afla|EED54530.1 afum|EAL90691.1 agla|OJJ80852.1 akaw|GAA88602.1 aluc|OJZ88510.1 anid|CBF88965.1 anig|EHA18216.1 aory|EIT78738.1 asyd|OJJ62798.1 ater|EAU39216.1 atub|OJI91095.1 aver|OJI96845.1 awen|OJJ35125.1 azon|OJJ49973.1

Group3111: abra|OJJ73555.1 acar|OOG00726.1 acla|EAW07473.1 afis|EAW22505.1 afla|EED54533.1 afum|EAL90688.1 agla|OJJ87536.1 akaw|GAA88598.1 aluc|OJZ88506.1 anid|CBF88970.1 anig|EHA18213.1 aory|EIT78812.1 asyd|OJJ62795.1 ater|EAU39210.1 atub|OJI91092.1 aver|OJI96842.1 awen|OJJ35130.1 azon|OJJ49969.1

Group3112: abra|OJJ73556.1 acar|OOG00728.1 acla|EAW07474.1 afis|EAW22506.1 afla|EED54534.1 afum|EAL90687.1 agla|OJJ87537.1 akaw|GAA88597.1 aluc|OJZ88505.1 anid|CBF88972.1 anig|EHA18212.1 aory|EIT78875.1 asyd|OJJ62794.1 ater|EAU39209.1 atub|OJI91091.1 aver|OJI96841.1 awen|OJJ35131.1 azon|OJJ49968.1

Group3113: abra|OJJ73560.1 acar|OOG00731.1 acla|EAW07477.1 afis|EAW22509.1 afla|EED54537.1 afum|EAL90684.1 agla|OJJ80867.1 akaw|GAA88594.1 aluc|OJZ88502.1 anid|CBF88977.1 anig|EHA18210.1 aory|EIT78879.1 asyd|OJJ62791.1 ater|EAU39205.1 atub|OJI91088.1 aver|OJI96838.1 awen|OJJ35134.1 azon|OJJ49964.1

Group3114: abra|OJJ73563.1 acar|OOG00746.1 acla|EAW07480.1 afis|EAW22512.1 afla|EED54540.1 afum|EAL90681.1 agla|OJJ80875.1 akaw|GAA88591.1 aluc|OJZ88498.1 anid|CBF88983.1 anig|EHA18207.1

aory|EIT78924.1 asyd|OJJ62788.1 ater|EAU39197.1 atub|OJI91085.1 aver|OJI96835.1 awen|OJJ35138.1 azon|OJJ49961.1

Group3115: abra|OJJ73566.1 acar|OOG00748.1 acla|EAW07482.1 afis|EAW22514.1 afla|EED54544.1 afum|EAL90679.1 agla|OJJ80873.1 akaw|GAA88589.1 aluc|OJZ88496.1 anid|CBF88984.1 anig|EHA18205.1 aory|EIT78876.1 asyd|OJJ62786.1 ater|EAU39195.1 atub|OJI91083.1 aver|OJI96833.1 awen|OJJ35140.1 azon|OJJ49959.1

Group3116: abra|OJJ73567.1 acar|OOG00745.1 acla|EAW07483.1 afis|EAW22515.1 afla|EED54545.1 afum|EAL90678.1 agla|OJJ80872.1 akaw|GAA88588.1 aluc|OJZ88495.1 anid|CBF88986.1 anig|EHA18204.1 aory|EIT78884.1 asyd|OJJ62785.1 ater|EAU39194.1 atub|OJI91082.1 aver|OJI96831.1 awen|OJJ35141.1 azon|OJJ49958.1

Group3117: abra|OJJ73568.1 acar|OOG00749.1 acla|EAW07484.1 afis|EAW22516.1 afla|EED54546.1 afum|EAL90677.1 agla|OJJ80877.1 akaw|GAA88587.1 aluc|OJZ88494.1 anid|CBF88988.1 anig|EHA18203.1 aory|EIT78756.1 asyd|OJJ62784.1 ater|EAU39193.1 atub|OJI91081.1 aver|OJI96830.1 awen|OJJ35142.1 azon|OJJ49957.1

Group3118: abra|OJJ73573.1 acar|OOG00754.1 acla|EAW07490.1 afis|EAW22523.1 afla|EED54548.1 afum|EAL90671.1 agla|OJJ80884.1 akaw|GAA88577.1 aluc|OJZ88480.1 anid|CBF89007.1 anig|EHA18196.1 aory|EIT78743.1 asyd|OJJ62776.1 ater|EAU39187.1 atub|OJI91074.1 aver|OJI96821.1 awen|OJJ35150.1 azon|OJJ49951.1

Group3119: abra|OJJ73574.1 acar|OOG00755.1 acla|EAW07491.1 afis|EAW22524.1 afla|EED54549.1 afum|EAL90670.1 agla|OJJ80885.1 akaw|GAA88576.1 aluc|OJZ88479.1 anid|CBF89009.1 anig|EHA18195.1 aory|EIT78837.1 asyd|OJJ62775.1 ater|EAU39186.1 atub|OJI91073.1 aver|OJI96820.1 awen|OJJ35151.1 azon|OJJ49950.1

Group3120: abra|OJJ73575.1 acar|OOG00756.1 acla|EAW07492.1 afis|EAW22525.1 afla|EED54553.1 afum|EAL90669.1 agla|OJJ80886.1 akaw|GAA88575.1 aluc|OJZ88477.1 anid|CBF89011.1 anig|EHA18194.1 aory|EIT78752.1 asyd|OJJ62774.1 ater|EAU39185.1 atub|OJI91072.1 aver|OJI96819.1 awen|OJJ35152.1 azon|OJJ49949.1

Group3121: abra|OJJ73578.1 acar|OOG00759.1 acla|EAW07496.1 afis|EAW22529.1 afla|EED54557.1 afum|EAL90665.1 agla|OJJ80890.1 akaw|GAA88573.1 aluc|OJZ88474.1 anid|CBF89014.1 anig|EHA18191.1 aory|EIT78796.1 asyd|OJJ62772.1 ater|EAU39182.1 atub|OJI91069.1 aver|OJI96817.1 awen|OJJ35156.1 azon|OJJ49945.1

Group3122: abra|OJJ73580.1 acar|OOG00761.1 acla|EAW07498.1 afis|EAW22531.1 afla|EED54559.1 afum|EAL90663.1 agla|OJJ80892.1 akaw|GAA88571.1 aluc|OJZ88472.1 anid|CBF89018.1 anig|EHA18189.1 aory|EIT78759.1 asyd|OJJ62770.1 ater|EAU39178.1 atub|OJI91067.1 aver|OJI96815.1 awen|OJJ35158.1 azon|OJJ49944.1

Group3123: abra|OJJ73581.1 acar|OOG00762.1 acla|EAW07499.1 afis|EAW22532.1 afla|EED54560.1 afum|EAL90662.1 agla|OJJ80893.1 akaw|GAA88570.1 aluc|OJZ88471.1 anid|CBF89020.1 anig|EHA18188.1 aory|EIT78827.1 asyd|OJJ62769.1 ater|EAU39177.1 atub|OJI91066.1 aver|OJI96814.1 awen|OJJ35159.1 azon|OJJ49943.1

Group3124: abra|OJJ73582.1 acar|OOG00763.1 acla|EAW07500.1 afis|EAW22533.1 afla|EED54561.1 afum|EAL90661.1 agla|OJJ80894.1 akaw|GAA88569.1 aluc|OJZ88470.1 anid|CBF89023.1 anig|EHA18187.1 aory|EIT78937.1 asyd|OJJ62768.1 ater|EAU39176.1 atub|OJI91065.1 aver|OJI96813.1 awen|OJJ35160.1 azon|OJJ49942.1

Group3125: abra|OJJ73583.1 acar|OOG00764.1 acla|EAW07501.1 afis|EAW22534.1 afla|EED54562.1 afum|EAL90660.1 agla|OJJ80895.1 akaw|GAA88568.1 aluc|OJZ88469.1 anid|CBF89025.1 anig|EHA18186.1 aory|EIT78784.1 asyd|OJJ62767.1 ater|EAU39175.1 atub|OJI91064.1 aver|OJI96812.1 awen|OJJ35161.1 azon|OJJ49941.1

Group3126: abra|OJJ73585.1 acar|OOG00766.1 acla|EAW07503.1 afis|EAW22536.1 afla|EED54564.1 afum|EAL90658.1 agla|OJJ80897.1 akaw|GAA88566.1 aluc|OJZ88467.1 anid|CBF89029.1 anig|EHA18184.1 aory|EIT78903.1 asyd|OJJ62765.1 ater|EAU39173.1 atub|OJI91062.1 aver|OJI96810.1 awen|OJJ35163.1 azon|OJJ49940.1

Group3127: abra|OJJ73586.1 acar|OOG00767.1 acla|EAW07504.1 afis|EAW22537.1 afla|EED54565.1 afum|EAL90657.1 agla|OJJ80898.1 akaw|GAA88565.1 aluc|OJZ88466.1 anid|CBF89031.1 anig|EHA18183.1 aory|EIT78873.1 asyd|OJJ62764.1 ater|EAU39172.1 atub|OJI91061.1 aver|OJI96809.1 awen|OJJ35165.1 azon|OJJ49939.1

Group3128: abra|OJJ73589.1 acar|OOG00771.1 acla|EAW07507.1 afis|EAW22539.1 afla|EED54568.1 afum|EAL90655.1 agla|OJJ80899.1 akaw|GAA88563.1 aluc|OJZ88461.1 anid|CBF89036.1 anig|EHA18181.1 aory|EIT78782.1 asyd|OJJ62762.1 ater|EAU39170.1 atub|OJI91059.1 aver|OJI96805.1 awen|OJJ35168.1 azon|OJJ49938.1

Group3129: abra|OJJ73592.1 acar|OOG00777.1 acla|EAW07511.1 afis|EAW22542.1 afla|EED54573.1 afum|EAL90652.1 agla|OJJ80904.1 akaw|GAA88560.1 aluc|OJZ88458.1 anid|CBF89040.1 anig|EHA18178.1 aory|EIT78776.1 asyd|OJJ62760.1 ater|EAU39166.1 atub|OJI91056.1 aver|OJI96803.1 awen|OJJ35171.1 azon|OJJ49936.1

Group3130: abra|OJJ73596.1 acar|OOG00782.1 akaw|GAA88556.1 aluc|OJZ88454.1 anig|EHA18175.1 atub|OJI91052.1 abra|OJJ74453.1 acar|OOF95299.1 afla|EED49262.1 akaw|GAA87347.1 aluc|OJZ89667.1 anig|EHA26904.1 aory|EIT82192.1 asyd|OJJ62435.1 ater|EAU29288.1 atub|OJI85133.1 aver|OJI96571.1 aver|OJI98027.1

Group3131: abra|OJJ73598.1 acar|OOG00784.1 acla|EAW07516.1 afis|EAW22546.1 afla|EED54578.1 afum|EAL90649.1 agla|OJJ80908.1 akaw|GAA88554.1 aluc|OJZ88452.1 anid|CBF89049.1 anig|EHA18173.1 aory|EIT78955.1 asyd|OJJ62755.1 ater|EAU39159.1 atub|OJI91050.1 aver|OJI96797.1 awen|OJJ35177.1 azon|OJJ49931.1

Group3132: abra|OJJ73599.1 acar|OOG00785.1 acla|EAW07517.1 afis|EAW22547.1 afla|EED54579.1 afum|EAL90648.1 agla|OJJ80909.1 akaw|GAA88553.1 aluc|OJZ88451.1 anid|CBF89051.1 anig|EHA18172.1 aory|EIT78753.1 asyd|OJJ62754.1 ater|EAU39158.1 atub|OJI91049.1 aver|OJI96796.1 awen|OJJ35178.1 azon|OJJ49930.1

Group3133: abra|OJJ73601.1 acar|OOG00786.1 acla|EAW07519.1 afis|EAW22549.1 afla|EED54582.1 afum|EAL90646.1 agla|OJJ80911.1 akaw|GAA88551.1 aluc|OJZ88447.1 anid|CBF89057.1 anig|EHA18170.1 aory|EIT78932.1 asyd|OJJ62752.1 ater|EAU39155.1 atub|OJI91046.1 aver|OJI96794.1 awen|OJJ35180.1 azon|OJJ49927.1

Group3134: abra|OJJ73606.1 acar|OOG00790.1 acla|EAW07523.1 afis|EAW22553.1 afla|EED54585.1 afum|EAL90642.1 agla|OJJ80915.1 akaw|GAA88547.1 aluc|OJZ88443.1 anid|CBF89062.1 anig|EHA18167.1 aory|EIT78786.1 asyd|OJJ62748.1 ater|EAU39151.1 atub|OJI91041.1 aver|OJI96788.1 awen|OJJ35184.1 azon|OJJ49924.1

Group3135: abra|OJJ73607.1 acar|OOG00791.1 acla|EAW07524.1 afis|EAW22554.1 afla|EED54586.1 afum|EAL90641.1 agla|OJJ80916.1 akaw|GAA88546.1 aluc|OJZ88442.1 anid|CBF89063.1 anig|EHA18166.1 aory|EIT78927.1 asyd|OJJ62747.1 ater|EAU39150.1 atub|OJI91040.1 aver|OJI96787.1 awen|OJJ35185.1 azon|OJJ49923.1

Group3136: abra|OJJ73609.1 acar|OOG00793.1 acla|EAW07526.1 afis|EAW22556.1 afla|EED54587.1 afum|EAL90639.1 agla|OJJ80917.1 akaw|GAA88544.1 aluc|OJZ88440.1 anid|CBF89065.1 anig|EHA18164.1 aory|EIT78856.1 asyd|OJJ62745.1 ater|EAU39148.1 atub|OJI91038.1 aver|OJI96785.1 awen|OJJ35187.1 azon|OJJ49921.1

Group3137: abra|OJJ73611.1 acar|OOG00794.1 acla|EAW07528.1 afis|EAW22558.1 afla|EED54589.1 afum|EAL90637.1 agla|OJJ80918.1 akaw|GAA88542.1 aluc|OJZ88439.1 anid|CBF89071.1 anig|EHA18162.1 aory|EIT78846.1 asyd|OJJ62743.1 ater|EAU39146.1 atub|OJI91036.1 aver|OJI96784.1 awen|OJJ35188.1 azon|OJJ49918.1

Group3138: abra|OJJ73612.1 acar|OOG00795.1 acla|EAW07529.1 afis|EAW22559.1 afla|EED54590.1 afum|EAL90636.1 agla|OJJ80919.1 akaw|GAA88541.1 aluc|OJZ88437.1 anid|CBF88377.1 anig|EHA18161.1 aory|EIT78952.1 asyd|OJJ63734.1 ater|EAU39145.1 atub|OJI91034.1 aver|OJI97206.1 awen|OJJ35189.1 azon|OJJ49917.1

Group3139: abra|OJJ73616.1 acar|OOG00800.1 acla|EAW07531.1 afis|EAW22562.1 afla|EED54592.1 afum|EAL90633.1 agla|OJJ80921.1 akaw|GAA88537.1 aluc|OJZ88433.1 anid|CBF88314.1 anig|EHA18157.1 aory|EIT78915.1 asyd|OJJ63700.1 ater|EAU39142.1 atub|OJI91030.1 aver|OJI97239.1 awen|OJJ35191.1 azon|OJJ49915.1

Group3140: abra|OJJ73617.1 acar|OOG00799.1 acla|EAW07532.1 afis|EAW22561.1 afla|EED54596.1 afum|EAL90634.1 agla|OJJ80922.1 akaw|GAA88536.1 aluc|OJZ88432.1 anid|CBF88316.1 anig|EHA18156.1 aory|EIT78857.1 asyd|OJJ63701.1 ater|EAU39141.1 atub|OJI91029.1 aver|OJI97238.1 awen|OJJ35192.1 azon|OJJ49914.1

Group3141: abra|OJJ73618.1 acar|OOG00801.1 acla|EAW07533.1 afis|EAW22567.1 afla|EED54597.1 afum|EAL90631.1 agla|OJJ80923.1 akaw|GAA88535.1 aluc|OJZ88431.1 anid|CBF88318.1 anig|EHA18155.1 aory|EIT78761.1 asyd|OJJ63702.1 ater|EAU39140.1 atub|OJI91028.1 aver|OJI97237.1 awen|OJJ35193.1 azon|OJJ49913.1

Group3142: abra|OJJ73619.1 acar|OOG00802.1 acla|EAW07534.1 afis|EAW22568.1 afla|EED54598.1 afum|EAL90630.1 agla|OJJ80924.1 akaw|GAA88534.1 aluc|OJZ88430.1 anid|CBF88320.1 anig|EHA18154.1 aory|EIT78762.1 asyd|OJJ63703.1 ater|EAU39139.1 atub|OJI91027.1 aver|OJI97236.1 awen|OJJ35194.1 azon|OJJ49912.1

Group3143: abra|OJJ73621.1 acar|OOG00805.1 acla|EAW07536.1 afis|EAW22570.1 afla|EED54600.1 afum|EAL90628.1 agla|OJJ80926.1 akaw|GAA88532.1 aluc|OJZ88427.1 anid|CBF88323.1 anig|EHA18152.1

aory|EIT78841.1 asyd|OJJ63705.1 ater|EAU39137.1 atub|OJI91025.1 aver|OJI97234.1 awen|OJJ35196.1 azon|OJJ49911.1

Group3144: abra|OJJ73622.1 acar|OOG00804.1 acla|EAW07537.1 afis|EAW22571.1 afla|EED54601.1 afum|EAL90627.1 agla|OJJ80928.1 akaw|GAA88531.1 aluc|OJZ88426.1 anid|CBF88325.1 anig|EHA18151.1 aory|EIT78890.1 asyd|OJJ63706.1 ater|EAU39136.1 atub|OJI91024.1 aver|OJI97233.1 awen|OJJ35198.1 azon|OJJ49910.1

Group3145: abra|OJJ73624.1 acar|OOG00807.1 acla|EAW07539.1 afis|EAW22573.1 afla|EED54602.1 afum|EAL90625.1 agla|OJJ80930.1 akaw|GAA88529.1 aluc|OJZ88424.1 anid|CBF88328.1 anig|EHA18149.1 aory|EIT78905.1 asyd|OJJ63708.1 ater|EAU39134.1 atub|OJI91020.1 aver|OJI97230.1 awen|OJJ35200.1 azon|OJJ49907.1

Group3146: abra|OJJ73627.1 acar|OOG00809.1 acla|EAW07541.1 afis|EAW22575.1 afla|EED54604.1 afum|EAL90623.1 agla|OJJ80931.1 akaw|GAA88526.1 aluc|OJZ88422.1 anid|CBF88332.1 anig|EHA18147.1 aory|EIT78895.1 asyd|OJJ63711.1 ater|EAU39129.1 atub|OJI91018.1 aver|OJI97228.1 awen|OJJ35203.1 azon|OJJ49905.1

Group3147: abra|OJJ73632.1 acar|OOG00814.1 acla|EAW07546.1 afis|EAW22580.1 afla|EED54625.1 afum|EAL90619.2 agla|OJJ80940.1 akaw|GAA88521.1 aluc|OJZ88417.1 anid|CBF88341.1 anig|EHA18143.1 aory|EIT78840.1 asyd|OJJ63716.1 ater|EAU39124.1 atub|OJI91013.1 aver|OJI97224.1 awen|OJJ35208.1 azon|OJJ49901.1

Group3148: abra|OJJ73633.1 acar|OOG00815.1 acla|EAW07547.1 afis|EAW22581.1 afla|EED54626.1 afum|EAL90618.1 agla|OJJ80941.1 akaw|GAA88520.1 aluc|OJZ88416.1 anid|CBF88343.1 anig|EHA18142.1 aory|EIT78811.1 asyd|OJJ63717.1 ater|EAU39123.1 atub|OJI91012.1 aver|OJI97223.1 awen|OJJ35209.1 azon|OJJ49900.1

Group3149: abra|OJJ73637.1 acar|OOG00817.1 acla|EAW07549.1 afis|EAW22583.1 afla|EED54628.1 afum|EAL90616.1 agla|OJJ80943.1 akaw|GAA88518.1 aluc|OJZ88413.1 anid|CBF88349.1 anig|EHA18140.1 aory|EIT78801.1 asyd|OJJ63719.1 ater|EAU39121.1 atub|OJI91009.1 aver|OJI97221.1 awen|OJJ35211.1 azon|OJJ49898.1

Group3150: abra|OJJ73638.1 acar|OOG00818.1 acla|EAW07550.1 afis|EAW22584.1 afla|EED54629.1 afum|EAL90615.1 agla|OJJ80944.1 akaw|GAA88517.1 aluc|OJZ88412.1 anid|CBF88350.1 anig|EHA18139.1 aory|EIT78791.1 asyd|OJJ63720.1 ater|EAU39120.1 atub|OJI91007.1 aver|OJI97220.1 awen|OJJ35212.1 azon|OJJ49897.1

Group3151: abra|OJJ73640.1 acar|OOG00820.1 acla|EAW07553.1 afis|EAW22587.1 afla|EED54633.1 afum|EAL90612.1 agla|OJJ80947.1 akaw|GAA88515.1 aluc|OJZ88410.1 anid|CBF88356.1 anig|EHA18137.1 aory|EIT78894.1 asyd|OJJ63723.1 ater|EAU39113.1 atub|OJI91006.1 aver|OJI97217.1 awen|OJJ35215.1 azon|OJJ49894.1

Group3152: abra|OJJ73641.1 acar|OOG00822.1 acla|EAW07554.1 afis|EAW22588.1 afla|EED54634.1 afum|EAL90611.1 agla|OJJ80948.1 akaw|GAA88514.1 aluc|OJZ88409.1 anid|CBF88358.1 anig|EHA18136.1 aory|EIT78862.1 asyd|OJJ63724.1 ater|EAU39112.1 atub|OJI91005.1 aver|OJI97216.1 awen|OJJ35216.1 azon|OJJ49893.1

Group3153: abra|OJJ73643.1 acar|OOG00824.1 acla|EAW07556.1 afis|EAW22590.1 afla|EED54636.1 afum|EAL90609.1 agla|OJJ80950.1 akaw|GAA88512.1 aluc|OJZ88407.1 anid|CBF88360.1 anig|EHA18134.1 aory|EIT78889.1 asyd|OJJ63726.1 ater|EAU39119.1 atub|OJI91003.1 aver|OJI97214.1 awen|OJJ35218.1 azon|OJJ49890.1

Group3154: abra|OJJ73644.1 acar|OOG00825.1 acla|EAW07557.1 afis|EAW22591.1 afla|EED54641.1 afum|EAL90608.1 agla|OJJ80951.1 akaw|GAA88511.1 aluc|OJZ88406.1 anid|CBF88364.1 anig|EHA18133.1 aory|EIT78790.1 asyd|OJJ63727.1 ater|EAU39111.1 atub|OJI91002.1 aver|OJI97213.1 awen|OJJ35219.1 azon|OJJ49889.1

Group3155: abra|OJJ73645.1 acar|OOG00826.1 acla|EAW07558.1 afis|EAW22592.1 afla|EED54642.1 afum|EAL90607.1 agla|OJJ80952.1 akaw|GAA88510.1 aluc|OJZ88405.1 anid|CBF88365.1 anig|EHA18132.1 aory|EIT78880.1 asyd|OJJ63728.1 ater|EAU39110.1 atub|OJI91001.1 aver|OJI97212.1 awen|OJJ35220.1 azon|OJJ49888.1

Group3156: abra|OJJ73647.1 acar|OOG00829.1 acla|EAW07560.1 afis|EAW22594.1 afla|EED54644.1 afum|EAL90605.1 agla|OJJ80954.1 akaw|GAA88508.1 aluc|OJZ88403.1 anid|CBF88369.1 anig|EHA18131.1 aory|EIT78768.1 asyd|OJJ63730.1 ater|EAU39109.1 atub|OJI90999.1 aver|OJI97210.1 awen|OJJ35222.1 azon|OJJ49886.1

Group3157: abra|OJJ73654.1 acar|OOG00832.1 acla|EAW07564.1 afis|EAW22598.1 afla|EED54720.1 afum|EAL90600.1 agla|OJJ80966.1 akaw|GAA88504.1 aluc|OJZ88399.1 anid|CBF88312.1 anig|EHA18127.1 aory|EIT78821.1 asyd|OJJ63698.1 ater|EAU39105.1 atub|OJI90994.1 aver|OJI97241.1 awen|OJJ35226.1 azon|OJJ49884.1

Group3158: abra|OJJ73655.1 acar|OOG00833.1 acla|EAW07565.1 afis|EAW22599.1 afla|EED54719.1 afum|EAL90599.1 agla|OJJ80967.1 akaw|GAA88503.1 aluc|OJZ88398.1 anid|CBF88310.1 anig|EHA18126.1 aory|EIT78916.1 asyd|OJJ63697.1 ater|EAU39104.1 atub|OJI90993.1 aver|OJI97242.1 awen|OJJ35228.1 azon|OJJ49883.1

Group3159: abra|OJJ73656.1 acar|OOG00834.1 acla|EAW07566.1 afis|EAW22600.1 afla|EED54718.1 afum|EAL90598.1 agla|OJJ80968.1 akaw|GAA88502.1 aluc|OJZ88397.1 anid|CBF88308.1 anig|EHA18125.1 aory|EIT78770.1 asyd|OJJ63696.1 ater|EAU39103.1 atub|OJI90992.1 aver|OJI97243.1 awen|OJJ35229.1 azon|OJJ49882.1

Group3160: abra|OJJ73657.1 acar|OOG00835.1 acla|EAW07567.1 afis|EAW22601.1 afla|EED54717.1 afum|EAL90597.1 agla|OJJ80971.1 akaw|GAA88501.1 aluc|OJZ88396.1 anid|CBF88256.1 anig|EHA18124.1 aory|EIT78942.1 asyd|OJJ63672.1 ater|EAU39102.1 atub|OJI90991.1 aver|OJI97266.1 awen|OJJ35230.1 azon|OJJ49881.1

Group3161: abra|OJJ73658.1 acar|OOG00836.1 acla|EAW07568.1 afis|EAW22602.1 afla|EED54716.1 afum|EAL90596.1 agla|OJJ80972.1 akaw|GAA88500.1 aluc|OJZ88395.1 anid|CBF88257.1 anig|EHA18123.1 aory|EIT78742.1 asyd|OJJ63673.1 ater|EAU39101.1 atub|OJI90990.1 aver|OJI97265.1 awen|OJJ35231.1 azon|OJJ49880.1

Group3162: abra|OJJ73665.1 acar|OOG00843.1 acla|EAW07572.1 afis|EAW22606.1 afla|EED49514.1 afum|EAL90592.1 agla|OJJ80975.1 akaw|GAA88491.1 aluc|OJZ88387.1 anid|CBF88263.1 anig|EHA18114.1 aory|EIT82633.1 asyd|OJJ63677.1 ater|EAU39097.1 atub|OJI90982.1 aver|OJI97262.1 awen|OJJ35234.1 azon|OJJ49877.1

Group3163: abra|OJJ73667.1 acar|OOG00844.1 acla|EAW07573.1 afis|EAW22607.1 afla|EED49805.1 afum|EAL90591.1 agla|OJJ80977.1 akaw|GAA88490.1 aluc|OJZ88385.1 anid|CBF88265.1 anig|EHA18113.1 aory|EIT77683.1 asyd|OJJ63678.1 ater|EAU39096.1 atub|OJI90980.1 aver|OJI97261.1 awen|OJJ35235.1 azon|OJJ49876.1

Group3164: abra|OJJ73668.1 acar|OOG00845.1 acla|EAW07575.1 afis|EAW22609.1 afla|EED49807.1 afum|EAL90589.1 agla|OJJ81009.1 akaw|GAA88489.1 aluc|OJZ88384.1 anid|CBF88269.1 anig|EHA18112.1 aory|EIT77641.1 asyd|OJJ63680.1 ater|EAU39095.1 atub|OJI90979.1 aver|OJI97259.1 awen|OJJ35237.1 azon|OJJ49870.1

Group3165: abra|OJJ73671.1 acar|OOG00846.1 acla|EAW07583.1 afis|EAW22611.1 afla|EED49809.1 afum|EAL90587.1 agla|OJJ81005.1 akaw|GAA88486.1 aluc|OJZ88382.1 anid|CBF88281.1 anig|EHA18109.1 aory|EIT77633.1 asyd|OJJ63684.1 ater|EAU39092.1 atub|OJI90976.1 aver|OJI97255.1 awen|OJJ35239.1 azon|OJJ49868.1

Group3166: abra|OJJ73672.1 acar|OOG00847.1 acla|EAW07582.1 afis|EAW22612.1 afla|EED49810.1 afum|EAL90586.1 agla|OJJ81004.1 akaw|GAA88485.1 aluc|OJZ88381.1 anid|CBF88283.1 anig|EHA18108.1 aory|EIT77693.1 asyd|OJJ63685.1 ater|EAU39091.1 atub|OJI90975.1 aver|OJI97254.1 awen|OJJ35240.1 azon|OJJ49867.1

Group3167: abra|OJJ73673.1 acar|OOG00848.1 acla|EAW07581.1 afis|EAW22613.1 afla|EED49811.1 afum|EAL90585.1 agla|OJJ81003.1 akaw|GAA88484.1 aluc|OJZ88379.1 anid|CBF88287.1 anig|EHA18106.1 aory|EIT77666.1 asyd|OJJ63686.1 ater|EAU39090.1 atub|OJI90973.1 aver|OJI97253.1 awen|OJJ35241.1 azon|OJJ49866.1

Group3168: abra|OJJ73681.1 acar|OOG00853.1 acla|EAW07596.1 afis|EAW22627.1 afla|EED49827.1 afum|EAL90569.1 agla|OJJ80994.1 akaw|GAA88478.1 aluc|OJZ88368.1 anid|CBF88241.1 anig|EHA18101.1 aory|EIT77681.1 asyd|OJJ63662.1 ater|EAU39082.1 atub|OJI90966.1 aver|OJI97272.1 awen|OJJ35248.1 azon|OJJ49858.1

Group3169: abra|OJJ73684.1 acar|OOG00854.1 acla|EAW07599.1 afis|EAW22630.1 afla|EED49879.1 afum|EAL90565.1 agla|OJJ80990.1 akaw|GAA88475.1 aluc|OJZ88365.1 anid|CBF88236.1 anig|EHA18097.1 aory|EIT72464.1 asyd|OJJ63659.1 ater|EAU39078.1 atub|OJI90963.1 aver|OJI97275.1 awen|OJJ35251.1 azon|OJJ49855.1

Group3170: abra|OJJ73688.1 acar|OOG00860.1 acla|EAW07603.1 afis|EAW22634.1 afla|EED49884.1 afum|EAL90559.1 agla|OJJ80985.1 akaw|GAA88471.1 aluc|OJZ88361.1 anid|CBF88229.1 anig|EHA18093.1 aory|EIT72459.1 asyd|OJJ63655.1 ater|EAU39074.1 atub|OJI90957.1 aver|OJI97279.1 awen|OJJ35256.1 azon|OJJ49851.1

Group3171: abra|OJJ73689.1 acar|OOG00861.1 acla|EAW07604.1 afis|EAW22635.1 afum|EAL90558.2 akaw|GAA88470.1 aluc|OJZ88360.1 anid|CBF88227.1 anig|EHA18092.1 asyd|OJJ63654.1 ater|EAU39071.1 atub|OJI90956.1 aver|OJI97280.1 awen|OJJ35257.1 azon|OJJ49850.1 afla|EED49885.1 aory|EIT72467.1 agla|OJJ80984.1

Group3172: abra|OJJ73692.1 acar|OOG00863.1 acla|EAW07607.1 afis|EAW22637.1 afla|EED49888.1 afum|EAL90555.1 agla|OJJ80981.1 akaw|GAA88468.1 aluc|OJZ88357.1 anid|CBF88221.1 anig|EHA18090.1

aory|EIT72465.1 asyd|OJJ63652.1 ater|EAU39069.1 atub|OJI90952.1 aver|OJI97282.1 awen|OJJ35260.1 azon|OJJ49847.1

Group3173: abra|OJJ73724.1 acar|OOG00893.1 acla|EAW07613.1 afis|EAW22643.1 afla|EED49912.1 afum|EAL90549.2 agla|OJJ81025.1 akaw|GAA88438.1 aluc|OJZ88326.1 anid|CBF88175.1 anig|EHA18062.1 aory|EIT72277.1 asyd|OJJ63625.1 ater|EAU39064.1 atub|OJI90921.1 aver|OJI97311.1 awen|OJJ35281.1 azon|OJJ49845.1

Group3174: abra|OJJ73731.1 acar|OOG00897.1 acla|EAW07616.1 afis|EAW22645.1 afla|EED49914.1 afum|EAL90547.1 agla|OJJ81028.1 akaw|GAA88432.1 aluc|OJZ88319.1 anid|CBF88179.1 anig|EHA18055.1 aory|EIT72268.1 asyd|OJJ63627.1 ater|EAU39062.1 atub|OJI90914.1 aver|OJI97309.1 awen|OJJ35278.1 azon|OJJ49841.1

Group3175: abra|OJJ73733.1 acar|OOG00898.1 acla|EAW07617.1 afis|EAW22646.1 afla|EED49915.1 afum|EAL90546.1 agla|OJJ81029.1 akaw|GAA88431.1 aluc|OJZ88318.1 anid|CBF88180.1 anig|EHA18054.1 aory|EIT72267.1 asyd|OJJ63628.1 ater|EAU39061.1 atub|OJI90913.1 aver|OJI97308.1 awen|OJJ35277.1 azon|OJJ49840.1

Group3176: abra|OJJ73734.1 acar|OOG00899.1 acla|EAW07618.1 afis|EAW22647.1 afla|EED49916.1 afum|EAL90545.1 agla|OJJ81030.1 akaw|GAA88430.1 aluc|OJZ88317.1 anid|CBF88182.1 anig|EHA18053.1 aory|EIT72271.1 asyd|OJJ63629.1 ater|EAU39060.1 atub|OJI90912.1 aver|OJI97307.1 awen|OJJ35276.1 azon|OJJ49839.1

Group3177: abra|OJJ73735.1 acar|OOG00900.1 acla|EAW07619.1 afis|EAW22648.1 afla|EED49917.1 afum|EAL90544.1 agla|OJJ81031.1 akaw|GAA88429.1 aluc|OJZ88316.1 anid|CBF88184.1 anig|EHA18052.1 aory|EIT72269.1 asyd|OJJ63630.1 ater|EAU39059.1 atub|OJI90911.1 aver|OJI97306.1 awen|OJJ35275.1 azon|OJJ49832.1

Group3178: abra|OJJ73736.1 acar|OOG00901.1 acla|EAW07620.1 afis|EAW22649.1 afla|EED49918.1 afum|EAL90543.1 agla|OJJ81032.1 akaw|GAA88428.1 aluc|OJZ88315.1 anid|CBF88186.1 anig|EHA18051.1 aory|EIT72278.1 asyd|OJJ63631.1 ater|EAU39058.1 atub|OJI90910.1 aver|OJI97305.1 awen|OJJ35274.1 azon|OJJ49833.1

Group3179: abra|OJJ73739.1 acar|OOG00902.1 acla|EAW07633.1 afis|EAW22653.1 afla|EED49921.1 afum|EAL90539.1 agla|OJJ81036.1 akaw|GAA88425.1 aluc|OJZ88312.1 anid|CBF88197.1 anig|EHA18049.1 aory|EIT72265.1 asyd|OJJ63634.1 ater|EAU39053.1 atub|OJI90907.1 aver|OJI97302.1 awen|OJJ35270.1 azon|OJJ49837.1

Group3180: abra|OJJ73741.1 acar|OOG00904.1 acla|EAW07635.1 afis|EAW22655.1 afla|EED49923.1 afum|EAL90537.1 agla|OJJ81038.1 akaw|GAA88424.1 aluc|OJZ88311.1 anid|CBF88201.1 anig|EHA18048.1 aory|EIT72272.1 asyd|OJJ63637.1 ater|EAU39051.1 atub|OJI90906.1 aver|OJI97299.1 awen|OJJ35268.1 azon|OJJ49831.1

Group3181: abra|OJJ73744.1 acar|OOG00908.1 acla|EAW07640.1 afis|EAW22660.1 afla|EED49930.1 afum|EAL90532.1 agla|OJJ81041.1 akaw|GAA88421.1 aluc|OJZ88308.1 anid|CBF88218.1 anig|EHA18045.1 aory|EIT72279.1 asyd|OJJ63649.1 ater|EAU39048.1 atub|OJI90903.1 aver|OJI97284.1 awen|OJJ35265.1 azon|OJJ49828.1

Group3182: abra|OJJ73745.1 acar|OOG00909.1 acla|EAW07639.1 afis|EAW22659.1 afla|EED49931.1 afum|EAL90533.1 agla|OJJ81042.1 akaw|GAA88420.1 aluc|OJZ88307.1 anid|CBF88215.1 anig|EHA18044.1 aory|EIT72275.1 asyd|OJJ63648.1 ater|EAU39049.1 atub|OJI90902.1 aver|OJI97285.1 awen|OJJ35283.1 azon|OJJ49827.1

Group3183: abra|OJJ73751.1 acar|OOG00914.1 acla|EAW07642.1 afis|EAW22662.1 afla|EED49935.1 afum|EAL90530.1 agla|OJJ79062.1 akaw|GAA88416.1 aluc|OJZ88301.1 anid|CBF88168.1 anig|EHA18040.1 aory|EIT83423.1 asyd|OJJ63622.1 ater|EAU39046.1 atub|OJI90896.1 aver|OJI97315.1 awen|OJJ35287.1 azon|OJJ49826.1

Group3184: abra|OJJ73752.1 acar|OOG00915.1 acla|EAW07643.1 afis|EAW22663.1 afla|EED49937.1 afum|EAL90529.1 akaw|GAA88415.1 aluc|OJZ88300.1 anid|CBF88166.1 anig|EHA18039.1 aory|EIT83433.1 asyd|OJJ63621.1 ater|EAU39045.1 atub|OJI90894.1 aver|OJI97316.1 awen|OJJ35288.1 azon|OJJ49825.1 agla|OJJ79061.1

Group3185: abra|OJJ73753.1 acar|OOG00916.1 acla|EAW07644.1 afis|EAW22664.1 afla|EED49938.1 afum|EAL90528.1 agla|OJJ79060.1 akaw|GAA88414.1 aluc|OJZ88298.1 anid|CBF88164.1 anig|EHA18038.1 aory|EIT83434.1 asyd|OJJ63620.1 ater|EAU39044.1 atub|OJI90893.1 aver|OJI97318.1 awen|OJJ35290.1 azon|OJJ49824.1

Group3186: abra|OJJ73756.1 acar|OOG00918.1 acla|EAW07646.1 afis|EAW22666.1 afla|EED49940.1 afum|EAL90526.1 agla|OJJ79058.1 akaw|GAA88412.1 aluc|OJZ88296.1 anid|CBF88161.1 anig|EHA18036.1 aory|EIT83451.1 asyd|OJJ63619.1 ater|EAU39042.1 atub|OJI90892.1 aver|OJI97320.1 awen|OJJ35292.1 azon|OJJ49822.1

Group3187: abra|OJJ73757.1 acar|OOG00919.1 acla|EAW07647.1 afis|EAW22667.1 afla|EED49941.1 afum|EAL90525.1 agla|OJJ79057.1 akaw|GAA88411.1 aluc|OJZ88295.1 anid|CBF88159.1 anig|EHA18035.1 aory|EIT83456.1 asyd|OJJ63618.1 ater|EAU39041.1 atub|OJI90891.1 aver|OJI97321.1 awen|OJJ35293.1 azon|OJJ49821.1

Group3188: abra|OJJ73758.1 acar|OOG00920.1 acla|EAW07648.1 afis|EAW22668.1 afla|EED49942.1 afum|EAL90524.1 agla|OJJ79056.1 akaw|GAA88410.1 aluc|OJZ88294.1 anid|CBF88157.1 anig|EHA18034.1 aory|EIT83429.1 asyd|OJJ63617.1 ater|EAU39040.1 atub|OJI90890.1 aver|OJI97322.1 awen|OJJ35294.1 azon|OJJ49820.1

Group3189: abra|OJJ73766.1 acar|OOG00925.1 acla|EAW07652.1 afis|EAW22670.1 afla|EED49947.1 afum|EAL90522.1 agla|OJJ79049.1 akaw|GAA88408.1 aluc|OJZ88290.1 anid|CBF88155.1 anig|EHA18031.1 aory|EIT83455.1 asyd|OJJ63608.1 ater|EAU39038.1 atub|OJI90887.1 aver|OJI97329.1 awen|OJJ35298.1 azon|OJJ49817.1

Group3190: abra|OJJ73768.1 acar|OOG00927.1 acla|EAW07653.1 afis|EAW22671.1 afla|EED49949.1 afum|EAL90521.1 agla|OJJ79048.1 akaw|GAA88407.1 aluc|OJZ88289.1 anid|CBF88151.1 anig|EHA18030.1 aory|EIT83435.1 asyd|OJJ63607.1 ater|EAU39037.1 atub|OJI90886.1 aver|OJI97330.1 awen|OJJ35299.1 azon|OJJ49816.1

Group3191: abra|OJJ73769.1 acar|OOG00926.1 acla|EAW07654.1 afis|EAW22672.1 afla|EED49951.1 afum|EAL90520.1 agla|OJJ79046.1 akaw|GAA88406.1 aluc|OJZ88288.1 anid|CBF88145.1 anig|EHA18029.1 aory|EIT83454.1 asyd|OJJ63604.1 ater|EAU39036.1 atub|OJI90885.1 aver|OJI97332.1 awen|OJJ35300.1 azon|OJJ49815.1

Group3192: abra|OJJ73776.1 acar|OOG00933.1 acla|EAW07661.1 afis|EAW22678.1 afla|EED49959.1 afum|EAL90514.1 agla|OJJ79038.1 akaw|GAA88400.1 aluc|OJZ88278.1 anid|CBF88130.1 anig|EHA18023.1 aory|EIT83426.1 asyd|OJJ63588.1 ater|EAU39031.1 atub|OJI90878.1 aver|OJI97345.1 awen|OJJ35306.1 azon|OJJ49809.1

Group3193: abra|OJJ73778.1 acar|OOG00934.1 acla|EAW07662.1 afis|EAW22679.1 afla|EED49960.1 afum|EAL90513.1 agla|OJJ79037.1 akaw|GAA88399.1 aluc|OJZ88277.1 anid|CBF88128.1 anig|EHA18022.1 aory|EIT83445.1 asyd|OJJ63587.1 ater|EAU39030.1 atub|OJI90877.1 aver|OJI97347.1 awen|OJJ35307.1 azon|OJJ49806.1

Group3194: abra|OJJ73781.1 acar|OOG00937.1 acla|EAW07666.1 afis|EAW22683.1 afla|EED49967.1 afum|EAL90509.1 agla|OJJ79035.1 akaw|GAA88396.1 aluc|OJZ88274.1 anid|CBF88119.1 anig|EHA18019.1 aory|EIT83437.1 asyd|OJJ63583.1 ater|EAU39026.1 atub|OJI90874.1 aver|OJI97351.1 awen|OJJ35317.1 azon|OJJ49799.1

Group3195: abra|OJJ73783.1 acar|OOG00939.1 acla|EAW07668.1 afis|EAW22685.1 afla|EED49969.1 afum|EAL90508.1 agla|OJJ79032.1 akaw|GAA88394.1 aluc|OJZ88272.1 anid|CBF88111.1 anig|EHA18017.1 aory|EIT83432.1 asyd|OJJ63580.1 ater|EAU39019.1 atub|OJI90872.1 aver|OJI97356.1 awen|OJJ35318.1 azon|OJJ49798.1

Group3196: abra|OJJ73784.1 acar|OOG00940.1 acla|EAW07671.1 afis|EAW22686.1 afla|EED49971.1 afum|EAL90507.1 agla|OJJ79031.1 akaw|GAA88393.1 aluc|OJZ88271.1 anid|CBF88109.1 anig|EHA18016.1 aory|EIT74696.1 asyd|OJJ63578.1 ater|EAU39021.1 atub|OJI90871.1 aver|OJI97357.1 awen|OJJ35320.1 azon|OJJ49796.1

Group3197: abra|OJJ73785.1 acar|OOG00941.1 acla|EAW07672.1 afis|EAW22687.1 afla|EED49972.1 afum|EAL90506.1 agla|OJJ79029.1 akaw|GAA88392.1 aluc|OJZ88270.1 anid|CBF88107.1 anig|EHA18014.1 aory|EIT74698.1 asyd|OJJ63577.1 ater|EAU39022.1 atub|OJI90870.1 aver|OJI97359.1 awen|OJJ35321.1 azon|OJJ49794.1

Group3198: abra|OJJ73786.1 acar|OOG00942.1 acla|EAW07673.1 afis|EAW22688.1 afla|EED49973.1 afum|EAL90505.1 agla|OJJ79028.1 akaw|GAA88391.1 aluc|OJZ88269.1 anid|CBF88106.1 anig|EHA18013.1 aory|EIT74642.1 asyd|OJJ63576.1 ater|EAU39023.1 atub|OJI90869.1 aver|OJI97360.1 awen|OJJ35322.1 azon|OJJ49793.1

Group3199: abra|OJJ73787.1 acar|OOF90345.1 acla|EAW07674.1 afis|EAW22689.1 afla|EED49974.1 afum|EAL90504.1 agla|OJJ79024.1 akaw|GAA88390.1 aluc|OJZ88267.1 anid|CBF88104.1 anig|EHA18012.1 aory|EIT74677.1 asyd|OJJ63575.1 ater|EAU39024.1 atub|OJI90867.1 aver|OJI97361.1 awen|OJJ35323.1 azon|OJJ49792.1

Group3200: abra|OJJ73791.1 acar|OOF90347.1 acla|EAW07680.1 afis|EAW22695.1 afla|EED49977.1 afum|EAL90498.1 agla|OJJ79021.1 akaw|GAA89410.1 aluc|OJZ88263.1 anid|CBF88098.1 anig|EHA18009.1 aory|EIT74603.1 asyd|OJJ63570.1 ater|EAU39015.1 atub|OJI90862.1 aver|OJI97364.1 awen|OJJ35325.1 azon|OJJ49789.1

Group3201: abra|OJJ73801.1 acar|OOG00957.1 acla|EAW07683.1 afis|EAW22698.1 afla|EED49984.1 afum|EAL90494.1 agla|OJJ79012.1 akaw|GAA89421.1 aluc|OJZ88252.1 anid|CBF88087.1 anig|EHA17994.1

aory|EIT74667.1 asyd|OJJ63563.1 ater|EAU39008.1 atub|OJI90850.1 aver|OJI97370.1 awen|OJJ35334.1 azon|OJJ49783.1

Group3202: abra|OJJ73804.1 acar|OOG00959.1 acla|EAW07676.1 afis|EAW22691.1 afla|EED49982.1 afum|EAL90501.1 agla|OJJ79016.1 akaw|GAA89423.1 aluc|OJZ88249.1 anid|CBF88090.1 anig|EHA17992.1 aory|EIT74721.1 asyd|OJJ63565.1 ater|EAU39010.1 atub|OJI90848.1 aver|OJI97368.1 awen|OJJ35333.1 azon|OJJ49785.1

Group3203: abra|OJJ73805.1 acar|OOG00960.1 acla|EAW07677.1 afis|EAW22692.1 afla|EED49980.1 afum|EBA27409.1 agla|OJJ79018.1 akaw|GAA89425.1 aluc|OJZ88247.1 anid|CBF88095.1 anig|EHA17991.1 aory|EIT74734.1 asyd|OJJ63566.1 ater|EAU39012.1 atub|OJI90847.1 aver|OJI97367.1 awen|OJJ35332.1 azon|OJJ49786.1

Group3204: abra|OJJ73806.1 acar|OOG00961.1 acla|EAW07694.1 afis|EAW22709.1 afla|EED49996.1 afum|EAL90483.1 agla|OJJ79001.1 akaw|GAA89426.1 aluc|OJZ88246.1 anid|CBF88064.1 anig|EHA17990.1 aory|EIT74673.1 asyd|OJJ63549.1 ater|EAU38992.1 atub|OJI90846.1 aver|OJI97385.1 awen|OJJ35346.1 azon|OJJ49771.1

Group3205: abra|OJJ73807.1 acar|OOG00963.1 acla|EAW07695.1 afis|EAW22712.1 afla|EED49997.1 afum|EAL90482.1 agla|OJJ79000.1 akaw|GAA89427.1 aluc|OJZ88243.1 anid|CBF88062.1 anig|EHA17989.1 aory|EIT74610.1 asyd|OJJ63548.1 ater|EAU38993.1 atub|OJI90845.1 aver|OJI97386.1 awen|OJJ35347.1 azon|OJJ49770.1

Group3206: abra|OJJ73816.1 acar|OOG00970.1 acla|EAW07703.1 afis|EAW22721.1 afla|EED50006.1 afum|EAL90474.1 agla|OJJ83682.1 akaw|GAA89435.1 aluc|OJZ88234.1 anid|CBF88036.1 anig|EHA17980.1 aory|EIT74720.1 asyd|OJJ63535.1 ater|EAU38983.1 atub|OJI90836.1 aver|OJI97397.1 awen|OJJ35415.1 azon|OJJ49760.1

Group3207: abra|OJJ73821.1 acar|OOG00973.1 acla|EAW07710.1 afis|EAW22726.1 afla|EED50012.1 afum|EAL90469.1 agla|OJJ83676.1 akaw|GAA89440.1 aluc|OJZ88227.1 anid|CBF88028.1 anig|EHA17975.1 aory|EIT74707.1 asyd|OJJ63529.1 ater|EAU38978.1 atub|OJI90828.1 aver|OJI97402.1 awen|OJJ35421.1 azon|OJJ49755.1

Group3208: abra|OJJ73830.1 acar|OOG00981.1 acla|EAW07725.1 afis|EAW22757.1 afla|EED50018.1 afum|EAL90442.2 agla|OJJ83666.1 akaw|GAA89448.1 aluc|OJZ88219.1 anid|CBF88015.1 anig|EHA17966.1 aory|EIT74656.1 asyd|OJJ63519.1 ater|EAU38962.1 atub|OJI90819.1 aver|OJI97411.1 awen|OJJ35429.1 azon|OJJ49747.1

Group3209: abra|OJJ73831.1 acar|OOG00982.1 acla|EAW07726.1 afis|EAW22758.1 afla|EED50019.1 afum|EAL90441.1 agla|OJJ83662.1 akaw|GAA89449.1 aluc|OJZ88218.1 anig|EHA17965.1 aory|EIT74708.1 asyd|OJJ63518.1 ater|EAU38961.1 atub|OJI90818.1 aver|OJI97412.1 awen|OJJ35430.1 azon|OJJ49746.1 aory|EIT74649.1

Group3210: abra|OJJ73832.1 acar|OOG00983.1 acla|EAW07727.1 afis|EAW22759.1 afla|EED50020.1 afum|EAL90440.1 agla|OJJ83661.1 akaw|GAA89450.1 aluc|OJZ88217.1 anid|CBF88013.1 anig|EHA17964.1 aory|EIT74627.1 asyd|OJJ63517.1 ater|EAU38960.1 atub|OJI90817.1 aver|OJI97413.1 awen|OJJ35431.1 azon|OJJ49745.1

Group3211: abra|OJJ73833.1 acar|OOG00984.1 acla|EAW07728.1 afis|EAW22760.1 afla|EED50021.1 afum|EAL90439.1 agla|OJJ83660.1 akaw|GAA89451.1 aluc|OJZ88216.1 anid|CBF88011.1 anig|EHA17963.1 aory|EIT74690.1 asyd|OJJ63516.1 ater|EAU38959.1 atub|OJI90816.1 aver|OJI97414.1 awen|OJJ35433.1 azon|OJJ49744.1

Group3212: abra|OJJ73834.1 acar|OOG00985.1 acla|EAW07729.1 afis|EAW22761.1 afla|EED50022.1 afum|EAL90438.1 agla|OJJ83659.1 akaw|GAA89452.1 aluc|OJZ88215.1 anid|CBF88009.1 anig|EHA17962.1 aory|EIT74638.1 asyd|OJJ63515.1 ater|EAU38958.1 atub|OJI90815.1 aver|OJI97415.1 awen|OJJ35434.1 azon|OJJ49743.1

Group3213: abra|OJJ73836.1 acar|OOG00987.1 acla|EAW07731.1 afis|EAW22763.1 afla|EED50025.1 afum|EAL90436.2 agla|OJJ83657.1 akaw|GAA89454.1 aluc|OJZ88213.1 anid|CBF88005.1 anig|EHA17961.1 aory|EIT74679.1 asyd|OJJ63513.1 ater|EAU38956.1 atub|OJI90813.1 aver|OJI97417.1 awen|OJJ35436.1 azon|OJJ49741.1

Group3214: abra|OJJ73837.1 acar|OOG00988.1 acla|EAW07732.1 afis|EAW22764.1 afla|EED50026.1 afum|EAL90435.1 agla|OJJ83656.1 akaw|GAA89455.1 aluc|OJZ88212.1 anid|CBF88003.1 anig|EHA17960.1 aory|EIT74726.1 asyd|OJJ63509.1 ater|EAU38953.1 atub|OJI90812.1 aver|OJI97421.1 awen|OJJ38969.1 azon|OJJ49740.1

Group3215: abra|OJJ73838.1 acar|OOG00989.1 acla|EAW07733.1 afis|EAW22765.1 afla|EED50027.1 afum|EAL90434.1 agla|OJJ83655.1 akaw|GAA89456.1 aluc|OJZ88211.1 anid|CBF88001.1 anig|EHA17958.1 aory|EIT74637.1 asyd|OJJ63508.1 ater|EAU38952.1 atub|OJI90811.1 aver|OJI97422.1 awen|OJJ38968.1 azon|OJJ49738.1

Group3216: abra|OJJ73841.1 acar|OOG00992.1 acla|EAW07736.1 afis|EAW22772.1 afla|EED50031.1 afum|EAL90426.1 agla|OJJ83602.1 akaw|GAA89459.1 aluc|OJZ88207.1 anid|CBF87996.1 anig|EHA17955.1 aory|EIT74633.1 asyd|OJJ63505.1 ater|EAU38941.1 atub|OJI90808.1 aver|OJI97425.1 awen|OJJ38965.1 azon|OJJ49736.1

Group3217: abra|OJJ73842.1 acar|OOG00993.1 acla|EAW07737.1 afis|EAW22773.1 afla|EED50032.1 afum|EAL90425.1 agla|OJJ83603.1 akaw|GAA89460.1 aluc|OJZ88206.1 anid|CBF87994.1 anig|EHA17954.1 aory|EIT74732.1 asyd|OJJ63504.1 ater|EAU38940.1 atub|OJI90807.1 aver|OJI97426.1 awen|OJJ38964.1 azon|OJJ49735.1

Group3218: abra|OJJ73843.1 acar|OOG00994.1 acla|EAW07738.1 afis|EAW22774.1 afla|EED50033.1 afum|EAL90424.1 agla|OJJ83604.1 akaw|GAA89461.1 aluc|OJZ88205.1 anid|CBF87992.1 anig|EHA17953.1 aory|EIT74665.1 asyd|OJJ63503.1 ater|EAU38939.1 atub|OJI90806.1 aver|OJI97427.1 awen|OJJ38963.1 azon|OJJ49734.1

Group3219: abra|OJJ73844.1 acar|OOG00995.1 acla|EAW07739.1 afis|EAW22775.1 afla|EED50034.1 afum|EAL90423.1 agla|OJJ83605.1 akaw|GAA89462.1 aluc|OJZ88204.1 anid|CBF87991.1 anig|EHA17952.1 aory|EIT74686.1 asyd|OJJ63502.1 ater|EAU38938.1 atub|OJI90805.1 aver|OJI97428.1 awen|OJJ38962.1 azon|OJJ49733.1

Group3220: abra|OJJ73846.1 acar|OOG00997.1 acla|EAW07741.1 afis|EAW22777.1 afla|EED50037.1 afum|EAL90421.1 agla|OJJ83628.1 akaw|GAA89464.1 aluc|OJZ88202.1 anid|CBF87987.1 anig|EHA17950.1 aory|EIT74713.1 asyd|OJJ63500.1 ater|EAU38937.1 atub|OJI90802.1 aver|OJI97430.1 awen|OJJ38960.1 azon|OJJ49731.1

Group3221: abra|OJJ73848.1 acar|OOG00998.1 acla|EAW07742.1 afis|EAW22778.1 afla|EED50038.1 afum|EAL90420.1 agla|OJJ83632.1 akaw|GAA89465.1 aluc|OJZ88201.1 anid|CBF87983.1 anig|EHA17949.1 aory|EIT74616.1 asyd|OJJ63497.1 ater|EAU38936.1 atub|OJI90799.1 aver|OJI97432.1 awen|OJJ38959.1 azon|OJJ49730.1

Group3222: abra|OJJ73850.1 acar|OOG01000.1 acla|EAW07743.1 afis|EAW22780.1 afla|EED50041.1 afum|EAL90419.1 agla|OJJ83633.1 akaw|GAA89467.1 aluc|OJZ88199.1 anid|CBF87970.1 anig|EHA17947.1 aory|EIT74714.1 asyd|OJJ63494.1 ater|EAU38932.1 atub|OJI90797.1 aver|OJI97435.1 awen|OJJ38957.1 azon|OJJ49729.1

Group3223: abra|OJJ73851.1 acar|OOG01001.1 acla|EAW07745.1 afis|EAW22782.1 afla|EED50044.1 afum|EAL90417.1 agla|OJJ83644.1 akaw|GAA89468.1 aluc|OJZ88197.1 anid|CBF87972.1 anig|EHA17946.1 aory|EIT74643.1 asyd|OJJ63493.1 ater|EAU38930.1 atub|OJI90796.1 aver|OJI97436.1 awen|OJJ38955.1 azon|OJJ49727.1

Group3224: abra|OJJ73854.1 acar|OOG01002.1 acla|EAW07746.1 afis|EAW22783.1 afla|EED50045.1 afum|EAL90416.1 agla|OJJ83653.1 akaw|GAA89470.1 aluc|OJZ88194.1 anid|CBF87974.1 anig|EHA17945.1 aory|EIT74701.1 asyd|OJJ63492.1 ater|EAU38926.1 atub|OJI90793.1 aver|OJI97437.1 awen|OJJ38954.1 azon|OJJ49726.1

Group3225: abra|OJJ73861.1 acar|OOG01007.1 acla|EAW07751.1 afis|EAW22788.1 afla|EED50052.1 afum|EAL90411.1 agla|OJJ83588.1 akaw|GAA89481.1 aluc|OJZ88182.1 anid|CBF87963.1 anig|EHA17934.1 aory|EIT74670.1 asyd|OJJ63484.1 ater|EAU38922.1 atub|OJI90781.1 aver|OJI97445.1 awen|OJJ38946.1 azon|OJJ49718.1

Group3226: abra|OJJ73864.1 acar|OOF89914.1 acla|EAW07754.1 afis|EAW22791.1 afla|EED50058.1 afum|EAL90408.1 agla|OJJ83567.1 akaw|GAA89478.1 aluc|OJZ88185.1 anid|CBF87957.1 anig|EHA17936.1 aory|EIT74695.1 asyd|OJJ63481.1 ater|EAU38920.1 atub|OJI90784.1 aver|OJI97449.1 awen|OJJ38943.1 azon|OJJ49716.1

Group3227: abra|OJJ73867.1 acar|OOF89912.1 acla|EAW07755.1 afis|EAW22792.1 afla|EED50059.1 afum|EAL90407.2 agla|OJJ83568.1 akaw|GAA89475.1 aluc|OJZ88188.1 anid|CBF87955.1 anig|EHA17939.1 aory|EIT74728.1 asyd|OJJ63480.1 ater|EAU38919.1 atub|OJI90787.1 aver|OJI97450.1 awen|OJJ38941.1 azon|OJJ49715.1

Group3228: abra|OJJ73869.1 acar|OOG01012.1 acla|EAW07757.1 afis|EAW22794.1 afla|EED50062.1 afum|EAL90405.1 agla|OJJ83570.1 akaw|GAA89473.1 aluc|OJZ88190.1 anid|CBF87952.1 anig|EHA17941.1 aory|EIT74606.1 asyd|OJJ63478.1 ater|EAU38917.1 atub|OJI90789.1 aver|OJI97452.1 awen|OJJ38939.1 azon|OJJ49713.1

Group3229: abra|OJJ73874.1 acar|OOG01014.1 acla|EAW07759.1 afis|EAW22795.1 afla|EED50063.1 afum|EAL90404.1 agla|OJJ83571.1 akaw|GAA89486.1 aluc|OJZ88176.1 anid|CBF87950.1 anig|EHA17928.1 aory|EIT74699.1 asyd|OJJ63477.1 ater|EAU38915.1 atub|OJI90774.1 aver|OJI97453.1 awen|OJJ38938.1 azon|OJJ49712.1

Group3230: abra|OJJ73876.1 acar|OOG01016.1 acla|EAW07761.1 afis|EAW22797.1 afla|EED50066.1 afum|EAL90402.1 agla|OJJ83574.1 akaw|GAA89488.1 aluc|OJZ88173.1 anid|CBF87946.1 anig|EHA17926.1

aory|EIT74703.1 asyd|OJJ63475.1 ater|EAU38913.1 atub|OJI90771.1 aver|OJI97455.1 awen|OJJ38932.1 azon|OJJ49708.1

Group3231: abra|OJJ73881.1 acar|OOG01020.1 acla|EAW07763.1 afis|EAW22799.1 afla|EED50069.1 afum|EAL90400.1 agla|OJJ83576.1 akaw|GAA89492.1 aluc|OJZ88169.1 anid|CBF87936.1 anig|EHA17923.1 aory|EIT74640.1 asyd|OJJ63466.1 ater|EAU38910.1 atub|OJI90767.1 aver|OJI97466.1 awen|OJJ38930.1 azon|OJJ49706.1

Group3232: abra|OJJ73882.1 acar|OOG01021.1 acla|EAW07764.1 afis|EAW22800.1 afla|EED50070.1 afum|EAL90399.1 agla|OJJ83577.1 akaw|GAA89493.1 aluc|OJZ88168.1 anid|CBF87934.1 anig|EHA17922.1 aory|EIT74735.1 asyd|OJJ63465.1 ater|EAU38909.1 atub|OJI90766.1 aver|OJI97467.1 awen|OJJ38929.1 azon|OJJ49705.1

Group3233: abra|OJJ73883.1 acar|OOG01022.1 acla|EAW07765.1 afis|EAW22801.1 afla|EED50071.1 afum|EAL90398.1 agla|OJJ83578.1 akaw|GAA89494.1 aluc|OJZ88167.1 anid|CBF87932.1 anig|EHA17921.1 aory|EIT74619.1 asyd|OJJ63464.1 ater|EAU38908.1 atub|OJI90765.1 aver|OJI97468.1 awen|OJJ38928.1 azon|OJJ49704.1

Group3234: abra|OJJ73884.1 acar|OOG01023.1 acla|EAW07766.1 afis|EAW22802.1 afla|EED50072.1 afum|EAL90397.1 agla|OJJ83579.1 akaw|GAA89495.1 aluc|OJZ88166.1 anid|CBF87930.1 anig|EHA17920.1 aory|EIT74719.1 asyd|OJJ63463.1 ater|EAU38907.1 atub|OJI90764.1 aver|OJI97469.1 awen|OJJ38927.1 azon|OJJ49703.1

Group3235: abra|OJJ73886.1 acar|OOG01024.1 acla|EAW07768.1 afis|EAW22804.1 afla|EED50074.1 afum|EAL90395.1 agla|OJJ83581.1 akaw|GAA89497.1 aluc|OJZ88164.1 anid|CBF87926.1 anig|EHA17918.1 aory|EIT74660.1 asyd|OJJ63461.1 ater|EAU38905.1 atub|OJI90762.1 aver|OJI97471.1 awen|OJJ38924.1 azon|OJJ49701.1

Group3236: abra|OJJ73889.1 acar|OOG01027.1 acla|EAW07773.1 afis|EAW22809.1 afla|EED50078.1 afum|EAL90392.1 agla|OJJ83584.1 akaw|GAA89501.1 aluc|OJZ88160.1 anid|CBF87904.1 anig|EHA17915.1 aory|EIT74748.1 asyd|OJJ63457.1 ater|EAU38902.1 atub|OJI90759.1 aver|OJI97475.1 awen|OJJ38921.1 azon|OJJ49698.1

Group3237: abra|OJJ73890.1 acar|OOG01030.1 acla|EAW07774.1 afis|EAW22810.1 afla|EED50079.1 afum|EAL90391.1 agla|OJJ83585.1 akaw|GAA89502.1 aluc|OJZ88159.1 anid|CBF87906.1 anig|EHA17914.1 aory|EIT74744.1 asyd|OJJ63456.1 ater|EAU38901.1 atub|OJI90758.1 aver|OJI97476.1 awen|OJJ38920.1 azon|OJJ49697.1

Group3238: abra|OJJ73891.1 acar|OOG01029.1 acla|EAW07775.1 afis|EAW22811.1 afla|EED50080.1 afum|EAL90390.1 agla|OJJ83566.1 akaw|GAA89503.1 aluc|OJZ88158.1 anid|CBF87908.1 anig|EHA17913.1 aory|EIT74652.1 asyd|OJJ63455.1 ater|EAU38900.1 atub|OJI90757.1 aver|OJI97477.1 awen|OJJ38915.1 azon|OJJ49696.1

Group3239: abra|OJJ73897.1 acar|OOG01034.1 acla|EAW07778.1 afis|EAW22815.1 afla|EED50088.1 afum|EAL90386.1 agla|OJJ83527.1 akaw|GAA89507.1 aluc|OJZ88149.1 anid|CBF87915.1 anig|EHA17910.1 aory|EIT74671.1 asyd|OJJ63448.1 ater|EAU38896.1 atub|OJI90752.1 aver|OJI97486.1 awen|OJJ38913.1 azon|OJJ49693.1

Group3240: abra|OJJ73901.1 acar|OOG01036.1 acla|EAW07780.1 afis|EAW22816.1 afla|EED50089.1 afum|EAL90385.1 agla|OJJ83524.1 akaw|GAA89508.1 aluc|OJZ88145.1 anid|CBF87918.1 anig|EHA17908.1 aory|EIT74605.1 asyd|OJJ63447.1 ater|EAU38895.1 atub|OJI90749.1 aver|OJI97489.1 awen|OJJ38911.1 azon|OJJ49692.1

Group3241: abra|OJJ73902.1 acar|OOG01037.1 acla|EAW07781.1 afis|EAW22817.1 afla|EED50090.1 afum|EAL90384.1 agla|OJJ83521.1 akaw|GAA89509.1 aluc|OJZ88144.1 anid|CBF87920.1 anig|EHA17907.1 aory|EIT74746.1 asyd|OJJ63446.1 ater|EAU38894.1 atub|OJI90748.1 aver|OJI97490.1 awen|OJJ38910.1 azon|OJJ49691.1

Group3242: abra|OJJ73904.1 acar|OOG01038.1 acla|EAW07783.1 afis|EAW22819.1 afla|EED50092.1 afum|EAL90382.1 agla|OJJ83519.1 akaw|GAA89511.1 aluc|OJZ88142.1 anid|CBF87899.1 anig|EHA17905.1 aory|EIT74751.1 asyd|OJJ63444.1 ater|EAU38891.1 atub|OJI90746.1 aver|OJI97492.1 awen|OJJ38908.1 azon|OJJ49689.1

Group3243: abra|OJJ73908.1 acar|OOG01041.1 acla|EAW07786.1 afis|EAW22822.1 afla|EED50097.1 afum|EAL90378.2 agla|OJJ83516.1 akaw|GAA89515.1 aluc|OJZ88139.1 anid|CBF87894.1 anig|EHA17903.1 aory|EIT74681.1 asyd|OJJ63441.1 ater|EAU38888.1 atub|OJI90742.1 aver|OJI97495.1 awen|OJJ38905.1 azon|OJJ49686.1

Group3244: abra|OJJ73909.1 acar|OOG01042.1 acla|EAW07787.1 afis|EAW22823.1 afla|EED50098.1 afum|EAL90377.1 agla|OJJ83515.1 akaw|GAA89516.1 aluc|OJZ88138.1 anid|CBF87892.1 anig|EHA17902.1 aory|EIT74745.1 asyd|OJJ63440.1 ater|EAU38887.1 atub|OJI90741.1 aver|OJI97496.1 awen|OJJ38904.1 azon|OJJ49685.1

Group3245: abra|OJJ73910.1 acar|OOG01043.1 acla|EAW07788.1 afis|EAW22824.1 afla|EED50099.1 afum|EAL90376.1 agla|OJJ83514.1 akaw|GAA89517.1 aluc|OJZ88137.1 anid|CBF87890.1 anig|EHA17901.1 aory|EIT74620.1 asyd|OJJ63439.1 ater|EAU38886.1 atub|OJI90740.1 aver|OJI97497.1 awen|OJJ38903.1 azon|OJJ49684.1

Group3246: abra|OJJ73912.1 acar|OOG01044.1 acla|EAW07789.1 afis|EAW22825.1 afla|EED50100.1 afum|EAL90375.2 agla|OJJ83513.1 akaw|GAA89518.1 aluc|OJZ88136.1 anid|CBF87888.1 anig|EHA17899.1 aory|EIT74685.1 asyd|OJJ63438.1 ater|EAU38885.1 atub|OJI90739.1 aver|OJI97499.1 awen|OJJ38902.1 azon|OJJ49683.1

Group3247: abra|OJJ73915.1 acar|OOG01047.1 acla|EAW07791.1 afis|EAW22829.1 afla|EED50104.1 afum|EAL90373.2 agla|OJJ83510.1 akaw|GAA89521.1 aluc|OJZ88133.1 anid|CBF87886.1 anig|EHA17897.1 aory|EIT74692.1 asyd|OJJ63437.1 ater|EAU38882.1 atub|OJI90737.1 aver|OJI97500.1 awen|OJJ38899.1 azon|OJJ49680.1

Group3248: abra|OJJ73916.1 acar|OOG01048.1 acla|EAW07792.1 afis|EAW22830.1 afla|EED50105.1 afum|EAL90372.2 agla|OJJ83509.1 akaw|GAA89522.1 aluc|OJZ88131.1 anid|CBF87883.1 anig|EHA17896.1 aory|EIT74639.1 asyd|OJJ63435.1 ater|EAU38881.1 atub|OJI90736.1 aver|OJI97502.1 awen|OJJ38898.1 azon|OJJ49679.1

Group3249: abra|OJJ73920.1 acar|OOG01051.1 acla|EAW07797.1 afis|EAW22834.1 afla|EED50109.1 afum|EAL90368.1 agla|OJJ83505.1 akaw|GAA89527.1 aluc|OJZ88126.1 anid|CBF87876.1 anig|EHA17892.1 aory|EIT74609.1 asyd|OJJ63432.1 ater|EAU38875.1 atub|OJI90733.1 aver|OJI97505.1 awen|OJJ38890.1 azon|OJJ49676.1

Group3250: abra|OJJ73923.1 acar|OOG01056.1 acla|EAW07800.1 afis|EAW22837.1 afla|EED50112.1 afum|EAL90365.2 agla|OJJ83504.1 akaw|GAA89530.1 aluc|OJZ88120.1 anid|CBF87867.1 anig|EHA17889.1 aory|EIT74684.1 asyd|OJJ63428.1 ater|EAU38873.1 atub|OJI90730.1 aver|OJI97509.1 awen|OJJ38887.1 azon|OJJ49675.1

Group3251: abra|OJJ73924.1 acar|OOG01057.1 acla|EAW07801.1 afis|EAW22838.1 afla|EED50113.1 afum|EAL90364.2 agla|OJJ83503.1 akaw|GAA89531.1 aluc|OJZ88119.1 anid|CBF87865.1 anig|EHA17888.1 aory|EIT74621.1 asyd|OJJ63427.1 ater|EAU38872.1 atub|OJI90729.1 aver|OJI97510.1 awen|OJJ38885.1 azon|OJJ49674.1

Group3252: abra|OJJ73926.1 acar|OOG01058.1 acla|EAW07803.1 afis|EAW22840.1 afla|EED50115.1 afum|EAL90363.1 agla|OJJ83501.1 akaw|GAA89533.1 aluc|OJZ88117.1 anid|CBF87861.1 anig|EHA17887.1 aory|EIT74702.1 asyd|OJJ63425.1 ater|EAU38870.1 atub|OJI90727.1 aver|OJI97512.1 awen|OJJ38883.1 azon|OJJ49672.1

Group3253: abra|OJJ73927.1 acar|OOG01060.1 acla|EAW07804.1 afis|EAW22841.1 afla|EED50116.1 afum|EAL90362.2 agla|OJJ83500.1 akaw|GAA89534.1 aluc|OJZ88116.1 anid|CBF87860.1 anig|EHA17886.1 aory|EIT74608.1 asyd|OJJ63424.1 ater|EAU38869.1 atub|OJI90725.1 aver|OJI97513.1 awen|OJJ38882.1 azon|OJJ49671.1

Group3254: abra|OJJ73928.1 acar|OOG01061.1 acla|EAW07805.1 afis|EAW22842.1 afla|EED50117.1 afum|EAL90361.1 agla|OJJ83496.1 akaw|GAA89535.1 aluc|OJZ88115.1 anid|CBF87858.1 anig|EHA17885.1 aory|EIT74650.1 asyd|OJJ63423.1 ater|EAU38868.1 atub|OJI90724.1 aver|OJI97514.1 awen|OJJ38881.1 azon|OJJ49670.1

Group3255: abra|OJJ73930.1 acar|OOG01062.1 acla|EAW07808.1 afis|EAW22845.1 afla|EED50120.1 afum|EAL90357.1 agla|OJJ83492.1 akaw|GAA89538.1 aluc|OJZ88112.1 anid|CBF87846.1 anig|EHA17882.1 aory|EIT74689.1 asyd|OJJ63421.1 ater|EAU38865.1 atub|OJI90720.1 aver|OJI97517.1 awen|OJJ38878.1 azon|OJJ49667.1

Group3256: abra|OJJ73931.1 acar|OOG01063.1 acla|EAW07809.1 afis|EAW22846.1 afla|EED50121.1 afum|EAL90356.1 agla|OJJ83485.1 akaw|GAA89539.1 aluc|OJZ88111.1 anid|CBF87848.1 anig|EHA17881.1 aory|EIT74731.1 asyd|OJJ63420.1 ater|EAU38864.1 atub|OJI90719.1 aver|OJI97518.1 awen|OJJ38877.1 azon|OJJ49666.1

Group3257: abra|OJJ73938.1 acar|OOG01069.1 afis|EAW22850.1 afla|EED50125.1 afum|EAL90350.1 agla|OJJ83473.1 akaw|GAA89543.1 aluc|OJZ88106.1 anid|CBF87855.1 anig|EHA17875.1 aory|EIT74705.1 asyd|OJJ63416.1 ater|EAU38859.1 atub|OJI90714.1 aver|OJI97522.1 awen|OJJ38871.1 azon|OJJ49663.1 acla|EAW07814.1

Group3258: abra|OJJ73939.1 acar|OOG01068.1 acla|EAW07816.1 afis|EAW22851.1 afla|EED50126.1 afum|EAL90349.1 agla|OJJ83471.1 akaw|GAA89544.1 aluc|OJZ88105.1 anid|CBF87841.1 anig|EHA17874.1 aory|EIT74661.1 asyd|OJJ63414.1 ater|EAU38858.1 atub|OJI90713.1 aver|OJI97524.1 awen|OJJ38867.1 azon|OJJ49661.1

Group3259: abra|OJJ73944.1 acar|OOG01072.1 acla|EAW07817.1 afis|EAW22852.1 afla|EED50132.1 afum|EAL90348.1 agla|OJJ83469.1 akaw|GAA89549.1 aluc|OJZ88100.1 anid|CBF87839.1 anig|EHA17869.1

aory|EIT74662.1 asyd|OJJ63413.1 ater|EAU38857.1 atub|OJI90708.1 aver|OJI97525.1 awen|OJJ38866.1 azon|OJJ49660.1

Group3260: abra|OJJ73945.1 acar|OOG01073.1 acla|EAW07818.1 afis|EAW22853.1 afla|EED50133.1 afum|EAL90347.1 agla|OJJ83468.1 akaw|GAA89550.1 aluc|OJZ88099.1 anid|CBF87838.1 anig|EHA17868.1 aory|EIT74757.1 asyd|OJJ63412.1 ater|EAU38856.1 atub|OJI90707.1 aver|OJI97526.1 awen|OJJ38865.1 azon|OJJ49659.1

Group3261: abra|OJJ73946.1 acar|OOG01074.1 acla|EAW07819.1 afis|EAW22854.1 afla|EED50134.1 afum|EAL90346.1 agla|OJJ83467.1 akaw|GAA89551.1 aluc|OJZ88098.1 anid|CBF87836.1 anig|EHA17867.1 aory|EIT74630.1 asyd|OJJ63410.1 atub|OJI90706.1 aver|OJI97527.1 awen|OJJ38864.1 azon|OJJ49658.1 ater|EAU38807.1

Group3262: abra|OJJ73949.1 acar|OOG01077.1 acla|EAW07823.1 afis|EAW22859.1 afla|EED50137.1 afum|EAL90341.1 agla|OJJ83465.1 akaw|GAA89554.1 aluc|OJZ88095.1 anid|CBF87833.1 anig|EHA17865.1 aory|EIT74739.1 asyd|OJJ63408.1 ater|EAU38809.1 atub|OJI90703.1 aver|OJI97529.1 awen|OJJ38861.1 azon|OJJ49655.1

Group3263: abra|OJJ73952.1 acar|OOG01080.1 acla|EAW07826.1 afis|EAW22862.1 afla|EED50146.1 afum|EAL90338.1 agla|OJJ83462.1 akaw|GAA89557.1 aluc|OJZ88092.1 anid|CBF87832.1 anig|EHA17862.1 aory|EIT74749.1 asyd|OJJ63407.1 ater|EAU38816.1 atub|OJI90700.1 aver|OJI97530.1 awen|OJJ38858.1 azon|OJJ49656.1

Group3264: abra|OJJ73959.1 acar|OOG01086.1 acla|EAW07828.1 afis|EAW22864.1 afla|EED50148.1 afum|EAL90336.1 agla|OJJ83460.1 akaw|GAA89562.1 aluc|OJZ88086.1 anid|CBF87829.1 anig|EHA17857.1 aory|EIT74755.1 asyd|OJJ63405.1 ater|EAU38824.1 atub|OJI90695.1 aver|OJI97531.1 awen|OJJ38856.1 azon|OJJ49654.1

Group3265: abra|OJJ73961.1 acar|OOG01087.1 acla|EAW07830.1 afis|EAW22866.1 afla|EED50150.1 afum|EAL90333.1 agla|OJJ83458.1 akaw|GAA89564.1 aluc|OJZ88084.1 anid|CBF87827.1 anig|EHA17855.1 aory|EIT74659.1 asyd|OJJ63403.1 ater|EAU38827.1 atub|OJI90693.1 aver|OJI97533.1 awen|OJJ38854.1 azon|OJJ49652.1

Group3266: abra|OJJ73962.1 acar|OOG01089.1 acla|EAW07831.1 afis|EAW22867.1 afla|EED50151.1 afum|EAL90332.1 agla|OJJ83457.1 akaw|GAA89565.1 aluc|OJZ88083.1 anid|CBF87826.1 anig|EHA17854.1 aory|EIT74722.1 asyd|OJJ63402.1 ater|EAU38828.1 atub|OJI90692.1 aver|OJI97534.1 awen|OJJ38853.1 azon|OJJ49651.1

Group3267: abra|OJJ73964.1 acar|OOG01090.1 acla|EAW07834.1 afis|EAW22871.1 afla|EED50154.1 afum|EAL90327.1 agla|OJJ83452.1 akaw|GAA89567.1 aluc|OJZ88080.1 anid|CBF87822.1 anig|EHA17852.1 aory|EIT76938.1 asyd|OJJ63400.1 ater|EAU38835.1 atub|OJI90689.1 aver|OJI97537.1 awen|OJJ38851.1 azon|OJJ49647.1

Group3268: abra|OJJ73967.1 acar|OOG01093.1 acla|EAW07837.1 afis|EAW22874.1 afla|EED50157.1 afum|EAL90324.1 agla|OJJ83443.1 akaw|GAA89570.1 aluc|OJZ88077.1 anid|CBF87816.1 anig|EHA17849.1 aory|EIT76898.1 asyd|OJJ63395.1 ater|EAU38838.1 atub|OJI90686.1 aver|OJI97541.1 awen|OJJ38848.1 azon|OJJ49644.1

Group3269: abra|OJJ73971.1 acar|OOG01097.1 acla|EAW07842.1 afis|EAW22878.1 afla|EED50183.1 afum|EAL90321.1 agla|OJJ83446.1 akaw|GAA89574.1 aluc|OJZ88073.1 anid|CBF87811.1 anig|EHA17846.1 aory|EIT77380.1 asyd|OJJ63390.1 ater|EAU38842.1 atub|OJI90682.1 aver|OJI97545.1 awen|OJJ38844.1 azon|OJJ49641.1

Group3270: abra|OJJ73976.1 acar|OOG01103.1 acla|EAW07847.1 afis|EAW22883.1 afla|EED50176.1 afum|EAL90316.1 agla|OJJ83441.1 akaw|GAA89579.1 aluc|OJZ88068.1 anid|CBF87803.1 anig|EHA17841.1 aory|EIT72282.1 asyd|OJJ63384.1 ater|EAU38846.1 atub|OJI90677.1 aver|OJI97551.1 awen|OJJ38839.1 azon|OJJ49636.1

Group3271: abra|OJJ73978.1 acar|OOG01104.1 acla|EAW07849.1 afis|EAW22885.1 afla|EED50174.1 afum|EAL90313.1 agla|OJJ83438.1 akaw|GAA89581.1 aluc|OJZ88066.1 anid|CBF87801.1 anig|EHA17839.1 aory|EIT72285.1 asyd|OJJ63382.1 ater|EAU35978.1 atub|OJI90675.1 aver|OJI97553.1 awen|OJJ38837.1 azon|OJJ49634.1

Group3272: abra|OJJ73980.1 acar|OOG01107.1 acla|EAW07851.1 afis|EAW22887.1 afla|EED50172.1 afum|EAL90311.1 agla|OJJ83436.1 akaw|GAA89583.1 aluc|OJZ88064.1 anid|CBF87798.1 anig|EHA17837.1 aory|EIT72283.1 asyd|OJJ63380.1 ater|EAU35985.1 atub|OJI90673.1 aver|OJI97555.1 awen|OJJ38835.1 azon|OJJ49632.1

Group3273: abra|OJJ73982.1 acar|OOG01109.1 acla|EAW07854.1 afis|EAW22890.1 afla|EED54859.1 afum|EAL90308.1 agla|OJJ83420.1 akaw|GAA89586.1 aluc|OJZ88062.1 anid|CBF87791.1 anig|EHA17835.1 aory|EIT79448.1 asyd|OJJ63377.1 ater|EAU35982.1 atub|OJI90671.1 aver|OJI97569.1 awen|OJJ38831.1 azon|OJJ49630.1

Group3274: abra|OJJ73983.1 acar|OOG01110.1 acla|EAW07855.1 afis|EAW22891.1 afla|EED54860.1 afum|EAL90307.1 agla|OJJ83407.1 akaw|GAA89587.1 aluc|OJZ88061.1 anid|CBF87790.1 anig|EHA17834.1 aory|EIT79392.1 asyd|OJJ63376.1 ater|EAU36000.1 atub|OJI90670.1 aver|OJI97570.1 awen|OJJ38830.1 azon|OJJ49629.1

Group3275: abra|OJJ73986.1 acar|OOG01111.1 acla|EAW07857.1 afis|EAW22893.1 afla|EED54862.1 afum|EAL90305.2 agla|OJJ83410.1 akaw|GAA87469.1 aluc|OJZ88059.1 anid|CBF87783.1 anig|EHA17831.1 aory|EIT79335.1 asyd|OJJ63374.1 ater|EAU35990.1 atub|OJI90667.1 aver|OJI97572.1 awen|OJJ38827.1 azon|OJJ49627.1

Group3276: abra|OJJ73989.1 acar|OOG01115.1 acla|EAW07860.1 afis|EAW22895.1 afla|EED54864.1 afum|EAL90303.1 agla|OJJ83415.1 akaw|GAA87472.1 aluc|OJZ88056.1 anid|CBF74977.1 anig|EHA17828.1 aory|EIT79412.1 asyd|OJJ60090.1 ater|EAU35987.1 atub|OJI90664.1 aver|OJJ04177.1 awen|OJJ38823.1 azon|OJJ49622.1

Group3277: abra|OJJ73991.1 acar|OOG01117.1 acla|EAW07862.1 afis|EAW22897.1 afla|EED54868.1 afum|EAL90301.1 agla|OJJ83417.1 akaw|GAA87474.1 aluc|OJZ88054.1 anid|CBF87775.1 anig|EHA17826.1 aory|EIT79359.1 asyd|OJJ63369.1 ater|EAU35981.1 atub|OJI90662.1 aver|OJI97577.1 awen|OJJ38820.1 azon|OJJ49620.1

Group3278: abra|OJJ73992.1 acar|OOG01119.1 acla|EAW07863.1 afis|EAW22898.1 afla|EED54869.1 afum|EAL90299.1 agla|OJJ83393.1 akaw|GAA87475.1 aluc|OJZ88053.1 anid|CBF87773.1 anig|EHA17825.1 asyd|OJJ63368.1 ater|EAU36001.1 atub|OJI90661.1 aver|OJI97578.1 awen|OJJ38732.1 azon|OJJ49619.1 agla|OJJ81800.1

Group3279: abra|OJJ73993.1 acar|OOG01118.1 acla|EAW07864.1 afis|EAW22899.1 afla|EED54870.1 afum|EAL90298.1 agla|OJJ83392.1 akaw|GAA87476.1 aluc|OJZ88052.1 anid|CBF87771.1 anig|EHA17824.1 aory|EIT79377.1 asyd|OJJ63367.1 ater|EAU36002.1 atub|OJI90660.1 aver|OJI97579.1 awen|OJJ38731.1 azon|OJJ49618.1

Group3280: abra|OJJ74000.1 acar|OOG01121.1 acla|EAW07866.1 afis|EAW22901.1 afla|EED54872.1 afum|EAL90296.1 agla|OJJ83389.1 akaw|GAA87482.1 aluc|OJZ88043.1 anid|CBF87767.1 anig|EHA17816.1 aory|EIT79383.1 asyd|OJJ63365.1 ater|EAU36006.1 atub|OJI90650.1 aver|OJI97582.1 awen|OJJ38729.1 azon|OJJ49615.1

Group3281: abra|OJJ74002.1 acar|OOG01122.1 acla|EAW07867.1 afis|EAW22902.1 afla|EED54873.1 afum|EAL90295.1 agla|OJJ83388.1 akaw|GAA87483.1 aluc|OJZ88042.1 anid|CBF87766.1 anig|EHA17815.1 aory|EIT79350.1 asyd|OJJ63364.1 ater|EAU36007.1 atub|OJI90649.1 aver|OJI97583.1 awen|OJJ38728.1 azon|OJJ49614.1

Group3282: abra|OJJ74003.1 acar|OOG01124.1 acla|EAW07868.1 afis|EAW22903.1 afla|EED54874.1 afum|EAL90294.1 agla|OJJ83387.1 akaw|GAA87484.1 aluc|OJZ88041.1 anid|CBF87764.1 anig|EHA17814.1 aory|EIT79433.1 asyd|OJJ63363.1 ater|EAU36010.1 atub|OJI90648.1 aver|OJI97584.1 awen|OJJ38727.1 azon|OJJ49613.1

Group3283: abra|OJJ74016.1 acar|OOG01137.1 acla|EAW07870.1 afis|EAW22905.1 afla|EED54897.1 afum|EAL90292.1 agla|OJJ83384.1 akaw|GAA87498.1 aluc|OJZ88024.1 anid|CBF87713.1 anig|EHA17801.1 aory|EIT80309.1 asyd|OJJ63338.1 ater|EAU36034.1 atub|OJI90632.1 aver|OJI97623.1 awen|OJJ38725.1 azon|OJJ49609.1

Group3284: abra|OJJ74017.1 acar|OOG01138.1 acla|EAW07871.1 afis|EAW22906.1 afla|EED54898.1 afum|EAL90291.1 agla|OJJ83383.1 akaw|GAA87499.1 aluc|OJZ88022.1 anid|CBF87712.1 anig|EHA17800.1 aory|EIT80293.1 asyd|OJJ63337.1 ater|EAU36033.1 atub|OJI90631.1 aver|OJI97625.1 awen|OJJ38724.1 azon|OJJ49608.1

Group3285: abra|OJJ74020.1 acar|OOG01139.1 acla|EAW07874.1 afis|EAW22909.1 afla|EED54901.1 afum|EAL90288.1 agla|OJJ83380.1 akaw|GAA87502.1 aluc|OJZ88019.1 anid|CBF87705.1 anig|EHA17798.1 aory|EIT80370.1 asyd|OJJ63332.1 ater|EAU36030.1 atub|OJI90628.1 aver|OJI97630.1 awen|OJJ38656.1 azon|OJJ49605.1

Group3286: abra|OJJ74021.1 acar|OOG01140.1 acla|EAW07875.1 afis|EAW22910.1 afla|EED54902.1 afum|EAL90287.1 agla|OJJ83379.1 akaw|GAA87503.1 aluc|OJZ88018.1 anid|CBF87703.1 anig|EHA17797.1 aory|EIT80365.1 asyd|OJJ63331.1 ater|EAU36029.1 atub|OJI90627.1 aver|OJI97631.1 awen|OJJ38655.1 azon|OJJ49604.1

Group3287: abra|OJJ74022.1 acar|OOG01141.1 acla|EAW07876.1 afis|EAW22911.1 afla|EED54904.1 afum|EAL90286.1 agla|OJJ83378.1 akaw|GAA87505.1 aluc|OJZ88015.1 anid|CBF87694.1 anig|EHA17796.1 aory|EIT80313.1 asyd|OJJ63326.1 ater|EAU36028.1 atub|OJI90625.1 aver|OJI97636.1 awen|OJJ38654.1 azon|OJJ49603.1

Group3288: abra|OJJ74023.1 acar|OOG01142.1 acla|EAW07877.1 afis|EAW22912.1 afla|EED54905.1 afum|EAL90285.1 agla|OJJ83377.1 akaw|GAA87504.1 aluc|OJZ88017.1 anid|CBF87695.1 anig|EHA17795.1

aory|EIT80317.1 asyd|OJJ63327.1 ater|EAU36027.1 atub|OJI90626.1 aver|OJI97635.1 awen|OJJ38653.1 azon|OJJ49602.1

Group3289: abra|OJJ74024.1 acar|OOG01143.1 acla|EAW07878.1 afis|EAW22913.1 afla|EED54906.1 afum|EAL90284.1 agla|OJJ83376.1 akaw|GAA87506.1 aluc|OJZ88014.1 anid|CBF87699.1 anig|EHA17794.1 aory|EIT80324.1 asyd|OJJ63329.1 ater|EAU36026.1 atub|OJI90623.1 aver|OJI97633.1 awen|OJJ38652.1 azon|OJJ49601.1

Group3290: abra|OJJ74027.1 acar|OOG01146.1 acla|EAW07879.1 afis|EAW22914.1 afla|EED54907.1 afum|EAL90283.1 agla|OJJ83375.1 akaw|GAA87510.1 aluc|OJZ88010.1 anid|CBF87697.1 anig|EHA17790.1 aory|EIT80315.1 asyd|OJJ63328.1 ater|EAU30708.1 atub|OJI90619.1 aver|OJI97634.1 awen|OJJ38651.1 azon|OJJ49599.1

Group3291: abra|OJJ74029.1 acar|OOG01147.1 acla|EAW07880.1 afis|EAW22915.1 afla|EED54911.1 afum|EAL90282.1 agla|OJJ83373.1 akaw|GAA87512.1 aluc|OJZ88008.1 anid|CBF87690.1 anig|EHA17788.1 aory|EIT80322.1 asyd|OJJ63325.1 ater|EAU30707.1 atub|OJI90617.1 aver|OJI97637.1 awen|OJJ38649.1 azon|OJJ49595.1

Group3292: abra|OJJ74032.1 acar|OOG01151.1 acla|EAW07883.1 afis|EAW22918.1 afla|EED54915.1 afum|EAL90279.1 agla|OJJ83371.1 akaw|GAA87516.1 aluc|OJZ88005.1 anid|CBF87687.1 anig|EHA17786.1 aory|EIT80289.1 asyd|OJJ63320.1 ater|EAU30705.1 atub|OJI90613.1 aver|OJI97639.1 awen|OJJ38646.1 azon|OJJ49588.1

Group3293: abra|OJJ74033.1 acar|OOG01152.1 acla|EAW07884.1 afis|EAW22919.1 afla|EED54916.1 afum|EAL90278.1 agla|OJJ83369.1 akaw|GAA87517.1 aluc|OJZ88004.1 anid|CBF87685.1 anig|EHA17785.1 aory|EIT80336.1 asyd|OJJ63319.1 ater|EAU30704.1 atub|OJI90612.1 aver|OJI97640.1 awen|OJJ38645.1 azon|OJJ49587.1

Group3294: abra|OJJ74035.1 acla|EAW07885.1 afis|EAW22920.1 afla|EED54917.1 afum|EAL90277.1 agla|OJJ83368.1 akaw|GAA87518.1 anid|CBF87683.1 aory|EIT80285.1 asyd|OJJ63318.1 ater|EAU30703.1 aver|OJI97641.1 awen|OJJ38644.1 azon|OJJ49586.1 acar|OOG01153.1 aluc|OJZ88003.1 anig|EHA17784.1 atub|OJI90611.1

Group3295: abra|OJJ74036.1 acar|OOG01155.1 acla|EAW07886.1 afis|EAW22921.1 afla|EED54918.1 afum|EAL90276.1 agla|OJJ83367.1 akaw|GAA87519.1 aluc|OJZ88000.1 anid|CBF87681.1 anig|EHA17782.1 aory|EIT80300.1 asyd|OJJ63317.1 ater|EAU30702.1 atub|OJI90609.1 aver|OJI97642.1 awen|OJJ38643.1 azon|OJJ49585.1

Group3296: abra|OJJ74037.1 acar|OOG01156.1 acla|EAW07887.1 afis|EAW22922.1 afla|EED54919.1 afum|EAL90275.1 agla|OJJ83366.1 akaw|GAA87520.1 aluc|OJZ87999.1 anid|CBF87680.1 anig|EHA17781.1 aory|EIT80390.1 asyd|OJJ63316.1 ater|EAU30701.1 atub|OJI90608.1 aver|OJI97643.1 awen|OJJ38642.1 azon|OJJ49584.1

Group3297: abra|OJJ74038.1 acar|OOG01157.1 acla|EAW07888.1 afis|EAW22923.1 afla|EED54920.1 afum|EAL90274.1 agla|OJJ83365.1 akaw|GAA87521.1 aluc|OJZ87998.1 anid|CBF87678.1 anig|EHA17780.1 aory|EIT80297.1 asyd|OJJ63315.1 ater|EAU30700.1 atub|OJI90607.1 aver|OJI97644.1 awen|OJJ38641.1 azon|OJJ49583.1

Group3298: abra|OJJ74039.1 acar|OOG01158.1 acla|EAW07889.1 afis|EAW22924.1 afla|EED54921.1 afum|EAL90273.1 agla|OJJ83364.1 akaw|GAA87522.1 aluc|OJZ87997.1 anid|CBF87676.1 anig|EHA17779.1 aory|EIT80292.1 asyd|OJJ63314.1 ater|EAU30699.1 atub|OJI90606.1 aver|OJI97645.1 awen|OJJ38640.1 azon|OJJ49582.1

Group3299: abra|OJJ74040.1 acar|OOG01159.1 acla|EAW07890.1 afis|EAW22925.1 afla|EED54922.1 afum|EAL90272.1 agla|OJJ83363.1 akaw|GAA87523.1 aluc|OJZ87996.1 anid|CBF87674.1 anig|EHA17778.1 aory|EIT80341.1 asyd|OJJ63313.1 ater|EAU30698.1 atub|OJI90605.1 aver|OJI97646.1 awen|OJJ38639.1 azon|OJJ49581.1

Group3300: abra|OJJ74043.1 acar|OOG01161.1 acla|EAW07892.1 afis|EAW22927.1 afla|EED54924.1 afum|EAL90270.1 agla|OJJ83361.1 akaw|GAA87525.1 aluc|OJZ87994.1 anid|CBF87671.1 anig|EHA17776.1 aory|EIT80287.1 asyd|OJJ63311.1 ater|EAU30696.1 atub|OJI90603.1 aver|OJI97648.1 awen|OJJ38637.1 azon|OJJ49579.1

Group3301: abra|OJJ74045.1 acar|OOG01164.1 acla|EAW07895.1 afis|EAW22930.1 afla|EED58123.1 afum|EAL90267.1 agla|OJJ83358.1 akaw|GAA87528.1 aluc|OJZ87990.1 anid|CBF87667.1 anig|EHA17773.1 aory|EIT79163.1 asyd|OJJ63308.1 ater|EAU30694.1 atub|OJI90600.1 aver|OJI97651.1 awen|OJJ38634.1 azon|OJJ49576.1

Group3302: abra|OJJ74048.1 acar|OOF89921.1 acla|EAW07897.1 afis|EAW22932.1 afla|EED58126.1 afum|EAL90264.1 agla|OJJ83355.1 akaw|GAA87530.1 aluc|OJZ87987.1 anid|CBF87662.1 anig|EHA17770.1 aory|EIT79157.1 asyd|OJJ63305.1 ater|EAU30692.1 atub|OJI90597.1 aver|OJI97654.1 awen|OJJ38631.1 azon|OJJ49573.1

Group3303: abra|OJJ74049.1 acar|OOF89922.1 acla|EAW07898.1 afis|EAW22934.1 afla|EED58127.1 afum|EAL90262.1 agla|OJJ83354.1 akaw|GAA87531.1 aluc|OJZ87986.1 anid|CBF87661.1 anig|EHA17769.1 aory|EIT79153.1 asyd|OJJ63304.1 ater|EAU30691.1 atub|OJI90596.1 aver|OJI97655.1 awen|OJJ38630.1 azon|OJJ49572.1

Group3304: abra|OJJ74050.1 acar|OOF89923.1 acla|EAW07899.1 afis|EAW22935.1 afla|EED58128.1 afum|EAL90261.1 agla|OJJ83353.1 akaw|GAA87532.1 aluc|OJZ87985.1 anid|CBF87659.1 anig|EHA17768.1 aory|EIT79137.1 asyd|OJJ63303.1 ater|EAU30690.1 atub|OJI90595.1 aver|OJI97656.1 awen|OJJ38629.1 azon|OJJ49571.1

Group3305: abra|OJJ74053.1 acar|OOG01166.1 acla|EAW07901.1 afis|EAW22938.1 afla|EED58131.1 afum|EAL90258.1 agla|OJJ83351.1 akaw|GAA87535.1 aluc|OJZ87982.1 anid|CBF87654.1 anig|EHA17765.1 aory|EIT79120.1 asyd|OJJ63299.1 ater|EAU30687.1 atub|OJI90592.1 aver|OJI97659.1 awen|OJJ38626.1 azon|OJJ49568.1

Group3306: abra|OJJ74054.1 acar|OOG01167.1 acla|EAW07902.1 afis|EAW22939.1 afla|EED58133.1 afum|EAL90257.1 agla|OJJ83349.1 akaw|GAA87536.1 aluc|OJZ87981.1 anid|CBF87652.1 anig|EHA17764.1 aory|EIT79138.1 asyd|OJJ63298.1 ater|EAU30686.1 atub|OJI90591.1 aver|OJI97660.1 awen|OJJ38625.1 azon|OJJ49567.1

Group3307: abra|OJJ74056.1 acar|OOG01169.1 acla|EAW07903.1 afis|EAW22940.1 afla|EED58154.1 afum|EAL90256.1 agla|OJJ83348.1 akaw|GAA87538.1 aluc|OJZ87979.1 anid|CBF87650.1 anig|EHA17762.1 aory|EIT79084.1 asyd|OJJ63297.1 ater|EAU30684.1 atub|OJI90589.1 aver|OJI97661.1 awen|OJJ38624.1 azon|OJJ49565.1

Group3308: abra|OJJ74060.1 acar|OOG01172.1 acla|EAW07905.1 afis|EAW22943.1 afla|EED58153.1 afum|EAL90254.1 agla|OJJ83345.1 akaw|GAA87541.1 aluc|OJZ87976.1 anid|CBF87646.1 anig|EHA17760.1 aory|EIT79073.1 asyd|OJJ63295.1 ater|EAU30681.1 atub|OJI90586.1 aver|OJI97663.1 awen|OJJ38621.1 azon|OJJ49562.1

Group3309: abra|OJJ74061.1 acar|OOG01173.1 acla|EAW07906.1 afis|EAW22944.1 afla|EED58151.1 afum|EAL90253.1 agla|OJJ83344.1 akaw|GAA87542.1 aluc|OJZ87975.1 anid|CBF87644.1 anig|EHA17759.1 aory|EIT79080.1 asyd|OJJ63294.1 ater|EAU30680.1 atub|OJI90585.1 aver|OJI97664.1 awen|OJJ38620.1 azon|OJJ49561.1

Group3310: abra|OJJ74062.1 acar|OOG01175.1 acla|EAW07907.1 afis|EAW22945.1 afla|EED58150.1 afum|EAL90252.1 agla|OJJ83343.1 akaw|GAA87543.1 aluc|OJZ87974.1 anid|CBF87642.1 anig|EHA17758.1 aory|EIT79097.1 asyd|OJJ63293.1 ater|EAU30679.1 atub|OJI90584.1 aver|OJI97665.1 awen|OJJ38619.1 azon|OJJ49560.1

Group3311: abra|OJJ74063.1 acar|OOG01176.1 acla|EAW07908.1 afis|EAW22946.1 afla|EED58149.1 afum|EAL90251.1 agla|OJJ83342.1 akaw|GAA87544.1 aluc|OJZ87973.1 anid|CBF87640.1 anig|EHA17757.1 aory|EIT79094.1 asyd|OJJ63292.1 ater|EAU30678.1 atub|OJI90583.1 aver|OJI97666.1 awen|OJJ38618.1 azon|OJJ49559.1

Group3312: abra|OJJ74064.1 acar|OOG01174.1 acla|EAW07909.1 afis|EAW22947.1 afla|EED58148.1 afum|EAL90250.1 agla|OJJ83341.1 akaw|GAA87545.1 aluc|OJZ87972.1 anid|CBF87639.1 anig|EHA17756.1 aory|EIT79089.1 asyd|OJJ63291.1 ater|EAU30677.1 atub|OJI90582.1 aver|OJI97667.1 awen|OJJ38617.1 azon|OJJ49558.1

Group3313: abra|OJJ74065.1 acar|OOG01177.1 acla|EAW07910.1 afis|EAW22948.1 afla|EED58147.1 afum|EAL90249.1 agla|OJJ83339.1 akaw|GAA87546.1 aluc|OJZ87971.1 anid|CBF87637.1 anig|EHA17755.1 aory|EIT79088.1 asyd|OJJ63290.1 ater|EAU30676.1 atub|OJI90581.1 aver|OJI97668.1 awen|OJJ38616.1 azon|OJJ49557.1

Group3314: abra|OJJ74066.1 acar|OOG01179.1 acla|EAW07911.1 afis|EAW22949.1 afla|EED58146.1 afum|EAL90248.1 agla|OJJ83338.1 akaw|GAA87547.1 aluc|OJZ87970.1 anid|CBF87636.1 anig|EHA17754.1 aory|EIT79070.1 asyd|OJJ63289.1 ater|EAU30675.1 atub|OJI90580.1 aver|OJI97669.1 awen|OJJ38615.1 azon|OJJ49556.1

Group3315: abra|OJJ74067.1 acar|OOG01178.1 acla|EAW07912.1 afis|EAW22950.1 afla|EED58145.1 afum|EAL90247.1 agla|OJJ83337.1 akaw|GAA87548.1 aluc|OJZ87969.1 anid|CBF87634.1 anig|EHA17753.1 aory|EIT79067.1 asyd|OJJ63288.1 ater|EAU30674.1 atub|OJI90579.1 aver|OJI97670.1 awen|OJJ38614.1 azon|OJJ49555.1

Group3316: abra|OJJ74068.1 acar|OOG01180.1 acla|EAW07913.1 afis|EAW22951.1 afla|EED58144.1 afum|EAL90246.1 agla|OJJ83336.1 akaw|GAA87549.1 aluc|OJZ87968.1 anid|CBF87632.1 anig|EHA17752.1 aory|EIT79092.1 asyd|OJJ63287.1 ater|EAU30673.1 atub|OJI90578.1 aver|OJI97671.1 awen|OJJ38613.1 azon|OJJ49554.1

Group3317: abra|OJJ74069.1 acar|OOG01181.1 acla|EAW07914.1 afis|EAW22952.1 afla|EED58143.1 afum|EAL90245.1 agla|OJJ83335.1 akaw|GAA87550.1 aluc|OJZ87967.1 anid|CBF87630.1 anig|EHA17751.1

aory|EIT79082.1 asyd|OJJ63286.1 ater|EAU30672.1 atub|OJI90577.1 aver|OJI97672.1 awen|OJJ38612.1 azon|OJJ49553.1

Group3318: abra|OJJ74070.1 acar|OOG01182.1 acla|EAW07915.1 afis|EAW22953.1 afla|EED58142.1 afum|EAL90244.1 agla|OJJ83334.1 akaw|GAA87551.1 aluc|OJZ87966.1 anid|CBF87629.1 anig|EHA17750.1 aory|EIT79065.1 asyd|OJJ63285.1 ater|EAU30671.1 atub|OJI90576.1 aver|OJI97673.1 awen|OJJ38611.1 azon|OJJ49552.1

Group3319: abra|OJJ74071.1 acar|OOG01183.1 acla|EAW07916.1 afis|EAW22954.1 afla|EED58141.1 afum|EAL90243.1 agla|OJJ83333.1 akaw|GAA87552.1 aluc|OJZ87965.1 anid|CBF87627.1 anig|EHA17749.1 aory|EIT79103.1 asyd|OJJ63284.1 ater|EAU30670.1 atub|OJI90575.1 aver|OJI97674.1 awen|OJJ38610.1 azon|OJJ49551.1

Group3320: abra|OJJ74072.1 acar|OOG01184.1 acla|EAW07917.1 afis|EAW22955.1 afla|EED58140.1 afum|EAL90242.1 agla|OJJ83332.1 akaw|GAA87553.1 aluc|OJZ87964.1 anid|CBF87625.1 anig|EHA17748.1 aory|EIT79149.1 asyd|OJJ63283.1 ater|EAU30669.1 atub|OJI90574.1 aver|OJI97675.1 awen|OJJ38608.1 azon|OJJ49550.1

Group3321: abra|OJJ74075.1 acar|OOG01187.1 acla|EAW07920.1 afis|EAW22958.1 afla|EED58138.1 afum|EAL90239.1 agla|OJJ83330.1 akaw|GAA87556.1 aluc|OJZ87961.1 anid|CBF87624.1 anig|EHA17745.1 aory|EIT79160.1 asyd|OJJ63281.1 ater|EAU30667.1 atub|OJI90571.1 aver|OJI97677.1 awen|OJJ38605.1 azon|OJJ49547.1

Group3322: abra|OJJ74076.1 acar|OOG01188.1 acla|EAW07921.1 afis|EAW22959.1 afla|EED58137.1 afum|EAL90238.1 agla|OJJ83329.1 akaw|GAA87557.1 aluc|OJZ87960.1 anid|CBF87622.1 anig|EHA17744.1 aory|EIT79140.1 asyd|OJJ63280.1 ater|EAU30666.1 atub|OJI90570.1 aver|OJI97678.1 awen|OJJ38604.1 azon|OJJ49546.1

Group3323: abra|OJJ74078.1 acar|OOG01190.1 acla|EAW07923.1 afis|EAW22961.1 afla|EED58135.1 afum|EAL90236.1 agla|OJJ83327.1 akaw|GAA87559.1 aluc|OJZ87957.1 anid|CBF87621.1 anig|EHA17742.1 aory|EIT79121.1 asyd|OJJ63278.1 ater|EAU30664.1 atub|OJI90568.1 aver|OJI97680.1 awen|OJJ38602.1 azon|OJJ49544.1

Group3324: abra|OJJ74079.1 acar|OOG01191.1 acla|EAW07924.1 afis|EAW22962.1 afla|EED58155.1 afum|EAL90235.1 agla|OJJ83326.1 akaw|GAA87560.1 aluc|OJZ87956.1 anid|CBF87619.1 anig|EHA17741.1 aory|EIT79083.1 asyd|OJJ63277.1 ater|EAU30663.1 atub|OJI90567.1 aver|OJI97681.1 awen|OJJ38601.1 azon|OJJ49543.1

Group3325: abra|OJJ74080.1 acar|OOG01192.1 acla|EAW07925.1 afis|EAW22963.1 afla|EED58156.1 afum|EAL90234.1 agla|OJJ83324.1 akaw|GAA87561.1 aluc|OJZ87955.1 anid|CBF87617.1 anig|EHA17740.1 aory|EIT79101.1 asyd|OJJ63276.1 ater|EAU30662.1 atub|OJI90566.1 aver|OJI97682.1 awen|OJJ38600.1 azon|OJJ49542.1

Group3326: abra|OJJ74081.1 acar|OOG01193.1 acla|EAW07926.1 afis|EAW22964.1 afla|EED58157.1 afum|EAL90233.1 agla|OJJ83323.1 akaw|GAA87562.1 aluc|OJZ87954.1 anid|CBF87615.1 anig|EHA17739.1 aory|EIT79095.1 asyd|OJJ63275.1 ater|EAU30661.1 atub|OJI90565.1 aver|OJI97683.1 awen|OJJ38599.1 azon|OJJ49541.1

Group3327: abra|OJJ74082.1 acar|OOG01194.1 acla|EAW07927.1 afis|EAW22965.1 afla|EED58158.1 afum|EAL90232.1 agla|OJJ83322.1 akaw|GAA87563.1 aluc|OJZ87953.1 anid|CBF87613.1 anig|EHA17738.1 aory|EIT79104.1 asyd|OJJ63274.1 ater|EAU30653.1 atub|OJI90564.1 aver|OJI97684.1 awen|OJJ38598.1 azon|OJJ49540.1

Group3328: abra|OJJ74084.1 acar|OOG01196.1 acla|EAW07928.1 afis|EAW22966.1 afla|EED58160.1 afum|EAL90231.1 agla|OJJ83321.1 akaw|GAA87564.1 aluc|OJZ87951.1 anid|CBF87612.1 anig|EHA17737.1 aory|EIT79091.1 asyd|OJJ63272.1 ater|EAU30654.1 atub|OJI90563.1 aver|OJI97685.1 awen|OJJ38597.1 azon|OJJ49538.1

Group3329: abra|OJJ74086.1 acar|OOG01197.1 acla|EAW07930.1 afis|EAW22968.1 afla|EED58161.1 afum|EAL90229.1 agla|OJJ83319.1 akaw|GAA87566.1 aluc|OJZ87949.1 anid|CBF87609.1 anig|EHA17735.1 aory|EIT79100.1 asyd|OJJ63270.1 ater|EAU30659.1 atub|OJI90560.1 aver|OJI97687.1 awen|OJJ38595.1 azon|OJJ49537.1

Group3330: abra|OJJ74089.1 acar|OOG01198.1 acla|EAW07932.1 afis|EAW22970.1 afla|EED58163.1 afum|EAL90227.2 agla|OJJ83317.1 akaw|GAA87568.1 aluc|OJZ87947.1 anid|CBF74548.1 anig|EHA17733.1 aory|EIT79078.1 asyd|OJJ57686.1 ater|EAU30652.1 atub|OJI90558.1 aver|OJJ05492.1 awen|OJJ38593.1 azon|OJJ49536.1

Group3331: abra|OJJ74090.1 acar|OOG01199.1 acla|EAW07933.1 afis|EAW22971.1 afla|EED58164.1 afum|EAL90226.1 agla|OJJ83316.1 akaw|GAA87569.1 aluc|OJZ87946.1 anid|CBF87607.1 anig|EHA17732.1 aory|EIT79076.1 asyd|OJJ63269.1 ater|EAU30651.1 atub|OJI90557.1 aver|OJI97688.1 awen|OJJ38591.1 azon|OJJ49535.1

Group3332: abra|OJJ74091.1 acar|OOG01200.1 acla|EAW07934.1 afis|EAW22972.1 afla|EED58166.1 afum|EAL90225.1 agla|OJJ83315.1 akaw|GAA87570.1 aluc|OJZ87945.1 anid|CBF87606.1 anig|EHA17731.1 aory|EIT79075.1 asyd|OJJ63268.1 ater|EAU30650.1 atub|OJI90556.1 aver|OJI97689.1 awen|OJJ38590.1 azon|OJJ49533.1

Group3333: abra|OJJ74093.1 acar|OOG01202.1 acla|EAW07936.1 afis|EAW22974.1 afla|EED58168.1 afum|EAL90223.1 agla|OJJ83313.1 akaw|GAA87572.1 aluc|OJZ87943.1 anid|CBF87602.1 anig|EHA17730.1 aory|EIT79071.1 asyd|OJJ63266.1 ater|EAU30647.1 atub|OJI90554.1 aver|OJI97691.1 awen|OJJ38589.1 azon|OJJ49532.1

Group3334: abra|OJJ74094.1 acar|OOG01204.1 acla|EAW07937.1 afis|EAW22975.1 afla|EED58169.1 afum|EAL90222.1 agla|OJJ83312.1 akaw|GAA87573.1 aluc|OJZ87942.1 anid|CBF87600.1 anig|EHA17729.1 aory|EIT79096.1 asyd|OJJ63265.1 ater|EAU30646.1 atub|OJI90553.1 aver|OJI97692.1 awen|OJJ38588.1 azon|OJJ49531.1

Group3335: abra|OJJ74095.1 acar|OOG01203.1 acla|EAW07938.1 afis|EAW22976.1 afla|EED58170.1 afum|EAL90221.1 agla|OJJ83311.1 akaw|GAA87574.1 aluc|OJZ87941.1 anid|CBF87598.1 anig|EHA17728.1 aory|EIT79085.1 asyd|OJJ63264.1 ater|EAU30645.1 atub|OJI90552.1 aver|OJI97693.1 awen|OJJ38587.1 azon|OJJ49530.1

Group3336: abra|OJJ74096.1 acar|OOG01205.1 acla|EAW07940.1 afis|EAW22978.1 afla|EED58171.1 afum|EAL90219.1 agla|OJJ83310.1 akaw|GAA87575.1 aluc|OJZ87940.1 anid|CBF87596.1 anig|EHA17727.1 aory|EIT79081.1 asyd|OJJ63262.1 ater|EAU30644.1 atub|OJI90551.1 aver|OJI97695.1 awen|OJJ38586.1 azon|OJJ49528.1

Group3337: abra|OJJ74100.1 acar|OOG01209.1 acla|EAW07943.1 afis|EAW22981.1 afla|EED58174.1 afum|EAL90216.1 agla|OJJ83306.1 akaw|GAA87578.1 aluc|OJZ87936.1 anid|CBF87589.1 anig|EHA17724.1 aory|EIT79069.1 asyd|OJJ63258.1 ater|EAU30640.1 atub|OJI90548.1 aver|OJI97699.1 awen|OJJ38582.1 azon|OJJ49524.1

Group3338: abra|OJJ74102.1 acar|OOG01211.1 acla|EAW07945.1 afis|EAW22983.1 afla|EED58176.1 afum|EAL90215.1 agla|OJJ83304.1 akaw|GAA87580.1 aluc|OJZ87934.1 anid|CBF87597.1 anig|EHA17723.1 aory|EIT79086.1 asyd|OJJ63263.1 ater|EAU30638.1 atub|OJI90546.1 aver|OJI97694.1 awen|OJJ38580.1 azon|OJJ49522.1

Group3339: abra|OJJ74103.1 acar|OOG01213.1 acla|EAW07946.1 afis|EAW22985.1 afla|EED58177.1 afum|EAL90214.1 agla|OJJ83302.1 akaw|GAA87581.1 aluc|OJZ87933.1 anid|CBF87584.1 anig|EHA17721.1 aory|EIT79072.1 asyd|OJJ63255.1 ater|EAU30637.1 atub|OJI90545.1 aver|OJI97702.1 awen|OJJ38579.1 azon|OJJ49520.1

Group3340: abra|OJJ74109.1 acar|OOG01217.1 acla|EAW07953.1 afis|EAW22990.1 afla|EED58182.1 afum|EAL90210.1 agla|OJJ83296.1 akaw|GAA87585.1 aluc|OJZ87925.1 anid|CBF87579.1 anig|EHA17717.1 aory|EIT79105.1 asyd|OJJ63251.1 ater|EAU30632.1 atub|OJI90537.1 aver|OJI97707.1 awen|OJJ38574.1 azon|OJJ49515.1

Group3341: abra|OJJ74116.1 acar|OOG01220.1 acla|EAW07958.1 afis|EAW22993.1 afla|EED58186.1 afum|EAL90208.1 agla|OJJ83291.1 akaw|GAA87589.1 aluc|OJZ87918.1 anid|CBF87570.1 anig|EHA17712.1 aory|EIT79074.1 asyd|OJJ63244.1 ater|EAU30627.1 atub|OJI90529.1 aver|OJI97712.1 awen|OJJ38568.1 azon|OJJ44834.1

Group3342: abra|OJJ74118.1 acar|OOG01223.1 acla|EAW07960.1 afis|EAW22995.1 afla|EED58188.1 afum|EAL90204.1 agla|OJJ83289.1 akaw|GAA87591.1 aluc|OJZ87914.1 anid|CBF87567.1 anig|EHA17708.1 aory|EIT79079.1 asyd|OJJ63242.1 ater|EAU30623.1 atub|OJI90526.1 aver|OJI97714.1 awen|OJJ38566.1 azon|OJJ50435.1

Group3343: abra|OJJ74159.1 acar|OOF95567.1 afum|EAL91052.1 akaw|GAA89725.1 aluc|OJZ80658.1 anid|CBF80314.1 anig|EHA27174.1 asyd|OJJ58662.1 atub|OJI87513.1 aver|OJJ01268.1 azon|OJJ43927.1 acar|OOF96898.1 akaw|GAA89030.1 aluc|OJZ80574.1 anig|EHA25625.1 atub|OJI88358.1 afis|EAW22138.1 azon|OJJ42294.1

Group3344: abra|OJJ74170.1 acar|OOF95550.1 acla|EAW07066.1 afis|EAW19330.1 afla|EED56451.1 afum|EAL94047.1 agla|OJJ84974.1 akaw|GAA86846.1 aluc|OJZ89965.1 anid|CBF78050.1 anig|EHA27165.1 asyd|OJJ55213.1 ater|EAU36486.1 atub|OJI84840.1 aver|OJJ07002.1 awen|OJJ31742.1 azon|OJJ42370.1 asyd|OJJ57144.1

Group3345: abra|OJJ74188.1 acar|OOF95533.1 acla|EAW07238.1 afis|EAW22261.1 afla|EED57649.1 afum|EAL90919.1 agla|OJJ87796.1 akaw|GAA86829.1 aluc|OJZ89947.1 anid|CBF88508.1 anig|EHA27148.1 aory|EIT79898.1 asyd|OJJ63801.1 ater|EAU34268.1 atub|OJI84859.1 aver|OJI97138.1 awen|OJJ34833.1 azon|OJJ45713.1

Group3346: abra|OJJ74189.1 acar|OOF95532.1 acla|EAW07241.1 afis|EAW22262.1 afla|EED57651.1 afum|EAL90918.1 agla|OJJ87794.1 akaw|GAA86828.1 aluc|OJZ89946.1 anid|CBF88511.1 anig|EHA27145.1

aory|EIT79914.1 asyd|OJJ63803.1 ater|EAU34269.1 atub|OJI84860.1 aver|OJI97135.1 awen|OJJ34835.1 azon|OJJ45712.1

Group3347: abra|OJJ74190.1 acar|OOF95531.1 acla|EAW07242.1 afis|EAW22263.1 afla|EED57655.1 afum|EAL90917.1 agla|OJJ87792.1 akaw|GAA86827.1 aluc|OJZ89945.1 anid|CBF88513.1 anig|EHA27144.1 aory|EIT79925.1 asyd|OJJ63804.1 ater|EAU34273.1 atub|OJI84861.1 aver|OJI97134.1 awen|OJJ34836.1 azon|OJJ45711.1

Group3348: abra|OJJ74193.1 acar|OOF95529.1 acla|EAW07245.1 afis|EAW22266.1 afla|EED57658.1 afum|EAL90914.1 agla|OJJ87789.1 akaw|GAA86824.1 aluc|OJZ89942.1 anid|CBF88519.1 anig|EHA27141.1 aory|EIT79939.1 asyd|OJJ63807.1 ater|EAU34277.1 atub|OJI84864.1 aver|OJI97131.1 awen|OJJ34838.1 azon|OJJ45708.1

Group3349: abra|OJJ74194.1 acar|OOF95527.1 acla|EAW07247.1 afis|EAW22268.1 afla|EED57660.1 afum|EAL90912.1 agla|OJJ87787.1 akaw|GAA86823.1 aluc|OJZ89941.1 anid|CBF88523.1 anig|EHA27140.1 aory|EIT79938.1 asyd|OJJ63808.1 ater|EAU34279.1 atub|OJI84865.1 aver|OJI97129.1 awen|OJJ34841.1 azon|OJJ45706.1

Group3350: abra|OJJ74195.1 acar|OOF95526.1 acla|EAW07248.1 afis|EAW22269.1 afla|EED57661.1 afum|EAL90911.1 agla|OJJ87786.1 akaw|GAA86822.1 aluc|OJZ89939.1 anid|CBF88527.1 anig|EHA27139.1 aory|EIT79909.1 asyd|OJJ63810.1 ater|EAU34280.1 atub|OJI84866.1 aver|OJI97127.1 awen|OJJ34842.1 azon|OJJ45705.1

Group3351: abra|OJJ74196.1 acar|OOF95524.1 acla|EAW07249.1 afis|EAW22270.1 afla|EED57662.1 afum|EAL90910.1 agla|OJJ87785.1 akaw|GAA86821.1 aluc|OJZ89938.1 anid|CBF88528.1 anig|EHA27138.1 aory|EIT79867.1 asyd|OJJ63811.1 ater|EAU34281.1 atub|OJI84868.1 aver|OJI97126.1 awen|OJJ34843.1 azon|OJJ45704.1

Group3352: abra|OJJ74197.1 acar|OOF95523.1 acla|EAW07250.1 afis|EAW22271.1 afla|EED57663.1 afum|EAL90909.1 agla|OJJ87784.1 akaw|GAA86820.1 aluc|OJZ89937.1 anid|CBF88530.1 anig|EHA27137.1 aory|EIT79903.1 asyd|OJJ63812.1 ater|EAU34282.1 atub|OJI84869.1 aver|OJI97125.1 awen|OJJ34844.1 azon|OJJ45703.1

Group3353: abra|OJJ74198.1 acar|OOF95521.1 acla|EAW07252.1 afis|EAW22273.1 afla|EED57665.1 afum|EAL90908.1 agla|OJJ87781.1 akaw|GAA86819.1 aluc|OJZ89936.1 anid|CBF88536.1 anig|EHA27136.1 aory|EIT79943.1 asyd|OJJ63820.1 ater|EAU34283.1 atub|OJI84870.1 aver|OJI97118.1 awen|OJJ34846.1 azon|OJJ45702.1

Group3354: abra|OJJ74199.1 acar|OOF95520.1 acla|EAW07253.1 afis|EAW22274.1 afla|EED57666.1 afum|EAL90907.1 agla|OJJ87780.1 akaw|GAA86818.1 aluc|OJZ89935.1 anid|CBF88537.1 anig|EHA27135.1 aory|EIT79937.1 asyd|OJJ63821.1 ater|EAU34284.1 atub|OJI84871.1 aver|OJI97117.1 awen|OJJ34847.1 azon|OJJ45700.1

Group3355: abra|OJJ74201.1 acar|OOF95519.1 acla|EAW07254.1 afis|EAW22275.1 afla|EED57667.1 afum|EAL90906.1 agla|OJJ87779.1 akaw|GAA86817.1 aluc|OJZ89934.1 anid|CBF88539.1 anig|EHA27134.1 aory|EIT79875.1 asyd|OJJ63822.1 ater|EAU34285.1 atub|OJI84872.1 aver|OJI97116.1 awen|OJJ34848.1 azon|OJJ45699.1

Group3356: abra|OJJ74202.1 acar|OOF95518.1 afla|EED57668.1 afum|EAL90905.1 agla|OJJ87777.1 akaw|GAA86816.1 aluc|OJZ89933.1 anid|CBF88541.1 anig|EHA27133.1 aory|EIT79884.1 ater|EAU34286.1 atub|OJI84873.1 aver|OJI97115.1 azon|OJJ45698.1 acla|EAW07255.1 afis|EAW22276.1 asyd|OJJ63823.1 awen|OJJ34849.1

Group3357: abra|OJJ74203.1 acar|OOF95517.1 acla|EAW07256.1 afis|EAW22277.1 afla|EED57669.1 afum|EAL90904.1 agla|OJJ87776.1 akaw|GAA86815.1 aluc|OJZ89932.1 anid|CBF88543.1 anig|EHA27132.1 aory|EIT79926.1 asyd|OJJ63824.1 ater|EAU34287.1 atub|OJI84874.1 aver|OJI97114.1 awen|OJJ34850.1 azon|OJJ45697.1

Group3358: abra|OJJ74204.1 acar|OOF95516.1 acla|EAW07257.1 afis|EAW22278.1 afla|EED57671.1 afum|EAL90903.1 agla|OJJ87775.1 akaw|GAA86814.1 aluc|OJZ89931.1 anid|CBF88545.1 anig|EHA27131.1 aory|EIT79927.1 asyd|OJJ63825.1 ater|EAU34288.1 atub|OJI84875.1 aver|OJI97113.1 awen|OJJ34851.1 azon|OJJ45696.1

Group3359: abra|OJJ74206.1 acar|OOF95514.1 acla|EAW07259.1 afis|EAW22280.1 afla|EED57677.1 afum|EAL90901.1 agla|OJJ87773.1 akaw|GAA86812.1 aluc|OJZ89929.1 anid|CBF88548.1 anig|EHA27129.1 aory|EIT79874.1 asyd|OJJ63827.1 ater|EAU34290.1 atub|OJI84877.1 aver|OJI97111.1 awen|OJJ34853.1 azon|OJJ45677.1

Group3360: abra|OJJ74207.1 acar|OOF95513.1 acla|EAW07260.1 afis|EAW22281.1 afla|EED57678.1 afum|EAL90900.1 agla|OJJ87768.1 akaw|GAA86811.1 aluc|OJZ89928.1 anid|CBF88550.1 anig|EHA27128.1 aory|EIT79890.1 asyd|OJJ63828.1 ater|EAU34291.1 atub|OJI84878.1 aver|OJI97110.1 awen|OJJ34854.1 azon|OJJ45676.1

Group3361: abra|OJJ74208.1 acar|OOF95512.1 acla|EAW07261.1 afis|EAW22282.1 afla|EED57679.1 afum|EAL90899.1 agla|OJJ87767.1 akaw|GAA86810.1 aluc|OJZ89926.1 anid|CBF88552.1 anig|EHA27127.1 aory|EIT79907.1 asyd|OJJ63829.1 ater|EAU34292.1 atub|OJI84879.1 aver|OJI97109.1 awen|OJJ34855.1 azon|OJJ45674.1

Group3362: abra|OJJ74209.1 acar|OOF95511.1 acla|EAW07262.1 afis|EAW22283.1 afla|EED57680.1 afum|EAL90898.1 agla|OJJ87766.1 akaw|GAA86809.1 aluc|OJZ89925.1 anid|CBF88554.1 anig|EHA27126.1 aory|EIT79921.1 asyd|OJJ63830.1 ater|EAU34293.1 atub|OJI84880.1 aver|OJI97108.1 awen|OJJ34856.1 azon|OJJ45673.1

Group3363: abra|OJJ74210.1 acar|OOF95510.1 acla|EAW07263.1 afis|EAW22284.1 afla|EED57682.1 afum|EAL90897.1 agla|OJJ87764.1 akaw|GAA86808.1 aluc|OJZ89924.1 anid|CBF88556.1 anig|EHA27124.1 aory|EIT79901.1 asyd|OJJ63831.1 ater|EAU34296.1 atub|OJI84881.1 aver|OJI97107.1 awen|OJJ34857.1 azon|OJJ45671.1

Group3364: abra|OJJ74211.1 acar|OOF95509.1 acla|EAW07264.1 afis|EAW22285.1 afla|EED57685.1 afum|EAL90896.1 agla|OJJ87763.1 akaw|GAA86807.1 aluc|OJZ89923.1 anid|CBF88558.1 anig|EHA27123.1 aory|EIT79887.1 asyd|OJJ63832.1 ater|EAU34297.1 atub|OJI84882.1 aver|OJI97106.1 awen|OJJ34858.1 azon|OJJ45670.1

Group3365: abra|OJJ74218.1 acar|OOF95503.1 acla|EAW07273.1 afis|EAW22294.1 afla|EED57690.1 afum|EAL90887.1 agla|OJJ87758.1 akaw|GAA86800.1 aluc|OJZ89916.1 anid|CBF88569.1 anig|EHA27116.1 aory|EIT79904.1 asyd|OJJ63837.1 ater|EAU34303.1 atub|OJI84889.1 aver|OJI97100.1 awen|OJJ34864.1 azon|OJJ45665.1

Group3366: abra|OJJ74220.1 acar|OOF90433.1 acla|EAW07284.1 afis|EAW22315.1 afla|EED57695.1 afum|EAL90875.1 agla|OJJ87735.1 akaw|GAA86798.1 aluc|OJZ89914.1 anid|CBF88577.1 anig|EHA27114.1 aory|EIT79936.1 asyd|OJJ63841.1 ater|EAU34308.1 atub|OJI84891.1 aver|OJI97096.1 awen|OJJ34868.1 azon|OJJ50045.1

Group3367: abra|OJJ74223.1 acar|OOF90434.1 acla|EAW07282.1 afis|EAW22313.1 afla|EED57697.1 afum|EAL90877.1 agla|OJJ87737.1 akaw|GAA86796.1 aluc|OJZ89911.1 anid|CBF88580.1 anig|EHA27111.1 aory|EIT79934.1 asyd|OJJ63843.1 ater|EAU34310.1 atub|OJI84895.1 aver|OJI97094.1 awen|OJJ34872.1 azon|OJJ50047.1

Group3368: abra|OJJ74224.1 acar|OOF90435.1 acla|EAW07281.1 afis|EAW22312.1 afla|EED57698.1 afum|EAL90878.1 agla|OJJ87738.1 akaw|GAA86795.1 aluc|OJZ89910.1 anid|CBF88582.1 anig|EHA27110.1 aory|EIT79916.1 asyd|OJJ63844.1 ater|EAU34311.1 atub|OJI84896.1 aver|OJI97093.1 awen|OJJ34873.1 azon|OJJ50048.1

Group3369: abra|OJJ74225.1 acar|OOF95502.1 acla|EAW07280.1 afis|EAW22311.1 afla|EED57701.1 afum|EAL90879.1 agla|OJJ87739.1 akaw|GAA86794.1 aluc|OJZ89908.1 anid|CBF88584.1 anig|EHA27108.1 aory|EIT79931.1 asyd|OJJ63846.1 ater|EAU34313.1 atub|OJI84898.1 aver|OJI97090.1 awen|OJJ34876.1 azon|OJJ50049.1

Group3370: abra|OJJ74228.1 acar|OOF95498.1 acla|EAW07276.1 afis|EAW22297.1 afla|EED57705.1 afum|EAL90884.1 agla|OJJ87748.1 akaw|GAA86791.1 aluc|OJZ89905.1 anid|CBF88592.1 anig|EHA27105.1 aory|EIT79881.1 asyd|OJJ63850.1 ater|EAU34317.1 atub|OJI84901.1 aver|OJI97087.1 awen|OJJ34881.1 azon|OJJ50050.1

Group3371: abra|OJJ74231.1 acar|OOF95495.1 acla|EAW07286.1 afis|EAW22319.1 afla|EED57710.1 afum|EAL90871.1 agla|OJJ87745.1 akaw|GAA86788.1 aluc|OJZ89902.1 anid|CBF88599.1 anig|EHA27102.1 aory|EIT79923.1 asyd|OJJ63854.1 ater|EAU34320.1 atub|OJI84905.1 aver|OJI97084.1 awen|OJJ34885.1 azon|OJJ50053.1

Group3372: abra|OJJ74237.1 acar|OOF95492.1 acla|EAW07288.1 afis|EAW22321.1 afla|EED57718.1 afum|EAL90870.1 agla|OJJ87749.1 akaw|GAA86783.1 aluc|OJZ89895.1 anid|CBF88602.1 anig|EHA27094.1 aory|EIT79929.1 asyd|OJJ63855.1 ater|EAU34321.1 atub|OJI84912.1 aver|OJI97083.1 awen|OJJ34888.1 azon|OJJ50055.1

Group3373: abra|OJJ74242.1 acar|OOF95488.1 acla|EAW07291.1 afis|EAW22325.1 afla|EED57723.1 afum|EAL90867.1 agla|OJJ87721.1 akaw|GAA86778.1 aluc|OJZ89890.1 anid|CBF88609.1 anig|EHA27089.1 aory|EIT79878.1 asyd|OJJ63859.1 ater|EAU34324.1 atub|OJI84917.1 aver|OJI97080.1 awen|OJJ34896.1 azon|OJJ50065.1

Group3374: abra|OJJ74246.1 acar|OOF95484.1 acla|EAW07298.1 afis|EAW22347.1 afla|EED57735.1 afum|EAL90845.1 agla|OJJ87687.1 akaw|GAA86774.1 aluc|OJZ89886.1 anid|CBF88623.1 anig|EHA27085.1 aory|EIT79913.1 asyd|OJJ63868.1 ater|EAU39483.1 atub|OJI84921.1 aver|OJI97071.1 awen|OJJ34928.1 azon|OJJ50070.1

Group3375: abra|OJJ74249.1 acar|OOF95479.1 acla|EAW07293.1 afis|EAW22327.1 afla|EED57726.1 afum|EAL90865.1 agla|OJJ87723.1 akaw|GAA86772.1 aluc|OJZ89883.1 anid|CBF88613.1 anig|EHA27082.1

aory|EIT79866.1 asyd|OJJ63861.1 ater|EAU34326.1 atub|OJI84924.1 aver|OJI97078.1 awen|OJJ34894.1 azon|OJJ50063.1

Group3376: abra|OJJ74250.1 acar|OOF95480.1 acla|EAW07294.1 afis|EAW22328.1 afla|EED57727.1 afum|EAL90864.1 agla|OJJ87724.1 akaw|GAA86771.1 aluc|OJZ89882.1 anid|CBF88615.1 anig|EHA27081.1 aory|EIT79906.1 asyd|OJJ63862.1 ater|EAU34327.1 atub|OJI84925.1 aver|OJI97077.1 awen|OJJ34893.1 azon|OJJ50062.1

Group3377: abra|OJJ74252.1 acar|OOF95477.1 acla|EAW07296.1 afis|EAW22330.1 afla|EED57729.1 afum|EAL90862.1 agla|OJJ87752.1 akaw|GAA86769.1 aluc|OJZ89880.1 anid|CBF88619.1 anig|EHA27079.1 aory|EIT79894.1 asyd|OJJ63864.1 ater|EAU34331.1 atub|OJI84927.1 aver|OJI97075.1 awen|OJJ34891.1 azon|OJJ50060.1

Group3378: abra|OJJ74256.1 acar|OOF95472.1 acla|EAW07302.1 afis|EAW22343.1 afla|EED57738.1 afum|EAL90848.1 agla|OJJ87691.1 akaw|GAA86765.1 aluc|OJZ89876.1 anid|CBF88627.1 anig|EHA27075.1 aory|EIT79915.1 asyd|OJJ63870.1 ater|EAU39481.1 atub|OJI84931.1 aver|OJI97069.1 awen|OJJ34922.1 azon|OJJ50072.1

Group3379: abra|OJJ74257.1 acar|OOF95471.1 acla|EAW07303.1 afis|EAW22342.1 afla|EED57739.1 afum|EAL90849.1 agla|OJJ87692.1 akaw|GAA86764.1 aluc|OJZ89875.1 anid|CBF88628.1 anig|EHA27074.1 aory|EIT79892.1 asyd|OJJ63871.1 ater|EAU39480.1 atub|OJI84932.1 aver|OJI97068.1 awen|OJJ34921.1 azon|OJJ50073.1

Group3380: abra|OJJ74260.1 acar|OOF89900.1 acla|EAW07305.1 afis|EAW22340.1 afla|EED57741.1 afum|EAL90852.1 agla|OJJ87698.1 akaw|GAA86761.1 aluc|OJZ89871.1 anid|CBF88632.1 anig|EHA27071.1 aory|EIT79893.1 asyd|OJJ63874.1 ater|EAU39478.1 atub|OJI84936.1 aver|OJI97064.1 awen|OJJ34919.1 azon|OJJ50075.1

Group3381: abra|OJJ74261.1 acar|OOF89901.1 acla|EAW07306.1 afis|EAW22339.1 afla|EED57742.1 afum|EAL90853.1 agla|OJJ87699.1 akaw|GAA86760.1 aluc|OJZ89870.1 anid|CBF88634.1 anig|EHA27070.1 aory|EIT79922.1 asyd|OJJ63875.1 ater|EAU39477.1 atub|OJI84937.1 aver|OJI97063.1 awen|OJJ34918.1 azon|OJJ50076.1

Group3382: abra|OJJ74264.1 acar|OOF95466.1 acla|EAW07308.1 afis|EAW22338.1 afla|EED57743.1 afum|EAL90854.1 agla|OJJ87700.1 akaw|GAA86759.1 aluc|OJZ89869.1 anid|CBF88636.1 anig|EHA27069.1 aory|EIT81083.1 asyd|OJJ63877.1 ater|EAU39475.1 atub|OJI84939.1 aver|OJI97062.1 awen|OJJ34916.1 azon|OJJ50079.1

Group3383: abra|OJJ74265.1 acar|OOF95465.1 acla|EAW07309.1 afis|EAW22337.1 afla|EED57744.1 afum|EAL90855.1 agla|OJJ87701.1 akaw|GAA86758.1 aluc|OJZ89868.1 anid|CBF88638.1 anig|EHA27068.1 aory|EIT81047.1 asyd|OJJ63878.1 ater|EAU39474.1 atub|OJI84940.1 aver|OJI97061.1 awen|OJJ34915.1 azon|OJJ45569.1

Group3384: abra|OJJ74266.1 acar|OOF95464.1 acla|EAW07310.1 afis|EAW22336.1 afla|EED57746.1 afum|EAL90856.1 agla|OJJ87702.1 akaw|GAA86757.1 aluc|OJZ89867.1 anid|CBF88640.1 anig|EHA27067.1 aory|EIT81092.1 asyd|OJJ63879.1 ater|EAU39473.1 atub|OJI84941.1 aver|OJI97060.1 awen|OJJ34914.1 azon|OJJ45570.1

Group3385: abra|OJJ74267.1 acar|OOF89814.1 acla|EAW07311.1 afis|EAW22335.1 afla|EED57747.1 afum|EAL90857.2 agla|OJJ87715.1 akaw|GAA86756.1 aluc|OJZ89866.1 anid|CBF88641.1 anig|EHA27066.1 aory|EIT81022.1 asyd|OJJ63880.1 ater|EAU39472.1 atub|OJI84942.1 aver|OJI97059.1 awen|OJJ34913.1 azon|OJJ45571.1

Group3386: abra|OJJ74270.1 acar|OOF95460.1 acla|EAW07314.1 afis|EAW22332.1 afla|EED57751.1 afum|EAL90860.1 agla|OJJ87712.1 akaw|GAA86753.1 aluc|OJZ89863.1 anid|CBF88647.1 anig|EHA27063.1 aory|EIT81116.1 asyd|OJJ63882.1 ater|EAU39461.1 atub|OJI84945.1 aver|OJI97057.1 awen|OJJ34910.1 azon|OJJ45575.1

Group3387: abra|OJJ74272.1 acar|OOF95458.1 acla|EAW07317.1 afis|EAW22349.1 afla|EED57753.1 afum|EAL90843.1 agla|OJJ87710.1 akaw|GAA86751.1 aluc|OJZ89861.1 anid|CBF88651.1 anig|EHA27061.1 aory|EIT81051.1 asyd|OJJ63885.1 ater|EAU39459.1 atub|OJI84947.1 aver|OJI97055.1 awen|OJJ34905.1 azon|OJJ45577.1

Group3388: abra|OJJ74276.1 acar|OOF95455.1 acla|EAW07320.1 afis|EAW22352.1 afla|EED57756.1 afum|EAL90840.1 agla|OJJ87707.1 akaw|GAA86748.1 aluc|OJZ89857.1 anid|CBF88661.1 anig|EHA27058.1 aory|EIT81039.1 asyd|OJJ63889.1 ater|EAU39488.1 atub|OJI84949.1 aver|OJI97050.1 awen|OJJ34903.1 azon|OJJ45579.1

Group3389: abra|OJJ74277.1 acar|OOF95454.1 acla|EAW07321.1 afis|EAW22353.1 afla|EED57759.1 afum|EAL90839.1 agla|OJJ87716.1 akaw|GAA86747.1 aluc|OJZ89856.1 anid|CBF88662.1 anig|EHA27057.1 aory|EIT81042.1 asyd|OJJ63890.1 ater|EAU39487.1 atub|OJI84950.1 aver|OJI97049.1 awen|OJJ34902.1 azon|OJJ45580.1

Group3390: abra|OJJ74278.1 acar|OOF95453.1 acla|EAW07322.1 afis|EAW22354.1 afla|EED57760.1 afum|EAL90838.1 agla|OJJ87717.1 akaw|GAA86746.1 aluc|OJZ89855.1 anid|CBF88664.1 anig|EHA27056.1 aory|EIT81028.1 asyd|OJJ63891.1 ater|EAU39486.1 atub|OJI84951.1 aver|OJI97048.1 awen|OJJ34901.1 azon|OJJ45581.1

Group3391: abra|OJJ74280.1 acar|OOF95451.1 acla|EAW07324.1 afis|EAW22356.1 afla|EED57762.1 afum|EAL90835.1 agla|OJJ87598.1 akaw|GAA86744.1 aluc|OJZ89853.1 anid|CBF88668.1 anig|EHA27054.1 aory|EIT81074.1 asyd|OJJ63893.1 ater|EAU39490.1 atub|OJI84953.1 aver|OJI97046.1 awen|OJJ34930.1 azon|OJJ45583.1

Group3392: abra|OJJ74281.1 acar|OOF95450.1 acla|EAW07325.1 afis|EAW22357.1 afla|EED57763.1 afum|EAL90834.1 agla|OJJ87682.1 akaw|GAA86743.1 aluc|OJZ89852.1 anid|CBF88672.1 anig|EHA27053.1 aory|EIT81043.1 asyd|OJJ63895.1 ater|EAU39491.1 atub|OJI84954.1 aver|OJI97044.1 awen|OJJ34931.1 azon|OJJ45584.1

Group3393: abra|OJJ74282.1 acar|OOF95449.1 acla|EAW07326.1 afis|EAW22358.1 afla|EED57764.1 afum|EAL90833.1 agla|OJJ87683.1 akaw|GAA86742.1 aluc|OJZ89851.1 anid|CBF88674.1 anig|EHA27052.1 aory|EIT81053.1 asyd|OJJ63896.1 ater|EAU39492.1 atub|OJI84955.1 aver|OJI97043.1 awen|OJJ34932.1 azon|OJJ45585.1

Group3394: abra|OJJ74283.1 acar|OOF95447.1 acla|EAW07327.1 afis|EAW22359.1 afla|EED57765.1 afum|EAL90832.1 agla|OJJ87684.1 akaw|GAA86741.1 aluc|OJZ89850.1 anid|CBF88676.1 anig|EHA27051.1 aory|EIT81093.1 asyd|OJJ63897.1 ater|EAU39493.1 atub|OJI84956.1 aver|OJI97042.1 awen|OJJ34933.1 azon|OJJ45586.1

Group3395: abra|OJJ74290.1 acar|OOF95441.1 acla|EAW07334.1 afis|EAW22366.1 afla|EED57774.1 afum|EBA27415.1 agla|OJJ87624.1 akaw|GAA86734.1 aluc|OJZ89844.1 anid|CBF88692.1 anig|EHA27045.1 aory|EIT81109.1 asyd|OJJ63910.1 ater|EAU39501.1 atub|OJI84964.1 aver|OJI97031.1 awen|OJJ34940.1 azon|OJJ45592.1

Group3396: abra|OJJ74291.1 acar|OOF95440.1 acla|EAW07335.1 afis|EAW22367.1 afla|EED57775.1 afum|EAL90825.1 agla|OJJ87622.1 akaw|GAA86733.1 aluc|OJZ89843.1 anid|CBF88694.1 anig|EHA27044.1 aory|EIT81063.1 asyd|OJJ63911.1 ater|EAU39502.1 atub|OJI84965.1 aver|OJI97030.1 awen|OJJ34941.1 azon|OJJ45593.1

Group3397: abra|OJJ74292.1 acar|OOF95439.1 acla|EAW07336.1 afis|EAW22368.1 afla|EED57776.1 afum|EAL90824.1 akaw|GAA86732.1 aluc|OJZ89842.1 anid|CBF88695.1 anig|EHA27043.1 aory|EIT81045.1 asyd|OJJ63913.1 ater|EAU39503.1 atub|OJI84966.1 aver|OJI97029.1 awen|OJJ34942.1 azon|OJJ45594.1 agla|OJJ87621.1

Group3398: abra|OJJ74293.1 acar|OOF95438.1 acla|EAW07337.1 afis|EAW22369.1 afla|EED57777.1 afum|EAL90823.1 agla|OJJ87620.1 akaw|GAA86731.1 aluc|OJZ89841.1 anid|CBF88697.1 anig|EHA27042.1 aory|EIT81034.1 asyd|OJJ63914.1 ater|EAU39504.1 atub|OJI84967.1 aver|OJI97028.1 awen|OJJ34944.1 azon|OJJ45595.1

Group3399: abra|OJJ74294.1 acar|OOF95437.1 acla|EAW07339.1 afis|EAW22370.1 afla|EED57778.1 afum|EAL90822.2 agla|OJJ87619.1 akaw|GAA86730.1 aluc|OJZ89840.1 anid|CBF88699.1 anig|EHA27041.1 aory|EIT81037.1 asyd|OJJ63915.1 ater|EAU39505.1 atub|OJI84968.1 aver|OJI97027.1 awen|OJJ34945.1 azon|OJJ45596.1

Group3400: abra|OJJ74295.1 acar|OOF95436.1 acla|EAW07340.1 afis|EAW22372.1 afla|EED57779.1 afum|EAL90820.2 agla|OJJ87617.1 akaw|GAA86729.1 aluc|OJZ89839.1 anid|CBF88703.1 anig|EHA27040.1 aory|EIT81082.1 asyd|OJJ63919.1 ater|EAU39506.1 atub|OJI84969.1 aver|OJI97024.1 awen|OJJ34947.1 azon|OJJ45597.1

Group3401: abra|OJJ74296.1 acar|OOF95435.1 acla|EAW07341.1 afis|EAW22373.1 afla|EED57780.1 afum|EAL90819.1 agla|OJJ87626.1 akaw|GAA86728.1 aluc|OJZ89838.1 anid|CBF88705.1 anig|EHA27039.1 aory|EIT81056.1 asyd|OJJ63920.1 ater|EAU39507.1 atub|OJI84970.1 aver|OJI97023.1 awen|OJJ34948.1 azon|OJJ45598.1

Group3402: abra|OJJ74297.1 acar|OOF95434.1 acla|EAW07342.1 afis|EAW22374.1 afla|EED57781.1 afum|EAL90818.1 agla|OJJ87627.1 akaw|GAA86727.1 aluc|OJZ89837.1 anid|CBF88706.1 anig|EHA27038.1 aory|EIT81052.1 asyd|OJJ63923.1 ater|EAU39508.1 atub|OJI84971.1 aver|OJI97022.1 awen|OJJ34949.1 azon|OJJ45599.1

Group3403: abra|OJJ74298.1 acar|OOF95433.1 acla|EAW07343.1 afis|EAW22375.1 afum|EAL90817.1 agla|OJJ87628.1 akaw|GAA86726.1 aluc|OJZ89835.1 anid|CBF88710.1 anig|EHA27037.1 aory|EIT81071.1 asyd|OJJ63924.1 ater|EAU39509.1 atub|OJI84972.1 aver|OJI97020.1 awen|OJJ34950.1 azon|OJJ45600.1 afla|EED57783.1

Group3404: abra|OJJ74299.1 acar|OOF95431.1 acla|EAW07344.1 afis|EAW22376.1 afla|EED57784.1 afum|EAL90816.1 agla|OJJ87629.1 akaw|GAA86725.1 aluc|OJZ89834.1 anid|CBF88712.1 anig|EHA27036.1

aory|EIT81070.1 asyd|OJJ63925.1 ater|EAU39510.1 atub|OJI84973.1 aver|OJI97019.1 awen|OJJ34951.1 azon|OJJ45601.1

Group3405: abra|OJJ74301.1 acar|OOF95429.1 acla|EAW07346.1 afis|EAW22378.1 afla|EED57786.1 afum|EAL90814.1 agla|OJJ87631.1 akaw|GAA86724.1 aluc|OJZ89832.1 anid|CBF88714.1 anig|EHA27034.1 aory|EIT81069.1 asyd|OJJ63926.1 ater|EAU39512.1 atub|OJI84974.1 aver|OJI97018.1 awen|OJJ34952.1 azon|OJJ45602.1

Group3406: abra|OJJ74302.1 acar|OOF95428.1 acla|EAW07349.1 afis|EAW22379.1 afla|EED57787.1 afum|EAL90812.1 agla|OJJ87635.1 akaw|GAA86723.1 aluc|OJZ89830.1 anid|CBF88716.1 anig|EHA27033.1 aory|EIT81084.1 asyd|OJJ63927.1 ater|EAU39513.1 atub|OJI84975.1 aver|OJI97017.1 awen|OJJ34953.1 azon|OJJ45603.1

Group3407: abra|OJJ74304.1 acar|OOF95424.1 acla|EAW07352.1 afis|EAW22381.1 afla|EED57790.1 afum|EAL90810.1 agla|OJJ87640.1 akaw|GAA86720.1 aluc|OJZ89824.1 anid|CBF88718.1 anig|EHA27031.1 aory|EIT81089.1 asyd|OJJ63928.1 ater|EAU39515.1 atub|OJI84982.1 aver|OJI97016.1 awen|OJJ34956.1 azon|OJJ45608.1

Group3408: abra|OJJ74305.1 acar|OOF95423.1 acla|EAW07353.1 afis|EAW22382.1 afla|EED56040.1 afum|EAL90809.1 agla|OJJ87641.1 akaw|GAA86719.1 aluc|OJZ89822.1 anid|CBF88721.1 anig|EHA27030.1 aory|EIT76582.1 asyd|OJJ63929.1 ater|EAU39516.1 atub|OJI84983.1 aver|OJI97014.1 awen|OJJ34957.1 azon|OJJ45609.1

Group3409: abra|OJJ74306.1 acar|OOF95422.1 acla|EAW07354.1 afis|EAW22383.1 afla|EED56041.1 afum|EAL90808.1 agla|OJJ87642.1 akaw|GAA86718.1 aluc|OJZ89821.1 anid|CBF88723.1 anig|EHA27029.1 aory|EIT76563.1 asyd|OJJ63930.1 ater|EAU39517.1 atub|OJI84984.1 aver|OJI97013.1 awen|OJJ34958.1 azon|OJJ45610.1

Group3410: abra|OJJ74307.1 acar|OOF95420.1 acla|EAW07355.1 afis|EAW22384.1 afla|EED56044.1 afum|EAL90806.1 agla|OJJ87644.1 akaw|GAA86717.1 aluc|OJZ89820.1 anid|CBF88725.1 anig|EHA27028.1 aory|EIT76594.1 asyd|OJJ63931.1 ater|EAU39337.1 atub|OJI84985.1 aver|OJI97012.1 awen|OJJ34959.1 azon|OJJ45611.1

Group3411: abra|OJJ74310.1 acar|OOF95419.1 acla|EAW07357.1 afis|EAW22386.1 afla|EED56046.1 afum|EAL90804.1 agla|OJJ87647.1 akaw|GAA86714.1 aluc|OJZ89816.1 anid|CBF88730.1 anig|EHA27025.1 aory|EIT76610.1 asyd|OJJ63934.1 ater|EAU39335.1 atub|OJI84989.1 aver|OJI97008.1 awen|OJJ34961.1 azon|OJJ45613.1

Group3412: abra|OJJ74311.1 acar|OOF95417.1 acla|EAW07358.1 afis|EAW22387.1 afla|EED56047.1 afum|EAL90803.1 agla|OJJ87648.1 akaw|GAA86713.1 aluc|OJZ89815.1 anid|CBF88732.1 anig|EHA27024.1 aory|EIT76591.1 asyd|OJJ63935.1 ater|EAU39334.1 atub|OJI84990.1 aver|OJI97007.1 awen|OJJ34962.1 azon|OJJ45614.1

Group3413: abra|OJJ74315.1 acar|OOF95412.1 acla|EAW07366.1 afis|EAW22393.1 afla|EED56056.1 afum|EAL90798.1 agla|OJJ87655.1 akaw|GAA86710.1 aluc|OJZ89811.1 anid|CBF88743.1 anig|EHA27021.1 aory|EIT76567.1 asyd|OJJ63954.1 ater|EAU39325.1 atub|OJI84994.1 aver|OJI96987.1 awen|OJJ34970.1 azon|OJJ45619.1

Group3414: abra|OJJ74316.1 acar|OOF95411.1 acla|EAW07367.1 afis|EAW22394.1 afla|EED56057.1 afum|EAL90797.1 agla|OJJ87657.1 akaw|GAA86709.1 aluc|OJZ89810.1 anid|CBF88745.1 anig|EHA27020.1 aory|EIT76865.1 asyd|OJJ63955.1 ater|EAU39324.1 atub|OJI84995.1 aver|OJI96986.1 awen|OJJ34971.1 azon|OJJ45620.1

Group3415: abra|OJJ74320.1 acar|OOF95409.1 acla|EAW07372.1 afis|EAW22397.1 afla|EED56061.1 afum|EAL90794.1 agla|OJJ87660.1 akaw|GAA86706.1 aluc|OJZ89808.1 anid|CBF88750.1 anig|EHA27018.1 aory|EIT76888.1 asyd|OJJ63958.1 ater|EAU39320.1 atub|OJI84998.1 aver|OJI96983.1 awen|OJJ34975.1 azon|OJJ45624.1

Group3416: abra|OJJ74325.1 acar|OOF95406.1 acla|EAW07377.1 afis|EAW22402.1 afla|EED56066.1 afum|EAL90789.1 agla|OJJ87665.1 akaw|GAA86701.1 aluc|OJZ89803.1 anid|CBF88765.1 anig|EHA27016.1 aory|EIT76819.1 asyd|OJJ63981.1 ater|EAU39306.1 atub|OJI85003.1 aver|OJI96972.1 awen|OJJ34981.1 azon|OJJ45629.1

Group3417: abra|OJJ74326.1 acar|OOF95405.1 acla|EAW07378.1 afis|EAW22403.1 afla|EED56067.1 afum|EAL90788.1 agla|OJJ87666.1 akaw|GAA86700.1 aluc|OJZ89802.1 anid|CBF88763.1 anig|EHA27015.1 aory|EIT76544.1 asyd|OJJ63982.1 ater|EAU39305.1 atub|OJI85004.1 aver|OJI96971.1 awen|OJJ34982.1 azon|OJJ45631.1

Group3418: abra|OJJ74329.1 acar|OOF95402.1 acla|EAW07379.1 afis|EAW22404.1 afla|EED56068.1 afum|EAL90787.1 agla|OJJ87667.1 akaw|GAA86699.1 aluc|OJZ89801.1 anid|CBF88785.1 anig|EHA27014.1 aory|EIT76781.1 asyd|OJJ62903.1 ater|EAU39304.1 atub|OJI85005.1 aver|OJI96957.1 awen|OJJ34983.1 azon|OJJ45633.1

Group3419: abra|OJJ74336.1 acar|OOF95391.1 acla|EAW07386.1 afis|EAW22411.1 afla|EED52491.1 afum|EAL90781.1 agla|OJJ87678.1 akaw|GAA86693.1 aluc|OJZ89794.1 anid|CBF88796.1 anig|EHA27007.1 aory|EIT82814.1 asyd|OJJ62897.1 ater|EAU39294.1 atub|OJI85012.1 aver|OJI96950.1 awen|OJJ34992.1 azon|OJJ45635.1

Group3420: abra|OJJ74340.1 acar|OOF89991.1 acla|EAW07391.1 afis|EAW22417.1 afla|EED52495.1 afum|EAL90775.1 agla|OJJ87599.1 akaw|GAA86690.1 aluc|OJZ89791.1 anid|CBF88802.1 anig|EHA27004.1 aory|EIT82790.1 asyd|OJJ62895.1 ater|EAU39289.1 atub|OJI85015.1 aver|OJI96947.1 awen|OJJ34996.1 azon|OJJ45639.1

Group3421: abra|OJJ74341.1 acar|OOF95389.1 acla|EAW07392.1 afis|EAW22418.1 afla|EED52496.1 afum|EAL90774.1 agla|OJJ87600.1 akaw|GAA86689.1 aluc|OJZ89790.1 anid|CBF88804.1 anig|EHA27003.1 aory|EIT82800.1 asyd|OJJ62894.1 ater|EAU39288.1 atub|OJI85016.1 aver|OJI96946.1 awen|OJJ34997.1 azon|OJJ45646.1

Group3422: abra|OJJ74343.1 acar|OOF95387.1 acla|EAW07397.1 afis|EAW22423.1 afla|EED52501.1 afum|EAL90768.2 agla|OJJ87605.1 akaw|GAA86687.1 aluc|OJZ89788.1 anid|CBF88813.1 anig|EHA27002.1 aory|EIT82762.1 asyd|OJJ62889.1 ater|EAU39284.1 atub|OJI85018.1 aver|OJI96941.1 awen|OJJ35002.1 azon|OJJ45642.1

Group3423: abra|OJJ74344.1 acar|OOF95386.1 acla|EAW07398.1 afis|EAW22424.1 afla|EED52502.1 afum|EAL90767.1 agla|OJJ87606.1 akaw|GAA86686.1 aluc|OJZ89787.1 anid|CBF75162.1 anig|EHA27001.1 aory|EIT82765.1 asyd|OJJ59961.1 ater|EAU39283.1 atub|OJI85019.1 aver|OJJ02329.1 awen|OJJ35003.1 azon|OJJ45641.1

Group3424: abra|OJJ74347.1 acar|OOF95382.1 acla|EAW07401.1 afis|EAW22427.1 afla|EED52505.1 afum|EAL90764.1 agla|OJJ87596.1 akaw|GAA86683.1 aluc|OJZ89783.1 anid|CBF88819.1 anig|EHA26998.1 aory|EIT82749.1 asyd|OJJ62886.1 ater|EAU39280.1 atub|OJI85022.1 aver|OJI96938.1 awen|OJJ35006.1 azon|OJJ45648.1

Group3425: abra|OJJ74348.1 acar|OOF95380.1 acla|EAW07402.1 afis|EAW22428.1 afla|EED52506.1 afum|EAL90763.1 agla|OJJ87595.1 akaw|GAA86682.1 aluc|OJZ89782.1 anid|CBF88822.1 anig|EHA26997.1 aory|EIT82757.1 asyd|OJJ62885.1 ater|EAU39279.1 atub|OJI85024.1 aver|OJI96937.1 awen|OJJ35007.1 azon|OJJ45649.1

Group3426: abra|OJJ74352.1 acar|OOF95378.1 acla|EAW07405.1 afis|EAW22431.1 afla|EED52510.1 afum|EAL90760.1 agla|OJJ87592.1 akaw|GAA86679.1 aluc|OJZ89776.1 anid|CBF88828.1 anig|EHA26994.1 aory|EIT82794.1 asyd|OJJ62882.1 ater|EAU39276.1 atub|OJI85027.1 aver|OJI96934.1 awen|OJJ35010.1 azon|OJJ45651.1

Group3427: abra|OJJ74353.1 acar|OOF95377.1 acla|EAW07406.1 afis|EAW22432.1 afla|EED52513.1 afum|EAL90759.1 agla|OJJ87590.1 akaw|GAA86678.1 aluc|OJZ89775.1 anid|CBF88832.1 anig|EHA26993.1 aory|EIT82776.1 asyd|OJJ62880.1 ater|EAU39275.1 atub|OJI85028.1 aver|OJI96932.1 awen|OJJ35011.1 azon|OJJ45652.1

Group3428: abra|OJJ74355.1 acar|OOF95375.1 acla|EAW07409.1 afis|EAW22435.1 afla|EED52515.1 afum|EAL90756.1 agla|OJJ87587.1 akaw|GAA86676.1 aluc|OJZ89773.1 anid|CBF88835.1 anig|EHA26991.1 aory|EIT82785.1 asyd|OJJ62878.1 ater|EAU39273.1 atub|OJI85030.1 aver|OJI96930.1 awen|OJJ35013.1 azon|OJJ45654.1

Group3429: abra|OJJ74359.1 acar|OOF95370.1 acla|EAW07410.1 afis|EAW22436.1 afla|EED54446.1 afum|EAL90755.1 agla|OJJ87586.1 akaw|GAA86672.1 aluc|OJZ89769.1 anid|CBF88882.1 anig|EHA26989.1 asyd|OJJ62877.1 ater|EAU39272.1 atub|OJI85034.1 aver|OJI96929.1 awen|OJJ35014.1 azon|OJJ45656.1 aory|EIT81137.1

Group3430: abra|OJJ74361.1 acar|OOF95368.1 acla|EAW07413.1 afis|EAW22439.1 afla|EED54449.1 afum|EAL90752.1 agla|OJJ87584.1 akaw|GAA86669.1 aluc|OJZ89766.1 anid|CBF88862.1 anig|EHA26986.1 aory|EIT81135.1 asyd|OJJ62874.1 ater|EAU39269.1 atub|OJI85036.1 aver|OJI96927.1 awen|OJJ35016.1 azon|OJJ45661.1

Group3431: abra|OJJ74364.1 acar|OOF95366.1 acla|EAW07414.1 afis|EAW22441.1 afla|EED54457.1 afum|EAL90750.1 agla|OJJ87583.1 akaw|GAA86666.1 aluc|OJZ89763.1 anid|CBF88845.1 anig|EHA26983.1 aory|EIT81148.1 asyd|OJJ62867.1 ater|EAU39263.1 atub|OJI85039.1 aver|OJI96920.1 awen|OJJ35018.1 azon|OJJ50043.1

Group3432: abra|OJJ74386.1 acar|OOF95352.1 acla|EAW07427.1 afis|EAW22453.1 afla|EED54469.1 afum|EAL90738.1 agla|OJJ87570.1 akaw|GAA87287.1 aluc|OJZ89732.1 anid|CBF88931.1 anig|EHA26958.1 aory|EIT81158.1 asyd|OJJ62850.1 ater|EAU39253.1 atub|OJI85069.1 aver|OJI96903.1 awen|OJJ35078.1 azon|OJJ50029.1

Group3433: abra|OJJ74395.1 acar|OOF95341.1 acla|EAW06819.1 afis|EAW16367.1 afla|EED53469.1 afum|EAL88992.2 agla|OJJ78588.1 akaw|GAA87293.1 aluc|OJZ89725.1 anid|CBF89350.1 anig|EHA26953.1

aory|EIT75003.1 asyd|OJJ62502.1 ater|EAU34081.1 atub|OJI85075.1 aver|OJI96623.1 awen|OJJ37772.1 azon|OJJ45083.1

Group3434: abra|OJJ74414.1 acar|OOF95329.1 acla|EAW06828.1 afis|EAW16378.1 afla|EED53458.1 afum|EAL89003.1 agla|OJJ78596.1 akaw|GAA87305.1 aluc|OJZ89712.1 anid|CBF88042.1 anig|EHA26943.1 aory|EIT75071.1 asyd|OJJ63543.1 ater|EAU34074.1 atub|OJI85090.1 aver|OJI97391.1 awen|OJJ37912.1 azon|OJJ45091.1

Group3435: abra|OJJ74417.1 acar|OOF95326.1 acla|EAW06832.1 afis|EAW16382.1 afla|EED53452.1 afum|EAL89007.1 agla|OJJ88205.1 akaw|GAA87308.1 aluc|OJZ89709.1 anid|CBF89369.1 anig|EHA26940.1 aory|EIT75129.1 asyd|OJJ62475.1 ater|EAU34070.1 atub|OJI85093.1 aver|OJI96611.1 awen|OJJ37915.1 azon|OJJ46177.1

Group3436: abra|OJJ74438.1 acar|OOF95308.1 acla|EAW12299.1 afis|EAW23841.1 afla|EED49145.1 afum|EAL87272.1 agla|OJJ79053.1 akaw|GAA87331.1 aluc|OJZ89683.1 anid|CBF89626.1 anig|EHA26919.1 aory|EIT78517.1 asyd|OJJ62298.1 ater|EAU29595.1 atub|OJI85117.1 aver|OJI96406.1 awen|OJJ35935.1 azon|OJJ49112.1

Group3437: abra|OJJ74456.1 acar|OOF95294.1 acla|EAW07237.1 afis|EAW22260.1 afla|EED57647.1 afum|EAL90920.1 agla|OJJ87795.1 akaw|GAA87350.1 aluc|OJZ89664.1 anid|CBF88509.1 anig|EHA26901.1 aory|EIT79886.1 asyd|OJJ63802.1 ater|EAU34263.1 atub|OJI85136.1 aver|OJI97136.1 awen|OJJ34832.1 azon|OJJ45714.1

Group3438: abra|OJJ74457.1 acar|OOF95293.1 acla|EAW07236.1 afis|EAW22259.1 afla|EED57646.1 afum|EAL90921.1 agla|OJJ87799.1 akaw|GAA87351.1 aluc|OJZ89663.1 anid|CBF88506.1 anig|EHA26900.1 aory|EIT79868.1 asyd|OJJ63799.1 ater|EAU34262.1 atub|OJI85137.1 aver|OJI97139.1 awen|OJJ34831.1 azon|OJJ45715.1

Group3439: abra|OJJ74463.1 acar|OOF95288.1 acla|EAW07235.1 afis|EAW22256.1 afla|EED57643.1 afum|EAL90922.1 agla|OJJ87800.1 akaw|GAA87355.1 aluc|OJZ89659.1 anid|CBF88504.1 anig|EHA26896.1 aory|EIT79873.1 asyd|OJJ63798.1 ater|EAU34257.1 atub|OJI85141.1 aver|OJI97140.1 awen|OJJ34828.1 azon|OJJ45720.1

Group3440: abra|OJJ74464.1 acar|OOF95286.1 acla|EAW07234.1 afis|EAW22255.1 afla|EED57641.1 afum|EAL90923.1 agla|OJJ87801.1 akaw|GAA87356.1 aluc|OJZ89658.1 anid|CBF88502.1 anig|EHA26895.1 aory|EIT82944.1 asyd|OJJ63796.1 ater|EAU34256.1 atub|OJI85143.1 aver|OJI97141.1 awen|OJJ34827.1 azon|OJJ45722.1

Group3441: abra|OJJ74466.1 acar|OOF95281.1 acla|EAW07231.1 afis|EAW22245.1 afla|EED57638.1 afum|EAL90948.1 agla|OJJ87805.1 akaw|GAA87359.1 aluc|OJZ89654.1 anid|CBF88496.1 anig|EHA26892.1 aory|EIT82948.1 asyd|OJJ63793.1 ater|EAU34253.1 atub|OJI85150.1 aver|OJI97144.1 awen|OJJ34823.1 azon|OJJ45727.1

Group3442: abra|OJJ74467.1 acar|OOF95280.1 acla|EAW07230.1 afis|EAW22244.1 afla|EED57637.1 afum|EAL90949.1 agla|OJJ87807.1 akaw|GAA87360.1 aluc|OJZ89653.1 anid|CBF88493.1 anig|EHA26891.1 aory|EIT82977.1 asyd|OJJ63792.1 ater|EAU34252.1 atub|OJI85151.1 aver|OJI97145.1 awen|OJJ34822.1 azon|OJJ45728.1

Group3443: abra|OJJ74468.1 acar|OOF95279.1 acla|EAW07229.1 afis|EAW22243.1 afla|EED57636.1 afum|EAL90950.1 agla|OJJ87808.1 akaw|GAA87361.1 aluc|OJZ89652.1 anid|CBF88491.1 anig|EHA26890.1 aory|EIT82978.1 asyd|OJJ63791.1 ater|EAU34251.1 atub|OJI85152.1 aver|OJI97146.1 awen|OJJ34821.1 azon|OJJ45729.1

Group3444: abra|OJJ74473.1 acar|OOF95271.1 acla|EAW07224.1 afis|EAW22238.1 afla|EED57627.1 afum|EAL90955.1 agla|OJJ87813.1 akaw|GAA87367.1 aluc|OJZ89645.1 anid|CBF88481.1 anig|EHA26884.1 aory|EIT82957.1 asyd|OJJ63785.1 ater|EAU34245.1 atub|OJI85158.1 aver|OJI97152.1 awen|OJJ34816.1 azon|OJJ45733.1

Group3445: abra|OJJ74477.1 acar|OOF95269.1 acla|EAW07223.1 afis|EAW22237.1 afla|EED57626.1 afum|EAL90956.1 agla|OJJ87814.1 akaw|GAA87370.1 aluc|OJZ89642.1 anid|CBF88479.1 anig|EHA26881.1 aory|EIT82989.1 asyd|OJJ63784.1 ater|EAU34244.1 atub|OJI85161.1 aver|OJI97153.1 awen|OJJ34815.1 azon|OJJ45734.1

Group3446: abra|OJJ74478.1 acar|OOF95268.1 acla|EAW07222.1 afis|EAW22236.1 afla|EED57625.1 afum|EAL90957.1 agla|OJJ87815.1 akaw|GAA87371.1 aluc|OJZ89641.1 anid|CBF88477.1 anig|EHA26880.1 aory|EIT82990.1 asyd|OJJ63783.1 ater|EAU34243.1 atub|OJI85162.1 aver|OJI97154.1 awen|OJJ34814.1 azon|OJJ45735.1

Group3447: abra|OJJ74481.1 acar|OOF95267.1 acla|EAW07220.1 afis|EAW22234.1 afla|EED57623.1 afum|EAL90959.1 agla|OJJ87817.1 akaw|GAA87374.1 aluc|OJZ89638.1 anid|CBF88474.1 anig|EHA26879.1 aory|EIT82951.1 asyd|OJJ63782.1 ater|EAU34237.1 atub|OJI85165.1 aver|OJI97155.1 awen|OJJ34813.1 azon|OJJ45736.1

Group3448: abra|OJJ74484.1 acar|OOF95266.1 acla|EAW07217.1 afis|EAW22231.1 afla|EED57620.1 afum|EAL90962.1 agla|OJJ87820.1 akaw|GAA87377.1 aluc|OJZ89635.1 anid|CBF88469.1 anig|EHA26877.1 aory|EIT82992.1 asyd|OJJ63780.1 ater|EAU34234.1 atub|OJI85168.1 aver|OJI97157.1 awen|OJJ34811.1 azon|OJJ45738.1

Group3449: abra|OJJ74489.1 acar|OOF95260.1 acla|EAW07206.1 afis|EAW22218.1 afla|EED57611.1 afum|EAL90976.1 agla|OJJ87826.1 akaw|GAA87383.1 aluc|OJZ89629.1 anid|CBF88456.1 anig|EHA26872.1 aory|EIT82939.1 asyd|OJJ63771.1 ater|EAU34228.1 atub|OJI85173.1 aver|OJI97167.1 awen|OJJ34804.1 azon|OJJ45749.1

Group3450: abra|OJJ74490.1 acar|OOF95259.1 acla|EAW07204.1 afis|EAW22216.1 afla|EED57610.1 afum|EAL90978.1 agla|OJJ87830.1 akaw|GAA87384.1 aluc|OJZ89628.1 anid|CBF88454.1 anig|EHA26871.1 aory|EIT82985.1 asyd|OJJ63770.1 ater|EAU34224.1 atub|OJI85174.1 aver|OJI97168.1 awen|OJJ34801.1 azon|OJJ45748.1

Group3451: abra|OJJ74491.1 acar|OOF95258.1 acla|EAW07203.1 afis|EAW22215.1 afla|EED57609.1 afum|EAL90979.1 agla|OJJ87829.1 akaw|GAA87385.1 aluc|OJZ89627.1 anid|CBF88452.1 anig|EHA26870.1 aory|EIT82971.1 asyd|OJJ63769.1 ater|EAU34223.1 atub|OJI85175.1 aver|OJI97169.1 awen|OJJ34800.1 azon|OJJ45747.1

Group3452: abra|OJJ74493.1 acar|OOF95256.1 acla|EAW07201.1 afis|EAW22213.1 afla|EED57607.1 afum|EAL90981.2 agla|OJJ87831.1 akaw|GAA87386.1 aluc|OJZ89625.1 anid|CBF88448.1 anig|EHA26869.1 aory|EIT82952.1 asyd|OJJ63768.1 ater|EAU34222.1 atub|OJI85177.1 aver|OJI97170.1 awen|OJJ34798.1 azon|OJJ45745.1

Group3453: abra|OJJ74494.1 acar|OOF95255.1 acla|EAW07200.1 afis|EAW22212.1 afla|EED57606.1 afum|EAL90982.1 agla|OJJ87832.1 akaw|GAA87387.1 aluc|OJZ89624.1 anid|CBF88446.1 anig|EHA26868.1 aory|EIT82953.1 asyd|OJJ63767.1 ater|EAU34221.1 atub|OJI85178.1 aver|OJI97171.1 awen|OJJ34797.1 azon|OJJ45744.1

Group3454: abra|OJJ74496.1 acar|OOF95254.1 acla|EAW07197.1 afis|EAW22209.1 afla|EED57603.1 afum|EAL90985.1 agla|OJJ87835.1 akaw|GAA87390.1 aluc|OJZ89622.1 anig|EHA26866.1 asyd|OJJ63765.1 ater|EAU34219.1 atub|OJI85180.1 aver|OJI97174.1 awen|OJJ34794.1 anid|CBF88442.1 aory|EIT82943.1 azon|OJJ45750.1

Group3455: abra|OJJ74497.1 acar|OOF95253.1 acla|EAW07195.1 afis|EAW22207.1 afla|EED57602.1 afum|EAL90986.2 agla|OJJ87837.1 akaw|GAA87392.1 aluc|OJZ89621.1 anid|CBF88439.1 anig|EHA26865.1 aory|EIT82955.1 asyd|OJJ63764.1 ater|EAU34218.1 atub|OJI85181.1 aver|OJI97176.1 awen|OJJ34793.1 azon|OJJ45753.1

Group3456: abra|OJJ74498.1 acar|OOF95252.1 acla|EAW13756.1 afis|EAW19810.1 afla|EED57601.1 afum|EAL85082.1 agla|OJJ87838.1 akaw|GAA87393.1 aluc|OJZ89620.1 anid|CBF88437.1 anig|EHA26864.1 aory|EIT82959.1 asyd|OJJ63763.1 ater|EAU34217.1 atub|OJI85182.1 aver|OJI97177.1 awen|OJJ34792.1 azon|OJJ45754.1

Group3457: abra|OJJ74503.1 acar|OOF95241.1 acla|EAW07189.1 afis|EAW22202.1 afla|EED57593.1 afum|EAL90990.1 agla|OJJ87842.1 akaw|GAA87397.1 aluc|OJZ89615.1 anid|CBF88430.1 anig|EHA26860.1 aory|EIT82963.1 asyd|OJJ63760.1 ater|EAU34212.1 atub|OJI85188.1 aver|OJI97180.1 awen|OJJ34787.1 azon|OJJ45771.1

Group3458: abra|OJJ74506.1 acar|OOF89839.1 acla|EAW07192.1 afis|EAW22205.1 afla|EED57588.1 afum|EAL90988.1 agla|OJJ87845.1 akaw|GAA88823.1 aluc|OJZ89611.1 anid|CBF88417.1 anig|EHA26856.1 aory|EIT82968.1 asyd|OJJ63756.1 ater|EAU34209.1 atub|OJI85192.1 aver|OJI97184.1 awen|OJJ34786.1 azon|OJJ45766.1

Group3459: abra|OJJ74508.1 acar|OOF95234.1 acla|EAW07184.1 afis|EAW22197.1 afla|EED57584.1 afum|EAL90994.1 agla|OJJ87848.1 akaw|GAA82091.1 aluc|OJZ92089.1 anid|CBF88409.1 anig|EHA26852.1 aory|EIT82932.1 asyd|OJJ63754.1 ater|EAU34207.1 atub|OJI86519.1 aver|OJI97187.1 awen|OJJ34780.1 azon|OJJ45761.1

Group3460: abra|OJJ74513.1 acar|OOF95230.1 acla|EAW07177.1 afis|EAW22191.1 afla|EED57575.1 afum|EAL91000.1 agla|OJJ87856.1 akaw|GAA82086.1 aluc|OJZ92094.1 anid|CBF88400.1 anig|EHA26847.1 aory|EIT82946.1 asyd|OJJ63748.1 ater|EAU34199.1 atub|OJI86527.1 aver|OJI97192.1 awen|OJJ34771.1 azon|OJJ45775.1

Group3461: abra|OJJ74519.1 acar|OOF95229.1 acla|EAW07174.1 afis|EAW22188.1 afla|EED57572.1 afum|EAL91002.1 agla|OJJ87861.1 akaw|GAA82081.1 aluc|OJZ92098.1 anid|CBF88395.1 anig|EHA26844.1 aory|EIT82984.1 asyd|OJJ63745.1 ater|EAU34195.1 atub|OJI86531.1 aver|OJI97195.1 awen|OJJ34768.1 azon|OJJ45778.1

Group3462: abra|OJJ74523.1 acar|OOF95226.1 acla|EAW07170.1 afis|EAW22185.1 afla|EED57563.1 afum|EAL91004.1 agla|OJJ87870.1 akaw|GAA82078.1 aluc|OJZ92102.1 anid|CBF88391.1 anig|EHA26841.1

aory|EIT81827.1 asyd|OJJ63742.1 ater|EAU34193.1 atub|OJI86536.1 aver|OJI97198.1 awen|OJJ34765.1 azon|OJJ45780.1

Group3463: abra|OJJ74524.1 acar|OOF95225.1 acla|EAW07169.1 afis|EAW22184.1 afla|EED57562.1 afum|EAL91005.1 agla|OJJ87869.1 akaw|GAA82077.1 aluc|OJZ92104.1 anid|CBF75419.1 anig|EHA26840.1 aory|EIT81846.1 asyd|OJJ59792.1 ater|EAU34188.1 atub|OJI86538.1 aver|OJJ02495.1 awen|OJJ34764.1 azon|OJJ45781.1

Group3464: abra|OJJ74527.1 acar|OOF95223.1 acla|EAW07164.1 afis|EAW22179.1 afla|EED57558.1 afum|EAL91010.1 agla|OJJ87876.1 akaw|GAA82074.1 aluc|OJZ92108.1 anid|CBF88384.1 anig|EHA26837.1 aory|EIT81836.1 asyd|OJJ63738.1 ater|EAU34184.1 atub|OJI86542.1 aver|OJI97202.1 awen|OJJ34760.1 azon|OJJ45791.1

Group3465: abra|OJJ74532.1 acar|OOF95220.1 acla|EAW07160.1 afis|EAW22175.1 afla|EED57554.1 afum|EAL91014.1 agla|OJJ87879.1 akaw|GAA82068.1 aluc|OJZ92114.1 anid|CBF89077.1 anig|EHA26831.1 aory|EIT81832.1 asyd|OJJ62739.1 ater|EAU34180.1 atub|OJI86547.1 aver|OJI96782.1 awen|OJJ39689.1 azon|OJJ45808.1

Group3466: abra|OJJ74533.1 acar|OOF95219.1 acla|EAW07159.1 afis|EAW22174.1 afla|EED57553.1 afum|EAL91015.1 agla|OJJ87880.1 akaw|GAA82067.1 aluc|OJZ92115.1 anid|CBF89079.1 anig|EHA26830.1 aory|EIT81853.1 asyd|OJJ62738.1 ater|EAU34179.1 atub|OJI86548.1 aver|OJI96781.1 awen|OJJ34749.1 azon|OJJ45810.1

Group3467: abra|OJJ74546.1 acar|OOF95210.1 acla|EAW07152.1 afis|EAW22166.1 afla|EED57536.1 afum|EAL91025.1 agla|OJJ87891.1 akaw|GAA82052.1 aluc|OJZ92128.1 anid|CBF89087.1 anig|EHA26820.1 aory|EIT81873.1 asyd|OJJ62734.1 ater|EAU34173.1 atub|OJI86560.1 aver|OJI96777.1 awen|OJJ34740.1 azon|OJJ45819.1

Group3468: abra|OJJ74548.1 acar|OOF95209.1 acla|EAW07150.1 afis|EAW22164.1 afla|EED57535.1 afum|EAL91027.1 agla|OJJ87893.1 akaw|GAA82050.1 aluc|OJZ92130.1 anid|CBF89090.1 anig|EHA26818.1 aory|EIT81851.1 asyd|OJJ62732.1 ater|EAU34171.1 atub|OJI86562.1 aver|OJI96775.1 awen|OJJ34738.1 azon|OJJ45821.1

Group3469: abra|OJJ74549.1 acar|OOF95208.1 acla|EAW07149.1 afis|EAW22163.1 afla|EED57534.1 afum|EAL91028.1 agla|OJJ87894.1 akaw|GAA82049.1 aluc|OJZ92131.1 anid|CBF89094.1 anig|EHA26817.1 aory|EIT81858.1 asyd|OJJ62731.1 ater|EAU34166.1 atub|OJI86563.1 aver|OJI96774.1 awen|OJJ34737.1 azon|OJJ45822.1

Group3470: abra|OJJ74553.1 acar|OOF95202.1 acla|EAW07145.1 afis|EAW22159.1 afla|EED57531.1 afum|EAL91031.1 agla|OJJ87905.1 akaw|GAA82044.1 aluc|OJZ92136.1 anid|CBF89101.1 anig|EHA26813.1 aory|EIT81829.1 asyd|OJJ62727.1 ater|EAU34162.1 atub|OJI86568.1 aver|OJI96770.1 awen|OJJ34733.1 azon|OJJ45826.1

Group3471: abra|OJJ74569.1 acar|OOF95192.1 acla|EAW07140.1 afis|EAW22153.1 afla|EED53614.1 afum|EAL91037.1 agla|OJJ87909.1 akaw|GAA82036.1 aluc|OJZ92147.1 anid|CBF89105.1 anig|EHA26802.1 aory|EIT75057.1 asyd|OJJ62725.1 ater|EAU34157.1 atub|OJI86575.1 aver|OJI96768.1 awen|OJJ34729.1 azon|OJJ45834.1

Group3472: abra|OJJ74571.1 acar|OOF95191.1 acla|EAW07138.1 afis|EAW22151.1 afla|EED53613.1 afum|EAL91039.1 agla|OJJ87911.1 akaw|GAA82034.1 aluc|OJZ92149.1 anid|CBF89108.1 anig|EHA26800.1 aory|EIT75195.1 asyd|OJJ62667.1 ater|EAU34156.1 atub|OJI86577.1 aver|OJI96765.1 awen|OJJ34727.1 azon|OJJ45835.1

Group3473: abra|OJJ74573.1 acar|OOF95189.1 acla|EAW07136.1 afis|EAW22149.1 afla|EED53610.1 afum|EAL91041.1 agla|OJJ87914.1 akaw|GAA82032.1 aluc|OJZ92151.1 anid|CBF89112.1 anig|EHA26798.1 aory|EIT74840.1 asyd|OJJ62665.1 ater|EAU34155.1 atub|OJI86579.1 aver|OJI96763.1 awen|OJJ34725.1 azon|OJJ45839.1

Group3474: abra|OJJ74577.1 acar|OOF95187.1 acla|EAW07133.1 afis|EAW22146.1 afla|EED45848.1 afum|EAL91044.1 agla|OJJ87922.1 akaw|GAA82028.1 aluc|OJZ92155.1 anid|CBF78365.1 anig|EHA26794.1 aory|EIT75625.1 asyd|OJJ57324.1 ater|EAU34149.1 atub|OJI86583.1 aver|OJJ05836.1 awen|OJJ34721.1 azon|OJJ46642.1

Group3475: abra|OJJ74578.1 acar|OOF95186.1 acla|EAW07132.1 afis|EAW22145.1 afla|EED45849.1 afum|EAL91045.1 agla|OJJ87923.1 akaw|GAA82027.1 aluc|OJZ92156.1 anid|CBF78367.1 anig|EHA26793.1 aory|EIT75756.1 asyd|OJJ57325.1 ater|EAU34148.1 atub|OJI86584.1 aver|OJJ05835.1 awen|OJJ34720.1 azon|OJJ46643.1

Group3476: abra|OJJ74581.1 acar|OOF95183.1 acla|EAW07131.1 afis|EAW22142.1 afla|EED53568.1 afum|EAL91048.2 agla|OJJ87932.1 akaw|GAA82024.1 aluc|OJZ92159.1 anid|CBF89555.1 anig|EHA26790.1 aory|EIT75148.1 asyd|OJJ62347.1 ater|EAU31752.1 atub|OJI86587.1 aver|OJI96453.1 awen|OJJ34717.1 azon|OJJ45846.1

Group3477: abra|OJJ74584.1 acar|OOF95182.1 acla|EAW10794.1 afis|EAW24782.1 afla|EED47203.1 afum|EAL89421.1 agla|OJJ82878.1 akaw|GAA82021.1 aluc|OJZ92162.1 anid|CBF81444.1 anig|EHA26787.1 aory|EIT79171.1 asyd|OJJ59430.1 ater|EAU36526.1 atub|OJI86590.1 aver|OJJ02841.1 awen|OJJ40368.1 azon|OJJ49217.1

Group3478: abra|OJJ74585.1 acar|OOF95181.1 acla|EAW10793.1 afis|EAW24783.1 afla|EED45489.1 afum|EAL89422.1 akaw|GAA82020.1 aluc|OJZ92163.1 anid|CBF81449.1 anig|EHA26786.1 aory|EIT79242.1 asyd|OJJ59432.1 atub|OJI86592.1 aver|OJJ02839.1 awen|OJJ36318.1 azon|OJJ43590.1 aory|EIT75730.1 afla|EED47202.1

Group3479: abra|OJJ74588.1 acar|OOF95178.1 acla|EAW10790.1 afis|EAW24786.1 afla|EED47199.1 afum|EAL89425.1 agla|OJJ82884.1 akaw|GAA82017.1 aluc|OJZ92165.1 anid|CBF81455.1 anig|EHA26784.1 aory|EIT79195.1 asyd|OJJ59435.1 ater|EAU36523.1 atub|OJI86595.1 aver|OJJ02836.1 awen|OJJ40370.1 azon|OJJ49220.1

Group3480: abra|OJJ74589.1 acar|OOF95177.1 acla|EAW10789.1 afis|EAW24787.1 afla|EED47197.1 afum|EAL89426.1 agla|OJJ82883.1 akaw|GAA82016.1 aluc|OJZ92167.1 anid|CBF81457.1 anig|EHA26782.1 aory|EIT79260.1 asyd|OJJ59437.1 ater|EAU36522.1 atub|OJI86598.1 aver|OJJ02834.1 awen|OJJ40371.1 azon|OJJ49221.1

Group3481: abra|OJJ74594.1 acar|OOF95175.1 acla|EAW10788.1 afis|EAW24791.1 afla|EED47195.1 afum|EAL89433.1 agla|OJJ82881.1 akaw|GAA82014.1 aluc|OJZ92169.1 anid|CBF81460.1 anig|EHA26780.1 aory|EIT79181.1 asyd|OJJ59439.1 ater|EAU36521.1 atub|OJI86601.1 aver|OJJ02833.1 awen|OJJ40372.1 azon|OJJ49223.1

Group3482: abra|OJJ74595.1 acar|OOF95174.1 acla|EAW10786.1 afis|EAW24792.1 afla|EED47194.1 afum|EAL89434.1 agla|OJJ82879.1 akaw|GAA82013.1 aluc|OJZ92170.1 anid|CBF81464.1 anig|EHA26779.1 aory|EIT79314.1 asyd|OJJ59442.1 ater|EAU36516.1 atub|OJI86602.1 aver|OJJ02831.1 awen|OJJ40373.1 azon|OJJ49224.1

Group3483: abra|OJJ74629.1 acla|EAW10752.1 afis|EAW24821.1 afla|EED47154.1 afum|EAL89465.1 agla|OJJ82919.1 akaw|GAA81979.1 aluc|OJZ92211.1 anid|CBF85870.1 anig|EHA26747.1 aory|EIT79196.1 asyd|OJJ64372.1 ater|EAU36484.1 atub|OJI86640.1 aver|OJI98468.1 awen|OJJ40407.1 azon|OJJ49256.1 afla|EED47155.1

Group3484: abra|OJJ74705.1 acar|OOF89903.1 acla|EAW10692.1 afis|EAW24878.1 afla|EED47088.1 afum|EAL89528.1 agla|OJJ82100.1 akaw|GAA81912.1 aluc|OJZ92285.1 anid|CBF75756.1 anig|EHA26683.1 aory|EIT72477.1 asyd|OJJ57526.1 ater|EAU36423.1 atub|OJI86716.1 aver|OJJ05652.1 awen|OJJ40468.1 azon|OJJ49332.1

Group3485: abra|OJJ74707.1 acar|OOF89905.1 acla|EAW10691.1 afis|EAW24879.1 afla|EED47087.1 afum|EAL89529.1 agla|OJJ82101.1 akaw|GAA81910.1 aluc|OJZ92288.1 anid|CBF75754.1 anig|EHA26681.1 aory|EIT72507.1 asyd|OJJ57525.1 ater|EAU36422.1 atub|OJI86718.1 aver|OJJ05653.1 awen|OJJ40469.1 azon|OJJ49333.1

Group3486: abra|OJJ74708.1 acar|OOF95718.1 acla|EAW10690.1 afis|EAW24880.1 afla|EED47086.1 afum|EAL89530.1 agla|OJJ82102.1 akaw|GAA81909.1 aluc|OJZ92289.1 anid|CBF75752.1 anig|EHA26680.1 aory|EIT72478.1 asyd|OJJ57524.1 ater|EAU36421.1 atub|OJI86719.1 aver|OJJ05654.1 awen|OJJ40470.1 azon|OJJ49334.1

Group3487: abra|OJJ74709.1 acar|OOF95719.1 acla|EAW10689.1 afis|EAW24881.1 afla|EED47085.1 afum|EAL89531.1 agla|OJJ82103.1 akaw|GAA81908.1 aluc|OJZ92290.1 anid|CBF75750.1 anig|EHA26679.1 aory|EIT72499.1 asyd|OJJ57523.1 ater|EAU36420.1 atub|OJI86720.1 aver|OJJ05655.1 awen|OJJ40471.1 azon|OJJ49335.1

Group3488: abra|OJJ74711.1 acar|OOF95722.1 acla|EAW10687.1 afis|EAW24883.1 afla|EED47083.1 afum|EAL89533.1 agla|OJJ82105.1 akaw|GAA81906.1 aluc|OJZ92292.1 anid|CBF75747.1 anig|EHA26677.1 aory|EIT72512.1 asyd|OJJ57521.1 ater|EAU36418.1 atub|OJI86722.1 aver|OJJ05657.1 awen|OJJ40473.1 azon|OJJ49337.1

Group3489: abra|OJJ74714.1 acar|OOF95723.1 acla|EAW10685.1 afis|EAW24885.1 afla|EED47081.1 afum|EAL89535.2 agla|OJJ82108.1 akaw|GAA81904.1 aluc|OJZ92294.1 anid|CBF75617.1 anig|EHA26675.1 aory|EIT72482.1 asyd|OJJ57452.1 ater|EAU36416.1 atub|OJI86724.1 aver|OJJ05726.1 awen|OJJ40475.1 azon|OJJ49339.1

Group3490: abra|OJJ74715.1 acar|OOF95724.1 acla|EAW10684.1 afis|EAW24886.1 afla|EED47080.1 afum|EAL89537.1 agla|OJJ82109.1 akaw|GAA81903.1 aluc|OJZ92295.1 anid|CBF75619.1 anig|EHA26674.1 aory|EIT72515.1 asyd|OJJ57453.1 ater|EAU36415.1 atub|OJI86725.1 aver|OJJ05725.1 awen|OJJ40476.1 azon|OJJ49340.1

Group3491: abra|OJJ74717.1 acar|OOF95728.1 acla|EAW10681.1 afis|EAW24889.1 afla|EED46965.1 afum|EAL89540.1 agla|OJJ82112.1 akaw|GAA81899.1 aluc|OJZ92299.1 anid|CBF75625.1 anig|EHA26672.1

aory|EIT72622.1 asyd|OJJ57456.1 ater|EAU36412.1 atub|OJI86728.1 aver|OJJ05723.1 awen|OJJ40479.1 azon|OJJ49343.1

Group3492: abra|OJJ74722.1 acar|OOF95731.1 acla|EAW10675.1 afis|EAW24891.1 afla|EED46970.1 afum|EAL89544.1 agla|OJJ82115.1 akaw|GAA81894.1 aluc|OJZ92306.1 anid|CBF75630.1 anig|EHA26662.1 aory|EIT72703.1 asyd|OJJ57459.1 ater|EAU36407.1 atub|OJI86736.1 aver|OJJ05721.1 awen|OJJ40484.1 azon|OJJ49346.1

Group3493: abra|OJJ74723.1 acar|OOF95732.1 acla|EAW10674.1 afis|EAW24892.1 afla|EED46971.1 afum|EAL89545.1 agla|OJJ82116.1 akaw|GAA81893.1 aluc|OJZ92307.1 anid|CBF75632.1 anig|EHA26661.1 aory|EIT72525.1 asyd|OJJ57460.1 ater|EAU36406.1 atub|OJI86737.1 aver|OJJ05720.1 awen|OJJ40485.1 azon|OJJ49347.1

Group3494: abra|OJJ74724.1 acar|OOF95733.1 acla|EAW10673.1 afis|EAW24893.1 afla|EED46972.1 afum|EAL89546.1 agla|OJJ82117.1 akaw|GAA81892.1 aluc|OJZ92308.1 anid|CBF75634.1 anig|EHA26660.1 aory|EIT72626.1 asyd|OJJ57461.1 ater|EAU36405.1 atub|OJI86738.1 aver|OJJ05719.1 awen|OJJ40486.1 azon|OJJ49348.1

Group3495: abra|OJJ74727.1 acar|OOF90030.1 acla|EAW10671.1 afis|EAW24895.1 afla|EED46975.1 afum|EAL89549.1 agla|OJJ82119.1 akaw|GAA81890.1 aluc|OJZ92311.1 anid|CBF75657.1 anig|EHA26658.1 aory|EIT72598.1 asyd|OJJ57463.1 ater|EAU36403.1 atub|OJI86740.1 aver|OJJ05717.1 awen|OJJ40488.1 azon|OJJ49351.1

Group3496: abra|OJJ74735.1 acar|OOF95734.1 acla|EAW10667.1 afis|EAW24899.1 afla|EED46978.1 afum|EAL89553.1 agla|OJJ82124.1 akaw|GAA83674.1 aluc|OJZ92317.1 anid|CBF75649.1 anig|EHA26654.1 aory|EIT72632.1 asyd|OJJ57470.1 ater|EAU36400.1 atub|OJI86746.1 aver|OJJ05712.1 awen|OJJ40493.1 azon|OJJ49355.1

Group3497: abra|OJJ74736.1 acar|OOF95735.1 acla|EAW10666.1 afis|EAW24900.1 afla|EED46980.1 afum|EAL89554.1 agla|OJJ82125.1 akaw|GAA83673.1 aluc|OJZ92318.1 anid|CBF75648.1 anig|EHA26653.1 aory|EIT72726.1 asyd|OJJ57472.1 ater|EAU36399.1 atub|OJI86747.1 aver|OJJ05711.1 awen|OJJ40494.1 azon|OJJ49356.1

Group3498: abra|OJJ74738.1 acar|OOF95736.1 acla|EAW10665.1 afis|EAW24901.1 afla|EED46981.1 afum|EAL89555.1 agla|OJJ82126.1 akaw|GAA83672.1 aluc|OJZ92319.1 anid|CBF75646.1 anig|EHA26651.1 aory|EIT72528.1 asyd|OJJ57473.1 ater|EAU36398.1 atub|OJI86748.1 aver|OJJ05710.1 awen|OJJ40495.1 azon|OJJ49357.1

Group3499: abra|OJJ74739.1 acar|OOF95737.1 acla|EAW10664.1 afis|EAW24902.1 afla|EED46982.1 afum|EAL89556.1 agla|OJJ82127.1 akaw|GAA83671.1 aluc|OJZ92320.1 anid|CBF75644.1 anig|EHA26650.1 aory|EIT72548.1 asyd|OJJ57474.1 ater|EAU36397.1 atub|OJI86749.1 aver|OJJ05707.1 awen|OJJ40496.1 azon|OJJ49358.1

Group3500: abra|OJJ74749.1 acar|OOF95742.1 acla|EAW10661.1 afis|EAW24906.1 afla|EED46989.1 afum|EAL89559.1 agla|OJJ82130.1 akaw|GAA83662.1 aluc|OJZ92330.1 anid|CBF75661.1 anig|EHA26641.1 aory|EIT72764.1 asyd|OJJ57478.1 ater|EAU36392.1 atub|OJI86759.1 aver|OJJ05703.1 awen|OJJ40499.1 azon|OJJ49361.1

Group3501: abra|OJJ74750.1 acar|OOF95743.1 acla|EAW10660.1 afis|EAW24907.1 afla|EED46991.1 afum|EAL89560.1 agla|OJJ82131.1 akaw|GAA83661.1 aluc|OJZ92331.1 anid|CBF75665.1 anig|EHA26640.1 aory|EIT72700.1 asyd|OJJ57479.1 ater|EAU36391.1 atub|OJI86760.1 aver|OJJ05702.1 awen|OJJ40500.1 azon|OJJ49362.1

Group3502: abra|OJJ74751.1 acar|OOF95744.1 acla|EAW10659.1 afis|EAW24908.1 afla|EED46992.1 afum|EAL89561.1 agla|OJJ82132.1 akaw|GAA83660.1 aluc|OJZ92332.1 anid|CBF75667.1 anig|EHA26639.1 aory|EIT72681.1 asyd|OJJ57480.1 ater|EAU36390.1 atub|OJI86761.1 aver|OJJ05701.1 awen|OJJ40501.1 azon|OJJ49363.1

Group3503: abra|OJJ74752.1 acar|OOF95745.1 acla|EAW10658.1 afis|EAW24909.1 afla|EED46994.1 afum|EAL89562.1 agla|OJJ82133.1 akaw|GAA83659.1 aluc|OJZ92333.1 anid|CBF75669.1 anig|EHA26638.1 aory|EIT72691.1 asyd|OJJ57481.1 ater|EAU36389.1 atub|OJI86762.1 aver|OJJ05700.1 awen|OJJ40503.1 azon|OJJ49364.1

Group3504: abra|OJJ74754.1 acar|OOF95747.1 acla|EAW10656.1 afis|EAW24911.1 afla|EED46996.1 afum|EAL89565.1 agla|OJJ82135.1 akaw|GAA83657.1 aluc|OJZ92335.1 anid|CBF75671.1 anig|EHA26636.1 aory|EIT72517.1 asyd|OJJ57483.1 ater|EAU36387.1 atub|OJI86764.1 aver|OJJ05698.1 awen|OJJ40506.1 azon|OJJ49366.1

Group3505: abra|OJJ74756.1 acar|OOF95749.1 acla|EAW10655.1 afis|EAW24912.1 afla|EED46997.1 afum|EAL89566.2 agla|OJJ82141.1 akaw|GAA83655.1 aluc|OJZ92338.1 anid|CBF75673.1 anig|EHA26634.1 aory|EIT72758.1 asyd|OJJ57484.1 ater|EAU36386.1 atub|OJI86766.1 aver|OJJ05697.1 awen|OJJ40508.1 azon|OJJ49367.1

Group3506: abra|OJJ74758.1 acar|OOF95751.1 acla|EAW10653.1 afis|EAW24914.1 afla|EED46999.1 afum|EAL89568.1 agla|OJJ82142.1 akaw|GAA83653.1 aluc|OJZ92342.1 anid|CBF75677.1 anig|EHA26632.1 aory|EIT72541.1 asyd|OJJ57486.1 ater|EAU36384.1 atub|OJI86768.1 aver|OJJ05694.1 awen|OJJ40510.1 azon|OJJ49369.1

Group3507: abra|OJJ74759.1 acar|OOF95752.1 acla|EAW10652.1 afis|EAW24915.1 afla|EED47000.1 afum|EAL89569.1 agla|OJJ82143.1 akaw|GAA83652.1 aluc|OJZ92343.1 anid|CBF75679.1 anig|EHA26631.1 aory|EIT72710.1 asyd|OJJ57487.1 ater|EAU36383.1 atub|OJI86769.1 aver|OJJ05693.1 awen|OJJ40511.1 azon|OJJ49370.1

Group3508: abra|OJJ74760.1 acar|OOF95753.1 acla|EAW10651.1 afis|EAW24916.1 afla|EED47001.1 afum|EAL89571.1 agla|OJJ82144.1 akaw|GAA83651.1 aluc|OJZ92344.1 anid|CBF75681.1 anig|EHA26630.1 aory|EIT72621.1 asyd|OJJ57488.1 ater|EAU36382.1 atub|OJI86770.1 aver|OJJ05692.1 awen|OJJ40512.1 azon|OJJ49371.1

Group3509: abra|OJJ74762.1 acar|OOF95755.1 acla|EAW10649.1 afis|EAW24918.1 afla|EED47003.1 afum|EAL89573.1 agla|OJJ82147.1 akaw|GAA83649.1 aluc|OJZ92346.1 anid|CBF75687.1 anig|EHA26628.1 aory|EIT72583.1 asyd|OJJ57490.1 ater|EAU36380.1 atub|OJI86773.1 aver|OJJ05690.1 awen|OJJ40515.1 azon|OJJ49374.1

Group3510: abra|OJJ74763.1 acar|OOF95756.1 acla|EAW10647.1 afis|EAW24919.1 afla|EED47004.1 afum|EAL89574.1 agla|OJJ82148.1 akaw|GAA83648.1 aluc|OJZ92347.1 anid|CBF75689.1 anig|EHA26627.1 aory|EIT72654.1 asyd|OJJ57491.1 ater|EAU36379.1 atub|OJI86774.1 aver|OJJ05689.1 awen|OJJ40516.1 azon|OJJ49375.1

Group3511: abra|OJJ74766.1 acar|OOF95758.1 acla|EAW10645.1 afis|EAW24921.1 afla|EED55552.1 afum|EAL89576.1 agla|OJJ82150.1 akaw|GAA83646.1 aluc|OJZ92349.1 anid|CBF75693.1 anig|EHA26625.1 aory|EIT82689.1 asyd|OJJ57493.1 ater|EAU35653.1 atub|OJI86777.1 aver|OJJ05687.1 awen|OJJ40518.1 azon|OJJ49377.1

Group3512: abra|OJJ74767.1 acar|OOF95759.1 acla|EAW10644.1 afis|EAW24922.1 afla|EED55551.1 afum|EAL89577.2 agla|OJJ82151.1 akaw|GAA83645.1 aluc|OJZ92350.1 anid|CBF75695.1 anig|EHA26624.1 aory|EIT82687.1 asyd|OJJ57494.1 ater|EAU35654.1 atub|OJI86778.1 aver|OJJ05686.1 awen|OJJ40519.1 azon|OJJ49378.1

Group3513: abra|OJJ74772.1 acar|OOF95762.1 acla|EAW10639.1 afis|EAW24927.1 afla|EED55546.1 afum|EAL89583.1 agla|OJJ82155.1 akaw|GAA83640.1 aluc|OJZ92353.1 anid|CBF75704.1 anig|EHA26621.1 aory|EIT82681.1 asyd|OJJ57497.1 ater|EAU35657.1 atub|OJI86783.1 aver|OJJ05681.1 awen|OJJ40523.1 azon|OJJ49383.1

Group3514: abra|OJJ74774.1 acar|OOF95763.1 acla|EAW10637.1 afis|EAW24929.1 afla|EED55544.1 afum|EAL89585.1 agla|OJJ82157.1 akaw|GAA83638.1 aluc|OJZ92355.1 anid|CBF75708.1 anig|EHA26619.1 aory|EIT82710.1 asyd|OJJ57499.1 ater|EAU35659.1 atub|OJI86785.1 aver|OJJ05679.1 awen|OJJ40525.1 azon|OJJ49385.1

Group3515: abra|OJJ74775.1 acar|OOF95764.1 acla|EAW10636.1 afis|EAW24930.1 afla|EED55543.1 afum|EAL89586.1 agla|OJJ82158.1 akaw|GAA83637.1 aluc|OJZ92356.1 anid|CBF75710.1 anig|EHA26618.1 aory|EIT82674.1 asyd|OJJ57500.1 ater|EAU35660.1 atub|OJI86786.1 aver|OJJ05678.1 awen|OJJ40526.1 azon|OJJ49386.1

Group3516: abra|OJJ74776.1 acar|OOF95765.1 acla|EAW10635.1 afis|EAW24931.1 afla|EED55542.1 afum|EAL89587.1 agla|OJJ82159.1 akaw|GAA83636.1 aluc|OJZ92357.1 anid|CBF75712.1 anig|EHA26617.1 aory|EIT82695.1 asyd|OJJ57502.1 ater|EAU35661.1 atub|OJI86787.1 aver|OJJ05676.1 awen|OJJ40527.1 azon|OJJ49387.1

Group3517: abra|OJJ74779.1 acar|OOF95769.1 acla|EAW10631.1 afis|EAW24936.1 afla|EED55536.1 afum|EAL89593.1 agla|OJJ82163.1 akaw|GAA83633.1 aluc|OJZ92360.1 anid|CBF75723.1 anig|EHA26613.1 aory|EIT80806.1 asyd|OJJ57506.1 ater|EAU35665.1 atub|OJI86790.1 aver|OJJ05672.1 awen|OJJ40531.1 azon|OJJ49391.1

Group3518: abra|OJJ74780.1 acar|OOF95771.1 acla|EAW10630.1 afis|EAW24937.1 afla|EED55535.1 afum|EAL89594.1 agla|OJJ82164.1 akaw|GAA83632.1 aluc|OJZ92361.1 anid|CBF75725.1 anig|EHA26612.1 aory|EIT80813.1 asyd|OJJ57507.1 ater|EAU35666.1 atub|OJI86791.1 aver|OJJ05671.1 awen|OJJ40532.1 azon|OJJ49392.1

Group3519: abra|OJJ74781.1 acar|OOF95772.1 acla|EAW10629.1 afis|EAW24938.1 afla|EED55534.1 afum|EAL89595.1 agla|OJJ82165.1 akaw|GAA83631.1 aluc|OJZ92362.1 anid|CBF75727.1 anig|EHA26611.1 aory|EIT80903.1 asyd|OJJ57508.1 ater|EAU35667.1 atub|OJI86792.1 aver|OJJ05670.1 awen|OJJ40533.1 azon|OJJ49393.1

Group3520: abra|OJJ74784.1 acar|OOF95774.1 acla|EAW10627.1 afis|EAW24940.1 afla|EED55525.1 afum|EAL89597.1 agla|OJJ82167.1 akaw|GAA83629.1 aluc|OJZ92364.1 anid|CBF75735.1 anig|EHA26609.1

aory|EIT80775.1 asyd|OJJ57512.1 ater|EAU35669.1 atub|OJI86794.1 aver|OJJ05667.1 awen|OJJ40536.1 azon|OJJ49395.1

Group3521: abra|OJJ74786.1 acar|OOF95776.1 acla|EAW10625.1 afis|EAW24942.1 afla|EED55523.1 afum|EAL89599.1 agla|OJJ82169.1 akaw|GAA83627.1 aluc|OJZ92366.1 anid|CBF75741.1 anig|EHA26607.1 aory|EIT80928.1 asyd|OJJ57517.1 ater|EAU35674.1 atub|OJI86796.1 aver|OJJ05662.1 awen|OJJ40537.1 azon|OJJ49397.1

Group3522: abra|OJJ74788.1 acar|OOF95777.1 acla|EAW10623.1 afis|EAW24944.1 afla|EED55521.1 afum|EAL89601.1 agla|OJJ82171.1 akaw|GAA83625.1 aluc|OJZ92368.1 anid|CBF75743.1 anig|EHA26605.1 aory|EIT80857.1 asyd|OJJ57519.1 ater|EAU35675.1 atub|OJI86798.1 aver|OJJ05659.1 awen|OJJ40539.1 azon|OJJ49399.1

Group3523: abra|OJJ74789.1 acar|OOF95779.1 acla|EAW10622.1 afis|EAW24945.1 afla|EED55520.1 afum|EAL89602.1 agla|OJJ82173.1 akaw|GAA83624.1 aluc|OJZ92369.1 anid|CBF75615.1 anig|EHA26604.1 aory|EIT80768.1 asyd|OJJ57451.1 ater|EAU35676.1 atub|OJI86799.1 aver|OJJ05727.1 awen|OJJ40540.1 azon|OJJ49400.1

Group3524: abra|OJJ74790.1 acar|OOF95780.1 acla|EAW10621.1 afis|EAW24946.1 afla|EED55519.1 afum|EAL89603.1 agla|OJJ82174.1 akaw|GAA83623.1 aluc|OJZ92370.1 anid|CBF81729.1 anig|EHA26603.1 aory|EIT80898.1 asyd|OJJ59591.1 ater|EAU35678.1 atub|OJI86800.1 aver|OJJ02678.1 awen|OJJ40541.1 azon|OJJ49401.1

Group3525: abra|OJJ74791.1 acar|OOF95781.1 acla|EAW10620.1 afis|EAW24947.1 afla|EED55518.1 afum|EAL89604.1 agla|OJJ82175.1 akaw|GAA83622.1 aluc|OJZ92371.1 anid|CBF81728.1 anig|EHA26602.1 aory|EIT80815.1 asyd|OJJ59590.1 ater|EAU35679.1 atub|OJI86801.1 aver|OJJ02679.1 awen|OJJ40542.1 azon|OJJ49402.1

Group3526: abra|OJJ74792.1 acar|OOF95782.1 acla|EAW10619.1 afis|EAW24948.1 afla|EED55517.1 afum|EAL89605.1 agla|OJJ82176.1 akaw|GAA83621.1 aluc|OJZ92372.1 anid|CBF81726.1 anig|EHA26601.1 aory|EIT80913.1 asyd|OJJ59589.1 ater|EAU35680.1 atub|OJI86802.1 aver|OJJ02680.1 awen|OJJ40543.1 azon|OJJ49403.1

Group3527: abra|OJJ74797.1 acar|OOF95789.1 acla|EAW10609.1 afis|EAW24958.1 afla|EED55505.1 afum|EAL89616.1 agla|OJJ82189.1 akaw|GAA83615.1 aluc|OJZ92378.1 anid|CBF81663.1 anig|EHA26595.1 aory|EIT80867.1 asyd|OJJ59551.1 ater|EAU35697.1 atub|OJI86808.1 aver|OJJ02717.1 awen|OJJ40552.1 azon|OJJ49411.1

Group3528: abra|OJJ74802.1 acar|OOF95795.1 acla|EAW10614.1 afis|EAW24953.1 afla|EED55511.1 afum|EAL89610.1 agla|OJJ82184.1 akaw|GAA83611.1 aluc|OJZ92383.1 anid|CBF81717.1 anig|EHA26592.1 aory|EIT80894.1 asyd|OJJ59584.1 ater|EAU35685.1 atub|OJI86812.1 aver|OJJ02686.1 awen|OJJ40548.1 azon|OJJ49408.1

Group3529: abra|OJJ74804.1 acar|OOF95796.1 acla|EAW10616.1 afis|EAW24951.1 afla|EED55513.1 afum|EAL89608.1 agla|OJJ82187.1 akaw|GAA83609.1 aluc|OJZ92385.1 anid|CBF81721.1 anig|EHA26590.1 aory|EIT80836.1 asyd|OJJ59586.1 atub|OJI86814.1 aver|OJJ02683.1 awen|OJJ40546.1 azon|OJJ49406.1 ater|EAU35683.1

Group3530: abra|OJJ74805.1 acar|OOF95797.1 acla|EAW10617.1 afis|EAW24950.1 afla|EED55514.1 afum|EAL89607.1 agla|OJJ82179.1 akaw|GAA83608.1 aluc|OJZ92386.1 anid|CBF81722.1 anig|EHA26589.1 aory|EIT80921.1 asyd|OJJ59587.1 ater|EAU35682.1 atub|OJI86815.1 aver|OJJ02682.1 awen|OJJ40545.1 azon|OJJ49405.1

Group3531: abra|OJJ74806.1 acar|OOF95798.1 acla|EAW10618.1 afis|EAW24949.1 afla|EED55516.1 afum|EAL89606.1 agla|OJJ82177.1 akaw|GAA83606.1 aluc|OJZ92387.1 anid|CBF81724.1 anig|EHA26588.1 aory|EIT80820.1 asyd|OJJ59588.1 ater|EAU35681.1 atub|OJI86817.1 aver|OJJ02681.1 awen|OJJ40544.1 azon|OJJ49404.1

Group3532: abra|OJJ74808.1 acar|OOF95800.1 acla|EAW10606.1 afis|EAW24961.1 afla|EED55501.1 afum|EAL89619.1 agla|OJJ82196.1 akaw|GAA83604.1 aluc|OJZ92389.1 anid|CBF81658.1 anig|EHA26585.1 aory|EIT80766.1 asyd|OJJ59548.1 ater|EAU35694.1 atub|OJI86819.1 aver|OJJ02720.1 awen|OJJ40554.1 azon|OJJ49416.1

Group3533: abra|OJJ74830.1 acar|OOF95817.1 acla|EAW11373.1 afis|EAW18122.1 afla|EED58048.1 afum|EAL87480.1 agla|OJJ87016.1 akaw|GAA83590.1 aluc|OJZ92407.1 anid|CBF75358.1 anig|EHA26558.1 aory|EIT79720.1 asyd|OJJ59851.1 ater|EAU29689.1 atub|OJI86835.1 aver|OJJ02436.1 awen|OJJ35524.1 azon|OJJ46961.1

Group3534: abra|OJJ74831.1 acar|OOF95819.1 acla|EAW11372.1 afis|EAW18123.1 afla|EED58049.1 afum|EAL87481.1 agla|OJJ87017.1 akaw|GAA83589.1 aluc|OJZ92408.1 anid|CBF80348.1 anig|EHA26557.1 aory|EIT79773.1 asyd|OJJ54832.1 ater|EAU29688.1 atub|OJI86836.1 aver|OJJ07329.1 awen|OJJ35525.1 azon|OJJ46959.1

Group3535: abra|OJJ74840.1 acar|OOF95843.1 acla|EAW09369.1 afis|EAW21766.1 afla|EED46332.1 afum|EAL92796.1 agla|OJJ84385.1 akaw|GAA83581.1 aluc|OJZ92416.1 anid|CBF83756.1 anig|EHA26550.1 aory|EIT81192.1 asyd|OJJ60860.1 ater|EAU38449.1 atub|OJI86846.1 aver|OJJ00185.1 awen|OJJ34049.1 azon|OJJ48182.1

Group3536: abra|OJJ74841.1 acar|OOF94168.1 acla|EAW08544.1 afis|EAW23581.1 afla|EED57489.1 afum|EAL87884.1 agla|OJJ86848.1 akaw|GAA83580.1 aluc|OJZ92417.1 anid|CBF89912.1 anig|EHA26549.1 aory|EIT81889.1 asyd|OJJ62129.1 ater|EAU35371.1 atub|OJI86847.1 aver|OJI96238.1 awen|OJJ39909.1 azon|OJJ45885.1

Group3537: abra|OJJ74847.1 acar|OOF95848.1 acla|EAW08541.1 afis|EAW23578.1 afla|EED57477.1 afum|EAL87887.1 agla|OJJ86843.1 akaw|GAA83575.1 aluc|OJZ92422.1 anid|CBF89598.1 anig|EHA26545.1 aory|EIT80110.1 asyd|OJJ62311.1 ater|EAU35366.1 atub|OJI86852.1 aver|OJI96419.1 awen|OJJ39906.1 azon|OJJ45883.1

Group3538: abra|OJJ74848.1 acar|OOF95849.1 acla|EAW08540.1 afis|EAW23577.1 afla|EED57476.1 afum|EAL87888.1 agla|OJJ86838.1 akaw|GAA83574.1 aluc|OJZ92423.1 anid|CBF89600.1 anig|EHA26544.1 aory|EIT80147.1 asyd|OJJ62310.1 ater|EAU35367.1 atub|OJI86853.1 aver|OJI96418.1 awen|OJJ39905.1 azon|OJJ45882.1

Group3539: abra|OJJ74852.1 acar|OOF95851.1 acla|EAW08535.1 afis|EAW23573.1 afla|EED50962.1 afum|EAL87892.2 agla|OJJ86833.1 akaw|GAA83570.1 aluc|OJZ92428.1 anid|CBF89610.1 anig|EHA26539.1 aory|EIT77015.1 asyd|OJJ62305.1 ater|EAU35343.1 atub|OJI86858.1 aver|OJI96413.1 awen|OJJ39901.1 azon|OJJ42316.1

Group3540: abra|OJJ74854.1 acar|OOF95854.1 acla|EAW08531.1 afis|EAW23570.1 afla|EED57463.1 afum|EAL87897.2 agla|OJJ86831.1 akaw|GAA83568.1 aluc|OJZ92430.1 anid|CBF89617.1 anig|EHA26536.1 aory|EIT80101.1 asyd|OJJ62302.1 ater|EAU35346.1 atub|OJI86860.1 aver|OJI96411.1 awen|OJJ39899.1 azon|OJJ45876.1

Group3541: abra|OJJ74856.1 acar|OOF95857.1 acla|EAW08528.1 afis|EAW23568.1 afla|EED57459.1 afum|EAL87899.1 agla|OJJ86829.1 akaw|GAA83566.1 aluc|OJZ92432.1 anid|CBF89621.1 anig|EHA26534.1 aory|EIT80063.1 asyd|OJJ62300.1 ater|EAU35349.1 atub|OJI86862.1 aver|OJI96408.1 awen|OJJ39896.1 azon|OJJ45874.1

Group3542: abra|OJJ74859.1 acar|OOF95859.1 acla|EAW08521.1 afis|EAW23559.1 afla|EED57454.1 afum|EAL87907.1 agla|OJJ86825.1 akaw|GAA83561.1 aluc|OJZ92437.1 anid|CBF89634.1 anig|EHA26531.1 aory|EIT80133.1 asyd|OJJ62295.1 ater|EAU35342.1 atub|OJI86866.1 aver|OJI96402.1 awen|OJJ39893.1 azon|OJJ43217.1

Group3543: abra|OJJ74860.1 acar|OOF95860.1 acla|EAW08520.1 afis|EAW23558.1 afla|EED57453.1 afum|EAL87908.1 agla|OJJ86826.1 akaw|GAA83560.1 aluc|OJZ92438.1 anid|CBF89636.1 anig|EHA26530.1 aory|EIT80071.1 asyd|OJJ62294.1 ater|EAU35341.1 atub|OJI86867.1 aver|OJI96401.1 awen|OJJ39891.1 azon|OJJ43218.1

Group3544: abra|OJJ74862.1 acar|OOF95862.1 acla|EAW08519.1 afis|EAW23557.1 afla|EED57452.1 afum|EAL87909.1 agla|OJJ86819.1 akaw|GAA83558.1 aluc|OJZ92440.1 anid|CBF89638.1 anig|EHA26527.1 aory|EIT80146.1 asyd|OJJ62292.1 ater|EAU35340.1 atub|OJI86869.1 aver|OJI96400.1 awen|OJJ39888.1 azon|OJJ43219.1

Group3545: abra|OJJ74867.1 acar|OOF95868.1 acla|EAW08513.1 afis|EAW23551.1 afla|EED57446.1 afum|EAL87915.1 agla|OJJ86807.1 akaw|GAA83553.1 aluc|OJZ92445.1 anid|CBF89646.1 anig|EHA26523.1 aory|EIT80111.1 asyd|OJJ62286.1 ater|EAU35335.1 atub|OJI86873.1 aver|OJI96394.1 awen|OJJ39882.1 azon|OJJ43223.1

Group3546: abra|OJJ74869.1 acar|OOF95870.1 acla|EAW08511.1 afis|EAW23549.1 afla|EED57444.1 afum|EAL87917.2 agla|OJJ86805.1 akaw|GAA83551.1 aluc|OJZ92447.1 anid|CBF89650.1 anig|EHA26521.1 aory|EIT80076.1 asyd|OJJ62284.1 ater|EAU35334.1 atub|OJI86875.1 aver|OJI96392.1 awen|OJJ39880.1 azon|OJJ43224.1

Group3547: abra|OJJ74872.1 acar|OOF95873.1 acla|EAW08508.1 afis|EAW23546.1 afla|EED57441.1 afum|EAL87920.1 agla|OJJ86800.1 akaw|GAA83548.1 aluc|OJZ92450.1 anid|CBF89654.1 anig|EHA26518.1 aory|EIT80061.1 asyd|OJJ62282.1 ater|EAU35331.1 atub|OJI86878.1 aver|OJI96390.1 awen|OJJ39877.1 azon|OJJ43226.1

Group3548: abra|OJJ74874.1 acar|OOF95875.1 acla|EAW08498.1 afis|EAW23543.1 afla|EED57440.1 afum|EAL87923.2 agla|OJJ86797.1 akaw|GAA83546.1 aluc|OJZ92452.1 anid|CBF89657.1 anig|EHA26516.1 aory|EIT80096.1 asyd|OJJ62279.1 ater|EAU35329.1 atub|OJI86880.1 aver|OJI96388.1 awen|OJJ39875.1 azon|OJJ43227.1

Group3549: abra|OJJ74877.1 acar|OOF95880.1 acla|EAW08493.1 afis|EAW23539.1 afla|EED57437.1 afum|EAL87927.1 agla|OJJ86794.1 akaw|GAA83543.1 aluc|OJZ92454.1 anid|CBF89665.1 anig|EHA26513.1

aory|EIT80117.1 asyd|OJJ62275.1 ater|EAU35325.1 atub|OJI86883.1 aver|OJI96384.1 awen|OJJ39870.1 azon|OJJ43231.1

Group3550: abra|OJJ74879.1 acar|OOF95882.1 acla|EAW08491.1 afis|EAW23537.1 afla|EED57435.1 afum|EAL87929.1 agla|OJJ86792.1 akaw|GAA83541.1 aluc|OJZ92456.1 anid|CBF89668.1 anig|EHA26511.1 aory|EIT80166.1 asyd|OJJ62273.1 ater|EAU35323.1 atub|OJI86885.1 aver|OJI96382.1 awen|OJJ39867.1 azon|OJJ43233.1

Group3551: abra|OJJ74881.1 acar|OOF95884.1 acla|EAW08489.1 afis|EAW23535.1 afla|EED57432.1 afum|EAL87931.1 agla|OJJ86790.1 akaw|GAA83539.1 aluc|OJZ92458.1 anid|CBF89672.1 anig|EHA26509.1 aory|EIT80162.1 asyd|OJJ62271.1 ater|EAU35321.1 atub|OJI86887.1 aver|OJI96380.1 awen|OJJ39865.1 azon|OJJ43235.1

Group3552: abra|OJJ74882.1 acar|OOF95886.1 acla|EAW08487.1 afis|EAW23533.1 afla|EED57430.1 afum|EAL87933.1 agla|OJJ86782.1 akaw|GAA83537.1 aluc|OJZ92459.1 anid|CBF89678.1 anig|EHA26508.1 aory|EIT80077.1 asyd|OJJ62269.1 ater|EAU35319.1 atub|OJI86888.1 aver|OJI96378.1 awen|OJJ39862.1 azon|OJJ43236.1

Group3553: abra|OJJ74884.1 akaw|GAA83535.1 aluc|OJZ92461.1 anig|EHA26506.1 atub|OJI86890.1 acar|OOF95889.1 acla|EAW08483.1 afis|EAW23529.1 afla|EED57428.1 afum|EAL87937.1 agla|OJJ86778.1 anid|CBF89683.1 asyd|OJJ62266.1 ater|EAU35317.1 aver|OJI96375.1 awen|OJJ39858.1 azon|OJJ43239.1 aory|EIT80152.1

Group3554: abra|OJJ74888.1 acar|OOF95893.1 acla|EAW08479.1 afis|EAW23525.1 afla|EED57424.1 afum|EAL87940.1 agla|OJJ86774.1 akaw|GAA83531.1 aluc|OJZ92465.1 anid|CBF89691.1 anig|EHA26502.1 aory|EIT80080.1 asyd|OJJ62262.1 ater|EAU35313.1 atub|OJI86894.1 aver|OJI96371.1 awen|OJJ39854.1 azon|OJJ43245.1

Group3555: abra|OJJ74889.1 acar|OOF95892.1 acla|EAW08478.1 afis|EAW23524.1 afla|EED50993.1 afum|EAL87941.1 agla|OJJ86773.1 akaw|GAA83530.1 aluc|OJZ92466.1 anid|CBF76066.1 anig|EHA26501.1 aory|EIT76998.1 asyd|OJJ60396.1 ater|EAU31002.1 atub|OJI86895.1 aver|OJI99725.1 awen|OJJ39853.1 azon|OJJ50688.1

Group3556: abra|OJJ74897.1 acar|OOF95897.1 acla|EAW08475.1 afis|EAW23521.1 afum|EAL87943.1 agla|OJJ86768.1 akaw|GAA83524.1 aluc|OJZ92471.1 anig|EHA26497.1 ater|EAU35309.1 atub|OJI86901.1 awen|OJJ39846.1 afla|EED57421.1 aory|EIT80149.1 aver|OJI96367.1 azon|OJJ43248.1 anid|CBF89696.1 asyd|OJJ62258.1

Group3557: abra|OJJ74901.1 acar|OOF95900.1 acla|EAW08472.1 afis|EAW23518.1 afla|EED57418.1 afum|EAL87946.1 agla|OJJ86765.1 akaw|GAA83521.1 aluc|OJZ92474.1 anid|CBF89702.1 anig|EHA26495.1 aory|EIT80062.1 asyd|OJJ62254.1 atub|OJI86905.1 aver|OJI96364.1 awen|OJJ39843.1 azon|OJJ43250.1 ater|EAU35306.1

Group3558: abra|OJJ74905.1 acar|OOF90301.1 acla|EAW08467.1 afis|EAW23514.1 afla|EED57413.1 afum|EAL87949.1 agla|OJJ86762.1 akaw|GAA83517.1 aluc|OJZ92478.1 anig|EHA26492.1 aory|EIT80167.1 asyd|OJJ59973.1 ater|EAU35303.1 atub|OJI86909.1 aver|OJJ02318.1 awen|OJJ39839.1 azon|OJJ43253.1 anid|CBF75148.1

Group3559: abra|OJJ74907.1 acar|OOF90299.1 acla|EAW08465.1 afis|EAW23510.1 afla|EED57410.1 afum|EAL87951.1 agla|OJJ86760.1 akaw|GAA83515.1 aluc|OJZ92480.1 anid|CBF89718.1 anig|EHA26490.1 aory|EIT80124.1 asyd|OJJ62245.1 ater|EAU35298.1 atub|OJI86911.1 aver|OJI96355.1 awen|OJJ39836.1 azon|OJJ43256.1

Group3560: abra|OJJ74909.1 acar|OOF90294.1 acla|EAW08463.1 afis|EAW23508.1 afla|EED57408.1 afum|EAL87953.1 agla|OJJ86758.1 akaw|GAA83513.1 aluc|OJZ92482.1 anid|CBF89714.1 anig|EHA26489.1 aory|EIT80104.1 asyd|OJJ62247.1 ater|EAU35296.1 atub|OJI86913.1 aver|OJI96357.1 awen|OJJ39834.1 azon|OJJ43258.1

Group3561: abra|OJJ74910.1 acar|OOF90297.1 acla|EAW08462.1 afis|EAW23507.1 afla|EED57407.1 afum|EAL87954.1 agla|OJJ86757.1 akaw|GAA83512.1 aluc|OJZ92483.1 anid|CBF89712.1 anig|EHA26488.1 aory|EIT80089.1 asyd|OJJ62248.1 ater|EAU35295.1 atub|OJI86914.1 aver|OJI96358.1 awen|OJJ39833.1 azon|OJJ43259.1

Group3562: abra|OJJ74916.1 acar|OOF95908.1 acla|EAW08454.1 afis|EAW23501.1 afla|EED57399.1 afum|EAL87961.1 agla|OJJ86746.1 akaw|GAA83506.1 aluc|OJZ92488.1 anid|CBF89733.1 anig|EHA26484.1 aory|EIT80085.1 asyd|OJJ62233.1 ater|EAU35288.1 atub|OJI86921.1 aver|OJI96342.1 awen|OJJ39824.1 azon|OJJ47948.1

Group3563: abra|OJJ74919.1 acar|OOF95910.1 afis|EAW23497.1 afla|EED57397.1 afum|EAL87966.1 agla|OJJ86744.1 akaw|GAA83504.1 aluc|OJZ92490.1 anid|CBF89737.1 anig|EHA26482.1 aory|EIT80105.1 asyd|OJJ62231.1 ater|EAU35286.1 atub|OJI86923.1 aver|OJI96340.1 awen|OJJ39822.1 azon|OJJ47946.1 acla|EAW08452.1

Group3564: abra|OJJ74922.1 acar|OOF95912.1 acla|EAW08449.1 afis|EAW23494.1 afla|EED57394.1 afum|EAL87969.1 agla|OJJ86741.1 akaw|GAA83501.1 aluc|OJZ92493.1 anid|CBF89742.1 anig|EHA26479.1 aory|EIT80159.1 asyd|OJJ62228.1 ater|EAU35283.1 atub|OJI86926.1 aver|OJI96337.1 awen|OJJ39819.1 azon|OJJ47943.1

Group3565: abra|OJJ74923.1 acar|OOF95914.1 acla|EAW08448.1 afis|EAW23493.1 afla|EED57393.1 afum|EAL87970.1 agla|OJJ86740.1 akaw|GAA83500.1 aluc|OJZ92494.1 anid|CBF89744.1 anig|EHA26478.1 aory|EIT80132.1 asyd|OJJ62227.1 ater|EAU35282.1 atub|OJI86927.1 aver|OJI96336.1 awen|OJJ39818.1 azon|OJJ47941.1

Group3566: abra|OJJ74924.1 acar|OOF95915.1 acla|EAW08447.1 afis|EAW23492.1 afla|EED57391.1 afum|EAL87971.1 agla|OJJ86737.1 akaw|GAA83499.1 aluc|OJZ92495.1 anid|CBF89746.1 anig|EHA26477.1 aory|EIT80165.1 asyd|OJJ62225.1 ater|EAU35279.1 atub|OJI86928.1 aver|OJI96334.1 awen|OJJ39816.1 azon|OJJ47939.1

Group3567: abra|OJJ74925.1 acar|OOF95916.1 acla|EAW08446.1 afis|EAW23491.1 afla|EED57390.1 afum|EAL87972.1 agla|OJJ86736.1 akaw|GAA83498.1 aluc|OJZ92496.1 anid|CBF89747.1 anig|EHA26476.1 aory|EIT80058.1 asyd|OJJ62224.1 ater|EAU35278.1 atub|OJI86929.1 aver|OJI96333.1 awen|OJJ39815.1 azon|OJJ47938.1

Group3568: abra|OJJ74928.1 acar|OOF95917.1 acla|EAW08442.1 afis|EAW23487.1 afla|EED57386.1 afum|EAL87976.1 agla|OJJ86732.1 akaw|GAA83494.1 aluc|OJZ92499.1 anid|CBF89755.1 anig|EHA26473.1 aory|EIT80123.1 asyd|OJJ62220.1 ater|EAU35274.1 atub|OJI86932.1 aver|OJI96330.1 awen|OJJ39810.1 azon|OJJ47934.1

Group3569: abra|OJJ74930.1 acar|OOF95920.1 acla|EAW08439.1 afis|EAW23484.1 afla|EED57379.1 afum|EAL87979.1 agla|OJJ86720.1 akaw|GAA83491.1 aluc|OJZ92502.1 anid|CBF89760.1 anig|EHA26471.1 aory|EIT80056.1 asyd|OJJ62217.1 ater|EAU35271.1 atub|OJI86934.1 aver|OJI96328.1 awen|OJJ39808.1 azon|OJJ47929.1

Group3570: abra|OJJ74931.1 acar|OOF95921.1 acla|EAW08438.1 afis|EAW23483.1 afla|EED57378.1 afum|EAL87980.1 agla|OJJ86719.1 akaw|GAA83490.1 aluc|OJZ92503.1 anid|CBF89762.1 anig|EHA26470.1 aory|EIT80160.1 asyd|OJJ62216.1 ater|EAU36899.1 atub|OJI86935.1 aver|OJI96327.1 awen|OJJ39807.1 azon|OJJ47928.1

Group3571: abra|OJJ74932.1 acar|OOF95922.1 acla|EAW08437.1 afis|EAW23482.1 afla|EED57377.1 afum|EAL87982.1 agla|OJJ86718.1 akaw|GAA83488.1 aluc|OJZ92505.1 anid|CBF89764.1 anig|EHA26468.1 aory|EIT80168.1 asyd|OJJ62215.1 ater|EAU36900.1 atub|OJI86937.1 aver|OJI96326.1 awen|OJJ39806.1 azon|OJJ47927.1

Group3572: abra|OJJ74935.1 acar|OOF97377.1 acla|EAW08436.1 afis|EAW23481.1 afla|EED57376.1 afum|EAL87983.1 agla|OJJ86717.1 akaw|GAA83485.1 aluc|OJZ92508.1 anid|CBF89766.1 anig|EHA26465.1 aory|EIT80075.1 asyd|OJJ62214.1 ater|EAU36901.1 atub|OJI86941.1 aver|OJI96325.1 awen|OJJ39805.1 azon|OJJ47926.1

Group3573: abra|OJJ74938.1 acar|OOF97376.1 acla|EAW08433.1 afis|EAW23476.1 afla|EED57372.1 afum|EAL87989.2 agla|OJJ86715.1 akaw|GAA83482.1 aluc|OJZ92513.1 anid|CBF89771.1 anig|EHA26462.1 aory|EIT80114.1 asyd|OJJ62212.1 ater|EAU36904.1 atub|OJI86944.1 aver|OJI96323.1 awen|OJJ39802.1 azon|OJJ47923.1

Group3574: abra|OJJ74939.1 acar|OOF97375.1 acla|EAW08431.1 afis|EAW23474.1 afla|EED57370.1 afum|EAL87991.2 agla|OJJ86713.1 akaw|GAA83481.1 aluc|OJZ92514.1 anid|CBF89775.1 anig|EHA26461.1 aory|EIT80156.1 asyd|OJJ62209.1 ater|EAU36906.1 atub|OJI86945.1 aver|OJI96321.1 awen|OJJ39800.1 azon|OJJ47922.1

Group3575: abra|OJJ74941.1 acar|OOF97374.1 acla|EAW08429.1 afis|EAW23473.1 afla|EED57369.1 afum|EAL87993.1 agla|OJJ86712.1 akaw|GAA83480.1 aluc|OJZ92515.1 anid|CBF89777.1 anig|EHA26460.1 aory|EIT80066.1 asyd|OJJ62208.1 ater|EAU36907.1 atub|OJI86948.1 aver|OJI96320.1 awen|OJJ39799.1 azon|OJJ47920.1

Group3576: abra|OJJ74943.1 acar|OOF97372.1 acla|EAW08427.1 afis|EAW23471.1 afla|EED57367.1 afum|EAL87995.1 agla|OJJ86710.1 akaw|GAA83478.1 aluc|OJZ92517.1 anid|CBF89782.1 anig|EHA26458.1 aory|EIT80170.1 asyd|OJJ62205.1 ater|EAU36909.1 atub|OJI86950.1 aver|OJI96317.1 awen|OJJ39796.1 azon|OJJ47918.1

Group3577: abra|OJJ74945.1 acar|OOF97371.1 acla|EAW08426.1 afis|EAW23442.1 afla|EED57366.1 afum|EAL88021.1 agla|OJJ86709.1 akaw|GAA83477.1 aluc|OJZ92518.1 anid|CBF89784.1 anig|EHA26457.1 aory|EIT80053.1 asyd|OJJ62204.1 ater|EAU36910.1 atub|OJI86952.1 aver|OJI96316.1 awen|OJJ39790.1 azon|OJJ47917.1

Group3578: abra|OJJ74946.1 acar|OOF97370.1 acla|EAW08425.1 afis|EAW23444.1 afla|EED57365.1 afum|EAL88020.1 agla|OJJ86707.1 akaw|GAA83476.1 aluc|OJZ92519.1 anid|CBF89786.1 anig|EHA26456.1

aory|EIT80115.1 asyd|OJJ62203.1 ater|EAU36911.1 atub|OJI86953.1 aver|OJI96315.1 awen|OJJ39786.1 azon|OJJ47916.1

Group3579: abra|OJJ74947.1 acar|OOF97369.1 acla|EAW08424.1 afis|EAW23445.1 afla|EED57364.1 afum|EAL88019.1 agla|OJJ86706.1 akaw|GAA83475.1 aluc|OJZ92520.1 anid|CBF89788.1 anig|EHA26455.1 aory|EIT80057.1 asyd|OJJ62202.1 ater|EAU36912.1 atub|OJI86954.1 aver|OJI96314.1 awen|OJJ39785.1 azon|OJJ47915.1

Group3580: abra|OJJ74949.1 acar|OOF97367.1 acla|EAW08422.1 afis|EAW23447.1 afla|EED57362.1 afum|EAL88018.1 agla|OJJ86702.1 akaw|GAA83473.1 aluc|OJZ92522.1 anid|CBF89791.1 anig|EHA26453.1 aory|EIT80100.1 asyd|OJJ62200.1 ater|EAU36913.1 atub|OJI86956.1 aver|OJI96312.1 awen|OJJ39782.1 azon|OJJ47913.1

Group3581: abra|OJJ74952.1 acar|OOF97362.1 acla|EAW08421.1 afis|EAW23448.1 afla|EED57359.1 afum|EAL88017.1 agla|OJJ86701.1 akaw|GAA83471.1 aluc|OJZ92524.1 anid|CBF89793.1 anig|EHA26450.1 aory|EIT80142.1 asyd|OJJ62199.1 ater|EAU36914.1 atub|OJI86959.1 aver|OJI96311.1 awen|OJJ39781.1 azon|OJJ47912.1

Group3582: abra|OJJ74953.1 acar|OOF97360.1 acla|EAW08420.1 afis|EAW23449.1 afla|EED57358.1 afum|EBA27332.1 agla|OJJ86700.1 akaw|GAA83470.1 aluc|OJZ92525.1 anid|CBF89795.1 anig|EHA26449.1 aory|EIT80082.1 asyd|OJJ62198.1 ater|EAU36916.1 atub|OJI86961.1 awen|OJJ39780.1 azon|OJJ47911.1 aver|OJI96310.1

Group3583: abra|OJJ74954.1 acar|OOF97359.1 acla|EAW08418.1 afis|EAW23451.1 afla|EED57352.1 afum|EAL88015.1 agla|OJJ86699.1 akaw|GAA83469.1 aluc|OJZ92526.1 anid|CBF89799.1 anig|EHA26448.1 aory|EIT80134.1 asyd|OJJ62196.1 ater|EAU36917.1 atub|OJI86962.1 aver|OJI96308.1 awen|OJJ39778.1 azon|OJJ47908.1

Group3584: abra|OJJ74956.1 acar|OOF97357.1 acla|EAW08416.1 afis|EAW23453.1 afla|EED57350.1 afum|EAL88013.1 agla|OJJ86697.1 akaw|GAA83467.1 aluc|OJZ92528.1 anid|CBF89802.1 anig|EHA26447.1 aory|EIT80106.1 asyd|OJJ62194.1 ater|EAU36919.1 atub|OJI86964.1 aver|OJI96306.1 awen|OJJ39776.1 azon|OJJ47906.1

Group3585: abra|OJJ74958.1 acar|OOF97355.1 acla|EAW08413.1 afis|EAW23456.1 afla|EED57345.1 afum|EBA27330.1 akaw|GAA83464.1 aluc|OJZ92530.1 anid|CBF89808.1 anig|EHA26444.1 aory|EIT80113.1 asyd|OJJ62191.1 ater|EAU36922.1 atub|OJI86967.1 aver|OJI96301.1 awen|OJJ39773.1 azon|OJJ47902.1 agla|OJJ86693.1

Group3586: abra|OJJ74959.1 acar|OOF97354.1 acla|EAW08412.1 afis|EAW23457.1 afla|EED57344.1 afum|EBA27329.1 agla|OJJ86692.1 akaw|GAA83463.1 aluc|OJZ92531.1 anid|CBF89810.1 anig|EHA26443.1 aory|EIT80054.1 asyd|OJJ62190.1 ater|EAU36923.1 atub|OJI86968.1 aver|OJI96300.1 awen|OJJ39772.1 azon|OJJ47903.1

Group3587: abra|OJJ74965.1 acar|OOF97349.1 acla|EAW08409.1 afis|EAW23460.1 afla|EED57340.1 afum|EAL88006.1 agla|OJJ86689.1 akaw|GAA83461.1 aluc|OJZ92535.1 anid|CBF89820.1 anig|EHA26440.1 aory|EIT74102.1 asyd|OJJ62187.1 ater|EAU37473.1 atub|OJI86971.1 aver|OJI96296.1 awen|OJJ39770.1 azon|OJJ47900.1

Group3588: abra|OJJ74966.1 acar|OOF97348.1 acla|EAW08408.1 afis|EAW23461.1 afla|EED57339.1 afum|EAL88005.1 agla|OJJ86688.1 akaw|GAA83460.1 aluc|OJZ92536.1 anid|CBF89850.1 anig|EHA26439.1 aory|EIT74042.1 asyd|OJJ62186.1 ater|EAU37474.1 atub|OJI86972.1 aver|OJI96295.1 awen|OJJ39769.1 azon|OJJ47899.1

Group3589: abra|OJJ74967.1 acar|OOF97346.1 acla|EAW08407.1 afis|EAW23462.1 afla|EED57338.1 afum|EAL88004.1 agla|OJJ86687.1 akaw|GAA83459.1 aluc|OJZ92537.1 anid|CBF89848.1 anig|EHA26438.1 aory|EIT74136.1 asyd|OJJ62185.1 ater|EAU37475.1 atub|OJI86973.1 aver|OJI96294.1 awen|OJJ39768.1 azon|OJJ47898.1

Group3590: abra|OJJ74969.1 acar|OOF97345.1 acla|EAW08405.1 afis|EAW23464.1 afla|EED57335.1 afum|EAL88002.1 agla|OJJ86685.1 akaw|GAA83457.1 aluc|OJZ92539.1 anid|CBF89843.1 anig|EHA26437.1 aory|EIT74036.1 asyd|OJJ62183.1 ater|EAU37480.1 atub|OJI86975.1 aver|OJI96292.1 awen|OJJ39766.1 azon|OJJ47896.1

Group3591: abra|OJJ74975.1 acar|OOF97344.1 acla|EAW08404.1 afis|EAW23465.1 afla|EED57334.1 afum|EAL88001.1 agla|OJJ86684.1 akaw|GAA83456.1 aluc|OJZ92540.1 anid|CBF89841.1 anig|EHA26436.1 aory|EIT73981.1 asyd|OJJ62182.1 ater|EAU37481.1 atub|OJI86976.1 aver|OJI96291.1 awen|OJJ39765.1 azon|OJJ47895.1

Group3592: abra|OJJ74976.1 acar|OOF97343.1 acla|EAW08403.1 afis|EAW23466.1 afla|EED57333.1 afum|EAL88000.1 agla|OJJ86683.1 akaw|GAA83455.1 aluc|OJZ92541.1 anid|CBF89839.1 anig|EHA26435.1 aory|EIT74149.1 asyd|OJJ62181.1 ater|EAU37482.1 atub|OJI86977.1 aver|OJI96290.1 awen|OJJ39764.1 azon|OJJ47894.1

Group3593: abra|OJJ74980.1 acar|OOF97340.1 acla|EAW08400.1 afis|EAW23469.1 afla|EED57331.1 afum|EAL87997.1 agla|OJJ86680.1 akaw|GAA83452.1 aluc|OJZ92544.1 anid|CBF89833.1 anig|EHA26432.1 aory|EIT74122.1 asyd|OJJ62178.1 ater|EAU37485.1 atub|OJI86980.1 aver|OJI96287.1 awen|OJJ39760.1 azon|OJJ47889.1

Group3594: abra|OJJ74981.1 acar|OOF97339.1 acla|EAW08399.1 afis|EAW23470.1 afla|EED57330.1 afum|EAL87996.1 agla|OJJ86679.1 akaw|GAA83451.1 aluc|OJZ92545.1 anid|CBF89831.1 anig|EHA26431.1 aory|EIT74162.1 asyd|OJJ62177.1 ater|EAU37486.1 atub|OJI86981.1 aver|OJI96286.1 awen|OJJ39759.1 azon|OJJ47888.1

Group3595: abra|OJJ74983.1 acar|OOF97338.1 acla|EAW08397.1 afis|EAW23440.1 afla|EED57329.1 afum|EAL88022.1 agla|OJJ86677.1 akaw|GAA83449.1 aluc|OJZ92547.1 anid|CBF89828.1 anig|EHA26429.1 asyd|OJJ62169.1 ater|EAU37487.1 atub|OJI86983.1 aver|OJI96281.1 awen|OJJ39757.1 azon|OJJ47885.1 anid|CBF89830.1

Group3596: abra|OJJ74988.1 acar|OOF97333.1 acla|EAW08394.1 afis|EAW23437.1 afla|EED57327.1 afum|EAL88025.1 agla|OJJ86674.1 akaw|GAA83446.1 aluc|OJZ92550.1 anid|CBF89852.1 anig|EHA26427.1 aory|EIT74016.1 asyd|OJJ62176.1 ater|EAU37489.1 atub|OJI86987.1 aver|OJI96285.1 awen|OJJ39754.1 azon|OJJ47881.1

Group3597: abra|OJJ74991.1 acar|OOF97329.1 acla|EAW08392.1 afis|EAW23423.1 afla|EED57322.1 afum|EAL88033.1 agla|OJJ86673.1 akaw|GAA83444.1 aluc|OJZ92555.1 anid|CBF89824.1 anig|EHA26424.1 aory|EIT74106.1 asyd|OJJ62171.1 ater|EAU37494.1 atub|OJI86991.1 aver|OJI96283.1 awen|OJJ39753.1 azon|OJJ47880.1

Group3598: abra|OJJ74993.1 acar|OOF97326.1 acla|EAW08390.1 afis|EAW23419.1 afla|EED57321.1 afum|EAL88050.1 agla|OJJ86644.1 akaw|GAA83442.1 aluc|OJZ92559.1 anid|CBF89854.1 anig|EHA26421.1 aory|EIT74073.1 asyd|OJJ62161.1 ater|EAU37503.1 atub|OJI86995.1 aver|OJI96271.1 awen|OJJ39752.1 azon|OJJ47878.1

Group3599: abra|OJJ74996.1 acar|OOF97324.1 acla|EAW08388.1 afis|EAW23417.1 afla|EED57318.1 afum|EAL88052.2 agla|OJJ86634.1 akaw|GAA88620.1 aluc|OJZ92562.1 anid|CBF89861.1 anig|EHA26419.1 aory|EIT73973.1 asyd|OJJ62158.1 ater|EAU37505.1 atub|OJI86998.1 aver|OJI96268.1 awen|OJJ39750.1 azon|OJJ47876.1

Group3600: abra|OJJ74997.1 acar|OOF97323.1 acla|EAW08387.1 afis|EAW23416.1 afla|EED57317.1 afum|EAL88053.2 agla|OJJ86633.1 akaw|GAA88621.1 aluc|OJZ92563.1 anid|CBF89862.1 anig|EHA26418.1 aory|EIT74065.1 asyd|OJJ62157.1 ater|EAU37506.1 atub|OJI86999.1 aver|OJI96267.1 awen|OJJ39749.1 azon|OJJ47875.1

Group3601: abra|OJJ74999.1 acar|OOF97322.1 acla|EAW08386.1 afis|EAW23415.1 afla|EED57316.1 afum|EAL88054.1 agla|OJJ86632.1 akaw|GAA88622.1 aluc|OJZ92564.1 anid|CBF89864.1 anig|EHA26417.1 aory|EIT73976.1 asyd|OJJ62156.1 ater|EAU37507.1 atub|OJI87000.1 aver|OJI96266.1 awen|OJJ39748.1 azon|OJJ47874.1

Group3602: abra|OJJ75000.1 acar|OOF97321.1 acla|EAW08385.1 afis|EAW23414.1 afla|EED57315.1 afum|EAL88055.1 agla|OJJ86631.1 akaw|GAA88623.1 aluc|OJZ92565.1 anid|CBF89865.1 anig|EHA26416.1 aory|EIT74071.1 asyd|OJJ62155.1 ater|EAU37508.1 atub|OJI87001.1 aver|OJI96265.1 awen|OJJ39747.1 azon|OJJ47873.1

Group3603: abra|OJJ75002.1 acar|OOF90037.1 acla|EAW08383.1 afis|EAW23412.1 afla|EED57313.1 afum|EAL88057.1 agla|OJJ86629.1 akaw|GAA88625.1 aluc|OJZ92567.1 anid|CBF89869.1 anig|EHA26414.1 aory|EIT74133.1 asyd|OJJ62153.1 ater|EAU37510.1 atub|OJI87003.1 aver|OJI96263.1 awen|OJJ39745.1 azon|OJJ47871.1

Group3604: abra|OJJ75003.1 acar|OOF90036.1 acla|EAW08382.1 afis|EAW23411.1 afla|EED57312.1 afum|EAL88058.1 agla|OJJ86628.1 akaw|GAA88626.1 aluc|OJZ92568.1 anid|CBF89871.1 anig|EHA26413.1 aory|EIT74029.1 asyd|OJJ62152.1 ater|EAU37511.1 atub|OJI87004.1 aver|OJI96262.1 awen|OJJ39744.1 azon|OJJ47870.1

Group3605: abra|OJJ75004.1 acar|OOF90035.1 acla|EAW08381.1 afis|EAW23410.1 afla|EED57310.1 afum|EAL88060.1 agla|OJJ86627.1 akaw|GAA88627.1 aluc|OJZ92569.1 anig|EHA26412.1 aory|EIT74104.1 asyd|OJJ62151.1 ater|EAU37516.1 atub|OJI87005.1 aver|OJI96261.1 awen|OJJ39743.1 azon|OJJ47869.1 agla|OJJ86626.1

Group3606: abra|OJJ75005.1 acar|OOF97320.1 acla|EAW08380.1 afis|EAW23409.1 afla|EED57309.1 afum|EAL88061.2 agla|OJJ86625.1 akaw|GAA88628.1 aluc|OJZ92570.1 anid|CBF89873.1 anig|EHA26411.1 aory|EIT74087.1 asyd|OJJ62150.1 ater|EAU37515.1 atub|OJI87006.1 aver|OJI96260.1 awen|OJJ39742.1 azon|OJJ47868.1

Group3607: abra|OJJ75006.1 acar|OOF97319.1 acla|EAW08379.1 afis|EAW23408.1 afla|EED57308.1 afum|EAL88062.1 agla|OJJ86624.1 akaw|GAA88629.1 aluc|OJZ92571.1 anid|CBF89874.1 anig|EHA26410.1

aory|EIT74117.1 asyd|OJJ62149.1 ater|EAU37512.1 atub|OJI87007.1 aver|OJI96259.1 awen|OJJ39735.1 azon|OJJ47867.1

Group3608: abra|OJJ75007.1 acar|OOF97318.1 acla|EAW08378.1 afis|EAW23407.1 afla|EED57307.1 afum|EAL88063.2 agla|OJJ86623.1 akaw|GAA88630.1 aluc|OJZ92572.1 anid|CBF89876.1 anig|EHA26409.1 aory|EIT74022.1 asyd|OJJ62148.1 ater|EAU37513.1 atub|OJI87008.1 aver|OJI96258.1 awen|OJJ39734.1 azon|OJJ47866.1

Group3609: abra|OJJ75009.1 acar|OOF97317.1 acla|EAW08204.1 afis|EAW23238.1 afla|EED55718.1 afum|EAL88244.1 agla|OJJ86622.1 akaw|GAA88632.1 aluc|OJZ92574.1 anid|CBF89882.1 anig|EHA26407.1 aory|EIT83207.1 asyd|OJJ62146.1 ater|EAU37519.1 atub|OJI87010.1 aver|OJI96256.1 awen|OJJ39732.1 azon|OJJ47864.1

Group3610: abra|OJJ75010.1 acar|OOF97316.1 acla|EAW08205.1 afis|EAW23239.1 afla|EED55719.1 afum|EAL88243.1 agla|OJJ86620.1 akaw|GAA88633.1 aluc|OJZ92575.1 anid|CBF89884.1 anig|EHA26406.1 aory|EIT83209.1 asyd|OJJ62145.1 ater|EAU37520.1 atub|OJI87011.1 aver|OJI96255.1 awen|OJJ39731.1 azon|OJJ47863.1

Group3611: abra|OJJ75011.1 acar|OOF97315.1 acla|EAW08206.1 afis|EAW23240.1 afla|EED55720.1 afum|EAL88242.1 agla|OJJ86619.1 akaw|GAA88634.1 aluc|OJZ92576.1 anid|CBF89885.1 anig|EHA26405.1 aory|EIT83217.1 asyd|OJJ62144.1 ater|EAU37521.1 atub|OJI87012.1 aver|OJI96254.1 awen|OJJ39730.1 azon|OJJ47862.1

Group3612: abra|OJJ75012.1 acar|OOF97314.1 acla|EAW08207.1 afis|EAW23241.1 afla|EED55722.1 afum|EAL88241.1 agla|OJJ86618.1 akaw|GAA88635.1 aluc|OJZ92577.1 anid|CBF89887.1 anig|EHA26404.1 aory|EIT83231.1 asyd|OJJ62143.1 ater|EAU37522.1 atub|OJI87013.1 aver|OJI96253.1 awen|OJJ39729.1 azon|OJJ47861.1

Group3613: abra|OJJ75014.1 acar|OOF97312.1 acla|EAW08209.1 afis|EAW23243.1 afla|EED55724.1 afum|EAL88239.1 agla|OJJ86617.1 akaw|GAA88637.1 aluc|OJZ92579.1 anid|CBF89891.1 anig|EHA26402.1 aory|EIT83225.1 asyd|OJJ62140.1 ater|EAU37524.1 atub|OJI87015.1 aver|OJI96250.1 awen|OJJ39727.1 azon|OJJ47858.1

Group3614: abra|OJJ75022.1 acar|OOF89975.1 acla|EAW08214.1 afis|EAW23248.1 afla|EED55729.1 afum|EAL88233.1 agla|OJJ86609.1 akaw|GAA88642.1 aluc|OJZ92589.1 anid|CBF84108.1 anig|EHA26397.1 aory|EIT83221.1 asyd|OJJ57101.1 ater|EAU37530.1 atub|OJI87021.1 aver|OJJ03198.1 awen|OJJ39716.1 azon|OJJ47847.1

Group3615: abra|OJJ75025.1 acar|OOF97307.1 acla|EAW08217.1 afis|EAW23251.1 afla|EED55732.1 afum|EAL88230.1 agla|OJJ86603.1 akaw|GAA88645.1 aluc|OJZ92592.1 anid|CBF84103.1 anig|EHA26394.1 aory|EIT83210.1 asyd|OJJ57097.1 ater|EAU37549.1 atub|OJI87025.1 aver|OJJ03202.1 awen|OJJ39713.1 azon|OJJ47844.1

Group3616: abra|OJJ75026.1 acar|OOF97306.1 acla|EAW08218.1 afis|EAW23252.1 afla|EED55733.1 afum|EAL88229.1 agla|OJJ86602.1 akaw|GAA88646.1 aluc|OJZ92593.1 anid|CBF84101.1 anig|EHA26393.1 aory|EIT83216.1 asyd|OJJ57096.1 ater|EAU37548.1 atub|OJI87027.1 aver|OJJ03203.1 awen|OJJ39712.1 azon|OJJ47843.1

Group3617: abra|OJJ75027.1 acar|OOF97305.1 acla|EAW08219.1 afis|EAW23253.1 afla|EED55734.1 afum|EAL88228.1 agla|OJJ86601.1 akaw|GAA88647.1 aluc|OJZ92594.1 anid|CBF84100.1 anig|EHA26392.1 aory|EIT83226.1 asyd|OJJ57095.1 ater|EAU37547.1 atub|OJI87028.1 aver|OJJ03204.1 awen|OJJ39711.1 azon|OJJ47842.1

Group3618: abra|OJJ75030.1 acar|OOF97303.1 acla|EAW08221.1 afis|EAW23255.1 afla|EED55736.1 afum|EAL88226.1 agla|OJJ86599.1 akaw|GAA88649.1 aluc|OJZ92597.1 anid|CBF84096.1 anig|EHA26390.1 aory|EIT83204.1 asyd|OJJ57093.1 ater|EAU37544.1 atub|OJI87031.1 aver|OJJ03206.1 awen|OJJ39709.1 azon|OJJ47840.1

Group3619: abra|OJJ75031.1 acar|OOF97302.1 acla|EAW08222.1 afis|EAW23256.1 afla|EED55737.1 afum|EAL88225.1 agla|OJJ86598.1 akaw|GAA88650.1 aluc|OJZ92598.1 anid|CBF84092.1 anig|EHA26389.1 aory|EIT83229.1 asyd|OJJ57092.1 ater|EAU37543.1 atub|OJI87032.1 aver|OJJ03208.1 awen|OJJ39708.1 azon|OJJ47839.1

Group3620: abra|OJJ75032.1 acar|OOF97301.1 acla|EAW08223.1 afis|EAW23257.1 afla|EED55738.1 afum|EAL88224.1 agla|OJJ86597.1 akaw|GAA88651.1 aluc|OJZ92599.1 anid|CBF84094.1 anig|EHA26388.1 aory|EIT83202.1 asyd|OJJ57091.1 ater|EAU37542.1 atub|OJI87034.1 aver|OJJ03209.1 awen|OJJ39707.1 azon|OJJ47838.1

Group3621: abra|OJJ75033.1 acar|OOF97300.1 acla|EAW08224.1 afis|EAW23258.1 afla|EED55739.1 afum|EAL88223.1 agla|OJJ86596.1 akaw|GAA88652.1 aluc|OJZ92600.1 anid|CBF84116.1 anig|EHA26387.1 aory|EIT83200.1 asyd|OJJ57105.1 ater|EAU37541.1 atub|OJI87035.1 aver|OJJ03194.1 awen|OJJ39706.1 azon|OJJ47837.1

Group3622: abra|OJJ75034.1 acar|OOF97299.1 acla|EAW08225.1 afis|EAW23259.1 afla|EED55740.1 afum|EAL88222.1 agla|OJJ86595.1 akaw|GAA88653.1 aluc|OJZ92602.1 anid|CBF84117.1 anig|EHA26386.1 aory|EIT83218.1 asyd|OJJ57106.1 ater|EAU37540.1 atub|OJI87036.1 aver|OJJ03193.1 awen|OJJ39705.1 azon|OJJ47835.1

Group3623: abra|OJJ75035.1 acar|OOF97298.1 acla|EAW08226.1 afis|EAW23260.1 afla|EED55741.1 afum|EAL88221.1 agla|OJJ86594.1 akaw|GAA88654.1 aluc|OJZ92603.1 anid|CBF84119.1 anig|EHA26385.1 aory|EIT83223.1 asyd|OJJ57107.1 ater|EAU37539.1 atub|OJI87037.1 aver|OJJ03192.1 awen|OJJ39704.1 azon|OJJ47834.1

Group3624: abra|OJJ75036.1 acar|OOF97297.1 acla|EAW08231.1 afis|EAW23265.1 afla|EED55746.1 afum|EAL88215.1 agla|OJJ86587.1 akaw|GAA88655.1 aluc|OJZ92604.1 anid|CBF84128.1 anig|EHA26384.1 aory|EIT83198.1 asyd|OJJ57112.1 ater|EAU37536.1 atub|OJI87038.1 aver|OJJ03187.1 awen|OJJ39699.1 azon|OJJ47829.1

Group3625: abra|OJJ75038.1 acar|OOF97294.1 acla|EAW08229.1 afis|EAW23263.1 afla|EED55744.1 afum|EAL88217.1 agla|OJJ86589.1 akaw|GAA88657.1 aluc|OJZ92606.1 anid|CBF84124.1 anig|EHA26382.1 aory|EIT83214.1 asyd|OJJ57110.1 ater|EAU37534.1 atub|OJI87040.1 aver|OJJ03189.1 awen|OJJ39701.1 azon|OJJ47831.1

Group3626: abra|OJJ75039.1 acar|OOF97293.1 acla|EAW08227.1 afis|EAW23261.1 afla|EED55742.1 afum|EAL88219.1 agla|OJJ86591.1 akaw|GAA88658.1 aluc|OJZ92607.1 anid|CBF84123.1 anig|EHA26381.1 aory|EIT83212.1 asyd|OJJ57109.1 ater|EAU37538.1 atub|OJI87041.1 aver|OJJ03190.1 awen|OJJ39703.1 azon|OJJ47833.1

Group3627: abra|OJJ75040.1 acar|OOF97292.1 acla|EAW08228.1 afis|EAW23262.1 afla|EED55743.1 afum|EAL88218.1 agla|OJJ86590.1 akaw|GAA88659.1 aluc|OJZ92608.1 anid|CBF84121.1 anig|EHA26380.1 aory|EIT83233.1 asyd|OJJ57108.1 ater|EAU37537.1 atub|OJI87042.1 aver|OJJ03191.1 awen|OJJ39702.1 azon|OJJ47832.1

Group3628: abra|OJJ75046.1 acar|OOF89831.1 acla|EAW08235.1 afis|EAW23269.1 afla|EED55710.1 afum|EAL88210.2 agla|OJJ86583.1 akaw|GAA88663.1 aluc|OJZ92612.1 anid|CBF89550.1 anig|EHA26375.1 aory|EIT83205.1 asyd|OJJ62375.1 ater|EAU37553.1 atub|OJI87046.1 aver|OJI96509.1 awen|OJJ39692.1 azon|OJJ47825.1

Group3629: abra|OJJ75047.1 acar|OOF97288.1 acla|EAW08236.1 afis|EAW23270.1 afla|EED55709.1 afum|EAL88209.1 agla|OJJ86582.1 akaw|GAA88664.1 aluc|OJZ92613.1 anid|CBF89548.1 anig|EHA26374.1 aory|EIT83232.1 asyd|OJJ62376.1 ater|EAU37554.1 atub|OJI87047.1 aver|OJI96510.1 awen|OJJ39691.1 azon|OJJ47824.1

Group3630: abra|OJJ75048.1 acar|OOF97286.1 acla|EAW08237.1 afis|EAW23271.1 afla|EED55706.1 afum|EAL88208.1 agla|OJJ86578.1 akaw|GAA88665.1 aluc|OJZ92615.1 anid|CBF89546.1 anig|EHA26373.1 aory|EIT83206.1 asyd|OJJ62377.1 ater|EAU37556.1 atub|OJI87049.1 aver|OJI96511.1 awen|OJJ39687.1 azon|OJJ47823.1

Group3631: abra|OJJ75052.1 acar|OOF97284.1 acla|EAW08238.1 afis|EAW23272.1 afla|EED55705.1 afum|EAL88207.1 agla|OJJ86577.1 akaw|GAA88666.1 aluc|OJZ92618.1 anid|CBF89544.1 anig|EHA26370.1 aory|EIT77089.1 asyd|OJJ62379.1 ater|EAU37557.1 atub|OJI87052.1 aver|OJI96513.1 awen|OJJ39685.1 azon|OJJ47822.1

Group3632: abra|OJJ75056.1 acar|OOF97281.1 acla|EAW08241.1 afis|EAW23275.1 afla|EED55703.1 afum|EAL88204.1 agla|OJJ86573.1 akaw|GAA88669.1 aluc|OJZ92622.1 anid|CBF89539.1 anig|EHA26367.1 aory|EIT77072.1 asyd|OJJ62382.1 ater|EAU37560.1 atub|OJI87055.1 aver|OJI96516.1 awen|OJJ39682.1 azon|OJJ47819.1

Group3633: abra|OJJ75057.1 acar|OOF97279.1 acla|EAW08242.1 afis|EAW23276.1 afla|EED55701.1 afum|EAL88203.1 agla|OJJ86571.1 akaw|GAA88670.1 aluc|OJZ92626.1 anid|CBF89533.1 anig|EHA26364.1 aory|EIT77071.1 asyd|OJJ62384.1 ater|EAU37561.1 atub|OJI87056.1 aver|OJI96518.1 awen|OJJ39680.1 azon|OJJ47818.1

Group3634: abra|OJJ75064.1 acar|OOF97270.1 acla|EAW08245.1 afis|EAW23279.1 afla|EED55695.1 afum|EAL88200.1 agla|OJJ86566.1 akaw|GAA88676.1 aluc|OJZ92630.1 anid|CBF89527.1 anig|EHA26360.1 aory|EIT77073.1 asyd|OJJ62388.1 ater|EAU37592.1 atub|OJI87061.1 aver|OJI96521.1 awen|OJJ39676.1 azon|OJJ47816.1

Group3635: abra|OJJ75066.1 acar|OOF97268.1 acla|EAW08247.1 afis|EAW23281.1 afla|EED55694.1 afum|EAL88198.2 agla|OJJ86565.1 akaw|GAA88678.1 aluc|OJZ92632.1 anid|CBF89525.1 anig|EHA26359.1 aory|EIT77087.1 asyd|OJJ62389.1 ater|EAU37591.1 atub|OJI87063.1 aver|OJI96522.1 awen|OJJ39675.1 azon|OJJ47805.1

Group3636: abra|OJJ75069.1 acar|OOF97264.1 acla|EAW08250.1 afis|EAW23284.1 afla|EED55691.1 afum|EAL88195.1 agla|OJJ86562.1 akaw|GAA88681.1 aluc|OJZ92635.1 anid|CBF89520.1 anig|EHA26356.1

aory|EIT77082.1 asyd|OJJ62392.1 ater|EAU37588.1 atub|OJI87066.1 aver|OJI96525.1 awen|OJJ39672.1 azon|OJJ47802.1

Group3637: abra|OJJ75070.1 acar|OOF97265.1 acla|EAW08251.1 afis|EAW23285.1 afla|EED55690.1 afum|EAL88194.1 agla|OJJ86561.1 akaw|GAA88682.1 aluc|OJZ92636.1 anid|CBF89518.1 anig|EHA26355.1 aory|EIT77093.1 asyd|OJJ62393.1 ater|EAU37587.1 atub|OJI87067.1 aver|OJI96526.1 awen|OJJ39671.1 azon|OJJ47801.1

Group3638: abra|OJJ75077.1 acar|OOF97256.1 acla|EAW08287.1 afis|EAW23320.1 afla|EED55678.1 afum|EAL88156.2 agla|OJJ86545.1 akaw|GAA88690.1 aluc|OJZ92643.1 anid|CBF89504.1 anig|EHA26349.1 aory|EIT77070.1 asyd|OJJ62400.1 ater|EAU37581.1 atub|OJI87074.1 aver|OJI96533.1 awen|OJJ39665.1 azon|OJJ47793.1

Group3639: abra|OJJ75080.1 acar|OOF97253.1 acla|EAW08284.1 afis|EAW23317.1 afla|EED55676.1 afum|EBA27338.1 agla|OJJ86542.1 akaw|GAA88693.1 aluc|OJZ92646.1 anid|CBF89487.1 anig|EHA26346.1 aory|EIT77097.1 asyd|OJJ62404.1 ater|EAU35544.1 atub|OJI87077.1 aver|OJI96537.1 awen|OJJ39662.1 azon|OJJ47791.1

Group3640: abra|OJJ75081.1 acar|OOF97252.1 acla|EAW08283.1 afis|EAW23316.1 afla|EED55675.1 afum|EAL88161.2 agla|OJJ86541.1 akaw|GAA88694.1 aluc|OJZ92647.1 anid|CBF89489.1 anig|EHA26345.1 aory|EIT77095.1 asyd|OJJ62405.1 ater|EAU35545.1 atub|OJI87078.1 aver|OJI96538.1 awen|OJJ39661.1 azon|OJJ47789.1

Group3641: abra|OJJ75082.1 acar|OOF97251.1 acla|EAW08282.1 afis|EAW23315.1 afla|EED55674.1 afum|EAL88162.1 agla|OJJ86540.1 akaw|GAA88695.1 aluc|OJZ92648.1 anid|CBF89491.1 anig|EHA26344.1 aory|EIT77078.1 asyd|OJJ62406.1 ater|EAU35546.1 atub|OJI87079.1 aver|OJI96539.1 awen|OJJ39660.1 azon|OJJ47788.1

Group3642: abra|OJJ75084.1 acar|OOF97249.1 acla|EAW08280.1 afis|EAW23313.1 afla|EED55672.1 afum|EAL88164.1 agla|OJJ86538.1 akaw|GAA88697.1 aluc|OJZ92650.1 anid|CBF89495.1 anig|EHA26342.1 aory|EIT77083.1 asyd|OJJ62408.1 ater|EAU35548.1 atub|OJI87081.1 aver|OJI96541.1 awen|OJJ39658.1 azon|OJJ47786.1

Group3643: abra|OJJ75087.1 acar|OOF97246.1 acla|EAW08277.1 afis|EAW23310.1 afla|EED55665.1 afum|EAL88167.1 akaw|GAA88699.1 aluc|OJZ92653.1 anid|CBF89485.1 anig|EHA26341.1 asyd|OJJ62412.1 ater|EAU35551.1 atub|OJI87084.1 aver|OJI96546.1 awen|OJJ39655.1 azon|OJJ47783.1 agla|OJJ86535.1 aory|EIT81799.1

Group3644: abra|OJJ75089.1 acar|OOF97244.1 acla|EAW08275.1 afis|EAW23308.1 afla|EED55663.1 afum|EAL88169.1 agla|OJJ86533.1 akaw|GAA88701.1 aluc|OJZ92655.1 anid|CBF89482.1 anig|EHA26339.1 aory|EIT81804.1 asyd|OJJ62413.1 ater|EAU35553.1 atub|OJI87086.1 aver|OJI96548.1 awen|OJJ39653.1 azon|OJJ47781.1

Group3645: abra|OJJ75090.1 acar|OOF97243.1 acla|EAW08274.1 afis|EAW23307.1 afla|EED55662.1 afum|EAL88170.1 agla|OJJ86532.1 akaw|GAA88702.1 aluc|OJZ92656.1 anid|CBF89480.1 anig|EHA26338.1 aory|EIT81820.1 asyd|OJJ62414.1 ater|EAU35554.1 atub|OJI87087.1 aver|OJI96549.1 awen|OJJ39652.1 azon|OJJ47780.1

Group3646: abra|OJJ75091.1 acar|OOF97241.1 acla|EAW08273.1 afis|EAW23306.1 afla|EED55661.1 afum|EAL88171.1 agla|OJJ86531.1 akaw|GAA88703.1 aluc|OJZ92657.1 anid|CBF89478.1 anig|EHA26337.1 aory|EIT81795.1 asyd|OJJ62415.1 ater|EAU35569.1 atub|OJI87088.1 aver|OJI96550.1 awen|OJJ39590.1 azon|OJJ47779.1

Group3647: abra|OJJ75093.1 acar|OOF97239.1 acla|EAW08271.1 afis|EAW23304.1 afla|EED55659.1 afum|EAL88173.1 agla|OJJ86529.1 akaw|GAA88705.1 anid|CBF89474.1 anig|EHA26335.1 aory|EIT81791.1 asyd|OJJ62418.1 ater|EAU35568.1 atub|OJI87090.1 aver|OJI96553.1 awen|OJJ39586.1 azon|OJJ47766.1 aluc|OJZ92659.1

Group3648: abra|OJJ75094.1 acar|OOF97238.1 acla|EAW08270.1 afis|EAW23303.1 afla|EED55656.1 afum|EAL88175.1 agla|OJJ86528.1 akaw|GAA88706.1 aluc|OJZ92660.1 anid|CBF89472.1 anig|EHA26334.1 aory|EIT81787.1 asyd|OJJ62419.1 ater|EAU35567.1 atub|OJI87091.1 aver|OJI96555.1 awen|OJJ39585.1 azon|OJJ47767.1

Group3649: abra|OJJ75096.1 acar|OOF97236.1 acla|EAW08268.1 afis|EAW23301.1 afla|EED55653.1 afum|EAL88178.1 agla|OJJ86507.1 akaw|GAA88708.1 aluc|OJZ92662.1 anid|CBF89468.1 anig|EHA26332.1 aory|EIT81800.1 asyd|OJJ62421.1 ater|EAU35565.1 atub|OJI87093.1 aver|OJI96556.1 awen|OJJ39583.1 azon|OJJ47769.1

Group3650: abra|OJJ75102.1 acar|OOF97232.1 acla|EAW08266.1 afis|EAW23299.1 afla|EED55649.1 afum|EAL88180.1 agla|OJJ86511.1 akaw|GAA88711.1 aluc|OJZ92666.1 anid|CBF89465.1 anig|EHA26329.1 aory|EIT81776.1 asyd|OJJ62424.1 ater|EAU35561.1 atub|OJI87097.1 aver|OJI96561.1 awen|OJJ39581.1 azon|OJJ47771.1

Group3651: abra|OJJ75103.1 acar|OOF97231.1 acla|EAW08265.1 afis|EAW23298.1 afla|EED55648.1 afum|EAL88181.2 agla|OJJ86513.1 akaw|GAA88712.1 aluc|OJZ92668.1 anid|CBF89463.1 anig|EHA26328.1 aory|EIT81789.1 asyd|OJJ62425.1 ater|EAU35559.1 atub|OJI87098.1 aver|OJI96562.1 awen|OJJ39579.1 azon|OJJ47772.1

Group3652: abra|OJJ75106.1 acar|OOF97228.1 acla|EAW08262.1 afis|EAW23295.1 afla|EED55645.1 afum|EAL88183.1 agla|OJJ86516.1 akaw|GAA88715.1 aluc|OJZ92671.1 anid|CBF89459.1 anig|EHA26326.1 aory|EIT81783.1 asyd|OJJ62427.1 ater|EAU35558.1 atub|OJI87100.1 aver|OJI96564.1 awen|OJJ39576.1 azon|OJJ47775.1

Group3653: abra|OJJ75107.1 acar|OOF97227.1 acla|EAW08261.1 afis|EAW23294.1 afla|EED55644.1 afum|EAL88184.1 agla|OJJ86517.1 akaw|GAA88716.1 aluc|OJZ92672.1 anid|CBF89457.1 anig|EHA26325.1 aory|EIT81790.1 asyd|OJJ62428.1 ater|EAU35557.1 atub|OJI87101.1 aver|OJI96565.1 awen|OJJ39575.1 azon|OJJ47776.1

Group3654: abra|OJJ75109.1 acar|OOF97223.1 acla|EAW08315.1 afis|EAW23325.1 afla|EED55641.1 afum|EAL88154.1 agla|OJJ86520.1 akaw|GAA88718.1 aluc|OJZ92675.1 anid|CBF89453.1 anig|EHA26323.1 aory|EIT81818.1 asyd|OJJ62431.1 ater|EAU35573.1 atub|OJI87103.1 aver|OJI96567.1 awen|OJJ39569.1 azon|OJJ47762.1

Group3655: abra|OJJ75114.1 acar|OOF97217.1 acla|EAW08309.1 afis|EAW23330.1 afla|EED55637.1 afum|EAL88149.2 agla|OJJ86525.1 akaw|GAA88724.1 aluc|OJZ92681.1 anid|CBF89441.1 anig|EHA26318.1 aory|EIT81798.1 asyd|OJJ62439.1 ater|EAU35577.1 atub|OJI87109.1 aver|OJI96575.1 awen|OJJ39564.1 azon|OJJ47757.1

Group3656: abra|OJJ75115.1 acar|OOF97216.1 acla|EAW08308.1 afis|EAW23331.1 afla|EED55636.1 afum|EAL88148.1 agla|OJJ86503.1 akaw|GAA88725.1 aluc|OJZ92682.1 anid|CBF89439.1 anig|EHA26317.1 aory|EIT81819.1 asyd|OJJ62440.1 ater|EAU35578.1 atub|OJI87110.1 aver|OJI96576.1 awen|OJJ39563.1 azon|OJJ47756.1

Group3657: abra|OJJ75120.1 acar|OOF97213.1 acla|EAW08306.1 afis|EAW23332.1 afla|EED55634.1 afum|EAL88146.2 agla|OJJ86500.1 akaw|GAA88729.1 aluc|OJZ92687.1 anid|CBF89437.1 anig|EHA26313.1 aory|EIT81801.1 asyd|OJJ62442.1 ater|EAU35580.1 atub|OJI87115.1 aver|OJI96578.1 awen|OJJ39561.1 azon|OJJ47754.1

Group3658: abra|OJJ75122.1 acar|OOF97212.1 acla|EAW08305.1 afis|EAW23334.1 afla|EED55632.1 afum|EAL88144.1 agla|OJJ86499.1 akaw|GAA88730.1 aluc|OJZ92688.1 anid|CBF89435.1 anig|EHA26312.1 aory|EIT81824.1 asyd|OJJ62443.1 ater|EAU35581.1 atub|OJI87116.1 aver|OJI96579.1 awen|OJJ39560.1 azon|OJJ47753.1

Group3659: abra|OJJ75124.1 acar|OOF97210.1 acla|EAW08304.1 afis|EAW23336.1 afla|EED55630.1 afum|EAL88142.1 agla|OJJ86494.1 akaw|GAA88732.1 aluc|OJZ92690.1 anid|CBF89432.1 anig|EHA26310.1 aory|EIT81797.1 asyd|OJJ62444.1 ater|EAU35582.1 atub|OJI87118.1 aver|OJI96581.1 awen|OJJ39558.1 azon|OJJ47749.1

Group3660: abra|OJJ75126.1 acar|OOF97209.1 acla|EAW08303.1 afis|EAW23337.1 afla|EED55629.1 afum|EAL88141.1 agla|OJJ86493.1 akaw|GAA88733.1 aluc|OJZ92691.1 anid|CBF89430.1 anig|EHA26309.1 aory|EIT81788.1 asyd|OJJ62445.1 ater|EAU35583.1 atub|OJI87119.1 aver|OJI96582.1 awen|OJJ39557.1 azon|OJJ47750.1

Group3661: abra|OJJ75129.1 acar|OOF97207.1 acla|EAW08300.1 afis|EAW23340.1 afla|EED55626.1 afum|EAL88138.1 agla|OJJ86490.1 akaw|GAA88736.1 aluc|OJZ92694.1 anid|CBF89425.1 anig|EHA26306.1 aory|EIT81810.1 asyd|OJJ62448.1 ater|EAU35585.1 atub|OJI87122.1 aver|OJI96585.1 awen|OJJ39554.1 azon|OJJ47748.1

Group3662: abra|OJJ75130.1 acar|OOF97205.1 acla|EAW08298.1 afis|EAW23342.1 afla|EED55624.1 afum|EAL88136.2 agla|OJJ86486.1 akaw|GAA88737.1 aluc|OJZ92695.1 anid|CBF89422.1 anig|EHA26304.1 aory|EIT81811.1 asyd|OJJ62451.1 ater|EAU35586.1 atub|OJI87123.1 aver|OJI96586.1 awen|OJJ39552.1 azon|OJJ47747.1

Group3663: abra|OJJ75131.1 acar|OOF97204.1 acla|EAW08297.1 afis|EAW23343.1 afla|EED55623.1 afum|EAL88135.1 agla|OJJ86485.1 akaw|GAA88738.1 aluc|OJZ92696.1 anid|CBF89420.1 anig|EHA26302.1 aory|EIT81805.1 asyd|OJJ62452.1 ater|EAU35587.1 atub|OJI87124.1 aver|OJI96587.1 awen|OJJ39550.1 azon|OJJ47746.1

Group3664: abra|OJJ75133.1 acar|OOF97203.1 acla|EAW08294.1 afis|EAW23345.1 afla|EED55621.1 afum|EAL88132.1 agla|OJJ86482.1 akaw|GAA88740.1 aluc|OJZ92699.1 anid|CBF89412.1 anig|EHA26300.1 aory|EIT81825.1 asyd|OJJ62455.1 ater|EAU35589.1 atub|OJI87127.1 aver|OJI96590.1 awen|OJJ39547.1 azon|OJJ47744.1

Group3665: abra|OJJ75134.1 acar|OOF97202.1 acla|EAW08293.1 afis|EAW23346.1 afla|EED55620.1 afum|EAL88130.1 agla|OJJ86481.1 akaw|GAA88741.1 aluc|OJZ92700.1 anid|CBF74918.1 anig|EHA26299.1

aory|EIT81822.1 asyd|OJJ60098.1 ater|EAU35590.1 atub|OJI87128.1 aver|OJJ04164.1 awen|OJJ39545.1 azon|OJJ47743.1

Group3666: abra|OJJ75136.1 acar|OOF97201.1 acla|EAW08292.1 afis|EAW23347.1 afla|EED55619.1 afum|EAL88129.1 agla|OJJ86480.1 akaw|GAA88742.1 aluc|OJZ92702.1 anid|CBF74916.1 anig|EHA26298.1 aory|EIT81794.1 asyd|OJJ60099.1 ater|EAU35591.1 atub|OJI87129.1 aver|OJJ04163.1 awen|OJJ39544.1 azon|OJJ47742.1

Group3667: abra|OJJ75146.1 acar|OOF97192.1 acla|EAW08319.1 afis|EAW23354.1 afla|EED55604.1 afum|EAL88122.2 agla|OJJ86471.1 akaw|GAA88751.1 aluc|OJZ92711.1 anid|CBF74886.1 anig|EHA26288.1 aory|EIT82711.1 asyd|OJJ60107.1 ater|EAU35606.1 atub|OJI87137.1 aver|OJJ04155.1 awen|OJJ39532.1 azon|OJJ47729.1

Group3668: abra|OJJ75149.1 acar|OOF97188.1 acla|EAW08323.1 afis|EAW23358.1 afla|EED55608.1 afum|EAL88118.1 agla|OJJ86468.1 akaw|GAA88755.1 aluc|OJZ92715.1 anid|CBF74882.1 anig|EHA26285.1 aory|EIT81782.1 asyd|OJJ60110.1 ater|EAU35609.1 atub|OJI87141.1 aver|OJJ04152.1 awen|OJJ39529.1 azon|OJJ47725.1

Group3669: abra|OJJ75150.1 acar|OOF97187.1 acla|EAW08324.1 afis|EAW23359.1 afla|EED55610.1 afum|EBA27336.1 agla|OJJ86467.1 akaw|GAA88756.1 aluc|OJZ92716.1 anid|CBF74880.1 anig|EHA26283.1 aory|EIT81806.1 asyd|OJJ60111.1 ater|EAU35610.1 atub|OJI87142.1 aver|OJJ04151.1 awen|OJJ39528.1 azon|OJJ47724.1

Group3670: abra|OJJ75152.1 acar|OOF97185.1 acla|EAW08327.1 afis|EAW23362.1 afla|EED55596.1 afum|EAL88112.1 agla|OJJ86464.1 akaw|GAA88757.1 aluc|OJZ92720.1 anid|CBF74837.1 anig|EHA26280.1 aory|EIT82684.1 asyd|OJJ60134.1 ater|EAU35613.1 atub|OJI87145.1 aver|OJJ04128.1 awen|OJJ39525.1 azon|OJJ47721.1

Group3671: abra|OJJ75153.1 acar|OOF97184.1 acla|EAW08328.1 afis|EAW23363.1 afla|EED55595.1 afum|EBA27334.1 agla|OJJ86463.1 akaw|GAA88758.1 aluc|OJZ92721.1 anid|CBF74835.1 anig|EHA26279.1 aory|EIT82682.1 asyd|OJJ60135.1 ater|EAU35614.1 atub|OJI87146.1 aver|OJJ04127.1 awen|OJJ39524.1 azon|OJJ47720.1

Group3672: abra|OJJ75157.1 acar|OOF97182.1 acla|EAW08332.1 afis|EAW23367.1 afla|EED55590.1 afum|EAL88106.1 agla|OJJ86457.1 akaw|GAA88763.1 aluc|OJZ92727.1 anid|CBF74823.1 anig|EHA26274.1 aory|EIT82667.1 asyd|OJJ60140.1 ater|EAU35618.1 atub|OJI87151.1 aver|OJJ04122.1 awen|OJJ39520.1 azon|OJJ47716.1

Group3673: abra|OJJ75159.1 acar|OOF97179.1 acla|EAW08335.1 afis|EAW23370.1 afla|EED55587.1 afum|EAL88103.2 agla|OJJ86453.1 akaw|GAA88770.1 aluc|OJZ92729.1 anid|CBF74817.1 anig|EHA26272.1 aory|EIT82666.1 asyd|OJJ60143.1 ater|EAU35621.1 atub|OJI87153.1 aver|OJJ04120.1 awen|OJJ39513.1 azon|OJJ47712.1

Group3674: abra|OJJ75160.1 acar|OOF97178.1 acla|EAW08336.1 afis|EAW23371.1 afla|EED55586.1 afum|EAL88101.1 agla|OJJ86452.1 akaw|GAA88771.1 aluc|OJZ92730.1 anid|CBF74815.1 anig|EHA26271.1 aory|EIT82693.1 asyd|OJJ60144.1 ater|EAU35622.1 atub|OJI87154.1 aver|OJJ04119.1 awen|OJJ39512.1 azon|OJJ47711.1

Group3675: abra|OJJ75162.1 acar|OOF97176.1 acla|EAW08338.1 afis|EAW23373.1 afla|EED55585.1 afum|EAL88099.1 agla|OJJ86450.1 akaw|GAA88773.1 aluc|OJZ92732.1 anid|CBF74811.1 anig|EHA26269.1 aory|EIT82709.1 asyd|OJJ60146.1 ater|EAU35624.1 atub|OJI87156.1 aver|OJJ04117.1 awen|OJJ39510.1 azon|OJJ47709.1

Group3676: abra|OJJ75165.1 acar|OOF97173.1 acla|EAW08339.1 afis|EAW23374.1 afla|EED55584.1 afum|EAL88097.1 agla|OJJ86449.1 akaw|GAA88776.1 aluc|OJZ92736.1 anid|CBF74839.1 anig|EHA26266.1 aory|EIT82678.1 asyd|OJJ60133.1 ater|EAU35628.1 atub|OJI87160.1 aver|OJJ04129.1 awen|OJJ39509.1 azon|OJJ47708.1

Group3677: abra|OJJ75166.1 acar|OOF97172.1 acla|EAW08340.1 afis|EAW23375.1 afla|EED55583.1 afum|EAL88096.1 agla|OJJ86448.1 akaw|GAA88777.1 aluc|OJZ92737.1 anid|CBF74841.1 anig|EHA26265.1 aory|EIT82676.1 asyd|OJJ60132.1 ater|EAU35629.1 atub|OJI87161.1 aver|OJJ04130.1 awen|OJJ39508.1 azon|OJJ47707.1

Group3678: abra|OJJ75168.1 acar|OOF94606.1 acla|EAW08132.1 afis|EAW23169.1 afla|EED46952.1 afum|EAL88315.1 agla|OJJ89680.1 akaw|GAA88778.1 aluc|OJZ92738.1 anid|CBF74480.1 anig|EHA26264.1 aory|EIT72535.1 asyd|OJJ57708.1 ater|EAU36298.1 atub|OJI87163.1 aver|OJJ05456.1 awen|OJJ36614.1 azon|OJJ47599.1

Group3679: abra|OJJ75172.1 acar|OOF94604.1 acla|EAW13004.1 afis|EAW17664.1 afla|EED50493.1 afum|EAL91478.1 agla|OJJ79537.1 akaw|GAA88779.1 aluc|OJZ92740.1 anid|CBF90052.1 anig|EHA26263.1 aory|EIT82027.1 asyd|OJJ62053.1 ater|EAU31989.1 atub|OJI87165.1 aver|OJI96159.1 awen|OJJ37177.1 azon|OJJ50877.1

Group3680: abra|OJJ75177.1 acar|OOF94602.1 acla|EAW13001.1 afis|EAW17667.1 afla|EED50488.1 afum|EAL91481.1 agla|OJJ79539.1 akaw|GAA88782.1 aluc|OJZ92743.1 anid|CBF90043.1 anig|EHA26260.1 aory|EIT82124.1 asyd|OJJ62058.1 ater|EAU31980.1 atub|OJI87168.1 aver|OJI96164.1 awen|OJJ37174.1 azon|OJJ50873.1

Group3681: abra|OJJ75178.1 acar|OOF94601.1 acla|EAW13000.1 afis|EAW17668.1 afla|EED50487.1 afum|EAL91482.1 agla|OJJ79540.1 akaw|GAA88783.1 aluc|OJZ92744.1 anid|CBF90041.1 anig|EHA26259.1 aory|EIT82171.1 asyd|OJJ62059.1 ater|EAU31979.1 atub|OJI87169.1 aver|OJI96165.1 awen|OJJ37173.1 azon|OJJ50872.1

Group3682: abra|OJJ75179.1 acar|OOF94600.1 acla|EAW12999.1 afis|EAW17669.1 afla|EED50486.1 afum|EAL91483.1 agla|OJJ79541.1 akaw|GAA88784.1 aluc|OJZ92745.1 anid|CBF90039.1 anig|EHA26258.1 aory|EIT82129.1 asyd|OJJ62060.1 ater|EAU31978.1 atub|OJI87170.1 aver|OJI96166.1 awen|OJJ37172.1 azon|OJJ50871.1

Group3683: abra|OJJ75182.1 acar|OOF94598.1 acla|EAW12995.1 afis|EAW17671.1 afla|EED50482.1 afum|EAL91485.1 agla|OJJ79543.1 akaw|GAA88786.1 aluc|OJZ92746.1 anid|CBF90036.1 anig|EHA26257.1 aory|EIT82081.1 asyd|OJJ62061.1 ater|EAU31976.1 atub|OJI87173.1 aver|OJI96167.1 awen|OJJ37171.1 azon|OJJ50867.1

Group3684: abra|OJJ75186.1 acar|OOF94596.1 acla|EAW12991.1 afis|EAW17674.1 afla|EED50477.1 afum|EAL91487.1 agla|OJJ79515.1 akaw|GAA88788.1 aluc|OJZ92749.1 anid|CBF90029.1 anig|EHA26254.1 aory|EIT82056.1 asyd|OJJ62064.1 ater|EAU31972.1 atub|OJI87175.1 aver|OJI96170.1 awen|OJJ37167.1 azon|OJJ50863.1

Group3685: abra|OJJ75189.1 acar|OOF94594.1 acla|EAW12990.1 afis|EAW17675.1 afla|EED50476.1 afum|EAL91488.1 agla|OJJ79514.1 akaw|GAA88790.1 aluc|OJZ92752.1 anid|CBF90024.1 anig|EHA26252.1 aory|EIT82044.1 asyd|OJJ62067.1 ater|EAU31971.1 atub|OJI87177.1 aver|OJI96173.1 awen|OJJ37165.1 azon|OJJ50862.1

Group3686: abra|OJJ75190.1 acar|OOF94593.1 acla|EAW12989.1 afis|EAW17676.1 afla|EED50475.1 afum|EAL91489.1 agla|OJJ79500.1 akaw|GAA88791.1 aluc|OJZ92754.1 anid|CBF90022.1 anig|EHA26251.1 aory|EIT82153.1 asyd|OJJ62068.1 ater|EAU31970.1 atub|OJI87179.1 aver|OJI96174.1 awen|OJJ37164.1 azon|OJJ50860.1

Group3687: abra|OJJ75198.1 acar|OOF94588.1 acla|EAW12983.1 afis|EAW17681.1 afla|EED50470.1 afum|EAL91495.1 agla|OJJ79499.1 akaw|GAA91750.1 aluc|OJZ92761.1 anid|CBF90009.1 anig|EHA26245.1 aory|EIT82018.1 asyd|OJJ62073.1 ater|EAU31944.1 atub|OJI87205.1 aver|OJI96180.1 awen|OJJ37158.1 azon|OJJ50856.1

Group3688: abra|OJJ75199.1 acar|OOF94589.1 acla|EAW12982.1 afis|EAW17682.1 afla|EED50469.1 afum|EAL91496.1 agla|OJJ79498.1 akaw|GAA91751.1 aluc|OJZ92762.1 anid|CBF90007.1 anig|EHA26244.1 aory|EIT82107.1 asyd|OJJ62074.1 ater|EAU31945.1 atub|OJI87206.1 aver|OJI96181.1 awen|OJJ37157.1 azon|OJJ50855.1

Group3689: abra|OJJ75201.1 acar|OOF94587.1 acla|EAW12981.1 afis|EAW17683.1 afla|EED50468.1 afum|EAL91497.1 agla|OJJ79496.1 akaw|GAA91752.1 aluc|OJZ92763.1 anid|CBF90005.1 anig|EHA26243.1 aory|EIT82022.1 asyd|OJJ62075.1 ater|EAU31946.1 atub|OJI87207.1 aver|OJI96182.1 awen|OJJ37156.1 azon|OJJ50854.1

Group3690: abra|OJJ75203.1 acar|OOF94585.1 acla|EAW12979.1 afis|EAW17685.1 afla|EED50466.1 afum|EAL91499.1 agla|OJJ79494.1 akaw|GAA91754.1 aluc|OJZ92766.1 anid|CBF90002.1 anig|EHA26241.1 aory|EIT82013.1 asyd|OJJ62078.1 ater|EAU31948.1 atub|OJI87210.1 aver|OJI96185.1 awen|OJJ37154.1 azon|OJJ50852.1

Group3691: abra|OJJ75206.1 acar|OOF94583.1 acla|EAW12977.1 afis|EAW17687.1 afla|EED50464.1 afum|EAL91501.1 agla|OJJ79492.1 akaw|GAA91756.1 aluc|OJZ92768.1 anid|CBF89987.1 anig|EHA26239.1 asyd|OJJ56120.1 ater|EAU31950.1 atub|OJI87212.1 aver|OJJ05061.1 awen|OJJ37152.1 asyd|OJJ62085.1 aver|OJI96192.1

Group3692: abra|OJJ75207.1 acar|OOF94582.1 acla|EAW12976.1 afis|EAW17688.1 afla|EED50463.1 afum|EAL91502.1 agla|OJJ79491.1 akaw|GAA91757.1 aluc|OJZ92769.1 anid|CBF89985.1 anig|EHA26238.1 aory|EIT82159.1 asyd|OJJ62086.1 ater|EAU31951.1 atub|OJI87213.1 aver|OJI96193.1 awen|OJJ37151.1 azon|OJJ50848.1

Group3693: abra|OJJ75208.1 acar|OOF94581.1 acla|EAW12975.1 afis|EAW17689.1 afla|EED50455.1 afum|EAL91503.1 agla|OJJ79490.1 akaw|GAA91758.1 aluc|OJZ92770.1 anid|CBF89984.1 anig|EHA26237.1 aory|EIT82021.1 asyd|OJJ62087.1 ater|EAU31952.1 atub|OJI87214.1 aver|OJI96194.1 awen|OJJ37147.1 azon|OJJ50847.1

Group3694: abra|OJJ75212.1 acar|OOF94572.1 acla|EAW13091.1 afis|EAW17572.1 afla|EED49249.1 afum|EAL91386.1 agla|OJJ81386.1 akaw|GAA91762.1 aluc|OJZ92776.1 anid|CBF90252.1 anig|EHA26234.1

aory|EIT82180.1 asyd|OJJ61946.1 ater|EAU37009.1 atub|OJI87218.1 aver|OJI96038.1 awen|OJJ37275.1 azon|OJJ50975.1

Group3695: abra|OJJ75213.1 acar|OOF94571.1 acla|EAW13092.1 afis|EAW17571.1 afla|EED49250.1 afum|EBA27432.1 agla|OJJ81387.1 akaw|GAA91763.1 aluc|OJZ92777.1 anid|CBF90254.1 anig|EHA26233.1 aory|EIT82196.1 asyd|OJJ61945.1 ater|EAU37008.1 atub|OJI87219.1 aver|OJI96037.1 awen|OJJ37278.1 azon|OJJ50976.1

Group3696: abra|OJJ75214.1 acar|OOF94570.1 acla|EAW13093.1 afis|EAW17570.1 afla|EED49251.1 afum|EAL91383.1 agla|OJJ81390.1 akaw|GAA91764.1 aluc|OJZ92778.1 anid|CBF90256.1 anig|EHA26232.1 aory|EIT82218.1 asyd|OJJ61944.1 ater|EAU37007.1 atub|OJI87220.1 aver|OJI96036.1 awen|OJJ37281.1 azon|OJJ50977.1

Group3697: abra|OJJ75215.1 acar|OOF94569.1 acla|EAW13094.1 afis|EAW17569.1 afla|EED49252.1 afum|EAL91382.1 agla|OJJ81395.1 akaw|GAA91765.1 aluc|OJZ92779.1 anid|CBF90258.1 anig|EHA26231.1 aory|EIT82201.1 asyd|OJJ61943.1 ater|EAU37006.1 atub|OJI87221.1 awen|OJJ37282.1 azon|OJJ50978.1 aver|OJI96035.1

Group3698: abra|OJJ75227.1 acar|OOF94562.1 acla|EAW12254.1 afis|EAW16982.1 afla|EED48301.1 afum|EAL86509.1 agla|OJJ79789.1 akaw|GAA91773.1 aluc|OJZ92788.1 anid|CBF86682.1 anig|EHA26219.1 aory|EIT76292.1 asyd|OJJ64757.1 ater|EAU31794.1 atub|OJI87231.1 aver|OJI98859.1 awen|OJJ35994.1 azon|OJJ50263.1

Group3699: abra|OJJ75272.1 acar|OOF94531.1 acla|EAW12236.1 afis|EAW16991.1 afla|EED48964.1 afum|EAL86515.1 agla|OJJ81058.1 akaw|GAA91810.1 aluc|OJZ92832.1 anid|CBF87436.1 anig|EHA26177.1 asyd|OJJ59098.1 ater|EAU31737.1 atub|OJI87289.1 aver|OJJ01689.1 awen|OJJ35979.1 azon|OJJ50286.1 aory|EIT80025.1

Group3700: abra|OJJ75274.1 acar|OOF94529.1 acla|EAW12235.1 afis|EAW16993.1 afla|EED48956.1 afum|EAL86517.1 agla|OJJ79819.1 akaw|GAA91812.1 aluc|OJZ92834.1 anid|CBF87472.1 anig|EHA26175.1 aory|EIT80048.1 asyd|OJJ52939.1 ater|EAU31735.1 atub|OJI87291.1 aver|OJJ08906.1 awen|OJJ35982.1 azon|OJJ50283.1

Group3701: abra|OJJ75279.1 afis|EAW25673.1 afla|EED49857.1 akaw|GAA91817.1 aluc|OJZ92840.1 anid|CBF82114.1 anig|EHA26171.1 aory|EIT83389.1 asyd|OJJ54289.1 ater|EAU31659.1 atub|OJI87296.1 aver|OJJ05243.1 awen|OJJ40812.1 afla|EED48617.1 ater|EAU33458.1 aory|EIT73274.1 ater|EAU35345.1 ater|EAU35445.1

Group3702: abra|OJJ75284.1 acar|OOF94523.1 acla|EAW12234.1 afis|EAW16994.1 afla|EED48957.1 afum|EAL86518.2 agla|OJJ79820.1 akaw|GAA91822.1 aluc|OJZ92845.1 anid|CBF87470.1 anig|EHA26165.1 aory|EIT79980.1 asyd|OJJ52938.1 ater|EAU31734.1 atub|OJI87301.1 aver|OJJ08905.1 awen|OJJ35983.1 azon|OJJ50282.1

Group3703: abra|OJJ75307.1 acar|OOF94522.1 acla|EAW12233.1 afis|EAW16995.1 afla|EED48958.1 afum|EAL86519.2 agla|OJJ79822.1 akaw|GAA91991.1 aluc|OJZ92861.1 anid|CBF87468.1 anig|EHA26104.1 aory|EIT79977.1 asyd|OJJ52937.1 ater|EAU31733.1 atub|OJI87319.1 aver|OJJ08904.1 awen|OJJ35984.1 azon|OJJ50281.1

Group3704: abra|OJJ75309.1 acar|OOF94520.1 acla|EAW12231.1 afis|EAW16997.1 afla|EED48962.1 afum|EAL86522.2 agla|OJJ79824.1 akaw|GAA91993.1 aluc|OJZ92863.1 anid|CBF87465.1 anig|EHA26102.1 aory|EIT79990.1 asyd|OJJ52935.1 ater|EAU31731.1 atub|OJI87321.1 aver|OJJ08902.1 awen|OJJ35986.1 azon|OJJ42560.1

Group3705: abra|OJJ75310.1 acar|OOF94519.1 acla|EAW12230.1 afis|EAW16998.1 afla|EED48952.1 afum|EAL86523.1 agla|OJJ79825.1 akaw|GAA91994.1 aluc|OJZ92864.1 anid|CBF87476.1 anig|EHA26101.1 aory|EIT79981.1 asyd|OJJ52941.1 ater|EAU31729.1 atub|OJI87322.1 aver|OJJ08908.1 awen|OJJ35987.1 azon|OJJ50276.1

Group3706: abra|OJJ75314.1 acar|OOF94515.1 acla|EAW10816.1 afis|EAW24758.1 afla|EED56728.1 afum|EAL89400.1 agla|OJJ78831.1 akaw|GAA91998.1 aluc|OJZ92868.1 anid|CBF89399.1 anig|EHA26097.1 aory|EIT73833.1 asyd|OJJ62458.1 ater|EAU34827.1 atub|OJI87326.1 aver|OJI96593.1 awen|OJJ33340.1 azon|OJJ42602.1

Group3707: abra|OJJ75322.1 acar|OOF94504.1 acla|EAW12347.1 afis|EAW16741.1 afla|EED48943.1 afum|EAL86316.1 akaw|GAA92017.1 aluc|OJZ92883.1 anid|CBF87481.1 anig|EHA26077.1 aory|EIT80047.1 asyd|OJJ64592.1 ater|EAU31723.1 atub|OJI87342.1 aver|OJI98689.1 awen|OJJ34684.1 azon|OJJ47373.1 anid|CBF75776.1

Group3708: abra|OJJ75342.1 acar|OOF89833.1 afla|EED48926.1 agla|OJJ81063.1 akaw|GAA92035.1 aluc|OJZ92905.1 anid|CBF87500.1 anig|EHA26058.1 atub|OJI87364.1 aver|OJJ08935.1 awen|OJJ35965.1 azon|OJJ50308.1 acla|EAW12182.1 afis|EAW17116.1 afum|EAL86578.1 asyd|OJJ52969.1 ater|EAU36171.1 aory|EIT80020.1

Group3709: abra|OJJ75368.1 acar|OOF94179.1 afla|EED48878.1 agla|OJJ83621.1 akaw|GAA92062.1 aluc|OJZ92937.1 anig|EHA26035.1 aory|EIT81224.1 asyd|OJJ57425.1 ater|EAU36149.1 atub|OJI87397.1 aver|OJJ04755.1 awen|OJJ31289.1 azon|OJJ50359.1 asyd|OJJ55807.1 awen|OJJ38783.1 azon|OJJ42402.1 awen|OJJ29661.1

Group3710: abra|OJJ75381.1 acar|OOF94466.1 acla|EAW12111.1 afis|EAW17274.1 afla|EED48845.1 afum|EAL86692.1 agla|OJJ81078.1 akaw|GAA92743.1 aluc|OJZ92951.1 anid|CBF87538.1 anig|EHA26024.1 aory|EIT81278.1 asyd|OJJ52990.1 ater|EAU34713.1 atub|OJI87410.1 aver|OJJ08959.1 awen|OJJ35760.1 azon|OJJ45524.1

Group3711: abra|OJJ75383.1 acar|OOF94464.1 acla|EAW12109.1 afis|EAW17276.1 afla|EED48843.1 afum|EAL86694.1 agla|OJJ81080.1 akaw|GAA92745.1 aluc|OJZ92953.1 anid|CBF87542.1 anig|EHA26022.1 aory|EIT81280.1 asyd|OJJ52992.1 ater|EAU34711.1 atub|OJI87412.1 aver|OJJ08961.1 awen|OJJ35758.1 azon|OJJ45526.1

Group3712: abra|OJJ75384.1 acar|OOF94463.1 acla|EAW12108.1 afis|EAW17277.1 afla|EED48842.1 afum|EAL86695.1 agla|OJJ81081.1 akaw|GAA92746.1 aluc|OJZ92954.1 anid|CBF87544.1 anig|EHA26021.1 aory|EIT81283.1 asyd|OJJ52993.1 ater|EAU34710.1 atub|OJI87413.1 aver|OJJ08962.1 awen|OJJ35757.1 azon|OJJ45527.1

Group3713: abra|OJJ75385.1 acar|OOF94461.1 acla|EAW12107.1 afis|EAW17278.1 afla|EED48841.1 afum|EAL86696.1 agla|OJJ81082.1 akaw|GAA92747.1 aluc|OJZ92955.1 anid|CBF87545.1 anig|EHA26020.1 aory|EIT81279.1 asyd|OJJ52994.1 ater|EAU34709.1 atub|OJI87414.1 aver|OJJ08963.1 awen|OJJ35756.1 azon|OJJ45528.1

Group3714: abra|OJJ75391.1 acar|OOF94456.1 acla|EAW12101.1 afis|EAW17287.1 afla|EED48833.1 afum|EAL86705.1 agla|OJJ81090.1 akaw|GAA92754.1 aluc|OJZ92962.1 anid|CBF87556.1 anig|EHA26013.1 aory|EIT81270.1 asyd|OJJ53001.1 ater|EAU34702.1 atub|OJI87420.1 aver|OJJ08970.1 awen|OJJ35750.1 azon|OJJ45536.1

Group3715: abra|OJJ75394.1 acar|OOF94455.1 acla|EAW12099.1 afis|EAW17285.1 afla|EED48831.1 afum|EAL86703.1 agla|OJJ81092.1 akaw|GAA92756.1 aluc|OJZ92965.1 anid|CBF87558.1 anig|EHA26010.1 aory|EIT81288.1 asyd|OJJ53003.1 ater|EAU34700.1 atub|OJI87423.1 aver|OJJ08971.1 awen|OJJ35749.1 azon|OJJ45542.1

Group3716: abra|OJJ75396.1 acar|OOF94453.1 acla|EAW12097.1 afis|EAW17289.1 afla|EED48829.1 afum|EAL86709.1 agla|OJJ81094.1 akaw|GAA92758.1 aluc|OJZ92968.1 anid|CBF81166.1 anig|EHA26008.1 aory|EIT81272.1 asyd|OJJ59287.1 ater|EAU34699.1 atub|OJI87425.1 aver|OJJ02988.1 awen|OJJ35747.1 azon|OJJ45540.1

Group3717: abra|OJJ75397.1 acar|OOF94451.1 acla|EAW12096.1 afis|EAW17290.1 afla|EED48828.1 afum|EAL86710.1 agla|OJJ81095.1 akaw|GAA92759.1 aluc|OJZ92969.1 anid|CBF74295.1 anig|EHA26007.1 aory|EIT81266.1 asyd|OJJ51909.1 ater|EAU34698.1 atub|OJI87426.1 aver|OJJ05350.1 awen|OJJ35746.1 azon|OJJ45539.1

Group3718: abra|OJJ75399.1 acar|OOF90746.1 acla|EAW13772.1 afis|EAW18360.1 afla|EED48826.1 afum|EAL87724.1 agla|OJJ88025.1 akaw|GAA84178.1 aluc|OJZ88851.1 anid|CBF79349.1 anig|EHA23652.1 aory|EIT72979.1 asyd|OJJ57798.1 ater|EAU34692.1 atub|OJI85939.1 aver|OJJ04589.1 awen|OJJ35744.1 azon|OJJ49497.1

Group3719: abra|OJJ75400.1 acar|OOF90747.1 acla|EAW13773.1 afis|EAW18359.1 afla|EED48825.1 afum|EAL87723.1 agla|OJJ88026.1 akaw|GAA84179.1 aluc|OJZ88852.1 anid|CBF79351.1 anig|EHA23651.1 aory|EIT73008.1 asyd|OJJ57799.1 ater|EAU34691.1 atub|OJI85938.1 aver|OJJ04588.1 awen|OJJ35743.1 azon|OJJ42023.1

Group3720: abra|OJJ75401.1 acar|OOF90748.1 acla|EAW08507.1 afis|EAW23545.1 afla|EED48824.1 afum|EAL87921.2 agla|OJJ79762.1 akaw|GAA84180.1 aluc|OJZ88853.1 anid|CBF78889.1 anig|EHA23650.1 aory|EIT73018.1 asyd|OJJ64596.1 ater|EAU31465.1 atub|OJI85937.1 aver|OJI98693.1 awen|OJJ36969.1 anig|EHA19941.1

Group3721: abra|OJJ75402.1 acar|OOF90749.1 acla|EAW13774.1 afis|EAW18358.1 afla|EED48823.1 afum|EAL87722.1 agla|OJJ88027.1 akaw|GAA84181.1 aluc|OJZ88854.1 anid|CBF79353.1 anig|EHA23649.1 aory|EIT72976.1 asyd|OJJ57800.1 ater|EAU34690.1 atub|OJI85936.1 aver|OJJ04587.1 awen|OJJ35742.1 azon|OJJ43326.1

Group3722: abra|OJJ75404.1 acar|OOF90751.1 acla|EAW13776.1 afis|EAW18356.1 afum|EAL87719.1 agla|OJJ88029.1 akaw|GAA84182.1 aluc|OJZ88856.1 anid|CBF79356.1 anig|EHA23647.1 aory|EIT72985.1 asyd|OJJ57802.1 ater|EAU34689.1 atub|OJI85934.1 aver|OJJ04585.1 awen|OJJ35740.1 azon|OJJ43323.1 afla|EED48821.1

Group3723: abra|OJJ75408.1 acar|OOF90754.1 acla|EAW13781.1 afis|EAW18351.1 afla|EED48816.1 afum|EAL87714.1 agla|OJJ88033.1 akaw|GAA84184.1 aluc|OJZ88859.1 anid|CBF79365.1 anig|EHA23644.1

aory|EIT72998.1 asyd|OJJ57807.1 ater|EAU34685.1 atub|OJI85931.1 aver|OJJ04581.1 awen|OJJ35735.1 azon|OJJ43321.1

Group3724: abra|OJJ75409.1 acar|OOF90755.1 acla|EAW13780.1 afis|EAW18352.1 afla|EED48817.1 afum|EAL87715.1 agla|OJJ88032.1 akaw|GAA84185.1 aluc|OJZ88860.1 anid|CBF79364.1 anig|EHA23643.1 aory|EIT73011.1 asyd|OJJ57806.1 ater|EAU34686.1 atub|OJI85930.1 aver|OJJ04582.1 awen|OJJ35736.1 azon|OJJ43322.1

Group3725: abra|OJJ75410.1 acar|OOF90756.1 acla|EAW13782.1 afis|EAW18350.1 afla|EED48815.1 afum|EAL87713.1 agla|OJJ88034.1 akaw|GAA84186.1 aluc|OJZ88861.1 anid|CBF79367.1 anig|EHA23642.1 aory|EIT73016.1 asyd|OJJ57808.1 ater|EAU34684.1 atub|OJI85929.1 aver|OJJ04580.1 awen|OJJ35734.1 azon|OJJ43319.1

Group3726: abra|OJJ75413.1 acar|OOF90757.1 acla|EAW13785.1 afis|EAW18347.1 afla|EED48812.1 afum|EAL87711.1 agla|OJJ88037.1 akaw|GAA84188.1 aluc|OJZ88864.1 anid|CBF79386.1 anig|EHA23639.1 aory|EIT73027.1 asyd|OJJ57811.1 ater|EAU34682.1 atub|OJI85926.1 aver|OJJ04577.1 awen|OJJ35731.1 azon|OJJ43318.1

Group3727: abra|OJJ75414.1 acar|OOF90760.1 acla|EAW13786.1 afis|EAW18346.1 afla|EED48811.1 afum|EAL87710.1 agla|OJJ88038.1 akaw|GAA84189.1 aluc|OJZ88865.1 anid|CBF79384.1 anig|EHA23638.1 aory|EIT73037.1 asyd|OJJ57812.1 ater|EAU34681.1 atub|OJI85925.1 aver|OJJ04576.1 awen|OJJ35730.1 azon|OJJ43317.1

Group3728: abra|OJJ75415.1 acar|OOF90759.1 acla|EAW13787.1 afis|EAW18345.1 afla|EED48810.1 afum|EAL87709.1 agla|OJJ88039.1 akaw|GAA84190.1 aluc|OJZ88866.1 anid|CBF79382.1 anig|EHA23637.1 aory|EIT72992.1 asyd|OJJ57813.1 ater|EAU34680.1 atub|OJI85924.1 aver|OJJ04575.1 awen|OJJ35729.1 azon|OJJ43316.1

Group3729: abra|OJJ75417.1 acar|OOF90762.1 acla|EAW13789.1 afis|EAW18343.1 afla|EED48808.1 afum|EAL87706.1 agla|OJJ88041.1 akaw|GAA84192.1 aluc|OJZ88868.1 anid|CBF79378.1 anig|EHA23635.1 aory|EIT73022.1 asyd|OJJ57815.1 ater|EAU34678.1 atub|OJI85922.1 aver|OJJ04573.1 awen|OJJ35727.1 azon|OJJ43315.1

Group3730: abra|OJJ75421.1 acar|OOF90765.1 acla|EAW13793.1 afis|EAW18339.1 afla|EED48803.1 afum|EAL87702.1 agla|OJJ88045.1 akaw|GAA84196.1 aluc|OJZ88873.1 anid|CBF79387.1 anig|EHA23632.1 aory|EIT73019.1 asyd|OJJ57819.1 ater|EAU34674.1 atub|OJI85918.1 aver|OJJ04572.1 awen|OJJ35722.1 azon|OJJ43312.1

Group3731: abra|OJJ75425.1 acar|OOF90770.1 acla|EAW13795.1 afis|EAW18337.1 afla|EED48801.1 afum|EAL87701.1 agla|OJJ88047.1 akaw|GAA84200.1 aluc|OJZ88877.1 anid|CBF79391.1 anig|EHA23628.1 aory|EIT73024.1 asyd|OJJ57821.1 ater|EAU33350.1 atub|OJI85914.1 aver|OJJ04570.1 awen|OJJ35720.1 azon|OJJ43310.1

Group3732: abra|OJJ75426.1 acar|OOF90771.1 acla|EAW13796.1 afis|EAW18336.1 afla|EED48800.1 afum|EAL87700.1 agla|OJJ88049.1 akaw|GAA84201.1 aluc|OJZ88878.1 anid|CBF79395.1 anig|EHA23627.1 aory|EIT73005.1 asyd|OJJ57825.1 ater|EAU33349.1 atub|OJI85913.1 aver|OJJ04567.1 awen|OJJ30717.1 azon|OJJ43309.1

Group3733: abra|OJJ75427.1 acar|OOF90773.1 acla|EAW13798.1 afis|EAW18334.1 afla|EED48798.1 afum|EAL87699.1 agla|OJJ88051.1 akaw|GAA84203.1 aluc|OJZ88880.1 anid|CBF79393.1 anig|EHA23626.1 aory|EIT73035.1 asyd|OJJ57822.1 ater|EAU33347.1 atub|OJI85912.1 aver|OJJ04569.1 awen|OJJ30715.1 azon|OJJ43307.1

Group3734: abra|OJJ75428.1 acar|OOF90774.1 acla|EAW11898.1 afis|EAW19508.1 afla|EED56888.1 afum|EAL86986.1 agla|OJJ88052.1 akaw|GAA84204.1 aluc|OJZ88881.1 anid|CBF82976.1 anig|EHA23625.1 aory|EIT77571.1 asyd|OJJ61561.1 ater|EAU33346.1 atub|OJI85911.1 aver|OJJ00884.1 awen|OJJ30714.1 azon|OJJ45886.1

Group3735: abra|OJJ75429.1 acar|OOF90775.1 acla|EAW13799.1 afis|EAW18333.1 afla|EED48797.1 afum|EAL87698.1 agla|OJJ88053.1 akaw|GAA84205.1 aluc|OJZ88882.1 anid|CBF79397.1 anig|EHA23624.1 aory|EIT73004.1 asyd|OJJ57826.1 ater|EAU33345.1 atub|OJI85910.1 aver|OJJ04566.1 awen|OJJ30713.1 azon|OJJ43306.1

Group3736: abra|OJJ75432.1 acar|OOF90777.1 acla|EAW13804.1 afis|EAW18327.1 afla|EED48793.1 afum|EAL87693.1 agla|OJJ88056.1 akaw|GAA84208.1 aluc|OJZ88885.1 anid|CBF79406.1 anig|EHA23621.1 aory|EIT72996.1 asyd|OJJ57829.1 ater|EAU33336.1 atub|OJI85905.1 aver|OJJ04563.1 awen|OJJ30711.1 azon|OJJ43303.1

Group3737: abra|OJJ75433.1 acar|OOF90778.1 acla|EAW13805.1 afis|EAW18325.1 afla|EED48792.1 afum|EAL87691.1 agla|OJJ88066.1 akaw|GAA84209.1 aluc|OJZ88886.1 anid|CBF79408.1 anig|EHA23620.1 aory|EIT72982.1 asyd|OJJ57830.1 ater|EAU33334.1 atub|OJI85904.1 aver|OJJ04562.1 awen|OJJ30707.1 azon|OJJ43302.1

Group3738: abra|OJJ75435.1 acar|OOF90779.1 acla|EAW13827.1 afis|EAW18304.1 afla|EED48773.1 afum|EAL87670.1 agla|OJJ88119.1 akaw|GAA84210.1 aluc|OJZ88888.1 anid|CBF79447.1 anig|EHA23619.1 aory|EIT73020.1 asyd|OJJ57851.1 ater|EAU33311.1 atub|OJI85901.1 aver|OJJ04541.1 awen|OJJ30678.1 azon|OJJ43279.1

Group3739: abra|OJJ75436.1 acar|OOF90780.1 acla|EAW13826.1 afis|EAW18305.1 afla|EED48774.1 afum|EAL87671.1 agla|OJJ88118.1 akaw|GAA84211.1 aluc|OJZ88889.1 anid|CBF79445.1 anig|EHA23618.1 aory|EIT73030.1 asyd|OJJ57850.1 ater|EAU33312.1 atub|OJI85900.1 aver|OJJ04542.1 awen|OJJ30679.1 azon|OJJ43280.1

Group3740: abra|OJJ75438.1 acar|OOF90783.1 acla|EAW12400.1 afis|EAW16848.1 afla|EED51031.1 afum|EAL86394.1 agla|OJJ81486.1 akaw|GAA84213.1 aluc|OJZ88891.1 anid|CBF81652.1 anig|EHA23615.1 aory|EIT77036.1 asyd|OJJ59545.1 ater|EAU30241.1 atub|OJI85898.1 aver|OJJ02722.1 awen|OJJ30081.1 azon|OJJ44766.1

Group3741: abra|OJJ75442.1 acar|OOF90787.1 acla|EAW13821.1 afis|EAW18310.1 afla|EED48779.1 afum|EAL87676.1 agla|OJJ88079.1 akaw|GAA84217.1 aluc|OJZ88895.1 anid|CBF79434.1 anig|EHA23611.1 aory|EIT73023.1 asyd|OJJ57844.1 ater|EAU33320.1 atub|OJI85894.1 aver|OJJ04548.1 awen|OJJ30685.1 azon|OJJ43284.1

Group3742: abra|OJJ75443.1 acar|OOF90788.1 acla|EAW13820.1 afis|EAW18311.1 afla|EED48780.1 afum|EAL87677.1 agla|OJJ88080.1 akaw|GAA84218.1 aluc|OJZ88896.1 anid|CBF79432.1 anig|EHA23610.1 aory|EIT72999.1 asyd|OJJ57845.1 ater|EAU33321.1 atub|OJI85893.1 aver|OJJ04547.1 awen|OJJ30686.1 azon|OJJ43285.1

Group3743: abra|OJJ75445.1 acar|OOF90789.1 acla|EAW13818.1 afis|EAW18312.1 afla|EED48781.1 afum|EAL87679.1 agla|OJJ88081.1 akaw|GAA84220.1 aluc|OJZ88900.1 anid|CBF79430.1 anig|EHA23608.1 aory|EIT73002.1 asyd|OJJ57841.1 ater|EAU33322.1 atub|OJI85891.1 aver|OJJ04549.1 awen|OJJ30688.1 azon|OJJ43286.1

Group3744: abra|OJJ75446.1 acar|OOF90790.1 acla|EAW13817.1 afis|EAW18313.1 afla|EED48782.1 afum|EAL87680.1 agla|OJJ88082.1 akaw|GAA84221.1 aluc|OJZ88901.1 anid|CBF79428.1 anig|EHA23607.1 aory|EIT73015.1 asyd|OJJ57840.1 ater|EAU33323.1 atub|OJI85890.1 aver|OJJ04550.1 awen|OJJ30689.1 azon|OJJ43287.1

Group3745: abra|OJJ75447.1 acar|OOF90791.1 acla|EAW13816.1 afis|EAW18314.1 afla|EED48783.1 afum|EAL87681.1 agla|OJJ88083.1 akaw|GAA84222.1 aluc|OJZ88902.1 anid|CBF79427.1 anig|EHA23606.1 aory|EIT73014.1 asyd|OJJ57839.1 ater|EAU33324.1 atub|OJI85889.1 aver|OJJ04551.1 awen|OJJ30690.1 azon|OJJ43288.1

Group3746: abra|OJJ75448.1 acar|OOF90792.1 acla|EAW13815.1 afis|EAW18315.1 afla|EED48784.1 afum|EAL87682.1 agla|OJJ88084.1 akaw|GAA84223.1 aluc|OJZ88903.1 anid|CBF79423.1 anig|EHA23605.1 aory|EIT72980.1 asyd|OJJ57837.1 ater|EAU33325.1 atub|OJI85888.1 aver|OJJ04553.1 awen|OJJ30691.1 azon|OJJ43289.1

Group3747: abra|OJJ75450.1 acar|OOF90794.1 acla|EAW13813.1 afis|EAW18317.1 afla|EED48785.1 afum|EAL87684.1 agla|OJJ88101.1 akaw|GAA84225.1 aluc|OJZ88905.1 anid|CBF79421.1 anig|EHA23603.1 aory|EIT73000.1 asyd|OJJ57836.1 ater|EAU33326.1 atub|OJI85886.1 aver|OJJ04554.1 awen|OJJ30699.1 azon|OJJ43292.1

Group3748: abra|OJJ75452.1 acar|OOF90796.1 acla|EAW13811.1 afis|EAW18319.1 afla|EED48787.1 afum|EAL87686.1 agla|OJJ88073.1 akaw|GAA84227.1 aluc|OJZ88907.1 anid|CBF87726.1 anig|EHA23601.1 aory|EIT72981.1 asyd|OJJ63342.1 ater|EAU33329.1 atub|OJI85884.1 aver|OJI97619.1 awen|OJJ30701.1 azon|OJJ50771.1

Group3749: abra|OJJ75454.1 acar|OOF90798.1 acla|EAW13809.1 afis|EAW18321.1 afla|EED48789.1 afum|EAL87688.1 agla|OJJ88070.1 akaw|GAA84229.1 aluc|OJZ88909.1 anid|CBF79416.1 anig|EHA23599.1 aory|EIT73021.1 asyd|OJJ57833.1 ater|EAU33331.1 atub|OJI85882.1 aver|OJJ04557.1 awen|OJJ30703.1 azon|OJJ43296.1

Group3750: abra|OJJ75457.1 acar|OOF90802.1 acla|EAW13808.1 afis|EAW18322.1 afla|EED48790.1 afum|EAL87689.2 agla|OJJ88069.1 akaw|GAA84232.1 aluc|OJZ88913.1 anid|CBF79414.1 anig|EHA23597.1 aory|EIT73033.1 asyd|OJJ57832.1 ater|EAU33332.1 atub|OJI85878.1 aver|OJJ04558.1 awen|OJJ30704.1 azon|OJJ43297.1

Group3751: abra|OJJ75460.1 acar|OOF90279.1 acla|EAW13828.1 afis|EAW18303.1 afla|EED48772.1 afum|EAL87669.1 agla|OJJ88120.1 akaw|GAA84234.1 aluc|OJZ88917.1 anid|CBF79449.1 anig|EHA23595.1 aory|EIT72974.1 asyd|OJJ57852.1 ater|EAU33310.1 atub|OJI85875.1 aver|OJJ04540.1 awen|OJJ30677.1 azon|OJJ43278.1

Group3752: abra|OJJ75463.1 acar|OOF90277.1 acla|EAW13830.1 afis|EAW18301.1 afla|EED48770.1 afum|EAL87667.2 agla|OJJ88122.1 akaw|GAA84236.1 aluc|OJZ88919.1 anid|CBF79480.1 anig|EHA23593.1

aory|EIT72978.1 asyd|OJJ57871.1 ater|EAU33309.1 atub|OJI85873.1 aver|OJJ04523.1 awen|OJJ30676.1 azon|OJJ43276.1

Group3753: abra|OJJ75465.1 acar|OOF90275.1 acla|EAW13832.1 afis|EAW18299.1 afla|EED48768.1 afum|EAL87665.1 agla|OJJ88124.1 akaw|GAA84238.1 aluc|OJZ88921.1 anid|CBF79476.1 anig|EHA23591.1 aory|EIT73017.1 asyd|OJJ57869.1 ater|EAU33307.1 atub|OJI85871.1 aver|OJJ04525.1 awen|OJJ30673.1 azon|OJJ43274.1

Group3754: abra|OJJ75466.1 acar|OOF90803.1 acla|EAW13833.1 afis|EAW18298.1 afla|EED48767.1 afum|EAL87664.2 agla|OJJ88125.1 akaw|GAA84239.1 aluc|OJZ88922.1 anid|CBF79472.1 anig|EHA23590.1 aory|EIT73036.1 asyd|OJJ57867.1 ater|EAU33306.1 atub|OJI85870.1 aver|OJJ04526.1 awen|OJJ30672.1 azon|OJJ43272.1

Group3755: abra|OJJ75468.1 acar|OOF90805.1 acla|EAW13834.1 afis|EAW18297.1 afla|EED48763.1 afum|EAL87663.1 agla|OJJ88126.1 akaw|GAA84240.1 aluc|OJZ88925.1 anid|CBF79470.1 anig|EHA23588.1 aory|EIT73013.1 asyd|OJJ57865.1 ater|EAU33305.1 atub|OJI85867.1 aver|OJJ04528.1 awen|OJJ30671.1 azon|OJJ43271.1

Group3756: abra|OJJ75472.1 acar|OOF90809.1 acla|EAW13837.1 afis|EAW18294.1 afla|EED48761.1 afum|EAL87660.1 agla|OJJ88129.1 akaw|GAA84243.1 aluc|OJZ88931.1 anid|CBF79465.1 anig|EHA23585.1 aory|EIT73038.1 asyd|OJJ57861.1 ater|EAU33303.1 atub|OJI85861.1 aver|OJJ04531.1 awen|OJJ30669.1 azon|OJJ43268.1

Group3757: abra|OJJ75473.1 acar|OOF90810.1 acla|EAW13838.1 afis|EAW18293.1 afla|EED48760.1 afum|EAL87659.1 agla|OJJ88132.1 akaw|GAA84244.1 aluc|OJZ88932.1 anid|CBF79463.1 anig|EHA23584.1 aory|EIT72991.1 asyd|OJJ57860.1 ater|EAU33302.1 atub|OJI85860.1 aver|OJJ04532.1 awen|OJJ30668.1 azon|OJJ43267.1

Group3758: abra|OJJ75474.1 acar|OOF90811.1 acla|EAW13839.1 afis|EAW18292.1 afla|EED48759.1 afum|EAL87658.1 agla|OJJ88133.1 akaw|GAA84245.1 aluc|OJZ88933.1 anid|CBF79461.1 anig|EHA23583.1 aory|EIT72984.1 asyd|OJJ57859.1 ater|EAU33301.1 atub|OJI85859.1 aver|OJJ04534.1 awen|OJJ30667.1 azon|OJJ43265.1

Group3759: abra|OJJ75476.1 acar|OOF90812.1 acla|EAW13841.1 afis|EAW18291.1 afla|EED48758.1 afum|EAL87656.1 agla|OJJ88134.1 akaw|GAA84246.1 aluc|OJZ88935.1 anid|CBF79459.1 anig|EHA23582.1 aory|EIT73034.1 asyd|OJJ57858.1 ater|EAU33299.1 atub|OJI85857.1 aver|OJJ04535.1 awen|OJJ30666.1 azon|OJJ43263.1

Group3760: abra|OJJ75480.1 acar|OOF90816.1 acla|EAW11214.1 afis|EAW18288.1 afla|EED48756.1 afum|EAL87654.2 agla|OJJ88136.1 akaw|GAA84249.1 aluc|OJZ88939.1 anid|CBF79454.1 anig|EHA23579.1 aory|EIT73128.1 asyd|OJJ57856.1 ater|EAU33296.1 atub|OJI85851.1 aver|OJJ04538.1 awen|OJJ30661.1 azon|OJJ47954.1

Group3761: abra|OJJ75486.1 acar|OOF90818.1 acla|EAW11217.1 afis|EAW18286.1 afla|EED48753.1 afum|EAL87652.1 agla|OJJ88138.1 akaw|GAA84252.1 aluc|OJZ88945.1 anid|CBF79450.1 anig|EHA23576.1 aory|EIT73238.1 asyd|OJJ57853.1 ater|EAU33292.1 atub|OJI85844.1 aver|OJJ04539.1 awen|OJJ30658.1 azon|OJJ47956.1

Group3762: abra|OJJ75487.1 acar|OOF90820.1 acla|EAW11218.1 afis|EAW18285.1 afla|EED48752.1 afum|EAL87651.1 agla|OJJ88139.1 akaw|GAA84253.1 aluc|OJZ88946.1 anid|CBF79482.1 anig|EHA23575.1 aory|EIT73219.1 asyd|OJJ57872.1 ater|EAU33291.1 atub|OJI85843.1 aver|OJJ04522.1 awen|OJJ30657.1 azon|OJJ47957.1

Group3763: abra|OJJ75488.1 acar|OOF90819.1 acla|EAW11219.1 afis|EAW18284.1 afla|EED48751.1 afum|EAL87650.1 agla|OJJ88140.1 akaw|GAA84254.1 aluc|OJZ88947.1 anid|CBF79483.1 anig|EHA23574.1 aory|EIT73141.1 asyd|OJJ57873.1 ater|EAU33290.1 atub|OJI85842.1 aver|OJJ04521.1 awen|OJJ30655.1 azon|OJJ47959.1

Group3764: abra|OJJ75491.1 acar|OOF90823.1 acla|EAW11223.1 afis|EAW18280.1 afla|EED48747.1 afum|EAL87646.1 agla|OJJ88146.1 akaw|GAA84256.1 aluc|OJZ88950.1 anid|CBF79491.1 anig|EHA23571.1 aory|EIT73260.1 asyd|OJJ57876.1 ater|EAU33286.1 atub|OJI85839.1 aver|OJJ04518.1 awen|OJJ30652.1 azon|OJJ47963.1

Group3765: abra|OJJ75493.1 acar|OOF90826.1 acla|EAW11226.1 afis|EAW18277.1 afla|EED48744.1 afum|EAL87643.1 agla|OJJ88149.1 akaw|GAA84259.1 aluc|OJZ88952.1 anid|CBF79496.1 anig|EHA23568.1 aory|EIT73150.1 asyd|OJJ57878.1 ater|EAU33283.1 atub|OJI85837.1 aver|OJJ04516.1 awen|OJJ30650.1 azon|OJJ47965.1

Group3766: abra|OJJ75494.1 acar|OOF90827.1 acla|EAW09288.1 afis|EAW21859.1 afla|EED46451.1 afum|EAL92872.1 agla|OJJ84276.1 akaw|GAA84260.1 aluc|OJZ88953.1 anid|CBF76751.1 anig|EHA23567.1 aory|EIT72375.1 asyd|OJJ61260.1 ater|EAU30858.1 atub|OJI85836.1 aver|OJJ00583.1 awen|OJJ34143.1 azon|OJJ43483.1

Group3767: abra|OJJ75495.1 acar|OOF90828.1 acla|EAW09289.1 afis|EAW21858.1 afla|EED46452.1 afum|EAL92871.1 agla|OJJ84277.1 akaw|GAA84261.1 aluc|OJZ88954.1 anid|CBF76749.1 anig|EHA23566.1 aory|EIT72324.1 asyd|OJJ61259.1 ater|EAU30857.1 atub|OJI85835.1 aver|OJJ00582.1 awen|OJJ34142.1 azon|OJJ43484.1

Group3768: abra|OJJ75496.1 acar|OOF90829.1 acla|EAW09290.1 afis|EAW21857.1 afla|EED46453.1 afum|EAL92870.1 agla|OJJ84278.1 akaw|GAA84262.1 aluc|OJZ88955.1 anid|CBF76747.1 anig|EHA23565.1 aory|EIT72382.1 asyd|OJJ61258.1 ater|EAU30856.1 atub|OJI85834.1 aver|OJJ00581.1 awen|OJJ34141.1 azon|OJJ43485.1

Group3769: abra|OJJ75497.1 acar|OOF90830.1 acla|EAW09291.1 afis|EAW21856.1 afla|EED46454.1 afum|EAL92869.1 agla|OJJ84279.1 akaw|GAA84263.1 aluc|OJZ88956.1 anid|CBF76745.1 aory|EIT72446.1 asyd|OJJ61257.1 ater|EAU30855.1 atub|OJI85833.1 aver|OJJ00580.1 awen|OJJ34140.1 azon|OJJ43486.1 anig|EHA23564.1

Group3770: abra|OJJ75506.1 acar|OOF89956.1 acla|EAW09296.1 afis|EAW21851.1 afla|EED46459.1 afum|EAL92864.1 agla|OJJ84284.1 akaw|GAA84269.1 aluc|OJZ88962.1 anid|CBF76735.1 anig|EHA23559.1 aory|EIT72419.1 asyd|OJJ61252.1 ater|EAU30850.1 atub|OJI85828.1 aver|OJJ00576.1 awen|OJJ34134.1 azon|OJJ44210.1

Group3771: abra|OJJ75507.1 acar|OOF89955.1 acla|EAW09297.1 afis|EAW21850.1 afla|EED46460.1 afum|EAL92863.2 agla|OJJ84285.1 akaw|GAA84270.1 aluc|OJZ88963.1 anid|CBF76733.1 anig|EHA23558.1 aory|EIT72388.1 asyd|OJJ61251.1 ater|EAU30849.1 atub|OJI85827.1 aver|OJJ00575.1 awen|OJJ34133.1 azon|OJJ46409.1

Group3772: abra|OJJ75511.1 acar|OOF90837.1 acla|EAW09299.1 afis|EAW21848.1 afum|EAL92861.2 agla|OJJ84287.1 akaw|GAA84275.1 aluc|OJZ88968.1 anid|CBF76727.1 anig|EHA23555.1 asyd|OJJ61248.1 ater|EAU30847.1 atub|OJI85822.1 aver|OJJ00572.1 awen|OJJ34131.1 azon|OJJ44212.1 afla|EED46464.1 aory|EIT72335.1

Group3773: abra|OJJ75512.1 acar|OOF90839.1 acla|EAW09300.1 afis|EAW21847.1 afla|EED46468.1 afum|EAL92860.1 agla|OJJ84288.1 akaw|GAA84276.1 aluc|OJZ88969.1 anid|CBF76725.1 anig|EHA23554.1 aory|EIT72394.1 asyd|OJJ61246.1 ater|EAU30846.1 atub|OJI85821.1 aver|OJJ00570.1 awen|OJJ34130.1 azon|OJJ44213.1

Group3774: abra|OJJ75518.1 acar|OOF90845.1 acla|EAW09302.1 afis|EAW21844.1 afla|EED46526.1 afum|EAL92856.1 agla|OJJ84292.1 akaw|GAA84282.1 aluc|OJZ88975.1 anid|CBF76717.1 anig|EHA23550.1 aory|EIT72426.1 asyd|OJJ61242.1 ater|EAU38521.1 atub|OJI85814.1 aver|OJJ00565.1 awen|OJJ34128.1 azon|OJJ44215.1

Group3775: abra|OJJ75521.1 acar|OOF90848.1 acla|EAW09308.1 afis|EAW21837.1 afla|EED46538.1 afum|EAL92851.1 agla|OJJ84305.1 akaw|GAA84285.1 aluc|OJZ88978.1 anid|CBF76701.1 anig|EHA23547.1 aory|EIT72362.1 asyd|OJJ61233.1 ater|EAU38512.1 atub|OJI85811.1 aver|OJJ00556.1 awen|OJJ34113.1 azon|OJJ44227.1

Group3776: abra|OJJ75523.1 acar|OOF90850.1 acla|EAW09310.1 afis|EAW21835.1 afla|EED46540.1 afum|EAL92850.1 agla|OJJ84309.1 akaw|GAA84286.1 aluc|OJZ88980.1 anid|CBF76699.1 anig|EHA23546.1 aory|EIT72364.1 asyd|OJJ61232.1 ater|EAU38510.1 atub|OJI85809.1 aver|OJJ00555.1 awen|OJJ34109.1 azon|OJJ44232.1

Group3777: abra|OJJ75527.1 acar|OOF90854.1 acla|EAW09313.1 afis|EAW21832.1 afla|EED46544.1 afum|EAL92847.1 agla|OJJ84312.1 akaw|GAA84289.1 aluc|OJZ88983.1 anid|CBF76694.1 anig|EHA23543.1 aory|EIT77942.1 asyd|OJJ61229.1 ater|EAU38508.1 atub|OJI85806.1 aver|OJJ00552.1 awen|OJJ34105.1 azon|OJJ44236.1

Group3778: abra|OJJ75528.1 acar|OOF90855.1 acla|EAW09314.1 afis|EAW21831.1 afla|EED46545.1 afum|EAL92846.1 agla|OJJ84313.1 akaw|GAA84290.1 aluc|OJZ88984.1 anid|CBF76680.1 anig|EHA23542.1 aory|EIT77881.1 asyd|OJJ61223.1 ater|EAU38507.1 atub|OJI85804.1 aver|OJJ00546.1 awen|OJJ34104.1 azon|OJJ44238.1

Group3779: abra|OJJ75539.1 acar|OOF90858.1 acla|EAW09315.1 afis|EAW21830.1 afla|EED46546.1 afum|EAL92845.1 agla|OJJ84314.1 akaw|GAA84293.1 aluc|OJZ88987.1 anid|CBF76676.1 anig|EHA23539.1 aory|EIT77971.1 asyd|OJJ61221.1 ater|EAU38506.1 atub|OJI85801.1 aver|OJJ00545.1 awen|OJJ34103.1 azon|OJJ44239.1

Group3780: abra|OJJ75544.1 acar|OOF90864.1 acla|EAW09316.1 afis|EAW21829.1 afla|EED46547.1 afum|EAL92844.1 agla|OJJ84315.1 akaw|GAA84300.1 aluc|OJZ88994.1 anid|CBF76674.1 anig|EHA23533.1 aory|EIT77834.1 asyd|OJJ61220.1 ater|EAU38505.1 atub|OJI85794.1 aver|OJJ00544.1 awen|OJJ34102.1 azon|OJJ44242.1

Group3781: abra|OJJ75545.1 acar|OOF90866.1 acla|EAW09317.1 afis|EAW21828.1 afla|EED46548.1 afum|EAL92843.1 agla|OJJ84316.1 akaw|GAA84301.1 aluc|OJZ88996.1 anid|CBF76670.1 anig|EHA23532.1

aory|EIT77919.1 asyd|OJJ61219.1 ater|EAU38504.1 atub|OJI85793.1 aver|OJJ00543.1 awen|OJJ34101.1 azon|OJJ44243.1

Group3782: abra|OJJ75548.1 acar|OOF90868.1 acla|EAW09319.1 afis|EAW21826.1 afla|EED46553.1 afum|EAL92841.1 agla|OJJ84318.1 akaw|GAA84303.1 aluc|OJZ88998.1 anid|CBF76666.1 anig|EHA23530.1 aory|EIT77928.1 asyd|OJJ61217.1 ater|EAU38502.1 atub|OJI85791.1 aver|OJJ00541.1 awen|OJJ34099.1 azon|OJJ44245.1

Group3783: abra|OJJ75552.1 acar|OOF90870.1 acla|EAW09324.1 afis|EAW21821.1 afla|EED46558.1 afum|EAL92837.1 agla|OJJ84321.1 akaw|GAA84307.1 aluc|OJZ89002.1 anid|CBF76660.1 anig|EHA23526.1 aory|EIT77966.1 asyd|OJJ61212.1 ater|EAU38495.1 atub|OJI85787.1 aver|OJJ00537.1 awen|OJJ34095.1 azon|OJJ44249.1

Group3784: abra|OJJ75553.1 acar|OOF90872.1 acla|EAW09355.1 afis|EAW21820.1 afla|EED46559.1 afum|EAL92836.1 agla|OJJ84322.1 akaw|GAA84308.1 aluc|OJZ89003.1 anid|CBF76658.1 anig|EHA23525.1 aory|EIT77877.1 asyd|OJJ61211.1 ater|EAU38494.1 atub|OJI85786.1 aver|OJJ00535.1 awen|OJJ34094.1 azon|OJJ44252.1

Group3785: abra|OJJ75554.1 acar|OOF90873.1 acla|EAW09354.1 afis|EAW21819.1 afla|EED46564.1 afum|EAL92835.1 agla|OJJ84327.1 akaw|GAA84309.1 aluc|OJZ89004.1 anid|CBF76654.1 anig|EHA23524.1 aory|EIT77873.1 asyd|OJJ61210.1 ater|EAU38490.1 atub|OJI85785.1 aver|OJJ00534.1 awen|OJJ34093.1 azon|OJJ44253.1

Group3786: abra|OJJ75574.1 acar|OOF90890.1 acla|EAW09346.1 afis|EAW21799.1 afla|EED46590.1 afum|EAL92824.2 agla|OJJ84324.1 akaw|GAA84325.1 aluc|OJZ89022.1 anid|CBF76633.1 anig|EHA23509.1 aory|EIT77934.1 asyd|OJJ61197.1 ater|EAU38481.1 atub|OJI85767.1 aver|OJJ00521.1 awen|OJJ34086.1 azon|OJJ44256.1

Group3787: abra|OJJ75575.1 acar|OOF90893.1 acla|EAW09343.1 afis|EAW21797.1 afla|EED46598.1 afum|EAL92822.2 agla|OJJ84331.1 akaw|GAA84327.1 aluc|OJZ89024.1 anid|CBF76629.1 anig|EHA23508.1 aory|EIT77937.1 asyd|OJJ61196.1 ater|EAU38479.1 atub|OJI85766.1 aver|OJJ00520.1 awen|OJJ34082.1 azon|OJJ44260.1

Group3788: akaw|GAA84326.1 aluc|OJZ89023.1 acar|OOF90891.1 acla|EAW09345.1 afis|EAW21798.1 afla|EED46597.1 afum|EAL92823.2 agla|OJJ84323.1 aory|EIT77884.1 awen|OJJ34085.1 acar|OOF90892.1 acla|EAW09344.1 agla|OJJ84329.1 anid|CBF76631.1 aory|EIT77853.1 ater|EAU38480.1 awen|OJJ34084.1 azon|OJJ44257.1

Group3789: abra|OJJ75579.1 acar|OOF90896.1 acla|EAW09340.1 afis|EAW21795.1 afla|EED46600.1 afum|EAL92820.1 agla|OJJ84333.1 akaw|GAA84331.1 aluc|OJZ89030.1 anid|CBF76625.1 anig|EHA23504.1 aory|EIT77931.1 asyd|OJJ61194.1 ater|EAU38477.1 atub|OJI85761.1 aver|OJJ00518.1 azon|OJJ44262.1 awen|OJJ34079.1

Group3790: abra|OJJ75581.1 acar|OOF90899.1 acla|EAW09337.1 afis|EAW21791.1 afla|EED46604.1 afum|EAL92816.1 agla|OJJ88519.1 akaw|GAA84333.1 aluc|OJZ89032.1 anid|CBF76623.1 anig|EHA23502.1 aory|EIT77970.1 asyd|OJJ61193.1 ater|EAU38475.1 atub|OJI85758.1 aver|OJJ00517.1 awen|OJJ37963.1 azon|OJJ46824.1

Group3791: abra|OJJ75592.1 acar|OOG00236.1 acla|EAW09330.1 afis|EAW21786.1 afla|EED46610.1 afum|EAL92813.1 agla|OJJ84337.1 akaw|GAA84344.1 aluc|OJZ89044.1 anid|CBF76617.1 anig|EHA23491.1 aory|EIT77835.1 asyd|OJJ61189.1 ater|EAU38469.1 atub|OJI85743.1 aver|OJJ00513.1 awen|OJJ34072.1 azon|OJJ48155.1

Group3792: abra|OJJ75602.1 acar|OOG00235.1 acla|EAW09327.1 afis|EAW21784.1 afla|EED46611.1 afum|EAL92811.1 agla|OJJ84339.1 akaw|GAA84352.1 aluc|OJZ89052.1 anid|CBF76613.1 anig|EHA23486.1 aory|EIT77805.1 asyd|OJJ61188.1 ater|EAU38467.1 atub|OJI85735.1 aver|OJJ00511.1 awen|OJJ34071.1 azon|OJJ48157.1

Group3793: abra|OJJ75605.1 acar|OOG00232.1 acla|EAW09367.1 afis|EAW21770.1 afla|EED46330.1 afum|EAL92799.1 agla|OJJ84378.1 akaw|GAA84356.1 aluc|OJZ89058.1 anid|CBF83758.1 anig|EHA23484.1 aory|EIT81243.1 asyd|OJJ60861.1 ater|EAU38448.1 atub|OJI85728.1 aver|OJJ00187.1 awen|OJJ34057.1 azon|OJJ48175.1

Group3794: abra|OJJ75610.1 acar|OOG00228.1 acla|EAW09372.1 afis|EAW21763.1 afla|EED46336.1 afum|EAL92793.1 agla|OJJ84388.1 akaw|GAA84360.1 aluc|OJZ89062.1 anid|CBF83751.1 anig|EHA23476.1 aory|EIT81185.1 asyd|OJJ60857.1 ater|EAU38452.1 atub|OJI85724.1 aver|OJJ00181.1 awen|OJJ34046.1 azon|OJJ48185.1

Group3795: abra|OJJ75611.1 acar|OOG00227.1 acla|EAW09373.1 afis|EAW21762.1 afla|EED46337.1 afum|EAL92792.1 agla|OJJ84389.1 akaw|GAA84361.1 aluc|OJZ89063.1 anid|CBF83749.1 anig|EHA23475.1 aory|EIT81205.1 asyd|OJJ60856.1 ater|EAU38453.1 atub|OJI85723.1 aver|OJJ00180.1 awen|OJJ34045.1 azon|OJJ48186.1

Group3796: abra|OJJ75615.1 acar|OOG00215.1 acla|EAW09363.1 afis|EAW21774.1 afla|EED46340.1 afum|EAL92803.1 agla|OJJ84350.1 akaw|GAA84364.1 aluc|OJZ89068.1 anid|CBF76597.1 anig|EHA23468.1 aory|EIT81181.1 asyd|OJJ61178.1 ater|EAU38464.1 atub|OJI85720.1 aver|OJJ00504.1 awen|OJJ34062.1 azon|OJJ48170.1

Group3797: abra|OJJ75625.1 acar|OOG00211.1 acla|EAW09360.1 afis|EAW21777.1 afla|EED46344.1 afum|EAL92806.1 agla|OJJ84342.1 akaw|GAA84372.1 aluc|OJZ89076.1 anid|CBF76603.1 anig|EHA23461.1 aory|EIT81165.1 asyd|OJJ61182.1 ater|EAU38462.1 atub|OJI85709.1 aver|OJJ00507.1 awen|OJJ34066.1 azon|OJJ48165.1

Group3798: abra|OJJ75628.1 acar|OOG00210.1 acla|EAW09465.1 afis|EAW21671.1 afla|EED57935.1 afum|EAL92706.1 agla|OJJ79745.1 akaw|GAA84375.1 aluc|OJZ89079.1 anid|CBF83585.1 anig|EHA23458.1 aory|EIT82333.1 asyd|OJJ60779.1 ater|EAU38355.1 atub|OJI85706.1 aver|OJJ00112.1 awen|OJJ33935.1 azon|OJJ48642.1

Group3799: abra|OJJ75630.1 acar|OOG00205.1 acla|EAW09463.1 afis|EAW21674.1 afla|EED57936.1 afum|EAL92708.1 agla|OJJ79747.1 akaw|GAA84377.1 aluc|OJZ89081.1 anid|CBF83589.1 anig|EHA23456.1 aory|EIT82341.1 asyd|OJJ60776.1 ater|EAU38357.1 atub|OJI85703.1 aver|OJJ00110.1 awen|OJJ33940.1 azon|OJJ48629.1

Group3800: abra|OJJ75633.1 acar|OOG00202.1 acla|EAW09461.1 agla|OJJ79927.1 akaw|GAA84380.1 aluc|OJZ89084.1 anig|EHA23453.1 asyd|OJJ60773.1 ater|EAU38359.1 atub|OJI85700.1 aver|OJJ00107.1 awen|OJJ33943.1 afla|EED57941.1 anid|CBF83591.1 afis|EAW21676.1 afum|EAL92710.1 aory|EIT82327.1 azon|OJJ48627.1

Group3801: abra|OJJ75634.1 acar|OOG00201.1 acla|EAW09460.1 afis|EAW21677.1 afla|EED57942.1 afum|EAL92711.1 agla|OJJ79926.1 akaw|GAA84381.1 aluc|OJZ89085.1 anid|CBF83593.1 anig|EHA23452.1 aory|EIT82349.1 asyd|OJJ60772.1 ater|EAU38360.1 atub|OJI85699.1 aver|OJJ00106.1 awen|OJJ33944.1 azon|OJJ48626.1

Group3802: abra|OJJ75636.1 acar|OOG00200.1 acla|EAW09459.1 afis|EAW21678.1 afla|EED57943.1 afum|EAL92712.1 agla|OJJ79925.1 akaw|GAA84382.1 aluc|OJZ89086.1 anid|CBF83595.1 anig|EHA23451.1 aory|EIT82316.1 asyd|OJJ60771.1 ater|EAU38361.1 atub|OJI85698.1 aver|OJJ00105.1 awen|OJJ33945.1 azon|OJJ48625.1

Group3803: abra|OJJ75637.1 acar|OOG00199.1 acla|EAW09458.1 afis|EAW21679.1 afla|EED57944.1 afum|EAL92713.1 agla|OJJ79924.1 akaw|GAA84383.1 aluc|OJZ89087.1 anid|CBF83597.1 anig|EHA23450.1 aory|EIT82325.1 asyd|OJJ60770.1 ater|EAU38362.1 atub|OJI85697.1 aver|OJJ00104.1 awen|OJJ33946.1 azon|OJJ48624.1

Group3804: abra|OJJ75639.1 acar|OOG00198.1 acla|EAW09457.1 afis|EAW21680.1 afla|EED57945.1 afum|EAL92714.1 agla|OJJ79923.1 akaw|GAA84384.1 aluc|OJZ89088.1 anid|CBF83598.1 anig|EHA23449.1 aory|EIT82335.1 asyd|OJJ60769.1 ater|EAU38363.1 atub|OJI85694.1 aver|OJJ00103.1 awen|OJJ33947.1 azon|OJJ48623.1

Group3805: abra|OJJ75642.1 acar|OOG00196.1 acla|EAW09456.1 afis|EAW21681.1 afla|EED57946.1 afum|EAL92715.1 agla|OJJ79921.1 akaw|GAA84386.1 aluc|OJZ89091.1 anid|CBF83600.1 anig|EHA23447.1 aory|EIT82326.1 asyd|OJJ60768.1 ater|EAU38364.1 atub|OJI85692.1 aver|OJJ00102.1 awen|OJJ33948.1 azon|OJJ48622.1

Group3806: abra|OJJ75644.1 acar|OOG00195.1 acla|EAW09454.1 afis|EAW21683.1 afla|EED57950.1 afum|EAL92717.1 agla|OJJ79918.1 akaw|GAA84389.1 aluc|OJZ89093.1 anid|CBF83605.1 anig|EHA23444.1 aory|EIT82322.1 asyd|OJJ60766.1 ater|EAU38367.1 atub|OJI85688.1 aver|OJJ00100.1 awen|OJJ33950.1 azon|OJJ48620.1

Group3807: abra|OJJ75645.1 acar|OOG00194.1 acla|EAW09453.1 afis|EAW21684.1 afla|EED57951.1 afum|EAL92718.2 agla|OJJ79917.1 akaw|GAA84390.1 aluc|OJZ89094.1 anid|CBF83607.1 anig|EHA23443.1 aory|EIT82348.1 asyd|OJJ60765.1 ater|EAU38368.1 atub|OJI85687.1 aver|OJJ00099.1 awen|OJJ33951.1 azon|OJJ48619.1

Group3808: abra|OJJ75650.1 acar|OOG00190.1 acla|EAW09452.1 afis|EAW21685.1 afla|EED57956.1 afum|EAL92719.1 agla|OJJ79914.1 akaw|GAA84393.1 aluc|OJZ89098.1 anid|CBF83611.1 anig|EHA23440.1 aory|EIT79731.1 asyd|OJJ60762.1 ater|EAU38371.1 atub|OJI85683.1 aver|OJJ00096.1 awen|OJJ33955.1 azon|OJJ48615.1

Group3809: abra|OJJ75651.1 acar|OOG00189.1 acla|EAW09449.1 afis|EAW21687.1 afla|EED57957.1 afum|EAL92721.2 agla|OJJ79913.1 akaw|GAA84394.1 aluc|OJZ89099.1 anid|CBF83612.1 anig|EHA23439.1 aory|EIT79765.1 asyd|OJJ60761.1 ater|EAU38372.1 atub|OJI85682.1 aver|OJJ00095.1 awen|OJJ33956.1 azon|OJJ48612.1

Group3810: abra|OJJ75652.1 acar|OOG00188.1 acla|EAW09448.1 afis|EAW21688.1 afla|EED57958.1 afum|EAL92722.1 agla|OJJ79912.1 akaw|GAA84395.1 aluc|OJZ89100.1 anid|CBF83614.1 anig|EHA23438.1

aory|EIT79734.1 asyd|OJJ60760.1 ater|EAU38373.1 atub|OJI85681.1 aver|OJJ00094.1 awen|OJJ33957.1 azon|OJJ48611.1

Group3811: abra|OJJ75654.1 acar|OOG00187.1 acla|EAW09446.1 afis|EAW21690.1 afla|EED57960.1 afum|EAL92724.1 agla|OJJ79910.1 akaw|GAA84397.1 aluc|OJZ89103.1 anid|CBF83617.1 anig|EHA23436.1 aory|EIT79781.1 asyd|OJJ60758.1 ater|EAU38375.1 atub|OJI85678.1 aver|OJJ00092.1 awen|OJJ33959.1 azon|OJJ48609.1

Group3812: abra|OJJ75656.1 acar|OOG00186.1 acla|EAW09444.1 afis|EAW21692.1 afla|EED57962.1 afum|EAL92726.1 agla|OJJ79907.1 akaw|GAA84399.1 aluc|OJZ89106.1 anid|CBF83621.1 anig|EHA23434.1 aory|EIT79782.1 asyd|OJJ60756.1 ater|EAU38377.1 atub|OJI85676.1 aver|OJJ00090.1 awen|OJJ33961.1 azon|OJJ48607.1

Group3813: abra|OJJ75657.1 acar|OOG00185.1 acla|EAW09442.1 afis|EAW21694.1 afla|EED57963.1 afum|EAL92728.1 agla|OJJ79905.1 akaw|GAA84400.1 aluc|OJZ89107.1 anid|CBF83623.1 anig|EHA23433.1 aory|EIT79708.1 asyd|OJJ60755.1 ater|EAU38379.1 atub|OJI85675.1 aver|OJJ00089.1 awen|OJJ33963.1 azon|OJJ48605.1

Group3814: abra|OJJ75658.1 acar|OOG00184.1 acla|EAW09441.1 afis|EAW21695.1 afla|EED57964.1 afum|EAL92729.1 agla|OJJ79904.1 akaw|GAA84401.1 aluc|OJZ89109.1 anid|CBF83624.1 anig|EHA23432.1 aory|EIT79788.1 asyd|OJJ60754.1 ater|EAU38380.1 atub|OJI85674.1 aver|OJJ00088.1 awen|OJJ33964.1 azon|OJJ48603.1

Group3815: abra|OJJ75662.1 acar|OOG00181.1 acla|EAW09439.1 afis|EAW21697.1 afla|EED57966.1 afum|EAL92731.1 agla|OJJ79901.1 akaw|GAA84403.1 aluc|OJZ89113.1 anid|CBF83628.1 anig|EHA23430.1 aory|EIT79705.1 asyd|OJJ60752.1 ater|EAU38382.1 atub|OJI85671.1 aver|OJJ00085.1 awen|OJJ33966.1 azon|OJJ48600.1

Group3816: abra|OJJ75664.1 acar|OOG00180.1 acla|EAW09437.1 afis|EAW21699.1 afla|EED57968.1 afum|EAL92733.1 agla|OJJ79899.1 akaw|GAA84405.1 aluc|OJZ89115.1 anid|CBF83632.1 anig|EHA23428.1 aory|EIT79768.1 asyd|OJJ60750.1 ater|EAU38384.1 atub|OJI85669.1 aver|OJJ00083.1 awen|OJJ33968.1 azon|OJJ48598.1

Group3817: abra|OJJ75665.1 acar|OOG00179.1 acla|EAW09436.1 afis|EAW21700.1 afla|EED57969.1 afum|EAL92734.1 agla|OJJ79898.1 akaw|GAA84406.1 aluc|OJZ89116.1 anid|CBF83633.1 anig|EHA23427.1 aory|EIT79802.1 asyd|OJJ60782.1 ater|EAU38385.1 atub|OJI85668.1 aver|OJJ00115.1 awen|OJJ33970.1 azon|OJJ48589.1

Group3818: abra|OJJ75667.1 acar|OOG00177.1 acla|EAW09434.1 afis|EAW21702.1 afla|EED57971.1 afum|EAL92736.1 agla|OJJ79896.1 akaw|GAA84408.1 aluc|OJZ89118.1 anid|CBF83639.1 anig|EHA23425.1 aory|EIT79763.1 asyd|OJJ60785.1 ater|EAU38387.1 atub|OJI85666.1 aver|OJJ00118.1 awen|OJJ33972.1 azon|OJJ48587.1

Group3819: abra|OJJ75668.1 acar|OOG00176.1 acla|EAW09433.1 afis|EAW21703.1 afla|EED57972.1 afum|EAL92737.1 agla|OJJ84485.1 akaw|GAA84409.1 aluc|OJZ89119.1 anid|CBF83649.1 anig|EHA23424.1 aory|EIT79804.1 asyd|OJJ60792.1 ater|EAU38388.1 atub|OJI85665.1 aver|OJJ00125.1 awen|OJJ33973.1 azon|OJJ48586.1

Group3820: abra|OJJ75670.1 acar|OOG00175.1 acla|EAW09432.1 afis|EAW21704.1 afla|EED57973.1 afum|EAL92738.1 agla|OJJ79888.1 akaw|GAA84410.1 aluc|OJZ89120.1 anid|CBF83648.1 anig|EHA23423.1 aory|EIT79806.1 asyd|OJJ60791.1 ater|EAU38389.1 atub|OJI85664.1 aver|OJJ00124.1 awen|OJJ33975.1 azon|OJJ48581.1

Group3821: abra|OJJ75672.1 acar|OOG00173.1 acla|EAW09425.1 afis|EAW21711.1 afla|EED57975.1 afum|EAL92746.1 agla|OJJ79890.1 akaw|GAA84413.1 aluc|OJZ89122.1 anid|CBF83646.1 anig|EHA23421.1 aory|EIT79737.1 asyd|OJJ60789.1 ater|EAU38391.1 atub|OJI85662.1 aver|OJJ00122.1 awen|OJJ33977.1 azon|OJJ48583.1

Group3822: abra|OJJ75673.1 acar|OOG00172.1 acla|EAW09426.1 afis|EAW21710.1 afla|EED57976.1 afum|EAL92745.1 agla|OJJ79891.1 akaw|GAA84414.1 aluc|OJZ89123.1 anid|CBF83644.1 anig|EHA23420.1 aory|EIT79753.1 asyd|OJJ60788.1 ater|EAU38392.1 atub|OJI85661.1 aver|OJJ00121.1 awen|OJJ33980.1 azon|OJJ48575.1

Group3823: abra|OJJ75674.1 acar|OOG00171.1 acla|EAW09427.1 afis|EAW21709.1 afla|EED57977.1 afum|EAL92744.1 agla|OJJ79892.1 akaw|GAA84415.1 aluc|OJZ89124.1 anid|CBF83642.1 anig|EHA23419.1 aory|EIT79730.1 asyd|OJJ60787.1 ater|EAU38393.1 atub|OJI85660.1 aver|OJJ00120.1 awen|OJJ33981.1 azon|OJJ48574.1

Group3824: abra|OJJ75675.1 acar|OOG00170.1 acla|EAW09428.1 afis|EAW21708.1 afla|EED57978.1 afum|EAL92743.1 agla|OJJ79893.1 akaw|GAA84416.1 aluc|OJZ89125.1 anid|CBF83641.1 anig|EHA23418.1 aory|EIT79718.1 asyd|OJJ60786.1 ater|EAU38394.1 atub|OJI85659.1 aver|OJJ00119.1 awen|OJJ33982.1 azon|OJJ48573.1

Group3825: abra|OJJ75677.1 acar|OOG00169.1 acla|EAW09423.1 afis|EAW21712.1 afla|EED57980.1 afum|EAL92747.1 agla|OJJ79895.1 akaw|GAA84418.1 aluc|OJZ89127.1 anid|CBF83653.1 anig|EHA23416.1 aory|EIT79727.1 asyd|OJJ60794.1 ater|EAU38396.1 atub|OJI85657.1 aver|OJJ00127.1 awen|OJJ33984.1 azon|OJJ48570.1

Group3826: abra|OJJ75679.1 acar|OOG00168.1 acla|EAW09424.1 afis|EAW21713.1 afla|EED57981.1 afum|EAL92748.1 agla|OJJ79886.1 akaw|GAA84419.1 aluc|OJZ89129.1 anid|CBF83655.1 anig|EHA23415.1 aory|EIT79767.1 asyd|OJJ60795.1 ater|EAU38398.1 atub|OJI85656.1 aver|OJJ00128.1 awen|OJJ33985.1 azon|OJJ48568.1

Group3827: abra|OJJ75680.1 acar|OOG00166.1 acla|EAW09422.1 afis|EAW21714.1 afla|EED57984.1 afum|EAL92749.1 agla|OJJ79885.1 akaw|GAA84420.1 aluc|OJZ89130.1 anid|CBF83657.1 anig|EHA23414.1 aory|EIT79791.1 asyd|OJJ60796.1 ater|EAU38399.1 atub|OJI85655.1 aver|OJJ00129.1 awen|OJJ33986.1 azon|OJJ48567.1

Group3828: abra|OJJ75682.1 acla|EAW09420.1 afis|EAW21716.1 afla|EED57986.1 afum|EAL92750.1 agla|OJJ79882.1 agla|OJJ79883.1 akaw|GAA84422.1 aluc|OJZ89133.1 anid|CBF83658.1 anig|EHA23412.1 aory|EIT79755.1 asyd|OJJ60798.1 ater|EAU38401.1 atub|OJI85653.1 aver|OJJ00131.1 awen|OJJ33988.1 azon|OJJ48565.1

Group3829: abra|OJJ75686.1 acar|OOG00164.1 acla|EAW09419.1 afis|EAW21717.1 afla|EED57987.1 afum|EAL92751.1 agla|OJJ79880.1 akaw|GAA84424.1 aluc|OJZ89137.1 anid|CBF83708.1 anig|EHA23410.1 aory|EIT79780.1 asyd|OJJ60831.1 ater|EAU38402.1 atub|OJI85648.1 aver|OJJ00156.1 awen|OJJ33990.1 azon|OJJ48563.1

Group3830: abra|OJJ75687.1 acar|OOG00163.1 acla|EAW09417.1 afis|EAW21719.1 afla|EED57990.1 afum|EAL92752.1 agla|OJJ79865.1 akaw|GAA84425.1 aluc|OJZ89138.1 anid|CBF83710.1 anig|EHA23407.1 aory|EIT79712.1 asyd|OJJ60833.1 ater|EAU38403.1 atub|OJI85647.1 aver|OJJ00157.1 awen|OJJ33992.1 azon|OJJ48561.1

Group3831: abra|OJJ75688.1 acar|OOG00162.1 acla|EAW09416.1 afis|EAW21720.1 afla|EED57989.1 afum|EAL92753.1 agla|OJJ79864.1 akaw|GAA84426.1 aluc|OJZ89139.1 anid|CBF83711.1 anig|EHA23406.1 aory|EIT79747.1 asyd|OJJ60834.1 ater|EAU38404.1 atub|OJI85646.1 aver|OJJ00158.1 awen|OJJ33993.1 azon|OJJ48560.1

Group3832: abra|OJJ75689.1 acar|OOG00161.1 acla|EAW09415.1 afis|EAW21721.1 afla|EED57988.1 afum|EAL92754.1 agla|OJJ79863.1 akaw|GAA84427.1 aluc|OJZ89140.1 anid|CBF83713.1 anig|EHA23405.1 aory|EIT79743.1 asyd|OJJ60835.1 ater|EAU38405.1 atub|OJI85645.1 aver|OJJ00159.1 awen|OJJ33994.1 azon|OJJ48559.1

Group3833: abra|OJJ75692.1 acar|OOG00160.1 acla|EAW09414.1 afis|EAW21722.1 afla|EED57991.1 afum|EAL92755.1 agla|OJJ79862.1 akaw|GAA84429.1 aluc|OJZ89143.1 anid|CBF83715.1 anig|EHA23404.1 aory|EIT79797.1 asyd|OJJ60836.1 ater|EAU38406.1 atub|OJI85644.1 aver|OJJ00160.1 awen|OJJ33995.1 azon|OJJ48558.1

Group3834: abra|OJJ75696.1 acar|OOG00158.1 acla|EAW09412.1 afis|EAW21724.1 afla|EED57993.1 afum|EAL92757.1 agla|OJJ79860.1 akaw|GAA84432.1 aluc|OJZ89147.1 anid|CBF83718.1 anig|EHA23401.1 aory|EIT79706.1 asyd|OJJ60838.1 ater|EAU38408.1 atub|OJI85641.1 aver|OJJ00162.1 awen|OJJ33997.1 azon|OJJ48555.1

Group3835: abra|OJJ75698.1 acar|OOG00155.1 acla|EAW09410.1 afis|EAW21726.1 afla|EED57995.1 afum|EAL92759.1 agla|OJJ79858.1 akaw|GAA84434.1 aluc|OJZ89149.1 anid|CBF83722.1 anig|EHA23399.1 aory|EIT79805.1 asyd|OJJ60840.1 ater|EAU38410.1 atub|OJI85639.1 aver|OJJ00164.1 awen|OJJ33999.1 azon|OJJ48553.1

Group3836: abra|OJJ75701.1 acar|OOG00153.1 acla|EAW09407.1 afis|EAW21729.1 afla|EED57998.1 afum|EAL92762.1 agla|OJJ79855.1 akaw|GAA84437.1 aluc|OJZ89152.1 anid|CBF83727.1 anig|EHA23397.1 aory|EIT79787.1 asyd|OJJ60843.1 ater|EAU38413.1 atub|OJI85636.1 aver|OJJ00168.1 awen|OJJ34002.1 azon|OJJ48550.1

Group3837: abra|OJJ75707.1 acar|OOG00151.1 acla|EAW09404.1 afis|EAW21732.1 afla|EED58001.1 afum|EAL92765.1 agla|OJJ79853.1 akaw|GAA84442.1 aluc|OJZ89158.1 anid|CBF83732.1 anig|EHA23395.1 aory|EIT79716.1 asyd|OJJ60846.1 ater|EAU38415.1 atub|OJI85632.1 aver|OJJ00171.1 awen|OJJ34005.1 azon|OJJ48546.1

Group3838: abra|OJJ75709.1 acar|OOG00149.1 acla|EAW09403.1 afis|EAW21733.1 afla|EED58002.1 afum|EAL92766.1 agla|OJJ79850.1 akaw|GAA84443.1 aluc|OJZ89160.1 anid|CBF83734.1 anig|EHA23394.1 aory|EIT79783.1 asyd|OJJ60848.1 ater|EAU38416.1 atub|OJI85630.1 aver|OJJ00174.1 awen|OJJ34006.1 azon|OJJ48545.1

Group3839: abra|OJJ75710.1 acar|OOG00150.1 acla|EAW09402.1 afis|EAW21734.1 afla|EED58006.1 afum|EAL92767.1 agla|OJJ79849.1 akaw|GAA84444.1 aluc|OJZ89161.1 anid|CBF83736.1 anig|EHA23393.1

aory|EIT79710.1 asyd|OJJ60850.1 ater|EAU36017.1 atub|OJI85629.1 aver|OJJ00172.1 awen|OJJ34007.1 azon|OJJ48544.1

Group3840: abra|OJJ75716.1 acar|OOG00143.1 acla|EAW09397.1 afis|EAW21737.1 afla|EED58011.1 afum|EAL92771.1 agla|OJJ84451.1 akaw|GAA84450.1 aluc|OJZ89168.1 anid|CBF83703.1 anig|EHA23387.1 aory|EIT79723.1 asyd|OJJ60826.1 ater|EAU38419.1 atub|OJI85623.1 aver|OJJ00152.1 awen|OJJ34019.1 azon|OJJ48539.1

Group3841: abra|OJJ75719.1 acar|OOG00137.1 acla|EAW09375.1 afis|EAW21760.1 afla|EED58018.1 afum|EAL92790.2 agla|OJJ84390.1 akaw|GAA84455.1 aluc|OJZ89171.1 anid|CBF83695.1 anig|EHA23384.1 aory|EIT79793.1 asyd|OJJ60824.1 ater|EAU38430.1 atub|OJI85619.1 aver|OJJ00150.1 awen|OJJ34044.1 azon|OJJ48187.1

Group3842: abra|OJJ75722.1 acar|OOG00136.1 acla|EAW09378.1 afis|EAW21757.1 afla|EED58020.1 afum|EAL92787.1 agla|OJJ84393.1 akaw|GAA84458.1 aluc|OJZ89174.1 anid|CBF83682.1 anig|EHA23381.1 aory|EIT79724.1 asyd|OJJ60821.1 ater|EAU38433.1 atub|OJI85616.1 aver|OJJ00147.1 awen|OJJ34041.1 azon|OJJ48198.1

Group3843: abra|OJJ75723.1 acar|OOG00135.1 acla|EAW09379.1 afis|EAW21756.1 afla|EED58022.1 afum|EAL92786.1 agla|OJJ84394.1 akaw|GAA84459.1 aluc|OJZ89175.1 anid|CBF83684.1 anig|EHA23380.1 asyd|OJJ60820.1 ater|EAU38434.1 atub|OJI85615.1 aver|OJJ00146.1 awen|OJJ34040.1 azon|OJJ48197.1 aory|EIT79736.1

Group3844: abra|OJJ75724.1 acar|OOG00134.1 acla|EAW09380.1 afis|EAW21755.1 afla|EED58023.1 afum|EAL92785.1 agla|OJJ84398.1 akaw|GAA84460.1 aluc|OJZ89176.1 anid|CBF83685.1 anig|EHA23379.1 aory|EIT79744.1 asyd|OJJ60819.1 ater|EAU38435.1 atub|OJI85613.1 aver|OJJ00145.1 awen|OJJ34039.1 azon|OJJ48196.1

Group3845: abra|OJJ75726.1 acar|OOG00132.1 acla|EAW09383.1 afis|EAW21752.1 afla|EED58026.1 afum|EAL92782.1 agla|OJJ84401.1 akaw|GAA84462.1 aluc|OJZ89178.1 anid|CBF83693.1 anig|EHA23377.1 aory|EIT79749.1 asyd|OJJ60816.1 ater|EAU38438.1 atub|OJI85611.1 aver|OJJ00142.1 awen|OJJ34035.1 azon|OJJ48207.1

Group3846: abra|OJJ75728.1 acar|OOF90230.1 acla|EAW08986.1 afis|EAW24686.1 afla|EED54452.1 afum|EAL89327.1 akaw|GAA84463.1 aluc|OJZ89180.1 anid|CBF83674.1 anig|EHA23375.1 aory|EIT76299.1 asyd|OJJ60813.1 ater|EAU39265.1 atub|OJI85609.1 aver|OJJ00140.1 awen|OJJ29740.1 afla|EED48344.1 ater|EAU31420.1

Group3847: abra|OJJ75732.1 acar|OOF89927.1 acla|EAW14909.1 afis|EAW24430.1 afla|EED55243.1 afum|EAL86098.1 agla|OJJ79244.1 akaw|GAA84465.1 aluc|OJZ89183.1 anid|CBF73775.1 anig|EHA23373.1 aory|EIT83557.1 asyd|OJJ56594.1 ater|EAU30015.1 atub|OJI85606.1 aver|OJJ06131.1 awen|OJJ32772.1 azon|OJJ44003.1

Group3848: abra|OJJ75734.1 acar|OOF89928.1 acla|EAW14908.1 afis|EAW24429.1 afla|EED55249.1 afum|EAL86097.1 agla|OJJ79242.1 akaw|GAA84467.1 aluc|OJZ89185.1 anid|CBF73777.1 anig|EHA23372.1 aory|EIT83561.1 asyd|OJJ56596.1 ater|EAU30016.1 atub|OJI85604.1 aver|OJJ06129.1 awen|OJJ32774.1 azon|OJJ44004.1

Group3849: abra|OJJ75735.1 acar|OOF89929.1 acla|EAW14907.1 afis|EAW24428.1 afla|EED55250.1 afum|EAL86096.1 agla|OJJ79241.1 akaw|GAA84468.1 aluc|OJZ89186.1 anid|CBF73778.1 anig|EHA23371.1 aory|EIT83631.1 asyd|OJJ56597.1 ater|EAU30017.1 atub|OJI85603.1 aver|OJJ06128.1 awen|OJJ32775.1 azon|OJJ44005.1

Group3850: abra|OJJ75737.1 acar|OOF89931.1 acla|EAW14904.1 afis|EAW24425.1 afla|EED55254.1 afum|EAL86093.1 agla|OJJ79239.1 akaw|GAA84470.1 aluc|OJZ89189.1 anid|CBF73782.1 anig|EHA23368.1 aory|EIT83565.1 asyd|OJJ56599.1 ater|EAU30020.1 atub|OJI85601.1 aver|OJJ06126.1 awen|OJJ32781.1 azon|OJJ44007.1

Group3851: abra|OJJ75738.1 acar|OOG00131.1 acla|EAW14903.1 afis|EAW24424.1 afla|EED55258.1 afum|EAL86092.1 agla|OJJ79238.1 akaw|GAA84471.1 aluc|OJZ89191.1 anid|CBF73784.1 anig|EHA23367.1 aory|EIT83539.1 asyd|OJJ56600.1 ater|EAU30021.1 atub|OJI85599.1 aver|OJJ06125.1 awen|OJJ32780.1 azon|OJJ44009.1

Group3852: abra|OJJ75784.1 acar|OOG00104.1 acla|EAW14898.1 afis|EAW24420.1 afla|EED47748.1 afum|EAL86089.1 agla|OJJ79224.1 akaw|GAA84500.1 aluc|OJZ89227.1 anid|CBF73792.1 anig|EHA23335.1 aory|EIT77177.1 asyd|OJJ56604.1 ater|EAU30307.1 atub|OJI85565.1 aver|OJJ06119.1 awen|OJJ32789.1 azon|OJJ44022.1

Group3853: abra|OJJ75785.1 acar|OOG00103.1 acla|EAW14897.1 afis|EAW24419.1 afla|EED47749.1 afum|EAL86088.1 agla|OJJ79226.1 akaw|GAA84501.1 aluc|OJZ89229.1 anid|CBF73795.1 anig|EHA23334.1 aory|EIT77122.1 asyd|OJJ56605.1 ater|EAU30308.1 atub|OJI85564.1 aver|OJJ06118.1 awen|OJJ32790.1 azon|OJJ44021.1

Group3854: abra|OJJ75786.1 acar|OOG00102.1 acla|EAW14896.1 afis|EAW24418.1 afla|EED47750.1 afum|EAL86087.1 agla|OJJ79227.1 akaw|GAA84502.1 aluc|OJZ89230.1 anid|CBF73797.1 anig|EHA23333.1 aory|EIT77133.1 asyd|OJJ56607.1 ater|EAU30309.1 atub|OJI85563.1 aver|OJJ06117.1 awen|OJJ32791.1 azon|OJJ44020.1

Group3855: abra|OJJ75787.1 acar|OOG00101.1 acla|EAW14895.1 afis|EAW24417.1 afla|EED47751.1 afum|EAL86086.1 agla|OJJ79228.1 akaw|GAA84503.1 aluc|OJZ89231.1 anid|CBF73799.1 anig|EHA23332.1 aory|EIT77182.1 asyd|OJJ56608.1 ater|EAU30310.1 atub|OJI85562.1 aver|OJJ06116.1 awen|OJJ32792.1 azon|OJJ44019.1

Group3856: abra|OJJ75788.1 acar|OOG00099.1 acla|EAW14893.1 afis|EAW24415.1 afla|EED47766.1 afum|EAL86084.1 agla|OJJ79218.1 akaw|GAA84504.1 aluc|OJZ89232.1 anid|CBF73803.1 anig|EHA23331.1 aory|EIT77193.1 asyd|OJJ56615.1 ater|EAU30314.1 atub|OJI85561.1 aver|OJJ06109.1 awen|OJJ32794.1 azon|OJJ49041.1

Group3857: abra|OJJ75789.1 acar|OOG00098.1 acla|EAW14892.1 afis|EAW24414.1 afla|EED47765.1 afum|EAL86083.1 agla|OJJ79217.1 akaw|GAA84505.1 aluc|OJZ89233.1 anid|CBF73747.1 anig|EHA23330.1 aory|EIT77149.1 asyd|OJJ56622.1 ater|EAU30315.1 atub|OJI85560.1 aver|OJJ06104.1 awen|OJJ32795.1 azon|OJJ49038.1

Group3858: abra|OJJ75792.1 acar|OOG00086.1 acla|EAW14889.1 afis|EAW24411.1 afum|EAL86080.1 agla|OJJ79214.1 akaw|GAA84511.1 aluc|OJZ89240.1 anid|CBF73741.1 anig|EHA23327.1 aory|EIT77168.1 asyd|OJJ56625.1 ater|EAU30318.1 atub|OJI85558.1 aver|OJJ06101.1 awen|OJJ32799.1 azon|OJJ49033.1 afla|EED47762.1

Group3859: abra|OJJ75796.1 acar|OOG00085.1 acla|EAW14888.1 afis|EAW24410.1 afla|EED47761.1 afum|EAL86079.1 agla|OJJ79213.1 akaw|GAA84514.1 aluc|OJZ89243.1 anid|CBF73739.1 anig|EHA23322.1 aory|EIT77110.1 asyd|OJJ56626.1 ater|EAU30319.1 atub|OJI85552.1 aver|OJJ06100.1 awen|OJJ32800.1 azon|OJJ49032.1

Group3860: abra|OJJ75804.1 acar|OOG00071.1 acla|EAW14879.1 afis|EAW24396.1 afla|EED47814.1 afum|EAL86068.1 agla|OJJ78876.1 akaw|GAA84520.1 aluc|OJZ89252.1 anid|CBF73724.1 anig|EHA23315.1 aory|EIT77191.1 asyd|OJJ56638.1 ater|EAU30339.1 atub|OJI85546.1 aver|OJJ06088.1 awen|OJJ32809.1 azon|OJJ49030.1

Group3861: abra|OJJ75805.1 acar|OOG00070.1 acla|EAW14878.1 afis|EAW24395.1 afla|EED47815.1 afum|EAL86067.1 agla|OJJ78875.1 akaw|GAA84521.1 aluc|OJZ89253.1 anid|CBF73720.1 anig|EHA23314.1 aory|EIT77139.1 asyd|OJJ56639.1 ater|EAU30340.1 atub|OJI85545.1 aver|OJJ06087.1 awen|OJJ32810.1 azon|OJJ49031.1

Group3862: abra|OJJ75808.1 acar|OOG00068.1 acla|EAW14864.1 afis|EAW24392.1 afla|EED47818.1 afum|EAL86064.1 agla|OJJ78872.1 akaw|GAA84523.1 aluc|OJZ89256.1 anid|CBF73713.1 anig|EHA23312.1 aory|EIT77112.1 asyd|OJJ56642.1 ater|EAU30342.1 atub|OJI85543.1 aver|OJJ06084.1 awen|OJJ32812.1 azon|OJJ49020.1

Group3863: abra|OJJ75810.1 acar|OOG00067.1 acla|EAW14863.1 afis|EAW24391.1 afla|EED47819.1 afum|EAL86063.1 agla|OJJ78871.1 akaw|GAA84524.1 aluc|OJZ89257.1 anid|CBF73711.1 anig|EHA23311.1 aory|EIT77158.1 asyd|OJJ56643.1 ater|EAU30343.1 atub|OJI85542.1 aver|OJJ06083.1 awen|OJJ32813.1 azon|OJJ49019.1

Group3864: abra|OJJ75813.1 acar|OOG00062.1 acla|EAW14860.1 afis|EAW24388.1 afla|EED47821.1 afum|EAL86060.1 agla|OJJ83284.1 akaw|GAA84527.1 aluc|OJZ89260.1 anid|CBF73705.1 anig|EHA23308.1 aory|EIT77124.1 asyd|OJJ56646.1 ater|EAU30346.1 atub|OJI85539.1 aver|OJJ06080.1 awen|OJJ32817.1 azon|OJJ49015.1

Group3865: abra|OJJ75814.1 acar|OOG00061.1 acla|EAW14859.1 afis|EAW24387.1 afla|EED47822.1 afum|EAL86059.1 agla|OJJ83283.1 akaw|GAA84528.1 aluc|OJZ89261.1 anid|CBF73703.1 anig|EHA23307.1 aory|EIT77142.1 asyd|OJJ56647.1 ater|EAU30347.1 atub|OJI85538.1 aver|OJJ06079.1 awen|OJJ32818.1 azon|OJJ49014.1

Group3866: abra|OJJ75817.1 acar|OOG00058.1 acla|EAW14858.1 afis|EAW24385.1 afla|EED47824.1 afum|EAL86057.1 agla|OJJ83281.1 akaw|GAA84531.1 aluc|OJZ89264.1 anid|CBF73697.1 anig|EHA23304.1 aory|EIT77137.1 asyd|OJJ56650.1 ater|EAU30350.1 atub|OJI85535.1 aver|OJJ06075.1 awen|OJJ32825.1 azon|OJJ49011.1

Group3867: abra|OJJ75821.1 acar|OOG00055.1 acla|EAW14854.1 afis|EAW24381.1 afla|EED47829.1 afum|EAL86053.1 agla|OJJ83269.1 akaw|GAA84534.1 aluc|OJZ89267.1 anid|CBF73667.1 anig|EHA23301.1 aory|EIT77192.1 asyd|OJJ56666.1 ater|EAU30354.1 atub|OJI85532.1 aver|OJJ06057.1 awen|OJJ32831.1 azon|OJJ49005.1

Group3868: abra|OJJ75822.1 acar|OOG00054.1 acla|EAW14853.1 afis|EAW24380.1 afla|EED47830.1 afum|EAL86052.1 agla|OJJ83268.1 akaw|GAA84535.1 aluc|OJZ89268.1 anid|CBF73669.1 anig|EHA23300.1

aory|EIT77144.1 asyd|OJJ56665.1 ater|EAU30355.1 atub|OJI85531.1 aver|OJJ06058.1 awen|OJJ32832.1 azon|OJJ49004.1

Group3869: abra|OJJ75828.1 acar|OOG00052.1 acla|EAW14850.1 afis|EAW24377.1 afla|EED47834.1 afum|EAL86049.1 agla|OJJ83263.1 akaw|GAA84538.1 aluc|OJZ89271.1 anid|CBF73674.1 anig|EHA23298.1 aory|EIT77135.1 asyd|OJJ56662.1 ater|EAU30359.1 atub|OJI85528.1 aver|OJJ06062.1 awen|OJJ32835.1 azon|OJJ49000.1

Group3870: abra|OJJ75829.1 acar|OOG00051.1 acla|EAW14849.1 afis|EAW24376.1 afla|EED47835.1 afum|EAL86048.1 agla|OJJ83262.1 akaw|GAA84539.1 aluc|OJZ89272.1 anid|CBF73680.1 anig|EHA23297.1 aory|EIT77202.1 asyd|OJJ56659.1 ater|EAU30360.1 atub|OJI85527.1 aver|OJJ06066.1 awen|OJJ32836.1 azon|OJJ48998.1

Group3871: abra|OJJ75831.1 acar|OOG00049.1 acla|EAW14847.1 afis|EAW24374.1 afla|EED47837.1 afum|EAL86046.1 agla|OJJ83260.1 akaw|GAA84541.1 aluc|OJZ89274.1 anid|CBF73682.1 anig|EHA23295.1 aory|EIT77185.1 asyd|OJJ56657.1 ater|EAU30362.1 atub|OJI85525.1 aver|OJJ06068.1 awen|OJJ32838.1 azon|OJJ48996.1

Group3872: abra|OJJ75832.1 acar|OOG00048.1 acla|EAW14846.1 afis|EAW24373.1 afla|EED47838.1 afum|EAL86045.1 agla|OJJ83259.1 akaw|GAA84542.1 aluc|OJZ89275.1 anid|CBF73684.1 anig|EHA23294.1 aory|EIT77136.1 asyd|OJJ56656.1 ater|EAU30363.1 atub|OJI85524.1 aver|OJJ06069.1 awen|OJJ32839.1 azon|OJJ48995.1

Group3873: abra|OJJ75843.1 acar|OOG00045.1 acla|EAW14844.1 afis|EAW24371.1 afla|EED47846.1 afum|EAL86043.1 agla|OJJ83256.1 akaw|GAA84553.1 aluc|OJZ89285.1 anid|CBF73688.1 anig|EHA23281.1 aory|EIT77207.1 asyd|OJJ56654.1 ater|EAU30368.1 atub|OJI85514.1 aver|OJJ06071.1 awen|OJJ32844.1 azon|OJJ46816.1

Group3874: abra|OJJ75851.1 acar|OOG00042.1 acla|EAW14835.1 afis|EAW24364.1 afla|EED47849.1 afum|EBA27254.1 agla|OJJ83249.1 akaw|GAA84556.1 aluc|OJZ89288.1 anid|CBF73935.1 anig|EHA23273.1 aory|EIT77237.1 asyd|OJJ53672.1 ater|EAU30373.1 atub|OJI85511.1 aver|OJJ07779.1 awen|OJJ32869.1 azon|OJJ48991.1

Group3875: abra|OJJ75854.1 acar|OOG00041.1 acla|EAW14833.1 afis|EAW24362.1 afla|EED47851.1 afum|EAL86032.1 agla|OJJ83247.1 akaw|GAA84559.1 aluc|OJZ89291.1 anid|CBF73928.1 anig|EHA23271.1 aory|EIT77246.1 asyd|OJJ53677.1 ater|EAU30376.1 atub|OJI85508.1 aver|OJJ07793.1 awen|OJJ32865.1 azon|OJJ48989.1

Group3876: abra|OJJ75855.1 acar|OOG00040.1 acla|EAW14832.1 afis|EAW24361.1 afla|EED47852.1 afum|EAL86031.1 agla|OJJ83246.1 akaw|GAA84560.1 aluc|OJZ89293.1 anid|CBF73926.1 anig|EHA23270.1 aory|EIT77210.1 asyd|OJJ53678.1 ater|EAU30377.1 atub|OJI85507.1 aver|OJJ07794.1 awen|OJJ32864.1 azon|OJJ48988.1

Group3877: abra|OJJ75856.1 acar|OOG00038.1 acla|EAW14831.1 afis|EAW24360.1 afla|EED47853.1 afum|EAL86030.1 agla|OJJ83245.1 akaw|GAA84561.1 aluc|OJZ89294.1 anid|CBF73924.1 anig|EHA23269.1 aory|EIT77236.1 asyd|OJJ53679.1 ater|EAU30378.1 atub|OJI85506.1 aver|OJJ07795.1 awen|OJJ32863.1 azon|OJJ48987.1

Group3878: abra|OJJ75858.1 acar|OOG00037.1 acla|EAW14829.1 afis|EAW24358.1 afla|EED47856.1 afum|EAL86028.1 agla|OJJ83242.1 akaw|GAA84563.1 aluc|OJZ89296.1 anid|CBF73921.1 anig|EHA23267.1 aory|EIT77221.1 asyd|OJJ53681.1 ater|EAU30381.1 atub|OJI85503.1 aver|OJJ07797.1 awen|OJJ32861.1 azon|OJJ48985.1

Group3879: abra|OJJ75864.1 acar|OOG00032.1 acla|EAW14824.1 afis|EAW24353.1 afla|EED47910.1 afum|EAL86024.2 agla|OJJ83233.1 akaw|GAA84570.1 aluc|OJZ89306.1 anid|CBF73944.1 anig|EHA23260.1 aory|EIT83380.1 asyd|OJJ53698.1 ater|EAU30386.1 atub|OJI85493.1 aver|OJJ07816.1 awen|OJJ32872.1 azon|OJJ48973.1

Group3880: abra|OJJ75865.1 acar|OOG00029.1 acla|EAW09926.1 afis|EAW21174.1 afla|EED54761.1 afum|EAL92194.1 agla|OJJ80004.1 akaw|GAA84571.1 aluc|OJZ89307.1 anid|CBF83297.1 anig|EHA23259.1 aory|EIT79426.1 asyd|OJJ60558.1 ater|EAU35841.1 atub|OJI85491.1 aver|OJI99884.1 awen|OJJ36513.1 azon|OJJ44389.1

Group3881: abra|OJJ75867.1 acar|OOG00027.1 acla|EAW09928.1 afis|EAW21176.1 afla|EED54763.1 afum|EAL92196.1 agla|OJJ80001.1 akaw|GAA84573.1 aluc|OJZ89309.1 anid|CBF83306.1 anig|EHA23257.1 aory|EIT79399.1 asyd|OJJ60555.1 ater|EAU35843.1 atub|OJI85490.1 aver|OJI99881.1 awen|OJJ36515.1 azon|OJJ44391.1

Group3882: abra|OJJ75873.1 acar|OOG00022.1 acla|EAW09932.1 afis|EAW21179.1 afla|EED54769.1 afum|EAL92199.1 agla|OJJ79993.1 akaw|GAA84576.1 aluc|OJZ89317.1 anid|CBF83303.1 anig|EHA23254.1 aory|EIT79449.1 asyd|OJJ60552.1 ater|EAU35845.1 atub|OJI85480.1 aver|OJI99878.1 awen|OJJ36518.1 azon|OJJ44393.1

Group3883: abra|OJJ75875.1 acar|OOG00020.1 acla|EAW09935.1 afis|EAW21180.1 afla|EED54785.1 afum|EAL92201.1 agla|OJJ79997.1 akaw|GAA84577.1 aluc|OJZ89319.1 anid|CBF83301.1 anig|EHA23253.1 aory|EIT79362.1 asyd|OJJ60551.1 ater|EAU35846.1 atub|OJI85479.1 aver|OJI99877.1 awen|OJJ36519.1 azon|OJJ44395.1

Group3884: abra|OJJ75876.1 acar|OOG00017.1 acla|EAW09938.1 afis|EAW21183.1 afla|EED54776.1 afum|EAL92204.1 agla|OJJ79987.1 akaw|GAA84578.1 aluc|OJZ89320.1 anid|CBF83312.1 anig|EHA23252.1 aory|EIT79349.1 asyd|OJJ60548.1 ater|EAU35849.1 atub|OJI85478.1 aver|OJI99874.1 awen|OJJ36522.1 azon|OJJ44398.1

Group3885: abra|OJJ75879.1 acar|OOG00014.1 acla|EAW09940.1 afis|EAW21185.1 afla|EED54774.1 afum|EAL92207.1 agla|OJJ79982.1 akaw|GAA84582.1 aluc|OJZ89325.1 anid|CBF83315.1 anig|EHA23249.1 aory|EIT79429.1 asyd|OJJ60547.1 ater|EAU35850.1 atub|OJI85474.1 aver|OJI99873.1 awen|OJJ36526.1 azon|OJJ44399.1

Group3886: abra|OJJ75880.1 acar|OOG00013.1 acla|EAW09942.1 afis|EAW21187.1 afla|EED54773.1 afum|EAL92209.1 agla|OJJ79979.1 akaw|GAA84583.1 aluc|OJZ89326.1 anid|CBF83319.1 anig|EHA23248.1 aory|EIT79437.1 asyd|OJJ60546.1 ater|EAU35851.1 atub|OJI85473.1 aver|OJI99872.1 awen|OJJ36529.1 azon|OJJ44401.1

Group3887: abra|OJJ75884.1 acar|OOF90248.1 acla|EAW09944.1 afis|EAW21222.1 afla|EED54784.1 afum|EBA27454.1 agla|OJJ78510.1 akaw|GAA84592.1 aluc|OJZ89337.1 anid|CBF83295.1 anig|EHA23244.1 aory|EIT79434.1 asyd|OJJ60560.1 ater|EAU35852.1 atub|OJI85462.1 aver|OJI99886.1 awen|OJJ36530.1 azon|OJJ44404.1

Group3888: abra|OJJ75886.1 acar|OOF90247.1 acla|EAW09945.1 afis|EAW21223.1 afla|EED54783.1 afum|EAL92236.1 agla|OJJ78507.1 akaw|GAA84593.1 aluc|OJZ89338.1 anid|CBF83293.1 anig|EHA23243.1 aory|EIT79443.1 asyd|OJJ60561.1 ater|EAU35853.1 atub|OJI85461.1 aver|OJI99887.1 awen|OJJ36531.1 azon|OJJ44405.1

Group3889: abra|OJJ75888.1 acar|OOF90245.1 acla|EAW09946.1 afis|EAW21224.1 afla|EED54782.1 afum|EAL92237.1 agla|OJJ78506.1 akaw|GAA84594.1 aluc|OJZ89340.1 anid|CBF83291.1 anig|EHA23242.1 aory|EIT79409.1 asyd|OJJ60562.1 ater|EAU35854.1 atub|OJI85459.1 aver|OJI99888.1 awen|OJJ36532.1 azon|OJJ44406.1

Group3890: abra|OJJ75889.1 acar|OOF90244.1 acla|EAW09948.1 afis|EAW21226.1 afla|EED54779.1 afum|EBA27456.1 agla|OJJ78504.1 akaw|GAA84596.1 aluc|OJZ89341.1 anid|CBF83289.1 anig|EHA23241.1 aory|EIT79431.1 asyd|OJJ60563.1 ater|EAU35857.1 atub|OJI85458.1 aver|OJI99890.1 awen|OJJ36534.1 azon|OJJ44408.1

Group3891: abra|OJJ75890.1 acar|OOG00012.1 acla|EAW09898.1 afis|EAW21228.1 afla|EED54786.1 afum|EAL92240.1 agla|OJJ78502.1 akaw|GAA84597.1 aluc|OJZ89342.1 anid|CBF83280.1 anig|EHA23240.1 aory|EIT79332.1 asyd|OJJ60568.1 ater|EAU35859.1 atub|OJI85457.1 aver|OJI99893.1 awen|OJJ36537.1 azon|OJJ44351.1

Group3892: abra|OJJ75891.1 acar|OOG00011.1 acla|EAW09897.1 afis|EAW21229.1 afla|EED54787.1 afum|EAL92241.2 agla|OJJ78501.1 akaw|GAA84598.1 aluc|OJZ89343.1 anid|CBF83278.1 anig|EHA23239.1 aory|EIT79422.1 asyd|OJJ60569.1 ater|EAU35860.1 atub|OJI85456.1 aver|OJI99894.1 awen|OJJ36538.1 azon|OJJ44349.1

Group3893: abra|OJJ75892.1 acar|OOG00010.1 acla|EAW09896.1 afis|EAW21230.1 afla|EED54788.1 afum|EAL92242.1 agla|OJJ78492.1 akaw|GAA84599.1 aluc|OJZ89344.1 anid|CBF83277.1 anig|EHA23238.1 aory|EIT79453.1 asyd|OJJ60570.1 ater|EAU35863.1 atub|OJI85455.1 aver|OJI99895.1 awen|OJJ36539.1 azon|OJJ44348.1

Group3894: abra|OJJ75897.1 acar|OOG00007.1 acla|EAW09894.1 afis|EAW21232.1 afla|EED54793.1 afum|EAL92244.1 agla|OJJ78494.1 akaw|GAA84603.1 aluc|OJZ89347.1 anid|CBF83273.1 anig|EHA23234.1 aory|EIT79342.1 asyd|OJJ60573.1 ater|EAU35865.1 atub|OJI85451.1 aver|OJI99898.1 awen|OJJ36542.1 azon|OJJ44346.1

Group3895: abra|OJJ75898.1 acar|OOG00006.1 acla|EAW09893.1 afis|EAW21233.1 afla|EED54794.1 afum|EAL92245.1 agla|OJJ78495.1 akaw|GAA84604.1 aluc|OJZ89348.1 anid|CBF83271.1 anig|EHA23233.1 aory|EIT79361.1 asyd|OJJ60574.1 ater|EAU35866.1 atub|OJI85450.1 aver|OJI99899.1 awen|OJJ36543.1 azon|OJJ44345.1

Group3896: abra|OJJ75899.1 acar|OOG00005.1 acla|EAW09892.1 afis|EAW21234.1 afla|EED54795.1 afum|EAL92246.1 agla|OJJ78496.1 akaw|GAA84605.1 aluc|OJZ89349.1 anid|CBF83270.1 anig|EHA23232.1 aory|EIT79442.1 asyd|OJJ60575.1 ater|EAU35867.1 atub|OJI85449.1 aver|OJI99900.1 awen|OJJ36544.1 azon|OJJ44344.1

Group3897: abra|OJJ75902.1 acar|OOG00002.1 acla|EAW09888.1 afis|EAW21238.1 afla|EED54798.1 afum|EAL92250.1 agla|OJJ78500.1 akaw|GAA84608.1 aluc|OJZ89352.1 anid|CBF83266.1 anig|EHA23229.1

aory|EIT79394.1 asyd|OJJ60579.1 ater|EAU35879.1 atub|OJI85446.1 aver|OJI99904.1 awen|OJJ36549.1 azon|OJJ44340.1

Group3898: abra|OJJ75903.1 acar|OOG00001.1 acla|EAW09887.1 afis|EAW21239.1 afla|EED54799.1 afum|EAL92252.1 agla|OJJ78491.1 akaw|GAA84609.1 aluc|OJZ89353.1 anid|CBF83264.1 anig|EHA23228.1 aory|EIT79376.1 asyd|OJJ60580.1 ater|EAU35880.1 atub|OJI85445.1 aver|OJI99905.1 awen|OJJ36550.1 azon|OJJ44339.1

Group3899: abra|OJJ75904.1 acar|OOF99999.1 acla|EAW09886.1 afis|EAW21240.1 afla|EED54800.1 afum|EAL92253.1 agla|OJJ78490.1 akaw|GAA84610.1 aluc|OJZ89354.1 anid|CBF83262.1 anig|EHA23227.1 aory|EIT79341.1 asyd|OJJ60581.1 ater|EAU35881.1 atub|OJI85444.1 aver|OJI99906.1 awen|OJJ36551.1 azon|OJJ44338.1

Group3900: abra|OJJ75905.1 acar|OOF99998.1 acla|EAW09885.1 afis|EAW21241.1 afla|EED54802.1 afum|EAL92254.1 agla|OJJ78489.1 akaw|GAA84611.1 aluc|OJZ89355.1 anid|CBF83259.1 anig|EHA23226.1 aory|EIT79395.1 asyd|OJJ60584.1 ater|EAU35882.1 atub|OJI85443.1 aver|OJI99907.1 awen|OJJ36552.1 azon|OJJ44337.1

Group3901: abra|OJJ75906.1 acar|OOF99997.1 acla|EAW09883.1 afis|EAW21243.1 afla|EED54803.1 afum|EAL92256.1 agla|OJJ78487.1 akaw|GAA84612.1 aluc|OJZ89356.1 anid|CBF83255.1 anig|EHA23225.1 aory|EIT79348.1 asyd|OJJ60587.1 ater|EAU35883.1 atub|OJI85442.1 aver|OJI99910.1 awen|OJJ36554.1 azon|OJJ44335.1

Group3902: abra|OJJ75910.1 acar|OOF99993.1 acla|EAW09879.1 afis|EAW21247.1 afla|EED54807.1 afum|EAL92260.1 agla|OJJ87509.1 akaw|GAA84616.1 aluc|OJZ89360.1 anid|CBF83240.1 anig|EHA23221.1 aory|EIT79440.1 asyd|OJJ60598.1 ater|EAU35887.1 atub|OJI85438.1 aver|OJI99921.1 awen|OJJ36560.1 azon|OJJ44331.1

Group3903: abra|OJJ75911.1 acar|OOF99992.1 acla|EAW09878.1 afis|EAW21248.1 afla|EED54808.1 afum|EAL92261.1 agla|OJJ87508.1 akaw|GAA84617.1 aluc|OJZ89361.1 anid|CBF83242.1 anig|EHA23220.1 aory|EIT79438.1 asyd|OJJ60597.1 ater|EAU35888.1 atub|OJI85437.1 aver|OJI99920.1 awen|OJJ36561.1 azon|OJJ44327.1

Group3904: abra|OJJ75914.1 acar|OOF99989.1 acla|EAW09876.1 afis|EAW21250.1 afla|EED54810.1 afum|EAL92263.1 agla|OJJ87501.1 akaw|GAA84619.1 aluc|OJZ89363.1 anid|CBF83245.1 anig|EHA23219.1 aory|EIT79398.1 asyd|OJJ60594.1 ater|EAU35890.1 atub|OJI85435.1 aver|OJI99918.1 awen|OJJ36564.1 azon|OJJ44324.1

Group3905: abra|OJJ75916.1 acar|OOF90029.1 acla|EAW09874.1 afis|EAW21252.1 afla|EED54812.1 afum|EAL92266.1 agla|OJJ87495.1 akaw|GAA84621.1 aluc|OJZ89365.1 anid|CBF83247.1 anig|EHA23217.1 aory|EIT79364.1 asyd|OJJ60592.1 ater|EAU35892.1 atub|OJI85433.1 aver|OJI99916.1 awen|OJJ36567.1 azon|OJJ44321.1

Group3906: abra|OJJ75919.1 acar|OOF90028.1 acla|EAW09873.1 afis|EAW21253.1 afla|EED54815.1 afum|EAL92268.1 agla|OJJ87497.1 akaw|GAA84622.1 aluc|OJZ89368.1 anid|CBF83249.1 anig|EHA23215.1 aory|EIT79356.1 asyd|OJJ60590.1 ater|EAU35893.1 atub|OJI85430.1 aver|OJI99913.1 awen|OJJ36568.1 azon|OJJ44319.1

Group3907: abra|OJJ75920.1 acar|OOF99987.1 acla|EAW09872.1 afis|EAW21254.1 afla|EED54816.1 afum|EAL92269.1 agla|OJJ87498.1 akaw|GAA84623.1 aluc|OJZ89369.1 anid|CBF83251.1 anig|EHA23214.1 aory|EIT79384.1 asyd|OJJ60589.1 ater|EAU35894.1 atub|OJI85429.1 aver|OJI99912.1 awen|OJJ36569.1 azon|OJJ44318.1

Group3908: abra|OJJ75922.1 acar|OOF99986.1 acla|EAW09866.1 afis|EAW21259.1 afla|EED54818.1 afum|EAL92274.1 agla|OJJ87491.1 akaw|GAA84626.1 aluc|OJZ89372.1 anid|CBF83325.1 anig|EHA23212.1 aory|EIT79360.1 asyd|OJJ60601.1 ater|EAU35896.1 atub|OJI85426.1 aver|OJI99924.1 awen|OJJ36574.1 azon|OJJ44315.1

Group3909: abra|OJJ75924.1 acar|OOF99984.1 acla|EAW09865.1 afis|EAW21260.1 afla|EED54819.1 afum|EAL92275.1 agla|OJJ87490.1 akaw|GAA84627.1 aluc|OJZ89373.1 anid|CBF83329.1 anig|EHA23210.1 aory|EIT79435.1 asyd|OJJ60605.1 ater|EAU35897.1 atub|OJI85425.1 aver|OJI99929.1 awen|OJJ36575.1 azon|OJJ44314.1

Group3910: abra|OJJ75931.1 acar|OOF99979.1 acla|EAW09862.1 afis|EAW21264.1 afla|EED54823.1 afum|EAL92279.1 agla|OJJ87471.1 akaw|GAA84632.1 aluc|OJZ89378.1 anid|CBF83340.1 anig|EHA23205.1 aory|EIT79396.1 asyd|OJJ60609.1 ater|EAU35901.1 atub|OJI85420.1 aver|OJI99933.1 awen|OJJ36581.1 azon|OJJ44309.1

Group3911: abra|OJJ75932.1 acar|OOF99978.1 acla|EAW09861.1 afis|EAW21265.1 afla|EED54824.1 afum|EAL92280.1 agla|OJJ87472.1 akaw|GAA84633.1 aluc|OJZ89379.1 anid|CBF83342.1 anig|EHA23204.1 aory|EIT79447.1 asyd|OJJ60610.1 ater|EAU35902.1 atub|OJI85419.1 aver|OJI99934.1 awen|OJJ36582.1 azon|OJJ44308.1

Group3912: abra|OJJ75934.1 acar|OOF99976.1 acla|EAW09859.1 afis|EAW21267.1 afla|EED54826.1 afum|EAL92282.1 agla|OJJ87474.1 akaw|GAA84635.1 aluc|OJZ89381.1 anid|CBF83345.1 anig|EHA23202.1 aory|EIT79337.1 asyd|OJJ60612.1 ater|EAU35903.1 atub|OJI85417.1 aver|OJI99936.1 awen|OJJ36584.1 azon|OJJ47371.1

Group3913: abra|OJJ75935.1 acar|OOF99975.1 acla|EAW09858.1 afis|EAW21268.1 afla|EED54827.1 afum|EAL92283.1 agla|OJJ87475.1 akaw|GAA84636.1 aluc|OJZ89382.1 anid|CBF83347.1 anig|EHA23201.1 aory|EIT79343.1 asyd|OJJ60613.1 ater|EAU35904.1 atub|OJI85416.1 aver|OJI99937.1 awen|OJJ39424.1 azon|OJJ44306.1

Group3914: abra|OJJ75936.1 acar|OOF99974.1 acla|EAW09857.1 afis|EAW21269.1 afla|EED54829.1 afum|EBA27457.1 agla|OJJ87476.1 akaw|GAA84637.1 aluc|OJZ89383.1 anid|CBF83349.1 anig|EHA23200.1 aory|EIT79419.1 asyd|OJJ60614.1 ater|EAU35905.1 atub|OJI85415.1 aver|OJI99938.1 awen|OJJ39423.1 azon|OJJ44305.1

Group3915: abra|OJJ75942.1 acar|OOF99968.1 acla|EAW09851.1 afis|EAW21284.1 afla|EED54835.1 afum|EAL92301.1 agla|OJJ87479.1 akaw|GAA84644.1 aluc|OJZ89389.1 anid|CBF83363.1 anig|EHA23195.1 aory|EIT79451.1 asyd|OJJ60620.1 ater|EAU35913.1 atub|OJI85408.1 aver|OJI99944.1 awen|OJJ39415.1 azon|OJJ44298.1

Group3916: abra|OJJ75944.1 acar|OOF99967.1 acla|EAW09849.1 afis|EAW21286.1 afla|EED54837.1 afum|EAL92303.1 agla|OJJ87468.1 akaw|GAA84646.1 aluc|OJZ89391.1 anid|CBF83367.1 anig|EHA23193.1 aory|EIT79347.1 asyd|OJJ60622.1 ater|EAU35918.1 atub|OJI85406.1 aver|OJI99946.1 awen|OJJ39413.1 azon|OJJ44296.1

Group3917: abra|OJJ75949.1 acar|OOF99964.1 acla|EAW09847.1 afis|EAW21288.1 afla|EED54839.1 afum|EAL92305.1 agla|OJJ87466.1 akaw|GAA84652.1 aluc|OJZ89398.1 anid|CBF83371.1 anig|EHA23189.1 aory|EIT79388.1 asyd|OJJ60624.1 ater|EAU35920.1 atub|OJI85399.1 aver|OJI99948.1 awen|OJJ39411.1 azon|OJJ44294.1

Group3918: abra|OJJ75950.1 acar|OOF99962.1 acla|EAW09846.1 afis|EAW21289.1 afla|EED54840.1 afum|EAL92306.1 agla|OJJ87464.1 akaw|GAA84653.1 aluc|OJZ89399.1 anid|CBF83372.1 anig|EHA23188.1 aory|EIT79351.1 asyd|OJJ60625.1 ater|EAU35921.1 atub|OJI85398.1 aver|OJI99949.1 awen|OJJ39410.1 azon|OJJ44293.1

Group3919: abra|OJJ75951.1 acar|OOF99961.1 acla|EAW09845.1 afis|EAW21290.1 afla|EED54842.1 afum|EAL92307.1 agla|OJJ87463.1 akaw|GAA84654.1 aluc|OJZ89400.1 anid|CBF85245.1 anig|EHA23187.1 aory|EIT79441.1 asyd|OJJ62975.1 ater|EAU35922.1 atub|OJI85397.1 aver|OJI97994.1 awen|OJJ35914.1 azon|OJJ44292.1

Group3920: abra|OJJ75952.1 acar|OOF99960.1 acla|EAW09844.1 afis|EAW21291.1 afla|EED54843.1 afum|EAL92308.1 agla|OJJ87462.1 akaw|GAA84655.1 aluc|OJZ89401.1 anid|CBF83374.1 anig|EHA23186.1 aory|EIT79331.1 asyd|OJJ60626.1 ater|EAU35923.1 atub|OJI85396.1 aver|OJI99950.1 awen|OJJ39408.1 azon|OJJ44291.1

Group3921: abra|OJJ75958.1 acar|OOF99956.1 acla|EAW09840.1 afis|EAW21295.1 afla|EED54846.1 afum|EAL92312.1 agla|OJJ87444.1 akaw|GAA84659.1 aluc|OJZ89406.1 anid|CBF83381.1 anig|EHA23181.1 aory|EIT79417.1 asyd|OJJ60629.1 ater|EAU35928.1 atub|OJI85391.1 aver|OJI99953.1 awen|OJJ39404.1 azon|OJJ44288.1

Group3922: abra|OJJ75959.1 acar|OOF99955.1 acla|EAW09839.1 afis|EAW21296.1 afla|EED54847.1 afum|EAL92313.1 agla|OJJ87443.1 akaw|GAA84660.1 aluc|OJZ89407.1 anid|CBF83383.1 anig|EHA23180.1 aory|EIT79405.1 asyd|OJJ60630.1 ater|EAU35929.1 atub|OJI85388.1 aver|OJI99954.1 awen|OJJ39403.1 azon|OJJ44287.1

Group3923: abra|OJJ75963.1 acar|OOF90049.1 acla|EAW09834.1 afis|EAW21299.1 afla|EED54851.1 afum|EAL92316.1 agla|OJJ87439.1 akaw|GAA84663.1 aluc|OJZ89411.1 anid|CBF83388.1 anig|EHA23176.1 aory|EIT79436.1 asyd|OJJ60634.1 ater|EAU35932.1 atub|OJI85385.1 aver|OJI99957.1 awen|OJJ39399.1 azon|OJJ44283.1

Group3924: abra|OJJ75965.1 acar|OOF90050.1 acla|EAW09833.1 afis|EAW21300.1 afla|EED54852.1 afum|EAL92317.1 agla|OJJ87438.1 akaw|GAA84664.1 aluc|OJZ89412.1 anid|CBF83392.1 anig|EHA23175.1 aory|EIT79390.1 asyd|OJJ60635.1 ater|EAU35933.1 atub|OJI85384.1 aver|OJI99958.1 awen|OJJ39398.1 azon|OJJ44282.1

Group3925: abra|OJJ75966.1 acar|OOF90052.1 acla|EAW09832.1 afis|EAW21301.1 afla|EED57261.1 afum|EAL92318.1 agla|OJJ87436.1 akaw|GAA84665.1 aluc|OJZ89413.1 anid|CBF83394.1 anig|EHA23174.1 aory|EIT73984.1 asyd|OJJ60636.1 ater|EAU35934.1 atub|OJI85382.1 aver|OJI99959.1 awen|OJJ39397.1 azon|OJJ44281.1

Group3926: abra|OJJ75973.1 acar|OOF99946.1 acla|EAW09823.1 afis|EAW21310.1 afla|EED57266.1 afum|EAL92326.1 agla|OJJ87432.1 akaw|GAA88983.1 aluc|OJZ89420.1 anid|CBF83404.1 anig|EHA23167.1

aory|EIT74110.1 asyd|OJJ60640.1 ater|EAU35940.1 atub|OJI85373.1 aver|OJI99961.1 awen|OJJ39390.1 azon|OJJ44278.1

Group3927: abra|OJJ75977.1 acar|OOF99943.1 acla|EAW09821.1 afis|EAW21313.1 afla|EED57270.1 afum|EAL92330.1 agla|OJJ87429.1 akaw|GAA88980.1 aluc|OJZ89423.1 anid|CBF83407.1 anig|EHA23164.1 aory|EIT74037.1 asyd|OJJ60647.1 ater|EAU35942.1 atub|OJI85370.1 aver|OJI99965.1 awen|OJJ39386.1 azon|OJJ44275.1

Group3928: abra|OJJ75978.1 acar|OOF99941.1 acla|EAW09820.1 afis|EAW21314.1 afla|EED57271.1 afum|EAL92331.1 agla|OJJ87426.1 akaw|GAA88979.1 aluc|OJZ89424.1 anid|CBF83408.1 anig|EHA23163.1 aory|EIT74076.1 asyd|OJJ60648.1 ater|EAU35943.1 atub|OJI85369.1 aver|OJI99968.1 awen|OJJ39383.1 azon|OJJ44273.1

Group3929: abra|OJJ75980.1 acar|OOF99939.1 acla|EAW09819.1 afis|EAW21315.1 afla|EED57273.1 afum|EAL92332.1 agla|OJJ87424.1 akaw|GAA88977.1 aluc|OJZ89426.1 anid|CBF83412.1 anig|EHA23160.1 aory|EIT74008.1 asyd|OJJ60650.1 ater|EAU35944.1 atub|OJI85367.1 aver|OJI99970.1 awen|OJJ39381.1 azon|OJJ44271.1

Group3930: abra|OJJ75982.1 acar|OOF99937.1 acla|EAW09817.1 afis|EAW21317.1 afla|EED57275.1 afum|EAL92334.1 agla|OJJ87422.1 akaw|GAA88975.1 aluc|OJZ89428.1 anid|CBF83416.1 anig|EHA23159.1 aory|EIT74079.1 asyd|OJJ60652.1 ater|EAU35945.1 atub|OJI85365.1 aver|OJI99972.1 awen|OJJ39379.1 azon|OJJ44269.1

Group3931: abra|OJJ75983.1 acar|OOF99936.1 acla|EAW09816.1 afis|EAW21318.1 afla|EED57276.1 afum|EAL92335.1 agla|OJJ87421.1 akaw|GAA88974.1 aluc|OJZ89429.1 anid|CBF83417.1 anig|EHA23158.1 aory|EIT74114.1 asyd|OJJ60653.1 ater|EAU35946.1 atub|OJI85364.1 aver|OJI99973.1 awen|OJJ39378.1 azon|OJJ44268.1

Group3932: abra|OJJ75984.1 acar|OOF99935.1 acla|EAW09815.1 afis|EAW21319.1 afla|EED57277.1 afum|EAL92336.1 agla|OJJ87420.1 akaw|GAA88973.1 aluc|OJZ89430.1 anid|CBF83419.1 anig|EHA23157.1 aory|EIT74055.1 asyd|OJJ60654.1 ater|EAU35947.1 atub|OJI85363.1 aver|OJI99974.1 awen|OJJ39377.1 azon|OJJ44267.1

Group3933: abra|OJJ75987.1 acar|OOF99934.1 acla|EAW09813.1 afis|EAW21321.1 afla|EED57279.1 afum|EAL92338.1 agla|OJJ87416.1 akaw|GAA88971.1 aluc|OJZ89432.1 anid|CBF83421.1 anig|EHA23155.1 aory|EIT74031.1 asyd|OJJ60655.1 ater|EAU35948.1 atub|OJI85361.1 aver|OJI99976.1 awen|OJJ39375.1 azon|OJJ44265.1

Group3934: abra|OJJ75988.1 acar|OOF99933.1 acla|EAW09812.1 afis|EAW21323.1 afla|EED57280.1 afum|EAL92340.1 agla|OJJ87415.1 akaw|GAA88970.1 aluc|OJZ89433.1 anid|CBF83423.1 anig|EHA23154.1 aory|EIT74009.1 asyd|OJJ60656.1 ater|EAU35949.1 atub|OJI85360.1 aver|OJI99977.1 awen|OJJ39374.1 azon|OJJ44264.1

Group3935: abra|OJJ75995.1 acar|OOF99930.1 acla|EAW09810.1 afis|EAW21325.1 afla|EED57283.1 afum|EAL92343.1 agla|OJJ87413.1 akaw|GAA88964.1 aluc|OJZ89441.1 anid|CBF83426.1 anig|EHA23145.1 aory|EIT74153.1 asyd|OJJ60658.1 ater|EAU35951.1 atub|OJI85352.1 aver|OJI99981.1 awen|OJJ39372.1 azon|OJJ51712.1

Group3936: abra|OJJ76003.1 acar|OOF99921.1 acla|EAW09802.1 afis|EAW21334.1 afla|EED57292.1 afum|EAL92352.1 agla|OJJ87399.1 akaw|GAA88957.1 aluc|OJZ89448.1 anid|CBF76279.1 anig|EHA23138.1 aory|EIT74027.1 asyd|OJJ60988.1 ater|EAU35959.1 atub|OJI85345.1 aver|OJJ00308.1 awen|OJJ39364.1 azon|OJJ51703.1

Group3937: abra|OJJ76004.1 acar|OOF99922.1 acla|EAW09801.1 afis|EAW21335.1 afla|EED57293.1 afum|EAL92353.1 agla|OJJ87398.1 akaw|GAA88956.1 aluc|OJZ89449.1 anid|CBF76277.1 anig|EHA23137.1 aory|EIT73971.1 asyd|OJJ60987.1 ater|EAU35960.1 atub|OJI85344.1 aver|OJJ00307.1 awen|OJJ39363.1 azon|OJJ51702.1

Group3938: abra|OJJ76005.1 acar|OOF99920.1 acla|EAW09800.1 afis|EAW21336.1 afla|EED57294.1 afum|EAL92354.1 agla|OJJ82541.1 akaw|GAA88955.1 aluc|OJZ89451.1 anid|CBF83881.1 anig|EHA23136.1 aory|EIT74083.1 asyd|OJJ60940.1 ater|EAU35961.1 atub|OJI85343.1 aver|OJJ00266.1 awen|OJJ39362.1 azon|OJJ51701.1

Group3939: abra|OJJ76007.1 acar|OOF99919.1 acla|EAW09798.1 afis|EAW21338.1 afla|EED57296.1 afum|EAL92356.1 agla|OJJ87396.1 akaw|GAA88953.1 aluc|OJZ89453.1 anid|CBF83884.1 anig|EHA23134.1 aory|EIT74155.1 asyd|OJJ60942.1 ater|EAU35963.1 atub|OJI85341.1 aver|OJJ00268.1 awen|OJJ39360.1 azon|OJJ51699.1

Group3940: abra|OJJ76008.1 acar|OOF99917.1 acla|EAW09797.1 afis|EAW21339.1 afla|EED55754.1 afum|EBA27459.1 agla|OJJ87395.1 akaw|GAA88952.1 aluc|OJZ89454.1 anid|CBF83886.1 anig|EHA23133.1 aory|EIT76828.1 asyd|OJJ60943.1 ater|EAU35964.1 atub|OJI85340.1 aver|OJJ00269.1 awen|OJJ39359.1 azon|OJJ51698.1

Group3941: abra|OJJ76010.1 acar|OOF99914.1 acla|EAW09795.1 afis|EAW21341.1 afla|EED55756.1 afum|EAL92358.1 agla|OJJ87391.1 akaw|GAA88950.1 aluc|OJZ89456.1 anid|CBF76183.1 anig|EHA23131.1 aory|EIT76602.1 asyd|OJJ60962.1 ater|EAU35965.1 atub|OJI85338.1 aver|OJJ00287.1 awen|OJJ39356.1 azon|OJJ51697.1

Group3942: abra|OJJ76023.1 acar|OOF99901.1 acla|EAW09785.1 afis|EAW21353.1 afla|EED55768.1 afum|EAL92368.1 agla|OJJ87381.1 akaw|GAA88934.1 aluc|OJZ89473.1 anid|CBF76255.1 anig|EHA23118.1 aory|EIT76752.1 asyd|OJJ60965.1 ater|EAU35974.1 atub|OJI85325.1 aver|OJJ00290.1 awen|OJJ39347.1 azon|OJJ51691.1

Group3943: abra|OJJ76025.1 acar|OOF99902.1 acla|EAW09784.1 afis|EAW21354.1 afla|EED55769.1 afum|EAL92369.1 agla|OJJ87375.1 akaw|GAA88933.1 aluc|OJZ89475.1 anid|CBF76253.1 anig|EHA23117.1 aory|EIT76709.1 asyd|OJJ60966.1 ater|EAU35204.1 atub|OJI85324.1 aver|OJJ00291.1 awen|OJJ39346.1 azon|OJJ51690.1

Group3944: abra|OJJ76035.1 acar|OOF99893.1 acla|EAW09768.1 afis|EAW21370.1 afla|EED55788.1 afum|EAL92387.1 agla|OJJ85492.1 akaw|GAA88929.1 aluc|OJZ89490.1 anid|CBF83869.1 anig|EHA23108.1 aory|EIT76876.1 asyd|OJJ60933.1 ater|EAU35186.1 atub|OJI85313.1 aver|OJJ00259.1 awen|OJJ39327.1 azon|OJJ51672.1

Group3945: abra|OJJ76037.1 acar|OOF99892.1 acla|EAW09769.1 afis|EAW21369.1 afla|EED55786.1 afum|EAL92386.1 agla|OJJ85493.1 akaw|GAA88928.1 aluc|OJZ89492.1 anid|CBF83867.1 anig|EHA23107.1 aory|EIT76715.1 asyd|OJJ60932.1 ater|EAU35187.1 atub|OJI85311.1 aver|OJJ00258.1 awen|OJJ39328.1 azon|OJJ51673.1

Group3946: abra|OJJ76039.1 acar|OOF99890.1 acla|EAW09770.1 afis|EAW21368.1 afla|EED55782.1 afum|EAL92385.2 agla|OJJ85494.1 akaw|GAA88926.1 aluc|OJZ89495.1 anid|CBF83874.1 anig|EHA23105.1 aory|EIT76864.1 asyd|OJJ60937.1 ater|EAU35189.1 atub|OJI85308.1 aver|OJJ00263.1 awen|OJJ39329.1 azon|OJJ51674.1

Group3947: abra|OJJ76040.1 acar|OOF99889.1 acla|EAW09771.1 afis|EAW21367.1 afla|EED55781.1 afum|EAL92384.1 agla|OJJ85495.1 akaw|GAA88925.1 aluc|OJZ89496.1 anid|CBF83872.1 anig|EHA23104.1 aory|EIT76541.1 asyd|OJJ60936.1 ater|EAU35190.1 atub|OJI85307.1 aver|OJJ00262.1 awen|OJJ39330.1 azon|OJJ51675.1

Group3948: abra|OJJ76044.1 acar|OOF99886.1 acla|EAW09774.1 afis|EAW21364.1 afla|EED55778.1 afum|EAL92380.1 agla|OJJ85485.1 akaw|GAA88921.1 aluc|OJZ89500.1 anid|CBF83875.1 anig|EHA23101.1 aory|EIT76770.1 asyd|OJJ60938.1 ater|EAU35196.1 atub|OJI85303.1 aver|OJJ00264.1 awen|OJJ39334.1 azon|OJJ51679.1

Group3949: abra|OJJ76046.1 acar|OOF99884.1 acla|EAW09777.1 afis|EAW21361.1 afla|EED55775.1 afum|EAL92377.1 agla|OJJ85480.1 akaw|GAA88919.1 aluc|OJZ89502.1 anid|CBF76267.1 anig|EHA23098.1 aory|EIT76605.1 asyd|OJJ60973.1 ater|EAU35199.1 atub|OJI85301.1 aver|OJJ00298.1 awen|OJJ39337.1 azon|OJJ51682.1

Group3950: abra|OJJ76048.1 acar|OOF99882.1 acla|EAW09779.1 afis|EAW21359.1 afla|EED55774.1 afum|EAL92375.1 agla|OJJ85478.1 akaw|GAA88917.1 aluc|OJZ89504.1 anid|CBF76265.1 anig|EHA23096.1 aory|EIT76598.1 asyd|OJJ60972.1 ater|EAU35200.1 atub|OJI85299.1 aver|OJJ00297.1 awen|OJJ39339.1 azon|OJJ51684.1

Group3951: abra|OJJ76056.1 acar|OOF99874.1 acla|EAW09762.1 afis|EAW21375.1 afla|EED55793.1 afum|EAL92392.1 agla|OJJ85508.1 akaw|GAA88910.1 aluc|OJZ89511.1 anid|CBF83860.1 anig|EHA23088.1 aory|EIT76584.1 asyd|OJJ60926.1 ater|EAU35182.1 atub|OJI85292.1 aver|OJJ00252.1 awen|OJJ39322.1 azon|OJJ51667.1

Group3952: abra|OJJ76057.1 acar|OOF99873.1 acla|EAW09763.1 afis|EAW21374.1 afla|EED55792.1 afum|EAL92391.1 agla|OJJ85500.1 akaw|GAA88909.1 aluc|OJZ89512.1 anid|CBF83862.1 anig|EHA23087.1 aory|EIT76633.1 asyd|OJJ60927.1 ater|EAU35183.1 atub|OJI85291.1 aver|OJJ00253.1 awen|OJJ39323.1 azon|OJJ51668.1

Group3953: abra|OJJ76058.1 acar|OOF99872.1 acla|EAW09760.1 afis|EAW21381.1 afla|EED55795.1 afum|EAL92396.1 agla|OJJ85505.1 akaw|GAA88908.1 aluc|OJZ89513.1 anid|CBF83856.1 anig|EHA23086.1 aory|EIT76717.1 asyd|OJJ60924.1 ater|EAU35180.1 atub|OJI85290.1 aver|OJJ00250.1 awen|OJJ39320.1 azon|OJJ51665.1

Group3954: abra|OJJ76059.1 acar|OOF99871.1 acla|EAW09759.1 afis|EAW21382.1 afla|EED55796.1 afum|EAL92397.1 agla|OJJ85504.1 akaw|GAA88907.1 aluc|OJZ89514.1 anid|CBF83854.1 anig|EHA23085.1 aory|EIT76703.1 asyd|OJJ60923.1 ater|EAU35179.1 atub|OJI85289.1 aver|OJJ00249.1 awen|OJJ39319.1 azon|OJJ51664.1

Group3955: abra|OJJ76061.1 acar|OOF99870.1 acla|EAW09757.1 afis|EAW21384.1 afla|EED55798.1 afum|EAL92399.1 agla|OJJ85509.1 akaw|GAA88905.1 aluc|OJZ89516.1 anid|CBF83847.1 anig|EHA23083.1

aory|EIT76813.1 asyd|OJJ60919.1 ater|EAU35178.1 atub|OJI85287.1 aver|OJJ00245.1 awen|OJJ39317.1 azon|OJJ51530.1

Group3956: abra|OJJ76062.1 acar|OOF99869.1 acla|EAW09756.1 afis|EAW21385.1 afla|EED55799.1 afum|EAL92400.1 agla|OJJ85510.1 akaw|GAA88904.1 aluc|OJZ89517.1 anid|CBF83845.1 anig|EHA23082.1 aory|EIT76797.1 asyd|OJJ60918.1 ater|EAU35171.1 atub|OJI85286.1 aver|OJJ00244.1 awen|OJJ39316.1 azon|OJJ51531.1

Group3957: abra|OJJ76064.1 acar|OOF99865.1 acla|EAW09754.1 afis|EAW21387.1 afla|EED55801.1 afum|EAL92402.1 agla|OJJ85512.1 akaw|GAA88902.1 aluc|OJZ89519.1 anid|CBF83842.1 anig|EHA23081.1 aory|EIT76678.1 asyd|OJJ60916.1 ater|EAU35173.1 atub|OJI85284.1 aver|OJJ00243.1 awen|OJJ39314.1 azon|OJJ51532.1

Group3958: abra|OJJ76069.1 acar|OOF99862.1 acla|EAW09751.1 afis|EAW21391.1 afla|EED55805.1 afum|EAL92405.1 agla|OJJ85515.1 akaw|GAA88898.1 aluc|OJZ89523.1 anid|CBF83838.1 anig|EHA23078.1 aory|EIT76825.1 asyd|OJJ60913.1 ater|EAU35176.1 atub|OJI85280.1 aver|OJJ00240.1 awen|OJJ39312.1 azon|OJJ51538.1

Group3959: abra|OJJ76071.1 acar|OOF99857.1 acla|EAW09748.1 afis|EAW21394.1 afla|EED55811.1 afum|EAL92408.1 agla|OJJ85518.1 akaw|GAA88896.1 aluc|OJZ89525.1 anid|CBF83826.1 anig|EHA23076.1 aory|EIT76796.1 asyd|OJJ60904.1 ater|EAU35162.1 atub|OJI85278.1 aver|OJJ00230.1 awen|OJJ39310.1 azon|OJJ51542.1

Group3960: abra|OJJ76079.1 acar|OOF99846.1 acla|EAW09742.1 afis|EAW21400.1 afla|EED55821.1 afum|EAL92414.1 agla|OJJ85525.1 akaw|GAA88887.1 aluc|OJZ89536.1 anid|CBF83829.1 anig|EHA23068.1 aory|EIT76664.1 asyd|OJJ60906.1 ater|EAU35150.1 atub|OJI85270.1 aver|OJJ00232.1 awen|OJJ39303.1 azon|OJJ51549.1

Group3961: abra|OJJ76080.1 acar|OOF99845.1 acla|EAW09741.1 afis|EAW21401.1 afla|EED55822.1 afum|EAL92415.1 agla|OJJ85528.1 akaw|GAA88886.1 aluc|OJZ89538.1 anid|CBF83820.1 anig|EHA23067.1 aory|EIT76768.1 asyd|OJJ60901.1 ater|EAU35149.1 atub|OJI85269.1 aver|OJJ00227.1 awen|OJJ39302.1 azon|OJJ51550.1

Group3962: abra|OJJ76082.1 acar|OOF99844.1 acla|EAW09739.1 afis|EAW21402.1 afla|EED55824.1 afum|EAL92416.1 agla|OJJ85530.1 akaw|GAA88884.1 aluc|OJZ89540.1 anid|CBF83819.1 anig|EHA23066.1 aory|EIT76826.1 asyd|OJJ60900.1 ater|EAU29588.1 atub|OJI85267.1 aver|OJJ00225.1 awen|OJJ30539.1 azon|OJJ51552.1

Group3963: abra|OJJ76083.1 acar|OOF99843.1 acla|EAW09738.1 afis|EAW21403.1 afla|EED55825.1 afum|EAL92417.1 agla|OJJ85531.1 akaw|GAA88883.1 aluc|OJZ89541.1 anid|CBF83817.1 anig|EHA23065.1 aory|EIT76589.1 asyd|OJJ60899.1 ater|EAU35147.1 atub|OJI85266.1 aver|OJJ00224.1 awen|OJJ39300.1 azon|OJJ51553.1

Group3964: abra|OJJ76084.1 acar|OOF99842.1 acla|EAW09737.1 afis|EAW21404.1 afla|EED55826.1 afum|EAL92418.1 agla|OJJ85533.1 akaw|GAA88882.1 aluc|OJZ89542.1 anid|CBF83815.1 anig|EHA23064.1 aory|EIT76656.1 asyd|OJJ60898.1 ater|EAU35146.1 atub|OJI85265.1 aver|OJJ00223.1 awen|OJJ39299.1 azon|OJJ51554.1

Group3965: abra|OJJ76088.1 acar|OOF99837.1 acla|EAW09734.1 afis|EAW21407.1 afla|EED55829.1 afum|EAL92421.1 agla|OJJ85537.1 akaw|GAA88879.1 aluc|OJZ89545.1 anid|CBF83810.1 anig|EHA23061.1 aory|EIT76811.1 asyd|OJJ60895.1 ater|EAU35141.1 atub|OJI85261.1 aver|OJJ00220.1 awen|OJJ39294.1 azon|OJJ51560.1

Group3966: abra|OJJ76090.1 acar|OOF99835.1 acla|EAW09731.1 afla|EED55831.1 agla|OJJ85541.1 akaw|GAA88877.1 aluc|OJZ89547.1 anig|EHA23059.1 aory|EIT76755.1 atub|OJI85259.1 awen|OJJ39291.1 azon|OJJ51563.1 afis|EAW21410.1 afum|EAL92424.1 anid|CBF83805.1 asyd|OJJ60892.1 ater|EAU35136.1 aver|OJJ00216.1

Group3967: abra|OJJ76092.1 acar|OOF99833.1 acla|EAW09730.1 afis|EAW21411.1 afla|EED55834.1 afum|EAL92425.1 agla|OJJ85542.1 akaw|GAA88875.1 aluc|OJZ89548.1 anid|CBF83803.1 anig|EHA23058.1 aory|EIT76568.1 asyd|OJJ60891.1 ater|EAU35135.1 atub|OJI85258.1 aver|OJJ00215.1 awen|OJJ39290.1 azon|OJJ51564.1

Group3968: abra|OJJ76095.1 acar|OOF99829.1 acla|EAW09723.1 afis|EAW21418.1 afla|EED55843.1 afum|EAL92433.1 agla|OJJ85551.1 akaw|GAA88871.1 aluc|OJZ89551.1 anid|CBF83792.1 anig|EHA23055.1 aory|EIT76877.1 asyd|OJJ60881.1 ater|EAU35131.1 atub|OJI85254.1 aver|OJJ00207.1 awen|OJJ39285.1 azon|OJJ51569.1

Group3969: abra|OJJ76096.1 acar|OOF99828.1 acla|EAW09722.1 afis|EAW21419.1 afla|EED55844.1 afum|EAL92434.1 agla|OJJ85553.1 akaw|GAA88870.1 aluc|OJZ89552.1 anid|CBF83790.1 anig|EHA23054.1 aory|EIT76586.1 asyd|OJJ60880.1 ater|EAU35130.1 atub|OJI85253.1 aver|OJJ00206.1 awen|OJJ39284.1 azon|OJJ51570.1

Group3970: abra|OJJ76098.1 acar|OOF99827.1 acla|EAW09720.1 afis|EAW21421.1 afla|EED55846.1 afum|EAL92436.1 agla|OJJ85557.1 akaw|GAA88868.1 aluc|OJZ89554.1 anid|CBF83787.1 anig|EHA23052.1 aory|EIT76632.1 asyd|OJJ60878.1 ater|EAU35128.1 atub|OJI85251.1 aver|OJJ00203.1 awen|OJJ39280.1 azon|OJJ51572.1

Group3971: abra|OJJ76099.1 acar|OOF99825.1 acla|EAW09719.1 afis|EAW21422.1 afla|EED55847.1 afum|EAL92437.1 agla|OJJ85558.1 akaw|GAA88867.1 aluc|OJZ89555.1 anid|CBF83785.1 anig|EHA23051.1 aory|EIT76561.1 asyd|OJJ60877.1 ater|EAU35127.1 atub|OJI85249.1 aver|OJJ00202.1 awen|OJJ39279.1 azon|OJJ51573.1

Group3972: abra|OJJ76100.1 acar|OOF99824.1 acla|EAW09718.1 afis|EAW21423.1 afla|EED55848.1 afum|EAL92438.1 agla|OJJ85559.1 akaw|GAA88866.1 aluc|OJZ89556.1 anid|CBF83783.1 anig|EHA23050.1 aory|EIT76818.1 asyd|OJJ60876.1 ater|EAU35126.1 atub|OJI85248.1 aver|OJJ00201.1 awen|OJJ39278.1 azon|OJJ51574.1

Group3973: abra|OJJ76102.1 acar|OOF99823.1 acla|EAW09716.1 afis|EAW21425.1 afla|EED55850.1 afum|EAL92440.1 agla|OJJ85562.1 akaw|GAA88864.1 aluc|OJZ89558.1 anid|CBF83781.1 anig|EHA23048.1 aory|EIT76846.1 asyd|OJJ60874.1 ater|EAU35124.1 atub|OJI85246.1 aver|OJJ00199.1 awen|OJJ39276.1 azon|OJJ51576.1

Group3974: abra|OJJ76103.1 acar|OOF99822.1 acla|EAW09715.1 afis|EAW21426.1 afla|EED55853.1 afum|EAL92441.1 agla|OJJ85563.1 akaw|GAA88863.1 aluc|OJZ89559.1 anid|CBF83780.1 anig|EHA23047.1 aory|EIT76704.1 asyd|OJJ60873.1 ater|EAU35123.1 atub|OJI85245.1 aver|OJJ00198.1 awen|OJJ39275.1 azon|OJJ51577.1

Group3975: abra|OJJ76104.1 acar|OOF99821.1 acla|EAW09714.1 afis|EAW21427.1 afla|EED55854.1 afum|EAL92442.1 agla|OJJ85564.1 akaw|GAA88862.1 aluc|OJZ89560.1 anid|CBF83778.1 anig|EHA23046.1 aory|EIT76600.1 asyd|OJJ60871.1 ater|EAU35121.1 atub|OJI85244.1 aver|OJJ00197.1 awen|OJJ39274.1 azon|OJJ51579.1

Group3976: abra|OJJ76107.1 acar|OOF99819.1 acla|EAW09713.1 afis|EAW21428.1 afla|EED55855.1 afum|EAL92443.1 agla|OJJ85566.1 akaw|GAA88859.1 aluc|OJZ89563.1 anid|CBF83776.1 anig|EHA23043.1 aory|EIT76883.1 asyd|OJJ60870.1 ater|EAU35120.1 atub|OJI85241.1 aver|OJJ00195.1 awen|OJJ39273.1 azon|OJJ51582.1

Group3977: abra|OJJ76108.1 acar|OOF99818.1 acla|EAW09712.1 afis|EAW21429.1 afla|EED55856.1 afum|EAL92444.1 agla|OJJ85567.1 akaw|GAA88858.1 aluc|OJZ89564.1 anid|CBF83774.1 anig|EHA23042.1 aory|EIT76794.1 asyd|OJJ60869.1 ater|EAU35119.1 atub|OJI85240.1 aver|OJJ00194.1 awen|OJJ39271.1 azon|OJJ51583.1

Group3978: abra|OJJ76109.1 acar|OOF99817.1 acla|EAW09711.1 afis|EAW21430.1 afla|EED55857.1 afum|EAL92445.1 agla|OJJ85568.1 akaw|GAA88857.1 aluc|OJZ89565.1 anid|CBF83772.1 anig|EHA23041.1 aory|EIT76635.1 asyd|OJJ60868.1 ater|EAU35118.1 atub|OJI85239.1 aver|OJJ00193.1 awen|OJJ39270.1 azon|OJJ51584.1

Group3979: abra|OJJ76111.1 acar|OOF99815.1 acla|EAW09709.1 afis|EAW21431.1 afla|EED55859.1 afum|EAL92446.2 agla|OJJ85572.1 akaw|GAA88856.1 aluc|OJZ89567.1 anid|CBF83769.1 anig|EHA23040.1 aory|EIT76616.1 asyd|OJJ60865.1 ater|EAU35117.1 atub|OJI85237.1 aver|OJJ00191.1 awen|OJJ39269.1 azon|OJJ51587.1

Group3980: abra|OJJ76112.1 acar|OOF99813.1 acla|EAW09708.1 afis|EAW21432.1 afla|EED55860.1 afum|EAL92447.1 agla|OJJ85573.1 akaw|GAA88855.1 aluc|OJZ89568.1 anid|CBF83767.1 anig|EHA23039.1 aory|EIT76829.1 asyd|OJJ60864.1 ater|EAU35116.1 atub|OJI85236.1 aver|OJJ00190.1 awen|OJJ39267.1 azon|OJJ51588.1

Group3981: abra|OJJ76113.1 acar|OOF99812.1 acla|EAW09707.1 afis|EAW21433.1 afla|EED55862.1 afum|EAL92448.2 agla|OJJ85574.1 akaw|GAA88854.1 aluc|OJZ89569.1 anid|CBF83765.1 anig|EHA23037.1 aory|EIT76807.1 asyd|OJJ60863.1 ater|EAU35111.1 atub|OJI85234.1 aver|OJJ00189.1 awen|OJJ39266.1 azon|OJJ51589.1

Group3982: abra|OJJ76114.1 acar|OOF99811.1 acla|EAW09706.1 afis|EAW21434.1 afla|EED55863.1 afum|EAL92449.1 agla|OJJ85575.1 akaw|GAA88853.1 aluc|OJZ89571.1 anid|CBF76591.1 anig|EHA23036.1 aory|EIT76762.1 asyd|OJJ61175.1 ater|EAU35110.1 atub|OJI85233.1 aver|OJJ00501.1 awen|OJJ39265.1 azon|OJJ51590.1

Group3983: abra|OJJ76116.1 acar|OOF99810.1 acla|EAW09704.1 afis|EAW21436.1 afla|EED55864.1 afum|EAL92451.1 agla|OJJ85579.1 akaw|GAA88851.1 aluc|OJZ89573.1 anid|CBF76587.1 anig|EHA23034.1 aory|EIT76708.1 asyd|OJJ61173.1 ater|EAU35109.1 atub|OJI85231.1 aver|OJJ00499.1 awen|OJJ39263.1 azon|OJJ51592.1

Group3984: abra|OJJ76118.1 acar|OOF99807.1 acla|EAW09701.1 afis|EAW21438.1 afla|EED55866.1 afum|EAL92453.2 agla|OJJ85583.1 akaw|GAA88849.1 aluc|OJZ89575.1 anid|CBF76583.1 anig|EHA23032.1

aory|EIT76853.1 asyd|OJJ61171.1 ater|EAU35107.1 atub|OJI85229.1 aver|OJJ00496.1 awen|OJJ39260.1 azon|OJJ51595.1

Group3985: abra|OJJ76120.1 acar|OOF99805.1 acla|EAW09699.1 afis|EAW21440.1 afla|EED55868.1 afum|EAL92456.1 agla|OJJ85587.1 akaw|GAA88847.1 aluc|OJZ89577.1 anid|CBF76579.1 anig|EHA23031.1 aory|EIT76773.1 asyd|OJJ61169.1 ater|EAU35105.1 atub|OJI85227.1 aver|OJJ00494.1 awen|OJJ39258.1 azon|OJJ51598.1

Group3986: abra|OJJ76121.1 acar|OOF99804.1 acla|EAW09698.1 afis|EAW21441.1 afla|EED55869.1 afum|EAL92458.2 agla|OJJ85588.1 akaw|GAA88846.1 aluc|OJZ89578.1 anid|CBF76577.1 anig|EHA23030.1 aory|EIT76534.1 asyd|OJJ61168.1 ater|EAU35104.1 atub|OJI85226.1 aver|OJJ00493.1 awen|OJJ39257.1 azon|OJJ51599.1

Group3987: abra|OJJ76122.1 acar|OOF99803.1 acla|EAW09697.1 afis|EAW21442.1 afla|EED55870.1 afum|EAL92460.1 agla|OJJ85589.1 akaw|GAA88845.1 aluc|OJZ89579.1 anid|CBF76575.1 anig|EHA23029.1 aory|EIT76841.1 asyd|OJJ61167.1 ater|EAU35103.1 atub|OJI85225.1 aver|OJJ00492.1 awen|OJJ39256.1 azon|OJJ51600.1

Group3988: abra|OJJ76123.1 acar|OOF99802.1 acla|EAW09696.1 afis|EAW21443.1 afla|EED55871.1 afum|EAL92461.2 agla|OJJ85590.1 akaw|GAA88844.1 aluc|OJZ89580.1 anid|CBF76573.1 anig|EHA23028.1 aory|EIT76804.1 asyd|OJJ61166.1 ater|EAU35102.1 atub|OJI85224.1 aver|OJJ00491.1 awen|OJJ39255.1 azon|OJJ51601.1

Group3989: abra|OJJ76125.1 acar|OOF99800.1 acla|EAW09694.1 afis|EAW21445.1 afla|EED55873.1 afum|EAL92464.2 agla|OJJ85593.1 akaw|GAA88842.1 aluc|OJZ89583.1 anid|CBF76569.1 anig|EHA23026.1 aory|EIT76681.1 asyd|OJJ61164.1 ater|EAU35099.1 atub|OJI85220.1 aver|OJJ00488.1 awen|OJJ39253.1 azon|OJJ51602.1

Group3990: abra|OJJ76126.1 acar|OOF99798.1 acla|EAW09693.1 afis|EAW21446.1 afla|EED55874.1 afum|EAL92465.2 agla|OJJ85594.1 akaw|GAA88841.1 aluc|OJZ89587.1 anid|CBF76565.1 anig|EHA23024.1 aory|EIT76673.1 asyd|OJJ61162.1 ater|EAU35097.1 atub|OJI85216.1 aver|OJJ00486.1 awen|OJJ39252.1 azon|OJJ51603.1

Group3991: abra|OJJ76127.1 acar|OOF99796.1 acla|EAW09692.1 afis|EAW21447.1 afla|EED55876.1 afum|EAL92466.1 agla|OJJ85595.1 akaw|GAA88840.1 aluc|OJZ89588.1 anid|CBF76563.1 anig|EHA23023.1 aory|EIT76593.1 asyd|OJJ61161.1 ater|EAU35096.1 atub|OJI85214.1 aver|OJJ00483.1 awen|OJJ39251.1 azon|OJJ51604.1

Group3992: abra|OJJ76128.1 acar|OOF99795.1 acla|EAW09691.1 afis|EAW21448.1 afla|EED55877.1 afum|EAL92467.1 agla|OJJ85596.1 akaw|GAA88839.1 aluc|OJZ89589.1 anid|CBF76559.1 anig|EHA23022.1 aory|EIT76881.1 asyd|OJJ61159.1 ater|EAU35095.1 atub|OJI85213.1 aver|OJJ00481.1 awen|OJJ39250.1 azon|OJJ51605.1

Group3993: abra|OJJ76129.1 acar|OOF99794.1 acla|EAW09690.1 afis|EAW21449.1 afla|EED55878.1 afum|EAL92468.1 agla|OJJ85597.1 akaw|GAA88838.1 aluc|OJZ89590.1 anid|CBF76557.1 anig|EHA23021.1 aory|EIT76638.1 asyd|OJJ61158.1 ater|EAU35094.1 atub|OJI85212.1 aver|OJJ00480.1 awen|OJJ39249.1 azon|OJJ51606.1

Group3994: abra|OJJ76133.1 acar|OOF99790.1 acla|EAW09685.1 afis|EAW21454.1 afla|EED55884.1 afum|EAL92473.1 agla|OJJ85602.1 akaw|GAA88834.1 aluc|OJZ89594.1 anid|CBF76548.1 anig|EHA23017.1 aory|EIT76723.1 asyd|OJJ61155.1 ater|EAU35081.1 atub|OJI85208.1 aver|OJJ00477.1 awen|OJJ39246.1 azon|OJJ51609.1

Group3995: abra|OJJ76138.1 acar|OOF99786.1 acla|EAW09681.1 afis|EAW21458.1 afla|EED55888.1 afum|EAL92477.1 agla|OJJ85605.1 akaw|GAA88830.1 aluc|OJZ89601.1 anid|CBF76540.1 anig|EHA23014.1 aory|EIT76702.1 asyd|OJJ61151.1 ater|EAU35077.1 atub|OJI85203.1 aver|OJJ00473.1 awen|OJJ39242.1 azon|OJJ51613.1

Group3996: abra|OJJ76140.1 acar|OOF99785.1 acla|EAW09679.1 afis|EAW21460.1 afla|EED55891.1 afum|EAL92479.1 agla|OJJ85608.1 akaw|GAA88828.1 aluc|OJZ89603.1 anid|CBF76536.1 anig|EHA23012.1 aory|EIT76836.1 asyd|OJJ61149.1 ater|EAU35075.1 atub|OJI85201.1 aver|OJJ00471.1 awen|OJJ39239.1 azon|OJJ51615.1

Group3997: abra|OJJ76141.1 acar|OOF99784.1 acla|EAW09664.1 afis|EAW21473.1 afla|EED55908.1 afum|EAL92492.1 agla|OJJ85622.1 akaw|GAA88826.1 aluc|OJZ89605.1 anid|CBF76513.1 anig|EHA23011.1 aory|EIT76607.1 asyd|OJJ61136.1 ater|EAU35060.1 atub|OJI85198.1 aver|OJJ00458.1 awen|OJJ39224.1 azon|OJJ51630.1

Group3998: abra|OJJ76142.1 acar|OOF99783.1 acla|EAW09665.1 afis|EAW21472.1 afla|EED55902.1 afum|EAL92491.1 agla|OJJ85621.1 akaw|GAA88825.1 aluc|OJZ89606.1 anid|CBF76515.1 anig|EHA23010.1 aory|EIT76799.1 asyd|OJJ61137.1 ater|EAU35061.1 atub|OJI85197.1 aver|OJJ00459.1 awen|OJJ39225.1 azon|OJJ51629.1

Group3999: abra|OJJ76143.1 acar|OOF99782.1 acla|EAW09666.1 afis|EAW21471.1 afla|EED55904.1 afum|EAL92490.1 agla|OJJ85620.1 akaw|GAA88824.1 aluc|OJZ89607.1 anid|CBF76517.1 anig|EHA23009.1 aory|EIT76648.1 asyd|OJJ61138.1 ater|EAU35062.1 atub|OJI85196.1 aver|OJJ00460.1 awen|OJJ39227.1 azon|OJJ51628.1

Group4000: abra|OJJ76145.1 acar|OOF99781.1 acla|EAW09668.1 afis|EAW21469.1 afla|EED55907.1 afum|EAL92488.1 agla|OJJ85618.1 akaw|GAA82094.1 aluc|OJZ92086.1 anid|CBF76521.1 anig|EHA23008.1 aory|EIT76871.1 asyd|OJJ61140.1 ater|EAU35064.1 atub|OJI86515.1 aver|OJJ00462.1 awen|OJJ39229.1 azon|OJJ51626.1

Group4001: abra|OJJ76146.1 acar|OOF99780.1 acla|EAW09671.1 afis|EAW21467.1 afla|EED55900.1 afum|EAL92487.1 agla|OJJ85617.1 akaw|GAA82095.1 aluc|OJZ92084.1 anid|CBF76523.1 anig|EHA23007.1 aory|EIT76694.1 asyd|OJJ61141.1 ater|EAU35066.1 atub|OJI86514.1 aver|OJJ00463.1 awen|OJJ39230.1 azon|OJJ51625.1

Group4002: abra|OJJ76150.1 acar|OOF99777.1 acla|EAW09675.1 afis|EAW21464.1 afla|EED55896.1 afum|EAL92484.1 agla|OJJ85612.1 akaw|GAA82099.1 aluc|OJZ92079.1 anid|CBF76529.1 anig|EHA23005.1 aory|EIT76592.1 asyd|OJJ61145.1 ater|EAU35071.1 atub|OJI86509.1 aver|OJJ00467.1 awen|OJJ39235.1 azon|OJJ51620.1

Group4003: abra|OJJ76151.1 acar|OOF89898.1 acla|EAW09676.1 afis|EAW21463.1 afla|EED55895.1 afum|EAL92483.1 agla|OJJ85611.1 akaw|GAA82100.1 aluc|OJZ92078.1 anid|CBF76530.1 anig|EHA23004.1 aory|EIT76817.1 asyd|OJJ61146.1 ater|EAU35072.1 atub|OJI86508.1 aver|OJJ00468.1 awen|OJJ39236.1 azon|OJJ51619.1

Group4004: abra|OJJ76152.1 acar|OOF89899.1 acla|EAW09677.1 afis|EAW21462.1 afla|EED55894.1 afum|EAL92481.1 agla|OJJ85610.1 akaw|GAA82101.1 aluc|OJZ92077.1 anid|CBF76532.1 anig|EHA23003.1 aory|EIT76700.1 asyd|OJJ61147.1 ater|EAU35073.1 atub|OJI86507.1 aver|OJJ00469.1 awen|OJJ39237.1 azon|OJJ51618.1

Group4005: abra|OJJ76157.1 acar|OOF99775.1 acla|EAW09661.1 afis|EAW21478.1 afla|EED55910.1 afum|EAL92498.1 agla|OJJ85630.1 akaw|GAA82105.1 aluc|OJZ92071.1 anid|CBF76508.1 anig|EHA23000.1 aory|EIT76734.1 asyd|OJJ61133.1 ater|EAU35058.1 atub|OJI86503.1 aver|OJJ00454.1 awen|OJJ39221.1 azon|OJJ51635.1

Group4006: abra|OJJ76160.1 acar|OOF99772.1 acla|EAW09658.1 afis|EAW21481.1 afla|EED55913.1 afum|EAL92501.1 agla|OJJ85635.1 akaw|GAA82107.1 aluc|OJZ92069.1 anid|CBF76502.1 anig|EHA22997.1 aory|EIT76621.1 asyd|OJJ61130.1 ater|EAU35056.1 atub|OJI86500.1 aver|OJJ00451.1 awen|OJJ39216.1 azon|OJJ51638.1

Group4007: abra|OJJ76166.1 acar|OOF99771.1 acla|EAW09657.1 afis|EAW21482.1 afla|EED55914.1 afum|EAL92502.1 agla|OJJ85636.1 akaw|GAA82114.1 aluc|OJZ92062.1 anid|CBF76498.1 anig|EHA22991.1 aory|EIT76649.1 asyd|OJJ61128.1 ater|EAU35055.1 atub|OJI86494.1 aver|OJJ00449.1 awen|OJJ39215.1 azon|OJJ51640.1

Group4008: abra|OJJ76172.1 acar|OOF99767.1 acla|EAW09653.1 afis|EAW21486.1 afla|EED55919.1 afum|EAL92506.1 agla|OJJ85645.1 akaw|GAA82118.1 aluc|OJZ92058.1 anid|CBF76490.1 anig|EHA22987.1 aory|EIT76891.1 asyd|OJJ61124.1 ater|EAU35051.1 atub|OJI86490.1 aver|OJJ00445.1 awen|OJJ39211.1 azon|OJJ51643.1

Group4009: abra|OJJ76174.1 acar|OOF99766.1 acla|EAW09652.1 afis|EAW21488.1 afla|EED55923.1 afum|EAL92507.1 agla|OJJ85648.1 akaw|GAA82119.1 aluc|OJZ92056.1 anid|CBF76488.1 anig|EHA22985.1 aory|EIT76645.1 asyd|OJJ61123.1 ater|EAU35050.1 atub|OJI86487.1 aver|OJJ00441.1 awen|OJJ39207.1 azon|OJJ51645.1

Group4010: abra|OJJ76175.1 acar|OOF99765.1 acla|EAW09651.1 afis|EAW21489.1 afla|EED55924.1 afum|EAL92508.1 agla|OJJ85649.1 akaw|GAA82120.1 aluc|OJZ92055.1 anid|CBF76486.1 anig|EHA22984.1 aory|EIT76531.1 asyd|OJJ61122.1 ater|EAU35049.1 atub|OJI86486.1 aver|OJJ00440.1 awen|OJJ39206.1 azon|OJJ51646.1

Group4011: abra|OJJ76176.1 acar|OOF99764.1 acla|EAW09650.1 afis|EAW21490.1 afla|EED55925.1 afum|EAL92509.1 agla|OJJ85652.1 akaw|GAA82121.1 aluc|OJZ92054.1 anid|CBF76484.1 anig|EHA22983.1 aory|EIT76596.1 asyd|OJJ61121.1 ater|EAU35048.1 atub|OJI86485.1 aver|OJJ00439.1 awen|OJJ39205.1 azon|OJJ51647.1

Group4012: abra|OJJ76177.1 acar|OOF99762.1 acla|EAW09649.1 afis|EAW21491.1 afla|EED55926.1 afum|EAL92510.1 agla|OJJ85653.1 akaw|GAA82122.1 aluc|OJZ92053.1 anid|CBF76482.1 anig|EHA22982.1 aory|EIT76879.1 asyd|OJJ61120.1 ater|EAU35047.1 atub|OJI86484.1 aver|OJJ00438.1 awen|OJJ39204.1 azon|OJJ51648.1

Group4013: abra|OJJ76178.1 acar|OOF99763.1 acla|EAW09648.1 afis|EAW21492.1 afla|EED55927.1 afum|EAL92511.1 agla|OJJ85654.1 akaw|GAA82123.1 aluc|OJZ92052.1 anid|CBF76480.1 anig|EHA22981.1

aory|EIT76815.1 asyd|OJJ61119.1 ater|EAU35046.1 atub|OJI86483.1 aver|OJJ00437.1 awen|OJJ39203.1 azon|OJJ51649.1

Group4014: abra|OJJ76180.1 acar|OOF99760.1 acla|EAW09646.1 afis|EAW21494.1 afla|EED55929.1 afum|EAL92513.1 agla|OJJ85656.1 akaw|GAA82125.1 aluc|OJZ92049.1 anid|CBF76476.1 anig|EHA22979.1 aory|EIT76689.1 asyd|OJJ61117.1 ater|EAU35044.1 atub|OJI86481.1 aver|OJJ00435.1 awen|OJJ39201.1 azon|OJJ51526.1

Group4015: abra|OJJ76181.1 acar|OOF99759.1 acla|EAW09645.1 afis|EAW21495.1 afla|EED55930.1 afum|EAL92514.1 agla|OJJ85657.1 akaw|GAA82126.1 aluc|OJZ92048.1 anid|CBF76474.1 anig|EHA22978.1 aory|EIT76659.1 asyd|OJJ61116.1 ater|EAU35043.1 atub|OJI86480.1 aver|OJJ00434.1 awen|OJJ39200.1 azon|OJJ51525.1

Group4016: abra|OJJ76182.1 acar|OOF99758.1 acla|EAW09644.1 afis|EAW21496.1 afla|EED55931.1 afum|EAL92515.1 agla|OJJ85658.1 akaw|GAA82127.1 aluc|OJZ92047.1 anid|CBF76472.1 anig|EHA22977.1 aory|EIT76569.1 asyd|OJJ61115.1 ater|EAU35042.1 atub|OJI86479.1 aver|OJJ00433.1 awen|OJJ39199.1 azon|OJJ51524.1

Group4017: abra|OJJ76183.1 acar|OOF99757.1 acla|EAW09643.1 afis|EAW21497.1 afla|EED55933.1 afum|EAL92516.1 agla|OJJ85660.1 akaw|GAA82128.1 aluc|OJZ92046.1 anid|CBF76468.1 anig|EHA22976.1 aory|EIT76844.1 asyd|OJJ61114.1 ater|EAU35041.1 atub|OJI86478.1 aver|OJJ00430.1 awen|OJJ39198.1 azon|OJJ51523.1

Group4018: abra|OJJ76187.1 acar|OOF99755.1 acla|EAW09640.1 afis|EAW21500.1 afla|EED55936.1 afum|EAL92519.1 agla|OJJ85663.1 akaw|GAA82131.1 aluc|OJZ92042.1 anid|CBF76459.1 anig|EHA22974.1 aory|EIT76547.1 asyd|OJJ61111.1 ater|EAU35038.1 atub|OJI86475.1 aver|OJJ00427.1 awen|OJJ39195.1 azon|OJJ51522.1

Group4019: abra|OJJ76188.1 acar|OOF99754.1 acla|EAW09639.1 afis|EAW21501.1 afla|EED55937.1 afum|EAL92520.1 agla|OJJ85664.1 akaw|GAA82132.1 aluc|OJZ92041.1 anid|CBF76457.1 anig|EHA22973.1 aory|EIT76679.1 asyd|OJJ61110.1 ater|EAU35037.1 atub|OJI86474.1 aver|OJJ00426.1 awen|OJJ39194.1 azon|OJJ51519.1

Group4020: abra|OJJ76191.1 acar|OOF99751.1 acla|EAW13914.1 afis|EAW18625.1 afla|EED52717.1 afum|EAL93337.1 agla|OJJ81242.1 akaw|GAA82134.1 aluc|OJZ92038.1 anid|CBF70486.1 anig|EHA22970.1 aory|EIT78208.1 asyd|OJJ58585.1 ater|EAU34925.1 atub|OJI86472.1 aver|OJJ01189.1 awen|OJJ32090.1 azon|OJJ46667.1

Group4021: abra|OJJ76192.1 acar|OOF99752.1 acla|EAW13915.1 afis|EAW18626.1 afla|EED52716.1 afum|EAL93338.1 agla|OJJ81246.1 akaw|GAA82135.1 aluc|OJZ92037.1 anid|CBF70484.1 anig|EHA22969.1 aory|EIT78180.1 asyd|OJJ58586.1 ater|EAU34926.1 atub|OJI86470.1 aver|OJJ01190.1 awen|OJJ32092.1 azon|OJJ46669.1

Group4022: abra|OJJ76193.1 acar|OOF99750.1 acla|EAW13916.1 afis|EAW18627.1 afla|EED52715.1 afum|EAL93339.1 agla|OJJ81245.1 akaw|GAA82136.1 aluc|OJZ92036.1 anid|CBF70478.1 anig|EHA22968.1 aory|EIT78215.1 asyd|OJJ58587.1 ater|EAU38285.1 atub|OJI86469.1 aver|OJJ01191.1 awen|OJJ32093.1 azon|OJJ46670.1

Group4023: abra|OJJ76194.1 acar|OOF99749.1 acla|EAW13917.1 afis|EAW18628.1 afla|EED52714.1 afum|EAL93340.1 agla|OJJ81244.1 akaw|GAA82137.1 aluc|OJZ92035.1 anid|CBF70480.1 anig|EHA22967.1 aory|EIT78226.1 asyd|OJJ58588.1 ater|EAU38284.1 atub|OJI86468.1 aver|OJJ01192.1 awen|OJJ32094.1 azon|OJJ46671.1

Group4024: abra|OJJ76197.1 acar|OOF99747.1 acla|EAW13918.1 afis|EAW18629.1 afla|EED52712.1 afum|EAL93341.1 agla|OJJ81251.1 akaw|GAA82139.1 aluc|OJZ92032.1 anid|CBF70482.1 anig|EHA22965.1 aory|EIT78174.1 asyd|OJJ58592.1 ater|EAU38282.1 atub|OJI86465.1 aver|OJJ01194.1 awen|OJJ32096.1 azon|OJJ46675.1

Group4025: abra|OJJ76200.1 acar|OOF99744.1 acla|EAW13921.1 afis|EAW18632.1 afla|EED52709.1 afum|EAL93345.1 agla|OJJ82445.1 akaw|GAA82143.1 aluc|OJZ92028.1 anid|CBF70472.1 anig|EHA22962.1 aory|EIT74199.1 asyd|OJJ58595.1 ater|EAU38279.1 atub|OJI86462.1 aver|OJJ01197.1 awen|OJJ32099.1 azon|OJJ46681.1

Group4026: abra|OJJ76201.1 acar|OOF99743.1 acla|EAW13922.1 afis|EAW18633.1 afla|EED52708.1 afum|EAL93346.1 agla|OJJ82444.1 akaw|GAA82144.1 aluc|OJZ92027.1 anid|CBF70470.1 anig|EHA22961.1 aory|EIT74198.1 asyd|OJJ58596.1 ater|EAU38278.1 atub|OJI86461.1 aver|OJJ01198.1 awen|OJJ32100.1 azon|OJJ46682.1

Group4027: abra|OJJ76202.1 acar|OOF99742.1 acla|EAW13923.1 afis|EAW18634.1 afla|EED52707.1 afum|EAL93347.1 agla|OJJ82443.1 akaw|GAA82145.1 aluc|OJZ92026.1 anid|CBF70468.1 anig|EHA22960.1 aory|EIT74275.1 asyd|OJJ58597.1 ater|EAU38277.1 atub|OJI86460.1 aver|OJJ01199.1 awen|OJJ32101.1 azon|OJJ46683.1

Group4028: abra|OJJ76204.1 acar|OOF99741.1 acla|EAW13925.1 afis|EAW18636.1 afla|EED52705.1 afum|EAL93349.1 akaw|GAA82147.1 aluc|OJZ92024.1 anid|CBF70461.1 anig|EHA22958.1 aory|EIT74316.1 asyd|OJJ58600.1 atub|OJI86458.1 aver|OJJ01202.1 awen|OJJ32103.1 azon|OJJ46685.1 agla|OJJ82439.1 ater|EAU38275.1

Group4029: abra|OJJ76210.1 acar|OOF99739.1 acla|EAW13927.1 afis|EAW18638.1 afla|EED52702.1 afum|EAL93351.1 agla|OJJ82436.1 akaw|GAA82150.1 aluc|OJZ92018.1 anid|CBF70453.1 anig|EHA22956.1 aory|EIT74209.1 asyd|OJJ58604.1 ater|EAU38273.1 atub|OJI86453.1 aver|OJJ01207.1 awen|OJJ32106.1 azon|OJJ46687.1

Group4030: abra|OJJ76213.1 acar|OOF99737.1 acla|EAW13929.1 afis|EAW18640.1 afla|EED52700.1 afum|EAL93354.2 agla|OJJ82434.1 akaw|GAA82152.1 aluc|OJZ92016.1 anid|CBF70449.1 anig|EHA22955.1 aory|EIT74286.1 asyd|OJJ58606.1 ater|EAU38271.1 atub|OJI86451.1 aver|OJJ01210.1 awen|OJJ32108.1 azon|OJJ46689.1

Group4031: abra|OJJ76214.1 acar|OOF99736.1 acla|EAW13930.1 afis|EAW18641.1 afla|EED52699.1 afum|EAL93355.1 agla|OJJ82433.1 akaw|GAA82153.1 aluc|OJZ92015.1 anid|CBF70447.1 anig|EHA22954.1 aory|EIT74167.1 asyd|OJJ58607.1 ater|EAU38270.1 atub|OJI86450.1 aver|OJJ01211.1 awen|OJJ32109.1 azon|OJJ46690.1

Group4032: abra|OJJ76215.1 acar|OOF99735.1 acla|EAW13931.1 afis|EAW18642.1 afla|EED52698.1 afum|EAL93356.1 agla|OJJ82432.1 akaw|GAA82154.1 aluc|OJZ92014.1 anid|CBF70445.1 anig|EHA22953.1 aory|EIT74249.1 asyd|OJJ58608.1 ater|EAU38269.1 atub|OJI86449.1 aver|OJJ01212.1 awen|OJJ32110.1 azon|OJJ46691.1

Group4033: abra|OJJ76220.1 acar|OOF99731.1 acla|EAW13935.1 afis|EAW18646.1 afla|EED52696.1 afum|EAL93361.1 agla|OJJ82429.1 akaw|GAA82158.1 aluc|OJZ92010.1 anid|CBF70437.1 anig|EHA22950.1 aory|EIT74305.1 asyd|OJJ58611.1 ater|EAU38265.1 atub|OJI86445.1 aver|OJJ01217.1 awen|OJJ32114.1 azon|OJJ46695.1

Group4034: abra|OJJ76225.1 acar|OOF99472.1 acla|EAW14365.1 afis|EAW18915.1 afla|EED50899.1 afum|EBA27498.1 agla|OJJ81529.1 akaw|GAA82161.1 aluc|OJZ92006.1 anid|CBF69691.1 anig|EHA22946.1 aory|EIT78987.1 asyd|OJJ59055.1 ater|EAU38082.1 atub|OJI86441.1 aver|OJJ01649.1 awen|OJJ31508.1 azon|OJJ47227.1

Group4035: abra|OJJ76226.1 acar|OOF99473.1 acla|EAW14366.1 afis|EAW18916.1 afla|EED50900.1 afum|EAL93647.1 agla|OJJ81530.1 akaw|GAA82162.1 aluc|OJZ92005.1 anid|CBF69689.1 anig|EHA22945.1 aory|EIT79047.1 asyd|OJJ59056.1 ater|EAU38083.1 atub|OJI86440.1 aver|OJJ01650.1 awen|OJJ31509.1 azon|OJJ47228.1

Group4036: abra|OJJ76227.1 acar|OOF99474.1 acla|EAW14367.1 afis|EAW18917.1 afla|EED50901.1 afum|EAL93648.1 agla|OJJ81531.1 akaw|GAA82164.1 aluc|OJZ92002.1 anid|CBF69688.1 anig|EHA22943.1 aory|EIT79014.1 asyd|OJJ59057.1 ater|EAU38084.1 atub|OJI86438.1 aver|OJJ01651.1 awen|OJJ31510.1 azon|OJJ47229.1

Group4037: abra|OJJ76228.1 acar|OOF99475.1 acla|EAW14368.1 afis|EAW18918.1 afla|EED50902.1 afum|EAL93649.2 agla|OJJ81532.1 akaw|GAA82165.1 aluc|OJZ92001.1 anid|CBF69686.1 anig|EHA22942.1 aory|EIT78973.1 asyd|OJJ59058.1 ater|EAU38085.1 atub|OJI86437.1 aver|OJJ01652.1 awen|OJJ31511.1 azon|OJJ47230.1

Group4038: abra|OJJ76242.1 acar|OOF99487.1 acla|EAW14197.1 afis|EAW18925.1 afla|EED50914.1 afum|EBA27500.1 agla|OJJ85252.1 akaw|GAA82175.1 aluc|OJZ91989.1 anid|CBF81679.1 anig|EHA22936.1 aory|EIT79011.1 asyd|OJJ59560.1 ater|EAU38100.1 atub|OJI86424.1 aver|OJJ02708.1 awen|OJJ31519.1 azon|OJJ47235.1

Group4039: abra|OJJ76245.1 acar|OOF99491.1 acla|EAW14362.1 afis|EAW18927.1 afla|EED50920.1 afum|EAL93658.1 agla|OJJ85256.1 akaw|GAA82178.1 aluc|OJZ91986.1 anid|CBF69671.1 anig|EHA22932.1 aory|EIT78966.1 asyd|OJJ59063.1 ater|EAU38106.1 atub|OJI86421.1 aver|OJJ01657.1 awen|OJJ31521.1 azon|OJJ47237.1

Group4040: abra|OJJ76246.1 acar|OOF99492.1 acla|EAW14361.1 afis|EAW18928.1 afla|EED50921.1 afum|EAL93659.1 agla|OJJ85257.1 akaw|GAA82179.1 aluc|OJZ91985.1 anid|CBF69669.1 anig|EHA22931.1 aory|EIT79013.1 asyd|OJJ59062.1 ater|EAU38107.1 atub|OJI86420.1 aver|OJJ01656.1 awen|OJJ31522.1 azon|OJJ47238.1

Group4041: abra|OJJ76248.1 acar|OOF89844.1 acla|EAW14359.1 afis|EAW18931.1 afla|EED50936.1 afum|EAL93663.1 agla|OJJ85260.1 akaw|GAA82181.1 aluc|OJZ91982.1 anid|CBF86796.1 anig|EHA22929.1 aory|EIT78972.1 asyd|OJJ64670.1 ater|EAU38110.1 atub|OJI86418.1 aver|OJI98772.1 awen|OJJ31525.1 azon|OJJ47240.1

Group4042: abra|OJJ76250.1 acar|OOF99495.1 acla|EAW14357.1 afis|EAW18933.1 afla|EED50938.1 afum|EAL93666.2 agla|OJJ85262.1 akaw|GAA82183.1 aluc|OJZ91977.1 anid|CBF86800.1 anig|EHA22927.1

aory|EIT77014.1 asyd|OJJ64668.1 ater|EAU38112.1 atub|OJI86416.1 aver|OJI98770.1 awen|OJJ31527.1 azon|OJJ47242.1

Group4043: abra|OJJ76251.1 acar|OOF99496.1 acla|EAW14356.1 afis|EAW18934.1 afla|EED50939.1 afum|EAL93667.2 agla|OJJ85263.1 akaw|GAA82184.1 aluc|OJZ91976.1 anid|CBF86802.1 anig|EHA22926.1 aory|EIT77060.1 asyd|OJJ64667.1 ater|EAU38113.1 atub|OJI86415.1 aver|OJI98769.1 awen|OJJ31528.1 azon|OJJ47243.1

Group4044: abra|OJJ76252.1 acar|OOF99497.1 acla|EAW14355.1 afis|EAW18935.1 afla|EED50940.1 afum|EAL93668.2 agla|OJJ85264.1 akaw|GAA82185.1 aluc|OJZ91975.1 anid|CBF86804.1 anig|EHA22925.1 aory|EIT76996.1 asyd|OJJ64666.1 ater|EAU38114.1 atub|OJI86414.1 aver|OJI98768.1 awen|OJJ31529.1 azon|OJJ47244.1

Group4045: abra|OJJ76253.1 acar|OOF99498.1 acla|EAW14354.1 afis|EAW18936.1 afla|EED50941.1 afum|EAL93669.2 agla|OJJ85265.1 akaw|GAA82186.1 aluc|OJZ91974.1 anid|CBF86805.1 anig|EHA22924.1 aory|EIT77063.1 asyd|OJJ64665.1 ater|EAU38115.1 atub|OJI86413.1 aver|OJI98767.1 awen|OJJ31530.1 azon|OJJ47245.1

Group4046: abra|OJJ76254.1 acar|OOF99499.1 acla|EAW14353.1 afis|EAW18937.1 afla|EED50942.1 afum|EAL93670.1 agla|OJJ85270.1 akaw|GAA82188.1 aluc|OJZ91972.1 anid|CBF86807.1 anig|EHA22918.1 aory|EIT77016.1 asyd|OJJ64664.1 ater|EAU38116.1 atub|OJI86411.1 aver|OJI98766.1 awen|OJJ31532.1 azon|OJJ47246.1

Group4047: abra|OJJ76256.1 acar|OOF99501.1 acla|EAW14350.1 afis|EAW18940.1 afla|EED50966.1 afum|EAL93673.1 agla|OJJ85271.1 akaw|GAA82190.1 aluc|OJZ91970.1 anid|CBF86811.1 anig|EHA22920.1 aory|EIT76988.1 asyd|OJJ64661.1 ater|EAU38119.1 atub|OJI86409.1 aver|OJI98763.1 awen|OJJ31534.1 azon|OJJ47250.1

Group4048: abra|OJJ76259.1 acar|OOF99504.1 acla|EAW14351.1 afis|EAW18939.1 afla|EED50969.1 afum|EAL93672.1 agla|OJJ85268.1 akaw|GAA82192.1 aluc|OJZ91964.1 anid|CBF86815.1 anig|EHA22923.1 aory|EIT77055.1 asyd|OJJ64658.1 ater|EAU38120.1 atub|OJI86407.1 aver|OJI98761.1 awen|OJJ31535.1 azon|OJJ47249.1

Group4049: abra|OJJ76262.1 acar|OOF99506.1 acla|EAW14346.1 afis|EAW18944.1 afla|EED50978.1 afum|EAL93677.1 agla|OJJ85278.1 akaw|GAA82194.1 aluc|OJZ91960.1 anid|CBF86822.1 anig|EHA22916.1 aory|EIT77009.1 asyd|OJJ64655.1 ater|EAU38124.1 atub|OJI86404.1 aver|OJI98757.1 awen|OJJ31540.1 azon|OJJ47257.1

Group4050: abra|OJJ76266.1 afla|EED50981.1 afum|EAL93681.2 anid|CBF86829.1 aory|EIT76942.1 asyd|OJJ64651.1 atub|OJI86400.1 aver|OJI98752.1 acar|OOF99509.1 acla|EAW14343.1 afis|EAW18947.1 agla|OJJ85308.1 akaw|GAA82198.1 aluc|OJZ91954.1 anig|EHA22913.1 ater|EAU38127.1 awen|OJJ31543.1 azon|OJJ47260.1

Group4051: abra|OJJ76267.1 acar|OOF99511.1 acla|EAW14341.1 afis|EAW18949.1 afla|EED50985.1 afum|EAL93683.1 agla|OJJ85320.1 akaw|GAA82200.1 aluc|OJZ91953.1 anid|CBF86833.1 anig|EHA22912.1 aory|EIT77049.1 asyd|OJJ64649.1 ater|EAU38135.1 atub|OJI86398.1 aver|OJI98749.1 awen|OJJ31547.1 azon|OJJ47263.1

Group4052: abra|OJJ76269.1 acar|OOF99517.1 acla|EAW12851.1 afis|EAW18014.1 afla|EED51880.1 afum|EAL91826.1 agla|OJJ85368.1 akaw|GAA82202.1 aluc|OJZ91951.1 anid|CBF80042.1 anig|EHA22910.1 aory|EIT81710.1 asyd|OJJ58276.1 ater|EAU31530.1 atub|OJI86396.1 aver|OJJ02148.1 awen|OJJ41526.1 azon|OJJ48247.1

Group4053: abra|OJJ76288.1 acar|OOF99529.1 acla|EAW14328.1 afis|EAW18960.1 afla|EED51006.1 afum|EAL93693.1 agla|OJJ85342.1 akaw|GAA82217.1 aluc|OJZ91934.1 anid|CBF69600.1 anig|EHA22894.1 aory|EIT77056.1 asyd|OJJ59135.1 ater|EAU38143.1 atub|OJI86376.1 aver|OJJ01724.1 awen|OJJ31559.1 azon|OJJ47276.1

Group4054: abra|OJJ76290.1 acar|OOF99535.1 acla|EAW14325.1 afis|EAW18963.1 afla|EED51010.1 afum|EAL93695.1 agla|OJJ85344.1 akaw|GAA82219.1 aluc|OJZ91932.1 anid|CBF69605.1 anig|EHA22893.1 aory|EIT76961.1 asyd|OJJ59132.1 ater|EAU38148.1 atub|OJI86374.1 aver|OJJ01721.1 awen|OJJ31562.1 azon|OJJ47283.1

Group4055: abra|OJJ76292.1 acar|OOF99537.1 acla|EAW14324.1 afis|EAW18964.1 afla|EED51011.1 afum|EAL93696.1 agla|OJJ85346.1 akaw|GAA82220.1 aluc|OJZ91930.1 anid|CBF69607.1 anig|EHA22892.1 aory|EIT77048.1 asyd|OJJ59130.1 atub|OJI86373.1 aver|OJJ01720.1 awen|OJJ31564.1 azon|OJJ47282.1 ater|EAU38149.1

Group4056: abra|OJJ76296.1 acar|OOF99542.1 acla|EAW14322.1 afis|EAW18966.1 afla|EED51032.1 afum|EAL93698.2 agla|OJJ85352.1 akaw|GAA82223.1 aluc|OJZ91926.1 anid|CBF69611.1 anig|EHA22889.1 aory|EIT76971.1 asyd|OJJ59128.1 ater|EAU38151.1 atub|OJI86370.1 aver|OJJ01716.1 awen|OJJ31566.1 azon|OJJ47284.1

Group4057: abra|OJJ76297.1 acar|OOF99545.1 acla|EAW14321.1 afis|EAW18967.1 afla|EED51033.1 afum|EAL93699.1 agla|OJJ85353.1 akaw|GAA82224.1 aluc|OJZ91925.1 anid|CBF69613.1 anig|EHA22888.1 aory|EIT76951.1 asyd|OJJ59127.1 ater|EAU38153.1 atub|OJI86369.1 aver|OJJ01715.1 awen|OJJ31567.1 azon|OJJ47285.1

Group4058: abra|OJJ76299.1 acar|OOF99546.1 acla|EAW14319.1 afis|EAW18969.1 afla|EED51035.1 afum|EAL93702.1 agla|OJJ85355.1 akaw|GAA82226.1 aluc|OJZ91923.1 anid|CBF69617.1 anig|EHA22887.1 aory|EIT77006.1 asyd|OJJ59124.1 ater|EAU38155.1 atub|OJI86367.1 aver|OJJ01713.1 awen|OJJ31569.1 azon|OJJ47287.1

Group4059: abra|OJJ76300.1 acar|OOF99544.1 acla|EAW14318.1 afis|EAW18970.1 afla|EED51036.1 afum|EAL93703.1 agla|OJJ85356.1 akaw|GAA82227.1 aluc|OJZ91922.1 anid|CBF69619.1 anig|EHA22886.1 aory|EIT77012.1 asyd|OJJ59123.1 ater|EAU38156.1 atub|OJI86366.1 aver|OJJ01712.1 awen|OJJ31570.1 azon|OJJ47288.1

Group4060: abra|OJJ76301.1 acar|OOF99547.1 acla|EAW14317.1 afis|EAW18971.1 afla|EED51037.1 afum|EAL93704.1 agla|OJJ85357.1 akaw|GAA82228.1 aluc|OJZ91921.1 anid|CBF69621.1 anig|EHA22885.1 aory|EIT77040.1 asyd|OJJ59122.1 ater|EAU38157.1 atub|OJI86365.1 aver|OJJ01711.1 awen|OJJ31571.1 azon|OJJ47289.1

Group4061: abra|OJJ76302.1 acar|OOF99548.1 acla|EAW14316.1 afis|EAW18972.1 afla|EED51038.1 afum|EAL93705.1 agla|OJJ85360.1 akaw|GAA82232.1 aluc|OJZ91916.1 anid|CBF69627.1 anig|EHA22884.1 aory|EIT77044.1 asyd|OJJ59119.1 ater|EAU38158.1 atub|OJI86364.1 aver|OJJ01708.1 awen|OJJ31572.1 azon|OJJ47291.1

Group4062: abra|OJJ76303.1 acar|OOF99549.1 acla|EAW14315.1 afis|EAW18973.1 afla|EED51039.1 afum|EAL93706.1 agla|OJJ85361.1 akaw|GAA82233.1 aluc|OJZ91914.1 anid|CBF69629.1 anig|EHA22883.1 aory|EIT77038.1 asyd|OJJ59117.1 ater|EAU38159.1 atub|OJI86363.1 aver|OJJ01707.1 awen|OJJ31573.1 azon|OJJ47290.1

Group4063: abra|OJJ76307.1 acar|OOF99552.1 acla|EAW14108.1 afis|EAW18810.1 afla|EED50762.1 afum|EAL93546.1 agla|OJJ80543.1 akaw|GAA82238.1 aluc|OJZ91908.1 anid|CBF69793.1 anig|EHA22882.1 aory|EIT78035.1 asyd|OJJ58963.1 ater|EAU37962.1 atub|OJI86358.1 aver|OJJ01557.1 awen|OJJ32385.1 azon|OJJ47146.1

Group4064: abra|OJJ76314.1 acar|OOF99566.1 acla|EAW14101.1 afis|EAW18802.1 afla|EED50774.1 afum|EAL93538.1 agla|OJJ80532.1 akaw|GAA82245.1 aluc|OJZ91900.1 anid|CBF69784.1 anig|EHA22879.1 aory|EIT77990.1 asyd|OJJ58960.1 ater|EAU37940.1 atub|OJI86353.1 aver|OJJ01554.1 awen|OJJ32376.1 azon|OJJ47139.1

Group4065: abra|OJJ76323.1 acar|OOF99575.1 acla|EAW14092.1 afis|EAW18798.1 afla|EED50791.1 afum|EAL93534.1 agla|OJJ80527.1 akaw|GAA82252.1 aluc|OJZ91892.1 anid|CBF69777.1 anig|EHA22871.1 aory|EIT78043.1 asyd|OJJ58956.1 ater|EAU37930.1 atub|OJI86344.1 aver|OJJ01550.1 awen|OJJ32372.1 azon|OJJ47135.1

Group4066: abra|OJJ76325.1 acar|OOF99577.1 acla|EAW14090.1 afis|EAW18796.1 afla|EED50793.1 afum|EAL93532.1 agla|OJJ80525.1 akaw|GAA82254.1 aluc|OJZ91890.1 anid|CBF69773.1 anig|EHA22870.1 aory|EIT78128.1 asyd|OJJ58954.1 ater|EAU37927.1 atub|OJI86342.1 aver|OJJ01548.1 awen|OJJ32370.1 azon|OJJ47133.1

Group4067: abra|OJJ76326.1 acar|OOF99578.1 afis|EAW18795.1 afla|EED50794.1 afum|EAL93531.1 agla|OJJ80524.1 akaw|GAA82255.1 aluc|OJZ91889.1 anid|CBF69771.1 anig|EHA22869.1 aory|EIT78000.1 asyd|OJJ58953.1 ater|EAU37926.1 atub|OJI86341.1 aver|OJJ01547.1 awen|OJJ32369.1 azon|OJJ47132.1 acla|EAW14089.1

Group4068: abra|OJJ76327.1 acar|OOF99580.1 acla|EAW14088.1 afis|EAW18794.1 afla|EED50795.1 afum|EAL93530.1 agla|OJJ80523.1 akaw|GAA82256.1 aluc|OJZ91888.1 anid|CBF69769.1 anig|EHA22868.1 aory|EIT78079.1 asyd|OJJ58952.1 ater|EAU37925.1 atub|OJI86340.1 aver|OJJ01546.1 awen|OJJ32367.1 azon|OJJ47131.1

Group4069: abra|OJJ76332.1 acar|OOF99584.1 acla|EAW14083.1 afis|EAW18790.1 afla|EED50748.1 afum|EAL93525.1 agla|OJJ80519.1 akaw|GAA82260.1 aluc|OJZ91881.1 anid|CBF69761.1 anig|EHA22859.1 aory|EIT78056.1 asyd|OJJ58947.1 ater|EAU37921.1 atub|OJI86332.1 aver|OJJ01542.1 awen|OJJ32363.1 azon|OJJ47126.1

Group4070: abra|OJJ76334.1 acar|OOF99587.1 acla|EAW14079.1 afis|EAW18786.1 afla|EED50745.1 afum|EAL93519.1 agla|OJJ80509.1 akaw|GAA82262.1 aluc|OJZ91878.1 anid|CBF69752.1 anig|EHA22856.1 aory|EIT78009.1 asyd|OJJ58941.1 ater|EAU37918.1 atub|OJI86329.1 aver|OJJ01537.1 awen|OJJ32360.1 azon|OJJ47123.1

Group4071: abra|OJJ76336.1 acar|OOF99588.1 acla|EAW14078.1 afis|EAW18785.1 afla|EED50744.1 afum|EAL93517.2 agla|OJJ80507.1 akaw|GAA82264.1 aluc|OJZ91876.1 anid|CBF69748.1 anig|EHA22855.1

aory|EIT78137.1 asyd|OJJ58938.1 ater|EAU37917.1 atub|OJI86327.1 aver|OJJ01534.1 awen|OJJ32358.1 azon|OJJ47121.1

Group4072: abra|OJJ76337.1 acar|OOF99589.1 acla|EAW14077.1 afis|EAW18784.1 afla|EED50743.1 afum|EAL93516.2 agla|OJJ80506.1 akaw|GAA82265.1 aluc|OJZ91875.1 anid|CBF69746.1 anig|EHA22854.1 aory|EIT78027.1 asyd|OJJ58937.1 ater|EAU37916.1 atub|OJI86326.1 aver|OJJ01533.1 awen|OJJ32357.1 azon|OJJ47120.1

Group4073: abra|OJJ76338.1 acar|OOF99590.1 acla|EAW14076.1 afis|EAW18783.1 afla|EED50742.1 afum|EAL93515.1 agla|OJJ80504.1 akaw|GAA82266.1 aluc|OJZ91874.1 anid|CBF69744.1 anig|EHA22853.1 aory|EIT78012.1 asyd|OJJ58936.1 ater|EAU37915.1 atub|OJI86325.1 aver|OJJ01532.1 awen|OJJ32356.1 azon|OJJ47119.1

Group4074: abra|OJJ76340.1 acar|OOF99591.1 acla|EAW14075.1 afis|EAW18782.1 afla|EED50741.1 afum|EAL93514.1 agla|OJJ80502.1 akaw|GAA82267.1 aluc|OJZ91873.1 anid|CBF69742.1 anig|EHA22852.1 aory|EIT78041.1 asyd|OJJ58935.1 ater|EAU37914.1 atub|OJI86324.1 aver|OJJ01531.1 awen|OJJ32354.1 azon|OJJ47118.1

Group4075: abra|OJJ76342.1 acar|OOF99593.1 acla|EAW14073.1 afis|EAW18780.1 afla|EED50739.1 afum|EAL93512.1 agla|OJJ80500.1 akaw|GAA82269.1 aluc|OJZ91871.1 anid|CBF69738.1 anig|EHA22850.1 aory|EIT77981.1 asyd|OJJ58933.1 ater|EAU37912.1 atub|OJI86322.1 aver|OJJ01529.1 awen|OJJ32352.1 azon|OJJ47116.1

Group4076: abra|OJJ76344.1 acar|OOF99595.1 acla|EAW14071.1 afis|EAW18778.1 afla|EED50737.1 afum|EAL93510.1 agla|OJJ80495.1 akaw|GAA82271.1 aluc|OJZ91869.1 anid|CBF69734.1 anig|EHA22849.1 aory|EIT78123.1 asyd|OJJ58931.1 ater|EAU37910.1 atub|OJI86320.1 aver|OJJ01527.1 awen|OJJ32349.1 azon|OJJ47114.1

Group4077: abra|OJJ76345.1 acar|OOF99596.1 acla|EAW14070.1 afis|EAW18777.1 afla|EED50736.1 afum|EAL93509.1 agla|OJJ80494.1 akaw|GAA82272.1 aluc|OJZ91868.1 anid|CBF69732.1 anig|EHA22848.1 aory|EIT78071.1 asyd|OJJ58930.1 ater|EAU37909.1 atub|OJI86319.1 aver|OJJ01526.1 awen|OJJ32348.1 azon|OJJ47113.1

Group4078: abra|OJJ76346.1 acar|OOF99597.1 acla|EAW14069.1 afis|EAW18776.1 afla|EED50735.1 afum|EAL93508.1 agla|OJJ80493.1 akaw|GAA82273.1 aluc|OJZ91867.1 anid|CBF69730.1 anig|EHA22847.1 aory|EIT78130.1 asyd|OJJ58929.1 ater|EAU37908.1 atub|OJI86318.1 aver|OJJ01525.1 awen|OJJ32347.1 azon|OJJ47112.1

Group4079: abra|OJJ76347.1 acar|OOF99598.1 acla|EAW14068.1 afis|EAW18775.1 afla|EED50734.1 afum|EAL93507.1 agla|OJJ80491.1 akaw|GAA82274.1 aluc|OJZ91866.1 anid|CBF69728.1 anig|EHA22846.1 aory|EIT78021.1 asyd|OJJ58928.1 ater|EAU37906.1 atub|OJI86317.1 aver|OJJ01524.1 awen|OJJ32346.1 azon|OJJ47111.1

Group4080: abra|OJJ76349.1 acar|OOF99600.1 acla|EAW14066.1 afis|EAW18773.1 afla|EED50732.1 afum|EAL93505.1 agla|OJJ80489.1 akaw|GAA82276.1 aluc|OJZ91864.1 anid|CBF69724.1 anig|EHA22844.1 aory|EIT78029.1 asyd|OJJ58927.1 ater|EAU37904.1 atub|OJI86315.1 aver|OJJ01523.1 awen|OJJ32344.1 azon|OJJ47110.1

Group4081: abra|OJJ76351.1 acar|OOF99603.1 acla|EAW14064.1 afis|EAW18771.1 afla|EED50730.1 afum|EAL93502.1 agla|OJJ80467.1 akaw|GAA82278.1 aluc|OJZ91862.1 anid|CBF69934.1 anig|EHA22842.1 aory|EIT78074.1 asyd|OJJ58922.1 ater|EAU37902.1 atub|OJI86314.1 aver|OJJ01519.1 awen|OJJ32340.1 azon|OJJ47106.1

Group4082: abra|OJJ76353.1 acar|OOF99604.1 acla|EAW14062.1 afis|EAW18769.1 afla|EED50728.1 afum|EAL93500.2 agla|OJJ82285.1 akaw|GAA82279.1 aluc|OJZ91860.1 anid|CBF69938.1 anig|EHA22840.1 aory|EIT78133.1 asyd|OJJ58920.1 ater|EAU37900.1 atub|OJI86311.1 aver|OJJ01517.1 awen|OJJ32338.1 azon|OJJ47103.1

Group4083: abra|OJJ76354.1 acar|OOF99605.1 acla|EAW14061.1 afis|EAW18768.1 afla|EED50727.1 afum|EAL93499.1 agla|OJJ82284.1 akaw|GAA82280.1 aluc|OJZ91859.1 anid|CBF69940.1 anig|EHA22839.1 aory|EIT78038.1 asyd|OJJ58919.1 ater|EAU37899.1 atub|OJI86310.1 aver|OJJ01516.1 awen|OJJ32337.1 azon|OJJ47102.1

Group4084: abra|OJJ76356.1 acar|OOF99607.1 acla|EAW14059.1 afis|EAW18765.1 afla|EED50725.1 afum|EAL93497.1 agla|OJJ80456.1 akaw|GAA82282.1 aluc|OJZ91857.1 anid|CBF69944.1 anig|EHA22837.1 aory|EIT78144.1 asyd|OJJ58917.1 ater|EAU37895.1 atub|OJI86308.1 aver|OJJ01514.1 awen|OJJ32335.1 azon|OJJ47101.1

Group4085: abra|OJJ76358.1 acar|OOF99609.1 acla|EAW14058.1 afis|EAW18764.1 afla|EED50724.1 afum|EAL93496.2 agla|OJJ80457.1 akaw|GAA82284.1 aluc|OJZ91855.1 anid|CBF69945.1 anig|EHA22836.1 aory|EIT78031.1 asyd|OJJ58915.1 ater|EAU37894.1 atub|OJI86306.1 aver|OJJ01512.1 awen|OJJ32333.1 azon|OJJ47099.1

Group4086: abra|OJJ76359.1 acar|OOF99610.1 acla|EAW14057.1 afis|EAW18763.1 afla|EED50723.1 afum|EAL93495.1 agla|OJJ80458.1 akaw|GAA82285.1 aluc|OJZ91854.1 anid|CBF69947.1 anig|EHA22835.1 aory|EIT78108.1 asyd|OJJ58914.1 ater|EAU37893.1 atub|OJI86305.1 aver|OJJ01511.1 awen|OJJ32332.1 azon|OJJ47098.1

Group4087: abra|OJJ76362.1 acar|OOF99613.1 acla|EAW14054.1 afis|EAW18760.1 afla|EED50720.1 afum|EAL93491.1 agla|OJJ82333.1 akaw|GAA82288.1 aluc|OJZ91850.1 anid|CBF69953.1 anig|EHA22832.1 aory|EIT78011.1 asyd|OJJ58911.1 ater|EAU37891.1 atub|OJI86302.1 aver|OJJ01508.1 awen|OJJ32330.1 azon|OJJ47095.1

Group4088: abra|OJJ76363.1 acar|OOF99614.1 acla|EAW14053.1 afis|EAW18759.1 afla|EED50719.1 afum|EAL93489.1 agla|OJJ82332.1 akaw|GAA82289.1 aluc|OJZ91849.1 anid|CBF69955.1 anig|EHA22831.1 aory|EIT78126.1 asyd|OJJ58910.1 ater|EAU37928.1 atub|OJI86301.1 aver|OJJ01507.1 awen|OJJ32329.1 azon|OJJ47094.1

Group4089: abra|OJJ76364.1 acar|OOF99615.1 acla|EAW14052.1 afis|EAW18758.1 afla|EED50718.1 afum|EAL93488.2 agla|OJJ82331.1 akaw|GAA82290.1 aluc|OJZ91848.1 anid|CBF69957.1 anig|EHA22830.1 aory|EIT78109.1 asyd|OJJ58909.1 ater|EAU37890.1 atub|OJI86300.1 aver|OJJ01506.1 awen|OJJ32328.1 azon|OJJ47093.1

Group4090: abra|OJJ76366.1 acar|OOF89987.1 acla|EAW14050.1 afis|EAW18756.1 afla|EED50716.1 afum|EAL93484.1 agla|OJJ82324.1 akaw|GAA82292.1 aluc|OJZ91846.1 anid|CBF69963.1 anig|EHA22828.1 aory|EIT78115.1 asyd|OJJ58907.1 ater|EAU37887.1 atub|OJI86298.1 aver|OJJ01504.1 awen|OJJ32325.1 azon|OJJ47092.1

Group4091: abra|OJJ76368.1 acar|OOF99617.1 acla|EAW14048.1 afis|EAW18754.1 afla|EED50714.1 afum|EAL93481.1 agla|OJJ82322.1 akaw|GAA82294.1 aluc|OJZ91844.1 anid|CBF69971.1 anig|EHA22827.1 aory|EIT77980.1 asyd|OJJ58901.1 ater|EAU37885.1 atub|OJI86296.1 aver|OJJ01501.1 awen|OJJ32323.1 azon|OJJ47090.1

Group4092: abra|OJJ76370.1 acar|OOF99619.1 acla|EAW14047.1 afis|EAW18753.1 afla|EED50712.1 afum|EAL93480.1 agla|OJJ82321.1 akaw|GAA82296.1 aluc|OJZ91842.1 anid|CBF69973.1 anig|EHA22825.1 aory|EIT78076.1 asyd|OJJ58900.1 ater|EAU37884.1 atub|OJI86294.1 aver|OJJ01500.1 awen|OJJ32322.1 azon|OJJ47089.1

Group4093: abra|OJJ76378.1 acar|OOF99623.1 acla|EAW14044.1 afis|EAW18750.1 afla|EED50709.1 afum|EAL93477.1 agla|OJJ82317.1 akaw|GAA82299.1 aluc|OJZ91836.1 anid|CBF69979.1 anig|EHA22823.1 aory|EIT78032.1 asyd|OJJ58896.1 ater|EAU37879.1 atub|OJI86290.1 aver|OJJ01497.1 awen|OJJ32317.1 azon|OJJ47085.1

Group4094: abra|OJJ76379.1 acar|OOF99624.1 acla|EAW14043.1 afis|EAW18749.1 afla|EED50708.1 afum|EAL93475.1 agla|OJJ82316.1 akaw|GAA82300.1 aluc|OJZ91835.1 anid|CBF69983.1 anig|EHA22822.1 aory|EIT77996.1 asyd|OJJ58895.1 ater|EAU37878.1 atub|OJI86289.1 aver|OJJ01496.1 awen|OJJ32316.1 azon|OJJ47084.1

Group4095: abra|OJJ76381.1 acar|OOF89925.1 acla|EAW14041.1 afis|EAW18747.1 afla|EED50706.1 afum|EAL93473.1 agla|OJJ82314.1 akaw|GAA82302.1 aluc|OJZ91833.1 anid|CBF69987.1 anig|EHA22820.1 aory|EIT78116.1 asyd|OJJ58893.1 ater|EAU37876.1 atub|OJI86287.1 aver|OJJ01494.1 awen|OJJ32313.1 azon|OJJ47082.1

Group4096: abra|OJJ76382.1 acar|OOF89924.1 acla|EAW14040.1 afis|EAW18746.1 afla|EED50705.1 afum|EAL93472.1 agla|OJJ82313.1 akaw|GAA82303.1 aluc|OJZ91832.1 anid|CBF69992.1 anig|EHA22819.1 aory|EIT78094.1 asyd|OJJ58891.1 ater|EAU37875.1 atub|OJI86286.1 aver|OJJ01492.1 awen|OJJ32312.1 azon|OJJ42712.1

Group4097: abra|OJJ76387.1 acar|OOF99628.1 acla|EAW14036.1 afis|EAW18742.1 afla|EED50698.1 afum|EAL93468.1 agla|OJJ82308.1 akaw|GAA82307.1 aluc|OJZ91826.1 anid|CBF69998.1 anig|EHA22815.1 aory|EIT78061.1 asyd|OJJ58887.1 ater|EAU37869.1 atub|OJI86281.1 aver|OJJ01488.1 awen|OJJ32307.1 azon|OJJ42719.1

Group4098: abra|OJJ76389.1 acar|OOF99630.1 acla|EAW14035.1 afis|EAW18741.1 afla|EED50696.1 afum|EAL93467.1 agla|OJJ82307.1 akaw|GAA82308.1 aluc|OJZ91824.1 anid|CBF70000.1 anig|EHA22814.1 aory|EIT78051.1 asyd|OJJ58886.1 ater|EAU37864.1 atub|OJI86279.1 aver|OJJ01487.1 awen|OJJ32306.1 azon|OJJ42720.1

Group4099: abra|OJJ76390.1 acar|OOF99631.1 acla|EAW14034.1 afis|EAW18740.1 afla|EED50695.1 afum|EAL93466.1 agla|OJJ82306.1 akaw|GAA82309.1 aluc|OJZ91823.1 anid|CBF70002.1 anig|EHA22813.1 aory|EIT78073.1 asyd|OJJ58884.1 ater|EAU37863.1 atub|OJI86278.1 aver|OJJ01486.1 awen|OJJ32305.1 azon|OJJ42721.1

Group4100: abra|OJJ76392.1 acar|OOF99632.1 acla|EAW14032.1 afis|EAW18738.1 afla|EED50693.1 afum|EAL93464.1 agla|OJJ82304.1 akaw|GAA82311.1 aluc|OJZ91821.1 anid|CBF70006.1 anig|EHA22811.1

aory|EIT78112.1 asyd|OJJ58882.1 ater|EAU37861.1 atub|OJI86276.1 aver|OJJ01484.1 awen|OJJ32302.1 azon|OJJ42723.1

Group4101: abra|OJJ76393.1 acar|OOF99633.1 acla|EAW14031.1 afis|EAW18737.1 afla|EED50692.1 afum|EAL93462.1 agla|OJJ82303.1 akaw|GAA82312.1 aluc|OJZ91820.1 anid|CBF70008.1 anig|EHA22810.1 aory|EIT78068.1 asyd|OJJ58881.1 ater|EAU37860.1 atub|OJI86275.1 aver|OJJ01483.1 awen|OJJ32301.1 azon|OJJ42724.1

Group4102: abra|OJJ76395.1 acar|OOF99635.1 acla|EAW14029.1 afis|EAW18735.1 afla|EED50690.1 afum|EAL93460.1 agla|OJJ82301.1 akaw|GAA82314.1 aluc|OJZ91818.1 anid|CBF70012.1 anig|EHA22808.1 aory|EIT78132.1 asyd|OJJ58879.1 ater|EAU37858.1 atub|OJI86273.1 aver|OJJ01481.1 awen|OJJ32298.1 azon|OJJ42734.1

Group4103: abra|OJJ76396.1 acar|OOF99637.1 acla|EAW14027.1 afis|EAW18733.1 afla|EED50688.1 afum|EAL93458.1 agla|OJJ82299.1 akaw|GAA82316.1 aluc|OJZ91816.1 anid|CBF70014.1 anig|EHA22807.1 aory|EIT78099.1 asyd|OJJ58878.1 ater|EAU37848.1 atub|OJI86272.1 aver|OJJ01480.1 awen|OJJ32296.1 azon|OJJ42736.1

Group4104: abra|OJJ76404.1 acar|OOF99647.1 acla|EAW14017.1 afis|EAW18725.1 afla|EED50682.1 afum|EAL93450.1 agla|OJJ80475.1 akaw|GAA82324.1 aluc|OJZ91808.1 anid|CBF70021.1 anig|EHA22801.1 aory|EIT77978.1 asyd|OJJ58874.1 ater|EAU38179.1 atub|OJI86263.1 aver|OJJ01477.1 awen|OJJ32229.1 azon|OJJ46783.1

Group4105: abra|OJJ76406.1 acar|OOF99649.1 acla|EAW14019.1 afis|EAW18723.1 afla|EED50680.1 afum|EAL93448.1 agla|OJJ80477.1 akaw|GAA82326.1 aluc|OJZ91806.1 anid|CBF70688.1 anig|EHA22799.1 aory|EIT78149.1 asyd|OJJ58468.1 ater|EAU38180.1 atub|OJI86261.1 aver|OJJ01067.1 awen|OJJ32231.1 azon|OJJ46781.1

Group4106: abra|OJJ76409.1 acar|OOF99651.1 acla|EAW14015.1 afis|EAW18721.1 afla|EED50678.1 afum|EAL93445.1 agla|OJJ82334.1 akaw|GAA82328.1 aluc|OJZ91802.1 anid|CBF70681.1 anig|EHA22797.1 aory|EIT78078.1 asyd|OJJ58470.1 ater|EAU38183.1 atub|OJI86257.1 aver|OJJ01071.1 awen|OJJ32227.1 azon|OJJ46778.1

Group4107: abra|OJJ76411.1 acar|OOF99652.1 acla|EAW14014.1 afis|EAW18720.1 afla|EED50677.1 afum|EAL93444.1 agla|OJJ82335.1 akaw|GAA82329.1 aluc|OJZ91801.1 anid|CBF70679.1 anig|EHA22796.1 aory|EIT78134.1 asyd|OJJ58471.1 ater|EAU38184.1 atub|OJI86256.1 aver|OJJ01072.1 awen|OJJ32226.1 azon|OJJ46777.1

Group4108: abra|OJJ76417.1 acar|OOF99659.1 acla|EAW14009.1 afis|EAW18715.1 afla|EED50669.1 afum|EAL93440.1 agla|OJJ82344.1 akaw|GAA82332.1 aluc|OJZ91795.1 anid|CBF70670.1 anig|EHA22792.1 aory|EIT78070.1 asyd|OJJ58476.1 ater|EAU38192.1 atub|OJI86251.1 aver|OJJ01079.1 awen|OJJ32202.1 azon|OJJ46771.1

Group4109: abra|OJJ76418.1 acar|OOF99660.1 acla|EAW14008.1 afis|EAW18714.1 afla|EED50667.1 afum|EAL93439.1 agla|OJJ82345.1 akaw|GAA82333.1 aluc|OJZ91794.1 anid|CBF70668.1 anig|EHA22791.1 aory|EIT78037.1 asyd|OJJ58477.1 ater|EAU38193.1 atub|OJI86250.1 aver|OJJ01080.1 awen|OJJ32201.1 azon|OJJ46770.1

Group4110: abra|OJJ76419.1 acar|OOF99661.1 acla|EAW14007.1 afis|EAW18713.1 afla|EED50666.1 afum|EAL93438.1 agla|OJJ82346.1 akaw|GAA82334.1 aluc|OJZ91793.1 anid|CBF70666.1 anig|EHA22790.1 aory|EIT78088.1 asyd|OJJ58478.1 ater|EAU38194.1 atub|OJI86249.1 aver|OJJ01081.1 awen|OJJ32200.1 azon|OJJ46769.1

Group4111: abra|OJJ76420.1 acar|OOF99662.1 acla|EAW14006.1 afis|EAW18712.1 afla|EED50665.1 afum|EAL93437.1 agla|OJJ82347.1 akaw|GAA82335.1 aluc|OJZ91792.1 anid|CBF70665.1 anig|EHA22789.1 aory|EIT77998.1 asyd|OJJ58480.1 ater|EAU38195.1 atub|OJI86248.1 aver|OJJ01082.1 awen|OJJ32199.1 azon|OJJ46768.1

Group4112: abra|OJJ76422.1 acar|OOF99665.1 acla|EAW14003.1 afis|EAW18711.1 afla|EED50662.1 afum|EAL93434.1 agla|OJJ82348.1 akaw|GAA82337.1 aluc|OJZ91790.1 anid|CBF70661.1 anig|EHA22787.1 asyd|OJJ58481.1 ater|EAU38197.1 atub|OJI86246.1 aver|OJJ01083.1 awen|OJJ32197.1 azon|OJJ46766.1 aory|EIT77986.1

Group4113: abra|OJJ76423.1 acar|OOF99667.1 acla|EAW14002.1 afis|EAW18710.1 afla|EED50660.1 afum|EAL93433.1 agla|OJJ82349.1 akaw|GAA82338.1 aluc|OJZ91789.1 anid|CBF70657.1 anig|EHA22786.1 aory|EIT78084.1 asyd|OJJ58482.1 ater|EAU38198.1 atub|OJI86245.1 aver|OJJ01085.1 awen|OJJ32196.1 azon|OJJ46765.1

Group4114: abra|OJJ76429.1 acar|OOF99673.1 acla|EAW13996.1 afis|EAW18704.1 afla|EED50653.1 afum|EAL93427.2 agla|OJJ82354.1 akaw|GAA82344.1 aluc|OJZ91783.1 anid|CBF70646.1 anig|EHA22780.1 aory|EIT78100.1 asyd|OJJ58488.1 ater|EAU38202.1 atub|OJI86239.1 aver|OJJ01091.1 awen|OJJ32187.1 azon|OJJ46760.1

Group4115: abra|OJJ76430.1 acar|OOF99674.1 acla|EAW13995.1 afis|EAW18703.1 afla|EED50652.1 afum|EAL93426.1 agla|OJJ82355.1 akaw|GAA82345.1 aluc|OJZ91782.1 anid|CBF70644.1 anig|EHA22779.1 aory|EIT78020.1 asyd|OJJ58489.1 ater|EAU38203.1 atub|OJI86238.1 aver|OJJ01092.1 awen|OJJ32186.1 azon|OJJ46761.1

Group4116: abra|OJJ76431.1 acar|OOF99675.1 acla|EAW13994.1 afis|EAW18702.1 afla|EED50651.1 afum|EAL93425.1 agla|OJJ82359.1 akaw|GAA82346.1 aluc|OJZ91781.1 anid|CBF70642.1 anig|EHA22778.1 aory|EIT78030.1 asyd|OJJ58490.1 ater|EAU38204.1 atub|OJI86237.1 aver|OJJ01093.1 awen|OJJ32185.1 azon|OJJ46762.1

Group4117: abra|OJJ76433.1 acar|OOF99677.1 acla|EAW13991.1 afis|EAW18699.1 afla|EED50647.1 afum|EAL93421.1 agla|OJJ82362.1 akaw|GAA82348.1 aluc|OJZ91779.1 anid|CBF70635.1 anig|EHA22776.1 aory|EIT78093.1 asyd|OJJ58494.1 ater|EAU38207.1 atub|OJI86235.1 aver|OJJ01097.1 awen|OJJ32181.1 azon|OJJ46756.1

Group4118: abra|OJJ76434.1 acar|OOF99679.1 acla|EAW13989.1 afis|EAW18697.1 afla|EED50645.1 afum|EAL93419.1 agla|OJJ82371.1 akaw|GAA82349.1 aluc|OJZ91778.1 anid|CBF70631.1 anig|EHA22775.1 aory|EIT78142.1 asyd|OJJ58496.1 ater|EAU38209.1 atub|OJI86234.1 aver|OJJ01099.1 awen|OJJ32168.1 azon|OJJ46752.1

Group4119: abra|OJJ76440.1 acar|OOF99685.1 acla|EAW13984.1 afis|EAW18693.1 afla|EED50642.1 afum|EAL93414.1 agla|OJJ82368.1 akaw|GAA82354.1 aluc|OJZ91769.1 anid|CBF70618.1 anig|EHA22769.1 aory|EIT78025.1 asyd|OJJ58503.1 ater|EAU38215.1 atub|OJI86227.1 aver|OJJ01104.1 awen|OJJ32164.1 azon|OJJ46745.1

Group4120: abra|OJJ76441.1 acar|OOF99686.1 acla|EAW13983.1 afis|EAW18692.1 afla|EED50641.1 afum|EAL93413.2 agla|OJJ82366.1 akaw|GAA82355.1 aluc|OJZ91768.1 anid|CBF70620.1 anig|EHA22768.1 aory|EIT78095.1 asyd|OJJ58502.1 ater|EAU38216.1 atub|OJI86226.1 aver|OJJ01103.1 awen|OJJ32163.1 azon|OJJ46746.1

Group4121: abra|OJJ76446.1 acar|OOF99689.1 acla|EAW13982.1 afis|EAW18691.1 afla|EED50638.1 afum|EAL93412.1 agla|OJJ82381.1 akaw|GAA82358.1 aluc|OJZ91764.1 anid|CBF70624.1 anig|EHA22765.1 aory|EIT78085.1 asyd|OJJ58499.1 ater|EAU38217.1 atub|OJI86222.1 aver|OJJ01102.1 awen|OJJ32162.1 azon|OJJ46748.1

Group4122: abra|OJJ76448.1 acar|OOF99690.1 acla|EAW13980.1 afis|EAW18689.1 afla|EED50637.1 afum|EAL93410.1 agla|OJJ82383.1 akaw|GAA82360.1 aluc|OJZ91762.1 anid|CBF70616.1 anig|EHA22763.1 aory|EIT78124.1 asyd|OJJ58506.1 ater|EAU38219.1 atub|OJI86220.1 aver|OJJ01107.1 awen|OJJ32160.1 azon|OJJ46744.1

Group4123: abra|OJJ76449.1 acar|OOF99692.1 acla|EAW13979.1 afis|EAW18688.1 afla|EED50636.1 afum|EAL93409.1 agla|OJJ82384.1 akaw|GAA82361.1 aluc|OJZ91761.1 anid|CBF70612.1 anig|EHA22762.1 aory|EIT78146.1 asyd|OJJ58510.1 ater|EAU38220.1 atub|OJI86219.1 aver|OJJ01111.1 awen|OJJ32159.1 azon|OJJ46743.1

Group4124: abra|OJJ76455.1 acar|OOF99696.1 acla|EAW13975.1 afis|EAW18684.1 afla|EED50622.1 afum|EAL93405.1 agla|OJJ82387.1 akaw|GAA82367.1 aluc|OJZ91755.1 anid|CBF70607.1 anig|EHA22758.1 aory|EIT78141.1 asyd|OJJ58514.1 ater|EAU38226.1 atub|OJI86213.1 aver|OJJ01115.1 awen|OJJ32155.1 azon|OJJ46739.1

Group4125: abra|OJJ76456.1 acar|OOF99695.1 acla|EAW13974.1 afis|EAW18683.1 afla|EED50627.1 afum|EAL93404.1 agla|OJJ82392.1 akaw|GAA82368.1 aluc|OJZ91754.1 anid|CBF70605.1 anig|EHA22757.1 aory|EIT77985.1 asyd|OJJ58515.1 ater|EAU38227.1 atub|OJI86212.1 aver|OJJ01116.1 awen|OJJ32154.1 azon|OJJ46738.1

Group4126: abra|OJJ76460.1 acar|OOF99699.1 acla|EAW13970.1 afis|EAW18679.1 afla|EED50631.1 afum|EAL93399.1 agla|OJJ82396.1 akaw|GAA82373.1 aluc|OJZ91749.1 anid|CBF70596.1 anig|EHA22752.1 aory|EIT78016.1 asyd|OJJ58519.1 ater|EAU38232.1 atub|OJI86207.1 aver|OJJ01120.1 awen|OJJ32151.1 azon|OJJ46735.1

Group4127: abra|OJJ76464.1 acar|OOF99702.1 acla|EAW13967.1 afis|EAW18676.1 afla|EED50634.1 afum|EAL93396.1 agla|OJJ82400.1 akaw|GAA82376.1 aluc|OJZ91746.1 anid|CBF70590.1 anig|EHA22749.1 aory|EIT78033.1 asyd|OJJ58522.1 ater|EAU38234.1 atub|OJI86204.1 aver|OJJ01123.1 awen|OJJ32148.1 azon|OJJ46732.1

Group4128: abra|OJJ76467.1 acar|OOF99705.1 acla|EAW13964.1 afis|EAW18673.1 afla|EED50626.1 afum|EAL93393.2 agla|OJJ82402.1 akaw|GAA82378.1 aluc|OJZ91743.1 anid|CBF70584.1 anig|EHA22746.1 aory|EIT78060.1 asyd|OJJ58525.1 ater|EAU38237.1 atub|OJI86201.1 aver|OJJ01126.1 awen|OJJ32145.1 azon|OJJ46729.1

Group4129: abra|OJJ76468.1 acar|OOF99706.1 acla|EAW13963.1 afis|EAW18672.1 afla|EED50621.1 afum|EAL93391.1 agla|OJJ82403.1 akaw|GAA82379.1 aluc|OJZ91742.1 anid|CBF70583.1 anig|EHA22745.1

aory|EIT78045.1 asyd|OJJ58526.1 ater|EAU38238.1 atub|OJI86200.1 aver|OJJ01127.1 awen|OJJ32144.1 azon|OJJ46728.1

Group4130: abra|OJJ76472.1 acar|OOF99709.1 acla|EAW13961.1 afis|EAW18670.1 afla|EED52658.1 afum|EAL93388.1 agla|OJJ82405.1 akaw|GAA82384.1 aluc|OJZ91737.1 anid|CBF70579.1 anig|EHA22743.1 aory|EIT74274.1 asyd|OJJ58528.1 ater|EAU38241.1 atub|OJI86195.1 aver|OJJ01130.1 awen|OJJ32142.1 azon|OJJ46723.1

Group4131: abra|OJJ76479.1 acar|OOF90192.1 acla|EAW13956.1 afis|EAW18666.1 afla|EED52663.1 afum|EAL93384.1 agla|OJJ82411.1 akaw|GAA82392.1 aluc|OJZ91728.1 anid|CBF70569.1 anig|EHA22735.1 aory|EIT74303.1 asyd|OJJ58533.1 ater|EAU38246.1 atub|OJI86186.1 aver|OJJ01135.1 awen|OJJ32134.1 azon|OJJ46717.1

Group4132: acar|OOF90191.1 aluc|OJZ91727.1 anig|EHA22734.1 atub|OJI86185.1 abra|OJJ76481.1 acla|EAW13954.1 afis|EAW18664.1 afla|EED52665.1 afum|EAL93382.1 agla|OJJ82412.1 akaw|GAA82394.1 anid|CBF70567.1 aory|EIT74310.1 asyd|OJJ58535.1 ater|EAU38248.1 aver|OJJ01136.1 awen|OJJ32132.1 azon|OJJ46715.1

Group4133: abra|OJJ76482.1 acar|OOF90190.1 acla|EAW13953.1 afis|EAW18663.1 afla|EED52666.1 afum|EBA27491.1 agla|OJJ82413.1 akaw|GAA82395.1 aluc|OJZ91726.1 anid|CBF70565.1 anig|EHA22733.1 aory|EIT74219.1 asyd|OJJ58536.1 ater|EAU38249.1 atub|OJI86184.1 aver|OJJ01137.1 awen|OJJ32131.1 azon|OJJ46714.1

Group4134: abra|OJJ76483.1 acar|OOF99714.1 acla|EAW13951.1 afis|EAW18661.1 afla|EED52669.1 afum|EAL93380.1 agla|OJJ82416.1 akaw|GAA82397.1 aluc|OJZ91725.1 anid|CBF70557.1 anig|EHA22732.1 aory|EIT74184.1 asyd|OJJ58538.1 ater|EAU38251.1 atub|OJI86182.1 aver|OJJ01138.1 awen|OJJ32128.1 azon|OJJ46711.1

Group4135: abra|OJJ76484.1 acar|OOF99715.1 acla|EAW13950.1 afis|EAW18659.1 afla|EED52670.1 afum|EAL93377.1 agla|OJJ82417.1 akaw|GAA82398.1 aluc|OJZ91724.1 anid|CBF70556.1 anig|EHA22731.1 aory|EIT74290.1 asyd|OJJ58539.1 ater|EAU38252.1 atub|OJI86181.1 aver|OJJ01139.1 awen|OJJ32127.1 azon|OJJ46710.1

Group4136: abra|OJJ76485.1 acar|OOF99718.1 acla|EAW13948.1 afis|EAW18658.1 afla|EED52672.1 afum|EAL93375.1 agla|OJJ82418.1 akaw|GAA82401.1 aluc|OJZ91721.1 anid|CBF70552.1 anig|EHA22728.1 aory|EIT74194.1 asyd|OJJ58540.1 ater|EAU38254.1 atub|OJI86178.1 aver|OJJ01141.1 awen|OJJ32125.1 azon|OJJ46709.1

Group4137: abra|OJJ76487.1 acar|OOF99720.1 acla|EAW13946.1 afis|EAW18656.1 afla|EED52674.1 afum|EAL93372.1 agla|OJJ82419.1 akaw|GAA82403.1 aluc|OJZ91718.1 anid|CBF70546.1 anig|EHA22726.1 aory|EIT74203.1 asyd|OJJ58543.1 ater|EAU38256.1 atub|OJI86175.1 aver|OJJ01143.1 awen|OJJ32123.1 azon|OJJ46708.1

Group4138: abra|OJJ76501.1 acar|OOF99466.1 acla|EAW14195.1 afis|EAW18911.1 afla|EED50838.1 afum|EAL93642.1 agla|OJJ81515.1 akaw|GAA82417.1 aluc|OJZ91701.1 anid|CBF69699.1 anig|EHA22715.1 aory|EIT79018.1 asyd|OJJ59050.1 ater|EAU38078.1 atub|OJI86160.1 aver|OJJ01646.1 awen|OJJ31504.1 azon|OJJ47224.1

Group4139: abra|OJJ76513.1 acar|OOF99459.1 acla|EAW14187.1 afis|EAW18902.1 afla|EED50846.1 afum|EAL93634.1 agla|OJJ81507.1 akaw|GAA82426.1 aluc|OJZ91692.1 anid|CBF69923.1 anig|EHA22705.1 aory|EIT79043.1 asyd|OJJ59039.1 ater|EAU38072.1 atub|OJI86145.1 aver|OJJ01634.1 awen|OJJ32459.1 azon|OJJ47217.1

Group4140: abra|OJJ76516.1 acar|OOF99457.1 acla|EAW14186.1 afis|EAW18901.1 afla|EED50847.1 afum|EAL93633.1 agla|OJJ81506.1 akaw|GAA82428.1 aluc|OJZ91690.1 anid|CBF69921.1 anig|EHA22703.1 aory|EIT79032.1 asyd|OJJ58987.1 ater|EAU38069.1 atub|OJI86143.1 aver|OJJ01629.1 awen|OJJ32458.1 azon|OJJ47216.1

Group4141: abra|OJJ76520.1 acar|OOF99453.1 acla|EAW14181.1 afis|EAW18896.1 afla|EED50867.1 afum|EAL93628.1 agla|OJJ81502.1 akaw|GAA82433.1 aluc|OJZ91688.1 anid|CBF69913.1 anig|EHA22700.1 aory|EIT79004.1 asyd|OJJ58992.1 ater|EAU38066.1 atub|OJI86139.1 aver|OJJ01623.1 awen|OJJ32454.1 azon|OJJ47211.1

Group4142: abra|OJJ76524.1 acar|OOF99445.1 acla|EAW14173.1 afis|EAW18887.1 afla|EED50878.1 afum|EAL93620.1 agla|OJJ81493.1 akaw|GAA82437.1 aluc|OJZ91684.1 anid|CBF69898.1 anig|EHA22696.1 aory|EIT79044.1 asyd|OJJ59000.1 ater|EAU38059.1 atub|OJI86136.1 aver|OJJ01615.1 awen|OJJ32445.1 azon|OJJ47203.1

Group4143: abra|OJJ76528.1 acar|OOF99447.1 acla|EAW14175.1 afis|EAW18889.1 afla|EED50874.1 afum|EAL93622.1 agla|OJJ81495.1 akaw|GAA82439.1 aluc|OJZ91677.1 anid|CBF69902.1 anig|EHA22694.1 aory|EIT78988.1 asyd|OJJ58998.1 ater|EAU38061.1 atub|OJI86132.1 aver|OJJ01617.1 awen|OJJ32448.1 azon|OJJ47206.1

Group4144: abra|OJJ76530.1 acar|OOF99443.1 acla|EAW14171.1 afis|EAW18885.1 afla|EED50880.1 afum|EAL93618.1 agla|OJJ80620.1 akaw|GAA82442.1 aluc|OJZ91674.1 anid|CBF69894.1 anig|EHA22692.1 aory|EIT78957.1 asyd|OJJ59002.1 ater|EAU38057.1 atub|OJI86129.1 aver|OJJ01613.1 awen|OJJ32443.1 azon|OJJ47201.1

Group4145: abra|OJJ76534.1 acar|OOF99440.1 acla|EAW14169.1 afis|EAW18883.1 afla|EED50882.1 afum|EAL93616.1 agla|OJJ80618.1 akaw|GAA82445.1 aluc|OJZ91671.1 anid|CBF69892.1 anig|EHA22689.1 aory|EIT79022.1 asyd|OJJ59005.1 ater|EAU38055.1 atub|OJI86126.1 aver|OJJ01611.1 awen|OJJ32440.1 azon|OJJ47199.1

Group4146: abra|OJJ76535.1 acar|OOF99439.1 acla|EAW14168.1 afis|EAW18882.1 afla|EED50883.1 afum|EAL93614.1 agla|OJJ80617.1 akaw|GAA82446.1 aluc|OJZ91670.1 anid|CBF70466.1 anig|EHA22688.1 aory|EIT78979.1 asyd|OJJ58598.1 ater|EAU38054.1 atub|OJI86125.1 aver|OJJ01200.1 awen|OJJ32441.1 azon|OJJ47198.1

Group4147: abra|OJJ76536.1 acar|OOF99432.1 acla|EAW14162.1 afis|EAW18875.1 afla|EED50892.1 afum|EAL93607.1 agla|OJJ80612.1 akaw|GAA82447.1 aluc|OJZ91669.1 anid|CBF69869.1 anig|EHA22687.1 aory|EIT79035.1 asyd|OJJ59019.1 ater|EAU38040.1 atub|OJI86124.1 aver|OJJ01597.1 awen|OJJ32434.1 azon|OJJ47192.1

Group4148: abra|OJJ76537.1 acar|OOF99433.1 acla|EAW14163.1 afis|EAW18876.1 afla|EED50891.1 afum|EAL93608.1 agla|OJJ80613.1 akaw|GAA82448.1 aluc|OJZ91667.1 anid|CBF69873.1 anig|EHA22686.1 aory|EIT79049.1 asyd|OJJ59018.1 ater|EAU38041.1 atub|OJI86122.1 aver|OJJ01598.1 awen|OJJ32435.1 azon|OJJ47191.1

Group4149: abra|OJJ76540.1 acar|OOF99435.1 acla|EAW14165.1 afis|EAW18878.1 afla|EED50888.1 afum|EAL93610.1 agla|OJJ80615.1 akaw|GAA82450.1 aluc|OJZ91663.1 anid|CBF69877.1 anig|EHA22683.1 aory|EIT79008.1 asyd|OJJ59015.1 ater|EAU38043.1 atub|OJI86120.1 aver|OJJ01600.1 awen|OJJ32437.1 azon|OJJ47196.1

Group4150: abra|OJJ76542.1 acar|OOF99437.1 acla|EAW12334.1 afis|EAW16888.1 afla|EED49676.1 afum|EAL86426.1 agla|OJJ79936.1 akaw|GAA82451.1 aluc|OJZ91661.1 anid|CBF84569.1 anig|EHA22681.1 aory|EIT80968.1 asyd|OJJ53441.1 ater|EAU39143.1 atub|OJI86117.1 aver|OJJ08310.1 awen|OJJ29756.1 azon|OJJ43187.1

Group4151: abra|OJJ76543.1 acar|OOF99431.1 acla|EAW14161.1 afis|EAW18874.1 afla|EED50893.1 afum|EAL93606.1 agla|OJJ80611.1 akaw|GAA82452.1 aluc|OJZ91660.1 anid|CBF69867.1 anig|EHA22680.1 aory|EIT79007.1 asyd|OJJ59020.1 ater|EAU38039.1 atub|OJI86116.1 aver|OJJ01596.1 awen|OJJ32433.1 azon|OJJ47193.1

Group4152: abra|OJJ76545.1 acar|OOF99429.1 acla|EAW14159.1 afis|EAW18872.1 afla|EED50895.1 afum|EAL93604.1 agla|OJJ80609.1 akaw|GAA82454.1 aluc|OJZ91658.1 anid|CBF69863.1 anig|EHA22678.1 aory|EIT79057.1 asyd|OJJ59023.1 ater|EAU38036.1 atub|OJI86114.1 aver|OJJ01594.1 awen|OJJ32431.1 azon|OJJ47188.1

Group4153: abra|OJJ76546.1 acar|OOF99428.1 acla|EAW14158.1 afis|EAW18871.1 afla|EED50831.1 afum|EAL93603.1 agla|OJJ80595.1 akaw|GAA82455.1 aluc|OJZ91657.1 anid|CBF69852.1 anig|EHA22677.1 aory|EIT79042.1 asyd|OJJ59028.1 ater|EAU38035.1 atub|OJI86112.1 aver|OJJ01588.1 awen|OJJ32430.1 azon|OJJ47185.1

Group4154: abra|OJJ76547.1 acar|OOF99427.1 acla|EAW14157.1 afis|EAW18870.1 afla|EED50832.1 afum|EAL93602.1 agla|OJJ80596.1 akaw|GAA82456.1 aluc|OJZ91656.1 anid|CBF69854.1 anig|EHA22676.1 aory|EIT79054.1 asyd|OJJ59027.1 ater|EAU38034.1 atub|OJI86111.1 aver|OJJ01589.1 awen|OJJ32429.1 azon|OJJ47184.1

Group4155: abra|OJJ76549.1 acar|OOF99426.1 acla|EAW14156.1 afis|EAW18869.1 afla|EED50833.1 afum|EAL93601.1 agla|OJJ80600.1 akaw|GAA82457.1 aluc|OJZ91654.1 anid|CBF69856.1 anig|EHA22675.1 aory|EIT78961.1 asyd|OJJ59026.1 ater|EAU38033.1 atub|OJI86108.1 aver|OJJ01591.1 awen|OJJ32428.1 azon|OJJ47183.1

Group4156: abra|OJJ76550.1 acar|OOF89946.1 acla|EAW14155.1 afis|EAW18868.1 afla|EED50834.1 afum|EAL93600.1 agla|OJJ80601.1 akaw|GAA82458.1 aluc|OJZ91653.1 anid|CBF69857.1 anig|EHA22674.1 aory|EIT78982.1 asyd|OJJ59025.1 ater|EAU38032.1 atub|OJI86107.1 aver|OJJ01592.1 awen|OJJ32427.1 azon|OJJ47182.1

Group4157: abra|OJJ76551.1 acar|OOF89945.1 acla|EAW14153.1 afis|EAW18866.1 afla|EED50897.1 afum|EAL93598.1 agla|OJJ80603.1 akaw|GAA82459.1 aluc|OJZ91652.1 anid|CBF69861.1 anig|EHA22673.1 aory|EIT79016.1 asyd|OJJ59024.1 ater|EAU38030.1 atub|OJI86106.1 aver|OJJ01593.1 awen|OJJ32425.1 azon|OJJ47180.1

Group4158: abra|OJJ76554.1 acar|OOF99424.1 acla|EAW14150.1 afis|EAW18863.1 afla|EED50830.1 afum|EAL93595.1 agla|OJJ80586.1 akaw|GAA82461.1 aluc|OJZ91650.1 anid|CBF69848.1 anig|EHA22671.1

aory|EIT79053.1 asyd|OJJ59030.1 ater|EAU38025.1 atub|OJI86104.1 aver|OJJ01586.1 awen|OJJ32419.1 azon|OJJ47176.1

Group4159: abra|OJJ76558.1 acar|OOF99421.1 acla|EAW14147.1 afis|EAW18860.1 afla|EED50827.1 afum|EAL93592.1 agla|OJJ80583.1 akaw|GAA82464.1 aluc|OJZ91647.1 anid|CBF69842.1 anig|EHA22668.1 aory|EIT78975.1 asyd|OJJ59033.1 ater|EAU38023.1 atub|OJI86101.1 aver|OJJ01583.1 awen|OJJ32416.1 azon|OJJ47173.1

Group4160: abra|OJJ76563.1 acar|OOF99419.1 acla|EAW14144.1 afis|EAW18857.1 afla|EED50824.1 afum|EAL93589.2 agla|OJJ80579.1 akaw|GAA82467.1 aluc|OJZ91643.1 anid|CBF69834.1 anig|EHA22665.1 aory|EIT78963.1 asyd|OJJ58985.1 ater|EAU38020.1 atub|OJI86097.1 aver|OJJ01580.1 awen|OJJ32413.1 azon|OJJ47170.1

Group4161: abra|OJJ76565.1 acar|OOF99418.1 acla|EAW14143.1 afis|EAW18856.1 afla|EED50823.1 afum|EAL93587.1 akaw|GAA82469.1 aluc|OJZ91641.1 anid|CBF69831.1 anig|EHA22664.1 aory|EIT79024.1 asyd|OJJ58983.1 ater|EAU38019.1 atub|OJI86095.1 aver|OJJ01578.1 awen|OJJ32411.1 azon|OJJ47168.1 agla|OJJ80577.1

Group4162: abra|OJJ76566.1 acar|OOF99355.1 acla|EAW14142.1 afis|EAW18855.1 afla|EED50822.1 afum|EAL93586.1 agla|OJJ80574.1 akaw|GAA82470.1 aluc|OJZ91640.1 anid|CBF69829.1 anig|EHA22663.1 aory|EIT78993.1 asyd|OJJ58981.1 ater|EAU38013.1 atub|OJI86094.1 aver|OJJ01576.1 awen|OJJ32410.1 azon|OJJ47167.1

Group4163: abra|OJJ76567.1 acar|OOF99417.1 acla|EAW14141.1 afis|EAW18854.1 afla|EED50821.1 afum|EAL93585.1 agla|OJJ80573.1 akaw|GAA82471.1 aluc|OJZ91639.1 anid|CBF69827.1 anig|EHA22662.1 aory|EIT78968.1 asyd|OJJ58980.1 ater|EAU38012.1 atub|OJI86093.1 aver|OJJ01575.1 awen|OJJ32409.1 azon|OJJ47166.1

Group4164: abra|OJJ76569.1 acar|OOF99416.1 acla|EAW14139.1 afis|EAW18852.1 afla|EED50819.1 afum|EAL93583.1 agla|OJJ80571.1 akaw|GAA82473.1 aluc|OJZ91636.1 anid|CBF89917.1 anig|EHA22660.1 aory|EIT79009.1 asyd|OJJ62126.1 ater|EAU38005.1 atub|OJI86089.1 aver|OJI96234.1 awen|OJJ32406.1 azon|OJJ47164.1

Group4165: abra|OJJ76570.1 acar|OOF99415.1 acla|EAW14138.1 afis|EAW18851.1 afla|EED50818.1 afum|EAL93582.1 agla|OJJ80570.1 akaw|GAA82474.1 aluc|OJZ91635.1 anid|CBF69823.1 anig|EHA22659.1 aory|EIT78969.1 asyd|OJJ58978.1 ater|EAU38004.1 atub|OJI86088.1 aver|OJJ01573.1 awen|OJJ32405.1 azon|OJJ47163.1

Group4166: abra|OJJ76571.1 acar|OOF99414.1 acla|EAW14137.1 afis|EAW18850.1 afla|EED50817.1 afum|EAL93581.1 agla|OJJ80569.1 akaw|GAA82475.1 aluc|OJZ91634.1 anid|CBF69821.1 anig|EHA22658.1 aory|EIT79031.1 asyd|OJJ58977.1 ater|EAU38003.1 atub|OJI86087.1 aver|OJJ01572.1 awen|OJJ32404.1 azon|OJJ47162.1

Group4167: abra|OJJ76572.1 acar|OOF99413.1 acla|EAW14136.1 afis|EAW18849.1 afla|EED50816.1 afum|EAL93580.1 agla|OJJ80568.1 akaw|GAA82476.1 aluc|OJZ91633.1 anid|CBF69819.1 anig|EHA22657.1 aory|EIT78989.1 asyd|OJJ58976.1 ater|EAU38002.1 atub|OJI86086.1 aver|OJJ01571.1 awen|OJJ32403.1 azon|OJJ47161.1

Group4168: abra|OJJ76573.1 acar|OOF99412.1 acla|EAW14134.1 afis|EAW18847.1 afla|EED50814.1 afum|EAL93579.1 agla|OJJ80567.1 akaw|GAA82478.1 aluc|OJZ91632.1 anid|CBF69817.1 anig|EHA22656.1 aory|EIT79029.1 asyd|OJJ58975.1 ater|EAU38001.1 atub|OJI86085.1 aver|OJJ01570.1 awen|OJJ32402.1 azon|OJJ47160.1

Group4169: abra|OJJ76576.1 acar|OOF99408.1 acla|EAW14117.1 afis|EAW18819.1 afla|EED50808.1 afum|EAL93555.1 agla|OJJ80554.1 akaw|GAA82481.1 aluc|OJZ91629.1 anid|CBF69809.1 anig|EHA22653.1 aory|EIT78964.1 asyd|OJJ58971.1 ater|EAU37998.1 atub|OJI86082.1 aver|OJJ01566.1 awen|OJJ32395.1 azon|OJJ47157.1

Group4170: abra|OJJ76577.1 acar|OOF99407.1 acla|EAW14116.1 afis|EAW18818.1 afla|EED50809.1 afum|EAL93554.1 agla|OJJ80552.1 akaw|GAA82482.1 aluc|OJZ91628.1 anid|CBF69811.1 anig|EHA22652.1 aory|EIT78995.1 asyd|OJJ58972.1 ater|EAU37997.1 atub|OJI86081.1 aver|OJJ01567.1 awen|OJJ32394.1 azon|OJJ47156.1

Group4171: abra|OJJ76579.1 acar|OOF99405.1 acla|EAW14114.1 afis|EAW18816.1 afla|EED50804.1 afum|EAL93552.1 agla|OJJ80549.1 akaw|GAA82484.1 aluc|OJZ91626.1 anid|CBF69815.1 anig|EHA22650.1 aory|EIT78104.1 asyd|OJJ58974.1 ater|EAU37991.1 atub|OJI86079.1 aver|OJJ01569.1 awen|OJJ32392.1 azon|OJJ47153.1

Group4172: abra|OJJ76581.1 acar|OOF99404.1 acla|EAW14113.1 afis|EAW18815.1 afla|EED50802.1 afum|EAL93551.1 agla|OJJ80548.1 akaw|GAA82485.1 aluc|OJZ91624.1 anid|CBF69803.1 anig|EHA22649.1 aory|EIT77988.1 asyd|OJJ58968.1 ater|EAU37982.1 atub|OJI86077.1 aver|OJJ01563.1 awen|OJJ32391.1 azon|OJJ47152.1

Group4173: abra|OJJ76582.1 acar|OOF99403.1 acla|EAW14112.1 afis|EAW18814.1 afla|EED50800.1 afum|EAL93550.2 agla|OJJ80547.1 akaw|GAA82486.1 aluc|OJZ91623.1 anid|CBF69801.1 anig|EHA22648.1 aory|EIT78081.1 asyd|OJJ58967.1 ater|EAU37981.1 atub|OJI86076.1 aver|OJJ01562.1 awen|OJJ32390.1 azon|OJJ47151.1

Group4174: abra|OJJ76588.1 acar|OOF99398.1 acla|EAW14313.1 afis|EAW18974.1 afla|EED53230.1 afum|EAL93708.1 agla|OJJ82930.1 akaw|GAA82492.1 aluc|OJZ91618.1 anid|CBF69639.1 anig|EHA22645.1 aory|EIT74874.1 asyd|OJJ59084.1 ater|EAU38166.1 atub|OJI86071.1 aver|OJJ01677.1 awen|OJJ31576.1 azon|OJJ47293.1

Group4175: abra|OJJ76590.1 acar|OOF99397.1 acla|EAW14311.1 afis|EAW18975.1 afla|EED53222.1 afum|EAL93710.1 agla|OJJ82932.1 akaw|GAA82494.1 aluc|OJZ91616.1 anid|CBF69642.1 anig|EHA22644.1 aory|EIT74886.1 asyd|OJJ59081.1 ater|EAU38164.1 atub|OJI86069.1 aver|OJJ01674.1 awen|OJJ31580.1 azon|OJJ47295.1

Group4176: abra|OJJ76591.1 acar|OOF99396.1 acla|EAW14310.1 afis|EAW18976.1 afla|EED53221.1 afum|EAL93711.2 agla|OJJ82933.1 akaw|GAA82495.1 aluc|OJZ91615.1 anid|CBF69644.1 anig|EHA22643.1 aory|EIT74842.1 asyd|OJJ59080.1 ater|EAU38163.1 atub|OJI86068.1 aver|OJJ01673.1 awen|OJJ31581.1 azon|OJJ47296.1

Group4177: abra|OJJ76594.1 acar|OOF99393.1 acla|EAW14307.1 afis|EAW18978.1 afla|EED53192.1 afum|EAL93714.1 agla|OJJ82936.1 akaw|GAA82498.1 aluc|OJZ91612.1 anid|CBF69649.1 anig|EHA22641.1 aory|EIT75152.1 asyd|OJJ59077.1 ater|EAU38169.1 atub|OJI86065.1 aver|OJJ01670.1 awen|OJJ31584.1 azon|OJJ47299.1

Group4178: abra|OJJ76601.1 acar|OOF99390.1 acla|EAW14305.1 afis|EAW18979.1 afla|EED53188.1 afum|EAL93715.1 agla|OJJ82941.1 akaw|GAA82504.1 aluc|OJZ91607.1 anid|CBF69652.1 anig|EHA22636.1 aory|EIT74907.1 asyd|OJJ59076.1 ater|EAU37838.1 atub|OJI86059.1 aver|OJJ01669.1 awen|OJJ31587.1 azon|OJJ47303.1

Group4179: abra|OJJ76605.1 acar|OOF99386.1 acla|EAW14301.1 afis|EAW18983.1 afla|EED53161.1 afum|EAL93719.1 agla|OJJ82958.1 akaw|GAA82509.1 aluc|OJZ91603.1 anid|CBF69659.1 anig|EHA22632.1 aory|EIT74944.1 asyd|OJJ59072.1 ater|EAU37822.1 atub|OJI86055.1 aver|OJJ01665.1 awen|OJJ31600.1 azon|OJJ47307.1

Group4180: abra|OJJ76606.1 acar|OOF99384.1 acla|EAW14300.1 afis|EAW18984.1 afla|EED53162.1 afum|EAL93720.1 agla|OJJ82959.1 akaw|GAA82510.1 aluc|OJZ91602.1 anid|CBF69661.1 anig|EHA22631.1 aory|EIT74985.1 asyd|OJJ59071.1 ater|EAU37821.1 atub|OJI86054.1 aver|OJJ01664.1 awen|OJJ31604.1 azon|OJJ47308.1

Group4181: abra|OJJ76609.1 acar|OOF99383.1 acla|EAW14297.1 afis|EAW18986.1 afla|EED53164.1 afum|EAL93722.1 agla|OJJ82964.1 akaw|GAA82512.1 aluc|OJZ91599.1 anid|CBF69665.1 anig|EHA22629.1 aory|EIT75120.1 asyd|OJJ59069.1 ater|EAU37819.1 atub|OJI86050.1 aver|OJJ01662.1 awen|OJJ31606.1 azon|OJJ47310.1

Group4182: abra|OJJ76624.1 acar|OOF99374.1 acla|EAW14295.1 afis|EAW18988.1 afla|EED53173.1 afum|EAL93724.1 agla|OJJ82975.1 akaw|GAA92833.1 aluc|OJZ91588.1 anid|CBF86791.1 anig|EHA22612.1 aory|EIT75118.1 asyd|OJJ64674.1 ater|EAU37816.1 atub|OJI86037.1 aver|OJI98775.1 awen|OJJ31609.1 azon|OJJ47312.1

Group4183: abra|OJJ76631.1 acar|OOF99366.1 acla|EAW08624.1 afis|EAW15808.1 afla|EED53158.1 afum|EAL85543.1 agla|OJJ81974.1 akaw|GAA92825.1 aluc|OJZ91580.1 anid|CBF86787.1 anig|EHA22607.1 aory|EIT75169.1 asyd|OJJ64676.1 ater|EAU37812.1 atub|OJI86029.1 aver|OJI98777.1 awen|OJJ38519.1 azon|OJJ50132.1

Group4184: abra|OJJ76632.1 acar|OOF99365.1 acla|EAW14288.1 afis|EAW18995.1 afla|EED53157.1 afum|EAL93730.1 agla|OJJ83149.1 akaw|GAA92824.1 aluc|OJZ91578.1 anid|CBF86656.1 anig|EHA22605.1 aory|EIT74972.1 asyd|OJJ64788.1 atub|OJI86028.1 aver|OJJ07926.1 awen|OJJ29936.1 aver|OJI98879.1 aver|OJI98900.1

Group4185: abra|OJJ76658.1 acar|OOF99343.1 acla|EAW14620.1 afis|EAW19358.1 afla|EED52157.1 afum|EAL94078.1 akaw|GAA92998.1 aluc|OJZ91547.1 anid|CBF73438.1 anig|EHA22582.1 aory|EIT74377.1 asyd|OJJ52447.1 ater|EAU36409.1 atub|OJI86000.1 aver|OJJ08112.1 azon|OJJ42907.1 azon|OJJ42486.1 aver|OJJ03969.1

Group4186: abra|OJJ76671.1 acar|OOF99339.1 acla|EAW09095.1 afis|EAW25689.1 afum|EAL84358.1 agla|OJJ78884.1 akaw|GAA92521.1 aluc|OJZ91535.1 anig|EHA22572.1 asyd|OJJ55501.1 atub|OJI85989.1 aver|OJJ06579.1 awen|OJJ38060.1 azon|OJJ45319.1 acar|OOG00611.1 anid|CBF80316.1 asyd|OJJ57586.1 aver|OJJ05596.1

Group4187: abra|OJJ76753.1 acar|OOF91249.1 acla|EAW13307.1 afis|EAW20421.1 afla|EED44850.1 afum|EAL84629.1 agla|OJJ84904.1 akaw|GAA87271.1 aluc|OJZ84156.1 anid|CBF79342.1 anig|EHA27518.1

aory|EIT77742.1 asyd|OJJ55796.1 ater|EAU37735.1 atub|OJI89928.1 aver|OJJ06885.1 awen|OJJ33114.1 azon|OJJ46330.1

Group4188: abra|OJJ76756.1 acar|OOF91251.1 acla|EAW13310.1 afis|EAW20424.1 afla|EED44846.1 afum|EAL84624.2 agla|OJJ84901.1 akaw|GAA87269.1 aluc|OJZ84159.1 anid|CBF79340.1 anig|EHA27521.1 aory|EIT77736.1 asyd|OJJ55792.1 ater|EAU37733.1 atub|OJI89925.1 aver|OJJ06882.1 awen|OJJ33117.1 azon|OJJ46333.1

Group4189: abra|OJJ76758.1 acar|OOF91253.1 acla|EAW13312.1 afis|EAW20426.1 afla|EED44842.1 afum|EAL84623.1 agla|OJJ84899.1 akaw|GAA87268.1 aluc|OJZ84161.1 anid|CBF79338.1 anig|EHA27522.1 aory|EIT77746.1 asyd|OJJ55790.1 ater|EAU37731.1 atub|OJI89923.1 aver|OJJ06880.1 awen|OJJ33119.1 azon|OJJ46335.1

Group4190: abra|OJJ76760.1 acar|OOF91255.1 acla|EAW13314.1 afis|EAW20428.1 afla|EED44840.1 afum|EAL84622.1 agla|OJJ84897.1 akaw|GAA87266.1 aluc|OJZ84163.1 anid|CBF79334.1 anig|EHA27524.1 aory|EIT77728.1 asyd|OJJ55788.1 ater|EAU37729.1 atub|OJI89921.1 aver|OJJ06878.1 awen|OJJ33121.1 azon|OJJ46337.1

Group4191: abra|OJJ76761.1 acar|OOF91256.1 acla|EAW13315.1 afis|EAW20429.1 afla|EED44839.1 afum|EAL84621.1 agla|OJJ84896.1 akaw|GAA87265.1 aluc|OJZ84164.1 anid|CBF79332.1 anig|EHA27525.1 aory|EIT77744.1 asyd|OJJ55787.1 ater|EAU37728.1 atub|OJI89920.1 aver|OJJ06877.1 awen|OJJ33122.1 azon|OJJ46338.1

Group4192: abra|OJJ76762.1 acar|OOF91257.1 acla|EAW13316.1 afis|EAW20430.1 afla|EED44838.1 afum|EAL84620.1 agla|OJJ84895.1 akaw|GAA87264.1 aluc|OJZ84165.1 anid|CBF79330.1 anig|EHA27526.1 aory|EIT77755.1 asyd|OJJ55786.1 ater|EAU37727.1 atub|OJI89919.1 aver|OJJ06876.1 awen|OJJ33123.1 azon|OJJ46339.1

Group4193: abra|OJJ76764.1 acar|OOF91259.1 acla|EAW13318.1 afis|EAW20432.1 afla|EED44836.1 afum|EAL84619.1 agla|OJJ84893.1 akaw|GAA87262.1 aluc|OJZ84168.1 anid|CBF79325.1 anig|EHA27528.1 aory|EIT77720.1 asyd|OJJ55784.1 ater|EAU37725.1 atub|OJI89916.1 aver|OJJ06874.1 awen|OJJ33125.1 azon|OJJ46341.1

Group4194: abra|OJJ76766.1 acar|OOF91260.1 acla|EAW13320.1 afis|EAW20435.1 afla|EED44835.1 afum|EAL84617.1 agla|OJJ84891.1 akaw|GAA87260.1 aluc|OJZ84169.1 anid|CBF79321.1 anig|EHA27530.1 aory|EIT77727.1 asyd|OJJ55783.1 ater|EAU37723.1 atub|OJI89915.1 aver|OJJ06872.1 awen|OJJ33126.1 azon|OJJ46342.1

Group4195: abra|OJJ76768.1 acar|OOF91262.1 acla|EAW13322.1 afis|EAW20437.1 afla|EED44833.1 afum|EAL84615.1 agla|OJJ84886.1 akaw|GAA87258.1 aluc|OJZ84171.1 anid|CBF79317.1 anig|EHA27532.1 aory|EIT77715.1 asyd|OJJ55781.1 ater|EAU37721.1 atub|OJI89913.1 aver|OJJ06870.1 awen|OJJ33128.1 azon|OJJ46344.1

Group4196: abra|OJJ76769.1 acar|OOF91263.1 acla|EAW13323.1 afis|EAW20438.1 afla|EED44832.1 afum|EAL84614.1 agla|OJJ84885.1 akaw|GAA87257.1 aluc|OJZ84172.1 anid|CBF79312.1 anig|EHA27533.1 aory|EIT77757.1 asyd|OJJ55780.1 ater|EAU37720.1 atub|OJI89912.1 aver|OJJ06869.1 awen|OJJ33129.1 azon|OJJ46345.1

Group4197: abra|OJJ76774.1 acar|OOF91269.1 acla|EAW13331.1 afis|EAW20446.1 afla|EED44824.1 afum|EAL84607.1 agla|OJJ84875.1 akaw|GAA87250.1 aluc|OJZ84178.1 anid|CBF79292.1 anig|EHA27537.1 aory|EIT77756.1 asyd|OJJ55775.1 ater|EAU37713.1 atub|OJI89905.1 aver|OJJ06864.1 awen|OJJ33136.1 azon|OJJ46349.1

Group4198: abra|OJJ76776.1 acar|OOF91271.1 acla|EAW13335.1 afis|EAW20450.1 afla|EED44818.1 afum|EAL84603.1 agla|OJJ84872.1 akaw|GAA87248.1 aluc|OJZ84180.1 anid|CBF79297.1 anig|EHA27540.1 aory|EIT77719.1 asyd|OJJ55773.1 ater|EAU37709.1 atub|OJI89903.1 aver|OJJ06860.1 awen|OJJ33140.1 azon|OJJ46351.1

Group4199: abra|OJJ76779.1 acar|OOF91275.1 acla|EAW13338.1 afis|EAW20453.1 afla|EED44815.1 afum|EAL84600.1 agla|OJJ84869.1 akaw|GAA87245.1 aluc|OJZ84183.1 anid|CBF79284.1 anig|EHA27543.1 aory|EIT77717.1 asyd|OJJ55770.1 ater|EAU37706.1 atub|OJI89900.1 aver|OJJ06856.1 awen|OJJ33143.1 azon|OJJ46354.1

Group4200: abra|OJJ76780.1 acar|OOF91276.1 acla|EAW13339.1 afis|EAW20454.1 afla|EED44814.1 afum|EAL84599.1 agla|OJJ84868.1 akaw|GAA87244.1 aluc|OJZ84184.1 anid|CBF79286.1 anig|EHA27544.1 aory|EIT77725.1 asyd|OJJ55769.1 ater|EAU37705.1 atub|OJI89899.1 aver|OJJ06855.1 awen|OJJ33144.1 azon|OJJ46355.1

Group4201: abra|OJJ76784.1 acar|OOF91279.1 acla|EAW13342.1 afis|EAW20457.1 afla|EED44810.1 afum|EAL84595.1 agla|OJJ84862.1 akaw|GAA87241.1 aluc|OJZ84187.1 anid|CBF79283.1 anig|EHA27547.1 aory|EIT77730.1 asyd|OJJ55766.1 ater|EAU37703.1 atub|OJI89896.1 aver|OJJ06852.1 awen|OJJ33149.1 azon|OJJ46359.1

Group4202: abra|OJJ76787.1 acar|OOF91281.1 acla|EAW13345.1 afis|EAW20460.1 afla|EED44808.1 afum|EAL84593.1 agla|OJJ84858.1 akaw|GAA87238.1 aluc|OJZ84189.1 anid|CBF79277.1 anig|EHA27549.1 aory|EIT77754.1 asyd|OJJ55763.1 ater|EAU37700.1 atub|OJI89893.1 aver|OJJ06849.1 awen|OJJ33153.1 azon|OJJ46362.1

Group4203: abra|OJJ76790.1 acar|OOF91284.1 acla|EAW13348.1 afis|EAW20463.1 afla|EED44805.1 afum|EAL84590.1 agla|OJJ84854.1 akaw|GAA87235.1 aluc|OJZ84193.1 anid|CBF79267.1 anig|EHA27552.1 aory|EIT77734.1 asyd|OJJ55760.1 ater|EAU37738.1 atub|OJI89890.1 aver|OJJ06845.1 awen|OJJ33157.1 azon|OJJ46366.1

Group4204: abra|OJJ76792.1 acar|OOF91283.1 acla|EAW13351.1 afis|EAW20465.1 afla|EED44802.1 afum|EAL84588.1 agla|OJJ84852.1 akaw|GAA87233.1 aluc|OJZ84196.1 anid|CBF79269.1 anig|EHA27553.1 aory|EIT77749.1 asyd|OJJ55759.1 ater|EAU37698.1 atub|OJI89888.1 aver|OJJ06844.1 awen|OJJ33159.1 azon|OJJ46367.1

Group4205: abra|OJJ76793.1 acar|OOF91286.1 acla|EAW13352.1 afis|EAW20466.1 afla|EED44801.1 afum|EAL84586.2 agla|OJJ84851.1 akaw|GAA87232.1 aluc|OJZ84197.1 anid|CBF79271.1 anig|EHA27554.1 aory|EIT77739.1 asyd|OJJ55758.1 ater|EAU37697.1 atub|OJI89887.1 aver|OJJ06843.1 awen|OJJ33160.1 azon|OJJ46368.1

Group4206: abra|OJJ76796.1 acar|OOF91289.1 acla|EAW13355.1 afis|EAW20469.1 afla|EED44799.1 afum|EAL84583.1 agla|OJJ84849.1 akaw|GAA87227.1 aluc|OJZ84203.1 anid|CBF79257.1 anig|EHA27557.1 aory|EIT77745.1 asyd|OJJ55748.1 ater|EAU37694.1 atub|OJI89881.1 aver|OJJ06835.1 awen|OJJ33163.1 azon|OJJ46370.1

Group4207: abra|OJJ76798.1 acar|OOF91291.1 acla|EAW13357.1 afis|EAW20471.1 afla|EED44797.1 afum|EAL84582.1 agla|OJJ84847.1 akaw|GAA87225.1 aluc|OJZ84205.1 anid|CBF79258.1 anig|EHA27558.1 aory|EIT77743.1 asyd|OJJ55750.1 ater|EAU37691.1 atub|OJI89879.1 aver|OJJ06837.1 awen|OJJ33165.1 azon|OJJ46372.1

Group4208: abra|OJJ76799.1 acar|OOF91292.1 acla|EAW13358.1 afis|EAW20472.1 afla|EED44796.1 afum|EAL84580.2 agla|OJJ84846.1 akaw|GAA87224.1 aluc|OJZ84206.1 anid|CBF79260.1 anig|EHA27559.1 aory|EIT77751.1 asyd|OJJ55751.1 ater|EAU37690.1 atub|OJI89878.1 aver|OJJ06838.1 awen|OJJ33166.1 azon|OJJ46373.1

Group4209: abra|OJJ76800.1 acar|OOF91294.1 acla|EAW13359.1 afis|EAW20473.1 afla|EED44794.1 afum|EAL84579.1 agla|OJJ84845.1 akaw|GAA87223.1 aluc|OJZ84207.1 anid|CBF79262.1 anig|EHA27561.1 aory|EIT77790.1 asyd|OJJ55754.1 ater|EAU37689.1 atub|OJI89877.1 aver|OJJ06839.1 awen|OJJ33167.1 azon|OJJ46375.1

Group4210: abra|OJJ76802.1 acar|OOF91296.1 acla|EAW13361.1 afis|EAW20475.1 afla|EED44793.1 afum|EAL84578.1 agla|OJJ84842.1 akaw|GAA87221.1 aluc|OJZ84209.1 anid|CBF79255.1 anig|EHA27563.1 aory|EIT77775.1 asyd|OJJ55747.1 ater|EAU37687.1 atub|OJI89874.1 aver|OJJ06834.1 awen|OJJ33170.1 azon|OJJ46377.1

Group4211: abra|OJJ76806.1 acar|OOF91300.1 acla|EAW13365.1 afis|EAW20479.1 afla|EED44789.1 afum|EAL84575.1 agla|OJJ84831.1 akaw|GAA87217.1 aluc|OJZ84213.1 anid|CBF79213.1 anig|EHA27566.1 aory|EIT77792.1 asyd|OJJ55738.1 ater|EAU29558.1 atub|OJI89870.1 aver|OJJ06824.1 awen|OJJ33174.1 azon|OJJ46381.1

Group4212: abra|OJJ76809.1 acar|OOF91305.1 acla|EAW13370.1 afis|EAW20484.1 afla|EED44784.1 afum|EAL84571.1 agla|OJJ84813.1 akaw|GAA87212.1 aluc|OJZ84218.1 anid|CBF79235.1 anig|EHA27569.1 aory|EIT77759.1 asyd|OJJ55742.1 ater|EAU29562.1 atub|OJI89866.1 aver|OJJ06829.1 awen|OJJ33185.1 azon|OJJ46384.1

Group4213: abra|OJJ76810.1 acar|OOF91306.1 acla|EAW13371.1 afis|EAW20485.1 afla|EED44782.1 afum|EAL84570.1 agla|OJJ84822.1 akaw|GAA87211.1 aluc|OJZ84221.1 anid|CBF79237.1 anig|EHA27570.1 aory|EIT77768.1 asyd|OJJ55743.1 ater|EAU29563.1 atub|OJI89865.1 aver|OJJ06830.1 awen|OJJ33186.1 azon|OJJ46388.1

Group4214: abra|OJJ76811.1 acar|OOF91307.1 acla|EAW13372.1 afis|EAW20486.1 afla|EED44781.1 afum|EAL84569.1 agla|OJJ84825.1 akaw|GAA87210.1 aluc|OJZ84222.1 anid|CBF79215.1 anig|EHA27571.1 aory|EIT77774.1 asyd|OJJ55737.1 ater|EAU29564.1 atub|OJI89864.1 aver|OJJ06823.1 awen|OJJ33187.1 azon|OJJ46389.1

Group4215: abra|OJJ76813.1 acar|OOF91309.1 acla|EAW13374.1 afis|EAW20488.1 afla|EED44776.1 afum|EAL84567.1 agla|OJJ84826.1 akaw|GAA87208.1 aluc|OJZ84224.1 anid|CBF79219.1 anig|EHA27573.1 aory|EIT77760.1 asyd|OJJ55736.1 ater|EAU29566.1 atub|OJI89861.1 aver|OJJ06822.1 awen|OJJ33188.1 azon|OJJ46390.1

Group4216: abra|OJJ76819.1 acar|OOF91315.1 acla|EAW13379.1 afis|EAW20493.1 afla|EED44737.1 afum|EAL84561.1 agla|OJJ84833.1 akaw|GAA87203.1 aluc|OJZ84229.1 anid|CBF79226.1 anig|EHA27578.1

aory|EIT77797.1 asyd|OJJ55732.1 ater|EAU29569.1 atub|OJI89855.1 aver|OJJ06818.1 awen|OJJ33193.1 azon|OJJ46403.1

Group4217: abra|OJJ76821.1 acar|OOF91316.1 acla|EAW13381.1 afis|EAW20495.1 afla|EED44771.1 afum|EAL84559.1 agla|OJJ84807.1 akaw|GAA87201.1 aluc|OJZ84232.1 anid|CBF79222.1 anig|EHA27579.1 aory|EIT77762.1 asyd|OJJ55734.1 ater|EAU37189.1 atub|OJI89853.1 aver|OJJ06820.1 awen|OJJ33195.1 azon|OJJ46401.1

Group4218: abra|OJJ76823.1 acar|OOF91319.1 acla|EAW13383.1 afis|EAW20497.1 afla|EED44739.1 afum|EAL84557.1 agla|OJJ84805.1 akaw|GAA87199.1 aluc|OJZ84234.1 anid|CBF79208.1 anig|EHA27581.1 aory|EIT77796.1 asyd|OJJ55729.1 ater|EAU37191.1 atub|OJI89851.1 aver|OJJ06815.1 awen|OJJ33197.1 azon|OJJ46399.1

Group4219: abra|OJJ76827.1 acar|OOF91321.1 acla|EAW13387.1 afis|EAW20501.1 afla|EED44744.1 afum|EAL84554.1 agla|OJJ84797.1 akaw|GAA87195.1 aluc|OJZ84239.1 anid|CBF79200.1 anig|EHA27586.1 aory|EIT77773.1 asyd|OJJ55725.1 ater|EAU37195.1 atub|OJI89844.1 aver|OJJ06811.1 awen|OJJ33202.1 azon|OJJ46396.1

Group4220: abra|OJJ76830.1 acar|OOF91324.1 acla|EAW13392.1 afis|EAW20504.1 afla|EED44747.1 afum|EAL84552.1 agla|OJJ84800.1 akaw|GAA87192.1 aluc|OJZ84242.1 anid|CBF79195.1 anig|EHA27587.1 aory|EIT77784.1 asyd|OJJ55722.1 ater|EAU37199.1 atub|OJI89841.1 aver|OJJ06808.1 awen|OJJ33205.1 azon|OJJ46478.1

Group4221: abra|OJJ76832.1 acar|OOF91326.1 afis|EAW20506.1 afum|EAL84550.1 akaw|GAA87190.1 aluc|OJZ84244.1 anig|EHA27589.1 asyd|OJJ55720.1 ater|EAU37200.1 atub|OJI89839.1 aver|OJJ06806.1 awen|OJJ33207.1 acla|EAW13395.1 afla|EED52484.1 aory|EIT82804.1 afla|EED52486.1 anid|CBF79191.1 aory|EIT82815.1

Group4222: abra|OJJ76834.1 acar|OOF91327.1 acla|EAW13396.1 afis|EAW20507.1 afla|EED52448.1 afum|EAL84549.1 agla|OJJ84784.1 akaw|GAA87189.1 aluc|OJZ84245.1 anid|CBF79193.1 anig|EHA27590.1 aory|EIT82772.1 asyd|OJJ55719.1 ater|EAU37201.1 atub|OJI89838.1 aver|OJJ06805.1 awen|OJJ33208.1 azon|OJJ46481.1

Group4223: abra|OJJ76835.1 acar|OOF91328.1 acla|EAW13397.1 afis|EAW20508.1 afla|EED52449.1 afum|EAL84548.1 agla|OJJ84785.1 akaw|GAA87188.1 aluc|OJZ84246.1 anid|CBF79190.1 anig|EHA27591.1 aory|EIT82778.1 asyd|OJJ55718.1 ater|EAU37202.1 atub|OJI89837.1 aver|OJJ06804.1 awen|OJJ33209.1 azon|OJJ46482.1

Group4224: abra|OJJ76836.1 acar|OOF90072.1 acla|EAW13398.1 afis|EAW20509.1 afla|EED52450.1 afum|EAL84547.1 agla|OJJ84786.1 akaw|GAA87187.1 aluc|OJZ84247.1 anid|CBF79189.1 anig|EHA27592.1 aory|EIT82759.1 asyd|OJJ55717.1 ater|EAU37203.1 atub|OJI89836.1 aver|OJJ06803.1 awen|OJJ33210.1 azon|OJJ46483.1

Group4225: abra|OJJ76838.1 acar|OOF91330.1 acla|EAW13400.1 afis|EAW20511.1 afla|EED52452.1 afum|EAL84545.2 agla|OJJ84788.1 akaw|GAA87185.1 aluc|OJZ84249.1 anid|CBF79186.1 anig|EHA27594.1 aory|EIT82799.1 asyd|OJJ55715.1 ater|EAU37205.1 atub|OJI89834.1 aver|OJJ06801.1 awen|OJJ33213.1 azon|OJJ46485.1

Group4226: abra|OJJ76839.1 acar|OOF91331.1 acla|EAW13401.1 afis|EAW20512.1 afla|EED52453.1 afum|EBA27192.1 agla|OJJ84789.1 akaw|GAA87184.1 aluc|OJZ84250.1 anid|CBF79184.1 anig|EHA27595.1 aory|EIT82753.1 asyd|OJJ55714.1 ater|EAU37208.1 atub|OJI89833.1 aver|OJJ06800.1 awen|OJJ33214.1 azon|OJJ46486.1

Group4227: abra|OJJ76840.1 acar|OOF91333.1 acla|EAW13403.1 afis|EAW20514.1 afla|EED52468.1 afum|EAL84543.2 agla|OJJ84776.1 akaw|GAA87182.1 aluc|OJZ84251.1 anid|CBF79055.1 anig|EHA27596.1 aory|EIT82784.1 asyd|OJJ55710.1 ater|EAU37187.1 atub|OJI89832.1 aver|OJJ06796.1 awen|OJJ33220.1 azon|OJJ46488.1

Group4228: abra|OJJ76843.1 acar|OOF89969.1 acla|EAW13406.1 afis|EAW20517.1 afla|EED52458.1 afum|EAL84540.1 agla|OJJ84769.1 akaw|GAA87179.1 aluc|OJZ84254.1 anid|CBF88058.1 anig|EHA27599.1 aory|EIT82793.1 asyd|OJJ63541.1 ater|EAU37184.1 atub|OJI89829.1 aver|OJI97392.1 awen|OJJ33223.1 azon|OJJ46553.1

Group4229: abra|OJJ76844.1 acar|OOF89968.1 acla|EAW13407.1 afis|EAW20518.1 afla|EED52457.1 afum|EAL84539.1 agla|OJJ84768.1 akaw|GAA87177.1 aluc|OJZ84256.1 anid|CBF79061.1 anig|EHA27600.1 aory|EIT82769.1 asyd|OJJ55707.1 ater|EAU37183.1 atub|OJI89827.1 aver|OJJ06793.1 awen|OJJ33224.1 azon|OJJ46581.1

Group4230: abra|OJJ76845.1 acar|OOF91335.1 acla|EAW13408.1 afis|EAW20519.1 afla|EED52455.1 afum|EAL84538.1 agla|OJJ84767.1 akaw|GAA87176.1 aluc|OJZ84257.1 anid|CBF79067.1 anig|EHA27601.1 aory|EIT82798.1 asyd|OJJ55706.1 ater|EAU37182.1 atub|OJI89826.1 aver|OJJ06792.1 awen|OJJ33225.1 azon|OJJ46580.1

Group4231: abra|OJJ76847.1 acar|OOF91337.1 acla|EAW13410.1 afis|EAW20521.1 afla|EED52470.1 afum|EAL84536.1 agla|OJJ84765.1 akaw|GAA87174.1 aluc|OJZ84259.1 anid|CBF79070.1 anig|EHA27602.1 aory|EIT82770.1 asyd|OJJ55704.1 ater|EAU37167.1 atub|OJI89824.1 aver|OJJ06790.1 awen|OJJ33227.1 azon|OJJ46577.1

Group4232: abra|OJJ76849.1 acar|OOF91338.1 acla|EAW13412.1 afis|EAW20523.1 afla|EED52472.1 afum|EAL84534.1 agla|OJJ84740.1 akaw|GAA87172.1 aluc|OJZ84261.1 anid|CBF79074.1 anig|EHA27604.1 aory|EIT82821.1 asyd|OJJ55702.1 ater|EAU37165.1 atub|OJI89822.1 aver|OJJ06787.1 awen|OJJ33229.1 azon|OJJ46575.1

Group4233: abra|OJJ76851.1 acar|OOF91339.1 acla|EAW13413.1 afis|EAW20524.1 afla|EED52473.1 afum|EAL84533.1 agla|OJJ84742.1 akaw|GAA87171.1 aluc|OJZ84262.1 anid|CBF79076.1 anig|EHA27605.1 aory|EIT82758.1 asyd|OJJ55701.1 ater|EAU37164.1 atub|OJI89821.1 aver|OJJ06786.1 awen|OJJ33230.1 azon|OJJ46574.1

Group4234: abra|OJJ76852.1 acar|OOF91340.1 acla|EAW13414.1 afis|EAW20525.1 afla|EED52474.1 afum|EAL84532.1 agla|OJJ84743.1 akaw|GAA87170.1 aluc|OJZ84264.1 anid|CBF79078.1 anig|EHA27606.1 aory|EIT82817.1 asyd|OJJ55699.1 ater|EAU37163.1 atub|OJI89819.1 aver|OJJ06784.1 awen|OJJ33231.1 azon|OJJ46573.1

Group4235: abra|OJJ76855.1 acar|OOF91341.1 acla|EAW13416.1 afis|EAW20527.1 afla|EED52476.1 afum|EAL84530.1 agla|OJJ84745.1 akaw|GAA87167.1 aluc|OJZ84267.1 anid|CBF79081.1 anig|EHA27609.1 aory|EIT82771.1 asyd|OJJ55696.1 ater|EAU37161.1 atub|OJI89816.1 aver|OJJ06781.1 awen|OJJ33234.1 azon|OJJ46570.1

Group4236: abra|OJJ76857.1 acar|OOF91343.1 acla|EAW13418.1 afis|EAW20529.1 afla|EED52478.1 afum|EAL84528.1 agla|OJJ84706.1 akaw|GAA87165.1 aluc|OJZ84269.1 anid|CBF79006.1 anig|EHA27611.1 aory|EIT82803.1 asyd|OJJ55691.1 ater|EAU37159.1 atub|OJI89814.1 aver|OJJ06776.1 awen|OJJ33236.1 azon|OJJ50608.1

Group4237: abra|OJJ76903.1 acar|OOF91351.1 acla|EAW13425.1 afis|EAW20536.1 afla|EED52442.1 afum|EAL84522.1 agla|OJJ84695.1 akaw|GAA87135.1 aluc|OJZ84302.1 anid|CBF78995.1 anig|EHA27638.1 aory|EIT82797.1 asyd|OJJ55685.1 ater|EAU37143.1 atub|OJI89766.1 aver|OJJ06769.1 awen|OJJ33243.1 azon|OJJ50622.1

Group4238: abra|OJJ76916.1 acar|OOF91363.1 acla|EAW13428.1 afis|EAW20544.1 afla|EED52424.1 afum|EAL84514.1 agla|OJJ88753.1 akaw|GAA87122.1 aluc|OJZ84316.1 anid|CBF78988.1 anig|EHA27651.1 aory|EIT83288.1 asyd|OJJ55682.1 ater|EAU37137.1 atub|OJI89752.1 aver|OJJ06766.1 awen|OJJ33252.1 azon|OJJ50638.1

Group4239: abra|OJJ76917.1 acar|OOF91364.1 acla|EAW13429.1 afis|EAW20545.1 afla|EED52423.1 afum|EBA27188.1 agla|OJJ88752.1 akaw|GAA87121.1 aluc|OJZ84317.1 anid|CBF78986.1 anig|EHA27652.1 aory|EIT83281.1 asyd|OJJ55681.1 ater|EAU37136.1 atub|OJI89751.1 aver|OJJ06765.1 awen|OJJ33253.1 azon|OJJ50637.1

Group4240: abra|OJJ76918.1 acar|OOF91365.1 acla|EAW13430.1 afis|EAW20546.1 afla|EED52422.1 afum|EAL84513.1 agla|OJJ88751.1 akaw|GAA87120.1 aluc|OJZ84318.1 anid|CBF78985.1 anig|EHA27653.1 aory|EIT83315.1 asyd|OJJ55680.1 ater|EAU37135.1 atub|OJI89750.1 aver|OJJ06763.1 awen|OJJ33254.1 azon|OJJ50636.1

Group4241: abra|OJJ76919.1 acar|OOF91366.1 acla|EAW13431.1 afis|EAW20547.1 afla|EED52421.1 afum|EAL84512.1 agla|OJJ88750.1 akaw|GAA87119.1 aluc|OJZ84319.1 anid|CBF78983.1 anig|EHA27654.1 aory|EIT83284.1 asyd|OJJ55679.1 ater|EAU37134.1 atub|OJI89749.1 aver|OJJ06762.1 awen|OJJ33255.1 azon|OJJ50635.1

Group4242: abra|OJJ76923.1 acar|OOF91370.1 acla|EAW13432.1 afis|EAW20548.1 afla|EED52408.1 afum|EAL84511.1 agla|OJJ88739.1 akaw|GAA87115.1 aluc|OJZ84323.1 anid|CBF78977.1 anig|EHA27658.1 aory|EIT83309.1 asyd|OJJ55675.1 ater|EAU37129.1 atub|OJI89745.1 aver|OJJ06758.1 awen|OJJ33260.1 aver|OJJ07651.1

Group4243: abra|OJJ76937.1 acar|OOF91386.1 acla|EAW12145.1 afis|EAW17130.1 afla|EED55089.1 afum|EAL86590.1 agla|OJJ82232.1 akaw|GAA87102.1 aluc|OJZ84336.1 anid|CBF80327.1 anig|EHA27670.1 aory|EIT81595.1 asyd|OJJ54844.1 ater|EAU31746.1 atub|OJI89732.1 aver|OJJ07318.1 awen|OJJ31282.1 awen|OJJ38657.1

Group4244: abra|OJJ76950.1 acar|OOF91396.1 acla|EAW13470.1 afis|EAW20582.1 afla|EED52277.1 afum|EAL84482.1 agla|OJJ88700.1 akaw|GAA87091.1 aluc|OJZ84347.1 anid|CBF78946.1 anig|EHA27683.1 aory|EIT74415.1 asyd|OJJ55651.1 ater|EAU37100.1 atub|OJI89721.1 aver|OJJ06736.1 awen|OJJ33287.1 azon|OJJ50663.1

Group4245: abra|OJJ76954.1 acar|OOF91398.1 acla|EAW13476.1 afis|EAW20583.1 afla|EED52245.1 afum|EBA27180.1 agla|OJJ88686.1 akaw|GAA87088.1 aluc|OJZ84350.1 anid|CBF78930.1 anig|EHA27687.1

aory|EIT74502.1 asyd|OJJ55642.1 ater|EAU37097.1 atub|OJI89717.1 aver|OJJ06725.1 awen|OJJ33300.1 azon|OJJ50671.1

Group4246: abra|OJJ76970.1 acar|OOF91415.1 acla|EAW13495.1 afis|EAW20595.1 afla|EED52229.1 afum|EAL84466.1 agla|OJJ88661.1 akaw|GAA87074.1 aluc|OJZ84366.1 anid|CBF78911.1 anig|EHA27701.1 aory|EIT74324.1 asyd|OJJ55615.1 ater|EAU35241.1 atub|OJI89702.1 aver|OJJ06690.1 awen|OJJ33385.1 azon|OJJ50691.1

Group4247: abra|OJJ76971.1 acar|OOF91416.1 acla|EAW13496.1 afis|EAW20596.1 afla|EED52228.1 afum|EAL84465.1 agla|OJJ88660.1 akaw|GAA87073.1 aluc|OJZ84367.1 anid|CBF78909.1 anig|EHA27702.1 aory|EIT74434.1 asyd|OJJ55614.1 ater|EAU35242.1 atub|OJI89701.1 aver|OJJ06689.1 awen|OJJ33388.1 azon|OJJ50692.1

Group4248: abra|OJJ76972.1 acar|OOF91417.1 acla|EAW13498.1 afis|EAW20597.1 afla|EED52213.1 afum|EAL84464.1 agla|OJJ88659.1 akaw|GAA87072.1 aluc|OJZ84368.1 anid|CBF78908.1 anig|EHA27703.1 aory|EIT74430.1 asyd|OJJ55612.1 ater|EAU35243.1 atub|OJI89700.1 aver|OJJ06687.1 awen|OJJ33389.1 azon|OJJ50693.1

Group4249: abra|OJJ76976.1 acar|OOF91419.1 acla|EAW13502.1 afis|EAW20601.1 afla|EED47729.1 afum|EAL84460.1 agla|OJJ88650.1 akaw|GAA87069.1 aluc|OJZ84372.1 anid|CBF75136.1 anig|EHA27707.1 aory|EIT77171.1 asyd|OJJ59976.1 ater|EAU35246.1 atub|OJI89695.1 aver|OJJ02315.1 awen|OJJ33393.1 azon|OJJ50697.1

Group4250: abra|OJJ76977.1 acar|OOF91422.1 acla|EAW13504.1 afis|EAW20603.1 afla|EED47732.1 afum|EAL84458.1 agla|OJJ88646.1 akaw|GAA87068.1 aluc|OJZ84374.1 anid|CBF86930.1 anig|EHA27708.1 aory|EIT77153.1 asyd|OJJ55600.1 ater|EAU35248.1 atub|OJI89694.1 aver|OJJ06680.1 awen|OJJ33395.1 azon|OJJ50698.1

Group4251: abra|OJJ76978.1 acar|OOF91424.1 acla|EAW13505.1 afis|EAW20604.1 afla|EED47733.1 afum|EAL84457.1 agla|OJJ88645.1 akaw|GAA87067.1 aluc|OJZ84376.1 anid|CBF86928.1 anig|EHA27709.1 aory|EIT77125.1 asyd|OJJ55601.1 ater|EAU35250.1 atub|OJI89693.1 aver|OJJ06681.1 awen|OJJ33396.1 azon|OJJ50394.1

Group4252: abra|OJJ76981.1 acar|OOF91427.1 acla|EAW12964.1 afis|EAW17699.1 afla|EED45692.1 afum|EAL91514.1 agla|OJJ79475.1 akaw|GAA87064.1 aluc|OJZ84381.1 anid|CBF89963.1 anig|EHA27712.1 aory|EIT75518.1 asyd|OJJ62104.1 ater|EAU31915.1 atub|OJI89690.1 aver|OJI96210.1 awen|OJJ37134.1 azon|OJJ50834.1

Group4253: abra|OJJ76984.1 acar|OOF91429.1 acla|EAW12966.1 afis|EAW17697.1 afla|EED50445.1 afum|EAL91511.1 agla|OJJ79478.1 akaw|GAA87062.1 aluc|OJZ84383.1 anid|CBF89965.1 anig|EHA27714.1 aory|EIT82014.1 asyd|OJJ62100.1 ater|EAU31918.1 atub|OJI89688.1 aver|OJI96206.1 awen|OJJ37137.1 azon|OJJ50836.1

Group4254: abra|OJJ76985.1 acar|OOF91430.1 acla|EAW12967.1 afis|EAW17696.1 afla|EED50446.1 afum|EAL91510.1 agla|OJJ79479.1 akaw|GAA87061.1 aluc|OJZ84384.1 anid|CBF89966.1 anig|EHA27715.1 aory|EIT82098.1 asyd|OJJ62099.1 ater|EAU31919.1 atub|OJI89687.1 aver|OJI96205.1 awen|OJJ37139.1 azon|OJJ50839.1

Group4255: abra|OJJ76986.1 acar|OOF92639.1 acla|EAW12968.1 afis|EAW17695.1 afla|EED50447.1 afum|EAL91509.1 agla|OJJ79483.1 akaw|GAA87060.1 aluc|OJZ84385.1 anid|CBF89968.1 anig|EHA27716.1 aory|EIT82034.1 asyd|OJJ62098.1 ater|EAU31920.1 atub|OJI89686.1 aver|OJI96204.1 awen|OJJ37140.1 azon|OJJ50840.1

Group4256: abra|OJJ76987.1 acar|OOF91431.1 acla|EAW12969.1 afis|EAW17694.1 afla|EED50448.1 afum|EAL91508.1 agla|OJJ79485.1 akaw|GAA87059.1 aluc|OJZ84386.1 anid|CBF89970.1 anig|EHA27717.1 aory|EIT82134.1 asyd|OJJ62097.1 ater|EAU31961.1 atub|OJI89685.1 aver|OJI96203.1 awen|OJJ37141.1 azon|OJJ50841.1

Group4257: abra|OJJ76993.1 acar|OOF91438.1 acla|EAW13455.1 afis|EAW20563.1 afum|EAL84498.1 agla|OJJ88728.1 akaw|GAA87053.1 aluc|OJZ84393.1 anid|CBF78959.1 anig|EHA27722.1 aory|EIT74501.1 asyd|OJJ55658.1 ater|EAU37112.1 atub|OJI89678.1 aver|OJJ06742.1 awen|OJJ33269.1 azon|OJJ50641.1 afla|EED52341.1

Group4258: abra|OJJ76996.1 acar|OOF91443.1 acla|EAW13460.1 afis|EAW20566.1 afla|EED52324.1 afum|EAL84495.1 agla|OJJ88727.1 akaw|GAA87050.1 aluc|OJZ84396.1 anid|CBF78935.1 anig|EHA27726.1 aory|EIT74477.1 asyd|OJJ55644.1 ater|EAU37111.1 atub|OJI89676.1 aver|OJJ06728.1 awen|OJJ33271.1 azon|OJJ50646.1

Group4259: abra|OJJ77016.1 acar|OOF95265.1 acla|EAW07210.1 afis|EAW22222.1 afla|EED57616.1 afum|EAL90972.1 agla|OJJ87822.1 akaw|GAA87034.1 aluc|OJZ84418.1 anid|CBF88465.1 anig|EHA27740.1 aory|EIT82988.1 asyd|OJJ63777.1 ater|EAU34232.1 atub|OJI89656.1 aver|OJI97160.1 awen|OJJ34809.1 azon|OJJ45741.1

Group4260: abra|OJJ77024.1 acar|OOF91461.1 acla|EAW12057.1 afis|EAW19698.1 afla|EED54134.1 afum|EAL86829.1 agla|OJJ80810.1 akaw|GAA87028.1 aluc|OJZ84424.1 anid|CBF81131.1 anig|EHA27746.1 aory|EIT76042.1 asyd|OJJ56500.1 ater|EAU32171.1 atub|OJI89649.1 aver|OJJ06219.1 awen|OJJ41934.1 azon|OJJ48232.1

Group4261: abra|OJJ77043.1 acar|OOF90273.1 acla|EAW13554.1 afis|EAW20631.1 afla|EED49762.1 afum|EAL84436.1 agla|OJJ80447.1 akaw|GAA87010.1 aluc|OJZ84445.1 anid|CBF83219.1 anig|EHA27763.1 aory|EIT77598.1 asyd|OJJ55568.1 ater|EAU32166.1 atub|OJI89629.1 aver|OJJ06647.1 awen|OJJ33600.1 azon|OJJ50722.1

Group4262: abra|OJJ77049.1 acar|OOF92938.1 acla|EAW13549.1 afis|EAW20628.1 afla|EED49778.1 afum|EAL84442.1 agla|OJJ88595.1 akaw|GAA87004.1 aluc|OJZ84453.1 anid|CBF83230.1 anig|EHA27768.1 aory|EIT77696.1 asyd|OJJ55582.1 ater|EAU32163.1 atub|OJI89623.1 aver|OJJ06661.1 awen|OJJ33461.1 azon|OJJ50717.1

Group4263: abra|OJJ77050.1 acar|OOF92939.1 acla|EAW13544.1 afis|EAW20616.1 afla|EED54198.1 afum|EAL84445.1 agla|OJJ88594.1 akaw|GAA87003.1 aluc|OJZ84454.1 anid|CBF86935.1 anig|EHA27769.1 aory|EIT75987.1 asyd|OJJ55584.1 ater|EAU35262.1 atub|OJI89621.1 aver|OJJ06663.1 awen|OJJ33460.1 azon|OJJ50716.1

Group4264: abra|OJJ77064.1 acar|OOF92948.1 acla|EAW13511.1 afis|EAW20609.1 afla|EED54207.1 afum|EAL84451.1 agla|OJJ88630.1 akaw|GAA86988.1 aluc|OJZ84472.1 anid|CBF86951.1 anig|EHA27781.1 aory|EIT76020.1 asyd|OJJ55594.1 ater|EAU35257.1 atub|OJI89606.1 aver|OJJ06673.1 awen|OJJ33452.1 azon|OJJ50709.1

Group4265: abra|OJJ77068.1 acar|OOF92951.1 acla|EAW13508.1 afis|EAW20606.1 afla|EED47725.1 afum|EAL84454.1 agla|OJJ88635.1 akaw|GAA86985.1 aluc|OJZ84476.1 anid|CBF86946.1 anig|EHA27784.1 aory|EIT77120.1 asyd|OJJ55591.1 ater|EAU35260.1 atub|OJI89603.1 aver|OJJ06670.1 awen|OJJ33449.1 azon|OJJ50706.1

Group4266: abra|OJJ77076.1 acar|OOF92958.1 acla|EAW11204.1 afis|EAW15934.1 afla|EED54927.1 afum|EAL88570.1 agla|OJJ89373.1 akaw|GAA86977.1 aluc|OJZ84486.1 anid|CBF86351.1 anig|EHA27790.1 aory|EIT80330.1 asyd|OJJ64980.1 ater|EAU35519.1 atub|OJI89594.1 aver|OJI99102.1 awen|OJJ35645.1 azon|OJJ46891.1

Group4267: abra|OJJ77079.1 acar|OOF92961.1 acla|EAW12041.1 afis|EAW19690.1 afla|EED57010.1 afum|EAL86836.1 agla|OJJ80804.1 akaw|GAA86974.1 aluc|OJZ84489.1 anid|CBF75140.1 anig|EHA27793.1 aory|EIT82303.1 asyd|OJJ59969.1 atub|OJI89591.1 aver|OJJ02322.1 awen|OJJ41917.1 azon|OJJ42689.1 ater|EAU32856.1

Group4268: abra|OJJ77090.1 acar|OOF92974.1 acla|EAW08632.1 afis|EAW15799.1 afla|EED51246.1 afum|EAL85551.1 agla|OJJ81963.1 akaw|GAA86965.1 aluc|OJZ84499.1 anid|CBF70921.1 anig|EHA27799.1 aory|EIT83184.1 asyd|OJJ52607.1 ater|EAU32343.1 atub|OJI89581.1 aver|OJJ04634.1 awen|OJJ38507.1 azon|OJJ43744.1

Group4269: abra|OJJ77091.1 acar|OOF92976.1 acla|EAW10011.1 afis|EAW21045.1 afla|EED51625.1 afum|EAL92081.1 agla|OJJ82363.1 akaw|GAA86964.1 aluc|OJZ84500.1 anid|CBF87097.1 anig|EHA27800.1 aory|EIT80549.1 asyd|OJJ60392.1 ater|EAU36817.1 atub|OJI89580.1 aver|OJI99721.1 awen|OJJ32179.1 azon|OJJ46755.1

Group4270: abra|OJJ77094.1 acar|OOF92977.1 acla|EAW10918.1 afis|EAW20940.1 afla|EED50295.1 afum|EAL84694.1 agla|OJJ83947.1 akaw|GAA86961.1 aluc|OJZ84503.1 anid|CBF82521.1 anig|EHA27803.1 aory|EIT77252.1 asyd|OJJ54070.1 ater|EAU38585.1 atub|OJI89577.1 aver|OJJ08672.1 awen|OJJ34377.1 azon|OJJ48111.1

Group4271: abra|OJJ77095.1 acar|OOF92978.1 acla|EAW10919.1 afis|EAW20939.1 afla|EED50294.1 afum|EAL84695.1 agla|OJJ83948.1 akaw|GAA86960.1 aluc|OJZ84504.1 anid|CBF82523.1 anig|EHA27804.1 aory|EIT77268.1 asyd|OJJ54071.1 ater|EAU38584.1 atub|OJI89576.1 aver|OJJ08671.1 awen|OJJ34378.1 azon|OJJ48112.1

Group4272: abra|OJJ77106.1 acar|OOF92991.1 acla|EAW11608.1 afis|EAW23840.1 afla|EED52551.1 afum|EAL87271.1 agla|OJJ82489.1 akaw|GAA86950.1 aluc|OJZ84515.1 anid|CBF77077.1 anig|EHA27813.1 aory|EIT74221.1 asyd|OJJ56983.1 ater|EAU33507.1 atub|OJI89566.1 aver|OJJ03304.1 awen|OJJ33810.1 azon|OJJ48724.1

Group4273: abra|OJJ77111.1 acar|OOF93001.1 acla|EAW07270.1 afis|EAW22291.1 afla|EED57693.1 afum|EAL90890.1 agla|OJJ87755.1 akaw|GAA90817.1 aluc|OJZ84520.1 anid|CBF88575.1 anig|EHA27818.1 aory|EIT79896.1 asyd|OJJ63840.1 ater|EAU34306.1 atub|OJI89560.1 aver|OJI97097.1 awen|OJJ34867.1 azon|OJJ50044.1

Group4274: abra|OJJ77116.1 acar|OOF93005.1 acla|EAW12952.1 afis|EAW17704.1 afla|EED45696.1 afum|EAL91519.2 agla|OJJ81740.1 akaw|GAA90813.1 aluc|OJZ84526.1 anid|CBF89929.1 anig|EHA27823.1

aory|EIT75367.1 asyd|OJJ62119.1 ater|EAU31912.1 atub|OJI89554.1 aver|OJI96226.1 awen|OJJ37131.1 azon|OJJ50830.1

Group4275: abra|OJJ77120.1 acar|OOF93007.1 acla|EAW12932.1 afis|EAW17709.1 afla|EED45711.1 afum|EAL91524.2 agla|OJJ81738.1 akaw|GAA90809.1 aluc|OJZ84530.1 anid|CBF89906.1 anig|EHA27827.1 aory|EIT75650.1 asyd|OJJ62132.1 ater|EAU31903.1 atub|OJI89550.1 aver|OJI96241.1 awen|OJJ37127.1 azon|OJJ50826.1

Group4276: abra|OJJ77123.1 acar|OOF93009.1 acla|EAW12936.1 afis|EAW17712.1 afla|EED45714.1 afum|EAL91527.1 agla|OJJ81736.1 akaw|GAA90805.1 aluc|OJZ84533.1 anid|CBF76920.1 anig|EHA27831.1 aory|EIT75661.1 asyd|OJJ57072.1 ater|EAU31891.1 atub|OJI89547.1 aver|OJJ03226.1 awen|OJJ37124.1 azon|OJJ50823.1

Group4277: abra|OJJ77125.1 acar|OOF93011.1 acla|EAW12938.1 afis|EAW17714.1 afla|EED45716.1 afum|EAL91529.1 agla|OJJ81734.1 akaw|GAA90803.1 aluc|OJZ84535.1 anid|CBF76916.1 anig|EHA27833.1 aory|EIT75429.1 asyd|OJJ57079.1 ater|EAU31892.1 atub|OJI89545.1 aver|OJJ03224.1 awen|OJJ37122.1 azon|OJJ50821.1

Group4278: abra|OJJ77126.1 acar|OOF93012.1 acla|EAW12939.1 afis|EAW17715.1 afla|EED45717.1 afum|EAL91530.1 agla|OJJ81733.1 akaw|GAA90802.1 aluc|OJZ84536.1 anid|CBF76914.1 anig|EHA27834.1 aory|EIT75816.1 asyd|OJJ57080.1 ater|EAU31893.1 atub|OJI89544.1 aver|OJJ03223.1 awen|OJJ37121.1 azon|OJJ50820.1

Group4279: abra|OJJ77130.1 acar|OOF93018.1 acla|EAW12941.1 afis|EAW17717.1 afla|EED45719.1 afum|EAL91532.1 agla|OJJ81731.1 akaw|GAA90798.1 aluc|OJZ84540.1 anid|CBF84088.1 anig|EHA27838.1 aory|EIT75412.1 asyd|OJJ57082.1 ater|EAU31895.1 atub|OJI89540.1 aver|OJJ03220.1 awen|OJJ37119.1 azon|OJJ50818.1

Group4280: abra|OJJ77131.1 acar|OOF93019.1 acla|EAW12942.1 afis|EAW17718.1 afla|EED45720.1 afum|EAL91533.1 agla|OJJ81728.1 akaw|GAA90797.1 aluc|OJZ84541.1 anid|CBF76908.1 anig|EHA27839.1 aory|EIT75617.1 asyd|OJJ57083.1 ater|EAU31896.1 atub|OJI89539.1 aver|OJJ03219.1 awen|OJJ37118.1 azon|OJJ50817.1

Group4281: abra|OJJ77133.1 acar|OOF93021.1 acla|EAW12944.1 afis|EAW17720.1 afla|EED45722.1 afum|EAL91535.2 agla|OJJ81730.1 akaw|GAA90795.1 aluc|OJZ84543.1 anid|CBF76904.1 anig|EHA27841.1 aory|EIT75737.1 asyd|OJJ57085.1 ater|EAU31898.1 atub|OJI89537.1 aver|OJJ03217.1 awen|OJJ37116.1 azon|OJJ50815.1

Group4282: abra|OJJ77138.1 acar|OOF90283.1 acla|EAW12948.1 afis|EAW17723.1 afla|EED45725.1 afum|EAL91538.1 agla|OJJ81727.1 akaw|GAA90791.1 aluc|OJZ84549.1 anid|CBF86584.1 anig|EHA27845.1 aory|EIT75656.1 asyd|OJJ64839.1 ater|EAU31901.1 atub|OJI89531.1 aver|OJI98950.1 awen|OJJ37115.1 azon|OJJ50813.1

Group4283: abra|OJJ77139.1 acar|OOF90282.1 acla|EAW12950.1 afis|EAW17725.1 afla|EED45726.1 afum|EAL91540.1 agla|OJJ81726.1 akaw|GAA90790.1 aluc|OJZ84552.1 anid|CBF86583.1 anig|EHA27846.1 aory|EIT75496.1 asyd|OJJ64840.1 ater|EAU31881.1 atub|OJI89530.1 aver|OJI98951.1 awen|OJJ37113.1 azon|OJJ50812.1

Group4284: abra|OJJ77141.1 acar|OOF93027.1 acla|EAW12927.1 afis|EAW17728.1 afla|EED45728.1 afum|EAL91543.1 agla|OJJ81723.1 akaw|GAA90788.1 aluc|OJZ84555.1 anid|CBF86577.1 anig|EHA27849.1 aory|EIT75836.1 asyd|OJJ64842.1 ater|EAU31879.1 atub|OJI89527.1 aver|OJI98953.1 awen|OJJ37110.1 azon|OJJ50809.1

Group4285: abra|OJJ77142.1 acar|OOF93028.1 acla|EAW12926.1 afis|EAW17729.1 afla|EED45729.1 afum|EAL91544.1 agla|OJJ81722.1 akaw|GAA90787.1 aluc|OJZ84556.1 anid|CBF86575.1 anig|EHA27850.1 aory|EIT75504.1 asyd|OJJ64843.1 ater|EAU31878.1 atub|OJI89526.1 aver|OJI98954.1 awen|OJJ37109.1 azon|OJJ50808.1

Group4286: abra|OJJ77143.1 acar|OOF93029.1 acla|EAW12925.1 afis|EAW17730.1 afla|EED45730.1 afum|EAL91545.1 agla|OJJ81721.1 akaw|GAA90786.1 aluc|OJZ84558.1 anid|CBF86573.1 anig|EHA27851.1 aory|EIT75486.1 asyd|OJJ64844.1 ater|EAU31877.1 atub|OJI89525.1 aver|OJI98955.1 awen|OJJ37108.1 azon|OJJ50807.1

Group4287: abra|OJJ77144.1 acar|OOF93030.1 acla|EAW12924.1 afis|EAW17731.1 afla|EED45731.1 afum|EAL91546.1 agla|OJJ81720.1 akaw|GAA90785.1 aluc|OJZ84559.1 anid|CBF86588.1 anig|EHA27852.1 aory|EIT75623.1 asyd|OJJ64838.1 ater|EAU31876.1 atub|OJI89524.1 aver|OJI98948.1 awen|OJJ37107.1 azon|OJJ50806.1

Group4288: abra|OJJ77145.1 acar|OOF93031.1 acla|EAW12923.1 afis|EAW17732.1 afla|EED45732.1 afum|EAL91547.1 agla|OJJ81719.1 akaw|GAA90784.1 aluc|OJZ84560.1 anid|CBF86586.1 anig|EHA27853.1 aory|EIT75469.1 asyd|OJJ64837.1 ater|EAU31875.1 atub|OJI89523.1 aver|OJI98947.1 awen|OJJ37105.1 azon|OJJ50805.1

Group4289: abra|OJJ77146.1 acar|OOF93032.1 acla|EAW12922.1 afis|EAW17733.1 afla|EED45734.1 afum|EAL91548.1 agla|OJJ81718.1 akaw|GAA90783.1 aluc|OJZ84561.1 anid|CBF86597.1 anig|EHA27855.1 aory|EIT75719.1 asyd|OJJ64833.1 ater|EAU31874.1 atub|OJI89521.1 aver|OJI98946.1 awen|OJJ37104.1 azon|OJJ50804.1

Group4290: abra|OJJ77148.1 acar|OOF93034.1 acla|EAW12921.1 afis|EAW17734.1 afla|EED45735.1 afum|EAL91549.1 agla|OJJ81716.1 akaw|GAA90782.1 aluc|OJZ84562.1 anid|CBF86595.1 anig|EHA27856.1 aory|EIT75759.1 asyd|OJJ64832.1 ater|EAU31873.1 atub|OJI89520.1 aver|OJI98945.1 azon|OJJ50802.1 awen|OJJ37101.1

Group4291: abra|OJJ77149.1 acar|OOF93035.1 acla|EAW12920.1 afis|EAW17735.1 afla|EED45736.1 afum|EAL91550.1 agla|OJJ81715.1 akaw|GAA90781.1 aluc|OJZ84563.1 anid|CBF86593.1 anig|EHA27857.1 aory|EIT75423.1 asyd|OJJ64831.1 ater|EAU31872.1 atub|OJI89519.1 aver|OJI98944.1 awen|OJJ37100.1 azon|OJJ50801.1

Group4292: abra|OJJ77157.1 acar|OOF93044.1 acla|EAW12915.1 afis|EAW17745.1 afla|EED45745.1 afum|EAL91557.1 agla|OJJ81708.1 akaw|GAA90773.1 aluc|OJZ84571.1 anid|CBF86614.1 anig|EHA27866.1 aory|EIT75582.1 asyd|OJJ64818.1 ater|EAU31862.1 atub|OJI89511.1 aver|OJI98931.1 awen|OJJ37092.1 azon|OJJ50796.1

Group4293: abra|OJJ77164.1 acar|OOF93050.1 acla|EAW12910.1 afis|EAW17754.1 afla|EED45758.1 afum|EAL91567.1 agla|OJJ81699.1 akaw|GAA90768.1 aluc|OJZ84577.1 anid|CBF86624.1 anig|EHA27870.1 aory|EIT75538.1 asyd|OJJ64811.1 ater|EAU31854.1 atub|OJI89504.1 aver|OJI98924.1 awen|OJJ37082.1 azon|OJJ50787.1

Group4294: abra|OJJ77165.1 acar|OOF93054.1 acla|EAW12901.1 afis|EAW17764.1 afla|EED45766.1 afum|EAL91574.1 agla|OJJ81698.1 akaw|GAA90767.1 aluc|OJZ84578.1 anid|CBF86628.1 anig|EHA27871.1 aory|EIT75559.1 asyd|OJJ64809.1 ater|EAU31853.1 atub|OJI89503.1 aver|OJI98922.1 awen|OJJ37081.1 azon|OJJ50784.1

Group4295: abra|OJJ77166.1 acar|OOF93055.1 acla|EAW12900.1 afis|EAW17765.1 afla|EED45767.1 afum|EAL91575.1 agla|OJJ81697.1 akaw|GAA90766.1 aluc|OJZ84579.1 anid|CBF86630.1 anig|EHA27872.1 aory|EIT75709.1 asyd|OJJ64808.1 ater|EAU31852.1 atub|OJI89502.1 aver|OJI98921.1 awen|OJJ37080.1 azon|OJJ50783.1

Group4296: abra|OJJ77176.1 acar|OOF93064.1 acla|EAW13174.1 afis|EAW17464.1 afla|EED49356.1 afum|EAL91293.1 agla|OJJ80340.1 akaw|GAA90759.1 aluc|OJZ84587.1 anid|CBF71650.1 anig|EHA27880.1 aory|EIT79568.1 asyd|OJJ61840.1 ater|EAU33949.1 atub|OJI89491.1 aver|OJI95939.1 awen|OJJ37381.1 azon|OJJ51055.1

Group4297: abra|OJJ77177.1 acar|OOF93065.1 acla|EAW13175.1 afis|EAW17463.1 afla|EED49357.1 afum|EAL91292.1 agla|OJJ80342.1 akaw|GAA90758.1 aluc|OJZ84589.1 anid|CBF71651.1 anig|EHA27881.1 aory|EIT79484.1 asyd|OJJ61839.1 ater|EAU33948.1 atub|OJI89490.1 aver|OJI95938.1 awen|OJJ37382.1 azon|OJJ51056.1

Group4298: abra|OJJ77186.1 acar|OOF93074.1 acla|EAW13176.1 afis|EAW17462.1 afla|EED49360.1 afum|EAL91291.1 agla|OJJ80343.1 akaw|GAA90753.1 aluc|OJZ84595.1 anid|CBF71661.1 anig|EHA27884.1 aory|EIT79468.1 asyd|OJJ61834.1 ater|EAU33947.1 atub|OJI89484.1 aver|OJI95933.1 awen|OJJ37384.1 azon|OJJ51059.1

Group4299: abra|OJJ77188.1 acar|OOF92641.1 acla|EAW13178.1 afis|EAW17461.1 afla|EED49361.1 afum|EAL91289.1 agla|OJJ80344.1 akaw|GAA90752.1 aluc|OJZ84596.1 anid|CBF71663.1 anig|EHA27885.1 aory|EIT79490.1 asyd|OJJ61833.1 ater|EAU33946.1 atub|OJI89483.1 aver|OJI95932.1 awen|OJJ37385.1 azon|OJJ51060.1

Group4300: abra|OJJ77190.1 acar|OOF93076.1 acla|EAW13179.1 afis|EAW17460.1 afla|EED49362.1 afum|EAL91288.1 agla|OJJ80345.1 akaw|GAA90750.1 aluc|OJZ84598.1 anid|CBF71665.1 anig|EHA27887.1 aory|EIT79506.1 asyd|OJJ61832.1 ater|EAU33945.1 atub|OJI89481.1 aver|OJI95931.1 awen|OJJ37386.1 azon|OJJ51061.1

Group4301: abra|OJJ77193.1 acar|OOF93078.1 acla|EAW13183.1 afis|EAW17457.1 afla|EED49365.1 afum|EAL91285.1 agla|OJJ80354.1 akaw|GAA90748.1 aluc|OJZ84600.1 anid|CBF71668.1 anig|EHA27890.1 aory|EIT79576.1 asyd|OJJ61829.1 ater|EAU33944.1 atub|OJI89479.1 aver|OJI95929.1 awen|OJJ37390.1 azon|OJJ51063.1

Group4302: abra|OJJ77194.1 acar|OOF93079.1 acla|EAW13184.1 afis|EAW17456.1 afla|EED49366.1 afum|EAL91284.1 agla|OJJ80353.1 akaw|GAA90747.1 aluc|OJZ84601.1 anid|CBF71670.1 anig|EHA27891.1 aory|EIT79508.1 asyd|OJJ61828.1 ater|EAU33936.1 atub|OJI89478.1 aver|OJI95928.1 awen|OJJ37391.1 azon|OJJ51064.1

Group4303: abra|OJJ77195.1 acar|OOF93080.1 acla|EAW13186.1 afis|EAW17455.1 afla|EED49367.1 afum|EAL91283.1 agla|OJJ80352.1 akaw|GAA90746.1 aluc|OJZ84602.1 anid|CBF71671.1 anig|EHA27892.1

aory|EIT79573.1 asyd|OJJ61827.1 ater|EAU33935.1 atub|OJI89477.1 aver|OJI95927.1 awen|OJJ37392.1 azon|OJJ51065.1

Group4304: abra|OJJ77198.1 acar|OOF93081.1 acla|EAW13187.1 afis|EAW17454.1 afla|EED49368.1 afum|EAL91282.1 agla|OJJ80351.1 akaw|GAA90743.1 aluc|OJZ84605.1 anid|CBF71673.1 anig|EHA27894.1 aory|EIT79544.1 asyd|OJJ61826.1 ater|EAU33934.1 atub|OJI89474.1 aver|OJI95926.1 awen|OJJ37393.1 azon|OJJ51066.1

Group4305: abra|OJJ77201.1 acar|OOF93089.1 acla|EAW13192.1 afis|EAW17449.1 afla|EED49373.1 afum|EBA27429.1 akaw|GAA90737.1 aluc|OJZ84611.1 anid|CBF71685.1 anig|EHA27900.1 aory|EIT79520.1 asyd|OJJ61821.1 ater|EAU33929.1 atub|OJI89470.1 aver|OJI95921.1 awen|OJJ37397.1 azon|OJJ51070.1 agla|OJJ80368.1

Group4306: abra|OJJ77202.1 acar|OOF93090.1 acla|EAW13193.1 afis|EAW17448.1 afla|EED49374.1 afum|EAL91275.2 agla|OJJ80369.1 akaw|GAA90736.1 aluc|OJZ84612.1 anid|CBF71687.1 anig|EHA27901.1 aory|EIT79525.1 asyd|OJJ61820.1 ater|EAU33928.1 atub|OJI89469.1 aver|OJI95920.1 awen|OJJ37398.1 azon|OJJ51071.1

Group4307: abra|OJJ77204.1 acar|OOF93092.1 acla|EAW13196.1 akaw|GAA90734.1 aluc|OJZ84615.1 anid|CBF71688.1 anig|EHA27903.1 aory|EIT79511.1 asyd|OJJ61819.1 ater|EAU33925.1 atub|OJI89466.1 aver|OJI95919.1 awen|OJJ37400.1 afis|EAW17446.1 afla|EED49381.1 afum|EAL91272.1 agla|OJJ80371.1 azon|OJJ51074.1

Group4308: abra|OJJ77205.1 acar|OOF93094.1 acla|EAW13197.1 afis|EAW17445.1 afla|EED49382.1 afum|EAL91271.1 agla|OJJ80372.1 akaw|GAA90733.1 aluc|OJZ84616.1 anid|CBF71690.1 anig|EHA27904.1 aory|EIT79527.1 asyd|OJJ61818.1 ater|EAU33924.1 atub|OJI89465.1 aver|OJI95918.1 awen|OJJ37401.1 azon|OJJ51075.1

Group4309: abra|OJJ77206.1 acar|OOF93093.1 acla|EAW13198.1 afis|EAW17444.1 afla|EED49383.1 afum|EAL91270.1 agla|OJJ80373.1 akaw|GAA90732.1 aluc|OJZ84617.1 anid|CBF71692.1 anig|EHA27905.1 aory|EIT79499.1 asyd|OJJ61817.1 ater|EAU33923.1 atub|OJI89464.1 aver|OJI95917.1 awen|OJJ37402.1 azon|OJJ51076.1

Group4310: abra|OJJ77207.1 acar|OOF93095.1 acla|EAW13199.1 afis|EAW17443.1 afla|EED49384.1 afum|EAL91269.1 agla|OJJ80374.1 akaw|GAA90731.1 aluc|OJZ84618.1 anid|CBF71694.1 anig|EHA27906.1 aory|EIT79562.1 asyd|OJJ61816.1 ater|EAU33922.1 atub|OJI89463.1 aver|OJI95916.1 awen|OJJ37403.1 azon|OJJ51077.1

Group4311: abra|OJJ77209.1 acar|OOF93096.1 acla|EAW13201.1 afis|EAW17441.1 afla|EED49386.1 afum|EAL91267.1 agla|OJJ80376.1 akaw|GAA90729.1 aluc|OJZ84620.1 anid|CBF71698.1 anig|EHA27908.1 aory|EIT79558.1 asyd|OJJ61814.1 ater|EAU33920.1 atub|OJI89461.1 aver|OJI95913.1 awen|OJJ37405.1 azon|OJJ51079.1

Group4312: abra|OJJ77210.1 acar|OOF93098.1 acla|EAW13202.1 afis|EAW17440.1 afla|EED49387.1 afum|EAL91266.1 agla|OJJ80377.1 akaw|GAA90728.1 aluc|OJZ84621.1 anid|CBF71700.1 anig|EHA27909.1 aory|EIT79563.1 asyd|OJJ61813.1 ater|EAU33919.1 atub|OJI89460.1 aver|OJI95912.1 awen|OJJ37406.1 azon|OJJ51080.1

Group4313: abra|OJJ77211.1 acar|OOF93097.1 acla|EAW13203.1 afis|EAW17439.1 afla|EED49388.1 afum|EAL91265.1 agla|OJJ80378.1 akaw|GAA90727.1 aluc|OJZ84622.1 anid|CBF71702.1 anig|EHA27910.1 aory|EIT79503.1 asyd|OJJ61812.1 ater|EAU33918.1 atub|OJI89459.1 aver|OJI95911.1 awen|OJJ37407.1 azon|OJJ51081.1

Group4314: abra|OJJ77213.1 acar|OOF93099.1 acla|EAW13205.1 afis|EAW17437.1 afla|EED49390.1 afla|EED49391.1 afum|EAL91264.1 agla|OJJ80380.1 akaw|GAA90725.1 aluc|OJZ84624.1 anig|EHA27911.1 aory|EIT79569.1 asyd|OJJ61810.1 ater|EAU33916.1 atub|OJI89457.1 aver|OJI95909.1 awen|OJJ37409.1 azon|OJJ51083.1

Group4315: abra|OJJ77214.1 acar|OOF93101.1 acla|EAW13206.1 afis|EAW17436.1 afla|EED49392.1 afum|EAL91263.1 agla|OJJ80381.1 akaw|GAA90724.1 aluc|OJZ84625.1 anid|CBF71706.1 anig|EHA27912.1 aory|EIT79513.1 asyd|OJJ61809.1 ater|EAU33915.1 atub|OJI89456.1 aver|OJI95908.1 awen|OJJ37410.1 azon|OJJ51084.1

Group4316: abra|OJJ77215.1 acar|OOF93102.1 acla|EAW13207.1 afis|EAW17435.1 afla|EED49409.1 afum|EAL91262.1 agla|OJJ80383.1 akaw|GAA90723.1 aluc|OJZ84626.1 anid|CBF71708.1 anig|EHA27913.1 aory|EIT79458.1 asyd|OJJ61808.1 ater|EAU33914.1 atub|OJI89455.1 aver|OJI95907.1 awen|OJJ37411.1 azon|OJJ51085.1

Group4317: abra|OJJ77216.1 acar|OOF93103.1 acla|EAW13209.1 afis|EAW17432.1 afla|EED49408.1 afum|EAL91260.1 agla|OJJ80384.1 akaw|GAA90722.1 aluc|OJZ84627.1 anid|CBF71710.1 anig|EHA27914.1 aory|EIT79545.1 asyd|OJJ61807.1 ater|EAU33913.1 atub|OJI89452.1 aver|OJI95906.1 awen|OJJ37412.1 azon|OJJ51086.1

Group4318: abra|OJJ77220.1 acar|OOF93105.1 acla|EAW13210.1 afis|EAW17431.1 afla|EED49407.1 afum|EAL91259.1 agla|OJJ80387.1 akaw|GAA90719.1 aluc|OJZ84630.1 anid|CBF71712.1 anig|EHA27918.1 aory|EIT79560.1 asyd|OJJ61806.1 ater|EAU33912.1 atub|OJI89449.1 aver|OJI95905.1 awen|OJJ37413.1 azon|OJJ51087.1

Group4319: abra|OJJ77223.1 acar|OOF93108.1 acla|EAW13214.1 afis|EAW17428.1 afla|EED49402.1 afum|EAL91256.1 agla|OJJ80396.1 akaw|GAA90716.1 aluc|OJZ84633.1 anid|CBF71716.1 anig|EHA27921.1 aory|EIT79534.1 asyd|OJJ61804.1 ater|EAU33909.1 atub|OJI89446.1 aver|OJI95903.1 awen|OJJ37415.1 azon|OJJ51089.1

Group4320: abra|OJJ77225.1 acar|OOF93110.1 acla|EAW13216.1 afis|EAW17426.1 afla|EED49400.1 afum|EAL91254.1 agla|OJJ80398.1 akaw|GAA90714.1 aluc|OJZ84635.1 anid|CBF71720.1 anig|EHA27923.1 aory|EIT79524.1 asyd|OJJ61802.1 ater|EAU33908.1 atub|OJI89444.1 aver|OJI95901.1 awen|OJJ37417.1 azon|OJJ51091.1

Group4321: abra|OJJ77226.1 acar|OOF93111.1 acla|EAW13217.1 afis|EAW17425.1 afla|EED49399.1 afum|EAL91253.1 agla|OJJ80399.1 akaw|GAA90713.1 aluc|OJZ84636.1 anid|CBF71722.1 anig|EHA27924.1 aory|EIT79475.1 asyd|OJJ61801.1 ater|EAU33907.1 atub|OJI89443.1 aver|OJI95900.1 awen|OJJ37418.1 azon|OJJ51092.1

Group4322: abra|OJJ77230.1 acar|OOF93113.1 acla|EAW13219.1 afis|EAW17423.1 afla|EED49397.1 afum|EAL91251.1 agla|OJJ80401.1 akaw|GAA90711.1 aluc|OJZ84640.1 anid|CBF71729.1 anig|EHA27926.1 aory|EIT79564.1 asyd|OJJ61799.1 ater|EAU33905.1 atub|OJI89440.1 aver|OJI95898.1 awen|OJJ37421.1 azon|OJJ51094.1

Group4323: abra|OJJ77238.1 acar|OOF93119.1 acla|EAW13245.1 afis|EAW17391.1 afla|EED49477.1 afum|EAL91228.1 agla|OJJ80406.1 akaw|GAA90705.1 aluc|OJZ84648.1 anid|CBF71739.1 anig|EHA27932.1 aory|EIT79509.1 asyd|OJJ61792.1 ater|EAU33896.1 atub|OJI89432.1 aver|OJI95892.1 awen|OJJ37424.1 azon|OJJ51104.1

Group4324: abra|OJJ77240.1 acar|OOF93121.1 acla|EAW13247.1 afis|EAW17389.1 afla|EED49481.1 afum|EAL91225.1 agla|OJJ80408.1 akaw|GAA90703.1 aluc|OJZ84650.1 anid|CBF71743.1 anig|EHA27933.1 aory|EIT79553.1 asyd|OJJ61790.1 ater|EAU33894.1 atub|OJI89430.1 aver|OJI95890.1 awen|OJJ37426.1 azon|OJJ51102.1

Group4325: abra|OJJ77242.1 acar|OOF93123.1 acla|EAW14606.1 afis|EAW19335.1 afla|EED56869.1 afum|EAL94052.1 agla|OJJ79421.1 akaw|GAA90701.1 aluc|OJZ84652.1 anid|CBF82394.1 anig|EHA27935.1 aory|EIT77502.1 asyd|OJJ54156.1 atub|OJI89428.1 aver|OJJ08579.1 awen|OJJ38873.1 asyd|OJJ59232.1 aver|OJJ01842.1

Group4326: abra|OJJ77244.1 acar|OOF93125.1 acla|EAW13248.1 afis|EAW17388.1 afla|EED49482.1 afum|EAL91224.1 agla|OJJ80409.1 akaw|GAA90699.1 aluc|OJZ84654.1 anid|CBF71787.1 anig|EHA27937.1 aory|EIT79464.1 asyd|OJJ61746.1 ater|EAU33893.1 atub|OJI89426.1 aver|OJI95843.1 awen|OJJ37427.1 azon|OJJ51101.1

Group4327: abra|OJJ77247.1 acar|OOF90006.1 acla|EAW13252.1 afis|EAW17384.1 afla|EED49501.1 afum|EAL91219.1 agla|OJJ80419.1 akaw|GAA90695.1 aluc|OJZ84658.1 anid|CBF84196.1 anig|EHA27940.1 aory|EIT82601.1 asyd|OJJ57160.1 ater|EAU33890.1 atub|OJI89422.1 aver|OJJ03153.1 awen|OJJ33371.1 azon|OJJ51109.1

Group4328: abra|OJJ77250.1 acar|OOF90005.1 acla|EAW13253.1 afis|EAW17383.1 afla|EED49504.1 afum|EAL91218.1 agla|OJJ80420.1 akaw|GAA90693.1 aluc|OJZ84661.1 anid|CBF84198.1 anig|EHA27943.1 aory|EIT82610.1 asyd|OJJ57161.1 ater|EAU33880.1 atub|OJI89419.1 aver|OJJ03152.1 awen|OJJ37433.1 azon|OJJ51112.1

Group4329: abra|OJJ77260.1 acar|OOF93128.1 afla|EED49508.1 akaw|GAA90687.1 aluc|OJZ84667.1 anig|EHA27946.1 aory|EIT82587.1 asyd|OJJ57166.1 ater|EAU33878.1 atub|OJI89413.1 awen|OJJ37439.1 azon|OJJ51115.1 acla|EAW13256.1 afis|EAW17380.1 afum|EAL91215.1 agla|OJJ80422.1 anid|CBF84207.1 aver|OJJ03148.1

Group4330: abra|OJJ77261.1 acar|OOF93130.1 acla|EAW13257.1 afis|EAW17379.1 afla|EED49509.1 afum|EAL91214.1 agla|OJJ80423.1 akaw|GAA90686.1 aluc|OJZ84668.1 anid|CBF84209.1 anig|EHA27947.1 aory|EIT82656.1 asyd|OJJ57167.1 ater|EAU33877.1 atub|OJI89412.1 aver|OJJ03147.1 awen|OJJ37441.1 azon|OJJ51117.1

Group4331: abra|OJJ77262.1 acar|OOF93132.1 acla|EAW13258.1 afis|EAW17378.1 afla|EED49510.1 afum|EAL91213.1 agla|OJJ80424.1 akaw|GAA90685.1 aluc|OJZ84669.1 anid|CBF84210.1 anig|EHA27948.1 aory|EIT82657.1 asyd|OJJ57168.1 ater|EAU33876.1 atub|OJI89411.1 aver|OJJ03146.1 awen|OJJ37442.1 azon|OJJ51120.1

Group4332: abra|OJJ77263.1 acar|OOF93131.1 acla|EAW13259.1 afis|EAW17377.1 afla|EED49511.1 afum|EAL91212.1 agla|OJJ80425.1 akaw|GAA90684.1 aluc|OJZ84671.1 anid|CBF84212.1 anig|EHA27949.1

aory|EIT82654.1 asyd|OJJ57169.1 ater|EAU33875.1 atub|OJI89409.1 aver|OJJ03145.1 awen|OJJ37443.1 azon|OJJ51121.1

Group4333: abra|OJJ77275.1 acar|OOF93150.1 acla|EAW13275.1 afis|EAW17342.1 afla|EED49781.1 afum|EBA27424.1 agla|OJJ79342.1 akaw|GAA90674.1 aluc|OJZ84684.1 anid|CBF84217.1 anig|EHA27960.1 aory|EIT77695.1 asyd|OJJ57179.1 ater|EAU33855.1 atub|OJI89397.1 aver|OJJ03135.1 awen|OJJ29675.1 azon|OJJ45994.1

Group4334: abra|OJJ77289.1 acar|OOF93167.1 acla|EAW13173.1 afis|EAW17465.1 afla|EED49355.1 afum|EAL91294.1 agla|OJJ80339.1 akaw|GAA90660.1 aluc|OJZ84700.1 anid|CBF71646.1 anig|EHA27975.1 aory|EIT79489.1 asyd|OJJ61841.1 ater|EAU33950.1 atub|OJI89382.1 aver|OJI95941.1 awen|OJJ37380.1 azon|OJJ51054.1

Group4335: abra|OJJ77290.1 acar|OOF93168.1 acla|EAW13172.1 afis|EAW17466.1 afla|EED49354.1 afum|EAL91295.1 agla|OJJ80338.1 akaw|GAA90659.1 aluc|OJZ84701.1 anid|CBF71638.1 anig|EHA27976.1 aory|EIT79580.1 asyd|OJJ61844.1 ater|EAU33951.1 atub|OJI89381.1 aver|OJI95944.1 awen|OJJ37379.1 azon|OJJ51053.1

Group4336: abra|OJJ77291.1 acar|OOF93169.1 acla|EAW13171.1 afis|EAW17468.1 afla|EED49353.1 afum|EAL91297.1 agla|OJJ80336.1 akaw|GAA90658.1 aluc|OJZ84702.1 anid|CBF71640.1 anig|EHA27977.1 aory|EIT79590.1 asyd|OJJ61843.1 ater|EAU33952.1 atub|OJI89380.1 aver|OJI95943.1 awen|OJJ37378.1 azon|OJJ51052.1

Group4337: abra|OJJ77295.1 acar|OOF93172.1 acla|EAW13169.1 afis|EAW17470.1 afla|EED49348.1 afum|EAL91299.1 agla|OJJ78440.1 akaw|GAA91456.1 aluc|OJZ84707.1 anid|CBF71630.1 anig|EHA27981.1 aory|EIT79461.1 asyd|OJJ61849.1 ater|EAU33954.1 atub|OJI89376.1 aver|OJI95948.1 awen|OJJ37372.1 azon|OJJ51050.1

Group4338: abra|OJJ77297.1 acar|OOF93177.1 acla|EAW13167.1 afis|EAW17472.1 afla|EED49343.1 akaw|GAA91454.1 aluc|OJZ84709.1 anid|CBF71626.1 anig|EHA27983.1 asyd|OJJ61851.1 ater|EAU33956.1 atub|OJI89374.1 aver|OJI95950.1 afum|EAL91301.1 agla|OJJ81490.1 aory|EIT79471.1 awen|OJJ37370.1 azon|OJJ51049.1

Group4339: abra|OJJ77300.1 acar|OOF93179.1 acla|EAW13165.1 afis|EAW17474.1 afla|EED49340.1 afum|EAL91303.1 agla|OJJ81488.1 akaw|GAA91452.1 aluc|OJZ84712.1 anid|CBF71622.1 anig|EHA27985.1 aory|EIT79516.1 asyd|OJJ61854.1 ater|EAU33958.1 atub|OJI89371.1 aver|OJI95953.1 awen|OJJ37368.1 azon|OJJ51047.1

Group4340: abra|OJJ77301.1 acar|OOF93180.1 acla|EAW13164.1 afis|EAW17475.1 afla|EED49339.1 afum|EAL91304.1 agla|OJJ81487.1 akaw|GAA91451.1 aluc|OJZ84713.1 anid|CBF71620.1 anig|EHA27986.1 aory|EIT79565.1 asyd|OJJ61855.1 ater|EAU33959.1 atub|OJI89370.1 aver|OJI95954.1 awen|OJJ37367.1 azon|OJJ51046.1

Group4341: abra|OJJ77303.1 acar|OOF93183.1 acla|EAW13163.1 afis|EAW17477.1 afla|EED49337.1 afum|EAL91306.1 agla|OJJ81480.1 akaw|GAA91448.1 aluc|OJZ84716.1 anid|CBF71603.1 anig|EHA27988.1 aory|EIT79497.1 asyd|OJJ61857.1 ater|EAU33693.1 atub|OJI89367.1 aver|OJI95956.1 awen|OJJ37365.1 azon|OJJ51045.1

Group4342: abra|OJJ77304.1 acar|OOF93182.1 acla|EAW13162.1 afis|EAW17478.1 afla|EED49336.1 afum|EAL91307.1 agla|OJJ81479.1 akaw|GAA91447.1 aluc|OJZ84717.1 anid|CBF71605.1 anig|EHA27989.1 aory|EIT79535.1 asyd|OJJ61858.1 ater|EAU33694.1 atub|OJI89366.1 aver|OJI95957.1 awen|OJJ37364.1 azon|OJJ51044.1

Group4343: abra|OJJ77305.1 acar|OOF93184.1 acla|EAW13161.1 afis|EAW17479.1 afla|EED49335.1 afum|EAL91308.1 agla|OJJ81478.1 akaw|GAA91446.1 aluc|OJZ84718.1 anid|CBF71607.1 anig|EHA27990.1 aory|EIT79462.1 asyd|OJJ61859.1 ater|EAU33695.1 atub|OJI89365.1 aver|OJI95958.1 awen|OJJ37362.1 azon|OJJ51043.1

Group4344: abra|OJJ77306.1 acar|OOF93185.1 acla|EAW13160.1 afis|EAW17481.1 afla|EED49334.1 afum|EAL91309.1 agla|OJJ81477.1 akaw|GAA91445.1 aluc|OJZ84720.1 anid|CBF71609.1 anig|EHA27991.1 aory|EIT79526.1 asyd|OJJ61860.1 ater|EAU36927.1 atub|OJI89363.1 aver|OJI95959.1 awen|OJJ37357.1 azon|OJJ51041.1

Group4345: abra|OJJ77315.1 acar|OOF93191.1 acla|EAW13157.1 afis|EAW17489.1 afla|EED49327.1 afum|EAL91311.1 agla|OJJ81467.1 akaw|GAA91440.1 aluc|OJZ84725.1 anid|CBF71617.1 anig|EHA27999.1 aory|EIT79549.1 asyd|OJJ61864.1 ater|EAU36934.1 atub|OJI89357.1 aver|OJI95962.1 awen|OJJ37352.1 azon|OJJ51038.1

Group4346: abra|OJJ77318.1 acar|OOF93194.1 acla|EAW13154.1 afis|EAW17492.1 afla|EED49324.1 afum|EAL91314.1 agla|OJJ81464.1 akaw|GAA91438.1 aluc|OJZ84728.1 anid|CBF71599.1 anig|EHA28001.1 aory|EIT79543.1 asyd|OJJ61866.1 ater|EAU36936.1 atub|OJI89354.1 aver|OJI95964.1 awen|OJJ37349.1 azon|OJJ51035.1

Group4347: abra|OJJ77319.1 acar|OOF93195.1 acla|EAW13153.1 afis|EAW17493.1 afla|EED49323.1 afum|EAL91315.1 agla|OJJ81463.1 akaw|GAA91437.1 aluc|OJZ84729.1 anid|CBF71595.1 anig|EHA28002.1 aory|EIT79479.1 asyd|OJJ61868.1 ater|EAU36937.1 atub|OJI89353.1 aver|OJI95966.1 awen|OJJ37348.1 azon|OJJ51034.1

Group4348: abra|OJJ77320.1 acar|OOF93196.1 acla|EAW13152.1 afis|EAW17494.1 afla|EED49322.1 afum|EAL91316.1 agla|OJJ81460.1 akaw|GAA91436.1 aluc|OJZ84730.1 anid|CBF71589.1 anig|EHA28003.1 aory|EIT79493.1 asyd|OJJ61870.1 ater|EAU36938.1 atub|OJI89352.1 aver|OJI95968.1 awen|OJJ37347.1 azon|OJJ51033.1

Group4349: abra|OJJ77326.1 acar|OOF93202.1 acla|EAW13146.1 afis|EAW17518.1 afla|EED49315.1 afum|EAL91336.1 agla|OJJ81452.1 akaw|GAA91430.1 aluc|OJZ84736.1 anid|CBF71576.1 anig|EHA28009.1 aory|EIT82190.1 asyd|OJJ61877.1 ater|EAU36948.1 atub|OJI89345.1 aver|OJI95975.1 awen|OJJ37339.1 azon|OJJ51025.1

Group4350: abra|OJJ77328.1 acar|OOF93203.1 acla|EAW13145.1 afis|EAW17517.1 afla|EED49314.1 afum|EAL91335.1 agla|OJJ81451.1 akaw|GAA91429.1 aluc|OJZ84737.1 anid|CBF71574.1 anig|EHA28010.1 aory|EIT82219.1 asyd|OJJ61878.1 ater|EAU36949.1 atub|OJI89344.1 aver|OJI95976.1 awen|OJJ37338.1 azon|OJJ51024.1

Group4351: abra|OJJ77329.1 acar|OOF93204.1 acla|EAW13144.1 afis|EAW17516.1 afla|EED49313.1 afum|EAL91334.1 agla|OJJ81450.1 akaw|GAA91428.1 aluc|OJZ84738.1 anid|CBF71572.1 anig|EHA28011.1 aory|EIT82242.1 asyd|OJJ61879.1 ater|EAU36950.1 atub|OJI89343.1 aver|OJI95977.1 awen|OJJ37337.1 azon|OJJ51023.1

Group4352: abra|OJJ77334.1 acar|OOF89885.1 acla|EAW13138.1 afis|EAW17507.1 afla|EED49308.1 afum|EAL91329.1 agla|OJJ81445.1 akaw|GAA91424.1 aluc|OJZ84742.1 anid|CBF71533.1 anig|EHA28016.1 aory|EIT82210.1 asyd|OJJ61892.1 ater|EAU36953.1 atub|OJI89339.1 aver|OJI95988.1 awen|OJJ37332.1 azon|OJJ51019.1

Group4353: abra|OJJ77338.1 acar|OOF93209.1 acla|EAW13137.1 afis|EAW17506.1 afla|EED49306.1 afum|EAL91328.2 agla|OJJ81444.1 akaw|GAA91423.1 aluc|OJZ84747.1 anid|CBF71535.1 anig|EHA28017.1 aory|EIT82204.1 asyd|OJJ61889.1 ater|EAU36955.1 atub|OJI89335.1 aver|OJI95986.1 awen|OJJ37329.1 azon|OJJ51018.1

Group4354: abra|OJJ77339.1 acar|OOF93210.1 acla|EAW13136.1 afis|EAW17505.1 afla|EED49305.1 afum|EAL91327.1 agla|OJJ81443.1 akaw|GAA91422.1 aluc|OJZ84748.1 anid|CBF71537.1 anig|EHA28018.1 aory|EIT82207.1 asyd|OJJ61894.1 ater|EAU36956.1 atub|OJI89334.1 aver|OJI95990.1 awen|OJJ37328.1 azon|OJJ51017.1

Group4355: abra|OJJ77341.1 acar|OOF93211.1 acla|EAW13134.1 afis|EAW17503.1 afla|EED49301.1 afum|EBA27430.1 agla|OJJ81441.1 akaw|GAA91420.1 aluc|OJZ84750.1 anid|CBF71541.1 anig|EHA28020.1 aory|EIT82193.1 asyd|OJJ61896.1 ater|EAU36958.1 atub|OJI89332.1 aver|OJI95992.1 awen|OJJ37326.1 azon|OJJ51015.1

Group4356: abra|OJJ77342.1 acar|OOF93212.1 acla|EAW13133.1 afis|EAW17502.1 afla|EED49300.1 afum|EAL91325.1 agla|OJJ81440.1 akaw|GAA91419.1 aluc|OJZ84751.1 anid|CBF71543.1 anig|EHA28021.1 aory|EIT82195.1 asyd|OJJ61897.1 ater|EAU36959.1 atub|OJI89331.1 aver|OJI95993.1 awen|OJJ37325.1 azon|OJJ51014.1

Group4357: abra|OJJ77343.1 acar|OOF93213.1 acla|EAW13132.1 afis|EAW17501.1 afla|EED49299.1 afum|EAL91324.1 agla|OJJ81439.1 akaw|GAA91418.1 aluc|OJZ84753.1 anid|CBF85589.1 anig|EHA28022.1 aory|EIT82233.1 asyd|OJJ64171.1 ater|EAU36960.1 atub|OJI89330.1 aver|OJI98246.1 awen|OJJ37324.1 azon|OJJ51013.1

Group4358: abra|OJJ77344.1 acar|OOF93214.1 acla|EAW13131.1 afis|EAW17532.1 afla|EED49298.1 afum|EAL91346.1 agla|OJJ81438.1 akaw|GAA91417.1 aluc|OJZ84754.1 anid|CBF90335.1 anig|EHA28023.1 aory|EIT82183.1 asyd|OJJ61898.1 ater|EAU36961.1 atub|OJI89329.1 aver|OJI95994.1 awen|OJJ37320.1 azon|OJJ51012.1

Group4359: abra|OJJ77346.1 acar|OOF93216.1 acla|EAW13128.1 afis|EAW17535.1 afla|EED49294.1 afum|EAL91349.1 agla|OJJ81436.1 akaw|GAA91414.1 aluc|OJZ84757.1 anid|CBF90326.1 anig|EHA28026.1 aory|EIT82178.1 asyd|OJJ61907.1 ater|EAU36964.1 atub|OJI89327.1 aver|OJI96000.1 awen|OJJ37318.1 azon|OJJ51010.1

Group4360: abra|OJJ77347.1 acar|OOF93217.1 acla|EAW13127.1 afis|EAW17536.1 afla|EED49293.1 afum|EAL91350.1 agla|OJJ81433.1 akaw|GAA91413.1 aluc|OJZ84758.1 anid|CBF90324.1 anig|EHA28027.1 aory|EIT82228.1 asyd|OJJ61908.1 ater|EAU36965.1 atub|OJI89326.1 aver|OJI96001.1 awen|OJJ37317.1 azon|OJJ51009.1

Group4361: abra|OJJ77348.1 acar|OOF93218.1 acla|EAW13126.1 afis|EAW17537.1 afla|EED49290.1 afum|EAL91351.1 agla|OJJ81432.1 akaw|GAA91412.1 aluc|OJZ84759.1 anid|CBF90322.1 anig|EHA28028.1

aory|EIT82241.1 asyd|OJJ61909.1 ater|EAU36967.1 atub|OJI89325.1 aver|OJI96002.1 awen|OJJ37315.1 azon|OJJ51007.1

Group4362: abra|OJJ77350.1 acar|OOF93220.1 acla|EAW13125.1 afis|EAW17539.1 afla|EED49288.1 afum|EAL91353.1 agla|OJJ81430.1 akaw|GAA91411.1 aluc|OJZ84761.1 anid|CBF90317.1 anig|EHA28030.1 aory|EIT82231.1 asyd|OJJ61911.1 ater|EAU36969.1 atub|OJI89323.1 aver|OJI96004.1 awen|OJJ37314.1 azon|OJJ51006.1

Group4363: abra|OJJ77353.1 acar|OOF93223.1 acla|EAW13121.1 afis|EAW17543.1 afla|EED49284.1 afum|EAL91357.1 agla|OJJ81426.1 akaw|GAA91408.1 aluc|OJZ84764.1 anid|CBF90308.1 anig|EHA28033.1 aory|EIT82220.1 asyd|OJJ61932.1 ater|EAU36973.1 atub|OJI89320.1 aver|OJI96024.1 awen|OJJ37310.1 azon|OJJ51002.1

Group4364: abra|OJJ77355.1 acar|OOF93225.1 acla|EAW13119.1 afis|EAW17545.1 afla|EED49282.1 afum|EAL91359.1 agla|OJJ81424.1 akaw|GAA91406.1 aluc|OJZ84765.1 anid|CBF90305.1 anig|EHA28034.1 aory|EIT82244.1 asyd|OJJ61930.1 ater|EAU36975.1 atub|OJI89319.1 aver|OJI96022.1 awen|OJJ37309.1 azon|OJJ51001.1

Group4365: abra|OJJ77356.1 acar|OOF93226.1 acla|EAW13118.1 afis|EAW17546.1 afla|EED49281.1 afum|EAL91360.1 agla|OJJ81423.1 akaw|GAA91405.1 aluc|OJZ84766.1 anid|CBF90303.1 anig|EHA28035.1 aory|EIT82243.1 asyd|OJJ61929.1 ater|EAU36976.1 atub|OJI89318.1 aver|OJI96021.1 awen|OJJ37308.1 azon|OJJ51000.1

Group4366: abra|OJJ77359.1 acar|OOF93228.1 acla|EAW13117.1 afis|EAW17547.1 afla|EED49278.1 afum|EAL91361.1 agla|OJJ81421.1 akaw|GAA91404.1 aluc|OJZ84767.1 anid|CBF90290.1 anig|EHA28037.1 aory|EIT82197.1 asyd|OJJ61921.1 ater|EAU36977.1 atub|OJI89315.1 aver|OJI96013.1 awen|OJJ37307.1 azon|OJJ50999.1

Group4367: abra|OJJ77361.1 acar|OOF93230.1 acla|EAW13115.1 afis|EAW17549.1 afla|EED49276.1 afum|EAL91363.1 agla|OJJ81419.1 akaw|GAA91402.1 aluc|OJZ84769.1 anid|CBF90294.1 anig|EHA28038.1 aory|EIT82240.1 asyd|OJJ61923.1 ater|EAU36985.1 atub|OJI89312.1 aver|OJI96015.1 awen|OJJ37305.1 azon|OJJ50997.1

Group4368: abra|OJJ77362.1 acar|OOF93231.1 acla|EAW13114.1 afis|EAW17550.1 afla|EED49275.1 afum|EAL91364.1 agla|OJJ81418.1 akaw|GAA91400.1 aluc|OJZ84771.1 anid|CBF90296.1 anig|EHA28039.1 aory|EIT82229.1 asyd|OJJ61924.1 ater|EAU36984.1 atub|OJI89310.1 aver|OJI96016.1 awen|OJJ37303.1 azon|OJJ50996.1

Group4369: abra|OJJ77365.1 acar|OOF93235.1 acla|EAW13111.1 afis|EAW17553.1 afla|EED49272.1 afum|EAL91367.1 agla|OJJ81415.1 akaw|GAA91397.1 aluc|OJZ84774.1 anid|CBF90299.1 anig|EHA28042.1 aory|EIT82208.1 asyd|OJJ61926.1 ater|EAU36981.1 atub|OJI89307.1 aver|OJI96018.1 awen|OJJ37300.1 azon|OJJ50993.1

Group4370: abra|OJJ77366.1 acar|OOF93236.1 acla|EAW13110.1 afis|EAW17554.1 afla|EED49271.1 afum|EAL91368.1 agla|OJJ81414.1 akaw|GAA91396.1 aluc|OJZ84775.1 anid|CBF90301.1 anig|EHA28043.1 aory|EIT82221.1 asyd|OJJ61927.1 ater|EAU36980.1 atub|OJI89306.1 aver|OJI96019.1 awen|OJJ37299.1 azon|OJJ50992.1

Group4371: abra|OJJ77368.1 acar|OOF93238.1 acla|EAW13108.1 afis|EAW17556.1 afla|EED49268.1 afum|EAL91370.1 agla|OJJ81412.1 akaw|GAA91394.1 aluc|OJZ84777.1 anid|CBF90282.1 anig|EHA28045.1 aory|EIT82203.1 asyd|OJJ61918.1 ater|EAU36988.1 atub|OJI89304.1 aver|OJI96010.1 awen|OJJ37297.1 azon|OJJ50990.1

Group4372: abra|OJJ77370.1 acar|OOF93241.1 acla|EAW13105.1 afis|EAW17559.1 afla|EED49265.1 afum|EAL91373.1 agla|OJJ81409.1 akaw|GAA91391.1 aluc|OJZ84780.1 anid|CBF90277.1 anig|EHA28047.1 aory|EIT82185.1 asyd|OJJ61933.1 ater|EAU36990.1 atub|OJI89302.1 aver|OJI96025.1 awen|OJJ37295.1 azon|OJJ50988.1

Group4373: abra|OJJ77371.1 acar|OOF93240.1 acla|EAW13104.1 afis|EAW17560.1 afla|EED49264.1 afum|EAL91374.1 agla|OJJ81408.1 akaw|GAA91390.1 aluc|OJZ84781.1 anid|CBF90275.1 anig|EHA28048.1 aory|EIT82187.1 asyd|OJJ61934.1 ater|EAU36991.1 atub|OJI89301.1 aver|OJI96026.1 awen|OJJ37294.1 azon|OJJ50987.1

Group4374: abra|OJJ77372.1 acar|OOF93242.1 acla|EAW13103.1 afis|EAW17561.1 afla|EED49263.1 afum|EAL91375.1 agla|OJJ81407.1 akaw|GAA91389.1 aluc|OJZ84782.1 anid|CBF90273.1 anig|EHA28049.1 aory|EIT82191.1 asyd|OJJ61935.1 ater|EAU36992.1 atub|OJI89300.1 aver|OJI96027.1 awen|OJJ37293.1 azon|OJJ50986.1

Group4375: abra|OJJ77373.1 acar|OOF93244.1 acla|EAW13102.1 afis|EAW17562.1 afla|EED49259.1 afum|EAL91376.1 agla|OJJ81406.1 akaw|GAA91387.1 aluc|OJZ84783.1 anid|CBF90271.1 anig|EHA28050.1 aory|EIT82238.1 asyd|OJJ61936.1 ater|EAU37000.1 atub|OJI89299.1 aver|OJI96028.1 awen|OJJ37292.1 azon|OJJ50985.1

Group4376: abra|OJJ77374.1 acar|OOF93243.1 acla|EAW13101.1 afis|EAW17563.1 afla|EED49258.1 afum|EAL91377.1 agla|OJJ81405.1 akaw|GAA91386.1 aluc|OJZ84784.1 anid|CBF90269.1 anig|EHA28051.1 aory|EIT82186.1 asyd|OJJ61937.1 ater|EAU37001.1 atub|OJI89298.1 aver|OJI96029.1 awen|OJJ37291.1 azon|OJJ50984.1

Group4377: abra|OJJ77375.1 acar|OOF93245.1 acla|EAW13100.1 afis|EAW17564.1 afla|EED49257.1 afum|EAL91378.1 agla|OJJ81402.1 akaw|GAA91385.1 aluc|OJZ84785.1 anid|CBF90267.1 anig|EHA28052.1 aory|EIT82246.1 asyd|OJJ61938.1 ater|EAU37002.1 atub|OJI89297.1 aver|OJI96030.1 awen|OJJ37290.1 azon|OJJ50983.1

Group4378: abra|OJJ77376.1 acar|OOF93246.1 acla|EAW13098.1 afis|EAW17565.1 afla|EED49256.1 afum|EAL91379.1 agla|OJJ81401.1 akaw|GAA91384.1 aluc|OJZ84786.1 anid|CBF90266.1 anig|EHA28053.1 aory|EIT82227.1 asyd|OJJ61939.1 ater|EAU37003.1 atub|OJI89296.1 aver|OJI96031.1 awen|OJJ37288.1 azon|OJJ50982.1

Group4379: abra|OJJ77378.1 acar|OOF93247.1 acla|EAW13096.1 afis|EAW17567.1 afla|EED49254.1 afum|EAL91381.1 agla|OJJ81398.1 akaw|GAA91382.1 aluc|OJZ84787.1 anid|CBF90262.1 anig|EHA28055.1 aory|EIT82184.1 asyd|OJJ61941.1 ater|EAU37004.1 atub|OJI89295.1 aver|OJI96033.1 awen|OJJ37286.1 azon|OJJ50980.1

Group4380: abra|OJJ77476.1 acar|OOF99249.1 acla|EAW06738.1 afis|EAW16288.1 afla|EED53265.1 afum|EAL88910.1 agla|OJJ88310.1 akaw|GAA84142.1 aluc|OJZ81062.1 anid|CBF86860.1 anig|EHA19306.1 aory|EIT75107.1 asyd|OJJ64634.1 ater|EAU37333.1 atub|OJI82523.1 aver|OJI98733.1 awen|OJJ37692.1 azon|OJJ44952.1

Group4381: abra|OJJ77477.1 acar|OOF99248.1 acla|EAW06739.1 afis|EAW16289.1 afla|EED53264.1 afum|EAL88911.1 agla|OJJ88311.1 akaw|GAA84141.1 aluc|OJZ81061.1 anid|CBF86858.1 anig|EHA19307.1 aory|EIT74825.1 asyd|OJJ64635.1 ater|EAU37334.1 atub|OJI82524.1 aver|OJI98734.1 awen|OJJ37691.1 azon|OJJ44951.1

Group4382: abra|OJJ77478.1 acar|OOF99247.1 acla|EAW06740.1 afis|EAW16290.1 afla|EED53263.1 afum|EAL88912.1 agla|OJJ88312.1 akaw|GAA84140.1 aluc|OJZ81060.1 anid|CBF86856.1 anig|EHA19308.1 aory|EIT75015.1 asyd|OJJ64636.1 ater|EAU37335.1 atub|OJI82525.1 aver|OJI98735.1 awen|OJJ37690.1 azon|OJJ44950.1

Group4383: abra|OJJ77479.1 acar|OOF99246.1 acla|EAW06741.1 afis|EAW16291.1 afla|EED53262.1 afum|EAL88913.1 agla|OJJ88313.1 akaw|GAA84139.1 aluc|OJZ81059.1 anid|CBF86854.1 anig|EHA19309.1 aory|EIT74947.1 asyd|OJJ64637.1 ater|EAU37336.1 atub|OJI82526.1 aver|OJI98736.1 awen|OJJ37689.1 azon|OJJ44949.1

Group4384: abra|OJJ77485.1 acar|OOF99240.1 acla|EAW06746.1 afis|EAW16296.1 afla|EED53258.1 afum|EAL88918.1 agla|OJJ88303.1 akaw|GAA84133.1 aluc|OJZ81051.1 anid|CBF86847.1 anig|EHA19316.1 aory|EIT75226.1 asyd|OJJ64642.1 ater|EAU29340.1 atub|OJI82533.1 aver|OJI98742.1 awen|OJJ37695.1 azon|OJJ44956.1

Group4385: abra|OJJ77489.1 acar|OOF99236.1 acla|EAW06751.1 afis|EAW16301.1 afla|EED53253.1 afum|EAL88923.1 agla|OJJ88293.1 akaw|GAA84130.1 aluc|OJZ81047.1 anid|CBF86841.1 anig|EHA19320.1 aory|EIT74998.1 asyd|OJJ64646.1 ater|EAU37346.1 atub|OJI82537.1 aver|OJI98746.1 awen|OJJ37700.1 azon|OJJ44961.1

Group4386: abra|OJJ77490.1 acar|OOF99235.1 acla|EAW06752.1 afis|EAW16302.1 afla|EED53252.1 afum|EAL88924.1 agla|OJJ88292.1 akaw|GAA84129.1 aluc|OJZ81046.1 anid|CBF86839.1 anig|EHA19321.1 aory|EIT75063.1 asyd|OJJ64647.1 ater|EAU37345.1 atub|OJI82538.1 aver|OJI98747.1 awen|OJJ37701.1 azon|OJJ44962.1

Group4387: abra|OJJ77493.1 acar|OOF99233.1 acla|EAW06754.1 afis|EAW16304.1 afla|EED53250.1 afum|EAL88926.1 agla|OJJ88290.1 akaw|GAA84127.1 aluc|OJZ81044.1 anid|CBF69582.1 anig|EHA19323.1 aory|EIT75043.1 asyd|OJJ59144.1 ater|EAU29332.1 atub|OJI82540.1 aver|OJJ01734.1 awen|OJJ37704.1 azon|OJJ44964.1

Group4388: abra|OJJ77494.1 acar|OOF99232.1 acla|EAW06755.1 afis|EAW16305.1 afla|EED53249.1 afum|EAL88927.1 agla|OJJ88267.1 akaw|GAA84126.1 aluc|OJZ81043.1 anid|CBF69583.1 anig|EHA19324.1 aory|EIT75153.1 asyd|OJJ59145.1 ater|EAU29333.1 atub|OJI82541.1 aver|OJJ01735.1 awen|OJJ37705.1 azon|OJJ44965.1

Group4389: abra|OJJ77495.1 acla|EAW06756.1 afis|EAW16306.1 afla|EED53248.1 afum|EAL88928.1 agla|OJJ88268.1 akaw|GAA84125.1 aluc|OJZ81042.1 anid|CBF69575.1 anig|EHA19325.1 aory|EIT75004.1 asyd|OJJ59149.1 ater|EAU29334.1 atub|OJI82542.1 aver|OJJ01739.1 awen|OJJ37706.1 azon|OJJ44966.1 aver|OJJ05343.1

Group4390: abra|OJJ77499.1 acar|OOF99227.1 acla|EAW06736.1 afis|EAW16287.1 afla|EED53269.1 afum|EAL88909.1 agla|OJJ88309.1 akaw|GAA84122.1 aluc|OJZ81039.1 anid|CBF86861.1 anig|EHA19328.1

aory|EIT74855.1 asyd|OJJ64633.1 ater|EAU37332.1 atub|OJI82545.1 aver|OJI98732.1 awen|OJJ37682.1 azon|OJJ44954.1

Group4391: abra|OJJ77502.1 acar|OOF99225.1 acla|EAW06734.1 afis|EAW16285.1 afla|EED53271.1 afum|EAL88906.1 agla|OJJ88306.1 akaw|GAA84119.1 aluc|OJZ81035.1 anid|CBF86869.1 anig|EHA19331.1 aory|EIT75236.1 asyd|OJJ64631.1 ater|EAU37329.1 atub|OJI82548.1 aver|OJI98729.1 awen|OJJ37679.1 azon|OJJ44945.1

Group4392: abra|OJJ77503.1 acar|OOF99224.1 acla|EAW06732.1 afis|EAW16284.1 afla|EED53272.1 afum|EAL88905.1 agla|OJJ88305.1 akaw|GAA84118.1 aluc|OJZ81034.1 anid|CBF86871.1 anig|EHA19332.1 aory|EIT75050.1 asyd|OJJ64630.1 ater|EAU37328.1 atub|OJI82549.1 aver|OJI98728.1 awen|OJJ37677.1 azon|OJJ44942.1

Group4393: abra|OJJ77504.1 acar|OOF99222.1 acla|EAW06730.1 afis|EAW16283.1 afla|EED53273.1 afum|EAL88904.1 agla|OJJ88318.1 akaw|GAA84116.1 aluc|OJZ81032.1 anid|CBF86876.1 anig|EHA19333.1 aory|EIT75123.1 asyd|OJJ64627.1 ater|EAU37327.1 atub|OJI82551.1 aver|OJI98725.1 awen|OJJ37675.1 azon|OJJ44940.1

Group4394: abra|OJJ77507.1 acar|OOF99219.1 acla|EAW06727.1 afis|EAW16280.1 afla|EED53276.1 afum|EAL88901.1 agla|OJJ88324.1 akaw|GAA84113.1 aluc|OJZ81029.1 anid|CBF86880.1 anig|EHA19336.1 aory|EIT74958.1 asyd|OJJ64626.1 ater|EAU37325.1 atub|OJI82554.1 aver|OJI98722.1 awen|OJJ37672.1 azon|OJJ44937.1

Group4395: abra|OJJ77508.1 acar|OOF99218.1 acla|EAW06726.1 afis|EAW16279.1 afla|EED53277.1 afum|EAL88900.1 agla|OJJ88323.1 akaw|GAA84112.1 aluc|OJZ81028.1 anid|CBF86882.1 anig|EHA19337.1 aory|EIT74848.1 asyd|OJJ64625.1 ater|EAU37324.1 atub|OJI82555.1 aver|OJI98721.1 awen|OJJ37671.1 azon|OJJ44936.1

Group4396: abra|OJJ77509.1 acar|OOF99217.1 acla|EAW06725.1 afis|EAW16278.1 afla|EED53279.1 afum|EAL88899.1 agla|OJJ88321.1 akaw|GAA84111.1 aluc|OJZ81027.1 anid|CBF86883.1 anig|EHA19338.1 aory|EIT74824.1 asyd|OJJ64624.1 ater|EAU37323.1 atub|OJI82556.1 aver|OJI98720.1 awen|OJJ37670.1 azon|OJJ44935.1

Group4397: abra|OJJ77510.1 acar|OOF99216.1 acla|EAW06724.1 afis|EAW16277.1 afla|EED53280.1 afum|EAL88898.1 agla|OJJ88328.1 akaw|GAA84110.1 aluc|OJZ81026.1 anid|CBF86885.1 anig|EHA19339.1 aory|EIT75207.1 asyd|OJJ64623.1 ater|EAU37322.1 atub|OJI82557.1 aver|OJI98719.1 awen|OJJ37669.1 azon|OJJ44934.1

Group4398: abra|OJJ77511.1 acar|OOF99214.1 acla|EAW06723.1 afis|EAW16276.1 afla|EED53281.1 afum|EAL88897.1 agla|OJJ88329.1 akaw|GAA84109.1 aluc|OJZ81025.1 anid|CBF86892.1 anig|EHA18669.1 aory|EIT75064.1 asyd|OJJ64622.1 ater|EAU37321.1 atub|OJI82558.1 aver|OJI98718.1 awen|OJJ37668.1 azon|OJJ44933.1

Group4399: abra|OJJ77513.1 acar|OOF99211.1 acla|EAW06722.1 afis|EAW16265.1 afla|EED53285.1 afum|EAL88886.1 agla|OJJ88330.1 akaw|GAA84107.1 aluc|OJZ81022.1 anid|CBF86894.1 anig|EHA19341.1 aory|EIT74876.1 asyd|OJJ64621.1 ater|EAU37438.1 atub|OJI82561.1 aver|OJI98717.1 awen|OJJ37667.1 azon|OJJ44930.1

Group4400: abra|OJJ77514.1 acar|OOF99210.1 acla|EAW06721.1 afis|EAW16266.1 afla|EED53284.1 afum|EAL88887.1 agla|OJJ88331.1 akaw|GAA84106.1 aluc|OJZ81021.1 anid|CBF86895.1 anig|EHA19342.1 aory|EIT75173.1 asyd|OJJ64620.1 ater|EAU37439.1 atub|OJI82562.1 aver|OJI98716.1 awen|OJJ37666.1 azon|OJJ44931.1

Group4401: abra|OJJ77516.1 acar|OOF99207.1 acla|EAW06720.1 afis|EAW16267.1 afla|EED53282.1 afum|EAL88888.1 agla|OJJ88333.1 akaw|GAA84104.1 aluc|OJZ81019.1 anid|CBF86897.1 anig|EHA19343.1 aory|EIT75090.1 asyd|OJJ64618.1 ater|EAU37440.1 atub|OJI82564.1 aver|OJI98714.1 awen|OJJ37664.1 azon|OJJ44923.1

Group4402: abra|OJJ77517.1 acar|OOF99208.1 acla|EAW06719.1 afis|EAW16268.1 afla|EED53288.1 afum|EAL88889.1 agla|OJJ88337.1 akaw|GAA84103.1 aluc|OJZ81018.1 anid|CBF86899.1 anig|EHA19344.1 aory|EIT74865.1 asyd|OJJ64617.1 ater|EAU37443.1 atub|OJI82565.1 aver|OJI98713.1 awen|OJJ37663.1 azon|OJJ44924.1

Group4403: abra|OJJ77520.1 acar|OOF99203.1 acla|EAW06715.1 afla|EED53293.1 afum|EAL88893.1 akaw|GAA84100.1 aluc|OJZ81016.1 anid|CBF86904.1 anig|EHA19346.1 aory|EIT74861.1 asyd|OJJ64613.1 atub|OJI82568.1 aver|OJI98709.1 awen|OJJ37656.1 azon|OJJ44929.1 afis|EAW16272.1 agla|OJJ88342.1 ater|EAU37447.1

Group4404: abra|OJJ77522.1 acar|OOF99202.1 acla|EAW06717.1 afis|EAW16270.1 afla|EED53295.1 afum|EAL88891.1 agla|OJJ88339.1 akaw|GAA84098.1 aluc|OJZ81014.1 anid|CBF86903.1 anig|EHA19347.1 aory|EIT74914.1 asyd|OJJ64615.1 ater|EAU37445.1 atub|OJI82570.1 aver|OJI98711.1 awen|OJJ37660.1 azon|OJJ44927.1

Group4405: abra|OJJ77524.1 acar|OOF99200.1 acla|EAW06718.1 afis|EAW16269.1 afla|EED53289.1 afum|EAL88890.1 agla|OJJ88338.1 akaw|GAA84095.1 aluc|OJZ81011.1 anid|CBF86901.1 anig|EHA19350.1 aory|EIT74922.1 asyd|OJJ64616.1 ater|EAU37444.1 atub|OJI82573.1 aver|OJI98712.1 awen|OJJ37661.1 azon|OJJ44926.1

Group4406: abra|OJJ77526.1 acar|OOF99198.1 acla|EAW06712.1 afis|EAW16275.1 afla|EED53297.1 afum|EAL88896.1 agla|OJJ88345.1 akaw|GAA84094.1 aluc|OJZ81009.1 anid|CBF86909.1 anig|EHA19351.1 aory|EIT75079.1 asyd|OJJ64610.1 ater|EAU37450.1 atub|OJI82575.1 aver|OJI98706.1 awen|OJJ37658.1 azon|OJJ44920.1

Group4407: abra|OJJ77535.1 acar|OOF99189.1 acla|EAW06704.1 afis|EAW16249.1 afla|EED53313.1 afum|EAL88877.1 agla|OJJ86260.1 akaw|GAA84085.1 aluc|OJZ81000.1 anid|CBF78891.1 anig|EHA19359.1 aory|EIT74965.1 asyd|OJJ64597.1 ater|EAU37466.1 atub|OJI82585.1 aver|OJI98694.1 awen|OJJ30206.1 azon|OJJ43931.1

Group4408: abra|OJJ77554.1 acar|OOF99173.1 acla|EAW06523.1 afis|EAW16239.1 afla|EED53338.1 afum|EAL88860.1 agla|OJJ86246.1 akaw|GAA84069.1 aluc|OJZ80983.1 anid|CBF78863.1 anig|EHA19377.1 aory|EIT75140.1 asyd|OJJ64574.1 ater|EAU33704.1 atub|OJI82603.1 aver|OJI98671.1 awen|OJJ30218.1 azon|OJJ43940.1

Group4409: abra|OJJ77557.1 acar|OOG00298.1 afis|EAW16398.1 afla|EED45629.1 afum|EAL89029.2 agla|OJJ82194.1 anid|CBF85652.1 anig|EHA19380.1 aory|EIT75689.1 asyd|OJJ54151.1 ater|EAU33011.1 aver|OJJ08588.1 awen|OJJ38072.1 aory|EIT76357.1 azon|OJJ46582.1 asyd|OJJ56065.1 aver|OJJ05008.1 awen|OJJ31109.1

Group4410: abra|OJJ77561.1 acar|OOF99169.1 acla|EAW06694.1 afis|EAW16072.1 afla|EED48591.1 afum|EAL88704.1 agla|OJJ86096.1 akaw|GAA84064.1 aluc|OJZ80977.1 anid|CBF75113.1 anig|EHA19382.1 aory|EIT73146.1 asyd|OJJ59998.1 ater|EAU33559.1 atub|OJI82608.1 aver|OJJ02293.1 awen|OJJ30362.1 azon|OJJ45381.1

Group4411: abra|OJJ77566.1 acar|OOF99164.1 acla|EAW06531.1 afis|EAW16064.1 afla|EED48578.1 afum|EAL88696.1 agla|OJJ86101.1 akaw|GAA84060.1 aluc|OJZ80972.1 anid|CBF75092.1 anig|EHA19387.1 aory|EIT73042.1 asyd|OJJ60011.1 ater|EAU33571.1 atub|OJI82613.1 aver|OJJ02274.1 awen|OJJ30375.1 azon|OJJ45391.1

Group4412: abra|OJJ77567.1 acar|OOF99162.1 acla|EAW06532.1 afis|EAW16063.1 afla|EED48577.1 afum|EAL88695.1 agla|OJJ86102.1 akaw|GAA84059.1 aluc|OJZ80971.1 anid|CBF75090.1 anig|EHA19388.1 aory|EIT73179.1 asyd|OJJ60012.1 ater|EAU33572.1 atub|OJI82614.1 aver|OJJ02273.1 awen|OJJ30376.1 azon|OJJ45392.1

Group4413: abra|OJJ77568.1 acar|OOF99163.1 acla|EAW06533.1 afis|EAW16062.1 afla|EED48576.1 afum|EAL88694.1 agla|OJJ86103.1 akaw|GAA84058.1 aluc|OJZ80970.1 anid|CBF75088.1 anig|EHA19389.1 aory|EIT73163.1 asyd|OJJ60013.1 ater|EAU33573.1 atub|OJI82615.1 aver|OJJ02272.1 awen|OJJ30377.1 azon|OJJ45393.1

Group4414: abra|OJJ77572.1 acar|OOF99157.1 acla|EAW06539.1 afis|EAW16056.1 afla|EED56442.1 afum|EAL88688.1 agla|OJJ86089.1 akaw|GAA84054.1 aluc|OJZ80966.1 anid|CBF75076.1 anig|EHA19392.1 aory|EIT72786.1 asyd|OJJ60016.1 ater|EAU33578.1 atub|OJI82620.1 aver|OJJ02269.1 awen|OJJ30383.1 azon|OJJ45400.1

Group4415: abra|OJJ77573.1 acar|OOF99156.1 acla|EAW06538.1 afis|EAW16057.1 afla|EED56441.1 afum|EAL88689.1 agla|OJJ86090.1 akaw|GAA84053.1 aluc|OJZ80965.1 anid|CBF75078.1 anig|EHA19393.1 aory|EIT72787.1 asyd|OJJ60017.1 ater|EAU33577.1 atub|OJI82621.1 aver|OJJ02268.1 awen|OJJ30382.1 azon|OJJ45399.1

Group4416: abra|OJJ77574.1 acar|OOF99155.1 acla|EAW06537.1 afis|EAW16058.1 afla|EED48572.1 afum|EAL88690.2 agla|OJJ86107.1 akaw|GAA84051.1 aluc|OJZ80963.1 anid|CBF75080.1 anig|EHA19394.1 aory|EIT73249.1 asyd|OJJ60018.1 ater|EAU33576.1 atub|OJI82623.1 aver|OJJ02267.1 awen|OJJ30381.1 azon|OJJ45398.1

Group4417: abra|OJJ77577.1 acar|OOF99150.1 acla|EAW06543.1 afis|EAW16052.1 afla|EED56446.1 afum|EAL88684.1 agla|OJJ86086.1 akaw|GAA84048.1 aluc|OJZ80959.1 anid|CBF75068.1 anig|EHA19397.1 aory|EIT72800.1 asyd|OJJ60022.1 ater|EAU33581.1 atub|OJI82626.1 aver|OJJ02262.1 awen|OJJ30390.1 azon|OJJ45402.1

Group4418: abra|OJJ77579.1 acar|OOF99148.1 acla|EAW06545.1 afis|EAW16049.1 afla|EED56438.1 afum|EAL88682.1 agla|OJJ86083.1 akaw|GAA84046.1 aluc|OJZ80956.1 anid|CBF75064.1 anig|EHA19399.1 aory|EIT72808.1 asyd|OJJ60024.1 ater|EAU33587.1 atub|OJI82628.1 aver|OJJ02260.1 awen|OJJ30394.1 azon|OJJ45404.1

Group4419: abra|OJJ77580.1 acar|OOF99147.1 acla|EAW06546.1 afis|EAW16048.1 afla|EED56437.1 afum|EAL88681.1 agla|OJJ86082.1 akaw|GAA84045.1 aluc|OJZ80955.1 anid|CBF75062.1 anig|EHA19400.1

aory|EIT72865.1 asyd|OJJ60025.1 ater|EAU33588.1 atub|OJI82629.1 aver|OJJ02259.1 awen|OJJ30396.1 azon|OJJ45405.1

Group4420: abra|OJJ77581.1 acar|OOF99146.1 acla|EAW06547.1 afis|EAW16047.1 afla|EED56436.1 afum|EAL88680.1 agla|OJJ86081.1 akaw|GAA84044.1 aluc|OJZ80954.1 anid|CBF75060.1 anig|EHA19401.1 aory|EIT72831.1 asyd|OJJ60026.1 ater|EAU33590.1 atub|OJI82630.1 aver|OJJ02258.1 awen|OJJ30398.1 azon|OJJ45406.1

Group4421: abra|OJJ77585.1 acar|OOF92640.1 acla|EAW06551.1 afis|EAW16044.1 afla|EED56433.1 afum|EAL88676.1 agla|OJJ86077.1 akaw|GAA84041.1 aluc|OJZ80950.1 anid|CBF75054.1 anig|EHA19404.1 aory|EIT72801.1 asyd|OJJ60029.1 ater|EAU33593.1 atub|OJI82633.1 aver|OJJ02255.1 awen|OJJ30402.1 azon|OJJ45410.1

Group4422: abra|OJJ77586.1 acar|OOF99144.1 acla|EAW06552.1 afis|EAW16043.1 afla|EED56432.1 afum|EAL88675.1 agla|OJJ86076.1 akaw|GAA84040.1 aluc|OJZ80949.1 anid|CBF75052.1 anig|EHA19405.1 aory|EIT72860.1 asyd|OJJ60030.1 ater|EAU33594.1 atub|OJI82634.1 aver|OJJ02254.1 awen|OJJ30403.1 azon|OJJ45411.1

Group4423: abra|OJJ77588.1 acar|OOF99143.1 acla|EAW06554.1 afis|EAW16041.1 afla|EED56430.1 afum|EAL88673.1 agla|OJJ86074.1 akaw|GAA84038.1 aluc|OJZ80947.1 anid|CBF75050.1 anig|EHA19407.1 aory|EIT72854.1 asyd|OJJ60032.1 ater|EAU33596.1 atub|OJI82636.1 aver|OJJ02252.1 awen|OJJ30405.1 azon|OJJ45413.1

Group4424: abra|OJJ77590.1 acar|OOF99141.1 acla|EAW06556.1 afis|EAW16039.1 afla|EED56427.1 afum|EAL88671.1 agla|OJJ86072.1 akaw|GAA84036.1 aluc|OJZ80945.1 anid|CBF75046.1 anig|EHA19409.1 aory|EIT72877.1 asyd|OJJ60034.1 ater|EAU33599.1 atub|OJI82638.1 aver|OJJ02250.1 awen|OJJ30407.1 azon|OJJ45415.1

Group4425: abra|OJJ77598.1 acar|OOF99128.1 acla|EAW06562.1 afis|EAW16034.1 afla|EED56420.1 afum|EAL88665.1 agla|OJJ86066.1 akaw|GAA84030.1 aluc|OJZ80935.1 anid|CBF75036.1 anig|EHA19415.1 aory|EIT72883.1 asyd|OJJ60049.1 ater|EAU33606.1 atub|OJI82645.1 aver|OJJ02240.1 awen|OJJ30413.1 azon|OJJ45421.1

Group4426: abra|OJJ77601.1 acar|OOF99126.1 acla|EAW06565.1 afis|EAW16031.1 afla|EED56413.1 afum|EAL88662.1 agla|OJJ86059.1 akaw|GAA84027.1 aluc|OJZ80932.1 anid|CBF75030.1 anig|EHA19418.1 aory|EIT72887.1 asyd|OJJ60052.1 ater|EAU33610.1 atub|OJI82648.1 aver|OJJ02237.1 awen|OJJ30417.1 azon|OJJ45424.1

Group4427: abra|OJJ77602.1 acar|OOF99124.1 acla|EAW06567.1 afis|EAW16029.1 afla|EED56412.1 afum|EAL88660.1 agla|OJJ86057.1 akaw|GAA84026.1 aluc|OJZ80931.1 anid|CBF75016.1 anig|EHA19419.1 aory|EIT72882.1 asyd|OJJ60070.1 ater|EAU33611.1 atub|OJI82649.1 aver|OJJ04192.1 awen|OJJ30419.1 azon|OJJ45426.1

Group4428: abra|OJJ77603.1 acar|OOF99123.1 acla|EAW06568.1 afis|EAW16028.1 afla|EED56411.1 afum|EAL88658.1 agla|OJJ86056.1 akaw|GAA84025.1 aluc|OJZ80930.1 anid|CBF75018.1 anig|EHA19420.1 aory|EIT72881.1 asyd|OJJ60069.1 ater|EAU33612.1 atub|OJI82650.1 aver|OJJ02226.1 awen|OJJ30420.1 azon|OJJ45427.1

Group4429: abra|OJJ77605.1 acar|OOF99122.1 acla|EAW06569.1 afis|EAW16027.1 afla|EED56410.1 afum|EAL88657.1 agla|OJJ86055.1 akaw|GAA84024.1 aluc|OJZ80928.1 anid|CBF75020.1 anig|EHA19421.1 aory|EIT83014.1 asyd|OJJ60068.1 ater|EAU33613.1 atub|OJI82651.1 aver|OJJ02227.1 awen|OJJ30421.1 azon|OJJ45428.1

Group4430: abra|OJJ77613.1 acar|OOF99115.1 acla|EAW06571.1 afis|EAW16025.1 afla|EED56406.1 afum|EAL88654.1 agla|OJJ86053.1 akaw|GAA84018.1 aluc|OJZ80921.1 anid|CBF75014.1 anig|EHA19426.1 aory|EIT83006.1 asyd|OJJ60057.1 ater|EAU33616.1 atub|OJI82659.1 aver|OJJ02232.1 awen|OJJ30432.1 azon|OJJ45431.1

Group4431: abra|OJJ77614.1 acar|OOF90324.1 acla|EAW11334.1 afis|EAW18161.1 afla|EED56332.1 afum|EAL87521.1 agla|OJJ85965.1 akaw|GAA84017.1 aluc|OJZ80920.1 anid|CBF85769.1 anig|EHA19427.1 aory|EIT73455.1 asyd|OJJ64306.1 ater|EAU33688.1 atub|OJI82660.1 aver|OJI98399.1 awen|OJJ30506.1 azon|OJJ45495.1

Group4432: abra|OJJ77615.1 acar|OOF90323.1 acla|EAW11335.1 afis|EAW18160.1 afla|EED56333.1 afum|EBA27314.1 agla|OJJ85966.1 akaw|GAA84016.1 aluc|OJZ80919.1 anid|CBF85772.1 anig|EHA19428.1 aory|EIT73306.1 asyd|OJJ64308.1 ater|EAU33687.1 atub|OJI82661.1 aver|OJI98400.1 awen|OJJ30505.1 azon|OJJ45494.1

Group4433: abra|OJJ77616.1 acar|OOF90325.1 acla|EAW11336.1 afis|EAW18159.1 afla|EED56335.1 afum|EAL87518.1 agla|OJJ85968.1 akaw|GAA84015.1 aluc|OJZ80918.1 anid|CBF85776.1 anig|EHA19429.1 aory|EIT73326.1 asyd|OJJ64310.1 ater|EAU33686.1 atub|OJI82662.1 aver|OJI98403.1 awen|OJJ30504.1 azon|OJJ45493.1

Group4434: abra|OJJ77617.1 acar|OOF90326.1 acla|EAW11337.1 afis|EAW18158.1 afla|EED56336.1 afum|EAL87517.1 agla|OJJ85969.1 akaw|GAA84014.1 aluc|OJZ80917.1 anid|CBF85777.1 anig|EHA19430.1 aory|EIT73425.1 asyd|OJJ64311.1 ater|EAU33685.1 atub|OJI82663.1 aver|OJI98404.1 awen|OJJ30503.1 azon|OJJ45492.1

Group4435: abra|OJJ77618.1 acar|OOF90327.1 acla|EAW11338.1 afis|EAW18157.1 afla|EED56337.1 afum|EAL87516.1 agla|OJJ85970.1 akaw|GAA84013.1 aluc|OJZ80916.1 anid|CBF85779.1 anig|EHA19431.1 aory|EIT73376.1 asyd|OJJ64312.1 ater|EAU33684.1 atub|OJI82664.1 aver|OJI98405.1 awen|OJJ30502.1 azon|OJJ45491.1

Group4436: abra|OJJ77619.1 acar|OOF90328.1 acla|EAW11339.1 afis|EAW18156.1 afla|EED56338.1 afum|EAL87515.1 agla|OJJ85971.1 akaw|GAA84012.1 aluc|OJZ80915.1 anid|CBF85781.1 anig|EHA19432.1 aory|EIT73330.1 asyd|OJJ64313.1 ater|EAU33683.1 atub|OJI82665.1 aver|OJI98406.1 awen|OJJ30500.1 azon|OJJ45490.1

Group4437: abra|OJJ77620.1 acar|OOF90329.1 acla|EAW11340.1 afis|EAW18155.1 afla|EED56339.1 afum|EAL87514.1 agla|OJJ85972.1 akaw|GAA84011.1 aluc|OJZ80914.1 anid|CBF85783.1 anig|EHA19433.1 aory|EIT73446.1 asyd|OJJ64314.1 ater|EAU33682.1 atub|OJI82666.1 aver|OJI98407.1 awen|OJJ30499.1 azon|OJJ45489.1

Group4438: abra|OJJ77621.1 acar|OOF99114.1 acla|EAW11341.1 afis|EAW18154.1 afla|EED56340.1 afum|EAL87513.1 agla|OJJ85973.1 akaw|GAA84010.1 aluc|OJZ80913.1 anid|CBF85784.1 anig|EHA19434.1 aory|EIT73328.1 asyd|OJJ64316.1 ater|EAU33681.1 atub|OJI82667.1 aver|OJI98409.1 awen|OJJ30498.1 azon|OJJ45488.1

Group4439: abra|OJJ77626.1 acar|OOF99112.1 acla|EAW11342.1 afis|EAW18152.1 afla|EED56342.1 afum|EAL87511.1 agla|OJJ85975.1 akaw|GAA84007.1 aluc|OJZ80905.1 anid|CBF85790.1 anig|EHA19436.1 aory|EIT73287.1 asyd|OJJ64325.1 ater|EAU33677.1 atub|OJI82672.1 aver|OJI98419.1 awen|OJJ30495.1 azon|OJJ45487.1

Group4440: abra|OJJ77629.1 acar|OOF99097.1 acla|EAW06480.1 afis|EAW15975.1 afla|EED56345.1 afum|EAL88607.1 agla|OJJ85980.1 akaw|GAA84005.1 aluc|OJZ80901.1 anid|CBF85795.1 anig|EHA19438.1 aory|EIT73353.1 asyd|OJJ64330.1 ater|EAU33676.1 atub|OJI82678.1 aver|OJI98423.1 awen|OJJ30494.1 azon|OJJ45485.1

Group4441: abra|OJJ77630.1 acar|OOF99098.1 acla|EAW06481.1 afis|EAW15976.1 afla|EED56346.1 afum|EAL88608.1 agla|OJJ85981.1 akaw|GAA84004.1 aluc|OJZ80900.1 anid|CBF85797.1 anig|EHA19439.1 aory|EIT73314.1 asyd|OJJ64331.1 ater|EAU33675.1 atub|OJI82679.1 aver|OJI98424.1 awen|OJJ30493.1 azon|OJJ45484.1

Group4442: abra|OJJ77631.1 acar|OOF99099.1 acla|EAW06482.1 afis|EAW15977.1 afla|EED56347.1 afum|EAL88609.1 agla|OJJ85982.1 akaw|GAA84003.1 aluc|OJZ80898.1 anid|CBF85799.1 anig|EHA19440.1 aory|EIT73397.1 asyd|OJJ64332.1 ater|EAU33674.1 atub|OJI82680.1 aver|OJI98425.1 awen|OJJ30492.1 azon|OJJ45483.1

Group4443: abra|OJJ77632.1 acar|OOF99100.1 acla|EAW06484.1 afis|EAW15979.1 afla|EED56349.1 afum|EAL88611.1 agla|OJJ85984.1 akaw|GAA84001.1 aluc|OJZ80897.1 anid|CBF85803.1 anig|EHA19442.1 aory|EIT73329.1 asyd|OJJ64334.1 ater|EAU33672.1 atub|OJI82681.1 aver|OJI98427.1 awen|OJJ30491.1 azon|OJJ45482.1

Group4444: abra|OJJ77637.1 acar|OOF99105.1 acla|EAW06487.1 afis|EAW15982.1 afla|EED56352.1 afum|EAL88614.1 agla|OJJ85985.1 akaw|GAA83997.1 aluc|OJZ80893.1 anid|CBF85806.1 anig|EHA19445.1 aory|EIT73461.1 asyd|OJJ64337.1 ater|EAU33670.1 atub|OJI82685.1 aver|OJI98430.1 awen|OJJ30488.1 azon|OJJ45479.1

Group4445: abra|OJJ77638.1 acar|OOF99106.1 acla|EAW06488.1 afis|EAW15983.1 afla|EED56353.1 afum|EAL88615.1 agla|OJJ85986.1 akaw|GAA83996.1 aluc|OJZ80892.1 anid|CBF85808.1 anig|EHA19446.1 aory|EIT73319.1 asyd|OJJ64339.1 ater|EAU33669.1 atub|OJI82686.1 aver|OJI98432.1 awen|OJJ30487.1 azon|OJJ45478.1

Group4446: abra|OJJ77640.1 acar|OOF99108.1 acla|EAW06490.1 afis|EAW15985.1 afla|EED56355.1 afum|EAL88617.1 agla|OJJ85988.1 akaw|GAA83994.1 aluc|OJZ80890.1 anid|CBF85812.1 anig|EHA19448.1 aory|EIT73389.1 asyd|OJJ64341.1 ater|EAU33667.1 atub|OJI82688.1 aver|OJI98434.1 awen|OJJ30485.1 azon|OJJ45476.1

Group4447: abra|OJJ77642.1 acar|OOF99094.1 acla|EAW06492.1 afis|EAW15988.1 afla|EED56360.1 afum|EAL88620.1 agla|OJJ85990.1 akaw|GAA83992.1 aluc|OJZ80888.1 anid|CBF85815.1 anig|EHA19450.1 aory|EIT73300.1 asyd|OJJ64342.1 ater|EAU33661.1 atub|OJI82690.1 aver|OJI98435.1 awen|OJJ30482.1 azon|OJJ45473.1

Group4448: abra|OJJ77644.1 acar|OOF99091.1 acla|EAW07074.1 afis|EAW20698.1 afla|EED46367.1 afum|EAL84873.1 akaw|GAA83990.1 aluc|OJZ80885.1 anid|CBF86210.1 anig|EHA19452.1 aory|EIT81200.1

asyd|OJJ64550.1 atub|OJI82692.1 aver|OJI98646.1 awen|OJJ36028.1 acar|OOF98893.1 afis|EAW20291.1 afum|EAL85423.1

Group4449: abra|OJJ77652.1 acar|OOF99084.1 acla|EAW06494.1 afis|EAW15990.1 afla|EED56362.1 afum|EAL88622.1 agla|OJJ85992.1 akaw|GAA83983.1 aluc|OJZ80877.1 anid|CBF85819.1 anig|EHA19459.1 aory|EIT73339.1 asyd|OJJ64343.1 ater|EAU33658.1 atub|OJI82700.1 aver|OJI98440.1 awen|OJJ30480.1 azon|OJJ45471.1

Group4450: abra|OJJ77654.1 acar|OOF99083.1 acla|EAW06496.1 afis|EAW15992.1 afla|EED56364.1 afum|EAL88624.2 agla|OJJ86003.1 akaw|GAA83981.1 aluc|OJZ80875.1 anid|CBF85823.1 anig|EHA19460.1 aory|EIT73404.1 asyd|OJJ64344.1 ater|EAU33657.1 atub|OJI82702.1 aver|OJI98442.1 awen|OJJ30478.1 azon|OJJ45470.1

Group4451: abra|OJJ77655.1 acar|OOF99081.1 acla|EAW06497.1 afis|EAW15993.1 afla|EED56365.1 afum|EAL88625.1 agla|OJJ86004.1 akaw|GAA83980.1 aluc|OJZ80874.1 anid|CBF85824.1 anig|EHA19462.1 aory|EIT73307.1 asyd|OJJ64345.1 ater|EAU33656.1 atub|OJI82704.1 aver|OJI98443.1 awen|OJJ30477.1 azon|OJJ45469.1

Group4452: abra|OJJ77656.1 acar|OOF99080.1 acla|EAW06498.1 afis|EAW15994.1 afla|EED56367.1 afum|EAL88626.1 agla|OJJ86008.1 akaw|GAA83979.1 aluc|OJZ80872.1 anid|CBF85828.1 anig|EHA19463.1 asyd|OJJ64348.1 ater|EAU33654.1 atub|OJI82705.1 aver|OJI98445.1 awen|OJJ30475.1 azon|OJJ45467.1 aory|EIT73315.1

Group4453: abra|OJJ77657.1 acar|OOF99079.1 acla|EAW06499.1 afis|EAW15995.1 afla|EED56369.1 afum|EAL88627.1 agla|OJJ86009.1 akaw|GAA83978.1 aluc|OJZ80871.1 anid|CBF85830.1 anig|EHA19464.1 aory|EIT73436.1 asyd|OJJ64349.1 ater|EAU33652.1 atub|OJI82706.1 aver|OJI98446.1 awen|OJJ30474.1 azon|OJJ45466.1

Group4454: abra|OJJ77659.1 acar|OOF99076.1 acla|EAW06501.1 afis|EAW15997.1 afla|EED56371.1 afum|EAL88629.1 agla|OJJ86017.1 akaw|GAA83976.1 aluc|OJZ80869.1 anid|CBF85832.1 anig|EHA19466.1 aory|EIT73442.1 asyd|OJJ64350.1 ater|EAU33649.1 atub|OJI82708.1 aver|OJI98447.1 awen|OJJ30472.1 azon|OJJ45464.1

Group4455: abra|OJJ77660.1 acar|OOF99075.1 acla|EAW06504.1 afis|EAW15998.1 afla|EED56372.1 afum|EAL88630.1 agla|OJJ86013.1 akaw|GAA83975.1 aluc|OJZ80868.1 anid|CBF85834.1 anig|EHA19467.1 aory|EIT73360.1 asyd|OJJ64351.1 ater|EAU33648.1 atub|OJI82709.1 aver|OJI98448.1 awen|OJJ30471.1 azon|OJJ45463.1

Group4456: abra|OJJ77661.1 acar|OOF99073.1 acla|EAW06598.1 afis|EAW16006.1 afla|EED56380.1 afum|EAL88638.2 agla|OJJ86025.1 akaw|GAA83974.1 aluc|OJZ80867.1 anid|CBF85868.1 anig|EHA19468.1 aory|EIT73448.1 asyd|OJJ64371.1 ater|EAU33637.1 atub|OJI82710.1 aver|OJI98467.1 awen|OJJ30461.1 azon|OJJ45451.1

Group4457: abra|OJJ77665.1 acar|OOF99070.1 acla|EAW06507.1 afis|EAW16002.1 afla|EED56376.1 afum|EAL88634.1 agla|OJJ86024.1 akaw|GAA83970.1 aluc|OJZ80863.1 anid|CBF85844.1 anig|EHA19472.1 aory|EIT73423.1 asyd|OJJ64356.1 ater|EAU33643.1 atub|OJI82714.1 aver|OJI98452.1 awen|OJJ30466.1 azon|OJJ45457.1

Group4458: abra|OJJ77667.1 acar|OOF99068.1 acla|EAW06506.1 afis|EAW16000.1 afla|EED56375.1 afum|EAL88633.1 agla|OJJ86020.1 akaw|GAA83962.1 aluc|OJZ80850.1 anid|CBF85840.1 anig|EHA19474.1 aory|EIT73310.1 asyd|OJJ64354.1 ater|EAU33645.1 atub|OJI82718.1 aver|OJI98450.1 awen|OJJ30469.1 azon|OJJ45461.1

Group4459: abra|OJJ77671.1 acar|OOF99066.1 acla|EAW06505.1 afis|EAW15999.1 afla|EED56374.1 afum|EAL88631.1 agla|OJJ86019.1 akaw|GAA83961.1 aluc|OJZ80848.1 anid|CBF85835.1 anig|EHA19475.1 aory|EIT73377.1 asyd|OJJ64352.1 ater|EAU33647.1 atub|OJI82719.1 aver|OJI98449.1 awen|OJJ30470.1 azon|OJJ45462.1

Group4460: abra|OJJ77680.1 acar|OOF99060.1 acla|EAW06591.1 afis|EAW16014.1 afla|EED56391.1 afum|EAL88643.1 agla|OJJ86040.1 akaw|GAA83954.1 aluc|OJZ80838.1 anid|CBF74562.1 anig|EHA19482.1 aory|EIT83009.1 asyd|OJJ57647.1 ater|EAU33630.1 atub|OJI82730.1 aver|OJJ05504.1 awen|OJJ30450.1 azon|OJJ45442.1

Group4461: abra|OJJ77681.1 acar|OOF99058.1 acla|EAW06592.1 afis|EAW16015.1 afla|EED56392.1 afum|EAL88644.1 agla|OJJ86041.1 akaw|GAA83952.1 aluc|OJZ80836.1 anid|CBF74560.1 anig|EHA19483.1 aory|EIT83002.1 asyd|OJJ57646.1 ater|EAU33629.1 atub|OJI82732.1 aver|OJJ05503.1 awen|OJJ30449.1 azon|OJJ45441.1

Group4462: abra|OJJ77682.1 acar|OOF99059.1 acla|EAW06594.1 afis|EAW16017.1 afla|EED56393.1 afum|EAL88646.1 agla|OJJ86042.1 akaw|GAA83951.1 aluc|OJZ80835.1 anid|CBF74558.1 anig|EHA19484.1 aory|EIT83003.1 asyd|OJJ57645.1 ater|EAU33628.1 atub|OJI82733.1 aver|OJJ05502.1 awen|OJJ30447.1 azon|OJJ45439.1

Group4463: abra|OJJ77683.1 acar|OOF99057.1 acla|EAW06595.1 afis|EAW16018.1 afla|EED56394.1 afum|EAL88647.1 agla|OJJ86045.1 akaw|GAA83950.1 aluc|OJZ80834.1 anid|CBF74555.1 anig|EHA19485.1 aory|EIT83010.1 asyd|OJJ57644.1 ater|EAU33627.1 atub|OJI82734.1 aver|OJJ05501.1 awen|OJJ30446.1 azon|OJJ45438.1

Group4464: abra|OJJ77684.1 acar|OOF99056.1 acla|EAW06596.1 afis|EAW16019.1 afla|EED56396.1 afum|EAL88648.1 agla|OJJ86046.1 akaw|GAA83949.1 aluc|OJZ80833.1 anid|CBF74553.1 anig|EHA19486.1 aory|EIT83017.1 asyd|OJJ57642.1 ater|EAU33626.1 atub|OJI82735.1 aver|OJJ05499.1 awen|OJJ30445.1 azon|OJJ45437.1

Group4465: abra|OJJ77690.1 acar|OOF99049.1 acla|EAW06575.1 afis|EAW16021.1 afla|EED56399.1 afum|EAL88650.1 agla|OJJ86048.1 akaw|GAA83944.1 aluc|OJZ80825.1 anid|CBF74538.1 anig|EHA19490.1 aory|EIT83016.1 asyd|OJJ57695.1 ater|EAU33619.1 atub|OJI82740.1 aver|OJJ05471.1 awen|OJJ30443.1 azon|OJJ45435.1

Group4466: abra|OJJ77691.1 acar|OOF99048.1 acla|EAW06574.1 afis|EAW16022.1 afla|EED56401.1 afum|EAL88651.1 agla|OJJ86051.1 akaw|GAA83939.1 aluc|OJZ80821.1 anid|CBF74540.1 anig|EHA19491.1 aory|EIT83005.1 asyd|OJJ57694.1 ater|EAU33618.1 atub|OJI82741.1 aver|OJJ05472.1 awen|OJJ30439.1 azon|OJJ45434.1

Group4467: abra|OJJ77695.1 acar|OOF99044.1 acla|EAW11326.1 afis|EAW18169.1 afla|EED56325.1 afum|EBA27315.1 agla|OJJ85948.1 akaw|GAA83935.1 aluc|OJZ90758.1 anid|CBF85744.1 anig|EHA19495.1 aory|EIT73371.1 asyd|OJJ64290.1 ater|EAU33835.1 atub|OJI82745.1 aver|OJI98383.1 awen|OJJ30516.1 azon|OJJ50197.1

Group4468: abra|OJJ77697.1 acar|OOF99043.1 acla|EAW11327.1 afis|EAW18168.1 afla|EED56326.1 afum|EAL87529.2 agla|OJJ85949.1 akaw|GAA83934.1 aluc|OJZ90761.1 anid|CBF85746.1 anig|EHA19496.1 aory|EIT73447.1 asyd|OJJ64291.1 ater|EAU33836.1 atub|OJI82748.1 aver|OJI98384.1 awen|OJJ30514.1 azon|OJJ50195.1

Group4469: abra|OJJ77699.1 acar|OOF99040.1 acla|EAW11324.1 afis|EAW18171.1 afla|EED56322.1 afum|EAL87533.1 agla|OJJ85935.1 akaw|GAA83931.1 aluc|OJZ90764.1 anid|CBF85742.1 anig|EHA19498.1 aory|EIT73405.1 asyd|OJJ64289.1 ater|EAU33833.1 atub|OJI82752.1 aver|OJI98382.1 awen|OJJ30518.1 azon|OJJ50199.1

Group4470: abra|OJJ77704.1 acar|OOF99036.1 acla|EAW11315.1 afis|EAW18183.1 afla|EED56304.1 afum|EAL87545.1 agla|OJJ85903.1 akaw|GAA83926.1 aluc|OJZ90769.1 anid|CBF85723.1 anig|EHA19504.1 aory|EIT73331.1 asyd|OJJ64277.1 ater|EAU33819.1 atub|OJI82757.1 aver|OJI98369.1 awen|OJJ30531.1 azon|OJJ50212.1

Group4471: abra|OJJ77708.1 acar|OOF99032.1 acla|EAW11317.1 afis|EAW18179.1 afla|EED56307.1 afum|EAL87540.1 agla|OJJ85910.1 akaw|GAA83922.1 aluc|OJZ90773.1 anid|CBF85729.1 anig|EHA19507.1 aory|EIT73380.1 asyd|OJJ64280.1 ater|EAU33821.1 atub|OJI82760.1 aver|OJI98372.1 awen|OJJ30527.1 azon|OJJ50208.1

Group4472: abra|OJJ77709.1 acar|OOF99031.1 acla|EAW11318.1 afis|EAW18178.1 afla|EED56306.1 afum|EAL87539.1 agla|OJJ85912.1 akaw|GAA83921.1 aluc|OJZ90775.1 anid|CBF85732.1 anig|EHA19508.1 aory|EIT73415.1 asyd|OJJ64283.1 ater|EAU33827.1 atub|OJI82761.1 aver|OJI98375.1 awen|OJJ30526.1 azon|OJJ50207.1

Group4473: abra|OJJ77710.1 acar|OOF99030.1 acla|EAW11319.1 afis|EAW18177.1 afla|EED56312.1 afum|EAL87538.1 agla|OJJ85913.1 akaw|GAA83920.1 aluc|OJZ90776.1 anid|CBF85733.1 anig|EHA19509.1 aory|EIT73345.1 asyd|OJJ64284.1 ater|EAU33828.1 atub|OJI82762.1 aver|OJI98376.1 awen|OJJ30525.1 azon|OJJ50206.1

Group4474: abra|OJJ77717.1 acar|OOF99022.1 acla|EAW11320.1 afis|EAW18176.1 afla|EED56313.1 afum|EAL87537.1 agla|OJJ85914.1 akaw|GAA83915.1 aluc|OJZ90782.1 anid|CBF85735.1 anig|EHA19515.1 aory|EIT73299.1 asyd|OJJ64285.1 ater|EAU33829.1 atub|OJI82769.1 aver|OJI98377.1 awen|OJJ30524.1 azon|OJJ50205.1

Group4475: abra|OJJ77718.1 acar|OOF99019.1 acla|EAW11321.1 afis|EAW18175.1 afla|EED56314.1 afum|EAL87536.1 agla|OJJ85915.1 akaw|GAA83914.1 aluc|OJZ90783.1 anid|CBF85737.1 anig|EHA19516.1 aory|EIT73381.1 asyd|OJJ64286.1 ater|EAU33830.1 atub|OJI82770.1 aver|OJI98378.1 awen|OJJ30523.1 azon|OJJ50204.1

Group4476: abra|OJJ77810.1 acar|OOF98957.1 acla|EAW11307.1 afis|EAW18194.1 afla|EED56284.1 afum|EBA27318.1 agla|OJJ85895.1 akaw|GAA83839.1 aluc|OJZ90877.1 anid|CBF86104.1 anig|EHA19597.1 aory|EIT73433.1 asyd|OJJ64482.1 ater|EAU33799.1 atub|OJI82874.1 aver|OJI98583.1 awen|OJJ30558.1 azon|OJJ45509.1

Group4477: abra|OJJ77811.1 acar|OOF98956.1 acla|EAW11306.1 afis|EAW18193.1 afla|EED56283.1 afum|EAL87559.1 agla|OJJ85893.1 akaw|GAA83838.1 aluc|OJZ90878.1 anid|CBF86108.1 anig|EHA19599.1

aory|EIT73325.1 asyd|OJJ64487.1 ater|EAU33798.1 atub|OJI82875.1 aver|OJI98588.1 awen|OJJ30559.1 azon|OJJ45508.1

Group4478: abra|OJJ77819.1 acar|OOF98949.1 acla|EAW11304.1 afis|EAW18191.1 afla|EED56281.1 afum|EAL87562.2 agla|OJJ85884.1 akaw|GAA83827.1 aluc|OJZ90891.1 anid|CBF86111.1 anig|EHA19610.1 aory|EIT73396.1 asyd|OJJ64489.1 ater|EAU33796.1 atub|OJI82885.1 aver|OJI98590.1 awen|OJJ30565.1 azon|OJJ45506.1

Group4479: abra|OJJ77821.1 acar|OOF98945.1 acla|EAW11303.1 afis|EAW18199.1 afla|EED56279.1 afum|EAL87564.1 agla|OJJ85883.1 akaw|GAA83824.1 aluc|OJZ90894.1 anid|CBF86115.1 anig|EHA19613.1 aory|EIT73401.1 asyd|OJJ64491.1 ater|EAU33795.1 atub|OJI82887.1 aver|OJI98593.1 awen|OJJ30567.1 azon|OJJ45505.1

Group4480: abra|OJJ77824.1 acar|OOF98944.1 acla|EAW11301.1 afis|EAW18201.1 afla|EED56277.1 afum|EAL87566.1 agla|OJJ85881.1 akaw|GAA83822.1 aluc|OJZ90897.1 anid|CBF86122.1 anig|EHA19615.1 aory|EIT73449.1 asyd|OJJ64496.1 ater|EAU33793.1 atub|OJI82889.1 aver|OJI98598.1 awen|OJJ30569.1 azon|OJJ45503.1

Group4481: abra|OJJ77826.1 acar|OOF98942.1 acla|EAW11299.1 afis|EAW18203.1 afla|EED56269.1 afum|EAL87568.1 agla|OJJ85879.1 akaw|GAA83820.1 aluc|OJZ90900.1 anid|CBF86119.1 anig|EHA19616.1 aory|EIT73413.1 asyd|OJJ64494.1 ater|EAU33786.1 atub|OJI82893.1 aver|OJI98596.1 awen|OJJ30571.1 azon|OJJ45501.1

Group4482: abra|OJJ77827.1 acar|OOF98941.1 acla|EAW11298.1 afis|EAW18204.1 afla|EED56268.1 afum|EAL87569.1 agla|OJJ85878.1 akaw|GAA83819.1 aluc|OJZ90901.1 anid|CBF86124.1 anig|EHA19617.1 aory|EIT73341.1 asyd|OJJ64497.1 ater|EAU33785.1 atub|OJI82894.1 aver|OJI98599.1 awen|OJJ30572.1 azon|OJJ45500.1

Group4483: abra|OJJ77830.1 acar|OOF98940.1 acla|EAW11296.1 afis|EAW18206.1 afla|EED56265.1 afum|EAL87571.1 agla|OJJ85876.1 akaw|GAA83817.1 aluc|OJZ90904.1 anid|CBF86132.1 anig|EHA19619.1 aory|EIT73429.1 asyd|OJJ64502.1 ater|EAU33783.1 atub|OJI82897.1 aver|OJI98601.1 awen|OJJ30578.1 azon|OJJ45498.1

Group4484: abra|OJJ77834.1 acar|OOF98937.1 acla|EAW11289.1 afis|EAW18210.1 afla|EED56260.1 afum|EAL87576.2 agla|OJJ85872.1 akaw|GAA83813.1 aluc|OJZ90908.1 anid|CBF86139.1 anig|EHA19623.1 aory|EIT73444.1 asyd|OJJ64506.1 ater|EAU33778.1 atub|OJI82901.1 aver|OJI98605.1 awen|OJJ30588.1 azon|OJJ50189.1

Group4485: abra|OJJ77841.1 acar|OOF98930.1 acla|EAW11282.1 afis|EAW18218.1 afla|EED56252.1 afum|EAL87584.1 agla|OJJ85864.1 akaw|GAA83806.1 aluc|OJZ90916.1 anid|CBF86148.1 anig|EHA19628.1 aory|EIT73348.1 asyd|OJJ64512.1 ater|EAU33773.1 atub|OJI82908.1 aver|OJI98610.1 awen|OJJ30596.1 azon|OJJ50172.1

Group4486: abra|OJJ77844.1 acar|OOF98928.1 acla|EAW11279.1 afis|EAW18221.1 afla|EED56249.1 afum|EAL87587.1 agla|OJJ85861.1 akaw|GAA83803.1 aluc|OJZ90919.1 anid|CBF86155.1 anig|EHA19631.1 aory|EIT73362.1 asyd|OJJ64515.1 ater|EAU33770.1 atub|OJI82911.1 aver|OJI98613.1 awen|OJJ30599.1 azon|OJJ50169.1

Group4487: abra|OJJ77845.1 acar|OOF98927.1 acla|EAW11278.1 afis|EAW18222.1 afla|EED56247.1 afum|EAL87588.1 agla|OJJ85860.1 akaw|GAA83802.1 aluc|OJZ90920.1 anid|CBF86153.1 anig|EHA19632.1 aory|EIT73414.1 asyd|OJJ64516.1 ater|EAU33769.1 atub|OJI82912.1 aver|OJI98614.1 awen|OJJ30600.1 azon|OJJ50168.1

Group4488: abra|OJJ77847.1 acar|OOF98925.1 acla|EAW11276.1 afis|EAW18224.1 afla|EED56245.1 afum|EAL87590.1 agla|OJJ85858.1 akaw|GAA83800.1 aluc|OJZ90922.1 anid|CBF86159.1 anig|EHA19634.1 aory|EIT73297.1 asyd|OJJ64518.1 ater|EAU33767.1 atub|OJI82914.1 aver|OJI98616.1 awen|OJJ30602.1 azon|OJJ50166.1

Group4489: abra|OJJ77851.1 acar|OOF98922.1 acla|EAW11275.1 afis|EAW18225.1 afla|EED56244.1 afum|EAL87591.1 agla|OJJ85857.1 akaw|GAA83797.1 aluc|OJZ90925.1 anid|CBF86161.1 anig|EHA19637.1 aory|EIT73369.1 asyd|OJJ64519.1 ater|EAU33766.1 atub|OJI82917.1 aver|OJI98617.1 awen|OJJ30603.1 azon|OJJ50165.1

Group4490: abra|OJJ77852.1 acar|OOF98921.1 acla|EAW11274.1 afis|EAW18226.1 afla|EED56243.1 afum|EAL87593.2 agla|OJJ85856.1 akaw|GAA83796.1 aluc|OJZ90926.1 anid|CBF86162.1 anig|EHA19638.1 aory|EIT73293.1 asyd|OJJ64520.1 ater|EAU33765.1 atub|OJI82918.1 aver|OJI98618.1 awen|OJJ30604.1 azon|OJJ50164.1

Group4491: abra|OJJ77853.1 acar|OOF98920.1 acla|EAW11273.1 afis|EAW18227.1 afla|EED56242.1 afum|EAL87594.1 agla|OJJ85855.1 akaw|GAA83795.1 aluc|OJZ90927.1 anid|CBF86164.1 anig|EHA19639.1 aory|EIT73313.1 asyd|OJJ64521.1 ater|EAU33764.1 atub|OJI82919.1 aver|OJI98619.1 awen|OJJ30605.1 azon|OJJ50163.1

Group4492: abra|OJJ77854.1 acar|OOF98919.1 acla|EAW11272.1 afis|EAW18228.1 afla|EED56241.1 afum|EAL87595.1 agla|OJJ85854.1 akaw|GAA83794.1 aluc|OJZ90928.1 anid|CBF86166.1 anig|EHA19640.1 aory|EIT73375.1 asyd|OJJ64522.1 ater|EAU33763.1 atub|OJI82920.1 aver|OJI98620.1 awen|OJJ30606.1 azon|OJJ50162.1

Group4493: abra|OJJ77857.1 acar|OOF98917.1 acla|EAW11269.1 afis|EAW18230.1 afla|EED56238.1 afum|EAL87597.1 agla|OJJ85844.1 akaw|GAA83792.1 aluc|OJZ90931.1 anid|CBF86172.1 anig|EHA19642.1 aory|EIT73450.1 asyd|OJJ64525.1 ater|EAU33761.1 atub|OJI82923.1 aver|OJI98623.1 awen|OJJ30609.1 azon|OJJ50159.1

Group4494: abra|OJJ77862.1 acar|OOF98911.1 acla|EAW11264.1 afis|EAW18234.1 afla|EED56232.1 afum|EBA27319.1 agla|OJJ85850.1 akaw|GAA83787.1 aluc|OJZ90937.1 anid|CBF86182.1 anig|EHA19645.1 aory|EIT73318.1 asyd|OJJ64530.1 ater|EAU33752.1 atub|OJI82928.1 aver|OJI98628.1 awen|OJJ30614.1 azon|OJJ48003.1

Group4495: abra|OJJ77864.1 acar|OOF98909.1 acla|EAW11262.1 afis|EAW18236.1 afla|EED56230.1 afum|EAL87603.1 agla|OJJ85838.1 akaw|GAA83785.1 aluc|OJZ90939.1 anid|CBF86185.1 anig|EHA19647.1 aory|EIT73458.1 asyd|OJJ64532.1 ater|EAU33749.1 atub|OJI82930.1 aver|OJI98630.1 awen|OJJ30616.1 azon|OJJ48001.1

Group4496: abra|OJJ77865.1 acar|OOF98907.1 acla|EAW11261.1 afis|EAW18237.1 afla|EED56229.1 afum|EAL87604.2 agla|OJJ85837.1 akaw|GAA83784.1 aluc|OJZ90940.1 anid|CBF86187.1 anig|EHA19648.1 aory|EIT73322.1 asyd|OJJ64534.1 ater|EAU33748.1 atub|OJI82931.1 aver|OJI98631.1 awen|OJJ30617.1 azon|OJJ48000.1

Group4497: abra|OJJ77866.1 acar|OOF98906.1 acla|EAW11259.1 afis|EAW18239.1 afla|EED56227.1 afum|EBA27321.1 agla|OJJ85831.1 akaw|GAA83782.1 aluc|OJZ90942.1 anid|CBF86191.1 anig|EHA19650.1 aory|EIT73427.1 asyd|OJJ64536.1 ater|EAU33744.1 atub|OJI82933.1 aver|OJI98633.1 awen|OJJ30619.1 azon|OJJ47998.1

Group4498: abra|OJJ77876.1 acar|OOF98895.1 acla|EAW06509.1 afis|EAW16222.1 afla|EED50188.1 afum|EAL88850.1 agla|OJJ86234.1 akaw|GAA83776.1 aluc|OJZ90951.1 anid|CBF86216.1 anig|EHA19657.1 aory|EIT77385.1 asyd|OJJ64558.1 ater|EAU33719.1 atub|OJI82942.1 aver|OJI98655.1 awen|OJJ30248.1 azon|OJJ43951.1

Group4499: abra|OJJ77878.1 acar|OOF98894.1 acla|EAW06602.1 afis|EAW16221.1 afla|EED50189.1 afum|EAL88849.1 agla|OJJ86233.1 akaw|GAA83775.1 aluc|OJZ90952.1 anid|CBF86215.1 anig|EHA19658.1 aory|EIT77381.1 asyd|OJJ64557.1 ater|EAU33720.1 atub|OJI82943.1 aver|OJI98654.1 awen|OJJ30250.1 azon|OJJ43952.1

Group4500: abra|OJJ77888.1 acar|OOF90363.1 afis|EAW17356.1 afla|EED49463.1 agla|OJJ82473.1 akaw|GAA82231.1 aluc|OJZ91917.1 anig|EHA19663.1 aory|EIT79554.1 asyd|OJJ56937.1 ater|EAU32565.1 aver|OJJ05741.1 awen|OJJ32672.1 aory|EIT75760.1 asyd|OJJ57437.1 awen|OJJ31967.1 afla|EED45648.1 asyd|OJJ57600.1

Group4501: abra|OJJ77890.1 acar|OOF98890.1 acla|EAW11245.1 afis|EAW18255.1 afla|EED56196.1 afum|EAL87618.1 agla|OJJ88189.1 akaw|GAA83768.1 aluc|OJZ90960.1 anid|CBF86206.1 anig|EHA19665.1 aory|EIT73437.1 asyd|OJJ64546.1 ater|EAU33727.1 atub|OJI82951.1 aver|OJI98642.1 awen|OJJ30629.1 azon|OJJ46567.1

Group4502: abra|OJJ77893.1 acar|OOF98889.1 acla|EAW11246.1 afis|EAW18254.1 afla|EED56198.1 afum|EAL87617.1 agla|OJJ88190.1 akaw|GAA83767.1 aluc|OJZ90963.1 anid|CBF86205.1 anig|EHA19667.1 aory|EIT73303.1 asyd|OJJ64545.1 ater|EAU33726.1 atub|OJI82954.1 aver|OJI98641.1 awen|OJJ30628.1 azon|OJJ47987.1

Group4503: abra|OJJ77897.1 acar|OOF98887.1 acla|EAW11248.1 afis|EAW18249.1 afla|EED56215.1 afum|EAL87614.1 agla|OJJ85821.1 akaw|GAA83765.1 aluc|OJZ90967.1 anid|CBF86200.1 anig|EHA19669.1 aory|EIT73361.1 asyd|OJJ64542.1 ater|EAU33731.1 atub|OJI82957.1 aver|OJI98638.1 awen|OJJ30625.1 azon|OJJ47993.1

Group4504: abra|OJJ77898.1 acar|OOF98886.1 acla|EAW11249.1 afis|EAW18248.1 afla|EED56216.1 afum|EAL87613.1 agla|OJJ85822.1 akaw|GAA83764.1 aluc|OJZ90968.1 anid|CBF86198.1 anig|EHA19670.1 aory|EIT73459.1 asyd|OJJ64541.1 ater|EAU33732.1 atub|OJI82958.1 aver|OJI98637.1 awen|OJJ30624.1 azon|OJJ47994.1

Group4505: abra|OJJ77902.1 acar|OOF98881.1 acla|EAW06512.1 afis|EAW16230.1 afla|EED53351.1 afum|EAL88853.1 agla|OJJ86245.1 akaw|GAA83760.1 aluc|OJZ90971.1 anid|CBF86223.1 anig|EHA19673.1 aory|EIT75232.1 asyd|OJJ64563.1 ater|EAU33714.1 atub|OJI82962.1 aver|OJI98660.1 awen|OJJ30232.1 azon|OJJ43949.1

Group4506: abra|OJJ77903.1 acar|OOF98879.1 acla|EAW06513.1 afis|EAW16231.1 afla|EED53350.1 afum|EAL88854.1 agla|OJJ86244.1 akaw|GAA83759.1 aluc|OJZ90972.1 anid|CBF86224.1 anig|EHA19674.1

aory|EIT74926.1 asyd|OJJ64564.1 ater|EAU33713.1 atub|OJI82963.1 aver|OJI98661.1 awen|OJJ30231.1 azon|OJJ43948.1

Group4507: abra|OJJ77910.1 acar|OOF98875.1 acla|EAW06518.1 afis|EAW16236.1 afla|EED53345.1 afum|EAL88857.1 agla|OJJ86239.1 akaw|GAA83749.1 aluc|OJZ90982.1 anid|CBF86232.1 anig|EHA19687.1 aory|EIT74884.1 asyd|OJJ64566.1 ater|EAU33708.1 atub|OJI82973.1 aver|OJI98663.1 awen|OJJ30224.1 azon|OJJ43944.1

Group4508: anig|EHA19693.1 abra|OJJ77921.1 acar|OOF98864.1 acla|EAW06687.1 afis|EAW16078.1 afla|EED48598.1 afum|EAL88710.1 agla|OJJ86111.1 akaw|GAA83742.1 aluc|OJZ90989.1 anid|CBF75119.1 aory|EIT73211.1 asyd|OJJ59993.1 ater|EAU33144.1 atub|OJI82980.1 aver|OJJ02298.1 awen|OJJ30355.1 azon|OJJ45377.1

Group4509: abra|OJJ77936.1 acar|OOF98872.1 acla|EAW06678.1 afis|EAW16087.1 afla|EED48608.1 afum|EAL88719.1 agla|OJJ86119.1 akaw|GAA83727.1 aluc|OJZ91004.1 anid|CBF75125.1 anig|EHA19709.1 aory|EIT73213.1 asyd|OJJ59984.1 ater|EAU33157.1 atub|OJI82995.1 aver|OJJ02307.1 awen|OJJ30344.1 azon|OJJ45369.1

Group4510: abra|OJJ77937.1 acar|OOF98873.1 acla|EAW06677.1 afis|EAW16088.1 afla|EED48609.1 afum|EAL88720.1 agla|OJJ86120.1 akaw|GAA83726.1 aluc|OJZ91005.1 anid|CBF75127.1 anig|EHA19710.1 aory|EIT73207.1 asyd|OJJ59983.1 ater|EAU33159.1 atub|OJI82996.1 aver|OJJ02308.1 awen|OJJ30343.1 azon|OJJ45368.1

Group4511: abra|OJJ77968.1 acar|OOF98847.1 acla|EAW06672.1 afis|EAW16093.1 afla|EED48629.1 afum|EAL88725.1 akaw|GAA83694.1 aluc|OJZ91038.1 anid|CBF85460.1 anig|EHA19737.1 aory|EIT73172.1 asyd|OJJ64101.1 ater|EAU33165.1 atub|OJI83034.1 aver|OJI98184.1 awen|OJJ30337.1 azon|OJJ45367.1 agla|OJJ86128.1

Group4512: abra|OJJ77971.1 acar|OOF98843.1 acla|EAW06670.1 afis|EAW16095.1 afla|EED48631.1 afum|EAL88727.1 agla|OJJ86126.1 akaw|GAA83691.1 aluc|OJZ91041.1 anid|CBF85455.1 anig|EHA19740.1 aory|EIT73094.1 asyd|OJJ64098.1 ater|EAU33171.1 atub|OJI83037.1 aver|OJI98181.1 awen|OJJ30335.1 azon|OJJ45363.1

Group4513: abra|OJJ77972.1 acar|OOF98842.1 acla|EAW06667.1 afis|EAW16098.1 afla|EED48633.1 afum|EAL88730.1 agla|OJJ86124.1 akaw|GAA83689.1 aluc|OJZ91043.1 anid|CBF85451.1 anig|EHA19742.1 aory|EIT73279.1 asyd|OJJ64096.1 ater|EAU33173.1 atub|OJI83039.1 aver|OJI98179.1 awen|OJJ30333.1 azon|OJJ45362.1

Group4514: abra|OJJ77973.1 acar|OOF98841.1 acla|EAW06665.1 afis|EAW16100.1 afla|EED48635.1 afum|EAL88732.1 agla|OJJ86122.1 anid|CBF85447.1 anig|EHA19743.1 aory|EIT73131.1 asyd|OJJ64095.1 ater|EAU33174.1 aver|OJI98177.1 awen|OJJ30332.1 azon|OJJ45360.1 akaw|GAA83687.1 aluc|OJZ91045.1 atub|OJI83041.1

Group4515: abra|OJJ77988.1 acar|OOF98835.1 acla|EAW06661.1 afis|EAW16104.1 afla|EED48640.1 afum|EAL88736.1 agla|OJJ86133.1 akaw|GAA85992.1 aluc|OJZ91063.1 anid|CBF85480.1 anig|EHA19767.1 aory|EIT73283.1 asyd|OJJ64108.1 ater|EAU33183.1 atub|OJI83059.1 aver|OJI98190.1 awen|OJJ30328.1 azon|OJJ45345.1

Group4516: abra|OJJ77989.1 acar|OOF98834.1 acla|EAW06660.1 afis|EAW16105.1 afla|EED48641.1 afum|EAL88737.1 agla|OJJ86134.1 akaw|GAA85991.1 aluc|OJZ91064.1 anid|CBF85482.1 anig|EHA19768.1 aory|EIT73197.1 asyd|OJJ64109.1 ater|EAU33184.1 atub|OJI83060.1 aver|OJI98191.1 awen|OJJ30327.1 azon|OJJ45344.1

Group4517: abra|OJJ77990.1 acar|OOF98833.1 acla|EAW06659.1 afis|EAW16106.1 afla|EED48642.1 afum|EAL88738.1 agla|OJJ86135.1 akaw|GAA85990.1 aluc|OJZ91066.1 anid|CBF85484.1 anig|EHA19769.1 aory|EIT73122.1 asyd|OJJ64110.1 ater|EAU33185.1 atub|OJI83061.1 aver|OJI98192.1 awen|OJJ30325.1 azon|OJJ45343.1

Group4518: abra|OJJ77991.1 acar|OOF98831.1 acla|EAW06658.1 afis|EAW16107.1 afla|EED48643.1 afum|EAL88739.2 agla|OJJ86136.1 akaw|GAA85989.1 aluc|OJZ91067.1 anid|CBF85486.1 anig|EHA19770.1 aory|EIT73085.1 asyd|OJJ64111.1 ater|EAU33190.1 atub|OJI83062.1 aver|OJI98193.1 awen|OJJ30324.1 azon|OJJ45342.1

Group4519: abra|OJJ77992.1 acar|OOF98830.1 acla|EAW06657.1 afis|EAW16108.1 afla|EED48644.1 afum|EAL88740.1 agla|OJJ86137.1 akaw|GAA85988.1 aluc|OJZ91068.1 anid|CBF85487.1 anig|EHA19771.1 aory|EIT73264.1 asyd|OJJ64112.1 ater|EAU33191.1 atub|OJI83063.1 aver|OJI98194.1 awen|OJJ30323.1 azon|OJJ45341.1

Group4520: abra|OJJ77993.1 acar|OOF98829.1 acla|EAW06655.1 afis|EAW16110.1 afla|EED48650.1 afum|EAL88741.1 agla|OJJ86145.1 akaw|GAA85987.1 aluc|OJZ91069.1 anid|CBF85492.1 anig|EHA19772.1 aory|EIT73074.1 asyd|OJJ64113.1 ater|EAU33193.1 atub|OJI83064.1 aver|OJI98196.1 awen|OJJ30320.1 azon|OJJ45339.1

Group4521: abra|OJJ77994.1 acar|OOF98828.1 acla|EAW06654.1 afis|EAW16111.1 afla|EED48651.1 afum|EAL88742.1 agla|OJJ86146.1 akaw|GAA85986.1 aluc|OJZ91070.1 anid|CBF85494.1 anig|EHA19773.1 aory|EIT73265.1 asyd|OJJ64114.1 ater|EAU33194.1 atub|OJI83065.1 aver|OJI98197.1 awen|OJJ30319.1 azon|OJJ45338.1

Group4522: abra|OJJ77995.1 acar|OOF98827.1 acla|EAW06653.1 afis|EAW16112.1 afla|EED48652.1 afum|EAL88743.1 agla|OJJ86151.1 akaw|GAA85985.1 aluc|OJZ91071.1 anid|CBF85496.1 anig|EHA19774.1 aory|EIT73201.1 asyd|OJJ64115.1 ater|EAU33195.1 atub|OJI83066.1 aver|OJI98198.1 awen|OJJ30318.1 azon|OJJ45337.1

Group4523: abra|OJJ78001.1 acar|OOF98821.1 acla|EAW06649.1 afis|EAW16116.1 afla|EED48657.1 afum|EAL88747.1 agla|OJJ86155.1 akaw|GAA85982.1 aluc|OJZ91077.1 anid|CBF85505.1 anig|EHA19778.1 aory|EIT73053.1 asyd|OJJ64119.1 ater|EAU33199.1 atub|OJI83072.1 aver|OJI98202.1 awen|OJJ30314.1 azon|OJJ45331.1

Group4524: abra|OJJ78004.1 acar|OOF98819.1 acla|EAW06646.1 afis|EAW16119.1 afla|EED48660.1 afum|EAL88750.1 agla|OJJ86160.1 akaw|GAA85979.1 aluc|OJZ91080.1 anid|CBF85543.1 anig|EHA19781.1 aory|EIT73098.1 asyd|OJJ64143.1 ater|EAU33202.1 atub|OJI83075.1 aver|OJI98224.1 awen|OJJ30311.1 azon|OJJ45328.1

Group4525: abra|OJJ78005.1 acar|OOF98818.1 acla|EAW06645.1 afis|EAW16120.1 afla|EED48661.1 afum|EAL88751.1 agla|OJJ86161.1 akaw|GAA85978.1 aluc|OJZ91081.1 anid|CBF85541.1 anig|EHA19782.1 aory|EIT73067.1 asyd|OJJ64142.1 ater|EAU33203.1 atub|OJI83077.1 aver|OJI98223.1 awen|OJJ30310.1 azon|OJJ45327.1

Group4526: abra|OJJ78008.1 acar|OOF98815.1 acla|EAW06642.1 afis|EAW16123.1 afla|EED48663.1 afum|EAL88753.1 agla|OJJ86164.1 akaw|GAA85975.1 aluc|OJZ91084.1 anid|CBF85534.1 anig|EHA19785.1 aory|EIT73192.1 asyd|OJJ64136.1 ater|EAU33206.1 atub|OJI83080.1 aver|OJI98219.1 awen|OJJ30307.1 azon|OJJ45310.1

Group4527: abra|OJJ78010.1 acar|OOF98813.1 acla|EAW06639.1 afis|EAW16126.1 afla|EED48666.1 afum|EAL88756.1 agla|OJJ86168.1 akaw|GAA85973.1 aluc|OJZ91086.1 anid|CBF85529.1 anig|EHA19787.1 aory|EIT73204.1 asyd|OJJ64133.1 ater|EAU33209.1 atub|OJI83082.1 aver|OJI98216.1 awen|OJJ30303.1 azon|OJJ45304.1

Group4528: abra|OJJ78011.1 acar|OOF98812.1 acla|EAW06638.1 afis|EAW16127.1 afla|EED48667.1 afum|EAL88757.1 agla|OJJ86169.1 akaw|GAA85972.1 aluc|OJZ91087.1 anid|CBF85527.1 anig|EHA19788.1 aory|EIT73187.1 asyd|OJJ64132.1 ater|EAU33210.1 atub|OJI83083.1 aver|OJI98215.1 awen|OJJ30302.1 azon|OJJ45303.1

Group4529: abra|OJJ78012.1 acar|OOF98811.1 acla|EAW06637.1 afis|EAW16128.1 afla|EED48668.1 afum|EAL88758.1 agla|OJJ86174.1 akaw|GAA85971.1 aluc|OJZ91088.1 anid|CBF85525.1 anig|EHA19789.1 aory|EIT73230.1 asyd|OJJ64131.1 ater|EAU33211.1 atub|OJI83085.1 aver|OJI98214.1 awen|OJJ30301.1 azon|OJJ45302.1

Group4530: abra|OJJ78013.1 acar|OOF98810.1 acla|EAW06635.1 afis|EAW16130.1 afla|EED48670.1 afum|EAL88760.1 agla|OJJ86175.1 akaw|GAA85970.1 aluc|OJZ91089.1 anid|CBF85523.1 anig|EHA19790.1 aory|EIT73154.1 asyd|OJJ64130.1 ater|EAU33212.1 atub|OJI83086.1 aver|OJI98213.1 awen|OJJ30299.1 azon|OJJ43987.1

Group4531: abra|OJJ78014.1 acar|OOF98809.1 acla|EAW06634.1 afis|EAW16131.1 afla|EED48669.1 afum|EAL88761.1 agla|OJJ86176.1 akaw|GAA85969.1 aluc|OJZ91090.1 anid|CBF85521.1 anig|EHA19791.1 aory|EIT73222.1 asyd|OJJ64128.1 ater|EAU33213.1 atub|OJI83087.1 aver|OJI98211.1 awen|OJJ30298.1 azon|OJJ43986.1

Group4532: abra|OJJ78018.1 acar|OOF98805.1 acla|EAW06629.1 afis|EAW16136.1 afla|EED48676.1 afum|EAL88766.2 agla|OJJ86181.1 akaw|GAA85965.1 aluc|OJZ91094.1 anid|CBF85512.1 anig|EHA19795.1 aory|EIT73051.1 asyd|OJJ64123.1 ater|EAU33218.1 atub|OJI83091.1 aver|OJI98206.1 awen|OJJ30292.1 azon|OJJ43981.1

Group4533: abra|OJJ78021.1 acar|OOF98803.1 acla|EAW06627.1 afis|EAW16138.1 afla|EED48678.1 afum|EAL88767.1 agla|OJJ86183.1 akaw|GAA85964.1 aluc|OJZ91096.1 anid|CBF85545.1 anig|EHA19797.1 aory|EIT73188.1 asyd|OJJ64144.1 ater|EAU33219.1 atub|OJI83094.1 aver|OJI98225.1 awen|OJJ30290.1 azon|OJJ43979.1

Group4534: abra|OJJ78023.1 acar|OOF98802.1 acla|EAW06626.1 afis|EAW16139.1 afla|EED48679.1 afum|EAL88768.1 agla|OJJ86185.1 akaw|GAA85963.1 aluc|OJZ91098.1 anid|CBF85546.1 anig|EHA19798.1 aory|EIT73152.1 asyd|OJJ64148.1 ater|EAU33220.1 atub|OJI83095.1 aver|OJI98228.1 awen|OJJ30289.1 azon|OJJ43978.1

Group4535: abra|OJJ78024.1 acar|OOF98801.1 acla|EAW06625.1 afis|EAW16140.1 afla|EED48681.1 afum|EAL88769.2 agla|OJJ86189.1 akaw|GAA85962.1 aluc|OJZ91099.1 anid|CBF85548.1 anig|EHA19799.1

aory|EIT73077.1 asyd|OJJ64149.1 ater|EAU33225.1 atub|OJI83096.1 aver|OJI98229.1 awen|OJJ30288.1 azon|OJJ43977.1

Group4536: abra|OJJ78026.1 acar|OOF98799.1 acla|EAW06623.1 afis|EAW16142.1 afla|EED48683.1 afum|EAL88771.1 agla|OJJ86190.1 akaw|GAA85960.1 aluc|OJZ91101.1 anid|CBF85551.1 anig|EHA19801.1 aory|EIT73175.1 asyd|OJJ64151.1 ater|EAU33227.1 atub|OJI83098.1 aver|OJI98231.1 awen|OJJ30286.1 azon|OJJ43975.1

Group4537: abra|OJJ78027.1 acar|OOF98796.1 acla|EAW06622.1 afis|EAW16143.1 afla|EED48684.1 afum|EAL88772.1 agla|OJJ86191.1 akaw|GAA85959.1 aluc|OJZ91102.1 anid|CBF79592.1 anig|EHA19802.1 aory|EIT73194.1 asyd|OJJ57983.1 ater|EAU33228.1 atub|OJI83099.1 aver|OJJ04418.1 awen|OJJ30285.1 azon|OJJ43974.1

Group4538: abra|OJJ78060.1 acar|OOF98793.1 acla|EAW13353.1 afis|EAW20467.1 afla|EED47863.1 afum|EAL84585.1 akaw|GAA85932.1 aluc|OJZ91131.1 anid|CBF79265.1 anig|EHA19830.1 aory|EIT77212.1 asyd|OJJ55757.1 atub|OJI83142.1 aver|OJJ06842.1 awen|OJJ33161.1 azon|OJJ51457.1 acar|OOF91287.1 awen|OJJ32661.1

Group4539: abra|OJJ78062.1 acar|OOF98792.1 acla|EAW06621.1 afis|EAW16144.1 afla|EED48686.1 afum|EAL88773.1 agla|OJJ86192.1 akaw|GAA85931.1 aluc|OJZ91133.1 anid|CBF79590.1 anig|EHA19832.1 aory|EIT73255.1 asyd|OJJ57981.1 ater|EAU33229.1 atub|OJI83143.1 aver|OJJ04419.1 awen|OJJ30283.1 azon|OJJ43973.1

Group4540: abra|OJJ78073.1 acar|OOF98790.1 acla|EAW15356.1 afis|EAW20356.1 afla|EED46131.1 afum|EAL85473.1 agla|OJJ81763.1 akaw|GAA85921.1 aluc|OJZ91146.1 anid|CBF74973.1 anig|EHA19844.1 aory|EIT76365.1 asyd|OJJ60088.1 ater|EAU34035.1 atub|OJI83160.1 aver|OJJ04175.1 awen|OJJ38942.1 azon|OJJ45059.1

Group4541: abra|OJJ78074.1 acar|OOF98789.1 afis|EAW16145.1 afla|EED48687.1 agla|OJJ86198.1 akaw|GAA85920.1 aluc|OJZ91147.1 anid|CBF79583.1 anig|EHA19845.1 aory|EIT73058.1 asyd|OJJ57977.1 ater|EAU33230.1 atub|OJI83161.1 aver|OJJ04423.1 awen|OJJ30281.1 azon|OJJ43972.1 acla|EAW06620.1 afum|EAL88774.1

Group4542: abra|OJJ78078.1 acar|OOF98786.1 acla|EAW06611.1 afis|EAW16204.1 afla|EED48697.1 afum|EAL88832.1 agla|OJJ86215.1 akaw|GAA85916.1 aluc|OJZ91150.1 anid|CBF79563.1 anig|EHA19848.1 aory|EIT73216.1 asyd|OJJ57913.1 ater|EAU33246.1 atub|OJI83164.1 aver|OJJ04480.1 awen|OJJ30268.1 azon|OJJ43964.1

Group4543: abra|OJJ78083.1 acar|OOF98783.1 acla|EAW06613.1 afis|EAW16177.1 afla|EED54377.1 afum|EAL88810.1 agla|OJJ86213.1 akaw|GAA85909.1 aluc|OJZ91157.1 anid|CBF79566.1 anig|EHA19853.1 aory|EIT75854.1 asyd|OJJ57917.1 ater|EAU33243.1 atub|OJI83172.1 aver|OJJ04474.1 awen|OJJ30272.1 azon|OJJ43966.1

Group4544: abra|OJJ78105.1 acar|OOF98760.1 acla|EAW06617.1 afis|EAW16148.1 afla|EED48691.1 afum|EBA27357.1 agla|OJJ86200.1 akaw|GAA85900.1 aluc|OJZ91169.1 anig|EHA19864.1 aory|EIT73263.1 ater|EAU33231.1 atub|OJI83184.1 awen|OJJ30276.1 asyd|OJJ57971.1 aver|OJJ04431.1 anig|EHA19863.1 aver|OJJ04432.1

Group4545: abra|OJJ78110.1 acar|OOF98756.1 acla|EAW06606.1 afis|EAW16217.1 afla|EED48711.1 afum|EAL88844.1 agla|OJJ86227.1 akaw|GAA85896.1 aluc|OJZ91175.1 anid|CBF79538.1 anig|EHA19867.1 aory|EIT73084.1 asyd|OJJ57899.1 ater|EAU33261.1 atub|OJI83188.1 aver|OJJ04496.1 awen|OJJ30255.1 azon|OJJ43955.1

Group4546: abra|OJJ78112.1 acar|OOF98755.1 acla|EAW06605.1 afis|EAW16218.1 afla|EED48712.1 afum|EAL88846.1 agla|OJJ86225.1 akaw|GAA85895.1 aluc|OJZ91177.1 anid|CBF79536.1 anig|EHA19868.1 aory|EIT73123.1 asyd|OJJ57898.1 ater|EAU33262.1 atub|OJI83189.1 aver|OJJ04497.1 awen|OJJ30254.1 azon|OJJ43957.1

Group4547: abra|OJJ78113.1 acar|OOF98753.1 acla|EAW07002.1 afis|EAW16619.1 afla|EED48441.1 afum|EAL89218.1 akaw|GAA85894.1 aluc|OJZ91178.1 anig|EHA19869.1 aory|EIT81677.1 asyd|OJJ60003.1 ater|EAU30738.1 atub|OJI83190.1 aver|OJJ02288.1 awen|OJJ30079.1 azon|OJJ42185.1 aory|EIT74880.1 awen|OJJ37785.1

Group4548: abra|OJJ78189.1 acar|OOF98670.1 acla|EAW06798.1 afis|EAW16347.1 afla|EED51752.1 afum|EAL88972.1 agla|OJJ88241.1 akaw|GAA85840.1 aluc|OJZ91260.1 anid|CBF89212.1 anig|EHA19948.1 aory|EIT80660.1 asyd|OJJ62523.1 ater|EAU36051.1 atub|OJI83249.1 aver|OJI96642.1 awen|OJJ37750.1 azon|OJJ47558.1

Group4549: abra|OJJ78192.1 acar|OOF98665.1 afum|EAL85073.2 akaw|GAA85837.1 aluc|OJZ91263.1 anid|CBF87346.1 asyd|OJJ56202.1 atub|OJI83252.1 aver|OJJ06457.1 acar|OOF90595.1 afis|EAW19818.1 afla|EED53224.1 aory|EIT75036.1 aver|OJJ05129.1 azon|OJJ42171.1 afla|EED53225.1 anig|EHA19951.1 anid|CBF75950.1

Group4550: abra|OJJ78206.1 acar|OOF98648.1 acla|EAW11242.1 afis|EAW18261.1 afla|EED48720.1 afum|EAL87625.1 agla|OJJ88165.1 akaw|GAA85820.1 aluc|OJZ91279.1 anid|CBF79529.1 anig|EHA19964.1 aory|EIT73088.1 asyd|OJJ57894.1 ater|EAU33269.1 atub|OJI83269.1 aver|OJJ04501.1 awen|OJJ30635.1 azon|OJJ47983.1

Group4551: abra|OJJ78213.1 acar|OOF98642.1 acla|EAW11237.1 afis|EAW18268.1 afla|EED48733.1 afum|EAL87633.1 agla|OJJ88159.1 akaw|GAA85814.1 aluc|OJZ91285.1 anid|CBF79516.1 anig|EHA19970.1 aory|EIT73048.1 asyd|OJJ57888.1 ater|EAU33275.1 atub|OJI83275.1 aver|OJJ04506.1 awen|OJJ30642.1 azon|OJJ47973.1

Group4552: abra|OJJ78216.1 acar|OOF98640.1 acla|EAW11234.1 afis|EAW18271.1 afla|EED48737.1 afum|EAL87636.1 agla|OJJ88156.1 akaw|GAA85811.1 aluc|OJZ91288.1 anid|CBF79510.1 anig|EHA19973.1 aory|EIT73125.1 asyd|OJJ57885.1 ater|EAU33277.1 atub|OJI83278.1 aver|OJJ04509.1 awen|OJJ30644.1 azon|OJJ47971.1

Group4553: abra|OJJ78217.1 acar|OOF98639.1 acla|EAW11232.1 afis|EAW18272.1 afla|EED48738.1 afum|EAL87637.1 agla|OJJ88155.1 akaw|GAA85807.1 aluc|OJZ91291.1 anid|CBF79508.1 anig|EHA19974.1 aory|EIT73235.1 asyd|OJJ57884.1 ater|EAU33278.1 atub|OJI83282.1 aver|OJJ04510.1 awen|OJJ30645.1 azon|OJJ47970.1

Group4554: abra|OJJ78218.1 acar|OOF98638.1 acla|EAW11231.1 afis|EAW18273.1 afla|EED48739.1 afum|EAL87638.1 agla|OJJ88154.1 akaw|GAA85806.1 aluc|OJZ91292.1 anid|CBF79507.1 anig|EHA19975.1 aory|EIT73224.1 asyd|OJJ57883.1 ater|EAU33279.1 atub|OJI83283.1 aver|OJJ04511.1 awen|OJJ30646.1 azon|OJJ47969.1

Group4555: abra|OJJ78220.1 acar|OOF98636.1 acla|EAW11229.1 afis|EAW18274.1 afla|EED48741.1 afum|EAL87640.1 agla|OJJ88152.1 akaw|GAA85805.1 aluc|OJZ91294.1 anid|CBF79503.1 anig|EHA19977.1 aory|EIT73281.1 asyd|OJJ57881.1 ater|EAU33280.1 atub|OJI83285.1 aver|OJJ04513.1 awen|OJJ30647.1 azon|OJJ47968.1

Group4556: abra|OJJ78221.1 acar|OOF98635.1 acla|EAW11228.1 afis|EAW18275.1 afla|EED48742.1 afum|EAL87641.1 agla|OJJ88151.1 akaw|GAA85804.1 aluc|OJZ91296.1 anid|CBF79499.1 anig|EHA19978.1 aory|EIT73223.1 asyd|OJJ57880.1 ater|EAU33281.1 atub|OJI83286.1 aver|OJJ04514.1 awen|OJJ30648.1 azon|OJJ47967.1

Group4557: abra|OJJ78226.1 acar|OOF98632.1 acla|EAW09284.1 afis|EAW21863.1 afla|EED46448.1 afum|EAL92877.2 agla|OJJ84270.1 akaw|GAA85799.1 aluc|OJZ91301.1 anid|CBF76755.1 anig|EHA19983.1 aory|EIT72377.1 asyd|OJJ61262.1 ater|EAU30861.1 atub|OJI83292.1 aver|OJJ00588.1 awen|OJJ34153.1 azon|OJJ43475.1

Group4558: abra|OJJ78229.1 acar|OOF98630.1 acla|EAW09280.1 afis|EAW21867.1 afla|EED46442.1 afum|EAL92881.1 agla|OJJ84268.1 akaw|GAA85795.1 aluc|OJZ91304.1 anid|CBF76763.1 anig|EHA19986.1 aory|EIT72329.1 asyd|OJJ61265.1 ater|EAU30864.1 atub|OJI83295.1 aver|OJJ00592.1 awen|OJJ34156.1 azon|OJJ43473.1

Group4559: abra|OJJ78230.1 acar|OOF98629.1 acla|EAW09279.1 afis|EAW21868.1 afla|EED46441.1 afum|EAL92882.1 agla|OJJ84267.1 akaw|GAA85794.1 aluc|OJZ91305.1 anid|CBF76765.1 anig|EHA19987.1 aory|EIT72366.1 asyd|OJJ61266.1 ater|EAU30866.1 atub|OJI83296.1 aver|OJJ00593.1 awen|OJJ34157.1 azon|OJJ43472.1

Group4560: abra|OJJ78231.1 acar|OOF98628.1 acla|EAW09278.1 afis|EAW21869.1 afla|EED46440.1 afum|EAL92883.1 agla|OJJ84264.1 akaw|GAA85793.1 aluc|OJZ91306.1 anid|CBF76767.1 anig|EHA19988.1 aory|EIT72428.1 asyd|OJJ61267.1 ater|EAU30867.1 atub|OJI83297.1 aver|OJJ00594.1 awen|OJJ34158.1 azon|OJJ43471.1

Group4561: abra|OJJ78233.1 acar|OOF98627.1 acla|EAW09277.1 afis|EAW21870.1 afla|EED46438.1 afum|EAL92884.1 agla|OJJ84262.1 akaw|GAA85792.1 aluc|OJZ91307.1 anid|CBF76769.1 anig|EHA19991.1 aory|EIT72393.1 asyd|OJJ61268.1 ater|EAU30868.1 atub|OJI83300.1 aver|OJJ00597.1 awen|OJJ34159.1 azon|OJJ43470.1

Group4562: abra|OJJ78234.1 acar|OOF98626.1 acla|EAW09276.1 afis|EAW21871.1 afla|EED46437.1 afum|EAL92885.1 agla|OJJ84261.1 akaw|GAA85791.1 aluc|OJZ91308.1 anid|CBF76771.1 anig|EHA19992.1 aory|EIT72319.1 asyd|OJJ61269.1 ater|EAU30869.1 atub|OJI83301.1 aver|OJJ00598.1 awen|OJJ34160.1 azon|OJJ43469.1

Group4563: abra|OJJ78237.1 acar|OOF98623.1 acla|EAW09273.1 afis|EAW21875.1 afla|EED46433.1 afum|EAL92889.2 agla|OJJ84257.1 akaw|GAA85788.1 aluc|OJZ91311.1 anid|CBF76776.1 anig|EHA19995.1 aory|EIT72389.1 asyd|OJJ61273.1 ater|EAU30874.1 atub|OJI83304.1 aver|OJJ00601.1 awen|OJJ34164.1 azon|OJJ43466.1

Group4564: abra|OJJ78238.1 acar|OOF98622.1 acla|EAW09272.1 afis|EAW21876.1 afla|EED46432.1 afum|EAL92890.1 agla|OJJ84256.1 akaw|GAA85787.1 aluc|OJZ91312.1 anid|CBF76778.1 anig|EHA19996.1

aory|EIT72368.1 asyd|OJJ61274.1 ater|EAU30875.1 atub|OJI83305.1 aver|OJJ00602.1 awen|OJJ34165.1 azon|OJJ43465.1

Group4565: abra|OJJ78239.1 acar|OOF98621.1 acla|EAW09271.1 afis|EAW21877.1 afla|EED46431.1 afum|EAL92891.2 agla|OJJ84255.1 akaw|GAA85786.1 aluc|OJZ91313.1 anid|CBF76780.1 anig|EHA19997.1 aory|EIT72420.1 asyd|OJJ61275.1 ater|EAU30876.1 atub|OJI83306.1 aver|OJJ00603.1 awen|OJJ34166.1 azon|OJJ43464.1

Group4566: abra|OJJ78241.1 acar|OOF98619.1 acla|EAW09269.1 afis|EAW21879.1 afla|EED46429.1 afum|EAL92893.1 agla|OJJ84253.1 akaw|GAA85783.1 aluc|OJZ91315.1 anid|CBF76784.1 anig|EHA19998.1 aory|EIT72355.1 asyd|OJJ61277.1 ater|EAU30877.1 atub|OJI83309.1 aver|OJJ00605.1 awen|OJJ34168.1 azon|OJJ43462.1

Group4567: abra|OJJ78242.1 acar|OOF98618.1 acla|EAW09268.1 afis|EAW21880.1 afla|EED46428.1 afum|EAL92894.2 agla|OJJ84252.1 akaw|GAA85782.1 aluc|OJZ91316.1 anid|CBF76786.1 anig|EHA19999.1 aory|EIT72330.1 asyd|OJJ61278.1 ater|EAU30878.1 atub|OJI83310.1 aver|OJJ00606.1 awen|OJJ34169.1 azon|OJJ43461.1

Group4568: abra|OJJ78244.1 acar|OOF98616.1 acla|EAW09265.1 afis|EAW21883.1 afla|EED46424.1 afum|EAL92897.1 agla|OJJ84251.1 akaw|GAA85780.1 aluc|OJZ91318.1 anid|CBF76788.1 anig|EHA20001.1 aory|EIT72383.1 asyd|OJJ61280.1 ater|EAU30881.1 atub|OJI83312.1 aver|OJJ00607.1 awen|OJJ34172.1 azon|OJJ43460.1

Group4569: abra|OJJ78247.1 acar|OOF98615.1 acla|EAW09263.1 afis|EAW21884.1 afla|EED46423.1 afum|EAL92898.1 agla|OJJ84249.1 akaw|GAA85779.1 aluc|OJZ91321.1 anid|CBF76790.1 anig|EHA20003.1 aory|EIT72443.1 asyd|OJJ61281.1 ater|EAU30882.1 atub|OJI83314.1 aver|OJJ00608.1 awen|OJJ34173.1 azon|OJJ43459.1

Group4570: abra|OJJ78249.1 acar|OOF98614.1 acla|EAW09262.1 afis|EAW21885.1 afla|EED46422.1 afum|EAL92899.1 agla|OJJ84248.1 akaw|GAA85778.1 aluc|OJZ91322.1 anid|CBF76792.1 anig|EHA20004.1 aory|EIT72353.1 asyd|OJJ61282.1 ater|EAU30883.1 atub|OJI83315.1 aver|OJJ00609.1 awen|OJJ34174.1 azon|OJJ43458.1

Group4571: abra|OJJ78250.1 acar|OOF98612.1 acla|EAW09261.1 afis|EAW21886.1 afla|EED46421.1 afum|EAL92900.1 agla|OJJ84247.1 akaw|GAA85777.1 aluc|OJZ91323.1 anid|CBF76794.1 anig|EHA20005.1 aory|EIT72442.1 asyd|OJJ61283.1 ater|EAU30884.1 atub|OJI83316.1 aver|OJJ00610.1 awen|OJJ34175.1 azon|OJJ43457.1

Group4572: abra|OJJ78252.1 acar|OOF98611.1 acla|EAW09259.1 afis|EAW21888.1 afla|EED46419.1 afum|EAL92902.1 agla|OJJ84245.1 akaw|GAA85775.1 aluc|OJZ91325.1 anid|CBF76798.1 anig|EHA20007.1 aory|EIT72406.1 asyd|OJJ61285.1 ater|EAU30885.1 atub|OJI83318.1 aver|OJJ00611.1 awen|OJJ34177.1 azon|OJJ43455.1

Group4573: abra|OJJ78253.1 acar|OOF98610.1 acla|EAW09258.1 afis|EAW21889.1 afla|EED46418.1 afum|EAL92903.1 agla|OJJ84244.1 akaw|GAA85774.1 aluc|OJZ91326.1 anid|CBF76800.1 anig|EHA20008.1 aory|EIT72432.1 asyd|OJJ61286.1 ater|EAU30886.1 atub|OJI83319.1 aver|OJJ00612.1 awen|OJJ34178.1 azon|OJJ43454.1

Group4574: abra|OJJ78255.1 acar|OOF98607.1 acla|EAW09254.1 afis|EAW21893.1 afla|EED46415.1 afum|EAL92907.1 agla|OJJ84239.1 akaw|GAA85771.1 aluc|OJZ91329.1 anid|CBF76806.1 anig|EHA20011.1 aory|EIT72313.1 asyd|OJJ61289.1 ater|EAU30889.1 atub|OJI83322.1 aver|OJJ00615.1 awen|OJJ34182.1 azon|OJJ43450.1

Group4575: abra|OJJ78256.1 acar|OOF98606.1 acla|EAW09253.1 afis|EAW21894.1 afla|EED46414.1 afum|EAL92908.1 agla|OJJ84238.1 akaw|GAA85770.1 aluc|OJZ91330.1 anid|CBF76808.1 anig|EHA20012.1 aory|EIT72378.1 asyd|OJJ61290.1 ater|EAU30890.1 atub|OJI83323.1 aver|OJJ00616.1 awen|OJJ34183.1 azon|OJJ43449.1

Group4576: abra|OJJ78260.1 acar|OOF98600.1 acla|EAW09250.1 afis|EAW21897.1 afla|EED46411.1 afum|EAL92910.1 agla|OJJ84227.1 akaw|GAA85765.1 aluc|OJZ91335.1 anid|CBF76814.1 anig|EHA20016.1 aory|EIT72440.1 asyd|OJJ61293.1 ater|EAU30892.1 atub|OJI83327.1 aver|OJJ00619.1 awen|OJJ34186.1 azon|OJJ43447.1

Group4577: abra|OJJ78261.1 acar|OOF98599.1 acla|EAW09249.1 afis|EAW21898.1 afla|EED46408.1 afum|EAL92911.2 agla|OJJ84228.1 akaw|GAA85764.1 aluc|OJZ91336.1 anid|CBF76816.1 anig|EHA20017.1 aory|EIT72346.1 asyd|OJJ61294.1 ater|EAU30893.1 atub|OJI83328.1 aver|OJJ00620.1 awen|OJJ34187.1 azon|OJJ43446.1

Group4578: abra|OJJ78263.1 acar|OOF98598.1 acla|EAW09248.1 afis|EAW21899.1 afla|EED46407.1 afum|EAL92912.1 agla|OJJ84229.1 akaw|GAA85762.1 aluc|OJZ91338.1 anid|CBF76818.1 anig|EHA20018.1 aory|EIT72328.1 asyd|OJJ61295.1 ater|EAU30894.1 atub|OJI83330.1 aver|OJJ00621.1 awen|OJJ34188.1 azon|OJJ43445.1

Group4579: abra|OJJ78264.1 acar|OOF98597.1 acla|EAW09247.1 afis|EAW21900.1 afla|EED46406.1 afum|EAL92913.1 agla|OJJ84230.1 akaw|GAA85761.1 aluc|OJZ91339.1 anid|CBF76820.1 anig|EHA20019.1 aory|EIT72352.1 asyd|OJJ61296.1 ater|EAU30895.1 atub|OJI83331.1 aver|OJJ00622.1 awen|OJJ34189.1 azon|OJJ43444.1

Group4580: abra|OJJ78266.1 acar|OOF98596.1 acla|EAW09246.1 afis|EAW21901.1 afla|EED46405.1 afum|EAL92914.1 agla|OJJ84231.1 akaw|GAA85760.1 aluc|OJZ91340.1 anid|CBF76822.1 anig|EHA20020.1 aory|EIT72444.1 asyd|OJJ61297.1 ater|EAU30896.1 atub|OJI83332.1 aver|OJJ00623.1 awen|OJJ34190.1 azon|OJJ43443.1

Group4581: abra|OJJ78267.1 acar|OOF98595.1 acla|EAW09245.1 afis|EAW21902.1 afla|EED46403.1 afum|EAL92915.1 agla|OJJ84232.1 akaw|GAA85759.1 aluc|OJZ91341.1 anid|CBF76824.1 anig|EHA20021.1 aory|EIT72339.1 asyd|OJJ61298.1 ater|EAU30897.1 atub|OJI83333.1 aver|OJJ00624.1 awen|OJJ34191.1 azon|OJJ43442.1

Group4582: abra|OJJ78268.1 acar|OOF98594.1 acla|EAW09244.1 afis|EAW21903.1 afla|EED46402.1 afum|EAL92916.1 agla|OJJ84233.1 akaw|GAA85758.1 aluc|OJZ91342.1 anid|CBF76826.1 anig|EHA20022.1 aory|EIT72439.1 asyd|OJJ61299.1 ater|EAU30898.1 atub|OJI83334.1 aver|OJJ00625.1 awen|OJJ34192.1 azon|OJJ43441.1

Group4583: abra|OJJ78275.1 acar|OOF98592.1 acla|EAW09239.1 afis|EAW21909.1 afla|EED46641.1 afum|EAL92921.1 agla|OJJ84222.1 akaw|GAA85751.1 aluc|OJZ91350.1 anid|CBF76837.1 anig|EHA20028.1 aory|EIT77855.1 asyd|OJJ61305.1 ater|EAU30909.1 atub|OJI83342.1 aver|OJJ00630.1 awen|OJJ34199.1 azon|OJJ43435.1

Group4584: abra|OJJ78277.1 acar|OOF98590.1 acla|EAW09237.1 afis|EAW21911.1 afla|EED46639.1 afum|EAL92923.1 agla|OJJ84220.1 akaw|GAA85749.1 aluc|OJZ91352.1 anid|CBF76842.1 anig|EHA20030.1 aory|EIT77915.1 asyd|OJJ61307.1 ater|EAU30911.1 atub|OJI83344.1 aver|OJJ00632.1 awen|OJJ34201.1 azon|OJJ43433.1

Group4585: abra|OJJ78278.1 acar|OOF98588.1 acla|EAW09236.1 afis|EAW21912.1 afla|EED46638.1 afum|EAL92924.1 agla|OJJ84219.1 akaw|GAA85748.1 aluc|OJZ91353.1 anid|CBF76844.1 anig|EHA20031.1 aory|EIT77829.1 asyd|OJJ61308.1 ater|EAU30912.1 atub|OJI83345.1 aver|OJJ00633.1 awen|OJJ34202.1 azon|OJJ43432.1

Group4586: abra|OJJ78280.1 acar|OOF98589.1 acla|EAW09235.1 afis|EAW21913.1 afla|EED46637.1 afum|EAL92925.1 agla|OJJ84210.1 akaw|GAA85747.1 aluc|OJZ91354.1 anid|CBF76846.1 anig|EHA20032.1 aory|EIT77951.1 asyd|OJJ61309.1 ater|EAU30914.1 atub|OJI83347.1 aver|OJJ00634.1 awen|OJJ34203.1 azon|OJJ43431.1

Group4587: abra|OJJ78282.1 acar|OOF98586.1 acla|EAW09233.1 afis|EAW21915.1 afla|EED46634.1 afum|EAL92927.1 agla|OJJ84208.1 akaw|GAA85745.1 aluc|OJZ91356.1 anid|CBF76850.1 anig|EHA20034.1 aory|EIT77867.1 asyd|OJJ61311.1 ater|EAU30916.1 atub|OJI83349.1 aver|OJJ00636.1 awen|OJJ34205.1 azon|OJJ43429.1

Group4588: abra|OJJ78284.1 acar|OOF98584.1 acla|EAW09231.1 afis|EAW21917.1 afla|EED46632.1 afum|EAL92929.1 agla|OJJ84206.1 akaw|GAA85743.1 aluc|OJZ91358.1 anid|CBF76854.1 anig|EHA20036.1 aory|EIT77843.1 asyd|OJJ61313.1 ater|EAU30919.1 atub|OJI83351.1 aver|OJJ00638.1 awen|OJJ34207.1 azon|OJJ43427.1

Group4589: abra|OJJ78285.1 acar|OOF98583.1 acla|EAW09230.1 afis|EAW21918.1 afum|EAL92930.1 agla|OJJ84205.1 akaw|GAA85742.1 aluc|OJZ91359.1 anid|CBF76856.1 anig|EHA20037.1 asyd|OJJ61314.1 ater|EAU30920.1 atub|OJI83352.1 aver|OJJ00639.1 awen|OJJ34208.1 azon|OJJ43426.1 afla|EED46631.1 aory|EIT77816.1

Group4590: abra|OJJ78287.1 acar|OOF98581.1 acla|EAW09228.1 afis|EAW21920.1 afla|EED46630.1 afum|EAL92932.1 agla|OJJ84203.1 akaw|GAA85740.1 aluc|OJZ91361.1 anid|CBF76860.1 anig|EHA20039.1 aory|EIT77823.1 asyd|OJJ61316.1 ater|EAU30925.1 atub|OJI83354.1 aver|OJJ00641.1 awen|OJJ34210.1 azon|OJJ43424.1

Group4591: abra|OJJ78288.1 acar|OOF98579.1 acla|EAW09227.1 afis|EAW21921.1 afla|EED46629.1 afum|EAL92934.1 agla|OJJ84202.1 akaw|GAA85739.1 aluc|OJZ91362.1 anid|CBF76862.1 anig|EHA20040.1 aory|EIT77918.1 asyd|OJJ61317.1 ater|EAU30926.1 atub|OJI83355.1 aver|OJJ00642.1 awen|OJJ34211.1 azon|OJJ43423.1

Group4592: abra|OJJ78289.1 acar|OOF98580.1 acla|EAW09226.1 afis|EAW21922.1 afla|EED46628.1 afum|EAL92935.1 agla|OJJ84201.1 akaw|GAA85738.1 aluc|OJZ91363.1 anid|CBF76864.1 anig|EHA20041.1 aory|EIT77831.1 asyd|OJJ61318.1 ater|EAU30927.1 atub|OJI83356.1 aver|OJJ00643.1 awen|OJJ34212.1 azon|OJJ43422.1

Group4593: abra|OJJ78290.1 acar|OOF98578.1 acla|EAW09225.1 afis|EAW21923.1 afla|EED46627.1 afum|EAL92936.1 agla|OJJ84200.1 akaw|GAA85737.1 aluc|OJZ91364.1 anid|CBF76866.1 anig|EHA20042.1

aory|EIT77896.1 asyd|OJJ61319.1 ater|EAU30928.1 atub|OJI83357.1 aver|OJJ00644.1 awen|OJJ34213.1 azon|OJJ43421.1

Group4594: abra|OJJ78292.1 acar|OOF98576.1 afis|EAW21924.1 afla|EED46615.1 afum|EAL92937.1 akaw|GAA85735.1 aluc|OJZ91366.1 anig|EHA20044.1 aory|EIT77890.1 ater|EAU30929.1 atub|OJI83359.1 aver|OJJ00647.1 azon|OJJ43420.1 acla|EAW09224.1 agla|OJJ84199.1 anid|CBF76872.1 asyd|OJJ61322.1 awen|OJJ34214.1

Group4595: abra|OJJ78293.1 acar|OOF98575.1 acla|EAW09223.1 afis|EAW21925.1 afla|EED46616.1 afum|EAL92938.1 agla|OJJ84198.1 akaw|GAA85734.1 aluc|OJZ91367.1 anid|CBF76882.1 anig|EHA20045.1 aory|EIT77869.1 asyd|OJJ61327.1 ater|EAU30930.1 atub|OJI83360.1 aver|OJJ00652.1 awen|OJJ34215.1 azon|OJJ43419.1

Group4596: abra|OJJ78298.1 acar|OOF98571.1 acla|EAW09218.1 afis|EAW21930.1 afla|EED46623.1 afum|EAL92942.1 agla|OJJ84191.1 akaw|GAA85729.1 aluc|OJZ91372.1 anid|CBF76884.1 anig|EHA20050.1 aory|EIT77852.1 asyd|OJJ61328.1 ater|EAU30936.1 atub|OJI83365.1 aver|OJJ00654.1 awen|OJJ34220.1 azon|OJJ43414.1

Group4597: abra|OJJ78299.1 acar|OOF98567.1 acla|EAW09216.1 afis|EAW21931.1 afla|EED46626.1 afum|EAL92943.2 agla|OJJ84190.1 akaw|GAA85727.1 aluc|OJZ91376.1 anid|CBF76885.1 anig|EHA20051.1 aory|EIT77815.1 asyd|OJJ61330.1 ater|EAU30937.1 atub|OJI83366.1 aver|OJJ00655.1 awen|OJJ34222.1 azon|OJJ43413.1

Group4598: abra|OJJ78301.1 acar|OOF98565.1 acla|EAW09214.1 afis|EAW21933.1 afla|EED46353.1 afum|EAL92946.1 agla|OJJ84187.1 akaw|GAA85725.1 aluc|OJZ91378.1 anid|CBF76887.1 anig|EHA20053.1 aory|EIT81186.1 asyd|OJJ61333.1 ater|EAU30938.1 atub|OJI83371.1 aver|OJJ00657.1 awen|OJJ34224.1 azon|OJJ43411.1

Group4599: abra|OJJ78302.1 acar|OOF98564.1 acla|EAW09213.1 afis|EAW21934.1 afla|EED46354.1 afum|EAL92947.1 agla|OJJ84186.1 akaw|GAA85724.1 aluc|OJZ91379.1 anid|CBF76889.1 anig|EHA20054.1 aory|EIT81211.1 asyd|OJJ61334.1 ater|EAU30939.1 atub|OJI83372.1 aver|OJJ00658.1 awen|OJJ34225.1 azon|OJJ43410.1

Group4600: abra|OJJ78308.1 acar|OOF98558.1 acla|EAW09212.1 afis|EAW21935.1 afla|EED46652.1 afum|EAL92948.2 agla|OJJ84185.1 akaw|GAA85721.1 aluc|OJZ91383.1 anid|CBF76899.1 anig|EHA20060.1 aory|EIT77952.1 asyd|OJJ61338.1 ater|EAU30941.1 atub|OJI83376.1 aver|OJJ00662.1 awen|OJJ34226.1 azon|OJJ43409.1

Group4601: abra|OJJ78319.1 acar|OOF98556.1 acla|EAW09206.1 afis|EAW21941.1 afla|EED46658.1 afum|EAL92955.1 agla|OJJ84182.1 akaw|GAA85711.1 aluc|OJZ91392.1 anid|CBF84078.1 anig|EHA20070.1 aory|EIT77820.1 asyd|OJJ61342.1 ater|EAU30946.1 atub|OJI83385.1 aver|OJJ00666.1 awen|OJJ34232.1 azon|OJJ43403.1

Group4602: abra|OJJ78323.1 acar|OOF98551.1 acla|EAW09202.1 afis|EAW21945.1 afla|EED46662.1 afum|EAL92959.1 agla|OJJ84179.1 akaw|GAA85705.1 aluc|OJZ91398.1 anid|CBF84050.1 anig|EHA20075.1 aory|EIT77962.1 asyd|OJJ61343.1 ater|EAU30993.1 atub|OJI83390.1 aver|OJJ00667.1 awen|OJJ34236.1 azon|OJJ43400.1

Group4603: abra|OJJ78325.1 acar|OOF98550.1 acla|EAW09201.1 afis|EAW21946.1 afla|EED46663.1 afum|EAL92960.1 agla|OJJ84174.1 akaw|GAA85704.1 aluc|OJZ91399.1 anid|CBF84059.1 anig|EHA20076.1 aory|EIT77808.1 asyd|OJJ61345.1 ater|EAU30994.1 atub|OJI83391.1 aver|OJJ00670.1 awen|OJJ34237.1 azon|OJJ43399.1

Group4604: abra|OJJ78327.1 acar|OOF98548.1 acla|EAW09199.1 afis|EAW21948.1 afla|EED46666.1 afum|EAL92962.1 agla|OJJ84172.1 akaw|GAA85702.1 aluc|OJZ91401.1 anid|CBF84062.1 anig|EHA20078.1 aory|EIT77953.1 asyd|OJJ61347.1 ater|EAU30996.1 atub|OJI83393.1 aver|OJJ00674.1 awen|OJJ34240.1 azon|OJJ43397.1

Group4605: abra|OJJ78328.1 acar|OOF98547.1 acla|EAW09198.1 afis|EAW21949.1 afum|EAL92963.1 agla|OJJ84171.1 akaw|GAA85701.1 aluc|OJZ91402.1 anid|CBF84064.1 anig|EHA20079.1 aory|EIT77849.1 asyd|OJJ61348.1 ater|EAU30997.1 atub|OJI83395.1 aver|OJJ00675.1 awen|OJJ34241.1 azon|OJJ43396.1 afla|EED46667.1

Group4606: abra|OJJ78329.1 acar|OOF98546.1 acla|EAW09197.1 afis|EAW21950.1 afla|EED46668.1 afum|EAL92964.1 agla|OJJ84170.1 akaw|GAA85700.1 aluc|OJZ91403.1 anid|CBF84066.1 anig|EHA20080.1 aory|EIT77926.1 asyd|OJJ61349.1 ater|EAU30998.1 atub|OJI83396.1 aver|OJJ00676.1 awen|OJJ34242.1 azon|OJJ43395.1

Group4607: abra|OJJ78334.1 acar|OOF98544.1 acla|EAW09194.1 afis|EAW21952.1 afla|EED46669.1 afum|EAL92966.1 agla|OJJ84159.1 akaw|GAA85696.1 aluc|OJZ91407.1 anid|CBF84069.1 anig|EHA20083.1 aory|EIT77916.1 asyd|OJJ61351.1 ater|EAU31000.1 atub|OJI83400.1 aver|OJJ00678.1 awen|OJJ34244.1 azon|OJJ43394.1

Group4608: abra|OJJ78337.1 acar|OOF98543.1 acla|EAW09193.1 afis|EAW21953.1 afla|EED46671.1 afum|EAL92967.1 agla|OJJ84160.1 akaw|GAA85695.1 aluc|OJZ91409.1 anid|CBF84070.1 anig|EHA20084.1 aory|EIT77807.1 asyd|OJJ61352.1 ater|EAU31001.1 atub|OJI83403.1 aver|OJJ00679.1 awen|OJJ34245.1 azon|OJJ43393.1

Group4609: abra|OJJ78339.1 acar|OOF98541.1 acla|EAW09192.1 afis|EAW21954.1 afla|EED46672.1 afum|EAL92968.1 agla|OJJ84161.1 akaw|GAA85694.1 aluc|OJZ91411.1 anid|CBF84074.1 anig|EHA20085.1 aory|EIT77806.1 asyd|OJJ61353.1 ater|EAU31060.1 atub|OJI83404.1 aver|OJJ00680.1 awen|OJJ34246.1 azon|OJJ43392.1

Group4610: abra|OJJ78345.1 acar|OOF90269.1 acla|EAW09189.1 afis|EAW21957.1 afla|EED46676.1 afum|EAL92971.1 agla|OJJ84164.1 akaw|GAA85688.1 aluc|OJZ91418.1 anid|CBF84037.1 anig|EHA20090.1 aory|EIT77863.1 asyd|OJJ61362.1 ater|EAU31063.1 atub|OJI83412.1 aver|OJJ00689.1 awen|OJJ34249.1 azon|OJJ43389.1

Group4611: abra|OJJ78346.1 acar|OOF90268.1 acla|EAW09188.1 afis|EAW21958.1 afla|EED46678.1 afum|EAL92972.1 agla|OJJ84158.1 akaw|GAA85687.1 aluc|OJZ91420.1 anid|CBF84041.1 anig|EHA20091.1 aory|EIT77880.1 asyd|OJJ61360.1 ater|EAU31078.1 atub|OJI83413.1 aver|OJJ00687.1 awen|OJJ34251.1 azon|OJJ43387.1

Group4612: abra|OJJ78347.1 acar|OOF90267.1 acla|EAW09187.1 afis|EAW21959.1 afla|EED46679.1 afum|EAL92973.1 agla|OJJ84157.1 akaw|GAA85686.1 aluc|OJZ91421.1 anid|CBF84043.1 anig|EHA20092.1 aory|EIT77956.1 asyd|OJJ61359.1 ater|EAU31079.1 atub|OJI83414.1 aver|OJJ00686.1 awen|OJJ34252.1 azon|OJJ43386.1

Group4613: abra|OJJ78351.1 acar|OOF98536.1 acla|EAW09184.1 afis|EAW21962.1 afla|EED46683.1 afum|EAL92976.1 agla|OJJ84141.1 akaw|GAA85683.1 aluc|OJZ91425.1 anid|CBF82742.1 anig|EHA20095.1 aory|EIT77876.1 asyd|OJJ61622.1 ater|EAU31082.1 atub|OJI83418.1 aver|OJJ00946.1 awen|OJJ34256.1 azon|OJJ43383.1

Group4614: abra|OJJ78353.1 acar|OOF98535.1 acla|EAW09182.1 afis|EAW21964.1 afla|EED46685.1 afum|EAL92978.2 agla|OJJ84139.1 akaw|GAA85681.1 aluc|OJZ91428.1 anid|CBF82738.1 anig|EHA20097.1 aory|EIT77960.1 asyd|OJJ61624.1 ater|EAU31084.1 atub|OJI83421.1 aver|OJJ00949.1 awen|OJJ34258.1 azon|OJJ43381.1

Group4615: abra|OJJ78354.1 acar|OOF98534.1 acla|EAW09181.1 afis|EAW21965.1 afla|EED46686.1 afum|EAL92979.1 agla|OJJ84138.1 akaw|GAA85680.1 aluc|OJZ91429.1 anid|CBF82736.1 anig|EHA20098.1 aory|EIT77854.1 asyd|OJJ61625.1 ater|EAU31085.1 atub|OJI83422.1 aver|OJJ00950.1 awen|OJJ34259.1 azon|OJJ43378.1

Group4616: abra|OJJ78356.1 acar|OOF98533.1 acla|EAW09178.1 afis|EAW21968.1 afla|EED46689.1 afum|EAL92982.1 agla|OJJ84131.1 akaw|GAA85677.1 aluc|OJZ91431.1 anid|CBF82731.1 anig|EHA20100.1 aory|EIT77885.1 asyd|OJJ61628.1 ater|EAU31088.1 atub|OJI83424.1 aver|OJJ00952.1 awen|OJJ34261.1 azon|OJJ43376.1

Group4617: abra|OJJ78363.1 acar|OOF98524.1 acla|EAW09172.1 afis|EAW21974.1 afla|EED46697.1 afum|EAL92989.1 agla|OJJ84128.1 akaw|GAA85670.1 aluc|OJZ91438.1 anid|CBF82716.1 anig|EHA20103.1 aory|EIT77925.1 asyd|OJJ61637.1 ater|EAU31092.1 atub|OJI83432.1 aver|OJJ00960.1 awen|OJJ34267.1 azon|OJJ43369.1

Group4618: abra|OJJ78365.1 acar|OOF98521.1 acla|EAW09170.1 afis|EAW21976.1 afla|EED46699.1 afum|EAL92991.1 agla|OJJ84125.1 akaw|GAA85668.1 aluc|OJZ91442.1 anid|CBF82720.1 anig|EHA20106.1 aory|EIT77848.1 asyd|OJJ61635.1 ater|EAU31094.1 atub|OJI83436.1 aver|OJJ00958.1 awen|OJJ34269.1 azon|OJJ43368.1

Group4619: abra|OJJ78366.1 acar|OOF98520.1 acla|EAW09169.1 afis|EAW21977.1 afla|EED46700.1 afum|EAL92992.1 agla|OJJ84124.1 akaw|GAA85667.1 aluc|OJZ91443.1 anid|CBF82722.1 anig|EHA20107.1 aory|EIT77809.1 asyd|OJJ61634.1 ater|EAU31095.1 atub|OJI83437.1 aver|OJJ00957.1 awen|OJJ34270.1 azon|OJJ43366.1

Group4620: abra|OJJ78368.1 acar|OOF98518.1 acla|EAW09167.1 afis|EAW21979.1 afla|EED46702.1 afum|EAL92994.1 agla|OJJ84120.1 akaw|GAA85665.1 aluc|OJZ91445.1 anid|CBF82711.1 anig|EHA20109.1 aory|EIT77847.1 asyd|OJJ61640.1 ater|EAU31097.1 atub|OJI83439.1 aver|OJJ00963.1 awen|OJJ34272.1 azon|OJJ43364.1

Group4621: abra|OJJ78369.1 acar|OOF90130.1 acla|EAW09166.1 afis|EAW21980.1 afla|EED46704.1 afum|EAL92995.1 agla|OJJ84119.1 akaw|GAA85664.1 aluc|OJZ91446.1 anid|CBF82709.1 anig|EHA20110.1 aory|EIT77845.1 asyd|OJJ61641.1 ater|EAU31100.1 atub|OJI83440.1 aver|OJJ00964.1 awen|OJJ34274.1 azon|OJJ43363.1

Group4622: abra|OJJ78371.1 acar|OOF90128.1 acla|EAW09164.1 afis|EAW21982.1 afla|EED46706.1 afum|EAL92997.1 agla|OJJ84118.1 akaw|GAA85662.1 aluc|OJZ91448.1 anid|CBF82705.1 anig|EHA20112.1

aory|EIT77874.1 asyd|OJJ61643.1 ater|EAU31101.1 atub|OJI83441.1 aver|OJJ00965.1 awen|OJJ34276.1 azon|OJJ43361.1

Group4623: abra|OJJ78372.1 acar|OOF90125.1 acla|EAW09163.1 afis|EAW21983.1 afla|EED46707.1 afum|EAL92998.1 agla|OJJ84115.1 akaw|GAA85661.1 aluc|OJZ91449.1 anid|CBF82703.1 anig|EHA20113.1 aory|EIT77859.1 asyd|OJJ61644.1 ater|EAU31102.1 atub|OJI83442.1 aver|OJJ00966.1 awen|OJJ34280.1 azon|OJJ43360.1

Group4624: abra|OJJ78374.1 acar|OOF98515.1 acla|EAW09161.1 afis|EAW21985.1 afla|EED46709.1 afum|EAL93000.1 agla|OJJ84113.1 akaw|GAA85659.1 aluc|OJZ91451.1 anid|CBF82698.1 anig|EHA20115.1 aory|EIT77948.1 asyd|OJJ61646.1 ater|EAU31104.1 atub|OJI83444.1 aver|OJJ00968.1 awen|OJJ34282.1 azon|OJJ43358.1

Group4625: abra|OJJ78375.1 acar|OOF98514.1 acla|EAW09160.1 afis|EAW21986.1 afla|EED46710.1 afum|EAL93002.1 agla|OJJ84112.1 akaw|GAA85658.1 aluc|OJZ91452.1 anid|CBF82696.1 anig|EHA20116.1 aory|EIT77888.1 asyd|OJJ61647.1 ater|EAU31106.1 atub|OJI83445.1 aver|OJJ00969.1 awen|OJJ34283.1 azon|OJJ43357.1

Group4626: abra|OJJ78378.1 acar|OOF98510.1 acla|EAW09155.1 afis|EAW21991.1 afla|EED46715.1 afum|EAL93006.2 agla|OJJ84107.1 akaw|GAA85654.1 aluc|OJZ91457.1 anid|CBF82687.1 anig|EHA20120.1 aory|EIT77887.1 asyd|OJJ61652.1 ater|EAU31110.1 atub|OJI83450.1 aver|OJJ00973.1 awen|OJJ34287.1 azon|OJJ43351.1

Group4627: abra|OJJ78380.1 acar|OOF98508.1 acla|EAW09153.1 afis|EAW21993.1 afla|EED46717.1 afum|EAL93008.1 agla|OJJ84104.1 akaw|GAA85652.1 aluc|OJZ91459.1 anid|CBF82683.1 anig|EHA20122.1 aory|EIT77842.1 asyd|OJJ61655.1 ater|EAU31112.1 atub|OJI83452.1 aver|OJJ00976.1 awen|OJJ34289.1 azon|OJJ43349.1

Group4628: abra|OJJ78381.1 acar|OOF98507.1 acla|EAW09152.1 afis|EAW21994.1 afla|EED46718.1 afum|EAL93009.1 agla|OJJ84103.1 akaw|GAA85651.1 aluc|OJZ91460.1 anid|CBF82681.1 anig|EHA20123.1 aory|EIT77902.1 asyd|OJJ61656.1 ater|EAU31113.1 atub|OJI83453.1 aver|OJJ00977.1 awen|OJJ34290.1 azon|OJJ43348.1

Group4629: abra|OJJ78382.1 acar|OOF98506.1 acla|EAW09151.1 afis|EAW21995.1 afla|EED46719.1 afum|EAL93010.1 agla|OJJ84102.1 akaw|GAA85650.1 aluc|OJZ91461.1 anid|CBF82679.1 anig|EHA20124.1 aory|EIT77906.1 asyd|OJJ61657.1 ater|EAU31114.1 atub|OJI83454.1 aver|OJJ00978.1 awen|OJJ34291.1 azon|OJJ43347.1

Group4630: abra|OJJ78383.1 acar|OOF98505.1 acla|EAW09150.1 afis|EAW21996.1 afla|EED46720.1 afum|EAL93011.1 agla|OJJ84101.1 akaw|GAA85649.1 aluc|OJZ91462.1 anid|CBF82677.1 anig|EHA20125.1 aory|EIT77812.1 asyd|OJJ61658.1 ater|EAU31115.1 atub|OJI83455.1 aver|OJJ00979.1 awen|OJJ34292.1 azon|OJJ43346.1

Group4631: abra|OJJ78384.1 acar|OOF98504.1 acla|EAW09149.1 afis|EAW21997.1 afla|EED46721.1 afum|EAL93012.1 agla|OJJ84100.1 akaw|GAA85648.1 aluc|OJZ91463.1 anid|CBF82675.1 anig|EHA20126.1 aory|EIT77875.1 asyd|OJJ61659.1 ater|EAU31116.1 atub|OJI83456.1 aver|OJJ00980.1 awen|OJJ34293.1 azon|OJJ43345.1

Group4632: abra|OJJ78385.1 acar|OOF98503.1 acla|EAW09148.1 afis|EAW21998.1 afla|EED46722.1 afum|EAL93013.1 agla|OJJ84099.1 akaw|GAA85647.1 aluc|OJZ91464.1 anid|CBF82674.1 anig|EHA20127.1 aory|EIT77891.1 asyd|OJJ61660.1 ater|EAU31117.1 atub|OJI83457.1 aver|OJJ00981.1 awen|OJJ34294.1 azon|OJJ43344.1

Group4633: abra|OJJ78386.1 acar|OOF98502.1 acla|EAW09147.1 afis|EAW21999.1 afla|EED46723.1 afum|EAL93014.1 agla|OJJ84098.1 akaw|GAA85646.1 aluc|OJZ91465.1 anid|CBF82672.1 anig|EHA20128.1 aory|EIT77907.1 asyd|OJJ61661.1 ater|EAU31118.1 atub|OJI83458.1 aver|OJJ00982.1 awen|OJJ34295.1 azon|OJJ43343.1

Group4634: abra|OJJ78388.1 acar|OOF98500.1 acla|EAW09145.1 afis|EAW22002.1 afla|EED46727.1 afum|EAL93017.1 agla|OJJ84096.1 akaw|GAA85644.1 aluc|OJZ91467.1 anid|CBF82666.1 anig|EHA20129.1 aory|EIT77844.1 asyd|OJJ61664.1 ater|EAU31121.1 atub|OJI83460.1 aver|OJJ00985.1 awen|OJJ34297.1 azon|OJJ43341.1

Group4635: acar|OOG00510.1 afis|EAW25745.1 akaw|GAA87910.1 aluc|OJZ88780.1 anid|CBF84274.1 anig|EHA18474.1 aory|EIT75519.1 asyd|OJJ57228.1 ater|EAU32770.1 atub|OJI91352.1 aver|OJI97599.1 awen|OJJ32621.1 afis|EAW24516.1 afla|EED47665.1 aver|OJJ03061.1 aluc|OJZ85622.1 aory|EIT80232.1 atub|OJI79415.1

Group4636: acla|EAW13541.1 afis|EAW20614.1 afla|EED54203.1 afum|EAL84447.1 agla|OJJ88592.1 akaw|GAA87001.1 aluc|OJZ84456.1 anid|CBF86940.1 anig|EHA27771.1 aory|EIT75953.1 asyd|OJJ55587.1 atub|OJI89619.1 aver|OJJ06666.1 awen|OJJ33458.1 azon|OJJ50714.1 ater|EAU35253.1 asyd|OJJ55586.1 aver|OJJ06665.1

Group4637: abra|OJJ65435.1 abra|OJJ65508.1 abra|OJJ65821.1 asyd|OJJ51932.1 asyd|OJJ52321.1 asyd|OJJ51963.1 asyd|OJJ51982.1 awen|OJJ29548.1 awen|OJJ29621.1 akaw|GAA93288.1 afis|EAW20669.1 agla|OJJ83544.1 aluc|OJZ79579.1 anig|EHA17693.1 ater|EAU30109.1 akaw|GAA93088.1 aluc|OJZ80408.1

Group4638: abra|OJJ65486.1 abra|OJJ65539.1 akaw|GAA92968.1 anig|EHA20201.1 afla|EED53240.1 agla|OJJ82729.1 anid|CBF82095.1 aory|EIT74878.1 aver|OJJ05216.1 awen|OJJ30575.1 abra|OJJ75529.1 acar|OOF90856.1 akaw|GAA84291.1 aluc|OJZ88985.1 anig|EHA23541.1 atub|OJI85803.1 ater|EAU39592.1

Group4639: abra|OJJ65671.1 abra|OJJ69720.1 acar|OOF92570.1 acar|OOG00879.1 afla|EED52525.1 afla|EED53603.1 akaw|GAA90173.1 akaw|GAA93196.1 aory|EIT74948.1 aory|EIT82807.1 acla|EAW11895.1 afum|EAL84364.1 aluc|OJZ83177.1 anig|EHA28251.1 atub|OJI87760.1 awen|OJJ36063.1 azon|OJJ42163.1

Group4640: abra|OJJ68069.1 abra|OJJ68262.1 akaw|GAA83203.1 aluc|OJZ83704.1 anig|EHA18854.1 atub|OJI90076.1 acar|OOF97541.1 afla|EED45998.1 agla|OJJ80130.1 anid|CBF90047.1 aory|EIT75615.1 asyd|OJJ62055.1 ater|EAU38008.1 aver|OJI96161.1 awen|OJJ30952.1 anid|CBF79559.1 awen|OJJ36502.1

Group4641: acar|OOF90194.1 acar|OOF90196.1 abra|OJJ67673.1 afis|EAW24512.1 afum|EBA27260.1 akaw|GAA89189.1 aluc|OJZ81642.1 anid|CBF75855.1 asyd|OJJ62592.1 ater|EAU37120.1 atub|OJI88540.1 aver|OJJ04434.1 awen|OJJ36101.1 afis|EAW15651.1 aluc|OJZ79951.1 aluc|OJZ80155.1 asyd|OJJ57969.1

Group4642: acar|OOF90453.1 acar|OOF90460.1 abra|OJJ71883.1 acla|EAW09638.1 afis|EAW21502.1 afla|EED55938.1 afum|EAL92521.1 agla|OJJ85665.1 akaw|GAA85033.1 aluc|OJZ86621.1 anig|EHA22045.1 asyd|OJJ61108.1 ater|EAU35036.1 atub|OJI83904.1 aver|OJJ00425.1 awen|OJJ39193.1 azon|OJJ51518.1

Group4643: acar|OOF92292.1 acar|OOF92333.1 abra|OJJ67509.1 acla|EAW06263.1 afis|EAW18369.1 afum|EAL93044.1 agla|OJJ84934.1 akaw|GAA86330.1 aluc|OJZ84855.1 anid|CBF70810.1 anig|EHA25088.1 aory|EIT77442.1 asyd|OJJ58407.1 atub|OJI80741.1 aver|OJJ01008.1 awen|OJJ31780.1 azon|OJJ42022.1

Group4644: acar|OOF92294.1 acar|OOF92335.1 abra|OJJ67506.1 acla|EAW06265.1 afis|EAW18371.1 afla|EED53035.1 afum|EAL93046.1 akaw|GAA86332.1 aluc|OJZ84857.1 anid|CBF70806.1 anig|EHA25090.1 aory|EIT77400.1 asyd|OJJ58409.1 ater|EAU32241.1 atub|OJI80739.1 aver|OJJ01011.1 azon|OJJ50444.1

Group4645: acar|OOF92315.1 acar|OOF92357.1 abra|OJJ67484.1 acla|EAW06290.1 afis|EAW18396.1 afla|EED53012.1 afum|EAL93072.1 agla|OJJ84967.1 akaw|GAA86355.1 aluc|OJZ84884.1 anid|CBF70760.1 anig|EHA25113.1 aory|EIT77413.1 asyd|OJJ58432.1 atub|OJI80716.1 aver|OJJ01033.1 azon|OJJ50471.1

Group4646: acar|OOF93666.1 acar|OOF93750.1 abra|OJJ66646.1 acla|EAW11789.1 afis|EAW19399.1 afum|EAL87095.1 agla|OJJ81110.1 akaw|GAA89599.1 aluc|OJZ80803.1 anid|CBF74334.1 anig|EHA27231.1 aory|EIT83139.1 asyd|OJJ51907.1 ater|EAU36125.1 atub|OJI87651.1 awen|OJJ36794.1 azon|OJJ43005.1

Group4647: acar|OOF93683.1 acar|OOF93768.1 abra|OJJ66668.1 acla|EAW11809.1 afis|EAW19419.1 afla|EED50376.1 afum|EAL87075.1 akaw|GAA89617.1 aluc|OJZ80779.1 anid|CBF74367.1 anig|EHA27250.1 aory|EIT83147.1 ater|EAU34725.1 atub|OJI87631.1 aver|OJJ05381.1 awen|OJJ36817.1 azon|OJJ42985.1

Group4648: acar|OOF93688.1 acar|OOF93773.1 abra|OJJ66674.1 acla|EAW11814.1 afis|EAW19424.1 afla|EED50382.1 afum|EAL87070.1 agla|OJJ81136.1 akaw|GAA89622.1 aluc|OJZ80774.1 anig|EHA27256.1 aory|EIT83125.1 asyd|OJJ57779.1 ater|EAU34728.1 atub|OJI87626.1 aver|OJJ05386.1 awen|OJJ36822.1

Group4649: acar|OOF93689.1 acar|OOF93774.1 abra|OJJ66676.1 acla|EAW11815.1 afis|EAW19426.1 afum|EAL87068.1 agla|OJJ81138.1 akaw|GAA89624.1 aluc|OJZ80772.1 anig|EHA27258.1 aory|EIT83128.1 asyd|OJJ57777.1 ater|EAU34730.1 atub|OJI87624.1 aver|OJJ05388.1 awen|OJJ36824.1 azon|OJJ42491.1

Group4650: acar|OOF93695.1 acar|OOF93780.1 abra|OJJ66686.1 acla|EAW11821.1 afis|EAW19432.1 afla|EED56637.1 agla|OJJ81144.1 akaw|GAA89630.1 aluc|OJZ80763.1 anig|EHA27264.1 aory|EIT79667.1 asyd|OJJ59390.1 atub|OJI87617.1 aver|OJJ02882.1 awen|OJJ36830.1 azon|OJJ42973.1 anid|CBF81378.1

Group4651: acar|OOF93698.1 acar|OOF93783.1 abra|OJJ66689.1 acla|EAW11824.1 afis|EAW19435.1 afla|EED56640.1 afum|EAL87060.1 agla|OJJ81147.1 akaw|GAA89633.1 aluc|OJZ80760.1 anig|EHA27267.1 aory|EIT79665.1 asyd|OJJ59394.1 ater|EAU34737.1 atub|OJI87614.1 aver|OJJ02878.1 azon|OJJ42970.1

Group4652: acar|OOF93704.1 acar|OOF93789.1 abra|OJJ66696.1 afla|EED56650.1 afum|EAL87053.1 agla|OJJ81156.1 akaw|GAA89640.1 aluc|OJZ80749.1 anid|CBF81397.1 anig|EHA27273.1 aory|EIT79652.1 asyd|OJJ59402.1 ater|EAU34744.1 atub|OJI87606.1 aver|OJJ02871.1 awen|OJJ36843.1 azon|OJJ42957.1

Group4653: acar|OOF93713.1 acar|OOF93799.1 abra|OJJ66711.1 acla|EAW11842.1 afis|EAW19453.1 afla|EED56661.1 afum|EAL87043.1 agla|OJJ81168.1 akaw|GAA89653.1 aluc|OJZ80736.1 anid|CBF81418.1 asyd|OJJ59413.1 ater|EAU34755.1 atub|OJI87593.1 aver|OJJ02860.1 awen|OJJ36854.1 azon|OJJ42946.1

Group4654: acar|OOF93722.1 acar|OOF93809.1 abra|OJJ66720.1 acla|EAW11851.1 afis|EAW19462.1 afla|EED56673.1 afum|EAL87034.1 agla|OJJ81179.1 akaw|GAA89662.1 aluc|OJZ80727.1 anid|CBF75493.1 asyd|OJJ59751.1 ater|EAU34765.1 atub|OJI87583.1 aver|OJJ02536.1 awen|OJJ36864.1 azon|OJJ42937.1

Group4655: acar|OOF94041.1 acar|OOF94111.1 abra|OJJ66400.1 afla|EED49553.1 akaw|GAA90268.1 aluc|OJZ83073.1 anig|EHA28137.1 aory|EIT82620.1 asyd|OJJ61454.1 atub|OJI87873.1 aver|OJJ00790.1 awen|OJJ39517.1 abra|OJJ69534.1 agla|OJJ79509.1 akaw|GAA88384.1 anig|EHA28392.1 acar|OOF92662.1

Group4656: acar|OOF94050.1 acar|OOF94120.1 abra|OJJ66395.1 acla|EAW12087.1 afla|EED49519.1 akaw|GAA90260.1 aluc|OJZ83080.1 anid|CBF73558.1 anig|EHA28143.1 aory|EIT82588.1 asyd|OJJ56699.1 atub|OJI87865.1 aver|OJJ06026.1 awen|OJJ40206.1 azon|OJJ47473.1 awen|OJJ35025.1 afis|EAW19730.1

Group4657: acar|OOF94052.1 acar|OOF94122.1 abra|OJJ66392.1 acla|EAW13709.1 afis|EAW19881.1 afla|EED46102.1 afum|EAL85017.1 akaw|GAA90258.1 aluc|OJZ83082.1 anid|CBF75188.1 anig|EHA28145.1 aory|EIT76369.1 asyd|OJJ59947.1 ater|EAU35603.1 atub|OJI87863.1 aver|OJJ02342.1 azon|OJJ51816.1

Group4658: acar|OOF94076.1 acar|OOF94146.1 abra|OJJ66026.1 acla|EAW07129.1 afis|EAW22139.1 anid|CBF89247.1 atub|OJI87440.1 awen|OJJ40945.1 azon|OJJ42435.1 abra|OJJ67016.1 afum|EAL91051.1 aver|OJJ03311.1 abra|OJJ67290.1 akaw|GAA86523.1 aluc|OJZ85069.1 anig|EHA25283.1 atub|OJI80529.1

Group4659: acar|OOF95088.1 acar|OOF95154.1 abra|OJJ74616.1 acla|EAW10766.1 afis|EAW24810.1 afla|EED47173.1 afum|EAL89453.2 agla|OJJ82905.1 akaw|GAA81990.1 aluc|OJZ92199.1 anig|EHA26758.1 aory|EIT79174.1 asyd|OJJ64383.1 atub|OJI86626.1 aver|OJI98479.1 awen|OJJ40394.1 azon|OJJ49247.1

Group4660: acar|OOF95105.1 acar|OOF95171.1 abra|OJJ74599.1 acla|EAW10780.1 afis|EAW24794.1 afla|EED46333.1 afum|EAL89436.1 agla|OJJ79801.1 akaw|GAA82009.1 aluc|OJZ92178.1 anig|EHA26775.1 aory|EIT81232.1 asyd|OJJ59471.1 ater|EAU36511.1 atub|OJI86606.1 awen|OJJ40379.1 aver|OJJ02803.1

Group4661: acar|OOF96237.1 acar|OOF96274.1 afum|EAL85171.1 afum|EAL87775.1 abra|OJJ71521.1 afis|EAW20050.1 afla|EED45330.1 agla|OJJ88910.1 akaw|GAA89396.1 aluc|OJZ86260.1 anid|CBF84802.1 anig|EHA21606.1 aory|EIT73757.1 ater|EAU38660.1 atub|OJI84283.1 awen|OJJ31808.1 azon|OJJ46314.1

Group4662: acar|OOF96905.1 acar|OOF96940.1 abra|OJJ70805.1 acla|EAW11119.1 afis|EAW15849.1 afum|EBA27349.1 agla|OJJ87260.1 akaw|GAA90120.1 aluc|OJZ84151.1 anid|CBF81170.1 anig|EHA19255.1 asyd|OJJ59289.1 ater|EAU31664.1 atub|OJI90523.1 aver|OJJ02986.1 awen|OJJ36779.1 azon|OJJ43020.1

Group4663: acar|OOF96910.1 acar|OOF96944.1 abra|OJJ70800.1 acla|EAW11124.1 afis|EAW15853.1 afla|EED56520.1 afum|EAL88489.1 agla|OJJ87255.1 akaw|GAA90116.1 aluc|OJZ84145.1 anid|CBF81179.1 asyd|OJJ59294.1 ater|EAU31667.1 atub|OJI90519.1 aver|OJJ02981.1 awen|OJJ36775.1 azon|OJJ47423.1

Group4664: acar|OOF96925.1 acar|OOF96960.1 abra|OJJ70784.1 acla|EAW11136.1 afis|EAW15868.1 afum|EAL88502.1 agla|OJJ87240.1 akaw|GAA90102.1 aluc|OJZ84130.1 anid|CBF81207.1 anig|EHA19238.1 asyd|OJJ59308.1 ater|EAU31683.1 atub|OJI90504.1 aver|OJJ02967.1 awen|OJJ29537.1 azon|OJJ47437.1

Group4665: acar|OOF96926.1 acar|OOF96961.1 abra|OJJ70782.1 afla|EED56506.1 agla|OJJ87239.1 akaw|GAA90101.1 aluc|OJZ84129.1 anid|CBF81209.1 anig|EHA19237.1 aory|EIT72792.1 asyd|OJJ59309.1 atub|OJI90503.1 aver|OJJ02966.1 awen|OJJ36761.1 azon|OJJ47438.1 acla|EAW11137.1 awen|OJJ29536.1

Group4666: acar|OOF97656.1 acar|OOF97703.1 abra|OJJ71501.1 acla|EAW10047.1 afla|EED44713.1 afum|EBA27399.1 agla|OJJ81159.1 akaw|GAA82519.1 aluc|OJZ87046.1 anig|EHA20319.1 aory|EIT83457.1 asyd|OJJ53008.1 ater|EAU33378.1 atub|OJI81589.1 aver|OJJ03835.1 awen|OJJ41279.1 azon|OJJ49502.1

Group4667: acar|OOF98479.1 acar|OOF98492.1 abra|OJJ78395.1 acla|EAW09137.1 afis|EAW22010.1 afla|EED46734.1 afum|EAL93025.1 agla|OJJ84088.1 akaw|GAA85636.1 aluc|OJZ91474.1 anig|EHA20136.1 asyd|OJJ61672.1 atub|OJI83467.1 awen|OJJ34305.1 azon|OJJ43332.1 anid|CBF82651.1 aver|OJJ00993.1

Group4668: acar|OOF99310.1 acar|OOF99311.1 acla|EAW09382.1 afis|EAW21753.1 afla|EED58025.1 afum|EAL92783.1 agla|OJJ84400.1 anid|CBF87161.1 aory|EIT79777.1 asyd|OJJ58846.1 ater|EAU38437.1 aver|OJJ01447.1 awen|OJJ34037.1 azon|OJJ45317.1 anid|CBF83691.1 asyd|OJJ60817.1 aver|OJJ00143.1

Group4669: acar|OOG00248.1 acar|OOG00266.1 abra|OJJ67542.1 acla|EAW13608.1 afis|EAW20002.1 afla|EED45812.1 afum|EAL84917.1 akaw|GAA88117.1 aluc|OJZ81390.1 anid|CBF78636.1 anig|EHA18683.1 aory|EIT75677.1 asyd|OJJ65124.1 ater|EAU37945.1 atub|OJI91595.1 aver|OJI99252.1 azon|OJJ47329.1

Group4670: afla|EED49436.1 afla|EED53000.1 aory|EIT77416.1 aory|EIT79505.1 abra|OJJ67468.1 acar|OOF92366.1 afis|EAW18409.1 afum|EAL93089.1 agla|OJJ84976.1 akaw|GAA86369.1 aluc|OJZ84898.1 anid|CBF70739.1 anig|EHA25125.1 asyd|OJJ58440.1 atub|OJI80700.1 aver|OJJ01040.1 awen|OJJ31740.1

Group4671: agla|OJJ81704.1 agla|OJJ85950.1 anig|EHA24273.1 anig|EHA24275.1 asyd|OJJ53801.1 asyd|OJJ63907.1 awen|OJJ37085.1 awen|OJJ39694.1 acla|EAW12912.1 afis|EAW17749.1 afla|EED55100.1 afum|EAL91563.1 anid|CBF80577.1 ater|EAU31857.1 aver|OJI97034.1 azon|OJJ44442.1 aory|EIT81664.1

Group4672: agla|OJJ84696.1 agla|OJJ88299.1 abra|OJJ65609.1 afla|EED54385.1 aory|EIT75847.1 abra|OJJ76926.1 acar|OOF91371.1 afla|EED46863.1 akaw|GAA87114.1 aluc|OJZ84324.1 anid|CBF74601.1 anig|EHA27659.1 aory|EIT72704.1 asyd|OJJ61216.1 atub|OJI89744.1 aver|OJJ00540.1 anig|EHA18441.1

Group4673: akaw|GAA82959.1 akaw|GAA85567.1 aluc|OJZ83501.1 aluc|OJZ87540.1 anig|EHA18316.1 anig|EHA24335.1 atub|OJI81107.1 atub|OJI82395.1 abra|OJJ72993.1 acar|OOF96169.1 afla|EED49853.1 agla|OJJ88852.1 anid|CBF73514.1 aory|EIT83415.1 asyd|OJJ61476.1 aver|OJJ00818.1 awen|OJJ31873.1

Group4674: akaw|GAA87889.1 akaw|GAA92892.1 asyd|OJJ52060.1 asyd|OJJ56757.1 abra|OJJ66170.1 afla|EED52364.1 afum|EAL88779.1 agla|OJJ83554.1 anid|CBF75918.1 anig|EHA26110.1 asyd|OJJ52025.1 azon|OJJ42071.1 acla|EAW11462.1 agla|OJJ81341.1 aluc|OJZ79991.1 acla|EAW11473.1 afis|EAW17037.1

Group4675: anid|CBF79042.1 anid|CBF89822.1 abra|OJJ74989.1 acar|OOF97331.1 acla|EAW08393.1 afis|EAW23436.1 afla|EED57325.1 afum|EAL88026.1 akaw|GAA83445.1 aluc|OJZ92551.1 anig|EHA26426.1 aory|EIT74067.1 asyd|OJJ62175.1 ater|EAU37490.1 atub|OJI86989.1 aver|OJI96284.1 azon|OJJ44145.1

Group4676: anid|CBF83201.1 anid|CBF86907.1 asyd|OJJ55555.1 asyd|OJJ58381.1 asyd|OJJ64611.1 ater|EAU29240.1 ater|EAU35231.1 ater|EAU39358.1 aver|OJI98707.1 aver|OJJ06635.1 aver|OJJ06637.1 azon|OJJ46028.1 afis|EAW17327.1 afla|EED54401.1 afum|EAL91164.1 agla|OJJ79869.1 aory|EIT75849.1

Group4677: asyd|OJJ53784.1 asyd|OJJ56688.1 abra|OJJ67840.1 acar|OOF92016.1 acla|EAW12885.1 afis|EAW17792.1 afla|EED53092.1 agla|OJJ85532.1 akaw|GAA91846.1 aluc|OJZ82052.1 anid|CBF73574.1 anig|EHA22207.1 aory|EIT74818.1 ater|EAU38815.1 atub|OJI84610.1 aver|OJJ06035.1 awen|OJJ35973.1

Group4678: asyd|OJJ54404.1 asyd|OJJ54753.1 asyd|OJJ64139.1 abra|OJJ65870.1 afla|EED51457.1 agla|OJJ85481.1 akaw|GAA92276.1 aluc|OJZ90170.1 aory|EIT80565.1 ater|EAU33973.1 atub|OJI81598.1 aver|OJI99607.1 awen|OJJ36266.1 asyd|OJJ58123.1 aver|OJJ08845.1 anid|CBF71445.1 asyd|OJJ56514.1

Group4679: asyd|OJJ54407.1 asyd|OJJ56037.1 abra|OJJ69897.1 acar|OOF93416.1 acla|EAW08923.1 afis|EAW15470.1 agla|OJJ80150.1 akaw|GAA87753.1 aluc|OJZ85386.1 anid|CBF78232.1 anig|EHA25570.1 aory|EIT72697.1 ater|EAU32658.1 atub|OJI80184.1 aver|OJJ04983.1 awen|OJJ38120.1 afum|EAL85852.1

Group4680: asyd|OJJ54582.1 asyd|OJJ59187.1 awen|OJJ38449.1 awen|OJJ38819.1 abra|OJJ67702.1 akaw|GAA92941.1 abra|OJJ77957.1 afis|EAW20204.1 afla|EED45854.1 aluc|OJZ83508.1 anig|EHA19712.1 aory|EIT75426.1 ater|EAU35724.1 atub|OJI91308.1 azon|OJJ45690.1 ater|EAU31781.1 agla|OJJ83813.1

Group4681: asyd|OJJ58287.1 asyd|OJJ62495.1 abra|OJJ68999.1 acar|OOF92711.1 afis|EAW18027.1 afla|EED51868.1 afum|EAL91838.1 agla|OJJ85400.1 akaw|GAA90518.1 aluc|OJZ85983.1 anid|CBF80061.1 anig|EHA24623.1 ater|EAU31541.1 atub|OJI79773.1 aver|OJJ02137.1 awen|OJJ41512.1 azon|OJJ48928.1

Group4682: ater|EAU30451.1 ater|EAU31200.1 abra|OJJ68616.1 acar|OOF92127.1 acla|EAW12608.1 afis|EAW18076.1 afla|EED51468.1 afum|EAL91888.1 akaw|GAA92905.1 aluc|OJZ81938.1 anid|CBF86402.1 anig|EHA22310.1 aory|EIT80658.1 asyd|OJJ64950.1 atub|OJI84491.1 aver|OJI99072.1 awen|OJJ41465.1

Group4683: ater|EAU30817.1 ater|EAU34589.1 abra|OJJ67750.1 acar|OOF90955.1 acla|EAW10989.1 afis|EAW20871.1 afla|EED45553.1 afum|EAL84763.1 akaw|GAA92441.1 aluc|OJZ81579.1 anid|CBF82846.1 anig|EHA25860.1 asyd|OJJ61591.1 atub|OJI88602.1 aver|OJJ00913.1 awen|OJJ34490.1 azon|OJJ51767.1

Group4684: atub|OJI81487.1 atub|OJI81722.1 abra|OJJ71387.1 acar|OOF97781.1 afis|EAW25058.1 afum|EAL89726.1 agla|OJJ79169.1 akaw|GAA82607.1 aluc|OJZ87153.1 anig|EHA20412.1 aver|OJJ05909.1 awen|OJJ33722.1 azon|OJJ46843.1 acar|OOF98188.1 aluc|OJZ90300.1 anig|EHA23788.1 akaw|GAA86066.1

Group4685: awen|OJJ30759.1 awen|OJJ31722.1 abra|OJJ67450.1 acar|OOF90340.1 acla|EAW06314.1 afla|EED52985.1 agla|OJJ84993.1 akaw|GAA86380.1 aluc|OJZ84914.1 anid|CBF70692.1 anig|EHA25135.1 aory|EIT78708.1 asyd|OJJ58465.1 ater|EAU29844.1 atub|OJI80684.1 aver|OJJ01064.1 azon|OJJ50485.1

Group4686: awen|OJJ30765.1 awen|OJJ31716.1 abra|OJJ67445.1 acar|OOF92382.1 acla|EAW06318.1 afis|EAW18432.1 afum|EAL93113.1 agla|OJJ84998.1 akaw|GAA86384.1 aluc|OJZ84918.1 anid|CBF70027.1 anig|EHA25138.1 asyd|OJJ58871.1 ater|EAU29848.1 atub|OJI80680.1 aver|OJJ01473.1 azon|OJJ50480.1

Group4687: awen|OJJ30766.1 awen|OJJ31715.1 abra|OJJ67444.1 acar|OOF92383.1 acla|EAW06319.1 afis|EAW18433.1 afum|EAL93114.1 agla|OJJ84999.1 akaw|GAA86385.1 aluc|OJZ84920.1 anid|CBF70029.1 anig|EHA25139.1 asyd|OJJ58870.1 ater|EAU29849.1 atub|OJI80679.1 aver|OJJ01472.1 azon|OJJ50479.1

Group4688: azon|OJJ42401.1 azon|OJJ48129.1 abra|OJJ68278.1 acar|OOG00615.1 afla|EED57500.1 akaw|GAA85996.1 aluc|OJZ83693.1 anig|EHA19762.1 aory|EIT81886.1 asyd|OJJ62122.1 ater|EAU34294.1 atub|OJI90065.1 aver|OJI96230.1 acar|OOF96457.1 aluc|OJZ91059.1 atub|OJI83055.1 azon|OJJ47352.1

Group4689: azon|OJJ47108.1 azon|OJJ47396.1 abra|OJJ67013.1 acar|OOF93607.1 acla|EAW15355.1 afis|EAW20357.1 afum|EAL85474.1 agla|OJJ80638.1 akaw|GAA92160.1 aluc|OJZ85567.1 anig|EHA24951.1 aory|EIT76052.1 asyd|OJJ64186.1 ater|EAU31045.1 atub|OJI79362.1 aver|OJI98263.1 awen|OJJ33014.1

Group4690: azon|OJJ50112.1 azon|OJJ50416.1 abra|OJJ68033.1 acar|OOF91810.1 acla|EAW06966.1 afla|EED55046.1 afum|EAL86259.2 agla|OJJ83082.1 akaw|GAA90919.1 aluc|OJZ82907.1 anig|EHA21330.1 aory|EIT81961.1 asyd|OJJ53715.1 ater|EAU36650.1 atub|OJI89133.1 aver|OJJ07841.1 awen|OJJ33892.1

Group4691: azon|OJJ50963.1 azon|OJJ51217.1 abra|OJJ67715.1 afis|EAW16634.1 anig|EHA27628.1 asyd|OJJ65143.1 aver|OJI99311.1 abra|OJJ68588.1 afis|EAW25637.1 afum|EBA27174.1 aver|OJJ02989.1 awen|OJJ33493.1 akaw|GAA85860.1 aluc|OJZ91219.1 ater|EAU36712.1 aver|OJJ06469.1 ater|EAU33968.1

Group4692: acla|EAW12782.1 awen|OJJ36316.1 acar|OOF94301.1 afis|EAW24051.1 afla|EED49757.1 afum|EAL87858.1 agla|OJJ80393.1 akaw|GAA87492.1 aluc|OJZ88032.1 anig|EHA17806.1 aory|EIT77642.1 asyd|OJJ56970.1 atub|OJI90638.1 aver|OJJ03318.1 afis|EAW23607.1 awen|OJJ41819.1 asyd|OJJ51929.1

Group4693: abra|OJJ65584.1 acar|OOF94381.1 acla|EAW15324.1 afis|EAW24551.1 afla|EED49772.1 afum|EAL91136.1 anid|CBF84508.1 aory|EIT77709.1 asyd|OJJ53518.1 ater|EAU38754.1 aver|OJJ08233.1 awen|OJJ37782.1 azon|OJJ42493.1 aver|OJJ07928.1 awen|OJJ31444.1 acla|EAW15019.1 afum|EAL86216.1

Group4694: abra|OJJ65640.1 afla|EED47428.1 aluc|OJZ91197.1 anid|CBF82230.1 aory|EIT81901.1 ater|EAU35410.1 awen|OJJ40835.1 abra|OJJ65888.1 acar|OOF94950.1 acla|EAW12250.1 agla|OJJ79071.1 akaw|GAA86016.1 aluc|OJZ90244.1 anig|EHA23739.1 asyd|OJJ57337.1 atub|OJI81672.1 aver|OJJ05825.1

Group4695: abra|OJJ65778.1 agla|OJJ79806.1 aluc|OJZ82947.1 atub|OJI89179.1 aver|OJJ07886.1 awen|OJJ35386.1 abra|OJJ69436.1 acla|EAW11436.1 afis|EAW23987.1 afla|EED52190.1 afum|EAL87416.1 aluc|OJZ82272.1 anid|CBF82043.1 asyd|OJJ54224.1 atub|OJI88167.1 aver|OJJ05177.1 afla|EED57914.1

Group4696: abra|OJJ65915.1 acar|OOF94966.1 afis|EAW19972.1 afla|EED50462.1 afum|EAL84938.1 agla|OJJ80064.1 akaw|GAA92317.1 aluc|OJZ90215.1 anid|CBF84460.1 anig|EHA23713.1 aory|EIT82029.1 asyd|OJJ53547.1 ater|EAU31470.1 atub|OJI81646.1 aver|OJJ08206.1 awen|OJJ36012.1 azon|OJJ46810.1

Group4697: abra|OJJ65924.1 acar|OOF94985.1 afis|EAW19706.1 afla|EED57047.1 afum|EAL86820.1 agla|OJJ78913.1 akaw|GAA92309.1 aluc|OJZ90207.1 anid|CBF90284.1 anig|EHA23704.1 aory|EIT82250.1 asyd|OJJ61919.1 ater|EAU32843.1 atub|OJI81636.1 aver|OJI96011.1 awen|OJJ29912.1 acla|EAW12063.1

Group4698: abra|OJJ65938.1 acar|OOF91905.1 acla|EAW12331.1 afis|EAW16891.1 afla|EED46396.1 afum|EAL86429.1 agla|OJJ79945.1 akaw|GAA92298.1 aluc|OJZ90195.1 anid|CBF78493.1 anig|EHA23694.1 aory|EIT72430.1 asyd|OJJ65325.1 ater|EAU30223.1 atub|OJI81622.1 aver|OJI99470.1 awen|OJJ29747.1

Group4699: abra|OJJ65941.1 acar|OOF95011.1 acla|EAW09018.1 afis|EAW19837.1 afla|EED47289.1 afum|EAL85104.1 akaw|GAA92294.1 aluc|OJZ90191.1 anid|CBF84686.1 anig|EHA23690.1 aory|EIT79198.1 asyd|OJJ53334.1 atub|OJI81618.1 aver|OJJ08249.1 aver|OJJ08410.1 acla|EAW13741.1 asyd|OJJ53496.1

Group4700: abra|OJJ65962.1 acar|OOF92912.1 acla|EAW13581.1 afis|EAW20020.1 afla|EED46028.1 afum|EAL84897.1 agla|OJJ88014.1 akaw|GAA90837.1 aluc|OJZ82996.1 anig|EHA21411.1 aory|EIT75634.1 asyd|OJJ53300.1 ater|EAU29382.1 atub|OJI89231.1 aver|OJJ08441.1 awen|OJJ30050.1 azon|OJJ43697.1

Group4701: abra|OJJ65966.1 acar|OOF92909.1 acla|EAW13585.1 afla|EED46023.1 afum|EAL84901.1 agla|OJJ87990.1 akaw|GAA90841.1 aluc|OJZ82992.1 anid|CBF84726.1 anig|EHA21406.1 aory|EIT75803.1 asyd|OJJ53306.1 ater|EAU29377.1 atub|OJI89226.1 aver|OJJ08435.1 awen|OJJ30034.1 azon|OJJ43683.1

Group4702: abra|OJJ65974.1 acar|OOF92898.1 acla|EAW13595.1 afis|EAW20009.1 afla|EED46015.1 afum|EAL84909.1 agla|OJJ87974.1 akaw|GAA90848.1 aluc|OJZ82985.1 anid|CBF84710.1 anig|EHA21399.1 aory|EIT75417.1 asyd|OJJ53323.1 ater|EAU29351.1 atub|OJI89219.1 aver|OJJ08420.1 awen|OJJ30022.1

Group4703: abra|OJJ65975.1 acar|OOF92897.1 acla|EAW13596.1 afis|EAW20008.1 afla|EED46014.1 afum|EAL84910.1 agla|OJJ87973.1 akaw|GAA90849.1 aluc|OJZ82984.1 anid|CBF84708.1 anig|EHA21398.1 aory|EIT75605.1 asyd|OJJ53322.1 ater|EAU29350.1 atub|OJI89218.1 aver|OJJ08421.1 awen|OJJ30021.1

Group4704: abra|OJJ65988.1 acar|OOF92882.1 afis|EAW19525.1 afla|EED49677.1 agla|OJJ87965.1 akaw|GAA90862.1 aluc|OJZ82971.1 anid|CBF78949.1 anig|EHA21387.1 aory|EIT80940.1 asyd|OJJ55654.1 ater|EAU29343.1 atub|OJI89204.1 aver|OJJ06738.1 awen|OJJ30015.1 azon|OJJ43645.1 anig|EHA21388.1

Group4705: abra|OJJ66002.1 acar|OOF92096.1 acla|EAW13639.1 afis|EAW19989.1 afla|EED45931.1 afum|EAL84926.1 agla|OJJ78858.1 akaw|GAA90874.1 aluc|OJZ82958.1 anid|CBF84689.1 anig|EHA21373.1 aory|EIT75732.1 asyd|OJJ53331.1 atub|OJI89190.1 aver|OJJ08413.1 awen|OJJ30007.1 azon|OJJ43636.1

Group4706: abra|OJJ66003.1 acar|OOF92095.1 acla|EAW13640.1 afis|EAW19990.1 afla|EED45930.1 afum|EAL84925.1 agla|OJJ79320.1 akaw|GAA90875.1 aluc|OJZ82957.1 anid|CBF82047.1 anig|EHA21372.1 aory|EIT75591.1 ater|EAU32162.1 atub|OJI89189.1 awen|OJJ30006.1 asyd|OJJ54230.1 aver|OJJ05183.1

Group4707: abra|OJJ66126.1 afis|EAW15452.1 afla|EED45675.1 anid|CBF89338.1 aory|EIT75361.1 abra|OJJ77946.1 akaw|GAA83717.1 aluc|OJZ91016.1 anid|CBF71846.1 asyd|OJJ59800.1 ater|EAU29638.1 atub|OJI83007.1 afum|EAL84194.2 anid|CBF80232.1 afis|EAW22090.1 aver|OJJ02490.1 azon|OJJ46524.1

Group4708: abra|OJJ66193.1 akaw|GAA89095.1 anig|EHA25697.1 atub|OJI88434.1 abra|OJJ77442.1 acla|EAW11799.1 afis|EAW19409.1 afla|EED54078.1 afum|EAL87085.1 agla|OJJ81123.1 anid|CBF74355.1 aory|EIT76033.1 asyd|OJJ57792.1 ater|EAU35399.1 aver|OJJ05372.1 awen|OJJ36805.1 ater|EAU34716.1

Group4709: abra|OJJ66245.1 acar|OOF91971.1 acla|EAW08549.1 afis|EAW23590.1 afla|EED51817.1 afum|EAL87876.1 agla|OJJ79051.1 akaw|GAA89046.1 aluc|OJZ80550.1 anid|CBF74981.1 anig|EHA25646.1 aory|EIT81750.1 asyd|OJJ54563.1 ater|EAU37769.1 atub|OJI88381.1 aver|OJJ07593.1 awen|OJJ35894.1

Group4710: abra|OJJ66282.1 acla|EAW10347.1 afis|EAW20261.1 anig|EHA23662.1 ater|EAU32983.1 atub|OJI89941.1 abra|OJJ70126.1 afla|EED49622.1 akaw|GAA86644.1 aluc|OJZ85201.1 anig|EHA25407.1 asyd|OJJ54176.1 atub|OJI80400.1 aver|OJJ08563.1 azon|OJJ42583.1 afla|EED54705.1 aory|EIT78877.1

Group4711: abra|OJJ66374.1 acla|EAW12222.1 afis|EAW20296.1 afla|EED51865.1 agla|OJJ84731.1 akaw|GAA90242.1 aluc|OJZ83098.1 anig|EHA28162.1 aory|EIT81762.1 asyd|OJJ64162.1 ater|EAU31056.1 atub|OJI87838.1 aver|OJI98239.1 awen|OJJ35379.1 anig|EHA25212.1 aluc|OJZ87036.1 atub|OJI83479.1

Group4712: abra|OJJ66475.1 afla|EED46093.1 akaw|GAA92461.1 aluc|OJZ81556.1 anid|CBF82161.1 anig|EHA25878.1 aory|EIT74924.1 asyd|OJJ54339.1 ater|EAU37565.1 atub|OJI88629.1 aver|OJJ05291.1 awen|OJJ39935.1 aory|EIT76371.1 afla|EED53488.1 asyd|OJJ56980.1 ater|EAU30836.1 aver|OJJ03307.1

Group4713: abra|OJJ66482.1 acar|OOF90959.1 acla|EAW10987.1 afis|EAW20873.1 afla|EED45555.1 afum|EAL84761.1 agla|OJJ83980.1 akaw|GAA92467.1 aluc|OJZ81552.1 anid|CBF82849.1 anig|EHA25882.1 aory|EIT75793.1 asyd|OJJ61593.1 atub|OJI88633.1 aver|OJJ00915.1 awen|OJJ34487.1 azon|OJJ51764.1

Group4714: abra|OJJ66487.1 acar|OOF90963.1 acla|EAW10982.1 afis|EAW20877.1 afla|EED45560.1 afum|EAL84755.1 agla|OJJ83987.1 akaw|GAA92470.1 aluc|OJZ81548.1 anid|CBF82859.1 anig|EHA25887.1 asyd|OJJ61596.1 ater|EAU30812.1 atub|OJI88636.1 aver|OJJ00920.1 awen|OJJ34482.1 azon|OJJ51759.1

Group4715: abra|OJJ66520.1 acar|OOF90974.1 acla|EAW10970.1 afis|EAW20890.1 afla|EED45585.1 afum|EAL84743.1 agla|OJJ83998.1 akaw|GAA91192.1 aluc|OJZ81511.1 anig|EHA25909.1 aory|EIT75393.1 asyd|OJJ61611.1 ater|EAU30829.1 atub|OJI88660.1 aver|OJJ00934.1 awen|OJJ34457.1 azon|OJJ51744.1

Group4716: abra|OJJ66521.1 acar|OOF90975.1 acla|EAW10969.1 afis|EAW20891.1 afla|EED45586.1 afum|EAL84742.1 agla|OJJ83999.1 akaw|GAA91191.1 aluc|OJZ81512.1 anig|EHA25910.1 aory|EIT75451.1 asyd|OJJ61612.1 ater|EAU30830.1 atub|OJI88661.1 aver|OJJ00935.1 awen|OJJ34456.1 azon|OJJ51743.1

Group4717: abra|OJJ66526.1 acar|OOF90979.1 acla|EAW10964.1 afis|EAW20896.1 afla|EED50245.1 afum|EAL84737.2 agla|OJJ84003.1 akaw|GAA91187.1 aluc|OJZ81516.1 anid|CBF82464.1 anig|EHA25914.1 asyd|OJJ54117.1 ater|EAU38530.1 atub|OJI88665.1 aver|OJJ08619.1 awen|OJJ34452.1 azon|OJJ51740.1

Group4718: abra|OJJ66547.1 acar|OOF89822.1 acla|EAW10944.1 afis|EAW20914.1 afla|EED50271.1 afum|EAL84720.1 agla|OJJ84038.1 akaw|GAA91172.1 aluc|OJZ81500.1 anid|CBF82481.1 anig|EHA25930.1 aory|EIT77321.1 asyd|OJJ54102.1 atub|OJI88681.1 aver|OJJ08633.1 awen|OJJ34407.1 azon|OJJ48149.1

Group4719: asyd|OJJ54108.1 abra|OJJ66555.1 acar|OOF91000.1 acla|EAW10939.1 afis|EAW20922.1 afla|EED50276.1 afum|EAL84712.1 agla|OJJ83967.1 akaw|GAA91160.1 aluc|OJZ81488.1 anid|CBF82447.1 aory|EIT77271.1 ater|EAU38569.1 atub|OJI88692.1 aver|OJJ08626.1 awen|OJJ34398.1 azon|OJJ48134.1

Group4720: abra|OJJ66561.1 acar|OOF91002.1 acla|EAW10936.1 afis|EAW20925.1 afla|EED50280.1 afum|EAL84709.1 agla|OJJ83962.1 akaw|GAA91156.1 aluc|OJZ81484.1 anid|CBF82499.1 anig|EHA25943.1 aory|EIT77329.1 asyd|OJJ54089.1 atub|OJI88697.1 aver|OJJ08649.1 awen|OJJ34393.1 azon|OJJ48124.1

Group4721: abra|OJJ66567.1 acar|OOF91008.1 acla|EAW10928.1 afis|EAW20933.1 afla|EED50287.1 afum|EAL84701.1 akaw|GAA91150.1 aluc|OJZ81477.1 anid|CBF82506.1 anig|EHA25949.1 aory|EIT77334.1 asyd|OJJ54079.1 ater|EAU38578.1 atub|OJI88702.1 aver|OJJ08659.1 awen|OJJ34385.1 azon|OJJ48116.1

Group4722: abra|OJJ66575.1 acar|OOF91016.1 acla|EAW10915.1 afis|EAW20943.1 afla|EED50297.1 afum|EAL84691.2 agla|OJJ83944.1 akaw|GAA91143.1 aluc|OJZ81469.1 anid|CBF82538.1 anig|EHA25958.1 asyd|OJJ54067.1 ater|EAU38587.1 atub|OJI88709.1 aver|OJJ08675.1 awen|OJJ34373.1 azon|OJJ48109.1

Group4723: abra|OJJ66586.1 acar|OOF91023.1 acla|EAW10905.1 afis|EAW20953.1 afla|EED50309.1 afum|EAL84681.2 agla|OJJ83927.1 akaw|GAA91131.1 aluc|OJZ81460.1 anid|CBF82556.1 anig|EHA25969.1 asyd|OJJ54057.1 ater|EAU38604.1 atub|OJI88719.1 aver|OJJ08684.1 awen|OJJ34363.1 azon|OJJ48099.1

Group4724: abra|OJJ66599.1 acla|EAW10893.1 afis|EAW20965.1 afla|EED50319.1 afum|EAL84668.1 agla|OJJ84042.1 akaw|GAA91120.1 aluc|OJZ81448.1 anid|CBF82576.1 anig|EHA25978.1 aory|EIT77287.1 asyd|OJJ54044.1 ater|EAU38615.1 atub|OJI88731.1 aver|OJJ08698.1 awen|OJJ34350.1 azon|OJJ48087.1

Group4725: abra|OJJ66611.1 acar|OOF91037.1 acla|EAW10885.1 afis|EAW20973.1 afla|EED50328.1 afum|EAL84660.1 agla|OJJ84051.1 akaw|GAA91112.1 aluc|OJZ81438.1 anid|CBF82591.1 anig|EHA25986.1 asyd|OJJ54034.1 ater|EAU38623.1 atub|OJI88743.1 aver|OJJ08706.1 awen|OJJ34340.1 azon|OJJ48080.1

Group4726: abra|OJJ66614.1 acla|EAW10882.1 afis|EAW20976.1 afla|EED50331.1 afum|EAL84657.1 agla|OJJ84054.1 akaw|GAA91109.1 aluc|OJZ81435.1 anid|CBF82596.1 anig|EHA25989.1 aory|EIT77310.1 asyd|OJJ54031.1 ater|EAU38626.1 atub|OJI88746.1 aver|OJJ08709.1 awen|OJJ34336.1 azon|OJJ48077.1

Group4727: abra|OJJ66618.1 acar|OOF91041.1 acla|EAW10879.1 afis|EAW20979.1 afla|EED50334.1 afum|EAL84654.1 agla|OJJ84056.1 akaw|GAA91107.1 aluc|OJZ81431.1 anid|CBF82600.1 anig|EHA25991.1 aory|EIT77351.1 asyd|OJJ54029.1 atub|OJI88749.1 aver|OJJ08711.1 awen|OJJ34333.1 azon|OJJ48075.1

Group4728: abra|OJJ66620.1 acar|OOF91043.1 acla|EAW10877.1 afis|EAW20981.1 afla|EED50336.1 afum|EAL84649.2 agla|OJJ84058.1 akaw|GAA91105.1 aluc|OJZ81429.1 anid|CBF82602.1 anig|EHA25993.1 aory|EIT77328.1 asyd|OJJ54028.1 ater|EAU38629.1 atub|OJI88751.1 aver|OJJ08712.1 awen|OJJ34331.1

Group4729: abra|OJJ66625.1 acar|OOF91049.1 acla|EAW10871.1 afis|EAW20987.1 afla|EED50342.1 afum|EAL84643.1 agla|OJJ84066.1 akaw|GAA91100.1 aluc|OJZ81423.1 anid|CBF82609.1 anig|EHA25998.1 aory|EIT77368.1 asyd|OJJ54024.1 ater|EAU38636.1 atub|OJI88758.1 aver|OJJ08716.1 awen|OJJ34323.1

Group4730: abra|OJJ66659.1 acla|EAW11801.1 afis|EAW19411.1 afla|EED50366.1 afum|EAL87083.1 agla|OJJ81125.1 akaw|GAA89609.1 aluc|OJZ80789.1 anid|CBF74351.1 anig|EHA27241.1 aory|EIT83143.1 asyd|OJJ57790.1 ater|EAU34717.1 atub|OJI87640.1 aver|OJJ05374.1 awen|OJJ36807.1 azon|OJJ42993.1

Group4731: abra|OJJ66692.1 acla|EAW11827.1 afis|EAW19439.1 afla|EED56646.1 afum|EAL87057.1 agla|OJJ81151.1 akaw|GAA89636.1 aluc|OJZ80754.1 anid|CBF81391.1 anig|EHA27270.1 aory|EIT79642.1 asyd|OJJ59398.1 ater|EAU34740.1 atub|OJI87610.1 aver|OJJ02874.1 awen|OJJ36836.1 azon|OJJ42962.1

Group4732: abra|OJJ66705.1 acla|EAW11836.1 afis|EAW19447.1 afla|EED56655.1 afum|EAL87048.1 agla|OJJ81162.1 akaw|GAA89647.1 aluc|OJZ80742.1 anid|CBF81407.1 anig|EHA27280.1 aory|EIT79638.1 asyd|OJJ59407.1 ater|EAU34749.1 atub|OJI87599.1 aver|OJJ02866.1 awen|OJJ36848.1 azon|OJJ42952.1

Group4733: abra|OJJ66707.1 acla|EAW11838.1 afis|EAW19449.1 afla|EED56657.1 afum|EAL87047.1 akaw|GAA89649.1 aluc|OJZ80740.1 anid|CBF81410.1 anig|EHA27282.1 aory|EIT79686.1 asyd|OJJ59409.1 ater|EAU34751.1 atub|OJI87597.1 aver|OJJ02864.1 awen|OJJ36850.1 azon|OJJ42950.1 agla|OJJ81164.1

Group4734: abra|OJJ66724.1 acar|OOF93812.1 acla|EAW11854.1 afis|EAW19465.1 afla|EED56676.1 afum|EAL87031.2 agla|OJJ81183.1 akaw|GAA89665.1 aluc|OJZ80724.1 anid|CBF75488.1 anig|EHA27296.1 asyd|OJJ59755.1 ater|EAU34775.1 atub|OJI87580.1 aver|OJJ02533.1 awen|OJJ36868.1 azon|OJJ42934.1

Group4735: abra|OJJ66756.1 acar|OOF93837.1 acla|EAW11882.1 afis|EAW19493.1 afla|EED56712.1 afum|EAL87001.1 agla|OJJ81223.1 akaw|GAA89690.1 aluc|OJZ80695.1 anid|CBF75448.1 anig|EHA27321.1 aory|EIT73873.1 asyd|OJJ59776.1 atub|OJI87554.1 aver|OJJ02510.1 awen|OJJ36909.1 azon|OJJ42906.1

Group4736: abra|OJJ66763.1 acar|OOF93844.1 acla|EAW11889.1 afis|EAW19501.1 afla|EED56721.1 afum|EAL86992.2 agla|OJJ81672.1 akaw|GAA89699.1 aluc|OJZ80686.1 anid|CBF75440.1 anig|EHA27328.1 asyd|OJJ59782.1 ater|EAU34816.1 atub|OJI87544.1 aver|OJJ02506.1 awen|OJJ36916.1 azon|OJJ42901.1

Group4737: abra|OJJ66766.1 acar|OOF93847.1 acla|EAW11892.1 afis|EAW19503.1 afla|EED56726.1 afum|EAL86990.1 akaw|GAA89703.1 aluc|OJZ80681.1 anid|CBF75427.1 anig|EHA27331.1 aory|EIT73853.1 asyd|OJJ59788.1 ater|EAU34818.1 atub|OJI87540.1 aver|OJJ02500.1 awen|OJJ36922.1 azon|OJJ43879.1

Group4738: abra|OJJ66797.1 acar|OOF93875.1 acla|EAW14976.1 afis|EAW24501.1 afla|EED45009.1 afum|EAL86171.1 akaw|GAA86862.1 aluc|OJZ89986.1 anid|CBF82363.1 anig|EHA27357.1 aory|EIT73712.1 asyd|OJJ53691.1 ater|EAU36993.1 atub|OJI84824.1 aver|OJJ03933.1 awen|OJJ33356.1 azon|OJJ42743.1

Group4739: abra|OJJ66827.1 acar|OOF91605.1 acla|EAW14410.1 afis|EAW19118.1 afla|EED54296.1 afum|EAL93850.1 agla|OJJ80433.1 akaw|GAA90463.1 aluc|OJZ85760.1 anid|CBF79726.1 anig|EHA24796.1 aory|EIT76076.1 asyd|OJJ58074.1 atub|OJI79545.1 aver|OJJ04333.1 awen|OJJ37449.1 azon|OJJ51126.1

Group4740: abra|OJJ66837.1 acar|OOF91615.1 acla|EAW14421.1 afis|EAW19128.1 afla|EED54286.1 afum|EAL93860.1 agla|OJJ88598.1 akaw|GAA90474.1 aluc|OJZ85750.1 anid|CBF79741.1 anig|EHA24806.1 aory|EIT76040.1 asyd|OJJ58085.1 ater|EAU37395.1 aver|OJJ04323.1 awen|OJJ37461.1 azon|OJJ51139.1

Group4741: abra|OJJ66867.1 acar|OOF91645.1 acla|EAW14443.1 afis|EAW19151.1 afla|EED54254.1 afum|EAL93882.1 agla|OJJ88571.1 akaw|GAA91541.1 aluc|OJZ85715.1 anid|CBF86303.1 anig|EHA24829.1 aory|EIT76113.1 asyd|OJJ65007.1 atub|OJI79505.1 aver|OJI99126.1 awen|OJJ37504.1 azon|OJJ51169.1

Group4742: abra|OJJ66868.1 acar|OOF91644.1 acla|EAW14444.1 afis|EAW19152.1 afla|EED54253.1 afum|EAL93883.1 agla|OJJ88570.1 akaw|GAA91540.1 aluc|OJZ85714.1 anid|CBF86301.1 anig|EHA24830.1 aory|EIT76066.1 asyd|OJJ65008.1 atub|OJI79504.1 aver|OJI99127.1 awen|OJJ37505.1 azon|OJJ51171.1

Group4743: abra|OJJ66882.1 acar|OOF91656.1 acla|EAW14456.1 afis|EAW19164.1 afla|EED54233.1 afum|EAL93894.1 agla|OJJ88560.1 akaw|GAA91528.1 aluc|OJZ85701.1 anig|EHA24842.1 aory|EIT75922.1 asyd|OJJ65019.1 ater|EAU37309.1 atub|OJI79491.1 aver|OJI99137.1 awen|OJJ37517.1 azon|OJJ51184.1

Group4744: abra|OJJ66892.1 acla|EAW14464.1 afis|EAW19172.1 afla|EED54223.1 afum|EAL93902.1 agla|OJJ88552.1 akaw|GAA91520.1 aluc|OJZ85690.1 anid|CBF86268.1 anig|EHA24851.1 aory|EIT76131.1 asyd|OJJ65027.1 ater|EAU37301.1 atub|OJI79480.1 aver|OJI99146.1 awen|OJJ37525.1 azon|OJJ51191.1

Group4745: abra|OJJ66901.1 acla|EAW14472.1 afis|EAW19181.1 afla|EED54214.1 afum|EAL93911.1 agla|OJJ88542.1 akaw|GAA91511.1 aluc|OJZ85681.1 anid|CBF86255.1 anig|EHA24860.1 aory|EIT76103.1 asyd|OJJ65037.1 ater|EAU37292.1 atub|OJI79471.1 aver|OJI99156.1 awen|OJJ37535.1 azon|OJJ51201.1

Group4746: abra|OJJ66922.1 acar|OOF91689.1 acla|EAW14488.1 afis|EAW19202.1 afla|EED47710.1 afum|EAL93932.1 agla|OJJ88510.1 akaw|GAA91497.1 aluc|OJZ85658.1 anid|CBF78552.1 anig|EHA24875.1 asyd|OJJ65058.1 ater|EAU37278.1 atub|OJI79450.1 aver|OJI99183.1 awen|OJJ37555.1 azon|OJJ51216.1

Group4747: abra|OJJ66925.1 acar|OOF91693.1 acla|EAW14491.1 afis|EAW19205.1 afla|EED47707.1 afum|EAL93935.1 agla|OJJ88506.1 akaw|GAA91492.1 aluc|OJZ85653.1 anid|CBF78559.1 anig|EHA24879.1 aory|EIT80229.1 asyd|OJJ65062.1 ater|EAU37267.1 atub|OJI79445.1 aver|OJI99186.1 awen|OJJ37558.1

Group4748: abra|OJJ66941.1 acar|OOF91707.1 acla|EAW14499.1 afis|EAW19213.1 afla|EED47687.1 afum|EAL93946.1 agla|OJJ88483.1 akaw|GAA91480.1 aluc|OJZ85641.1 anid|CBF78587.1 anig|EHA24893.1 aory|EIT80252.1 asyd|OJJ65082.1 atub|OJI79432.1 aver|OJI99207.1 awen|OJJ37570.1 azon|OJJ51230.1

Group4749: abra|OJJ66943.1 acar|OOF91709.1 acla|EAW14502.1 afis|EAW19216.1 afla|EED47684.1 afum|EAL93949.1 agla|OJJ88481.1 akaw|GAA91477.1 aluc|OJZ85639.1 anid|CBF78593.1 anig|EHA24895.1 asyd|OJJ65085.1 ater|EAU37252.1 atub|OJI79430.1 aver|OJI99210.1 awen|OJJ37573.1 azon|OJJ51232.1

Group4750: acla|EAW15049.1 afis|EAW24612.1 afla|EED56125.1 afum|EAL86242.1 aory|EIT75528.1 asyd|OJJ61495.1 ater|EAU36605.1 aver|OJJ07466.1 awen|OJJ35562.1 acar|OOF93879.1 akaw|GAA86866.1 aluc|OJZ89989.1 anig|EHA27359.1 atub|OJI84820.1 afis|EAW22020.1 afla|EED45483.1 awen|OJJ30949.1

Group4751: ater|EAU37911.1 abra|OJJ76343.1 acar|OOF99594.1 acla|EAW14072.1 afis|EAW18779.1 afla|EED50738.1 afum|EAL93511.1 agla|OJJ80499.1 akaw|GAA82270.1 aluc|OJZ91870.1 anid|CBF69736.1 aory|EIT78057.1 asyd|OJJ58932.1 atub|OJI86321.1 aver|OJJ01528.1 awen|OJJ32350.1 azon|OJJ47115.1

Group4752: abra|OJJ67071.1 acla|EAW06987.1 afis|EAW16601.1 afla|EED45020.1 afum|EAL89197.1 agla|OJJ82651.1 akaw|GAA91084.1 aluc|OJZ87901.1 anid|CBF84000.1 anig|EHA20197.1 aory|EIT73543.1 asyd|OJJ64729.1 ater|EAU36681.1 atub|OJI80750.1 aver|OJI98833.1 awen|OJJ40205.1 azon|OJJ49072.1

Group4753: abra|OJJ67141.1 acar|OOF98415.1 afis|EAW15396.1 afla|EED47554.1 agla|OJJ83078.1 akaw|GAA91028.1 aluc|OJZ87831.1 anid|CBF80436.1 anig|EHA20310.1 aory|EIT74522.1 asyd|OJJ54780.1 ater|EAU29457.1 atub|OJI80810.1 aver|OJJ07385.1 awen|OJJ38130.1 agla|OJJ86276.1 aver|OJJ05913.1

Group4754: abra|OJJ67165.1 acar|OOF98396.1 acla|EAW14563.1 afis|EAW19276.1 afum|EAL94002.1 agla|OJJ88397.1 akaw|GAA91009.1 aluc|OJZ87810.1 anid|CBF80662.1 anig|EHA20292.1 aory|EIT80262.1 asyd|OJJ54608.1 ater|EAU38851.1 atub|OJI80835.1 aver|OJJ07548.1 awen|OJJ37636.1 azon|OJJ44884.1

Group4755: abra|OJJ67166.1 acar|OOF98395.1 acla|EAW14562.1 afis|EAW19275.1 afum|EAL94001.1 agla|OJJ88398.1 akaw|GAA91008.1 aluc|OJZ87809.1 anid|CBF78749.1 anig|EHA20291.1 aory|EIT80174.1 asyd|OJJ65226.1 ater|EAU29489.1 atub|OJI80836.1 aver|OJI99376.1 awen|OJJ37635.1 azon|OJJ44885.1

Group4756: abra|OJJ67170.1 acar|OOF98393.1 acla|EAW14559.1 afis|EAW19272.1 afla|EED47604.1 afum|EAL93998.1 agla|OJJ88411.1 akaw|GAA91004.1 aluc|OJZ87805.1 anid|CBF78725.1 anig|EHA20287.1 aory|EIT80192.1 asyd|OJJ65224.1 ater|EAU29492.1 atub|OJI80840.1 aver|OJI99373.1 awen|OJJ37633.1

Group4757: abra|OJJ67184.1 acla|EAW14539.1 afis|EAW19257.1 afla|EED47619.1 afum|EAL93984.1 agla|OJJ88438.1 akaw|GAA90992.1 aluc|OJZ87791.1 anid|CBF78675.1 anig|EHA20272.1 aory|EIT80266.1 asyd|OJJ65207.1 ater|EAU29518.1 atub|OJI80856.1 aver|OJI99356.1 awen|OJJ37618.1 azon|OJJ47007.1

Group4758: abra|OJJ67185.1 acar|OOF98378.1 acla|EAW14538.1 afis|EAW19254.1 afla|EED47620.1 afum|EAL93983.1 agla|OJJ88441.1 akaw|GAA90991.1 aluc|OJZ87789.1 anid|CBF78673.1 anig|EHA20271.1 asyd|OJJ65206.1 ater|EAU29513.1 atub|OJI80857.1 aver|OJI99355.1 awen|OJJ37617.1 azon|OJJ47006.1

Group4759: abra|OJJ67194.1 acar|OOF98370.1 acla|EAW14530.1 afis|EAW19248.1 afla|EED47630.1 afum|EAL93977.1 agla|OJJ88449.1 akaw|GAA90984.1 aluc|OJZ87781.1 anid|CBF78660.1 aory|EIT80194.1 asyd|OJJ65198.1 ater|EAU29502.1 atub|OJI80866.1 aver|OJI99347.1 awen|OJJ37605.1 azon|OJJ51258.1

Group4760: abra|OJJ67196.1 acar|OOF98368.1 acla|EAW14528.1 afis|EAW19246.1 afla|EED47632.1 afum|EAL93975.1 agla|OJJ88451.1 akaw|GAA90982.1 aluc|OJZ87779.1 anid|CBF78656.1 anig|EHA20261.1 aory|EIT80195.1 asyd|OJJ65196.1 atub|OJI80868.1 aver|OJI99345.1 awen|OJJ37603.1 azon|OJJ51256.1

Group4761: abra|OJJ67198.1 acar|OOF98367.1 acla|EAW14527.1 afis|EAW19245.1 afum|EAL93974.1 agla|OJJ88452.1 akaw|GAA90981.1 aluc|OJZ87776.1 anid|CBF85414.1 anig|EHA20260.1 aory|EIT80249.1 asyd|OJJ64070.1 ater|EAU36216.1 atub|OJI80869.1 aver|OJI98157.1 awen|OJJ37602.1 azon|OJJ50356.1

Group4762: abra|OJJ67214.1 acar|OOF98344.1 acla|EAW14514.1 afis|EAW19232.1 afla|EED47656.1 afum|EAL93961.1 agla|OJJ88474.1 akaw|GAA90963.1 aluc|OJZ87759.1 anid|CBF78621.1 anig|EHA20244.1 asyd|OJJ65162.1 ater|EAU37231.1 atub|OJI80885.1 aver|OJI99313.1 awen|OJJ37589.1 azon|OJJ51243.1

Group4763: agla|OJJ81007.1 abra|OJJ73670.1 acla|EAW07584.1 afis|EAW22610.1 afla|EED49808.1 afum|EAL90588.1 akaw|GAA88487.1 aluc|OJZ88383.1 anid|CBF88276.1 anig|EHA18110.1 aory|EIT77703.1 asyd|OJJ63682.1 ater|EAU39094.1 atub|OJI90977.1 awen|OJJ35238.1 azon|OJJ49869.1 aver|OJI97256.1

Group4764: abra|OJJ67253.1 acar|OOF98308.1 acla|EAW10399.1 afis|EAW25178.1 afla|EED53636.1 afum|EAL89852.1 agla|OJJ81059.1 akaw|GAA91670.1 aluc|OJZ87725.1 anid|CBF85243.1 aory|EIT75074.1 asyd|OJJ62971.1 ater|EAU34337.1 atub|OJI80918.1 aver|OJI97996.1 awen|OJJ33660.1 azon|OJJ42611.1

Group4765: abra|OJJ67256.1 acar|OOF98307.1 acla|EAW10397.1 afis|EAW25180.1 afla|EED48197.1 afum|EAL89854.1 agla|OJJ84746.1 akaw|GAA91671.1 aluc|OJZ87723.1 anig|EHA20217.1 aory|EIT74879.1 asyd|OJJ62969.1 atub|OJI80919.1 aver|OJI97998.1 azon|OJJ47985.1 aluc|OJZ87724.1 aory|EIT72909.1

Group4766: abra|OJJ67270.1 acar|OOF98295.1 acla|EAW10415.1 afis|EAW25154.1 afla|EED55279.1 afum|EAL89829.2 akaw|GAA91685.1 aluc|OJZ87709.1 anid|CBF85640.1 anig|EHA20891.1 aory|EIT83616.1 asyd|OJJ64240.1 ater|EAU39433.1 atub|OJI80933.1 aver|OJI98333.1 awen|OJJ40784.1 azon|OJJ46869.1

Group4767: abra|OJJ67317.1 acar|OOF89802.1 acla|EAW06419.1 afis|EAW18523.1 afum|EAL93218.1 agla|OJJ85135.1 akaw|GAA86501.1 aluc|OJZ85043.1 anid|CBF70217.1 anig|EHA25261.1 aory|EIT78635.1 asyd|OJJ58776.1 ater|EAU30045.1 atub|OJI80554.1 aver|OJJ01385.1 awen|OJJ31075.1 azon|OJJ50584.1

Group4768: abra|OJJ67318.1 acar|OOF92497.1 acla|EAW06418.1 afis|EAW18522.1 afum|EAL93217.1 agla|OJJ85134.1 akaw|GAA86500.1 aluc|OJZ85042.1 anid|CBF70221.1 anig|EHA25260.1 aory|EIT78625.1 asyd|OJJ58775.1 ater|EAU30046.1 atub|OJI80555.1 aver|OJJ01384.1 awen|OJJ31074.1 azon|OJJ50583.1

Group4769: abra|OJJ67348.1 acar|OOF92467.1 acla|EAW06388.1 afis|EAW18494.1 afla|EED52910.1 afum|EAL93184.2 agla|OJJ85089.1 akaw|GAA86470.1 aluc|OJZ85011.1 anid|CBF70166.1 anig|EHA25232.1 asyd|OJJ58798.1 ater|EAU29920.1 atub|OJI80586.1 aver|OJJ01402.1 awen|OJJ31047.1 azon|OJJ50558.1

Group4770: abra|OJJ67349.1 acla|EAW06387.1 afis|EAW18493.1 afla|EED52911.1 afum|EAL93183.1 agla|OJJ85088.1 akaw|GAA86469.1 aluc|OJZ85010.1 anid|CBF70164.1 anig|EHA25231.1 aory|EIT78601.1 asyd|OJJ58799.1 ater|EAU29919.1 atub|OJI80587.1 aver|OJJ01403.1 awen|OJJ31046.1 azon|OJJ50557.1

Group4771: abra|OJJ67361.1 acar|OOF92461.1 acla|EAW06382.1 afis|EAW18488.1 afla|EED52919.1 afum|EAL93178.1 agla|OJJ85084.1 akaw|GAA86459.1 aluc|OJZ85002.1 anid|CBF70153.1 anig|EHA25219.1 aory|EIT78648.1 asyd|OJJ58805.1 ater|EAU29913.1 atub|OJI80597.1 aver|OJJ01408.1 awen|OJJ31042.1

Group4772: abra|OJJ67384.1 acar|OOF92429.1 acla|EAW06375.1 afis|EAW18481.1 afla|EED52928.1 afum|EAL93170.1 agla|OJJ85064.1 akaw|GAA86440.1 aluc|OJZ84982.1 anig|EHA25195.1 aory|EIT78719.1 asyd|OJJ58811.1 ater|EAU29904.1 atub|OJI80619.1 aver|OJJ01414.1 awen|OJJ31039.1 azon|OJJ50546.1

Group4773: abra|OJJ67390.1 acla|EAW06371.1 afis|EAW18477.1 afla|EED52933.1 afum|EAL93165.1 agla|OJJ85069.1 akaw|GAA86435.1 aluc|OJZ84977.1 anid|CBF70134.1 anig|EHA25190.1 aory|EIT78710.1 asyd|OJJ58815.1 ater|EAU29894.1 atub|OJI80624.1 aver|OJJ01418.1 awen|OJJ31030.1 azon|OJJ50543.1

Group4774: abra|OJJ67402.1 acar|OOF92414.1 acla|EAW06356.1 afis|EAW18467.1 afla|EED52943.1 afum|EAL93153.1 agla|OJJ85052.1 akaw|GAA86425.1 aluc|OJZ84966.1 anid|CBF70098.1 anig|EHA25179.1 asyd|OJJ58833.1 ater|EAU29884.1 atub|OJI80634.1 aver|OJJ01436.1 awen|OJJ31019.1 azon|OJJ50533.1

Group4775: abra|OJJ67407.1 acar|OOF92409.1 acla|EAW06351.1 afis|EAW18463.1 afla|EED52946.1 afum|EAL93149.1 agla|OJJ85047.1 akaw|GAA86420.1 aluc|OJZ84961.1 anig|EHA25175.1 aory|EIT78711.1 asyd|OJJ58839.1 ater|EAU29881.1 atub|OJI80639.1 aver|OJJ01441.1 awen|OJJ31015.1 azon|OJJ50528.1

Group4776: abra|OJJ67408.1 acar|OOF92408.1 acla|EAW06350.1 afis|EAW18462.1 afum|EAL93148.2 agla|OJJ85046.1 akaw|GAA86419.1 aluc|OJZ84960.1 anid|CBF70091.1 anig|EHA25174.1 aory|EIT78697.1 asyd|OJJ58840.1 ater|EAU29880.1 atub|OJI80640.1 aver|OJJ01442.1 awen|OJJ31014.1 azon|OJJ50527.1

Group4777: abra|OJJ67427.1 acla|EAW06337.1 afis|EAW18449.1 afla|EED52959.1 afum|EAL93134.1 agla|OJJ85027.1 akaw|GAA86403.1 aluc|OJZ84940.1 anid|CBF70066.1 anig|EHA25157.1 aory|EIT78716.1 asyd|OJJ58849.1 ater|EAU29867.1 atub|OJI80660.1 aver|OJJ01450.1 awen|OJJ31001.1 azon|OJJ50515.1

Group4778: abra|OJJ67430.1 acar|OOF92396.1 acla|EAW06334.1 afis|EAW18446.1 afla|EED52965.1 afum|EAL93131.1 agla|OJJ85023.1 akaw|GAA86400.1 aluc|OJZ84936.1 anig|EHA25154.1 aory|EIT78687.1 asyd|OJJ56611.1 ater|EAU29865.1 atub|OJI80663.1 aver|OJJ06112.1 awen|OJJ30998.1 azon|OJJ50512.1

Group4779: abra|OJJ67436.1 acar|OOF92392.1 acla|EAW06326.1 afis|EAW18440.1 afla|EED52973.1 afum|EAL93122.1 agla|OJJ85005.1 akaw|GAA86394.1 aluc|OJZ84928.1 anid|CBF70051.1 anig|EHA25147.1 aory|EIT78715.1 asyd|OJJ58860.1 ater|EAU36110.1 atub|OJI80671.1 aver|OJJ01461.1 awen|OJJ30988.1

Group4780: abra|OJJ67464.1 acar|OOF92368.1 acla|EAW06304.1 afis|EAW18413.1 afum|EAL93093.2 agla|OJJ84980.1 akaw|GAA86372.1 aluc|OJZ84901.1 anid|CBF70733.1 anig|EHA25127.1 aory|EIT77394.1 asyd|OJJ58443.1 ater|EAU29828.1 atub|OJI80697.1 aver|OJJ01043.1 awen|OJJ31737.1 azon|OJJ50496.1

Group4781: abra|OJJ67573.1 acar|OOF91947.1 acla|EAW14347.1 afis|EAW18943.1 afla|EED50976.1 afum|EAL93676.1 agla|OJJ83697.1 akaw|GAA89102.1 anid|CBF79785.1 anig|EHA25704.1 aory|EIT77067.1 asyd|OJJ58115.1 ater|EAU33169.1 atub|OJI88441.1 aver|OJJ04290.1 awen|OJJ30692.1 azon|OJJ47252.1

Group4782: abra|OJJ67611.1 acar|OOF91958.1 afla|EED45464.1 akaw|GAA89128.1 aluc|OJZ81710.1 anid|CBF87139.1 anig|EHA19582.1 aory|EIT82377.1 asyd|OJJ60382.1 ater|EAU30789.1 atub|OJI88474.1 aver|OJI99711.1 awen|OJJ41962.1 azon|OJJ48033.1 anid|CBF87137.1 ater|EAU29990.1 azon|OJJ46093.1

Group4783: abra|OJJ67612.1 acar|OOF91957.1 acla|EAW11005.1 afis|EAW20848.1 afum|EAL84785.1 agla|OJJ83719.1 akaw|GAA89129.1 aluc|OJZ81709.1 anid|CBF82870.1 anig|EHA25729.1 aory|EIT82455.1 asyd|OJJ61571.1 ater|EAU30790.1 atub|OJI88475.1 aver|OJJ00891.1 awen|OJJ34515.1 azon|OJJ51801.1

Group4784: abra|OJJ67727.1 acar|OOF89959.1 acla|EAW11003.1 afis|EAW20853.1 afum|EAL84779.1 agla|OJJ83722.1 akaw|GAA89221.1 aluc|OJZ81607.1 anid|CBF82773.1 anig|EHA25838.1 aory|EIT82434.1 asyd|OJJ61572.1 ater|EAU30792.1 atub|OJI88578.1 aver|OJJ00892.1 awen|OJJ34520.1 azon|OJJ51799.1

Group4785: abra|OJJ67740.1 acar|OOF90939.1 acla|EAW11002.1 afis|EAW20858.1 afla|EED45450.1 afum|EAL84777.1 agla|OJJ83739.1 akaw|GAA92431.1 aluc|OJZ81591.1 anid|CBF82824.1 anig|EHA25850.1 aory|EIT82442.1 asyd|OJJ61579.1 ater|EAU30801.1 atub|OJI88592.1 aver|OJJ00901.1 awen|OJJ34524.1

Group4786: abra|OJJ67741.1 acar|OOF90942.1 acla|EAW11001.1 afis|EAW20860.1 afla|EED45446.1 afum|EAL84775.2 agla|OJJ83746.1 akaw|GAA92432.1 aluc|OJZ81589.1 anid|CBF84701.1 anig|EHA25851.1 aory|EIT82460.1 asyd|OJJ53318.1 ater|EAU30803.1 atub|OJI88593.1 aver|OJJ08425.1 awen|OJJ34510.1

Group4787: abra|OJJ67745.1 acar|OOF90946.1 acla|EAW10994.1 afis|EAW20866.1 afla|EED45545.1 afum|EAL84768.1 agla|OJJ83974.1 akaw|GAA92436.1 aluc|OJZ81584.1 anid|CBF82836.1 anig|EHA25855.1 aory|EIT75798.1 asyd|OJJ61586.1 ater|EAU30822.1 atub|OJI88597.1 aver|OJJ00908.1 azon|OJJ51775.1

Group4788: abra|OJJ67856.1 acar|OOF92835.1 acla|EAW11744.1 afis|EAW23777.1 afla|EED51577.1 afum|EAL87211.1 agla|OJJ78928.1 akaw|GAA91862.1 aluc|OJZ82036.1 anid|CBF79830.1 anig|EHA22220.1 aory|EIT80612.1 asyd|OJJ58143.1 atub|OJI84591.1 aver|OJJ04260.1 awen|OJJ33700.1 azon|OJJ42841.1

Group4789: abra|OJJ67858.1 acar|OOF92838.1 acla|EAW11740.1 afis|EAW23773.1 afum|EAL87207.1 agla|OJJ78925.1 akaw|GAA91865.1 aluc|OJZ82034.1 anid|CBF79824.1 anig|EHA22222.1 aory|EIT80706.1 asyd|OJJ58142.1 ater|EAU31317.1 atub|OJI84589.1 aver|OJJ04262.1 awen|OJJ33702.1 azon|OJJ42839.1

Group4790: abra|OJJ67862.1 acar|OOF92842.1 acla|EAW11736.1 afis|EAW23769.1 afla|EED51570.1 afum|EAL87203.1 agla|OJJ78951.1 akaw|GAA91869.1 aluc|OJZ82030.1 anid|CBF79819.1 anig|EHA22226.1 asyd|OJJ58139.1 ater|EAU31313.1 atub|OJI84584.1 aver|OJJ04266.1 awen|OJJ33706.1 azon|OJJ42834.1

Group4791: abra|OJJ67868.1 acar|OOF92850.1 acla|EAW11729.1 afis|EAW23761.1 afla|EED51548.1 afum|EBA27305.1 agla|OJJ78961.1 akaw|GAA91875.1 aluc|OJZ82024.1 anid|CBF79789.1 anig|EHA22231.1 aory|EIT80682.1 asyd|OJJ58124.1 atub|OJI84577.1 aver|OJJ04282.1 awen|OJJ33686.1 azon|OJJ42828.1

Group4792: acar|OOF90067.1 abra|OJJ73116.1 acla|EAW15001.1 afla|EED51114.1 agla|OJJ82961.1 akaw|GAA88053.1 aluc|OJZ81315.1 anid|CBF80452.1 anig|EHA18619.1 aory|EIT81295.1 asyd|OJJ54765.1 atub|OJI91518.1 aver|OJJ07402.1 awen|OJJ40911.1 afla|EED45581.1 agla|OJJ78689.1 aory|EIT75557.1

Group4793: abra|OJJ67907.1 acar|OOF92871.1 acla|EAW11701.1 afis|EAW23722.1 afla|EED51506.1 afum|EAL87162.1 agla|OJJ78977.1 akaw|GAA91912.1 aluc|OJZ81980.1 anid|CBF85571.1 anig|EHA22274.1 aory|EIT80734.1 asyd|OJJ58128.1 atub|OJI84532.1 aver|OJJ04277.1 afis|EAW24685.1 afum|EAL89325.1

Group4794: abra|OJJ67912.1 acar|OOF92764.1 acla|EAW12797.1 afis|EAW17960.1 afla|EED51947.1 afum|EAL91771.1 agla|OJJ80208.1 akaw|GAA91284.1 aluc|OJZ82785.1 anid|CBF79952.1 aory|EIT81471.1 asyd|OJJ58217.1 ater|EAU31449.1 atub|OJI89017.1 aver|OJJ02205.1 awen|OJJ41581.1 azon|OJJ48294.1

Group4795: abra|OJJ67914.1 acar|OOF92765.1 acla|EAW12795.1 afis|EAW17958.1 afum|EAL91769.1 agla|OJJ80206.1 akaw|GAA91282.1 aluc|OJZ82787.1 anid|CBF79949.1 anig|EHA21228.1 aory|EIT81485.1 asyd|OJJ58215.1 ater|EAU31447.1 atub|OJI89019.1 aver|OJJ02207.1 awen|OJJ41584.1 azon|OJJ48296.1

Group4796: abra|OJJ67916.1 acar|OOF92768.1 acla|EAW12793.1 afis|EAW17956.1 afla|EED51951.1 afum|EAL91767.1 agla|OJJ80203.1 akaw|GAA91280.1 aluc|OJZ82789.1 anid|CBF79945.1 anig|EHA21230.1 aory|EIT81421.1 asyd|OJJ58213.1 ater|EAU31444.1 aver|OJJ02209.1 awen|OJJ41586.1 azon|OJJ48301.1

Group4797: abra|OJJ67924.1 acla|EAW12786.1 afis|EAW17949.1 afla|EED51958.1 afum|EAL91759.1 agla|OJJ80196.1 akaw|GAA91273.1 aluc|OJZ82796.1 anid|CBF79927.1 anig|EHA21237.1 aory|EIT81486.1 asyd|OJJ58203.1 ater|EAU31432.1 atub|OJI89027.1 aver|OJJ04203.1 awen|OJJ41594.1 azon|OJJ48309.1

Group4798: abra|OJJ67932.1 acar|OOF92782.1 afis|EAW23834.1 afla|EED53910.1 afum|EAL87265.1 agla|OJJ82500.1 akaw|GAA91265.1 aluc|OJZ82804.1 anid|CBF79896.1 anig|EHA21245.1 aory|EIT78489.1 asyd|OJJ58183.1 ater|EAU31393.1 atub|OJI89035.1 aver|OJJ04222.1 awen|OJJ33792.1 azon|OJJ48752.1

Group4799: abra|OJJ67933.1 acar|OOF92783.1 afis|EAW20400.1 afla|EED51970.1 afum|EAL85499.2 agla|OJJ82479.1 akaw|GAA91264.1 aluc|OJZ82805.1 anid|CBF79898.1 anig|EHA21246.1 aory|EIT81412.1 asyd|OJJ58184.1 ater|EAU31392.1 atub|OJI89036.1 aver|OJJ04221.1 awen|OJJ33791.1 azon|OJJ48751.1

Group4800: abra|OJJ67935.1 acar|OOF92784.1 afis|EAW23832.1 afla|EED51973.1 akaw|GAA91261.1 aluc|OJZ82807.1 aory|EIT81418.1 asyd|OJJ58186.1 ater|EAU31390.1 atub|OJI89038.1 awen|OJJ33790.1 azon|OJJ48756.1 acla|EAW11624.1 afum|EAL87264.2 agla|OJJ82480.1 aver|OJJ04219.1 anid|CBF79902.1

Group4801: abra|OJJ67941.1 acar|OOF92790.1 acla|EAW11631.1 afis|EAW23825.1 afla|EED51981.1 agla|OJJ82487.1 akaw|GAA91255.1 aluc|OJZ82813.1 anig|EHA21253.1 aory|EIT81429.1 asyd|OJJ57257.1 ater|EAU31380.1 atub|OJI89044.1 aver|OJJ05344.1 awen|OJJ33783.1 azon|OJJ42885.1 ater|EAU29233.1

Group4802: abra|OJJ67948.1 acar|OOF92797.1 acla|EAW11638.1 afis|EAW23819.1 afla|EED51986.1 afum|EAL87254.1 agla|OJJ82465.1 akaw|GAA91249.1 aluc|OJZ82821.1 anid|CBF79892.1 anig|EHA21260.1 asyd|OJJ58178.1 ater|EAU31374.1 atub|OJI89050.1 aver|OJJ04226.1 awen|OJJ33776.1 azon|OJJ42881.1

Group4803: abra|OJJ67949.1 acar|OOF92796.1 acla|EAW11639.1 afis|EAW23818.1 afla|EED51987.1 afum|EAL87253.1 agla|OJJ82466.1 akaw|GAA91248.1 aluc|OJZ82822.1 anid|CBF79891.1 aory|EIT81399.1 asyd|OJJ58177.1 ater|EAU31375.1 atub|OJI89051.1 aver|OJJ04227.1 awen|OJJ33775.1 azon|OJJ42880.1

Group4804: abra|OJJ67950.1 acla|EAW11640.1 afla|EED51988.1 afum|EAL87252.1 agla|OJJ82463.1 akaw|GAA91247.1 aluc|OJZ82823.1 anid|CBF79889.1 anig|EHA21261.1 asyd|OJJ58176.1 ater|EAU31372.1 atub|OJI89052.1 aver|OJJ04228.1 awen|OJJ33774.1 azon|OJJ42878.1 afis|EAW23817.1 aory|EIT81468.1

Group4805: abra|OJJ67953.1 acla|EAW11642.1 afis|EAW23815.1 afla|EED51990.1 afum|EAL87250.1 agla|OJJ82456.1 akaw|GAA91244.1 aluc|OJZ82826.1 anid|CBF79885.1 anig|EHA21263.1 aory|EIT81442.1 asyd|OJJ58174.1 ater|EAU31370.1 atub|OJI89056.1 aver|OJJ04230.1 awen|OJJ33772.1 azon|OJJ42875.1

Group4806: abra|OJJ67956.1 acar|OOF92801.1 acla|EAW11645.1 afis|EAW23811.1 afla|EED52004.1 afum|EAL87246.1 agla|OJJ82451.1 akaw|GAA91240.1 aluc|OJZ82829.1 anid|CBF79877.1 anig|EHA21267.1 asyd|OJJ58170.1 ater|EAU31364.1 atub|OJI89059.1 aver|OJJ04233.1 awen|OJJ33767.1 azon|OJJ42871.1

Group4807: abra|OJJ67959.1 acar|OOF92805.1 acla|EAW11649.1 afis|EAW23807.1 afla|EED52008.1 afum|EAL87241.1 agla|OJJ82447.1 akaw|GAA91237.1 aluc|OJZ82832.1 anid|CBF79870.1 anig|EHA21271.1 asyd|OJJ58166.1 ater|EAU31359.1 atub|OJI89062.1 aver|OJJ04236.1 awen|OJJ33764.1 azon|OJJ42867.1

Group4808: abra|OJJ67965.1 acar|OOF92811.1 acla|EAW11655.1 afis|EAW23798.1 afum|EAL87233.1 agla|OJJ78846.1 akaw|GAA91232.1 aluc|OJZ82838.1 anid|CBF79861.1 anig|EHA21275.1 aory|EIT80592.1 asyd|OJJ58161.1 ater|EAU31336.1 atub|OJI89068.1 aver|OJJ04241.1 awen|OJJ33757.1 azon|OJJ42863.1

Group4809: abra|OJJ67967.1 acar|OOF92812.1 acla|EAW11656.1 afis|EAW23797.1 afla|EED51616.1 afum|EAL87231.2 agla|OJJ78845.1 akaw|GAA91231.1 aluc|OJZ82839.1 anid|CBF79860.1 anig|EHA21276.1 asyd|OJJ58160.1 ater|EAU31335.1 atub|OJI89069.1 aver|OJJ04242.1 awen|OJJ33755.1 azon|OJJ42862.1

Group4810: abra|OJJ67987.1 acar|OOF92029.1 acla|EAW12429.1 afis|EAW16736.1 afla|EED47284.1 afum|EAL86311.1 akaw|GAA91215.1 aluc|OJZ82856.1 anid|CBF73583.1 anig|EHA21291.1 aory|EIT79240.1 asyd|OJJ56684.1 ater|EAU29285.1 atub|OJI89086.1 aver|OJJ06039.1 awen|OJJ39986.1 azon|OJJ43671.1

Group4811: abra|OJJ68043.1 acar|OOF90405.1 acla|EAW13685.1 afis|EAW19921.1 afla|EED45891.1 afum|EAL84988.1 akaw|GAA90910.1 aluc|OJZ82916.1 anig|EHA21339.1 aory|EIT75607.1 atub|OJI89142.1 aver|OJJ07332.1 awen|OJJ29957.1 agla|OJJ79335.1 anig|EHA23723.1 atub|OJI82465.1 azon|OJJ42323.1

Group4812: abra|OJJ68119.1 acar|OOF93913.1 acla|EAW09088.1 afis|EAW25680.1 afla|EED51102.1 afum|EAL84350.1 agla|OJJ78750.1 akaw|GAA86900.1 aluc|OJZ90023.1 anid|CBF83938.1 anig|EHA27399.1 aory|EIT81294.1 asyd|OJJ61451.1 ater|EAU29935.1 atub|OJI84784.1 aver|OJJ00786.1 awen|OJJ35396.1

Group4813: abra|OJJ68122.1 acar|OOF93916.1 acla|EAW12863.1 afis|EAW17812.1 afla|EED45597.1 afum|EAL91621.1 agla|OJJ81684.1 akaw|GAA86903.1 aluc|OJZ90026.1 anid|CBF80724.1 anig|EHA27402.1 aory|EIT75545.1 asyd|OJJ54569.1 ater|EAU30289.1 atub|OJI84781.1 aver|OJJ07585.1 awen|OJJ36929.1

Group4814: abra|OJJ68124.1 acar|OOF93918.1 acla|EAW12868.1 afis|EAW17816.1 afla|EED45600.1 afum|EAL91626.1 agla|OJJ81688.1 akaw|GAA86905.1 aluc|OJZ90028.1 anid|CBF80746.1 anig|EHA27404.1 aory|EIT75820.1 asyd|OJJ54540.1 ater|EAU30286.1 atub|OJI84779.1 aver|OJJ07641.1 awen|OJJ36933.1

Group4815: abra|OJJ68159.1 acla|EAW08944.1 akaw|GAA83283.1 aluc|OJZ83793.1 anig|EHA18929.1 atub|OJI90161.1 awen|OJJ34623.1 abra|OJJ68167.1 acar|OOF93405.1 akaw|GAA83276.1 aluc|OJZ83782.1 anig|EHA22297.1 aory|EIT74145.1 asyd|OJJ63613.1 ater|EAU35823.1 atub|OJI90152.1 afum|EAL91589.1

Group4816: abra|OJJ68187.1 acar|OOF97480.1 acla|EAW13068.1 afis|EAW17596.1 afla|EED49224.1 afum|EAL91409.1 agla|OJJ81334.1 akaw|GAA83258.1 aluc|OJZ83764.1 anid|CBF90205.1 anig|EHA18905.1 aory|EIT83157.1 asyd|OJJ61975.1 ater|EAU37034.1 atub|OJI90135.1 aver|OJI96068.1 awen|OJJ37250.1

Group4817: abra|OJJ68191.1 acar|OOF97487.1 acla|EAW13073.1 afis|EAW17591.1 afla|EED49231.1 afum|EAL91404.1 agla|OJJ81358.1 akaw|GAA83254.1 aluc|OJZ83755.1 anid|CBF90214.1 anig|EHA18902.1 aory|EIT83155.1 ater|EAU37028.1 atub|OJI90130.1 aver|OJI96061.1 awen|OJJ37255.1 azon|OJJ50950.1

Group4818: abra|OJJ68192.1 acla|EAW13074.1 afis|EAW17590.1 afla|EED49232.1 afum|EAL91403.1 agla|OJJ81359.1 akaw|GAA83253.1 aluc|OJZ83754.1 anid|CBF90216.1 anig|EHA18901.1 aory|EIT83156.1 asyd|OJJ61968.1 ater|EAU37027.1 atub|OJI90129.1 aver|OJI96060.1 awen|OJJ37256.1 azon|OJJ50951.1

Group4819: abra|OJJ68195.1 acla|EAW13078.1 afis|EAW17586.1 afla|EED49236.1 afum|EAL91399.1 agla|OJJ81366.1 akaw|GAA83250.1 aluc|OJZ83751.1 anid|CBF90223.1 anig|EHA18898.1 aory|EIT83150.1 asyd|OJJ61964.1 ater|EAU37021.1 atub|OJI90126.1 aver|OJI96056.1 awen|OJJ37260.1 azon|OJJ50955.1

Group4820: abra|OJJ68214.1 acar|OOF97497.1 acla|EAW12971.1 afis|EAW17693.1 afla|EED50451.1 afum|EAL91507.1 agla|OJJ79486.1 akaw|GAA83232.1 aluc|OJZ83732.1 anid|CBF89976.1 anig|EHA18881.1 asyd|OJJ62092.1 ater|EAU31959.1 atub|OJI90109.1 aver|OJI96198.1 awen|OJJ37143.1 azon|OJJ50843.1

Group4821: abra|OJJ68216.1 acar|OOF97500.1 acla|EAW12280.1 afis|EAW16960.1 afum|EAL86488.2 agla|OJJ85938.1 akaw|GAA83230.1 aluc|OJZ83730.1 anid|CBF78343.1 anig|EHA18879.1 aory|EIT77597.1 asyd|OJJ56130.1 ater|EAU34038.1 atub|OJI90107.1 aver|OJJ05071.1 awen|OJJ35391.1 azon|OJJ49853.1

Group4822: abra|OJJ68225.1 acar|OOF97522.1 afis|EAW23333.1 afum|EAL88145.1 akaw|GAA83218.1 aluc|OJZ83720.1 anid|CBF84160.1 anig|EHA18869.1 asyd|OJJ57124.1 ater|EAU33402.1 atub|OJI90094.1 aver|OJJ03180.1 awen|OJJ38697.1 azon|OJJ47302.1 anid|CBF78048.1 atub|OJI86339.1 awen|OJJ40872.1

Group4823: abra|OJJ68270.1 acar|OOF97549.1 acla|EAW07692.1 afis|EAW22707.1 afla|EED49993.1 afum|EAL90485.1 agla|OJJ79003.1 akaw|GAA83199.1 aluc|OJZ83700.1 anid|CBF88067.1 anig|EHA18851.1 aory|EIT74717.1 asyd|OJJ63551.1 atub|OJI90072.1 aver|OJI97383.1 awen|OJJ35344.1 azon|OJJ49773.1

Group4824: abra|OJJ68273.1 acar|OOF97553.1 acla|EAW09070.1 afis|EAW25638.1 afla|EED55165.1 afum|EAL84325.1 agla|OJJ78891.1 akaw|GAA83196.1 aluc|OJZ83698.1 anid|CBF83138.1 anig|EHA18849.1 aory|EIT81612.1 asyd|OJJ55426.1 ater|EAU32709.1 atub|OJI90069.1 aver|OJJ06505.1 awen|OJJ36108.1

Group4825: abra|OJJ68286.1 acar|OOF90109.1 afis|EAW22105.1 afla|EED55143.1 afum|EAL91086.1 agla|OJJ82697.1 akaw|GAA83186.1 aluc|OJZ83685.1 anid|CBF84478.1 anig|EHA18839.1 aory|EIT81640.1 asyd|OJJ53529.1 ater|EAU32729.1 atub|OJI90056.1 aver|OJJ08221.1 awen|OJJ40237.1 azon|OJJ48799.1

Group4826: abra|OJJ68304.1 acar|OOF96684.1 acla|EAW12949.1 afis|EAW17724.1 afla|EED55147.1 afum|EAL91539.1 agla|OJJ81577.1 akaw|GAA83170.1 aluc|OJZ83669.1 anid|CBF83920.1 anig|EHA18824.1 aory|EIT81584.1 asyd|OJJ55430.1 ater|EAU32724.1 atub|OJI90040.1 aver|OJJ06509.1 awen|OJJ31293.1

Group4827: abra|OJJ68343.1 acar|OOF97641.1 acla|EAW08568.1 afis|EAW23614.1 afla|EED49585.1 afum|EAL87848.1 akaw|GAA92110.1 aluc|OJZ83615.1 anid|CBF76101.1 anig|EHA18772.1 asyd|OJJ59936.1 ater|EAU33073.1 atub|OJI89988.1 aver|OJJ02351.1 awen|OJJ39632.1 azon|OJJ50314.1 anid|CBF75197.1

Group4828: abra|OJJ68358.1 acar|OOF99377.1 afla|EED48863.1 akaw|GAA92838.1 aluc|OJZ91595.1 anig|EHA22625.1 asyd|OJJ53448.1 aver|OJJ08304.1 abra|OJJ78138.1 afla|EED53385.1 aluc|OJZ82076.1 aory|EIT75247.1 awen|OJJ29928.1 anig|EHA21837.1 aory|EIT81210.1 awen|OJJ31481.1 awen|OJJ29599.1

Group4829: abra|OJJ68384.1 acla|EAW12359.1 afis|EAW16783.1 afla|EED47253.1 afum|EAL86354.1 akaw|GAA92561.1 aluc|OJZ90107.1 anid|CBF80601.1 anig|EHA20963.1 aory|EIT79232.1 ater|EAU30268.1 atub|OJI84695.1 awen|OJJ29849.1 azon|OJJ43199.1 agla|OJJ79765.1 asyd|OJJ54699.1 aver|OJJ07475.1

Group4830: abra|OJJ68411.1 acar|OOF93966.1 acla|EAW13740.1 afis|EAW19836.1 afum|EAL85056.1 agla|OJJ86861.1 akaw|GAA92539.1 aluc|OJZ90087.1 anid|CBF81579.1 anig|EHA20987.1 aory|EIT79584.1 asyd|OJJ57609.1 atub|OJI84719.1 aver|OJJ05575.1 awen|OJJ31663.1 azon|OJJ45931.1 awen|OJJ39923.1

Group4831: abra|OJJ68416.1 acar|OOF93924.1 acla|EAW13751.1 afis|EAW19814.1 afla|EED45417.1 afum|EAL85077.1 agla|OJJ83693.1 akaw|GAA92532.1 aluc|OJZ90081.1 anid|CBF82421.1 anig|EHA20992.1 aory|EIT82441.1 asyd|OJJ54129.1 ater|EAU30780.1 atub|OJI84725.1 aver|OJJ08607.1 awen|OJJ34554.1

Group4832: abra|OJJ68446.1 acar|OOF93949.1 acla|EAW13754.1 afis|EAW19812.1 afla|EED45411.1 afum|EAL85079.1 agla|OJJ83694.1 akaw|GAA86928.1 aluc|OJZ90052.1 anid|CBF82391.1 anig|EHA21018.1 aory|EIT82394.1 asyd|OJJ54153.1 ater|EAU30741.1 atub|OJI84755.1 aver|OJJ08587.1 awen|OJJ34559.1

Group4833: abra|OJJ68461.1 acar|OOF95945.1 acla|EAW11549.1 afis|EAW23904.1 afla|EED52639.1 agla|OJJ79371.1 akaw|GAA92383.1 aluc|OJZ83242.1 anid|CBF77137.1 anig|EHA21097.1 asyd|OJJ56915.1 ater|EAU32333.1 atub|OJI87694.1 aver|OJJ03377.1 awen|OJJ39045.1 afum|EAL87332.1 azon|OJJ51374.1

Group4834: abra|OJJ68469.1 acar|OOF90193.1 acla|EAW13957.1 afis|EAW18667.1 afla|EED52662.1 afum|EAL93385.1 agla|OJJ82410.1 akaw|GAA92390.1 aluc|OJZ83250.1 anid|CBF70571.1 anig|EHA21103.1 asyd|OJJ58532.1 ater|EAU38245.1 atub|OJI87687.1 aver|OJJ01134.1 awen|OJJ32135.1 azon|OJJ46718.1

Group4835: abra|OJJ68473.1 acar|OOF94210.1 acla|EAW14616.1 afis|EAW19354.1 afla|EED46571.1 afum|EAL85102.1 akaw|GAA92394.1 aluc|OJZ83254.1 anid|CBF76650.1 anig|EHA21108.1 aory|EIT77929.1 asyd|OJJ61207.1 ater|EAU38488.1 atub|OJI87683.1 aver|OJJ00531.1 ater|EAU36160.1 afum|EAL94074.2

Group4836: abra|OJJ68481.1 acar|OOF91471.1 acla|EAW13584.1 afis|EAW20017.1 afla|EED46025.1 afum|EAL84900.1 agla|OJJ87994.1 akaw|GAA92405.1 aluc|OJZ83266.1 anid|CBF84729.1 anig|EHA21119.1 aory|EIT75379.1 asyd|OJJ53304.1 atub|OJI87671.1 aver|OJJ08437.1 awen|OJJ30036.1 azon|OJJ43680.1

Group4837: abra|OJJ68488.1 acar|OOF95813.1 acla|EAW13752.1 afis|EAW19815.1 afla|EED45414.1 afum|EAL85076.1 agla|OJJ83692.1 akaw|GAA92783.1 aluc|OJZ82700.1 anid|CBF82393.1 anig|EHA21156.1 aory|EIT82424.1 asyd|OJJ54157.1 ater|EAU30740.1 atub|OJI88932.1 aver|OJJ08580.1 awen|OJJ36130.1

Group4838: abra|OJJ68489.1 acar|OOF99836.1 acla|EAW09732.1 afis|EAW21409.1 afum|EAL92423.1 agla|OJJ85539.1 akaw|GAA92781.1 aluc|OJZ82702.1 anid|CBF83806.1 anig|EHA21157.1 aory|EIT76599.1 asyd|OJJ60893.1 ater|EAU35137.1 atub|OJI88935.1 aver|OJJ00217.1 awen|OJJ39292.1 azon|OJJ51562.1

Group4839: abra|OJJ68491.1 acar|OOF92638.1 acla|EAW08749.1 afis|EAW15687.1 afla|EED51367.1 afum|EAL85661.1 agla|OJJ81835.1 akaw|GAA92779.1 aluc|OJZ82704.1 anid|CBF71140.1 anig|EHA21159.1 aory|EIT80679.1 asyd|OJJ55835.1 atub|OJI88937.1 aver|OJJ04777.1 awen|OJJ38378.1 azon|OJJ43862.1

Group4840: abra|OJJ68506.1 acar|OOF95814.1 acla|EAW12116.1 afis|EAW17267.1 afla|EED48852.1 afum|EAL86684.1 agla|OJJ81072.1 akaw|GAA92767.1 aluc|OJZ82718.1 anig|EHA21172.1 aory|EIT81274.1 asyd|OJJ59981.1 atub|OJI88952.1 aver|OJJ02310.1 awen|OJJ35766.1 azon|OJJ45516.1 ater|EAU36132.1

Group4841: abra|OJJ68509.1 acar|OOF92671.1 acla|EAW07689.1 afis|EAW22704.1 afla|EED49990.1 afum|EAL90488.1 agla|OJJ79005.1 akaw|GAA92762.1 aluc|OJZ82723.1 anid|CBF88072.1 anig|EHA21177.1 aory|EIT74747.1 asyd|OJJ63554.1 atub|OJI88957.1 aver|OJI97380.1 awen|OJJ35341.1 azon|OJJ49777.1

Group4842: abra|OJJ68513.1 acar|OOF95416.1 acla|EAW07359.1 afis|EAW22388.1 afla|EED56048.1 afum|EAL90802.1 akaw|GAA91333.1 aluc|OJZ82728.1 anid|CBF88736.1 anig|EHA21181.1 aory|EIT76661.1 asyd|OJJ63936.1 ater|EAU39333.1 atub|OJI88961.1 aver|OJI97006.1 awen|OJJ34963.1 azon|OJJ45615.1

Group4843: abra|OJJ68520.1 acar|OOF92725.1 acla|EAW12844.1 afis|EAW18007.1 afla|EED51888.1 afum|EAL91819.1 agla|OJJ80267.1 akaw|GAA91329.1 aluc|OJZ82734.1 anid|CBF80028.1 anig|EHA21185.1 asyd|OJJ58268.1 ater|EAU31523.1 atub|OJI88967.1 aver|OJJ02156.1 awen|OJJ41534.1 azon|OJJ48253.1

Group4844: abra|OJJ68523.1 acar|OOF92728.1 acla|EAW12842.1 afis|EAW18005.1 afla|EED51890.1 afum|EAL91817.1 agla|OJJ80271.1 akaw|GAA91326.1 aluc|OJZ82737.1 anid|CBF81140.1 anig|EHA21188.1 aory|EIT82920.1 asyd|OJJ56508.1 ater|EAU31517.1 atub|OJI88972.1 aver|OJJ06211.1 awen|OJJ41537.1

Group4845: abra|OJJ68534.1 acla|EAW12824.1 afis|EAW17989.1 afla|EED51914.1 afum|EAL91798.1 agla|OJJ80249.1 akaw|GAA91317.1 aluc|OJZ82747.1 anid|CBF79981.1 anig|EHA21197.1 aory|EIT81368.1 asyd|OJJ58246.1 ater|EAU31499.1 atub|OJI88981.1 aver|OJJ02180.1 awen|OJJ41557.1 azon|OJJ48272.1

Group4846: abra|OJJ68543.1 acar|OOF92741.1 acla|EAW12831.1 afis|EAW17996.1 afla|EED51905.1 afum|EAL91806.2 akaw|GAA91310.1 aluc|OJZ82757.1 anid|CBF80013.1 anig|EHA21204.1 aory|EIT76178.1 asyd|OJJ58258.1 ater|EAU31508.1 atub|OJI88992.1 aver|OJJ02165.1 awen|OJJ41547.1 azon|OJJ48265.1

Group4847: abra|OJJ68550.1 acar|OOF92746.1 acla|EAW12817.1 afis|EAW17982.1 afla|EED51923.1 afum|EAL91791.1 agla|OJJ80239.1 akaw|GAA91303.1 aluc|OJZ82767.1 anid|CBF79994.1 anig|EHA21210.1 aory|EIT81466.1 asyd|OJJ58237.1 ater|EAU31492.1 aver|OJJ02188.1 awen|OJJ41563.1 azon|OJJ48278.1

Group4848: abra|OJJ68553.1 acar|OOF92748.1 acla|EAW12814.1 afis|EAW17979.1 afla|EED51926.1 afum|EAL91788.1 agla|OJJ80236.1 akaw|GAA91300.1 aluc|OJZ82770.1 anid|CBF79978.1 anig|EHA21212.1 aory|EIT81426.1 asyd|OJJ58233.1 atub|OJI89001.1 aver|OJJ02191.1 awen|OJJ41566.1 azon|OJJ44835.1

Group4849: abra|OJJ68558.1 acar|OOF92753.1 acla|EAW12807.1 afis|EAW17973.1 afla|EED51931.1 afum|EAL91782.1 agla|OJJ80231.1 akaw|GAA91295.1 anid|CBF79970.1 anig|EHA21217.1 aory|EIT81474.1 asyd|OJJ58228.1 ater|EAU31485.1 atub|OJI89006.1 aver|OJJ02196.1 awen|OJJ41571.1 azon|OJJ48284.1

Group4850: abra|OJJ68563.1 acar|OOF92756.1 acla|EAW12803.1 afis|EAW17969.1 afla|EED51936.1 afum|EAL91778.1 agla|OJJ80227.1 akaw|GAA91291.1 aluc|OJZ82778.1 anid|CBF79966.1 anig|EHA21221.1 aory|EIT81395.1 asyd|OJJ58226.1 ater|EAU31479.1 atub|OJI89011.1 awen|OJJ41575.1 azon|OJJ48288.1

Group4851: abra|OJJ68566.1 acar|OOF92762.1 acla|EAW12799.1 afis|EAW17965.1 afla|EED51945.1 afum|EBA27440.1 agla|OJJ80210.1 akaw|GAA91287.1 aluc|OJZ82781.1 anid|CBF79956.1 anig|EHA21224.1 aory|EIT81435.1 asyd|OJJ58219.1 atub|OJI89014.1 aver|OJJ02203.1 awen|OJJ41579.1 azon|OJJ48292.1

Group4852: abra|OJJ68569.1 acar|OOF98183.1 afla|EED57921.1 afum|EAL86294.1 agla|OJJ78785.1 aluc|OJZ81967.1 anid|CBF90243.1 anig|EHA22280.1 aory|EIT82365.1 asyd|OJJ61952.1 atub|OJI84518.1 aver|OJI96043.1 awen|OJJ35925.1 azon|OJJ42130.1 azon|OJJ47732.1 acar|OOF91241.1 aver|OJI99323.1

Group4853: abra|OJJ68600.1 acar|OOF92110.1 acla|EAW12639.1 afis|EAW18046.1 afla|EED51500.1 afum|EAL91858.1 agla|OJJ85417.1 akaw|GAA91924.1 aluc|OJZ81956.1 anid|CBF79777.1 anig|EHA22291.1 asyd|OJJ58120.1 ater|EAU31277.1 atub|OJI84507.1 aver|OJJ04285.1 awen|OJJ41488.1 azon|OJJ48218.1

Group4854: abra|OJJ68635.1 acar|OOF92139.1 acla|EAW12617.1 afis|EAW18067.1 afla|EED51451.1 afum|EBA27444.1 agla|OJJ87355.1 akaw|GAA89774.1 aluc|OJZ81920.1 anid|CBF86418.1 aory|EIT80649.1 asyd|OJJ64940.1 ater|EAU30461.1 atub|OJI84472.1 aver|OJI99062.1 awen|OJJ41454.1 azon|OJJ48912.1

Group4855: abra|OJJ68651.1 acla|EAW12627.1 afis|EAW18057.1 afla|EED51425.1 afum|EBA27441.1 agla|OJJ87329.1 akaw|GAA89791.1 aluc|OJZ81900.1 anid|CBF86437.1 anig|EHA22343.1 aory|EIT80763.1 asyd|OJJ64930.1 ater|EAU30477.1 atub|OJI84453.1 aver|OJI99049.1 awen|OJJ41440.1 azon|OJJ48902.1

Group4856: abra|OJJ68674.1 acar|OOF92175.1 acla|EAW12574.1 afis|EAW24074.1 afla|EED51399.1 afum|EAL91930.2 agla|OJJ87143.1 akaw|GAA89827.1 aluc|OJZ81861.1 anid|CBF86469.1 aory|EIT80583.1 asyd|OJJ64907.1 ater|EAU30496.1 atub|OJI84428.1 aver|OJI99029.1 awen|OJJ41411.1 azon|OJJ48881.1

Group4857: abra|OJJ68675.1 acar|OOF92176.1 acla|EAW12573.1 afis|EAW24075.1 afla|EED51398.1 afum|EBA27446.1 agla|OJJ87144.1 akaw|GAA89828.1 aluc|OJZ81860.1 anid|CBF86471.1 anig|EHA22365.1 aory|EIT80528.1 asyd|OJJ64906.1 atub|OJI84427.1 aver|OJI99028.1 awen|OJJ41412.1 azon|OJJ48880.1

Group4858: abra|OJJ68683.1 acar|OOF92185.1 acla|EAW12564.1 afis|EAW24086.1 afla|EED51387.1 afum|EAL91941.1 agla|OJJ87156.1 akaw|GAA89836.1 aluc|OJZ81850.1 anid|CBF86487.1 anig|EHA22372.1 asyd|OJJ64897.1 ater|EAU30509.1 atub|OJI84419.1 aver|OJI99018.1 awen|OJJ41403.1 azon|OJJ48869.1

Group4859: abra|OJJ68688.1 acar|OOF92191.1 acla|EAW12557.1 afis|EAW24093.1 afum|EAL91948.1 agla|OJJ87167.1 akaw|GAA89843.1 aluc|OJZ81845.1 anid|CBF86496.1 anig|EHA22377.1 aory|EIT72686.1 asyd|OJJ64892.1 ater|EAU30518.1 atub|OJI84414.1 aver|OJI99013.1 awen|OJJ41396.1 azon|OJJ48864.1

Group4860: abra|OJJ68699.1 acar|OOF92203.1 acla|EAW12545.1 afis|EAW24105.1 afla|EED46828.1 afum|EAL91959.1 agla|OJJ87180.1 akaw|GAA89854.1 aluc|OJZ81832.1 anid|CBF86515.1 anig|EHA22389.1 asyd|OJJ64879.1 ater|EAU30533.1 atub|OJI84401.1 aver|OJI98993.1 awen|OJJ41383.1 azon|OJJ48852.1

Group4861: abra|OJJ68700.1 acar|OOF92204.1 acla|EAW12544.1 afis|EAW24106.1 afla|EED46827.1 afum|EAL91960.1 akaw|GAA89855.1 aluc|OJZ81831.1 anid|CBF86516.1 anig|EHA22390.1 aory|EIT72643.1 asyd|OJJ64878.1 ater|EAU30534.1 atub|OJI84400.1 aver|OJI98992.1 awen|OJJ41382.1 azon|OJJ48851.1

Group4862: abra|OJJ68705.1 acar|OOF92207.1 acla|EAW12540.1 afis|EAW24109.1 afla|EED46823.1 afum|EAL91963.1 agla|OJJ87184.1 akaw|GAA89859.1 aluc|OJZ81825.1 anid|CBF86522.1 anig|EHA22393.1 asyd|OJJ64875.1 ater|EAU30537.1 atub|OJI84395.1 aver|OJI98988.1 awen|OJJ41379.1 azon|OJJ48847.1

Group4863: abra|OJJ68711.1 acla|EAW12534.1 afis|EAW24115.1 afla|EED46818.1 afum|EAL91969.1 agla|OJJ87192.1 akaw|GAA89865.1 aluc|OJZ81816.1 anid|CBF86533.1 anig|EHA22399.1 aory|EIT72744.1 asyd|OJJ64869.1 ater|EAU30543.1 atub|OJI84389.1 aver|OJI98981.1 awen|OJJ41373.1 azon|OJJ48841.1

Group4864: abra|OJJ68720.1 acar|OOF92219.1 acla|EAW12527.1 afis|EAW24122.1 afum|EAL91978.1 agla|OJJ87197.1 akaw|GAA89871.1 aluc|OJZ81809.1 anid|CBF86548.1 anig|EHA22408.1 aory|EIT72689.1 asyd|OJJ64859.1 ater|EAU30550.1 atub|OJI84381.1 aver|OJI98971.1 awen|OJJ41367.1 azon|OJJ48834.1

Group4865: abra|OJJ68740.1 acla|EAW12507.1 afis|EAW24139.1 afla|EED46795.1 afum|EAL91994.1 agla|OJJ87217.1 akaw|GAA89888.1 aluc|OJZ81787.1 anid|CBF77940.1 anig|EHA22428.1 aory|EIT72678.1 asyd|OJJ55284.1 ater|EAU30568.1 atub|OJI84361.1 aver|OJJ06944.1 awen|OJJ41350.1 azon|OJJ48813.1

Group4866: abra|OJJ68754.1 acar|OOF92241.1 acla|EAW12496.1 afis|EAW24149.1 afla|EED46782.1 afum|EAL92004.1 agla|OJJ87229.1 akaw|GAA89899.1 aluc|OJZ81772.1 anid|CBF77917.1 anig|EHA22444.1 aory|EIT72767.1 asyd|OJJ55296.1 ater|EAU30581.1 atub|OJI84345.1 aver|OJJ06930.1 awen|OJJ41329.1

Group4867: abra|OJJ68760.1 acar|OOF92247.1 acla|EAW12491.1 afis|EAW24154.1 afum|EAL92009.1 agla|OJJ89529.1 akaw|GAA89904.1 aluc|OJZ81766.1 anid|CBF77906.1 anig|EHA22448.1 aory|EIT72713.1 asyd|OJJ55300.1 ater|EAU30586.1 atub|OJI84338.1 aver|OJJ06925.1 awen|OJJ41323.1 azon|OJJ48796.1

Group4868: abra|OJJ68771.1 acar|OOF92256.1 acla|EAW12481.1 afis|EAW24164.1 afla|EED46769.1 afum|EAL92021.1 agla|OJJ89516.1 akaw|GAA89914.1 aluc|OJZ81755.1 anid|CBF77886.1 anig|EHA22457.1 asyd|OJJ55310.1 ater|EAU30595.1 atub|OJI84326.1 aver|OJJ06916.1 awen|OJJ41313.1 azon|OJJ48787.1

Group4869: abra|OJJ68772.1 acar|OOF92257.1 acla|EAW12480.1 afis|EAW24165.1 afla|EED46768.1 afum|EAL92022.1 agla|OJJ89515.1 akaw|GAA89915.1 aluc|OJZ81754.1 anid|CBF77885.1 anig|EHA22458.1 asyd|OJJ55311.1 ater|EAU30596.1 atub|OJI84325.1 aver|OJJ06915.1 awen|OJJ41312.1 azon|OJJ48786.1

Group4870: abra|OJJ68773.1 acar|OOF92258.1 acla|EAW12479.1 afis|EAW24166.1 afla|EED46767.1 afum|EAL92023.1 agla|OJJ89514.1 akaw|GAA89916.1 aluc|OJZ81753.1 anid|CBF77883.1 anig|EHA22459.1 asyd|OJJ55312.1 ater|EAU30597.1 atub|OJI84324.1 aver|OJJ06914.1 awen|OJJ41311.1 azon|OJJ48785.1

Group4871: abra|OJJ68775.1 acar|OOF92262.1 acla|EAW12476.1 afis|EAW24167.1 afla|EED46765.1 afum|EAL92025.1 agla|OJJ89511.1 akaw|GAA89919.1 aluc|OJZ81751.1 anid|CBF77876.1 anig|EHA22461.1 asyd|OJJ55314.1 ater|EAU30600.1 atub|OJI84322.1 aver|OJJ06912.1 awen|OJJ41309.1 azon|OJJ48782.1

Group4872: abra|OJJ68776.1 acar|OOF92260.1 acla|EAW12475.1 afis|EAW24168.1 afla|EED46764.1 afum|EAL92026.1 agla|OJJ89510.1 akaw|GAA89920.1 aluc|OJZ81750.1 anid|CBF77874.1 anig|EHA22462.1 asyd|OJJ55315.1 ater|EAU30601.1 atub|OJI84321.1 aver|OJJ06911.1 awen|OJJ41308.1 azon|OJJ48781.1

Group4873: abra|OJJ68779.1 acar|OOF92265.1 acla|EAW12472.1 afis|EAW24169.1 afla|EED46761.1 afum|EAL92031.1 agla|OJJ89507.1 akaw|GAA89923.1 aluc|OJZ81747.1 anid|CBF77869.1 anig|EHA22465.1 asyd|OJJ55319.1 ater|EAU30604.1 atub|OJI84318.1 aver|OJJ06909.1 awen|OJJ41304.1 azon|OJJ48778.1

Group4874: abra|OJJ68782.1 acar|OOF92267.1 acla|EAW12470.1 afis|EAW24171.1 afla|EED46758.1 afum|EAL92033.1 agla|OJJ89504.1 akaw|GAA89926.1 aluc|OJZ81744.1 anid|CBF77863.1 anig|EHA22468.1 asyd|OJJ55322.1 ater|EAU30606.1 atub|OJI84315.1 aver|OJJ06906.1 awen|OJJ41301.1 azon|OJJ48775.1

Group4875: abra|OJJ68786.1 acar|OOF92270.1 acla|EAW12468.1 afis|EAW24173.1 afla|EED46755.1 afum|EAL92035.1 agla|OJJ89501.1 akaw|GAA89928.1 aluc|OJZ81742.1 anid|CBF77859.1 anig|EHA22470.1 asyd|OJJ55324.1 ater|EAU30608.1 atub|OJI84313.1 aver|OJJ06904.1 awen|OJJ41297.1 azon|OJJ48773.1

Group4876: abra|OJJ68789.1 acla|EAW12464.1 afis|EAW24177.1 afla|EED46752.1 afum|EAL92039.1 agla|OJJ89497.1 akaw|GAA89931.1 aluc|OJZ81738.1 anid|CBF77851.1 anig|EHA22474.1 aory|EIT72262.1 asyd|OJJ55328.1 ater|EAU30612.1 atub|OJI84309.1 aver|OJJ06900.1 awen|OJJ41292.1 azon|OJJ48766.1

Group4877: abra|OJJ68792.1 acar|OOF92276.1 acla|EAW12461.1 afis|EAW24180.1 afla|EED46750.1 afum|EAL92041.1 agla|OJJ89494.1 aluc|OJZ81735.1 anid|CBF77846.1 anig|EHA22477.1 aory|EIT72256.1 asyd|OJJ55331.1 ater|EAU30614.1 atub|OJI84306.1 aver|OJJ06897.1 awen|OJJ41289.1 azon|OJJ48763.1

Group4878: abra|OJJ68795.1 acar|OOF92279.1 acla|EAW14659.1 afis|EAW24184.1 afla|EED46747.1 afum|EAL92044.1 agla|OJJ89491.1 aluc|OJZ81732.1 anid|CBF77840.1 anig|EHA22480.1 aory|EIT72264.1 asyd|OJJ55334.1 ater|EAU30617.1 atub|OJI84303.1 aver|OJJ06894.1 awen|OJJ41286.1 azon|OJJ42002.1

Group4879: abra|OJJ68799.1 acar|OOF92283.1 acla|EAW14654.1 afis|EAW24187.1 afla|EED46744.1 afum|EAL92047.1 agla|OJJ89487.1 akaw|GAA89938.1 aluc|OJZ81728.1 anid|CBF77833.1 aory|EIT72263.1 asyd|OJJ55338.1 ater|EAU30621.1 atub|OJI84299.1 aver|OJJ06889.1 awen|OJJ41282.1 azon|OJJ43706.1

Group4880: abra|OJJ68803.1 acar|OOF91597.1 acla|EAW14402.1 afis|EAW19109.1 afla|EED54306.1 afum|EAL93840.1 agla|OJJ84497.1 akaw|GAA90450.1 aluc|OJZ85774.1 anid|CBF79695.1 aory|EIT76105.1 asyd|OJJ58049.1 ater|EAU37367.1 atub|OJI79559.1 aver|OJJ04354.1 awen|OJJ32506.1 azon|OJJ46032.1

Group4881: abra|OJJ68849.1 acar|OOF90540.1 acla|EAW15272.1 afis|EAW20242.1 afla|EED57103.1 afum|EAL85359.1 agla|OJJ84630.1 akaw|GAA90400.1 aluc|OJZ85829.1 anid|CBF85142.1 anig|EHA24732.1 asyd|OJJ63037.1 ater|EAU32034.1 atub|OJI79617.1 aver|OJI97930.1 awen|OJJ31237.1 azon|OJJ46159.1

Group4882: abra|OJJ68857.1 acar|OOF90531.1 acla|EAW15254.1 afis|EAW20228.1 afla|EED57126.1 afum|EAL85345.2 agla|OJJ84634.1 akaw|GAA90396.1 aluc|OJZ85835.1 anid|CBF86775.1 anig|EHA24728.1 aory|EIT74040.1 asyd|OJJ64690.1 ater|EAU32030.1 atub|OJI79623.1 aver|OJI98793.1 awen|OJJ31231.1

Group4883: abra|OJJ68858.1 acla|EAW15251.1 afis|EAW20224.1 afla|EED57127.1 afum|EAL85339.1 agla|OJJ84637.1 akaw|GAA90395.1 aluc|OJZ85836.1 anid|CBF85118.1 anig|EHA24727.1 aory|EIT74129.1 asyd|OJJ63046.1 ater|EAU32029.1 atub|OJI79624.1 aver|OJI97917.1 awen|OJJ31230.1 azon|OJJ46153.1

Group4884: abra|OJJ68873.1 acar|OOF89966.1 acla|EAW15247.1 afis|EAW20213.1 afla|EED57131.1 afum|EAL85330.1 agla|OJJ84641.1 akaw|GAA90382.1 aluc|OJZ85849.1 anid|CBF85109.1 anig|EHA28097.1 aory|EIT74159.1 asyd|OJJ63052.1 ater|EAU32023.1 aver|OJI97911.1 awen|OJJ31219.1 azon|OJJ46138.1

Group4885: abra|OJJ68875.1 acla|EAW15245.1 afis|EAW20212.1 afla|EED57132.1 afum|EAL85329.1 agla|OJJ84642.1 akaw|GAA90380.1 aluc|OJZ85851.1 anid|CBF85107.1 anig|EHA28096.1 aory|EIT73979.1 asyd|OJJ63054.1 ater|EAU32021.1 atub|OJI79638.1 aver|OJI97909.1 awen|OJJ31217.1 azon|OJJ46140.1

Group4886: acar|OOF92273.1 acla|EAW12465.1 afis|EAW24176.1 afla|EED46753.1 afum|EAL92038.1 agla|OJJ89498.1 akaw|GAA89930.1 aluc|OJZ81739.1 anid|CBF77853.1 anig|EHA22473.1 aory|EIT72253.1 asyd|OJJ55327.1 ater|EAU30611.1 atub|OJI84310.1 aver|OJJ06901.1 awen|OJJ41294.1 azon|OJJ48770.1

Group4887: abra|OJJ68877.1 acar|OOF90522.1 acla|EAW15243.1 afis|EAW20210.1 afla|EED57133.1 afum|EAL85327.1 agla|OJJ84654.1 akaw|GAA90378.1 aluc|OJZ85853.1 anid|CBF85102.1 aory|EIT73969.1 asyd|OJJ63056.1 ater|EAU32020.1 atub|OJI79640.1 aver|OJI97906.1 awen|OJJ31215.1 azon|OJJ46145.1

Group4888: abra|OJJ68878.1 acar|OOF97048.1 acla|EAW07071.1 afis|EAW25746.1 afum|EBA27245.1 asyd|OJJ52302.1 ater|EAU38964.1 azon|OJJ44440.1 abra|OJJ69169.1 acla|EAW08352.1 akaw|GAA86294.1 aluc|OJZ86156.1 anid|CBF89377.1 anig|EHA24469.1 atub|OJI79948.1 awen|OJJ37486.1 acla|EAW12600.1

Group4889: abra|OJJ68916.1 acar|OOF91504.1 acla|EAW14257.1 afis|EAW19023.1 afla|EED56625.1 afum|EAL93754.1 agla|OJJ84609.1 akaw|GAA90333.1 aluc|OJZ85899.1 anid|CBF85582.1 anig|EHA24693.1 aory|EIT79644.1 asyd|OJJ64175.1 ater|EAU32065.1 atub|OJI79684.1 aver|OJI98250.1 awen|OJJ31398.1

Group4890: abra|OJJ68924.1 acar|OOF91510.1 acla|EAW13235.1 afis|EAW16374.1 afla|EED45428.1 afum|EAL88998.1 akaw|GAA90324.1 aluc|OJZ85909.1 anid|CBF83908.1 anig|EHA24683.1 aory|EIT82385.1 asyd|OJJ55419.1 atub|OJI79693.1 aver|OJJ06496.1 awen|OJJ31350.1 azon|OJJ46550.1 acar|OOF92458.1

Group4891: abra|OJJ68934.1 acar|OOF91517.1 acla|EAW14244.1 afis|EAW19036.1 afla|EED50390.1 afum|EAL93765.2 agla|OJJ84585.1 akaw|GAA90315.1 aluc|OJZ85919.1 anid|CBF79603.1 anig|EHA24673.1 asyd|OJJ57992.1 ater|EAU32075.1 atub|OJI79711.1 aver|OJJ04410.1 awen|OJJ31465.1 azon|OJJ46167.1

Group4892: abra|OJJ68955.1 acar|OOF91533.1 acla|EAW14231.1 afis|EAW19050.1 afla|EED54442.1 afum|EAL93779.1 agla|OJJ84556.1 akaw|GAA90483.1 aluc|OJZ85941.1 anig|EHA24658.1 aory|EIT81131.1 asyd|OJJ58006.1 ater|EAU32088.1 atub|OJI79729.1 aver|OJJ04394.1 awen|OJJ31479.1 azon|OJJ46192.1

Group4893: abra|OJJ68977.1 acar|OOF91552.1 acla|EAW14211.1 afis|EAW19070.1 afla|EED54435.1 afum|EAL93798.1 agla|OJJ84535.1 akaw|GAA90502.1 aluc|OJZ85965.1 anid|CBF79660.1 anig|EHA24639.1 asyd|OJJ58026.1 ater|EAU32118.1 atub|OJI79753.1 aver|OJJ04372.1 awen|OJJ31502.1 azon|OJJ46217.1

Group4894: abra|OJJ68983.1 acla|EAW14208.1 afis|EAW19073.1 afla|EED54438.1 afum|EAL93801.1 agla|OJJ84530.1 akaw|GAA90505.1 aluc|OJZ85969.1 anid|CBF79665.1 anig|EHA24636.1 aory|EIT75855.1 asyd|OJJ58030.1 ater|EAU32123.1 atub|OJI79757.1 aver|OJJ04369.1 awen|OJJ32467.1 azon|OJJ46239.1

Group4895: abra|OJJ68988.1 acar|OOF91559.1 acla|EAW14203.1 afis|EAW19079.1 afla|EED54337.1 afum|EAL93807.1 agla|OJJ84527.1 akaw|GAA90509.1 aluc|OJZ85974.1 anid|CBF79668.1 aory|EIT76081.1 asyd|OJJ58033.1 ater|EAU32125.1 atub|OJI79763.1 aver|OJJ04366.1 awen|OJJ32471.1 azon|OJJ46235.1

Group4896: abra|OJJ69001.1 acar|OOF92713.1 acla|EAW12656.1 afis|EAW18024.1 afla|EED51870.1 afum|EAL91836.1 agla|OJJ85376.1 akaw|GAA90520.1 aluc|OJZ85985.1 anid|CBF80058.1 aory|EIT81751.1 asyd|OJJ58285.1 ater|EAU31538.1 atub|OJI79775.1 aver|OJJ02139.1 awen|OJJ41516.1 azon|OJJ48239.1

Group4897: abra|OJJ69003.1 acar|OOF92716.1 acla|EAW12856.1 afis|EAW18019.1 afla|EED51872.1 afum|EAL91831.1 agla|OJJ85373.1 akaw|GAA90521.1 aluc|OJZ85989.1 anid|CBF80054.1 anig|EHA24621.1 asyd|OJJ58281.1 ater|EAU31535.1 atub|OJI79778.1 aver|OJJ02142.1 awen|OJJ41520.1 azon|OJJ48242.1

Group4898: abra|OJJ69007.1 acla|EAW12852.1 afis|EAW18015.1 afla|EED51879.1 afum|EAL91827.1 agla|OJJ85367.1 akaw|GAA90524.1 aluc|OJZ85992.1 anid|CBF80046.1 anig|EHA24618.1 aory|EIT81690.1 asyd|OJJ58277.1 ater|EAU31531.1 atub|OJI79781.1 aver|OJJ02147.1 awen|OJJ41525.1 azon|OJJ48246.1

Group4899: abra|OJJ69009.1 acar|OOF92720.1 acla|EAW12849.1 afis|EAW18012.1 afla|EED51884.1 afum|EAL91824.1 agla|OJJ80284.1 akaw|GAA90526.1 aluc|OJZ85995.1 anid|CBF80039.1 anig|EHA24615.1 asyd|OJJ58274.1 ater|EAU31528.1 atub|OJI79783.1 aver|OJJ02150.1 awen|OJJ41529.1 azon|OJJ48249.1

Group4900: abra|OJJ69033.1 acar|OOF92651.1 acla|EAW08890.1 afis|EAW15499.1 afla|EED49014.1 afum|EAL85824.1 agla|OJJ80177.1 akaw|GAA90548.1 aluc|OJZ86018.1 anig|EHA24593.1 aory|EIT82509.1 asyd|OJJ56012.1 ater|EAU32620.1 atub|OJI79807.1 aver|OJJ04957.1 awen|OJJ38158.1 azon|OJJ43131.1

Group4901: abra|OJJ69035.1 acar|OOF92650.1 acla|EAW14479.1 afis|EAW19188.1 afla|EED47723.1 afum|EAL93919.1 agla|OJJ88533.1 akaw|GAA90550.1 aluc|OJZ86020.1 anid|CBF86244.1 anig|EHA24591.1 aory|EIT77180.1 asyd|OJJ65045.1 atub|OJI79809.1 aver|OJI99166.1 awen|OJJ37546.1 azon|OJJ51209.1

Group4902: abra|OJJ69040.1 acar|OOF92967.1 acla|EAW11878.1 afis|EAW19489.1 afla|EED56707.1 afum|EAL87005.1 agla|OJJ81219.1 akaw|GAA90555.1 aluc|OJZ86025.1 anid|CBF75454.1 anig|EHA24586.1 aory|EIT73897.1 asyd|OJJ59773.1 atub|OJI79814.1 aver|OJJ02513.1 awen|OJJ36905.1 azon|OJJ42910.1

Group4903: abra|OJJ69056.1 acar|OOF91227.1 acla|EAW08758.1 afis|EAW15675.1 afla|EED46855.1 afum|EAL85672.1 agla|OJJ81822.1 akaw|GAA90586.1 aluc|OJZ86044.1 anid|CBF71159.1 anig|EHA24567.1 asyd|OJJ55843.1 ater|EAU32485.1 atub|OJI79841.1 aver|OJJ04786.1 awen|OJJ38353.1 azon|OJJ43867.1

Group4904: abra|OJJ69073.1 acar|OOF91233.1 acla|EAW08769.1 afis|EAW15661.1 afla|EED46884.1 afum|EAL85687.1 agla|OJJ81783.1 akaw|GAA90604.1 aluc|OJZ86062.1 anid|CBF71184.1 anig|EHA24551.1 aory|EIT72716.1 asyd|OJJ55860.1 atub|OJI79859.1 aver|OJJ04801.1 awen|OJJ38289.1 azon|OJJ43026.1

Group4905: abra|OJJ69086.1 acar|OOF91207.1 acla|EAW08746.1 afis|EAW15690.1 afum|EAL85658.1 agla|OJJ81837.1 akaw|GAA90615.1 aluc|OJZ86074.1 anid|CBF71135.1 anig|EHA24541.1 aory|EIT80497.1 asyd|OJJ55832.1 ater|EAU32464.1 atub|OJI79871.1 aver|OJJ04774.1 awen|OJJ38381.1 azon|OJJ43856.1

Group4906: abra|OJJ69087.1 acar|OOF91206.1 acla|EAW08745.1 afis|EAW15691.1 afla|EED51363.1 afum|EAL85657.1 agla|OJJ81838.1 akaw|GAA90616.1 aluc|OJZ86075.1 anid|CBF71133.1 anig|EHA24540.1 aory|EIT80567.1 asyd|OJJ55831.1 atub|OJI79872.1 aver|OJJ04773.1 awen|OJJ38382.1 azon|OJJ43855.1

Group4907: abra|OJJ69088.1 acar|OOF91205.1 acla|EAW08744.1 afis|EAW15692.1 afla|EED51362.1 afum|EAL85656.1 agla|OJJ81839.1 akaw|GAA90617.1 aluc|OJZ86076.1 anid|CBF71131.1 anig|EHA24539.1 asyd|OJJ55830.1 ater|EAU32463.1 atub|OJI79873.1 aver|OJJ04772.1 awen|OJJ38383.1 azon|OJJ43854.1

Group4908: abra|OJJ69095.1 acar|OOF91199.1 acla|EAW08736.1 afis|EAW15700.1 afla|EED51351.1 afum|EAL85648.1 agla|OJJ81848.1 akaw|GAA90623.1 aluc|OJZ86085.1 anid|CBF71125.1 anig|EHA24531.1 asyd|OJJ55824.1 ater|EAU32448.1 atub|OJI79881.1 aver|OJJ04766.1 awen|OJJ38392.1 azon|OJJ43846.1

Group4909: abra|OJJ69103.1 acla|EAW08730.1 afis|EAW15704.1 afla|EED51346.1 afum|EAL85643.1 agla|OJJ81854.1 akaw|GAA90628.1 aluc|OJZ86091.1 anid|CBF71102.1 anig|EHA24526.1 aory|EIT80416.1 asyd|OJJ52708.1 ater|EAU32439.1 atub|OJI79887.1 aver|OJJ04737.1 awen|OJJ38401.1 azon|OJJ43842.1

Group4910: abra|OJJ69105.1 acar|OOF91179.1 acla|EAW08727.1 afis|EAW15735.1 afla|EED51344.1 afum|EBA27236.1 akaw|GAA90630.1 aluc|OJZ86093.1 anid|CBF83999.1 anig|EHA24524.1 aory|EIT80417.1 asyd|OJJ61395.1 ater|EAU32437.1 atub|OJI79889.1 aver|OJJ00724.1 awen|OJJ38403.1 azon|OJJ43840.1

Group4911: abra|OJJ69106.1 acar|OOF91180.1 acla|EAW08726.1 afis|EAW15734.1 afla|EED51343.1 afum|EBA27237.1 agla|OJJ81859.1 akaw|GAA90631.1 aluc|OJZ86094.1 anid|CBF71098.1 aory|EIT81543.1 asyd|OJJ52706.1 ater|EAU32436.1 atub|OJI79890.1 aver|OJJ04735.1 awen|OJJ38404.1 azon|OJJ43838.1

Group4912: abra|OJJ69107.1 acla|EAW08725.1 afis|EAW15733.1 afla|EED51342.1 afum|EAL85614.1 agla|OJJ81860.1 akaw|GAA90632.1 aluc|OJZ86095.1 anid|CBF71096.1 anig|EHA24523.1 aory|EIT81551.1 asyd|OJJ52705.1 ater|EAU32435.1 atub|OJI79891.1 aver|OJJ04734.1 awen|OJJ38405.1 azon|OJJ43837.1

Group4913: abra|OJJ69109.1 acar|OOF91182.1 acla|EAW08723.1 afis|EAW15731.1 afla|EED51340.1 afum|EAL85616.1 agla|OJJ81862.1 akaw|GAA90634.1 aluc|OJZ86097.1 anid|CBF71092.1 anig|EHA24521.1 asyd|OJJ52702.1 ater|EAU32433.1 atub|OJI79893.1 aver|OJJ04730.1 awen|OJJ38407.1 azon|OJJ43835.1

Group4914: abra|OJJ69114.1 acar|OOF91187.1 acla|EAW08718.1 afis|EAW15729.1 afla|EED51334.1 afum|EAL85618.1 agla|OJJ81868.1 akaw|GAA90639.1 aluc|OJZ86103.1 anid|CBF71083.1 anig|EHA24516.1 asyd|OJJ52697.1 ater|EAU32429.1 atub|OJI79898.1 aver|OJJ04725.1 awen|OJJ38413.1 azon|OJJ43829.1

Group4915: abra|OJJ69118.1 acar|OOF91190.1 acla|EAW08714.1 afis|EAW15722.1 afla|EED51330.1 afum|EAL85625.2 agla|OJJ81872.1 akaw|GAA90643.1 aluc|OJZ86107.1 anid|CBF71077.1 anig|EHA23837.1 asyd|OJJ52693.1 ater|EAU32425.1 atub|OJI79902.1 aver|OJJ04721.1 awen|OJJ38417.1 azon|OJJ43825.1

Group4916: abra|OJJ69124.1 acar|OOF91176.1 acla|EAW08708.1 afis|EAW15716.1 afla|EED51324.1 afum|EAL85632.1 agla|OJJ81880.1 akaw|GAA90649.1 aluc|OJZ86113.1 anid|CBF71066.1 anig|EHA24508.1 aory|EIT81491.1 asyd|OJJ52686.1 atub|OJI79908.1 aver|OJJ04714.1 awen|OJJ38424.1 azon|OJJ43819.1

Group4917: abra|OJJ69146.1 acar|OOF91157.1 acla|EAW08668.1 afis|EAW15756.1 afla|EED51301.1 afum|EAL85593.1 agla|OJJ81907.1 akaw|GAA86312.1 aluc|OJZ86132.1 anid|CBF71029.1 anig|EHA24489.1 aory|EIT81548.1 asyd|OJJ52662.1 ater|EAU32399.1 atub|OJI79928.1 aver|OJJ04689.1 awen|OJJ38448.1

Group4918: abra|OJJ69153.1 acar|OOF91150.1 acla|EAW08670.1 afis|EAW15755.1 afla|EED51299.1 afum|EAL85594.1 agla|OJJ81908.1 akaw|GAA86309.1 aluc|OJZ86140.1 anid|CBF71031.1 anig|EHA24484.1 asyd|OJJ52663.1 ater|EAU32398.1 atub|OJI79932.1 aver|OJJ04690.1 awen|OJJ38452.1 azon|OJJ43752.1

Group4919: abra|OJJ69160.1 acar|OOF91143.1 acla|EAW08677.1 afis|EAW15747.1 afla|EED51292.1 agla|OJJ81916.1 akaw|GAA86302.1 aluc|OJZ86147.1 anid|CBF71011.1 anig|EHA24477.1 aory|EIT81495.1 asyd|OJJ52652.1 ater|EAU32391.1 atub|OJI79939.1 aver|OJJ04681.1 awen|OJJ38459.1 azon|OJJ43760.1

Group4920: abra|OJJ69185.1 acar|OOF91127.1 acla|EAW08661.1 afis|EAW15770.1 afla|EED51277.1 afum|EAL85578.1 agla|OJJ81934.1 akaw|GAA86280.1 aluc|OJZ86171.1 anid|CBF70981.1 anig|EHA24454.1 asyd|OJJ52635.1 ater|EAU32371.1 atub|OJI79963.1 aver|OJJ04663.1 awen|OJJ38478.1 azon|OJJ43778.1

Group4921: abra|OJJ69190.1 acar|OOF91123.1 acla|EAW08655.1 afis|EAW15776.1 afla|EED51270.1 afum|EAL85573.1 agla|OJJ81939.1 akaw|GAA86274.1 aluc|OJZ86176.1 anig|EHA24448.1 aory|EIT81527.1 asyd|OJJ52630.1 ater|EAU32365.1 atub|OJI79968.1 aver|OJJ04658.1 awen|OJJ38483.1 azon|OJJ43784.1

Group4922: abra|OJJ69202.1 acar|OOF91114.1 acla|EAW08645.1 afis|EAW15786.1 afum|EAL85564.1 agla|OJJ81949.1 akaw|GAA92214.1 aluc|OJZ86186.1 anid|CBF70956.1 anig|EHA24438.1 aory|EIT81494.1 asyd|OJJ52620.1 ater|EAU32355.1 atub|OJI79978.1 aver|OJJ04647.1 awen|OJJ38493.1 azon|OJJ43795.1

Group4923: abra|OJJ69203.1 acar|OOF91113.1 acla|EAW08644.1 afis|EAW15787.1 afla|EED51258.1 afum|EAL85563.1 agla|OJJ81950.1 akaw|GAA92215.1 aluc|OJZ86187.1 anid|CBF70958.1 anig|EHA24437.1 asyd|OJJ52619.1 ater|EAU32354.1 atub|OJI79979.1 aver|OJJ04646.1 awen|OJJ38495.1 azon|OJJ43796.1

Group4924: abra|OJJ69223.1 acar|OOF91097.1 acla|EAW08623.1 afis|EAW15809.1 afum|EAL85542.1 agla|OJJ81975.1 akaw|GAA92231.1 aluc|OJZ86203.1 anid|CBF70907.1 anig|EHA24419.1 aory|EIT83175.1 asyd|OJJ52598.1 ater|EAU33439.1 atub|OJI79997.1 aver|OJJ04625.1 awen|OJJ38520.1 azon|OJJ43735.1

Group4925: abra|OJJ69225.1 acar|OOF91095.1 afis|EAW15811.1 afum|EAL85540.1 agla|OJJ81977.1 akaw|GAA92233.1 aluc|OJZ86205.1 anid|CBF75271.1 aory|EIT83193.1 asyd|OJJ55915.1 ater|EAU33621.1 atub|OJI79999.1 aver|OJJ04855.1 awen|OJJ38522.1 abra|OJJ70041.1 anig|EHA18723.1 afla|EED51236.1

Group4926: abra|OJJ69233.1 acar|OOF91087.1 acla|EAW08616.1 afis|EAW15818.1 afla|EED51229.1 afum|EAL85533.1 agla|OJJ81982.1 akaw|GAA92241.1 aluc|OJZ86213.1 anid|CBF70893.1 anig|EHA24413.1 asyd|OJJ52592.1 ater|EAU33430.1 atub|OJI80008.1 aver|OJJ04619.1 awen|OJJ38532.1 azon|OJJ43730.1

Group4927: abra|OJJ69245.1 acar|OOF91075.1 acla|EAW08604.1 afis|EAW15829.1 afla|EED51216.1 afum|EAL85522.2 agla|OJJ81993.1 akaw|GAA92253.1 aluc|OJZ86226.1 anid|CBF70867.1 anig|EHA24402.1 aory|EIT80404.1 asyd|OJJ52579.1 atub|OJI80019.1 aver|OJJ04607.1 awen|OJJ38543.1 azon|OJJ43720.1

Group4928: abra|OJJ69266.1 acar|OOF89860.1 acla|EAW14656.1 afis|EAW24190.1 afum|EAL85863.1 agla|OJJ79689.1 akaw|GAA88142.1 aluc|OJZ82095.1 anid|CBF80855.1 anig|EHA28634.1 aory|EIT82483.1 asyd|OJJ56290.1 ater|EAU31141.1 atub|OJI88342.1 aver|OJJ06427.1 awen|OJJ33109.1 azon|OJJ44840.1

Group4929: abra|OJJ69272.1 acar|OOF94449.1 acla|EAW14661.1 afis|EAW24192.1 afla|EED48079.1 agla|OJJ79687.1 akaw|GAA88146.1 aluc|OJZ82099.1 anid|CBF74310.1 anig|EHA28630.1 aory|EIT82480.1 asyd|OJJ54006.1 ater|EAU31143.1 atub|OJI88338.1 aver|OJJ08101.1 awen|OJJ33107.1 azon|OJJ44841.1

Group4930: abra|OJJ69303.1 acla|EAW14689.1 afis|EAW24220.1 afla|EED48051.1 afum|EAL85889.1 agla|OJJ79650.1 akaw|GAA88172.1 aluc|OJZ82131.1 anid|CBF74246.1 anig|EHA28600.1 aory|EIT83327.1 asyd|OJJ53974.1 ater|EAU31170.1 atub|OJI88305.1 aver|OJJ08073.1 awen|OJJ33076.1 azon|OJJ44870.1

Group4931: abra|OJJ69304.1 acar|OOF94422.1 acla|EAW14690.1 afis|EAW24221.1 afum|EAL85890.1 agla|OJJ79649.1 akaw|GAA88173.1 aluc|OJZ82132.1 anid|CBF74244.1 anig|EHA28599.1 aory|EIT83331.1 asyd|OJJ53973.1 ater|EAU31171.1 atub|OJI88304.1 aver|OJJ08072.1 awen|OJJ33075.1 azon|OJJ44871.1

Group4932: abra|OJJ69317.1 acar|OOF90304.1 acla|EAW14731.1 afis|EAW24264.1 afla|EED48013.1 afum|EAL85932.1 agla|OJJ79608.1 akaw|GAA88186.1 aluc|OJZ82146.1 anid|CBF74177.1 anig|EHA28584.1 aory|EIT82900.1 asyd|OJJ53942.1 atub|OJI88290.1 aver|OJJ08039.1 awen|OJJ33037.1 azon|OJJ48482.1

Group4933: abra|OJJ69320.1 acar|OOF94410.1 acla|EAW14725.1 afis|EAW24261.1 afla|EED48017.1 afum|EAL85929.1 agla|OJJ79611.1 akaw|GAA88189.1 aluc|OJZ82149.1 anid|CBF74187.1 anig|EHA28581.1 asyd|OJJ53948.1 ater|EAU31206.1 atub|OJI88286.1 aver|OJJ08044.1 awen|OJJ33041.1 azon|OJJ48595.1

Group4934: abra|OJJ69324.1 acar|OOF94403.1 acla|EAW14716.1 afis|EAW24251.1 afla|EED48022.1 afum|EAL85920.2 agla|OJJ79622.1 akaw|GAA88195.1 aluc|OJZ82156.1 anid|CBF74197.1 anig|EHA28576.1 asyd|OJJ53951.1 ater|EAU31192.1 atub|OJI88281.1 aver|OJJ08047.1 awen|OJJ33048.1 azon|OJJ48471.1

Group4935: abra|OJJ69325.1 acar|OOF94402.1 acla|EAW14715.1 afis|EAW24250.1 afla|EED48023.1 afum|EAL85919.1 agla|OJJ79623.1 akaw|GAA88196.1 aluc|OJZ82157.1 anid|CBF74199.1 anig|EHA28575.1 asyd|OJJ53952.1 ater|EAU31191.1 atub|OJI88280.1 aver|OJJ08048.1 awen|OJJ33062.1 azon|OJJ48470.1

Group4936: abra|OJJ69326.1 acar|OOF94400.1 acla|EAW14714.1 afis|EAW24249.1 afla|EED48024.1 afum|EAL85917.2 agla|OJJ79624.1 akaw|GAA88197.1 aluc|OJZ82158.1 anid|CBF74201.1 anig|EHA28574.1 asyd|OJJ53953.1 ater|EAU31190.1 atub|OJI88279.1 aver|OJJ08049.1 awen|OJJ33061.1 azon|OJJ48469.1

Group4937: abra|OJJ69339.1 acla|EAW14707.1 afis|EAW24238.1 afla|EED48033.1 afum|EAL85906.1 agla|OJJ79633.1 akaw|GAA88208.1 aluc|OJZ82171.1 anid|CBF74215.1 anig|EHA28562.1 aory|EIT83347.1 asyd|OJJ53959.1 ater|EAU31184.1 atub|OJI88266.1 aver|OJJ08055.1 awen|OJJ33052.1 azon|OJJ48463.1

Group4938: abra|OJJ69347.1 acar|OOF94377.1 acla|EAW14699.1 afis|EAW24230.1 afla|EED48041.1 afum|EAL85899.2 agla|OJJ79641.1 akaw|GAA88216.1 aluc|OJZ82179.1 anid|CBF74226.1 anig|EHA28555.1 aory|EIT83348.1 asyd|OJJ53966.1 atub|OJI88258.1 aver|OJJ08062.1 awen|OJJ33067.1 azon|OJJ48456.1

Group4939: abra|OJJ69356.1 acar|OOF94368.1 acla|EAW14747.1 afis|EAW24278.1 afla|EED47998.1 afum|EAL85947.1 agla|OJJ79584.1 akaw|GAA88226.1 aluc|OJZ82188.1 anid|CBF74153.1 anig|EHA28547.1 aory|EIT82913.1 asyd|OJJ53924.1 atub|OJI88247.1 aver|OJJ08022.1 awen|OJJ33019.1 azon|OJJ48498.1

Group4940: abra|OJJ69358.1 acar|OOF94366.1 acla|EAW14749.1 afis|EAW24280.1 afla|EED47996.1 afum|EAL85949.1 agla|OJJ80634.1 akaw|GAA88228.1 aluc|OJZ82190.1 anid|CBF74149.1 anig|EHA28545.1 aory|EIT82910.1 asyd|OJJ53922.1 atub|OJI88245.1 aver|OJJ08020.1 awen|OJJ33017.1 azon|OJJ48500.1

Group4941: abra|OJJ69364.1 acar|OOF94361.1 acla|EAW14754.1 afis|EAW24285.1 afla|EED47991.1 afum|EAL85954.1 agla|OJJ83147.1 akaw|GAA88233.1 aluc|OJZ82198.1 anid|CBF74133.1 aory|EIT79826.1 asyd|OJJ53914.1 ater|EAU31236.1 atub|OJI88238.1 aver|OJJ08012.1 awen|OJJ33011.1 azon|OJJ48504.1

Group4942: abra|OJJ69365.1 acar|OOF94360.1 acla|EAW14755.1 afis|EAW24286.1 afla|EED47990.1 afum|EAL85955.1 agla|OJJ83150.1 akaw|GAA88234.1 aluc|OJZ82199.1 anig|EHA28540.1 aory|EIT79821.1 asyd|OJJ53913.1 ater|EAU31237.1 atub|OJI88237.1 aver|OJJ08011.1 awen|OJJ33010.1 azon|OJJ48505.1

Group4943: abra|OJJ69372.1 acar|OOF94356.1 acla|EAW14762.1 afis|EAW24293.1 afum|EAL85961.1 agla|OJJ83158.1 akaw|GAA88241.1 aluc|OJZ82206.1 anid|CBF74120.1 anig|EHA28533.1 aory|EIT79846.1 asyd|OJJ53906.1 ater|EAU31243.1 atub|OJI88230.1 aver|OJJ08004.1 awen|OJJ33003.1 azon|OJJ48512.1

Group4944: abra|OJJ69379.1 acar|OOF94346.1 acla|EAW14767.1 afis|EAW24298.1 afla|EED47974.1 afum|EAL85966.1 agla|OJJ83165.1 akaw|GAA88247.1 aluc|OJZ82214.1 anid|CBF74112.1 anig|EHA28526.1 asyd|OJJ53902.1 ater|EAU31249.1 atub|OJI88221.1 aver|OJJ07999.1 awen|OJJ32994.1 azon|OJJ48521.1

Group4945: abra|OJJ69392.1 acla|EAW14777.1 afis|EAW24308.1 afum|EAL85976.1 agla|OJJ83177.1 akaw|GAA88260.1 aluc|OJZ82229.1 anid|CBF74089.1 anig|EHA28510.1 aory|EIT79815.1 asyd|OJJ53886.1 ater|EAU31261.1 atub|OJI88208.1 aver|OJJ07984.1 awen|OJJ32984.1 azon|OJJ48933.1 afla|EED47961.1

Group4946: abra|OJJ69395.1 acar|OOF94334.1 afis|EAW24310.1 afla|EED47957.1 akaw|GAA88262.1 aluc|OJZ82231.1 anig|EHA28508.1 aory|EIT79855.1 asyd|OJJ53884.1 ater|EAU31263.1 atub|OJI88206.1 aver|OJJ07982.1 awen|OJJ32982.1 azon|OJJ48935.1 acla|EAW14779.1 afum|EBA27252.1 anid|CBF74085.1

Group4947: abra|OJJ69399.1 acar|OOF94331.1 acla|EAW14781.1 afis|EAW24312.1 afla|EED47955.1 afum|EAL85980.1 agla|OJJ83180.1 akaw|GAA88265.1 aluc|OJZ82235.1 anig|EHA28506.1 aory|EIT79818.1 asyd|OJJ53881.1 ater|EAU31265.1 atub|OJI88203.1 aver|OJJ07979.1 awen|OJJ32980.1 azon|OJJ48937.1

Group4948: abra|OJJ69403.1 acar|OOF94328.1 acla|EAW14785.1 afis|EAW24316.1 afla|EED47951.1 afum|EAL85984.1 agla|OJJ83184.1 akaw|GAA88268.1 aluc|OJZ82239.1 anid|CBF74076.1 anig|EHA28502.1 asyd|OJJ53877.1 ater|EAU31268.1 atub|OJI88199.1 aver|OJJ07975.1 awen|OJJ32976.1 azon|OJJ48941.1

Group4949: abra|OJJ69406.1 acla|EAW14788.1 afis|EAW24319.1 afla|EED47948.1 afum|EAL85989.1 agla|OJJ83187.1 akaw|GAA88271.1 aluc|OJZ82243.1 anig|EHA28499.1 ater|EAU31272.1 atub|OJI88195.1 aver|OJJ07972.1 awen|OJJ32973.1 anid|CBF74071.1 asyd|OJJ53874.1 azon|OJJ48944.1 aory|EIT79838.1

Group4950: abra|OJJ69416.1 acar|OOF94316.1 acla|EAW14797.1 afis|EAW24328.1 afla|EED47938.1 afum|EAL85998.1 agla|OJJ83195.1 akaw|GAA88284.1 aluc|OJZ82253.1 anig|EHA28488.1 aory|EIT79811.1 asyd|OJJ53866.1 ater|EAU30425.1 atub|OJI88184.1 aver|OJJ07964.1 awen|OJJ32966.1 azon|OJJ48951.1

Group4951: abra|OJJ69438.1 acar|OOF94302.1 acla|EAW14810.1 afis|EAW24339.1 afla|EED47924.1 afum|EAL86010.1 agla|OJJ83227.1 aluc|OJZ82274.1 anid|CBF74023.1 anig|EHA28471.1 aory|EIT83369.1 asyd|OJJ53843.1 ater|EAU30409.1 atub|OJI88165.1 aver|OJJ07946.1 awen|OJJ32883.1 azon|OJJ48972.1

Group4952: abra|OJJ69444.1 acar|OOF94295.1 acla|EAW11741.1 afis|EAW23774.1 afla|EED57694.1 afum|EAL87208.1 akaw|GAA88374.1 aluc|OJZ82280.1 anid|CBF69507.1 anig|EHA28466.1 aory|EIT79917.1 asyd|OJJ59220.1 ater|EAU31469.1 atub|OJI88159.1 aver|OJJ01831.1 awen|OJJ40003.1 azon|OJJ49871.1

Group4953: abra|OJJ69454.1 acar|OOF94287.1 acla|EAW14818.1 afis|EAW24348.1 afla|EED47915.1 afum|EAL86018.1 agla|OJJ83236.1 akaw|GAA88358.1 aluc|OJZ82297.1 anid|CBF74015.1 anig|EHA28458.1 aory|EIT83377.1 asyd|OJJ53839.1 atub|OJI88142.1 aver|OJJ07942.1 awen|OJJ32876.1 ater|EAU32154.1

Group4954: abra|OJJ69461.1 acar|OOF94279.1 acla|EAW09923.1 afis|EAW21171.1 afla|EED54736.1 afum|EBA27452.1 agla|OJJ80009.1 akaw|GAA88351.1 aluc|OJZ82305.1 anid|CBF83234.1 anig|EHA28451.1 asyd|OJJ60542.1 ater|EAU35838.1 atub|OJI88133.1 aver|OJI99869.1 awen|OJJ36509.1 azon|OJJ44380.1

Group4955: abra|OJJ69466.1 acar|OOF94276.1 acla|EAW09921.1 afis|EAW21169.1 afla|EED54734.1 afum|EAL92188.2 agla|OJJ80011.1 akaw|GAA88347.1 aluc|OJZ82309.1 anig|EHA28448.1 aory|EIT78858.1 asyd|OJJ60540.1 ater|EAU35836.1 atub|OJI88130.1 aver|OJI99867.1 awen|OJJ36506.1 azon|OJJ44378.1

Group4956: abra|OJJ69479.1 acla|EAW09910.1 afis|EAW21159.1 afla|EED54650.1 afum|EAL92177.1 agla|OJJ80029.1 akaw|GAA88337.1 aluc|OJZ82321.1 anid|CBF86979.1 anig|EHA28437.1 aory|EIT78788.1 asyd|OJJ60526.1 ater|EAU35821.1 atub|OJI88119.1 aver|OJI99854.1 awen|OJJ36491.1 azon|OJJ44357.1

Group4957: abra|OJJ69480.1 acar|OOF94264.1 acla|EAW09909.1 afis|EAW21158.1 afum|EAL92176.1 agla|OJJ80030.1 akaw|GAA88336.1 aluc|OJZ82322.1 anid|CBF86981.1 anig|EHA28436.1 aory|EIT78777.1 asyd|OJJ60525.1 ater|EAU35820.1 atub|OJI88118.1 aver|OJI99853.1 awen|OJJ36490.1 azon|OJJ44358.1

Group4958: abra|OJJ69493.1 acar|OOF94252.1 acla|EAW09901.1 afis|EAW21149.1 afla|EED54663.1 afum|EAL92168.1 agla|OJJ80052.1 akaw|GAA88327.1 aluc|OJZ82333.1 anid|CBF75142.1 anig|EHA28427.1 aory|EIT78868.1 asyd|OJJ59970.1 ater|EAU36737.1 atub|OJI88107.1 aver|OJJ02321.1 awen|OJJ36476.1

Group4959: abra|OJJ69503.1 acar|OOF94248.1 afla|EED54665.1 akaw|GAA88320.1 aluc|OJZ82342.1 anig|EHA28418.1 aory|EIT78849.1 atub|OJI88098.1 awen|OJJ36473.1 acla|EAW09899.1 afis|EAW21147.1 afum|EAL92166.1 agla|OJJ80054.1 anid|CBF83168.1 asyd|OJJ55503.1 aver|OJJ06583.1 ater|EAU39451.1

Group4960: abra|OJJ69504.1 acla|EAW09963.1 afis|EAW21132.1 afla|EED51810.1 afum|EAL92150.1 agla|OJJ80085.1 akaw|GAA88319.1 aluc|OJZ82343.1 anig|EHA28417.1 aory|EIT81769.1 asyd|OJJ60462.1 ater|EAU36775.1 atub|OJI88097.1 aver|OJI99782.1 awen|OJJ36455.1 anid|CBF87017.1 azon|OJJ44421.1

Group4961: abra|OJJ69512.1 acar|OOF94241.1 acla|EAW09956.1 afis|EAW21140.1 afla|EED46143.1 afum|EAL92159.1 agla|OJJ80067.1 akaw|GAA88312.1 aluc|OJZ82350.1 anid|CBF87005.1 anig|EHA28410.1 aory|EIT76500.1 asyd|OJJ60469.1 atub|OJI88090.1 aver|OJI99789.1 awen|OJJ36464.1 azon|OJJ44414.1

Group4962: abra|OJJ69530.1 acar|OOF92669.1 acla|EAW07395.1 afis|EAW22421.1 afla|EED52499.1 afum|EAL90771.1 agla|OJJ87603.1 akaw|GAA88381.1 aluc|OJZ82369.1 anid|CBF88809.1 anig|EHA28396.1 aory|EIT82809.1 asyd|OJJ62891.1 atub|OJI88073.1 aver|OJI96942.1 awen|OJJ35000.1 azon|OJJ45643.1

Group4963: abra|OJJ69551.1 acar|OOF95793.1 acla|EAW10612.1 afis|EAW24955.1 afla|EED55509.1 afum|EAL89613.1 agla|OJJ82182.1 akaw|GAA91563.1 aluc|OJZ82381.1 anid|CBF81670.1 anig|EHA28384.1 aory|EIT80837.1 asyd|OJJ59554.1 ater|EAU35702.1 atub|OJI88060.1 aver|OJJ02714.1 awen|OJJ40550.1

Group4964: abra|OJJ69578.1 acar|OOF91577.1 acla|EAW14392.1 afis|EAW19099.1 afla|EED54333.1 afum|EAL93831.1 agla|OJJ84502.1 akaw|GAA91574.1 aluc|OJZ82396.1 anid|CBF79698.1 aory|EIT75935.1 asyd|OJJ58052.1 ater|EAU32146.1 atub|OJI88046.1 aver|OJJ04352.1 awen|OJJ32492.1 azon|OJJ46039.1

Group4965: abra|OJJ69579.1 acar|OOF91575.1 acla|EAW14389.1 afis|EAW19096.1 afla|EED54329.1 afum|EBA27504.1 agla|OJJ84505.1 akaw|GAA91575.1 aluc|OJZ82397.1 anid|CBF79704.1 asyd|OJJ58054.1 ater|EAU32144.1 atub|OJI88045.1 aver|OJJ04349.1 awen|OJJ32489.1 azon|OJJ46042.1 aory|EIT76045.1

Group4966: abra|OJJ69596.1 acar|OOF91563.1 acla|EAW14374.1 afis|EAW19082.1 afla|EED54314.1 afum|EAL93811.1 agla|OJJ84524.1 akaw|GAA91588.1 aluc|OJZ82411.1 anid|CBF79670.1 anig|EHA28356.1 aory|EIT75963.1 asyd|OJJ58035.1 ater|EAU32129.1 atub|OJI88030.1 aver|OJJ04364.1 awen|OJJ32477.1

Group4967: abra|OJJ69613.1 acar|OOF92693.1 acla|EAW11766.1 afis|EAW23684.1 afla|EED51826.1 afum|EAL87129.1 agla|OJJ86862.1 akaw|GAA91597.1 aluc|OJZ82421.1 anid|CBF80086.1 anig|EHA28348.1 aory|EIT81697.1 asyd|OJJ58305.1 ater|EAU31583.1 atub|OJI88020.1 aver|OJJ02107.1 azon|OJJ45792.1

Group4968: abra|OJJ69651.1 acar|OOG00403.1 acla|EAW06766.1 afis|EAW16316.1 afla|EED53565.1 afum|EAL88939.1 agla|OJJ88277.1 akaw|GAA91628.1 aluc|OJZ82455.1 anid|CBF89144.1 anig|EHA28320.1 aory|EIT75184.1 asyd|OJJ62570.1 ater|EAU34141.1 atub|OJI87983.1 aver|OJI96694.1 awen|OJJ37714.1

Group4969: abra|OJJ69653.1 acar|OOF93647.1 acla|EAW14511.1 afis|EAW19229.1 afla|EED47661.1 agla|OJJ88470.1 akaw|GAA91632.1 aluc|OJZ82459.1 anid|CBF78612.1 anig|EHA28316.1 asyd|OJJ65179.1 ater|EAU37234.1 atub|OJI87979.1 aver|OJI99332.1 awen|OJJ37587.1 azon|OJJ51241.1 aory|EIT80209.1

Group4970: abra|OJJ69678.1 acar|OOF93500.1 acla|EAW08823.1 afis|EAW15574.1 afla|EED49083.1 afum|EAL85760.1 agla|OJJ85178.1 akaw|GAA88803.1 aluc|OJZ82486.1 anid|CBF78103.1 anig|EHA28288.1 aory|EIT82543.1 asyd|OJJ55894.1 ater|EAU32557.1 atub|OJI87946.1 aver|OJJ04839.1 awen|OJJ38234.1

Group4971: abra|OJJ69700.1 acar|OOF90941.1 afis|EAW20859.1 afla|EED45448.1 afum|EAL84776.1 agla|OJJ83745.1 akaw|GAA88817.1 aluc|OJZ82501.1 anid|CBF84702.1 anig|EHA28268.1 aory|EIT82427.1 asyd|OJJ53319.1 ater|EAU30802.1 atub|OJI87930.1 aver|OJJ08424.1 awen|OJJ34514.1 azon|OJJ45199.1

Group4972: abra|OJJ69708.1 acar|OOF94202.1 acla|EAW09975.1 afis|EAW21114.1 afla|EED51765.1 afum|EAL92137.1 agla|OJJ80097.1 akaw|GAA90157.1 aluc|OJZ83190.1 anid|CBF87034.1 anig|EHA28260.1 aory|EIT80039.1 asyd|OJJ60453.1 ater|EAU36785.1 atub|OJI87749.1 aver|OJI99771.1 awen|OJJ36444.1

Group4973: abra|OJJ69711.1 acar|OOF94201.1 acla|EAW09974.1 afis|EAW21115.1 afla|EED51766.1 afum|EAL92138.1 agla|OJJ80096.1 akaw|GAA90161.1 aluc|OJZ83186.1 anid|CBF87032.1 anig|EHA28258.1 asyd|OJJ60454.1 ater|EAU36784.1 atub|OJI87753.1 aver|OJI99772.1 awen|OJJ36445.1 azon|OJJ44431.1

Group4974: abra|OJJ69731.1 acar|OOF94190.1 acla|EAW11683.1 afla|EED51759.1 agla|OJJ86787.1 akaw|GAA90184.1 aluc|OJZ83165.1 anid|CBF73433.1 anig|EHA28229.1 aory|EIT80571.1 atub|OJI87772.1 aver|OJJ08109.1 awen|OJJ38074.1 azon|OJJ42488.1 acar|OOF93383.1 afum|EAL86339.2 afis|EAW16771.1

Group4975: abra|OJJ69741.1 acar|OOF94180.1 acla|EAW09983.1 afis|EAW21109.1 afla|EED51713.1 afum|EAL92131.1 akaw|GAA90193.1 aluc|OJZ83155.1 anid|CBF80441.1 anig|EHA28221.1 aory|EIT80457.1 asyd|OJJ54783.1 ater|EAU36790.1 atub|OJI87782.1 aver|OJJ07383.1 awen|OJJ36437.1 azon|OJJ45175.1

Group4976: abra|OJJ69743.1 acar|OOG00883.1 acla|EAW07453.1 afis|EAW22485.1 afla|EED56932.1 afum|EAL89359.2 akaw|GAA90196.1 aluc|OJZ83152.1 anig|EHA28218.1 aory|EIT77590.1 ater|EAU39229.1 atub|OJI87785.1 awen|OJJ35104.1 afla|EED47459.1 afum|EAL90708.1 aory|EIT79961.1 afla|EED56933.1

Group4977: abra|OJJ69747.1 acar|OOF94175.1 acla|EAW09986.1 afis|EAW21107.1 afla|EED51688.1 afum|EAL92128.1 agla|OJJ80109.1 akaw|GAA90199.1 aluc|OJZ83149.1 anig|EHA28215.1 aory|EIT80537.1 asyd|OJJ60447.1 ater|EAU36793.1 atub|OJI87788.1 aver|OJI99765.1 awen|OJJ36368.1 azon|OJJ51535.1

Group4978: abra|OJJ69772.1 agla|OJJ89535.1 akaw|GAA87882.1 aluc|OJZ85527.1 anig|EHA25058.1 asyd|OJJ59178.1 atub|OJI80041.1 aver|OJJ01798.1 abra|OJJ76053.1 afla|EED48186.1 akaw|GAA88912.1 aluc|OJZ89509.1 anig|EHA23091.1 aory|EIT72940.1 ater|EAU37178.1 atub|OJI85294.1 awen|OJJ39948.1

Group4979: acar|OOF93589.1 anig|EHA25040.1 atub|OJI80057.1 abra|OJJ69791.1 acla|EAW11764.1 afis|EAW23687.1 afla|EED57653.1 afum|EAL87132.1 agla|OJJ78642.1 akaw|GAA87867.1 aluc|OJZ85508.1 anid|CBF84203.1 aory|EIT79945.1 asyd|OJJ57163.1 ater|EAU34272.1 aver|OJJ03150.1 awen|OJJ33314.1

Group4980: abra|OJJ69803.1 acar|OOF93573.1 acla|EAW11066.1 afis|EAW20808.1 afla|EED51436.1 afum|EAL84820.1 akaw|GAA87856.1 aluc|OJZ85498.1 anid|CBF80564.1 anig|EHA25031.1 aory|EIT80626.1 asyd|OJJ54681.1 ater|EAU30269.1 atub|OJI80067.1 aver|OJJ07487.1 awen|OJJ31034.1 azon|OJJ42532.1

Group4981: abra|OJJ69845.1 acar|OOF93368.1 acla|EAW08980.1 afis|EAW15429.1 afla|EED55190.1 afum|EAL84214.1 agla|OJJ80141.1 akaw|GAA87814.1 aluc|OJZ85449.1 anid|CBF78299.1 anig|EHA24992.1 aory|EIT83595.1 asyd|OJJ56101.1 ater|EAU32702.1 atub|OJI80118.1 aver|OJJ05044.1 awen|OJJ38049.1

Group4982: anid|CBF87945.1 abra|OJJ77908.1 acla|EAW13240.1 afis|EAW17394.1 afla|EED55526.1 afum|EAL91232.1 agla|OJJ86275.1 akaw|GAA83751.1 aluc|OJZ90980.1 anig|EHA19685.1 aory|EIT80850.1 asyd|OJJ63474.1 ater|EAU38912.1 atub|OJI82971.1 aver|OJI97456.1 awen|OJJ32549.1 azon|OJJ47335.1

Group4983: abra|OJJ69868.1 acar|OOF90336.1 acla|EAW15316.1 afis|EAW20318.1 afla|EED57091.1 afum|EAL85446.1 akaw|GAA87788.1 aluc|OJZ85423.1 anig|EHA25602.1 aory|EIT82311.1 asyd|OJJ56070.1 ater|EAU32680.1 atub|OJI80152.1 aver|OJJ05013.1 awen|OJJ31345.1 asyd|OJJ52769.1 aver|OJJ08752.1

Group4984: abra|OJJ69874.1 acar|OOF93387.1 acla|EAW08929.1 afis|EAW15467.1 afla|EED46919.1 afum|EAL85855.1 agla|OJJ80148.1 akaw|GAA87783.1 aluc|OJZ85418.1 anid|CBF78284.1 anig|EHA25597.1 aory|EIT72664.1 asyd|OJJ56069.1 ater|EAU32661.1 atub|OJI80158.1 aver|OJJ05012.1 awen|OJJ38113.1

Group4985: abra|OJJ69904.1 acar|OOF97941.1 acla|EAW08919.1 afis|EAW15475.1 afla|EED47464.1 afum|EAL85846.1 agla|OJJ89051.1 akaw|GAA87748.1 aluc|OJZ85381.1 anid|CBF86603.1 anig|EHA25564.1 aory|EIT79959.1 asyd|OJJ64826.1 ater|EAU32653.1 atub|OJI80189.1 aver|OJI98939.1 awen|OJJ36394.1

Group4986: abra|OJJ69912.1 acar|OOF90389.1 acla|EAW08916.1 afis|EAW15478.1 afum|EAL85843.1 agla|OJJ78432.1 akaw|GAA87745.1 aluc|OJZ85376.1 anid|CBF78223.1 anig|EHA25559.1 aory|EIT72649.1 asyd|OJJ56031.1 ater|EAU32649.1 atub|OJI80193.1 aver|OJJ04978.1 awen|OJJ38135.1 azon|OJJ43156.1

Group4987: abra|OJJ69915.1 acar|OOF93427.1 acla|EAW08913.1 afis|EAW15482.1 afla|EED46931.1 afum|EAL85840.1 agla|OJJ80161.1 akaw|GAA87741.1 aluc|OJZ85373.1 anid|CBF78219.1 anig|EHA25557.1 aory|EIT72658.1 asyd|OJJ56027.1 atub|OJI80197.1 aver|OJJ04974.1 awen|OJJ38141.1 azon|OJJ43149.1

Group4988: abra|OJJ69919.1 acar|OOF93431.1 acla|EAW08910.1 afis|EAW15486.1 afla|EED46941.1 afum|EAL85837.1 agla|OJJ80164.1 akaw|GAA87737.1 aluc|OJZ85369.1 anig|EHA25552.1 aory|EIT72770.1 asyd|OJJ56024.1 ater|EAU32646.1 atub|OJI80201.1 aver|OJJ04971.1 awen|OJJ38143.1 azon|OJJ43145.1

Group4989: abra|OJJ69924.1 acar|OOF93435.1 acla|EAW08906.1 afis|EAW15490.1 afla|EED49004.1 afum|EAL85833.1 agla|OJJ80167.1 akaw|GAA87733.1 aluc|OJZ85364.1 anig|EHA25547.1 aory|EIT82535.1 asyd|OJJ56021.1 ater|EAU32629.1 atub|OJI80206.1 aver|OJJ04968.1 awen|OJJ38146.1 azon|OJJ43141.1

Group4990: abra|OJJ69926.1 acar|OOF93437.1 acla|EAW08904.1 afis|EAW15492.1 afla|EED49007.1 afum|EAL85831.1 agla|OJJ80169.1 akaw|GAA87731.1 aluc|OJZ85362.1 anig|EHA25545.1 aory|EIT82513.1 asyd|OJJ56019.1 ater|EAU32627.1 atub|OJI80209.1 aver|OJJ04966.1 awen|OJJ38148.1 azon|OJJ43139.1

Group4991: abra|OJJ69927.1 acar|OOF93438.1 acla|EAW08903.1 afis|EAW15493.1 afla|EED49008.1 afum|EAL85830.1 agla|OJJ80170.1 akaw|GAA87730.1 aluc|OJZ85361.1 anig|EHA25544.1 aory|EIT82514.1 asyd|OJJ56018.1 ater|EAU32626.1 atub|OJI80210.1 aver|OJJ04964.1 awen|OJJ38149.1 azon|OJJ43138.1

Group4992: abra|OJJ69928.1 acar|OOF93439.1 acla|EAW08902.1 afis|EAW15494.1 afla|EED49009.1 afum|EAL85829.1 agla|OJJ80171.1 akaw|GAA87729.1 aluc|OJZ85359.1 anig|EHA25543.1 aory|EIT82523.1 asyd|OJJ56017.1 ater|EAU32625.1 atub|OJI80212.1 aver|OJJ04963.1 awen|OJJ38150.1 azon|OJJ43137.1

Group4993: abra|OJJ69929.1 acar|OOF93440.1 acla|EAW08901.1 afis|EAW15495.1 afla|EED49010.1 afum|EAL85828.1 agla|OJJ80173.1 akaw|GAA87728.1 aluc|OJZ85358.1 anig|EHA25542.1 aory|EIT82519.1 asyd|OJJ56016.1 ater|EAU32624.1 atub|OJI80213.1 aver|OJJ04962.1 awen|OJJ38151.1 azon|OJJ43136.1

Group4994: abra|OJJ69930.1 acar|OOF93441.1 acla|EAW08893.1 afis|EAW15496.1 afla|EED49011.1 afum|EAL85827.1 agla|OJJ80175.1 akaw|GAA87727.1 aluc|OJZ85357.1 anig|EHA25541.1 aory|EIT82520.1 asyd|OJJ56015.1 ater|EAU32623.1 atub|OJI80214.1 aver|OJJ04961.1 awen|OJJ38154.1 azon|OJJ43135.1

Group4995: abra|OJJ69934.1 acar|OOF93443.1 acla|EAW08892.1 afis|EAW15497.1 afla|EED49012.1 afum|EAL85826.1 agla|OJJ80176.1 akaw|GAA87725.1 aluc|OJZ85353.1 anig|EHA25539.1 aory|EIT82506.1 asyd|OJJ56014.1 ater|EAU32622.1 atub|OJI80217.1 aver|OJJ04959.1 awen|OJJ38156.1 azon|OJJ43134.1

Group4996: abra|OJJ69937.1 acar|OOF93446.1 acla|EAW08886.1 afis|EAW15503.1 afla|EED49016.1 afum|EAL85821.1 agla|OJJ80180.1 akaw|GAA87722.1 aluc|OJZ85350.1 anig|EHA25537.1 aory|EIT82554.1 asyd|OJJ56008.1 ater|EAU32615.1 atub|OJI80220.1 aver|OJJ04953.1 awen|OJJ38163.1 azon|OJJ43124.1

Group4997: abra|OJJ69938.1 acar|OOF93447.1 acla|EAW08885.1 afis|EAW15504.1 afla|EED49017.1 afum|EAL85820.1 agla|OJJ80181.1 akaw|GAA87721.1 aluc|OJZ85349.1 anig|EHA25536.1 aory|EIT82559.1 asyd|OJJ56007.1 ater|EAU32614.1 atub|OJI80221.1 aver|OJJ04952.1 awen|OJJ38164.1 azon|OJJ43125.1

Group4998: abra|OJJ69939.1 acar|OOF93448.1 acla|EAW08884.1 afis|EAW15505.1 afla|EED49018.1 afum|EAL85819.1 agla|OJJ80182.1 akaw|GAA87720.1 aluc|OJZ85348.1 anig|EHA25535.1 aory|EIT82495.1 asyd|OJJ56006.1 ater|EAU32613.1 atub|OJI80222.1 aver|OJJ04951.1 awen|OJJ38165.1 azon|OJJ43126.1

Group4999: abra|OJJ69944.1 acar|OOF93453.1 acla|EAW08879.1 afis|EAW15516.1 afla|EED49022.1 afum|EAL85813.1 agla|OJJ80187.1 akaw|GAA87716.1 aluc|OJZ85343.1 anig|EHA25531.1 aory|EIT82503.1 asyd|OJJ56002.1 ater|EAU32608.1 atub|OJI80228.1 aver|OJJ04948.1 awen|OJJ38177.1 azon|OJJ43120.1

Group5000: abra|OJJ69945.1 acar|OOF93454.1 acla|EAW08878.1 afis|EAW15517.1 afla|EED49023.1 afum|EAL85812.1 agla|OJJ85228.1 akaw|GAA87715.1 aluc|OJZ85342.1 anig|EHA25530.1 aory|EIT82489.1 asyd|OJJ56001.1 ater|EAU32607.1 atub|OJI80229.1 aver|OJJ04947.1 awen|OJJ38178.1 azon|OJJ43119.1

Group5001: abra|OJJ69946.1 acar|OOF93455.1 acla|EAW08877.1 afis|EAW15518.1 afla|EED49025.1 afum|EAL85811.1 agla|OJJ85227.1 akaw|GAA87714.1 aluc|OJZ85341.1 anig|EHA25529.1 aory|EIT82492.1 asyd|OJJ56000.1 ater|EAU32606.1 atub|OJI80230.1 aver|OJJ04946.1 awen|OJJ38179.1 azon|OJJ43118.1

Group5002: abra|OJJ69949.1 acar|OOF93457.1 acla|EAW08875.1 afis|EAW15520.1 afla|EED49027.1 afum|EAL85809.1 agla|OJJ85226.1 akaw|GAA87711.1 aluc|OJZ85338.1 anig|EHA25526.1 aory|EIT82544.1 asyd|OJJ55997.1 ater|EAU32604.1 atub|OJI80233.1 aver|OJJ04944.1 awen|OJJ38181.1 azon|OJJ43112.1

Group5003: abra|OJJ69957.1 acar|OOF93467.1 acla|EAW08865.1 afis|EAW15529.1 afla|EED49040.1 afum|EAL85799.1 akaw|GAA87703.1 aluc|OJZ85325.1 anid|CBF78208.1 anig|EHA25515.1 aory|EIT82500.1 asyd|OJJ55987.1 ater|EAU32596.1 atub|OJI80247.1 aver|OJJ04935.1 awen|OJJ38193.1 azon|OJJ43101.1

Group5004: abra|OJJ69966.1 acla|EAW08838.1 afis|EAW15562.1 afla|EED49068.1 afum|EAL85770.1 agla|OJJ85189.1 akaw|GAA87693.1 aluc|OJZ85316.1 anid|CBF78123.1 anig|EHA25509.1 aory|EIT82555.1 asyd|OJJ55904.1 ater|EAU32570.1 atub|OJI80256.1 aver|OJJ04849.1 awen|OJJ38222.1 azon|OJJ43073.1

Group5005: abra|OJJ69969.1 acar|OOF93474.1 acla|EAW08840.1 afis|EAW15561.1 afla|EED49065.1 afum|EAL85771.1 agla|OJJ85193.1 akaw|GAA87692.1 aluc|OJZ85313.1 anig|EHA25508.1 aory|EIT82526.1 asyd|OJJ55911.1 ater|EAU32571.1 atub|OJI80258.1 aver|OJJ04851.1 awen|OJJ38219.1 azon|OJJ43074.1

Group5006: abra|OJJ69970.1 acar|OOF93476.1 acla|EAW08841.1 afis|EAW15560.1 afla|EED49064.1 afum|EAL85772.1 agla|OJJ85194.1 akaw|GAA87691.1 aluc|OJZ85312.1 anig|EHA25507.1 aory|EIT82521.1 asyd|OJJ55912.1 ater|EAU32572.1 atub|OJI80259.1 aver|OJJ04852.1 awen|OJJ38218.1 azon|OJJ43075.1

Group5007: abra|OJJ69972.1 acar|OOF93477.1 acla|EAW08843.1 afis|EAW15558.1 afla|EED49062.1 afum|EAL85774.1 agla|OJJ85196.1 akaw|GAA87689.1 aluc|OJZ85310.1 anid|CBF78161.1 anig|EHA25505.1 asyd|OJJ55962.1 ater|EAU32576.1 atub|OJI80261.1 aver|OJJ04907.1 awen|OJJ38216.1 azon|OJJ43079.1

Group5008: abra|OJJ69980.1 acla|EAW08850.1 afis|EAW15546.1 afum|EAL85786.1 agla|OJJ85206.1 akaw|GAA87681.1 aluc|OJZ85300.1 anid|CBF78178.1 anig|EHA25497.1 aory|EIT82561.1 asyd|OJJ55971.1 ater|EAU32581.1 atub|OJI80270.1 aver|OJJ04918.1 awen|OJJ38207.1 azon|OJJ43088.1 afla|EED49053.1

Group5009: abra|OJJ69988.1 acar|OOF93493.1 acla|EAW08859.1 afis|EAW15535.1 afla|EED49045.1 afum|EAL85793.1 agla|OJJ85212.1 akaw|GAA87673.1 aluc|OJZ85291.1 anid|CBF78195.1 anig|EHA25489.1 aory|EIT82566.1 asyd|OJJ55980.1 ater|EAU32590.1 atub|OJI80279.1 aver|OJJ04928.1 awen|OJJ38199.1

Group5010: abra|OJJ69998.1 acar|OOF93505.1 acla|EAW08821.1 afis|EAW15576.1 afla|EED49085.1 afum|EAL85758.1 agla|OJJ85176.1 akaw|GAA87663.1 aluc|OJZ85281.1 anig|EHA25478.1 aory|EIT82569.1 asyd|OJJ55892.1 ater|EAU32555.1 atub|OJI80289.1 aver|OJJ04837.1 awen|OJJ38236.1 azon|OJJ46061.1

Group5011: abra|OJJ70002.1 acar|OOF93508.1 acla|EAW08817.1 afis|EAW15580.1 afla|EED49092.1 afum|EAL85754.1 agla|OJJ85172.1 akaw|GAA87659.1 aluc|OJZ85277.1 anid|CBF78098.1 anig|EHA25474.1 asyd|OJJ55888.1 ater|EAU32550.1 atub|OJI80293.1 aver|OJJ04832.1 awen|OJJ38242.1 azon|OJJ43056.1

Group5012: abra|OJJ70005.1 acar|OOF93510.1 acla|EAW07277.1 afis|EAW22308.1 afla|EED57704.1 afum|EAL90882.1 agla|OJJ87742.1 akaw|GAA87656.1 aluc|OJZ85274.1 anid|CBF88590.1 anig|EHA25472.1 aory|EIT79905.1 asyd|OJJ63849.1 ater|EAU34316.1 atub|OJI80296.1 aver|OJI97088.1 awen|OJJ34879.1

Group5013: abra|OJJ70010.1 acar|OOF93516.1 acla|EAW08814.1 afis|EAW15583.1 afla|EED49087.1 afum|EAL85751.1 agla|OJJ85168.1 akaw|GAA87652.1 aluc|OJZ85269.1 anid|CBF78058.1 anig|EHA25468.1 aory|EIT82488.1 ater|EAU32546.1 atub|OJI80302.1 aver|OJJ04812.1 awen|OJJ38248.1 azon|OJJ43028.1

Group5014: abra|OJJ70012.1 acar|OOF93518.1 acla|EAW08812.1 afis|EAW15585.1 afla|EED54444.1 afum|EAL85747.1 agla|OJJ85164.1 akaw|GAA87650.1 aluc|OJZ85266.1 anid|CBF78061.1 anig|EHA25465.1 asyd|OJJ55869.1 ater|EAU32544.1 atub|OJI80304.1 aver|OJJ04814.1 awen|OJJ38251.1 azon|OJJ43031.1

Group5015: abra|OJJ70137.1 acar|OOF93594.1 acla|EAW13744.1 afis|EAW19842.1 afla|EED46533.1 afum|EAL85053.1 akaw|GAA86628.1 aluc|OJZ85183.1 anid|CBF76707.1 anig|EHA25395.1 aory|EIT72392.1 asyd|OJJ61237.1 ater|EAU38516.1 atub|OJI80414.1 aver|OJJ00560.1 awen|OJJ34118.1 azon|OJJ49476.1

Group5016: abra|OJJ70140.1 acar|OOF93597.1 acla|EAW13733.1 afis|EAW19843.1 afla|EED46534.1 afum|EAL85052.1 akaw|GAA86625.1 aluc|OJZ85180.1 anid|CBF76705.1 anig|EHA25392.1 aory|EIT72326.1 asyd|OJJ61236.1 ater|EAU38515.1 atub|OJI80417.1 aver|OJJ00559.1 awen|OJJ34117.1 azon|OJJ49477.1

Group5017: abra|OJJ70193.1 acar|OOF92582.1 acla|EAW06999.1 afis|EAW16614.1 afla|EED49260.1 afum|EAL89222.2 agla|OJJ82625.1 akaw|GAA86582.1 aluc|OJZ85133.1 anid|CBF86355.1 anig|EHA25352.1 asyd|OJJ57360.1 ater|EAU36698.1 atub|OJI80460.1 aver|OJJ05806.1 awen|OJJ40191.1 afla|EED44998.1

Group5018: abra|OJJ70232.1 acar|OOF93866.1 afla|EED56154.1 afum|EAL90745.1 agla|OJJ88092.1 akaw|GAA86552.1 aluc|OJZ85100.1 anig|EHA26109.1 asyd|OJJ59762.1 ater|EAU37896.1 aver|OJJ02526.1 awen|OJJ39210.1 azon|OJJ50036.1 anig|EHA25322.1 acla|EAW07420.1 afis|EAW22446.1 awen|OJJ35071.1

Group5019: abra|OJJ70304.1 acar|OOF91757.1 acla|EAW12288.1 afis|EAW25658.1 afla|EED45243.1 afum|EAL86477.2 agla|OJJ79117.1 akaw|GAA93018.1 aluc|OJZ82567.1 anid|CBF80474.1 anig|EHA27507.1 aory|EIT73558.1 asyd|OJJ54715.1 ater|EAU30190.1 atub|OJI88811.1 aver|OJJ07452.1 afis|EAW16950.1

Group5020: abra|OJJ70306.1 acar|OOF91761.1 acla|EAW12289.1 afis|EAW16947.1 afla|EED45244.1 afum|EAL86474.1 agla|OJJ79960.1 akaw|GAA93016.1 aluc|OJZ82569.1 anid|CBF80495.1 anig|EHA27505.1 aory|EIT73662.1 asyd|OJJ54729.1 ater|EAU30192.1 atub|OJI88813.1 aver|OJJ07439.1 awen|OJJ29711.1

Group5021: abra|OJJ70329.1 acar|OOF91784.1 acla|EAW12320.1 afis|EAW16901.1 afla|EED45272.1 afum|EAL86436.1 agla|OJJ79952.1 akaw|GAA92584.1 aluc|OJZ82598.1 anid|CBF80772.1 anig|EHA27479.1 aory|EIT73781.1 asyd|OJJ54500.1 ater|EAU30216.1 atub|OJI88834.1 aver|OJJ07680.1 awen|OJJ29730.1

Group5022: abra|OJJ70366.1 acar|OOF91906.1 acla|EAW12330.1 afis|EAW16893.1 afum|EAL86431.1 agla|OJJ79946.1 akaw|GAA91985.1 aluc|OJZ82628.1 anid|CBF80796.1 anig|EHA27449.1 aory|EIT75240.1 asyd|OJJ54490.1 ater|EAU30222.1 atub|OJI88861.1 aver|OJJ07689.1 awen|OJJ29744.1 azon|OJJ42525.1

Group5023: abra|OJJ70382.1 acar|OOF91889.1 afis|EAW16883.1 afla|EED47346.1 afum|EAL86419.1 agla|OJJ79750.1 akaw|GAA91974.1 aluc|OJZ82641.1 anid|CBF80677.1 anig|EHA27429.1 aory|EIT82848.1 asyd|OJJ54589.1 ater|EAU30232.1 atub|OJI88874.1 aver|OJJ07564.1 awen|OJJ29766.1 azon|OJJ50379.1

Group5024: abra|OJJ70471.1 acar|OOF97454.1 acla|EAW13043.1 afis|EAW17624.1 afla|EED49193.1 afum|EAL91438.1 agla|OJJ81279.1 akaw|GAA83320.1 aluc|OJZ83815.1 anid|CBF90154.1 anig|EHA18960.1 asyd|OJJ61999.1 ater|EAU37058.1 atub|OJI90182.1 aver|OJI96091.1 awen|OJJ37220.1 azon|OJJ50923.1

Group5025: abra|OJJ70486.1 acar|OOF97440.1 acla|EAW13030.1 afis|EAW17637.1 afla|EED50527.1 afum|EAL91452.1 agla|OJJ81257.1 akaw|GAA83334.1 aluc|OJZ83830.1 anid|CBF90130.1 anig|EHA18973.1 asyd|OJJ62016.1 ater|EAU37073.1 atub|OJI90197.1 aver|OJI96107.1 awen|OJJ37207.1 azon|OJJ50906.1

Group5026: abra|OJJ70494.1 acar|OOF97433.1 acla|EAW13023.1 afis|EAW17644.1 afla|EED50516.1 agla|OJJ79560.1 akaw|GAA83342.1 aluc|OJZ83838.1 anid|CBF90116.1 anig|EHA18981.1 aory|EIT82026.1 asyd|OJJ62023.1 ater|EAU37080.1 atub|OJI90205.1 aver|OJI96114.1 awen|OJJ37201.1 azon|OJJ50900.1

Group5027: abra|OJJ70501.1 acar|OOF97426.1 acla|EAW08173.1 afis|EAW23209.1 afla|EED47042.1 afum|EAL88275.1 agla|OJJ78736.1 akaw|GAA83349.1 aluc|OJZ83846.1 anig|EHA18988.1 aory|EIT72737.1 asyd|OJJ60187.1 ater|EAU36337.1 atub|OJI90212.1 aver|OJJ04077.1 awen|OJJ39442.1 azon|OJJ47644.1

Group5028: abra|OJJ70503.1 acar|OOF97423.1 acla|EAW08172.1 afis|EAW23208.1 afla|EED47046.1 afum|EBA27341.1 agla|OJJ78738.1 akaw|GAA83351.1 aluc|OJZ83848.1 anig|EHA18990.1 aory|EIT72619.1 asyd|OJJ60188.1 ater|EAU36333.1 atub|OJI90214.1 aver|OJJ04076.1 awen|OJJ39441.1 azon|OJJ47642.1

Group5029: abra|OJJ70511.1 acar|OOF97418.1 acla|EAW08164.1 afis|EAW23202.1 afla|EED47052.1 afum|EAL88281.1 agla|OJJ78746.1 akaw|GAA83358.1 aluc|OJZ83856.1 anid|CBF74726.1 anig|EHA18997.1 asyd|OJJ60196.1 ater|EAU36330.1 atub|OJI90223.1 aver|OJJ04069.1 awen|OJJ39432.1 azon|OJJ47636.1

Group5030: abra|OJJ70523.1 acla|EAW08152.1 afis|EAW23187.1 afla|EED47066.1 afum|EAL88296.1 agla|OJJ86362.1 akaw|GAA83370.1 aluc|OJZ83868.1 anid|CBF74683.1 anig|EHA19009.1 aory|EIT72474.1 asyd|OJJ60221.1 ater|EAU36318.1 atub|OJI90236.1 aver|OJJ04047.1 awen|OJJ36589.1 azon|OJJ47617.1

Group5031: abra|OJJ70527.1 acar|OOF97404.1 acla|EAW08140.1 afis|EAW23176.1 afla|EED46943.1 afum|EAL88307.2 agla|OJJ89688.1 akaw|GAA83372.1 aluc|OJZ83870.1 anid|CBF74500.1 anig|EHA19011.1 aory|EIT72572.1 asyd|OJJ57721.1 atub|OJI90238.1 aver|OJJ05447.1 awen|OJJ36602.1 azon|OJJ47606.1

Group5032: abra|OJJ70533.1 acar|OOF97399.1 afis|EAW16751.1 afla|EED47075.1 agla|OJJ86369.1 akaw|GAA83378.1 aluc|OJZ83876.1 anig|EHA19016.1 aory|EIT72481.1 atub|OJI90244.1 awen|OJJ36596.1 azon|OJJ50257.1 afis|EAW23182.1 acla|EAW12439.1 acla|EAW12440.1 afum|EBA27268.1 afum|EAL88300.1

Group5033: abra|OJJ70547.1 acla|EAW08134.1 afis|EAW23170.1 afla|EED46951.1 afum|EAL88314.1 agla|OJJ89681.1 akaw|GAA83391.1 aluc|OJZ83891.1 anid|CBF74482.1 anig|EHA19027.1 asyd|OJJ57710.1 ater|EAU36299.1 atub|OJI90258.1 aver|OJJ05455.1 awen|OJJ36613.1 azon|OJJ47600.1 agla|OJJ89682.1

Group5034: abra|OJJ70551.1 acar|OOF97169.1 acla|EAW08343.1 afis|EAW23378.1 afla|EED55579.1 afum|EAL88093.1 agla|OJJ86445.1 akaw|GAA83395.1 aluc|OJZ83894.1 anid|CBF74847.1 anig|EHA19030.1 asyd|OJJ60129.1 atub|OJI90262.1 aver|OJJ04132.1 awen|OJJ39506.1 azon|OJJ47705.1 ater|EAU35632.1

Group5035: abra|OJJ70553.1 acar|OOF97166.1 acla|EAW08348.1 afis|EAW23382.1 afla|EED55569.1 afum|EAL88090.1 agla|OJJ86441.1 akaw|GAA83398.1 aluc|OJZ83896.1 anig|EHA19032.1 aory|EIT82669.1 ater|EAU35635.1 atub|OJI90265.1 awen|OJJ39502.1 azon|OJJ47702.1 afum|EAL88089.1 anig|EHA19033.1

Group5036: abra|OJJ70562.1 acar|OOF97157.1 acla|EAW08361.1 afis|EAW23392.1 afla|EED55565.1 afum|EAL88078.1 agla|OJJ86425.1 akaw|GAA83407.1 aluc|OJZ83906.1 anid|CBF74862.1 anig|EHA19041.1 asyd|OJJ60122.1 ater|EAU35645.1 atub|OJI90277.1 aver|OJJ04140.1 awen|OJJ39487.1 azon|OJJ47690.1

Group5037: abra|OJJ70586.1 acar|OOF97134.1 acla|EAW08195.1 afis|EAW23229.1 afla|EED47021.1 afum|EAL88254.1 agla|OJJ86386.1 akaw|GAA83429.1 aluc|OJZ83930.1 anid|CBF74773.1 anig|EHA19062.1 asyd|OJJ60166.1 ater|EAU36358.1 atub|OJI90302.1 aver|OJJ04098.1 awen|OJJ39464.1 azon|OJJ47664.1

Group5038: abra|OJJ70587.1 acar|OOF97133.1 acla|EAW08194.1 afis|EAW23228.1 afla|EED47022.1 afum|EAL88255.1 agla|OJJ86385.1 akaw|GAA83430.1 aluc|OJZ83931.1 anid|CBF74771.1 anig|EHA19063.1 asyd|OJJ60167.1 ater|EAU36357.1 atub|OJI90303.1 aver|OJJ04097.1 awen|OJJ39463.1 azon|OJJ47663.1

Group5039: abra|OJJ70590.1 acar|OOF97129.1 acla|EAW08191.1 afis|EAW23225.1 afla|EED47025.1 afum|EAL88258.1 agla|OJJ86382.1 akaw|GAA83434.1 aluc|OJZ83935.1 anid|CBF74765.1 anig|EHA19066.1 aory|EIT72720.1 asyd|OJJ60170.1 atub|OJI90307.1 aver|OJJ04094.1 awen|OJJ39460.1 azon|OJJ47659.1

Group5040: abra|OJJ70591.1 afla|EED47026.1 afum|EAL88260.1 agla|OJJ86381.1 akaw|GAA83435.1 aluc|OJZ83937.1 anid|CBF74763.1 aory|EIT72526.1 azon|OJJ47658.1 awen|OJJ39459.1 acla|EAW08190.1 afis|EAW23224.1 anig|EHA19067.1 asyd|OJJ60171.1 ater|EAU36355.1 atub|OJI90309.1 aver|OJJ04093.1

Group5041: abra|OJJ70597.1 acar|OOF97122.1 afis|EAW23218.1 afla|EED47033.1 afum|EAL88266.1 agla|OJJ86374.1 akaw|GAA83441.1 aluc|OJZ83943.1 anid|CBF74754.1 anig|EHA19072.1 aory|EIT72711.1 asyd|OJJ60177.1 ater|EAU36349.1 atub|OJI90315.1 aver|OJJ04087.1 awen|OJJ39452.1 azon|OJJ47652.1

Group5042: abra|OJJ70601.1 acar|OOF97119.1 acla|EAW08181.1 afis|EAW23216.1 afum|EAL88268.1 agla|OJJ86372.1 akaw|GAA89941.1 aluc|OJZ83946.1 anid|CBF74747.1 anig|EHA19075.1 aory|EIT72596.1 asyd|OJJ60179.1 ater|EAU36346.1 atub|OJI90317.1 aver|OJJ04083.1 awen|OJJ39450.1 azon|OJJ47650.1

Group5043: abra|OJJ70614.1 acar|OOF97107.1 acla|EAW13013.1 afis|EAW17654.1 afla|EED50504.1 afum|EAL91469.1 agla|OJJ79550.1 akaw|GAA89953.1 aluc|OJZ83960.1 anig|EHA19088.1 aory|EIT82141.1 atub|OJI90332.1 aver|OJI96149.1 awen|OJJ37191.1 azon|OJJ50890.1 anid|CBF90070.1 asyd|OJJ62043.1

Group5044: abra|OJJ70615.1 acar|OOF97106.1 acla|EAW13012.1 afis|EAW17655.1 afla|EED50503.1 afum|EAL91470.1 akaw|GAA89954.1 aluc|OJZ83961.1 anid|CBF90068.1 anig|EHA19089.1 aory|EIT82061.1 asyd|OJJ62044.1 ater|EAU31998.1 atub|OJI90333.1 aver|OJI96150.1 awen|OJJ37190.1 azon|OJJ48749.1

Group5045: abra|OJJ70631.1 acar|OOF97092.1 acla|EAW08125.1 afis|EAW23162.1 afla|EED46959.1 afum|EAL88321.1 agla|OJJ89673.1 akaw|GAA89970.1 aluc|OJZ83979.1 anid|CBF74465.1 anig|EHA19106.1 aory|EIT72687.1 asyd|OJJ57699.1 atub|OJI90352.1 aver|OJJ05465.1 awen|OJJ36621.1 azon|OJJ47592.1

Group5046: abra|OJJ70636.1 acar|OOF97078.1 acla|EAW08120.1 afis|EAW23131.1 afla|EED48997.1 afum|EAL88350.2 agla|OJJ89668.1 akaw|GAA89974.1 aluc|OJZ83984.1 anid|CBF74460.1 aory|EIT80038.1 asyd|OJJ57680.1 ater|EAU36284.1 atub|OJI90355.1 aver|OJJ05485.1 awen|OJJ36629.1 azon|OJJ47588.1

Group5047: abra|OJJ70638.1 acar|OOF97079.1 acla|EAW08119.1 afis|EAW23132.1 afla|EED48996.1 afum|EAL88349.1 agla|OJJ89667.1 aluc|OJZ83985.1 anid|CBF74458.1 anig|EHA19111.1 aory|EIT80017.1 asyd|OJJ57678.1 ater|EAU36285.1 atub|OJI90356.1 aver|OJJ05484.1 awen|OJJ36628.1 azon|OJJ47587.1

Group5048: abra|OJJ70652.1 acla|EAW08107.1 afis|EAW23144.1 afla|EED48507.1 afum|EAL88337.2 agla|OJJ89658.1 akaw|GAA89988.1 aluc|OJZ84002.1 anid|CBF74437.1 anig|EHA19124.1 aory|EIT73158.1 asyd|OJJ57746.1 ater|EAU36267.1 atub|OJI90371.1 aver|OJJ05421.1 awen|OJJ36639.1 azon|OJJ47576.1

Group5049: abra|OJJ70655.1 acla|EAW08104.1 afis|EAW23147.1 afla|EED48510.1 afum|EAL88334.1 agla|OJJ89655.1 akaw|GAA89991.1 aluc|OJZ84005.1 anid|CBF74432.1 anig|EHA19127.1 aory|EIT73217.1 asyd|OJJ57749.1 ater|EAU36264.1 atub|OJI90374.1 aver|OJJ05418.1 awen|OJJ36642.1 azon|OJJ47572.1

Group5050: abra|OJJ70658.1 acar|OOF97064.1 acla|EAW08101.1 afis|EAW23150.1 afla|EED48513.1 afum|EAL88331.1 agla|OJJ89652.1 akaw|GAA89994.1 aluc|OJZ84008.1 anid|CBF74426.1 anig|EHA19130.1 asyd|OJJ57752.1 ater|EAU36262.1 atub|OJI90377.1 aver|OJJ05415.1 awen|OJJ36646.1 azon|OJJ47569.1

Group5051: abra|OJJ70687.1 acar|OOF97040.1 afis|EAW23113.1 afla|EED53331.1 afum|EAL88366.1 agla|OJJ89627.1 akaw|GAA90019.1 aluc|OJZ84035.1 anid|CBF76022.1 anig|EHA19156.1 aory|EIT75261.1 asyd|OJJ52449.1 ater|EAU36241.1 atub|OJI90407.1 aver|OJJ03967.1 awen|OJJ36668.1 azon|OJJ47531.1

Group5052: abra|OJJ70688.1 acar|OOF97039.1 afis|EAW23112.1 afla|EED53332.1 afum|EAL88367.1 agla|OJJ89629.1 akaw|GAA90020.1 aluc|OJZ84036.1 anid|CBF76020.1 anig|EHA19157.1 aory|EIT75265.1 asyd|OJJ52448.1 ater|EAU36240.1 atub|OJI90408.1 aver|OJJ03968.1 awen|OJJ36669.1 azon|OJJ47530.1

Group5053: abra|OJJ70690.1 acla|EAW08074.1 afis|EAW23110.1 afla|EED48538.1 afum|EAL88369.1 agla|OJJ89631.1 akaw|GAA90022.1 aluc|OJZ84038.1 anid|CBF74387.1 anig|EHA19159.1 aory|EIT73045.1 asyd|OJJ57772.1 ater|EAU36238.1 atub|OJI90410.1 aver|OJJ05394.1 awen|OJJ36671.1 azon|OJJ47528.1

Group5054: abra|OJJ70692.1 acla|EAW08072.1 afis|EAW23108.1 afla|EED48540.1 afum|EAL88371.1 agla|OJJ89633.1 akaw|GAA90023.1 aluc|OJZ84039.1 anid|CBF81372.1 anig|EHA19161.1 aory|EIT73071.1 asyd|OJJ59387.1 ater|EAU36236.1 atub|OJI90411.1 aver|OJJ02886.1 awen|OJJ36673.1 azon|OJJ47526.1

Group5055: abra|OJJ70695.1 acar|OOF97034.1 acla|EAW08070.1 afis|EAW23106.1 afla|EED48542.1 afum|EAL88373.1 agla|OJJ89637.1 akaw|GAA90026.1 aluc|OJZ84042.1 anid|CBF81369.1 anig|EHA19164.1 asyd|OJJ59385.1 ater|EAU36234.1 atub|OJI90414.1 aver|OJJ02888.1 awen|OJJ36675.1 azon|OJJ47523.1

Group5056: abra|OJJ70703.1 acla|EAW08062.1 afis|EAW23098.1 afla|EED48546.1 afum|EAL88380.2 agla|OJJ89610.1 akaw|GAA90034.1 aluc|OJZ84050.1 anid|CBF81354.1 anig|EHA19170.1 aory|EIT73119.1 asyd|OJJ59378.1 ater|EAU36231.1 atub|OJI90421.1 aver|OJJ02895.1 awen|OJJ36683.1 azon|OJJ47514.1

Group5057: abra|OJJ70712.1 acar|OOF97024.1 acla|EAW08052.1 afis|EAW23088.1 afla|EED48560.1 afum|EAL88391.1 agla|OJJ89598.1 akaw|GAA90043.1 aluc|OJZ84059.1 anid|CBF81137.1 anig|EHA19179.1 aory|EIT73115.1 asyd|OJJ56502.1 atub|OJI90430.1 aver|OJJ06217.1 awen|OJJ36696.1 azon|OJJ47501.1

Group5058: abra|OJJ70713.1 acar|OOF97023.1 acla|EAW08050.1 afis|EAW23086.1 afla|EED48562.1 afum|EAL88393.1 agla|OJJ89596.1 akaw|GAA90044.1 aluc|OJZ84060.1 anid|CBF81341.1 anig|EHA19180.1 asyd|OJJ59370.1 ater|EAU36210.1 atub|OJI90431.1 aver|OJJ02904.1 awen|OJJ36698.1 azon|OJJ47499.1

Group5059: abra|OJJ70724.1 acar|OOF97014.1 acla|EAW08041.1 afis|EAW23076.1 afum|EAL88403.1 agla|OJJ89583.1 akaw|GAA90053.1 aluc|OJZ84069.1 anid|CBF81322.1 anig|EHA19189.1 aory|EIT73284.1 asyd|OJJ59360.1 ater|EAU36200.1 atub|OJI90442.1 aver|OJJ02914.1 awen|OJJ36708.1 azon|OJJ47491.1

Group5060: abra|OJJ70732.1 acar|OOF97006.1 acla|EAW11181.1 afis|EAW15912.1 afla|EED56456.1 afum|EAL88547.2 agla|OJJ89576.1 akaw|GAA90060.1 aluc|OJZ84077.1 anid|CBF81303.1 anig|EHA19196.1 aory|EIT72846.1 asyd|OJJ59351.1 atub|OJI90449.1 aver|OJJ02923.1 awen|OJJ36718.1 azon|OJJ47477.1

Group5061: abra|OJJ70739.1 acla|EAW11177.1 afis|EAW15908.1 afla|EED56460.1 afum|EAL88543.1 agla|OJJ89572.1 akaw|GAA90064.1 aluc|OJZ84084.1 anid|CBF81261.1 anig|EHA19200.1 aory|EIT72862.1 asyd|OJJ59331.1 ater|EAU36187.1 atub|OJI90454.1 aver|OJJ02943.1 awen|OJJ36722.1 azon|OJJ47481.1

Group5062: abra|OJJ70749.1 acar|OOF96996.1 acla|EAW11172.1 afis|EAW15903.1 afla|EED56467.1 afum|EAL88538.1 agla|OJJ89563.1 akaw|GAA90069.1 aluc|OJZ84093.1 anid|CBF81278.1 anig|EHA19206.1 aory|EIT72833.1 asyd|OJJ59336.1 atub|OJI90465.1 aver|OJJ02938.1 awen|OJJ36728.1 azon|OJJ47469.1

Group5063: abra|OJJ70750.1 acar|OOF96995.1 acla|EAW11171.1 afis|EAW15902.1 afla|EED56468.1 afum|EAL88537.1 agla|OJJ89562.1 akaw|GAA90070.1 aluc|OJZ84094.1 anid|CBF81279.1 anig|EHA19207.1 aory|EIT72855.1 asyd|OJJ59337.1 atub|OJI90466.1 aver|OJJ02937.1 awen|OJJ36729.1 azon|OJJ47468.1

Group5064: abra|OJJ70757.1 acar|OOF96987.1 acla|EAW11164.1 afis|EAW15895.1 afum|EAL88530.1 agla|OJJ89549.1 akaw|GAA90077.1 aluc|OJZ84102.1 anid|CBF81292.1 anig|EHA19214.1 aory|EIT72812.1 asyd|OJJ59345.1 ater|EAU36174.1 atub|OJI90475.1 aver|OJJ02929.1 awen|OJJ36736.1 azon|OJJ47461.1

Group5065: abra|OJJ70761.1 acar|OOF96983.1 acla|EAW11160.1 afis|EAW15891.1 afla|EED56481.1 afum|EAL88526.2 agla|OJJ89531.1 akaw|GAA90081.1 aluc|OJZ84106.1 anid|CBF81258.1 anig|EHA19218.1 aory|EIT72809.1 asyd|OJJ59329.1 ater|EAU31713.1 atub|OJI90480.1 aver|OJJ02945.1 azon|OJJ47458.1

Group5066: abra|OJJ70763.1 afis|EAW15889.1 afla|EED56483.1 akaw|GAA90083.1 aluc|OJZ84108.1 anid|CBF81254.1 anig|EHA19220.1 aory|EIT72840.1 asyd|OJJ59327.1 atub|OJI90482.1 aver|OJJ02947.1 awen|OJJ36741.1 azon|OJJ47456.1 acla|EAW11158.1 afum|EAL88523.1 agla|OJJ89533.1 ater|EAU31711.1

Group5067: abra|OJJ70764.1 acar|OOF96981.1 acla|EAW11157.1 afis|EAW15888.1 afla|EED56484.1 afum|EAL88522.2 akaw|GAA90084.1 aluc|OJZ84109.1 anid|CBF81252.1 anig|EHA19221.1 aory|EIT72836.1 asyd|OJJ59326.1 ater|EAU31710.1 atub|OJI90483.1 aver|OJJ02948.1 awen|OJJ36742.1 azon|OJJ47455.1

Group5068: abra|OJJ70772.1 acar|OOF96974.1 acla|EAW11150.1 afis|EAW15881.1 afla|EED56491.1 afum|EAL88515.1 agla|OJJ89540.1 akaw|GAA90091.1 aluc|OJZ84118.1 anid|CBF81236.1 anig|EHA19227.1 asyd|OJJ59318.1 ater|EAU31701.1 atub|OJI90491.1 aver|OJJ02955.1 awen|OJJ36750.1 azon|OJJ47449.1

Group5069: abra|OJJ70810.1 acar|OOF98292.1 acla|EAW10419.1 afis|EAW25150.1 afla|EED55285.1 afum|EAL89825.1 agla|OJJ89005.1 akaw|GAA91690.1 aluc|OJZ87704.1 anid|CBF85670.1 anig|EHA20887.1 asyd|OJJ64248.1 ater|EAU39418.1 atub|OJI80939.1 aver|OJI98339.1 awen|OJJ40779.1 azon|OJJ46873.1

Group5070: abra|OJJ70813.1 acar|OOF98289.1 acla|EAW10422.1 afis|EAW25147.1 afla|EED55291.1 afum|EAL89822.1 agla|OJJ89010.1 akaw|GAA91693.1 aluc|OJZ87701.1 anid|CBF85675.1 aory|EIT83562.1 asyd|OJJ64252.1 ater|EAU39421.1 atub|OJI80942.1 aver|OJI98342.1 awen|OJJ40772.1 azon|OJJ46876.1

Group5071: abra|OJJ70814.1 acar|OOF98288.1 acla|EAW10423.1 afis|EAW25146.1 afla|EED55292.1 afum|EAL89821.1 agla|OJJ89009.1 akaw|GAA91694.1 aluc|OJZ87700.1 anid|CBF85677.1 aory|EIT83581.1 asyd|OJJ64253.1 ater|EAU39422.1 atub|OJI80943.1 aver|OJI98343.1 awen|OJJ40771.1 azon|OJJ46877.1

Group5072: abra|OJJ70835.1 acar|OOF98279.1 acla|EAW10430.1 afis|EAW25139.1 afla|EED55306.1 afum|EAL89811.1 agla|OJJ89028.1 akaw|GAA91703.1 aluc|OJZ87687.1 anid|CBF86097.1 anig|EHA20876.1 aory|EIT83522.1 asyd|OJJ64475.1 atub|OJI80953.1 aver|OJI98579.1 awen|OJJ40763.1 azon|OJJ44128.1

Group5073: abra|OJJ70847.1 acar|OOF98268.1 acla|EAW10438.1 afis|EAW25129.1 afla|EED55315.1 afum|EAL89800.1 agla|OJJ89037.1 akaw|GAA91715.1 aluc|OJZ87674.1 anid|CBF85699.1 anig|EHA20864.1 asyd|OJJ64264.1 ater|EAU39402.1 atub|OJI80969.1 aver|OJI98356.1 awen|OJJ40754.1 azon|OJJ44118.1

Group5074: abra|OJJ70849.1 acar|OOF98267.1 acla|EAW10440.1 afis|EAW25127.1 afum|EAL89798.1 agla|OJJ89043.1 akaw|GAA91717.1 aluc|OJZ87671.1 anid|CBF85695.1 anig|EHA20862.1 aory|EIT83507.1 asyd|OJJ64262.1 ater|EAU39400.1 atub|OJI80971.1 aver|OJI98354.1 awen|OJJ40752.1 azon|OJJ44116.1

Group5075: abra|OJJ70863.1 acar|OOF98253.1 acla|EAW10455.1 afis|EAW25112.1 afla|EED55330.1 afum|EAL89783.2 agla|OJJ89069.1 akaw|GAA91730.1 aluc|OJZ87657.1 anid|CBF86075.1 aory|EIT76201.1 asyd|OJJ64462.1 ater|EAU39382.1 atub|OJI80986.1 aver|OJI98565.1 awen|OJJ40729.1 azon|OJJ44101.1

Group5076: abra|OJJ70868.1 acar|OOF90122.1 acla|EAW10460.1 afis|EAW25107.1 afum|EAL89778.1 agla|OJJ89080.1 akaw|GAA91735.1 aluc|OJZ87651.1 anid|CBF86066.1 anig|EHA20845.1 aory|EIT76240.1 asyd|OJJ64457.1 ater|EAU39375.1 atub|OJI80992.1 aver|OJI98560.1 awen|OJJ40725.1 azon|OJJ44097.1

Group5077: abra|OJJ70870.1 acar|OOF98249.1 acla|EAW10461.1 afis|EAW25106.1 afla|EED55338.1 afum|EAL89777.1 agla|OJJ89073.1 akaw|GAA91736.1 aluc|OJZ87650.1 anid|CBF86025.1 anig|EHA20844.1 asyd|OJJ64455.1 ater|EAU39374.1 atub|OJI80993.1 aver|OJI98558.1 awen|OJJ40719.1 azon|OJJ44094.1

Group5078: abra|OJJ70877.1 acar|OOF98243.1 acla|EAW10466.1 afis|EAW25101.1 afla|EED55344.1 afum|EAL89772.1 agla|OJJ85449.1 akaw|GAA91740.1 aluc|OJZ87645.1 anid|CBF86017.1 anig|EHA20838.1 asyd|OJJ64451.1 ater|EAU39368.1 atub|OJI81000.1 aver|OJI98554.1 awen|OJJ40715.1 azon|OJJ44092.1

Group5079: abra|OJJ70881.1 acar|OOF98240.1 acla|EAW10469.1 afis|EAW25098.1 afla|EED55348.1 afum|EAL89769.1 akaw|GAA91743.1 aluc|OJZ87642.1 anid|CBF86012.1 anig|EHA20835.1 aory|EIT80829.1 asyd|OJJ64448.1 ater|EAU39356.1 atub|OJI81003.1 aver|OJI98551.1 awen|OJJ40712.1 azon|OJJ44089.1

Group5080: abra|OJJ70891.1 acar|OOF90216.1 acla|EAW10473.1 afis|EAW25094.1 afum|EAL89765.2 agla|OJJ89083.1 akaw|GAA87407.1 aluc|OJZ87632.1 anid|CBF86003.1 anig|EHA20828.1 aory|EIT80821.1 asyd|OJJ64444.1 ater|EAU39352.1 atub|OJI81017.1 aver|OJI98546.1 awen|OJJ40708.1 azon|OJJ44084.1

Group5081: abra|OJJ70893.1 acla|EAW10475.1 afis|EAW25092.1 afla|EED55355.1 afum|EAL89762.2 agla|OJJ89085.1 akaw|GAA87409.1 aluc|OJZ87630.1 anid|CBF85999.1 anig|EHA20826.1 aory|EIT80902.1 asyd|OJJ64442.1 ater|EAU39350.1 atub|OJI81019.1 aver|OJI98544.1 awen|OJJ40706.1 azon|OJJ44082.1

Group5082: abra|OJJ70905.1 acar|OOF98215.1 acla|EAW10483.1 afis|EAW25084.1 afla|EED55366.1 afum|EAL89753.1 agla|OJJ89101.1 akaw|GAA87419.1 aluc|OJZ87619.1 anid|CBF85986.1 anig|EHA20817.1 aory|EIT80910.1 asyd|OJJ64433.1 ater|EAU39340.1 atub|OJI81030.1 aver|OJI98534.1 awen|OJJ40696.1

Group5083: abra|OJJ70907.1 acar|OOF98213.1 acla|EAW10485.1 afis|EAW25082.1 afla|EED55368.1 afum|EAL89751.1 agla|OJJ89103.1 akaw|GAA87421.1 aluc|OJZ87617.1 anid|CBF85983.1 anig|EHA20815.1 aory|EIT80932.1 asyd|OJJ64431.1 atub|OJI81032.1 aver|OJI98532.1 awen|OJJ40694.1 azon|OJJ44071.1

Group5084: abra|OJJ70919.1 acar|OOF98205.1 acla|EAW10493.1 afis|EAW25074.1 afla|EED55379.1 afum|EAL89743.1 agla|OJJ89118.1 akaw|GAA83021.1 aluc|OJZ87607.1 anig|EHA20805.1 aory|EIT80877.1 asyd|OJJ64422.1 ater|EAU39551.1 atub|OJI81041.1 aver|OJI98521.1 awen|OJJ40685.1 azon|OJJ44062.1

Group5085: abra|OJJ70924.1 acla|EAW13722.1 afis|EAW21022.1 afla|EED51437.1 afum|EAL92061.1 akaw|GAA83016.1 aluc|OJZ87602.1 anid|CBF87343.1 anig|EHA20801.1 asyd|OJJ52825.1 ater|EAU33971.1 atub|OJI81046.1 aver|OJJ08801.1 ater|EAU36221.1 aver|OJJ04859.1 aory|EIT80432.1 ater|EAU31577.1

Group5086: abra|OJJ70932.1 acla|EAW10498.1 afis|EAW25069.1 afla|EED55385.1 afum|EAL89738.1 agla|OJJ89125.1 akaw|GAA83010.1 aluc|OJZ87595.1 anid|CBF85956.1 anig|EHA20795.1 aory|EIT80780.1 asyd|OJJ64417.1 ater|EAU35816.1 atub|OJI81052.1 aver|OJI98516.1 awen|OJJ40676.1 azon|OJJ44057.1

Group5087: abra|OJJ70935.1 acar|OOF89983.1 acla|EAW10501.1 afis|EAW25066.1 afla|EED55388.1 afum|EAL89735.2 agla|OJJ89127.1 aluc|OJZ87592.1 anid|CBF85951.1 anig|EHA20793.1 aory|EIT80771.1 asyd|OJJ64415.1 ater|EAU35813.1 atub|OJI81055.1 aver|OJI98514.1 awen|OJJ40674.1 azon|OJJ44055.1

Group5088: abra|OJJ70938.1 acla|EAW10503.1 afis|EAW25064.1 afla|EED55390.1 afum|EAL89733.1 agla|OJJ89129.1 akaw|GAA83006.1 aluc|OJZ87590.1 anid|CBF85945.1 anig|EHA20791.1 aory|EIT80842.1 asyd|OJJ64413.1 ater|EAU35811.1 atub|OJI81058.1 aver|OJI98512.1 awen|OJJ40671.1 azon|OJJ44052.1

Group5089: abra|OJJ70943.1 acar|OOF98189.1 acla|EAW10508.1 afis|EAW25059.1 afla|EED55394.1 afum|EAL89727.1 agla|OJJ89136.1 akaw|GAA83001.1 aluc|OJZ87585.1 anid|CBF85938.1 anig|EHA20786.1 asyd|OJJ64409.1 ater|EAU35807.1 atub|OJI81063.1 aver|OJI98507.1 awen|OJJ40666.1 azon|OJJ44047.1

Group5090: abra|OJJ70956.1 acar|OOF98174.1 acla|EAW10517.1 afis|EAW25049.1 afla|EED55402.1 afum|EAL89717.1 agla|OJJ89143.1 akaw|GAA82991.1 aluc|OJZ87576.1 anid|CBF85921.1 anig|EHA20778.1 aory|EIT80931.1 asyd|OJJ64403.1 ater|EAU35799.1 atub|OJI81073.1 aver|OJI98499.1 azon|OJJ44038.1

Group5091: abra|OJJ70962.1 acla|EAW10523.1 afis|EAW25043.1 afla|EED55409.1 afum|EAL89711.1 agla|OJJ89151.1 akaw|GAA82985.1 aluc|OJZ87570.1 anig|EHA20772.1 aory|EIT80767.1 asyd|OJJ64397.1 ater|EAU35793.1 atub|OJI81079.1 aver|OJI98493.1 awen|OJJ40648.1 azon|OJJ44033.1 anid|CBF85910.1

Group5092: abra|OJJ70970.1 acar|OOF98160.1 acla|EAW10530.1 afis|EAW24997.1 akaw|GAA82978.1 aluc|OJZ87562.1 anid|CBF85899.1 anig|EHA20764.1 asyd|OJJ64391.1 ater|EAU35787.1 atub|OJI81085.1 awen|OJJ40642.1 azon|OJJ44027.1 afla|EED55417.1 afum|EAL89704.1 aory|EIT80790.1 aver|OJI98486.1

Group5093: abra|OJJ70994.1 acar|OOF90032.1 acla|EAW10549.1 afis|EAW25017.1 afla|EED55440.1 afum|EAL89678.1 agla|OJJ89185.1 akaw|GAA82954.1 aluc|OJZ87534.1 anid|CBF81534.1 anig|EHA20743.1 asyd|OJJ59484.1 ater|EAU35769.1 atub|OJI81115.1 aver|OJJ02790.1 awen|OJJ40623.1 azon|OJJ49471.1

Group5094: abra|OJJ70995.1 acar|OOF90033.1 acla|EAW10551.1 afis|EAW25018.1 afla|EED55441.1 afum|EAL89677.1 agla|OJJ89186.1 akaw|GAA82953.1 aluc|OJZ87533.1 anid|CBF81536.1 anig|EHA20742.1 asyd|OJJ59487.1 ater|EAU35768.1 atub|OJI81116.1 aver|OJJ02787.1 awen|OJJ40622.1 azon|OJJ49470.1

Group5095: abra|OJJ70997.1 acla|EAW10553.1 afis|EAW25020.1 afla|EED55443.1 afum|EAL89675.1 agla|OJJ89188.1 akaw|GAA82951.1 aluc|OJZ87530.1 anid|CBF81540.1 anig|EHA20740.1 aory|EIT80782.1 asyd|OJJ59489.1 ater|EAU35766.1 atub|OJI81118.1 aver|OJJ02785.1 awen|OJJ40620.1 azon|OJJ49468.1

Group5096: abra|OJJ71012.1 acar|OOF98131.1 acla|EAW10563.1 afis|EAW25030.1 afla|EED55453.1 afum|EAL89665.1 agla|OJJ79154.1 akaw|GAA82942.1 aluc|OJZ87519.1 anid|CBF81562.1 anig|EHA20730.1 aory|EIT80874.1 asyd|OJJ59501.1 ater|EAU35757.1 atub|OJI81130.1 aver|OJJ02773.1 awen|OJJ40609.1

Group5097: abra|OJJ71020.1 acar|OOF98120.1 acla|EAW10569.1 afis|EAW25035.1 afla|EED55461.1 afum|EAL89659.2 agla|OJJ82256.1 akaw|GAA82935.1 aluc|OJZ87512.1 anid|CBF81576.1 anig|EHA20724.1 asyd|OJJ59508.1 ater|EAU35748.1 atub|OJI81139.1 aver|OJJ02766.1 awen|OJJ40603.1 azon|OJJ49453.1

Group5098: abra|OJJ71026.1 acar|OOF98115.1 acla|EAW10576.1 afis|EAW24991.1 afla|EED55468.1 afum|EAL89651.1 akaw|GAA82929.1 aluc|OJZ87505.1 anid|CBF81594.1 anig|EHA20718.1 aory|EIT80848.1 asyd|OJJ59516.1 ater|EAU35742.1 atub|OJI81145.1 aver|OJJ02757.1 awen|OJJ40596.1 azon|OJJ49447.1

Group5099: abra|OJJ71039.1 acar|OOF98099.1 acla|EAW10588.1 afis|EAW24980.1 afla|EED55479.1 afum|EAL89640.1 agla|OJJ79137.1 akaw|GAA82917.1 aluc|OJZ87489.1 anid|CBF81610.1 anig|EHA20709.1 asyd|OJJ59524.1 ater|EAU35732.1 atub|OJI81159.1 aver|OJJ02748.1 awen|OJJ40577.1 azon|OJJ49432.1

Group5100: abra|OJJ71046.1 acar|OOF98093.1 acla|EAW10594.1 afis|EAW24974.1 afum|EAL89634.1 agla|OJJ82240.1 akaw|GAA82910.1 aluc|OJZ87480.1 anid|CBF81632.1 anig|EHA20702.1 aory|EIT80851.1 asyd|OJJ59532.1 ater|EAU35722.1 atub|OJI81168.1 aver|OJJ02738.1 awen|OJJ40571.1 azon|OJJ49428.1

Group5101: abra|OJJ71058.1 acla|EAW10601.1 afis|EAW24967.1 afla|EED55493.1 afum|EAL89625.2 agla|OJJ82221.1 akaw|GAA82901.1 aluc|OJZ87470.1 anid|CBF81645.1 anig|EHA20692.1 aory|EIT80912.1 asyd|OJJ59539.1 ater|EAU35710.1 atub|OJI81178.1 aver|OJJ02729.1 awen|OJJ40560.1 azon|OJJ49421.1

Group5102: abra|OJJ71060.1 acar|OOF98081.1 acla|EAW10599.1 afis|EAW24969.1 afla|EED55492.1 afum|EAL89628.1 agla|OJJ82235.1 akaw|GAA82899.1 aluc|OJZ87468.1 anid|CBF81641.1 anig|EHA20690.1 asyd|OJJ59537.1 ater|EAU35714.1 atub|OJI81180.1 aver|OJJ02732.1 awen|OJJ40563.1 azon|OJJ49423.1

Group5103: abra|OJJ71065.1 acar|OOF98080.1 afis|EAW17397.1 afla|EED54883.1 afum|EAL91235.1 agla|OJJ83972.1 akaw|GAA82897.1 aluc|OJZ87466.1 anid|CBF88997.1 anig|EHA20688.1 aory|EIT79374.1 asyd|OJJ62782.1 ater|EAU29507.1 atub|OJI81182.1 aver|OJI96827.1 awen|OJJ36326.1 acla|EAW13233.1

Group5104: abra|OJJ71068.1 acar|OOF98077.1 acla|EAW13231.1 afis|EAW17399.1 afla|EED54882.1 afum|EAL91237.1 agla|OJJ83970.1 akaw|GAA82894.1 aluc|OJZ87463.1 anid|CBF83002.1 anig|EHA20685.1 aory|EIT79368.1 asyd|OJJ61518.1 ater|EAU29509.1 atub|OJI81185.1 aver|OJJ00854.1 awen|OJJ36327.1

Group5105: abra|OJJ71113.1 acar|OOF98037.1 acla|EAW10379.1 afis|EAW25214.1 afla|EED53668.1 afum|EAL89875.1 akaw|GAA82856.1 aluc|OJZ87421.1 anid|CBF85252.1 aory|EIT75344.1 asyd|OJJ62922.1 atub|OJI81227.1 aver|OJI98067.1 awen|OJJ40876.1 azon|OJJ46811.1 agla|OJJ88947.1 ater|EAU34371.1

Group5106: abra|OJJ71127.1 acla|EAW10371.1 afis|EAW25221.1 afla|EED53685.1 afum|EAL89882.1 agla|OJJ79188.1 akaw|GAA82847.1 aluc|OJZ87408.1 anid|CBF85260.1 anig|EHA20638.1 aory|EIT75267.1 asyd|OJJ62912.1 ater|EAU34380.1 atub|OJI81238.1 aver|OJI98077.1 awen|OJJ40887.1 azon|OJJ44516.1

Group5107: abra|OJJ71131.1 acar|OOF98021.1 acla|EAW10366.1 afis|EAW25226.1 afla|EED53670.1 afum|EAL89887.1 agla|OJJ79182.1 akaw|GAA82843.1 aluc|OJZ87404.1 anid|CBF85267.1 anig|EHA20633.1 aory|EIT75270.1 asyd|OJJ62907.1 atub|OJI81242.1 aver|OJI98081.1 awen|OJJ40891.1 azon|OJJ44520.1

Group5108: abra|OJJ71144.1 acar|OOF98012.1 acla|EAW10362.1 afis|EAW25231.1 afla|EED53677.1 afum|EAL89892.1 agla|OJJ79176.1 akaw|GAA82833.1 aluc|OJZ87394.1 anid|CBF85331.1 anig|EHA20624.1 aory|EIT75332.1 asyd|OJJ64019.1 atub|OJI81252.1 aver|OJI98105.1 awen|OJJ40897.1 azon|OJJ44524.1

Group5109: abra|OJJ71149.1 acla|EAW10353.1 afis|EAW25241.1 afla|EED53703.1 afum|EAL89903.1 agla|OJJ89207.1 akaw|GAA82828.1 aluc|OJZ87389.1 anid|CBF85318.1 anig|EHA20618.1 aory|EIT75342.1 asyd|OJJ64010.1 ater|EAU34366.1 atub|OJI81258.1 aver|OJI98098.1 awen|OJJ40954.1 azon|OJJ44539.1

Group5110: abra|OJJ71152.1 acar|OOF98005.1 acla|EAW10345.1 afis|EAW25244.1 afla|EED53699.1 afum|EAL89906.1 agla|OJJ89212.1 akaw|GAA82825.1 aluc|OJZ87386.1 anig|EHA20615.1 aory|EIT75333.1 asyd|OJJ64013.1 ater|EAU34359.1 atub|OJI81261.1 aver|OJI98101.1 awen|OJJ40957.1 azon|OJJ44536.1

Group5111: abra|OJJ71168.1 acla|EAW10334.1 afis|EAW25258.1 afla|EED53723.1 afum|EAL89919.1 agla|OJJ89229.1 akaw|GAA82813.1 aluc|OJZ87373.1 anid|CBF85358.1 anig|EHA20606.1 aory|EIT75289.1 asyd|OJJ64038.1 ater|EAU34399.1 atub|OJI81273.1 aver|OJI98122.1 awen|OJJ40971.1 azon|OJJ44557.1

Group5112: abra|OJJ71176.1 acar|OOF97990.1 acla|EAW10329.1 afis|EAW25265.1 afla|EED53729.1 afum|EAL89928.1 akaw|GAA82808.1 aluc|OJZ87367.1 anid|CBF85371.1 anig|EHA20601.1 aory|EIT75310.1 asyd|OJJ64046.1 ater|EAU34405.1 atub|OJI81278.1 aver|OJI98131.1 awen|OJJ40977.1 azon|OJJ44565.1

Group5113: abra|OJJ71191.1 acar|OOF97976.1 acla|EAW10313.1 afis|EAW25279.1 afla|EED53746.1 afum|EAL89942.1 agla|OJJ89253.1 akaw|GAA82794.1 aluc|OJZ87353.1 anid|CBF85416.1 anig|EHA20587.1 aory|EIT75305.1 asyd|OJJ64071.1 atub|OJI81292.1 aver|OJI98158.1 awen|OJJ40993.1 azon|OJJ44582.1

Group5114: abra|OJJ71196.1 acar|OOF97971.1 acla|EAW10306.1 afis|EAW25286.1 afum|EAL89949.1 agla|OJJ89259.1 akaw|GAA82787.1 aluc|OJZ87346.1 anid|CBF85427.1 anig|EHA20583.1 aory|EIT78443.1 asyd|OJJ64081.1 ater|EAU34427.1 atub|OJI81298.1 aver|OJI98167.1 awen|OJJ40998.1 azon|OJJ44588.1

Group5115: abra|OJJ71219.1 akaw|GAA82767.1 aluc|OJZ87326.1 anig|EHA20563.1 atub|OJI81319.1 abra|OJJ75210.1 acar|OOF94578.1 acla|EAW15349.1 afla|EED53880.1 aory|EIT80547.1 asyd|OJJ52896.1 aver|OJJ08867.1 awen|OJJ32543.1 aluc|OJZ90682.1 afla|EED51735.1 akaw|GAA85321.1 aory|EIT78423.1

Group5116: abra|OJJ71225.1 acar|OOF97954.1 acla|EAW10297.1 afis|EAW25295.1 afum|EAL89958.1 agla|OJJ89271.1 akaw|GAA82764.1 aluc|OJZ87322.1 anig|EHA20556.1 ater|EAU34436.1 atub|OJI81323.1 aver|OJI98153.1 azon|OJJ44600.1 afla|EED53761.1 anid|CBF85407.1 asyd|OJJ64066.1 awen|OJJ41009.1

Group5117: abra|OJJ71226.1 acar|OOF97953.1 acla|EAW10296.1 afis|EAW25296.1 afla|EED53762.1 afum|EAL89959.2 agla|OJJ89272.1 akaw|GAA82763.1 aluc|OJZ87321.1 anig|EHA20555.1 aory|EIT78357.1 asyd|OJJ64065.1 ater|EAU34437.1 atub|OJI81324.1 aver|OJI98152.1 awen|OJJ41010.1 azon|OJJ44599.1

Group5118: abra|OJJ71232.1 acla|EAW13299.1 afis|EAW17306.1 afla|EED54153.1 afum|EAL91142.1 agla|OJJ82653.1 akaw|GAA82756.1 aluc|OJZ87313.1 anid|CBF78484.1 anig|EHA20549.1 aory|EIT76008.1 asyd|OJJ65333.1 ater|EAU33840.1 atub|OJI81332.1 aver|OJI99478.1 awen|OJJ34703.1 azon|OJJ50430.1

Group5119: abra|OJJ71249.1 acla|EAW10286.1 afis|EAW25306.1 afla|EED53784.1 afum|EAL89969.1 agla|OJJ89285.1 akaw|GAA82738.1 aluc|OJZ87293.1 anid|CBF75294.1 anig|EHA20534.1 aory|EIT78475.1 asyd|OJJ59885.1 ater|EAU34454.1 atub|OJI81347.1 aver|OJJ02401.1 awen|OJJ41020.1 azon|OJJ44607.1

Group5120: abra|OJJ71260.1 afla|EED53793.1 acla|EAW10276.1 afis|EAW25318.1 afum|EAL89979.1 aory|EIT78464.1 asyd|OJJ59894.1 ater|EAU34463.1 awen|OJJ41045.1 azon|OJJ44618.1 agla|OJJ83856.1 akaw|GAA82729.1 aluc|OJZ87283.1 anid|CBF75279.1 anig|EHA20525.1 atub|OJI81358.1 aver|OJJ02392.1

Group5121: abra|OJJ71282.1 acar|OOF97890.1 acla|EAW10265.1 afis|EAW25329.1 afla|EED53817.1 afum|EAL89990.1 agla|OJJ83879.1 akaw|GAA82706.1 aluc|OJZ87261.1 anid|CBF75230.1 anig|EHA20504.1 asyd|OJJ59916.1 ater|EAU34482.1 atub|OJI81381.1 aver|OJJ02370.1 awen|OJJ41024.1 azon|OJJ44641.1

Group5122: abra|OJJ71283.1 acar|OOF97889.1 acla|EAW10266.1 afis|EAW25328.1 afla|EED53818.1 afum|EAL89989.1 agla|OJJ83880.1 akaw|GAA82705.1 aluc|OJZ87260.1 anid|CBF75228.1 aory|EIT78476.1 asyd|OJJ59918.1 ater|EAU34484.1 atub|OJI81382.1 aver|OJJ02368.1 awen|OJJ41023.1 azon|OJJ44642.1

Group5123: abra|OJJ71284.1 acar|OOF97888.1 acla|EAW10252.1 afis|EAW25340.1 afla|EED53821.1 afum|EAL90003.1 agla|OJJ83882.1 akaw|GAA82704.1 aluc|OJZ87259.1 anid|CBF75215.1 anig|EHA20503.1 asyd|OJJ59930.1 ater|EAU34486.1 atub|OJI81383.1 aver|OJJ02356.1 awen|OJJ41059.1 azon|OJJ44643.1

Group5124: abra|OJJ71298.1 acar|OOF97872.1 acla|EAW10240.1 afis|EAW25352.1 afla|EED53833.1 agla|OJJ83900.1 akaw|GAA82691.1 aluc|OJZ87245.1 anid|CBF77436.1 anig|EHA20490.1 aory|EIT78382.1 asyd|OJJ53224.1 ater|EAU34500.1 atub|OJI81397.1 aver|OJJ03628.1 awen|OJJ41075.1 azon|OJJ44657.1

Group5125: abra|OJJ71314.1 acar|OOF97858.1 acla|EAW10225.1 afis|EAW25367.1 afla|EED53848.1 afum|EAL90027.1 agla|OJJ83754.1 akaw|GAA82677.1 aluc|OJZ87230.1 anid|CBF77448.1 anig|EHA20477.1 asyd|OJJ53209.1 ater|EAU34512.1 atub|OJI81412.1 aver|OJJ03641.1 awen|OJJ41093.1 azon|OJJ44672.1

Group5126: abra|OJJ71318.1 acla|EAW10221.1 afis|EAW25371.1 afla|EED53852.1 afum|EAL90031.1 agla|OJJ83758.1 akaw|GAA82673.1 aluc|OJZ87226.1 anid|CBF77440.1 anig|EHA20473.1 aory|EIT78361.1 asyd|OJJ53205.1 ater|EAU34516.1 atub|OJI81416.1 aver|OJJ03645.1 awen|OJJ41097.1 azon|OJJ44676.1

Group5127: abra|OJJ71319.1 acla|EAW10220.1 afis|EAW25372.1 afla|EED53853.1 afum|EAL90032.1 agla|OJJ83759.1 akaw|GAA82672.1 aluc|OJZ87225.1 anid|CBF77473.1 anig|EHA20472.1 aory|EIT78430.1 asyd|OJJ53204.1 ater|EAU34517.1 atub|OJI81417.1 aver|OJJ03646.1 awen|OJJ41098.1 azon|OJJ44677.1

Group5128: abra|OJJ71331.1 acar|OOF97844.1 acla|EAW10209.1 afis|EAW25386.1 afla|EED53863.1 afum|EAL90045.1 agla|OJJ83787.1 akaw|GAA82661.1 aluc|OJZ87211.1 anid|CBF77494.1 anig|EHA20462.1 asyd|OJJ53193.1 ater|EAU34533.1 atub|OJI81428.1 aver|OJJ03657.1 awen|OJJ41113.1 azon|OJJ44689.1

Group5129: abra|OJJ71332.1 acar|OOF97843.1 acla|EAW10208.1 afis|EAW25387.1 afla|EED53864.1 afum|EAL90046.2 agla|OJJ83788.1 akaw|GAA82660.1 aluc|OJZ87210.1 anid|CBF77496.1 aory|EIT78422.1 asyd|OJJ53192.1 ater|EAU34534.1 atub|OJI81429.1 aver|OJJ03658.1 awen|OJJ41114.1 azon|OJJ44690.1

Group5130: abra|OJJ71335.1 acar|OOF97837.1 acla|EAW10204.1 afis|EAW25391.1 afla|EED53868.1 afum|EAL90050.1 agla|OJJ87029.1 akaw|GAA82657.1 aluc|OJZ87207.1 anid|CBF77504.1 aory|EIT78375.1 asyd|OJJ53188.1 ater|EAU34539.1 atub|OJI81432.1 aver|OJJ03662.1 awen|OJJ41118.1 azon|OJJ44694.1

Group5131: abra|OJJ71348.1 acar|OOF97822.1 acla|EAW10190.1 afis|EAW25405.1 afla|EED53884.1 afum|EAL90064.1 agla|OJJ87046.1 akaw|GAA82644.1 aluc|OJZ87191.1 anid|CBF77534.1 anig|EHA20448.1 asyd|OJJ53175.1 ater|EAU34556.1 atub|OJI81447.1 aver|OJJ03677.1 awen|OJJ41132.1 azon|OJJ44706.1

Group5132: abra|OJJ71350.1 acar|OOF97820.1 acla|EAW10188.1 afis|EAW25407.1 afla|EED53886.1 afum|EAL90066.1 agla|OJJ87048.1 akaw|GAA82642.1 aluc|OJZ87189.1 anid|CBF77538.1 anig|EHA20446.1 asyd|OJJ53173.1 ater|EAU34558.1 atub|OJI81449.1 aver|OJJ03679.1 awen|OJJ41134.1 azon|OJJ44708.1

Group5133: abra|OJJ71369.1 acar|OOF97803.1 acla|EAW10169.1 afis|EAW25425.1 afum|EAL90084.1 agla|OJJ87067.1 akaw|GAA82625.1 aluc|OJZ87171.1 anid|CBF77555.1 anig|EHA20429.1 aory|EIT78498.1 asyd|OJJ53154.1 ater|EAU34579.1 atub|OJI81467.1 aver|OJJ03697.1 awen|OJJ41153.1 azon|OJJ44732.1

Group5134: abra|OJJ71375.1 acar|OOF97793.1 afis|EAW25432.1 afum|EAL90090.1 agla|OJJ87077.1 akaw|GAA82619.1 aluc|OJZ87165.1 anig|EHA20424.1 aory|EIT78287.1 atub|OJI81474.1 aver|OJJ03704.1 awen|OJJ41161.1 azon|OJJ44739.1 acla|EAW10162.1 afla|EED53920.1 asyd|OJJ53147.1 anid|CBF77542.1

Group5135: abra|OJJ71377.1 acla|EAW10147.1 afis|EAW25446.1 afla|EED53930.1 afum|EAL90104.1 agla|OJJ87088.1 akaw|GAA82617.1 aluc|OJZ87163.1 anid|CBF77608.1 anig|EHA20422.1 aory|EIT78346.1 asyd|OJJ53136.1 ater|EAU34614.1 atub|OJI81476.1 aver|OJJ03716.1 awen|OJJ41170.1 azon|OJJ44749.1

Group5136: abra|OJJ71381.1 acar|OOF97788.1 acla|EAW10156.1 afis|EAW25438.1 afla|EED53926.1 afum|EAL90096.1 agla|OJJ87083.1 akaw|GAA82613.1 aluc|OJZ87159.1 anid|CBF77598.1 anig|EHA20418.1 asyd|OJJ53141.1 ater|EAU34610.1 atub|OJI81481.1 aver|OJJ03711.1 awen|OJJ41166.1 azon|OJJ44744.1

Group5137: abra|OJJ71385.1 acar|OOF97784.1 acla|EAW10160.1 afis|EAW25434.1 afla|EED53922.1 afum|EAL90092.1 agla|OJJ87079.1 akaw|GAA82609.1 aluc|OJZ87155.1 anig|EHA20414.1 aory|EIT78289.1 asyd|OJJ53145.1 ater|EAU34606.1 atub|OJI81485.1 aver|OJJ03707.1 awen|OJJ41163.1 azon|OJJ44741.1

Group5138: abra|OJJ71388.1 acar|OOF97780.1 acla|EAW10149.1 afis|EAW25444.1 afla|EED53932.1 afum|EAL90102.1 agla|OJJ87091.1 akaw|GAA82606.1 aluc|OJZ87152.1 anid|CBF77612.1 anig|EHA20411.1 aory|EIT78343.1 asyd|OJJ53134.1 atub|OJI81488.1 aver|OJJ03718.1 awen|OJJ41175.1 azon|OJJ44751.1

Group5139: abra|OJJ71396.1 acar|OOF97773.1 acla|EAW10143.1 afis|EAW25450.1 afla|EED53937.1 afum|EAL90109.2 agla|OJJ87102.1 akaw|GAA82601.1 aluc|OJZ87146.1 anid|CBF77627.1 anig|EHA20405.1 asyd|OJJ53129.1 ater|EAU34620.1 atub|OJI81495.1 aver|OJJ03725.1 awen|OJJ41181.1 azon|OJJ44757.1

Group5140: abra|OJJ71422.1 acla|EAW10112.1 afis|EAW25473.1 afla|EED53965.1 afum|EAL90136.2 agla|OJJ87121.1 akaw|GAA82580.1 aluc|OJZ87116.1 anid|CBF77676.1 anig|EHA20382.1 aory|EIT78353.1 asyd|OJJ53090.1 ater|EAU34642.1 atub|OJI81524.1 aver|OJJ03756.1 awen|OJJ41208.1 azon|OJJ44794.1

Group5141: abra|OJJ71433.1 acar|OOF90400.1 acla|EAW10105.1 afis|EAW25482.1 afla|EED53980.1 afum|EAL90147.1 agla|OJJ87130.1 akaw|GAA82570.1 aluc|OJZ87105.1 anid|CBF77699.1 anig|EHA20373.1 asyd|OJJ53071.1 ater|EAU34653.1 atub|OJI81533.1 aver|OJJ03774.1 awen|OJJ41223.1 azon|OJJ44784.1

Group5142: abra|OJJ71445.1 acla|EAW10101.1 afis|EAW25486.1 afla|EED53984.1 afum|EAL90151.1 agla|OJJ87134.1 akaw|GAA82564.1 aluc|OJZ87096.1 anid|CBF77715.1 anig|EHA20367.1 aory|EIT78340.1 asyd|OJJ53066.1 ater|EAU34657.1 atub|OJI81542.1 aver|OJJ03779.1 awen|OJJ41227.1 azon|OJJ44780.1

Group5143: abra|OJJ71457.1 acla|EAW10092.1 afis|EAW25495.1 afla|EED53995.1 afum|EAL90161.1 agla|OJJ87310.1 akaw|GAA82554.1 aluc|OJZ87086.1 anid|CBF77747.1 anig|EHA20356.1 aory|EIT78281.1 asyd|OJJ53048.1 ater|EAU34666.1 atub|OJI81552.1 aver|OJJ03797.1 awen|OJJ41237.1 azon|OJJ44803.1

Group5144: abra|OJJ71482.1 acla|EAW10067.1 afis|EAW25518.1 afla|EED54019.1 afum|EAL90183.1 agla|OJJ87285.1 akaw|GAA82537.1 aluc|OJZ87064.1 anid|CBF77790.1 anig|EHA20337.1 aory|EIT78295.1 asyd|OJJ53027.1 ater|EAU33364.1 atub|OJI81571.1 aver|OJJ03817.1 awen|OJJ41261.1 azon|OJJ44826.1

Group5145: abra|OJJ71539.1 acla|EAW15094.1 afis|EAW20063.1 afla|EED45314.1 afum|EAL85183.1 agla|OJJ88896.1 akaw|GAA89382.1 aluc|OJZ86277.1 anid|CBF84830.1 anig|EHA21594.1 aory|EIT73621.1 asyd|OJJ63210.1 ater|EAU38672.1 atub|OJI84262.1 aver|OJI97745.1 awen|OJJ31824.1 azon|OJJ46301.1

Group5146: abra|OJJ71546.1 acla|EAW15100.1 afis|EAW20072.1 afla|EED45307.1 afum|EAL85190.1 agla|OJJ88888.1 akaw|GAA89376.1 aluc|OJZ86284.1 anid|CBF84847.1 anig|EHA21587.1 aory|EIT73619.1 asyd|OJJ63203.1 ater|EAU38679.1 atub|OJI84254.1 aver|OJI97752.1 awen|OJJ31831.1 azon|OJJ46293.1

Group5147: abra|OJJ71547.1 acla|EAW15101.1 afis|EAW20073.1 afla|EED45306.1 afum|EAL85191.1 agla|OJJ88887.1 akaw|GAA89375.1 aluc|OJZ86285.1 anid|CBF84849.1 anig|EHA21586.1 aory|EIT73795.1 asyd|OJJ63202.1 ater|EAU38681.1 atub|OJI84253.1 aver|OJI97753.1 awen|OJJ31832.1 azon|OJJ46292.1

Group5148: abra|OJJ71552.1 acla|EAW15106.1 afis|EAW20078.1 afla|EED45297.1 afum|EAL85196.1 agla|OJJ88882.1 akaw|GAA89370.1 aluc|OJZ86290.1 anid|CBF84862.1 anig|EHA21581.1 aory|EIT73726.1 asyd|OJJ63197.1 ater|EAU38686.1 atub|OJI84248.1 aver|OJI97758.1 awen|OJJ31837.1 azon|OJJ46286.1

Group5149: abra|OJJ71557.1 acla|EAW15110.1 afis|EAW20081.1 afla|EED45293.1 afum|EAL85200.1 agla|OJJ88878.1 akaw|GAA89366.1 aluc|OJZ86295.1 anid|CBF84868.1 anig|EHA21577.1 aory|EIT73574.1 asyd|OJJ63193.1 ater|EAU38689.1 atub|OJI84242.1 aver|OJI97762.1 awen|OJJ31841.1 azon|OJJ46282.1

Group5150: abra|OJJ71559.1 acar|OOF90419.1 acla|EAW15111.1 afis|EAW20082.1 afla|EED45292.1 afum|EAL85201.1 agla|OJJ88876.1 akaw|GAA89365.1 aluc|OJZ86296.1 anid|CBF84870.1 anig|EHA21576.1 asyd|OJJ63192.1 ater|EAU38690.1 atub|OJI84241.1 aver|OJI97764.1 awen|OJJ31842.1 azon|OJJ46281.1

Group5151: abra|OJJ71564.1 acar|OOF96300.1 acla|EAW15116.1 afis|EAW20087.1 afla|EED45286.1 afum|EAL85206.1 akaw|GAA89360.1 aluc|OJZ86301.1 anid|CBF84879.1 anig|EHA21571.1 aory|EIT73732.1 asyd|OJJ63187.1 ater|EAU38696.1 atub|OJI84235.1 aver|OJI97769.1 awen|OJJ31847.1 azon|OJJ46276.1

Group5152: abra|OJJ71571.1 acla|EAW15120.1 afis|EAW20090.1 afla|EED45280.1 afum|EAL85209.1 agla|OJJ88866.1 akaw|GAA89354.1 aluc|OJZ86308.1 anid|CBF84884.1 anig|EHA21565.1 aory|EIT73541.1 asyd|OJJ63183.1 ater|EAU38699.1 atub|OJI84228.1 aver|OJI97772.1 awen|OJJ31852.1 azon|OJJ46272.1

Group5153: abra|OJJ71572.1 acar|OOF96304.1 acla|EAW15121.1 afis|EAW20091.1 afla|EED45279.1 afum|EAL85210.1 agla|OJJ88863.1 akaw|GAA89352.1 aluc|OJZ86309.1 anid|CBF84886.1 anig|EHA21564.1 asyd|OJJ63181.1 ater|EAU38700.1 atub|OJI84226.1 aver|OJI97773.1 awen|OJJ31854.1 azon|OJJ46271.1

Group5154: abra|OJJ71578.1 acar|OOF96308.1 acla|EAW15125.1 afis|EAW20095.1 afla|EED45275.1 afum|EAL85213.1 agla|OJJ88859.1 akaw|GAA89346.1 aluc|OJZ86315.1 anid|CBF84891.1 anig|EHA21560.1 aory|EIT73717.1 asyd|OJJ63178.1 atub|OJI84220.1 aver|OJI97776.1 awen|OJJ31858.1 azon|OJJ46267.1

Group5155: abra|OJJ71583.1 acar|OOF96312.1 acla|EAW15128.1 afis|EAW20098.1 afla|EED53373.1 afum|EAL85216.1 agla|OJJ88856.1 akaw|GAA89340.1 aluc|OJZ86323.1 anid|CBF84899.1 anig|EHA21555.1 aory|EIT74942.1 asyd|OJJ63174.1 atub|OJI84211.1 aver|OJI97780.1 awen|OJJ31866.1 azon|OJJ46264.1

Group5156: abra|OJJ71589.1 acar|OOF96318.1 acla|EAW15134.1 afis|EAW20106.1 afum|EAL85223.1 agla|OJJ88847.1 akaw|GAA89334.1 aluc|OJZ86331.1 anid|CBF84912.1 anig|EHA21549.1 aory|EIT74895.1 asyd|OJJ63167.1 ater|EAU38725.1 atub|OJI84205.1 aver|OJI97787.1 awen|OJJ31878.1 azon|OJJ46256.1

Group5157: abra|OJJ71591.1 acla|EAW15137.1 afis|EAW20109.1 afla|EED53363.1 afum|EAL85226.1 akaw|GAA89331.1 aluc|OJZ86334.1 anid|CBF84917.1 anig|EHA21547.1 aory|EIT75266.1 asyd|OJJ63165.1 ater|EAU38727.1 atub|OJI84201.1 aver|OJI97790.1 awen|OJJ31880.1 azon|OJJ46254.1 agla|OJJ88843.1

Group5158: abra|OJJ71598.1 acar|OOF96325.1 acla|EAW15146.1 afis|EAW20114.1 afla|EED53359.1 afum|EAL85231.1 agla|OJJ88837.1 akaw|GAA89326.1 aluc|OJZ86339.1 anid|CBF84928.1 anig|EHA21542.1 aory|EIT74994.1 asyd|OJJ63160.1 ater|EAU38731.1 atub|OJI84196.1 aver|OJI97797.1 awen|OJJ31886.1

Group5159: abra|OJJ71607.1 acla|EAW15149.1 afis|EAW20117.1 afla|EED53354.1 afum|EAL85234.1 agla|OJJ88832.1 akaw|GAA89321.1 aluc|OJZ86349.1 anid|CBF84936.1 anig|EHA21537.1 aory|EIT75234.1 asyd|OJJ63157.1 ater|EAU38735.1 atub|OJI84185.1 aver|OJI97799.1 awen|OJJ31887.1 azon|OJJ46247.1

Group5160: abra|OJJ71616.1 acar|OOF96332.1 acla|EAW15158.1 afis|EAW20126.1 afla|EED57250.1 afum|EAL85244.1 agla|OJJ88821.1 akaw|GAA89311.1 aluc|OJZ86357.1 anid|CBF84950.1 anig|EHA21533.1 aory|EIT74017.1 asyd|OJJ63146.1 atub|OJI84177.1 aver|OJI97808.1 awen|OJJ31895.1 azon|OJJ46241.1

Group5161: abra|OJJ71634.1 acar|OOF91865.1 afis|EAW20144.1 afla|EED57204.1 afum|EAL85259.1 agla|OJJ88788.1 akaw|GAA89292.1 aluc|OJZ86377.1 anid|CBF78474.1 anig|EHA21517.1 aory|EIT74128.1 asyd|OJJ58349.1 ater|EAU38761.1 atub|OJI84157.1 aver|OJJ02058.1 awen|OJJ32015.1 azon|OJJ46072.1

Group5162: abra|OJJ71636.1 acar|OOF96353.1 acla|EAW15176.1 afis|EAW20146.1 afla|EED57203.1 afum|EAL85261.1 agla|OJJ88787.1 akaw|GAA89290.1 aluc|OJZ86379.1 anig|EHA21515.1 aory|EIT74050.1 asyd|OJJ63129.1 ater|EAU38762.1 atub|OJI84155.1 aver|OJI97826.1 awen|OJJ32020.1 azon|OJJ46070.1

Group5163: abra|OJJ71638.1 acar|OOF96355.1 acla|EAW15178.1 afis|EAW20148.1 afla|EED57201.1 afum|EAL85263.1 agla|OJJ88785.1 akaw|GAA89288.1 aluc|OJZ86381.1 anid|CBF84982.1 aory|EIT74108.1 asyd|OJJ63126.1 ater|EAU38764.1 atub|OJI84152.1 aver|OJI97829.1 awen|OJJ32022.1 azon|OJJ46068.1

Group5164: abra|OJJ71650.1 acla|EAW15187.1 afis|EAW20157.1 afla|EED57193.1 afum|EAL85273.1 agla|OJJ88773.1 akaw|GAA89279.1 aluc|OJZ86394.1 anid|CBF85000.1 anig|EHA21504.1 aory|EIT74099.1 asyd|OJJ63117.1 ater|EAU38771.1 atub|OJI84139.1 aver|OJI97842.1 awen|OJJ32051.1 azon|OJJ46084.1

Group5165: abra|OJJ71651.1 acar|OOF96364.1 acla|EAW15188.1 afla|EED57190.1 afum|EAL85274.1 agla|OJJ88772.1 akaw|GAA89278.1 aluc|OJZ86395.1 anid|CBF85002.1 anig|EHA21503.1 asyd|OJJ63116.1 ater|EAU38772.1 atub|OJI84138.1 aver|OJI97843.1 awen|OJJ32052.1 azon|OJJ46085.1 agla|OJJ79968.1

Group5166: abra|OJJ71652.1 acar|OOF96365.1 acla|EAW15189.1 afis|EAW20159.1 afla|EED57189.1 afum|EAL85276.1 akaw|GAA89277.1 aluc|OJZ86398.1 anid|CBF85004.1 anig|EHA21502.1 aory|EIT74024.1 asyd|OJJ63115.1 ater|EAU38775.1 atub|OJI84137.1 aver|OJI97844.1 awen|OJJ32053.1 azon|OJJ46086.1

Group5167: abra|OJJ71653.1 akaw|GAA89276.1 aluc|OJZ86399.1 anig|EHA21501.1 ater|EAU38776.1 atub|OJI84136.1 awen|OJJ32054.1 acar|OOF96367.1 acla|EAW15191.1 afis|EAW20160.1 afla|EED57188.1 afum|EAL85277.1 anid|CBF85006.1 asyd|OJJ63114.1 aver|OJI97845.1 agla|OJJ88771.1 azon|OJJ46087.1

Group5168: abra|OJJ71658.1 acar|OOF96371.1 acla|EAW15195.1 afis|EAW20164.1 afla|EED57185.1 afum|EAL85282.1 agla|OJJ88766.1 akaw|GAA89272.1 aluc|OJZ86403.1 anid|CBF85013.1 anig|EHA21497.1 aory|EIT74112.1 asyd|OJJ63110.1 ater|EAU38795.1 atub|OJI84132.1 aver|OJI97850.1 awen|OJJ32059.1

Group5169: abra|OJJ71664.1 acar|OOF96374.1 acla|EAW15200.1 afis|EAW20166.1 afla|EED57182.1 afum|EAL85284.1 agla|OJJ88763.1 akaw|GAA89269.1 aluc|OJZ86406.1 anid|CBF85020.1 anig|EHA21493.1 asyd|OJJ63106.1 ater|EAU38798.1 atub|OJI84125.1 aver|OJI97855.1 awen|OJJ32063.1 azon|OJJ46096.1

Group5170: abra|OJJ71680.1 acar|OOF96391.1 acla|EAW15219.1 afis|EAW20184.1 afla|EED57162.1 afum|EAL85302.2 agla|OJJ84682.1 akaw|GAA89252.1 aluc|OJZ86422.1 anig|EHA21477.1 aory|EIT74012.1 asyd|OJJ63090.1 ater|EAU35220.1 atub|OJI84109.1 aver|OJI97872.1 awen|OJJ31183.1 azon|OJJ46114.1

Group5171: abra|OJJ71681.1 acar|OOF96392.1 acla|EAW15220.1 afis|EAW20185.1 afla|EED57163.1 afum|EAL85304.2 agla|OJJ84681.1 akaw|GAA89251.1 aluc|OJZ86423.1 anid|CBF85052.1 anig|EHA21475.1 asyd|OJJ63089.1 ater|EAU35221.1 atub|OJI84108.1 aver|OJI97873.1 awen|OJJ31185.1 azon|OJJ46115.1

Group5172: abra|OJJ71694.1 acar|OOF96401.1 acla|EAW15231.1 afis|EAW20195.1 afla|EED57149.1 afum|EAL85315.1 agla|OJJ84670.1 akaw|GAA89241.1 aluc|OJZ86438.1 anid|CBF85071.1 anig|EHA21465.1 aory|EIT73968.1 asyd|OJJ63079.1 ater|EAU32003.1 atub|OJI84092.1 aver|OJI97885.1 awen|OJJ31201.1

Group5173: abra|OJJ71695.1 acar|OOF96403.1 acla|EAW15232.1 afis|EAW20196.1 afla|EED57148.1 afum|EAL85316.1 agla|OJJ84669.1 akaw|GAA89240.1 aluc|OJZ86439.1 anid|CBF85073.1 anig|EHA21464.1 aory|EIT74078.1 asyd|OJJ63078.1 ater|EAU32004.1 atub|OJI84090.1 aver|OJI97886.1 awen|OJJ31202.1

Group5174: abra|OJJ71698.1 acar|OOF96406.1 acla|EAW15234.1 afis|EAW20198.1 afla|EED57146.1 afum|EAL85318.1 agla|OJJ84667.1 akaw|GAA89237.1 aluc|OJZ86441.1 anid|CBF85080.1 anig|EHA21460.1 aory|EIT74082.1 asyd|OJJ63075.1 atub|OJI84087.1 aver|OJI97889.1 awen|OJJ31204.1 azon|OJJ46129.1

Group5175: abra|OJJ71712.1 acar|OOF96417.1 acla|EAW06436.1 afis|EAW18541.1 afla|EED52854.1 afum|EAL93241.1 agla|OJJ81573.1 akaw|GAA85177.1 aluc|OJZ86456.1 anid|CBF70194.1 anig|EHA21451.1 aory|EIT78642.1 asyd|OJJ58789.1 ater|EAU30024.1 atub|OJI84071.1 aver|OJJ01396.1 awen|OJJ31097.1

Group5176: abra|OJJ71724.1 acar|OOF96428.1 acla|EAW10028.1 afla|EED54155.1 akaw|GAA85166.1 aluc|OJZ86470.1 anig|EHA21440.1 aory|EIT81946.1 asyd|OJJ53835.1 ater|EAU30727.1 atub|OJI84059.1 aver|OJI98821.1 awen|OJJ38368.1 azon|OJJ45250.1 afla|EED55034.1 asyd|OJJ64718.1 ater|EAU37147.1

Group5177: acla|EAW09043.1 afis|EAW25210.1 aory|EIT73738.1 akaw|GAA86660.1 aluc|OJZ89756.1 atub|OJI85045.1 awen|OJJ36425.1 azon|OJJ48013.1 acla|EAW09045.1 afla|EED45079.1 anid|CBF79126.1 asyd|OJJ53761.1 ater|EAU35542.1 afum|EAL89871.1 asyd|OJJ57939.1 aory|EIT73659.1 agla|OJJ78860.1

Group5178: abra|OJJ71744.1 acar|OOF96435.1 acla|EAW06448.1 afis|EAW18552.1 afum|EAL93252.1 agla|OJJ81604.1 akaw|GAA85158.1 aluc|OJZ86479.1 anid|CBF70279.1 anig|EHA22163.1 aory|EIT78165.1 asyd|OJJ58742.1 ater|EAU34840.1 atub|OJI84051.1 aver|OJJ01352.1 awen|OJJ31113.1 azon|OJJ46589.1

Group5179: abra|OJJ71802.1 acar|OOF96482.1 acla|EAW13843.1 afis|EAW18557.1 afla|EED52809.1 afum|EAL93257.1 agla|OJJ81602.1 akaw|GAA85108.1 aluc|OJZ86532.1 anid|CBF70284.1 anig|EHA22114.1 aory|EIT78228.1 asyd|OJJ58738.1 ater|EAU34846.1 atub|OJI83989.1 aver|OJJ01349.1 awen|OJJ31115.1

Group5180: abra|OJJ71807.1 acar|OOF96485.1 acla|EAW13846.1 afis|EAW18560.1 afla|EED52800.1 afum|EAL93260.1 agla|OJJ81599.1 akaw|GAA85105.1 aluc|OJZ86537.1 anid|CBF70290.1 anig|EHA22111.1 aory|EIT78240.1 asyd|OJJ58734.1 atub|OJI83986.1 aver|OJJ01346.1 awen|OJJ31118.1 azon|OJJ46591.1

Group5181: abra|OJJ71832.1 acar|OOF96508.1 acla|EAW13867.1 afis|EAW18579.1 afla|EED52769.1 afum|EAL93282.1 akaw|GAA85079.1 aluc|OJZ86564.1 anid|CBF70356.1 anig|EHA22087.1 aory|EIT78246.1 asyd|OJJ58656.1 ater|EAU34872.1 atub|OJI83957.1 aver|OJJ01264.1 awen|OJJ31138.1 azon|OJJ46618.1

Group5182: abra|OJJ71847.1 acar|OOF96516.1 acla|EAW13878.1 afis|EAW18590.1 afla|EED52756.1 afum|EAL93297.1 agla|OJJ81642.1 akaw|GAA85070.1 aluc|OJZ86580.1 anid|CBF70377.1 aory|EIT78203.1 asyd|OJJ58645.1 ater|EAU34883.1 atub|OJI83942.1 aver|OJJ01251.1 awen|OJJ31153.1 azon|OJJ46628.1

Group5183: abra|OJJ71848.1 acar|OOF96517.1 acla|EAW13879.1 afis|EAW18591.1 afum|EAL93298.1 agla|OJJ81643.1 akaw|GAA85069.1 aluc|OJZ86581.1 anid|CBF70379.1 anig|EHA22077.1 aory|EIT78204.1 asyd|OJJ58644.1 ater|EAU34884.1 atub|OJI83941.1 aver|OJJ01250.1 awen|OJJ31154.1 azon|OJJ46629.1

Group5184: abra|OJJ71852.1 acar|OOF96522.1 acla|EAW13884.1 afis|EAW18596.1 afla|EED52751.1 afum|EAL93304.2 agla|OJJ81649.1 akaw|GAA85065.1 aluc|OJZ86586.1 anid|CBF70393.1 anig|EHA22074.1 asyd|OJJ58635.1 ater|EAU34890.1 atub|OJI83937.1 aver|OJJ01241.1 awen|OJJ31159.1 azon|OJJ46635.1

Group5185: abra|OJJ71863.1 acla|EAW13893.1 afis|EAW18605.1 afla|EED52741.1 afum|EAL93314.1 agla|OJJ81657.1 akaw|GAA85053.1 aluc|OJZ86600.1 anid|CBF70410.1 anig|EHA22063.1 aory|EIT78173.1 asyd|OJJ58626.1 ater|EAU34900.1 atub|OJI83923.1 aver|OJJ01232.1 awen|OJJ31171.1 azon|OJJ46648.1

Group5186: abra|OJJ71867.1 acar|OOF96537.1 acla|EAW13897.1 afis|EAW18609.1 afla|EED52737.1 afum|EAL93319.1 agla|OJJ81664.1 akaw|GAA85048.1 aluc|OJZ86604.1 anid|CBF70418.1 anig|EHA22059.1 aory|EIT78182.1 asyd|OJJ58621.1 ater|EAU34907.1 atub|OJI83919.1 aver|OJJ01228.1 awen|OJJ32070.1

Group5187: abra|OJJ71872.1 acar|OOF96543.1 acla|EAW13902.1 afis|EAW18614.1 afla|EED52732.1 afum|EAL93324.1 agla|OJJ81230.1 akaw|GAA85043.1 aluc|OJZ86611.1 anid|CBF70430.1 anig|EHA22055.1 asyd|OJJ58615.1 ater|EAU34911.1 atub|OJI83915.1 aver|OJJ01222.1 awen|OJJ32075.1 azon|OJJ46655.1

Group5188: abra|OJJ71877.1 aluc|OJZ86615.1 atub|OJI83911.1 acla|EAW13906.1 afla|EED52727.1 afis|EAW18618.1 anig|EHA22051.1 afum|EAL93328.1 anid|CBF70503.1 aory|EIT78247.1 agla|OJJ81226.1 akaw|GAA85039.1 asyd|OJJ58578.1 ater|EAU34918.1 aver|OJJ01182.1 awen|OJJ32080.1 azon|OJJ46659.1

Group5189: abra|OJJ71893.1 acar|OOF96553.1 acla|EAW09629.1 afis|EAW21511.1 afla|EED55949.1 afum|EAL92531.1 agla|OJJ85675.1 akaw|GAA85025.1 aluc|OJZ86632.1 anid|CBF76437.1 anig|EHA22037.1 aory|EIT76814.1 asyd|OJJ61099.1 atub|OJI83896.1 aver|OJJ00414.1 awen|OJJ39183.1 azon|OJJ51510.1

Group5190: abra|OJJ71894.1 acar|OOF96555.1 acla|EAW09628.1 afis|EAW21512.1 afla|EED55950.1 afum|EAL92532.1 agla|OJJ85676.1 akaw|GAA85024.1 aluc|OJZ86633.1 anid|CBF76435.1 anig|EHA22036.1 asyd|OJJ61097.1 ater|EAU35028.1 atub|OJI83895.1 aver|OJJ00412.1 awen|OJJ39182.1 azon|OJJ51511.1

Group5191: abra|OJJ71897.1 acar|OOF96558.1 acla|EAW09625.1 afis|EAW21515.1 afla|EED55953.1 afum|EAL92535.1 agla|OJJ85679.1 akaw|GAA85021.1 aluc|OJZ86636.1 anid|CBF76429.1 anig|EHA22033.1 asyd|OJJ61094.1 ater|EAU35025.1 atub|OJI83892.1 aver|OJJ00409.1 awen|OJJ39179.1 azon|OJJ51517.1

Group5192: abra|OJJ71904.1 acar|OOF96563.1 acla|EAW09619.1 afis|EAW21520.1 afla|EED55958.1 afum|EAL92541.1 agla|OJJ85683.1 akaw|GAA85016.1 aluc|OJZ86641.1 anid|CBF76417.1 anig|EHA22028.1 aory|EIT76716.1 asyd|OJJ61089.1 atub|OJI83886.1 aver|OJJ00404.1 awen|OJJ39173.1 azon|OJJ51500.1

Group5193: abra|OJJ71906.1 acla|EAW09617.1 afis|EAW21522.1 afla|EED55960.1 afum|EAL92543.1 agla|OJJ85685.1 akaw|GAA85014.1 aluc|OJZ86643.1 anid|CBF76414.1 anig|EHA22026.1 aory|EIT76548.1 asyd|OJJ61087.1 ater|EAU35019.1 atub|OJI83884.1 aver|OJJ00402.1 awen|OJJ39171.1 azon|OJJ51502.1

Group5194: abra|OJJ71910.1 afis|EAW21525.1 afla|EED55963.1 afum|EAL92546.1 agla|OJJ85688.1 akaw|GAA85010.1 aluc|OJZ86647.1 anid|CBF76408.1 anig|EHA22023.1 aory|EIT76627.1 asyd|OJJ61084.1 ater|EAU35016.1 atub|OJI83880.1 aver|OJJ00399.1 awen|OJJ39168.1 azon|OJJ51497.1 acla|EAW09614.1

Group5195: abra|OJJ71919.1 acar|OOF96576.1 acla|EAW09611.1 afis|EAW21528.1 afla|EED55976.1 afum|EAL92549.1 agla|OJJ85692.1 akaw|GAA85003.1 aluc|OJZ86656.1 anid|CBF76402.1 anig|EHA22015.1 asyd|OJJ61080.1 ater|EAU35013.1 atub|OJI83871.1 aver|OJJ00396.1 awen|OJJ39165.1 azon|OJJ51494.1

Group5196: abra|OJJ71925.1 acar|OOF96582.1 acla|EAW09603.1 afis|EAW21534.1 afla|EED55984.1 afum|EBA27462.1 akaw|GAA84997.1 aluc|OJZ86662.1 anid|CBF76390.1 anig|EHA22009.1 aory|EIT76780.1 asyd|OJJ61072.1 ater|EAU35005.1 atub|OJI83863.1 aver|OJJ00389.1 awen|OJJ39158.1 azon|OJJ51488.1

Group5197: abra|OJJ71941.1 acar|OOF96597.1 acla|EAW09589.1 afis|EAW21549.1 afum|EAL92569.1 agla|OJJ85727.1 akaw|GAA84984.1 aluc|OJZ86677.1 anid|CBF76360.1 anig|EHA21996.1 aory|EIT76795.1 asyd|OJJ61058.1 ater|EAU34990.1 atub|OJI83850.1 aver|OJJ00375.1 awen|OJJ39141.1 azon|OJJ51466.1

Group5198: abra|OJJ71948.1 acla|EAW09586.1 afis|EAW21552.1 afla|EED56004.1 afum|EAL92572.2 agla|OJJ85730.1 akaw|GAA84979.1 aluc|OJZ86683.1 anid|CBF76354.1 anig|EHA21990.1 aory|EIT76847.1 asyd|OJJ61055.1 ater|EAU34986.1 atub|OJI83844.1 aver|OJJ00372.1 awen|OJJ39138.1 azon|OJJ51471.1

Group5199: abra|OJJ71949.1 acla|EAW09585.1 afis|EAW21553.1 afla|EED56005.1 afum|EAL92573.1 agla|OJJ85733.1 akaw|GAA84978.1 aluc|OJZ86684.1 anid|CBF76352.1 anig|EHA21989.1 aory|EIT76552.1 asyd|OJJ61054.1 ater|EAU34981.1 atub|OJI83843.1 aver|OJJ00371.1 awen|OJJ39137.1 azon|OJJ51472.1

Group5200: abra|OJJ71954.1 acla|EAW09582.1 afis|EAW21555.1 afla|EED56007.1 afum|EBA27465.1 agla|OJJ85734.1 akaw|GAA84974.1 aluc|OJZ86688.1 anid|CBF76348.1 aory|EIT76671.1 asyd|OJJ61052.1 ater|EAU34979.1 atub|OJI83837.1 aver|OJJ00368.1 awen|OJJ39135.1 azon|OJJ51463.1 anig|EHA21985.1

Group5201: abra|OJJ71955.1 acar|OOF96617.1 acla|EAW09580.1 afis|EAW21557.1 afla|EED56009.1 afum|EAL92576.1 agla|OJJ85735.1 akaw|GAA84972.1 aluc|OJZ86690.1 anig|EHA21983.1 aory|EIT76882.1 asyd|OJJ61050.1 ater|EAU34977.1 atub|OJI83834.1 aver|OJJ00366.1 awen|OJJ39133.1 azon|OJJ51461.1

Group5202: abra|OJJ71964.1 acar|OOF89892.1 acla|EAW09572.1 afis|EAW21564.1 afla|EED56020.1 afum|EAL92583.1 agla|OJJ85749.1 akaw|GAA84964.1 aluc|OJZ86700.1 anid|CBF76332.1 anig|EHA21976.1 asyd|OJJ61037.1 ater|EAU34968.1 atub|OJI83826.1 aver|OJJ00355.1 awen|OJJ39122.1 azon|OJJ51452.1

Group5203: abra|OJJ71981.1 acar|OOF96632.1 acla|EAW09558.1 afis|EAW21579.1 afla|EED57796.1 afum|EAL92597.1 agla|OJJ85760.1 akaw|GAA84948.1 aluc|OJZ86718.1 anid|CBF76296.1 anig|EHA21961.1 aory|EIT81066.1 asyd|OJJ61003.1 atub|OJI83811.1 aver|OJJ00321.1 awen|OJJ39102.1 azon|OJJ51432.1

Group5204: abra|OJJ71982.1 acar|OOF96633.1 acla|EAW09557.1 afis|EAW21580.1 afla|EED57797.1 afum|EAL92598.1 agla|OJJ85759.1 akaw|GAA84947.1 aluc|OJZ86719.1 anid|CBF76298.1 anig|EHA21960.1 asyd|OJJ61002.1 ater|EAU34954.1 atub|OJI83810.1 awen|OJJ39101.1 azon|OJJ51433.1 aver|OJJ00320.1

Group5205: abra|OJJ71984.1 acar|OOF96636.1 acla|EAW09554.1 afis|EAW21582.1 afla|EED57802.1 afum|EAL92600.1 agla|OJJ85755.1 akaw|GAA84945.1 aluc|OJZ86721.1 anig|EHA21958.1 aory|EIT81095.1 asyd|OJJ60998.1 ater|EAU34951.1 atub|OJI83807.1 aver|OJJ00318.1 awen|OJJ39099.1 azon|OJJ51435.1

Group5206: abra|OJJ71985.1 acar|OOF96639.1 acla|EAW09553.1 afis|EAW21583.1 afla|EED57804.1 afum|EAL92601.2 agla|OJJ85773.1 akaw|GAA84944.1 aluc|OJZ86722.1 anid|CBF76291.1 anig|EHA21957.1 asyd|OJJ60997.1 ater|EAU34948.1 atub|OJI83806.1 aver|OJJ00317.1 awen|OJJ39098.1 azon|OJJ51436.1

Group5207: abra|OJJ71986.1 acar|OOF96640.1 acla|EAW09552.1 afis|EAW21584.1 afla|EED57805.1 afum|EAL92602.1 agla|OJJ85776.1 akaw|GAA84943.1 aluc|OJZ86723.1 anig|EHA21956.1 aory|EIT81057.1 asyd|OJJ60996.1 ater|EAU34947.1 atub|OJI83805.1 aver|OJJ00316.1 awen|OJJ39097.1 azon|OJJ51430.1

Group5208: abra|OJJ71989.1 acar|OOF96643.1 acla|EAW09546.1 afis|EAW21587.1 afla|EED57808.1 afum|EAL92605.1 agla|OJJ85780.1 akaw|GAA84940.1 aluc|OJZ86726.1 anig|EHA21953.1 aory|EIT81079.1 asyd|OJJ60993.1 ater|EAU34944.1 atub|OJI83802.1 aver|OJJ00313.1 awen|OJJ39094.1 azon|OJJ51427.1

Group5209: abra|OJJ72011.1 acla|EAW09530.1 afis|EAW21602.1 afla|EED57830.1 afum|EAL92621.1 agla|OJJ85798.1 akaw|GAA84922.1 aluc|OJZ86747.1 anid|CBF83464.1 anig|EHA21932.1 aory|EIT81055.1 asyd|OJJ60685.1 ater|EAU38293.1 atub|OJI83780.1 aver|OJJ00010.1 awen|OJJ39079.1 azon|OJJ51411.1

Group5210: abra|OJJ72013.1 acar|OOF96662.1 acla|EAW09527.1 afis|EAW21605.1 afla|EED57832.1 afum|EAL92624.1 agla|OJJ85800.1 akaw|GAA84920.1 aluc|OJZ86749.1 anid|CBF83467.1 anig|EHA21930.1 asyd|OJJ60687.1 ater|EAU38294.1 atub|OJI83778.1 aver|OJJ00012.1 awen|OJJ39077.1 azon|OJJ51409.1

Group5211: abra|OJJ72023.1 acar|OOF96672.1 acla|EAW09518.1 afis|EAW21614.1 afla|EED57845.1 afum|EAL92648.1 agla|OJJ78625.1 akaw|GAA84909.1 aluc|OJZ86760.1 anid|CBF83487.1 anig|EHA21919.1 asyd|OJJ60699.1 ater|EAU38306.1 atub|OJI83767.1 aver|OJJ00025.1 awen|OJJ39066.1 azon|OJJ51397.1

Group5212: abra|OJJ72025.1 acar|OOF96674.1 acla|EAW09514.1 afis|EAW21618.1 afla|EED57847.1 afum|EAL92654.1 agla|OJJ78627.1 akaw|GAA84907.1 aluc|OJZ86762.1 anid|CBF83491.1 anig|EHA21917.1 asyd|OJJ60701.1 ater|EAU38308.1 atub|OJI83765.1 aver|OJJ00027.1 awen|OJJ39064.1 azon|OJJ51395.1

Group5213: abra|OJJ72051.1 acar|OOF96708.1 acla|EAW09496.1 afis|EAW21640.1 afla|EED57903.1 afum|EAL92674.1 agla|OJJ79707.1 akaw|GAA84883.1 aluc|OJZ86788.1 anid|CBF83525.1 aory|EIT82339.1 asyd|OJJ60717.1 ater|EAU38325.1 atub|OJI83731.1 aver|OJJ00050.1 awen|OJJ33897.1 azon|OJJ48663.1

Group5214: abra|OJJ72053.1 acar|OOF96710.1 acla|EAW09494.1 afis|EAW21642.1 afla|EED57901.1 afum|EAL92676.1 agla|OJJ79709.1 akaw|GAA84881.1 aluc|OJZ86790.1 anig|EHA21893.1 aory|EIT82345.1 asyd|OJJ60719.1 ater|EAU38327.1 atub|OJI83729.1 aver|OJJ00052.1 awen|OJJ33899.1 azon|OJJ48661.1

Group5215: abra|OJJ72059.1 acar|OOF89883.1 acla|EAW09490.1 afis|EAW21646.1 afla|EED48456.1 afum|EAL92680.1 agla|OJJ79714.1 akaw|GAA84876.1 aluc|OJZ86795.1 anid|CBF83533.1 anig|EHA21888.1 asyd|OJJ60723.1 ater|EAU32953.1 atub|OJI83723.1 aver|OJJ00056.1 awen|OJJ33904.1 azon|OJJ48657.1

Group5216: abra|OJJ72080.1 acar|OOF96728.1 acla|EAW09487.1 afis|EAW21649.1 afla|EED57920.1 afum|EAL92683.1 agla|OJJ79729.1 aluc|OJZ86834.1 anid|CBF83557.1 anig|EHA21801.1 aory|EIT82340.1 asyd|OJJ60737.1 ater|EAU38344.1 atub|OJI83684.1 aver|OJJ00069.1 awen|OJJ33918.1 azon|OJJ48644.1

Group5217: abra|OJJ72107.1 acar|OOF96750.1 acla|EAW09384.1 afis|EAW21751.1 afla|EED58027.1 afum|EBA27473.1 agla|OJJ84423.1 akaw|GAA84815.1 aluc|OJZ86862.1 anid|CBF83669.1 aory|EIT79785.1 asyd|OJJ60809.1 ater|EAU38439.1 atub|OJI83658.1 aver|OJJ00137.1 awen|OJJ34032.1 azon|OJJ48205.1

Group5218: abra|OJJ72114.1 acar|OOF96754.1 acla|EAW14915.1 afis|EAW24437.1 afla|EED55242.1 afum|EAL86105.1 agla|OJJ78776.1 akaw|GAA84808.1 aluc|OJZ86869.1 anid|CBF73764.1 anig|EHA21772.1 aory|EIT83629.1 asyd|OJJ56589.1 atub|OJI83650.1 aver|OJJ06136.1 awen|OJJ32762.1 azon|OJJ43997.1

Group5219: abra|OJJ72122.1 acar|OOF96762.1 acla|EAW14922.1 afis|EAW24444.1 afla|EED55233.1 afum|EAL86113.1 agla|OJJ80289.1 akaw|GAA84801.1 aluc|OJZ86879.1 anid|CBF73751.1 anig|EHA21765.1 aory|EIT83604.1 asyd|OJJ56581.1 ater|EAU30003.1 atub|OJI83640.1 aver|OJJ06143.1 awen|OJJ32755.1

Group5220: abra|OJJ72142.1 acar|OOF96780.1 acla|EAW14936.1 afis|EAW24451.1 afum|EAL86120.1 agla|OJJ80297.1 akaw|GAA84781.1 aluc|OJZ86899.1 anid|CBF73817.1 anig|EHA21743.1 aory|EIT83518.1 asyd|OJJ56573.1 ater|EAU29995.1 atub|OJI83620.1 aver|OJJ06150.1 awen|OJJ32746.1 azon|OJJ45298.1

Group5221: abra|OJJ72151.1 afis|EAW24461.1 afla|EED55212.1 agla|OJJ80309.1 akaw|GAA84773.1 aluc|OJZ86910.1 anid|CBF73833.1 anig|EHA21735.1 asyd|OJJ56559.1 atub|OJI83612.1 awen|OJJ32735.1 azon|OJJ45286.1 acla|EAW14946.1 afla|EED55213.1 afum|EAL86131.1 aory|EIT83597.1 ater|EAU29981.1

Group5222: abra|OJJ72192.1 afis|EAW21198.1 afum|EAL90047.1 akaw|GAA84741.1 aluc|OJZ86943.1 anig|EHA21701.1 aory|EIT78387.1 asyd|OJJ53191.1 atub|OJI83579.1 aver|OJI97565.1 awen|OJJ41115.1 azon|OJJ44691.1 acla|EAW10207.1 afis|EAW25388.1 afla|EED53865.1 ater|EAU34535.1 aver|OJJ03659.1

Group5223: abra|OJJ72201.1 acar|OOF96821.1 acla|EAW14948.1 afis|EAW24463.1 afla|EED55203.1 afum|EAL86134.1 agla|OJJ80321.1 akaw|GAA84735.1 aluc|OJZ86951.1 anid|CBF73839.1 anig|EHA21696.1 aory|EIT83590.1 asyd|OJJ56556.1 ater|EAU29977.1 atub|OJI83572.1 aver|OJJ06164.1 awen|OJJ32728.1

Group5224: abra|OJJ72208.1 acar|OOF96829.1 acla|EAW14968.1 afis|EAW24491.1 afum|EAL86160.1 agla|OJJ80328.1 akaw|GAA84726.1 aluc|OJZ86960.1 anid|CBF73841.1 anig|EHA21688.1 aory|EIT78779.1 asyd|OJJ56555.1 ater|EAU29943.1 atub|OJI83563.1 aver|OJJ06165.1 awen|OJJ32704.1 azon|OJJ45096.1

Group5225: abra|OJJ72253.1 acar|OOF96884.1 acla|EAW12203.1 afis|EAW17077.1 afla|EED52139.1 agla|OJJ81367.1 akaw|GAA84684.1 aluc|OJZ87004.1 anid|CBF79753.1 anig|EHA21642.1 aory|EIT74390.1 asyd|OJJ58098.1 ater|EAU30779.1 atub|OJI83515.1 aver|OJJ04306.1 awen|OJJ36962.1 azon|OJJ45322.1

Group5226: abra|OJJ72287.1 acar|OOF94936.1 acla|EAW12015.1 afis|EAW19654.1 afla|EED57006.1 afum|EAL86868.1 agla|OJJ80800.1 akaw|GAA86039.1 aluc|OJZ90266.1 anid|CBF71365.1 anig|EHA23761.1 asyd|OJJ55082.1 ater|EAU32874.1 atub|OJI81691.1 aver|OJJ07125.1 awen|OJJ41906.1 azon|OJJ42696.1

Group5227: abra|OJJ72289.1 acar|OOF94934.1 acla|EAW12013.1 afis|EAW19653.1 afla|EED57004.1 afum|EAL86869.1 agla|OJJ80799.1 akaw|GAA86041.1 aluc|OJZ90269.1 anid|CBF71363.1 anig|EHA23762.1 asyd|OJJ55083.1 ater|EAU32875.1 atub|OJI81694.1 aver|OJJ07124.1 awen|OJJ41904.1 azon|OJJ42693.1

Group5228: abra|OJJ72291.1 acar|OOF94929.1 acla|EAW12011.1 afis|EAW19651.1 afla|EED57002.1 afum|EAL86871.1 agla|OJJ80797.1 akaw|GAA86043.1 aluc|OJZ90272.1 anid|CBF71359.1 anig|EHA23764.1 asyd|OJJ55085.1 ater|EAU32877.1 atub|OJI81696.1 aver|OJJ07122.1 awen|OJJ41902.1 azon|OJJ42698.1

Group5229: abra|OJJ72292.1 acar|OOF94932.1 acla|EAW12010.1 afis|EAW19650.1 afla|EED57001.1 agla|OJJ80796.1 akaw|GAA86044.1 aluc|OJZ90273.1 anid|CBF71357.1 anig|EHA23765.1 aory|EIT77584.1 asyd|OJJ55086.1 ater|EAU32878.1 atub|OJI81697.1 aver|OJJ07121.1 awen|OJJ41901.1 azon|OJJ42699.1

Group5230: abra|OJJ72293.1 acar|OOF94930.1 acla|EAW12009.1 afis|EAW19649.1 afla|EED57000.1 afum|EAL86873.1 agla|OJJ80791.1 akaw|GAA86045.1 aluc|OJZ90274.1 anig|EHA23766.1 aory|EIT77489.1 asyd|OJJ55087.1 ater|EAU32879.1 atub|OJI81698.1 aver|OJJ07120.1 awen|OJJ41900.1 azon|OJJ42700.1

Group5231: abra|OJJ72294.1 acar|OOF94931.1 acla|EAW12008.1 afis|EAW19648.1 afla|EED56999.1 afum|EAL86874.1 agla|OJJ80790.1 akaw|GAA86046.1 aluc|OJZ90275.1 anid|CBF71356.1 anig|EHA23767.1 asyd|OJJ55088.1 ater|EAU32880.1 atub|OJI81699.1 aver|OJJ07119.1 awen|OJJ41899.1 azon|OJJ42701.1

Group5232: abra|OJJ72300.1 acar|OOF94925.1 acla|EAW12004.1 afis|EAW19644.1 afla|EED56995.1 afum|EAL86878.1 agla|OJJ80786.1 akaw|GAA86051.1 aluc|OJZ90281.1 anid|CBF71348.1 anig|EHA23771.1 aory|EIT77548.1 asyd|OJJ55093.1 atub|OJI81705.1 aver|OJJ07115.1 awen|OJJ41895.1 azon|OJJ42705.1

Group5233: abra|OJJ72308.1 acar|OOF94905.1 acla|EAW13442.1 afis|EAW20849.1 afum|EAL84784.1 aluc|OJZ90289.1 anid|CBF89941.1 anig|EHA23779.1 asyd|OJJ61203.1 ater|EAU29955.1 atub|OJI81713.1 aver|OJJ00527.1 awen|OJJ41681.1 azon|OJJ50849.1 anig|EHA21820.1 asyd|OJJ61389.1 aver|OJJ00717.1

Group5234: abra|OJJ72315.1 acar|OOF94895.1 afis|EAW19532.1 afla|EED48961.1 afum|EAL86974.1 akaw|GAA86062.1 aluc|OJZ90296.1 anig|EHA23785.1 aory|EIT79991.1 ater|EAU31962.1 atub|OJI81720.1 awen|OJJ35408.1 azon|OJJ49084.1 afla|EED56029.1 aory|EIT76743.1 atub|OJI89169.1 awen|OJJ35688.1

Group5235: abra|OJJ72326.1 acar|OOF94888.1 acla|EAW11997.1 afis|EAW19637.1 afla|EED56989.1 afum|EAL86885.1 agla|OJJ80779.1 akaw|GAA86076.1 aluc|OJZ90311.1 anid|CBF71337.1 anig|EHA23798.1 asyd|OJJ55102.1 ater|EAU32891.1 atub|OJI81736.1 aver|OJJ07105.1 awen|OJJ41882.1 azon|OJJ47080.1

Group5236: abra|OJJ72329.1 acar|OOF94885.1 acla|EAW11994.1 afis|EAW19634.1 afla|EED56985.1 afum|EAL86888.1 agla|OJJ86606.1 akaw|GAA86079.1 aluc|OJZ90314.1 anid|CBF71331.1 anig|EHA23801.1 aory|EIT77448.1 asyd|OJJ55105.1 ater|EAU32894.1 atub|OJI81739.1 aver|OJJ07102.1 awen|OJJ41879.1

Group5237: abra|OJJ72350.1 acar|OOF94868.1 acla|EAW11976.1 afis|EAW19610.1 afla|EED56970.1 afum|EAL86912.1 agla|OJJ80760.1 akaw|GAA86097.1 aluc|OJZ90329.1 anid|CBF71304.1 anig|EHA23813.1 asyd|OJJ55122.1 ater|EAU32911.1 atub|OJI81756.1 aver|OJJ07081.1 awen|OJJ41860.1 azon|OJJ47054.1

Group5238: abra|OJJ72357.1 acar|OOF94861.1 acla|EAW11962.1 afis|EAW19599.1 afla|EED56955.1 afum|EAL86919.1 agla|OJJ80741.1 akaw|GAA86104.1 aluc|OJZ90336.1 anid|CBF71289.1 anig|EHA23820.1 aory|EIT77466.1 ater|EAU32926.1 atub|OJI81763.1 aver|OJJ07075.1 awen|OJJ41842.1 azon|OJJ44889.1

Group5239: abra|OJJ72378.1 acar|OOF94839.1 acla|EAW11956.1 afis|EAW19593.1 afla|EED56949.1 afum|EAL86925.1 akaw|GAA86126.1 aluc|OJZ90365.1 anid|CBF71271.1 anig|EHA23840.1 aory|EIT77564.1 asyd|OJJ55148.1 ater|EAU32931.1 atub|OJI81792.1 aver|OJJ07059.1 awen|OJJ41810.1 azon|OJJ47022.1

Group5240: abra|OJJ72383.1 acar|OOF90287.1 acla|EAW11952.1 afis|EAW19589.1 afla|EED56943.1 afum|EAL86929.1 agla|OJJ80729.1 akaw|GAA86132.1 aluc|OJZ90370.1 anig|EHA23845.1 aory|EIT77526.1 asyd|OJJ55143.1 ater|EAU32937.1 atub|OJI81796.1 aver|OJJ07064.1 awen|OJJ41804.1 azon|OJJ47027.1

Group5241: abra|OJJ72386.1 acar|OOF90292.1 acla|EAW11950.1 afis|EAW19587.1 afla|EED56915.1 afum|EAL86931.1 agla|OJJ80724.1 akaw|GAA86135.1 aluc|OJZ90373.1 anig|EHA23847.1 aory|EIT77573.1 asyd|OJJ55152.1 ater|EAU32961.1 atub|OJI81799.1 aver|OJJ07055.1 awen|OJJ41801.1 azon|OJJ47030.1

Group5242: abra|OJJ72398.1 acla|EAW11942.1 afis|EAW19578.1 afla|EED56803.1 afum|EAL86940.2 agla|OJJ80707.1 akaw|GAA86143.1 aluc|OJZ90383.1 anid|CBF71237.1 anig|EHA23855.1 aory|EIT73849.1 asyd|OJJ55164.1 ater|EAU32977.1 atub|OJI81809.1 aver|OJJ07043.1 awen|OJJ41791.1 azon|OJJ47042.1

Group5243: abra|OJJ72407.1 acar|OOF94822.1 acla|EAW11935.1 afis|EAW19570.1 afla|EED56813.1 agla|OJJ80698.1 akaw|GAA86150.1 aluc|OJZ90391.1 anid|CBF71224.1 anig|EHA23861.1 aory|EIT73940.1 asyd|OJJ55172.1 ater|EAU32993.1 atub|OJI81820.1 aver|OJJ07035.1 awen|OJJ41783.1 azon|OJJ44895.1

Group5244: abra|OJJ72432.1 acla|EAW12664.1 afis|EAW17828.1 afla|EED46170.1 afum|EAL91638.1 agla|OJJ80649.1 akaw|GAA86172.1 aluc|OJZ90415.1 anid|CBF77999.1 anig|EHA23881.1 aory|EIT76464.1 asyd|OJJ55248.1 ater|EAU33032.1 atub|OJI81848.1 aver|OJJ06975.1 awen|OJJ41758.1 azon|OJJ48449.1

Group5245: abra|OJJ72433.1 acar|OOF94802.1 acla|EAW12665.1 afis|EAW17829.1 afla|EED46171.1 afum|EAL91639.1 agla|OJJ80650.1 akaw|GAA86173.1 aluc|OJZ90416.1 anid|CBF77998.1 anig|EHA23884.1 aory|EIT76389.1 asyd|OJJ55249.1 ater|EAU33033.1 atub|OJI81849.1 aver|OJJ06974.1 azon|OJJ48450.1

Group5246: abra|OJJ72436.1 acar|OOF94799.1 acla|EAW12669.1 afis|EAW17833.1 afla|EED46175.1 afum|EAL91643.1 akaw|GAA86176.1 aluc|OJZ90420.1 anid|CBF77990.1 anig|EHA23890.1 aory|EIT76472.1 asyd|OJJ55254.1 ater|EAU33036.1 atub|OJI81852.1 aver|OJJ06970.1 awen|OJJ41748.1 azon|OJJ48441.1

Group5247: abra|OJJ72438.1 acla|EAW12671.1 afis|EAW17834.1 afla|EED46179.1 afum|EAL91644.1 agla|OJJ83132.1 akaw|GAA86177.1 aluc|OJZ90422.1 anid|CBF77988.1 anig|EHA23891.1 aory|EIT76441.1 asyd|OJJ55255.1 ater|EAU33038.1 atub|OJI81854.1 aver|OJJ06969.1 awen|OJJ41742.1 azon|OJJ48438.1

Group5248: abra|OJJ72441.1 acar|OOF94796.1 acla|EAW12674.1 afis|EAW17837.1 afla|EED46182.1 afum|EAL91647.1 agla|OJJ83129.1 akaw|GAA86180.1 aluc|OJZ90425.1 anid|CBF77983.1 anig|EHA23894.1 aory|EIT76412.1 asyd|OJJ55258.1 ater|EAU33041.1 atub|OJI81857.1 awen|OJJ41739.1 azon|OJJ48435.1

Group5249: abra|OJJ72447.1 acar|OOF94789.1 acla|EAW12680.1 afis|EAW17843.1 afla|EED46188.1 afum|EAL91652.1 agla|OJJ83124.1 akaw|GAA86185.1 aluc|OJZ90431.1 anid|CBF77971.1 anig|EHA23900.1 aory|EIT76400.1 asyd|OJJ55264.1 ater|EAU33045.1 atub|OJI81863.1 awen|OJJ41733.1 azon|OJJ48430.1

Group5250: abra|OJJ72452.1 acar|OOF94784.1 acla|EAW12686.1 afis|EAW17849.1 afla|EED46202.1 afum|EAL91658.1 agla|OJJ83120.1 akaw|GAA86189.1 aluc|OJZ90437.1 anig|EHA23904.1 aory|EIT76465.1 asyd|OJJ55271.1 ater|EAU33051.1 atub|OJI81868.1 aver|OJJ06956.1 awen|OJJ41724.1 azon|OJJ48422.1

Group5251: abra|OJJ72463.1 acar|OOF94776.1 acla|EAW12695.1 afis|EAW17859.1 afla|EED46214.1 afum|EAL91667.1 agla|OJJ83110.1 akaw|GAA86199.1 aluc|OJZ90450.1 anid|CBF77945.1 anig|EHA23913.1 aory|EIT76391.1 asyd|OJJ55280.1 atub|OJI81878.1 aver|OJJ06947.1 awen|OJJ41713.1 azon|OJJ48408.1

Group5252: abra|OJJ72472.1 acar|OOF94763.1 acla|EAW12705.1 afis|EAW17868.1 afum|EAL91676.1 agla|OJJ83088.1 akaw|GAA86210.1 aluc|OJZ90459.1 anid|CBF89895.1 anig|EHA23921.1 aory|EIT76489.1 asyd|OJJ62138.1 ater|EAU33086.1 atub|OJI81886.1 aver|OJI96248.1 awen|OJJ41703.1 azon|OJJ48399.1

Group5253: abra|OJJ72473.1 acar|OOF94762.1 acla|EAW12706.1 afis|EAW17869.1 afla|EED46237.1 afum|EAL91677.1 agla|OJJ83089.1 akaw|GAA86211.1 aluc|OJZ90460.1 anid|CBF89897.1 anig|EHA23922.1 asyd|OJJ62137.1 ater|EAU33087.1 atub|OJI81887.1 aver|OJI96247.1 awen|OJJ41702.1 azon|OJJ48398.1

Group5254: abra|OJJ72477.1 acar|OOF94759.1 acla|EAW12708.1 afis|EAW17871.1 afum|EAL91679.1 agla|OJJ83091.1 akaw|GAA86213.1 aluc|OJZ90463.1 anid|CBF89900.1 anig|EHA23924.1 aory|EIT76455.1 asyd|OJJ62135.1 ater|EAU33089.1 atub|OJI81889.1 aver|OJI96245.1 awen|OJJ41699.1 azon|OJJ48396.1

Group5255: abra|OJJ72479.1 acar|OOF94757.1 acla|EAW12710.1 afis|EAW17873.1 afla|EED46240.1 afum|EAL91681.1 agla|OJJ83075.1 akaw|GAA86215.1 aluc|OJZ90467.1 anid|CBF89932.1 anig|EHA23926.1 aory|EIT76355.1 asyd|OJJ62115.1 ater|EAU33091.1 atub|OJI81891.1 aver|OJI96222.1 awen|OJJ41695.1

Group5256: abra|OJJ72486.1 acar|OOF94749.1 afis|EAW17880.1 afla|EED51757.1 afum|EAL91688.1 akaw|GAA86221.1 aluc|OJZ90473.1 anid|CBF88138.1 anig|EHA23932.1 aory|EIT80744.1 asyd|OJJ63602.1 ater|EAU33100.1 atub|OJI81897.1 aver|OJJ05546.1 awen|OJJ36319.1 ater|EAU29427.1 aver|OJI97334.1

Group5257: abra|OJJ72491.1 acar|OOF94746.1 acla|EAW12721.1 afis|EAW17884.1 afla|EED46271.1 afum|EAL91692.1 agla|OJJ83063.1 akaw|GAA86225.1 aluc|OJZ90477.1 anid|CBF77031.1 anig|EHA23936.1 asyd|OJJ57010.1 ater|EAU33104.1 atub|OJI81902.1 aver|OJJ03277.1 awen|OJJ41675.1 azon|OJJ48390.1

Group5258: abra|OJJ72504.1 acla|EAW12728.1 afis|EAW17891.1 afla|EED46283.1 afum|EAL91700.1 agla|OJJ83052.1 akaw|GAA86232.1 aluc|OJZ90487.1 anid|CBF77012.1 anig|EHA23945.1 aory|EIT76351.1 asyd|OJJ57018.1 ater|EAU33111.1 atub|OJI81911.1 aver|OJJ03271.1 awen|OJJ41662.1 azon|OJJ48381.1

Group5259: abra|OJJ72506.1 acar|OOF94736.1 acla|EAW12729.1 afis|EAW17892.1 afla|EED46285.1 afum|EAL91701.1 agla|OJJ83050.1 akaw|GAA86234.1 aluc|OJZ90489.1 anid|CBF77008.1 anig|EHA23947.1 asyd|OJJ57020.1 ater|EAU33113.1 atub|OJI81913.1 aver|OJJ03269.1 awen|OJJ41660.1 azon|OJJ48380.1

Group5260: abra|OJJ72533.1 acar|OOF94720.1 acla|EAW12740.1 afis|EAW17903.1 afla|EED46295.1 afum|EAL91711.1 agla|OJJ83040.1 akaw|GAA86255.1 aluc|OJZ90512.1 anid|CBF76988.1 aory|EIT76329.1 ater|EAU33122.1 atub|OJI81937.1 aver|OJJ03258.1 awen|OJJ41647.1 asyd|OJJ57034.1 azon|OJJ48369.1

Group5261: abra|OJJ72536.1 acla|EAW12743.1 afis|EAW17906.1 afla|EED46300.1 afum|EAL91715.1 agla|OJJ83036.1 akaw|GAA86259.1 aluc|OJZ90517.1 anid|CBF76983.1 anig|EHA23971.1 aory|EIT76325.1 asyd|OJJ57036.1 ater|EAU33124.1 atub|OJI81941.1 aver|OJJ03256.1 awen|OJJ41644.1 azon|OJJ48365.1

Group5262: abra|OJJ72546.1 acar|OOF94707.1 acla|EAW12750.1 afis|EAW17914.1 afla|EED46317.1 afum|EAL91724.1 agla|OJJ83029.1 akaw|GAA85179.1 aluc|OJZ90533.1 anid|CBF76973.1 anig|EHA23982.1 asyd|OJJ57046.1 ater|EAU33135.1 atub|OJI81954.1 aver|OJJ03249.1 awen|OJJ41633.1 azon|OJJ48358.1

Group5263: abra|OJJ72563.1 acar|OOF94691.1 acla|EAW12763.1 afis|EAW17925.1 afum|EAL91736.1 agla|OJJ83017.1 akaw|GAA85192.1 aluc|OJZ90547.1 anid|CBF76944.1 anig|EHA23995.1 aory|EIT81166.1 asyd|OJJ57061.1 ater|EAU33547.1 atub|OJI81969.1 aver|OJJ03237.1 awen|OJJ41621.1 azon|OJJ48344.1

Group5264: abra|OJJ72572.1 acar|OOF99282.1 akaw|GAA85198.1 aluc|OJZ90553.1 anig|EHA24000.1 asyd|OJJ56262.1 atub|OJI81976.1 abra|OJJ78185.1 acla|EAW08831.1 anig|EHA19946.1 ater|EAU38338.1 aver|OJJ08473.1 acar|OOF96678.1 afla|EED51444.1 agla|OJJ88347.1 aory|EIT80756.1 ater|EAU37977.1

Group5265: abra|OJJ72579.1 acar|OOF94673.1 afla|EED56580.1 agla|OJJ88581.1 akaw|GAA85205.1 aluc|OJZ90560.1 anig|EHA24006.1 aory|EIT79653.1 asyd|OJJ60421.1 ater|EAU30076.1 atub|OJI81983.1 aver|OJJ01323.1 awen|OJJ33716.1 azon|OJJ42313.1 asyd|OJJ58710.1 awen|OJJ40585.1 akaw|GAA87620.1

Group5266: abra|OJJ72586.1 acar|OOF94663.1 acla|EAW12772.1 afis|EAW17935.1 afum|EAL91746.1 agla|OJJ82997.1 akaw|GAA85212.1 aluc|OJZ90567.1 anid|CBF77040.1 anig|EHA24013.1 aory|EIT78504.1 asyd|OJJ57001.1 ater|EAU33535.1 atub|OJI81990.1 aver|OJJ03286.1 awen|OJJ41611.1 azon|OJJ48330.1

Group5267: abra|OJJ72587.1 acar|OOF94664.1 acla|EAW12773.1 afis|EAW17936.1 afla|EED49106.1 agla|OJJ82996.1 akaw|GAA85213.1 aluc|OJZ90568.1 anid|CBF77042.1 aory|EIT78510.1 asyd|OJJ57000.1 ater|EAU33534.1 atub|OJI81991.1 aver|OJJ03287.1 awen|OJJ41610.1 azon|OJJ48329.1 afum|EAL91747.1

Group5268: abra|OJJ72590.1 acar|OOF94660.1 acla|EAW12776.1 afis|EAW17939.1 afla|EED49102.1 afum|EAL91750.1 agla|OJJ82993.1 akaw|GAA85216.1 aluc|OJZ90571.1 anid|CBF77047.1 anig|EHA24016.1 asyd|OJJ56997.1 ater|EAU33529.1 atub|OJI81994.1 aver|OJJ03290.1 awen|OJJ41606.1 azon|OJJ48324.1

Group5269: abra|OJJ72615.1 acla|EAW07008.1 afis|EAW16624.1 afla|EED44980.1 afum|EAL89213.1 agla|OJJ82624.1 akaw|GAA85240.1 aluc|OJZ90597.1 anid|CBF83188.1 anig|EHA24036.1 aory|EIT73566.1 asyd|OJJ55547.1 ater|EAU36728.1 atub|OJI82021.1 aver|OJJ06624.1 awen|OJJ36245.1 azon|OJJ42444.1

Group5270: abra|OJJ72621.1 acar|OOF90202.1 acla|EAW11603.1 afis|EAW23855.1 afla|EED49096.1 afum|EAL87283.1 agla|OJJ82506.1 akaw|GAA85242.1 aluc|OJZ90599.1 anid|CBF77246.1 aory|EIT82505.1 asyd|OJJ56811.1 ater|EAU33500.1 atub|OJI82023.1 aver|OJJ03524.1 awen|OJJ33816.1 azon|OJJ48722.1

Group5271: abra|OJJ72624.1 acar|OOF94640.1 acla|EAW11600.1 afis|EAW23857.1 afla|EED52582.1 afum|EAL87285.1 agla|OJJ82511.1 akaw|GAA85244.1 aluc|OJZ90601.1 anid|CBF77238.1 aory|EIT74177.1 asyd|OJJ56813.1 ater|EAU33494.1 atub|OJI82025.1 aver|OJJ03522.1 awen|OJJ33818.1 azon|OJJ48717.1

Group5272: abra|OJJ72631.1 acar|OOF94634.1 afis|EAW23863.1 afla|EED52587.1 afum|EAL87291.1 agla|OJJ82529.1 akaw|GAA85251.1 aluc|OJZ90608.1 anid|CBF77226.1 anig|EHA24046.1 aory|EIT74239.1 asyd|OJJ56818.1 ater|EAU33488.1 atub|OJI82032.1 aver|OJJ03517.1 awen|OJJ33823.1 azon|OJJ48712.1

Group5273: abra|OJJ72634.1 acar|OOF93305.1 afis|EAW18838.1 afla|EED47562.1 afum|EAL93572.1 akaw|GAA85255.1 aluc|OJZ90613.1 anid|CBF74711.1 anig|EHA24051.1 aory|EIT74513.1 asyd|OJJ59476.1 ater|EAU38813.1 atub|OJI82039.1 aver|OJJ02797.1 aver|OJJ08238.1 awen|OJJ32594.1 asyd|OJJ53508.1

Group5274: abra|OJJ72637.1 acla|EAW11587.1 afis|EAW23866.1 afla|EED52594.1 afum|EAL87294.1 agla|OJJ82535.1 akaw|GAA85258.1 aluc|OJZ90616.1 anid|CBF77211.1 anig|EHA24053.1 aory|EIT74298.1 asyd|OJJ56826.1 ater|EAU33476.1 atub|OJI82042.1 aver|OJJ03509.1 awen|OJJ33842.1 azon|OJJ48707.1

Group5275: abra|OJJ72638.1 acla|EAW11586.1 afis|EAW23867.1 afla|EED52595.1 afum|EAL87295.1 agla|OJJ82536.1 akaw|GAA85259.1 aluc|OJZ90617.1 anid|CBF77213.1 anig|EHA24054.1 aory|EIT74234.1 asyd|OJJ56825.1 ater|EAU33477.1 atub|OJI82043.1 aver|OJJ03510.1 awen|OJJ33843.1 azon|OJJ48704.1

Group5276: abra|OJJ72639.1 acar|OOF94632.1 acla|EAW11585.1 afis|EAW23868.1 afla|EED52596.1 afum|EAL87296.1 agla|OJJ82537.1 akaw|GAA85260.1 aluc|OJZ90618.1 anid|CBF77215.1 anig|EHA24055.1 aory|EIT74172.1 asyd|OJJ56824.1 atub|OJI82044.1 aver|OJJ03511.1 awen|OJJ33844.1 azon|OJJ48703.1

Group5277: abra|OJJ72649.1 acar|OOF94621.1 acla|EAW11579.1 afis|EAW23875.1 afla|EED52603.1 afum|EAL87303.1 agla|OJJ82545.1 akaw|GAA85270.1 aluc|OJZ90628.1 anid|CBF77207.1 anig|EHA24064.1 asyd|OJJ56830.1 ater|EAU33471.1 atub|OJI82054.1 aver|OJJ03506.1 awen|OJJ33851.1 azon|OJJ48697.1

Group5278: abra|OJJ72651.1 acla|EAW11578.1 afis|EAW23876.1 afla|EED52604.1 afum|EAL87304.1 agla|OJJ82546.1 akaw|GAA85271.1 aluc|OJZ90629.1 anid|CBF77205.1 anig|EHA24065.1 aory|EIT74232.1 asyd|OJJ56831.1 ater|EAU33470.1 atub|OJI82055.1 aver|OJJ03505.1 awen|OJJ33852.1 azon|OJJ48696.1

Group5279: abra|OJJ72656.1 acar|OOF94618.1 acla|EAW11519.1 afis|EAW23934.1 afla|EED50584.1 afum|EAL87359.1 agla|OJJ79414.1 akaw|GAA85278.1 aluc|OJZ90637.1 anid|CBF77271.1 anig|EHA24072.1 asyd|OJJ56798.1 ater|EAU32303.1 atub|OJI82065.1 aver|OJJ03536.1 awen|OJJ39021.1 azon|OJJ51346.1

Group5280: abra|OJJ72660.1 acar|OOF94613.1 acla|EAW11524.1 afis|EAW23929.1 afla|EED50589.1 afum|EAL87354.1 agla|OJJ79408.1 akaw|GAA85283.1 aluc|OJZ90641.1 anid|CBF77279.1 anig|EHA24076.1 aory|EIT82068.1 asyd|OJJ56794.1 atub|OJI82069.1 aver|OJJ03540.1 awen|OJJ39025.1 azon|OJJ51351.1

Group5281: abra|OJJ72674.1 acar|OOF97383.1 acla|EAW11536.1 afis|EAW23918.1 afla|EED50607.1 afum|EAL87344.1 agla|OJJ79387.1 akaw|GAA85293.1 aluc|OJZ90652.1 anid|CBF77094.1 anig|EHA24087.1 asyd|OJJ56956.1 ater|EAU32317.1 atub|OJI82081.1 aver|OJJ03329.1 awen|OJJ39035.1 azon|OJJ51363.1

Group5282: abra|OJJ72718.1 acar|OOF95950.1 acla|EAW11554.1 afis|EAW23899.1 afla|EED52633.1 afum|EAL87326.1 agla|OJJ78641.1 aluc|OJZ90687.1 anid|CBF77146.1 anig|EHA24115.1 aory|EIT74241.1 asyd|OJJ56911.1 ater|EAU32338.1 atub|OJI82112.1 aver|OJJ03382.1 awen|OJJ39049.1 azon|OJJ51381.1

Group5283: abra|OJJ72724.1 acar|OOF95957.1 acla|EAW11561.1 afis|EAW23892.1 afla|EED52622.1 afum|EAL87320.1 agla|OJJ78632.1 akaw|GAA85330.1 aluc|OJZ90693.1 anid|CBF77169.1 aory|EIT74282.1 asyd|OJJ56852.1 ater|EAU33450.1 atub|OJI82118.1 aver|OJJ03467.1 awen|OJJ39057.1 azon|OJJ51388.1

Group5284: abra|OJJ72726.1 acar|OOF95959.1 acla|EAW11563.1 afis|EAW23890.1 afla|EED52620.1 afum|EAL87318.1 agla|OJJ78630.1 akaw|GAA85332.1 aluc|OJZ90695.1 anid|CBF77173.1 anig|EHA24120.1 asyd|OJJ56849.1 ater|EAU33453.1 atub|OJI82121.1 aver|OJJ03490.1 awen|OJJ39059.1 azon|OJJ51390.1

Group5285: abra|OJJ72731.1 acla|EAW11566.1 afis|EAW23887.1 afla|EED52616.1 afum|EAL87314.2 agla|OJJ82574.1 akaw|GAA85337.1 aluc|OJZ90700.1 anid|CBF77179.1 anig|EHA24126.1 aory|EIT74278.1 asyd|OJJ56845.1 ater|EAU33459.1 atub|OJI82126.1 aver|OJJ03493.1 awen|OJJ33869.1 azon|OJJ48685.1

Group5286: abra|OJJ72734.1 acar|OOF95964.1 acla|EAW11569.1 afis|EAW23884.1 afla|EED52613.1 afum|EAL87312.1 agla|OJJ82571.1 akaw|GAA85340.1 aluc|OJZ90702.1 anid|CBF77185.1 anig|EHA24128.1 aory|EIT74192.1 asyd|OJJ56842.1 ater|EAU33462.1 atub|OJI82129.1 aver|OJJ03496.1 awen|OJJ33866.1

Group5287: abra|OJJ72740.1 acar|OOF95968.1 acla|EAW11575.1 afis|EAW23879.1 afla|EED52609.1 afum|EAL87307.1 agla|OJJ82569.1 akaw|GAA85350.1 aluc|OJZ90712.1 anig|EHA24132.1 aory|EIT74197.1 asyd|OJJ56837.1 ater|EAU33466.1 atub|OJI82139.1 aver|OJJ03500.1 awen|OJJ33860.1 azon|OJJ48692.1

Group5288: abra|OJJ72752.1 acar|OOF95975.1 acla|EAW11511.1 afis|EAW23942.1 afla|EED50574.1 afum|EAL87367.1 agla|OJJ79430.1 akaw|GAA85364.1 aluc|OJZ90726.1 anid|CBF77258.1 anig|EHA24144.1 asyd|OJJ56805.1 ater|EAU32291.1 atub|OJI82154.1 aver|OJJ03529.1 awen|OJJ39013.1 azon|OJJ51338.1

Group5289: abra|OJJ72758.1 acar|OOF95979.1 acla|EAW11506.1 afis|EAW23946.1 afum|EAL87372.1 agla|OJJ79439.1 akaw|GAA85368.1 aluc|OJZ90732.1 anid|CBF77288.1 anig|EHA24148.1 aory|EIT82036.1 asyd|OJJ56789.1 ater|EAU32288.1 atub|OJI82158.1 aver|OJJ03547.1 awen|OJJ39005.1 azon|OJJ51334.1

Group5290: abra|OJJ72759.1 acla|EAW11505.1 afis|EAW23947.1 afla|EED50569.1 afum|EAL87373.1 agla|OJJ79440.1 akaw|GAA85369.1 aluc|OJZ90733.1 anid|CBF77286.1 anig|EHA24149.1 aory|EIT82150.1 asyd|OJJ56790.1 ater|EAU32287.1 atub|OJI82159.1 aver|OJJ03546.1 awen|OJJ39004.1 azon|OJJ51333.1

Group5291: abra|OJJ72760.1 acla|EAW11504.1 afis|EAW23948.1 afla|EED50568.1 afum|EAL87374.1 agla|OJJ79441.1 akaw|GAA85370.1 aluc|OJZ90734.1 anid|CBF77285.1 anig|EHA24150.1 aory|EIT82140.1 asyd|OJJ56791.1 ater|EAU29804.1 atub|OJI82161.1 aver|OJJ03543.1 awen|OJJ39002.1 azon|OJJ51332.1

Group5292: abra|OJJ72771.1 acar|OOF95990.1 acla|EAW11494.1 afis|EAW23958.1 afla|EED50554.1 afum|EAL87386.1 agla|OJJ86881.1 akaw|GAA85380.1 aluc|OJZ90748.1 anid|CBF77310.1 aory|EIT82130.1 asyd|OJJ56775.1 ater|EAU29794.1 atub|OJI82174.1 aver|OJJ03558.1 awen|OJJ38989.1 azon|OJJ51321.1

Group5293: abra|OJJ72777.1 acar|OOF95995.1 acla|EAW11489.1 afis|EAW23964.1 afla|EED50546.1 afum|EAL87394.1 agla|OJJ86893.1 akaw|GAA85386.1 aluc|OJZ90755.1 anid|CBF77318.1 aory|EIT82079.1 asyd|OJJ56771.1 ater|EAU29788.1 atub|OJI82179.1 aver|OJJ03562.1 awen|OJJ38981.1 azon|OJJ51315.1

Group5294: abra|OJJ72787.1 acar|OOF96004.1 acla|EAW11482.1 afis|EAW23971.1 afla|EED50537.1 afum|EAL87401.1 agla|OJJ86900.1 akaw|GAA85394.1 aluc|OJZ83285.1 anid|CBF77333.1 anig|EHA24171.1 asyd|OJJ53277.1 ater|EAU29781.1 atub|OJI82188.1 aver|OJJ03571.1 awen|OJJ38972.1 azon|OJJ51307.1

Group5295: abra|OJJ72790.1 acar|OOF90124.1 acla|EAW11481.1 afis|EAW23972.1 afla|EED50536.1 afum|EAL87402.1 aluc|OJZ83287.1 anid|CBF77337.1 anig|EHA24172.1 aory|EIT82163.1 ater|EAU29780.1 atub|OJI82189.1 aver|OJJ03573.1 awen|OJJ38971.1 azon|OJJ51306.1 agla|OJJ86901.1 asyd|OJJ53276.1

Group5296: abra|OJJ72793.1 acla|EAW11479.1 afis|EAW23974.1 afla|EED50534.1 afum|EAL87404.1 agla|OJJ86903.1 akaw|GAA85396.1 aluc|OJZ83289.1 anid|CBF77341.1 anig|EHA24174.1 aory|EIT82105.1 asyd|OJJ53274.1 ater|EAU29778.1 atub|OJI82191.1 aver|OJJ03575.1 awen|OJJ35439.1 azon|OJJ51304.1

Group5297: abra|OJJ72799.1 acar|OOF96007.1 acla|EAW11445.1 afis|EAW23980.1 afla|EED49179.1 afum|EAL87410.1 agla|OJJ86912.1 akaw|GAA85402.1 aluc|OJZ83297.1 anid|CBF77354.1 anig|EHA24180.1 aory|EIT78569.1 asyd|OJJ53269.1 atub|OJI82197.1 awen|OJJ35448.1 azon|OJJ51298.1 aver|OJJ03582.1

Group5298: abra|OJJ72802.1 acar|OOF96010.1 acla|EAW11442.1 afis|EAW23983.1 afla|EED49176.1 afum|EBA27313.1 agla|OJJ86917.1 akaw|GAA85405.1 aluc|OJZ83300.1 anid|CBF77360.1 aory|EIT78516.1 asyd|OJJ53266.1 ater|EAU29768.1 atub|OJI82200.1 aver|OJJ03585.1 awen|OJJ35451.1 azon|OJJ51292.1

Group5299: abra|OJJ72806.1 afla|EED49171.1 akaw|GAA85409.1 aluc|OJZ83304.1 anig|EHA24187.1 aory|EIT78556.1 ater|EAU29763.1 atub|OJI82204.1 awen|OJJ35455.1 acla|EAW11435.1 afis|EAW23988.1 afum|EAL87417.2 agla|OJJ86921.1 anid|CBF77366.1 asyd|OJJ53263.1 aver|OJJ03588.1 azon|OJJ51288.1

Group5300: abra|OJJ72816.1 acar|OOF96022.1 acla|EAW11424.1 afis|EAW23997.1 agla|OJJ86931.1 akaw|GAA85417.1 aluc|OJZ83318.1 anid|CBF77381.1 anig|EHA24196.1 aory|EIT78578.1 asyd|OJJ53252.1 ater|EAU29752.1 atub|OJI82214.1 aver|OJJ03600.1 awen|OJJ35468.1 azon|OJJ51276.1 afum|EAL87427.1

Group5301: abra|OJJ72845.1 acar|OOF96046.1 acla|EAW11398.1 afis|EAW24023.1 afla|EED49131.1 afum|EAL87454.1 agla|OJJ86956.1 akaw|GAA85446.1 aluc|OJZ83353.1 anid|CBF75302.1 anig|EHA24219.1 asyd|OJJ59880.1 ater|EAU29723.1 atub|OJI82245.1 aver|OJJ02406.1 awen|OJJ35496.1 azon|OJJ46993.1

Group5302: abra|OJJ72846.1 acar|OOF96047.1 acla|EAW11397.1 afis|EAW24024.1 afla|EED49130.1 afum|EAL87455.1 agla|OJJ86957.1 akaw|GAA85447.1 aluc|OJZ83354.1 anid|CBF75304.1 anig|EHA24220.1 asyd|OJJ59879.1 ater|EAU29722.1 atub|OJI82246.1 aver|OJJ02407.1 awen|OJJ35497.1 azon|OJJ46992.1

Group5303: abra|OJJ72849.1 acar|OOF96050.1 acla|EAW11394.1 afis|EAW24027.1 afla|EED49127.1 afum|EAL87458.1 agla|OJJ86960.1 akaw|GAA85450.1 aluc|OJZ83358.1 anid|CBF75308.1 anig|EHA24223.1 asyd|OJJ59875.1 ater|EAU29717.1 atub|OJI82249.1 aver|OJJ02411.1 awen|OJJ35500.1 azon|OJJ46989.1

Group5304: abra|OJJ72851.1 acla|EAW11392.1 afis|EAW24029.1 afla|EED49125.1 afum|EAL87460.1 agla|OJJ86963.1 akaw|GAA85452.1 aluc|OJZ83360.1 anid|CBF75312.1 anig|EHA24225.1 aory|EIT78520.1 asyd|OJJ59873.1 ater|EAU29715.1 atub|OJI82251.1 aver|OJJ02414.1 awen|OJJ35502.1 azon|OJJ46987.1

Group5305: abra|OJJ72864.1 acar|OOF96063.1 acla|EAW11381.1 afis|EAW18115.1 afla|EED58040.1 afum|EAL87472.1 agla|OJJ86981.1 akaw|GAA85463.1 aluc|OJZ83377.1 anid|CBF75338.1 anig|EHA24236.1 asyd|OJJ59860.1 ater|EAU29702.1 atub|OJI82263.1 aver|OJJ02427.1 awen|OJJ35514.1 azon|OJJ46969.1

Group5306: abra|OJJ72867.1 acar|OOF96066.1 acla|EAW11378.1 afis|EAW18117.1 afla|EED58042.1 afum|EAL87475.1 agla|OJJ86979.1 akaw|GAA85466.1 aluc|OJZ83380.1 anid|CBF75342.1 anig|EHA24239.1 asyd|OJJ59858.1 ater|EAU29700.1 atub|OJI82266.1 aver|OJJ02429.1 awen|OJJ35516.1 azon|OJJ46968.1

Group5307: abra|OJJ72868.1 acar|OOF96067.1 acla|EAW11377.1 afis|EAW18118.1 afla|EED58043.1 afum|EAL87476.1 agla|OJJ86978.1 akaw|GAA85467.1 aluc|OJZ83381.1 anid|CBF75344.1 anig|EHA24240.1 aory|EIT79729.1 asyd|OJJ59857.1 ater|EAU29699.1 atub|OJI82267.1 aver|OJJ02430.1 awen|OJJ35517.1

Group5308: abra|OJJ72897.1 acla|EAW11354.1 afis|EAW18140.1 afla|EED58066.1 afum|EAL87497.1 agla|OJJ83825.1 akaw|GAA85484.1 aluc|OJZ83406.1 anid|CBF75390.1 anig|EHA24257.1 aory|EIT79738.1 asyd|OJJ59835.1 ater|EAU35463.1 atub|OJI82295.1 aver|OJJ02453.1 awen|OJJ35582.1 azon|OJJ46940.1

Group5309: abra|OJJ72908.1 acar|OOF96091.1 acla|EAW11345.1 afis|EAW18149.1 afla|EED58077.1 afum|EAL87507.1 agla|OJJ83834.1 akaw|GAA85493.1 aluc|OJZ83418.1 anid|CBF81112.1 anig|EHA24267.1 asyd|OJJ56484.1 ater|EAU35471.1 atub|OJI82305.1 aver|OJJ06235.1 awen|OJJ35592.1 azon|OJJ46914.1

Group5310: abra|OJJ72925.1 acar|OOF96106.1 acla|EAW06464.1 afis|EAW15959.1 afla|EED58099.1 afum|EAL88592.1 agla|OJJ89353.1 akaw|GAA85508.1 aluc|OJZ83435.1 aory|EIT79128.1 ater|EAU35497.1 atub|OJI82326.1 aver|OJJ06259.1 awen|OJJ35614.1 azon|OJJ46909.1 anid|CBF81076.1 asyd|OJJ56462.1

Group5311: abra|OJJ72926.1 acar|OOF96107.1 acla|EAW06463.1 afis|EAW15958.1 afla|EED58100.1 afum|EAL88591.1 agla|OJJ89354.1 akaw|GAA85509.1 aluc|OJZ83436.1 anid|CBF81074.1 anig|EHA24284.1 aory|EIT79118.1 asyd|OJJ56461.1 atub|OJI82327.1 aver|OJJ06260.1 awen|OJJ35615.1 azon|OJJ46908.1

Group5312: abra|OJJ72943.1 acla|EAW11208.1 afis|EAW15941.1 afla|EED58117.1 afum|EAL88575.1 agla|OJJ89369.1 akaw|GAA85525.1 aluc|OJZ83453.1 anig|EHA24299.1 aory|EIT79113.1 asyd|OJJ56347.1 ater|EAU35515.1 atub|OJI82345.1 aver|OJJ06366.1 azon|OJJ46894.1 anid|CBF80960.1 awen|OJJ35641.1

Group5313: abra|OJJ72944.1 acar|OOF96125.1 acla|EAW11207.1 afis|EAW15937.1 afla|EED58118.1 afum|EAL88573.1 agla|OJJ89370.1 akaw|GAA85526.1 aluc|OJZ83454.1 anid|CBF80962.1 anig|EHA24300.1 aory|EIT79136.1 asyd|OJJ56348.1 atub|OJI82346.1 aver|OJJ06365.1 awen|OJJ35642.1 azon|OJJ46893.1

Group5314: abra|OJJ72951.1 acar|OOF96133.1 acla|EAW11197.1 afis|EAW15927.1 afla|EED54934.1 afum|EAL88563.1 agla|OJJ89378.1 akaw|GAA85532.1 aluc|OJZ83461.1 anid|CBF81022.1 aory|EIT80375.1 asyd|OJJ56384.1 ater|EAU35528.1 atub|OJI82353.1 aver|OJJ06328.1 awen|OJJ35653.1 azon|OJJ46885.1

Group5315: abra|OJJ72955.1 acar|OOF96136.1 acla|EAW11200.1 afis|EAW15930.1 afla|EED54931.1 afum|EAL88566.1 agla|OJJ89376.1 akaw|GAA85536.1 aluc|OJZ83464.1 anid|CBF81028.1 anig|EHA24308.1 asyd|OJJ56387.1 ater|EAU35522.1 atub|OJI82357.1 aver|OJJ06326.1 awen|OJJ35650.1 azon|OJJ46888.1

Group5316: abra|OJJ72964.1 acar|OOF96146.1 acla|EAW11190.1 afis|EAW15921.1 afla|EED54940.1 afum|EAL88557.2 agla|OJJ89389.1 akaw|GAA85542.1 aluc|OJZ83473.1 anid|CBF81005.1 aory|EIT80301.1 asyd|OJJ56378.1 ater|EAU35535.1 atub|OJI82363.1 aver|OJJ06335.1 awen|OJJ35660.1 azon|OJJ46880.1

Group5317: abra|OJJ72995.1 acar|OOF96172.1 acla|EAW08017.1 afis|EAW23051.1 afum|EAL88429.1 agla|OJJ89428.1 akaw|GAA85569.1 aluc|OJZ83503.1 anid|CBF89694.1 anig|EHA24337.1 aory|EIT80308.1 asyd|OJJ56447.1 ater|EAU29597.1 atub|OJI82397.1 aver|OJJ06275.1 awen|OJJ35687.1 azon|OJJ50729.1

Group5318: abra|OJJ73005.1 acla|EAW08008.1 afis|EAW23043.1 afla|EED54975.1 afum|EAL88438.1 agla|OJJ89435.1 akaw|GAA85583.1 aluc|OJZ83520.1 anid|CBF80940.1 anig|EHA24343.1 aory|EIT80357.1 asyd|OJJ56336.1 ater|EAU29604.1 atub|OJI82409.1 aver|OJJ06377.1 awen|OJJ35697.1 azon|OJJ44161.1

Group5319: abra|OJJ73007.1 acar|OOF96179.1 acla|EAW08006.1 afis|EAW23041.1 afla|EED54977.1 afum|EAL88440.1 agla|OJJ89437.1 akaw|GAA85585.1 aluc|OJZ83522.1 anid|CBF80937.1 anig|EHA24345.1 aory|EIT80307.1 asyd|OJJ56313.1 atub|OJI82411.1 aver|OJJ06401.1 awen|OJJ35701.1 azon|OJJ44163.1

Group5320: abra|OJJ73011.1 acar|OOF96184.1 acla|EAW08001.1 afis|EAW23036.1 afla|EED54981.1 afum|EAL88445.1 agla|OJJ89441.1 akaw|GAA85590.1 aluc|OJZ83526.1 anid|CBF80927.1 anig|EHA24349.1 asyd|OJJ56317.1 ater|EAU29622.1 atub|OJI82415.1 aver|OJJ06397.1 awen|OJJ35705.1 azon|OJJ44167.1

Group5321: abra|OJJ73015.1 acla|EAW07998.1 afis|EAW23033.1 afla|EED54983.1 afum|EAL88448.1 akaw|GAA85593.1 aluc|OJZ83530.1 anid|CBF80922.1 anig|EHA24352.1 aory|EIT80306.1 asyd|OJJ56321.1 ater|EAU29624.1 atub|OJI82420.1 awen|OJJ35712.1 azon|OJJ44170.1 aver|OJJ06392.1 agla|OJJ89444.1

Group5322: abra|OJJ73017.1 acar|OOF96187.1 acla|EAW07996.1 afis|EAW23031.1 afum|EAL88451.2 agla|OJJ89446.1 akaw|GAA85595.1 aluc|OJZ83532.1 anid|CBF80912.1 anig|EHA24354.1 asyd|OJJ56327.1 ater|EAU29626.1 atub|OJI82422.1 awen|OJJ35714.1 afla|EED54985.1 aory|EIT80319.1 aver|OJJ06386.1

Group5323: abra|OJJ73026.1 acar|OOF96193.1 acla|EAW07990.1 afis|EAW23025.1 afla|EED54990.1 afum|EAL88457.1 agla|OJJ89454.1 akaw|GAA85600.1 aluc|OJZ83538.1 anid|CBF80909.1 anig|EHA24359.1 aory|EIT80325.1 asyd|OJJ56330.1 ater|EAU29650.1 atub|OJI82429.1 awen|OJJ30718.1 azon|OJJ44179.1

Group5324: abra|OJJ73047.1 acla|EAW07972.1 afis|EAW23007.1 afla|EED48101.1 afum|EAL88473.1 agla|OJJ80223.1 akaw|GAA85618.1 aluc|OJZ83558.1 anid|CBF80877.1 anig|EHA24376.1 aory|EIT76233.1 asyd|OJJ56301.1 ater|EAU29666.1 atub|OJI82448.1 aver|OJJ06416.1 awen|OJJ31316.1 azon|OJJ50668.1

Group5325: abra|OJJ73054.1 acar|OOF96216.1 acla|EAW07966.1 afis|EAW23001.1 afla|EED55014.1 afum|EAL88479.2 agla|OJJ89478.1 aluc|OJZ83565.1 anid|CBF80866.1 anig|EHA24382.1 aory|EIT80356.1 asyd|OJJ56295.1 ater|EAU29672.1 atub|OJI82455.1 aver|OJJ06422.1 awen|OJJ30740.1 azon|OJJ44206.1

Group5326: abra|OJJ73057.1 acar|OOF96219.1 acla|EAW07964.1 afis|EAW22999.1 afla|EED55016.1 afum|EAL88481.1 agla|OJJ89480.1 akaw|GAA85625.1 aluc|OJZ83568.1 anid|CBF80862.1 anig|EHA24384.1 aory|EIT80386.1 asyd|OJJ56293.1 atub|OJI82457.1 aver|OJJ06424.1 awen|OJJ30742.1 azon|OJJ44203.1

Group5327: abra|OJJ73083.1 acar|OOG00317.1 acla|EAW12281.1 afis|EAW19608.1 afla|EED57664.1 afum|EAL89356.1 akaw|GAA88077.1 aluc|OJZ81340.1 anid|CBF88728.1 anig|EHA18646.1 aory|EIT79918.1 asyd|OJJ63933.1 ater|EAU35193.1 atub|OJI91545.1 aver|OJI97009.1 awen|OJJ31943.1 azon|OJJ50754.1

Group5328: abra|OJJ73108.1 acar|OOG00341.1 acla|EAW07041.1 afis|EAW16668.1 afla|EED44908.1 afum|EAL89272.1 agla|OJJ78701.1 akaw|GAA88058.1 aluc|OJZ81320.1 anid|CBF82081.1 anig|EHA18626.1 aory|EIT73518.1 asyd|OJJ54255.1 atub|OJI91523.1 aver|OJJ05208.1 awen|OJJ40045.1 azon|OJJ50124.1

Group5329: abra|OJJ73144.1 acar|OOG00356.1 acla|EAW07020.1 afis|EAW16646.1 afla|EED44954.1 afum|EAL89250.1 agla|OJJ82597.1 akaw|GAA88028.1 aluc|OJZ81287.1 anid|CBF82056.1 anig|EHA18586.1 asyd|OJJ54237.1 ater|EAU39571.1 atub|OJI91494.1 aver|OJJ05189.1 awen|OJJ40068.1 azon|OJJ45556.1

Group5330: abra|OJJ73150.1 acar|OOG00362.1 acla|EAW07027.1 afis|EAW16653.1 afla|EED44947.1 afum|EAL89257.1 agla|OJJ78686.1 akaw|GAA88022.1 aluc|OJZ81280.1 anig|EHA18580.1 aory|EIT73514.1 asyd|OJJ54243.1 ater|EAU39577.1 atub|OJI91488.1 aver|OJJ05195.1 awen|OJJ40061.1 azon|OJJ45566.1

Group5331: abra|OJJ73157.1 acar|OOG00370.1 acla|EAW07031.1 afis|EAW16657.1 afla|EED44942.1 afum|EAL89261.1 agla|OJJ78715.1 akaw|GAA88015.1 aluc|OJZ81273.1 anig|EHA18573.1 aory|EIT73488.1 asyd|OJJ54247.1 ater|EAU39580.1 atub|OJI91480.1 aver|OJJ05201.1 awen|OJJ40057.1 azon|OJJ50085.1

Group5332: abra|OJJ73169.1 acla|EAW10795.1 afis|EAW24781.1 afla|EED47204.1 afum|EAL89420.1 agla|OJJ82877.1 akaw|GAA88002.1 aluc|OJZ81258.1 anid|CBF81440.1 anig|EHA18565.1 aory|EIT79259.1 asyd|OJJ59429.1 atub|OJI91454.1 aver|OJJ02842.1 awen|OJJ40367.1 azon|OJJ49216.1 ater|EAU36527.1

Group5333: abra|OJJ73174.1 acar|OOG00414.1 acla|EAW06857.1 afis|EAW16474.1 afla|EED47209.1 afum|EAL89068.1 agla|OJJ82872.1 akaw|GAA87997.1 aluc|OJZ81253.1 anid|CBF81433.1 anig|EHA18561.1 asyd|OJJ59423.1 ater|EAU36568.1 atub|OJI91449.1 aver|OJJ02847.1 awen|OJJ40362.1 azon|OJJ49210.1

Group5334: abra|OJJ73182.1 acar|OOG00419.1 acla|EAW06860.1 afis|EAW16477.1 afla|EED47214.1 afum|EAL89071.1 agla|OJJ82868.1 akaw|GAA87993.1 aluc|OJZ81247.1 anid|CBF75497.1 anig|EHA18556.1 aory|EIT79205.1 asyd|OJJ59749.1 atub|OJI91443.1 aver|OJJ02539.1 awen|OJJ40358.1 azon|OJJ49207.1

Group5335: abra|OJJ73188.1 acar|OOG00423.1 acla|EAW06864.1 afis|EAW16481.1 afla|EED47218.1 afum|EAL89075.1 agla|OJJ82864.1 akaw|GAA87989.1 aluc|OJZ81243.1 anig|EHA18552.1 aory|EIT79179.1 asyd|OJJ59745.1 ater|EAU36557.1 atub|OJI91439.1 aver|OJJ02543.1 awen|OJJ40353.1 azon|OJJ49203.1

Group5336: abra|OJJ73192.1 acar|OOG00427.1 acla|EAW06868.1 afis|EAW16485.1 afla|EED47225.1 afum|EAL89079.1 agla|OJJ82859.1 akaw|GAA87985.1 aluc|OJZ81237.1 anid|CBF75518.1 anig|EHA18548.1 asyd|OJJ59737.1 ater|EAU36553.1 atub|OJI91435.1 aver|OJJ02550.1 awen|OJJ40344.1 azon|OJJ49199.1

Group5337: abra|OJJ73197.1 acar|OOG00436.1 acla|EAW06874.1 afis|EAW16491.1 afla|EED47232.1 afum|EAL89085.1 agla|OJJ82830.1 akaw|GAA87980.1 aluc|OJZ81232.1 anid|CBF75524.1 anig|EHA18543.1 asyd|OJJ59734.1 ater|EAU36546.1 atub|OJI91429.1 aver|OJJ02553.1 awen|OJJ40338.1 azon|OJJ49192.1

Group5338: abra|OJJ73198.1 acar|OOG00437.1 acla|EAW06875.1 afis|EAW16492.1 afla|EED47233.1 afum|EAL89086.1 agla|OJJ82831.1 akaw|GAA87979.1 aluc|OJZ81231.1 anid|CBF75525.1 anig|EHA18542.1 aory|EIT79252.1 asyd|OJJ59733.1 atub|OJI91428.1 aver|OJJ02554.1 awen|OJJ40337.1 azon|OJJ49191.1

Group5339: abra|OJJ73201.1 acar|OOG00440.1 acla|EAW06878.1 afis|EAW16495.1 afum|EAL89089.1 agla|OJJ82834.1 akaw|GAA87976.1 aluc|OJZ81228.1 anid|CBF75608.1 anig|EHA18539.1 aory|EIT79203.1 asyd|OJJ59728.1 ater|EAU36544.1 atub|OJI91425.1 aver|OJJ02558.1 awen|OJJ40334.1 azon|OJJ49188.1

Group5340: abra|OJJ73214.1 acar|OOG00451.1 acla|EAW06889.1 afis|EAW16506.1 afla|EED56172.1 afum|EAL89101.1 agla|OJJ82819.1 akaw|GAA87965.1 aluc|OJZ81214.1 anid|CBF75596.1 anig|EHA18528.1 aory|EIT76809.1 asyd|OJJ59717.1 atub|OJI91409.1 aver|OJJ02569.1 awen|OJJ40319.1 azon|OJJ49177.1

Group5341: abra|OJJ73221.1 acar|OOG00459.1 acla|EAW06896.1 afis|EAW16513.1 afla|EED56160.1 afum|EAL89108.1 agla|OJJ82809.1 akaw|GAA87957.1 aluc|OJZ81206.1 anid|CBF75564.1 anig|EHA18521.1 asyd|OJJ59707.1 ater|EAU36579.1 atub|OJI91402.1 aver|OJJ02578.1 awen|OJJ40311.1 azon|OJJ49171.1

Group5342: abra|OJJ73243.1 acar|OOG00482.1 acla|EAW06917.1 afis|EAW16534.1 afla|EED56135.1 afum|EAL89131.1 akaw|GAA87937.1 aluc|OJZ81185.1 anid|CBF81738.1 anig|EHA18502.1 aory|EIT76790.1 asyd|OJJ59595.1 ater|EAU36597.1 atub|OJI91381.1 aver|OJJ02673.1 awen|OJJ40288.1 azon|OJJ49149.1

Group5343: abra|OJJ73267.1 acar|OOG00507.1 acla|EAW06944.1 afis|EAW16560.1 afla|EED56103.1 afum|EAL89158.1 agla|OJJ82764.1 akaw|GAA87915.1 aluc|OJZ81162.1 anid|CBF81804.1 anig|EHA18479.1 aory|EIT76866.1 asyd|OJJ59643.1 ater|EAU36626.1 atub|OJI91357.1 aver|OJJ02639.1 awen|OJJ40258.1

Group5344: abra|OJJ73272.1 acar|OOG00512.1 acla|EAW06949.1 afis|EAW16564.1 afla|EED56099.1 afum|EAL89161.1 agla|OJJ82757.1 akaw|GAA87908.1 aluc|OJZ88777.1 anid|CBF81811.1 anig|EHA18471.1 aory|EIT76849.1 asyd|OJJ59647.1 atub|OJI91349.1 aver|OJJ02635.1 awen|OJJ40254.1 azon|OJJ49116.1

Group5345: abra|OJJ73282.1 acar|OOG00518.1 acla|EAW06957.1 afis|EAW16572.1 afla|EED56092.1 afum|EAL89169.2 agla|OJJ82704.1 akaw|GAA92723.1 aluc|OJZ88767.1 anig|EHA18463.1 aory|EIT76859.1 asyd|OJJ59658.1 ater|EAU36640.1 atub|OJI91340.1 aver|OJJ02624.1 awen|OJJ40246.1 azon|OJJ49107.1

Group5346: abra|OJJ73285.1 acar|OOG00521.1 acla|EAW06960.1 afis|EAW16575.1 afla|EED56089.1 afum|EBA27368.1 agla|OJJ82702.1 akaw|GAA92720.1 aluc|OJZ88764.1 anig|EHA18460.1 aory|EIT76647.1 asyd|OJJ59661.1 atub|OJI91337.1 aver|OJJ02621.1 awen|OJJ40243.1 azon|OJJ49104.1 anid|CBF81831.1

Group5347: abra|OJJ73298.1 afis|EAW19827.1 afla|EED51781.1 afum|EAL85065.1 akaw|GAA92708.1 aluc|OJZ88750.1 anig|EHA18452.1 asyd|OJJ59177.1 atub|OJI91321.1 aver|OJJ01796.1 awen|OJJ34692.1 azon|OJJ49039.1 agla|OJJ79503.1 acar|OOF93317.1 asyd|OJJ62364.1 aver|OJI96498.1 awen|OJJ30127.1

Group5348: abra|OJJ73323.1 acla|EAW15335.1 afis|EAW18094.1 afla|EED54259.1 afum|EAL91902.1 agla|OJJ78550.1 akaw|GAA86996.1 aluc|OJZ84463.1 anid|CBF75962.1 anig|EHA18708.1 aory|EIT76021.1 ater|EAU36995.1 atub|OJI89614.1 aver|OJJ00568.1 awen|OJJ32938.1 azon|OJJ42340.1 anid|CBF75964.1

Group5349: abra|OJJ73354.1 acla|EAW15249.1 afis|EAW20217.1 afla|EED44916.1 afum|EAL85332.1 agla|OJJ84639.1 akaw|GAA83130.1 aluc|OJZ88686.1 anid|CBF78511.1 anig|EHA18366.1 aory|EIT73493.1 asyd|OJJ65310.1 atub|OJI91266.1 aver|OJI99450.1 awen|OJJ31229.1 afla|EED57128.1 aory|EIT74120.1

Group5350: acla|EAW13469.1 agla|OJJ88694.1 abra|OJJ76945.1 acar|OOF91395.1 afla|EED52278.1 afum|EAL86634.1 akaw|GAA87095.1 aluc|OJZ84343.1 anid|CBF71790.1 anig|EHA27679.1 aory|EIT74372.1 asyd|OJJ61714.1 ater|EAU37102.1 atub|OJI89725.1 aver|OJI95809.1 awen|OJJ33286.1 azon|OJJ50664.1

Group5351: abra|OJJ73356.1 afla|EED45197.1 agla|OJJ82670.1 akaw|GAA83128.1 aluc|OJZ88684.1 anid|CBF81892.1 aory|EIT73648.1 asyd|OJJ59679.1 ater|EAU36665.1 atub|OJI91264.1 aver|OJJ02609.1 awen|OJJ40226.1 azon|OJJ49091.1 acla|EAW06975.1 afis|EAW16589.1 afum|EAL89184.1 anig|EHA18364.1

Group5352: abra|OJJ73363.1 acar|OOG00570.1 acla|EAW06983.1 afis|EAW16597.1 afla|EED45016.1 afum|EAL89193.1 agla|OJJ82657.1 akaw|GAA83121.1 aluc|OJZ88677.1 anid|CBF81904.1 aory|EIT73770.1 asyd|OJJ59687.1 ater|EAU36676.1 atub|OJI91258.1 aver|OJJ02600.1 awen|OJJ40211.1 azon|OJJ49077.1

Group5353: abra|OJJ73373.1 acar|OOG00578.1 acla|EAW06991.1 afis|EAW16605.1 afla|EED45024.1 afum|EAL89202.2 agla|OJJ82648.1 akaw|GAA83113.1 aluc|OJZ88668.1 anid|CBF81912.1 aory|EIT73735.1 asyd|OJJ59693.1 ater|EAU36687.1 atub|OJI91250.1 aver|OJJ02594.1 awen|OJJ40201.1 azon|OJJ49066.1

Group5354: abra|OJJ73377.1 acar|OOG00580.1 acla|EAW06990.1 afis|EAW16604.1 afla|EED45023.1 afum|EAL89200.1 akaw|GAA83112.1 aluc|OJZ88666.1 anid|CBF81911.1 anig|EHA18351.1 aory|EIT73764.1 asyd|OJJ59692.1 ater|EAU36686.1 atub|OJI91249.1 aver|OJJ02595.1 awen|OJJ40202.1 azon|OJJ49070.1

Group5355: abra|OJJ73379.1 acla|EAW06993.1 afis|EAW16608.1 afla|EED45026.1 afum|EAL89204.1 agla|OJJ82645.1 akaw|GAA83110.1 aluc|OJZ88664.1 anid|CBF81966.1 anig|EHA18349.1 aory|EIT73734.1 asyd|OJJ57354.1 ater|EAU36692.1 atub|OJI91246.1 aver|OJJ05811.1 awen|OJJ40199.1 azon|OJJ49068.1

Group5356: abra|OJJ73380.1 acar|OOF90416.1 acla|EAW06996.1 afis|EAW16610.1 afla|EED45002.1 afum|EAL89206.1 agla|OJJ82636.1 akaw|GAA83109.1 aluc|OJZ88663.1 anid|CBF81964.1 aory|EIT73646.1 asyd|OJJ57357.1 ater|EAU36695.1 atub|OJI91244.1 aver|OJJ05809.1 awen|OJJ40196.1 azon|OJJ49063.1

Group5357: abra|OJJ73386.1 acar|OOG00588.1 acla|EAW07010.1 afis|EAW16629.1 afla|EED44985.1 afum|EAL89224.2 agla|OJJ82614.1 akaw|GAA83103.1 aluc|OJZ88651.1 anid|CBF81970.1 anig|EHA18342.1 aory|EIT73790.1 asyd|OJJ57346.1 ater|EAU36722.1 atub|OJI91238.1 aver|OJJ05819.1 azon|OJJ44014.1

Group5358: abra|OJJ73387.1 acar|OOG00589.1 acla|EAW07001.1 afis|EAW16616.1 afla|EED44988.1 afum|EAL89220.1 agla|OJJ82631.1 akaw|GAA83102.1 aluc|OJZ88650.1 anid|CBF81968.1 anig|EHA18341.1 aory|EIT73725.1 asyd|OJJ57348.1 ater|EAU36704.1 atub|OJI91237.1 aver|OJJ05817.1 awen|OJJ40182.1

Group5359: abra|OJJ73404.1 acar|OOG00602.1 acla|EAW07011.1 afis|EAW16631.1 afla|EED44967.1 afum|EAL89227.1 agla|OJJ82618.1 akaw|GAA83086.1 aluc|OJZ88633.1 anid|CBF82012.1 anig|EHA18323.1 aory|EIT73524.1 asyd|OJJ56189.1 atub|OJI91217.1 aver|OJJ05117.1 awen|OJJ40078.1 azon|OJJ49493.1

Group5360: abra|OJJ73441.1 acar|OOG00647.1 acla|EAW06776.1 afis|EAW16327.1 afla|EED53554.1 afum|EAL88951.1 agla|OJJ88265.1 aluc|OJZ88599.1 anid|CBF89166.1 anig|EHA18293.1 aory|EIT74850.1 asyd|OJJ62544.1 ater|EAU34130.1 atub|OJI91184.1 aver|OJI96671.1 awen|OJJ37728.1 azon|OJJ44987.1

Group5361: abra|OJJ73444.1 acla|EAW06779.1 afis|EAW16330.1 afla|EED53548.1 afum|EAL88954.1 agla|OJJ88262.1 akaw|GAA83050.1 aluc|OJZ88596.1 anid|CBF89171.1 anig|EHA18290.1 aory|EIT75161.1 asyd|OJJ62541.1 ater|EAU34128.1 atub|OJI91181.1 aver|OJI96668.1 awen|OJJ37731.1 azon|OJJ44989.1

Group5362: abra|OJJ73457.1 acar|OOG00658.1 acla|EAW06784.1 afis|EAW16334.1 afla|EED53542.1 afum|EAL88959.1 agla|OJJ88258.1 akaw|GAA83041.1 aluc|OJZ88586.1 anid|CBF89190.1 aory|EIT75196.1 asyd|OJJ62535.1 ater|EAU34124.1 atub|OJI91172.1 aver|OJI96660.1 awen|OJJ37735.1 azon|OJJ44994.1

Group5363: abra|OJJ73459.1 acar|OOG00660.1 acla|EAW06786.1 afla|EED53540.1 afum|EAL88961.1 agla|OJJ88253.1 akaw|GAA83039.1 aluc|OJZ88584.1 anid|CBF89194.1 anig|EHA18279.1 aory|EIT74903.1 asyd|OJJ62533.1 ater|EAU34122.1 atub|OJI91170.1 aver|OJI96658.1 awen|OJJ37738.1 azon|OJJ44999.1

Group5364: abra|OJJ73461.1 acar|OOG00662.1 acla|EAW06788.1 afis|EAW16337.1 afla|EED53538.1 afum|EAL88963.1 agla|OJJ88255.1 akaw|GAA83037.1 aluc|OJZ88582.1 anid|CBF89197.1 aory|EIT75002.1 asyd|OJJ62531.1 ater|EAU34120.1 atub|OJI91168.1 aver|OJI96656.1 awen|OJJ37740.1 azon|OJJ45001.1

Group5365: abra|OJJ73499.1 acar|OOG00691.1 acla|EAW07433.1 afis|EAW22459.1 afla|EED54485.1 afum|EAL90733.1 agla|OJJ87564.1 akaw|GAA87448.1 aluc|OJZ88549.1 anid|CBF88916.1 anig|EHA18247.1 asyd|OJJ62843.1 ater|EAU39246.1 atub|OJI91134.1 aver|OJI96887.1 awen|OJJ35084.1 azon|OJJ50023.1

Group5366: abra|OJJ73501.1 acar|OOG00689.1 acla|EAW07434.1 afis|EAW22460.1 afla|EED54484.1 afum|EAL90731.1 agla|OJJ87562.1 akaw|GAA87449.1 aluc|OJZ88546.1 anid|CBF88914.1 anig|EHA18246.1 asyd|OJJ62842.1 ater|EAU39245.1 atub|OJI91131.1 aver|OJI96886.1 awen|OJJ35086.1 azon|OJJ50021.1

Group5367: abra|OJJ73504.1 acar|OOG00687.1 acla|EAW07437.1 afis|EAW22463.1 afla|EED54481.1 afum|EAL90728.1 agla|OJJ87560.1 aluc|OJZ88544.1 anid|CBF88909.1 anig|EHA18244.1 aory|EIT81141.1 asyd|OJJ62840.1 ater|EAU39243.1 atub|OJI91129.1 aver|OJI96883.1 awen|OJJ35089.1 azon|OJJ50019.1

Group5368: abra|OJJ73505.1 acla|EAW07438.1 afis|EAW22465.1 afla|EED54496.1 afum|EAL90726.1 agla|OJJ87559.1 akaw|GAA87452.1 aluc|OJZ88543.1 anid|CBF88907.1 anig|EHA18243.1 aory|EIT78760.1 asyd|OJJ62839.1 ater|EAU39242.1 atub|OJI91128.1 aver|OJI96882.1 awen|OJJ35090.1 azon|OJJ50018.1

Group5369: abra|OJJ73511.1 acla|EAW07444.1 afis|EAW22471.1 afla|EED54503.1 afum|EAL90720.1 agla|OJJ87553.1 akaw|GAA87459.1 aluc|OJZ88536.1 anid|CBF88898.1 anig|EHA18237.1 aory|EIT78805.1 asyd|OJJ62833.1 atub|OJI91121.1 aver|OJI96876.1 awen|OJJ35097.1 azon|OJJ50012.1 ater|EAU39237.1

Group5370: abra|OJJ73514.1 acar|OOG00702.1 acla|EAW07448.1 afis|EAW22475.1 afla|EED54507.1 afum|EAL90716.1 agla|OJJ87549.1 akaw|GAA87463.1 aluc|OJZ88533.1 anid|CBF88890.1 anig|EHA18234.1 asyd|OJJ62830.1 ater|EAU39234.1 atub|OJI91118.1 aver|OJI96873.1 awen|OJJ35100.1 azon|OJJ50008.1

Group5371: abra|OJJ73517.1 acar|OOG00707.1 acla|EAW07452.1 afis|EAW22484.1 afla|EED54510.1 afum|EAL90710.1 agla|OJJ87542.1 akaw|GAA87466.1 aluc|OJZ88530.1 anid|CBF88885.1 anig|EHA18231.1 asyd|OJJ62825.1 ater|EAU39233.1 atub|OJI91115.1 aver|OJI96870.1 awen|OJJ35103.1 azon|OJJ50004.1

Group5372: abra|OJJ73553.1 acar|OOG00725.1 acla|EAW07471.1 afis|EAW22504.1 afla|EED54531.1 afum|EAL90689.1 agla|OJJ80845.1 akaw|GAA88600.1 aluc|OJZ88508.1 anid|CBF88968.1 anig|EHA18215.1 asyd|OJJ62796.1 ater|EAU39214.1 atub|OJI91094.1 aver|OJI96843.1 awen|OJJ35128.1 azon|OJJ49972.1

Group5373: abra|OJJ73558.1 acar|OOG00729.1 acla|EAW07475.1 afis|EAW22507.1 afla|EED54535.1 afum|EAL90686.1 agla|OJJ87538.1 akaw|GAA88596.1 aluc|OJZ88504.1 anid|CBF88973.1 anig|EHA18211.1 asyd|OJJ62793.1 ater|EAU39206.1 atub|OJI91090.1 aver|OJI96840.1 awen|OJJ35132.1 azon|OJJ49966.1

Group5374: abra|OJJ73569.1 acar|OOG00751.1 acla|EAW07486.1 afis|EAW22519.1 afum|EAL90674.1 agla|OJJ80879.1 akaw|GAA88586.1 aluc|OJZ88491.1 anid|CBF89002.1 anig|EHA18200.1 aory|EIT78853.1 asyd|OJJ62780.1 ater|EAU39190.1 atub|OJI91078.1 aver|OJI96825.1 awen|OJJ35146.1 azon|OJJ49954.1

Group5375: abra|OJJ73577.1 acar|OOG00758.1 acla|EAW07495.1 afis|EAW22528.1 afla|EED54556.1 afum|EAL90666.2 agla|OJJ80888.1 aluc|OJZ88475.1 anid|CBF89012.1 anig|EHA18192.1 aory|EIT78898.1 asyd|OJJ62773.1 ater|EAU39183.1 atub|OJI91070.1 aver|OJI96818.1 awen|OJJ35155.1 azon|OJJ49946.1

Group5376: abra|OJJ73579.1 acar|OOG00760.1 acla|EAW07497.1 afis|EAW22530.1 afla|EED54558.1 afum|EAL90664.1 agla|OJJ80891.1 akaw|GAA88572.1 aluc|OJZ88473.1 anid|CBF89016.1 anig|EHA18190.1 aory|EIT81483.1 asyd|OJJ62771.1 ater|EAU39179.1 atub|OJI91068.1 aver|OJI96816.1 awen|OJJ35157.1

Group5377: abra|OJJ73584.1 acar|OOG00765.1 acla|EAW07502.1 afis|EAW22535.1 afla|EED54563.1 afum|EAL90659.1 agla|OJJ80896.1 akaw|GAA88567.1 aluc|OJZ88468.1 anid|CBF89027.1 anig|EHA18185.1 aory|EIT78861.1 asyd|OJJ62766.1 ater|EAU39174.1 atub|OJI91063.1 aver|OJI96811.1 awen|OJJ35162.1

Group5378: abra|OJJ73595.1 acar|OOG00780.1 acla|EAW07514.1 afis|EAW22545.1 afla|EED54576.1 afum|EAL90650.1 agla|OJJ80907.1 akaw|GAA88557.1 aluc|OJZ88455.1 anid|CBF89045.1 anig|EHA18176.1 asyd|OJJ62757.1 ater|EAU39163.1 atub|OJI91053.1 aver|OJI96799.1 awen|OJJ35174.1 azon|OJJ49933.1

Group5379: abra|OJJ73600.1 acla|EAW07518.1 afis|EAW22548.1 afla|EED54580.1 afum|EAL90647.1 agla|OJJ80910.1 akaw|GAA88552.1 aluc|OJZ88450.1 anid|CBF89053.1 anig|EHA18171.1 aory|EIT78826.1 asyd|OJJ62753.1 ater|EAU39157.1 atub|OJI91048.1 aver|OJI96795.1 awen|OJJ35179.1 azon|OJJ49929.1

Group5380: abra|OJJ73613.1 acar|OOG00796.1 afla|EED54591.1 agla|OJJ80920.1 akaw|GAA88540.1 aluc|OJZ88436.1 anid|CBF88375.1 anig|EHA18160.1 aory|EIT78809.1 atub|OJI91033.1 aver|OJI97207.1 azon|OJJ49916.1 afis|EAW22560.1 afum|EAL90635.1 awen|OJJ35190.1 acla|EAW07530.1 ater|EAU39144.1

Group5381: abra|OJJ73620.1 acar|OOG00803.1 acla|EAW07535.1 afis|EAW22569.1 afla|EED54599.1 afum|EAL90629.1 agla|OJJ80925.1 akaw|GAA88533.1 aluc|OJZ88429.1 anid|CBF88321.1 anig|EHA18153.1 aory|EIT78845.1 asyd|OJJ63704.1 ater|EAU39138.1 atub|OJI91026.1 aver|OJI97235.1 awen|OJJ35195.1

Group5382: abra|OJJ73623.1 acar|OOG00806.1 acla|EAW07538.1 afis|EAW22572.1 afum|EAL90626.1 agla|OJJ80929.1 akaw|GAA88530.1 aluc|OJZ88425.1 anid|CBF88327.1 anig|EHA18150.1 aory|EIT78792.1 asyd|OJJ63707.1 ater|EAU39135.1 atub|OJI91023.1 aver|OJI97232.1 awen|OJJ35199.1 azon|OJJ49909.1

Group5383: abra|OJJ73626.1 acar|OOG00808.1 acla|EAW07540.1 afis|EAW22574.1 afla|EED54603.1 afum|EAL90624.2 akaw|GAA88527.1 aluc|OJZ88423.1 anid|CBF88330.1 anig|EHA18148.1 aory|EIT78769.1 asyd|OJJ63710.1 ater|EAU39130.1 atub|OJI91019.1 aver|OJI97229.1 awen|OJJ35202.1 azon|OJJ49906.1

Group5384: abra|OJJ73628.1 acla|EAW07542.1 afis|EAW22576.1 afla|EED54615.1 afum|EAL90622.2 agla|OJJ80936.1 akaw|GAA88525.1 aluc|OJZ88421.1 anid|CBF88334.1 anig|EHA18146.1 aory|EIT78740.1 asyd|OJJ63712.1 ater|EAU39128.1 atub|OJI91017.1 aver|OJI97227.1 awen|OJJ35204.1 azon|OJJ49904.1

Group5385: abra|OJJ73630.1 acar|OOG00812.1 acla|EAW07544.1 afis|EAW22578.1 afla|EED54621.1 afum|EAL90620.2 agla|OJJ80938.1 akaw|GAA88523.1 aluc|OJZ88419.1 anid|CBF88337.1 anig|EHA18145.1 aory|EIT78944.1 asyd|OJJ63714.1 ater|EAU39126.1 atub|OJI91015.1 awen|OJJ35206.1 azon|OJJ49902.1

Group5386: abra|OJJ73642.1 acar|OOG00823.1 acla|EAW07555.1 afis|EAW22589.1 afla|EED54635.1 afum|EAL90610.1 agla|OJJ80949.1 akaw|GAA88513.1 aluc|OJZ88408.1 anid|CBF88362.1 anig|EHA18135.1 asyd|OJJ63725.1 ater|EAU39118.1 atub|OJI91004.1 aver|OJI97215.1 awen|OJJ35217.1 azon|OJJ49892.1

Group5387: abra|OJJ73664.1 acla|EAW07571.1 afis|EAW22605.1 afla|EED49515.1 afum|EAL90593.1 agla|OJJ80973.1 akaw|GAA88492.1 aluc|OJZ88388.1 anid|CBF88261.1 anig|EHA18115.1 aory|EIT82604.1 asyd|OJJ63675.1 ater|EAU39099.1 atub|OJI90983.1 aver|OJI97263.1 awen|OJJ35233.1 azon|OJJ49878.1

Group5388: abra|OJJ73677.1 acla|EAW07578.1 afis|EAW22616.1 afla|EED49824.1 afum|EAL90581.1 agla|OJJ80998.1 akaw|GAA88482.1 aluc|OJZ88372.1 anid|CBF88249.1 anig|EHA18105.1 aory|EIT77667.1 asyd|OJJ63668.1 ater|EAU39085.1 atub|OJI90970.1 aver|OJI97269.1 awen|OJJ35243.1 azon|OJJ49862.1

Group5389: abra|OJJ73679.1 acar|OOG00850.1 acla|EAW07594.1 afis|EAW22625.1 afla|EED49826.1 afum|EAL90571.1 agla|OJJ80996.1 akaw|GAA88480.1 aluc|OJZ88370.1 anid|CBF88245.1 anig|EHA18103.1 aory|EIT77671.1 asyd|OJJ63665.1 ater|EAU39084.1 atub|OJI90968.1 aver|OJI97270.1 awen|OJJ35245.1

Group5390: abra|OJJ73682.1 acar|OOG00852.1 acla|EAW07597.1 afis|EAW22628.1 afla|EED49877.1 afum|EAL90568.1 agla|OJJ80993.1 akaw|GAA88477.1 aluc|OJZ88367.1 anid|CBF88239.1 anig|EHA18100.1 asyd|OJJ63661.1 ater|EAU39080.1 atub|OJI90965.1 aver|OJI97273.1 awen|OJJ35249.1 azon|OJJ49857.1

Group5391: abra|OJJ73683.1 afis|EAW22629.1 afla|EED49878.1 afum|EAL90566.1 agla|OJJ80991.1 akaw|GAA88476.1 aluc|OJZ88366.1 anid|CBF88238.1 aory|EIT72447.1 asyd|OJJ63660.1 ater|EAU39079.1 atub|OJI90964.1 aver|OJI97274.1 awen|OJJ35250.1 azon|OJJ49856.1 acla|EAW07598.1 anig|EHA18098.1

Group5392: abra|OJJ73687.1 acar|OOG00859.1 afis|EAW22633.1 afla|EED49883.1 afum|EAL90560.1 agla|OJJ80986.1 akaw|GAA88472.1 aluc|OJZ88362.1 anid|CBF88230.1 anig|EHA18094.1 aory|EIT72448.1 asyd|OJJ63656.1 ater|EAU39075.1 atub|OJI90958.1 aver|OJI97278.1 awen|OJJ35255.1 acla|EAW07602.1

Group5393: abra|OJJ73691.1 acla|EAW07606.1 afis|EAW22636.1 afla|EED49887.1 afum|EAL90556.2 agla|OJJ80982.1 akaw|GAA88469.1 aluc|OJZ88358.1 anid|CBF88223.1 anig|EHA18091.1 aory|EIT72468.1 asyd|OJJ63653.1 ater|EAU39070.1 atub|OJI90953.1 aver|OJI97281.1 awen|OJJ35259.1 azon|OJJ49848.1

Group5394: abra|OJJ73721.1 acar|OOG00887.1 acla|EAW12122.1 afis|EAW17258.1 afla|EED45960.1 afum|EAL86675.1 agla|OJJ83737.1 akaw|GAA88441.1 aluc|OJZ88329.1 anid|CBF88172.1 anig|EHA18064.1 aory|EIT75555.1 asyd|OJJ63623.1 ater|EAU32152.1 atub|OJI90924.1 aver|OJI97313.1 awen|OJJ37054.1

Group5395: abra|OJJ73730.1 acar|OOG00895.1 acla|EAW07614.1 afis|EAW22644.1 afla|EED49913.1 afum|EAL90548.1 agla|OJJ81026.1 aluc|OJZ88320.1 anid|CBF88177.1 anig|EHA18056.1 aory|EIT72270.1 asyd|OJJ63626.1 ater|EAU39063.1 atub|OJI90915.1 aver|OJI97310.1 awen|OJJ35280.1 azon|OJJ49844.1

Group5396: abra|OJJ73738.1 acar|OOG00903.1 acla|EAW07622.1 afis|EAW22651.1 afla|EED49920.1 afum|EAL90541.1 agla|OJJ81034.1 akaw|GAA88426.1 aluc|OJZ88313.1 anid|CBF88192.1 anig|EHA18050.1 asyd|OJJ63633.1 ater|EAU39056.1 atub|OJI90908.1 aver|OJI97303.1 awen|OJJ35272.1 azon|OJJ49835.1

Group5397: abra|OJJ73765.1 acla|EAW07651.1 afis|EAW22669.1 afla|EED49946.1 afum|EAL90523.1 agla|OJJ79052.1 akaw|GAA88409.1 aluc|OJZ88291.1 anid|CBF88153.1 anig|EHA18032.1 aory|EIT83448.1 asyd|OJJ63614.1 ater|EAU39039.1 atub|OJI90888.1 aver|OJI97324.1 awen|OJJ35297.1 azon|OJJ49819.1

Group5398: abra|OJJ73772.1 acar|OOG00930.1 acla|EAW07656.1 afis|EAW22674.1 afla|EED49955.1 afum|EAL90518.1 agla|OJJ79043.1 akaw|GAA88404.1 aluc|OJZ88284.1 anid|CBF88144.1 anig|EHA18027.1 aory|EIT83450.1 asyd|OJJ63600.1 atub|OJI90882.1 aver|OJI97337.1 awen|OJJ35303.1 azon|OJJ49813.1

Group5399: abra|OJJ73780.1 acar|OOG00936.1 acla|EAW07664.1 afis|EAW22681.1 afla|EED49962.1 afum|EAL90511.1 akaw|GAA88397.1 aluc|OJZ88275.1 anid|CBF88122.1 anig|EHA18020.1 aory|EIT83441.1 asyd|OJJ63586.1 ater|EAU39028.1 atub|OJI90875.1 aver|OJI97348.1 awen|OJJ35309.1 azon|OJJ49802.1

Group5400: abra|OJJ73796.1 acar|OOG00951.1 acla|EAW07687.1 afis|EAW22702.1 afum|EAL90490.1 agla|OJJ79007.1 akaw|GAA89414.1 aluc|OJZ88259.1 anid|CBF88076.1 anig|EHA18004.1 aory|EIT74750.1 asyd|OJJ63556.1 ater|EAU39000.1 atub|OJI90858.1 aver|OJI97378.1 awen|OJJ35339.1 azon|OJJ49779.1

Group5401: abra|OJJ73799.1 acar|OOG00948.1 acla|EAW07693.1 afis|EAW22708.1 afla|EED49994.1 afum|EAL90484.1 agla|OJJ79002.1 akaw|GAA89417.1 aluc|OJZ88256.1 anid|CBF88066.1 aory|EIT74634.1 asyd|OJJ63550.1 ater|EAU38994.1 atub|OJI90854.1 aver|OJI97384.1 awen|OJJ35345.1 azon|OJJ49772.1

Group5402: abra|OJJ73814.1 acla|EAW07701.1 afis|EAW22718.1 afla|EED50004.1 afum|EAL90476.1 agla|OJJ83685.1 akaw|GAA89433.1 aluc|OJZ88236.1 anid|CBF88048.1 anig|EHA17983.1 aory|EIT74712.1 asyd|OJJ63538.1 ater|EAU38985.1 atub|OJI90838.1 aver|OJI97395.1 awen|OJJ35412.1 azon|OJJ49764.1

Group5403: abra|OJJ73815.1 acar|OOG00969.1 acla|EAW07702.1 afis|EAW22719.1 afla|EED50005.1 afum|EAL90475.1 agla|OJJ83684.1 akaw|GAA89434.1 aluc|OJZ88235.1 anid|CBF88040.1 anig|EHA17982.1 asyd|OJJ63537.1 ater|EAU38984.1 atub|OJI90837.1 aver|OJI97396.1 awen|OJJ35413.1 azon|OJJ49763.1

Group5404: abra|OJJ73817.1 acar|OOG00971.1 acla|EAW07706.1 afis|EAW22723.1 afla|EED50008.1 afum|EAL90473.1 agla|OJJ83680.1 akaw|GAA89436.1 aluc|OJZ88233.1 anid|CBF88032.1 anig|EHA17979.1 aory|EIT74623.1 asyd|OJJ63533.1 ater|EAU38982.1 atub|OJI90834.1 aver|OJI97399.1 awen|OJJ35417.1

Group5405: abra|OJJ73818.1 acla|EAW07707.1 afis|EAW22724.1 afla|EED50009.1 afum|EAL90472.1 agla|OJJ83679.1 akaw|GAA89437.1 aluc|OJZ88232.1 anid|CBF88031.1 anig|EHA17978.1 aory|EIT74693.1 ater|EAU38981.1 atub|OJI90833.1 awen|OJJ35418.1 azon|OJJ49759.1 asyd|OJJ63532.1 aver|OJI97400.1

Group5406: abra|OJJ73820.1 acla|EAW07709.1 afis|EAW22725.1 afla|EED50011.1 afum|EAL90471.1 agla|OJJ83677.1 akaw|GAA89439.1 aluc|OJZ88229.1 anid|CBF88029.1 anig|EHA17976.1 aory|EIT74724.1 asyd|OJJ63531.1 ater|EAU38979.1 atub|OJI90831.1 aver|OJI97401.1 awen|OJJ35420.1 azon|OJJ49757.1

Group5407: abra|OJJ73826.1 acar|OOG00978.1 acla|EAW07721.1 afis|EAW22753.1 afum|EAL90446.2 agla|OJJ83672.1 akaw|GAA89444.1 aluc|OJZ88223.1 anid|CBF88020.1 anig|EHA17970.1 aory|EIT74674.1 asyd|OJJ63523.1 ater|EAU38967.1 atub|OJI90823.1 aver|OJI97407.1 awen|OJJ35425.1 azon|OJJ49751.1

Group5408: abra|OJJ73828.1 acar|OOG00980.1 acla|EAW07723.1 afis|EAW22755.1 afla|EED50016.1 afum|EAL90444.2 agla|OJJ83670.1 akaw|GAA89446.1 aluc|OJZ88221.1 anig|EHA17968.1 aory|EIT74646.1 asyd|OJJ63521.1 ater|EAU38965.1 atub|OJI90821.1 aver|OJI97409.1 awen|OJJ35427.1 azon|OJJ49749.1

Group5409: abra|OJJ73829.1 acla|EAW07724.1 afis|EAW22756.1 afla|EED50017.1 afum|EAL90443.1 agla|OJJ83667.1 akaw|GAA89447.1 aluc|OJZ88220.1 anid|CBF88017.1 aory|EIT74727.1 asyd|OJJ63520.1 ater|EAU38963.1 atub|OJI90820.1 aver|OJI97410.1 awen|OJJ35428.1 azon|OJJ49748.1 anig|EHA17967.1

Group5410: abra|OJJ73845.1 acar|OOG00996.1 acla|EAW07740.1 afis|EAW22776.1 afla|EED50036.1 afum|EAL90422.1 agla|OJJ83612.1 akaw|GAA89463.1 aluc|OJZ88203.1 anid|CBF87989.1 anig|EHA17951.1 aory|EIT74654.1 asyd|OJJ63501.1 atub|OJI90803.1 aver|OJI97429.1 awen|OJJ38961.1 azon|OJJ49732.1

Group5411: abra|OJJ73858.1 acla|EAW07749.1 afis|EAW22786.1 afla|EED50049.1 afum|EAL90413.1 agla|OJJ83591.1 akaw|GAA89483.1 aluc|OJZ88179.1 anid|CBF87967.1 anig|EHA17931.1 aory|EIT74754.1 asyd|OJJ63487.1 ater|EAU38924.1 atub|OJI90778.1 aver|OJI97442.1 awen|OJJ38949.1 azon|OJJ49721.1

Group5412: abra|OJJ73860.1 acar|OOG01008.1 acla|EAW07750.1 afis|EAW22787.1 afla|EED50051.1 afum|EAL90412.1 agla|OJJ83589.1 akaw|GAA89482.1 aluc|OJZ88181.1 anid|CBF87965.1 anig|EHA17933.1 asyd|OJJ63485.1 ater|EAU38923.1 atub|OJI90780.1 aver|OJI97444.1 awen|OJJ38947.1 azon|OJJ49719.1

Group5413: abra|OJJ73863.1 acar|OOF89915.1 acla|EAW07753.1 afis|EAW22790.1 afla|EED50057.1 afum|EAL90409.1 agla|OJJ83586.1 akaw|GAA89479.1 aluc|OJZ88184.1 anid|CBF87959.1 anig|EHA17935.1 aory|EIT74648.1 asyd|OJJ63482.1 atub|OJI90783.1 aver|OJI97448.1 awen|OJJ38944.1 azon|OJJ49717.1

Group5414: abra|OJJ73875.1 acla|EAW07760.1 afis|EAW22796.1 afla|EED50065.1 afum|EAL90403.1 agla|OJJ83573.1 akaw|GAA89487.1 aluc|OJZ88174.1 anid|CBF87948.1 anig|EHA17927.1 aory|EIT74629.1 asyd|OJJ63476.1 ater|EAU38914.1 atub|OJI90772.1 aver|OJI97454.1 awen|OJJ38933.1 azon|OJJ49709.1

Group5415: abra|OJJ73879.1 acar|OOG01019.1 acla|EAW07762.1 afis|EAW22798.1 afla|EED50068.1 afum|EAL90401.1 agla|OJJ83575.1 akaw|GAA89491.1 aluc|OJZ88170.1 anid|CBF87937.1 anig|EHA17924.1 aory|EIT74758.1 asyd|OJJ63467.1 atub|OJI90768.1 aver|OJI97465.1 awen|OJJ38931.1 azon|OJJ49707.1

Group5416: abra|OJJ73885.1 acla|EAW07767.1 afis|EAW22803.1 afla|EED50073.1 afum|EAL90396.1 agla|OJJ83580.1 akaw|GAA89496.1 aluc|OJZ88165.1 anid|CBF87928.1 anig|EHA17919.1 aory|EIT74628.1 asyd|OJJ63462.1 ater|EAU38906.1 atub|OJI90763.1 aver|OJI97470.1 awen|OJJ38925.1 azon|OJJ49702.1

Group5417: abra|OJJ73895.1 acla|EAW07777.1 afis|EAW22813.1 afla|EED50086.1 afum|EAL90388.2 agla|OJJ83556.1 akaw|GAA89505.1 aluc|OJZ88152.1 anid|CBF87911.1 anig|EHA17912.1 aory|EIT74666.1 asyd|OJJ63450.1 ater|EAU38897.1 atub|OJI90754.1 aver|OJI97484.1 awen|OJJ38918.1 azon|OJJ49695.1

Group5418: abra|OJJ73903.1 acla|EAW07782.1 afis|EAW22818.1 afla|EED50091.1 afum|EAL90383.1 akaw|GAA89510.1 aluc|OJZ88143.1 anig|EHA17906.1 aory|EIT74676.1 ater|EAU38892.1 atub|OJI90747.1 awen|OJJ38909.1 azon|OJJ49690.1 anid|CBF87900.1 asyd|OJJ63445.1 aver|OJI97491.1 agla|OJJ83520.1

Group5419: abra|OJJ73914.1 acar|OOG01046.1 acla|EAW07790.1 afis|EAW22826.1 afla|EED50102.1 afum|EAL90374.2 agla|OJJ83511.1 akaw|GAA89520.1 aluc|OJZ88134.1 anid|CBF87885.1 anig|EHA17898.1 aory|EIT74725.1 asyd|OJJ63436.1 ater|EAU38883.1 aver|OJI97501.1 awen|OJJ38900.1 azon|OJJ49681.1

Group5420: abra|OJJ73929.1 acla|EAW07806.1 afis|EAW22843.1 afla|EED50118.1 afum|EAL90360.1 agla|OJJ83495.1 akaw|GAA89536.1 aluc|OJZ88114.1 anid|CBF87856.1 anig|EHA17884.1 aory|EIT74743.1 asyd|OJJ63422.1 ater|EAU38867.1 atub|OJI90723.1 aver|OJI97515.1 awen|OJJ38880.1 azon|OJJ49669.1

Group5421: abra|OJJ73950.1 acar|OOG01078.1 acla|EAW07825.1 afis|EAW22861.1 afla|EED50144.1 afum|EAL90339.1 agla|OJJ83463.1 akaw|GAA89555.1 aluc|OJZ88094.1 anid|CBF84626.1 anig|EHA17864.1 aory|EIT74618.1 asyd|OJJ60002.1 atub|OJI90702.1 aver|OJJ02289.1 awen|OJJ38859.1 agla|OJJ78756.1

Group5422: abra|OJJ73951.1 acar|OOG01079.1 acla|EAW07824.1 afis|EAW22860.1 afla|EED50141.1 afum|EAL90340.1 agla|OJJ83464.1 akaw|GAA89556.1 aluc|OJZ88093.1 anid|CBF84628.1 anig|EHA17863.1 aory|EIT74753.1 asyd|OJJ60001.1 atub|OJI90701.1 aver|OJJ02290.1 awen|OJJ38860.1 agla|OJJ78752.1

Group5423: abra|OJJ73955.1 acar|OOG01082.1 acla|EAW14167.1 afis|EAW18881.1 afla|EED50885.1 afum|EAL93613.1 agla|OJJ88981.1 akaw|GAA89560.1 aluc|OJZ88088.1 anid|CBF80630.1 anig|EHA17859.1 aory|EIT77121.1 asyd|OJJ54635.1 ater|EAU38049.1 atub|OJI90697.1 aver|OJJ07521.1 afla|EED47797.1

Group5424: abra|OJJ73958.1 acar|OOG01085.1 acla|EAW07827.1 afis|EAW22863.1 afla|EED50147.1 afum|EAL90337.1 agla|OJJ83461.1 akaw|GAA89561.1 aluc|OJZ88087.1 anid|CBF87830.1 anig|EHA17858.1 aory|EIT74683.1 asyd|OJJ63406.1 ater|EAU38823.1 atub|OJI90696.1 awen|OJJ38857.1 azon|OJJ49657.1

Group5425: abra|OJJ73960.1 acar|OOG01088.1 acla|EAW07829.1 afis|EAW22865.1 afla|EED50149.1 afum|EAL90334.1 agla|OJJ83459.1 akaw|GAA89563.1 aluc|OJZ88085.1 anig|EHA17856.1 aory|EIT74687.1 asyd|OJJ63404.1 ater|EAU38825.1 atub|OJI90694.1 aver|OJI97532.1 awen|OJJ38855.1 azon|OJJ49653.1

Group5426: abra|OJJ73965.1 acar|OOG01091.1 acla|EAW07835.1 afis|EAW22872.1 afla|EED50155.1 afum|EAL90326.1 agla|OJJ83451.1 akaw|GAA89568.1 aluc|OJZ88079.1 anid|CBF87819.1 anig|EHA17851.1 asyd|OJJ63398.1 ater|EAU38836.1 atub|OJI90688.1 aver|OJI97538.1 awen|OJJ38850.1 azon|OJJ49646.1

Group5427: abra|OJJ73966.1 acar|OOG01092.1 acla|EAW07836.1 afis|EAW22873.1 afla|EED50156.1 afum|EAL90325.2 agla|OJJ83450.1 akaw|GAA89569.1 aluc|OJZ88078.1 anid|CBF87817.1 anig|EHA17850.1 asyd|OJJ63396.1 ater|EAU38837.1 atub|OJI90687.1 aver|OJI97540.1 awen|OJJ38849.1 azon|OJJ49645.1

Group5428: abra|OJJ73981.1 acar|OOG01108.1 acla|EAW07853.1 afis|EAW22889.1 afla|EED50170.1 afum|EAL90309.1 agla|OJJ83421.1 akaw|GAA89585.1 aluc|OJZ88063.1 anid|CBF87794.1 anig|EHA17836.1 asyd|OJJ63379.1 ater|EAU35984.1 atub|OJI90672.1 aver|OJI97556.1 awen|OJJ38833.1 azon|OJJ49631.1

Group5429: abra|OJJ73984.1 acla|EAW07856.1 afis|EAW22892.1 afla|EED54861.1 afum|EAL90306.1 agla|OJJ83409.1 akaw|GAA89588.1 aluc|OJZ88060.1 anid|CBF87788.1 anig|EHA17833.1 aory|EIT79389.1 asyd|OJJ63375.1 ater|EAU35991.1 atub|OJI90668.1 aver|OJI97571.1 awen|OJJ38828.1 azon|OJJ49626.1

Group5430: abra|OJJ74014.1 acla|EAW07869.1 afis|EAW22904.1 afla|EED54896.1 afum|EAL90293.1 agla|OJJ83386.1 akaw|GAA87497.1 aluc|OJZ88025.1 anid|CBF87715.1 anig|EHA17802.1 aory|EIT79432.1 asyd|OJJ63339.1 ater|EAU36035.1 atub|OJI90633.1 aver|OJI97622.1 awen|OJJ38726.1 azon|OJJ49612.1

Group5431: abra|OJJ74042.1 acar|OOG01160.1 acla|EAW07891.1 afis|EAW22926.1 afla|EED54923.1 afum|EAL90271.1 agla|OJJ83362.1 akaw|GAA87524.1 aluc|OJZ87995.1 anid|CBF87673.1 anig|EHA17777.1 asyd|OJJ63312.1 ater|EAU30697.1 atub|OJI90604.1 aver|OJI97647.1 awen|OJJ38638.1 azon|OJJ49580.1

Group5432: abra|OJJ74046.1 acar|OOG01163.1 acla|EAW07896.1 afis|EAW22931.1 afla|EED58124.1 afum|EAL90266.1 agla|OJJ83357.1 akaw|GAA87529.1 aluc|OJZ87989.1 anid|CBF87666.1 anig|EHA17772.1 asyd|OJJ63307.1 ater|EAU30693.1 atub|OJI90599.1 aver|OJI97652.1 awen|OJJ38633.1 azon|OJJ49575.1

Group5433: abra|OJJ74051.1 acar|OOF89807.1 acla|EAW07900.1 afis|EAW22936.1 afla|EED58129.1 afum|EAL90260.1 agla|OJJ83352.1 akaw|GAA87533.1 aluc|OJZ87984.1 anid|CBF87657.1 anig|EHA17767.1 aory|EIT79148.1 asyd|OJJ63302.1 atub|OJI90594.1 aver|OJI97657.1 awen|OJJ38628.1 azon|OJJ49570.1

Group5434: abra|OJJ74074.1 acar|OOG01186.1 acla|EAW07919.1 afis|EAW22957.1 afla|EED58139.1 afum|EAL90240.1 agla|OJJ83331.1 akaw|GAA87555.1 aluc|OJZ87962.1 anig|EHA17746.1 aory|EIT79129.1 asyd|OJJ63282.1 ater|EAU30668.1 atub|OJI90572.1 aver|OJI97676.1 awen|OJJ38606.1 azon|OJJ49548.1

Group5435: abra|OJJ74077.1 acar|OOG01189.1 acla|EAW07922.1 afis|EAW22960.1 afla|EED58136.1 afum|EAL90237.1 agla|OJJ83328.1 akaw|GAA87558.1 aluc|OJZ87959.1 anig|EHA17743.1 aory|EIT79122.1 asyd|OJJ63279.1 ater|EAU30665.1 atub|OJI90569.1 aver|OJI97679.1 awen|OJJ38603.1 azon|OJJ49545.1

Group5436: abra|OJJ74098.1 acar|OOG01207.1 acla|EAW07941.1 afis|EAW22979.1 afla|EED58172.1 agla|OJJ83308.1 akaw|GAA87576.1 aluc|OJZ87938.1 anid|CBF87592.1 anig|EHA17726.1 aory|EIT79098.1 asyd|OJJ63260.1 ater|EAU30642.1 atub|OJI90550.1 aver|OJI97697.1 awen|OJJ38584.1 azon|OJJ49526.1

Group5437: abra|OJJ74099.1 acar|OOG01208.1 acla|EAW07942.1 afis|EAW22980.1 afla|EED58173.1 afum|EAL90217.1 agla|OJJ83307.1 akaw|GAA87577.1 aluc|OJZ87937.1 anid|CBF87590.1 anig|EHA17725.1 asyd|OJJ63259.1 ater|EAU30641.1 atub|OJI90549.1 aver|OJI97698.1 awen|OJJ38583.1 azon|OJJ49525.1

Group5438: abra|OJJ74105.1 acla|EAW07950.1 afis|EAW22987.1 afla|EED58179.1 afum|EAL90212.1 agla|OJJ83300.1 akaw|GAA87583.1 aluc|OJZ87931.1 anid|CBF87581.1 anig|EHA17720.1 aory|EIT79066.1 asyd|OJJ63253.1 ater|EAU30636.1 atub|OJI90543.1 aver|OJI97705.1 awen|OJJ38577.1 azon|OJJ49519.1

Group5439: abra|OJJ74106.1 acar|OOG01215.1 acla|EAW07951.1 afis|EAW22988.1 afla|EED58180.1 afum|EAL90211.1 agla|OJJ83298.1 aluc|OJZ87930.1 anid|CBF75423.1 anig|EHA17719.1 aory|EIT79102.1 asyd|OJJ59794.1 ater|EAU30635.1 atub|OJI90542.1 aver|OJJ02493.1 awen|OJJ38576.1 azon|OJJ49518.1

Group5440: abra|OJJ74111.1 acar|OOF89999.1 acla|EAW07954.1 afis|EAW22991.1 afla|EED58183.1 afum|EAL90209.1 agla|OJJ83295.1 akaw|GAA87586.1 aluc|OJZ87924.1 anid|CBF87578.1 anig|EHA17716.1 aory|EIT79107.1 asyd|OJJ63249.1 ater|EAU30631.1 atub|OJI90536.1 aver|OJI97708.1 awen|OJJ38573.1

Group5441: abra|OJJ74114.1 acar|OOG01219.1 acla|EAW07956.1 afis|EAW22992.1 afla|EED58184.1 afum|EBA27400.1 agla|OJJ83293.1 akaw|GAA87588.1 aluc|OJZ87920.1 anid|CBF87574.1 anig|EHA17714.1 asyd|OJJ63246.1 ater|EAU30628.1 atub|OJI90531.1 aver|OJI97710.1 awen|OJJ38570.1 azon|OJJ42000.1

Group5442: abra|OJJ74131.1 acar|OOF95603.1 acla|EAW11690.1 afis|EAW23706.1 afla|EED56137.1 afum|EAL87150.1 agla|OJJ78978.1 akaw|GAA89755.1 aluc|OJZ80628.1 anid|CBF84383.1 anig|EHA27203.1 aory|EIT76571.1 asyd|OJJ58682.1 ater|EAU36843.1 atub|OJI87484.1 aver|OJJ01294.1 awen|OJJ30914.1

Group5443: abra|OJJ74154.1 acar|OOF95571.1 acla|EAW12278.1 afis|EAW16962.1 afla|EED54137.1 afum|EAL86489.1 agla|OJJ88175.1 akaw|GAA89729.1 aluc|OJZ80654.1 anid|CBF89392.1 anig|EHA27178.1 aory|EIT75949.1 asyd|OJJ62463.1 ater|EAU34050.1 atub|OJI87509.1 aver|OJI96598.1 awen|OJJ40679.1

Group5444: abra|OJJ74174.1 acar|OOF95554.1 acla|EAW10815.1 afis|EAW24759.1 afla|EED48096.1 afum|EAL89401.1 agla|OJJ79193.1 akaw|GAA86842.1 aluc|OJZ89961.1 anid|CBF89394.1 anig|EHA27161.1 aory|EIT76187.1 asyd|OJJ62462.1 ater|EAU34013.1 atub|OJI84844.1 aver|OJI96597.1 awen|OJJ30980.1

Group5445: abra|OJJ74205.1 acar|OOF95515.1 acla|EAW07258.1 afis|EAW22279.1 afla|EED57672.1 afum|EAL90902.1 agla|OJJ87774.1 akaw|GAA86813.1 aluc|OJZ89930.1 anid|CBF88546.1 anig|EHA27130.1 asyd|OJJ63826.1 ater|EAU34289.1 atub|OJI84876.1 aver|OJI97112.1 awen|OJJ34852.1 azon|OJJ45695.1

Group5446: abra|OJJ74216.1 acar|OOF95505.1 acla|EAW07268.1 afis|EAW22289.1 afla|EED57688.1 afum|EAL90892.1 agla|OJJ87760.1 akaw|GAA86802.1 aluc|OJZ89918.1 anid|CBF88566.1 anig|EHA27118.1 asyd|OJJ63836.1 ater|EAU34301.1 atub|OJI84887.1 aver|OJI97101.1 awen|OJJ34862.1 azon|OJJ45666.1

Group5447: abra|OJJ74219.1 acar|OOF90432.1 acla|EAW07272.1 afis|EAW22293.1 afla|EED57691.1 afum|EAL90888.1 agla|OJJ87757.1 akaw|GAA86799.1 aluc|OJZ89915.1 anid|CBF88571.1 anig|EHA27115.1 asyd|OJJ63838.1 ater|EAU34304.1 atub|OJI84890.1 aver|OJI97099.1 awen|OJJ34865.1 azon|OJJ45664.1

Group5448: abra|OJJ74222.1 acla|EAW07283.1 afis|EAW22314.1 afla|EED57696.1 afum|EAL90876.1 agla|OJJ87736.1 akaw|GAA86797.1 aluc|OJZ89912.1 anid|CBF88579.1 anig|EHA27112.1 aory|EIT79872.1 asyd|OJJ63842.1 ater|EAU34309.1 atub|OJI84893.1 aver|OJI97095.1 awen|OJJ34870.1 azon|OJJ50046.1

Group5449: abra|OJJ74227.1 acar|OOF95499.1 acla|EAW07278.1 afis|EAW22309.1 afla|EED57703.1 afum|EAL90881.1 agla|OJJ87741.1 akaw|GAA86792.1 aluc|OJZ89906.1 anid|CBF88588.1 anig|EHA27106.1 aory|EIT79911.1 asyd|OJJ63847.1 ater|EAU34315.1 atub|OJI84900.1 aver|OJI97089.1 awen|OJJ34878.1

Group5450: abra|OJJ74230.1 acar|OOF95496.1 acla|EAW07274.1 afis|EAW22295.1 afla|EED57708.1 afum|EAL90886.1 agla|OJJ87747.1 akaw|GAA86789.1 aluc|OJZ89903.1 anid|CBF88596.1 anig|EHA27103.1 asyd|OJJ63852.1 ater|EAU34319.1 atub|OJI84903.1 aver|OJI97085.1 awen|OJJ34883.1 azon|OJJ50051.1

Group5451: abra|OJJ74238.1 acar|OOF95493.1 acla|EAW07289.1 afis|EAW22322.1 afla|EED57719.1 afum|EAL90869.1 agla|OJJ87750.1 akaw|GAA86782.1 aluc|OJZ89894.1 anid|CBF88603.1 anig|EHA27093.1 aory|EIT79885.1 asyd|OJJ63856.1 atub|OJI84913.1 aver|OJI97082.1 awen|OJJ34889.1 azon|OJJ50056.1

Group5452: abra|OJJ74243.1 acar|OOF95487.1 acla|EAW07292.1 afis|EAW22326.1 afla|EED57725.1 afum|EAL90866.1 agla|OJJ87722.1 akaw|GAA86777.1 aluc|OJZ89889.1 anid|CBF88611.1 anig|EHA27088.1 aory|EIT79882.1 asyd|OJJ63860.1 ater|EAU34325.1 atub|OJI84918.1 aver|OJI97079.1 awen|OJJ34895.1

Group5453: abra|OJJ74247.1 acar|OOF95482.1 acla|EAW07301.1 afis|EAW22344.1 afla|EED57737.1 afum|EAL90847.1 agla|OJJ87690.1 aluc|OJZ89885.1 anid|CBF88625.1 anig|EHA27084.1 aory|EIT79933.1 asyd|OJJ63869.1 ater|EAU39482.1 atub|OJI84922.1 aver|OJI97070.1 awen|OJJ34923.1 azon|OJJ50071.1

Group5454: abra|OJJ74251.1 acar|OOF95478.1 acla|EAW07295.1 afis|EAW22329.1 afla|EED57728.1 afum|EAL90863.1 agla|OJJ87754.1 akaw|GAA86770.1 aluc|OJZ89881.1 anid|CBF88617.1 anig|EHA27080.1 aory|EIT79902.1 asyd|OJJ63863.1 atub|OJI84926.1 aver|OJI97076.1 awen|OJJ34892.1 azon|OJJ50061.1

Group5455: abra|OJJ74253.1 acar|OOF95476.1 acla|EAW07297.1 afis|EAW22348.1 afla|EED57730.1 afum|EAL90844.1 agla|OJJ87751.1 akaw|GAA86768.1 aluc|OJZ89879.1 anid|CBF88621.1 anig|EHA27078.1 aory|EIT79888.1 asyd|OJJ63865.1 ater|EAU39456.1 atub|OJI84928.1 aver|OJI97074.1 azon|OJJ50059.1

Group5456: abra|OJJ74259.1 acla|EAW07304.1 afis|EAW22341.1 afla|EED57740.1 afum|EAL90851.1 agla|OJJ87694.1 akaw|GAA86762.1 aluc|OJZ89872.1 anid|CBF88630.1 anig|EHA27072.1 aory|EIT79897.1 asyd|OJJ63873.1 ater|EAU39479.1 atub|OJI84935.1 aver|OJI97065.1 awen|OJJ34920.1 azon|OJJ50074.1

Group5457: abra|OJJ74287.1 acar|OOF95444.1 acla|EAW07331.1 afis|EAW22363.1 afla|EED57771.1 afum|EAL90828.1 agla|OJJ87612.1 akaw|GAA86737.1 aluc|OJZ89846.1 anig|EHA27047.1 aory|EIT81068.1 asyd|OJJ63905.1 ater|EAU39497.1 atub|OJI84960.1 aver|OJI97036.1 awen|OJJ34937.1 azon|OJJ45589.1

Group5458: abra|OJJ74288.1 acar|OOF95443.1 acla|EAW07332.1 afis|EAW22364.1 afla|EED57772.1 afum|EAL90827.1 akaw|GAA86735.1 aluc|OJZ89845.1 anid|CBF88688.1 anig|EHA27046.1 aory|EIT81112.1 asyd|OJJ63906.1 ater|EAU39498.1 atub|OJI84961.1 aver|OJI97035.1 awen|OJJ34938.1 azon|OJJ45590.1

Group5459: abra|OJJ74308.1 acla|EAW07356.1 afis|EAW22385.1 afla|EED56045.1 afum|EAL90805.1 agla|OJJ87646.1 akaw|GAA86716.1 aluc|OJZ89819.1 anid|CBF88727.1 anig|EHA27027.1 aory|EIT76587.1 asyd|OJJ63932.1 ater|EAU39336.1 atub|OJI84987.1 aver|OJI97011.1 awen|OJJ34960.1 azon|OJJ45612.1

Group5460: abra|OJJ74313.1 acar|OOF95415.1 acla|EAW07364.1 afis|EAW22391.1 afla|EED56054.1 afum|EBA27413.1 agla|OJJ87653.1 aluc|OJZ89813.1 anid|CBF88739.1 anig|EHA27023.1 aory|EIT76735.1 asyd|OJJ63952.1 ater|EAU39327.1 atub|OJI84992.1 aver|OJI96989.1 awen|OJJ34968.1 azon|OJJ45617.1

Group5461: abra|OJJ74321.1 acla|EAW07373.1 afis|EAW22398.1 afla|EED56062.1 afum|EAL90793.1 agla|OJJ87661.1 akaw|GAA86705.1 aluc|OJZ89807.1 anid|CBF88752.1 anig|EHA27017.1 aory|EIT76767.1 asyd|OJJ63959.1 ater|EAU39315.1 atub|OJI84999.1 aver|OJI96982.1 awen|OJJ34976.1 azon|OJJ45625.1

Group5462: abra|OJJ74323.1 acar|OOF95407.1 acla|EAW07375.1 afis|EAW22400.1 afla|EED56064.1 afum|EAL90791.1 agla|OJJ87663.1 akaw|GAA86703.1 aluc|OJZ89805.1 anid|CBF88768.1 aory|EIT76742.1 asyd|OJJ63979.1 ater|EAU39307.1 atub|OJI85001.1 aver|OJI96974.1 awen|OJJ34979.1 azon|OJJ45627.1

Group5463: abra|OJJ74330.1 acar|OOF95401.1 acla|EAW07381.1 afis|EAW22406.1 afla|EED56070.1 afum|EAL90786.1 agla|OJJ87669.1 akaw|GAA86698.1 aluc|OJZ89799.1 anid|CBF88789.1 anig|EHA27012.1 asyd|OJJ62901.1 ater|EAU39303.1 atub|OJI85006.1 aver|OJI96955.1 awen|OJJ34985.1 azon|OJJ42755.1

Group5464: abra|OJJ74334.1 acar|OOF95398.1 acla|EAW07384.1 afis|EAW22409.1 afla|EED52489.1 afum|EAL90783.1 agla|OJJ87677.1 akaw|GAA86695.1 aluc|OJZ89795.1 anid|CBF82129.1 anig|EHA27009.1 aory|EIT82766.1 asyd|OJJ54297.1 ater|EAU39299.1 atub|OJI85010.1 aver|OJJ05252.1 awen|OJJ34989.1

Group5465: abra|OJJ74350.1 acar|OOF95379.1 acla|EAW07404.1 afis|EAW22430.1 afla|EED52508.1 agla|OJJ87593.1 akaw|GAA86680.1 aluc|OJZ89780.1 anid|CBF88826.1 anig|EHA26995.1 asyd|OJJ62883.1 ater|EAU39277.1 atub|OJI85026.1 aver|OJI96935.1 awen|OJJ35009.1 azon|OJJ45650.1 afum|EAL90761.1

Group5466: abra|OJJ74354.1 acar|OOF95376.1 acla|EAW07407.1 afis|EAW22433.1 afla|EED52514.1 afum|EAL90758.1 agla|OJJ87588.1 akaw|GAA86677.1 aluc|OJZ89774.1 anid|CBF88834.1 anig|EHA26992.1 asyd|OJJ62879.1 ater|EAU39274.1 atub|OJI85029.1 aver|OJI96931.1 awen|OJJ35012.1 azon|OJJ45653.1

Group5467: abra|OJJ74360.1 acla|EAW07411.1 afis|EAW22437.1 afla|EED54447.1 afum|EAL90754.1 agla|OJJ87585.1 akaw|GAA86671.1 aluc|OJZ89768.1 anid|CBF88866.1 anig|EHA26988.1 aory|EIT81146.1 asyd|OJJ62876.1 ater|EAU39271.1 atub|OJI85035.1 aver|OJI96928.1 awen|OJJ35015.1 azon|OJJ45660.1

Group5468: abra|OJJ74380.1 acar|OOF95357.1 acla|EAW07421.1 afis|EAW22447.1 afla|EED54461.1 afum|EAL90744.1 agla|OJJ87577.1 akaw|GAA87280.1 aluc|OJZ89740.1 anid|CBF88852.1 anig|EHA26964.1 asyd|OJJ62863.1 ater|EAU39260.1 atub|OJI85061.1 aver|OJI96916.1 awen|OJJ35072.1 azon|OJJ50035.1

Group5469: abra|OJJ74381.1 acar|OOF95356.1 acla|EAW07422.1 afis|EAW22448.1 afla|EED54463.1 afum|EAL90743.1 agla|OJJ87576.1 akaw|GAA87281.1 aluc|OJZ89739.1 anid|CBF88854.1 anig|EHA26963.1 asyd|OJJ62862.1 ater|EAU39259.1 atub|OJI85062.1 aver|OJI96915.1 awen|OJJ35073.1 azon|OJJ50034.1

Group5470: abra|OJJ74382.1 acar|OOF95355.1 acla|EAW07425.1 afis|EAW22451.1 afla|EED54464.1 afum|EAL90740.1 akaw|GAA87283.1 aluc|OJZ89736.1 anid|CBF88856.1 anig|EHA26962.1 aory|EIT81161.1 asyd|OJJ62870.1 ater|EAU39258.1 atub|OJI85065.1 aver|OJI96923.1 awen|OJJ35074.1 azon|OJJ50033.1

Group5471: abra|OJJ74385.1 afis|EAW22452.1 agla|OJJ87571.1 akaw|GAA87286.1 aluc|OJZ89733.1 anid|CBF88929.1 anig|EHA26959.1 ater|EAU39254.1 atub|OJI85068.1 aver|OJI96904.1 awen|OJJ35077.1 azon|OJJ50030.1 acla|EAW07426.1 afum|EAL90739.1 asyd|OJJ62851.1 afla|EED54468.1 aory|EIT81152.1

Group5472: abra|OJJ74390.1 acar|OOF95346.1 acla|EAW06816.1 afis|EAW16365.1 afum|EAL88990.1 agla|OJJ78586.1 akaw|GAA87290.1 aluc|OJZ89729.1 anid|CBF89345.1 anig|EHA26956.1 aory|EIT78416.1 asyd|OJJ62506.1 ater|EAU34087.1 atub|OJI85072.1 aver|OJI96626.1 awen|OJJ37769.1 afla|EED53778.1

Group5473: abra|OJJ74394.1 acla|EAW06818.1 afis|EAW16366.1 afla|EED53470.1 afum|EAL88991.1 agla|OJJ78587.1 akaw|GAA87292.1 aluc|OJZ89726.1 anid|CBF89348.1 anig|EHA26954.1 aory|EIT75141.1 asyd|OJJ62503.1 ater|EAU34082.1 atub|OJI85074.1 aver|OJI96624.1 awen|OJJ37771.1 azon|OJJ45082.1

Group5474: abra|OJJ74398.1 acar|OOF95339.1 acla|EAW06821.1 afis|EAW16369.1 afla|EED53466.1 afum|EAL88994.1 agla|OJJ78590.1 akaw|GAA87296.1 aluc|OJZ89722.1 anid|CBF89353.1 anig|EHA26950.1 aory|EIT74909.1 atub|OJI85078.1 aver|OJJ03881.1 awen|OJJ37906.1 azon|OJJ45084.1 aver|OJI96621.1

Group5475: abra|OJJ74409.1 acar|OOF95333.1 acla|EAW06823.1 afis|EAW16371.1 afla|EED53464.1 afum|EAL88996.1 agla|OJJ78592.1 akaw|GAA87302.1 aluc|OJZ89716.1 anid|CBF89355.1 aory|EIT75209.1 asyd|OJJ62500.1 ater|EAU34078.1 atub|OJI85086.1 aver|OJI96620.1 awen|OJJ37908.1 azon|OJJ45086.1

Group5476: abra|OJJ74413.1 acar|OOF95330.1 acla|EAW06825.1 afis|EAW16376.1 afla|EED53461.1 afum|EAL89001.1 agla|OJJ78594.1 akaw|GAA87304.1 aluc|OJZ89713.1 anid|CBF84606.1 anig|EHA26944.1 aory|EIT74960.1 asyd|OJJ53373.1 ater|EAU34076.1 atub|OJI85089.1 aver|OJJ08371.1 awen|OJJ37910.1

Group5477: abra|OJJ74419.1 acar|OOF95321.1 acla|EAW06835.1 afis|EAW16383.1 afla|EED53449.1 afum|EAL89011.1 agla|OJJ88210.1 akaw|GAA87313.1 aluc|OJZ89704.1 anid|CBF69425.1 anig|EHA26938.1 aory|EIT75059.1 asyd|OJJ54913.1 ater|EAU34063.1 atub|OJI85095.1 aver|OJJ07290.1 awen|OJJ37921.1

Group5478: abra|OJJ74424.1 acar|OOF95316.1 acla|EAW06836.1 afis|EAW16384.1 afla|EED54095.1 afum|EAL89012.1 akaw|GAA87317.1 aluc|OJZ89700.1 anid|CBF74710.1 anig|EHA26934.1 aory|EIT75993.1 asyd|OJJ60207.1 ater|EAU39463.1 atub|OJI85101.1 aver|OJJ04060.1 awen|OJJ37924.1 azon|OJJ45093.1

Group5479: abra|OJJ74470.1 acar|OOF95277.1 acla|EAW07227.1 afis|EAW22241.1 afla|EED57634.1 afum|EAL90952.1 agla|OJJ87810.1 akaw|GAA87363.1 aluc|OJZ89650.1 anid|CBF88487.1 anig|EHA26888.1 aory|EIT82970.1 asyd|OJJ63789.1 ater|EAU34248.1 atub|OJI85154.1 aver|OJI97149.1 awen|OJJ34819.1

Group5480: abra|OJJ74483.1 acla|EAW07218.1 afis|EAW22232.1 afla|EED57621.1 afum|EAL90961.1 agla|OJJ87819.1 akaw|GAA87376.1 aluc|OJZ89636.1 anid|CBF88470.1 anig|EHA26878.1 aory|EIT82969.1 asyd|OJJ63781.1 ater|EAU34235.1 atub|OJI85167.1 aver|OJI97156.1 awen|OJJ34812.1 azon|OJJ45737.1

Group5481: abra|OJJ74485.1 acar|OOF95264.1 acla|EAW09107.1 afis|EAW25711.1 afla|EED54262.1 afum|EAL84384.1 agla|OJJ84991.1 akaw|GAA87378.1 aluc|OJZ89634.1 anid|CBF88463.1 anig|EHA26876.1 aory|EIT76037.1 asyd|OJJ63776.1 ater|EAU37423.1 atub|OJI85169.1 aver|OJI97161.1 awen|OJJ38524.1

Group5482: abra|OJJ74501.1 acar|OOF95243.1 acla|EAW07185.1 afis|EAW22198.1 afla|EED57599.1 afum|EBA27419.1 akaw|GAA87395.1 aluc|OJZ89617.1 anid|CBF88433.1 anig|EHA26862.1 aory|EIT82949.1 asyd|OJJ63761.1 ater|EAU34216.1 atub|OJI85186.1 aver|OJI97179.1 awen|OJJ34791.1 azon|OJJ45755.1

Group5483: abra|OJJ74504.1 acar|OOF95240.1 acla|EAW07190.1 afis|EAW22203.1 afla|EED57592.1 afum|EAL90989.1 agla|OJJ87843.1 akaw|GAA87399.1 aluc|OJZ89613.1 anig|EHA26859.1 aory|EIT82935.1 asyd|OJJ63759.1 ater|EAU34211.1 atub|OJI85189.1 aver|OJI97181.1 awen|OJJ32000.1 azon|OJJ50253.1

Group5484: abra|OJJ74507.1 acla|EAW07194.1 afis|EAW22206.1 afla|EED57585.1 afum|EAL90987.2 agla|OJJ87847.1 akaw|GAA82092.1 aluc|OJZ92088.1 anid|CBF88411.1 anig|EHA26853.1 aory|EIT82967.1 asyd|OJJ63755.1 ater|EAU34208.1 atub|OJI86517.1 aver|OJI97186.1 awen|OJJ34781.1 azon|OJJ45763.1

Group5485: abra|OJJ74515.1 acla|EAW07176.1 afis|EAW22190.1 afla|EED57574.1 afum|EAL91001.1 agla|OJJ87857.1 akaw|GAA82084.1 aluc|OJZ92096.1 anid|CBF88398.1 anig|EHA26845.1 aory|EIT82936.1 asyd|OJJ63747.1 ater|EAU34198.1 atub|OJI86529.1 aver|OJI97193.1 awen|OJJ34770.1 azon|OJJ45776.1

Group5486: abra|OJJ74525.1 acla|EAW07166.1 afis|EAW22181.1 afla|EED57560.1 afum|EAL91008.1 agla|OJJ87874.1 akaw|GAA82076.1 aluc|OJZ92105.1 anid|CBF88387.1 anig|EHA26839.1 aory|EIT81854.1 asyd|OJJ63740.1 ater|EAU34186.1 atub|OJI86539.1 aver|OJI97200.1 awen|OJJ34762.1 azon|OJJ45788.1

Group5487: abra|OJJ74526.1 acla|EAW07165.1 afis|EAW22180.1 afla|EED57559.1 afum|EAL91009.1 agla|OJJ87875.1 akaw|GAA82075.1 aluc|OJZ92107.1 anid|CBF88385.1 anig|EHA26838.1 aory|EIT81828.1 asyd|OJJ63739.1 ater|EAU34185.1 atub|OJI86541.1 aver|OJI97201.1 awen|OJJ34761.1 azon|OJJ45789.1

Group5488: abra|OJJ74550.1 acla|EAW07148.1 afis|EAW22162.1 afla|EED57533.1 afum|EAL91029.1 agla|OJJ87902.1 akaw|GAA82047.1 aluc|OJZ92133.1 anid|CBF89096.1 anig|EHA26816.1 aory|EIT81833.1 asyd|OJJ62730.1 ater|EAU34164.1 atub|OJI86565.1 aver|OJI96773.1 awen|OJJ34736.1 azon|OJJ45823.1

Group5489: abra|OJJ74551.1 acar|OOF95204.1 acla|EAW07147.1 afis|EAW22161.1 afla|EED57532.1 afum|EAL91030.1 agla|OJJ87903.1 akaw|GAA82046.1 aluc|OJZ92134.1 anid|CBF89097.1 anig|EHA26815.1 aory|EIT81868.1 asyd|OJJ62729.1 atub|OJI86566.1 aver|OJI96772.1 awen|OJJ34735.1 azon|OJJ45824.1

Group5490: abra|OJJ74572.1 acar|OOF95190.1 acla|EAW07137.1 afis|EAW22150.1 afla|EED53612.1 afum|EAL91040.1 agla|OJJ87912.1 akaw|GAA82033.1 aluc|OJZ92150.1 anid|CBF89110.1 anig|EHA26799.1 aory|EIT75168.1 asyd|OJJ62666.1 atub|OJI86578.1 aver|OJI96764.1 awen|OJJ34726.1 azon|OJJ45836.1

Group5491: abra|OJJ74574.1 acar|OOF95188.1 acla|EAW07135.1 afis|EAW22148.1 afla|EED53609.1 afum|EAL91042.1 agla|OJJ87913.1 akaw|GAA82031.1 aluc|OJZ92152.1 anid|CBF87368.1 anig|EHA26797.1 aory|EIT74984.1 asyd|OJJ52838.1 ater|EAU34154.1 atub|OJI86580.1 aver|OJJ08809.1 awen|OJJ34724.1

Group5492: abra|OJJ74580.1 acar|OOF95184.1 afis|EAW22143.1 afla|EED53569.1 afum|EAL91047.1 agla|OJJ87931.1 akaw|GAA82025.1 aluc|OJZ92158.1 anid|CBF78371.1 anig|EHA26791.1 aory|EIT75000.1 asyd|OJJ57327.1 ater|EAU34146.1 atub|OJI86586.1 aver|OJJ05833.1 awen|OJJ34718.1 azon|OJJ46644.1

Group5493: abra|OJJ74625.1 acla|EAW10757.1 afis|EAW24817.1 afla|EED47165.1 afum|EAL89461.1 agla|OJJ82916.1 akaw|GAA81983.1 aluc|OJZ92207.1 anid|CBF85877.1 anig|EHA26751.1 aory|EIT79320.1 asyd|OJJ64376.1 ater|EAU36489.1 atub|OJI86636.1 aver|OJI98472.1 awen|OJJ40403.1 azon|OJJ49252.1

Group5494: abra|OJJ74638.1 acla|EAW10750.1 afis|EAW24824.1 afla|EED47151.1 afum|EAL89468.1 agla|OJJ82012.1 akaw|GAA81973.1 aluc|OJZ92217.1 anid|CBF85855.1 anig|EHA26741.1 aory|EIT79315.1 asyd|OJJ64365.1 ater|EAU36481.1 atub|OJI86648.1 aver|OJI98460.1 awen|OJJ40410.1 azon|OJJ49259.1

Group5495: abra|OJJ74642.1 acla|EAW10746.1 afis|EAW24828.1 afla|EED47147.1 agla|OJJ82018.1 akaw|GAA81969.1 aluc|OJZ92222.1 anid|CBF85863.1 anig|EHA26736.1 aory|EIT79265.1 asyd|OJJ64369.1 ater|EAU36477.1 atub|OJI86652.1 aver|OJI98465.1 awen|OJJ40414.1 afum|EAL89473.1 azon|OJJ49264.1

Group5496: abra|OJJ74710.1 acar|OOF95720.1 acla|EAW10688.1 afis|EAW24882.1 afla|EED47084.1 afum|EAL89532.1 agla|OJJ82104.1 akaw|GAA81907.1 aluc|OJZ92291.1 anid|CBF75748.1 anig|EHA26678.1 asyd|OJJ57522.1 ater|EAU36419.1 atub|OJI86721.1 aver|OJJ05656.1 awen|OJJ40472.1 azon|OJJ49336.1

Group5497: abra|OJJ74716.1 acar|OOF95725.1 acla|EAW10683.1 afis|EAW24887.1 afum|EAL89538.2 agla|OJJ82110.1 akaw|GAA81902.1 aluc|OJZ92296.1 anid|CBF75621.1 anig|EHA26673.1 aory|EIT72493.1 asyd|OJJ57454.1 ater|EAU36413.1 atub|OJI86727.1 aver|OJJ05724.1 awen|OJJ40477.1 azon|OJJ49341.1

Group5498: abra|OJJ74728.1 acar|OOF90031.1 acla|EAW10670.1 afis|EAW24896.1 afum|EAL89550.1 agla|OJJ82120.1 akaw|GAA81889.1 aluc|OJZ92312.1 anid|CBF75655.1 anig|EHA26657.1 aory|EIT72662.1 asyd|OJJ57464.1 ater|EAU36402.1 atub|OJI86741.1 aver|OJJ05716.1 awen|OJJ40489.1 azon|OJJ49352.1

Group5499: abra|OJJ74748.1 acla|EAW10662.1 afis|EAW24905.1 afla|EED46988.1 afum|EAL89558.1 agla|OJJ82129.1 akaw|GAA83663.1 aluc|OJZ92329.1 anid|CBF75659.1 anig|EHA26642.1 asyd|OJJ57477.1 ater|EAU36393.1 atub|OJI86758.1 aver|OJJ05704.1 awen|OJJ40498.1 azon|OJJ49360.1 aory|EIT72554.1

Group5500: abra|OJJ74753.1 acar|OOF95746.1 acla|EAW10657.1 afis|EAW24910.1 afla|EED46995.1 afum|EAL89564.1 agla|OJJ82134.1 akaw|GAA83658.1 aluc|OJZ92334.1 anig|EHA26637.1 aory|EIT72538.1 asyd|OJJ57482.1 ater|EAU36388.1 atub|OJI86763.1 aver|OJJ05699.1 awen|OJJ40505.1 azon|OJJ49365.1

Group5501: akaw|GAA83616.1 aluc|OJZ92377.1 acar|OOF95790.1 acla|EAW10610.1 afis|EAW24957.1 afla|EED55506.1 afum|EAL89615.2 agla|OJJ82188.1 anid|CBF81665.1 anig|EHA26596.1 aory|EIT80801.1 asyd|OJJ59552.1 ater|EAU35698.1 atub|OJI86807.1 aver|OJJ02716.1 awen|OJJ40551.1 azon|OJJ49410.1

Group5502: abra|OJJ74801.1 acar|OOF95794.1 acla|EAW10613.1 afis|EAW24954.1 afla|EED55510.1 afum|EAL89611.1 agla|OJJ82183.1 akaw|GAA83612.1 aluc|OJZ92382.1 anid|CBF81715.1 anig|EHA26593.1 aory|EIT80800.1 asyd|OJJ59583.1 ater|EAU35704.1 atub|OJI86811.1 awen|OJJ40549.1 azon|OJJ49409.1

Group5503: abra|OJJ74803.1 acla|EAW10615.1 afis|EAW24952.1 afla|EED55512.1 afum|EAL89609.1 agla|OJJ82185.1 akaw|GAA83610.1 aluc|OJZ92384.1 anid|CBF81719.1 anig|EHA26591.1 aory|EIT80787.1 asyd|OJJ59585.1 ater|EAU35684.1 atub|OJI86813.1 aver|OJJ02685.1 awen|OJJ40547.1 azon|OJJ49407.1

Group5504: abra|OJJ74832.1 acar|OOF95820.1 acla|EAW11371.1 afis|EAW18124.1 afla|EED58050.1 afum|EAL87482.1 akaw|GAA83588.1 aluc|OJZ92409.1 anid|CBF75360.1 anig|EHA26556.1 aory|EIT79713.1 asyd|OJJ59850.1 atub|OJI86837.1 aver|OJJ02437.1 awen|OJJ35526.1 azon|OJJ46958.1 agla|OJJ87018.1

Group5505: abra|OJJ74845.1 acar|OOF95846.1 acla|EAW08543.1 afis|EAW23580.1 afla|EED57485.1 afum|EAL87885.1 agla|OJJ86846.1 akaw|GAA83577.1 aluc|OJZ92420.1 anig|EHA26546.1 aory|EIT81877.1 asyd|OJJ62315.1 ater|EAU35364.1 atub|OJI86850.1 aver|OJI96422.1 awen|OJJ39908.1 azon|OJJ50702.1

Group5506: abra|OJJ74846.1 acar|OOF95847.1 acla|EAW08542.1 afis|EAW23579.1 afla|EED57478.1 afum|EAL87886.1 agla|OJJ86844.1 akaw|GAA83576.1 aluc|OJZ92421.1 anid|CBF89596.1 aory|EIT80108.1 asyd|OJJ62312.1 ater|EAU35365.1 atub|OJI86851.1 aver|OJI96420.1 awen|OJJ39907.1 azon|OJJ45884.1

Group5507: abra|OJJ74851.1 acar|OOF95852.1 acla|EAW08536.1 afis|EAW23574.1 afla|EED57465.1 afum|EAL87891.1 agla|OJJ86834.1 akaw|GAA83571.1 aluc|OJZ92427.1 anid|CBF89608.1 anig|EHA26540.1 aory|EIT80065.1 asyd|OJJ62306.1 atub|OJI86857.1 aver|OJI96414.1 awen|OJJ39902.1 azon|OJJ45878.1

Group5508: abra|OJJ74871.1 acar|OOF95872.1 acla|EAW08509.1 afis|EAW23547.1 afla|EED57442.1 afum|EAL87919.1 akaw|GAA83549.1 aluc|OJZ92449.1 anid|CBF89652.1 anig|EHA26519.1 aory|EIT80070.1 asyd|OJJ62283.1 ater|EAU35332.1 atub|OJI86877.1 aver|OJI96391.1 awen|OJJ39878.1 azon|OJJ43225.1

Group5509: abra|OJJ74878.1 acar|OOF95881.1 acla|EAW08492.1 afis|EAW23538.1 afla|EED57436.1 afum|EAL87928.1 agla|OJJ86793.1 akaw|GAA83542.1 aluc|OJZ92455.1 anid|CBF89666.1 anig|EHA26512.1 asyd|OJJ62274.1 ater|EAU35324.1 atub|OJI86884.1 aver|OJI96383.1 awen|OJJ39869.1 azon|OJJ43232.1

Group5510: abra|OJJ74885.1 acar|OOF95890.1 acla|EAW08482.1 afis|EAW23528.1 afla|EED57427.1 afum|EBA27327.1 akaw|GAA83534.1 aluc|OJZ92462.1 anig|EHA26505.1 aory|EIT80107.1 asyd|OJJ62265.1 ater|EAU35316.1 atub|OJI86891.1 aver|OJI96374.1 awen|OJJ39857.1 azon|OJJ43241.1 agla|OJJ86777.1

Group5511: abra|OJJ74887.1 acar|OOF95891.1 acla|EAW08480.1 afis|EAW23526.1 afla|EED57425.1 afum|EAL87939.1 agla|OJJ86775.1 akaw|GAA83532.1 aluc|OJZ92464.1 anid|CBF89689.1 anig|EHA26503.1 asyd|OJJ62263.1 ater|EAU35314.1 atub|OJI86893.1 aver|OJI96372.1 awen|OJJ39855.1 azon|OJJ43246.1

Group5512: abra|OJJ74899.1 acar|OOF95899.1 acla|EAW08473.1 afis|EAW23519.1 afla|EED57419.1 afum|EAL87945.2 agla|OJJ86766.1 akaw|GAA83522.1 aluc|OJZ92473.1 anid|CBF89700.1 anig|EHA26496.1 aory|EIT80138.1 asyd|OJJ62256.1 ater|EAU35307.1 atub|OJI86904.1 aver|OJI96365.1 awen|OJJ39844.1

Group5513: abra|OJJ74906.1 acar|OOF90300.1 acla|EAW08466.1 afis|EAW23513.1 afla|EED57411.1 afum|EAL87950.1 agla|OJJ86761.1 akaw|GAA83516.1 aluc|OJZ92479.1 anid|CBF89707.1 anig|EHA26491.1 asyd|OJJ62251.1 ater|EAU35302.1 atub|OJI86910.1 aver|OJI96361.1 awen|OJJ39837.1 azon|OJJ43254.1

Group5514: abra|OJJ74908.1 acar|OOF90298.1 acla|EAW08464.1 afis|EAW23509.1 afla|EED57409.1 afum|EAL87952.1 agla|OJJ86759.1 akaw|GAA83514.1 aluc|OJZ92481.1 aory|EIT80131.1 asyd|OJJ62246.1 ater|EAU35297.1 atub|OJI86912.1 aver|OJI96356.1 awen|OJJ39835.1 azon|OJJ43257.1 anid|CBF89716.1

Group5515: abra|OJJ74911.1 acar|OOF90295.1 acla|EAW08461.1 afis|EAW23506.1 afla|EED57406.1 afum|EAL87955.1 agla|OJJ86756.1 akaw|GAA83511.1 aluc|OJZ92484.1 anid|CBF89710.1 aory|EIT80118.1 asyd|OJJ62249.1 ater|EAU35294.1 atub|OJI86915.1 aver|OJI96359.1 awen|OJJ39832.1 azon|OJJ43260.1

Group5516: abra|OJJ74912.1 acar|OOF90296.1 acla|EAW08460.1 afis|EAW23505.1 afla|EED57405.1 afum|EAL87956.1 agla|OJJ86752.1 akaw|GAA83510.1 aluc|OJZ92485.1 anid|CBF89721.1 aory|EIT80097.1 asyd|OJJ62237.1 ater|EAU35293.1 atub|OJI86916.1 aver|OJI96346.1 awen|OJJ39831.1 azon|OJJ46407.1

Group5517: abra|OJJ74921.1 acar|OOF95913.1 acla|EAW08450.1 afis|EAW23495.1 afla|EED57395.1 afum|EAL87968.1 agla|OJJ86742.1 akaw|GAA83502.1 aluc|OJZ92492.1 anid|CBF89740.1 anig|EHA26480.1 asyd|OJJ62229.1 ater|EAU35284.1 atub|OJI86925.1 aver|OJI96338.1 awen|OJJ39820.1 azon|OJJ47944.1

Group5518: abra|OJJ74926.1 acla|EAW08443.1 afis|EAW23488.1 afla|EED57389.1 afum|EAL87975.1 agla|OJJ86735.1 akaw|GAA83497.1 aluc|OJZ92497.1 anid|CBF89749.1 anig|EHA26475.1 aory|EIT80081.1 asyd|OJJ62222.1 ater|EAU35277.1 atub|OJI86930.1 aver|OJI96332.1 awen|OJJ39811.1 azon|OJJ47937.1

Group5519: abra|OJJ74929.1 acar|OOF95919.1 acla|EAW08440.1 afis|EAW23485.1 afla|EED57384.1 afum|EAL87978.2 akaw|GAA83492.1 aluc|OJZ92500.1 anid|CBF89759.1 anig|EHA26472.1 aory|EIT80161.1 asyd|OJJ62219.1 ater|EAU35273.1 atub|OJI86933.1 aver|OJI96329.1 awen|OJJ39809.1 azon|OJJ47931.1

Group5520: abra|OJJ74937.1 acla|EAW08434.1 afis|EAW23477.1 afla|EED57374.1 afum|EAL87988.1 agla|OJJ86716.1 akaw|GAA83483.1 aluc|OJZ92512.1 anid|CBF89769.1 anig|EHA26463.1 aory|EIT80086.1 asyd|OJJ62213.1 ater|EAU36903.1 atub|OJI86943.1 aver|OJI96324.1 awen|OJJ39803.1 azon|OJJ47924.1

Group5521: abra|OJJ74968.1 acar|OOF97347.1 acla|EAW08406.1 afis|EAW23463.1 afla|EED57337.1 afum|EAL88003.1 agla|OJJ86686.1 akaw|GAA83458.1 aluc|OJZ92538.1 anid|CBF89845.1 aory|EIT74160.1 asyd|OJJ62184.1 ater|EAU37478.1 atub|OJI86974.1 aver|OJI96293.1 awen|OJJ39767.1 azon|OJJ47897.1

Group5522: abra|OJJ74978.1 acar|OOF97342.1 acla|EAW08402.1 afis|EAW23467.1 afum|EAL87999.1 agla|OJJ86681.1 akaw|GAA83454.1 aluc|OJZ92542.1 anid|CBF89837.1 anig|EHA26434.1 aory|EIT74085.1 asyd|OJJ62180.1 ater|EAU37483.1 atub|OJI86978.1 aver|OJI96289.1 awen|OJJ39762.1 azon|OJJ47891.1

Group5523: abra|OJJ75001.1 acar|OOF90038.1 acla|EAW08384.1 afis|EAW23413.1 afla|EED57314.1 afum|EAL88056.1 agla|OJJ86630.1 akaw|GAA88624.1 aluc|OJZ92566.1 anid|CBF89867.1 anig|EHA26415.1 asyd|OJJ62154.1 ater|EAU37509.1 atub|OJI87002.1 aver|OJI96264.1 awen|OJJ39746.1 azon|OJJ47872.1

Group5524: abra|OJJ75013.1 acar|OOF97313.1 acla|EAW08208.1 afis|EAW23242.1 afla|EED55723.1 afum|EAL88240.1 akaw|GAA88636.1 aluc|OJZ92578.1 anid|CBF89889.1 anig|EHA26403.1 aory|EIT83219.1 asyd|OJJ62141.1 ater|EAU37523.1 atub|OJI87014.1 aver|OJI96251.1 awen|OJJ39728.1 azon|OJJ47859.1

Group5525: abra|OJJ75016.1 acar|OOF97310.1 acla|EAW08211.1 afis|EAW23245.1 afla|EED55726.1 afum|EAL88237.1 agla|OJJ86615.1 akaw|GAA88639.1 aluc|OJZ92582.1 anid|CBF84114.1 anig|EHA26400.1 aory|EIT83230.1 asyd|OJJ57104.1 ater|EAU37526.1 atub|OJI87017.1 aver|OJJ03195.1 awen|OJJ39725.1

Group5526: abra|OJJ75021.1 acar|OOF89973.1 acla|EAW08213.1 afis|EAW23247.1 afla|EED55728.1 afum|EAL88234.2 akaw|GAA88641.1 aluc|OJZ92588.1 anid|CBF84110.1 anig|EHA26398.1 aory|EIT83211.1 asyd|OJJ57102.1 ater|EAU37529.1 atub|OJI87020.1 aver|OJJ03197.1 awen|OJJ39722.1 azon|OJJ47848.1

Group5527: abra|OJJ75023.1 acar|OOF97309.1 acla|EAW08215.1 afis|EAW23249.1 afla|EED55730.1 afum|EAL88232.2 agla|OJJ86607.1 akaw|GAA88643.1 aluc|OJZ92590.1 anid|CBF84107.1 anig|EHA26396.1 asyd|OJJ57099.1 ater|EAU37551.1 atub|OJI87022.1 aver|OJJ03200.1 awen|OJJ39715.1 azon|OJJ47846.1

Group5528: abra|OJJ75024.1 acla|EAW08216.1 afis|EAW23250.1 afla|EED55731.1 afum|EAL88231.2 agla|OJJ86604.1 akaw|GAA88644.1 aluc|OJZ92591.1 anid|CBF84105.1 anig|EHA26395.1 aory|EIT83234.1 asyd|OJJ57098.1 ater|EAU37550.1 atub|OJI87024.1 aver|OJJ03201.1 awen|OJJ39714.1 azon|OJJ47845.1

Group5529: abra|OJJ75029.1 acar|OOF97304.1 acla|EAW08220.1 afis|EAW23254.1 afla|EED55735.1 afum|EAL88227.1 agla|OJJ86600.1 akaw|GAA88648.1 aluc|OJZ92596.1 anid|CBF84098.1 anig|EHA26391.1 asyd|OJJ57094.1 ater|EAU37545.1 atub|OJI87030.1 aver|OJJ03205.1 awen|OJJ39710.1 azon|OJJ47841.1

Group5530: abra|OJJ75037.1 acar|OOF97295.1 acla|EAW08230.1 afis|EAW23264.1 afla|EED55745.1 afum|EAL88216.1 agla|OJJ86588.1 akaw|GAA88656.1 aluc|OJZ92605.1 anig|EHA26383.1 aory|EIT83208.1 asyd|OJJ57111.1 ater|EAU37535.1 atub|OJI87039.1 aver|OJJ03188.1 awen|OJJ39700.1 azon|OJJ47830.1

Group5531: abra|OJJ75045.1 acar|OOF89832.1 acla|EAW08234.1 afis|EAW23268.1 afla|EED55714.1 afum|EAL88211.1 agla|OJJ86584.1 akaw|GAA88662.1 aluc|OJZ92611.1 anid|CBF89552.1 anig|EHA26376.1 asyd|OJJ62374.1 ater|EAU37531.1 atub|OJI87045.1 aver|OJI96508.1 awen|OJJ39693.1 azon|OJJ47826.1

Group5532: abra|OJJ75055.1 acar|OOF97282.1 acla|EAW08240.1 afis|EAW23274.1 afla|EED55704.1 afum|EAL88205.1 agla|OJJ86574.1 akaw|GAA88668.1 aluc|OJZ92620.1 anid|CBF89540.1 anig|EHA26368.1 asyd|OJJ62381.1 ater|EAU37559.1 atub|OJI87054.1 aver|OJI96515.1 awen|OJJ39683.1 azon|OJJ47820.1

Group5533: abra|OJJ75058.1 acar|OOF97276.1 acla|EAW08243.1 afis|EAW23277.1 afla|EED55699.1 afum|EAL88202.1 agla|OJJ86569.1 akaw|GAA88671.1 aluc|OJZ92627.1 anid|CBF89532.1 anig|EHA26363.1 asyd|OJJ62385.1 ater|EAU37562.1 atub|OJI87057.1 aver|OJI96519.1 awen|OJJ39679.1 azon|OJJ47817.1

Group5534: abra|OJJ75067.1 acar|OOF97267.1 acla|EAW08248.1 afis|EAW23282.1 afla|EED55693.1 afum|EAL88197.2 agla|OJJ86564.1 akaw|GAA88679.1 aluc|OJZ92633.1 anid|CBF89523.1 anig|EHA26358.1 asyd|OJJ62390.1 ater|EAU37590.1 atub|OJI87064.1 aver|OJI96523.1 awen|OJJ39674.1 azon|OJJ47804.1

Group5535: abra|OJJ75068.1 acar|OOF97266.1 acla|EAW08249.1 afis|EAW23283.1 afla|EED55692.1 afum|EAL88196.2 agla|OJJ86563.1 akaw|GAA88680.1 aluc|OJZ92634.1 anid|CBF89522.1 anig|EHA26357.1 asyd|OJJ62391.1 ater|EAU37589.1 atub|OJI87065.1 aver|OJI96524.1 awen|OJJ39673.1 azon|OJJ47803.1

Group5536: abra|OJJ75073.1 acar|OOF97260.1 acla|EAW08255.1 afis|EAW23289.1 afla|EED55685.1 afum|EAL88190.1 agla|OJJ86556.1 akaw|GAA88686.1 aluc|OJZ92639.1 anid|CBF89511.1 anig|EHA26352.1 aory|EIT77081.1 asyd|OJJ62396.1 ater|EAU37584.1 atub|OJI87070.1 aver|OJI96530.1 awen|OJJ39668.1

Group5537: abra|OJJ75078.1 acar|OOF97255.1 acla|EAW08286.1 afis|EAW23319.1 afla|EED55677.1 afum|EAL88157.1 agla|OJJ86544.1 akaw|GAA88691.1 aluc|OJZ92644.1 anid|CBF89502.1 anig|EHA26348.1 asyd|OJJ62401.1 ater|EAU37580.1 atub|OJI87075.1 aver|OJI96534.1 awen|OJJ39664.1 azon|OJJ47794.1

Group5538: abra|OJJ75079.1 acar|OOF97254.1 acla|EAW08285.1 afis|EAW23318.1 afum|EAL88158.1 agla|OJJ86543.1 akaw|GAA88692.1 aluc|OJZ92645.1 anid|CBF89500.1 anig|EHA26347.1 aory|EIT77080.1 asyd|OJJ62402.1 ater|EAU37579.1 atub|OJI87076.1 aver|OJI96535.1 awen|OJJ39663.1 azon|OJJ47795.1

Group5539: abra|OJJ75083.1 acar|OOF97250.1 acla|EAW08281.1 afis|EAW23314.1 afla|EED55673.1 afum|EAL88163.1 agla|OJJ86539.1 akaw|GAA88696.1 aluc|OJZ92649.1 anid|CBF89493.1 anig|EHA26343.1 asyd|OJJ62407.1 ater|EAU35547.1 atub|OJI87080.1 aver|OJI96540.1 awen|OJJ39659.1 azon|OJJ47787.1

Group5540: abra|OJJ75088.1 acar|OOF97245.1 acla|EAW08276.1 afis|EAW23309.1 afla|EED55664.1 afum|EAL88168.1 agla|OJJ86534.1 akaw|GAA88700.1 aluc|OJZ92654.1 anid|CBF89483.1 anig|EHA26340.1 aory|EIT81812.1 ater|EAU35552.1 atub|OJI87085.1 aver|OJI96547.1 awen|OJJ39654.1 azon|OJJ47782.1

Group5541: abra|OJJ75092.1 acar|OOF97240.1 acla|EAW08272.1 afis|EAW23305.1 afla|EED55660.1 afum|EAL88172.1 agla|OJJ86530.1 akaw|GAA88704.1 aluc|OJZ92658.1 anid|CBF89476.1 anig|EHA26336.1 aory|EIT81785.1 asyd|OJJ62417.1 atub|OJI87089.1 aver|OJI96552.1 awen|OJJ39587.1 azon|OJJ47778.1

Group5542: abra|OJJ75095.1 acar|OOF97237.1 acla|EAW08269.1 afis|EAW23302.1 afla|EED55654.1 afum|EAL88177.1 agla|OJJ86527.1 akaw|GAA88707.1 aluc|OJZ92661.1 anid|CBF89470.1 anig|EHA26333.1 aory|EIT81821.1 asyd|OJJ62420.1 ater|EAU35566.1 atub|OJI87092.1 awen|OJJ39584.1 azon|OJJ47768.1

Group5543: abra|OJJ75100.1 acar|OOF97234.1 acla|EAW08267.1 afis|EAW23300.1 afla|EED55652.1 afum|EAL88179.1 agla|OJJ86509.1 akaw|GAA88709.1 aluc|OJZ92664.1 anid|CBF89467.1 anig|EHA26330.1 asyd|OJJ62422.1 ater|EAU35563.1 atub|OJI87095.1 aver|OJI96559.1 awen|OJJ39582.1 azon|OJJ47770.1

Group5544: abra|OJJ75105.1 acar|OOF97229.1 acla|EAW08263.1 afis|EAW23296.1 afla|EED55646.1 afum|EAL88182.1 agla|OJJ86515.1 akaw|GAA88714.1 aluc|OJZ92670.1 anid|CBF89461.1 anig|EHA26327.1 aory|EIT81781.1 asyd|OJJ62426.1 atub|OJI87099.1 aver|OJI96563.1 awen|OJJ39577.1 azon|OJJ47774.1

Group5545: abra|OJJ75108.1 acar|OOF97225.1 acla|EAW08260.1 afis|EAW23293.1 afla|EED55643.1 afum|EAL88186.1 agla|OJJ86518.1 akaw|GAA88717.1 aluc|OJZ92674.1 anid|CBF89455.1 anig|EHA26324.1 asyd|OJJ62430.1 ater|EAU35556.1 atub|OJI87102.1 aver|OJI96566.1 awen|OJJ39574.1 azon|OJJ47764.1

Group5546: abra|OJJ75112.1 acar|OOF97219.1 acla|EAW08312.1 afis|EAW23328.1 afla|EED55639.1 afum|EAL88151.1 agla|OJJ86523.1 akaw|GAA88722.1 aluc|OJZ92679.1 anid|CBF89444.1 anig|EHA26320.1 asyd|OJJ62437.1 ater|EAU35575.1 atub|OJI87107.1 aver|OJI96573.1 awen|OJJ39566.1 azon|OJJ47759.1

Group5547: abra|OJJ75113.1 acar|OOF97218.1 acla|EAW08311.1 afis|EAW23329.1 afla|EED55638.1 afum|EAL88150.1 agla|OJJ86524.1 akaw|GAA88723.1 aluc|OJZ92680.1 anid|CBF89443.1 anig|EHA26319.1 asyd|OJJ62438.1 ater|EAU35576.1 atub|OJI87108.1 awen|OJJ39565.1 azon|OJJ47758.1 aver|OJI96574.1

Group5548: abra|OJJ75128.1 akaw|GAA88735.1 anig|EHA26307.1 atub|OJI87121.1 asyd|OJJ62447.1 aver|OJI96584.1 acla|EAW08301.1 afla|EED55627.1 afum|EAL88139.1 aory|EIT81809.1 afis|EAW23339.1 anid|CBF89427.1 ater|EAU35584.1 agla|OJJ86491.1 awen|OJJ39555.1 azon|OJJ47752.1 aluc|OJZ92693.1

Group5549: abra|OJJ75132.1 acla|EAW08295.1 afis|EAW23344.1 afla|EED55622.1 afum|EAL88133.1 agla|OJJ86483.1 akaw|GAA88739.1 aluc|OJZ92698.1 anid|CBF89414.1 anig|EHA26301.1 aory|EIT81796.1 asyd|OJJ62454.1 ater|EAU35588.1 atub|OJI87126.1 aver|OJI96589.1 awen|OJJ39548.1 azon|OJJ47745.1

Group5550: abra|OJJ75151.1 acar|OOF97186.1 acla|EAW08326.1 afis|EAW23361.1 afla|EED55597.1 afum|EAL88113.1 agla|OJJ86465.1 aluc|OJZ92719.1 anid|CBF74877.1 anig|EHA26281.1 aory|EIT82694.1 asyd|OJJ60113.1 ater|EAU35612.1 atub|OJI87144.1 aver|OJJ04149.1 awen|OJJ39526.1 azon|OJJ47722.1

Group5551: abra|OJJ75154.1 acla|EAW08329.1 afis|EAW23364.1 afla|EED55594.1 afum|EAL88109.1 agla|OJJ86462.1 akaw|GAA88759.1 aluc|OJZ92722.1 anid|CBF74833.1 anig|EHA26278.1 aory|EIT82699.1 asyd|OJJ60136.1 ater|EAU35615.1 atub|OJI87147.1 aver|OJJ04126.1 awen|OJJ39523.1 azon|OJJ47719.1

Group5552: abra|OJJ75176.1 acar|OOF94603.1 acla|EAW13002.1 afis|EAW17666.1 afla|EED50489.1 afum|EAL91480.2 akaw|GAA88781.1 aluc|OJZ92742.1 anid|CBF90045.1 anig|EHA26261.1 aory|EIT82041.1 asyd|OJJ62057.1 ater|EAU31987.1 atub|OJI87167.1 aver|OJI96163.1 awen|OJJ37175.1 azon|OJJ50874.1

Group5553: abra|OJJ75191.1 acar|OOF94592.1 acla|EAW12988.1 afis|EAW17677.1 afla|EED50474.1 afum|EAL91490.1 agla|OJJ79501.1 akaw|GAA88792.1 aluc|OJZ92755.1 anid|CBF90020.1 anig|EHA26250.1 asyd|OJJ62069.1 ater|EAU31969.1 atub|OJI87181.1 aver|OJI96175.1 awen|OJJ37163.1 azon|OJJ50859.1

Group5554: abra|OJJ75192.1 acar|OOF94591.1 acla|EAW12987.1 afis|EAW17678.1 afla|EED50472.1 afum|EAL91491.1 agla|OJJ79502.1 akaw|GAA91745.1 aluc|OJZ92756.1 anid|CBF90017.1 anig|EHA26249.1 asyd|OJJ62071.1 ater|EAU31968.1 atub|OJI87183.1 aver|OJI96176.1 awen|OJJ37161.1 azon|OJJ50858.1

Group5555: abra|OJJ75202.1 acar|OOF94586.1 acla|EAW12980.1 afis|EAW17684.1 afla|EED50467.1 afum|EAL91498.1 agla|OJJ79495.1 akaw|GAA91753.1 aluc|OJZ92765.1 anid|CBF90004.1 anig|EHA26242.1 asyd|OJJ62077.1 ater|EAU31947.1 atub|OJI87209.1 aver|OJI96184.1 awen|OJJ37155.1 azon|OJJ50853.1

Group5556: abra|OJJ75204.1 acar|OOF94584.1 acla|EAW12978.1 afis|EAW17686.1 afla|EED50465.1 afum|EAL91500.1 agla|OJJ79493.1 akaw|GAA91755.1 aluc|OJZ92767.1 anid|CBF89989.1 anig|EHA26240.1 asyd|OJJ62084.1 ater|EAU31949.1 atub|OJI87211.1 aver|OJI96191.1 awen|OJJ37153.1 azon|OJJ50851.1

Group5557: abra|OJJ75216.1 acar|OOF94568.1 acla|EAW13095.1 afis|EAW17568.1 afla|EED49253.1 afum|EBA27431.1 agla|OJJ81397.1 akaw|GAA91766.1 aluc|OJZ92780.1 anid|CBF90260.1 anig|EHA26230.1 asyd|OJJ61942.1 ater|EAU37005.1 atub|OJI87222.1 aver|OJI96034.1 awen|OJJ37283.1 azon|OJJ50979.1

Group5558: abra|OJJ75243.1 acar|OOF94550.1 acla|EAW13700.1 afis|EAW19906.1 afla|EED45982.1 afum|EAL85001.1 akaw|GAA91786.1 aluc|OJZ92805.1 anid|CBF78482.1 anig|EHA26204.1 aory|EIT75561.1 asyd|OJJ65334.1 ater|EAU31595.1 atub|OJI87248.1 aver|OJI99479.1 awen|OJJ30802.1 azon|OJJ47379.1

Group5559: awen|OJJ30208.1 abra|OJJ77534.1 acar|OOF99192.1 afis|EAW16248.1 afla|EED53312.1 afum|EAL88876.1 agla|OJJ86258.1 akaw|GAA84087.1 aluc|OJZ81002.1 anid|CBF78877.1 anig|EHA19358.1 aory|EIT74927.1 asyd|OJJ64593.1 ater|EAU37458.1 atub|OJI82583.1 aver|OJI98690.1 azon|OJJ43932.1

Group5560: abra|OJJ75273.1 acar|OOF94530.1 afis|EAW16992.1 afla|EED48963.1 afum|EAL86516.1 akaw|GAA91811.1 aluc|OJZ92833.1 anid|CBF77694.1 anig|EHA26176.1 aory|EIT80034.1 asyd|OJJ53081.1 ater|EAU31736.1 atub|OJI87290.1 aver|OJJ03766.1 awen|OJJ35980.1 azon|OJJ50285.1 agla|OJJ80492.1

Group5561: abra|OJJ75340.1 acar|OOF89855.1 acla|EAW07715.1 afum|EAL86289.1 akaw|GAA92033.1 aluc|OJZ92903.1 anid|CBF81870.1 anig|EHA26060.1 asyd|OJJ53488.1 atub|OJI87362.1 aver|OJJ00349.1 awen|OJJ40854.1 azon|OJJ43191.1 azon|OJJ50965.1 ater|EAU31350.1 azon|OJJ42649.1 azon|OJJ45030.1

Group5562: abra|OJJ75378.1 acar|OOF94469.1 acla|EAW12114.1 afis|EAW17271.1 afla|EED48848.1 afum|EAL86688.1 agla|OJJ81075.1 akaw|GAA92740.1 aluc|OJZ92948.1 anid|CBF87533.1 aory|EIT81268.1 asyd|OJJ52987.1 ater|EAU36135.1 atub|OJI87407.1 aver|OJJ08956.1 awen|OJJ35763.1 azon|OJJ45521.1

Group5563: abra|OJJ75379.1 acar|OOF94468.1 acla|EAW12113.1 afis|EAW17272.1 afla|EED48847.1 afum|EAL86689.1 agla|OJJ81076.1 akaw|GAA92741.1 aluc|OJZ92949.1 anid|CBF87535.1 aory|EIT81287.1 asyd|OJJ52988.1 ater|EAU36136.1 atub|OJI87408.1 aver|OJJ08957.1 awen|OJJ35762.1 azon|OJJ45522.1

Group5564: abra|OJJ75380.1 acar|OOF94467.1 acla|EAW12112.1 afis|EAW17273.1 afla|EED48846.1 afum|EAL86691.1 agla|OJJ81077.1 akaw|GAA92742.1 aluc|OJZ92950.1 anid|CBF87537.1 aory|EIT81267.1 asyd|OJJ52989.1 ater|EAU34714.1 atub|OJI87409.1 aver|OJJ08958.1 awen|OJJ35761.1 azon|OJJ45523.1

Group5565: abra|OJJ75386.1 acar|OOF94462.1 acla|EAW12106.1 afis|EAW17279.1 afla|EED48840.1 afum|EAL86697.1 agla|OJJ81083.1 akaw|GAA92748.1 aluc|OJZ92956.1 anid|CBF87547.1 anig|EHA26019.1 asyd|OJJ52995.1 ater|EAU34708.1 atub|OJI87415.1 aver|OJJ08964.1 awen|OJJ35755.1 azon|OJJ45529.1

Group5566: abra|OJJ75387.1 acar|OOF94459.1 acla|EAW12105.1 afis|EAW17281.1 afla|EED48838.1 afum|EAL86699.1 agla|OJJ81084.1 akaw|GAA92750.1 aluc|OJZ92958.1 anid|CBF87551.1 anig|EHA26017.1 asyd|OJJ52997.1 ater|EAU34707.1 atub|OJI87416.1 aver|OJJ08966.1 awen|OJJ35754.1 azon|OJJ45530.1

Group5567: abra|OJJ75403.1 acar|OOF90750.1 acla|EAW13775.1 afis|EAW18357.1 afla|EED48822.1 afum|EAL87721.1 agla|OJJ88028.1 aluc|OJZ88855.1 anid|CBF79354.1 anig|EHA23648.1 aory|EIT73028.1 asyd|OJJ57801.1 atub|OJI85935.1 aver|OJJ04586.1 awen|OJJ35741.1 azon|OJJ43325.1 afum|EAL87720.1

Group5568: abra|OJJ75407.1 acar|OOF90752.1 acla|EAW13779.1 afis|EAW18353.1 afla|EED48818.1 afum|EAL87716.1 agla|OJJ88031.1 aluc|OJZ88858.1 anid|CBF79362.1 anig|EHA23645.1 aory|EIT73012.1 asyd|OJJ57805.1 ater|EAU34687.1 atub|OJI85932.1 aver|OJJ04583.1 awen|OJJ35737.1 azon|OJJ50439.1

Group5569: abra|OJJ75416.1 acar|OOF90761.1 acla|EAW13788.1 afis|EAW18344.1 afla|EED48809.1 afum|EAL87708.1 agla|OJJ88040.1 akaw|GAA84191.1 aluc|OJZ88867.1 anid|CBF79380.1 anig|EHA23636.1 aory|EIT73003.1 asyd|OJJ57814.1 ater|EAU34679.1 atub|OJI85923.1 aver|OJJ04574.1 awen|OJJ35728.1

Group5570: abra|OJJ75418.1 acar|OOF90763.1 acla|EAW13790.1 afis|EAW18342.1 afla|EED48807.1 afum|EAL87705.1 agla|OJJ88042.1 akaw|GAA84193.1 aluc|OJZ88869.1 anid|CBF79376.1 anig|EHA23634.1 aory|EIT73006.1 asyd|OJJ57816.1 ater|EAU34677.1 atub|OJI85921.1 awen|OJJ35726.1 azon|OJJ43314.1

Group5571: abra|OJJ75422.1 acar|OOF90767.1 acla|EAW13794.1 afis|EAW18338.1 afla|EED48802.1 agla|OJJ88046.1 akaw|GAA84197.1 aluc|OJZ88874.1 anid|CBF79389.1 anig|EHA23631.1 aory|EIT72987.1 asyd|OJJ57820.1 ater|EAU34673.1 atub|OJI85917.1 aver|OJJ04571.1 awen|OJJ35721.1 azon|OJJ43311.1

Group5572: abra|OJJ75437.1 acar|OOF90781.1 acla|EAW13825.1 afis|EAW18306.1 afla|EED48775.1 afum|EAL87672.1 agla|OJJ88117.1 akaw|GAA84212.1 aluc|OJZ88890.1 anid|CBF79443.1 anig|EHA23617.1 aory|EIT72975.1 asyd|OJJ57849.1 ater|EAU33316.1 atub|OJI85899.1 aver|OJJ04543.1 awen|OJJ30680.1

Group5573: abra|OJJ75440.1 acar|OOF90784.1 acla|EAW13823.1 afis|EAW18308.1 afum|EAL87674.1 agla|OJJ88074.1 akaw|GAA84215.1 aluc|OJZ88893.1 anid|CBF79440.1 anig|EHA23613.1 aory|EIT72977.1 asyd|OJJ57847.1 ater|EAU33318.1 atub|OJI85896.1 aver|OJJ04545.1 awen|OJJ30682.1 azon|OJJ43282.1

Group5574: abra|OJJ75451.1 acar|OOF90795.1 acla|EAW13812.1 afis|EAW18318.1 afla|EED48786.1 afum|EAL87685.1 agla|OJJ88100.1 akaw|GAA84226.1 aluc|OJZ88906.1 anid|CBF79419.1 anig|EHA23602.1 asyd|OJJ57835.1 ater|EAU33328.1 atub|OJI85885.1 aver|OJJ04555.1 awen|OJJ30700.1 azon|OJJ43293.1

Group5575: abra|OJJ75453.1 acar|OOF90797.1 acla|EAW13810.1 afis|EAW18320.1 afla|EED48788.1 agla|OJJ88071.1 akaw|GAA84228.1 aluc|OJZ88908.1 anid|CBF79417.1 anig|EHA23600.1 aory|EIT73031.1 asyd|OJJ57834.1 ater|EAU33330.1 atub|OJI85883.1 aver|OJJ04556.1 awen|OJJ30702.1 azon|OJJ43294.1

Group5576: abra|OJJ75458.1 acar|OOF90280.1 acla|EAW13807.1 afis|EAW18323.1 afla|EED48791.1 afum|EAL87690.2 agla|OJJ88068.1 akaw|GAA84233.1 aluc|OJZ88915.1 anid|CBF79412.1 anig|EHA23596.1 asyd|OJJ57831.1 ater|EAU33333.1 atub|OJI85877.1 aver|OJJ04559.1 awen|OJJ30705.1 azon|OJJ43299.1

Group5577: abra|OJJ75469.1 acar|OOF90804.1 acla|EAW13835.1 afis|EAW18296.1 afla|EED48762.1 afum|EAL87662.1 agla|OJJ88127.1 akaw|GAA84241.1 aluc|OJZ88926.1 anid|CBF79468.1 anig|EHA23587.1 asyd|OJJ57864.1 ater|EAU33304.1 atub|OJI85866.1 aver|OJJ04529.1 awen|OJJ30670.1 azon|OJJ43270.1

Group5578: abra|OJJ75489.1 acar|OOF90821.1 acla|EAW11220.1 afis|EAW18283.1 afla|EED48750.1 afum|EAL87649.1 agla|OJJ88144.1 aluc|OJZ88948.1 anid|CBF79485.1 anig|EHA23573.1 aory|EIT73079.1 asyd|OJJ57874.1 ater|EAU33289.1 atub|OJI85841.1 aver|OJJ04520.1 awen|OJJ30654.1 azon|OJJ47960.1

Group5579: abra|OJJ75524.1 acar|OOF90851.1 acla|EAW09311.1 afis|EAW21834.1 afla|EED46541.1 afum|EAL92849.1 agla|OJJ84310.1 akaw|GAA84287.1 aluc|OJZ88981.1 anig|EHA23545.1 aory|EIT77894.1 asyd|OJJ61231.1 ater|EAU38509.1 atub|OJI85808.1 aver|OJJ00554.1 awen|OJJ34107.1 azon|OJJ44234.1

Group5580: abra|OJJ75547.1 acar|OOF90867.1 acla|EAW09318.1 afis|EAW21827.1 afla|EED46549.1 afum|EAL92842.1 agla|OJJ84317.1 akaw|GAA84302.1 aluc|OJZ88997.1 anid|CBF76668.1 anig|EHA23531.1 asyd|OJJ61218.1 ater|EAU38503.1 atub|OJI85792.1 aver|OJJ00542.1 awen|OJJ34100.1 azon|OJJ44244.1

Group5581: abra|OJJ75550.1 acar|OOF90869.1 afis|EAW21823.1 afla|EED46555.1 afum|EAL92839.1 agla|OJJ84319.1 akaw|GAA84305.1 aluc|OJZ89000.1 anid|CBF86661.1 anig|EHA23528.1 aory|EIT77858.1 asyd|OJJ61620.1 ater|EAU38497.1 atub|OJI85789.1 aver|OJJ00944.1 awen|OJJ34097.1 azon|OJJ44247.1

Group5582: abra|OJJ75551.1 acar|OOF90871.1 acla|EAW09323.1 afis|EAW21822.1 afla|EED46557.1 afum|EAL92838.1 agla|OJJ84320.1 akaw|GAA84306.1 aluc|OJZ89001.1 anid|CBF76662.1 anig|EHA23527.1 asyd|OJJ61214.1 ater|EAU38496.1 atub|OJI85788.1 aver|OJJ00538.1 awen|OJJ34096.1 azon|OJJ44248.1

Group5583: abra|OJJ75580.1 acar|OOF90898.1 acla|EAW09338.1 afis|EAW21793.1 afla|EED46606.1 afum|EAL92818.1 agla|OJJ83098.1 akaw|GAA84332.1 aluc|OJZ89031.1 anid|CBF76621.1 anig|EHA23503.1 aory|EIT77822.1 asyd|OJJ61191.1 ater|EAU38472.1 atub|OJI85760.1 aver|OJJ00515.1 awen|OJJ34077.1

Group5584: abra|OJJ75591.1 acar|OOG00237.1 acla|EAW09331.1 afis|EAW21787.1 afla|EED46609.1 afum|EAL92814.1 agla|OJJ84336.1 akaw|GAA84343.1 aluc|OJZ89043.1 anid|CBF76619.1 anig|EHA23492.1 asyd|OJJ61190.1 ater|EAU38470.1 atub|OJI85744.1 aver|OJJ00514.1 awen|OJJ34073.1 azon|OJJ48156.1

Group5585: abra|OJJ75604.1 acar|OOG00233.1 acla|EAW09366.1 afis|EAW21771.1 afla|EED46329.1 afum|EAL92800.1 agla|OJJ84379.1 akaw|GAA84355.1 aluc|OJZ89056.1 anid|CBF83760.1 anig|EHA23485.1 asyd|OJJ60862.1 ater|EAU38447.1 atub|OJI85731.1 aver|OJJ00188.1 awen|OJJ34058.1 azon|OJJ48174.1

Group5586: abra|OJJ75613.1 acar|OOG00223.1 acla|EAW09365.1 afis|EAW21772.1 afla|EED46338.1 afum|EAL92801.1 agla|OJJ84352.1 akaw|GAA84362.1 aluc|OJZ89066.1 anid|CBF76593.1 anig|EHA23470.1 aory|EIT81227.1 asyd|OJJ61176.1 atub|OJI85722.1 aver|OJJ00502.1 awen|OJJ34059.1 azon|OJJ48173.1

Group5587: abra|OJJ75617.1 acla|EAW09361.1 afis|EAW21776.1 afla|EED46342.1 afum|EAL92805.1 agla|OJJ84348.1 akaw|GAA84366.1 aluc|OJZ89070.1 anid|CBF76601.1 anig|EHA23466.1 aory|EIT81196.1 asyd|OJJ61181.1 ater|EAU38463.1 atub|OJI85718.1 aver|OJJ00506.1 awen|OJJ34064.1 azon|OJJ48168.1

Group5588: abra|OJJ75623.1 acar|OOG00219.1 acla|EAW09358.1 afis|EAW21779.1 afla|EED46346.1 afum|EAL92808.1 akaw|GAA84370.1 aluc|OJZ89075.1 anid|CBF76605.1 anig|EHA23463.1 aory|EIT81237.1 asyd|OJJ61184.1 ater|EAU38461.1 atub|OJI85711.1 aver|OJJ00508.1 awen|OJJ34068.1 azon|OJJ48161.1

Group5589: abra|OJJ75627.1 acla|EAW09466.1 afis|EAW21670.1 afla|EED57934.1 afum|EAL92705.1 agla|OJJ79744.1 akaw|GAA84374.1 aluc|OJZ89078.1 anid|CBF83584.1 anig|EHA23459.1 aory|EIT82346.1 asyd|OJJ60780.1 ater|EAU38354.1 atub|OJI85707.1 aver|OJJ00113.1 awen|OJJ33934.1 azon|OJJ48641.1

Group5590: abra|OJJ75629.1 acar|OOG00206.1 acla|EAW09464.1 afis|EAW21672.1 afum|EAL92707.1 agla|OJJ79746.1 akaw|GAA84376.1 aluc|OJZ89080.1 anid|CBF83587.1 anig|EHA23457.1 aory|EIT82330.1 asyd|OJJ60778.1 ater|EAU38356.1 atub|OJI85705.1 aver|OJJ00111.1 awen|OJJ33937.1 azon|OJJ48630.1

Group5591: abra|OJJ75643.1 acla|EAW09455.1 afis|EAW21682.1 afla|EED57949.1 afum|EAL92716.2 agla|OJJ79919.1 akaw|GAA84388.1 aluc|OJZ89092.1 anid|CBF83604.1 anig|EHA23445.1 aory|EIT82329.1 asyd|OJJ60767.1 ater|EAU38366.1 atub|OJI85689.1 aver|OJJ00101.1 awen|OJJ33949.1 azon|OJJ48621.1

Group5592: abra|OJJ75653.1 acla|EAW09447.1 afis|EAW21689.1 afla|EED57959.1 afum|EAL92723.2 agla|OJJ79911.1 akaw|GAA84396.1 aluc|OJZ89102.1 anid|CBF83616.1 anig|EHA23437.1 aory|EIT79725.1 asyd|OJJ60759.1 ater|EAU38374.1 atub|OJI85679.1 aver|OJJ00093.1 awen|OJJ33958.1 azon|OJJ48610.1

Group5593: abra|OJJ75655.1 acla|EAW09445.1 afis|EAW21691.1 afla|EED57961.1 afum|EAL92725.1 agla|OJJ79909.1 akaw|GAA84398.1 aluc|OJZ89104.1 anid|CBF83619.1 anig|EHA23435.1 aory|EIT79711.1 asyd|OJJ60757.1 ater|EAU38376.1 atub|OJI85677.1 aver|OJJ00091.1 awen|OJJ33960.1 azon|OJJ48608.1

Group5594: abra|OJJ75660.1 acar|OOG00183.1 acla|EAW09440.1 afis|EAW21696.1 afla|EED57965.1 afum|EAL92730.1 agla|OJJ79903.1 akaw|GAA84402.1 aluc|OJZ89110.1 anid|CBF83626.1 anig|EHA23431.1 asyd|OJJ60753.1 ater|EAU38381.1 atub|OJI85673.1 aver|OJJ00087.1 awen|OJJ33965.1 azon|OJJ48604.1

Group5595: abra|OJJ75663.1 acar|OOG00182.1 acla|EAW09438.1 afis|EAW21698.1 afla|EED57967.1 afum|EAL92732.1 agla|OJJ79900.1 akaw|GAA84404.1 aluc|OJZ89114.1 anid|CBF83630.1 anig|EHA23429.1 asyd|OJJ60751.1 ater|EAU38383.1 atub|OJI85670.1 aver|OJJ00084.1 awen|OJJ33967.1 azon|OJJ48599.1

Group5596: abra|OJJ75666.1 acar|OOG00178.1 acla|EAW09435.1 afis|EAW21701.1 afla|EED57970.1 afum|EAL92735.1 agla|OJJ79897.1 akaw|GAA84407.1 aluc|OJZ89117.1 anid|CBF83635.1 anig|EHA23426.1 asyd|OJJ60783.1 ater|EAU38386.1 atub|OJI85667.1 aver|OJJ00116.1 awen|OJJ33971.1 azon|OJJ48588.1

Group5597: abra|OJJ75676.1 acla|EAW09429.1 afis|EAW21707.1 afla|EED57979.1 afum|EAL92742.1 agla|OJJ79894.1 akaw|GAA84417.1 aluc|OJZ89126.1 anid|CBF83651.1 anig|EHA23417.1 aory|EIT79779.1 asyd|OJJ60793.1 ater|EAU38395.1 atub|OJI85658.1 aver|OJJ00126.1 awen|OJJ33983.1 azon|OJJ48576.1

Group5598: abra|OJJ75681.1 acar|OOG00167.1 acla|EAW09421.1 afis|EAW21715.1 afla|EED57985.1 afum|EBA27470.1 agla|OJJ79884.1 akaw|GAA84421.1 aluc|OJZ89132.1 anig|EHA23413.1 aory|EIT79799.1 asyd|OJJ60797.1 ater|EAU38400.1 atub|OJI85654.1 aver|OJJ00130.1 awen|OJJ33987.1 azon|OJJ48566.1

Group5599: abra|OJJ75693.1 acar|OOG00159.1 acla|EAW07961.1 afis|EAW22996.1 afla|EED49502.1 agla|OJJ83288.1 akaw|GAA84430.1 aluc|OJZ89145.1 anid|CBF80648.1 anig|EHA23403.1 aory|EIT82603.1 asyd|OJJ54616.1 ater|EAU29307.1 atub|OJI85643.1 aver|OJJ07543.1 awen|OJJ38565.1 azon|OJJ43626.1

Group5600: abra|OJJ75725.1 acar|OOG00133.1 acla|EAW09381.1 afis|EAW21754.1 afla|EED58024.1 afum|EAL92784.1 agla|OJJ84399.1 akaw|GAA84461.1 aluc|OJZ89177.1 anid|CBF83689.1 anig|EHA23378.1 asyd|OJJ60818.1 ater|EAU38436.1 atub|OJI85612.1 aver|OJJ00144.1 awen|OJJ34038.1 azon|OJJ48208.1

Group5601: abra|OJJ75782.1 acla|EAW14900.1 afis|EAW24422.1 afla|EED47746.1 afum|EAL86090.1 agla|OJJ79222.1 akaw|GAA84499.1 aluc|OJZ89224.1 anid|CBF73788.1 anig|EHA23337.1 aory|EIT77143.1 asyd|OJJ56602.1 ater|EAU30306.1 atub|OJI85567.1 aver|OJJ06122.1 awen|OJJ32787.1 azon|OJJ49043.1

Group5602: abra|OJJ75803.1 acar|OOG00073.1 acla|EAW14880.1 afis|EAW24398.1 afla|EED47813.1 afum|EAL86069.1 agla|OJJ78879.1 akaw|GAA84519.1 aluc|OJZ89251.1 anid|CBF73726.1 anig|EHA23316.1 aory|EIT77134.1 asyd|OJJ56635.1 atub|OJI85547.1 aver|OJJ06091.1 awen|OJJ32807.1 azon|OJJ49029.1

Group5603: abra|OJJ75818.1 acar|OOG00057.1 acla|EAW14857.1 afis|EAW24384.1 afla|EED47825.1 afum|EAL86056.1 akaw|GAA84532.1 aluc|OJZ89265.1 anid|CBF73699.1 anig|EHA23303.1 aory|EIT77173.1 asyd|OJJ56649.1 ater|EAU30351.1 atub|OJI85534.1 aver|OJJ06077.1 awen|OJJ32827.1 azon|OJJ49010.1

Group5604: abra|OJJ75826.1 acar|OOG00053.1 acla|EAW14851.1 afis|EAW24378.1 afla|EED47833.1 afum|EAL86050.1 agla|OJJ83264.1 akaw|GAA84537.1 aluc|OJZ89270.1 anid|CBF73672.1 anig|EHA23299.1 aory|EIT77204.1 asyd|OJJ56663.1 atub|OJI85529.1 aver|OJJ06061.1 awen|OJJ32834.1 azon|OJJ49001.1

Group5605: abra|OJJ75830.1 acar|OOG00050.1 acla|EAW14848.1 afis|EAW24375.1 afla|EED47836.1 afum|EAL86047.1 agla|OJJ83261.1 akaw|GAA84540.1 aluc|OJZ89273.1 anig|EHA23296.1 aory|EIT77176.1 asyd|OJJ56658.1 ater|EAU30361.1 atub|OJI85526.1 aver|OJJ06067.1 awen|OJJ32837.1 azon|OJJ48997.1

Group5606: abra|OJJ75842.1 acar|OOG00046.1 acla|EAW14845.1 afis|EAW24372.1 afla|EED47842.1 afum|EAL86044.1 agla|OJJ83257.1 akaw|GAA84552.1 aluc|OJZ89284.1 anid|CBF73686.1 anig|EHA23282.1 aory|EIT77226.1 asyd|OJJ56655.1 ater|EAU30365.1 atub|OJI85515.1 aver|OJJ06070.1 awen|OJJ32842.1

Group5607: abra|OJJ75853.1 acla|EAW14834.1 afis|EAW24363.1 afla|EED47850.1 afum|EAL86034.1 agla|OJJ83248.1 akaw|GAA84558.1 aluc|OJZ89290.1 anid|CBF73930.1 anig|EHA23272.1 aory|EIT77222.1 asyd|OJJ53676.1 ater|EAU30375.1 atub|OJI85509.1 aver|OJJ07792.1 awen|OJJ32868.1 azon|OJJ48990.1

Group5608: abra|OJJ75857.1 acar|OOG00039.1 acla|EAW14830.1 afis|EAW24359.1 afla|EED47854.1 afum|EAL86029.1 agla|OJJ83244.1 akaw|GAA84562.1 aluc|OJZ89295.1 anid|CBF73923.1 anig|EHA23268.1 asyd|OJJ53680.1 ater|EAU30379.1 atub|OJI85505.1 aver|OJJ07796.1 awen|OJJ32862.1 azon|OJJ48986.1

Group5609: abra|OJJ75861.1 acar|OOG00035.1 acla|EAW14828.1 afis|EAW24357.1 afla|EED47857.1 afum|EAL86027.1 agla|OJJ83239.1 akaw|GAA84566.1 aluc|OJZ89299.1 anid|CBF73919.1 anig|EHA23264.1 aory|EIT77243.1 asyd|OJJ53682.1 ater|EAU30382.1 atub|OJI85499.1 aver|OJJ07799.1 awen|OJJ32860.1

Group5610: abra|OJJ75862.1 acar|OOG00034.1 acla|EAW14827.1 afis|EAW24356.1 afla|EED47906.1 afum|EAL86026.2 akaw|GAA84568.1 aluc|OJZ89303.1 anid|CBF73945.1 anig|EHA23263.1 aory|EIT83365.1 asyd|OJJ53697.1 ater|EAU30390.1 atub|OJI85496.1 aver|OJJ07813.1 awen|OJJ32870.1 azon|OJJ48983.1

Group5611: abra|OJJ75868.1 acar|OOG00026.1 acla|EAW09929.1 afis|EAW21177.1 afla|EED54764.1 afum|EAL92197.1 agla|OJJ80000.1 akaw|GAA84574.1 aluc|OJZ89310.1 anid|CBF83304.1 anig|EHA23256.1 aory|EIT79450.1 asyd|OJJ60554.1 atub|OJI85489.1 aver|OJI99880.1 awen|OJJ36516.1 azon|OJJ44392.1

Group5612: abra|OJJ75894.1 acar|OOG00009.1 acla|EAW09895.1 afis|EAW21231.1 afum|EAL92243.1 agla|OJJ78493.1 akaw|GAA84600.1 aluc|OJZ89345.1 anid|CBF83275.1 anig|EHA23237.1 aory|EIT79428.1 asyd|OJJ60571.1 ater|EAU35864.1 atub|OJI85454.1 aver|OJI99896.1 awen|OJJ36541.1 azon|OJJ44347.1

Group5613: abra|OJJ75901.1 acar|OOG00004.1 acla|EAW09889.1 afis|EAW21237.1 afla|EED54797.1 afum|EAL92249.1 agla|OJJ78499.1 akaw|GAA84607.1 aluc|OJZ89351.1 anid|CBF83268.1 anig|EHA23230.1 asyd|OJJ60577.1 ater|EAU35878.1 atub|OJI85447.1 aver|OJI99903.1 awen|OJJ36548.1 azon|OJJ44341.1

Group5614: abra|OJJ75913.1 acar|OOF99991.1 afla|EED54809.1 akaw|GAA84618.1 aluc|OJZ89362.1 aory|EIT79401.1 ater|EAU35889.1 atub|OJI85436.1 awen|OJJ36562.1 azon|OJJ44326.1 acla|EAW09877.1 afis|EAW21249.1 agla|OJJ87507.1 anid|CBF83244.1 asyd|OJJ60596.1 aver|OJI99919.1 afum|EAL92262.1

Group5615: abra|OJJ75930.1 acar|OOF99980.1 acla|EAW09863.1 afis|EAW21263.1 afla|EED54822.1 afum|EAL92278.1 akaw|GAA84631.1 aluc|OJZ89377.1 anid|CBF83338.1 anig|EHA23206.1 aory|EIT79385.1 asyd|OJJ60608.1 ater|EAU35900.1 atub|OJI85421.1 aver|OJI99932.1 awen|OJJ36580.1 azon|OJJ44310.1

Group5616: abra|OJJ75943.1 acla|EAW09850.1 afis|EAW21285.1 afla|EED54836.1 afum|EAL92302.1 agla|OJJ87469.1 akaw|GAA84645.1 aluc|OJZ89390.1 anid|CBF83365.1 anig|EHA23194.1 aory|EIT79369.1 asyd|OJJ60621.1 ater|EAU35917.1 atub|OJI85407.1 aver|OJI99945.1 awen|OJJ39414.1 azon|OJJ44297.1

Group5617: abra|OJJ75999.1 acar|OOF99925.1 acla|EAW09805.1 afis|EAW21331.1 afla|EED57289.1 afum|EAL92349.1 agla|OJJ87408.1 akaw|GAA88960.1 aluc|OJZ89445.1 anid|CBF83433.1 anig|EHA23141.1 asyd|OJJ60663.1 ater|EAU35957.1 atub|OJI85348.1 aver|OJI99986.1 awen|OJJ39367.1 azon|OJJ51707.1

Group5618: abra|OJJ76002.1 acar|OOF99923.1 acla|EAW09803.1 afis|EAW21333.1 afla|EED57291.1 afum|EAL92351.1 agla|OJJ87401.1 akaw|GAA88958.1 aluc|OJZ89447.1 anid|CBF83437.1 anig|EHA23139.1 aory|EIT73967.1 asyd|OJJ60665.1 atub|OJI85346.1 aver|OJI99988.1 awen|OJJ39365.1 azon|OJJ51704.1

Group5619: abra|OJJ76006.1 acar|OOF99918.1 acla|EAW09799.1 afis|EAW21337.1 afla|EED57295.1 afum|EAL92355.2 agla|OJJ87397.1 akaw|GAA88954.1 aluc|OJZ89452.1 anid|CBF83883.1 anig|EHA23135.1 asyd|OJJ60941.1 ater|EAU35962.1 atub|OJI85342.1 aver|OJJ00267.1 awen|OJJ39361.1 azon|OJJ51700.1

Group5620: abra|OJJ76050.1 acar|OOF99881.1 acla|EAW09781.1 afis|EAW21357.1 afla|EED55772.1 afum|EAL92372.1 agla|OJJ87379.1 akaw|GAA88915.1 aluc|OJZ89506.1 anid|CBF76261.1 anig|EHA23094.1 aory|EIT76646.1 asyd|OJJ60970.1 atub|OJI85297.1 aver|OJJ00295.1 awen|OJJ39341.1 azon|OJJ51687.1

Group5621: abra|OJJ76052.1 acar|OOF99875.1 acla|EAW09765.1 afis|EAW21372.1 afla|EED55790.1 afum|EAL92389.1 agla|OJJ85497.1 akaw|GAA88913.1 aluc|OJZ89508.1 anid|CBF83865.1 anig|EHA23092.1 aory|EIT76543.1 asyd|OJJ60931.1 atub|OJI85295.1 aver|OJJ00257.1 awen|OJJ39325.1 azon|OJJ51670.1

Group5622: abra|OJJ76054.1 acla|EAW09764.1 afis|EAW21373.1 afla|EED55791.1 afum|EAL92390.1 agla|OJJ85498.1 akaw|GAA88911.1 aluc|OJZ89510.1 anid|CBF83863.1 anig|EHA23090.1 aory|EIT76741.1 asyd|OJJ60929.1 ater|EAU35184.1 atub|OJI85293.1 aver|OJJ00255.1 awen|OJJ39324.1 azon|OJJ51669.1

Group5623: abra|OJJ76075.1 acar|OOF99850.1 acla|EAW09746.1 afis|EAW21396.1 afla|EED55815.1 afum|EAL92410.1 agla|OJJ85521.1 akaw|GAA88892.1 aluc|OJZ89531.1 anid|CBF83835.1 anig|EHA23071.1 aory|EIT76634.1 asyd|OJJ60911.1 atub|OJI85274.1 aver|OJJ00238.1 awen|OJJ39307.1 azon|OJJ51547.1

Group5624: abra|OJJ76085.1 acar|OOF99840.1 acla|EAW09736.1 afis|EAW21405.1 afla|EED55827.1 afum|EAL92419.1 agla|OJJ85534.1 akaw|GAA88881.1 aluc|OJZ89543.1 anid|CBF83813.1 anig|EHA23063.1 asyd|OJJ60897.1 ater|EAU35145.1 atub|OJI85264.1 aver|OJJ00222.1 awen|OJJ39298.1 azon|OJJ51555.1

Group5625: abra|OJJ76086.1 acar|OOF99838.1 acla|EAW09735.1 afis|EAW21406.1 afla|EED55828.1 afum|EAL92420.1 agla|OJJ85535.1 akaw|GAA88880.1 aluc|OJZ89544.1 anid|CBF83812.1 anig|EHA23062.1 asyd|OJJ60896.1 ater|EAU35142.1 atub|OJI85262.1 aver|OJJ00221.1 awen|OJJ39297.1 azon|OJJ51558.1

Group5626: abra|OJJ76089.1 acla|EAW09733.1 afis|EAW21408.1 afla|EED55830.1 afum|EAL92422.1 agla|OJJ85538.1 akaw|GAA88878.1 aluc|OJZ89546.1 anid|CBF83808.1 anig|EHA23060.1 aory|EIT76672.1 asyd|OJJ60894.1 ater|EAU35140.1 atub|OJI85260.1 aver|OJJ00219.1 awen|OJJ39293.1 azon|OJJ51561.1

Group5627: abra|OJJ76093.1 acar|OOF99832.1 acla|EAW09726.1 afis|EAW21412.1 afla|EED55837.1 afum|EAL92426.1 agla|OJJ85545.1 akaw|GAA88874.1 aluc|OJZ89549.1 anid|CBF83796.1 anig|EHA23057.1 aory|EIT76701.1 asyd|OJJ60886.1 atub|OJI85257.1 aver|OJJ00210.1 awen|OJJ39287.1 azon|OJJ51567.1

Group5628: abra|OJJ76097.1 acla|EAW09721.1 afis|EAW21420.1 afla|EED55845.1 afum|EAL92435.1 agla|OJJ85554.1 akaw|GAA88869.1 aluc|OJZ89553.1 anid|CBF83789.1 anig|EHA23053.1 aory|EIT76872.1 asyd|OJJ60879.1 ater|EAU35129.1 atub|OJI85252.1 aver|OJJ00205.1 awen|OJJ39283.1 azon|OJJ51571.1

Group5629: abra|OJJ76101.1 acar|OOF99826.1 acla|EAW09717.1 afis|EAW21424.1 afla|EED55849.1 afum|EAL92439.1 agla|OJJ85561.1 akaw|GAA88865.1 aluc|OJZ89557.1 anig|EHA23049.1 aory|EIT76560.1 asyd|OJJ60875.1 ater|EAU35125.1 atub|OJI85247.1 aver|OJJ00200.1 awen|OJJ39277.1 azon|OJJ51575.1

Group5630: abra|OJJ76119.1 acar|OOF99806.1 acla|EAW09700.1 afis|EAW21439.1 afla|EED55867.1 afum|EAL92455.1 agla|OJJ85584.1 akaw|GAA88848.1 aluc|OJZ89576.1 anid|CBF76581.1 aory|EIT76843.1 asyd|OJJ61170.1 ater|EAU35106.1 atub|OJI85228.1 aver|OJJ00495.1 awen|OJJ39259.1 azon|OJJ51596.1

Group5631: abra|OJJ76135.1 acar|OOF99788.1 acla|EAW09683.1 afis|EAW21456.1 afla|EED55886.1 afum|EAL92475.1 agla|OJJ85604.1 akaw|GAA88832.1 aluc|OJZ89598.1 anid|CBF76544.1 anig|EHA23015.1 asyd|OJJ61153.1 ater|EAU35079.1 atub|OJI85206.1 aver|OJJ00475.1 awen|OJJ39244.1 azon|OJJ51611.1

Group5632: abra|OJJ76139.1 acla|EAW09680.1 afis|EAW21459.1 afla|EED55890.1 afum|EAL92478.1 agla|OJJ85607.1 akaw|GAA88829.1 aluc|OJZ89602.1 anid|CBF76538.1 anig|EHA23013.1 aory|EIT76890.1 asyd|OJJ61150.1 ater|EAU35076.1 atub|OJI85202.1 aver|OJJ00472.1 awen|OJJ39240.1 azon|OJJ51614.1

Group5633: abra|OJJ76147.1 acar|OOF99779.1 acla|EAW09672.1 afis|EAW21466.1 afla|EED55899.1 afum|EAL92486.1 agla|OJJ85616.1 akaw|GAA82097.1 aluc|OJZ92082.1 anid|CBF76525.1 aory|EIT76748.1 asyd|OJJ61142.1 atub|OJI86512.1 aver|OJJ00464.1 awen|OJJ39231.1 ater|EAU35067.1 azon|OJJ51624.1

Group5634: abra|OJJ76159.1 acar|OOF99773.1 acla|EAW09659.1 afis|EAW21480.1 afla|EED55912.1 afum|EAL92500.1 agla|OJJ85634.1 akaw|GAA92403.1 aluc|OJZ83264.1 anid|CBF76504.1 anig|EHA21117.1 asyd|OJJ61131.1 ater|EAU35057.1 atub|OJI87673.1 aver|OJJ00452.1 awen|OJJ39217.1 azon|OJJ51637.1

Group5635: abra|OJJ76184.1 acla|EAW09642.1 afis|EAW21498.1 afla|EED55934.1 afum|EAL92517.1 agla|OJJ85661.1 akaw|GAA82129.1 aluc|OJZ92045.1 anid|CBF76464.1 anig|EHA22975.1 aory|EIT76682.1 asyd|OJJ61113.1 ater|EAU35040.1 atub|OJI86477.1 aver|OJJ00429.1 awen|OJJ39197.1 azon|OJJ51520.1

Group5636: abra|OJJ76186.1 acar|OOF99756.1 acla|EAW09641.1 afis|EAW21499.1 afla|EED55935.1 afum|EAL92518.1 agla|OJJ85662.1 akaw|GAA82130.1 aluc|OJZ92043.1 anid|CBF76460.1 aory|EIT76668.1 asyd|OJJ61112.1 ater|EAU35039.1 atub|OJI86476.1 aver|OJJ00428.1 awen|OJJ39196.1 azon|OJJ51521.1

Group5637: abra|OJJ76198.1 acar|OOF99746.1 acla|EAW13919.1 afis|EAW18630.1 afla|EED52710.1 afum|EAL93342.1 agla|OJJ81255.1 akaw|GAA82140.1 aluc|OJZ92031.1 anid|CBF70476.1 anig|EHA22964.1 asyd|OJJ58593.1 ater|EAU38281.1 atub|OJI86464.1 aver|OJJ01195.1 awen|OJJ32097.1 azon|OJJ46676.1

Group5638: abra|OJJ76199.1 acar|OOF99745.1 acla|EAW13920.1 afis|EAW18631.1 afla|EED52711.1 afum|EAL93343.2 agla|OJJ82446.1 akaw|GAA82142.1 aluc|OJZ92029.1 anid|CBF70475.1 anig|EHA22963.1 asyd|OJJ58594.1 ater|EAU38280.1 atub|OJI86463.1 aver|OJJ01196.1 awen|OJJ32098.1 azon|OJJ46678.1

Group5639: abra|OJJ76203.1 acla|EAW13924.1 afis|EAW18635.1 afla|EED52706.1 afum|EAL93348.2 agla|OJJ82441.1 akaw|GAA82146.1 aluc|OJZ92025.1 anid|CBF70464.1 anig|EHA22959.1 aory|EIT74268.1 asyd|OJJ58599.1 ater|EAU38276.1 atub|OJI86459.1 aver|OJJ01201.1 awen|OJJ32102.1 azon|OJJ46684.1

Group5640: abra|OJJ76205.1 acla|EAW13926.1 afis|EAW18637.1 afla|EED52704.1 afum|EAL93350.1 agla|OJJ82438.1 akaw|GAA82148.1 aluc|OJZ92023.1 anid|CBF70459.1 anig|EHA22957.1 aory|EIT74180.1 asyd|OJJ58601.1 ater|EAU38274.1 atub|OJI86456.1 aver|OJJ01203.1 awen|OJJ32105.1 azon|OJJ46686.1

Group5641: abra|OJJ76216.1 acar|OOF99734.1 acla|EAW13932.1 afis|EAW18643.1 afla|EED52693.1 afum|EAL93357.1 agla|OJJ82431.1 akaw|GAA82155.1 aluc|OJZ92013.1 anid|CBF70443.1 anig|EHA22952.1 asyd|OJJ58609.1 ater|EAU38268.1 atub|OJI86448.1 aver|OJJ01214.1 awen|OJJ32111.1 azon|OJJ46692.1

Group5642: abra|OJJ76218.1 acar|OOF99733.1 acla|EAW13933.1 afis|EAW18644.1 afla|EED52694.1 afum|EAL93359.1 agla|OJJ82430.1 akaw|GAA82156.1 aluc|OJZ92012.1 anid|CBF70441.1 anig|EHA22951.1 asyd|OJJ58610.1 ater|EAU38267.1 atub|OJI86447.1 aver|OJJ01215.1 awen|OJJ32112.1 azon|OJJ46693.1

Group5643: abra|OJJ76224.1 acar|OOF99471.1 acla|EAW14364.1 afis|EAW18914.1 afla|EED50898.1 afum|EBA27497.1 agla|OJJ81518.1 akaw|GAA82160.1 aluc|OJZ92007.1 anid|CBF69693.1 aory|EIT79056.1 asyd|OJJ59054.1 ater|EAU38081.1 atub|OJI86442.1 aver|OJJ01648.1 awen|OJJ31507.1 azon|OJJ47226.1

Group5644: abra|OJJ76235.1 acla|EAW14372.1 afis|EAW18921.1 afla|EED50907.1 afum|EAL93652.1 agla|OJJ81522.1 akaw|GAA82170.1 aluc|OJZ91995.1 anid|CBF69680.1 anig|EHA22940.1 aory|EIT78977.1 asyd|OJJ59061.1 ater|EAU38094.1 atub|OJI86430.1 aver|OJJ01655.1 awen|OJJ31514.1 azon|OJJ47232.1

Group5645: abra|OJJ76236.1 acar|OOF99480.1 acla|EAW14373.1 afis|EAW18922.1 afla|EED50908.1 afum|EAL93653.1 agla|OJJ81521.1 akaw|GAA82171.1 aluc|OJZ91994.1 anid|CBF69678.1 anig|EHA22939.1 asyd|OJJ59066.1 ater|EAU38095.1 atub|OJI86429.1 aver|OJJ01660.1 awen|OJJ31515.1 azon|OJJ47233.1

Group5646: abra|OJJ76247.1 acar|OOF99493.1 acla|EAW14360.1 afis|EAW18930.1 afla|EED50923.1 afum|EAL93661.1 agla|OJJ85259.1 akaw|GAA82180.1 aluc|OJZ91984.1 anig|EHA22930.1 aory|EIT78976.1 asyd|OJJ64673.1 atub|OJI86419.1 aver|OJI98774.1 awen|OJJ31524.1 azon|OJJ47239.1 anid|CBF86792.1

Group5647: abra|OJJ76249.1 acar|OOF99494.1 acla|EAW14358.1 afis|EAW18932.1 afla|EED50937.1 afum|EAL93664.1 agla|OJJ85261.1 akaw|GAA82182.1 aluc|OJZ91981.1 anid|CBF86798.1 anig|EHA22928.1 asyd|OJJ64669.1 ater|EAU38111.1 atub|OJI86417.1 aver|OJI98771.1 awen|OJJ31526.1 azon|OJJ47241.1

Group5648: abra|OJJ76255.1 acar|OOF99500.1 acla|EAW14352.1 afis|EAW18938.1 afla|EED50943.1 afum|EAL93671.1 agla|OJJ85269.1 akaw|GAA82189.1 aluc|OJZ91971.1 anid|CBF86809.1 anig|EHA22919.1 asyd|OJJ64663.1 ater|EAU38117.1 atub|OJI86410.1 aver|OJI98765.1 awen|OJJ31533.1 azon|OJJ47247.1

Group5649: abra|OJJ76261.1 acar|OOF99505.1 afis|EAW18942.1 afla|EED50975.1 afum|EAL93675.1 akaw|GAA82193.1 aluc|OJZ91961.1 anid|CBF86818.1 anig|EHA22917.1 aory|EIT76997.1 asyd|OJJ64656.1 ater|EAU38123.1 atub|OJI86405.1 aver|OJI98759.1 awen|OJJ31539.1 azon|OJJ47251.1 acla|EAW14349.1

Group5650: abra|OJJ76263.1 acar|OOF99507.1 acla|EAW14345.1 afis|EAW18945.1 afla|EED50979.1 afum|EAL93678.1 agla|OJJ85272.1 akaw|GAA82195.1 aluc|OJZ91959.1 anid|CBF86824.1 anig|EHA22915.1 aory|EIT76970.1 asyd|OJJ64653.1 ater|EAU38125.1 atub|OJI86402.1 aver|OJI98755.1 awen|OJJ31541.1

Group5651: abra|OJJ76265.1 acar|OOF99508.1 acla|EAW14344.1 afis|EAW18946.1 afla|EED50982.1 afum|EAL93680.1 akaw|GAA82197.1 aluc|OJZ91956.1 anid|CBF86827.1 anig|EHA22914.1 aory|EIT77065.1 asyd|OJJ64652.1 ater|EAU38128.1 atub|OJI86401.1 aver|OJI98753.1 awen|OJJ31544.1 azon|OJJ47261.1

Group5652: abra|OJJ76273.1 acar|OOF99521.1 acla|EAW14339.1 afis|EAW18951.1 afla|EED50988.1 afum|EAL93686.1 agla|OJJ85322.1 akaw|GAA82206.1 aluc|OJZ91947.1 anid|CBF69580.1 anig|EHA22906.1 aory|EIT77062.1 asyd|OJJ59143.1 ater|EAU38130.1 atub|OJI86392.1 aver|OJJ01733.1 azon|OJJ47265.1

Group5653: abra|OJJ76280.1 acar|OOF99525.1 acla|EAW14335.1 afis|EAW18954.1 afla|EED51001.1 agla|OJJ85329.1 akaw|GAA82212.1 aluc|OJZ91940.1 anid|CBF69592.1 anig|EHA22899.1 aory|EIT77031.1 asyd|OJJ59140.1 ater|EAU38138.1 atub|OJI86382.1 aver|OJJ01729.1 awen|OJJ31552.1 azon|OJJ47269.1

Group5654: abra|OJJ76284.1 acar|OOF99526.1 acla|EAW14332.1 afis|EAW18956.1 afla|EED51003.1 agla|OJJ85334.1 akaw|GAA82213.1 aluc|OJZ91937.1 anid|CBF69594.1 anig|EHA22897.1 asyd|OJJ59139.1 ater|EAU38139.1 atub|OJI86379.1 aver|OJJ01728.1 awen|OJJ31555.1 azon|OJJ47272.1 afum|EAL93690.1

Group5655: abra|OJJ76285.1 acar|OOF99528.1 acla|EAW14331.1 afis|EAW18957.1 afla|EED51004.1 agla|OJJ85335.1 akaw|GAA82214.1 aluc|OJZ91936.1 anid|CBF69596.1 anig|EHA22896.1 aory|EIT77061.1 asyd|OJJ59137.1 ater|EAU38140.1 atub|OJI86378.1 aver|OJJ01727.1 awen|OJJ31556.1 azon|OJJ47273.1

Group5656: abra|OJJ76315.1 acla|EAW14100.1 afis|EAW18801.1 afla|EED50775.1 agla|OJJ80531.1 akaw|GAA82246.1 aluc|OJZ91899.1 anid|CBF69782.1 anig|EHA22878.1 aory|EIT78048.1 asyd|OJJ58959.1 ater|EAU37933.1 atub|OJI86352.1 aver|OJJ01553.1 awen|OJJ32375.1 afum|EAL93537.1 azon|OJJ47138.1

Group5657: abra|OJJ76319.1 acar|OOF99572.1 acla|EAW14098.1 afis|EAW18800.1 afla|EED50776.1 afum|EAL93536.1 agla|OJJ80529.1 akaw|GAA82249.1 aluc|OJZ91895.1 anid|CBF69780.1 anig|EHA22874.1 asyd|OJJ58958.1 ater|EAU37932.1 atub|OJI86348.1 aver|OJJ01552.1 awen|OJJ32374.1 azon|OJJ47137.1

Group5658: abra|OJJ76321.1 acar|OOF99574.1 acla|EAW14093.1 afis|EAW18799.1 afla|EED50790.1 afum|EAL93535.1 akaw|GAA82250.1 aluc|OJZ91894.1 anid|CBF69778.1 anig|EHA22873.1 aory|EIT78129.1 asyd|OJJ58957.1 ater|EAU37931.1 atub|OJI86346.1 aver|OJJ01551.1 awen|OJJ32373.1 azon|OJJ47136.1

Group5659: abra|OJJ76324.1 acar|OOF99576.1 acla|EAW14091.1 afis|EAW18797.1 afla|EED50792.1 afum|EAL93533.1 agla|OJJ80526.1 akaw|GAA82253.1 aluc|OJZ91891.1 anid|CBF69775.1 aory|EIT78127.1 asyd|OJJ58955.1 ater|EAU37929.1 atub|OJI86343.1 aver|OJJ01549.1 awen|OJJ32371.1 azon|OJJ47134.1

Group5660: abra|OJJ76330.1 acar|OOF99582.1 acla|EAW14085.1 afis|EAW18792.1 afla|EED50750.1 afum|EAL93527.1 agla|OJJ80521.1 aluc|OJZ91883.1 anid|CBF69765.1 anig|EHA22861.1 aory|EIT77994.1 asyd|OJJ58949.1 ater|EAU37923.1 atub|OJI86334.1 aver|OJJ01544.1 awen|OJJ32365.1 azon|OJJ47129.1

Group5661: abra|OJJ76331.1 acar|OOF99583.1 acla|EAW14084.1 afis|EAW18791.1 afla|EED50749.1 afum|EAL93526.1 agla|OJJ80520.1 akaw|GAA82259.1 aluc|OJZ91882.1 anid|CBF69763.1 anig|EHA22860.1 asyd|OJJ58948.1 ater|EAU37922.1 atub|OJI86333.1 aver|OJJ01543.1 awen|OJJ32364.1 azon|OJJ47127.1

Group5662: abra|OJJ76333.1 acar|OOF99585.1 acla|EAW14082.1 afis|EAW18789.1 afla|EED50747.1 afum|EAL93523.1 agla|OJJ80518.1 aluc|OJZ91880.1 anid|CBF69759.1 anig|EHA22858.1 aory|EIT78001.1 asyd|OJJ58945.1 ater|EAU37920.1 atub|OJI86331.1 aver|OJJ01541.1 awen|OJJ32362.1 azon|OJJ47125.1

Group5663: abra|OJJ76341.1 acar|OOF99592.1 acla|EAW14074.1 afis|EAW18781.1 afla|EED50740.1 afum|EAL93513.1 agla|OJJ80501.1 akaw|GAA82268.1 aluc|OJZ91872.1 anid|CBF69740.1 anig|EHA22851.1 asyd|OJJ58934.1 ater|EAU37913.1 atub|OJI86323.1 aver|OJJ01530.1 awen|OJJ32353.1 azon|OJJ47117.1

Group5664: abra|OJJ76352.1 acar|OOF99602.1 acla|EAW14063.1 afis|EAW18770.1 afum|EAL93501.2 agla|OJJ80472.1 aluc|OJZ91861.1 anid|CBF69936.1 anig|EHA22841.1 aory|EIT78053.1 asyd|OJJ58921.1 ater|EAU37901.1 atub|OJI86312.1 aver|OJJ01518.1 awen|OJJ32339.1 azon|OJJ47105.1 afla|EED50729.1

Group5665: abra|OJJ76361.1 acar|OOF99612.1 acla|EAW14055.1 afis|EAW18761.1 afla|EED50721.1 afum|EAL93492.2 agla|OJJ80460.1 akaw|GAA82287.1 aluc|OJZ91851.1 anid|CBF69951.1 anig|EHA22833.1 asyd|OJJ58912.1 ater|EAU37892.1 atub|OJI86303.1 aver|OJJ01509.1 awen|OJJ32331.1 azon|OJJ47096.1

Group5666: abra|OJJ76365.1 acar|OOF99616.1 acla|EAW14051.1 afis|EAW18757.1 afla|EED50717.1 afum|EAL93486.1 agla|OJJ82325.1 akaw|GAA82291.1 aluc|OJZ91847.1 anid|CBF69959.1 anig|EHA22829.1 aory|EIT78006.1 asyd|OJJ58908.1 ater|EAU37889.1 atub|OJI86299.1 aver|OJJ01505.1 awen|OJJ32327.1

Group5667: abra|OJJ76367.1 acar|OOF89986.1 acla|EAW14049.1 afis|EAW18755.1 afla|EED50715.1 afum|EAL93482.1 akaw|GAA82293.1 aluc|OJZ91845.1 aory|EIT78024.1 asyd|OJJ58906.1 atub|OJI86297.1 aver|OJJ01503.1 awen|OJJ32324.1 azon|OJJ47091.1 anid|CBF69965.1 ater|EAU37886.1 agla|OJJ82323.1

Group5668: abra|OJJ76372.1 acar|OOF99620.1 acla|EAW14046.1 afis|EAW18752.1 afla|EED50711.1 afum|EAL93479.1 agla|OJJ82320.1 akaw|GAA82297.1 aluc|OJZ91841.1 anid|CBF69975.1 aory|EIT77974.1 asyd|OJJ58899.1 ater|EAU37883.1 atub|OJI86293.1 aver|OJJ01499.1 awen|OJJ32321.1 azon|OJJ47088.1

Group5669: abra|OJJ76373.1 acar|OOF99621.1 acla|EAW14045.1 afis|EAW18751.1 afla|EED50710.1 afum|EAL93478.1 agla|OJJ82319.1 akaw|GAA82298.1 aluc|OJZ91840.1 anig|EHA22824.1 aory|EIT78136.1 asyd|OJJ58898.1 ater|EAU37882.1 atub|OJI86292.1 aver|OJJ01498.1 awen|OJJ32320.1 azon|OJJ47087.1

Group5670: abra|OJJ76380.1 acar|OOF89926.1 acla|EAW14042.1 afis|EAW18748.1 afla|EED50707.1 afum|EAL93474.1 agla|OJJ82315.1 akaw|GAA82301.1 aluc|OJZ91834.1 anid|CBF69985.1 anig|EHA22821.1 asyd|OJJ58894.1 ater|EAU37877.1 atub|OJI86288.1 aver|OJJ01495.1 awen|OJJ32315.1 azon|OJJ47083.1

Group5671: abra|OJJ76385.1 acar|OOF99626.1 acla|EAW14039.1 afis|EAW18745.1 afla|EED50701.1 afum|EAL93471.1 agla|OJJ82310.1 akaw|GAA82304.1 aluc|OJZ91830.1 anid|CBF69991.1 anig|EHA22817.1 aory|EIT78013.1 asyd|OJJ58890.1 atub|OJI86284.1 aver|OJJ01490.1 awen|OJJ32310.1 azon|OJJ42714.1

Group5672: abra|OJJ76391.1 acla|EAW14033.1 afis|EAW18739.1 afla|EED50694.1 afum|EAL93465.1 agla|OJJ82305.1 akaw|GAA82310.1 aluc|OJZ91822.1 anid|CBF70004.1 anig|EHA22812.1 aory|EIT78125.1 asyd|OJJ58883.1 ater|EAU37862.1 atub|OJI86277.1 aver|OJJ01485.1 awen|OJJ32304.1 azon|OJJ42722.1

Group5673: abra|OJJ76394.1 acar|OOF99634.1 acla|EAW14030.1 afis|EAW18736.1 afla|EED50691.1 afum|EAL93461.1 agla|OJJ82302.1 akaw|GAA82313.1 aluc|OJZ91819.1 anid|CBF70010.1 anig|EHA22809.1 asyd|OJJ58880.1 ater|EAU37859.1 atub|OJI86274.1 aver|OJJ01482.1 awen|OJJ32300.1 azon|OJJ42733.1

Group5674: abra|OJJ76403.1 acar|OOF99646.1 acla|EAW14016.1 afis|EAW18726.1 afla|EED50683.1 afum|EAL93451.1 agla|OJJ80473.1 akaw|GAA82323.1 aluc|OJZ91809.1 anid|CBF70019.1 anig|EHA22802.1 asyd|OJJ58876.1 ater|EAU38178.1 atub|OJI86264.1 aver|OJJ01478.1 awen|OJJ32228.1 azon|OJJ46784.1

Group5675: abra|OJJ76405.1 acar|OOF99648.1 acla|EAW14018.1 afis|EAW18724.1 afla|EED50681.1 afum|EAL93449.1 agla|OJJ80476.1 akaw|GAA82325.1 aluc|OJZ91807.1 anid|CBF70686.1 anig|EHA22800.1 aory|EIT78066.1 asyd|OJJ58467.1 atub|OJI86262.1 aver|OJJ01066.1 awen|OJJ32230.1 azon|OJJ46782.1

Group5676: abra|OJJ76416.1 acar|OOF99656.1 acla|EAW14011.1 afis|EAW18717.1 afla|EED50671.1 afum|EBA27493.1 agla|OJJ82342.1 akaw|GAA82331.1 aluc|OJZ91797.1 anid|CBF70674.1 anig|EHA22793.1 asyd|OJJ58474.1 ater|EAU38188.1 atub|OJI86253.1 aver|OJJ01076.1 awen|OJJ32211.1 azon|OJJ46772.1

Group5677: abra|OJJ76425.1 acar|OOF99669.1 acla|EAW14000.1 afis|EAW18708.1 afla|EED50657.1 afum|EAL93431.1 agla|OJJ82350.1 akaw|GAA82340.1 aluc|OJZ91787.1 anid|CBF70652.1 anig|EHA22784.1 aory|EIT78022.1 asyd|OJJ58484.1 ater|EAU38200.1 atub|OJI86243.1 aver|OJJ01087.1 awen|OJJ32191.1

Group5678: abra|OJJ76426.1 acar|OOF99670.1 acla|EAW13999.1 afis|EAW18707.1 afla|EED50656.1 afum|EAL93430.1 agla|OJJ82351.1 akaw|GAA82341.1 aluc|OJZ91786.1 anid|CBF70650.1 anig|EHA22783.1 aory|EIT78121.1 asyd|OJJ58485.1 ater|EAU38201.1 atub|OJI86242.1 aver|OJJ01088.1 awen|OJJ32190.1

Group5679: abra|OJJ76427.1 acar|OOF99672.1 acla|EAW13998.1 afis|EAW18706.1 afla|EED50655.1 afum|EAL93429.1 agla|OJJ82352.1 akaw|GAA82342.1 aluc|OJZ91785.1 anid|CBF70649.1 anig|EHA22782.1 aory|EIT78135.1 asyd|OJJ58486.1 atub|OJI86241.1 aver|OJJ01089.1 awen|OJJ32189.1 azon|OJJ46758.1

Group5680: abra|OJJ76437.1 acla|EAW13986.1 afis|EAW18694.1 afla|EED50643.1 afum|EAL93416.1 agla|OJJ82369.1 akaw|GAA82352.1 aluc|OJZ91773.1 anid|CBF70626.1 anig|EHA22773.1 aory|EIT78019.1 asyd|OJJ58498.1 ater|EAU38212.1 atub|OJI86230.1 aver|OJJ01101.1 awen|OJJ32165.1 azon|OJJ46750.1

Group5681: abra|OJJ76450.1 acar|OOF99693.1 acla|EAW13978.1 afis|EAW18687.1 afla|EED50635.1 afum|EAL93408.2 agla|OJJ82385.1 akaw|GAA82362.1 aluc|OJZ91760.1 anid|CBF70611.1 aory|EIT78005.1 asyd|OJJ58511.1 ater|EAU38221.1 atub|OJI86218.1 aver|OJJ01112.1 awen|OJJ32158.1 azon|OJJ46742.1

Group5682: abra|OJJ76454.1 acla|EAW13976.1 afis|EAW18685.1 afla|EED50623.1 afum|EAL93406.1 agla|OJJ82386.1 akaw|GAA82366.1 aluc|OJZ91756.1 anid|CBF70609.1 anig|EHA22759.1 aory|EIT78050.1 asyd|OJJ58513.1 ater|EAU38223.1 atub|OJI86214.1 aver|OJJ01114.1 awen|OJJ32156.1 azon|OJJ46740.1

Group5683: abra|OJJ76458.1 acar|OOF99698.1 acla|EAW13971.1 afis|EAW18680.1 afla|EED50630.1 afum|EAL93401.1 agla|OJJ82395.1 akaw|GAA82371.1 aluc|OJZ91751.1 anid|CBF70599.1 anig|EHA22754.1 asyd|OJJ58518.1 ater|EAU38229.1 atub|OJI86209.1 aver|OJJ01119.1 awen|OJJ32152.1 azon|OJJ46736.1

Group5684: abra|OJJ76461.1 acar|OOF99700.1 acla|EAW13969.1 afis|EAW18678.1 afla|EED50632.1 agla|OJJ82398.1 akaw|GAA82374.1 aluc|OJZ91748.1 anid|CBF70594.1 anig|EHA22751.1 aory|EIT78090.1 asyd|OJJ58520.1 ater|EAU38233.1 atub|OJI86206.1 aver|OJJ01121.1 awen|OJJ32150.1 azon|OJJ46734.1

Group5685: abra|OJJ76463.1 acar|OOF99701.1 acla|EAW13968.1 afis|EAW18677.1 afla|EED50633.1 afum|EAL93397.1 agla|OJJ82399.1 akaw|GAA82375.1 aluc|OJZ91747.1 anid|CBF70592.1 anig|EHA22750.1 aory|EIT78096.1 asyd|OJJ58521.1 atub|OJI86205.1 aver|OJJ01122.1 awen|OJJ32149.1 azon|OJJ46733.1

Group5686: abra|OJJ76473.1 acar|OOF99710.1 acla|EAW13960.1 afis|EAW18669.1 afla|EED52659.1 afum|EAL93387.1 agla|OJJ82406.1 akaw|GAA82385.1 aluc|OJZ91736.1 anid|CBF70577.1 anig|EHA22742.1 asyd|OJJ58529.1 ater|EAU38242.1 atub|OJI86194.1 aver|OJJ01131.1 awen|OJJ32141.1 azon|OJJ46722.1

Group5687: abra|OJJ76490.1 acla|EAW13945.1 afis|EAW18654.1 afla|EED52682.1 afum|EAL93370.1 agla|OJJ82421.1 akaw|GAA82405.1 aluc|OJZ91715.1 anid|CBF70542.1 anig|EHA22725.1 aory|EIT74201.1 asyd|OJJ58548.1 ater|EAU38258.1 atub|OJI86173.1 aver|OJJ01146.1 awen|OJJ32121.1 azon|OJJ46707.1

Group5688: abra|OJJ76492.1 acar|OOF99724.1 acla|EAW13940.1 afis|EAW18651.1 afla|EED52688.1 afum|EAL93367.1 agla|OJJ82423.1 akaw|GAA82408.1 aluc|OJZ91712.1 anid|CBF70537.1 anig|EHA22723.1 asyd|OJJ58550.1 ater|EAU38261.1 atub|OJI86171.1 aver|OJJ01148.1 awen|OJJ32119.1 azon|OJJ46705.1

Group5689: abra|OJJ76494.1 acar|OOF99726.1 acla|EAW13938.1 afis|EAW18649.1 afla|EED52690.1 afum|EAL93365.1 agla|OJJ82425.1 akaw|GAA82410.1 aluc|OJZ91710.1 anid|CBF70535.1 anig|EHA22721.1 aory|EIT74193.1 asyd|OJJ58552.1 atub|OJI86169.1 aver|OJJ01150.1 awen|OJJ32117.1 azon|OJJ46703.1

Group5690: azon|OJJ47218.1 abra|OJJ76512.1 acar|OOF99460.1 acla|EAW14188.1 afis|EAW18903.1 afla|EED50845.1 afum|EAL93635.1 agla|OJJ81508.1 akaw|GAA82425.1 aluc|OJZ91693.1 anid|CBF69928.1 anig|EHA22706.1 asyd|OJJ59040.1 ater|EAU38071.1 atub|OJI86147.1 aver|OJJ01635.1 awen|OJJ32460.1

Group5691: abra|OJJ76522.1 acar|OOF99451.1 acla|EAW14179.1 afis|EAW18894.1 afla|EED50869.1 afum|EAL93626.1 agla|OJJ81500.1 akaw|GAA82435.1 aluc|OJZ91686.1 anid|CBF69910.1 anig|EHA22698.1 aory|EIT78965.1 asyd|OJJ58994.1 ater|EAU38065.1 atub|OJI86138.1 awen|OJJ32452.1 azon|OJJ47210.1

Group5692: abra|OJJ76532.1 acar|OOF99441.1 acla|EAW14170.1 afis|EAW18884.1 afla|EED50881.1 afum|EAL93617.1 agla|OJJ80619.1 akaw|GAA82444.1 aluc|OJZ91672.1 anig|EHA22690.1 aory|EIT78974.1 asyd|OJJ59003.1 ater|EAU38056.1 atub|OJI86127.1 aver|OJJ01612.1 azon|OJJ47200.1 awen|OJJ32439.1

Group5693: abra|OJJ76544.1 acar|OOF99430.1 acla|EAW14160.1 afis|EAW18873.1 afla|EED50894.1 afum|EAL93605.1 agla|OJJ80610.1 akaw|GAA82453.1 aluc|OJZ91659.1 anig|EHA22679.1 aory|EIT78984.1 asyd|OJJ59022.1 ater|EAU38037.1 atub|OJI86115.1 aver|OJJ01595.1 awen|OJJ32432.1 azon|OJJ47190.1

Group5694: abra|OJJ76553.1 acar|OOF89944.1 acla|EAW14152.1 afis|EAW18865.1 afla|EED50896.1 afum|EAL93597.1 agla|OJJ80587.1 akaw|GAA82460.1 aluc|OJZ91651.1 anid|CBF69850.1 anig|EHA22672.1 aory|EIT79012.1 asyd|OJJ59029.1 atub|OJI86105.1 aver|OJJ01587.1 awen|OJJ32424.1 azon|OJJ47179.1

Group5695: abra|OJJ76556.1 acar|OOF99423.1 acla|EAW14149.1 afis|EAW18862.1 afla|EED50829.1 afum|EAL93594.1 agla|OJJ80585.1 akaw|GAA82462.1 aluc|OJZ91649.1 anid|CBF69846.1 anig|EHA22670.1 asyd|OJJ59031.1 ater|EAU38024.1 atub|OJI86103.1 aver|OJJ01585.1 awen|OJJ32418.1 azon|OJJ47175.1

Group5696: abra|OJJ76559.1 acla|EAW14146.1 afis|EAW18859.1 afla|EED50826.1 afum|EAL93591.1 agla|OJJ80582.1 akaw|GAA82465.1 aluc|OJZ91646.1 anid|CBF69840.1 anig|EHA22667.1 aory|EIT78992.1 asyd|OJJ59034.1 ater|EAU38022.1 atub|OJI86100.1 aver|OJJ01582.1 awen|OJJ32415.1 azon|OJJ47172.1

Group5697: abra|OJJ76560.1 acla|EAW14145.1 afis|EAW18858.1 afla|EED50825.1 afum|EAL93590.2 agla|OJJ80580.1 akaw|GAA82466.1 aluc|OJZ91644.1 anid|CBF69838.1 anig|EHA22666.1 aory|EIT78960.1 asyd|OJJ59035.1 ater|EAU38021.1 atub|OJI86099.1 aver|OJJ01581.1 awen|OJJ32414.1 azon|OJJ47171.1

Group5698: abra|OJJ76568.1 afis|EAW18853.1 afum|EAL93584.1 agla|OJJ80572.1 akaw|GAA82472.1 aluc|OJZ91638.1 anid|CBF69825.1 anig|EHA22661.1 asyd|OJJ58979.1 atub|OJI86091.1 aver|OJJ01574.1 awen|OJJ32407.1 azon|OJJ47165.1 acla|EAW14140.1 afla|EED50820.1 aory|EIT79040.1 ater|EAU38009.1

Group5699: abra|OJJ76574.1 acar|OOF99410.1 afis|EAW18821.1 afla|EED50805.1 afum|EAL93558.1 agla|OJJ80565.1 akaw|GAA82479.1 aluc|OJZ91631.1 anid|CBF82810.1 anig|EHA22654.1 aory|EIT78039.1 asyd|OJJ60269.1 ater|EAU38000.1 atub|OJI86084.1 aver|OJJ03985.1 awen|OJJ32399.1 azon|OJJ47159.1

Group5700: abra|OJJ76575.1 acar|OOF99409.1 acla|EAW14118.1 afis|EAW18820.1 afla|EED50807.1 afum|EAL93557.1 agla|OJJ80555.1 akaw|GAA82480.1 aluc|OJZ91630.1 anid|CBF69807.1 aory|EIT79005.1 asyd|OJJ58970.1 ater|EAU37999.1 atub|OJI86083.1 aver|OJJ01565.1 awen|OJJ32396.1 azon|OJJ47158.1

Group5701: abra|OJJ76578.1 acar|OOF99406.1 acla|EAW14115.1 afis|EAW18817.1 afla|EED50803.1 afum|EAL93553.1 agla|OJJ80550.1 akaw|GAA82483.1 aluc|OJZ91627.1 anid|CBF69813.1 anig|EHA22651.1 asyd|OJJ58973.1 ater|EAU37992.1 atub|OJI86080.1 aver|OJJ01568.1 awen|OJJ32393.1 azon|OJJ47154.1

Group5702: abra|OJJ76602.1 acar|OOF99389.1 acla|EAW14302.1 afis|EAW18982.1 afla|EED52436.1 afum|EAL93718.1 agla|OJJ82953.1 akaw|GAA82505.1 aluc|OJZ91606.1 anid|CBF69657.1 anig|EHA22635.1 asyd|OJJ59073.1 ater|EAU37829.1 atub|OJI86058.1 aver|OJJ01666.1 awen|OJJ31597.1 azon|OJJ47306.1

Group5703: abra|OJJ76607.1 acla|EAW14299.1 afis|EAW18985.1 afla|EED53163.1 afum|EAL93721.1 agla|OJJ82960.1 akaw|GAA82511.1 aluc|OJZ91600.1 anid|CBF69663.1 anig|EHA22630.1 aory|EIT75021.1 asyd|OJJ59070.1 ater|EAU37820.1 atub|OJI86051.1 aver|OJJ01663.1 awen|OJJ31605.1 azon|OJJ47309.1

Group5704: abra|OJJ76610.1 agla|OJJ82965.1 akaw|GAA82513.1 aluc|OJZ91598.1 anig|EHA22628.1 atub|OJI86049.1 acla|EAW14296.1 afis|EAW18987.1 afla|EED53166.1 afum|EAL93723.1 anid|CBF69667.1 aory|EIT75116.1 asyd|OJJ59068.1 ater|EAU37818.1 aver|OJJ01661.1 awen|OJJ31607.1 azon|OJJ47311.1

Group5705: abra|OJJ76625.1 acar|OOF99373.1 acla|EAW14294.1 afis|EAW18989.1 afla|EED53174.1 afum|EAL93725.1 agla|OJJ82976.1 akaw|GAA92832.1 aluc|OJZ91587.1 anid|CBF75890.1 anig|EHA22611.1 aory|EIT75047.1 asyd|OJJ60261.1 atub|OJI86036.1 aver|OJJ03992.1 awen|OJJ31610.1 anid|CBF82973.1

Group5706: abra|OJJ76683.1 acar|OOF99328.1 acla|EAW12908.1 afis|EAW17784.1 afla|EED57245.1 afum|EAL91592.1 agla|OJJ78673.1 akaw|GAA92511.1 aluc|OJZ91524.1 anid|CBF82109.1 aory|EIT74118.1 asyd|OJJ54283.1 ater|EAU37365.1 atub|OJI85979.1 aver|OJJ05237.1 awen|OJJ33522.1 azon|OJJ50362.1

Group5707: abra|OJJ76685.1 acar|OOF99326.1 acla|EAW07047.1 afis|EAW16672.1 afla|EED52488.1 afum|EAL89277.1 agla|OJJ78694.1 akaw|GAA92509.1 aluc|OJZ91522.1 anig|EHA22559.1 aory|EIT82788.1 asyd|OJJ56612.1 ater|EAU33886.1 atub|OJI85977.1 aver|OJJ06111.1 awen|OJJ40037.1 azon|OJJ45870.1

Group5708: abra|OJJ76694.1 acla|EAW11086.1 afis|EAW20772.1 afla|EED45937.1 afum|EAL84847.1 akaw|GAA92502.1 aluc|OJZ91515.1 anid|CBF71393.1 anig|EHA22553.1 aory|EIT75761.1 asyd|OJJ55030.1 ater|EAU36978.1 atub|OJI85967.1 aver|OJJ07173.1 awen|OJJ29779.1 anid|CBF69396.1 awen|OJJ41548.1

Group5709: abra|OJJ76751.1 acar|OOF91247.1 acla|EAW13770.1 afis|EAW20419.1 afum|EAL84630.1 agla|OJJ84906.1 akaw|GAA87273.1 aluc|OJZ84154.1 anid|CBF79344.1 aory|EIT77737.1 asyd|OJJ55798.1 ater|EAU37737.1 atub|OJI89930.1 aver|OJI98814.1 awen|OJJ33112.1 azon|OJJ45537.1 agla|OJJ83727.1

Group5710: abra|OJJ76754.1 acar|OOF91250.1 acla|EAW13308.1 afis|EAW20422.1 afla|EED44849.1 afum|EAL84628.1 agla|OJJ84903.1 akaw|GAA87270.1 aluc|OJZ84157.1 anig|EHA27519.1 aory|EIT77733.1 asyd|OJJ55795.1 ater|EAU37734.1 atub|OJI89927.1 aver|OJJ06884.1 awen|OJJ33115.1 azon|OJJ46331.1

Group5711: abra|OJJ76763.1 acar|OOF91258.1 acla|EAW13317.1 afis|EAW20431.1 afla|EED44837.1 agla|OJJ84894.1 akaw|GAA87263.1 aluc|OJZ84166.1 anid|CBF79329.1 anig|EHA27527.1 aory|EIT77758.1 asyd|OJJ55785.1 atub|OJI89918.1 aver|OJJ06875.1 awen|OJJ33124.1 azon|OJJ46340.1 ater|EAU37726.1

Group5712: abra|OJJ76770.1 acar|OOF91265.1 acla|EAW13324.1 afis|EAW20439.1 afla|EED44831.1 afum|EAL84613.1 agla|OJJ84884.1 akaw|GAA87256.1 aluc|OJZ84174.1 anid|CBF79313.1 anig|EHA27534.1 aory|EIT77752.1 asyd|OJJ55779.1 ater|EAU37719.1 atub|OJI89911.1 aver|OJJ06867.1 awen|OJJ33130.1

Group5713: abra|OJJ76773.1 acar|OOF91268.1 acla|EAW13330.1 afis|EAW20445.1 afla|EED44825.1 afum|EBA27197.1 agla|OJJ84876.1 akaw|GAA87251.1 aluc|OJZ84177.1 anid|CBF79305.1 anig|EHA27536.1 aory|EIT77731.1 ater|EAU37714.1 atub|OJI89906.1 aver|OJJ06865.1 awen|OJJ33135.1 azon|OJJ46348.1

Group5714: abra|OJJ76777.1 acar|OOF91273.1 acla|EAW13336.1 afis|EAW20451.1 afla|EED44817.1 afum|EAL84602.1 agla|OJJ84871.1 akaw|GAA87247.1 aluc|OJZ84181.1 anig|EHA27541.1 aory|EIT77722.1 asyd|OJJ52097.1 ater|EAU37708.1 atub|OJI89902.1 aver|OJJ03020.1 awen|OJJ33141.1 azon|OJJ46352.1

Group5715: abra|OJJ76782.1 acla|EAW13340.1 afis|EAW20455.1 afla|EED44813.1 afum|EBA27196.1 agla|OJJ84867.1 akaw|GAA87243.1 aluc|OJZ84185.1 anid|CBF79288.1 anig|EHA27545.1 aory|EIT77735.1 asyd|OJJ55768.1 ater|EAU37704.1 atub|OJI89898.1 aver|OJJ06854.1 awen|OJJ33145.1 azon|OJJ46357.1

Group5716: abra|OJJ76783.1 acar|OOF91277.1 acla|EAW13341.1 afis|EAW20456.1 afla|EED44812.1 afum|EAL84596.1 agla|OJJ84866.1 akaw|GAA87242.1 aluc|OJZ84186.1 anid|CBF79290.1 anig|EHA27546.1 aory|EIT77713.1 asyd|OJJ55767.1 atub|OJI89897.1 aver|OJJ06853.1 awen|OJJ33146.1 azon|OJJ46358.1

Group5717: abra|OJJ76788.1 acla|EAW13346.1 afis|EAW20461.1 afla|EED44807.1 afum|EAL84592.1 agla|OJJ84856.1 akaw|GAA87237.1 aluc|OJZ84190.1 anid|CBF79275.1 anig|EHA27550.1 aory|EIT77750.1 asyd|OJJ55762.1 ater|EAU37699.1 atub|OJI89892.1 aver|OJJ06847.1 awen|OJJ33154.1 azon|OJJ46363.1

Group5718: abra|OJJ76789.1 acar|OOF91282.1 acla|EAW13347.1 afis|EAW20462.1 afla|EED44806.1 afum|EAL84591.1 agla|OJJ84855.1 akaw|GAA87236.1 aluc|OJZ84191.1 anid|CBF79273.1 anig|EHA27551.1 aory|EIT77738.1 asyd|OJJ55761.1 atub|OJI89891.1 aver|OJJ06846.1 awen|OJJ33155.1 azon|OJJ46364.1

Group5719: abra|OJJ76805.1 acla|EAW13364.1 afis|EAW20478.1 afla|EED44790.1 afum|EAL84576.1 agla|OJJ84839.1 akaw|GAA87218.1 aluc|OJZ84212.1 anid|CBF79250.1 anig|EHA27565.1 aory|EIT77766.1 asyd|OJJ55744.1 ater|EAU29556.1 atub|OJI89871.1 aver|OJJ06831.1 awen|OJJ33173.1 azon|OJJ46380.1

Group5720: abra|OJJ76820.1 acar|OOF91314.1 acla|EAW13380.1 afis|EAW20494.1 afla|EED44770.1 afum|EAL84560.2 agla|OJJ84808.1 akaw|GAA87202.1 aluc|OJZ84230.1 anid|CBF79224.1 aory|EIT77791.1 asyd|OJJ55733.1 ater|EAU29570.1 atub|OJI89854.1 aver|OJJ06819.1 awen|OJJ33194.1 azon|OJJ46402.1

Group5721: abra|OJJ76841.1 acar|OOF91334.1 acla|EAW13404.1 afis|EAW20515.1 afla|EED52467.1 afum|EAL84542.1 agla|OJJ84775.1 akaw|GAA87181.1 aluc|OJZ84252.1 anid|CBF79057.1 anig|EHA27597.1 aory|EIT82760.1 asyd|OJJ55709.1 ater|EAU37186.1 atub|OJI89831.1 aver|OJJ06795.1 awen|OJJ33221.1

Group5722: abra|OJJ76846.1 acar|OOF91336.1 acla|EAW13409.1 afis|EAW20520.1 afla|EED52469.1 afum|EAL84537.1 agla|OJJ84766.1 akaw|GAA87175.1 aluc|OJZ84258.1 anid|CBF79069.1 aory|EIT82811.1 asyd|OJJ55705.1 ater|EAU37168.1 atub|OJI89825.1 aver|OJJ06791.1 awen|OJJ33226.1 azon|OJJ46578.1

Group5723: abra|OJJ76848.1 acla|EAW13411.1 afis|EAW20522.1 afla|EED52471.1 afum|EAL84535.1 agla|OJJ84739.1 akaw|GAA87173.1 aluc|OJZ84260.1 anid|CBF79072.1 anig|EHA27603.1 aory|EIT82780.1 asyd|OJJ55703.1 ater|EAU37166.1 atub|OJI89823.1 aver|OJJ06789.1 awen|OJJ33228.1 azon|OJJ46576.1

Group5724: abra|OJJ76853.1 acla|EAW13415.1 afis|EAW20526.1 afla|EED52475.1 afum|EAL84531.1 agla|OJJ84744.1 akaw|GAA87169.1 aluc|OJZ84265.1 anid|CBF79080.1 anig|EHA27607.1 aory|EIT82795.1 asyd|OJJ55698.1 ater|EAU37162.1 atub|OJI89818.1 aver|OJJ06783.1 awen|OJJ33232.1 azon|OJJ46572.1

Group5725: abra|OJJ76858.1 acar|OOF91346.1 acla|EAW13419.1 afis|EAW20530.1 afla|EED52480.1 afum|EAL84527.1 agla|OJJ84704.1 akaw|GAA87164.1 aluc|OJZ84270.1 anid|CBF79004.1 anig|EHA27612.1 aory|EIT82752.1 asyd|OJJ55690.1 ater|EAU37155.1 atub|OJI89813.1 aver|OJJ06774.1 awen|OJJ33237.1

Group5726: abra|OJJ76902.1 acar|OOF91350.1 acla|EAW13423.1 afis|EAW20535.1 afla|EED54118.1 afum|EAL84523.1 akaw|GAA87136.1 aluc|OJZ84301.1 anid|CBF78997.1 anig|EHA27637.1 aory|EIT76011.1 asyd|OJJ55686.1 ater|EAU37148.1 atub|OJI89767.1 aver|OJJ06770.1 awen|OJJ33242.1 azon|OJJ50621.1

Group5727: abra|OJJ76915.1 acar|OOF91362.1 afis|EAW20543.1 afla|EED52430.1 afum|EAL84515.1 agla|OJJ88755.1 akaw|GAA87123.1 aluc|OJZ84315.1 anid|CBF78989.1 anig|EHA27650.1 aory|EIT83314.1 asyd|OJJ55683.1 ater|EAU37138.1 atub|OJI89753.1 aver|OJJ06767.1 awen|OJJ33250.1 azon|OJJ42414.1

Group5728: abra|OJJ76974.1 acar|OOF91418.1 acla|EAW13500.1 afis|EAW20599.1 afla|EED52205.1 afum|EAL84462.1 agla|OJJ88656.1 akaw|GAA87071.1 aluc|OJZ84370.1 anid|CBF78906.1 anig|EHA27705.1 asyd|OJJ55610.1 ater|EAU35244.1 atub|OJI89697.1 aver|OJJ06685.1 awen|OJJ33391.1 azon|OJJ50695.1

Group5729: abra|OJJ76975.1 acar|OOF91420.1 acla|EAW13501.1 afis|EAW20600.1 afla|EED47728.1 afum|EAL84461.1 agla|OJJ88657.1 akaw|GAA87070.1 aluc|OJZ84371.1 anid|CBF86933.1 anig|EHA27706.1 aory|EIT77127.1 asyd|OJJ55598.1 atub|OJI89696.1 aver|OJJ06678.1 awen|OJJ33392.1 azon|OJJ50696.1

Group5730: abra|OJJ76999.1 acar|OOF91445.1 acla|EAW13462.1 afis|EAW20568.1 afla|EED52322.1 afum|EAL84494.1 agla|OJJ88725.1 akaw|GAA87049.1 aluc|OJZ84397.1 anid|CBF87400.1 anig|EHA27728.1 aory|EIT74352.1 asyd|OJJ52861.1 ater|EAU37115.1 atub|OJI89675.1 aver|OJJ08839.1 awen|OJJ33272.1

Group5731: abra|OJJ77006.1 acar|OOF91451.1 afis|EAW25624.1 afla|EED54122.1 afum|EAL84312.1 agla|OJJ78904.1 akaw|GAA87042.1 aluc|OJZ84405.1 anid|CBF83175.1 anig|EHA27735.1 aory|EIT76161.1 asyd|OJJ55524.1 ater|EAU32184.1 atub|OJI89667.1 aver|OJJ06602.1 awen|OJJ36043.1 azon|OJJ42823.1

Group5732: abra|OJJ77035.1 acla|EAW06829.1 afis|EAW16379.1 afla|EED54135.1 afum|EAL89004.1 agla|OJJ88206.1 akaw|GAA87016.1 aluc|OJZ84438.1 anid|CBF81133.1 anig|EHA27757.1 aory|EIT76057.1 asyd|OJJ56501.1 ater|EAU32642.1 atub|OJI89635.1 aver|OJJ06218.1 awen|OJJ37913.1 azon|OJJ45092.1

Group5733: abra|OJJ77047.1 acar|OOF92936.1 acla|EAW13551.1 afis|EAW20626.1 afla|EED49776.1 afum|EAL84440.1 agla|OJJ80442.1 akaw|GAA87006.1 aluc|OJZ84451.1 anid|CBF83226.1 anig|EHA27766.1 aory|EIT77694.1 asyd|OJJ55580.1 atub|OJI89625.1 aver|OJJ06659.1 awen|OJJ33463.1 azon|OJJ50719.1

Group5734: abra|OJJ77051.1 acar|OOF92940.1 acla|EAW13542.1 afis|EAW20615.1 afla|EED54202.1 afum|EAL84446.1 agla|OJJ88593.1 akaw|GAA87002.1 aluc|OJZ84455.1 anid|CBF86938.1 anig|EHA27770.1 asyd|OJJ55588.1 ater|EAU35252.1 atub|OJI89620.1 aver|OJJ06667.1 awen|OJJ33459.1 azon|OJJ45125.1

Group5735: abra|OJJ77052.1 acar|OOF92941.1 acla|EAW13540.1 afis|EAW20613.1 afum|EAL84448.1 agla|OJJ88622.1 akaw|GAA87000.1 aluc|OJZ84457.1 anid|CBF86943.1 anig|EHA27772.1 aory|EIT76152.1 asyd|OJJ55585.1 ater|EAU35254.1 atub|OJI89618.1 aver|OJJ06664.1 awen|OJJ33457.1 azon|OJJ50713.1

Group5736: abra|OJJ77062.1 acla|EAW13513.1 afis|EAW20611.1 afla|EED54206.1 afum|EAL84450.1 agla|OJJ88628.1 akaw|GAA86990.1 aluc|OJZ84470.1 anid|CBF86957.1 anig|EHA27780.1 aory|EIT75918.1 asyd|OJJ55596.1 ater|EAU35256.1 atub|OJI89608.1 aver|OJJ06675.1 awen|OJJ33454.1 azon|OJJ50711.1

Group5737: abra|OJJ77066.1 acar|OOF92950.1 acla|EAW13509.1 afis|EAW20607.1 afla|EED54208.1 afum|EAL84453.1 agla|OJJ88632.1 akaw|GAA86986.1 aluc|OJZ84475.1 anid|CBF86948.1 anig|EHA27783.1 asyd|OJJ55592.1 ater|EAU35259.1 atub|OJI89604.1 aver|OJJ06671.1 awen|OJJ33450.1 azon|OJJ50707.1

Group5738: abra|OJJ77070.1 acla|EAW09727.1 afis|EAW21413.1 afla|EED55836.1 afum|EAL92427.1 agla|OJJ85544.1 akaw|GAA86983.1 aluc|OJZ84478.1 anid|CBF83797.1 anig|EHA27786.1 aory|EIT76757.1 asyd|OJJ60887.1 ater|EAU35133.1 atub|OJI89601.1 aver|OJJ00211.1 awen|OJJ39288.1 azon|OJJ51566.1

Group5739: abra|OJJ77071.1 acar|OOF92953.1 acla|EAW09728.1 afis|EAW21414.1 afla|EED55835.1 afum|EAL92428.1 agla|OJJ85543.1 akaw|GAA86982.1 aluc|OJZ84479.1 anig|EHA27787.1 aory|EIT76805.1 asyd|OJJ60888.1 ater|EAU35134.1 atub|OJI89600.1 aver|OJJ00212.1 awen|OJJ39289.1 azon|OJJ51565.1

Group5740: abra|OJJ77074.1 acar|OOF92955.1 acla|EAW12062.1 afis|EAW19705.1 afla|EED57046.1 afum|EAL86821.1 agla|OJJ78914.1 akaw|GAA86979.1 aluc|OJZ84484.1 anid|CBF71407.1 anig|EHA27788.1 aory|EIT82262.1 asyd|OJJ55014.1 ater|EAU32844.1 atub|OJI89596.1 aver|OJJ07197.1 awen|OJJ29913.1

Group5741: abra|OJJ77080.1 acar|OOF92963.1 acla|EAW06701.1 afis|EAW16246.1 afla|EED53307.1 afum|EAL88874.1 agla|OJJ86255.1 akaw|GAA86973.1 aluc|OJZ84490.1 anid|CBF78902.1 anig|EHA27794.1 aory|EIT75199.1 asyd|OJJ64604.1 ater|EAU37456.1 atub|OJI89589.1 aver|OJI98701.1 awen|OJJ30210.1

Group5742: abra|OJJ77087.1 acar|OOF92971.1 afis|EAW21842.1 afla|EED46528.1 afum|EAL92854.1 agla|OJJ84295.1 akaw|GAA86968.1 aluc|OJZ84496.1 anid|CBF76713.1 anig|EHA27798.1 aory|EIT72340.1 asyd|OJJ61240.1 ater|EAU38519.1 atub|OJI89584.1 aver|OJJ00563.1 awen|OJJ34126.1 azon|OJJ44217.1

Group5743: abra|OJJ77088.1 acar|OOF92972.1 acla|EAW10185.1 afis|EAW25410.1 afla|EED53889.1 afum|EAL90069.1 agla|OJJ87051.1 akaw|GAA86967.1 aluc|OJZ84497.1 anid|CBF77586.1 aory|EIT78478.1 asyd|OJJ53170.1 ater|EAU34561.1 atub|OJI89583.1 aver|OJJ03682.1 awen|OJJ41137.1 azon|OJJ44716.1

Group5744: abra|OJJ77089.1 acar|OOF92973.1 acla|EAW08631.1 afis|EAW15800.1 afla|EED51245.1 afum|EAL85550.2 agla|OJJ81967.1 akaw|GAA86966.1 aluc|OJZ84498.1 anid|CBF70920.1 aory|EIT83182.1 asyd|OJJ52606.1 ater|EAU32342.1 atub|OJI89582.1 aver|OJJ04633.1 awen|OJJ38509.1 azon|OJJ43743.1

Group5745: abra|OJJ77099.1 acar|OOF92981.1 afis|EAW16895.1 afla|EED53380.1 afum|EAL86432.1 agla|OJJ79067.1 akaw|GAA86957.1 aluc|OJZ84507.1 anid|CBF80785.1 anig|EHA27806.1 aory|EIT75077.1 asyd|OJJ54496.1 ater|EAU30219.1 atub|OJI89573.1 aver|OJJ07684.1 awen|OJJ29738.1 agla|OJJ79950.1

Group5746: abra|OJJ77115.1 acar|OOF93004.1 acla|EAW12953.1 afis|EAW17703.1 afla|EED45697.1 afum|EAL91518.1 agla|OJJ79466.1 akaw|GAA90814.1 aluc|OJZ84525.1 anid|CBF89930.1 anig|EHA27822.1 aory|EIT75749.1 asyd|OJJ62118.1 ater|EAU31911.1 atub|OJI89555.1 aver|OJI96225.1 awen|OJJ37132.1

Group5747: abra|OJJ77117.1 acar|OOF93234.1 acla|EAW12934.1 afis|EAW17706.1 afla|EED45708.1 afum|EAL91520.2 akaw|GAA90812.1 aluc|OJZ84527.1 anig|EHA27824.1 aory|EIT75608.1 asyd|OJJ62133.1 ater|EAU31913.1 atub|OJI89553.1 aver|OJI96242.1 awen|OJJ37130.1 azon|OJJ50829.1 afis|EAW17705.1

Group5748: abra|OJJ77119.1 acla|EAW12931.1 afis|EAW17708.1 afla|EED45710.1 afum|EAL91522.2 agla|OJJ81739.1 akaw|GAA90810.1 aluc|OJZ84529.1 anid|CBF89908.1 anig|EHA27826.1 aory|EIT75384.1 asyd|OJJ62131.1 ater|EAU31904.1 atub|OJI89551.1 aver|OJI96240.1 awen|OJJ37128.1 azon|OJJ50827.1

Group5749: abra|OJJ77124.1 acar|OOF93010.1 acla|EAW12937.1 afis|EAW17713.1 afla|EED45715.1 afum|EAL91528.1 agla|OJJ81735.1 akaw|GAA90804.1 aluc|OJZ84534.1 anid|CBF76918.1 anig|EHA27832.1 aory|EIT75535.1 asyd|OJJ57073.1 atub|OJI89546.1 aver|OJJ03225.1 awen|OJJ37123.1 azon|OJJ50822.1

Group5750: abra|OJJ77129.1 acla|EAW12940.1 afis|EAW17716.1 afla|EED45718.1 afum|EAL91531.1 agla|OJJ81732.1 akaw|GAA90799.1 aluc|OJZ84539.1 anid|CBF84090.1 anig|EHA27837.1 aory|EIT75537.1 asyd|OJJ57081.1 ater|EAU31894.1 atub|OJI89541.1 aver|OJJ03221.1 awen|OJJ37120.1 azon|OJJ50819.1

Group5751: abra|OJJ77132.1 acar|OOF93020.1 acla|EAW12943.1 afis|EAW17719.1 afla|EED45721.1 afum|EAL91534.2 agla|OJJ81729.1 akaw|GAA90796.1 aluc|OJZ84542.1 anid|CBF76906.1 anig|EHA27840.1 asyd|OJJ57084.1 ater|EAU31897.1 atub|OJI89538.1 aver|OJJ03218.1 awen|OJJ37117.1 azon|OJJ50816.1

Group5752: abra|OJJ77137.1 acar|OOF90284.1 afis|EAW16801.1 afla|EED47170.1 afum|EAL84245.1 agla|OJJ88968.1 akaw|GAA90792.1 aluc|OJZ84548.1 anid|CBF78601.1 anig|EHA27844.1 asyd|OJJ59267.1 atub|OJI89532.1 aver|OJJ01879.1 awen|OJJ32033.1 acla|EAW12954.1 afis|EAW15402.1 aory|EIT79292.1

Group5753: abra|OJJ77151.1 acar|OOF93037.1 acla|EAW12918.1 afis|EAW17737.1 afla|EED45738.1 afum|EAL91552.1 akaw|GAA90779.1 aluc|OJZ84565.1 anid|CBF86590.1 anig|EHA27859.1 aory|EIT75449.1 asyd|OJJ64829.1 ater|EAU31870.1 atub|OJI89517.1 aver|OJI98942.1 awen|OJJ37098.1 azon|OJJ50799.1

Group5754: abra|OJJ77169.1 acar|OOF93060.1 acla|EAW12897.1 afis|EAW17772.1 afla|EED45786.1 afum|EAL91581.1 agla|OJJ81695.1 akaw|GAA90764.1 aluc|OJZ84582.1 anig|EHA27875.1 aory|EIT75547.1 ater|EAU31841.1 atub|OJI89498.1 awen|OJJ37074.1 acla|EAW15270.1 afis|EAW20239.1 afum|EAL85357.1

Group5755: abra|OJJ77208.1 acla|EAW13200.1 afis|EAW17442.1 afla|EED49385.1 afum|EAL91268.1 agla|OJJ80375.1 akaw|GAA90730.1 aluc|OJZ84619.1 anid|CBF71696.1 anig|EHA27907.1 aory|EIT79456.1 asyd|OJJ61815.1 ater|EAU33921.1 atub|OJI89462.1 aver|OJI95914.1 awen|OJJ37404.1 azon|OJJ51078.1

Group5756: abra|OJJ77224.1 acar|OOF93109.1 acla|EAW13215.1 afis|EAW17427.1 afla|EED49401.1 afum|EAL91255.1 agla|OJJ80397.1 akaw|GAA90715.1 aluc|OJZ84634.1 anid|CBF71718.1 anig|EHA27922.1 aory|EIT79578.1 asyd|OJJ61803.1 atub|OJI89445.1 aver|OJI95902.1 awen|OJJ37416.1 azon|OJJ51090.1

Group5757: abra|OJJ77227.1 acar|OOF93112.1 acla|EAW13218.1 afis|EAW17424.1 afla|EED49398.1 afum|EAL91252.1 agla|OJJ80400.1 akaw|GAA90712.1 aluc|OJZ84637.1 anid|CBF71724.1 anig|EHA27925.1 asyd|OJJ61800.1 ater|EAU33906.1 atub|OJI89442.1 aver|OJI95899.1 awen|OJJ37419.1 azon|OJJ51093.1

Group5758: abra|OJJ77237.1 acar|OOF93118.1 acla|EAW13244.1 afis|EAW17392.1 afla|EED49476.1 afum|EAL91230.1 agla|OJJ80402.1 akaw|GAA90706.1 aluc|OJZ84647.1 anid|CBF71737.1 anig|EHA27931.1 asyd|OJJ61793.1 ater|EAU33897.1 atub|OJI89433.1 aver|OJI95893.1 awen|OJJ37423.1 azon|OJJ51106.1

Group5759: abra|OJJ77239.1 acar|OOF93120.1 acla|EAW13246.1 afis|EAW17390.1 afla|EED49478.1 afum|EAL91226.1 agla|OJJ80407.1 akaw|GAA90704.1 aluc|OJZ84649.1 anid|CBF71741.1 aory|EIT79473.1 asyd|OJJ61791.1 ater|EAU33895.1 atub|OJI89431.1 aver|OJI95891.1 awen|OJJ37425.1 azon|OJJ51103.1

Group5760: abra|OJJ77245.1 acla|EAW13249.1 afis|EAW17387.1 afla|EED49496.1 afum|EAL91221.2 agla|OJJ80412.1 akaw|GAA90697.1 aluc|OJZ84656.1 anid|CBF84185.1 anig|EHA27938.1 aory|EIT82598.1 asyd|OJJ57142.1 ater|EAU33892.1 atub|OJI89424.1 aver|OJJ03162.1 awen|OJJ37429.1 azon|OJJ51100.1

Group5761: abra|OJJ77246.1 acar|OOF93127.1 acla|EAW13250.1 afis|EAW17386.1 afla|EED49497.1 afum|EBA27427.1 agla|OJJ80416.1 akaw|GAA90696.1 aluc|OJZ84657.1 anid|CBF84181.1 anig|EHA27939.1 aory|EIT82594.1 asyd|OJJ57158.1 atub|OJI89423.1 aver|OJJ03155.1 awen|OJJ37430.1 azon|OJJ51107.1

Group5762: abra|OJJ77251.1 acar|OOF90004.1 acla|EAW13254.1 afis|EAW17382.1 afla|EED49505.1 afum|EAL91217.1 akaw|GAA90692.1 aluc|OJZ84662.1 anid|CBF84200.1 anig|EHA27944.1 aory|EIT82574.1 asyd|OJJ57162.1 ater|EAU33879.1 atub|OJI89418.1 aver|OJJ03151.1 awen|OJJ37435.1 azon|OJJ51113.1

Group5763: abra|OJJ77259.1 acar|OOF90003.1 acla|EAW13255.1 afis|EAW17381.1 afla|EED49507.1 afum|EAL91216.1 agla|OJJ80421.1 akaw|GAA90688.1 aluc|OJZ84666.1 anid|CBF84205.1 anig|EHA27945.1 aory|EIT82652.1 asyd|OJJ57165.1 atub|OJI89415.1 aver|OJJ03149.1 awen|OJJ37437.1 azon|OJJ51114.1

Group5764: abra|OJJ77276.1 acar|OOF93151.1 acla|EAW13276.1 afis|EAW17341.1 afla|EED49780.1 afum|EAL91186.1 agla|OJJ79343.1 akaw|GAA90673.1 aluc|OJZ84686.1 anid|CBF84219.1 anig|EHA27961.1 aory|EIT77670.1 asyd|OJJ57177.1 atub|OJI89396.1 aver|OJJ03137.1 awen|OJJ36045.1 azon|OJJ45995.1

Group5765: abra|OJJ77280.1 acar|OOF93154.1 acla|EAW13279.1 afis|EAW17338.1 afla|EED51043.1 afum|EAL91182.1 agla|OJJ79348.1 akaw|GAA90669.1 aluc|OJZ84690.1 anig|EHA27965.1 aory|EIT76982.1 asyd|OJJ57176.1 ater|EAU33848.1 atub|OJI89392.1 aver|OJJ03138.1 awen|OJJ40817.1 azon|OJJ50225.1

Group5766: abra|OJJ77294.1 acar|OOF93171.1 acla|EAW13170.1 afis|EAW17469.1 afla|EED49347.1 afum|EAL91298.1 agla|OJJ78441.1 akaw|GAA91457.1 aluc|OJZ84706.1 anid|CBF71628.1 anig|EHA27980.1 aory|EIT79559.1 asyd|OJJ61848.1 ater|EAU33953.1 atub|OJI89377.1 aver|OJI95947.1 awen|OJJ37373.1

Group5767: abra|OJJ77296.1 acar|OOF93173.1 acla|EAW13168.1 afis|EAW17471.1 afla|EED49349.1 afum|EAL91300.1 agla|OJJ81491.1 akaw|GAA91455.1 aluc|OJZ84708.1 anid|CBF71632.1 anig|EHA27982.1 aory|EIT79572.1 asyd|OJJ61850.1 ater|EAU33955.1 atub|OJI89375.1 aver|OJI95949.1 awen|OJJ37371.1

Group5768: abra|OJJ77299.1 acar|OOF93178.1 acla|EAW13166.1 afis|EAW17473.1 afla|EED49342.1 afum|EAL91302.1 agla|OJJ81489.1 akaw|GAA91453.1 aluc|OJZ84711.1 anid|CBF71624.1 anig|EHA27984.1 asyd|OJJ61852.1 ater|EAU33957.1 atub|OJI89372.1 aver|OJI95952.1 awen|OJJ37369.1 azon|OJJ51048.1

Group5769: abra|OJJ77302.1 acar|OOF93181.1 afis|EAW17476.1 afla|EED49338.1 afum|EAL91305.1 agla|OJJ81485.1 akaw|GAA91449.1 aluc|OJZ84715.1 anid|CBF71601.1 anig|EHA27987.1 aory|EIT79492.1 asyd|OJJ61856.1 ater|EAU33692.1 atub|OJI89368.1 aver|OJI95955.1 awen|OJJ37366.1 azon|OJJ45454.1

Group5770: abra|OJJ77316.1 acar|OOF93192.1 acla|EAW13156.1 afis|EAW17490.1 afla|EED49326.1 afum|EAL91312.1 agla|OJJ81466.1 aluc|OJZ84726.1 anid|CBF71619.1 anig|EHA28000.1 aory|EIT79591.1 asyd|OJJ61865.1 ater|EAU36935.1 atub|OJI89356.1 aver|OJI95963.1 awen|OJJ37351.1 azon|OJJ51037.1

Group5771: abra|OJJ77323.1 acar|OOF93199.1 afis|EAW17497.1 afla|EED49319.1 afum|EAL91320.1 agla|OJJ81458.1 akaw|GAA91433.1 aluc|OJZ84733.1 anid|CBF71584.1 anig|EHA28006.1 asyd|OJJ61873.1 ater|EAU36941.1 atub|OJI89348.1 aver|OJI95971.1 awen|OJJ37344.1 azon|OJJ51030.1 ater|EAU38793.1

Group5772: abra|OJJ77324.1 acar|OOF93200.1 acla|EAW13148.1 afis|EAW17499.1 afla|EED49317.1 afum|EAL91322.1 agla|OJJ81454.1 akaw|GAA91432.1 aluc|OJZ84734.1 anid|CBF71580.1 anig|EHA28007.1 asyd|OJJ61875.1 ater|EAU36944.1 atub|OJI89347.1 aver|OJI95973.1 awen|OJJ37342.1 azon|OJJ51027.1

Group5773: abra|OJJ77330.1 acar|OOF93205.1 acla|EAW13143.1 afis|EAW17515.1 afla|EED49312.1 afum|EAL91333.1 agla|OJJ81449.1 akaw|GAA91427.1 aluc|OJZ84739.1 anid|CBF71570.1 anig|EHA28012.1 aory|EIT82234.1 asyd|OJJ61880.1 atub|OJI89342.1 aver|OJI95978.1 awen|OJJ37336.1 azon|OJJ51022.1

Group5774: abra|OJJ77332.1 acar|OOF93206.1 acla|EAW13142.1 afis|EAW17514.1 afla|EED49311.1 afum|EAL91332.1 agla|OJJ81448.1 akaw|GAA91426.1 aluc|OJZ84740.1 anid|CBF73599.1 anig|EHA28013.1 aory|EIT82247.1 asyd|OJJ56681.1 ater|EAU36951.1 atub|OJI89341.1 aver|OJJ06042.1 awen|OJJ37335.1

Group5775: abra|OJJ77340.1 acla|EAW13135.1 afis|EAW17504.1 afla|EED49304.1 afum|EAL91326.1 agla|OJJ81442.1 akaw|GAA91421.1 aluc|OJZ84749.1 anid|CBF71539.1 anig|EHA28019.1 aory|EIT82224.1 asyd|OJJ61895.1 ater|EAU36957.1 atub|OJI89333.1 aver|OJI95991.1 awen|OJJ37327.1 azon|OJJ51016.1

Group5776: abra|OJJ77360.1 acar|OOF93229.1 acla|EAW13116.1 afis|EAW17548.1 afla|EED49277.1 afum|EAL91362.1 agla|OJJ81420.1 akaw|GAA91403.1 aluc|OJZ84768.1 anid|CBF90292.1 aory|EIT82223.1 asyd|OJJ61922.1 ater|EAU36987.1 atub|OJI89314.1 aver|OJI96014.1 awen|OJJ37306.1 azon|OJJ50998.1

Group5777: abra|OJJ77389.1 acar|OOF93258.1 acla|EAW08796.1 afis|EAW15599.1 afla|EED45975.1 afum|EAL85733.1 akaw|GAA91372.1 aluc|OJZ84797.1 anid|CBF78871.1 anig|EHA28064.1 aory|EIT75420.1 asyd|OJJ64587.1 ater|EAU30348.1 atub|OJI89285.1 aver|OJI98684.1 awen|OJJ37986.1 azon|OJJ43887.1

Group5778: abra|OJJ77395.1 acar|OOF93263.1 afis|EAW17805.1 afla|EED45800.1 afum|EAL91613.1 agla|OJJ78527.1 akaw|GAA91367.1 aluc|OJZ84803.1 anid|CBF86646.1 anig|EHA28068.1 aory|EIT75621.1 asyd|OJJ64799.1 ater|EAU31836.1 atub|OJI89280.1 aver|OJI98913.1 awen|OJJ41514.1 agla|OJJ80100.1

Group5779: abra|OJJ77466.1 acar|OOF99268.1 acla|EAW07126.1 afis|EAW22136.1 afla|EED54175.1 afum|EAL91054.1 akaw|GAA84151.1 aluc|OJZ81072.1 anid|CBF69552.1 anig|EHA19297.1 aory|EIT75896.1 asyd|OJJ59166.1 ater|EAU29320.1 atub|OJI82514.1 aver|OJJ01756.1 awen|OJJ34713.1 aory|EIT76090.1

Group5780: abra|OJJ77469.1 acar|OOF99255.1 acla|EAW06765.1 afis|EAW16315.1 afla|EED54179.1 afum|EAL88938.1 agla|OJJ88273.1 akaw|GAA84148.1 aluc|OJZ81069.1 anid|CBF69555.1 anig|EHA19300.1 aory|EIT75912.1 asyd|OJJ59164.1 ater|EAU29325.1 atub|OJI82517.1 aver|OJJ01754.1 awen|OJJ37713.1

Group5781: abra|OJJ77470.1 acar|OOF99254.1 acla|EAW06762.1 afis|EAW16312.1 afla|EED54183.1 afum|EAL88934.1 agla|OJJ88271.1 akaw|GAA84147.1 aluc|OJZ81068.1 anig|EHA19301.1 aory|EIT76029.1 asyd|OJJ59157.1 ater|EAU29330.1 atub|OJI82518.1 aver|OJJ01747.1 awen|OJJ37712.1 azon|OJJ44971.1

Group5782: abra|OJJ77472.1 acar|OOF99252.1 acla|EAW06759.1 afis|EAW16309.1 afla|EED54185.1 afum|EAL88931.1 agla|OJJ86289.1 akaw|GAA84145.1 aluc|OJZ81065.1 anid|CBF69562.1 anig|EHA19303.1 aory|EIT75996.1 asyd|OJJ59155.1 ater|EAU29338.1 atub|OJI82520.1 aver|OJJ01745.1 awen|OJJ37708.1

Group5783: abra|OJJ77473.1 acar|OOF99251.1 acla|EAW06758.1 afis|EAW16308.1 afla|EED53241.1 afum|EAL88930.1 agla|OJJ86288.1 akaw|GAA84144.1 aluc|OJZ81064.1 anid|CBF69564.1 anig|EHA19304.1 aory|EIT75223.1 asyd|OJJ59154.1 ater|EAU29337.1 atub|OJI82521.1 aver|OJJ01744.1 awen|OJJ37707.1

Group5784: abra|OJJ77481.1 acar|OOF99244.1 acla|EAW06743.1 afis|EAW16293.1 afum|EAL88915.1 agla|OJJ88315.1 akaw|GAA84137.1 aluc|OJZ81056.1 anid|CBF86850.1 anig|EHA19311.1 aory|EIT75158.1 asyd|OJJ64639.1 ater|EAU37337.1 atub|OJI82528.1 aver|OJI98738.1 awen|OJJ37687.1 azon|OJJ44947.1

Group5785: abra|OJJ77483.1 acar|OOF99243.1 acla|EAW06744.1 afis|EAW16294.1 afla|EED53260.1 afum|EAL88916.1 agla|OJJ88316.1 akaw|GAA84136.1 aluc|OJZ81054.1 anid|CBF86848.1 anig|EHA19313.1 aory|EIT74888.1 asyd|OJJ64640.1 atub|OJI82530.1 aver|OJI98740.1 awen|OJJ37685.1 azon|OJJ44946.1

Group5786: abra|OJJ77484.1 acar|OOF99241.1 acla|EAW06747.1 afis|EAW16297.1 afla|EED53257.1 afum|EAL88919.1 agla|OJJ88298.1 akaw|GAA84134.1 aluc|OJZ81052.1 anid|CBF86845.1 anig|EHA19315.1 asyd|OJJ64643.1 ater|EAU29339.1 atub|OJI82532.1 aver|OJI98743.1 awen|OJJ37696.1 azon|OJJ44957.1

Group5787: abra|OJJ77487.1 acar|OOF99238.1 acla|EAW06749.1 afis|EAW16299.1 afla|EED53255.1 afum|EAL88921.1 agla|OJJ88296.1 akaw|GAA84132.1 aluc|OJZ81049.1 anig|EHA19318.1 aory|EIT75108.1 asyd|OJJ64645.1 ater|EAU37341.1 atub|OJI82535.1 aver|OJI98745.1 awen|OJJ37698.1 azon|OJJ44959.1

Group5788: abra|OJJ77519.1 acar|OOF99205.1 acla|EAW06713.1 afis|EAW16274.1 afla|EED53296.1 afum|EAL88895.1 agla|OJJ78577.1 akaw|GAA84102.1 aluc|OJZ81017.1 anid|CBF78275.1 anig|EHA19345.1 aory|EIT75252.1 asyd|OJJ56075.1 ater|EAU37449.1 atub|OJI82566.1 aver|OJJ05018.1 azon|OJJ50409.1

Group5789: abra|OJJ77531.1 acla|EAW06706.1 afis|EAW16263.1 afla|EED53301.1 afum|EAL88884.1 agla|OJJ88379.1 akaw|GAA84089.1 aluc|OJZ81004.1 anid|CBF86916.1 anig|EHA19356.1 aory|EIT75165.1 asyd|OJJ64606.1 ater|EAU37452.1 atub|OJI82581.1 aver|OJI98703.1 awen|OJJ37653.1 azon|OJJ44917.1

Group5790: abra|OJJ77536.1 acar|OOF99190.1 acla|EAW06711.1 afis|EAW16257.1 afla|EED53323.1 afum|EAL88878.1 akaw|GAA84084.1 aluc|OJZ80998.1 anid|CBF78893.1 anig|EHA19362.1 aory|EIT75253.1 asyd|OJJ64598.1 ater|EAU37468.1 atub|OJI82586.1 aver|OJI98695.1 awen|OJJ30202.1 azon|OJJ50681.1

Group5791: abra|OJJ77546.1 acar|OOF99177.1 acla|EAW06708.1 afis|EAW16261.1 afla|EED53325.1 afum|EAL88882.1 agla|OJJ88382.1 akaw|GAA84077.1 aluc|OJZ80991.1 anid|CBF78870.1 anig|EHA19371.1 asyd|OJJ64588.1 ater|EAU33700.1 atub|OJI82594.1 aver|OJI98685.1 awen|OJJ37651.1 azon|OJJ44915.1

Group5792: abra|OJJ77547.1 acar|OOF99176.1 acla|EAW06707.1 afis|EAW16262.1 afla|EED53326.1 afum|EAL88883.1 agla|OJJ88381.1 akaw|GAA84076.1 aluc|OJZ80990.1 anid|CBF78868.1 aory|EIT74871.1 asyd|OJJ64589.1 ater|EAU33701.1 atub|OJI82595.1 aver|OJI98686.1 awen|OJJ37652.1 azon|OJJ44916.1

Group5793: abra|OJJ77549.1 acar|OOF95414.1 acla|EAW08956.1 afla|EED49536.1 afum|EAL88030.1 agla|OJJ78723.1 akaw|GAA84074.1 aluc|OJZ80988.1 anid|CBF70524.1 anig|EHA19372.1 asyd|OJJ57565.1 atub|OJI82598.1 aver|OJJ05614.1 awen|OJJ32036.1 azon|OJJ46060.1 asyd|OJJ56222.1 aver|OJJ05154.1

Group5794: abra|OJJ77558.1 acar|OOG00297.1 afis|EAW16397.1 afla|EED51760.1 afum|EAL89030.2 agla|OJJ88351.1 akaw|GAA84066.1 aluc|OJZ80980.1 anid|CBF85653.1 anig|EHA19381.1 aory|EIT80739.1 asyd|OJJ62365.1 ater|EAU33012.1 atub|OJI82606.1 aver|OJI96499.1 awen|OJJ30119.1 agla|OJJ82192.1

Group5795: abra|OJJ77562.1 acla|EAW06695.1 afis|EAW16071.1 afla|EED48590.1 afum|EAL88703.2 agla|OJJ86097.1 akaw|GAA84063.1 aluc|OJZ80976.1 anid|CBF75105.1 anig|EHA19383.1 aory|EIT73151.1 asyd|OJJ59999.1 ater|EAU33560.1 atub|OJI82609.1 aver|OJJ02292.1 awen|OJJ30363.1 azon|OJJ45382.1

Group5796: abra|OJJ77565.1 acar|OOF99165.1 acla|EAW06530.1 afis|EAW16065.1 afla|EED48586.1 afum|EAL88697.1 agla|OJJ86100.1 akaw|GAA84061.1 aluc|OJZ80973.1 anid|CBF75096.1 anig|EHA19386.1 asyd|OJJ60008.1 ater|EAU33568.1 atub|OJI82612.1 aver|OJJ02277.1 awen|OJJ30370.1 azon|OJJ45390.1

Group5797: abra|OJJ77570.1 acla|EAW06535.1 afis|EAW16060.1 afla|EED48574.1 afum|EAL88693.1 akaw|GAA84056.1 aluc|OJZ80968.1 anid|CBF75084.1 anig|EHA19390.1 aory|EIT73112.1 asyd|OJJ60015.1 atub|OJI82618.1 aver|OJJ02270.1 awen|OJJ30379.1 azon|OJJ45395.1 agla|OJJ86105.1 ater|EAU33574.1

Group5798: abra|OJJ77571.1 acar|OOF99158.1 acla|EAW06540.1 afis|EAW16055.1 afla|EED56443.1 afum|EAL88687.1 agla|OJJ86088.1 akaw|GAA84055.1 aluc|OJZ80967.1 anid|CBF75074.1 anig|EHA19391.1 aory|EIT72848.1 asyd|OJJ60020.1 ater|EAU33579.1 atub|OJI82619.1 aver|OJJ02265.1 awen|OJJ30384.1

Group5799: abra|OJJ77578.1 acar|OOF99149.1 acla|EAW06544.1 afis|EAW16050.1 afla|EED56439.1 afum|EAL88683.1 agla|OJJ86084.1 akaw|GAA84047.1 aluc|OJZ80957.1 anid|CBF75066.1 anig|EHA19398.1 asyd|OJJ60023.1 atub|OJI82627.1 aver|OJJ02261.1 awen|OJJ30393.1 azon|OJJ45403.1 ater|EAU33586.1

Group5800: abra|OJJ77594.1 acla|EAW06558.1 afis|EAW16038.1 afla|EED56426.1 afum|EAL88669.1 agla|OJJ86062.1 akaw|GAA84034.1 aluc|OJZ80941.1 anid|CBF75038.1 anig|EHA19412.1 aory|EIT72878.1 asyd|OJJ60036.1 ater|EAU33602.1 atub|OJI82641.1 aver|OJJ02249.1 awen|OJJ30408.1 azon|OJJ45416.1

Group5801: abra|OJJ77595.1 acar|OOF99132.1 acla|EAW06559.1 afis|EAW16037.1 afla|EED56425.1 afum|EBA27355.1 agla|OJJ86063.1 akaw|GAA84033.1 aluc|OJZ80940.1 anid|CBF75042.1 aory|EIT72875.1 asyd|OJJ60037.1 ater|EAU33603.1 atub|OJI82642.1 aver|OJJ02248.1 awen|OJJ30409.1 azon|OJJ45417.1

Group5802: abra|OJJ77596.1 acar|OOF99130.1 acla|EAW06560.1 afis|EAW16036.1 afla|EED56424.1 afum|EBA27354.1 agla|OJJ86064.1 akaw|GAA84032.1 aluc|OJZ80939.1 anid|CBF75040.1 anig|EHA19413.1 asyd|OJJ60038.1 ater|EAU33604.1 atub|OJI82643.1 aver|OJJ02247.1 awen|OJJ30410.1 azon|OJJ45419.1

Group5803: abra|OJJ77600.1 acar|OOF99125.1 acla|EAW06564.1 afis|EAW16032.1 afla|EED56414.1 afum|EAL88663.1 agla|OJJ86060.1 akaw|GAA84028.1 aluc|OJZ80933.1 anid|CBF75032.1 anig|EHA19417.1 asyd|OJJ60051.1 ater|EAU33609.1 atub|OJI82647.1 aver|OJJ02238.1 awen|OJJ30415.1 azon|OJJ45423.1

Group5804: abra|OJJ77647.1 acar|OOF99089.1 acla|EAW06493.1 afis|EAW15989.1 afla|EED56361.1 afum|EAL88621.1 agla|OJJ85991.1 akaw|GAA83988.1 aluc|OJZ80882.1 anid|CBF85817.1 anig|EHA19454.1 aory|EIT73312.1 ater|EAU33660.1 atub|OJI82695.1 aver|OJI98436.1 awen|OJJ30481.1 azon|OJJ45472.1

Group5805: abra|OJJ77649.1 acar|OOF99087.1 akaw|GAA83986.1 aluc|OJZ80880.1 anig|EHA19456.1 atub|OJI82697.1 aver|OJI98438.1 abra|OJJ78132.1 afis|EAW20412.1 afla|EED52849.1 afum|EAL89232.1 aluc|OJZ91194.1 aory|EIT78655.1 ater|EAU29526.1 atub|OJI83213.1 awen|OJJ38076.1 anid|CBF87263.1

Group5806: abra|OJJ77662.1 acar|OOF99072.1 acla|EAW06599.1 afis|EAW16005.1 afla|EED56379.1 afum|EAL88637.2 agla|OJJ86029.1 akaw|GAA83973.1 aluc|OJZ80866.1 anid|CBF85850.1 anig|EHA19469.1 asyd|OJJ64361.1 ater|EAU33638.1 atub|OJI82711.1 aver|OJI98457.1 awen|OJJ30462.1 azon|OJJ45452.1

Group5807: abra|OJJ77663.1 acla|EAW06600.1 afis|EAW16004.1 afum|EAL88636.1 agla|OJJ86028.1 akaw|GAA83972.1 aluc|OJZ80865.1 anid|CBF85848.1 anig|EHA19470.1 aory|EIT73333.1 asyd|OJJ64358.1 ater|EAU33639.1 atub|OJI82712.1 aver|OJI98454.1 awen|OJJ30463.1 azon|OJJ45453.1 afla|EED56378.1

Group5808: abra|OJJ77676.1 acla|EAW06588.1 afis|EAW16011.1 afla|EED56388.1 afum|EAL88641.1 agla|OJJ86036.1 akaw|GAA83956.1 aluc|OJZ80843.1 anid|CBF74567.1 anig|EHA19480.1 aory|EIT83019.1 asyd|OJJ57650.1 ater|EAU33632.1 atub|OJI82724.1 aver|OJJ05509.1 awen|OJJ30454.1 azon|OJJ45445.1

Group5809: abra|OJJ77677.1 acla|EAW06589.1 afis|EAW16012.1 afla|EED56389.1 afum|EAL88642.1 agla|OJJ86037.1 akaw|GAA83955.1 aluc|OJZ80842.1 anid|CBF74566.1 anig|EHA19481.1 aory|EIT83001.1 asyd|OJJ57649.1 ater|EAU33631.1 atub|OJI82725.1 aver|OJJ05508.1 awen|OJJ30453.1 azon|OJJ45444.1

Group5810: abra|OJJ77692.1 acar|OOF99047.1 acla|EAW06573.1 afis|EAW16023.1 afla|EED56402.1 afum|EAL88652.1 agla|OJJ86052.1 akaw|GAA83938.1 aluc|OJZ80820.1 anid|CBF75011.1 anig|EHA19492.1 aory|EIT82997.1 asyd|OJJ60055.1 atub|OJI82742.1 aver|OJJ02234.1 awen|OJJ30438.1 azon|OJJ45433.1

Group5811: abra|OJJ77698.1 acar|OOF99042.1 acla|EAW11328.1 afis|EAW18167.1 afla|EED56328.1 afum|EAL87527.2 agla|OJJ85954.1 akaw|GAA83932.1 aluc|OJZ90763.1 anid|CBF85748.1 anig|EHA19497.1 aory|EIT73301.1 asyd|OJJ64294.1 ater|EAU33837.1 atub|OJI82751.1 aver|OJI98386.1 awen|OJJ30513.1

Group5812: abra|OJJ77705.1 acar|OOF99035.1 acla|EAW11316.1 afis|EAW18180.1 afla|EED56308.1 afum|EAL87541.1 akaw|GAA83925.1 aluc|OJZ90770.1 anid|CBF85727.1 anig|EHA19505.1 aory|EIT73291.1 asyd|OJJ64279.1 ater|EAU33820.1 atub|OJI82758.1 aver|OJI98371.1 awen|OJJ30528.1 azon|OJJ50210.1

Group5813: abra|OJJ77739.1 acar|OOF98987.1 afis|EAW21097.1 afla|EED49614.1 afum|EAL92120.1 agla|OJJ80115.1 akaw|GAA83890.1 aluc|OJZ90805.1 anid|CBF84486.1 anig|EHA19532.1 aory|EIT80984.1 asyd|OJJ53525.1 ater|EAU37778.1 atub|OJI82794.1 aver|OJJ08225.1 awen|OJJ36023.1 acla|EAW09993.1

Group5814: abra|OJJ77772.1 acar|OOF98962.1 acla|EAW11312.1 afis|EAW18186.1 afla|EED56292.1 afum|EAL87549.1 agla|OJJ85898.1 akaw|GAA83874.1 aluc|OJZ90822.1 anid|CBF85717.1 anig|EHA19548.1 aory|EIT73285.1 asyd|OJJ64274.1 ater|EAU33808.1 atub|OJI82810.1 aver|OJI98366.1 awen|OJJ30541.1

Group5815: abra|OJJ77773.1 acar|OOF98964.1 acla|EAW08571.1 afis|EAW23629.1 afla|EED46113.1 afum|EAL87831.1 akaw|GAA83873.1 aluc|OJZ90825.1 anid|CBF84280.1 anig|EHA19549.1 aory|EIT76433.1 asyd|OJJ57232.1 ater|EAU38029.1 atub|OJI82812.1 aver|OJJ03057.1 awen|OJJ36003.1 azon|OJJ51529.1

Group5816: abra|OJJ77774.1 acar|OOF98965.1 acla|EAW08572.1 afis|EAW23630.1 afla|EED46115.1 afum|EBA27325.1 akaw|GAA83872.1 aluc|OJZ90826.1 anid|CBF84282.1 anig|EHA19550.1 aory|EIT76372.1 asyd|OJJ57233.1 ater|EAU38028.1 atub|OJI82813.1 aver|OJJ03056.1 awen|OJJ36002.1 azon|OJJ51527.1

Group5817: abra|OJJ77776.1 acar|OOF98966.1 acla|EAW11313.1 afis|EAW18184.1 afla|EED56301.1 afum|EBA27317.1 agla|OJJ85900.1 akaw|GAA83871.1 aluc|OJZ90829.1 anid|CBF85715.1 anig|EHA19552.1 aory|EIT73399.1 asyd|OJJ64273.1 ater|EAU33816.1 atub|OJI82815.1 aver|OJI98365.1 awen|OJJ30532.1

Group5818: abra|OJJ77780.1 acar|OOF98970.1 afla|EED53399.1 akaw|GAA83867.1 aluc|OJZ90833.1 anid|CBF80617.1 anig|EHA19556.1 aory|EIT74939.1 asyd|OJJ54627.1 atub|OJI82819.1 aver|OJJ07529.1 awen|OJJ34502.1 afla|EED44856.1 aory|EIT82733.1 aver|OJI99413.1 anid|CBF78813.1 asyd|OJJ65262.1

Group5819: abra|OJJ77823.1 acar|OOF98946.1 acla|EAW11302.1 afis|EAW18200.1 afla|EED56278.1 afum|EAL87565.1 agla|OJJ85882.1 akaw|GAA83823.1 aluc|OJZ90895.1 anid|CBF86117.1 anig|EHA19614.1 aory|EIT73354.1 ater|EAU33794.1 atub|OJI82888.1 aver|OJI98595.1 awen|OJJ30568.1 azon|OJJ45504.1

Group5820: abra|OJJ77829.1 acla|EAW11297.1 afis|EAW18205.1 afla|EED56267.1 afum|EAL87570.1 agla|OJJ85877.1 akaw|GAA83818.1 aluc|OJZ90903.1 anid|CBF86130.1 anig|EHA19618.1 aory|EIT73457.1 asyd|OJJ64500.1 ater|EAU33784.1 atub|OJI82896.1 aver|OJI98600.1 awen|OJJ30576.1 azon|OJJ45499.1

Group5821: abra|OJJ77832.1 acar|OOF98938.1 acla|EAW11294.1 afis|EAW18208.1 afla|EED56262.1 afum|EAL87574.1 agla|OJJ85874.1 akaw|GAA83815.1 aluc|OJZ90906.1 anid|CBF86135.1 anig|EHA19621.1 asyd|OJJ64504.1 ater|EAU33780.1 atub|OJI82899.1 aver|OJI98603.1 awen|OJJ30580.1 azon|OJJ50191.1

Group5822: abra|OJJ77833.1 acla|EAW11293.1 afis|EAW18209.1 afla|EED56261.1 afum|EAL87575.2 agla|OJJ85873.1 akaw|GAA83814.1 aluc|OJZ90907.1 anid|CBF86137.1 anig|EHA19622.1 aory|EIT73295.1 asyd|OJJ64505.1 ater|EAU33779.1 atub|OJI82900.1 aver|OJI98604.1 awen|OJJ30581.1 azon|OJJ50190.1

Group5823: abra|OJJ77835.1 acar|OOF98936.1 acla|EAW11288.1 afis|EAW18211.1 afum|EAL87577.1 agla|OJJ85871.1 akaw|GAA83812.1 aluc|OJZ90909.1 anid|CBF86140.1 anig|EHA19624.1 aory|EIT73309.1 asyd|OJJ64507.1 ater|EAU33777.1 atub|OJI82902.1 aver|OJI98606.1 awen|OJJ30589.1 azon|OJJ50187.1

Group5824: abra|OJJ77842.1 acla|EAW11281.1 afis|EAW18219.1 afla|EED56251.1 afum|EAL87585.1 agla|OJJ85863.1 akaw|GAA83805.1 aluc|OJZ90917.1 anid|CBF86150.1 anig|EHA19629.1 aory|EIT73410.1 asyd|OJJ64513.1 ater|EAU33772.1 atub|OJI82909.1 aver|OJI98611.1 awen|OJJ30597.1 azon|OJJ50171.1

Group5825: abra|OJJ77843.1 acar|OOF98929.1 acla|EAW11280.1 afis|EAW18220.1 afla|EED56250.1 afum|EAL87586.1 agla|OJJ85862.1 akaw|GAA83804.1 aluc|OJZ90918.1 anid|CBF86157.1 anig|EHA19630.1 asyd|OJJ64514.1 ater|EAU33771.1 atub|OJI82910.1 aver|OJI98612.1 awen|OJJ30598.1 azon|OJJ50170.1

Group5826: abra|OJJ77856.1 acar|OOF98918.1 acla|EAW11270.1 afis|EAW18229.1 afla|EED56239.1 afum|EAL87596.1 agla|OJJ85852.1 akaw|GAA83793.1 aluc|OJZ90930.1 anid|CBF86170.1 anig|EHA19641.1 asyd|OJJ64524.1 ater|EAU33762.1 atub|OJI82922.1 aver|OJI98622.1 awen|OJJ30608.1 azon|OJJ50160.1

Group5827: abra|OJJ77863.1 acar|OOF98910.1 acla|EAW11263.1 afis|EAW18235.1 afla|EED56231.1 afum|EAL87602.2 agla|OJJ85839.1 akaw|GAA83786.1 aluc|OJZ90938.1 anid|CBF86184.1 anig|EHA19646.1 asyd|OJJ64531.1 ater|EAU33750.1 atub|OJI82929.1 aver|OJI98629.1 awen|OJJ30615.1 azon|OJJ48002.1

Group5828: abra|OJJ77909.1 acar|OOF98876.1 acla|EAW06517.1 afis|EAW16235.1 afla|EED53346.1 agla|OJJ86240.1 akaw|GAA83750.1 aluc|OJZ90981.1 anid|CBF86229.1 anig|EHA19686.1 aory|EIT75244.1 asyd|OJJ64565.1 ater|EAU33709.1 atub|OJI82972.1 aver|OJI98662.1 awen|OJJ30225.1 azon|OJJ43945.1

Group5829: abra|OJJ77922.1 acla|EAW06686.1 afis|EAW16079.1 afla|EED48599.1 afum|EAL88711.1 agla|OJJ86112.1 akaw|GAA83741.1 aluc|OJZ90990.1 anid|CBF75121.1 anig|EHA19694.1 aory|EIT73245.1 asyd|OJJ59992.1 ater|EAU33145.1 atub|OJI82981.1 aver|OJJ02299.1 awen|OJJ30354.1 azon|OJJ45376.1

Group5830: abra|OJJ77927.1 acar|OOF98865.1 acla|EAW06685.1 afis|EAW16080.1 afla|EED48600.1 afum|EAL88712.1 agla|OJJ86113.1 akaw|GAA83737.1 aluc|OJZ90995.1 anig|EHA19702.1 aory|EIT73241.1 asyd|OJJ59991.1 ater|EAU33146.1 atub|OJI82985.1 aver|OJJ02300.1 awen|OJJ30353.1 azon|OJJ45375.1

Group5831: abra|OJJ77928.1 acar|OOF98866.1 acla|EAW06684.1 afis|EAW16081.1 afla|EED48601.1 afum|EAL88713.1 agla|OJJ86114.1 akaw|GAA83736.1 aluc|OJZ90996.1 anig|EHA19703.1 aory|EIT73043.1 asyd|OJJ59990.1 ater|EAU33147.1 atub|OJI82986.1 aver|OJJ02301.1 awen|OJJ30352.1 azon|OJJ45374.1

Group5832: abra|OJJ77931.1 acar|OOF98868.1 acla|EAW06683.1 afis|EAW16082.1 afla|EED48602.1 afum|EAL88714.1 agla|OJJ86116.1 akaw|GAA83732.1 aluc|OJZ90999.1 anig|EHA19706.1 aory|EIT73073.1 asyd|OJJ59989.1 ater|EAU33148.1 atub|OJI82989.1 aver|OJJ02302.1 awen|OJJ30351.1 azon|OJJ45373.1

Group5833: abra|OJJ77932.1 acar|OOF98869.1 acla|EAW06682.1 afis|EAW16083.1 afla|EED48603.1 afum|EAL88715.1 agla|OJJ86115.1 akaw|GAA83731.1 aluc|OJZ91000.1 anig|EHA19707.1 aory|EIT73153.1 asyd|OJJ59988.1 ater|EAU33149.1 atub|OJI82990.1 aver|OJJ02303.1 awen|OJJ30350.1 azon|OJJ45372.1

Group5834: abra|OJJ77934.1 acla|EAW06680.1 afis|EAW16085.1 afla|EED48605.1 afum|EAL88717.1 agla|OJJ86117.1 akaw|GAA83729.1 aluc|OJZ91002.1 anid|CBF75123.1 anig|EHA19708.1 aory|EIT73254.1 asyd|OJJ59986.1 ater|EAU33151.1 atub|OJI82992.1 aver|OJJ02305.1 awen|OJJ30347.1 azon|OJJ45371.1

Group5835: abra|OJJ77953.1 afla|EED49771.1 agla|OJJ89120.1 akaw|GAA83710.1 aluc|OJZ91024.1 anid|CBF87975.1 anig|EHA19724.1 aory|EIT77708.1 asyd|OJJ63489.1 ater|EAU33503.1 atub|OJI83016.1 aver|OJI97440.1 awen|OJJ30074.1 azon|OJJ42166.1 atub|OJI83019.1 agla|OJJ84720.1 agla|OJJ88987.1

Group5836: abra|OJJ77986.1 acar|OOF98837.1 acla|EAW06662.1 afis|EAW16103.1 afla|EED48638.1 afum|EAL88735.1 akaw|GAA85994.1 aluc|OJZ91061.1 anid|CBF85479.1 anig|EHA19764.1 aory|EIT73148.1 asyd|OJJ64107.1 ater|EAU33180.1 atub|OJI83057.1 aver|OJI98189.1 awen|OJJ30329.1 azon|OJJ45346.1

Group5837: abra|OJJ77996.1 acar|OOF98825.1 acla|EAW06652.1 afis|EAW16113.1 afla|EED48653.1 afum|EAL88744.1 agla|OJJ86152.1 akaw|GAA85984.1 aluc|OJZ91072.1 anid|CBF85498.1 anig|EHA19775.1 aory|EIT73209.1 asyd|OJJ64116.1 ater|EAU33196.1 atub|OJI83067.1 aver|OJI98199.1 awen|OJJ30317.1

Group5838: abra|OJJ78002.1 acla|EAW06648.1 afis|EAW16117.1 afla|EED48658.1 afum|EAL88748.1 agla|OJJ86156.1 akaw|GAA85981.1 aluc|OJZ91078.1 anid|CBF85507.1 anig|EHA19779.1 aory|EIT73114.1 asyd|OJJ64120.1 ater|EAU33200.1 atub|OJI83073.1 aver|OJI98203.1 awen|OJJ30313.1 azon|OJJ45330.1

Group5839: abra|OJJ78003.1 acar|OOF98820.1 acla|EAW06647.1 afis|EAW16118.1 afla|EED48659.1 afum|EAL88749.1 agla|OJJ86159.1 akaw|GAA85980.1 aluc|OJZ91079.1 anid|CBF85509.1 anig|EHA19780.1 asyd|OJJ64121.1 ater|EAU33201.1 atub|OJI83074.1 aver|OJI98204.1 awen|OJJ30312.1 azon|OJJ45329.1

Group5840: abra|OJJ78006.1 acar|OOF98816.1 acla|EAW06644.1 afis|EAW16121.1 afum|EAL88752.1 agla|OJJ86162.1 akaw|GAA85977.1 aluc|OJZ91082.1 anid|CBF85539.1 anig|EHA19783.1 aory|EIT73243.1 asyd|OJJ64141.1 ater|EAU33204.1 atub|OJI83078.1 aver|OJI98222.1 awen|OJJ30309.1 azon|OJJ45312.1

Group5841: abra|OJJ78009.1 acar|OOF98814.1 acla|EAW06640.1 afis|EAW16125.1 afla|EED48665.1 afum|EAL88755.1 agla|OJJ86167.1 akaw|GAA85974.1 aluc|OJZ91085.1 anid|CBF85530.1 anig|EHA19786.1 asyd|OJJ64134.1 ater|EAU33208.1 atub|OJI83081.1 aver|OJI98217.1 awen|OJJ30304.1 azon|OJJ45305.1

Group5842: abra|OJJ78016.1 acar|OOF98808.1 acla|EAW06632.1 afis|EAW16133.1 afla|EED48672.1 afum|EAL88763.1 agla|OJJ86178.1 akaw|GAA85967.1 aluc|OJZ91092.1 anid|CBF85518.1 anig|EHA19793.1 asyd|OJJ64126.1 ater|EAU33215.1 atub|OJI83089.1 aver|OJI98209.1 awen|OJJ30295.1 azon|OJJ43985.1

Group5843: abra|OJJ78017.1 acar|OOF98806.1 acla|EAW06630.1 afis|EAW16135.1 afla|EED48674.1 afum|EAL88765.1 agla|OJJ86180.1 akaw|GAA85966.1 aluc|OJZ91093.1 anid|CBF85514.1 anig|EHA19794.1 asyd|OJJ64124.1 ater|EAU33217.1 atub|OJI83090.1 aver|OJI98207.1 awen|OJJ30293.1 azon|OJJ43982.1

Group5844: abra|OJJ78025.1 acar|OOF98800.1 acla|EAW06624.1 afis|EAW16141.1 afla|EED48682.1 afum|EAL88770.2 akaw|GAA85961.1 aluc|OJZ91100.1 anid|CBF85550.1 anig|EHA19800.1 aory|EIT73246.1 asyd|OJJ64150.1 ater|EAU33226.1 atub|OJI83097.1 aver|OJI98230.1 awen|OJJ30287.1 azon|OJJ43976.1

Group5845: abra|OJJ78056.1 afis|EAW20703.1 afla|EED51737.1 afum|EAL84878.1 agla|OJJ85717.1 akaw|GAA85936.1 aluc|OJZ91126.1 anid|CBF70385.1 anig|EHA19827.1 aory|EIT80428.1 asyd|OJJ58639.1 atub|OJI83137.1 aver|OJJ01246.1 asyd|OJJ56271.1 aver|OJJ08494.1 anid|CBF79145.1 atub|OJI79837.1

Group5846: afla|EED48481.1 ater|EAU29541.1 acar|OOF98852.1 acla|EAW09008.1 akaw|GAA93107.1 aluc|OJZ80597.1 anig|EHA22185.1 asyd|OJJ64213.1 awen|OJJ29864.1 anid|CBF80392.1 aory|EIT73087.1 asyd|OJJ54790.1 aver|OJJ07373.1 azon|OJJ43202.1 afis|EAW17320.1 aver|OJJ06502.1 anid|CBF89303.1

Group5847: abra|OJJ78077.1 acla|EAW06610.1 afis|EAW16205.1 afla|EED48698.1 agla|OJJ86216.1 akaw|GAA85917.1 aluc|OJZ91149.1 anid|CBF79565.1 anig|EHA19847.1 aory|EIT73041.1 asyd|OJJ57914.1 ater|EAU33247.1 atub|OJI83163.1 aver|OJJ04479.1 awen|OJJ30267.1 azon|OJJ43963.1 afum|EAL88833.1

Group5848: abra|OJJ78095.1 acar|OOF98778.1 acla|EAW11017.1 afis|EAW20831.1 afla|EED57883.1 afum|EAL84801.1 akaw|GAA85903.1 aluc|OJZ91164.1 anid|CBF69443.1 anig|EHA19859.1 aory|EIT83273.1 asyd|OJJ52396.1 ater|EAU34241.1 atub|OJI83180.1 aver|OJJ01927.1 azon|OJJ48593.1 azon|OJJ42147.1

Group5849: abra|OJJ78102.1 acar|OOF90265.1 acla|EAW06615.1 afis|EAW16152.1 afla|EED48694.1 agla|OJJ86204.1 akaw|GAA85901.1 aluc|OJZ91167.1 anid|CBF79575.1 anig|EHA19860.1 aory|EIT73256.1 asyd|OJJ57908.1 ater|EAU33236.1 atub|OJI83182.1 aver|OJJ04484.1 awen|OJJ30273.1 azon|OJJ43967.1

Group5850: abra|OJJ78123.1 acar|OOF97938.1 acla|EAW13664.1 afis|EAW24511.1 afla|EED51090.1 afum|EBA27259.1 akaw|GAA85885.1 aluc|OJZ91185.1 anid|CBF83112.1 anig|EHA19879.1 asyd|OJJ54499.1 ater|EAU31566.1 atub|OJI83204.1 aver|OJJ07681.1 awen|OJJ38687.1 acar|OOF98765.1 anid|CBF80779.1

Group5851: abra|OJJ78208.1 acar|OOF98646.1 afis|EAW18263.1 afla|EED48722.1 afum|EAL87627.1 agla|OJJ88163.1 akaw|GAA85818.1 aluc|OJZ91281.1 anid|CBF79532.1 anig|EHA19966.1 aory|EIT73066.1 asyd|OJJ57896.1 ater|EAU29421.1 atub|OJI83271.1 aver|OJJ04499.1 awen|OJJ30637.1 azon|OJJ47979.1

Group5852: abra|OJJ78222.1 acar|OOF98634.1 acla|EAW11227.1 afis|EAW18276.1 afla|EED48743.1 afum|EAL87642.1 agla|OJJ88150.1 akaw|GAA85803.1 aluc|OJZ91297.1 anid|CBF79497.1 anig|EHA19979.1 asyd|OJJ57879.1 ater|EAU33282.1 atub|OJI83287.1 aver|OJJ04515.1 awen|OJJ30649.1 azon|OJJ47966.1

Group5853: abra|OJJ78227.1 acla|EAW09282.1 afis|EAW21865.1 afla|EED46446.1 afum|EAL92879.1 agla|OJJ84272.1 akaw|GAA85797.1 aluc|OJZ91302.1 anid|CBF76757.1 anig|EHA19984.1 aory|EIT72438.1 asyd|OJJ61263.1 ater|EAU30862.1 atub|OJI83293.1 aver|OJJ00589.1 awen|OJJ34151.1 azon|OJJ43478.1

Group5854: abra|OJJ78271.1 acar|OOF98593.1 acla|EAW09242.1 afis|EAW21905.1 afla|EED46646.1 afum|EBA27480.1 agla|OJJ84225.1 akaw|GAA85755.1 aluc|OJZ91345.1 anid|CBF76830.1 anig|EHA20024.1 asyd|OJJ61301.1 ater|EAU30900.1 atub|OJI83337.1 aver|OJJ00627.1 awen|OJJ34195.1 azon|OJJ43439.1

Group5855: abra|OJJ78273.1 acla|EAW09241.1 afis|EAW21907.1 afum|EAL92919.2 akaw|GAA85753.1 aluc|OJZ91347.1 anig|EHA20026.1 asyd|OJJ61303.1 ater|EAU30901.1 atub|OJI83339.1 aver|OJJ00628.1 awen|OJJ34197.1 afla|EED46644.1 agla|OJJ84224.1 aory|EIT77882.1 anid|CBF76833.1 azon|OJJ43437.1

Group5856: abra|OJJ78274.1 acla|EAW09240.1 afis|EAW21908.1 afla|EED46643.1 afum|EAL92920.1 agla|OJJ84223.1 akaw|GAA85752.1 aluc|OJZ91349.1 anid|CBF76835.1 anig|EHA20027.1 aory|EIT77824.1 asyd|OJJ61304.1 ater|EAU30908.1 atub|OJI83340.1 aver|OJJ00629.1 awen|OJJ34198.1 azon|OJJ43436.1

Group5857: abra|OJJ78276.1 acar|OOF98591.1 acla|EAW09238.1 afis|EAW21910.1 afla|EED46640.1 afum|EAL92922.2 agla|OJJ84221.1 akaw|GAA85750.1 aluc|OJZ91351.1 anid|CBF76839.1 anig|EHA20029.1 asyd|OJJ61306.1 ater|EAU30910.1 atub|OJI83343.1 aver|OJJ00631.1 awen|OJJ34200.1 azon|OJJ43434.1

Group5858: abra|OJJ78281.1 acar|OOF98587.1 acla|EAW09234.1 afis|EAW21914.1 afla|EED46636.1 afum|EBA27482.1 agla|OJJ84209.1 akaw|GAA85746.1 aluc|OJZ91355.1 anid|CBF76848.1 anig|EHA20033.1 asyd|OJJ61310.1 ater|EAU30915.1 atub|OJI83348.1 aver|OJJ00635.1 awen|OJJ34204.1 azon|OJJ43430.1

Group5859: abra|OJJ78286.1 acar|OOF98582.1 acla|EAW09229.1 afis|EAW21919.1 afum|EAL92931.1 agla|OJJ84204.1 akaw|GAA85741.1 aluc|OJZ91360.1 anid|CBF76858.1 anig|EHA20038.1 aory|EIT77872.1 asyd|OJJ61315.1 ater|EAU30924.1 atub|OJI83353.1 aver|OJJ00640.1 awen|OJJ34209.1 azon|OJJ43425.1

Group5860: abra|OJJ78295.1 acla|EAW09221.1 afis|EAW21927.1 afla|EED46620.1 afum|EAL92940.1 agla|OJJ84194.1 akaw|GAA85732.1 aluc|OJZ91369.1 anid|CBF76878.1 anig|EHA20047.1 aory|EIT77851.1 asyd|OJJ61325.1 ater|EAU30933.1 atub|OJI83362.1 aver|OJJ00650.1 awen|OJJ34217.1 azon|OJJ43417.1

Group5861: abra|OJJ78296.1 acar|OOF98572.1 acla|EAW09220.1 afis|EAW21928.1 afla|EED46621.1 afum|EAL92941.1 agla|OJJ84193.1 akaw|GAA85731.1 aluc|OJZ91370.1 anid|CBF76876.1 anig|EHA20048.1 asyd|OJJ61324.1 ater|EAU30934.1 atub|OJI83363.1 aver|OJJ00649.1 awen|OJJ34218.1 azon|OJJ43416.1

Group5862: abra|OJJ78306.1 acar|OOF98560.1 acla|EAW09210.1 afis|EAW21937.1 afla|EED46654.1 afum|EAL92950.2 akaw|GAA85719.1 aluc|OJZ91385.1 anid|CBF76897.1 anig|EHA20058.1 aory|EIT77879.1 asyd|OJJ61337.1 ater|EAU30943.1 atub|OJI83378.1 aver|OJJ00661.1 awen|OJJ34228.1 azon|OJJ43407.1

Group5863: abra|OJJ78315.1 acar|OOF90168.1 acla|EAW12152.1 afis|EAW17176.1 afla|EED54162.1 afum|EAL86606.1 akaw|GAA85715.1 aluc|OJZ91388.1 anid|CBF76640.1 anig|EHA20066.1 aory|EIT76137.1 asyd|OJJ61201.1 ater|EAU35069.1 atub|OJI83381.1 aver|OJJ00526.1 awen|OJJ37819.1 azon|OJJ42628.1

Group5864: abra|OJJ78318.1 acla|EAW09207.1 afis|EAW21940.1 afla|EED46657.1 afum|EAL92953.1 agla|OJJ84183.1 akaw|GAA85712.1 aluc|OJZ91391.1 anid|CBF84079.1 anig|EHA20069.1 aory|EIT77958.1 asyd|OJJ61341.1 ater|EAU30945.1 atub|OJI83384.1 aver|OJJ00665.1 awen|OJJ34231.1 azon|OJJ43404.1

Group5865: abra|OJJ78322.1 acar|OOF98552.1 acla|EAW09203.1 afis|EAW21944.1 afla|EED46661.1 afum|EAL92958.1 agla|OJJ84180.1 akaw|GAA85706.1 aluc|OJZ91397.1 anid|CBF84051.1 anig|EHA20074.1 aory|EIT77927.1 ater|EAU30992.1 atub|OJI83389.1 aver|OJJ00668.1 awen|OJJ34235.1 azon|OJJ43401.1

Group5866: abra|OJJ78326.1 afla|EED46664.1 acar|OOF98549.1 akaw|GAA85703.1 aluc|OJZ91400.1 anid|CBF84060.1 anig|EHA20077.1 asyd|OJJ61346.1 atub|OJI83392.1 aver|OJJ00671.1 azon|OJJ43398.1 acla|EAW09200.1 afis|EAW21947.1 afum|EAL92961.1 ater|EAU30995.1 agla|OJJ84173.1 awen|OJJ34238.1

Group5867: abra|OJJ78344.1 acar|OOF90270.1 acla|EAW09190.1 afis|EAW21956.1 afla|EED46675.1 afum|EAL92970.2 agla|OJJ84163.1 akaw|GAA85689.1 aluc|OJZ91417.1 anid|CBF84036.1 anig|EHA20089.1 aory|EIT77945.1 asyd|OJJ61363.1 atub|OJI83411.1 aver|OJJ00690.1 awen|OJJ34248.1 azon|OJJ43390.1

Group5868: abra|OJJ78349.1 afis|EAW21961.1 afla|EED46682.1 afum|EAL92975.1 agla|OJJ84155.1 akaw|GAA85684.1 aluc|OJZ91424.1 anid|CBF84046.1 anig|EHA20094.1 aory|EIT77949.1 asyd|OJJ61357.1 ater|EAU31081.1 atub|OJI83417.1 aver|OJJ00684.1 awen|OJJ34255.1 acla|EAW09185.1 azon|OJJ43384.1

Group5869: abra|OJJ78352.1 acla|EAW09183.1 afis|EAW21963.1 afla|EED46684.1 afum|EAL92977.1 agla|OJJ84140.1 akaw|GAA85682.1 aluc|OJZ91427.1 anid|CBF82740.1 anig|EHA20096.1 aory|EIT77811.1 asyd|OJJ61623.1 ater|EAU31083.1 atub|OJI83420.1 aver|OJJ00948.1 awen|OJJ34257.1 azon|OJJ43382.1

Group5870: aver|OJJ00953.1 abra|OJJ78358.1 acar|OOF98528.1 acla|EAW09176.1 afis|EAW21970.1 afla|EED46692.1 afum|EAL92984.1 agla|OJJ84133.1 akaw|GAA85675.1 aluc|OJZ91433.1 anid|CBF82727.1 aory|EIT77959.1 asyd|OJJ61630.1 ater|EAU31090.1 atub|OJI83426.1 awen|OJJ34263.1 azon|OJJ43374.1

Group5871: abra|OJJ78367.1 acla|EAW09168.1 afis|EAW21978.1 afla|EED46701.1 afum|EAL92993.1 agla|OJJ84122.1 akaw|GAA85666.1 aluc|OJZ91444.1 anid|CBF82712.1 anig|EHA20108.1 aory|EIT77893.1 asyd|OJJ61639.1 ater|EAU31096.1 atub|OJI83438.1 aver|OJJ00962.1 awen|OJJ34271.1 azon|OJJ43365.1

Group5872: abra|OJJ78373.1 acar|OOF98516.1 acla|EAW09162.1 afis|EAW21984.1 afla|EED46708.1 afum|EAL92999.1 agla|OJJ84114.1 akaw|GAA85660.1 aluc|OJZ91450.1 anid|CBF82701.1 anig|EHA20114.1 asyd|OJJ61645.1 ater|EAU31103.1 atub|OJI83443.1 aver|OJJ00967.1 awen|OJJ34281.1 azon|OJJ43359.1

Group5873: abra|OJJ78376.1 acar|OOF98512.1 acla|EAW09158.1 afis|EAW21988.1 afla|EED46712.1 afum|EAL93004.1 agla|OJJ84110.1 akaw|GAA85657.1 aluc|OJZ91454.1 anid|CBF82692.1 anig|EHA20117.1 aory|EIT77910.1 asyd|OJJ61649.1 atub|OJI83447.1 aver|OJJ00970.1 awen|OJJ34284.1 azon|OJJ43354.1

Group5874: abra|OJJ78377.1 acla|EAW09157.1 afis|EAW21989.1 afla|EED46713.1 afum|EAL93005.2 agla|OJJ84109.1 akaw|GAA85656.1 aluc|OJZ91455.1 anid|CBF82690.1 anig|EHA20118.1 aory|EIT77905.1 asyd|OJJ61650.1 ater|EAU31108.1 atub|OJI83448.1 aver|OJJ00971.1 awen|OJJ34285.1 azon|OJJ43353.1

Group5875: abra|OJJ78379.1 acar|OOF98509.1 acla|EAW09154.1 afis|EAW21992.1 afla|EED46716.1 afum|EAL93007.1 agla|OJJ84105.1 akaw|GAA85653.1 aluc|OJZ91458.1 anid|CBF82685.1 anig|EHA20121.1 asyd|OJJ61654.1 ater|EAU31111.1 atub|OJI83451.1 aver|OJJ00975.1 awen|OJJ34288.1 azon|OJJ43350.1

Group5876: abra|OJJ78389.1 acar|OOF98499.1 acla|EAW09144.1 afis|EAW22003.1 afla|EED46728.1 afum|EAL93018.1 agla|OJJ84095.1 akaw|GAA85643.1 aluc|OJZ91468.1 anid|CBF82664.1 anig|EHA20130.1 asyd|OJJ61665.1 ater|EAU31122.1 atub|OJI83461.1 aver|OJJ00986.1 awen|OJJ34298.1 azon|OJJ43340.1

Group5877: acar|OOF90682.1 acla|EAW14843.1 afis|EAW24370.1 afla|EED49848.1 afum|EAL86042.1 anid|CBF87190.1 awen|OJJ30943.1 acar|OOG00289.1 acla|EAW12454.1 afla|EED51634.1 anid|CBF84176.1 aory|EIT80587.1 asyd|OJJ57129.1 aver|OJJ03175.1 azon|OJJ46526.1 awen|OJJ32845.1 awen|OJJ36074.1

Group5878: acar|OOF91612.1 acla|EAW14418.1 afis|EAW19125.1 afla|EED54289.1 afum|EAL93857.1 agla|OJJ80440.1 akaw|GAA90471.1 aluc|OJZ85753.1 anid|CBF79735.1 anig|EHA24803.1 aory|EIT76095.1 asyd|OJJ58082.1 ater|EAU37392.1 atub|OJI79537.1 aver|OJJ04326.1 awen|OJJ37456.1 azon|OJJ51135.1

Group5879: acar|OOF93414.1 acla|EAW08926.1 afis|EAW15468.1 afla|EED46920.1 afum|EAL85854.1 agla|OJJ80149.1 akaw|GAA87755.1 aluc|OJZ85388.1 anid|CBF78258.1 anig|EHA25572.1 aory|EIT72527.1 asyd|OJJ56063.1 ater|EAU32660.1 atub|OJI80182.1 aver|OJJ05006.1 awen|OJJ38118.1 azon|OJJ51809.1

Group5880: acar|OOF98511.1 acla|EAW09156.1 afis|EAW21990.1 afla|EED46714.1 afum|EBA27484.1 agla|OJJ84108.1 akaw|GAA85655.1 aluc|OJZ91456.1 anid|CBF82688.1 anig|EHA20119.1 aory|EIT77857.1 asyd|OJJ61651.1 ater|EAU31109.1 atub|OJI83449.1 aver|OJJ00972.1 awen|OJJ34286.1 azon|OJJ43352.1

Group5881: acar|OOG01162.1 acla|EAW07893.1 afis|EAW22928.1 afla|EED58121.1 afum|EAL90269.1 agla|OJJ83360.1 akaw|GAA87526.1 aluc|OJZ87993.1 anid|CBF87669.1 anig|EHA17775.1 aory|EIT79165.1 asyd|OJJ63310.1 ater|EAU30695.1 atub|OJI90602.1 aver|OJI97649.1 awen|OJJ38636.1 azon|OJJ49578.1

Group5882: acla|EAW09522.1 afis|EAW21610.1 afla|EED57839.1 afum|EAL92644.1 agla|OJJ85805.1 akaw|GAA84915.1 aluc|OJZ86754.1 anid|CBF83478.1 anig|EHA21925.1 aory|EIT81104.1 asyd|OJJ60693.1 ater|EAU38299.1 aver|OJJ00018.1 awen|OJJ39072.1 azon|OJJ51403.1 azon|OJJ51404.1 aver|OJJ00019.1

Group5883: acla|EAW14913.1 afis|EAW24434.1 afla|EED45083.1 afum|EAL86102.1 akaw|GAA88070.1 aluc|OJZ81335.1 aory|EIT73616.1 asyd|OJJ56673.1 ater|EAU29576.1 atub|OJI91538.1 aver|OJJ06051.1 awen|OJJ38123.1 azon|OJJ48557.1 agla|OJJ87162.1 ater|EAU29898.1 awen|OJJ30898.1 aory|EIT74968.1

Group5884: abra|OJJ66154.1 abra|OJJ75559.1 asyd|OJJ58905.1 asyd|OJJ59701.1 aver|OJJ01502.1 aver|OJJ02586.1 atub|OJI85780.1 awen|OJJ38282.1 acar|OOF90877.1 agla|OJJ88520.1 akaw|GAA84314.1 aluc|OJZ89009.1 anid|CBF69967.1 anig|EHA23519.1 aory|EIT75405.1 ater|EAU30290.1

Group5885: abra|OJJ67080.1 abra|OJJ68912.1 acar|OOF91499.1 afis|EAW20326.1 afla|EED57092.1 agla|OJJ78923.1 akaw|GAA90337.1 aluc|OJZ85895.1 anid|CBF81708.1 anig|EHA24697.1 aory|EIT82304.1 asyd|OJJ59577.1 ater|EAU32053.1 atub|OJI79680.1 aver|OJJ02692.1 awen|OJJ38052.1

Group5886: acar|OOF91513.1 acar|OOF98315.1 abra|OJJ67236.1 afis|EAW24732.1 afla|EED46087.1 afum|EAL89379.1 agla|OJJ86786.1 akaw|GAA91660.1 aluc|OJZ87739.1 anid|CBF69482.1 anig|EHA20231.1 aory|EIT76445.1 asyd|OJJ59273.1 atub|OJI80901.1 aver|OJJ01886.1 awen|OJJ40220.1

Group5887: acar|OOF92290.1 acar|OOF92331.1 abra|OJJ67511.1 acla|EAW06261.1 afla|EED57038.1 afum|EAL93042.1 agla|OJJ84932.1 aluc|OJZ84853.1 anid|CBF70812.1 aory|EIT77427.1 asyd|OJJ58406.1 ater|EAU32237.1 atub|OJI80743.1 aver|OJJ01007.1 awen|OJJ31782.1 asyd|OJJ61026.1

Group5888: acar|OOF92605.1 acar|OOF92626.1 abra|OJJ69658.1 acla|EAW13777.1 afis|EAW18355.1 afla|EED48820.1 akaw|GAA91637.1 aluc|OJZ82464.1 anid|CBF79358.1 anig|EHA28311.1 aory|EIT73001.1 asyd|OJJ57803.1 ater|EAU34688.1 atub|OJI87974.1 awen|OJJ35739.1 azon|OJJ43324.1

Group5889: acar|OOF92826.1 acar|OOF98200.1 acla|EAW12158.1 acla|EAW13719.1 afis|EAW17190.1 afis|EAW21021.1 afum|EAL86616.2 afum|EAL92059.1 ater|EAU31621.1 ater|EAU36218.1 abra|OJJ70923.1 akaw|GAA83017.1 aluc|OJZ87603.1 anig|EHA20802.1 atub|OJI81045.1 aver|OJJ05979.1

Group5890: acar|OOF93694.1 acar|OOF93779.1 abra|OJJ66685.1 acla|EAW09000.1 afla|EED45351.1 agla|OJJ85436.1 akaw|GAA89629.1 aluc|OJZ80764.1 anid|CBF81376.1 anig|EHA27263.1 aory|EIT82467.1 asyd|OJJ59389.1 atub|OJI87618.1 aver|OJJ02883.1 awen|OJJ39926.1 azon|OJJ50230.1

Group5891: acar|OOF94035.1 acar|OOF94105.1 abra|OJJ66415.1 afis|EAW20234.1 afla|EED55132.1 afum|EAL85351.1 akaw|GAA90282.1 aluc|OJZ83058.1 anid|CBF73520.1 anig|EHA28124.1 aory|EIT81578.1 asyd|OJJ56746.1 ater|EAU37765.1 atub|OJI87888.1 aver|OJJ05984.1 azon|OJJ47372.1

Group5892: acar|OOF94043.1 acar|OOF94113.1 abra|OJJ66398.1 afis|EAW16735.1 afla|EED49556.1 afum|EAL86309.1 agla|OJJ79510.1 akaw|GAA90266.1 aluc|OJZ83075.1 anig|EHA28139.1 asyd|OJJ61452.1 atub|OJI87871.1 aver|OJJ00787.1 awen|OJJ39514.1 azon|OJJ51715.1 anid|CBF83128.1

Group5893: acar|OOF94086.1 acar|OOF94156.1 abra|OJJ66352.1 afis|EAW17344.1 afla|EED46461.1 afum|EAL91188.1 agla|OJJ88648.1 akaw|GAA90225.1 aluc|OJZ83116.1 anid|CBF82993.1 anig|EHA28179.1 aory|EIT72314.1 asyd|OJJ54261.1 atub|OJI87818.1 aver|OJJ05214.1 awen|OJJ40134.1

Group5894: acar|OOF94093.1 acar|OOF94163.1 abra|OJJ66344.1 acla|EAW09990.1 afis|EAW21102.1 afla|EED51684.1 afum|EAL92123.1 akaw|GAA90218.1 aluc|OJZ83123.1 anid|CBF86612.1 anig|EHA28187.1 aory|EIT80601.1 asyd|OJJ64820.1 ater|EAU36797.1 atub|OJI87809.1 aver|OJI98933.1

Group5895: acar|OOF94752.1 acar|OOF95317.1 abra|OJJ72483.1 acla|EAW12714.1 afis|EAW17876.1 afla|EED46294.1 afum|EAL91684.1 agla|OJJ83070.1 akaw|GAA86218.1 aluc|OJZ90470.1 anig|EHA23929.1 asyd|OJJ52333.1 ater|EAU33096.1 atub|OJI81894.1 aver|OJJ05550.1 awen|OJJ41688.1

Group5896: acar|OOF95617.1 acar|OOF95669.1 abra|OJJ67062.1 acla|EAW12224.1 afla|EED54494.1 agla|OJJ85767.1 akaw|GAA82365.1 aluc|OJZ91758.1 anid|CBF81941.1 anig|EHA18944.1 aory|EIT78917.1 asyd|OJJ56618.1 ater|EAU31797.1 atub|OJI86216.1 aver|OJJ04440.1 anig|EHA26144.1

Group5897: acar|OOF96227.1 acar|OOF96265.1 abra|OJJ71512.1 acla|EAW15069.1 afis|EAW20041.1 afum|EAL85161.2 agla|OJJ88921.1 aluc|OJZ86250.1 anid|CBF84782.1 anig|EHA21615.1 aory|EIT73556.1 asyd|OJJ63233.1 atub|OJI84292.1 aver|OJI97723.1 awen|OJJ31797.1 azon|OJJ41993.1

Group5898: acar|OOF96911.1 acar|OOF96946.1 abra|OJJ70799.1 afla|EED56519.1 agla|OJJ87254.1 akaw|GAA90115.1 aluc|OJZ84144.1 anid|CBF81181.1 anig|EHA19252.1 aory|EIT72828.1 asyd|OJJ59295.1 ater|EAU31668.1 atub|OJI90518.1 aver|OJJ02980.1 awen|OJJ36774.1 azon|OJJ47424.1

Group5899: acar|OOF97665.1 acar|OOF97712.1 abra|OJJ71492.1 acla|EAW10058.1 afis|EAW25526.1 afum|EAL90191.2 agla|OJJ87276.1 akaw|GAA82528.1 aluc|OJZ87055.1 anid|CBF77803.1 anig|EHA20328.1 aory|EIT78255.1 asyd|OJJ53018.1 atub|OJI81580.1 aver|OJJ03825.1 azon|OJJ49512.1

Group5900: acar|OOF98477.1 acar|OOF98490.1 abra|OJJ78397.1 acla|EAW09135.1 afla|EED46736.1 agla|OJJ84086.1 akaw|GAA85634.1 aluc|OJZ91476.1 anid|CBF82648.1 anig|EHA20138.1 asyd|OJJ61674.1 ater|EAU31133.1 atub|OJI83469.1 aver|OJJ00995.1 awen|OJJ34307.1 azon|OJJ43334.1

Group5901: acar|OOG00239.1 acar|OOG00258.1 abra|OJJ67535.1 acla|EAW11018.1 afis|EAW20832.1 afla|EED45130.1 afum|EAL84800.1 akaw|GAA88124.1 aluc|OJZ81397.1 anid|CBF80709.1 anig|EHA18689.1 aory|EIT73782.1 asyd|OJJ62619.1 atub|OJI91602.1 aver|OJI96745.1 azon|OJJ42683.1

Group5902: acar|OOG00251.1 acar|OOG00269.1 abra|OJJ67545.1 acla|EAW12867.1 afla|EED45590.1 agla|OJJ81687.1 akaw|GAA88115.1 aluc|OJZ81387.1 anid|CBF82370.1 anig|EHA18680.1 aory|EIT75758.1 asyd|OJJ54193.1 ater|EAU37669.1 atub|OJI91592.1 aver|OJJ08533.1 awen|OJJ35116.1

Group5903: afis|EAW17151.1 afis|EAW18536.1 agla|OJJ85246.1 agla|OJJ86248.1 abra|OJJ71801.1 afla|EED49211.1 akaw|GAA85109.1 aluc|OJZ86531.1 anid|CBF80198.1 anig|EHA22115.1 aory|EIT83151.1 asyd|OJJ62941.1 ater|EAU39447.1 atub|OJI83990.1 aver|OJI98049.1 awen|OJJ30934.1

Group5904: afis|EAW22768.1 afis|EAW24693.1 abra|OJJ73447.1 afla|EED45075.1 afum|EAL89335.1 agla|OJJ81339.1 akaw|GAA89025.1 aluc|OJZ80579.1 anid|CBF76030.1 anig|EHA25619.1 aory|EIT73722.1 asyd|OJJ52452.1 ater|EAU33314.1 atub|OJI88351.1 aver|OJJ03964.1 awen|OJJ38020.1

Group5905: afla|EED45997.1 afla|EED55181.1 aory|EIT75482.1 aory|EIT81589.1 asyd|OJJ55511.1 asyd|OJJ55533.1 asyd|OJJ57370.1 aver|OJJ05798.1 aver|OJJ06588.1 acla|EAW09058.1 afis|EAW25631.1 afum|EAL84319.1 anid|CBF83157.1 ater|EAU32719.1 awen|OJJ36051.1 aver|OJJ06611.1

Group5906: afla|EED49561.1 afla|EED49753.1 abra|OJJ67096.1 acar|OOF98458.1 acla|EAW08935.1 afis|EAW15463.1 afum|EAL85858.1 akaw|GAA91065.1 aluc|OJZ87878.1 anid|CBF74671.1 anig|EHA20182.1 aory|EIT82641.1 ater|EAU39081.1 atub|OJI80772.1 awen|OJJ37034.1 azon|OJJ44465.1

Group5907: afum|EAL86255.1 afum|EAL89177.1 abra|OJJ68037.1 acar|OOF91806.1 acla|EAW06968.1 afis|EAW16582.1 afla|EED55050.1 akaw|GAA90915.1 aluc|OJZ82911.1 anig|EHA21334.1 aory|EIT81979.1 asyd|OJJ53719.1 ater|EAU36654.1 atub|OJI89137.1 aver|OJJ07845.1 awen|OJJ33891.1

Group5908: agla|OJJ78990.1 agla|OJJ83688.1 abra|OJJ73812.1 acar|OOG00967.1 acla|EAW07699.1 afis|EAW22716.1 afla|EED50000.1 afum|EAL90478.1 akaw|GAA89431.1 aluc|OJZ88238.1 anid|CBF88051.1 anig|EHA17985.1 aory|EIT74675.1 atub|OJI90840.1 awen|OJJ35410.1 azon|OJJ49766.1

Group5909: agla|OJJ83138.1 agla|OJJ86319.1 abra|OJJ68823.1 afum|EAL86208.1 akaw|GAA90430.1 aluc|OJZ85799.1 anid|CBF78855.1 anig|EHA24758.1 asyd|OJJ55033.1 atub|OJI79584.1 aver|OJJ07170.1 awen|OJJ30830.1 afla|EED50787.1 aory|EIT77976.1 afla|EED57506.1 aory|EIT81869.1

Group5910: agla|OJJ85816.1 agla|OJJ85832.1 agla|OJJ85818.1 acar|OOF98908.1 acla|EAW11260.1 akaw|GAA83783.1 aluc|OJZ90941.1 anid|CBF86189.1 anig|EHA19649.1 asyd|OJJ64535.1 ater|EAU33747.1 atub|OJI82932.1 aver|OJI98632.1 awen|OJJ30618.1 azon|OJJ47999.1 afla|EED56228.1

Group5911: agla|OJJ86769.1 agla|OJJ86770.1 abra|OJJ74895.1 acar|OOF95896.1 acla|EAW08476.1 afis|EAW23522.1 afla|EED57422.1 akaw|GAA83525.1 aluc|OJZ92470.1 anig|EHA26498.1 aory|EIT80073.1 asyd|OJJ62259.1 atub|OJI86900.1 aver|OJI96368.1 awen|OJJ39847.1 afum|EAL87942.1

Group5912: anid|CBF71552.1 anid|CBF84169.1 acla|EAW08561.1 afis|EAW23603.1 afla|EED55065.1 afum|EAL87863.2 aory|EIT77141.1 asyd|OJJ56282.1 ater|EAU35362.1 aver|OJJ08479.1 awen|OJJ32575.1 acla|EAW13726.1 afla|EED50166.1 aory|EIT76902.1 afla|EED47738.1 anid|CBF71809.1

Group5913: aory|EIT75268.1 aory|EIT75646.1 abra|OJJ66105.1 acar|OOF93992.1 afis|EAW16816.1 afla|EED53712.1 afum|EAL86367.1 akaw|GAA92674.1 aluc|OJZ90129.1 anid|CBF78556.1 anig|EHA20942.1 asyd|OJJ65060.1 atub|OJI84667.1 azon|OJJ46860.1 acar|OOF91691.1 aory|EIT80433.1

Group5914: asyd|OJJ54204.1 asyd|OJJ54927.1 aver|OJJ01905.1 aver|OJJ07268.1 awen|OJJ35311.1 awen|OJJ36432.1 acla|EAW07665.1 afis|EAW22682.1 afla|EED49966.1 afum|EAL90510.1 agla|OJJ79757.1 anid|CBF88120.1 aory|EIT83446.1 asyd|OJJ63584.1 ater|EAU39027.1 aver|OJI97350.1

Group5915: asyd|OJJ54824.1 asyd|OJJ57889.1 ater|EAU33274.1 ater|EAU38853.1 acar|OOF98643.1 acla|EAW11238.1 afis|EAW18267.1 afum|EAL87632.1 anid|CBF79518.1 aory|EIT73252.1 atub|OJI88770.1 aver|OJJ04505.1 awen|OJJ30641.1 azon|OJJ47974.1 ater|EAU34238.1 aver|OJJ07336.1

Group5916: asyd|OJJ56712.1 asyd|OJJ57429.1 aver|OJJ05750.1 aver|OJJ06013.1 abra|OJJ67120.1 acar|OOF98431.1 acla|EAW11078.1 afla|EED52028.1 agla|OJJ86869.1 akaw|GAA91045.1 aluc|OJZ87855.1 anig|EHA20160.1 aory|EIT81420.1 atub|OJI80792.1 awen|OJJ33344.1 ater|EAU37270.1

Group5917: asyd|OJJ57667.1 asyd|OJJ64961.1 atub|OJI90353.1 abra|OJJ70633.1 acar|OOF97091.1 afis|EAW23159.1 afla|EED46961.1 afum|EAL88324.1 akaw|GAA89972.1 aluc|OJZ83981.1 anid|CBF86381.1 anig|EHA19108.1 aory|EIT72519.1 ater|EAU36292.1 aver|OJI99083.1 awen|OJJ36624.1

Group5918: asyd|OJJ60004.1 asyd|OJJ62600.1 awen|OJJ32651.1 awen|OJJ37094.1 abra|OJJ77155.1 acar|OOF93042.1 afla|EED45743.1 agla|OJJ81794.1 akaw|GAA90775.1 aluc|OJZ84569.1 anid|CBF79863.1 anig|EHA27864.1 aory|EIT75727.1 ater|EAU31865.1 atub|OJI89513.1 aver|OJJ02282.1

Group5919: asyd|OJJ60338.1 asyd|OJJ65170.1 abra|OJJ69828.1 acar|OOF93353.1 afum|EAL86293.1 agla|OJJ79198.1 akaw|GAA87832.1 aluc|OJZ85468.1 anid|CBF87111.1 anig|EHA25009.1 ater|EAU37987.1 atub|OJI80095.1 aver|OJI99322.1 awen|OJJ32858.1 azon|OJJ47733.1 azon|OJJ45214.1

Group5920: ater|EAU32685.1 ater|EAU32837.1 abra|OJJ69617.1 acar|OOF92689.1 akaw|GAA91601.1 aluc|OJZ82425.1 anid|CBF82164.1 anig|EHA28344.1 asyd|OJJ54340.1 atub|OJI88015.1 aver|OJJ05292.1 awen|OJJ36838.1 azon|OJJ43517.1 anig|EHA28204.1 akaw|GAA92799.1 aluc|OJZ88721.1

Group5921: awen|OJJ29955.1 awen|OJJ35055.1 afla|EED47573.1 abra|OJJ72994.1 acar|OOF96171.1 afis|EAW16182.1 afla|EED52079.1 akaw|GAA85568.1 aluc|OJZ83502.1 anig|EHA24336.1 aory|EIT76901.1 atub|OJI82396.1 awen|OJJ30004.1 afum|EAL88813.1 ater|EAU29301.1 aluc|OJZ80354.1

Group5922: awen|OJJ30767.1 awen|OJJ31714.1 abra|OJJ67443.1 acar|OOF92384.1 afis|EAW18434.1 afla|EED52981.1 afum|EAL93115.1 agla|OJJ85001.1 akaw|GAA86386.1 aluc|OJZ84921.1 anid|CBF70031.1 aory|EIT78703.1 asyd|OJJ58869.1 ater|EAU29850.1 atub|OJI80678.1 aver|OJJ01471.1

Group5923: awen|OJJ30908.1 awen|OJJ38004.1 abra|OJJ73700.1 acar|OOF90210.1 akaw|GAA88464.1 aluc|OJZ88353.1 anig|EHA18085.1 aory|EIT81772.1 atub|OJI90948.1 abra|OJJ75777.1 acar|OOF91917.1 acla|EAW07822.1 afis|EAW22858.1 afum|EAL90342.1 akaw|GAA87320.1 afla|EED51804.1

Group5924: awen|OJJ30944.1 awen|OJJ32848.1 abra|OJJ75187.1 acar|OOF94595.1 acla|EAW14838.1 afis|EAW24366.1 afla|EED46591.1 afum|EAL86038.2 agla|OJJ82520.1 akaw|GAA88789.1 aluc|OJZ92750.1 anid|CBF87187.1 anig|EHA26253.1 aory|EIT77903.1 atub|OJI87176.1 acar|OOF90681.1

Group5925: awen|OJJ32657.1 awen|OJJ33840.1 abra|OJJ66139.1 acar|OOF95826.1 acla|EAW11590.1 afla|EED52591.1 agla|OJJ88404.1 akaw|GAA92695.1 aluc|OJZ90153.1 anid|CBF83901.1 anig|EHA20916.1 aory|EIT73374.1 ater|EAU33482.1 atub|OJI84646.1 agla|OJJ82534.1 aory|EIT74204.1

Group5926: awen|OJJ33832.1 awen|OJJ36387.1 abra|OJJ72946.1 acar|OOF96127.1 afla|EED48176.1 akaw|GAA85528.1 aluc|OJZ83456.1 anid|CBF83118.1 anig|EHA24302.1 aory|EIT83024.1 asyd|OJJ53474.1 ater|EAU31957.1 atub|OJI82348.1 aver|OJJ03999.1 azon|OJJ42763.1 asyd|OJJ60256.1

Group5927: azon|OJJ42111.1 azon|OJJ45189.1 azon|OJJ44136.1 azon|OJJ49610.1 abra|OJJ73127.1 acar|OOG00350.1 afla|EED48144.1 akaw|GAA88037.1 aluc|OJZ81296.1 anig|EHA18604.1 aory|EIT76194.1 asyd|OJJ56200.1 ater|EAU39558.1 atub|OJI91505.1 aver|OJJ05127.1 awen|OJJ36253.1

Group5928: azon|OJJ44493.1 azon|OJJ50272.1 abra|OJJ71252.1 acar|OOF97918.1 acla|EAW10283.1 afis|EAW25309.1 akaw|GAA82735.1 aluc|OJZ87290.1 anig|EHA20531.1 atub|OJI81351.1 awen|OJJ41057.1 acar|OOG00279.1 akaw|GAA88103.1 aluc|OJZ81370.1 anig|EHA18670.1 atub|OJI91575.1

Group5929: acla|EAW11715.1 agla|OJJ78979.1 acar|OOF97633.1 akaw|GAA92095.1 aluc|OJZ83630.1 anig|EHA18783.1 aory|EIT80453.1 atub|OJI89999.1 afis|EAW23734.1 afla|EED51512.1 afum|EAL87172.1 anid|CBF87375.1 asyd|OJJ52839.1 ater|EAU31294.1 aver|OJJ08808.1 awen|OJJ33670.1

Group5930: abra|OJJ65588.1 acla|EAW13295.1 afis|EAW17315.1 asyd|OJJ60476.1 abra|OJJ68806.1 acar|OOF91595.1 afum|EAL91135.2 akaw|GAA90447.1 aluc|OJZ85778.1 anid|CBF74648.1 anig|EHA24776.1 ater|EAU39376.1 atub|OJI79563.1 aver|OJI99796.1 awen|OJJ35960.1 afum|EAL91151.1

Group5931: abra|OJJ65608.1 acar|OOF92455.1 acla|EAW12305.1 afla|EED51167.1 anid|CBF81654.1 aory|EIT81330.1 asyd|OJJ54344.1 ater|EAU31008.1 aver|OJJ05296.1 abra|OJJ77464.1 anig|EHA19296.1 asyd|OJJ56405.1 ater|EAU39560.1 aver|OJJ06307.1 anid|CBF80165.1 aver|OJI99520.1

Group5932: abra|OJJ65769.1 afum|EAL87801.2 anig|EHA21875.1 aver|OJI99576.1 awen|OJJ29597.1 acar|OOF90907.1 acla|EAW09702.1 agla|OJJ85451.1 anig|EHA21874.1 asyd|OJJ52319.1 acla|EAW12595.1 acla|EAW12596.1 aory|EIT75343.1 afis|EAW16225.1 afum|EAL87819.1 aory|EIT79546.1

Group5933: afis|EAW19924.1 acar|OOF93587.1 acla|EAW13682.1 abra|OJJ75101.1 acar|OOF97233.1 afla|EED55651.1 afum|EBA27214.1 agla|OJJ86510.1 akaw|GAA88710.1 aluc|OJZ92665.1 aory|EIT81813.1 asyd|OJJ62423.1 atub|OJI87096.1 aver|OJI96560.1 awen|OJJ39580.1 azon|OJJ50107.1

Group5934: afla|EED49755.1 afum|EAL84309.2 acar|OOF98690.1 acla|EAW13141.1 afis|EAW25621.1 agla|OJJ86173.1 aluc|OJZ88033.1 aory|EIT77706.1 asyd|OJJ56973.1 atub|OJI90639.1 aver|OJJ03316.1 awen|OJJ36298.1 afis|EAW17513.1 afla|EED52201.1 aluc|OJZ79945.1 aory|EIT74476.1

Group5935: abra|OJJ65917.1 acar|OOF94967.1 afla|EED50458.1 afum|EAL84940.1 agla|OJJ80059.1 akaw|GAA92315.1 aluc|OJZ90213.1 anid|CBF84461.1 anig|EHA23711.1 aory|EIT82170.1 asyd|OJJ53548.1 ater|EAU31471.1 atub|OJI81644.1 aver|OJJ08205.1 awen|OJJ31371.1 asyd|OJJ54265.1

Group5936: abra|OJJ65943.1 acar|OOF92930.1 acla|EAW13568.1 afis|EAW20034.1 afum|EAL84882.1 agla|OJJ88020.1 akaw|GAA90821.1 aluc|OJZ83014.1 anid|CBF84767.1 anig|EHA21429.1 asyd|OJJ53283.1 ater|EAU29400.1 atub|OJI89249.1 aver|OJJ08459.1 awen|OJJ30057.1 azon|OJJ46328.1

Group5937: abra|OJJ65947.1 acar|OOF92926.1 afla|EED46040.1 afum|EAL84886.1 agla|OJJ88017.1 akaw|GAA90825.1 aluc|OJZ83010.1 anid|CBF84758.1 anig|EHA21425.1 aory|EIT75670.1 asyd|OJJ53287.1 ater|EAU29394.1 atub|OJI89245.1 aver|OJJ08454.1 awen|OJJ30053.1 azon|OJJ43699.1

Group5938: abra|OJJ65955.1 acar|OOF92917.1 acla|EAW13576.1 afis|EAW20025.1 afla|EED46033.1 afum|EAL84892.1 agla|OJJ88005.1 akaw|GAA90832.1 aluc|OJZ83002.1 anid|CBF84747.1 anig|EHA21418.1 asyd|OJJ53294.1 ater|EAU29387.1 atub|OJI89236.1 aver|OJJ08446.1 azon|OJJ43692.1

Group5939: abra|OJJ65970.1 acar|OOF92903.1 acla|EAW13591.1 afla|EED46019.1 agla|OJJ87977.1 akaw|GAA90845.1 aluc|OJZ82988.1 anid|CBF87092.1 anig|EHA21403.1 aory|EIT75531.1 asyd|OJJ60402.1 atub|OJI89222.1 aver|OJI99731.1 awen|OJJ30026.1 afla|EED54119.1 awen|OJJ36029.1

Group5940: abra|OJJ66005.1 acar|OOF92092.1 acla|EAW13642.1 afis|EAW19993.1 afla|EED45929.1 afum|EAL84924.1 akaw|GAA90878.1 aluc|OJZ82955.1 anid|CBF84688.1 anig|EHA21370.1 aory|EIT75499.1 asyd|OJJ53332.1 ater|EAU35263.1 atub|OJI89187.1 aver|OJJ08412.1 azon|OJJ43627.1

Group5941: afis|EAW17530.1 agla|OJJ79940.1 anid|CBF76169.1 abra|OJJ69571.1 afla|EED57484.1 akaw|GAA92494.1 aluc|OJZ91506.1 anig|EHA28383.1 aory|EIT80112.1 asyd|OJJ64206.1 ater|EAU32458.1 atub|OJI82869.1 aver|OJI98294.1 awen|OJJ29896.1 agla|OJJ82855.1 agla|OJJ86642.1

Group5942: abra|OJJ66098.1 acar|OOF90927.1 acla|EAW14985.1 afis|EAW24507.1 afla|EED45890.1 afum|EAL86175.1 akaw|GAA93286.1 aluc|OJZ90158.1 anid|CBF89388.1 anig|EHA20914.1 aory|EIT75697.1 asyd|OJJ56086.1 ater|EAU35712.1 atub|OJI84641.1 aver|OJJ05029.1 awen|OJJ38109.1

Group5943: abra|OJJ66100.1 akaw|GAA92666.1 aluc|OJZ90120.1 anig|EHA20949.1 atub|OJI84675.1 abra|OJJ66420.1 acar|OOG00283.1 afla|EED54053.1 afum|EAL87184.1 anid|CBF86035.1 anig|EHA28120.1 aory|EIT76094.1 aver|OJJ07930.1 azon|OJJ48538.1 ater|EAU37533.1 awen|OJJ41204.1

Group5944: abra|OJJ66195.1 acar|OOF91936.1 acla|EAW12300.1 afis|EAW16934.1 afla|EED53214.1 afum|EAL86459.1 agla|OJJ86546.1 akaw|GAA89093.1 anid|CBF82438.1 anig|EHA25695.1 aory|EIT74921.1 asyd|OJJ54125.1 ater|EAU37364.1 atub|OJI88432.1 aver|OJJ08611.1 awen|OJJ30070.1

Group5945: abra|OJJ66233.1 acar|OOF90691.1 afla|EED47090.1 agla|OJJ84413.1 akaw|GAA89056.1 aluc|OJZ80538.1 anid|CBF85759.1 anig|EHA25657.1 aory|EIT72497.1 asyd|OJJ64301.1 atub|OJI88393.1 aver|OJJ07894.1 awen|OJJ31375.1 agla|OJJ80120.1 aver|OJI98394.1 awen|OJJ36365.1

Group5946: abra|OJJ66246.1 acar|OOF90704.1 afis|EAW24436.1 afla|EED55245.1 afum|EAL86104.1 agla|OJJ79251.1 akaw|GAA89044.1 aluc|OJZ80552.1 anid|CBF86218.1 anig|EHA25645.1 asyd|OJJ64561.1 ater|EAU30010.1 atub|OJI88379.1 aver|OJI98658.1 awen|OJJ32763.1 azon|OJJ42168.1

Group5947: abra|OJJ66261.1 acar|OOF90732.1 acla|EAW09306.1 afis|EAW21840.1 afla|EED46532.1 aluc|OJZ80569.1 anig|EHA25629.1 aory|EIT72369.1 asyd|OJJ59252.1 ater|EAU38517.1 atub|OJI88362.1 aver|OJJ01861.1 awen|OJJ34122.1 azon|OJJ48038.1 aory|EIT82363.1 afla|EED57910.1

Group5948: abra|OJJ66429.1 acar|OOF92895.1 afla|EED53122.1 akaw|GAA90297.1 aluc|OJZ83040.1 anid|CBF88837.1 anig|EHA28111.1 aory|EIT72885.1 asyd|OJJ62821.1 atub|OJI87909.1 azon|OJJ51632.1 akaw|GAA84751.1 aluc|OJZ86932.1 anig|EHA21714.1 atub|OJI83589.1 afla|EED56423.1

Group5949: abra|OJJ66471.1 afis|EAW16806.1 afla|EED53911.1 agla|OJJ89421.1 akaw|GAA92130.1 aluc|OJZ83592.1 anid|CBF80483.1 asyd|OJJ55016.1 ater|EAU31818.1 atub|OJI89964.1 aver|OJI99385.1 awen|OJJ37491.1 acar|OOF94407.1 awen|OJJ36143.1 anid|CBF78765.1 asyd|OJJ65236.1

Group5950: abra|OJJ66505.1 acar|OOF90969.1 acla|EAW10976.1 afis|EAW20883.1 afla|EED57731.1 afum|EAL84749.1 agla|OJJ85833.1 akaw|GAA91205.1 aluc|OJZ81540.1 anid|CBF85489.1 anig|EHA25896.1 aory|EIT79869.1 asyd|OJJ54961.1 atub|OJI88646.1 aver|OJJ07230.1 awen|OJJ34477.1

Group5951: abra|OJJ66533.1 acar|OOF90984.1 acla|EAW10959.1 afis|EAW20900.1 afla|EED50250.1 afum|EBA27203.1 agla|OJJ84013.1 akaw|GAA91182.1 aluc|OJZ81521.1 anig|EHA25919.1 asyd|OJJ54113.1 ater|EAU38533.1 atub|OJI88670.1 aver|OJJ08622.1 awen|OJJ34448.1 azon|OJJ51734.1

Group5952: abra|OJJ66537.1 acar|OOF90988.1 acla|EAW10954.1 afis|EAW20907.1 afla|EED50255.1 afum|EAL84726.1 agla|OJJ84023.1 akaw|GAA91178.1 aluc|OJZ81510.1 anig|EHA25923.1 aory|EIT77300.1 asyd|OJJ54099.1 ater|EAU38537.1 atub|OJI88674.1 aver|OJJ08639.1 awen|OJJ34444.1

Group5953: abra|OJJ66539.1 acar|OOF90992.1 acla|EAW10951.1 afis|EAW20910.1 afla|EED50258.1 afum|EAL84724.2 agla|OJJ84026.1 akaw|GAA91176.1 aluc|OJZ81505.1 anid|CBF82479.1 anig|EHA25925.1 aory|EIT77280.1 ater|EAU38540.1 atub|OJI88676.1 awen|OJJ34436.1 azon|OJJ51718.1

Group5954: abra|OJJ66549.1 acar|OOF89953.1 acla|EAW10943.1 afis|EAW20915.1 afla|EED48929.1 afum|EAL84719.1 agla|OJJ78905.1 akaw|GAA91174.1 aluc|OJZ81502.1 anid|CBF87874.1 anig|EHA25928.1 aory|EIT80003.1 asyd|OJJ65183.1 atub|OJI88679.1 aver|OJI99336.1 awen|OJJ30158.1

Group5955: abra|OJJ66560.1 acla|EAW10935.1 afis|EAW20926.1 afla|EED50279.1 afum|EAL84708.1 agla|OJJ83963.1 akaw|GAA91157.1 aluc|OJZ81485.1 anid|CBF82501.1 anig|EHA25942.1 aory|EIT77314.1 ater|EAU38572.1 atub|OJI88696.1 aver|OJJ08648.1 awen|OJJ34394.1 azon|OJJ48125.1

Group5956: abra|OJJ66571.1 acar|OOF91011.1 acla|EAW10922.1 afis|EAW20936.1 afum|EAL84698.1 agla|OJJ83950.1 akaw|GAA91147.1 aluc|OJZ81473.1 anid|CBF82528.1 anig|EHA25953.1 asyd|OJJ54074.1 ater|EAU38582.1 atub|OJI88705.1 aver|OJJ08665.1 awen|OJJ34381.1 azon|OJJ48113.1

Group5957: abra|OJJ66603.1 acar|OOF90115.1 acla|EAW10890.1 afis|EAW20968.1 afla|EED50322.1 afum|EBA27201.1 agla|OJJ84045.1 akaw|GAA91117.1 aluc|OJZ81443.1 anid|CBF82582.1 anig|EHA25981.1 aory|EIT77258.1 asyd|OJJ54040.1 ater|EAU38618.1 atub|OJI88735.1 awen|OJJ34347.1

Group5958: abra|OJJ66605.1 acar|OOF90113.1 acla|EAW10888.1 afis|EAW20970.1 afla|EED50324.1 afum|EAL84664.1 agla|OJJ84047.1 akaw|GAA91115.1 aluc|OJZ81441.1 anig|EHA25983.1 aory|EIT77253.1 asyd|OJJ54038.1 atub|OJI88737.1 aver|OJJ08702.1 awen|OJJ34345.1 azon|OJJ48082.1

Group5959: abra|OJJ66658.1 acar|OOF93675.1 acla|EAW11800.1 afis|EAW19410.1 afla|EED50365.1 afum|EAL87084.2 agla|OJJ81124.1 akaw|GAA89608.1 aluc|OJZ80790.1 anid|CBF74353.1 aory|EIT83118.1 asyd|OJJ57791.1 atub|OJI87641.1 aver|OJJ05373.1 awen|OJJ36806.1 acar|OOF93760.1

Group5960: abra|OJJ66762.1 acla|EAW11888.1 afis|EAW19500.1 afla|EED56719.1 afum|EBA27297.1 agla|OJJ81669.1 akaw|GAA89698.1 aluc|OJZ80687.1 anid|CBF75438.1 anig|EHA27327.1 asyd|OJJ59784.1 ater|EAU34815.1 atub|OJI87546.1 aver|OJJ02504.1 awen|OJJ36915.1 azon|OJJ42900.1

Group5961: abra|OJJ66768.1 acar|OOF93849.1 acla|EAW11605.1 afla|EED49852.1 agla|OJJ83400.1 akaw|GAA89705.1 aluc|OJZ80679.1 anid|CBF82408.1 anig|EHA27333.1 aory|EIT83408.1 asyd|OJJ54169.1 atub|OJI87538.1 aver|OJJ08570.1 awen|OJJ40864.1 azon|OJJ49073.1 awen|OJJ40207.1

Group5962: abra|OJJ66778.1 acar|OOF93857.1 afis|EAW16744.1 afla|EED45880.1 afum|EAL86319.2 agla|OJJ80280.1 akaw|GAA89713.1 aluc|OJZ80671.1 anid|CBF75770.1 anig|EHA27340.1 asyd|OJJ62106.1 ater|EAU36169.1 atub|OJI87525.1 aver|OJI96212.1 awen|OJJ29904.1 azon|OJJ42471.1

Group5963: abra|OJJ66785.1 acar|OOF96742.1 acla|EAW12028.1 afis|EAW19677.1 afum|EAL86845.1 akaw|GAA89719.1 aluc|OJZ80665.1 anid|CBF81955.1 anig|EHA27345.1 asyd|OJJ52953.1 atub|OJI87520.1 aver|OJJ08919.1 awen|OJJ37896.1 afla|EED47776.1 agla|OJJ81344.1 aory|EIT77196.1

Group5964: abra|OJJ66826.1 acla|EAW14408.1 afis|EAW19116.1 afla|EED54298.1 afum|EAL93848.2 agla|OJJ80431.1 akaw|GAA90462.1 aluc|OJZ85761.1 anid|CBF79730.1 anig|EHA24795.1 asyd|OJJ58072.1 ater|EAU37378.1 atub|OJI79546.1 aver|OJJ04335.1 awen|OJJ37446.1 azon|OJJ51124.1

Group5965: abra|OJJ66841.1 acar|OOF91618.1 acla|EAW14422.1 afis|EAW19129.1 afla|EED54285.1 afum|EAL93861.1 agla|OJJ88601.1 akaw|GAA90477.1 aluc|OJZ85743.1 anid|CBF79747.1 anig|EHA24809.1 asyd|OJJ58089.1 atub|OJI79528.1 aver|OJJ04311.1 awen|OJJ37462.1 azon|OJJ51140.1

Group5966: abra|OJJ66872.1 acar|OOF91648.1 acla|EAW14447.1 afis|EAW19155.1 afla|EED54247.1 afum|EAL93886.1 akaw|GAA91537.1 aluc|OJZ85709.1 anid|CBF86296.1 anig|EHA24833.1 aory|EIT76022.1 asyd|OJJ65011.1 ater|EAU37428.1 atub|OJI79500.1 aver|OJI99130.1 awen|OJJ37508.1

Group5967: abra|OJJ66877.1 acar|OOF91652.1 acla|EAW14452.1 afis|EAW19160.1 afla|EED54242.1 afum|EAL93890.2 akaw|GAA91532.1 aluc|OJZ85705.1 anid|CBF86287.1 anig|EHA24837.1 aory|EIT76015.1 asyd|OJJ65016.1 ater|EAU37433.1 atub|OJI79495.1 aver|OJI99134.1 awen|OJJ37513.1

Group5968: abra|OJJ66878.1 acar|OOF91653.1 acla|EAW14453.1 afis|EAW19161.1 afla|EED54241.1 afum|EAL93891.1 agla|OJJ88563.1 akaw|GAA91531.1 aluc|OJZ85704.1 anid|CBF75184.1 anig|EHA24838.1 aory|EIT76132.1 ater|EAU37319.1 atub|OJI79494.1 awen|OJJ37514.1 azon|OJJ51177.1

Group5969: abra|OJJ66959.1 acar|OOF93649.1 acla|EAW14961.1 afis|EAW24484.1 afla|EED52058.1 afum|EAL86153.1 agla|OJJ88957.1 akaw|GAA92202.1 aluc|OJZ85617.1 anig|EHA24909.1 aory|EIT81432.1 asyd|OJJ55928.1 ater|EAU37957.1 atub|OJI79410.1 aver|OJJ04878.1 azon|OJJ51805.1

Group5970: abra|OJJ66989.1 acar|OOF93623.1 acla|EAW12086.1 afis|EAW19729.1 afla|EED52168.1 afum|EAL86798.1 akaw|GAA92174.1 aluc|OJZ85584.1 anid|CBF87096.1 anig|EHA24937.1 aory|EIT74461.1 asyd|OJJ60395.1 atub|OJI79379.1 aver|OJI99724.1 awen|OJJ29795.1 azon|OJJ50255.1

Group5971: abra|OJJ67092.1 acar|OOF97365.1 acla|EAW12323.1 afis|EAW25701.1 afum|EAL84374.1 agla|OJJ86316.1 akaw|GAA91069.1 aluc|OJZ87883.1 anid|CBF82025.1 anig|EHA20186.1 asyd|OJJ58346.1 ater|EAU39413.1 atub|OJI80768.1 aver|OJJ02067.1 afis|EAW23748.1 anid|CBF80131.1

Group5972: abra|OJJ67100.1 acar|OOF98452.1 acla|EAW13713.1 afis|EAW20273.1 afla|EED48132.1 afum|EAL85407.1 agla|OJJ87505.1 akaw|GAA91061.1 aluc|OJZ87873.1 anid|CBF78240.1 anig|EHA20179.1 aory|EIT76216.1 ater|EAU31788.1 atub|OJI80776.1 awen|OJJ35880.1 afis|EAW23647.1

Group5973: abra|OJJ67109.1 acar|OOF98443.1 afis|EAW24483.1 afla|EED52545.1 afum|EAL86152.1 agla|OJJ85836.1 akaw|GAA91055.1 aluc|OJZ87866.1 anig|EHA20172.1 aory|EIT74263.1 ater|EAU30095.1 atub|OJI80782.1 awen|OJJ30838.1 ater|EAU33502.1 afum|EAL92050.2 asyd|OJJ52467.1

Group5974: abra|OJJ67132.1 acar|OOF99014.1 afla|EED53374.1 anid|CBF81678.1 aory|EIT74911.1 asyd|OJJ59559.1 aver|OJI98274.1 awen|OJJ38788.1 azon|OJJ42559.1 abra|OJJ70398.1 acar|OOF91875.1 akaw|GAA91960.1 aluc|OJZ82657.1 anig|EHA27416.1 atub|OJI88889.1 aver|OJJ02709.1

Group5975: abra|OJJ67156.1 acar|OOF98403.1 acla|EAW14569.1 afis|EAW19282.1 afla|EED47595.1 agla|OJJ88394.1 akaw|GAA91016.1 aluc|OJZ87817.1 anid|CBF78761.1 anig|EHA20298.1 aory|EIT80196.1 asyd|OJJ65234.1 atub|OJI80827.1 aver|OJI99383.1 awen|OJJ37643.1 azon|OJJ44907.1

Group5976: abra|OJJ67161.1 acar|OOF98399.1 afis|EAW19279.1 afla|EED47599.1 akaw|GAA91012.1 aluc|OJZ87813.1 anid|CBF78754.1 aory|EIT80238.1 asyd|OJJ65230.1 ater|EAU29484.1 atub|OJI80831.1 aver|OJI99379.1 awen|OJJ37639.1 acla|EAW14566.1 afum|EAL94005.1 azon|OJJ44880.1

Group5977: abra|OJJ67167.1 acar|OOF98394.1 acla|EAW14561.1 afis|EAW19274.1 afum|EAL94000.1 akaw|GAA91007.1 aluc|OJZ87808.1 anid|CBF78747.1 anig|EHA20290.1 asyd|OJJ65225.1 ater|EAU29490.1 atub|OJI80837.1 aver|OJI99375.1 afla|EED47602.1 agla|OJJ88399.1 aory|EIT80191.1

Group5978: abra|OJJ67230.1 acar|OOF98323.1 acla|EAW10403.1 afis|EAW25169.1 afla|EED57527.1 afum|EAL89842.1 agla|OJJ88986.1 akaw|GAA91655.1 aluc|OJZ87744.1 anid|CBF85618.1 asyd|OJJ64203.1 ater|EAU39453.1 atub|OJI80897.1 aver|OJI98292.1 awen|OJJ40820.1 azon|OJJ46864.1

Group5979: abra|OJJ67233.1 asyd|OJJ53601.1 aver|OJJ08147.1 awen|OJJ35947.1 azon|OJJ42412.1 abra|OJJ77556.1 acar|OOF99171.1 afla|EED46976.1 akaw|GAA84067.1 aluc|OJZ80981.1 anig|EHA19379.1 aory|EIT72531.1 ater|EAU33717.1 atub|OJI82605.1 ater|EAU31461.1 awen|OJJ34676.1

Group5980: abra|OJJ67269.1 acar|OOF98297.1 afis|EAW17110.1 afla|EED55274.1 afum|EAL86574.1 agla|OJJ88999.1 akaw|GAA91684.1 aluc|OJZ87710.1 anid|CBF85646.1 anig|EHA20204.1 aory|EIT83570.1 asyd|OJJ64237.1 ater|EAU39436.1 atub|OJI80932.1 aver|OJI98330.1 awen|OJJ40797.1

Group5981: abra|OJJ67271.1 acla|EAW10416.1 afis|EAW25153.1 afla|EED55280.1 afum|EAL89828.1 akaw|GAA91686.1 aluc|OJZ87708.1 anid|CBF85639.1 anig|EHA20890.1 aory|EIT83511.1 asyd|OJJ64241.1 atub|OJI80934.1 aver|OJI98334.1 awen|OJJ40782.1 azon|OJJ46870.1 agla|OJJ89001.1

Group5982: akaw|GAA86510.1 aluc|OJZ85054.1 anig|EHA25270.1 acar|OOF92509.1 acla|EAW06428.1 afis|EAW18532.1 afla|EED52868.1 agla|OJJ85145.1 anid|CBF70202.1 asyd|OJJ58785.1 ater|EAU30038.1 aver|OJJ01393.1 awen|OJJ31090.1 azon|OJJ50596.1 afum|EAL93230.1 aory|EIT78632.1

Group5983: abra|OJJ67307.1 acar|OOF92506.1 acla|EAW06426.1 afis|EAW18530.1 afla|EED52871.1 afum|EAL93227.1 agla|OJJ85143.1 akaw|GAA86508.1 aluc|OJZ85051.1 anid|CBF70206.1 anig|EHA25268.1 asyd|OJJ58783.1 atub|OJI80547.1 aver|OJJ01391.1 awen|OJJ31088.1 azon|OJJ50593.1

Group5984: abra|OJJ67312.1 acla|EAW06420.1 afis|EAW18524.1 afum|EAL93219.1 agla|OJJ85136.1 akaw|GAA86503.1 aluc|OJZ85046.1 anid|CBF70215.1 anig|EHA25263.1 aory|EIT78618.1 asyd|OJJ58777.1 ater|EAU30048.1 atub|OJI80552.1 aver|OJJ01386.1 awen|OJJ31078.1 azon|OJJ50586.1

Group5985: abra|OJJ67320.1 acar|OOF92494.1 acla|EAW06416.1 afis|EAW18520.1 afla|EED52878.1 afum|EAL93215.1 agla|OJJ85132.1 akaw|GAA86498.1 aluc|OJZ85040.1 anid|CBF70224.1 anig|EHA25258.1 aory|EIT78617.1 ater|EAU30050.1 atub|OJI80557.1 awen|OJJ31072.1 azon|OJJ50582.1

Group5986: abra|OJJ67333.1 acar|OOF92480.1 acla|EAW06401.1 afis|EAW18505.1 afla|EED52896.1 afum|EAL93199.1 agla|OJJ85117.1 akaw|GAA86483.1 aluc|OJZ85024.1 anid|CBF87122.1 anig|EHA25245.1 aory|EIT78621.1 asyd|OJJ60361.1 ater|EAU30066.1 atub|OJI80573.1 aver|OJI99698.1

Group5987: afla|EED54124.1 aory|EIT76004.1 abra|OJJ77008.1 afis|EAW25620.1 agla|OJJ78907.1 akaw|GAA87040.1 aluc|OJZ84407.1 anig|EHA27737.1 asyd|OJJ55522.1 ater|EAU32182.1 atub|OJI89665.1 aver|OJJ06600.1 awen|OJJ36039.1 azon|OJJ45113.1 acar|OOF91454.1 anid|CBF83171.1

Group5988: abra|OJJ67397.1 acar|OOF92420.1 acla|EAW06363.1 afis|EAW18472.1 afla|EED52938.1 afum|EAL93159.2 agla|OJJ85059.1 akaw|GAA86430.1 aluc|OJZ84972.1 anig|EHA25184.1 asyd|OJJ58828.1 ater|EAU29889.1 atub|OJI80629.1 aver|OJJ01431.1 awen|OJJ31024.1 azon|OJJ50538.1

Group5989: abra|OJJ67406.1 acar|OOF92410.1 acla|EAW06352.1 afis|EAW18464.1 afum|EAL93151.2 agla|OJJ85048.1 akaw|GAA86421.1 aluc|OJZ84962.1 anid|CBF70093.1 aory|EIT78734.1 asyd|OJJ58838.1 ater|EAU29882.1 atub|OJI80638.1 aver|OJJ01440.1 awen|OJJ31016.1 azon|OJJ50529.1

Group5990: abra|OJJ67428.1 acar|OOF92398.1 acla|EAW06336.1 afis|EAW18448.1 afla|EED52960.1 afum|EAL93133.2 agla|OJJ85025.1 akaw|GAA86402.1 aluc|OJZ84938.1 anid|CBF70068.1 anig|EHA25156.1 asyd|OJJ58848.1 atub|OJI80661.1 aver|OJJ01449.1 awen|OJJ31000.1 azon|OJJ50514.1

Group5991: abra|OJJ67532.1 acar|OOG01225.1 acla|EAW14583.1 afis|EAW20812.1 afla|EED55975.1 afum|EAL84816.1 akaw|GAA88126.1 anid|CBF77711.1 anig|EHA18691.1 asyd|OJJ53078.1 ater|EAU32035.1 atub|OJI91604.1 aver|OJJ03769.1 awen|OJJ32225.1 acla|EAW11063.1 awen|OJJ30947.1

Group5992: abra|OJJ67550.1 acar|OOF90250.1 acla|EAW13271.1 afis|EAW17346.1 afla|EED45503.1 afum|EAL91190.1 akaw|GAA88110.1 aluc|OJZ81380.1 anig|EHA18675.1 aory|EIT75626.1 asyd|OJJ56676.1 ater|EAU29418.1 atub|OJI91586.1 aver|OJJ06048.1 acar|OOF94391.1 ater|EAU35969.1

Group5993: abra|OJJ67617.1 afla|EED45202.1 akaw|GAA89134.1 aluc|OJZ81702.1 anig|EHA19895.1 aory|EIT73724.1 atub|OJI81348.1 abra|OJJ73693.1 acar|OOG00866.1 agla|OJJ78660.1 akaw|GAA88467.1 aluc|OJZ88356.1 anig|EHA18089.1 ater|EAU39318.1 atub|OJI90951.1 awen|OJJ33717.1

Group5994: abra|OJJ67624.1 acar|OOF91972.1 afis|EAW20775.1 afla|EED45948.1 afum|EAL84845.1 agla|OJJ79050.1 akaw|GAA89144.1 aluc|OJZ81692.1 anid|CBF74983.1 anig|EHA25752.1 aory|EIT75387.1 asyd|OJJ54562.1 atub|OJI88488.1 aver|OJJ07594.1 awen|OJJ35893.1 azon|OJJ42145.1

Group5995: abra|OJJ67669.1 acar|OOF90912.1 acla|EAW12248.1 afis|EAW16768.1 akaw|GAA89185.1 aluc|OJZ81646.1 anid|CBF84388.1 anig|EHA25797.1 asyd|OJJ54909.1 ater|EAU34859.1 atub|OJI88536.1 aver|OJJ07293.1 azon|OJJ43892.1 anid|CBF78140.1 asyd|OJJ53585.1 aver|OJJ08163.1

Group5996: abra|OJJ67692.1 acar|OOF97519.1 acla|EAW14628.1 afis|EAW17251.1 afla|EED53073.1 afum|EAL86670.1 akaw|GAA89201.1 aluc|OJZ81629.1 anid|CBF88038.1 anig|EHA25814.1 aory|EIT74791.1 asyd|OJJ58695.1 ater|EAU30968.1 atub|OJI88551.1 aver|OJJ01307.1 awen|OJJ32905.1

Group5997: abra|OJJ67729.1 acar|OOF90931.1 afis|EAW20573.1 afla|EED52328.1 afum|EAL84492.1 agla|OJJ88971.1 akaw|GAA89223.1 aluc|OJZ81605.1 anid|CBF74531.1 anig|EHA25840.1 aory|EIT74343.1 atub|OJI88580.1 aver|OJJ06519.1 awen|OJJ29984.1 asyd|OJJ55446.1 awen|OJJ36164.1

Group5998: abra|OJJ67829.1 acar|OOG00541.1 acla|EAW09505.1 afum|EAL84476.1 akaw|GAA91840.1 aluc|OJZ82058.1 anid|CBF78144.1 anig|EHA22199.1 ater|EAU32305.1 atub|OJI84617.1 aver|OJI99602.1 awen|OJJ31459.1 azon|OJJ43641.1 acar|OOF92586.1 azon|OJJ44381.1 aver|OJI99600.1

Group5999: abra|OJJ67857.1 acar|OOF92837.1 acla|EAW11743.1 afis|EAW23776.1 afla|EED51576.1 afum|EAL87210.1 agla|OJJ78927.1 akaw|GAA91863.1 aluc|OJZ82035.1 anid|CBF79828.1 anig|EHA22221.1 aory|EIT80538.1 atub|OJI84590.1 aver|OJJ04261.1 awen|OJJ33701.1 azon|OJJ42840.1

Group6000: abra|OJJ67860.1 acar|OOF92841.1 acla|EAW11738.1 afis|EAW23771.1 afla|EED51572.1 afum|EAL87205.1 agla|OJJ78948.1 akaw|GAA91867.1 aluc|OJZ82032.1 anig|EHA22224.1 aory|EIT80473.1 asyd|OJJ58140.1 ater|EAU31315.1 atub|OJI84587.1 aver|OJJ04265.1 awen|OJJ33704.1

Group6001: abra|OJJ67864.1 acar|OOF92844.1 afis|EAW23767.1 afla|EED51553.1 afum|EAL87201.1 agla|OJJ78955.1 akaw|GAA91871.1 aluc|OJZ82028.1 anid|CBF84704.1 anig|EHA22228.1 aory|EIT80560.1 asyd|OJJ53320.1 atub|OJI84582.1 aver|OJJ08423.1 awen|OJJ33696.1 azon|OJJ42545.1

Group6002: abra|OJJ67880.1 acar|OOF92856.1 acla|EAW11723.1 afis|EAW23746.1 afla|EED51538.1 afum|EAL87181.1 akaw|GAA91887.1 aluc|OJZ82010.1 anid|CBF87336.1 anig|EHA22247.1 aory|EIT80493.1 asyd|OJJ56363.1 atub|OJI84559.1 aver|OJJ06351.1 awen|OJJ33681.1 azon|OJJ44710.1

Group6003: abra|OJJ67882.1 acar|OOF92863.1 acla|EAW11720.1 afis|EAW23741.1 afla|EED51535.1 afum|EAL87178.1 agla|OJJ78964.1 akaw|GAA91889.1 aluc|OJZ82008.1 anid|CBF79813.1 anig|EHA22249.1 aory|EIT80693.1 ater|EAU31298.1 atub|OJI84557.1 awen|OJJ33678.1 azon|OJJ42819.1

Group6004: abra|OJJ67888.1 acla|EAW11717.1 afis|EAW23736.1 afla|EED51513.1 afum|EAL87174.1 agla|OJJ78970.1 akaw|GAA91894.1 aluc|OJZ82002.1 anid|CBF84245.1 anig|EHA22254.1 asyd|OJJ57198.1 ater|EAU31295.1 atub|OJI84552.1 aver|OJJ03093.1 awen|OJJ33671.1 azon|OJJ42814.1

Group6005: abra|OJJ67889.1 acar|OOF92859.1 afis|EAW23738.1 afla|EED51514.1 afum|EAL87175.1 agla|OJJ78967.1 akaw|GAA91895.1 aluc|OJZ82001.1 anid|CBF79809.1 anig|EHA22255.1 aory|EIT80532.1 asyd|OJJ58130.1 ater|EAU31296.1 atub|OJI84551.1 aver|OJJ04275.1 awen|OJJ33674.1

Group6006: abra|OJJ67890.1 acar|OOF92858.1 acla|EAW11060.1 afis|EAW20814.1 afla|EED47512.1 afum|EAL84814.1 akaw|GAA91896.1 aluc|OJZ82000.1 anid|CBF78358.1 anig|EHA22256.1 aory|EIT74562.1 asyd|OJJ56145.1 ater|EAU30844.1 atub|OJI84550.1 aver|OJJ05086.1 awen|OJJ35834.1

Group6007: ater|EAU35708.1 anid|CBF69522.1 afis|EAW21018.1 afla|EED56839.1 afum|EAL92056.1 agla|OJJ80564.1 aory|EIT77488.1 asyd|OJJ59206.1 aver|OJJ01817.1 awen|OJJ37826.1 afum|EAL85152.1 agla|OJJ88496.1 atub|OJI81729.1 akaw|GAA86070.1 aluc|OJZ90305.1 awen|OJJ31968.1

Group6008: acla|EAW12904.1 abra|OJJ78317.1 acar|OOF90170.1 akaw|GAA85713.1 aluc|OJZ91390.1 anid|CBF75868.1 anig|EHA20068.1 asyd|OJJ60241.1 atub|OJI83383.1 aver|OJJ04025.1 afis|EAW17776.1 afla|EED56744.1 afum|EAL91584.2 aory|EIT73880.1 ater|EAU31816.1 asyd|OJJ52432.1

Group6009: abra|OJJ67939.1 acar|OOF92788.1 acla|EAW11629.1 afis|EAW23827.1 afla|EED51979.1 afum|EAL87259.1 agla|OJJ82485.1 akaw|GAA91257.1 aluc|OJZ82811.1 anid|CBF79909.1 anig|EHA21250.1 aory|EIT81475.1 ater|EAU31382.1 atub|OJI89041.1 awen|OJJ33785.1 azon|OJJ42887.1

Group6010: abra|OJJ67972.1 acar|OOF92813.1 acla|EAW11762.1 afis|EAW23795.1 afla|EED51602.1 afum|EAL87229.2 agla|OJJ78844.1 akaw|GAA91230.1 aluc|OJZ82840.1 anid|CBF79856.1 anig|EHA21277.1 aory|EIT80530.1 atub|OJI89070.1 aver|OJJ04244.1 awen|OJJ33751.1 azon|OJJ42860.1

Group6011: abra|OJJ67984.1 acar|OOF92026.1 afis|EAW25692.1 afla|EED46905.1 afum|EAL84362.1 akaw|GAA91218.1 aluc|OJZ82852.1 anid|CBF78965.1 anig|EHA21288.1 aory|EIT72639.1 asyd|OJJ55667.1 ater|EAU35088.1 atub|OJI89082.1 aver|OJJ06748.1 awen|OJJ36205.1 azon|OJJ50743.1

Group6012: abra|OJJ68114.1 acar|OOF93908.1 afis|EAW15405.1 afla|EED44866.1 afum|EAL84241.1 agla|OJJ86207.1 akaw|GAA86895.1 aluc|OJZ90018.1 anid|CBF86936.1 anig|EHA27394.1 aory|EIT82714.1 asyd|OJJ55589.1 atub|OJI84789.1 aver|OJJ06668.1 awen|OJJ41765.1 agla|OJJ83696.1

Group6013: abra|OJJ68115.1 acar|OOF93909.1 afis|EAW16723.1 afla|EED45933.1 afum|EAL86297.2 agla|OJJ88975.1 akaw|GAA86896.1 aluc|OJZ90019.1 anig|EHA27395.1 aory|EIT75655.1 asyd|OJJ56047.1 atub|OJI84788.1 aver|OJJ04990.1 awen|OJJ36020.1 azon|OJJ45077.1 anid|CBF78249.1

Group6014: abra|OJJ68116.1 acar|OOF93910.1 afis|EAW16724.1 afla|EED45934.1 afum|EAL86298.1 agla|OJJ88976.1 akaw|GAA86897.1 aluc|OJZ90020.1 anid|CBF78251.1 anig|EHA27396.1 aory|EIT75493.1 asyd|OJJ56049.1 atub|OJI84787.1 aver|OJJ04992.1 acla|EAW13226.1 acar|OOF94484.1

Group6015: agla|OJJ87804.1 abra|OJJ74465.1 acla|EAW07232.1 afis|EAW22246.1 afum|EAL90947.1 akaw|GAA87358.1 aluc|OJZ89655.1 anid|CBF88498.1 anig|EHA26893.1 asyd|OJJ63794.1 ater|EAU34254.1 atub|OJI85149.1 aver|OJI97143.1 awen|OJJ34824.1 afla|EED57639.1 aory|EIT82987.1

Group6016: abra|OJJ68174.1 acar|OOF97472.1 acla|EAW13059.1 afis|EAW17606.1 afla|EED49216.1 afum|EAL91419.1 agla|OJJ81353.1 akaw|GAA83270.1 aluc|OJZ83776.1 anid|CBF90185.1 anig|EHA18915.1 asyd|OJJ61984.1 atub|OJI90146.1 aver|OJI96076.1 awen|OJJ37236.1 azon|OJJ50938.1

Group6017: abra|OJJ68180.1 acar|OOF97475.1 acla|EAW13062.1 afis|EAW17602.1 afla|EED49219.1 afum|EAL91415.1 akaw|GAA83266.1 aluc|OJZ83772.1 anig|EHA18912.1 aory|EIT83164.1 asyd|OJJ60509.1 ater|EAU37039.1 atub|OJI90143.1 aver|OJI99834.1 awen|OJJ37243.1 azon|OJJ50941.1

Group6018: abra|OJJ68188.1 acar|OOF97482.1 acla|EAW13070.1 afis|EAW17594.1 afla|EED49226.1 afum|EAL91407.1 akaw|GAA83257.1 aluc|OJZ83760.1 anid|CBF90208.1 anig|EHA18904.1 asyd|OJJ61971.1 ater|EAU37032.1 atub|OJI90133.1 aver|OJI96064.1 awen|OJJ37252.1 azon|OJJ50947.1

Group6019: abra|OJJ68189.1 acar|OOF97485.1 acla|EAW13071.1 afis|EAW17593.1 afla|EED49228.1 afum|EAL91406.1 agla|OJJ81332.1 akaw|GAA83256.1 aluc|OJZ83759.1 asyd|OJJ61970.1 ater|EAU37030.1 atub|OJI90132.1 aver|OJI96063.1 awen|OJJ37253.1 azon|OJJ50948.1 anid|CBF90210.1

Group6020: abra|OJJ68210.1 acar|OOF97494.1 acla|EAW12974.1 afis|EAW17690.1 afla|EED50454.1 afum|EAL91504.1 agla|OJJ79489.1 akaw|GAA83236.1 aluc|OJZ83735.1 anid|CBF89982.1 anig|EHA18884.1 asyd|OJJ62089.1 atub|OJI90113.1 aver|OJI96195.1 awen|OJJ37146.1 azon|OJJ50846.1

Group6021: abra|OJJ68215.1 acla|EAW12970.1 afla|EED50449.1 akaw|GAA83231.1 aluc|OJZ83731.1 anid|CBF89974.1 anig|EHA18880.1 aory|EIT82144.1 asyd|OJJ58334.1 ater|EAU29907.1 atub|OJI90108.1 aver|OJJ02076.1 awen|OJJ37142.1 agla|OJJ80592.1 asyd|OJJ62093.1 aver|OJI96199.1

Group6022: abra|OJJ68236.1 acla|EAW12223.1 afla|EED56299.1 anid|CBF87461.1 anig|EHA18314.1 aory|EIT73296.1 asyd|OJJ52778.1 ater|EAU35485.1 atub|OJI89935.1 aver|OJJ08340.1 awen|OJJ36976.1 anid|CBF87418.1 aver|OJJ03445.1 afla|EED56300.1 asyd|OJJ53411.1 aver|OJJ03428.1

Group6023: abra|OJJ68253.1 acar|OOF97532.1 afis|EAW24691.1 afla|EED46562.1 afum|EAL89332.1 akaw|GAA83208.1 aluc|OJZ83710.1 anid|CBF70720.1 anig|EHA18861.1 aory|EIT77828.1 asyd|OJJ61535.1 ater|EAU29994.1 atub|OJI90082.1 aver|OJJ00863.1 aver|OJJ07656.1 asyd|OJJ54532.1

Group6024: abra|OJJ68295.1 acar|OOF97562.1 acla|EAW08965.1 afis|EAW15446.1 afla|EED46465.1 afum|EAL84200.1 akaw|GAA83178.1 aluc|OJZ83677.1 anig|EHA18833.1 aory|EIT72318.1 atub|OJI90048.1 ater|EAU36715.1 asyd|OJJ53380.1 aver|OJJ08363.1 anid|CBF73480.1 anid|CBF87739.1

Group6025: abra|OJJ68325.1 acar|OOF97617.1 afla|EED57063.1 agla|OJJ78511.1 akaw|GAA92076.1 aluc|OJZ83648.1 anig|EHA18800.1 aory|EIT83411.1 asyd|OJJ56739.1 atub|OJI90017.1 awen|OJJ38714.1 asyd|OJJ52101.1 afla|EED49837.1 akaw|GAA93219.1 anid|CBF89124.1 anig|EHA22004.1

Group6026: abra|OJJ68348.1 acar|OOF94844.1 afis|EAW17519.1 afla|EED45478.1 afum|EAL91337.1 agla|OJJ78830.1 akaw|GAA92115.1 aluc|OJZ83610.1 anid|CBF80533.1 anig|EHA18769.1 aory|EIT75491.1 asyd|OJJ54654.1 ater|EAU36947.1 atub|OJI89983.1 aver|OJJ07507.1 awen|OJJ37340.1

Group6027: abra|OJJ68355.1 afla|EED53490.1 agla|OJJ82739.1 akaw|GAA92127.1 aluc|OJZ83595.1 anig|EHA18755.1 aory|EIT72952.1 asyd|OJJ56978.1 ater|EAU39005.1 atub|OJI89967.1 aver|OJJ03309.1 awen|OJJ36414.1 acar|OOF91796.1 afla|EED48206.1 ater|EAU30833.1 awen|OJJ37862.1

Group6028: abra|OJJ68383.1 acar|OOF93985.1 afis|EAW16782.1 afla|EED56189.1 afum|EAL86353.1 agla|OJJ79766.1 akaw|GAA92562.1 aluc|OJZ90109.1 anig|EHA20962.1 aory|EIT73337.1 asyd|OJJ59099.1 ater|EAU30270.1 atub|OJI84694.1 aver|OJJ01690.1 awen|OJJ29848.1 azon|OJJ50360.1

Group6029: abra|OJJ68391.1 afis|EAW19987.1 afum|EAL84928.1 akaw|GAA92556.1 aluc|OJZ90103.1 anig|EHA20967.1 aory|EIT82613.1 atub|OJI84702.1 acla|EAW13638.1 akaw|GAA92555.1 anid|CBF73971.1 anig|EHA20968.1 asyd|OJJ62368.1 aver|OJI96502.1 afla|EED53131.1 aory|EIT74881.1

Group6030: abra|OJJ68402.1 acar|OOF93970.1 afis|EAW17326.1 afla|EED54080.1 afum|EAL91163.1 akaw|GAA92545.1 aluc|OJZ90094.1 anid|CBF82407.1 anig|EHA20982.1 aory|EIT75924.1 asyd|OJJ54167.1 ater|EAU29239.1 atub|OJI84712.1 aver|OJJ08575.1 awen|OJJ29674.1 azon|OJJ46027.1

Group6031: abra|OJJ68447.1 acar|OOF93948.1 acla|EAW15354.1 afis|EAW20359.1 afla|EED47313.1 afum|EAL85476.1 akaw|GAA86927.1 aluc|OJZ90051.1 anid|CBF71653.1 anig|EHA21019.1 aory|EIT82832.1 asyd|OJJ61838.1 ater|EAU30275.1 atub|OJI84756.1 aver|OJI95937.1 azon|OJJ50770.1

Group6032: abra|OJJ68454.1 afla|EED56946.1 agla|OJJ87926.1 akaw|GAA86920.1 aluc|OJZ90044.1 anid|CBF70196.1 anig|EHA21027.1 aory|EIT77506.1 asyd|OJJ58788.1 ater|EAU30025.1 atub|OJI84763.1 aver|OJJ01395.1 awen|OJJ37051.1 azon|OJJ46448.1 afla|EED48364.1 agla|OJJ81574.1

Group6033: abra|OJJ68480.1 acar|OOF91469.1 acla|EAW06709.1 afis|EAW16259.1 afla|EED48914.1 afum|EAL88880.1 akaw|GAA92404.1 aluc|OJZ83265.1 anid|CBF80822.1 anig|EHA21118.1 asyd|OJJ54455.1 ater|EAU32190.1 atub|OJI87672.1 aver|OJJ07700.1 awen|OJJ38053.1 azon|OJJ43162.1

Group6034: abra|OJJ68493.1 acar|OOF90054.1 acla|EAW12143.1 afis|EAW17193.1 afum|EAL86621.1 agla|OJJ86661.1 akaw|GAA92777.1 aluc|OJZ82706.1 anid|CBF69881.1 anig|EHA21161.1 aory|EIT73563.1 asyd|OJJ59013.1 ater|EAU37674.1 atub|OJI88939.1 aver|OJJ01602.1 azon|OJJ42624.1

Group6035: abra|OJJ68547.1 afis|EAW17986.1 afla|EED51918.1 afum|EAL91795.1 akaw|GAA91307.1 aluc|OJZ82763.1 anid|CBF80002.1 anig|EHA21207.1 aory|EIT81366.1 asyd|OJJ58249.1 ater|EAU31496.1 atub|OJI88996.1 aver|OJJ02175.1 azon|OJJ48275.1 acla|EAW12821.1 awen|OJJ41560.1

Group6036: abra|OJJ68579.1 afla|EED56440.1 anid|CBF75415.1 aory|EIT72799.1 asyd|OJJ56522.1 atub|OJI91463.1 aver|OJJ06200.1 awen|OJJ37869.1 abra|OJJ68930.1 acar|OOF90673.1 afis|EAW19226.1 afum|EAL93957.1 akaw|GAA90321.1 aluc|OJZ85912.1 anig|EHA24676.1 atub|OJI79702.1

Group6037: anid|CBF78851.1 abra|OJJ74275.1 acar|OOF95456.1 acla|EAW07319.1 afis|EAW22351.1 afla|EED57755.1 afum|EAL90841.1 agla|OJJ87708.1 akaw|GAA86749.1 aluc|OJZ89858.1 anig|EHA27059.1 aory|EIT81036.1 asyd|OJJ63888.1 atub|OJI84948.1 aver|OJI97051.1 anid|CBF88659.1

Group6038: abra|OJJ68644.1 acar|OOF92147.1 acla|EAW07072.1 afis|EAW24963.1 afum|EAL89621.1 akaw|GAA89784.1 aluc|OJZ81907.1 anig|EHA22336.1 asyd|OJJ55374.1 ater|EAU30793.1 atub|OJI84460.1 aver|OJJ06453.1 awen|OJJ33779.1 azon|OJJ50607.1 anid|CBF82913.1 ater|EAU34596.1

Group6039: abra|OJJ68669.1 acar|OOF92169.1 acla|EAW12578.1 afis|EAW24070.1 afla|EED51405.1 afum|EAL91926.1 agla|OJJ87313.1 akaw|GAA89810.1 aluc|OJZ81880.1 anid|CBF86460.1 asyd|OJJ64913.1 ater|EAU30491.1 atub|OJI84434.1 aver|OJI99034.1 awen|OJJ41420.1 azon|OJJ48885.1

Group6040: abra|OJJ68671.1 acar|OOF92172.1 acla|EAW12576.1 afis|EAW24072.1 afum|EAL91928.1 agla|OJJ87141.1 akaw|GAA89824.1 aluc|OJZ81866.1 anid|CBF86464.1 anig|EHA22361.1 aory|EIT80514.1 asyd|OJJ64911.1 ater|EAU30493.1 atub|OJI84432.1 aver|OJI99032.1 awen|OJJ41408.1

Group6041: abra|OJJ68680.1 acar|OOF92182.1 acla|EAW12567.1 afis|EAW24083.1 afla|EED51390.1 afum|EAL91938.1 agla|OJJ87153.1 akaw|GAA89833.1 aluc|OJZ81853.1 anid|CBF86482.1 asyd|OJJ64900.1 ater|EAU30506.1 atub|OJI84422.1 aver|OJI99021.1 awen|OJJ41406.1 azon|OJJ48872.1

Group6042: abra|OJJ68737.1 acar|OOF92233.1 acla|EAW12510.1 afis|EAW24136.1 afum|EAL91991.1 agla|OJJ87213.1 akaw|GAA89885.1 aluc|OJZ81791.1 anid|CBF86571.1 anig|EHA22424.1 asyd|OJJ64846.1 ater|EAU30565.1 atub|OJI84364.1 aver|OJI98958.1 awen|OJJ41353.1 azon|OJJ48816.1

Group6043: abra|OJJ68764.1 acar|OOF92250.1 acla|EAW12487.1 afis|EAW24158.1 afum|EAL92014.2 agla|OJJ89523.1 akaw|GAA89908.1 aluc|OJZ81761.1 anid|CBF77898.1 anig|EHA22452.1 asyd|OJJ55304.1 ater|EAU30589.1 atub|OJI84333.1 aver|OJJ06922.1 awen|OJJ41320.1 azon|OJJ48793.1

Group6044: abra|OJJ68765.1 acar|OOF92251.1 acla|EAW12486.1 afis|EAW24159.1 afla|EED46774.1 afum|EAL92015.1 agla|OJJ89522.1 akaw|GAA89909.1 aluc|OJZ81760.1 anid|CBF77896.1 aory|EIT72588.1 asyd|OJJ55305.1 atub|OJI84332.1 aver|OJJ06921.1 awen|OJJ41319.1 azon|OJJ48792.1

Group6045: abra|OJJ68778.1 acar|OOF92264.1 acla|EAW12473.1 afla|EED46762.1 afum|EAL92028.1 agla|OJJ89508.1 akaw|GAA89922.1 aluc|OJZ81748.1 anid|CBF77870.1 anig|EHA22464.1 asyd|OJJ55318.1 ater|EAU30603.1 atub|OJI84319.1 aver|OJJ06910.1 awen|OJJ41305.1 azon|OJJ48779.1

Group6046: abra|OJJ68781.1 acar|OOF92268.1 acla|EAW12471.1 afla|EED46759.1 afum|EAL92032.1 agla|OJJ89505.1 akaw|GAA89925.1 aluc|OJZ81745.1 anid|CBF77865.1 anig|EHA22467.1 asyd|OJJ55321.1 ater|EAU30605.1 atub|OJI84316.1 aver|OJJ06907.1 awen|OJJ41302.1 azon|OJJ48776.1

Group6047: abra|OJJ68787.1 acar|OOF92271.1 acla|EAW12467.1 afis|EAW24174.1 afum|EAL92036.2 agla|OJJ89500.1 aluc|OJZ81741.1 anid|CBF77857.1 anig|EHA22471.1 aory|EIT72261.1 asyd|OJJ55325.1 ater|EAU30609.1 atub|OJI84312.1 aver|OJJ06903.1 awen|OJJ41296.1 azon|OJJ48772.1

Group6048: abra|OJJ68796.1 acar|OOF92280.1 acla|EAW14658.1 afis|EAW24185.1 afum|EAL92045.1 agla|OJJ89490.1 akaw|GAA89935.1 aluc|OJZ81731.1 anid|CBF77838.1 anig|EHA22481.1 asyd|OJJ55335.1 ater|EAU30618.1 atub|OJI84302.1 aver|OJJ06892.1 awen|OJJ41285.1 azon|OJJ48768.1

Group6049: acar|OOF97439.1 acla|EAW13029.1 afis|EAW17638.1 afum|EAL91453.1 agla|OJJ81258.1 anid|CBF90127.1 aory|EIT82094.1 asyd|OJJ62017.1 ater|EAU37074.1 aver|OJI96108.1 azon|OJJ50907.1 afla|EED50525.1 akaw|GAA83335.1 aluc|OJZ83831.1 anig|EHA18974.1 atub|OJI90198.1

Group6050: acla|EAW15285.1 afum|EAL85363.2 agla|OJJ84627.1 acar|OOF90548.1 afis|EAW20280.1 afla|EED57098.1 akaw|GAA90407.1 aluc|OJZ85821.1 anid|CBF85204.1 asyd|OJJ63028.1 atub|OJI79608.1 aver|OJI97940.1 awen|OJJ31242.1 azon|OJJ50674.1 acla|EAW15275.1 afis|EAW20245.1

Group6051: abra|OJJ68843.1 acar|OOF90546.1 acla|EAW11068.1 afis|EAW20806.1 afla|EED47530.1 afum|EAL84822.1 akaw|GAA90405.1 aluc|OJZ85823.1 anid|CBF85130.1 anig|EHA24738.1 aory|EIT74519.1 asyd|OJJ63043.1 ater|EAU37215.1 atub|OJI79610.1 aver|OJI97921.1 azon|OJJ44328.1

Group6052: abra|OJJ68845.1 acar|OOF90544.1 acla|EAW15280.1 afis|EAW20249.1 afla|EED57096.1 afum|EAL85367.1 akaw|GAA90403.1 aluc|OJZ85825.1 anid|CBF85206.1 anig|EHA24736.1 aory|EIT82292.1 asyd|OJJ63030.1 ater|EAU32040.1 atub|OJI79613.1 aver|OJI97938.1 azon|OJJ50397.1

Group6053: abra|OJJ68847.1 acla|EAW15274.1 afis|EAW20244.1 afla|EED57100.1 afum|EAL85361.1 agla|OJJ84629.1 aluc|OJZ85827.1 anid|CBF85138.1 anig|EHA24734.1 aory|EIT82275.1 asyd|OJJ63035.1 ater|EAU30255.1 atub|OJI79615.1 aver|OJI97932.1 awen|OJJ31241.1 akaw|GAA92649.1

Group6054: abra|OJJ68925.1 acar|OOF91511.1 acla|EAW14249.1 afis|EAW19031.1 afla|EED50384.1 afum|EAL93760.2 agla|OJJ84591.1 akaw|GAA90323.1 aluc|OJZ85910.1 anid|CBF79594.1 anig|EHA24682.1 asyd|OJJ57984.1 ater|EAU32070.1 atub|OJI79699.1 aver|OJJ04417.1 awen|OJJ31460.1

Group6055: abra|OJJ68928.1 afis|EAW16673.1 afum|EAL89278.1 akaw|GAA90319.1 aluc|OJZ85914.1 anig|EHA24678.1 atub|OJI79704.1 afla|EED48203.1 acar|OOF90674.1 agla|OJJ79205.1 anid|CBF83970.1 aory|EIT72893.1 asyd|OJJ61414.1 aver|OJJ00750.1 ater|EAU34353.1 awen|OJJ31870.1

Group6056: abra|OJJ68986.1 acar|OOF91557.1 afis|EAW19076.1 afla|EED54441.1 agla|OJJ83213.1 akaw|GAA90507.1 aluc|OJZ85972.1 anig|EHA24633.1 aory|EIT81163.1 asyd|OJJ58661.1 ater|EAU32124.1 atub|OJI79760.1 aver|OJJ01267.1 awen|OJJ32470.1 azon|OJJ46236.1 acar|OOF91848.1

Group6057: abra|OJJ69019.1 acar|OOF98449.1 afis|EAW20276.1 afla|EED53202.1 afum|EAL85410.1 agla|OJJ87502.1 akaw|GAA90536.1 aluc|OJZ86005.1 anid|CBF78245.1 anig|EHA24605.1 aory|EIT75242.1 asyd|OJJ56192.1 ater|EAU31765.1 atub|OJI79794.1 aver|OJJ05121.1 awen|OJJ35878.1

Group6058: acla|EAW14513.1 azon|OJJ44015.1 acar|OOF98345.1 afis|EAW19231.1 afla|EED47657.1 afum|EAL93960.1 akaw|GAA88443.1 aluc|OJZ88331.1 anid|CBF78616.1 anig|EHA18066.1 aory|EIT80280.1 asyd|OJJ65166.1 ater|EAU37232.1 atub|OJI90926.1 aver|OJI99318.1 afis|EAW25599.1

Group6059: abra|OJJ69042.1 acar|OOF89866.1 acla|EAW11011.1 afis|EAW16707.1 afla|EED53056.1 afum|EAL89298.1 akaw|GAA90571.1 aluc|OJZ86027.1 anid|CBF85134.1 anig|EHA24585.1 aory|EIT74798.1 asyd|OJJ63344.1 ater|EAU32201.1 atub|OJI79816.1 aver|OJI97617.1 azon|OJJ45979.1

Group6060: abra|OJJ69085.1 acla|EAW08747.1 afis|EAW15689.1 afla|EED51364.1 afum|EAL85659.1 akaw|GAA90614.1 aluc|OJZ86073.1 anid|CBF86339.1 anig|EHA24542.1 aory|EIT80722.1 asyd|OJJ64986.1 ater|EAU32465.1 atub|OJI79870.1 aver|OJI99106.1 awen|OJJ38380.1 azon|OJJ43864.1

Group6061: abra|OJJ69101.1 acar|OOF91195.1 acla|EAW08731.1 afis|EAW15705.1 afla|EED51347.1 afum|EAL85642.1 akaw|GAA90627.1 aluc|OJZ86090.1 anid|CBF71104.1 anig|EHA24527.1 asyd|OJJ52709.1 ater|EAU32440.1 atub|OJI79886.1 aver|OJJ04738.1 azon|OJJ43843.1 agla|OJJ81853.1

Group6062: abra|OJJ69112.1 acla|EAW08720.1 afum|EAL85620.1 agla|OJJ81866.1 akaw|GAA90637.1 aluc|OJZ86101.1 anid|CBF71087.1 anig|EHA24518.1 asyd|OJJ52699.1 aver|OJJ04727.1 awen|OJJ38411.1 azon|OJJ43831.1 acar|OOF91186.1 atub|OJI79896.1 afis|EAW15727.1 afla|EED51336.1

Group6063: abra|OJJ69117.1 acar|OOF91189.1 acla|EAW08715.1 afis|EAW15723.1 afla|EED51331.1 afum|EAL85624.1 agla|OJJ81871.1 akaw|GAA90642.1 aluc|OJZ86106.1 anig|EHA24514.1 asyd|OJJ52694.1 ater|EAU32426.1 atub|OJI79901.1 aver|OJJ04722.1 awen|OJJ38416.1 azon|OJJ43826.1

Group6064: abra|OJJ69129.1 acar|OOF91171.1 acla|EAW08703.1 afis|EAW15711.1 afla|EED51319.1 akaw|GAA90654.1 aluc|OJZ86118.1 anid|CBF71055.1 anig|EHA24503.1 aory|EIT81518.1 asyd|OJJ52677.1 ater|EAU32415.1 atub|OJI79914.1 aver|OJJ04703.1 awen|OJJ38429.1 azon|OJJ43814.1

Group6065: abra|OJJ69139.1 acla|EAW08695.1 afis|EAW15740.1 afla|EED51312.1 afum|EAL85608.1 agla|OJJ81901.1 akaw|GAA86318.1 aluc|OJZ86126.1 anig|EHA24495.1 aory|EIT81512.1 atub|OJI79922.1 awen|OJJ38439.1 azon|OJJ43806.1 anid|CBF71043.1 asyd|OJJ52670.1 aver|OJJ04696.1

Group6066: abra|OJJ69227.1 acar|OOF91093.1 acla|EAW08621.1 afis|EAW15812.1 afla|EED51234.1 afum|EAL85539.1 agla|OJJ81978.1 akaw|GAA92235.1 aluc|OJZ86207.1 anid|CBF75275.1 aory|EIT83185.1 asyd|OJJ55913.1 ater|EAU33442.1 atub|OJI80001.1 aver|OJJ04853.1 awen|OJJ38525.1

Group6067: abra|OJJ69258.1 acar|OOF91062.1 acla|EAW08591.1 afis|EAW15843.1 afla|EED51206.1 afum|EAL85508.1 agla|OJJ84914.1 akaw|GAA92264.1 aluc|OJZ86238.1 anid|CBF70841.1 asyd|OJJ52566.1 ater|EAU33392.1 atub|OJI80030.1 aver|OJJ04595.1 awen|OJJ38557.1 azon|OJJ43709.1

Group6068: abra|OJJ69270.1 acar|OOF94450.1 acla|EAW14660.1 afis|EAW24191.1 afum|EAL85864.1 agla|OJJ79688.1 akaw|GAA88145.1 aluc|OJZ82098.1 anid|CBF74308.1 anig|EHA28631.1 asyd|OJJ54007.1 ater|EAU31142.1 atub|OJI88339.1 aver|OJJ08102.1 awen|OJJ33108.1 azon|OJJ44839.1

Group6069: abra|OJJ69287.1 acar|OOF94435.1 acla|EAW14676.1 afis|EAW24207.1 afla|EED48064.1 afum|EAL85877.1 agla|OJJ79670.1 akaw|GAA88159.1 aluc|OJZ82116.1 anig|EHA28615.1 aory|EIT83334.1 asyd|OJJ53988.1 atub|OJI88321.1 aver|OJJ08086.1 awen|OJJ33091.1 azon|OJJ44857.1

Group6070: abra|OJJ69290.1 acar|OOF94432.1 acla|EAW14679.1 afis|EAW24210.1 afla|EED48061.1 agla|OJJ79656.1 akaw|GAA88162.1 aluc|OJZ82120.1 anid|CBF74265.1 anig|EHA28612.1 asyd|OJJ53985.1 ater|EAU31160.1 atub|OJI88318.1 aver|OJJ08083.1 awen|OJJ33086.1 azon|OJJ44860.1

Group6071: abra|OJJ69323.1 acar|OOF94405.1 acla|EAW14721.1 afis|EAW24257.1 afum|EAL85925.1 agla|OJJ79616.1 akaw|GAA88192.1 aluc|OJZ82153.1 anid|CBF74191.1 anig|EHA28578.1 asyd|OJJ53950.1 ater|EAU31203.1 atub|OJI88283.1 aver|OJJ08046.1 awen|OJJ33043.1 azon|OJJ48476.1

Group6072: abra|OJJ69327.1 acar|OOF94401.1 acla|EAW14713.1 afis|EAW24248.1 afla|EED48025.1 afum|EAL85916.1 agla|OJJ79625.1 akaw|GAA88198.1 aluc|OJZ82159.1 anig|EHA28573.1 asyd|OJJ53954.1 ater|EAU31189.1 atub|OJI88278.1 aver|OJJ08050.1 awen|OJJ33060.1 azon|OJJ48468.1

Group6073: abra|OJJ69344.1 acar|OOF94384.1 acla|EAW14702.1 afis|EAW24233.1 afla|EED48038.1 afum|EAL85902.1 agla|OJJ79638.1 akaw|GAA88213.1 aluc|OJZ82176.1 anig|EHA28558.1 aory|EIT83330.1 asyd|OJJ53963.1 ater|EAU31181.1 atub|OJI88261.1 aver|OJJ08059.1 azon|OJJ48459.1

Group6074: abra|OJJ69354.1 acar|OOF94371.1 acla|EAW14744.1 afis|EAW24275.1 afla|EED48001.1 afum|EAL85943.1 agla|OJJ79586.1 akaw|GAA88223.1 anig|EHA28549.1 aory|EIT82906.1 asyd|OJJ53926.1 ater|EAU31220.1 atub|OJI88250.1 aver|OJJ08025.1 awen|OJJ33022.1 azon|OJJ48495.1

Group6075: abra|OJJ69388.1 acar|OOF94338.1 acla|EAW14775.1 afis|EAW24306.1 afla|EED47964.1 afum|EAL85974.1 agla|OJJ83174.1 akaw|GAA88256.1 aluc|OJZ82222.1 anid|CBF74097.1 anig|EHA28518.1 asyd|OJJ53895.1 ater|EAU31259.1 atub|OJI88213.1 aver|OJJ07992.1 awen|OJJ32986.1

Group6076: abra|OJJ69431.1 acar|OOF94305.1 acla|EAW14807.1 afis|EAW24336.1 afla|EED47928.1 afum|EAL86006.1 akaw|GAA88296.1 aluc|OJZ82268.1 anid|CBF74032.1 anig|EHA28475.1 aory|EIT79835.1 asyd|OJJ53850.1 ater|EAU30412.1 atub|OJI88170.1 aver|OJJ07951.1 azon|OJJ48962.1

Group6077: abra|OJJ69451.1 acar|OOF94290.1 acla|EAW14815.1 afis|EAW24345.1 afla|EED47918.1 agla|OJJ83208.1 akaw|GAA88368.1 aluc|OJZ82287.1 anid|CBF74021.1 anig|EHA28461.1 aory|EIT83361.1 asyd|OJJ53842.1 ater|EAU30401.1 atub|OJI88152.1 aver|OJJ07945.1 awen|OJJ32954.1

Group6078: abra|OJJ69455.1 acar|OOF94286.1 acla|EAW07408.1 afis|EAW22434.1 afla|EED53608.1 afum|EAL90757.1 akaw|GAA88357.1 aluc|OJZ82298.1 anig|EHA28457.1 aory|EIT75075.1 asyd|OJJ57319.1 ater|EAU34153.1 atub|OJI88141.1 aver|OJJ05842.1 awen|OJJ36494.1 azon|OJJ44369.1

Group6079: abra|OJJ69500.1 aluc|OJZ82340.1 anig|EHA28420.1 atub|OJI88100.1 awen|OJJ30574.1 azon|OJJ45335.1 abra|OJJ78072.1 acar|OOF91543.1 acla|EAW15328.1 afum|EAL85362.2 aory|EIT73816.1 asyd|OJJ63986.1 aver|OJI96967.1 awen|OJJ40088.1 azon|OJJ46957.1 afis|EAW21468.1

Group6080: abra|OJJ69531.1 acar|OOF92664.1 afis|EAW18043.1 afla|EED51503.1 akaw|GAA88382.1 aluc|OJZ82370.1 anid|CBF75024.1 anig|EHA28394.1 asyd|OJJ60073.1 ater|EAU31283.1 atub|OJI88072.1 aver|OJJ04188.1 awen|OJJ41491.1 azon|OJJ48222.1 afum|EAL91854.1 aory|EIT80430.1

Group6081: akaw|GAA82187.1 abra|OJJ71154.1 acar|OOF93867.1 acla|EAW11932.1 afum|EAL92066.1 agla|OJJ88460.1 aluc|OJZ84300.1 asyd|OJJ52830.1 ater|EAU33249.1 atub|OJI89769.1 ater|EAU31455.1 acar|OOF98994.1 afis|EAW21129.1 aluc|OJZ91973.1 atub|OJI86412.1 akaw|GAA87137.1

Group6082: abra|OJJ69576.1 acar|OOF91579.1 acla|EAW14393.1 afis|EAW19101.1 afum|EAL93833.1 agla|OJJ84501.1 akaw|GAA91573.1 aluc|OJZ82394.1 anid|CBF79700.1 anig|EHA28371.1 aory|EIT75927.1 ater|EAU32147.1 atub|OJI88048.1 awen|OJJ32494.1 azon|OJJ46037.1 afla|EED54334.1

Group6083: abra|OJJ69597.1 acar|OOF91562.1 acla|EAW14201.1 afis|EAW19081.1 afla|EED54313.1 afum|EAL93810.1 akaw|GAA91589.1 aluc|OJZ82412.1 anid|CBF75163.1 anig|EHA28355.1 aory|EIT76019.1 asyd|OJJ59960.1 ater|EAU32128.1 atub|OJI88029.1 aver|OJJ02330.1 awen|OJJ32476.1

Group6084: abra|OJJ69674.1 afla|EED49568.1 anid|CBF90241.1 anig|EHA28292.1 aory|EIT82591.1 abra|OJJ69681.1 acar|OOF93865.1 afis|EAW19351.1 afum|EAL94070.1 akaw|GAA88806.1 aluc|OJZ82489.1 anig|EHA28285.1 ater|EAU37356.1 atub|OJI87943.1 awen|OJJ41942.1 atub|OJI85778.1

Group6085: abra|OJJ69689.1 acar|OOF93402.1 acla|EAW11116.1 afla|EED56732.1 agla|OJJ88353.1 akaw|GAA88813.1 aluc|OJZ82496.1 anid|CBF69500.1 anig|EHA28276.1 aory|EIT73912.1 asyd|OJJ59234.1 ater|EAU33166.1 atub|OJI87935.1 aver|OJJ01844.1 awen|OJJ29946.1 azon|OJJ50326.1

Group6086: abra|OJJ69739.1 acar|OOF94181.1 acla|EAW09980.1 afis|EAW21110.1 afla|EED51761.1 afum|EAL92132.1 akaw|GAA90192.1 aluc|OJZ83157.1 anid|CBF87039.1 anig|EHA28222.1 aory|EIT80519.1 asyd|OJJ60449.1 ater|EAU36788.1 atub|OJI87780.1 aver|OJI99767.1 azon|OJJ44437.1

Group6087: abra|OJJ69810.1 acar|OOF93567.1 afis|EAW25601.1 afla|EED57247.1 afum|EBA27172.1 akaw|GAA87850.1 aluc|OJZ85490.1 anid|CBF81957.1 anig|EHA25026.1 aory|EIT74113.1 asyd|OJJ54537.1 ater|EAU34250.1 atub|OJI80075.1 aver|OJJ07644.1 awen|OJJ35936.1 azon|OJJ43522.1

Group6088: abra|OJJ69818.1 acar|OOF93345.1 acla|EAW12083.1 afis|EAW19726.1 afla|EED51052.1 agla|OJJ78576.1 akaw|GAA87842.1 aluc|OJZ85478.1 anid|CBF78377.1 anig|EHA25019.1 aory|EIT76968.1 asyd|OJJ57329.1 ater|EAU33553.1 atub|OJI80085.1 aver|OJJ05832.1 awen|OJJ40132.1

Group6089: abra|OJJ69861.1 afis|EAW15433.1 afla|EED45625.1 afum|EAL84211.1 agla|OJJ86265.1 akaw|GAA87795.1 aluc|OJZ85430.1 anig|EHA25609.1 aory|EIT75726.1 asyd|OJJ64001.1 ater|EAU31307.1 atub|OJI80145.1 aver|OJI98090.1 awen|OJJ32639.1 aluc|OJZ83144.1 anig|EHA23287.1

Group6090: abra|OJJ69871.1 acar|OOF93385.1 acla|EAW08953.1 afis|EAW20571.1 afla|EED54199.1 afum|EBA27182.1 agla|OJJ80145.1 akaw|GAA87785.1 aluc|OJZ85420.1 anid|CBF78263.1 anig|EHA25599.1 asyd|OJJ56066.1 ater|EAU32669.1 atub|OJI80155.1 aver|OJJ05009.1 awen|OJJ38089.1

Group6091: abra|OJJ69922.1 acar|OOF93434.1 acla|EAW08907.1 afis|EAW15489.1 afla|EED49003.1 afum|EAL85834.1 akaw|GAA87734.1 aluc|OJZ85366.1 anig|EHA25548.1 asyd|OJJ56022.1 ater|EAU32631.1 atub|OJI80205.1 aver|OJJ04969.1 azon|OJJ43142.1 awen|OJJ38145.1 agla|OJJ80166.1

Group6092: abra|OJJ69936.1 acar|OOF93445.1 acla|EAW08889.1 afis|EAW15500.1 afum|EAL85823.1 agla|OJJ80178.1 akaw|GAA87723.1 aluc|OJZ85351.1 anig|EHA25538.1 aory|EIT82557.1 asyd|OJJ56011.1 ater|EAU32619.1 atub|OJI80219.1 aver|OJJ04956.1 awen|OJJ38159.1 azon|OJJ43130.1

Group6093: abra|OJJ69942.1 acar|OOF93452.1 acla|EAW08880.1 afis|EAW15515.1 afum|EAL85816.1 agla|OJJ80186.1 akaw|GAA87717.1 aluc|OJZ85344.1 anig|EHA25532.1 aory|EIT82510.1 asyd|OJJ56004.1 ater|EAU32609.1 atub|OJI80226.1 aver|OJJ04949.1 awen|OJJ38176.1 azon|OJJ43121.1

Group6094: abra|OJJ69950.1 acla|EAW08874.1 afis|EAW15522.1 afla|EED49028.1 afum|EBA27244.1 agla|OJJ85224.1 akaw|GAA87710.1 aluc|OJZ85336.1 anig|EHA25525.1 aory|EIT82532.1 asyd|OJJ55995.1 ater|EAU32603.1 atub|OJI80234.1 aver|OJJ04943.1 awen|OJJ38182.1 azon|OJJ43111.1

Group6095: abra|OJJ69954.1 acar|OOF93464.1 acla|EAW08868.1 afis|EAW15526.1 afla|EED49037.1 afum|EAL85802.1 agla|OJJ85220.1 akaw|GAA87705.1 aluc|OJZ85328.1 anid|CBF78213.1 aory|EIT82546.1 asyd|OJJ55990.1 atub|OJI80244.1 aver|OJJ04938.1 awen|OJJ38190.1 azon|OJJ43105.1

Group6096: abra|OJJ69956.1 acar|OOF93466.1 acla|EAW08866.1 afis|EAW15528.1 afla|EED49039.1 afum|EAL85800.1 agla|OJJ85218.1 aluc|OJZ85326.1 anid|CBF78210.1 anig|EHA25516.1 aory|EIT82502.1 asyd|OJJ55988.1 atub|OJI80246.1 aver|OJJ04936.1 awen|OJJ38192.1 azon|OJJ43102.1

Group6097: abra|OJJ69965.1 acar|OOF93473.1 acla|EAW08837.1 afis|EAW15563.1 afum|EAL85769.1 agla|OJJ85188.1 akaw|GAA87694.1 aluc|OJZ85318.1 anid|CBF78122.1 aory|EIT82485.1 asyd|OJJ55903.1 ater|EAU32569.1 atub|OJI80255.1 aver|OJJ04848.1 awen|OJJ38223.1 azon|OJJ43072.1

Group6098: abra|OJJ69977.1 acla|EAW08845.1 afis|EAW15555.1 afla|EED49059.1 afum|EAL85777.1 agla|OJJ85199.1 akaw|GAA87684.1 aluc|OJZ85303.1 anid|CBF78170.1 anig|EHA25500.1 aory|EIT82563.1 asyd|OJJ55966.1 ater|EAU32578.1 atub|OJI80267.1 aver|OJJ04913.1 awen|OJJ38213.1

Group6099: abra|OJJ69992.1 acar|OOF93498.1 acla|EAW08828.1 afis|EAW15569.1 afla|EED49079.1 afum|EAL85764.1 agla|OJJ85183.1 akaw|GAA87669.1 aluc|OJZ85287.1 anig|EHA25484.1 asyd|OJJ55898.1 ater|EAU32561.1 atub|OJI80283.1 aver|OJJ04843.1 awen|OJJ38228.1 azon|OJJ43067.1

Group6100: abra|OJJ69993.1 acar|OOF93499.1 acla|EAW08826.1 afis|EAW15571.1 afla|EED49080.1 afum|EAL85762.1 agla|OJJ85181.1 akaw|GAA87668.1 aluc|OJZ85286.1 anid|CBF74534.1 anig|EHA25483.1 aory|EIT82491.1 ater|EAU32560.1 atub|OJI80284.1 awen|OJJ38230.1 azon|OJJ43066.1

Group6101: abra|OJJ69994.1 acar|OOF93501.1 acla|EAW08825.1 afis|EAW15572.1 afla|EED49081.1 agla|OJJ85180.1 akaw|GAA87667.1 aluc|OJZ85285.1 anid|CBF78107.1 anig|EHA25482.1 aory|EIT82508.1 asyd|OJJ55896.1 ater|EAU32559.1 atub|OJI80285.1 aver|OJJ04841.1 awen|OJJ38231.1

Group6102: abra|OJJ69996.1 acar|OOF93503.1 acla|EAW08822.1 afis|EAW15575.1 afla|EED49084.1 afum|EAL85759.1 agla|OJJ85177.1 akaw|GAA87665.1 aluc|OJZ85283.1 anig|EHA25480.1 aory|EIT82540.1 asyd|OJJ55893.1 ater|EAU32556.1 atub|OJI80287.1 aver|OJJ04838.1 awen|OJJ38235.1

Group6103: abra|OJJ69999.1 acar|OOF93506.1 acla|EAW08820.1 afis|EAW15577.1 afla|EED49086.1 afum|EAL85757.1 agla|OJJ85175.1 akaw|GAA87662.1 aluc|OJZ85280.1 anig|EHA25477.1 aory|EIT82487.1 asyd|OJJ55891.1 ater|EAU32554.1 atub|OJI80290.1 aver|OJJ04836.1 awen|OJJ38238.1

Group6104: abra|OJJ70000.1 acla|EAW08819.1 afis|EAW15578.1 afla|EED49094.1 afum|EAL85756.1 agla|OJJ85174.1 akaw|GAA87661.1 aluc|OJZ85279.1 anid|CBF78102.1 anig|EHA25476.1 aory|EIT82512.1 asyd|OJJ55890.1 atub|OJI80291.1 aver|OJJ04835.1 awen|OJJ38240.1 azon|OJJ43059.1

Group6105: abra|OJJ70017.1 acar|OOF93523.1 acla|EAW08806.1 afis|EAW15590.1 afla|EED50413.1 afum|EAL85742.1 agla|OJJ85158.1 akaw|GAA87644.1 aluc|OJZ85259.1 anid|CBF78074.1 anig|EHA25460.1 asyd|OJJ55875.1 atub|OJI80309.1 aver|OJJ04820.1 awen|OJJ38263.1 azon|OJJ43022.1

Group6106: abra|OJJ70019.1 acar|OOF93525.1 afis|EAW15592.1 afla|EED50415.1 afum|EAL85740.1 akaw|GAA87642.1 aluc|OJZ85257.1 anid|CBF78078.1 anig|EHA25458.1 aory|EIT82135.1 asyd|OJJ55877.1 ater|EAU32536.1 atub|OJI80311.1 aver|OJJ04822.1 awen|OJJ38265.1 azon|OJJ43913.1

Group6107: abra|OJJ70033.1 afis|EAW17103.1 anig|EHA18718.1 atub|OJI79322.1 awen|OJJ29677.1 abra|OJJ72237.1 acar|OOF96859.1 afla|EED54714.1 akaw|GAA84702.1 aluc|OJZ86985.1 anid|CBF76384.1 anig|EHA21662.1 asyd|OJJ61069.1 ater|EAU29940.1 atub|OJI83534.1 aver|OJJ00385.1

Group6108: abra|OJJ70131.1 acar|OOF93325.1 acla|EAW09068.1 afis|EAW25640.1 afla|EED55189.1 afum|EAL84327.2 agla|OJJ80957.1 akaw|GAA86633.1 aluc|OJZ85190.1 anid|CBF86061.1 anig|EHA25404.1 aory|EIT83514.1 ater|EAU32713.1 atub|OJI80407.1 awen|OJJ36096.1 agla|OJJ78893.1

Group6109: abra|OJJ70178.1 acar|OOF91400.1 acla|EAW13479.1 afis|EAW20586.1 afum|EAL84479.1 agla|OJJ80828.1 akaw|GAA86595.1 aluc|OJZ85142.1 anig|EHA25358.1 asyd|OJJ55630.1 ater|EAU37093.1 atub|OJI80450.1 aver|OJJ03121.1 awen|OJJ33297.1 anig|EHA27684.1 aver|OJJ06707.1

Group6110: abra|OJJ70197.1 acar|OOF90057.1 acla|EAW14580.1 afis|EAW19314.1 afla|EED47568.1 afum|EAL94030.1 akaw|GAA86579.1 aluc|OJZ85130.1 anid|CBF78805.1 anig|EHA25349.1 aory|EIT74569.1 asyd|OJJ65257.1 ater|EAU29461.1 atub|OJI80463.1 aver|OJI99408.1 awen|OJJ35850.1

Group6111: abra|OJJ70293.1 acar|OOF91747.1 afis|EAW22073.1 afum|EAL91097.1 akaw|GAA92855.1 aluc|OJZ82558.1 anid|CBF82784.1 anig|EHA22520.1 aory|EIT73943.1 asyd|OJJ57122.1 ater|EAU30151.1 atub|OJI88800.1 aver|OJJ02365.1 awen|OJJ35472.1 azon|OJJ47361.1 awen|OJJ33482.1

Group6112: abra|OJJ70368.1 acar|OOF91902.1 afis|EAW16892.1 afla|EED48420.1 afum|EAL86430.2 agla|OJJ79761.1 akaw|GAA91983.1 aluc|OJZ82630.1 anig|EHA27447.1 aory|EIT83082.1 atub|OJI88863.1 awen|OJJ31356.1 azon|OJJ49611.1 afla|EED51051.1 aory|EIT77047.1 awen|OJJ35872.1

Group6113: abra|OJJ70384.1 acar|OOF91887.1 acla|EAW12345.1 afis|EAW16882.1 afum|EAL86418.1 agla|OJJ79752.1 akaw|GAA91973.1 aluc|OJZ82642.1 anid|CBF80675.1 anig|EHA27428.1 asyd|OJJ54588.1 ater|EAU30233.1 atub|OJI88875.1 aver|OJJ07566.1 awen|OJJ29769.1 azon|OJJ50378.1

Group6114: abra|OJJ70385.1 acar|OOF91886.1 acla|EAW12343.1 afis|EAW16879.1 afla|EED47344.1 afum|EAL86416.1 akaw|GAA91972.1 aluc|OJZ82643.1 anid|CBF80671.1 anig|EHA27427.1 aory|EIT82869.1 asyd|OJJ54586.1 ater|EAU30234.1 atub|OJI88876.1 aver|OJJ07568.1 awen|OJJ29771.1

Group6115: abra|OJJ70391.1 acar|OOF90062.1 acla|EAW12389.1 afis|EAW16836.1 afla|EED47326.1 afum|EAL86384.1 agla|OJJ89425.1 akaw|GAA91968.1 aluc|OJZ82648.1 anid|CBF74685.1 anig|EHA27424.1 asyd|OJJ60209.1 ater|EAU30243.1 atub|OJI88880.1 aver|OJJ04059.1 awen|OJJ35684.1

Group6116: abra|OJJ70472.1 acla|EAW13042.1 afis|EAW17625.1 afum|EAL91439.1 agla|OJJ81278.1 akaw|GAA83321.1 aluc|OJZ83816.1 anid|CBF90152.1 anig|EHA18961.1 aory|EIT78528.1 asyd|OJJ62000.1 ater|EAU37059.1 atub|OJI90183.1 aver|OJI96092.1 awen|OJJ37219.1 azon|OJJ50922.1

Group6117: abra|OJJ70506.1 afis|EAW19833.1 afla|EED46252.1 agla|OJJ88103.1 akaw|GAA83355.1 aluc|OJZ83851.1 anig|EHA18993.1 aory|EIT76504.1 asyd|OJJ56258.1 atub|OJI90217.1 aver|OJJ08506.1 awen|OJJ32240.1 afla|EED57615.1 afum|EAL85059.1 aory|EIT82986.1 ater|EAU31581.1

Group6118: abra|OJJ70520.1 acar|OOF97408.1 acla|EAW08155.1 afis|EAW23184.1 afla|EED47061.1 afum|EAL88299.1 akaw|GAA83367.1 aluc|OJZ83865.1 anid|CBF74700.1 anig|EHA19006.1 aory|EIT72501.1 asyd|OJJ60216.1 atub|OJI90233.1 aver|OJJ04052.1 awen|OJJ36586.1 azon|OJJ47621.1

Group6119: abra|OJJ70538.1 acar|OOF97395.1 acla|EAW08150.1 afis|EAW23189.1 afla|EED47068.1 afum|EAL88294.2 agla|OJJ86364.1 akaw|GAA83383.1 aluc|OJZ83882.1 anid|CBF74525.1 anig|EHA19020.1 asyd|OJJ57735.1 atub|OJI90250.1 aver|OJJ05432.1 awen|OJJ36591.1 azon|OJJ47615.1

Group6120: abra|OJJ70542.1 acar|OOF97391.1 acla|EAW08136.1 afis|EAW23172.1 afla|EED46947.1 afum|EAL88311.2 agla|OJJ89684.1 akaw|GAA83388.1 aluc|OJZ83887.1 anid|CBF74493.1 anig|EHA19024.1 aory|EIT72523.1 asyd|OJJ57717.1 atub|OJI90254.1 aver|OJJ05450.1 awen|OJJ36605.1

Group6121: abra|OJJ70573.1 acar|OOF97148.1 acla|EAW08372.1 afis|EAW23401.1 afla|EED47008.1 afum|EAL88069.1 akaw|GAA83417.1 aluc|OJZ83917.1 anid|CBF74797.1 anig|EHA19050.1 aory|EIT72568.1 asyd|OJJ60154.1 atub|OJI90287.1 aver|OJJ04110.1 awen|OJJ39477.1 azon|OJJ47680.1

Group6122: abra|OJJ70588.1 acar|OOF97132.1 acla|EAW08193.1 afis|EAW23227.1 afla|EED47023.1 afum|EAL88256.1 agla|OJJ86384.1 akaw|GAA83431.1 aluc|OJZ83932.1 anid|CBF74769.1 anig|EHA19064.1 asyd|OJJ60168.1 atub|OJI90304.1 aver|OJJ04096.1 awen|OJJ39462.1 azon|OJJ47662.1

Group6123: abra|OJJ70624.1 acla|EAW13006.1 afis|EAW17662.1 afla|EED50496.1 afum|EAL91476.2 agla|OJJ79531.1 akaw|GAA89962.1 aluc|OJZ83969.1 anid|CBF90058.1 anig|EHA19098.1 aory|EIT82078.1 ater|EAU31991.1 atub|OJI90342.1 aver|OJI96157.1 awen|OJJ37180.1 azon|OJJ50882.1

Group6124: abra|OJJ70645.1 acar|OOF97076.1 acla|EAW08115.1 afis|EAW23136.1 afla|EED48499.1 afum|EAL88345.2 agla|OJJ89664.1 akaw|GAA89981.1 aluc|OJZ83995.1 anid|CBF74452.1 asyd|OJJ57739.1 ater|EAU36281.1 atub|OJI90364.1 aver|OJJ05428.1 awen|OJJ36632.1 azon|OJJ47583.1

Group6125: abra|OJJ70649.1 acar|OOF97071.1 acla|EAW08110.1 afis|EAW23141.1 afum|EAL88340.1 agla|OJJ89660.1 akaw|GAA89985.1 aluc|OJZ83999.1 anid|CBF74442.1 aory|EIT73199.1 asyd|OJJ57743.1 ater|EAU36276.1 atub|OJI90368.1 aver|OJJ05424.1 awen|OJJ36637.1 azon|OJJ47579.1

Group6126: abra|OJJ70651.1 acar|OOF97069.1 acla|EAW08108.1 afis|EAW23143.1 afla|EED48506.1 afum|EAL88338.1 akaw|GAA89987.1 aluc|OJZ84001.1 anid|CBF74438.1 anig|EHA19123.1 aory|EIT73205.1 asyd|OJJ57745.1 ater|EAU36268.1 atub|OJI90370.1 aver|OJJ05422.1 azon|OJJ47577.1

Group6127: abra|OJJ70682.1 acar|OOF97045.1 acla|EAW13290.1 afis|EAW17319.1 afum|EAL91155.1 agla|OJJ89622.1 akaw|GAA90014.1 aluc|OJZ84030.1 anid|CBF86689.1 anig|EHA19151.1 asyd|OJJ64751.1 ater|EAU36247.1 atub|OJI90402.1 aver|OJI98852.1 awen|OJJ36663.1 acla|EAW08082.1

Group6128: abra|OJJ70683.1 acar|OOF97044.1 acla|EAW08079.1 afis|EAW23118.1 afum|EAL88362.1 agla|OJJ89623.1 akaw|GAA90015.1 aluc|OJZ84031.1 anid|CBF74395.1 anig|EHA19152.1 asyd|OJJ57768.1 ater|EAU36246.1 atub|OJI90403.1 aver|OJJ05398.1 awen|OJJ36664.1 azon|OJJ47535.1

Group6129: abra|OJJ70707.1 acar|OOF97030.1 acla|EAW08057.1 afis|EAW23093.1 afla|EED48555.1 afum|EAL88385.1 agla|OJJ89605.1 akaw|GAA90038.1 aluc|OJZ84054.1 anig|EHA19174.1 asyd|OJJ59375.1 ater|EAU36217.1 atub|OJI90425.1 aver|OJJ02898.1 awen|OJJ36689.1 azon|OJJ47508.1

Group6130: abra|OJJ70709.1 acar|OOF97027.1 acla|EAW08055.1 afis|EAW23091.1 afla|EED48557.1 afum|EAL88388.1 agla|OJJ89603.1 akaw|GAA90040.1 aluc|OJZ84056.1 anid|CBF81347.1 asyd|OJJ59373.1 ater|EAU36214.1 atub|OJI90427.1 aver|OJJ02901.1 awen|OJJ36691.1 azon|OJJ47505.1

Group6131: abra|OJJ70759.1 acar|OOF96985.1 acla|EAW11162.1 afis|EAW15893.1 afla|EED56478.1 afum|EAL88528.1 akaw|GAA90079.1 aluc|OJZ84104.1 anid|CBF81296.1 anig|EHA19216.1 aory|EIT72872.1 asyd|OJJ59347.1 ater|EAU36172.1 atub|OJI90478.1 aver|OJJ02927.1 awen|OJJ36738.1

Group6132: abra|OJJ70808.1 acar|OOF98294.1 acla|EAW10417.1 afis|EAW25152.1 afla|EED55281.1 afum|EAL89827.1 agla|OJJ89002.1 akaw|GAA91688.1 aluc|OJZ87706.1 anid|CBF89561.1 aory|EIT83566.1 ater|EAU39431.1 atub|OJI80937.1 aver|OJI96448.1 awen|OJJ40781.1 azon|OJJ46871.1

Group6133: abra|OJJ70811.1 acar|OOF98290.1 acla|EAW10420.1 afis|EAW25149.1 afla|EED55287.1 afum|EAL89824.1 agla|OJJ89006.1 akaw|GAA91691.1 aluc|OJZ87703.1 anid|CBF85672.1 anig|EHA20886.1 asyd|OJJ64249.1 atub|OJI80940.1 aver|OJI98340.1 awen|OJJ40777.1 azon|OJJ46874.1

Group6134: abra|OJJ70812.1 acar|OOF98286.1 acla|EAW10421.1 afis|EAW25148.1 afla|EED55288.1 afum|EAL89823.1 agla|OJJ89007.1 akaw|GAA91692.1 aluc|OJZ87702.1 anid|CBF85673.1 asyd|OJJ64251.1 ater|EAU39420.1 atub|OJI80941.1 aver|OJI98341.1 awen|OJJ40773.1 azon|OJJ46875.1

Group6135: abra|OJJ70819.1 acar|OOF98283.1 acla|EAW10426.1 afis|EAW25143.1 afla|EED55302.1 afum|EAL89815.2 agla|OJJ89024.1 akaw|GAA91698.1 aluc|OJZ87692.1 anid|CBF85685.1 anig|EHA20881.1 aory|EIT83611.1 ater|EAU39412.1 atub|OJI80948.1 aver|OJI98348.1 awen|OJJ40767.1

Group6136: abra|OJJ70836.1 acar|OOF98278.1 acla|EAW10431.1 afis|EAW25138.1 afla|EED55308.1 afum|EAL89809.2 akaw|GAA91704.1 aluc|OJZ87685.1 anid|CBF85712.1 anig|EHA20875.1 asyd|OJJ64271.1 ater|EAU39408.1 atub|OJI80958.1 aver|OJI98363.1 awen|OJJ40762.1 azon|OJJ44127.1

Group6137: abra|OJJ70838.1 acar|OOF98276.1 afis|EAW25136.1 afla|EED55310.1 afum|EAL89807.2 agla|OJJ89031.1 akaw|GAA91706.1 aluc|OJZ87683.1 anid|CBF85708.1 anig|EHA20873.1 aory|EIT83586.1 asyd|OJJ64269.1 atub|OJI80960.1 aver|OJI98361.1 awen|OJJ40760.1 azon|OJJ44124.1

Group6138: abra|OJJ70841.1 acar|OOF98273.1 acla|EAW13282.1 afis|EAW17335.1 afla|EED46106.1 afum|EAL91178.1 akaw|GAA91709.1 aluc|OJZ87680.1 anig|EHA20870.1 aory|EIT76506.1 asyd|OJJ56055.1 ater|EAU33889.1 atub|OJI80963.1 aver|OJJ04997.1 awen|OJJ33505.1 azon|OJJ50299.1

Group6139: acar|OOF98702.1 afis|EAW20762.1 afum|EAL84857.1 anid|CBF85637.1 aver|OJJ01161.1 azon|OJJ46003.1 acla|EAW11101.1 afla|EED53201.1 agla|OJJ86968.1 akaw|GAA92918.1 anig|EHA23722.1 aory|EIT75034.1 asyd|OJJ58562.1 ater|EAU29736.1 atub|OJI82462.1 awen|OJJ37494.1

Group6140: abra|OJJ70852.1 acar|OOF98264.1 acla|EAW10442.1 afis|EAW25125.1 afum|EAL89796.1 agla|OJJ89045.1 akaw|GAA91719.1 aluc|OJZ87669.1 anid|CBF85690.1 anig|EHA20860.1 asyd|OJJ64260.1 ater|EAU39398.1 atub|OJI80973.1 aver|OJI98352.1 awen|OJJ40743.1 azon|OJJ44114.1

Group6141: abra|OJJ70894.1 acar|OOF90218.1 acla|EAW10476.1 afis|EAW25091.1 afla|EED55356.1 afum|EAL89761.2 agla|OJJ89086.1 akaw|GAA87410.1 aluc|OJZ87629.1 anig|EHA20825.1 aory|EIT80828.1 asyd|OJJ64441.1 atub|OJI81020.1 aver|OJI98543.1 awen|OJJ40705.1 azon|OJJ44081.1

Group6142: abra|OJJ70900.1 acar|OOF98220.1 acla|EAW10478.1 afis|EAW25089.1 afla|EED55358.1 afum|EAL89758.2 agla|OJJ89088.1 akaw|GAA87415.1 aluc|OJZ87623.1 anid|CBF85995.1 anig|EHA20820.1 asyd|OJJ64439.1 atub|OJI81026.1 aver|OJI98540.1 awen|OJJ40702.1 azon|OJJ44079.1

Group6143: abra|OJJ70930.1 acar|OOF98197.1 acla|EAW10496.1 afis|EAW25071.1 afla|EED55383.1 afum|EAL89740.1 agla|OJJ89123.1 akaw|GAA83012.1 aluc|OJZ87597.1 anig|EHA20797.1 asyd|OJJ64419.1 ater|EAU39554.1 atub|OJI81050.1 aver|OJI98518.1 awen|OJJ40682.1 azon|OJJ44059.1

Group6144: abra|OJJ70936.1 acla|EAW10502.1 afis|EAW25065.1 afla|EED55389.1 afum|EAL89734.1 agla|OJJ89128.1 akaw|GAA83007.1 anid|CBF85947.1 anig|EHA20792.1 aory|EIT80920.1 asyd|OJJ64414.1 ater|EAU35812.1 atub|OJI81056.1 aver|OJI98513.1 awen|OJJ40673.1 azon|OJJ44053.1

Group6145: abra|OJJ70974.1 acar|OOF98157.1 acla|EAW10533.1 afis|EAW25000.1 afla|EED55420.1 afum|EAL89698.1 agla|OJJ89162.1 akaw|GAA82975.1 aluc|OJZ87558.1 anid|CBF81497.1 anig|EHA20761.1 asyd|OJJ59464.1 atub|OJI81089.1 aver|OJJ02810.1 awen|OJJ40639.1 azon|OJJ44023.1

Group6146: abra|OJJ70978.1 acar|OOF98152.1 afis|EAW25004.1 afla|EED55424.1 afum|EAL89694.1 agla|OJJ89167.1 akaw|GAA82971.1 aluc|OJZ87554.1 anid|CBF81483.1 anig|EHA20757.1 asyd|OJJ59455.1 ater|EAU35781.1 aver|OJJ02818.1 awen|OJJ40635.1 azon|OJJ49048.1 acla|EAW10537.1

Group6147: abra|OJJ71000.1 acar|OOF98140.1 afla|EED55446.1 akaw|GAA82948.1 aluc|OJZ87527.1 anig|EHA20737.1 aory|EIT80919.1 asyd|OJJ59492.1 atub|OJI81121.1 aver|OJJ02782.1 awen|OJJ40616.1 acla|EAW10557.1 afis|EAW25024.1 afum|EAL89671.1 anid|CBF81545.1 ater|EAU35763.1

Group6148: abra|OJJ71024.1 acar|OOF98116.1 acla|EAW10574.1 afis|EAW24993.1 afum|EAL89653.1 agla|OJJ82262.1 akaw|GAA82931.1 aluc|OJZ87507.1 anid|CBF81590.1 anig|EHA20720.1 asyd|OJJ59514.1 ater|EAU35744.1 atub|OJI81143.1 aver|OJJ02759.1 awen|OJJ40598.1 azon|OJJ49448.1

Group6149: abra|OJJ71025.1 acla|EAW10575.1 afis|EAW24992.1 afla|EED55467.1 afum|EAL89652.1 agla|OJJ82263.1 akaw|GAA82930.1 aluc|OJZ87506.1 anid|CBF81592.1 anig|EHA20719.1 aory|EIT80781.1 asyd|OJJ59515.1 ater|EAU35743.1 atub|OJI81144.1 awen|OJJ40597.1 aver|OJJ02758.1

Group6150: abra|OJJ71076.1 acar|OOF98068.1 acla|EAW11904.1 afis|EAW19816.1 afla|EED46069.1 afum|EAL85075.1 akaw|GAA82886.1 aluc|OJZ87454.1 anig|EHA20675.1 aory|EIT83499.1 atub|OJI81194.1 awen|OJJ34557.1 afis|EAW25580.1 afla|EED52141.1 afum|EAL84291.1 ater|EAU36155.1

Group6151: abra|OJJ71095.1 afla|EED48308.1 akaw|GAA90277.1 aluc|OJZ83064.1 anig|EHA28129.1 aory|EIT76315.1 atub|OJI87881.1 abra|OJJ71758.1 afis|EAW17369.1 afum|EAL91206.2 agla|OJJ79798.1 aluc|OJZ84837.1 anig|EHA25747.1 asyd|OJJ61751.1 aver|OJI95848.1 awen|OJJ37888.1

Group6152: abra|OJJ71098.1 acla|EAW14582.1 afis|EAW19316.1 afla|EED48318.1 afum|EAL94033.1 akaw|GAA83836.1 aluc|OJZ90881.1 anig|EHA28512.1 aory|EIT76307.1 atub|OJI83754.1 azon|OJJ46593.1 afis|EAW17411.1 afla|EED55973.1 aory|EIT76588.1 aver|OJJ06557.1 acla|EAW15051.1

Group6153: abra|OJJ71101.1 acar|OOF98042.1 afis|EAW25201.1 afum|EAL89864.1 akaw|GAA82867.1 aluc|OJZ87434.1 anig|EHA20656.1 atub|OJI81214.1 afla|EED53648.1 anid|CBF85217.1 aory|EIT75339.1 aver|OJI97967.1 awen|OJJ40836.1 acla|EAW10390.1 agla|OJJ88974.1 asyd|OJJ62999.1

Group6154: abra|OJJ71153.1 acar|OOF98004.1 acla|EAW10344.1 afis|EAW25245.1 afum|EAL89907.2 akaw|GAA82824.1 aluc|OJZ87385.1 anid|CBF85325.1 anig|EHA20614.1 asyd|OJJ64014.1 ater|EAU34358.1 atub|OJI81262.1 aver|OJI98102.1 awen|OJJ40958.1 azon|OJJ43036.1 anid|CBF82000.1

Group6155: abra|OJJ71159.1 acar|OOF98001.1 acla|EAW10340.1 afis|EAW25249.1 afla|EED53715.1 afum|EAL89911.1 agla|OJJ89221.1 akaw|GAA82819.1 aluc|OJZ87380.1 anid|CBF85303.1 anig|EHA20611.1 asyd|OJJ63997.1 atub|OJI81266.1 aver|OJI98086.1 awen|OJJ40963.1 azon|OJJ44544.1

Group6156: abra|OJJ71199.1 acar|OOF97969.1 afis|EAW20331.1 afla|EED48992.1 afum|EAL85457.1 agla|OJJ88970.1 akaw|GAA82784.1 aluc|OJZ87343.1 anid|CBF82369.1 anig|EHA20580.1 asyd|OJJ54197.1 ater|EAU37619.1 atub|OJI81301.1 aver|OJJ08527.1 awen|OJJ31947.1 akaw|GAA90166.1

Group6157: abra|OJJ71228.1 acar|OOF97951.1 acla|EAW10294.1 afis|EAW25298.1 afla|EED53764.1 afum|EAL89961.1 agla|OJJ89277.1 akaw|GAA82761.1 aluc|OJZ87319.1 anig|EHA20553.1 aory|EIT78432.1 ater|EAU34439.1 atub|OJI81326.1 aver|OJI98150.1 awen|OJJ41012.1 azon|OJJ44597.1

Group6158: abra|OJJ71247.1 acar|OOF97921.1 acla|EAW10288.1 afis|EAW25304.1 afum|EAL89967.2 agla|OJJ89283.1 akaw|GAA82740.1 aluc|OJZ87295.1 anid|CBF75298.1 anig|EHA20536.1 asyd|OJJ59883.1 ater|EAU34452.1 atub|OJI81345.1 aver|OJJ02403.1 awen|OJJ41018.1 azon|OJJ44605.1

Group6159: abra|OJJ71250.1 acla|EAW10285.1 afis|EAW25307.1 afum|EAL89970.1 agla|OJJ89286.1 akaw|GAA82737.1 aluc|OJZ87292.1 anid|CBF75293.1 anig|EHA20533.1 aory|EIT78358.1 asyd|OJJ59886.1 ater|EAU34455.1 atub|OJI81349.1 aver|OJJ02400.1 awen|OJJ41021.1 azon|OJJ44608.1

Group6160: abra|OJJ71313.1 acar|OOF97859.1 acla|EAW10226.1 afis|EAW25366.1 afla|EED53847.1 afum|EBA27393.1 akaw|GAA82678.1 aluc|OJZ87231.1 anid|CBF77450.1 aory|EIT78418.1 asyd|OJJ53210.1 atub|OJI81411.1 aver|OJJ03640.1 awen|OJJ41092.1 agla|OJJ83753.1 azon|OJJ44671.1

Group6161: abra|OJJ71323.1 acar|OOF97851.1 acla|EAW10216.1 afis|EAW25377.1 afla|EED53857.1 afum|EAL90036.1 agla|OJJ83766.1 akaw|GAA82668.1 aluc|OJZ87221.1 anid|CBF77481.1 aory|EIT78408.1 asyd|OJJ53200.1 ater|EAU34521.1 aver|OJJ03650.1 awen|OJJ41104.1 azon|OJJ44681.1

Group6162: abra|OJJ71338.1 acar|OOF97834.1 acla|EAW10201.1 afis|EAW25394.1 afla|EED53871.1 afum|EAL90053.1 agla|OJJ87032.1 akaw|GAA82654.1 aluc|OJZ87204.1 anid|CBF77510.1 anig|EHA20457.1 aory|EIT78385.1 ater|EAU34542.1 atub|OJI81436.1 awen|OJJ41120.1 azon|OJJ44695.1

Group6163: abra|OJJ71356.1 acar|OOF97814.1 acla|EAW10180.1 afis|EAW25414.1 afla|EED53893.1 afum|EAL90073.1 agla|OJJ87056.1 akaw|GAA82636.1 aluc|OJZ87183.1 asyd|OJJ53166.1 ater|EAU34565.1 atub|OJI81456.1 aver|OJJ03686.1 awen|OJJ41141.1 azon|OJJ44721.1 anid|CBF77578.1

Group6164: abra|OJJ71473.1 acla|EAW10077.1 afis|EAW25509.1 afla|EED54010.1 afum|EAL90175.2 agla|OJJ87296.1 akaw|GAA82543.1 aluc|OJZ87073.1 aory|EIT78294.1 asyd|OJJ53035.1 ater|EAU33356.1 atub|OJI81564.1 aver|OJJ03809.1 awen|OJJ41251.1 azon|OJJ44818.1 anid|CBF77773.1

Group6165: abra|OJJ71590.1 acar|OOF96319.1 acla|EAW15136.1 afis|EAW20108.1 afum|EAL85225.1 agla|OJJ88846.1 akaw|GAA89332.1 aluc|OJZ86333.1 anid|CBF84915.1 anig|EHA21548.1 asyd|OJJ63166.1 ater|EAU38726.1 atub|OJI84202.1 aver|OJI97789.1 awen|OJJ31879.1 azon|OJJ46255.1

Group6166: abra|OJJ71632.1 acar|OOF96349.1 acla|EAW15173.1 afis|EAW20142.1 afla|EED57207.1 afum|EAL85257.1 aluc|OJZ86375.1 anid|CBF84976.1 anig|EHA21519.1 aory|EIT74096.1 asyd|OJJ63131.1 ater|EAU38759.1 atub|OJI84159.1 aver|OJI97823.1 awen|OJJ32012.1 azon|OJJ46077.1

Group6167: abra|OJJ71642.1 acar|OOF96358.1 acla|EAW15182.1 afis|EAW20152.1 afum|EAL85267.1 agla|OJJ88782.1 akaw|GAA89284.1 aluc|OJZ86385.1 anid|CBF84989.1 anig|EHA21510.1 aory|EIT74139.1 asyd|OJJ63122.1 atub|OJI84148.1 aver|OJI97832.1 awen|OJJ32025.1 azon|OJJ46065.1

Group6168: abra|OJJ71703.1 acar|OOF96414.1 acla|EAW15238.1 afis|EAW20203.1 afla|EED57138.1 afum|EAL85323.1 agla|OJJ84662.1 aluc|OJZ86446.1 anid|CBF85089.1 anig|EHA21455.1 aory|EIT74000.1 asyd|OJJ63068.1 atub|OJI84082.1 aver|OJI97894.1 awen|OJJ31209.1 azon|OJJ46132.1

Group6169: abra|OJJ71814.1 acla|EAW13854.1 afis|EAW18568.1 afla|EED52783.1 afum|EAL93270.1 agla|OJJ81619.1 akaw|GAA85096.1 aluc|OJZ86547.1 anid|CBF70307.1 anig|EHA22103.1 aory|EIT78241.1 asyd|OJJ58725.1 atub|OJI83976.1 aver|OJJ01338.1 awen|OJJ31125.1 azon|OJJ46606.1

Group6170: abra|OJJ71841.1 acar|OOF96510.1 acla|EAW13872.1 afis|EAW18583.1 afum|EAL93288.1 agla|OJJ81635.1 akaw|GAA85076.1 aluc|OJZ86574.1 anid|CBF70364.1 anig|EHA22083.1 asyd|OJJ58651.1 ater|EAU34877.1 atub|OJI83949.1 aver|OJJ01257.1 awen|OJJ31147.1 azon|OJJ46622.1

Group6171: abra|OJJ71871.1 acar|OOF96542.1 acla|EAW13901.1 afis|EAW18613.1 afla|EED52731.1 afum|EAL93323.1 akaw|GAA85044.1 aluc|OJZ86610.1 anid|CBF70432.1 anig|EHA22056.1 aory|EIT78245.1 asyd|OJJ58614.1 ater|EAU34910.1 atub|OJI83916.1 aver|OJJ01221.1 awen|OJJ32074.1

Group6172: abra|OJJ71916.1 acar|OOF96573.1 afis|EAW20251.1 afla|EED46917.1 afum|EAL85369.1 agla|OJJ80821.1 akaw|GAA85006.1 aluc|OJZ86652.1 anid|CBF80720.1 anig|EHA22018.1 aory|EIT72602.1 asyd|OJJ54567.1 ater|EAU31463.1 atub|OJI83875.1 aver|OJJ07587.1 awen|OJJ33373.1

Group6173: abra|OJJ71926.1 acar|OOF96583.1 acla|EAW09602.1 afis|EAW21535.1 afla|EED55985.1 afum|EBA27463.1 agla|OJJ85698.1 akaw|GAA84996.1 aluc|OJZ86663.1 anid|CBF76388.1 asyd|OJJ61071.1 ater|EAU35004.1 atub|OJI83862.1 aver|OJJ00388.1 awen|OJJ39157.1 azon|OJJ51487.1

Group6174: abra|OJJ71930.1 acar|OOF96587.1 acla|EAW09599.1 afis|EAW21538.1 afla|EED55990.1 afum|EAL92558.2 agla|OJJ85702.1 akaw|GAA84993.1 aluc|OJZ86667.1 anid|CBF76380.1 anig|EHA22006.1 aory|EIT76800.1 asyd|OJJ61067.1 aver|OJJ00383.1 awen|OJJ39153.1 azon|OJJ51484.1

Group6175: abra|OJJ71936.1 acar|OOF96592.1 acla|EAW09594.1 afis|EAW21543.1 afla|EED55994.1 afum|EAL92563.1 agla|OJJ85706.1 akaw|GAA84988.1 aluc|OJZ86673.1 anid|CBF76372.1 anig|EHA22001.1 aory|EIT76857.1 ater|EAU34996.1 atub|OJI83855.1 awen|OJJ39147.1 azon|OJJ51483.1

Group6176: acar|OOF96616.1 anig|EHA21984.1 atub|OJI83835.1 acla|EAW09581.1 afis|EAW21556.1 afla|EED56008.1 afum|EBA27466.1 akaw|GAA84973.1 aluc|OJZ86689.1 anid|CBF76346.1 aory|EIT76788.1 asyd|OJJ61051.1 ater|EAU34978.1 aver|OJJ00367.1 awen|OJJ39134.1 azon|OJJ51462.1

Group6177: abra|OJJ72004.1 acla|EAW09535.1 afis|EAW21598.1 afla|EED57822.1 afum|EAL92617.1 agla|OJJ85793.1 akaw|GAA84928.1 aluc|OJZ86740.1 anid|CBF83455.1 anig|EHA21939.1 asyd|OJJ60678.1 ater|EAU38287.1 atub|OJI83788.1 aver|OJJ00001.1 awen|OJJ39084.1 azon|OJJ51416.1

Group6178: abra|OJJ72006.1 acar|OOF96656.1 afis|EAW17213.1 afla|EED56213.1 afum|EAL86640.1 akaw|GAA84927.1 aluc|OJZ86742.1 anid|CBF83257.1 anig|EHA21937.1 aory|EIT73317.1 asyd|OJJ60586.1 ater|EAU38288.1 atub|OJI83785.1 aver|OJI99909.1 awen|OJJ41450.1 azon|OJJ50967.1

Group6179: abra|OJJ72012.1 acar|OOF96661.1 acla|EAW09528.1 afis|EAW21604.1 afla|EED57831.1 afum|EAL92623.1 agla|OJJ85799.1 akaw|GAA84921.1 aluc|OJZ86748.1 anid|CBF83465.1 anig|EHA21931.1 asyd|OJJ60686.1 atub|OJI83779.1 aver|OJJ00011.1 awen|OJJ39078.1 azon|OJJ51410.1

Group6180: abra|OJJ72021.1 acar|OOF96670.1 acla|EAW09516.1 afis|EAW21616.1 afum|EAL92650.1 agla|OJJ78623.1 akaw|GAA84911.1 aluc|OJZ86758.1 anig|EHA21921.1 asyd|OJJ60697.1 atub|OJI83769.1 aver|OJJ00023.1 awen|OJJ39068.1 azon|OJJ51399.1 afla|EED57843.1 aory|EIT81106.1

Group6181: abra|OJJ72041.1 acar|OOF96700.1 afis|EAW21632.1 afla|EED57860.1 afum|EAL92665.1 akaw|GAA84892.1 aluc|OJZ86779.1 anig|EHA21903.1 aory|EIT81090.1 asyd|OJJ54353.1 ater|EAU38318.1 atub|OJI83741.1 aver|OJJ05307.1 awen|OJJ33881.1 azon|OJJ42153.1 afis|EAW19761.1

Group6182: abra|OJJ72047.1 acar|OOF96705.1 acla|EAW09499.1 afis|EAW21637.1 afla|EED57880.1 afum|EAL92671.1 agla|OJJ79702.1 akaw|GAA84887.1 aluc|OJZ86784.1 anid|CBF83519.1 anig|EHA21898.1 asyd|OJJ60715.1 atub|OJI83736.1 aver|OJJ00048.1 awen|OJJ33886.1 azon|OJJ48667.1

Group6183: abra|OJJ72066.1 acar|OOF96720.1 acla|EAW09481.1 afis|EAW21655.1 afla|EED57890.1 afum|EAL92690.1 akaw|GAA84868.1 aluc|OJZ86802.1 anid|CBF83546.1 anig|EHA21881.1 aory|EIT82739.1 asyd|OJJ60731.1 atub|OJI83716.1 aver|OJJ00063.1 awen|OJJ33912.1 azon|OJJ48649.1

Group6184: abra|OJJ72069.1 acar|OOF96725.1 acla|EAW09484.1 afis|EAW21652.1 afla|EED57885.1 afum|EBA27469.1 agla|OJJ79725.1 akaw|GAA84866.1 aluc|OJZ86804.1 anid|CBF83552.1 anig|EHA21878.1 aory|EIT83276.1 asyd|OJJ60734.1 atub|OJI83714.1 aver|OJJ00066.1 awen|OJJ33916.1

Group6185: abra|OJJ72088.1 acar|OOF96733.1 acla|EAW09469.1 afis|EAW21667.1 afla|EED57931.1 afum|EAL92702.1 agla|OJJ79738.1 akaw|GAA84832.1 aluc|OJZ86842.1 anid|CBF83571.1 anig|EHA21795.1 aory|EIT82338.1 asyd|OJJ60748.1 atub|OJI83676.1 aver|OJJ00080.1 azon|OJJ48638.1

Group6186: abra|OJJ72093.1 acar|OOF96738.1 acla|EAW09474.1 afis|EAW21663.1 afla|EED57926.1 afum|EAL92698.1 agla|OJJ79733.1 akaw|GAA84827.1 aluc|OJZ86848.1 anig|EHA21790.1 aory|EIT82353.1 asyd|OJJ60743.1 atub|OJI83671.1 aver|OJJ00075.1 awen|OJJ33925.1 azon|OJJ48633.1

Group6187: abra|OJJ72103.1 acar|OOF96747.1 acla|EAW09387.1 afis|EAW21747.1 afla|EED58031.1 afum|EBA27471.1 agla|OJJ84426.1 aluc|OJZ86858.1 anid|CBF83664.1 anig|EHA21781.1 aory|EIT79746.1 asyd|OJJ60805.1 atub|OJI83661.1 aver|OJJ00134.1 awen|OJJ34029.1 azon|OJJ48202.1

Group6188: abra|OJJ72121.1 acar|OOF96761.1 afla|EED55235.1 akaw|GAA84802.1 aluc|OJZ86878.1 anig|EHA21766.1 aory|EIT83535.1 asyd|OJJ52924.1 ater|EAU30004.1 atub|OJI83641.1 afis|EAW24443.1 afum|EAL86112.1 agla|OJJ80288.1 aver|OJJ06142.1 awen|OJJ32756.1 asyd|OJJ56582.1

Group6189: abra|OJJ72164.1 afis|EAW19708.1 afla|EED48321.1 afum|EBA27290.1 akaw|GAA84764.1 aluc|OJZ86919.1 anig|EHA21725.1 aory|EIT76289.1 asyd|OJJ52893.1 atub|OJI83602.1 aver|OJJ08864.1 awen|OJJ29680.1 azon|OJJ46785.1 acla|EAW07716.1 afum|EBA27266.1 awen|OJJ32267.1

Group6190: abra|OJJ72167.1 acar|OOF96800.1 acla|EAW12227.1 afis|EAW25720.1 afla|EED49350.1 afum|EBA27175.1 akaw|GAA84761.1 aluc|OJZ86922.1 anid|CBF79751.1 anig|EHA21722.1 asyd|OJJ58090.1 atub|OJI83599.1 aver|OJJ04310.1 awen|OJJ33494.1 azon|OJJ45920.1 ater|EAU38134.1

Group6191: abra|OJJ72170.1 acla|EAW10000.1 afis|EAW19663.1 afla|EED47278.1 afum|EAL86859.2 akaw|GAA84758.1 aluc|OJZ86925.1 anig|EHA21719.1 aory|EIT79218.1 ater|EAU37567.1 atub|OJI83596.1 aver|OJI99279.1 awen|OJJ30665.1 azon|OJJ49273.1 azon|OJJ46500.1 acla|EAW13620.1

Group6192: abra|OJJ72205.1 acar|OOF96825.1 afis|EAW17755.1 afla|EED47380.1 afum|EAL91568.1 agla|OJJ84453.1 akaw|GAA84729.1 aluc|OJZ86957.1 anid|CBF87748.1 anig|EHA21691.1 aory|EIT82863.1 asyd|OJJ53603.1 ater|EAU32573.1 atub|OJI83566.1 aver|OJJ08145.1 awen|OJJ40504.1

Group6193: abra|OJJ72214.1 acar|OOF96835.1 acla|EAW14971.1 afis|EAW24496.1 afla|EED51682.1 afum|EAL86166.1 akaw|GAA84720.1 aluc|OJZ86966.1 anid|CBF70528.1 anig|EHA21683.1 aory|EIT80642.1 asyd|OJJ65163.1 atub|OJI83556.1 aver|OJI99315.1 awen|OJJ32646.1 azon|OJJ47043.1

Group6194: abra|OJJ72226.1 acar|OOF96849.1 afis|EAW17406.1 afla|EED47644.1 afum|EAL91241.1 agla|OJJ81617.1 akaw|GAA84711.1 aluc|OJZ86976.1 anid|CBF78379.1 anig|EHA21672.1 aory|EIT80206.1 asyd|OJJ57330.1 ater|EAU30398.1 atub|OJI83544.1 aver|OJJ05831.1 awen|OJJ40137.1

Group6195: abra|OJJ72268.1 acar|OOF90352.1 afis|EAW19946.1 afla|EED46129.1 afum|EAL84969.1 agla|OJJ85570.1 akaw|GAA86023.1 aluc|OJZ90250.1 anid|CBF84247.1 anig|EHA23746.1 aory|EIT76403.1 asyd|OJJ57201.1 ater|EAU30028.1 atub|OJI81676.1 aver|OJJ03089.1 awen|OJJ31249.1

Group6196: abra|OJJ72330.1 acar|OOF94884.1 acla|EAW11993.1 afis|EAW19633.1 afla|EED56984.1 afum|EAL86889.1 akaw|GAA86080.1 aluc|OJZ90315.1 anid|CBF71329.1 anig|EHA23802.1 aory|EIT77577.1 asyd|OJJ55106.1 atub|OJI81740.1 aver|OJJ07100.1 awen|OJJ41877.1 azon|OJJ47076.1

Group6197: abra|OJJ72466.1 acla|EAW12698.1 afis|EAW17861.1 afla|EED46223.1 afum|EAL91669.2 agla|OJJ83105.1 akaw|GAA86202.1 aluc|OJZ90453.1 anid|CBF76903.1 anig|EHA23915.1 aory|EIT76388.1 ater|EAU33066.1 atub|OJI81880.1 aver|OJJ03216.1 awen|OJJ41710.1 azon|OJJ48406.1

Group6198: abra|OJJ72508.1 acar|OOF94734.1 acla|EAW12731.1 afis|EAW17894.1 afla|EED46268.1 afum|EBA27438.1 agla|OJJ83048.1 akaw|GAA86236.1 aluc|OJZ90491.1 anig|EHA23949.1 aory|EIT76520.1 asyd|OJJ57022.1 atub|OJI81915.1 aver|OJJ03267.1 awen|OJJ41658.1 azon|OJJ48378.1

Group6199: abra|OJJ72540.1 acar|OOF94713.1 acla|EAW14602.1 afis|EAW17909.1 afla|EED46311.1 afum|EAL91718.1 agla|OJJ80664.1 akaw|GAA86264.1 aluc|OJZ90524.1 anig|EHA23976.1 aory|EIT76346.1 ater|EAU33129.1 atub|OJI81946.1 aver|OJJ07237.1 awen|OJJ41640.1 azon|OJJ48362.1

Group6200: abra|OJJ72547.1 acar|OOF94705.1 acla|EAW12752.1 afis|EAW17915.1 afum|EAL91725.2 agla|OJJ83028.1 akaw|GAA85180.1 aluc|OJZ90535.1 anid|CBF76977.1 anig|EHA23983.1 ater|EAU33137.1 atub|OJI81956.1 aver|OJJ03246.1 awen|OJJ41630.1 azon|OJJ48355.1 asyd|OJJ57050.1

Group6201: abra|OJJ72552.1 acar|OOF94701.1 acla|EAW12757.1 afis|EAW17920.1 afum|EAL91731.1 agla|OJJ83022.1 akaw|GAA85184.1 aluc|OJZ90539.1 anid|CBF76953.1 anig|EHA23987.1 aory|EIT81208.1 asyd|OJJ57055.1 atub|OJI81961.1 aver|OJJ03242.1 awen|OJJ41626.1 azon|OJJ48349.1

Group6202: abra|OJJ72601.1 acar|OOF94652.1 acla|EAW11616.1 afis|EAW23850.1 afla|EED52567.1 afum|EAL87278.2 agla|OJJ82493.1 akaw|GAA85229.1 aluc|OJZ90584.1 anid|CBF77065.1 aory|EIT74171.1 asyd|OJJ56988.1 ater|EAU33517.1 atub|OJI82007.1 aver|OJJ03299.1 azon|OJJ48739.1

Group6203: abra|OJJ72602.1 acar|OOF94651.1 acla|EAW11606.1 afis|EAW23854.1 afla|EED52554.1 afum|EAL87282.1 akaw|GAA85230.1 aluc|OJZ90585.1 anid|CBF77073.1 anig|EHA24025.1 aory|EIT74267.1 asyd|OJJ56985.1 atub|OJI82008.1 aver|OJJ03302.1 awen|OJJ33808.1 azon|OJJ48738.1

Group6204: abra|OJJ72628.1 acar|OOF94636.1 acla|EAW12131.1 afis|EAW17256.1 afla|EED49528.1 afum|EAL86674.1 akaw|GAA85248.1 aluc|OJZ90605.1 anid|CBF75430.1 anig|EHA24042.1 aory|EIT82589.1 asyd|OJJ59786.1 ater|EAU39597.1 atub|OJI82029.1 aver|OJJ02502.1 azon|OJJ50767.1

Group6205: abra|OJJ72645.1 acar|OOF94624.1 afis|EAW23873.1 afla|EED52601.1 afum|EAL87301.1 agla|OJJ82543.1 akaw|GAA85266.1 aluc|OJZ90624.1 anid|CBF84519.1 anig|EHA24061.1 aory|EIT74214.1 asyd|OJJ60279.1 ater|EAU33473.1 atub|OJI82050.1 aver|OJJ03977.1 awen|OJJ33849.1

Group6206: abra|OJJ72681.1 acar|OOF95930.1 akaw|GAA85299.1 aluc|OJZ90659.1 anig|EHA24094.1 atub|OJI82088.1 agla|OJJ82475.1 awen|OJJ36222.1 acar|OOG00299.1 aluc|OJZ82670.1 anig|EHA22183.1 aory|EIT79504.1 asyd|OJJ64226.1 atub|OJI88905.1 aver|OJI98319.1 akaw|GAA91948.1

Group6207: abra|OJJ72723.1 acar|OOF95956.1 acla|EAW11560.1 afis|EAW23893.1 afla|EED52627.1 afum|EAL87321.1 agla|OJJ78633.1 akaw|GAA85329.1 aluc|OJZ90692.1 aory|EIT74264.1 asyd|OJJ56853.1 ater|EAU33449.1 atub|OJI82117.1 aver|OJJ03468.1 awen|OJJ39056.1 azon|OJJ51387.1

Group6208: abra|OJJ72732.1 acar|OOF95962.1 acla|EAW11567.1 afis|EAW23886.1 afla|EED52615.1 afum|EBA27308.1 agla|OJJ82573.1 akaw|GAA85338.1 anid|CBF77181.1 aory|EIT74191.1 asyd|OJJ56844.1 ater|EAU33460.1 atub|OJI82127.1 aver|OJJ03494.1 awen|OJJ33868.1 azon|OJJ48686.1

Group6209: abra|OJJ72762.1 acar|OOF95981.1 acla|EAW11502.1 afis|EAW23950.1 afla|EED50566.1 afum|EAL87376.1 akaw|GAA85372.1 aluc|OJZ90737.1 anid|CBF77294.1 anig|EHA24152.1 aory|EIT82082.1 ater|EAU29802.1 atub|OJI82164.1 aver|OJJ03550.1 awen|OJJ38998.1 azon|OJJ51330.1

Group6210: abra|OJJ72796.1 acar|OOF89881.1 acla|EAW11476.1 afis|EAW23977.1 afla|EED49184.1 afum|EAL87407.1 agla|OJJ86907.1 akaw|GAA85399.1 aluc|OJZ83293.1 anid|CBF77347.1 anig|EHA24177.1 aory|EIT78540.1 ater|EAU29775.1 atub|OJI82194.1 aver|OJJ03578.1 awen|OJJ35444.1

Group6211: abra|OJJ72809.1 acar|OOF96016.1 acla|EAW11431.1 afis|EAW23992.1 afum|EAL87422.1 agla|OJJ86925.1 akaw|GAA85412.1 aluc|OJZ83307.1 anid|CBF77371.1 anig|EHA24190.1 asyd|OJJ53259.1 ater|EAU29759.1 atub|OJI82207.1 aver|OJJ03592.1 awen|OJJ35461.1 azon|OJJ51282.1

Group6212: abra|OJJ72817.1 acla|EAW11423.1 afis|EAW23998.1 afla|EED49163.1 afum|EAL87428.1 akaw|GAA85418.1 aluc|OJZ83319.1 anig|EHA24197.1 aory|EIT78511.1 asyd|OJJ53251.1 ater|EAU29751.1 atub|OJI82216.1 aver|OJJ03601.1 awen|OJJ35469.1 agla|OJJ86932.1 anid|CBF77383.1

Group6213: abra|OJJ72819.1 afla|EED49161.1 akaw|GAA85420.1 aluc|OJZ83321.1 anig|EHA24199.1 asyd|OJJ53249.1 ater|EAU29749.1 atub|OJI82218.1 awen|OJJ35473.1 azon|OJJ51273.1 acla|EAW11420.1 afis|EAW24002.1 afum|EAL87432.1 agla|OJJ86934.1 aver|OJJ03603.1 anid|CBF77388.1

Group6214: abra|OJJ72888.1 acar|OOF96078.1 acla|EAW11362.1 afla|EED58057.1 agla|OJJ83816.1 akaw|GAA85475.1 aluc|OJZ83396.1 anid|CBF75374.1 anig|EHA24249.1 aory|EIT79739.1 asyd|OJJ59845.1 ater|EAU35455.1 atub|OJI82282.1 aver|OJJ02444.1 awen|OJJ35573.1 azon|OJJ46948.1

Group6215: abra|OJJ72899.1 afis|EAW18142.1 agla|OJJ83827.1 akaw|GAA85486.1 aluc|OJZ83408.1 anig|EHA24259.1 aver|OJJ06228.1 awen|OJJ35584.1 acla|EAW11352.1 afum|EAL87499.1 afla|EED58068.1 aory|EIT79790.1 anid|CBF81127.1 asyd|OJJ56490.1 ater|EAU35465.1 azon|OJJ46938.1

Group6216: abra|OJJ72916.1 acar|OOF96098.1 afis|EAW15969.1 afla|EED58088.1 afum|EAL88602.1 akaw|GAA85499.1 aluc|OJZ83426.1 anid|CBF81096.1 anig|EHA24278.1 aory|EIT79150.1 asyd|OJJ56473.1 atub|OJI82312.1 aver|OJJ06245.1 awen|OJJ35604.1 acla|EAW06475.1 ater|EAU35479.1

Group6217: abra|OJJ72918.1 acar|OOF96100.1 acla|EAW06473.1 afis|EAW15967.1 afla|EED58090.1 afum|EAL88600.1 akaw|GAA85501.1 aluc|OJZ83428.1 anid|CBF81092.1 anig|EHA24280.1 asyd|OJJ56469.1 atub|OJI82316.1 aver|OJJ06250.1 awen|OJJ35606.1 azon|OJJ46926.1 ater|EAU35481.1

Group6218: abra|OJJ72919.1 acar|OOF96101.1 acla|EAW06472.1 afis|EAW15966.1 afum|EAL88599.1 agla|OJJ89345.1 akaw|GAA85502.1 aluc|OJZ83429.1 anid|CBF81091.1 aory|EIT79159.1 asyd|OJJ56468.1 ater|EAU35482.1 atub|OJI82317.1 aver|OJJ06251.1 awen|OJJ35607.1 azon|OJJ46927.1

Group6219: abra|OJJ72924.1 acar|OOF96105.1 acla|EAW13672.1 afis|EAW17353.1 afum|EAL91195.1 akaw|GAA85507.1 aluc|OJZ83434.1 anid|CBF80731.1 anig|EHA24283.1 asyd|OJJ55183.1 ater|EAU32472.1 atub|OJI82325.1 aver|OJJ07572.1 awen|OJJ32564.1 acar|OOF94912.1 aver|OJJ07024.1

Group6220: abra|OJJ72928.1 acar|OOF96109.1 acla|EAW06461.1 afis|EAW15957.1 afum|EAL88590.1 agla|OJJ89356.1 akaw|GAA85510.1 aluc|OJZ83438.1 anid|CBF81070.1 anig|EHA24285.1 asyd|OJJ56459.1 ater|EAU35499.1 atub|OJI82329.1 aver|OJJ06263.1 awen|OJJ35617.1 azon|OJJ46906.1

Group6221: abra|OJJ72939.1 acar|OOF96124.1 afis|EAW15944.1 afla|EED58114.1 afum|EAL88578.2 akaw|GAA85522.1 aluc|OJZ83450.1 anid|CBF80955.1 anig|EHA24297.1 aory|EIT79115.1 asyd|OJJ56344.1 ater|EAU35512.1 atub|OJI82342.1 aver|OJJ06369.1 awen|OJJ35638.1 azon|OJJ46897.1

Group6222: abra|OJJ72941.1 acla|EAW11210.1 afis|EAW15943.1 afla|EED58115.1 afum|EAL88577.1 agla|OJJ89367.1 akaw|GAA85523.1 aluc|OJZ83451.1 anid|CBF80957.1 aory|EIT79131.1 asyd|OJJ56345.1 ater|EAU35513.1 atub|OJI82343.1 aver|OJJ06368.1 awen|OJJ35639.1 azon|OJJ46896.1

Group6223: abra|OJJ72957.1 acar|OOF96139.1 acla|EAW11202.1 afis|EAW15932.1 afla|EED54929.1 afum|EAL88568.1 akaw|GAA85538.1 aluc|OJZ83466.1 anid|CBF81031.1 anig|EHA24310.1 aory|EIT80333.1 asyd|OJJ56389.1 ater|EAU35520.1 atub|OJI82359.1 aver|OJJ06324.1 awen|OJJ35647.1

Group6224: abra|OJJ72977.1 acar|OOF96155.1 acla|EAW08032.1 afis|EAW23067.1 afla|EED54950.1 afum|EAL88412.1 agla|OJJ89402.1 akaw|GAA85552.1 aluc|OJZ83487.1 anid|CBF80983.1 anig|EHA24322.1 aory|EIT80376.1 ater|EAU37598.1 atub|OJI82379.1 awen|OJJ35668.1 azon|OJJ44139.1

Group6225: abra|OJJ73033.1 acla|EAW07984.1 afis|EAW23019.1 afla|EED54997.1 afum|EBA27346.1 agla|OJJ89460.1 akaw|GAA85606.1 aluc|OJZ83544.1 anid|CBF80898.1 anig|EHA24364.1 aory|EIT80344.1 asyd|OJJ56311.1 atub|OJI82436.1 aver|OJJ06403.1 awen|OJJ30724.1 azon|OJJ44185.1

Group6226: abra|OJJ73034.1 acar|OOF96199.1 acla|EAW07983.1 afis|EAW23018.1 afum|EBA27347.1 agla|OJJ89461.1 akaw|GAA85607.1 aluc|OJZ83545.1 anig|EHA24365.1 ater|EAU29656.1 atub|OJI82437.1 awen|OJJ30725.1 azon|OJJ44186.1 anid|CBF86993.1 aver|OJJ06407.1 asyd|OJJ56308.1

Group6227: abra|OJJ73043.1 acar|OOF96206.1 acla|EAW07976.1 afis|EAW23011.1 afla|EED55005.1 afum|EAL88470.1 agla|OJJ89469.1 akaw|GAA85614.1 aluc|OJZ83554.1 anid|CBF80883.1 anig|EHA24372.1 aory|EIT80373.1 asyd|OJJ56303.1 atub|OJI82445.1 aver|OJJ06412.1 azon|OJJ44193.1

Group6228: abra|OJJ73067.1 acar|OOG00301.1 afis|EAW16938.1 afla|EED57086.1 afum|EAL86463.1 agla|OJJ83600.1 akaw|GAA88093.1 aluc|OJZ81359.1 anid|CBF82093.1 anig|EHA18659.1 aory|EIT82299.1 asyd|OJJ54263.1 ater|EAU33842.1 atub|OJI91563.1 aver|OJJ05217.1 awen|OJJ32504.1

Group6229: abra|OJJ73109.1 acar|OOG00340.1 acla|EAW07040.1 afis|EAW16667.1 afla|EED44909.1 afum|EAL89271.2 agla|OJJ78702.1 akaw|GAA88057.1 aluc|OJZ81319.1 anig|EHA18625.1 aory|EIT73517.1 asyd|OJJ54658.1 ater|EAU39588.1 atub|OJI91522.1 awen|OJJ40046.1 azon|OJJ50125.1

Group6230: abra|OJJ73138.1 acar|OOG00351.1 acla|EAW11917.1 afis|EAW19544.1 afla|EED55716.1 afum|EAL86965.1 akaw|GAA88034.1 aluc|OJZ81293.1 anid|CBF86680.1 anig|EHA18592.1 aory|EIT83197.1 asyd|OJJ64777.1 ater|EAU30840.1 atub|OJI91502.1 aver|OJI98877.1 azon|OJJ42603.1

Group6231: abra|OJJ73200.1 acar|OOG00439.1 acla|EAW06877.1 afis|EAW16494.1 afla|EED47238.1 afum|EAL89088.1 agla|OJJ82833.1 akaw|GAA87977.1 aluc|OJZ81229.1 anid|CBF75613.1 anig|EHA18540.1 asyd|OJJ59731.1 atub|OJI91426.1 aver|OJJ02556.1 awen|OJJ40335.1 azon|OJJ49189.1

Group6232: abra|OJJ73211.1 acar|OOG00450.1 acla|EAW06887.1 afis|EAW16504.1 afla|EED56174.1 agla|OJJ82817.1 akaw|GAA87967.1 aluc|OJZ81216.1 anid|CBF75593.1 anig|EHA18530.1 aory|EIT76644.1 asyd|OJJ59719.1 atub|OJI91413.1 aver|OJJ02567.1 awen|OJJ40322.1 azon|OJJ49179.1

Group6233: abra|OJJ73215.1 acar|OOG00452.1 acla|EAW06890.1 afis|EAW16507.1 afla|EED56171.1 afum|EAL89102.1 agla|OJJ82821.1 akaw|GAA87964.1 aluc|OJZ81213.1 anig|EHA18527.1 asyd|OJJ59715.1 ater|EAU36533.1 atub|OJI91408.1 aver|OJJ02571.1 awen|OJJ40318.1 azon|OJJ49176.1

Group6234: abra|OJJ73217.1 acar|OOG00454.1 afis|EAW19888.1 afla|EED56169.1 afum|EAL85010.1 agla|OJJ82813.1 akaw|GAA87962.1 aluc|OJZ81211.1 anid|CBF75574.1 anig|EHA18525.1 aory|EIT76660.1 asyd|OJJ59712.1 ater|EAU36574.1 atub|OJI91406.1 aver|OJJ02574.1 awen|OJJ40316.1

Group6235: abra|OJJ73244.1 acar|OOG00483.1 acla|EAW06918.1 afis|EAW16535.1 afum|EAL89132.1 agla|OJJ82787.1 akaw|GAA87936.1 aluc|OJZ81184.1 anid|CBF81739.1 anig|EHA18501.1 asyd|OJJ59596.1 ater|EAU36598.1 atub|OJI91380.1 aver|OJJ02672.1 awen|OJJ40287.1 azon|OJJ49148.1

Group6236: abra|OJJ73250.1 acar|OOG00492.1 acla|EAW06927.1 afis|EAW16542.1 afla|EED56126.1 afum|EAL89139.1 agla|OJJ82781.1 akaw|GAA87930.1 aluc|OJZ81177.1 anid|CBF81750.1 asyd|OJJ59602.1 ater|EAU36604.1 atub|OJI91373.1 aver|OJJ02666.1 awen|OJJ40278.1 azon|OJJ49140.1

Group6237: abra|OJJ73255.1 acar|OOG00497.1 acla|EAW06932.1 afis|EAW16547.1 afla|EED56117.1 afum|EAL89145.1 agla|OJJ82776.1 akaw|GAA87926.1 aluc|OJZ81173.1 anid|CBF81759.1 anig|EHA18491.1 aory|EIT76577.1 ater|EAU36610.1 atub|OJI91368.1 awen|OJJ40270.1 azon|OJJ49134.1

Group6238: abra|OJJ73271.1 acar|OOG00511.1 acla|EAW06947.1 afis|EAW16562.1 afla|EED56100.1 akaw|GAA87909.1 aluc|OJZ88778.1 anid|CBF81809.1 anig|EHA18472.1 aory|EIT76542.1 asyd|OJJ59646.1 ater|EAU36629.1 atub|OJI91350.1 aver|OJJ02636.1 awen|OJJ40255.1 azon|OJJ49118.1

Group6239: abra|OJJ73384.1 acar|OOG00585.1 acar|OOG00586.1 afla|EED51029.1 akaw|GAA83104.1 aluc|OJZ88652.1 anid|CBF84515.1 anig|EHA18343.1 anig|EHA18344.1 asyd|OJJ53501.1 ater|EAU33844.1 atub|OJI91239.1 aver|OJJ08244.1 awen|OJJ35851.1 azon|OJJ50133.1 afla|EED46379.1

Group6240: abra|OJJ73391.1 acar|OOG00593.1 acla|EAW14262.1 afla|EED45223.1 agla|OJJ86281.1 akaw|GAA83098.1 aluc|OJZ88645.1 anid|CBF84157.1 anig|EHA18336.1 aory|EIT73559.1 asyd|OJJ57115.1 ater|EAU30181.1 atub|OJI91232.1 aver|OJJ03185.1 awen|OJJ35938.1 azon|OJJ50415.1

Group6241: abra|OJJ73431.1 acar|OOG00638.1 acla|EAW06770.1 afis|EAW16321.1 afum|EBA27364.1 agla|OJJ88283.1 akaw|GAA83061.1 aluc|OJZ88610.1 anid|CBF89155.1 anig|EHA18303.1 asyd|OJJ62564.1 ater|EAU34136.1 atub|OJI91195.1 aver|OJI96688.1 awen|OJJ37721.1 azon|OJJ44980.1

Group6242: abra|OJJ73443.1 acar|OOG00649.1 acla|EAW06778.1 afis|EAW16329.1 afla|EED53549.1 afum|EAL88953.1 agla|OJJ88263.1 akaw|GAA83051.1 aluc|OJZ88597.1 anid|CBF89169.1 anig|EHA18291.1 aory|EIT74883.1 asyd|OJJ62542.1 atub|OJI91182.1 aver|OJI96667.1 awen|OJJ37730.1

Group6243: abra|OJJ73480.1 acla|EAW06804.1 afis|EAW16353.1 afum|EAL88978.1 agla|OJJ88250.1 akaw|GAA87430.1 aluc|OJZ88567.1 anid|CBF88378.1 anig|EHA18264.1 aory|EIT75259.1 asyd|OJJ63735.1 ater|EAU34108.1 atub|OJI91153.1 aver|OJI97205.1 awen|OJJ37757.1 azon|OJJ45017.1

Group6244: abra|OJJ73483.1 acar|OOG00677.1 acla|EAW06806.1 afis|EAW16355.1 afum|EAL88980.1 agla|OJJ88238.1 akaw|GAA87432.1 aluc|OJZ88565.1 anid|CBF89228.1 anig|EHA18262.1 asyd|OJJ62516.1 ater|EAU34105.1 atub|OJI91151.1 aver|OJI96635.1 awen|OJJ37759.1 azon|OJJ45057.1

Group6245: abra|OJJ73498.1 acar|OOG00692.1 acla|EAW07432.1 afis|EAW22458.1 afla|EED54486.1 agla|OJJ87566.1 akaw|GAA87447.1 aluc|OJZ88550.1 anid|CBF88918.1 anig|EHA18248.1 asyd|OJJ62844.1 ater|EAU39247.1 atub|OJI91135.1 aver|OJI96888.1 awen|OJJ35083.1 azon|OJJ50024.1

Group6246: abra|OJJ73509.1 acla|EAW07442.1 afis|EAW22469.1 afla|EED54501.1 afum|EAL90722.1 agla|OJJ87555.1 akaw|GAA87457.1 aluc|OJZ88538.1 anig|EHA18239.1 aory|EIT78948.1 asyd|OJJ62835.1 ater|EAU39239.1 atub|OJI91123.1 aver|OJI96878.1 awen|OJJ35095.1 azon|OJJ50014.1

Group6247: abra|OJJ73559.1 acar|OOG00730.1 acla|EAW07476.1 afis|EAW22508.1 afla|EED54536.1 afum|EAL90685.1 agla|OJJ87539.1 akaw|GAA88595.1 aluc|OJZ88503.1 anid|CBF88975.1 aory|EIT78907.1 asyd|OJJ62792.1 atub|OJI91089.1 aver|OJI96839.1 awen|OJJ35133.1 azon|OJJ49965.1

Group6248: abra|OJJ73565.1 acar|OOG00747.1 acla|EAW07481.1 afis|EAW22513.1 afla|EED54543.1 afum|EAL90680.1 agla|OJJ80874.1 akaw|GAA88590.1 aluc|OJZ88497.1 anig|EHA18206.1 asyd|OJJ62787.1 ater|EAU39196.1 atub|OJI91084.1 aver|OJI96834.1 awen|OJJ35139.1 azon|OJJ49960.1

Group6249: abra|OJJ73571.1 acar|OOG00753.1 acla|EAW07489.1 afis|EAW22522.1 afum|EAL90672.1 agla|OJJ80883.1 akaw|GAA88584.1 aluc|OJZ88489.1 anid|CBF89006.1 anig|EHA18198.1 asyd|OJJ62777.1 ater|EAU39188.1 atub|OJI91076.1 aver|OJI96822.1 awen|OJJ35149.1 azon|OJJ49952.1

Group6250: abra|OJJ73593.1 acar|OOG00778.1 acla|EAW07512.1 afis|EAW22543.1 afla|EED54574.1 afum|EBA27412.1 agla|OJJ80905.1 akaw|GAA88559.1 aluc|OJZ88457.1 anid|CBF89042.1 asyd|OJJ62759.1 ater|EAU39165.1 atub|OJI91055.1 aver|OJI96802.1 awen|OJJ35172.1 azon|OJJ49935.1

Group6251: abra|OJJ73594.1 acar|OOG00779.1 acla|EAW07513.1 afis|EAW22544.1 afum|EAL90651.1 agla|OJJ80906.1 akaw|GAA88558.1 aluc|OJZ88456.1 anid|CBF89044.1 anig|EHA18177.1 asyd|OJJ62758.1 ater|EAU39164.1 atub|OJI91054.1 aver|OJI96800.1 awen|OJJ35173.1 azon|OJJ49934.1

Group6252: abra|OJJ73603.1 acar|OOG00787.1 afis|EAW22550.1 afum|EAL90645.1 agla|OJJ80912.1 akaw|GAA88550.1 aluc|OJZ88446.1 anid|CBF89058.1 anig|EHA18169.1 asyd|OJJ62751.1 ater|EAU39154.1 atub|OJI91044.1 aver|OJI96793.1 awen|OJJ35181.1 azon|OJJ49926.1 acla|EAW07520.1

Group6253: abra|OJJ73629.1 acar|OOG00811.1 acla|EAW07543.1 afis|EAW22577.1 afla|EED54618.1 afum|EAL90621.1 agla|OJJ80937.1 akaw|GAA88524.1 aluc|OJZ88420.1 anid|CBF88336.1 asyd|OJJ63713.1 ater|EAU39127.1 atub|OJI91016.1 aver|OJI97226.1 awen|OJJ35205.1 azon|OJJ49903.1

Group6254: abra|OJJ73631.1 acar|OOG00813.1 acla|EAW07545.1 afis|EAW22579.1 afla|EED54624.1 afum|EBA27411.1 agla|OJJ80939.1 akaw|GAA88522.1 aluc|OJZ88418.1 anid|CBF88339.1 anig|EHA18144.1 asyd|OJJ63715.1 ater|EAU39125.1 atub|OJI91014.1 aver|OJI97225.1 awen|OJJ35207.1

Group6255: abra|OJJ73639.1 acar|OOG00821.1 acla|EAW07552.1 afis|EAW22586.1 afla|EED54631.1 afum|EAL90613.1 agla|OJJ80946.1 akaw|GAA88516.1 aluc|OJZ88411.1 anid|CBF88354.1 anig|EHA18138.1 asyd|OJJ63722.1 aver|OJI97218.1 awen|OJJ35214.1 azon|OJJ49895.1 afla|EED54632.1

Group6256: abra|OJJ73648.1 acar|OOG00828.1 acla|EAW07561.1 afis|EAW22595.1 afla|EED54645.1 afum|EAL90604.1 akaw|GAA88507.1 aluc|OJZ88402.1 anid|CBF88371.1 anig|EHA18130.1 aory|EIT78751.1 asyd|OJJ63731.1 ater|EAU39108.1 atub|OJI90998.1 aver|OJI97209.1 awen|OJJ35223.1

Group6257: abra|OJJ73659.1 acar|OOG00837.1 acla|EAW07569.1 afis|EAW22603.1 afla|EED49516.1 afum|EAL90595.1 akaw|GAA88499.1 aluc|OJZ88394.1 anid|CBF88259.1 anig|EHA18122.1 aory|EIT82614.1 ater|EAU39100.1 atub|OJI90989.1 aver|OJI97264.1 awen|OJJ35232.1 azon|OJJ49879.1

Group6258: abra|OJJ73678.1 acar|OOG00851.1 acla|EAW07577.1 afis|EAW22617.1 afla|EED49825.1 afum|EAL90580.1 agla|OJJ80997.1 akaw|GAA88481.1 aluc|OJZ88371.1 anid|CBF88247.1 anig|EHA18104.1 aory|EIT77617.1 asyd|OJJ63666.1 atub|OJI90969.1 awen|OJJ35244.1 azon|OJJ49860.1

Group6259: abra|OJJ73680.1 acla|EAW07595.1 afis|EAW22626.1 afum|EAL90570.1 akaw|GAA88479.1 aluc|OJZ88369.1 anig|EHA18102.1 aory|EIT77592.1 ater|EAU39083.1 atub|OJI90967.1 awen|OJJ35247.1 azon|OJJ49859.1 agla|OJJ80995.1 anid|CBF88243.1 asyd|OJJ63663.1 aver|OJI97271.1

Group6260: abra|OJJ73742.1 acar|OOG00905.1 acla|EAW07636.1 afis|EAW22656.1 afla|EED49924.1 afum|EAL90536.1 agla|OJJ81039.1 akaw|GAA88423.1 aluc|OJZ88310.1 anid|CBF88203.1 anig|EHA18047.1 asyd|OJJ63638.1 atub|OJI90905.1 aver|OJI97298.1 awen|OJJ35267.1 azon|OJJ49830.1

Group6261: abra|OJJ73782.1 acar|OOG00938.1 acla|EAW07667.1 afis|EAW22684.1 afla|EED49968.1 agla|OJJ79034.1 akaw|GAA88395.1 aluc|OJZ88273.1 anid|CBF88115.1 anig|EHA18018.1 asyd|OJJ63581.1 ater|EAU39018.1 atub|OJI90873.1 aver|OJI97355.1 awen|OJJ35312.1 azon|OJJ49801.1

Group6262: abra|OJJ73789.1 acar|OOF90346.1 acla|EAW07682.1 afis|EAW22697.1 afla|EED49975.1 afum|EAL90495.1 agla|OJJ79023.1 akaw|GAA89408.1 aluc|OJZ88265.1 anid|CBF88102.1 anig|EHA18011.1 asyd|OJJ63574.1 ater|EAU39025.1 atub|OJI90864.1 aver|OJI97362.1 azon|OJJ49791.1

Group6263: abra|OJJ73819.1 acar|OOG00972.1 acla|EAW07708.1 afla|EED50010.1 agla|OJJ83678.1 akaw|GAA89438.1 aluc|OJZ88230.1 anid|CBF78043.1 anig|EHA17977.1 aory|EIT74631.1 asyd|OJJ55216.1 ater|EAU38980.1 atub|OJI90832.1 aver|OJJ07000.1 awen|OJJ35419.1 azon|OJJ49758.1

Group6264: abra|OJJ73825.1 acar|OOG00977.1 acla|EAW07720.1 afis|EAW22752.1 afum|EAL90447.1 agla|OJJ83673.1 akaw|GAA89443.1 aluc|OJZ88224.1 anid|CBF88022.1 anig|EHA17971.1 asyd|OJJ63524.1 ater|EAU38968.1 atub|OJI90824.1 aver|OJI97406.1 awen|OJJ35424.1 azon|OJJ49752.1

Group6265: abra|OJJ73839.1 acar|OOG00990.1 acla|EAW07734.1 afis|EAW22766.1 afla|EED50028.1 afum|EAL90433.1 agla|OJJ83654.1 akaw|GAA89457.1 aluc|OJZ88210.1 anid|CBF87999.1 anig|EHA17957.1 asyd|OJJ63507.1 ater|EAU38951.1 aver|OJI97423.1 awen|OJJ38967.1 azon|OJJ49737.1

Group6266: abra|OJJ73862.1 acar|OOG01010.1 acla|EAW07752.1 afis|EAW22789.1 afla|EED50054.1 afum|EAL90410.2 agla|OJJ83587.1 akaw|GAA89480.1 aluc|OJZ88183.1 anid|CBF87961.1 aory|EIT74718.1 asyd|OJJ63483.1 ater|EAU38921.1 atub|OJI90782.1 aver|OJI97447.1 awen|OJJ38945.1

Group6267: abra|OJJ73888.1 acar|OOG01028.1 afis|EAW22808.1 afla|EED50077.1 afum|EBA27406.1 agla|OJJ83583.1 akaw|GAA89500.1 aluc|OJZ88161.1 anid|CBF87902.1 aory|EIT74657.1 asyd|OJJ63458.1 atub|OJI90760.1 aver|OJI97474.1 awen|OJJ38922.1 azon|OJJ49699.1 acla|EAW07772.1

Group6268: abra|OJJ73892.1 acar|OOG01031.1 acla|EAW07776.1 afis|EAW22812.1 afla|EED50081.1 agla|OJJ83560.1 akaw|GAA89504.1 aluc|OJZ88157.1 anid|CBF87909.1 aory|EIT74614.1 asyd|OJJ63454.1 ater|EAU38899.1 atub|OJI90756.1 aver|OJI97478.1 awen|OJJ38916.1 afum|EAL90389.1

Group6269: abra|OJJ73907.1 acar|OOG01040.1 acla|EAW07785.1 afis|EAW22821.1 afla|EED50096.1 afum|EAL90379.2 agla|OJJ83517.1 akaw|GAA89514.1 aluc|OJZ88140.1 anid|CBF87895.1 asyd|OJJ63442.1 ater|EAU38889.1 atub|OJI90743.1 aver|OJI97494.1 awen|OJJ38906.1 azon|OJJ49687.1

Group6270: abra|OJJ73919.1 acar|OOG01050.1 acla|EAW07795.1 afis|EAW22832.1 afum|EAL90370.1 agla|OJJ83507.1 akaw|GAA89525.1 aluc|OJZ88127.1 anid|CBF87879.1 anig|EHA17893.1 asyd|OJJ63433.1 ater|EAU38877.1 atub|OJI90734.1 aver|OJI97504.1 awen|OJJ38892.1 azon|OJJ49677.1

Group6271: abra|OJJ73925.1 acar|OOG01059.1 acla|EAW07802.1 afis|EAW22839.1 afla|EED50114.1 afum|EBA27404.1 agla|OJJ83502.1 akaw|GAA89532.1 aluc|OJZ88118.1 anid|CBF87863.1 asyd|OJJ63426.1 ater|EAU38871.1 atub|OJI90728.1 aver|OJI97511.1 awen|OJJ38884.1 azon|OJJ49673.1

Group6272: abra|OJJ73933.1 acar|OOG01065.1 acla|EAW07811.1 afla|EED50123.1 akaw|GAA89541.1 aluc|OJZ88109.1 anid|CBF87849.1 anig|EHA17879.1 ater|EAU38862.1 atub|OJI90717.1 aver|OJI97519.1 awen|OJJ38875.1 afis|EAW22848.1 afum|EAL90354.1 asyd|OJJ63419.1 azon|OJJ49664.1

Group6273: abra|OJJ73969.1 acar|OOG01095.1 acla|EAW07840.1 afis|EAW22876.1 afla|EED50162.1 afum|EAL90323.1 agla|OJJ83444.1 akaw|GAA89572.1 aluc|OJZ88075.1 aory|EIT76932.1 asyd|OJJ63392.1 ater|EAU38840.1 atub|OJI90684.1 aver|OJI97544.1 awen|OJJ38845.1 azon|OJJ49642.1

Group6274: abra|OJJ73973.1 acar|OOG01100.1 acla|EAW07844.1 afis|EAW22880.1 afla|EED50180.1 afum|EAL90319.1 akaw|GAA89576.1 aluc|OJZ88071.1 anid|CBF87808.1 anig|EHA17844.1 asyd|OJJ63388.1 ater|EAU38844.1 atub|OJI90680.1 aver|OJI97547.1 awen|OJJ38842.1 azon|OJJ49639.1

Group6275: abra|OJJ73975.1 acar|OOG01102.1 acla|EAW07846.1 afis|EAW22882.1 afla|EED50178.1 afum|EAL90317.1 akaw|GAA89578.1 aluc|OJZ88069.1 anid|CBF87805.1 anig|EHA17842.1 aory|EIT77389.1 atub|OJI90678.1 aver|OJI97550.1 awen|OJJ38840.1 agla|OJJ83442.1 azon|OJJ49637.1

Group6276: abra|OJJ73977.1 acar|OOG01105.1 acla|EAW07848.1 afis|EAW22884.1 afla|EED50175.1 afum|EAL90314.1 agla|OJJ83439.1 akaw|GAA89580.1 aluc|OJZ88067.1 anig|EHA17840.1 asyd|OJJ63383.1 ater|EAU35977.1 atub|OJI90676.1 aver|OJI97552.1 awen|OJJ38838.1 azon|OJJ49635.1

Group6277: abra|OJJ73987.1 acar|OOG01112.1 acla|EAW07858.1 afla|EED54863.1 afum|EBA27402.1 agla|OJJ83411.1 akaw|GAA87470.1 aluc|OJZ88058.1 anid|CBF87780.1 anig|EHA17830.1 asyd|OJJ63372.1 ater|EAU35989.1 atub|OJI90666.1 aver|OJI97574.1 awen|OJJ38826.1 azon|OJJ49628.1

Group6278: abra|OJJ73988.1 acar|OOG01113.1 acla|EAW07859.1 afis|EAW22894.1 afum|EBA27401.1 agla|OJJ83412.1 akaw|GAA87471.1 aluc|OJZ88057.1 anid|CBF87778.1 anig|EHA17829.1 asyd|OJJ63371.1 ater|EAU35988.1 atub|OJI90665.1 aver|OJI97575.1 awen|OJJ38825.1 azon|OJJ49624.1

Group6279: abra|OJJ73997.1 acar|OOG01120.1 acla|EAW07865.1 afis|EAW22900.1 afla|EED54871.1 afum|EAL90297.2 agla|OJJ83391.1 akaw|GAA87481.1 aluc|OJZ88045.1 anid|CBF87769.1 anig|EHA17817.1 asyd|OJJ63366.1 atub|OJI90653.1 aver|OJI97580.1 awen|OJJ38730.1 azon|OJJ49617.1

Group6280: abra|OJJ74018.1 acla|EAW07872.1 afis|EAW22907.1 afla|EED54899.1 afum|EAL90290.1 agla|OJJ83382.1 akaw|GAA87500.1 aluc|OJZ88021.1 anid|CBF87710.1 aory|EIT80302.1 asyd|OJJ63336.1 ater|EAU36031.1 atub|OJI90630.1 aver|OJI97626.1 awen|OJJ38723.1 azon|OJJ49607.1

Group6281: abra|OJJ74019.1 acla|EAW07873.1 afis|EAW22908.1 afla|EED54900.1 afum|EAL90289.1 agla|OJJ83381.1 akaw|GAA87501.1 aluc|OJZ88020.1 anid|CBF87708.1 anig|EHA17799.1 aory|EIT80354.1 asyd|OJJ63335.1 atub|OJI90629.1 aver|OJI97627.1 awen|OJJ38722.1 azon|OJJ49606.1

Group6282: abra|OJJ74052.1 acar|OOG01165.1 afis|EAW22937.1 afla|EED58130.1 afum|EAL90259.1 akaw|GAA87534.1 aluc|OJZ87983.1 anid|CBF87656.1 anig|EHA17766.1 aory|EIT79132.1 asyd|OJJ63300.1 ater|EAU30688.1 atub|OJI90593.1 aver|OJI97658.1 awen|OJJ38627.1 azon|OJJ49569.1

Group6283: abra|OJJ74073.1 acar|OOG01185.1 acla|EAW07918.1 afis|EAW22956.1 afla|EED52146.1 afum|EAL90241.1 akaw|GAA87554.1 aluc|OJZ87963.1 anid|CBF74656.1 anig|EHA17747.1 aory|EIT74328.1 asyd|OJJ56119.1 atub|OJI90573.1 aver|OJJ05060.1 azon|OJJ49549.1 asyd|OJJ54385.1

Group6284: abra|OJJ74092.1 acar|OOG01201.1 acla|EAW07935.1 afis|EAW22973.1 afla|EED58167.1 afum|EAL90224.1 agla|OJJ83314.1 akaw|GAA87571.1 aluc|OJZ87944.1 anid|CBF87604.1 aory|EIT79099.1 asyd|OJJ63267.1 ater|EAU30649.1 atub|OJI90555.1 aver|OJI97690.1 azon|OJJ49534.1

Group6285: abra|OJJ74101.1 acar|OOG01210.1 acla|EAW07944.1 afis|EAW22982.1 afla|EED58175.1 agla|OJJ83305.1 akaw|GAA87579.1 aluc|OJZ87935.1 anid|CBF87587.1 aory|EIT79087.1 asyd|OJJ63257.1 ater|EAU30639.1 atub|OJI90547.1 aver|OJI97700.1 awen|OJJ38581.1 azon|OJJ49523.1

Group6286: abra|OJJ74158.1 acar|OOF95568.1 acla|EAW11004.1 afla|EED49739.1 agla|OJJ83720.1 akaw|GAA89726.1 aluc|OJZ80657.1 anid|CBF89386.1 anig|EHA27175.1 aory|EIT77622.1 asyd|OJJ62471.1 ater|EAU34048.1 atub|OJI87512.1 aver|OJI96606.1 awen|OJJ34519.1 azon|OJJ51800.1

Group6287: abra|OJJ74171.1 acar|OOF95551.1 acla|EAW14600.1 afis|EAW19331.1 afla|EED56724.1 afum|EAL94048.2 agla|OJJ81675.1 akaw|GAA86845.1 aluc|OJZ89964.1 anig|EHA27164.1 ater|EAU34817.1 atub|OJI84841.1 awen|OJJ36921.1 acla|EAW11891.1 afis|EAW19502.1 afum|EAL86991.2

Group6288: abra|OJJ74183.1 acar|OOF95539.1 acla|EAW14230.1 afis|EAW19051.1 afla|EED52862.1 afum|EAL93780.1 akaw|GAA86834.1 aluc|OJZ89953.1 anid|CBF81033.1 anig|EHA27154.1 aory|EIT75883.1 asyd|OJJ58008.1 atub|OJI84853.1 aver|OJJ04392.1 awen|OJJ31480.1 afla|EED54405.1

Group6289: abra|OJJ74229.1 acar|OOF95497.1 acla|EAW07275.1 afis|EAW22296.1 afla|EED57706.1 afum|EAL90885.1 akaw|GAA86790.1 aluc|OJZ89904.1 anid|CBF88594.1 anig|EHA27104.1 aory|EIT79877.1 asyd|OJJ63851.1 ater|EAU34318.1 atub|OJI84902.1 aver|OJI97086.1 awen|OJJ34882.1

Group6290: abra|OJJ74241.1 acar|OOF95489.1 acla|EAW07290.1 afis|EAW22324.1 afla|EED57722.1 afum|EAL90868.1 agla|OJJ87720.1 akaw|GAA86779.1 aluc|OJZ89891.1 anid|CBF88607.1 anig|EHA27090.1 asyd|OJJ63858.1 atub|OJI84916.1 aver|OJI97081.1 awen|OJJ34899.1 azon|OJJ50066.1

Group6291: abra|OJJ74255.1 acar|OOF95473.1 acla|EAW07157.1 afis|EAW22172.1 afla|EED57544.1 afum|EAL91017.1 akaw|GAA86766.1 aluc|OJZ89877.1 anid|CBF89083.1 anig|EHA27076.1 aory|EIT81850.1 asyd|OJJ62736.1 ater|EAU34177.1 atub|OJI84930.1 aver|OJI96779.1 azon|OJJ45812.1

Group6292: abra|OJJ74286.1 acar|OOF95445.1 acla|EAW07330.1 afis|EAW22362.1 afla|EED57770.1 afum|EAL90829.1 agla|OJJ87611.1 akaw|GAA86738.1 aluc|OJZ89847.1 anig|EHA27048.1 asyd|OJJ63903.1 ater|EAU39496.1 atub|OJI84959.1 aver|OJI97037.1 awen|OJJ34936.1 azon|OJJ45588.1

Group6293: abra|OJJ74312.1 acla|EAW07363.1 afis|EAW22390.1 afla|EED56053.1 afum|EBA27414.1 agla|OJJ87652.1 akaw|GAA86712.1 aluc|OJZ89814.1 ater|EAU39328.1 atub|OJI84991.1 awen|OJJ34967.1 azon|OJJ45616.1 acla|EAW07362.1 afla|EED56052.1 aory|EIT76640.1 awen|OJJ34966.1

Group6294: abra|OJJ74322.1 acla|EAW07374.1 afis|EAW22399.1 afla|EED56063.1 afum|EAL90792.1 agla|OJJ87662.1 akaw|GAA86704.1 aluc|OJZ89806.1 anid|CBF88754.1 aory|EIT76775.1 asyd|OJJ63960.1 ater|EAU39314.1 atub|OJI85000.1 aver|OJI96981.1 awen|OJJ34978.1 azon|OJJ45626.1

Group6295: abra|OJJ74331.1 acar|OOF95400.1 acla|EAW07382.1 afis|EAW22407.1 afla|EED56071.1 afum|EAL90785.1 agla|OJJ87670.1 akaw|GAA86697.1 aluc|OJZ89798.1 anid|CBF88791.1 anig|EHA27011.1 asyd|OJJ62899.1 ater|EAU39301.1 atub|OJI85007.1 aver|OJI96953.1 awen|OJJ34987.1

Group6296: abra|OJJ74342.1 acar|OOF95388.1 acla|EAW07393.1 afis|EAW22419.1 afla|EED52497.1 afum|EAL90773.1 agla|OJJ87601.1 akaw|GAA86688.1 aluc|OJZ89789.1 anid|CBF88806.1 asyd|OJJ62893.1 ater|EAU39287.1 atub|OJI85017.1 aver|OJI96945.1 awen|OJJ34998.1 azon|OJJ45645.1

Group6297: abra|OJJ74387.1 acla|EAW07428.1 afis|EAW22454.1 afla|EED54476.1 afum|EAL90737.1 akaw|GAA87288.1 aluc|OJZ89731.1 anid|CBF88933.1 anig|EHA26957.1 aory|EIT81126.1 asyd|OJJ62849.1 ater|EAU39252.1 atub|OJI85070.1 aver|OJI96902.1 awen|OJJ35079.1 azon|OJJ50028.1

Group6298: abra|OJJ74446.1 akaw|GAA87338.1 aluc|OJZ89676.1 anig|EHA26911.1 asyd|OJJ62990.1 ater|EAU30284.1 atub|OJI85124.1 aver|OJI97978.1 awen|OJJ30984.1 abra|OJJ76869.1 afis|EAW16916.1 afla|EED51653.1 aluc|OJZ84280.1 anig|EHA27625.1 aory|EIT80516.1 atub|OJI89801.1

Group6299: abra|OJJ74449.1 acar|OOF95302.1 afis|EAW17050.1 afla|EED54125.1 agla|OJJ78792.1 akaw|GAA87340.1 aluc|OJZ89674.1 anid|CBF78431.1 anig|EHA26909.1 aory|EIT76080.1 asyd|OJJ52557.1 ater|EAU32136.1 atub|OJI85126.1 aver|OJJ03843.1 awen|OJJ39931.1 azon|OJJ42221.1

Group6300: abra|OJJ74510.1 acla|EAW07181.1 afis|EAW22194.1 afum|EAL90997.1 agla|OJJ87854.1 akaw|GAA82089.1 aluc|OJZ92091.1 anid|CBF88406.1 anig|EHA26850.1 aory|EIT82934.1 asyd|OJJ63751.1 ater|EAU34203.1 atub|OJI86522.1 aver|OJI97189.1 awen|OJJ34775.1 azon|OJJ45759.1

Group6301: abra|OJJ74545.1 acla|EAW07153.1 afis|EAW22167.1 afla|EED57537.1 afum|EAL91024.1 agla|OJJ87890.1 akaw|GAA82053.1 aluc|OJZ92127.1 anid|CBF89085.1 anig|EHA26821.1 asyd|OJJ62735.1 ater|EAU34174.1 atub|OJI86559.1 aver|OJI96778.1 awen|OJJ34741.1 azon|OJJ45818.1

Group6302: abra|OJJ74547.1 acar|OOF95211.1 acla|EAW07151.1 afis|EAW22165.1 afum|EAL91026.1 agla|OJJ87892.1 akaw|GAA82051.1 aluc|OJZ92129.1 anid|CBF89088.1 anig|EHA26819.1 asyd|OJJ62733.1 ater|EAU34172.1 atub|OJI86561.1 aver|OJI96776.1 awen|OJJ34739.1 azon|OJJ45820.1

Group6303: abra|OJJ74558.1 acar|OOF95198.1 acla|EAW07144.1 afis|EAW22156.1 afla|EED53616.1 afum|EAL91034.1 akaw|GAA82039.1 aluc|OJZ92140.1 anid|CBF89103.1 anig|EHA26809.1 aory|EIT74954.1 asyd|OJJ62726.1 atub|OJI86572.1 aver|OJI96769.1 awen|OJJ34730.1 azon|OJJ45829.1

Group6304: abra|OJJ74579.1 acar|OOF95185.1 afis|EAW22144.1 afla|EED45850.1 afum|EAL91046.1 agla|OJJ87930.1 akaw|GAA82026.1 aluc|OJZ92157.1 anid|CBF78369.1 anig|EHA26792.1 asyd|OJJ57326.1 ater|EAU34147.1 atub|OJI86585.1 aver|OJJ05834.1 awen|OJJ34719.1 azon|OJJ46645.1

Group6305: abra|OJJ74656.1 afla|EED47426.1 afla|EED51743.1 agla|OJJ78810.1 akaw|GAA81954.1 aluc|OJZ92234.1 anig|EHA26720.1 anig|EHA26721.1 aory|EIT80436.1 aory|EIT81913.1 asyd|OJJ62822.1 atub|OJI86667.1 aver|OJI96866.1 azon|OJJ43656.1 asyd|OJJ62824.1 aver|OJI96868.1

Group6306: abra|OJJ74726.1 afum|EAL89548.1 akaw|GAA81891.1 aluc|OJZ92310.1 anig|EHA26659.1 atub|OJI86739.1 azon|OJJ49350.1 acla|EAW10672.1 afis|EAW24894.1 afla|EED46974.1 agla|OJJ82118.1 aory|EIT72672.1 ater|EAU36404.1 anid|CBF75636.1 asyd|OJJ57462.1 aver|OJJ05718.1

Group6307: abra|OJJ74741.1 acla|EAW10663.1 afis|EAW24904.1 afla|EED46984.1 afum|EAL89557.1 akaw|GAA83669.1 aluc|OJZ92322.1 anid|CBF75640.1 anig|EHA26647.1 aory|EIT72733.1 asyd|OJJ57476.1 ater|EAU36395.1 atub|OJI86751.1 aver|OJJ05705.1 awen|OJJ40497.1 azon|OJJ49359.1

Group6308: abra|OJJ74773.1 acla|EAW10638.1 afis|EAW24928.1 afla|EED55545.1 afum|EAL89584.2 agla|OJJ82156.1 akaw|GAA83639.1 aluc|OJZ92354.1 anid|CBF75706.1 anig|EHA26620.1 asyd|OJJ57498.1 ater|EAU35658.1 atub|OJI86784.1 aver|OJJ05680.1 awen|OJJ40524.1 azon|OJJ49384.1

Group6309: abra|OJJ74782.1 acar|OOF95773.1 acla|EAW10628.1 afis|EAW24939.1 afla|EED55533.1 afum|EAL89596.1 agla|OJJ82166.1 akaw|GAA83630.1 aluc|OJZ92363.1 anid|CBF75729.1 anig|EHA26610.1 asyd|OJJ57509.1 atub|OJI86793.1 aver|OJJ05669.1 awen|OJJ40534.1 azon|OJJ49394.1

Group6310: abra|OJJ74798.1 acar|OOF95788.1 acla|EAW10608.1 afis|EAW24959.1 afla|EED55504.1 afum|EAL89617.2 agla|OJJ82190.1 akaw|GAA83614.1 aluc|OJZ92379.1 anid|CBF81662.1 aory|EIT80863.1 asyd|OJJ59550.1 ater|EAU35696.1 atub|OJI86809.1 aver|OJJ02718.1 azon|OJJ49412.1

Group6311: abra|OJJ74843.1 acar|OOF95844.1 acla|EAW08547.1 afis|EAW23584.1 afla|EED57491.1 afum|EAL87880.1 akaw|GAA83579.1 aluc|OJZ92418.1 anid|CBF89589.1 anig|EHA26548.1 aory|EIT81885.1 asyd|OJJ62316.1 ater|EAU35370.1 atub|OJI86848.1 aver|OJI96424.1 awen|OJJ39911.1

Group6312: abra|OJJ74853.1 acar|OOF95853.1 acla|EAW08534.1 afis|EAW23572.1 afla|EED57464.1 afum|EAL87894.1 agla|OJJ86832.1 akaw|GAA83569.1 aluc|OJZ92429.1 anid|CBF89616.1 anig|EHA26537.1 aory|EIT80120.1 asyd|OJJ62303.1 atub|OJI86859.1 awen|OJJ39900.1 azon|OJJ45877.1

Group6313: asyd|OJJ62301.1 ater|EAU35348.1 afla|EED57460.1 agla|OJJ79115.1 akaw|GAA84856.1 aluc|OJZ86815.1 anid|CBF89623.1 aory|EIT80095.1 ater|EAU33845.1 atub|OJI83702.1 aver|OJI96409.1 aver|OJJ08705.1 awen|OJJ33633.1 asyd|OJJ55383.1 anid|CBF82589.1 asyd|OJJ54035.1

Group6314: abra|OJJ74857.1 acla|EAW08527.1 afis|EAW23567.1 afla|EED57458.1 afum|EAL87900.1 agla|OJJ86828.1 akaw|GAA83565.1 aluc|OJZ92433.1 anid|CBF89625.1 anig|EHA26533.1 aory|EIT80060.1 asyd|OJJ62299.1 ater|EAU35350.1 atub|OJI86863.1 aver|OJI96407.1 awen|OJJ39895.1

Group6315: abra|OJJ74858.1 acar|OOF95858.1 acla|EAW08525.1 afis|EAW23565.1 afla|EED57457.1 afum|EAL87903.1 agla|OJJ86827.1 akaw|GAA83562.1 aluc|OJZ92436.1 anid|CBF89630.1 anig|EHA26532.1 asyd|OJJ62297.1 ater|EAU35351.1 atub|OJI86865.1 aver|OJI96405.1 awen|OJJ39894.1

Group6316: abra|OJJ74865.1 acar|OOF95867.1 acla|EAW08514.1 afis|EAW23552.1 afla|EED57447.1 afum|EAL87914.2 agla|OJJ86808.1 aluc|OJZ92443.1 anid|CBF89645.1 anig|EHA26524.1 aory|EIT80099.1 asyd|OJJ62287.1 atub|OJI86872.1 aver|OJI96395.1 awen|OJJ39883.1 azon|OJJ43222.1

Group6317: abra|OJJ74883.1 acar|OOF95887.1 acla|EAW08486.1 afis|EAW23532.1 afla|EED57429.1 agla|OJJ86781.1 akaw|GAA83536.1 aluc|OJZ92460.1 anid|CBF89680.1 anig|EHA26507.1 aory|EIT80137.1 asyd|OJJ62268.1 atub|OJI86889.1 aver|OJI96377.1 awen|OJJ39861.1 azon|OJJ43237.1

Group6318: abra|OJJ74898.1 acar|OOF95898.1 acla|EAW08474.1 afis|EAW23520.1 afla|EED57420.1 afum|EAL87944.1 akaw|GAA83523.1 aluc|OJZ92472.1 aory|EIT80084.1 asyd|OJJ62257.1 ater|EAU35308.1 atub|OJI86903.1 awen|OJJ39845.1 aver|OJI96366.1 azon|OJJ43249.1 agla|OJJ86767.1

Group6319: abra|OJJ74927.1 acar|OOF95918.1 acla|EAW08444.1 afis|EAW23489.1 afla|EED57388.1 afum|EAL87974.1 agla|OJJ86734.1 akaw|GAA83496.1 aluc|OJZ92498.1 anid|CBF89751.1 anig|EHA26474.1 asyd|OJJ62221.1 ater|EAU35276.1 atub|OJI86931.1 aver|OJI96331.1 azon|OJJ47936.1

Group6320: abra|OJJ74955.1 acar|OOF97358.1 acla|EAW08417.1 afis|EAW23452.1 afla|EED57351.1 afum|EAL88014.1 agla|OJJ86698.1 akaw|GAA83468.1 aluc|OJZ92527.1 anid|CBF89801.1 asyd|OJJ62195.1 ater|EAU36918.1 atub|OJI86963.1 aver|OJI96307.1 awen|OJJ39777.1 azon|OJJ47907.1

Group6321: abra|OJJ74995.1 acar|OOF97325.1 acla|EAW08389.1 afis|EAW23418.1 afla|EED57319.1 afum|EAL88051.2 agla|OJJ86635.1 akaw|GAA88619.1 aluc|OJZ92561.1 anid|CBF89859.1 asyd|OJJ62159.1 ater|EAU37504.1 atub|OJI86997.1 aver|OJI96269.1 awen|OJJ39751.1 azon|OJJ47877.1

Group6322: abra|OJJ75044.1 acar|OOF97289.1 acla|EAW08233.1 afis|EAW23267.1 afla|EED55715.1 afum|EAL88212.1 akaw|GAA88661.1 aluc|OJZ92610.1 anid|CBF87389.1 anig|EHA26377.1 aory|EIT83222.1 asyd|OJJ52845.1 atub|OJI87044.1 aver|OJJ08822.1 awen|OJJ39695.1 azon|OJJ47827.1

Group6323: abra|OJJ75072.1 acar|OOF97262.1 acla|EAW08253.1 afis|EAW23287.1 afla|EED55687.1 afum|EAL88192.1 agla|OJJ86560.1 akaw|GAA88684.1 aluc|OJZ92638.1 anig|EHA26353.1 asyd|OJJ62395.1 atub|OJI87069.1 aver|OJI96528.1 awen|OJJ39669.1 azon|OJJ47799.1 anid|CBF89514.1

Group6324: abra|OJJ75076.1 acar|OOF97257.1 acla|EAW08259.1 afis|EAW23292.1 afla|EED55679.1 afum|EAL88187.1 agla|OJJ86547.1 akaw|GAA88689.1 aluc|OJZ92642.1 anid|CBF89506.1 anig|EHA26350.1 asyd|OJJ62399.1 atub|OJI87073.1 aver|OJI96532.1 awen|OJJ39666.1 azon|OJJ47792.1

Group6325: abra|OJJ75155.1 acla|EAW08330.1 afis|EAW23365.1 afla|EED55593.1 afum|EAL88108.1 agla|OJJ86459.1 akaw|GAA88760.1 aluc|OJZ92723.1 anid|CBF74831.1 anig|EHA26277.1 asyd|OJJ60137.1 ater|EAU35616.1 atub|OJI87148.1 aver|OJJ04125.1 awen|OJJ39522.1 azon|OJJ47718.1

Group6326: abra|OJJ75183.1 acar|OOF94597.1 afis|EAW17673.1 afla|EED50480.1 afum|EBA27437.1 agla|OJJ79519.1 akaw|GAA88787.1 aluc|OJZ92747.1 anid|CBF90032.1 anig|EHA26256.1 aory|EIT82045.1 asyd|OJJ62062.1 ater|EAU31974.1 atub|OJI87174.1 aver|OJI96168.1 awen|OJJ37169.1

Group6327: abra|OJJ75231.1 acar|OOF94561.1 acla|EAW12435.1 afis|EAW16745.1 afla|EED48386.1 afum|EAL86320.1 agla|OJJ81565.1 akaw|GAA91775.1 aluc|OJZ92790.1 anig|EHA26215.1 aory|EIT83073.1 asyd|OJJ58458.1 atub|OJI87234.1 aver|OJJ01058.1 awen|OJJ31577.1 azon|OJJ50252.1

Group6328: abra|OJJ75239.1 acla|EAW13711.1 afis|EAW19879.1 afum|EAL85019.1 akaw|GAA91782.1 aluc|OJZ92801.1 anid|CBF76173.1 anig|EHA26208.1 asyd|OJJ63077.1 ater|EAU35155.1 atub|OJI87244.1 aver|OJI97887.1 azon|OJJ42808.1 afis|EAW19980.1 afum|EAL84935.1 anid|CBF85075.1

Group6329: abra|OJJ75282.1 acar|OOF94526.1 acla|EAW09348.1 afis|EAW21803.1 afla|EED46584.1 afum|EAL92829.2 akaw|GAA91820.1 aluc|OJZ92843.1 anid|CBF73984.1 anig|EHA26168.1 aory|EIT77871.1 asyd|OJJ52386.1 ater|EAU31475.1 atub|OJI87299.1 aver|OJJ01937.1 azon|OJJ42848.1

Group6330: abra|OJJ75311.1 acar|OOF94518.1 acla|EAW12229.1 afis|EAW16999.1 afla|EED48951.1 afum|EAL86524.1 agla|OJJ79826.1 akaw|GAA91995.1 aluc|OJZ92865.1 anig|EHA26100.1 aory|EIT80050.1 asyd|OJJ52942.1 atub|OJI87323.1 aver|OJJ08909.1 awen|OJJ35988.1 azon|OJJ50275.1

Group6331: abra|OJJ75369.1 acar|OOF89961.1 acla|EAW12121.1 afis|EAW17259.1 afla|EED48877.1 afum|EAL86676.1 agla|OJJ81068.1 akaw|GAA92063.1 aluc|OJZ92938.1 anig|EHA26034.1 aory|EIT81198.1 asyd|OJJ54367.1 ater|EAU36150.1 atub|OJI87398.1 aver|OJJ05320.1 awen|OJJ35774.1

Group6332: abra|OJJ75376.1 afis|EAW17268.1 afla|EED48851.1 afum|EAL86685.1 agla|OJJ81073.1 akaw|GAA92738.1 aluc|OJZ92946.1 anid|CBF87529.1 anig|EHA26026.1 aory|EIT81264.1 asyd|OJJ52985.1 ater|EAU36133.1 atub|OJI87405.1 aver|OJJ08954.1 awen|OJJ35765.1 azon|OJJ45517.1

Group6333: abra|OJJ75377.1 afis|EAW17269.1 afla|EED48849.1 afum|EAL86686.1 agla|OJJ81074.1 akaw|GAA92739.1 aluc|OJZ92947.1 anid|CBF87531.1 anig|EHA26025.1 aory|EIT81289.1 asyd|OJJ52986.1 ater|EAU36134.1 atub|OJI87406.1 aver|OJJ08955.1 awen|OJJ35764.1 azon|OJJ45518.1

Group6334: abra|OJJ75382.1 acar|OOF94465.1 acla|EAW12110.1 afis|EAW17275.1 afla|EED48844.1 afum|EAL86693.1 agla|OJJ81079.1 akaw|GAA92744.1 aluc|OJZ92952.1 anid|CBF87540.1 anig|EHA26023.1 asyd|OJJ52991.1 atub|OJI87411.1 aver|OJJ08960.1 awen|OJJ35759.1 azon|OJJ45525.1

Group6335: abra|OJJ75390.1 acla|EAW12102.1 afis|EAW17284.1 afla|EED48835.1 afum|EAL86702.1 agla|OJJ81087.1 akaw|GAA92753.1 aluc|OJZ92961.1 anig|EHA26014.1 aory|EIT81290.1 asyd|OJJ53000.1 ater|EAU34704.1 atub|OJI87419.1 aver|OJJ08969.1 awen|OJJ35751.1 azon|OJJ45533.1

Group6336: abra|OJJ75398.1 acar|OOF94452.1 afis|EAW17291.1 afla|EED48827.1 agla|OJJ81096.1 akaw|GAA92760.1 aluc|OJZ92970.1 anid|CBF74293.1 anig|EHA26006.1 aory|EIT81276.1 asyd|OJJ51910.1 ater|EAU34697.1 atub|OJI87427.1 aver|OJJ05351.1 awen|OJJ35745.1 azon|OJJ45538.1

Group6337: abra|OJJ75405.1 acar|OOF90753.1 acla|EAW13778.1 afis|EAW18354.1 agla|OJJ88030.1 akaw|GAA84183.1 aluc|OJZ88857.1 anid|CBF79360.1 anig|EHA23646.1 aory|EIT72997.1 asyd|OJJ57804.1 atub|OJI85933.1 aver|OJJ04584.1 awen|OJJ35738.1 azon|OJJ50440.1 afla|EED48819.1

Group6338: abra|OJJ75420.1 acar|OOF90766.1 acla|EAW13792.1 afis|EAW18340.1 afla|EED48805.1 afum|EAL87703.1 agla|OJJ88044.1 akaw|GAA84195.1 aluc|OJZ88872.1 anid|CBF79373.1 aory|EIT72983.1 asyd|OJJ57818.1 ater|EAU34675.1 atub|OJI85919.1 awen|OJJ35724.1 azon|OJJ43313.1

Group6339: abra|OJJ75490.1 acar|OOF90822.1 afis|EAW18282.1 afla|EED48749.1 afum|EAL87648.1 agla|OJJ88145.1 aluc|OJZ88949.1 anid|CBF79487.1 anig|EHA23572.1 aory|EIT73144.1 asyd|OJJ57875.1 atub|OJI85840.1 aver|OJJ04519.1 awen|OJJ30653.1 azon|OJJ47962.1 acla|EAW11221.1

Group6340: abra|OJJ75498.1 acar|OOF90831.1 acla|EAW09292.1 afis|EAW21855.1 afum|EAL92868.1 agla|OJJ84280.1 akaw|GAA84264.1 aluc|OJZ88957.1 anid|CBF76743.1 anig|EHA23563.1 asyd|OJJ61256.1 ater|EAU30854.1 atub|OJI85832.1 aver|OJJ00579.1 awen|OJJ34139.1 azon|OJJ43487.1

Group6341: abra|OJJ75499.1 acar|OOF90832.1 acla|EAW09293.1 afla|EED46455.1 agla|OJJ84281.1 akaw|GAA84265.1 aluc|OJZ88958.1 anid|CBF76741.1 anig|EHA23562.1 aory|EIT72343.1 asyd|OJJ61255.1 ater|EAU30853.1 atub|OJI85831.1 aver|OJJ00578.1 awen|OJJ34138.1 afis|EAW21854.1

Group6342: abra|OJJ75525.1 acar|OOF90852.1 acla|EAW09312.1 afis|EAW21833.1 afla|EED46542.1 afum|EAL92848.1 agla|OJJ84311.1 akaw|GAA84288.1 aluc|OJZ88982.1 anid|CBF76696.1 anig|EHA23544.1 asyd|OJJ61230.1 atub|OJI85807.1 aver|OJJ00553.1 awen|OJJ34106.1 azon|OJJ44235.1

Group6343: abra|OJJ75555.1 acar|OOF90874.1 acla|EAW09353.1 afis|EAW21818.1 afla|EED46565.1 afum|EAL92834.1 agla|OJJ84326.1 akaw|GAA84310.1 aluc|OJZ89005.1 anid|CBF76656.1 anig|EHA23523.1 aory|EIT77933.1 asyd|OJJ61209.1 atub|OJI85784.1 aver|OJJ00533.1 awen|OJJ34092.1

Group6344: abra|OJJ75606.1 acar|OOG00231.1 acla|EAW09368.1 afis|EAW21769.1 afla|EED46331.1 afum|EAL92798.1 akaw|GAA84357.1 aluc|OJZ89059.1 anid|CBF69640.1 anig|EHA23483.1 aory|EIT81239.1 asyd|OJJ59082.1 atub|OJI85727.1 aver|OJJ01675.1 awen|OJJ30662.1 azon|OJJ48176.1

Group6345: abra|OJJ75614.1 afla|EED46339.1 agla|OJJ84351.1 akaw|GAA84363.1 aluc|OJZ89067.1 anig|EHA23469.1 aory|EIT81226.1 asyd|OJJ61177.1 atub|OJI85721.1 aver|OJJ00503.1 awen|OJJ34060.1 azon|OJJ48172.1 acla|EAW09364.1 afis|EAW21773.1 afum|EAL92802.1 anid|CBF76595.1

Group6346: abra|OJJ75616.1 acar|OOG00216.1 acla|EAW09362.1 afis|EAW21775.1 afla|EED46341.1 afum|EAL92804.1 agla|OJJ84349.1 akaw|GAA84365.1 aluc|OJZ89069.1 anid|CBF76599.1 anig|EHA23467.1 asyd|OJJ61180.1 atub|OJI85719.1 aver|OJJ00505.1 awen|OJJ34063.1 azon|OJJ48169.1

Group6347: abra|OJJ75626.1 acla|EAW12341.1 afis|EAW16873.1 afla|EED53975.1 afum|EAL86411.1 akaw|GAA84373.1 aluc|OJZ89077.1 anid|CBF78447.1 anig|EHA23460.1 aory|EIT78316.1 asyd|OJJ59237.1 atub|OJI85708.1 aver|OJJ01847.1 awen|OJJ38677.1 azon|OJJ45244.1 azon|OJJ45813.1

Group6348: abra|OJJ75671.1 acar|OOG00174.1 acla|EAW09431.1 afis|EAW21705.1 afla|EED57974.1 afum|EAL92739.1 agla|OJJ79889.1 akaw|GAA84411.1 aluc|OJZ89121.1 anig|EHA23422.1 aory|EIT79776.1 asyd|OJJ60790.1 atub|OJI85663.1 aver|OJJ00123.1 awen|OJJ33976.1 azon|OJJ48582.1

Group6349: abra|OJJ75697.1 acar|OOG00156.1 acla|EAW09411.1 afis|EAW21725.1 afla|EED57994.1 afum|EAL92758.1 agla|OJJ79859.1 akaw|GAA84433.1 aluc|OJZ89148.1 anid|CBF83720.1 anig|EHA23400.1 asyd|OJJ60839.1 atub|OJI85640.1 aver|OJJ00163.1 awen|OJJ33998.1 azon|OJJ48554.1

Group6350: abra|OJJ75797.1 acar|OOG00077.1 acla|EAW08955.1 afis|EAW15456.1 afla|EED48146.1 afum|EAL84189.1 akaw|GAA84515.1 aluc|OJZ89246.1 anid|CBF73694.1 anig|EHA23320.1 aory|EIT76198.1 asyd|OJJ56653.1 ater|EAU35490.1 atub|OJI85551.1 aver|OJJ06072.1 awen|OJJ32596.1

Group6351: abra|OJJ75802.1 acar|OOG00074.1 acla|EAW14883.1 afis|EAW24402.1 afla|EED47811.1 afum|EAL86073.1 akaw|GAA84518.1 aluc|OJZ89250.1 anid|CBF73733.1 anig|EHA23317.1 aory|EIT77114.1 asyd|OJJ56631.1 ater|EAU30337.1 atub|OJI85548.1 aver|OJJ06095.1 azon|OJJ43213.1

Group6352: abra|OJJ75806.1 acar|OOG00069.1 acla|EAW14877.1 afis|EAW24394.1 afla|EED47816.1 afum|EAL86066.1 akaw|GAA84522.1 aluc|OJZ89254.1 anid|CBF73719.1 anig|EHA23313.1 aory|EIT77200.1 asyd|OJJ56640.1 ater|EAU30341.1 atub|OJI85544.1 aver|OJJ06086.1 awen|OJJ32811.1

Group6353: abra|OJJ75820.1 acar|OOG00056.1 acla|EAW14855.1 afis|EAW24382.1 afla|EED47828.1 afum|EAL86054.1 agla|OJJ83270.1 akaw|GAA84533.1 aluc|OJZ89266.1 anid|CBF73665.1 anig|EHA23302.1 asyd|OJJ56667.1 atub|OJI85533.1 aver|OJJ06056.1 awen|OJJ32830.1 azon|OJJ49006.1

Group6354: abra|OJJ75866.1 acar|OOG00028.1 acla|EAW09927.1 afis|EAW21175.1 afla|EED54762.1 afum|EAL92195.1 agla|OJJ80002.1 akaw|GAA84572.1 aluc|OJZ89308.1 anid|CBF83299.1 anig|EHA23258.1 asyd|OJJ60557.1 ater|EAU35842.1 aver|OJI99883.1 awen|OJJ36514.1 azon|OJJ44390.1

Group6355: abra|OJJ75929.1 afis|EAW21262.1 afla|EED54821.1 agla|OJJ87470.1 akaw|GAA84630.1 aluc|OJZ89376.1 anig|EHA23207.1 aory|EIT79410.1 asyd|OJJ60607.1 ater|EAU35899.1 atub|OJI85422.1 afum|EAL92277.1 anid|CBF83336.1 aver|OJI99927.1 awen|OJJ36579.1 azon|OJJ44311.1

Group6356: abra|OJJ75941.1 acar|OOF99969.1 acla|EAW09852.1 afis|EAW21283.1 afla|EED54834.1 afum|EAL92300.2 akaw|GAA84643.1 aluc|OJZ89388.1 anid|CBF83361.1 anig|EHA23196.1 asyd|OJJ60619.1 ater|EAU35912.1 atub|OJI85409.1 aver|OJI99943.1 awen|OJJ39416.1 azon|OJJ44299.1

Group6357: abra|OJJ76017.1 acar|OOF99909.1 acla|EAW09789.1 afis|EAW21347.1 afla|EED55762.1 afum|EAL92364.1 agla|OJJ87382.1 akaw|GAA88943.1 aluc|OJZ89464.1 anid|CBF76197.1 anig|EHA23124.1 asyd|OJJ60959.1 ater|EAU35970.1 atub|OJI85331.1 aver|OJJ00284.1 azon|OJJ51692.1

Group6358: abra|OJJ76021.1 acar|OOF99904.1 acla|EAW08988.1 afis|EAW15423.1 afla|EED55766.1 afum|EAL84221.1 akaw|GAA88939.1 aluc|OJZ89468.1 anid|CBF76201.1 anig|EHA23120.1 aory|EIT76861.1 asyd|OJJ60956.1 ater|EAU35972.1 atub|OJI85327.1 aver|OJJ00283.1 azon|OJJ45285.1

Group6359: abra|OJJ76022.1 acar|OOF99903.1 acla|EAW09786.1 afis|EAW21352.1 afla|EED55767.1 afum|EAL92367.1 akaw|GAA88935.1 aluc|OJZ89472.1 anid|CBF76203.1 anig|EHA23119.1 aory|EIT76835.1 asyd|OJJ60952.1 ater|EAU35973.1 atub|OJI85326.1 aver|OJJ00279.1 awen|OJJ39348.1

Group6360: abra|OJJ76045.1 acar|OOF99885.1 acla|EAW09776.1 afis|EAW21362.1 afla|EED55776.1 afum|EAL92378.1 akaw|GAA88920.1 aluc|OJZ89501.1 anid|CBF83879.1 anig|EHA23099.1 asyd|OJJ60939.1 ater|EAU35198.1 atub|OJI85302.1 aver|OJJ00265.1 awen|OJJ39336.1 azon|OJJ51681.1

Group6361: abra|OJJ76047.1 acar|OOF99883.1 acla|EAW09778.1 afis|EAW21360.1 afla|EED46576.1 afum|EAL92376.1 agla|OJJ85479.1 akaw|GAA88918.1 aluc|OJZ89503.1 anid|CBF86925.1 anig|EHA23097.1 asyd|OJJ52230.1 ater|EAU34094.1 atub|OJI85300.1 azon|OJJ51683.1 acla|EAW09086.1

Group6362: abra|OJJ76051.1 acar|OOF99877.1 acla|EAW09767.1 afis|EAW21371.1 afla|EED55789.1 afum|EAL92388.1 agla|OJJ85491.1 akaw|GAA88914.1 aluc|OJZ89507.1 anid|CBF83870.1 anig|EHA23093.1 asyd|OJJ60934.1 atub|OJI85296.1 aver|OJJ00260.1 awen|OJJ39326.1 azon|OJJ51671.1

Group6363: abra|OJJ76074.1 acla|EAW09747.1 afis|EAW21395.1 afla|EED55814.1 afum|EAL92409.1 agla|OJJ85520.1 akaw|GAA88893.1 aluc|OJZ89530.1 anid|CBF83837.1 anig|EHA23072.1 asyd|OJJ60912.1 atub|OJI85275.1 aver|OJJ00239.1 azon|OJJ51545.1 aory|EIT76625.1 ater|EAU35161.1

Group6364: abra|OJJ76105.1 acar|OOF99820.1 acla|EAW06763.1 afis|EAW16313.1 afla|EED54182.1 afum|EAL88935.1 akaw|GAA88861.1 aluc|OJZ89561.1 anid|CBF69558.1 anig|EHA23045.1 aory|EIT76112.1 asyd|OJJ59162.1 ater|EAU29329.1 atub|OJI85243.1 aver|OJJ01752.1 awen|OJJ36250.1

Group6365: abra|OJJ76137.1 acar|OOF99787.1 acla|EAW09682.1 afis|EAW21457.1 afla|EED55887.1 afum|EAL92476.1 akaw|GAA88831.1 aluc|OJZ89600.1 anid|CBF76542.1 aory|EIT76595.1 asyd|OJJ61152.1 ater|EAU35078.1 atub|OJI85204.1 aver|OJJ00474.1 awen|OJJ39243.1 azon|OJJ51612.1

Group6366: abra|OJJ76144.1 acla|EAW09667.1 afis|EAW21470.1 afla|EED55906.1 afum|EAL92489.1 agla|OJJ85619.1 akaw|GAA82093.1 aluc|OJZ92087.1 anid|CBF76519.1 aory|EIT76732.1 asyd|OJJ61139.1 ater|EAU35063.1 atub|OJI86516.1 aver|OJJ00461.1 awen|OJJ39228.1 azon|OJJ51627.1

Group6367: aory|EIT76874.1 abra|OJJ76156.1 acla|EAW09662.1 afis|EAW21477.1 afla|EED55909.1 afum|EAL92497.1 agla|OJJ85626.1 akaw|GAA82104.1 aluc|OJZ92072.1 anid|CBF76509.1 aory|EIT76651.1 asyd|OJJ61134.1 atub|OJI86504.1 aver|OJJ00455.1 azon|OJJ51634.1 agla|OJJ85625.1

Group6368: abra|OJJ76233.1 acar|OOF99479.1 acla|EAW14369.1 afis|EAW18919.1 afla|EED50903.1 afum|EAL93650.2 akaw|GAA82168.1 aluc|OJZ91998.1 anid|CBF69684.1 anig|EHA22941.1 aory|EIT79023.1 asyd|OJJ59059.1 ater|EAU38086.1 atub|OJI86433.1 aver|OJJ01653.1 awen|OJJ31512.1

Group6369: abra|OJJ76268.1 acla|EAW14340.1 afis|EAW18950.1 afum|EAL93684.1 agla|OJJ85321.1 akaw|GAA82201.1 aluc|OJZ91952.1 anid|CBF86831.1 anig|EHA22911.1 aory|EIT77032.1 asyd|OJJ64650.1 ater|EAU38136.1 atub|OJI86397.1 aver|OJI98750.1 awen|OJJ31548.1 azon|OJJ47264.1

Group6370: abra|OJJ76275.1 acar|OOF99523.1 acla|EAW14337.1 afis|EAW18952.1 afla|EED50990.1 afum|EAL93688.1 agla|OJJ85323.1 aluc|OJZ91945.1 anid|CBF69589.1 anig|EHA22904.1 aory|EIT76952.1 asyd|OJJ59141.1 ater|EAU38137.1 atub|OJI86390.1 aver|OJJ01731.1 azon|OJJ47268.1

Group6371: abra|OJJ76298.1 acar|OOF99543.1 acla|EAW14320.1 afis|EAW18968.1 afla|EED51034.1 afum|EAL93700.2 agla|OJJ85354.1 akaw|GAA82225.1 aluc|OJZ91924.1 anid|CBF69615.1 asyd|OJJ59126.1 ater|EAU38154.1 atub|OJI86368.1 aver|OJJ01714.1 awen|OJJ31568.1 azon|OJJ47286.1

Group6372: abra|OJJ76350.1 acar|OOF99601.1 acla|EAW14065.1 afis|EAW18772.1 afla|EED50731.1 afum|EAL93504.1 agla|OJJ80488.1 akaw|GAA82277.1 aluc|OJZ91863.1 anid|CBF69932.1 anig|EHA22843.1 aory|EIT78147.1 asyd|OJJ58923.1 ater|EAU37903.1 awen|OJJ32341.1 azon|OJJ47109.1

Group6373: abra|OJJ76360.1 acar|OOF99611.1 acla|EAW14056.1 afis|EAW18762.1 afla|EED50722.1 afum|EAL93494.1 agla|OJJ80459.1 akaw|GAA82286.1 aluc|OJZ91853.1 anid|CBF69949.1 anig|EHA22834.1 aory|EIT78002.1 asyd|OJJ58913.1 atub|OJI86304.1 aver|OJJ01510.1 azon|OJJ47097.1

Group6374: abra|OJJ76414.1 acar|OOF99654.1 acla|EAW14012.1 afis|EAW18718.1 afla|EED50672.1 afum|EAL93442.2 agla|OJJ82340.1 anid|CBF70675.1 anig|EHA22794.1 aory|EIT77982.1 asyd|OJJ58473.1 atub|OJI86254.1 aver|OJJ01074.1 awen|OJJ32222.1 azon|OJJ46774.1 afla|EED50673.1

Group6375: abra|OJJ76428.1 acar|OOF99671.1 acla|EAW13997.1 afis|EAW18705.1 afla|EED50654.1 afum|EAL93428.1 agla|OJJ82353.1 akaw|GAA82343.1 aluc|OJZ91784.1 anid|CBF70647.1 anig|EHA22781.1 asyd|OJJ58487.1 atub|OJI86240.1 aver|OJJ01090.1 awen|OJJ32188.1 azon|OJJ46759.1

Group6376: abra|OJJ76435.1 acar|OOF99680.1 acla|EAW13988.1 afis|EAW18696.1 afla|EED50644.1 afum|EAL93418.1 agla|OJJ82370.1 akaw|GAA82350.1 aluc|OJZ91777.1 anid|CBF70629.1 asyd|OJJ58497.1 ater|EAU38210.1 atub|OJI86233.1 aver|OJJ01100.1 awen|OJJ32167.1 azon|OJJ46751.1

Group6377: abra|OJJ76478.1 acar|OOF99713.1 afum|EAL94102.1 agla|OJJ82409.1 akaw|GAA82391.1 aluc|OJZ91729.1 anid|CBF70573.1 anig|EHA22736.1 aory|EIT74210.1 asyd|OJJ58531.1 ater|EAU38244.1 atub|OJI86187.1 aver|OJJ01133.1 awen|OJJ32136.1 azon|OJJ46719.1 acar|OOF91475.1

Group6378: abra|OJJ76486.1 acar|OOF99719.1 acla|EAW13947.1 afis|EAW18657.1 afla|EED52673.1 afum|EAL93374.1 akaw|GAA82402.1 aluc|OJZ91720.1 anid|CBF70550.1 anig|EHA22727.1 aory|EIT74222.1 asyd|OJJ58541.1 ater|EAU38255.1 atub|OJI86177.1 aver|OJJ01142.1 awen|OJJ32124.1

Group6379: abra|OJJ76493.1 acar|OOF99725.1 acla|EAW13939.1 afis|EAW18650.1 afla|EED52689.1 afum|EAL93366.1 agla|OJJ82424.1 akaw|GAA82409.1 aluc|OJZ91711.1 anig|EHA22722.1 asyd|OJJ58551.1 ater|EAU38262.1 atub|OJI86170.1 aver|OJJ01149.1 awen|OJJ32118.1 azon|OJJ46704.1

Group6380: abra|OJJ76511.1 acla|EAW14189.1 afis|EAW18904.1 afla|EED50844.1 afum|EAL93636.1 agla|OJJ81509.1 akaw|GAA82424.1 aluc|OJZ91694.1 anid|CBF69717.1 anig|EHA22707.1 aory|EIT79060.1 asyd|OJJ59045.1 ater|EAU38074.1 atub|OJI86148.1 aver|OJJ01641.1 awen|OJJ32461.1

Group6381: abra|OJJ76527.1 acar|OOF99448.1 acla|EAW14174.1 afis|EAW18888.1 afla|EED50875.1 afum|EAL93621.1 agla|OJJ81494.1 aluc|OJZ91678.1 anid|CBF69900.1 anig|EHA22695.1 aory|EIT78967.1 asyd|OJJ58999.1 ater|EAU38060.1 atub|OJI86133.1 awen|OJJ32446.1 azon|OJJ47205.1

Group6382: abra|OJJ76539.1 acar|OOF99434.1 acla|EAW14164.1 afis|EAW18877.1 afla|EED50889.1 afum|EAL93609.1 agla|OJJ80614.1 akaw|GAA82449.1 aluc|OJZ91664.1 anig|EHA22684.1 asyd|OJJ59016.1 ater|EAU38042.1 atub|OJI86121.1 aver|OJJ01599.1 awen|OJJ32436.1 azon|OJJ47194.1

Group6383: abra|OJJ76541.1 acar|OOF99436.1 acla|EAW14166.1 afis|EAW18879.1 afla|EED50887.1 afum|EAL93611.1 agla|OJJ80616.1 aluc|OJZ91662.1 anid|CBF69879.1 anig|EHA22682.1 aory|EIT78990.1 asyd|OJJ59014.1 atub|OJI86119.1 aver|OJJ01601.1 awen|OJJ32438.1 azon|OJJ47197.1

Group6384: abra|OJJ76557.1 acar|OOF99422.1 acla|EAW14148.1 afis|EAW18861.1 afla|EED50828.1 afum|EAL93593.2 agla|OJJ80584.1 akaw|GAA82463.1 aluc|OJZ91648.1 anid|CBF69844.1 asyd|OJJ59032.1 atub|OJI86102.1 aver|OJJ01584.1 awen|OJJ32417.1 azon|OJJ47174.1 anig|EHA22669.1

Group6385: abra|OJJ76585.1 acar|OOF99401.1 acla|EAW14110.1 afis|EAW18812.1 afla|EED50797.1 afum|EAL93548.1 agla|OJJ80545.1 akaw|GAA82488.1 aluc|OJZ91621.1 anid|CBF69797.1 asyd|OJJ58965.1 ater|EAU37979.1 atub|OJI86074.1 aver|OJJ01559.1 awen|OJJ32388.1 azon|OJJ47148.1

Group6386: abra|OJJ76626.1 acar|OOF99372.1 afis|EAW18990.1 afla|EED53175.1 afum|EAL93726.1 agla|OJJ85977.1 akaw|GAA92831.1 aluc|OJZ91586.1 anig|EHA22610.1 aory|EIT75103.1 asyd|OJJ53667.1 ater|EAU37640.1 atub|OJI86035.1 awen|OJJ31611.1 asyd|OJJ57923.1 aver|OJJ04469.1

Group6387: abra|OJJ76628.1 acar|OOF99370.1 acla|EAW07509.1 afla|EED47282.1 agla|OJJ86876.1 akaw|GAA92829.1 aluc|OJZ91584.1 anig|EHA22609.1 aory|EIT79279.1 asyd|OJJ56229.1 ater|EAU33542.1 atub|OJI86033.1 aver|OJJ05164.1 awen|OJJ37547.1 azon|OJJ42292.1 agla|OJJ81403.1

Group6388: abra|OJJ76633.1 acla|EAW14286.1 afis|EAW18996.1 afla|EED53146.1 afum|EAL93731.1 agla|OJJ82984.1 akaw|GAA92823.1 aluc|OJZ91577.1 anid|CBF86736.1 anig|EHA22604.1 aory|EIT75154.1 asyd|OJJ64693.1 atub|OJI86026.1 aver|OJI98797.1 awen|OJJ31617.1 azon|OJJ47319.1

Group6389: abra|OJJ76642.1 acar|OOF99354.1 afis|EAW19003.1 afla|EED53118.1 afum|EAL93737.1 agla|OJJ85231.1 akaw|GAA92816.1 aluc|OJZ91567.1 anid|CBF83913.1 anig|EHA22596.1 asyd|OJJ55494.1 ater|EAU37794.1 atub|OJI86017.1 aver|OJJ06572.1 awen|OJJ31632.1 azon|OJJ47332.1

Group6390: abra|OJJ76651.1 acar|OOF99351.1 afis|EAW19013.1 afla|EED53115.1 afum|EAL93747.2 agla|OJJ81764.1 akaw|GAA92991.1 aluc|OJZ91556.1 anid|CBF86756.1 anig|EHA22590.1 aory|EIT74783.1 asyd|OJJ64709.1 ater|EAU37788.1 atub|OJI86008.1 aver|OJI98811.1 awen|OJJ31650.1

Group6391: abra|OJJ76661.1 acar|OOF99342.1 acla|EAW11020.1 afis|EAW20833.1 afla|EED50430.1 afum|EAL84799.1 agla|OJJ80142.1 akaw|GAA93002.1 aluc|OJZ91541.1 anig|EHA22578.1 aory|EIT81448.1 atub|OJI85995.1 awen|OJJ36031.1 azon|OJJ45265.1 afla|EED52050.1 aory|EIT82047.1

Group6392: abra|OJJ76759.1 acar|OOF91254.1 acla|EAW13313.1 afis|EAW20427.1 afum|EBA27200.1 agla|OJJ84898.1 akaw|GAA87267.1 aluc|OJZ84162.1 anid|CBF79336.1 anig|EHA27523.1 asyd|OJJ55789.1 ater|EAU37730.1 atub|OJI89922.1 aver|OJJ06879.1 awen|OJJ33120.1 azon|OJJ46336.1

Group6393: abra|OJJ76775.1 acar|OOF91272.1 acla|EAW13334.1 afis|EAW20449.1 afla|EED44819.1 agla|OJJ84873.1 akaw|GAA87249.1 aluc|OJZ84179.1 anid|CBF79299.1 anig|EHA27539.1 asyd|OJJ55774.1 ater|EAU37710.1 atub|OJI89904.1 aver|OJJ06861.1 awen|OJJ33139.1 azon|OJJ46350.1

Group6394: abra|OJJ76778.1 acar|OOF91274.1 acla|EAW13337.1 afis|EAW20452.1 afum|EAL84601.1 agla|OJJ84870.1 akaw|GAA87246.1 aluc|OJZ84182.1 anid|CBF79303.1 anig|EHA27542.1 asyd|OJJ55771.1 ater|EAU37707.1 atub|OJI89901.1 aver|OJJ06857.1 awen|OJJ33142.1 azon|OJJ46353.1

Group6395: abra|OJJ76785.1 acar|OOF91278.1 acla|EAW13343.1 afis|EAW20458.1 afum|EBA27195.1 agla|OJJ84861.1 akaw|GAA87240.1 aluc|OJZ84188.1 anid|CBF79281.1 anig|EHA27548.1 asyd|OJJ55765.1 ater|EAU37702.1 atub|OJI89895.1 aver|OJJ06851.1 awen|OJJ33150.1 azon|OJJ46360.1

Group6396: abra|OJJ76786.1 acar|OOF91280.1 acla|EAW13344.1 afis|EAW20459.1 afla|EED44809.1 afum|EAL84594.2 agla|OJJ84860.1 akaw|GAA87239.1 anid|CBF79279.1 aory|EIT77712.1 asyd|OJJ55764.1 ater|EAU37701.1 atub|OJI89894.1 aver|OJJ06850.1 awen|OJJ33151.1 azon|OJJ46361.1

Group6397: abra|OJJ76801.1 acar|OOF91295.1 acla|EAW13360.1 afis|EAW20474.1 afum|EBA27193.1 agla|OJJ84844.1 akaw|GAA87222.1 aluc|OJZ84208.1 anid|CBF79264.1 anig|EHA27562.1 asyd|OJJ55755.1 ater|EAU37688.1 atub|OJI89876.1 aver|OJJ06840.1 awen|OJJ33168.1 azon|OJJ46376.1

Group6398: abra|OJJ76803.1 acar|OOF91297.1 acla|EAW13362.1 afis|EAW20476.1 afum|EAL84577.1 agla|OJJ84841.1 akaw|GAA87220.1 aluc|OJZ84210.1 anid|CBF79253.1 anig|EHA27564.1 asyd|OJJ55746.1 ater|EAU29554.1 atub|OJI89873.1 aver|OJJ06833.1 awen|OJJ33171.1 azon|OJJ46378.1

Group6399: abra|OJJ76829.1 acar|OOF91323.1 acla|EAW13390.1 afis|EAW20503.1 afla|EED44746.1 afum|EAL84553.1 agla|OJJ84799.1 akaw|GAA87193.1 aluc|OJZ84241.1 anid|CBF79197.1 aory|EIT77770.1 asyd|OJJ55723.1 atub|OJI89842.1 aver|OJJ06809.1 awen|OJJ33204.1 azon|OJJ46394.1

Group6400: abra|OJJ76920.1 acar|OOF91367.1 akaw|GAA87118.1 aluc|OJZ84320.1 anig|EHA27655.1 atub|OJI89748.1 awen|OJJ33256.1 afis|EAW20100.1 afum|EAL85217.1 ater|EAU39419.1 azon|OJJ45280.1 afla|EED55286.1 aory|EIT83559.1 asyd|OJJ53769.1 aver|OJJ07877.1 awen|OJJ40778.1

Group6401: abra|OJJ76933.1 acar|OOF91382.1 acla|EAW12082.1 afis|EAW19723.1 afla|EED52289.1 afum|EAL86803.1 agla|OJJ79589.1 akaw|GAA87106.1 aluc|OJZ84332.1 anid|CBF78536.1 anig|EHA27666.1 asyd|OJJ65296.1 ater|EAU33142.1 atub|OJI89737.1 aver|OJI99436.1 awen|OJJ31939.1

Group6402: abra|OJJ77065.1 acar|OOF92949.1 acla|EAW13510.1 afis|EAW20608.1 afum|EAL84452.1 agla|OJJ88631.1 akaw|GAA86987.1 aluc|OJZ84473.1 anid|CBF86949.1 anig|EHA27782.1 asyd|OJJ55593.1 ater|EAU35258.1 atub|OJI89605.1 aver|OJJ06672.1 awen|OJJ33451.1 azon|OJJ50708.1

Group6403: abra|OJJ77073.1 acar|OOF96453.1 acla|EAW11108.1 afla|EED48433.1 afum|EAL85153.1 agla|OJJ81759.1 akaw|GAA86980.1 aluc|OJZ84483.1 aory|EIT77054.1 asyd|OJJ62356.1 atub|OJI89597.1 aver|OJI99662.1 awen|OJJ40115.1 awen|OJJ32899.1 afla|EED51078.1 agla|OJJ80487.1

Group6404: abra|OJJ77075.1 acar|OOF92956.1 acla|EAW11000.1 afis|EAW20861.1 afla|EED45445.1 afum|EAL84774.1 agla|OJJ83748.1 akaw|GAA86978.1 aluc|OJZ84485.1 anid|CBF82829.1 anig|EHA27789.1 asyd|OJJ61581.1 ater|EAU30804.1 atub|OJI89595.1 aver|OJJ00903.1 awen|OJJ34509.1

Group6405: abra|OJJ77081.1 acar|OOF92965.1 afis|EAW23835.1 afla|EED51967.1 afum|EAL87266.1 agla|OJJ82499.1 akaw|GAA86972.1 aluc|OJZ84491.1 anid|CBF74533.1 anig|EHA27795.1 aory|EIT81457.1 asyd|OJJ57683.1 atub|OJI89588.1 aver|OJJ05489.1 awen|OJJ33795.1 azon|OJJ48747.1

Group6406: abra|OJJ77140.1 acar|OOF90281.1 acla|EAW12951.1 afis|EAW17726.1 afla|EED45727.1 afum|EAL91541.1 agla|OJJ81725.1 akaw|GAA90789.1 aluc|OJZ84553.1 anid|CBF86581.1 anig|EHA27847.1 aory|EIT75454.1 ater|EAU31880.1 atub|OJI89529.1 awen|OJJ37112.1 azon|OJJ50811.1

Group6407: abra|OJJ77156.1 acar|OOF93043.1 acla|EAW12916.1 afis|EAW17744.1 afla|EED45744.1 afum|EAL91556.1 agla|OJJ81709.1 akaw|GAA90774.1 aluc|OJZ84570.1 anig|EHA27865.1 asyd|OJJ64819.1 ater|EAU31864.1 atub|OJI89512.1 aver|OJI98932.1 awen|OJJ37093.1 azon|OJJ50797.1

Group6408: abra|OJJ77189.1 acar|OOF93075.1 afis|EAW20801.1 afla|EED49544.1 afum|EAL84828.1 akaw|GAA90751.1 aluc|OJZ84597.1 anid|CBF71376.1 anig|EHA27886.1 aory|EIT82575.1 asyd|OJJ55076.1 ater|EAU32871.1 atub|OJI89482.1 aver|OJJ07132.1 awen|OJJ41913.1 azon|OJJ47357.1

Group6409: abra|OJJ77212.1 acar|OOF93100.1 acla|EAW13204.1 afis|EAW17438.1 afla|EED49389.1 agla|OJJ80379.1 akaw|GAA90726.1 aluc|OJZ84623.1 anid|CBF71703.1 aory|EIT79556.1 asyd|OJJ61811.1 ater|EAU33917.1 atub|OJI89458.1 aver|OJI95910.1 awen|OJJ37408.1 azon|OJJ51082.1

Group6410: abra|OJJ77222.1 acar|OOF93107.1 acla|EAW13213.1 afis|EAW17429.1 afum|EAL91257.1 agla|OJJ80389.1 akaw|GAA90717.1 aluc|OJZ84632.1 anid|CBF71714.1 anig|EHA27920.1 asyd|OJJ61805.1 ater|EAU33910.1 atub|OJI89447.1 aver|OJI95904.1 awen|OJJ37414.1 azon|OJJ51088.1

Group6411: abra|OJJ77267.1 acar|OOF93136.1 acla|EAW13261.1 afis|EAW17364.1 afla|EED49803.1 afum|EAL91201.1 akaw|GAA90681.1 aluc|OJZ84674.1 anid|CBF78904.1 anig|EHA27952.1 aory|EIT77591.1 asyd|OJJ64605.1 ater|EAU33870.1 atub|OJI89406.1 aver|OJI98702.1 awen|OJJ38423.1

Group6412: abra|OJJ77269.1 acar|OOF93138.1 afis|EAW17307.1 afla|EED49712.1 agla|OJJ88114.1 akaw|GAA90679.1 aluc|OJZ84676.1 anid|CBF76018.1 anig|EHA27954.1 asyd|OJJ52446.1 ater|EAU34226.1 atub|OJI89404.1 aver|OJJ03970.1 awen|OJJ29857.1 azon|OJJ43039.1 atub|OJI83508.1

Group6413: abra|OJJ77317.1 acar|OOF93193.1 acla|EAW13155.1 afis|EAW17491.1 afla|EED49325.1 afum|EAL91313.1 agla|OJJ81465.1 akaw|GAA91439.1 aluc|OJZ84727.1 anid|CBF71597.1 aory|EIT79496.1 asyd|OJJ61867.1 atub|OJI89355.1 aver|OJI95965.1 awen|OJJ37350.1 azon|OJJ51036.1

Group6414: abra|OJJ77321.1 acar|OOF93197.1 acla|EAW13151.1 afis|EAW17495.1 afum|EAL91317.1 agla|OJJ81461.1 akaw|GAA91435.1 aluc|OJZ84731.1 anid|CBF71587.1 anig|EHA28004.1 asyd|OJJ61871.1 ater|EAU36939.1 atub|OJI89351.1 aver|OJI95969.1 awen|OJJ37346.1 azon|OJJ51032.1

Group6415: abra|OJJ77351.1 acar|OOF95303.1 acla|EAW07415.1 afis|EAW19197.1 afum|EAL93929.2 akaw|GAA91410.1 aluc|OJZ84762.1 anig|EHA28031.1 asyd|OJJ57245.1 ater|EAU35862.1 atub|OJI89322.1 aver|OJJ02853.1 azon|OJJ42365.1 aver|OJJ02229.1 afis|EAW15631.1 asyd|OJJ60061.1

Group6416: abra|OJJ77367.1 acar|OOF93237.1 acla|EAW13109.1 afis|EAW17555.1 afum|EAL91369.1 agla|OJJ81413.1 akaw|GAA91395.1 aluc|OJZ84776.1 anig|EHA28044.1 aory|EIT82194.1 asyd|OJJ61928.1 ater|EAU36979.1 atub|OJI89305.1 aver|OJI96020.1 awen|OJJ37298.1 azon|OJJ50991.1

Group6417: abra|OJJ77396.1 acar|OOF93292.1 afis|EAW17804.1 afla|EED45798.1 afum|EAL91612.1 agla|OJJ80101.1 akaw|GAA91366.1 aluc|OJZ84804.1 anid|CBF86648.1 anig|EHA28069.1 aory|EIT75488.1 asyd|OJJ64800.1 ater|EAU31835.1 atub|OJI89279.1 aver|OJI98914.1 awen|OJJ31649.1

Group6418: abra|OJJ77407.1 acar|OOF93273.1 acla|EAW15350.1 afis|EAW20363.1 afla|EED48150.1 afum|EAL85479.1 akaw|GAA91358.1 aluc|OJZ84814.1 anid|CBF82412.1 anig|EHA28078.1 aory|EIT76248.1 asyd|OJJ54171.1 ater|EAU30296.1 atub|OJI89271.1 aver|OJJ08568.1 awen|OJJ33329.1

Group6419: abra|OJJ77409.1 acar|OOF93276.1 afla|EED54609.1 agla|OJJ87938.1 akaw|GAA91355.1 aluc|OJZ84817.1 aory|EIT80035.1 atub|OJI89268.1 aver|OJI98815.1 awen|OJJ36312.1 asyd|OJJ64712.1 aver|OJJ08404.1 ater|EAU39115.1 afla|EED48935.1 awen|OJJ34699.1 asyd|OJJ53341.1

Group6420: abra|OJJ77454.1 acar|OOF99272.1 acla|EAW07123.1 afis|EAW22133.1 afum|EAL91057.2 agla|OJJ87941.1 akaw|GAA84154.1 aluc|OJZ81075.1 anid|CBF69544.1 anig|EHA19287.1 asyd|OJJ59170.1 ater|EAU29319.1 atub|OJI82510.1 aver|OJJ01760.1 awen|OJJ34708.1 azon|OJJ45856.1

Group6421: abra|OJJ77505.1 acar|OOF99221.1 acla|EAW06729.1 afis|EAW16282.1 afla|EED53274.1 afum|EAL88903.1 agla|OJJ88319.1 akaw|GAA84115.1 aluc|OJZ81031.1 anig|EHA19334.1 aory|EIT75058.1 ater|EAU37326.1 atub|OJI82552.1 aver|OJI98724.1 awen|OJJ37674.1 azon|OJJ44939.1

Group6422: abra|OJJ77506.1 acar|OOF99220.1 acla|EAW06728.1 afis|EAW16281.1 afla|EED53275.1 afum|EAL88902.1 agla|OJJ88320.1 akaw|GAA84114.1 aluc|OJZ81030.1 anid|CBF86878.1 anig|EHA19335.1 aory|EIT75011.1 atub|OJI82553.1 aver|OJI98723.1 awen|OJJ37673.1 azon|OJJ44938.1

Group6423: abra|OJJ77537.1 agla|OJJ86263.1 akaw|GAA84083.1 aluc|OJZ80997.1 anig|EHA19363.1 ater|EAU37470.1 atub|OJI82587.1 awen|OJJ30196.1 acla|EAW06710.1 afis|EAW16258.1 afum|EAL88879.1 aory|EIT75045.1 asyd|OJJ64599.1 aver|OJI98696.1 azon|OJJ43930.1 anid|CBF78895.1

Group6424: abra|OJJ77538.1 acar|OOF99191.1 afis|EAW16260.1 afla|EED53324.1 afum|EAL88881.1 agla|OJJ86264.1 akaw|GAA91934.1 aluc|OJZ82688.1 anid|CBF78897.1 anig|EHA19364.1 aory|EIT75144.1 asyd|OJJ64601.1 ater|EAU33702.1 atub|OJI88920.1 aver|OJI98698.1 awen|OJJ30188.1

Group6425: abra|OJJ77553.1 acar|OOF99174.1 afis|EAW15596.1 afum|EBA27240.1 agla|OJJ81010.1 akaw|GAA84070.1 aluc|OJZ80984.1 anid|CBF85308.1 anig|EHA19376.1 asyd|OJJ64002.1 atub|OJI82602.1 aver|OJI98091.1 awen|OJJ30880.1 azon|OJJ44352.1 acla|EAW08801.1 afla|EED54704.1

Group6426: abra|OJJ77583.1 acla|EAW06550.1 afis|EAW16045.1 afla|EED56434.1 afum|EAL88677.1 akaw|GAA84042.1 aluc|OJZ80951.1 anid|CBF75056.1 anig|EHA19403.1 aory|EIT72842.1 asyd|OJJ60028.1 ater|EAU33592.1 atub|OJI82632.1 aver|OJJ02256.1 awen|OJJ30401.1 azon|OJJ45409.1

Group6427: abra|OJJ77587.1 acla|EAW06553.1 afis|EAW16042.1 afla|EED56431.1 afum|EAL88674.1 agla|OJJ86075.1 akaw|GAA84039.1 aluc|OJZ80948.1 anig|EHA19406.1 aory|EIT72832.1 asyd|OJJ60031.1 ater|EAU33595.1 atub|OJI82635.1 aver|OJJ02253.1 awen|OJJ30404.1 azon|OJJ45412.1

Group6428: abra|OJJ77589.1 acar|OOF99142.1 acla|EAW06555.1 afis|EAW16040.1 afla|EED56429.1 afum|EAL88672.1 agla|OJJ86071.1 akaw|GAA84037.1 aluc|OJZ80946.1 anid|CBF75048.1 anig|EHA19408.1 asyd|OJJ60033.1 atub|OJI82637.1 aver|OJJ02251.1 awen|OJJ30406.1 azon|OJJ45414.1

Group6429: abra|OJJ77628.1 acar|OOF99096.1 afis|EAW22624.1 afla|EED53780.1 afum|EAL90572.1 agla|OJJ86605.1 akaw|GAA84006.1 aluc|OJZ80902.1 anid|CBF84273.1 anig|EHA19437.1 aory|EIT78403.1 asyd|OJJ57224.1 ater|EAU30257.1 atub|OJI82677.1 aver|OJJ03063.1 awen|OJJ35803.1

Group6430: abra|OJJ77633.1 acar|OOF99101.1 acla|EAW06485.1 afis|EAW15980.1 afla|EED56350.1 afum|EAL88612.1 agla|OJJ85983.1 akaw|GAA84000.1 aluc|OJZ80896.1 anid|CBF85805.1 anig|EHA19443.1 asyd|OJJ64335.1 atub|OJI82682.1 aver|OJI98428.1 awen|OJJ30490.1 azon|OJJ45481.1

Group6431: abra|OJJ77723.1 acar|OOF99017.1 acla|EAW11323.1 afis|EAW18172.1 afla|EED56320.1 afum|EAL87534.1 agla|OJJ85934.1 akaw|GAA83912.1 aluc|OJZ90785.1 anid|CBF85741.1 anig|EHA19518.1 aory|EIT73350.1 ater|EAU33832.1 atub|OJI82772.1 awen|OJJ30519.1 azon|OJJ50200.1

Group6432: abra|OJJ77820.1 acar|OOF98948.1 afis|EAW18198.1 afla|EED56280.1 afum|EAL87563.1 akaw|GAA83826.1 aluc|OJZ90892.1 anid|CBF73614.1 anig|EHA19611.1 aory|EIT73308.1 asyd|OJJ61700.1 atub|OJI82886.1 aver|OJI95786.1 awen|OJJ30566.1 azon|OJJ42707.1 acar|OOF97585.1

Group6433: abra|OJJ77825.1 acar|OOF98943.1 acla|EAW11300.1 afis|EAW18202.1 afla|EED56276.1 afum|EAL87567.1 agla|OJJ85880.1 akaw|GAA83821.1 aluc|OJZ90899.1 anid|CBF86120.1 asyd|OJJ64495.1 ater|EAU33792.1 atub|OJI82892.1 aver|OJI98597.1 awen|OJJ30570.1 azon|OJJ45502.1

Group6434: abra|OJJ77836.1 acar|OOF98935.1 acla|EAW11287.1 afis|EAW18212.1 afla|EED56259.1 afum|EAL87578.1 agla|OJJ85870.1 akaw|GAA83811.1 aluc|OJZ90910.1 anid|CBF86142.1 anig|EHA19625.1 aory|EIT73408.1 asyd|OJJ64508.1 atub|OJI82903.1 awen|OJJ30590.1 azon|OJJ50186.1

Group6435: abra|OJJ77837.1 acar|OOF98934.1 afis|EAW18214.1 afla|EED56257.1 agla|OJJ85868.1 akaw|GAA83810.1 anig|EHA19626.1 aory|EIT73324.1 asyd|OJJ64509.1 ater|EAU33775.1 atub|OJI82904.1 aver|OJI98607.1 awen|OJJ30592.1 acla|EAW11285.1 afum|EAL87580.1 aluc|OJZ90911.1

Group6436: abra|OJJ77860.1 acar|OOF98913.1 acla|EAW11266.1 afis|EAW18232.1 afla|EED56234.1 afum|EAL87600.1 agla|OJJ85848.1 akaw|GAA83789.1 aluc|OJZ90934.1 anid|CBF86178.1 anig|EHA19644.1 asyd|OJJ64528.1 ater|EAU33754.1 atub|OJI82926.1 aver|OJI98626.1 awen|OJJ30612.1

Group6437: azon|OJJ48004.1 abra|OJJ77861.1 acla|EAW11265.1 afis|EAW18233.1 afla|EED56233.1 afum|EAL87601.2 agla|OJJ85849.1 akaw|GAA83788.1 aluc|OJZ90936.1 anid|CBF86180.1 aory|EIT73454.1 ater|EAU33753.1 atub|OJI82927.1 awen|OJJ30613.1 asyd|OJJ64529.1 aver|OJI98627.1

Group6438: abra|OJJ77894.1 acla|EAW10030.1 afis|EAW18250.1 afla|EED56211.1 afum|EAL87615.1 agla|OJJ85820.1 akaw|GAA83766.1 aluc|OJZ90964.1 anid|CBF86201.1 anig|EHA19668.1 aory|EIT73417.1 asyd|OJJ64543.1 atub|OJI82955.1 aver|OJI98639.1 awen|OJJ30626.1 azon|OJJ47992.1

Group6439: abra|OJJ77899.1 acar|OOF98885.1 acla|EAW11250.1 afis|EAW18247.1 afla|EED56219.1 afum|EAL87612.1 agla|OJJ85823.1 akaw|GAA83763.1 anid|CBF86196.1 aory|EIT73320.1 asyd|OJJ64540.1 ater|EAU33740.1 atub|OJI82959.1 aver|OJI98636.1 awen|OJJ30622.1 azon|OJJ47995.1

Group6440: abra|OJJ77911.1 acla|EAW06519.1 afis|EAW16237.1 afla|EED53343.1 afum|EAL88858.1 agla|OJJ86238.1 akaw|GAA83748.1 aluc|OJZ90983.1 anid|CBF81925.1 anig|EHA19688.1 asyd|OJJ57424.1 atub|OJI82974.1 aver|OJJ05753.1 awen|OJJ30223.1 azon|OJJ43943.1 ater|EAU37269.1

Group6441: abra|OJJ77920.1 acla|EAW06688.1 afis|EAW16077.1 afla|EED48597.1 afum|EAL88709.1 agla|OJJ86110.1 akaw|GAA83743.1 aluc|OJZ90988.1 anid|CBF75117.1 aory|EIT73124.1 asyd|OJJ59994.1 ater|EAU33143.1 atub|OJI82979.1 aver|OJJ02297.1 awen|OJJ30356.1 azon|OJJ45378.1

Group6442: abra|OJJ77925.1 afla|EED48702.1 akaw|GAA83739.1 aluc|OJZ90992.1 anid|CBF75954.1 anig|EHA19700.1 aory|EIT73185.1 asyd|OJJ59612.1 atub|OJI82983.1 aver|OJJ02657.1 awen|OJJ32885.1 azon|OJJ46805.1 acar|OOG00945.1 anid|CBF78736.1 asyd|OJJ59810.1 aver|OJJ02478.1

Group6443: abra|OJJ77997.1 acar|OOF98826.1 acla|EAW06651.1 afis|EAW16114.1 afla|EED48654.1 akaw|GAA85983.1 aluc|OJZ91073.1 anid|CBF85499.1 anig|EHA19776.1 aory|EIT73269.1 asyd|OJJ64117.1 ater|EAU33197.1 atub|OJI83068.1 aver|OJI98200.1 awen|OJJ30316.1 azon|OJJ45334.1

Group6444: abra|OJJ78007.1 acar|OOF98817.1 acla|EAW06643.1 afis|EAW16122.1 afla|EED48662.1 agla|OJJ86163.1 akaw|GAA85976.1 aluc|OJZ91083.1 anid|CBF73662.1 anig|EHA19784.1 asyd|OJJ56669.1 ater|EAU33205.1 atub|OJI83079.1 aver|OJJ06053.1 awen|OJJ30308.1 azon|OJJ45311.1

Group6445: abra|OJJ78020.1 acar|OOF98804.1 acla|EAW06628.1 afis|EAW16137.1 afla|EED48677.1 afum|EBA27356.1 agla|OJJ86182.1 aluc|OJZ91095.1 anid|CBF85511.1 anig|EHA19796.1 aory|EIT73251.1 asyd|OJJ64122.1 atub|OJI83093.1 aver|OJI98205.1 awen|OJJ30291.1 azon|OJJ43980.1

Group6446: abra|OJJ78084.1 acar|OOF98782.1 afis|EAW16178.1 afla|EED48695.1 afum|EAL88811.1 agla|OJJ86212.1 akaw|GAA85908.1 aluc|OJZ91158.1 anid|CBF79572.1 anig|EHA19854.1 aory|EIT73135.1 asyd|OJJ57911.1 ater|EAU33238.1 atub|OJI83173.1 aver|OJJ04481.1 awen|OJJ30271.1

Group6447: abra|OJJ78107.1 acar|OOF98758.1 afis|EAW16214.1 afla|EED48708.1 afum|EAL88841.1 agla|OJJ86221.1 akaw|GAA85899.1 aluc|OJZ91172.1 anid|CBF86724.1 anig|EHA19865.1 aory|EIT73189.1 asyd|OJJ64714.1 ater|EAU33254.1 atub|OJI83186.1 aver|OJI98817.1 awen|OJJ30263.1

Group6448: abra|OJJ78243.1 acar|OOF98617.1 acla|EAW09267.1 afis|EAW21881.1 afla|EED46427.1 afum|EAL92895.1 akaw|GAA85781.1 aluc|OJZ91317.1 anid|CBF80463.1 anig|EHA20000.1 aory|EIT72327.1 asyd|OJJ52555.1 ater|EAU30879.1 atub|OJI83311.1 aver|OJJ03846.1 awen|OJJ34170.1

Group6449: abra|OJJ78251.1 acar|OOF98613.1 acla|EAW09260.1 afis|EAW21887.1 afla|EED46420.1 afum|EAL92901.1 agla|OJJ84246.1 akaw|GAA85776.1 aluc|OJZ91324.1 anid|CBF76796.1 anig|EHA20006.1 aory|EIT72399.1 asyd|OJJ61284.1 atub|OJI83317.1 awen|OJJ34176.1 azon|OJJ43456.1

Group6450: abra|OJJ78269.1 acla|EAW09243.1 afis|EAW21904.1 afla|EED46648.1 afum|EAL92917.1 agla|OJJ84226.1 akaw|GAA85757.1 aluc|OJZ91343.1 anid|CBF76828.1 anig|EHA20023.1 aory|EIT77920.1 asyd|OJJ61300.1 atub|OJI83335.1 aver|OJJ00626.1 awen|OJJ34193.1 azon|OJJ43440.1

Group6451: abra|OJJ78297.1 acar|OOF98570.1 acla|EAW09219.1 afis|EAW21929.1 afla|EED46622.1 agla|OJJ84192.1 akaw|GAA85730.1 aluc|OJZ91371.1 anid|CBF76874.1 anig|EHA20049.1 asyd|OJJ61323.1 ater|EAU30935.1 atub|OJI83364.1 aver|OJJ00648.1 awen|OJJ34219.1 azon|OJJ43415.1

Group6452: abra|OJJ78331.1 acar|OOF98545.1 acla|EAW09196.1 afis|EAW21951.1 afum|EAL92965.1 agla|OJJ84169.1 akaw|GAA85699.1 aluc|OJZ91404.1 anid|CBF84067.1 anig|EHA20081.1 aory|EIT77813.1 asyd|OJJ61350.1 ater|EAU30999.1 atub|OJI83397.1 aver|OJJ00677.1 awen|OJJ34243.1

Group6453: abra|OJJ78348.1 acla|EAW09186.1 afis|EAW21960.1 afla|EED46680.1 afum|EAL92974.1 agla|OJJ84156.1 akaw|GAA85685.1 aluc|OJZ91422.1 anid|CBF84045.1 anig|EHA20093.1 aory|EIT77833.1 asyd|OJJ61358.1 atub|OJI83415.1 aver|OJJ00685.1 awen|OJJ34253.1 azon|OJJ43385.1

Group6454: abra|OJJ78357.1 acla|EAW09177.1 afis|EAW21969.1 afla|EED46691.1 afum|EAL92983.1 agla|OJJ84132.1 akaw|GAA85676.1 aluc|OJZ91432.1 anid|CBF82729.1 anig|EHA20101.1 aory|EIT77810.1 asyd|OJJ61629.1 ater|EAU31089.1 atub|OJI83425.1 awen|OJJ34262.1 azon|OJJ43375.1

Group6455: acar|OOF99242.1 acla|EAW06748.1 afis|EAW16298.1 afla|EED53256.1 afum|EAL88920.1 agla|OJJ88297.1 akaw|GAA84135.1 aluc|OJZ81053.1 anid|CBF86843.1 anig|EHA19314.1 asyd|OJJ64644.1 ater|EAU37342.1 atub|OJI82531.1 aver|OJI98744.1 awen|OJJ37697.1 azon|OJJ44958.1

Group6456: acar|OOF99586.1 acla|EAW14080.1 afis|EAW18787.1 afla|EED50746.1 afum|EAL93520.1 agla|OJJ80517.1 akaw|GAA82261.1 aluc|OJZ91879.1 anid|CBF69758.1 anig|EHA22857.1 asyd|OJJ58944.1 ater|EAU37919.1 atub|OJI86330.1 aver|OJJ01540.1 awen|OJJ32361.1 azon|OJJ47124.1

Group6457: acar|OOF99738.1 acla|EAW13928.1 afis|EAW18639.1 afla|EED52701.1 afum|EBA27489.1 agla|OJJ82435.1 akaw|GAA82151.1 aluc|OJZ92017.1 anid|CBF70451.1 aory|EIT74270.1 asyd|OJJ58605.1 ater|EAU38272.1 atub|OJI86452.1 aver|OJJ01209.1 awen|OJJ32107.1 azon|OJJ46688.1

Group6458: acar|OOG01106.1 acla|EAW07850.1 afis|EAW22886.1 afla|EED50173.1 afum|EAL90312.1 agla|OJJ83437.1 akaw|GAA89582.1 aluc|OJZ88065.1 anid|CBF87800.1 anig|EHA17838.1 aory|EIT72281.1 asyd|OJJ63381.1 ater|EAU35979.1 aver|OJI97554.1 awen|OJJ38836.1 azon|OJJ49633.1

Group6459: acla|EAW06283.1 afis|EAW18389.1 afla|EED53018.1 afum|EAL93064.1 agla|OJJ84952.1 akaw|GAA86349.1 aluc|OJZ84876.1 anid|CBF70774.1 aory|EIT77404.1 asyd|OJJ58426.1 ater|EAU32262.1 atub|OJI80722.1 aver|OJJ01027.1 awen|OJJ31762.1 azon|OJJ50462.1 azon|OJJ50463.1

Group6460: acla|EAW12582.1 afis|EAW24066.1 afla|EED51408.1 afum|EAL91921.1 agla|OJJ87348.1 akaw|GAA89808.1 aluc|OJZ81883.1 anid|CBF86455.1 anig|EHA22357.1 aory|EIT80581.1 asyd|OJJ64916.1 ater|EAU30489.1 atub|OJI84436.1 aver|OJI99036.1 awen|OJJ41423.1 azon|OJJ48888.1

Group6461: acla|EAW13089.1 afis|EAW17575.1 afla|EED49246.1 afum|EAL91388.1 agla|OJJ81382.1 akaw|GAA83238.1 aluc|OJZ83737.1 anid|CBF90248.1 anig|EHA18886.1 aory|EIT82188.1 asyd|OJJ61949.1 ater|EAU37011.1 atub|OJI90115.1 awen|OJJ37272.1 azon|OJJ50972.1 agla|OJJ81383.1

Group6462: abra|OJJ65466.1 abra|OJJ65661.1 abra|OJJ65708.1 akaw|GAA92342.1 akaw|GAA93087.1 aluc|OJZ79597.1 aluc|OJZ80086.1 aluc|OJZ80127.1 akaw|GAA92648.1 afis|EAW16765.1 asyd|OJJ52281.1 anig|EHA17694.1 afis|EAW22033.1 afum|EAL87734.1 aluc|OJZ80411.1

Group6463: abra|OJJ66441.1 abra|OJJ73349.1 acar|OOF91488.1 acar|OOG00561.1 aver|OJJ00756.1 aver|OJJ01680.1 acla|EAW08966.1 afla|EED56166.1 agla|OJJ89199.1 aluc|OJZ88690.1 anig|EHA18371.1 aory|EIT76753.1 asyd|OJJ59090.1 atub|OJI91270.1 ater|EAU31722.1

Group6464: abra|OJJ68104.1 abra|OJJ72188.1 akaw|GAA84745.1 akaw|GAA86888.1 aluc|OJZ86939.1 aluc|OJZ90012.1 atub|OJI83583.1 atub|OJI84795.1 acar|OOF96814.1 afla|EED48887.1 anig|EHA21705.1 anig|EHA27387.1 aory|EIT81215.1 awen|OJJ34661.1 anid|CBF79549.1

Group6465: abra|OJJ68870.1 abra|OJJ74824.1 afis|EAW17098.1 afla|EED49690.1 afum|EAL86572.1 akaw|GAA90384.1 aluc|OJZ85847.1 anig|EHA18384.1 aory|EIT80999.1 ater|EAU34158.1 atub|OJI79635.1 aver|OJJ03074.1 awen|OJJ38320.1 azon|OJJ43674.1 anid|CBF84320.1

Group6466: abra|OJJ69717.1 abra|OJJ74254.1 aluc|OJZ83180.1 aluc|OJZ89878.1 anig|EHA27077.1 anig|EHA28254.1 atub|OJI84929.1 atub|OJI87757.1 acar|OOF95474.1 afis|EAW17024.1 akaw|GAA86767.1 akaw|GAA90170.1 anid|CBF76231.1 asyd|OJJ53104.1 aver|OJJ03748.1

Group6467: abra|OJJ71057.1 abra|OJJ75794.1 awen|OJJ29799.1 awen|OJJ30198.1 awen|OJJ33368.1 awen|OJJ34646.1 acar|OOG00490.1 acla|EAW09127.1 afla|EED46258.1 agla|OJJ88373.1 anig|EHA23324.1 aory|EIT76517.1 ater|EAU29446.1 atub|OJI85555.1 acla|EAW13548.1

Group6468: acar|OOF90492.1 acar|OOF90500.1 abra|OJJ65600.1 abra|OJJ72519.1 afum|EAL85383.2 agla|OJJ80044.1 akaw|GAA92639.1 aluc|OJZ90500.1 abra|OJJ77804.1 agla|OJJ87320.1 ater|EAU37954.1 atub|OJI84682.1 awen|OJJ39106.1 acar|OOF98428.1 agla|OJJ79450.1

Group6469: acar|OOF92285.1 acar|OOF92326.1 abra|OJJ67517.1 acla|EAW06256.1 afis|EAW18363.1 afla|EED53042.1 afum|EAL93036.1 akaw|GAA86323.1 aluc|OJZ84847.1 anid|CBF70823.1 aory|EIT77435.1 asyd|OJJ58400.1 ater|EAU32231.1 aver|OJJ01001.1 azon|OJJ50436.1

Group6470: acar|OOF92321.1 acar|OOF92363.1 abra|OJJ67477.1 acla|EAW06295.1 afis|EAW18401.1 afla|EED53005.1 afum|EAL93079.1 agla|OJJ84972.1 akaw|GAA86361.1 anid|CBF70747.1 aory|EIT77410.1 ater|EAU29817.1 atub|OJI80710.1 awen|OJJ31748.1 azon|OJJ50506.1

Group6471: acar|OOF93680.1 acar|OOF93766.1 abra|OJJ66665.1 acla|EAW11806.1 afis|EAW19416.1 afla|EED50372.1 afum|EAL87078.1 akaw|GAA89614.1 aluc|OJZ80782.1 anig|EHA27247.1 aory|EIT83137.1 ater|EAU34722.1 atub|OJI87634.1 awen|OJJ36813.1 azon|OJJ42988.1

Group6472: acar|OOF93740.1 acar|OOF93830.1 abra|OJJ66745.1 acla|EAW11873.1 afis|EAW19484.1 afla|EED56700.1 afum|EAL87011.1 agla|OJJ81209.1 akaw|GAA89682.1 aluc|OJZ80703.1 anid|CBF75462.1 aory|EIT73938.1 ater|EAU34797.1 aver|OJJ02519.1 awen|OJJ36896.1

Group6473: acar|OOF93973.1 acar|OOF94546.1 akaw|GAA91790.1 akaw|GAA92547.1 aluc|OJZ90096.1 aluc|OJZ92810.1 abra|OJJ68400.1 afla|EED51092.1 anig|EHA20980.1 anig|EHA26198.1 aory|EIT81996.1 atub|OJI84710.1 atub|OJI87267.1 abra|OJJ75251.1 azon|OJJ42265.1

Group6474: acar|OOF94044.1 acar|OOF94114.1 abra|OJJ66397.1 acla|EAW13678.1 afis|EAW19927.1 afum|EAL84982.2 akaw|GAA90265.1 aluc|OJZ83076.1 anid|CBF88380.1 anig|EHA28140.1 asyd|OJJ63736.1 ater|EAU32192.1 atub|OJI87870.1 aver|OJI97204.1 awen|OJJ32244.1

Group6475: acar|OOF95659.1 acar|OOF95710.1 abra|OJJ74700.1 acla|EAW10699.1 afis|EAW24871.1 afla|EED47111.1 afum|EAL89521.1 agla|OJJ82092.1 akaw|GAA81918.1 aluc|OJZ92278.1 anid|CBF75796.1 anig|EHA26686.1 aory|EIT79276.1 atub|OJI86711.1 awen|OJJ40462.1

Group6476: acar|OOF96225.1 acar|OOF96263.1 abra|OJJ71510.1 acla|EAW15068.1 afis|EAW20040.1 afla|EED45340.1 afum|EAL85160.1 agla|OJJ88923.1 aluc|OJZ86248.1 anid|CBF84780.1 anig|EHA21616.1 asyd|OJJ63235.1 atub|OJI84294.1 aver|OJI97721.1 awen|OJJ31795.1

Group6477: acar|OOF96230.1 acar|OOF96268.1 abra|OJJ71514.1 acla|EAW15071.1 afis|EAW20043.1 afum|EAL85163.1 agla|OJJ88919.1 akaw|GAA89402.1 aluc|OJZ86252.1 anid|CBF84788.1 anig|EHA21613.1 asyd|OJJ63231.1 aver|OJI97725.1 awen|OJJ31799.1 azon|OJJ46320.1

Group6478: acar|OOF96238.1 acar|OOF96275.1 abra|OJJ71522.1 acla|EAW15079.1 afis|EAW20051.1 afla|EED45329.1 agla|OJJ88909.1 akaw|GAA89395.1 aluc|OJZ86261.1 anid|CBF84804.1 asyd|OJJ63223.1 atub|OJI84282.1 aver|OJI97732.1 awen|OJJ31809.1 azon|OJJ46313.1

Group6479: acar|OOF96258.1 acar|OOF96297.1 abra|OJJ71553.1 acla|EAW15107.1 afum|EAL85197.1 agla|OJJ88881.1 akaw|GAA89369.1 aluc|OJZ86291.1 anid|CBF84860.1 anig|EHA21580.1 asyd|OJJ63196.1 atub|OJI84247.1 aver|OJI97759.1 awen|OJJ31838.1 azon|OJJ46285.1

Group6480: acar|OOF96904.1 acar|OOF96939.1 abra|OJJ70806.1 acla|EAW11118.1 afis|EAW15848.1 afum|EAL88484.1 agla|OJJ87261.1 aluc|OJZ84152.1 anid|CBF81168.1 anig|EHA19256.1 asyd|OJJ59288.1 atub|OJI90524.1 aver|OJJ02987.1 awen|OJJ36780.1 azon|OJJ43019.1

Group6481: acar|OOF96907.1 acar|OOF96941.1 abra|OJJ70804.1 acla|EAW11120.1 afum|EAL88485.1 agla|OJJ87259.1 aluc|OJZ84150.1 anid|CBF81172.1 anig|EHA19254.1 aory|EIT72856.1 asyd|OJJ59290.1 atub|OJI90522.1 aver|OJJ02985.1 awen|OJJ36778.1 azon|OJJ47420.1

Group6482: acar|OOG00256.1 acar|OOG01224.1 abra|OJJ67533.1 afis|EAW16788.1 afla|EED52192.1 afum|EAL86358.1 agla|OJJ84235.1 akaw|GAA88125.1 aluc|OJZ81398.1 anig|EHA18690.1 aory|EIT74382.1 ater|EAU37169.1 atub|OJI91603.1 aver|OJJ05093.1 awen|OJJ31292.1

Group6483: afla|EED46070.1 afla|EED48411.1 agla|OJJ79282.1 agla|OJJ86339.1 aory|EIT83061.1 aory|EIT83495.1 ater|EAU32179.1 ater|EAU38631.1 awen|OJJ36035.1 awen|OJJ38326.1 afis|EAW20219.1 afum|EAL85334.1 anid|CBF82403.1 asyd|OJJ54162.1 aver|OJJ08586.1

Group6484: afla|EED46467.1 afla|EED50409.1 abra|OJJ68297.1 acar|OOF97564.1 acla|EAW08963.1 afis|EAW15444.1 afum|EAL84203.1 akaw|GAA83176.1 aluc|OJZ83675.1 anig|EHA18831.1 aory|EIT72404.1 atub|OJI90046.1 awen|OJJ37059.1 azon|OJJ50216.1 aory|EIT82156.1

Group6485: afla|EED49603.1 ater|EAU30205.1 awen|OJJ41173.1 abra|OJJ68053.1 afis|EAW20390.1 akaw|GAA90902.1 aluc|OJZ82924.1 anig|EHA21347.1 atub|OJI89153.1 afis|EAW20394.1 atub|OJI84686.1 agla|OJJ84018.1 anid|CBF76038.1 asyd|OJJ52457.1 aver|OJJ03956.1

Group6486: afla|EED52142.1 afla|EED54470.1 akaw|GAA85879.1 akaw|GAA91949.1 aory|EIT74353.1 aory|EIT81140.1 awen|OJJ31330.1 awen|OJJ35028.1 abra|OJJ78128.1 afis|EAW17305.1 agla|OJJ82856.1 aluc|OJZ91190.1 anig|EHA19886.1 ater|EAU30276.1 atub|OJI83209.1

Group6487: agla|OJJ78553.1 agla|OJJ79588.1 aory|EIT75808.1 aory|EIT77002.1 abra|OJJ69870.1 acar|OOF90338.1 afla|EED50977.1 akaw|GAA87786.1 aluc|OJZ85421.1 anig|EHA25600.1 atub|OJI80154.1 awen|OJJ34596.1 asyd|OJJ52487.1 afla|EED45617.1 aver|OJJ03917.1

Group6488: agla|OJJ80656.1 agla|OJJ88730.1 abra|OJJ76929.1 acar|OOF91375.1 afis|EAW20552.1 afla|EED52399.1 afum|EBA27185.1 akaw|GAA87111.1 aluc|OJZ84327.1 anid|CBF78979.1 aory|EIT83293.1 asyd|OJJ55676.1 atub|OJI89741.1 aver|OJJ06759.1 azon|OJJ50639.1

Group6489: agla|OJJ86274.1 agla|OJJ86315.1 abra|OJJ66362.1 acar|OOF91802.1 acla|EAW09126.1 afis|EAW19349.1 afla|EED54384.1 afum|EAL94069.1 akaw|GAA90235.1 aluc|OJZ83106.1 anig|EHA28169.1 aory|EIT75885.1 ater|EAU34937.1 atub|OJI87828.1 awen|OJJ41007.1

Group6490: agla|OJJ87321.1 agla|OJJ88357.1 ater|EAU31741.1 ater|EAU34065.1 abra|OJJ67564.1 acar|OOG00282.1 afla|EED46126.1 akaw|GAA88101.1 aluc|OJZ81368.1 anig|EHA18667.1 aory|EIT76425.1 asyd|OJJ62818.1 atub|OJI91573.1 aver|OJI96862.1 awen|OJJ32677.1

Group6491: akaw|GAA86876.1 akaw|GAA91587.1 aluc|OJZ82410.1 aluc|OJZ90000.1 abra|OJJ68091.1 acar|OOF94969.1 anig|EHA27375.1 asyd|OJJ56861.1 ater|EAU37871.1 atub|OJI84808.1 aver|OJJ01691.1 afla|EED47244.1 anid|CBF77165.1 atub|OJI88031.1 asyd|OJJ59100.1

Group6492: akaw|GAA89006.1 akaw|GAA92332.1 akaw|GAA93318.1 akaw|GAA93178.1 aluc|OJZ79789.1 aluc|OJZ80052.1 aluc|OJZ80373.1 aluc|OJZ80417.1 aluc|OJZ80292.1 acla|EAW11465.1 agla|OJJ88685.1 anig|EHA17704.1 ater|EAU30126.1 agla|OJJ78803.1 akaw|GAA93248.1

Group6493: akaw|GAA91886.1 akaw|GAA92573.1 aluc|OJZ79607.1 aluc|OJZ82011.1 aluc|OJZ82584.1 anig|EHA22246.1 anig|EHA27489.1 azon|OJJ42325.1 azon|OJJ43504.1 azon|OJJ47559.1 abra|OJJ67879.1 acar|OOF92855.1 agla|OJJ81435.1 atub|OJI84560.1 awen|OJJ39344.1

Group6494: anig|EHA20955.1 ater|EAU32742.1 aver|OJJ08257.1 awen|OJJ32580.1 aluc|OJZ92195.1 acar|OOF93601.1 awen|OJJ33281.1 afis|EAW19749.1 afis|EAW17510.1 afla|EED47902.1 akaw|GAA81994.1 aory|EIT77215.1 asyd|OJJ61760.1 atub|OJI87954.1 afum|EAL85149.1

Group6495: anig|EHA21432.1 anig|EHA24295.1 abra|OJJ70060.1 agla|OJJ82971.1 anid|CBF88858.1 asyd|OJJ58687.1 ater|EAU39309.1 aver|OJJ01298.1 azon|OJJ43584.1 abra|OJJ71728.1 akaw|GAA85165.1 aluc|OJZ86471.1 atub|OJI84058.1 awen|OJJ36417.1 asyd|OJJ52531.1

Group6496: anig|EHA22579.1 anig|EHA26969.1 aver|OJI95836.1 aver|OJJ07608.1 acla|EAW07588.1 afis|EAW24582.1 afum|EAL91344.1 agla|OJJ84347.1 akaw|GAA93001.1 aluc|OJZ91542.1 anid|CBF71545.1 asyd|OJJ61739.1 ater|EAU37849.1 atub|OJI85996.1 azon|OJJ42379.1

Group6497: asyd|OJJ51930.1 asyd|OJJ54851.1 asyd|OJJ59010.1 aver|OJJ01605.1 aver|OJJ01789.1 abra|OJJ77438.1 acla|EAW13587.1 afis|EAW20015.1 afum|EAL84903.1 anid|CBF82909.1 anig|EHA26145.1 aory|EIT80686.1 ater|EAU29375.1 awen|OJJ30140.1 afla|EED51517.1

Group6498: asyd|OJJ55952.1 asyd|OJJ56682.1 abra|OJJ66112.1 acar|OOF94007.1 afis|EAW20771.1 afla|EED51742.1 afum|EAL84848.1 akaw|GAA92676.1 aluc|OJZ90131.1 anid|CBF78155.1 anig|EHA20940.1 aory|EIT80531.1 ater|EAU30761.1 atub|OJI84665.1 aver|OJJ04898.1

Group6499: asyd|OJJ58820.1 asyd|OJJ65390.1 abra|OJJ76257.1 acar|OOF99502.1 afla|EED50967.1 agla|OJJ85266.1 akaw|GAA82191.1 aluc|OJZ91969.1 anid|CBF70114.1 anig|EHA22921.1 aory|EIT76962.1 ater|EAU39425.1 atub|OJI86408.1 aver|OJJ01424.1 awen|OJJ31537.1

Group6500: aver|OJJ07741.1 aver|OJJ07747.1 abra|OJJ75456.1 acar|OOF90801.1 acla|EAW14623.1 afla|EED51516.1 afum|EAL91245.1 akaw|GAA84231.1 aluc|OJZ88912.1 anid|CBF78441.1 anig|EHA23598.1 aory|EIT80439.1 ater|EAU30772.1 atub|OJI85879.1 awen|OJJ35595.1

Group6501: awen|OJJ30745.1 awen|OJJ31736.1 abra|OJJ67463.1 acar|OOF92369.1 acla|EAW06305.1 afis|EAW18414.1 afla|EED52995.1 afum|EAL93095.1 aluc|OJZ84902.1 anid|CBF70732.1 aory|EIT77392.1 asyd|OJJ58444.1 atub|OJI80696.1 aver|OJJ01044.1 azon|OJJ50495.1

Group6502: awen|OJJ30746.1 awen|OJJ31735.1 abra|OJJ67462.1 acar|OOF92370.1 acla|EAW06306.1 afis|EAW18415.1 afla|EED52994.1 afum|EAL93097.1 akaw|GAA86373.1 aluc|OJZ84903.1 anig|EHA25128.1 ater|EAU29829.1 atub|OJI80695.1 agla|OJJ84981.1 aory|EIT77412.1

Group6503: awen|OJJ30780.1 awen|OJJ31701.1 abra|OJJ74544.1 acar|OOF95214.1 afis|EAW18924.1 afum|EBA27499.1 agla|OJJ86350.1 akaw|GAA82054.1 aluc|OJZ92126.1 anig|EHA26822.1 aory|EIT79041.1 atub|OJI86558.1 agla|OJJ85251.1 afla|EED50911.1 ater|EAU38099.1

Group6504: awen|OJJ32533.1 awen|OJJ36341.1 abra|OJJ65873.1 acar|OOF92851.1 acla|EAW08930.1 afla|EED51543.1 akaw|GAA92671.1 aluc|OJZ90126.1 anig|EHA20945.1 aory|EIT80758.1 ater|EAU31302.1 atub|OJI84670.1 azon|OJJ42135.1 acla|EAW09998.1 ater|EAU36341.1

Group6505: awen|OJJ33603.1 awen|OJJ38321.1 abra|OJJ65909.1 acar|OOF90094.1 afis|EAW16174.1 afum|EAL88806.1 akaw|GAA92323.1 aluc|OJZ90221.1 anid|CBF70181.1 anig|EHA23719.1 ater|EAU32517.1 atub|OJI81653.1 aver|OJJ03072.1 afla|EED53604.1 aory|EIT75012.1

Group6506: awen|OJJ36041.1 awen|OJJ37946.1 abra|OJJ74165.1 acar|OOF95557.1 afla|EED49699.1 agla|OJJ88220.1 akaw|GAA86851.1 aluc|OJZ89971.1 anid|CBF82398.1 anig|EHA27169.1 aory|EIT77647.1 asyd|OJJ54159.1 ater|EAU34030.1 atub|OJI84836.1 aver|OJJ08583.1

Group6507: awen|OJJ36383.1 awen|OJJ37566.1 abra|OJJ69745.1 acar|OOF94177.1 acla|EAW09984.1 afis|EAW20374.1 afla|EED51712.1 afum|EAL85488.1 agla|OJJ79481.1 akaw|GAA90197.1 aluc|OJZ83151.1 anig|EHA28217.1 aory|EIT80472.1 ater|EAU36791.1 atub|OJI87786.1

Group6508: azon|OJJ43178.1 azon|OJJ45073.1 azon|OJJ50086.1 azon|OJJ48050.1 azon|OJJ45124.1 anid|CBF87498.1 afla|EED47862.1 aory|EIT77241.1 asyd|OJJ52968.1 aver|OJJ08934.1 awen|OJJ36411.1 aluc|OJZ87024.1 atub|OJI83494.1 ater|EAU34445.1 aver|OJI98277.1

Group6509: azon|OJJ43505.1 azon|OJJ43545.1 acar|OOF90108.1 acla|EAW11253.1 afis|EAW18244.1 afla|EED55184.1 anid|CBF84910.1 aory|EIT81586.1 asyd|OJJ52868.1 aver|OJJ01275.1 afum|EAL87609.1 anid|CBF85151.1 asyd|OJJ65141.1 aver|OJJ08849.1 asyd|OJJ58666.1

Group6510: azon|OJJ45159.1 azon|OJJ47882.1 abra|OJJ71798.1 acar|OOF96478.1 afis|EAW18326.1 afla|EED51024.1 afum|EAL87692.1 agla|OJJ87481.1 akaw|GAA85112.1 aluc|OJZ86528.1 anig|EHA22118.1 aory|EIT77030.1 ater|EAU33335.1 atub|OJI83993.1 awen|OJJ36577.1

Group6511: abra|OJJ65517.1 afla|EED56699.1 abra|OJJ66119.1 acar|OOF94011.1 afis|EAW22074.1 afum|EAL91096.1 akaw|GAA92681.1 aluc|OJZ90136.1 anid|CBF82786.1 anig|EHA20930.1 asyd|OJJ57365.1 ater|EAU30144.1 atub|OJI84660.1 aver|OJJ05802.1 azon|OJJ47362.1

Group6512: abra|OJJ65776.1 akaw|GAA90886.1 aluc|OJZ82945.1 anig|EHA21363.1 atub|OJI89177.1 abra|OJJ66450.1 afis|EAW15392.1 anid|CBF76217.1 abra|OJJ72699.1 afla|EED45915.1 agla|OJJ79095.1 aory|EIT75728.1 asyd|OJJ54672.1 aver|OJJ07498.1 awen|OJJ30437.1

Group6513: abra|OJJ65905.1 acar|OOF90096.1 afis|EAW24555.1 afla|EED53606.1 afum|EAL89025.2 agla|OJJ88982.1 akaw|GAA92326.1 aluc|OJZ90224.1 anid|CBF86676.1 anig|EHA23725.1 aory|EIT75186.1 ater|EAU29646.1 atub|OJI81657.1 aver|OJJ01776.1 aver|OJJ03073.1

Group6514: abra|OJJ65923.1 acar|OOF94984.1 afis|EAW21278.1 afla|EED53912.1 afum|EAL92293.2 agla|OJJ79460.1 akaw|GAA92310.1 aluc|OJZ90208.1 anig|EHA23705.1 aory|EIT78500.1 ater|EAU32848.1 atub|OJI81638.1 awen|OJJ41848.1 agla|OJJ82035.1 ater|EAU31598.1

Group6515: abra|OJJ65973.1 acar|OOF92899.1 afis|EAW20010.1 afla|EED46016.1 afum|EAL84908.1 agla|OJJ87975.1 akaw|GAA90847.1 aluc|OJZ82986.1 anid|CBF84711.1 anig|EHA21400.1 asyd|OJJ53324.1 ater|EAU36307.1 atub|OJI89220.1 aver|OJJ08419.1 awen|OJJ30023.1

Group6516: abra|OJJ66055.1 acla|EAW09050.1 afla|EED54364.1 agla|OJJ85926.1 akaw|GAA93125.1 aluc|OJZ81109.1 anid|CBF80067.1 anig|EHA19260.1 aory|EIT75951.1 asyd|OJJ52547.1 ater|EAU29733.1 atub|OJI82470.1 aver|OJJ03857.1 awen|OJJ35036.1 awen|OJJ41945.1

Group6517: abra|OJJ66076.1 acar|OOF99298.1 afis|EAW19550.1 afum|EAL86959.1 akaw|GAA84170.1 aluc|OJZ81095.1 anig|EHA19276.1 ater|EAU31596.1 atub|OJI82487.1 abra|OJJ74744.1 afla|EED45803.1 agla|OJJ84420.1 anig|EHA26644.1 aory|EIT75509.1 awen|OJJ37997.1

Group6518: abra|OJJ66152.1 aory|EIT75964.1 ater|EAU35158.1 abra|OJJ69079.1 acar|OOF91212.1 afis|EAW15681.1 afum|EAL85667.1 akaw|GAA90608.1 aluc|OJZ86067.1 anig|EHA24547.1 asyd|OJJ59217.1 ater|EAU32481.1 atub|OJI79863.1 aver|OJJ01827.1 awen|OJJ38371.1

Group6519: abra|OJJ66216.1 acar|OOF90654.1 afis|EAW17334.1 afla|EED44865.1 afum|EAL91177.1 agla|OJJ81057.1 akaw|GAA89070.1 aluc|OJZ80522.1 anid|CBF87474.1 anig|EHA25673.1 aory|EIT82732.1 asyd|OJJ52940.1 atub|OJI88408.1 aver|OJJ08907.1 awen|OJJ35981.1

Group6520: abra|OJJ66232.1 acar|OOF90690.1 afis|EAW19974.1 afla|EED48919.1 agla|OJJ84412.1 akaw|GAA89057.1 aluc|OJZ80537.1 anig|EHA25658.1 aory|EIT81204.1 asyd|OJJ64302.1 atub|OJI88394.1 aver|OJI98395.1 awen|OJJ40795.1 azon|OJJ51819.1 agla|OJJ88997.1

Group6521: abra|OJJ66371.1 acar|OOF90878.1 acla|EAW12441.1 afis|EAW16752.1 afum|EAL86324.1 akaw|GAA90240.1 aluc|OJZ83100.1 anig|EHA28164.1 asyd|OJJ52537.1 ater|EAU35858.1 atub|OJI87835.1 aver|OJJ03866.1 awen|OJJ40150.1 awen|OJJ36212.1 ater|EAU35156.1

Group6522: abra|OJJ66401.1 acla|EAW11050.1 afis|EAW17509.1 afla|EED49633.1 afum|EAL88818.2 akaw|GAA90269.1 aluc|OJZ83072.1 anig|EHA28136.1 aory|EIT81977.1 ater|EAU36748.1 atub|OJI87874.1 azon|OJJ45944.1 aory|EIT80973.1 afis|EAW16187.1 afla|EED55032.1

Group6523: afis|EAW24579.1 acar|OOF93376.1 acla|EAW11104.1 afum|EBA27360.1 agla|OJJ80466.1 akaw|GAA91092.1 aluc|OJZ87912.1 anid|CBF87522.1 asyd|OJJ54643.1 ater|EAU33006.1 aver|OJJ07515.1 acla|EAW07585.1 aory|EIT79515.1 ater|EAU37171.1 anid|CBF80610.1

Group6524: abra|OJJ66411.1 afis|EAW19775.1 afla|EED45892.1 afum|EAL85120.1 agla|OJJ89264.1 akaw|GAA90276.1 aluc|OJZ83065.1 anid|CBF83011.1 anig|EHA28130.1 aory|EIT75647.1 asyd|OJJ56089.1 ater|EAU36326.1 aver|OJJ05032.1 azon|OJJ48537.1 aory|EIT75470.1

Group6525: abra|OJJ66535.1 acar|OOF90985.1 acla|EAW10956.1 afis|EAW20902.1 afum|EAL84729.2 agla|OJJ84019.1 akaw|GAA91180.1 aluc|OJZ81523.1 anid|CBF82466.1 anig|EHA25921.1 asyd|OJJ54097.1 ater|EAU38535.1 atub|OJI88672.1 aver|OJJ08642.1 awen|OJJ34446.1

Group6526: afis|EAW25736.1 agla|OJJ84593.1 awen|OJJ40855.1 agla|OJJ80566.1 acla|EAW07659.1 afla|EED45769.1 akaw|GAA84843.1 aluc|OJZ86828.1 anig|EHA21809.1 aory|EIT75398.1 asyd|OJJ57996.1 atub|OJI83690.1 aver|OJJ04406.1 azon|OJJ44444.1 aory|EIT75446.1

Group6527: abra|OJJ66574.1 acar|OOF91015.1 acla|EAW10916.1 afis|EAW20942.1 afum|EAL84692.1 agla|OJJ83945.1 akaw|GAA91144.1 aluc|OJZ81470.1 anid|CBF82536.1 anig|EHA25957.1 asyd|OJJ54068.1 atub|OJI88708.1 aver|OJJ08674.1 awen|OJJ34374.1 azon|OJJ48110.1

Group6528: abra|OJJ66631.1 acar|OOF91055.1 acla|EAW10865.1 afis|EAW20993.1 afum|EAL84635.1 agla|OJJ84071.1 aluc|OJZ81418.1 anid|CBF82613.1 anig|EHA26003.1 asyd|OJJ54016.1 ater|EAU38642.1 atub|OJI88763.1 aver|OJJ08725.1 awen|OJJ34317.1 azon|OJJ43331.1

Group6529: abra|OJJ66638.1 akaw|GAA89591.1 aluc|OJZ80811.1 anig|EHA27223.1 atub|OJI87659.1 acla|EAW11781.1 afis|EAW19391.1 afla|EED50349.1 afum|EAL87103.1 agla|OJJ81102.1 anid|CBF74287.1 ater|EAU36054.1 aver|OJJ05355.1 azon|OJJ43014.1 awen|OJJ36784.1

Group6530: abra|OJJ66670.1 acla|EAW11811.1 afis|EAW19421.1 afla|EED50378.1 afum|EAL87073.1 agla|OJJ81133.1 akaw|GAA89619.1 aluc|OJZ80777.1 anid|CBF74371.1 anig|EHA27252.1 asyd|OJJ57782.1 atub|OJI87629.1 aver|OJJ05383.1 awen|OJJ36819.1 azon|OJJ42983.1

Group6531: abra|OJJ66677.1 acla|EAW11816.1 afis|EAW19427.1 afla|EED54172.1 afum|EAL87067.1 agla|OJJ81139.1 aluc|OJZ80771.1 anid|CBF74380.1 aory|EIT76134.1 asyd|OJJ57776.1 atub|OJI87623.1 aver|OJJ05389.1 awen|OJJ36825.1 azon|OJJ42978.1 afis|EAW20339.1

Group6532: abra|OJJ66738.1 acar|OOF93824.1 afis|EAW19476.1 afum|EAL87020.1 agla|OJJ81199.1 akaw|GAA89676.1 aluc|OJZ80710.1 anig|EHA27307.1 ater|EAU34788.1 atub|OJI87568.1 acla|EAW11865.1 afla|EED56684.1 asyd|OJJ58316.1 aver|OJJ02095.1 awen|OJJ36881.1

Group6533: abra|OJJ66808.1 afla|EED53419.1 agla|OJJ85967.1 anid|CBF75004.1 anig|EHA24972.1 aory|EIT73383.1 asyd|OJJ57602.1 ater|EAU33737.1 atub|OJI79334.1 aver|OJJ05582.1 awen|OJJ32727.1 aory|EIT75066.1 afla|EED56295.1 atub|OJI89931.1 aver|OJJ03885.1

Group6534: abra|OJJ66863.1 acar|OOF91639.1 acla|EAW14439.1 afis|EAW19147.1 afla|EED54266.1 afum|EAL93878.1 agla|OJJ88577.1 akaw|GAA91545.1 aluc|OJZ85720.1 anig|EHA24825.1 asyd|OJJ65003.1 atub|OJI79509.1 aver|OJI99122.1 awen|OJJ37500.1 azon|OJJ51165.1

Group6535: abra|OJJ66898.1 acar|OOF91670.1 afis|EAW19178.1 afum|EAL93908.1 agla|OJJ88545.1 akaw|GAA91514.1 aluc|OJZ85684.1 anig|EHA24857.1 aory|EIT75923.1 asyd|OJJ65034.1 ater|EAU37295.1 atub|OJI79474.1 aver|OJI99153.1 awen|OJJ37532.1 azon|OJJ51198.1

Group6536: abra|OJJ66936.1 acar|OOF91702.1 acla|EAW15006.1 afla|EED47701.1 akaw|GAA91484.1 aluc|OJZ85646.1 anid|CBF78584.1 anig|EHA24889.1 asyd|OJJ65080.1 ater|EAU37257.1 atub|OJI79437.1 aver|OJI99205.1 awen|OJJ38054.1 azon|OJJ50617.1 agla|OJJ86875.1

Group6537: abra|OJJ66949.1 acar|OOF91715.1 akaw|GAA91471.1 aluc|OJZ85631.1 anig|EHA24900.1 atub|OJI79421.1 abra|OJJ68287.1 afla|EED57470.1 akaw|GAA83185.1 aluc|OJZ83684.1 anig|EHA18838.1 aory|EIT80135.1 ater|EAU37317.1 atub|OJI90055.1 azon|OJJ45779.1

Group6538: abra|OJJ66981.1 acar|OOF93630.1 acla|EAW15027.1 afis|EAW24557.1 afla|EED52331.1 afum|EAL84825.1 akaw|GAA92183.1 aluc|OJZ85593.1 anid|CBF89361.1 anig|EHA24929.1 aory|EIT74464.1 asyd|OJJ54577.1 atub|OJI79389.1 aver|OJJ07577.1 afis|EAW20803.1

Group6539: abra|OJJ66982.1 acla|EAW14291.1 afis|EAW24717.1 afla|EED52778.1 agla|OJJ82980.1 akaw|GAA92182.1 aluc|OJZ85592.1 anid|CBF78277.1 asyd|OJJ52419.1 atub|OJI79388.1 aver|OJJ03364.1 awen|OJJ40136.1 azon|OJJ47314.1 afis|EAW18992.1 awen|OJJ31614.1

Group6540: abra|OJJ66983.1 acar|OOF93629.1 acla|EAW06325.1 afis|EAW18439.1 afla|EED52974.1 afum|EAL93121.2 akaw|GAA92181.1 aluc|OJZ85591.1 anid|CBF71469.1 anig|EHA24930.1 aory|EIT78681.1 asyd|OJJ54946.1 ater|EAU37196.1 atub|OJI79387.1 aver|OJJ07249.1

Group6541: abra|OJJ66984.1 acar|OOF93628.1 afis|EAW15610.1 afla|EED44877.1 afum|EAL85724.1 agla|OJJ89447.1 akaw|GAA92180.1 aluc|OJZ85590.1 anid|CBF79935.1 anig|EHA24931.1 aory|EIT83254.1 asyd|OJJ58208.1 atub|OJI79385.1 aver|OJJ02214.1 awen|OJJ35715.1

Group6542: abra|OJJ67004.1 acar|OOF93610.1 acla|EAW12184.1 afis|EAW17119.1 afla|EED48925.1 afum|EAL86581.1 akaw|GAA92163.1 aluc|OJZ85570.1 anid|CBF79544.1 anig|EHA24948.1 aory|EIT80049.1 ater|EAU38948.1 atub|OJI79365.1 awen|OJJ36385.1 azon|OJJ42810.1

Group6543: acla|EAW14794.1 afla|EED52852.1 afum|EAL85995.1 anig|EHA18938.1 asyd|OJJ53869.1 acar|OOF94319.1 afis|EAW24325.1 afla|EED47941.1 agla|OJJ83192.1 anid|CBF74058.1 aory|EIT79825.1 ater|EAU30428.1 aver|OJJ07967.1 awen|OJJ32969.1 azon|OJJ48948.1

Group6544: abra|OJJ67102.1 acar|OOF98450.1 afis|EAW20275.1 afla|EED48127.1 afum|EAL85409.1 agla|OJJ87506.1 akaw|GAA91059.1 aluc|OJZ87871.1 anid|CBF78244.1 anig|EHA20177.1 aory|EIT76225.1 asyd|OJJ56195.1 ater|EAU31784.1 atub|OJI80778.1 aver|OJJ05122.1

Group6545: abra|OJJ67106.1 acar|OOF98445.1 acla|EAW14949.1 afis|EAW24466.1 afla|EED55055.1 afum|EAL86137.2 agla|OJJ84375.1 akaw|GAA91057.1 aluc|OJZ87868.1 anid|CBF78857.1 anig|EHA20174.1 aory|EIT81943.1 asyd|OJJ64569.1 atub|OJI80780.1 aver|OJI98666.1

Group6546: abra|OJJ67113.1 acar|OOF98439.1 afis|EAW20325.1 afla|EED47501.1 afum|EAL85450.1 akaw|GAA91051.1 aluc|OJZ87862.1 anid|CBF82806.1 anig|EHA20168.1 aory|EIT74564.1 asyd|OJJ65274.1 atub|OJI80786.1 aver|OJI99426.1 awen|OJJ38373.1 azon|OJJ47727.1

Group6547: abra|OJJ67117.1 acar|OOF98435.1 afla|EED57082.1 agla|OJJ86026.1 akaw|GAA91048.1 aluc|OJZ87858.1 anid|CBF84004.1 anig|EHA20164.1 aory|EIT82248.1 asyd|OJJ63809.1 ater|EAU31388.1 atub|OJI80789.1 aver|OJI97128.1 awen|OJJ31653.1 azon|OJJ42556.1

Group6548: abra|OJJ67125.1 acar|OOF98427.1 acla|EAW13223.1 afum|EAL92064.1 akaw|GAA91042.1 aluc|OJZ87852.1 anig|EHA20155.1 aory|EIT78866.1 ater|EAU39256.1 atub|OJI80796.1 awen|OJJ33217.1 azon|OJJ50336.1 afis|EAW19498.1 afla|EED54513.1 agla|OJJ84619.1

Group6549: abra|OJJ67127.1 acar|OOF98426.1 afis|EAW19342.1 afla|EED47538.1 agla|OJJ79330.1 akaw|GAA91041.1 aluc|OJZ87851.1 anid|CBF80125.1 anig|EHA20154.1 aory|EIT74588.1 asyd|OJJ58343.1 ater|EAU29444.1 atub|OJI80797.1 awen|OJJ35789.1 azon|OJJ42318.1

Group6550: afis|EAW20743.1 acar|OOF95023.1 afla|EED53084.1 akaw|GAA88129.1 aluc|OJZ81401.1 anig|EHA18694.1 aory|EIT74814.1 asyd|OJJ52801.1 atub|OJI91607.1 aver|OJJ08778.1 azon|OJJ42879.1 afla|EED47484.1 asyd|OJJ61688.1 afla|EED48103.1 aory|EIT76221.1

Group6551: abra|OJJ67146.1 acar|OOF98409.1 afis|EAW19303.1 afla|EED47579.1 afum|EAL94021.1 akaw|GAA91024.1 aluc|OJZ87827.1 anid|CBF83911.1 anig|EHA20306.1 aory|EIT74541.1 asyd|OJJ55496.1 ater|EAU29465.1 atub|OJI80814.1 aver|OJJ06574.1 awen|OJJ30170.1

Group6552: abra|OJJ67171.1 acla|EAW14553.1 afis|EAW19270.1 afum|EBA27510.1 agla|OJJ88428.1 akaw|GAA91003.1 aluc|OJZ87804.1 anid|CBF78701.1 anig|EHA20286.1 asyd|OJJ65221.1 ater|EAU29550.1 atub|OJI80843.1 aver|OJI99370.1 awen|OJJ37630.1 azon|OJJ47018.1

Group6553: abra|OJJ67174.1 acar|OOF98389.1 acla|EAW14549.1 afis|EAW19266.1 akaw|GAA91000.1 aluc|OJZ87801.1 anid|CBF78693.1 anig|EHA20282.1 asyd|OJJ65217.1 ater|EAU29547.1 atub|OJI80846.1 aver|OJI99367.1 awen|OJJ37627.1 azon|OJJ50093.1 afum|EAL93993.1

Group6554: abra|OJJ67177.1 acar|OOF98387.1 acla|EAW14546.1 afla|EED47611.1 agla|OJJ88422.1 aluc|OJZ87798.1 anid|CBF78686.1 anig|EHA20279.1 aory|EIT80173.1 asyd|OJJ65213.1 ater|EAU29544.1 atub|OJI80849.1 aver|OJI99363.1 awen|OJJ37624.1 azon|OJJ47013.1

Group6555: abra|OJJ67210.1 acar|OOF98349.1 acla|EAW08556.1 afis|EAW23598.1 afla|EED55270.1 afum|EAL87868.1 agla|OJJ86857.1 akaw|GAA90967.1 aluc|OJZ87763.1 anig|EHA20248.1 aory|EIT83544.1 ater|EAU35358.1 atub|OJI80881.1 awen|OJJ39920.1 azon|OJJ45894.1

Group6556: abra|OJJ67227.1 acar|OOF98325.1 acla|EAW10407.1 afis|EAW25161.1 afla|EED53621.1 afum|EAL89836.1 agla|OJJ86387.1 akaw|GAA91653.1 aluc|OJZ87747.1 anig|EHA20236.1 aory|EIT75179.1 ater|EAU34332.1 atub|OJI80895.1 awen|OJJ36214.1 azon|OJJ43928.1

Group6557: abra|OJJ67259.1 acar|OOF98304.1 acla|EAW10394.1 afis|EAW25183.1 afla|EED53641.1 afum|EAL89857.1 akaw|GAA91674.1 aluc|OJZ87720.1 anid|CBF73876.1 anig|EHA20214.1 aory|EIT75349.1 asyd|OJJ54775.1 ater|EAU34340.1 atub|OJI80922.1 aver|OJJ07391.1

Group6558: abra|OJJ67285.1 acar|OOF93926.1 afla|EED51076.1 afum|EAL91134.1 akaw|GAA86945.1 aluc|OJZ90070.1 anid|CBF87380.1 anig|EHA25286.1 aory|EIT77018.1 asyd|OJJ56243.1 atub|OJI84737.1 aver|OJJ05166.1 anig|EHA21003.1 afla|EED53132.1 aory|EIT74829.1

Group6559: abra|OJJ67302.1 acar|OOF92512.1 acla|EAW06435.1 afis|EAW20289.1 afla|EED52855.1 afum|EAL85420.1 akaw|GAA86514.1 aluc|OJZ85059.1 anid|CBF70198.1 anig|EHA25274.1 aory|EIT78646.1 asyd|OJJ58787.1 ater|EAU30034.1 atub|OJI80539.1 aver|OJJ01394.1

Group6560: abra|OJJ67309.1 acar|OOF92503.1 acla|EAW06423.1 afis|EAW18527.1 afla|EED52874.1 afum|EBA27487.1 akaw|GAA86506.1 aluc|OJZ85049.1 anid|CBF70209.1 anig|EHA25266.1 aory|EIT78614.1 asyd|OJJ58780.1 ater|EAU30042.1 atub|OJI80549.1 azon|OJJ50141.1

Group6561: abra|OJJ67338.1 acar|OOF92475.1 acla|EAW06396.1 afis|EAW18500.1 afum|EAL93191.2 agla|OJJ85127.1 akaw|GAA86478.1 aluc|OJZ85019.1 anid|CBF70255.1 anig|EHA25240.1 asyd|OJJ58756.1 atub|OJI80578.1 aver|OJJ01364.1 awen|OJJ31054.1 azon|OJJ50562.1

Group6562: abra|OJJ67447.1 acar|OOF90342.1 acla|EAW06316.1 afis|EAW18430.1 afla|EED52983.1 afum|EAL93110.2 agla|OJJ84996.1 akaw|GAA86382.1 aluc|OJZ84916.1 anid|CBF70023.1 asyd|OJJ58873.1 ater|EAU29846.1 atub|OJI80682.1 aver|OJJ01476.1 awen|OJJ30763.1

Group6563: abra|OJJ67466.1 acar|OOF92367.1 afis|EAW18411.1 afla|EED53001.1 afum|EAL93091.1 agla|OJJ84977.1 akaw|GAA86370.1 aluc|OJZ84899.1 anid|CBF70737.1 anig|EHA25126.1 aory|EIT77393.1 asyd|OJJ58441.1 atub|OJI80699.1 aver|OJJ01041.1 awen|OJJ31739.1

Group6564: abra|OJJ67473.1 afis|EAW18406.1 afla|EED53002.1 afum|EAL93086.1 agla|OJJ84957.1 aluc|OJZ84892.1 anid|CBF69756.1 aory|EIT77429.1 atub|OJI80706.1 aver|OJJ01539.1 awen|OJJ31744.1 azon|OJJ50504.1 acla|EAW06299.1 akaw|GAA86364.1 asyd|OJJ58943.1

Group6565: anid|CBF80146.1 aory|EIT81308.1 acla|EAW14607.1 afis|EAW19336.1 afla|EED51165.1 afum|EAL94053.1 agla|OJJ79422.1 anid|CBF90105.1 asyd|OJJ59231.1 aver|OJJ01841.1 afis|EAW24674.1 asyd|OJJ57397.1 aver|OJI96118.1 asyd|OJJ62027.1 aver|OJI99534.1

Group6566: afis|EAW15462.1 azon|OJJ42091.1 acar|OOF90313.1 afla|EED51800.1 aluc|OJZ83718.1 anid|CBF83023.1 anig|EHA25927.1 asyd|OJJ61031.1 ater|EAU33407.1 awen|OJJ30427.1 akaw|GAA83216.1 aory|EIT81712.1 asyd|OJJ57906.1 awen|OJJ29893.1 agla|OJJ84402.1

Group6567: abra|OJJ67599.1 acar|OOF91951.1 acla|EAW12420.1 afla|EED49737.1 agla|OJJ86654.1 akaw|GAA89118.1 aluc|OJZ81721.1 anid|CBF83048.1 anig|EHA25718.1 aory|EIT77628.1 asyd|OJJ64554.1 ater|EAU34047.1 atub|OJI88460.1 aver|OJI98651.1 ater|EAU36466.1

Group6568: abra|OJJ67693.1 acar|OOF97520.1 acla|EAW14630.1 afis|EAW17253.1 afla|EED53071.1 afum|EAL86672.2 akaw|GAA89202.1 aluc|OJZ81628.1 anig|EHA25815.1 asyd|OJJ55210.1 ater|EAU30970.1 atub|OJI88552.1 aver|OJJ07007.1 awen|OJJ32906.1 anid|CBF78046.1

Group6569: abra|OJJ67903.1 acar|OOF92875.1 acla|EAW11718.1 afis|EAW23739.1 afla|EED51515.1 afum|EAL87176.1 akaw|GAA91908.1 aluc|OJZ81985.1 anid|CBF79811.1 anig|EHA22270.1 asyd|OJJ58131.1 ater|EAU31297.1 atub|OJI84536.1 aver|OJJ04274.1 awen|OJJ33675.1

Group6570: abra|OJJ67905.1 acar|OOF92873.1 afis|EAW23709.1 afum|EAL87153.1 akaw|GAA91910.1 aluc|OJZ81982.1 anig|EHA22272.1 asyd|OJJ54985.1 ater|EAU31291.1 atub|OJI84534.1 aver|OJJ07216.1 awen|OJJ35904.1 afla|EED45764.1 afum|EBA27287.1 aory|EIT75534.1

Group6571: abra|OJJ67906.1 acar|OOF92872.1 acla|EAW11693.1 afis|EAW23710.1 afla|EED45765.1 afum|EBA27286.1 akaw|GAA91911.1 aluc|OJZ81981.1 anig|EHA22273.1 aory|EIT75397.1 asyd|OJJ54986.1 atub|OJI84533.1 aver|OJJ07215.1 awen|OJJ35905.1 ater|EAU31290.1

Group6572: abra|OJJ67927.1 acar|OOF92778.1 acla|EAW12783.1 afis|EAW17945.1 afla|EED51960.1 afum|EAL91756.1 agla|OJJ82988.1 akaw|GAA91270.1 aluc|OJZ82799.1 aory|EIT81460.1 ater|EAU31430.1 atub|OJI89030.1 aver|OJJ04204.1 awen|OJJ41597.1 azon|OJJ48313.1

Group6573: abra|OJJ67937.1 acar|OOF92786.1 acla|EAW11626.1 afis|EAW23830.1 afla|EED51975.1 afum|EAL87262.1 agla|OJJ82482.1 akaw|GAA91259.1 aluc|OJZ82809.1 anid|CBF79906.1 aory|EIT81481.1 asyd|OJJ58188.1 aver|OJJ04217.1 awen|OJJ33788.1 azon|OJJ48758.1

Group6574: abra|OJJ67961.1 acar|OOF92807.1 afis|EAW23804.1 afla|EED52025.1 afum|EAL87238.1 agla|OJJ81899.1 akaw|GAA91236.1 aluc|OJZ82834.1 anig|EHA21272.1 aory|EIT81419.1 asyd|OJJ55237.1 ater|EAU31357.1 atub|OJI89064.1 aver|OJJ06986.1 awen|OJJ33761.1

Group6575: abra|OJJ67963.1 acar|OOF92809.1 acla|EAW11653.1 afla|EED51618.1 akaw|GAA91234.1 aluc|OJZ82836.1 anid|CBF79867.1 anig|EHA21274.1 aory|EIT80710.1 asyd|OJJ58165.1 ater|EAU31352.1 atub|OJI89066.1 aver|OJJ04237.1 awen|OJJ33759.1 azon|OJJ42865.1

Group6576: abra|OJJ67964.1 acar|OOF92810.1 acla|EAW11654.1 afis|EAW23799.1 afla|EED51617.1 afum|EAL87234.2 agla|OJJ78847.1 akaw|GAA91233.1 aluc|OJZ82837.1 anid|CBF79865.1 asyd|OJJ58164.1 ater|EAU31351.1 atub|OJI89067.1 aver|OJJ04238.1 awen|OJJ33758.1

Group6577: abra|OJJ67999.1 acar|OOF92042.1 afla|EED47248.1 agla|OJJ88174.1 akaw|GAA90951.1 aluc|OJZ82873.1 anid|CBF85192.1 anig|EHA21299.1 aory|EIT79211.1 asyd|OJJ64571.1 atub|OJI89097.1 aver|OJJ04298.1 awen|OJJ39963.1 azon|OJJ46818.1 azon|OJJ46410.1

Group6578: abra|OJJ68031.1 acar|OOF92076.1 acla|EAW11089.1 afla|EED56027.1 agla|OJJ79962.1 akaw|GAA90921.1 aluc|OJZ82905.1 anig|EHA21328.1 aory|EIT76559.1 asyd|OJJ53443.1 ater|EAU36890.1 atub|OJI89131.1 aver|OJJ08308.1 awen|OJJ33579.1 azon|OJJ43620.1

Group6579: abra|OJJ68032.1 acar|OOF92077.1 acla|EAW11090.1 afla|EED45881.1 agla|OJJ79961.1 akaw|GAA90920.1 aluc|OJZ82906.1 anig|EHA21329.1 aory|EIT75827.1 asyd|OJJ53444.1 ater|EAU36889.1 atub|OJI89132.1 aver|OJJ08307.1 awen|OJJ33580.1 azon|OJJ43619.1

Group6580: afis|EAW19754.1 afum|EAL85143.1 acla|EAW13626.1 afis|EAW21093.1 ater|EAU29533.1 afum|EAL92116.1 agla|OJJ78720.1 aluc|OJZ85485.1 anid|CBF87240.1 anig|EHA26573.1 asyd|OJJ53804.1 atub|OJI80079.1 aver|OJJ07432.1 afis|EAW25734.1 anid|CBF82281.1

Group6581: abra|OJJ68107.1 acar|OOF93907.1 acla|EAW12450.1 afis|EAW16754.1 afla|EED55088.1 afum|EAL86326.1 akaw|GAA86889.1 aluc|OJZ90013.1 anid|CBF73885.1 anig|EHA27388.1 asyd|OJJ53657.1 ater|EAU33724.1 atub|OJI84794.1 aver|OJJ07765.1 akaw|GAA86890.1

Group6582: abra|OJJ68147.1 acar|OOF90644.1 acla|EAW12084.1 afis|EAW25676.1 afla|EED47641.1 agla|OJJ79768.1 akaw|GAA92372.1 aluc|OJZ83230.1 anid|CBF78373.1 anig|EHA21088.1 aory|EIT80233.1 asyd|OJJ57328.1 atub|OJI87706.1 afum|EAL84346.1 agla|OJJ87493.1

Group6583: abra|OJJ68282.1 acar|OOF97556.1 afla|EED56846.1 agla|OJJ86179.1 akaw|GAA83190.1 aluc|OJZ83689.1 anid|CBF89594.1 anig|EHA18843.1 aory|EIT77522.1 asyd|OJJ62314.1 ater|EAU39319.1 atub|OJI90061.1 aver|OJI96421.1 awen|OJJ38769.1 azon|OJJ49590.1

Group6584: abra|OJJ68335.1 acar|OOF97602.1 afla|EED57057.1 agla|OJJ89061.1 akaw|GAA92088.1 aluc|OJZ83635.1 anid|CBF71747.1 anig|EHA18788.1 aory|EIT82312.1 asyd|OJJ55947.1 atub|OJI90004.1 aver|OJI98883.1 awen|OJJ41747.1 aver|OJI95777.1 agla|OJJ80132.1

Group6585: abra|OJJ68340.1 acar|OOF89851.1 acla|EAW14396.1 afis|EAW19103.1 afla|EED55063.1 akaw|GAA92107.1 aluc|OJZ83618.1 anid|CBF84412.1 anig|EHA18776.1 aory|EIT81966.1 asyd|OJJ53561.1 atub|OJI89991.1 aver|OJJ08191.1 afum|EAL93835.1 aory|EIT81968.1

Group6586: abra|OJJ68352.1 acar|OOF90073.1 acla|EAW12404.1 afis|EAW16863.1 afla|EED56821.1 afum|EBA27272.1 agla|OJJ85254.1 aluc|OJZ83602.1 anid|CBF88782.1 aory|EIT73928.1 asyd|OJJ63993.1 ater|EAU33019.1 atub|OJI89974.1 awen|OJJ35930.1 azon|OJJ45220.1

Group6587: abra|OJJ68382.1 acar|OOF93986.1 acla|EAW12358.1 afis|EAW16781.1 afum|EAL86352.1 agla|OJJ79769.1 akaw|GAA92565.1 aluc|OJZ90113.1 anig|EHA20961.1 ater|EAU30271.1 atub|OJI84693.1 awen|OJJ38253.1 acla|EAW08810.1 agla|OJJ85167.1 ater|EAU32542.1

Group6588: abra|OJJ68410.1 acar|OOF93967.1 afla|EED55615.1 agla|OJJ88415.1 akaw|GAA92540.1 aluc|OJZ90088.1 anig|EHA20986.1 aory|EIT81814.1 asyd|OJJ58699.1 ater|EAU32552.1 atub|OJI84718.1 aver|OJJ01311.1 awen|OJJ33712.1 anig|EHA20906.1 awen|OJJ40592.1

Group6589: abra|OJJ68412.1 acar|OOF93965.1 afla|EED45887.1 agla|OJJ80674.1 akaw|GAA92538.1 aluc|OJZ90086.1 anig|EHA20988.1 aory|EIT75567.1 atub|OJI84720.1 awen|OJJ31664.1 azon|OJJ45178.1 anid|CBF75880.1 asyd|OJJ60249.1 aver|OJJ04008.1 awen|OJJ35875.1

Group6590: abra|OJJ68424.1 acar|OOF93929.1 acla|EAW09508.1 afis|EAW21625.1 afum|EAL92661.1 agla|OJJ79692.1 akaw|GAA86948.1 aluc|OJZ90073.1 anid|CBF82427.1 anig|EHA21000.1 asyd|OJJ54127.1 ater|EAU30754.1 atub|OJI84733.1 aver|OJJ08609.1 awen|OJJ33875.1

Group6591: abra|OJJ68425.1 acar|OOF93928.1 afis|EAW23752.1 akaw|GAA86947.1 aluc|OJZ90072.1 anig|EHA21001.1 aory|EIT82579.1 atub|OJI84734.1 awen|OJJ38776.1 azon|OJJ45873.1 afla|EED49559.1 afum|EAL87189.1 anid|CBF85105.1 asyd|OJJ63055.1 aver|OJI97908.1

Group6592: abra|OJJ68441.1 acar|OOF93954.1 afis|EAW19802.1 afla|EED52096.1 agla|OJJ87951.1 akaw|GAA86931.1 aluc|OJZ90056.1 anid|CBF82420.1 anig|EHA21015.1 aory|EIT76908.1 asyd|OJJ54149.1 ater|EAU30747.1 atub|OJI84751.1 aver|OJJ08592.1 awen|OJJ29693.1

Group6593: abra|OJJ68470.1 acar|OOF90640.1 acla|EAW07154.1 afis|EAW22168.1 afla|EED54171.1 afum|EAL91023.1 agla|OJJ87886.1 akaw|GAA92391.1 aluc|OJZ83251.1 anid|CBF73660.1 anig|EHA21104.1 asyd|OJJ56670.1 ater|EAU30397.1 atub|OJI87686.1 aver|OJJ06052.1

Group6594: abra|OJJ68495.1 acar|OOF98750.1 acla|EAW12141.1 afis|EAW17195.1 afla|EED45177.1 afum|EAL86623.1 agla|OJJ86659.1 akaw|GAA92775.1 aluc|OJZ82708.1 anig|EHA21163.1 aory|EIT73805.1 asyd|OJJ59008.1 atub|OJI88941.1 anid|CBF69889.1 aver|OJJ01607.1

Group6595: abra|OJJ68661.1 acar|OOF92163.1 acla|EAW12592.1 afis|EAW24061.1 afla|EED51413.1 afum|EAL91916.1 agla|OJJ87343.1 akaw|GAA89801.1 aluc|OJZ81890.1 anid|CBF86447.1 aory|EIT80456.1 asyd|OJJ64921.1 atub|OJI84443.1 awen|OJJ41429.1 azon|OJJ43871.1

Group6596: abra|OJJ68676.1 acar|OOF92178.1 afis|EAW24077.1 afla|EED51396.1 akaw|GAA89829.1 aluc|OJZ81858.1 anid|CBF86474.1 anig|EHA22366.1 aory|EIT80737.1 asyd|OJJ64904.1 atub|OJI84426.1 aver|OJI99026.1 awen|OJJ41415.1 afum|EAL91932.1 ater|EAU30498.1

Group6597: abra|OJJ68694.1 acar|OOF92198.1 acla|EAW12550.1 agla|OJJ87175.1 akaw|GAA89849.1 aluc|OJZ81837.1 anig|EHA22384.1 asyd|OJJ64884.1 ater|EAU30528.1 atub|OJI84406.1 aver|OJI98998.1 awen|OJJ41388.1 azon|OJJ48857.1 afla|EED46833.1 aory|EIT72625.1

Group6598: abra|OJJ68721.1 acar|OOF92220.1 acla|EAW12525.1 afis|EAW24123.1 afla|EED46811.1 afum|EAL91979.1 agla|OJJ87198.1 aluc|OJZ81808.1 anid|CBF86549.1 anig|EHA22409.1 asyd|OJJ64858.1 atub|OJI84380.1 aver|OJI98970.1 awen|OJJ41366.1 azon|OJJ48833.1

Group6599: abra|OJJ68741.1 acar|OOF90305.1 afis|EAW24140.1 afla|EED46794.1 afum|EAL91995.1 agla|OJJ87219.1 aluc|OJZ81785.1 anid|CBF77938.1 anig|EHA22430.1 asyd|OJJ55285.1 ater|EAU30569.1 atub|OJI84359.1 aver|OJJ06943.1 awen|OJJ41348.1 azon|OJJ48812.1

Group6600: abra|OJJ68798.1 acar|OOF92282.1 acla|EAW14655.1 afla|EED46745.1 agla|OJJ89488.1 akaw|GAA89937.1 aluc|OJZ81729.1 anid|CBF77835.1 anig|EHA22483.1 asyd|OJJ55337.1 ater|EAU30620.1 atub|OJI84300.1 aver|OJJ06890.1 awen|OJJ41283.1 azon|OJJ43705.1

Group6601: abra|OJJ68804.1 acar|OOF91596.1 acla|EAW14400.1 afis|EAW19107.1 afla|EED54303.1 afum|EAL93838.1 akaw|GAA90449.1 aluc|OJZ85775.1 anig|EHA24779.1 aory|EIT75984.1 asyd|OJJ58067.1 ater|EAU29824.1 atub|OJI79560.1 aver|OJJ04340.1 awen|OJJ32508.1

Group6602: abra|OJJ68867.1 akaw|GAA90386.1 aluc|OJZ85845.1 anig|EHA24719.1 ater|EAU29643.1 atub|OJI79633.1 awen|OJJ32556.1 azon|OJJ50747.1 acla|EAW13300.1 afis|EAW16173.1 afla|EED45260.1 afum|EAL88805.1 aory|EIT73657.1 ater|EAU34150.1 awen|OJJ36062.1

Group6603: abra|OJJ68920.1 acar|OOF90184.1 acla|EAW14255.1 afis|EAW19025.1 afla|EED56630.1 afum|EAL93756.1 akaw|GAA90329.1 aluc|OJZ85904.1 anid|CBF85588.1 anig|EHA24689.1 aory|EIT79677.1 asyd|OJJ64172.1 atub|OJI79689.1 aver|OJI98247.1 azon|OJJ44373.1

Group6604: abra|OJJ68921.1 acar|OOF90182.1 akaw|GAA90328.1 aluc|OJZ85905.1 anig|EHA24688.1 atub|OJI79690.1 abra|OJJ76678.1 acar|OOF99330.1 anid|CBF74091.1 anig|EHA22563.1 asyd|OJJ53891.1 atub|OJI87198.1 aver|OJJ07989.1 awen|OJJ33629.1 awen|OJJ36547.1

Group6605: abra|OJJ68958.1 acar|OOF91536.1 acla|EAW14227.1 afis|EAW19054.1 afla|EED54412.1 afum|EAL93783.1 agla|OJJ84553.1 akaw|GAA90486.1 aluc|OJZ85944.1 anig|EHA24655.1 asyd|OJJ58011.1 atub|OJI79732.1 aver|OJJ04389.1 awen|OJJ31485.1 azon|OJJ46195.1

Group6606: abra|OJJ68985.1 acar|OOF91556.1 acla|EAW14206.1 afis|EAW19075.1 afla|EED54440.1 afum|EAL93803.1 agla|OJJ84528.1 aluc|OJZ85971.1 anig|EHA24634.1 asyd|OJJ58032.1 ater|EAU32121.1 atub|OJI79759.1 aver|OJJ04367.1 awen|OJJ32469.1 azon|OJJ46237.1

Group6607: abra|OJJ69055.1 acar|OOF91235.1 acla|EAW08771.1 afis|EAW15652.1 afla|EED46894.1 afum|EAL85698.2 agla|OJJ81781.1 akaw|GAA90585.1 aluc|OJZ86043.1 anid|CBF71188.1 asyd|OJJ55858.1 atub|OJI79839.1 aver|OJJ04799.1 awen|OJJ38288.1 azon|OJJ47472.1

Group6608: abra|OJJ69070.1 acar|OOF95825.1 afis|EAW21326.1 akaw|GAA90601.1 aluc|OJZ86059.1 atub|OJI79856.1 azon|OJJ46525.1 acla|EAW11918.1 agla|OJJ82478.1 anig|EHA21144.1 akaw|GAA89042.1 azon|OJJ45151.1 afum|EAL92344.1 azon|OJJ47740.1 azon|OJJ50420.1

Group6609: abra|OJJ69115.1 afla|EED51333.1 afum|EAL85622.1 agla|OJJ81869.1 akaw|GAA90640.1 aluc|OJZ86104.1 anid|CBF71081.1 asyd|OJJ52696.1 ater|EAU32428.1 atub|OJI79899.1 aver|OJJ04724.1 awen|OJJ38414.1 azon|OJJ43828.1 acla|EAW08717.1 afis|EAW15725.1

Group6610: abra|OJJ69130.1 acar|OOF91170.1 acla|EAW08702.1 afis|EAW15710.1 afla|EED51318.1 afum|EAL85637.1 agla|OJJ81891.1 akaw|GAA90655.1 aluc|OJZ86119.1 anig|EHA24502.1 asyd|OJJ52676.1 atub|OJI79915.1 aver|OJJ04702.1 awen|OJJ38430.1 azon|OJJ43813.1

Group6611: abra|OJJ69140.1 acla|EAW08694.1 afis|EAW15741.1 afla|EED51311.1 afum|EAL85607.1 agla|OJJ81902.1 akaw|GAA86317.1 anid|CBF71041.1 anig|EHA24494.1 aory|EIT81509.1 asyd|OJJ52669.1 ater|EAU32404.1 aver|OJJ04695.1 awen|OJJ38440.1 azon|OJJ43805.1

Group6612: abra|OJJ69253.1 acar|OOF91068.1 acla|EAW08596.1 afis|EAW15837.1 afla|EED51209.1 afum|EAL85514.1 agla|OJJ82001.1 anid|CBF70852.1 anig|EHA24396.1 asyd|OJJ52572.1 ater|EAU33411.1 atub|OJI80027.1 aver|OJJ04601.1 awen|OJJ38549.1 azon|OJJ43712.1

Group6613: abra|OJJ69254.1 acar|OOF91067.1 acla|EAW08595.1 afis|EAW15838.1 afum|EAL85513.1 agla|OJJ82002.1 akaw|GAA92260.1 aluc|OJZ86233.1 anid|CBF70850.1 anig|EHA24395.1 aory|EIT80407.1 asyd|OJJ52571.1 ater|EAU33397.1 aver|OJJ04600.1 awen|OJJ38551.1

Group6614: abra|OJJ69255.1 acar|OOF91066.1 acla|EAW08594.1 afis|EAW15840.1 afum|EAL85511.2 agla|OJJ84919.1 akaw|GAA92261.1 aluc|OJZ86234.1 anid|CBF70846.1 anig|EHA24393.1 aory|EIT80411.1 asyd|OJJ52569.1 ater|EAU33395.1 aver|OJJ04598.1 awen|OJJ38553.1

Group6615: abra|OJJ69273.1 acar|OOF94448.1 acla|EAW14662.1 afis|EAW24193.1 agla|OJJ79686.1 akaw|GAA88147.1 aluc|OJZ82101.1 anid|CBF74311.1 anig|EHA28629.1 asyd|OJJ54005.1 ater|EAU31144.1 atub|OJI88335.1 aver|OJJ08100.1 awen|OJJ33106.1 azon|OJJ44842.1

Group6616: abra|OJJ69280.1 acar|OOF94442.1 acla|EAW14669.1 afis|EAW24200.1 afum|EBA27247.1 agla|OJJ79677.1 aluc|OJZ82108.1 anid|CBF74325.1 anig|EHA28622.1 asyd|OJJ53995.1 ater|EAU31151.1 atub|OJI88328.1 aver|OJJ08093.1 awen|OJJ33098.1 azon|OJJ44849.1

Group6617: abra|OJJ69306.1 acar|OOF94421.1 acla|EAW14692.1 afis|EAW24223.1 afla|EED48049.1 afum|EBA27249.1 akaw|GAA88175.1 aluc|OJZ82135.1 anid|CBF74237.1 anig|EHA28596.1 aory|EIT83337.1 atub|OJI88302.1 aver|OJJ08069.1 awen|OJJ33073.1 azon|OJJ44874.1

Group6618: abra|OJJ69309.1 acar|OOF94417.1 acla|EAW14696.1 afis|EAW24227.1 afla|EED48044.1 afum|EAL85896.1 agla|OJJ79644.1 akaw|GAA88178.1 aluc|OJZ82138.1 anig|EHA28593.1 ater|EAU31176.1 atub|OJI88299.1 aver|OJJ08065.1 awen|OJJ33069.1 azon|OJJ48453.1

Group6619: abra|OJJ69318.1 afis|EAW24263.1 afla|EED48014.1 afum|EAL85931.1 agla|OJJ79609.1 akaw|GAA88187.1 aluc|OJZ82147.1 anig|EHA28583.1 aory|EIT82899.1 atub|OJI88289.1 awen|OJJ33038.1 acla|EAW14730.1 anid|CBF74178.1 ater|EAU31208.1 azon|OJJ48481.1

Group6620: abra|OJJ69321.1 acar|OOF94409.1 acla|EAW14724.1 afis|EAW24260.1 afum|EAL85928.1 agla|OJJ79612.1 akaw|GAA88190.1 aluc|OJZ82150.1 anid|CBF74189.1 anig|EHA28580.1 asyd|OJJ53949.1 atub|OJI88285.1 aver|OJJ08045.1 awen|OJJ33042.1 azon|OJJ48596.1

Group6621: abra|OJJ69322.1 acar|OOF99033.1 afis|EAW22828.1 afla|EED45971.1 agla|OJJ81579.1 akaw|GAA88191.1 aluc|OJZ82152.1 anid|CBF69421.1 anig|EHA28579.1 aory|EIT75572.1 asyd|OJJ60158.1 ater|EAU32277.1 atub|OJI88284.1 awen|OJJ29835.1 aluc|OJZ79516.1

Group6622: aory|EIT82917.1 acar|OOF94404.1 acla|EAW14717.1 afis|EAW24252.1 afla|EED48021.1 afum|EAL85921.2 agla|OJJ79621.1 akaw|GAA88194.1 aluc|OJZ82155.1 anid|CBF74195.1 anig|EHA28577.1 ater|EAU31195.1 atub|OJI88282.1 awen|OJJ33047.1 azon|OJJ48472.1

Group6623: abra|OJJ69340.1 acar|OOF94388.1 acla|EAW14706.1 afis|EAW24237.1 afla|EED48034.1 agla|OJJ79635.1 akaw|GAA88209.1 aluc|OJZ82172.1 anid|CBF74217.1 anig|EHA28561.1 asyd|OJJ53960.1 atub|OJI88265.1 aver|OJJ08056.1 awen|OJJ33051.1 azon|OJJ48462.1

Group6624: abra|OJJ69352.1 acar|OOF94374.1 acla|EAW14741.1 afis|EAW24272.1 afla|EED48005.1 afum|EAL85940.2 agla|OJJ79593.1 aluc|OJZ82184.1 anid|CBF74162.1 anig|EHA28551.1 aory|EIT82911.1 asyd|OJJ53932.1 ater|EAU31217.1 aver|OJJ08030.1 awen|OJJ33027.1

Group6625: abra|OJJ69368.1 acar|OOF89838.1 acla|EAW14758.1 afis|EAW24289.1 afla|EED47987.1 afum|EBA27251.1 agla|OJJ83154.1 akaw|GAA88237.1 aluc|OJZ82202.1 anig|EHA28537.1 asyd|OJJ53910.1 atub|OJI88234.1 aver|OJJ08008.1 awen|OJJ33007.1 azon|OJJ48508.1

Group6626: abra|OJJ69389.1 acar|OOF94337.1 acla|EAW14776.1 afis|EAW24307.1 afla|EED47963.1 afum|EAL85975.1 agla|OJJ83175.1 akaw|GAA88257.1 aluc|OJZ82224.1 anid|CBF74095.1 anig|EHA28516.1 asyd|OJJ53893.1 atub|OJI88211.1 aver|OJJ07991.1 awen|OJJ32985.1

Group6627: abra|OJJ69393.1 acar|OOF94335.1 acla|EAW14778.1 afis|EAW24309.1 afum|EAL85977.2 akaw|GAA88261.1 aluc|OJZ82230.1 anid|CBF74087.1 anig|EHA28509.1 asyd|OJJ53885.1 ater|EAU31262.1 atub|OJI88207.1 aver|OJJ07983.1 awen|OJJ32983.1 azon|OJJ48934.1

Group6628: abra|OJJ69430.1 acar|OOF94307.1 acla|EAW14806.1 afis|EAW24335.1 afla|EED47929.1 afum|EAL86005.1 agla|OJJ83204.1 anid|CBF74034.1 anig|EHA28476.1 aory|EIT79858.1 ater|EAU30413.1 atub|OJI88171.1 aver|OJJ07952.1 awen|OJJ32959.1 azon|OJJ48956.1

Group6629: abra|OJJ69443.1 acar|OOF94296.1 afis|EAW17312.1 afum|EAL91147.1 akaw|GAA88375.1 aluc|OJZ82279.1 anid|CBF84390.1 anig|EHA28467.1 asyd|OJJ53584.1 atub|OJI88160.1 aver|OJJ08164.1 azon|OJJ43891.1 asyd|OJJ64146.1 aver|OJI98227.1 azon|OJJ42534.1

Group6630: abra|OJJ69509.1 afis|EAW21139.1 afla|EED46137.1 afum|EAL92158.2 agla|OJJ80079.1 akaw|GAA88315.1 aluc|OJZ82347.1 anid|CBF86998.1 anig|EHA28413.1 aory|EIT76469.1 asyd|OJJ60466.1 atub|OJI88093.1 aver|OJI99786.1 awen|OJJ36460.1 azon|OJJ44417.1

Group6631: abra|OJJ69550.1 acar|OOF95792.1 acla|EAW10611.1 afis|EAW24956.1 afla|EED55508.1 afum|EAL89614.2 akaw|GAA91562.1 aluc|OJZ82380.1 anid|CBF81669.1 anig|EHA28385.1 aory|EIT80914.1 asyd|OJJ59553.1 ater|EAU35701.1 atub|OJI88061.1 aver|OJJ02715.1

Group6632: abra|OJJ69611.1 acar|OOF92696.1 agla|OJJ83949.1 akaw|GAA91595.1 aluc|OJZ82419.1 anid|CBF82527.1 anig|EHA28350.1 asyd|OJJ54073.1 atub|OJI88022.1 aver|OJJ08666.1 awen|OJJ34380.1 acar|OOF91012.1 acla|EAW10921.1 afis|EAW20937.1 afum|EAL84697.1

Group6633: abra|OJJ69661.1 afis|EAW23620.1 afla|EED47586.1 afum|EAL87842.1 akaw|GAA91629.1 aluc|OJZ82456.1 anid|CBF78710.1 anig|EHA28319.1 aory|EIT74583.1 asyd|OJJ55361.1 ater|EAU35232.1 atub|OJI87982.1 aver|OJJ08471.1 awen|OJJ30195.1 azon|OJJ42645.1

Group6634: abra|OJJ69673.1 acar|OOF95528.1 acla|EAW07246.1 afis|EAW22267.1 afla|EED57659.1 afum|EAL90913.1 agla|OJJ87788.1 akaw|GAA88797.1 aluc|OJZ82480.1 anig|EHA28295.1 aory|EIT79924.1 atub|OJI87958.1 awen|OJJ34840.1 azon|OJJ45707.1 ater|EAU34278.1

Group6635: abra|OJJ69725.1 afis|EAW20757.1 afla|EED47461.1 afla|EED57110.1 afum|EAL84862.1 agla|OJJ82213.1 akaw|GAA90178.1 aluc|OJZ83172.1 anig|EHA28246.1 aory|EIT74047.1 ater|EAU34036.1 atub|OJI87766.1 awen|OJJ35686.1 azon|OJJ47342.1 aory|EIT79962.1

Group6636: afla|EED45788.1 aory|EIT75775.1 abra|OJJ77174.1 acar|OOF93062.1 akaw|GAA90760.1 aluc|OJZ84586.1 anig|EHA27879.1 atub|OJI89494.1 afis|EAW17778.1 ater|EAU31838.1 acar|OOF90557.1 agla|OJJ88651.1 asyd|OJJ63643.1 aver|OJI97293.1 afum|EAL91587.1

Group6637: abra|OJJ69787.1 acar|OOF93591.1 akaw|GAA87870.1 aluc|OJZ85511.1 anig|EHA25043.1 atub|OJI80055.1 awen|OJJ37150.1 abra|OJJ78040.1 acla|EAW10012.1 afis|EAW20263.1 afla|EED51626.1 aory|EIT80659.1 ater|EAU36816.1 afis|EAW21044.1 afum|EBA27447.1

Group6638: abra|OJJ69811.1 acar|OOF93566.1 afis|EAW23600.1 afla|EED53575.1 afum|EAL87867.1 agla|OJJ86859.1 akaw|GAA87846.1 aluc|OJZ85484.1 anig|EHA25023.1 aory|EIT74887.1 asyd|OJJ64227.1 ater|EAU35359.1 atub|OJI80080.1 aver|OJI98320.1 awen|OJJ39925.1

Group6639: abra|OJJ69890.1 acar|OOF93411.1 acla|EAW10351.1 akaw|GAA87760.1 aluc|OJZ85394.1 anid|CBF88252.1 anig|EHA25578.1 asyd|OJJ54130.1 ater|EAU38931.1 atub|OJI80177.1 aver|OJJ08605.1 aory|EIT83383.1 asyd|OJJ63670.1 aver|OJI97267.1 anid|CBF81833.1

Group6640: abra|OJJ69896.1 acar|OOF93415.1 acla|EAW08924.1 afis|EAW15469.1 afla|EED46921.1 afum|EAL85853.1 akaw|GAA87754.1 aluc|OJZ85387.1 anid|CBF85148.1 anig|EHA25571.1 asyd|OJJ56064.1 ater|EAU32659.1 atub|OJI80183.1 aver|OJJ05007.1 awen|OJJ38119.1

Group6641: abra|OJJ69907.1 acar|OOF93424.1 acla|EAW08917.1 afis|EAW15477.1 afla|EED46938.1 afum|EAL85844.1 agla|OJJ80156.1 akaw|GAA87746.1 aluc|OJZ85379.1 anig|EHA25562.1 aory|EIT72647.1 ater|EAU32650.1 atub|OJI80191.1 awen|OJJ38134.1 azon|OJJ43158.1

Group6642: abra|OJJ69920.1 acar|OOF93432.1 acla|EAW08911.1 afis|EAW15484.1 afla|EED46940.1 afum|EAL85838.1 agla|OJJ80163.1 akaw|GAA87736.1 aluc|OJZ85368.1 anig|EHA25551.1 aory|EIT72589.1 asyd|OJJ56025.1 atub|OJI80202.1 aver|OJJ04972.1 azon|OJJ43144.1

Group6643: abra|OJJ69935.1 acar|OOF93444.1 acla|EAW08891.1 afis|EAW15498.1 afla|EED49013.1 afum|EAL85825.1 akaw|GAA87724.1 aluc|OJZ85352.1 aory|EIT82542.1 asyd|OJJ56013.1 ater|EAU32621.1 atub|OJI80218.1 aver|OJJ04958.1 awen|OJJ38157.1 azon|OJJ43132.1

Group6644: abra|OJJ69948.1 acla|EAW08876.1 afis|EAW15519.1 afla|EED49026.1 agla|OJJ78534.1 akaw|GAA87712.1 aluc|OJZ85339.1 anig|EHA25527.1 aory|EIT82538.1 asyd|OJJ55999.1 ater|EAU32605.1 atub|OJI80232.1 aver|OJJ04945.1 awen|OJJ38180.1 azon|OJJ42826.1

Group6645: abra|OJJ69964.1 acar|OOF93472.1 acla|EAW08836.1 afis|EAW15564.1 afla|EED49073.1 afum|EBA27241.1 agla|OJJ85187.1 akaw|GAA87695.1 aluc|OJZ85319.1 anid|CBF78120.1 asyd|OJJ55902.1 atub|OJI80254.1 aver|OJJ04847.1 awen|OJJ38224.1 azon|OJJ43071.1

Group6646: abra|OJJ69976.1 acar|OOF93482.1 afla|EED49058.1 afum|EAL85778.1 agla|OJJ85200.1 akaw|GAA87685.1 aluc|OJZ85305.1 anid|CBF78169.1 anig|EHA25501.1 asyd|OJJ55967.1 ater|EAU32579.1 atub|OJI80266.1 aver|OJJ04914.1 awen|OJJ38211.1 azon|OJJ43085.1

Group6647: abra|OJJ69986.1 acar|OOF93492.1 acla|EAW08857.1 afis|EAW15539.1 afla|EED57487.1 afum|EAL85791.1 akaw|GAA87675.1 aluc|OJZ85293.1 anig|EHA25491.1 aory|EIT81876.1 asyd|OJJ55978.1 ater|EAU32588.1 atub|OJI80277.1 aver|OJJ04926.1 acla|EAW10022.1

Group6648: abra|OJJ70043.1 acar|OOF93538.1 acla|EAW09891.1 agla|OJJ78497.1 akaw|GAA87632.1 aluc|OJZ85248.1 anig|EHA25448.1 ater|EAU35868.1 atub|OJI80321.1 afis|EAW21235.1 afla|EED54796.1 afum|EAL92247.1 aory|EIT79408.1 awen|OJJ36546.1 azon|OJJ44343.1

Group6649: abra|OJJ70076.1 acar|OOF93555.1 acla|EAW08783.1 afla|EED48901.1 agla|OJJ81676.1 akaw|GAA87607.1 aluc|OJZ85227.1 anid|CBF79625.1 anig|EHA25431.1 aory|EIT81256.1 asyd|OJJ58007.1 ater|EAU38523.1 atub|OJI80353.1 aver|OJJ04393.1 awen|OJJ37011.1

Group6650: abra|OJJ70082.1 acar|OOF94970.1 acla|EAW11425.1 afis|EAW20383.1 afla|EED45579.1 agla|OJJ83251.1 akaw|GAA87605.1 aluc|OJZ85225.1 anig|EHA25429.1 aory|EIT75740.1 asyd|OJJ54468.1 ater|EAU32964.1 atub|OJI80356.1 aver|OJJ07781.1 awen|OJJ30094.1

Group6651: abra|OJJ70087.1 acla|EAW12309.1 afis|EAW21036.1 afum|EAL92075.1 abra|OJJ74144.1 acar|OOF95585.1 afla|EED51068.1 akaw|GAA89740.1 aluc|OJZ80644.1 anig|EHA27188.1 aory|EIT77034.1 atub|OJI87499.1 awen|OJJ30201.1 azon|OJJ45080.1 atub|OJI89254.1

Group6652: abra|OJJ70139.1 acar|OOF90638.1 acla|EAW14614.1 afis|EAW19344.1 afla|EED56898.1 afum|EAL94063.1 agla|OJJ79571.1 akaw|GAA86626.1 aluc|OJZ85181.1 anid|CBF80127.1 anig|EHA25393.1 aory|EIT77543.1 asyd|OJJ55351.1 atub|OJI80416.1 awen|OJJ30884.1

Group6653: abra|OJJ70142.1 acar|OOF90637.1 acla|EAW13599.1 afis|EAW20006.1 afla|EED46483.1 afum|EAL84912.1 akaw|GAA86623.1 aluc|OJZ85178.1 anid|CBF87145.1 anig|EHA25390.1 aory|EIT72437.1 asyd|OJJ60300.1 atub|OJI80419.1 aver|OJI99640.1 azon|OJJ49479.1

Group6654: abra|OJJ70146.1 afla|EED52391.1 akaw|GAA90562.1 aory|EIT83299.1 awen|OJJ35566.1 abra|OJJ71760.1 acar|OOF90583.1 akaw|GAA92836.1 aluc|OJZ84840.1 aluc|OJZ91593.1 anig|EHA22617.1 anig|EHA25750.1 asyd|OJJ53405.1 atub|OJI86044.1 aver|OJJ04196.1

Group6655: abra|OJJ70203.1 acar|OOF92505.1 acla|EAW06425.1 afis|EAW18529.1 afla|EED52872.1 afum|EAL93226.1 akaw|GAA86574.1 aluc|OJZ85124.1 anig|EHA25343.1 aory|EIT78674.1 asyd|OJJ58782.1 ater|EAU30040.1 atub|OJI80469.1 aver|OJJ01390.1 awen|OJJ31087.1

Group6656: abra|OJJ70217.1 acar|OOF92563.1 agla|OJJ80299.1 akaw|GAA86560.1 aluc|OJZ85109.1 anid|CBF71762.1 anig|EHA25330.1 aory|EIT79760.1 asyd|OJJ61725.1 atub|OJI80484.1 aver|OJI95823.1 awen|OJJ32745.1 afla|EED53095.1 awen|OJJ36310.1 afla|EED57954.1

Group6657: abra|OJJ70234.1 acar|OOF93308.1 afla|EED54309.1 agla|OJJ84752.1 akaw|GAA86550.1 aluc|OJZ85098.1 anid|CBF90396.1 anig|EHA26106.1 asyd|OJJ56223.1 ater|EAU35998.1 atub|OJI80500.1 aver|OJJ05155.1 awen|OJJ37024.1 azon|OJJ42474.1 azon|OJJ43935.1

Group6658: abra|OJJ70323.1 acar|OOF91779.1 acla|EAW12317.1 afis|EAW16905.1 afla|EED45263.1 akaw|GAA92579.1 aluc|OJZ82592.1 anid|CBF80776.1 anig|EHA27483.1 asyd|OJJ54504.1 ater|EAU30212.1 atub|OJI88827.1 aver|OJJ07678.1 awen|OJJ29715.1 aory|EIT73794.1

Group6659: abra|OJJ70363.1 acla|EAW12635.1 afis|EAW24543.1 afla|EED46220.1 afum|EAL86205.2 akaw|GAA91988.1 aluc|OJZ82625.1 anig|EHA27451.1 aory|EIT76522.1 asyd|OJJ61035.1 atub|OJI88859.1 aver|OJJ00353.1 acla|EAW12634.1 afum|EAL86203.1 akaw|GAA91989.1

Group6660: abra|OJJ70369.1 acar|OOF91903.1 afla|EED47349.1 agla|OJJ79929.1 akaw|GAA91982.1 aluc|OJZ82631.1 anid|CBF80680.1 anig|EHA27446.1 aory|EIT82890.1 asyd|OJJ54591.1 ater|EAU30231.1 atub|OJI88864.1 aver|OJJ07561.1 awen|OJJ29764.1 azon|OJJ50383.1

Group6661: abra|OJJ70421.1 acar|OOF91767.1 afla|EED46000.1 agla|OJJ85947.1 akaw|GAA91936.1 aluc|OJZ82686.1 anid|CBF78384.1 anig|EHA21054.1 aory|EIT75811.1 asyd|OJJ62599.1 ater|EAU35269.1 atub|OJI88918.1 aver|OJI96744.1 awen|OJJ29995.1 ater|EAU32499.1

Group6662: abra|OJJ70436.1 acar|OOF91846.1 acla|EAW12327.1 afis|EAW18106.1 afla|EED49485.1 afum|EBA27494.1 akaw|GAA90144.1 aluc|OJZ83203.1 anid|CBF90397.1 anig|EHA21065.1 asyd|OJJ56616.1 ater|EAU35997.1 atub|OJI87734.1 aver|OJJ06108.1 awen|OJJ38309.1

Group6663: abra|OJJ70439.1 acar|OOF91840.1 acla|EAW12226.1 afla|EED45473.1 akaw|GAA90141.1 aluc|OJZ83206.1 anid|CBF82356.1 anig|EHA21067.1 aory|EIT75713.1 asyd|OJJ54211.1 ater|EAU39363.1 atub|OJI87731.1 aver|OJJ08520.1 awen|OJJ30577.1 azon|OJJ49489.1

Group6664: abra|OJJ70460.1 acar|OOF98860.1 afis|EAW24540.1 afla|EED52291.1 agla|OJJ88705.1 akaw|GAA83306.1 aluc|OJZ83802.1 anig|EHA18950.1 aory|EIT82851.1 asyd|OJJ52928.1 atub|OJI90170.1 aver|OJJ08896.1 awen|OJJ41936.1 afla|EED47306.1 aory|EIT74327.1

Group6665: abra|OJJ70474.1 acar|OOF97451.1 acla|EAW13040.1 afis|EAW17627.1 afla|EED49191.1 afum|EAL91441.1 akaw|GAA83323.1 aluc|OJZ83818.1 anid|CBF90148.1 anig|EHA18963.1 aory|EIT78538.1 ater|EAU37061.1 atub|OJI90185.1 awen|OJJ37217.1 azon|OJJ50920.1

Group6666: abra|OJJ70496.1 acar|OOF97430.1 acla|EAW13021.1 afum|EAL91461.1 akaw|GAA83344.1 aluc|OJZ83841.1 anig|EHA18983.1 asyd|OJJ62025.1 ater|EAU37082.1 atub|OJI90207.1 awen|OJJ37199.1 azon|OJJ50898.1 afis|EAW17646.1 anid|CBF90112.1 aver|OJI96116.1

Group6667: abra|OJJ70505.1 acla|EAW08169.1 afis|EAW23205.1 afla|EED47048.1 afum|EAL88278.1 agla|OJJ78740.1 akaw|GAA83354.1 aluc|OJZ83850.1 anig|EHA18992.1 aory|EIT73768.1 asyd|OJJ60190.1 ater|EAU36335.1 atub|OJI90216.1 aver|OJJ04074.1 awen|OJJ39437.1

Group6668: abra|OJJ70517.1 acar|OOF97411.1 acla|EAW08158.1 afis|EAW23196.1 afla|EED47058.1 afum|EAL88287.1 agla|OJJ86333.1 akaw|GAA83364.1 aluc|OJZ83862.1 anid|CBF74708.1 asyd|OJJ60220.1 atub|OJI90230.1 aver|OJJ04048.1 awen|OJJ39426.1 azon|OJJ47627.1

Group6669: abra|OJJ70535.1 acla|EAW08147.1 afis|EAW23192.1 afla|EED47071.1 afum|EAL88291.1 akaw|GAA83380.1 aluc|OJZ83879.1 anid|CBF74520.1 anig|EHA19017.1 aory|EIT72510.1 atub|OJI90246.1 aver|OJJ05435.1 azon|OJJ47613.1 ater|EAU36316.1 awen|OJJ36594.1

Group6670: abra|OJJ70558.1 acla|EAW08356.1 afis|EAW23387.1 afla|EED55575.1 afum|EAL88084.1 agla|OJJ86431.1 akaw|GAA83403.1 aluc|OJZ83901.1 anid|CBF74871.1 asyd|OJJ60116.1 ater|EAU35641.1 atub|OJI90271.1 aver|OJJ04146.1 awen|OJJ39494.1 azon|OJJ47695.1

Group6671: abra|OJJ70565.1 acla|EAW08363.1 afis|EAW23394.1 afla|EED55563.1 afum|EAL88076.1 agla|OJJ86421.1 akaw|GAA83409.1 aluc|OJZ83908.1 anid|CBF74858.1 asyd|OJJ60124.1 ater|EAU35647.1 atub|OJI90279.1 aver|OJJ04138.1 awen|OJJ39485.1 azon|OJJ47688.1

Group6672: abra|OJJ70603.1 acar|OOF97118.1 acla|EAW08179.1 afis|EAW23214.1 afum|EBA27339.1 agla|OJJ78730.1 akaw|GAA89943.1 aluc|OJZ83948.1 anid|CBF74743.1 anig|EHA19077.1 aory|EIT72723.1 asyd|OJJ60181.1 ater|EAU36344.1 atub|OJI90319.1 awen|OJJ39448.1

Group6673: abra|OJJ70610.1 acar|OOF97112.1 acla|EAW13017.1 afis|EAW17650.1 agla|OJJ79554.1 akaw|GAA89949.1 aluc|OJZ83956.1 anig|EHA19083.1 aory|EIT82154.1 asyd|OJJ62038.1 ater|EAU37086.1 atub|OJI90326.1 aver|OJI96144.1 awen|OJJ37195.1 azon|OJJ50894.1

Group6674: abra|OJJ70632.1 acla|EAW08124.1 afis|EAW23161.1 afla|EED46960.1 afum|EAL88322.1 agla|OJJ89672.1 akaw|GAA89971.1 aluc|OJZ83980.1 anid|CBF74463.1 anig|EHA19107.1 aory|EIT72750.1 asyd|OJJ57698.1 aver|OJJ05466.1 awen|OJJ36622.1 azon|OJJ47591.1

Group6675: abra|OJJ70676.1 acar|OOF97051.1 afla|EED48529.1 agla|OJJ89615.1 akaw|GAA90008.1 aluc|OJZ84024.1 anid|CBF89219.1 anig|EHA19145.1 aory|EIT73177.1 asyd|OJJ62525.1 ater|EAU30325.1 atub|OJI90396.1 aver|OJI96648.1 awen|OJJ36657.1 azon|OJJ47541.1

Group6676: abra|OJJ70699.1 acar|OOF90047.1 acla|EAW08066.1 afis|EAW23102.1 afla|EED48550.1 agla|OJJ89614.1 akaw|GAA90030.1 aluc|OJZ84045.1 anid|CBF81361.1 asyd|OJJ59382.1 ater|EAU36226.1 atub|OJI90417.1 aver|OJJ02891.1 awen|OJJ36679.1 azon|OJJ47518.1

Group6677: abra|OJJ70730.1 acar|OOF97008.1 acla|EAW11183.1 afis|EAW15914.1 afum|EAL88549.1 agla|OJJ89578.1 akaw|GAA90058.1 aluc|OJZ84075.1 anid|CBF81307.1 anig|EHA19194.1 asyd|OJJ59353.1 atub|OJI90447.1 aver|OJJ02921.1 awen|OJJ36716.1 azon|OJJ47475.1

Group6678: abra|OJJ70765.1 acar|OOF96980.1 acla|EAW13067.1 afis|EAW17597.1 afla|EED49891.1 afum|EBA27433.1 agla|OJJ86461.1 akaw|GAA90085.1 aluc|OJZ84111.1 aory|EIT72454.1 ater|EAU36530.1 atub|OJI90484.1 awen|OJJ35909.1 acla|EAW13492.1 agla|OJJ84144.1

Group6679: awen|OJJ34621.1 abra|OJJ75538.1 acla|EAW08949.1 afum|EAL91102.1 akaw|GAA91958.1 aluc|OJZ82660.1 anig|EHA27434.1 asyd|OJJ60065.1 ater|EAU29412.1 atub|OJI88892.1 aver|OJI99005.1 acla|EAW07628.1 akaw|GAA83289.1 anig|EHA18934.1 acla|EAW13518.1

Group6680: abra|OJJ70846.1 acar|OOF98269.1 acla|EAW10437.1 afis|EAW25130.1 afum|EAL89801.1 agla|OJJ89036.1 akaw|GAA91714.1 aluc|OJZ87675.1 anid|CBF85701.1 anig|EHA20865.1 asyd|OJJ64265.1 atub|OJI80968.1 aver|OJI98357.1 awen|OJJ40755.1 azon|OJJ44119.1

Group6681: abra|OJJ70866.1 acar|OOF98250.1 acla|EAW10458.1 afis|EAW25109.1 afla|EED55333.1 afum|EAL89780.2 akaw|GAA91733.1 aluc|OJZ87653.1 anid|CBF86070.1 anig|EHA20847.1 asyd|OJJ64459.1 ater|EAU39379.1 atub|OJI80989.1 aver|OJI98562.1 azon|OJJ45099.1

Group6682: abra|OJJ70965.1 acar|OOF98165.1 acla|EAW10526.1 afis|EAW25040.1 afla|EED55412.1 afum|EAL89708.1 akaw|GAA82982.1 aluc|OJZ87567.1 anid|CBF85905.1 anig|EHA20769.1 aory|EIT80904.1 asyd|OJJ64394.1 ater|EAU35790.1 aver|OJI98489.1 azon|OJJ44030.1

Group6683: abra|OJJ71015.1 acar|OOF98125.1 acla|EAW10565.1 afla|EED55456.1 afum|EAL89663.1 agla|OJJ82243.1 akaw|GAA82940.1 aluc|OJZ87517.1 anig|EHA20728.1 aory|EIT80924.1 asyd|OJJ59503.1 ater|EAU33531.1 atub|OJI81133.1 aver|OJJ02771.1 azon|OJJ49457.1

Group6684: abra|OJJ71027.1 acar|OOF98112.1 acla|EAW10578.1 afis|EAW24989.1 afum|EAL89649.1 akaw|GAA82928.1 aluc|OJZ87503.1 anid|CBF81597.1 asyd|OJJ59518.1 ater|EAU35740.1 atub|OJI81146.1 aver|OJJ02755.1 awen|OJJ40594.1 azon|OJJ49445.1 agla|OJJ82268.1

Group6685: abra|OJJ71066.1 acar|OOF98079.1 acla|EAW13232.1 afis|EAW17398.1 afla|EED54884.1 afum|EAL91236.1 agla|OJJ83971.1 akaw|GAA82896.1 aluc|OJZ87465.1 anid|CBF89000.1 anig|EHA20687.1 aory|EIT79355.1 ater|EAU29508.1 atub|OJI81183.1 azon|OJJ47028.1

Group6686: abra|OJJ71081.1 acar|OOF98058.1 acla|EAW10411.1 afis|EAW25157.1 afla|EED57496.1 afum|EAL89832.2 akaw|GAA82880.1 aluc|OJZ87448.1 anid|CBF85655.1 anig|EHA20671.1 aory|EIT81882.1 asyd|OJJ64231.1 atub|OJI81201.1 awen|OJJ36557.1 azon|OJJ50305.1

Group6687: abra|OJJ71096.1 atub|OJI83751.1 aver|OJJ02223.1 abra|OJJ71756.1 afla|EED48306.1 aluc|OJZ84835.1 anig|EHA25745.1 awen|OJJ37891.1 abra|OJJ75283.1 acar|OOF94524.1 akaw|GAA91821.1 aluc|OJZ92844.1 anig|EHA26166.1 atub|OJI87300.1 azon|OJJ43899.1

Group6688: abra|OJJ71138.1 afis|EAW25230.1 afla|EED53674.1 afum|EAL89891.1 agla|OJJ79178.1 akaw|GAA82836.1 aluc|OJZ87397.1 anig|EHA20627.1 aory|EIT75330.1 atub|OJI81249.1 aver|OJI98108.1 anid|CBF85336.1 asyd|OJJ64022.1 ater|EAU34388.1 awen|OJJ40895.1

Group6689: abra|OJJ71160.1 acla|EAW10338.1 afis|EAW25251.1 afla|EED53718.1 afum|EAL89913.1 akaw|GAA82818.1 aluc|OJZ87379.1 anid|CBF85343.1 asyd|OJJ64026.1 ater|EAU34354.1 atub|OJI81267.1 aver|OJI98112.1 azon|OJJ44549.1 agla|OJJ89224.1 awen|OJJ40965.1

Group6690: abra|OJJ71180.1 acar|OOF97985.1 acla|EAW10324.1 afis|EAW25268.1 afla|EED53733.1 afum|EAL89931.1 agla|OJJ89238.1 akaw|GAA82805.1 aluc|OJZ87363.1 anid|CBF85378.1 anig|EHA20598.1 aory|EIT75272.1 ater|EAU34411.1 atub|OJI81282.1 awen|OJJ40982.1

Group6691: abra|OJJ71186.1 acla|EAW10317.1 afis|EAW25275.1 afla|EED53740.1 afum|EAL89938.1 agla|OJJ89249.1 akaw|GAA82799.1 anid|CBF85391.1 anig|EHA20592.1 aory|EIT75341.1 asyd|OJJ64057.1 ater|EAU34418.1 aver|OJI98142.1 awen|OJJ40989.1 azon|OJJ44576.1

Group6692: ater|EAU36088.1 abra|OJJ74682.1 akaw|GAA81932.1 aluc|OJZ92259.1 atub|OJI86693.1 anid|CBF82907.1 afis|EAW19769.1 afla|EED49601.1 afum|EBA27220.1 akaw|GAA82112.1 aluc|OJZ92064.1 anig|EHA22525.1 aver|OJJ03005.1 azon|OJJ50039.1 aory|EIT80948.1

Group6693: acar|OOF97935.1 abra|OJJ74828.1 afla|EED51414.1 akaw|GAA83591.1 anig|EHA26560.1 ater|EAU38714.1 atub|OJI86833.1 abra|OJJ74829.1 acla|EAW09105.1 aluc|OJZ92406.1 anig|EHA26559.1 acar|OOF95816.1 agla|OJJ89327.1 asyd|OJJ52422.1 anig|EHA26216.1

Group6694: abra|OJJ71254.1 acar|OOF97917.1 afla|EED53786.1 agla|OJJ83847.1 akaw|GAA82734.1 aluc|OJZ87289.1 anid|CBF75289.1 anig|EHA20530.1 aory|EIT78457.1 asyd|OJJ59888.1 ater|EAU34457.1 atub|OJI81352.1 aver|OJJ02398.1 awen|OJJ41056.1 azon|OJJ44615.1

Group6695: abra|OJJ71309.1 acla|EAW10231.1 afis|EAW25361.1 afla|EED53843.1 afum|EAL90023.1 agla|OJJ83911.1 akaw|GAA82682.1 aluc|OJZ87235.1 anid|CBF77457.1 anig|EHA20482.1 asyd|OJJ53215.1 atub|OJI81407.1 aver|OJJ03636.1 awen|OJJ41087.1 azon|OJJ44667.1

Group6696: abra|OJJ71344.1 acar|OOF97828.1 acla|EAW10195.1 afis|EAW25400.1 afla|EED53878.1 afum|EAL90059.1 agla|OJJ87040.1 akaw|GAA82648.1 aluc|OJZ87197.1 anid|CBF77522.1 aory|EIT78462.1 ater|EAU34551.1 atub|OJI81442.1 awen|OJJ41127.1 azon|OJJ44701.1

Group6697: abra|OJJ71406.1 acla|EAW10129.1 afis|EAW25458.1 afum|EAL90118.2 aluc|OJZ87136.1 anig|EHA20395.1 asyd|OJJ53121.1 atub|OJI81506.1 afla|EED53947.1 agla|OJJ87111.1 anid|CBF77643.1 aory|EIT78342.1 ater|EAU34628.1 aver|OJJ03733.1 awen|OJJ41190.1

Group6698: abra|OJJ71484.1 acla|EAW10066.1 afis|EAW25519.1 afla|EED54021.1 afum|EAL90184.1 agla|OJJ87283.1 akaw|GAA82536.1 aluc|OJZ87063.1 anid|CBF77792.1 anig|EHA20336.1 asyd|OJJ53026.1 ater|EAU33365.1 atub|OJI81572.1 awen|OJJ41262.1 azon|OJJ44827.1

Group6699: abra|OJJ71579.1 acar|OOF96309.1 acla|EAW15126.1 afis|EAW20096.1 afla|EED45274.1 agla|OJJ88858.1 akaw|GAA89345.1 aluc|OJZ86316.1 anid|CBF84890.1 anig|EHA21559.1 asyd|OJJ63179.1 atub|OJI84219.1 aver|OJI97775.1 awen|OJJ31859.1 azon|OJJ46266.1

Group6700: acar|OOF90408.1 acla|EAW15147.1 afla|EED53356.1 akaw|GAA89323.1 aluc|OJZ86347.1 anig|EHA21539.1 aory|EIT74898.1 ater|EAU38733.1 atub|OJI84187.1 afis|EAW20115.1 afum|EAL85232.1 agla|OJJ88834.1 anid|CBF84932.1 asyd|OJJ63159.1 azon|OJJ46249.1

Group6701: abra|OJJ71648.1 acar|OOF96362.1 afla|EED55106.1 agla|OJJ88187.1 akaw|GAA89281.1 aluc|OJZ86392.1 anig|EHA21506.1 ater|EAU31825.1 atub|OJI84141.1 awen|OJJ33397.1 azon|OJJ46081.1 abra|OJJ75690.1 asyd|OJJ52523.1 aver|OJJ06741.1 aory|EIT81590.1

Group6702: abra|OJJ71660.1 acar|OOF96372.1 acla|EAW15197.1 afla|EED57184.1 agla|OJJ88765.1 akaw|GAA89271.1 aluc|OJZ86404.1 anig|EHA21496.1 asyd|OJJ63108.1 ater|EAU38796.1 atub|OJI84129.1 aver|OJI97853.1 awen|OJJ32060.1 azon|OJJ46094.1 aory|EIT73999.1

Group6703: abra|OJJ71684.1 acar|OOF96394.1 acla|EAW15222.1 afis|EAW20187.1 afla|EED57159.1 afum|EAL85306.1 agla|OJJ84679.1 akaw|GAA89249.1 aluc|OJZ86426.1 anig|EHA21473.1 asyd|OJJ63087.1 atub|OJI84105.1 aver|OJI97875.1 awen|OJJ31187.1 azon|OJJ46117.1

Group6704: abra|OJJ71704.1 acar|OOF90838.1 acla|EAW11767.1 afis|EAW23682.1 akaw|GAA92698.1 aluc|OJZ90156.1 anig|EHA24839.1 aory|EIT78101.1 asyd|OJJ52062.1 ater|EAU29953.1 aver|OJI99826.1 awen|OJJ29989.1 azon|OJJ42782.1 asyd|OJJ60502.1 afla|EED50753.1

Group6705: abra|OJJ71721.1 acar|OOF96426.1 acla|EAW06443.1 afla|EED52827.1 afum|EAL93248.1 agla|OJJ81593.1 akaw|GAA85169.1 aluc|OJZ86467.1 anig|EHA21443.1 aory|EIT78233.1 ater|EAU34837.1 atub|OJI84062.1 awen|OJJ31106.1 azon|OJJ46555.1 afis|EAW18548.1

Group6706: abra|OJJ71731.1 acar|OOF97589.1 afis|EAW22619.1 afla|EED54115.1 afum|EAL90577.1 agla|OJJ78574.1 anid|CBF82374.1 aory|EIT76007.1 asyd|OJJ57664.1 ater|EAU34345.1 aver|OJJ05526.1 awen|OJJ31955.1 aory|EIT72769.1 afla|EED46969.1 agla|OJJ78787.1

Group6707: abra|OJJ71767.1 acar|OOF90207.1 afla|EED45608.1 akaw|GAA85140.1 aluc|OJZ86497.1 anig|EHA22147.1 aory|EIT75590.1 asyd|OJJ53487.1 atub|OJI84029.1 aver|OJJ01718.1 aluc|OJZ80178.1 acar|OOF94848.1 akaw|GAA93285.1 aluc|OJZ90159.1 asyd|OJJ65407.1

Group6708: abra|OJJ71790.1 acar|OOF96470.1 afla|EED56596.1 agla|OJJ83771.1 akaw|GAA85120.1 aluc|OJZ86520.1 anid|CBF82149.1 anig|EHA22126.1 aory|EIT77664.1 asyd|OJJ54326.1 atub|OJI84002.1 aver|OJJ05280.1 awen|OJJ30240.1 afla|EED49716.1 awen|OJJ29855.1

Group6709: abra|OJJ71796.1 acar|OOF96476.1 acla|EAW12078.1 afis|EAW19719.1 afla|EED52814.1 afum|EAL86807.2 akaw|GAA85114.1 aluc|OJZ86526.1 anid|CBF78803.1 anig|EHA22120.1 aory|EIT78244.1 asyd|OJJ55182.1 ater|EAU34845.1 atub|OJI83996.1 aver|OJJ07025.1

Group6710: abra|OJJ71808.1 acar|OOF96486.1 acla|EAW13847.1 afis|EAW18561.1 afla|EED52799.1 afum|EAL93262.1 agla|OJJ81598.1 akaw|GAA85104.1 aluc|OJZ86538.1 anid|CBF70292.1 aory|EIT78167.1 asyd|OJJ58733.1 atub|OJI83985.1 aver|OJJ01345.1 awen|OJJ31119.1

Group6711: abra|OJJ71815.1 acla|EAW13855.1 afis|EAW18569.1 afla|EED52782.1 afum|EAL93271.1 agla|OJJ81620.1 akaw|GAA85095.1 anid|CBF70309.1 anig|EHA22102.1 aory|EIT78229.1 asyd|OJJ58724.1 atub|OJI83975.1 aver|OJJ01337.1 awen|OJJ31126.1 azon|OJJ46607.1

Group6712: abra|OJJ71856.1 acar|OOF96526.1 acla|EAW13888.1 afis|EAW18600.1 afum|EAL93308.1 akaw|GAA85061.1 aluc|OJZ86590.1 anid|CBF70401.1 anig|EHA22070.1 asyd|OJJ58631.1 ater|EAU34894.1 atub|OJI83933.1 aver|OJJ01237.1 awen|OJJ31165.1 azon|OJJ46639.1

Group6713: abra|OJJ71892.1 acar|OOF96554.1 acla|EAW09630.1 afis|EAW21510.1 afla|EED55948.1 afum|EAL92530.1 agla|OJJ85674.1 akaw|GAA85026.1 aluc|OJZ86631.1 anid|CBF76439.1 asyd|OJJ61100.1 atub|OJI83897.1 aver|OJJ00415.1 awen|OJJ39184.1 azon|OJJ51509.1

Group6714: abra|OJJ71962.1 afis|EAW21563.1 afla|EED56017.1 afum|EAL92582.1 agla|OJJ85743.1 akaw|GAA84965.1 anid|CBF76334.1 anig|EHA21977.1 aory|EIT76724.1 asyd|OJJ61043.1 ater|EAU34970.1 aver|OJJ00359.1 awen|OJJ39124.1 azon|OJJ51454.1 acla|EAW09574.1

Group6715: akaw|GAA84877.1 abra|OJJ72058.1 acar|OOF89882.1 aluc|OJZ86794.1 atub|OJI83724.1 acar|OOF89761.1 afla|EED57898.1 anig|EHA21889.1 asyd|OJJ60722.1 acla|EAW09491.1 afis|EAW21645.1 afum|EAL92679.1 ater|EAU38330.1 agla|OJJ84562.1 anid|CBF83532.1

Group6716: abra|OJJ72060.1 acar|OOF89884.1 afla|EED57897.1 agla|OJJ79717.1 akaw|GAA84875.1 anid|CBF83535.1 anig|EHA21887.1 aory|EIT82737.1 asyd|OJJ60724.1 ater|EAU38331.1 atub|OJI83722.1 aver|OJJ00057.1 awen|OJJ33905.1 azon|OJJ48656.1 aluc|OJZ86796.1

Group6717: abra|OJJ72128.1 acar|OOF96772.1 afla|EED46870.1 afum|EAL90429.1 agla|OJJ86322.1 akaw|GAA84795.1 aluc|OJZ86885.1 anid|CBF77734.1 anig|EHA21759.1 aory|EIT72614.1 asyd|OJJ53056.1 ater|EAU33000.1 atub|OJI83634.1 aver|OJJ03788.1 awen|OJJ32568.1

Group6718: abra|OJJ72191.1 acar|OOF96817.1 afla|EED48888.1 agla|OJJ85283.1 akaw|GAA84742.1 aluc|OJZ86942.1 anig|EHA21702.1 aory|EIT81212.1 asyd|OJJ57933.1 atub|OJI83580.1 aver|OJJ04464.1 awen|OJJ34665.1 azon|OJJ50423.1 anid|CBF79551.1 awen|OJJ36918.1

Group6719: abra|OJJ72202.1 acar|OOF96822.1 afis|EAW24464.1 afla|EED55202.1 afum|EAL86135.1 agla|OJJ80322.1 akaw|GAA84734.1 aluc|OJZ86952.1 anig|EHA21695.1 aory|EIT83510.1 asyd|OJJ57432.1 ater|EAU29970.1 atub|OJI83571.1 aver|OJJ05748.1 awen|OJJ32726.1

Group6720: abra|OJJ72210.1 acla|EAW09560.1 afis|EAW21188.1 afla|EED53237.1 afum|EAL92210.1 akaw|GAA84724.1 aluc|OJZ86962.1 anig|EHA21686.1 aory|EIT75124.1 asyd|OJJ56246.1 ater|EAU34395.1 atub|OJI83561.1 aver|OJJ05169.1 azon|OJJ49087.1 anid|CBF82034.1

Group6721: abra|OJJ72250.1 acar|OOF96881.1 acla|EAW14644.1 afis|EAW24569.1 afla|EED50216.1 agla|OJJ79377.1 akaw|GAA84687.1 aluc|OJZ87001.1 anid|CBF71507.1 anig|EHA21645.1 aory|EIT77275.1 asyd|OJJ54920.1 ater|EAU36842.1 atub|OJI83518.1 afla|EED50215.1

Group6722: abra|OJJ72325.1 acar|OOF94889.1 acla|EAW11998.1 afis|EAW19638.1 afum|EAL86884.1 agla|OJJ80780.1 akaw|GAA86075.1 aluc|OJZ90310.1 anid|CBF71339.1 anig|EHA23797.1 asyd|OJJ55100.1 atub|OJI81735.1 aver|OJJ07106.1 awen|OJJ41883.1 azon|OJJ47081.1

Group6723: abra|OJJ72356.1 acar|OOF97624.1 acla|EAW11960.1 afis|EAW19597.1 afla|EED56953.1 afum|EAL86921.1 agla|OJJ80739.1 akaw|GAA86103.1 aluc|OJZ90335.1 anig|EHA23819.1 asyd|OJJ54506.1 ater|EAU32928.1 atub|OJI81762.1 aver|OJJ07676.1 awen|OJJ41845.1

Group6724: abra|OJJ72358.1 acar|OOF94860.1 agla|OJJ81678.1 akaw|GAA86105.1 aluc|OJZ90337.1 anid|CBF87127.1 anig|EHA23821.1 asyd|OJJ60367.1 ater|EAU31885.1 atub|OJI81764.1 aver|OJJ01794.1 awen|OJJ33424.1 aver|OJI99704.1 acla|EAW15010.1 anid|CBF85560.1

Group6725: abra|OJJ72364.1 acar|OOF94858.1 afis|EAW19111.1 afla|EED52118.1 afum|EAL93842.2 agla|OJJ84490.1 akaw|GAA86115.1 aluc|OJZ90351.1 anid|CBF89579.1 anig|EHA23835.1 asyd|OJJ52177.1 ater|EAU37369.1 atub|OJI81778.1 aver|OJI95854.1 awen|OJJ32509.1

Group6726: abra|OJJ72385.1 acar|OOF90291.1 acla|EAW11951.1 afis|EAW19588.1 afum|EAL86930.1 agla|OJJ80725.1 akaw|GAA86134.1 aluc|OJZ90372.1 anig|EHA23846.1 asyd|OJJ55151.1 ater|EAU32960.1 atub|OJI81798.1 aver|OJJ07056.1 awen|OJJ41802.1 azon|OJJ47029.1

Group6727: abra|OJJ72403.1 acar|OOF94825.1 acla|EAW11938.1 afis|EAW19573.1 afla|EED56810.1 afum|EAL86944.1 agla|OJJ80701.1 akaw|GAA86147.1 aluc|OJZ90388.1 anid|CBF71230.1 asyd|OJJ55168.1 atub|OJI81816.1 aver|OJJ07038.1 awen|OJJ41786.1 azon|OJJ44892.1

Group6728: abra|OJJ72411.1 acar|OOF94818.1 acla|EAW11930.1 afis|EAW19564.1 afum|EAL86948.1 akaw|GAA86154.1 aluc|OJZ90395.1 anid|CBF78022.1 anig|EHA23864.1 asyd|OJJ55234.1 ater|EAU32990.1 atub|OJI81825.1 aver|OJJ06989.1 awen|OJJ41779.1 azon|OJJ44898.1

Group6729: abra|OJJ72416.1 acar|OOF94813.1 acla|EAW12249.1 afis|EAW17524.1 afla|EED45961.1 afum|EAL91339.2 akaw|GAA86159.1 aluc|OJZ90400.1 anid|CBF73870.1 anig|EHA23870.1 asyd|OJJ55449.1 atub|OJI81831.1 aver|OJJ06522.1 awen|OJJ41196.1 azon|OJJ46820.1

Group6730: abra|OJJ72420.1 acar|OOF90369.1 acla|EAW09083.1 afis|EAW25670.1 afla|EED46155.1 afum|EAL84342.1 akaw|GAA86162.1 aluc|OJZ90403.1 anid|CBF69527.1 anig|EHA23871.1 aory|EIT76456.1 asyd|OJJ59184.1 ater|EAU33005.1 atub|OJI81834.1 ater|EAU31774.1

Group6731: abra|OJJ72442.1 acar|OOF94794.1 acla|EAW12675.1 afis|EAW17838.1 afla|EED46183.1 afum|EAL91648.1 akaw|GAA86181.1 aluc|OJZ90426.1 anid|CBF77981.1 anig|EHA23896.1 aory|EIT76436.1 asyd|OJJ55259.1 atub|OJI81858.1 aver|OJJ06966.1 awen|OJJ41738.1

Group6732: abra|OJJ72464.1 acar|OOF94774.1 acla|EAW12697.1 afis|EAW17860.1 afla|EED54666.1 afum|EAL91668.1 agla|OJJ83107.1 akaw|GAA86201.1 aluc|OJZ90452.1 anig|EHA23914.1 aory|EIT78803.1 ater|EAU33065.1 atub|OJI81879.1 awen|OJJ41712.1 azon|OJJ48407.1

Group6733: abra|OJJ72485.1 acar|OOF94750.1 acla|EAW12716.1 afis|EAW17878.1 afum|EAL91686.1 agla|OJJ83068.1 akaw|GAA86220.1 aluc|OJZ90472.1 anid|CBF82881.1 anig|EHA23931.1 asyd|OJJ55065.1 atub|OJI81896.1 aver|OJJ05548.1 awen|OJJ41686.1 aver|OJJ07146.1

Group6734: abra|OJJ72534.1 acar|OOF94718.1 acla|EAW12741.1 afis|EAW17904.1 afla|EED46296.1 afum|EAL91712.1 agla|OJJ83038.1 akaw|GAA86256.1 aluc|OJZ90513.1 anid|CBF76987.1 anig|EHA23969.1 aory|EIT76331.1 atub|OJI81938.1 awen|OJJ41646.1 azon|OJJ45694.1

Group6735: abra|OJJ72541.1 acla|EAW12746.1 afis|EAW17910.1 afla|EED46312.1 afum|EAL91719.2 agla|OJJ83033.1 akaw|GAA86265.1 aluc|OJZ90525.1 anid|CBF76965.1 anig|EHA23977.1 aory|EIT76322.1 asyd|OJJ57039.1 atub|OJI81947.1 awen|OJJ41639.1 azon|OJJ48361.1

Group6736: abra|OJJ72548.1 afla|EED46318.1 agla|OJJ83027.1 aluc|OJZ90536.1 anig|EHA23984.1 aory|EIT81238.1 atub|OJI81957.1 awen|OJJ41629.1 azon|OJJ48354.1 acla|EAW12753.1 afis|EAW17916.1 afum|EAL91726.1 anid|CBF76975.1 asyd|OJJ57049.1 aver|OJJ03247.1

Group6737: abra|OJJ72553.1 acar|OOF94699.1 acla|EAW12758.1 agla|OJJ83021.1 akaw|GAA85185.1 aluc|OJZ90540.1 anig|EHA23988.1 aory|EIT81219.1 asyd|OJJ57056.1 ater|EAU33551.1 atub|OJI81962.1 awen|OJJ41625.1 azon|OJJ48348.1 afla|EED46323.1 ater|EAU30195.1

Group6738: abra|OJJ72594.1 acla|EAW12779.1 afis|EAW17942.1 afum|EAL91753.1 agla|OJJ82990.1 akaw|GAA85222.1 aluc|OJZ90576.1 anid|CBF77051.1 anig|EHA24020.1 asyd|OJJ56995.1 ater|EAU33522.1 atub|OJI82000.1 aver|OJJ03292.1 awen|OJJ41601.1 azon|OJJ48321.1

Group6739: abra|OJJ72807.1 acar|OOF96013.1 acla|EAW11434.1 afla|EED49170.1 afum|EAL87418.1 agla|OJJ86922.1 akaw|GAA85410.1 aluc|OJZ83305.1 anig|EHA24188.1 asyd|OJJ53262.1 ater|EAU29762.1 atub|OJI82205.1 aver|OJJ03589.1 awen|OJJ35456.1 azon|OJJ51287.1

Group6740: abra|OJJ72821.1 acar|OOF96026.1 acla|EAW11418.1 afis|EAW24004.1 afla|EED49157.1 afum|EAL87434.1 akaw|GAA85422.1 aluc|OJZ83323.1 anid|CBF77392.1 anig|EHA24201.1 aory|EIT78585.1 asyd|OJJ53246.1 ater|EAU29747.1 atub|OJI82220.1 awen|OJJ35475.1

Group6741: abra|OJJ72889.1 acar|OOF96079.1 afla|EED58058.1 agla|OJJ83817.1 akaw|GAA85476.1 aluc|OJZ83397.1 anid|CBF75382.1 anig|EHA24250.1 aory|EIT79702.1 asyd|OJJ59844.1 ater|EAU35456.1 atub|OJI82283.1 aver|OJJ02445.1 awen|OJJ35574.1 azon|OJJ46947.1

Group6742: abra|OJJ72981.1 acla|EAW08029.1 afis|EAW23063.1 afla|EED54954.1 afum|EAL88416.1 agla|OJJ89411.1 akaw|GAA85556.1 aluc|OJZ83491.1 anid|CBF80977.1 anig|EHA24326.1 aory|EIT80367.1 asyd|OJJ56356.1 atub|OJI82384.1 aver|OJJ06358.1 awen|OJJ35672.1

Group6743: abra|OJJ73000.1 acar|OOF89940.1 acla|EAW08013.1 afis|EAW23047.1 afla|EED54971.1 afum|EAL88433.1 akaw|GAA85574.1 aluc|OJZ83510.1 anid|CBF80947.1 aory|EIT80393.1 asyd|OJJ56340.1 ater|EAU29599.1 atub|OJI82403.1 aver|OJJ06373.1 awen|OJJ35692.1

Group6744: abra|OJJ73027.1 acar|OOF96194.1 acla|EAW07989.1 afis|EAW23024.1 afum|EAL88458.1 agla|OJJ89455.1 akaw|GAA85601.1 aluc|OJZ83539.1 anid|CBF80907.1 asyd|OJJ56331.1 atub|OJI82430.1 aver|OJJ06383.1 awen|OJJ30719.1 azon|OJJ44180.1 afla|EED54992.1

Group6745: abra|OJJ73073.1 acar|OOG00306.1 acla|EAW13615.1 afis|EAW19976.1 afla|EED45377.1 afum|EBA27213.1 akaw|GAA88088.1 aluc|OJZ81352.1 anig|EHA18654.1 aory|EIT82383.1 ater|EAU31775.1 atub|OJI91556.1 aver|OJJ02016.1 awen|OJJ29807.1 azon|OJJ42869.1

Group6746: abra|OJJ73141.1 acar|OOF92575.1 afis|EAW24505.1 afla|EED48167.1 afum|EAL86173.1 akaw|GAA88031.1 aluc|OJZ81290.1 anid|CBF76012.1 anig|EHA18589.1 aory|EIT83029.1 asyd|OJJ52436.1 atub|OJI91498.1 aver|OJJ03974.1 awen|OJJ31247.1 azon|OJJ47356.1

Group6747: abra|OJJ73146.1 acar|OOG00359.1 acla|EAW07022.1 afis|EAW16648.1 afum|EAL89252.1 agla|OJJ82594.1 akaw|GAA88026.1 aluc|OJZ81285.1 anid|CBF82059.1 anig|EHA18584.1 asyd|OJJ54239.1 atub|OJI91492.1 aver|OJJ05191.1 awen|OJJ40066.1 azon|OJJ45867.1

Group6748: abra|OJJ73155.1 acar|OOG00372.1 acla|EAW07029.1 afis|EAW16655.1 afla|EED44939.1 afum|EAL89259.1 akaw|GAA88017.1 aluc|OJZ81275.1 anid|CBF82069.1 anig|EHA18575.1 aory|EIT73497.1 asyd|OJJ54245.1 atub|OJI91483.1 aver|OJJ05197.1 awen|OJJ40059.1

Group6749: abra|OJJ73277.1 acla|EAW06952.1 afis|EAW16567.1 afla|EED56096.1 afum|EAL89164.1 agla|OJJ82754.1 akaw|GAA92728.1 aluc|OJZ88772.1 anid|CBF81820.1 aory|EIT76655.1 asyd|OJJ59653.1 ater|EAU36636.1 aver|OJJ02629.1 awen|OJJ40252.1 azon|OJJ49113.1

Group6750: abra|OJJ73327.1 acar|OOF97930.1 acla|EAW09072.1 afis|EAW25645.1 afla|EED52302.1 afum|EAL84331.1 akaw|GAA92791.1 aluc|OJZ88709.1 anid|CBF71476.1 anig|EHA18410.1 aory|EIT74506.1 asyd|OJJ54944.1 atub|OJI91295.1 aver|OJJ07251.1 azon|OJJ48164.1

Group6751: abra|OJJ73348.1 acar|OOG00562.1 acla|EAW12349.1 afis|EAW16770.1 afla|EED45246.1 afum|EAL86338.1 akaw|GAA83134.1 aluc|OJZ88691.1 anid|CBF87373.1 anig|EHA18372.1 aory|EIT73754.1 asyd|OJJ61421.1 atub|OJI91271.1 aver|OJJ00757.1 azon|OJJ43652.1

Group6752: abra|OJJ73364.1 acar|OOG00571.1 acla|EAW06984.1 afis|EAW16598.1 afla|EED45017.1 afum|EAL89194.1 agla|OJJ82656.1 akaw|GAA83120.1 aluc|OJZ88676.1 anig|EHA18358.1 aory|EIT73536.1 ater|EAU36677.1 atub|OJI91257.1 awen|OJJ40210.1 azon|OJJ49076.1

Group6753: abra|OJJ73430.1 acar|OOG00637.1 afis|EAW16320.1 afla|EED53561.1 agla|OJJ88281.1 akaw|GAA83062.1 aluc|OJZ88611.1 anid|CBF89153.1 anig|EHA18304.1 aory|EIT75018.1 asyd|OJJ62565.1 ater|EAU34137.1 atub|OJI91196.1 aver|OJI96689.1 azon|OJJ44979.1

Group6754: abra|OJJ73475.1 acar|OOG00672.1 acla|EAW06800.1 afis|EAW16350.1 afla|EED53522.1 afum|EAL88975.1 akaw|GAA83026.1 aluc|OJZ88572.1 anid|CBF89208.1 anig|EHA18270.1 asyd|OJJ62521.1 atub|OJI91158.1 aver|OJI96640.1 awen|OJJ37752.1 azon|OJJ45012.1

Group6755: abra|OJJ73506.1 acla|EAW07439.1 afis|EAW22466.1 afla|EED54498.1 afum|EAL90725.1 agla|OJJ87558.1 akaw|GAA87453.1 aluc|OJZ88542.1 anid|CBF88905.1 anig|EHA18242.1 asyd|OJJ62838.1 atub|OJI91127.1 aver|OJI96881.1 awen|OJJ35092.1 azon|OJJ50017.1

Group6756: abra|OJJ73536.1 acla|EAW07458.1 afis|EAW22489.1 afla|EED54518.1 afum|EAL90704.1 agla|OJJ80858.1 akaw|GAA88614.1 aluc|OJZ88524.1 anid|CBF88948.1 anig|EHA18226.1 aory|EIT78914.1 asyd|OJJ62812.1 atub|OJI91109.1 awen|OJJ35109.1 azon|OJJ49984.1

Group6757: abra|OJJ73546.1 acar|OOG00720.1 acla|EAW07465.1 afis|EAW22498.1 afla|EED54525.1 afum|EAL90695.1 akaw|GAA88606.1 aluc|OJZ88514.1 anid|CBF88958.1 anig|EHA18220.1 asyd|OJJ62806.1 ater|EAU39220.1 atub|OJI91099.1 aver|OJI96852.1 awen|OJJ35120.1

Group6758: abra|OJJ73570.1 afis|EAW22521.1 akaw|GAA88585.1 aluc|OJZ88490.1 anid|CBF89004.1 anig|EHA18199.1 asyd|OJJ62778.1 ater|EAU39189.1 atub|OJI91077.1 awen|OJJ35148.1 azon|OJJ49953.1 acla|EAW07488.1 agla|OJJ80882.1 aory|EIT78870.1 aver|OJI96823.1

Group6759: abra|OJJ73588.1 acar|OOG00770.1 acla|EAW07506.1 afis|EAW22538.1 afum|EAL90656.2 akaw|GAA88564.1 aluc|OJZ88462.1 anid|CBF89034.1 anig|EHA18182.1 aory|EIT78956.1 asyd|OJJ62763.1 ater|EAU39171.1 atub|OJI91060.1 aver|OJI96806.1 awen|OJJ35167.1

Group6760: abra|OJJ73674.1 afis|EAW22614.1 afla|EED49812.1 afum|EAL90584.1 akaw|GAA88483.1 aluc|OJZ88377.1 anid|CBF88254.1 aory|EIT77600.1 asyd|OJJ63671.1 ater|EAU39088.1 atub|OJI90972.1 acla|EAW07580.1 aluc|OJZ88378.1 awen|OJJ35242.1 azon|OJJ49865.1

Group6761: abra|OJJ73718.1 acar|OOG00880.1 acla|EAW07610.1 afis|EAW22640.1 afla|EED53497.1 afum|EAL90552.2 akaw|GAA88445.1 atub|OJI90928.1 aver|OJI99806.1 acla|EAW07609.1 afis|EAW22639.1 afla|EED53496.1 afum|EAL90553.2 anig|EHA18068.1 aory|EIT75156.1

Group6762: abra|OJJ73754.1 acar|OOG00917.1 acla|EAW07645.1 afis|EAW22665.1 afla|EED49939.1 afum|EAL90527.1 agla|OJJ79059.1 akaw|GAA88413.1 aluc|OJZ88297.1 anid|CBF88163.1 anig|EHA18037.1 aory|EIT83449.1 aver|OJI97319.1 awen|OJJ35291.1 azon|OJJ49823.1

Group6763: abra|OJJ73790.1 acla|EAW07681.1 akaw|GAA89409.1 aluc|OJZ88264.1 anig|EHA18010.1 ater|EAU39016.1 atub|OJI90863.1 aver|OJI97363.1 afis|EAW22696.1 afum|EAL90496.1 afla|EED49976.1 agla|OJJ79022.1 anid|CBF88100.1 aory|EIT74741.1 asyd|OJJ63571.1

Group6764: abra|OJJ73800.1 acar|OOG00943.1 acla|EAW07678.1 afis|EAW22693.1 agla|OJJ79019.1 akaw|GAA89420.1 aluc|OJZ88253.1 anig|EHA17995.1 asyd|OJJ63568.1 ater|EAU39013.1 atub|OJI90851.1 aver|OJI97366.1 awen|OJJ35331.1 afum|EBA27408.1 azon|OJJ50247.1

Group6765: abra|OJJ73808.1 acar|OOG00964.1 acla|EAW07696.1 afis|EAW22713.1 afla|EED49998.1 afum|EAL90481.1 agla|OJJ78999.1 akaw|GAA89428.1 aluc|OJZ88242.1 anig|EHA17988.1 asyd|OJJ63547.1 atub|OJI90844.1 aver|OJI97387.1 awen|OJJ35348.1 azon|OJJ49769.1

Group6766: abra|OJJ73835.1 afis|EAW22762.1 afla|EED50024.1 afum|EAL90437.1 agla|OJJ83658.1 akaw|GAA89453.1 aluc|OJZ88214.1 aory|EIT74697.1 asyd|OJJ63514.1 atub|OJI90814.1 aver|OJI97416.1 awen|OJJ35435.1 acar|OOG00986.1 azon|OJJ49742.1 anid|CBF88007.1

Group6767: abra|OJJ73917.1 acar|OOG01049.1 acla|EAW07793.1 afis|EAW22831.1 afla|EED50106.1 afum|EAL90371.2 agla|OJJ83508.1 akaw|GAA89523.1 aluc|OJZ88129.1 anid|CBF87881.1 anig|EHA17895.1 ater|EAU38880.1 atub|OJI90735.1 awen|OJJ38897.1 azon|OJJ49678.1

Group6768: abra|OJJ73932.1 acar|OOG01064.1 acla|EAW07810.1 afis|EAW22847.1 afla|EED50122.1 afum|EAL90355.1 agla|OJJ83477.1 akaw|GAA89540.1 aluc|OJZ88110.1 anig|EHA17880.1 aory|EIT74658.1 ater|EAU38863.1 atub|OJI90718.1 awen|OJJ38876.1 azon|OJJ49665.1

Group6769: abra|OJJ73947.1 acar|OOG01075.1 afis|EAW22855.1 afum|EAL90345.1 agla|OJJ83466.1 akaw|GAA89552.1 aluc|OJZ88097.1 anid|CBF87835.1 anig|EHA17866.1 aory|EIT74691.1 asyd|OJJ63409.1 ater|EAU38808.1 atub|OJI90705.1 aver|OJI97528.1 awen|OJJ38863.1

Group6770: abra|OJJ73968.1 acar|OOG01094.1 acla|EAW07839.1 afis|EAW22875.1 afla|EED50160.1 akaw|GAA89571.1 aluc|OJZ88076.1 anid|CBF87814.1 anig|EHA17848.1 aory|EIT76936.1 asyd|OJJ63393.1 ater|EAU38839.1 atub|OJI90685.1 aver|OJI97542.1 awen|OJJ38847.1

Group6771: abra|OJJ73970.1 acar|OOG01096.1 acla|EAW07841.1 afis|EAW22877.1 afla|EED50163.1 afum|EAL90322.1 agla|OJJ83445.1 akaw|GAA89573.1 aluc|OJZ88074.1 anid|CBF87813.1 anig|EHA17847.1 aory|EIT76903.1 asyd|OJJ63391.1 ater|EAU38841.1 atub|OJI90683.1

Group6772: abra|OJJ74113.1 acar|OOG01218.1 acla|EAW07955.1 agla|OJJ83294.1 akaw|GAA87587.1 aluc|OJZ87921.1 anid|CBF87576.1 anig|EHA17715.1 aory|EIT79068.1 asyd|OJJ63247.1 ater|EAU30629.1 atub|OJI90532.1 aver|OJI97709.1 awen|OJJ38571.1 azon|OJJ50433.1

Group6773: abra|OJJ74132.1 acla|EAW14127.1 afis|EAW18828.1 afla|EED55161.1 afum|EAL93562.1 agla|OJJ80864.1 akaw|GAA89754.1 aluc|OJZ80629.1 anig|EHA27202.1 aory|EIT81622.1 ater|EAU33514.1 atub|OJI87485.1 awen|OJJ41929.1 acar|OOF91236.1 agla|OJJ89620.1

Group6774: abra|OJJ74135.1 acar|OOF95596.1 afis|EAW21379.1 afla|EED45561.1 afum|EAL92394.1 akaw|GAA89751.1 aluc|OJZ80633.1 anid|CBF82348.1 anig|EHA27199.1 aory|EIT75532.1 asyd|OJJ55145.1 atub|OJI87488.1 aver|OJJ07062.1 awen|OJJ36399.1 awen|OJJ36158.1

Group6775: abra|OJJ74164.1 acar|OOF95558.1 acla|EAW06847.1 afis|EAW16468.1 afla|EED49700.1 akaw|GAA86852.1 aluc|OJZ89972.1 asyd|OJJ62459.1 ater|EAU34008.1 atub|OJI84835.1 aver|OJI96594.1 awen|OJJ33441.1 azon|OJJ45095.1 anid|CBF89397.1 agla|OJJ85937.1

Group6776: abra|OJJ74192.1 afla|EED57657.1 agla|OJJ87790.1 akaw|GAA86825.1 aluc|OJZ89943.1 anid|CBF88517.1 anig|EHA27142.1 aory|EIT79910.1 asyd|OJJ63806.1 ater|EAU34276.1 atub|OJI84863.1 aver|OJI97132.1 acla|EAW07244.1 afis|EAW22265.1 afum|EAL90915.1

Group6777: abra|OJJ74226.1 acar|OOF95500.1 acla|EAW07279.1 afis|EAW22310.1 afla|EED57702.1 afum|EAL90880.1 agla|OJJ87740.1 akaw|GAA86793.1 aluc|OJZ89907.1 anid|CBF88586.1 anig|EHA27107.1 aory|EIT79928.1 ater|EAU34314.1 atub|OJI84899.1 awen|OJJ34877.1

Group6778: abra|OJJ74324.1 acar|OOF95408.1 acla|EAW07376.1 afis|EAW22401.1 afla|EED56065.1 afum|EAL90790.1 agla|OJJ87664.1 akaw|GAA86702.1 aluc|OJZ89804.1 anid|CBF88767.1 asyd|OJJ63980.1 atub|OJI85002.1 aver|OJI96973.1 awen|OJJ34980.1 azon|OJJ45628.1

Group6779: abra|OJJ74428.1 acar|OOF95312.1 afla|EED54042.1 akaw|GAA87323.1 aluc|OJZ89692.1 anig|EHA26931.1 aory|EIT76115.1 asyd|OJJ54921.1 ater|EAU36078.1 atub|OJI85109.1 aver|OJJ07274.1 awen|OJJ32611.1 azon|OJJ49314.1 afla|EED54043.1 awen|OJJ32610.1

Group6780: abra|OJJ74432.1 acar|OOF95314.1 acla|EAW11099.1 afis|EAW20764.1 afla|EED55080.1 afum|EAL84855.1 agla|OJJ87704.1 akaw|GAA87326.1 aluc|OJZ89689.1 anid|CBF69548.1 anig|EHA26927.1 aory|EIT81650.1 asyd|OJJ59167.1 atub|OJI85112.1 aver|OJJ01757.1

Group6781: abra|OJJ74450.1 acar|OOF95301.1 afla|EED54126.1 agla|OJJ78793.1 akaw|GAA87341.1 aluc|OJZ89673.1 anid|CBF78429.1 anig|EHA26908.1 aory|EIT75911.1 asyd|OJJ52558.1 ater|EAU32137.1 atub|OJI85127.1 aver|OJJ03842.1 awen|OJJ39930.1 azon|OJJ42222.1

Group6782: abra|OJJ74471.1 acar|OOF95274.1 acla|EAW12406.1 afis|EAW16865.1 afla|EED51094.1 afum|EAL86404.1 agla|OJJ84377.1 akaw|GAA87364.1 aluc|OJZ89649.1 anig|EHA26887.1 aory|EIT81999.1 ater|EAU32022.1 atub|OJI85155.1 aver|OJJ07275.1 awen|OJJ38926.1

Group6783: abra|OJJ74511.1 acar|OOF95232.1 acla|EAW07180.1 afis|EAW22193.1 afum|EAL90998.1 akaw|GAA82088.1 aluc|OJZ92092.1 anig|EHA26849.1 asyd|OJJ63750.1 ater|EAU34201.1 atub|OJI86525.1 aver|OJI97190.1 awen|OJJ34774.1 azon|OJJ42148.1 awen|OJJ34608.1

Group6784: abra|OJJ74521.1 acar|OOF95227.1 acla|EAW07171.1 afis|EAW22186.1 afla|EED57565.1 agla|OJJ87864.1 akaw|GAA82079.1 aluc|OJZ92100.1 anid|CBF88393.1 anig|EHA26842.1 asyd|OJJ63743.1 ater|EAU34194.1 atub|OJI86533.1 aver|OJI97196.1 awen|OJJ34766.1

Group6785: abra|OJJ74537.1 acar|OOF95218.1 acla|EAW07156.1 afis|EAW22171.1 afla|EED57543.1 afum|EAL91018.1 agla|OJJ87884.1 akaw|GAA82064.1 aluc|OJZ92118.1 anig|EHA26826.1 ater|EAU34176.1 atub|OJI86551.1 awen|OJJ34747.1 azon|OJJ45815.1 aory|EIT81866.1

Group6786: abra|OJJ74570.1 acar|OOF95193.1 acla|EAW07139.1 afis|EAW22152.1 afum|EAL91038.1 agla|OJJ87910.1 akaw|GAA82035.1 aluc|OJZ92148.1 anid|CBF89106.1 anig|EHA26801.1 asyd|OJJ62724.1 atub|OJI86576.1 aver|OJI96767.1 awen|OJJ34728.1 azon|OJJ43283.1

Group6787: abra|OJJ74587.1 acar|OOF95179.1 acla|EAW10791.1 agla|OJJ82885.1 akaw|GAA82018.1 asyd|OJJ59434.1 atub|OJI86594.1 aver|OJJ02837.1 azon|OJJ49219.1 afla|EED47200.1 aory|EIT79256.1 ater|EAU36524.1 afis|EAW24785.1 afum|EAL89424.1 anid|CBF81453.1

Group6788: abra|OJJ74771.1 acar|OOF95761.1 acla|EAW10640.1 afis|EAW24926.1 afla|EED55547.1 afum|EAL89582.1 agla|OJJ82154.1 akaw|GAA83641.1 anid|CBF75703.1 aory|EIT82675.1 ater|EAU35656.1 atub|OJI86782.1 aver|OJJ05682.1 awen|OJJ40522.1 azon|OJJ49382.1

Group6789: abra|OJJ74778.1 acar|OOF95767.1 acla|EAW10633.1 afis|EAW24934.1 afla|EED55540.1 afum|EAL87766.1 agla|OJJ82161.1 akaw|GAA83634.1 aluc|OJZ92359.1 anig|EHA26614.1 aory|EIT80846.1 ater|EAU35663.1 atub|OJI86789.1 awen|OJJ40529.1 afum|EAL89591.1

Group6790: abra|OJJ74787.1 acar|OOF95778.1 acla|EAW10624.1 afis|EAW24943.1 afla|EED55522.1 afum|EAL89600.1 agla|OJJ82170.1 akaw|GAA83626.1 aluc|OJZ92367.1 anig|EHA26606.1 asyd|OJJ57518.1 atub|OJI86797.1 aver|OJJ05661.1 awen|OJJ40538.1 azon|OJJ49398.1

Group6791: abra|OJJ74807.1 acar|OOF95799.1 acla|EAW10857.1 afis|EAW20256.1 afla|EED53758.1 agla|OJJ85792.1 akaw|GAA83605.1 aluc|OJZ92388.1 anig|EHA26587.1 aory|EIT78405.1 asyd|OJJ53596.1 ater|EAU30254.1 atub|OJI86818.1 aver|OJJ08152.1 awen|OJJ40023.1

Group6792: abra|OJJ74849.1 acar|OOF95850.1 afla|EED57862.1 agla|OJJ87005.1 akaw|GAA83573.1 aluc|OJZ92425.1 anid|CBF89602.1 anig|EHA26542.1 aory|EIT81073.1 asyd|OJJ62309.1 ater|EAU33980.1 atub|OJI86855.1 aver|OJI96417.1 azon|OJJ45216.1 aver|OJJ03882.1

Group6793: abra|OJJ74855.1 acar|OOF95855.1 acla|EAW08529.1 afis|EAW23569.1 afla|EED57461.1 agla|OJJ86830.1 akaw|GAA83567.1 aluc|OJZ92431.1 anid|CBF89619.1 anig|EHA26535.1 atub|OJI86861.1 aver|OJI96410.1 awen|OJJ39897.1 azon|OJJ45875.1 afum|EAL87898.2

Group6794: abra|OJJ74948.1 acar|OOF97368.1 acla|EAW08423.1 afis|EAW23446.1 afla|EED57363.1 agla|OJJ86703.1 akaw|GAA83474.1 aluc|OJZ92521.1 anid|CBF89790.1 anig|EHA26454.1 asyd|OJJ62201.1 atub|OJI86955.1 aver|OJI96313.1 awen|OJJ39783.1 azon|OJJ47914.1

Group6795: abra|OJJ74985.1 acar|OOF97336.1 acla|EAW08396.1 afis|EAW23439.1 afla|EED57328.1 afum|EAL88023.1 agla|OJJ86676.1 akaw|GAA83448.1 aluc|OJZ92548.1 anid|CBF89826.1 asyd|OJJ62170.1 atub|OJI86985.1 aver|OJI96282.1 awen|OJJ39756.1 azon|OJJ47884.1

Group6796: abra|OJJ75015.1 acar|OOF97311.1 acla|EAW08210.1 afis|EAW23244.1 afla|EED55725.1 afum|EAL88238.1 agla|OJJ86616.1 akaw|GAA88638.1 aluc|OJZ92581.1 anig|EHA26401.1 aory|EIT83203.1 ater|EAU37525.1 atub|OJI87016.1 awen|OJJ39726.1 azon|OJJ47851.1

Group6797: abra|OJJ75111.1 acla|EAW08314.1 afis|EAW23326.1 afla|EED55640.1 afum|EAL88153.1 agla|OJJ86521.1 akaw|GAA88721.1 aluc|OJZ92678.1 anid|CBF89449.1 anig|EHA26321.1 aory|EIT81784.1 ater|EAU35574.1 atub|OJI87105.1 awen|OJJ39568.1 azon|OJJ47761.1

Group6798: abra|OJJ75141.1 acar|OOF97195.1 afis|EAW18092.1 afum|EAL91901.1 agla|OJJ79755.1 akaw|GAA88748.1 aluc|OJZ92708.1 asyd|OJJ62669.1 ater|EAU34821.1 atub|OJI87134.1 azon|OJJ42596.1 acla|EAW11029.1 anig|EHA26292.1 anid|CBF81859.1 ater|EAU29635.1

Group6799: abra|OJJ75147.1 acar|OOF97191.1 acla|EAW08320.1 afis|EAW23355.1 afla|EED55605.1 afum|EAL88121.1 agla|OJJ86470.1 akaw|GAA88752.1 aluc|OJZ92712.1 anig|EHA26287.1 atub|OJI87138.1 aver|OJJ04154.1 awen|OJJ39531.1 anid|CBF80815.1 asyd|OJJ51974.1

Group6800: abra|OJJ75161.1 acar|OOF97177.1 acla|EAW08337.1 afis|EAW23372.1 afum|EAL88100.1 akaw|GAA88772.1 aluc|OJZ92731.1 anid|CBF74813.1 anig|EHA26270.1 asyd|OJJ60145.1 ater|EAU35623.1 atub|OJI87155.1 aver|OJJ04118.1 awen|OJJ39511.1 azon|OJJ47710.1

Group6801: abra|OJJ75197.1 acar|OOF94590.1 acla|EAW12984.1 afis|EAW17680.1 afum|EAL91494.1 akaw|GAA91749.1 aluc|OJZ92760.1 anid|CBF90011.1 anig|EHA26246.1 aory|EIT82116.1 asyd|OJJ62072.1 ater|EAU31966.1 atub|OJI87204.1 aver|OJI96179.1 awen|OJJ37159.1

Group6802: abra|OJJ75223.1 acar|OOF94564.1 acla|EAW12257.1 afis|EAW16980.1 afla|EED48289.1 afum|EAL86507.1 akaw|GAA91771.1 aluc|OJZ92786.1 anid|CBF80361.1 anig|EHA26222.1 asyd|OJJ54822.1 ater|EAU37125.1 atub|OJI87229.1 aver|OJJ07339.1 awen|OJJ37844.1

Group6803: abra|OJJ75229.1 afis|EAW23610.1 afla|EED50212.1 afum|EAL87854.1 agla|OJJ78559.1 akaw|GAA91774.1 aluc|OJZ92789.1 anig|EHA26218.1 aory|EIT77297.1 asyd|OJJ53110.1 atub|OJI87233.1 aver|OJJ02078.1 awen|OJJ37793.1 agla|OJJ80608.1 agla|OJJ89254.1

Group6804: abra|OJJ75238.1 acar|OOF94554.1 acla|EAW15276.1 afis|EAW20246.1 afla|EED55538.1 afum|EAL87764.1 akaw|GAA91781.1 aluc|OJZ92800.1 anid|CBF78386.1 anig|EHA26209.1 aory|EIT80807.1 asyd|OJJ54698.1 atub|OJI87243.1 aver|OJJ07477.1 afum|EAL85364.1

Group6805: abra|OJJ75247.1 afis|EAW25208.1 afla|EED53658.1 afum|EAL89869.1 agla|OJJ88953.1 akaw|GAA91788.1 aluc|OJZ92807.1 anid|CBF89950.1 anig|EHA26201.1 aory|EIT75346.1 asyd|OJJ62107.1 ater|EAU34350.1 atub|OJI87257.1 aver|OJI96213.1 awen|OJJ40842.1

Group6806: abra|OJJ75281.1 acar|OOF94527.1 afis|EAW17073.1 afla|EED56842.1 afum|EAL86561.1 akaw|GAA91819.1 aluc|OJZ92842.1 anid|CBF83438.1 anig|EHA26169.1 aory|EIT77563.1 asyd|OJJ60666.1 ater|EAU31476.1 atub|OJI87298.1 aver|OJI99989.1 azon|OJJ42849.1

Group6807: abra|OJJ75291.1 afis|EAW23628.1 afum|EAL87832.1 akaw|GAA91829.1 aluc|OJZ92852.1 anid|CBF90359.1 anig|EHA26157.1 asyd|OJJ55367.1 ater|EAU36701.1 atub|OJI87308.1 aver|OJJ06446.1 azon|OJJ43549.1 afis|EAW15548.1 afum|EAL85784.1 agla|OJJ85203.1

Group6808: abra|OJJ75338.1 acar|OOF94493.1 afla|EED45104.1 agla|OJJ78434.1 akaw|GAA92031.1 aluc|OJZ92901.1 anid|CBF78846.1 anig|EHA26062.1 aory|EIT73740.1 asyd|OJJ65279.1 atub|OJI87359.1 aver|OJI99429.1 awen|OJJ32699.1 aory|EIT80021.1 awen|OJJ36391.1

Group6809: abra|OJJ75361.1 acar|OOF94477.1 acla|EAW08779.1 afis|EAW15622.1 afla|EED47178.1 afum|EAL85712.1 akaw|GAA92057.1 aluc|OJZ92929.1 anig|EHA26042.1 aory|EIT79311.1 asyd|OJJ61699.1 ater|EAU36501.1 atub|OJI87389.1 aver|OJI95785.1 awen|OJJ30946.1

Group6810: abra|OJJ75395.1 acar|OOF94454.1 acla|EAW12098.1 afis|EAW17288.1 afla|EED48830.1 agla|OJJ81093.1 akaw|GAA92757.1 aluc|OJZ92966.1 anid|CBF87565.1 anig|EHA26009.1 asyd|OJJ63241.1 atub|OJI87424.1 aver|OJI97715.1 awen|OJJ35748.1 azon|OJJ45541.1

Group6811: abra|OJJ75412.1 acla|EAW13784.1 afis|EAW18348.1 afla|EED48813.1 afum|EAL87712.2 agla|OJJ88036.1 akaw|GAA84187.1 aluc|OJZ88863.1 anid|CBF79371.1 anig|EHA23640.1 aory|EIT73007.1 asyd|OJJ57810.1 atub|OJI85927.1 aver|OJJ04578.1 awen|OJJ35732.1

Group6812: abra|OJJ75439.1 acla|EAW12403.1 afis|EAW16851.1 afla|EED45355.1 afum|EAL86398.1 agla|OJJ79793.1 akaw|GAA84214.1 aluc|OJZ88892.1 anig|EHA23614.1 ater|EAU34051.1 atub|OJI85897.1 awen|OJJ35814.1 azon|OJJ43496.1 anid|CBF73884.1 agla|OJJ88633.1

Group6813: abra|OJJ75485.1 acla|EAW11216.1 afis|EAW18287.1 afla|EED48754.1 afum|EAL87653.1 agla|OJJ88137.1 akaw|GAA84251.1 aluc|OJZ88943.1 anig|EHA23577.1 aory|EIT73136.1 asyd|OJJ57854.1 ater|EAU33293.1 atub|OJI85846.1 awen|OJJ30659.1 anid|CBF79452.1

Group6814: abra|OJJ75515.1 acar|OOF90842.1 acla|EAW08968.1 afis|EAW15447.1 afum|EAL84199.2 agla|OJJ86217.1 akaw|GAA84279.1 aluc|OJZ88972.1 anid|CBF76721.1 aory|EIT74529.1 asyd|OJJ61244.1 ater|EAU29468.1 atub|OJI85818.1 aver|OJJ00567.1 azon|OJJ43618.1

Group6815: abra|OJJ75564.1 acar|OOF90881.1 afis|EAW19373.1 afla|EED45373.1 afum|EAL94091.1 akaw|GAA84316.1 aluc|OJZ89012.1 anid|CBF80781.1 anig|EHA23518.1 aory|EIT82375.1 asyd|OJJ60289.1 ater|EAU30172.1 atub|OJI85777.1 aver|OJI99621.1 awen|OJJ36565.1

Group6816: abra|OJJ75589.1 acar|OOF90906.1 acla|EAW12146.1 afis|EAW20643.1 agla|OJJ89292.1 akaw|GAA84340.1 aluc|OJZ89040.1 anid|CBF83009.1 anig|EHA23495.1 aory|EIT76318.1 asyd|OJJ61506.1 atub|OJI85750.1 aver|OJJ00843.1 awen|OJJ39950.1 azon|OJJ45222.1

Group6817: acla|EAW12175.1 afla|EED47415.1 akaw|GAA87847.1 anid|CBF87351.1 anig|EHA25024.1 aory|EIT81919.1 asyd|OJJ60268.1 atub|OJI80077.1 aver|OJJ04868.1 acla|EAW14872.1 afis|EAW23641.1 aver|OJJ03986.1 asyd|OJJ55204.1 aver|OJJ07012.1 aluc|OJZ85487.1

Group6818: abra|OJJ75700.1 acar|OOG00154.1 afum|EAL92761.1 akaw|GAA84436.1 aluc|OJZ89151.1 aory|EIT79728.1 ater|EAU38412.1 atub|OJI85637.1 awen|OJJ34001.1 acla|EAW09408.1 afis|EAW21728.1 afla|EED57997.1 agla|OJJ79856.1 azon|OJJ48551.1 aory|EIT79784.1

Group6819: abra|OJJ75711.1 acar|OOG00148.1 acla|EAW09401.1 afis|EAW21735.1 afla|EED58007.1 afum|EAL92768.1 agla|OJJ79848.1 akaw|GAA84445.1 aluc|OJZ89162.1 anig|EHA23392.1 asyd|OJJ60849.1 atub|OJI85628.1 aver|OJJ00173.1 awen|OJJ34008.1 azon|OJJ48543.1

Group6820: abra|OJJ75739.1 acar|OOG00130.1 acla|EAW14901.1 afis|EAW24423.1 afla|EED55259.1 afum|EAL86091.1 akaw|GAA84472.1 aluc|OJZ89192.1 anid|CBF79768.1 anig|EHA23366.1 aory|EIT83547.1 asyd|OJJ58107.1 atub|OJI85598.1 aver|OJJ04297.1 awen|OJJ32785.1

Group6821: abra|OJJ75753.1 acar|OOG00962.1 afla|EED46568.1 afum|EAL86494.1 agla|OJJ83640.1 akaw|GAA84486.1 aluc|OJZ89207.1 anig|EHA23352.1 aory|EIT77923.1 asyd|OJJ54710.1 atub|OJI85585.1 aver|OJJ07456.1 awen|OJJ36177.1 afla|EED52361.1 awen|OJJ34573.1

Group6822: abra|OJJ75816.1 acar|OOG00059.1 afis|EAW24386.1 afla|EED47823.1 afum|EAL86058.1 agla|OJJ83282.1 akaw|GAA84530.1 aluc|OJZ89263.1 anig|EHA23305.1 aory|EIT77111.1 asyd|OJJ56648.1 atub|OJI85536.1 aver|OJJ06078.1 awen|OJJ32824.1 azon|OJJ49013.1

Group6823: abra|OJJ75915.1 acar|OOF99988.1 acla|EAW09875.1 afis|EAW21251.1 afla|EED54811.1 afum|EAL92264.1 agla|OJJ87494.1 akaw|GAA84620.1 aluc|OJZ89364.1 anig|EHA23218.1 aory|EIT79427.1 ater|EAU35891.1 atub|OJI85434.1 awen|OJJ36566.1 azon|OJJ44323.1

Group6824: abra|OJJ75981.1 acar|OOF99938.1 acla|EAW09818.1 afis|EAW21316.1 afla|EED57274.1 afum|EAL92333.1 agla|OJJ87423.1 akaw|GAA88976.1 aluc|OJZ89427.1 anid|CBF83414.1 asyd|OJJ60651.1 atub|OJI85366.1 aver|OJI99971.1 awen|OJJ39380.1 azon|OJJ44270.1

Group6825: abra|OJJ76016.1 acar|OOF99907.1 acla|EAW09788.1 afis|EAW21348.1 afla|EED55760.1 afum|EAL92365.1 akaw|GAA88944.1 aluc|OJZ89463.1 anid|CBF76195.1 anig|EHA23125.1 aory|EIT76680.1 asyd|OJJ60960.1 atub|OJI85332.1 aver|OJJ00285.1 awen|OJJ39351.1

Group6826: abra|OJJ76026.1 acar|OOF93547.1 afis|EAW17185.1 afla|EED57111.1 afum|EBA27281.1 agla|OJJ82967.1 akaw|GAA88932.1 aluc|OJZ89477.1 anid|CBF84193.1 anig|EHA23116.1 aory|EIT74116.1 asyd|OJJ54805.1 atub|OJI85323.1 aver|OJJ07356.1 awen|OJJ31334.1

Group6827: abra|OJJ76029.1 acar|OOF93544.1 afis|EAW17187.1 afla|EED45760.1 afum|EAL86613.2 agla|OJJ82966.1 aluc|OJZ89480.1 anig|EHA23113.1 aory|EIT75754.1 asyd|OJJ54804.1 atub|OJI85320.1 aver|OJJ07357.1 awen|OJJ40015.1 azon|OJJ47385.1 anid|CBF84194.1

Group6828: abra|OJJ76148.1 afis|EAW21465.1 afla|EED55898.1 akaw|GAA82098.1 aluc|OJZ92081.1 anig|EHA23006.1 aory|EIT76852.1 atub|OJI86511.1 awen|OJJ39233.1 acla|EAW09674.1 afum|EAL92485.1 ater|EAU35070.1 azon|OJJ51623.1 aver|OJJ00465.1 asyd|OJJ61144.1

Group6829: abra|OJJ76167.1 acar|OOF99770.1 acla|EAW09656.1 afis|EAW21483.1 afla|EED55915.1 afum|EAL92503.1 agla|OJJ85637.1 akaw|GAA82115.1 aluc|OJZ92061.1 anid|CBF76496.1 anig|EHA22990.1 asyd|OJJ61127.1 atub|OJI86493.1 aver|OJJ00448.1 awen|OJJ39214.1

Group6830: abra|OJJ76219.1 acar|OOF99732.1 acla|EAW13934.1 afis|EAW18645.1 afla|EED52695.1 afum|EAL93360.1 akaw|GAA82157.1 aluc|OJZ92011.1 anid|CBF70439.1 aory|EIT74225.1 ater|EAU38266.1 atub|OJI86446.1 aver|OJJ01216.1 awen|OJJ32113.1 azon|OJJ46694.1

Group6831: abra|OJJ76222.1 acar|OOF99729.1 acla|EAW13936.1 afis|EAW18647.1 afla|EED52697.1 afum|EAL93362.1 agla|OJJ82428.1 akaw|GAA82159.1 aluc|OJZ92009.1 anig|EHA22948.1 ater|EAU38264.1 atub|OJI86444.1 awen|OJJ32115.1 azon|OJJ46696.1 awen|OJJ33655.1

Group6832: abra|OJJ76234.1 acar|OOF99478.1 acla|EAW14370.1 afis|EAW18920.1 agla|OJJ81523.1 akaw|GAA82169.1 aluc|OJZ91997.1 anid|CBF69682.1 asyd|OJJ59060.1 ater|EAU38087.1 atub|OJI86432.1 aver|OJJ01654.1 awen|OJJ31513.1 azon|OJJ47231.1 afum|EAL93651.2

Group6833: abra|OJJ76317.1 acar|OOF99568.1 acla|EAW06524.1 afis|EAW16240.1 afla|EED48909.1 afum|EAL88861.1 akaw|GAA82247.1 aluc|OJZ91897.1 anig|EHA22876.1 aory|EIT81245.1 asyd|OJJ56319.1 atub|OJI86350.1 awen|OJJ30217.1 azon|OJJ50001.1 acar|OOF92999.1

Group6834: abra|OJJ76328.1 acar|OOF99581.1 acla|EAW14087.1 afis|EAW18793.1 afum|EAL93529.1 agla|OJJ80522.1 akaw|GAA82258.1 aluc|OJZ91886.1 anig|EHA22862.1 asyd|OJJ58951.1 atub|OJI86336.1 aver|OJJ01545.1 awen|OJJ32366.1 azon|OJJ47130.1 ater|EAU37924.1

Group6835: abra|OJJ76348.1 acar|OOF99599.1 acla|EAW14067.1 afis|EAW18774.1 afla|EED50733.1 afum|EAL93506.1 agla|OJJ80490.1 akaw|GAA82275.1 aluc|OJZ91865.1 anid|CBF69726.1 anig|EHA22845.1 aory|EIT77977.1 ater|EAU37905.1 atub|OJI86316.1 awen|OJJ32345.1

Group6836: abra|OJJ76355.1 acar|OOF99606.1 acla|EAW14060.1 afla|EED50726.1 akaw|GAA82281.1 aluc|OJZ91858.1 anid|CBF69942.1 anig|EHA22838.1 aory|EIT78007.1 asyd|OJJ58918.1 ater|EAU37897.1 atub|OJI86309.1 aver|OJJ01515.1 awen|OJJ35974.1 azon|OJJ45258.1

Group6837: abra|OJJ76413.1 acar|OOF99653.1 agla|OJJ82338.1 akaw|GAA82330.1 aluc|OJZ91799.1 anid|CBF73518.1 anig|EHA22795.1 aory|EIT78040.1 asyd|OJJ56747.1 atub|OJI86255.1 aver|OJJ05983.1 awen|OJJ36284.1 azon|OJJ42090.1 asyd|OJJ54494.1 aver|OJJ07686.1

Group6838: abra|OJJ76421.1 acar|OOF99663.1 acla|EAW14005.1 afla|EED50663.1 akaw|GAA82336.1 aluc|OJZ91791.1 anid|CBF78486.1 anig|EHA22788.1 aory|EIT78010.1 asyd|OJJ65332.1 ater|EAU38196.1 atub|OJI86247.1 aver|OJI99477.1 awen|OJJ32198.1 azon|OJJ42546.1

Group6839: abra|OJJ76502.1 acla|EAW14193.1 afis|EAW18908.1 afla|EED50839.1 afum|EAL93640.1 agla|OJJ81514.1 akaw|GAA82418.1 aluc|OJZ91700.1 anid|CBF69703.1 anig|EHA22714.1 asyd|OJJ59048.1 ater|EAU38077.1 atub|OJI86159.1 aver|OJJ01644.1 azon|OJJ47225.1

Group6840: abra|OJJ76518.1 acar|OOF99455.1 acla|EAW14183.1 afis|EAW18898.1 afla|EED50865.1 afum|EAL93630.1 agla|OJJ81504.1 akaw|GAA82431.1 anid|CBF69915.1 aory|EIT78985.1 asyd|OJJ58990.1 atub|OJI86141.1 aver|OJJ01625.1 awen|OJJ32456.1 azon|OJJ47213.1

Group6841: abra|OJJ76519.1 acar|OOF99454.1 acla|EAW14182.1 afis|EAW18897.1 afla|EED50866.1 afum|EAL93629.1 agla|OJJ81503.1 akaw|GAA82432.1 anig|EHA22701.1 aory|EIT78962.1 asyd|OJJ58991.1 atub|OJI86140.1 aver|OJJ01624.1 awen|OJJ32455.1 azon|OJJ47212.1

Group6842: abra|OJJ76521.1 acar|OOF99452.1 acla|EAW14180.1 afis|EAW18895.1 afla|EED50868.1 afum|EAL93627.1 agla|OJJ81501.1 akaw|GAA82434.1 aluc|OJZ91687.1 anid|CBF69911.1 anig|EHA22699.1 aory|EIT78971.1 asyd|OJJ58993.1 aver|OJJ01622.1 awen|OJJ32453.1

Group6843: abra|OJJ76599.1 acar|OOF99391.1 afla|EED53190.1 akaw|GAA82502.1 aluc|OJZ91609.1 anig|EHA22638.1 aory|EIT74987.1 asyd|OJJ54120.1 ater|EAU38170.1 atub|OJI86061.1 aver|OJI99295.1 awen|OJJ31585.1 afla|EED50956.1 aory|EIT77024.1 anid|CBF82431.1

Group6844: aory|EIT75135.1 acla|EAW14303.1 afis|EAW18981.1 afla|EED53184.1 afum|EAL93717.1 akaw|GAA82506.1 anid|CBF69655.1 anig|EHA22634.1 asyd|OJJ59074.1 ater|EAU37831.1 aver|OJJ01667.1 awen|OJJ31593.1 azon|OJJ47305.1 afla|EED48362.1 aory|EIT83093.1

Group6845: abra|OJJ76630.1 acar|OOF99367.1 acla|EAW14289.1 afis|EAW18994.1 afla|EED53159.1 afum|EAL93729.1 agla|OJJ80084.1 akaw|GAA92826.1 aluc|OJZ91581.1 anid|CBF86789.1 anig|EHA22608.1 ater|EAU37814.1 atub|OJI86030.1 awen|OJJ31616.1 aory|EIT74857.1

Group6846: abra|OJJ76637.1 acar|OOF99363.1 acla|EAW14278.1 afla|EED53143.1 akaw|GAA92821.1 aluc|OJZ91573.1 anid|CBF86762.1 anig|EHA22602.1 aory|EIT75028.1 asyd|OJJ64685.1 ater|EAU37807.1 atub|OJI86024.1 aver|OJI98787.1 awen|OJJ31624.1 azon|OJJ47322.1

Group6847: abra|OJJ76640.1 acar|OOF99357.1 afla|EED53120.1 agla|OJJ85229.1 akaw|GAA92818.1 aluc|OJZ91569.1 anid|CBF86738.1 anig|EHA22598.1 aory|EIT75177.1 asyd|OJJ64695.1 ater|EAU37797.1 atub|OJI86019.1 aver|OJI98800.1 awen|OJJ31630.1 azon|OJJ42456.1

Group6848: abra|OJJ76641.1 acar|OOF99356.1 acla|EAW14270.1 afla|EED53119.1 agla|OJJ85230.1 akaw|GAA92817.1 aluc|OJZ91568.1 anid|CBF86739.1 anig|EHA22597.1 aory|EIT75210.1 asyd|OJJ64696.1 ater|EAU37796.1 atub|OJI86018.1 aver|OJI98801.1 awen|OJJ31631.1

Group6849: abra|OJJ76691.1 acar|OOF99320.1 afis|EAW19661.1 afla|EED57014.1 afum|EAL86861.1 akaw|GAA92503.1 aluc|OJZ91516.1 anid|CBF71388.1 anig|EHA22554.1 aory|EIT82285.1 asyd|OJJ55068.1 ater|EAU32850.1 atub|OJI85971.1 aver|OJJ07143.1 awen|OJJ41920.1

Group6850: abra|OJJ76712.1 acar|OOF92524.1 acla|EAW14556.1 afis|EAW16941.1 afum|EBA27273.1 akaw|GAA83772.1 aluc|OJZ90956.1 anid|CBF84647.1 asyd|OJJ53357.1 ater|EAU33241.1 atub|OJI82946.1 aver|OJJ08384.1 azon|OJJ43514.1 aluc|OJZ80182.1 awen|OJJ36418.1

Group6851: abra|OJJ76715.1 acla|EAW07081.1 afis|EAW19676.1 afla|EED53626.1 afum|EAL86846.1 akaw|GAA92485.1 aluc|OJZ91497.1 anid|CBF86669.1 anig|EHA22537.1 aory|EIT74953.1 asyd|OJJ64775.1 ater|EAU38555.1 atub|OJI85950.1 aver|OJI98875.1 awen|OJJ30166.1

Group6852: abra|OJJ76825.1 acar|OOF91320.1 acla|EAW13385.1 afis|EAW20499.1 afla|EED44742.1 akaw|GAA87197.1 aluc|OJZ84236.1 anid|CBF79204.1 anig|EHA27583.1 aory|EIT77800.1 asyd|OJJ55727.1 ater|EAU37193.1 atub|OJI89849.1 aver|OJJ06813.1 awen|OJJ33200.1

Group6853: abra|OJJ76905.1 acar|OOF91353.1 afis|EAW20538.1 afla|EED52441.1 afum|EBA27189.1 akaw|GAA87133.1 aluc|OJZ84304.1 anig|EHA27640.1 aory|EIT82801.1 asyd|OJJ56118.1 ater|EAU37139.1 atub|OJI89764.1 aver|OJJ05059.1 awen|OJJ33245.1 azon|OJJ50624.1

Group6854: abra|OJJ76908.1 acar|OOF91359.1 afis|EAW20541.1 afla|EED52432.1 afum|EAL84517.2 akaw|GAA87130.1 aluc|OJZ84307.1 anig|EHA27643.1 aory|EIT83289.1 asyd|OJJ56121.1 ater|EAU37141.1 atub|OJI89761.1 aver|OJJ05062.1 awen|OJJ33248.1 azon|OJJ50627.1

Group6855: abra|OJJ76935.1 acar|OOF91384.1 afis|EAW20554.1 afla|EED53209.1 afum|EBA27184.1 akaw|GAA87104.1 aluc|OJZ84334.1 anid|CBF78972.1 anig|EHA27668.1 aory|EIT74844.1 asyd|OJJ55673.1 ater|EAU37122.1 atub|OJI89734.1 aver|OJJ06756.1 awen|OJJ33267.1

Group6856: abra|OJJ76953.1 acla|EAW13477.1 afis|EAW20584.1 afla|EED52254.1 akaw|GAA87089.1 aluc|OJZ84349.1 anig|EHA27686.1 aory|EIT74496.1 atub|OJI89719.1 afum|EBA27179.1 agla|OJJ88678.1 anid|CBF78923.1 ater|EAU37098.1 aver|OJJ06702.1 asyd|OJJ51988.1

Group6857: abra|OJJ76957.1 acar|OOF93291.1 afis|EAW19845.1 afla|EED54456.1 afum|EAL85049.1 agla|OJJ83721.1 akaw|GAA87086.1 aluc|OJZ84352.1 anid|CBF80472.1 anig|EHA27690.1 aory|EIT81149.1 asyd|OJJ54743.1 atub|OJI89714.1 aver|OJJ07421.1 awen|OJJ32032.1

Group6858: abra|OJJ76959.1 acar|OOF91404.1 acla|EAW11699.1 afis|EAW23715.1 afla|EED52264.1 afum|EAL87160.2 akaw|GAA87084.1 aluc|OJZ84354.1 anid|CBF82105.1 anig|EHA27692.1 asyd|OJJ54272.1 ater|EAU37092.1 atub|OJI89713.1 aver|OJJ05228.1 ater|EAU35157.1

Group6859: abra|OJJ76991.1 acar|OOF91435.1 acla|EAW13465.1 afis|EAW20577.1 afla|EED52312.1 afum|EAL84488.2 akaw|GAA87055.1 aluc|OJZ84391.1 anid|CBF78938.1 anig|EHA27720.1 aory|EIT74375.1 ater|EAU37108.1 atub|OJI89680.1 awen|OJJ33277.1 azon|OJJ50651.1

Group6860: abra|OJJ77014.1 acar|OOF91457.1 acla|EAW09004.1 afis|EAW25544.1 agla|OJJ78911.1 akaw|GAA87036.1 aluc|OJZ84414.1 anid|CBF83190.1 aory|EIT75992.1 asyd|OJJ55535.1 ater|EAU32180.1 atub|OJI89659.1 aver|OJJ06613.1 awen|OJJ36037.1 afla|EED54131.1

Group6861: abra|OJJ77046.1 acar|OOF89824.1 afla|EED45584.1 afum|EAL86588.1 agla|OJJ85383.1 akaw|GAA87007.1 aluc|OJZ84449.1 anig|EHA27765.1 aory|EIT75525.1 atub|OJI89626.1 awen|OJJ35839.1 afis|EAW17128.1 afla|EED47800.1 aory|EIT77186.1 awen|OJJ41963.1

Group6862: abra|OJJ77057.1 afis|EAW20755.1 afla|EED51084.1 afum|EAL84863.1 akaw|GAA86995.1 aluc|OJZ84464.1 anid|CBF87301.1 anig|EHA27776.1 asyd|OJJ55871.1 ater|EAU30141.1 atub|OJI89612.1 aver|OJJ04816.1 aory|EIT76985.1 awen|OJJ40179.1 anid|CBF78065.1

Group6863: abra|OJJ77082.1 acar|OOF92966.1 afla|EED51963.1 agla|OJJ88141.1 akaw|GAA86971.1 aluc|OJZ84492.1 anig|EHA27796.1 aory|EIT81436.1 asyd|OJJ61412.1 ater|EAU30963.1 atub|OJI89587.1 aver|OJJ00742.1 awen|OJJ33796.1 azon|OJJ48746.1 aory|EIT81482.1

Group6864: abra|OJJ77107.1 acar|OOF92992.1 acla|EAW13566.1 afis|EAW20035.1 agla|OJJ78423.1 aluc|OJZ84516.1 anid|CBF89780.1 anig|EHA27814.1 aory|EIT83237.1 asyd|OJJ62206.1 ater|EAU29401.1 atub|OJI89564.1 aver|OJI96318.1 awen|OJJ30058.1 azon|OJJ42029.1

Group6865: abra|OJJ77160.1 acar|OOF93046.1 afis|EAW24602.1 afla|EED56878.1 afum|EAL88037.1 aluc|OJZ84573.1 anig|EHA27867.1 aory|EIT77536.1 asyd|OJJ54816.1 atub|OJI89509.1 aver|OJJ07345.1 awen|OJJ37088.1 azon|OJJ50793.1 aver|OJI96492.1 anid|CBF80367.1

Group6866: abra|OJJ77161.1 acar|OOF93047.1 afis|EAW17748.1 afla|EED45754.1 afum|EAL91562.1 agla|OJJ81705.1 akaw|GAA90771.1 aluc|OJZ84574.1 anid|CBF86619.1 asyd|OJJ64814.1 ater|EAU31859.1 atub|OJI89508.1 aver|OJI98927.1 awen|OJJ37087.1 azon|OJJ50792.1

Group6867: abra|OJJ77268.1 acar|OOF93137.1 acla|EAW13270.1 afis|EAW17351.1 afum|EAL91194.1 akaw|GAA90680.1 aluc|OJZ84675.1 anid|CBF80095.1 anig|EHA27953.1 aory|EIT77668.1 asyd|OJJ58300.1 ater|EAU33857.1 atub|OJI89405.1 aver|OJJ02124.1 awen|OJJ33645.1

Group6868: abra|OJJ77322.1 acar|OOF93198.1 acla|EAW13150.1 afis|EAW17496.1 afla|EED49320.1 afum|EAL91319.1 agla|OJJ81462.1 akaw|GAA91434.1 aluc|OJZ84732.1 anig|EHA28005.1 aory|EIT79567.1 ater|EAU36940.1 atub|OJI89349.1 awen|OJJ37345.1 azon|OJJ51031.1

Group6869: abra|OJJ77349.1 acar|OOF93219.1 afla|EED49289.1 agla|OJJ81431.1 aluc|OJZ84760.1 anid|CBF90320.1 anig|EHA28029.1 asyd|OJJ61910.1 ater|EAU36968.1 atub|OJI89324.1 aver|OJI96003.1 awen|OJJ37316.1 azon|OJJ51008.1 afis|EAW17538.1 afum|EAL91352.1

Group6870: abra|OJJ77385.1 acar|OOF93254.1 acla|EAW11087.1 afis|EAW20768.1 afla|EED50190.1 afum|EAL84850.1 akaw|GAA91376.1 aluc|OJZ84793.1 anid|CBF73860.1 anig|EHA28061.1 asyd|OJJ56519.1 ater|EAU31813.1 atub|OJI89289.1 aver|OJJ06203.1 aory|EIT77383.1

Group6871: abra|OJJ77397.1 acar|OOF89919.1 acla|EAW12873.1 afis|EAW17803.1 afla|EED45797.1 afum|EAL91611.1 akaw|GAA91365.1 aluc|OJZ84805.1 anid|CBF86650.1 anig|EHA28070.1 aory|EIT75543.1 asyd|OJJ64801.1 ater|EAU31834.1 atub|OJI89278.1 aver|OJI98915.1

Group6872: abra|OJJ77411.1 acar|OOF93278.1 afis|EAW22076.1 afla|EED56696.1 afum|EAL91094.1 agla|OJJ81293.1 akaw|GAA91353.1 aluc|OJZ84820.1 anid|CBF84146.1 anig|EHA28082.1 aory|EIT73893.1 asyd|OJJ59789.1 atub|OJI89266.1 aver|OJJ02499.1 azon|OJJ47365.1

Group6873: abra|OJJ77429.1 afis|EAW24666.1 afla|EED51151.1 afum|EAL89343.1 agla|OJJ85716.1 akaw|GAA88066.1 aluc|OJZ81329.1 anid|CBF90090.1 anig|EHA18633.1 asyd|OJJ57398.1 ater|EAU31628.1 atub|OJI91532.1 afis|EAW24701.1 anid|CBF80153.1 aver|OJI99512.1

Group6874: abra|OJJ77480.1 acar|OOF99245.1 acla|EAW06742.1 afis|EAW16292.1 afla|EED53261.1 agla|OJJ88314.1 akaw|GAA84138.1 aluc|OJZ81058.1 anid|CBF86852.1 anig|EHA19310.1 asyd|OJJ64638.1 atub|OJI82527.1 aver|OJI98737.1 awen|OJJ37688.1 azon|OJJ44948.1

Group6875: abra|OJJ77486.1 acar|OOF99239.1 acla|EAW06745.1 afis|EAW16295.1 afla|EED53259.1 afum|EAL88917.1 agla|OJJ88304.1 aluc|OJZ81050.1 anig|EHA19317.1 aory|EIT75026.1 asyd|OJJ64641.1 atub|OJI82534.1 aver|OJI98741.1 awen|OJJ37694.1 azon|OJJ44955.1

Group6876: abra|OJJ77529.1 afis|EAW16243.1 afla|EED53304.1 afum|EAL88865.1 agla|OJJ86253.1 akaw|GAA84091.1 aluc|OJZ81006.1 anid|CBF78898.1 anig|EHA19354.1 asyd|OJJ64602.1 ater|EAU37453.1 atub|OJI82579.1 awen|OJJ30213.1 acla|EAW06698.1 aver|OJI98699.1

Group6877: abra|OJJ77530.1 acar|OOF99196.1 acla|EAW06699.1 afis|EAW16244.1 afla|EED53305.1 afum|EAL88872.1 akaw|GAA84090.1 aluc|OJZ81005.1 anid|CBF71516.1 anig|EHA19355.1 aory|EIT75076.1 atub|OJI82580.1 aver|OJJ07277.1 awen|OJJ30212.1 azon|OJJ43937.1

Group6878: abra|OJJ77545.1 acar|OOF99179.1 afla|EED56789.1 akaw|GAA84078.1 aluc|OJZ80992.1 anid|CBF71424.1 anig|EHA19369.1 aory|EIT73871.1 asyd|OJJ54994.1 ater|EAU38454.1 atub|OJI82593.1 aver|OJJ07207.1 awen|OJJ31995.1 azon|OJJ46050.1 ater|EAU32726.1

Group6879: abra|OJJ77559.1 acar|OOF99170.1 acla|EAW06692.1 afis|EAW16073.1 afla|EED48592.1 afum|EAL88705.1 agla|OJJ86095.1 akaw|GAA84065.1 aluc|OJZ80979.1 anid|CBF75111.1 asyd|OJJ59997.1 ater|EAU33558.1 atub|OJI82607.1 aver|OJJ02294.1 awen|OJJ30361.1

Group6880: abra|OJJ77575.1 acar|OOF99152.1 acla|EAW08860.1 afis|EAW15534.1 afla|EED54856.1 afum|EAL85794.1 agla|OJJ85213.1 aluc|OJZ80962.1 anid|CBF78197.1 asyd|OJJ55982.1 ater|EAU32591.1 atub|OJI82624.1 aver|OJJ04929.1 awen|OJJ38198.1 azon|OJJ45418.1

Group6881: abra|OJJ77673.1 acar|OOF99063.1 acla|EAW06582.1 afis|EAW16008.1 afla|EED56384.1 afum|EBA27353.1 akaw|GAA83959.1 aluc|OJZ80846.1 anid|CBF85865.1 anig|EHA19477.1 asyd|OJJ64370.1 ater|EAU33635.1 atub|OJI82721.1 aver|OJI98466.1 aory|EIT73411.1

Group6882: abra|OJJ77716.1 acar|OOF99023.1 afla|EED47308.1 agla|OJJ83229.1 akaw|GAA83916.1 aluc|OJZ90781.1 anid|CBF81696.1 anig|EHA19514.1 aory|EIT82843.1 asyd|OJJ59567.1 ater|EAU36653.1 atub|OJI82768.1 aver|OJJ02702.1 awen|OJJ29820.1 acar|OOF98222.1

Group6883: abra|OJJ77721.1 acar|OOF99018.1 afis|EAW18174.1 afla|EED56318.1 afum|EAL87535.1 agla|OJJ85916.1 akaw|GAA83913.1 aluc|OJZ90784.1 anig|EHA19517.1 aory|EIT73323.1 ater|EAU33831.1 atub|OJI82771.1 awen|OJJ30522.1 azon|OJJ50202.1 acla|EAW11322.1

Group6884: abra|OJJ77725.1 acar|OOF99015.1 afla|EED47524.1 akaw|GAA83910.1 aluc|OJZ90787.1 anid|CBF80342.1 anig|EHA19520.1 aory|EIT74520.1 asyd|OJJ54835.1 ater|EAU30843.1 atub|OJI82774.1 aver|OJJ07326.1 awen|OJJ30011.1 azon|OJJ43640.1 agla|OJJ79033.1

Group6885: abra|OJJ77727.1 acar|OOF99011.1 acla|EAW10005.1 afis|EAW21051.1 afla|EED46595.1 afum|EAL92087.1 akaw|GAA83909.1 aluc|OJZ90788.1 anig|EHA19521.1 aory|EIT77870.1 asyd|OJJ56448.1 atub|OJI82775.1 aver|OJJ06274.1 awen|OJJ36351.1 azon|OJJ49321.1

Group6886: abra|OJJ77777.1 acar|OOF98967.1 agla|OJJ85901.1 akaw|GAA83870.1 aluc|OJZ90830.1 anig|EHA19553.1 aory|EIT73292.1 atub|OJI82816.1 afla|EED56302.1 afum|EBA27316.1 acla|EAW11314.1 anid|CBF85714.1 asyd|OJJ64272.1 aver|OJI98364.1 azon|OJJ50213.1

Group6887: abra|OJJ77782.1 acar|OOF98971.1 afis|EAW24714.1 afla|EED54614.1 agla|OJJ87530.1 akaw|GAA83865.1 aluc|OJZ90835.1 anig|EHA19558.1 aory|EIT78767.1 asyd|OJJ53083.1 atub|OJI82821.1 aver|OJJ03763.1 awen|OJJ36610.1 anig|EHA26134.1 atub|OJI87458.1

Group6888: abra|OJJ77807.1 acar|OOF98960.1 afis|EAW18197.1 afla|EED56287.1 afum|EAL87554.1 agla|OJJ86043.1 akaw|GAA83842.1 aluc|OJZ90874.1 anig|EHA19594.1 aory|EIT73344.1 asyd|OJJ61784.1 ater|EAU33803.1 aver|OJI95884.1 awen|OJJ30554.1 azon|OJJ45512.1

Group6889: abra|OJJ77808.1 acar|OOF98959.1 acla|EAW11309.1 afis|EAW18196.1 afla|EED56289.1 afum|EAL87555.1 agla|OJJ85897.1 akaw|GAA83841.1 aluc|OJZ90875.1 anig|EHA19595.1 aory|EIT73382.1 asyd|OJJ64480.1 ater|EAU33802.1 awen|OJJ30556.1 azon|OJJ45511.1

Group6890: abra|OJJ77904.1 acar|OOF98880.1 acla|EAW06514.1 afis|EAW16232.1 afla|EED53349.1 afum|EAL88855.1 agla|OJJ86243.1 akaw|GAA83758.1 aluc|OJZ90973.1 anig|EHA19675.1 aory|EIT75117.1 ater|EAU33712.1 atub|OJI82964.1 awen|OJJ30228.1 azon|OJJ43947.1

Group6891: abra|OJJ77906.1 acar|OOF98877.1 acla|EAW06516.1 afis|EAW16234.1 afla|EED53347.1 agla|OJJ86241.1 akaw|GAA83756.1 aluc|OJZ90975.1 anid|CBF86227.1 anig|EHA19677.1 aory|EIT74992.1 ater|EAU33710.1 atub|OJI82966.1 awen|OJJ30226.1 azon|OJJ43946.1

Group6892: abra|OJJ77918.1 afla|EED51606.1 agla|OJJ82628.1 akaw|GAA92604.1 aluc|OJZ82623.1 anid|CBF73911.1 anig|EHA18403.1 aory|EIT80624.1 asyd|OJJ53670.1 atub|OJI88857.1 aver|OJJ07777.1 awen|OJJ40188.1 anig|EHA27455.1 asyd|OJJ53437.1 aver|OJJ08312.1

Group6893: abra|OJJ77955.1 afis|EAW19554.1 afla|EED45670.1 afum|EAL91021.2 anid|CBF80597.1 aory|EIT74504.1 asyd|OJJ53429.1 aver|OJJ05783.1 acla|EAW11909.1 afis|EAW16191.1 afla|EED52216.1 afum|EAL88822.1 azon|OJJ49842.1 aory|EIT75438.1 aver|OJJ08316.1

Group6894: abra|OJJ78000.1 acla|EAW06650.1 afis|EAW16115.1 afla|EED48656.1 afum|EAL88746.1 agla|OJJ86154.1 aluc|OJZ91076.1 anid|CBF85503.1 anig|EHA19777.1 asyd|OJJ64118.1 ater|EAU33198.1 atub|OJI83071.1 aver|OJI98201.1 awen|OJJ30315.1 azon|OJJ45332.1

Group6895: agla|OJJ81797.1 anid|CBF75762.1 ater|EAU29690.1 atub|OJI83003.1 azon|OJJ48592.1 acar|OOF91976.1 aluc|OJZ91012.1 anig|EHA19714.1 asyd|OJJ64671.1 aver|OJI98773.1 awen|OJJ33537.1 afla|EED52320.1 akaw|GAA91343.1 aluc|OJZ84830.1 anig|EHA25739.1

Group6896: abra|OJJ78307.1 acar|OOF98559.1 acla|EAW09211.1 afis|EAW21936.1 afla|EED46653.1 afum|EAL92949.2 agla|OJJ84184.1 akaw|GAA85720.1 aluc|OJZ91384.1 anid|CBF76895.1 anig|EHA20059.1 ater|EAU30942.1 atub|OJI83377.1 awen|OJJ34227.1 azon|OJJ43408.1

Group6897: abra|OJJ78359.1 acar|OOF98530.1 acla|EAW09175.1 afis|EAW21971.1 afla|EED46693.1 afum|EAL92986.1 agla|OJJ84134.1 akaw|GAA85674.1 aluc|OJZ91434.1 anid|CBF74905.1 aory|EIT77961.1 asyd|OJJ61631.1 atub|OJI83427.1 aver|OJJ00954.1 azon|OJJ43373.1

Group6898: abra|OJJ78360.1 acar|OOF98529.1 acla|EAW06848.1 afis|EAW16469.1 afla|EED46694.1 afum|EAL89063.1 agla|OJJ84135.1 akaw|GAA85673.1 aluc|OJZ91435.1 anid|CBF82725.1 aory|EIT77825.1 asyd|OJJ61632.1 atub|OJI83428.1 aver|OJJ00955.1 azon|OJJ43372.1

Group6899: abra|OJJ78361.1 acar|OOF98526.1 acla|EAW09174.1 afis|EAW21972.1 afla|EED46695.1 afum|EAL92987.1 agla|OJJ84136.1 akaw|GAA85672.1 anid|CBF82723.1 aory|EIT77939.1 asyd|OJJ61633.1 atub|OJI83429.1 aver|OJJ00956.1 awen|OJJ34264.1 azon|OJJ43371.1

Group6900: acar|OOF90135.1 acla|EAW14633.1 afis|EAW23651.1 afla|EED52134.1 afum|EAL86791.1 agla|OJJ79804.1 aory|EIT74500.1 asyd|OJJ62551.1 ater|EAU32804.1 aver|OJJ07303.1 awen|OJJ40924.1 afis|EAW19738.1 afla|EED56525.1 aory|EIT82924.1 aver|OJI96674.1

Group6901: acar|OOF90407.1 acla|EAW15148.1 afis|EAW20116.1 afla|EED53355.1 afum|EAL85233.1 agla|OJJ88833.1 akaw|GAA89322.1 aluc|OJZ86348.1 anid|CBF84934.1 anig|EHA21538.1 aory|EIT74950.1 asyd|OJJ63158.1 ater|EAU38734.1 atub|OJI84186.1 azon|OJJ46248.1

Group6902: acar|OOF91304.1 acla|EAW13369.1 afis|EAW20483.1 afla|EED44785.1 agla|OJJ84812.1 akaw|GAA87213.1 aluc|OJZ84217.1 anid|CBF79233.1 aory|EIT77785.1 asyd|OJJ55741.1 ater|EAU29561.1 atub|OJI89867.1 aver|OJJ06827.1 awen|OJJ33181.1 azon|OJJ46387.1

Group6903: acar|OOF91572.1 acla|EAW14387.1 afis|EAW19093.1 afum|EAL93824.1 agla|OJJ84508.1 akaw|GAA91577.1 aluc|OJZ82399.1 anid|CBF79707.1 anig|EHA28366.1 asyd|OJJ58056.1 ater|EAU32142.1 atub|OJI88042.1 aver|OJJ04347.1 awen|OJJ32487.1 azon|OJJ46044.1

Group6904: acar|OOF91787.1 afis|EAW20744.1 afla|EED47485.1 akaw|GAA88128.1 aluc|OJZ81400.1 anig|EHA18693.1 aory|EIT79969.1 asyd|OJJ64559.1 atub|OJI91606.1 aver|OJI98656.1 azon|OJJ45978.1 afis|EAW17202.1 afum|EAL86286.2 anig|EHA21846.1 aver|OJJ03938.1

Group6905: acar|OOF94697.1 acla|EAW12761.1 afis|EAW17923.1 afum|EAL91734.1 akaw|GAA85188.1 aluc|OJZ90543.1 anid|CBF76947.1 anig|EHA23991.1 aory|EIT81229.1 asyd|OJJ57059.1 ater|EAU33548.1 atub|OJI81965.1 aver|OJJ03239.1 awen|OJJ41622.1 azon|OJJ48345.1

Group6906: acar|OOF94883.1 acla|EAW11991.1 afis|EAW19631.1 afla|EED56983.1 afum|EAL86891.1 agla|OJJ80776.1 akaw|GAA86083.1 anid|CBF71327.1 aory|EIT77443.1 asyd|OJJ55107.1 ater|EAU32895.1 atub|OJI81743.1 aver|OJJ07099.1 awen|OJJ41875.1 azon|OJJ47074.1

Group6907: acar|OOF95360.1 acla|EAW07419.1 afis|EAW22444.1 afla|EED54460.1 afum|EAL90747.1 akaw|GAA87277.1 aluc|OJZ89744.1 anid|CBF88850.1 anig|EHA26966.1 aory|EIT81138.1 atub|OJI85058.1 awen|OJJ35070.1 asyd|OJJ62864.1 aver|OJI96917.1 azon|OJJ50037.1

Group6908: acar|OOF98609.1 acla|EAW09257.1 afis|EAW21890.1 afum|EAL92904.1 agla|OJJ84243.1 akaw|GAA85773.1 aluc|OJZ91327.1 anid|CBF76802.1 anig|EHA20009.1 asyd|OJJ61287.1 ater|EAU30887.1 atub|OJI83320.1 aver|OJJ00613.1 awen|OJJ34179.1 azon|OJJ43453.1

Group6909: acar|OOF99563.1 acla|EAW14103.1 afis|EAW18804.1 afla|EED50772.1 afum|EAL93540.1 agla|OJJ80534.1 akaw|GAA82242.1 aluc|OJZ91903.1 anid|CBF88046.1 aory|EIT78114.1 asyd|OJJ63544.1 ater|EAU37944.1 aver|OJI97390.1 awen|OJJ32379.1 azon|OJJ47141.1

Group6910: acar|OOG00396.1 afla|EED47341.1 agla|OJJ88112.1 akaw|GAA84671.1 aluc|OJZ87018.1 aory|EIT82882.1 asyd|OJJ58096.1 atub|OJI83500.1 aver|OJJ08161.1 awen|OJJ29859.1 afla|EED49710.1 aory|EIT77634.1 asyd|OJJ53587.1 afla|EED51663.1 aory|EIT80543.1

Group6911: acla|EAW07807.1 afis|EAW22844.1 afla|EED50119.1 afum|EAL90359.1 agla|OJJ83494.1 akaw|GAA89537.1 aluc|OJZ88113.1 anid|CBF87844.1 anig|EHA17883.1 aory|EIT74663.1 ater|EAU38866.1 atub|OJI90722.1 aver|OJI97516.1 awen|OJJ38879.1 azon|OJJ49668.1

Group6912: acla|EAW12228.1 afis|EAW17000.1 afum|EAL86525.1 agla|OJJ84724.1 anid|CBF77120.1 anig|EHA25714.1 asyd|OJJ62718.1 ater|EAU38945.1 aver|OJI99732.1 awen|OJJ30973.1 afis|EAW20298.1 afum|EAL91117.1 agla|OJJ88388.1 afum|EAL85427.1 asyd|OJJ60403.1

Group6913: acla|EAW12410.1 afis|EAW16763.1 afum|EAL86334.1 agla|OJJ84649.1 akaw|GAA81985.1 aluc|OJZ92205.1 anid|CBF80579.1 aory|EIT73906.1 asyd|OJJ54683.1 ater|EAU29565.1 atub|OJI86634.1 aver|OJJ07486.1 awen|OJJ33303.1 asyd|OJJ54682.1 azon|OJJ46848.1

Group6914: acar|OOF90424.1 acar|OOF90428.1 afis|EAW16794.1 afum|EAL86363.1 asyd|OJJ61718.1 aver|OJI95812.1 awen|OJJ41828.1 afla|EED45973.1 agla|OJJ79963.1 aory|EIT75831.1 awen|OJJ33625.1 afla|EED49828.1 aory|EIT77643.1 anid|CBF71775.1

Group6915: acar|OOF90491.1 acar|OOF90501.1 abra|OJJ71085.1 afis|EAW25755.1 afla|EED49652.1 akaw|GAA82876.1 aluc|OJZ87443.1 anid|CBF75199.1 aory|EIT80942.1 asyd|OJJ51843.1 ater|EAU29905.1 atub|OJI81205.1 awen|OJJ38992.1 awen|OJJ29745.1

Group6916: acar|OOF92316.1 acar|OOF92358.1 abra|OJJ67483.1 afla|EED53011.1 akaw|GAA86356.1 aluc|OJZ84885.1 anid|CBF70758.1 anig|EHA25114.1 aory|EIT77440.1 asyd|OJJ58433.1 ater|EAU32283.1 atub|OJI80715.1 aver|OJJ01034.1 awen|OJJ31755.1

Group6917: acar|OOF93700.1 acar|OOF93786.1 abra|OJJ66691.1 afis|EAW19438.1 afum|EAL87058.1 akaw|GAA89635.1 aluc|OJZ80755.1 anid|CBF81389.1 anig|EHA27269.1 asyd|OJJ59397.1 ater|EAU34739.1 atub|OJI87611.1 aver|OJJ02875.1 awen|OJJ36835.1

Group6918: acar|OOF93725.1 acar|OOF93813.1 abra|OJJ66725.1 acla|EAW11855.1 afis|EAW19466.1 afum|EAL87030.1 agla|OJJ81184.1 akaw|GAA89666.1 aluc|OJZ80723.1 anig|EHA27297.1 ater|EAU34776.1 atub|OJI87579.1 awen|OJJ36870.1 azon|OJJ42933.1

Group6919: acar|OOF94033.1 acar|OOF94103.1 abra|OJJ66418.1 acla|EAW15257.1 afis|EAW20232.1 afla|EED55129.1 afum|EAL85349.1 akaw|GAA90284.1 aluc|OJZ83056.1 anig|EHA28122.1 aory|EIT81653.1 ater|EAU37763.1 atub|OJI87891.1 azon|OJJ47375.1

Group6920: acar|OOF94042.1 acar|OOF94112.1 abra|OJJ66399.1 afla|EED49555.1 agla|OJJ79508.1 akaw|GAA90267.1 aluc|OJZ83074.1 anig|EHA28138.1 aory|EIT82623.1 asyd|OJJ61453.1 atub|OJI87872.1 aver|OJJ00788.1 awen|OJJ39515.1 azon|OJJ45872.1

Group6921: acar|OOF94069.1 acar|OOF94139.1 abra|OJJ66353.1 acla|EAW10818.1 afis|EAW24755.1 afum|EAL89398.1 akaw|GAA90227.1 aluc|OJZ83114.1 anig|EHA28177.1 asyd|OJJ63866.1 ater|EAU33501.1 atub|OJI87820.1 aver|OJI97073.1 awen|OJJ33617.1

Group6922: acar|OOF94092.1 acar|OOF94162.1 abra|OJJ66345.1 afla|EED47093.1 agla|OJJ80117.1 akaw|GAA90219.1 aluc|OJZ83122.1 anid|CBF86770.1 anig|EHA28186.1 aory|EIT72479.1 atub|OJI87811.1 aver|OJI98790.1 awen|OJJ36362.1 asyd|OJJ64688.1

Group6923: acar|OOF95049.1 acar|OOF95116.1 abra|OJJ70453.1 afla|EED45438.1 akaw|GAA90130.1 aluc|OJZ83219.1 anid|CBF84672.1 anig|EHA21077.1 aory|EIT82469.1 asyd|OJJ53348.1 ater|EAU37647.1 atub|OJI87718.1 aver|OJJ08396.1 awen|OJJ38302.1

Group6924: acar|OOF95635.1 acar|OOF95686.1 abra|OJJ74672.1 akaw|GAA81941.1 aluc|OJZ92249.1 anid|CBF71781.1 anig|EHA26707.1 asyd|OJJ61744.1 atub|OJI86682.1 aver|OJI95841.1 acar|OOF91477.1 azon|OJJ50316.1 afla|EED55300.1 aory|EIT83599.1

Group6925: acar|OOF96226.1 acar|OOF96264.1 abra|OJJ71511.1 afla|EED45339.1 agla|OJJ88922.1 akaw|GAA89404.1 aluc|OJZ86249.1 aory|EIT73804.1 asyd|OJJ63234.1 ater|EAU38650.1 atub|OJI84293.1 aver|OJI97722.1 awen|OJJ31796.1 azon|OJJ45990.1

Group6926: acar|OOF96909.1 acar|OOF96945.1 abra|OJJ70801.1 acla|EAW11123.1 afis|EAW15852.1 afla|EED56521.1 afum|EAL88488.1 agla|OJJ87256.1 akaw|GAA90117.1 aluc|OJZ84146.1 anid|CBF81178.1 aory|EIT72788.1 asyd|OJJ59293.1 aver|OJJ02982.1

Group6927: acar|OOF96929.1 acar|OOF96964.1 abra|OJJ70779.1 acla|EAW11140.1 afis|EAW15871.1 afum|EAL88505.1 akaw|GAA90098.1 aluc|OJZ84126.1 anid|CBF81215.1 anig|EHA19234.1 aory|EIT72839.1 ater|EAU31688.1 atub|OJI90500.1 azon|OJJ47441.1

Group6928: acar|OOF97668.1 acar|OOF97716.1 abra|OJJ71488.1 acla|EAW10062.1 afum|EBA27396.1 agla|OJJ87279.1 akaw|GAA82532.1 aluc|OJZ87059.1 anig|EHA20332.1 asyd|OJJ53022.1 atub|OJI81576.1 aver|OJJ03821.1 awen|OJJ41266.1 azon|OJJ44831.1

Group6929: afis|EAW24052.1 afis|EAW25623.1 aluc|OJZ79946.1 aluc|OJZ85403.1 acar|OOF98689.1 afla|EED49756.1 afum|EAL84311.1 agla|OJJ86172.1 akaw|GAA87768.1 aory|EIT77610.1 asyd|OJJ56972.1 ater|EAU37499.1 atub|OJI80170.1 aver|OJJ03317.1

Group6930: afla|EED45951.1 afla|EED57818.1 aory|EIT75362.1 aory|EIT81086.1 acla|EAW09537.1 afis|EAW21595.1 afum|EAL92613.1 aluc|OJZ85107.1 ater|EAU34929.1 awen|OJJ36083.1 afla|EED45851.1 anig|EHA25328.1 afla|EED57817.1 akaw|GAA86559.1

Group6931: afla|EED48216.1 afla|EED52152.1 aory|EIT72949.1 aory|EIT74498.1 abra|OJJ75217.1 acar|OOF94567.1 acla|EAW15362.1 afis|EAW20350.1 afum|EAL85468.1 akaw|GAA91767.1 aluc|OJZ92781.1 anig|EHA26229.1 ater|EAU31806.1 atub|OJI87223.1

Group6932: afla|EED52305.1 afla|EED56023.1 aory|EIT74399.1 aory|EIT76551.1 acar|OOF91585.1 afis|EAW19560.1 afum|EAL86951.1 agla|OJJ86282.1 anid|CBF73581.1 asyd|OJJ56685.1 ater|EAU39598.1 aver|OJJ06038.1 awen|OJJ30876.1 azon|OJJ47317.1

Group6933: afum|EAL85412.1 afum|EAL91515.2 acar|OOG00333.1 acla|EAW12963.1 afis|EAW17700.1 asyd|OJJ57237.1 aver|OJJ03048.1 asyd|OJJ61526.1 ater|EAU36789.1 azon|OJJ49085.1 afum|EAL91513.1 anid|CBF80589.1 awen|OJJ36014.1 aver|OJJ00859.1

Group6934: agla|OJJ82009.1 agla|OJJ85461.1 acar|OOF93839.1 afla|EED50687.1 akaw|GAA89693.1 aluc|OJZ80691.1 anig|EHA27324.1 aory|EIT73601.1 atub|OJI87551.1 aver|OJJ03883.1 azon|OJJ46763.1 afla|EED45096.1 asyd|OJJ60303.1 awen|OJJ32713.1

Group6935: akaw|GAA92844.1 akaw|GAA92891.1 akaw|GAA93037.1 akaw|GAA93105.1 aluc|OJZ79646.1 aluc|OJZ80468.1 abra|OJJ65835.1 acar|OOF97962.1 afla|EED46520.1 aory|EIT72396.1 asyd|OJJ56740.1 ater|EAU34462.1 awen|OJJ36064.1 azon|OJJ45266.1

Group6936: aluc|OJZ80069.1 aluc|OJZ81388.1 abra|OJJ67544.1 acla|EAW12261.1 afla|EED48219.1 agla|OJJ82716.1 akaw|GAA88116.1 anid|CBF86667.1 anig|EHA18681.1 asyd|OJJ64774.1 ater|EAU31805.1 atub|OJI91593.1 aver|OJI98874.1 awen|OJJ30122.1

Group6937: aluc|OJZ80266.1 aluc|OJZ87735.1 anig|EHA17697.1 anig|EHA20226.1 abra|OJJ67240.1 acar|OOF89888.1 acla|EAW08784.1 afis|EAW25600.1 afum|EAL84305.1 agla|OJJ79165.1 akaw|GAA91665.1 aory|EIT81930.1 ater|EAU34041.1 atub|OJI80905.1

Group6938: anid|CBF77156.1 anid|CBF86820.1 asyd|OJJ56906.1 asyd|OJJ56933.1 aver|OJI95763.1 aver|OJJ03387.1 awen|OJJ36091.1 awen|OJJ39054.1 agla|OJJ78635.1 afla|EED48930.1 aory|EIT80041.1 ater|EAU36115.1 anig|EHA20661.1 azon|OJJ48055.1

Group6939: anid|CBF78784.1 anid|CBF86372.1 abra|OJJ67144.1 acar|OOF98410.1 afla|EED47581.1 afum|EAL84948.1 akaw|GAA91025.1 aluc|OJZ87828.1 anig|EHA20307.1 aory|EIT74572.1 asyd|OJJ65244.1 ater|EAU29469.1 atub|OJI80813.1 aver|OJI99396.1

Group6940: anid|CBF88140.1 anid|CBF90391.1 acar|OOF90187.1 afis|EAW20751.1 afla|EED52306.1 agla|OJJ83899.1 akaw|GAA91990.1 aluc|OJZ92860.1 anig|EHA26140.1 asyd|OJJ63603.1 ater|EAU29428.1 atub|OJI87318.1 aver|OJI97333.1 awen|OJJ29676.1

Group6941: asyd|OJJ52095.1 asyd|OJJ52431.1 abra|OJJ75939.1 acar|OOF99971.1 afla|EED45996.1 agla|OJJ81767.1 akaw|GAA84641.1 aluc|OJZ89386.1 anid|CBF86225.1 anig|EHA23198.1 aory|EIT75400.1 ater|EAU35909.1 atub|OJI85411.1 awen|OJJ39419.1

Group6942: asyd|OJJ52254.1 asyd|OJJ62108.1 abra|OJJ75248.1 afis|EAW25209.1 afum|EAL89870.1 agla|OJJ88954.1 akaw|GAA91789.1 aluc|OJZ92808.1 anid|CBF89952.1 anig|EHA26200.1 ater|EAU34351.1 atub|OJI87258.1 aver|OJI96214.1 awen|OJJ40843.1

Group6943: asyd|OJJ61017.1 asyd|OJJ61682.1 awen|OJJ32648.1 awen|OJJ37015.1 acar|OOF90107.1 acla|EAW10851.1 afis|EAW19664.1 afla|EED49410.1 anid|CBF84685.1 anig|EHA18841.1 aory|EIT79536.1 ater|EAU29364.1 atub|OJI90058.1 afla|EED49411.1

Group6944: ater|EAU29345.1 ater|EAU29346.1 abra|OJJ65751.1 afis|EAW17418.1 afum|EAL86499.2 akaw|GAA92847.1 aluc|OJZ88790.1 anid|CBF73627.1 anig|EHA26111.1 asyd|OJJ62709.1 aver|OJJ03130.1 awen|OJJ29578.1 anig|EHA18443.1 aluc|OJZ79962.1

Group6945: ater|EAU30324.1 ater|EAU34792.1 abra|OJJ66348.1 acar|OOF89957.1 afis|EAW17257.1 afla|EED47566.1 akaw|GAA90222.1 aluc|OJZ83119.1 anig|EHA28183.1 aory|EIT74539.1 asyd|OJJ56686.1 atub|OJI87814.1 aver|OJJ06037.1 awen|OJJ38070.1

Group6946: ater|EAU31648.1 ater|EAU32789.1 abra|OJJ74153.1 acar|OOF95572.1 afla|EED56734.1 agla|OJJ84116.1 akaw|GAA89730.1 aluc|OJZ80653.1 anig|EHA27179.1 aory|EIT73918.1 asyd|OJJ61411.1 atub|OJI87508.1 aver|OJJ00741.1 awen|OJJ34278.1

Group6947: atub|OJI81113.1 atub|OJI88409.1 abra|OJJ70992.1 afis|EAW21012.1 afla|EED53337.1 agla|OJJ82530.1 akaw|GAA89071.1 aluc|OJZ80521.1 anig|EHA20745.1 aory|EIT75239.1 ater|EAU30432.1 awen|OJJ35895.1 anig|EHA25674.1 aluc|OJZ87536.1

Group6948: aver|OJJ03412.1 aver|OJJ04807.1 abra|OJJ66799.1 acar|OOF93876.1 afla|EED55091.1 agla|OJJ78901.1 akaw|GAA86863.1 aluc|OJZ89987.1 anig|EHA27358.1 aory|EIT81582.1 ater|EAU29930.1 atub|OJI84823.1 awen|OJJ36049.1 azon|OJJ50226.1

Group6949: awen|OJJ29806.1 awen|OJJ37388.1 abra|OJJ68900.1 acar|OOF90515.1 acla|EAW12124.1 afis|EAW16876.1 akaw|GAA90351.1 aluc|OJZ85881.1 anig|EHA24705.1 atub|OJI79667.1 azon|OJJ45041.1 agla|OJJ78940.1 afum|EAL86413.1 asyd|OJJ54557.1

Group6950: awen|OJJ29925.1 awen|OJJ36429.1 abra|OJJ77388.1 acar|OOF93257.1 afla|EED51018.1 akaw|GAA91373.1 aluc|OJZ84796.1 anig|EHA28063.1 aory|EIT76995.1 asyd|OJJ57005.1 ater|EAU31809.1 atub|OJI89286.1 aver|OJJ03282.1 afla|EED48974.1

Group6951: awen|OJJ30251.1 awen|OJJ33419.1 azon|OJJ45688.1 azon|OJJ50381.1 acla|EAW15282.1 agla|OJJ81596.1 abra|OJJ77686.1 aluc|OJZ80830.1 aory|EIT76243.1 ater|EAU39391.1 atub|OJI82736.1 acla|EAW10823.1 afis|EAW19958.1 afla|EED55323.1

Group6952: awen|OJJ30777.1 awen|OJJ31704.1 abra|OJJ67680.1 acar|OOF90430.1 afis|EAW17170.1 afla|EED56039.1 afum|EAL86602.1 agla|OJJ87459.1 akaw|GAA89193.1 aluc|OJZ81638.1 anig|EHA25806.1 aory|EIT76820.1 atub|OJI88542.1 aver|OJJ03941.1

Group6953: awen|OJJ33374.1 awen|OJJ35476.1 awen|OJJ35903.1 acar|OOF90256.1 acla|EAW08587.1 akaw|GAA83752.1 aluc|OJZ90979.1 anid|CBF80129.1 anig|EHA19684.1 asyd|OJJ58345.1 atub|OJI83135.1 aver|OJJ02068.1 azon|OJJ45079.1 atub|OJI82970.1

Group6954: awen|OJJ36940.1 awen|OJJ40858.1 abra|OJJ68430.1 acar|OOF93960.1 afla|EED49405.1 agla|OJJ78444.1 akaw|GAA86941.1 aluc|OJZ90066.1 anid|CBF77083.1 anig|EHA21008.1 aory|EIT79528.1 asyd|OJJ56963.1 atub|OJI84741.1 aver|OJJ03323.1

Group6955: azon|OJJ42576.1 azon|OJJ45716.1 azon|OJJ45074.1 azon|OJJ43555.1 abra|OJJ66011.1 acar|OOF91994.1 acla|EAW11547.1 afis|EAW19001.1 afum|EAL87334.1 agla|OJJ84824.1 asyd|OJJ60039.1 aver|OJJ02246.1 awen|OJJ31270.1 afis|EAW23906.1

Group6956: azon|OJJ42644.1 azon|OJJ48760.1 azon|OJJ50425.1 azon|OJJ42305.1 abra|OJJ77949.1 agla|OJJ88431.1 akaw|GAA83715.1 aluc|OJZ91018.1 anig|EHA19720.1 atub|OJI88499.1 aluc|OJZ81682.1 afla|EED51049.1 awen|OJJ31287.1 afum|EAL86479.2

Group6957: azon|OJJ44233.1 azon|OJJ46858.1 abra|OJJ76476.1 afla|EED54050.1 akaw|GAA82388.1 aluc|OJZ91732.1 anig|EHA22739.1 aory|EIT76013.1 asyd|OJJ52398.1 ater|EAU29590.1 atub|OJI86191.1 aver|OJJ01925.1 afla|EED50664.1 aory|EIT78105.1

Group6958: azon|OJJ50342.1 azon|OJJ51822.1 abra|OJJ76238.1 acar|OOF99483.1 acla|EAW12382.1 afis|EAW22751.1 afla|EED50909.1 akaw|GAA82172.1 aluc|OJZ91993.1 anig|EHA22938.1 aory|EIT79036.1 ater|EAU38096.1 atub|OJI86428.1 awen|OJJ38781.1

Group6959: abra|OJJ65599.1 afla|EED45684.1 akaw|GAA92638.1 aluc|OJZ90501.1 aory|EIT75419.1 atub|OJI84074.1 azon|OJJ49035.1 abra|OJJ68604.1 acar|OOF94949.1 agla|OJJ80045.1 abra|OJJ68910.1 acar|OOF98557.1 ater|EAU35052.1 azon|OJJ50354.1

Group6960: abra|OJJ65686.1 acar|OOF93614.1 acla|EAW12339.1 afis|EAW15634.1 afla|EED44879.1 afum|EAL85703.1 anid|CBF82090.1 aory|EIT83252.1 asyd|OJJ54264.1 ater|EAU32954.1 atub|OJI83488.1 aver|OJJ05218.1 awen|OJJ31358.1 acla|EAW08773.1

Group6961: abra|OJJ65695.1 afla|EED56834.1 afum|EAL84264.1 aory|EIT73846.1 azon|OJJ47368.1 abra|OJJ68911.1 acar|OOF91496.1 acla|EAW12302.1 agla|OJJ88168.1 akaw|GAA90339.1 aluc|OJZ85893.1 anig|EHA24698.1 atub|OJI79678.1 awen|OJJ30630.1

Group6962: abra|OJJ65700.1 afla|EED56832.1 afum|EAL84270.1 agla|OJJ84578.1 anid|CBF84427.1 anig|EHA18819.1 aory|EIT73851.1 asyd|OJJ62606.1 ater|EAU33258.1 aver|OJJ07759.1 awen|OJJ30258.1 asyd|OJJ53640.1 aver|OJJ01166.1 anig|EHA23479.1

Group6963: abra|OJJ65812.1 acar|OOF94171.1 akaw|GAA92877.1 aluc|OJZ82532.1 anig|EHA22498.1 atub|OJI88777.1 abra|OJJ66511.1 afla|EED46594.1 aory|EIT77965.1 ater|EAU32617.1 abra|OJJ78186.1 acla|EAW13446.1 agla|OJJ79275.1 azon|OJJ45475.1

Group6964: abra|OJJ65859.1 acar|OOF95020.1 acla|EAW14586.1 afla|EED49636.1 agla|OJJ81812.1 akaw|GAA92286.1 aluc|OJZ90183.1 anid|CBF74599.1 anig|EHA23682.1 aory|EIT80958.1 ater|EAU32492.1 atub|OJI81610.1 azon|OJJ43053.1 acla|EAW11034.1

Group6965: abra|OJJ65880.1 acar|OOF95295.1 akaw|GAA88079.1 aluc|OJZ81342.1 anig|EHA18710.1 atub|OJI91547.1 azon|OJJ42627.1 aory|EIT74600.1 ater|EAU33327.1 anid|CBF88770.1 asyd|OJJ63983.1 aver|OJI96970.1 awen|OJJ35859.1 awen|OJJ30904.1

Group6966: abra|OJJ65892.1 afla|EED46587.1 agla|OJJ85291.1 akaw|GAA86013.1 aluc|OJZ90240.1 anig|EHA23737.1 aory|EIT77889.1 atub|OJI81669.1 awen|OJJ31278.1 akaw|GAA86638.1 aluc|OJZ85195.1 atub|OJI85130.1 akaw|GAA87344.1 aluc|OJZ89670.1

Group6967: abra|OJJ65894.1 acar|OOF94955.1 afla|EED53410.1 akaw|GAA86011.1 aluc|OJZ90238.1 aory|EIT74977.1 ater|EAU30301.1 atub|OJI81667.1 aver|OJJ06586.1 awen|OJJ32203.1 akaw|GAA85849.1 aluc|OJZ91250.1 ater|EAU29841.1 awen|OJJ37897.1

Group6968: abra|OJJ65918.1 afla|EED50457.1 afum|EAL84941.1 agla|OJJ80058.1 akaw|GAA92314.1 aluc|OJZ90212.1 anid|CBF84463.1 anig|EHA23710.1 aory|EIT82057.1 asyd|OJJ53549.1 ater|EAU31472.1 atub|OJI81643.1 aver|OJJ08204.1 awen|OJJ29952.1

Group6969: abra|OJJ65920.1 acar|OOF94981.1 afis|EAW22856.1 afum|EAL90344.1 akaw|GAA92312.1 aluc|OJZ90210.1 anig|EHA23708.1 atub|OJI81641.1 awen|OJJ30825.1 azon|OJJ43901.1 acar|OOF90237.1 anig|EHA19271.1 acla|EAW07820.1 azon|OJJ42247.1

Group6970: abra|OJJ65925.1 acar|OOF94986.1 acla|EAW12065.1 afla|EED57048.1 agla|OJJ78912.1 aluc|OJZ90206.1 anid|CBF71409.1 anig|EHA23703.1 asyd|OJJ55013.1 atub|OJI81635.1 aver|OJJ07198.1 awen|OJJ29906.1 acar|OOF94208.1 ater|EAU32841.1

Group6971: acla|EAW08976.1 afis|EAW15455.1 afla|EED54388.1 afum|EAL84190.1 aory|EIT75844.1 abra|OJJ68368.1 awen|OJJ40831.1 acar|OOF95006.1 akaw|GAA92295.1 aluc|OJZ90192.1 anig|EHA23691.1 ater|EAU37834.1 atub|OJI81619.1 azon|OJJ42817.1

Group6972: abra|OJJ65953.1 acar|OOF92919.1 afla|EED46035.1 agla|OJJ88003.1 akaw|GAA90830.1 aluc|OJZ83004.1 anig|EHA21420.1 aory|EIT75813.1 asyd|OJJ53292.1 ater|EAU29389.1 atub|OJI89238.1 aver|OJJ08448.1 awen|OJJ30044.1 azon|OJJ43690.1

Group6973: abra|OJJ65985.1 acar|OOF98728.1 afla|EED45376.1 agla|OJJ87971.1 akaw|GAA90858.1 aluc|OJZ82975.1 anid|CBF78951.1 anig|EHA21391.1 aory|EIT82468.1 asyd|OJJ55656.1 ater|EAU37435.1 atub|OJI89208.1 aver|OJJ06740.1 awen|OJJ30018.1

Group6974: abra|OJJ65989.1 acar|OOF92884.1 afis|EAW19526.1 afla|EED49679.1 agla|OJJ87964.1 akaw|GAA90863.1 aluc|OJZ82970.1 anig|EHA21386.1 aory|EIT81017.1 asyd|OJJ55655.1 atub|OJI89203.1 aver|OJJ06739.1 awen|OJJ30014.1 azon|OJJ43644.1

Group6975: abra|OJJ66004.1 acar|OOF92094.1 afla|EED50226.1 akaw|GAA90877.1 aluc|OJZ82956.1 anid|CBF80408.1 anig|EHA21371.1 aory|EIT76439.1 ater|EAU30955.1 atub|OJI89188.1 awen|OJJ34468.1 azon|OJJ45184.1 afla|EED46243.1 azon|OJJ45166.1

Group6976: abra|OJJ66113.1 acar|OOF94006.1 afis|EAW20770.1 afla|EED51741.1 afum|EAL84849.1 akaw|GAA92677.1 aluc|OJZ90132.1 anid|CBF78153.1 anig|EHA20939.1 aory|EIT80598.1 asyd|OJJ55951.1 ater|EAU30762.1 atub|OJI84664.1 aver|OJJ04897.1

Group6977: abra|OJJ66123.1 acar|OOF94013.1 acla|EAW12395.1 afis|EAW16843.1 akaw|GAA92684.1 aluc|OJZ90142.1 anid|CBF82118.1 anig|EHA20927.1 aory|EIT78834.1 asyd|OJJ54291.1 atub|OJI84657.1 aver|OJJ05247.1 awen|OJJ41473.1 afla|EED54594.1

Group6978: abra|OJJ66149.1 afis|EAW17097.1 afla|EED51083.1 agla|OJJ79833.1 akaw|GAA88362.1 aluc|OJZ82293.1 anig|EHA18736.1 aory|EIT77058.1 asyd|OJJ61752.1 ater|EAU38712.1 atub|OJI88146.1 aver|OJI95850.1 awen|OJJ29778.1 ater|EAU33991.1

Group6979: abra|OJJ66189.1 acar|OOF98736.1 afla|EED52248.1 aory|EIT74383.1 asyd|OJJ65286.1 awen|OJJ38315.1 abra|OJJ76496.1 afis|EAW16417.1 akaw|GAA82412.1 aluc|OJZ91707.1 anid|CBF80658.1 anig|EHA22719.1 atub|OJI86166.1 aver|OJJ08324.1

Group6980: abra|OJJ66252.1 acar|OOF90724.1 acla|EAW13652.1 afis|EAW19953.1 afla|EED57854.1 afum|EAL84962.1 akaw|GAA89035.1 aluc|OJZ80560.1 anig|EHA25637.1 aory|EIT81119.1 ater|EAU36872.1 atub|OJI88371.1 afla|EED45865.1 aory|EIT75804.1

Group6981: abra|OJJ66263.1 acar|OOF90737.1 afla|EED52626.1 agla|OJJ80715.1 akaw|GAA89032.1 aluc|OJZ80571.1 anid|CBF83203.1 anig|EHA25627.1 asyd|OJJ55553.1 ater|EAU29936.1 atub|OJI88360.1 aver|OJJ06630.1 awen|OJJ40108.1 azon|OJJ43920.1

Group6982: abra|OJJ66357.1 acla|EAW10007.1 afis|EAW21049.1 afla|EED51621.1 afum|EAL92085.1 akaw|GAA90231.1 aluc|OJZ83110.1 anid|CBF87102.1 anig|EHA28173.1 aory|EIT80670.1 asyd|OJJ60389.1 ater|EAU36821.1 atub|OJI87824.1 aver|OJI99718.1

Group6983: abra|OJJ66433.1 afla|EED49899.1 agla|OJJ84659.1 anid|CBF71844.1 aory|EIT72456.1 awen|OJJ35942.1 abra|OJJ70251.1 acar|OOF90243.1 akaw|GAA86540.1 aluc|OJZ85088.1 anig|EHA25307.1 atub|OJI80511.1 asyd|OJJ61490.1 aver|OJJ00831.1

Group6984: abra|OJJ66442.1 afla|EED53319.1 agla|OJJ85806.1 aory|EIT75251.1 asyd|OJJ59091.1 ater|EAU34526.1 aver|OJJ01681.1 abra|OJJ68019.1 acar|OOF92062.1 akaw|GAA90932.1 aluc|OJZ82893.1 anig|EHA21317.1 atub|OJI89117.1 asyd|OJJ56921.1

Group6985: abra|OJJ66457.1 akaw|GAA84342.1 aluc|OJZ89042.1 anid|CBF84580.1 atub|OJI85747.1 abra|OJJ70525.1 acar|OOF93024.1 aver|OJJ07638.1 awen|OJJ38692.1 azon|OJJ46468.1 abra|OJJ78045.1 asyd|OJJ54520.1 aver|OJJ05321.1 aver|OJJ04477.1

Group6986: abra|OJJ66484.1 acar|OOF90961.1 acla|EAW10985.1 afis|EAW20875.1 afla|EED45557.1 afum|EAL84759.1 agla|OJJ83982.1 anid|CBF82853.1 anig|EHA25884.1 aory|EIT75810.1 ater|EAU30814.1 aver|OJJ00917.1 awen|OJJ34485.1 azon|OJJ51762.1

Group6987: abra|OJJ66527.1 acla|EAW10963.1 afla|EED50246.1 afum|EAL84736.1 agla|OJJ84004.1 akaw|GAA91186.1 aluc|OJZ81517.1 anid|CBF82462.1 anig|EHA25915.1 aory|EIT72398.1 asyd|OJJ54116.1 atub|OJI88666.1 awen|OJJ34451.1 afla|EED46511.1

Group6988: abra|OJJ66623.1 acar|OOF91047.1 acla|EAW10873.1 afis|EAW20985.1 afum|EAL84645.1 agla|OJJ84064.1 anid|CBF82624.1 anig|EHA25996.1 aory|EIT77261.1 asyd|OJJ54021.1 ater|EAU38634.1 atub|OJI88755.1 aver|OJJ08719.1 azon|OJJ48070.1

Group6989: abra|OJJ66699.1 acar|OOF94909.1 acla|EAW10031.1 afis|EAW19514.1 afla|EED45381.1 afum|EAL86982.1 akaw|GAA89642.1 aluc|OJZ80747.1 anid|CBF87985.1 anig|EHA27275.1 aory|EIT82412.1 ater|EAU32110.1 atub|OJI79834.1 atub|OJI87604.1

Group6990: abra|OJJ66701.1 acar|OOF94910.1 acla|EAW10033.1 afis|EAW19513.1 afla|EED45382.1 afum|EAL86983.1 akaw|GAA89643.1 aluc|OJZ80746.1 anig|EHA27276.1 aory|EIT82444.1 asyd|OJJ65086.1 ater|EAU32109.1 atub|OJI87603.1 aver|OJI99212.1

Group6991: abra|OJJ66792.1 acar|OOF93869.1 acla|EAW12437.1 afis|EAW16746.1 afla|EED54713.1 afum|EAL86321.1 akaw|GAA86857.1 aluc|OJZ89980.1 anig|EHA27352.1 aory|EIT78785.1 asyd|OJJ52903.1 ater|EAU31716.1 atub|OJI84830.1 aver|OJJ08872.1

Group6992: abra|OJJ66844.1 acar|OOF91621.1 acla|EAW14423.1 afis|EAW19131.1 afla|EED54283.1 afum|EAL93863.1 agla|OJJ88603.1 akaw|GAA90480.1 aluc|OJZ85739.1 anig|EHA24812.1 aory|EIT76101.1 atub|OJI79524.1 awen|OJJ37466.1 azon|OJJ51144.1

Group6993: abra|OJJ66908.1 acar|OOF91678.1 afis|EAW19186.1 afum|EAL93917.1 akaw|GAA91506.1 aluc|OJZ85672.1 anig|EHA24867.1 atub|OJI79464.1 awen|OJJ37544.1 azon|OJJ51208.1 acla|EAW14477.1 afla|EED54209.1 aory|EIT75928.1 aver|OJI99164.1

Group6994: abra|OJJ66929.1 acar|OOF91697.1 afla|EED47704.1 agla|OJJ88503.1 akaw|GAA91488.1 aluc|OJZ85650.1 anid|CBF78565.1 anig|EHA24883.1 aory|EIT80210.1 asyd|OJJ65065.1 atub|OJI79441.1 aver|OJI99189.1 awen|OJJ37561.1 azon|OJJ51223.1

Group6995: abra|OJJ66963.1 acar|OOF93644.1 afla|EED55269.1 agla|OJJ86858.1 akaw|GAA92198.1 aluc|OJZ85613.1 anid|CBF76561.1 anig|EHA24913.1 aory|EIT83605.1 asyd|OJJ61160.1 ater|EAU39596.1 atub|OJI79406.1 aver|OJJ00482.1 awen|OJJ39921.1

Group6996: abra|OJJ66986.1 acar|OOF93626.1 afla|EED55059.1 afum|EAL87117.1 agla|OJJ78983.1 akaw|GAA92178.1 aluc|OJZ85588.1 anid|CBF90363.1 anig|EHA24933.1 aory|EIT81956.1 asyd|OJJ54789.1 atub|OJI79383.1 aver|OJJ07375.1 azon|OJJ42303.1

Group6997: abra|OJJ66995.1 acar|OOF93620.1 afla|EED52138.1 agla|OJJ81674.1 akaw|GAA92169.1 aluc|OJZ85577.1 anid|CBF80571.1 anig|EHA24942.1 aory|EIT74495.1 asyd|OJJ52851.1 atub|OJI79373.1 aver|OJJ08829.1 awen|OJJ36920.1 azon|OJJ42894.1

Group6998: abra|OJJ67035.1 afla|EED49393.1 akaw|GAA93115.1 aory|EIT79467.1 abra|OJJ67037.1 agla|OJJ80958.1 akaw|GAA93113.1 anig|EHA25076.1 abra|OJJ72134.1 akaw|GAA84788.1 aluc|OJZ86892.1 anig|EHA21751.1 atub|OJI83627.1 azon|OJJ42321.1

Group6999: abra|OJJ67099.1 acar|OOF98453.1 agla|OJJ81046.1 akaw|GAA91062.1 aluc|OJZ87874.1 anig|EHA20180.1 atub|OJI80775.1 awen|OJJ35991.1 abra|OJJ70413.1 afla|EED53421.1 akaw|GAA91943.1 aluc|OJZ82677.1 anig|EHA21046.1 aory|EIT75149.1

Group7000: abra|OJJ67114.1 acar|OOF98438.1 afis|EAW20324.1 afla|EED47500.1 afum|EAL85449.1 akaw|GAA91050.1 aluc|OJZ87860.1 anid|CBF82804.1 anig|EHA20167.1 aory|EIT74548.1 asyd|OJJ65273.1 atub|OJI80787.1 aver|OJI99425.1 awen|OJJ38372.1

Group7001: abra|OJJ67158.1 acar|OOF98401.1 afla|EED47597.1 agla|OJJ88392.1 akaw|GAA91014.1 aluc|OJZ87815.1 anid|CBF78758.1 anig|EHA20296.1 aory|EIT80282.1 asyd|OJJ65232.1 atub|OJI80829.1 aver|OJI99381.1 awen|OJJ37641.1 azon|OJJ44905.1

Group7002: abra|OJJ67228.1 acar|OOF98324.1 acla|EAW10404.1 afis|EAW25167.1 afum|EAL89840.1 akaw|GAA91654.1 aluc|OJZ87745.1 anid|CBF89142.1 anig|EHA20235.1 asyd|OJJ62573.1 ater|EAU39454.1 atub|OJI80896.1 aver|OJI96695.1 awen|OJJ40819.1

Group7003: abra|OJJ67237.1 acar|OOF98314.1 afis|EAW24733.1 afla|EED46088.1 afum|EAL89380.1 agla|OJJ86785.1 akaw|GAA91662.1 aluc|OJZ87738.1 anid|CBF69480.1 anig|EHA20230.1 asyd|OJJ59272.1 atub|OJI80902.1 aver|OJJ01885.1 awen|OJJ40221.1

Group7004: abra|OJJ67252.1 acla|EAW10398.1 afis|EAW25179.1 afla|EED53633.1 afum|EAL89853.2 agla|OJJ88988.1 akaw|GAA91669.1 aluc|OJZ87726.1 aory|EIT75085.1 asyd|OJJ62970.1 atub|OJI80917.1 aver|OJI97997.1 awen|OJJ40783.1 azon|OJJ48454.1

Group7005: abra|OJJ67261.1 afis|EAW21084.1 afum|EAL92106.1 agla|OJJ79289.1 akaw|GAA91676.1 aluc|OJZ87718.1 anid|CBF87308.1 anig|EHA20212.1 aory|EIT73510.1 asyd|OJJ61440.1 ater|EAU31935.1 atub|OJI80924.1 aver|OJJ00774.1 awen|OJJ35786.1

Group7006: abra|OJJ67264.1 afis|EAW21083.1 afla|EED44921.1 afum|EAL92104.2 agla|OJJ79285.1 akaw|GAA91680.1 aluc|OJZ87714.1 anig|EHA20208.1 aory|EIT73466.1 asyd|OJJ61441.1 ater|EAU31938.1 atub|OJI80928.1 aver|OJJ00776.1 awen|OJJ35783.1

Group7007: abra|OJJ67283.1 afum|EAL87109.1 agla|OJJ82023.1 akaw|GAA86526.1 aluc|OJZ85074.1 anid|CBF71657.1 anig|EHA25288.1 aory|EIT75950.1 asyd|OJJ61836.1 ater|EAU32861.1 atub|OJI80524.1 aver|OJI95935.1 awen|OJJ36118.1 afla|EED54339.1

Group7008: abra|OJJ67386.1 acla|EAW06373.1 afis|EAW18479.1 afla|EED52930.1 afum|EAL93168.1 agla|OJJ85067.1 akaw|GAA86438.1 anid|CBF70138.1 anig|EHA25193.1 aory|EIT78689.1 asyd|OJJ58813.1 atub|OJI80621.1 aver|OJJ01416.1 awen|OJJ31032.1

Group7009: abra|OJJ67465.1 acla|EAW06303.1 afis|EAW18412.1 afum|EAL93092.1 agla|OJJ84979.1 akaw|GAA86371.1 aluc|OJZ84900.1 anid|CBF70735.1 aory|EIT77434.1 asyd|OJJ58442.1 ater|EAU29827.1 atub|OJI80698.1 awen|OJJ31738.1 azon|OJJ50497.1

Group7010: abra|OJJ67567.1 acar|OOG00285.1 afla|EED48141.1 afum|EAL86219.1 agla|OJJ79350.1 akaw|GAA88098.1 aluc|OJZ81365.1 anig|EHA18665.1 aory|EIT76241.1 ater|EAU33061.1 atub|OJI91569.1 aver|OJJ07627.1 awen|OJJ39953.1 afis|EAW24554.1

Group7011: abra|OJJ67586.1 afla|EED46507.1 agla|OJJ80469.1 aory|EIT72341.1 abra|OJJ68809.1 acar|OOF90612.1 acla|EAW15298.1 afis|EAW25743.1 akaw|GAA90445.1 aluc|OJZ85781.1 anig|EHA24774.1 atub|OJI79566.1 awen|OJJ40650.1 azon|OJJ42396.1

Group7012: abra|OJJ67597.1 afis|EAW16583.1 afla|EED57872.1 afum|EAL89178.1 akaw|GAA89116.1 aluc|OJZ81723.1 anid|CBF69623.1 anig|EHA25716.1 aory|EIT83271.1 asyd|OJJ59121.1 ater|EAU36660.1 atub|OJI88458.1 aver|OJJ01710.1 awen|OJJ40233.1

Group7013: abra|OJJ67695.1 acla|EAW12035.1 afla|EED46151.1 akaw|GAA89205.1 aluc|OJZ81625.1 anid|CBF84623.1 anig|EHA25818.1 aory|EIT76443.1 ater|EAU32264.1 atub|OJI88555.1 awen|OJJ29707.1 azon|OJJ48894.1 asyd|OJJ56446.1 aver|OJJ06276.1

Group7014: abra|OJJ67700.1 acar|OOF99866.1 afla|EED48149.1 agla|OJJ78988.1 akaw|GAA89209.1 aluc|OJZ81620.1 anid|CBF81674.1 anig|EHA25823.1 aory|EIT76264.1 asyd|OJJ59556.1 atub|OJI88560.1 aver|OJJ02712.1 awen|OJJ33677.1 azon|OJJ43895.1

Group7015: abra|OJJ67726.1 acar|OOF89958.1 acla|EAW11900.1 afis|EAW19510.1 afla|EED54084.1 afum|EAL86985.1 akaw|GAA89220.1 aluc|OJZ81609.1 anid|CBF73808.1 anig|EHA25837.1 aory|EIT76030.1 asyd|OJJ56578.1 atub|OJI88577.1 azon|OJJ50137.1

Group7016: abra|OJJ67730.1 acar|OOF93977.1 afis|EAW25604.1 afla|EED55496.1 akaw|GAA89225.1 aluc|OJZ81601.1 anig|EHA25841.1 aory|EIT80776.1 ater|EAU31033.1 atub|OJI88581.1 awen|OJJ40146.1 acar|OOG00064.1 anid|CBF82319.1 ater|EAU32207.1

Group7017: abra|OJJ67737.1 acar|OOF90936.1 acla|EAW07080.1 afla|EED45455.1 akaw|GAA92428.1 aluc|OJZ81594.1 anid|CBF82819.1 anig|EHA25847.1 aory|EIT82388.1 asyd|OJJ61576.1 ater|EAU30798.1 atub|OJI88589.1 aver|OJJ00897.1 anid|CBF73492.1

Group7018: abra|OJJ67756.1 ater|EAU37754.1 atub|OJI84566.1 awen|OJJ36302.1 abra|OJJ68832.1 acar|OOF91920.1 afla|EED49279.1 agla|OJJ78834.1 akaw|GAA90420.1 aluc|OJZ85810.1 aory|EIT82230.1 asyd|OJJ52481.1 atub|OJI79596.1 aver|OJJ03926.1

Group7019: abra|OJJ67806.1 acar|OOF97083.1 acla|EAW08938.1 afis|EAW24383.1 afla|EED47826.1 afum|EAL86055.1 agla|OJJ83271.1 aluc|OJZ90851.1 aory|EIT77203.1 asyd|OJJ60975.1 aver|OJJ00299.1 awen|OJJ32829.1 azon|OJJ49007.1 acar|OOF93187.1

Group7020: abra|OJJ67838.1 acar|OOF92012.1 afla|EED47865.1 akaw|GAA91844.1 aluc|OJZ82054.1 anid|CBF82519.1 anig|EHA22206.1 asyd|OJJ54086.1 atub|OJI84612.1 aver|OJJ08652.1 awen|OJJ34679.1 aory|EIT77225.1 agla|OJJ79694.1 aver|OJJ05932.1

Group7021: abra|OJJ67883.1 acar|OOF89797.1 acla|EAW11692.1 afis|EAW23708.1 afla|EED51508.1 afum|EAL87152.1 agla|OJJ83691.1 akaw|GAA91890.1 aluc|OJZ82007.1 anig|EHA22250.1 aory|EIT80464.1 ater|EAU30421.1 atub|OJI84556.1 awen|OJJ33667.1

Group7022: abra|OJJ67894.1 afla|EED50788.1 afum|EAL86207.1 agla|OJJ83137.1 akaw|GAA91899.1 aluc|OJZ81996.1 anid|CBF74829.1 anig|EHA22260.1 aory|EIT78072.1 asyd|OJJ55031.1 atub|OJI84546.1 aver|OJJ07172.1 awen|OJJ32933.1 anid|CBF78470.1

Group7023: abra|OJJ67901.1 acar|OOF90069.1 afla|EED47043.1 afum|EAL84243.1 agla|OJJ88180.1 akaw|GAA91905.1 aluc|OJZ81989.1 anid|CBF80370.1 anig|EHA22267.1 asyd|OJJ54814.1 atub|OJI84540.1 aver|OJJ07347.1 awen|OJJ35403.1 aory|EIT72724.1

Group7024: abra|OJJ68010.1 acar|OOF92052.1 acla|EAW12060.1 afis|EAW19701.1 afum|EAL86826.1 akaw|GAA90940.1 aluc|OJZ82884.1 anid|CBF83221.1 anig|EHA21310.1 asyd|OJJ55583.1 atub|OJI89109.1 aver|OJJ06662.1 awen|OJJ40821.1 azon|OJJ45544.1

Group7025: abra|OJJ68027.1 acar|OOF92070.1 acla|EAW13766.1 afis|EAW19792.1 afla|EED48126.1 afum|EAL85099.1 akaw|GAA90926.1 aluc|OJZ82900.1 anid|CBF80565.1 anig|EHA21325.1 aory|EIT76265.1 atub|OJI89126.1 awen|OJJ35357.1 azon|OJJ43533.1

Group7026: abra|OJJ68036.1 acar|OOF91807.1 acla|EAW06967.1 afla|EED55049.1 akaw|GAA90916.1 aluc|OJZ82910.1 anig|EHA21333.1 aory|EIT81960.1 asyd|OJJ53718.1 ater|EAU36652.1 atub|OJI89136.1 aver|OJJ07844.1 awen|OJJ33895.1 azon|OJJ45202.1

Group7027: abra|OJJ68057.1 acar|OOF92078.1 acla|EAW13674.1 afis|EAW19933.1 afla|EED45899.1 afum|EAL84976.2 agla|OJJ78944.1 akaw|GAA90899.1 aluc|OJZ82929.1 anig|EHA21349.1 asyd|OJJ53382.1 atub|OJI89158.1 aver|OJJ08361.1 awen|OJJ29750.1

Group7028: abra|OJJ68058.1 acar|OOF92079.1 afla|EED45900.1 akaw|GAA90898.1 aluc|OJZ82930.1 anig|EHA21350.1 aory|EIT75411.1 asyd|OJJ54997.1 atub|OJI89159.1 aver|OJJ07717.1 abra|OJJ68906.1 afis|EAW23720.1 anig|EHA21439.1 awen|OJJ38152.1

Group7029: abra|OJJ68059.1 acar|OOF92080.1 afla|EED45901.1 akaw|GAA90897.1 aluc|OJZ82931.1 anig|EHA21351.1 aory|EIT75786.1 atub|OJI89160.1 abra|OJJ68908.1 afis|EAW23717.1 agla|OJJ83729.1 anig|EHA21437.1 asyd|OJJ54998.1 aver|OJJ07718.1

Group7030: abra|OJJ68118.1 acar|OOF93912.1 akaw|GAA86899.1 aluc|OJZ90022.1 anig|EHA27398.1 atub|OJI84785.1 acla|EAW15013.1 afis|EAW20621.1 agla|OJJ83066.1 anid|CBF86716.1 asyd|OJJ64725.1 ater|EAU34581.1 aver|OJI98826.1 azon|OJJ42452.1

Group7031: abra|OJJ68121.1 acar|OOF93915.1 acla|EAW10812.1 afis|EAW24761.1 afum|EAL89403.2 agla|OJJ81683.1 akaw|GAA86902.1 aluc|OJZ90025.1 anig|EHA27401.1 asyd|OJJ53604.1 ater|EAU30291.1 atub|OJI84782.1 aver|OJJ08144.1 awen|OJJ36928.1

Group7032: abra|OJJ68134.1 acar|OOF93922.1 afis|EAW15619.1 afla|EED57117.1 afum|EAL85716.1 akaw|GAA86914.1 aluc|OJZ90037.1 anig|EHA27412.1 aory|EIT73963.1 asyd|OJJ54936.1 ater|EAU39396.1 atub|OJI84769.1 aver|OJJ07260.1 azon|OJJ45726.1

Group7033: abra|OJJ68164.1 acar|OOG00542.1 afis|EAW19094.1 afla|EED55028.1 afum|EAL93825.1 agla|OJJ84507.1 akaw|GAA83279.1 aluc|OJZ83789.1 anid|CBF81973.1 anig|EHA18926.1 aory|EIT81948.1 asyd|OJJ56411.1 atub|OJI90156.1 aver|OJJ06304.1

Group7034: abra|OJJ68166.1 acar|OOF95535.1 afum|EAL91588.1 akaw|GAA83277.1 aluc|OJZ83784.1 anig|EHA27152.1 aory|EIT81861.1 ater|EAU35824.1 atub|OJI90154.1 aluc|OJZ89951.1 atub|OJI84855.1 anig|EHA22298.1 akaw|GAA86832.1 aluc|OJZ83783.1

Group7035: acla|EAW12135.1 agla|OJJ78883.1 abra|OJJ68522.1 acar|OOF92727.1 afla|EED53957.1 akaw|GAA91327.1 aluc|OJZ82736.1 anig|EHA21187.1 aory|EIT78312.1 ater|EAU38557.1 atub|OJI88971.1 awen|OJJ32685.1 aver|OJJ01922.1 asyd|OJJ52399.1

Group7036: abra|OJJ68280.1 acar|OOF97554.1 afis|EAW19873.1 afla|EED57867.1 akaw|GAA83192.1 aluc|OJZ83691.1 anid|CBF83973.1 anig|EHA18845.1 asyd|OJJ61407.1 atub|OJI90063.1 aver|OJJ00737.1 awen|OJJ36018.1 acar|OOF98334.1 aory|EIT81088.1

Group7037: abra|OJJ68307.1 acar|OOF96680.1 afla|EED52176.1 agla|OJJ87681.1 akaw|GAA83167.1 aluc|OJZ83666.1 anig|EHA18822.1 aory|EIT74411.1 asyd|OJJ60085.1 ater|EAU34118.1 atub|OJI90037.1 aver|OJJ04171.1 awen|OJJ38013.1 anid|CBF78265.1

Group7038: abra|OJJ68336.1 acar|OOF97601.1 afla|EED57056.1 agla|OJJ79513.1 akaw|GAA92089.1 aluc|OJZ83634.1 anig|EHA18787.1 aory|EIT82279.1 asyd|OJJ55948.1 ater|EAU36072.1 atub|OJI90003.1 aver|OJI98882.1 awen|OJJ38045.1 azon|OJJ42407.1

Group7039: abra|OJJ68353.1 afla|EED45350.1 agla|OJJ88058.1 akaw|GAA92124.1 aluc|OJZ83598.1 anig|EHA18757.1 aory|EIT82458.1 asyd|OJJ56537.1 atub|OJI89970.1 aver|OJJ06185.1 awen|OJJ41255.1 agla|OJJ87293.1 asyd|OJJ56265.1 aver|OJJ08501.1

Group7040: abra|OJJ68379.1 acar|OOF93988.1 acla|EAW12354.1 afla|EED56191.1 afum|EAL86349.1 agla|OJJ79784.1 aluc|OJZ90111.1 anid|CBF80567.1 anig|EHA20959.1 asyd|OJJ54686.1 atub|OJI84691.1 aver|OJJ07483.1 awen|OJJ29950.1 azon|OJJ45007.1

Group7041: abra|OJJ68414.1 acar|OOF93964.1 afis|EAW16785.1 afla|EED54700.1 afum|EAL86355.1 akaw|GAA92534.1 aluc|OJZ90083.1 anig|EHA20990.1 aory|EIT78774.1 atub|OJI84723.1 aver|OJJ03097.1 awen|OJJ41510.1 asyd|OJJ57194.1 azon|OJJ43632.1

Group7042: abra|OJJ68450.1 acar|OOF93945.1 agla|OJJ80809.1 akaw|GAA86923.1 aluc|OJZ90047.1 anid|CBF87391.1 anig|EHA21023.1 aory|EIT76168.1 asyd|OJJ52856.1 atub|OJI84759.1 aver|OJJ08833.1 awen|OJJ35054.1 afla|EED51909.1 afla|EED51910.1

Group7043: agla|OJJ88993.1 awen|OJJ40800.1 azon|OJJ46866.1 acar|OOF98330.1 acla|EAW10410.1 afis|EAW25158.1 afla|EED57497.1 afum|EAL89833.1 anid|CBF85657.1 anig|EHA21109.1 aory|EIT81888.1 asyd|OJJ64230.1 ater|EAU39442.1 aver|OJI98323.1

Group7044: abra|OJJ68638.1 acar|OOF92142.1 afis|EAW17026.1 afla|EED55113.1 akaw|GAA89777.1 aluc|OJZ81917.1 anig|EHA22330.1 asyd|OJJ53597.1 atub|OJI84469.1 aver|OJJ08151.1 awen|OJJ33828.1 asyd|OJJ58568.1 aver|OJJ07899.1 awen|OJJ40156.1

Group7045: abra|OJJ68731.1 acar|OOF92229.1 acla|EAW12517.1 afis|EAW24130.1 afla|EED46802.1 afum|EAL91986.1 agla|OJJ87206.1 akaw|GAA89880.1 aluc|OJZ81799.1 anig|EHA22417.1 ater|EAU30560.1 atub|OJI84370.1 awen|OJJ41359.1 azon|OJJ48823.1

Group7046: abra|OJJ68751.1 acar|OOF92240.1 acla|EAW12500.1 agla|OJJ87225.1 akaw|GAA89895.1 aluc|OJZ81776.1 anid|CBF77925.1 anig|EHA22440.1 asyd|OJJ55293.1 ater|EAU30577.1 atub|OJI84349.1 aver|OJJ06933.1 awen|OJJ41334.1 azon|OJJ48805.1

Group7047: abra|OJJ68777.1 acar|OOF92263.1 afla|EED46763.1 afum|EAL92027.1 agla|OJJ89509.1 akaw|GAA89921.1 aluc|OJZ81749.1 anid|CBF77872.1 anig|EHA22463.1 asyd|OJJ55317.1 ater|EAU30602.1 atub|OJI84320.1 awen|OJJ41306.1 azon|OJJ48780.1

Group7048: abra|OJJ68780.1 acar|OOF92266.1 afis|EAW24170.1 afla|EED46760.1 agla|OJJ89506.1 akaw|GAA89924.1 aluc|OJZ81746.1 anid|CBF77867.1 anig|EHA22466.1 asyd|OJJ55320.1 atub|OJI84317.1 aver|OJJ06908.1 awen|OJJ41303.1 azon|OJJ48777.1

Group7049: abra|OJJ68834.1 afla|EED57089.1 afum|EAL85444.1 akaw|GAA90417.1 aluc|OJZ85813.1 anid|CBF85202.1 aory|EIT82267.1 asyd|OJJ63025.1 ater|EAU32052.1 atub|OJI79599.1 aver|OJI97943.1 acla|EAW15312.1 afis|EAW20316.1 anig|EHA24746.1

Group7050: acla|EAW11543.1 abra|OJJ72704.1 acar|OOF95938.1 afla|EED52653.1 afum|EAL87336.1 agla|OJJ84625.1 akaw|GAA85308.1 aluc|OJZ90668.1 anig|EHA24102.1 aory|EIT74238.1 asyd|OJJ56945.1 ater|EAU32326.1 atub|OJI82098.1 afis|EAW23908.1

Group7051: abra|OJJ68862.1 acar|OOF90527.1 akaw|GAA90391.1 aluc|OJZ85840.1 anig|EHA24724.1 asyd|OJJ59626.1 ater|EAU32026.1 atub|OJI79628.1 awen|OJJ31225.1 azon|OJJ44218.1 abra|OJJ75059.1 acar|OOF91774.1 agla|OJJ79767.1 akaw|GAA88672.1

Group7052: abra|OJJ68876.1 acar|OOF90524.1 acla|EAW15244.1 afis|EAW20211.1 afum|EAL85328.1 akaw|GAA90379.1 aluc|OJZ85852.1 anid|CBF85104.1 aory|EIT74109.1 atub|OJI79639.1 aver|OJI97907.1 awen|OJJ31216.1 azon|OJJ46144.1 agla|OJJ84653.1

Group7053: abra|OJJ68879.1 acar|OOF90520.1 akaw|GAA90377.1 aluc|OJZ85855.1 anig|EHA28095.1 aory|EIT73528.1 atub|OJI79641.1 awen|OJJ35628.1 azon|OJJ46809.1 afla|EED55102.1 afla|EED45285.1 akaw|GAA91034.1 aluc|OJZ87844.1 atub|OJI80804.1

Group7054: abra|OJJ68897.1 acar|OOF91485.1 afla|EED56747.1 akaw|GAA90354.1 aluc|OJZ85878.1 anid|CBF73541.1 anig|EHA24708.1 aory|EIT73948.1 asyd|OJJ56708.1 ater|EAU34585.1 atub|OJI79664.1 aver|OJJ06017.1 awen|OJJ34052.1 azon|OJJ48179.1

Group7055: abra|OJJ68898.1 afla|EED56746.1 agla|OJJ84383.1 aluc|OJZ85879.1 anid|CBF84142.1 anig|EHA24707.1 aory|EIT73921.1 asyd|OJJ56707.1 ater|EAU34586.1 atub|OJI79665.1 aver|OJJ06018.1 awen|OJJ34051.1 azon|OJJ48180.1 asyd|OJJ60979.1

Group7056: abra|OJJ68902.1 acar|OOF91490.1 afla|EED47316.1 agla|OJJ84016.1 akaw|GAA90349.1 aluc|OJZ85883.1 anid|CBF75663.1 anig|EHA24703.1 aory|EIT82838.1 asyd|OJJ63348.1 atub|OJI79669.1 aver|OJI97612.1 awen|OJJ29919.1 azon|OJJ44010.1

Group7057: abra|OJJ68904.1 acla|EAW12183.1 afis|EAW17117.1 akaw|GAA90347.1 aluc|OJZ85885.1 anig|EHA24701.1 asyd|OJJ52513.1 ater|EAU36170.1 atub|OJI79671.1 aver|OJJ05885.1 anid|CBF87502.1 asyd|OJJ52970.1 aver|OJJ08936.1 afum|EAL86579.1

Group7058: abra|OJJ68998.1 acar|OOF92710.1 acla|EAW12653.1 afla|EED51867.1 akaw|GAA90517.1 aluc|OJZ85982.1 anid|CBF81942.1 anig|EHA24624.1 aory|EIT81688.1 asyd|OJJ58288.1 ater|EAU31547.1 atub|OJI79772.1 aver|OJJ02136.1 azon|OJJ48231.1

Group7059: abra|OJJ69013.1 acla|EAW12344.1 afis|EAW16880.1 afla|EED47343.1 afum|EAL86417.1 akaw|GAA90530.1 aluc|OJZ85999.1 anid|CBF80673.1 anig|EHA24611.1 asyd|OJJ54587.1 atub|OJI79786.1 aver|OJJ07567.1 awen|OJJ29770.1 azon|OJJ46799.1

Group7060: abra|OJJ69015.1 acar|OOG00871.1 afis|EAW23588.1 afla|EED47660.1 akaw|GAA90532.1 aluc|OJZ86001.1 anid|CBF74928.1 anig|EHA24609.1 aory|EIT80264.1 asyd|OJJ63611.1 atub|OJI79788.1 aver|OJI97326.1 awen|OJJ37280.1 azon|OJJ42349.1

Group7061: abra|OJJ69016.1 acar|OOG00870.1 afis|EAW23587.1 afum|EAL87877.1 akaw|GAA90533.1 aluc|OJZ86002.1 anid|CBF74930.1 anig|EHA24608.1 aory|EIT80247.1 asyd|OJJ63612.1 atub|OJI79789.1 aver|OJI97325.1 awen|OJJ37279.1 azon|OJJ42350.1

Group7062: abra|OJJ69097.1 acar|OOF91198.1 afla|EED51350.1 afum|EAL85646.2 akaw|GAA90624.1 aluc|OJZ86087.1 anid|CBF71110.1 aory|EIT80714.1 asyd|OJJ52713.1 ater|EAU32444.1 atub|OJI79883.1 aver|OJJ04742.1 awen|OJJ38395.1 azon|OJJ43194.1

Group7063: abra|OJJ69207.1 acar|OOF91110.1 acla|EAW08640.1 afis|EAW15791.1 afla|EED51254.1 afum|EAL85559.1 agla|OJJ81954.1 anid|CBF70966.1 anig|EHA24432.1 aory|EIT81549.1 asyd|OJJ52615.1 aver|OJJ04642.1 awen|OJJ38499.1 azon|OJJ43800.1

Group7064: abra|OJJ69251.1 acar|OOF91070.1 acla|EAW08598.1 afis|EAW15835.1 afla|EED51211.1 afum|EAL85517.1 agla|OJJ81999.1 akaw|GAA92258.1 anid|CBF70856.1 aory|EIT80406.1 ater|EAU33413.1 atub|OJI80025.1 awen|OJJ38548.1 azon|OJJ43714.1

Group7065: abra|OJJ69267.1 acar|OOF89862.1 acla|EAW14652.1 afis|EAW24189.1 afla|EED48080.1 afum|EAL85862.1 agla|OJJ79690.1 aluc|OJZ82096.1 anig|EHA28633.1 asyd|OJJ54009.1 atub|OJI88341.1 aver|OJJ08104.1 awen|OJJ33110.1 azon|OJJ41998.1

Group7066: abra|OJJ69310.1 acar|OOF94416.1 acla|EAW14697.1 afis|EAW24228.1 afla|EED48043.1 afum|EAL85897.1 agla|OJJ79643.1 akaw|GAA88179.1 aluc|OJZ82139.1 anid|CBF74230.1 aory|EIT83344.1 ater|EAU31177.1 atub|OJI88298.1 aver|OJJ08064.1

Group7067: abra|OJJ69374.1 acar|OOF94353.1 acla|EAW14764.1 afis|EAW24295.1 afla|EED47981.1 afum|EAL85963.1 agla|OJJ83160.1 akaw|GAA88243.1 aluc|OJZ82209.1 anig|EHA28531.1 aory|EIT79813.1 ater|EAU31245.1 atub|OJI88227.1 aver|OJJ08002.1

Group7068: abra|OJJ69407.1 acar|OOF90930.1 acla|EAW07670.1 afis|EAW19533.1 afum|EAL92130.1 akaw|GAA88272.1 aluc|OJZ82244.1 anid|CBF78092.1 anig|EHA28498.1 asyd|OJJ56571.1 ater|EAU39311.1 atub|OJI88194.1 awen|OJJ35862.1 afum|EAL86973.1

Group7069: abra|OJJ69450.1 acar|OOF94291.1 acla|EAW14814.1 afis|EAW24344.1 afum|EAL86015.1 agla|OJJ83209.1 akaw|GAA88369.1 aluc|OJZ82286.1 anig|EHA28462.1 aory|EIT83368.1 asyd|OJJ53848.1 atub|OJI88153.1 awen|OJJ32953.1 azon|OJJ48967.1

Group7070: abra|OJJ69476.1 acla|EAW09912.1 afla|EED54648.1 afum|EBA27451.1 agla|OJJ80025.1 akaw|GAA88339.1 aluc|OJZ82319.1 anig|EHA28439.1 asyd|OJJ60529.1 atub|OJI88122.1 aver|OJI99856.1 awen|OJJ36493.1 azon|OJJ44368.1 aory|EIT78848.1

Group7071: abra|OJJ69494.1 acar|OOF94253.1 acla|EAW09900.1 afis|EAW21148.1 afla|EED54664.1 afum|EAL92167.1 agla|OJJ80053.1 akaw|GAA88326.1 aluc|OJZ82334.1 anig|EHA28426.1 aory|EIT78744.1 atub|OJI88106.1 awen|OJJ36475.1 asyd|OJJ55504.1

Group7072: abra|OJJ69521.1 acar|OOF94235.1 acla|EAW09967.1 afis|EAW21121.1 afla|EED51794.1 afum|EAL92144.1 agla|OJJ80090.1 aluc|OJZ82361.1 anid|CBF87023.1 asyd|OJJ60458.1 atub|OJI88081.1 aver|OJI99778.1 awen|OJJ36451.1 azon|OJJ44426.1

Group7073: afum|EAL90772.1 aory|EIT82768.1 acar|OOF92668.1 acla|EAW07394.1 afis|EAW22420.1 afla|EED52498.1 agla|OJJ87602.1 anid|CBF88808.1 anig|EHA28395.1 asyd|OJJ62892.1 ater|EAU39286.1 aver|OJI96943.1 awen|OJJ34999.1 azon|OJJ45644.1

Group7074: abra|OJJ69539.1 acar|OOF90591.1 afla|EED54770.1 agla|OJJ86497.1 akaw|GAA88387.1 aluc|OJZ82375.1 anid|CBF79837.1 anig|EHA28389.1 aory|EIT79393.1 asyd|OJJ58147.1 ater|EAU29309.1 atub|OJI88067.1 aver|OJJ04257.1 awen|OJJ29884.1

Group7075: abra|OJJ69559.1 acar|OOF91591.1 afla|EED54879.1 agla|OJJ83840.1 akaw|GAA91568.1 aluc|OJZ82386.1 anig|EHA28377.1 aory|EIT79415.1 asyd|OJJ64200.1 atub|OJI88054.1 aver|OJI98289.1 awen|OJJ40805.1 afis|EAW16972.1 azon|OJJ42646.1

Group7076: abra|OJJ69572.1 acar|OOG00214.1 afla|EED46072.1 agla|OJJ82290.1 akaw|GAA91569.1 aluc|OJZ82389.1 anig|EHA28374.1 asyd|OJJ53775.1 atub|OJI88052.1 aver|OJJ01985.1 acar|OOF94544.1 aory|EIT83492.1 asyd|OJJ55184.1 aver|OJJ07023.1

Group7077: abra|OJJ69625.1 acla|EAW12186.1 afis|EAW17122.1 afla|EED54750.1 afum|EAL86583.1 akaw|GAA91605.1 aluc|OJZ82430.1 anid|CBF75576.1 anig|EHA28341.1 asyd|OJJ59713.1 atub|OJI88011.1 aver|OJJ02573.1 awen|OJJ31950.1 azon|OJJ45909.1

Group7078: abra|OJJ69649.1 acar|OOF96862.1 afla|EED54712.1 agla|OJJ81815.1 akaw|GAA91626.1 aluc|OJZ82453.1 anid|CBF89149.1 anig|EHA28322.1 aory|EIT78933.1 asyd|OJJ62567.1 atub|OJI87985.1 aver|OJI96691.1 awen|OJJ36244.1 azon|OJJ49988.1

Group7079: abra|OJJ69715.1 acar|OOF96659.1 afis|EAW16198.1 afla|EED51070.1 anid|CBF84544.1 anig|EHA19586.1 aory|EIT76955.1 asyd|OJJ53470.1 ater|EAU37852.1 atub|OJI89167.1 aver|OJJ07493.1 azon|OJJ42561.1 aver|OJJ08286.1 asyd|OJJ61691.1

Group7080: abra|OJJ69738.1 acar|OOF94185.1 acla|EAW09979.1 afla|EED51762.1 afum|EAL92133.1 agla|OJJ80103.1 akaw|GAA90191.1 aluc|OJZ83158.1 anig|EHA28223.1 asyd|OJJ60450.1 atub|OJI87779.1 aver|OJI99768.1 awen|OJJ36439.1 azon|OJJ44434.1

Group7081: abra|OJJ69766.1 afla|EED45128.1 agla|OJJ87486.1 akaw|GAA87887.1 aluc|OJZ85532.1 anid|CBF74489.1 anig|EHA25063.1 aory|EIT73677.1 asyd|OJJ57715.1 atub|OJI80036.1 aver|OJJ05452.1 awen|OJJ37832.1 asyd|OJJ58686.1 aver|OJJ01297.1

Group7082: abra|OJJ69786.1 acar|OOF93251.1 afum|EAL89021.1 akaw|GAA87871.1 aluc|OJZ85512.1 anid|CBF74932.1 anig|EHA25044.1 asyd|OJJ57349.1 atub|OJI80054.1 aver|OJJ05816.1 awen|OJJ35828.1 afla|EED47405.1 aory|EIT81921.1 aver|OJJ08581.1

Group7083: abra|OJJ69792.1 acar|OOF93588.1 acla|EAW11765.1 afis|EAW23686.1 afum|EAL87131.1 agla|OJJ84604.1 akaw|GAA87866.1 aluc|OJZ85507.1 anig|EHA25039.1 aory|EIT79940.1 asyd|OJJ55413.1 atub|OJI80058.1 aver|OJJ06491.1 awen|OJJ33313.1

Group7084: abra|OJJ69830.1 acar|OOF93355.1 afla|EED52425.1 agla|OJJ79195.1 akaw|GAA87830.1 aluc|OJZ85466.1 anid|CBF85157.1 anig|EHA25007.1 asyd|OJJ54543.1 atub|OJI80097.1 awen|OJJ32854.1 azon|OJJ45921.1 aory|EIT83307.1 aver|OJJ07634.1

Group7085: abra|OJJ69852.1 acar|OOF95291.1 afla|EED55820.1 agla|OJJ88215.1 akaw|GAA83677.1 aluc|OJZ85441.1 anid|CBF71171.1 anig|EHA24764.1 aory|EIT76535.1 asyd|OJJ55851.1 atub|OJI80126.1 aver|OJJ04794.1 awen|OJJ29733.1 akaw|GAA87806.1

Group7086: abra|OJJ69886.1 acar|OOF93404.1 afla|EED49654.1 agla|OJJ79812.1 akaw|GAA87772.1 aluc|OJZ85407.1 anig|EHA25586.1 aory|EIT81008.1 asyd|OJJ59146.1 atub|OJI80169.1 aver|OJJ01736.1 awen|OJJ29886.1 azon|OJJ42186.1 anid|CBF69585.1

Group7087: abra|OJJ69921.1 acar|OOF93433.1 acla|EAW08908.1 afis|EAW15488.1 afla|EED49002.1 afum|EAL85835.1 akaw|GAA87735.1 aluc|OJZ85367.1 anig|EHA25549.1 asyd|OJJ56023.1 atub|OJI80204.1 awen|OJJ38144.1 azon|OJJ43143.1 agla|OJJ80165.1

Group7088: abra|OJJ70047.1 acar|OOF93539.1 afla|EED47674.1 agla|OJJ79818.1 akaw|GAA87628.1 aluc|OJZ85244.1 anid|CBF84436.1 anig|EHA25444.1 aory|EIT80186.1 ater|EAU34190.1 atub|OJI80327.1 aver|OJJ05338.1 asyd|OJJ53634.1 ater|EAU37246.1

Group7089: abra|OJJ70057.1 acar|OOG00094.1 afla|EED45413.1 agla|OJJ79846.1 akaw|GAA87617.1 aluc|OJZ85238.1 anid|CBF79014.1 anig|EHA25440.1 aory|EIT82433.1 ater|EAU29573.1 atub|OJI80337.1 aver|OJI98312.1 awen|OJJ38040.1 azon|OJJ50753.1

Group7090: abra|OJJ70077.1 acar|OOF93556.1 acla|EAW12089.1 afis|EAW19732.1 afla|EED50184.1 afum|EAL86796.1 akaw|GAA87606.1 aluc|OJZ85226.1 anid|CBF80340.1 anig|EHA25430.1 aory|EIT77387.1 asyd|OJJ54837.1 atub|OJI80354.1 aver|OJJ07324.1

Group7091: abra|OJJ70116.1 acar|OOF90680.1 afla|EED54140.1 agla|OJJ80717.1 akaw|GAA87594.1 aluc|OJZ85214.1 anid|CBF85149.1 anig|EHA25417.1 aory|EIT82640.1 asyd|OJJ58394.1 ater|EAU37768.1 atub|OJI80371.1 awen|OJJ29837.1 aory|EIT76075.1

Group7092: abra|OJJ70119.1 acar|OOF89936.1 acla|EAW12301.1 afis|EAW16936.1 afum|EAL86461.1 agla|OJJ79583.1 akaw|GAA87592.1 aluc|OJZ85212.1 anid|CBF80372.1 anig|EHA25415.1 asyd|OJJ54812.1 ater|EAU33526.1 atub|OJI80375.1 aver|OJJ07349.1

Group7093: abra|OJJ70129.1 acar|OOF90316.1 afla|EED49770.1 akaw|GAA86641.1 aluc|OJZ85198.1 anid|CBF87977.1 anig|EHA25405.1 aory|EIT77648.1 asyd|OJJ63490.1 ater|EAU30788.1 atub|OJI80403.1 aver|OJI97439.1 azon|OJJ42745.1 azon|OJJ42167.1

Group7094: abra|OJJ70188.1 acar|OOF93289.1 afla|EED57382.1 akaw|GAA86587.1 aluc|OJZ85134.1 anid|CBF74946.1 anig|EHA22865.1 asyd|OJJ63940.1 atub|OJI80458.1 aver|OJI97002.1 awen|OJJ40813.1 azon|OJJ47392.1 azon|OJJ42320.1 aory|EIT80088.1

Group7095: abra|OJJ70213.1 acar|OOF92567.1 acla|EAW13610.1 afis|EAW19996.1 afla|EED47541.1 afum|EAL84920.1 agla|OJJ80415.1 akaw|GAA86563.1 aluc|OJZ85113.1 anid|CBF84002.1 anig|EHA25333.1 asyd|OJJ64736.1 atub|OJI80481.1 aver|OJI98842.1

Group7096: abra|OJJ70279.1 acar|OOF91728.1 afla|EED45364.1 agla|OJJ84429.1 akaw|GAA92869.1 aluc|OJZ82542.1 anid|CBF85097.1 anig|EHA22510.1 aory|EIT82423.1 asyd|OJJ63060.1 ater|EAU36830.1 atub|OJI88786.1 aver|OJI97903.1 azon|OJJ42219.1

Group7097: abra|OJJ70335.1 acla|EAW11925.1 afla|EED48233.1 agla|OJJ83801.1 akaw|GAA92589.1 aluc|OJZ82605.1 anig|EHA27475.1 aory|EIT72899.1 asyd|OJJ65322.1 atub|OJI88840.1 aver|OJI98020.1 awen|OJJ36208.1 azon|OJJ42550.1 aver|OJI99467.1

Group7098: abra|OJJ70381.1 acar|OOF91890.1 afla|EED47347.1 agla|OJJ79748.1 akaw|GAA91975.1 aluc|OJZ82640.1 anid|CBF80678.1 anig|EHA27430.1 aory|EIT82858.1 asyd|OJJ54590.1 atub|OJI88873.1 aver|OJJ07563.1 awen|OJJ29765.1 azon|OJJ50380.1

Group7099: abra|OJJ70387.1 acar|OOF90064.1 acla|EAW12391.1 afis|EAW16838.1 afla|EED47315.1 agla|OJJ81677.1 akaw|GAA91970.1 aluc|OJZ82645.1 anid|CBF80643.1 asyd|OJJ54620.1 atub|OJI88878.1 aver|OJJ07538.1 awen|OJJ32207.1 azon|OJJ43044.1

Group7100: abra|OJJ70423.1 acar|OOF91769.1 afla|EED46003.1 agla|OJJ86987.1 akaw|GAA90154.1 aluc|OJZ83193.1 anid|CBF84634.1 aory|EIT75521.1 asyd|OJJ62597.1 ater|EAU35268.1 atub|OJI87745.1 aver|OJI96742.1 awen|OJJ29998.1 anig|EHA21057.1

Group7101: abra|OJJ70445.1 acar|OOF91811.1 asyd|OJJ62580.1 atub|OJI84005.1 aver|OJI96700.1 azon|OJJ45154.1 abra|OJJ73417.1 acar|OOF90624.1 akaw|GAA83074.1 aluc|OJZ85790.1 anig|EHA24770.1 atub|OJI79575.1 awen|OJJ32707.1 akaw|GAA90437.1

Group7102: abra|OJJ70497.1 acar|OOF97429.1 acla|EAW08177.1 afla|EED47038.1 agla|OJJ78732.1 akaw|GAA83345.1 aluc|OJZ83842.1 anig|EHA18984.1 aory|EIT72611.1 asyd|OJJ60183.1 atub|OJI90208.1 aver|OJJ04081.1 awen|OJJ39446.1 azon|OJJ47648.1

Group7103: abra|OJJ70516.1 acar|OOF97413.1 acla|EAW08159.1 afis|EAW23197.1 afla|EED47057.1 afum|EAL88286.1 akaw|GAA83363.1 aluc|OJZ83861.1 anig|EHA19002.1 aory|EIT72472.1 ater|EAU36324.1 atub|OJI90229.1 awen|OJJ39427.1 azon|OJJ47631.1

Group7104: abra|OJJ70528.1 acar|OOF97403.1 acla|EAW08141.1 afis|EAW23177.1 afum|EAL88306.1 agla|OJJ89689.1 akaw|GAA83373.1 aluc|OJZ83871.1 anid|CBF74511.1 anig|EHA19012.1 asyd|OJJ57729.1 ater|EAU36305.1 atub|OJI90239.1 aver|OJJ05440.1

Group7105: abra|OJJ70530.1 agla|OJJ89692.1 aluc|OJZ83873.1 acla|EAW08143.1 afis|EAW23179.1 afum|EAL88304.1 anid|CBF74514.1 anig|EHA19014.1 asyd|OJJ57731.1 atub|OJI90241.1 aver|OJJ05438.1 akaw|GAA83375.1 ater|EAU36313.1 awen|OJJ36599.1

Group7106: acar|OOF97127.1 acla|EAW08189.1 afis|EAW23223.1 afla|EED47027.1 afum|EAL88261.1 agla|OJJ86380.1 akaw|GAA83436.1 anid|CBF74761.1 aory|EIT72739.1 asyd|OJJ60172.1 ater|EAU36354.1 atub|OJI90310.1 aver|OJJ04092.1 azon|OJJ47657.1

Group7107: akaw|GAA89968.1 aluc|OJZ83977.1 atub|OJI90350.1 acar|OOF97094.1 acla|EAW08127.1 afis|EAW23164.1 afum|EBA27344.1 agla|OJJ89675.1 anig|EHA19104.1 aory|EIT72677.1 asyd|OJJ57703.1 aver|OJJ05461.1 awen|OJJ36619.1 azon|OJJ47594.1

Group7108: abra|OJJ70673.1 acla|EAW08091.1 afis|EAW23126.1 afla|EED48524.1 afum|EAL88354.1 agla|OJJ89642.1 akaw|GAA90005.1 aluc|OJZ84021.1 anig|EHA19142.1 asyd|OJJ57760.1 atub|OJI90392.1 aver|OJJ05406.1 awen|OJJ36654.1 azon|OJJ47550.1

Group7109: abra|OJJ70769.1 acar|OOF96975.1 acla|EAW11152.1 afis|EAW15883.1 afum|EAL88517.1 agla|OJJ89538.1 akaw|GAA90089.1 aluc|OJZ84115.1 anig|EHA19225.1 aory|EIT72829.1 atub|OJI90488.1 awen|OJJ36748.1 azon|OJJ47451.1 anid|CBF81239.1

Group7110: abra|OJJ70874.1 acar|OOF98246.1 acla|EAW10462.1 afis|EAW25105.1 afla|EED55341.1 afum|EAL89776.1 akaw|GAA91737.1 aluc|OJZ87648.1 anig|EHA20841.1 aory|EIT76196.1 ater|EAU39371.1 atub|OJI80997.1 awen|OJJ40720.1 azon|OJJ44096.1

Group7111: abra|OJJ70917.1 acar|OOF98207.1 acla|EAW10491.1 afis|EAW25076.1 afla|EED55377.1 afum|EAL89745.1 akaw|GAA87427.1 anid|CBF85966.1 aory|EIT80793.1 asyd|OJJ64424.1 ater|EAU39547.1 aver|OJI98523.1 awen|OJJ40687.1 azon|OJJ44064.1

Group7112: abra|OJJ70947.1 acar|OOF98184.1 afis|EAW25052.1 afla|EED55399.1 afum|EAL89720.1 akaw|GAA82996.1 aluc|OJZ87581.1 anid|CBF85927.1 anig|EHA20782.1 aory|EIT80860.1 ater|EAU35802.1 atub|OJI81067.1 aver|OJI98503.1 awen|OJJ40662.1

Group7113: abra|OJJ71004.1 afis|EAW19703.1 afla|EED48153.1 afum|EAL86823.1 agla|OJJ83778.1 akaw|GAA92514.1 aluc|OJZ91527.1 anig|EHA22564.1 aory|EIT76246.1 asyd|OJJ61200.1 ater|EAU32849.1 atub|OJI85981.1 aver|OJJ00525.1 awen|OJJ30441.1

Group7114: abra|OJJ71053.1 acar|OOF98086.1 acla|EAW10602.1 afis|EAW24966.1 afum|EAL89624.2 agla|OJJ82222.1 akaw|GAA82904.1 aluc|OJZ87474.1 anig|EHA20694.1 asyd|OJJ59540.1 atub|OJI81174.1 aver|OJJ02728.1 awen|OJJ40559.1 azon|OJJ49420.1

Group7115: abra|OJJ71079.1 acar|OOF98062.1 agla|OJJ79303.1 akaw|GAA82883.1 aluc|OJZ87451.1 anig|EHA20673.1 ater|EAU34335.1 atub|OJI81197.1 awen|OJJ35534.1 acar|OOF93359.1 atub|OJI85145.1 acar|OOF97611.1 aluc|OJZ80193.1 atub|OJI80137.1

Group7116: abra|OJJ71110.1 acar|OOF98039.1 afla|EED53100.1 agla|OJJ88955.1 akaw|GAA82859.1 aluc|OJZ87425.1 anid|CBF85602.1 anig|EHA20647.1 aory|EIT74809.1 asyd|OJJ64195.1 atub|OJI81224.1 aver|OJI98272.1 awen|OJJ32252.1 azon|OJJ50351.1

Group7117: abra|OJJ71193.1 acar|OOF97974.1 acla|EAW10311.1 afis|EAW25282.1 afla|EED53748.1 afum|EAL89945.1 agla|OJJ89256.1 akaw|GAA82790.1 aluc|OJZ87349.1 anig|EHA20585.1 ater|EAU34423.1 atub|OJI81294.1 awen|OJJ40995.1 azon|OJJ44585.1

Group7118: abra|OJJ71243.1 acar|OOF97925.1 acla|EAW14272.1 afis|EAW19002.1 afla|EED53771.1 afum|EAL93736.1 akaw|GAA82744.1 aluc|OJZ87299.1 anid|CBF85402.1 anig|EHA20540.1 aory|EIT78404.1 asyd|OJJ64062.1 atub|OJI81341.1 aver|OJI98148.1

Group7119: abra|OJJ71246.1 acar|OOF97922.1 afis|EAW25303.1 agla|OJJ89282.1 akaw|GAA82741.1 aluc|OJZ87296.1 anig|EHA20537.1 aory|EIT78446.1 asyd|OJJ59882.1 ater|EAU34451.1 atub|OJI81344.1 aver|OJJ02404.1 awen|OJJ41017.1 azon|OJJ44604.1

Group7120: abra|OJJ71275.1 acar|OOF97899.1 acla|EAW10255.1 afla|EED53808.1 agla|OJJ83866.1 akaw|GAA82715.1 aluc|OJZ87269.1 anid|CBF75244.1 aory|EIT78433.1 asyd|OJJ59909.1 atub|OJI81373.1 aver|OJJ02377.1 awen|OJJ41036.1 azon|OJJ44630.1

Group7121: abra|OJJ71336.1 acar|OOF97836.1 acla|EAW10203.1 afis|EAW25392.1 afla|EED53869.1 afum|EAL90051.1 akaw|GAA82656.1 aluc|OJZ87206.1 anid|CBF77506.1 asyd|OJJ53187.1 ater|EAU34540.1 atub|OJI81433.1 aver|OJJ03663.1 agla|OJJ87030.1

Group7122: abra|OJJ71462.1 acar|OOF97738.1 acla|EAW10089.1 afis|EAW25498.1 afla|EED53998.1 afum|EAL90164.1 agla|OJJ87306.1 aluc|OJZ87082.1 anid|CBF77753.1 anig|EHA20353.1 aory|EIT78313.1 asyd|OJJ53045.1 atub|OJI81555.1 azon|OJJ44806.1

Group7123: abra|OJJ71574.1 acla|EAW15308.1 agla|OJJ87100.1 akaw|GAA89350.1 aluc|OJZ86311.1 anid|CBF87247.1 anig|EHA21562.1 asyd|OJJ64556.1 ater|EAU34641.1 atub|OJI84224.1 aver|OJI98653.1 awen|OJJ38026.1 azon|OJJ42335.1 asyd|OJJ60372.1

Group7124: abra|OJJ71576.1 acar|OOF96307.1 acla|EAW15123.1 afis|EAW20093.1 afla|EED45277.1 agla|OJJ88861.1 akaw|GAA89348.1 aluc|OJZ86313.1 anid|CBF84895.1 asyd|OJJ63176.1 atub|OJI84222.1 aver|OJI97778.1 awen|OJJ31856.1 azon|OJJ46269.1

Group7125: abra|OJJ71654.1 acar|OOF96368.1 afum|EAL85278.1 agla|OJJ88770.1 akaw|GAA89275.1 aluc|OJZ86400.1 anid|CBF85007.1 anig|EHA21500.1 asyd|OJJ63113.1 ater|EAU38777.1 atub|OJI84135.1 aver|OJI97846.1 awen|OJJ32055.1 azon|OJJ46088.1

Group7126: abra|OJJ71738.1 afla|EED55107.1 agla|OJJ86611.1 akaw|GAA85151.1 aluc|OJZ86486.1 anig|EHA22179.1 aory|EIT81608.1 asyd|OJJ52525.1 ater|EAU37390.1 atub|OJI84043.1 aver|OJJ08763.1 awen|OJJ32705.1 atub|OJI89783.1 ater|EAU30096.1

Group7127: abra|OJJ71794.1 acar|OOF96474.1 afla|EED56589.1 agla|OJJ83768.1 akaw|GAA85116.1 aluc|OJZ86524.1 anig|EHA22122.1 aory|EIT79634.1 asyd|OJJ54321.1 atub|OJI83998.1 aver|OJJ05275.1 awen|OJJ30247.1 acar|OOF90387.1 asyd|OJJ58094.1

Group7128: abra|OJJ71831.1 acar|OOF96507.1 afla|EED51272.1 akaw|GAA85080.1 aluc|OJZ86563.1 anid|CBF89177.1 anig|EHA22088.1 aory|EIT81564.1 asyd|OJJ62538.1 ater|EAU34871.1 atub|OJI83958.1 aver|OJI96664.1 awen|OJJ31137.1 azon|OJJ46617.1

Group7129: abra|OJJ71849.1 acar|OOF96518.1 afis|EAW18592.1 afum|EAL93299.2 agla|OJJ81644.1 akaw|GAA85068.1 aluc|OJZ86582.1 asyd|OJJ58643.1 ater|EAU34886.1 atub|OJI83940.1 aver|OJJ01249.1 awen|OJJ31155.1 azon|OJJ46630.1 acla|EAW13880.1

Group7130: abra|OJJ71934.1 acla|EAW09596.1 afis|EAW21541.1 afla|EED55992.1 afum|EAL92561.1 akaw|GAA84990.1 aluc|OJZ86671.1 anid|CBF76376.1 aory|EIT76667.1 asyd|OJJ61065.1 ater|EAU34998.1 atub|OJI83857.1 awen|OJJ39150.1 azon|OJJ51480.1

Group7131: abra|OJJ71977.1 acar|OOF96629.1 afis|EAW21574.1 agla|OJJ85765.1 akaw|GAA84953.1 aluc|OJZ86714.1 anid|CBF76310.1 anig|EHA21964.1 aory|EIT81098.1 asyd|OJJ61008.1 ater|EAU34958.1 atub|OJI83815.1 aver|OJJ00326.1 awen|OJJ39107.1

Group7132: abra|OJJ72067.1 acla|EAW09482.1 afis|EAW21656.1 afla|EED57887.1 afum|EAL92691.1 agla|OJJ79723.1 anid|CBF83548.1 anig|EHA21880.1 aory|EIT83277.1 asyd|OJJ60732.1 ater|EAU38337.1 aver|OJJ00064.1 awen|OJJ33913.1 azon|OJJ48648.1

Group7133: abra|OJJ72079.1 acla|EAW09486.1 afis|EAW21650.1 afla|EED57919.1 afum|EAL92684.1 agla|OJJ79728.1 akaw|GAA84840.1 aluc|OJZ86833.1 anid|CBF83555.1 anig|EHA21802.1 aory|EIT82324.1 asyd|OJJ60736.1 atub|OJI83685.1 aver|OJJ00068.1

Group7134: abra|OJJ72102.1 acar|OOF97361.1 afla|EED49736.1 akaw|GAA84819.1 aluc|OJZ86857.1 anig|EHA21804.1 aory|EIT77704.1 asyd|OJJ58041.1 ater|EAU30969.1 atub|OJI83662.1 aver|OJJ05862.1 awen|OJJ40097.1 awen|OJJ35799.1 aver|OJJ04360.1

Group7135: abra|OJJ72133.1 acar|OOF96769.1 afla|EED46874.1 afum|EAL90427.1 agla|OJJ86320.1 akaw|GAA84789.1 aluc|OJZ86891.1 anid|CBF77739.1 anig|EHA21752.1 aory|EIT72613.1 asyd|OJJ53053.1 atub|OJI83628.1 aver|OJJ03791.1 awen|OJJ32566.1

Group7136: abra|OJJ72189.1 acar|OOF96815.1 acla|EAW14722.1 afis|EAW24258.1 afla|EED48885.1 afum|EAL85926.2 akaw|GAA84744.1 aluc|OJZ86940.1 anig|EHA21704.1 ater|EAU31204.1 atub|OJI83582.1 awen|OJJ34659.1 azon|OJJ48477.1 afla|EED48019.1

Group7137: abra|OJJ72224.1 acar|OOF96847.1 afla|EED52265.1 agla|OJJ82580.1 akaw|GAA84713.1 aluc|OJZ86974.1 anig|EHA21674.1 aory|EIT74358.1 asyd|OJJ56431.1 atub|OJI83547.1 aver|OJJ06289.1 awen|OJJ31658.1 azon|OJJ45108.1 awen|OJJ35906.1

Group7138: abra|OJJ72238.1 acar|OOF96860.1 akaw|GAA84701.1 aluc|OJZ86986.1 anid|CBF73875.1 anig|EHA21661.1 aory|EIT80235.1 asyd|OJJ56550.1 atub|OJI83533.1 aver|OJJ06172.1 awen|OJJ34533.1 azon|OJJ43185.1 asyd|OJJ57935.1 aver|OJJ04459.1

Group7139: abra|OJJ72251.1 acar|OOF96882.1 afis|EAW24570.1 afla|EED45430.1 afum|EAL86233.1 agla|OJJ79374.1 akaw|GAA84686.1 aluc|OJZ87002.1 anid|CBF71505.1 anig|EHA21644.1 aory|EIT82437.1 ater|EAU30385.1 atub|OJI83517.1 awen|OJJ36198.1

Group7140: abra|OJJ72260.1 acar|OOF94178.1 acla|EAW11896.1 afis|EAW25664.1 afla|EED46456.1 akaw|GAA84670.1 aluc|OJZ87019.1 anig|EHA21636.1 aory|EIT74483.1 asyd|OJJ55671.1 atub|OJI83499.1 aver|OJJ06751.1 azon|OJJ42663.1 aory|EIT72349.1

Group7141: abra|OJJ72272.1 acla|EAW12458.1 afis|EAW20690.1 afum|EAL86236.1 agla|OJJ85124.1 akaw|GAA86027.1 aluc|OJZ90254.1 anig|EHA23749.1 aory|EIT81681.1 ater|EAU30625.1 atub|OJI81680.1 aver|OJJ00345.1 awen|OJJ35811.1 awen|OJJ30155.1

Group7142: abra|OJJ72281.1 acar|OOF94940.1 afis|EAW20799.1 afla|EED49546.1 agla|OJJ85013.1 akaw|GAA86033.1 aluc|OJZ90260.1 anid|CBF71373.1 anig|EHA23756.1 asyd|OJJ55078.1 atub|OJI81686.1 aver|OJJ07130.1 awen|OJJ41911.1 azon|OJJ47359.1

Group7143: abra|OJJ72354.1 acar|OOF94864.1 acla|EAW11971.1 afis|EAW19615.1 afla|EED55110.1 afum|EAL86907.1 akaw|GAA86101.1 aluc|OJZ90333.1 anig|EHA23817.1 aory|EIT81605.1 asyd|OJJ58818.1 atub|OJI81760.1 aver|OJJ01422.1 agla|OJJ80756.1

Group7144: abra|OJJ72409.1 acar|OOF94820.1 acla|EAW11931.1 afis|EAW19568.1 afla|EED56815.1 agla|OJJ80693.1 akaw|GAA86152.1 aluc|OJZ90393.1 anid|CBF78031.1 asyd|OJJ55226.1 atub|OJI81822.1 aver|OJJ06995.1 awen|OJJ41781.1 azon|OJJ44897.1

Group7145: abra|OJJ72444.1 acar|OOF94792.1 acla|EAW12677.1 afis|EAW17840.1 afla|EED46185.1 agla|OJJ83127.1 aluc|OJZ90428.1 anid|CBF77977.1 aory|EIT76356.1 asyd|OJJ55261.1 atub|OJI81860.1 aver|OJJ06964.1 awen|OJJ41736.1 azon|OJJ48433.1

Group7146: abra|OJJ72453.1 acar|OOF94783.1 acla|EAW12688.1 afla|EED46205.1 akaw|GAA86190.1 aluc|OJZ90438.1 anig|EHA23905.1 ater|EAU33053.1 atub|OJI81869.1 awen|OJJ41722.1 azon|OJJ48420.1 agla|OJJ83118.1 aory|EIT76525.1 afis|EAW17851.1

Group7147: abra|OJJ72487.1 acar|OOF94748.1 afla|EED46287.1 akaw|GAA86222.1 aluc|OJZ90474.1 anig|EHA23933.1 ater|EAU33101.1 atub|OJI81899.1 anid|CBF77037.1 asyd|OJJ57004.1 acla|EAW12718.1 afis|EAW17881.1 afum|EAL91689.2 aory|EIT76320.1

Group7148: abra|OJJ72505.1 acar|OOF94737.1 afla|EED46284.1 agla|OJJ83051.1 akaw|GAA86233.1 aluc|OJZ90488.1 anid|CBF77010.1 anig|EHA23946.1 aory|EIT76344.1 asyd|OJJ57019.1 ater|EAU33112.1 atub|OJI81912.1 aver|OJJ03270.1 awen|OJJ41661.1

Group7149: abra|OJJ72530.1 acar|OOF94723.1 afla|EED53072.1 akaw|GAA86252.1 aluc|OJZ90509.1 anig|EHA23966.1 aory|EIT74797.1 asyd|OJJ64691.1 atub|OJI81934.1 aver|OJI98796.1 awen|OJJ38084.1 acar|OOF96878.1 awen|OJJ29969.1 azon|OJJ43170.1

Group7150: abra|OJJ72576.1 acar|OOF94683.1 afla|EED56584.1 agla|OJJ88579.1 akaw|GAA85202.1 aluc|OJZ90557.1 anig|EHA24004.1 aory|EIT79670.1 asyd|OJJ58700.1 ater|EAU30078.1 atub|OJI81980.1 aver|OJJ01312.1 awen|OJJ40589.1 azon|OJJ42311.1

Group7151: abra|OJJ72580.1 acar|OOF94672.1 afla|EED56581.1 agla|OJJ79967.1 akaw|GAA85206.1 aluc|OJZ90561.1 anid|CBF70336.1 anig|EHA24007.1 aory|EIT79650.1 ater|EAU37872.1 atub|OJI81984.1 aver|OJJ01322.1 awen|OJJ40586.1 azon|OJJ42307.1

Group7152: abra|OJJ72591.1 acar|OOF94659.1 afis|EAW17940.1 afla|EED49101.1 afum|EAL91751.1 agla|OJJ82992.1 akaw|GAA85217.1 aluc|OJZ90572.1 anig|EHA24017.1 aory|EIT82515.1 atub|OJI81995.1 awen|OJJ41605.1 acla|EAW12777.1 azon|OJJ48319.1

Group7153: abra|OJJ72606.1 acla|EAW11607.1 afis|EAW23839.1 afla|EED52552.1 afum|EAL87270.1 akaw|GAA85233.1 aluc|OJZ90589.1 anid|CBF77075.1 anig|EHA24028.1 atub|OJI82012.1 azon|OJJ48725.1 asyd|OJJ56984.1 aver|OJJ03303.1 awen|OJJ33809.1

Group7154: abra|OJJ72632.1 acla|EAW15264.1 afis|EAW20533.1 afla|EED49494.1 agla|OJJ83528.1 akaw|GAA85252.1 aluc|OJZ90609.1 ater|EAU30517.1 atub|OJI82034.1 aver|OJI99707.1 awen|OJJ40340.1 ater|EAU38189.1 aory|EIT82593.1 ater|EAU30522.1

Group7155: abra|OJJ72686.1 acar|OOF95935.1 afis|EAW24408.1 afum|EAL86077.1 akaw|GAA85304.1 aluc|OJZ90664.1 anig|EHA24099.1 aory|EIT79027.1 asyd|OJJ53367.1 ater|EAU31789.1 atub|OJI82093.1 aver|OJJ08380.1 afla|EED50925.1 anid|CBF80323.1

Group7156: abra|OJJ72713.1 acar|OOF93166.1 afis|EAW20784.1 afla|EED44940.1 agla|OJJ85770.1 akaw|GAA85314.1 aluc|OJZ90675.1 anig|EHA24110.1 aory|EIT73511.1 asyd|OJJ60270.1 ater|EAU39593.1 atub|OJI82108.1 aver|OJJ03981.1 awen|OJJ30824.1

Group7157: abra|OJJ72720.1 acar|OOF95953.1 acla|EAW11556.1 afis|EAW23897.1 afla|EED52631.1 afum|EBA27310.1 agla|OJJ78639.1 aluc|OJZ90689.1 anid|CBF77150.1 asyd|OJJ56909.1 atub|OJI82114.1 aver|OJJ03384.1 awen|OJJ39051.1 azon|OJJ51383.1

Group7158: abra|OJJ72878.1 acar|OOF96072.1 acla|EAW11367.1 afis|EAW18128.1 afla|EED58052.1 agla|OJJ87020.1 aluc|OJZ83389.1 anid|CBF75364.1 asyd|OJJ59849.1 ater|EAU35449.1 atub|OJI82275.1 aver|OJJ02439.1 awen|OJJ35556.1 azon|OJJ46954.1

Group7159: abra|OJJ72948.1 acar|OOF96128.1 acla|EAW11194.1 afis|EAW15924.1 afum|EBA27352.1 agla|OJJ89381.1 akaw|GAA85529.1 aluc|OJZ83458.1 anid|CBF81017.1 asyd|OJJ56381.1 atub|OJI82350.1 aver|OJJ06332.1 awen|OJJ35656.1 azon|OJJ46883.1

Group7160: abra|OJJ73044.1 acar|OOF96207.1 acla|EAW07975.1 afis|EAW23010.1 afum|EBA27348.1 agla|OJJ89470.1 akaw|GAA85615.1 aluc|OJZ83555.1 anig|EHA24373.1 ater|EAU29663.1 atub|OJI82446.1 aver|OJJ06413.1 awen|OJJ30732.1 azon|OJJ44194.1

Group7161: abra|OJJ73063.1 acar|OOG00287.1 acla|EAW13236.1 afis|EAW17395.1 afla|EED56758.1 afum|EAL91233.1 akaw|GAA88096.1 aluc|OJZ81363.1 anig|EHA18663.1 aory|EIT73957.1 asyd|OJJ55047.1 ater|EAU34829.1 atub|OJI91567.1 aver|OJJ07157.1

Group7162: abra|OJJ73103.1 acar|OOG00343.1 acla|EAW07042.1 afis|EAW16669.1 afum|EAL89273.1 akaw|GAA88062.1 aluc|OJZ81325.1 anid|CBF82087.1 anig|EHA18630.1 asyd|OJJ54258.1 ater|EAU39590.1 atub|OJI91527.1 aver|OJJ05211.1 awen|OJJ40040.1

Group7163: ater|EAU34264.1 abra|OJJ74172.1 acar|OOF95552.1 afla|EED45879.1 akaw|GAA86844.1 aluc|OJZ89963.1 anig|EHA27163.1 aory|EIT76254.1 asyd|OJJ56505.1 atub|OJI84842.1 aver|OJJ06214.1 awen|OJJ33199.1 aory|EIT75447.1 afla|EED48137.1

Group7164: abra|OJJ73195.1 acar|OOG00432.1 acla|EAW06872.1 afis|EAW16489.1 afla|EED47230.1 afum|EAL89083.1 agla|OJJ82845.1 akaw|GAA87982.1 aluc|OJZ81234.1 anig|EHA18545.1 aory|EIT79329.1 atub|OJI91431.1 awen|OJJ40341.1 azon|OJJ49195.1

Group7165: abra|OJJ73225.1 acar|OOG00463.1 acla|EAW06900.1 afis|EAW16517.1 afla|EED56156.1 afum|EAL89112.1 agla|OJJ82804.1 aluc|OJZ81202.1 anid|CBF75556.1 asyd|OJJ59703.1 atub|OJI91397.1 aver|OJJ02584.1 awen|OJJ40306.1 azon|OJJ49167.1

Group7166: abra|OJJ73233.1 acla|EAW06908.1 afis|EAW16525.1 afla|EED56146.1 afum|EAL89120.1 akaw|GAA87947.1 aluc|OJZ81195.1 anid|CBF75539.1 anig|EHA18510.1 aory|EIT76683.1 atub|OJI91390.1 aver|OJJ05733.1 awen|OJJ40298.1 azon|OJJ49160.1

Group7167: abra|OJJ73260.1 acla|EAW06937.1 afis|EAW16552.1 afla|EED56110.1 afum|EAL89150.1 agla|OJJ82771.1 aluc|OJZ81168.1 anig|EHA18486.1 aory|EIT76885.1 asyd|OJJ59614.1 atub|OJI91363.1 aver|OJJ02655.1 awen|OJJ40265.1 azon|OJJ49129.1

Group7168: abra|OJJ73279.1 acla|EAW13748.1 afis|EAW19822.1 afla|EED46509.1 afum|EBA27218.1 akaw|GAA92726.1 aluc|OJZ88770.1 anid|CBF87169.1 anig|EHA18466.1 asyd|OJJ60291.1 atub|OJI91343.1 aver|OJI99631.1 azon|OJJ43888.1 aory|EIT72336.1

Group7169: abra|OJJ73286.1 acla|EAW06961.1 afis|EAW16576.1 afla|EED56088.1 afum|EBA27369.1 agla|OJJ82701.1 akaw|GAA92719.1 anid|CBF81874.1 aory|EIT76789.1 asyd|OJJ59664.1 ater|EAU36644.1 atub|OJI91336.1 aver|OJJ02619.1 awen|OJJ40242.1

Group7170: afum|EAL93420.1 ater|EAU38208.1 azon|OJJ43129.1 acar|OOF94198.1 acla|EAW13990.1 afis|EAW18698.1 afla|EED50646.1 agla|OJJ82379.1 aluc|OJZ80605.1 anid|CBF70633.1 aory|EIT78017.1 asyd|OJJ58495.1 aver|OJJ01098.1 awen|OJJ32172.1

Group7171: abra|OJJ73335.1 afis|EAW18089.1 afum|EAL92289.2 agla|OJJ85996.1 akaw|GAA83150.1 aluc|OJZ88706.1 anid|CBF76810.1 anig|EHA18392.1 ater|EAU31602.1 atub|OJI91288.1 awen|OJJ41847.1 azon|OJJ45143.1 afis|EAW21274.1 azon|OJJ42731.1

Group7172: abra|OJJ73403.1 acar|OOG00601.1 afis|EAW16630.1 afum|EAL89226.2 akaw|GAA83087.1 aluc|OJZ88634.1 anig|EHA18324.1 asyd|OJJ52757.1 asyd|OJJ56187.1 ater|EAU36753.1 atub|OJI91218.1 aver|OJJ05114.1 aver|OJJ08739.1 awen|OJJ40079.1

Group7173: abra|OJJ73448.1 afis|EAW24694.1 afla|EED45074.1 afum|EAL89336.1 agla|OJJ81337.1 akaw|GAA89026.1 aluc|OJZ80578.1 anid|CBF76026.1 anig|EHA25620.1 asyd|OJJ52451.1 ater|EAU33315.1 atub|OJI88352.1 aver|OJJ03965.1 awen|OJJ38021.1

Group7174: abra|OJJ73454.1 acar|OOG00655.1 acla|EAW15267.1 afis|EAW20387.1 afla|EED55194.1 afum|EAL89010.1 akaw|GAA83043.1 aluc|OJZ88588.1 anig|EHA18283.1 aory|EIT83592.1 asyd|OJJ60076.1 atub|OJI91174.1 awen|OJJ37580.1 azon|OJJ45162.1

Group7175: abra|OJJ73495.1 acar|OOG00695.1 acla|EAW07429.1 afis|EAW22455.1 afla|EED54489.1 afum|EAL90736.1 agla|OJJ87569.1 akaw|GAA87444.1 aluc|OJZ88553.1 anig|EHA18251.1 ater|EAU39250.1 atub|OJI91138.1 awen|OJJ35080.1 azon|OJJ50027.1

Group7176: abra|OJJ73532.1 acar|OOF99639.1 agla|OJJ87024.1 akaw|GAA83969.1 aluc|OJZ80861.1 anig|EHA21807.1 atub|OJI82716.1 abra|OJJ75769.1 afla|EED54892.1 aory|EIT79386.1 asyd|OJJ58675.1 atub|OJI87189.1 aver|OJJ01287.1 awen|OJJ30923.1

Group7177: abra|OJJ73543.1 acar|OOG00718.1 acla|EAW07463.1 afis|EAW22494.1 afla|EED54523.1 afum|EAL90699.1 akaw|GAA88608.1 aluc|OJZ88516.1 anid|CBF88957.1 anig|EHA18222.1 ater|EAU39221.1 atub|OJI91101.1 awen|OJJ35118.1 azon|OJJ49977.1

Group7178: abra|OJJ73551.1 acla|EAW07470.1 afis|EAW22503.1 afum|EAL90690.1 akaw|GAA88601.1 anid|CBF88967.1 aory|EIT78772.1 asyd|OJJ62797.1 ater|EAU39215.1 aver|OJI96844.1 awen|OJJ35127.1 azon|OJJ49971.1 abra|OJJ73552.1 aluc|OJZ88509.1

Group7179: abra|OJJ73604.1 acar|OOG00788.1 acla|EAW07521.1 afis|EAW22551.1 afla|EED54583.1 agla|OJJ80913.1 akaw|GAA88549.1 aluc|OJZ88445.1 aory|EIT78825.1 atub|OJI91043.1 aver|OJI96792.1 awen|OJJ35182.1 azon|OJJ49925.1 ater|EAU39153.1

Group7180: abra|OJJ73608.1 acar|OOG00792.1 acla|EAW07525.1 afis|EAW22555.1 afum|EAL90640.1 akaw|GAA88545.1 aluc|OJZ88441.1 anig|EHA18165.1 asyd|OJJ62746.1 ater|EAU39149.1 atub|OJI91039.1 aver|OJI96786.1 awen|OJJ35186.1 azon|OJJ49922.1

Group7181: abra|OJJ73646.1 acar|OOG00827.1 acla|EAW07559.1 afis|EAW22593.1 afla|EED54643.1 afum|EAL90606.1 agla|OJJ80953.1 akaw|GAA88509.1 aluc|OJZ88404.1 asyd|OJJ63729.1 atub|OJI91000.1 aver|OJI97211.1 awen|OJJ35221.1 azon|OJJ49887.1

Group7182: abra|OJJ73727.1 acla|EAW07612.1 afis|EAW22642.1 afla|EED49911.1 afum|EAL90550.2 agla|OJJ81014.1 akaw|GAA88436.1 aluc|OJZ88324.1 anid|CBF88174.1 anig|EHA18060.1 ater|EAU39065.1 atub|OJI90919.1 awen|OJJ35282.1 azon|OJJ49846.1

Group7183: abra|OJJ73750.1 acar|OOG00912.1 acla|EAW07638.1 afis|EAW22658.1 afla|EED49932.1 afum|EAL90534.1 akaw|GAA88417.1 aluc|OJZ88302.1 anig|EHA18041.1 aory|EIT83439.1 asyd|OJJ63624.1 atub|OJI90897.1 aver|OJI97312.1 awen|OJJ35286.1

Group7184: abra|OJJ73865.1 acar|OOF89913.1 acla|EAW13618.1 afis|EAW19962.1 afum|EAL84943.1 agla|OJJ78582.1 akaw|GAA89477.1 aluc|OJZ88186.1 anid|CBF76002.1 anig|EHA17937.1 asyd|OJJ60264.1 atub|OJI90785.1 azon|OJJ50331.1 aver|OJJ03989.1

Group7185: abra|OJJ73896.1 acar|OOG01033.1 afis|EAW22814.1 afla|EED50087.1 afum|EAL90387.1 akaw|GAA89506.1 aluc|OJZ88150.1 anid|CBF87913.1 anig|EHA17911.1 aory|EIT74635.1 asyd|OJJ63449.1 atub|OJI90753.1 aver|OJI97485.1 awen|OJJ38914.1

Group7186: abra|OJJ73996.1 acar|OOF94226.1 afis|EAW25671.1 afla|EED54637.1 akaw|GAA87478.1 aluc|OJZ88048.1 anid|CBF82311.1 anig|EHA17820.1 aory|EIT76476.1 atub|OJI90656.1 afum|EAL84343.1 ater|EAU36071.1 afla|EED46248.1 aory|EIT78781.1

Group7187: abra|OJJ74005.1 acar|OOG01126.1 afis|EAW16956.1 afum|EAL86483.2 agla|OJJ83456.1 akaw|GAA87486.1 aluc|OJZ88039.1 anig|EHA17812.1 asyd|OJJ52815.1 ater|EAU31916.1 atub|OJI90646.1 aver|OJJ08794.1 awen|OJJ30845.1 azon|OJJ44441.1

Group7188: abra|OJJ74044.1 acla|EAW07894.1 afis|EAW22929.1 afum|EAL90268.1 agla|OJJ83359.1 aluc|OJZ87991.1 aory|EIT79123.1 asyd|OJJ63309.1 atub|OJI90601.1 aver|OJI97650.1 awen|OJJ38635.1 azon|OJJ49577.1 anig|EHA17774.1 akaw|GAA87527.1

Group7189: abra|OJJ74059.1 acar|OOG01171.1 acla|EAW07904.1 afis|EAW22942.1 agla|OJJ83346.1 akaw|GAA87540.1 aluc|OJZ87977.1 anid|CBF87647.1 asyd|OJJ63296.1 ater|EAU30682.1 atub|OJI90587.1 aver|OJI97662.1 awen|OJJ38622.1 azon|OJJ49563.1

Group7190: abra|OJJ74088.1 acla|EAW07931.1 afis|EAW22969.1 afla|EED58162.1 afum|EAL90228.1 agla|OJJ83318.1 akaw|GAA87567.1 aluc|OJZ87948.1 anid|CBF74907.1 anig|EHA17734.1 aory|EIT79093.1 ater|EAU30660.1 atub|OJI90559.1 awen|OJJ38594.1

Group7191: abra|OJJ74104.1 acar|OOG01214.1 acla|EAW07947.1 afis|EAW22986.1 afla|EED58178.1 afum|EAL90213.1 agla|OJJ83301.1 akaw|GAA87582.1 aluc|OJZ87932.1 anid|CBF87583.1 asyd|OJJ63254.1 atub|OJI90544.1 aver|OJI97703.1 awen|OJJ38578.1

Group7192: abra|OJJ74139.1 acar|OOF95591.1 afla|EED47376.1 agla|OJJ78697.1 akaw|GAA89746.1 aluc|OJZ80638.1 anid|CBF85351.1 anig|EHA27195.1 aory|EIT82868.1 asyd|OJJ64034.1 ater|EAU32379.1 atub|OJI87493.1 aver|OJI98119.1 awen|OJJ40039.1

Group7193: abra|OJJ74142.1 acar|OOF95588.1 afla|EED47374.1 agla|OJJ78695.1 akaw|GAA89743.1 aluc|OJZ80641.1 anid|CBF85353.1 anig|EHA27191.1 aory|EIT82855.1 asyd|OJJ64035.1 ater|EAU32381.1 atub|OJI87496.1 aver|OJI98120.1 awen|OJJ40038.1

Group7194: abra|OJJ74150.1 acar|OOF95579.1 afla|EED51809.1 agla|OJJ84495.1 akaw|GAA89733.1 aluc|OJZ80650.1 anid|CBF81689.1 anig|EHA27182.1 aory|EIT81747.1 asyd|OJJ52210.1 atub|OJI87505.1 awen|OJJ35888.1 azon|OJJ51659.1 acar|OOF98447.1

Group7195: abra|OJJ74166.1 acar|OOF89849.1 acla|EAW06846.1 afis|EAW16466.1 afla|EED49697.1 afum|EAL89062.1 akaw|GAA86849.1 aluc|OJZ89970.1 anig|EHA27168.1 aory|EIT77629.1 asyd|OJJ52107.1 ater|EAU34012.1 atub|OJI84837.1 aver|OJI95751.1

Group7196: abra|OJJ74177.1 acar|OOF95547.1 afla|EED45245.1 akaw|GAA86839.1 aluc|OJZ89958.1 anid|CBF85438.1 anig|EHA27158.1 aory|EIT73807.1 asyd|OJJ64089.1 ater|EAU34330.1 atub|OJI84848.1 aver|OJI98173.1 awen|OJJ38749.1 acar|OOF95244.1

Group7197: abra|OJJ74309.1 acar|OOF95418.1 afla|EED46915.1 akaw|GAA86715.1 aluc|OJZ89817.1 anid|CBF80697.1 anig|EHA27026.1 aory|EIT72567.1 asyd|OJJ54566.1 ater|EAU36826.1 atub|OJI84988.1 aver|OJJ07588.1 awen|OJJ33372.1 ater|EAU35085.1

Group7198: abra|OJJ74362.1 afis|EAW22440.1 afla|EED54450.1 afum|EAL90751.1 akaw|GAA86668.1 aluc|OJZ89765.1 anid|CBF88841.1 anig|EHA26985.1 aory|EIT81159.1 asyd|OJJ62869.1 ater|EAU39268.1 atub|OJI85037.1 aver|OJI96922.1 awen|OJJ35017.1

Group7199: abra|OJJ74420.1 acar|OOF95320.1 afla|EED53450.1 agla|OJJ88212.1 akaw|GAA87314.1 aluc|OJZ89703.1 anid|CBF69427.1 anig|EHA26937.1 aory|EIT74891.1 asyd|OJJ54914.1 ater|EAU34062.1 atub|OJI85096.1 aver|OJJ07289.1 awen|OJJ37923.1

Group7200: abra|OJJ74426.1 acar|OOF95315.1 acla|EAW10813.1 afis|EAW24760.1 afla|EED53441.1 afum|EAL89402.1 agla|OJJ88213.1 akaw|GAA87322.1 aluc|OJZ89694.1 anig|EHA26932.1 aory|EIT75145.1 ater|EAU34060.1 atub|OJI85108.1 awen|OJJ37926.1

Group7201: abra|OJJ74434.1 acar|OOF95310.1 acla|EAW08987.1 afis|EAW15425.1 afum|EAL84219.1 agla|OJJ82707.1 akaw|GAA87328.1 aluc|OJZ89687.1 anid|CBF85295.1 anig|EHA26925.1 asyd|OJJ54705.1 atub|OJI85114.1 aver|OJJ07467.1 awen|OJJ37917.1

Group7202: abra|OJJ74495.1 acla|EAW07199.1 afis|EAW22211.1 afla|EED57605.1 afum|EAL90983.2 agla|OJJ87833.1 akaw|GAA87388.1 aluc|OJZ89623.1 aory|EIT82941.1 asyd|OJJ63766.1 atub|OJI85179.1 aver|OJI97172.1 awen|OJJ34796.1 azon|OJJ45752.1

Group7203: abra|OJJ74502.1 acar|OOF95242.1 acla|EAW07187.1 afis|EAW22200.1 afla|EED57596.1 afum|EAL90992.1 agla|OJJ87841.1 akaw|GAA87396.1 aluc|OJZ89616.1 anig|EHA26861.1 aory|EIT82965.1 ater|EAU34214.1 atub|OJI85187.1 awen|OJJ34789.1

Group7204: abra|OJJ74505.1 acar|OOF95239.1 acla|EAW07191.1 afis|EAW22204.1 afla|EED57590.1 agla|OJJ83744.1 akaw|GAA87400.1 aluc|OJZ89612.1 anid|CBF88424.1 anig|EHA26858.1 aory|EIT82947.1 asyd|OJJ63758.1 atub|OJI85190.1 aver|OJI97182.1

Group7205: abra|OJJ74552.1 acar|OOF95203.1 acla|EAW07146.1 afis|EAW22160.1 agla|OJJ87904.1 akaw|GAA82045.1 aluc|OJZ92135.1 anid|CBF89099.1 anig|EHA26814.1 asyd|OJJ62728.1 atub|OJI86567.1 aver|OJI96771.1 awen|OJJ34734.1 azon|OJJ45825.1

Group7206: abra|OJJ74554.1 acar|OOF95201.1 afis|EAW22158.1 afla|EED57530.1 afum|EAL91032.1 agla|OJJ87906.1 akaw|GAA82043.1 aluc|OJZ92137.1 anig|EHA26812.1 aory|EIT81838.1 ater|EAU34161.1 atub|OJI86569.1 awen|OJJ34732.1 azon|OJJ45827.1

Group7207: abra|OJJ74555.1 acar|OOF95200.1 afis|EAW22157.1 afla|EED57529.1 afum|EAL91033.1 agla|OJJ87907.1 akaw|GAA82041.1 aluc|OJZ92138.1 anig|EHA26811.1 aory|EIT81843.1 ater|EAU34160.1 atub|OJI86570.1 awen|OJJ34731.1 azon|OJJ45828.1

Group7208: abra|OJJ74576.1 acla|EAW07134.1 afis|EAW22147.1 afum|EAL91043.1 akaw|GAA82029.1 aluc|OJZ92154.1 anig|EHA26795.1 asyd|OJJ52409.1 atub|OJI86582.1 aver|OJI98308.1 awen|OJJ34723.1 azon|OJJ45255.1 aver|OJJ08301.1 asyd|OJJ62652.1

Group7209: abra|OJJ74655.1 acla|EAW10734.1 afis|EAW24841.1 afla|EED47136.1 afum|EAL89487.1 agla|OJJ82031.1 akaw|GAA81955.1 anid|CBF74632.1 aory|EIT79284.1 asyd|OJJ57582.1 ater|EAU36463.1 atub|OJI86666.1 awen|OJJ40426.1 azon|OJJ49286.1

Group7210: abra|OJJ74676.1 acla|EAW10721.1 afis|EAW24852.1 afla|EED47119.1 afum|EAL89499.1 aluc|OJZ92253.1 anid|CBF75833.1 aory|EIT79237.1 asyd|OJJ57554.1 ater|EAU36451.1 atub|OJI86686.1 aver|OJJ05624.1 azon|OJJ49298.1 awen|OJJ40437.1

Group7211: abra|OJJ74718.1 akaw|GAA81898.1 aluc|OJZ92301.1 anig|EHA26670.1 ater|EAU36411.1 atub|OJI86730.1 awen|OJJ40480.1 azon|OJJ49344.1 acla|EAW10680.1 afis|EAW24890.1 afla|EED46967.1 afum|EAL89541.2 agla|OJJ82113.1 aory|EIT72699.1

Group7212: abra|OJJ74768.1 acla|EAW10643.1 afis|EAW24923.1 afla|EED55550.1 afum|EAL89578.2 agla|OJJ82152.1 akaw|GAA83644.1 aluc|OJZ92351.1 anig|EHA26623.1 aory|EIT82697.1 asyd|OJJ57495.1 atub|OJI86779.1 aver|OJJ05685.1 azon|OJJ49379.1

Group7213: abra|OJJ74844.1 acar|OOF95845.1 acla|EAW08545.1 afis|EAW23582.1 afum|EAL87882.1 agla|OJJ86849.1 akaw|GAA83578.1 aluc|OJZ92419.1 anid|CBF89591.1 anig|EHA26547.1 atub|OJI86849.1 aver|OJI96423.1 azon|OJJ45887.1 awen|OJJ39910.1

Group7214: abra|OJJ74868.1 acla|EAW08512.1 afis|EAW23550.1 afla|EED57445.1 afum|EAL87916.1 agla|OJJ86806.1 akaw|GAA83552.1 aluc|OJZ92446.1 anid|CBF89648.1 anig|EHA26522.1 asyd|OJJ62285.1 atub|OJI86874.1 aver|OJI96393.1 awen|OJJ39881.1

Group7215: abra|OJJ74870.1 acar|OOF95871.1 acla|EAW08510.1 afis|EAW23548.1 afla|EED57443.1 afum|EAL87918.2 agla|OJJ86801.1 akaw|GAA83550.1 aluc|OJZ92448.1 anig|EHA26520.1 aory|EIT80059.1 ater|EAU35333.1 atub|OJI86876.1 awen|OJJ39879.1

Group7216: abra|OJJ74894.1 acar|OOF95895.1 acla|EAW08477.1 agla|OJJ86771.1 akaw|GAA83526.1 aluc|OJZ92469.1 anid|CBF89693.1 anig|EHA26499.1 asyd|OJJ62260.1 ater|EAU35310.1 atub|OJI86899.1 aver|OJI96369.1 awen|OJJ39848.1 azon|OJJ43242.1

Group7217: abra|OJJ74936.1 acar|OOF97378.1 acla|EAW08435.1 afis|EAW23480.1 afla|EED57375.1 afum|EAL87984.1 akaw|GAA83484.1 aluc|OJZ92509.1 anig|EHA26464.1 aory|EIT80103.1 ater|EAU36902.1 atub|OJI86942.1 awen|OJJ39804.1 azon|OJJ47925.1

Group7218: abra|OJJ74987.1 acar|OOF97334.1 acla|EAW08395.1 afis|EAW23438.1 afum|EAL88024.1 agla|OJJ86675.1 akaw|GAA83447.1 aluc|OJZ92549.1 anig|EHA26428.1 aory|EIT74111.1 ater|EAU37488.1 atub|OJI86986.1 awen|OJJ39755.1 azon|OJJ47883.1

Group7219: abra|OJJ75085.1 acar|OOF97248.1 acla|EAW08279.1 afis|EAW23312.1 afla|EED55667.1 afum|EAL88165.1 agla|OJJ86537.1 aluc|OJZ92651.1 anid|CBF89496.1 asyd|OJJ62409.1 atub|OJI87082.1 aver|OJI96542.1 awen|OJJ39657.1 azon|OJJ47785.1

Group7220: abra|OJJ75123.1 acar|OOF97211.1 afis|EAW23335.1 afla|EED55631.1 afum|EAL88143.1 agla|OJJ86498.1 akaw|GAA88731.1 aluc|OJZ92689.1 anid|CBF89433.1 anig|EHA26311.1 aory|EIT81777.1 atub|OJI87117.1 aver|OJI96580.1 awen|OJJ39559.1

Group7221: abra|OJJ75233.1 acar|OOF94559.1 afla|EED54622.1 agla|OJJ88701.1 akaw|GAA91776.1 aluc|OJZ92793.1 anid|CBF71788.1 anig|EHA26214.1 aory|EIT78816.1 asyd|OJJ61716.1 ater|EAU36016.1 atub|OJI87236.1 aver|OJI95810.1 awen|OJJ33290.1

Group7222: abra|OJJ75268.1 acar|OOF94535.1 afla|EED48989.1 agla|OJJ81047.1 akaw|GAA91806.1 aluc|OJZ92828.1 anid|CBF71428.1 anig|EHA26181.1 aory|EIT80042.1 asyd|OJJ54996.1 ater|EAU31740.1 atub|OJI87285.1 aver|OJJ07205.1 awen|OJJ35992.1

Group7223: abra|OJJ75275.1 acar|OOF97939.1 acla|EAW15333.1 afis|EAW20371.1 afla|EED53113.1 afum|EAL85485.1 akaw|GAA91815.1 aluc|OJZ92835.1 anig|EHA26172.1 aory|EIT74765.1 asyd|OJJ56415.1 atub|OJI87293.1 aver|OJJ06300.1 awen|OJJ36523.1

Group7224: abra|OJJ75323.1 acar|OOF94501.1 afla|EED56848.1 agla|OJJ83738.1 akaw|GAA92018.1 aluc|OJZ92884.1 anid|CBF81531.1 anig|EHA26076.1 aory|EIT77520.1 asyd|OJJ59486.1 ater|EAU33512.1 atub|OJI87343.1 aver|OJJ02788.1 awen|OJJ38739.1

Group7225: abra|OJJ75334.1 acar|OOF94499.1 afis|EAW24598.1 agla|OJJ80218.1 akaw|GAA92028.1 aluc|OJZ92898.1 anid|CBF71503.1 anig|EHA26065.1 ater|EAU33062.1 atub|OJI87356.1 aver|OJJ07276.1 awen|OJJ33328.1 agla|OJJ84487.1 aver|OJI96489.1

Group7226: abra|OJJ75339.1 acar|OOF94492.1 acla|EAW12073.1 afis|EAW19714.1 afla|EED56203.1 afum|EAL86813.1 akaw|GAA92032.1 aluc|OJZ92902.1 anig|EHA26061.1 aory|EIT73373.1 asyd|OJJ55950.1 awen|OJJ29967.1 azon|OJJ42680.1 aver|OJJ04895.1

Group7227: abra|OJJ75350.1 acar|OOF94481.1 acla|EAW12386.1 afla|EED47329.1 agla|OJJ80355.1 akaw|GAA92042.1 aluc|OJZ92912.1 anid|CBF80633.1 anig|EHA26053.1 aory|EIT82840.1 asyd|OJJ54622.1 ater|EAU30238.1 atub|OJI87372.1 aver|OJJ07536.1

Group7228: abra|OJJ75372.1 acar|OOF89963.1 acla|EAW08075.1 afis|EAW20377.1 akaw|GAA92065.1 aluc|OJZ92940.1 anid|CBF82140.1 anig|EHA26032.1 asyd|OJJ54320.1 atub|OJI87400.1 aver|OJJ05274.1 awen|OJJ31303.1 afum|EAL85491.1 ater|EAU29315.1

Group7229: azon|OJJ43281.1 acar|OOF90782.1 acla|EAW13824.1 afis|EAW18307.1 afla|EED48776.1 afum|EAL87673.1 agla|OJJ88116.1 anid|CBF79442.1 anig|EHA23616.1 aory|EIT72990.1 asyd|OJJ57848.1 ater|EAU33317.1 aver|OJJ04544.1 awen|OJJ30681.1

Group7230: abra|OJJ75514.1 acar|OOF90841.1 afis|EAW25707.1 afla|EED46469.1 afum|EAL84380.1 agla|OJJ79319.1 akaw|GAA84278.1 aluc|OJZ88971.1 anid|CBF71220.1 anig|EHA23553.1 asyd|OJJ58670.1 atub|OJI85819.1 aver|OJJ01281.1 awen|OJJ30879.1

Group7231: abra|OJJ75569.1 acar|OOF90887.1 afla|EED45967.1 agla|OJJ80728.1 akaw|GAA84320.1 aluc|OJZ89017.1 anid|CBF88701.1 anig|EHA23513.1 aory|EIT75698.1 asyd|OJJ63916.1 atub|OJI85772.1 aver|OJI97026.1 awen|OJJ35840.1 azon|OJJ42079.1

Group7232: abra|OJJ75624.1 acar|OOG00212.1 acla|EAW09359.1 afis|EAW21778.1 afla|EED46345.1 afum|EAL92807.1 agla|OJJ84343.1 akaw|GAA84371.1 anig|EHA23462.1 aory|EIT81182.1 asyd|OJJ61183.1 atub|OJI85710.1 awen|OJJ34067.1 azon|OJJ48160.1

Group7233: acar|OOF93294.1 acla|EAW08080.1 afis|EAW20391.1 akaw|GAA85129.1 aluc|OJZ86511.1 anig|EHA20898.1 asyd|OJJ56867.1 atub|OJI84016.1 aver|OJJ03459.1 awen|OJJ35976.1 afla|EED52282.1 anig|EHA22136.1 aory|EIT74357.1 awen|OJJ36061.1

Group7234: abra|OJJ75699.1 acla|EAW09409.1 afis|EAW21727.1 afla|EED57996.1 afum|EAL92760.1 akaw|GAA84435.1 aluc|OJZ89150.1 anid|CBF83723.1 anig|EHA23398.1 aory|EIT79751.1 ater|EAU38411.1 atub|OJI85638.1 awen|OJJ34000.1 azon|OJJ48552.1

Group7235: abra|OJJ75705.1 agla|OJJ79854.1 aluc|OJZ89156.1 anig|EHA23396.1 ater|EAU38414.1 atub|OJI85635.1 acla|EAW09405.1 afis|EAW21731.1 afum|EAL92764.1 aver|OJJ00170.1 awen|OJJ34004.1 azon|OJJ48548.1 akaw|GAA84440.1 asyd|OJJ60845.1

Group7236: abra|OJJ75783.1 acar|OOG00105.1 acla|EAW14899.1 afis|EAW24421.1 afla|EED47747.1 agla|OJJ79223.1 aluc|OJZ89226.1 anid|CBF73790.1 anig|EHA23336.1 asyd|OJJ56603.1 atub|OJI85566.1 aver|OJJ06120.1 awen|OJJ32788.1 azon|OJJ49044.1

Group7237: abra|OJJ75833.1 acar|OOG00047.1 agla|OJJ83258.1 akaw|GAA84543.1 aluc|OJZ89276.1 anig|EHA23293.1 ater|EAU29962.1 atub|OJI85523.1 afla|EED53211.1 aory|EIT74885.1 asyd|OJJ64552.1 aver|OJJ06998.1 awen|OJJ31383.1 asyd|OJJ55222.1

Group7238: abra|OJJ75845.1 acla|EAW14839.1 afla|EED47848.1 afum|EBA27255.1 akaw|GAA84554.1 aluc|OJZ89286.1 anig|EHA23280.1 aory|EIT77214.1 asyd|OJJ53663.1 atub|OJI85513.1 aver|OJJ07770.1 awen|OJJ32847.1 afis|EAW24367.1 anid|CBF73890.1

Group7239: abra|OJJ75900.1 acar|OOG00003.1 acla|EAW09890.1 afis|EAW21236.1 afum|EAL92248.1 agla|OJJ78498.1 akaw|GAA84606.1 aluc|OJZ89350.1 anig|EHA23231.1 asyd|OJJ60576.1 ater|EAU35877.1 atub|OJI85448.1 aver|OJI99902.1 azon|OJJ44342.1

Group7240: abra|OJJ75998.1 acar|OOF99926.1 acla|EAW09806.1 afis|EAW21330.1 afla|EED57288.1 afum|EAL92348.1 agla|OJJ87409.1 aluc|OJZ89444.1 asyd|OJJ60662.1 ater|EAU35956.1 atub|OJI85349.1 aver|OJI99985.1 awen|OJJ39368.1 azon|OJJ51708.1

Group7241: abra|OJJ76043.1 acar|OOF99887.1 acla|EAW09773.1 afis|EAW21365.1 afla|EED55779.1 afum|EAL92381.1 akaw|GAA88923.1 aluc|OJZ89498.1 anig|EHA23102.1 aory|EIT76574.1 ater|EAU35192.1 atub|OJI85305.1 awen|OJJ39332.1 azon|OJJ51677.1

Group7242: abra|OJJ76063.1 acar|OOF99868.1 acla|EAW09755.1 afis|EAW21386.1 afla|EED55800.1 afum|EAL92401.1 agla|OJJ85511.1 akaw|GAA88903.1 aluc|OJZ89518.1 anid|CBF83844.1 asyd|OJJ60917.1 ater|EAU35172.1 atub|OJI85285.1 awen|OJJ39315.1

Group7243: abra|OJJ76115.1 afis|EAW21435.1 afum|EAL92450.1 agla|OJJ85577.1 akaw|GAA88852.1 aluc|OJZ89572.1 anig|EHA23035.1 asyd|OJJ61174.1 atub|OJI85232.1 aver|OJJ00500.1 awen|OJJ39264.1 azon|OJJ51591.1 acla|EAW09705.1 anid|CBF76589.1

Group7244: abra|OJJ76439.1 acar|OOF99684.1 acla|EAW13985.1 afis|EAW24719.1 afum|EAL89368.2 aluc|OJZ91770.1 anid|CBF87159.1 anig|EHA22770.1 aory|EIT77975.1 asyd|OJJ58845.1 ater|EAU38214.1 atub|OJI86228.1 aver|OJJ01446.1 azon|OJJ45350.1

Group7245: abra|OJJ76447.1 acar|OOF99691.1 acla|EAW13981.1 afis|EAW18690.1 afum|EAL93411.2 agla|OJJ82382.1 akaw|GAA82359.1 aluc|OJZ91763.1 anig|EHA22764.1 asyd|OJJ58505.1 atub|OJI86221.1 aver|OJJ01106.1 awen|OJJ32161.1 azon|OJJ46747.1

Group7246: abra|OJJ76489.1 acar|OOF99721.1 afis|EAW18655.1 afla|EED52676.1 afum|EAL93371.1 akaw|GAA82404.1 aluc|OJZ91716.1 anid|CBF70544.1 aory|EIT74257.1 asyd|OJJ58547.1 ater|EAU38257.1 atub|OJI86174.1 aver|OJJ01145.1 awen|OJJ32122.1

Group7247: abra|OJJ76648.1 acar|OOF99353.1 afis|EAW19011.1 afla|EED56274.1 afum|EAL93745.2 akaw|GAA92989.1 aluc|OJZ91560.1 anid|CBF86743.1 anig|EHA22592.1 ater|EAU37791.1 atub|OJI86012.1 aver|OJI96703.1 asyd|OJJ64698.1 aver|OJI98803.1

Group7248: abra|OJJ76724.1 acla|EAW11290.1 afla|EED45040.1 afum|EAL86531.2 akaw|GAA92477.1 aluc|OJZ91487.1 anid|CBF78525.1 anig|EHA22531.1 aory|EIT73672.1 ater|EAU32966.1 atub|OJI85942.1 aver|OJI99446.1 awen|OJJ40910.1 azon|OJJ42296.1

Group7249: abra|OJJ76752.1 acar|OOF91248.1 acla|EAW13771.1 afis|EAW20420.1 agla|OJJ84905.1 akaw|GAA87272.1 aluc|OJZ84155.1 anig|EHA27517.1 aory|EIT77740.1 asyd|OJJ55797.1 ater|EAU37736.1 atub|OJI89929.1 aver|OJJ06886.1 awen|OJJ33113.1

Group7250: abra|OJJ76771.1 acla|EAW13325.1 afis|EAW20440.1 afla|EED44830.1 afum|EAL84612.1 akaw|GAA87255.1 aluc|OJZ84175.1 anid|CBF79315.1 aory|EIT77724.1 asyd|OJJ55778.1 ater|EAU37718.1 atub|OJI89910.1 awen|OJJ33131.1 azon|OJJ46346.1

Group7251: abra|OJJ76797.1 acar|OOF91290.1 acla|EAW13356.1 afis|EAW20470.1 afla|EED44798.1 afum|EBA27194.1 agla|OJJ84848.1 akaw|GAA87226.1 aluc|OJZ84204.1 asyd|OJJ55749.1 atub|OJI89880.1 aver|OJJ06836.1 awen|OJJ33164.1 azon|OJJ46371.1

Group7252: abra|OJJ76828.1 acar|OOF91322.1 acla|EAW13389.1 afis|EAW20502.1 afla|EED44745.1 agla|OJJ84798.1 akaw|GAA87194.1 aluc|OJZ84240.1 anid|CBF79199.1 asyd|OJJ55724.1 atub|OJI89843.1 aver|OJJ06810.1 awen|OJJ33203.1 azon|OJJ46395.1

Group7253: abra|OJJ76904.1 acar|OOF91352.1 acla|EAW13426.1 afis|EAW20537.1 afum|EBA27190.1 agla|OJJ84694.1 akaw|GAA87134.1 aluc|OJZ84303.1 anid|CBF78993.1 anig|EHA27639.1 ater|EAU37142.1 atub|OJI89765.1 awen|OJJ33244.1 azon|OJJ50623.1

Group7254: abra|OJJ76909.1 acar|OOF91358.1 afis|EAW20542.1 afla|EED52431.1 afum|EAL84516.1 akaw|GAA87129.1 aluc|OJZ84308.1 anig|EHA27644.1 aory|EIT83301.1 asyd|OJJ56122.1 atub|OJI89760.1 aver|OJJ05063.1 awen|OJJ33249.1 azon|OJJ50628.1

Group7255: abra|OJJ76910.1 acar|OOF91360.1 afis|EAW24763.1 afla|EED56327.1 agla|OJJ82591.1 akaw|GAA87128.1 aluc|OJZ84309.1 anig|EHA27645.1 aory|EIT73407.1 asyd|OJJ54749.1 ater|EAU39055.1 atub|OJI89759.1 aver|OJJ07418.1 awen|OJJ33554.1

Group7256: abra|OJJ76928.1 acar|OOF91374.1 afis|EAW16739.1 agla|OJJ82711.1 akaw|GAA87112.1 aluc|OJZ84326.1 anid|CBF78016.1 anig|EHA27661.1 asyd|OJJ55236.1 ater|EAU30321.1 atub|OJI89742.1 aver|OJJ06987.1 awen|OJJ30974.1 azon|OJJ51732.1

Group7257: abra|OJJ77000.1 acar|OOF98680.1 acla|EAW13452.1 afis|EAW16435.1 afla|EED47786.1 agla|OJJ89569.1 akaw|GAA87048.1 aluc|OJZ84398.1 anig|EHA27729.1 aory|EIT77178.1 ater|EAU38162.1 atub|OJI89674.1 awen|OJJ32553.1 acar|OOF90078.1

Group7258: abra|OJJ77112.1 acar|OOF93002.1 acla|EAW07059.1 afis|EAW16683.1 afla|EED45706.1 afum|EAL89287.1 akaw|GAA90816.1 aluc|OJZ84521.1 anig|EHA27819.1 aory|EIT75620.1 ater|EAU31914.1 atub|OJI89558.1 azon|OJJ48992.1 ater|EAU31030.1

Group7259: abra|OJJ77153.1 acar|OOF93039.1 afla|EED45740.1 agla|OJJ80497.1 akaw|GAA90777.1 aluc|OJZ84567.1 anid|CBF86599.1 anig|EHA27861.1 aory|EIT75632.1 asyd|OJJ64828.1 atub|OJI89515.1 aver|OJI98941.1 awen|OJJ33824.1 azon|OJJ44402.1

Group7260: abra|OJJ77171.1 afis|EAW15676.1 afla|EED51771.1 afum|EAL85671.1 akaw|GAA90762.1 aluc|OJZ84584.1 anig|EHA27877.1 aory|EIT81698.1 ater|EAU34097.1 atub|OJI89496.1 asyd|OJJ63636.1 aver|OJI97300.1 azon|OJJ43868.1 anid|CBF69536.1

Group7261: abra|OJJ77266.1 acar|OOF93134.1 acla|EAW13260.1 afis|EAW17366.1 afla|EED49804.1 aluc|OJZ84673.1 anid|CBF84213.1 anig|EHA27951.1 asyd|OJJ57171.1 ater|EAU33871.1 atub|OJI89407.1 aver|OJJ03144.1 awen|OJJ37444.1 azon|OJJ51123.1

Group7262: abra|OJJ77382.1 acar|OOF93250.1 afla|EED55717.1 afum|EAL87849.1 agla|OJJ87591.1 akaw|GAA91379.1 aluc|OJZ84790.1 anig|EHA28058.1 aory|EIT83227.1 asyd|OJJ59183.1 atub|OJI89292.1 aver|OJJ01804.1 awen|OJJ30872.1 acar|OOF95534.1

Group7263: abra|OJJ77444.1 acla|EAW15000.1 afis|EAW24524.1 afla|EED54711.1 afum|EAL86189.1 agla|OJJ79087.1 anid|CBF80443.1 asyd|OJJ54784.1 ater|EAU29961.1 aver|OJJ04453.1 awen|OJJ36201.1 asyd|OJJ57943.1 asyd|OJJ60276.1 aver|OJJ07382.1

Group7264: abra|OJJ77488.1 acar|OOF99237.1 acla|EAW06750.1 afis|EAW16300.1 afla|EED53254.1 afum|EAL88922.1 akaw|GAA84131.1 aluc|OJZ81048.1 anig|EHA19319.1 aory|EIT75243.1 ater|EAU37343.1 atub|OJI82536.1 awen|OJJ37699.1 azon|OJJ44960.1

Group7265: abra|OJJ77501.1 agla|OJJ88307.1 akaw|GAA84120.1 aluc|OJZ81036.1 anid|CBF86867.1 anig|EHA19330.1 asyd|OJJ64632.1 atub|OJI82547.1 aver|OJI98730.1 acla|EAW06735.1 afis|EAW16286.1 afum|EAL88908.1 awen|OJJ37680.1 azon|OJJ44944.1

Group7266: abra|OJJ77521.1 afum|EAL88892.1 akaw|GAA84099.1 aluc|OJZ81015.1 ater|EAU37446.1 atub|OJI82569.1 awen|OJJ37659.1 azon|OJJ44928.1 acla|EAW06716.1 afis|EAW16271.1 afla|EED53294.1 asyd|OJJ64614.1 aver|OJI98710.1 agla|OJJ88340.1

Group7267: abra|OJJ77569.1 acar|OOF99161.1 acla|EAW06534.1 afis|EAW16061.1 afla|EED48575.1 agla|OJJ86104.1 akaw|GAA84057.1 aluc|OJZ80969.1 anid|CBF75086.1 asyd|OJJ60014.1 atub|OJI82617.1 aver|OJJ02271.1 awen|OJJ30378.1 azon|OJJ45394.1

Group7268: abra|OJJ77635.1 acar|OOF99102.1 afla|EED51641.1 akaw|GAA83999.1 aluc|OJZ80895.1 anid|CBF89719.1 anig|EHA19444.1 asyd|OJJ62244.1 ater|EAU37813.1 atub|OJI82683.1 aver|OJJ01807.1 awen|OJJ31252.1 aver|OJI96354.1 asyd|OJJ59190.1

Group7269: abra|OJJ77666.1 acar|OOF99069.1 aluc|OJZ80862.1 ater|EAU33644.1 atub|OJI82715.1 aver|OJI98451.1 awen|OJJ30468.1 azon|OJJ45459.1 agla|OJJ86021.1 anid|CBF85842.1 aory|EIT73356.1 asyd|OJJ64355.1 afis|EAW16001.1 azon|OJJ45460.1

Group7270: abra|OJJ77688.1 acar|OOF99055.1 acla|EAW06576.1 afis|EAW16020.1 afum|EAL88649.1 agla|OJJ86047.1 akaw|GAA83948.1 aluc|OJZ80829.1 anid|CBF74536.1 anig|EHA19487.1 ater|EAU33624.1 atub|OJI82737.1 awen|OJJ30444.1 azon|OJJ45436.1

Group7271: abra|OJJ77700.1 acar|OOF99041.1 afla|EED53096.1 agla|OJJ84143.1 akaw|GAA83930.1 aluc|OJZ90765.1 anig|EHA19499.1 aory|EIT74802.1 atub|OJI82753.1 aver|OJI99804.1 awen|OJJ33497.1 azon|OJJ42269.1 afla|EED46279.1 aory|EIT76324.1

Group7272: abra|OJJ77786.1 afla|EED55125.1 afum|EAL84256.1 agla|OJJ83665.1 akaw|GAA83861.1 aluc|OJZ90841.1 anid|CBF84135.1 anig|EHA19562.1 aory|EIT81666.1 asyd|OJJ55632.1 ater|EAU39626.1 atub|OJI82828.1 aver|OJJ06709.1 awen|OJJ40274.1

Group7273: abra|OJJ77787.1 afla|EED55126.1 afum|EAL84255.1 agla|OJJ83664.1 akaw|GAA83860.1 aluc|OJZ90842.1 anid|CBF84137.1 anig|EHA19563.1 aory|EIT81624.1 asyd|OJJ55631.1 ater|EAU39625.1 atub|OJI82829.1 aver|OJJ06708.1 awen|OJJ40273.1

Group7274: abra|OJJ77816.1 acar|OOF98954.1 afis|EAW23617.1 afla|EED53805.1 afum|EAL87845.1 akaw|GAA83830.1 aluc|OJZ90888.1 anid|CBF75250.1 anig|EHA19607.1 aory|EIT78460.1 ater|EAU34476.1 atub|OJI82882.1 aver|OJJ04189.1 awen|OJJ39947.1

Group7275: abra|OJJ77858.1 acar|OOF98916.1 acla|EAW11268.1 afla|EED56237.1 afum|EAL87598.1 agla|OJJ85845.1 akaw|GAA83791.1 aluc|OJZ90932.1 anid|CBF86175.1 asyd|OJJ64526.1 atub|OJI82924.1 aver|OJI98624.1 awen|OJJ30610.1 azon|OJJ50158.1

Group7276: abra|OJJ77867.1 acar|OOF98905.1 afis|EAW18240.1 afla|EED56226.1 agla|OJJ85826.1 akaw|GAA83781.1 aluc|OJZ90943.1 anig|EHA19651.1 asyd|OJJ64537.1 ater|EAU33743.1 atub|OJI82934.1 aver|OJI98634.1 awen|OJJ30620.1 azon|OJJ47997.1

Group7277: abra|OJJ77939.1 afla|EED54357.1 aluc|OJZ90948.1 anig|EHA20678.1 aory|EIT75969.1 asyd|OJJ54282.1 atub|OJI82949.1 aver|OJJ05236.1 awen|OJJ40815.1 azon|OJJ42433.1 atub|OJI81191.1 agla|OJJ83742.1 aory|EIT77972.1 anig|EHA19718.1

Group7278: akaw|GAA83690.1 aluc|OJZ91042.1 anig|EHA19741.1 atub|OJI83038.1 acar|OOF98844.1 acla|EAW06668.1 afis|EAW16097.1 afla|EED48632.1 afum|EAL88729.1 agla|OJJ86125.1 anid|CBF85453.1 asyd|OJJ64097.1 aver|OJI98180.1 awen|OJJ30334.1

Group7279: abra|OJJ77975.1 acar|OOF91245.1 acla|EAW12898.1 afla|EED49851.1 akaw|GAA83685.1 aluc|OJZ91049.1 anid|CBF80320.1 anig|EHA19745.1 aory|EIT83397.1 asyd|OJJ52366.1 atub|OJI83044.1 aver|OJJ01981.1 awen|OJJ37000.1 acar|OOF97583.1

Group7280: abra|OJJ78076.1 acar|OOF98787.1 afla|EED48706.1 agla|OJJ86228.1 akaw|GAA85918.1 anid|CBF79570.1 asyd|OJJ57915.1 ater|EAU33251.1 aver|OJJ04476.1 awen|OJJ30266.1 azon|OJJ43961.1 acla|EAW06609.1 afis|EAW16206.1 afum|EAL88834.1

Group7281: abra|OJJ78104.1 acar|OOF98761.1 acla|EAW06616.1 afis|EAW16149.1 afla|EED48692.1 agla|OJJ86202.1 aluc|OJZ91168.1 anid|CBF79579.1 anig|EHA19862.1 asyd|OJJ57970.1 atub|OJI83183.1 aver|OJJ04433.1 awen|OJJ30274.1 azon|OJJ43968.1

Group7282: abra|OJJ78124.1 acar|OOF97937.1 acla|EAW12352.1 afis|EAW16777.1 afla|EED49599.1 afum|EAL86346.1 akaw|GAA85884.1 aluc|OJZ91186.1 anid|CBF73562.1 anig|EHA19880.1 aory|EIT79312.1 ater|EAU31567.1 atub|OJI83205.1 azon|OJJ45860.1

Group7283: abra|OJJ78126.1 acar|OOF94027.1 afla|EED51361.1 akaw|GAA85882.1 aluc|OJZ91188.1 anid|CBF75425.1 anig|EHA19884.1 aory|EIT80444.1 asyd|OJJ59790.1 ater|EAU30521.1 atub|OJI83207.1 aver|OJJ02498.1 awen|OJJ33364.1 azon|OJJ42494.1

Group7284: abra|OJJ78144.1 afis|EAW19675.1 afla|EED55112.1 akaw|GAA92008.1 aluc|OJZ92873.1 anid|CBF79033.1 anig|EHA28256.1 aory|EIT81601.1 ater|EAU33098.1 atub|OJI87331.1 awen|OJJ30818.1 acar|OOF90918.1 aluc|OJZ83576.1 atub|OJI89250.1

Group7285: abra|OJJ78219.1 acar|OOF98637.1 acla|EAW11230.1 afla|EED48740.1 afum|EAL87639.1 agla|OJJ88153.1 aluc|OJZ91293.1 anid|CBF79505.1 anig|EHA19976.1 aory|EIT73180.1 asyd|OJJ57882.1 atub|OJI83284.1 aver|OJJ04512.1 azon|OJJ47961.1

Group7286: abra|OJJ78240.1 acar|OOF98620.1 acla|EAW09270.1 afis|EAW21878.1 afla|EED46430.1 afum|EAL92892.1 agla|OJJ84254.1 akaw|GAA85785.1 aluc|OJZ91314.1 asyd|OJJ61276.1 atub|OJI83308.1 aver|OJJ00604.1 awen|OJJ34167.1 azon|OJJ43463.1

Group7287: abra|OJJ78333.1 acar|OOF98231.1 acla|EAW06320.1 afis|EAW16713.1 afla|EED53311.1 afum|EAL85780.1 akaw|GAA85698.1 aluc|OJZ91405.1 anid|CBF79053.1 anig|EHA20082.1 aory|EIT75151.1 ater|EAU34393.1 atub|OJI83399.1 awen|OJJ34746.1

Group7288: abra|OJJ78340.1 acar|OOF98540.1 acla|EAW09191.1 afis|EAW21955.1 afla|EED46673.1 afum|EAL92969.1 agla|OJJ84162.1 akaw|GAA85693.1 aluc|OJZ91412.1 anid|CBF84076.1 ater|EAU31061.1 atub|OJI83406.1 awen|OJJ34247.1 azon|OJJ43391.1

Group7289: abra|OJJ78370.1 acar|OOF90129.1 acla|EAW09165.1 afis|EAW21981.1 afum|EAL92996.1 akaw|GAA85663.1 aluc|OJZ91447.1 anid|CBF82707.1 anig|EHA20111.1 aory|EIT77895.1 asyd|OJJ61642.1 awen|OJJ34275.1 azon|OJJ43362.1 afla|EED46705.1

Group7290: acar|OOF90254.1 acla|EAW12853.1 afis|EAW18016.1 afla|EED51878.1 afum|EAL91828.1 agla|OJJ85370.1 anid|CBF80048.1 anig|EHA27366.1 aory|EIT81738.1 asyd|OJJ58278.1 ater|EAU31532.1 aver|OJJ02145.1 awen|OJJ41523.1 azon|OJJ48245.1

Group7291: acar|OOF90348.1 acla|EAW07679.1 afis|EAW22694.1 afla|EED49978.1 afum|EAL90499.1 agla|OJJ79020.1 anid|CBF88096.1 anig|EHA18008.1 aory|EIT74752.1 asyd|OJJ63569.1 ater|EAU39014.1 aver|OJI97365.1 awen|OJJ35326.1 azon|OJJ49788.1

Group7292: acar|OOF93126.1 acla|EAW11421.1 afis|EAW24001.1 afla|EED49483.1 afum|EAL87431.1 agla|OJJ88142.1 akaw|GAA90698.1 aluc|OJZ84655.1 anid|CBF71785.1 aory|EIT79531.1 asyd|OJJ61745.1 ater|EAU34696.1 awen|OJJ40502.1 azon|OJJ50776.1

Group7293: acar|OOF93495.1 anig|EHA25487.1 azon|OJJ43097.1 acla|EAW08862.1 afis|EAW15532.1 afla|EED49043.1 afum|EAL85796.2 agla|OJJ85215.1 anid|CBF78200.1 asyd|OJJ55984.1 ater|EAU32593.1 aver|OJJ04931.1 awen|OJJ38196.1 aory|EIT82545.1

Group7294: acar|OOF95324.1 acla|EAW09283.1 afis|EAW21864.1 afla|EED46447.1 afum|EAL92878.2 agla|OJJ84271.1 akaw|GAA87310.1 aluc|OJZ89707.1 anid|CBF77195.1 asyd|OJJ56835.1 ater|EAU32981.1 aver|OJJ03502.1 awen|OJJ34152.1 azon|OJJ43476.1

Group7295: acar|OOF96104.1 acla|EAW06468.1 afis|EAW15963.1 afla|EED58094.1 afum|EAL88596.1 agla|OJJ89349.1 akaw|GAA85505.1 anid|CBF81083.1 aory|EIT79119.1 ater|EAU35491.1 atub|OJI82322.1 aver|OJJ06255.1 awen|OJJ35611.1 azon|OJJ46933.1

Group7296: acar|OOF96168.1 acla|EAW08019.1 afis|EAW23053.1 afla|EED54965.1 afum|EAL88427.1 agla|OJJ89423.1 akaw|GAA85566.1 aluc|OJZ83500.1 anid|CBF81045.1 aory|EIT80366.1 asyd|OJJ56440.1 ater|EAU37611.1 atub|OJI82394.1 azon|OJJ44151.1

Group7297: acar|OOF96379.1 acla|EAW15207.1 afis|EAW20172.1 afla|EED57176.1 afum|EAL85290.1 agla|OJJ88758.1 aluc|OJZ86410.1 anid|CBF85030.1 anig|EHA21489.1 asyd|OJJ63102.1 ater|EAU35208.1 atub|OJI84121.1 aver|OJI97860.1 azon|OJJ46100.1

Group7298: afis|EAW20697.1 agla|OJJ86296.1 azon|OJJ47338.1 afla|EED45521.1 asyd|OJJ61710.1 afum|EAL84871.1 aluc|OJZ83608.1 anig|EHA18767.1 aory|EIT75681.1 atub|OJI89981.1 aver|OJI95806.1 awen|OJJ33573.1 akaw|GAA92117.1 awen|OJJ31973.1

Group7299: acla|EAW09761.1 afis|EAW21376.1 afla|EED55794.1 afum|EAL92393.1 agla|OJJ78767.1 akaw|GAA83966.1 aluc|OJZ80855.1 anid|CBF83858.1 aory|EIT76823.1 asyd|OJJ60925.1 ater|EAU35181.1 aver|OJJ00251.1 awen|OJJ39321.1 azon|OJJ51666.1

Group7300: acla|EAW12193.1 afis|EAW17089.1 afla|EED53701.1 afum|EAL86567.1 akaw|GAA84584.1 anid|CBF78067.1 aory|EIT75279.1 atub|OJI84548.1 awen|OJJ33470.1 anig|EHA23247.1 aluc|OJZ81998.1 awen|OJJ30116.1 asyd|OJJ52156.1 ater|EAU35602.1

Group7301: acla|EAW12242.1 afla|EED52536.1 afis|EAW21476.1 afis|EAW23696.1 agla|OJJ87419.1 aluc|OJZ80170.1 aory|EIT74591.1 ater|EAU33757.1 awen|OJJ30150.1 afla|EED47559.1 afum|EAL92496.1 aluc|OJZ90162.1 anig|EHA21660.1 atub|OJI84639.1

Group7302: acla|EAW15159.1 afis|EAW20127.1 afla|EED57224.1 afum|EAL85245.1 akaw|GAA90165.1 aluc|OJZ83182.1 anid|CBF84954.1 anig|EHA18394.1 aory|EIT74093.1 asyd|OJJ63144.1 ater|EAU38746.1 aver|OJI97810.1 awen|OJJ37487.1 azon|OJJ46221.1

Group7303: afis|EAW18365.1 afum|EAL93039.1 agla|OJJ84928.1 akaw|GAA86328.1 aluc|OJZ84852.1 anid|CBF70819.1 anig|EHA25086.1 aory|EIT77425.1 asyd|OJJ58402.1 ater|EAU32233.1 atub|OJI80744.1 aver|OJJ01003.1 awen|OJJ31786.1 azon|OJJ50441.1

Group7304: abra|OJJ65625.1 abra|OJJ65711.1 aluc|OJZ79679.1 aluc|OJZ81150.1 acla|EAW15091.1 agla|OJJ85723.1 akaw|GAA92894.1 afis|EAW21070.1 afla|EED46368.1 anig|EHA20203.1 atub|OJI80392.1 akaw|GAA93306.1 aluc|OJZ88793.1

Group7305: abra|OJJ66410.1 abra|OJJ67969.1 agla|OJJ86641.1 agla|OJJ87771.1 ater|EAU32456.1 ater|EAU32813.1 acla|EAW15054.1 afis|EAW17011.1 aver|OJJ06537.1 azon|OJJ42141.1 afla|EED55206.1 aory|EIT83552.1 awen|OJJ37013.1

Group7306: abra|OJJ74542.1 abra|OJJ77038.1 aluc|OJZ90093.1 aluc|OJZ92122.1 atub|OJI84713.1 atub|OJI86555.1 awen|OJJ30779.1 awen|OJJ31702.1 acar|OOF95216.1 akaw|GAA82060.1 ater|EAU31041.1 azon|OJJ45316.1 anid|CBF73564.1

Group7307: acar|OOF90466.1 acar|OOG00946.1 abra|OJJ70070.1 acla|EAW12442.1 afla|EED47625.1 akaw|GAA87881.1 aluc|OJZ85523.1 aory|EIT74251.1 ater|EAU38526.1 anig|EHA25055.1 aory|EIT80234.1 atub|OJI80044.1 afla|EED52535.1

Group7308: acar|OOF94032.1 acar|OOF94102.1 abra|OJJ66421.1 acla|EAW08781.1 afis|EAW15620.1 afum|EAL85715.1 akaw|GAA90286.1 aluc|OJZ83054.1 anig|EHA28119.1 asyd|OJJ56692.1 atub|OJI87893.1 aver|OJJ06032.1 azon|OJJ51149.1

Group7309: acar|OOF94036.1 acar|OOF94106.1 abra|OJJ66414.1 afis|EAW20231.1 afla|EED55133.1 afum|EAL85348.1 akaw|GAA90281.1 aluc|OJZ83059.1 anid|CBF73522.1 anig|EHA28125.1 aory|EIT81631.1 atub|OJI87887.1 azon|OJJ47377.1

Group7310: acar|OOF94084.1 acar|OOF94154.1 abra|OJJ71261.1 acla|EAW12246.1 afla|EED53196.1 akaw|GAA82728.1 aluc|OJZ87282.1 anig|EHA20524.1 aory|EIT74916.1 ater|EAU30957.1 atub|OJI81359.1 azon|OJJ45974.1 anid|CBF80413.1

Group7311: acar|OOF94085.1 acar|OOF94155.1 abra|OJJ69769.1 afis|EAW23675.1 afla|EED49539.1 afum|EAL87118.1 akaw|GAA87884.1 aluc|OJZ85529.1 anid|CBF80787.1 anig|EHA25060.1 aory|EIT82650.1 asyd|OJJ65400.1 atub|OJI80039.1

Group7312: acar|OOF95050.1 acar|OOF95117.1 abra|OJJ68392.1 afla|EED46845.1 agla|OJJ89056.1 anig|EHA20969.1 aory|EIT72552.1 asyd|OJJ52726.1 aver|OJJ04488.1 awen|OJJ34415.1 asyd|OJJ62241.1 aver|OJJ07602.1 aver|OJI96350.1

Group7313: acar|OOF95071.1 acar|OOF95137.1 abra|OJJ74644.1 acla|EAW10744.1 afis|EAW24830.1 afla|EED47145.1 afum|EAL89475.1 akaw|GAA81967.1 anid|CBF74574.1 anig|EHA26734.1 ater|EAU36475.1 awen|OJJ40417.1 azon|OJJ49275.1

Group7314: acar|OOF95072.1 acar|OOF95138.1 abra|OJJ74643.1 acla|EAW10745.1 afis|EAW24829.1 afla|EED47146.1 afum|EAL89474.1 akaw|GAA81968.1 anid|CBF74572.1 anig|EHA26735.1 ater|EAU36476.1 awen|OJJ40415.1 azon|OJJ49270.1

Group7315: acar|OOF95082.1 acar|OOF95147.1 abra|OJJ74627.1 acla|EAW10755.1 afis|EAW24819.1 afum|EAL89463.1 akaw|GAA81981.1 aluc|OJZ92209.1 anig|EHA26749.1 ater|EAU36487.1 atub|OJI86638.1 awen|OJJ40405.1 azon|OJJ49254.1

Group7316: acar|OOF95623.1 acar|OOF95675.1 awen|OJJ31312.1 awen|OJJ34512.1 asyd|OJJ63066.1 aver|OJI97896.1 acla|EAW10139.1 agla|OJJ82667.1 aory|EIT75881.1 azon|OJJ47348.1 afla|EED54390.1 anid|CBF85091.1 aory|EIT73682.1

Group7317: acar|OOF96410.1 acar|OOF96792.1 acar|OOF99159.1 afla|EED45799.1 afla|EED54109.1 abra|OJJ72157.1 afis|EAW20301.1 agla|OJJ79843.1 akaw|GAA84769.1 aluc|OJZ86914.1 anig|EHA21730.1 atub|OJI83607.1 azon|OJJ51777.1

Group7318: acar|OOF97653.1 acar|OOF97700.1 abra|OJJ71504.1 afla|EED44717.1 akaw|GAA82516.1 aluc|OJZ87043.1 anig|EHA20316.1 aory|EIT83460.1 asyd|OJJ53005.1 ater|EAU33381.1 atub|OJI81592.1 aver|OJJ03838.1 azon|OJJ49498.1

Group7319: acar|OOF98697.1 acar|OOG00709.1 aory|EIT76303.1 aory|EIT81949.1 abra|OJJ65604.1 afla|EED55021.1 anig|EHA19920.1 atub|OJI83232.1 akaw|GAA85865.1 aluc|OJZ91214.1 agla|OJJ82283.1 afla|EED48305.1 aory|EIT81992.1

Group7320: acar|OOG00238.1 acar|OOG00255.1 abra|OJJ70094.1 acla|EAW13718.1 afis|EAW17168.1 akaw|GAA87601.1 aluc|OJZ85221.1 anig|EHA25425.1 ater|EAU34027.1 atub|OJI80362.1 aver|OJJ05978.1 acar|OOF92822.1 ater|EAU36219.1

Group7321: acar|OOG00240.1 acar|OOG00259.1 abra|OJJ75763.1 afis|EAW16465.1 afum|EAL89061.1 akaw|GAA86037.1 aluc|OJZ90264.1 anig|EHA23760.1 asyd|OJJ54942.1 ater|EAU31651.1 atub|OJI81690.1 aver|OJJ07253.1 awen|OJJ32194.1

Group7322: afis|EAW16721.1 afis|EAW19720.1 abra|OJJ71190.1 acar|OOF97977.1 acla|EAW12079.1 afla|EED52128.1 afum|EAL86806.1 akaw|GAA82795.1 aluc|OJZ87354.1 anig|EHA20588.1 aory|EIT74423.1 atub|OJI81291.1 acar|OOF95351.1

Group7323: afla|EED49302.1 afla|EED51803.1 abra|OJJ73154.1 acar|OOG00366.1 afis|EAW23609.1 afum|EAL87855.2 akaw|GAA88018.1 aluc|OJZ81276.1 anig|EHA18576.1 aory|EIT81692.1 ater|EAU31012.1 atub|OJI91484.1 aory|EIT82217.1

Group7324: afla|EED52443.1 afla|EED55159.1 aory|EIT81613.1 aory|EIT82756.1 anid|CBF78247.1 abra|OJJ66501.1 afis|EAW19736.1 afum|EAL86793.1 agla|OJJ89295.1 asyd|OJJ64194.1 ater|EAU29805.1 aver|OJI98271.1 awen|OJJ36110.1

Group7325: afum|EAL87804.2 afum|EBA27481.1 abra|OJJ65723.1 acla|EAW15117.1 akaw|GAA92609.1 aluc|OJZ80248.1 anid|CBF77053.1 anig|EHA21829.1 azon|OJJ50041.1 atub|OJI79447.1 acar|OOF90669.1 akaw|GAA93131.1 aluc|OJZ85655.1

Group7326: afum|EAL90932.1 afum|EAL91125.1 abra|OJJ66324.1 afis|EAW20675.1 afla|EED52353.1 akaw|GAA87893.1 anid|CBF84441.1 anig|EHA21855.1 asyd|OJJ52040.1 aver|OJJ03116.1 azon|OJJ42061.1 afis|EAW20715.1 afum|EAL87749.2

Group7327: agla|OJJ78572.1 agla|OJJ83839.1 abra|OJJ72314.1 acar|OOF94896.1 akaw|GAA86061.1 aluc|OJZ90295.1 anig|EHA23784.1 asyd|OJJ52860.1 ater|EAU38830.1 atub|OJI81719.1 azon|OJJ46699.1 aver|OJJ08838.1 awen|OJJ33752.1

Group7328: agla|OJJ86078.1 agla|OJJ83488.1 agla|OJJ83489.1 agla|OJJ83490.1 agla|OJJ87769.1 agla|OJJ86007.1 agla|OJJ79776.1 agla|OJJ79777.1 agla|OJJ83486.1 agla|OJJ79778.1 agla|OJJ82186.1 agla|OJJ83487.1 agla|OJJ87770.1

Group7329: agla|OJJ81016.1 agla|OJJ82741.1 awen|OJJ36089.1 awen|OJJ37947.1 abra|OJJ75237.1 acar|OOF94555.1 afla|EED47785.1 akaw|GAA91780.1 aluc|OJZ92798.1 anig|EHA26210.1 aory|EIT77101.1 ater|EAU37358.1 atub|OJI87242.1

Group7330: agla|OJJ81590.1 agla|OJJ85374.1 acla|EAW12658.1 afis|EAW18021.1 afla|EED51871.1 afum|EAL91833.1 anid|CBF80056.1 aory|EIT81742.1 asyd|OJJ58282.1 ater|EAU31536.1 aver|OJJ02141.1 awen|OJJ41519.1 azon|OJJ48241.1

Group7331: agla|OJJ83535.1 agla|OJJ86812.1 abra|OJJ68865.1 akaw|GAA90388.1 aluc|OJZ85843.1 anig|EHA24721.1 asyd|OJJ56360.1 atub|OJI79631.1 aver|OJJ06354.1 abra|OJJ70353.1 aver|OJJ03414.1 awen|OJJ41927.1 aluc|OJZ80609.1

Group7332: agla|OJJ83636.1 agla|OJJ88102.1 anid|CBF69578.1 anid|CBF70072.1 anid|CBF78172.1 anid|CBF89925.1 afla|EED52413.1 akaw|GAA92412.1 aluc|OJZ83270.1 aory|EIT83305.1 aver|OJJ02701.1 awen|OJJ33609.1 agla|OJJ79698.1

Group7333: agla|OJJ83706.1 agla|OJJ85925.1 awen|OJJ29699.1 awen|OJJ41943.1 acla|EAW13758.1 afis|EAW19809.1 afla|EED53501.1 afum|EAL85083.1 anid|CBF84506.1 aory|EIT74925.1 asyd|OJJ53519.1 ater|EAU29283.1 aver|OJJ08231.1

Group7334: agla|OJJ86205.1 agla|OJJ86206.1 abra|OJJ67776.1 afis|EAW15655.1 afla|EED53126.1 afum|EAL86481.2 aory|EIT75150.1 asyd|OJJ59174.1 ater|EAU32443.1 aver|OJJ01763.1 awen|OJJ33264.1 azon|OJJ44220.1 afis|EAW16954.1

Group7335: agla|OJJ89526.1 agla|OJJ89527.1 agla|OJJ89528.1 aver|OJI97616.1 aver|OJJ07740.1 acla|EAW07124.1 afis|EAW22134.1 afum|EAL91056.1 ater|EAU31467.1 awen|OJJ38813.1 asyd|OJJ54663.1 azon|OJJ47921.1 aver|OJJ07501.1

Group7336: akaw|GAA93154.1 akaw|GAA93176.1 akaw|GAA93247.1 aluc|OJZ79787.1 aluc|OJZ80418.1 acla|EAW11464.1 afis|EAW20659.1 anig|EHA17702.1 anid|CBF87230.1 asyd|OJJ52330.1 aluc|OJZ79978.1 anig|EHA21850.1 asyd|OJJ52139.1

Group7337: aluc|OJZ80039.1 aluc|OJZ88966.1 abra|OJJ75510.1 acar|OOF90836.1 afla|EED49953.1 agla|OJJ82554.1 akaw|GAA84273.1 anig|EHA23556.1 aory|EIT83425.1 asyd|OJJ56398.1 ater|EAU39565.1 atub|OJI85824.1 aver|OJJ06314.1

Group7338: aluc|OJZ89236.1 aluc|OJZ91106.1 abra|OJJ78029.1 acar|OOG00096.1 afis|EAW15604.1 afla|EED45424.1 afum|EAL85729.1 akaw|GAA85956.1 anig|EHA19804.1 aory|EIT82389.1 atub|OJI83103.1 aver|OJI99882.1 akaw|GAA84507.1

Group7339: anid|CBF69416.1 anid|CBF87272.1 abra|OJJ69759.1 afla|EED52105.1 afum|EAL88029.1 aory|EIT76922.1 asyd|OJJ55659.1 ater|EAU29407.1 awen|OJJ35887.1 afis|EAW17758.1 aver|OJJ06743.1 anid|CBF78961.1 ater|EAU31006.1

Group7340: aory|EIT73388.1 aory|EIT77973.1 acar|OOF99554.1 acla|EAW14107.1 afis|EAW18808.1 afla|EED50769.1 afla|EED56309.1 afum|EAL93544.1 asyd|OJJ63757.1 ater|EAU37959.1 aver|OJI97183.1 awen|OJJ32383.1 anid|CBF88422.1

Group7341: asyd|OJJ53576.1 asyd|OJJ62638.1 aver|OJI99291.1 aver|OJJ08181.1 abra|OJJ69683.1 acar|OOF93998.1 afla|EED52270.1 akaw|GAA88807.1 aluc|OJZ82490.1 anig|EHA28283.1 aory|EIT74510.1 atub|OJI87942.1 awen|OJJ38314.1

Group7342: asyd|OJJ54571.1 asyd|OJJ55108.1 aver|OJJ07098.1 aver|OJJ07583.1 acla|EAW14964.1 afis|EAW24487.1 afla|EED52186.1 afum|EAL86156.1 agla|OJJ83084.1 anid|CBF71325.1 aory|EIT74354.1 ater|EAU31591.1 awen|OJJ31454.1

Group7343: asyd|OJJ56962.1 asyd|OJJ64221.1 aver|OJJ03324.1 aver|OJJ07424.1 aver|OJJ05559.1 aver|OJJ06598.1 acar|OOF90702.1 afla|EED56542.1 anid|CBF77081.1 aory|EIT79693.1 ater|EAU32038.1 awen|OJJ34782.1 asyd|OJJ55520.1

Group7344: ater|EAU29369.1 ater|EAU32785.1 ater|EAU30091.1 ater|EAU32227.1 ater|EAU30716.1 ater|EAU36853.1 ater|EAU36854.1 ater|EAU32787.1 agla|OJJ79772.1 agla|OJJ86298.1 akaw|GAA93358.1 aluc|OJZ88799.1 anid|CBF80312.1

Group7345: ater|EAU29454.1 ater|EAU37672.1 abra|OJJ67139.1 acar|OOF98416.1 afla|EED47552.1 agla|OJJ78797.1 akaw|GAA91030.1 aluc|OJZ87833.1 anig|EHA20312.1 atub|OJI80808.1 awen|OJJ30096.1 azon|OJJ45318.1 aory|EIT74563.1

Group7346: atub|OJI80394.1 atub|OJI81725.1 abra|OJJ72317.1 acar|OOF95833.1 agla|OJJ79170.1 akaw|GAA86652.1 aluc|OJZ85210.1 anig|EHA25413.1 asyd|OJJ60280.1 aver|OJJ05907.1 awen|OJJ32563.1 afla|EED48202.1 aory|EIT72944.1

Group7347: aver|OJJ05171.1 aver|OJJ07902.1 abra|OJJ75300.1 afla|EED48251.1 agla|OJJ87358.1 akaw|GAA91833.1 aluc|OJZ92855.1 anid|CBF82036.1 anig|EHA26151.1 aory|EIT72920.1 asyd|OJJ54219.1 ater|EAU29252.1 atub|OJI87312.1

Group7348: awen|OJJ30458.1 awen|OJJ31969.1 awen|OJJ40902.1 awen|OJJ34825.1 abra|OJJ70664.1 akaw|GAA90000.1 aluc|OJZ84014.1 anig|EHA19137.1 atub|OJI90385.1 aver|OJJ06098.1 asyd|OJJ56628.1 acla|EAW12208.1 asyd|OJJ62336.1

Group7349: awen|OJJ30752.1 awen|OJJ31729.1 acla|EAW06309.1 afis|EAW18418.1 afla|EED45520.1 afum|EAL93102.1 agla|OJJ84986.1 anid|CBF70712.1 aory|EIT75484.1 asyd|OJJ58450.1 ater|EAU29834.1 aver|OJJ01051.1 azon|OJJ42190.1

Group7350: awen|OJJ30792.1 awen|OJJ31689.1 abra|OJJ66774.1 acar|OOF93852.1 afla|EED47254.1 akaw|GAA89709.1 aluc|OJZ80675.1 anig|EHA27338.1 aory|EIT79327.1 ater|EAU31717.1 atub|OJI87534.1 aver|OJI99829.1 agla|OJJ84418.1

Group7351: awen|OJJ31094.1 awen|OJJ32622.1 abra|OJJ77287.1 akaw|GAA90662.1 aluc|OJZ84698.1 anig|EHA27973.1 asyd|OJJ57282.1 ater|EAU31772.1 atub|OJI89384.1 aver|OJJ05874.1 afla|EED45792.1 aory|EIT75819.1 ater|EAU31829.1

Group7352: awen|OJJ38691.1 awen|OJJ38773.1 abra|OJJ77880.1 acar|OOF96530.1 afis|EAW22770.1 afum|EAL89320.2 agla|OJJ87882.1 akaw|GAA83773.1 aluc|OJZ90955.1 anig|EHA19660.1 asyd|OJJ57693.1 ater|EAU31721.1 atub|OJI82945.1

Group7353: azon|OJJ42557.1 azon|OJJ47248.1 abra|OJJ70067.1 agla|OJJ83497.1 akaw|GAA90340.1 aluc|OJZ85892.1 anig|EHA25435.1 asyd|OJJ63451.1 ater|EAU37388.1 atub|OJI79677.1 aver|OJJ03764.1 awen|OJJ34783.1 asyd|OJJ60829.1

Group7354: azon|OJJ46490.1 azon|OJJ47685.1 abra|OJJ67091.1 akaw|GAA91070.1 aluc|OJZ87884.1 anig|EHA20187.1 atub|OJI80767.1 abra|OJJ68144.1 acar|OOF89989.1 akaw|GAA92374.1 aluc|OJZ83232.1 anig|EHA21090.1 atub|OJI87704.1

Group7355: afis|EAW22095.1 abra|OJJ66781.1 afla|EED49683.1 agla|OJJ89077.1 akaw|GAA89716.1 aluc|OJZ80668.1 anig|EHA27343.1 aory|EIT80939.1 ater|EAU32526.1 awen|OJJ32823.1 azon|OJJ46503.1 aver|OJJ03199.1 acar|OOF96366.1

Group7356: abra|OJJ65645.1 acla|EAW06925.1 afum|EAL89009.2 agla|OJJ85284.1 awen|OJJ35901.1 abra|OJJ67674.1 afis|EAW15380.1 akaw|GAA83879.1 aluc|OJZ90817.1 anig|EHA25871.1 awen|OJJ31453.1 acar|OOF89911.1 afis|EAW20386.1

Group7357: abra|OJJ65698.1 afla|EED56837.1 afum|EAL84268.1 agla|OJJ84577.1 aluc|OJZ88718.1 anid|CBF85277.1 anig|EHA18817.1 aory|EIT74597.1 ater|EAU33257.1 aver|OJJ07758.1 awen|OJJ30260.1 asyd|OJJ61767.1 anig|EHA28201.1

Group7358: afis|EAW15627.1 afla|EED45354.1 aory|EIT82475.1 abra|OJJ70088.1 acla|EAW12308.1 afis|EAW21035.1 anig|EHA22484.1 atub|OJI89253.1 awen|OJJ30200.1 afum|EAL85707.1 akaw|GAA92131.1 anig|EHA26723.1 atub|OJI89962.1

Group7359: abra|OJJ65850.1 afis|EAW19899.1 asyd|OJJ52738.1 ater|EAU30137.1 aver|OJJ02994.1 abra|OJJ69836.1 acar|OOF89897.1 afla|EED50801.1 akaw|GAA87824.1 aluc|OJZ85460.1 anig|EHA25002.1 aory|EIT78046.1 atub|OJI80103.1

Group7360: abra|OJJ65851.1 acar|OOF95012.1 acla|EAW12070.1 afla|EED49952.1 agla|OJJ82555.1 akaw|GAA92293.1 aluc|OJZ90190.1 anig|EHA23689.1 aory|EIT83442.1 asyd|OJJ56407.1 atub|OJI81617.1 aver|OJJ06305.1 azon|OJJ46786.1

Group7361: abra|OJJ65862.1 acar|OOF95033.1 afla|EED46490.1 agla|OJJ78843.1 akaw|GAA92283.1 aluc|OJZ90180.1 anig|EHA23679.1 aory|EIT72435.1 asyd|OJJ56860.1 ater|EAU32957.1 atub|OJI81607.1 aver|OJJ03489.1 awen|OJJ34662.1

Group7362: abra|OJJ65916.1 afla|EED50461.1 afum|EAL84939.1 agla|OJJ80063.1 akaw|GAA92316.1 aluc|OJZ90214.1 anid|CBF86991.1 anig|EHA23712.1 aory|EIT82100.1 atub|OJI81645.1 aver|OJJ05232.1 awen|OJJ29951.1 asyd|OJJ54279.1

Group7363: abra|OJJ65932.1 acar|OOF94997.1 afla|EED54753.1 akaw|GAA92303.1 aluc|OJZ90200.1 anid|CBF75866.1 anig|EHA23697.1 aory|EIT78794.1 asyd|OJJ60224.1 ater|EAU34019.1 atub|OJI81628.1 aver|OJJ04043.1 awen|OJJ36434.1

Group7364: abra|OJJ65965.1 acar|OOF92908.1 afla|EED46024.1 agla|OJJ87992.1 akaw|GAA90840.1 aluc|OJZ82993.1 anig|EHA21407.1 aory|EIT75431.1 asyd|OJJ53305.1 atub|OJI89227.1 aver|OJJ08436.1 awen|OJJ30035.1 azon|OJJ43682.1

Group7365: abra|OJJ66010.1 afis|EAW16373.1 afla|EED57913.1 afum|EAL88997.1 agla|OJJ79810.1 aluc|OJZ82951.1 anig|EHA21366.1 aory|EIT82362.1 asyd|OJJ53105.1 atub|OJI89183.1 aver|OJJ03747.1 awen|OJJ35383.1 ater|EAU36730.1

Group7366: abra|OJJ66064.1 ater|EAU36630.1 abra|OJJ69774.1 acar|OOF91483.1 acla|EAW13724.1 afis|EAW19867.1 akaw|GAA87879.1 aluc|OJZ85520.1 anig|EHA25053.1 asyd|OJJ58291.1 atub|OJI80046.1 aver|OJJ02133.1 azon|OJJ45966.1

Group7367: abra|OJJ66081.1 acar|OOF99288.1 afla|EED46900.1 agla|OJJ79338.1 akaw|GAA84165.1 aluc|OJZ81090.1 anid|CBF70514.1 anig|EHA19280.1 aory|EIT72730.1 asyd|OJJ64106.1 atub|OJI82494.1 aver|OJI98188.1 awen|OJJ33665.1

Group7368: abra|OJJ66161.1 acla|EAW09047.1 afla|EED49666.1 aory|EIT81013.1 asyd|OJJ61205.1 aver|OJJ00529.1 awen|OJJ32034.1 abra|OJJ68395.1 akaw|GAA92553.1 aluc|OJZ90101.1 anig|EHA20971.1 atub|OJI84705.1 aver|OJJ04319.1

Group7369: abra|OJJ66162.1 afla|EED52446.1 agla|OJJ82380.1 akaw|GAA92448.1 aluc|OJZ81570.1 aory|EIT82786.1 asyd|OJJ54191.1 ater|EAU32218.1 atub|OJI88615.1 aver|OJJ01809.1 awen|OJJ30882.1 aver|OJJ03360.1 asyd|OJJ59193.1

Group7370: abra|OJJ66212.1 acar|OOF90648.1 afis|EAW16638.1 afla|EED55072.1 afum|EAL89240.1 akaw|GAA89075.1 aluc|OJZ80516.1 anig|EHA25677.1 aory|EIT81974.1 asyd|OJJ52112.1 ater|EAU37747.1 atub|OJI88414.1 azon|OJJ51721.1

Group7371: abra|OJJ66214.1 acar|OOF90647.1 afis|EAW16637.1 afla|EED55073.1 afum|EAL89239.1 akaw|GAA89074.1 aluc|OJZ80517.1 anig|EHA25676.1 aory|EIT81988.1 asyd|OJJ52113.1 ater|EAU37746.1 atub|OJI88413.1 azon|OJJ51722.1

Group7372: abra|OJJ66215.1 acar|OOF89826.1 afis|EAW16636.1 afla|EED55074.1 afum|EAL89238.1 akaw|GAA89073.1 aluc|OJZ80518.1 anig|EHA25675.1 aory|EIT81954.1 asyd|OJJ52114.1 ater|EAU37745.1 atub|OJI88412.1 azon|OJJ51723.1

Group7373: abra|OJJ66218.1 acar|OOF91502.1 afla|EED53203.1 agla|OJJ79257.1 akaw|GAA89068.1 aluc|OJZ80524.1 anid|CBF80522.1 aory|EIT75054.1 asyd|OJJ64214.1 atub|OJI88406.1 aver|OJI99460.1 awen|OJJ36184.1 aver|OJJ05265.1

Group7374: abra|OJJ66235.1 acar|OOF90693.1 agla|OJJ84415.1 akaw|GAA89054.1 aluc|OJZ80540.1 anid|CBF85755.1 anig|EHA25655.1 aory|EIT81171.1 asyd|OJJ64298.1 atub|OJI88391.1 aver|OJJ07892.1 awen|OJJ31374.1 aver|OJI98391.1

Group7375: abra|OJJ66249.1 acar|OOF90707.1 acla|EAW07084.1 afis|EAW20351.1 afla|EED49660.1 afum|EAL85469.1 akaw|GAA89039.1 aluc|OJZ80556.1 anid|CBF83083.1 anig|EHA25641.1 aory|EIT80972.1 atub|OJI88375.1 azon|OJJ43479.1

Group7376: abra|OJJ66250.1 acar|OOF90721.1 afis|EAW19305.1 afla|EED47578.1 afum|EAL94023.1 akaw|GAA89037.1 aluc|OJZ80558.1 anid|CBF78788.1 anig|EHA25639.1 aory|EIT74579.1 asyd|OJJ65245.1 atub|OJI88373.1 aver|OJI99397.1

Group7377: azon|OJJ42793.1 abra|OJJ70220.1 akaw|GAA92418.1 aluc|OJZ81490.1 anig|EHA25329.1 ater|EAU32956.1 atub|OJI80486.1 awen|OJJ36220.1 abra|OJJ74462.1 agla|OJJ86653.1 atub|OJI88690.1 akaw|GAA91162.1 anig|EHA25939.1

Group7378: abra|OJJ66438.1 acar|OOF90289.1 acla|EAW12207.1 afis|EAW17055.1 afla|EED56939.1 afum|EAL86552.2 agla|OJJ84864.1 akaw|GAA90300.1 aluc|OJZ83037.1 asyd|OJJ55135.1 ater|EAU32959.1 aver|OJJ07068.1 awen|OJJ37361.1

Group7379: akaw|GAA88006.1 aluc|OJZ81264.1 afis|EAW20651.1 afla|EED50436.1 afum|EAL84421.1 agla|OJJ88364.1 aory|EIT82011.1 asyd|OJJ53617.1 atub|OJI91464.1 aver|OJJ05937.1 awen|OJJ33663.1 ater|EAU33965.1 awen|OJJ30582.1

Group7380: acla|EAW11663.1 abra|OJJ76866.1 afla|EED51650.1 akaw|GAA87157.1 aluc|OJZ84277.1 anig|EHA27622.1 aory|EIT80664.1 asyd|OJJ62987.1 ater|EAU30283.1 atub|OJI89804.1 aver|OJI97981.1 awen|OJJ30983.1 awen|OJJ38752.1

Group7381: abra|OJJ66470.1 afis|EAW22767.1 afla|EED52194.1 anid|CBF74556.1 aory|EIT74441.1 asyd|OJJ57640.1 ater|EAU37828.1 aver|OJJ05498.1 awen|OJJ37843.1 azon|OJJ45252.1 anid|CBF87339.1 asyd|OJJ52804.1 aver|OJJ08784.1

Group7382: abra|OJJ66572.1 acar|OOF91013.1 acla|EAW10920.1 afis|EAW20938.1 afla|EED50293.1 afum|EAL84696.1 akaw|GAA91146.1 aluc|OJZ81472.1 anid|CBF82525.1 anig|EHA25955.1 aory|EIT77251.1 ater|EAU38583.1 atub|OJI88706.1

Group7383: abra|OJJ66578.1 afla|EED50300.1 akaw|GAA91140.1 aluc|OJZ81466.1 anig|EHA25961.1 atub|OJI88712.1 azon|OJJ48106.1 acla|EAW10912.1 afis|EAW20946.1 afum|EAL84688.1 aory|EIT77333.1 awen|OJJ34370.1 agla|OJJ83940.1

Group7384: abra|OJJ66633.1 acar|OOF91056.1 acla|EAW10862.1 afis|EAW20996.1 afum|EAL84632.1 agla|OJJ84074.1 aluc|OJZ81415.1 anid|CBF82635.1 asyd|OJJ54014.1 atub|OJI88765.1 aver|OJJ08727.1 awen|OJJ34314.1 azon|OJJ48059.1

Group7385: abra|OJJ66761.1 akaw|GAA89697.1 aluc|OJZ80688.1 atub|OJI87547.1 acar|OOF93843.1 afis|EAW19499.1 afla|EED56718.1 agla|OJJ81668.1 anid|CBF75436.1 ater|EAU34814.1 azon|OJJ42899.1 acla|EAW11887.1 afum|EBA27298.1

Group7386: abra|OJJ66769.1 acar|OOF93850.1 akaw|GAA89706.1 aluc|OJZ80678.1 anig|EHA27334.1 ater|EAU35185.1 atub|OJI87537.1 anid|CBF82286.1 abra|OJJ70434.1 afla|EED55096.1 aory|EIT81580.1 asyd|OJJ62603.1 afis|EAW20378.1

Group7387: abra|OJJ66777.1 acar|OOF93856.1 acla|EAW10822.1 afis|EAW24750.1 afla|EED55054.1 afum|EAL89395.2 akaw|GAA89712.1 aluc|OJZ80672.1 anig|EHA27339.1 aory|EIT81989.1 atub|OJI87526.1 aver|OJI95888.1 awen|OJJ33476.1

Group7388: abra|OJJ66845.1 acar|OOF91623.1 acla|EAW14424.1 afla|EED54282.1 agla|OJJ88606.1 akaw|GAA91560.1 aluc|OJZ85738.1 aory|EIT75991.1 asyd|OJJ64987.1 atub|OJI79523.1 aver|OJI99107.1 awen|OJJ37467.1 azon|OJJ51145.1

Group7389: abra|OJJ66852.1 acar|OOF91628.1 acla|EAW14430.1 afis|EAW19137.1 afum|EAL93869.1 agla|OJJ88616.1 akaw|GAA91554.1 aluc|OJZ85732.1 anid|CBF86325.1 aory|EIT76124.1 atub|OJI79517.1 awen|OJJ37474.1 azon|OJJ51153.1

Group7390: abra|OJJ66916.1 acar|OOF91683.1 acla|EAW14483.1 afis|EAW19192.1 afum|EAL93923.2 agla|OJJ88517.1 akaw|GAA91502.1 aluc|OJZ85663.1 anid|CBF86237.1 aory|EIT80237.1 ater|EAU37283.1 atub|OJI79457.1 aver|OJI99172.1

Group7391: abra|OJJ66917.1 acar|OOF91684.1 acla|EAW14484.1 afis|EAW19193.1 afum|EAL93924.1 agla|OJJ88516.1 akaw|GAA91501.1 aluc|OJZ85662.1 anid|CBF86236.1 anig|EHA24871.1 ater|EAU37282.1 atub|OJI79456.1 aver|OJI99173.1

Group7392: abra|OJJ66967.1 acar|OOF93642.1 acla|EAW08557.1 afla|EED55267.1 agla|OJJ86860.1 akaw|GAA92196.1 aluc|OJZ85611.1 anig|EHA24915.1 aory|EIT83620.1 ater|EAU35360.1 atub|OJI79404.1 awen|OJJ39924.1 azon|OJJ45896.1

Group7393: abra|OJJ66974.1 acar|OOF93639.1 acla|EAW12274.1 afis|EAW16448.1 afla|EED54345.1 agla|OJJ80869.1 akaw|GAA92191.1 aluc|OJZ85603.1 anig|EHA24921.1 aory|EIT75903.1 ater|EAU32187.1 atub|OJI79399.1 afla|EED52036.1

Group7394: abra|OJJ66993.1 acar|OOF93621.1 acla|EAW10840.1 afis|EAW24725.1 afla|EED45253.1 afum|EBA27376.1 akaw|GAA92171.1 aluc|OJZ85579.1 anid|CBF78503.1 anig|EHA24940.1 aory|EIT73778.1 atub|OJI79375.1 aver|OJI99472.1

Group7395: abra|OJJ67027.1 acar|OOF94678.1 afis|EAW17090.1 afum|EAL86568.1 akaw|GAA92147.1 aluc|OJZ85552.1 anig|EHA24961.1 asyd|OJJ53614.1 atub|OJI79353.1 aver|OJJ08323.1 awen|OJJ37029.1 akaw|GAA83299.1 asyd|OJJ53423.1

Group7396: abra|OJJ67068.1 acar|OOF90569.1 afis|EAW22303.1 agla|OJJ82727.1 anid|CBF85188.1 aory|EIT78616.1 atub|OJI83264.1 aver|OJJ05782.1 awen|OJJ31408.1 anig|EHA19961.1 akaw|GAA85826.1 aluc|OJZ91275.1 awen|OJJ39637.1

Group7397: afis|EAW16375.1 abra|OJJ69960.1 acar|OOF99006.1 akaw|GAA87699.1 aluc|OJZ85323.1 anig|EHA25513.1 atub|OJI80250.1 azon|OJJ45136.1 acla|EAW06824.1 afum|EAL88999.1 agla|OJJ78593.1 awen|OJJ37909.1 azon|OJJ45089.1

Group7398: abra|OJJ67134.1 acar|OOF95941.1 akaw|GAA91035.1 aluc|OJZ87845.1 atub|OJI80803.1 azon|OJJ46808.1 abra|OJJ68592.1 anid|CBF80418.1 asyd|OJJ59279.1 ater|EAU35716.1 aver|OJJ07407.1 anid|CBF69456.1 ater|EAU35440.1

Group7399: abra|OJJ67181.1 acar|OOF98381.1 acla|EAW14542.1 afis|EAW19260.1 afla|EED47615.1 afum|EAL93987.1 agla|OJJ88435.1 akaw|GAA90995.1 aluc|OJZ87794.1 anig|EHA20275.1 ater|EAU29515.1 atub|OJI80853.1 aver|OJI99359.1

Group7400: abra|OJJ67206.1 acar|OOF98355.1 afis|EAW19234.1 akaw|GAA90971.1 aluc|OJZ87768.1 anid|CBF78627.1 anig|EHA20252.1 asyd|OJJ65184.1 ater|EAU37229.1 atub|OJI80877.1 aver|OJI99337.1 awen|OJJ37591.1 acla|EAW14516.1

Group7401: abra|OJJ67248.1 acar|OOG00373.1 acla|EAW14979.1 afum|EAL92019.1 aluc|OJZ87729.1 anig|EHA20221.1 asyd|OJJ61733.1 ater|EAU34169.1 atub|OJI80911.1 aver|OJI95827.1 awen|OJJ29626.1 azon|OJJ45147.1 acla|EAW13306.1

Group7402: abra|OJJ67260.1 afis|EAW21086.1 afum|EAL92109.1 agla|OJJ79283.1 akaw|GAA91675.1 aluc|OJZ87719.1 anig|EHA20213.1 aory|EIT73486.1 asyd|OJJ61438.1 ater|EAU31934.1 atub|OJI80923.1 aver|OJJ00772.1 awen|OJJ35787.1

Group7403: abra|OJJ67344.1 acar|OOF92470.1 acla|EAW12129.1 afis|EAW17250.1 afum|EAL86669.1 akaw|GAA86473.1 aluc|OJZ85014.1 anid|CBF76006.1 anig|EHA25235.1 asyd|OJJ60262.1 ater|EAU36012.1 atub|OJI80583.1 aver|OJJ03991.1

Group7404: abra|OJJ67356.1 acar|OOF92465.1 afla|EED53299.1 agla|OJJ87954.1 akaw|GAA86465.1 aluc|OJZ85006.1 anid|CBF70183.1 anig|EHA25224.1 aory|EIT74866.1 asyd|OJJ55012.1 atub|OJI80592.1 aver|OJJ07200.1 awen|OJJ33477.1

Group7405: abra|OJJ67425.1 acar|OOF89875.1 acla|EAW06339.1 afis|EAW18451.1 afla|EED52957.1 afum|EAL93136.1 agla|OJJ85029.1 aluc|OJZ84942.1 anig|EHA25159.1 asyd|OJJ58851.1 atub|OJI80658.1 aver|OJJ01452.1 awen|OJJ31003.1

Group7406: azon|OJJ50368.1 afis|EAW24609.1 afla|EED56494.1 afum|EAL86239.1 akaw|GAA90756.1 anid|CBF81501.1 anig|EHA25067.1 ater|EAU39132.1 atub|OJI89487.1 akaw|GAA89017.1 aluc|OJZ84591.1 asyd|OJJ56397.1 aver|OJJ06316.1

Group7407: abra|OJJ67577.1 akaw|GAA89106.1 atub|OJI88445.1 abra|OJJ73994.1 acar|OOF96468.1 acla|EAW07117.1 akaw|GAA87477.1 aluc|OJZ88050.1 anig|EHA17822.1 asyd|OJJ59175.1 atub|OJI90658.1 awen|OJJ38785.1 azon|OJJ45128.1

Group7408: abra|OJJ67621.1 acar|OOF91969.1 afla|EED47432.1 akaw|GAA89141.1 aluc|OJZ81695.1 anid|CBF82186.1 anig|EHA25736.1 aory|EIT81905.1 asyd|OJJ52189.1 ater|EAU29296.1 atub|OJI88485.1 aver|OJJ05957.1 awen|OJJ36974.1

Group7409: abra|OJJ67623.1 acar|OOF98678.1 afla|EED47424.1 akaw|GAA89143.1 aluc|OJZ81693.1 anid|CBF82192.1 anig|EHA25738.1 aory|EIT81894.1 asyd|OJJ52187.1 ater|EAU29298.1 atub|OJI88487.1 aver|OJJ05958.1 awen|OJJ36972.1

Group7410: abra|OJJ67649.1 acar|OOF91984.1 afla|EED52575.1 agla|OJJ84703.1 akaw|GAA89165.1 aluc|OJZ81668.1 anig|EHA25775.1 aory|EIT74170.1 asyd|OJJ53747.1 atub|OJI88515.1 aver|OJJ07862.1 awen|OJJ39959.1 agla|OJJ84647.1

Group7411: abra|OJJ67653.1 acar|OOF91985.1 afla|EED57547.1 agla|OJJ83873.1 akaw|GAA89169.1 aluc|OJZ81664.1 anid|CBF80430.1 anig|EHA25779.1 asyd|OJJ52823.1 ater|EAU29530.1 atub|OJI88519.1 aver|OJJ08799.1 aver|OJJ07624.1

Group7412: abra|OJJ67794.1 acar|OOF95393.1 afis|EAW17791.1 afla|EED47240.1 afum|EAL91600.1 agla|OJJ84168.1 anid|CBF75217.1 asyd|OJJ53463.1 aver|OJJ08292.1 awen|OJJ32923.1 anig|EHA18418.1 aory|EIT78082.1 ater|EAU29258.1

Group7413: abra|OJJ67849.1 acar|OOF92830.1 acla|EAW11750.1 afis|EAW23783.1 afla|EED51592.1 afum|EAL87217.2 agla|OJJ78935.1 aluc|OJZ82042.1 asyd|OJJ58151.1 atub|OJI84601.1 awen|OJJ33728.1 azon|OJJ42854.1 aver|OJJ04253.1

Group7414: abra|OJJ67884.1 acar|OOF92862.1 acla|EAW11691.1 afis|EAW23707.1 afla|EED51510.1 afum|EAL87151.1 akaw|GAA91891.1 aluc|OJZ82005.1 anig|EHA22251.1 aory|EIT80627.1 ater|EAU30420.1 atub|OJI84555.1 awen|OJJ33668.1

Group7415: abra|OJJ67887.1 acar|OOF92860.1 acla|EAW11714.1 afis|EAW23733.1 afla|EED51511.1 afum|EAL87171.1 akaw|GAA91893.1 aluc|OJZ82003.1 anig|EHA22253.1 aory|EIT80635.1 ater|EAU31293.1 atub|OJI84553.1 awen|OJJ33669.1

Group7416: abra|OJJ67898.1 acar|OOF90066.1 afis|EAW22155.1 afla|EED52135.1 afum|EAL91035.1 agla|OJJ88704.1 akaw|GAA91902.1 aluc|OJZ81992.1 anig|EHA22264.1 aory|EIT74401.1 ater|EAU38340.1 aver|OJJ03874.1 awen|OJJ29943.1

Group7417: abra|OJJ67900.1 acar|OOF90068.1 afla|EED47044.1 afum|EAL84242.1 agla|OJJ88179.1 akaw|GAA91904.1 aluc|OJZ81990.1 anid|CBF80368.1 anig|EHA22266.1 aory|EIT72634.1 asyd|OJJ54815.1 atub|OJI84541.1 aver|OJJ07346.1

Group7418: abra|OJJ67910.1 acar|OOF98178.1 afla|EED46477.1 akaw|GAA91913.1 aluc|OJZ81979.1 anid|CBF90240.1 anig|EHA22275.1 aory|EIT72386.1 asyd|OJJ61956.1 atub|OJI84531.1 aver|OJI96048.1 awen|OJJ35928.1 azon|OJJ42126.1

Group7419: abra|OJJ67930.1 acar|OOF92780.1 akaw|GAA91267.1 aluc|OJZ82802.1 anig|EHA21243.1 aory|EIT73390.1 asyd|OJJ61413.1 ater|EAU30954.1 atub|OJI89033.1 aver|OJJ00743.1 awen|OJJ33794.1 azon|OJJ48748.1 afla|EED56214.1

Group7420: abra|OJJ67945.1 acar|OOF92794.1 acla|EAW11635.1 afis|EAW23822.1 afla|EED51984.1 agla|OJJ80056.1 akaw|GAA91251.1 aluc|OJZ82817.1 anig|EHA21257.1 aory|EIT81424.1 ater|EAU31377.1 atub|OJI89048.1 azon|OJJ42246.1

Group7421: abra|OJJ67985.1 acar|OOF92027.1 afis|EAW25693.1 afla|EED46904.1 afum|EAL84363.1 akaw|GAA91217.1 aluc|OJZ82853.1 anig|EHA21289.1 aory|EIT72721.1 asyd|OJJ55668.1 ater|EAU35089.1 atub|OJI89083.1 aver|OJJ06749.1

Group7422: abra|OJJ68000.1 acar|OOF92043.1 akaw|GAA90950.1 aluc|OJZ82874.1 anid|CBF81948.1 anig|EHA21300.1 asyd|OJJ57368.1 atub|OJI89098.1 aver|OJJ05800.1 awen|OJJ40803.1 acar|OOF90715.1 afla|EED45646.1 aory|EIT75683.1

Group7423: abra|OJJ68011.1 acar|OOF92053.1 acla|EAW13750.1 afis|EAW19813.1 afla|EED47561.1 afum|EAL85078.1 akaw|GAA90939.1 aluc|OJZ82885.1 anig|EHA21311.1 aory|EIT79581.1 atub|OJI89110.1 awen|OJJ35921.1 azon|OJJ50657.1

Group7424: abra|OJJ68035.1 acar|OOF91808.1 afla|EED55048.1 afum|EAL86258.1 akaw|GAA90917.1 aluc|OJZ82909.1 anig|EHA21332.1 aory|EIT81969.1 asyd|OJJ53717.1 atub|OJI89135.1 aver|OJJ07843.1 awen|OJJ33894.1 azon|OJJ42784.1

Group7425: abra|OJJ68063.1 acar|OOF92084.1 acla|EAW13646.1 afis|EAW19959.1 afla|EED45907.1 afum|EAL84956.1 agla|OJJ79326.1 akaw|GAA90895.1 aluc|OJZ82934.1 anig|EHA21354.1 atub|OJI89163.1 awen|OJJ29990.1 azon|OJJ43625.1

Group7426: abra|OJJ68085.1 acar|OOF93883.1 acla|EAW09009.1 afis|EAW25588.1 afla|EED48864.1 afum|EAL84295.2 akaw|GAA86870.1 aluc|OJZ89994.1 aory|EIT81170.1 asyd|OJJ61551.1 atub|OJI84814.1 aver|OJJ00876.1 anid|CBF82996.1

Group7427: abra|OJJ68090.1 acar|OOF93894.1 agla|OJJ81061.1 akaw|GAA86875.1 aluc|OJZ89999.1 anid|CBF85662.1 anig|EHA27374.1 ater|EAU29581.1 atub|OJI84809.1 awen|OJJ29840.1 agla|OJJ88676.1 asyd|OJJ64243.1 aver|OJI98335.1

Group7428: abra|OJJ68094.1 acar|OOF93897.1 afla|EED45939.1 akaw|GAA86878.1 aluc|OJZ90002.1 anid|CBF79119.1 anig|EHA27377.1 aory|EIT75636.1 atub|OJI84806.1 aver|OJI96964.1 anid|CBF73549.1 asyd|OJJ63990.1 atub|OJI85466.1

Group7429: abra|OJJ68112.1 afis|EAW19856.1 afla|EED57112.1 afum|EAL85039.1 akaw|GAA86894.1 aluc|OJZ90017.1 anid|CBF78829.1 anig|EHA27393.1 aory|EIT83382.1 asyd|OJJ54898.1 atub|OJI84790.1 aver|OJJ03939.1 aory|EIT73989.1

Group7430: abra|OJJ68148.1 acar|OOF93943.1 afis|EAW22414.1 afla|EED54477.1 afum|EAL90778.1 aluc|OJZ83583.1 anid|CBF88923.1 aory|EIT81162.1 asyd|OJJ62847.1 ater|EAU39251.1 atub|OJI89957.1 aver|OJI96891.1 anig|EHA27993.1

Group7431: afum|EAL91395.1 acar|OOF90118.1 acla|EAW13082.1 afis|EAW17582.1 agla|OJJ81371.1 akaw|GAA83246.1 anid|CBF90231.1 anig|EHA18894.1 aory|EIT82200.1 asyd|OJJ61960.1 ater|EAU37017.1 aver|OJI96052.1 awen|OJJ37266.1

Group7432: abra|OJJ68206.1 afis|EAW17573.1 afla|EED49248.1 agla|OJJ81385.1 akaw|GAA83240.1 aluc|OJZ83739.1 anig|EHA18888.1 asyd|OJJ61947.1 atub|OJI90117.1 aver|OJI96039.1 awen|OJJ37274.1 azon|OJJ50974.1 acar|OOF97492.1

Group7433: abra|OJJ68258.1 acar|OOF97535.1 acla|EAW09080.1 akaw|GAA83204.1 aluc|OJZ83706.1 anig|EHA18857.1 ater|EAU29580.1 atub|OJI90078.1 awen|OJJ37962.1 azon|OJJ51745.1 afis|EAW16406.1 afla|EED45131.1 aory|EIT73565.1

Group7434: abra|OJJ68316.1 acar|OOF97576.1 afla|EED55061.1 akaw|GAA83157.1 aluc|OJZ83657.1 anid|CBF78294.1 anig|EHA18809.1 aory|EIT81958.1 asyd|OJJ56107.1 ater|EAU32200.1 atub|OJI90027.1 aver|OJJ05050.1 azon|OJJ46804.1

Group7435: abra|OJJ68342.1 acar|OOF97640.1 afla|EED46510.1 agla|OJJ78917.1 akaw|GAA92109.1 aluc|OJZ83616.1 anid|CBF83851.1 anig|EHA18773.1 aory|EIT72415.1 asyd|OJJ60921.1 atub|OJI89989.1 aver|OJJ00247.1 awen|OJJ33611.1

Group7436: abra|OJJ68344.1 acar|OOF97642.1 afla|EED45993.1 akaw|GAA92111.1 aluc|OJZ83614.1 anid|CBF88055.1 anig|EHA18771.1 aory|EIT75448.1 ater|EAU31718.1 atub|OJI89987.1 awen|OJJ40161.1 azon|OJJ47958.1 awen|OJJ30656.1

Group7437: aory|EIT75783.1 acar|OOF90738.1 acla|EAW09089.1 afis|EAW25681.1 afla|EED45592.1 afum|EAL84351.1 agla|OJJ85285.1 anid|CBF82743.1 asyd|OJJ61619.1 atub|OJI91328.1 aver|OJJ00943.1 awen|OJJ35900.1 azon|OJJ42101.1

Group7438: abra|OJJ68438.1 acar|OOF93956.1 afla|EED52097.1 akaw|GAA86934.1 aluc|OJZ90059.1 anid|CBF82381.1 anig|EHA21012.1 aory|EIT76912.1 asyd|OJJ54137.1 ater|EAU30749.1 atub|OJI84748.1 aver|OJJ08599.1 awen|OJJ29692.1

Group7439: abra|OJJ68442.1 acar|OOF93952.1 acla|EAW13760.1 afis|EAW19804.1 afla|EED45407.1 afum|EAL85089.1 agla|OJJ83221.1 akaw|GAA86930.1 aluc|OJZ90055.1 anig|EHA21016.1 aory|EIT82409.1 ater|EAU30745.1 atub|OJI84752.1

Group7440: abra|OJJ68448.1 acla|EAW13158.1 afis|EAW17488.1 afla|EED55784.1 agla|OJJ81468.1 akaw|GAA86926.1 aluc|OJZ90050.1 anig|EHA21020.1 awen|OJJ37354.1 azon|OJJ51039.1 anid|CBF71615.1 asyd|OJJ61863.1 aver|OJI95961.1

Group7441: abra|OJJ68603.1 acla|EAW12589.1 afis|EAW18088.1 afum|EAL91900.1 akaw|GAA92917.1 anid|CBF79766.1 anig|EHA22296.1 aory|EIT80666.1 asyd|OJJ58104.1 ater|EAU30434.1 atub|OJI84504.1 aver|OJJ04300.1 azon|OJJ48215.1

Group7442: abra|OJJ68646.1 acar|OOF92148.1 afla|EED51431.1 agla|OJJ87324.1 akaw|GAA89786.1 aluc|OJZ81905.1 anid|CBF86428.1 anig|EHA22338.1 asyd|OJJ64935.1 ater|EAU30469.1 atub|OJI84458.1 aver|OJI99056.1 awen|OJJ41448.1

Group7443: abra|OJJ68791.1 acar|OOF92275.1 acla|EAW12462.1 afis|EAW24179.1 agla|OJJ89495.1 aluc|OJZ81737.1 anid|CBF77847.1 anig|EHA22476.1 asyd|OJJ55330.1 atub|OJI84307.1 aver|OJJ06898.1 awen|OJJ41290.1 azon|OJJ48764.1

Group7444: abra|OJJ68805.1 afis|EAW17767.1 afla|EED44891.1 afum|EAL91576.1 akaw|GAA90448.1 aluc|OJZ85777.1 anid|CBF74647.1 anig|EHA24777.1 aory|EIT83241.1 asyd|OJJ54533.1 atub|OJI79562.1 aver|OJJ07655.1 awen|OJJ30892.1

Group7445: abra|OJJ68850.1 acar|OOF90538.1 acla|EAW15261.1 agla|OJJ87632.1 aluc|OJZ85830.1 anid|CBF74588.1 asyd|OJJ54708.1 atub|OJI79618.1 aver|OJJ07458.1 awen|OJJ31236.1 azon|OJJ46158.1 acar|OOF90618.1 agla|OJJ79329.1

Group7446: abra|OJJ68893.1 afis|EAW20640.1 afla|EED47502.1 agla|OJJ82459.1 akaw|GAA90358.1 aluc|OJZ85874.1 anid|CBF82442.1 anig|EHA24714.1 asyd|OJJ54122.1 atub|OJI79660.1 aver|OJJ08615.1 awen|OJJ37240.1 anig|EHA18313.1

Group7447: abra|OJJ68894.1 afla|EED56749.1 akaw|GAA90356.1 aluc|OJZ85876.1 anid|CBF73545.1 anig|EHA24711.1 aory|EIT73862.1 asyd|OJJ56710.1 ater|EAU34583.1 atub|OJI79662.1 aver|OJJ06015.1 awen|OJJ34055.1 azon|OJJ48177.1

Group7448: abra|OJJ68895.1 afla|EED56748.1 akaw|GAA90355.1 aluc|OJZ85877.1 anid|CBF73543.1 anig|EHA24710.1 aory|EIT73900.1 asyd|OJJ56709.1 ater|EAU34584.1 atub|OJI79663.1 aver|OJJ06016.1 awen|OJJ34054.1 azon|OJJ48178.1

Group7449: abra|OJJ69060.1 acar|OOF91223.1 afla|EED46858.1 akaw|GAA90590.1 aluc|OJZ86048.1 anid|CBF71167.1 anig|EHA24563.1 aory|EIT72620.1 asyd|OJJ55847.1 ater|EAU32488.1 atub|OJI79845.1 aver|OJJ04790.1 awen|OJJ38349.1

Group7450: abra|OJJ69077.1 acar|OOF91214.1 afis|EAW15679.1 afum|EAL85669.1 akaw|GAA90606.1 aluc|OJZ86065.1 anig|EHA24549.1 asyd|OJJ59219.1 ater|EAU32483.1 atub|OJI79861.1 aver|OJJ01829.1 awen|OJJ38356.1 azon|OJJ48602.1

Group7451: abra|OJJ69170.1 acar|OOF91134.1 afla|EED55555.1 akaw|GAA86293.1 aluc|OJZ86157.1 anig|EHA24467.1 aory|EIT82688.1 asyd|OJJ61354.1 ater|EAU32678.1 atub|OJI79949.1 aver|OJJ00681.1 awen|OJJ38094.1 azon|OJJ42538.1

Group7452: abra|OJJ69172.1 acar|OOF91132.1 afla|EED55557.1 akaw|GAA86291.1 aluc|OJZ86159.1 anig|EHA24465.1 aory|EIT82661.1 asyd|OJJ61356.1 ater|EAU32676.1 atub|OJI79951.1 aver|OJJ00683.1 awen|OJJ38096.1 azon|OJJ42540.1

Group7453: abra|OJJ69265.1 acar|OOF91058.1 acla|EAW13565.1 afla|EED51202.1 agla|OJJ84910.1 aluc|OJZ86243.1 anid|CBF70831.1 aory|EIT80408.1 asyd|OJJ52562.1 atub|OJI80035.1 aver|OJJ04590.1 awen|OJJ38563.1 azon|OJJ48767.1

Group7454: aory|EIT82901.1 acar|OOF94369.1 acla|EAW14746.1 afis|EAW24277.1 afla|EED47999.1 afum|EAL85946.1 akaw|GAA88225.1 anid|CBF74155.1 ater|EAU31227.1 atub|OJI88248.1 aver|OJJ08023.1 awen|OJJ33020.1 azon|OJJ48497.1

Group7455: abra|OJJ69382.1 akaw|GAA88250.1 asyd|OJJ53900.1 awen|OJJ32992.1 azon|OJJ48524.1 acla|EAW14769.1 afis|EAW24300.1 afum|EAL85968.1 aver|OJJ07997.1 ater|EAU31251.1 afla|EED47972.1 agla|OJJ83168.1 anid|CBF74106.1

Group7456: abra|OJJ69391.1 acar|OOF94336.1 afla|EED47867.1 akaw|GAA88259.1 aluc|OJZ82228.1 anig|EHA28511.1 aory|EIT77240.1 ater|EAU31260.1 atub|OJI88209.1 aver|OJJ07919.1 awen|OJJ36230.1 asyd|OJJ53819.1 awen|OJJ34075.1

Group7457: abra|OJJ69401.1 acla|EAW14783.1 afis|EAW24314.1 afla|EED47953.1 afum|EAL85982.1 agla|OJJ83182.1 aluc|OJZ82237.1 anig|EHA28504.1 asyd|OJJ53879.1 atub|OJI88201.1 aver|OJJ07977.1 awen|OJJ32978.1 azon|OJJ48939.1

Group7458: abra|OJJ69465.1 acar|OOF94277.1 acla|EAW07122.1 afis|EAW22132.1 afum|EAL91058.1 akaw|GAA88348.1 aluc|OJZ82308.1 anig|EHA28449.1 ater|EAU34982.1 atub|OJI88131.1 afla|EED51690.1 aory|EIT80487.1 awen|OJJ32637.1

Group7459: abra|OJJ69491.1 acar|OOF94250.1 akaw|GAA88329.1 aluc|OJZ82331.1 anig|EHA28429.1 atub|OJI88109.1 abra|OJJ71732.1 afis|EAW20342.1 afum|EAL85462.1 asyd|OJJ53414.1 aver|OJJ08337.1 awen|OJJ38124.1 azon|OJJ43050.1

Group7460: abra|OJJ69499.1 acar|OOF94258.1 acla|EAW09953.1 afis|EAW21143.1 afla|EED54671.1 afum|EAL92162.2 agla|OJJ80073.1 akaw|GAA88322.1 aluc|OJZ82339.1 anig|EHA28421.1 aory|EIT78899.1 atub|OJI88101.1 awen|OJJ36472.1

Group7461: abra|OJJ69515.1 acar|OOF95345.1 afla|EED56902.1 akaw|GAA88309.1 aluc|OJZ82353.1 anig|EHA28407.1 aory|EIT77545.1 asyd|OJJ60238.1 ater|EAU38102.1 atub|OJI88087.1 aver|OJJ04027.1 azon|OJJ51783.1 acar|OOG00884.1

Group7462: acla|EAW11726.1 abra|OJJ74800.1 afis|EAW23751.1 afla|EED50581.1 afum|EAL87188.1 akaw|GAA83613.1 aluc|OJZ92381.1 anid|CBF85466.1 asyd|OJJ64103.1 atub|OJI86810.1 aver|OJI98186.1 agla|OJJ87545.1 azon|OJJ47330.1

Group7463: abra|OJJ69609.1 acar|OOF92698.1 acla|EAW11684.1 agla|OJJ84464.1 akaw|GAA91594.1 aluc|OJZ82417.1 anig|EHA28351.1 aory|EIT81722.1 asyd|OJJ60324.1 ater|EAU31564.1 atub|OJI88024.1 aver|OJI99672.1 awen|OJJ33649.1

Group7464: abra|OJJ69612.1 acar|OOF92694.1 afis|EAW20306.1 afla|EED56731.1 akaw|GAA91596.1 aluc|OJZ82420.1 anid|CBF71186.1 anig|EHA28349.1 aory|EIT73861.1 asyd|OJJ55859.1 atub|OJI88021.1 aver|OJJ04800.1 azon|OJJ45630.1

Group7465: abra|OJJ69680.1 acar|OOG00315.1 acla|EAW14998.1 afla|EED44880.1 afum|EAL86187.1 agla|OJJ85918.1 akaw|GAA88805.1 aluc|OJZ82488.1 anig|EHA28286.1 ater|EAU39594.1 atub|OJI87944.1 afis|EAW24522.1 asyd|OJJ56165.1

Group7466: abra|OJJ69733.1 acar|OOF94188.1 acla|EAW11686.1 afla|EED48937.1 akaw|GAA90186.1 aluc|OJZ83163.1 anig|EHA28227.1 aory|EIT80012.1 asyd|OJJ65340.1 ater|EAU39367.1 atub|OJI87774.1 aver|OJI99484.1 azon|OJJ47813.1

Group7467: abra|OJJ69751.1 acar|OOF97950.1 afis|EAW20379.1 agla|OJJ82717.1 akaw|GAA90202.1 aluc|OJZ83146.1 anig|EHA28210.1 asyd|OJJ56934.1 atub|OJI87791.1 aver|OJJ03350.1 awen|OJJ31442.1 azon|OJJ43563.1 awen|OJJ36135.1

Group7468: abra|OJJ69768.1 acar|OOF90889.1 afla|EED45806.1 agla|OJJ78514.1 akaw|GAA87885.1 aluc|OJZ85530.1 anig|EHA25061.1 aory|EIT75772.1 asyd|OJJ60227.1 atub|OJI80038.1 aver|OJJ04040.1 awen|OJJ33640.1 azon|OJJ47399.1

Group7469: abra|OJJ69831.1 acar|OOF93356.1 acla|EAW12369.1 agla|OJJ79196.1 akaw|GAA87829.1 aluc|OJZ85465.1 anig|EHA25006.1 asyd|OJJ57216.1 ater|EAU31943.1 atub|OJI80098.1 aver|OJI98298.1 awen|OJJ32853.1 anid|CBF77164.1

Group7470: abra|OJJ69894.1 acar|OOG00773.1 afla|EED54570.1 agla|OJJ83056.1 anid|CBF71733.1 aory|EIT78739.1 asyd|OJJ61797.1 ater|EAU39169.1 aver|OJI95896.1 awen|OJJ41668.1 azon|OJJ43876.1 acla|EAW08580.1 aory|EIT78892.1

Group7471: abra|OJJ69917.1 acar|OOF93429.1 acla|EAW08909.1 afis|EAW15487.1 afla|EED49001.1 afum|EAL85836.1 akaw|GAA87739.1 aluc|OJZ85371.1 anig|EHA25555.1 asyd|OJJ52102.1 ater|EAU32634.1 atub|OJI80199.1 aver|OJI97592.1

Group7472: abra|OJJ69981.1 acar|OOF93485.1 acla|EAW08851.1 afis|EAW15545.1 afla|EED49052.1 afum|EBA27242.1 aluc|OJZ85299.1 anid|CBF78180.1 asyd|OJJ55972.1 atub|OJI80271.1 aver|OJJ04919.1 awen|OJJ38206.1 azon|OJJ43089.1

Group7473: abra|OJJ70009.1 acar|OOF93515.1 afis|EAW25679.1 afla|EED46059.1 akaw|GAA87653.1 aluc|OJZ85270.1 anig|EHA25469.1 aory|EIT83484.1 ater|EAU37660.1 atub|OJI80301.1 awen|OJJ39964.1 afum|EAL84349.1 aver|OJI99713.1

Group7474: abra|OJJ70021.1 acar|OOF93527.1 acla|EAW08797.1 afis|EAW15600.1 afla|EED50424.1 agla|OJJ81556.1 akaw|GAA87640.1 aluc|OJZ85255.1 anig|EHA25456.1 aory|EIT82104.1 atub|OJI80313.1 awen|OJJ38277.1 azon|OJJ43047.1

Group7475: abra|OJJ70114.1 afla|EED55193.1 aory|EIT83589.1 asyd|OJJ58806.1 aver|OJJ05475.1 azon|OJJ45163.1 abra|OJJ73453.1 akaw|GAA83044.1 aluc|OJZ88589.1 anig|EHA18284.1 atub|OJI91175.1 aver|OJJ01409.1 awen|OJJ37581.1

Group7476: abra|OJJ70158.1 acar|OOF93318.1 afla|EED45634.1 afum|EAL91080.1 akaw|GAA86605.1 aluc|OJZ85155.1 anid|CBF80294.1 anig|EHA25371.1 aory|EIT75694.1 asyd|OJJ60439.1 atub|OJI80432.1 aver|OJJ02283.1 awen|OJJ29853.1

Group7477: abra|OJJ70179.1 acla|EAW13224.1 afis|EAW16396.1 afla|EED47676.1 agla|OJJ86670.1 akaw|GAA86593.1 aluc|OJZ85140.1 anig|EHA25356.1 aory|EIT80220.1 asyd|OJJ54662.1 ater|EAU38225.1 atub|OJI80452.1 aver|OJJ03122.1

Group7478: abra|OJJ70233.1 acar|OOF93309.1 afla|EED54310.1 agla|OJJ84751.1 akaw|GAA86551.1 aluc|OJZ85099.1 anig|EHA25321.1 aory|EIT76032.1 asyd|OJJ56224.1 ater|EAU35999.1 atub|OJI80499.1 aver|OJJ05156.1 azon|OJJ45165.1

Group7479: abra|OJJ70244.1 acar|OOF92521.1 akaw|GAA86543.1 aluc|OJZ85092.1 anid|CBF79134.1 anig|EHA25312.1 asyd|OJJ65410.1 atub|OJI80507.1 aver|OJJ07739.1 azon|OJJ45127.1 aver|OJJ01719.1 awen|OJJ40032.1 asyd|OJJ61775.1

Group7480: afum|EAL86040.1 acla|EAW14841.1 afis|EAW24368.1 afla|EED51156.1 anid|CBF82288.1 aory|EIT81291.1 ater|EAU32941.1 aver|OJJ08259.1 azon|OJJ45963.1 afla|EED45148.1 aory|EIT73814.1 afla|EED45171.1 ater|EAU37757.1

Group7481: abra|OJJ70269.1 acla|EAW06604.1 afis|EAW25674.1 afla|EED56544.1 afum|EAL88847.1 agla|OJJ86231.1 anid|CBF82350.1 aory|EIT79694.1 asyd|OJJ52249.1 ater|EAU34095.1 awen|OJJ30253.1 azon|OJJ43959.1 afis|EAW16219.1

Group7482: abra|OJJ70274.1 acar|OOF98672.1 agla|OJJ82824.1 akaw|GAA92875.1 aluc|OJZ82536.1 anid|CBF78791.1 anig|EHA22504.1 aory|EIT79189.1 asyd|OJJ55020.1 atub|OJI88780.1 aver|OJJ07185.1 awen|OJJ32615.1 afla|EED47288.1

Group7483: abra|OJJ70294.1 acar|OOF91748.1 afis|EAW22072.1 afla|EED56698.1 afum|EAL91098.2 akaw|GAA93025.1 aluc|OJZ82559.1 anid|CBF82782.1 aory|EIT73867.1 asyd|OJJ52973.1 ater|EAU30150.1 atub|OJI88801.1 awen|OJJ35471.1

Group7484: abra|OJJ70296.1 acar|OOF91750.1 acla|EAW06429.1 afis|EAW18534.1 afla|EED52865.1 afum|EAL93232.1 agla|OJJ81549.1 akaw|GAA93024.1 aluc|OJZ82561.1 anig|EHA22522.1 atub|OJI88803.1 awen|OJJ31092.1 azon|OJJ50601.1

Group7485: abra|OJJ70339.1 acar|OOF92847.1 afla|EED48270.1 agla|OJJ88108.1 akaw|GAA92592.1 aluc|OJZ82609.1 anid|CBF89179.1 anig|EHA27473.1 aory|EIT72938.1 asyd|OJJ65382.1 atub|OJI88844.1 aver|OJI99552.1 awen|OJJ37855.1

Group7486: awen|OJJ33342.1 abra|OJJ77817.1 acar|OOF98951.1 acla|EAW10282.1 afis|EAW25312.1 agla|OJJ78832.1 akaw|GAA83829.1 aluc|OJZ90889.1 anig|EHA19608.1 atub|OJI82883.1 awen|OJJ33341.1 afum|EAL89973.1 agla|OJJ78833.1

Group7487: abra|OJJ70424.1 acar|OOF91770.1 afla|EED46005.1 agla|OJJ86984.1 akaw|GAA90153.1 aluc|OJZ83194.1 anig|EHA21058.1 asyd|OJJ62596.1 ater|EAU35267.1 atub|OJI87744.1 aver|OJI96741.1 awen|OJJ29999.1 afla|EED46004.1

Group7488: abra|OJJ70430.1 afis|EAW16965.1 afum|EAL86492.1 akaw|GAA90150.1 aluc|OJZ83198.1 anig|EHA21061.1 atub|OJI87740.1 aver|OJJ05858.1 abra|OJJ76669.1 afla|EED53956.1 agla|OJJ80055.1 aory|EIT78323.1 acar|OOF95783.1

Group7489: abra|OJJ70548.1 acar|OOF95010.1 afis|EAW22097.1 afum|EAL89915.1 akaw|GAA83392.1 aluc|OJZ83892.1 anig|EHA19028.1 asyd|OJJ61427.1 ater|EAU29365.1 atub|OJI90259.1 aver|OJJ00763.1 azon|OJJ45988.1 afis|EAW16226.1

Group7490: abra|OJJ70623.1 acar|OOF97100.1 afis|EAW17661.1 afum|EBA27435.1 agla|OJJ79530.1 akaw|GAA89961.1 aluc|OJZ83968.1 anig|EHA19097.1 asyd|OJJ62051.1 atub|OJI90341.1 aver|OJI96156.1 awen|OJJ37181.1 azon|OJJ50883.1

Group7491: abra|OJJ70634.1 acar|OOF97090.1 afis|EAW23158.1 afla|EED46962.1 afum|EAL88325.1 akaw|GAA89973.1 aluc|OJZ83982.1 anig|EHA19109.1 aory|EIT72715.1 asyd|OJJ56067.1 atub|OJI90354.1 aver|OJJ05010.1 awen|OJJ36625.1

Group7492: abra|OJJ70643.1 acar|OOF97077.1 acla|EAW08116.1 afis|EAW23135.1 afla|EED48498.1 afum|EAL88346.1 akaw|GAA89980.1 aluc|OJZ83994.1 anid|CBF74453.1 anig|EHA19118.1 asyd|OJJ57675.1 atub|OJI90363.1 awen|OJJ36631.1

Group7493: abra|OJJ70925.1 acar|OOF98199.1 acla|EAW12449.1 afis|EAW21019.1 afum|EAL92057.1 akaw|GAA83015.1 aluc|OJZ87601.1 anig|EHA20800.1 aory|EIT74202.1 atub|OJI81047.1 asyd|OJJ55695.1 aver|OJJ06780.1 afla|EED52543.1

Group7494: abra|OJJ70969.1 acar|OOF98161.1 acla|EAW10529.1 afis|EAW24996.1 afla|EED55416.1 afum|EAL89705.1 agla|OJJ89156.1 akaw|GAA82979.1 aluc|OJZ87563.1 anig|EHA20765.1 aory|EIT80769.1 atub|OJI81084.1 awen|OJJ40643.1

Group7495: abra|OJJ71005.1 afis|EAW19704.1 afla|EED48154.1 afum|EAL86822.1 akaw|GAA92515.1 aluc|OJZ91528.1 anig|EHA22565.1 aory|EIT76190.1 asyd|OJJ61199.1 ater|EAU32845.1 atub|OJI85982.1 aver|OJJ00524.1 awen|OJJ30442.1

Group7496: ater|EAU29316.1 abra|OJJ73238.1 acar|OOG00475.1 afla|EED56142.1 akaw|GAA87943.1 aluc|OJZ81190.1 anid|CBF75531.1 anig|EHA18506.1 aory|EIT76793.1 atub|OJI91386.1 aver|OJJ05729.1 awen|OJJ40294.1 azon|OJJ49156.1

Group7497: abra|OJJ71155.1 acar|OOF98003.1 acla|EAW10343.1 afis|EAW25246.1 afla|EED53709.1 afum|EAL89908.1 agla|OJJ89213.1 akaw|GAA82823.1 aluc|OJZ87384.1 aory|EIT75300.1 ater|EAU34357.1 awen|OJJ40960.1 azon|OJJ44530.1

Group7498: abra|OJJ71229.1 acar|OOF97942.1 afla|EED46060.1 agla|OJJ82198.1 akaw|GAA82759.1 aluc|OJZ87318.1 anig|EHA20552.1 aory|EIT83482.1 atub|OJI81327.1 awen|OJJ36263.1 azon|OJJ46516.1 anig|EHA19259.1 atub|OJI82512.1

Group7499: abra|OJJ71230.1 acar|OOF97943.1 afla|EED56275.1 agla|OJJ81775.1 akaw|GAA82758.1 aluc|OJZ87317.1 anig|EHA20551.1 aory|EIT73346.1 asyd|OJJ62583.1 ater|EAU33790.1 atub|OJI81328.1 aver|OJI96702.1 awen|OJJ31637.1

Group7500: abra|OJJ71236.1 acar|OOF97947.1 afla|EED56271.1 agla|OJJ81771.1 akaw|GAA82752.1 aluc|OJZ87309.1 anig|EHA20545.1 aory|EIT73402.1 asyd|OJJ56498.1 ater|EAU33787.1 atub|OJI81336.1 aver|OJJ06221.1 awen|OJJ31643.1

Group7501: abra|OJJ71281.1 acla|EAW10264.1 afum|EAL89991.1 agla|OJJ83878.1 akaw|GAA82707.1 aluc|OJZ87262.1 anid|CBF75232.1 asyd|OJJ59915.1 atub|OJI81380.1 azon|OJJ44640.1 afla|EED53816.1 aory|EIT78417.1 aver|OJJ02371.1

Group7502: abra|OJJ71326.1 acar|OOF97848.1 acla|EAW10213.1 afis|EAW25380.1 afum|EBA27394.1 agla|OJJ83773.1 akaw|GAA82665.1 aluc|OJZ87217.1 anig|EHA20466.1 atub|OJI81423.1 awen|OJJ41107.1 azon|OJJ44684.1 afla|EED49572.1

Group7503: abra|OJJ71373.1 acar|OOF97795.1 acla|EAW10164.1 agla|OJJ87075.1 akaw|GAA82621.1 aluc|OJZ87167.1 anid|CBF77545.1 asyd|OJJ53149.1 atub|OJI81472.1 aver|OJJ03702.1 awen|OJJ41159.1 ater|EAU34601.1 azon|OJJ44737.1

Group7504: afla|EED55188.1 acla|EAW08982.1 afis|EAW15426.1 afum|EAL84218.1 agla|OJJ80139.1 anid|CBF75421.1 aory|EIT83577.1 asyd|OJJ59793.1 ater|EAU32703.1 aver|OJJ02494.1 awen|OJJ33347.1 aver|OJI98309.1 asyd|OJJ52408.1

Group7505: abra|OJJ71544.1 acla|EAW15098.1 afis|EAW20070.1 afum|EAL85188.1 agla|OJJ88890.1 akaw|GAA89378.1 aluc|OJZ86282.1 anig|EHA21589.1 atub|OJI84256.1 awen|OJJ31829.1 azon|OJJ46295.1 afla|EED45309.1 aory|EIT73730.1

Group7506: abra|OJJ71610.1 acla|EAW15154.1 afis|EAW20122.1 afla|EED57254.1 afum|EAL85240.1 aluc|OJZ86354.1 atub|OJI84183.1 azon|OJJ46245.1 agla|OJJ88827.1 anid|CBF84943.1 asyd|OJJ63150.1 aver|OJI97804.1 awen|OJJ31891.1

Group7507: abra|OJJ71702.1 afis|EAW20202.1 afla|EED57142.1 afum|EAL85322.1 akaw|GAA89233.1 aluc|OJZ86445.1 anid|CBF85087.1 anig|EHA21456.1 aory|EIT74074.1 asyd|OJJ63069.1 atub|OJI84083.1 aver|OJI97893.1 awen|OJJ31208.1

Group7508: abra|OJJ71740.1 acar|OOF96431.1 afla|EED56963.1 agla|OJJ80108.1 akaw|GAA85154.1 aluc|OJZ86483.1 anid|CBF78820.1 anig|EHA22176.1 aory|EIT77505.1 ater|EAU36025.1 atub|OJI84046.1 aver|OJI99417.1 azon|OJJ46020.1

Group7509: abra|OJJ71757.1 acar|OOF98108.1 afis|EAW17370.1 afla|EED53905.1 afum|EAL91207.2 agla|OJJ79799.1 aluc|OJZ84836.1 anig|EHA25746.1 aory|EIT78481.1 asyd|OJJ61750.1 ater|EAU32672.1 aver|OJI95847.1 awen|OJJ37892.1

Group7510: abra|OJJ71771.1 acar|OOF96444.1 afla|EED47495.1 akaw|GAA85136.1 aluc|OJZ86501.1 anid|CBF70338.1 anig|EHA22143.1 aory|EIT74589.1 asyd|OJJ58708.1 ater|EAU33072.1 atub|OJI84025.1 aver|OJJ01321.1 awen|OJJ31942.1

Group7511: abra|OJJ72082.1 afla|EED56585.1 agla|OJJ82914.1 akaw|GAA84838.1 aluc|OJZ86836.1 anid|CBF70326.1 aory|EIT79601.1 asyd|OJJ58715.1 ater|EAU30077.1 atub|OJI83682.1 aver|OJJ01328.1 awen|OJJ40591.1 azon|OJJ42312.1

Group7512: abra|OJJ72132.1 acar|OOF96770.1 afla|EED46872.1 afum|EAL90428.1 agla|OJJ86321.1 akaw|GAA84790.1 aluc|OJZ86890.1 anig|EHA21753.1 aory|EIT72539.1 asyd|OJJ53055.1 atub|OJI83629.1 aver|OJJ03789.1 awen|OJJ32567.1

Group7513: afis|EAW19362.1 afum|EAL94082.1 anid|CBF69511.1 asyd|OJJ59215.1 aver|OJJ01825.1 acla|EAW14624.1 afla|EED49733.1 aory|EIT77677.1 ater|EAU29441.1 awen|OJJ30083.1 azon|OJJ48994.1 afla|EED47546.1 aory|EIT74554.1

Group7514: abra|OJJ72193.1 acar|OOF94917.1 afla|EED45443.1 agla|OJJ79307.1 akaw|GAA84740.1 aluc|OJZ86944.1 anig|EHA21700.1 aory|EIT82473.1 asyd|OJJ59729.1 atub|OJI83578.1 aver|OJI97567.1 awen|OJJ36157.1 aver|OJJ02557.1

Group7515: abra|OJJ72203.1 acar|OOF96823.1 afis|EAW24465.1 afla|EED55201.1 afum|EAL86136.1 agla|OJJ80323.1 akaw|GAA84733.1 aluc|OJZ86953.1 anig|EHA21694.1 aory|EIT83578.1 ater|EAU29969.1 atub|OJI83570.1 awen|OJJ32725.1

Group7516: abra|OJJ72213.1 acar|OOF96834.1 afis|EAW23666.1 afla|EED54474.1 afum|EAL87111.1 akaw|GAA84721.1 aluc|OJZ86965.1 anid|CBF75940.1 anig|EHA21684.1 aory|EIT81139.1 asyd|OJJ59159.1 atub|OJI83557.1 aver|OJJ01749.1

Group7517: abra|OJJ72255.1 acar|OOF96886.1 acla|EAW13649.1 afis|EAW19957.1 afla|EED55750.1 akaw|GAA84682.1 aluc|OJZ87006.1 anig|EHA21640.1 aory|EIT78227.1 atub|OJI83513.1 awen|OJJ39613.1 aory|EIT76733.1 afla|EED52821.1

Group7518: abra|OJJ72304.1 acar|OOF94914.1 afis|EAW17159.1 afum|EAL86596.1 agla|OJJ82674.1 akaw|GAA86055.1 aluc|OJZ90285.1 anid|CBF81939.1 anig|EHA23775.1 asyd|OJJ57378.1 atub|OJI81709.1 aver|OJJ05780.1 awen|OJJ32539.1

Group7519: abra|OJJ72334.1 acar|OOF94881.1 acla|EAW11988.1 afis|EAW19628.1 afla|EED56980.1 afum|EBA27292.1 agla|OJJ80774.1 akaw|GAA86086.1 aory|EIT77534.1 asyd|OJJ55110.1 ater|EAU32898.1 aver|OJJ07096.1 azon|OJJ47070.1

Group7520: abra|OJJ72341.1 acar|OOF94876.1 afis|EAW19623.1 afla|EED56975.1 afum|EBA27294.1 akaw|GAA86091.1 aluc|OJZ90320.1 anig|EHA23806.1 asyd|OJJ55115.1 ater|EAU32901.1 aver|OJJ07091.1 awen|OJJ41870.1 azon|OJJ47063.1

Group7521: abra|OJJ72342.1 acar|OOF94875.1 afis|EAW19622.1 afla|EED56974.1 afum|EAL86899.2 aluc|OJZ90321.1 anid|CBF71312.1 anig|EHA23807.1 asyd|OJJ55116.1 ater|EAU32904.1 aver|OJJ07090.1 awen|OJJ41869.1 azon|OJJ47062.1

Group7522: abra|OJJ72359.1 acar|OOF94859.1 akaw|GAA86106.1 aluc|OJZ90338.1 anid|CBF87130.1 anig|EHA23822.1 asyd|OJJ60366.1 ater|EAU31884.1 atub|OJI81765.1 aver|OJI99703.1 awen|OJJ33425.1 azon|OJJ45034.1 aory|EIT77542.1

Group7523: abra|OJJ72451.1 acar|OOF94785.1 afla|EED46200.1 akaw|GAA86188.1 aluc|OJZ90436.1 anig|EHA23903.1 asyd|OJJ55270.1 ater|EAU33050.1 atub|OJI81867.1 aver|OJJ06957.1 agla|OJJ83121.1 aory|EIT76366.1 anid|CBF77962.1

Group7524: abra|OJJ72557.1 acar|OOF94696.1 acla|EAW12762.1 afis|EAW17924.1 afla|EED46326.1 afum|EAL91735.2 agla|OJJ83018.1 akaw|GAA85190.1 aluc|OJZ90545.1 anid|CBF76945.1 asyd|OJJ57060.1 atub|OJI81967.1 aver|OJJ03238.1

Group7525: abra|OJJ72717.1 acar|OOF95951.1 acla|EAW11553.1 afis|EAW23900.1 afla|EED52634.1 afum|EAL87327.1 akaw|GAA85325.1 aluc|OJZ90686.1 anid|CBF77144.1 anig|EHA24114.1 aory|EIT74229.1 aver|OJJ03381.1 azon|OJJ51379.1

Group7526: abra|OJJ72923.1 acla|EAW06465.1 afis|EAW15960.1 afla|EED58098.1 afum|EAL88593.1 agla|OJJ89352.1 akaw|GAA85506.1 aluc|OJZ83433.1 anid|CBF81077.1 aory|EIT79151.1 ater|EAU35496.1 atub|OJI82323.1 azon|OJJ46910.1

Group7527: acla|EAW06450.1 agla|OJJ89366.1 anig|EHA24296.1 acar|OOF96122.1 afis|EAW15947.1 afla|EED58112.1 afum|EAL88580.1 anid|CBF81050.1 aory|EIT79145.1 asyd|OJJ56449.1 ater|EAU35511.1 aver|OJJ06273.1 awen|OJJ35637.1

Group7528: abra|OJJ72965.1 acar|OOF96147.1 afla|EED54941.1 akaw|GAA85544.1 aluc|OJZ83475.1 aory|EIT80320.1 asyd|OJJ56377.1 ater|EAU35536.1 aver|OJJ06336.1 agla|OJJ89390.1 anig|EHA24313.1 atub|OJI82365.1 afis|EAW15920.1

Group7529: abra|OJJ73004.1 acar|OOF96177.1 acla|EAW08009.1 afis|EAW23044.1 afum|EAL88437.1 agla|OJJ89434.1 akaw|GAA85582.1 aluc|OJZ83518.1 anid|CBF80942.1 asyd|OJJ56337.1 aver|OJJ06376.1 awen|OJJ35696.1 azon|OJJ44160.1

Group7530: abra|OJJ73079.1 acar|OOG00314.1 acla|EAW12286.1 afis|EAW16952.1 afum|EAL86480.1 akaw|GAA88083.1 aluc|OJZ81347.1 anid|CBF82879.1 anig|EHA18650.1 asyd|OJJ62954.1 atub|OJI91551.1 aver|OJI98030.1 azon|OJJ46840.1

Group7531: abra|OJJ73133.1 acar|OOF98128.1 acla|EAW13624.1 afis|EAW21095.1 afum|EAL92118.1 akaw|GAA92663.1 aluc|OJZ90117.1 anig|EHA19697.1 ater|EAU31342.1 atub|OJI84680.1 awen|OJJ30105.1 acar|OOF98459.1 acla|EAW08506.1

Group7532: abra|OJJ73140.1 acar|OOF92576.1 afis|EAW25132.1 afla|EED48164.1 afum|EAL89803.1 agla|OJJ82671.1 akaw|GAA88032.1 aluc|OJZ81291.1 anid|CBF76014.1 anig|EHA18590.1 asyd|OJJ52435.1 atub|OJI91499.1 aver|OJJ03975.1

Group7533: abra|OJJ73171.1 acar|OOG00412.1 acla|EAW10797.1 afla|EED47206.1 agla|OJJ82875.1 akaw|GAA88000.1 aluc|OJZ81256.1 aory|EIT79227.1 asyd|OJJ59427.1 atub|OJI91452.1 aver|OJJ02844.1 awen|OJJ40365.1 azon|OJJ49214.1

Group7534: abra|OJJ73241.1 acar|OOG00480.1 acla|EAW06914.1 afis|EAW16531.1 afla|EED56139.1 afum|EAL89128.1 agla|OJJ82789.1 akaw|GAA87940.1 anid|CBF81733.1 asyd|OJJ59593.1 aver|OJJ02675.1 awen|OJJ40290.1 azon|OJJ49152.1

Group7535: abra|OJJ73261.1 acla|EAW06938.1 afis|EAW16553.1 afla|EED56109.1 afum|EAL89151.2 agla|OJJ82770.1 akaw|GAA87921.1 anid|CBF81793.1 anig|EHA18485.1 aory|EIT76758.1 ater|EAU36618.1 aver|OJJ02645.1 azon|OJJ49128.1

Group7536: abra|OJJ73276.1 acla|EAW06951.1 afis|EAW16566.1 afla|EED56097.1 afum|EAL89163.1 agla|OJJ82755.1 akaw|GAA92729.1 aluc|OJZ88773.1 anid|CBF81819.1 asyd|OJJ59652.1 ater|EAU36635.1 aver|OJJ02630.1 azon|OJJ49114.1

Group7537: abra|OJJ73336.1 aluc|OJZ88704.1 atub|OJI91285.1 akaw|GAA83147.1 ater|EAU39536.1 abra|OJJ78151.1 acar|OOF98715.1 afis|EAW16455.1 afla|EED56916.1 afum|EAL89050.1 aluc|OJZ91203.1 anig|EHA19908.1 aory|EIT77472.1

Group7538: abra|OJJ73389.1 acar|OOG00591.1 akaw|GAA83100.1 aluc|OJZ88648.1 anig|EHA18339.1 aory|EIT76038.1 asyd|OJJ54922.1 ater|EAU36079.1 atub|OJI91235.1 aver|OJJ07273.1 awen|OJJ32623.1 azon|OJJ49315.1 afla|EED54040.1

Group7539: abra|OJJ73411.1 acar|OOG00608.1 acla|EAW12385.1 afis|EAW16832.1 afum|EAL86380.1 akaw|GAA83080.1 aluc|OJZ88626.1 anid|CBF87378.1 anig|EHA18318.1 asyd|OJJ52841.1 atub|OJI91211.1 aver|OJJ08812.1 azon|OJJ48153.1

Group7540: abra|OJJ73416.1 acar|OOG00620.1 acla|EAW07038.1 afis|EAW16664.1 afla|EED44897.1 afum|EAL89268.1 agla|OJJ78705.1 akaw|GAA83075.1 anid|CBF82080.1 aory|EIT83244.1 ater|EAU39586.1 awen|OJJ40049.1 azon|OJJ50119.1

Group7541: abra|OJJ73433.1 acar|OOG00640.1 aluc|OJZ88608.1 anig|EHA18301.1 aory|EIT74830.1 ater|EAU31397.1 atub|OJI91192.1 aver|OJI99557.1 awen|OJJ33326.1 afla|EED53596.1 afla|EED53597.1 anid|CBF77720.1 ater|EAU36690.1

Group7542: abra|OJJ73436.1 acar|OOG00375.1 afis|EAW24840.1 afum|EAL89486.1 akaw|GAA83057.1 aluc|OJZ88605.1 anid|CBF83578.1 anig|EHA18298.1 asyd|OJJ56751.1 ater|EAU32493.1 atub|OJI91189.1 aver|OJJ05974.1 azon|OJJ45683.1

Group7543: abra|OJJ73809.1 acar|OOG00966.1 acla|EAW07698.1 afis|EAW22715.1 afla|EED49999.1 afum|EAL90479.1 akaw|GAA89429.1 aluc|OJZ88241.1 aory|EIT74738.1 ater|EAU38990.1 atub|OJI90843.1 awen|OJJ35409.1 azon|OJJ49767.1

Group7544: abra|OJJ73810.1 acla|EAW09052.1 afis|EAW25628.1 afla|EED56730.1 afum|EAL84316.1 aluc|OJZ88240.1 anig|EHA17987.1 asyd|OJJ59451.1 ater|EAU33004.1 atub|OJI90842.1 aver|OJJ02821.1 awen|OJJ30199.1 azon|OJJ50244.1

Group7545: abra|OJJ73956.1 acar|OOG01083.1 acla|EAW13229.1 afla|EED57673.1 aory|EIT79891.1 awen|OJJ30246.1 abra|OJJ74367.1 agla|OJJ80513.1 akaw|GAA82002.1 aluc|OJZ92185.1 atub|OJI86614.1 atub|OJI83979.1 agla|OJJ79591.1

Group7546: abra|OJJ74047.1 acar|OOF89920.1 afla|EED58125.1 afum|EAL90265.1 agla|OJJ83356.1 aluc|OJZ87988.1 anid|CBF87664.1 anig|EHA17771.1 asyd|OJJ63306.1 atub|OJI90598.1 aver|OJI97653.1 awen|OJJ38632.1 azon|OJJ49574.1

Group7547: abra|OJJ74058.1 acar|OOG01170.1 afis|EAW22941.1 afum|EAL90255.1 agla|OJJ83347.1 akaw|GAA87539.1 aluc|OJZ87978.1 anid|CBF87649.1 anig|EHA17761.1 ater|EAU30683.1 atub|OJI90588.1 awen|OJJ38623.1 azon|OJJ49564.1

Group7548: abra|OJJ74127.1 acar|OOF95609.1 afla|EED45392.1 agla|OJJ88369.1 akaw|GAA89757.1 aluc|OJZ80626.1 anig|EHA27206.1 aory|EIT82418.1 asyd|OJJ57318.1 ater|EAU33964.1 atub|OJI87481.1 aver|OJJ05843.1 azon|OJJ51096.1

Group7549: abra|OJJ74186.1 afla|EED46892.1 agla|OJJ81679.1 akaw|GAA86830.1 aluc|OJZ89949.1 anig|EHA27150.1 aory|EIT72565.1 asyd|OJJ65128.1 ater|EAU38482.1 atub|OJI84858.1 aver|OJI99256.1 awen|OJJ33921.1 anig|EHA22243.1

Group7550: acla|EAW13277.1 afum|EAL91185.1 anid|CBF80804.1 abra|OJJ77277.1 afis|EAW17340.1 afla|EED54195.1 akaw|GAA90672.1 aluc|OJZ84687.1 anig|EHA27962.1 asyd|OJJ54486.1 atub|OJI89395.1 aver|OJJ07693.1 agla|OJJ84306.1

Group7551: abra|OJJ74217.1 acar|OOF95504.1 acla|EAW07269.1 afis|EAW22290.1 afla|EED57689.1 agla|OJJ87759.1 akaw|GAA86801.1 aluc|OJZ89917.1 anid|CBF88567.1 anig|EHA27117.1 ater|EAU34302.1 atub|OJI84888.1 awen|OJJ34863.1

Group7552: abra|OJJ74431.1 acar|OOF95313.1 acla|EAW12293.1 afla|EED56193.1 akaw|GAA87325.1 aluc|OJZ89690.1 anid|CBF74694.1 anig|EHA26928.1 aory|EIT73441.1 asyd|OJJ60213.1 atub|OJI85111.1 aver|OJJ04055.1 azon|OJJ50250.1

Group7553: abra|OJJ74435.1 acar|OOF95309.1 afis|EAW20279.1 afla|EED48229.1 afum|EAL85414.1 akaw|GAA87329.1 aluc|OJZ89686.1 anig|EHA26924.1 ater|EAU36999.1 atub|OJI85115.1 aver|OJJ07464.1 awen|OJJ35547.1 ater|EAU34774.1

Group7554: abra|OJJ74460.1 acar|OOF95290.1 afis|EAW22258.1 afla|EED57645.1 akaw|GAA87353.1 aluc|OJZ89661.1 anig|EHA26898.1 aory|EIT79876.1 asyd|OJJ55350.1 ater|EAU36273.1 atub|OJI85139.1 awen|OJJ34830.1 azon|OJJ45717.1

Group7555: abra|OJJ74461.1 acar|OOF95289.1 afla|EED57644.1 akaw|GAA87354.1 aluc|OJZ89660.1 anig|EHA26897.1 aory|EIT79870.1 asyd|OJJ55349.1 ater|EAU36274.1 atub|OJI85140.1 awen|OJJ34829.1 azon|OJJ45719.1 afis|EAW22257.1

Group7556: abra|OJJ74492.1 acar|OOF95257.1 acla|EAW07202.1 afis|EAW22214.1 afla|EED57608.1 afum|EAL90980.2 agla|OJJ87828.1 aluc|OJZ89626.1 anid|CBF88450.1 aory|EIT82974.1 atub|OJI85176.1 awen|OJJ34799.1 azon|OJJ45746.1

Group7557: aory|EIT82929.1 azon|OJJ45751.1 acla|EAW07198.1 afis|EAW22210.1 afla|EED57604.1 afum|EAL90984.2 agla|OJJ87834.1 akaw|GAA87389.1 anid|CBF88444.1 anig|EHA26867.1 ater|EAU34220.1 aver|OJI97173.1 awen|OJJ34795.1

Group7558: abra|OJJ74530.1 acla|EAW07162.1 afis|EAW22177.1 afla|EED57556.1 afum|EAL91012.1 akaw|GAA82070.1 aluc|OJZ92112.1 anid|CBF89073.1 asyd|OJJ62741.1 ater|EAU34182.1 atub|OJI86545.1 awen|OJJ34754.1 azon|OJJ45804.1

Group7559: abra|OJJ74743.1 acar|OOF95739.1 afis|EAW19551.1 afla|EED45805.1 afum|EAL86958.1 agla|OJJ84419.1 akaw|GAA83667.1 aluc|OJZ92325.1 anig|EHA26645.1 aory|EIT75693.1 atub|OJI86753.1 awen|OJJ37996.1 azon|OJJ46002.1

Group7560: abra|OJJ74822.1 acar|OOF92577.1 afis|EAW16433.1 afla|EED53098.1 agla|OJJ80040.1 akaw|GAA86590.1 aluc|OJZ85137.1 anig|EHA18382.1 ater|EAU29269.1 atub|OJI80455.1 aver|OJJ01774.1 awen|OJJ34672.1 anig|EHA25369.1

Group7561: abra|OJJ75138.1 acar|OOF97198.1 afis|EAW23351.1 afla|EED55612.1 afum|EAL88125.2 agla|OJJ86475.1 akaw|GAA88744.1 aluc|OJZ92704.1 anig|EHA26295.1 asyd|OJJ60101.1 atub|OJI87131.1 awen|OJJ39538.1 azon|OJJ47735.1

Group7562: acla|EAW13657.1 afis|EAW19940.1 afum|EAL84972.1 anid|CBF84259.1 anid|CBF71413.1 agla|OJJ80510.1 asyd|OJJ57205.1 atub|OJI87195.1 aver|OJJ03084.1 awen|OJJ29828.1 azon|OJJ44623.1 asyd|OJJ58059.1 aver|OJJ04344.1

Group7563: abra|OJJ75254.1 acar|OOF89846.1 afla|EED57153.1 akaw|GAA91794.1 aluc|OJZ92813.1 anid|CBF69546.1 anig|EHA26196.1 aory|EIT74141.1 asyd|OJJ59173.1 ater|EAU31999.1 atub|OJI87270.1 aver|OJJ01762.1 azon|OJJ45568.1

Group7564: abra|OJJ75258.1 acar|OOF93396.1 afla|EED47370.1 agla|OJJ87649.1 akaw|GAA92056.1 aluc|OJZ92928.1 anig|EHA26192.1 aory|EIT82871.1 atub|OJI87388.1 azon|OJJ46464.1 aluc|OJZ92815.1 atub|OJI87273.1 akaw|GAA91796.1

Group7565: abra|OJJ75262.1 acar|OOF94540.1 afla|EED56573.1 akaw|GAA91800.1 aluc|OJZ92820.1 anid|CBF80336.1 anig|EHA26188.1 aory|EIT79623.1 asyd|OJJ54840.1 atub|OJI87278.1 aver|OJJ07321.1 awen|OJJ33405.1 azon|OJJ46004.1

Group7566: asyd|OJJ63976.1 abra|OJJ75356.1 afis|EAW23649.1 afla|EED48391.1 akaw|GAA92052.1 aluc|OJZ92922.1 anig|EHA26047.1 aory|EIT83060.1 atub|OJI87383.1 aver|OJJ05182.1 afum|EAL87725.1 aver|OJI96978.1 asyd|OJJ54229.1

Group7567: abra|OJJ75346.1 acar|OOF94490.1 afis|EAW17118.1 afla|EED55671.1 afum|EAL86580.1 akaw|GAA92039.1 aluc|OJZ92908.1 anig|EHA26056.1 aory|EIT77096.1 asyd|OJJ52959.1 ater|EAU35549.1 atub|OJI87368.1 awen|OJJ38032.1

Group7568: abra|OJJ75375.1 acar|OOF91864.1 afla|EED48856.1 akaw|GAA92737.1 aluc|OJZ92945.1 anid|CBF78476.1 anig|EHA26027.1 aory|EIT81275.1 asyd|OJJ58350.1 ater|EAU36129.1 atub|OJI87404.1 aver|OJJ02057.1 awen|OJJ35768.1

Group7569: abra|OJJ75411.1 acar|OOF90758.1 acla|EAW13783.1 afis|EAW18349.1 afla|EED48814.1 aluc|OJZ88862.1 anid|CBF79369.1 anig|EHA23641.1 asyd|OJJ57809.1 ater|EAU34683.1 aver|OJJ04579.1 awen|OJJ35733.1 atub|OJI85928.1

Group7570: abra|OJJ75431.1 acla|EAW13803.1 afis|EAW18328.1 afum|EAL87694.1 agla|OJJ88055.1 anid|CBF79404.1 anig|EHA23622.1 asyd|OJJ57828.1 ater|EAU33337.1 atub|OJI85906.1 aver|OJJ04564.1 azon|OJJ43304.1 afis|EAW18329.1

Group7571: abra|OJJ75631.1 acar|OOG00203.1 acla|EAW09462.1 afis|EAW21675.1 afla|EED57938.1 afum|EAL92709.1 akaw|GAA84378.1 aluc|OJZ89082.1 anig|EHA23455.1 aory|EIT82351.1 ater|EAU38358.1 atub|OJI85702.1 awen|OJJ33941.1

Group7572: abra|OJJ75646.1 acar|OOG00193.1 afla|EED57952.1 agla|OJJ79916.1 akaw|GAA84391.1 aluc|OJZ89095.1 anid|CBF83609.1 anig|EHA23442.1 atub|OJI85686.1 awen|OJJ33952.1 asyd|OJJ60764.1 aver|OJJ00098.1 azon|OJJ48618.1

Group7573: abra|OJJ75702.1 acar|OOG00152.1 acla|EAW09406.1 afis|EAW21730.1 afla|EED57999.1 afum|EAL92763.1 akaw|GAA84438.1 aluc|OJZ89153.1 anid|CBF83728.1 asyd|OJJ60844.1 aver|OJJ00169.1 awen|OJJ34003.1 azon|OJJ48549.1

Group7574: abra|OJJ75754.1 acar|OOG00108.1 afla|EED51163.1 akaw|GAA84488.1 aluc|OJZ89209.1 anid|CBF81244.1 anig|EHA23350.1 aory|EIT81300.1 asyd|OJJ59322.1 atub|OJI85583.1 aver|OJJ02951.1 aver|OJI99531.1 anid|CBF80149.1

Group7575: abra|OJJ75801.1 acar|OOG00075.1 acla|EAW14884.1 afis|EAW24403.1 afla|EED47810.1 afum|EAL86074.1 akaw|GAA84517.1 aluc|OJZ89249.1 anig|EHA23318.1 aory|EIT77119.1 asyd|OJJ54139.1 atub|OJI85549.1 aver|OJJ08597.1

Group7576: abra|OJJ75815.1 acar|OOG00060.1 afis|EAW20376.1 afla|EED47802.1 afum|EAL85490.1 akaw|GAA84529.1 aluc|OJZ89262.1 anid|CBF83573.1 anig|EHA23306.1 asyd|OJJ52178.1 atub|OJI85537.1 aver|OJJ05967.1 awen|OJJ32819.1

Group7577: abra|OJJ75850.1 acar|OOG00043.1 afis|EAW19556.1 afum|EAL86955.2 akaw|GAA84555.1 aluc|OJZ89287.1 anid|CBF73904.1 anig|EHA23274.1 asyd|OJJ53665.1 ater|EAU30372.1 atub|OJI85512.1 aver|OJJ07772.1 awen|OJJ32850.1

Group7578: abra|OJJ75921.1 afis|EAW21258.1 afla|EED54817.1 afum|EAL92273.1 akaw|GAA84625.1 aluc|OJZ89371.1 anig|EHA23213.1 aory|EIT79444.1 ater|EAU35895.1 atub|OJI85428.1 acla|EAW09867.1 aver|OJI99926.1 anid|CBF83327.1

Group7579: acla|EAW15320.1 afis|EAW16157.1 afla|EED46084.1 agla|OJJ81262.1 anid|CBF81446.1 aory|EIT76480.1 asyd|OJJ59431.1 ater|EAU32998.1 aver|OJJ02840.1 awen|OJJ39814.1 afla|EED45361.1 aory|EIT82435.1 awen|OJJ39864.1

Group7580: abra|OJJ75946.1 acar|OOF99966.1 acla|EAW12303.1 afis|EAW16937.1 afla|EED45589.1 afum|EAL86462.1 agla|OJJ78888.1 akaw|GAA84648.1 aluc|OJZ89394.1 anig|EHA23191.1 aory|EIT75565.1 atub|OJI85403.1 awen|OJJ36295.1

Group7581: abra|OJJ75974.1 acar|OOF99945.1 afis|EAW21311.1 afum|EAL92327.1 agla|OJJ87431.1 akaw|GAA88982.1 aluc|OJZ89421.1 anid|CBF83402.1 anig|EHA23166.1 asyd|OJJ60641.1 atub|OJI85372.1 awen|OJJ39389.1 azon|OJJ44277.1

Group7582: abra|OJJ75986.1 acla|EAW09814.1 afis|EAW21320.1 afla|EED57278.1 afum|EAL92337.1 akaw|GAA88972.1 aluc|OJZ89431.1 anig|EHA23156.1 aory|EIT74148.1 atub|OJI85362.1 aver|OJI99975.1 awen|OJJ39376.1 azon|OJJ44266.1

Group7583: abra|OJJ76018.1 acar|OOF99908.1 afla|EED46077.1 akaw|GAA88942.1 aluc|OJZ89465.1 anid|CBF73507.1 anig|EHA23123.1 aory|EIT76501.1 asyd|OJJ55811.1 atub|OJI85330.1 aver|OJJ04759.1 awen|OJJ32715.1 azon|OJJ50219.1

Group7584: abra|OJJ76042.1 acar|OOF99888.1 acla|EAW09772.1 afis|EAW21366.1 afla|EED55780.1 afum|EAL92382.2 akaw|GAA88924.1 aluc|OJZ89497.1 anig|EHA23103.1 aory|EIT76663.1 atub|OJI85306.1 awen|OJJ39331.1 azon|OJJ51676.1

Group7585: abra|OJJ76239.1 acar|OOF99484.1 acla|EAW14199.1 afum|EAL93654.1 agla|OJJ81520.1 akaw|GAA82173.1 aluc|OJZ91992.1 anid|CBF69672.1 ater|EAU38097.1 atub|OJI86427.1 aver|OJJ01659.1 awen|OJJ31516.1 afla|EED46399.1

Group7586: abra|OJJ76276.1 acar|OOF99524.1 acla|EAW14336.1 afis|EAW18953.1 afum|EAL93689.2 akaw|GAA82208.1 aluc|OJZ91944.1 atub|OJI86387.1 awen|OJJ31550.1 agla|OJJ85328.1 afla|EED50991.1 aory|EIT76990.1 azon|OJJ47266.1

Group7587: abra|OJJ76278.1 acla|EAW07089.1 afum|EAL89047.1 akaw|GAA82210.1 aluc|OJZ91942.1 anig|EHA22901.1 atub|OJI86385.1 aver|OJJ03910.1 aory|EIT75549.1 acla|EAW07088.1 ater|EAU35312.1 afis|EAW16447.1 afla|EED45668.1

Group7588: abra|OJJ76311.1 acar|OOF99553.1 afla|EED50765.1 agla|OJJ80542.1 akaw|GAA82239.1 aluc|OJZ91907.1 aory|EIT77995.1 ater|EAU37960.1 atub|OJI86356.1 awen|OJJ32384.1 azon|OJJ47144.1 afis|EAW18809.1 afum|EAL93545.1

Group7589: acar|OOF99564.1 afum|EAL93539.1 akaw|GAA82244.1 aluc|OJZ91901.1 awen|OJJ32378.1 acla|EAW14102.1 afis|EAW18803.1 afla|EED50773.1 agla|OJJ80533.1 anid|CBF69786.1 aory|EIT78059.1 ater|EAU37941.1 azon|OJJ47140.1

Group7590: abra|OJJ76436.1 acar|OOF99681.1 acla|EAW13987.1 afis|EAW18695.1 afum|EAL93417.1 akaw|GAA82351.1 aluc|OJZ91776.1 anid|CBF70628.1 anig|EHA22774.1 aory|EIT78143.1 ater|EAU38211.1 atub|OJI86232.1 awen|OJJ32166.1

Group7591: abra|OJJ76459.1 acla|EAW12026.1 afis|EAW20067.1 afum|EAL88796.1 akaw|GAA82372.1 aluc|OJZ91750.1 anig|EHA22753.1 asyd|OJJ54642.1 atub|OJI86208.1 aver|OJJ07516.1 awen|OJJ38106.1 afis|EAW20719.1 afum|EAL85185.1

Group7592: abra|OJJ76477.1 acar|OOF99712.1 acla|EAW10942.1 afis|EAW20918.1 afla|EED52661.1 agla|OJJ82408.1 akaw|GAA82389.1 aluc|OJZ91731.1 anig|EHA22738.1 asyd|OJJ54405.1 atub|OJI86188.1 awen|OJJ32137.1 awen|OJJ34403.1

Group7593: akaw|GAA82400.1 aluc|OJZ91722.1 anig|EHA22729.1 atub|OJI86179.1 acar|OOF99717.1 acla|EAW13949.1 afis|EAW18660.1 afla|EED52671.1 afum|EAL93376.1 anid|CBF70554.1 aory|EIT74287.1 aver|OJJ01140.1 awen|OJJ32126.1

Group7594: abra|OJJ76508.1 acar|OOF99463.1 acla|EAW14190.1 afis|EAW18905.1 afum|EAL93637.1 agla|OJJ81510.1 akaw|GAA82421.1 aluc|OJZ91696.1 anig|EHA22709.1 atub|OJI86151.1 aver|OJJ01640.1 awen|OJJ32462.1 azon|OJJ47219.1

Group7595: abra|OJJ76620.1 acar|OOF92933.1 afis|EAW19772.1 afum|EAL85128.1 agla|OJJ81606.1 akaw|GAA85576.1 aluc|OJZ83512.1 anid|CBF75779.1 ater|EAU35166.1 atub|OJI85467.1 aver|OJJ07924.1 afis|EAW23639.1 anid|CBF79117.1

Group7596: abra|OJJ76638.1 acar|OOF99361.1 acla|EAW14274.1 afis|EAW19000.1 afla|EED53125.1 afum|EAL93735.1 agla|OJJ85234.1 akaw|GAA92820.1 aluc|OJZ91571.1 anig|EHA22600.1 aory|EIT75061.1 ater|EAU37804.1 atub|OJI86021.1

Group7597: abra|OJJ76689.1 acar|OOF99322.1 agla|OJJ87953.1 akaw|GAA92505.1 aluc|OJZ91518.1 anid|CBF84585.1 anig|EHA22556.1 asyd|OJJ63694.1 atub|OJI85973.1 aver|OJI97245.1 awen|OJJ33478.1 asyd|OJJ55011.1 aver|OJJ07201.1

Group7598: abra|OJJ76699.1 acar|OOF96791.1 afla|EED53493.1 akaw|GAA92500.1 aluc|OJZ91513.1 anig|EHA22552.1 aory|EIT78829.1 atub|OJI85965.1 azon|OJJ42675.1 akaw|GAA89228.1 aluc|OJZ86452.1 atub|OJI84078.1 aory|EIT75025.1

Group7599: abra|OJJ76733.1 afis|EAW17071.1 afla|EED49616.1 afum|EAL86559.2 agla|OJJ79086.1 aluc|OJZ80163.1 aory|EIT83018.1 asyd|OJJ58392.1 aver|OJJ07901.1 awen|OJJ32614.1 asyd|OJJ60948.1 aver|OJJ00275.1 aory|EIT80981.1

Group7600: abra|OJJ76757.1 acar|OOF91252.1 acla|EAW13311.1 afis|EAW20425.1 afla|EED44843.1 agla|OJJ84900.1 aluc|OJZ84160.1 aory|EIT77741.1 asyd|OJJ55791.1 atub|OJI89924.1 aver|OJJ06881.1 awen|OJJ33118.1 azon|OJJ46334.1

Group7601: asyd|OJJ55776.1 acar|OOF91267.1 acla|EAW13328.1 afis|EAW20443.1 afla|EED44827.1 afum|EBA27199.1 agla|OJJ84882.1 akaw|GAA87252.1 anid|CBF79306.1 aory|EIT77753.1 ater|EAU37716.1 atub|OJI89907.1 awen|OJJ33133.1

Group7602: abra|OJJ76804.1 acar|OOF91298.1 acla|EAW13363.1 afis|EAW20477.1 agla|OJJ84840.1 akaw|GAA87219.1 aluc|OJZ84211.1 anid|CBF79251.1 asyd|OJJ55745.1 atub|OJI89872.1 aver|OJJ06832.1 awen|OJJ33172.1 azon|OJJ46379.1

Group7603: abra|OJJ76826.1 akaw|GAA87196.1 aluc|OJZ84238.1 anig|EHA27585.1 atub|OJI89845.1 acla|EAW13386.1 ater|EAU37194.1 aver|OJJ06812.1 azon|OJJ46397.1 afis|EAW20500.1 afla|EED44743.1 afum|EAL84555.1 awen|OJJ33201.1

Group7604: anid|CBF80489.1 acla|EAW11604.1 afis|EAW17175.1 afum|EAL86605.1 agla|OJJ89217.1 aluc|OJZ83018.1 azon|OJJ50115.1 acar|OOF89864.1 azon|OJJ45906.1 afis|EAW16802.1 anid|CBF74922.1 asyd|OJJ60080.1 aver|OJJ04167.1

Group7605: abra|OJJ76932.1 acar|OOF91376.1 acla|EAW09285.1 afis|EAW21862.1 afla|EED57684.1 afum|EBA27477.1 agla|OJJ84273.1 akaw|GAA87107.1 aluc|OJZ84331.1 anig|EHA27665.1 aory|EIT79941.1 atub|OJI89738.1 awen|OJJ34147.1

Group7606: abra|OJJ76958.1 acar|OOF91403.1 acla|EAW13480.1 afis|EAW20588.1 afla|EED52262.1 afum|EAL84473.1 akaw|GAA87085.1 aluc|OJZ84353.1 anig|EHA27691.1 ater|EAU37091.1 awen|OJJ33296.1 afla|EED52261.1 azon|OJJ50676.1

Group7607: abra|OJJ76973.1 acla|EAW13499.1 afis|EAW20598.1 afla|EED52206.1 afum|EAL84463.1 agla|OJJ88655.1 aluc|OJZ84369.1 anig|EHA27704.1 asyd|OJJ55611.1 atub|OJI89699.1 aver|OJJ06686.1 awen|OJJ33390.1 azon|OJJ50694.1

Group7608: abra|OJJ77003.1 acar|OOF91449.1 acla|EAW09012.1 afis|EAW25625.1 afla|EED54120.1 afum|EAL84313.2 akaw|GAA87045.1 aluc|OJZ84402.1 anig|EHA27732.1 aory|EIT75973.1 ater|EAU32186.1 atub|OJI89670.1 awen|OJJ36358.1

Group7609: abra|OJJ77063.1 acar|OOF92947.1 acla|EAW13512.1 afis|EAW20610.1 agla|OJJ88629.1 akaw|GAA86989.1 aluc|OJZ84471.1 anid|CBF86955.1 asyd|OJJ55595.1 atub|OJI89607.1 aver|OJJ06674.1 awen|OJJ33453.1 azon|OJJ50710.1

Group7610: abra|OJJ77108.1 acar|OOF92993.1 afis|EAW21597.1 afla|EED57821.1 afum|EAL92616.1 agla|OJJ85791.1 akaw|GAA90820.1 aluc|OJZ84517.1 anig|EHA27815.1 aory|EIT81081.1 ater|EAU38286.1 atub|OJI89563.1 awen|OJJ39085.1

Group7611: abra|OJJ77135.1 acar|OOF93026.1 acla|EAW12946.1 afis|EAW17721.1 afla|EED45723.1 akaw|GAA90794.1 aluc|OJZ84545.1 anig|EHA27842.1 aory|EIT75556.1 ater|EAU31899.1 atub|OJI89535.1 afum|EAL91536.1 awen|OJJ35791.1

Group7612: abra|OJJ77241.1 acar|OOF93122.1 afla|EED49532.1 agla|OJJ88644.1 akaw|GAA90702.1 aluc|OJZ84651.1 anid|CBF74979.1 anig|EHA27934.1 aory|EIT82638.1 asyd|OJJ56620.1 atub|OJI89429.1 aver|OJJ06106.1 awen|OJJ33583.1

Group7613: abra|OJJ77311.1 aluc|OJZ84723.1 anig|EHA27997.1 atub|OJI89360.1 acar|OOF91476.1 afis|EAW17113.1 afla|EED51657.1 akaw|GAA91442.1 aluc|OJZ88724.1 acar|OOG00671.1 anig|EHA18429.1 ater|EAU37965.1 akaw|GAA86060.1

Group7614: abra|OJJ77377.1 acla|EAW13097.1 afis|EAW17566.1 afla|EED49255.1 agla|OJJ81400.1 akaw|GAA91383.1 anid|CBF90264.1 anig|EHA28054.1 aory|EIT82177.1 asyd|OJJ61940.1 aver|OJI96032.1 awen|OJJ37287.1 azon|OJJ50981.1

Group7615: abra|OJJ77394.1 acar|OOF93262.1 afla|EED47334.1 akaw|GAA91368.1 aluc|OJZ84802.1 anid|CBF81560.1 anig|EHA28067.1 aory|EIT82889.1 asyd|OJJ59500.1 ater|EAU38091.1 atub|OJI89281.1 aver|OJJ02774.1 awen|OJJ32599.1

Group7616: afis|EAW16273.1 agla|OJJ88343.1 ater|EAU37448.1 acar|OOF99204.1 acla|EAW06714.1 afla|EED53292.1 afum|EAL88894.1 akaw|GAA84101.1 anid|CBF86906.1 aory|EIT74828.1 asyd|OJJ64612.1 atub|OJI82567.1 aver|OJI98708.1

Group7617: ater|EAU34098.1 acla|EAW11545.1 afis|EAW16825.1 afum|EAL86534.1 agla|OJJ84964.1 akaw|GAA85253.1 aluc|OJZ90611.1 anig|EHA24049.1 awen|OJJ34434.1 azon|OJJ51296.1 afla|EED52804.1 atub|OJI82033.1 aver|OJJ03112.1

Group7618: abra|OJJ77582.1 acar|OOF99145.1 afis|EAW16046.1 afla|EED56435.1 agla|OJJ86079.1 akaw|GAA84043.1 anid|CBF75058.1 aory|EIT72830.1 asyd|OJJ60027.1 ater|EAU33591.1 aver|OJJ02257.1 awen|OJJ30399.1 azon|OJJ45407.1

Group7619: abra|OJJ77609.1 acar|OOF99120.1 afla|EED45043.1 agla|OJJ88528.1 akaw|GAA84020.1 aluc|OJZ80924.1 anid|CBF83741.1 aory|EIT73540.1 atub|OJI82654.1 aver|OJJ08840.1 awen|OJJ38696.1 azon|OJJ47380.1 aluc|OJZ79623.1

Group7620: abra|OJJ77653.1 afis|EAW15991.1 afla|EED56363.1 afum|EAL88623.1 agla|OJJ86002.1 akaw|GAA83982.1 aluc|OJZ80876.1 anid|CBF85821.1 aory|EIT73432.1 atub|OJI82701.1 aver|OJI98441.1 awen|OJJ30479.1 acla|EAW06495.1

Group7621: abra|OJJ77678.1 acar|OOF90214.1 acla|EAW06590.1 afis|EAW16013.1 afla|EED56390.1 agla|OJJ86038.1 aluc|OJZ80841.1 anid|CBF74564.1 asyd|OJJ57648.1 atub|OJI82726.1 aver|OJJ05507.1 awen|OJJ30452.1 azon|OJJ45443.1

Group7622: abra|OJJ77746.1 afis|EAW25749.1 afla|EED56205.1 agla|OJJ78981.1 akaw|GAA83888.1 aluc|OJZ90807.1 anig|EHA19535.1 aory|EIT73302.1 asyd|OJJ52526.1 atub|OJI82796.1 aver|OJJ03446.1 afis|EAW16451.1 afla|EED56206.1

Group7623: abra|OJJ77788.1 afla|EED55127.1 afum|EAL84254.1 akaw|GAA83859.1 aluc|OJZ90843.1 anig|EHA19564.1 aory|EIT81611.1 asyd|OJJ55633.1 ater|EAU39624.1 atub|OJI82830.1 aver|OJJ06710.1 awen|OJJ40272.1 anid|CBF84133.1

Group7624: abra|OJJ77884.1 acar|OOF89934.1 acla|EAW12221.1 afla|EED55364.1 agla|OJJ80663.1 akaw|GAA83770.1 aluc|OJZ90958.1 anig|EHA19661.1 aory|EIT80852.1 ater|EAU30465.1 atub|OJI82948.1 awen|OJJ38337.1 azon|OJJ45939.1

Group7625: abra|OJJ77956.1 afis|EAW17321.1 anid|CBF80599.1 aory|EIT74761.1 asyd|OJJ61777.1 ater|EAU35011.1 aver|OJJ06501.1 afla|EED48336.1 afla|EED56881.1 afum|EAL91157.1 anid|CBF84289.1 aory|EIT77553.1 aver|OJJ05784.1

Group7626: abra|OJJ77966.1 afla|EED48626.1 akaw|GAA83697.1 aluc|OJZ91036.1 anig|EHA19735.1 aory|EIT73214.1 ater|EAU33162.1 atub|OJI83032.1 acla|EAW15160.1 agla|OJJ80060.1 awen|OJJ31982.1 afis|EAW20129.1 azon|OJJ45558.1

Group7627: abra|OJJ77985.1 acar|OOF96458.1 acla|EAW14397.1 afla|EED57498.1 akaw|GAA85995.1 aluc|OJZ91060.1 anig|EHA19763.1 aory|EIT81880.1 asyd|OJJ62123.1 ater|EAU34295.1 atub|OJI83056.1 aver|OJI96231.1 azon|OJJ50091.1

Group7628: abra|OJJ78183.1 acar|OOF90019.1 afla|EED52129.1 agla|OJJ80837.1 akaw|GAA85852.1 aluc|OJZ91247.1 anid|CBF78478.1 anig|EHA19945.1 aory|EIT74481.1 asyd|OJJ65351.1 atub|OJI83244.1 aver|OJI99499.1 awen|OJJ36155.1

Group7629: abra|OJJ78225.1 acar|OOF98633.1 afis|EAW21861.1 afla|EED46449.1 afum|EAL92874.1 akaw|GAA85800.1 aluc|OJZ91300.1 anig|EHA19982.1 aory|EIT72436.1 ater|EAU30860.1 atub|OJI83291.1 azon|OJJ43480.1 acla|EAW09286.1

Group7630: ater|EAU31107.1 acar|OOF98513.1 acla|EAW09159.1 afis|EAW21987.1 afla|EED46711.1 afum|EAL93003.1 agla|OJJ84111.1 aluc|OJZ91453.1 anid|CBF82694.1 aory|EIT77827.1 asyd|OJJ61648.1 atub|OJI83446.1 azon|OJJ43356.1

Group7631: aory|EIT82553.1 acla|EAW08882.1 afis|EAW15513.1 afla|EED49020.1 afum|EBA27246.1 agla|OJJ80184.1 aluc|OJZ85346.1 asyd|OJJ56005.1 ater|EAU32611.1 atub|OJI80224.1 aver|OJJ04950.1 awen|OJJ38167.1 azon|OJJ43127.1

Group7632: acar|OOF91385.1 akaw|GAA87103.1 aluc|OJZ84335.1 anig|EHA27669.1 atub|OJI89733.1 afla|EED51040.1 agla|OJJ89266.1 aory|EIT77026.1 ater|EAU38160.1 aver|OJJ07298.1 awen|OJJ35815.1 aory|EIT77224.1 ater|EAU29251.1

Group7633: acar|OOF92448.1 afla|EED52678.1 agla|OJJ89361.1 akaw|GAA87160.1 aluc|OJZ84274.1 anid|CBF74993.1 anig|EHA27618.1 aory|EIT74189.1 asyd|OJJ52340.1 atub|OJI89807.1 aver|OJJ07183.1 awen|OJJ32627.1 asyd|OJJ55023.1

Group7634: acar|OOF93488.1 acla|EAW08854.1 afis|EAW15542.1 afum|EBA27243.1 akaw|GAA87678.1 aluc|OJZ85296.1 anid|CBF78185.1 anig|EHA25494.1 asyd|OJJ55975.1 atub|OJI80274.1 aver|OJJ04923.1 awen|OJJ38203.1 azon|OJJ43092.1

Group7635: acar|OOF93828.1 acla|EAW11871.1 afis|EAW19482.1 afla|EED56692.1 afum|EAL87013.2 akaw|GAA89680.1 anid|CBF75466.1 aory|EIT73954.1 asyd|OJJ59766.1 ater|EAU34794.1 atub|OJI87563.1 awen|OJJ36889.1 aver|OJJ02521.1

Group7636: acar|OOF93947.1 acla|EAW10402.1 afis|EAW25172.1 afum|EAL89846.1 agla|OJJ85939.1 akaw|GAA86925.1 aluc|OJZ90049.1 anid|CBF80114.1 anig|EHA21021.1 asyd|OJJ58328.1 ater|EAU32737.1 atub|OJI84757.1 aver|OJJ02083.1

Group7637: acar|OOF93976.1 acla|EAW15056.1 afis|EAW20267.1 afla|EED46404.1 afum|EAL87186.2 anid|CBF80632.1 aory|EIT72320.1 asyd|OJJ54649.1 ater|EAU36376.1 aver|OJJ07513.1 awen|OJJ33496.1 azon|OJJ45798.1 agla|OJJ88715.1

Group7638: acar|OOF96492.1 acla|EAW13852.1 afis|EAW18566.1 afla|EED52787.1 afum|EAL93268.1 agla|OJJ81615.1 akaw|GAA85098.1 anid|CBF70303.1 anig|EHA22105.1 aory|EIT78212.1 asyd|OJJ58727.1 aver|OJJ01340.1 afla|EED52788.1

Group7639: acar|OOF96721.1 acla|EAW09480.1 afla|EED57891.1 akaw|GAA84869.1 aory|EIT82734.1 aver|OJJ00062.1 afis|EAW21654.1 agla|OJJ79722.1 asyd|OJJ60730.1 ater|EAU38336.1 awen|OJJ33911.1 anid|CBF83544.1 azon|OJJ48650.1

Group7640: acar|OOF97569.1 acla|EAW08582.1 afla|EED45084.1 agla|OJJ85248.1 akaw|GAA93165.1 aluc|OJZ83024.1 anid|CBF80424.1 asyd|OJJ60255.1 ater|EAU32189.1 aver|OJI96217.1 awen|OJJ38283.1 ater|EAU30502.1 aver|OJJ04002.1

Group7641: acar|OOF98032.1 acla|EAW10377.1 afis|EAW25216.1 afla|EED53691.1 afum|EAL89877.1 agla|OJJ79191.1 akaw|GAA82852.1 anid|CBF85256.1 aory|EIT75334.1 ater|EAU34373.1 aver|OJI98069.1 awen|OJJ40879.1 azon|OJJ44507.1

Group7642: acar|OOF99981.1 acla|EAW09864.1 afis|EAW21261.1 afla|EED54820.1 afum|EAL92276.1 anid|CBF83335.1 aory|EIT79334.1 asyd|OJJ60606.1 ater|EAU35898.1 aver|OJI99928.1 awen|OJJ36578.1 azon|OJJ44312.1 agla|OJJ87480.1

Group7643: acar|OOG00100.1 afis|EAW17241.1 afla|EED53344.1 agla|OJJ87208.1 anid|CBF81011.1 anig|EHA18742.1 aory|EIT75023.1 ater|EAU30922.1 atub|OJI86839.1 aver|OJI95797.1 awen|OJJ40918.1 azon|OJJ42282.1 azon|OJJ42283.1

Group7644: acar|OOG00456.1 acla|EAW06893.1 afis|EAW16510.1 afla|EED56163.1 afum|EAL89105.1 akaw|GAA87960.1 aluc|OJZ81209.1 anid|CBF75570.1 aory|EIT76557.1 asyd|OJJ59710.1 ater|EAU36576.1 awen|OJJ40314.1 azon|OJJ49174.1

Group7645: acar|OOG00632.1 aluc|OJZ88347.1 atub|OJI90943.1 acla|EAW13661.1 afla|EED53471.1 acla|EAW13662.1 afis|EAW20293.1 afla|EED53472.1 aory|EIT75166.1 aver|OJI98795.1 asyd|OJJ57227.1 anig|EHA18080.1 asyd|OJJ60232.1

Group7646: acla|EAW10694.1 afis|EAW24876.1 afla|EED47116.1 afum|EAL89526.1 agla|OJJ82097.1 akaw|GAA81915.1 aluc|OJZ92283.1 anid|CBF75760.1 aory|EIT79321.1 asyd|OJJ57535.1 ater|EAU36425.1 atub|OJI86714.1 azon|OJJ49329.1

Group7647: acla|EAW10856.1 afla|EED55753.1 agla|OJJ88738.1 akaw|GAA86558.1 aluc|OJZ85106.1 aory|EIT77602.1 asyd|OJJ60982.1 ater|EAU29511.1 awen|OJJ36084.1 afla|EED45950.1 afla|EED49752.1 aory|EIT76787.1 awen|OJJ30847.1

Group7648: acla|EAW11041.1 afis|EAW16184.1 afla|EED49624.1 afum|EAL88815.1 akaw|GAA89230.1 aluc|OJZ86450.1 anid|CBF74597.1 aory|EIT80992.1 atub|OJI84076.1 azon|OJJ45953.1 azon|OJJ45948.1 afla|EED53494.1 aory|EIT75178.1

Group7649: acla|EAW12225.1 afis|EAW15409.1 afla|EED45427.1 afum|EAL84237.1 agla|OJJ85886.1 anid|CBF82111.1 aory|EIT82438.1 asyd|OJJ54285.1 ater|EAU36066.1 atub|OJI89934.1 aver|OJJ05239.1 afla|EED56298.1 aory|EIT73343.1

Group7650: acla|EAW15253.1 afis|EAW20226.1 afla|EED51673.1 afum|EAL86331.1 agla|OJJ79162.1 aory|EIT80656.1 asyd|OJJ53352.1 ater|EAU29654.1 aver|OJJ08391.1 awen|OJJ30859.1 azon|OJJ42818.1 afis|EAW16759.1 ater|EAU36806.1

Group7651: agla|OJJ80803.1 asyd|OJJ53436.1 aver|OJJ08313.1 awen|OJJ41916.1 afis|EAW19695.1 afla|EED57009.1 afum|EAL86832.1 anid|CBF80586.1 aory|EIT82270.1 aver|OJJ07164.1 anid|CBF71380.1 ater|EAU32858.1 asyd|OJJ55043.1

Group7652: afis|EAW16829.1 afla|EED45150.1 afum|EAL86377.1 aory|EIT73769.1 asyd|OJJ57563.1 aver|OJJ05615.1 awen|OJJ40310.1 agla|OJJ84439.1 anid|CBF77116.1 asyd|OJJ53724.1 ater|EAU32940.1 awen|OJJ33485.1 aver|OJI98279.1

Group7653: afla|EED52180.1 anid|CBF80450.1 aory|EIT74338.1 ater|EAU32198.1 afla|EED57326.1 anid|CBF79044.1 aory|EIT74098.1 asyd|OJJ54776.1 aver|OJJ07390.1 azon|OJJ42451.1 ater|EAU37845.1 azon|OJJ45999.1 ater|EAU39619.1

Group7654: abra|OJJ65689.1 abra|OJJ75757.1 acar|OOF96842.1 afla|EED53478.1 akaw|GAA84489.1 aluc|OJZ89210.1 anig|EHA23348.1 aory|EIT75235.1 asyd|OJJ58455.1 atub|OJI85582.1 aver|OJJ01055.1 awen|OJJ30911.1

Group7655: abra|OJJ67660.1 abra|OJJ69554.1 afla|EED52538.1 akaw|GAA89175.1 aluc|OJZ81658.1 anig|EHA25787.1 asyd|OJJ62553.1 ater|EAU37830.1 atub|OJI88525.1 awen|OJJ31283.1 aver|OJI96679.1 awen|OJJ38789.1

Group7656: abra|OJJ67877.1 abra|OJJ74185.1 aluc|OJZ82012.1 aluc|OJZ89950.1 afis|EAW21800.1 afum|EAL92825.2 akaw|GAA91885.1 anid|CBF78714.1 atub|OJI84561.1 akaw|GAA86831.1 anig|EHA27151.1 atub|OJI84856.1

Group7657: acar|OOF90155.1 acar|OOF90156.1 abra|OJJ68813.1 akaw|GAA90441.1 aluc|OJZ85786.1 anig|EHA24772.1 asyd|OJJ56211.1 atub|OJI79570.1 aver|OJJ05140.1 afla|EED46079.1 aory|EIT76405.1 awen|OJJ31260.1

Group7658: acar|OOF94056.1 acar|OOF94126.1 abra|OJJ66387.1 afla|EED55698.1 akaw|GAA90253.1 aluc|OJZ83087.1 anig|EHA28149.1 aory|EIT77086.1 ater|EAU36287.1 atub|OJI87857.1 awen|OJJ33623.1 azon|OJJ50003.1

Group7659: acar|OOF94071.1 acar|OOF94141.1 afla|EED52544.1 agla|OJJ79026.1 anid|CBF89913.1 awen|OJJ30839.1 akaw|GAA90226.1 aluc|OJZ83115.1 anig|EHA28178.1 asyd|OJJ62128.1 atub|OJI87819.1 aver|OJI96236.1

Group7660: acar|OOF94091.1 acar|OOF94161.1 abra|OJJ66346.1 afla|EED47091.1 agla|OJJ80119.1 akaw|GAA90220.1 aluc|OJZ83121.1 anid|CBF86771.1 anig|EHA28185.1 atub|OJI87812.1 awen|OJJ36364.1 asyd|OJJ64687.1

Group7661: acar|OOF95047.1 acar|OOF95114.1 abra|OJJ68050.1 afla|EED47436.1 akaw|GAA90905.1 aluc|OJZ82921.1 anid|CBF76686.1 anig|EHA21344.1 aory|EIT81904.1 asyd|OJJ61225.1 aver|OJJ00548.1 awen|OJJ33879.1

Group7662: acar|OOF95094.1 acar|OOF95160.1 abra|OJJ74609.1 afla|EED47182.1 agla|OJJ82898.1 akaw|GAA81999.1 aluc|OJZ92189.1 asyd|OJJ59462.1 atub|OJI86618.1 aver|OJJ02812.1 awen|OJJ40387.1 azon|OJJ49237.1

Group7663: acar|OOF95624.1 acar|OOF95676.1 afis|EAW22113.1 afla|EED49706.1 afum|EAL91072.1 agla|OJJ78653.1 anid|CBF79167.1 anig|EHA26949.1 asyd|OJJ54536.1 aver|OJJ07297.1 awen|OJJ38661.1 azon|OJJ44489.1

Group7664: acar|OOF95642.1 acar|OOF95693.1 abra|OJJ74680.1 acla|EAW10715.1 agla|OJJ82070.1 akaw|GAA81934.1 aluc|OJZ92257.1 asyd|OJJ57551.1 atub|OJI86691.1 aver|OJJ05627.1 awen|OJJ40444.1 azon|OJJ44350.1

Group7665: acar|OOF95664.1 acar|OOF95715.1 abra|OJJ76653.1 afla|EED51631.1 akaw|GAA92993.1 aluc|OJZ91552.1 anig|EHA22587.1 asyd|OJJ53928.1 ater|EAU35677.1 atub|OJI86005.1 aver|OJJ08027.1 awen|OJJ34602.1

Group7666: acar|OOF96221.1 acar|OOF96259.1 abra|OJJ71505.1 afum|EBA27222.1 agla|OJJ78424.1 aluc|OJZ86244.1 anid|CBF84771.1 asyd|OJJ63240.1 atub|OJI84298.1 aver|OJI97716.1 awen|OJJ31790.1 azon|OJJ46324.1

Group7667: acar|OOF96920.1 acar|OOF96955.1 abra|OJJ70790.1 afis|EAW15864.1 agla|OJJ87245.1 akaw|GAA90106.1 aluc|OJZ84135.1 anid|CBF81198.1 anig|EHA19243.1 ater|EAU31677.1 atub|OJI90509.1 azon|OJJ47432.1

Group7668: acar|OOF96923.1 acar|OOF96958.1 abra|OJJ70787.1 agla|OJJ87242.1 aluc|OJZ84132.1 anid|CBF81204.1 anig|EHA19240.1 asyd|OJJ59306.1 atub|OJI90506.1 aver|OJJ02969.1 awen|OJJ36763.1 azon|OJJ47435.1

Group7669: acar|OOF97676.1 acar|OOF97724.1 anig|EHA20341.1 acla|EAW10074.1 afis|EAW25512.1 afla|EED54013.1 afum|EBA27395.1 aory|EIT78283.1 asyd|OJJ53032.1 aver|OJJ03812.1 awen|OJJ41254.1 azon|OJJ49623.1

Group7670: acar|OOG00249.1 acar|OOG00267.1 abra|OJJ67543.1 afla|EED44855.1 aluc|OJZ81389.1 anid|CBF78816.1 anig|EHA18682.1 aory|EIT82716.1 asyd|OJJ65264.1 atub|OJI91594.1 aver|OJI99415.1 awen|OJJ34504.1

Group7671: afis|EAW17068.1 afis|EAW18107.1 afis|EAW22051.1 afis|EAW24035.1 afis|EAW25562.1 afis|EAW22070.1 afis|EAW22021.1 afis|EAW22711.1 afis|EAW25561.1 afis|EAW22022.1 anid|CBF71258.1 anid|CBF84328.1

Group7672: afla|EED44759.1 afla|EED52734.1 aory|EIT77776.1 aory|EIT78252.1 abra|OJJ65869.1 afis|EAW20794.1 akaw|GAA92277.1 aluc|OJZ90171.1 anid|CBF80606.1 asyd|OJJ54706.1 atub|OJI81599.1 aver|OJJ07463.1

Group7673: afla|EED45192.1 afla|EED56934.1 afum|EAL89358.1 afum|EAL91840.1 aory|EIT73583.1 aory|EIT77457.1 afis|EAW18029.1 agla|OJJ79482.1 agla|OJJ86419.1 asyd|OJJ60254.1 ater|EAU31810.1 aver|OJJ04003.1

Group7674: afla|EED52293.1 afla|EED55253.1 aory|EIT74395.1 acar|OOF92107.1 akaw|GAA82888.1 anig|EHA20677.1 ater|EAU34227.1 atub|OJI81192.1 awen|OJJ36197.1 aory|EIT83537.1 ater|EAU30380.1 ater|EAU39011.1

Group7675: agla|OJJ78541.1 agla|OJJ79072.1 agla|OJJ85470.1 agla|OJJ80123.1 agla|OJJ84570.1 aluc|OJZ79503.1 aluc|OJZ79504.1 aluc|OJZ79505.1 aluc|OJZ79506.1 aluc|OJZ80207.1 anid|CBF81124.1 anid|CBF80690.1

Group7676: agla|OJJ78953.1 agla|OJJ86170.1 aver|OJJ01157.1 aver|OJJ02588.1 acar|OOF93635.1 asyd|OJJ59699.1 ater|EAU29902.1 awen|OJJ33469.1 azon|OJJ44552.1 agla|OJJ79879.1 asyd|OJJ58558.1 awen|OJJ39793.1

Group7677: agla|OJJ83762.1 agla|OJJ88099.1 afla|EED45943.1 anid|CBF76181.1 aory|EIT75503.1 asyd|OJJ54376.1 aver|OJJ05330.1 afla|EED53128.1 aory|EIT75245.1 ater|EAU38147.1 awen|OJJ35602.1 azon|OJJ46533.1

Group7678: agla|OJJ85586.1 agla|OJJ87986.1 agla|OJJ87987.1 abra|OJJ65967.1 acar|OOF92907.1 acla|EAW13588.1 akaw|GAA90842.1 aluc|OJZ82991.1 asyd|OJJ53311.1 atub|OJI89225.1 aver|OJJ08432.1 awen|OJJ30030.1

Group7679: agla|OJJ87968.1 agla|OJJ87970.1 acla|EAW06333.1 aory|EIT73259.1 aver|OJJ03126.1 awen|OJJ40054.1 acar|OOF96480.1 aluc|OJZ80027.1 asyd|OJJ61373.1 azon|OJJ42338.1 azon|OJJ44456.1 aver|OJJ00700.1

Group7680: agla|OJJ88217.1 abra|OJJ66972.1 akaw|GAA92192.1 aluc|OJZ85607.1 anig|EHA24919.1 atub|OJI79400.1 acar|OOF89937.1 acla|EAW15004.1 asyd|OJJ53754.1 aver|OJJ04037.1 agla|OJJ80331.1 ater|EAU29897.1

Group7681: akaw|GAA86475.1 akaw|GAA88273.1 aluc|OJZ82245.1 aluc|OJZ85016.1 anig|EHA25237.1 anig|EHA28497.1 abra|OJJ69408.1 acar|OOF94324.1 atub|OJI88193.1 azon|OJJ45519.1 acar|OOF92472.1 atub|OJI80581.1

Group7682: akaw|GAA92631.1 akaw|GAA93217.1 akaw|GAA93206.1 acla|EAW11094.1 afla|EED47301.1 aluc|OJZ79896.1 aory|EIT82879.1 acar|OOF98664.1 agla|OJJ87887.1 atub|OJI86643.1 anig|EHA20977.1 aluc|OJZ79813.1

Group7683: akaw|GAA93035.1 akaw|GAA93188.1 acla|EAW08081.1 anid|CBF76131.1 acla|EAW13564.1 aory|EIT74602.1 afla|EED44723.1 agla|OJJ83536.1 aory|EIT74601.1 asyd|OJJ54400.1 ater|EAU29677.1 agla|OJJ79573.1

Group7684: aluc|OJZ79517.1 aluc|OJZ82151.1 abra|OJJ77707.1 afis|EAW22827.1 agla|OJJ81580.1 akaw|GAA83923.1 awen|OJJ29836.1 azon|OJJ49065.1 ater|EAU39006.1 agla|OJJ80694.1 anid|CBF69419.1 aluc|OJZ90772.1

Group7685: anid|CBF74471.1 anid|CBF83141.1 acar|OOF90730.1 acla|EAW11661.1 agla|OJJ80959.1 aluc|OJZ80568.1 anid|CBF69458.1 anig|EHA25630.1 asyd|OJJ57702.1 atub|OJI88363.1 aver|OJJ05462.1 ater|EAU32818.1

Group7686: anig|EHA25380.1 anig|EHA27101.1 atub|OJI80429.1 atub|OJI84906.1 abra|OJJ73396.1 acar|OOF95494.1 afla|EED48296.1 aory|EIT76312.1 ater|EAU37658.1 abra|OJJ74232.1 akaw|GAA86787.1 aluc|OJZ89901.1

Group7687: aory|EIT75329.1 aory|EIT81312.1 acla|EAW10356.1 afis|EAW25238.1 afla|EED51106.1 afum|EAL89899.1 anid|CBF71827.1 asyd|OJJ54971.1 ater|EAU34363.1 aver|OJJ07220.1 awen|OJJ40951.1 afla|EED53705.1

Group7688: asyd|OJJ52874.1 asyd|OJJ56093.1 aver|OJJ05036.1 aver|OJJ05544.1 acla|EAW06675.1 afis|EAW16090.1 afla|EED56822.1 afum|EAL88722.1 agla|OJJ89210.1 anid|CBF78311.1 aory|EIT73863.1 ater|EAU32515.1

Group7689: asyd|OJJ53582.1 asyd|OJJ53655.1 ater|EAU29680.1 ater|EAU38473.1 abra|OJJ70110.1 afla|EED52397.1 anid|CBF89322.1 anig|EHA24974.1 aory|EIT83285.1 atub|OJI87919.1 awen|OJJ30809.1 ater|EAU30258.1

Group7690: asyd|OJJ56530.1 asyd|OJJ59832.1 aver|OJJ02456.1 aver|OJJ06193.1 abra|OJJ67083.1 afis|EAW17161.1 afla|EED45643.1 afum|EAL86598.1 akaw|GAA91076.1 aluc|OJZ87891.1 aory|EIT75802.1 atub|OJI80761.1

Group7691: ater|EAU34014.1 ater|EAU37263.1 abra|OJJ68840.1 acar|OOF90550.1 afla|EED48419.1 afum|EAL89284.1 akaw|GAA90409.1 aluc|OJZ85819.1 anig|EHA24741.1 aory|EIT83107.1 atub|OJI79605.1 awen|OJJ31448.1

Group7692: ater|EAU35408.1 ater|EAU37500.1 abra|OJJ66201.1 afis|EAW19378.1 afla|EED45776.1 afum|EAL94092.1 akaw|GAA89083.1 aluc|OJZ80507.1 anid|CBF84662.1 anig|EHA25686.1 aory|EIT75685.1 atub|OJI88423.1

Group7693: aver|OJI98033.1 aver|OJI98035.1 abra|OJJ73322.1 anig|EHA18329.1 asyd|OJJ62081.1 atub|OJI88678.1 awen|OJJ38257.1 atub|OJI85498.1 akaw|GAA84567.1 aluc|OJZ89300.1 asyd|OJJ65301.1 aver|OJI96189.1

Group7694: aver|OJJ06494.1 aver|OJJ07722.1 abra|OJJ68455.1 acar|OOF93942.1 acla|EAW14590.1 akaw|GAA86919.1 aluc|OJZ90042.1 anid|CBF83906.1 anig|EHA21028.1 asyd|OJJ55417.1 atub|OJI84764.1 aluc|OJZ92924.1

Group7695: awen|OJJ30775.1 awen|OJJ31706.1 acla|EAW12283.1 afla|EED45674.1 afum|EAL84398.1 akaw|GAA92930.1 aluc|OJZ82523.1 aory|EIT75753.1 afis|EAW25725.1 anid|CBF83904.1 asyd|OJJ55415.1 aver|OJJ06492.1

Group7696: awen|OJJ30786.1 awen|OJJ31695.1 abra|OJJ68903.1 afla|EED51119.1 akaw|GAA90348.1 aluc|OJZ85884.1 anig|EHA24702.1 aory|EIT81361.1 asyd|OJJ57238.1 atub|OJI79670.1 aver|OJJ07748.1 azon|OJJ42366.1

Group7697: awen|OJJ33531.1 awen|OJJ38755.1 abra|OJJ67826.1 acla|EAW11667.1 afla|EED44780.1 agla|OJJ83419.1 anig|EHA22196.1 aory|EIT77765.1 asyd|OJJ60486.1 ater|EAU32819.1 aver|OJI99812.1 azon|OJJ45348.1

Group7698: awen|OJJ34695.1 awen|OJJ35397.1 abra|OJJ76901.1 acar|OOF94976.1 afla|EED57113.1 akaw|GAA87138.1 aluc|OJZ84299.1 anid|CBF73949.1 anig|EHA27636.1 aory|EIT73972.1 ater|EAU30247.1 atub|OJI89770.1

Group7699: azon|OJJ42584.1 azon|OJJ45682.1 azon|OJJ50353.1 abra|OJJ68641.1 akaw|GAA89779.1 aluc|OJZ81913.1 anig|EHA22332.1 atub|OJI84465.1 afis|EAW15860.1 abra|OJJ77938.1 acar|OOF95217.1 azon|OJJ45691.1

Group7700: azon|OJJ44016.1 azon|OJJ44483.1 afis|EAW16698.1 afum|EAL89295.1 anig|EHA21621.1 aory|EIT82617.1 asyd|OJJ61736.1 ater|EAU35154.1 afla|EED49533.1 aver|OJI95829.1 afla|EED54073.1 aory|EIT75934.1

Group7701: azon|OJJ45223.1 azon|OJJ50375.1 azon|OJJ48030.1 afis|EAW20692.1 afum|EAL86274.1 afis|EAW24588.1 agla|OJJ82377.1 asyd|OJJ59621.1 aver|OJI96482.1 awen|OJJ38307.1 awen|OJJ33829.1 agla|OJJ86208.1

Group7702: ater|EAU29224.1 ater|EAU39627.1 ater|EAU37739.1 ater|EAU29404.1 ater|EAU36851.1 ater|EAU32783.1 ater|EAU36852.1 ater|EAU34004.1 ater|EAU34003.1 ater|EAU34001.1 ater|EAU31662.1 ater|EAU32784.1

Group7703: abra|OJJ65456.1 acla|EAW12622.1 afis|EAW18062.1 afla|EED51430.1 afum|EAL91873.1 akaw|GAA92603.1 aluc|OJZ82621.1 anid|CBF70082.1 aory|EIT80753.1 atub|OJI84571.1 aver|OJI99055.1 awen|OJJ39990.1

Group7704: abra|OJJ65521.1 acar|OOG00382.1 acla|EAW11461.1 afla|EED47679.1 akaw|GAA92935.1 awen|OJJ32784.1 abra|OJJ75276.1 akaw|GAA91816.1 aluc|OJZ92838.1 aory|EIT80248.1 atub|OJI87294.1 acla|EAW11448.1

Group7705: abra|OJJ65591.1 acla|EAW15024.1 afum|EAL91140.1 aluc|OJZ87727.1 anid|CBF70726.1 anig|EHA20219.1 asyd|OJJ55956.1 ater|EAU37181.1 atub|OJI80916.1 aver|OJJ04902.1 awen|OJJ30889.1 atub|OJI80915.1

Group7706: abra|OJJ65632.1 asyd|OJJ64077.1 ater|EAU35401.1 aver|OJI98165.1 abra|OJJ74155.1 acar|OOF98984.1 akaw|GAA89728.1 aluc|OJZ80655.1 anig|EHA27177.1 ater|EAU34546.1 atub|OJI87510.1 awen|OJJ35908.1

Group7707: abra|OJJ65685.1 afla|EED44898.1 akaw|GAA85808.1 aory|EIT83245.1 atub|OJI83487.1 abra|OJJ66513.1 acla|EAW08894.1 akaw|GAA91198.1 aluc|OJZ81533.1 anig|EHA25903.1 atub|OJI88654.1 atub|OJI83281.1

Group7708: abra|OJJ65696.1 afla|EED56835.1 afum|EAL84265.1 agla|OJJ84576.1 anig|EHA18816.1 aory|EIT73898.1 ater|EAU33255.1 aver|OJJ07756.1 awen|OJJ30262.1 azon|OJJ43501.1 afla|EED54691.1 aory|EIT78815.1

Group7709: abra|OJJ65740.1 acar|OOF98411.1 acla|EAW14577.1 agla|OJJ86269.1 akaw|GAA91461.1 aluc|OJZ81130.1 anid|CBF78778.1 asyd|OJJ65242.1 awen|OJJ30186.1 azon|OJJ44976.1 atub|OJI84622.1 aver|OJI99393.1

Group7710: abra|OJJ65744.1 afis|EAW25302.1 afla|EED53781.1 akaw|GAA93269.1 aory|EIT78412.1 asyd|OJJ53534.1 ater|EAU30712.1 aver|OJJ07518.1 awen|OJJ31268.1 azon|OJJ43189.1 acar|OOF92061.1 asyd|OJJ54638.1

Group7711: abra|OJJ65774.1 acla|EAW15144.1 afis|EAW22742.1 afum|EAL84238.1 akaw|GAA82879.1 aluc|OJZ87447.1 anig|EHA20670.1 ater|EAU36809.1 atub|OJI81202.1 aver|OJI99435.1 azon|OJJ50701.1 afum|EAL90453.1

Group7712: abra|OJJ65779.1 afis|EAW17148.1 afla|EED51030.1 agla|OJJ79807.1 akaw|GAA90884.1 aluc|OJZ82948.1 anig|EHA21365.1 aory|EIT77069.1 asyd|OJJ53779.1 atub|OJI89180.1 aver|OJJ07887.1 awen|OJJ35385.1

Group7713: abra|OJJ65797.1 acar|OOF89998.1 afla|EED46363.1 agla|OJJ88225.1 anig|EHA22244.1 aory|EIT81235.1 abra|OJJ70155.1 akaw|GAA85143.1 aluc|OJZ86494.1 anig|EHA22150.1 atub|OJI84032.1 anid|CBF74180.1

Group7714: abra|OJJ65810.1 acar|OOF96445.1 afla|EED47494.1 akaw|GAA92878.1 aluc|OJZ82530.1 anid|CBF70340.1 anig|EHA22496.1 aory|EIT74547.1 asyd|OJJ58707.1 ater|EAU33071.1 atub|OJI88775.1 aver|OJJ01320.1

Group7715: abra|OJJ65898.1 acar|OOF97277.1 akaw|GAA86006.1 aluc|OJZ90232.1 anig|EHA23733.1 atub|OJI81662.1 abra|OJJ66271.1 afla|EED52054.1 aory|EIT81394.1 ater|EAU30259.1 agla|OJJ80348.1 agla|OJJ80349.1

Group7716: abra|OJJ65904.1 acar|OOF93264.1 afla|EED56024.1 afum|EAL84406.1 akaw|GAA92327.1 aluc|OJZ90225.1 anid|CBF74692.1 anig|EHA23726.1 asyd|OJJ60212.1 atub|OJI81658.1 aver|OJJ04056.1 awen|OJJ30875.1

Group7717: abra|OJJ65921.1 acar|OOF94982.1 afla|EED46586.1 agla|OJJ85290.1 akaw|GAA87345.1 aluc|OJZ89669.1 anig|EHA23707.1 aory|EIT77838.1 atub|OJI81640.1 awen|OJJ31277.1 akaw|GAA92311.1 atub|OJI85131.1

Group7718: abra|OJJ65929.1 acar|OOF94990.1 afla|EED50955.1 agla|OJJ86022.1 akaw|GAA92307.1 aluc|OJZ90203.1 anig|EHA23700.1 aory|EIT76963.1 asyd|OJJ62113.1 atub|OJI81632.1 aver|OJI99296.1 awen|OJJ30467.1

Group7719: abra|OJJ65933.1 acar|OOF94998.1 afla|EED54754.1 akaw|GAA92302.1 aluc|OJZ90199.1 anid|CBF75864.1 aory|EIT78874.1 asyd|OJJ60226.1 ater|EAU34018.1 atub|OJI81627.1 aver|OJJ04041.1 awen|OJJ36435.1

Group7720: agla|OJJ88524.1 awen|OJJ30076.1 abra|OJJ75971.1 acar|OOF99949.1 afla|EED54790.1 akaw|GAA88985.1 aluc|OJZ89418.1 anig|EHA23170.1 aory|EIT79391.1 asyd|OJJ52192.1 atub|OJI85376.1 aver|OJJ05954.1

Group7721: abra|OJJ66060.1 afla|EED49908.1 akaw|GAA93130.1 aluc|OJZ81103.1 anid|CBF84615.1 anig|EHA19266.1 aory|EIT72451.1 asyd|OJJ55638.1 ater|EAU30987.1 atub|OJI82476.1 aver|OJJ06721.1 awen|OJJ33567.1

Group7722: afla|EED52065.1 azon|OJJ51778.1 abra|OJJ71647.1 acar|OOF96361.1 akaw|GAA89282.1 aluc|OJZ86391.1 anig|EHA21507.1 asyd|OJJ63120.1 ater|EAU36529.1 atub|OJI84142.1 aver|OJI97838.1 awen|OJJ32029.1

Group7723: abra|OJJ66253.1 acar|OOF90723.1 acla|EAW13651.1 afis|EAW19954.1 afla|EED45864.1 afum|EAL84961.1 akaw|GAA89034.1 aluc|OJZ80561.1 anig|EHA25636.1 aory|EIT75666.1 ater|EAU36873.1 atub|OJI88370.1

Group7724: abra|OJJ66265.1 acar|OOF97513.1 afla|EED48579.1 agla|OJJ82945.1 akaw|GAA89029.1 aluc|OJZ80575.1 anig|EHA25623.1 aory|EIT73107.1 asyd|OJJ58036.1 atub|OJI88356.1 aver|OJJ04363.1 awen|OJJ32355.1

Group7725: afla|EED49619.1 afum|EAL89790.1 aory|EIT78191.1 afis|EAW25119.1 awen|OJJ40738.1 acla|EAW11037.1 aory|EIT80982.1 akaw|GAA84863.1 aluc|OJZ86808.1 atub|OJI83709.1 awen|OJJ33430.1 afla|EED52806.1

Group7726: abra|OJJ66293.1 afis|EAW15415.1 abra|OJJ68852.1 acar|OOF98662.1 aluc|OJZ91912.1 anid|CBF75772.1 asyd|OJJ62871.1 aver|OJI96924.1 azon|OJJ48282.1 azon|OJJ50411.1 akaw|GAA82235.1 atub|OJI86362.1

Group7727: abra|OJJ66363.1 afum|EAL84456.1 agla|OJJ85514.1 aluc|OJZ83105.1 anig|EHA28168.1 asyd|OJJ62939.1 atub|OJI87829.1 aver|OJI98052.1 awen|OJJ37289.1 azon|OJJ42066.1 agla|OJJ81972.1 atub|OJI87830.1

Group7728: abra|OJJ66439.1 acar|OOF90290.1 afis|EAW17056.1 afla|EED56940.1 afum|EBA27278.1 agla|OJJ84863.1 akaw|GAA90301.1 aluc|OJZ83036.1 asyd|OJJ55134.1 aver|OJJ07070.1 awen|OJJ37360.1 aver|OJJ07069.1

Group7729: abra|OJJ66480.1 afla|EED46095.1 akaw|GAA92463.1 aluc|OJZ81554.1 anid|CBF82153.1 anig|EHA25880.1 aory|EIT76478.1 asyd|OJJ54330.1 ater|EAU37662.1 atub|OJI88631.1 aver|OJJ05284.1 awen|OJJ41955.1

Group7730: abra|OJJ66498.1 acla|EAW11079.1 afla|EED48918.1 agla|OJJ78855.1 aory|EIT81209.1 asyd|OJJ61762.1 ater|EAU32698.1 aver|OJI95861.1 azon|OJJ44219.1 anid|CBF85162.1 asyd|OJJ54546.1 aver|OJJ07631.1

Group7731: abra|OJJ66565.1 acla|EAW10931.1 afis|EAW20930.1 afla|EED50284.1 afum|EAL84704.1 agla|OJJ83958.1 aluc|OJZ81480.1 anid|CBF82512.1 asyd|OJJ54082.1 aver|OJJ08656.1 awen|OJJ34388.1 azon|OJJ48120.1

Group7732: abra|OJJ66626.1 acar|OOF91050.1 akaw|GAA91099.1 aluc|OJZ81422.1 anig|EHA25999.1 asyd|OJJ54025.1 ater|EAU38637.1 atub|OJI88759.1 awen|OJJ34322.1 azon|OJJ48068.1 anid|CBF82607.1 aver|OJJ08715.1

Group7733: abra|OJJ66634.1 acar|OOF91057.1 acla|EAW10861.1 afis|EAW20997.1 afum|EAL84631.1 akaw|GAA91094.1 aluc|OJZ81413.1 anid|CBF82631.1 anig|EHA26005.1 ater|EAU38645.1 atub|OJI88766.1 aver|OJJ08729.1

Group7734: ater|EAU39538.1 acla|EAW12270.1 akaw|GAA93168.1 aluc|OJZ83027.1 asyd|OJJ54186.1 afla|EED51446.1 agla|OJJ78762.1 aory|EIT80520.1 asyd|OJJ56240.1 aver|OJJ07428.1 aory|EIT81292.1 aory|EIT74777.1

Group7735: abra|OJJ66800.1 acar|OOF89950.1 afla|EED50558.1 agla|OJJ86487.1 akaw|GAA86864.1 aluc|OJZ89988.1 aory|EIT82148.1 ater|EAU38752.1 atub|OJI84822.1 awen|OJJ33316.1 azon|OJJ42587.1 awen|OJJ35519.1

Group7736: abra|OJJ66807.1 acar|OOF89865.1 afla|EED45504.1 afum|EAL86260.1 anid|CBF74948.1 anig|EHA22864.1 asyd|OJJ63939.1 atub|OJI79335.1 aver|OJI97003.1 awen|OJJ31669.1 azon|OJJ47393.1 aory|EIT75746.1

Group7737: abra|OJJ66823.1 akaw|GAA90458.1 aluc|OJZ85766.1 anig|EHA24790.1 ater|EAU31569.1 atub|OJI79550.1 azon|OJJ45133.1 aory|EIT74723.1 afla|EED50055.1 afla|EED50056.1 awen|OJJ31438.1 awen|OJJ40138.1

Group7738: abra|OJJ66848.1 acla|EAW14427.1 afis|EAW19134.1 afum|EAL93866.1 agla|OJJ88609.1 akaw|GAA91557.1 aluc|OJZ85735.1 anid|CBF86334.1 anig|EHA24815.1 atub|OJI79520.1 awen|OJJ37470.1 azon|OJJ51148.1

Group7739: abra|OJJ66884.1 acar|OOF91658.1 afla|EED54231.1 akaw|GAA91526.1 aluc|OJZ85698.1 anig|EHA24844.1 asyd|OJJ65021.1 ater|EAU37307.1 atub|OJI79488.1 aver|OJI99140.1 awen|OJJ37519.1 anid|CBF86278.1

Group7740: abra|OJJ66947.1 akaw|GAA91473.1 aluc|OJZ85633.1 anig|EHA24898.1 asyd|OJJ63595.1 atub|OJI79423.1 aver|OJI97341.1 azon|OJJ50918.1 abra|OJJ76875.1 afla|EED49864.1 aory|EIT83407.1 ater|EAU38550.1

Group7741: azon|OJJ46492.1 abra|OJJ72303.1 acar|OOF94916.1 afla|EED44929.1 akaw|GAA86054.1 aluc|OJZ90284.1 anid|CBF81770.1 anig|EHA23774.1 asyd|OJJ59613.1 atub|OJI81708.1 aver|OJJ02656.1 aory|EIT73476.1

Group7742: abra|OJJ67079.1 afis|EAW17160.1 afla|EED45644.1 afum|EAL86597.1 akaw|GAA91077.1 aluc|OJZ87892.1 aory|EIT75452.1 asyd|OJJ59830.1 atub|OJI80760.1 aver|OJJ02457.1 asyd|OJJ56531.1 aver|OJJ06192.1

Group7743: abra|OJJ67085.1 afis|EAW17162.1 afla|EED45642.1 afum|EAL86599.1 akaw|GAA91074.1 aluc|OJZ87889.1 aory|EIT75477.1 asyd|OJJ59824.1 atub|OJI80763.1 aver|OJJ02463.1 asyd|OJJ56529.1 aver|OJJ06194.1

Group7744: abra|OJJ67088.1 acar|OOF98739.1 afla|EED52556.1 akaw|GAA91073.1 aluc|OJZ87887.1 anig|EHA20190.1 aory|EIT74195.1 atub|OJI80764.1 aver|OJJ02171.1 awen|OJJ39932.1 akaw|GAA93120.1 agla|OJJ80749.1

Group7745: abra|OJJ67121.1 acar|OOF98430.1 afla|EED49575.1 akaw|GAA91044.1 aluc|OJZ87854.1 anid|CBF75960.1 anig|EHA20159.1 aory|EIT82634.1 ater|EAU35486.1 atub|OJI80793.1 aver|OJI98019.1 azon|OJJ42389.1

Group7746: abra|OJJ67122.1 acar|OOF98429.1 afla|EED52834.1 akaw|GAA91043.1 aluc|OJZ87853.1 anig|EHA20158.1 aory|EIT78628.1 ater|EAU30047.1 atub|OJI80794.1 aver|OJJ05221.1 aory|EIT78659.1 aver|OJJ05222.1

Group7747: abra|OJJ67216.1 acar|OOF98342.1 afla|EED48099.1 aluc|OJZ87756.1 anid|CBF73467.1 anig|EHA20242.1 aory|EIT76260.1 asyd|OJJ59932.1 atub|OJI80887.1 aver|OJJ02355.1 akaw|GAA90961.1 anid|CBF75223.1

Group7748: abra|OJJ67235.1 acar|OOF98317.1 acla|EAW14733.1 akaw|GAA91659.1 aluc|OJZ87740.1 anig|EHA20232.1 atub|OJI80900.1 ater|EAU32758.1 agla|OJJ80474.1 anid|CBF87447.1 aory|EIT82827.1 ater|EAU33881.1

Group7749: abra|OJJ67262.1 afla|EED44923.1 agla|OJJ79287.1 akaw|GAA91678.1 aluc|OJZ87716.1 anig|EHA20210.1 aory|EIT73516.1 ater|EAU31936.1 atub|OJI80926.1 aver|OJJ00775.1 awen|OJJ35785.1 afum|EAL92105.1

Group7750: abra|OJJ67288.1 acla|EAW11926.1 asyd|OJJ65353.1 aver|OJJ08392.1 abra|OJJ67574.1 acar|OOF91948.1 afla|EED49682.1 akaw|GAA89103.1 anig|EHA25705.1 aory|EIT81004.1 atub|OJI88442.1 aver|OJJ05142.1

Group7751: abra|OJJ67292.1 acar|OOF92519.1 afla|EED45659.1 agla|OJJ79146.1 akaw|GAA86521.1 aluc|OJZ85067.1 anid|CBF81706.1 anig|EHA25281.1 aory|EIT75796.1 asyd|OJJ59568.1 atub|OJI80531.1 aver|OJJ02699.1

Group7752: abra|OJJ67295.1 acar|OOF92517.1 afla|EED45655.1 agla|OJJ79149.1 akaw|GAA86518.1 aluc|OJZ85064.1 anid|CBF81700.1 anig|EHA25278.1 aory|EIT75769.1 asyd|OJJ59571.1 atub|OJI80534.1 aver|OJJ02696.1

Group7753: abra|OJJ67469.1 acar|OOF92365.1 afis|EAW18408.1 afla|EED52999.1 afum|EAL93088.1 agla|OJJ84975.1 akaw|GAA86367.1 aluc|OJZ84896.1 anig|EHA25124.1 aory|EIT77441.1 atub|OJI80702.1 awen|OJJ31741.1

Group7754: abra|OJJ67527.1 afla|EED45121.1 akaw|GAA88135.1 aluc|OJZ81406.1 anig|EHA18700.1 aory|EIT73612.1 ater|EAU38026.1 atub|OJI91613.1 azon|OJJ46856.1 abra|OJJ71709.1 afis|EAW16733.1 aluc|OJZ82080.1

Group7755: abra|OJJ67547.1 akaw|GAA88113.1 aluc|OJZ81385.1 anig|EHA18678.1 atub|OJI91590.1 awen|OJJ30837.1 acar|OOF91839.1 afum|EAL86145.1 afis|EAW24472.1 afla|EED51419.1 aory|EIT80675.1 azon|OJJ46868.1

Group7756: abra|OJJ67600.1 acar|OOF91953.1 afla|EED57077.1 akaw|GAA89119.1 aluc|OJZ81720.1 anig|EHA25719.1 ater|EAU36925.1 atub|OJI88461.1 awen|OJJ41650.1 acar|OOF99338.1 aory|EIT82257.1 atub|OJI85988.1

Group7757: abra|OJJ67610.1 akaw|GAA89127.1 aluc|OJZ81711.1 anig|EHA25728.1 atub|OJI88473.1 abra|OJJ77181.1 acar|OOF91800.1 afla|EED47268.1 agla|OJJ84568.1 aory|EIT79210.1 asyd|OJJ64972.1 aver|OJI99096.1

Group7758: abra|OJJ67628.1 afis|EAW22230.1 afla|EED48945.1 afum|EAL90963.1 agla|OJJ86822.1 akaw|GAA89148.1 aluc|OJZ81688.1 anig|EHA25755.1 aory|EIT80023.1 ater|EAU30273.1 atub|OJI88491.1 awen|OJJ33560.1

Group7759: abra|OJJ67668.1 acla|EAW12018.1 afis|EAW19659.1 afum|EAL86864.1 akaw|GAA89184.1 aluc|OJZ81647.1 anig|EHA25796.1 asyd|OJJ56442.1 atub|OJI88535.1 aver|OJJ06279.1 awen|OJJ40941.1 azon|OJJ47339.1

Group7760: abra|OJJ67689.1 acar|OOF97598.1 akaw|GAA89200.1 aluc|OJZ81630.1 anig|EHA25813.1 aory|EIT72450.1 asyd|OJJ62817.1 ater|EAU33823.1 atub|OJI88550.1 aver|OJI96861.1 awen|OJJ35797.1 afla|EED49893.1

Group7761: asyd|OJJ56367.1 abra|OJJ74235.1 akaw|GAA86785.1 aluc|OJZ89896.1 anig|EHA27095.1 ater|EAU37996.1 atub|OJI84911.1 azon|OJJ43210.1 afla|EED56906.1 aory|EIT77585.1 ater|EAU37207.1 aver|OJJ06347.1

Group7762: abra|OJJ67736.1 acar|OOF90933.1 afla|EED45459.1 akaw|GAA92426.1 aluc|OJZ81596.1 anid|CBF82814.1 anig|EHA25845.1 aory|EIT82392.1 ater|EAU30795.1 atub|OJI88586.1 aver|OJJ00894.1 acla|EAW15012.1

Group7763: abra|OJJ67778.1 anig|EHA26129.1 awen|OJJ36195.1 abra|OJJ74149.1 acar|OOF95576.1 agla|OJJ78610.1 akaw|GAA89734.1 aluc|OJZ80649.1 anig|EHA27183.1 asyd|OJJ61973.1 atub|OJI87504.1 aver|OJI96066.1

Group7764: abra|OJJ67832.1 acar|OOG00391.1 afis|EAW16691.1 afla|EED54749.1 akaw|GAA91841.1 aluc|OJZ82057.1 anig|EHA22202.1 atub|OJI84616.1 azon|OJJ42293.1 acar|OOG00607.1 acla|EAW07057.1 ater|EAU38378.1

Group7765: abra|OJJ67837.1 acar|OOF92011.1 afla|EED45645.1 akaw|GAA91842.1 aluc|OJZ82055.1 anid|CBF82517.1 anig|EHA22205.1 aory|EIT75527.1 asyd|OJJ54085.1 atub|OJI84614.1 aver|OJJ08653.1 awen|OJJ34680.1

Group7766: aory|EIT80699.1 acla|EAW11754.1 afis|EAW23787.1 afla|EED51596.1 afum|EAL87221.1 agla|OJJ78814.1 akaw|GAA91852.1 anid|CBF79849.1 awen|OJJ40590.1 azon|OJJ42846.1 asyd|OJJ58714.1 aver|OJJ01327.1

Group7767: abra|OJJ67875.1 acla|EAW09035.1 afis|EAW22733.1 afum|EAL90462.1 agla|OJJ78521.1 akaw|GAA91883.1 aluc|OJZ82014.1 anid|CBF84340.1 anig|EHA22242.1 ater|EAU38784.1 atub|OJI84563.1 ater|EAU39160.1

Group7768: abra|OJJ67893.1 acar|OOF92867.1 afla|EED51694.1 akaw|GAA91898.1 aluc|OJZ81997.1 anid|CBF84531.1 anig|EHA22259.1 aory|EIT80449.1 atub|OJI84547.1 awen|OJJ36322.1 asyd|OJJ53499.1 aver|OJJ08246.1

Group7769: abra|OJJ67947.1 acar|OOF92795.1 acla|EAW11634.1 afis|EAW23823.1 afla|EED51985.1 akaw|GAA91250.1 aluc|OJZ82819.1 anig|EHA21259.1 aory|EIT81415.1 ater|EAU31378.1 atub|OJI89049.1 awen|OJJ31423.1

Group7770: abra|OJJ68013.1 acar|OOF92055.1 akaw|GAA90938.1 aluc|OJZ82886.1 anid|CBF87409.1 anig|EHA21312.1 asyd|OJJ55062.1 atub|OJI89111.1 awen|OJJ31349.1 azon|OJJ50427.1 aver|OJJ07149.1 anid|CBF82887.1

Group7771: abra|OJJ68034.1 acar|OOF91809.1 afla|EED55047.1 afum|EAL86257.1 akaw|GAA90918.1 aluc|OJZ82908.1 anig|EHA21331.1 asyd|OJJ53716.1 ater|EAU36651.1 atub|OJI89134.1 aver|OJJ07842.1 awen|OJJ33893.1

Group7772: abra|OJJ68044.1 acar|OOF90406.1 acla|EAW13675.1 afis|EAW19932.1 afla|EED52285.1 afum|EAL84977.2 akaw|GAA90909.1 aluc|OJZ82917.1 anig|EHA21340.1 aory|EIT74359.1 atub|OJI89143.1 awen|OJJ31365.1

Group7773: abra|OJJ68082.1 acar|OOF93881.1 afla|EED51805.1 akaw|GAA86868.1 aluc|OJZ89992.1 anid|CBF71451.1 anig|EHA27363.1 aory|EIT81765.1 asyd|OJJ54952.1 atub|OJI84817.1 aver|OJJ07244.1 azon|OJJ45036.1

Group7774: abra|OJJ68097.1 acar|OOF93900.1 akaw|GAA86882.1 aluc|OJZ90005.1 anid|CBF78037.1 anig|EHA27380.1 asyd|OJJ55223.1 ater|EAU39549.1 atub|OJI84803.1 akaw|GAA86881.1 anid|CBF78039.1 atub|OJI84802.1

Group7775: abra|OJJ68130.1 acar|OOF90084.1 afla|EED45616.1 agla|OJJ85560.1 akaw|GAA86910.1 aluc|OJZ90033.1 anid|CBF80537.1 anig|EHA27409.1 asyd|OJJ54656.1 atub|OJI84773.1 aver|OJJ07506.1 awen|OJJ36982.1

Group7776: abra|OJJ68183.1 acar|OOF97478.1 afum|EAL91413.1 agla|OJJ81346.1 akaw|GAA83263.1 aluc|OJZ83769.1 anig|EHA18909.1 asyd|OJJ61979.1 atub|OJI90140.1 aver|OJI96071.1 awen|OJJ37246.1 azon|OJJ50944.1

Group7777: abra|OJJ68226.1 acar|OOF97523.1 acla|EAW14601.1 afla|EED46365.1 agla|OJJ79978.1 akaw|GAA83217.1 aluc|OJZ83719.1 anig|EHA18868.1 aory|EIT81222.1 asyd|OJJ57588.1 atub|OJI90093.1 aver|OJJ05594.1

Group7778: abra|OJJ68252.1 acar|OOF97531.1 afla|EED44935.1 afum|EAL91141.1 akaw|GAA83209.1 aluc|OJZ83711.1 anid|CBF74645.1 anig|EHA18862.1 aory|EIT73520.1 asyd|OJJ55958.1 atub|OJI90084.1 aver|OJJ04904.1

Group7779: abra|OJJ68255.1 acla|EAW08789.1 agla|OJJ84445.1 akaw|GAA83206.1 aluc|OJZ83708.1 anid|CBF71209.1 anig|EHA18859.1 asyd|OJJ54941.1 atub|OJI90080.1 awen|OJJ32016.1 azon|OJJ51812.1 agla|OJJ84446.1

Group7780: abra|OJJ68271.1 acar|OOF97550.1 akaw|GAA83198.1 awen|OJJ30297.1 abra|OJJ78223.1 acla|EAW12024.1 akaw|GAA85802.1 aluc|OJZ91298.1 anig|EHA19980.1 asyd|OJJ53516.1 atub|OJI83288.1 aver|OJJ08235.1

Group7781: ater|EAU30250.1 abra|OJJ74423.1 afla|EED53650.1 agla|OJJ88972.1 akaw|GAA87316.1 aluc|OJZ89701.1 anig|EHA26935.1 aory|EIT75315.1 atub|OJI85100.1 awen|OJJ40838.1 atub|OJI80442.1 aver|OJI95754.1

Group7782: abra|OJJ68319.1 acar|OOF93891.1 afla|EED52687.1 agla|OJJ85435.1 akaw|GAA83154.1 aluc|OJZ83654.1 anig|EHA18805.1 aory|EIT74301.1 asyd|OJJ63047.1 atub|OJI90023.1 aver|OJI97916.1 awen|OJJ39602.1

Group7783: abra|OJJ68321.1 acar|OOF97580.1 afla|EED52685.1 agla|OJJ85434.1 akaw|GAA83152.1 aluc|OJZ83652.1 anig|EHA18803.1 aory|EIT74311.1 atub|OJI90021.1 awen|OJJ39604.1 aver|OJJ04463.1 azon|OJJ50222.1

Group7784: abra|OJJ68354.1 acar|OOF96867.1 afla|EED53489.1 aluc|OJZ83596.1 anig|EHA18756.1 aory|EIT74860.1 asyd|OJJ56979.1 ater|EAU32939.1 atub|OJI89968.1 aver|OJJ03308.1 awen|OJJ36413.1 akaw|GAA92126.1

Group7785: abra|OJJ68356.1 afla|EED49517.1 akaw|GAA92128.1 aluc|OJZ83594.1 anig|EHA18754.1 aory|EIT75080.1 asyd|OJJ56977.1 ater|EAU39004.1 atub|OJI89966.1 aver|OJJ03310.1 awen|OJJ36415.1 afla|EED53491.1

Group7786: abra|OJJ68420.1 afla|EED50221.1 akaw|GAA92528.1 aluc|OJZ90077.1 anig|EHA20996.1 aory|EIT77318.1 ater|EAU34260.1 atub|OJI84729.1 agla|OJJ78584.1 asyd|OJJ59203.1 awen|OJJ36121.1 aver|OJJ01815.1

Group7787: abra|OJJ68498.1 acla|EAW14276.1 afis|EAW18998.1 afum|EAL93733.1 agla|OJJ81777.1 aluc|OJZ82711.1 anig|EHA21166.1 asyd|OJJ64683.1 atub|OJI88944.1 aver|OJI98785.1 awen|OJJ38286.1 azon|OJJ50677.1

Group7788: ater|EAU34701.1 acla|EAW12100.1 afis|EAW17286.1 afla|EED48832.1 afum|EAL86704.1 agla|OJJ81091.1 akaw|GAA92755.1 aluc|OJZ92964.1 anig|EHA26011.1 aory|EIT81284.1 atub|OJI87422.1 azon|OJJ45535.1

Group7789: abra|OJJ68733.1 acla|EAW12515.1 afis|EAW24132.1 agla|OJJ87209.1 aluc|OJZ81797.1 anid|CBF86564.1 anig|EHA22419.1 asyd|OJJ64850.1 atub|OJI84368.1 aver|OJI98962.1 awen|OJJ41357.1 azon|OJJ48821.1

Group7790: acar|OOF92243.1 akaw|GAA89898.1 anig|EHA22443.1 acla|EAW12497.1 afis|EAW24148.1 afla|EED46785.1 afum|EAL92003.1 agla|OJJ87228.1 anid|CBF77919.1 aory|EIT72557.1 ater|EAU30580.1 awen|OJJ41331.1

Group7791: abra|OJJ68836.1 acar|OOF90554.1 acla|EAW15291.1 afis|EAW20302.1 afum|EAL85432.1 akaw|GAA90415.1 aluc|OJZ85815.1 anid|CBF85210.1 asyd|OJJ63032.1 ater|EAU32048.1 atub|OJI79601.1 aver|OJI97936.1

Group7792: abra|OJJ68846.1 acar|OOF90543.1 afis|EAW20250.1 afla|EED57095.1 afum|EAL85368.1 akaw|GAA90402.1 aluc|OJZ85826.1 anig|EHA24735.1 aory|EIT82289.1 ater|EAU32041.1 atub|OJI79614.1 awen|OJJ31254.1

Group7793: abra|OJJ68899.1 afla|EED56741.1 akaw|GAA90352.1 aluc|OJZ85880.1 anig|EHA24706.1 aory|EIT73870.1 asyd|OJJ56704.1 ater|EAU34587.1 atub|OJI79666.1 aver|OJJ06021.1 awen|OJJ34050.1 azon|OJJ48181.1

Group7794: abra|OJJ68922.1 acar|OOF90181.1 agla|OJJ87041.1 akaw|GAA90326.1 aluc|OJZ85907.1 anig|EHA24685.1 asyd|OJJ65138.1 ater|EAU36408.1 atub|OJI79691.1 aver|OJJ03960.1 awen|OJJ37940.1 anig|EHA24687.1

Group7795: abra|OJJ69096.1 acla|EAW08735.1 afis|EAW15701.1 afum|EAL85647.2 aluc|OJZ86086.1 anid|CBF71112.1 anig|EHA24530.1 asyd|OJJ52714.1 atub|OJI79882.1 aver|OJJ04743.1 azon|OJJ43193.1 awen|OJJ38394.1

Group7796: abra|OJJ69141.1 acar|OOF91161.1 agla|OJJ81903.1 akaw|GAA86316.1 aluc|OJZ86127.1 anid|CBF71040.1 anig|EHA24493.1 asyd|OJJ52668.1 atub|OJI79923.1 aver|OJJ04694.1 awen|OJJ38441.1 azon|OJJ43804.1

Group7797: abra|OJJ69221.1 acla|EAW08625.1 afis|EAW15807.1 afla|EED51239.1 afum|EAL85544.1 akaw|GAA92229.1 asyd|OJJ52600.1 ater|EAU33437.1 atub|OJI79995.1 awen|OJJ38517.1 anig|EHA24421.1 aory|EIT83183.1

Group7798: abra|OJJ69226.1 acar|OOF91094.1 afla|EED51235.1 akaw|GAA92234.1 aluc|OJZ86206.1 anid|CBF75273.1 aory|EIT83173.1 asyd|OJJ55914.1 ater|EAU33441.1 atub|OJI80000.1 aver|OJJ04854.1 awen|OJJ38523.1

Group7799: abra|OJJ69261.1 acar|OOF91061.1 afis|EAW15845.1 afum|EAL85506.2 akaw|GAA92266.1 aluc|OJZ86240.1 anid|CBF70837.1 anig|EHA24389.1 aory|EIT80402.1 aver|OJJ04593.1 awen|OJJ38560.1 azon|OJJ43707.1

Group7800: abra|OJJ69331.1 acar|OOF94396.1 acla|EAW14709.1 afis|EAW24244.1 afum|EAL85912.1 akaw|GAA88202.1 aluc|OJZ82163.1 anid|CBF74212.1 anig|EHA28569.1 atub|OJI88274.1 awen|OJJ33055.1 afla|EED48030.1

Group7801: abra|OJJ69390.1 acar|OOF98420.1 afla|EED47296.1 agla|OJJ87839.1 akaw|GAA88258.1 aluc|OJZ82225.1 anig|EHA20735.1 aory|EIT79184.1 atub|OJI88210.1 awen|OJJ38322.1 azon|OJJ46846.1 acar|OOF98138.1

Group7802: akaw|GAA88303.1 acar|OOF94232.1 acla|EAW09970.1 afis|EAW21118.1 afla|EED51791.1 afum|EAL92142.2 agla|OJJ80093.1 anid|CBF87027.1 aory|EIT81687.1 ater|EAU36781.1 aver|OJI99775.1 awen|OJJ36448.1

Group7803: abra|OJJ69540.1 acar|OOF90592.1 afla|EED54771.1 agla|OJJ86495.1 akaw|GAA88388.1 aluc|OJZ82376.1 anid|CBF79835.1 anig|EHA28388.1 asyd|OJJ58146.1 atub|OJI88066.1 aver|OJJ04258.1 awen|OJJ29883.1

Group7804: abra|OJJ69556.1 afla|EED56588.1 agla|OJJ83853.1 akaw|GAA91565.1 aluc|OJZ82383.1 anig|EHA28380.1 aory|EIT79666.1 asyd|OJJ53860.1 atub|OJI88057.1 aver|OJJ07960.1 awen|OJJ39006.1 acar|OOF92424.1

Group7805: abra|OJJ69558.1 acar|OOF91592.1 afla|EED54880.1 akaw|GAA91567.1 aluc|OJZ82385.1 anid|CBF85622.1 anig|EHA28378.1 aory|EIT79358.1 asyd|OJJ64201.1 atub|OJI88055.1 aver|OJI98290.1 awen|OJJ40806.1

Group7806: abra|OJJ69608.1 acar|OOF92699.1 afis|EAW23703.1 afla|EED51841.1 agla|OJJ88173.1 akaw|GAA91593.1 aluc|OJZ82416.1 anig|EHA28352.1 aory|EIT81731.1 ater|EAU31563.1 atub|OJI88025.1 awen|OJJ33654.1

Group7807: abra|OJJ69643.1 acar|OOF95564.1 acla|EAW10801.1 afis|EAW24776.1 afum|EBA27382.1 agla|OJJ88224.1 akaw|GAA91621.1 aluc|OJZ82447.1 anig|EHA28326.1 atub|OJI87992.1 awen|OJJ37936.1 azon|OJJ51806.1

Group7808: abra|OJJ69644.1 acar|OOF95563.1 acla|EAW10804.1 afis|EAW24774.1 afla|EED51027.1 akaw|GAA91622.1 aluc|OJZ82448.1 anid|CBF89919.1 anig|EHA28325.1 aory|EIT77043.1 atub|OJI87991.1 afum|EAL89415.1

Group7809: abra|OJJ69645.1 acar|OOF96863.1 agla|OJJ83837.1 akaw|GAA91623.1 aluc|OJZ82449.1 anid|CBF70110.1 anig|EHA28323.1 asyd|OJJ58822.1 atub|OJI87990.1 aver|OJJ01426.1 awen|OJJ35596.1 azon|OJJ46917.1

Group7810: abra|OJJ69685.1 acar|OOF93997.1 afis|EAW20308.1 agla|OJJ86261.1 akaw|GAA88809.1 aluc|OJZ82492.1 anig|EHA28280.1 aory|EIT79333.1 asyd|OJJ53574.1 atub|OJI87940.1 aver|OJJ08183.1 awen|OJJ31264.1

Group7811: abra|OJJ69800.1 acar|OOF93577.1 afis|EAW24740.1 afum|EBA27380.1 akaw|GAA87859.1 aluc|OJZ85500.1 anid|CBF80398.1 anig|EHA25033.1 asyd|OJJ54760.1 atub|OJI80065.1 aver|OJJ07410.1 azon|OJJ45020.1

Group7812: abra|OJJ69813.1 acar|OOF93564.1 afla|EED53287.1 akaw|GAA87844.1 aluc|OJZ85482.1 anig|EHA25022.1 aory|EIT74928.1 atub|OJI80082.1 abra|OJJ72703.1 asyd|OJJ54670.1 aver|OJJ07500.1 azon|OJJ44532.1

Group7813: abra|OJJ69858.1 acar|OOF90331.1 afla|EED56644.1 akaw|GAA87798.1 aluc|OJZ85433.1 anig|EHA24978.1 aory|EIT81635.1 asyd|OJJ57658.1 atub|OJI80135.1 aver|OJJ05519.1 afla|EED55141.1 anid|CBF83114.1

Group7814: abra|OJJ69866.1 acar|OOF90334.1 acla|EAW15314.1 akaw|GAA87790.1 aluc|OJZ85425.1 anid|CBF89416.1 anig|EHA25604.1 asyd|OJJ52768.1 atub|OJI80150.1 aver|OJJ08751.1 awen|OJJ38713.1 awen|OJJ31346.1

Group7815: abra|OJJ69893.1 acar|OOF93413.1 acla|EAW09497.1 afis|EAW21639.1 afum|EAL92673.1 akaw|GAA87757.1 aluc|OJZ85390.1 anid|CBF83523.1 anig|EHA25574.1 ater|EAU37628.1 atub|OJI80180.1 ater|EAU38324.1

Group7816: abra|OJJ69900.1 acar|OOF93418.1 afis|EAW15472.1 akaw|GAA87751.1 aluc|OJZ85384.1 anid|CBF78230.1 anig|EHA25568.1 asyd|OJJ56035.1 atub|OJI80186.1 aver|OJJ04981.1 azon|OJJ43161.1 afum|EAL85849.1

Group7817: abra|OJJ69973.1 acar|OOF93478.1 afis|EAW15557.1 agla|OJJ85197.1 akaw|GAA87688.1 aluc|OJZ85308.1 anig|EHA25504.1 asyd|OJJ55963.1 atub|OJI80263.1 aver|OJJ04909.1 awen|OJJ38215.1 azon|OJJ43080.1

Group7818: abra|OJJ69997.1 acar|OOF93504.1 agla|OJJ81896.1 akaw|GAA87664.1 aluc|OJZ85282.1 anid|CBF75974.1 anig|EHA25479.1 aory|EIT80225.1 atub|OJI80288.1 aver|OJJ05474.1 awen|OJJ37818.1 asyd|OJJ57690.1

Group7819: abra|OJJ70108.1 acar|OOG00113.1 afis|EAW25649.1 agla|OJJ81746.1 akaw|GAA92808.1 aluc|OJZ79628.1 aluc|OJZ88731.1 anig|EHA26920.1 aory|EIT75909.1 azon|OJJ50376.1 afla|EED54160.1 anig|EHA19939.1

Group7820: abra|OJJ70120.1 acar|OOF89935.1 afla|EED46964.1 akaw|GAA86653.1 aluc|OJZ85211.1 anid|CBF86364.1 anig|EHA25414.1 aory|EIT72580.1 asyd|OJJ64968.1 atub|OJI80393.1 aver|OJI99092.1 awen|OJJ35043.1

Group7821: abra|OJJ70171.1 acla|EAW13458.1 afis|EAW19015.1 afla|EED53215.1 afum|EBA27502.1 anid|CBF82209.1 anig|EHA25802.1 aory|EIT75022.1 asyd|OJJ57406.1 ater|EAU31285.1 aver|OJI99546.1 awen|OJJ32946.1

Group7822: abra|OJJ70172.1 acla|EAW13459.1 afis|EAW19016.1 afla|EED53216.1 afum|EAL93749.1 anid|CBF83507.1 anig|EHA25803.1 aory|EIT75109.1 asyd|OJJ52785.1 ater|EAU31286.1 aver|OJI99547.1 awen|OJJ32945.1

Group7823: abra|OJJ70226.1 acla|EAW15360.1 afis|EAW17375.1 afum|EAL91211.1 akaw|GAA86556.1 aluc|OJZ85104.1 anid|CBF84300.1 anig|EHA25325.1 asyd|OJJ58561.1 atub|OJI80493.1 aver|OJJ01160.1 azon|OJJ46423.1

Group7824: abra|OJJ70250.1 acar|OOF92547.1 acla|EAW08787.1 afis|EAW15616.1 afum|EAL85719.1 akaw|GAA86541.1 aluc|OJZ85089.1 anid|CBF78513.1 anig|EHA25309.1 ater|EAU31808.1 atub|OJI80510.1 awen|OJJ33723.1

Group7825: abra|OJJ70286.1 acar|OOF91739.1 acla|EAW09934.1 afla|EED50211.1 akaw|GAA92862.1 aluc|OJZ82551.1 anig|EHA22515.1 aory|EIT77364.1 ater|EAU30374.1 atub|OJI88792.1 awen|OJJ38796.1 azon|OJJ45868.1

Group7826: abra|OJJ70290.1 acar|OOF91743.1 afla|EED46566.1 akaw|GAA92858.1 aluc|OJZ82555.1 anig|EHA22519.1 aory|EIT77917.1 asyd|OJJ54711.1 atub|OJI88797.1 afis|EAW17781.1 aver|OJJ07455.1 afla|EED46567.1

Group7827: acla|EAW14390.1 acar|OOF91576.1 afis|EAW19097.1 afla|EED54331.1 afum|EAL93829.1 agla|OJJ84504.1 aory|EIT76031.1 asyd|OJJ58053.1 ater|EAU32145.1 aver|OJJ04350.1 awen|OJJ32490.1 anid|CBF79702.1

Group7828: abra|OJJ70309.1 acar|OOF91764.1 akaw|GAA93013.1 aluc|OJZ82572.1 anid|CBF80500.1 anig|EHA27502.1 asyd|OJJ61524.1 ater|EAU30199.1 atub|OJI88816.1 aver|OJJ00856.1 awen|OJJ36272.1 azon|OJJ50648.1

Group7829: abra|OJJ70320.1 acar|OOF91766.1 afla|EED56893.1 akaw|GAA92576.1 aluc|OJZ82589.1 anid|CBF73568.1 anig|EHA27486.1 asyd|OJJ56696.1 atub|OJI88824.1 aver|OJJ06028.1 awen|OJJ34442.1 aory|EIT77499.1

Group7830: abra|OJJ70327.1 acar|OOF91782.1 acla|EAW12321.1 afis|EAW25727.1 afum|EBA27177.1 agla|OJJ89276.1 akaw|GAA92582.1 aluc|OJZ82596.1 anid|CBF86212.1 atub|OJI88831.1 awen|OJJ37148.1 azon|OJJ42554.1

Group7831: abra|OJJ70351.1 afla|EED52083.1 afum|EAL91020.1 agla|OJJ79299.1 akaw|GAA83851.1 aluc|OJZ90864.1 anig|EHA19577.1 aory|EIT76900.1 atub|OJI82862.1 aver|OJJ03454.1 awen|OJJ40164.1 azon|OJJ42391.1

Group7832: abra|OJJ70371.1 acar|OOF91899.1 afis|EAW19877.1 afla|EED45164.1 afum|EAL85020.1 akaw|GAA91979.1 aluc|OJZ82635.1 anig|EHA27444.1 aory|EIT73622.1 ater|EAU29244.1 atub|OJI88866.1 azon|OJJ42477.1

Group7833: abra|OJJ70412.1 acar|OOF91861.1 akaw|GAA91944.1 aluc|OJZ82676.1 anig|EHA21044.1 anig|EHA21045.1 atub|OJI88911.1 acar|OOF91860.1 atub|OJI88912.1 ater|EAU31053.1 anid|CBF87094.1 aver|OJJ03744.1

Group7834: abra|OJJ70417.1 acar|OOF91854.1 afla|EED54715.1 agla|OJJ79104.1 akaw|GAA91940.1 aluc|OJZ82682.1 anig|EHA21050.1 aory|EIT78950.1 asyd|OJJ60504.1 atub|OJI88914.1 aver|OJI99828.1 awen|OJJ32604.1

Group7835: abra|OJJ70420.1 acar|OOF91849.1 afis|EAW19696.1 afla|EED47623.1 afum|EAL86831.1 akaw|GAA91937.1 aluc|OJZ82685.1 anid|CBF86383.1 anig|EHA21053.1 aory|EIT80179.1 ater|EAU32952.1 atub|OJI88917.1

Group7836: abra|OJJ70422.1 acar|OOF91768.1 afla|EED46001.1 akaw|GAA91935.1 aluc|OJZ82687.1 anig|EHA21055.1 aory|EIT75442.1 asyd|OJJ62598.1 atub|OJI88919.1 aver|OJI96743.1 awen|OJJ29996.1 azon|OJJ45158.1

Group7837: abra|OJJ70425.1 acar|OOF91771.1 afla|EED46006.1 agla|OJJ86983.1 akaw|GAA90152.1 aluc|OJZ83196.1 anig|EHA21059.1 asyd|OJJ62595.1 ater|EAU35266.1 atub|OJI87743.1 aver|OJI96740.1 awen|OJJ30000.1

Group7838: abra|OJJ70442.1 acar|OOF93068.1 afla|EED46067.1 akaw|GAA90138.1 aluc|OJZ83208.1 anig|EHA21069.1 aory|EIT83478.1 asyd|OJJ62584.1 ater|EAU31776.1 atub|OJI87729.1 aver|OJI96706.1 awen|OJJ39649.1

Group7839: abra|OJJ70842.1 acla|EAW13283.1 afla|EED48191.1 agla|OJJ82047.1 akaw|GAA91710.1 aluc|OJZ87679.1 anig|EHA20869.1 aory|EIT72946.1 asyd|OJJ52438.1 ater|EAU34819.1 atub|OJI80964.1 awen|OJJ36923.1

Group7840: abra|OJJ70887.1 acar|OOF98226.1 acla|EAW10810.1 afla|EED55841.1 akaw|GAA87405.1 aluc|OJZ87635.1 anig|EHA20832.1 aory|EIT76887.1 atub|OJI81013.1 agla|OJJ86415.1 awen|OJJ37992.1 azon|OJJ51825.1

Group7841: abra|OJJ70904.1 acar|OOF98216.1 afla|EED45869.1 akaw|GAA87418.1 aluc|OJZ87620.1 anid|CBF84155.1 anig|EHA20818.1 asyd|OJJ57116.1 atub|OJI81029.1 aver|OJJ03184.1 awen|OJJ35402.1 azon|OJJ43192.1

Group7842: abra|OJJ70939.1 acar|OOF98194.1 acla|EAW10504.1 afis|EAW25063.1 afum|EAL89732.1 agla|OJJ89132.1 akaw|GAA83005.1 aluc|OJZ87589.1 anig|EHA20790.1 atub|OJI81059.1 awen|OJJ40670.1 azon|OJJ44051.1

Group7843: abra|OJJ70993.1 afis|EAW23803.1 afum|EAL87237.1 agla|OJJ84592.1 akaw|GAA82955.1 aluc|OJZ87535.1 anig|EHA20744.1 atub|OJI81114.1 awen|OJJ33418.1 akaw|GAA89072.1 aluc|OJZ80520.1 atub|OJI88410.1

Group7844: abra|OJJ71094.1 afla|EED46486.1 agla|OJJ79797.1 abra|OJJ76614.1 afis|EAW17368.1 afum|EAL91205.2 aluc|OJZ84838.1 anig|EHA22619.1 asyd|OJJ53392.1 ater|EAU32673.1 aver|OJI95849.1 awen|OJJ33526.1

Group7845: abra|OJJ71116.1 acar|OOF98034.1 acla|EAW13262.1 agla|OJJ80844.1 akaw|GAA82854.1 aluc|OJZ87419.1 anig|EHA20643.1 ater|EAU31348.1 atub|OJI81229.1 awen|OJJ34859.1 azon|OJJ45669.1 ater|EAU37564.1

Group7846: afis|EAW21014.1 afum|EAL92052.1 azon|OJJ49135.1 afla|EED56757.1 akaw|GAA84860.1 aluc|OJZ86811.1 anid|CBF86706.1 aory|EIT73891.1 asyd|OJJ64739.1 atub|OJI83706.1 aver|OJI98843.1 anig|EHA21811.1

Group7847: abra|OJJ71164.1 afla|EED57983.1 akaw|GAA82815.1 aluc|OJZ87376.1 anid|CBF82377.1 anig|EHA20608.1 aory|EIT79735.1 asyd|OJJ54196.1 ater|EAU39203.1 atub|OJI81270.1 aver|OJJ08530.1 aluc|OJZ82082.1

Group7848: abra|OJJ71195.1 acar|OOF97973.1 afla|EED53750.1 akaw|GAA82788.1 aluc|OJZ87347.1 aory|EIT78458.1 atub|OJI81296.1 ater|EAU34426.1 afis|EAW25284.1 awen|OJJ40997.1 azon|OJJ44587.1 agla|OJJ89258.1

Group7849: abra|OJJ71330.1 afis|EAW25385.1 afla|EED53862.1 afum|EAL90044.1 agla|OJJ83781.1 akaw|GAA82662.1 aluc|OJZ87212.1 anig|EHA20463.1 aory|EIT78452.1 ater|EAU34532.1 atub|OJI81427.1 awen|OJJ41112.1

Group7850: abra|OJJ71581.1 akaw|GAA89343.1 aluc|OJZ86318.1 anid|CBF71811.1 anig|EHA21557.1 aory|EIT73902.1 asyd|OJJ53778.1 ater|EAU38707.1 atub|OJI84217.1 aver|OJJ07883.1 awen|OJJ31863.1 azon|OJJ45160.1

Group7851: acla|EAW12315.1 agla|OJJ80335.1 azon|OJJ51051.1 afis|EAW16906.1 afla|EED56571.1 afum|EAL86443.2 aory|EIT79619.1 asyd|OJJ54823.1 aver|OJJ07338.1 awen|OJJ37376.1 anid|CBF80359.1 agla|OJJ79458.1

Group7852: acar|OOF96344.1 akaw|GAA89299.1 acla|EAW15167.1 afis|EAW20136.1 afla|EED57216.1 afum|EBA27226.1 anid|CBF84967.1 aory|EIT74069.1 asyd|OJJ63137.1 ater|EAU38757.1 aver|OJI97817.1 awen|OJJ32005.1

Group7853: abra|OJJ71713.1 acar|OOF96418.1 acla|EAW06437.1 afla|EED52843.1 agla|OJJ81583.1 akaw|GAA85176.1 aluc|OJZ86457.1 anig|EHA21450.1 aory|EIT78624.1 ater|EAU30023.1 atub|OJI84070.1 awen|OJJ31098.1

Group7854: abra|OJJ71737.1 agla|OJJ86803.1 akaw|GAA85152.1 aluc|OJZ86485.1 anig|EHA22178.1 aory|EIT77234.1 asyd|OJJ52527.1 ater|EAU30949.1 atub|OJI84044.1 aver|OJJ08764.1 aluc|OJZ85544.1 aver|OJI99258.1

Group7855: abra|OJJ71777.1 acar|OOF96451.1 afla|EED52267.1 akaw|GAA85131.1 aluc|OJZ86507.1 anig|EHA22138.1 aory|EIT74460.1 asyd|OJJ65341.1 ater|EAU37665.1 atub|OJI84020.1 aver|OJI99487.1 awen|OJJ31285.1

Group7856: afum|EAL86238.1 afis|EAW24608.1 afla|EED56495.1 agla|OJJ80104.1 akaw|GAA89018.1 aluc|OJZ80586.1 anig|EHA25066.1 aory|EIT72853.1 asyd|OJJ55480.1 ater|EAU39133.1 aver|OJJ06559.1 awen|OJJ35866.1

Group7857: abra|OJJ71913.1 acar|OOF96571.1 agla|OJJ82336.1 akaw|GAA85008.1 aluc|OJZ86650.1 anid|CBF73516.1 anig|EHA22020.1 asyd|OJJ54492.1 atub|OJI83877.1 aver|OJJ07687.1 awen|OJJ36285.1 anid|CBF80792.1

Group7858: abra|OJJ71963.1 acar|OOF96608.1 acla|EAW09573.1 afla|EED56018.1 agla|OJJ85744.1 aluc|OJZ86699.1 asyd|OJJ61042.1 ater|EAU34969.1 atub|OJI83827.1 aver|OJJ00358.1 awen|OJJ39123.1 azon|OJJ51453.1

Group7859: abra|OJJ72026.1 acar|OOF96675.1 afla|EED45205.1 akaw|GAA84906.1 aluc|OJZ86763.1 anid|CBF75720.1 anig|EHA21916.1 aory|EIT73679.1 asyd|OJJ57504.1 atub|OJI83764.1 aver|OJJ05674.1 awen|OJJ37803.1

Group7860: abra|OJJ72027.1 acar|OOF96676.1 afla|EED45204.1 akaw|GAA84905.1 aluc|OJZ86764.1 anid|CBF75722.1 anig|EHA21915.1 aory|EIT73567.1 asyd|OJJ57505.1 atub|OJI83763.1 aver|OJJ05673.1 awen|OJJ37804.1

Group7861: abra|OJJ72070.1 acar|OOF90293.1 afis|EAW17757.1 afum|EAL85397.1 akaw|GAA84864.1 aluc|OJZ86807.1 anid|CBF82213.1 anig|EHA21876.1 asyd|OJJ54441.1 ater|EAU30971.1 atub|OJI83710.1 acla|EAW13297.1

Group7862: abra|OJJ72127.1 acar|OOF96767.1 afis|EAW21627.1 afum|EAL92662.1 akaw|GAA84796.1 aluc|OJZ86884.1 anid|CBF76682.1 anig|EHA21760.1 asyd|OJJ61224.1 atub|OJI83635.1 aver|OJJ00547.1 awen|OJJ33878.1

Group7863: abra|OJJ72177.1 acar|OOF99533.1 afla|EED54144.1 aory|EIT76039.1 asyd|OJJ61371.1 aver|OJI99825.1 abra|OJJ76995.1 agla|OJJ86188.1 akaw|GAA87051.1 anig|EHA27725.1 awen|OJJ34432.1 aver|OJJ00698.1

Group7864: abra|OJJ72190.1 acar|OOF96816.1 afla|EED48889.1 agla|OJJ85282.1 akaw|GAA84743.1 aluc|OJZ86941.1 anig|EHA21703.1 aory|EIT81228.1 asyd|OJJ57932.1 atub|OJI83581.1 aver|OJJ04465.1 awen|OJJ34658.1

Group7865: abra|OJJ72228.1 acar|OOF96851.1 afla|EED48168.1 akaw|GAA84709.1 aluc|OJZ86978.1 anid|CBF76010.1 aory|EIT83028.1 asyd|OJJ52437.1 atub|OJI83542.1 aver|OJJ03973.1 awen|OJJ31248.1 azon|OJJ43481.1

Group7866: abra|OJJ72231.1 acar|OOF96854.1 afla|EED50944.1 akaw|GAA84706.1 aluc|OJZ86981.1 anid|CBF74713.1 anig|EHA21668.1 aory|EIT76980.1 asyd|OJJ60206.1 atub|OJI83539.1 aver|OJJ04061.1 awen|OJJ35937.1

Group7867: abra|OJJ72242.1 acar|OOF96868.1 afla|EED50952.1 akaw|GAA84697.1 aluc|OJZ86990.1 anig|EHA21655.1 aory|EIT76950.1 asyd|OJJ62640.1 ater|EAU39427.1 atub|OJI83529.1 aver|OJI99288.1 awen|OJJ34409.1

Group7868: abra|OJJ72243.1 acar|OOF96869.1 afla|EED50953.1 akaw|GAA84696.1 aluc|OJZ86991.1 anig|EHA21654.1 aory|EIT76992.1 asyd|OJJ62641.1 atub|OJI83528.1 aver|OJI99287.1 awen|OJJ34410.1 azon|OJJ50347.1

Group7869: abra|OJJ72263.1 acar|OOF96895.1 acla|EAW12115.1 afis|EAW17270.1 afla|EED56563.1 afum|EAL86687.1 akaw|GAA84668.1 aluc|OJZ87021.1 aory|EIT79621.1 ater|EAU37128.1 atub|OJI83497.1 azon|OJJ45520.1

Group7870: abra|OJJ72270.1 afis|EAW19948.1 afum|EAL84967.1 akaw|GAA86025.1 aluc|OJZ90252.1 anid|CBF84251.1 anig|EHA23748.1 asyd|OJJ57199.1 ater|EAU30030.1 atub|OJI81678.1 aver|OJJ03091.1 awen|OJJ31253.1

Group7871: abra|OJJ72274.1 acar|OOF94944.1 agla|OJJ80826.1 akaw|GAA86029.1 aluc|OJZ90256.1 anid|CBF71384.1 anig|EHA23751.1 asyd|OJJ55066.1 atub|OJI81682.1 aver|OJJ07145.1 azon|OJJ48317.1 agla|OJJ78691.1

Group7872: abra|OJJ72286.1 acla|EAW12016.1 afis|EAW19655.1 afla|EED57007.1 afum|EAL86867.1 agla|OJJ80801.1 akaw|GAA86038.1 anid|CBF71367.1 aory|EIT82254.1 ater|EAU32865.1 awen|OJJ41907.1 azon|OJJ42692.1

Group7873: abra|OJJ72569.1 afla|EED57598.1 agla|OJJ88512.1 anid|CBF83005.1 aory|EIT82930.1 asyd|OJJ61505.1 aver|OJJ00842.1 awen|OJJ34790.1 acla|EAW07186.1 afis|EAW22199.1 afum|EBA27418.1 ater|EAU34215.1

Group7874: abra|OJJ72921.1 acla|EAW06470.1 afla|EED58092.1 agla|OJJ89347.1 aluc|OJZ83431.1 anid|CBF81087.1 aory|EIT79162.1 asyd|OJJ56466.1 atub|OJI82319.1 aver|OJJ06253.1 awen|OJJ35609.1 azon|OJJ46929.1

Group7875: abra|OJJ73092.1 acar|OOF92447.1 acla|EAW13486.1 afla|EED52090.1 afum|EAL92097.1 agla|OJJ81318.1 akaw|GAA88068.1 aluc|OJZ81333.1 anid|CBF78401.1 anig|EHA18637.1 ater|EAU32753.1 atub|OJI91536.1

Group7876: anig|EHA18495.1 aory|EIT76893.1 acar|OOG00493.1 acla|EAW06928.1 afis|EAW16543.1 afla|EED56122.1 afum|EAL89141.1 agla|OJJ82780.1 anid|CBF81752.1 asyd|OJJ59603.1 ater|EAU36606.1 awen|OJJ40277.1

Group7877: abra|OJJ73329.1 afla|EED48232.1 aluc|OJZ91199.1 anig|EHA19899.1 aory|EIT72888.1 asyd|OJJ57942.1 ater|EAU37460.1 atub|OJI83216.1 aver|OJJ04454.1 afis|EAW25728.1 aver|OJJ01758.1 awen|OJJ30936.1

Group7878: abra|OJJ73413.1 acar|OOG00610.1 afis|EAW19923.1 afla|EED51718.1 akaw|GAA83078.1 aluc|OJZ88624.1 aory|EIT80668.1 atub|OJI91209.1 aver|OJJ01286.1 azon|OJJ49481.1 acla|EAW13683.1 afum|EBA27215.1

Group7879: abra|OJJ73472.1 acla|EAW06796.1 akaw|GAA83029.1 aluc|OJZ88575.1 atub|OJI91161.1 azon|OJJ45009.1 afis|EAW16345.1 afla|EED53525.1 afum|EAL88970.1 agla|OJJ88243.1 aver|OJI96644.1 awen|OJJ37748.1

Group7880: abra|OJJ73478.1 acar|OOG00278.1 afis|EAW17177.1 afla|EED54490.1 akaw|GAA83023.1 aluc|OJZ88569.1 anid|CBF73633.1 anig|EHA18266.1 aory|EIT72967.1 atub|OJI91155.1 awen|OJJ29932.1 aory|EIT78797.1

Group7881: abra|OJJ73490.1 acla|EAW06815.1 afis|EAW16364.1 afla|EED53505.1 afum|EAL88989.1 akaw|GAA87439.1 aluc|OJZ88558.1 anid|CBF89243.1 aory|EIT75052.1 atub|OJI91143.1 awen|OJJ37767.1 azon|OJJ45659.1

Group7882: abra|OJJ73492.1 acar|OOF96693.1 akaw|GAA87441.1 aluc|OJZ88556.1 anig|EHA18254.1 asyd|OJJ61392.1 atub|OJI91141.1 aver|OJJ00722.1 azon|OJJ51393.1 aory|EIT73591.1 asyd|OJJ56722.1 aver|OJJ06003.1

Group7883: abra|OJJ73590.1 acar|OOG00772.1 acla|EAW07508.1 afis|EAW22540.1 afla|EED54569.1 afum|EAL90654.2 akaw|GAA88562.1 aluc|OJZ88460.1 anig|EHA18180.1 aory|EIT78746.1 atub|OJI91058.1 awen|OJJ35169.1

Group7884: abra|OJJ73610.1 acla|EAW07527.1 afis|EAW22557.1 afla|EED54588.1 afum|EAL90638.1 akaw|GAA88543.1 anid|CBF89067.1 anig|EHA18163.1 aory|EIT78831.1 ater|EAU39147.1 azon|OJJ49920.1 atub|OJI91037.1

Group7885: abra|OJJ73694.1 acar|OOG00559.1 acla|EAW12597.1 akaw|GAA88466.1 aluc|OJZ88355.1 anig|EHA21867.1 ater|EAU30061.1 atub|OJI90950.1 awen|OJJ32890.1 azon|OJJ42142.1 akaw|GAA93227.1 azon|OJJ49026.1

Group7886: abra|OJJ73707.1 acla|EAW14997.1 afla|EED48402.1 agla|OJJ84463.1 akaw|GAA88456.1 aluc|OJZ88344.1 anig|EHA18077.1 aory|EIT83053.1 asyd|OJJ60325.1 atub|OJI90940.1 aver|OJI99673.1 awen|OJJ33650.1

Group7887: abra|OJJ73708.1 acla|EAW14996.1 afla|EED48403.1 agla|OJJ84462.1 akaw|GAA88455.1 aluc|OJZ88343.1 anig|EHA18076.1 aory|EIT83112.1 asyd|OJJ60326.1 atub|OJI90939.1 aver|OJI99674.1 awen|OJJ33651.1

Group7888: awen|OJJ41209.1 acla|EAW10113.1 afis|EAW25474.1 afla|EED53966.1 afum|EAL90137.1 agla|OJJ78864.1 anid|CBF77662.1 aory|EIT78351.1 asyd|OJJ53101.1 ater|EAU34645.1 aver|OJJ03752.1 azon|OJJ42088.1

Group7889: abra|OJJ73770.1 acar|OOG00928.1 aluc|OJZ88286.1 anig|EHA18028.1 atub|OJI90884.1 abra|OJJ73771.1 akaw|GAA88405.1 atub|OJI90883.1 afla|EED49954.1 aory|EIT83424.1 anid|CBF88142.1 awen|OJJ35301.1

Group7890: abra|OJJ73779.1 acar|OOG00935.1 acla|EAW07663.1 afis|EAW22680.1 afla|EED49961.1 agla|OJJ79036.1 akaw|GAA88398.1 aluc|OJZ88276.1 anig|EHA18021.1 ater|EAU39029.1 atub|OJI90876.1 awen|OJJ35308.1

Group7891: anid|CBF69705.1 aory|EIT74413.1 aver|OJJ01642.1 abra|OJJ78198.1 acar|OOF98657.1 afla|EED52184.1 akaw|GAA85832.1 aluc|OJZ91269.1 anig|EHA19957.1 asyd|OJJ59046.1 atub|OJI83258.1 acar|OOF93983.1

Group7892: abra|OJJ74025.1 acar|OOF99121.1 akaw|GAA87507.1 aluc|OJZ88013.1 anig|EHA17793.1 ater|EAU38038.1 atub|OJI90622.1 aver|OJJ04441.1 afla|EED57597.1 aory|EIT82950.1 agla|OJJ87362.1 ater|EAU35731.1

Group7893: abra|OJJ74030.1 acar|OOG01149.1 afla|EED54913.1 agla|OJJ83372.1 akaw|GAA87514.1 aluc|OJZ88007.1 anig|EHA17787.1 aory|EIT80377.1 ater|EAU30706.1 atub|OJI90615.1 awen|OJJ38647.1 azon|OJJ49593.1

Group7894: abra|OJJ74031.1 akaw|GAA87515.1 aluc|OJZ88006.1 atub|OJI90614.1 acar|OOG01150.1 afis|EAW22917.1 acla|EAW07882.1 afum|EAL90280.1 asyd|OJJ63323.1 azon|OJJ49592.1 aver|OJI97638.1 anid|CBF87688.1

Group7895: abra|OJJ74115.1 acar|OOG01221.1 acla|EAW07957.1 afla|EED58185.1 agla|OJJ83292.1 aluc|OJZ87919.1 anig|EHA17713.1 aory|EIT79077.1 atub|OJI90530.1 aver|OJI97711.1 awen|OJJ38569.1 azon|OJJ44833.1

Group7896: abra|OJJ74140.1 acar|OOF95590.1 afla|EED47375.1 akaw|GAA89745.1 aluc|OJZ80639.1 anig|EHA27194.1 aory|EIT82875.1 ater|EAU32380.1 atub|OJI87494.1 awen|OJJ29701.1 aver|OJJ06010.1 asyd|OJJ56715.1

Group7897: abra|OJJ74145.1 acar|OOF95583.1 afla|EED51486.1 akaw|GAA89739.1 aluc|OJZ80645.1 anid|CBF78620.1 anig|EHA27187.1 asyd|OJJ65164.1 atub|OJI87500.1 aver|OJI99316.1 awen|OJJ31240.1 azon|OJJ43380.1

Group7898: abra|OJJ74147.1 acar|OOF95581.1 afla|EED51487.1 akaw|GAA89738.1 aluc|OJZ80646.1 anid|CBF78618.1 aory|EIT80636.1 asyd|OJJ65165.1 atub|OJI87501.1 aver|OJI99317.1 awen|OJJ31239.1 azon|OJJ43379.1

Group7899: akaw|GAA86670.1 anig|EHA26987.1 acar|OOF95369.1 acla|EAW07412.1 afis|EAW22438.1 afla|EED54448.1 afum|EAL90753.1 aluc|OJZ89767.1 anid|CBF88864.1 aory|EIT81124.1 asyd|OJJ62875.1 ater|EAU39270.1

Group7900: abra|OJJ74415.1 acar|OOF95328.1 acla|EAW06830.1 afis|EAW19937.1 afum|EAL89005.1 akaw|GAA87306.1 aluc|OJZ89711.1 anid|CBF87793.1 anig|EHA26942.1 ater|EAU34072.1 atub|OJI85091.1 afis|EAW16380.1

Group7901: abra|OJJ74421.1 acar|OOF95319.1 acla|EAW12039.1 afis|EAW19688.1 afum|EAL86838.1 akaw|GAA87315.1 aluc|OJZ89702.1 asyd|OJJ65223.1 atub|OJI85097.1 aver|OJI99371.1 awen|OJJ34664.1 afis|EAW19686.1

Group7902: abra|OJJ74429.1 afis|EAW20220.1 afla|EED46887.1 akaw|GAA87324.1 aluc|OJZ89691.1 anig|EHA26930.1 aory|EIT72563.1 asyd|OJJ56971.1 ater|EAU29632.1 atub|OJI85110.1 aver|OJJ05883.1 awen|OJJ33420.1

Group7903: abra|OJJ74436.1 agla|OJJ85389.1 akaw|GAA87330.1 aluc|OJZ89685.1 anig|EHA26923.1 asyd|OJJ52854.1 ater|EAU32806.1 atub|OJI85116.1 aver|OJI95983.1 awen|OJJ40908.1 aluc|OJZ79624.1 asyd|OJJ61885.1

Group7904: abra|OJJ74482.1 acla|EAW07219.1 afis|EAW22233.1 afla|EED57622.1 afum|EAL90960.1 agla|OJJ87818.1 akaw|GAA87375.1 aluc|OJZ89637.1 anid|CBF88472.1 aory|EIT82961.1 ater|EAU34236.1 atub|OJI85166.1

Group7905: abra|OJJ74695.1 acla|EAW10703.1 afis|EAW24867.1 afum|EAL89517.2 agla|OJJ82085.1 aluc|OJZ92272.1 asyd|OJJ57538.1 ater|EAU36433.1 atub|OJI86707.1 aver|OJJ05640.1 awen|OJJ40458.1 azon|OJJ49317.1

Group7906: abra|OJJ74740.1 afla|EED46983.1 agla|OJJ82128.1 akaw|GAA83670.1 aluc|OJZ92321.1 anid|CBF75642.1 anig|EHA26648.1 aory|EIT72629.1 asyd|OJJ57475.1 ater|EAU36396.1 atub|OJI86750.1 aver|OJJ05706.1

Group7907: anig|EHA26586.1 awen|OJJ40553.1 acla|EAW10607.1 afis|EAW24960.1 afla|EED55503.1 afum|EAL89618.1 agla|OJJ82191.1 anid|CBF81660.1 asyd|OJJ59549.1 ater|EAU35695.1 aver|OJJ02719.1 azon|OJJ49413.1

Group7908: abra|OJJ74836.1 acar|OOF95827.1 afis|EAW23586.1 afla|EED55273.1 afum|EAL87878.1 agla|OJJ86850.1 akaw|GAA83584.1 aluc|OJZ92413.1 anig|EHA26552.1 aory|EIT83520.1 atub|OJI86843.1 awen|OJJ39913.1

Group7909: abra|OJJ74875.1 acar|OOF95876.1 acla|EAW08496.1 afis|EAW23542.1 afum|EAL87924.1 agla|OJJ86796.1 anid|CBF89659.1 asyd|OJJ62278.1 ater|EAU35328.1 aver|OJI96387.1 awen|OJJ39873.1 azon|OJJ43228.1

Group7910: azon|OJJ47798.1 acar|OOF97261.1 acla|EAW08254.1 afis|EAW23288.1 afla|EED55686.1 afum|EAL88191.1 agla|OJJ86559.1 akaw|GAA88685.1 anid|CBF89513.1 aory|EIT77077.1 ater|EAU37585.1 aver|OJI96529.1

Group7911: abra|OJJ75180.1 acar|OOF94599.1 acla|EAW12996.1 afis|EAW17670.1 afla|EED50484.1 afum|EAL91484.1 agla|OJJ79542.1 akaw|GAA88785.1 anid|CBF90038.1 ater|EAU31977.1 atub|OJI87171.1 azon|OJJ50868.1

Group7912: abra|OJJ75245.1 acar|OOF90093.1 acla|EAW14871.1 afis|EAW23642.1 akaw|GAA85578.1 aluc|OJZ83514.1 anig|EHA26202.1 aory|EIT76993.1 ater|EAU35168.1 atub|OJI82405.1 awen|OJJ34614.1 azon|OJJ43969.1

Group7913: abra|OJJ75287.1 afis|EAW23626.1 afla|EED48184.1 afum|EAL87834.1 akaw|GAA91825.1 aluc|OJZ92848.1 anid|CBF76056.1 anig|EHA26161.1 asyd|OJJ55369.1 atub|OJI87304.1 aver|OJJ06448.1 aory|EIT72928.1

Group7914: anid|CBF80524.1 azon|OJJ46018.1 afis|EAW19545.1 afla|EED52109.1 afum|EAL86964.1 aory|EIT76933.1 asyd|OJJ53786.1 aver|OJI99284.1 asyd|OJJ65287.1 aver|OJJ01919.1 afla|EED56861.1 aory|EIT77556.1

Group7915: abra|OJJ75315.1 acar|OOF94513.1 afis|EAW19846.1 afla|EED48944.1 akaw|GAA91999.1 aluc|OJZ92869.1 anid|CBF82998.1 anig|EHA26096.1 aory|EIT80043.1 asyd|OJJ61547.1 atub|OJI87327.1 aver|OJJ00874.1

Group7916: abra|OJJ75345.1 afis|EAW19992.1 afla|EED51077.1 agla|OJJ86027.1 akaw|GAA92038.1 aluc|OJZ92907.1 asyd|OJJ58162.1 atub|OJI87367.1 aver|OJJ04240.1 awen|OJJ30003.1 azon|OJJ43628.1 aluc|OJZ88788.1

Group7917: aory|EIT81269.1 acar|OOF94460.1 afis|EAW17280.1 afla|EED48839.1 afum|EAL86698.1 agla|OJJ78405.1 akaw|GAA92749.1 aluc|OJZ92957.1 anid|CBF87549.1 anig|EHA26018.1 asyd|OJJ52996.1 aver|OJJ08965.1

Group7918: abra|OJJ75423.1 acar|OOF90768.1 agla|OJJ83093.1 akaw|GAA84198.1 aluc|OJZ88875.1 anig|EHA23630.1 ater|EAU33729.1 atub|OJI85916.1 aver|OJI98829.1 awen|OJJ41696.1 azon|OJJ48394.1 ater|EAU32644.1

Group7919: abra|OJJ75461.1 acar|OOF90278.1 acla|EAW13829.1 afis|EAW18302.1 afla|EED48771.1 afum|EAL87668.2 akaw|GAA84235.1 aluc|OJZ88918.1 anig|EHA23594.1 aory|EIT72986.1 atub|OJI85874.1 azon|OJJ43277.1

Group7920: abra|OJJ75715.1 acar|OOG00144.1 afla|EED58010.1 agla|OJJ84452.1 akaw|GAA84449.1 aluc|OJZ89166.1 anig|EHA23388.1 asyd|OJJ62645.1 ater|EAU38418.1 atub|OJI85624.1 awen|OJJ34018.1 azon|OJJ48540.1

Group7921: abra|OJJ75749.1 afla|EED52149.1 akaw|GAA84481.1 aluc|OJZ89201.1 anig|EHA23356.1 atub|OJI85589.1 awen|OJJ32797.1 azon|OJJ48883.1 acar|OOF90579.1 afis|EAW25381.1 aory|EIT74322.1 ater|EAU38539.1

Group7922: abra|OJJ75759.1 acar|OOF93580.1 acla|EAW08560.1 afis|EAW23602.1 afla|EED52608.1 afum|EAL87865.1 anid|CBF81651.1 aory|EIT74254.1 asyd|OJJ59543.1 ater|EAU30757.1 aver|OJJ02725.1 azon|OJJ42334.1

Group7923: abra|OJJ75766.1 asyd|OJJ58683.1 ater|EAU29471.1 atub|OJI87192.1 aver|OJJ01295.1 awen|OJJ30927.1 abra|OJJ76680.1 afla|EED48159.1 anig|EHA22561.1 aory|EIT83026.1 asyd|OJJ53890.1 aver|OJJ07988.1

Group7924: abra|OJJ75780.1 acar|OOF93583.1 afis|EAW17231.1 afla|EED54397.1 agla|OJJ86976.1 akaw|GAA84497.1 aluc|OJZ89222.1 anig|EHA23340.1 aory|EIT75880.1 asyd|OJJ53739.1 atub|OJI85570.1 aver|OJJ07898.1

Group7925: abra|OJJ75877.1 acar|OOG00016.1 acla|EAW07593.1 afis|EAW20265.1 afla|EED52337.1 afum|EAL85373.1 akaw|GAA84579.1 aluc|OJZ89321.1 anig|EHA23251.1 aory|EIT74435.1 atub|OJI85477.1 azon|OJJ42382.1

Group7926: abra|OJJ75968.1 acar|OOF99951.1 acla|EAW09830.1 afis|EAW21303.1 afla|EED57263.1 afum|EAL92320.1 akaw|GAA88988.1 aluc|OJZ89415.1 anig|EHA23172.1 aory|EIT73970.1 atub|OJI85380.1 awen|OJJ39395.1

Group7927: abra|OJJ75989.1 afla|EED52372.1 afum|EAL85021.1 akaw|GAA88969.1 aluc|OJZ89434.1 anid|CBF80378.1 anig|EHA23153.1 aory|EIT83634.1 asyd|OJJ54809.1 ater|EAU32113.1 atub|OJI85359.1 aver|OJJ07352.1

Group7928: abra|OJJ75991.1 afla|EED52368.1 afum|EAL85023.1 akaw|GAA88966.1 aluc|OJZ89437.1 anig|EHA23151.1 aory|EIT82368.1 asyd|OJJ54806.1 ater|EAU32111.1 atub|OJI85356.1 aver|OJJ07355.1 asyd|OJJ61705.1

Group7929: abra|OJJ76230.1 afis|EAW24532.1 afla|EED47409.1 afum|EAL86198.1 agla|OJJ86404.1 anig|EHA22167.1 asyd|OJJ54668.1 aver|OJJ02021.1 abra|OJJ77428.1 afla|EED53645.1 aory|EIT75298.1 awen|OJJ29985.1

Group7930: abra|OJJ76274.1 acar|OOF99522.1 acla|EAW14338.1 afla|EED50989.1 akaw|GAA82207.1 aluc|OJZ91946.1 anig|EHA22905.1 aory|EIT76975.1 asyd|OJJ59142.1 atub|OJI86391.1 aver|OJJ01732.1 awen|OJJ31549.1

Group7931: abra|OJJ76295.1 acar|OOF99539.1 afis|EAW19942.1 afla|EED53217.1 afum|EAL84971.1 akaw|GAA82222.1 aluc|OJZ91927.1 anid|CBF77661.1 anig|EHA22890.1 aory|EIT75072.1 atub|OJI86371.1 awen|OJJ40850.1

Group7932: abra|OJJ76312.1 acar|OOF99560.1 acla|EAW08993.1 afis|EAW15412.1 afla|EED46401.1 afum|EAL84234.1 akaw|GAA82240.1 aluc|OJZ91905.1 anig|EHA22881.1 aory|EIT72391.1 ater|EAU31771.1 atub|OJI86355.1

Group7933: aory|EIT77993.1 acar|OOF99657.1 acla|EAW14010.1 afis|EAW18716.1 afum|EBA27492.1 agla|OJJ82343.1 anid|CBF70672.1 asyd|OJJ58475.1 ater|EAU38191.1 aver|OJJ01077.1 awen|OJJ32210.1 afla|EED50670.1

Group7934: abra|OJJ76564.1 acar|OOF99420.1 afum|EAL93588.1 agla|OJJ80578.1 akaw|GAA82468.1 aluc|OJZ91642.1 anid|CBF69832.1 asyd|OJJ58984.1 atub|OJI86096.1 aver|OJJ01579.1 azon|OJJ47169.1 awen|OJJ32412.1

Group7935: abra|OJJ76622.1 acar|OOF92931.1 afla|EED52230.1 anid|CBF69490.1 aory|EIT74471.1 asyd|OJJ59239.1 ater|EAU37442.1 aver|OJJ01849.1 acar|OOG00740.1 afis|EAW23428.1 ater|EAU30841.1 azon|OJJ49327.1

Group7936: abra|OJJ76639.1 acar|OOF99358.1 afla|EED53121.1 agla|OJJ80190.1 akaw|GAA92819.1 aluc|OJZ91570.1 anid|CBF84681.1 anig|EHA22599.1 aory|EIT74934.1 atub|OJI86020.1 awen|OJJ31626.1 ater|EAU37855.1

Group7937: abra|OJJ76656.1 acar|OOF99346.1 afla|EED53112.1 akaw|GAA92996.1 aluc|OJZ91549.1 anid|CBF86727.1 anig|EHA22584.1 aory|EIT74803.1 ater|EAU37782.1 atub|OJI86002.1 awen|OJJ38069.1 azon|OJJ43903.1

Group7938: abra|OJJ76701.1 acar|OOF99317.1 afla|EED51717.1 agla|OJJ85387.1 akaw|GAA92498.1 aluc|OJZ91510.1 anid|CBF80099.1 anig|EHA22549.1 asyd|OJJ58298.1 ater|EAU34017.1 atub|OJI85962.1 aver|OJJ02126.1

Group7939: abra|OJJ76722.1 acar|OOG00872.1 afla|EED51531.1 agla|OJJ89413.1 akaw|GAA92479.1 aluc|OJZ91489.1 anig|EHA22533.1 aory|EIT80628.1 asyd|OJJ56396.1 atub|OJI85944.1 aver|OJJ06317.1 awen|OJJ35675.1

Group7940: afis|EAW20448.1 anig|EHA27538.1 acar|OOF91270.1 acla|EAW13333.1 afla|EED44822.1 afum|EAL84604.1 agla|OJJ84874.1 anid|CBF79301.1 aory|EIT77747.1 ater|EAU37711.1 aver|OJJ06862.1 awen|OJJ33138.1

Group7941: abra|OJJ76812.1 acar|OOF91308.1 acla|EAW13373.1 afis|EAW20487.1 afla|EED44777.1 afum|EAL84568.1 akaw|GAA87209.1 aluc|OJZ84223.1 anid|CBF79217.1 anig|EHA27572.1 aory|EIT77789.1 atub|OJI89863.1

Group7942: abra|OJJ76887.1 afla|EED47433.1 agla|OJJ84734.1 akaw|GAA91072.1 aluc|OJZ87886.1 anid|CBF71511.1 anig|EHA26975.1 aory|EIT81892.1 asyd|OJJ54924.1 atub|OJI80765.1 aver|OJJ07271.1 awen|OJJ32682.1

Group7943: afum|EAL85122.1 ater|EAU37570.1 anid|CBF90348.1 afla|EED51127.1 agla|OJJ81313.1 aory|EIT81315.1 asyd|OJJ55577.1 aver|OJJ06656.1 afla|EED52426.1 aory|EIT83303.1 azon|OJJ42156.1 anid|CBF76191.1

Group7944: abra|OJJ76930.1 acla|EAW13435.1 afis|EAW20551.1 afum|EBA27186.1 akaw|GAA87110.1 aluc|OJZ84328.1 anig|EHA27662.1 aory|EIT83280.1 atub|OJI89740.1 awen|OJJ33266.1 azon|OJJ50630.1 ater|EAU31069.1

Group7945: abra|OJJ76961.1 acar|OOF91406.1 afla|EED48151.1 akaw|GAA87082.1 aluc|OJZ84356.1 aory|EIT76227.1 asyd|OJJ55022.1 ater|EAU33244.1 atub|OJI89711.1 aver|OJJ07184.1 awen|OJJ37978.1 azon|OJJ42641.1

Group7946: abra|OJJ77058.1 acar|OOF95835.1 acla|EAW14557.1 akaw|GAA86994.1 aluc|OJZ84465.1 anid|CBF70126.1 anig|EHA27777.1 asyd|OJJ59808.1 ater|EAU32470.1 atub|OJI89611.1 awen|OJJ30216.1 aory|EIT77220.1

Group7947: abra|OJJ77182.1 acar|OOF91801.1 afla|EED47269.1 agla|OJJ84566.1 aory|EIT79233.1 asyd|OJJ64975.1 aver|OJI99097.1 awen|OJJ37796.1 azon|OJJ45194.1 acar|OOG00110.1 afla|EED47897.1 aory|EIT77231.1

Group7948: abra|OJJ77203.1 acar|OOF93091.1 acla|EAW13195.1 afis|EAW17447.1 afum|EAL91274.1 agla|OJJ80370.1 akaw|GAA90735.1 aluc|OJZ84613.1 anig|EHA27902.1 atub|OJI89468.1 awen|OJJ37399.1 azon|OJJ51072.1

Group7949: abra|OJJ77243.1 acar|OOF93124.1 acla|EAW14605.1 afis|EAW19334.1 afum|EAL94051.2 agla|OJJ79420.1 akaw|GAA90700.1 aluc|OJZ84653.1 anig|EHA27936.1 asyd|OJJ59233.1 atub|OJI89427.1 aver|OJJ01843.1

Group7950: abra|OJJ77354.1 acar|OOF93224.1 acla|EAW13120.1 afis|EAW17544.1 afla|EED49283.1 afum|EAL91358.1 agla|OJJ81425.1 akaw|GAA91407.1 anid|CBF90306.1 asyd|OJJ61931.1 ater|EAU36974.1 aver|OJI96023.1

Group7951: abra|OJJ77437.1 acar|OOF92528.1 afla|EED54491.1 agla|OJJ82053.1 anig|EHA26146.1 aory|EIT78832.1 ater|EAU39533.1 aver|OJJ01790.1 awen|OJJ31427.1 aory|EIT76911.1 afla|EED52077.1 awen|OJJ30426.1

Group7952: abra|OJJ77460.1 acar|OOF97635.1 afla|EED49592.1 akaw|GAA92097.1 aluc|OJZ83628.1 anig|EHA19293.1 aory|EIT82658.1 asyd|OJJ56400.1 ater|EAU39564.1 atub|OJI89997.1 aver|OJJ06312.1 anig|EHA18781.1

Group7953: abra|OJJ77497.1 acar|OOF99229.1 afis|EAW25461.1 afla|EED53950.1 afum|EAL90124.1 akaw|GAA84124.1 aluc|OJZ81041.1 anig|EHA19326.1 aory|EIT78317.1 atub|OJI82543.1 awen|OJJ41197.1 azon|OJJ42903.1

Group7954: abra|OJJ77515.1 acar|OOF99209.1 afla|EED53283.1 agla|OJJ88332.1 akaw|GAA84105.1 aluc|OJZ81020.1 aory|EIT75185.1 asyd|OJJ64619.1 atub|OJI82563.1 aver|OJI98715.1 awen|OJJ37665.1 azon|OJJ44932.1

Group7955: abra|OJJ77713.1 acar|OOF99027.1 agla|OJJ85978.1 akaw|GAA83919.1 aluc|OJZ90778.1 anid|CBF81695.1 anig|EHA19511.1 asyd|OJJ59564.1 atub|OJI82765.1 aver|OJJ02705.1 awen|OJJ41676.1 aory|EIT83272.1

Group7956: abra|OJJ77756.1 acar|OOF98694.1 akaw|GAA83883.1 aluc|OJZ90813.1 anig|EHA19541.1 asyd|OJJ61417.1 ater|EAU31760.1 atub|OJI82803.1 awen|OJJ32585.1 azon|OJJ46428.1 asyd|OJJ53798.1 aver|OJI96898.1

Group7957: abra|OJJ77758.1 acar|OOF98977.1 akaw|GAA83881.1 aluc|OJZ90815.1 anig|EHA19543.1 asyd|OJJ62928.1 atub|OJI82805.1 aver|OJI98062.1 awen|OJJ39984.1 acar|OOG00864.1 anid|CBF80410.1 ater|EAU30956.1

Group7958: abra|OJJ77759.1 acar|OOF98978.1 acla|EAW12027.1 afis|EAW15413.1 afla|EED45861.1 afum|EAL84233.1 akaw|GAA83880.1 aluc|OJZ90816.1 anig|EHA19544.1 aory|EIT75386.1 atub|OJI82806.1 azon|OJJ50242.1

Group7959: abra|OJJ77799.1 afla|EED54466.1 akaw|GAA83849.1 aluc|OJZ90866.1 anid|CBF88117.1 anig|EHA19583.1 aory|EIT81145.1 atub|OJI82864.1 aver|OJJ02027.1 afla|EED48243.1 aory|EIT72921.1 aver|OJI97353.1

Group7960: abra|OJJ77855.1 acla|EAW11271.1 afla|EED56240.1 agla|OJJ85853.1 aluc|OJZ90929.1 anid|CBF86168.1 aory|EIT73289.1 asyd|OJJ64523.1 atub|OJI82921.1 aver|OJI98621.1 awen|OJJ30607.1 azon|OJJ50161.1

Group7961: abra|OJJ77964.1 acar|OOF95818.1 afis|EAW24661.1 afla|EED47737.1 afum|EAL87778.1 akaw|GAA83701.1 aluc|OJZ91031.1 anig|EHA19732.1 asyd|OJJ52847.1 ater|EAU38131.1 atub|OJI83027.1 aver|OJJ08825.1

Group7962: akaw|GAA83688.1 aluc|OJZ91044.1 atub|OJI83040.1 acla|EAW06666.1 afis|EAW16099.1 afla|EED48634.1 afum|EAL88731.1 agla|OJJ86123.1 anid|CBF85449.1 aory|EIT73226.1 aver|OJI98178.1 azon|OJJ45361.1

Group7963: abra|OJJ78057.1 afis|EAW20701.1 afum|EAL84876.1 akaw|GAA85935.1 aluc|OJZ91127.1 anid|CBF70383.1 anig|EHA19828.1 asyd|OJJ56273.1 atub|OJI83138.1 aver|OJJ08492.1 asyd|OJJ58640.1 aver|OJJ01247.1

Group7964: abra|OJJ78075.1 acar|OOF98788.1 afis|EAW16208.1 afla|EED48707.1 afum|EAL88835.1 agla|OJJ86223.1 akaw|GAA85919.1 aluc|OJZ91148.1 anig|EHA19846.1 ater|EAU33252.1 atub|OJI83162.1 awen|OJJ30265.1

Group7965: abra|OJJ78082.1 acar|OOF98784.1 afla|EED55363.1 akaw|GAA85910.1 aluc|OJZ91156.1 anid|CBF73594.1 anig|EHA19852.1 aory|EIT80908.1 asyd|OJJ56690.1 atub|OJI83171.1 aver|OJJ06033.1 aver|OJJ06041.1

Group7966: abra|OJJ78115.1 acla|EAW09077.1 afla|EED45412.1 akaw|GAA85892.1 aluc|OJZ91180.1 anid|CBF73728.1 anig|EHA19871.1 aory|EIT82408.1 asyd|OJJ56636.1 atub|OJI83193.1 aver|OJJ06090.1 aver|OJJ01921.1

Group7967: abra|OJJ78199.1 acar|OOF98656.1 afum|EAL87798.1 akaw|GAA85831.1 aluc|OJZ91270.1 ater|EAU33020.1 atub|OJI83259.1 awen|OJJ37932.1 azon|OJJ41995.1 akaw|GAA85830.1 agla|OJJ86067.1 ater|EAU32286.1

Group7968: abra|OJJ78272.1 afis|EAW21906.1 afla|EED46645.1 afum|EAL92918.2 akaw|GAA85754.1 aluc|OJZ91346.1 anid|CBF76832.1 anig|EHA20025.1 asyd|OJJ61302.1 atub|OJI83338.1 awen|OJJ34196.1 azon|OJJ43438.1

Group7969: abra|OJJ78303.1 acar|OOF98563.1 afla|EED46355.1 akaw|GAA85723.1 aluc|OJZ91380.1 anid|CBF82931.1 anig|EHA20055.1 asyd|OJJ52921.1 ater|EAU30948.1 atub|OJI83373.1 aver|OJJ08890.1 azon|OJJ45182.1

Group7970: abra|OJJ78305.1 acar|OOF98561.1 afis|EAW22060.1 afum|EAL92951.2 agla|OJJ78429.1 akaw|GAA85718.1 anid|CBF76901.1 asyd|OJJ61339.1 aver|OJJ00663.1 awen|OJJ34229.1 azon|OJJ43406.1 acla|EAW09209.1

Group7971: acar|OOF89791.1 acla|EAW13952.1 afis|EAW18662.1 afla|EED52667.1 afum|EBA27490.1 akaw|GAA82396.1 anid|CBF70563.1 aory|EIT74292.1 ater|EAU38250.1 atub|OJI86183.1 awen|OJJ32129.1 azon|OJJ46713.1

Group7972: acar|OOF89948.1 acla|EAW11544.1 afla|EED52802.1 akaw|GAA89086.1 aluc|OJZ80505.1 anig|EHA25688.1 aory|EIT78187.1 ater|EAU30248.1 atub|OJI88425.1 aver|OJJ07309.1 awen|OJJ35833.1 ater|EAU34435.1

Group7973: acla|EAW14252.1 afis|EAW19029.1 afla|EED52649.1 afum|EAL93759.1 agla|OJJ85922.1 anid|CBF71834.1 aory|EIT74276.1 asyd|OJJ59935.1 aver|OJJ02352.1 awen|OJJ37063.1 azon|OJJ48237.1 anid|CBF85566.1

Group7974: acar|OOF90772.1 acla|EAW13797.1 afis|EAW18335.1 afla|EED48799.1 agla|OJJ88050.1 akaw|GAA84202.1 aluc|OJZ88879.1 asyd|OJJ57824.1 ater|EAU33348.1 aver|OJJ04568.1 awen|OJJ30716.1 azon|OJJ43308.1

Group7975: acar|OOF90991.1 acla|EAW10952.1 afla|EED50257.1 afum|EBA27202.1 agla|OJJ84025.1 aluc|OJZ81507.1 anid|CBF82473.1 asyd|OJJ54101.1 aver|OJJ08637.1 awen|OJJ34437.1 azon|OJJ51719.1 afis|EAW20909.1

Group7976: acar|OOF91836.1 afla|EED54607.1 akaw|GAA86592.1 aluc|OJZ85139.1 anig|EHA25355.1 aory|EIT78758.1 asyd|OJJ57415.1 ater|EAU35411.1 atub|OJI80453.1 aver|OJJ03415.1 awen|OJJ32893.1 aver|OJJ05762.1

Group7977: ater|EAU32979.1 acla|EAW14129.1 afis|EAW18824.1 afla|EED47307.1 akaw|GAA91939.1 aluc|OJZ82683.1 anid|CBF86781.1 anig|EHA21051.1 aory|EIT82866.1 atub|OJI88915.1 afla|EED52171.1 aory|EIT74365.1

Group7978: acar|OOF92154.1 acla|EAW12629.1 afis|EAW18055.1 afla|EED51422.1 afum|EAL91866.1 agla|OJJ87331.1 akaw|GAA89793.1 aluc|OJZ81898.1 anid|CBF86442.1 aory|EIT80614.1 atub|OJI84451.1 awen|OJJ41438.1

Group7979: acar|OOF92760.1 afis|EAW17966.1 afla|EED51943.1 awen|OJJ41578.1 acla|EAW12800.1 afum|EAL91775.1 aory|EIT81377.1 azon|OJJ48291.1 asyd|OJJ58220.1 aver|OJJ02202.1 agla|OJJ80212.1 anid|CBF79958.1

Group7980: acar|OOF93461.1 acla|EAW08871.1 afis|EAW15523.1 afla|EED49032.1 afum|EAL85805.1 agla|OJJ85223.1 aory|EIT82496.1 asyd|OJJ55994.1 ater|EAU32600.1 aver|OJJ04941.1 awen|OJJ38187.1 azon|OJJ43108.1

Group7981: acar|OOF94545.1 acla|EAW07361.1 afla|EED56050.1 afum|EAL86532.1 agla|OJJ82292.1 aory|EIT76842.1 asyd|OJJ52087.1 ater|EAU39329.1 aver|OJJ03477.1 awen|OJJ35551.1 asyd|OJJ63951.1 aver|OJI96990.1

Group7982: afla|EED57687.1 anid|CBF88564.1 awen|OJJ34861.1 acla|EAW07267.1 afis|EAW22288.1 afum|EAL90893.1 agla|OJJ87761.1 aory|EIT79935.1 asyd|OJJ63835.1 ater|EAU34300.1 aver|OJI97102.1 azon|OJJ45667.1

Group7983: acar|OOF95561.1 acla|EAW10799.1 afis|EAW24778.1 afla|EED49723.1 afum|EAL89418.1 agla|OJJ88222.1 anid|CBF73605.1 aory|EIT77687.1 asyd|OJJ56677.1 ater|EAU34010.1 aver|OJJ06045.1 anid|CBF82799.1

Group7984: acar|OOF95562.1 acla|EAW10800.1 afis|EAW24777.1 afla|EED49724.1 afum|EBA27383.1 agla|OJJ88223.1 anid|CBF73603.1 aory|EIT77625.1 asyd|OJJ56679.1 ater|EAU34011.1 aver|OJJ06044.1 afla|EED45748.1

Group7985: acar|OOF95838.1 acla|EAW10413.1 afis|EAW25156.1 afla|EED57495.1 afum|EAL89831.1 agla|OJJ88995.1 anid|CBF85650.1 aory|EIT81875.1 ater|EAU39439.1 awen|OJJ40799.1 asyd|OJJ64233.1 aver|OJI98325.1

Group7986: acar|OOF95865.1 acla|EAW08516.1 afis|EAW23554.1 afla|EED57449.1 afum|EAL87912.2 agla|OJJ86814.1 akaw|GAA83555.1 aory|EIT80121.1 asyd|OJJ62289.1 ater|EAU35337.1 aver|OJI96397.1 awen|OJJ39885.1

Group7987: acar|OOF96832.1 afla|EED54673.1 agla|OJJ80327.1 aory|EIT78946.1 ater|EAU29967.1 awen|OJJ31988.1 azon|OJJ45264.1 anid|CBF80727.1 asyd|OJJ54769.1 aver|OJJ07398.1 ater|EAU33083.1 ater|EAU37146.1

Group7988: acar|OOF97167.1 acla|EAW08345.1 afis|EAW23380.1 afla|EED55577.1 afum|EAL88091.1 akaw|GAA83397.1 anid|CBF74851.1 aory|EIT82700.1 ater|EAU35634.1 atub|OJI90264.1 aver|OJJ04134.1 azon|OJJ47703.1

Group7989: acar|OOF97593.1 acla|EAW14960.1 afis|EAW24480.1 afla|EED56129.1 afum|EAL86151.2 anid|CBF86922.1 aory|EIT76863.1 asyd|OJJ58105.1 aver|OJJ04299.1 awen|OJJ33343.1 aory|EIT76601.1 azon|OJJ43602.1

Group7990: acar|OOF98901.1 afis|EAW24712.1 afla|EED45533.1 agla|OJJ78671.1 anig|EHA28381.1 atub|OJI88058.1 awen|OJJ32909.1 azon|OJJ42264.1 afla|EED44881.1 aory|EIT75725.1 awen|OJJ29789.1 aory|EIT83255.1

Group7991: acar|OOF99636.1 acla|EAW14028.1 afis|EAW18734.1 afla|EED50689.1 afum|EAL93459.1 agla|OJJ82300.1 akaw|GAA82315.1 aluc|OJZ91817.1 aory|EIT78042.1 ater|EAU37857.1 awen|OJJ32297.1 azon|OJJ42735.1

Group7992: acar|OOG00142.1 acla|EAW09395.1 afis|EAW21738.1 afla|EED58013.1 afum|EAL92772.1 agla|OJJ84450.1 akaw|GAA84451.1 anid|CBF83701.1 aory|EIT79800.1 atub|OJI85622.1 awen|OJJ34020.1 azon|OJJ48536.1

Group7993: acar|OOG01216.1 acla|EAW07952.1 afis|EAW22989.1 afla|EED58181.1 akaw|GAA87584.1 aluc|OJZ87927.1 asyd|OJJ63252.1 ater|EAU30634.1 atub|OJI90539.1 aver|OJI97706.1 azon|OJJ49517.1 agla|OJJ83297.1

Group7994: acla|EAW07158.1 afis|EAW22173.1 afla|EED57552.1 afum|EAL91016.1 agla|OJJ87881.1 anid|CBF89081.1 aory|EIT81826.1 asyd|OJJ62737.1 ater|EAU34178.1 awen|OJJ34748.1 azon|OJJ45811.1 aver|OJI96780.1

Group7995: acla|EAW07360.1 afis|EAW22389.1 afla|EED56049.1 afum|EAL90801.1 agla|OJJ87651.1 anid|CBF88738.1 aory|EIT76759.1 asyd|OJJ63950.1 ater|EAU39330.1 aver|OJI96991.1 awen|OJJ34964.1 azon|OJJ50264.1

Group7996: acla|EAW08122.1 afis|EAW23157.1 afla|EED47076.1 afum|EAL88326.1 agla|OJJ85093.1 anid|CBF74461.1 aory|EIT72513.1 asyd|OJJ57697.1 ater|EAU36291.1 aver|OJJ05467.1 awen|OJJ36626.1 azon|OJJ47585.1

Group7997: acla|EAW08748.1 afis|EAW15688.1 afla|EED51365.1 afum|EAL85660.1 agla|OJJ81836.1 aluc|OJZ86072.1 anid|CBF71138.1 asyd|OJJ55834.1 atub|OJI79869.1 aver|OJJ04776.1 awen|OJJ38379.1 azon|OJJ43863.1

Group7998: acla|EAW08996.1 afis|EAW22252.1 anid|CBF81866.1 asyd|OJJ65275.1 ater|EAU32670.1 aver|OJI99427.1 acla|EAW12059.1 afis|EAW19700.1 afla|EED51442.1 aory|EIT80577.1 ater|EAU31759.1 anid|CBF70179.1

Group7999: acla|EAW09011.1 afis|EAW25590.1 afla|EED54127.1 afum|EAL84298.2 agla|OJJ84434.1 anid|CBF69467.1 aory|EIT78941.1 asyd|OJJ55527.1 ater|EAU31693.1 aver|OJJ06605.1 awen|OJJ36339.1 afla|EED54744.1

Group8000: acla|EAW09811.1 afis|EAW21324.1 afla|EED57282.1 afum|EAL92341.1 agla|OJJ87414.1 anid|CBF83425.1 aory|EIT74105.1 asyd|OJJ60657.1 ater|EAU35950.1 aver|OJI99980.1 awen|OJJ39373.1 azon|OJJ51714.1

Group8001: acla|EAW10049.1 afis|EAW25535.1 afla|EED44711.1 afum|EAL90199.1 agla|OJJ87268.1 anid|CBF75629.1 aory|EIT83459.1 asyd|OJJ57458.1 ater|EAU31795.1 aver|OJJ05722.1 awen|OJJ41277.1 azon|OJJ49500.1

Group8002: acla|EAW10841.1 afis|EAW24724.1 afla|EED45494.1 afum|EAL89373.1 agla|OJJ89011.1 anid|CBF78354.1 aory|EIT75594.1 asyd|OJJ56144.1 ater|EAU37770.1 aver|OJJ05084.1 awen|OJJ37709.1 azon|OJJ44968.1

Group8003: acla|EAW10847.1 afis|EAW20281.1 afla|EED45816.1 afum|EAL85415.1 agla|OJJ80880.1 aory|EIT75459.1 asyd|OJJ54574.1 ater|EAU34007.1 aver|OJJ07580.1 awen|OJJ30005.1 afis|EAW16940.1 afum|EAL86467.1

Group8004: acla|EAW10948.1 afis|EAW20912.1 afla|EED50260.1 afum|EAL84722.1 agla|OJJ84030.1 anid|CBF82197.1 aory|EIT77315.1 asyd|OJJ52184.1 ater|EAU38560.1 aver|OJJ05961.1 awen|OJJ34426.1 ater|EAU29294.1

Group8005: acla|EAW11333.1 afis|EAW18162.1 afla|EED56331.1 afum|EAL87522.1 agla|OJJ85964.1 anid|CBF85767.1 aory|EIT73363.1 asyd|OJJ64305.1 ater|EAU33689.1 aver|OJI98398.1 awen|OJJ30507.1 azon|OJJ45496.1

Group8006: acla|EAW11518.1 afis|EAW23935.1 afla|EED50582.1 afum|EAL87360.1 agla|OJJ79415.1 anid|CBF77269.1 asyd|OJJ56799.1 ater|EAU32302.1 aver|OJJ03535.1 awen|OJJ39020.1 azon|OJJ50744.1 azon|OJJ51345.1

Group8007: acla|EAW12899.1 afis|EAW17751.1 afla|EED45756.1 afum|EAL91564.1 agla|OJJ81703.1 anid|CBF86620.1 aory|EIT75826.1 asyd|OJJ64813.1 ater|EAU31856.1 aver|OJI98926.1 awen|OJJ37084.1 azon|OJJ43544.1

Group8008: acla|EAW14594.1 afis|EAW19323.1 afla|EED47556.1 afum|EAL94041.1 aory|EIT74540.1 asyd|OJJ54476.1 ater|EAU29459.1 aver|OJJ01973.1 azon|OJJ45105.1 afla|EED57235.1 aory|EIT74026.1 asyd|OJJ52376.1

Group8009: acla|EAW14894.1 afis|EAW24416.1 afla|EED47752.1 afum|EAL86085.2 agla|OJJ79229.1 anid|CBF73801.1 aory|EIT77104.1 asyd|OJJ56609.1 ater|EAU30313.1 aver|OJJ06115.1 awen|OJJ32793.1 azon|OJJ49042.1

Group8010: acla|EAW14994.1 afis|EAW24520.1 afla|EED54710.1 agla|OJJ80334.1 anid|CBF73856.1 aory|EIT78838.1 asyd|OJJ58690.1 ater|EAU34602.1 aver|OJJ01302.1 awen|OJJ32601.1 asyd|OJJ56518.1 aver|OJJ06204.1

Group8011: aluc|OJZ86495.1 anig|EHA22149.1 afla|EED46109.1 aory|EIT76485.1 asyd|OJJ61904.1 atub|OJI84031.1 akaw|GAA85142.1 anid|CBF85129.1 asyd|OJJ63354.1 aver|OJI97607.1 aver|OJI95997.1 azon|OJJ44491.1

Group8012: afis|EAW22248.1 afum|EAL90926.1 agla|OJJ84408.1 aver|OJJ02486.1 awen|OJJ33386.1 azon|OJJ50364.1 afla|EED53653.1 asyd|OJJ54692.1 awen|OJJ38675.1 azon|OJJ50078.1 agla|OJJ85930.1 asyd|OJJ59839.1

Group8013: afis|EAW24610.1 afla|EED56493.1 afum|EAL86240.1 agla|OJJ80105.1 akaw|GAA89016.1 aluc|OJZ80587.1 anig|EHA25068.1 aory|EIT72798.1 asyd|OJJ55481.1 ater|EAU39131.1 aver|OJJ06560.1 awen|OJJ35865.1

Group8014: afis|EAW25609.1 afla|EED53586.1 agla|OJJ85463.1 aory|EIT75042.1 asyd|OJJ55479.1 awen|OJJ36072.1 azon|OJJ49484.1 asyd|OJJ64006.1 anid|CBF78314.1 aory|EIT75715.1 agla|OJJ85466.1 awen|OJJ33321.1

Group8015: afla|EED54685.1 afum|EAL86542.1 agla|OJJ79565.1 akaw|GAA92700.1 aluc|OJZ88739.1 anig|EHA25057.1 aory|EIT78918.1 asyd|OJJ62354.1 atub|OJI80042.1 azon|OJJ42250.1 aluc|OJZ85526.1 awen|OJJ30177.1

Group8016: abra|OJJ65592.1 abra|OJJ67250.1 acla|EAW15023.1 anid|CBF70728.1 asyd|OJJ55955.1 afum|EAL91139.1 anig|EHA20220.1 ater|EAU37180.1 atub|OJI80914.1 aver|OJJ04901.1 awen|OJJ30891.1

Group8017: acar|OOF92309.1 acar|OOF92350.1 anig|EHA25105.1 acla|EAW06280.1 afis|EAW18386.1 afum|EAL93061.1 asyd|OJJ58428.1 ater|EAU32255.1 aver|OJJ01030.1 awen|OJJ31765.1 azon|OJJ50459.1

Group8018: acar|OOF94057.1 acar|OOF94127.1 abra|OJJ66385.1 afla|EED45638.1 akaw|GAA90251.1 aluc|OJZ83089.1 anig|EHA28151.1 aory|EIT75777.1 atub|OJI87855.1 awen|OJJ39958.1 azon|OJJ42768.1

Group8019: acar|OOF94070.1 acar|OOF94140.1 abra|OJJ66354.1 afla|EED46178.1 agla|OJJ83133.1 akaw|GAA90228.1 aluc|OJZ83113.1 anig|EHA28176.1 aory|EIT76384.1 atub|OJI87821.1 awen|OJJ41743.1

Group8020: acar|OOF94073.1 acar|OOF94143.1 abra|OJJ75211.1 afis|EAW20315.1 afum|EAL85441.1 atub|OJI83214.1 awen|OJJ36678.1 aluc|OJZ82088.1 anig|EHA19893.1 ater|EAU35396.1 azon|OJJ46792.1

Group8021: acar|OOF95046.1 acar|OOF95113.1 abra|OJJ68048.1 akaw|GAA90908.1 aluc|OJZ82918.1 anig|EHA21342.1 aory|EIT81634.1 asyd|OJJ60513.1 atub|OJI89144.1 aver|OJI99838.1 afla|EED55168.1

Group8022: acar|OOF95621.1 acar|OOF95673.1 acar|OOF93549.1 afis|EAW17184.1 afum|EBA27280.1 awen|OJJ38774.1 azon|OJJ48414.1 akaw|GAA92706.1 aluc|OJZ88746.1 anig|EHA18450.1 atub|OJI91318.1

Group8023: afis|EAW15399.1 afis|EAW23606.1 awen|OJJ38137.1 awen|OJJ39933.1 awen|OJJ41505.1 acla|EAW08564.1 afla|EED52204.1 afum|EAL87859.1 aory|EIT73522.1 acla|EAW14639.1 afla|EED44903.1

Group8024: afla|EED51189.1 afla|EED52082.1 abra|OJJ66811.1 atub|OJI79331.1 awen|OJJ36115.1 aory|EIT76913.1 aver|OJI96134.1 anid|CBF79164.1 asyd|OJJ62635.1 agla|OJJ80814.1 ater|EAU33810.1

Group8025: agla|OJJ78650.1 agla|OJJ80358.1 abra|OJJ75947.1 acar|OOF99965.1 akaw|GAA84649.1 aluc|OJZ89395.1 asyd|OJJ63512.1 atub|OJI85402.1 aver|OJI97418.1 azon|OJJ42102.1 aory|EIT77229.1

Group8026: agla|OJJ78651.1 agla|OJJ80356.1 acar|OOF99963.1 akaw|GAA84650.1 aluc|OJZ89396.1 anid|CBF73405.1 asyd|OJJ63511.1 aver|OJI97419.1 azon|OJJ42103.1 anig|EHA23190.1 atub|OJI85401.1

Group8027: agla|OJJ78657.1 agla|OJJ87534.1 abra|OJJ77783.1 acar|OOF98972.1 akaw|GAA83864.1 aluc|OJZ90836.1 anig|EHA19559.1 ater|EAU37643.1 atub|OJI82822.1 aver|OJJ06546.1 awen|OJJ36608.1

Group8028: agla|OJJ78722.1 agla|OJJ83804.1 acla|EAW09078.1 afla|EED49822.1 akaw|GAA93232.1 aluc|OJZ79644.1 aory|EIT77660.1 asyd|OJJ62958.1 ater|EAU37158.1 aver|OJI98009.1 awen|OJJ38685.1

Group8029: agla|OJJ79874.1 agla|OJJ86310.1 agla|OJJ85501.1 agla|OJJ89554.1 agla|OJJ85929.1 acar|OOF93326.1 acla|EAW11906.1 afla|EED48385.1 aory|EIT83083.1 awen|OJJ37791.1 awen|OJJ32674.1

Group8030: akaw|GAA86109.1 akaw|GAA88997.1 aluc|OJZ79582.1 aluc|OJZ80398.1 aluc|OJZ90341.1 abra|OJJ72360.1 anig|EHA23824.1 aory|EIT82048.1 ater|EAU31484.1 atub|OJI81768.1 afla|EED50485.1

Group8031: akaw|GAA89003.1 akaw|GAA93082.1 akaw|GAA93160.1 akaw|GAA93259.1 aluc|OJZ79724.1 aluc|OJZ79865.1 aluc|OJZ80228.1 aluc|OJZ80385.1 aluc|OJZ79934.1 aluc|OJZ80214.1 asyd|OJJ51955.1

Group8032: anid|CBF70122.1 anid|CBF76213.1 abra|OJJ66449.1 afis|EAW15394.1 afla|EED47388.1 agla|OJJ79143.1 aory|EIT82877.1 asyd|OJJ56277.1 aver|OJJ05920.1 awen|OJJ29900.1 aver|OJJ08487.1

Group8033: anid|CBF74934.1 anid|CBF79106.1 asyd|OJJ53787.1 asyd|OJJ57613.1 aver|OJI99285.1 aver|OJJ05572.1 afis|EAW19546.1 afla|EED56860.1 afum|EAL86963.1 aory|EIT77497.1 azon|OJJ46017.1

Group8034: anid|CBF75968.1 anid|CBF80729.1 asyd|OJJ52926.1 asyd|OJJ63977.1 aver|OJI96976.1 aver|OJJ08894.1 afla|EED49562.1 aory|EIT76523.1 ater|EAU37179.1 awen|OJJ35537.1 ater|EAU39437.1

Group8035: asyd|OJJ51879.1 asyd|OJJ57331.1 acla|EAW06840.1 afis|EAW16461.1 afum|EAL89057.2 anid|CBF78381.1 ater|EAU30974.1 aver|OJJ05830.1 afis|EAW20328.1 atub|OJI84624.1 aluc|OJZ80015.1

Group8036: asyd|OJJ52429.1 asyd|OJJ57204.1 aver|OJJ03085.1 aver|OJJ03899.1 aver|OJJ05542.1 acla|EAW11074.1 anid|CBF84256.1 aory|EIT73975.1 ater|EAU33926.1 azon|OJJ43046.1 afla|EED57248.1

Group8037: asyd|OJJ53788.1 asyd|OJJ57624.1 anid|CBF74936.1 azon|OJJ46019.1 afis|EAW19547.1 afla|EED52110.1 afum|EAL86962.1 aory|EIT77454.1 aver|OJJ05562.1 aver|OJI99286.1 afla|EED56862.1

Group8038: asyd|OJJ56931.1 asyd|OJJ62711.1 abra|OJJ74539.1 afum|EAL86341.1 akaw|GAA82063.1 aluc|OJZ92119.1 anig|EHA26825.1 atub|OJI86552.1 awen|OJJ40807.1 afla|EED49332.1 aluc|OJZ79685.1

Group8039: asyd|OJJ58567.1 asyd|OJJ59622.1 awen|OJJ33825.1 awen|OJJ40155.1 abra|OJJ69773.1 acar|OOF91484.1 afis|EAW17033.1 akaw|GAA87880.1 aluc|OJZ85522.1 atub|OJI80045.1 aver|OJJ04806.1

Group8040: asyd|OJJ59816.1 asyd|OJJ61416.1 aver|OJJ00752.1 aver|OJJ02474.1 abra|OJJ77757.1 agla|OJJ79448.1 akaw|GAA83882.1 aluc|OJZ90814.1 anig|EHA19542.1 atub|OJI82804.1 awen|OJJ32586.1

Group8041: ater|EAU29577.1 ater|EAU34057.1 acar|OOF94502.1 afis|EAW17046.1 afla|EED53439.1 anid|CBF69407.1 aory|EIT75055.1 asyd|OJJ62008.1 aver|OJI96100.1 aver|OJJ06555.1 awen|OJJ29783.1

Group8042: awen|OJJ29560.1 awen|OJJ31897.1 abra|OJJ67105.1 acar|OOF93562.1 afis|EAW21629.1 afla|EED54261.1 akaw|GAA86631.1 aluc|OJZ85186.1 aory|EIT76082.1 atub|OJI80409.1 azon|OJJ43599.1

Group8043: awen|OJJ29962.1 awen|OJJ39646.1 afis|EAW19737.1 afum|EAL86792.1 aory|EIT75574.1 afla|EED45894.1 agla|OJJ79333.1 anid|CBF85600.1 asyd|OJJ64193.1 ater|EAU36894.1 aver|OJI98270.1

Group8044: awen|OJJ29988.1 awen|OJJ35841.1 abra|OJJ68829.1 acar|OOF94025.1 akaw|GAA90423.1 aluc|OJZ85806.1 anig|EHA24749.1 asyd|OJJ54678.1 atub|OJI79592.1 aver|OJJ07489.1 aory|EIT77344.1

Group8045: awen|OJJ30092.1 awen|OJJ30120.1 awen|OJJ30095.1 abra|OJJ71734.1 acar|OOF90208.1 akaw|GAA85148.1 aluc|OJZ86489.1 anig|EHA22182.1 agla|OJJ83273.1 atub|OJI84040.1 azon|OJJ44998.1

Group8046: awen|OJJ30282.1 awen|OJJ31244.1 awen|OJJ31538.1 abra|OJJ77948.1 acla|EAW13702.1 agla|OJJ88430.1 akaw|GAA83716.1 aluc|OJZ91017.1 anig|EHA19719.1 ater|EAU35598.1 atub|OJI83008.1

Group8047: awen|OJJ30387.1 awen|OJJ37881.1 abra|OJJ66211.1 acar|OOF90649.1 afla|EED47533.1 akaw|GAA89076.1 aluc|OJZ80515.1 anig|EHA25678.1 aory|EIT74542.1 atub|OJI88415.1 azon|OJJ45139.1

Group8048: awen|OJJ30794.1 awen|OJJ31687.1 afis|EAW20856.1 afla|EED45802.1 afum|EAL84778.1 aluc|OJZ90879.1 anig|EHA19600.1 aory|EIT75421.1 atub|OJI82876.1 awen|OJJ40932.1 azon|OJJ42352.1

Group8049: awen|OJJ31392.1 awen|OJJ37043.1 abra|OJJ78202.1 acar|OOF98652.1 afla|EED55373.1 akaw|GAA85827.1 aluc|OJZ91273.1 anig|EHA19960.1 aory|EIT80865.1 ater|EAU31198.1 atub|OJI83262.1

Group8050: awen|OJJ31446.1 awen|OJJ35962.1 abra|OJJ76659.1 afis|EAW16761.1 afum|EAL86332.1 akaw|GAA92999.1 aluc|OJZ91544.1 anig|EHA22581.1 asyd|OJJ53611.1 atub|OJI85998.1 aver|OJJ08138.1

Group8051: awen|OJJ32912.1 awen|OJJ37496.1 awen|OJJ35377.1 azon|OJJ43561.1 azon|OJJ46698.1 azon|OJJ46168.1 awen|OJJ30279.1 azon|OJJ42825.1 ater|EAU36022.1 ater|EAU32855.1 awen|OJJ33312.1

Group8052: awen|OJJ35361.1 awen|OJJ36277.1 afla|EED51192.1 agla|OJJ82229.1 anid|CBF86353.1 asyd|OJJ60252.1 ater|EAU32101.1 aver|OJJ04005.1 afla|EED54550.1 aory|EIT78943.1 asyd|OJJ55219.1

Group8053: awen|OJJ41678.1 awen|OJJ41680.1 acar|OOF94904.1 acla|EAW13441.1 afis|EAW16680.1 agla|OJJ83769.1 asyd|OJJ61388.1 atub|OJI81715.1 aver|OJJ00716.1 acla|EAW07054.1 azon|OJJ46174.1

Group8054: azon|OJJ42012.1 azon|OJJ47279.1 acla|EAW14327.1 afis|EAW18961.1 afla|EED51007.1 anid|CBF69602.1 aory|EIT76960.1 asyd|OJJ59134.1 aver|OJJ01723.1 awen|OJJ31560.1 ater|EAU38144.1

Group8055: azon|OJJ45098.1 azon|OJJ50418.1 abra|OJJ67372.1 acar|OOF92439.1 akaw|GAA86450.1 aluc|OJZ84993.1 anig|EHA25205.1 asyd|OJJ64319.1 ater|EAU32175.1 atub|OJI80607.1 aver|OJI98412.1

Group8056: azon|OJJ45957.1 azon|OJJ45961.1 acla|EAW13611.1 afla|EED53482.1 akaw|GAA81993.1 aluc|OJZ92196.1 anig|EHA20151.1 aory|EIT75106.1 asyd|OJJ61761.1 atub|OJI87951.1 agla|OJJ80452.1

Group8057: abra|OJJ65488.1 acar|OOF90656.1 afla|EED47478.1 afum|EAL85424.1 akaw|GAA88140.1 anig|EHA18428.1 aory|EIT79975.1 atub|OJI91618.1 aver|OJJ06578.1 asyd|OJJ55500.1 azon|OJJ42390.1

Group8058: abra|OJJ65684.1 acar|OOF90739.1 acla|EAW11008.1 afis|EAW20846.1 afla|EED55713.1 afum|EAL84787.1 akaw|GAA92270.1 aluc|OJZ90164.1 anid|CBF80207.1 aory|EIT83199.1 ater|EAU37552.1

Group8059: abra|OJJ65762.1 acar|OOF93287.1 afis|EAW15656.1 afla|EED45011.1 afum|EAL86262.1 aluc|OJZ80174.1 anig|EHA22866.1 aory|EIT73797.1 asyd|OJJ63942.1 aver|OJI97000.1 azon|OJJ47390.1

Group8060: abra|OJJ65780.1 afis|EAW16372.1 agla|OJJ79808.1 akaw|GAA90883.1 aluc|OJZ82949.1 anid|CBF77672.1 asyd|OJJ53107.1 atub|OJI89181.1 aver|OJJ03745.1 awen|OJJ35384.1 awen|OJJ37007.1

Group8061: abra|OJJ65791.1 acla|EAW07096.1 afis|EAW22107.1 afla|EED47860.1 afum|EAL91077.1 anid|CBF73938.1 anig|EHA18730.1 aory|EIT77219.1 asyd|OJJ53809.1 ater|EAU29312.1 aver|OJJ07909.1

Group8062: abra|OJJ65792.1 acla|EAW07097.1 afis|EAW22108.1 afla|EED54169.1 afum|EAL91076.2 anid|CBF73940.1 anig|EHA18731.1 aory|EIT76083.1 asyd|OJJ53808.1 ater|EAU29313.1 aver|OJJ07908.1

Group8063: acla|EAW06486.1 afis|EAW15981.1 afum|EAL88613.1 awen|OJJ30489.1 azon|OJJ45480.1 acar|OOF99104.1 afla|EED56351.1 aory|EIT73357.1 asyd|OJJ64336.1 ater|EAU33671.1 aver|OJI98429.1

Group8064: abra|OJJ65834.1 acar|OOF97961.1 afla|EED52116.1 agla|OJJ84459.1 akaw|GAA92845.1 aluc|OJZ88787.1 anid|CBF81710.1 asyd|OJJ52026.1 ater|EAU31337.1 aver|OJJ03366.1 awen|OJJ32285.1

Group8065: abra|OJJ65844.1 acla|EAW11454.1 afis|EAW16227.1 afla|EED53064.1 agla|OJJ79219.1 akaw|GAA92840.1 aluc|OJZ88786.1 anig|EHA22172.1 aory|EIT74759.1 aver|OJJ03132.1 akaw|GAA93198.1

Group8066: abra|OJJ65856.1 acar|OOF95016.1 akaw|GAA92289.1 aluc|OJZ90186.1 anig|EHA23685.1 atub|OJI81613.1 abra|OJJ66187.1 afis|EAW16418.1 asyd|OJJ65285.1 aver|OJI99510.1 asyd|OJJ65361.1

Group8067: abra|OJJ65860.1 acar|OOF95025.1 afla|EED46398.1 akaw|GAA92285.1 aluc|OJZ90182.1 anig|EHA23681.1 aory|EIT72358.1 atub|OJI81609.1 awen|OJJ37988.1 azon|OJJ47980.1 ater|EAU31822.1

Group8068: abra|OJJ65881.1 acar|OOF98858.1 afla|EED49468.1 akaw|GAA83746.1 aluc|OJZ90985.1 anid|CBF71213.1 anig|EHA19690.1 asyd|OJJ55198.1 atub|OJI82976.1 aver|OJJ07016.1 awen|OJJ35945.1

Group8069: abra|OJJ65996.1 afla|EED53500.1 agla|OJJ85432.1 akaw|GAA90868.1 aluc|OJZ82965.1 anig|EHA21381.1 aory|EIT74951.1 ater|EAU29732.1 atub|OJI89197.1 awen|OJJ41944.1 awen|OJJ37715.1

Group8070: abra|OJJ66079.1 acar|OOF99289.1 afla|EED46899.1 agla|OJJ79339.1 akaw|GAA84166.1 aluc|OJZ81091.1 anid|CBF70516.1 anig|EHA19279.1 aory|EIT72524.1 atub|OJI82493.1 awen|OJJ33666.1

Group8071: abra|OJJ66083.1 acar|OOF99285.1 akaw|GAA84162.1 aluc|OJZ81087.1 anig|EHA19283.1 asyd|OJJ56886.1 atub|OJI82497.1 awen|OJJ30430.1 azon|OJJ44589.1 acar|OOF94971.1 atub|OJI89657.1

Group8072: acar|OOF99278.1 asyd|OJJ59182.1 aver|OJJ01803.1 abra|OJJ76647.1 akaw|GAA92988.1 aluc|OJZ91562.1 anig|EHA22593.1 atub|OJI86013.1 anid|CBF69529.1 azon|OJJ43247.1 awen|OJJ29873.1

Group8073: abra|OJJ66131.1 acar|OOF94019.1 afla|EED50864.1 akaw|GAA92688.1 aluc|OJZ90146.1 anid|CBF71758.1 anig|EHA20924.1 aory|EIT78958.1 asyd|OJJ61737.1 atub|OJI84653.1 aver|OJI95830.1

Group8074: abra|OJJ66133.1 acar|OOF94020.1 afis|EAW15420.1 afla|EED51825.1 afum|EBA27170.1 akaw|GAA92690.1 aluc|OJZ90148.1 anig|EHA20922.1 aory|EIT81711.1 ater|EAU36667.1 atub|OJI84651.1

Group8075: abra|OJJ66136.1 acar|OOF94024.1 afla|EED45757.1 akaw|GAA92693.1 aluc|OJZ90151.1 anig|EHA20919.1 aory|EIT75460.1 atub|OJI84648.1 acla|EAW12911.1 afis|EAW17753.1 afum|EAL91566.1

Group8076: abra|OJJ66157.1 asyd|OJJ54351.1 atub|OJI83132.1 aver|OJJ05304.1 awen|OJJ34587.1 abra|OJJ68081.1 acar|OOF93880.1 akaw|GAA86867.1 aluc|OJZ89990.1 anig|EHA27360.1 atub|OJI84819.1

Group8077: afis|EAW24547.1 abra|OJJ77744.1 acla|EAW12412.1 afis|EAW20266.1 afla|EED47769.1 afum|EAL86211.1 anig|EHA25377.1 aory|EIT77164.1 asyd|OJJ64794.1 aver|OJI98908.1 ater|EAU29413.1

Group8078: acla|EAW11746.1 agla|OJJ78930.1 anid|CBF79834.1 asyd|OJJ58145.1 awen|OJJ33698.1 acar|OOF92833.1 afis|EAW23779.1 afla|EED51581.1 afum|EAL87213.1 aory|EIT80483.1 ater|EAU31320.1

Group8079: abra|OJJ66202.1 afla|EED51058.1 akaw|GAA87479.1 aluc|OJZ88047.1 anig|EHA17819.1 ater|EAU29612.1 atub|OJI90655.1 aver|OJJ05901.1 awen|OJJ36099.1 azon|OJJ43914.1 aory|EIT76948.1

Group8080: abra|OJJ66234.1 acar|OOF90692.1 agla|OJJ84414.1 akaw|GAA89055.1 aluc|OJZ80539.1 anid|CBF85757.1 anig|EHA25656.1 asyd|OJJ64299.1 atub|OJI88392.1 aver|OJJ07893.1 awen|OJJ34688.1

Group8081: abra|OJJ66238.1 acar|OOF90700.1 afis|EAW16784.1 akaw|GAA89051.1 aluc|OJZ80543.1 anig|EHA25652.1 asyd|OJJ57311.1 atub|OJI88388.1 aver|OJJ05851.1 awen|OJJ29850.1 azon|OJJ45022.1

Group8082: azon|OJJ42357.1 abra|OJJ68265.1 acar|OOF97544.1 agla|OJJ88624.1 akaw|GAA83200.1 aluc|OJZ83701.1 anid|CBF87287.1 anig|EHA18852.1 asyd|OJJ56366.1 atub|OJI90073.1 aver|OJJ06348.1

Group8083: abra|OJJ66268.1 afis|EAW15637.1 afla|EED54380.1 afum|EAL85702.1 agla|OJJ88682.1 anig|EHA18763.1 aory|EIT75861.1 asyd|OJJ52551.1 aver|OJI98881.1 awen|OJJ32688.1 aver|OJJ03849.1

Group8084: abra|OJJ66269.1 afis|EAW15638.1 afum|EAL85701.2 anig|EHA18762.1 aver|OJI98880.1 abra|OJJ76725.1 acar|OOF91357.1 akaw|GAA92476.1 aluc|OJZ91486.1 anig|EHA22530.1 atub|OJI85940.1

Group8085: abra|OJJ66297.1 acar|OOG00129.1 afis|EAW24606.1 akaw|GAA89856.1 aluc|OJZ81830.1 asyd|OJJ52051.1 ater|EAU33800.1 atub|OJI84399.1 aver|OJJ04478.1 awen|OJJ31949.1 azon|OJJ42430.1

Group8086: abra|OJJ66473.1 afis|EAW17243.1 afum|EAL86663.2 akaw|GAA92459.1 aluc|OJZ81558.1 anid|CBF75986.1 anig|EHA25876.1 asyd|OJJ53456.1 atub|OJI88627.1 aver|OJJ08299.1 asyd|OJJ53457.1

Group8087: abra|OJJ66479.1 afla|EED46094.1 akaw|GAA92462.1 aluc|OJZ81555.1 anid|CBF82152.1 anig|EHA25879.1 aory|EIT76390.1 asyd|OJJ54329.1 atub|OJI88630.1 aver|OJJ05283.1 awen|OJJ41956.1

Group8088: abra|OJJ66486.1 acla|EAW10983.1 afla|EED45559.1 afum|EAL84756.1 agla|OJJ83984.1 aluc|OJZ81549.1 anid|CBF82857.1 anig|EHA25886.1 aver|OJJ00919.1 awen|OJJ34483.1 azon|OJJ51760.1

Group8089: abra|OJJ66545.1 acar|OOF90997.1 afla|EED50269.1 akaw|GAA91170.1 aluc|OJZ81498.1 anig|EHA25932.1 aory|EIT77367.1 ater|EAU38565.1 atub|OJI88683.1 aluc|OJZ80365.1 anig|EHA20400.1

Group8090: afla|EED50292.1 aory|EIT77326.1 acla|EAW10923.1 afis|EAW20935.1 afum|EAL84699.1 anid|CBF82530.1 anig|EHA25952.1 asyd|OJJ54075.1 ater|EAU38581.1 aver|OJJ08664.1 azon|OJJ50774.1

Group8091: abra|OJJ66758.1 acar|OOF93840.1 agla|OJJ85834.1 akaw|GAA89694.1 aluc|OJZ80690.1 anig|EHA27325.1 aory|EIT75473.1 asyd|OJJ54962.1 ater|EAU30488.1 atub|OJI87550.1 aver|OJJ07229.1

Group8092: abra|OJJ66764.1 acar|OOF93845.1 afla|EED56723.1 akaw|GAA89700.1 aluc|OJZ80685.1 anid|CBF69518.1 anig|EHA27329.1 aory|EIT73881.1 asyd|OJJ59211.1 atub|OJI87543.1 aver|OJJ01821.1

Group8093: abra|OJJ66770.1 acar|OOF93851.1 afis|EAW15609.1 afla|EED56864.1 akaw|GAA89707.1 aluc|OJZ80677.1 anig|EHA27335.1 aory|EIT77549.1 atub|OJI87536.1 awen|OJJ35552.1 acar|OOF93886.1

Group8094: abra|OJJ66779.1 acar|OOF93858.1 afla|EED49902.1 agla|OJJ81856.1 akaw|GAA89714.1 aluc|OJZ80670.1 anig|EHA27341.1 aory|EIT72458.1 ater|EAU32527.1 atub|OJI87524.1 awen|OJJ32044.1

Group8095: abra|OJJ66786.1 acar|OOF93860.1 afla|EED50228.1 akaw|GAA89720.1 aluc|OJZ80664.1 anig|EHA27346.1 aory|EIT77264.1 asyd|OJJ58370.1 atub|OJI87519.1 aver|OJJ02036.1 awen|OJJ31989.1

Group8096: abra|OJJ66787.1 acar|OOF93861.1 afla|EED50229.1 akaw|GAA89721.1 aluc|OJZ80663.1 anig|EHA27347.1 aory|EIT77320.1 asyd|OJJ58369.1 atub|OJI87518.1 aver|OJJ02037.1 awen|OJJ31990.1

Group8097: acar|OOF91635.1 afla|EED54270.1 akaw|GAA91549.1 aory|EIT76006.1 acla|EAW14435.1 afis|EAW19143.1 afum|EAL93874.1 anid|CBF86316.1 asyd|OJJ64998.1 ater|EAU37415.1 aver|OJI99118.1

Group8098: abra|OJJ66954.1 acar|OOF93654.1 agla|OJJ79211.1 akaw|GAA92207.1 aluc|OJZ85624.1 anig|EHA24904.1 asyd|OJJ65095.1 ater|EAU37244.1 atub|OJI79417.1 aver|OJI99219.1 awen|OJJ30185.1

Group8099: abra|OJJ66968.1 acar|OOF93636.1 afla|EED47522.1 akaw|GAA92195.1 aluc|OJZ85610.1 anig|EHA24916.1 aory|EIT74560.1 asyd|OJJ62820.1 ater|EAU31758.1 atub|OJI79403.1 aver|OJI96865.1

Group8100: abra|OJJ66971.1 acar|OOG00221.1 afla|EED56781.1 akaw|GAA92193.1 aluc|OJZ85608.1 anig|EHA24918.1 aory|EIT73837.1 atub|OJI79401.1 asyd|OJJ57306.1 aver|OJI99612.1 aory|EIT79995.1

Group8101: abra|OJJ67001.1 acar|OOF93613.1 afla|EED45809.1 akaw|GAA92164.1 aluc|OJZ85571.1 anid|CBF83990.1 anig|EHA24947.1 aory|EIT75678.1 atub|OJI79367.1 aver|OJI96468.1 awen|OJJ37853.1

Group8102: abra|OJJ67005.1 acar|OOF93609.1 afla|EED49662.1 akaw|GAA92162.1 aluc|OJZ85569.1 anid|CBF87113.1 anig|EHA24949.1 asyd|OJJ60328.1 atub|OJI79364.1 aver|OJI99676.1 awen|OJJ37893.1

Group8103: afla|EED51775.1 awen|OJJ35033.1 abra|OJJ74169.1 acar|OOF95556.1 akaw|GAA86847.1 aluc|OJZ89968.1 anid|CBF77079.1 anig|EHA27166.1 asyd|OJJ56967.1 atub|OJI84839.1 aver|OJJ03320.1

Group8104: abra|OJJ67211.1 acar|OOF98348.1 akaw|GAA90966.1 aluc|OJZ87762.1 anid|CBF89581.1 anig|EHA20247.1 asyd|OJJ62332.1 atub|OJI80882.1 aver|OJI96438.1 awen|OJJ39919.1 ater|EAU35357.1

Group8105: abra|OJJ67232.1 acar|OOF98319.1 afla|EED45990.1 agla|OJJ89178.1 akaw|GAA91657.1 aluc|OJZ87742.1 anig|EHA20234.1 aory|EIT75662.1 asyd|OJJ64982.1 atub|OJI80899.1 awen|OJJ33648.1

Group8106: abra|OJJ67263.1 afis|EAW21082.1 afla|EED44922.1 afum|EAL92103.1 agla|OJJ79286.1 akaw|GAA91679.1 aluc|OJZ87715.1 anig|EHA20209.1 atub|OJI80927.1 aver|OJJ00777.1 awen|OJJ35784.1

Group8107: abra|OJJ67291.1 acar|OOF92522.1 akaw|GAA86522.1 aluc|OJZ85068.1 anid|CBF81693.1 anig|EHA25282.1 aory|EIT75508.1 atub|OJI80530.1 afla|EED45685.1 asyd|OJJ59563.1 agla|OJJ79145.1

Group8108: abra|OJJ67293.1 acar|OOF92518.1 afla|EED45658.1 agla|OJJ79147.1 akaw|GAA86520.1 aluc|OJZ85066.1 anid|CBF81704.1 anig|EHA25280.1 asyd|OJJ59569.1 atub|OJI80532.1 aver|OJJ02698.1

Group8109: abra|OJJ67294.1 acar|OOF92516.1 agla|OJJ79148.1 akaw|GAA86519.1 aluc|OJZ85065.1 anid|CBF81702.1 anig|EHA25279.1 aory|EIT75593.1 asyd|OJJ59572.1 atub|OJI80533.1 aver|OJJ02695.1

Group8110: abra|OJJ67314.1 afla|EED46371.1 aory|EIT81658.1 asyd|OJJ59224.1 ater|EAU32714.1 aver|OJJ01834.1 awen|OJJ36094.1 azon|OJJ50751.1 acar|OOF91718.1 atub|OJI89682.1 afla|EED55175.1

Group8111: abra|OJJ67383.1 acar|OOF92430.1 afla|EED52927.1 agla|OJJ84468.1 akaw|GAA86441.1 aluc|OJZ84983.1 anig|EHA25196.1 aory|EIT78683.1 ater|EAU29906.1 atub|OJI80617.1 awen|OJJ31909.1

Group8112: aluc|OJZ89562.1 anid|CBF69557.1 awen|OJJ36249.1 acla|EAW06764.1 ater|EAU29328.1 atub|OJI85242.1 afla|EED54180.1 akaw|GAA88860.1 anig|EHA23044.1 asyd|OJJ59163.1 aver|OJJ01753.1

Group8113: abra|OJJ67493.1 acla|EAW06277.1 afis|EAW18383.1 afla|EED53023.1 afum|EAL93058.1 agla|OJJ84947.1 akaw|GAA86345.1 aluc|OJZ84871.1 anig|EHA25102.1 aory|EIT77432.1 ater|EAU32252.1

Group8114: abra|OJJ67555.1 acar|OOF90044.1 afla|EED45512.1 akaw|GAA88109.1 aluc|OJZ81379.1 anig|EHA18674.1 aory|EIT75413.1 atub|OJI91585.1 azon|OJJ45314.1 acar|OOF94222.1 agla|OJJ86352.1

Group8115: abra|OJJ67559.1 acar|OOG00273.1 afla|EED53442.1 akaw|GAA88107.1 aluc|OJZ81374.1 anig|EHA18672.1 aory|EIT74821.1 asyd|OJJ53831.1 ater|EAU37629.1 atub|OJI91580.1 aver|OJJ07935.1

Group8116: abra|OJJ67570.1 acar|OOF91942.1 afla|EED45401.1 agla|OJJ83699.1 akaw|GAA89100.1 anig|EHA25702.1 aory|EIT82457.1 asyd|OJJ58150.1 atub|OJI88438.1 aver|OJJ04254.1 awen|OJJ30694.1

Group8117: abra|OJJ67598.1 acar|OOF91949.1 afla|EED47298.1 akaw|GAA89117.1 aluc|OJZ81722.1 anig|EHA25717.1 aory|EIT82872.1 asyd|OJJ63585.1 atub|OJI88459.1 aver|OJI97349.1 awen|OJJ33436.1

Group8118: abra|OJJ67638.1 afla|EED46470.1 afum|EAL86600.1 akaw|GAA89156.1 aluc|OJZ81678.1 anig|EHA25763.1 ater|EAU35976.1 atub|OJI88503.1 awen|OJJ40906.1 aory|EIT72325.1 azon|OJJ48532.1

Group8119: abra|OJJ67675.1 afis|EAW15381.1 afla|EED48117.1 akaw|GAA83878.1 aluc|OJZ90818.1 anid|CBF83147.1 anig|EHA25870.1 aory|EIT76237.1 asyd|OJJ55490.1 aver|OJJ06568.1 awen|OJJ31452.1

Group8120: abra|OJJ67766.1 acar|OOF98574.1 afla|EED47855.1 agla|OJJ84197.1 akaw|GAA89764.1 aluc|OJZ80619.1 anig|EHA27213.1 ater|EAU30931.1 atub|OJI87473.1 afla|EED46617.1 anig|EHA27212.1

Group8121: abra|OJJ67777.1 afla|EED51686.1 agla|OJJ78473.1 ater|EAU39528.1 azon|OJJ42638.1 abra|OJJ68156.1 afla|EED47515.1 akaw|GAA90125.1 aluc|OJZ83225.1 anig|EHA21082.1 atub|OJI87711.1

Group8122: awen|OJJ32655.1 abra|OJJ77418.1 afla|EED53244.1 akaw|GAA91337.1 aluc|OJZ84842.1 asyd|OJJ54864.1 atub|OJI89256.1 aver|OJJ07914.1 aory|EIT75212.1 atub|OJI89978.1 asyd|OJJ53813.1

Group8123: abra|OJJ67897.1 agla|OJJ84454.1 akaw|GAA91901.1 aluc|OJZ81993.1 anig|EHA22263.1 asyd|OJJ64966.1 atub|OJI84543.1 ater|EAU29594.1 afis|EAW20706.1 awen|OJJ34083.1 aver|OJI99089.1

Group8124: abra|OJJ67911.1 afis|EAW16401.1 afum|EAL89026.1 agla|OJJ83731.1 aluc|OJZ81975.1 anid|CBF86678.1 anig|EHA22278.1 atub|OJI84526.1 aver|OJJ05880.1 awen|OJJ29891.1 aver|OJJ01777.1

Group8125: abra|OJJ67931.1 acar|OOF92781.1 afis|EAW20401.1 afum|EAL85500.2 akaw|GAA91266.1 aluc|OJZ82803.1 anig|EHA21244.1 ater|EAU31394.1 atub|OJI89034.1 awen|OJJ33793.1 azon|OJJ48753.1

Group8126: awen|OJJ31422.1 anig|EHA19747.1 atub|OJI85733.1 acar|OOF91999.1 acla|EAW13942.1 akaw|GAA83683.1 anid|CBF76229.1 acla|EAW13943.1 atub|OJI83046.1 akaw|GAA84353.1 aluc|OJZ89054.1

Group8127: abra|OJJ68038.1 acar|OOF91805.1 afis|EAW24633.1 afla|EED55051.1 akaw|GAA90914.1 aluc|OJZ82912.1 anig|EHA21335.1 aory|EIT81965.1 asyd|OJJ53720.1 ater|EAU36658.1 atub|OJI89138.1

Group8128: abra|OJJ68108.1 afla|EED53403.1 akaw|GAA86891.1 aluc|OJZ90014.1 anid|CBF80624.1 anig|EHA27389.1 aory|EIT75164.1 asyd|OJJ54633.1 atub|OJI84793.1 aver|OJJ07523.1 awen|OJJ33178.1

Group8129: abra|OJJ68109.1 afla|EED53402.1 akaw|GAA86892.1 aluc|OJZ90015.1 anid|CBF80626.1 anig|EHA27390.1 aory|EIT75119.1 asyd|OJJ54632.1 atub|OJI84792.1 aver|OJJ07524.1 awen|OJJ33177.1

Group8130: abra|OJJ68160.1 akaw|GAA83282.1 aluc|OJZ83791.1 anig|EHA18928.1 atub|OJI90159.1 azon|OJJ43666.1 abra|OJJ78087.1 ater|EAU37599.1 acar|OOF96465.1 anid|CBF86357.1 akaw|GAA83281.1

Group8131: abra|OJJ68288.1 acla|EAW12903.1 afis|EAW17775.1 afla|EED56743.1 afum|EAL91583.1 akaw|GAA83183.1 aluc|OJZ83683.1 anig|EHA18837.1 aory|EIT73838.1 atub|OJI90054.1 aver|OJI99931.1

Group8132: abra|OJJ68296.1 acar|OOF97563.1 acla|EAW08964.1 afis|EAW15445.1 afla|EED46466.1 afum|EAL84202.1 akaw|GAA83177.1 aluc|OJZ83676.1 anig|EHA18832.1 aory|EIT72322.1 atub|OJI90047.1

Group8133: abra|OJJ68298.1 acar|OOF97573.1 aluc|OJZ83674.1 anig|EHA18830.1 atub|OJI90045.1 abra|OJJ77761.1 acla|EAW12360.1 afum|EAL91922.1 azon|OJJ49864.1 afla|EED51123.1 aory|EIT81326.1

Group8134: abra|OJJ68320.1 acar|OOF97581.1 afla|EED52686.1 akaw|GAA83153.1 aluc|OJZ83653.1 anig|EHA18804.1 aory|EIT74233.1 asyd|OJJ59949.1 atub|OJI90022.1 aver|OJJ02340.1 awen|OJJ39603.1

Group8135: abra|OJJ68323.1 afla|EED49521.1 akaw|GAA92074.1 aluc|OJZ83650.1 anid|CBF78523.1 anig|EHA18801.1 aory|EIT82597.1 atub|OJI90019.1 aver|OJI99447.1 awen|OJJ39149.1 azon|OJJ51481.1

Group8136: abra|OJJ68332.1 acar|OOF97607.1 afis|EAW25713.1 afla|EED46086.1 afum|EAL84385.1 agla|OJJ84491.1 akaw|GAA92084.1 aluc|OJZ83640.1 anig|EHA18793.1 aory|EIT76410.1 atub|OJI90009.1

Group8137: acla|EAW08804.1 afis|EAW15593.1 afum|EAL85739.2 ater|EAU32534.1 afla|EED54741.1 agla|OJJ85154.1 anid|CBF78081.1 aory|EIT78817.1 asyd|OJJ55879.1 aver|OJJ04824.1 awen|OJJ38269.1

Group8138: abra|OJJ68378.1 acar|OOF93989.1 acla|EAW14618.1 afis|EAW19356.1 afum|EAL94076.1 akaw|GAA92566.1 aluc|OJZ90114.1 anig|EHA20954.1 atub|OJI84685.1 azon|OJJ43675.1 azon|OJJ51821.1

Group8139: abra|OJJ68427.1 acar|OOF93936.1 agla|OJJ86941.1 akaw|GAA86944.1 aluc|OJZ90069.1 anig|EHA21005.1 atub|OJI84738.1 awen|OJJ33587.1 acar|OOF93935.1 anig|EHA21004.1 awen|OJJ37600.1

Group8140: abra|OJJ68457.1 acar|OOF93940.1 acla|EAW11092.1 afis|EAW20765.1 afla|EED50227.1 afum|EAL84854.1 akaw|GAA86917.1 aluc|OJZ90040.1 anig|EHA21031.1 ater|EAU32754.1 atub|OJI84766.1

Group8141: abra|OJJ68516.1 acar|OOF95565.1 afis|EAW24773.1 afla|EED49729.1 aluc|OJZ82730.1 anid|CBF89921.1 asyd|OJJ62124.1 atub|OJI88963.1 aver|OJI96232.1 awen|OJJ37933.1 azon|OJJ43610.1

Group8142: abra|OJJ68577.1 acar|OOF92103.1 afla|EED49944.1 akaw|GAA91916.1 aluc|OJZ81964.1 anid|CBF71749.1 anig|EHA22283.1 aory|EIT83427.1 asyd|OJJ61731.1 atub|OJI84515.1 aver|OJI95825.1

Group8143: anid|CBF73475.1 acar|OOF91851.1 afis|EAW25678.1 afla|EED52546.1 afum|EAL84348.1 agla|OJJ86553.1 aory|EIT74174.1 asyd|OJJ54791.1 aver|OJJ07372.1 awen|OJJ29895.1 azon|OJJ43601.1

Group8144: abra|OJJ68822.1 acar|OOF99136.1 agla|OJJ82564.1 anig|EHA24759.1 aory|EIT74227.1 atub|OJI79583.1 awen|OJJ36433.1 afis|EAW20257.1 acla|EAW12444.1 asyd|OJJ55238.1 aver|OJJ06985.1

Group8145: abra|OJJ68826.1 akaw|GAA90427.1 aluc|OJZ85802.1 anig|EHA24754.1 asyd|OJJ55693.1 ater|EAU37617.1 atub|OJI79587.1 aver|OJJ06778.1 abra|OJJ71062.1 acla|EAW11659.1 azon|OJJ45881.1

Group8146: abra|OJJ68853.1 acar|OOF98661.1 anid|CBF75774.1 asyd|OJJ62872.1 aver|OJI96925.1 azon|OJJ48281.1 abra|OJJ75195.1 akaw|GAA91747.1 aluc|OJZ92758.1 anig|EHA26248.1 atub|OJI87202.1

Group8147: abra|OJJ68861.1 acar|OOF90528.1 afla|EED56733.1 agla|OJJ81558.1 akaw|GAA90392.1 aluc|OJZ85839.1 anig|EHA24725.1 aory|EIT73860.1 atub|OJI79627.1 awen|OJJ31227.1 azon|OJJ46150.1

Group8148: abra|OJJ69011.1 acar|OOF92722.1 acla|EAW12847.1 afis|EAW18010.1 afum|EAL91822.1 agla|OJJ80276.1 akaw|GAA90528.1 aluc|OJZ85997.1 anig|EHA24613.1 aory|EIT81701.1 ater|EAU31526.1

Group8149: abra|OJJ69022.1 acar|OOF89908.1 aluc|OJZ86008.1 anid|CBF88670.1 anig|EHA24602.1 aory|EIT76676.1 asyd|OJJ63894.1 atub|OJI79797.1 aver|OJI97045.1 awen|OJJ30870.1 azon|OJJ42016.1

Group8150: abra|OJJ69023.1 acar|OOF91421.1 afis|EAW23608.1 afla|EED51808.1 afum|EAL87857.1 akaw|GAA90540.1 aluc|OJZ86010.1 aory|EIT81716.1 atub|OJI79799.1 anig|EHA22502.1 acla|EAW08565.1

Group8151: abra|OJJ69044.1 acar|OOF98654.1 afla|EED46057.1 akaw|GAA90573.1 aluc|OJZ86029.1 anig|EHA24583.1 aory|EIT83476.1 asyd|OJJ53816.1 atub|OJI79818.1 aver|OJJ06538.1 awen|OJJ38012.1

Group8152: abra|OJJ69246.1 acar|OOF91074.1 acla|EAW08603.1 afis|EAW15830.1 afum|EBA27234.1 agla|OJJ81994.1 aluc|OJZ86227.1 anid|CBF70865.1 asyd|OJJ52578.1 atub|OJI80020.1 azon|OJJ43719.1

Group8153: abra|OJJ69333.1 anig|EHA28567.1 abra|OJJ72542.1 acar|OOG00675.1 afla|EED53624.1 akaw|GAA86266.1 aluc|OJZ90526.1 aory|EIT74957.1 atub|OJI81948.1 awen|OJJ31579.1 agla|OJJ89571.1

Group8154: abra|OJJ69335.1 acar|OOF94394.1 afla|EED45505.1 akaw|GAA88205.1 aluc|OJZ82167.1 anig|EHA28565.1 aory|EIT75585.1 ater|EAU29420.1 atub|OJI88270.1 acar|OOF90008.1 awen|OJJ35535.1

Group8155: abra|OJJ69341.1 acar|OOF94387.1 acla|EAW14705.1 afis|EAW24236.1 afla|EED48035.1 afum|EAL85905.1 akaw|GAA88210.1 aluc|OJZ82173.1 aory|EIT83333.1 atub|OJI88264.1 awen|OJJ33050.1

Group8156: abra|OJJ69433.1 acar|OOF94304.1 afis|EAW23589.1 afla|EED49524.1 akaw|GAA88297.1 aluc|OJZ82269.1 anig|EHA28474.1 aory|EIT82611.1 atub|OJI88169.1 afla|EED48207.1 aory|EIT72964.1

Group8157: ater|EAU33216.1 azon|OJJ43984.1 acla|EAW06631.1 afis|EAW16134.1 afla|EED48673.1 afum|EAL88764.1 anid|CBF85516.1 aory|EIT73100.1 asyd|OJJ64125.1 aver|OJI98208.1 awen|OJJ30294.1

Group8158: abra|OJJ69561.1 afis|EAW24399.1 afum|EAL86070.1 agla|OJJ78881.1 anid|CBF73731.1 anig|EHA28376.1 asyd|OJJ56632.1 ater|EAU30338.1 aver|OJJ06094.1 awen|OJJ32806.1 azon|OJJ49028.1

Group8159: abra|OJJ69574.1 acar|OOF91582.1 acla|EAW14399.1 afla|EED45439.1 akaw|GAA91571.1 aluc|OJZ82391.1 aory|EIT82459.1 ater|EAU32011.1 atub|OJI88050.1 awen|OJJ32499.1 azon|OJJ46033.1

Group8160: abra|OJJ69599.1 acar|OOF93069.1 atub|OJI83025.1 awen|OJJ38317.1 abra|OJJ71063.1 acla|EAW11660.1 anid|CBF71513.1 asyd|OJJ57389.1 ater|EAU36458.1 aver|OJJ07279.1 acar|OOF97615.1

Group8161: abra|OJJ69662.1 acar|OOF90603.1 afla|EED54757.1 agla|OJJ85393.1 akaw|GAA91630.1 aluc|OJZ82457.1 anig|EHA28318.1 aory|EIT79455.1 atub|OJI87981.1 awen|OJJ40351.1 ater|EAU32696.1

Group8162: abra|OJJ69692.1 afla|EED48235.1 agla|OJJ86810.1 aluc|OJZ80602.1 anid|CBF84032.1 aory|EIT80596.1 asyd|OJJ61368.1 aver|OJJ00695.1 awen|OJJ38793.1 afla|EED51647.1 agla|OJJ83217.1

Group8163: abra|OJJ69693.1 afla|EED48234.1 agla|OJJ86809.1 agla|OJJ88731.1 aluc|OJZ80606.1 anid|CBF84034.1 aory|EIT72897.1 asyd|OJJ61369.1 aver|OJJ00696.1 awen|OJJ41926.1 agla|OJJ87453.1

Group8164: abra|OJJ69697.1 afla|EED48379.1 akaw|GAA88937.1 aluc|OJZ89470.1 anig|EHA28272.1 aory|EIT80938.1 asyd|OJJ63605.1 atub|OJI88346.1 aver|OJI97331.1 aory|EIT83092.1 agla|OJJ86353.1

Group8165: abra|OJJ69704.1 acar|OOF90537.1 afis|EAW19198.1 afum|EAL93930.1 anid|CBF88879.1 anig|EHA28262.1 asyd|OJJ57244.1 ater|EAU35861.1 aver|OJJ02855.1 azon|OJJ42364.1 acla|EAW07626.1

Group8166: acar|OOF99053.1 azon|OJJ49414.1 acla|EAW09549.1 akaw|GAA83945.1 aluc|OJZ80826.1 anig|EHA19489.1 asyd|OJJ58668.1 atub|OJI82739.1 aver|OJJ07956.1 asyd|OJJ53316.1 agla|OJJ88733.1

Group8167: abra|OJJ69790.1 acla|EAW11672.1 afis|EAW23688.1 afla|EED57654.1 afum|EAL87134.1 agla|OJJ79366.1 akaw|GAA87868.1 aluc|OJZ85509.1 anid|CBF84202.1 aory|EIT79864.1 awen|OJJ33315.1

Group8168: abra|OJJ69812.1 acar|OOF93565.1 akaw|GAA87845.1 aluc|OJZ85483.1 anid|CBF71117.1 asyd|OJJ52715.1 atub|OJI80081.1 aver|OJJ04744.1 afla|EED50974.1 aory|EIT77027.1 ater|EAU38133.1

Group8169: abra|OJJ69832.1 acar|OOF93357.1 afla|EED52428.1 agla|OJJ79199.1 akaw|GAA87828.1 aluc|OJZ85464.1 aory|EIT81265.1 atub|OJI80099.1 awen|OJJ32852.1 afla|EED48859.1 aory|EIT83300.1

Group8170: abra|OJJ69838.1 acar|OOF89869.1 afla|EED51496.1 akaw|GAA87822.1 aluc|OJZ85458.1 anig|EHA25000.1 aory|EIT80655.1 asyd|OJJ56208.1 ater|EAU30100.1 atub|OJI80105.1 aver|OJJ04864.1

Group8171: abra|OJJ69865.1 acar|OOF90333.1 acla|EAW15318.1 akaw|GAA87791.1 aluc|OJZ85426.1 anig|EHA25605.1 asyd|OJJ52771.1 ater|EAU32683.1 atub|OJI80149.1 aver|OJJ08754.1 awen|OJJ31348.1

Group8172: abra|OJJ69867.1 acar|OOF90335.1 acla|EAW15315.1 akaw|GAA87789.1 aluc|OJZ85424.1 anig|EHA25603.1 asyd|OJJ52770.1 ater|EAU32681.1 atub|OJI80151.1 aver|OJJ08753.1 awen|OJJ31344.1

Group8173: abra|OJJ69881.1 acar|OOF93394.1 acla|EAW08288.1 afis|EAW19380.1 akaw|GAA87777.1 aluc|OJZ85412.1 anid|CBF84571.1 anig|EHA25591.1 asyd|OJJ53387.1 atub|OJI80165.1 aver|OJJ08356.1

Group8174: abra|OJJ69885.1 acar|OOF93403.1 agla|OJJ79811.1 akaw|GAA87773.1 aluc|OJZ85408.1 anid|CBF69587.1 anig|EHA25587.1 asyd|OJJ59147.1 atub|OJI80168.1 aver|OJJ01737.1 awen|OJJ29887.1

Group8175: asyd|OJJ52395.1 aver|OJJ01928.1 afla|EED53341.1 agla|OJJ87119.1 akaw|GAA92123.1 aluc|OJZ83599.1 anig|EHA18758.1 aory|EIT75010.1 atub|OJI89971.1 awen|OJJ35024.1 awen|OJJ31259.1

Group8176: agla|OJJ88720.1 anig|EHA27459.1 akaw|GAA92601.1 aluc|OJZ82618.1 anid|CBF86707.1 asyd|OJJ54372.1 atub|OJI88853.1 aver|OJJ08239.1 awen|OJJ40863.1 azon|OJJ43553.1 asyd|OJJ53507.1

Group8177: abra|OJJ69895.1 acar|OOG00774.1 afla|EED54571.1 agla|OJJ83057.1 anid|CBF71735.1 aory|EIT78904.1 asyd|OJJ61796.1 ater|EAU39168.1 aver|OJI95895.1 awen|OJJ41669.1 azon|OJJ43877.1

Group8178: aory|EIT77582.1 acla|EAW12037.1 anid|CBF71549.1 asyd|OJJ62469.1 ater|EAU29265.1 aver|OJI96604.1 awen|OJJ32576.1 azon|OJJ50293.1 afla|EED56859.1 asyd|OJJ58382.1 aver|OJJ08477.1

Group8179: afla|EED49439.1 afis|EAW24663.1 anid|CBF80281.1 aory|EIT79585.1 asyd|OJJ62362.1 ater|EAU35138.1 aver|OJI96496.1 awen|OJJ33814.1 asyd|OJJ55554.1 aver|OJJ00802.1 ater|EAU33898.1

Group8180: abra|OJJ70020.1 acar|OOF93526.1 akaw|GAA87641.1 aluc|OJZ85256.1 anig|EHA25457.1 aory|EIT79606.1 asyd|OJJ64929.1 atub|OJI80312.1 aver|OJI99048.1 awen|OJJ37945.1 afla|EED56645.1

Group8181: abra|OJJ70049.1 afla|EED49529.1 akaw|GAA87626.1 aluc|OJZ85242.1 anid|CBF87406.1 anig|EHA25442.1 aory|EIT82586.1 asyd|OJJ52889.1 atub|OJI80329.1 aver|OJJ08859.1 awen|OJJ35375.1

Group8182: afis|EAW17081.1 afum|EBA27279.1 abra|OJJ70068.1 acar|OOG00526.1 afla|EED57140.1 akaw|GAA90341.1 aluc|OJZ85891.1 aory|EIT73992.1 ater|EAU37389.1 atub|OJI79676.1 aver|OJI98349.1

Group8183: abra|OJJ70115.1 acar|OOF91821.1 afis|EAW24597.1 afla|EED47458.1 akaw|GAA89324.1 aluc|OJZ86344.1 atub|OJI84190.1 aver|OJI96488.1 abra|OJJ70245.1 agla|OJJ84792.1 awen|OJJ34669.1

Group8184: abra|OJJ70174.1 afis|EAW18190.1 afla|EED45821.1 akaw|GAA86596.1 aluc|OJZ85143.1 aory|EIT75436.1 asyd|OJJ55822.1 atub|OJI80448.1 ater|EAU33791.1 aver|OJJ04764.1 ater|EAU29322.1

Group8185: abra|OJJ70177.1 afis|EAW16395.1 afla|EED48109.1 agla|OJJ86671.1 akaw|GAA86594.1 aluc|OJZ85141.1 anig|EHA25357.1 aory|EIT76191.1 asyd|OJJ57218.1 atub|OJI80451.1 aver|OJJ03123.1

Group8186: abra|OJJ70215.1 aluc|OJZ85111.1 azon|OJJ51439.1 abra|OJJ78119.1 acar|OOF90504.1 agla|OJJ80751.1 akaw|GAA85889.1 anig|EHA19875.1 asyd|OJJ61549.1 atub|OJI83200.1 aver|OJJ06187.1

Group8187: abra|OJJ70228.1 acla|EAW12153.1 afis|EAW17178.1 afla|EED49680.1 afum|EAL86608.1 akaw|GAA93010.1 aluc|OJZ82575.1 anid|CBF78875.1 aory|EIT80947.1 ater|EAU29536.1 awen|OJJ31952.1

Group8188: abra|OJJ70257.1 acar|OOF90565.1 acla|EAW13529.1 afla|EED44860.1 agla|OJJ89546.1 akaw|GAA89224.1 aluc|OJZ81604.1 anig|EHA25306.1 aory|EIT82728.1 aver|OJJ08166.1 acar|OOF95349.1

Group8189: abra|OJJ70283.1 acar|OOF91735.1 afla|EED45375.1 akaw|GAA92866.1 aluc|OJZ82548.1 anig|EHA22512.1 aory|EIT82391.1 atub|OJI88789.1 afis|EAW19537.1 afum|EAL86969.2 azon|OJJ45081.1

Group8190: abra|OJJ70308.1 acar|OOF91763.1 akaw|GAA93014.1 aluc|OJZ82571.1 anid|CBF80499.1 anig|EHA27503.1 asyd|OJJ61522.1 ater|EAU30198.1 atub|OJI88815.1 awen|OJJ36271.1 azon|OJJ50649.1

Group8191: abra|OJJ70316.1 acar|OOF91908.1 afla|EED54367.1 agla|OJJ84148.1 akaw|GAA92571.1 aluc|OJZ82582.1 anid|CBF71562.1 aory|EIT75696.1 asyd|OJJ54215.1 aver|OJJ08516.1 afla|EED45496.1

Group8192: abra|OJJ70332.1 acar|OOF91792.1 afla|EED45228.1 akaw|GAA92587.1 aluc|OJZ82602.1 anig|EHA27477.1 aory|EIT73736.1 asyd|OJJ56051.1 atub|OJI88838.1 awen|OJJ32249.1 acla|EAW08765.1

Group8193: abra|OJJ70333.1 acar|OOF91795.1 afla|EED51848.1 agla|OJJ78674.1 akaw|GAA92588.1 aluc|OJZ82603.1 aory|EIT81682.1 asyd|OJJ52960.1 ater|EAU29425.1 atub|OJI88839.1 awen|OJJ33656.1

Group8194: abra|OJJ70343.1 acar|OOF91923.1 acla|EAW08792.1 agla|OJJ79972.1 akaw|GAA92765.1 aluc|OJZ82720.1 anig|EHA21174.1 aory|EIT83363.1 atub|OJI88954.1 awen|OJJ35056.1 acar|OOF96900.1

Group8195: abra|OJJ70364.1 acla|EAW12636.1 afis|EAW24544.1 afla|EED46221.1 akaw|GAA91987.1 aluc|OJZ82626.1 aory|EIT76430.1 asyd|OJJ61036.1 aver|OJJ00354.1 awen|OJJ35634.1 azon|OJJ42138.1

Group8196: abra|OJJ70365.1 acar|OOF92537.1 acla|EAW07832.1 afis|EAW22869.1 afum|EAL90330.2 akaw|GAA91986.1 aluc|OJZ82627.1 anig|EHA27450.1 atub|OJI88860.1 awen|OJJ31366.1 azon|OJJ49480.1

Group8197: abra|OJJ70375.1 acar|OOF91894.1 acla|EAW12335.1 afis|EAW16887.1 afla|EED47355.1 afum|EAL86425.2 agla|OJJ79934.1 anig|EHA27433.1 ater|EAU30226.1 awen|OJJ29757.1 azon|OJJ42519.1

Group8198: abra|OJJ70502.1 acar|OOF97424.1 afla|EED47045.1 agla|OJJ78737.1 akaw|GAA83350.1 aluc|OJZ83847.1 anig|EHA18989.1 aory|EIT72605.1 ater|EAU36336.1 atub|OJI90213.1 azon|OJJ47643.1

Group8199: abra|OJJ70641.1 acar|OOF97086.1 akaw|GAA89977.1 aluc|OJZ83988.1 anid|CBF86684.1 anig|EHA19114.1 asyd|OJJ64755.1 ater|EAU36164.1 atub|OJI90360.1 aver|OJI98857.1 afla|EED52060.1

Group8200: abra|OJJ70820.1 acar|OOF98282.1 afla|EED56895.1 akaw|GAA91699.1 aluc|OJZ87691.1 anig|EHA20880.1 aory|EIT77507.1 ater|EAU33221.1 atub|OJI80949.1 awen|OJJ41833.1 azon|OJJ50734.1

Group8201: afis|EAW16807.1 afla|EED49870.1 abra|OJJ77980.1 akaw|GAA86000.1 aluc|OJZ91055.1 anig|EHA19758.1 asyd|OJJ54859.1 atub|OJI83051.1 acar|OOF94408.1 anid|CBF80481.1 aory|EIT83410.1

Group8202: abra|OJJ71108.1 acar|OOF90225.1 afis|EAW16704.1 afla|EED54349.1 akaw|GAA82861.1 aluc|OJZ87427.1 anig|EHA20649.1 aory|EIT76100.1 asyd|OJJ53361.1 atub|OJI81222.1 aver|OJJ07647.1

Group8203: abra|OJJ71223.1 acar|OOF96454.1 afla|EED48464.1 akaw|GAA82770.1 aluc|OJZ87329.1 anig|EHA20567.1 aory|EIT76276.1 atub|OJI81315.1 awen|OJJ35993.1 agla|OJJ81048.1 azon|OJJ50273.1

Group8204: abra|OJJ71327.1 acar|OOF97847.1 acla|EAW10212.1 afis|EAW25382.1 agla|OJJ83774.1 aluc|OJZ87216.1 asyd|OJJ53196.1 ater|EAU34528.1 aver|OJJ03654.1 awen|OJJ41108.1 azon|OJJ44686.1

Group8205: abra|OJJ71398.1 acla|EAW10136.1 afla|EED53939.1 agla|OJJ87104.1 akaw|GAA82599.1 aluc|OJZ87144.1 asyd|OJJ53127.1 atub|OJI81497.1 awen|OJJ41183.1 azon|OJJ44759.1 aver|OJJ03727.1

Group8206: abra|OJJ71541.1 acar|OOF96288.1 afla|EED45312.1 aluc|OJZ86279.1 anid|CBF84827.1 anig|EHA21592.1 asyd|OJJ63208.1 atub|OJI84260.1 aver|OJI97747.1 awen|OJJ31826.1 azon|OJJ46298.1

Group8207: abra|OJJ71582.1 afla|EED56708.1 akaw|GAA89342.1 aluc|OJZ86319.1 anig|EHA21556.1 asyd|OJJ53777.1 ater|EAU38708.1 atub|OJI84216.1 aver|OJJ07882.1 awen|OJJ31864.1 aory|EIT73841.1

Group8208: akaw|GAA89304.1 acla|EAW15162.1 afis|EAW20131.1 afla|EED57221.1 afum|EAL85247.1 anid|CBF84958.1 aory|EIT74080.1 asyd|OJJ63142.1 aver|OJI97812.1 awen|OJJ31997.1 azon|OJJ46052.1

Group8209: abra|OJJ71783.1 acar|OOF96462.1 afla|EED45772.1 afla|EED56598.1 akaw|GAA85127.1 aluc|OJZ86513.1 aory|EIT75471.1 aory|EIT79648.1 ater|EAU37978.1 atub|OJI84011.1 awen|OJJ35890.1

Group8210: abra|OJJ71791.1 acar|OOF96471.1 afla|EED56593.1 akaw|GAA85119.1 aluc|OJZ86521.1 anid|CBF82145.1 anig|EHA22125.1 asyd|OJJ54324.1 atub|OJI84001.1 aver|OJJ05278.1 awen|OJJ30243.1

Group8211: abra|OJJ71979.1 acar|OOF96630.1 agla|OJJ85763.1 akaw|GAA84951.1 aluc|OJZ86716.1 anid|CBF76308.1 asyd|OJJ61007.1 atub|OJI83813.1 aver|OJJ00325.1 awen|OJJ39104.1 azon|OJJ51437.1

Group8212: abra|OJJ72055.1 acar|OOF96712.1 akaw|GAA84879.1 aluc|OJZ86792.1 anig|EHA21891.1 asyd|OJJ62344.1 atub|OJI83727.1 aver|OJI96449.1 azon|OJJ44925.1 aory|EIT81929.1 afla|EED52243.1

Group8213: abra|OJJ72135.1 acar|OOF96774.1 afis|EAW18427.1 afla|EED55067.1 afum|EAL93108.1 akaw|GAA84787.1 aluc|OJZ86893.1 anig|EHA21750.1 aory|EIT81983.1 atub|OJI83626.1 awen|OJJ34806.1

Group8214: abra|OJJ72137.1 acar|OOF96775.1 afis|EAW18423.1 agla|OJJ82205.1 akaw|GAA84785.1 aluc|OJZ86895.1 anid|CBF77131.1 anig|EHA21748.1 asyd|OJJ56942.1 atub|OJI83624.1 aver|OJJ03342.1

Group8215: abra|OJJ72162.1 acar|OOF96797.1 akaw|GAA84766.1 aluc|OJZ86917.1 anid|CBF87753.1 anig|EHA21727.1 asyd|OJJ56864.1 atub|OJI83604.1 aver|OJJ06179.1 akaw|GAA89021.1 aluc|OJZ80583.1

Group8216: abra|OJJ72168.1 acar|OOF96802.1 agla|OJJ81392.1 akaw|GAA84760.1 aluc|OJZ86923.1 anig|EHA21721.1 asyd|OJJ64156.1 atub|OJI83598.1 aver|OJI98234.1 awen|OJJ41507.1 azon|OJJ43660.1

Group8217: abra|OJJ72173.1 acar|OOF96138.1 afis|EAW17528.1 afla|EED53696.1 akaw|GAA84755.1 aluc|OJZ86928.1 aory|EIT75271.1 atub|OJI83593.1 acar|OOF90362.1 akaw|GAA90582.1 aluc|OJZ86040.1

Group8218: abra|OJJ72206.1 acar|OOF96826.1 agla|OJJ79955.1 akaw|GAA84728.1 aluc|OJZ86958.1 anig|EHA21690.1 aory|EIT81912.1 atub|OJI83565.1 awen|OJJ32668.1 afla|EED47449.1 asyd|OJJ52212.1

Group8219: afis|EAW18893.1 afla|EED50870.1 afum|EAL93625.1 agla|OJJ81499.1 anid|CBF69908.1 aory|EIT78999.1 awen|OJJ32451.1 aver|OJJ01621.1 acla|EAW14178.1 asyd|OJJ58995.1 ater|EAU38064.1

Group8220: abra|OJJ72227.1 acar|OOF96850.1 afla|EED56073.1 agla|OJJ78578.1 akaw|GAA84710.1 aluc|OJZ86977.1 anid|CBF75000.1 anig|EHA21671.1 aory|EIT76545.1 ater|EAU36082.1 atub|OJI83543.1

Group8221: abra|OJJ72273.1 acar|OOF94945.1 agla|OJJ80824.1 akaw|GAA86028.1 aluc|OJZ90255.1 anid|CBF71386.1 anig|EHA23750.1 asyd|OJJ55067.1 atub|OJI81681.1 aver|OJJ07144.1 azon|OJJ46419.1

Group8222: abra|OJJ72492.1 acar|OOF94980.1 agla|OJJ86015.1 akaw|GAA86226.1 aluc|OJZ90478.1 anig|EHA23937.1 ater|EAU31292.1 atub|OJI81903.1 awen|OJJ29973.1 azon|OJJ42026.1 azon|OJJ49012.1

Group8223: abra|OJJ72566.1 acar|OOF94688.1 afis|EAW17929.1 agla|OJJ83004.1 akaw|GAA85193.1 aluc|OJZ90548.1 anig|EHA23996.1 ater|EAU33541.1 atub|OJI81970.1 awen|OJJ41617.1 afum|EAL91740.1

Group8224: abra|OJJ72611.1 acla|EAW09029.1 afis|EAW22738.1 afla|EED49794.1 afum|EAL90457.1 agla|OJJ78515.1 akaw|GAA85237.1 aluc|OJZ90593.1 anig|EHA24032.1 aory|EIT77614.1 atub|OJI82016.1

Group8225: abra|OJJ72633.1 acar|OOF93306.1 afis|EAW20380.1 afla|EED53181.1 afum|EAL85493.1 akaw|GAA85254.1 aluc|OJZ90612.1 anig|EHA24050.1 aory|EIT74849.1 ater|EAU30709.1 atub|OJI82038.1

Group8226: abra|OJJ72744.1 aory|EIT82466.1 azon|OJJ45796.1 afla|EED45368.1 acar|OOG00576.1 afis|EAW21380.1 ater|EAU30220.1 atub|OJI86518.1 awen|OJJ35063.1 awen|OJJ41701.1 agla|OJJ82531.1

Group8227: abra|OJJ72774.1 acla|EAW11492.1 afis|EAW23961.1 afla|EED50549.1 afum|EAL87390.1 agla|OJJ86886.1 akaw|GAA85383.1 aluc|OJZ90751.1 ater|EAU29791.1 awen|OJJ38986.1 azon|OJJ51318.1

Group8228: abra|OJJ72829.1 afla|EED44984.1 aory|EIT73701.1 ater|EAU31052.1 awen|OJJ40615.1 azon|OJJ42824.1 atub|OJI82790.1 azon|OJJ42226.1 atub|OJI81974.1 akaw|GAA83894.1 aluc|OJZ90801.1

Group8229: abra|OJJ72875.1 acla|EAW11368.1 afis|EAW18127.1 afum|EAL87485.1 akaw|GAA85471.1 aluc|OJZ83387.1 anid|CBF75362.1 aory|EIT79714.1 ater|EAU35450.1 atub|OJI82273.1 afla|EED58051.1

Group8230: azon|OJJ50214.1 acla|EAW11062.1 afis|EAW20813.1 afla|EED57085.1 afum|EAL84815.1 anid|CBF84024.1 aory|EIT82273.1 asyd|OJJ61374.1 ater|EAU32258.1 aver|OJJ00701.1 awen|OJJ32660.1

Group8231: afum|EAL88569.1 agla|OJJ89374.1 acla|EAW11203.1 afis|EAW15933.1 afla|EED54928.1 anid|CBF86349.1 aory|EIT80384.1 aver|OJI99101.1 awen|OJJ35646.1 azon|OJJ46890.1 asyd|OJJ64979.1

Group8232: abra|OJJ72966.1 acar|OOF96148.1 afis|EAW15919.1 afla|EED54942.1 akaw|GAA85545.1 aluc|OJZ83477.1 anig|EHA24314.1 aory|EIT80351.1 asyd|OJJ56375.1 atub|OJI82366.1 aver|OJJ06337.1

Group8233: abra|OJJ72978.1 acar|OOF96157.1 afis|EAW23066.1 afla|EED54951.1 agla|OJJ89408.1 akaw|GAA85553.1 aluc|OJZ83488.1 anig|EHA24323.1 atub|OJI82380.1 awen|OJJ35669.1 afum|EAL88413.1

Group8234: abra|OJJ73072.1 acar|OOG00305.1 afis|EAW19681.1 afla|EED52420.1 afum|EAL86841.1 agla|OJJ85236.1 akaw|GAA88089.1 aluc|OJZ81353.1 aory|EIT83313.1 ater|EAU29957.1 atub|OJI91557.1

Group8235: abra|OJJ73093.1 acar|OOG00325.1 akaw|GAA88067.1 aluc|OJZ81332.1 anid|CBF78568.1 anig|EHA18636.1 aory|EIT82629.1 asyd|OJJ65068.1 atub|OJI91535.1 aver|OJI99192.1 afla|EED49586.1

Group8236: abra|OJJ73120.1 afla|EED47382.1 akaw|GAA88046.1 aluc|OJZ81307.1 anid|CBF76271.1 anig|EHA18614.1 aory|EIT82854.1 asyd|OJJ60977.1 atub|OJI91515.1 aver|OJJ00301.1 awen|OJJ40827.1

Group8237: abra|OJJ73178.1 acla|EAW08985.1 afis|EAW24687.1 afla|EED48345.1 afum|EAL89328.1 aory|EIT76280.1 asyd|OJJ55424.1 ater|EAU31418.1 aver|OJJ06503.1 awen|OJJ35368.1 anid|CBF83132.1

Group8238: abra|OJJ73317.1 acar|OOF98708.1 afla|EED48338.1 akaw|GAA92803.1 aluc|OJZ88727.1 anig|EHA18432.1 asyd|OJJ52013.1 atub|OJI91305.1 aver|OJJ07818.1 azon|OJJ50290.1 aory|EIT76306.1

Group8239: akaw|GAA91200.1 anid|CBF85440.1 afla|EED54857.1 agla|OJJ88276.1 aluc|OJZ81535.1 anig|EHA25901.1 aory|EIT79406.1 asyd|OJJ64090.1 ater|EAU36997.1 atub|OJI88652.1 aver|OJI98174.1

Group8240: abra|OJJ73350.1 acar|OOG00560.1 afla|EED46514.1 akaw|GAA83133.1 aluc|OJZ88689.1 anig|EHA18370.1 aory|EIT72405.1 asyd|OJJ61420.1 atub|OJI91269.1 aver|OJJ00755.1 awen|OJJ31683.1

Group8241: azon|OJJ45838.1 afla|EED56851.1 agla|OJJ80512.1 akaw|GAA83095.1 aluc|OJZ90880.1 aory|EIT77511.1 atub|OJI83753.1 awen|OJJ32547.1 aluc|OJZ88642.1 atub|OJI91227.1 agla|OJJ81564.1

Group8242: abra|OJJ73434.1 acar|OOG00641.1 afla|EED53595.1 akaw|GAA83059.1 aluc|OJZ88607.1 anid|CBF77718.1 anig|EHA18300.1 aory|EIT75104.1 ater|EAU36689.1 atub|OJI91191.1 aver|OJI99558.1

Group8243: ater|EAU39218.1 acla|EAW07467.1 afis|EAW22500.1 afla|EED54528.1 afum|EAL90693.1 agla|OJJ80850.1 aory|EIT78819.1 asyd|OJJ62801.1 aver|OJI96848.1 awen|OJJ35123.1 anid|CBF88963.1

Group8244: abra|OJJ73720.1 acar|OOF95372.1 agla|OJJ81470.1 akaw|GAA88442.1 aluc|OJZ88330.1 anid|CBF86465.1 anig|EHA18065.1 asyd|OJJ64909.1 ater|EAU37936.1 atub|OJI90925.1 aver|OJI99031.1

Group8245: abra|OJJ73729.1 acar|OOG00896.1 acla|EAW11756.1 afis|EAW23789.1 afla|EED53093.1 afum|EAL87223.2 aluc|OJZ88321.1 anig|EHA18057.1 atub|OJI90916.1 awen|OJJ29816.1 azon|OJJ42845.1

Group8246: abra|OJJ73859.1 acar|OOG01006.1 afla|EED50050.1 agla|OJJ83590.1 aluc|OJZ88180.1 anig|EHA17932.1 asyd|OJJ63486.1 atub|OJI90779.1 aver|OJI97443.1 awen|OJJ38948.1 azon|OJJ49720.1

Group8247: abra|OJJ74006.1 acar|OOG01128.1 afla|EED47788.1 agla|OJJ79634.1 akaw|GAA87487.1 aluc|OJZ88038.1 anig|EHA17811.1 aory|EIT77140.1 asyd|OJJ56545.1 atub|OJI90645.1 aory|EIT78695.1

Group8248: abra|OJJ74167.1 acar|OOF89850.1 afla|EED55901.1 agla|OJJ87090.1 akaw|GAA86848.1 aluc|OJZ89969.1 anig|EHA27167.1 aory|EIT76783.1 atub|OJI84838.1 awen|OJJ36033.1 azon|OJJ48006.1

Group8249: abra|OJJ74300.1 acar|OOF95430.1 acla|EAW07345.1 afis|EAW22377.1 afum|EAL90815.1 agla|OJJ87630.1 aluc|OJZ89833.1 anid|CBF89405.1 anig|EHA27035.1 ater|EAU39511.1 azon|OJJ42609.1

Group8250: abra|OJJ74370.1 afla|EED45946.1 akaw|GAA86661.1 aluc|OJZ89758.1 anid|CBF84510.1 anig|EHA26978.1 aory|EIT75597.1 asyd|OJJ53515.1 atub|OJI85042.1 aver|OJJ08236.1 awen|OJJ40866.1

Group8251: abra|OJJ74410.1 acar|OOF95332.1 afla|EED53463.1 agla|OJJ85405.1 anid|CBF89357.1 anig|EHA26945.1 aory|EIT75147.1 asyd|OJJ62499.1 ater|EAU34077.1 aver|OJI96619.1 azon|OJJ46508.1

Group8252: abra|OJJ74447.1 acar|OOF95548.1 acla|EAW10184.1 akaw|GAA87339.1 aluc|OJZ89675.1 anig|EHA26910.1 aory|EIT82309.1 ater|EAU34842.1 atub|OJI85125.1 acar|OOF98747.1 afla|EED57083.1

Group8253: abra|OJJ74479.1 acla|EAW07221.1 afis|EAW22235.1 afla|EED57624.1 afum|EAL90958.1 agla|OJJ87816.1 akaw|GAA87372.1 aluc|OJZ89640.1 anid|CBF88475.1 aory|EIT82938.1 atub|OJI85163.1

Group8254: abra|OJJ74561.1 afla|EED48714.1 agla|OJJ82643.1 aluc|OJZ92143.1 anid|CBF84656.1 anig|EHA26806.1 aory|EIT73078.1 asyd|OJJ54383.1 ater|EAU33265.1 atub|OJI86157.1 awen|OJJ36280.1

Group8255: abra|OJJ74562.1 afla|EED48715.1 agla|OJJ82642.1 aluc|OJZ92142.1 anid|CBF84658.1 anig|EHA26807.1 aory|EIT73277.1 asyd|OJJ54382.1 ater|EAU33266.1 atub|OJI86156.1 awen|OJJ36281.1

Group8256: abra|OJJ74583.1 acar|OOF90134.1 afla|EED45976.1 akaw|GAA82022.1 aluc|OJZ92161.1 anig|EHA26788.1 aory|EIT75498.1 ater|EAU38427.1 atub|OJI86589.1 azon|OJJ50619.1 aory|EIT75794.1

Group8257: abra|OJJ74821.1 afla|EED45370.1 agla|OJJ80041.1 akaw|GAA88360.1 aluc|OJZ82295.1 anig|EHA18381.1 aory|EIT82386.1 ater|EAU29268.1 atub|OJI88144.1 aver|OJJ01773.1 awen|OJJ30072.1

Group8258: abra|OJJ74827.1 acar|OOF95812.1 afla|EED50886.1 akaw|GAA83594.1 aluc|OJZ92402.1 anig|EHA26564.1 aory|EIT79033.1 ater|EAU32477.1 atub|OJI86830.1 awen|OJJ35400.1 azon|OJJ50135.1

Group8259: abra|OJJ74838.1 acar|OOF95842.1 afla|EED50992.1 agla|OJJ84176.1 akaw|GAA83582.1 aluc|OJZ92415.1 aory|EIT77064.1 ater|EAU37156.1 atub|OJI86845.1 awen|OJJ33797.1 azon|OJJ50128.1

Group8260: abra|OJJ75104.1 acar|OOF97230.1 acla|EAW08264.1 afis|EAW23297.1 afla|EED55647.1 agla|OJJ86514.1 akaw|GAA88713.1 aluc|OJZ92669.1 aory|EIT81817.1 awen|OJJ39578.1 azon|OJJ47773.1

Group8261: abra|OJJ75116.1 akaw|GAA88726.1 aluc|OJZ92683.1 atub|OJI87111.1 acla|EAW08307.1 asyd|OJJ62441.1 aver|OJI96577.1 awen|OJJ39562.1 agla|OJJ86501.1 ater|EAU35579.1 anid|CBF89438.1

Group8262: abra|OJJ75164.1 acar|OOF97174.1 akaw|GAA88775.1 aluc|OJZ92735.1 anid|CBF71797.1 anig|EHA26267.1 ater|EAU35627.1 atub|OJI87159.1 agla|OJJ78945.1 asyd|OJJ64766.1 aver|OJI98866.1

Group8263: abra|OJJ75286.1 acar|OOF90101.1 afla|EED45609.1 akaw|GAA91824.1 aluc|OJZ92847.1 anig|EHA26163.1 aory|EIT75599.1 atub|OJI87303.1 asyd|OJJ55353.1 aver|OJJ06442.1 anig|EHA26162.1

Group8264: abra|OJJ75288.1 afis|EAW23622.1 afla|EED48180.1 afum|EAL87838.2 akaw|GAA91826.1 aluc|OJZ92849.1 anig|EHA26160.1 aory|EIT83036.1 asyd|OJJ55365.1 atub|OJI87305.1 aver|OJJ08468.1

Group8265: abra|OJJ75289.1 afis|EAW23623.1 afla|EED48179.1 afum|EAL87837.1 akaw|GAA91827.1 aluc|OJZ92850.1 anig|EHA26159.1 aory|EIT83030.1 asyd|OJJ55366.1 atub|OJI87306.1 aver|OJJ08467.1

Group8266: abra|OJJ75290.1 afis|EAW23624.1 afla|EED48178.1 afum|EAL87836.1 akaw|GAA91828.1 aluc|OJZ92851.1 anig|EHA26158.1 aory|EIT83027.1 asyd|OJJ55364.1 atub|OJI87307.1 aver|OJJ08469.1

Group8267: abra|OJJ75306.1 acar|OOF99290.1 acla|EAW09100.1 akaw|GAA91470.1 aluc|OJZ85630.1 anig|EHA21435.1 asyd|OJJ57294.1 atub|OJI79425.1 aver|OJJ05861.1 awen|OJJ35050.1 azon|OJJ44457.1

Group8268: abra|OJJ75313.1 acar|OOF94516.1 afla|EED46217.1 agla|OJJ88346.1 akaw|GAA91997.1 aluc|OJZ92867.1 anig|EHA26098.1 aory|EIT76449.1 asyd|OJJ56261.1 atub|OJI87325.1 awen|OJJ32218.1

Group8269: abra|OJJ75357.1 acar|OOF94480.1 afis|EAW15481.1 akaw|GAA92053.1 aluc|OJZ92923.1 anig|EHA26046.1 atub|OJI87384.1 azon|OJJ43150.1 afum|EAL85841.1 agla|OJJ80160.1 awen|OJJ38139.1

Group8270: abra|OJJ75455.1 acar|OOF90800.1 acla|EAW14622.1 afum|EBA27428.1 akaw|GAA84230.1 aluc|OJZ88911.1 anid|CBF78443.1 ater|EAU30773.1 atub|OJI85880.1 aver|OJI99496.1 afis|EAW17413.1

Group8271: akaw|GAA84242.1 aluc|OJZ88927.1 aory|EIT72988.1 acar|OOF90806.1 acla|EAW13836.1 afis|EAW18295.1 afum|EAL87661.2 agla|OJJ88128.1 anig|EHA23586.1 asyd|OJJ57863.1 azon|OJJ43269.1

Group8272: abra|OJJ75477.1 acar|OOF90813.1 afis|EAW18290.1 afum|EAL87655.1 akaw|GAA84247.1 aluc|OJZ88936.1 anig|EHA23581.1 atub|OJI85856.1 awen|OJJ30663.1 azon|OJJ43262.1 anid|CBF79458.1

Group8273: abra|OJJ75508.1 acla|EAW09298.1 afis|EAW21849.1 afla|EED46463.1 afum|EAL92862.1 akaw|GAA84271.1 aluc|OJZ88964.1 anig|EHA23557.1 aory|EIT72403.1 ater|EAU29685.1 atub|OJI85825.1

Group8274: abra|OJJ75513.1 acar|OOF90840.1 acla|EAW09301.1 afis|EAW21846.1 afum|EAL92858.1 agla|OJJ84289.1 akaw|GAA84277.1 aluc|OJZ88970.1 atub|OJI85820.1 awen|OJJ34129.1 azon|OJJ44214.1

Group8275: abra|OJJ75517.1 acar|OOF90844.1 afla|EED46524.1 akaw|GAA84281.1 aluc|OJZ88974.1 anid|CBF76719.1 anig|EHA23551.1 aory|EIT72337.1 asyd|OJJ61243.1 atub|OJI85816.1 aver|OJJ00566.1

Group8276: abra|OJJ75549.1 acla|EAW09320.1 afis|EAW21825.1 afla|EED46554.1 akaw|GAA84304.1 aluc|OJZ88999.1 anig|EHA23529.1 ater|EAU30086.1 atub|OJI85790.1 awen|OJJ34098.1 azon|OJJ44246.1

Group8277: acla|EAW06528.1 afis|EAW16067.1 afum|EAL88699.1 acar|OOF99167.1 afla|EED48588.1 anid|CBF75100.1 asyd|OJJ60006.1 ater|EAU33562.1 aver|OJJ02280.1 awen|OJJ30368.1 azon|OJJ45387.1

Group8278: abra|OJJ75684.1 acar|OOG00165.1 agla|OJJ88407.1 akaw|GAA84423.1 aluc|OJZ89134.1 anig|EHA23411.1 asyd|OJJ55072.1 atub|OJI85651.1 awen|OJJ33989.1 agla|OJJ79455.1 awen|OJJ32650.1

Group8279: abra|OJJ75718.1 acar|OOG00138.1 acla|EAW09374.1 afla|EED58017.1 akaw|GAA84454.1 aluc|OJZ89170.1 anig|EHA23385.1 aory|EIT79703.1 atub|OJI85620.1 afis|EAW21761.1 afum|EAL92791.1

Group8280: abra|OJJ75740.1 acar|OOG00121.1 akaw|GAA84473.1 aluc|OJZ89193.1 anid|CBF84717.1 anig|EHA23365.1 asyd|OJJ57258.1 ater|EAU31798.1 atub|OJI85597.1 awen|OJJ32786.1 aory|EIT83626.1

Group8281: abra|OJJ75748.1 acar|OOG00552.1 afla|EED49480.1 agla|OJJ86504.1 akaw|GAA84008.1 aluc|OJZ80908.1 aory|EIT79477.1 atub|OJI82670.1 awen|OJJ38033.1 azon|OJJ43113.1 awen|OJJ35327.1

Group8282: akaw|GAA84581.1 aluc|OJZ89324.1 acla|EAW09939.1 afis|EAW21184.1 afla|EED54775.1 afum|EAL92205.1 agla|OJJ79985.1 anid|CBF83314.1 aory|EIT79372.1 atub|OJI85475.1 awen|OJJ36525.1

Group8283: akaw|GAA84595.1 acar|OOF90246.1 acla|EAW09947.1 afis|EAW21225.1 afla|EED54781.1 afum|EBA27455.1 agla|OJJ78505.1 ater|EAU35855.1 aver|OJI99889.1 awen|OJJ36533.1 azon|OJJ44407.1

Group8284: abra|OJJ75895.1 afla|EED54792.1 agla|OJJ85404.1 akaw|GAA84601.1 anig|EHA23236.1 aory|EIT79367.1 asyd|OJJ60572.1 atub|OJI85453.1 awen|OJJ36126.1 acar|OOF90505.1 agla|OJJ80750.1

Group8285: abra|OJJ76009.1 acar|OOF99916.1 acla|EAW09796.1 afis|EAW21340.1 afla|EED55755.1 afum|EBA27460.1 akaw|GAA88951.1 aluc|OJZ89455.1 anig|EHA23132.1 aory|EIT76666.1 atub|OJI85339.1

Group8286: abra|OJJ76011.1 acar|OOF99912.1 afis|EAW21350.1 afla|EED55757.1 akaw|GAA88949.1 aluc|OJZ89457.1 anig|EHA23130.1 asyd|OJJ54361.1 atub|OJI85337.1 aver|OJJ05314.1 awen|OJJ36188.1

Group8287: abra|OJJ76012.1 afis|EAW21349.1 afla|EED55758.1 akaw|GAA88948.1 aluc|OJZ89458.1 anig|EHA23129.1 aory|EIT76833.1 asyd|OJJ54360.1 atub|OJI85336.1 aver|OJJ05313.1 awen|OJJ36189.1

Group8288: abra|OJJ76028.1 acar|OOF93546.1 afla|EED45762.1 aluc|OJZ89479.1 anig|EHA23114.1 atub|OJI85321.1 azon|OJJ47387.1 aory|EIT75395.1 agla|OJJ82968.1 akaw|GAA88930.1 aory|EIT75791.1

Group8289: abra|OJJ76078.1 acar|OOF99847.1 afla|EED54036.1 akaw|GAA88888.1 aluc|OJZ89534.1 anig|EHA23069.1 asyd|OJJ62433.1 ater|EAU35152.1 atub|OJI85271.1 aver|OJI96569.1 awen|OJJ33339.1

Group8290: abra|OJJ76081.1 acla|EAW09740.1 afla|EED55823.1 agla|OJJ85529.1 akaw|GAA88885.1 aluc|OJZ89539.1 ater|EAU35148.1 atub|OJI85268.1 aver|OJJ00226.1 awen|OJJ39301.1 azon|OJJ51551.1

Group8291: acar|OOF99510.1 akaw|GAA82199.1 atub|OJI86399.1 acla|EAW14342.1 afis|EAW18948.1 afla|EED50980.1 afum|EAL93682.1 agla|OJJ85307.1 anid|CBF86835.1 aory|EIT77021.1 awen|OJJ31542.1

Group8292: awen|OJJ31561.1 afum|EAL93694.1 acla|EAW14326.1 afis|EAW18962.1 afla|EED51009.1 anid|CBF69604.1 aory|EIT77037.1 asyd|OJJ59133.1 ater|EAU38145.1 aver|OJJ01722.1 azon|OJJ47280.1

Group8293: abra|OJJ76480.1 acla|EAW13955.1 afis|EAW18665.1 afla|EED52664.1 afum|EAL93383.1 akaw|GAA82393.1 aory|EIT74237.1 asyd|OJJ58534.1 ater|EAU38247.1 awen|OJJ32133.1 azon|OJJ46716.1

Group8294: abra|OJJ76679.1 acar|OOF99329.1 afla|EED48158.1 akaw|GAA92513.1 aluc|OJZ91526.1 anid|CBF78397.1 anig|EHA22562.1 aory|EIT83025.1 atub|OJI85980.1 aver|OJJ05531.1 aver|OJJ05532.1

Group8295: abra|OJJ76684.1 acar|OOF99327.1 afla|EED52487.1 akaw|GAA92510.1 aluc|OJZ91523.1 anig|EHA22560.1 aory|EIT82751.1 asyd|OJJ56613.1 ater|EAU33885.1 atub|OJI85978.1 aver|OJJ06110.1

Group8296: agla|OJJ79332.1 acla|EAW13679.1 afis|EAW19926.1 afla|EED45895.1 afum|EAL84983.1 anid|CBF84574.1 aory|EIT75374.1 asyd|OJJ53385.1 ater|EAU36895.1 aver|OJJ08357.1 awen|OJJ29968.1

Group8297: abra|OJJ76690.1 acar|OOF99321.1 acla|EAW15334.1 agla|OJJ80666.1 akaw|GAA92504.1 aluc|OJZ91517.1 anid|CBF84583.1 anig|EHA22555.1 asyd|OJJ63695.1 atub|OJI85972.1 aver|OJI97244.1

Group8298: abra|OJJ76714.1 acar|OOF97649.1 afis|EAW20852.1 akaw|GAA92486.1 aluc|OJZ91498.1 anid|CBF87489.1 ater|EAU30419.1 atub|OJI85951.1 awen|OJJ38285.1 acar|OOF98045.1 ater|EAU30791.1

Group8299: abra|OJJ76765.1 acla|EAW13319.1 afis|EAW20433.1 afum|EAL84618.1 agla|OJJ84892.1 akaw|GAA87261.1 anid|CBF79323.1 anig|EHA27529.1 aory|EIT77721.1 ater|EAU37724.1 aver|OJJ06873.1

Group8300: abra|OJJ76791.1 acar|OOF91285.1 acla|EAW13350.1 afis|EAW20464.1 afla|EED44803.1 afum|EAL84589.1 akaw|GAA87234.1 aluc|OJZ84195.1 atub|OJI89889.1 awen|OJJ33158.1 azon|OJJ46365.1

Group8301: abra|OJJ76868.1 afla|EED51652.1 akaw|GAA87155.1 aluc|OJZ84279.1 anig|EHA27624.1 aory|EIT80421.1 asyd|OJJ62989.1 ater|EAU30282.1 atub|OJI89802.1 aver|OJI97979.1 awen|OJJ30985.1

Group8302: abra|OJJ76898.1 afla|EED44884.1 agla|OJJ86439.1 aluc|OJZ84291.1 anig|EHA27631.1 aory|EIT83263.1 atub|OJI89778.1 aver|OJJ05827.1 awen|OJJ40828.1 asyd|OJJ57334.1 akaw|GAA87146.1

Group8303: abra|OJJ76899.1 agla|OJJ86438.1 akaw|GAA87145.1 aluc|OJZ84292.1 anig|EHA27632.1 asyd|OJJ57333.1 atub|OJI89777.1 aver|OJJ05828.1 awen|OJJ40830.1 afla|EED44883.1 aory|EIT83257.1

Group8304: abra|OJJ76906.1 acar|OOF91354.1 afis|EAW20539.1 afla|EED52440.1 afum|EAL84519.2 akaw|GAA87132.1 aluc|OJZ84305.1 anig|EHA27641.1 aory|EIT82775.1 atub|OJI89763.1 azon|OJJ50625.1

Group8305: ater|EAU30470.1 acla|EAW14578.1 afis|EAW19304.1 afla|EED50043.1 afum|EAL94022.1 aory|EIT74632.1 asyd|OJJ57611.1 aver|OJJ05574.1 awen|OJJ39678.1 azon|OJJ50361.1 agla|OJJ80391.1

Group8306: abra|OJJ77030.1 acar|OOF91468.1 afla|EED49751.1 akaw|GAA87021.1 aluc|OJZ84433.1 anid|CBF83217.1 anig|EHA27752.1 aory|EIT77601.1 atub|OJI89640.1 awen|OJJ33607.1 azon|OJJ50728.1

Group8307: acar|OOF97159.1 acla|EAW08359.1 afis|EAW23390.1 afla|EED55567.1 afum|EAL88081.1 anid|CBF74866.1 aory|EIT82701.1 asyd|OJJ60120.1 ater|EAU35643.1 awen|OJJ39490.1 azon|OJJ47692.1

Group8308: abra|OJJ77248.1 acar|OOF92495.1 akaw|GAA90694.1 aluc|OJZ84659.1 anig|EHA27941.1 asyd|OJJ52058.1 atub|OJI89421.1 aver|OJJ06036.1 azon|OJJ47624.1 acla|EAW11446.1 asyd|OJJ56687.1

Group8309: abra|OJJ77285.1 acar|OOF93162.1 agla|OJJ82983.1 akaw|GAA90664.1 aluc|OJZ84695.1 anid|CBF84517.1 anig|EHA27971.1 asyd|OJJ64780.1 atub|OJI89387.1 aver|OJI98888.1 azon|OJJ44996.1

Group8310: abra|OJJ77408.1 akaw|GAA91357.1 aluc|OJZ84815.1 anig|EHA28079.1 aory|EIT78902.1 asyd|OJJ54172.1 ater|EAU37354.1 atub|OJI89270.1 aver|OJJ08567.1 awen|OJJ39965.1 acar|OOF93274.1

Group8311: abra|OJJ77457.1 acar|OOF97634.1 afla|EED49593.1 akaw|GAA92096.1 aluc|OJZ83629.1 anig|EHA18782.1 aory|EIT82624.1 asyd|OJJ56403.1 ater|EAU39562.1 atub|OJI89998.1 aver|OJJ06309.1

Group8312: afis|EAW17047.1 afla|EED53437.1 aory|EIT74988.1 asyd|OJJ62007.1 aver|OJJ06553.1 awen|OJJ29782.1 acar|OOF90140.1 anid|CBF69408.1 ater|EAU34055.1 aver|OJI96099.1 ater|EAU29578.1

Group8313: abra|OJJ77636.1 acar|OOF99103.1 afla|EED51640.1 akaw|GAA83998.1 aluc|OJZ80894.1 aory|EIT80550.1 asyd|OJJ59189.1 atub|OJI82684.1 aver|OJJ01806.1 awen|OJJ32278.1 azon|OJJ50432.1

Group8314: abra|OJJ77734.1 akaw|GAA83898.1 aluc|OJZ90797.1 anig|EHA19529.1 atub|OJI82786.1 abra|OJJ78097.1 acar|OOF98316.1 aluc|OJZ81971.1 atub|OJI79697.1 awen|OJJ36261.1 azon|OJJ51725.1

Group8315: abra|OJJ77751.1 agla|OJJ83614.1 akaw|GAA85927.1 aluc|OJZ91138.1 ater|EAU32665.1 atub|OJI82801.1 azon|OJJ46415.1 acla|EAW08896.1 atub|OJI83149.1 agla|OJJ86354.1 azon|OJJ48579.1

Group8316: atub|OJI84529.1 abra|OJJ77962.1 afla|EED56643.1 aluc|OJZ91030.1 anig|EHA19729.1 awen|OJJ35884.1 azon|OJJ46497.1 acar|OOF97936.1 aory|EIT74808.1 atub|OJI83023.1 aluc|OJZ81976.1

Group8317: abra|OJJ77778.1 acar|OOF98968.1 afla|EED53401.1 akaw|GAA83869.1 aluc|OJZ90831.1 anid|CBF80622.1 anig|EHA19554.1 aory|EIT74913.1 asyd|OJJ54630.1 atub|OJI82817.1 aver|OJJ07526.1

Group8318: abra|OJJ77779.1 acar|OOF98969.1 afla|EED53400.1 akaw|GAA83868.1 aluc|OJZ90832.1 anid|CBF80619.1 anig|EHA19555.1 aory|EIT75211.1 asyd|OJJ54628.1 atub|OJI82818.1 aver|OJJ07528.1

Group8319: abra|OJJ77838.1 acar|OOF98933.1 afis|EAW18215.1 afla|EED56256.1 afum|EAL87581.1 agla|OJJ85867.1 akaw|GAA83809.1 aluc|OJZ90912.1 atub|OJI82905.1 awen|OJJ30593.1 azon|OJJ50173.1

Group8320: abra|OJJ77913.1 afla|EED51612.1 agla|OJJ82630.1 anid|CBF73910.1 anig|EHA18405.1 aory|EIT80557.1 asyd|OJJ53669.1 ater|EAU37638.1 aver|OJJ07776.1 awen|OJJ40186.1 akaw|GAA92003.1

Group8321: abra|OJJ77959.1 acla|EAW14094.1 afis|EAW21811.1 afum|EBA27363.1 agla|OJJ86302.1 akaw|GAA83703.1 aluc|OJZ91028.1 anig|EHA19725.1 aory|EIT73767.1 ater|EAU30721.1 atub|OJI83021.1

Group8322: afla|EED45124.1 agla|OJJ86301.1 ater|EAU30312.1 acar|OOF99948.1 afis|EAW18020.1 akaw|GAA88984.1 aluc|OJZ89419.1 anig|EHA23169.1 aory|EIT73544.1 atub|OJI85375.1 afum|EAL91832.1

Group8323: abra|OJJ78051.1 afla|EED53546.1 akaw|GAA85940.1 aluc|OJZ91122.1 anid|CBF76245.1 anig|EHA19822.1 aory|EIT75016.1 asyd|OJJ65129.1 atub|OJI83127.1 aver|OJI99257.1 awen|OJJ31402.1

Group8324: abra|OJJ78059.1 acla|EAW06393.1 afis|EAW20702.1 afum|EAL84877.1 akaw|GAA85933.1 aluc|OJZ91129.1 anig|EHA19829.1 asyd|OJJ56272.1 atub|OJI83140.1 aver|OJJ08493.1 azon|OJJ46854.1

Group8325: acla|EAW11747.1 afum|EAL87214.1 agla|OJJ78931.1 ater|EAU31321.1 awen|OJJ33697.1 acar|OOF98955.1 afis|EAW23780.1 afla|EED51584.1 aory|EIT80450.1 asyd|OJJ54457.1 aver|OJJ07699.1

Group8326: anid|CBF87734.1 acar|OOF90599.1 akaw|GAA91564.1 aluc|OJZ82382.1 anig|EHA28382.1 asyd|OJJ52703.1 atub|OJI88059.1 aver|OJJ04732.1 azon|OJJ42290.1 afla|EED57160.1 aory|EIT74157.1

Group8327: abra|OJJ78174.1 acar|OOF90378.1 afla|EED45140.1 akaw|GAA85858.1 aluc|OJZ91223.1 anig|EHA19929.1 aory|EIT73743.1 asyd|OJJ58338.1 ater|EAU30391.1 atub|OJI83237.1 aver|OJJ02073.1

Group8328: afla|EED47360.1 aory|EIT82857.1 acar|OOF97636.1 akaw|GAA92098.1 aluc|OJZ83627.1 anig|EHA18780.1 asyd|OJJ53730.1 atub|OJI89996.1 aver|OJI99816.1 awen|OJJ33530.1 asyd|OJJ60490.1

Group8329: azon|OJJ49722.1 acla|EAW07748.1 afis|EAW22785.1 afla|EED45210.1 afum|EAL90414.1 agla|OJJ83592.1 anid|CBF69506.1 aory|EIT73745.1 asyd|OJJ59222.1 aver|OJJ01832.1 awen|OJJ30113.1

Group8330: acar|OOF89777.1 acla|EAW12928.1 agla|OJJ81724.1 aluc|OJZ84554.1 anid|CBF86579.1 anig|EHA27848.1 asyd|OJJ64841.1 atub|OJI89528.1 aver|OJI98952.1 awen|OJJ37111.1 azon|OJJ50810.1

Group8331: acar|OOF91032.1 acla|EAW10894.1 afis|EAW20964.1 afum|EAL84669.1 agla|OJJ84041.1 anid|CBF82574.1 asyd|OJJ54045.1 ater|EAU38614.1 aver|OJJ08697.1 awen|OJJ34351.1 azon|OJJ48088.1

Group8332: afla|EED52198.1 aory|EIT77473.1 aver|OJJ08834.1 azon|OJJ43562.1 anid|CBF87393.1 awen|OJJ31363.1 aory|EIT74499.1 ater|EAU33233.1 awen|OJJ30836.1 afla|EED56918.1 asyd|OJJ52857.1

Group8333: anid|CBF86967.1 asyd|OJJ60533.1 acla|EAW10016.1 afis|EAW21038.1 afla|EED52178.1 afum|EAL92077.2 agla|OJJ88274.1 aory|EIT74334.1 ater|EAU31077.1 awen|OJJ37569.1 aver|OJI99860.1

Group8334: acar|OOF91788.1 acla|EAW12355.1 afis|EAW16779.1 afum|EAL86350.1 akaw|GAA92920.1 anig|EHA23721.1 asyd|OJJ52440.1 ater|EAU29739.1 atub|OJI82463.1 azon|OJJ42771.1 asyd|OJJ54744.1

Group8335: acar|OOF91924.1 afis|EAW23806.1 afla|EED52017.1 afum|EAL87239.1 agla|OJJ78849.1 aory|EIT81443.1 asyd|OJJ52831.1 ater|EAU31358.1 aver|OJJ08806.1 awen|OJJ33763.1 azon|OJJ50679.1

Group8336: afis|EAW22729.1 afum|EAL90466.1 acla|EAW15033.1 agla|OJJ83540.1 akaw|GAA91348.1 aluc|OJZ84826.1 anig|EHA28084.1 asyd|OJJ59208.1 ater|EAU32746.1 atub|OJI89261.1 aver|OJJ01819.1

Group8337: acar|OOF92531.1 acla|EAW10014.1 afla|EED45877.1 anid|CBF84190.1 aory|EIT75644.1 asyd|OJJ57134.1 ater|EAU39449.1 aver|OJJ00721.1 awen|OJJ30878.1 azon|OJJ42801.1 aver|OJJ03169.1

Group8338: acar|OOF92815.1 afla|EED48582.1 afum|EAL87228.2 anig|EHA21279.1 asyd|OJJ62860.1 aver|OJI96913.1 awen|OJJ38043.1 akaw|GAA92524.1 aluc|OJZ91538.1 anig|EHA22575.1 atub|OJI85992.1

Group8339: acar|OOF94008.1 afis|EAW19783.1 afla|EED44763.1 aory|EIT78409.1 asyd|OJJ52912.1 ater|EAU36519.1 aver|OJJ08881.1 awen|OJJ33437.1 afis|EAW17062.1 aory|EIT83012.1 aory|EIT77779.1

Group8340: acar|OOF94668.1 asyd|OJJ53759.1 ater|EAU37276.1 aver|OJJ07870.1 azon|OJJ48021.1 afla|EED53078.1 agla|OJJ80576.1 atub|OJI88869.1 anid|CBF87274.1 asyd|OJJ64167.1 aver|OJJ08275.1

Group8341: acar|OOF95273.1 akaw|GAA87365.1 aluc|OJZ89648.1 anid|CBF88485.1 anig|EHA26886.1 asyd|OJJ63788.1 ater|EAU34247.1 atub|OJI85156.1 aver|OJI97150.1 awen|OJJ34818.1 azon|OJJ45731.1

Group8342: acar|OOF95727.1 acla|EAW10682.1 afis|EAW24888.1 afum|EAL89539.2 agla|OJJ82111.1 akaw|GAA81900.1 aluc|OJZ92298.1 anid|CBF75623.1 asyd|OJJ57455.1 awen|OJJ40478.1 azon|OJJ49342.1

Group8343: acar|OOF95805.1 acla|EAW08505.1 afis|EAW21090.1 afla|EED54674.1 akaw|GAA83598.1 aluc|OJZ92396.1 anig|EHA26570.1 aory|EIT78919.1 atub|OJI86826.1 awen|OJJ30106.1 acla|EAW13629.1

Group8344: afla|EED54299.1 acla|EAW14407.1 afis|EAW19115.1 afum|EAL93847.2 agla|OJJ80430.1 aluc|OJZ85762.1 anid|CBF79732.1 anig|EHA24794.1 asyd|OJJ58071.1 aver|OJJ04336.1 awen|OJJ37445.1

Group8345: acar|OOF97206.1 acla|EAW08299.1 afis|EAW23341.1 afla|EED55625.1 afum|EAL88137.2 agla|OJJ86489.1 anid|CBF89423.1 anig|EHA26305.1 aory|EIT81815.1 asyd|OJJ62450.1 awen|OJJ39553.1

Group8346: acar|OOF97220.1 acla|EAW08313.1 afis|EAW23327.1 afum|EAL88152.1 agla|OJJ86522.1 atub|OJI87106.1 awen|OJJ39567.1 azon|OJJ47760.1 asyd|OJJ62436.1 aver|OJI96572.1 anid|CBF89448.1

Group8347: acar|OOF98466.1 afis|EAW16909.1 afla|EED46359.1 afum|EAL86445.1 anig|EHA25226.1 aory|EIT81189.1 ater|EAU33597.1 atub|OJI82375.1 awen|OJJ39792.1 awen|OJJ35924.1 awen|OJJ38690.1

Group8348: acar|OOF98603.1 acla|EAW06636.1 afis|EAW16129.1 afla|EED45888.1 afum|EAL88759.1 anid|CBF80467.1 aory|EIT75433.1 asyd|OJJ54773.1 ater|EAU33904.1 aver|OJJ07393.1 awen|OJJ30300.1

Group8349: awen|OJJ39409.1 acla|EAW12397.1 afis|EAW16845.1 afla|EED53680.1 afum|EAL86391.1 agla|OJJ87798.1 anid|CBF73978.1 aory|EIT75322.1 ater|EAU30295.1 asyd|OJJ53745.1 aver|OJJ07859.1

Group8350: acla|EAW06378.1 afis|EAW18484.1 afla|EED52924.1 afum|EBA27485.1 agla|OJJ85073.1 anid|CBF70143.1 aory|EIT78661.1 asyd|OJJ58810.1 aver|OJJ01413.1 awen|OJJ31036.1 azon|OJJ50549.1

Group8351: acla|EAW06430.1 afis|EAW18535.1 afla|EED55531.1 afum|EAL93233.1 agla|OJJ81548.1 anid|CBF82130.1 aory|EIT80923.1 asyd|OJJ54310.1 aver|OJJ05258.1 awen|OJJ31093.1 azon|OJJ46171.1

Group8352: acla|EAW08558.1 afis|EAW23601.1 afla|EED47740.1 afum|EAL87866.1 agla|OJJ86868.1 anid|CBF89576.1 aory|EIT77169.1 asyd|OJJ62334.1 ater|EAU35361.1 aver|OJI96440.1 awen|OJJ39927.1

Group8353: acla|EAW09916.1 afis|EAW21164.1 afla|EED54728.1 afum|EAL92183.1 agla|OJJ78619.1 anid|CBF86965.1 aory|EIT78741.1 asyd|OJJ60534.1 ater|EAU35831.1 aver|OJI99861.1 awen|OJJ36498.1

Group8354: acla|EAW10114.1 afis|EAW25475.1 afla|EED53967.1 afum|EAL90138.2 agla|OJJ78866.1 anid|CBF77680.1 asyd|OJJ53089.1 ater|EAU34643.1 aver|OJJ03757.1 awen|OJJ41211.1 azon|OJJ42087.1

Group8355: acla|EAW10193.1 afis|EAW25402.1 afla|EED53881.1 afum|EAL90061.1 agla|OJJ87043.1 aory|EIT78364.1 asyd|OJJ53178.1 ater|EAU34553.1 aver|OJJ03674.1 awen|OJJ41129.1 azon|OJJ44703.1

Group8356: acla|EAW10414.1 afis|EAW25155.1 afla|EED57494.1 afum|EAL89830.1 agla|OJJ82513.1 anid|CBF85648.1 aory|EIT81887.1 asyd|OJJ64235.1 ater|EAU39438.1 aver|OJI98327.1 awen|OJJ40798.1

Group8357: acla|EAW10866.1 afis|EAW20992.1 afla|EED50346.1 afum|EAL84637.1 agla|OJJ84070.1 anid|CBF82615.1 aory|EIT77358.1 asyd|OJJ54017.1 ater|EAU38641.1 aver|OJJ08724.1 awen|OJJ34318.1

Group8358: acla|EAW10947.1 afla|EED52376.1 aluc|OJZ88751.1 anig|EHA18453.1 atub|OJI91322.1 aver|OJJ06237.1 awen|OJJ40913.1 asyd|OJJ58113.1 aver|OJJ04292.1 anid|CBF83892.1 asyd|OJJ53372.1

Group8359: acla|EAW11687.1 afis|EAW23704.1 afla|EED53411.1 afum|EAL87148.1 anid|CBF78352.1 aory|EIT74993.1 asyd|OJJ56143.1 ater|EAU30088.1 aver|OJJ05083.1 afis|EAW16210.1 afum|EAL88837.1

Group8360: acla|EAW11707.1 afis|EAW23724.1 afla|EED50513.1 akaw|GAA85949.1 aluc|OJZ91113.1 anig|EHA19811.1 aory|EIT82001.1 asyd|OJJ55358.1 ater|EAU31558.1 atub|OJI83111.1 aory|EIT82067.1

Group8361: acla|EAW12296.1 afis|EAW16927.1 afum|EAL86453.1 anid|CBF82135.1 anig|EHA21632.1 asyd|OJJ54314.1 ater|EAU32987.1 aver|OJJ05268.1 azon|OJJ42780.1 aluc|OJZ87026.1 atub|OJI83492.1

Group8362: acla|EAW13190.1 afis|EAW17451.1 afla|EED49371.1 afum|EAL91279.1 agla|OJJ80361.1 akaw|GAA90741.1 aluc|OJZ84607.1 anid|CBF71681.1 aory|EIT79502.1 ater|EAU33931.1 awen|OJJ37395.1

Group8363: acla|EAW13643.1 afis|EAW19994.1 afla|EED45928.1 afum|EAL84923.1 agla|OJJ88076.1 anid|CBF84677.1 aory|EIT75618.1 asyd|OJJ53343.1 ater|EAU35264.1 aver|OJJ08402.1 awen|OJJ30002.1

Group8364: acla|EAW13656.1 afis|EAW19936.1 afla|EED45905.1 afum|EAL84974.1 aory|EIT75425.1 asyd|OJJ53376.1 ater|EAU30903.1 aver|OJJ08367.1 awen|OJJ29979.1 anid|CBF70076.1 aory|EIT75908.1

Group8365: acla|EAW13696.1 afis|EAW19908.1 afum|EAL84998.1 akaw|GAA91346.1 aluc|OJZ84827.1 anid|CBF83988.1 anig|EHA28085.1 asyd|OJJ61405.1 atub|OJI89260.1 aver|OJJ00735.1 akaw|GAA91347.1

Group8366: acla|EAW14275.1 afis|EAW18999.1 afla|EED53127.1 afum|EAL93734.1 anid|CBF86777.1 aory|EIT75125.1 asyd|OJJ64684.1 ater|EAU37805.1 aver|OJI98786.1 awen|OJJ29914.1 azon|OJJ47177.1

Group8367: acla|EAW14596.1 afis|EAW19325.1 afla|EED57234.1 afum|EAL94043.1 aory|EIT74546.1 asyd|OJJ52375.1 ater|EAU29458.1 aver|OJJ01974.1 azon|OJJ45104.1 aory|EIT74049.1 afla|EED47555.1

Group8368: acla|EAW14597.1 afis|EAW19326.1 afla|EED52550.1 afum|EBA27512.1 aory|EIT74035.1 asyd|OJJ52374.1 ater|EAU33506.1 aver|OJJ01975.1 awen|OJJ30098.1 azon|OJJ45103.1 afla|EED57233.1

Group8369: acla|EAW14974.1 afis|EAW22748.1 afla|EED54370.1 agla|OJJ79726.1 akaw|GAA91089.1 aluc|OJZ87906.1 aory|EIT76018.1 asyd|OJJ55933.1 aver|OJJ04882.1 awen|OJJ36946.1 aory|EIT80427.1

Group8370: acla|EAW15067.1 afis|EAW20039.1 afla|EED45341.1 afum|EBA27223.1 agla|OJJ88924.1 anid|CBF84779.1 aory|EIT73728.1 asyd|OJJ63236.1 ater|EAU38652.1 aver|OJI97720.1 awen|OJJ31794.1

Group8371: afis|EAW15647.1 afum|EAL87785.1 anid|CBF85170.1 asyd|OJJ60246.1 ater|EAU30252.1 aver|OJJ04015.1 afis|EAW21202.1 afum|EAL92212.1 ater|EAU31233.1 awen|OJJ32937.1 azon|OJJ45229.1

Group8372: afis|EAW17266.1 afum|EAL86683.1 agla|OJJ81071.1 anid|CBF87526.1 aory|EIT81263.1 asyd|OJJ52983.1 ater|EAU36130.1 aver|OJJ08952.1 azon|OJJ45515.1 afla|EED48854.1 awen|OJJ35767.1

Group8373: afis|EAW18533.1 afla|EED52867.1 afum|EAL93231.1 agla|OJJ85147.1 anid|CBF75405.1 aory|EIT78596.1 asyd|OJJ59813.1 ater|EAU30035.1 aver|OJJ02475.1 awen|OJJ31091.1 azon|OJJ50597.1

Group8374: afla|EED56626.1 aver|OJJ07661.1 afla|EED52559.1 akaw|GAA88922.1 aluc|OJZ89499.1 anid|CBF86953.1 aory|EIT74185.1 atub|OJI85304.1 agla|OJJ82849.1 asyd|OJJ63468.1 aver|OJI97464.1

Group8375: afis|EAW24571.1 afla|EED53136.1 afum|EAL86234.1 anid|CBF83980.1 aory|EIT75246.1 asyd|OJJ57973.1 ater|EAU31428.1 aver|OJJ04428.1 azon|OJJ42716.1 asyd|OJJ61404.1 aver|OJJ00734.1

Group8376: afla|EED56549.1 agla|OJJ78761.1 anid|CBF84227.1 aory|EIT79610.1 asyd|OJJ63008.1 ater|EAU33859.1 aver|OJI97952.1 awen|OJJ34569.1 asyd|OJJ62574.1 asyd|OJJ57181.1 aver|OJJ03109.1

Group8377: abra|OJJ65483.1 abra|OJJ65634.1 afla|EED55265.1 agla|OJJ79068.1 anid|CBF82221.1 aory|EIT83579.1 asyd|OJJ55371.1 ater|EAU35403.1 aver|OJJ06450.1 awen|OJJ36001.1

Group8378: abra|OJJ65628.1 abra|OJJ65709.1 aluc|OJZ80119.1 aluc|OJZ88802.1 asyd|OJJ51944.1 asyd|OJJ52124.1 akaw|GAA93005.1 abra|OJJ66180.1 afis|EAW20677.1 acla|EAW11466.1

Group8379: abra|OJJ65666.1 abra|OJJ70428.1 anid|CBF81161.1 anid|CBF82242.1 anid|CBF87226.1 akaw|GAA93194.1 aluc|OJZ81153.1 atub|OJI88768.1 asyd|OJJ57260.1 aver|OJJ03036.1

Group8380: acar|OOF91916.1 acar|OOF98336.1 abra|OJJ70254.1 afla|EED44862.1 anig|EHA25303.1 aory|EIT82731.1 asyd|OJJ55178.1 aver|OJJ07029.1 awen|OJJ35530.1 asyd|OJJ62626.1

Group8381: acar|OOF92322.1 acar|OOF92364.1 abra|OJJ67476.1 afis|EAW18403.1 afum|EAL93081.1 asyd|OJJ58697.1 ater|EAU29820.1 atub|OJI80709.1 aver|OJJ01309.1 awen|OJJ31747.1

Group8382: acar|OOF94055.1 acar|OOF94125.1 abra|OJJ66389.1 afla|EED52182.1 akaw|GAA90255.1 aluc|OJZ83085.1 aory|EIT74406.1 ater|EAU31726.1 atub|OJI87859.1 azon|OJJ45262.1

Group8383: acar|OOF94067.1 acar|OOF94137.1 awen|OJJ31275.1 abra|OJJ66355.1 afla|EED47477.1 akaw|GAA90229.1 aluc|OJZ83112.1 anig|EHA28175.1 aory|EIT79963.1 atub|OJI87822.1

Group8384: acar|OOF95040.1 acar|OOF95107.1 acar|OOF90589.1 azon|OJJ45969.1 acla|EAW11067.1 afla|EED54161.1 anig|EHA19940.1 aory|EIT75957.1 atub|OJI79304.1 asyd|OJJ57430.1

Group8385: acar|OOF95078.1 acar|OOF95143.1 abra|OJJ74634.1 akaw|GAA81975.1 aluc|OJZ92214.1 anig|EHA26743.1 asyd|OJJ62239.1 atub|OJI86645.1 aver|OJI96348.1 awen|OJJ37607.1

Group8386: acar|OOF95096.1 acar|OOF95161.1 abra|OJJ74607.1 afla|EED49964.1 akaw|GAA82001.1 aluc|OJZ92187.1 anig|EHA26766.1 aory|EIT83447.1 atub|OJI86616.1 azon|OJJ47334.1

Group8387: acar|OOF95097.1 acar|OOF95163.1 abra|OJJ74606.1 afla|EED49963.1 aluc|OJZ92186.1 anig|EHA26767.1 aory|EIT83431.1 atub|OJI86615.1 azon|OJJ47333.1 azon|OJJ47301.1

Group8388: acar|OOF95620.1 acar|OOF95672.1 afla|EED56570.1 aluc|OJZ88747.1 anig|EHA18451.1 aory|EIT79617.1 asyd|OJJ61712.1 ater|EAU29928.1 atub|OJI91319.1 aver|OJI95807.1

Group8389: acar|OOF95652.1 acar|OOF95703.1 abra|OJJ74692.1 afla|EED47157.1 aluc|OJZ92269.1 aory|EIT79192.1 ater|EAU38044.1 atub|OJI86703.1 agla|OJJ78428.1 aluc|OJZ83217.1

Group8390: acar|OOF95655.1 acar|OOF95706.1 abra|OJJ76469.1 afla|EED46484.1 agla|OJJ87727.1 akaw|GAA82381.1 aluc|OJZ91740.1 atub|OJI86198.1 awen|OJJ31394.1 aory|EIT72384.1

Group8391: acar|OOF96446.1 acar|OOG00865.1 abra|OJJ70058.1 afla|EED54159.1 akaw|GAA87616.1 aluc|OJZ85237.1 aory|EIT76003.1 asyd|OJJ54511.1 atub|OJI80338.1 aver|OJJ07672.1

Group8392: acar|OOG00241.1 acar|OOG00260.1 abra|OJJ67536.1 akaw|GAA88123.1 aluc|OJZ81396.1 anig|EHA18688.1 asyd|OJJ53704.1 atub|OJI91601.1 aver|OJJ07828.1 azon|OJJ43560.1

Group8393: afla|EED45918.1 afla|EED45920.1 abra|OJJ69032.1 acar|OOF92652.1 afis|EAW19901.1 akaw|GAA90547.1 aluc|OJZ86017.1 anig|EHA24594.1 atub|OJI79806.1 abra|OJJ74401.1

Group8394: afum|EAL86719.2 afum|EAL92228.1 akaw|GAA92886.1 akaw|GAA92946.1 acla|EAW11456.1 aluc|OJZ87839.1 anid|CBF87743.1 ater|EAU32044.1 awen|OJJ40122.1 azon|OJJ45924.1

Group8395: agla|OJJ78529.1 agla|OJJ84460.1 afis|EAW19320.1 afla|EED48990.1 afum|EAL94038.1 aory|EIT80001.1 asyd|OJJ64749.1 ater|EAU31743.1 aver|OJI98850.1 awen|OJJ31889.1

Group8396: agla|OJJ86080.1 agla|OJJ78838.1 abra|OJJ72688.1 acar|OOF95937.1 akaw|GAA85306.1 aluc|OJZ90666.1 anig|EHA24101.1 atub|OJI82095.1 awen|OJJ31985.1 afum|EAL86718.1

Group8397: agla|OJJ79716.1 agla|OJJ88991.1 agla|OJJ82731.1 azon|OJJ47655.1 afis|EAW20587.1 afis|EAW21873.1 afum|EAL88434.1 ater|EAU34446.1 atub|OJI86118.1 anid|CBF83577.1

Group8398: agla|OJJ81519.1 agla|OJJ81528.1 abra|OJJ70448.1 afla|EED51815.1 akaw|GAA90133.1 aluc|OJZ83214.1 anid|CBF73692.1 anig|EHA21074.1 aory|EIT81708.1 atub|OJI87722.1

Group8399: agla|OJJ82390.1 agla|OJJ88105.1 awen|OJJ29753.1 awen|OJJ38325.1 acla|EAW15022.1 afla|EED48979.1 agla|OJJ87354.1 aory|EIT79986.1 aver|OJJ04712.1 awen|OJJ29941.1

Group8400: agla|OJJ84307.1 agla|OJJ84308.1 abra|OJJ75522.1 acar|OOF90849.1 acla|EAW09309.1 afis|EAW21836.1 afla|EED46539.1 atub|OJI85810.1 awen|OJJ34112.1 azon|OJJ44231.1

Group8401: agla|OJJ87455.1 agla|OJJ89093.1 abra|OJJ76700.1 acar|OOF99318.1 afla|EED48317.1 akaw|GAA92499.1 aluc|OJZ91512.1 anig|EHA22550.1 aory|EIT76304.1 atub|OJI85964.1

Group8402: akaw|GAA83188.1 akaw|GAA90127.1 aluc|OJZ83222.1 aluc|OJZ83687.1 atub|OJI87714.1 atub|OJI90059.1 abra|OJJ68284.1 acar|OOF94475.1 anig|EHA21080.1 azon|OJJ45238.1

Group8403: akaw|GAA88991.1 akaw|GAA93233.1 abra|OJJ65743.1 acar|OOG00878.1 acla|EAW11894.1 afla|EED52522.1 aluc|OJZ79643.1 aver|OJI98825.1 awen|OJJ36066.1 azon|OJJ47413.1

Group8404: akaw|GAA92025.1 akaw|GAA92119.1 aluc|OJZ83604.1 aluc|OJZ92893.1 atub|OJI87351.1 atub|OJI89976.1 abra|OJJ68351.1 acar|OOF90144.1 anig|EHA18764.1 anig|EHA26070.1

Group8405: aluc|OJZ79988.1 aluc|OJZ84722.1 abra|OJJ77310.1 akaw|GAA91443.1 anig|EHA27996.1 atub|OJI89361.1 acar|OOF91981.1 afis|EAW17114.1 afla|EED51656.1 acar|OOF99569.1

Group8406: anid|CBF89276.1 anid|CBF89323.1 awen|OJJ36372.1 acar|OOF93282.1 anig|EHA24975.1 ater|EAU34904.1 atub|OJI80323.1 awen|OJJ30808.1 agla|OJJ82139.1 aver|OJJ03024.1

Group8407: anig|EHA21148.1 anig|EHA28208.1 abra|OJJ69753.1 acar|OOG00377.1 acla|EAW13607.1 akaw|GAA90204.1 atub|OJI87793.1 awen|OJJ37499.1 acar|OOF95027.1 asyd|OJJ58374.1

Group8408: asyd|OJJ52165.1 asyd|OJJ53371.1 asyd|OJJ61758.1 aver|OJI95856.1 aver|OJI99567.1 aver|OJJ08373.1 agla|OJJ87524.1 anid|CBF84630.1 anig|EHA21030.1 anid|CBF88873.1

Group8409: asyd|OJJ56881.1 asyd|OJJ59210.1 abra|OJJ66476.1 afis|EAW17352.1 afla|EED51997.1 akaw|GAA91463.1 aluc|OJZ82068.1 aory|EIT81417.1 ater|EAU33008.1 atub|OJI84625.1

Group8410: asyd|OJJ57202.1 asyd|OJJ60345.1 aver|OJI99686.1 aver|OJJ03087.1 afis|EAW17363.1 afla|EED49455.1 afum|EAL91200.1 anid|CBF84252.1 aory|EIT76139.1 ater|EAU33868.1

Group8411: ater|EAU29225.1 ater|EAU29370.1 ater|EAU31354.1 afis|EAW22028.1 agla|OJJ79773.1 aluc|OJZ81912.1 anid|CBF80838.1 anig|EHA21043.1 awen|OJJ38846.1 agla|OJJ79276.1

Group8412: ater|EAU29238.1 ater|EAU31399.1 acar|OOF99900.1 acla|EAW07173.1 afla|EED57571.1 agla|OJJ87862.1 anid|CBF84514.1 aory|EIT82976.1 asyd|OJJ53502.1 aver|OJJ08243.1

Group8413: ater|EAU29963.1 ater|EAU36059.1 afis|EAW18832.1 afla|EED53422.1 afum|EAL93566.1 anid|CBF79052.1 asyd|OJJ64204.1 aver|OJI98293.1 awen|OJJ41494.1 anid|CBF85615.1

Group8414: ater|EAU30556.1 ater|EAU35249.1 afis|EAW24045.1 abra|OJJ73943.1 afum|EBA27445.1 akaw|GAA89548.1 aluc|OJZ88101.1 anig|EHA17870.1 atub|OJI90709.1 azon|OJJ43062.1

Group8415: ater|EAU31622.1 ater|EAU34595.1 aver|OJI99627.1 aver|OJJ07368.1 abra|OJJ68596.1 acar|OOF92450.1 afis|EAW24695.1 afum|EAL89337.1 anid|CBF69453.1 ater|EAU32761.1

Group8416: ater|EAU31627.1 ater|EAU34593.1 azon|OJJ45985.1 anig|EHA23409.1 awen|OJJ31932.1 acar|OOF99005.1 afis|EAW24700.1 afla|EED53388.1 afum|EAL89342.1 aory|EIT75160.1

Group8417: aver|OJJ03450.1 aver|OJJ06658.1 afla|EED51125.1 agla|OJJ81315.1 anid|CBF90344.1 aory|EIT81329.1 asyd|OJJ55579.1 ater|EAU37571.1 asyd|OJJ62613.1 asyd|OJJ62615.1

Group8418: awen|OJJ30159.1 awen|OJJ36055.1 abra|OJJ68149.1 acar|OOF95997.1 afla|EED45662.1 akaw|GAA92371.1 aluc|OJZ83229.1 anig|EHA21086.1 ater|EAU36032.1 atub|OJI87707.1

Group8419: awen|OJJ30762.1 awen|OJJ31719.1 abra|OJJ69691.1 afla|EED48236.1 agla|OJJ84995.1 aluc|OJZ80608.1 anid|CBF84030.1 aory|EIT72891.1 asyd|OJJ61367.1 aver|OJJ00694.1

Group8420: awen|OJJ30776.1 awen|OJJ31705.1 afis|EAW17200.1 afum|EAL86627.2 ater|EAU37835.1 afla|EED45211.1 anid|CBF78548.1 aory|EIT73675.1 asyd|OJJ65056.1 aver|OJI99179.1

Group8421: awen|OJJ30959.1 awen|OJJ36540.1 awen|OJJ39737.1 awen|OJJ33713.1 acla|EAW11064.1 afis|EAW20810.1 afum|EAL84818.1 anid|CBF87080.1 asyd|OJJ60416.1 azon|OJJ42504.1

Group8422: awen|OJJ30979.1 awen|OJJ35555.1 abra|OJJ76316.1 acar|OOF99567.1 aluc|OJZ91898.1 anig|EHA22877.1 asyd|OJJ63041.1 atub|OJI86351.1 aver|OJI97927.1 afis|EAW16162.1

Group8423: awen|OJJ32041.1 awen|OJJ33427.1 abra|OJJ72830.1 acar|OOF96034.1 akaw|GAA85429.1 aluc|OJZ83331.1 anig|EHA24208.1 atub|OJI82227.1 afis|EAW19852.1 agla|OJJ79014.1

Group8424: azon|OJJ42161.1 abra|OJJ66115.1 afis|EAW21210.1 afla|EED50949.1 afum|EAL92221.1 aory|EIT77051.1 asyd|OJJ57281.1 ater|EAU36270.1 aver|OJJ05875.1 awen|OJJ37951.1

Group8425: azon|OJJ44008.1 azon|OJJ48147.1 agla|OJJ82609.1 awen|OJJ40073.1 afla|EED57859.1 aluc|OJZ80321.1 aory|EIT81094.1 asyd|OJJ60285.1 aver|OJI99617.1 asyd|OJJ54869.1

Group8426: aluc|OJZ79479.1 aluc|OJZ80339.1 aluc|OJZ80254.1 aluc|OJZ79612.1 aluc|OJZ79697.1 aluc|OJZ79554.1 aluc|OJZ80167.1 aluc|OJZ79747.1 aluc|OJZ79877.1 aluc|OJZ79901.1

Group8427: abra|OJJ65437.1 akaw|GAA92646.1 abra|OJJ66323.1 afla|EED47879.1 afum|EAL85394.1 anid|CBF82965.1 asyd|OJJ62692.1 ater|EAU30107.1 awen|OJJ29581.1 akaw|GAA87900.1

Group8428: abra|OJJ65515.1 acla|EAW07045.1 afla|EED44910.1 aory|EIT73468.1 atub|OJI82843.1 awen|OJJ32271.1 afla|EED44911.1 azon|OJJ50129.1 awen|OJJ35039.1 azon|OJJ42748.1

Group8429: abra|OJJ65697.1 afla|EED56836.1 afum|EAL84267.1 agla|OJJ84575.1 anig|EHA18815.1 aory|EIT73939.1 asyd|OJJ62604.1 ater|EAU33256.1 aver|OJJ07757.1 awen|OJJ30261.1

Group8430: abra|OJJ65701.1 afla|EED56831.1 afum|EAL84271.2 anid|CBF83516.1 anig|EHA18820.1 aory|EIT73884.1 asyd|OJJ60706.1 ater|EAU38173.1 aver|OJJ00032.1 ater|EAU38313.1

Group8431: afum|EAL88787.1 anig|EHA26113.1 asyd|OJJ52039.1 afum|EAL88783.1 azon|OJJ42060.1 aluc|OJZ80012.1 atub|OJI87462.1 aluc|OJZ79617.1 anig|EHA26116.1 aluc|OJZ80024.1

Group8432: afla|EED52381.1 abra|OJJ67028.1 acar|OOF91717.1 akaw|GAA83905.1 aluc|OJZ92124.1 anig|EHA24970.1 atub|OJI82779.1 atub|OJI86556.1 akaw|GAA82056.1 aver|OJJ04874.1

Group8433: abra|OJJ65861.1 afla|EED51768.1 akaw|GAA92284.1 aluc|OJZ90181.1 anig|EHA23680.1 aory|EIT80033.1 asyd|OJJ54550.1 atub|OJI81608.1 aver|OJJ07598.1 awen|OJJ29824.1

Group8434: abra|OJJ65895.1 afla|EED56889.1 akaw|GAA86010.1 aluc|OJZ90236.1 anig|EHA23736.1 aory|EIT77462.1 ater|EAU30299.1 atub|OJI81666.1 aver|OJJ06587.1 awen|OJJ35825.1

Group8435: abra|OJJ65968.1 acar|OOF92906.1 akaw|GAA90843.1 aluc|OJZ82990.1 anig|EHA21405.1 asyd|OJJ53325.1 ater|EAU29362.1 atub|OJI89224.1 aver|OJJ08418.1 awen|OJJ29772.1

Group8436: abra|OJJ66059.1 afla|EED49906.1 akaw|GAA93129.1 aluc|OJZ81104.1 anig|EHA19265.1 aory|EIT72470.1 asyd|OJJ55639.1 atub|OJI82475.1 aver|OJJ06722.1 awen|OJJ33566.1

Group8437: abra|OJJ66063.1 acar|OOF89827.1 afla|EED56764.1 akaw|GAA84175.1 aluc|OJZ81100.1 aory|EIT73917.1 atub|OJI82480.1 abra|OJJ71214.1 anig|EHA20557.1 acar|OOF93979.1

Group8438: abra|OJJ66070.1 acla|EAW12247.1 akaw|GAA84174.1 aluc|OJZ81099.1 anig|EHA19272.1 atub|OJI82481.1 abra|OJJ74890.1 akaw|GAA83529.1 aluc|OJZ92467.1 atub|OJI86896.1

Group8439: abra|OJJ66104.1 afis|EAW16817.1 afla|EED45840.1 afum|EAL86368.2 akaw|GAA92673.1 aluc|OJZ90128.1 anig|EHA20943.1 aory|EIT75700.1 atub|OJI84668.1 afla|EED45839.1

Group8440: abra|OJJ66114.1 afis|EAW21211.1 afla|EED50948.1 afum|EAL92222.1 aory|EIT77000.1 asyd|OJJ57280.1 ater|EAU36269.1 aver|OJJ05876.1 awen|OJJ37952.1 azon|OJJ42160.1

Group8441: abra|OJJ66120.1 acar|OOF94012.1 akaw|GAA92682.1 aluc|OJZ90137.1 anid|CBF82787.1 anig|EHA20929.1 asyd|OJJ57366.1 ater|EAU30145.1 atub|OJI84659.1 azon|OJJ47363.1

Group8442: abra|OJJ66166.1 afis|EAW18034.1 afum|EAL91845.1 anid|CBF84528.1 aory|EIT76120.1 acla|EAW12651.1 afla|EED54072.1 asyd|OJJ62164.1 aver|OJI96275.1 awen|OJJ40173.1

Group8443: awen|OJJ37693.1 acar|OOF91998.1 afis|EAW23726.1 afla|EED53317.1 agla|OJJ86824.1 aory|EIT75193.1 ater|EAU36803.1 awen|OJJ33348.1 azon|OJJ45552.1 awen|OJJ39890.1

Group8444: abra|OJJ66204.1 acar|OOF91929.1 afla|EED50169.1 akaw|GAA89081.1 aluc|OJZ80510.1 anig|EHA25683.1 aory|EIT72280.1 asyd|OJJ61320.1 atub|OJI88420.1 aver|OJJ00645.1

Group8445: abra|OJJ66231.1 acar|OOF90689.1 agla|OJJ84411.1 akaw|GAA89058.1 aluc|OJZ80536.1 anig|EHA25659.1 asyd|OJJ64300.1 atub|OJI88395.1 aver|OJI98393.1 awen|OJJ34687.1

Group8446: abra|OJJ66236.1 acar|OOF90697.1 agla|OJJ84417.1 akaw|GAA89053.1 aluc|OJZ80541.1 anig|EHA25654.1 asyd|OJJ53791.1 atub|OJI88390.1 aver|OJJ07889.1 awen|OJJ31377.1

Group8447: abra|OJJ66262.1 acar|OOF90733.1 acla|EAW09305.1 aluc|OJZ80570.1 anig|EHA25628.1 asyd|OJJ59251.1 atub|OJI88361.1 aver|OJJ01860.1 awen|OJJ34123.1 azon|OJJ48039.1

Group8448: acla|EAW14333.1 awen|OJJ31553.1 afis|EAW18955.1 agla|OJJ88792.1 anig|EHA18744.1 ater|EAU30923.1 aver|OJI95799.1 azon|OJJ47271.1 aory|EIT74943.1 agla|OJJ85332.1

Group8449: abra|OJJ66286.1 acla|EAW07167.1 afis|EAW22182.1 afla|EED57561.1 afum|EAL91007.1 anig|EHA23666.1 aory|EIT81852.1 ater|EAU34187.1 atub|OJI89937.1 awen|OJJ34763.1

Group8450: abra|OJJ66333.1 acar|OOF96844.1 agla|OJJ83531.1 akaw|GAA90210.1 aluc|OJZ83135.1 anig|EHA28194.1 aory|EIT77484.1 atub|OJI87799.1 azon|OJJ47327.1 asyd|OJJ56279.1

Group8451: abra|OJJ66339.1 afla|EED53167.1 akaw|GAA90213.1 aluc|OJZ83130.1 anig|EHA28190.1 aory|EIT75194.1 atub|OJI87805.1 awen|OJJ35030.1 aver|OJJ01854.1 anid|CBF69485.1

Group8452: abra|OJJ66375.1 afla|EED54381.1 akaw|GAA90243.1 aluc|OJZ83097.1 anig|EHA28161.1 aory|EIT75879.1 asyd|OJJ61448.1 ater|EAU38988.1 atub|OJI87839.1 awen|OJJ32931.1

Group8453: abra|OJJ66377.1 acar|OOF98137.1 afum|EAL94060.1 akaw|GAA90245.1 aluc|OJZ83095.1 anig|EHA28159.1 aory|EIT78381.1 atub|OJI87841.1 azon|OJJ42527.1 agla|OJJ86818.1

Group8454: afis|EAW19341.1 abra|OJJ67128.1 acar|OOF98425.1 acla|EAW14612.1 afla|EED47545.1 akaw|GAA91040.1 aluc|OJZ87850.1 anig|EHA20153.1 atub|OJI80798.1 aory|EIT74552.1

Group8455: abra|OJJ66386.1 afla|EED45823.1 akaw|GAA90252.1 aluc|OJZ83088.1 anig|EHA28150.1 aory|EIT75601.1 asyd|OJJ55225.1 ater|EAU36363.1 atub|OJI87856.1 aver|OJJ06996.1

Group8456: abra|OJJ66474.1 afis|EAW17242.1 afum|EBA27283.1 akaw|GAA92460.1 aluc|OJZ81557.1 anid|CBF75984.1 anig|EHA25877.1 asyd|OJJ53455.1 atub|OJI88628.1 aver|OJJ08300.1

Group8457: abra|OJJ66524.1 acar|OOF90977.1 akaw|GAA91189.1 aluc|OJZ81514.1 anig|EHA25912.1 atub|OJI88663.1 anid|CBF87384.1 afla|EED53133.1 afum|EAL91114.2 aory|EIT75039.1

Group8458: abra|OJJ66791.1 afis|EAW20403.1 akaw|GAA86856.1 aluc|OJZ89978.1 anig|EHA27351.1 asyd|OJJ52424.1 atub|OJI84831.1 aver|OJJ08460.1 awen|OJJ35381.1 aver|OJI96708.1

Group8459: abra|OJJ66794.1 acar|OOF93871.1 afla|EED44982.1 akaw|GAA86859.1 aluc|OJZ89982.1 anig|EHA27354.1 aory|EIT73545.1 atub|OJI84828.1 azon|OJJ50414.1 aver|OJJ05859.1

Group8460: abra|OJJ66818.1 acar|OOF91601.1 akaw|GAA90456.1 aluc|OJZ85768.1 anig|EHA24785.1 aory|EIT76062.1 asyd|OJJ58068.1 ater|EAU37371.1 atub|OJI79553.1 aver|OJJ04339.1

Group8461: afla|EED48048.1 ater|EAU31173.1 agla|OJJ79647.1 asyd|OJJ53970.1 anid|CBF74235.1 awen|OJJ33072.1 acla|EAW14693.1 afis|EAW24224.1 afum|EBA27250.1 aver|OJJ08068.1

Group8462: abra|OJJ66996.1 acar|OOF93619.1 afla|EED52137.1 agla|OJJ81673.1 akaw|GAA92168.1 aluc|OJZ85576.1 anig|EHA24943.1 aory|EIT74409.1 atub|OJI79372.1 awen|OJJ36919.1

Group8463: agla|OJJ89337.1 abra|OJJ67011.1 afla|EED53268.1 akaw|GAA85914.1 aluc|OJZ91152.1 aory|EIT74894.1 atub|OJI83166.1 aluc|OJZ87778.1 awen|OJJ33695.1 anig|EHA22294.1

Group8464: abra|OJJ67012.1 acar|OOF93608.1 akaw|GAA92161.1 aluc|OJZ85568.1 anig|EHA24950.1 asyd|OJJ53490.1 ater|EAU31044.1 atub|OJI79363.1 aver|OJJ03352.1 azon|OJJ47107.1

Group8465: abra|OJJ67015.1 afla|EED52172.1 akaw|GAA92158.1 aluc|OJZ85565.1 anid|CBF82918.1 anig|EHA24953.1 aory|EIT74453.1 asyd|OJJ57303.1 atub|OJI79359.1 aver|OJJ05856.1

Group8466: abra|OJJ67115.1 acar|OOF98437.1 acla|EAW11657.1 agla|OJJ83143.1 anid|CBF89632.1 anig|EHA20166.1 asyd|OJJ61474.1 ater|EAU31281.1 aver|OJJ00815.1 awen|OJJ36970.1

Group8467: abra|OJJ67168.1 afla|EED47603.1 akaw|GAA91006.1 aluc|OJZ87807.1 anig|EHA20289.1 atub|OJI80838.1 aory|EIT80219.1 azon|OJJ41965.1 azon|OJJ44887.1 aver|OJI99374.1

Group8468: abra|OJJ67224.1 acar|OOF98328.1 akaw|GAA91651.1 aluc|OJZ87750.1 anig|EHA20238.1 asyd|OJJ60284.1 ater|EAU39445.1 atub|OJI80893.1 aver|OJI99616.1 afum|EAL86282.1

Group8469: abra|OJJ67266.1 acar|OOF98300.1 agla|OJJ81020.1 akaw|GAA91682.1 aluc|OJZ87712.1 ater|EAU37170.1 atub|OJI80930.1 afla|EED45389.1 aory|EIT82395.1 ater|EAU36037.1

Group8470: abra|OJJ67281.1 afla|EED49573.1 akaw|GAA86528.1 aluc|OJZ85076.1 anid|CBF83018.1 anig|EHA25291.1 aory|EIT82642.1 asyd|OJJ61509.1 atub|OJI80522.1 aver|OJJ00846.1

Group8471: abra|OJJ67351.1 acar|OOF93557.1 akaw|GAA86467.1 aluc|OJZ85008.1 anig|EHA25229.1 atub|OJI80589.1 awen|OJJ32046.1 azon|OJJ43962.1 acar|OOF98888.1 azon|OJJ50778.1

Group8472: agla|OJJ82649.1 azon|OJJ50009.1 acla|EAW06989.1 afis|EAW16603.1 afla|EED45022.1 afum|EAL89199.1 anid|CBF78453.1 aory|EIT73688.1 ater|EAU36684.1 awen|OJJ40203.1

Group8473: abra|OJJ67370.1 afla|EED49792.1 akaw|GAA93172.1 aluc|OJZ83031.1 anig|EHA28107.1 aory|EIT77626.1 asyd|OJJ56702.1 ater|EAU31039.1 atub|OJI87917.1 aver|OJJ06023.1

Group8474: abra|OJJ67413.1 acla|EAW07113.1 akaw|GAA86414.1 aluc|OJZ84955.1 asyd|OJJ65312.1 ater|EAU33566.1 atub|OJI80645.1 aver|OJI99394.1 acar|OOF94725.1 ater|EAU31127.1

Group8475: abra|OJJ67528.1 acar|OOF99206.1 afla|EED50912.1 akaw|GAA88134.1 aluc|OJZ81405.1 anig|EHA18699.1 aory|EIT79025.1 asyd|OJJ54793.1 atub|OJI91612.1 aver|OJJ02014.1

Group8476: ater|EAU34364.1 awen|OJJ40952.1 acla|EAW10355.1 afis|EAW25240.1 afla|EED53706.1 afum|EAL89901.1 aory|EIT75314.1 asyd|OJJ64007.1 aver|OJI98094.1 azon|OJJ44542.1

Group8477: abra|OJJ67560.1 acar|OOG00274.1 afla|EED53443.1 akaw|GAA88106.1 aluc|OJZ81373.1 asyd|OJJ53830.1 atub|OJI91579.1 aver|OJJ07934.1 azon|OJJ50340.1 aory|EIT75217.1

Group8478: abra|OJJ67580.1 acla|EAW13490.1 afla|EED49873.1 akaw|GAA85343.1 aluc|OJZ90705.1 anig|EHA24578.1 aory|EIT83405.1 asyd|OJJ57242.1 atub|OJI82132.1 azon|OJJ42362.1

Group8479: abra|OJJ67631.1 acar|OOF91978.1 akaw|GAA89151.1 aluc|OJZ81685.1 anig|EHA25758.1 aory|EIT77909.1 atub|OJI88496.1 azon|OJJ45916.1 acla|EAW09333.1 afla|EED46608.1

Group8480: abra|OJJ67636.1 acar|OOG00390.1 afis|EAW21768.1 akaw|GAA89155.1 aluc|OJZ81680.1 anig|EHA25762.1 asyd|OJJ57934.1 atub|OJI88501.1 azon|OJJ42789.1 awen|OJJ38051.1

Group8481: abra|OJJ67654.1 acar|OOF91986.1 akaw|GAA89170.1 aluc|OJZ81663.1 anid|CBF82092.1 anig|EHA25780.1 asyd|OJJ54268.1 atub|OJI88520.1 aver|OJJ05220.1 azon|OJJ51658.1

Group8482: abra|OJJ67664.1 afla|EED45195.1 akaw|GAA89180.1 aluc|OJZ81653.1 anid|CBF83091.1 aory|EIT73614.1 atub|OJI88529.1 abra|OJJ73455.1 acar|OOG00657.1 anig|EHA18282.1

Group8483: abra|OJJ67684.1 acar|OOG00732.1 akaw|GAA89196.1 aluc|OJZ81634.1 anid|CBF73535.1 anig|EHA25809.1 asyd|OJJ56720.1 ater|EAU30905.1 atub|OJI88546.1 aver|OJJ06005.1

Group8484: abra|OJJ67688.1 afis|EAW18096.1 agla|OJJ79878.1 akaw|GAA89199.1 aluc|OJZ81631.1 anig|EHA25812.1 asyd|OJJ53374.1 ater|EAU33707.1 atub|OJI88549.1 aver|OJJ08370.1

Group8485: abra|OJJ67701.1 akaw|GAA89210.1 anig|EHA25824.1 atub|OJI88561.1 afla|EED49815.1 abra|OJJ68462.1 acar|OOF90074.1 aory|EIT77612.1 awen|OJJ32891.1 aluc|OJZ81619.1

Group8486: afum|EAL89281.1 abra|OJJ68146.1 acar|OOF91873.1 afis|EAW16676.1 afum|EAL91340.1 agla|OJJ89194.1 anig|EHA21124.1 awen|OJJ32336.1 acar|OOF95856.1 afis|EAW17525.1

Group8487: abra|OJJ67708.1 acla|EAW12291.1 afis|EAW16946.1 afum|EAL86473.1 akaw|GAA89217.1 aluc|OJZ81612.1 anid|CBF82181.1 anig|EHA25828.1 ater|EAU31755.1 atub|OJI88573.1

Group8488: abra|OJJ67731.1 acla|EAW13296.1 afis|EAW17314.1 afla|EED51565.1 afum|EBA27288.1 anid|CBF70701.1 asyd|OJJ58459.1 aver|OJJ01059.1 azon|OJJ43027.1 afum|EAL91150.1

Group8489: abra|OJJ67734.1 acar|OOF98896.1 agla|OJJ78426.1 ater|EAU39365.1 aver|OJJ05244.1 awen|OJJ35802.1 agla|OJJ85936.1 asyd|OJJ56536.1 aver|OJJ07603.1 aver|OJJ06186.1

Group8490: acla|EAW07641.1 afis|EAW22661.1 afum|EAL90531.1 anid|CBF88219.1 asyd|OJJ63651.1 aver|OJI97283.1 acar|OOG00907.1 afla|EED49926.1 aory|EIT72276.1 ater|EAU39047.1

Group8491: abra|OJJ67818.1 afis|EAW20294.1 afum|EAL86773.1 anid|CBF82005.1 anig|EHA20932.1 ater|EAU38015.1 aver|OJJ07617.1 abra|OJJ72559.1 anid|CBF79160.1 awen|OJJ36345.1

Group8492: abra|OJJ67823.1 acar|OOG00540.1 acla|EAW09506.1 akaw|GAA91838.1 aluc|OJZ82060.1 anig|EHA22192.1 atub|OJI84619.1 aver|OJI99601.1 anid|CBF78142.1 asyd|OJJ55556.1

Group8493: abra|OJJ67830.1 anig|EHA22200.1 abra|OJJ68154.1 acla|EAW11666.1 afla|EED47517.1 aory|EIT74525.1 ater|EAU32820.1 awen|OJJ38754.1 afla|EED53909.1 aory|EIT78479.1

Group8494: akaw|GAA91864.1 ater|EAU31318.1 acar|OOF92836.1 acla|EAW11742.1 afis|EAW23775.1 afla|EED51575.1 afum|EAL87209.1 agla|OJJ78926.1 anid|CBF79826.1 aory|EIT80582.1

Group8495: abra|OJJ67952.1 afla|EED52892.1 agla|OJJ83937.1 akaw|GAA91245.1 aluc|OJZ82825.1 aory|EIT78643.1 asyd|OJJ54510.1 atub|OJI89054.1 aver|OJJ07673.1 awen|OJJ33890.1

Group8496: ater|EAU32068.1 acar|OOF95577.1 afla|EED48704.1 akaw|GAA86621.1 aluc|OJZ85176.1 anid|CBF78739.1 anig|EHA23743.1 aory|EIT73057.1 acar|OOF98774.1 atub|OJI79791.1

Group8497: abra|OJJ67996.1 acar|OOF92039.1 afis|EAW16855.1 afla|EED48418.1 akaw|GAA90953.1 aluc|OJZ82871.1 aory|EIT83059.1 atub|OJI89094.1 awen|OJJ31329.1 azon|OJJ42617.1

Group8498: ater|EAU33696.1 abra|OJJ77550.1 afla|EED45583.1 akaw|GAA84073.1 aluc|OJZ80987.1 anig|EHA19373.1 aory|EIT75671.1 atub|OJI82599.1 aver|OJI98668.1 awen|OJJ30215.1

Group8499: abra|OJJ68095.1 acar|OOF93898.1 afla|EED45837.1 akaw|GAA86879.1 aluc|OJZ90003.1 anid|CBF86409.1 anig|EHA27378.1 asyd|OJJ64945.1 atub|OJI84805.1 aver|OJI99067.1

Group8500: abra|OJJ68100.1 acar|OOF93903.1 afla|EED47252.1 akaw|GAA86885.1 aluc|OJZ90008.1 anig|EHA27383.1 aory|EIT79239.1 atub|OJI84799.1 azon|OJJ44487.1 awen|OJJ35121.1

Group8501: acar|OOF95726.1 afis|EAW16737.1 afum|EAL86312.1 anid|CBF75876.1 acla|EAW12430.1 agla|OJJ82712.1 akaw|GAA81901.1 aluc|OJZ92297.1 asyd|OJJ60231.1 aver|OJJ04035.1

Group8502: abra|OJJ68135.1 acar|OOG00330.1 afla|EED45836.1 akaw|GAA86915.1 aluc|OJZ90038.1 anig|EHA27413.1 aory|EIT75832.1 ater|EAU38812.1 atub|OJI84768.1 aver|OJJ07591.1

Group8503: abra|OJJ68157.1 acla|EAW11669.1 afla|EED47514.1 akaw|GAA90126.1 aluc|OJZ83224.1 anig|EHA21081.1 aory|EIT74585.1 atub|OJI87712.1 awen|OJJ38758.1 azon|OJJ42175.1

Group8504: abra|OJJ68163.1 acar|OOF90232.1 afla|EED54707.1 afum|EAL84366.1 aory|EIT78936.1 asyd|OJJ52027.1 ater|EAU29956.1 aver|OJJ06576.1 azon|OJJ43611.1 asyd|OJJ55498.1

Group8505: abra|OJJ68256.1 acar|OOF92459.1 afla|EED57066.1 agla|OJJ80955.1 akaw|GAA92929.1 aluc|OJZ82522.1 aory|EIT82301.1 ater|EAU32759.1 atub|OJI88771.1 ater|EAU31618.1

Group8506: abra|OJJ68263.1 acar|OOF97540.1 afla|EED56315.1 akaw|GAA83202.1 aluc|OJZ83703.1 aory|EIT73351.1 asyd|OJJ52858.1 atub|OJI90075.1 aver|OJJ08835.1 ater|EAU38457.1

Group8507: afis|EAW19797.1 afla|EED55700.1 afum|EAL85093.1 acar|OOF99879.1 acla|EAW13763.1 agla|OJJ80903.1 aory|EIT77079.1 awen|OJJ32634.1 azon|OJJ45126.1 anig|EHA18746.1

Group8508: abra|OJJ68283.1 acar|OOF97557.1 afla|EED56847.1 akaw|GAA83189.1 aluc|OJZ83688.1 anig|EHA18842.1 aory|EIT77513.1 atub|OJI90060.1 awen|OJJ40736.1 ater|EAU39316.1

Group8509: abra|OJJ68303.1 afla|EED48106.1 akaw|GAA83171.1 aluc|OJZ83670.1 anig|EHA18825.1 aory|EIT76195.1 asyd|OJJ56506.1 atub|OJI90041.1 aver|OJJ06213.1 aory|EIT76252.1

Group8510: acla|EAW15043.1 afis|EAW24566.1 afum|EAL86229.1 anid|CBF82903.1 aory|EIT72962.1 asyd|OJJ63345.1 afla|EED48256.1 aluc|OJZ81377.1 atub|OJI91583.1 aver|OJI97615.1

Group8511: abra|OJJ68326.1 acar|OOF90097.1 akaw|GAA92077.1 aluc|OJZ83647.1 anig|EHA18799.1 aory|EIT83111.1 ater|EAU35876.1 atub|OJI90016.1 awen|OJJ29972.1 azon|OJJ50258.1

Group8512: abra|OJJ68333.1 acar|OOF97605.1 akaw|GAA92085.1 aluc|OJZ83638.1 anid|CBF87144.1 anig|EHA18790.1 atub|OJI90007.1 azon|OJJ50659.1 asyd|OJJ60298.1 aver|OJI99639.1

Group8513: afla|EED49557.1 abra|OJJ77650.1 acar|OOF99086.1 acla|EAW12090.1 akaw|GAA83985.1 aluc|OJZ80879.1 anig|EHA19457.1 atub|OJI82698.1 aver|OJI98439.1 aory|EIT82606.1

Group8514: acla|EAW14966.1 afis|EAW24489.1 afum|EAL86158.1 afla|EED52188.1 agla|OJJ83083.1 aory|EIT74403.1 asyd|OJJ54573.1 ater|EAU31589.1 aver|OJJ07581.1 awen|OJJ31458.1

Group8515: abra|OJJ68437.1 acar|OOF94352.1 acla|EAW06852.1 afum|EAL89861.2 agla|OJJ82172.1 asyd|OJJ52065.1 ater|EAU32989.1 atub|OJI84747.1 azon|OJJ48405.1 aluc|OJZ90060.1

Group8516: acar|OOF97878.1 agla|OJJ79108.1 awen|OJJ32534.1 acla|EAW14865.1 afla|EED45241.1 aory|EIT73666.1 asyd|OJJ65108.1 aver|OJI99235.1 asyd|OJJ55396.1 aver|OJJ08261.1

Group8517: abra|OJJ68585.1 acar|OOF91918.1 afla|EED46153.1 afum|EAL85013.1 agla|OJJ79202.1 aory|EIT76463.1 atub|OJI91457.1 awen|OJJ29959.1 aver|OJJ07751.1 ater|EAU34352.1

Group8518: abra|OJJ68597.1 afis|EAW16795.1 afla|EED51091.1 afum|EAL86364.1 agla|OJJ84441.1 akaw|GAA91921.1 aluc|OJZ81959.1 anig|EHA22288.1 aory|EIT81998.1 atub|OJI84510.1

Group8519: abra|OJJ68645.1 acar|OOF90075.1 afla|EED54068.1 akaw|GAA89785.1 aluc|OJZ81906.1 anig|EHA22337.1 aory|EIT75989.1 atub|OJI84459.1 azon|OJJ50287.1 azon|OJJ50765.1

Group8520: agla|OJJ87145.1 afis|EAW24076.1 afla|EED51397.1 afum|EAL91931.2 aory|EIT80467.1 awen|OJJ41414.1 azon|OJJ48878.1 anid|CBF86472.1 ater|EAU30497.1 aver|OJI99027.1

Group8521: abra|OJJ68830.1 afla|EED57507.1 afum|EAL86206.2 agla|OJJ83139.1 aluc|OJZ85808.1 anid|CBF78853.1 asyd|OJJ55032.1 atub|OJI79594.1 aver|OJJ07171.1 awen|OJJ30829.1

Group8522: agla|OJJ81776.1 acla|EAW14268.1 afis|EAW19010.1 afum|EAL93744.1 anid|CBF86741.1 aory|EIT74782.1 asyd|OJJ64697.1 ater|EAU37792.1 aver|OJI98802.1 awen|OJJ31636.1

Group8523: abra|OJJ68890.1 afla|EED46933.1 agla|OJJ83414.1 akaw|GAA90366.1 aluc|OJZ85866.1 aory|EIT72536.1 asyd|OJJ52059.1 ater|EAU33158.1 atub|OJI79652.1 aver|OJI99473.1

Group8524: abra|OJJ69017.1 acar|OOF91642.1 afla|EED51547.1 akaw|GAA90534.1 aluc|OJZ86003.1 anig|EHA24607.1 aory|EIT80625.1 asyd|OJJ60637.1 ater|EAU38822.1 atub|OJI79790.1

Group8525: abra|OJJ69142.1 acar|OOF91160.1 afis|EAW15742.1 afum|EAL85606.1 akaw|GAA86315.1 aluc|OJZ86128.1 asyd|OJJ52667.1 ater|EAU32403.1 atub|OJI79924.1 awen|OJJ38442.1

Group8526: abra|OJJ69541.1 acar|OOF90593.1 afla|EED45994.1 akaw|GAA88389.1 aluc|OJZ82377.1 anig|EHA28387.1 aory|EIT75564.1 atub|OJI88065.1 awen|OJJ30869.1 azon|OJJ42099.1

Group8527: abra|OJJ69567.1 afum|EAL86533.1 asyd|OJJ53776.1 aver|OJJ07192.1 afla|EED49428.1 agla|OJJ82291.1 afla|EED49427.1 aver|OJJ03476.1 aory|EIT79583.1 asyd|OJJ52086.1

Group8528: abra|OJJ69606.1 akaw|GAA87142.1 aluc|OJZ84295.1 aory|EIT81683.1 asyd|OJJ65403.1 atub|OJI89774.1 aver|OJI95768.1 awen|OJJ30863.1 afla|EED51836.1 awen|OJJ38681.1

Group8529: abra|OJJ69614.1 acar|OOF92692.1 afla|EED51844.1 akaw|GAA91598.1 aluc|OJZ82422.1 anig|EHA28347.1 aory|EIT81748.1 ater|EAU36721.1 atub|OJI88019.1 awen|OJJ33375.1

Group8530: abra|OJJ69618.1 acar|OOF92688.1 afis|EAW19835.1 afum|EAL85057.1 akaw|GAA91602.1 aluc|OJZ82427.1 atub|OJI88014.1 azon|OJJ43516.1 awen|OJJ38281.1 ater|EAU37434.1

Group8531: abra|OJJ69619.1 acar|OOF92687.1 akaw|GAA91603.1 aluc|OJZ82428.1 anig|EHA28343.1 asyd|OJJ54527.1 atub|OJI88013.1 aver|OJJ03006.1 awen|OJJ36839.1 azon|OJJ43515.1

Group8532: abra|OJJ69624.1 akaw|GAA92927.1 aluc|OJZ82518.1 anid|CBF81979.1 asyd|OJJ56153.1 atub|OJI88774.1 aver|OJJ05092.1 afla|EED56560.1 anig|EHA20901.1 awen|OJJ33377.1

Group8533: abra|OJJ69633.1 agla|OJJ89114.1 akaw|GAA91613.1 aory|EIT79306.1 asyd|OJJ54598.1 ater|EAU30447.1 atub|OJI88001.1 aver|OJJ07554.1 azon|OJJ50114.1 afla|EED47295.1

Group8534: abra|OJJ69660.1 acar|OOF99512.1 afis|EAW20409.1 afum|EAL94101.1 akaw|GAA91640.1 aluc|OJZ82467.1 anig|EHA28308.1 atub|OJI87971.1 awen|OJJ32178.1 afum|EAL89235.1

Group8535: abra|OJJ69679.1 acar|OOF92106.1 afla|EED56765.1 akaw|GAA88804.1 aluc|OJZ82487.1 anig|EHA28287.1 ater|EAU32563.1 atub|OJI87945.1 aver|OJI96617.1 ater|EAU33068.1

Group8536: abra|OJJ69684.1 acar|OOF93995.1 afla|EED52268.1 akaw|GAA88808.1 aluc|OJZ82491.1 anig|EHA28281.1 asyd|OJJ62639.1 atub|OJI87941.1 aver|OJI99290.1 awen|OJJ38313.1

Group8537: abra|OJJ69709.1 acar|OOF94203.1 akaw|GAA90158.1 aluc|OJZ83189.1 anig|EHA28259.1 atub|OJI87750.1 aver|OJJ07620.1 awen|OJJ36347.1 acar|OOF91992.1 anig|EHA20935.1

Group8538: anid|CBF84263.1 asyd|OJJ57213.1 aver|OJJ03076.1 azon|OJJ45849.1 agla|OJJ78889.1 acla|EAW09099.1 afis|EAW25695.1 afum|EAL84369.1 ater|EAU31571.1 awen|OJJ29700.1

Group8539: abra|OJJ69749.1 acar|OOF99050.1 afis|EAW19964.1 afum|EAL84945.2 akaw|GAA90200.1 aluc|OJZ83148.1 anig|EHA28213.1 asyd|OJJ52906.1 atub|OJI87789.1 awen|OJJ35861.1

Group8540: abra|OJJ69797.1 acar|OOF92534.1 akaw|GAA87862.1 aluc|OJZ85503.1 atub|OJI80062.1 aver|OJJ02051.1 ater|EAU33076.1 anid|CBF82435.1 asyd|OJJ58356.1 azon|OJJ42230.1

Group8541: abra|OJJ69802.1 acar|OOF93575.1 afis|EAW24738.1 afum|EAL89385.1 akaw|GAA87857.1 aluc|OJZ85499.1 anig|EHA25032.1 atub|OJI80066.1 asyd|OJJ55919.1 aver|OJJ04872.1

Group8542: abra|OJJ69842.1 acar|OOF93365.1 akaw|GAA87818.1 aluc|OJZ85452.1 anig|EHA24996.1 atub|OJI80112.1 akaw|GAA87817.1 atub|OJI80114.1 anig|EHA24995.1 aver|OJI95871.1

Group8543: abra|OJJ69844.1 acar|OOF93367.1 akaw|GAA87815.1 aluc|OJZ85450.1 anig|EHA24993.1 asyd|OJJ61770.1 ater|EAU31226.1 atub|OJI80117.1 aver|OJI95872.1 anid|CBF84353.1

Group8544: afla|EED53085.1 aory|EIT74813.1 ater|EAU38424.1 awen|OJJ32258.1 acla|EAW09120.1 atub|OJI80560.1 aver|OJJ05803.1 asyd|OJJ57364.1 anid|CBF82277.1 asyd|OJJ54899.1

Group8545: afum|EAL90594.1 acla|EAW07570.1 afis|EAW22604.1 afla|EED53103.1 anid|CBF73873.1 aory|EIT74774.1 asyd|OJJ56549.1 ater|EAU31548.1 aver|OJJ06173.1 azon|OJJ44953.1

Group8546: acla|EAW15171.1 afis|EAW20140.1 afla|EED47645.1 afum|EAL85255.1 asyd|OJJ52227.1 acar|OOG00545.1 agla|OJJ88796.1 anid|CBF78273.1 aory|EIT80255.1 awen|OJJ32009.1

Group8547: abra|OJJ70095.1 acar|OOF97934.1 afis|EAW17169.1 akaw|GAA87600.1 aluc|OJZ85220.1 anig|EHA25424.1 asyd|OJJ61499.1 ater|EAU34028.1 atub|OJI80363.1 aver|OJJ00836.1

Group8548: abra|OJJ70109.1 acar|OOF92006.1 afis|EAW25650.1 afla|EED46124.1 agla|OJJ81747.1 akaw|GAA92807.1 aluc|OJZ88730.1 anig|EHA18434.1 aver|OJJ05888.1 azon|OJJ50374.1

Group8549: aory|EIT80953.1 asyd|OJJ52682.1 akaw|GAA92001.1 aluc|OJZ92872.1 atub|OJI87330.1 acar|OOF98683.1 acla|EAW08794.1 aver|OJJ08668.1 azon|OJJ45038.1 acla|EAW11039.1

Group8550: asyd|OJJ56560.1 abra|OJJ77608.1 acar|OOF91822.1 afla|EED52319.1 akaw|GAA84021.1 aluc|OJZ80925.1 anig|EHA19422.1 atub|OJI82653.1 azon|OJJ44042.1 aver|OJJ06161.1

Group8551: abra|OJJ70205.1 acar|OOF92574.1 akaw|GAA86572.1 aluc|OJZ85122.1 anig|EHA25342.1 atub|OJI80471.1 awen|OJJ36218.1 abra|OJJ76882.1 agla|OJJ83401.1 asyd|OJJ56898.1

Group8552: abra|OJJ70225.1 acar|OOF95982.1 afla|EED48241.1 akaw|GAA92567.1 aluc|OJZ82577.1 anig|EHA25326.1 atub|OJI80492.1 azon|OJJ42530.1 azon|OJJ42788.1 anid|CBF88083.1

Group8553: abra|OJJ70238.1 acar|OOF92556.1 akaw|GAA86547.1 aluc|OJZ85095.1 anig|EHA25317.1 atub|OJI80503.1 aver|OJI96728.1 abra|OJJ71438.1 agla|OJJ82297.1 awen|OJJ41773.1

Group8554: abra|OJJ70239.1 acar|OOF92555.1 afla|EED47781.1 akaw|GAA86546.1 aluc|OJZ85094.1 anig|EHA25316.1 aory|EIT77105.1 asyd|OJJ55212.1 atub|OJI80504.1 aver|OJJ07003.1

Group8555: abra|OJJ70260.1 acar|OOF90564.1 afis|EAW15398.1 afum|EAL84250.1 agla|OJJ89320.1 akaw|GAA86537.1 aluc|OJZ85084.1 anig|EHA25299.1 ater|EAU32205.1 atub|OJI80514.1

Group8556: abra|OJJ70313.1 afis|EAW17105.1 afla|EED47134.1 akaw|GAA92569.1 aluc|OJZ82579.1 anig|EHA27492.1 ater|EAU29939.1 atub|OJI88820.1 awen|OJJ37982.1 aory|EIT79243.1

Group8557: abra|OJJ70340.1 afla|EED48271.1 agla|OJJ88107.1 akaw|GAA92593.1 aluc|OJZ82610.1 anig|EHA27472.1 aory|EIT72917.1 asyd|OJJ65381.1 atub|OJI88845.1 aver|OJI99551.1

Group8558: abra|OJJ70348.1 afla|EED48623.1 akaw|GAA92600.1 aluc|OJZ82617.1 anig|EHA27460.1 aory|EIT73276.1 asyd|OJJ60739.1 atub|OJI88852.1 aver|OJJ00071.1 awen|OJJ33353.1

Group8559: abra|OJJ70349.1 afla|EED51524.1 agla|OJJ88978.1 akaw|GAA83850.1 aluc|OJZ90865.1 anig|EHA19578.1 aory|EIT80574.1 atub|OJI82863.1 aver|OJJ03457.1 awen|OJJ38030.1

Group8560: abra|OJJ70372.1 acar|OOF91897.1 afla|EED47359.1 akaw|GAA91978.1 aluc|OJZ82636.1 anig|EHA27443.1 ater|EAU30229.1 atub|OJI88867.1 awen|OJJ29760.1 agla|OJJ79931.1

Group8561: abra|OJJ70386.1 acar|OOF91885.1 akaw|GAA91971.1 aluc|OJZ82644.1 anig|EHA27426.1 asyd|OJJ56716.1 atub|OJI88877.1 aver|OJJ06009.1 awen|OJJ29702.1 azon|OJJ50134.1

Group8562: abra|OJJ70401.1 acar|OOF98650.1 acla|EAW07591.1 afum|EAL91131.1 akaw|GAA87027.1 aluc|OJZ80469.1 ater|EAU34897.1 atub|OJI89648.1 azon|OJJ43571.1 aluc|OJZ84425.1

Group8563: abra|OJJ70414.1 acar|OOF91858.1 afla|EED57576.1 akaw|GAA91942.1 aluc|OJZ82678.1 anig|EHA21047.1 aory|EIT82964.1 asyd|OJJ65358.1 aver|OJI99507.1 awen|OJJ40094.1

Group8564: abra|OJJ70426.1 afis|EAW20641.1 afum|EAL84429.1 agla|OJJ84763.1 akaw|GAA90151.1 aluc|OJZ83197.1 anig|EHA21060.1 ater|EAU32980.1 atub|OJI87741.1 aver|OJI99688.1

Group8565: abra|OJJ70437.1 acar|OOF91845.1 afla|EED45396.1 agla|OJJ78985.1 akaw|GAA90143.1 aluc|OJZ83204.1 anig|EHA21066.1 aory|EIT82379.1 atub|OJI87733.1 azon|OJJ48044.1

Group8566: abra|OJJ70457.1 acar|OOF95615.1 acla|EAW12032.1 afla|EED56210.1 akaw|GAA83303.1 aluc|OJZ83798.1 anig|EHA18946.1 aory|EIT73342.1 atub|OJI90167.1 azon|OJJ43168.1

Group8567: abra|OJJ70461.1 akaw|GAA83307.1 aluc|OJZ83804.1 anig|EHA18951.1 atub|OJI90171.1 abra|OJJ77762.1 afis|EAW16209.1 afum|EAL88836.1 asyd|OJJ57955.1 ater|EAU38498.1

Group8568: abra|OJJ70480.1 afis|EAW16858.1 afla|EED48435.1 afum|EAL86401.1 aory|EIT81680.1 asyd|OJJ55447.1 aver|OJJ06520.1 awen|OJJ35635.1 asyd|OJJ61385.1 aver|OJJ00713.1

Group8569: abra|OJJ70691.1 acar|OOF97037.1 acla|EAW08073.1 afis|EAW23109.1 afla|EED48539.1 afum|EAL88370.1 agla|OJJ89632.1 anig|EHA19160.1 ater|EAU36237.1 awen|OJJ36672.1

Group8570: abra|OJJ70934.1 acla|EAW10500.1 afis|EAW25067.1 afla|EED55387.1 afum|EAL89736.1 akaw|GAA83008.1 aluc|OJZ87593.1 anid|CBF85953.1 ater|EAU35814.1 atub|OJI81054.1

Group8571: abra|OJJ71003.1 afis|EAW25683.1 afla|EED55158.1 agla|OJJ78887.1 anid|CBF83931.1 anig|EHA18440.1 aory|EIT81575.1 asyd|OJJ55436.1 ater|EAU32720.1 aver|OJI96710.1

Group8572: abra|OJJ71234.1 acar|OOF97945.1 afla|EED45827.1 akaw|GAA82753.1 aluc|OJZ87310.1 anig|EHA20546.1 aory|EIT75462.1 atub|OJI81335.1 awen|OJJ40448.1 azon|OJJ45934.1

Group8573: abra|OJJ71235.1 acar|OOG00345.1 acla|EAW13487.1 afla|EED45437.1 akaw|GAA88047.1 aluc|OJZ81306.1 aory|EIT82398.1 atub|OJI91513.1 azon|OJJ47990.1 atub|OJI91514.1

Group8574: abra|OJJ71612.1 acar|OOF96328.1 afla|EED53953.1 agla|OJJ85548.1 akaw|GAA89315.1 anid|CBF87163.1 aory|EIT78260.1 atub|OJI84181.1 awen|OJJ29931.1 anid|CBF79656.1

Group8575: abra|OJJ71613.1 acar|OOF96329.1 afis|EAW19672.1 afla|EED46495.1 akaw|GAA89314.1 aory|EIT72331.1 atub|OJI84180.1 aory|EIT72401.1 afis|EAW19673.1 afla|EED46496.1

Group8576: abra|OJJ71635.1 acar|OOF96352.1 afla|EED54063.1 akaw|GAA89291.1 aluc|OJZ86378.1 anig|EHA21516.1 aory|EIT76025.1 atub|OJI84156.1 azon|OJJ50320.1 acar|OOF92013.1

Group8577: abra|OJJ71707.1 afis|EAW24690.1 afla|EED49331.1 afum|EAL89331.1 anig|EHA22157.1 aory|EIT79538.1 awen|OJJ35380.1 azon|OJJ50279.1 afla|EED52321.1 aory|EIT74480.1

Group8578: abra|OJJ71743.1 acar|OOF96434.1 akaw|GAA85157.1 aluc|OJZ86480.1 asyd|OJJ58743.1 atub|OJI84050.1 aver|OJJ01353.1 awen|OJJ31112.1 azon|OJJ46588.1 acla|EAW06447.1

Group8579: abra|OJJ71747.1 afla|EED48442.1 agla|OJJ79876.1 akaw|GAA85161.1 aluc|OJZ86475.1 anig|EHA22160.1 aory|EIT81672.1 ater|EAU35870.1 atub|OJI84054.1 awen|OJJ37784.1

Group8580: abra|OJJ71779.1 acar|OOF90183.1 afis|EAW17531.1 afum|EAL93576.2 agla|OJJ80575.1 akaw|GAA85130.1 aluc|OJZ86510.1 anig|EHA22137.1 awen|OJJ34616.1 azon|OJJ49597.1

Group8581: abra|OJJ71792.1 acar|OOF96472.1 afla|EED56592.1 akaw|GAA85118.1 aluc|OJZ86522.1 anig|EHA22124.1 asyd|OJJ54323.1 atub|OJI84000.1 aver|OJJ05277.1 awen|OJJ30244.1

Group8582: abra|OJJ71826.1 acar|OOF96503.1 akaw|GAA85084.1 aluc|OJZ86558.1 anid|CBF85400.1 anig|EHA22091.1 asyd|OJJ64061.1 ater|EAU39384.1 atub|OJI83964.1 aver|OJI98147.1

Group8583: abra|OJJ71839.1 acar|OOF90147.1 acla|EAW13870.1 afla|EED52765.1 afum|EAL93285.1 agla|OJJ81633.1 aluc|OJZ86572.1 atub|OJI83951.1 awen|OJJ31145.1 azon|OJJ46620.1

Group8584: abra|OJJ71914.1 acar|OOF96572.1 agla|OJJ82337.1 akaw|GAA85007.1 aluc|OJZ86651.1 anig|EHA22019.1 asyd|OJJ54493.1 ater|EAU33699.1 atub|OJI83876.1 anid|CBF80791.1

Group8585: abra|OJJ71932.1 acar|OOF96589.1 acla|EAW07948.1 afis|EAW19524.1 akaw|GAA84991.1 aluc|OJZ86669.1 ater|EAU39264.1 atub|OJI83858.1 awen|OJJ29589.1 azon|OJJ45027.1

Group8586: abra|OJJ71965.1 acar|OOF89891.1 afla|EED56021.1 akaw|GAA84963.1 aluc|OJZ86701.1 anig|EHA21975.1 aory|EIT76827.1 atub|OJI83825.1 awen|OJJ39121.1 azon|OJJ51451.1

Group8587: abra|OJJ72076.1 akaw|GAA84841.1 aluc|OJZ86830.1 anig|EHA21805.1 atub|OJI83688.1 abra|OJJ76924.1 acar|OOF91616.1 afla|EED50450.1 aory|EIT82050.1 aver|OJI98301.1

Group8588: abra|OJJ72084.1 acar|OOF96729.1 akaw|GAA84836.1 aluc|OJZ86838.1 anid|CBF69410.1 anig|EHA21799.1 asyd|OJJ63819.1 atub|OJI83680.1 aver|OJI97119.1 azon|OJJ51797.1

Group8589: agla|OJJ84829.1 ater|EAU38680.1 afla|EED53378.1 aory|EIT75163.1 awen|OJJ37899.1 afla|EED55748.1 aory|EIT83201.1 asyd|OJJ52198.1 aver|OJI98022.1 ater|EAU37793.1

Group8590: abra|OJJ72109.1 acar|OOF90021.1 akaw|GAA84813.1 aluc|OJZ86864.1 anig|EHA21777.1 aory|EIT75415.1 atub|OJI83656.1 awen|OJJ37056.1 afla|EED45957.1 afla|EED45958.1

Group8591: abra|OJJ72136.1 afis|EAW18424.1 agla|OJJ82206.1 akaw|GAA84786.1 aluc|OJZ86894.1 anid|CBF77133.1 anig|EHA21749.1 asyd|OJJ56944.1 atub|OJI83625.1 aver|OJJ03340.1

Group8592: abra|OJJ72153.1 acar|OOF93888.1 acla|EAW06696.1 akaw|GAA83563.1 aluc|OJZ92435.1 asyd|OJJ57413.1 ater|EAU37618.1 afla|EED49526.1 atub|OJI86864.1 awen|OJJ40937.1

Group8593: abra|OJJ72179.1 afis|EAW15945.1 afum|EAL88579.1 abra|OJJ72619.1 acar|OOF90209.1 akaw|GAA92011.1 aluc|OJZ92876.1 anig|EHA26083.1 atub|OJI87336.1 azon|OJJ43898.1

Group8594: agla|OJJ82503.1 asyd|OJJ52341.1 azon|OJJ42372.1 acar|OOF93295.1 aluc|OJZ84275.1 anig|EHA27619.1 asyd|OJJ65289.1 atub|OJI89806.1 akaw|GAA90169.1 asyd|OJJ57146.1

Group8595: abra|OJJ72215.1 acar|OOF96836.1 afis|EAW23701.1 afum|EAL87142.1 agla|OJJ79039.1 akaw|GAA84719.1 aluc|OJZ86967.1 atub|OJI83555.1 acla|EAW06801.1 awen|OJJ37753.1

Group8596: abra|OJJ72216.1 acar|OOF96838.1 agla|OJJ89329.1 aluc|OJZ86968.1 anig|EHA21682.1 asyd|OJJ54158.1 atub|OJI83554.1 aver|OJJ08582.1 awen|OJJ29881.1 acar|OOG00379.1

Group8597: abra|OJJ72248.1 acar|OOF96877.1 afla|EED48185.1 agla|OJJ84382.1 akaw|GAA84690.1 aluc|OJZ86998.1 anig|EHA21648.1 atub|OJI83521.1 awen|OJJ32733.1 azon|OJJ46152.1

Group8598: aory|EIT75256.1 akaw|GAA88127.1 asyd|OJJ62937.1 aver|OJI98054.1 asyd|OJJ61690.1 afis|EAW15654.1 anig|EHA18692.1 atub|OJI91605.1 awen|OJJ31668.1 aluc|OJZ81399.1

Group8599: abra|OJJ72371.1 acar|OOF94853.1 afis|EAW19750.1 afla|EED56959.1 afum|EAL85147.2 akaw|GAA86121.1 aluc|OJZ90360.1 anig|EHA23831.1 aory|EIT77558.1 atub|OJI81787.1

Group8600: aver|OJJ06961.1 acla|EAW12681.1 afis|EAW17844.1 afla|EED46191.1 afum|EAL91653.1 anid|CBF77969.1 aory|EIT76376.1 asyd|OJJ55265.1 ater|EAU33046.1 azon|OJJ48429.1

Group8601: abra|OJJ72529.1 acar|OOF94724.1 afla|EED49525.1 agla|OJJ86479.1 akaw|GAA86251.1 aluc|OJZ90508.1 anig|EHA23965.1 aory|EIT82605.1 asyd|OJJ54907.1 atub|OJI81932.1

Group8602: abra|OJJ72630.1 afla|EED52588.1 anig|EHA24045.1 aory|EIT74244.1 asyd|OJJ56819.1 aver|OJJ03516.1 akaw|GAA85250.1 aluc|OJZ90607.1 atub|OJI82031.1 anid|CBF77224.1

Group8603: abra|OJJ72692.1 acar|OOF95537.1 afis|EAW19896.1 afla|EED51786.1 afum|EAL85005.1 agla|OJJ86286.1 aory|EIT81709.1 asyd|OJJ52354.1 aver|OJI98306.1 aory|EIT81770.1

Group8604: abra|OJJ72852.1 acar|OOF98952.1 afla|EED47911.1 akaw|GAA85453.1 aluc|OJZ83361.1 anig|EHA24226.1 aory|EIT83379.1 atub|OJI82252.1 azon|OJJ42496.1 afum|EAL85699.2

Group8605: abra|OJJ72901.1 acar|OOF89982.1 afis|EAW18144.1 afum|EAL87501.1 akaw|GAA85488.1 aluc|OJZ83410.1 anig|EHA24261.1 ater|EAU35466.1 atub|OJI82298.1 awen|OJJ29667.1

Group8606: abra|OJJ73021.1 acar|OOF96189.1 acla|EAW07994.1 afis|EAW23029.1 agla|OJJ89450.1 aluc|OJZ83534.1 asyd|OJJ56325.1 atub|OJI82425.1 aver|OJJ06388.1 azon|OJJ44175.1

Group8607: abra|OJJ73075.1 acar|OOG00310.1 agla|OJJ83255.1 akaw|GAA88086.1 aluc|OJZ81350.1 anig|EHA18652.1 atub|OJI91554.1 awen|OJJ40004.1 azon|OJJ50412.1 awen|OJJ37994.1

Group8608: abra|OJJ73086.1 acar|OOG00319.1 afla|EED57717.1 akaw|GAA88074.1 anig|EHA18643.1 atub|OJI91542.1 aver|OJJ06545.1 awen|OJJ34511.1 aory|EIT79944.1 agla|OJJ80281.1

Group8609: abra|OJJ73115.1 afla|EED47395.1 akaw|GAA88054.1 aluc|OJZ81316.1 anig|EHA18620.1 aory|EIT77357.1 ater|EAU38561.1 atub|OJI91519.1 acla|EAW13710.1 ater|EAU34083.1

Group8610: abra|OJJ73300.1 acar|OOF91872.1 afis|EAW20652.1 akaw|GAA92705.1 aluc|OJZ88745.1 anig|EHA18449.1 asyd|OJJ60305.1 atub|OJI91317.1 aver|OJI99646.1 awen|OJJ34595.1

Group8611: abra|OJJ73351.1 acar|OOF98071.1 afla|EED55527.1 agla|OJJ85523.1 akaw|GAA83132.1 aluc|OJZ88688.1 anig|EHA18368.1 aory|EIT80909.1 atub|OJI91268.1 awen|OJJ40292.1

Group8612: abra|OJJ73360.1 acar|OOF96593.1 acla|EAW09766.1 akaw|GAA83124.1 aluc|OJZ88680.1 anig|EHA18361.1 aory|EIT76747.1 atub|OJI91261.1 awen|OJJ34492.1 azon|OJJ43956.1

Group8613: abra|OJJ73390.1 acar|OOG00592.1 afla|EED45860.1 agla|OJJ87868.1 akaw|GAA83099.1 aluc|OJZ88646.1 anig|EHA18337.1 aory|EIT75773.1 ater|EAU35441.1 atub|OJI91233.1

Group8614: abra|OJJ73426.1 acar|OOG00630.1 agla|OJJ78773.1 akaw|GAA83066.1 aluc|OJZ88617.1 anig|EHA18308.1 atub|OJI91202.1 awen|OJJ34076.1 azon|OJJ42202.1 atub|OJI84004.1

Group8615: abra|OJJ73428.1 acar|OOG00634.1 afis|EAW24000.1 afum|EAL87430.1 akaw|GAA83064.1 aluc|OJZ88615.1 anid|CBF89126.1 anig|EHA18306.1 ater|EAU37866.1 atub|OJI91200.1

Group8616: abra|OJJ73661.1 acar|OOG00841.1 afla|EED48352.1 akaw|GAA88495.1 aluc|OJZ88391.1 anig|EHA18118.1 atub|OJI90986.1 awen|OJJ38333.1 acla|EAW07714.1 afum|EAL86290.2

Group8617: abra|OJJ73666.1 afla|EED52813.1 agla|OJJ81786.1 aluc|OJZ88386.1 aory|EIT78202.1 asyd|OJJ60492.1 ater|EAU34949.1 atub|OJI90981.1 afla|EED51528.1 agla|OJJ85745.1

Group8618: abra|OJJ73690.1 acar|OOG00862.1 acla|EAW07605.1 afla|EED49886.1 agla|OJJ80983.1 aluc|OJZ88359.1 anid|CBF88225.1 atub|OJI90955.1 awen|OJJ35258.1 azon|OJJ49849.1

Group8619: abra|OJJ73726.1 acar|OOF93330.1 afla|EED56886.1 akaw|GAA88437.1 aluc|OJZ88325.1 anig|EHA18061.1 aory|EIT77470.1 ater|EAU34163.1 atub|OJI90920.1 azon|OJJ49300.1

Group8620: acar|OOG01154.1 aluc|OJZ88002.1 anig|EHA17783.1 atub|OJI90610.1 anid|CBF73534.1 asyd|OJJ62477.1 aver|OJI96614.1 afla|EED50986.1 aory|EIT76966.1 aver|OJJ03878.1

Group8621: abra|OJJ74148.1 acar|OOF95580.1 agla|OJJ78612.1 akaw|GAA89735.1 aluc|OJZ80648.1 anig|EHA27184.1 asyd|OJJ61972.1 atub|OJI87503.1 aver|OJI96065.1 awen|OJJ31296.1

Group8622: abra|OJJ74213.1 acar|OOF95507.1 akaw|GAA86805.1 aluc|OJZ89921.1 anig|EHA27121.1 asyd|OJJ54348.1 atub|OJI84884.1 aver|OJJ05300.1 afla|EED52429.1 aory|EIT83291.1

Group8623: abra|OJJ74233.1 acla|EAW07287.1 afis|EAW22320.1 agla|OJJ87744.1 akaw|GAA86786.1 aluc|OJZ89900.1 anig|EHA27100.1 atub|OJI84907.1 awen|OJJ34886.1 azon|OJJ50054.1

Group8624: agla|OJJ87659.1 acla|EAW07371.1 afis|EAW22396.1 afum|EAL90795.1 awen|OJJ34974.1 afla|EED56060.1 aory|EIT76623.1 asyd|OJJ63957.1 anid|CBF88748.1 aver|OJI96984.1

Group8625: acla|EAW07418.1 afis|EAW22443.1 afla|EED54459.1 aory|EIT81156.1 ater|EAU39261.1 azon|OJJ50038.1 aver|OJI96918.1 awen|OJJ35069.1 asyd|OJJ62865.1 anid|CBF88848.1

Group8626: afla|EED52294.1 afla|EED48293.1 aory|EIT76302.1 aluc|OJZ85149.1 anig|EHA25386.1 aory|EIT74332.1 ater|EAU37316.1 atub|OJI80423.1 akaw|GAA86601.1 aory|EIT74450.1

Group8627: abra|OJJ74459.1 acar|OOF95292.1 afla|EED44901.1 akaw|GAA87352.1 aluc|OJZ89662.1 anig|EHA26899.1 asyd|OJJ59445.1 atub|OJI85138.1 aver|OJJ02827.1 aory|EIT73487.1

Group8628: abra|OJJ74557.1 acar|OOF98302.1 acla|EAW12252.1 afis|EAW16984.1 afla|EED54813.1 afum|EBA27275.1 agla|OJJ82691.1 aory|EIT79339.1 ater|EAU32971.1 azon|OJJ44320.1

Group8629: abra|OJJ75060.1 acar|OOF97273.1 acla|EAW08244.1 afis|EAW23278.1 afum|EAL88201.1 agla|OJJ86568.1 akaw|GAA88673.1 aory|EIT77076.1 asyd|OJJ62386.1 ater|EAU37563.1

Group8630: abra|OJJ75143.1 akaw|GAA88749.1 aluc|OJZ92709.1 anid|CBF74888.1 anig|EHA26290.1 asyd|OJJ60106.1 ater|EAU35605.1 atub|OJI87135.1 aver|OJJ04156.1 awen|OJJ39533.1

Group8631: abra|OJJ75163.1 acar|OOF97175.1 akaw|GAA88774.1 anig|EHA26268.1 atub|OJI87157.1 agla|OJJ78946.1 ater|EAU35626.1 aluc|OJZ92733.1 aluc|OJZ92734.1 atub|OJI87158.1

Group8632: abra|OJJ75271.1 afla|EED48967.1 agla|OJJ84613.1 akaw|GAA91809.1 aluc|OJZ92831.1 anig|EHA26178.1 asyd|OJJ52915.1 atub|OJI87288.1 aver|OJJ08885.1 awen|OJJ35978.1

Group8633: abra|OJJ75349.1 acar|OOF94487.1 afis|EAW24559.1 akaw|GAA92041.1 aluc|OJZ92911.1 anig|EHA26054.1 asyd|OJJ55939.1 atub|OJI87371.1 aver|OJJ04888.1 azon|OJJ43548.1

Group8634: abra|OJJ75355.1 acar|OOF94486.1 akaw|GAA92051.1 aluc|OJZ92921.1 anid|CBF80201.1 anig|EHA26048.1 asyd|OJJ57901.1 atub|OJI87382.1 aver|OJJ04493.1 ater|EAU29641.1

Group8635: abra|OJJ75367.1 afla|EED55374.1 agla|OJJ87991.1 akaw|GAA92060.1 aluc|OJZ92936.1 anig|EHA26036.1 aory|EIT80789.1 ater|EAU29840.1 atub|OJI87396.1 aluc|OJZ92933.1

Group8636: abra|OJJ75557.1 acar|OOF90875.1 agla|OJJ88523.1 akaw|GAA84312.1 aluc|OJZ89007.1 anig|EHA23521.1 aory|EIT75455.1 asyd|OJJ59698.1 atub|OJI85782.1 aver|OJJ02589.1

Group8637: abra|OJJ75566.1 acar|OOF90883.1 afla|EED45935.1 akaw|GAA84318.1 aluc|OJZ89014.1 anig|EHA23516.1 aory|EIT75515.1 atub|OJI85775.1 awen|OJJ35902.1 azon|OJJ42467.1

Group8638: abra|OJJ75596.1 acar|OOF93959.1 acla|EAW11912.1 afla|EED52181.1 agla|OJJ79875.1 akaw|GAA84346.1 aluc|OJZ89047.1 anig|EHA23489.1 aory|EIT74345.1 atub|OJI85740.1

Group8639: abra|OJJ75603.1 acar|OOG00234.1 acla|EAW09326.1 afis|EAW21783.1 afla|EED46612.1 afum|EBA27474.1 akaw|GAA84354.1 aluc|OJZ89055.1 aory|EIT77866.1 atub|OJI85732.1

Group8640: abra|OJJ75727.1 acar|OOF90229.1 aluc|OJZ89179.1 anid|CBF83676.1 anig|EHA23376.1 aory|EIT81144.1 asyd|OJJ60814.1 atub|OJI85610.1 aver|OJJ00141.1 awen|OJJ29739.1

Group8641: abra|OJJ75742.1 acar|OOF91837.1 afla|EED48726.1 akaw|GAA84475.1 aluc|OJZ89195.1 anig|EHA23363.1 asyd|OJJ56157.1 atub|OJI85595.1 aver|OJJ05097.1 awen|OJJ40001.1

Group8642: abra|OJJ75979.1 acar|OOF99940.1 akaw|GAA88978.1 aluc|OJZ89425.1 anig|EHA23161.1 asyd|OJJ60649.1 atub|OJI85368.1 aver|OJI99969.1 awen|OJJ39382.1 anid|CBF83410.1

Group8643: abra|OJJ75990.1 afla|EED52371.1 akaw|GAA88967.1 aluc|OJZ89436.1 anid|CBF84666.1 aory|EIT82366.1 asyd|OJJ54807.1 ater|EAU32112.1 atub|OJI85357.1 aver|OJJ07354.1

Group8644: abra|OJJ76013.1 afla|EED55759.1 akaw|GAA88947.1 aluc|OJZ89459.1 anig|EHA23128.1 aory|EIT76777.1 asyd|OJJ54358.1 atub|OJI85335.1 aver|OJJ05312.1 awen|OJJ36190.1

Group8645: abra|OJJ76027.1 acar|OOF93545.1 afis|EAW17186.1 afum|EAL86612.1 akaw|GAA88931.1 aluc|OJZ89478.1 anig|EHA23115.1 atub|OJI85322.1 awen|OJJ31339.1 azon|OJJ42796.1

Group8646: abra|OJJ76034.1 acar|OOF99895.1 agla|OJJ87634.1 anig|EHA23109.1 asyd|OJJ58373.1 atub|OJI85316.1 aver|OJJ05952.1 afla|EED51787.1 aory|EIT81775.1 aluc|OJZ89486.1

Group8647: abra|OJJ76060.1 afla|EED55797.1 akaw|GAA88906.1 aluc|OJZ89515.1 anig|EHA23084.1 atub|OJI85288.1 agla|OJJ85503.1 aory|EIT76562.1 awen|OJJ39318.1 aver|OJJ00248.1

Group8648: abra|OJJ76065.1 acar|OOF99864.1 afis|EAW21389.1 agla|OJJ87676.1 akaw|GAA88901.1 aluc|OJZ89520.1 asyd|OJJ60915.1 atub|OJI85283.1 aver|OJJ00242.1 awen|OJJ39313.1

Group8649: abra|OJJ76244.1 acar|OOF99490.1 agla|OJJ89206.1 akaw|GAA82177.1 aluc|OJZ91987.1 anig|EHA22933.1 atub|OJI86422.1 azon|OJJ42359.1 anig|EHA18806.1 atub|OJI90024.1

Group8650: abra|OJJ76335.1 agla|OJJ80508.1 akaw|GAA82263.1 aluc|OJZ91877.1 anid|CBF69750.1 asyd|OJJ58939.1 atub|OJI86328.1 aver|OJJ01535.1 awen|OJJ32359.1 azon|OJJ47122.1

Group8651: abra|OJJ76398.1 acar|OOF99641.1 afla|EED50686.1 agla|OJJ82293.1 akaw|GAA82318.1 aluc|OJZ91814.1 anig|EHA22805.1 aory|EIT78087.1 atub|OJI86270.1 awen|OJJ32275.1

Group8652: agla|OJJ85516.1 acla|EAW09750.1 afis|EAW21392.1 afum|EAL92406.1 anid|CBF83827.1 asyd|OJJ60905.1 ater|EAU35175.1 aver|OJJ00231.1 azon|OJJ51539.1 afla|EED55807.1

Group8653: abra|OJJ76514.1 acar|OOF99458.1 afla|EED48156.1 agla|OJJ84830.1 akaw|GAA82427.1 aluc|OJZ91691.1 anig|EHA22704.1 atub|OJI86144.1 awen|OJJ36353.1 azon|OJJ44584.1

Group8654: abra|OJJ76603.1 acar|OOF99388.1 afla|EED46909.1 akaw|GAA82507.1 aluc|OJZ91605.1 anig|EHA22633.1 aory|EIT72679.1 ater|EAU34984.1 atub|OJI86057.1 azon|OJJ42097.1

Group8655: abra|OJJ76627.1 acar|OOF99371.1 acla|EAW14293.1 agla|OJJ82978.1 aluc|OJZ91585.1 asyd|OJJ64675.1 atub|OJI86034.1 aver|OJI98776.1 awen|OJJ31612.1 azon|OJJ43048.1

Group8656: abra|OJJ76695.1 acar|OOF99119.1 afis|EAW22868.1 afum|EAL90331.1 anid|CBF83074.1 asyd|OJJ65378.1 atub|OJI85966.1 aver|OJI99548.1 acar|OOF96679.1 awen|OJJ32261.1

Group8657: abra|OJJ76719.1 acar|OOF97650.1 afla|EED47261.1 agla|OJJ86864.1 akaw|GAA92481.1 aluc|OJZ91493.1 anig|EHA22535.1 aory|EIT79288.1 atub|OJI85948.1 awen|OJJ38036.1

Group8658: abra|OJJ76795.1 afis|EAW21126.1 afla|EED48222.1 akaw|GAA87230.1 aluc|OJZ84199.1 anig|EHA27556.1 aory|EIT72926.1 asyd|OJJ52878.1 ater|EAU37153.1 atub|OJI89885.1

Group8659: abra|OJJ76867.1 afla|EED51651.1 akaw|GAA87156.1 aluc|OJZ84278.1 anig|EHA27623.1 aory|EIT80683.1 asyd|OJJ62988.1 atub|OJI89803.1 aver|OJI97980.1 awen|OJJ30981.1

Group8660: abra|OJJ76877.1 afis|EAW16808.1 afla|EED49867.1 agla|OJJ85994.1 anid|CBF80479.1 asyd|OJJ65373.1 ater|EAU38552.1 aver|OJI99540.1 ater|EAU35087.1 aory|EIT83392.1

Group8661: abra|OJJ76896.1 afla|EED52108.1 akaw|GAA87148.1 aluc|OJZ84289.1 anig|EHA27629.1 aory|EIT76926.1 asyd|OJJ54792.1 aver|OJJ02013.1 azon|OJJ46450.1 agla|OJJ78807.1

Group8662: abra|OJJ76900.1 acar|OOF98423.1 afis|EAW24058.1 akaw|GAA87140.1 aluc|OJZ84297.1 anig|EHA27633.1 atub|OJI89772.1 akaw|GAA88827.1 aluc|OJZ89604.1 atub|OJI85199.1

Group8663: abra|OJJ76938.1 acla|EAW13467.1 afum|EAL84485.2 akaw|GAA87101.1 aluc|OJZ84337.1 anig|EHA27671.1 atub|OJI89731.1 acar|OOF91388.1 afis|EAW20579.1 azon|OJJ50655.1

Group8664: aver|OJJ01956.1 abra|OJJ77104.1 acar|OOF92989.1 afla|EED45184.1 akaw|GAA86952.1 aluc|OJZ84513.1 anig|EHA27811.1 aory|EIT73680.1 atub|OJI89568.1 azon|OJJ42742.1

Group8665: abra|OJJ76967.1 acar|OOF91411.1 afla|EED52237.1 agla|OJJ88667.1 akaw|GAA87077.1 aluc|OJZ84363.1 anig|EHA27698.1 aory|EIT74351.1 atub|OJI89705.1 awen|OJJ33309.1

Group8666: abra|OJJ77004.1 acar|OOF91452.1 akaw|GAA87044.1 aluc|OJZ84403.1 anid|CBF83173.1 anig|EHA27733.1 asyd|OJJ55525.1 atub|OJI89669.1 aver|OJJ06603.1 awen|OJJ36046.1

Group8667: abra|OJJ77021.1 acar|OOF91459.1 akaw|GAA87031.1 aluc|OJZ84421.1 anig|EHA27743.1 asyd|OJJ64322.1 ater|EAU32174.1 atub|OJI89652.1 aver|OJI98415.1 azon|OJJ50419.1

Group8668: abra|OJJ77093.1 acar|OOG00597.1 akaw|GAA86962.1 aluc|OJZ84502.1 anig|EHA27802.1 atub|OJI89578.1 abra|OJJ78139.1 afla|EED47404.1 aluc|OJZ82075.1 aory|EIT81902.1

Group8669: acla|EAW11185.1 afis|EAW15916.1 afum|EAL88551.2 afla|EED56452.1 anid|CBF81311.1 aory|EIT72826.1 asyd|OJJ59355.1 ater|EAU36194.1 aver|OJJ02919.1 awen|OJJ36714.1

Group8670: ater|EAU33891.1 acla|EAW13251.1 afis|EAW17385.1 afla|EED49498.1 afum|EBA27426.1 agla|OJJ80417.1 aory|EIT82619.1 asyd|OJJ52535.1 aver|OJJ03868.1 awen|OJJ37431.1

Group8671: abra|OJJ77264.1 acar|OOF93133.1 afla|EED52247.1 agla|OJJ83705.1 akaw|GAA90683.1 aluc|OJZ84672.1 anig|EHA27950.1 aory|EIT74408.1 asyd|OJJ60410.1 atub|OJI89408.1

Group8672: abra|OJJ77274.1 acar|OOF93142.1 acla|EAW13263.1 afla|EED49798.1 akaw|GAA90677.1 aluc|OJZ84680.1 anid|CBF84234.1 anig|EHA27958.1 aory|EIT77679.1 atub|OJI89400.1

Group8673: abra|OJJ77284.1 acar|OOF93161.1 akaw|GAA90665.1 aluc|OJZ84694.1 anig|EHA27970.1 asyd|OJJ64781.1 atub|OJI89388.1 aver|OJI98889.1 anid|CBF80818.1 agla|OJJ82982.1

Group8674: abra|OJJ77312.1 acar|OOF99913.1 agla|OJJ80696.1 akaw|GAA91441.1 aluc|OJZ84724.1 anig|EHA27998.1 asyd|OJJ55558.1 atub|OJI89359.1 aver|OJJ06639.1 azon|OJJ45680.1

Group8675: abra|OJJ77414.1 acar|OOF93280.1 afla|EED45194.1 agla|OJJ84823.1 akaw|GAA91351.1 aluc|OJZ84822.1 aory|EIT73534.1 atub|OJI89264.1 awen|OJJ31927.1 azon|OJJ50738.1

Group8676: abra|OJJ77451.1 acar|OOF99275.1 afla|EED54168.1 akaw|GAA84157.1 aluc|OJZ81078.1 aory|EIT75983.1 asyd|OJJ59171.1 atub|OJI82506.1 aver|OJJ01761.1 awen|OJJ37964.1

Group8677: abra|OJJ77491.1 acar|OOF99234.1 afla|EED53251.1 akaw|GAA84128.1 aluc|OJZ81045.1 anig|EHA19322.1 aory|EIT74841.1 atub|OJI82539.1 awen|OJJ37702.1 agla|OJJ88291.1

Group8678: afis|EAW17045.1 afla|EED53440.1 anid|CBF69405.1 asyd|OJJ62010.1 aver|OJJ06554.1 awen|OJJ29786.1 acar|OOF94503.1 aory|EIT75180.1 ater|EAU34058.1 aver|OJI96102.1

Group8679: anig|EHA19441.1 acla|EAW06483.1 afis|EAW15978.1 afum|EAL88610.1 akaw|GAA84002.1 anid|CBF85801.1 aory|EIT73421.1 asyd|OJJ64333.1 ater|EAU33673.1 aver|OJI98426.1

Group8680: abra|OJJ77735.1 agla|OJJ79109.1 akaw|GAA83897.1 aluc|OJZ90798.1 anig|EHA19530.1 atub|OJI82787.1 awen|OJJ36260.1 azon|OJJ51724.1 aver|OJJ08264.1 asyd|OJJ55393.1

Group8681: abra|OJJ77768.1 asyd|OJJ58669.1 abra|OJJ78086.1 acar|OOF99707.1 afla|EED47274.1 atub|OJI86197.1 aver|OJJ01280.1 awen|OJJ38935.1 agla|OJJ79396.1 anig|EHA27634.1

Group8682: abra|OJJ77784.1 acar|OOF98975.1 akaw|GAA83863.1 aluc|OJZ90837.1 anig|EHA19560.1 asyd|OJJ56882.1 atub|OJI82824.1 aory|EIT77282.1 afla|EED50218.1 awen|OJJ33550.1

Group8683: abra|OJJ77875.1 afla|EED45771.1 akaw|GAA83777.1 aluc|OJZ90950.1 aory|EIT75664.1 asyd|OJJ63559.1 atub|OJI82941.1 aver|OJI97373.1 awen|OJJ40359.1 agla|OJJ79938.1

Group8684: abra|OJJ77917.1 afla|EED51607.1 agla|OJJ82627.1 anid|CBF73913.1 anig|EHA18402.1 aory|EIT80700.1 asyd|OJJ53671.1 ater|EAU37636.1 aver|OJJ07778.1 awen|OJJ40189.1

Group8685: abra|OJJ77945.1 acar|OOF91772.1 akaw|GAA83718.1 aluc|OJZ91015.1 anid|CBF71854.1 anig|EHA19717.1 asyd|OJJ59801.1 atub|OJI83006.1 aver|OJJ02489.1 asyd|OJJ53758.1

Group8686: abra|OJJ77952.1 afla|EED55614.1 agla|OJJ89121.1 akaw|GAA83711.1 aluc|OJZ91023.1 anid|CBF90342.1 anig|EHA19723.1 aory|EIT81816.1 atub|OJI83015.1 awen|OJJ30075.1

Group8687: abra|OJJ78050.1 acar|OOF95285.1 akaw|GAA85941.1 aluc|OJZ91121.1 anid|CBF86028.1 anig|EHA19821.1 asyd|OJJ56423.1 atub|OJI83126.1 awen|OJJ40096.1 azon|OJJ42792.1

Group8688: abra|OJJ78052.1 acar|OOF93015.1 afla|EED56742.1 anid|CBF87750.1 anig|EHA19823.1 aory|EIT73934.1 asyd|OJJ64215.1 ater|EAU35699.1 awen|OJJ40180.1 anig|EHA23883.1

Group8689: abra|OJJ78064.1 acar|OOF98791.1 akaw|GAA85930.1 aluc|OJZ91135.1 anid|CBF79585.1 anig|EHA19833.1 asyd|OJJ57978.1 atub|OJI83145.1 aver|OJJ04422.1 azon|OJJ45148.1

Group8690: abra|OJJ78117.1 acar|OOF94026.1 afis|EAW20619.1 akaw|GAA85890.1 aluc|OJZ91182.1 anig|EHA19873.1 asyd|OJJ56225.1 atub|OJI83195.1 aver|OJJ05157.1 azon|OJJ42619.1

Group8691: abra|OJJ78125.1 acar|OOF90392.1 acla|EAW12156.1 afis|EAW17188.1 afum|EAL86614.1 akaw|GAA85883.1 aluc|OJZ91187.1 anig|EHA19883.1 atub|OJI83206.1 azon|OJJ50182.1

Group8692: abra|OJJ78127.1 acla|EAW07051.1 akaw|GAA85881.1 aluc|OJZ91189.1 anid|CBF74942.1 atub|OJI83208.1 azon|OJJ50040.1 akaw|GAA85880.1 azon|OJJ42019.1 azon|OJJ42018.1

Group8693: abra|OJJ78130.1 acar|OOF92539.1 afis|EAW20650.1 afla|EED50437.1 afum|EAL84423.1 akaw|GAA85877.1 aluc|OJZ91192.1 anig|EHA19888.1 aory|EIT82127.1 atub|OJI83211.1

Group8694: abra|OJJ78152.1 acar|OOF98714.1 afla|EED45419.1 akaw|GAA85874.1 aluc|OJZ91204.1 anig|EHA19909.1 aory|EIT82384.1 atub|OJI83221.1 aver|OJJ03359.1 anig|EHA25893.1

Group8695: abra|OJJ78164.1 acar|OOF92532.1 afla|EED56896.1 agla|OJJ82607.1 akaw|GAA85861.1 aluc|OJZ91218.1 anig|EHA19924.1 ater|EAU36290.1 atub|OJI83236.1 awen|OJJ39943.1

Group8696: aory|EIT72935.1 abra|OJJ78224.1 acla|EAW12025.1 agla|OJJ81172.1 akaw|GAA85801.1 aluc|OJZ91299.1 anig|EHA19981.1 asyd|OJJ53517.1 atub|OJI83289.1 aver|OJJ08234.1

Group8697: abra|OJJ78188.1 acar|OOF98671.1 afla|EED45661.1 akaw|GAA85841.1 aluc|OJZ91259.1 anig|EHA19947.1 aory|EIT75485.1 ater|EAU30768.1 atub|OJI83248.1 awen|OJJ37792.1

Group8698: abra|OJJ78262.1 acar|OOF93149.1 akaw|GAA85763.1 aluc|OJZ91337.1 atub|OJI83329.1 awen|OJJ29559.1 azon|OJJ51295.1 asyd|OJJ52299.1 agla|OJJ83668.1 awen|OJJ34751.1

Group8699: abra|OJJ78304.1 acar|OOF98562.1 afis|EAW22061.1 afum|EAL92952.2 agla|OJJ81051.1 akaw|GAA85717.1 asyd|OJJ61340.1 aver|OJJ00664.1 awen|OJJ34230.1 azon|OJJ43405.1

Group8700: asyd|OJJ61336.1 aver|OJJ00660.1 acla|EAW12155.1 afis|EAW17179.1 afla|EED47299.1 afum|EAL86609.1 anid|CBF76893.1 aory|EIT82830.1 awen|OJJ33780.1 azon|OJJ45040.1

Group8701: acar|OOF90205.1 afis|EAW19850.1 afum|EAL85043.1 agla|OJJ81560.1 anig|EHA26561.1 asyd|OJJ54751.1 aver|OJJ07416.1 awen|OJJ33517.1 aver|OJJ07191.1 azon|OJJ42417.1

Group8702: aver|OJJ02169.1 acar|OOF94543.1 akaw|GAA91797.1 aluc|OJZ92817.1 anig|EHA26191.1 atub|OJI87275.1 afla|EED51755.1 aory|EIT80500.1 agla|OJJ82987.1 asyd|OJJ58254.1

Group8703: acar|OOF90534.1 afla|EED57937.1 agla|OJJ84631.1 anid|CBF85124.1 aory|EIT82350.1 ater|EAU32033.1 aver|OJI97920.1 awen|OJJ31234.1 azon|OJJ46157.1 awen|OJJ33711.1

Group8704: acar|OOF90645.1 acla|EAW12240.1 afis|EAW16987.1 afum|EBA27277.1 agla|OJJ79129.1 awen|OJJ34463.1 azon|OJJ51753.1 afla|EED53772.1 aory|EIT78465.1 awen|OJJ35911.1

Group8705: anid|CBF80416.1 afla|EED46245.1 akaw|GAA85952.1 aluc|OJZ91110.1 aory|EIT76424.1 ater|EAU30959.1 atub|OJI83749.1 azon|OJJ42267.1 anig|EHA19807.1 atub|OJI83107.1

Group8706: acar|OOF90940.1 acla|EAW12081.1 afis|EAW19722.1 afum|EAL86804.2 akaw|GAA88819.1 aluc|OJZ82503.1 anig|EHA28267.1 atub|OJI87928.1 awen|OJJ34530.1 azon|OJJ45114.1

Group8707: acar|OOF91332.1 acla|EAW13402.1 afis|EAW20513.1 afla|EED52454.1 afum|EBA27191.1 akaw|GAA87183.1 anid|CBF79182.1 aory|EIT82743.1 ater|EAU37209.1 awen|OJJ33215.1

Group8708: acar|OOF91786.1 afis|EAW20741.1 akaw|GAA88131.1 aluc|OJZ81403.1 anig|EHA18696.1 aory|EIT79960.1 asyd|OJJ64560.1 atub|OJI91609.1 aver|OJI98657.1 asyd|OJJ61686.1

Group8709: acar|OOF92587.1 afla|EED45910.1 agla|OJJ88075.1 anid|CBF84632.1 aory|EIT75466.1 ater|EAU35265.1 aver|OJJ08374.1 awen|OJJ30001.1 azon|OJJ45970.1 asyd|OJJ53370.1

Group8710: acar|OOF92804.1 acla|EAW11648.1 afis|EAW23808.1 afla|EED52007.1 afum|EAL87242.1 anid|CBF79872.1 anig|EHA21270.1 aory|EIT81430.1 asyd|OJJ58167.1 ater|EAU31361.1

Group8711: acar|OOF93611.1 acla|EAW10001.1 ater|EAU37568.1 aver|OJJ06315.1 awen|OJJ32924.1 azon|OJJ46499.1 acar|OOF99541.1 afis|EAW20618.1 ater|EAU33869.1 aory|EIT77091.1

Group8712: acar|OOF93841.1 acla|EAW11884.1 afis|EAW19496.1 afla|EED56715.1 afum|EAL86998.1 agla|OJJ81662.1 akaw|GAA89695.1 anid|CBF75432.1 ater|EAU34808.1 awen|OJJ36913.1

Group8713: acar|OOF93874.1 afla|EED45884.1 aory|EIT75492.1 asyd|OJJ54701.1 aver|OJJ07472.1 acla|EAW12125.1 afis|EAW17223.1 afum|EAL86645.1 agla|OJJ81066.1 awen|OJJ35778.1

Group8714: acar|OOF93938.1 acla|EAW07102.1 afis|EAW22117.1 afla|EED45472.1 afum|EAL91069.1 aory|EIT75606.1 asyd|OJJ54719.1 aver|OJJ07449.1 awen|OJJ34553.1 azon|OJJ50399.1

Group8715: acar|OOF94234.1 acla|EAW09969.1 afis|EAW21119.1 afla|EED51792.1 afum|EAL92143.1 agla|OJJ80092.1 akaw|GAA88304.1 aory|EIT81684.1 aver|OJI99776.1 awen|OJJ36449.1

Group8716: awen|OJJ35852.1 agla|OJJ82563.1 akaw|GAA91338.1 aluc|OJZ84841.1 anig|EHA18765.1 ater|EAU37118.1 atub|OJI89257.1 azon|OJJ42300.1 aluc|OJZ83606.1 atub|OJI89979.1

Group8717: acar|OOF95373.1 agla|OJJ85923.1 akaw|GAA92957.1 asyd|OJJ57074.1 afis|EAW24593.1 anid|CBF80221.1 akaw|GAA93201.1 aluc|OJZ79704.1 aver|OJJ07710.1 anid|CBF81144.1

Group8718: acar|OOF95885.1 acla|EAW08488.1 afis|EAW23534.1 afla|EED57431.1 afum|EAL87932.1 agla|OJJ86783.1 akaw|GAA83538.1 anid|CBF89676.1 aory|EIT80079.1 ater|EAU35320.1

Group8719: acar|OOF96231.1 acla|EAW15073.1 afum|EAL85165.1 agla|OJJ88917.1 aluc|OJZ86254.1 anid|CBF84791.1 asyd|OJJ63229.1 atub|OJI84289.1 awen|OJJ31801.1 azon|OJJ46322.1

Group8720: acar|OOF96801.1 afis|EAW15657.1 afla|EED45409.1 afum|EAL85689.1 agla|OJJ81801.1 aory|EIT82452.1 asyd|OJJ54185.1 awen|OJJ33737.1 anid|CBF84452.1 asyd|OJJ64256.1

Group8721: acar|OOF96972.1 acla|EAW11148.1 afis|EAW15879.1 afla|EED56496.1 afum|EAL88513.1 akaw|GAA90093.1 aluc|OJZ84120.1 anid|CBF81232.1 aory|EIT72850.1 ater|EAU31700.1

Group8722: acar|OOF97384.1 afla|EED48121.1 akaw|GAA85866.1 aluc|OJZ91213.1 anid|CBF80007.1 anig|EHA19919.1 asyd|OJJ58252.1 atub|OJI83231.1 aver|OJJ02172.1 awen|OJJ31654.1

Group8723: afis|EAW20839.1 afum|EAL84794.2 acla|EAW11025.1 afla|EED51814.1 agla|OJJ80752.1 anid|CBF84523.1 asyd|OJJ62168.1 aver|OJI96280.1 awen|OJJ40169.1 ater|EAU31956.1

Group8724: acar|OOF98730.1 akaw|GAA87770.1 aluc|OJZ85405.1 anig|EHA25584.1 aory|EIT81183.1 asyd|OJJ65052.1 aver|OJI98891.1 awen|OJJ37890.1 afla|EED48912.1 aver|OJI99175.1

Group8725: acar|OOF98764.1 afis|EAW19773.1 afla|EED46883.1 afum|EAL85127.1 agla|OJJ83933.1 anig|EHA19874.1 aory|EIT72745.1 asyd|OJJ52082.1 ater|EAU29584.1 aver|OJJ03021.1

Group8726: acar|OOF99188.1 agla|OJJ86259.1 anid|CBF78881.1 asyd|OJJ64595.1 ater|EAU31457.1 atub|OJI80365.1 aver|OJI98692.1 awen|OJJ30207.1 azon|OJJ50373.1 aver|OJJ05528.1

Group8727: acar|OOF99223.1 acla|EAW06731.1 akaw|GAA84117.1 aluc|OJZ81033.1 anid|CBF86874.1 asyd|OJJ64628.1 atub|OJI82550.1 aver|OJI98726.1 awen|OJJ37676.1 azon|OJJ44941.1

Group8728: acar|OOF99348.1 afla|EED53111.1 agla|OJJ88749.1 aory|EIT74779.1 awen|OJJ36022.1 afla|EED49457.1 aory|EIT79530.1 ater|EAU31594.1 asyd|OJJ55202.1 aver|OJJ07014.1

Group8729: acar|OOF99513.1 afis|EAW23668.1 afla|EED50426.1 afum|EAL87113.1 anid|CBF75936.1 aory|EIT82003.1 awen|OJJ39942.1 azon|OJJ42345.1 afla|EED54472.1 aory|EIT81132.1

Group8730: acar|OOG01148.1 acla|EAW07881.1 afis|EAW22916.1 afla|EED54912.1 afum|EAL90281.1 akaw|GAA87513.1 anid|CBF87692.1 aory|EIT80372.1 atub|OJI90616.1 awen|OJJ38648.1

Group8731: acla|EAW06293.1 afis|EAW18399.1 afum|EAL93076.2 ater|EAU29815.1 awen|OJJ31751.1 agla|OJJ84970.1 azon|OJJ50473.1 anid|CBF70753.1 asyd|OJJ58437.1 aver|OJJ01037.1

Group8732: acla|EAW06854.1 afis|EAW16471.1 afla|EED49701.1 afum|EAL89065.1 anid|CBF89396.1 aory|EIT77699.1 asyd|OJJ62460.1 ater|EAU34009.1 aver|OJI96595.1 awen|OJJ34632.1

Group8733: acla|EAW06986.1 afis|EAW16600.1 afla|EED45019.1 afum|EAL89196.1 agla|OJJ82652.1 asyd|OJJ59689.1 aver|OJJ02598.1 awen|OJJ40208.1 azon|OJJ49074.1 anid|CBF81907.1

Group8734: acla|EAW07380.1 afis|EAW22405.1 afla|EED56069.1 agla|OJJ87668.1 aluc|OJZ89800.1 anid|CBF88787.1 asyd|OJJ62902.1 aver|OJI96956.1 awen|OJJ34984.1 azon|OJJ42756.1

Group8735: acla|EAW07974.1 afis|EAW23009.1 afla|EED55006.1 afum|EAL88471.1 agla|OJJ89471.1 akaw|GAA85616.1 anid|CBF80881.1 aory|EIT80305.1 ater|EAU29664.1 azon|OJJ44195.1

Group8736: acla|EAW08445.1 afis|EAW23490.1 afla|EED57387.1 afum|EAL87973.2 agla|OJJ86733.1 akaw|GAA83495.1 anid|CBF89753.1 aory|EIT80140.1 ater|EAU35275.1 azon|OJJ47935.1

Group8737: acla|EAW08887.1 afis|EAW15502.1 afla|EED49015.1 afum|EAL85822.1 aory|EIT82541.1 asyd|OJJ56009.1 ater|EAU32616.1 aver|OJJ04954.1 awen|OJJ38162.1 azon|OJJ43123.1

Group8738: acla|EAW08972.1 afis|EAW15451.1 afla|EED54387.1 afum|EAL84195.2 agla|OJJ87010.1 anid|CBF86652.1 aory|EIT75869.1 asyd|OJJ64798.1 ater|EAU31837.1 aver|OJI98912.1

Group8739: acla|EAW09111.1 afis|EAW23756.1 afla|EED52087.1 afum|EAL87192.1 anid|CBF78719.1 aory|EIT76934.1 asyd|OJJ55648.1 ater|EAU29642.1 aver|OJJ06733.1 ater|EAU33187.1

Group8740: acla|EAW10097.1 afis|EAW25490.1 afum|EAL90155.1 akaw|GAA82560.1 anid|CBF77730.1 anig|EHA20363.1 asyd|OJJ53058.1 ater|EAU34661.1 aver|OJJ03786.1 awen|OJJ41231.1

Group8741: acla|EAW10292.1 afis|EAW25299.1 afla|EED53773.1 afum|EAL89962.2 agla|OJJ89279.1 anid|CBF75191.1 asyd|OJJ59939.1 ater|EAU34447.1 aver|OJJ02348.1 awen|OJJ41014.1

Group8742: acla|EAW10544.1 afis|EAW25012.1 afum|EAL89684.1 agla|OJJ89176.1 anid|CBF81521.1 asyd|OJJ59477.1 ater|EAU35774.1 aver|OJJ02796.1 awen|OJJ40628.1 azon|OJJ49055.1

Group8743: acla|EAW10900.1 afis|EAW20958.1 afla|EED50314.1 afum|EAL84675.1 agla|OJJ83919.1 akaw|GAA91126.1 anid|CBF82565.1 ater|EAU38609.1 aver|OJJ08691.1 azon|OJJ48094.1

Group8744: acla|EAW10967.1 afis|EAW20893.1 afla|EED45588.1 afum|EAL84740.1 anid|CBF82749.1 aory|EIT75638.1 asyd|OJJ61614.1 ater|EAU30832.1 aver|OJJ00937.1 awen|OJJ32625.1

Group8745: acla|EAW11075.1 afis|EAW20792.1 afla|EED47369.1 afum|EAL84834.1 aory|EIT82870.1 asyd|OJJ57140.1 aver|OJJ03165.1 anid|CBF84183.1 anid|CBF90380.1 ater|EAU31730.1

Group8746: acla|EAW11186.1 afis|EAW15917.1 afum|EAL88552.1 anid|CBF81313.1 aory|EIT72813.1 asyd|OJJ59356.1 ater|EAU36195.1 aver|OJJ02918.1 awen|OJJ36713.1 azon|OJJ47486.1

Group8747: acla|EAW12353.1 afis|EAW16778.1 afla|EED56192.1 afum|EAL86348.2 anid|CBF80569.1 aory|EIT73352.1 asyd|OJJ54685.1 ater|EAU30274.1 aver|OJJ07484.1 awen|OJJ29949.1

Group8748: acla|EAW12379.1 afis|EAW16827.1 afla|EED57472.1 afum|EAL86376.1 anid|CBF87493.1 aory|EIT80051.1 asyd|OJJ52965.1 ater|EAU31804.1 aver|OJJ08931.1 afis|EAW16725.1

Group8749: acla|EAW12914.1 afis|EAW17746.1 afla|EED45746.1 afum|EAL91558.1 aory|EIT75371.1 asyd|OJJ64817.1 ater|EAU31861.1 aver|OJI98930.1 awen|OJJ37091.1 azon|OJJ50795.1

Group8750: acla|EAW12985.1 afis|EAW17679.1 afla|EED50471.1 afum|EAL91493.1 anid|CBF90013.1 aory|EIT82122.1 ater|EAU31967.1 aver|OJI96178.1 awen|OJJ37160.1 azon|OJJ50857.1

Group8751: acla|EAW13483.1 afis|EAW20591.1 afla|EED52236.1 afum|EAL84470.1 anid|CBF86917.1 aory|EIT74432.1 asyd|OJJ55622.1 ater|EAU35230.1 aver|OJJ06697.1 awen|OJJ33310.1

Group8752: acla|EAW13598.1 afla|EED50431.1 anid|CBF84555.1 aory|EIT82020.1 asyd|OJJ53465.1 aver|OJJ08290.1 awen|OJJ30897.1 azon|OJJ43647.1 agla|OJJ84354.1 anid|CBF84554.1

Group8753: acla|EAW13736.1 afis|EAW19834.1 afla|EED52179.1 afum|EAL85058.1 anid|CBF80447.1 aory|EIT74473.1 asyd|OJJ54757.1 ater|EAU32197.1 aver|OJJ07413.1 azon|OJJ47556.1

Group8754: acla|EAW14245.1 afis|EAW19035.1 afla|EED57302.1 afum|EAL93764.2 agla|OJJ89326.1 aory|EIT78186.1 ater|EAU35143.1 awen|OJJ30855.1 aory|EIT74123.1 afla|EED52797.1

Group8755: acla|EAW14579.1 afis|EAW19313.1 afla|EED53657.1 afum|EAL94029.1 anid|CBF79587.1 aory|EIT75352.1 asyd|OJJ57979.1 ater|EAU32519.1 aver|OJJ04421.1 awen|OJJ40846.1

Group8756: acla|EAW14604.1 afis|EAW19333.1 afum|EAL94050.2 agla|OJJ79419.1 aory|EIT79822.1 asyd|OJJ59228.1 aver|OJJ01838.1 awen|OJJ32996.1 azon|OJJ48518.1 afla|EED47976.1

Group8757: acla|EAW14720.1 afis|EAW24256.1 afla|EED48020.1 afum|EAL85924.1 akaw|GAA88193.1 anid|CBF74193.1 aory|EIT82908.1 ater|EAU31197.1 awen|OJJ33044.1 azon|OJJ48475.1

Group8758: acla|EAW14970.1 afis|EAW24493.1 afla|EED54699.1 afum|EAL86162.1 agla|OJJ84580.1 anid|CBF73845.1 asyd|OJJ56554.1 ater|EAU29946.1 aver|OJJ06167.1 awen|OJJ32702.1

Group8759: acla|EAW15153.1 afis|EAW20121.1 afla|EED57255.1 afum|EAL85239.1 anid|CBF84941.1 aory|EIT74015.1 asyd|OJJ63151.1 ater|EAU38739.1 aver|OJI97803.1 awen|OJJ31890.1

Group8760: acla|EAW15170.1 afis|EAW20139.1 afla|EED47779.1 afum|EAL85254.1 anid|CBF84971.1 aory|EIT77146.1 asyd|OJJ63134.1 aver|OJI97820.1 awen|OJJ32008.1 azon|OJJ42033.1

Group8761: afis|EAW15432.1 afla|EED45624.1 agla|OJJ86266.1 aluc|OJZ83141.1 anig|EHA23289.1 aory|EIT75551.1 asyd|OJJ64000.1 ater|EAU31306.1 aver|OJI98089.1 awen|OJJ32640.1

Group8762: afis|EAW16966.1 agla|OJJ83931.1 anid|CBF85598.1 aory|EIT72785.1 asyd|OJJ65318.1 atub|OJI87260.1 aver|OJI99463.1 awen|OJJ36296.1 afum|EAL91082.1 anid|CBF80298.1

Group8763: afis|EAW17042.1 afla|EED44725.1 asyd|OJJ54278.1 aver|OJI99266.1 awen|OJJ40175.1 azon|OJJ42360.1 akaw|GAA92656.1 anid|CBF69573.1 anig|EHA19539.1 aluc|OJZ81155.1

Group8764: afis|EAW18182.1 afum|EAL87544.1 anid|CBF85725.1 asyd|OJJ64278.1 aver|OJI98370.1 awen|OJJ30530.1 aory|EIT82861.1 asyd|OJJ52193.1 afla|EED47321.1 aver|OJJ05953.1

Group8765: afis|EAW18188.1 afla|EED55528.1 afum|EAL87551.1 anid|CBF86098.1 aory|EIT80825.1 asyd|OJJ64478.1 ater|EAU33806.1 aver|OJI98580.1 awen|OJJ30551.1 azon|OJJ45514.1

Group8766: afis|EAW21085.1 afla|EED44924.1 afum|EAL92107.2 akaw|GAA91677.1 aluc|OJZ87717.1 anig|EHA20211.1 aory|EIT73525.1 asyd|OJJ61439.1 atub|OJI80925.1 aver|OJJ00773.1

Group8767: afis|EAW25641.1 afla|EED51109.1 afum|EAL84328.2 akaw|GAA92605.1 aluc|OJZ82624.1 aory|EIT81346.1 asyd|OJJ54684.1 ater|EAU31799.1 atub|OJI88858.1 aver|OJJ07485.1

Group8768: afla|EED48329.1 agla|OJJ78800.1 akaw|GAA83720.1 aluc|OJZ91013.1 anig|EHA19715.1 asyd|OJJ55448.1 ater|EAU32774.1 atub|OJI83004.1 aver|OJJ06521.1 awen|OJJ33538.1

Group8769: afla|EED51054.1 agla|OJJ85437.1 anid|CBF84153.1 aory|EIT76946.1 asyd|OJJ58763.1 ater|EAU38829.1 aver|OJJ01373.1 awen|OJJ37828.1 asyd|OJJ61432.1 aver|OJJ00767.1

Group8770: afla|EED57392.1 agla|OJJ86738.1 aory|EIT80083.1 asyd|OJJ62243.1 ater|EAU35281.1 aver|OJI96353.1 azon|OJJ47940.1 anid|CBF81868.1 ater|EAU32329.1 azon|OJJ45101.1

Group8771: abra|OJJ65451.1 abra|OJJ65502.1 abra|OJJ65814.1 awen|OJJ29571.1 awen|OJJ29595.1 awen|OJJ29635.1 awen|OJJ29618.1 akaw|GAA93238.1 ater|EAU30120.1

Group8772: abra|OJJ65548.1 abra|OJJ65554.1 afis|EAW24053.1 afum|EAL91907.2 akaw|GAA93142.1 asyd|OJJ56982.1 aver|OJJ03305.1 anid|CBF82344.1 akaw|GAA93186.1

Group8773: abra|OJJ65560.1 abra|OJJ65734.1 akaw|GAA93244.1 akaw|GAA93292.1 awen|OJJ29572.1 awen|OJJ29636.1 awen|OJJ29617.1 awen|OJJ29594.1 ater|EAU30121.1

Group8774: abra|OJJ65830.1 abra|OJJ68080.1 anig|EHA21142.1 anig|EHA21847.1 aluc|OJZ79866.1 aory|EIT74448.1 afla|EED52212.1 akaw|GAA93352.1 ater|EAU37381.1

Group8775: abra|OJJ67069.1 abra|OJJ72180.1 abra|OJJ72620.1 anig|EHA18937.1 anig|EHA26082.1 agla|OJJ82519.1 akaw|GAA92012.1 aluc|OJZ92877.1 atub|OJI87337.1

Group8776: acar|OOF90251.1 acar|OOF94223.1 abra|OJJ67549.1 afla|EED45499.1 akaw|GAA88111.1 aluc|OJZ81381.1 anig|EHA18676.1 atub|OJI91587.1 azon|OJJ45315.1

Group8777: acar|OOF90262.1 acar|OOF90263.1 abra|OJJ68046.1 acar|OOF98217.1 aluc|OJZ83950.1 anig|EHA19079.1 abra|OJJ78103.1 anig|EHA19861.1 awen|OJJ29577.1

Group8778: acar|OOF90382.1 acar|OOF90384.1 abra|OJJ66512.1 akaw|GAA91199.1 aluc|OJZ81534.1 anig|EHA25902.1 ater|EAU37133.1 atub|OJI88653.1 awen|OJJ31634.1

Group8779: acar|OOF93360.1 acar|OOF97612.1 agla|OJJ79302.1 agla|OJJ83534.1 atub|OJI80136.1 atub|OJI85146.1 abra|OJJ70356.1 aluc|OJZ80196.1 awen|OJJ35533.1

Group8780: acar|OOF94039.1 acar|OOF94109.1 abra|OJJ66412.1 akaw|GAA90278.1 aluc|OJZ83063.1 anig|EHA28128.1 atub|OJI87882.1 aver|OJJ07280.1 azon|OJJ43651.1

Group8781: acar|OOF94053.1 acar|OOF94123.1 azon|OJJ43612.1 azon|OJJ49265.1 abra|OJJ66391.1 akaw|GAA90257.1 aluc|OJZ83083.1 anig|EHA28146.1 atub|OJI87862.1

Group8782: acar|OOF94077.1 acar|OOF94147.1 afis|EAW16983.1 afum|EAL86510.1 anid|CBF88830.1 asyd|OJJ62881.1 aver|OJI96933.1 afla|EED48302.1 aory|EIT76296.1

Group8783: acar|OOF95051.1 acar|OOF95118.1 abra|OJJ68393.1 afla|EED46846.1 afum|EAL92231.1 anig|EHA20970.1 aory|EIT72743.1 aver|OJJ04487.1 awen|OJJ34413.1

Group8784: acar|OOF95348.1 acar|OOG00095.1 abra|OJJ70253.1 afla|EED44863.1 akaw|GAA86538.1 aluc|OJZ85086.1 anig|EHA25302.1 aory|EIT82718.1 atub|OJI80513.1

Group8785: acar|OOF99307.1 acar|OOF99312.1 akaw|GAA92346.1 akaw|GAA93056.1 abra|OJJ71069.1 acla|EAW13473.1 anid|CBF73463.1 azon|OJJ44546.1 abra|OJJ72523.1

Group8786: acar|OOG00247.1 acar|OOG00265.1 abra|OJJ67541.1 akaw|GAA88118.1 aluc|OJZ81391.1 anig|EHA18684.1 asyd|OJJ64470.1 atub|OJI91596.1 aver|OJI98574.1

Group8787: acla|EAW12164.1 acla|EAW14981.1 abra|OJJ67663.1 akaw|GAA89178.1 aluc|OJZ81655.1 anig|EHA25790.1 asyd|OJJ59826.1 atub|OJI88528.1 aver|OJJ02461.1

Group8788: afis|EAW23659.1 afis|EAW25762.1 asyd|OJJ52423.1 asyd|OJJ60340.1 aver|OJI99683.1 afum|EAL89841.1 anid|CBF89329.1 afis|EAW25168.1 aory|EIT74325.1

Group8789: afla|EED50425.1 afla|EED54473.1 aory|EIT81160.1 aory|EIT82025.1 azon|OJJ42343.1 acar|OOF99515.1 afis|EAW23667.1 afum|EAL87112.1 anid|CBF75938.1

Group8790: afum|EAL90934.1 afum|EAL91122.1 afis|EAW20714.1 afla|EED52356.1 agla|OJJ78986.1 akaw|GAA87891.1 anig|EHA21853.1 asyd|OJJ52038.1 azon|OJJ42064.1

Group8791: afum|EAL90946.1 afum|EAL91115.2 agla|OJJ79315.1 agla|OJJ83717.1 asyd|OJJ62706.1 asyd|OJJ62717.1 afis|EAW15512.1 aluc|OJZ80028.1 anig|EHA20894.1

Group8792: agla|OJJ79258.1 agla|OJJ83426.1 agla|OJJ81210.1 anid|CBF83975.1 anid|CBF83977.1 abra|OJJ69175.1 aluc|OJZ86162.1 anig|EHA24462.1 atub|OJI79954.1

Group8793: agla|OJJ79506.1 agla|OJJ85828.1 agla|OJJ83718.1 awen|OJJ36133.1 abra|OJJ72502.1 akaw|GAA92424.1 aluc|OJZ81598.1 anig|EHA25843.1 atub|OJI88584.1

Group8794: akaw|GAA83148.1 akaw|GAA92140.1 atub|OJI79312.1 atub|OJI91286.1 aluc|OJZ88705.1 anid|CBF87325.1 asyd|OJJ59111.1 aver|OJJ01703.1 anig|EHA18389.1

Group8795: akaw|GAA83292.1 akaw|GAA85869.1 aluc|OJZ83794.1 aluc|OJZ91210.1 atub|OJI83228.1 atub|OJI90164.1 abra|OJJ67070.1 acar|OOF96438.1 anig|EHA18936.1

Group8796: aluc|OJZ79493.1 aluc|OJZ80465.1 aluc|OJZ79841.1 aluc|OJZ79687.1 aluc|OJZ80286.1 aluc|OJZ79880.1 agla|OJJ78477.1 asyd|OJJ52216.1 aver|OJJ07337.1

Group8797: aluc|OJZ79785.1 aluc|OJZ80377.1 aluc|OJZ80035.1 aluc|OJZ80422.1 aluc|OJZ79853.1 aluc|OJZ79976.1 asyd|OJJ51957.1 asyd|OJJ52282.1 asyd|OJJ52136.1

Group8798: aluc|OJZ80106.1 aluc|OJZ86035.1 abra|OJJ67578.1 afla|EED56236.1 akaw|GAA90577.1 anig|EHA24580.1 aory|EIT73434.1 atub|OJI79822.1 aver|OJI98015.1

Group8799: aluc|OJZ80144.1 aluc|OJZ89697.1 afis|EAW25568.1 acar|OOF90908.1 agla|OJJ87888.1 anig|EHA26933.1 asyd|OJJ55925.1 atub|OJI85106.1 awen|OJJ41481.1

Group8800: aluc|OJZ80441.1 aluc|OJZ90711.1 abra|OJJ72739.1 acar|OOG00406.1 agla|OJJ85079.1 akaw|GAA85349.1 anig|EHA24131.1 atub|OJI82138.1 awen|OJJ33532.1

Group8801: anid|CBF74938.1 anid|CBF79108.1 asyd|OJJ53785.1 asyd|OJJ57623.1 aver|OJI99283.1 aver|OJJ05563.1 afis|EAW19548.1 afum|EAL86961.1 azon|OJJ46016.1

Group8802: anid|CBF76143.1 anid|CBF80990.1 agla|OJJ84359.1 akaw|GAA81951.1 aluc|OJZ92237.1 anig|EHA26717.1 asyd|OJJ54883.1 atub|OJI86671.1 awen|OJJ32017.1

Group8803: anig|EHA21625.1 anig|EHA27497.1 abra|OJJ70050.1 akaw|GAA85317.1 aluc|OJZ90678.1 azon|OJJ42543.1 anid|CBF88936.1 asyd|OJJ62856.1 aver|OJI96908.1

Group8804: aory|EIT82280.1 aory|EIT83563.1 abra|OJJ65852.1 acar|OOF95013.1 afla|EED57052.1 akaw|GAA92292.1 aluc|OJZ90189.1 anig|EHA23688.1 atub|OJI81616.1

Group8805: asyd|OJJ52061.1 asyd|OJJ57195.1 acla|EAW14822.1 afla|EED48428.1 afum|EAL90942.1 aory|EIT81675.1 ater|EAU37469.1 aver|OJJ03096.1 awen|OJJ32279.1

Group8806: asyd|OJJ52779.1 asyd|OJJ53480.1 asyd|OJJ54408.1 afis|EAW19800.1 afum|EAL85090.1 ater|EAU33009.1 aver|OJJ08758.1 awen|OJJ33979.1 azon|OJJ48572.1

Group8807: asyd|OJJ54218.1 asyd|OJJ57155.1 asyd|OJJ62487.1 aver|OJJ03157.1 aver|OJJ05170.1 ater|EAU33942.1 afla|EED57475.1 aory|EIT80078.1 awen|OJJ40016.1

Group8808: asyd|OJJ54422.1 asyd|OJJ55849.1 aver|OJI96304.1 aver|OJJ04792.1 ater|EAU32489.1 afla|EED46861.1 anid|CBF76241.1 aory|EIT72566.1 awen|OJJ36354.1

Group8809: asyd|OJJ54727.1 asyd|OJJ58679.1 asyd|OJJ55034.1 abra|OJJ66493.1 afla|EED44961.1 aver|OJJ05224.1 afis|EAW24634.1 ater|EAU35914.1 ater|EAU32225.1

Group8810: asyd|OJJ54886.1 asyd|OJJ54987.1 asyd|OJJ55459.1 aver|OJJ05933.1 aver|OJJ07214.1 afla|EED50434.1 anid|CBF71415.1 aory|EIT82120.1 ater|EAU32832.1

Group8811: asyd|OJJ55385.1 asyd|OJJ60334.1 asyd|OJJ59628.1 agla|OJJ79113.1 afis|EAW17031.1 aluc|OJZ79600.1 aver|OJI99679.1 awen|OJJ36175.1 aver|OJJ08272.1

Group8812: asyd|OJJ59241.1 asyd|OJJ60243.1 acla|EAW07120.1 afla|EED45694.1 agla|OJJ85747.1 aory|EIT75416.1 azon|OJJ44490.1 aver|OJJ04022.1 azon|OJJ45725.1

Group8813: ater|EAU30364.1 ater|EAU31308.1 awen|OJJ32840.1 awen|OJJ36219.1 agla|OJJ89215.1 anid|CBF79787.1 aory|EIT80645.1 asyd|OJJ58108.1 aver|OJJ04296.1

Group8814: ater|EAU31139.1 ater|EAU36855.1 afis|EAW16718.1 afis|EAW20349.1 afis|EAW22038.1 akaw|GAA87762.1 anid|CBF74296.1 atub|OJI80175.1 anig|EHA24807.1

Group8815: ater|EAU38677.1 ater|EAU39213.1 afis|EAW25163.1 afum|EAL89837.1 anid|CBF85826.1 asyd|OJJ64346.1 aver|OJI98444.1 awen|OJJ33748.1 afla|EED53476.1

Group8816: aver|OJI99583.1 aver|OJI99593.1 abra|OJJ69735.1 acar|OOF94186.1 akaw|GAA90188.1 aluc|OJZ83161.1 anig|EHA28225.1 atub|OJI87776.1 ater|EAU34770.1

Group8817: aver|OJJ01780.1 aver|OJJ01786.1 acla|EAW07213.1 afis|EAW22226.1 afla|EED49534.1 afum|EAL90968.1 aory|EIT82651.1 afis|EAW16699.1 afum|EAL89296.1

Group8818: awen|OJJ29775.1 awen|OJJ31332.1 afis|EAW23426.1 akaw|GAA92715.1 aluc|OJZ88761.1 anig|EHA18457.1 ater|EAU36827.1 atub|OJI91331.1 azon|OJJ48299.1

Group8819: awen|OJJ30862.1 awen|OJJ38682.1 afla|EED51835.1 akaw|GAA87144.1 aluc|OJZ84293.1 aory|EIT81768.1 atub|OJI89776.1 aver|OJI95767.1 aver|OJI95765.1

Group8820: awen|OJJ33516.1 awen|OJJ39640.1 awen|OJJ39595.1 abra|OJJ72040.1 acar|OOF99380.1 afis|EAW16612.1 akaw|GAA84894.1 aluc|OJZ86777.1 atub|OJI83743.1

Group8821: awen|OJJ34552.1 awen|OJJ36961.1 acla|EAW11716.1 afis|EAW23735.1 afla|EED51561.1 afum|EAL87173.1 aory|EIT80459.1 afla|EED51560.1 aory|EIT80688.1

Group8822: azon|OJJ42004.1 azon|OJJ45156.1 abra|OJJ77523.1 acar|OOF99201.1 acla|EAW06693.1 afis|EAW16922.1 akaw|GAA84097.1 aluc|OJZ81013.1 anig|EHA19348.1

Group8823: azon|OJJ45307.1 azon|OJJ49529.1 acla|EAW07939.1 afis|EAW22977.1 afum|EAL90220.2 anid|CBF82133.1 asyd|OJJ54313.1 ater|EAU32988.1 aver|OJJ05267.1

Group8824: azon|OJJ47349.1 azon|OJJ50598.1 acla|EAW14194.1 afis|EAW18909.1 afum|EAL93641.1 anid|CBF69701.1 asyd|OJJ59049.1 aver|OJJ01645.1 awen|OJJ32465.1

Group8825: azon|OJJ48993.1 azon|OJJ50417.1 azon|OJJ50113.1 abra|OJJ66237.1 acar|OOF90699.1 akaw|GAA89052.1 aluc|OJZ80542.1 anig|EHA25653.1 atub|OJI88389.1

Group8826: agla|OJJ78461.1 agla|OJJ81492.1 agla|OJJ83286.1 agla|OJJ83546.1 agla|OJJ89323.1 agla|OJJ88335.1 agla|OJJ86194.1 agla|OJJ83142.1 agla|OJJ89324.1

Group8827: abra|OJJ65540.1 afla|EED54088.1 akaw|GAA92411.1 anig|EHA21122.1 aory|EIT75899.1 asyd|OJJ58564.1 aver|OJJ01164.1 azon|OJJ42799.1 acar|OOG00222.1

Group8828: abra|OJJ65636.1 afla|EED55263.1 agla|OJJ79070.1 anid|CBF82224.1 aory|EIT83528.1 asyd|OJJ55372.1 ater|EAU35405.1 aver|OJJ06451.1 awen|OJJ35999.1

Group8829: abra|OJJ65637.1 agla|OJJ79069.1 aluc|OJZ91198.1 anid|CBF82226.1 asyd|OJJ55370.1 ater|EAU35406.1 aver|OJJ05826.1 awen|OJJ36000.1 asyd|OJJ57336.1

Group8830: abra|OJJ65651.1 ater|EAU36882.1 anid|CBF85860.1 afla|EED47059.1 agla|OJJ88059.1 aory|EIT72485.1 asyd|OJJ56267.1 aver|OJJ08500.1 awen|OJJ30710.1

Group8831: abra|OJJ65775.1 acar|OOF92087.1 afla|EED54748.1 akaw|GAA90887.1 aluc|OJZ82944.1 anig|EHA21362.1 aory|EIT78754.1 atub|OJI89176.1 azon|OJJ45774.1

Group8832: abra|OJJ65802.1 afum|EAL85708.1 akaw|GAA92132.1 aluc|OJZ83591.1 anig|EHA26724.1 atub|OJI89961.1 afla|EED45353.1 aory|EIT82378.1 afis|EAW15626.1

Group8833: abra|OJJ65803.1 afis|EAW15625.1 afla|EED45352.1 afum|EAL85709.1 akaw|GAA92133.1 aluc|OJZ83590.1 anig|EHA26725.1 aory|EIT82463.1 atub|OJI89960.1

Group8834: abra|OJJ65806.1 afis|EAW20221.1 afla|EED53057.1 afum|EAL85335.1 akaw|GAA92934.1 aluc|OJZ82527.1 anig|EHA22493.1 aory|EIT74775.1 awen|OJJ30065.1

Group8835: abra|OJJ65809.1 afla|EED47456.1 akaw|GAA89023.1 aluc|OJZ80581.1 anig|EHA18752.1 aory|EIT79972.1 atub|OJI88349.1 awen|OJJ37830.1 azon|OJJ50431.1

Group8836: abra|OJJ65868.1 afla|EED52735.1 akaw|GAA92278.1 aluc|OJZ90172.1 anid|CBF80608.1 anig|EHA23671.1 asyd|OJJ54707.1 atub|OJI81600.1 aver|OJJ07462.1

Group8837: abra|OJJ65871.1 acar|OOF99185.1 afla|EED50203.1 akaw|GAA92272.1 aluc|OJZ90166.1 anig|EHA23670.1 aory|EIT77365.1 atub|OJI81594.1 aory|EIT77269.1

Group8838: abra|OJJ65872.1 acla|EAW08931.1 anig|EHA20944.1 ater|EAU36342.1 atub|OJI84669.1 aluc|OJZ90127.1 awen|OJJ36342.1 afla|EED51542.1 akaw|GAA92672.1

Group8839: abra|OJJ65884.1 acar|OOF94539.1 afis|EAW20560.1 afla|EED54350.1 afum|EAL84501.1 anid|CBF84309.1 aory|EIT75919.1 ater|EAU37622.1 azon|OJJ50732.1

Group8840: abra|OJJ65885.1 agla|OJJ82870.1 akaw|GAA89040.1 aluc|OJZ80555.1 anig|EHA25642.1 asyd|OJJ55009.1 atub|OJI88376.1 aver|OJI99608.1 awen|OJJ36097.1

Group8841: abra|OJJ65899.1 agla|OJJ85281.1 akaw|GAA86005.1 aluc|OJZ90231.1 anig|EHA23732.1 aory|EIT81180.1 atub|OJI81661.1 awen|OJJ41745.1 azon|OJJ45155.1

Group8842: abra|OJJ65901.1 akaw|GAA92329.1 aluc|OJZ90227.1 anig|EHA23728.1 atub|OJI81660.1 aver|OJJ05777.1 awen|OJJ38787.1 azon|OJJ50184.1 asyd|OJJ57381.1

Group8843: abra|OJJ65902.1 acar|OOF93266.1 afum|EAL84408.1 akaw|GAA92328.1 aluc|OJZ90226.1 anig|EHA23727.1 aory|EIT76803.1 atub|OJI81659.1 awen|OJJ30873.1

Group8844: abra|OJJ65934.1 acar|OOF95000.1 afla|EED48874.1 akaw|GAA92301.1 aluc|OJZ90198.1 anig|EHA23696.1 aory|EIT77788.1 atub|OJI81626.1 aver|OJJ05898.1

Group8845: agla|OJJ82750.1 acla|EAW11255.1 afla|EED56223.1 aory|EIT73393.1 asyd|OJJ63640.1 ater|EAU33741.1 aver|OJI97296.1 awen|OJJ39937.1 azon|OJJ44919.1

Group8846: awen|OJJ32624.1 acar|OOF92097.1 afla|EED47378.1 agla|OJJ88252.1 anig|EHA21374.1 aory|EIT82894.1 asyd|OJJ57558.1 ater|EAU36897.1 aver|OJJ05620.1

Group8847: abra|OJJ66022.1 afis|EAW17084.1 afla|EED56776.1 akaw|GAA83857.1 anid|CBF88194.1 aory|EIT73959.1 asyd|OJJ61423.1 atub|OJI82846.1 aver|OJJ05745.1

Group8848: abra|OJJ66023.1 afis|EAW17083.1 afla|EED56775.1 anid|CBF88196.1 aory|EIT73910.1 asyd|OJJ61422.1 atub|OJI82847.1 aver|OJJ05746.1 akaw|GAA83856.1

Group8849: abra|OJJ66028.1 acar|OOF90023.1 afla|EED46902.1 anig|EHA22566.1 aory|EIT72748.1 asyd|OJJ52492.1 atub|OJI87438.1 aver|OJJ02991.1 aver|OJJ03418.1

Group8850: abra|OJJ66085.1 acar|OOF99279.1 agla|OJJ84602.1 akaw|GAA84160.1 aluc|OJZ81083.1 asyd|OJJ62094.1 atub|OJI82501.1 aver|OJI96200.1 awen|OJJ33594.1

Group8851: abra|OJJ66087.1 acar|OOF99277.1 afla|EED51470.1 agla|OJJ84709.1 asyd|OJJ59181.1 atub|OJI82503.1 aver|OJJ01802.1 awen|OJJ37966.1 aory|EIT80465.1

Group8852: abra|OJJ66093.1 aluc|OJZ81265.1 anid|CBF78286.1 asyd|OJJ55817.1 atub|OJI91465.1 afla|EED50997.1 akaw|GAA88007.1 asyd|OJJ53628.1 aver|OJJ05945.1

Group8853: abra|OJJ66103.1 acar|OOF90360.1 acla|EAW15357.1 afis|EAW20355.1 afum|EAL85472.1 akaw|GAA92670.1 aluc|OJZ90125.1 anig|EHA20946.1 atub|OJI84671.1

Group8854: abra|OJJ66106.1 afla|EED56755.1 akaw|GAA92675.1 aluc|OJZ90130.1 anig|EHA20941.1 aory|EIT73895.1 asyd|OJJ57251.1 atub|OJI84666.1 aver|OJJ03043.1

Group8855: abra|OJJ66118.1 acar|OOF94010.1 afla|EED52122.1 akaw|GAA92680.1 aluc|OJZ90135.1 anig|EHA20931.1 aory|EIT74381.1 atub|OJI84661.1 aver|OJI96544.1

Group8856: abra|OJJ66127.1 afis|EAW24043.1 afla|EED54372.1 afum|EAL84419.1 akaw|GAA92686.1 aluc|OJZ90144.1 aory|EIT76043.1 ater|EAU31255.1 atub|OJI84655.1

Group8857: abra|OJJ66155.1 acar|OOF92445.1 afla|EED52224.1 awen|OJJ31085.1 akaw|GAA86454.1 aluc|OJZ84996.1 anig|EHA25209.1 atub|OJI80602.1 aory|EIT74340.1

Group8858: afis|EAW20704.1 aluc|OJZ79614.1 aory|EIT83020.1 afum|EAL89354.1 agla|OJJ84888.1 akaw|GAA92657.1 afla|EED52365.1 agla|OJJ88573.1 asyd|OJJ52005.1

Group8859: abra|OJJ66190.1 acar|OOF90355.1 afis|EAW25757.1 afla|EED45251.1 anig|EHA25699.1 aory|EIT73746.1 ater|EAU33160.1 atub|OJI88436.1 azon|OJJ51136.1

Group8860: abra|OJJ66220.1 acar|OOF90663.1 acla|EAW14125.1 afum|EAL93564.1 akaw|GAA89066.1 aluc|OJZ80526.1 anig|EHA25669.1 atub|OJI88404.1 afis|EAW18830.1

Group8861: abra|OJJ66229.1 acar|OOF90687.1 akaw|GAA89060.1 aluc|OJZ80534.1 anig|EHA25661.1 asyd|OJJ64303.1 atub|OJI88397.1 aver|OJI98396.1 awen|OJJ31376.1

Group8862: abra|OJJ66270.1 acla|EAW12150.1 afis|EAW19364.1 akaw|GAA84341.1 aluc|OJZ89041.1 atub|OJI85749.1 awen|OJJ31987.1 azon|OJJ45320.1 afum|EAL94083.1

Group8863: abra|OJJ66330.1 afla|EED47462.1 akaw|GAA90207.1 aluc|OJZ83138.1 anid|CBF74204.1 anig|EHA28197.1 aory|EIT79966.1 atub|OJI87796.1 awen|OJJ31027.1

Group8864: abra|OJJ66340.1 acar|OOF98712.1 afla|EED54087.1 akaw|GAA90214.1 aluc|OJZ83124.1 anig|EHA28189.1 atub|OJI87808.1 agla|OJJ87383.1 aluc|OJZ83129.1

Group8865: awen|OJJ36570.1 acla|EAW09871.1 afis|EAW21255.1 afla|EED53205.1 afum|EAL92270.1 anid|CBF87366.1 aory|EIT75204.1 asyd|OJJ52837.1 aver|OJJ08810.1

Group8866: abra|OJJ66419.1 acla|EAW08780.1 afis|EAW15621.1 afum|EAL85713.2 akaw|GAA90285.1 aluc|OJZ83055.1 anig|EHA28121.1 atub|OJI87892.1 azon|OJJ51150.1

Group8867: abra|OJJ66431.1 acar|OOF93322.1 akaw|GAA90298.1 aluc|OJZ83039.1 atub|OJI87910.1 asyd|OJJ53812.1 aver|OJJ07913.1 acar|OOF93086.1 agla|OJJ84363.1

Group8868: abra|OJJ66440.1 acar|OOF91486.1 afla|EED47532.1 agla|OJJ85807.1 aory|EIT74584.1 asyd|OJJ59093.1 ater|EAU32970.1 aver|OJJ01683.1 awen|OJJ41964.1

Group8869: aver|OJJ08488.1 abra|OJJ66453.1 afis|EAW15393.1 afla|EED47384.1 agla|OJJ79142.1 anid|CBF76215.1 aory|EIT82853.1 asyd|OJJ56276.1 aver|OJJ05915.1

Group8870: abra|OJJ66464.1 afla|EED53775.1 aory|EIT78428.1 abra|OJJ75242.1 acar|OOF94551.1 akaw|GAA91785.1 aluc|OJZ92804.1 anig|EHA26205.1 atub|OJI87247.1

Group8871: abra|OJJ66541.1 afla|EED46585.1 agla|OJJ85289.1 akaw|GAA91166.1 aluc|OJZ81494.1 anig|EHA25936.1 aory|EIT77950.1 atub|OJI88687.1 awen|OJJ31276.1

Group8872: abra|OJJ66747.1 acar|OOF93831.1 afla|EED56702.1 akaw|GAA89683.1 aluc|OJZ80702.1 anig|EHA27314.1 aory|EIT73908.1 atub|OJI87561.1 azon|OJJ42916.1

Group8873: abra|OJJ66748.1 acla|EAW13730.1 akaw|GAA89684.1 aluc|OJZ80701.1 anig|EHA27315.1 atub|OJI87560.1 afla|EED56703.1 ater|EAU37677.1 awen|OJJ36900.1

Group8874: afla|EED51157.1 aory|EIT81303.1 abra|OJJ72182.1 azon|OJJ44533.1 akaw|GAA92010.1 aluc|OJZ92875.1 anig|EHA26084.1 atub|OJI87335.1 aver|OJI99513.1

Group8875: abra|OJJ66950.1 akaw|GAA89085.1 aluc|OJZ80506.1 anid|CBF85596.1 anig|EHA25687.1 asyd|OJJ61738.1 atub|OJI88424.1 aver|OJI95835.1 akaw|GAA89084.1

Group8876: abra|OJJ66979.1 acar|OOF93632.1 akaw|GAA92185.1 aluc|OJZ85595.1 anig|EHA24927.1 atub|OJI79391.1 awen|OJJ33815.1 afla|EED52199.1 aory|EIT74405.1

Group8877: abra|OJJ66985.1 acar|OOF93627.1 afla|EED53106.1 akaw|GAA92179.1 aluc|OJZ85589.1 anig|EHA24932.1 aory|EIT74805.1 atub|OJI79384.1 awen|OJJ33628.1

Group8878: abra|OJJ67043.1 acar|OOF91820.1 anig|EHA18600.1 abra|OJJ67565.1 akaw|GAA88100.1 aluc|OJZ81367.1 anig|EHA18666.1 atub|OJI91571.1 afla|EED51381.1

Group8879: abra|OJJ67089.1 acar|OOF97130.1 acla|EAW06853.1 agla|OJJ84328.1 abra|OJJ67529.1 akaw|GAA88133.1 aluc|OJZ81404.1 atub|OJI91611.1 aver|OJI99294.1

Group8880: abra|OJJ67111.1 acar|OOF98441.1 akaw|GAA91053.1 aluc|OJZ87864.1 anig|EHA20170.1 asyd|OJJ54576.1 atub|OJI80784.1 aver|OJJ07578.1 azon|OJJ45866.1

Group8881: abra|OJJ67143.1 acar|OOF98412.1 afis|EAW18022.1 afum|EAL91834.1 akaw|GAA91026.1 aluc|OJZ87829.1 anig|EHA20308.1 atub|OJI80812.1 awen|OJJ32548.1

Group8882: abra|OJJ67249.1 acla|EAW10029.1 afis|EAW20707.1 afum|EAL86558.1 agla|OJJ79456.1 akaw|GAA91668.1 aluc|OJZ87728.1 atub|OJI80912.1 awen|OJJ36438.1

Group8883: abra|OJJ67275.1 acar|OOF95205.1 afla|EED55153.1 agla|OJJ79074.1 akaw|GAA92071.1 aluc|OJZ81158.1 anid|CBF81933.1 aory|EIT81614.1 aver|OJJ01872.1

Group8884: abra|OJJ67305.1 acar|OOF92508.1 afla|EED45063.1 akaw|GAA86511.1 aluc|OJZ85056.1 anig|EHA25272.1 aory|EIT73803.1 ater|EAU36095.1 atub|OJI80544.1

Group8885: abra|OJJ67371.1 acar|OOF92440.1 akaw|GAA86451.1 aluc|OJZ84994.1 asyd|OJJ64318.1 ater|EAU32176.1 atub|OJI80606.1 aver|OJI98411.1 azon|OJJ50421.1

Group8886: abra|OJJ67373.1 acar|OOF92438.1 afla|EED46879.1 akaw|GAA86449.1 aluc|OJZ84992.1 anig|EHA25204.1 aory|EIT72545.1 atub|OJI80608.1 azon|OJJ42630.1

Group8887: abra|OJJ67389.1 acar|OOF92426.1 akaw|GAA86436.1 aluc|OJZ84979.1 anig|EHA25191.1 atub|OJI80623.1 aver|OJJ00924.1 afis|EAW15660.1 afis|EAW18259.1

Group8888: aory|EIT78684.1 asyd|OJJ58835.1 aver|OJJ01438.1 acar|OOF92411.1 awen|OJJ31017.1 azon|OJJ50530.1 acla|EAW06353.1 agla|OJJ85049.1 asyd|OJJ58837.1

Group8889: asyd|OJJ52824.1 aver|OJJ08800.1 abra|OJJ77425.1 afla|EED48705.1 akaw|GAA86622.1 aluc|OJZ85177.1 anid|CBF78741.1 aory|EIT73120.1 ater|EAU33969.1

Group8890: aluc|OJZ87017.1 afla|EED49711.1 agla|OJJ88115.1 akaw|GAA84672.1 anid|CBF84384.1 asyd|OJJ53588.1 atub|OJI83501.1 aver|OJJ08160.1 awen|OJJ29858.1

Group8891: agla|OJJ80602.1 awen|OJJ32426.1 acla|EAW14154.1 afis|EAW18867.1 afla|EED54191.1 afum|EAL93599.1 asyd|OJJ53475.1 ater|EAU38031.1 aver|OJJ08282.1

Group8892: abra|OJJ67639.1 afum|EAL86601.1 akaw|GAA89157.1 aluc|OJZ81677.1 anig|EHA25764.1 asyd|OJJ56417.1 ater|EAU35975.1 atub|OJI88504.1 awen|OJJ40907.1

Group8893: abra|OJJ67642.1 acar|OOG00328.1 acla|EAW11927.1 agla|OJJ80933.1 akaw|GAA89162.1 aluc|OJZ81672.1 anig|EHA25768.1 atub|OJI88509.1 ater|EAU32792.1

Group8894: abra|OJJ67655.1 acar|OOF91987.1 agla|OJJ86443.1 akaw|GAA89171.1 aluc|OJZ81662.1 anig|EHA25781.1 asyd|OJJ62557.1 atub|OJI88521.1 aver|OJI96682.1

Group8895: abra|OJJ67661.1 afla|EED45941.1 akaw|GAA89176.1 aluc|OJZ81657.1 anig|EHA25788.1 ater|EAU36101.1 atub|OJI88526.1 awen|OJJ40653.1 aory|EIT75369.1

Group8896: abra|OJJ67681.1 acla|EAW10034.1 afis|EAW19874.1 afum|EAL85025.1 aluc|OJZ81637.1 anig|EHA25807.1 ater|EAU30906.1 atub|OJI88543.1 akaw|GAA89194.1

Group8897: abra|OJJ67696.1 acar|OOF90176.1 afla|EED54114.1 akaw|GAA89206.1 aluc|OJZ81624.1 anig|EHA25819.1 aory|EIT76104.1 atub|OJI88556.1 awen|OJJ37736.1

Group8898: afla|EED56802.1 aluc|OJZ81576.1 azon|OJJ43211.1 aory|EIT73865.1 asyd|OJJ56359.1 ater|EAU37206.1 atub|OJI88605.1 aver|OJJ06355.1 akaw|GAA92443.1

Group8899: abra|OJJ67719.1 acar|OOF92594.1 anig|EHA25832.1 atub|OJI88570.1 abra|OJJ70189.1 akaw|GAA86586.1 anid|CBF81815.1 azon|OJJ43538.1 aver|OJI95867.1

Group8900: abra|OJJ67770.1 acla|EAW09030.1 akaw|GAA92456.1 aluc|OJZ81561.1 anig|EHA25872.1 atub|OJI88623.1 acar|OOF98687.1 ater|EAU37843.1 afum|EAL84353.1

Group8901: abra|OJJ67773.1 agla|OJJ85341.1 akaw|GAA82754.1 aluc|OJZ87311.1 anid|CBF76237.1 asyd|OJJ61546.1 atub|OJI81334.1 aver|OJJ00873.1 awen|OJJ32606.1

Group8902: abra|OJJ67814.1 afla|EED49432.1 agla|OJJ85427.1 akaw|GAA91086.1 aluc|OJZ87903.1 aver|OJJ00270.1 awen|OJJ29686.1 agla|OJJ84211.1 asyd|OJJ60944.1

Group8903: anid|CBF78806.1 acla|EAW14611.1 afis|EAW19340.1 afla|EED47548.1 afum|EAL94059.1 aory|EIT74531.1 asyd|OJJ65258.1 ater|EAU29442.1 aver|OJI99409.1

Group8904: abra|OJJ67878.1 akaw|GAA83771.1 aluc|OJZ90957.1 anig|EHA22245.1 asyd|OJJ55666.1 ater|EAU37359.1 atub|OJI82947.1 afla|EED44755.1 aory|EIT77771.1

Group8905: abra|OJJ67992.1 acar|OOF92034.1 afla|EED45378.1 akaw|GAA90956.1 aluc|OJZ82866.1 anig|EHA21296.1 aory|EIT82449.1 atub|OJI89091.1 awen|OJJ36168.1

Group8906: abra|OJJ68003.1 acar|OOF92045.1 akaw|GAA90947.1 aluc|OJZ82877.1 anig|EHA21303.1 ater|EAU31764.1 atub|OJI89101.1 azon|OJJ48316.1 agla|OJJ78692.1

Group8907: abra|OJJ68049.1 acar|OOF98060.1 afla|EED53913.1 akaw|GAA90907.1 aluc|OJZ82919.1 anig|EHA21343.1 ater|EAU33484.1 atub|OJI89145.1 azon|OJJ50310.1

Group8908: abra|OJJ68093.1 acar|OOF93895.1 akaw|GAA86877.1 aluc|OJZ90001.1 anid|CBF77160.1 anig|EHA27376.1 asyd|OJJ64792.1 atub|OJI84807.1 aver|OJI98904.1

Group8909: abra|OJJ68098.1 acar|OOF93901.1 akaw|GAA86883.1 aluc|OJZ90006.1 anig|EHA27381.1 ater|EAU31284.1 atub|OJI84801.1 awen|OJJ34742.1 azon|OJJ45817.1

Group8910: abra|OJJ68101.1 acar|OOF93904.1 agla|OJJ79459.1 aluc|OJZ90009.1 anid|CBF84395.1 anig|EHA27384.1 atub|OJI84798.1 azon|OJJ43872.1 agla|OJJ82036.1

Group8911: abra|OJJ68102.1 acar|OOF93905.1 akaw|GAA86886.1 aluc|OJZ90010.1 anig|EHA27385.1 atub|OJI84797.1 awen|OJJ36287.1 agla|OJJ82040.1 azon|OJJ43873.1

Group8912: abra|OJJ68127.1 afla|EED46561.1 agla|OJJ79758.1 akaw|GAA86908.1 aluc|OJZ90031.1 anig|EHA27407.1 ater|EAU32450.1 atub|OJI84776.1 aory|EIT77840.1

Group8913: ater|EAU37042.1 afis|EAW17607.1 afum|EAL91420.1 agla|OJJ81354.1 anid|CBF90183.1 anig|EHA18916.1 asyd|OJJ61985.1 aver|OJI96077.1 awen|OJJ37235.1

Group8914: afis|EAW19918.1 afum|EAL84990.1 ater|EAU32636.1 afla|EED46192.1 agla|OJJ82676.1 akaw|GAA90167.1 aory|EIT76440.1 awen|OJJ36109.1 azon|OJJ50724.1

Group8915: abra|OJJ68232.1 afla|EED46472.1 agla|OJJ86621.1 aluc|OJZ85539.1 aory|EIT73944.1 asyd|OJJ55531.1 aver|OJJ06606.1 aory|EIT72409.1 afla|EED56760.1

Group8916: abra|OJJ68261.1 ater|EAU38186.1 abra|OJJ71565.1 acla|EAW08364.1 afis|EAW20354.1 akaw|GAA89359.1 aluc|OJZ86302.1 anig|EHA21570.1 atub|OJI84234.1

Group8917: abra|OJJ68300.1 acar|OOF89816.1 akaw|GAA83174.1 anid|CBF78847.1 anig|EHA18828.1 asyd|OJJ56504.1 aver|OJJ06215.1 asyd|OJJ64567.1 aver|OJI98664.1

Group8918: abra|OJJ68308.1 acar|OOF94000.1 akaw|GAA83166.1 aluc|OJZ83665.1 anig|EHA18821.1 asyd|OJJ57949.1 atub|OJI90036.1 azon|OJJ42302.1 azon|OJJ42657.1

Group8919: abra|OJJ68331.1 acar|OOF97608.1 acla|EAW11059.1 agla|OJJ82217.1 akaw|GAA92083.1 aluc|OJZ83641.1 anig|EHA18794.1 atub|OJI90010.1 awen|OJJ30930.1

Group8920: abra|OJJ68349.1 acar|OOF94843.1 afla|EED45477.1 akaw|GAA92116.1 aluc|OJZ83609.1 anid|CBF80535.1 anig|EHA18768.1 asyd|OJJ54655.1 atub|OJI89982.1

Group8921: abra|OJJ68436.1 afla|EED56857.1 akaw|GAA86935.1 aluc|OJZ90061.1 anig|EHA19937.1 aory|EIT77557.1 asyd|OJJ55051.1 atub|OJI84746.1 aver|OJI99561.1

Group8922: abra|OJJ68443.1 afis|EAW20783.1 afla|EED47899.1 afum|EAL84839.2 akaw|GAA85957.1 aluc|OJZ91104.1 atub|OJI83102.1 awen|OJJ29871.1 aluc|OJZ91105.1

Group8923: abra|OJJ68560.1 awen|OJJ41573.1 acla|EAW12805.1 afum|EAL91780.1 akaw|GAA91293.1 aluc|OJZ82776.1 anig|EHA21219.1 atub|OJI89008.1 azon|OJJ48286.1

Group8924: abra|OJJ68584.1 acla|EAW14987.1 atub|OJI91458.1 abra|OJJ76613.1 afla|EED48309.1 anig|EHA22622.1 aory|EIT76294.1 asyd|OJJ53394.1 aver|OJJ02224.1

Group8925: abra|OJJ68810.1 acar|OOF90613.1 afla|EED51121.1 akaw|GAA90444.1 aluc|OJZ85782.1 anid|CBF71803.1 anig|EHA24773.1 aory|EIT81320.1 atub|OJI79567.1

Group8926: abra|OJJ68812.1 akaw|GAA90442.1 aluc|OJZ85785.1 anid|CBF87323.1 ater|EAU30074.1 atub|OJI79569.1 awen|OJJ40584.1 akaw|GAA92139.1 atub|OJI79311.1

Group8927: abra|OJJ68824.1 akaw|GAA90428.1 aluc|OJZ85801.1 anid|CBF76151.1 anig|EHA24756.1 asyd|OJJ55694.1 ater|EAU37616.1 atub|OJI79586.1 aver|OJJ06779.1

Group8928: abra|OJJ68825.1 acar|OOF97565.1 acla|EAW12169.1 afis|EAW25751.1 anig|EHA24755.1 asyd|OJJ53428.1 ater|EAU36731.1 aver|OJJ03483.1 aver|OJJ08317.1

Group8929: abra|OJJ68874.1 acar|OOF89967.1 acla|EAW15246.1 akaw|GAA90381.1 aluc|OJZ85850.1 asyd|OJJ63053.1 awen|OJJ31218.1 azon|OJJ46139.1 aver|OJI97910.1

Group8930: abra|OJJ68880.1 agla|OJJ80106.1 akaw|GAA90376.1 aluc|OJZ85856.1 anig|EHA28094.1 asyd|OJJ54912.1 ater|EAU31014.1 atub|OJI79642.1 aver|OJJ07291.1

Group8931: abra|OJJ68901.1 acar|OOF90516.1 agla|OJJ85340.1 akaw|GAA90350.1 aluc|OJZ85882.1 anig|EHA24704.1 atub|OJI79668.1 azon|OJJ45042.1 afis|EAW17222.1

Group8932: abra|OJJ69018.1 acar|OOF94255.1 afla|EED45435.1 akaw|GAA90535.1 aluc|OJZ86004.1 anig|EHA24606.1 aory|EIT82428.1 atub|OJI79793.1 azon|OJJ49774.1

Group8933: abra|OJJ69041.1 acar|OOF90577.1 akaw|GAA90570.1 aluc|OJZ86026.1 asyd|OJJ59199.1 ater|EAU31579.1 atub|OJI79815.1 aver|OJI98873.1 asyd|OJJ64773.1

Group8934: abra|OJJ69050.1 acla|EAW09006.1 afis|EAW25586.1 anid|CBF80548.1 asyd|OJJ61364.1 ater|EAU37357.1 atub|OJI79827.1 aver|OJJ00691.1 asyd|OJJ54680.1

Group8935: abra|OJJ69334.1 acar|OOF94393.1 acla|EAW13648.1 akaw|GAA88204.1 aluc|OJZ82166.1 anig|EHA28566.1 atub|OJI88271.1 afum|EAL84835.1 afis|EAW15537.1

Group8936: afum|EBA27265.1 agla|OJJ83228.1 aory|EIT74428.1 ater|EAU30410.1 awen|OJJ32881.1 afum|EAL86734.1 aluc|OJZ79925.1 afum|EAL87784.1 anig|EHA21823.1

Group8937: abra|OJJ69445.1 akaw|GAA88373.1 aluc|OJZ82281.1 atub|OJI88158.1 azon|OJJ49268.1 acar|OOG00594.1 awen|OJJ33653.1 agla|OJJ84838.1 ater|EAU36550.1

Group8938: abra|OJJ69538.1 acla|EAW13735.1 afla|EED53088.1 agla|OJJ85439.1 akaw|GAA88386.1 aluc|OJZ82374.1 anig|EHA28390.1 atub|OJI88068.1 aory|EIT74780.1

Group8939: abra|OJJ69623.1 afis|EAW16749.1 akaw|GAA92928.1 aluc|OJZ82519.1 anid|CBF81981.1 asyd|OJJ56152.1 atub|OJI88773.1 aver|OJJ05091.1 azon|OJJ46546.1

Group8940: abra|OJJ69719.1 afla|EED55123.1 akaw|GAA90172.1 aluc|OJZ83178.1 anid|CBF84456.1 anig|EHA28252.1 aory|EIT81596.1 ater|EAU34461.1 atub|OJI87759.1

Group8941: abra|OJJ69723.1 acar|OOF94193.1 akaw|GAA90176.1 aluc|OJZ83174.1 anig|EHA28248.1 aory|EIT74486.1 atub|OJI87763.1 azon|OJJ42480.1 afla|EED52161.1

Group8942: abra|OJJ69742.1 acar|OOF98455.1 akaw|GAA90195.1 aluc|OJZ83153.1 anig|EHA28219.1 atub|OJI87784.1 azon|OJJ50964.1 agla|OJJ84791.1 aory|EIT82297.1

Group8943: acla|EAW09548.1 aver|OJJ01277.1 awen|OJJ35779.1 azon|OJJ49415.1 asyd|OJJ58667.1 acar|OOF99052.1 akaw|GAA83946.1 aluc|OJZ80827.1 anig|EHA19488.1

Group8944: abra|OJJ69754.1 acar|OOG00378.1 agla|OJJ78965.1 akaw|GAA90205.1 aluc|OJZ83140.1 anig|EHA21149.1 asyd|OJJ56043.1 atub|OJI87794.1 anig|EHA28207.1

Group8945: abra|OJJ69770.1 afis|EAW19884.1 afla|EED49540.1 afum|EAL85015.1 akaw|GAA87883.1 aluc|OJZ85528.1 anig|EHA25059.1 aory|EIT82582.1 atub|OJI80040.1

Group8946: abra|OJJ69783.1 afis|EAW20781.1 afla|EED47442.1 akaw|GAA87873.1 aluc|OJZ85514.1 anig|EHA25047.1 aory|EIT81907.1 ater|EAU36877.1 atub|OJI80052.1

Group8947: abra|OJJ69825.1 acar|OOF93350.1 afla|EED45808.1 akaw|GAA87835.1 aluc|OJZ85471.1 anig|EHA25012.1 aory|EIT75743.1 atub|OJI80092.1 abra|OJJ78054.1

Group8948: afis|EAW17801.1 afum|EAL91609.1 acla|EAW12875.1 afla|EED45793.1 aory|EIT75679.1 ater|EAU31830.1 asyd|OJJ62341.1 awen|OJJ35263.1 azon|OJJ46433.1

Group8949: abra|OJJ69855.1 acar|OOF97521.1 akaw|GAA87801.1 aluc|OJZ85436.1 anig|EHA24982.1 asyd|OJJ52920.1 atub|OJI80132.1 aver|OJJ08889.1 awen|OJJ38306.1

Group8950: abra|OJJ69875.1 acar|OOF93389.1 akaw|GAA87781.1 aluc|OJZ85416.1 anig|EHA25595.1 asyd|OJJ52367.1 ater|EAU36015.1 atub|OJI80160.1 aver|OJJ01980.1

Group8951: abra|OJJ69878.1 acar|OOF93393.1 agla|OJJ86558.1 akaw|GAA87779.1 aluc|OJZ85414.1 anig|EHA25593.1 asyd|OJJ61435.1 atub|OJI80163.1 aver|OJJ00769.1

Group8952: abra|OJJ70030.1 afis|EAW17108.1 aluc|OJZ80263.1 anig|EHA18715.1 asyd|OJJ54267.1 ater|EAU29704.1 atub|OJI79319.1 aver|OJJ05219.1 awen|OJJ31369.1

Group8953: afis|EAW16686.1 afla|EED45678.1 afis|EAW19302.1 agla|OJJ87730.1 asyd|OJJ57614.1 aver|OJJ05571.1 awen|OJJ32519.1 akaw|GAA85268.1 aluc|OJZ90626.1

Group8954: acar|OOF90922.1 abra|OJJ74950.1 agla|OJJ88326.1 akaw|GAA83472.1 aluc|OJZ92523.1 anig|EHA26452.1 atub|OJI86957.1 awen|OJJ39641.1 acar|OOF95614.1

Group8955: abra|OJJ70097.1 afis|EAW23566.1 afla|EED56882.1 afum|EAL87902.1 anid|CBF71436.1 aory|EIT77475.1 asyd|OJJ57197.1 aver|OJJ03094.1 acar|OOF95926.1

Group8956: abra|OJJ70152.1 afla|EED52836.1 anid|CBF90331.1 aory|EIT78651.1 asyd|OJJ60244.1 aver|OJJ04017.1 awen|OJJ40936.1 asyd|OJJ61905.1 aver|OJI95998.1

Group8957: abra|OJJ70162.1 acar|OOF94978.1 akaw|GAA86609.1 aluc|OJZ85160.1 ater|EAU37188.1 atub|OJI80436.1 awen|OJJ40818.1 azon|OJJ43906.1 awen|OJJ38140.1

Group8958: abra|OJJ70168.1 acar|OOF92520.1 akaw|GAA86616.1 aluc|OJZ85168.1 anid|CBF81950.1 anig|EHA25365.1 ater|EAU31386.1 atub|OJI80445.1 awen|OJJ32777.1

Group8959: agla|OJJ83231.1 acla|EAW10881.1 afis|EAW20977.1 afla|EED57501.1 afum|EAL84656.1 aory|EIT81862.1 asyd|OJJ52800.1 aver|OJJ08777.1 awen|OJJ34383.1

Group8960: afum|EAL93660.1 awen|OJJ31523.1 afis|EAW18929.1 ater|EAU38108.1 aver|OJJ07470.1 afla|EED50922.1 agla|OJJ85258.1 aory|EIT79050.1 asyd|OJJ54703.1

Group8961: abra|OJJ70240.1 agla|OJJ79705.1 akaw|GAA89191.1 aluc|OJZ81640.1 anig|EHA25315.1 asyd|OJJ54752.1 atub|OJI84631.1 awen|OJJ37884.1 awen|OJJ37480.1

Group8962: abra|OJJ70270.1 afis|EAW22229.1 anig|EHA22501.1 asyd|OJJ57193.1 ater|EAU37173.1 atub|OJI88778.1 aver|OJJ04287.1 aver|OJJ03098.1 asyd|OJJ58118.1

Group8963: abra|OJJ70276.1 acar|OOF98674.1 akaw|GAA92873.1 aluc|OJZ82538.1 anig|EHA22506.1 asyd|OJJ55018.1 atub|OJI88782.1 aver|OJJ07187.1 awen|OJJ32618.1

Group8964: abra|OJJ70288.1 acar|OOF91740.1 afla|EED48872.1 akaw|GAA92860.1 aluc|OJZ82553.1 anig|EHA22516.1 aory|EIT81240.1 atub|OJI88794.1 awen|OJJ38798.1

Group8965: abra|OJJ70305.1 acar|OOF91758.1 acla|EAW15279.1 afla|EED56373.1 akaw|GAA93017.1 aluc|OJZ82568.1 anig|EHA27506.1 aory|EIT73336.1 atub|OJI88812.1

Group8966: abra|OJJ70307.1 acar|OOF91762.1 akaw|GAA93015.1 aluc|OJZ82570.1 anid|CBF80497.1 anig|EHA27504.1 ater|EAU30197.1 atub|OJI88814.1 awen|OJJ36273.1

Group8967: abra|OJJ70337.1 agla|OJJ81765.1 akaw|GAA92590.1 aluc|OJZ82607.1 atub|OJI88842.1 anid|CBF73505.1 aver|OJJ04761.1 asyd|OJJ55813.1 azon|OJJ45029.1

Group8968: abra|OJJ70360.1 acar|OOF90575.1 acla|EAW11010.1 afis|EAW20624.1 agla|OJJ78535.1 akaw|GAA87152.1 aluc|OJZ84283.1 anid|CBF71200.1 atub|OJI89798.1

Group8969: abra|OJJ70410.1 afis|EAW17944.1 afum|EAL91755.1 agla|OJJ80394.1 aluc|OJZ82674.1 anig|EHA21040.1 asyd|OJJ57668.1 atub|OJI88909.1 awen|OJJ41817.1

Group8970: abra|OJJ70419.1 acar|OOF91850.1 akaw|GAA91938.1 aluc|OJZ82684.1 anig|EHA21052.1 atub|OJI88916.1 aver|OJJ04011.1 azon|OJJ43523.1 azon|OJJ42214.1

Group8971: abra|OJJ70435.1 afla|EED49380.1 akaw|GAA90145.1 aluc|OJZ83202.1 anig|EHA21064.1 atub|OJI87735.1 aver|OJI97289.1 awen|OJJ41820.1 asyd|OJJ63645.1

Group8972: abra|OJJ70605.1 acar|OOG01013.1 akaw|GAA89945.1 aluc|OJZ83951.1 aory|EIT74963.1 atub|OJI90322.1 awen|OJJ40125.1 abra|OJJ73870.1 ater|EAU36764.1

Group8973: anid|CBF79805.1 asyd|OJJ58127.1 aver|OJJ04278.1 acla|EAW11700.1 afis|EAW23721.1 afla|EED51507.1 afum|EAL87161.1 ater|EAU31289.1 agla|OJJ79866.1

Group8974: abra|OJJ70921.1 acar|OOF98202.1 afis|EAW21023.1 afum|EAL92062.1 akaw|GAA83019.1 aluc|OJZ87605.1 anig|EHA20804.1 atub|OJI81043.1 ater|EAU31617.1

Group8975: abra|OJJ70949.1 acar|OOF98181.1 afla|EED47498.1 agla|OJJ79400.1 akaw|GAA82995.1 aluc|OJZ87580.1 aory|EIT74595.1 atub|OJI81068.1 awen|OJJ35926.1

Group8976: abra|OJJ71055.1 agla|OJJ80677.1 awen|OJJ30197.1 abra|OJJ75795.1 acla|EAW09128.1 anig|EHA23323.1 ater|EAU39612.1 atub|OJI85554.1 awen|OJJ29800.1

Group8977: abra|OJJ71067.1 acar|OOF98078.1 akaw|GAA82895.1 aluc|OJZ87464.1 anig|EHA20686.1 atub|OJI81184.1 azon|OJJ50343.1 acla|EAW13536.1 afis|EAW16450.1

Group8978: abra|OJJ71072.1 acar|OOF98074.1 afla|EED52241.1 akaw|GAA82890.1 aluc|OJZ87459.1 anig|EHA20681.1 aory|EIT74433.1 atub|OJI81189.1 awen|OJJ30451.1

Group8979: abra|OJJ71086.1 acar|OOF98051.1 afla|EED55098.1 akaw|GAA82875.1 aluc|OJZ87442.1 anig|EHA20667.1 aory|EIT81606.1 atub|OJI81206.1 azon|OJJ42080.1

Group8980: abra|OJJ71171.1 afla|EED53726.1 afum|EBA27390.1 agla|OJJ89232.1 aluc|OJZ87370.1 anid|CBF85364.1 asyd|OJJ64041.1 azon|OJJ44561.1 afis|EAW25261.1

Group8981: abra|OJJ71200.1 akaw|GAA82783.1 aluc|OJZ87342.1 anig|EHA20579.1 asyd|OJJ60978.1 ater|EAU38955.1 atub|OJI81302.1 aver|OJI99490.1 afis|EAW16709.1

Group8982: abra|OJJ71209.1 acar|OOF97958.1 afla|EED54680.1 akaw|GAA82775.1 aluc|OJZ87334.1 anig|EHA20570.1 asyd|OJJ65152.1 ater|EAU36929.1 atub|OJI81310.1

Group8983: abra|OJJ71218.1 akaw|GAA82766.1 aluc|OJZ87325.1 anid|CBF87231.1 anig|EHA20562.1 asyd|OJJ60478.1 ater|EAU39529.1 atub|OJI81320.1 aver|OJI99798.1

Group8984: abra|OJJ71239.1 afis|EAW16931.1 afum|EAL90573.1 aluc|OJZ87306.1 awen|OJJ31868.1 azon|OJJ44471.1 awen|OJJ31396.1 afis|EAW22622.1 azon|OJJ46440.1

Group8985: abra|OJJ71292.1 acar|OOF97879.1 akaw|GAA82697.1 aluc|OJZ87251.1 anig|EHA20496.1 asyd|OJJ53542.1 ater|EAU38010.1 atub|OJI81391.1 aver|OJJ08211.1

Group8986: abra|OJJ71471.1 acla|EAW10079.1 afis|EAW25507.1 afla|EED54007.1 afum|EAL90173.2 anid|CBF77770.1 anig|EHA20344.1 aory|EIT78341.1 atub|OJI81562.1

Group8987: abra|OJJ71620.1 acar|OOF96336.1 akaw|GAA89308.1 aluc|OJZ86361.1 anig|EHA21530.1 asyd|OJJ52377.1 atub|OJI84173.1 aver|OJJ01970.1 azon|OJJ46014.1

Group8988: abra|OJJ71718.1 acar|OOF96424.1 afis|EAW18544.1 afum|EAL93244.1 akaw|GAA85172.1 aluc|OJZ86464.1 anig|EHA21446.1 atub|OJI84065.1 awen|OJJ31102.1

Group8989: aluc|OJZ84831.1 acar|OOF90314.1 afis|EAW22621.1 afla|EED47487.1 afum|EAL90575.1 anig|EHA25740.1 ater|EAU34167.1 asyd|OJJ54194.1 aver|OJJ08532.1

Group8990: abra|OJJ71733.1 afla|EED57064.1 afum|EAL85781.1 aory|EIT78315.1 atub|OJI90938.1 awen|OJJ29930.1 afla|EED53954.1 akaw|GAA88454.1 aluc|OJZ88342.1

Group8991: abra|OJJ71748.1 acar|OOF90203.1 akaw|GAA85162.1 aluc|OJZ86474.1 anig|EHA22159.1 atub|OJI84055.1 acar|OOG00119.1 ater|EAU34843.1 awen|OJJ33439.1

Group8992: abra|OJJ71759.1 afla|EED52387.1 akaw|GAA90559.1 aluc|OJZ84839.1 anid|CBF78497.1 anig|EHA25749.1 aory|EIT83296.1 asyd|OJJ53404.1 aver|OJJ04197.1

Group8993: abra|OJJ71763.1 acar|OOF93296.1 agla|OJJ82502.1 akaw|GAA85145.1 aluc|OJZ86492.1 anig|EHA22156.1 asyd|OJJ65290.1 atub|OJI84035.1 awen|OJJ33417.1

Group8994: abra|OJJ71786.1 acar|OOF96463.1 afla|EED47808.1 akaw|GAA85125.1 aluc|OJZ86516.1 aory|EIT77128.1 atub|OJI84008.1 awen|OJJ29872.1 azon|OJJ45183.1

Group8995: abra|OJJ71907.1 acar|OOF96567.1 afla|EED55961.1 akaw|GAA85013.1 aluc|OJZ86644.1 anig|EHA22025.1 aory|EIT76850.1 ater|EAU35018.1 atub|OJI83883.1

Group8996: abra|OJJ71996.1 afla|EED51564.1 akaw|GAA83855.1 aluc|OJZ86734.1 anig|EHA21945.1 aory|EIT80511.1 atub|OJI83794.1 acla|EAW11233.1 aver|OJI96463.1

Group8997: abra|OJJ72034.1 ater|EAU38314.1 afla|EED57852.1 aory|EIT81102.1 awen|OJJ33876.1 akaw|GAA84898.1 aluc|OJZ86773.1 anig|EHA21908.1 atub|OJI83750.1

Group8998: anid|CBF85629.1 afla|EED49842.1 aluc|OJZ91052.1 aory|EIT83399.1 asyd|OJJ55532.1 atub|OJI83048.1 aver|OJJ02120.1 akaw|GAA83681.1 aver|OJJ06607.1

Group8999: abra|OJJ72074.1 afis|EAW16881.1 afla|EED47936.1 akaw|GAA90490.1 aluc|OJZ85948.1 anig|EHA24651.1 aory|EIT79857.1 ater|EAU37967.1 atub|OJI79737.1

Group9000: abra|OJJ72159.1 acar|OOF96793.1 akaw|GAA84768.1 aluc|OJZ86915.1 anid|CBF87087.1 anig|EHA21729.1 atub|OJI83606.1 aver|OJI99727.1 azon|OJJ45233.1

Group9001: abra|OJJ72169.1 afla|EED47276.1 akaw|GAA84759.1 aluc|OJZ86924.1 anig|EHA21720.1 aory|EIT79221.1 atub|OJI83597.1 aver|OJI99280.1 azon|OJJ49272.1

Group9002: abra|OJJ72171.1 afla|EED47279.1 akaw|GAA84757.1 aluc|OJZ86926.1 anig|EHA21718.1 aory|EIT79188.1 atub|OJI83595.1 aver|OJI99281.1 azon|OJJ49274.1

Group9003: abra|OJJ72211.1 acar|OOF96831.1 akaw|GAA84723.1 aluc|OJZ86963.1 anig|EHA21685.1 asyd|OJJ62585.1 ater|EAU39313.1 atub|OJI83560.1 awen|OJJ32710.1

Group9004: abra|OJJ72222.1 acar|OOF96845.1 afum|EAL89019.1 agla|OJJ84645.1 akaw|GAA84715.1 aluc|OJZ86972.1 anig|EHA21676.1 atub|OJI83549.1 aver|OJI99824.1

Group9005: abra|OJJ72247.1 acar|OOF96875.1 afla|EED45525.1 akaw|GAA84691.1 aluc|OJZ86996.1 anig|EHA21649.1 aory|EIT75809.1 atub|OJI83523.1 awen|OJJ38801.1

Group9006: abra|OJJ72249.1 acar|OOF96880.1 afis|EAW17082.1 akaw|GAA84688.1 aluc|OJZ87000.1 anig|EHA21646.1 ater|EAU32328.1 atub|OJI83519.1 awen|OJJ37062.1

Group9007: abra|OJJ72254.1 acar|OOF96885.1 afis|EAW25585.1 afla|EED51556.1 akaw|GAA84683.1 aluc|OJZ87005.1 anig|EHA21641.1 atub|OJI83514.1 anid|CBF73951.1

Group9008: abra|OJJ72259.1 acar|OOF96892.1 afis|EAW25190.1 afla|EED52816.1 akaw|GAA84679.1 aluc|OJZ87010.1 anig|EHA21637.1 aory|EIT78151.1 atub|OJI83509.1

Group9009: abra|OJJ72305.1 acar|OOF94913.1 agla|OJJ82675.1 akaw|GAA86056.1 aluc|OJZ90286.1 anid|CBF81930.1 anig|EHA23776.1 atub|OJI81710.1 awen|OJJ32540.1

Group9010: abra|OJJ72312.1 acar|OOF94898.1 afla|EED53607.1 aluc|OJZ90294.1 anig|EHA23783.1 aory|EIT75130.1 ater|EAU30388.1 atub|OJI81718.1 azon|OJJ42687.1

Group9011: abra|OJJ72556.1 afla|EED52281.1 akaw|GAA85189.1 aluc|OJZ90544.1 anig|EHA23992.1 aory|EIT74482.1 asyd|OJJ55179.1 atub|OJI81966.1 aver|OJJ07028.1

Group9012: abra|OJJ72614.1 akaw|GAA85239.1 aluc|OJZ90596.1 anid|CBF83186.1 anig|EHA24035.1 asyd|OJJ55548.1 atub|OJI82020.1 aver|OJJ06625.1 awen|OJJ36246.1

Group9013: abra|OJJ72636.1 afis|EAW20564.1 afla|EED49091.1 akaw|GAA85257.1 aluc|OJZ90615.1 aory|EIT82551.1 atub|OJI82041.1 awen|OJJ41931.1 afum|EAL84497.1

Group9014: abra|OJJ72702.1 acar|OOF92091.1 agla|OJJ79098.1 aluc|OJZ82954.1 asyd|OJJ54674.1 atub|OJI89186.1 aver|OJJ07495.1 awen|OJJ30434.1 akaw|GAA90879.1

Group9015: abra|OJJ72959.1 acar|OOF96141.1 afla|EED48496.1 akaw|GAA85539.1 aluc|OJZ83467.1 anig|EHA24311.1 aory|EIT73273.1 atub|OJI82361.1 aory|EIT73240.1

Group9016: abra|OJJ73001.1 acar|OOF96174.1 afla|EED54972.1 akaw|GAA85579.1 aluc|OJZ83515.1 aory|EIT80334.1 ater|EAU29598.1 atub|OJI82406.1 awen|OJJ35693.1

Group9017: abra|OJJ73066.1 acar|OOG00300.1 afla|EED46118.1 akaw|GAA88094.1 aluc|OJZ81360.1 anig|EHA18660.1 asyd|OJJ52257.1 atub|OJI91564.1 anig|EHA19258.1

Group9018: abra|OJJ73084.1 acar|OOG00318.1 agla|OJJ83103.1 akaw|GAA88076.1 aluc|OJZ81339.1 anid|CBF84304.1 anig|EHA18645.1 asyd|OJJ52229.1 atub|OJI91544.1

Group9019: abra|OJJ73090.1 acar|OOG00324.1 afla|EED48399.1 akaw|GAA88069.1 aluc|OJZ81334.1 anig|EHA18638.1 aory|EIT83056.1 atub|OJI91537.1 awen|OJJ37776.1

Group9020: azon|OJJ42284.1 abra|OJJ74440.1 acar|OOF95306.1 akaw|GAA87333.1 aluc|OJZ89681.1 anid|CBF90361.1 anig|EHA26917.1 atub|OJI85119.1 awen|OJJ34148.1

Group9021: abra|OJJ73136.1 afla|EED52276.1 akaw|GAA88036.1 aluc|OJZ81295.1 anig|EHA18593.1 aory|EIT74346.1 asyd|OJJ53764.1 atub|OJI91504.1 aver|OJJ07871.1

Group9022: abra|OJJ73162.1 acar|OOF93999.1 acla|EAW09982.1 afla|EED47463.1 akaw|GAA92475.1 aluc|OJZ91485.1 aory|EIT79946.1 awen|OJJ37807.1 azon|OJJ51118.1

Group9023: abra|OJJ73180.1 acar|OOG01004.1 aory|EIT81927.1 asyd|OJJ58379.1 aver|OJI95996.1 awen|OJJ29660.1 afla|EED47400.1 asyd|OJJ65394.1 asyd|OJJ61903.1

Group9024: abra|OJJ73273.1 acar|OOG00513.1 akaw|GAA92733.1 aluc|OJZ88776.1 anig|EHA18470.1 atub|OJI91348.1 agla|OJJ82552.1 afla|EED48991.1 awen|OJJ30164.1

Group9025: ater|EAU34865.1 acar|OOG00742.1 acla|EAW12163.1 afis|EAW24631.1 anid|CBF76034.1 asyd|OJJ52455.1 aver|OJJ03958.1 azon|OJJ46552.1 aory|EIT75408.1

Group9026: abra|OJJ73334.1 acla|EAW14727.1 afis|EAW18090.1 agla|OJJ85993.1 akaw|GAA83151.1 aluc|OJZ88707.1 anig|EHA18393.1 atub|OJI91289.1 azon|OJJ45141.1

Group9027: abra|OJJ73369.1 anig|EHA18353.1 abra|OJJ73370.1 acar|OOG00575.1 akaw|GAA83115.1 aluc|OJZ88670.1 atub|OJI91252.1 ater|EAU37399.1 awen|OJJ29987.1

Group9028: abra|OJJ73406.1 acar|OOG00603.1 afla|EED48700.1 akaw|GAA83085.1 aluc|OJZ88632.1 anid|CBF78732.1 anig|EHA18322.1 aory|EIT73282.1 atub|OJI91216.1

Group9029: abra|OJJ73538.1 acar|OOG00714.1 agla|OJJ80855.1 aluc|OJZ88521.1 asyd|OJJ62810.1 atub|OJI91106.1 aver|OJI96856.1 awen|OJJ35111.1 azon|OJJ49982.1

Group9030: abra|OJJ73554.1 acar|OOG00727.1 afis|EAW22445.1 afla|EED51935.1 afum|EAL90746.1 akaw|GAA88599.1 aluc|OJZ88507.1 anig|EHA18214.1 atub|OJI91093.1

Group9031: abra|OJJ73614.1 acar|OOG00797.1 acla|EAW14617.1 akaw|GAA88539.1 aluc|OJZ88435.1 anid|CBF85631.1 anig|EHA18159.1 ater|EAU33513.1 atub|OJI91032.1

Group9032: abra|OJJ73698.1 acar|OOF90621.1 afis|EAW17756.1 afla|EED46199.1 agla|OJJ87338.1 akaw|GAA93226.1 anig|EHA21868.1 azon|OJJ42143.1 azon|OJJ45205.1

Group9033: abra|OJJ73855.1 acar|OOF98203.1 afla|EED47131.1 agla|OJJ80857.1 akaw|GAA89471.1 aluc|OJZ88193.1 anig|EHA17944.1 aory|EIT79268.1 atub|OJI90791.1

Group9034: abra|OJJ73940.1 afis|EAW24046.1 akaw|GAA89545.1 aluc|OJZ88104.1 anig|EHA17873.1 ater|EAU30555.1 atub|OJI90712.1 afum|EAL91906.2 azon|OJJ43061.1

Group9035: afla|EED45229.1 aory|EIT73829.1 acar|OOF90596.1 afis|EAW24743.1 afum|EAL89389.1 anid|CBF84284.1 asyd|OJJ58313.1 aver|OJJ02098.1 azon|OJJ42092.1

Group9036: abra|OJJ74009.1 acar|OOG01132.1 akaw|GAA87490.1 aluc|OJZ88035.1 anig|EHA17808.1 asyd|OJJ59920.1 atub|OJI90641.1 aver|OJJ02366.1 anid|CBF75201.1

Group9037: abra|OJJ74097.1 acar|OOG01206.1 agla|OJJ83309.1 aluc|OJZ87939.1 asyd|OJJ63261.1 ater|EAU30643.1 aver|OJI97696.1 awen|OJJ38585.1 azon|OJJ49527.1

Group9038: abra|OJJ74130.1 acar|OOF95607.1 acla|EAW09987.1 afis|EAW21106.1 afum|EAL92127.1 akaw|GAA89756.1 aluc|OJZ80627.1 anig|EHA27204.1 atub|OJI87483.1

Group9039: abra|OJJ74162.1 acar|OOF95560.1 afla|EED49721.1 akaw|GAA86854.1 aluc|OJZ89974.1 anig|EHA27171.1 aory|EIT77653.1 atub|OJI84833.1 awen|OJJ30124.1

Group9040: abra|OJJ74239.1 acar|OOF95491.1 akaw|GAA86781.1 aluc|OJZ89893.1 anig|EHA27092.1 ater|EAU34322.1 atub|OJI84914.1 awen|OJJ34890.1 azon|OJJ50058.1

Group9041: abra|OJJ74244.1 afla|EED48142.1 akaw|GAA86776.1 aluc|OJZ89888.1 anig|EHA27087.1 aory|EIT76205.1 ater|EAU34831.1 atub|OJI84919.1 awen|OJJ39954.1

Group9042: abra|OJJ74248.1 acar|OOF95481.1 acla|EAW14927.1 afla|EED49821.1 akaw|GAA86773.1 aluc|OJZ89884.1 anig|EHA27083.1 aory|EIT77603.1 atub|OJI84923.1

Group9043: abra|OJJ74391.1 acar|OOF95344.1 afla|EED55060.1 agla|OJJ79760.1 aluc|OJZ89728.1 anig|EHA26955.1 aory|EIT81947.1 awen|OJJ33582.1 afis|EAW16388.1

Group9044: abra|OJJ74433.1 acar|OOF91974.1 akaw|GAA87327.1 aluc|OJZ89688.1 anig|EHA26926.1 atub|OJI85113.1 aory|EIT78644.1 acla|EAW09079.1 afla|EED52860.1

Group9045: abra|OJJ74443.1 akaw|GAA87336.1 aluc|OJZ89678.1 anig|EHA26914.1 atub|OJI85122.1 acla|EAW09670.1 azon|OJJ44353.1 afla|EED47742.1 aory|EIT77103.1

Group9046: abra|OJJ74455.1 acar|OOF95296.1 acla|EAW08566.1 afis|EAW23611.1 afla|EED44886.1 akaw|GAA87349.1 aluc|OJZ89665.1 anig|EHA26902.1 atub|OJI85135.1

Group9047: abra|OJJ74499.1 acar|OOF95251.1 afis|EAW21104.1 afum|EAL92125.1 akaw|GAA87394.1 aluc|OJZ89619.1 anid|CBF81975.1 anig|EHA26863.1 atub|OJI85183.1

Group9048: abra|OJJ74535.1 afla|EED49212.1 agla|OJJ79416.1 akaw|GAA82073.1 aluc|OJZ92109.1 anig|EHA26835.1 aory|EIT83158.1 aver|OJJ07284.1 atub|OJI86543.1

Group9049: abra|OJJ74563.1 afla|EED48717.1 agla|OJJ82641.1 aluc|OJZ92141.1 anid|CBF84660.1 asyd|OJJ54381.1 ater|EAU33267.1 atub|OJI86155.1 awen|OJJ36282.1

Group9050: abra|OJJ74590.1 acar|OOF95176.1 akaw|GAA82015.1 aluc|OJZ92168.1 anig|EHA26781.1 asyd|OJJ56081.1 ater|EAU38774.1 atub|OJI86599.1 aver|OJJ05024.1

Group9051: abra|OJJ74658.1 afla|EED47427.1 agla|OJJ78809.1 akaw|GAA81953.1 aluc|OJZ92235.1 anig|EHA26719.1 asyd|OJJ62823.1 atub|OJI86668.1 aver|OJI96867.1

Group9052: abra|OJJ74742.1 acar|OOF95738.1 akaw|GAA83668.1 aluc|OJZ92324.1 anig|EHA26646.1 atub|OJI86752.1 awen|OJJ35536.1 agla|OJJ80541.1 asyd|OJJ55949.1

Group9053: abra|OJJ74820.1 agla|OJJ82231.1 akaw|GAA83595.1 aluc|OJZ92399.1 anig|EHA26566.1 asyd|OJJ51849.1 awen|OJJ40025.1 azon|OJJ50369.1 awen|OJJ29651.1

Group9054: ater|EAU37514.1 azon|OJJ47865.1 acla|EAW08377.1 afis|EAW23406.1 afum|EAL88064.1 aver|OJI96257.1 awen|OJJ39733.1 anid|CBF89878.1 asyd|OJJ62147.1

Group9055: abra|OJJ75209.1 acar|OOF94580.1 akaw|GAA91759.1 aluc|OJZ92772.1 anid|CBF87348.1 anig|EHA26236.1 asyd|OJJ52817.1 atub|OJI87215.1 aver|OJJ08796.1

Group9056: abra|OJJ75240.1 acar|OOF94553.1 afla|EED56785.1 akaw|GAA91783.1 aluc|OJZ92802.1 anig|EHA26207.1 aory|EIT73960.1 atub|OJI87245.1 awen|OJJ35798.1

Group9057: abra|OJJ75257.1 acar|OOF93397.1 akaw|GAA91795.1 aluc|OJZ92814.1 anig|EHA26193.1 atub|OJI87272.1 azon|OJJ46463.1 ater|EAU34344.1 atub|OJI81733.1

Group9058: abra|OJJ75292.1 afis|EAW23627.1 afum|EAL87833.1 akaw|GAA91830.1 aluc|OJZ92853.1 anig|EHA26156.1 asyd|OJJ55368.1 atub|OJI87309.1 aver|OJJ06447.1

Group9059: abra|OJJ75293.1 afis|EAW23757.1 afla|EED45465.1 atub|OJI87310.1 acla|EAW12136.1 afla|EED47778.1 akaw|GAA91831.1 anid|CBF80310.1 anig|EHA26155.1

Group9060: abra|OJJ75317.1 acar|OOF94511.1 afis|EAW19724.1 akaw|GAA92013.1 aluc|OJZ92878.1 anid|CBF73560.1 anig|EHA26081.1 atub|OJI87338.1 azon|OJJ42858.1

Group9061: abra|OJJ75329.1 acar|OOF90143.1 afis|EAW16697.1 akaw|GAA92024.1 aluc|OJZ92892.1 anig|EHA26071.1 aory|EIT83095.1 atub|OJI87350.1 afum|EAL89294.1

Group9062: abra|OJJ75582.1 acar|OOF90900.1 afis|EAW21790.1 akaw|GAA84334.1 aluc|OJZ89033.1 anig|EHA23501.1 atub|OJI85757.1 aver|OJJ07240.1 acla|EAW09336.1

Group9063: abra|OJJ75632.1 acar|OOG00204.1 akaw|GAA84379.1 aluc|OJZ89083.1 anig|EHA23454.1 atub|OJI85701.1 agla|OJJ79928.1 awen|OJJ33942.1 azon|OJJ48628.1

Group9064: abra|OJJ75641.1 acar|OOG00197.1 afis|EAW21626.1 akaw|GAA84385.1 aluc|OJZ89090.1 anig|EHA23448.1 asyd|OJJ59505.1 atub|OJI85693.1 aver|OJJ02769.1

Group9065: abra|OJJ75714.1 acar|OOG00145.1 agla|OJJ88779.1 akaw|GAA84448.1 aluc|OJZ89165.1 anig|EHA23389.1 asyd|OJJ54601.1 atub|OJI85625.1 azon|OJJ48542.1

Group9066: abra|OJJ75743.1 acar|OOF91838.1 afla|EED48727.1 akaw|GAA84476.1 aluc|OJZ89196.1 anig|EHA23362.1 asyd|OJJ56158.1 atub|OJI85594.1 awen|OJJ40000.1

Group9067: abra|OJJ75744.1 akaw|GAA84477.1 aluc|OJZ89197.1 anig|EHA23361.1 aory|EIT73164.1 asyd|OJJ56159.1 atub|OJI85593.1 aver|OJJ05098.1 awen|OJJ39999.1

Group9068: abra|OJJ75878.1 acar|OOG00015.1 acla|EAW07592.1 afum|EAL85374.1 akaw|GAA84580.1 aluc|OJZ89322.1 anig|EHA23250.1 atub|OJI85476.1 azon|OJJ42381.1

Group9069: abra|OJJ75933.1 acar|OOF99977.1 afla|EED54825.1 akaw|GAA84634.1 aluc|OJZ89380.1 anig|EHA23203.1 aory|EIT79423.1 atub|OJI85418.1 agla|OJJ87473.1

Group9070: abra|OJJ76076.1 akaw|GAA88891.1 aluc|OJZ89532.1 atub|OJI85273.1 awen|OJJ39305.1 acla|EAW09745.1 afis|EAW21397.1 afum|EAL92411.1 agla|OJJ85522.1

Group9071: abra|OJJ76272.1 acar|OOF99520.1 akaw|GAA82205.1 aluc|OJZ91948.1 anig|EHA22907.1 asyd|OJJ58290.1 atub|OJI86393.1 aver|OJJ02134.1 azon|OJJ48315.1

Group9072: abra|OJJ76318.1 acar|OOF99570.1 afum|EAL84306.1 agla|OJJ84929.1 akaw|GAA82248.1 aluc|OJZ91896.1 anig|EHA22875.1 ater|EAU32234.1 atub|OJI86349.1

Group9073: abra|OJJ76604.1 acar|OOF97932.1 afla|EED46907.1 akaw|GAA82508.1 aluc|OJZ91604.1 ater|EAU34983.1 atub|OJI86056.1 azon|OJJ42096.1 acar|OOF99387.1

Group9074: abra|OJJ76871.1 afis|EAW16920.1 afla|EED51655.1 aluc|OJZ84282.1 anig|EHA27627.1 asyd|OJJ62994.1 atub|OJI89799.1 aver|OJI97974.1 awen|OJJ30987.1

Group9075: abra|OJJ76889.1 afis|EAW20742.1 afla|EED47483.1 akaw|GAA88130.1 aluc|OJZ81402.1 anig|EHA18695.1 aory|EIT79974.1 asyd|OJJ61687.1 atub|OJI91608.1

Group9076: asyd|OJJ55376.1 abra|OJJ76927.1 acar|OOF91373.1 akaw|GAA87113.1 aluc|OJZ84325.1 anig|EHA27660.1 atub|OJI89743.1 aver|OJJ06467.1 anid|CBF73479.1

Group9077: abra|OJJ76911.1 akaw|GAA87127.1 aluc|OJZ84311.1 anig|EHA27646.1 ater|EAU36008.1 atub|OJI89757.1 aver|OJJ05996.1 awen|OJJ37020.1 azon|OJJ42552.1

Group9078: abra|OJJ76914.1 acar|OOF91361.1 akaw|GAA87124.1 aluc|OJZ84314.1 anig|EHA27649.1 atub|OJI89754.1 awen|OJJ38290.1 azon|OJJ45871.1 agla|OJJ79984.1

Group9079: abra|OJJ76936.1 acar|OOF96837.1 acla|EAW14959.1 aory|EIT72660.1 ater|EAU29986.1 atub|OJI82860.1 awen|OJJ29878.1 acar|OOG00384.1 ater|EAU32857.1

Group9080: abra|OJJ76980.1 acar|OOF91426.1 afla|EED45952.1 akaw|GAA87065.1 aluc|OJZ84378.1 anig|EHA27711.1 atub|OJI89691.1 awen|OJJ36086.1 aory|EIT76167.1

Group9081: abra|OJJ76994.1 acar|OOF91441.1 acla|EAW10677.1 akaw|GAA87052.1 aluc|OJZ84394.1 anig|EHA27724.1 ater|EAU33601.1 atub|OJI89677.1 azon|OJJ42964.1

Group9082: abra|OJJ77039.1 acar|OOF90077.1 afla|EED52764.1 akaw|GAA87013.1 aluc|OJZ84441.1 anig|EHA27760.1 aory|EIT78251.1 atub|OJI89632.1 awen|OJJ35563.1

Group9083: abra|OJJ77054.1 acar|OOF92944.1 afla|EED51059.1 akaw|GAA86998.1 aluc|OJZ84459.1 anig|EHA27774.1 aory|EIT77022.1 ater|EAU31960.1 atub|OJI89616.1

Group9084: acla|EAW14398.1 afis|EAW19104.1 afla|EED52251.1 afum|EAL93836.1 agla|OJJ85946.1 ater|EAU34835.1 aver|OJJ00282.1 asyd|OJJ60955.1 aory|EIT74497.1

Group9085: abra|OJJ77101.1 acar|OOF92986.1 afla|EED45188.1 akaw|GAA86955.1 aluc|OJZ84509.1 anig|EHA27808.1 aory|EIT73729.1 atub|OJI89571.1 azon|OJJ42738.1

Group9086: abra|OJJ77102.1 acar|OOF92987.1 afla|EED45186.1 akaw|GAA86954.1 aluc|OJZ84511.1 anig|EHA27809.1 aory|EIT73793.1 atub|OJI89570.1 azon|OJJ42740.1

Group9087: abra|OJJ77113.1 acla|EAW10827.1 afla|EED45704.1 anig|EHA27820.1 aory|EIT75704.1 ater|EAU31906.1 atub|OJI89557.1 afis|EAW24737.1 afum|EAL89384.1

Group9088: abra|OJJ77184.1 acar|OOF93073.1 akaw|GAA90755.1 aluc|OJZ84592.1 anid|CBF73620.1 anig|EHA27882.1 atub|OJI89486.1 awen|OJJ31679.1 azon|OJJ47346.1

Group9089: abra|OJJ77196.1 akaw|GAA90745.1 aluc|OJZ84603.1 anig|EHA27893.1 aory|EIT76206.1 atub|OJI89476.1 aver|OJJ07906.1 awen|OJJ37284.1 afla|EED48097.1

Group9090: abra|OJJ77221.1 acar|OOF93106.1 acla|EAW13212.1 akaw|GAA90718.1 aluc|OJZ84631.1 anig|EHA27919.1 ater|EAU33911.1 atub|OJI89448.1 afla|EED49403.1

Group9091: afis|EAW19026.1 afum|EAL93757.1 ater|EAU36166.1 acla|EAW14254.1 afla|EED49653.1 anid|CBF87511.1 aory|EIT80949.1 asyd|OJJ52978.1 aver|OJJ08944.1

Group9092: abra|OJJ77256.1 anid|CBF85477.1 anig|EHA22319.1 asyd|OJJ54570.1 aver|OJJ07584.1 afla|EED54058.1 aory|EIT76142.1 ater|EAU35689.1 awen|OJJ36255.1

Group9093: abra|OJJ77406.1 acar|OOF93272.1 acla|EAW08567.1 afis|EAW23613.1 afum|EAL87850.1 akaw|GAA91359.1 aluc|OJZ84813.1 anig|EHA28077.1 atub|OJI89272.1

Group9094: abra|OJJ77417.1 afla|EED52415.1 agla|OJJ79834.1 akaw|GAA91344.1 aluc|OJZ84829.1 anig|EHA26668.1 aory|EIT74801.1 atub|OJI89258.1 anig|EHA26665.1

Group9095: afla|EED51174.1 aory|EIT81321.1 afis|EAW24706.1 afum|EAL89348.2 anid|CBF80158.1 asyd|OJJ57603.1 aver|OJJ05581.1 awen|OJJ40794.1 aver|OJI99516.1

Group9096: abra|OJJ77439.1 acar|OOF99276.1 afla|EED51469.1 agla|OJJ84708.1 aory|EIT80740.1 asyd|OJJ59180.1 atub|OJI82504.1 aver|OJJ01801.1 awen|OJJ37965.1

Group9097: acla|EAW11633.1 agla|OJJ82469.1 ater|EAU31379.1 afis|EAW23824.1 afla|EED51982.1 afum|EAL87257.1 aory|EIT81440.1 awen|OJJ33781.1 azon|OJJ42884.1

Group9098: aory|EIT82373.1 acar|OOF98767.1 afis|EAW24715.1 afla|EED52382.1 akaw|GAA90460.1 aluc|OJZ85764.1 anig|EHA24792.1 atub|OJI79548.1 afla|EED56739.1

Group9099: abra|OJJ77548.1 aory|EIT77201.1 acar|OOF97222.1 agla|OJJ82568.1 akaw|GAA84075.1 aluc|OJZ80989.1 atub|OJI82596.1 azon|OJJ49034.1 afla|EED47793.1

Group9100: abra|OJJ77610.1 acar|OOF99118.1 afum|EAL86782.1 aluc|OJZ80923.1 anig|EHA19423.1 asyd|OJJ52469.1 atub|OJI82656.1 aver|OJJ03946.1 awen|OJJ35351.1

Group9101: abra|OJJ77651.1 acar|OOF99085.1 akaw|GAA83984.1 aluc|OJZ80878.1 anig|EHA19458.1 asyd|OJJ60545.1 atub|OJI82699.1 aver|OJJ08474.1 aver|OJI99871.1

Group9102: abra|OJJ77769.1 agla|OJJ79715.1 afla|EED47272.1 akaw|GAA83280.1 anid|CBF86359.1 ater|EAU37600.1 azon|OJJ43665.1 atub|OJI90157.1 aluc|OJZ83790.1

Group9103: abra|OJJ77789.1 acla|EAW12265.1 afla|EED50778.1 aluc|OJZ90953.1 acla|EAW12370.1 afla|EED54249.1 agla|OJJ88405.1 aory|EIT78075.1 agla|OJJ88406.1

Group9104: abra|OJJ77896.1 aluc|OJZ90965.1 atub|OJI82956.1 awen|OJJ35885.1 azon|OJJ42025.1 acar|OOF97572.1 aluc|OJZ81977.1 anig|EHA19765.1 akaw|GAA93033.1

Group9105: abra|OJJ77951.1 afis|EAW19520.1 afla|EED55082.1 akaw|GAA83712.1 aluc|OJZ91022.1 anig|EHA19722.1 aory|EIT81637.1 ater|EAU38717.1 atub|OJI83014.1

Group9106: afla|EED53048.1 acar|OOF98857.1 akaw|GAA83747.1 aluc|OJZ90984.1 anid|CBF71211.1 anig|EHA19689.1 asyd|OJJ55199.1 atub|OJI82975.1 aver|OJJ07015.1

Group9107: abra|OJJ77960.1 acar|OOF91996.1 afis|EAW21810.1 afum|EAL88679.1 akaw|GAA83702.1 aluc|OJZ91029.1 atub|OJI83022.1 aver|OJJ02114.1 ater|EAU32564.1

Group9108: abra|OJJ78030.1 acar|OOF93442.1 afla|EED47542.1 akaw|GAA85955.1 aluc|OJZ91107.1 anig|EHA19805.1 aory|EIT74575.1 atub|OJI83104.1 azon|OJJ42777.1

Group9109: abra|OJJ78191.1 acar|OOF98666.1 afla|EED53445.1 agla|OJJ86666.1 akaw|GAA85838.1 aluc|OJZ91262.1 anig|EHA19950.1 atub|OJI83251.1 aory|EIT74872.1

Group9110: abra|OJJ78312.1 acar|OOF90132.1 akaw|GAA92407.1 aluc|OJZ91381.1 anig|EHA20063.1 asyd|OJJ61204.1 atub|OJI83374.1 aver|OJJ00528.1 akaw|GAA85722.1

Group9111: akaw|GAA85678.1 acla|EAW09179.1 afis|EAW21967.1 afla|EED46688.1 afum|EAL92981.1 agla|OJJ84130.1 anid|CBF82733.1 aory|EIT77924.1 ater|EAU31087.1

Group9112: acar|OOF89813.1 acla|EAW14632.1 afis|EAW23652.1 afla|EED52133.1 agla|OJJ79805.1 aory|EIT74392.1 ater|EAU32803.1 aver|OJJ07302.1 awen|OJJ40925.1

Group9113: acar|OOF90608.1 acla|EAW06503.1 afis|EAW16944.1 afum|EAL86471.1 akaw|GAA91561.1 aluc|OJZ82378.1 anid|CBF83060.1 atub|OJI88064.1 azon|OJJ42653.1

Group9114: acar|OOF90862.1 acla|EAW12398.1 afis|EAW16846.1 afum|EAL86392.1 asyd|OJJ54166.1 awen|OJJ36027.1 azon|OJJ47628.1 agla|OJJ78551.1 anid|CBF82405.1

Group9115: aver|OJI96460.1 acla|EAW13438.1 aory|EIT79267.1 awen|OJJ40042.1 afla|EED47281.1 anig|EHA25773.1 aory|EIT79322.1 atub|OJI88512.1 agla|OJJ78869.1

Group9116: acar|OOF91866.1 asyd|OJJ58348.1 aver|OJJ02065.1 acar|OOF97895.1 acla|EAW10259.1 afis|EAW25334.1 afla|EED53812.1 afum|EAL89997.1 akaw|GAA82711.1

Group9117: acar|OOF91944.1 afla|EED46098.1 agla|OJJ83701.1 akaw|GAA85823.1 aory|EIT76402.1 asyd|OJJ61027.1 awen|OJJ30695.1 akaw|GAA92465.1 afla|EED57040.1

Group9118: awen|OJJ33536.1 afla|EED48330.1 akaw|GAA83722.1 aluc|OJZ91011.1 anig|EHA19713.1 aory|EIT76309.1 asyd|OJJ60497.1 atub|OJI83002.1 azon|OJJ48591.1

Group9119: acar|OOF93058.1 afla|EED45785.1 afum|EAL91579.1 aory|EIT75464.1 ater|EAU31842.1 acla|EAW12894.1 afis|EAW17770.1 anid|CBF86635.1 awen|OJJ37073.1

Group9120: acar|OOF93391.1 agla|OJJ79771.1 anid|CBF87482.1 asyd|OJJ52946.1 ater|EAU36394.1 atub|OJI84520.1 aver|OJJ08913.1 awen|OJJ41683.1 azon|OJJ51829.1

Group9121: afla|EED46051.1 aory|EIT83485.1 afis|EAW23643.1 agla|OJJ82137.1 asyd|OJJ52372.1 ater|EAU35169.1 awen|OJJ39956.1 ater|EAU30990.1 awen|OJJ39961.1

Group9122: acar|OOF94769.1 acla|EAW12701.1 afis|EAW17864.1 afla|EED46227.1 afum|EAL91672.1 akaw|GAA86206.1 anid|CBF84083.1 aory|EIT76490.1 ater|EAU33079.1

Group9123: acar|OOF94900.1 acla|EAW14619.1 afis|EAW19357.1 afum|EAL94077.1 anid|CBF73428.1 asyd|OJJ58361.1 ater|EAU36410.1 aver|OJJ02045.1 aver|OJJ08106.1

Group9124: acar|OOF95037.1 afla|EED48165.1 aory|EIT83031.1 asyd|OJJ56127.1 aver|OJJ05069.1 acar|OOF96827.1 afis|EAW17236.1 afum|EAL86653.1 anid|CBF78337.1

Group9125: awen|OJJ36311.1 acla|EAW11091.1 afis|EAW20766.1 afum|EAL84853.1 asyd|OJJ64586.1 ater|EAU29633.1 aver|OJJ07575.1 asyd|OJJ54580.1 aver|OJJ05899.1

Group9126: acar|OOF95311.1 afla|EED49674.1 anid|CBF71486.1 aory|EIT80994.1 asyd|OJJ54940.1 awen|OJJ33541.1 asyd|OJJ58307.1 aver|OJJ02105.1 aver|OJJ07254.1

Group9127: acar|OOF95546.1 acla|EAW14598.1 afis|EAW19327.1 afla|EED48187.1 afum|EBA27513.1 agla|OJJ84438.1 anid|CBF82027.1 aory|EIT72925.1 awen|OJJ30151.1

Group9128: acla|EAW14391.1 afis|EAW19098.1 afla|EED54332.1 afum|EAL93830.1 aory|EIT76114.1 aver|OJJ04351.1 azon|OJJ46041.1 agla|OJJ84503.1 awen|OJJ32491.1

Group9129: acar|OOF96114.1 afis|EAW24504.1 afla|EED48395.1 anid|CBF89954.1 aory|EIT83057.1 asyd|OJJ62109.1 aver|OJI96215.1 azon|OJJ50120.1 afum|EAL86172.1

Group9130: acar|OOF96689.1 akaw|GAA85853.1 aluc|OJZ91246.1 anid|CBF83945.1 asyd|OJJ53664.1 atub|OJI83243.1 aver|OJJ07771.1 asyd|OJJ56721.1 aver|OJJ06004.1

Group9131: acar|OOF96855.1 acla|EAW14992.1 afis|EAW24518.1 afum|EAL86184.1 agla|OJJ81259.1 aory|EIT78854.1 ater|EAU32757.1 awen|OJJ32603.1 afla|EED54709.1

Group9132: aver|OJJ05436.1 acla|EAW08146.1 afis|EAW23193.1 afla|EED47072.1 afum|EAL88290.1 agla|OJJ86368.1 ater|EAU36317.1 awen|OJJ36595.1 anid|CBF74518.1

Group9133: acar|OOF97463.1 acla|EAW13052.1 afis|EAW17615.1 afla|EED49206.1 afum|EAL91428.1 akaw|GAA83310.1 anid|CBF90171.1 aory|EIT78545.1 ater|EAU37049.1

Group9134: acar|OOF97639.1 afla|EED55683.1 agla|OJJ80678.1 anid|CBF74954.1 anig|EHA18774.1 aory|EIT77090.1 asyd|OJJ60491.1 aver|OJI99817.1 awen|OJJ38071.1

Group9135: acar|OOF98602.1 afis|EAW21895.1 afla|EED46413.1 afum|EBA27478.1 agla|OJJ84237.1 aory|EIT72333.1 asyd|OJJ61291.1 aver|OJJ00617.1 awen|OJJ34184.1

Group9136: acar|OOF98772.1 acla|EAW06577.1 afis|EAW17131.1 afla|EED52292.1 afum|EAL86591.1 agla|OJJ85165.1 asyd|OJJ57691.1 awen|OJJ33404.1 agla|OJJ84600.1

Group9137: acar|OOF98849.1 acla|EAW06674.1 afis|EAW16091.1 afla|EED48627.1 afum|EAL88723.1 akaw|GAA83696.1 anid|CBF85464.1 aory|EIT73280.1 ater|EAU33163.1

Group9138: acar|OOF99007.1 afis|EAW23654.1 afla|EED45846.1 akaw|GAA91466.1 aluc|OJZ82071.1 asyd|OJJ52509.1 atub|OJI84628.1 aver|OJJ01856.1 asyd|OJJ63572.1

Group9139: agla|OJJ85921.1 acla|EAW14251.1 afis|EAW19030.1 afla|EED52648.1 anid|CBF85564.1 aory|EIT74253.1 asyd|OJJ59934.1 aver|OJJ02353.1 awen|OJJ37064.1

Group9140: acar|OOG00140.1 acla|EAW09393.1 afis|EAW21740.1 afla|EED58015.1 agla|OJJ84433.1 anid|CBF83697.1 aory|EIT79732.1 ater|EAU38421.1 awen|OJJ34022.1

Group9141: acar|OOG00207.1 afla|EED53210.1 agla|OJJ78700.1 anig|EHA26131.1 aory|EIT74945.1 atub|OJI82857.1 azon|OJJ50126.1 afla|EED44915.1 aory|EIT73506.1

Group9142: aory|EIT77690.1 afla|EED49709.1 agla|OJJ88111.1 akaw|GAA84676.1 aluc|OJZ87013.1 asyd|OJJ53592.1 atub|OJI83506.1 aver|OJJ08156.1 awen|OJJ29854.1

Group9143: acar|OOG00584.1 afla|EED47839.1 akaw|GAA83105.1 aluc|OJZ88653.1 anig|EHA18345.1 aory|EIT77165.1 atub|OJI91240.1 aver|OJI98273.1 azon|OJJ44492.1

Group9144: acar|OOG00781.1 acla|EAW07515.1 afla|EED54577.1 anid|CBF89047.1 aory|EIT78806.1 asyd|OJJ62756.1 aver|OJI96798.1 awen|OJJ35175.1 azon|OJJ49932.1

Group9145: acla|EAW06462.1 afla|EED58101.1 agla|OJJ89355.1 aory|EIT79126.1 asyd|OJJ56460.1 ater|EAU35498.1 aver|OJJ06261.1 awen|OJJ35616.1 azon|OJJ46907.1

Group9146: acla|EAW06697.1 afis|EAW16242.1 afum|EAL88864.1 anid|CBF69397.1 anid|CBF84546.1 asyd|OJJ53471.1 ater|EAU33900.1 aver|OJJ08285.1 awen|OJJ30214.1

Group9147: acla|EAW07056.1 afis|EAW16682.1 afla|EED47441.1 afum|EAL89286.1 anid|CBF89915.1 aory|EIT81914.1 asyd|OJJ62127.1 ater|EAU35872.1 aver|OJI96235.1

Group9148: acla|EAW07457.1 afis|EAW22488.1 afla|EED54517.1 afum|EAL90705.1 akaw|GAA88615.1 anid|CBF88946.1 aory|EIT78851.1 asyd|OJJ62813.1 ater|EAU39226.1

Group9149: acla|EAW07462.1 afis|EAW22493.1 afla|EED54522.1 afum|EAL90700.1 agla|OJJ80846.1 aory|EIT78808.1 ater|EAU39222.1 awen|OJJ35114.1 azon|OJJ49979.1

Group9150: acla|EAW07611.1 afis|EAW22641.1 afla|EED45986.1 afum|EAL90551.1 agla|OJJ84039.1 aory|EIT75586.1 aver|OJJ04317.1 awen|OJJ35264.1 awen|OJJ34412.1

Group9151: acla|EAW07704.1 afis|EAW22722.1 afla|EED50007.1 agla|OJJ83681.1 anid|CBF88034.1 asyd|OJJ63534.1 aver|OJI97398.1 awen|OJJ35416.1 aory|EIT74710.1

Group9152: acla|EAW07796.1 afis|EAW22833.1 afla|EED50108.1 afum|EAL90369.1 akaw|GAA89526.1 anid|CBF87878.1 aory|EIT74636.1 ater|EAU38876.1 awen|OJJ38891.1

Group9153: acla|EAW08692.1 anig|EHA24751.1 afis|EAW16421.1 afla|EED52252.1 agla|OJJ86219.1 aory|EIT74454.1 ater|EAU33982.1 awen|OJJ33627.1 ater|EAU29300.1

Group9154: acla|EAW08960.1 afis|EAW15434.1 afla|EED51779.1 afum|EAL84210.1 aluc|OJZ89488.1 aory|EIT81696.1 awen|OJJ29678.1 azon|OJJ50850.1 afis|EAW24647.1

Group9155: acla|EAW09082.1 afis|EAW25669.1 afla|EED46154.1 afum|EAL84341.1 anid|CBF69525.1 aory|EIT76492.1 asyd|OJJ59185.1 ater|EAU31773.1 awen|OJJ33422.1

Group9156: acla|EAW09106.1 afis|EAW25710.1 afla|EED51115.1 afum|EAL84383.1 anid|CBF83223.1 asyd|OJJ55569.1 ater|EAU33341.1 aver|OJJ06648.1 awen|OJJ30275.1

Group9157: acla|EAW09109.1 afis|EAW25716.1 afla|EED52044.1 afum|EAL84390.2 anid|CBF74827.1 aory|EIT81372.1 asyd|OJJ60138.1 ater|EAU34875.1 aver|OJJ04124.1

Group9158: acla|EAW09110.1 afis|EAW25715.1 afla|EED53391.1 afum|EAL84389.2 agla|OJJ84456.1 asyd|OJJ54848.1 ater|EAU31032.1 awen|OJJ34476.1 azon|OJJ42415.1

Group9159: acla|EAW09208.1 afis|EAW21939.1 afla|EED46656.1 agla|OJJ81052.1 aluc|OJZ91386.1 anid|CBF84081.1 anig|EHA20057.1 ater|EAU30944.1 atub|OJI83379.1

Group9160: acla|EAW09256.1 afis|EAW21891.1 afla|EED46417.1 afum|EAL92905.1 agla|OJJ84242.1 aory|EIT72332.1 ater|EAU30888.1 awen|OJJ34180.1 azon|OJJ43452.1

Group9161: acla|EAW09347.1 afis|EAW21802.1 afla|EED49375.1 afum|EAL92828.1 anid|CBF78346.1 aory|EIT79485.1 asyd|OJJ56139.1 aver|OJJ05078.1 awen|OJJ40847.1

Group9162: acla|EAW09623.1 afis|EAW21517.1 afla|EED55955.1 afum|EAL92537.1 anid|CBF76425.1 asyd|OJJ61092.1 ater|EAU35023.1 aver|OJJ00407.1 awen|OJJ39177.1

Group9163: acla|EAW09775.1 afis|EAW21363.1 afum|EAL92379.1 agla|OJJ85484.1 anid|CBF83877.1 anig|EHA23100.1 ater|EAU35197.1 awen|OJJ39335.1 azon|OJJ51680.1

Group9164: acla|EAW09903.1 afis|EAW21151.1 afla|EED57519.1 afum|EAL92170.1 anid|CBF71745.1 aory|EIT81840.1 asyd|OJJ61789.1 ater|EAU33856.1 aver|OJI95889.1

Group9165: acla|EAW09949.1 afis|EAW21227.1 afum|EAL92239.2 agla|OJJ78503.1 anid|CBF83284.1 asyd|OJJ60565.1 aver|OJI99891.1 awen|OJJ36535.1 azon|OJJ44410.1

Group9166: acla|EAW10098.1 afis|EAW25489.1 afum|EAL90154.1 asyd|OJJ53060.1 ater|EAU34660.1 aver|OJJ03784.1 azon|OJJ44797.1 agla|OJJ87137.1 anid|CBF77726.1

Group9167: acla|EAW10510.1 afis|EAW25057.1 afla|EED55395.1 afum|EAL89725.1 akaw|GAA83000.1 anid|CBF85935.1 aory|EIT80810.1 ater|EAU35806.1 azon|OJJ44046.1

Group9168: acla|EAW10849.1 afis|EAW20283.1 afla|EED51079.1 afum|EAL85417.1 agla|OJJ82442.1 aory|EIT81679.1 ater|EAU34005.1 afis|EAW24684.1 aory|EIT76981.1

Group9169: acla|EAW11370.1 afis|EAW18125.1 afum|EAL87483.1 agla|OJJ87019.1 awen|OJJ35527.1 azon|OJJ46956.1 anid|CBF86927.1 anig|EHA21087.1 asyd|OJJ52261.1

Group9170: acla|EAW11533.1 afis|EAW23921.1 afla|EED48136.1 afum|EBA27312.1 anid|CBF89559.1 aory|EIT76210.1 asyd|OJJ62343.1 aver|OJI96447.1 azon|OJJ45191.1

Group9171: acla|EAW12171.1 afis|EAW17191.1 afum|EAL86619.2 afla|EED46575.1 anid|CBF75737.1 aory|EIT77817.1 ater|EAU35875.1 asyd|OJJ57513.1 aver|OJJ05666.1

Group9172: acla|EAW12180.1 afis|EAW21257.1 afla|EED45193.1 afum|EAL92272.1 aory|EIT73658.1 asyd|OJJ56082.1 ater|EAU32949.1 aver|OJJ05025.1 aory|EIT75997.1

Group9173: acla|EAW12380.1 afis|EAW16826.1 afla|EED57473.1 afum|EAL86375.1 anid|CBF87495.1 aory|EIT80128.1 asyd|OJJ52966.1 ater|EAU31803.1 aver|OJJ08932.1

Group9174: anid|CBF85683.1 aver|OJI98347.1 afis|EAW16913.1 agla|OJJ80713.1 asyd|OJJ53486.1 ater|EAU37353.1 awen|OJJ35845.1 agla|OJJ89201.1 asyd|OJJ64257.1

Group9175: acla|EAW13309.1 agla|OJJ84902.1 aory|EIT77729.1 asyd|OJJ55794.1 atub|OJI89926.1 aver|OJJ06883.1 awen|OJJ33116.1 azon|OJJ46332.1 afla|EED44848.1

Group9176: acla|EAW13493.1 afis|EAW20593.1 afla|EED52232.1 afum|EAL84468.1 anid|CBF78913.1 aory|EIT74326.1 asyd|OJJ55616.1 ater|EAU35239.1 aver|OJJ06691.1

Group9177: afis|EAW19233.1 afum|EAL93962.1 agla|OJJ88475.1 aory|EIT80180.1 anid|CBF78623.1 asyd|OJJ65127.1 ater|EAU37230.1 aver|OJI99255.1 awen|OJJ37590.1

Group9178: acla|EAW13545.1 afis|EAW20617.1 afum|EAL84444.1 anid|CBF81468.1 asyd|OJJ59444.1 ater|EAU38172.1 aver|OJJ02829.1 afis|EAW23633.1 afum|EAL87827.2

Group9179: acla|EAW14792.1 afis|EAW24323.1 afla|EED47943.1 afum|EAL85993.1 akaw|GAA88280.1 anid|CBF74062.1 aory|EIT79830.1 ater|EAU30429.1 atub|OJI88188.1

Group9180: acla|EAW14876.1 afis|EAW24393.1 afla|EED47817.1 afum|EAL86065.1 agla|OJJ78873.1 anid|CBF73715.1 aory|EIT77108.1 asyd|OJJ56641.1 aver|OJJ06085.1

Group9181: acla|EAW15039.1 afis|EAW24563.1 afla|EED51162.1 afum|EAL86224.1 anid|CBF80178.1 aory|EIT81341.1 asyd|OJJ65362.1 aver|OJI99527.1 awen|OJJ34439.1

Group9182: acla|EAW15352.1 afis|EAW20361.1 afum|EAL84493.1 asyd|OJJ55542.1 ater|EAU38527.1 aver|OJJ06620.1 awen|OJJ33274.1 afis|EAW20569.1 ater|EAU37110.1

Group9183: acla|EAW15353.1 afis|EAW20360.1 afla|EED56798.1 afum|EAL85477.1 anid|CBF79110.1 aory|EIT73835.1 asyd|OJJ55045.1 aver|OJJ07162.1 awen|OJJ36891.1

Group9184: afis|EAW15407.1 afla|EED54363.1 aory|EIT75986.1 awen|OJJ40121.1 afis|EAW17367.1 afum|EAL91204.1 aver|OJJ08376.1 azon|OJJ42232.1 aver|OJJ02033.1

Group9185: afis|EAW15839.1 afum|EAL85512.1 agla|OJJ82003.1 anid|CBF70848.1 anig|EHA24394.1 asyd|OJJ52570.1 ater|EAU33396.1 aver|OJJ04599.1 awen|OJJ38552.1

Group9186: afis|EAW17022.1 akaw|GAA84315.1 aluc|OJZ89010.1 anig|EHA25070.1 atub|OJI85779.1 afis|EAW19370.1 afla|EED56074.1 afum|EAL94089.1 aory|EIT76714.1

Group9187: afis|EAW17614.1 afla|EED49205.1 afum|EAL91427.1 akaw|GAA83309.1 anid|CBF73974.1 aory|EIT78553.1 asyd|OJJ61247.1 ater|EAU36631.1 aver|OJJ00571.1

Group9188: afis|EAW19018.1 afum|EAL93750.1 agla|OJJ81482.1 anid|CBF73866.1 asyd|OJJ57950.1 ater|EAU36614.1 aver|OJI96129.1 agla|OJJ86909.1 aver|OJJ04449.1

Group9189: afis|EAW22745.1 agla|OJJ85233.1 akaw|GAA91090.1 aluc|OJZ87908.1 aory|EIT80680.1 aver|OJJ04886.1 awen|OJJ36950.1 afla|EED51629.1 asyd|OJJ55937.1

Group9190: afis|EAW23725.1 afla|EED54369.1 afum|EAL87164.1 anid|CBF80093.1 aory|EIT75901.1 asyd|OJJ58301.1 ater|EAU31559.1 aver|OJJ02123.1 awen|OJJ33662.1

Group9191: afis|EAW24589.1 afla|EED57467.1 agla|OJJ82376.1 anid|CBF82016.1 aory|EIT80068.1 asyd|OJJ60274.1 aver|OJI96483.1 awen|OJJ40075.1 aver|OJJ05119.1

Group9192: afis|EAW24710.1 afla|EED57577.1 afum|EAL89351.1 anid|CBF80539.1 aory|EIT82954.1 asyd|OJJ65359.1 ater|EAU30093.1 aver|OJI99508.1 awen|OJJ35407.1

Group9193: afis|EAW25705.1 afla|EED44735.1 afum|EAL84378.1 aory|EIT77803.1 asyd|OJJ63128.1 ater|EAU31742.1 aver|OJI97827.1 awen|OJJ34906.1 azon|OJJ44482.1

Group9194: afla|EED45154.1 anid|CBF84224.1 asyd|OJJ57183.1 ater|EAU33861.1 aver|OJJ03107.1 awen|OJJ34567.1 afla|EED56553.1 aory|EIT79604.1 ater|EAU37800.1

Group9195: aluc|OJZ85150.1 anig|EHA19891.1 aory|EIT76278.1 asyd|OJJ54433.1 awen|OJJ33529.1 akaw|GAA86602.1 anig|EHA25385.1 atub|OJI80424.1 ater|EAU37657.1

Group9196: afla|EED48373.1 akaw|GAA87139.1 aluc|OJZ84298.1 anid|CBF78786.1 anig|EHA27635.1 aory|EIT83076.1 asyd|OJJ52098.1 atub|OJI89771.1 aver|OJJ03019.1

Group9197: afla|EED51064.1 agla|OJJ86414.1 anid|CBF78517.1 aory|EIT77046.1 asyd|OJJ65309.1 aver|OJI99449.1 awen|OJJ33561.1 azon|OJJ50766.1 anid|CBF78515.1

Group9198: afla|EED54340.1 aory|EIT75900.1 awen|OJJ36117.1 afum|EAL87108.1 agla|OJJ82022.1 anid|CBF71659.1 asyd|OJJ61835.1 aver|OJI95934.1 ater|EAU32862.1

Group9199: afla|EED54365.1 agla|OJJ80330.1 akaw|GAA86635.1 aluc|OJZ85192.1 anid|CBF73847.1 aory|EIT76162.1 asyd|OJJ56552.1 ater|EAU29948.1 aver|OJJ06169.1

Group9200: afla|EED55116.1 anid|CBF75417.1 aory|EIT81627.1 asyd|OJJ59791.1 ater|EAU38524.1 aver|OJJ02496.1 awen|OJJ37900.1 asyd|OJJ61701.1 aver|OJI95790.1

Group9201: afla|EED57948.1 agla|OJJ79920.1 akaw|GAA84387.1 anid|CBF83602.1 anig|EHA23446.1 aory|EIT82352.1 ater|EAU38365.1 atub|OJI85690.1 aver|OJJ07833.1

Group9202: agla|OJJ82212.1 akaw|GAA90757.1 aluc|OJZ84590.1 asyd|OJJ60314.1 atub|OJI89488.1 aver|OJI99655.1 awen|OJJ37954.1 asyd|OJJ58903.1 ater|EAU33756.1

Group9203: akaw|GAA89007.1 aluc|OJZ80001.1 anig|EHA19810.1 asyd|OJJ51835.1 ater|EAU30105.1 awen|OJJ34752.1 azon|OJJ45153.1 aluc|OJZ86476.1 azon|OJJ51294.1

Group9204: abra|OJJ65506.1 abra|OJJ65818.1 awen|OJJ29551.1 awen|OJJ29615.1 awen|OJJ29592.1 awen|OJJ29638.1 awen|OJJ29574.1 akaw|GAA93240.1

Group9205: abra|OJJ65542.1 abra|OJJ67279.1 abra|OJJ65832.1 abra|OJJ78309.1 asyd|OJJ51968.1 asyd|OJJ51992.1 awen|OJJ29628.1 aver|OJI99571.1

Group9206: abra|OJJ65785.1 abra|OJJ68042.1 aluc|OJZ81112.1 aluc|OJZ82915.1 acar|OOF90404.1 anig|EHA21338.1 atub|OJI89141.1 akaw|GAA90911.1

Group9207: abra|OJJ67521.1 abra|OJJ70181.1 afis|EAW19751.1 afum|EAL85146.1 agla|OJJ80471.1 ater|EAU36092.1 azon|OJJ45960.1 acar|OOF95578.1

Group9208: abra|OJJ75992.1 abra|OJJ77978.1 anig|EHA19756.1 anig|EHA23150.1 akaw|GAA86002.1 aluc|OJZ91053.1 atub|OJI83049.1 acar|OOF98681.1

Group9209: acar|OOF94031.1 acar|OOF94101.1 abra|OJJ66422.1 agla|OJJ84436.1 akaw|GAA90287.1 aluc|OJZ83053.1 anig|EHA28118.1 atub|OJI87894.1

Group9210: acar|OOF94047.1 acar|OOF94117.1 aver|OJI96455.1 asyd|OJJ62360.1 awen|OJJ32291.1 akaw|GAA84177.1 aluc|OJZ81102.1 atub|OJI82478.1

Group9211: acar|OOF94054.1 acar|OOF94124.1 abra|OJJ66390.1 akaw|GAA90256.1 aluc|OJZ83084.1 anig|EHA28147.1 atub|OJI87860.1 azon|OJJ46015.1

Group9212: acar|OOF94089.1 acar|OOF94159.1 afla|EED51675.1 anid|CBF87054.1 aory|EIT80733.1 asyd|OJJ60441.1 ater|EAU36804.1 aver|OJI99757.1

Group9213: acar|OOF94795.1 acar|OOF95808.1 abra|OJJ69043.1 akaw|GAA90572.1 aluc|OJZ86028.1 anig|EHA24584.1 atub|OJI79817.1 azon|OJJ42399.1

Group9214: acar|OOF95042.1 acar|OOF95109.1 abra|OJJ74661.1 afis|EAW18425.1 akaw|GAA81950.1 aluc|OJZ92238.1 anig|EHA26716.1 atub|OJI86672.1

Group9215: acar|OOF95627.1 acar|OOF95678.1 acla|EAW10949.1 anid|CBF71795.1 asyd|OJJ57418.1 aver|OJI98867.1 awen|OJJ36985.1 asyd|OJJ64767.1

Group9216: afis|EAW15521.1 afis|EAW17759.1 acla|EAW15037.1 asyd|OJJ54438.1 afla|EED56155.1 aory|EIT76867.1 ater|EAU29606.1 afis|EAW25617.1

Group9217: afis|EAW17145.1 afis|EAW25652.1 anid|CBF71257.1 anid|CBF80826.1 agla|OJJ87916.1 asyd|OJJ57920.1 aver|OJJ04472.1 anid|CBF80832.1

Group9218: afis|EAW18099.1 afis|EAW20671.1 afis|EAW21807.1 afum|EAL85401.1 afum|EAL86716.1 akaw|GAA92340.1 akaw|GAA93212.1 awen|OJJ29632.1

Group9219: afla|EED50205.1 afla|EED52402.1 aory|EIT77283.1 aory|EIT83295.1 anid|CBF80133.1 asyd|OJJ57904.1 ater|EAU35873.1 aver|OJJ04490.1

Group9220: afum|EAL84367.1 afum|EAL86717.1 abra|OJJ68568.1 acla|EAW13720.1 afla|EED52526.1 aluc|OJZ81968.1 anig|EHA22279.1 atub|OJI84519.1

Group9221: afum|EAL90937.1 afum|EAL91119.1 acar|OOF98469.1 afis|EAW20300.1 agla|OJJ88439.1 asyd|OJJ62721.1 aver|OJJ01791.1 azon|OJJ49995.1

Group9222: agla|OJJ78667.1 agla|OJJ79271.1 agla|OJJ80298.1 asyd|OJJ60507.1 asyd|OJJ65157.1 acla|EAW13440.1 aver|OJI99833.1 awen|OJJ40852.1

Group9223: agla|OJJ78959.1 agla|OJJ85943.1 abra|OJJ78200.1 acar|OOF98655.1 akaw|GAA85829.1 aluc|OJZ91271.1 anig|EHA19958.1 atub|OJI83260.1

Group9224: agla|OJJ80410.1 agla|OJJ80411.1 abra|OJJ67601.1 acar|OOF91952.1 afla|EED47760.1 aory|EIT77150.1 asyd|OJJ59779.1 awen|OJJ36712.1

Group9225: agla|OJJ81527.1 abra|OJJ70446.1 acar|OOF92544.1 akaw|GAA90135.1 aluc|OJZ83212.1 anig|EHA21072.1 atub|OJI87725.1 azon|OJJ50644.1

Group9226: agla|OJJ82367.1 agla|OJJ87520.1 abra|OJJ76438.1 acar|OOF99683.1 akaw|GAA82353.1 aluc|OJZ91771.1 anig|EHA22771.1 atub|OJI86229.1

Group9227: agla|OJJ87623.1 agla|OJJ89294.1 asyd|OJJ52764.1 asyd|OJJ54087.1 aver|OJJ08651.1 aver|OJJ08747.1 afis|EAW19735.1 afum|EAL86794.1

Group9228: aluc|OJZ79779.1 aluc|OJZ80211.1 aluc|OJZ81116.1 aluc|OJZ80324.1 aluc|OJZ79800.1 aluc|OJZ88830.1 aluc|OJZ79861.1 akaw|GAA92961.1

Group9229: anid|CBF70488.1 anid|CBF84336.1 anid|CBF88734.1 anid|CBF89682.1 anid|CBF74751.1 anid|CBF74302.1 anid|CBF77827.1 anig|EHA26857.1

Group9230: anid|CBF73471.1 anid|CBF89307.1 abra|OJJ70827.1 acla|EAW09025.1 anig|EHA17996.1 afla|EED48483.1 aory|EIT73134.1 aory|EIT73198.1

Group9231: anig|EHA21298.1 anig|EHA21854.1 abra|OJJ67998.1 acar|OOF92040.1 akaw|GAA90952.1 aluc|OJZ82872.1 ater|EAU33480.1 atub|OJI89096.1

Group9232: asyd|OJJ54286.1 asyd|OJJ57635.1 aver|OJJ04437.1 aver|OJJ05240.1 aver|OJJ07873.1 anid|CBF74901.1 ater|EAU29682.1 anid|CBF87450.1

Group9233: asyd|OJJ55575.1 asyd|OJJ62614.1 afla|EED51129.1 agla|OJJ81311.1 anid|CBF90352.1 aory|EIT81299.1 ater|EAU37569.1 aver|OJJ06654.1

Group9234: asyd|OJJ56430.1 asyd|OJJ65104.1 acla|EAW14591.1 afis|EAW19321.1 afum|EAL94039.1 agla|OJJ82955.1 aver|OJJ06291.1 aver|OJI99231.1

Group9235: asyd|OJJ57392.1 asyd|OJJ61778.1 afis|EAW23425.1 afla|EED48337.1 afla|EED53049.1 aory|EIT76295.1 ater|EAU35010.1 anid|CBF84291.1

Group9236: ater|EAU31780.1 abra|OJJ75593.1 aluc|OJZ89045.1 anid|CBF81783.1 anig|EHA23490.1 atub|OJI85742.1 awen|OJJ33346.1 ater|EAU29819.1

Group9237: ater|EAU31681.1 ater|EAU32193.1 afis|EAW19665.1 afum|EAL86858.1 anid|CBF83924.1 asyd|OJJ55428.1 aver|OJJ06507.1 azon|OJJ50194.1

Group9238: ater|EAU33182.1 ater|EAU35438.1 abra|OJJ66403.1 akaw|GAA90271.1 aluc|OJZ83070.1 anid|CBF74241.1 anig|EHA28134.1 atub|OJI87876.1

Group9239: ater|EAU35500.1 ater|EAU37634.1 ater|EAU37635.1 abra|OJJ75366.1 afis|EAW22480.1 afla|EED54392.1 aory|EIT75858.1 agla|OJJ83599.1

Group9240: aver|OJI95756.1 aver|OJJ03474.1 aver|OJJ07017.1 afla|EED56874.1 aory|EIT77566.1 asyd|OJJ55196.1 ater|EAU37686.1 awen|OJJ40083.1

Group9241: awen|OJJ29742.1 awen|OJJ33472.1 awen|OJJ33471.1 awen|OJJ38762.1 awen|OJJ34539.1 awen|OJJ31077.1 awen|OJJ40128.1 acar|OOF90469.1

Group9242: awen|OJJ29863.1 awen|OJJ32233.1 awen|OJJ38934.1 abra|OJJ74437.1 ater|EAU38801.1 afla|EED57522.1 agla|OJJ86496.1 aory|EIT81865.1

Group9243: awen|OJJ30774.1 awen|OJJ31707.1 acla|EAW13759.1 afla|EED53053.1 anid|CBF75178.1 asyd|OJJ59953.1 ater|EAU30744.1 aver|OJJ02336.1

Group9244: awen|OJJ30781.1 awen|OJJ31700.1 abra|OJJ74543.1 acar|OOF95213.1 akaw|GAA82055.1 aluc|OJZ92125.1 anig|EHA26823.1 atub|OJI86557.1

Group9245: awen|OJJ30789.1 awen|OJJ31692.1 acla|EAW09118.1 ater|EAU36745.1 afis|EAW16193.1 afum|EAL88824.1 afla|EED49618.1 aory|EIT81011.1

Group9246: awen|OJJ31652.1 awen|OJJ39922.1 abra|OJJ74357.1 acar|OOF99852.1 afla|EED45403.1 akaw|GAA86673.1 aluc|OJZ89770.1 atub|OJI85033.1

Group9247: awen|OJJ32421.1 awen|OJJ38256.1 akaw|GAA83096.1 aluc|OJZ88643.1 abra|OJJ73393.1 anig|EHA18327.1 azon|OJJ51357.1 atub|OJI91228.1

Group9248: awen|OJJ33367.1 awen|OJJ34645.1 acla|EAW13546.1 afla|EED46259.1 agla|OJJ78469.1 aory|EIT76451.1 ater|EAU29447.1 agla|OJJ88372.1

Group9249: azon|OJJ42123.1 azon|OJJ42751.1 abra|OJJ72163.1 akaw|GAA84765.1 aluc|OJZ86918.1 anig|EHA21726.1 atub|OJI83603.1 awen|OJJ29679.1

Group9250: azon|OJJ42217.1 azon|OJJ42355.1 abra|OJJ77929.1 acar|OOF98867.1 akaw|GAA83734.1 aluc|OJZ90997.1 anig|EHA19705.1 atub|OJI82987.1

Group9251: azon|OJJ42380.1 azon|OJJ46456.1 abra|OJJ68370.1 acar|OOF95038.1 akaw|GAA92155.1 aluc|OJZ85562.1 anig|EHA25069.1 atub|OJI79342.1

Group9252: azon|OJJ42668.1 azon|OJJ49109.1 abra|OJJ75741.1 acar|OOG00122.1 akaw|GAA84474.1 aluc|OJZ89194.1 anig|EHA23364.1 atub|OJI85596.1

Group9253: azon|OJJ43541.1 azon|OJJ49998.1 afla|EED52563.1 anid|CBF79152.1 aory|EIT74188.1 asyd|OJJ55205.1 ater|EAU38341.1 aver|OJJ07011.1

Group9254: azon|OJJ43568.1 azon|OJJ45832.1 ater|EAU31965.1 awen|OJJ38329.1 acar|OOF90161.1 asyd|OJJ59615.1 afis|EAW22743.1 awen|OJJ29945.1

Group9255: azon|OJJ43587.1 azon|OJJ46445.1 abra|OJJ73314.1 acar|OOF98733.1 akaw|GAA92809.1 aluc|OJZ88732.1 anig|EHA18435.1 atub|OJI91307.1

Group9256: azon|OJJ43919.1 azon|OJJ50661.1 awen|OJJ35806.1 agla|OJJ87371.1 asyd|OJJ59396.1 agla|OJJ88048.1 aver|OJJ02876.1 anid|CBF79746.1

Group9257: azon|OJJ44126.1 azon|OJJ49997.1 acar|OOF93310.1 afis|EAW24585.1 afum|EAL88042.1 agla|OJJ80221.1 ater|EAU32202.1 azon|OJJ43540.1

Group9258: azon|OJJ45076.1 azon|OJJ50104.1 aory|EIT72347.1 afla|EED46498.1 aver|OJJ04023.1 aory|EIT72416.1 asyd|OJJ55510.1 ater|EAU32300.1

Group9259: azon|OJJ45115.1 azon|OJJ45686.1 azon|OJJ46496.1 azon|OJJ48981.1 azon|OJJ51110.1 azon|OJJ46510.1 aver|OJJ01292.1 asyd|OJJ58680.1

Group9260: azon|OJJ45169.1 azon|OJJ46865.1 acar|OOF96694.1 afis|EAW16437.1 afla|EED46603.1 aory|EIT77862.1 ater|EAU36666.1 awen|OJJ36274.1

Group9261: aluc|OJZ79663.1 aluc|OJZ80168.1 aluc|OJZ80047.1 aluc|OJZ80334.1 aluc|OJZ80488.1 aluc|OJZ80253.1 aluc|OJZ88813.1 aluc|OJZ79792.1

Group9262: abra|OJJ65421.1 afla|EED48896.1 akaw|GAA92642.1 anid|CBF87516.1 asyd|OJJ52980.1 ater|EAU36148.1 aver|OJJ08949.1 awen|OJJ35775.1

Group9263: abra|OJJ65633.1 anid|CBF82220.1 asyd|OJJ55373.1 ater|EAU35402.1 aver|OJJ06452.1 awen|OJJ31257.1 afla|EED54055.1 aory|EIT75907.1

Group9264: abra|OJJ65680.1 afis|EAW20903.1 anid|CBF82302.1 azon|OJJ47402.1 anig|EHA19604.1 akaw|GAA83833.1 aluc|OJZ90884.1 atub|OJI82879.1

Group9265: abra|OJJ65724.1 acar|OOG01114.1 afla|EED52284.1 agla|OJJ79398.1 asyd|OJJ52034.1 aver|OJI97995.1 azon|OJJ42308.1 asyd|OJJ62974.1

Group9266: abra|OJJ65788.1 acla|EAW10803.1 afis|EAW24775.1 afum|EAL89416.1 anid|CBF82868.1 anig|EHA18726.1 ater|EAU34032.1 awen|OJJ37934.1

Group9267: abra|OJJ65794.1 afla|EED52269.1 aory|EIT74396.1 asyd|OJJ65314.1 aver|OJI99455.1 azon|OJJ42715.1 acla|EAW14869.1 ater|EAU30136.1

Group9268: abra|OJJ65808.1 akaw|GAA92879.1 aluc|OJZ82529.1 anig|EHA22495.1 asyd|OJJ63153.1 atub|OJI88347.1 aver|OJI97802.1 asyd|OJJ63152.1

Group9269: afum|EAL85465.1 abra|OJJ77463.1 afis|EAW20345.1 akaw|GAA84153.1 aluc|OJZ81074.1 anig|EHA19295.1 atub|OJI82511.1 awen|OJJ41840.1

Group9270: abra|OJJ65863.1 acar|OOF95034.1 akaw|GAA92282.1 aluc|OJZ90179.1 anig|EHA23678.1 aory|EIT75041.1 atub|OJI81606.1 azon|OJJ51790.1

Group9271: abra|OJJ65889.1 acar|OOF94953.1 akaw|GAA86015.1 aluc|OJZ90243.1 asyd|OJJ65116.1 atub|OJI81671.1 aver|OJI99242.1 azon|OJJ47730.1

Group9272: abra|OJJ65914.1 acar|OOF94965.1 akaw|GAA92318.1 aluc|OJZ90216.1 anig|EHA23714.1 ater|EAU32851.1 atub|OJI81647.1 awen|OJJ35886.1

Group9273: ater|EAU31059.1 acla|EAW13302.1 aory|EIT74213.1 afis|EAW23645.1 afla|EED52624.1 anid|CBF84296.1 awen|OJJ29669.1 azon|OJJ42659.1

Group9274: abra|OJJ65926.1 acar|OOF94987.1 akaw|GAA92308.1 aluc|OJZ90205.1 anig|EHA23702.1 asyd|OJJ57266.1 atub|OJI81634.1 aver|OJJ03395.1

Group9275: abra|OJJ65936.1 acar|OOF95001.1 afla|EED46397.1 akaw|GAA92299.1 aluc|OJZ90196.1 aory|EIT72321.1 atub|OJI81623.1 azon|OJJ42446.1

Group9276: abra|OJJ65939.1 acar|OOF95002.1 afla|EED49788.1 akaw|GAA92297.1 aluc|OJZ90194.1 anig|EHA23693.1 aory|EIT77638.1 atub|OJI81621.1

Group9277: abra|OJJ66086.1 afla|EED54166.1 akaw|GAA84159.1 aluc|OJZ81082.1 atub|OJI82502.1 awen|OJJ33593.1 asyd|OJJ62095.1 aver|OJI96201.1

Group9278: afum|EAL92635.1 agla|OJJ87899.1 aluc|OJZ80198.1 anig|EHA26130.1 asyd|OJJ56528.1 ater|EAU29449.1 aver|OJJ06195.1 awen|OJJ31591.1

Group9279: abra|OJJ66107.1 afla|EED49582.1 anid|CBF85234.1 ater|EAU33589.1 aver|OJI97985.1 aory|EIT82612.1 asyd|OJJ62984.1 awen|OJJ39698.1

Group9280: akaw|GAA91918.1 asyd|OJJ62307.1 agla|OJJ87096.1 aluc|OJZ81962.1 anig|EHA22285.1 atub|OJI84513.1 awen|OJJ32716.1 aver|OJI96415.1

Group9281: abra|OJJ66122.1 acar|OOF90188.1 anid|CBF80738.1 anig|EHA22239.1 asyd|OJJ54542.1 aver|OJJ07639.1 azon|OJJ47323.1 azon|OJJ50406.1

Group9282: afla|EED49727.1 asyd|OJJ58706.1 anid|CBF84222.1 awen|OJJ32442.1 aory|EIT77632.1 asyd|OJJ63399.1 aver|OJJ01318.1 agla|OJJ83109.1

Group9283: aory|EIT77763.1 abra|OJJ72187.1 acar|OOF96812.1 afla|EED44769.1 akaw|GAA84746.1 aluc|OJZ86937.1 anig|EHA21707.1 atub|OJI83584.1

Group9284: abra|OJJ66135.1 acar|OOF94021.1 akaw|GAA92692.1 aluc|OJZ90150.1 anig|EHA20920.1 ater|EAU29938.1 atub|OJI84649.1 anid|CBF78933.1

Group9285: azon|OJJ42406.1 afla|EED52318.1 akaw|GAA83721.1 aory|EIT76283.1 agla|OJJ81796.1 aory|EIT74387.1 asyd|OJJ64672.1 anid|CBF75764.1

Group9286: abra|OJJ66227.1 acar|OOF90677.1 akaw|GAA89062.1 aluc|OJZ80532.1 anig|EHA25663.1 asyd|OJJ56693.1 atub|OJI88399.1 aver|OJJ06031.1

Group9287: abra|OJJ66242.1 acar|OOF98734.1 afis|EAW23512.1 abra|OJJ72743.1 akaw|GAA85354.1 aluc|OJZ90716.1 anig|EHA24135.1 atub|OJI82143.1

Group9288: anig|EHA24966.1 aory|EIT75035.1 aver|OJJ01836.1 afla|EED53155.1 akaw|GAA92153.1 aluc|OJZ85560.1 atub|OJI79345.1 asyd|OJJ59226.1

Group9289: asyd|OJJ58255.1 acar|OOF94380.1 akaw|GAA92136.1 aluc|OJZ83587.1 afla|EED57050.1 anig|EHA26575.1 aory|EIT82253.1 aver|OJJ02168.1

Group9290: abra|OJJ66332.1 acar|OOG00529.1 afla|EED56852.1 akaw|GAA90209.1 aluc|OJZ83136.1 anig|EHA28195.1 aory|EIT77516.1 atub|OJI87798.1

Group9291: abra|OJJ66376.1 afla|EED48192.1 akaw|GAA90244.1 aluc|OJZ83096.1 anig|EHA28160.1 aory|EIT72933.1 asyd|OJJ62482.1 atub|OJI87840.1

Group9292: afis|EAW19007.1 afla|EED49643.1 afum|EAL93741.1 aory|EIT80941.1 akaw|GAA90361.1 aluc|OJZ85871.1 atub|OJI79657.1 anig|EHA24712.1

Group9293: abra|OJJ66407.1 acla|EAW14868.1 afla|EED53097.1 akaw|GAA90275.1 aluc|OJZ83066.1 anig|EHA28131.1 aory|EIT74795.1 atub|OJI87880.1

Group9294: abra|OJJ66468.1 afla|EED45604.1 agla|OJJ85473.1 akaw|GAA92152.1 aluc|OJZ85559.1 aory|EIT75587.1 atub|OJI79346.1 awen|OJJ35057.1

Group9295: ater|EAU36796.1 acla|EAW09989.1 afis|EAW21103.1 afla|EED51685.1 afum|EAL92124.1 aory|EIT80523.1 awen|OJJ36359.1 azon|OJJ48034.1

Group9296: abra|OJJ66635.1 agla|OJJ81099.1 aluc|OJZ80814.1 anid|CBF74289.1 asyd|OJJ51913.1 atub|OJI87663.1 aver|OJJ05354.1 azon|OJJ43017.1

Group9297: abra|OJJ66783.1 acar|OOF98651.1 akaw|GAA89718.1 aluc|OJZ80666.1 aory|EIT77167.1 atub|OJI87521.1 afum|EAL86844.1 anig|EHA27344.1

Group9298: abra|OJJ66788.1 acar|OOF93862.1 akaw|GAA89722.1 aluc|OJZ80662.1 anig|EHA27348.1 atub|OJI87517.1 afla|EED45999.1 aory|EIT75755.1

Group9299: abra|OJJ66842.1 acar|OOF91619.1 akaw|GAA90478.1 aluc|OJZ85742.1 anig|EHA24810.1 ater|EAU37397.1 atub|OJI79527.1 awen|OJJ37463.1

Group9300: akaw|GAA91533.1 acla|EAW14451.1 afis|EAW19159.1 afla|EED54243.1 afum|EBA27507.1 anid|CBF86289.1 aory|EIT76061.1 ater|EAU37432.1

Group9301: afis|EAW17777.1 afum|EAL91586.1 afla|EED44889.1 aory|EIT83249.1 asyd|OJJ64750.1 ater|EAU31839.1 aver|OJI98851.1 awen|OJJ41518.1

Group9302: abra|OJJ66966.1 acar|OOF93643.1 afla|EED55268.1 akaw|GAA92197.1 aluc|OJZ85612.1 anig|EHA24914.1 aory|EIT83531.1 atub|OJI79405.1

Group9303: abra|OJJ66998.1 acar|OOF93618.1 akaw|GAA92167.1 aluc|OJZ85575.1 anig|EHA24944.1 atub|OJI79371.1 afum|EAL91341.2 afis|EAW25616.1

Group9304: abra|OJJ67024.1 akaw|GAA83296.1 anig|EHA24958.1 aory|EIT79521.1 asyd|OJJ52361.1 aver|OJI99599.1 aver|OJJ01987.1 agla|OJJ88611.1

Group9305: abra|OJJ67029.1 acar|OOF92968.1 acla|EAW10802.1 afla|EED49512.1 akaw|GAA82083.1 aluc|OJZ92097.1 aory|EIT82618.1 azon|OJJ47933.1

Group9306: abra|OJJ67030.1 acar|OOF91479.1 afis|EAW17354.1 akaw|GAA92150.1 aluc|OJZ85555.1 anig|EHA24964.1 atub|OJI79349.1 awen|OJJ36015.1

Group9307: abra|OJJ67093.1 afla|EED52013.1 akaw|GAA91068.1 aluc|OJZ87882.1 anig|EHA20185.1 aory|EIT81413.1 atub|OJI80769.1 aver|OJJ08326.1

Group9308: abra|OJJ67103.1 acar|OOF98446.1 acla|EAW07387.1 akaw|GAA91058.1 aluc|OJZ87870.1 anig|EHA20176.1 atub|OJI80779.1 azon|OJJ43579.1

Group9309: anig|EHA20161.1 acar|OOF98432.1 afla|EED55140.1 agla|OJJ78842.1 anid|CBF78818.1 asyd|OJJ65265.1 aver|OJI99416.1 awen|OJJ38064.1

Group9310: abra|OJJ67138.1 acar|OOF98418.1 agla|OJJ80244.1 akaw|GAA91031.1 aluc|OJZ87834.1 anig|EHA20313.1 atub|OJI80807.1 awen|OJJ36378.1

Group9311: abra|OJJ67207.1 acar|OOF98354.1 akaw|GAA90970.1 aluc|OJZ87767.1 anid|CBF89587.1 anig|EHA20251.1 asyd|OJJ62317.1 atub|OJI80878.1

Group9312: abra|OJJ67265.1 agla|OJJ81019.1 akaw|GAA91681.1 anig|EHA20207.1 atub|OJI80929.1 acar|OOF98301.1 aluc|OJZ87713.1 acla|EAW13208.1

Group9313: abra|OJJ67368.1 acar|OOF92442.1 akaw|GAA86456.1 aluc|OJZ84998.1 anig|EHA25207.1 aory|EIT74218.1 ater|EAU39602.1 atub|OJI80604.1

Group9314: abra|OJJ67414.1 akaw|GAA92400.1 aluc|OJZ83261.1 anig|EHA27888.1 atub|OJI87677.1 awen|OJJ38896.1 agla|OJJ88131.1 asyd|OJJ55566.1

Group9315: abra|OJJ67415.1 acar|OOF93051.1 afla|EED53045.1 agla|OJJ85100.1 anig|EHA25169.1 atub|OJI80649.1 azon|OJJ43513.1 awen|OJJ32479.1

Group9316: abra|OJJ67417.1 acar|OOF93053.1 akaw|GAA86411.1 aluc|OJZ84951.1 anig|EHA25167.1 atub|OJI80651.1 awen|OJJ38766.1 azon|OJJ43511.1

Group9317: abra|OJJ67448.1 acar|OOF99139.1 akaw|GAA89803.1 aluc|OJZ81888.1 ater|EAU35113.1 atub|OJI84441.1 azon|OJJ43519.1 afla|EED57539.1

Group9318: abra|OJJ67470.1 acar|OOF89933.1 acla|EAW06302.1 afla|EED52996.1 akaw|GAA86366.1 anig|EHA25123.1 aory|EIT77430.1 atub|OJI80703.1

Group9319: abra|OJJ67537.1 acar|OOG00242.1 akaw|GAA88122.1 aluc|OJZ81395.1 atub|OJI91600.1 aver|OJJ07829.1 azon|OJJ43559.1 asyd|OJJ53705.1

Group9320: abra|OJJ67563.1 acar|OOG00281.1 afla|EED55967.1 akaw|GAA88102.1 aluc|OJZ81369.1 anig|EHA18668.1 aory|EIT76720.1 atub|OJI91574.1

Group9321: abra|OJJ67595.1 afis|EAW20297.1 afum|EAL85426.1 agla|OJJ88387.1 ater|EAU38944.1 atub|OJI88456.1 anid|CBF77122.1 aver|OJI99733.1

Group9322: abra|OJJ67604.1 acar|OOF91954.1 akaw|GAA89121.1 aluc|OJZ81717.1 anig|EHA25722.1 asyd|OJJ61711.1 atub|OJI88465.1 aver|OJI98906.1

Group9323: abra|OJJ67635.1 acar|OOF96455.1 afla|EED48408.1 akaw|GAA89154.1 aluc|OJZ81681.1 anig|EHA25760.1 atub|OJI88500.1 agla|OJJ82592.1

Group9324: abra|OJJ67637.1 aluc|OJZ81679.1 anid|CBF87120.1 asyd|OJJ60359.1 atub|OJI88502.1 aver|OJI99697.1 awen|OJJ33616.1 ater|EAU29523.1

Group9325: abra|OJJ67679.1 akaw|GAA89192.1 aluc|OJZ81639.1 anig|EHA25805.1 asyd|OJJ53349.1 atub|OJI88541.1 aver|OJJ08395.1 acla|EAW12061.1

Group9326: abra|OJJ67682.1 afis|EAW19875.1 afum|EAL85024.1 akaw|GAA89195.1 aluc|OJZ81636.1 anig|EHA25808.1 atub|OJI88544.1 ater|EAU30907.1

Group9327: abra|OJJ67690.1 afum|EAL86724.1 aory|EIT72972.1 asyd|OJJ62707.1 ater|EAU32453.1 atub|OJI86665.1 awen|OJJ36179.1 anid|CBF78433.1

Group9328: abra|OJJ67704.1 acar|OOF98710.1 akaw|GAA89212.1 aluc|OJZ81617.1 anig|EHA25826.1 atub|OJI88563.1 azon|OJJ42195.1 aory|EIT75780.1

Group9329: abra|OJJ67716.1 afis|EAW19890.1 afla|EED46278.1 aory|EIT76343.1 asyd|OJJ60197.1 atub|OJI80366.1 aver|OJJ07195.1 aver|OJJ04068.1

Group9330: abra|OJJ67720.1 akaw|GAA92885.1 anig|EHA25833.1 atub|OJI88571.1 aver|OJI95869.1 abra|OJJ70190.1 ater|EAU31225.1 azon|OJJ43537.1

Group9331: abra|OJJ67786.1 acar|OOF90619.1 afla|EED51852.1 agla|OJJ78974.1 aory|EIT81700.1 asyd|OJJ60223.1 aver|OJJ04044.1 awen|OJJ33659.1

Group9332: abra|OJJ67843.1 acar|OOF92020.1 akaw|GAA91849.1 aluc|OJZ82049.1 anig|EHA22209.1 asyd|OJJ63642.1 atub|OJI84607.1 aver|OJI97294.1

Group9333: ater|EAU32015.1 acla|EAW11694.1 afis|EAW23711.1 afum|EAL87155.1 asyd|OJJ59161.1 aver|OJJ01751.1 awen|OJJ33504.1 azon|OJJ46437.1

Group9334: abra|OJJ67891.1 acar|OOF90315.1 agla|OJJ84835.1 akaw|GAA91897.1 aluc|OJZ81999.1 anig|EHA19897.1 ater|EAU39317.1 atub|OJI84549.1

Group9335: akaw|GAA91263.1 anig|EHA21247.1 afis|EAW20399.1 afla|EED51971.1 afum|EAL85498.1 anid|CBF79900.1 aory|EIT81375.1 azon|OJJ48750.1

Group9336: abra|OJJ67971.1 afis|EAW16715.1 afla|EED55256.1 agla|OJJ80337.1 aluc|OJZ80478.1 anig|EHA22003.1 aory|EIT83618.1 asyd|OJJ56138.1

Group9337: abra|OJJ68005.1 acar|OOF92047.1 akaw|GAA90945.1 aluc|OJZ82879.1 anig|EHA21305.1 asyd|OJJ64160.1 atub|OJI89103.1 aver|OJI98237.1

Group9338: abra|OJJ68008.1 acar|OOF92050.1 afum|EAL86543.1 akaw|GAA90942.1 aluc|OJZ82882.1 anig|EHA21308.1 atub|OJI89106.1 ater|EAU31761.1

Group9339: abra|OJJ68066.1 acar|OOF90622.1 acla|EAW15143.1 akaw|GAA90893.1 aluc|OJZ82936.1 atub|OJI89165.1 awen|OJJ33049.1 agla|OJJ82056.1

Group9340: agla|OJJ83641.1 asyd|OJJ56660.1 acla|EAW09999.1 afis|EAW21072.1 afla|EED47789.1 aory|EIT77187.1 aver|OJJ07406.1 aver|OJJ06064.1

Group9341: abra|OJJ68087.1 acar|OOF90255.1 akaw|GAA86872.1 aluc|OJZ89996.1 anig|EHA27371.1 asyd|OJJ56932.1 atub|OJI84812.1 azon|OJJ46836.1

Group9342: abra|OJJ68099.1 acar|OOF93902.1 akaw|GAA86884.1 aluc|OJZ90007.1 anig|EHA27382.1 atub|OJI84800.1 awen|OJJ37264.1 azon|OJJ46012.1

Group9343: abra|OJJ68136.1 acar|OOF93923.1 afis|EAW19309.1 afum|EAL94026.2 akaw|GAA86916.1 aluc|OJZ90039.1 anig|EHA27414.1 atub|OJI84767.1

Group9344: acar|OOF94576.1 aory|EIT75961.1 acla|EAW15344.1 afla|EED54355.1 asyd|OJJ56917.1 azon|OJJ46842.1 anid|CBF77240.1 aver|OJJ03375.1

Group9345: abra|OJJ68244.1 acar|OOG00394.1 afis|EAW18880.1 afla|EED49705.1 aory|EIT77710.1 atub|OJI84567.1 awen|OJJ32675.1 agla|OJJ85238.1

Group9346: abra|OJJ68246.1 acar|OOF97525.1 agla|OJJ83212.1 akaw|GAA83215.1 aluc|OJZ83717.1 anig|EHA18867.1 atub|OJI90092.1 awen|OJJ41816.1

Group9347: abra|OJJ68292.1 akaw|GAA83181.1 aluc|OJZ83681.1 anig|EHA18835.1 aory|EIT81661.1 atub|OJI90052.1 awen|OJJ41219.1 ater|EAU32731.1

Group9348: abra|OJJ68313.1 acar|OOF96695.1 afla|EED47754.1 akaw|GAA83160.1 aluc|OJZ83660.1 anig|EHA18812.1 atub|OJI90031.1 aver|OJJ07315.1

Group9349: abra|OJJ68341.1 acar|OOF89852.1 akaw|GAA92108.1 aluc|OJZ83617.1 anig|EHA18775.1 atub|OJI89990.1 awen|OJJ40124.1 azon|OJJ45193.1

Group9350: abra|OJJ68360.1 acar|OOF99379.1 afla|EED45109.1 akaw|GAA82514.1 aluc|OJZ91597.1 anig|EHA22627.1 aory|EIT73776.1 atub|OJI86048.1

Group9351: abra|OJJ68403.1 afla|EED48372.1 akaw|GAA92543.1 aluc|OJZ90091.1 anig|EHA20983.1 aory|EIT83048.1 ater|EAU33483.1 atub|OJI84714.1

Group9352: abra|OJJ68409.1 acar|OOF93968.1 akaw|GAA92541.1 aluc|OJZ90089.1 anig|EHA20985.1 atub|OJI84717.1 awen|OJJ33720.1 akaw|GAA87622.1

Group9353: afla|EED51305.1 afis|EAW20761.1 afum|EAL84858.1 agla|OJJ86655.1 anid|CBF71470.1 aory|EIT81498.1 asyd|OJJ54945.1 aver|OJJ07250.1

Group9354: abra|OJJ68591.1 afla|EED57758.1 akaw|GAA91920.1 aluc|OJZ81960.1 anig|EHA22287.1 aory|EIT81044.1 ater|EAU35718.1 atub|OJI84511.1

Group9355: afla|EED47362.1 aory|EIT82862.1 acar|OOF92454.1 ater|EAU32760.1 agla|OJJ86668.1 anid|CBF78928.1 asyd|OJJ55634.1 aver|OJJ06711.1

Group9356: abra|OJJ68595.1 afla|EED44779.1 agla|OJJ79463.1 anid|CBF69449.1 aory|EIT77767.1 ater|EAU35433.1 anid|CBF83149.1 atub|OJI80341.1

Group9357: abra|OJJ68868.1 afis|EAW16175.1 afla|EED51072.1 akaw|GAA90385.1 aluc|OJZ85846.1 anig|EHA24718.1 aory|EIT77053.1 atub|OJI79634.1

Group9358: abra|OJJ68905.1 afla|EED52081.1 afum|EAL91019.1 anig|EHA25685.1 aory|EIT76931.1 atub|OJI88422.1 aver|OJJ03455.1 awen|OJJ40167.1

Group9359: acla|EAW14232.1 afis|EAW19048.1 afum|EAL93777.1 aory|EIT82019.1 ater|EAU32087.1 awen|OJJ31478.1 agla|OJJ84557.1 afla|EED50407.1

Group9360: abra|OJJ69002.1 acar|OOF92715.1 agla|OJJ85375.1 aluc|OJZ85988.1 asyd|OJJ58283.1 aver|OJJ02140.1 awen|OJJ41517.1 azon|OJJ48240.1

Group9361: abra|OJJ69021.1 agla|OJJ87908.1 akaw|GAA90538.1 aluc|OJZ86007.1 anig|EHA24603.1 atub|OJI79796.1 agla|OJJ85711.1 ater|EAU29899.1

Group9362: abra|OJJ69027.1 akaw|GAA90543.1 aluc|OJZ86013.1 anid|CBF76221.1 anig|EHA24598.1 atub|OJI79802.1 awen|OJJ31674.1 afla|EED53396.1

Group9363: abra|OJJ69028.1 agla|OJJ79300.1 akaw|GAA90544.1 aluc|OJZ86014.1 anid|CBF76223.1 anig|EHA24597.1 atub|OJI79803.1 awen|OJJ31673.1

Group9364: abra|OJJ69174.1 acar|OOF91130.1 akaw|GAA86288.1 aluc|OJZ86161.1 anig|EHA24463.1 atub|OJI79953.1 azon|OJJ46834.1 acar|OOF93937.1

Group9365: abra|OJJ69348.1 anig|EHA18747.1 atub|OJI88256.1 awen|OJJ38174.1 azon|OJJ42137.1 anid|CBF83079.1 asyd|OJJ52796.1 aver|OJJ08774.1

Group9366: azon|OJJ48491.1 acar|OOF94373.1 agla|OJJ79592.1 anid|CBF74160.1 asyd|OJJ53930.1 ater|EAU31218.1 aver|OJJ08029.1 awen|OJJ33026.1

Group9367: abra|OJJ69425.1 acar|OOF94311.1 agla|OJJ79346.1 akaw|GAA88291.1 aluc|OJZ82262.1 anig|EHA28481.1 asyd|OJJ52459.1 atub|OJI88176.1

Group9368: abra|OJJ69427.1 acar|OOF94309.1 afla|EED52059.1 akaw|GAA88293.1 aluc|OJZ82264.1 anig|EHA28479.1 aory|EIT81422.1 atub|OJI88174.1

Group9369: abra|OJJ69428.1 acar|OOF94308.1 akaw|GAA88294.1 aluc|OJZ82265.1 anig|EHA28478.1 atub|OJI88173.1 awen|OJJ39638.1 azon|OJJ47765.1

Group9370: abra|OJJ69434.1 acar|OOF94303.1 afla|EED49523.1 akaw|GAA88298.1 aluc|OJZ82270.1 anig|EHA28473.1 aory|EIT82576.1 atub|OJI88168.1

Group9371: abra|OJJ69555.1 acla|EAW13604.1 abra|OJJ72384.1 acar|OOF90288.1 akaw|GAA86133.1 aluc|OJZ90371.1 atub|OJI81797.1 agla|OJJ87532.1

Group9372: abra|OJJ69563.1 aory|EIT79512.1 asyd|OJJ55176.1 aver|OJJ07031.1 awen|OJJ29814.1 acar|OOF98337.1 afla|EED49421.1 azon|OJJ42661.1

Group9373: abra|OJJ69566.1 aory|EIT79529.1 abra|OJJ73128.1 acar|OOF90604.1 acla|EAW11775.1 afla|EED49418.1 anig|EHA18702.1 azon|OJJ42289.1

Group9374: abra|OJJ69573.1 acar|OOF91581.1 acla|EAW14395.1 akaw|GAA91570.1 aluc|OJZ82390.1 anig|EHA28373.1 atub|OJI88051.1 awen|OJJ32496.1

Group9375: abra|OJJ69577.1 acar|OOF91578.1 afis|EAW19100.1 aluc|OJZ82395.1 aory|EIT76155.1 atub|OJI88047.1 awen|OJJ32493.1 azon|OJJ46038.1

Group9376: abra|OJJ69650.1 acar|OOF96861.1 akaw|GAA91627.1 aluc|OJZ82454.1 anig|EHA28321.1 atub|OJI87984.1 afis|EAW19668.1 afum|EAL86855.1

Group9377: abra|OJJ69726.1 acar|OOF93548.1 akaw|GAA90179.1 aluc|OJZ83171.1 anig|EHA28245.1 atub|OJI87767.1 azon|OJJ47343.1 aory|EIT75546.1

Group9378: abra|OJJ69750.1 afis|EAW19965.1 afum|EAL84946.1 akaw|GAA90201.1 aluc|OJZ83147.1 anig|EHA28211.1 atub|OJI87790.1 anig|EHA28212.1

Group9379: abra|OJJ69752.1 acar|OOG00536.1 akaw|GAA90203.1 aluc|OJZ83145.1 anig|EHA28209.1 atub|OJI87792.1 awen|OJJ31962.1 azon|OJJ42094.1

Group9380: abra|OJJ69755.1 afla|EED45418.1 akaw|GAA92452.1 aluc|OJZ81566.1 anig|EHA21150.1 aory|EIT82456.1 ater|EAU30037.1 atub|OJI88619.1

Group9381: abra|OJJ69782.1 afis|EAW19285.1 akaw|GAA87874.1 aluc|OJZ85515.1 anig|EHA25048.1 atub|OJI80051.1 awen|OJJ30810.1 afis|EAW18910.1

Group9382: abra|OJJ69785.1 acar|OOF97620.1 agla|OJJ85957.1 anig|EHA25045.1 aory|EIT74493.1 asyd|OJJ56178.1 awen|OJJ35829.1 azon|OJJ48193.1

Group9383: aory|EIT75364.1 acar|OOF93576.1 afis|EAW24739.1 afla|EED45537.1 afum|EBA27379.1 akaw|GAA87858.1 anid|CBF80396.1 azon|OJJ45019.1

Group9384: abra|OJJ69808.1 acar|OOF93569.1 afla|EED45863.1 akaw|GAA87852.1 aluc|OJZ85492.1 anig|EHA25027.1 aory|EIT75512.1 atub|OJI80073.1

Group9385: abra|OJJ69821.1 acar|OOF93342.1 afla|EED51605.1 akaw|GAA87839.1 aluc|OJZ85475.1 aory|EIT80578.1 atub|OJI80088.1 anig|EHA21834.1

Group9386: abra|OJJ69829.1 acar|OOF93354.1 agla|OJJ79194.1 akaw|GAA87831.1 aluc|OJZ85467.1 anig|EHA25008.1 atub|OJI80096.1 awen|OJJ32855.1

Group9387: abra|OJJ69851.1 acar|OOF93378.1 akaw|GAA87807.1 aluc|OJZ85442.1 anig|EHA24986.1 atub|OJI80125.1 afis|EAW21631.1 awen|OJJ40024.1

Group9388: abra|OJJ69880.1 acar|OOF93395.1 afla|EED46215.1 akaw|GAA87778.1 aluc|OJZ85413.1 anig|EHA25592.1 aory|EIT76375.1 atub|OJI80164.1

Group9389: abra|OJJ69882.1 acar|OOF93400.1 akaw|GAA87776.1 aluc|OJZ85411.1 anig|EHA25590.1 atub|OJI80166.1 awen|OJJ33524.1 azon|OJJ42492.1

Group9390: abra|OJJ70034.1 afis|EAW24625.1 afum|EAL86249.1 anid|CBF79092.1 anig|EHA18719.1 aver|OJJ08541.1 afis|EAW24621.1 aver|OJJ08544.1

Group9391: abra|OJJ70045.1 afis|EAW16416.1 agla|OJJ80726.1 akaw|GAA87629.1 aluc|OJZ85246.1 anig|EHA25446.1 asyd|OJJ57151.1 atub|OJI80325.1

Group9392: abra|OJJ70055.1 akaw|GAA85315.1 aluc|OJZ90676.1 aver|OJJ00039.1 afis|EAW24586.1 afum|EAL88040.1 anig|EHA27498.1 aver|OJJ08529.1

Group9393: abra|OJJ70065.1 acar|OOF93553.1 afla|EED46369.1 akaw|GAA87611.1 aluc|OJZ85232.1 anig|EHA25436.1 aory|EIT81169.1 atub|OJI80346.1

Group9394: anid|CBF80550.1 abra|OJJ76736.1 akaw|GAA90363.1 abra|OJJ76738.1 afis|EAW19005.1 afum|EAL93739.1 aluc|OJZ85869.1 atub|OJI79655.1

Group9395: abra|OJJ70086.1 acla|EAW12310.1 afis|EAW21037.1 afum|EAL92076.2 akaw|GAA92273.1 aluc|OJZ90167.1 atub|OJI89255.1 atub|OJI81595.1

Group9396: abra|OJJ70111.1 afla|EED51525.1 akaw|GAA83852.1 aluc|OJZ90863.1 anig|EHA19576.1 aory|EIT80609.1 atub|OJI82861.1 awen|OJJ40163.1

Group9397: abra|OJJ70117.1 akaw|GAA87593.1 aluc|OJZ85213.1 anid|CBF84298.1 anig|EHA25416.1 asyd|OJJ52179.1 atub|OJI80373.1 aver|OJJ05966.1

Group9398: afis|EAW24745.1 afla|EED48475.1 anid|CBF73382.1 aory|EIT73052.1 awen|OJJ29802.1 afum|EAL89391.1 ater|EAU37149.1 agla|OJJ78874.1

Group9399: abra|OJJ70148.1 afla|EED52388.1 akaw|GAA90560.1 aory|EIT83294.1 abra|OJJ71751.1 anig|EHA25741.1 asyd|OJJ53403.1 aver|OJJ04198.1

Group9400: abra|OJJ70161.1 akaw|GAA86608.1 aluc|OJZ85158.1 anig|EHA25374.1 atub|OJI80435.1 acar|OOG00107.1 agla|OJJ89298.1 awen|OJJ39599.1

Group9401: abra|OJJ70198.1 acla|EAW12119.1 afis|EAW17261.1 afum|EAL86678.1 akaw|GAA86578.1 aluc|OJZ85129.1 anig|EHA25348.1 atub|OJI80464.1

Group9402: abra|OJJ70200.1 acar|OOF92580.1 akaw|GAA93214.1 aluc|OJZ85127.1 anig|EHA25346.1 atub|OJI80466.1 awen|OJJ30063.1 akaw|GAA86576.1

Group9403: abra|OJJ70216.1 acar|OOF92564.1 akaw|GAA86561.1 aluc|OJZ85110.1 anig|EHA25331.1 asyd|OJJ61743.1 atub|OJI80483.1 aver|OJI95840.1

Group9404: abra|OJJ70222.1 akaw|GAA86557.1 aluc|OJZ85105.1 anig|EHA25327.1 atub|OJI80489.1 awen|OJJ38080.1 aver|OJI96677.1 aory|EIT76141.1

Group9405: abra|OJJ70249.1 acar|OOF92548.1 afla|EED50196.1 anig|EHA25310.1 aory|EIT77382.1 awen|OJJ34411.1 azon|OJJ46186.1 awen|OJJ30067.1

Group9406: abra|OJJ70255.1 acar|OOF93348.1 afla|EED55138.1 akaw|GAA86110.1 aluc|OJZ90342.1 anig|EHA25304.1 aory|EIT81616.1 atub|OJI81769.1

Group9407: abra|OJJ70261.1 afla|EED45530.1 akaw|GAA86536.1 aluc|OJZ85083.1 anig|EHA25297.1 aory|EIT75465.1 ater|EAU32500.1 atub|OJI80515.1

Group9408: abra|OJJ70277.1 akaw|GAA92872.1 aluc|OJZ82539.1 anig|EHA22507.1 asyd|OJJ55017.1 atub|OJI88783.1 aver|OJJ07188.1 awen|OJJ32617.1

Group9409: abra|OJJ70280.1 akaw|GAA92868.1 aluc|OJZ82544.1 atub|OJI88787.1 azon|OJJ45991.1 aver|OJJ01890.1 atub|OJI88354.1 awen|OJJ39639.1

Group9410: abra|OJJ70287.1 acar|OOF91738.1 afla|EED48873.1 akaw|GAA92861.1 aluc|OJZ82552.1 aory|EIT81179.1 atub|OJI88793.1 awen|OJJ38797.1

Group9411: anid|CBF69719.1 asyd|OJJ58925.1 acar|OOF98770.1 afis|EAW16870.1 afum|EAL91625.1 aver|OJJ01521.1 azon|OJJ45904.1 aver|OJI99598.1

Group9412: anid|CBF80471.1 afis|EAW17123.1 afla|EED46216.1 afum|EAL86584.1 aory|EIT76510.1 asyd|OJJ54771.1 ater|EAU35115.1 aver|OJJ07395.1

Group9413: abra|OJJ70302.1 acar|OOF91755.1 afla|EED45962.1 akaw|GAA93020.1 aluc|OJZ82565.1 anig|EHA27509.1 ater|EAU32148.1 atub|OJI88809.1

Group9414: abra|OJJ70303.1 acar|OOF91756.1 akaw|GAA93019.1 aluc|OJZ82566.1 anig|EHA27508.1 aory|EIT75475.1 atub|OJI88810.1 afla|EED45963.1

Group9415: abra|OJJ70325.1 acar|OOF91781.1 agla|OJJ87947.1 akaw|GAA92581.1 aluc|OJZ82594.1 anid|CBF89612.1 anig|EHA27481.1 atub|OJI88829.1

Group9416: abra|OJJ70330.1 acar|OOF91790.1 akaw|GAA92585.1 aluc|OJZ82599.1 anid|CBF87302.1 atub|OJI88835.1 aver|OJJ05911.1 afis|EAW15388.1

Group9417: abra|OJJ70331.1 acar|OOF91791.1 afis|EAW15387.1 akaw|GAA92586.1 aluc|OJZ82601.1 anig|EHA27478.1 atub|OJI88837.1 aver|OJJ05912.1

Group9418: abra|OJJ70347.1 akaw|GAA92599.1 aluc|OJZ82616.1 anig|EHA27461.1 asyd|OJJ60740.1 atub|OJI88851.1 aver|OJJ00072.1 awen|OJJ33352.1

Group9419: abra|OJJ70350.1 afis|EAW17019.1 afla|EED51526.1 akaw|GAA89818.1 aluc|OJZ81872.1 aory|EIT80752.1 aver|OJJ03456.1 awen|OJJ40165.1

Group9420: abra|OJJ70406.1 acar|OOF98106.1 afla|EED45215.1 akaw|GAA92455.1 aluc|OJZ81562.1 anig|EHA27370.1 aory|EIT73775.1 atub|OJI88622.1

Group9421: abra|OJJ70418.1 akaw|GAA88221.1 aluc|OJZ82185.1 anid|CBF69412.1 anig|EHA18391.1 asyd|OJJ63816.1 atub|OJI88252.1 aver|OJI97122.1

Group9422: acla|EAW11588.1 afis|EAW23865.1 afum|EAL87293.1 aory|EIT74285.1 acar|OOG00288.1 afla|EED52592.1 ater|EAU32689.1 atub|OJI87742.1

Group9423: acla|EAW06948.1 agla|OJJ89078.1 afis|EAW16563.1 afla|EED45794.1 afum|EAL89160.1 agla|OJJ82758.1 aory|EIT75797.1 azon|OJJ49117.1

Group9424: abra|OJJ70438.1 acar|OOF91841.1 afla|EED48140.1 akaw|GAA90142.1 aluc|OJZ83205.1 aory|EIT76222.1 atub|OJI87732.1 azon|OJJ51108.1

Group9425: abra|OJJ70922.1 acar|OOF98201.1 afis|EAW21020.1 afum|EAL92058.1 akaw|GAA83018.1 aluc|OJZ87604.1 anig|EHA20803.1 atub|OJI81044.1

Group9426: abra|OJJ71050.1 anig|EHA20698.1 abra|OJJ71051.1 akaw|GAA82906.1 aluc|OJZ87476.1 anig|EHA20697.1 atub|OJI81172.1 asyd|OJJ63592.1

Group9427: abra|OJJ71077.1 afla|EED46425.1 akaw|GAA82885.1 aluc|OJZ87453.1 ater|EAU30880.1 atub|OJI81195.1 awen|OJJ36404.1 awen|OJJ33727.1

Group9428: abra|OJJ71090.1 acar|OOF98049.1 agla|OJJ87148.1 akaw|GAA82871.1 aluc|OJZ87438.1 ater|EAU29809.1 atub|OJI81210.1 awen|OJJ35051.1

Group9429: abra|OJJ71099.1 akaw|GAA83835.1 aluc|OJZ90882.1 anig|EHA28513.1 aory|EIT76287.1 atub|OJI83755.1 afla|EED48319.1 azon|OJJ46592.1

Group9430: abra|OJJ71216.1 acar|OOF94574.1 acla|EAW15347.1 anig|EHA20560.1 atub|OJI87216.1 aver|OJJ03894.1 aluc|OJZ92773.1 akaw|GAA91760.1

Group9431: abra|OJJ71257.1 afla|EED53790.1 aluc|OJZ87286.1 asyd|OJJ59891.1 atub|OJI81355.1 aver|OJJ02395.1 awen|OJJ41049.1 azon|OJJ44611.1

Group9432: abra|OJJ71285.1 acar|OOF97887.1 agla|OJJ83883.1 aluc|OJZ87258.1 asyd|OJJ59929.1 atub|OJI81384.1 awen|OJJ41061.1 azon|OJJ44644.1

Group9433: abra|OJJ71736.1 akaw|GAA85150.1 aluc|OJZ86487.1 anig|EHA22180.1 aory|EIT73750.1 asyd|OJJ52524.1 atub|OJI84042.1 atub|OJI89784.1

Group9434: afla|EED55199.1 aory|EIT83630.1 afis|EAW17199.1 afum|EBA27282.1 anid|CBF80261.1 asyd|OJJ54174.1 ater|EAU36725.1 aver|OJJ08565.1

Group9435: abra|OJJ71772.1 acar|OOF90397.1 akaw|GAA85135.1 aluc|OJZ86502.1 anig|EHA22142.1 atub|OJI84024.1 awen|OJJ40723.1 azon|OJJ45843.1

Group9436: abra|OJJ71773.1 acar|OOF90396.1 akaw|GAA85134.1 aluc|OJZ86503.1 anig|EHA22141.1 atub|OJI84023.1 awen|OJJ40722.1 azon|OJJ45844.1

Group9437: abra|OJJ71774.1 acar|OOF90395.1 akaw|GAA85133.1 aluc|OJZ86504.1 anig|EHA22140.1 atub|OJI84022.1 awen|OJJ40721.1 azon|OJJ45845.1

Group9438: aory|EIT77623.1 afla|EED49715.1 akaw|GAA84675.1 aluc|OJZ87014.1 asyd|OJJ53591.1 atub|OJI83505.1 aver|OJJ08157.1 awen|OJJ29856.1

Group9439: abra|OJJ71799.1 acar|OOF96479.1 agla|OJJ82295.1 akaw|GAA85111.1 aluc|OJZ86529.1 anig|EHA22117.1 ater|EAU38224.1 atub|OJI83992.1

Group9440: abra|OJJ71911.1 acar|OOF96569.1 afla|EED51107.1 akaw|GAA85009.1 aluc|OJZ86648.1 anig|EHA22021.1 aory|EIT81335.1 atub|OJI83878.1

Group9441: abra|OJJ71953.1 acar|OOF96618.1 akaw|GAA84975.1 aluc|OJZ86687.1 anig|EHA21986.1 aory|EIT82635.1 atub|OJI83838.1 afla|EED49591.1

Group9442: abra|OJJ72073.1 acar|OOF99319.1 akaw|GAA84844.1 aluc|OJZ86827.1 anig|EHA21810.1 atub|OJI83691.1 aver|OJJ07659.1 azon|OJJ51731.1

Group9443: abra|OJJ72078.1 akaw|GAA83328.1 aluc|OJZ83824.1 anig|EHA18968.1 atub|OJI90191.1 aver|OJJ08590.1 afla|EED58003.1 aory|EIT79715.1

Group9444: abra|OJJ72098.1 acar|OOF96743.1 afis|EAW17296.1 akaw|GAA84823.1 aluc|OJZ86853.1 anig|EHA21786.1 atub|OJI83666.1 azon|OJJ42798.1

Group9445: abra|OJJ72155.1 acar|OOF96788.1 acla|EAW09342.1 akaw|GAA84772.1 aluc|OJZ86911.1 anig|EHA21733.1 atub|OJI83610.1 awen|OJJ38508.1

Group9446: abra|OJJ72160.1 acar|OOF96796.1 akaw|GAA84767.1 aluc|OJZ86916.1 anig|EHA21728.1 atub|OJI83605.1 awen|OJJ35674.1 azon|OJJ42753.1

Group9447: abra|OJJ72172.1 afla|EED53529.1 akaw|GAA84756.1 aluc|OJZ86927.1 asyd|OJJ60512.1 atub|OJI83594.1 aver|OJI99837.1 awen|OJJ36998.1

Group9448: abra|OJJ72225.1 acar|OOF96848.1 akaw|GAA84712.1 aluc|OJZ86975.1 anig|EHA21673.1 ater|EAU34020.1 atub|OJI83546.1 awen|OJJ36430.1

Group9449: abra|OJJ72235.1 acar|OOF96857.1 akaw|GAA84704.1 aluc|OJZ86983.1 anig|EHA21664.1 atub|OJI83536.1 awen|OJJ34545.1 azon|OJJ42224.1

Group9450: abra|OJJ72245.1 acar|OOF96872.1 akaw|GAA84693.1 aluc|OJZ86994.1 anig|EHA21651.1 atub|OJI83525.1 aver|OJJ02012.1 azon|OJJ47854.1

Group9451: abra|OJJ72246.1 acar|OOF96874.1 akaw|GAA84692.1 aluc|OJZ86995.1 anig|EHA21650.1 atub|OJI83524.1 aver|OJJ02010.1 azon|OJJ47856.1

Group9452: abra|OJJ72257.1 acar|OOF96889.1 akaw|GAA84680.1 aluc|OJZ87008.1 anig|EHA21638.1 aory|EIT78168.1 atub|OJI83511.1 afla|EED52819.1

Group9453: awen|OJJ39614.1 acar|OOF93144.1 afla|EED48639.1 agla|OJJ78980.1 aory|EIT76620.1 ater|EAU37752.1 aory|EIT73113.1 awen|OJJ38067.1

Group9454: abra|OJJ72262.1 afla|EED51740.1 agla|OJJ87797.1 abra|OJJ77983.1 akaw|GAA85997.1 aluc|OJZ91058.1 anig|EHA19761.1 atub|OJI83054.1

Group9455: abra|OJJ72301.1 acar|OOF94923.1 akaw|GAA86052.1 aluc|OJZ90282.1 anig|EHA23772.1 asyd|OJJ62082.1 atub|OJI81706.1 aver|OJI96190.1

Group9456: abra|OJJ72521.1 anid|CBF75174.1 afla|EED46473.1 akaw|GAA86068.1 aluc|OJZ90303.1 anig|EHA23790.1 aory|EIT72334.1 awen|OJJ30885.1

Group9457: abra|OJJ72525.1 acar|OOF99516.1 agla|OJJ88724.1 aluc|OJZ90505.1 anig|EHA23963.1 atub|OJI81929.1 awen|OJJ41812.1 azon|OJJ48045.1

Group9458: abra|OJJ72531.1 acar|OOF94722.1 akaw|GAA86253.1 aluc|OJZ90510.1 anid|CBF85175.1 anig|EHA23967.1 atub|OJI81935.1 awen|OJJ37800.1

Group9459: abra|OJJ72570.1 afla|EED53148.1 anid|CBF83014.1 aory|EIT74956.1 asyd|OJJ61508.1 ater|EAU37811.1 aver|OJI99644.1 aver|OJJ00845.1

Group9460: abra|OJJ72578.1 acar|OOF94676.1 akaw|GAA85204.1 aluc|OJZ90559.1 anid|CBF81860.1 anig|EHA24005.1 atub|OJI81982.1 azon|OJJ45930.1

Group9461: awen|OJJ30623.1 asyd|OJJ62304.1 ater|EAU31749.1 afla|EED45569.1 anid|CBF89614.1 aory|EIT75642.1 aver|OJI96412.1 azon|OJJ48028.1

Group9462: abra|OJJ72745.1 acar|OOF90853.1 acla|EAW12599.1 akaw|GAA85355.1 aluc|OJZ90717.1 anig|EHA24136.1 atub|OJI82144.1 acar|OOF90713.1

Group9463: aory|EIT77718.1 acla|EAW13329.1 afis|EAW20444.1 afla|EED44826.1 afum|EBA27198.1 agla|OJJ84877.1 ater|EAU37715.1 awen|OJJ33134.1

Group9464: ater|EAU35516.1 acla|EAW11206.1 afis|EAW15936.1 afla|EED58119.1 afum|EAL88572.1 anid|CBF80964.1 aory|EIT79154.1 awen|OJJ35643.1

Group9465: agla|OJJ82327.1 anig|EHA25324.1 asyd|OJJ54335.1 atub|OJI80495.1 awen|OJJ37033.1 awen|OJJ41692.1 akaw|GAA86555.1 aluc|OJZ85103.1

Group9466: abra|OJJ73077.1 acar|OOG00312.1 afla|EED53424.1 akaw|GAA88085.1 aluc|OJZ81349.1 anig|EHA18651.1 aory|EIT75086.1 atub|OJI91553.1

Group9467: abra|OJJ73089.1 acar|OOG00323.1 acla|EAW11115.1 afla|EED45467.1 akaw|GAA88071.1 aluc|OJZ81336.1 anig|EHA18640.1 atub|OJI91539.1

Group9468: abra|OJJ73106.1 agla|OJJ87928.1 anid|CBF87412.1 asyd|OJJ54090.1 aver|OJJ08647.1 azon|OJJ46446.1 asyd|OJJ52897.1 aver|OJJ08868.1

Group9469: abra|OJJ73126.1 acar|OOG00349.1 afla|EED53573.1 akaw|GAA88038.1 aluc|OJZ81297.1 anig|EHA18605.1 atub|OJI91506.1 anig|EHA18606.1

Group9470: abra|OJJ73152.1 acar|OOG00364.1 akaw|GAA88020.1 aluc|OJZ81278.1 anid|CBF86006.1 anig|EHA18578.1 atub|OJI91486.1 azon|OJJ51713.1

Group9471: abra|OJJ73328.1 afla|EED48231.1 aluc|OJZ91200.1 anig|EHA19900.1 aory|EIT72905.1 asyd|OJJ65323.1 atub|OJI83217.1 aver|OJI99468.1

Group9472: abra|OJJ73399.1 acar|OOG00599.1 akaw|GAA83091.1 aluc|OJZ88638.1 anig|EHA18331.1 atub|OJI91224.1 aver|OJJ06007.1 asyd|OJJ56718.1

Group9473: abra|OJJ73420.1 acar|OOG00623.1 acla|EAW13605.1 afis|EAW20247.1 afum|EAL85365.2 akaw|GAA83071.1 awen|OJJ37579.1 azon|OJJ48024.1

Group9474: abra|OJJ73427.1 acar|OOG00633.1 akaw|GAA83065.1 aluc|OJZ88616.1 anig|EHA18307.1 ater|EAU36880.1 atub|OJI91201.1 azon|OJJ43923.1

Group9475: abra|OJJ73435.1 acar|OOG00642.1 akaw|GAA83058.1 aluc|OJZ88606.1 anig|EHA18299.1 atub|OJI91190.1 aver|OJI96135.1 awen|OJJ33325.1

Group9476: abra|OJJ73450.1 acar|OOG00653.1 afla|EED51807.1 akaw|GAA83046.1 aluc|OJZ88591.1 anig|EHA18287.1 aory|EIT81691.1 atub|OJI91177.1

Group9477: acar|OOF90597.1 afis|EAW24744.1 afum|EAL89390.1 anid|CBF84285.1 asyd|OJJ58314.1 aver|OJJ02097.1 azon|OJJ42093.1 ater|EAU34259.1

Group9478: abra|OJJ73995.1 acar|OOF94225.1 afla|EED51679.1 aluc|OJZ88049.1 anig|EHA17821.1 aory|EIT77469.1 atub|OJI90657.1 aver|OJJ03001.1

Group9479: abra|OJJ74008.1 acar|OOG01130.1 akaw|GAA87489.1 aluc|OJZ88036.1 anig|EHA17809.1 asyd|OJJ61383.1 atub|OJI90643.1 aver|OJJ00710.1

Group9480: abra|OJJ74010.1 acar|OOG01131.1 akaw|GAA87491.1 aluc|OJZ88034.1 anig|EHA17807.1 asyd|OJJ61382.1 atub|OJI90640.1 aver|OJJ00709.1

Group9481: abra|OJJ74011.1 acar|OOG01133.1 afla|EED47557.1 akaw|GAA87493.1 aluc|OJZ88030.1 anig|EHA17805.1 atub|OJI90637.1 aory|EIT74565.1

Group9482: abra|OJJ74137.1 acar|OOF95593.1 akaw|GAA89748.1 aluc|OJZ80636.1 anig|EHA27197.1 ater|EAU39604.1 atub|OJI87491.1 awen|OJJ33586.1

Group9483: abra|OJJ74138.1 acar|OOF95592.1 akaw|GAA89747.1 aluc|OJZ80637.1 anig|EHA27196.1 ater|EAU39605.1 atub|OJI87492.1 awen|OJJ33585.1

Group9484: abra|OJJ74179.1 acar|OOF95542.1 afla|EED48879.1 akaw|GAA86836.1 aluc|OJZ89955.1 aory|EIT81178.1 atub|OJI84851.1 aver|OJJ03488.1

Group9485: abra|OJJ74365.1 acar|OOF95365.1 afla|EED48363.1 akaw|GAA86665.1 aluc|OJZ89762.1 anig|EHA26982.1 aory|EIT83079.1 atub|OJI85040.1

Group9486: abra|OJJ74369.1 acar|OOF95363.1 agla|OJJ89236.1 akaw|GAA86662.1 aluc|OJZ89759.1 anig|EHA26979.1 awen|OJJ40218.1 azon|OJJ49843.1

Group9487: abra|OJJ74372.1 acla|EAW07076.1 akaw|GAA86656.1 aluc|OJZ89750.1 anig|EHA26973.1 ater|EAU36709.1 atub|OJI85052.1 aver|OJJ06459.1

Group9488: abra|OJJ74379.1 acar|OOF95358.1 akaw|GAA87279.1 aluc|OJZ89741.1 anig|EHA26965.1 atub|OJI85060.1 awen|OJJ33358.1 azon|OJJ50315.1

Group9489: abra|OJJ74393.1 acar|OOF95342.1 akaw|GAA87291.1 aluc|OJZ89727.1 atub|OJI85073.1 aver|OJJ00246.1 agla|OJJ82677.1 asyd|OJJ60920.1

Group9490: aory|EIT79191.1 afum|EAL87119.1 afla|EED47271.1 agla|OJJ82620.1 ater|EAU32195.1 acla|EAW11773.1 afis|EAW23676.1 awen|OJJ29773.1

Group9491: abra|OJJ74559.1 acar|OOF93083.1 agla|OJJ87487.1 akaw|GAA82038.1 anig|EHA26808.1 atub|OJI86573.1 awen|OJJ41475.1 azon|OJJ43918.1

Group9492: abra|OJJ74747.1 acar|OOF95741.1 afla|EED49965.1 akaw|GAA83664.1 aluc|OJZ92327.1 anig|EHA26643.1 aory|EIT83428.1 atub|OJI86757.1

Group9493: abra|OJJ74873.1 acar|OOF95874.1 akaw|GAA83547.1 aluc|OJZ92451.1 anig|EHA26517.1 atub|OJI86879.1 awen|OJJ39876.1 asyd|OJJ62281.1

Group9494: akaw|GAA83493.1 agla|OJJ86731.1 acla|EAW08441.1 afis|EAW23486.1 afla|EED57385.1 afum|EAL87977.1 anid|CBF89757.1 azon|OJJ47932.1

Group9495: abra|OJJ74961.1 acar|OOF97353.1 acla|EAW08411.1 agla|OJJ86691.1 anid|CBF89817.1 asyd|OJJ62189.1 aver|OJI96298.1 azon|OJJ47904.1

Group9496: abra|OJJ75127.1 acar|OOF97208.1 akaw|GAA88734.1 aluc|OJZ92692.1 anid|CBF89429.1 anig|EHA26308.1 atub|OJI87120.1 azon|OJJ47751.1

Group9497: aver|OJJ03174.1 azon|OJJ46527.1 acar|OOG00290.1 acla|EAW12455.1 afla|EED51632.1 anid|CBF84178.1 aory|EIT80501.1 asyd|OJJ57130.1

Group9498: abra|OJJ75234.1 acar|OOF94558.1 akaw|GAA91777.1 aluc|OJZ92795.1 anig|EHA26213.1 atub|OJI87238.1 awen|OJJ39989.1 azon|OJJ45243.1

Group9499: abra|OJJ75249.1 acar|OOF91827.1 afla|EED51022.1 asyd|OJJ60674.1 atub|OJI87262.1 aver|OJI99997.1 awen|OJJ36156.1 acar|OOG00965.1

Group9500: abra|OJJ75260.1 acar|OOF94542.1 akaw|GAA91798.1 aluc|OJZ92818.1 anig|EHA26190.1 atub|OJI87276.1 azon|OJJ43908.1 awen|OJJ38761.1

Group9501: abra|OJJ75295.1 aluc|OJZ90845.1 anig|EHA25378.1 ater|EAU36044.1 atub|OJI82832.1 aver|OJJ06528.1 awen|OJJ40184.1 aver|OJI95774.1

Group9502: abra|OJJ75325.1 acar|OOF94645.1 acla|EAW12069.1 akaw|GAA92020.1 aluc|OJZ92888.1 anig|EHA26075.1 atub|OJI87346.1 awen|OJJ30536.1

Group9503: abra|OJJ75328.1 acar|OOF90142.1 afla|EED48381.1 akaw|GAA92023.1 aluc|OJZ92891.1 anig|EHA26072.1 aory|EIT83088.1 atub|OJI87349.1

Group9504: ater|EAU37650.1 acar|OOG00117.1 acla|EAW07061.1 afis|EAW16684.1 afla|EED48490.1 aory|EIT73090.1 azon|OJJ42235.1 azon|OJJ45218.1

Group9505: abra|OJJ75352.1 afis|EAW25185.1 afla|EED53819.1 akaw|GAA92045.1 aluc|OJZ92914.1 anig|EHA26051.1 aory|EIT78407.1 atub|OJI87374.1

Group9506: abra|OJJ75365.1 afis|EAW17962.1 afum|EAL91772.1 aluc|OJZ92935.1 anig|EHA26037.1 atub|OJI87395.1 atub|OJI80907.1 aver|OJJ01291.1

Group9507: abra|OJJ75535.1 afla|EED46910.1 akaw|GAA91955.1 aluc|OJZ82663.1 anig|EHA27437.1 aory|EIT72749.1 atub|OJI88896.1 azon|OJJ49040.1

Group9508: abra|OJJ75541.1 acar|OOF90860.1 akaw|GAA84295.1 aluc|OJZ88989.1 anig|EHA23537.1 atub|OJI85798.1 awen|OJJ35898.1 azon|OJJ43915.1

Group9509: abra|OJJ75567.1 acar|OOF90885.1 akaw|GAA84319.1 aluc|OJZ89015.1 anig|EHA23515.1 asyd|OJJ52898.1 ater|EAU31415.1 atub|OJI85774.1

Group9510: abra|OJJ75568.1 acar|OOF90886.1 afla|EED54163.1 agla|OJJ85899.1 aluc|OJZ89016.1 anig|EHA23514.1 aory|EIT75933.1 atub|OJI85773.1

Group9511: abra|OJJ75578.1 acar|OOF90895.1 acla|EAW13757.1 akaw|GAA84329.1 aluc|OJZ89028.1 anig|EHA23505.1 atub|OJI85763.1 azon|OJJ46062.1

Group9512: abra|OJJ75584.1 afla|EED55988.1 akaw|GAA84336.1 aluc|OJZ89035.1 anig|EHA23499.1 aory|EIT76706.1 atub|OJI85755.1 acla|EAW06995.1

Group9513: acla|EAW09370.1 afum|EAL92795.1 afis|EAW21765.1 afla|EED46334.1 agla|OJJ84386.1 aory|EIT81257.1 awen|OJJ34048.1 azon|OJJ48183.1

Group9514: abra|OJJ75704.1 afum|EAL89397.1 agla|OJJ83953.1 aluc|OJZ89155.1 ater|EAU31769.1 atub|OJI87801.1 awen|OJJ36182.1 azon|OJJ42007.1

Group9515: aory|EIT73848.1 ater|EAU30293.1 afis|EAW19505.1 afla|EED56796.1 anid|CBF83979.1 asyd|OJJ61406.1 aver|OJJ00736.1 afum|EAL86988.1

Group9516: abra|OJJ75883.1 afla|EED47845.1 akaw|GAA84586.1 aluc|OJZ89330.1 anig|EHA23245.1 aory|EIT77232.1 atub|OJI85470.1 awen|OJJ41919.1

Group9517: acla|EAW10263.1 afis|EAW25330.1 afum|EAL89992.1 afla|EED57803.1 aory|EIT81120.1 ater|EAU30152.1 aver|OJI98324.1 asyd|OJJ55359.1

Group9518: abra|OJJ76110.1 acar|OOF99816.1 acla|EAW09710.1 afum|EBA27461.1 aluc|OJZ89566.1 atub|OJI85238.1 azon|OJJ51585.1 asyd|OJJ60867.1

Group9519: abra|OJJ76149.1 acar|OOF99778.1 agla|OJJ85613.1 aluc|OJZ92080.1 atub|OJI86510.1 aver|OJJ00466.1 awen|OJJ39234.1 azon|OJJ51621.1

Group9520: acar|OOF99551.1 afla|EED53231.1 aory|EIT75189.1 afum|EAL93707.1 ater|EAU38165.1 awen|OJJ31575.1 acla|EAW14314.1 asyd|OJJ59085.1

Group9521: agla|OJJ79956.1 acla|EAW12319.1 afis|EAW16902.1 afla|EED45265.1 afum|EAL86437.1 aory|EIT73652.1 awen|OJJ29717.1 awen|OJJ29924.1

Group9522: abra|OJJ76357.1 acar|OOF99608.1 akaw|GAA82283.1 aluc|OJZ91856.1 atub|OJI86307.1 aver|OJJ01513.1 azon|OJJ47100.1 asyd|OJJ58916.1

Group9523: abra|OJJ76415.1 acar|OOF99655.1 agla|OJJ82341.1 aluc|OJZ91798.1 awen|OJJ32221.1 azon|OJJ46773.1 ater|EAU38187.1 aver|OJJ01075.1

Group9524: abra|OJJ76475.1 afis|EAW23592.1 afum|EAL87874.1 akaw|GAA82387.1 aluc|OJZ91733.1 anig|EHA22740.1 atub|OJI86192.1 awen|OJJ34755.1

Group9525: abra|OJJ76499.1 afla|EED56465.1 akaw|GAA86205.1 aory|EIT72845.1 asyd|OJJ60352.1 ater|EAU36868.1 afis|EAW23443.1 anid|CBF80766.1

Group9526: abra|OJJ76600.1 agla|OJJ79790.1 akaw|GAA82503.1 aluc|OJZ91608.1 anig|EHA22637.1 atub|OJI86060.1 awen|OJJ30242.1 azon|OJJ46833.1

Group9527: asyd|OJJ64031.1 abra|OJJ76646.1 akaw|GAA92814.1 aluc|OJZ91565.1 anig|EHA22594.1 atub|OJI86015.1 aver|OJI98116.1 asyd|OJJ56046.1

Group9528: abra|OJJ76675.1 acar|OOF99334.1 afis|EAW25184.1 afum|EAL89859.1 akaw|GAA92517.1 aluc|OJZ91530.1 anig|EHA22569.1 atub|OJI85984.1

Group9529: acla|EAW07062.1 azon|OJJ45052.1 afis|EAW16693.1 afla|EED45071.1 afum|EAL89291.1 aory|EIT73802.1 ater|EAU34152.1 aver|OJJ01782.1

Group9530: abra|OJJ76702.1 acar|OOF99316.1 agla|OJJ84121.1 akaw|GAA92497.1 aluc|OJZ91509.1 anid|CBF78146.1 anig|EHA22548.1 atub|OJI85961.1

Group9531: abra|OJJ76720.1 acar|OOF97651.1 afla|EED47263.1 akaw|GAA92480.1 aluc|OJZ91490.1 anig|EHA22534.1 aory|EIT79251.1 atub|OJI85947.1

Group9532: abra|OJJ76727.1 afla|EED56123.1 agla|OJJ86581.1 anig|EHA28102.1 aory|EIT76858.1 asyd|OJJ55398.1 ater|EAU30982.1 awen|OJJ30160.1

Group9533: abra|OJJ76734.1 afis|EAW21009.1 agla|OJJ82882.1 aluc|OJZ80162.1 asyd|OJJ58391.1 aver|OJJ07900.1 awen|OJJ32613.1 afis|EAW17034.1

Group9534: abra|OJJ76912.1 akaw|GAA87126.1 aluc|OJZ84312.1 anig|EHA27647.1 atub|OJI89756.1 aver|OJJ05995.1 awen|OJJ37021.1 agla|OJJ83564.1

Group9535: abra|OJJ76913.1 acar|OOF98741.1 akaw|GAA87125.1 aluc|OJZ84313.1 anig|EHA27648.1 atub|OJI89755.1 aver|OJJ05994.1 awen|OJJ37022.1

Group9536: abra|OJJ76988.1 acar|OOF91432.1 afla|EED46111.1 akaw|GAA87058.1 aluc|OJZ84388.1 anig|EHA27718.1 aory|EIT76518.1 atub|OJI89684.1

Group9537: abra|OJJ77005.1 acar|OOF91450.1 akaw|GAA87043.1 aluc|OJZ84404.1 anig|EHA27734.1 ater|EAU32185.1 atub|OJI89668.1 azon|OJJ50405.1

Group9538: afla|EED54149.1 aory|EIT76116.1 asyd|OJJ53602.1 aver|OJJ08146.1 akaw|GAA91860.1 aluc|OJZ82039.1 anig|EHA22218.1 atub|OJI84597.1

Group9539: awen|OJJ29670.1 acar|OOF98731.1 afla|EED48910.1 akaw|GAA87771.1 aluc|OJZ85406.1 anig|EHA25585.1 aver|OJJ08853.1 aver|OJI98890.1

Group9540: abra|OJJ77235.1 acar|OOF93117.1 akaw|GAA90708.1 aluc|OJZ84645.1 asyd|OJJ61794.1 ater|EAU33902.1 atub|OJI89435.1 aver|OJI95894.1

Group9541: abra|OJJ77420.1 acar|OOF90641.1 acla|EAW14626.1 akaw|GAA90295.1 aluc|OJZ83042.1 anig|EHA22538.1 atub|OJI87908.1 azon|OJJ45271.1

Group9542: abra|OJJ77422.1 afis|EAW24698.1 afla|EED51169.1 afum|EAL89340.1 anid|CBF90109.1 aory|EIT81319.1 ater|EAU31625.1 awen|OJJ40787.1

Group9543: abra|OJJ77431.1 afis|EAW24696.1 afum|EAL89338.1 anid|CBF90099.1 asyd|OJJ62030.1 ater|EAU31623.1 aver|OJI96121.1 awen|OJJ40792.1

Group9544: abra|OJJ77434.1 afis|EAW24699.1 afum|EAL89341.2 asyd|OJJ62029.1 aver|OJI96120.1 awen|OJJ40786.1 anid|CBF90101.1 ater|EAU31626.1

Group9545: abra|OJJ77435.1 acar|OOF97515.1 acla|EAW08777.1 afla|EED48086.1 afum|EAL86654.1 aory|EIT81472.1 azon|OJJ46405.1 afla|EED52055.1

Group9546: abra|OJJ77496.1 acar|OOF99231.1 agla|OJJ79563.1 aory|EIT78991.1 awen|OJJ36058.1 afla|EED50860.1 azon|OJJ45260.1 agla|OJJ79153.1

Group9547: aory|EIT73905.1 azon|OJJ46048.1 afla|EED56791.1 anid|CBF71422.1 asyd|OJJ54992.1 aver|OJJ07209.1 awen|OJJ31993.1 ater|EAU32727.1

Group9548: aory|EIT73792.1 agla|OJJ83250.1 awen|OJJ41102.1 anig|EHA20899.1 ater|EAU39541.1 atub|OJI80593.1 agla|OJJ79434.1 aluc|OJZ85005.1

Group9549: abra|OJJ77731.1 acar|OOF98995.1 afla|EED47651.1 akaw|GAA83904.1 aluc|OJZ90791.1 anig|EHA19525.1 aory|EIT80176.1 atub|OJI82780.1

Group9550: abra|OJJ77736.1 acar|OOF98985.1 afla|EED54278.1 akaw|GAA83893.1 aluc|OJZ90802.1 anig|EHA19531.1 atub|OJI82791.1 awen|OJJ36026.1

Group9551: abra|OJJ77749.1 acar|OOF90666.1 akaw|GAA83885.1 aluc|OJZ90810.1 anig|EHA19538.1 atub|OJI82799.1 azon|OJJ42324.1 akaw|GAA88768.1

Group9552: abra|OJJ77763.1 acar|OOF97506.1 acla|EAW12047.1 afis|EAW17016.1 akaw|GAA89815.1 aluc|OJZ81875.1 anig|EHA20909.1 acar|OOF89896.1

Group9553: abra|OJJ77794.1 acar|OOF90716.1 asyd|OJJ54626.1 aver|OJJ07530.1 awen|OJJ38009.1 abra|OJJ77797.1 agla|OJJ83011.1 anid|CBF80615.1

Group9554: abra|OJJ77812.1 acar|OOF93334.1 akaw|GAA83837.1 anid|CBF80740.1 anig|EHA19602.1 ater|EAU35344.1 atub|OJI82877.1 ater|EAU31073.1

Group9555: abra|OJJ77879.1 acar|OOF96529.1 afis|EAW22771.1 afum|EAL89321.1 akaw|GAA83774.1 aluc|OJZ90954.1 anig|EHA19659.1 atub|OJI82944.1

Group9556: aver|OJJ05263.1 awen|OJJ30807.1 azon|OJJ48046.1 afis|EAW24240.1 afum|EAL85908.1 atub|OJI87442.1 anig|EHA24976.1 anid|CBF89325.1

Group9557: abra|OJJ77977.1 acar|OOF93312.1 agla|OJJ88228.1 akaw|GAA89019.1 aluc|OJZ80585.1 anid|CBF87755.1 asyd|OJJ57292.1 aver|OJJ05864.1

Group9558: abra|OJJ78028.1 acar|OOF97287.1 akaw|GAA85958.1 aluc|OJZ91103.1 anig|EHA19803.1 ater|EAU31770.1 atub|OJI83100.1 azon|OJJ42009.1

Group9559: abra|OJJ78034.1 acar|OOF98463.1 agla|OJJ81557.1 akaw|GAA85948.1 aluc|OJZ91114.1 anig|EHA19813.1 atub|OJI83118.1 awen|OJJ35948.1

Group9560: abra|OJJ78101.1 acar|OOF98773.1 acla|EAW07103.1 akaw|GAA85902.1 aluc|OJZ91165.1 anig|EHA18602.1 atub|OJI83181.1 anid|CBF80283.1

Group9561: afis|EAW16147.1 azon|OJJ43970.1 acla|EAW06618.1 afla|EED48689.1 anid|CBF79581.1 aory|EIT73166.1 asyd|OJJ57972.1 aver|OJJ04429.1

Group9562: abra|OJJ78120.1 acar|OOF98723.1 akaw|GAA85888.1 aluc|OJZ91183.1 anig|EHA19876.1 asyd|OJJ54308.1 atub|OJI83201.1 aver|OJJ05254.1

Group9563: abra|OJJ78122.1 afla|EED49665.1 akaw|GAA85886.1 anig|EHA19878.1 aory|EIT74889.1 ater|EAU31568.1 atub|OJI83203.1 aory|EIT80979.1

Group9564: abra|OJJ78149.1 afis|EAW17181.1 aluc|OJZ91201.1 anig|EHA19906.1 asyd|OJJ61780.1 atub|OJI83218.1 aver|OJI95880.1 afum|EAL86610.1

Group9565: abra|OJJ78166.1 acla|EAW10842.1 afis|EAW24723.1 afla|EED45492.1 afum|EAL89372.1 agla|OJJ89012.1 anid|CBF78356.1 aory|EIT75360.1

Group9566: abra|OJJ78177.1 afis|EAW22124.1 akaw|GAA85856.1 aluc|OJZ91226.1 anid|CBF76068.1 anig|EHA19932.1 asyd|OJJ65399.1 atub|OJI83239.1

Group9567: abra|OJJ78195.1 acar|OOF98660.1 afla|EED52679.1 akaw|GAA85834.1 aluc|OJZ91266.1 anig|EHA19954.1 aory|EIT74230.1 atub|OJI83255.1

Group9568: abra|OJJ78196.1 acar|OOF98659.1 akaw|GAA85833.1 aluc|OJZ91267.1 anig|EHA19955.1 atub|OJI83256.1 awen|OJJ40833.1 azon|OJJ42579.1

Group9569: acar|OOF90271.1 acla|EAW13552.1 agla|OJJ80443.1 anid|CBF87441.1 aver|OJJ02466.1 asyd|OJJ59821.1 aory|EIT77635.1 awen|OJJ33464.1

Group9570: aver|OJI98259.1 acla|EAW10787.1 asyd|OJJ64181.1 acla|EAW12421.1 afla|EED52222.1 anid|CBF83046.1 aory|EIT74356.1 anid|CBF85573.1

Group9571: acar|OOF91813.1 agla|OJJ79114.1 asyd|OJJ55384.1 ater|EAU31022.1 aver|OJJ08273.1 awen|OJJ36173.1 asyd|OJJ54435.1 asyd|OJJ60332.1

Group9572: acar|OOF91835.1 afla|EED54608.1 akaw|GAA86591.1 aluc|OJZ85138.1 anig|EHA25354.1 aory|EIT78931.1 atub|OJI80454.1 awen|OJJ32894.1

Group9573: afum|EAL86182.1 agla|OJJ87465.1 azon|OJJ42591.1 afla|EED58055.1 aory|EIT79764.1 ater|EAU35453.1 awen|OJJ31869.1 awen|OJJ38824.1

Group9574: acar|OOF92005.1 afla|EED46125.1 agla|OJJ80835.1 aory|EIT76419.1 ater|EAU36061.1 aver|OJJ05887.1 awen|OJJ34649.1 asyd|OJJ59683.1

Group9575: acar|OOF92559.1 afla|EED47335.1 anid|CBF81558.1 aory|EIT82837.1 asyd|OJJ59499.1 ater|EAU38092.1 aver|OJJ02775.1 awen|OJJ32598.1

Group9576: acar|OOF93022.1 agla|OJJ84215.1 akaw|GAA89204.1 aluc|OJZ81626.1 anid|CBF84613.1 anig|EHA25817.1 atub|OJI88554.1 awen|OJJ29705.1

Group9577: acar|OOF93139.1 afla|EED45155.1 anid|CBF84225.1 aory|EIT73638.1 asyd|OJJ57182.1 ater|EAU33860.1 aver|OJJ03108.1 ater|EAU37802.1

Group9578: acar|OOF93252.1 akaw|GAA87876.1 aluc|OJZ85517.1 asyd|OJJ57352.1 atub|OJI80049.1 aver|OJJ05813.1 awen|OJJ35826.1 anig|EHA25050.1

Group9579: acar|OOF93301.1 afla|EED57510.1 agla|OJJ82926.1 aory|EIT81837.1 asyd|OJJ64583.1 aver|OJI98681.1 awen|OJJ36288.1 awen|OJJ29790.1

Group9580: acar|OOF93864.1 afla|EED47783.1 akaw|GAA86247.1 anid|CBF82004.1 anig|EHA23960.1 aory|EIT77175.1 atub|OJI81926.1 awen|OJJ36959.1

Group9581: acar|OOF94001.1 agla|OJJ79469.1 asyd|OJJ53706.1 aver|OJJ07830.1 awen|OJJ33637.1 afla|EED47078.1 aory|EIT72480.1 awen|OJJ36283.1

Group9582: acar|OOF94002.1 afla|EED45233.1 anid|CBF73947.1 aory|EIT73699.1 asyd|OJJ53707.1 aver|OJJ07831.1 awen|OJJ32558.1 afla|EED45232.1

Group9583: acar|OOF94505.1 afla|EED52367.1 akaw|GAA92890.1 aluc|OJZ80025.1 awen|OJJ36241.1 acla|EAW07590.1 azon|OJJ45227.1 akaw|GAA93085.1

Group9584: acar|OOF95224.1 acla|EAW07168.1 afis|EAW22183.1 afum|EAL91006.2 anid|CBF88389.1 asyd|OJJ63741.1 aver|OJI97199.1 azon|OJJ50295.1

Group9585: acar|OOF95323.1 acla|EAW06833.1 afla|EED53446.1 agla|OJJ88207.1 akaw|GAA87311.1 aluc|OJZ89706.1 aory|EIT74837.1 awen|OJJ37919.1

Group9586: acar|OOF95522.1 acla|EAW07251.1 afis|EAW22272.1 agla|OJJ87783.1 anid|CBF88532.1 asyd|OJJ63814.1 aver|OJI97124.1 awen|OJJ34845.1

Group9587: acar|OOF95545.1 aory|EIT82462.1 asyd|OJJ59655.1 atub|OJI85048.1 aver|OJJ02627.1 azon|OJJ46452.1 akaw|GAA86658.1 aluc|OJZ89754.1

Group9588: aory|EIT74997.1 ater|EAU29363.1 afla|EED48304.1 anig|EHA25873.1 aory|EIT76308.1 awen|OJJ35912.1 afla|EED53462.1 agla|OJJ80122.1

Group9589: acar|OOF96061.1 afla|EED45985.1 anid|CBF85237.1 aory|EIT75476.1 asyd|OJJ62982.1 ater|EAU33614.1 aver|OJI97987.1 azon|OJJ47328.1

Group9590: afis|EAW15418.1 afum|EAL84226.1 afla|EED50614.1 agla|OJJ80128.1 aory|EIT78054.1 asyd|OJJ64717.1 ater|EAU36021.1 aver|OJI98820.1

Group9591: agla|OJJ85407.1 acla|EAW12652.1 afis|EAW18032.1 afum|EAL91843.1 aory|EIT81764.1 asyd|OJJ58289.1 ater|EAU31549.1 aver|OJJ02135.1

Group9592: acar|OOF97516.1 acla|EAW12033.1 afis|EAW20284.1 afum|EBA27230.1 anid|CBF85595.1 asyd|OJJ64164.1 aver|OJI98242.1 azon|OJJ45219.1

Group9593: acar|OOF97537.1 afis|EAW20329.1 afla|EED52857.1 afum|EAL85456.1 anid|CBF86760.1 asyd|OJJ65101.1 aver|OJI99227.1 acla|EAW15055.1

Group9594: acar|OOF98716.1 afis|EAW24716.1 afla|EED46063.1 afum|EAL89364.1 anid|CBF87718.1 aory|EIT83486.1 asyd|OJJ60329.1 awen|OJJ32554.1

Group9595: acar|OOF98771.1 afis|EAW16869.1 afum|EAL86408.2 anid|CBF69720.1 asyd|OJJ58926.1 aver|OJJ01522.1 awen|OJJ36306.1 azon|OJJ45905.1

Group9596: acar|OOF99259.1 akaw|GAA91878.1 aluc|OJZ82021.1 anig|EHA22235.1 atub|OJI84573.1 acla|EAW11675.1 anid|CBF73436.1 aver|OJJ08111.1

Group9597: acar|OOG00292.1 acla|EAW12452.1 afla|EED51636.1 anid|CBF84174.1 aory|EIT80653.1 asyd|OJJ57128.1 aver|OJJ03176.1 azon|OJJ46529.1

Group9598: acar|OOG00544.1 acla|EAW11651.1 afis|EAW23802.1 afum|EAL87236.1 asyd|OJJ65160.1 ater|EAU31356.1 awen|OJJ35849.1 azon|OJJ42812.1

Group9599: ater|EAU39162.1 acla|EAW09026.1 afis|EAW22741.1 afla|EED49795.1 afum|EAL90454.1 agla|OJJ78547.1 aory|EIT77688.1 aver|OJI95793.1

Group9600: acar|OOG01212.1 afis|EAW22984.1 agla|OJJ83303.1 anid|CBF87585.1 anig|EHA17722.1 asyd|OJJ63256.1 aver|OJI97701.1 azon|OJJ49521.1

Group9601: acla|EAW06581.1 afis|EAW17092.1 afla|EED53428.1 afum|EAL86569.1 aory|EIT74920.1 ater|EAU31405.1 azon|OJJ50131.1 ater|EAU33863.1

Group9602: acla|EAW06797.1 afis|EAW16346.1 afum|EAL88971.1 anid|CBF89214.1 aory|EIT74969.1 ater|EAU34112.1 afla|EED53524.1 aver|OJI96643.1

Group9603: agla|OJJ83630.1 anid|CBF86700.1 ater|EAU33510.1 awen|OJJ34516.1 afla|EED54639.1 asyd|OJJ64742.1 aver|OJI98846.1 awen|OJJ40119.1

Group9604: acla|EAW07115.1 afis|EAW22127.1 afum|EAL91061.1 anid|CBF83959.1 asyd|OJJ61428.1 ater|EAU37214.1 aver|OJJ00764.1 azon|OJJ47383.1

Group9605: acla|EAW07188.1 afis|EAW22201.1 afla|EED57594.1 afum|EAL90991.1 agla|OJJ88513.1 aory|EIT82942.1 ater|EAU34213.1 awen|OJJ34788.1

Group9606: acla|EAW08018.1 afis|EAW23052.1 afum|EAL88428.2 agla|OJJ89424.1 aory|EIT80290.1 awen|OJJ35683.1 azon|OJJ44152.1 afla|EED54966.1

Group9607: acla|EAW08302.1 afis|EAW23338.1 afum|EAL88140.1 asyd|OJJ62446.1 afla|EED55628.1 awen|OJJ39556.1 agla|OJJ86492.1 aver|OJI96583.1

Group9608: acla|EAW08990.1 afla|EED57080.1 anid|CBF70705.1 asyd|OJJ58456.1 ater|EAU37969.1 aver|OJJ01056.1 agla|OJJ79340.1 aory|EIT82263.1

Group9609: acla|EAW09028.1 afis|EAW17359.1 afis|EAW22739.1 afum|EAL90456.1 agla|OJJ78545.1 ater|EAU39161.1 aver|OJI95794.1 afum|EAL91196.1

Group9610: acla|EAW09098.1 agla|OJJ78890.1 anid|CBF84261.1 ater|EAU31572.1 aver|OJJ03077.1 afis|EAW25694.1 afum|EAL84368.2 asyd|OJJ57211.1

Group9611: acla|EAW09123.1 afis|EAW19674.1 afla|EED45947.1 afum|EAL86847.2 agla|OJJ80970.1 aory|EIT75658.1 awen|OJJ32667.1 ater|EAU30256.1

Group9612: acla|EAW09566.1 afis|EAW21570.1 afum|EBA27468.1 agla|OJJ85772.1 anid|CBF76320.1 aory|EIT76631.1 asyd|OJJ61012.1 aver|OJJ00331.1

Group9613: acla|EAW09977.1 afis|EAW21112.1 afla|EED45988.1 afum|EAL92135.1 anid|CBF87179.1 aory|EIT75427.1 asyd|OJJ54077.1 aver|OJJ08661.1

Group9614: acla|EAW10253.1 afis|EAW25339.1 afla|EED53820.1 afum|EAL90002.1 agla|OJJ83881.1 aory|EIT78466.1 ater|EAU34485.1 awen|OJJ41058.1

Group9615: acla|EAW10396.1 afis|EAW25181.1 afla|EED53639.1 afum|EAL89855.1 anid|CBF69472.1 asyd|OJJ59266.1 ater|EAU34338.1 aver|OJJ01878.1

Group9616: acla|EAW10798.1 afis|EAW24779.1 agla|OJJ88221.1 anid|CBF83178.1 asyd|OJJ55540.1 aver|OJJ06618.1 awen|OJJ37941.1 afla|EED49722.1

Group9617: acla|EAW10843.1 afum|EAL89369.1 afis|EAW24720.1 afla|EED53233.1 agla|OJJ87995.1 asyd|OJJ64486.1 aver|OJI98587.1 azon|OJJ43679.1

Group9618: acla|EAW11085.1 afis|EAW20773.1 afum|EAL84846.1 anid|CBF90337.1 aory|EIT76641.1 asyd|OJJ59877.1 ater|EAU35825.1 aver|OJJ02409.1

Group9619: acla|EAW11222.1 afis|EAW18281.1 afla|EED48748.1 afum|EAL87647.1 akaw|GAA84255.1 anid|CBF79489.1 aory|EIT73200.1 ater|EAU33287.1

Group9620: acla|EAW11621.1 afis|EAW23837.1 afla|EED52573.1 afum|EAL87268.1 aory|EIT74212.1 asyd|OJJ56994.1 aver|OJJ03293.1 awen|OJJ33799.1

Group9621: acla|EAW11768.1 afis|EAW23681.1 afla|EED47434.1 afum|EAL87124.1 aory|EIT81922.1 asyd|OJJ55513.1 ater|EAU29290.1 aver|OJJ06590.1

Group9622: acla|EAW11868.1 afis|EAW19479.1 afla|EED56687.1 afum|EAL87017.1 aory|EIT73845.1 ater|EAU34791.1 awen|OJJ36884.1 azon|OJJ42922.1

Group9623: acla|EAW11974.1 afis|EAW19612.1 afla|EED56960.1 afum|EAL86910.1 aory|EIT77528.1 ater|EAU32913.1 awen|OJJ41858.1 azon|OJJ47052.1

Group9624: acla|EAW12130.1 afis|EAW17255.1 afla|EED48389.1 afum|EAL86673.2 akaw|GAA86865.1 anid|CBF82354.1 aory|EIT83091.1 atub|OJI84821.1

Group9625: acla|EAW12132.1 afis|EAW17219.1 afla|EED51385.1 afum|EAL86644.1 aluc|OJZ88710.1 aory|EIT80584.1 atub|OJI91297.1 azon|OJJ50154.1

Group9626: acla|EAW12262.1 afis|EAW16968.1 afum|EAL86496.1 anid|CBF83937.1 asyd|OJJ55433.1 aver|OJJ06510.1 awen|OJJ36436.1 azon|OJJ45263.1

Group9627: acla|EAW12340.1 afis|EAW15382.1 afla|EED56841.1 anid|CBF80189.1 aory|EIT77576.1 asyd|OJJ52070.1 ater|EAU31367.1 aver|OJJ03034.1

Group9628: acla|EAW12374.1 afis|EAW16820.1 afla|EED45874.1 afum|EBA27271.1 anid|CBF69524.1 asyd|OJJ59204.1 aver|OJJ01816.1 azon|OJJ50870.1

Group9629: acla|EAW12381.1 afis|EAW22413.1 afla|EED46912.1 afum|EAL92429.1 aory|EIT72559.1 awen|OJJ36112.1 azon|OJJ45692.1 afum|EAL90779.1

Group9630: acla|EAW12393.1 afis|EAW16841.1 afla|EED51875.1 afum|EAL86388.1 aory|EIT81706.1 asyd|OJJ63067.1 aver|OJI97895.1 azon|OJJ47714.1

Group9631: acla|EAW12559.1 afis|EAW24091.1 afla|EED46848.1 afum|EAL91946.2 agla|OJJ87165.1 akaw|GAA89841.1 ater|EAU30515.1 awen|OJJ41398.1

Group9632: acla|EAW12572.1 afis|EAW24078.1 afla|EED51395.1 afum|EAL91933.1 agla|OJJ87149.1 aory|EIT80743.1 ater|EAU30499.1 awen|OJJ41416.1

Group9633: acla|EAW12755.1 afis|EAW17918.1 afla|EED46321.1 afum|EAL91729.1 akaw|GAA85182.1 anid|CBF76957.1 aory|EIT81193.1 ater|EAU33141.1

Group9634: acla|EAW12864.1 afis|EAW17813.1 afum|EAL91622.1 anid|CBF80722.1 asyd|OJJ54568.1 ater|EAU30288.1 aver|OJJ07586.1 azon|OJJ44403.1

Group9635: acla|EAW12874.1 afis|EAW17802.1 afla|EED45796.1 afum|EAL91610.1 aory|EIT75648.1 ater|EAU31831.1 asyd|OJJ61386.1 aver|OJJ00714.1

Group9636: acla|EAW12994.1 afis|EAW17672.1 afla|EED50481.1 afum|EBA27436.1 agla|OJJ79544.1 aory|EIT82137.1 awen|OJJ37170.1 azon|OJJ50865.1

Group9637: acla|EAW13332.1 afis|EAW20447.1 afla|EED44823.1 afum|EAL84605.1 aory|EIT77748.1 ater|EAU37712.1 aver|OJJ06863.1 awen|OJJ33137.1

Group9638: acla|EAW13523.1 afla|EED57079.1 anid|CBF82203.1 asyd|OJJ52181.1 ater|EAU29291.1 aver|OJJ05964.1 awen|OJJ34429.1 aory|EIT82290.1

Group9639: acla|EAW13619.1 afis|EAW19979.1 afum|EBA27212.1 asyd|OJJ61567.1 ater|EAU37664.1 aver|OJJ00887.1 awen|OJJ36068.1 azon|OJJ50241.1

Group9640: awen|OJJ33831.1 afis|EAW16762.1 afla|EED48119.1 afum|EAL86333.1 ater|EAU29806.1 asyd|OJJ53532.1 aver|OJJ08218.1 azon|OJJ42616.1

Group9641: acla|EAW14271.1 afis|EAW22464.1 afla|EED49640.1 afum|EAL90727.1 agla|OJJ89360.1 anid|CBF82495.1 aory|EIT80935.1 awen|OJJ37069.1

Group9642: acla|EAW14382.1 afis|EAW19088.1 afla|EED54320.1 afum|EAL93818.1 agla|OJJ84516.1 akaw|GAA91582.1 anid|CBF79688.1 azon|OJJ46229.1

Group9643: acla|EAW14667.1 afis|EAW24198.1 afum|EAL85870.1 agla|OJJ79679.1 anid|CBF74321.1 asyd|OJJ53998.1 aver|OJJ08095.1 awen|OJJ33101.1

Group9644: acla|EAW14956.1 afis|EAW23763.1 afla|EED53475.1 afum|EAL87196.1 aory|EIT74970.1 asyd|OJJ57952.1 ater|EAU36896.1 aver|OJJ04445.1

Group9645: acla|EAW14958.1 afis|EAW24475.1 afla|EED52400.1 afum|EAL86147.1 agla|OJJ82598.1 aory|EIT83306.1 awen|OJJ30938.1 azon|OJJ50151.1

Group9646: acla|EAW14989.1 afis|EAW24510.1 afla|EED45970.1 afum|EAL86179.1 aluc|OJZ83019.1 anid|CBF73497.1 aory|EIT75818.1 ater|EAU30633.1

Group9647: acla|EAW15193.1 afis|EAW20162.1 afum|EAL85280.1 agla|OJJ88768.1 anid|CBF83902.1 asyd|OJJ55414.1 awen|OJJ32057.1 azon|OJJ46091.1

Group9648: acla|EAW15304.1 afis|EAW25739.1 agla|OJJ88185.1 anid|CBF69360.1 asyd|OJJ57312.1 aver|OJJ05850.1 awen|OJJ35540.1 atub|OJI88255.1

Group9649: acla|EAW15339.1 afis|EAW20370.1 afla|EED45212.1 afum|EAL85484.1 anid|CBF75990.1 aory|EIT73752.1 ater|EAU37784.1 awen|OJJ38806.1

Group9650: afis|EAW15613.1 afum|EAL85722.1 akaw|GAA83735.1 asyd|OJJ54690.1 ater|EAU30431.1 aver|OJJ07479.1 azon|OJJ42183.1 anig|EHA19704.1

Group9651: afis|EAW16671.1 afla|EED47560.1 afum|EAL89276.1 anid|CBF82085.1 aory|EIT74537.1 asyd|OJJ54259.1 ater|EAU34536.1 aver|OJJ05212.1

Group9652: afis|EAW17298.1 afla|EED49518.1 anid|CBF81687.1 asyd|OJJ59561.1 ater|EAU29436.1 aver|OJJ02707.1 aory|EIT82602.1 asyd|OJJ60042.1

Group9653: afis|EAW18030.1 afla|EED50491.1 afum|EAL91841.1 aluc|OJZ87315.1 aory|EIT82092.1 atub|OJI81330.1 aver|OJI97482.1 ater|EAU36807.1

Group9654: afis|EAW18104.1 afum|EAL89392.1 afis|EAW20338.1 afum|EAL85459.1 anid|CBF86695.1 asyd|OJJ64155.1 ater|EAU33491.1 aver|OJJ04471.1

Group9655: afis|EAW19776.1 afla|EED49440.1 afum|EAL85121.1 anid|CBF76189.1 aory|EIT79582.1 asyd|OJJ57342.1 ater|EAU39072.1 aver|OJJ05820.1

Group9656: afis|EAW19866.1 afla|EED48493.1 afum|EAL85030.1 aory|EIT73271.1 ater|EAU38716.1 awen|OJJ41888.1 asyd|OJJ62671.1 awen|OJJ37878.1

Group9657: afis|EAW19886.1 afla|EED57357.1 afum|EAL85011.1 anid|CBF82754.1 aory|EIT80143.1 asyd|OJJ61609.1 ater|EAU33759.1 aver|OJJ00932.1

Group9658: anid|CBF88868.1 afla|EED56605.1 aory|EIT79681.1 akaw|GAA93300.1 anig|EHA25611.1 asyd|OJJ54554.1 aver|OJI99480.1 awen|OJJ33512.1

Group9659: afis|EAW20319.1 afla|EED47508.1 afum|EAL85447.1 anid|CBF78532.1 aory|EIT74553.1 asyd|OJJ65298.1 aver|OJI99438.1 awen|OJJ36330.1

Group9660: afis|EAW20734.1 afla|EED51866.1 aluc|OJZ87038.1 anig|EHA23278.1 aory|EIT81721.1 asyd|OJJ54378.1 atub|OJI83477.1 anig|EHA25210.1

Group9661: afis|EAW21029.1 afla|EED55085.1 afum|EAL92067.2 agla|OJJ88640.1 anid|CBF84416.1 aory|EIT81592.1 asyd|OJJ53559.1 aver|OJJ08195.1

Group9662: afis|EAW21030.1 afla|EED55086.1 afum|EAL92069.1 agla|OJJ88643.1 anid|CBF84414.1 aory|EIT81594.1 asyd|OJJ53560.1 aver|OJJ08194.1

Group9663: afis|EAW21073.1 afum|EAL92096.1 ater|EAU36810.1 awen|OJJ36924.1 afis|EAW24619.1 afum|EAL86247.1 asyd|OJJ56920.1 aver|OJJ03372.1

Group9664: afis|EAW21128.1 afum|EAL88867.1 akaw|GAA87228.1 aluc|OJZ84201.1 anid|CBF73407.1 atub|OJI89883.1 afis|EAW24040.1 anig|EHA25772.1

Group9665: afis|EAW21628.1 afla|EED47435.1 afum|EAL92663.1 anid|CBF76690.1 aory|EIT81891.1 asyd|OJJ61227.1 aver|OJJ00550.1 awen|OJJ33877.1

Group9666: afis|EAW22304.1 afla|EED49500.1 agla|OJJ82728.1 anid|CBF85190.1 aory|EIT82581.1 ater|EAU38703.1 awen|OJJ31406.1 afum|EAL91593.1

Group9667: aver|OJJ08911.1 afla|EED45036.1 agla|OJJ81205.1 anid|CBF75956.1 aory|EIT73593.1 asyd|OJJ62489.1 awen|OJJ35943.1 asyd|OJJ52944.1

Group9668: anig|EHA19540.1 afla|EED48397.1 akaw|GAA85012.1 aluc|OJZ86645.1 aory|EIT83074.1 atub|OJI83882.1 asyd|OJJ55474.1 azon|OJJ42568.1

Group9669: afis|EAW23636.1 afla|EED50224.1 afum|EAL87824.1 anid|CBF70172.1 aory|EIT77338.1 asyd|OJJ58794.1 ater|EAU30758.1 aver|OJJ01399.1

Group9670: afis|EAW24452.1 afla|EED55222.1 afum|EAL86121.1 anid|CBF73818.1 aory|EIT83526.1 asyd|OJJ56572.1 ater|EAU29993.1 aver|OJJ06151.1

Group9671: agla|OJJ84650.1 anid|CBF83058.1 asyd|OJJ55508.1 ater|EAU32779.1 akaw|GAA83708.1 aluc|OJZ91027.1 atub|OJI83018.1 anid|CBF74142.1

Group9672: afis|EAW24527.1 afla|EED55266.1 afum|EAL86191.1 anid|CBF84640.1 aory|EIT83543.1 asyd|OJJ53358.1 ater|EAU37242.1 aver|OJJ08383.1

Group9673: afis|EAW24678.1 anid|CBF90088.1 asyd|OJJ57390.1 aver|OJI96124.1 afis|EAW24703.1 afum|EAL89345.1 ater|EAU31632.1 asyd|OJJ62034.1

Group9674: afis|EAW24722.1 afla|EED53236.1 afum|EAL89371.1 anid|CBF82176.1 aory|EIT74978.1 asyd|OJJ64483.1 aver|OJI98584.1 azon|OJJ43677.1

Group9675: afis|EAW25171.1 afla|EED51940.1 afum|EAL89844.1 asyd|OJJ56509.1 ater|EAU29519.1 aver|OJJ06210.1 awen|OJJ35896.1 aver|OJJ08252.1

Group9676: afis|EAW25627.1 afla|EED45969.1 afum|EAL84315.1 anid|CBF82423.1 aory|EIT75513.1 asyd|OJJ52964.1 ater|EAU36879.1 aver|OJJ08930.1

Group9677: afla|EED45122.1 aory|EIT73826.1 asyd|OJJ57416.1 awen|OJJ35842.1 afla|EED45123.1 agla|OJJ79344.1 aory|EIT75622.1 ater|EAU37751.1

Group9678: afla|EED46047.1 akaw|GAA92080.1 aluc|OJZ83644.1 aory|EIT83475.1 asyd|OJJ56880.1 ater|EAU30182.1 aver|OJJ03281.1 asyd|OJJ57006.1

Group9679: afla|EED47577.1 agla|OJJ80418.1 anid|CBF81533.1 aory|EIT74570.1 asyd|OJJ59485.1 aver|OJJ02789.1 awen|OJJ40853.1 agla|OJJ81823.1

Group9680: afla|EED47894.1 akaw|GAA86249.1 aluc|OJZ90504.1 anig|EHA23962.1 aory|EIT77238.1 atub|OJI81928.1 awen|OJJ31309.1 awen|OJJ38088.1

Group9681: afla|EED50239.1 agla|OJJ82251.1 aory|EIT77373.1 asyd|OJJ65272.1 aver|OJI99424.1 awen|OJJ35699.1 azon|OJJ43655.1 azon|OJJ50309.1

Group9682: afla|EED50965.1 aluc|OJZ80362.1 anid|CBF84013.1 aory|EIT77068.1 asyd|OJJ61380.1 aver|OJJ00707.1 awen|OJJ29826.1 awen|OJJ41774.1

Group9683: afla|EED51146.1 aory|EIT81353.1 asyd|OJJ56871.1 awen|OJJ30234.1 azon|OJJ46826.1 anid|CBF78574.1 asyd|OJJ65069.1 aver|OJI99194.1

Group9684: afla|EED51801.1 agla|OJJ78555.1 aory|EIT81703.1 asyd|OJJ57234.1 ater|EAU32272.1 aver|OJJ03051.1 awen|OJJ35819.1 azon|OJJ42502.1

Group9685: afla|EED52042.1 agla|OJJ82690.1 aory|EIT81445.1 asyd|OJJ61433.1 ater|EAU37779.1 aver|OJJ00768.1 asyd|OJJ52786.1 aver|OJJ08761.1

Group9686: afla|EED52175.1 akaw|GAA88461.1 aluc|OJZ88351.1 anig|EHA18083.1 aory|EIT74452.1 asyd|OJJ60084.1 atub|OJI90946.1 aver|OJJ04170.1

Group9687: afla|EED54201.1 afum|EAL85372.1 anid|CBF75978.1 aory|EIT76166.1 asyd|OJJ55418.1 ater|EAU32161.1 aver|OJJ06495.1 azon|OJJ50842.1

Group9688: afla|EED54742.1 anid|CBF78080.1 aory|EIT78954.1 asyd|OJJ55878.1 ater|EAU32535.1 aver|OJJ04823.1 awen|OJJ38266.1 aory|EIT78757.1

Group9689: afla|EED56558.1 agla|OJJ88413.1 akaw|GAA92923.1 aluc|OJZ82514.1 anig|EHA20904.1 asyd|OJJ57316.1 aver|OJJ05845.1 awen|OJJ33379.1

Group9690: afla|EED56567.1 aluc|OJZ88350.1 aory|EIT79594.1 asyd|OJJ56922.1 atub|OJI90945.1 aver|OJJ03371.1 azon|OJJ48671.1 asyd|OJJ62605.1

Group9691: afla|EED57356.1 agla|OJJ86314.1 anid|CBF74605.1 aory|EIT80139.1 asyd|OJJ58332.1 aver|OJJ01923.1 awen|OJJ34597.1 aver|OJJ02079.1

Group9692: afum|EAL92634.2 agla|OJJ87896.1 aluc|OJZ80194.1 asyd|OJJ52335.1 ater|EAU29451.1 aver|OJJ06197.1 awen|OJJ31628.1 asyd|OJJ56526.1

Group9693: agla|OJJ81745.1 akaw|GAA92806.1 aluc|OJZ88729.1 anig|EHA18332.1 atub|OJI91223.1 aver|OJI98066.1 akaw|GAA83090.1 asyd|OJJ62923.1

Group9694: agla|OJJ85513.1 anid|CBF87357.1 asyd|OJJ56542.1 ater|EAU36633.1 aver|OJJ05540.1 azon|OJJ51537.1 asyd|OJJ55375.1 aver|OJJ06461.1

Group9695: akaw|GAA89825.1 aluc|OJZ81863.1 anig|EHA22363.1 asyd|OJJ65112.1 atub|OJI84430.1 aver|OJI96896.1 awen|OJJ30085.1 asyd|OJJ53796.1

Group9696: abra|OJJ65444.1 abra|OJJ73318.1 akaw|GAA92802.1 aluc|OJZ88726.1 anig|EHA18431.1 atub|OJI91303.1 acar|OOF90376.1

Group9697: abra|OJJ65534.1 abra|OJJ65602.1 aluc|OJZ79709.1 aluc|OJZ79954.1 aluc|OJZ80151.1 awen|OJJ29544.1 awen|OJJ29650.1

Group9698: abra|OJJ65561.1 abra|OJJ65733.1 abra|OJJ65825.1 acla|EAW07109.1 afis|EAW22121.1 azon|OJJ50251.1 ater|EAU36018.1

Group9699: abra|OJJ70403.1 abra|OJJ74817.1 acar|OOF91878.1 aluc|OJZ92774.1 atub|OJI87217.1 akaw|GAA91761.1 anig|EHA26235.1

Group9700: acar|OOF90383.1 acar|OOF90385.1 afla|EED49707.1 asyd|OJJ54939.1 aver|OJJ07257.1 awen|OJJ30843.1 aver|OJJ08817.1

Group9701: acar|OOF90513.1 acar|OOF90514.1 abra|OJJ69552.1 akaw|GAA92501.1 aluc|OJZ91514.1 anig|EHA18709.1 atub|OJI85968.1

Group9702: acar|OOF92295.1 acar|OOF92336.1 abra|OJJ67505.1 akaw|GAA86333.1 aluc|OJZ84858.1 anig|EHA25091.1 atub|OJI80738.1

Group9703: acar|OOF94029.1 acar|OOF94099.1 abra|OJJ66424.1 akaw|GAA90288.1 aluc|OJZ83052.1 anig|EHA28116.1 atub|OJI87896.1

Group9704: acar|OOF94060.1 acar|OOF94130.1 abra|OJJ66012.1 acla|EAW12366.1 akaw|GAA92142.1 aluc|OJZ85547.1 atub|OJI79343.1

Group9705: acar|OOF95054.1 acar|OOF95121.1 abra|OJJ74659.1 akaw|GAA81952.1 aluc|OJZ92236.1 anig|EHA26718.1 atub|OJI86669.1

Group9706: acar|OOF95061.1 acar|OOF95128.1 abra|OJJ74654.1 akaw|GAA81956.1 aluc|OJZ92233.1 anig|EHA26722.1 atub|OJI86664.1

Group9707: acar|OOF95636.1 acar|OOF95687.1 abra|OJJ74673.1 akaw|GAA81940.1 aluc|OJZ92250.1 anig|EHA26706.1 atub|OJI86683.1

Group9708: acar|OOF98699.1 acar|OOF98700.1 abra|OJJ70124.1 akaw|GAA86645.1 aluc|OJZ85202.1 anig|EHA25408.1 atub|OJI80399.1

Group9709: acar|OOF99133.1 acar|OOG01134.1 abra|OJJ75543.1 akaw|GAA84299.1 aluc|OJZ88993.1 anig|EHA23534.1 atub|OJI85795.1

Group9710: acar|OOG00245.1 acar|OOG00263.1 abra|OJJ67539.1 akaw|GAA88120.1 aluc|OJZ81393.1 anig|EHA18686.1 atub|OJI91598.1

Group9711: acar|OOG00246.1 acar|OOG00264.1 abra|OJJ67540.1 akaw|GAA88119.1 aluc|OJZ81392.1 anig|EHA18685.1 atub|OJI91597.1

Group9712: afum|EAL85379.1 afum|EAL87817.1 aluc|OJZ79810.1 aluc|OJZ79957.1 afis|EAW19387.1 anig|EHA21845.1 awen|OJJ29569.1

Group9713: afum|EAL90933.1 afum|EAL91124.1 afla|EED52355.1 agla|OJJ83478.1 akaw|GAA87892.1 azon|OJJ42063.1 azon|OJJ42062.1

Group9714: agla|OJJ78566.1 agla|OJJ89693.1 acla|EAW08144.1 afis|EAW23180.1 afum|EAL88303.1 azon|OJJ47609.1 awen|OJJ36598.1

Group9715: agla|OJJ79308.1 agla|OJJ84499.1 afis|EAW19110.1 afla|EED57648.1 afum|EAL93841.2 aory|EIT79880.1 awen|OJJ32503.1

Group9716: agla|OJJ79364.1 agla|OJJ84921.1 agla|OJJ86637.1 agla|OJJ84922.1 agla|OJJ85121.1 anid|CBF80817.1 anig|EHA22300.1

Group9717: agla|OJJ80014.1 agla|OJJ80018.1 agla|OJJ80444.1 awen|OJJ38763.1 azon|OJJ51132.1 asyd|OJJ59795.1 awen|OJJ33433.1

Group9718: agla|OJJ80553.1 agla|OJJ85190.1 agla|OJJ86423.1 agla|OJJ84511.1 agla|OJJ84513.1 agla|OJJ87332.1 agla|OJJ87333.1

Group9719: agla|OJJ81317.1 agla|OJJ83542.1 agla|OJJ85999.1 afis|EAW20674.1 aluc|OJZ80140.1 anid|CBF84439.1 aory|EIT78799.1

Group9720: agla|OJJ83799.1 agla|OJJ86158.1 aluc|OJZ79693.1 aluc|OJZ79906.1 aluc|OJZ81126.1 asyd|OJJ52320.1 aver|OJJ05900.1

Group9721: akaw|GAA92341.1 akaw|GAA93146.1 akaw|GAA93220.1 akaw|GAA92653.1 akaw|GAA93096.1 akaw|GAA93271.1 aluc|OJZ80323.1

Group9722: akaw|GAA92359.1 akaw|GAA92361.1 abra|OJJ65460.1 agla|OJJ79296.1 asyd|OJJ52173.1 acla|EAW11447.1 afla|EED52346.1

Group9723: akaw|GAA92608.1 akaw|GAA92958.1 abra|OJJ65616.1 acla|EAW11449.1 afla|EED52349.1 abra|OJJ66313.1 aluc|OJZ80004.1

Group9724: akaw|GAA93274.1 akaw|GAA93310.1 aluc|OJZ79676.1 aluc|OJZ80120.1 aluc|OJZ81147.1 aluc|OJZ88801.1 abra|OJJ66179.1

Group9725: aluc|OJZ79625.1 aluc|OJZ90862.1 aluc|OJZ80185.1 abra|OJJ70106.1 agla|OJJ81743.1 akaw|GAA83853.1 anig|EHA26922.1

Group9726: aluc|OJZ79673.1 aluc|OJZ88804.1 asyd|OJJ52019.1 asyd|OJJ52273.1 abra|OJJ66183.1 akaw|GAA93191.1 akaw|GAA93344.1

Group9727: aluc|OJZ79696.1 aluc|OJZ88837.1 asyd|OJJ52056.1 asyd|OJJ52167.1 abra|OJJ65729.1 acar|OOF96902.1 afla|EED54492.1

Group9728: aluc|OJZ80014.1 aluc|OJZ80092.1 abra|OJJ66175.1 acar|OOF97139.1 acla|EAW08830.1 afis|EAW16714.1 anig|EHA26114.1

Group9729: anid|CBF79026.1 anid|CBF87275.1 acla|EAW14555.1 afis|EAW17237.1 afum|EAL86595.1 asyd|OJJ55940.1 aver|OJJ04889.1

Group9730: anid|CBF81400.1 anid|CBF83110.1 acla|EAW09041.1 afis|EAW15400.1 afla|EED45073.1 afum|EAL84248.1 aory|EIT73649.1

Group9731: anig|EHA19569.1 anig|EHA27836.1 abra|OJJ77128.1 acar|OOF93017.1 akaw|GAA90800.1 aluc|OJZ84538.1 atub|OJI89542.1

Group9732: anig|EHA21872.1 anig|EHA26121.1 abra|OJJ66306.1 akaw|GAA93149.1 aluc|OJZ91228.1 acar|OOF90507.1 aluc|OJZ80008.1

Group9733: asyd|OJJ51854.1 asyd|OJJ54878.1 afla|EED50242.1 akaw|GAA83269.1 anid|CBF88280.1 aory|EIT77311.1 aluc|OJZ87733.1

Group9734: asyd|OJJ52104.1 asyd|OJJ64009.1 aver|OJI97594.1 aver|OJI98097.1 anid|CBF85316.1 ater|EAU34365.1 azon|OJJ50183.1

Group9735: asyd|OJJ52161.1 asyd|OJJ52176.1 acla|EAW11072.1 afla|EED50551.1 aory|EIT82040.1 aver|OJJ00281.1 asyd|OJJ60953.1

Group9736: asyd|OJJ52271.1 asyd|OJJ52315.1 asyd|OJJ51920.1 asyd|OJJ51995.1 akaw|GAA89005.1 akaw|GAA92333.1 aluc|OJZ79473.1

Group9737: asyd|OJJ52888.1 asyd|OJJ55405.1 asyd|OJJ62611.1 asyd|OJJ63961.1 aver|OJJ03448.1 aver|OJJ06485.1 aver|OJJ07671.1

Group9738: asyd|OJJ54343.1 asyd|OJJ54528.1 aver|OJJ03007.1 aver|OJJ05295.1 akaw|GAA85810.1 aluc|OJZ91289.1 atub|OJI83279.1

Group9739: ater|EAU33181.1 ater|EAU35443.1 afla|EED45067.1 anid|CBF84552.1 aory|EIT73756.1 asyd|OJJ53482.1 aver|OJJ08277.1

Group9740: aver|OJI96131.1 aver|OJJ04892.1 acla|EAW15060.1 afum|EAL85111.2 asyd|OJJ55943.1 ater|EAU37825.1 agla|OJJ82585.1

Group9741: aver|OJJ06627.1 aver|OJJ07716.1 anid|CBF83182.1 agla|OJJ80382.1 ater|EAU36808.1 asyd|OJJ55550.1 awen|OJJ35782.1

Group9742: awen|OJJ30182.1 awen|OJJ30894.1 abra|OJJ74480.1 akaw|GAA87373.1 aluc|OJZ89639.1 atub|OJI85164.1 acar|OOF96768.1

Group9743: awen|OJJ30783.1 awen|OJJ31698.1 abra|OJJ67839.1 acar|OOF92014.1 akaw|GAA91845.1 aluc|OJZ82053.1 atub|OJI84611.1

Group9744: awen|OJJ30877.1 awen|OJJ33939.1 awen|OJJ33317.1 awen|OJJ30918.1 awen|OJJ31456.1 afis|EAW19028.1 afum|EAL93758.1

Group9745: awen|OJJ33688.1 abra|OJJ74729.1 acar|OOF92036.1 akaw|GAA81888.1 aluc|OJZ92313.1 anig|EHA26656.1 atub|OJI86742.1

Group9746: awen|OJJ31359.1 awen|OJJ38671.1 abra|OJJ68029.1 akaw|GAA90924.1 aluc|OJZ82902.1 atub|OJI89128.1 azon|OJJ46859.1

Group9747: awen|OJJ32700.1 awen|OJJ33320.1 awen|OJJ39607.1 awen|OJJ32701.1 afla|EED54688.1 afla|EED54689.1 aory|EIT78885.1

Group9748: awen|OJJ33465.1 awen|OJJ38085.1 afis|EAW20739.1 asyd|OJJ56939.1 aver|OJJ03346.1 afla|EED46860.1 aory|EIT72670.1

Group9749: awen|OJJ33556.1 awen|OJJ33557.1 asyd|OJJ52927.1 aver|OJJ08895.1 afla|EED50327.1 aory|EIT77355.1 afis|EAW16165.1

Group9750: awen|OJJ34707.1 awen|OJJ35048.1 acla|EAW12023.1 afis|EAW19832.1 afum|EAL85060.1 aver|OJJ08534.1 azon|OJJ50136.1

Group9751: azon|OJJ42218.1 azon|OJJ42501.1 azon|OJJ43570.1 acar|OOF90551.1 awen|OJJ41549.1 afis|EAW22623.1 agla|OJJ79471.1

Group9752: azon|OJJ50292.1 abra|OJJ69152.1 acar|OOF91151.1 akaw|GAA86310.1 aluc|OJZ86139.1 anig|EHA24485.1 atub|OJI79931.1

Group9753: azon|OJJ42472.1 azon|OJJ42785.1 azon|OJJ43529.1 azon|OJJ51058.1 azon|OJJ48053.1 azon|OJJ45122.1 azon|OJJ50325.1

Group9754: azon|OJJ42586.1 azon|OJJ45998.1 acar|OOF92160.1 akaw|GAA89799.1 aluc|OJZ81892.1 anig|EHA22350.1 atub|OJI84445.1

Group9755: azon|OJJ46005.1 azon|OJJ47953.1 acla|EAW08459.1 afis|EAW23504.1 afum|EAL87957.1 agla|OJJ86750.1 awen|OJJ39830.1

Group9756: agla|OJJ79522.1 agla|OJJ88709.1 agla|OJJ81045.1 agla|OJJ82034.1 agla|OJJ85366.1 agla|OJJ82928.1 agla|OJJ89310.1

Group9757: aluc|OJZ79560.1 aluc|OJZ80030.1 aluc|OJZ79858.1 aluc|OJZ88827.1 aluc|OJZ79782.1 aluc|OJZ80429.1 aluc|OJZ80241.1

Group9758: abra|OJJ65470.1 aluc|OJZ87040.1 anid|CBF76054.1 asyd|OJJ62366.1 atub|OJI83545.1 aver|OJI96500.1 awen|OJJ37045.1

Group9759: abra|OJJ65476.1 acar|OOF98953.1 afla|EED47796.1 ater|EAU31614.1 anid|CBF79154.1 asyd|OJJ62575.1 aver|OJI96697.1

Group9760: abra|OJJ65493.1 agla|OJJ85324.1 akaw|GAA91467.1 aluc|OJZ82072.1 anig|EHA18712.1 atub|OJI79314.1 aver|OJI98909.1

Group9761: abra|OJJ65547.1 acar|OOF97960.1 akaw|GAA93195.1 aluc|OJZ79942.1 atub|OJI87430.1 awen|OJJ29539.1 akaw|GAA93187.1

Group9762: abra|OJJ65607.1 acla|EAW12428.1 afis|EAW19528.1 afum|EAL86976.2 akaw|GAA93116.1 aluc|OJZ80588.1 azon|OJJ48014.1

Group9763: anid|CBF80257.1 asyd|OJJ54183.1 ater|EAU36723.1 aver|OJJ08557.1 acla|EAW13743.1 afis|EAW19839.1 afum|EAL85054.2

Group9764: awen|OJJ33738.1 acar|OOF91732.1 afla|EED52383.1 anig|EHA19836.1 aory|EIT83298.1 aluc|OJZ86320.1 atub|OJI84215.1

Group9765: abra|OJJ65683.1 acar|OOF95486.1 agla|OJJ78865.1 anid|CBF75920.1 ater|EAU30725.1 aver|OJJ08165.1 awen|OJJ34745.1

Group9766: afis|EAW25611.1 agla|OJJ88492.1 afla|EED53588.1 aory|EIT74897.1 aver|OJJ07160.1 asyd|OJJ62355.1 afum|EAL92214.1

Group9767: abra|OJJ65795.1 acar|OOF96896.1 afla|EED51402.1 aory|EIT80451.1 asyd|OJJ62238.1 aver|OJI96347.1 awen|OJJ37610.1

Group9768: abra|OJJ65807.1 afla|EED47496.1 aluc|OJZ82528.1 anig|EHA22494.1 aory|EIT74566.1 awen|OJJ33614.1 azon|OJJ42014.1

Group9769: abra|OJJ65837.1 akaw|GAA92980.1 aluc|OJZ79992.1 anig|EHA21147.1 akaw|GAA93036.1 ater|EAU35416.1 akaw|GAA93103.1

Group9770: abra|OJJ65867.1 aluc|OJZ90173.1 anig|EHA23672.1 atub|OJI81601.1 ater|EAU30769.1 awen|OJJ35394.1 azon|OJJ50615.1

Group9771: abra|OJJ65874.1 agla|OJJ84702.1 akaw|GAA92269.1 aluc|OJZ90163.1 anig|EHA23669.1 azon|OJJ45109.1 awen|OJJ38804.1

Group9772: abra|OJJ65882.1 akaw|GAA90453.1 aluc|OJZ85771.1 asyd|OJJ65100.1 atub|OJI79556.1 aver|OJI99226.1 ater|EAU36740.1

Group9773: abra|OJJ65891.1 acar|OOF94954.1 akaw|GAA86014.1 aluc|OJZ90241.1 anig|EHA23738.1 atub|OJI81670.1 awen|OJJ37827.1

Group9774: abra|OJJ65906.1 acar|OOF93399.1 akaw|GAA92325.1 aluc|OJZ90223.1 anig|EHA23724.1 ater|EAU34024.1 atub|OJI81656.1

Group9775: ater|EAU32518.1 akaw|GAA93171.1 aluc|OJZ83030.1 anig|EHA28106.1 asyd|OJJ52501.1 atub|OJI87918.1 aver|OJJ05539.1

Group9776: abra|OJJ65940.1 acar|OOF95005.1 agla|OJJ89023.1 akaw|GAA92296.1 aluc|OJZ90193.1 anig|EHA23692.1 atub|OJI81620.1

Group9777: abra|OJJ65950.1 acar|OOF92922.1 agla|OJJ87999.1 aluc|OJZ83007.1 atub|OJI89242.1 aver|OJJ08451.1 awen|OJJ30038.1

Group9778: abra|OJJ65977.1 acar|OOF92894.1 akaw|GAA90851.1 aluc|OJZ82982.1 anig|EHA21396.1 atub|OJI89216.1 awen|OJJ31295.1

Group9779: abra|OJJ65982.1 acar|OOF92891.1 akaw|GAA90855.1 aluc|OJZ82978.1 anig|EHA21393.1 atub|OJI89211.1 awen|OJJ35855.1

Group9780: abra|OJJ65997.1 akaw|GAA90869.1 aluc|OJZ82964.1 anig|EHA21380.1 ater|EAU33862.1 atub|OJI89196.1 anig|EHA24971.1

Group9781: asyd|OJJ62944.1 aver|OJI98046.1 afla|EED55278.1 aory|EIT83593.1 aver|OJJ07922.1 awen|OJJ33327.1 atub|OJI87200.1

Group9782: abra|OJJ66039.1 afla|EED49333.1 akaw|GAA86615.1 aluc|OJZ85167.1 aory|EIT79483.1 ater|EAU32795.1 atub|OJI80444.1

Group9783: abra|OJJ66074.1 afla|EED52684.1 akaw|GAA84172.1 aluc|OJZ81097.1 anig|EHA19274.1 aory|EIT74216.1 atub|OJI82485.1

Group9784: aver|OJI99624.1 acla|EAW13613.1 afla|EED53480.1 akaw|GAA88583.1 aluc|OJZ88487.1 anig|EHA20149.1 aory|EIT75250.1

Group9785: abra|OJJ66116.1 acar|OOF95021.1 akaw|GAA92678.1 aluc|OJZ90133.1 anig|EHA20938.1 asyd|OJJ52521.1 atub|OJI84663.1

Group9786: abra|OJJ66129.1 anig|EHA20925.1 abra|OJJ66130.1 akaw|GAA92687.1 aluc|OJZ90145.1 atub|OJI84654.1 awen|OJJ33721.1

Group9787: abra|OJJ66134.1 acar|OOF94022.1 afla|EED51824.1 akaw|GAA92691.1 aluc|OJZ90149.1 anig|EHA20921.1 atub|OJI84650.1

Group9788: abra|OJJ66141.1 akaw|GAA92696.1 aluc|OJZ90154.1 anig|EHA20915.1 asyd|OJJ64759.1 atub|OJI84645.1 azon|OJJ42386.1

Group9789: abra|OJJ66153.1 afla|EED45844.1 anid|CBF71482.1 asyd|OJJ60379.1 ater|EAU37971.1 aver|OJJ03873.1 awen|OJJ40104.1

Group9790: asyd|OJJ57411.1 ater|EAU35429.1 aver|OJJ05770.1 afis|EAW20691.1 afum|EAL86273.1 asyd|OJJ60275.1 ater|EAU31611.1

Group9791: abra|OJJ66167.1 afla|EED54668.1 akaw|GAA83593.1 aluc|OJZ92403.1 anig|EHA26563.1 aory|EIT78789.1 atub|OJI86831.1

Group9792: abra|OJJ66177.1 agla|OJJ85094.1 aluc|OJZ80013.1 anig|EHA26115.1 asyd|OJJ56758.1 atub|OJI87459.1 awen|OJJ38803.1

Group9793: abra|OJJ66197.1 acar|OOF95791.1 afla|EED53394.1 agla|OJJ81774.1 asyd|OJJ55465.1 awen|OJJ41831.1 awen|OJJ35538.1

Group9794: abra|OJJ66199.1 acar|OOF91931.1 afis|EAW16948.1 afum|EAL86475.1 akaw|GAA89090.1 anig|EHA25692.1 atub|OJI88429.1

Group9795: abra|OJJ66206.1 afis|EAW17408.1 afla|EED48166.1 agla|OJJ81618.1 anid|CBF76000.1 asyd|OJJ60265.1 ater|EAU37953.1

Group9796: abra|OJJ66208.1 acar|OOF90652.1 akaw|GAA89079.1 aluc|OJZ80512.1 anig|EHA25681.1 atub|OJI88418.1 azon|OJJ42563.1

Group9797: abra|OJJ66222.1 afis|EAW20825.1 akaw|GAA89064.1 aluc|OJZ80528.1 atub|OJI88402.1 awen|OJJ39957.1 afum|EAL84804.1

Group9798: abra|OJJ66241.1 afis|EAW16930.1 afla|EED53698.1 ater|EAU34192.1 azon|OJJ44470.1 agla|OJJ84652.1 aluc|OJZ80547.1

Group9799: abra|OJJ66278.1 acla|EAW15063.1 afla|EED45032.1 afum|EAL85114.1 anig|EHA23658.1 ater|EAU38973.1 atub|OJI89945.1

Group9800: abra|OJJ66279.1 acar|OOF96459.1 afis|EAW19507.1 afum|EAL86987.1 anig|EHA23659.1 ater|EAU36494.1 atub|OJI89944.1

Group9801: abra|OJJ66283.1 acar|OOF91927.1 acla|EAW10349.1 anig|EHA23663.1 ater|EAU38976.1 atub|OJI89940.1 ater|EAU38975.1

Group9802: abra|OJJ66328.1 afis|EAW17144.1 afla|EED47885.1 aluc|OJZ91241.1 anid|CBF82971.1 atub|OJI79303.1 afis|EAW20656.1

Group9803: afum|EAL90431.1 anid|CBF79083.1 afla|EED47305.1 aory|EIT82825.1 ater|EAU37854.1 aver|OJJ05552.1 awen|OJJ32867.1

Group9804: abra|OJJ66445.1 akaw|GAA86650.1 aluc|OJZ85208.1 anig|EHA26090.1 asyd|OJJ56744.1 aver|OJJ07916.1 azon|OJJ50084.1

Group9805: abra|OJJ66492.1 acar|OOF93283.1 akaw|GAA85441.1 aluc|OJZ83346.1 anig|EHA24979.1 ater|EAU34029.1 atub|OJI82240.1

Group9806: afis|EAW22298.1 afum|EAL90883.1 agla|OJJ86326.1 anid|CBF69368.1 asyd|OJJ53654.1 asyd|OJJ60387.1 ater|EAU36564.1

Group9807: acla|EAW11674.1 acar|OOF92982.1 afis|EAW23695.1 afla|EED51840.1 afum|EAL87139.1 aory|EIT81695.1 ater|EAU31565.1

Group9808: abra|OJJ66510.1 azon|OJJ51380.1 abra|OJJ75779.1 akaw|GAA84495.1 aluc|OJZ89219.1 anig|EHA23343.1 atub|OJI85573.1

Group9809: azon|OJJ46501.1 acla|EAW12906.1 afis|EAW17782.1 afla|EED56297.1 aory|EIT80563.1 atub|OJI89933.1 aory|EIT73403.1

Group9810: abra|OJJ66776.1 acar|OOF93855.1 afla|EED51424.1 akaw|GAA89711.1 aluc|OJZ80673.1 aory|EIT80482.1 atub|OJI87527.1

Group9811: abra|OJJ66798.1 afis|EAW22088.1 afla|EED57474.1 asyd|OJJ56428.1 atub|OJI89598.1 aver|OJJ03128.1 awen|OJJ31393.1

Group9812: abra|OJJ66810.1 afla|EED46150.1 agla|OJJ88619.1 aory|EIT76481.1 asyd|OJJ54552.1 atub|OJI79332.1 aver|OJI96980.1

Group9813: abra|OJJ66824.1 akaw|GAA90459.1 aluc|OJZ85765.1 anig|EHA24791.1 atub|OJI79549.1 acar|OOG00558.1 azon|OJJ49025.1

Group9814: abra|OJJ66935.1 acar|OOF94943.1 aluc|OJZ85389.1 anig|EHA24888.1 akaw|GAA87756.1 atub|OJI80181.1 anig|EHA25573.1

Group9815: abra|OJJ66946.1 akaw|GAA91474.1 aluc|OJZ85634.1 anig|EHA24897.1 asyd|OJJ63596.1 atub|OJI79424.1 aver|OJI97340.1

Group9816: abra|OJJ66969.1 afla|EED56783.1 akaw|GAA92194.1 aluc|OJZ85609.1 anig|EHA24917.1 aory|EIT73926.1 atub|OJI79402.1

Group9817: abra|OJJ66977.1 acar|OOF93640.1 akaw|GAA92187.1 aluc|OJZ85597.1 anig|EHA24925.1 atub|OJI79393.1 azon|OJJ46406.1

Group9818: abra|OJJ66978.1 acar|OOF93641.1 akaw|GAA92186.1 aluc|OJZ85596.1 anig|EHA24926.1 atub|OJI79392.1 azon|OJJ48711.1

Group9819: abra|OJJ67014.1 acar|OOF93606.1 akaw|GAA92159.1 aluc|OJZ85566.1 anig|EHA24952.1 atub|OJI79361.1 ater|EAU31042.1

Group9820: acla|EAW15303.1 afis|EAW17015.1 akaw|GAA89814.1 aluc|OJZ81876.1 acar|OOF92825.1 ater|EAU31620.1 afis|EAW25738.1

Group9821: abra|OJJ67026.1 acla|EAW11886.1 akaw|GAA83301.1 anid|CBF81862.1 ater|EAU31650.1 aver|OJI95821.1 ater|EAU30724.1

Group9822: abra|OJJ67034.1 afla|EED56204.1 akaw|GAA90579.1 aluc|OJZ86037.1 aory|EIT73386.1 ater|EAU36046.1 atub|OJI79824.1

Group9823: abra|OJJ67060.1 afla|EED48485.1 anid|CBF89311.1 anig|EHA22169.1 aory|EIT73239.1 asyd|OJJ59580.1 aver|OJJ02689.1

Group9824: abra|OJJ67066.1 acar|OOF96439.1 akaw|GAA83293.1 aluc|OJZ83795.1 anig|EHA18940.1 atub|OJI90165.1 azon|OJJ50262.1

Group9825: abra|OJJ67078.1 akaw|GAA91079.1 aluc|OJZ87894.1 anid|CBF70190.1 asyd|OJJ59829.1 atub|OJI80758.1 aver|OJJ02458.1

Group9826: abra|OJJ67130.1 acar|OOF99012.1 afla|EED53376.1 aory|EIT75221.1 asyd|OJJ59558.1 aver|OJJ02710.1 awen|OJJ39795.1

Group9827: abra|OJJ67155.1 acar|OOF91669.1 akaw|GAA91017.1 aluc|OJZ87818.1 anig|EHA20299.1 ater|EAU35207.1 atub|OJI80826.1

Group9828: abra|OJJ67217.1 acar|OOF93377.1 acla|EAW11102.1 akaw|GAA91093.1 aluc|OJZ87913.1 ater|EAU29431.1 atub|OJI79340.1

Group9829: asyd|OJJ58705.1 aver|OJJ01317.1 afla|EED56582.1 agla|OJJ79966.1 aory|EIT79628.1 awen|OJJ40587.1 azon|OJJ42309.1

Group9830: abra|OJJ67231.1 acar|OOF98322.1 agla|OJJ89179.1 akaw|GAA91656.1 aluc|OJZ87743.1 asyd|OJJ64981.1 atub|OJI80898.1

Group9831: anid|CBF78375.1 afis|EAW25697.1 afum|EAL84372.1 afla|EED56870.1 agla|OJJ79800.1 asyd|OJJ57320.1 aver|OJJ05841.1

Group9832: abra|OJJ67353.1 acar|OOF91852.1 afis|EAW25677.1 afum|EAL84347.1 anid|CBF73490.1 anig|EHA25228.1 asyd|OJJ54610.1

Group9833: abra|OJJ67367.1 acar|OOF92443.1 akaw|GAA86455.1 aluc|OJZ84997.1 anig|EHA25208.1 atub|OJI80603.1 azon|OJJ42413.1

Group9834: abra|OJJ67382.1 acar|OOF92431.1 akaw|GAA86442.1 aluc|OJZ84984.1 anig|EHA25197.1 atub|OJI80616.1 azon|OJJ45903.1

Group9835: abra|OJJ67557.1 acar|OOG00272.1 aluc|OJZ81376.1 aory|EIT74385.1 atub|OJI91582.1 awen|OJJ29920.1 agla|OJJ84145.1

Group9836: anid|CBF90356.1 ater|EAU36813.1 afla|EED51131.1 agla|OJJ81309.1 aory|EIT81314.1 asyd|OJJ55573.1 aver|OJJ06652.1

Group9837: ater|EAU34362.1 afla|EED56877.1 aory|EIT77518.1 asyd|OJJ54821.1 aver|OJJ07340.1 awen|OJJ37089.1 aver|OJJ08816.1

Group9838: abra|OJJ67581.1 afla|EED44748.1 akaw|GAA85344.1 aluc|OJZ90706.1 anig|EHA24577.1 atub|OJI82133.1 awen|OJJ36299.1

Group9839: abra|OJJ67589.1 akaw|GAA89108.1 anid|CBF84243.1 anig|EHA25708.1 asyd|OJJ57196.1 atub|OJI88449.1 aver|OJJ03095.1

Group9840: abra|OJJ67627.1 akaw|GAA89147.1 aluc|OJZ81689.1 anig|EHA25754.1 atub|OJI88490.1 ater|EAU30032.1 awen|OJJ36957.1

Group9841: abra|OJJ67640.1 akaw|GAA89158.1 aluc|OJZ81676.1 anid|CBF77412.1 anig|EHA25765.1 atub|OJI88505.1 azon|OJJ48531.1

Group9842: abra|OJJ67643.1 akaw|GAA93106.1 aluc|OJZ80598.1 anig|EHA25769.1 aory|EIT82302.1 asyd|OJJ55803.1 atub|OJI88510.1

Group9843: abra|OJJ67659.1 akaw|GAA89174.1 aluc|OJZ81659.1 atub|OJI88524.1 aver|OJI98977.1 awen|OJJ41960.1 anid|CBF70175.1

Group9844: abra|OJJ67683.1 aluc|OJZ81635.1 anid|CBF73537.1 asyd|OJJ56719.1 ater|EAU30904.1 atub|OJI88545.1 aver|OJJ06006.1

Group9845: abra|OJJ67706.1 afis|EAW23760.1 afum|EBA27303.1 anig|EHA21358.1 ater|EAU30777.1 atub|OJI88565.1 aver|OJI99276.1

Group9846: afla|EED50755.1 abra|OJJ73326.1 agla|OJJ84167.1 akaw|GAA92792.1 aluc|OJZ88712.1 anig|EHA18414.1 atub|OJI91299.1

Group9847: afla|EED48994.1 aory|EIT80027.1 anid|CBF87387.1 asyd|OJJ52843.1 acar|OOF96795.1 aver|OJJ08819.1 azon|OJJ51746.1

Group9848: abra|OJJ67805.1 acla|EAW13532.1 afis|EAW17482.1 aluc|OJZ90852.1 ater|EAU33223.1 atub|OJI82851.1 afum|EAL87841.1

Group9849: abra|OJJ67812.1 afis|EAW19367.1 afla|EED48199.1 afum|EAL94086.1 anid|CBF87297.1 aory|EIT72948.1 ater|EAU30774.1

Group9850: abra|OJJ67819.1 afum|EAL86777.1 anid|CBF79158.1 anig|EHA20933.1 ater|EAU38016.1 aver|OJJ07618.1 awen|OJJ36344.1

Group9851: abra|OJJ67876.1 acar|OOF92854.1 afis|EAW21801.1 afum|EAL92827.1 akaw|GAA91884.1 aluc|OJZ82013.1 atub|OJI84562.1

Group9852: asyd|OJJ56421.1 aver|OJJ08898.1 asyd|OJJ55463.1 afla|EED46393.1 aory|EIT72410.1 afum|EAL85341.1 azon|OJJ46466.1

Group9853: acla|EAW15309.1 afis|EAW20269.1 awen|OJJ39874.1 afla|EED52435.1 afum|EBA27229.1 aory|EIT83304.1 asyd|OJJ60803.1

Group9854: abra|OJJ67955.1 agla|OJJ82452.1 akaw|GAA91241.1 aluc|OJZ82828.1 anig|EHA21266.1 atub|OJI89058.1 acar|OOF92800.1

Group9855: abra|OJJ67979.1 acar|OOF92820.1 akaw|GAA91223.1 aluc|OJZ82847.1 anig|EHA21284.1 atub|OJI89077.1 awen|OJJ38660.1

Group9856: abra|OJJ67980.1 acar|OOF92022.1 akaw|GAA91222.1 aluc|OJZ82848.1 anig|EHA21285.1 atub|OJI89078.1 awen|OJJ30801.1

Group9857: abra|OJJ68004.1 acar|OOF92046.1 akaw|GAA90946.1 aluc|OJZ82878.1 anig|EHA21304.1 atub|OJI89102.1 acla|EAW10448.1

Group9858: abra|OJJ68006.1 acar|OOF92048.1 akaw|GAA90944.1 aluc|OJZ82880.1 anig|EHA21306.1 ater|EAU31763.1 atub|OJI89104.1

Group9859: abra|OJJ68017.1 acar|OOF92060.1 agla|OJJ78754.1 akaw|GAA90934.1 aluc|OJZ82890.1 anig|EHA21316.1 atub|OJI89115.1

Group9860: afis|EAW17410.1 afum|EAL91244.2 anid|CBF84542.1 asyd|OJJ53468.1 aver|OJJ08288.1 azon|OJJ50176.1 agla|OJJ79331.1

Group9861: abra|OJJ68105.1 acar|OOF93579.1 acla|EAW13241.1 agla|OJJ88425.1 asyd|OJJ61245.1 aver|OJI99635.1 agla|OJJ83824.1

Group9862: ater|EAU32652.1 abra|OJJ72593.1 acar|OOF97551.1 akaw|GAA85220.1 aluc|OJZ90574.1 atub|OJI81998.1 afla|EED47249.1

Group9863: abra|OJJ68289.1 acar|OOF97559.1 akaw|GAA83182.1 aluc|OJZ83682.1 anig|EHA18836.1 atub|OJI90053.1 awen|OJJ33733.1

Group9864: abra|OJJ68290.1 acla|EAW11095.1 awen|OJJ41508.1 abra|OJJ69501.1 agla|OJJ83611.1 ater|EAU32042.1 azon|OJJ47418.1

Group9865: abra|OJJ68293.1 acar|OOF97560.1 akaw|GAA83180.1 aluc|OJZ83680.1 anig|EHA18834.1 atub|OJI90051.1 awen|OJJ41482.1

Group9866: abra|OJJ68306.1 afis|EAW21032.1 agla|OJJ78583.1 akaw|GAA83168.1 aluc|OJZ83667.1 atub|OJI90038.1 asyd|OJJ52735.1

Group9867: abra|OJJ68310.1 acar|OOF96698.1 akaw|GAA83164.1 aluc|OJZ83663.1 atub|OJI90034.1 acla|EAW12363.1 azon|OJJ42237.1

Group9868: abra|OJJ68314.1 acar|OOF96690.1 akaw|GAA83159.1 aluc|OJZ83659.1 anig|EHA18811.1 atub|OJI90030.1 awen|OJJ30841.1

Group9869: abra|OJJ68315.1 acar|OOF97575.1 akaw|GAA83158.1 aluc|OJZ83658.1 anig|EHA18810.1 atub|OJI90028.1 azon|OJJ43591.1

Group9870: abra|OJJ68401.1 acar|OOF93972.1 akaw|GAA92546.1 aluc|OJZ90095.1 anig|EHA20981.1 atub|OJI84711.1 azon|OJJ49016.1

Group9871: abra|OJJ68429.1 acar|OOF93961.1 akaw|GAA86942.1 aluc|OJZ90067.1 anig|EHA21007.1 atub|OJI84740.1 awen|OJJ35052.1

Group9872: abra|OJJ68435.1 akaw|GAA86936.1 aluc|OJZ90062.1 anid|CBF80381.1 anig|EHA21011.1 atub|OJI84745.1 aver|OJJ05141.1

Group9873: abra|OJJ68581.1 asyd|OJJ52742.1 ater|EAU29645.1 atub|OJI91461.1 aver|OJJ02998.1 awen|OJJ38361.1 anid|CBF78423.1

Group9874: abra|OJJ68583.1 afla|EED54193.1 aory|EIT76123.1 asyd|OJJ52740.1 atub|OJI91459.1 aver|OJJ02997.1 awen|OJJ38358.1

Group9875: afis|EAW25655.1 afla|EED45054.1 agla|OJJ83647.1 anid|CBF78631.1 aory|EIT73702.1 asyd|OJJ65121.1 aver|OJI99249.1

Group9876: abra|OJJ68625.1 acar|OOF92134.1 akaw|GAA92896.1 aluc|OJZ81930.1 anig|EHA22317.1 atub|OJI84483.1 azon|OJJ42332.1

Group9877: abra|OJJ68639.1 acar|OOF92143.1 aluc|OJZ81916.1 anig|EHA22331.1 asyd|OJJ53598.1 atub|OJI84468.1 aver|OJJ08150.1

Group9878: afla|EED48267.1 agla|OJJ87541.1 aory|EIT72939.1 asyd|OJJ60930.1 atub|OJI80380.1 aver|OJJ00256.1 asyd|OJJ53338.1

Group9879: abra|OJJ68835.1 acar|OOF90555.1 akaw|GAA90416.1 aluc|OJZ85814.1 anig|EHA24745.1 atub|OJI79600.1 awen|OJJ35363.1

Group9880: abra|OJJ68838.1 afla|EED52962.1 akaw|GAA90412.1 aluc|OJZ85817.1 anig|EHA24744.1 atub|OJI79603.1 aory|EIT78717.1

Group9881: abra|OJJ68842.1 acar|OOF90547.1 akaw|GAA90406.1 aluc|OJZ85822.1 anig|EHA24739.1 atub|OJI79609.1 azon|OJJ44329.1

Group9882: abra|OJJ68891.1 acar|OOF97646.1 akaw|GAA90365.1 aluc|OJZ85867.1 anig|EHA28086.1 atub|OJI79653.1 awen|OJJ37241.1

Group9883: acla|EAW10335.1 afis|EAW25257.1 afum|EAL89918.1 asyd|OJJ52543.1 aver|OJJ03861.1 awen|OJJ40969.1 azon|OJJ44555.1

Group9884: abra|OJJ68931.1 acar|OOF91514.1 agla|OJJ84589.1 aver|OJJ04413.1 awen|OJJ31462.1 afla|EED50386.1 asyd|OJJ57989.1

Group9885: anid|CBF71418.1 afla|EED50433.1 aory|EIT82063.1 asyd|OJJ54989.1 ater|EAU32830.1 aver|OJJ07212.1 awen|OJJ29997.1

Group9886: abra|OJJ69020.1 akaw|GAA90537.1 aluc|OJZ86006.1 anig|EHA24604.1 atub|OJI79795.1 agla|OJJ85315.1 ater|EAU29900.1

Group9887: afla|EED53342.1 aory|EIT74834.1 akaw|GAA92122.1 aluc|OJZ83600.1 anig|EHA18759.1 atub|OJI89972.1 awen|OJJ35023.1

Group9888: abra|OJJ69059.1 acar|OOF91224.1 akaw|GAA90589.1 aluc|OJZ86047.1 anig|EHA24564.1 atub|OJI79844.1 awen|OJJ38350.1

Group9889: ater|EAU37437.1 afla|EED52297.1 akaw|GAA86600.1 aluc|OJZ85148.1 anig|EHA25387.1 aory|EIT74362.1 atub|OJI80422.1

Group9890: abra|OJJ69441.1 acar|OOF94298.1 acla|EAW11923.1 akaw|GAA88377.1 aluc|OJZ82277.1 anig|EHA28469.1 atub|OJI88162.1

Group9891: abra|OJJ69442.1 acar|OOF94297.1 akaw|GAA88376.1 aluc|OJZ82278.1 anig|EHA28468.1 atub|OJI88161.1 aver|OJI98014.1

Group9892: abra|OJJ69690.1 akaw|GAA88814.1 aluc|OJZ82497.1 anig|EHA28275.1 ater|EAU33152.1 atub|OJI87934.1 akaw|GAA92979.1

Group9893: abra|OJJ69760.1 ater|EAU29408.1 aory|EIT77531.1 akaw|GAA84865.1 aluc|OJZ86806.1 anig|EHA21877.1 atub|OJI83711.1

Group9894: abra|OJJ69793.1 acar|OOF93585.1 akaw|GAA87865.1 aluc|OJZ85506.1 anig|EHA25038.1 atub|OJI80059.1 awen|OJJ32206.1

Group9895: abra|OJJ69807.1 afis|EAW20366.1 afum|EAL85482.1 aluc|OJZ85494.1 ater|EAU29813.1 atub|OJI80071.1 awen|OJJ38363.1

Group9896: abra|OJJ69823.1 acar|OOF93340.1 akaw|GAA87837.1 aluc|OJZ85473.1 anig|EHA25014.1 atub|OJI80090.1 afla|EED56824.1

Group9897: abra|OJJ69826.1 acar|OOF93351.1 akaw|GAA87834.1 aluc|OJZ85470.1 anig|EHA25011.1 ater|EAU32700.1 atub|OJI80093.1

Group9898: anid|CBF70124.1 asyd|OJJ63947.1 aver|OJI96995.1 afla|EED53617.1 aory|EIT75095.1 atub|OJI83154.1 awen|OJJ29953.1

Group9899: abra|OJJ69835.1 akaw|GAA87825.1 aluc|OJZ85461.1 anig|EHA25003.1 atub|OJI80102.1 asyd|OJJ53551.1 aver|OJJ08203.1

Group9900: abra|OJJ69837.1 acar|OOF89895.1 akaw|GAA87823.1 aluc|OJZ85459.1 anig|EHA25001.1 asyd|OJJ53653.1 atub|OJI80104.1

Group9901: afis|EAW19691.1 acla|EAW12051.1 afla|EED48468.1 afum|EAL86835.1 aory|EIT76270.1 asyd|OJJ62961.1 aver|OJI98006.1

Group9902: abra|OJJ69892.1 agla|OJJ86388.1 akaw|GAA87758.1 aluc|OJZ85392.1 anig|EHA25576.1 asyd|OJJ54425.1 atub|OJI80179.1

Group9903: azon|OJJ46029.1 afis|EAW17328.1 afum|EAL91165.1 anid|CBF83207.1 asyd|OJJ55551.1 ater|EAU29241.1 aver|OJJ06628.1

Group9904: abra|OJJ69899.1 acar|OOF98467.1 afla|EED45550.1 aory|EIT75463.1 asyd|OJJ55060.1 ater|EAU37249.1 aver|OJJ07151.1

Group9905: agla|OJJ80662.1 acar|OOF98066.1 afis|EAW22082.1 afum|EAL85691.1 afla|EED49423.1 aory|EIT79507.1 azon|OJJ48138.1

Group9906: abra|OJJ69932.1 akaw|GAA87726.1 aluc|OJZ85354.1 anig|EHA25540.1 atub|OJI80216.1 aver|OJI95750.1 agla|OJJ88378.1

Group9907: aory|EIT73092.1 acar|OOF96082.1 acla|EAW12196.1 afla|EED48624.1 anid|CBF78879.1 asyd|OJJ64594.1 aver|OJI98691.1

Group9908: abra|OJJ70037.1 afis|EAW17095.1 akaw|GAA88364.1 aluc|OJZ82291.1 anig|EHA18734.1 ater|EAU38710.1 atub|OJI88148.1

Group9909: abra|OJJ70038.1 afis|EAW17094.1 aluc|OJZ82292.1 anig|EHA18735.1 ater|EAU38709.1 atub|OJI88147.1 akaw|GAA88363.1

Group9910: ater|EAU32025.1 akaw|GAA86452.1 aluc|OJZ84995.1 afla|EED46132.1 anig|EHA21904.1 afla|EED52089.1 atub|OJI80605.1

Group9911: abra|OJJ70051.1 afla|EED48110.1 aory|EIT76218.1 asyd|OJJ57159.1 ater|EAU37425.1 aver|OJJ03154.1 azon|OJJ42158.1

Group9912: abra|OJJ70059.1 aver|OJJ03870.1 azon|OJJ43583.1 aluc|OJZ83449.1 atub|OJI82341.1 awen|OJJ30957.1 akaw|GAA85521.1

Group9913: abra|OJJ70062.1 acar|OOF93550.1 akaw|GAA87614.1 aluc|OJZ85235.1 anig|EHA25439.1 ater|EAU29812.1 atub|OJI80343.1

Group9914: abra|OJJ70064.1 acar|OOF93552.1 afla|EED45173.1 akaw|GAA87612.1 aluc|OJZ85233.1 anig|EHA25437.1 atub|OJI80345.1

Group9915: azon|OJJ44018.1 acar|OOF95582.1 afla|EED53509.1 anig|EHA18388.1 aory|EIT74905.1 asyd|OJJ57341.1 aver|OJJ05821.1

Group9916: abra|OJJ70084.1 acar|OOF93328.1 akaw|GAA87603.1 aluc|OJZ85223.1 anig|EHA25427.1 atub|OJI80358.1 awen|OJJ32642.1

Group9917: abra|OJJ70089.1 acla|EAW12307.1 afis|EAW21034.1 afum|EAL92072.2 anig|EHA22485.1 atub|OJI89252.1 awen|OJJ39971.1

Group9918: abra|OJJ70104.1 akaw|GAA87596.1 aluc|OJZ85216.1 anig|EHA25419.1 asyd|OJJ58181.1 atub|OJI80369.1 azon|OJJ45181.1

Group9919: abra|OJJ70105.1 acar|OOF95839.1 akaw|GAA87595.1 aluc|OJZ85215.1 anig|EHA25418.1 ater|EAU30871.1 atub|OJI80370.1

Group9920: abra|OJJ70107.1 acar|OOG00112.1 agla|OJJ81742.1 akaw|GAA83854.1 aluc|OJZ90861.1 anig|EHA26921.1 aluc|OJZ79626.1

Group9921: afla|EED47795.1 aory|EIT77102.1 acla|EAW12020.1 anid|CBF81916.1 asyd|OJJ59696.1 aver|OJJ02591.1 awen|OJJ31360.1

Group9922: abra|OJJ70122.1 acar|OOF90917.1 agla|OJJ86726.1 akaw|GAA86647.1 aluc|OJZ85204.1 anig|EHA25409.1 atub|OJI80397.1

Group9923: abra|OJJ70147.1 afla|EED52390.1 akaw|GAA90561.1 anig|EHA25742.1 aory|EIT83292.1 asyd|OJJ53402.1 aver|OJJ04199.1

Group9924: abra|OJJ70160.1 acar|OOF93313.1 akaw|GAA86607.1 aluc|OJZ85157.1 anig|EHA25373.1 atub|OJI80434.1 awen|OJJ32182.1

Group9925: abra|OJJ70170.1 acar|OOG00402.1 akaw|GAA86618.1 aluc|OJZ85171.1 anid|CBF87114.1 atub|OJI80447.1 azon|OJJ47278.1

Group9926: abra|OJJ70196.1 acar|OOF90058.1 akaw|GAA86580.1 aluc|OJZ85131.1 anig|EHA25350.1 atub|OJI80462.1 awen|OJJ40107.1

Group9927: abra|OJJ70202.1 afis|EAW25730.1 aluc|OJZ85125.1 anig|EHA25344.1 ater|EAU30246.1 atub|OJI80468.1 awen|OJJ32208.1

Group9928: abra|OJJ70227.1 afla|EED52373.1 akaw|GAA93009.1 aluc|OJZ82576.1 anid|CBF73532.1 asyd|OJJ56723.1 aver|OJJ06002.1

Group9929: abra|OJJ70229.1 agla|OJJ88090.1 akaw|GAA86553.1 aluc|OJZ85101.1 anig|EHA25323.1 atub|OJI80496.1 awen|OJJ39209.1

Group9930: acla|EAW07216.1 afis|EAW19605.1 afla|EED48181.1 asyd|OJJ57100.1 ater|EAU36003.1 aver|OJJ08823.1 awen|OJJ37374.1

Group9931: abra|OJJ70281.1 acar|OOF91733.1 akaw|GAA92867.1 aluc|OJZ82546.1 asyd|OJJ62800.1 atub|OJI88788.1 aver|OJI96847.1

Group9932: aory|EIT81581.1 acla|EAW13266.1 afis|EAW17362.1 afum|EAL91199.1 asyd|OJJ57191.1 aver|OJJ03100.1 awen|OJJ29649.1

Group9933: abra|OJJ70404.1 afis|EAW19308.1 akaw|GAA91950.1 aluc|OJZ82668.1 atub|OJI88901.1 aver|OJJ03245.1 azon|OJJ42500.1

Group9934: abra|OJJ70407.1 afla|EED53381.1 akaw|GAA91947.1 aluc|OJZ82671.1 anig|EHA21038.1 aory|EIT75053.1 atub|OJI88907.1

Group9935: abra|OJJ70433.1 akaw|GAA90146.1 aluc|OJZ83201.1 anig|EHA21063.1 atub|OJI87736.1 aver|OJI96790.1 awen|OJJ41821.1

Group9936: abra|OJJ70465.1 acar|OOF97460.1 akaw|GAA83314.1 aluc|OJZ83808.1 atub|OJI90175.1 awen|OJJ37226.1 azon|OJJ50929.1

Group9937: acar|OOF97422.1 acla|EAW08171.1 afis|EAW23207.1 afum|EBA27342.1 akaw|GAA83352.1 aory|EIT72570.1 awen|OJJ39439.1

Group9938: abra|OJJ70663.1 afla|EED51846.1 akaw|GAA89999.1 aluc|OJZ84013.1 anig|EHA19136.1 atub|OJI90383.1 aory|EIT81693.1

Group9939: abra|OJJ70816.1 akaw|GAA91696.1 aluc|OJZ87696.1 anig|EHA20884.1 atub|OJI80946.1 azon|OJJ42050.1 ater|EAU37986.1

Group9940: abra|OJJ71041.1 acar|OOF97196.1 afla|EED56868.1 akaw|GAA82915.1 aluc|OJZ87487.1 aory|EIT77521.1 atub|OJI81161.1

Group9941: abra|OJJ71089.1 acar|OOF98048.1 akaw|GAA82872.1 aluc|OJZ87439.1 anig|EHA20664.1 atub|OJI81209.1 aver|OJI99591.1

Group9942: abra|OJJ71092.1 acar|OOF98046.1 akaw|GAA82869.1 aluc|OJZ87436.1 anig|EHA20662.1 atub|OJI81212.1 azon|OJJ43494.1

Group9943: abra|OJJ71109.1 acar|OOF98040.1 akaw|GAA82860.1 aluc|OJZ87426.1 anig|EHA20648.1 atub|OJI81223.1 azon|OJJ45898.1

Group9944: abra|OJJ71202.1 afis|EAW16710.1 akaw|GAA82782.1 aluc|OJZ87341.1 anig|EHA20578.1 atub|OJI81303.1 awen|OJJ38292.1

Group9945: abra|OJJ71238.1 akaw|GAA82750.1 aluc|OJZ87307.1 anig|EHA20543.1 atub|OJI81338.1 aver|OJI97600.1 agla|OJJ83108.1

Group9946: abra|OJJ71412.1 anid|CBF80562.1 anig|EHA19901.1 asyd|OJJ55495.1 ater|EAU30134.1 aver|OJJ06573.1 awen|OJJ39982.1

Group9947: afla|EED57198.1 awen|OJJ32024.1 afis|EAW20151.1 afum|EAL85266.1 ater|EAU38766.1 acla|EAW15181.1 azon|OJJ46066.1

Group9948: abra|OJJ71710.1 acar|OOF96416.1 akaw|GAA85178.1 aluc|OJZ86455.1 anig|EHA21452.1 atub|OJI84072.1 azon|OJJ46023.1

Group9949: abra|OJJ71746.1 acar|OOF96437.1 afla|EED49269.1 akaw|GAA85160.1 aluc|OJZ86477.1 aory|EIT82212.1 atub|OJI84053.1

Group9950: anid|CBF80176.1 acar|OOF92901.1 afla|EED51173.1 anig|EHA25376.1 aory|EIT81316.1 agla|OJJ88683.1 aver|OJI99526.1

Group9951: abra|OJJ71762.1 acar|OOF90731.1 afis|EAW20638.1 afla|EED45242.1 anid|CBF82414.1 aory|EIT73703.1 acla|EAW07348.1

Group9952: abra|OJJ71765.1 afis|EAW24474.1 afla|EED48647.1 anig|EHA22153.1 aory|EIT73075.1 asyd|OJJ52754.1 aver|OJJ07374.1

Group9953: abra|OJJ71766.1 acar|OOF90206.1 akaw|GAA85141.1 aluc|OJZ86496.1 anig|EHA22148.1 atub|OJI84030.1 azon|OJJ46960.1

Group9954: abra|OJJ71769.1 acar|OOF96442.1 akaw|GAA85138.1 aluc|OJZ86499.1 anig|EHA22145.1 atub|OJI84027.1 azon|OJJ44477.1

Group9955: ater|EAU34853.1 awen|OJJ31123.1 acla|EAW13851.1 afis|EAW18565.1 afum|EAL93267.2 agla|OJJ81614.1 azon|OJJ46604.1

Group9956: abra|OJJ71825.1 acar|OOF96502.1 akaw|GAA85085.1 aluc|OJZ86557.1 anig|EHA22092.1 ater|EAU36063.1 atub|OJI83965.1

Group9957: abra|OJJ71830.1 acar|OOF96506.1 akaw|GAA85081.1 aluc|OJZ86562.1 ater|EAU34870.1 atub|OJI83959.1 awen|OJJ31136.1

Group9958: abra|OJJ71915.1 acar|OOF92082.1 acla|EAW13304.1 atub|OJI89559.1 awen|OJJ31867.1 acar|OOF92446.1 azon|OJJ42304.1

Group9959: abra|OJJ71933.1 agla|OJJ79132.1 anid|CBF80476.1 asyd|OJJ54716.1 aver|OJJ07451.1 awen|OJJ37071.1 azon|OJJ50307.1

Group9960: abra|OJJ71939.1 aluc|OJZ85540.1 anig|EHA24968.1 ater|EAU35754.1 atub|OJI89782.1 aver|OJJ06472.1 asyd|OJJ65144.1

Group9961: abra|OJJ72030.1 agla|OJJ79437.1 akaw|GAA84901.1 aluc|OJZ86768.1 anig|EHA21911.1 atub|OJI83760.1 asyd|OJJ56786.1

Group9962: abra|OJJ72165.1 akaw|GAA84763.1 aluc|OJZ86920.1 anig|EHA21724.1 atub|OJI83601.1 awen|OJJ29681.1 acar|OOF96798.1

Group9963: abra|OJJ72166.1 acar|OOF96799.1 akaw|GAA84762.1 aluc|OJZ86921.1 anig|EHA21723.1 atub|OJI83600.1 azon|OJJ43589.1

Group9964: abra|OJJ72244.1 acar|OOF96870.1 akaw|GAA84695.1 aluc|OJZ86992.1 anig|EHA21653.1 atub|OJI83527.1 azon|OJJ46216.1

Group9965: abra|OJJ72271.1 afis|EAW20689.1 akaw|GAA86026.1 aluc|OJZ90253.1 atub|OJI81679.1 acla|EAW06361.1 awen|OJJ30154.1

Group9966: abra|OJJ72306.1 akaw|GAA86057.1 aluc|OJZ90287.1 anig|EHA23777.1 ater|EAU29587.1 atub|OJI81711.1 awen|OJJ30540.1

Group9967: abra|OJJ72310.1 aluc|OJZ90292.1 anid|CBF89252.1 anig|EHA23781.1 asyd|OJJ65158.1 ater|EAU39466.1 atub|OJI81716.1

Group9968: abra|OJJ72311.1 acar|OOF90706.1 agla|OJJ86229.1 aluc|OJZ90293.1 anig|EHA23782.1 atub|OJI81717.1 aver|OJJ06473.1

Group9969: abra|OJJ72361.1 acar|OOF93349.1 afla|EED57480.1 akaw|GAA86111.1 aluc|OJZ90343.1 aory|EIT80169.1 atub|OJI81770.1

Group9970: abra|OJJ72372.1 acar|OOF94854.1 akaw|GAA86122.1 aluc|OJZ90361.1 anig|EHA23830.1 atub|OJI81788.1 anig|EHA23825.1

Group9971: abra|OJJ72524.1 acar|OOF98685.1 agla|OJJ83808.1 akaw|GAA86248.1 aluc|OJZ90503.1 anig|EHA23961.1 atub|OJI81927.1

Group9972: abra|OJJ72532.1 acar|OOF94721.1 akaw|GAA86254.1 aluc|OJZ90511.1 anid|CBF85173.1 anig|EHA23968.1 atub|OJI81936.1

Group9973: abra|OJJ72537.1 aluc|OJZ90518.1 anig|EHA23973.1 atub|OJI81942.1 abra|OJJ77424.1 acar|OOF94714.1 aver|OJI99459.1

Group9974: abra|OJJ72616.1 afis|EAW15946.1 akaw|GAA92009.1 aluc|OJZ92874.1 anid|CBF81448.1 anig|EHA26085.1 atub|OJI87334.1

Group9975: abra|OJJ72709.1 afum|EAL85031.1 anig|EHA24107.1 atub|OJI82104.1 awen|OJJ41893.1 azon|OJJ42594.1 afis|EAW19865.1

Group9976: abra|OJJ72870.1 aluc|OJZ83383.1 anig|EHA24242.1 asyd|OJJ65315.1 atub|OJI82270.1 afum|EAL89186.1 agla|OJJ85840.1

Group9977: abra|OJJ73068.1 acar|OOG00303.1 akaw|GAA88092.1 aluc|OJZ81357.1 anig|EHA18657.1 atub|OJI91561.1 acla|EAW12432.1

Group9978: abra|OJJ73080.1 acar|OOF91965.1 akaw|GAA88082.1 aluc|OJZ81346.1 anig|EHA18649.1 atub|OJI91550.1 azon|OJJ42463.1

Group9979: abra|OJJ73082.1 acar|OOG00316.1 akaw|GAA88078.1 aluc|OJZ81341.1 anig|EHA18647.1 ater|EAU38109.1 atub|OJI91546.1

Group9980: abra|OJJ73087.1 afla|EED48882.1 akaw|GAA88073.1 aluc|OJZ81338.1 anig|EHA18642.1 aory|EIT81225.1 atub|OJI91541.1

Group9981: asyd|OJJ55509.1 acar|OOF97221.1 akaw|GAA83709.1 aluc|OJZ91026.1 ater|EAU32780.1 atub|OJI83017.1 awen|OJJ38712.1

Group9982: abra|OJJ73153.1 acar|OOG00365.1 akaw|GAA88019.1 aluc|OJZ81277.1 anig|EHA18577.1 atub|OJI91485.1 awen|OJJ29892.1

Group9983: abra|OJJ73301.1 acar|OOF91869.1 akaw|GAA92704.1 aluc|OJZ88744.1 anig|EHA18448.1 atub|OJI91316.1 azon|OJJ50500.1

Group9984: abra|OJJ73310.1 acar|OOF90736.1 akaw|GAA92812.1 aluc|OJZ88735.1 anig|EHA18438.1 ater|EAU30392.1 atub|OJI91311.1

Group9985: abra|OJJ73332.1 acar|OOF91493.1 agla|OJJ81113.1 akaw|GAA92789.1 anig|EHA18399.1 aory|EIT74059.1 afla|EED57240.1

Group9986: abra|OJJ73477.1 acar|OOF92543.1 acla|EAW12325.1 akaw|GAA83024.1 aluc|OJZ88570.1 anig|EHA18267.1 atub|OJI91156.1

Group9987: akaw|GAA87461.1 anid|CBF88894.1 acla|EAW07446.1 afis|EAW22473.1 afla|EED54505.1 afum|EAL90718.1 aory|EIT78749.1

Group9988: abra|OJJ73533.1 acar|OOF99640.1 agla|OJJ87023.1 akaw|GAA83963.1 aluc|OJZ80851.1 anig|EHA21808.1 atub|OJI82717.1

Group9989: abra|OJJ73540.1 acar|OOF92654.1 agla|OJJ85033.1 akaw|GAA88611.1 aluc|OJZ88519.1 anig|EHA21862.1 atub|OJI91104.1

Group9990: abra|OJJ73705.1 acar|OOG00869.1 akaw|GAA88458.1 aluc|OJZ88346.1 anig|EHA18079.1 ater|EAU30713.1 atub|OJI90942.1

Group9991: abra|OJJ73711.1 acla|EAW12409.1 akaw|GAA88452.1 aluc|OJZ88339.1 anig|EHA18074.1 atub|OJI90935.1 azon|OJJ50261.1

Group9992: abra|OJJ73715.1 acar|OOG00875.1 akaw|GAA88448.1 aluc|OJZ88335.1 anig|EHA18071.1 atub|OJI90931.1 azon|OJJ45232.1

Group9993: aory|EIT72286.1 acla|EAW07852.1 afis|EAW22888.1 afum|EAL90310.1 akaw|GAA89584.1 anid|CBF87796.1 awen|OJJ38834.1

Group9994: abra|OJJ74004.1 acar|OOG01125.1 akaw|GAA87485.1 aluc|OJZ88040.1 anig|EHA17813.1 atub|OJI90647.1 agla|OJJ83134.1

Group9995: abra|OJJ74013.1 acar|OOG01136.1 akaw|GAA87496.1 aluc|OJZ88026.1 anig|EHA17803.1 atub|OJI90634.1 aver|OJI96477.1

Group9996: acar|OOF90011.1 acla|EAW09033.1 afum|EAL90460.1 agla|OJJ78519.1 aory|EIT76954.1 afis|EAW22735.1 afla|EED51047.1

Group9997: aver|OJI95818.1 azon|OJJ42327.1 asyd|OJJ61723.1 afla|EED56597.1 aory|EIT79615.1 awen|OJJ30239.1 agla|OJJ80956.1

Group9998: abra|OJJ74120.1 afla|EED54374.1 agla|OJJ84790.1 akaw|GAA87149.1 aluc|OJZ84288.1 anig|EHA19566.1 aory|EIT75859.1

Group9999: abra|OJJ74133.1 acar|OOF95600.1 akaw|GAA89753.1 aluc|OJZ80630.1 anig|EHA27201.1 atub|OJI87486.1 azon|OJJ42575.1

Group10000: abra|OJJ74163.1 acar|OOF95559.1 akaw|GAA86853.1 aluc|OJZ89973.1 anig|EHA27170.1 atub|OJI84834.1 azon|OJJ50833.1

Group10001: abra|OJJ74184.1 acar|OOF95536.1 akaw|GAA86833.1 aluc|OJZ89952.1 anig|EHA27153.1 atub|OJI84854.1 azon|OJJ42445.1

Group10002: abra|OJJ74258.1 acar|OOF95470.1 agla|OJJ88793.1 akaw|GAA86763.1 aluc|OJZ89873.1 anig|EHA27073.1 atub|OJI84934.1

Group10003: abra|OJJ74303.1 atub|OJI84979.1 acar|OOF95427.1 afla|EED57788.1 akaw|GAA86722.1 aluc|OJZ89827.1 anid|CBF88720.1

Group10004: abra|OJJ74422.1 acar|OOF95318.1 anig|EHA26936.1 atub|OJI85099.1 afis|EAW19687.1 asyd|OJJ65222.1 awen|OJJ34663.1

Group10005: abra|OJJ74441.1 acar|OOF95305.1 afla|EED45620.1 akaw|GAA87334.1 aluc|OJZ89680.1 anig|EHA26916.1 atub|OJI85120.1

Group10006: abra|OJJ74442.1 acar|OOF95304.1 akaw|GAA87335.1 aluc|OJZ89679.1 anig|EHA26915.1 atub|OJI85121.1 acla|EAW12161.1

Group10007: afla|EED49714.1 aory|EIT77662.1 akaw|GAA84674.1 aluc|OJZ87015.1 asyd|OJJ53590.1 atub|OJI83504.1 aver|OJJ08158.1

Group10008: abra|OJJ74540.1 afla|EED51454.1 afum|EAL86342.1 akaw|GAA82062.1 aluc|OJZ92120.1 atub|OJI86553.1 aluc|OJZ79684.1

Group10009: ater|EAU33340.1 afis|EAW20286.1 akaw|GAA82048.1 aluc|OJZ92132.1 atub|OJI86564.1 awen|OJJ37823.1 azon|OJJ42240.1

Group10010: abra|OJJ74653.1 acar|OOG00867.1 akaw|GAA81957.1 aluc|OJZ92232.1 atub|OJI86662.1 awen|OJJ34777.1 afis|EAW22068.1

Group10011: abra|OJJ74691.1 afla|EED47158.1 aluc|OJZ92268.1 aory|EIT79287.1 ater|EAU38045.1 atub|OJI86702.1 aver|OJJ03743.1

Group10012: afla|EED46968.1 awen|OJJ40483.1 azon|OJJ49345.1 acla|EAW10676.1 afum|EAL89543.1 agla|OJJ82114.1 anid|CBF75627.1

Group10013: abra|OJJ74793.1 acar|OOF95784.1 akaw|GAA83620.1 aluc|OJZ92373.1 anig|EHA26600.1 atub|OJI86803.1 aver|OJJ02008.1

Group10014: abra|OJJ74816.1 acla|EAW12267.1 akaw|GAA83599.1 aluc|OJZ92395.1 anig|EHA26579.1 aver|OJJ05789.1 azon|OJJ42191.1

Group10015: abra|OJJ74818.1 acar|OOF95806.1 akaw|GAA83597.1 aluc|OJZ92397.1 anig|EHA26568.1 ater|EAU32633.1 atub|OJI86827.1

Group10016: abra|OJJ74825.1 acar|OOF99295.1 aluc|OJZ92400.1 atub|OJI86829.1 abra|OJJ74826.1 anig|EHA26565.1 aluc|OJZ92401.1

Group10017: abra|OJJ74934.1 acar|OOF95925.1 akaw|GAA83486.1 aluc|OJZ92507.1 anig|EHA26466.1 atub|OJI86940.1 aluc|OJZ80309.1

Group10018: abra|OJJ74979.1 acar|OOF97341.1 akaw|GAA83453.1 aluc|OJZ92543.1 anig|EHA26433.1 atub|OJI86979.1 awen|OJJ39761.1

Group10019: abra|OJJ74982.1 acar|OOF97337.1 akaw|GAA83450.1 aluc|OJZ92546.1 anig|EHA26430.1 atub|OJI86982.1 agla|OJJ86678.1

Group10020: abra|OJJ75117.1 acar|OOF97215.1 acla|EAW06578.1 akaw|GAA88727.1 aluc|OJZ92684.1 atub|OJI87112.1 azon|OJJ43664.1

Group10021: abra|OJJ75220.1 agla|OJJ83396.1 aluc|OJZ84409.1 asyd|OJJ55427.1 atub|OJI89662.1 aver|OJJ06506.1 akaw|GAA87038.1

Group10022: abra|OJJ75252.1 acar|OOF89847.1 akaw|GAA91791.1 aluc|OJZ92811.1 anig|EHA26197.1 atub|OJI87268.1 azon|OJJ44475.1

Group10023: abra|OJJ75319.1 acar|OOF94507.1 akaw|GAA92014.1 aluc|OJZ92880.1 anig|EHA26080.1 atub|OJI87339.1 azon|OJJ42604.1

Group10024: asyd|OJJ54676.1 afum|EAL86852.2 anid|CBF80541.1 awen|OJJ31938.1 afla|EED48443.1 aver|OJJ07491.1 azon|OJJ46598.1

Group10025: abra|OJJ75336.1 acar|OOF94494.1 akaw|GAA92030.1 aluc|OJZ92900.1 anig|EHA26063.1 atub|OJI87358.1 awen|OJJ36997.1

Group10026: abra|OJJ75344.1 acar|OOF94491.1 agla|OJJ84455.1 akaw|GAA92037.1 aluc|OJZ92906.1 anig|EHA26057.1 atub|OJI87366.1

Group10027: abra|OJJ75351.1 acar|OOF94483.1 akaw|GAA92043.1 aluc|OJZ92913.1 anig|EHA26052.1 atub|OJI87373.1 azon|OJJ45814.1

Group10028: abra|OJJ75353.1 acar|OOF94485.1 akaw|GAA92046.1 aluc|OJZ92915.1 anig|EHA26050.1 atub|OJI87375.1 awen|OJJ38721.1

Group10029: abra|OJJ75359.1 acar|OOF94479.1 akaw|GAA92054.1 aluc|OJZ92925.1 anig|EHA26044.1 atub|OJI87386.1 azon|OJJ43200.1

Group10030: abra|OJJ75362.1 acar|OOF89871.1 akaw|GAA92058.1 aluc|OJZ92931.1 anig|EHA26040.1 atub|OJI87391.1 azon|OJJ45869.1

Group10031: abra|OJJ75371.1 acar|OOF89962.1 akaw|GAA92064.1 aluc|OJZ92939.1 anig|EHA26033.1 atub|OJI87399.1 awen|OJJ31304.1

Group10032: abra|OJJ75424.1 akaw|GAA84199.1 aluc|OJZ88876.1 anig|EHA23629.1 ater|EAU33728.1 atub|OJI85915.1 awen|OJJ41697.1

Group10033: abra|OJJ75533.1 afla|EED46911.1 akaw|GAA91953.1 aluc|OJZ82665.1 anig|EHA27439.1 aory|EIT72640.1 atub|OJI88898.1

Group10034: abra|OJJ75536.1 akaw|GAA91956.1 aluc|OJZ82662.1 anig|EHA27436.1 atub|OJI88895.1 aver|OJI99010.1 azon|OJJ46514.1

Group10035: abra|OJJ75540.1 acar|OOF90859.1 akaw|GAA84294.1 aluc|OJZ88988.1 anig|EHA23538.1 atub|OJI85800.1 azon|OJJ43916.1

Group10036: abra|OJJ75556.1 acar|OOF90587.1 akaw|GAA84311.1 aluc|OJZ89006.1 anig|EHA23522.1 atub|OJI85783.1 azon|OJJ42073.1

Group10037: abra|OJJ75588.1 afla|EED55987.1 akaw|GAA84339.1 aluc|OJZ89039.1 anig|EHA23496.1 aory|EIT76606.1 atub|OJI85751.1

Group10038: afis|EAW19006.1 afum|EAL93740.1 awen|OJJ32644.1 acar|OOF93329.1 akaw|GAA90362.1 aluc|OJZ85870.1 atub|OJI79656.1

Group10039: afis|EAW19004.1 afum|EAL93738.1 anid|CBF82894.1 awen|OJJ32645.1 akaw|GAA90364.1 aluc|OJZ85868.1 atub|OJI79654.1

Group10040: abra|OJJ75765.1 afla|EED54893.1 aory|EIT79407.1 asyd|OJJ58671.1 atub|OJI87193.1 aver|OJJ01282.1 awen|OJJ30929.1

Group10041: abra|OJJ75926.1 acar|OOF99982.1 afla|EED46844.1 akaw|GAA84629.1 aluc|OJZ89375.1 anig|EHA23208.1 atub|OJI85423.1

Group10042: abra|OJJ76015.1 acar|OOF99910.1 akaw|GAA88945.1 aluc|OJZ89461.1 anig|EHA23127.1 atub|OJI85333.1 azon|OJJ50329.1

Group10043: abra|OJJ76020.1 acar|OOF99905.1 akaw|GAA88940.1 aluc|OJZ89467.1 anig|EHA23121.1 atub|OJI85328.1 aver|OJI99554.1

Group10044: abra|OJJ76055.1 afla|EED48993.1 anid|CBF86023.1 anig|EHA23089.1 asyd|OJJ64454.1 aver|OJI98557.1 awen|OJJ30147.1

Group10045: abra|OJJ76279.1 afla|EED48939.1 akaw|GAA82211.1 aluc|OJZ91941.1 anig|EHA22900.1 aory|EIT80029.1 atub|OJI86384.1

Group10046: abra|OJJ76470.1 acar|OOF96464.1 afla|EED47273.1 akaw|GAA82382.1 aluc|OJZ91739.1 aver|OJJ01279.1 awen|OJJ38936.1

Group10047: abra|OJJ76595.1 akaw|GAA82499.1 atub|OJI86064.1 acar|OOF90606.1 afla|EED53599.1 aory|EIT75174.1 anig|EHA22640.1

Group10048: afis|EAW19360.1 afum|EAL94080.1 acla|EAW14625.1 afla|EED49735.1 aory|EIT77607.1 asyd|OJJ62358.1 azon|OJJ45913.1

Group10049: abra|OJJ76654.1 acar|OOF99344.1 akaw|GAA92994.1 aluc|OJZ91551.1 anig|EHA22586.1 ater|EAU37780.1 atub|OJI86004.1

Group10050: abra|OJJ76657.1 acar|OOF99347.1 akaw|GAA92997.1 aluc|OJZ91548.1 anig|EHA22583.1 ater|EAU37783.1 atub|OJI86001.1

Group10051: ater|EAU32745.1 acla|EAW15034.1 afis|EAW22730.1 afum|EAL90465.1 anid|CBF69520.1 asyd|OJJ59207.1 aver|OJJ01818.1

Group10052: abra|OJJ76698.1 agla|OJJ78475.1 aory|EIT73159.1 azon|OJJ45928.1 anid|CBF87458.1 asyd|OJJ53409.1 aver|OJJ08342.1

Group10053: abra|OJJ76703.1 acar|OOF89910.1 akaw|GAA92496.1 aluc|OJZ91508.1 anig|EHA22547.1 atub|OJI85960.1 azon|OJJ51652.1

Group10054: abra|OJJ76728.1 acar|OOF90684.1 anig|EHA24276.1 abra|OJJ77450.1 afla|EED47460.1 aory|EIT79958.1 ater|EAU33901.1

Group10055: abra|OJJ76741.1 acar|OOF94681.1 afla|EED46914.1 akaw|GAA83302.1 anig|EHA24960.1 aory|EIT72665.1 azon|OJJ42574.1

Group10056: abra|OJJ76864.1 afla|EED47869.1 anig|EHA27616.1 aory|EIT77205.1 ater|EAU34267.1 atub|OJI89809.1 awen|OJJ38127.1

Group10057: abra|OJJ76865.1 afla|EED47868.1 anig|EHA27617.1 aory|EIT77216.1 ater|EAU34266.1 atub|OJI89808.1 awen|OJJ38128.1

Group10058: abra|OJJ76893.1 acar|OOF97193.1 agla|OJJ85292.1 anig|EHA25831.1 ater|EAU30174.1 awen|OJJ31279.1 azon|OJJ43581.1

Group10059: acla|EAW07063.1 afis|EAW16694.1 afla|EED45072.1 afum|EAL89292.1 aory|EIT73749.1 ater|EAU34151.1 azon|OJJ45051.1

Group10060: abra|OJJ76921.1 acar|OOF91368.1 akaw|GAA87117.1 aluc|OJZ84321.1 anig|EHA27656.1 atub|OJI89747.1 awen|OJJ33257.1

Group10061: abra|OJJ77023.1 afla|EED52861.1 akaw|GAA87029.1 aluc|OJZ84423.1 anig|EHA27745.1 aory|EIT78595.1 atub|OJI89650.1

Group10062: abra|OJJ77026.1 acar|OOF91464.1 akaw|GAA87025.1 aluc|OJZ84428.1 anig|EHA27747.1 atub|OJI89646.1 awen|OJJ30416.1

Group10063: abra|OJJ77034.1 acar|OOF91473.1 akaw|GAA87017.1 aluc|OJZ84437.1 anig|EHA27756.1 atub|OJI89636.1 ater|EAU29367.1

Group10064: abra|OJJ77053.1 acar|OOF92942.1 akaw|GAA86999.1 aluc|OJZ84458.1 anig|EHA27773.1 atub|OJI89617.1 awen|OJJ29905.1

Group10065: abra|OJJ77059.1 afla|EED52210.1 akaw|GAA86993.1 aluc|OJZ84466.1 anig|EHA27778.1 aory|EIT74394.1 atub|OJI89610.1

Group10066: agla|OJJ87921.1 afis|EAW19748.1 afla|EED45936.1 afum|EAL92049.1 aory|EIT75380.1 ater|EAU32158.1 awen|OJJ34722.1

Group10067: abra|OJJ77168.1 acar|OOF93059.1 acla|EAW12895.1 agla|OJJ81694.1 aluc|OJZ84581.1 anig|EHA27874.1 atub|OJI89499.1

Group10068: abra|OJJ77172.1 acar|OOF93061.1 acla|EAW13647.1 akaw|GAA90761.1 aluc|OJZ84585.1 anig|EHA27878.1 atub|OJI89495.1

Group10069: anig|EHA27454.1 atub|OJI89788.1 acla|EAW09119.1 asyd|OJJ54982.1 aory|EIT78998.1 aver|OJJ03925.1 afla|EED50917.1

Group10070: abra|OJJ77392.1 acar|OOF93260.1 akaw|GAA91369.1 aluc|OJZ84800.1 anig|EHA28066.1 atub|OJI89282.1 azon|OJJ50265.1

Group10071: abra|OJJ77415.1 acar|OOF98448.1 agla|OJJ78621.1 akaw|GAA91350.1 aluc|OJZ84824.1 atub|OJI89263.1 awen|OJJ41822.1

Group10072: abra|OJJ77456.1 afla|EED49594.1 anig|EHA19290.1 aory|EIT82609.1 asyd|OJJ56404.1 ater|EAU39561.1 aver|OJJ06308.1

Group10073: abra|OJJ77542.1 acar|OOF99182.1 akaw|GAA84080.1 aluc|OJZ80994.1 anig|EHA19367.1 ater|EAU38456.1 atub|OJI82590.1

Group10074: anid|CBF71420.1 azon|OJJ46047.1 afla|EED56792.1 asyd|OJJ54991.1 ater|EAU32728.1 aver|OJJ07210.1 awen|OJJ31992.1

Group10075: abra|OJJ77544.1 acar|OOF99181.1 akaw|GAA84079.1 aluc|OJZ80993.1 anig|EHA19368.1 ater|EAU38455.1 atub|OJI82591.1

Group10076: anid|CBF75958.1 afla|EED45676.1 aory|EIT75553.1 afla|EED52107.1 agla|OJJ78530.1 aory|EIT76907.1 ater|EAU36996.1

Group10077: abra|OJJ77719.1 acla|EAW08570.1 akaw|GAA83858.1 aluc|OJZ90846.1 aory|EIT77655.1 ater|EAU32756.1 atub|OJI82839.1

Group10078: abra|OJJ77730.1 acar|OOF99000.1 akaw|GAA83907.1 aluc|OJZ90790.1 anig|EHA19523.1 atub|OJI82778.1 awen|OJJ29956.1

Group10079: abra|OJJ77793.1 acar|OOF90717.1 agla|OJJ83012.1 asyd|OJJ58565.1 aver|OJJ02064.1 awen|OJJ38010.1 aver|OJJ01172.1

Group10080: abra|OJJ77814.1 acar|OOG01054.1 afis|EAW16953.1 aluc|OJZ88121.1 asyd|OJJ59176.1 aver|OJJ01795.1 awen|OJJ38388.1

Group10081: abra|OJJ77848.1 acar|OOF98924.1 acla|EAW14866.1 akaw|GAA83799.1 aluc|OJZ90923.1 anig|EHA19635.1 atub|OJI82915.1

Group10082: abra|OJJ77871.1 acar|OOF98900.1 akaw|GAA83779.1 aluc|OJZ90945.1 anig|EHA19653.1 atub|OJI82937.1 awen|OJJ30237.1

Group10083: abra|OJJ77950.1 afla|EED49859.1 aluc|OJZ91021.1 anig|EHA19721.1 aory|EIT83403.1 ater|EAU37436.1 atub|OJI83013.1

Group10084: abra|OJJ77970.1 acar|OOF98845.1 akaw|GAA83692.1 aluc|OJZ91040.1 anid|CBF85456.1 anig|EHA19739.1 atub|OJI83036.1

Group10085: abra|OJJ77987.1 akaw|GAA85993.1 aluc|OJZ91062.1 anig|EHA19766.1 atub|OJI83058.1 awen|OJJ33715.1 aory|EIT75619.1

Group10086: azon|OJJ43034.1 afis|EAW23758.1 afla|EED45475.1 aory|EIT75757.1 asyd|OJJ64547.1 ater|EAU37841.1 aver|OJI98643.1

Group10087: abra|OJJ78036.1 anig|EHA25166.1 afla|EED45652.1 atub|OJI80648.1 afla|EED52845.1 aory|EIT78592.1 ater|EAU33569.1

Group10088: abra|OJJ78046.1 akaw|GAA85942.1 aluc|OJZ91120.1 anid|CBF76243.1 anig|EHA19818.1 atub|OJI83124.1 awen|OJJ31401.1

Group10089: abra|OJJ78070.1 afis|EAW23479.1 anig|EHA19842.1 atub|OJI83159.1 aver|OJJ05976.1 acar|OOF92824.1 agla|OJJ82952.1

Group10090: abra|OJJ78096.1 afla|EED46926.1 akaw|GAA84547.1 aluc|OJZ81970.1 anig|EHA26143.1 aory|EIT72661.1 atub|OJI79698.1

Group10091: abra|OJJ78135.1 afla|EED47411.1 aluc|OJZ82085.1 aver|OJJ00303.1 awen|OJJ29926.1 afla|EED47410.1 aluc|OJZ82084.1

Group10092: abra|OJJ78136.1 afla|EED53386.1 aluc|OJZ82086.1 aory|EIT75084.1 asyd|OJJ52736.1 aver|OJJ02990.1 awen|OJJ29927.1

Group10093: abra|OJJ78157.1 afla|EED48913.1 akaw|GAA85868.1 aluc|OJZ91211.1 anig|EHA19916.1 aory|EIT81246.1 atub|OJI83229.1

Group10094: abra|OJJ78175.1 acar|OOF90380.1 aluc|OJZ91224.1 anig|EHA19930.1 asyd|OJJ57209.1 aver|OJJ03080.1 aory|EIT73676.1

Group10095: abra|OJJ78194.1 acar|OOF98663.1 akaw|GAA85835.1 aluc|OJZ91265.1 anid|CBF82953.1 anig|EHA19953.1 atub|OJI83254.1

Group10096: abra|OJJ78205.1 acar|OOF98649.1 akaw|GAA85822.1 aluc|OJZ91277.1 anig|EHA19963.1 atub|OJI83266.1 awen|OJJ29567.1

Group10097: abra|OJJ78311.1 acar|OOF90330.1 afis|EAW17201.1 afla|EED56925.1 afum|EAL86628.1 anig|EHA20062.1 ater|EAU29640.1

Group10098: abra|OJJ78314.1 akaw|GAA85716.1 aluc|OJZ91387.1 anig|EHA20065.1 asyd|OJJ56083.1 atub|OJI83380.1 aver|OJJ05026.1

Group10099: abra|OJJ78341.1 acar|OOF98539.1 akaw|GAA85692.1 aluc|OJZ91413.1 anig|EHA20086.1 ater|EAU31065.1 atub|OJI83407.1

Group10100: abra|OJJ78342.1 acar|OOF98538.1 agla|OJJ84605.1 akaw|GAA85691.1 aluc|OJZ91414.1 anig|EHA20087.1 atub|OJI83408.1

Group10101: afum|EAL86784.2 afis|EAW19742.1 afla|EED56524.1 agla|OJJ82488.1 aory|EIT82921.1 asyd|OJJ60342.1 aver|OJJ07300.1

Group10102: acar|OOF89843.1 acla|EAW10859.1 aory|EIT75121.1 ater|EAU29341.1 awen|OJJ36180.1 azon|OJJ44011.1 awen|OJJ40844.1

Group10103: acar|OOF90105.1 agla|OJJ84646.1 akaw|GAA87357.1 aluc|OJZ89657.1 anig|EHA26894.1 atub|OJI85144.1 awen|OJJ36224.1

Group10104: aver|OJJ01972.1 acla|EAW14595.1 afis|EAW19324.1 afla|EED57236.1 afum|EAL94042.1 asyd|OJJ54477.1 aory|EIT74119.1

Group10105: afla|EED44800.1 agla|OJJ84850.1 awen|OJJ33162.1 acla|EAW13354.1 afis|EAW20468.1 afum|EAL84584.1 ater|EAU37695.1

Group10106: acar|OOF91586.1 afla|EED45784.1 agla|OJJ85380.1 aory|EIT75424.1 asyd|OJJ59941.1 ater|EAU31846.1 aver|OJJ02346.1

Group10107: acar|OOF91587.1 afla|EED45783.1 agla|OJJ85381.1 aory|EIT75684.1 asyd|OJJ59943.1 ater|EAU31847.1 aver|OJJ02345.1

Group10108: acar|OOF91731.1 afla|EED52519.1 aluc|OJZ86321.1 anig|EHA19837.1 aory|EIT82774.1 atub|OJI84213.1 awen|OJJ33740.1

Group10109: anid|CBF84500.1 akaw|GAA92280.1 aluc|OJZ90177.1 anig|EHA23676.1 asyd|OJJ53505.1 atub|OJI81604.1 aver|OJJ08241.1

Group10110: acar|OOF91881.1 acla|EAW10805.1 afis|EAW24765.1 afla|EED51966.1 afum|EBA27381.1 aory|EIT81438.1 asyd|OJJ56523.1

Group10111: acar|OOF91983.1 afla|EED56296.1 aory|EIT73428.1 asyd|OJJ54284.1 atub|OJI89932.1 aver|OJJ05238.1 awen|OJJ37070.1

Group10112: acar|OOF92444.1 afis|EAW16803.1 agla|OJJ87317.1 ater|EAU32887.1 azon|OJJ49097.1 asyd|OJJ58339.1 aver|OJJ02072.1

Group10113: acar|OOF92546.1 afla|EED48227.1 akaw|GAA92149.1 aluc|OJZ85554.1 anig|EHA24963.1 aory|EIT72929.1 atub|OJI79351.1

Group10114: agla|OJJ84340.1 afis|EAW20668.1 afla|EED53061.1 anig|EHA17700.1 aory|EIT74773.1 asyd|OJJ60418.1 aver|OJI99742.1

Group10115: ater|EAU36151.1 acla|EAW12120.1 afis|EAW17260.1 afla|EED48875.1 afum|EAL86677.1 aory|EIT81221.1 awen|OJJ35773.1

Group10116: afis|EAW16921.1 afum|EAL86448.1 anid|CBF73423.1 afis|EAW16414.1 afum|EAL89038.1 asyd|OJJ54161.1 aver|OJJ08585.1

Group10117: acar|OOF93175.1 afla|EED47510.1 aory|EIT74549.1 asyd|OJJ63013.1 ater|EAU31932.1 aver|OJJ04461.1 azon|OJJ42258.1

Group10118: acar|OOF93327.1 afla|EED53182.1 anig|EHA26133.1 ater|EAU35986.1 aver|OJJ07601.1 awen|OJJ31083.1 awen|OJJ35808.1

Group10119: acar|OOF93361.1 afis|EAW20745.1 agla|OJJ85097.1 asyd|OJJ56493.1 atub|OJI82460.1 aver|OJI99444.1 afis|EAW16251.1

Group10120: aory|EIT73610.1 acla|EAW14096.1 afis|EAW21809.1 afum|EAL88829.2 agla|OJJ86299.1 anig|EHA19726.1 ater|EAU30722.1

Group10121: acar|OOF93384.1 agla|OJJ80146.1 aory|EIT72746.1 ater|EAU32667.1 asyd|OJJ61539.1 aver|OJJ00866.1 azon|OJJ43166.1

Group10122: acar|OOF93994.1 afla|EED46878.1 anid|CBF84499.1 aory|EIT72586.1 asyd|OJJ53528.1 aver|OJJ08222.1 azon|OJJ51729.1

Group10123: acar|OOF94573.1 acla|EAW15348.1 akaw|GAA82768.1 aluc|OJZ87327.1 anig|EHA19267.1 atub|OJI81318.1 aver|OJJ03895.1

Group10124: afum|EAL85105.2 acla|EAW09091.1 afis|EAW17508.1 aory|EIT76499.1 ater|EAU36366.1 afis|EAW19377.1 afis|EAW19789.1

Group10125: acla|EAW11968.1 afis|EAW19616.1 afla|EED56964.1 afum|EAL86906.2 aory|EIT77561.1 ater|EAU32916.1 awen|OJJ37898.1

Group10126: acar|OOF94907.1 aver|OJJ08790.1 awen|OJJ40865.1 afis|EAW24596.1 agla|OJJ80219.1 anig|EHA25389.1 aver|OJI96487.1

Group10127: acar|OOF94915.1 asyd|OJJ53586.1 acla|EAW14558.1 aver|OJJ08162.1 afla|EED45461.1 aory|EIT76816.1 afla|EED55783.1

Group10128: acar|OOF94995.1 akaw|GAA91649.1 aluc|OJZ87753.1 anig|EHA20240.1 ater|EAU39601.1 atub|OJI80891.1 awen|OJJ36100.1

Group10129: acar|OOF95587.1 akaw|GAA89742.1 aluc|OJZ80642.1 anig|EHA27190.1 atub|OJI87497.1 awen|OJJ37789.1 azon|OJJ43526.1

Group10130: acar|OOF95803.1 acla|EAW14957.1 afis|EAW17785.1 asyd|OJJ52954.1 aver|OJJ08920.1 acla|EAW09539.1 afla|EED52915.1

Group10131: acar|OOF96113.1 afis|EAW24503.1 afla|EED48396.1 aory|EIT83084.1 asyd|OJJ62110.1 aver|OJI96216.1 azon|OJJ50121.1

Group10132: acar|OOF96163.1 acla|EAW08024.1 afis|EAW23058.1 afum|EAL88421.1 akaw|GAA85561.1 anid|CBF81035.1 aory|EIT80328.1

Group10133: acar|OOF96565.1 acla|EAW11013.1 afis|EAW23732.1 anid|CBF78318.1 asyd|OJJ54556.1 aver|OJJ05038.1 asyd|OJJ56094.1

Group10134: akaw|GAA83443.1 aluc|OJZ92558.1 anig|EHA26422.1 atub|OJI86994.1 asyd|OJJ57626.1 aver|OJI99501.1 azon|OJJ43875.1

Group10135: ater|EAU34271.1 awen|OJJ40923.1 azon|OJJ45211.1 akaw|GAA92633.1 anid|CBF71813.1 asyd|OJJ52465.1 aver|OJJ03950.1

Group10136: acar|OOF96637.1 agla|OJJ81297.1 anid|CBF87941.1 asyd|OJJ57967.1 aver|OJJ04435.1 awen|OJJ33491.1 ater|EAU35869.1

Group10137: acar|OOF96871.1 akaw|GAA84694.1 azon|OJJ47852.1 aluc|OJZ86993.1 anig|EHA21652.1 atub|OJI83526.1 aver|OJJ02007.1

Group10138: acar|OOF97405.1 afis|EAW16894.1 akaw|GAA86524.1 aluc|OJZ85070.1 anig|EHA25284.1 atub|OJI80528.1 awen|OJJ29746.1

Group10139: acar|OOF98076.1 akaw|GAA82893.1 aluc|OJZ87462.1 anid|CBF86128.1 anig|EHA20684.1 ater|EAU32051.1 atub|OJI81186.1

Group10140: acar|OOF98088.1 acla|EAW10137.1 afis|EAW23635.1 afla|EED56572.1 afum|EAL87825.1 anid|CBF71566.1 aory|EIT79618.1

Group10141: acar|OOF98554.1 acla|EAW13534.1 afis|EAW17486.1 afum|EAL93436.2 anig|EHA20072.1 aory|EIT80022.1 atub|OJI83387.1

Group10142: acar|OOF98721.1 acla|EAW09869.1 afis|EAW19903.1 afum|EAL94096.1 agla|OJJ79564.1 ater|EAU37749.1 afum|EAL86548.1

Group10143: acar|OOF98729.1 akaw|GAA87769.1 aluc|OJZ85404.1 anig|EHA25583.1 asyd|OJJ52866.1 aver|OJJ08848.1 awen|OJJ40141.1

Group10144: acar|OOF98740.1 afis|EAW25597.1 afum|EAL84303.1 awen|OJJ40808.1 anid|CBF82031.1 asyd|OJJ53734.1 aver|OJJ07849.1

Group10145: acar|OOG00467.1 acla|EAW06903.1 afis|EAW16520.1 afla|EED56151.1 afum|EAL89115.1 anid|CBF75549.1 aory|EIT76578.1

Group10146: acla|EAW07006.1 afum|EBA27370.1 agla|OJJ82623.1 anid|CBF81986.1 asyd|OJJ56155.1 ater|EAU36735.1 aver|OJJ05095.1

Group10147: acla|EAW07118.1 afis|EAW17372.1 akaw|GAA90194.1 aluc|OJZ83154.1 anid|CBF76129.1 anig|EHA28220.1 atub|OJI87783.1

Group10148: acla|EAW08349.1 afla|EED55570.1 agla|OJJ83852.1 ater|EAU37951.1 aver|OJJ04135.1 anid|CBF74853.1 asyd|OJJ60127.1

Group10149: acla|EAW08532.1 afis|EAW23571.1 afla|EED54701.1 afum|EAL87895.2 aory|EIT78780.1 awen|OJJ41511.1 azon|OJJ48011.1

Group10150: acla|EAW08776.1 afis|EAW15630.1 afum|EAL85704.1 asyd|OJJ57953.1 ater|EAU31446.1 aver|OJJ04444.1 azon|OJJ49098.1

Group10151: acla|EAW08790.1 afis|EAW15612.1 afum|EAL85723.1 anid|CBF79048.1 ater|EAU37846.1 azon|OJJ42551.1 azon|OJJ42690.1

Group10152: acla|EAW08793.1 afis|EAW15633.1 afla|EED52683.1 aory|EIT74288.1 anid|CBF71036.1 asyd|OJJ52665.1 aver|OJJ04692.1

Group10153: acla|EAW08951.1 afis|EAW15459.1 afum|EAL85860.1 anid|CBF78360.1 asyd|OJJ57338.1 ater|EAU31585.1 aver|OJJ05824.1

Group10154: aory|EIT76507.1 afis|EAW15614.1 afla|EED46085.1 afum|EAL85721.1 agla|OJJ84457.1 ater|EAU37826.1 ater|EAU31985.1

Group10155: acla|EAW09328.1 afis|EAW21785.1 afum|EAL92812.1 aver|OJJ00512.1 anid|CBF76615.1 aory|EIT77901.1 ater|EAU38468.1

Group10156: acla|EAW10107.1 afis|EAW25480.1 afla|EED53978.1 afum|EAL90145.1 agla|OJJ87128.1 awen|OJJ41221.1 aory|EIT78329.1

Group10157: acla|EAW10391.1 afis|EAW25200.1 afla|EED53647.1 afum|EAL89863.2 aory|EIT75327.1 asyd|OJJ63000.1 aver|OJI97966.1

Group10158: asyd|OJJ62848.1 ater|EAU32016.1 aver|OJJ07736.1 agla|OJJ80505.1 azon|OJJ42028.1 agla|OJJ89600.1 awen|OJJ37495.1

Group10159: afla|EED48161.1 aory|EIT83035.1 ater|EAU32688.1 anid|CBF76016.1 asyd|OJJ52434.1 aver|OJJ03976.1 azon|OJJ45937.1

Group10160: acla|EAW11009.1 akaw|GAA85912.1 asyd|OJJ61457.1 ater|EAU32838.1 atub|OJI83169.1 aver|OJJ00795.1 aluc|OJZ91154.1

Group10161: acla|EAW11024.1 afis|EAW20838.1 anid|CBF82163.1 ater|EAU37264.1 afla|EED46649.1 afum|EAL84795.1 aory|EIT77935.1

Group10162: acla|EAW11038.1 afis|EAW16190.1 afla|EED49621.1 afum|EAL88821.2 aory|EIT80943.1 azon|OJJ45962.1 azon|OJJ45946.1

Group10163: ater|EAU36746.1 acla|EAW11049.1 afis|EAW16189.1 afla|EED49632.1 afum|EAL88820.2 aory|EIT80969.1 azon|OJJ45959.1

Group10164: acla|EAW11081.1 afis|EAW24664.1 afla|EED55932.1 afum|EAL89322.1 anid|CBF82329.1 aory|EIT76860.1 azon|OJJ45895.1

Group10165: acla|EAW11191.1 afla|EED54939.1 afum|EAL88558.2 akaw|GAA85541.1 aluc|OJZ83474.1 anid|CBF81003.1 ater|EAU34828.1

Group10166: acla|EAW11439.1 afis|EAW23986.1 afum|EAL87415.2 awen|OJJ35454.1 azon|OJJ51289.1 agla|OJJ86920.1 ater|EAU29765.1

Group10167: akaw|GAA92880.1 acla|EAW11450.1 afla|EED52348.1 agla|OJJ83278.1 aluc|OJZ80003.1 anig|EHA21870.1 awen|OJJ29564.1

Group10168: acla|EAW11711.1 afis|EAW23728.1 afla|EED52211.1 afum|EBA27302.1 anid|CBF85586.1 aory|EIT74336.1 ater|EAU32749.1

Group10169: acla|EAW11732.1 afis|EAW23764.1 afum|EAL87197.1 agla|OJJ78958.1 asyd|OJJ52071.1 aver|OJJ03032.1 azon|OJJ42829.1

Group10170: acla|EAW12042.1 afla|EED56293.1 aory|EIT73398.1 asyd|OJJ57620.1 ater|EAU37840.1 aver|OJJ05566.1 azon|OJJ42462.1

Group10171: acla|EAW12050.1 afla|EED56294.1 aory|EIT73305.1 asyd|OJJ57621.1 ater|EAU37839.1 aver|OJJ05565.1 azon|OJJ42461.1

Group10172: azon|OJJ50599.1 afum|EBA27289.1 akaw|GAA85221.1 aluc|OJZ90575.1 anig|EHA24019.1 atub|OJI81999.1 awen|OJJ36191.1

Group10173: acla|EAW12190.1 afla|EED52085.1 aory|EIT76919.1 asyd|OJJ55647.1 ater|EAU36876.1 aver|OJJ06732.1 ater|EAU33188.1

Group10174: asyd|OJJ54509.1 afis|EAW19961.1 afum|EAL84954.1 ater|EAU37380.1 awen|OJJ39610.1 afla|EED57051.1 aory|EIT82314.1

Group10175: acla|EAW12244.1 afis|EAW17146.1 asyd|OJJ59649.1 aver|OJJ02633.1 azon|OJJ48730.1 agla|OJJ85326.1 azon|OJJ42911.1

Group10176: afis|EAW17467.1 afla|EED54400.1 afum|EAL91296.1 aory|EIT75887.1 asyd|OJJ58076.1 ater|EAU35543.1 aver|OJJ04331.1

Group10177: acla|EAW12383.1 afla|EED54197.1 agla|OJJ78719.1 aory|EIT75932.1 awen|OJJ38699.1 ater|EAU33853.1 aver|OJI98299.1

Group10178: acla|EAW12425.1 afla|EED52218.1 akaw|GAA84849.1 aluc|OJZ86822.1 anid|CBF83038.1 aory|EIT74368.1 atub|OJI83695.1

Group10179: acla|EAW12446.1 afla|EED52537.1 aory|EIT74248.1 ater|EAU29648.1 acla|EAW15007.1 afis|EAW24535.1 afum|EAL86200.1

Group10180: acla|EAW12616.1 afis|EAW18068.1 afum|EAL91880.1 agla|OJJ81011.1 asyd|OJJ58329.1 aver|OJJ02082.1 awen|OJJ41455.1

Group10181: acla|EAW12724.1 afis|EAW17887.1 afla|EED46274.1 afum|EAL91696.1 agla|OJJ83060.1 awen|OJJ41672.1 azon|OJJ48387.1

Group10182: acla|EAW12816.1 afis|EAW17981.1 afla|EED51924.1 afum|EAL91790.1 agla|OJJ80238.1 aory|EIT81459.1 awen|OJJ41564.1

Group10183: afla|EED53694.1 aory|EIT75299.1 afis|EAW21378.1 agla|OJJ78598.1 anid|CBF85730.1 asyd|OJJ64282.1 aver|OJI98374.1

Group10184: acla|EAW12882.1 afis|EAW17795.1 afum|EAL91603.1 agla|OJJ84738.1 asyd|OJJ54200.1 ater|EAU36885.1 aver|OJJ08524.1

Group10185: afla|EED51128.1 anid|CBF90350.1 aory|EIT81350.1 agla|OJJ81312.1 asyd|OJJ55576.1 ater|EAU37573.1 aver|OJJ06655.1

Group10186: acla|EAW13211.1 afis|EAW17430.1 afla|EED47239.1 afum|EAL91258.1 agla|OJJ80131.1 aory|EIT79299.1 azon|OJJ47812.1

Group10187: acla|EAW13284.1 afla|EED48190.1 agla|OJJ82046.1 aory|EIT72947.1 asyd|OJJ56058.1 aver|OJJ05000.1 awen|OJJ33507.1

Group10188: acla|EAW13521.1 afis|EAW20747.1 afum|EAL87985.1 ater|EAU31793.1 awen|OJJ34473.1 afla|EED48969.1 aory|EIT79997.1

Group10189: acla|EAW13586.1 afum|EAL84902.1 agla|OJJ87989.1 asyd|OJJ52162.1 awen|OJJ30033.1 azon|OJJ43667.1 afis|EAW20016.1

Group10190: acla|EAW13617.1 afla|EED48924.1 agla|OJJ86967.1 aory|EIT80007.1 asyd|OJJ59205.1 awen|OJJ29736.1 azon|OJJ42772.1

Group10191: acla|EAW13673.1 afis|EAW19950.1 afla|EED54751.1 afum|EAL84965.1 aory|EIT78814.1 awen|OJJ31428.1 aory|EIT81673.1

Group10192: acla|EAW13676.1 afis|EAW19931.1 afla|EED48977.1 afum|EAL84978.1 anid|CBF76644.1 ater|EAU34786.1 awen|OJJ37835.1

Group10193: acla|EAW13727.1 afis|EAW20341.1 afla|EED50165.1 afum|EAL85463.1 anid|CBF79796.1 asyd|OJJ56280.1 aver|OJJ08480.1

Group10194: acla|EAW13731.1 afis|EAW19848.1 afum|EBA27217.1 anid|CBF82492.1 asyd|OJJ56116.1 ater|EAU39290.1 aver|OJJ05057.1

Group10195: acla|EAW13747.1 afis|EAW19823.1 afum|EAL85070.1 anid|CBF87170.1 asyd|OJJ60290.1 aver|OJI99630.1 azon|OJJ50616.1

Group10196: acla|EAW14554.1 afis|EAW19271.1 afum|EBA27511.1 agla|OJJ88410.1 ater|EAU29495.1 awen|OJJ37632.1 azon|OJJ47019.1

Group10197: acla|EAW14629.1 afis|EAW17252.1 afla|EED57076.1 afum|EAL86671.2 aory|EIT82261.1 asyd|OJJ56524.1 aver|OJJ06199.1

Group10198: acla|EAW14870.1 afla|EED56762.1 anid|CBF80292.1 aory|EIT73866.1 asyd|OJJ59194.1 ater|EAU31801.1 aver|OJJ01810.1

Group10199: acla|EAW14950.1 afis|EAW24467.1 afum|EAL86139.1 anid|CBF85607.1 asyd|OJJ64198.1 aver|OJI98287.1 azon|OJJ45772.1

Group10200: acla|EAW15017.1 afis|EAW24548.1 afum|EAL86212.1 asyd|OJJ56858.1 ater|EAU30730.1 aver|OJJ03471.1 awen|OJJ38777.1

Group10201: acla|EAW15325.1 afis|EAW20336.1 afla|EED53114.1 afum|EBA27233.1 aory|EIT74784.1 ater|EAU37787.1 awen|OJJ31651.1

Group10202: acla|EAW15366.1 afis|EAW17226.1 afla|EED51519.1 afum|EAL86648.1 aory|EIT80474.1 asyd|OJJ54522.1 aver|OJJ07662.1

Group10203: anid|CBF76052.1 afla|EED54194.1 aory|EIT76093.1 asyd|OJJ52529.1 ater|EAU33852.1 aver|OJJ03871.1 ater|EAU34638.1

Group10204: afis|EAW15553.1 afla|EED53618.1 aory|EIT74877.1 asyd|OJJ63945.1 atub|OJI83152.1 aver|OJI96997.1 awen|OJJ39626.1

Group10205: awen|OJJ41766.1 afla|EED48378.1 agla|OJJ86209.1 aory|EIT83086.1 asyd|OJJ64822.1 anid|CBF86608.1 aver|OJI98935.1

Group10206: afis|EAW16690.1 afum|EAL89290.1 asyd|OJJ52717.1 aver|OJJ04746.1 afis|EAW17132.1 afum|EAL86592.1 anid|CBF71114.1

Group10207: afis|EAW16758.1 afum|EAL86330.1 anid|CBF78505.1 asyd|OJJ56110.1 ater|EAU29655.1 aver|OJJ05053.1 afla|EED51667.1

Group10208: afis|EAW16923.1 afla|EED52030.1 afum|EAL86449.1 aory|EIT81450.1 asyd|OJJ54634.1 aver|OJJ07522.1 awen|OJJ29801.1

Group10209: akaw|GAA90159.1 asyd|OJJ64677.1 aluc|OJZ83188.1 azon|OJJ45179.1 afum|EAL91083.1 anid|CBF86785.1 aver|OJI98778.1

Group10210: afis|EAW17086.1 afla|EED54865.1 afum|EAL87816.2 anig|EHA22234.1 ater|EAU35234.1 atub|OJI84574.1 anig|EHA24788.1

Group10211: afis|EAW17329.1 afum|EAL91166.1 anid|CBF83206.1 asyd|OJJ55552.1 ater|EAU29242.1 aver|OJJ06629.1 azon|OJJ46030.1

Group10212: afis|EAW17361.1 afum|EAL91198.1 anid|CBF87332.1 asyd|OJJ56399.1 ater|EAU31360.1 aver|OJJ06313.1 anid|CBF76185.1

Group10213: afis|EAW17605.1 afum|EAL91418.1 akaw|GAA92445.1 aluc|OJZ81574.1 anig|EHA25863.1 atub|OJI88607.1 azon|OJJ44984.1

Group10214: afis|EAW18387.1 afum|EAL93062.1 anid|CBF70770.1 aver|OJJ01029.1 awen|OJJ31764.1 afla|EED53020.1 agla|OJJ84950.1

Group10215: afis|EAW19199.1 agla|OJJ86724.1 anid|CBF76193.1 asyd|OJJ53368.1 ater|EAU31800.1 aver|OJJ03315.1 aver|OJJ08379.1

Group10216: afis|EAW20335.1 afum|EBA27232.1 anid|CBF71679.1 asyd|OJJ61823.1 aver|OJI95923.1 akaw|GAA88745.1 aluc|OJZ92705.1

Group10217: afis|EAW20397.1 afla|EED57025.1 agla|OJJ84195.1 aory|EIT80002.1 asyd|OJJ64225.1 awen|OJJ37009.1 afla|EED48960.1

Group10218: afis|EAW21055.1 afla|EED54200.1 afum|EAL92091.1 aory|EIT76084.1 asyd|OJJ54691.1 ater|EAU32447.1 aver|OJJ07478.1

Group10219: afis|EAW21127.1 afla|EED48221.1 afum|EAL88871.1 akaw|GAA87229.1 aluc|OJZ84200.1 aory|EIT72951.1 atub|OJI89884.1

Group10220: afis|EAW21309.1 afum|EAL92325.1 anid|CBF83400.1 asyd|OJJ60643.1 ater|EAU35939.1 aver|OJI99962.1 azon|OJJ44279.1

Group10221: afis|EAW21792.1 afla|EED46605.1 afum|EAL92817.1 anid|CBF90027.1 aory|EIT77868.1 asyd|OJJ62065.1 aver|OJI96171.1

Group10222: afis|EAW22077.1 afla|EED56694.1 afum|EAL91093.2 anid|CBF84149.1 aory|EIT73955.1 asyd|OJJ57121.1 ater|EAU30147.1

Group10223: afis|EAW22098.1 afla|EED48272.1 afum|EAL91091.2 aory|EIT72934.1 asyd|OJJ54148.1 ater|EAU38943.1 aver|OJJ08593.1

Group10224: afis|EAW22223.1 afum|EAL90971.1 anid|CBF69392.1 asyd|OJJ56492.1 ater|EAU33092.1 aver|OJJ06226.1 azon|OJJ47605.1

Group10225: afis|EAW23625.1 afla|EED45750.1 afum|EAL87835.1 aory|EIT75598.1 asyd|OJJ55363.1 ater|EAU32046.1 aver|OJJ06445.1

Group10226: afis|EAW23685.1 afla|EED57652.1 afum|EAL87130.1 agla|OJJ78696.1 aory|EIT79919.1 ater|EAU34270.1 aver|OJI98899.1

Group10227: afis|EAW23849.1 afla|EED52566.1 afum|EAL87277.1 agla|OJJ82492.1 aory|EIT74262.1 ater|EAU33516.1 awen|OJJ33805.1

Group10228: afis|EAW23915.1 akaw|GAA85870.1 aluc|OJZ91208.1 awen|OJJ35046.1 azon|OJJ50613.1 anig|EHA19915.1 atub|OJI83224.1

Group10229: afla|EED49124.1 awen|OJJ36333.1 agla|OJJ86964.1 aver|OJJ02415.1 anid|CBF75314.1 asyd|OJJ59872.1 ater|EAU29714.1

Group10230: afis|EAW24477.1 anid|CBF69438.1 asyd|OJJ52390.1 aver|OJJ01933.1 afla|EED53176.1 agla|OJJ87867.1 aory|EIT75134.1

Group10231: afis|EAW24713.1 afla|EED54611.1 agla|OJJ87527.1 aory|EIT78802.1 asyd|OJJ53079.1 aver|OJJ03768.1 awen|OJJ36607.1

Group10232: afis|EAW25186.1 afla|EED47736.1 afum|EAL89860.2 ater|EAU35594.1 aver|OJJ08837.1 anid|CBF82361.1 aory|EIT77170.1

Group10233: afis|EAW25651.1 agla|OJJ81744.1 aluc|OJZ80187.1 asyd|OJJ62924.1 ater|EAU37898.1 aver|OJI98065.1 azon|OJJ50377.1

Group10234: afla|EED44854.1 anid|CBF78815.1 aory|EIT82719.1 aory|EIT82730.1 asyd|OJJ65263.1 aver|OJI99414.1 awen|OJJ34503.1

Group10235: afla|EED45044.1 agla|OJJ84353.1 aory|EIT73706.1 asyd|OJJ58384.1 ater|EAU37461.1 aver|OJJ02034.1 awen|OJJ39645.1

Group10236: afla|EED45300.1 akaw|GAA90425.1 aluc|OJZ85804.1 anig|EHA24750.1 aory|EIT73573.1 atub|OJI79590.1 ater|EAU30976.1

Group10237: afla|EED45562.1 aory|EIT75731.1 afla|EED57227.1 akaw|GAA87619.1 aluc|OJZ85240.1 atub|OJI80335.1 anig|EHA25441.1

Group10238: afla|EED45575.1 anid|CBF74994.1 aory|EIT75630.1 asyd|OJJ59242.1 aver|OJJ01852.1 ater|EAU37983.1 azon|OJJ45067.1

Group10239: afla|EED45886.1 agla|OJJ80673.1 anid|CBF75882.1 aory|EIT75675.1 asyd|OJJ60251.1 aver|OJJ04006.1 awen|OJJ35874.1

Group10240: afla|EED46046.1 anig|EHA27452.1 asyd|OJJ54976.1 atub|OJI89789.1 aver|OJJ03922.1 awen|OJJ32212.1 asyd|OJJ54984.1

Group10241: afla|EED46049.1 anid|CBF77118.1 aory|EIT83488.1 asyd|OJJ56878.1 ater|EAU39543.1 aory|EIT81916.1 atub|OJI81199.1

Group10242: afla|EED47345.1 agla|OJJ79751.1 aory|EIT82850.1 aver|OJJ07565.1 awen|OJJ29767.1 azon|OJJ49003.1 ater|EAU31777.1

Group10243: afla|EED47425.1 anid|CBF82193.1 aory|EIT81900.1 asyd|OJJ52186.1 ater|EAU29297.1 aver|OJJ05959.1 awen|OJJ36973.1

Group10244: afla|EED47469.1 agla|OJJ79971.1 anig|EHA25784.1 aory|EIT79970.1 ater|EAU31068.1 awen|OJJ35931.1 awen|OJJ34653.1

Group10245: afla|EED47870.1 anig|EHA27620.1 aory|EIT77230.1 asyd|OJJ61764.1 ater|EAU37245.1 aver|OJI95863.1 awen|OJJ39973.1

Group10246: afla|EED49454.1 agla|OJJ85294.1 anid|CBF80385.1 aory|EIT79463.1 asyd|OJJ54787.1 aver|OJJ07377.1 awen|OJJ31281.1

Group10247: afla|EED50207.1 anid|CBF82198.1 aory|EIT77343.1 asyd|OJJ52183.1 ater|EAU29293.1 aver|OJJ05962.1 awen|OJJ34427.1

Group10248: afla|EED50780.1 anid|CBF78322.1 aory|EIT78058.1 asyd|OJJ56096.1 aver|OJJ05040.1 ater|EAU36099.1 azon|OJJ42248.1

Group10249: afla|EED50796.1 aory|EIT78102.1 asyd|OJJ54742.1 ater|EAU31419.1 aver|OJJ07422.1 awen|OJJ32720.1 azon|OJJ51218.1

Group10250: afla|EED51075.1 anid|CBF87082.1 aory|EIT76983.1 asyd|OJJ60415.1 ater|EAU32194.1 aver|OJI99740.1 awen|OJJ38057.1

Group10251: afla|EED51527.1 agla|OJJ81393.1 asyd|OJJ54813.1 aver|OJJ07348.1 afla|EED53583.1 aory|EIT75065.1 awen|OJJ35388.1

Group10252: afla|EED51637.1 anid|CBF84495.1 aory|EIT80544.1 asyd|OJJ53526.1 ater|EAU35686.1 aver|OJJ08224.1 awen|OJJ32550.1

Group10253: afla|EED51695.1 akaw|GAA86245.1 anid|CBF82002.1 anig|EHA23959.1 aory|EIT80643.1 atub|OJI81925.1 akaw|GAA86246.1

Group10254: afla|EED52147.1 agla|OJJ88641.1 anid|CBF80186.1 asyd|OJJ52190.1 aver|OJJ05956.1 awen|OJJ29974.1 azon|OJJ45078.1

Group10255: afla|EED52227.1 anid|CBF82127.1 aory|EIT74449.1 asyd|OJJ54296.1 ater|EAU37212.1 aver|OJJ05251.1 awen|OJJ36070.1

Group10256: afla|EED52266.1 agla|OJJ82579.1 asyd|OJJ60517.1 aver|OJI99844.1 awen|OJJ35907.1 azon|OJJ45107.1 anid|CBF86988.1

Group10257: afla|EED52314.1 agla|OJJ87504.1 aory|EIT74376.1 asyd|OJJ54910.1 ater|EAU33815.1 aver|OJJ07292.1 awen|OJJ37811.1

Group10258: afla|EED52465.1 aluc|OJZ87035.1 anig|EHA25213.1 aory|EIT82789.1 atub|OJI83480.1 awen|OJJ30799.1 agla|OJJ84730.1

Group10259: afla|EED53550.1 agla|OJJ82963.1 aory|EIT74991.1 asyd|OJJ52687.1 aver|OJJ04715.1 agla|OJJ78662.1 awen|OJJ38323.1

Group10260: afla|EED54066.1 anid|CBF74924.1 aory|EIT76035.1 asyd|OJJ61408.1 ater|EAU31003.1 aver|OJJ00738.1 azon|OJJ42636.1

Group10261: afla|EED54378.1 aluc|OJZ87037.1 anig|EHA25211.1 aory|EIT75882.1 asyd|OJJ52554.1 atub|OJI83478.1 aver|OJJ03847.1

Group10262: afla|EED54888.1 akaw|GAA86659.1 aluc|OJZ89755.1 anig|EHA26977.1 aory|EIT79366.1 atub|OJI85047.1 aver|OJJ00799.1

Group10263: afla|EED55071.1 aory|EIT81945.1 ater|EAU37975.1 azon|OJJ50349.1 anid|CBF80573.1 asyd|OJJ52849.1 aver|OJJ08827.1

Group10264: afla|EED55764.1 anid|CBF87737.1 aory|EIT76801.1 asyd|OJJ65053.1 ater|EAU36714.1 aver|OJI99176.1 azon|OJJ47815.1

Group10265: afla|EED56551.1 anid|CBF80504.1 aory|EIT79636.1 asyd|OJJ54734.1 ater|EAU39450.1 aver|OJI97956.1 aver|OJJ07437.1

Group10266: afla|EED56557.1 agla|OJJ88412.1 akaw|GAA92924.1 aluc|OJZ82515.1 asyd|OJJ57317.1 aver|OJJ05844.1 awen|OJJ33378.1

Group10267: afla|EED56559.1 akaw|GAA92921.1 aluc|OJZ82510.1 anig|EHA20908.1 asyd|OJJ57314.1 aver|OJJ05847.1 awen|OJJ33376.1

Group10268: afla|EED56855.1 anig|EHA19935.1 aory|EIT77517.1 asyd|OJJ55049.1 aver|OJI99563.1 anig|EHA19934.1 aver|OJI99564.1

Group10269: afla|EED56903.1 agla|OJJ87523.1 aory|EIT77550.1 asyd|OJJ53514.1 ater|EAU30079.1 aver|OJJ08237.1 awen|OJJ40162.1

Group10270: afla|EED56921.1 akaw|GAA89749.1 aluc|OJZ80635.1 aory|EIT77565.1 atub|OJI87490.1 awen|OJJ32542.1 anid|CBF82770.1

Group10271: afla|EED57045.1 agla|OJJ82533.1 aory|EIT82268.1 asyd|OJJ52763.1 ater|EAU38773.1 aver|OJJ08746.1 awen|OJJ36237.1

Group10272: afla|EED57579.1 anid|CBF89418.1 aory|EIT82972.1 asyd|OJJ62453.1 ater|EAU36881.1 aver|OJI96588.1 awen|OJJ31389.1

Group10273: agla|OJJ78975.1 akaw|GAA83144.1 aluc|OJZ88701.1 anig|EHA18385.1 asyd|OJJ62632.1 atub|OJI91282.1 awen|OJJ32216.1

Group10274: agla|OJJ80901.1 akaw|GAA91207.1 aluc|OJZ81542.1 anig|EHA25892.1 azon|OJJ43255.1 ater|EAU36288.1 atub|OJI88644.1

Group10275: agla|OJJ82912.1 akaw|GAA84837.1 aluc|OJZ86837.1 asyd|OJJ56204.1 aver|OJJ05131.1 atub|OJI83681.1 anid|CBF75980.1

Group10276: agla|OJJ87897.1 aluc|OJZ80195.1 aory|EIT79523.1 asyd|OJJ52334.1 aver|OJJ05793.1 awen|OJJ31362.1 asyd|OJJ57375.1

Group10277: agla|OJJ88416.1 akaw|GAA92922.1 aluc|OJZ82513.1 anig|EHA20905.1 asyd|OJJ57313.1 aver|OJJ08123.1 awen|OJJ33718.1

Group10278: akaw|GAA91033.1 aluc|OJZ87843.1 asyd|OJJ54599.1 atub|OJI80805.1 aver|OJJ07553.1 awen|OJJ31645.1 asyd|OJJ57925.1

Group10279: asyd|OJJ59036.1 ater|EAU37517.1 atub|OJI79386.1 aver|OJJ01631.1 azon|OJJ49591.1 asyd|OJJ62323.1 aver|OJI96429.1

Group10280: abra|OJJ65452.1 abra|OJJ65570.1 abra|OJJ65527.1 akaw|GAA93119.1 akaw|GAA93278.1 aluc|OJZ80235.1

Group10281: abra|OJJ65509.1 abra|OJJ65822.1 abra|OJJ65731.1 awen|OJJ29546.1 awen|OJJ29607.1 akaw|GAA93289.1

Group10282: abra|OJJ68474.1 abra|OJJ69672.1 akaw|GAA88796.1 aluc|OJZ82479.1 anig|EHA28296.1 atub|OJI87959.1

Group10283: abra|OJJ73135.1 abra|OJJ78187.1 anid|CBF85171.1 asyd|OJJ60247.1 aver|OJJ04013.1 afis|EAW15646.1

Group10284: acar|OOF92657.1 acar|OOF97631.1 aluc|OJZ83632.1 anig|EHA18785.1 atub|OJI90001.1 akaw|GAA92093.1

Group10285: acar|OOF94040.1 acar|OOF94110.1 abra|OJJ66406.1 akaw|GAA90274.1 aluc|OJZ83067.1 atub|OJI87879.1

Group10286: acar|OOF95029.1 acar|OOF95877.1 akaw|GAA83684.1 aluc|OJZ91050.1 anig|EHA19746.1 atub|OJI83045.1

Group10287: acar|OOF95619.1 acar|OOF95671.1 abra|OJJ67064.1 akaw|GAA83294.1 aluc|OJZ83796.1 anig|EHA18942.1

Group10288: afum|EAL84881.1 afum|EAL87107.1 afum|EAL92048.1 afum|EAL94109.1 afum|EAL85503.1 afum|EAL91105.1

Group10289: afum|EAL86279.1 afum|EAL86759.1 afum|EAL86739.1 aluc|OJZ82090.1 anig|EHA21839.1 akaw|GAA93042.1

Group10290: agla|OJJ78751.1 agla|OJJ79298.1 agla|OJJ85892.1 agla|OJJ82208.1 agla|OJJ79871.1 aver|OJJ06000.1

Group10291: agla|OJJ78863.1 agla|OJJ86143.1 agla|OJJ84406.1 agla|OJJ84596.1 asyd|OJJ65342.1 aver|OJI99488.1

Group10292: agla|OJJ81525.1 agla|OJJ83613.1 afis|EAW25592.1 afum|EAL84299.2 anid|CBF82326.1 ater|EAU32211.1

Group10293: agla|OJJ83532.1 agla|OJJ83538.1 abra|OJJ70354.1 afla|EED48239.1 aluc|OJZ80604.1 aory|EIT72889.1

Group10294: agla|OJJ84147.1 agla|OJJ84853.1 asyd|OJJ58689.1 aver|OJJ01301.1 aory|EIT76069.1 awen|OJJ40081.1

Group10295: agla|OJJ85428.1 agla|OJJ86970.1 agla|OJJ82397.1 agla|OJJ85894.1 acla|EAW12314.1 aory|EIT77476.1

Group10296: agla|OJJ85647.1 agla|OJJ87448.1 anid|CBF81918.1 awen|OJJ31299.1 asyd|OJJ65137.1 aver|OJI99271.1

Group10297: akaw|GAA82059.1 akaw|GAA83940.1 aluc|OJZ80610.1 aluc|OJZ80822.1 abra|OJJ66437.1 afum|EBA27257.1

Group10298: akaw|GAA88999.1 akaw|GAA92353.1 akaw|GAA93062.1 aluc|OJZ80234.1 aluc|OJZ80393.1 aluc|OJZ80501.1

Group10299: akaw|GAA92615.1 akaw|GAA93181.1 aluc|OJZ80046.1 anig|EHA17706.1 atub|OJI87468.1 aluc|OJZ79875.1

Group10300: akaw|GAA92956.1 akaw|GAA93132.1 aluc|OJZ80249.1 aluc|OJZ80307.1 aluc|OJZ88842.1 abra|OJJ65722.1

Group10301: aluc|OJZ79549.1 aluc|OJZ79984.1 aluc|OJZ80077.1 aluc|OJZ79743.1 aluc|OJZ80258.1 akaw|GAA93216.1

Group10302: aluc|OJZ79574.1 aluc|OJZ88808.1 aluc|OJZ79754.1 aluc|OJZ80112.1 aluc|OJZ81138.1 aluc|OJZ79669.1

Group10303: aluc|OJZ79748.1 aluc|OJZ79793.1 abra|OJJ66061.1 acar|OOF99134.1 afla|EED57044.1 aory|EIT82260.1

Group10304: aluc|OJZ80108.1 aluc|OJZ86033.1 akaw|GAA90576.1 anid|CBF73579.1 anig|EHA24581.1 atub|OJI79821.1

Group10305: aluc|OJZ80121.1 aluc|OJZ88797.1 abra|OJJ65718.1 aver|OJJ03131.1 akaw|GAA92627.1 asyd|OJJ62710.1

Group10306: anid|CBF70120.1 anid|CBF76211.1 abra|OJJ66451.1 afis|EAW15395.1 aver|OJJ05919.1 aver|OJJ08486.1

Group10307: anid|CBF71194.1 anid|CBF71815.1 asyd|OJJ52514.1 aver|OJI98860.1 aver|OJI99668.1 asyd|OJJ64758.1

Group10308: anid|CBF75857.1 anid|CBF82342.1 agla|OJJ83798.1 asyd|OJJ56257.1 asyd|OJJ57579.1 aver|OJJ05601.1

Group10309: anid|CBF75872.1 anid|CBF81776.1 awen|OJJ38087.1 asyd|OJJ60236.1 aver|OJJ04029.1 asyd|OJJ54470.1

Group10310: anid|CBF76081.1 anid|CBF81135.1 anid|CBF80652.1 anid|CBF86730.1 afla|EED48083.1 asyd|OJJ53648.1

Group10311: anid|CBF78399.1 anid|CBF78480.1 afla|EED48094.1 asyd|OJJ65356.1 ater|EAU32693.1 aver|OJI99505.1

Group10312: aory|EIT76368.1 aory|EIT80996.1 acla|EAW12271.1 afla|EED46235.1 aluc|OJZ83582.1 afla|EED49611.1

Group10313: asyd|OJJ52245.1 asyd|OJJ52265.1 akaw|GAA92953.1 aluc|OJZ79529.1 ater|EAU35415.1 asyd|OJJ52287.1

Group10314: asyd|OJJ53369.1 asyd|OJJ64222.1 anid|CBF73868.1 ater|EAU31570.1 aver|OJJ08378.1 aver|OJI98305.1

Group10315: asyd|OJJ54675.1 asyd|OJJ55945.1 asyd|OJJ56883.1 aver|OJJ03439.1 aver|OJJ04894.1 aver|OJJ07628.1

Group10316: asyd|OJJ55469.1 asyd|OJJ62349.1 aver|OJI96456.1 aver|OJI99665.1 afis|EAW19744.1 anid|CBF85275.1

Group10317: asyd|OJJ59620.1 asyd|OJJ60333.1 afis|EAW17032.1 aluc|OJZ79602.1 aver|OJI99678.1 awen|OJJ36174.1

Group10318: ater|EAU29263.1 ater|EAU29264.1 anid|CBF80331.1 asyd|OJJ54841.1 aver|OJJ07320.1 asyd|OJJ61548.1

Group10319: ater|EAU29430.1 ater|EAU39347.1 anid|CBF80714.1 aory|EIT77684.1 asyd|OJJ58388.1 aver|OJJ02030.1

Group10320: ater|EAU30977.1 ater|EAU37123.1 abra|OJJ68828.1 akaw|GAA90424.1 aluc|OJZ85805.1 atub|OJI79591.1

Group10321: atub|OJI87842.1 abra|OJJ66373.1 acar|OOF91466.1 akaw|GAA90246.1 aluc|OJZ83094.1 anig|EHA28158.1

Group10322: aver|OJJ05186.1 aver|OJJ07923.1 afla|EED49650.1 anid|CBF82052.1 asyd|OJJ54234.1 ater|EAU35451.1

Group10323: awen|OJJ29631.1 awen|OJJ32583.1 abra|OJJ67274.1 akaw|GAA92072.1 aluc|OJZ81159.1 ater|EAU30099.1

Group10324: awen|OJJ30181.1 awen|OJJ35359.1 awen|OJJ33786.1 acar|OOF97382.1 afla|EED49773.1 aory|EIT77646.1

Group10325: awen|OJJ30229.1 awen|OJJ37834.1 awen|OJJ35460.1 aory|EIT82841.1 acar|OOF96655.1 afla|EED48288.1

Group10326: awen|OJJ30782.1 awen|OJJ31699.1 acar|OOF95212.1 afis|EAW21197.1 afla|EED51082.1 aory|EIT76947.1

Group10327: awen|OJJ31923.1 awen|OJJ31924.1 afis|EAW21057.1 afum|EAL92092.2 agla|OJJ86218.1 awen|OJJ31921.1

Group10328: awen|OJJ32578.1 awen|OJJ33915.1 awen|OJJ34575.1 afla|EED51615.1 agla|OJJ79230.1 aory|EIT80725.1

Group10329: awen|OJJ32587.1 awen|OJJ40143.1 anid|CBF82324.1 ater|EAU32210.1 afis|EAW25593.1 afum|EAL84300.1

Group10330: awen|OJJ35554.1 awen|OJJ38665.1 afla|EED45524.1 agla|OJJ79658.1 aory|EIT75717.1 agla|OJJ79657.1

Group10331: awen|OJJ36151.1 awen|OJJ39651.1 aory|EIT80995.1 afla|EED49642.1 aory|EIT80963.1 azon|OJJ45200.1

Group10332: azon|OJJ43902.1 azon|OJJ45176.1 aory|EIT76053.1 asyd|OJJ54879.1 awen|OJJ33642.1 awen|OJJ36384.1

Group10333: azon|OJJ46408.1 azon|OJJ46425.1 acla|EAW12201.1 agla|OJJ79017.1 awen|OJJ39828.1 agla|OJJ89184.1

Group10334: azon|OJJ47737.1 azon|OJJ47738.1 acla|EAW10002.1 asyd|OJJ52067.1 atub|OJI90794.1 aver|OJJ04279.1

Group10335: ater|EAU32799.1 ater|EAU32800.1 acla|EAW11704.1 afis|EAW25203.1 awen|OJJ40149.1 afis|EAW15641.1

Group10336: agla|OJJ79260.1 agla|OJJ83552.1 agla|OJJ80192.1 agla|OJJ86165.1 agla|OJJ83845.1 agla|OJJ81286.1

Group10337: agla|OJJ84815.1 agla|OJJ84818.1 agla|OJJ84817.1 agla|OJJ84819.1 agla|OJJ84820.1 agla|OJJ84821.1

Group10338: aluc|OJZ79561.1 aluc|OJZ79857.1 aluc|OJZ80400.1 aluc|OJZ80031.1 aluc|OJZ88825.1 aluc|OJZ80210.1

Group10339: aluc|OJZ79569.1 aluc|OJZ79651.1 aluc|OJZ79845.1 aluc|OJZ79805.1 aluc|OJZ80038.1 aluc|OJZ80181.1

Group10340: aluc|OJZ79594.1 aluc|OJZ79691.1 aluc|OJZ80032.1 aluc|OJZ79803.1 aluc|OJZ80428.1 aluc|OJZ80209.1

Group10341: aluc|OJZ79765.1 aluc|OJZ80131.1 aluc|OJZ79930.1 aluc|OJZ80443.1 aluc|OJZ81121.1 aluc|OJZ79913.1

Group10342: abra|OJJ65430.1 akaw|BAL04878.1 aluc|OJZ79468.1 aory|AFM82519.1 azon|OJJ41973.1 acar|OOF89757.1

Group10343: abra|OJJ65441.1 acar|OOF89781.1 akaw|BAL04886.1 aory|AFM82515.1 abra|OJJ74156.1 azon|OJJ41974.1

Group10344: abra|OJJ65471.1 acar|OOF91980.1 acla|EAW12245.1 afis|EAW16985.1 afum|EBA27276.1 aluc|OJZ87039.1

Group10345: abra|OJJ65578.1 acla|EAW08290.1 afis|EAW23352.1 afla|EED55611.1 afum|EAL88124.2 aory|EIT81778.1

Group10346: abra|OJJ65597.1 akaw|GAA92335.1 aluc|OJZ80458.1 anid|CBF78445.1 anig|EHA20202.1 asyd|OJJ52117.1

Group10347: abra|OJJ65605.1 asyd|OJJ52072.1 ater|EAU31024.1 aver|OJJ03031.1 azon|OJJ45965.1 awen|OJJ35369.1

Group10348: abra|OJJ65639.1 afla|EED47430.1 aluc|OJZ91196.1 anid|CBF82228.1 aory|EIT81917.1 ater|EAU35409.1

Group10349: abra|OJJ65720.1 afis|EAW17325.1 afum|EAL91162.1 agla|OJJ81285.1 anid|CBF82951.1 agla|OJJ86251.1

Group10350: asyd|OJJ61776.1 aver|OJI95879.1 azon|OJJ46057.1 afis|EAW19284.1 afum|EAL94010.1 ater|EAU34096.1

Group10351: acar|OOF92886.1 abra|OJJ65986.1 akaw|GAA90860.1 aluc|OJZ82973.1 atub|OJI89206.1 awen|OJJ29601.1

Group10352: awen|OJJ29547.1 afis|EAW22023.1 afum|EAL87807.1 akaw|GAA93360.1 anid|CBF75934.1 afum|EAL86764.1

Group10353: abra|OJJ65841.1 afla|EED44983.1 akaw|GAA92842.1 aory|EIT73608.1 asyd|OJJ51840.1 atub|OJI80376.1

Group10354: abra|OJJ65857.1 acar|OOF95017.1 akaw|GAA92288.1 aluc|OJZ90185.1 anig|EHA23684.1 atub|OJI81612.1

Group10355: abra|OJJ65879.1 anid|CBF70159.1 asyd|OJJ58802.1 ater|EAU36040.1 aver|OJJ01406.1 awen|OJJ40927.1

Group10356: agla|OJJ85712.1 anid|CBF82295.1 asyd|OJJ52730.1 afis|EAW17761.1 afum|EAL91571.1 ater|EAU32268.1

Group10357: abra|OJJ65907.1 acar|OOF90312.1 akaw|GAA92324.1 aluc|OJZ90222.1 anig|EHA23720.1 atub|OJI81655.1

Group10358: afis|EAW20774.1 acla|EAW11084.1 anid|CBF90339.1 asyd|OJJ59878.1 ater|EAU35826.1 aver|OJJ02408.1

Group10359: abra|OJJ65983.1 akaw|GAA90856.1 aluc|OJZ82977.1 atub|OJI89210.1 azon|OJJ50384.1 acar|OOF92890.1

Group10360: abra|OJJ66000.1 acar|OOF92100.1 akaw|GAA90872.1 aluc|OJZ82961.1 anig|EHA21377.1 atub|OJI89193.1

Group10361: abra|OJJ66042.1 afla|EED53086.1 afum|EAL85337.1 aory|EIT74810.1 atub|OJI87253.1 aver|OJJ08351.1

Group10362: abra|OJJ66048.1 afla|EED47798.1 anid|CBF85569.1 aory|EIT77152.1 asyd|OJJ61616.1 aver|OJJ00939.1

Group10363: abra|OJJ66062.1 akaw|GAA84176.1 aluc|OJZ81101.1 atub|OJI82479.1 abra|OJJ66068.1 anig|EHA19269.1

Group10364: abra|OJJ66075.1 afla|EED55023.1 anig|EHA22528.1 aory|EIT81973.1 asyd|OJJ54331.1 awen|OJJ41954.1

Group10365: abra|OJJ66077.1 acar|OOF99297.1 akaw|GAA84169.1 aluc|OJZ81094.1 anig|EHA19277.1 atub|OJI82489.1

Group10366: abra|OJJ66078.1 acar|OOF99294.1 akaw|GAA84168.1 aluc|OJZ81093.1 anig|EHA19278.1 atub|OJI82490.1

Group10367: abra|OJJ66084.1 acar|OOF99281.1 akaw|GAA84161.1 aluc|OJZ81084.1 anig|EHA19284.1 atub|OJI82500.1

Group10368: abra|OJJ66117.1 acar|OOF94009.1 akaw|GAA92679.1 aluc|OJZ90134.1 anig|EHA20937.1 atub|OJI84662.1

Group10369: abra|OJJ66147.1 aluc|OJZ87030.1 asyd|OJJ52810.1 ater|EAU32276.1 aver|OJJ08788.1 awen|OJJ32687.1

Group10370: anig|EHA22187.1 acar|OOF98855.1 agla|OJJ78699.1 aluc|OJZ80595.1 asyd|OJJ64212.1 awen|OJJ29866.1

Group10371: abra|OJJ66210.1 acar|OOF90650.1 akaw|GAA89077.1 aluc|OJZ80514.1 anig|EHA25679.1 atub|OJI88416.1

Group10372: abra|OJJ66240.1 akaw|GAA89049.1 aluc|OJZ80545.1 anig|EHA25650.1 ater|EAU37347.1 atub|OJI88386.1

Group10373: abra|OJJ66258.1 acar|OOF90728.1 aluc|OJZ80565.1 anig|EHA25633.1 atub|OJI88366.1 awen|OJJ38261.1

Group10374: abra|OJJ66264.1 acar|OOF90740.1 akaw|GAA89031.1 aluc|OJZ80572.1 anig|EHA25626.1 atub|OJI88359.1

Group10375: abra|OJJ66285.1 acar|OOF97578.1 afla|EED53302.1 anig|EHA23665.1 aory|EIT75229.1 atub|OJI89938.1

Group10376: afum|EAL86538.1 afis|EAW20678.1 agla|OJJ81546.1 anid|CBF84458.1 aory|EIT78065.1 asyd|OJJ52425.1

Group10377: acar|OOF93931.1 afis|EAW24668.1 akaw|GAA92101.1 aluc|OJZ83624.1 anig|EHA27516.1 atub|OJI87852.1

Group10378: abra|OJJ66338.1 aluc|OJZ83131.1 anig|EHA28191.1 atub|OJI87804.1 awen|OJJ38742.1 awen|OJJ35029.1

Group10379: abra|OJJ66372.1 acar|OOF90518.1 akaw|GAA90241.1 aluc|OJZ83099.1 anig|EHA28163.1 atub|OJI87836.1

Group10380: awen|OJJ32643.1 acar|OOF97524.1 afla|EED45828.1 anid|CBF82890.1 aory|EIT75667.1 ater|EAU29947.1

Group10381: asyd|OJJ54304.1 afis|EAW21272.1 afum|EAL92287.1 anid|CBF80705.1 ater|EAU31603.1 azon|OJJ45142.1

Group10382: afla|EED47337.1 awen|OJJ33857.1 anid|CBF81977.1 asyd|OJJ54442.1 aver|OJJ05822.1 asyd|OJJ57340.1

Group10383: abra|OJJ66434.1 acar|OOF93320.1 agla|OJJ84362.1 akaw|GAA90299.1 aluc|OJZ83038.1 atub|OJI87911.1

Group10384: azon|OJJ50390.1 afla|EED49675.1 anid|CBF80084.1 aory|EIT81002.1 asyd|OJJ58306.1 aver|OJJ02106.1

Group10385: abra|OJJ66447.1 afla|EED47381.1 aory|EIT82864.1 asyd|OJJ54739.1 aver|OJJ07426.1 awen|OJJ33682.1

Group10386: azon|OJJ46451.1 abra|OJJ74445.1 akaw|GAA87337.1 aluc|OJZ89677.1 anig|EHA26912.1 atub|OJI85123.1

Group10387: abra|OJJ66462.1 acar|OOF98735.1 asyd|OJJ63039.1 aver|OJI97928.1 azon|OJJ42626.1 abra|OJJ73164.1

Group10388: abra|OJJ66472.1 akaw|GAA92458.1 aluc|OJZ81559.1 anid|CBF75988.1 asyd|OJJ53458.1 atub|OJI88626.1

Group10389: abra|OJJ66490.1 acar|OOF90904.1 akaw|GAA91210.1 aluc|OJZ81545.1 anig|EHA25890.1 atub|OJI88639.1

Group10390: afla|EED45120.1 aluc|OJZ92859.1 aory|EIT73830.1 atub|OJI87317.1 awen|OJJ31226.1 akaw|GAA91837.1

Group10391: abra|OJJ66621.1 acar|OOF91044.1 akaw|GAA91104.1 aluc|OJZ81428.1 anig|EHA25994.1 atub|OJI88752.1

Group10392: abra|OJJ66765.1 acar|OOF93846.1 akaw|GAA89701.1 aluc|OJZ80683.1 anig|EHA27330.1 atub|OJI87542.1

Group10393: afum|EAL89363.1 akaw|GAA83432.1 atub|OJI90305.1 agla|OJJ87489.1 aluc|OJZ83933.1 asyd|OJJ64748.1

Group10394: abra|OJJ66782.1 acar|OOF93859.1 akaw|GAA89717.1 aluc|OJZ80667.1 atub|OJI87522.1 afis|EAW19290.1

Group10395: abra|OJJ66795.1 acar|OOF93872.1 akaw|GAA86860.1 aluc|OJZ89984.1 anig|EHA27355.1 atub|OJI84826.1

Group10396: abra|OJJ66801.1 acla|EAW14120.1 asyd|OJJ57638.1 ater|EAU37623.1 aver|OJJ01944.1 azon|OJJ44711.1

Group10397: abra|OJJ66817.1 acar|OOF91600.1 akaw|GAA90455.1 aluc|OJZ85769.1 atub|OJI79554.1 azon|OJJ50733.1

Group10398: abra|OJJ66838.1 akaw|GAA90475.1 aluc|OJZ85748.1 acar|OOF95831.1 ater|EAU35194.1 asyd|OJJ55923.1

Group10399: abra|OJJ66951.1 acar|OOF91716.1 akaw|GAA91469.1 aluc|OJZ85628.1 anig|EHA24901.1 atub|OJI79420.1

Group10400: abra|OJJ66961.1 acar|OOF93646.1 akaw|GAA92200.1 aluc|OJZ85615.1 anig|EHA24911.1 atub|OJI79408.1

Group10401: abra|OJJ66975.1 acar|OOF93638.1 akaw|GAA92190.1 aluc|OJZ85602.1 anig|EHA24922.1 atub|OJI79397.1

Group10402: afum|EBA27187.1 acla|EAW13433.1 afis|EAW20549.1 agla|OJJ88737.1 awen|OJJ33261.1 azon|OJJ50632.1

Group10403: abra|OJJ66999.1 acar|OOF93617.1 akaw|GAA92166.1 aluc|OJZ85574.1 anig|EHA24945.1 atub|OJI79369.1

Group10404: abra|OJJ67018.1 acar|OOF99009.1 afla|EED49608.1 anig|EHA18792.1 aory|EIT73526.1 afla|EED45163.1

Group10405: abra|OJJ67020.1 acar|OOG00388.1 akaw|GAA92143.1 aluc|OJZ85548.1 anig|EHA24954.1 atub|OJI79357.1

Group10406: abra|OJJ67021.1 acar|OOG00387.1 akaw|GAA92144.1 aluc|OJZ85549.1 anig|EHA24955.1 atub|OJI79356.1

Group10407: abra|OJJ67022.1 acar|OOG00386.1 akaw|GAA92145.1 aluc|OJZ85550.1 anig|EHA24956.1 atub|OJI79355.1

Group10408: abra|OJJ67023.1 acar|OOG00385.1 akaw|GAA92146.1 aluc|OJZ85551.1 anig|EHA24957.1 atub|OJI79354.1

Group10409: abra|OJJ67033.1 acar|OOF98677.1 akaw|GAA92156.1 aluc|OJZ85563.1 anig|EHA24969.1 atub|OJI79341.1

Group10410: abra|OJJ67041.1 afla|EED52067.1 agla|OJJ78564.1 anig|EHA18601.1 azon|OJJ42507.1 afla|EED53081.1

Group10411: abra|OJJ67044.1 afla|EED57120.1 aluc|OJZ83572.1 anig|EHA18599.1 azon|OJJ42511.1 azon|OJJ48982.1

Group10412: abra|OJJ67046.1 afla|EED52064.1 anig|EHA18597.1 aory|EIT72301.1 azon|OJJ42509.1 ater|EAU35195.1

Group10413: abra|OJJ67065.1 afla|EED45196.1 anid|CBF83087.1 anig|EHA18941.1 akaw|GAA89179.1 aluc|OJZ81654.1

Group10414: abra|OJJ67073.1 acar|OOF99306.1 akaw|GAA91082.1 aluc|OJZ87899.1 anig|EHA20196.1 atub|OJI80752.1

Group10415: abra|OJJ67076.1 acar|OOF99074.1 akaw|GAA92134.1 aluc|OJZ83589.1 anig|EHA22154.1 atub|OJI89955.1

Group10416: abra|OJJ67084.1 akaw|GAA91075.1 aluc|OJZ87890.1 asyd|OJJ59833.1 atub|OJI80762.1 aver|OJJ02455.1

Group10417: abra|OJJ67090.1 akaw|GAA91071.1 aluc|OJZ87885.1 anig|EHA20188.1 atub|OJI80766.1 awen|OJJ38312.1

Group10418: abra|OJJ67108.1 acar|OOF98444.1 akaw|GAA91056.1 aluc|OJZ87867.1 anig|EHA20173.1 atub|OJI80781.1

Group10419: abra|OJJ67110.1 acar|OOF98442.1 akaw|GAA91054.1 aluc|OJZ87865.1 anig|EHA20171.1 atub|OJI80783.1

Group10420: abra|OJJ67119.1 acar|OOF98433.1 akaw|GAA91046.1 aluc|OJZ87856.1 anig|EHA20162.1 atub|OJI80791.1

Group10421: afla|EED54178.1 acla|EAW08584.1 afis|EAW16400.1 afum|EBA27365.1 azon|OJJ47064.1 azon|OJJ42648.1

Group10422: abra|OJJ67222.1 acar|OOF92457.1 aluc|OJZ87754.1 anig|EHA20241.1 atub|OJI80890.1 awen|OJJ41933.1

Group10423: abra|OJJ67223.1 acar|OOF98329.1 akaw|GAA91650.1 aluc|OJZ87751.1 anig|EHA20239.1 atub|OJI80892.1

Group10424: agla|OJJ87686.1 afla|EED48226.1 aory|EIT72932.1 asyd|OJJ54915.1 ater|EAU30160.1 aver|OJJ07288.1

Group10425: abra|OJJ67267.1 akaw|GAA89063.1 aluc|OJZ80529.1 anig|EHA19570.1 atub|OJI88401.1 anig|EHA25666.1

Group10426: abra|OJJ67287.1 aluc|OJZ85071.1 atub|OJI80527.1 abra|OJJ67645.1 awen|OJJ35366.1 azon|OJJ45347.1

Group10427: abra|OJJ67316.1 acar|OOF92499.1 akaw|GAA86502.1 aluc|OJZ85045.1 anig|EHA25262.1 atub|OJI80553.1

Group10428: abra|OJJ67378.1 acar|OOF92434.1 akaw|GAA86445.1 aluc|OJZ84987.1 anig|EHA25200.1 atub|OJI80613.1

Group10429: afis|EAW21098.1 afum|EAL92121.1 acar|OOF91914.1 afla|EED50862.1 anig|EHA20145.1 aory|EIT79061.1

Group10430: awen|OJJ41051.1 afis|EAW16411.1 afum|EAL89035.1 anid|CBF73417.1 aver|OJJ03473.1 azon|OJJ44613.1

Group10431: abra|OJJ67525.1 agla|OJJ88365.1 akaw|GAA88138.1 aluc|OJZ81409.1 anig|EHA18707.1 atub|OJI91615.1

Group10432: agla|OJJ89427.1 afla|EED48105.1 aory|EIT76250.1 asyd|OJJ52803.1 aver|OJJ08780.1 awen|OJJ38062.1

Group10433: abra|OJJ67558.1 agla|OJJ86033.1 aluc|OJZ81375.1 atub|OJI91581.1 awen|OJJ30156.1 anid|CBF88274.1

Group10434: abra|OJJ67561.1 acar|OOG00277.1 akaw|GAA88105.1 aluc|OJZ81372.1 anig|EHA18671.1 atub|OJI91578.1

Group10435: abra|OJJ67582.1 akaw|GAA85347.1 aluc|OJZ90709.1 anid|CBF76179.1 anig|EHA24576.1 atub|OJI82136.1

Group10436: awen|OJJ31273.1 acla|EAW07079.1 acar|OOF90671.1 afla|EED52119.1 aory|EIT74478.1 anid|CBF89958.1

Group10437: abra|OJJ67656.1 acar|OOF91989.1 akaw|GAA89172.1 aluc|OJZ81661.1 anig|EHA25782.1 atub|OJI88522.1

Group10438: abra|OJJ67665.1 akaw|GAA89181.1 aluc|OJZ81651.1 anig|EHA25792.1 atub|OJI88531.1 azon|OJJ45276.1

Group10439: abra|OJJ67670.1 acar|OOF90914.1 akaw|GAA89186.1 aluc|OJZ81645.1 anig|EHA25798.1 atub|OJI88537.1

Group10440: abra|OJJ67671.1 acar|OOF90915.1 akaw|GAA89187.1 aluc|OJZ81644.1 anig|EHA25799.1 atub|OJI88538.1

Group10441: abra|OJJ67691.1 afis|EAW22025.1 afla|EED55372.1 afum|EAL85442.1 agla|OJJ86871.1 aory|EIT80897.1

Group10442: abra|OJJ67725.1 acar|OOF98717.1 akaw|GAA89219.1 aluc|OJZ81610.1 anig|EHA25836.1 atub|OJI88576.1

Group10443: abra|OJJ67757.1 acar|OOF98136.1 afla|EED56905.1 agla|OJJ84581.1 aory|EIT77493.1 afla|EED56904.1

Group10444: acla|EAW07632.1 anid|CBF79138.1 aory|EIT80510.1 aver|OJJ03897.1 afla|EED51734.1 ater|EAU33339.1

Group10445: abra|OJJ67780.1 afis|EAW15615.1 afla|EED45885.1 afum|EAL85720.1 aory|EIT75828.1 ater|EAU31882.1

Group10446: abra|OJJ67803.1 afla|EED53415.1 agla|OJJ84579.1 aluc|OJZ90854.1 aory|EIT75008.1 aver|OJJ07761.1

Group10447: abra|OJJ67815.1 acar|OOF94643.1 anid|CBF82006.1 anig|EHA20936.1 aory|EIT78893.1 ater|EAU38017.1

Group10448: abra|OJJ67816.1 acar|OOF91993.1 afum|EAL86778.1 ater|EAU38018.1 aver|OJJ07622.1 awen|OJJ36346.1

Group10449: afum|EAL85091.1 afis|EAW19799.1 agla|OJJ79579.1 anid|CBF80376.1 asyd|OJJ54811.1 aver|OJJ07350.1

Group10450: abra|OJJ67885.1 acar|OOF92861.1 akaw|GAA91892.1 aluc|OJZ82004.1 anig|EHA22252.1 atub|OJI84554.1

Group10451: abra|OJJ67902.1 acar|OOF89863.1 akaw|GAA91907.1 aluc|OJZ81987.1 anig|EHA22269.1 atub|OJI84538.1

Group10452: abra|OJJ67943.1 acar|OOF92792.1 akaw|GAA91253.1 aluc|OJZ82815.1 anig|EHA21255.1 atub|OJI89046.1

Group10453: abra|OJJ67982.1 acar|OOF92023.1 akaw|GAA91220.1 aluc|OJZ82850.1 anig|EHA21287.1 atub|OJI89080.1

Group10454: abra|OJJ67983.1 acar|OOF92024.1 akaw|GAA91219.1 aluc|OJZ82851.1 atub|OJI89081.1 anig|EHA23744.1

Group10455: abra|OJJ68007.1 acar|OOF92049.1 akaw|GAA90943.1 aluc|OJZ82881.1 anig|EHA21307.1 atub|OJI89105.1

Group10456: abra|OJJ68009.1 acar|OOF92051.1 akaw|GAA90941.1 aluc|OJZ82883.1 anig|EHA21309.1 atub|OJI89108.1

Group10457: abra|OJJ68014.1 acar|OOF92056.1 akaw|GAA90937.1 aluc|OJZ82887.1 anig|EHA21313.1 atub|OJI89112.1

Group10458: abra|OJJ68023.1 acar|OOF92067.1 akaw|GAA90928.1 aluc|OJZ82897.1 anig|EHA21321.1 atub|OJI89122.1

Group10459: abra|OJJ68025.1 akaw|GAA90927.1 aluc|OJZ82898.1 anig|EHA21322.1 atub|OJI89124.1 acar|OOF92068.1

Group10460: asyd|OJJ53538.1 acla|EAW12055.1 aluc|OJZ92004.1 atub|OJI86439.1 akaw|GAA82163.1 anig|EHA22944.1

Group10461: abra|OJJ68055.1 agla|OJJ87898.1 aluc|OJZ80197.1 asyd|OJJ57374.1 aver|OJJ05794.1 awen|OJJ31627.1

Group10462: abra|OJJ68132.1 acar|OOF90086.1 akaw|GAA86912.1 aluc|OJZ90035.1 atub|OJI84771.1 awen|OJJ39616.1

Group10463: abra|OJJ68155.1 afla|EED47516.1 akaw|GAA90124.1 aluc|OJZ83226.1 aory|EIT74590.1 atub|OJI87710.1

Group10464: abra|OJJ68161.1 agla|OJJ89558.1 akaw|GAA85356.1 aluc|OJZ90718.1 anig|EHA19728.1 atub|OJI82145.1

Group10465: abra|OJJ68162.1 akaw|GAA85357.1 aluc|OJZ90719.1 anig|EHA19727.1 atub|OJI82146.1 awen|OJJ37323.1

Group10466: aory|EIT79542.1 acar|OOF98704.1 anig|EHA20950.1 atub|OJI84676.1 azon|OJJ45278.1 awen|OJJ33319.1

Group10467: abra|OJJ68294.1 afla|EED46577.1 akaw|GAA83179.1 aluc|OJZ83678.1 asyd|OJJ52231.1 atub|OJI90049.1

Group10468: abra|OJJ68309.1 acar|OOF96699.1 akaw|GAA83165.1 aluc|OJZ83664.1 anig|EHA18814.1 atub|OJI90035.1

Group10469: abra|OJJ68312.1 acar|OOF96696.1 akaw|GAA83161.1 aluc|OJZ83661.1 anig|EHA18813.1 atub|OJI90032.1

Group10470: abra|OJJ68318.1 akaw|GAA83155.1 aluc|OJZ83655.1 anig|EHA18807.1 atub|OJI90025.1 azon|OJJ45026.1

Group10471: abra|OJJ68327.1 acar|OOF90098.1 akaw|GAA92078.1 aluc|OJZ83646.1 anig|EHA18798.1 atub|OJI90015.1

Group10472: abra|OJJ68328.1 acar|OOF97595.1 akaw|GAA92079.1 aluc|OJZ83645.1 anig|EHA18797.1 atub|OJI90014.1

Group10473: abra|OJJ68329.1 acar|OOF97594.1 akaw|GAA92081.1 aluc|OJZ83643.1 anig|EHA18796.1 atub|OJI90013.1

Group10474: abra|OJJ68330.1 acar|OOF97609.1 akaw|GAA92082.1 aluc|OJZ83642.1 anig|EHA18795.1 atub|OJI90012.1

Group10475: abra|OJJ68345.1 acar|OOF97644.1 akaw|GAA92112.1 aluc|OJZ83613.1 anig|EHA18770.1 atub|OJI89986.1

Group10476: abra|OJJ68374.1 akaw|GAA92665.1 aluc|OJZ90119.1 anig|EHA20952.1 atub|OJI84678.1 awen|OJJ31418.1

Group10477: abra|OJJ68394.1 afla|EED57178.1 akaw|GAA92554.1 aluc|OJZ90102.1 ater|EAU36573.1 atub|OJI84704.1

Group10478: abra|OJJ68413.1 afis|EAW15645.1 akaw|GAA92537.1 aluc|OJZ90085.1 anig|EHA20989.1 atub|OJI84721.1

Group10479: abra|OJJ68418.1 acar|OOF93925.1 akaw|GAA92530.1 aluc|OJZ90079.1 anig|EHA20994.1 atub|OJI84727.1

Group10480: anid|CBF73566.1 asyd|OJJ56697.1 aver|OJJ06027.1 acla|EAW14119.1 afis|EAW18827.1 afum|EAL93561.1

Group10481: abra|OJJ68576.1 acar|OOF92878.1 akaw|GAA91915.1 aluc|OJZ81965.1 anig|EHA22282.1 atub|OJI84516.1

Group10482: abra|OJJ68578.1 akaw|GAA91917.1 aluc|OJZ81963.1 anig|EHA22284.1 atub|OJI84514.1 acar|OOF92104.1

Group10483: abra|OJJ68640.1 agla|OJJ84965.1 akaw|GAA89778.1 aluc|OJZ81915.1 asyd|OJJ60398.1 atub|OJI84467.1

Group10484: abra|OJJ68808.1 acar|OOF90609.1 akaw|GAA90446.1 aluc|OJZ85780.1 anig|EHA24775.1 atub|OJI79565.1

Group10485: asyd|OJJ64662.1 aver|OJI97479.1 aver|OJI98764.1 anid|CBF84018.1 ater|EAU38121.1 awen|OJJ31107.1

Group10486: abra|OJJ68817.1 acar|OOF96411.1 akaw|GAA90435.1 aluc|OJZ85792.1 anig|EHA24768.1 atub|OJI79578.1

Group10487: abra|OJJ68820.1 akaw|GAA90431.1 aluc|OJZ85796.1 atub|OJI79582.1 awen|OJJ31951.1 azon|OJJ42671.1

Group10488: abra|OJJ68821.1 asyd|OJJ61025.1 awen|OJJ30915.1 afla|EED51704.1 aory|EIT80539.1 ater|EAU30961.1

Group10489: abra|OJJ68839.1 acar|OOF97928.1 akaw|GAA90410.1 aluc|OJZ85818.1 anig|EHA24743.1 atub|OJI79604.1

Group10490: abra|OJJ68848.1 acar|OOF90542.1 akaw|GAA90401.1 aluc|OJZ85828.1 anig|EHA24733.1 atub|OJI79616.1

Group10491: abra|OJJ68854.1 akaw|GAA90399.1 aluc|OJZ85832.1 anig|EHA24731.1 ater|EAU38595.1 atub|OJI79620.1

Group10492: abra|OJJ68869.1 anid|CBF79171.1 anig|EHA24717.1 asyd|OJJ56242.1 aver|OJJ05165.1 awen|OJJ36396.1

Group10493: abra|OJJ68881.1 acar|OOF90517.1 akaw|GAA90375.1 aluc|OJZ85857.1 anig|EHA28093.1 atub|OJI79643.1

Group10494: abra|OJJ68887.1 akaw|GAA90369.1 aluc|OJZ85863.1 anig|EHA28089.1 atub|OJI79649.1 awen|OJJ41446.1

Group10495: agla|OJJ85339.1 asyd|OJJ64783.1 aver|OJI98894.1 aluc|OJZ91594.1 atub|OJI86045.1 anig|EHA22623.1

Group10496: abra|OJJ68909.1 afis|EAW23719.1 anig|EHA21436.1 asyd|OJJ54999.1 aver|OJJ07719.1 awen|OJJ34233.1

Group10497: abra|OJJ68914.1 acar|OOF91501.1 akaw|GAA90335.1 aluc|OJZ85897.1 anig|EHA24695.1 atub|OJI79682.1

Group10498: abra|OJJ68995.1 acar|OOF92706.1 akaw|GAA90514.1 aluc|OJZ85979.1 anig|EHA24627.1 atub|OJI79769.1

Group10499: abra|OJJ69068.1 acar|OOF91230.1 akaw|GAA90599.1 aluc|OJZ86057.1 anig|EHA24555.1 atub|OJI79854.1

Group10500: abra|OJJ69135.1 acar|OOF91166.1 akaw|GAA86321.1 aluc|OJZ86123.1 anig|EHA24498.1 atub|OJI79919.1

Group10501: abra|OJJ69336.1 acar|OOF94390.1 akaw|GAA88206.1 aluc|OJZ82168.1 anig|EHA28564.1 atub|OJI88269.1

Group10502: abra|OJJ69502.1 acar|OOF94249.1 akaw|GAA88321.1 aluc|OJZ82341.1 anig|EHA28419.1 atub|OJI88099.1

Group10503: abra|OJJ69526.1 acar|OOF94215.1 akaw|GAA88299.1 aluc|OJZ82366.1 anig|EHA28399.1 atub|OJI88076.1

Group10504: aver|OJJ08860.1 acar|OOF91857.1 acla|EAW15093.1 afum|EAL91111.1 akaw|GAA87906.1 ater|EAU34633.1

Group10505: abra|OJJ69560.1 afis|EAW25594.1 awen|OJJ36408.1 acar|OOF91590.1 acla|EAW07717.1 ater|EAU29336.1

Group10506: abra|OJJ69626.1 acar|OOF92685.1 akaw|GAA91606.1 aluc|OJZ82431.1 anig|EHA28340.1 atub|OJI88010.1

Group10507: abra|OJJ69638.1 afum|EAL90965.1 akaw|GAA91618.1 aluc|OJZ82444.1 anig|EHA28329.1 atub|OJI87996.1

Group10508: abra|OJJ69641.1 acar|OOF96819.1 akaw|GAA91619.1 aluc|OJZ82445.1 anig|EHA28328.1 atub|OJI87995.1

Group10509: abra|OJJ69642.1 acar|OOF95888.1 akaw|GAA91620.1 aluc|OJZ82446.1 anig|EHA28327.1 atub|OJI87994.1

Group10510: abra|OJJ69656.1 acar|OOF98682.1 akaw|GAA91635.1 aluc|OJZ82462.1 anig|EHA28313.1 atub|OJI87976.1

Group10511: abra|OJJ69686.1 acar|OOF91754.1 akaw|GAA88810.1 aluc|OJZ82493.1 anig|EHA28279.1 atub|OJI87939.1

Group10512: abra|OJJ69694.1 afla|EED48240.1 anid|CBF84029.1 aory|EIT72961.1 asyd|OJJ61366.1 aver|OJJ00693.1

Group10513: abra|OJJ69714.1 acar|OOF96660.1 anig|EHA19585.1 atub|OJI89168.1 aver|OJJ07492.1 awen|OJJ30133.1

Group10514: abra|OJJ69724.1 acar|OOF94192.1 akaw|GAA90177.1 aluc|OJZ83173.1 anig|EHA28247.1 atub|OJI87765.1

Group10515: abra|OJJ69736.1 acar|OOF94184.1 akaw|GAA90189.1 aluc|OJZ83160.1 anig|EHA28224.1 atub|OJI87777.1

Group10516: abra|OJJ69771.1 afis|EAW25194.1 asyd|OJJ52879.1 ater|EAU31193.1 aver|OJJ07169.1 asyd|OJJ55037.1

Group10517: abra|OJJ69798.1 acar|OOF92533.1 akaw|GAA87861.1 aluc|OJZ85502.1 anig|EHA25035.1 atub|OJI80063.1

Group10518: abra|OJJ69820.1 acar|OOF93343.1 akaw|GAA87840.1 aluc|OJZ85476.1 anig|EHA25017.1 atub|OJI80087.1

Group10519: abra|OJJ69843.1 acar|OOF93366.1 akaw|GAA87816.1 aluc|OJZ85451.1 anig|EHA24994.1 atub|OJI80116.1

Group10520: abra|OJJ69849.1 acar|OOF93372.1 akaw|GAA87809.1 aluc|OJZ85444.1 anig|EHA24988.1 atub|OJI80123.1

Group10521: abra|OJJ69854.1 acar|OOF93380.1 akaw|GAA87804.1 aluc|OJZ85439.1 anig|EHA24984.1 atub|OJI80129.1

Group10522: abra|OJJ69876.1 akaw|GAA87780.1 aluc|OJZ85415.1 anig|EHA25594.1 atub|OJI80161.1 acla|EAW13220.1

Group10523: azon|OJJ48214.1 acar|OOF94715.1 akaw|GAA86261.1 aluc|OJZ90519.1 anig|EHA23972.1 atub|OJI81943.1

Group10524: abra|OJJ69918.1 acar|OOF93430.1 akaw|GAA87738.1 aluc|OJZ85370.1 anig|EHA25554.1 atub|OJI80200.1

Group10525: abra|OJJ69947.1 acar|OOF93456.1 akaw|GAA87713.1 aluc|OJZ85340.1 anig|EHA25528.1 atub|OJI80231.1

Group10526: abra|OJJ70007.1 acar|OOF93513.1 aluc|OJZ85272.1 ater|EAU32548.1 atub|OJI80299.1 awen|OJJ38245.1

Group10527: abra|OJJ70035.1 afis|EAW24623.1 afum|EAL86251.1 anid|CBF79088.1 anig|EHA18720.1 aver|OJJ08542.1

Group10528: abra|OJJ70056.1 acar|OOF93541.1 akaw|GAA87618.1 aluc|OJZ85239.1 anid|CBF83197.1 atub|OJI80336.1

Group10529: abra|OJJ70071.1 acar|OOF97445.1 akaw|GAA87608.1 aluc|OJZ85228.1 anig|EHA25432.1 atub|OJI80351.1

Group10530: abra|OJJ70073.1 acla|EAW12447.1 aory|EIT74228.1 acar|OOG00947.1 acla|EAW12166.1 anid|CBF75785.1

Group10531: abra|OJJ70090.1 acla|EAW12306.1 afis|EAW21033.1 afum|EAL92071.1 anig|EHA22486.1 atub|OJI89251.1

Group10532: abra|OJJ70103.1 akaw|GAA87597.1 aluc|OJZ85217.1 anig|EHA25421.1 atub|OJI80368.1 awen|OJJ39119.1

Group10533: abra|OJJ70136.1 acar|OOF93596.1 akaw|GAA86629.1 aluc|OJZ85184.1 anig|EHA25396.1 atub|OJI80413.1

Group10534: abra|OJJ70138.1 afis|EAW19841.1 akaw|GAA86627.1 aluc|OJZ85182.1 anig|EHA25394.1 atub|OJI80415.1

Group10535: abra|OJJ70166.1 acar|OOF94004.1 akaw|GAA86613.1 aluc|OJZ85164.1 anig|EHA25361.1 atub|OJI80440.1

Group10536: abra|OJJ70201.1 acar|OOF92579.1 akaw|GAA86575.1 aluc|OJZ85126.1 anig|EHA25345.1 atub|OJI80467.1

Group10537: abra|OJJ70210.1 acar|OOF90367.1 akaw|GAA86566.1 aluc|OJZ85117.1 anig|EHA25337.1 atub|OJI80476.1

Group10538: abra|OJJ70219.1 acar|OOF92560.1 aluc|OJZ85108.1 asyd|OJJ57681.1 atub|OJI80485.1 aver|OJJ05487.1

Group10539: abra|OJJ70221.1 afla|EED54371.1 aory|EIT75998.1 atub|OJI80487.1 agla|OJJ78677.1 awen|OJJ40934.1

Group10540: abra|OJJ70241.1 acar|OOF93542.1 agla|OJJ79704.1 akaw|GAA89190.1 aluc|OJZ81641.1 atub|OJI84630.1

Group10541: abra|OJJ70256.1 afla|EED44861.1 anig|EHA25305.1 aory|EIT82727.1 ater|EAU36571.1 anid|CBF69432.1

Group10542: abra|OJJ70284.1 acar|OOF91737.1 akaw|GAA92863.1 aluc|OJZ82550.1 anig|EHA22514.1 atub|OJI88791.1

Group10543: abra|OJJ70311.1 afis|EAW17295.1 akaw|GAA93012.1 aluc|OJZ82573.1 anig|EHA27500.1 atub|OJI88817.1

Group10544: abra|OJJ70322.1 acar|OOF91777.1 akaw|GAA92578.1 aluc|OJZ82591.1 anig|EHA27484.1 atub|OJI88826.1

Group10545: abra|OJJ70334.1 acar|OOF91812.1 akaw|GAA84587.1 aluc|OJZ89331.1 anig|EHA27476.1 atub|OJI85469.1

Group10546: abra|OJJ70338.1 akaw|GAA92591.1 aluc|OJZ82608.1 atub|OJI88843.1 aver|OJJ04185.1 awen|OJJ34534.1

Group10547: abra|OJJ70345.1 acar|OOF97499.1 akaw|GAA92595.1 aluc|OJZ82612.1 anig|EHA27466.1 atub|OJI88847.1

Group10548: abra|OJJ70399.1 acar|OOF91876.1 akaw|GAA91959.1 aluc|OJZ82658.1 anig|EHA27415.1 atub|OJI88890.1

Group10549: afla|EED52104.1 aory|EIT76899.1 ater|EAU33022.1 acar|OOF98874.1 awen|OJJ32047.1 acla|EAW13637.1

Group10550: abra|OJJ70411.1 acar|OOF90920.1 akaw|GAA91945.1 aluc|OJZ82675.1 anig|EHA21041.1 atub|OJI88910.1

Group10551: abra|OJJ70415.1 acar|OOF91855.1 akaw|GAA91941.1 aluc|OJZ82680.1 anig|EHA21048.1 atub|OJI88913.1

Group10552: akaw|GAA90155.1 anig|EHA21056.1 afla|EED46002.1 aluc|OJZ83192.1 aory|EIT75649.1 atub|OJI87746.1

Group10553: anid|CBF81919.1 asyd|OJJ65136.1 acar|OOF90359.1 agla|OJJ87447.1 aver|OJI99270.1 azon|OJJ46518.1

Group10554: abra|OJJ70443.1 akaw|GAA90137.1 aluc|OJZ83210.1 anig|EHA21070.1 atub|OJI87727.1 azon|OJJ50192.1

Group10555: abra|OJJ70447.1 acar|OOF92545.1 akaw|GAA90134.1 aluc|OJZ83213.1 anig|EHA21073.1 atub|OJI87724.1

Group10556: abra|OJJ70454.1 akaw|GAA90129.1 aluc|OJZ83220.1 anig|EHA21078.1 atub|OJI87717.1 awen|OJJ38301.1

Group10557: abra|OJJ70455.1 acar|OOG00546.1 akaw|GAA90128.1 aluc|OJZ83221.1 anig|EHA21079.1 atub|OJI87716.1

Group10558: abra|OJJ70459.1 afla|EED46518.1 aluc|OJZ83800.1 anig|EHA18948.1 aory|EIT72338.1 acar|OOG00889.1

Group10559: abra|OJJ70491.1 acar|OOF97434.1 aluc|OJZ83835.1 anig|EHA18978.1 atub|OJI90202.1 akaw|GAA83339.1

Group10560: abra|OJJ70524.1 akaw|GAA83371.1 aluc|OJZ83869.1 anig|EHA19010.1 atub|OJI90237.1 awen|OJJ30866.1

Group10561: abra|OJJ70693.1 agla|OJJ86502.1 akaw|GAA90024.1 aluc|OJZ84040.1 anig|EHA19162.1 atub|OJI90412.1

Group10562: abra|OJJ70829.1 afis|EAW18252.1 anig|EHA20879.1 aver|OJI99193.1 awen|OJJ37512.1 azon|OJJ48417.1

Group10563: awen|OJJ34594.1 asyd|OJJ52729.1 afis|EAW17760.1 afum|EAL91570.1 anid|CBF82297.1 ater|EAU32269.1

Group10564: abra|OJJ70948.1 aluc|OJZ89074.1 anid|CBF80794.1 asyd|OJJ54501.1 ater|EAU37594.1 atub|OJI85712.1

Group10565: abra|OJJ71010.1 acar|OOF98132.1 akaw|GAA82943.1 aluc|OJZ87520.1 anig|EHA20731.1 atub|OJI81128.1

Group10566: afis|EAW17009.1 afla|EED48915.1 afum|EAL86528.2 aory|EIT81230.1 azon|OJJ46210.1 awen|OJJ41834.1

Group10567: abra|OJJ71070.1 acar|OOF98075.1 akaw|GAA82892.1 aluc|OJZ87461.1 anig|EHA20683.1 atub|OJI81187.1

Group10568: abra|OJJ71078.1 acar|OOF98061.1 aluc|OJZ87452.1 anig|EHA20674.1 ater|EAU35427.1 atub|OJI81196.1

Group10569: abra|OJJ71080.1 acar|OOF98063.1 akaw|GAA82882.1 aluc|OJZ87450.1 anig|EHA20672.1 atub|OJI81198.1

Group10570: abra|OJJ71083.1 acar|OOF98055.1 akaw|GAA82878.1 aluc|OJZ87445.1 anig|EHA20669.1 atub|OJI81203.1

Group10571: acar|OOF93275.1 afla|EED56368.1 aory|EIT73394.1 akaw|GAA91356.1 aluc|OJZ84816.1 atub|OJI89269.1

Group10572: abra|OJJ71115.1 acar|OOF98035.1 akaw|GAA82855.1 aluc|OJZ87420.1 anig|EHA20644.1 atub|OJI81228.1

Group10573: abra|OJJ71148.1 afum|EAL89902.1 akaw|GAA82829.1 aluc|OJZ87390.1 anig|EHA20619.1 atub|OJI81257.1

Group10574: abra|OJJ71221.1 aluc|OJZ87328.1 anig|EHA20565.1 atub|OJI81317.1 awen|OJJ30958.1 akaw|GAA82769.1

Group10575: ater|EAU36915.1 acar|OOF90884.1 afis|EAW22125.1 asyd|OJJ55526.1 aver|OJJ06604.1 afum|EAL91063.1

Group10576: abra|OJJ71273.1 acar|OOF97901.1 akaw|GAA82717.1 aluc|OJZ87271.1 anig|EHA20512.1 atub|OJI81371.1

Group10577: abra|OJJ71410.1 acar|OOF97762.1 akaw|GAA82590.1 aluc|OJZ87130.1 anig|EHA20392.1 atub|OJI81510.1

Group10578: abra|OJJ71441.1 acar|OOF90398.1 akaw|GAA82566.1 aluc|OJZ87099.1 anig|EHA20369.1 atub|OJI81539.1

Group10579: abra|OJJ71575.1 akaw|GAA89349.1 aluc|OJZ86312.1 atub|OJI84223.1 asyd|OJJ64555.1 aver|OJI98652.1

Group10580: akaw|GAA89333.1 atub|OJI84204.1 acla|EAW15135.1 afis|EAW20107.1 afum|EAL85224.1 anid|CBF84913.1

Group10581: abra|OJJ71631.1 akaw|GAA89294.1 aluc|OJZ86373.1 anig|EHA25362.1 atub|OJI84161.1 atub|OJI80441.1

Group10582: abra|OJJ71641.1 acar|OOF96357.1 akaw|GAA89285.1 aluc|OJZ86384.1 anig|EHA21511.1 atub|OJI84149.1

Group10583: abra|OJJ71739.1 acar|OOF96430.1 akaw|GAA85153.1 aluc|OJZ86484.1 anig|EHA22177.1 atub|OJI84045.1

Group10584: abra|OJJ71782.1 akaw|GAA85128.1 aluc|OJZ86512.1 anid|CBF74969.1 anig|EHA22134.1 atub|OJI84015.1

Group10585: abra|OJJ71788.1 akaw|GAA85123.1 aluc|OJZ86518.1 asyd|OJJ62325.1 atub|OJI84006.1 aver|OJI96430.1

Group10586: abra|OJJ71789.1 acar|OOF96469.1 akaw|GAA85121.1 aluc|OJZ86519.1 anig|EHA22127.1 atub|OJI84003.1

Group10587: abra|OJJ71800.1 acar|OOF96481.1 akaw|GAA85110.1 aluc|OJZ86530.1 anig|EHA22116.1 atub|OJI83991.1

Group10588: abra|OJJ71812.1 acar|OOF96490.1 akaw|GAA85100.1 aluc|OJZ86543.1 anig|EHA22107.1 atub|OJI83980.1

Group10589: abra|OJJ71836.1 afla|EED52767.1 agla|OJJ81630.1 aluc|OJZ86567.1 asyd|OJJ58654.1 aver|OJJ01263.1

Group10590: abra|OJJ71859.1 acar|OOF96531.1 akaw|GAA85056.1 aluc|OJZ86596.1 anig|EHA22066.1 atub|OJI83927.1

Group10591: abra|OJJ71901.1 acar|OOF90471.1 akaw|GAA85019.1 aluc|OJZ86638.1 anig|EHA22031.1 atub|OJI83889.1

Group10592: afis|EAW19988.1 anid|CBF73969.1 asyd|OJJ62369.1 aver|OJI96503.1 afum|EAL84927.1 anig|EHA22195.1

Group10593: abra|OJJ72125.1 acar|OOF96765.1 akaw|GAA84798.1 aluc|OJZ86882.1 anig|EHA21762.1 atub|OJI83637.1

Group10594: ater|EAU31365.1 awen|OJJ34585.1 asyd|OJJ52932.1 anid|CBF87463.1 aver|OJJ08897.1 azon|OJJ42194.1

Group10595: abra|OJJ72156.1 acar|OOF96789.1 akaw|GAA84771.1 aluc|OJZ86912.1 anig|EHA21732.1 atub|OJI83609.1

Group10596: abra|OJJ72174.1 acar|OOF96805.1 akaw|GAA84752.1 aluc|OJZ86931.1 anig|EHA21717.1 atub|OJI83590.1

Group10597: abra|OJJ72175.1 acar|OOF96806.1 akaw|GAA84753.1 aluc|OJZ86930.1 anig|EHA21716.1 atub|OJI83591.1

Group10598: abra|OJJ72176.1 acar|OOF96807.1 akaw|GAA84754.1 aluc|OJZ86929.1 anig|EHA21715.1 atub|OJI83592.1

Group10599: abra|OJJ72178.1 akaw|GAA91085.1 aluc|OJZ87902.1 anid|CBF75982.1 anig|EHA20198.1 azon|OJJ50246.1

Group10600: abra|OJJ72184.1 akaw|GAA84750.1 aluc|OJZ86933.1 anig|EHA21713.1 asyd|OJJ57958.1 atub|OJI83588.1

Group10601: abra|OJJ72185.1 acar|OOF96809.1 akaw|GAA84749.1 aluc|OJZ86934.1 anig|EHA21710.1 atub|OJI83587.1

Group10602: abra|OJJ72186.1 acar|OOF96810.1 akaw|GAA84748.1 aluc|OJZ86935.1 anig|EHA21709.1 atub|OJI83586.1

Group10603: abra|OJJ72194.1 akaw|GAA84739.1 aluc|OJZ86945.1 anig|EHA21699.1 atub|OJI83577.1 aver|OJI97566.1

Group10604: abra|OJJ72204.1 akaw|GAA84732.1 aluc|OJZ86954.1 anig|EHA21693.1 atub|OJI83569.1 awen|OJJ29939.1

Group10605: abra|OJJ72221.1 acar|OOF96843.1 akaw|GAA84716.1 aluc|OJZ86971.1 anig|EHA21677.1 atub|OJI83550.1

Group10606: abra|OJJ72239.1 acar|OOF96864.1 akaw|GAA84700.1 aluc|OJZ86987.1 anig|EHA21659.1 atub|OJI83532.1

Group10607: abra|OJJ72240.1 acar|OOF96865.1 akaw|GAA84699.1 aluc|OJZ86988.1 anig|EHA21658.1 atub|OJI83531.1

Group10608: abra|OJJ72252.1 acar|OOF96883.1 akaw|GAA84685.1 aluc|OJZ87003.1 anig|EHA21643.1 atub|OJI83516.1

Group10609: abra|OJJ72256.1 acar|OOF96888.1 akaw|GAA84681.1 aluc|OJZ87007.1 anig|EHA21639.1 atub|OJI83512.1

Group10610: acla|EAW12258.1 agla|OJJ79787.1 afis|EAW16979.1 afum|EAL86506.1 awen|OJJ35996.1 azon|OJJ50268.1

Group10611: anid|CBF80595.1 afla|EED49584.1 aory|EIT82628.1 asyd|OJJ62163.1 ater|EAU31435.1 aver|OJI96273.1

Group10612: abra|OJJ72316.1 acar|OOF94894.1 akaw|GAA86063.1 aluc|OJZ90297.1 anig|EHA23786.1 atub|OJI81721.1

Group10613: abra|OJJ72367.1 acar|OOF94845.1 akaw|GAA86117.1 aluc|OJZ90355.1 atub|OJI81782.1 azon|OJJ45146.1

Group10614: acla|EAW08391.1 afum|EAL94103.2 afla|EED57628.1 agla|OJJ80817.1 aory|EIT82956.1 ater|EAU36097.1

Group10615: abra|OJJ72374.1 agla|OJJ80747.1 anig|EHA23827.1 asyd|OJJ55140.1 ater|EAU32918.1 atub|OJI81771.1

Group10616: abra|OJJ72435.1 acar|OOF94800.1 akaw|GAA86175.1 aluc|OJZ90418.1 anig|EHA23886.1 atub|OJI81851.1

Group10617: abra|OJJ72456.1 acar|OOF94782.1 akaw|GAA86192.1 aluc|OJZ90441.1 anig|EHA23906.1 atub|OJI81871.1

Group10618: abra|OJJ72501.1 acar|OOF94739.1 akaw|GAA86230.1 aluc|OJZ90484.1 anig|EHA23942.1 atub|OJI81909.1

Group10619: abra|OJJ72512.1 acar|OOF94731.1 akaw|GAA86239.1 aluc|OJZ90494.1 anig|EHA23953.1 atub|OJI81919.1

Group10620: abra|OJJ72574.1 acar|OOF94685.1 akaw|GAA85200.1 aluc|OJZ90555.1 anig|EHA24002.1 atub|OJI81978.1

Group10621: abra|OJJ72577.1 acar|OOF94682.1 agla|OJJ88580.1 akaw|GAA85203.1 aluc|OJZ90558.1 atub|OJI81981.1

Group10622: acla|EAW13708.1 acar|OOF94989.1 afla|EED47558.1 aluc|OJZ84462.1 aory|EIT74536.1 ater|EAU33405.1

Group10623: abra|OJJ72610.1 acar|OOF94644.1 akaw|GAA85236.1 aluc|OJZ90592.1 anig|EHA24031.1 atub|OJI82015.1

Group10624: abra|OJJ72617.1 akaw|GAA85241.1 aluc|OJZ90598.1 anig|EHA24037.1 atub|OJI82022.1 ater|EAU39349.1

Group10625: abra|OJJ72644.1 acar|OOF94626.1 akaw|GAA85265.1 aluc|OJZ90623.1 anig|EHA24060.1 atub|OJI82049.1

Group10626: abra|OJJ72671.1 acar|OOF97387.1 akaw|GAA85291.1 aluc|OJZ90649.1 anig|EHA24085.1 atub|OJI82079.1

Group10627: abra|OJJ72672.1 acar|OOF97386.1 akaw|GAA85292.1 aluc|OJZ90650.1 anig|EHA24086.1 atub|OJI82080.1

Group10628: abra|OJJ72676.1 acar|OOF97381.1 akaw|GAA85295.1 aluc|OJZ90654.1 anig|EHA24089.1 atub|OJI82083.1

Group10629: abra|OJJ72680.1 acar|OOF95929.1 akaw|GAA85298.1 aluc|OJZ90658.1 anig|EHA24093.1 atub|OJI82087.1

Group10630: asyd|OJJ58315.1 acar|OOF97087.1 akaw|GAA84853.1 aluc|OJZ86818.1 atub|OJI83699.1 aver|OJJ02096.1

Group10631: abra|OJJ72747.1 acar|OOF95970.1 akaw|GAA85359.1 aluc|OJZ90721.1 anig|EHA24139.1 atub|OJI82149.1

Group10632: abra|OJJ72893.1 acar|OOF96084.1 akaw|GAA85480.1 aluc|OJZ83402.1 anig|EHA24253.1 atub|OJI82289.1

Group10633: abra|OJJ72952.1 akaw|GAA85533.1 aluc|OJZ83462.1 anig|EHA24305.1 atub|OJI82354.1 anid|CBF77659.1

Group10634: abra|OJJ72961.1 acar|OOF96143.1 acla|EAW11188.1 afum|EAL88555.1 aluc|OJZ83469.1 anid|CBF81010.1

Group10635: abra|OJJ73069.1 acar|OOG00302.1 akaw|GAA88091.1 aluc|OJZ81356.1 anig|EHA18656.1 atub|OJI91560.1

Group10636: abra|OJJ73088.1 acar|OOG00320.1 akaw|GAA88072.1 aluc|OJZ81337.1 anig|EHA18641.1 atub|OJI91540.1

Group10637: abra|OJJ73137.1 akaw|GAA88035.1 aluc|OJZ81294.1 anig|EHA18603.1 atub|OJI91503.1 azon|OJJ47527.1

Group10638: abra|OJJ73166.1 acar|OOF89877.1 akaw|GAA88010.1 aluc|OJZ81268.1 anig|EHA18568.1 atub|OJI91474.1

Group10639: abra|OJJ73299.1 acla|EAW12038.1 afla|EED49379.1 akaw|GAA92707.1 aluc|OJZ88748.1 atub|OJI91320.1

Group10640: abra|OJJ73302.1 acar|OOF91870.1 akaw|GAA92703.1 aluc|OJZ88743.1 anig|EHA18447.1 atub|OJI91315.1

Group10641: abra|OJJ73311.1 acar|OOF90734.1 akaw|GAA92811.1 aluc|OJZ88734.1 anig|EHA18437.1 atub|OJI91310.1

Group10642: abra|OJJ73315.1 acar|OOF98706.1 akaw|GAA92805.1 aluc|OJZ88728.1 anig|EHA18433.1 atub|OJI91306.1

Group10643: abra|OJJ73316.1 acar|OOF98707.1 agla|OJJ85299.1 akaw|GAA92804.1 asyd|OJJ52015.1 aver|OJJ07820.1

Group10644: abra|OJJ73355.1 acar|OOG00564.1 akaw|GAA83129.1 aluc|OJZ88685.1 anig|EHA18365.1 atub|OJI91265.1

Group10645: abra|OJJ73398.1 akaw|GAA83089.1 aluc|OJZ88636.1 anig|EHA18334.1 atub|OJI91221.1 azon|OJJ42505.1

Group10646: abra|OJJ73429.1 acar|OOG00635.1 akaw|GAA83063.1 aluc|OJZ88614.1 anig|EHA18305.1 atub|OJI91199.1

Group10647: abra|OJJ73468.1 afla|EED55020.1 aory|EIT81957.1 asyd|OJJ62950.1 ater|EAU30736.1 aver|OJI98037.1

Group10648: abra|OJJ73494.1 acar|OOG00686.1 akaw|GAA87443.1 aluc|OJZ88554.1 anig|EHA18252.1 atub|OJI91139.1

Group10649: abra|OJJ73524.1 acar|OOF94525.1 afla|EED51013.1 anig|EHA18230.1 aory|EIT77005.1 ater|EAU38118.1

Group10650: abra|OJJ73662.1 akaw|GAA88494.1 aluc|OJZ88390.1 anig|EHA18117.1 atub|OJI90985.1 azon|OJJ46544.1

Group10651: abra|OJJ73663.1 akaw|GAA88493.1 aluc|OJZ88389.1 anig|EHA18116.1 atub|OJI90984.1 azon|OJJ50139.1

Group10652: abra|OJJ73695.1 anid|CBF87423.1 asyd|OJJ52904.1 aver|OJJ08874.1 afla|EED51191.1 anid|CBF87425.1

Group10653: abra|OJJ73702.1 acar|OOG00868.1 akaw|GAA88460.1 aluc|OJZ88349.1 anig|EHA18082.1 atub|OJI90944.1

Group10654: abra|OJJ73710.1 akaw|GAA88453.1 aluc|OJZ88340.1 anid|CBF76664.1 anig|EHA18075.1 atub|OJI90936.1

Group10655: abra|OJJ73723.1 acar|OOG00892.1 akaw|GAA88439.1 aluc|OJZ88327.1 anig|EHA18063.1 atub|OJI90922.1

Group10656: abra|OJJ73747.1 acar|OOG00911.1 akaw|GAA88419.1 aluc|OJZ88305.1 anig|EHA18043.1 atub|OJI90900.1

Group10657: abra|OJJ73748.1 acar|OOG00913.1 akaw|GAA88418.1 aluc|OJZ88304.1 anig|EHA18042.1 atub|OJI90899.1

Group10658: abra|OJJ73823.1 acar|OOG00975.1 akaw|GAA89442.1 aluc|OJZ88225.1 anig|EHA17973.1 atub|OJI90826.1

Group10659: abra|OJJ73852.1 afla|EED57230.1 anig|EHA19926.1 aory|EIT74060.1 asyd|OJJ55228.1 azon|OJJ42637.1

Group10660: abra|OJJ73873.1 acar|OOG01015.1 akaw|GAA89485.1 aluc|OJZ88177.1 anig|EHA17929.1 atub|OJI90775.1

Group10661: abra|OJJ73877.1 acar|OOG01017.1 akaw|GAA89489.1 aluc|OJZ88172.1 anig|EHA17925.1 atub|OJI90770.1

Group10662: abra|OJJ73941.1 acar|OOG01070.1 akaw|GAA89546.1 aluc|OJZ88103.1 anig|EHA17872.1 atub|OJI90711.1

Group10663: abra|OJJ74012.1 akaw|GAA87495.1 aluc|OJZ88027.1 anig|EHA17804.1 atub|OJI90635.1 aver|OJJ03919.1

Group10664: abra|OJJ74141.1 acar|OOF95589.1 akaw|GAA89744.1 aluc|OJZ80640.1 anig|EHA27192.1 atub|OJI87495.1

Group10665: abra|OJJ74175.1 acar|OOF95555.1 agla|OJJ85296.1 akaw|GAA86841.1 aluc|OJZ89960.1 atub|OJI84845.1

Group10666: abra|OJJ74176.1 akaw|GAA86840.1 aluc|OJZ89959.1 anig|EHA27159.1 atub|OJI84846.1 anig|EHA27160.1

Group10667: abra|OJJ74178.1 acar|OOF95543.1 aluc|OJZ89956.1 akaw|GAA86837.1 anig|EHA27156.1 atub|OJI84850.1

Group10668: abra|OJJ74182.1 acar|OOG00344.1 afla|EED47784.1 anig|EHA18613.1 aory|EIT77181.1 atub|OJI91314.1

Group10669: abra|OJJ74215.1 acar|OOF95506.1 akaw|GAA86803.1 aluc|OJZ89919.1 anig|EHA27119.1 atub|OJI84886.1

Group10670: abra|OJJ74221.1 afla|EED52530.1 aluc|OJZ89913.1 anig|EHA27113.1 aory|EIT74279.1 atub|OJI84892.1

Group10671: awen|OJJ37058.1 afis|EAW25698.1 afum|EAL84373.1 anid|CBF69711.1 asyd|OJJ59041.1 aver|OJJ01636.1

Group10672: abra|OJJ74319.1 akaw|GAA86707.1 aluc|OJZ89809.1 anig|EHA27019.1 atub|OJI84997.1 ater|EAU39321.1

Group10673: abra|OJJ74356.1 acar|OOF95374.1 akaw|GAA86675.1 aluc|OJZ89772.1 anig|EHA26990.1 atub|OJI85031.1

Group10674: abra|OJJ74368.1 akaw|GAA86663.1 aluc|OJZ89760.1 anig|EHA26980.1 aver|OJJ04193.1 awen|OJJ38112.1

Group10675: abra|OJJ74377.1 acar|OOF95361.1 akaw|GAA87276.1 aluc|OJZ89745.1 anig|EHA26967.1 atub|OJI85057.1

Group10676: abra|OJJ74404.1 acar|OOF95337.1 akaw|GAA87298.1 aluc|OJZ89720.1 anig|EHA26947.1 atub|OJI85080.1

Group10677: abra|OJJ74439.1 acar|OOF95307.1 akaw|GAA87332.1 aluc|OJZ89682.1 anig|EHA26918.1 atub|OJI85118.1

Group10678: abra|OJJ74451.1 acar|OOF95300.1 akaw|GAA87342.1 aluc|OJZ89672.1 anig|EHA26907.1 atub|OJI85128.1

Group10679: abra|OJJ74509.1 akaw|GAA82090.1 aluc|OJZ92090.1 anig|EHA26851.1 atub|OJI86520.1 acar|OOF95233.1

Group10680: abra|OJJ74520.1 akaw|GAA82080.1 aluc|OJZ92099.1 anig|EHA26843.1 atub|OJI86532.1 acar|OOF95228.1

Group10681: acla|EAW11015.1 afis|EAW20642.1 afla|EED53090.1 aory|EIT74794.1 afum|EBA27178.1 agla|OJJ84405.1

Group10682: aory|EIT82288.1 awen|OJJ35543.1 azon|OJJ42180.1 acla|EAW12959.1 afla|EED57072.1 ater|EAU32797.1

Group10683: abra|OJJ74721.1 acar|OOF91440.1 akaw|GAA81895.1 aluc|OJZ92305.1 anig|EHA26663.1 atub|OJI86735.1

Group10684: abra|OJJ74746.1 acar|OOF95740.1 afla|EED52026.1 akaw|GAA83665.1 aluc|OJZ92326.1 atub|OJI86756.1

Group10685: abra|OJJ74794.1 acar|OOF95785.1 akaw|GAA83619.1 aluc|OJZ92374.1 anig|EHA26599.1 atub|OJI86804.1

Group10686: abra|OJJ74795.1 acar|OOF95786.1 akaw|GAA83618.1 aluc|OJZ92375.1 anig|EHA26598.1 atub|OJI86805.1

Group10687: abra|OJJ74796.1 acar|OOF95787.1 akaw|GAA83617.1 aluc|OJZ92376.1 anig|EHA26597.1 atub|OJI86806.1

Group10688: afis|EAW23527.1 afum|EBA27328.1 acla|EAW08481.1 afla|EED57426.1 awen|OJJ39856.1 aory|EIT80094.1

Group10689: abra|OJJ74933.1 acar|OOF95924.1 akaw|GAA83487.1 aluc|OJZ92506.1 anig|EHA26467.1 atub|OJI86938.1

Group10690: abra|OJJ74972.1 afla|EED45924.1 anig|EHA21620.1 aver|OJJ07895.1 afum|EAL94095.1 ater|EAU36098.1

Group10691: abra|OJJ75065.1 acar|OOF97269.1 akaw|GAA88677.1 aluc|OJZ92631.1 ater|EAU39424.1 atub|OJI87062.1

Group10692: abra|OJJ75200.1 agla|OJJ79497.1 aluc|OJZ92764.1 asyd|OJJ62076.1 atub|OJI87208.1 aver|OJI96183.1

Group10693: abra|OJJ75235.1 akaw|GAA91778.1 aluc|OJZ92796.1 atub|OJI87239.1 anig|EHA26212.1 atub|OJI87240.1

Group10694: abra|OJJ75261.1 acar|OOF94541.1 akaw|GAA91799.1 aluc|OJZ92819.1 anig|EHA26189.1 atub|OJI87277.1

Group10695: abra|OJJ75267.1 acar|OOF94536.1 akaw|GAA91803.1 aluc|OJZ92825.1 anig|EHA26182.1 atub|OJI87282.1

Group10696: abra|OJJ75280.1 acar|OOF94528.1 akaw|GAA91818.1 aluc|OJZ92841.1 anig|EHA26170.1 atub|OJI87297.1

Group10697: abra|OJJ75301.1 acar|OOG00389.1 akaw|GAA91834.1 aluc|OJZ92856.1 anig|EHA26150.1 atub|OJI87313.1

Group10698: azon|OJJ44317.1 acla|EAW09870.1 afis|EAW21256.1 afum|EAL92271.1 agla|OJJ87499.1 awen|OJJ36571.1

Group10699: abra|OJJ75312.1 acar|OOF94517.1 akaw|GAA91996.1 aluc|OJZ92866.1 anig|EHA26099.1 atub|OJI87324.1

Group10700: abra|OJJ75324.1 acar|OOF90137.1 akaw|GAA92019.1 aluc|OJZ92886.1 atub|OJI87345.1 acla|EAW12068.1

Group10701: afla|EED48382.1 afis|EAW21075.1 afum|EAL92098.1 anid|CBF87052.1 aory|EIT83064.1 aver|OJI99759.1

Group10702: abra|OJJ75331.1 acar|OOF94496.1 akaw|GAA92026.1 aluc|OJZ92895.1 anig|EHA26068.1 atub|OJI87352.1

Group10703: abra|OJJ75333.1 acar|OOF94498.1 akaw|GAA92027.1 aluc|OJZ92897.1 anig|EHA26067.1 atub|OJI87355.1

Group10704: abra|OJJ75530.1 acar|OOF90857.1 akaw|GAA84292.1 aluc|OJZ88986.1 anig|EHA23540.1 atub|OJI85802.1

Group10705: abra|OJJ75531.1 akaw|GAA91951.1 aluc|OJZ82667.1 anig|EHA27441.1 atub|OJI88900.1 azon|OJJ43936.1

Group10706: abra|OJJ75565.1 acar|OOF90882.1 akaw|GAA84317.1 aluc|OJZ89013.1 anig|EHA23517.1 atub|OJI85776.1

Group10707: abra|OJJ75583.1 akaw|GAA84335.1 aluc|OJZ89034.1 anig|EHA23500.1 atub|OJI85756.1 acar|OOF90901.1

Group10708: abra|OJJ75599.1 afla|EED55137.1 akaw|GAA84350.1 aluc|OJZ89050.1 aory|EIT81597.1 atub|OJI85737.1

Group10709: afla|EED45366.1 agla|OJJ83652.1 aory|EIT82445.1 acar|OOG00334.1 afis|EAW22128.1 ater|EAU30031.1

Group10710: abra|OJJ75647.1 acar|OOG00191.1 akaw|GAA84392.1 aluc|OJZ89096.1 anig|EHA23441.1 atub|OJI85684.1

Group10711: akaw|GAA84412.1 acla|EAW09430.1 afis|EAW21706.1 afum|EAL92741.1 aory|EIT79758.1 ater|EAU38390.1

Group10712: acla|EAW08984.1 afum|EAL89329.1 ater|EAU31417.1 afla|EED48346.1 afis|EAW24688.1 aory|EIT76288.1

Group10713: abra|OJJ75751.1 akaw|GAA84484.1 aluc|OJZ89205.1 anig|EHA23354.1 atub|OJI85587.1 awen|OJJ36146.1

Group10714: abra|OJJ75755.1 acar|OOG00109.1 akaw|GAA84487.1 aluc|OJZ89208.1 anig|EHA23351.1 atub|OJI85584.1

Group10715: abra|OJJ75771.1 agla|OJJ79604.1 asyd|OJJ58676.1 atub|OJI87187.1 aver|OJJ01288.1 awen|OJJ30925.1

Group10716: abra|OJJ75775.1 akaw|GAA84493.1 aluc|OJZ89216.1 anid|CBF83116.1 anig|EHA23344.1 atub|OJI85576.1

Group10717: abra|OJJ75793.1 afum|EAL89022.1 akaw|GAA84512.1 aluc|OJZ89241.1 anig|EHA23326.1 atub|OJI85557.1

Group10718: abra|OJJ75859.1 acar|OOG00036.1 akaw|GAA84564.1 aluc|OJZ89297.1 anig|EHA23266.1 atub|OJI85501.1

Group10719: abra|OJJ75887.1 aluc|OJZ89339.1 acla|EAW12093.1 acar|OOF99323.1 atub|OJI85460.1 awen|OJJ37189.1

Group10720: abra|OJJ75925.1 acar|OOF99983.1 akaw|GAA84628.1 aluc|OJZ89374.1 anig|EHA23209.1 atub|OJI85424.1

Group10721: abra|OJJ75962.1 acar|OOF99953.1 akaw|GAA84662.1 aluc|OJZ89410.1 anig|EHA23177.1 atub|OJI85386.1

Group10722: abra|OJJ76124.1 acar|OOF99801.1 akaw|GAA88843.1 aluc|OJZ89581.1 anig|EHA23027.1 atub|OJI85223.1

Group10723: awen|OJJ39988.1 acar|OOF96879.1 akaw|GAA84689.1 aluc|OJZ86999.1 anig|EHA21647.1 atub|OJI83520.1

Group10724: abra|OJJ76369.1 akaw|GAA82295.1 aluc|OJZ91843.1 anig|EHA22826.1 atub|OJI86295.1 acar|OOF99618.1

Group10725: abra|OJJ76397.1 acar|OOF99638.1 akaw|GAA82317.1 aluc|OJZ91815.1 anig|EHA22806.1 atub|OJI86271.1

Group10726: abra|OJJ76596.1 akaw|GAA82500.1 aluc|OJZ91611.1 asyd|OJJ54847.1 atub|OJI86063.1 aver|OJJ07317.1

Group10727: abra|OJJ76611.1 acar|OOF99378.1 akaw|GAA82515.1 aluc|OJZ91596.1 anig|EHA22626.1 atub|OJI86047.1

Group10728: abra|OJJ76615.1 afis|EAW17371.1 afla|EED46485.1 afum|EAL91209.1 anig|EHA22618.1 ater|EAU32674.1

Group10729: abra|OJJ76616.1 aluc|OJZ91592.1 anid|CBF79101.1 ater|EAU31046.1 atub|OJI86043.1 azon|OJJ46457.1

Group10730: abra|OJJ76617.1 aluc|OJZ91591.1 anid|CBF79103.1 ater|EAU31047.1 atub|OJI86042.1 azon|OJJ42682.1

Group10731: abra|OJJ76655.1 acar|OOF99345.1 akaw|GAA92995.1 aluc|OJZ91550.1 anig|EHA22585.1 atub|OJI86003.1

Group10732: abra|OJJ76660.1 akaw|GAA93000.1 aluc|OJZ91543.1 anig|EHA22580.1 ater|EAU38719.1 atub|OJI85997.1

Group10733: abra|OJJ76682.1 asyd|OJJ52827.1 atub|OJI87914.1 aver|OJJ08803.1 akaw|GAA90303.1 aluc|OJZ83034.1

Group10734: abra|OJJ76706.1 akaw|GAA92492.1 aluc|OJZ91504.1 anid|CBF74590.1 anig|EHA22544.1 atub|OJI85957.1

Group10735: abra|OJJ76707.1 akaw|GAA92491.1 aluc|OJZ91503.1 anid|CBF74591.1 anig|EHA22543.1 atub|OJI85956.1

Group10736: abra|OJJ76708.1 akaw|GAA92490.1 aluc|OJZ91502.1 anid|CBF75584.1 anig|EHA22542.1 atub|OJI85955.1

Group10737: acla|EAW14593.1 afla|EED57237.1 aory|EIT74052.1 asyd|OJJ54482.1 ater|EAU32501.1 aver|OJJ01971.1

Group10738: abra|OJJ76710.1 akaw|GAA92488.1 aluc|OJZ91500.1 anig|EHA22540.1 asyd|OJJ53825.1 atub|OJI85953.1

Group10739: abra|OJJ76732.1 akaw|GAA92668.1 aluc|OJZ90122.1 anig|EHA20948.1 atub|OJI84673.1 azon|OJJ46853.1

Group10740: abra|OJJ76749.1 acar|OOF89995.1 akaw|GAA88936.1 aluc|OJZ89471.1 anig|EHA28271.1 aory|EIT81012.1

Group10741: abra|OJJ76750.1 acar|OOF89996.1 akaw|GAA88938.1 aluc|OJZ89469.1 anig|EHA28273.1 atub|OJI88343.1

Group10742: abra|OJJ76794.1 acar|OOF91288.1 akaw|GAA87231.1 aluc|OJZ84198.1 anig|EHA27555.1 atub|OJI89886.1

Group10743: abra|OJJ76876.1 agla|OJJ85995.1 asyd|OJJ65375.1 ater|EAU38551.1 aver|OJI99542.1 ater|EAU35086.1

Group10744: aver|OJI96465.1 anig|EHA24748.1 awen|OJJ41824.1 akaw|GAA90419.1 aluc|OJZ85811.1 atub|OJI79597.1

Group10745: abra|OJJ76960.1 acar|OOF91405.1 akaw|GAA87083.1 aluc|OJZ84355.1 anig|EHA27693.1 atub|OJI89712.1

Group10746: abra|OJJ76965.1 acar|OOF91409.1 akaw|GAA87079.1 aluc|OJZ84361.1 anig|EHA27696.1 atub|OJI89707.1

Group10747: abra|OJJ76966.1 acar|OOF91410.1 aluc|OJZ84362.1 anig|EHA27697.1 ater|EAU35236.1 atub|OJI89706.1

Group10748: abra|OJJ76983.1 acar|OOF91428.1 akaw|GAA87063.1 aluc|OJZ84382.1 anig|EHA27713.1 atub|OJI89689.1

Group10749: acla|EAW12965.1 afis|EAW17698.1 afum|EAL91512.1 awen|OJJ37135.1 agla|OJJ79477.1 azon|OJJ50835.1

Group10750: abra|OJJ76989.1 acar|OOF91433.1 akaw|GAA87057.1 aluc|OJZ84389.1 anig|EHA27719.1 atub|OJI89683.1

Group10751: abra|OJJ77029.1 akaw|GAA87022.1 aluc|OJZ84431.1 anig|EHA27751.1 atub|OJI89642.1 aver|OJJ08329.1

Group10752: abra|OJJ77118.1 acar|OOF93006.1 akaw|GAA90811.1 aluc|OJZ84528.1 anig|EHA27825.1 atub|OJI89552.1

Group10753: abra|OJJ77121.1 acar|OOF93008.1 akaw|GAA90808.1 aluc|OJZ84531.1 anig|EHA27828.1 atub|OJI89549.1

Group10754: abra|OJJ77162.1 akaw|GAA90770.1 aluc|OJZ84575.1 anig|EHA27868.1 atub|OJI89507.1 acar|OOF93048.1

Group10755: abra|OJJ77197.1 akaw|GAA90744.1 aluc|OJZ84604.1 aory|EIT80010.1 atub|OJI89475.1 awen|OJJ38743.1

Group10756: afla|EED45429.1 aluc|OJZ89737.1 aory|EIT82426.1 atub|OJI85064.1 akaw|GAA87282.1 aory|EIT77471.1

Group10757: abra|OJJ77249.1 akaw|GAA93200.1 aluc|OJZ84660.1 anig|EHA27942.1 atub|OJI89420.1 azon|OJJ47625.1

Group10758: abra|OJJ77252.1 anid|CBF71241.1 asyd|OJJ55161.1 ater|EAU32974.1 aver|OJJ07046.1 awen|OJJ41793.1

Group10759: abra|OJJ77283.1 acar|OOF93157.1 akaw|GAA90666.1 aluc|OJZ84693.1 anig|EHA27969.1 atub|OJI89389.1

Group10760: abra|OJJ77286.1 acar|OOF93163.1 akaw|GAA90663.1 aluc|OJZ84697.1 anig|EHA27972.1 atub|OJI89385.1

Group10761: abra|OJJ77293.1 acar|OOF93170.1 akaw|GAA91458.1 aluc|OJZ84705.1 anig|EHA27979.1 atub|OJI89378.1

Group10762: azon|OJJ44221.1 acla|EAW11080.1 agla|OJJ78856.1 asyd|OJJ61763.1 ater|EAU32699.1 aver|OJI95862.1

Group10763: abra|OJJ77314.1 acar|OOF93190.1 ater|EAU36933.1 atub|OJI89358.1 afla|EED49328.1 aory|EIT79488.1

Group10764: abra|OJJ77352.1 akaw|GAA91409.1 aluc|OJZ84763.1 anig|EHA28032.1 atub|OJI89321.1 awen|OJJ37311.1

Group10765: abra|OJJ77391.1 acar|OOF93259.1 akaw|GAA91370.1 aluc|OJZ84799.1 anig|EHA28065.1 atub|OJI89283.1

Group10766: abra|OJJ77399.1 acar|OOF89804.1 akaw|GAA91364.1 aluc|OJZ84806.1 anig|EHA28072.1 atub|OJI89277.1

Group10767: abra|OJJ77400.1 anig|EHA28073.1 abra|OJJ77402.1 akaw|GAA91363.1 aluc|OJZ84807.1 atub|OJI89276.1

Group10768: abra|OJJ77401.1 anig|EHA28074.1 abra|OJJ77403.1 akaw|GAA91362.1 aluc|OJZ84808.1 atub|OJI89275.1

Group10769: abra|OJJ77426.1 afis|EAW24704.1 afum|EAL89346.1 asyd|OJJ65317.1 ater|EAU31631.1 aver|OJI99457.1

Group10770: abra|OJJ77433.1 agla|OJJ82672.1 anid|CBF90094.1 asyd|OJJ62033.1 aver|OJI96123.1 afis|EAW24705.1

Group10771: afla|EED51170.1 anid|CBF78267.1 aory|EIT81331.1 aver|OJJ05022.1 afis|EAW24683.1 asyd|OJJ56079.1

Group10772: abra|OJJ77461.1 afla|EED49596.1 anig|EHA19294.1 asyd|OJJ56402.1 ater|EAU39563.1 aver|OJJ06310.1

Group10773: aory|EIT72890.1 afis|EAW19807.1 afla|EED48208.1 afum|EAL85085.1 ater|EAU30742.1 awen|OJJ34560.1

Group10774: atub|OJI89805.1 acar|OOF91775.1 aluc|OJZ84276.1 anig|EHA27621.1 awen|OJJ41855.1 azon|OJJ51657.1

Group10775: abra|OJJ77540.1 acar|OOF99186.1 akaw|GAA84082.1 aluc|OJZ80996.1 anig|EHA19365.1 atub|OJI82588.1

Group10776: abra|OJJ77541.1 acar|OOF99187.1 akaw|GAA84081.1 aluc|OJZ80995.1 anig|EHA19366.1 atub|OJI82589.1

Group10777: abra|OJJ77645.1 acar|OOF99090.1 akaw|GAA83989.1 aluc|OJZ80884.1 anig|EHA19453.1 atub|OJI82693.1

Group10778: anid|CBF87761.1 asyd|OJJ63917.1 afla|EED55599.1 aory|EIT82670.1 aver|OJI97025.1 awen|OJJ33421.1

Group10779: abra|OJJ77672.1 acar|OOF89815.1 akaw|GAA83960.1 aluc|OJZ80847.1 anig|EHA19476.1 atub|OJI82720.1

Group10780: abra|OJJ77679.1 afis|EAW19871.1 afla|EED56845.1 afum|EAL85026.1 aluc|OJZ79989.1 aory|EIT77461.1

Group10781: abra|OJJ77703.1 acar|OOF99037.1 akaw|GAA83927.1 aluc|OJZ90768.1 anig|EHA19503.1 atub|OJI82756.1

Group10782: abra|OJJ77706.1 acar|OOF99034.1 akaw|GAA83924.1 aluc|OJZ90771.1 anig|EHA19506.1 atub|OJI82759.1

Group10783: abra|OJJ77720.1 afla|EED52339.1 anig|EHA22491.1 aory|EIT74350.1 ater|EAU36727.1 afis|EAW18825.1

Group10784: abra|OJJ77728.1 acar|OOF99010.1 akaw|GAA83908.1 aluc|OJZ90789.1 anig|EHA19522.1 atub|OJI82777.1

Group10785: abra|OJJ77732.1 acar|OOF98991.1 akaw|GAA83903.1 aluc|OJZ90792.1 anig|EHA19526.1 atub|OJI82781.1

Group10786: abra|OJJ77733.1 acar|OOF98992.1 akaw|GAA83901.1 aluc|OJZ90794.1 anig|EHA19528.1 atub|OJI82783.1

Group10787: abra|OJJ77750.1 afis|EAW23114.1 afum|EAL92030.1 akaw|GAA83884.1 aluc|OJZ90811.1 atub|OJI82800.1

Group10788: abra|OJJ77770.1 agla|OJJ86554.1 akaw|GAA83876.1 aluc|OJZ90820.1 anig|EHA19546.1 atub|OJI82808.1

Group10789: abra|OJJ77785.1 acar|OOF98974.1 akaw|GAA83862.1 aluc|OJZ90838.1 anig|EHA19561.1 atub|OJI82825.1

Group10790: abra|OJJ77790.1 afla|EED54248.1 aory|EIT75978.1 awen|OJJ32560.1 agla|OJJ81250.1 agla|OJJ82010.1

Group10791: abra|OJJ77801.1 acar|OOF90928.1 akaw|GAA83848.1 aluc|OJZ90867.1 anig|EHA19587.1 atub|OJI82865.1

Group10792: abra|OJJ77803.1 acar|OOF90103.1 akaw|GAA83846.1 aluc|OJZ90869.1 anig|EHA19589.1 atub|OJI82867.1

Group10793: abra|OJJ77849.1 acar|OOF98923.1 akaw|GAA83798.1 aluc|OJZ90924.1 anig|EHA19636.1 atub|OJI82916.1

Group10794: afis|EAW25704.1 asyd|OJJ62981.1 ater|EAU33718.1 aver|OJI97988.1 afum|EAL84377.1 anid|CBF85239.1

Group10795: abra|OJJ77912.1 acar|OOF98859.1 akaw|GAA83745.1 aluc|OJZ90986.1 anig|EHA19691.1 atub|OJI82977.1

Group10796: abra|OJJ77963.1 acar|OOF92549.1 acla|EAW08133.1 agla|OJJ82391.1 anig|EHA19730.1 acla|EAW12841.1

Group10797: abra|OJJ78015.1 acar|OOF98807.1 akaw|GAA85968.1 aluc|OJZ91091.1 anig|EHA19792.1 atub|OJI83088.1

Group10798: abra|OJJ78035.1 acar|OOF98464.1 afis|EAW16432.1 akaw|GAA85947.1 aluc|OJZ91115.1 atub|OJI83119.1

Group10799: abra|OJJ78042.1 afis|EAW17174.1 aver|OJJ05322.1 awen|OJJ34517.1 acla|EAW12151.1 afum|EAL86604.2

Group10800: abra|OJJ78053.1 acla|EAW06391.1 akaw|GAA85938.1 aluc|OJZ91124.1 anig|EHA19824.1 atub|OJI83129.1

Group10801: agla|OJJ88216.1 afis|EAW16427.1 afla|EED51785.1 aory|EIT81717.1 awen|OJJ36320.1 aver|OJI98042.1

Group10802: abra|OJJ78066.1 akaw|GAA85924.1 aluc|OJZ91143.1 anig|EHA19839.1 atub|OJI83156.1 awen|OJJ32238.1

Group10803: abra|OJJ78068.1 afis|EAW23478.1 akaw|GAA85923.1 aluc|OJZ91144.1 anig|EHA19840.1 atub|OJI83157.1

Group10804: afla|EED56830.1 aory|EIT73894.1 anid|CBF87309.1 afis|EAW16917.1 asyd|OJJ62991.1 aver|OJI97977.1

Group10805: acar|OOF97898.1 anid|CBF76233.1 aluc|OJZ87268.1 anig|EHA20510.1 atub|OJI81374.1 akaw|GAA82714.1

Group10806: abra|OJJ78131.1 acar|OOF91246.1 akaw|GAA85876.1 aluc|OJZ91193.1 anig|EHA19889.1 atub|OJI83212.1

Group10807: abra|OJJ78134.1 aluc|OJZ91195.1 anig|EHA19894.1 atub|OJI83215.1 awen|OJJ29976.1 azon|OJJ50083.1

Group10808: abra|OJJ78162.1 afla|EED48082.1 akaw|GAA85863.1 aluc|OJZ91216.1 anig|EHA19922.1 atub|OJI83234.1

Group10809: abra|OJJ78168.1 asyd|OJJ59101.1 ater|EAU30168.1 aver|OJJ05743.1 asyd|OJJ57435.1 aver|OJJ01692.1

Group10810: anid|CBF81921.1 aver|OJI99269.1 awen|OJJ31300.1 acar|OOF90379.1 agla|OJJ87445.1 asyd|OJJ65135.1

Group10811: abra|OJJ78178.1 acar|OOF93332.1 akaw|GAA85855.1 aluc|OJZ91242.1 anig|EHA19942.1 atub|OJI83240.1

Group10812: abra|OJJ78182.1 acar|OOF92536.1 akaw|GAA85854.1 aluc|OJZ91245.1 anig|EHA19944.1 atub|OJI83242.1

Group10813: abra|OJJ78193.1 akaw|GAA85836.1 aluc|OJZ91264.1 anig|EHA19952.1 atub|OJI83253.1 aory|EIT81857.1

Group10814: abra|OJJ78201.1 acar|OOF98653.1 akaw|GAA85828.1 aluc|OJZ91272.1 anig|EHA19959.1 atub|OJI83261.1

Group10815: abra|OJJ78257.1 acar|OOF98605.1 akaw|GAA85769.1 aluc|OJZ91331.1 anig|EHA20013.1 atub|OJI83324.1

Group10816: abra|OJJ78291.1 acar|OOF98577.1 akaw|GAA85736.1 aluc|OJZ91365.1 anig|EHA20043.1 atub|OJI83358.1

Group10817: abra|OJJ78300.1 acar|OOF98566.1 akaw|GAA85726.1 aluc|OJZ91377.1 anig|EHA20052.1 atub|OJI83370.1

Group10818: abra|OJJ78310.1 anig|EHA20061.1 ater|EAU29639.1 afis|EAW19862.1 afum|EAL85033.1 awen|OJJ41889.1

Group10819: abra|OJJ78313.1 acar|OOF90131.1 agla|OJJ79272.1 akaw|GAA92408.1 anig|EHA20064.1 awen|OJJ33978.1

Group10820: abra|OJJ78320.1 acar|OOF98555.1 akaw|GAA85710.1 aluc|OJZ91393.1 anig|EHA20071.1 atub|OJI83386.1

Group10821: aory|EIT74091.1 ater|EAU32691.1 afis|EAW16871.1 afla|EED57119.1 afum|EAL86409.1 awen|OJJ38670.1

Group10822: anid|CBF82900.1 acla|EAW15045.1 asyd|OJJ63347.1 afla|EED57631.1 afum|EAL86231.1 aver|OJI97613.1

Group10823: aory|EIT81598.1 ater|EAU36165.1 afla|EED55135.1 anid|CBF85666.1 asyd|OJJ64246.1 aver|OJI98337.1

Group10824: acar|OOF90519.1 afla|EED53434.1 aory|EIT75046.1 asyd|OJJ63992.1 aver|OJI96962.1 awen|OJJ37883.1

Group10825: acar|OOF90568.1 akaw|GAA84846.1 aluc|OJZ86825.1 atub|OJI83693.1 aver|OJJ05791.1 ater|EAU31615.1

Group10826: acar|OOF90607.1 acla|EAW06502.1 afis|EAW16943.1 afum|EAL86470.1 anid|CBF83062.1 azon|OJJ42652.1

Group10827: acar|OOF90639.1 agla|OJJ79570.1 anid|CBF82096.1 asyd|OJJ54262.1 ater|EAU37952.1 awen|OJJ30883.1

Group10828: anid|CBF70282.1 afla|EED52810.1 aory|EIT78221.1 asyd|OJJ58740.1 aver|OJJ01350.1 azon|OJJ45701.1

Group10829: acar|OOF91474.1 afla|EED49607.1 aory|EIT80983.1 acla|EAW14635.1 afis|EAW22085.1 afum|EAL85694.1

Group10830: anid|CBF73401.1 asyd|OJJ62226.1 awen|OJJ39817.1 afis|EAW15643.1 ater|EAU35280.1 aver|OJI96335.1

Group10831: acar|OOF91710.1 afla|EED46364.1 akaw|GAA82958.1 aluc|OJZ87539.1 aory|EIT81251.1 atub|OJI81108.1

Group10832: acar|OOF91713.1 afla|EED46366.1 akaw|GAA82956.1 aluc|OJZ87537.1 aory|EIT81252.1 atub|OJI81110.1

Group10833: acar|OOF91933.1 agla|OJJ86357.1 asyd|OJJ61498.1 ater|EAU31269.1 aver|OJJ00835.1 afis|EAW15550.1

Group10834: acar|OOF92000.1 anig|EHA22204.1 awen|OJJ39936.1 aluc|OJZ82056.1 atub|OJI84615.1 anig|EHA22203.1

Group10835: azon|OJJ45295.1 afis|EAW16444.1 afum|EAL89045.1 anid|CBF83799.1 asyd|OJJ60889.1 aver|OJJ00213.1

Group10836: acar|OOF92395.1 afla|EED57546.1 aory|EIT81835.1 anid|CBF87345.1 asyd|OJJ52821.1 azon|OJJ50181.1

Group10837: afla|EED48899.1 aory|EIT81241.1 awen|OJJ31601.1 anid|CBF84022.1 asyd|OJJ61375.1 aver|OJJ00702.1

Group10838: acar|OOF92821.1 acla|EAW12157.1 afis|EAW17189.1 afum|EAL86615.1 ater|EAU36220.1 aver|OJJ05977.1

Group10839: acar|OOF93593.1 azon|OJJ49475.1 anid|CBF76709.1 asyd|OJJ61238.1 aver|OJJ00561.1 awen|OJJ34119.1

Group10840: ater|EAU35139.1 afis|EAW24662.1 afla|EED49438.1 aory|EIT79517.1 asyd|OJJ62363.1 aver|OJI96497.1

Group10841: acar|OOF93963.1 akaw|GAA92865.1 aluc|OJZ82549.1 anig|EHA22513.1 atub|OJI88790.1 azon|OJJ42553.1

Group10842: acar|OOF93969.1 acla|EAW13729.1 afis|EAW19857.1 afum|EAL85038.1 asyd|OJJ54928.1 azon|OJJ50684.1

Group10843: acar|OOF94016.1 ater|EAU31927.1 afis|EAW17059.1 afum|EAL86555.1 anid|CBF88302.1 azon|OJJ43508.1

Group10844: anid|CBF80757.1 acla|EAW15365.1 afis|EAW17225.1 afla|EED51522.1 afum|EAL86647.1 aory|EIT80644.1

Group10845: acar|OOF94259.1 agla|OJJ79983.1 asyd|OJJ63562.1 ater|EAU29647.1 aver|OJI97371.1 ater|EAU30278.1

Group10846: acar|OOF94674.1 afis|EAW25596.1 afum|EAL84302.1 anid|CBF82033.1 awen|OJJ40809.1 awen|OJJ40810.1

Group10847: acar|OOF94920.1 afla|EED53412.1 agla|OJJ79309.1 aory|EIT75202.1 asyd|OJJ59730.1 awen|OJJ36159.1

Group10848: afla|EED45497.1 aory|EIT75628.1 ater|EAU37348.1 acla|EAW12361.1 afis|EAW16790.1 afum|EBA27269.1

Group10849: azon|OJJ46838.1 ater|EAU38833.1 anid|CBF71821.1 asyd|OJJ52369.1 awen|OJJ37044.1 aver|OJJ01979.1

Group10850: acar|OOF95611.1 akaw|GAA89759.1 aluc|OJZ80624.1 anig|EHA21151.1 atub|OJI87478.1 aluc|OJZ85538.1

Group10851: acla|EAW14931.1 afla|EED55230.1 agla|OJJ80292.1 aory|EIT83524.1 awen|OJJ32752.1 ater|EAU30000.1

Group10852: aver|OJJ05286.1 afla|EED55024.1 anig|EHA22527.1 aory|EIT81971.1 asyd|OJJ54332.1 awen|OJJ41953.1

Group10853: acla|EAW11539.1 afis|EAW23912.1 afum|EAL87341.1 afla|EED50619.1 ater|EAU32320.1 aory|EIT78117.1

Group10854: acar|OOF98223.1 aory|EIT79698.1 asyd|OJJ57922.1 aver|OJJ04470.1 awen|OJJ29819.1 afla|EED56538.1

Group10855: acar|OOF98338.1 afla|EED49425.1 aory|EIT79491.1 asyd|OJJ55174.1 aver|OJJ07033.1 awen|OJJ41932.1

Group10856: acar|OOF98376.1 acla|EAW14535.1 akaw|GAA90989.1 aluc|OJZ87787.1 anig|EHA20269.1 atub|OJI80860.1

Group10857: azon|OJJ43412.1 acla|EAW09215.1 afis|EAW21932.1 afum|EAL92945.2 awen|OJJ34223.1 agla|OJJ84189.1

Group10858: acar|OOF98569.1 acla|EAW09217.1 afla|EED46624.1 akaw|GAA85728.1 aluc|OJZ91373.1 awen|OJJ34221.1

Group10859: afum|EAL91132.1 afla|EED52347.1 akaw|GAA91495.1 aluc|OJZ85656.1 atub|OJI79448.1 azon|OJJ43934.1

Group10860: acar|OOF98980.1 afla|EED51701.1 anid|CBF85292.1 aory|EIT80694.1 asyd|OJJ64962.1 aver|OJI99085.1

Group10861: acar|OOF99230.1 afis|EAW16459.1 afum|EAL89055.1 asyd|OJJ52560.1 azon|OJJ47381.1 awen|OJJ37788.1

Group10862: acar|OOF99269.1 acla|EAW10003.1 afum|EAL87166.1 akaw|GAA93034.1 aluc|OJZ86778.1 atub|OJI83742.1

Group10863: acar|OOF99359.1 afis|EAW15419.1 afum|EAL84225.1 agla|OJJ80129.1 asyd|OJJ64793.1 aver|OJI98907.1

Group10864: acar|OOF99534.1 afum|EAL92233.1 akaw|GAA91224.1 aluc|OJZ82846.1 atub|OJI89076.1 awen|OJJ30815.1

Group10865: acar|OOG00030.1 afis|EAW22476.1 akaw|GAA82110.1 asyd|OJJ64490.1 ater|EAU39429.1 anig|EHA23339.1

Group10866: acar|OOG00308.1 afla|EED45138.1 asyd|OJJ55192.1 ater|EAU31199.1 awen|OJJ32684.1 azon|OJJ43672.1

Group10867: acar|OOG00631.1 acla|EAW14122.1 afis|EAW18837.1 afum|EAL93571.1 agla|OJJ83237.1 awen|OJJ39741.1

Group10868: acla|EAW06433.1 afis|EAW18539.1 afla|EED45457.1 afum|EAL93237.1 aory|EIT82411.1 asyd|OJJ54410.1

Group10869: acla|EAW06527.1 afis|EAW16069.1 afum|EAL88701.2 agla|OJJ86099.1 awen|OJJ30366.1 azon|OJJ45385.1

Group10870: acla|EAW06980.1 afis|EAW16594.1 afum|EAL89190.1 anid|CBF81899.1 asyd|OJJ59684.1 aver|OJJ02603.1

Group10871: acla|EAW07016.1 afla|EED44965.1 agla|OJJ82605.1 aory|EIT73509.1 awen|OJJ40072.1 anid|CBF82042.1

Group10872: acla|EAW07046.1 afis|EAW16670.1 afum|EAL89275.1 asyd|OJJ64078.1 ater|EAU39591.1 aver|OJI98166.1

Group10873: acla|EAW07196.1 afis|EAW22208.1 afum|EBA27417.1 agla|OJJ87836.1 akaw|GAA87391.1 aver|OJI97175.1

Group10874: aory|EIT75409.1 afis|EAW21191.1 afla|EED45482.1 afum|EAL92211.1 asyd|OJJ61494.1 awen|OJJ30948.1

Group10875: acla|EAW07623.1 afis|EAW22652.1 afum|EAL90540.1 agla|OJJ81035.1 awen|OJJ35271.1 azon|OJJ49836.1

Group10876: acla|EAW08045.1 agla|OJJ89590.1 asyd|OJJ59365.1 aver|OJJ02909.1 awen|OJJ36703.1 anid|CBF81332.1

Group10877: acla|EAW08095.1 afla|EED47675.1 agla|OJJ82740.1 aory|EIT80240.1 ater|EAU31466.1 awen|OJJ32590.1

Group10878: acla|EAW08123.1 afis|EAW23160.1 afum|EAL88323.1 agla|OJJ89671.1 awen|OJJ36623.1 azon|OJJ47590.1

Group10879: acla|EAW08291.1 afis|EAW23349.1 afum|EAL88127.1 agla|OJJ86476.1 awen|OJJ39540.1 azon|OJJ47739.1

Group10880: acla|EAW08351.1 afis|EAW16824.1 afum|EAL90929.1 agla|OJJ83279.1 aluc|OJZ80477.1 afum|EAL90930.1

Group10881: acla|EAW08583.1 afla|EED51137.1 aory|EIT81310.1 asyd|OJJ61697.1 ater|EAU35444.1 aver|OJI95783.1

Group10882: acla|EAW08586.1 afis|EAW24711.1 afum|EAL89352.1 anid|CBF69534.1 asyd|OJJ55944.1 aver|OJJ04893.1

Group10883: acla|EAW09027.1 afis|EAW22740.1 afla|EED49801.1 afum|EAL90455.1 agla|OJJ78546.1 aory|EIT77594.1

Group10884: acla|EAW09031.1 afis|EAW22737.1 afla|EED49797.1 afum|EAL90458.2 agla|OJJ78516.1 aory|EIT77624.1

Group10885: acla|EAW09032.1 afis|EAW22736.1 afla|EED49799.1 afum|EAL90459.1 agla|OJJ78517.1 aory|EIT77686.1

Group10886: acla|EAW09060.1 afis|EAW25633.1 afla|EED55179.1 afum|EAL84321.1 aory|EIT81667.1 azon|OJJ50760.1

Group10887: acla|EAW09075.1 anid|CBF75396.1 asyd|OJJ65398.1 ater|EAU35625.1 aver|OJJ02470.1 asyd|OJJ59817.1

Group10888: acla|EAW09332.1 afis|EAW21788.1 agla|OJJ84335.1 aory|EIT77941.1 ater|EAU38471.1 awen|OJJ34074.1

Group10889: acla|EAW09334.1 afis|EAW21789.1 afum|EAL92815.1 ater|EAU35494.1 afla|EED44767.1 awen|OJJ31288.1

Group10890: acla|EAW09350.1 afis|EAW21815.1 afum|EAL92831.1 anid|CBF76635.1 asyd|OJJ61198.1 aver|OJJ00523.1

Group10891: acla|EAW09790.1 afis|EAW21346.1 afum|EAL92363.1 agla|OJJ87385.1 awen|OJJ39352.1 azon|OJJ51693.1

Group10892: acla|EAW09792.1 afis|EAW21344.1 afum|EAL92361.1 agla|OJJ87387.1 awen|OJJ39354.1 azon|OJJ51695.1

Group10893: acla|EAW09793.1 afis|EAW21343.1 afum|EAL92360.1 agla|OJJ87388.1 awen|OJJ39355.1 azon|OJJ51696.1

Group10894: afla|EED45049.1 afum|EBA27450.1 aory|EIT73758.1 awen|OJJ31602.1 anid|CBF80088.1 afis|EAW21153.1

Group10895: acla|EAW09941.1 afis|EAW21186.1 afum|EAL92208.1 agla|OJJ79981.1 awen|OJJ36527.1 azon|OJJ44400.1

Group10896: acla|EAW09943.1 afis|EAW21221.1 afum|EBA27453.1 asyd|OJJ57321.1 ater|EAU37400.1 aver|OJJ05840.1

Group10897: acla|EAW10023.1 aory|EIT81874.1 ater|EAU31048.1 aory|EIT81884.1 asyd|OJJ65355.1 aver|OJI99504.1

Group10898: acla|EAW10307.1 afla|EED45940.1 agla|OJJ79266.1 aory|EIT75741.1 asyd|OJJ64745.1 awen|OJJ38707.1

Group10899: acla|EAW10401.1 afis|EAW25177.1 afum|EAL89850.1 anid|CBF87304.1 asyd|OJJ61747.1 aver|OJI95844.1

Group10900: acla|EAW10409.1 afum|EAL85466.1 anid|CBF85659.1 asyd|OJJ64229.1 afis|EAW20346.1 aver|OJI98322.1

Group10901: acla|EAW10554.1 afis|EAW25021.1 afum|EAL89674.1 agla|OJJ89189.1 awen|OJJ40619.1 azon|OJJ49467.1

Group10902: acla|EAW10821.1 afis|EAW24751.1 afla|EED46134.1 aory|EIT76503.1 asyd|OJJ61398.1 aver|OJJ00727.1

Group10903: acla|EAW10836.1 afis|EAW24727.1 afum|EAL89375.2 agla|OJJ79449.1 awen|OJJ37857.1 agla|OJJ85244.1

Group10904: acla|EAW11031.1 afla|EED49639.1 afum|EAL88028.1 aory|EIT80989.1 azon|OJJ45952.1 aory|EIT80990.1

Group10905: acla|EAW11048.1 afis|EAW16188.1 afla|EED49631.1 afum|EAL88819.2 aory|EIT80944.1 azon|OJJ45945.1

Group10906: acla|EAW11369.1 afis|EAW18126.1 afum|EAL87484.1 ater|EAU29686.1 awen|OJJ35528.1 azon|OJJ46955.1

Group10907: acla|EAW11509.1 afis|EAW23944.1 afum|EAL87369.1 awen|OJJ39011.1 azon|OJJ51336.1 ater|EAU32290.1

Group10908: acla|EAW11697.1 afla|EED55120.1 aory|EIT81610.1 awen|OJJ31848.1 azon|OJJ46275.1 ater|EAU31406.1

Group10909: acla|EAW11774.1 akaw|GAA88136.1 aluc|OJZ81407.1 anig|EHA26066.1 atub|OJI91614.1 awen|OJJ31320.1

Group10910: acla|EAW11919.1 afis|EAW19552.1 afla|EED46516.1 afum|EAL86957.1 aory|EIT72359.1 azon|OJJ50150.1

Group10911: acla|EAW11922.1 afis|EAW18845.1 asyd|OJJ64701.1 aver|OJI98805.1 awen|OJJ35513.1 azon|OJJ51815.1

Group10912: asyd|OJJ57322.1 aver|OJJ05838.1 afla|EED51380.1 aory|EIT80730.1 awen|OJJ34572.1 azon|OJJ48189.1

Group10913: acla|EAW12133.1 afis|EAW17218.1 afla|EED51384.1 afum|EAL86643.2 aory|EIT80508.1 azon|OJJ50153.1

Group10914: acla|EAW12188.1 afis|EAW17126.1 afla|EED49687.1 afum|EAL86586.1 aory|EIT80962.1 awen|OJJ34565.1

Group10915: acla|EAW12284.1 afis|EAW16955.1 afum|EAL86482.2 agla|OJJ79123.1 asyd|OJJ55379.1 aver|OJJ08274.1

Group10916: asyd|OJJ53688.1 aver|OJJ07805.1 akaw|GAA92447.1 aluc|OJZ81571.1 anig|EHA25868.1 atub|OJI88614.1

Group10917: acla|EAW12368.1 afis|EAW16791.1 afla|EED54678.1 afum|EAL86360.1 aory|EIT78930.1 ater|EAU34475.1

Group10918: acla|EAW12436.1 afla|EED47340.1 aory|EIT82891.1 asyd|OJJ52819.1 ater|EAU36733.1 awen|OJJ31378.1

Group10919: anig|EHA22526.1 aver|OJJ04034.1 aver|OJJ08540.1 afla|EED52531.1 aory|EIT74259.1 awen|OJJ32262.1

Group10920: anid|CBF80374.1 afis|EAW19798.1 afum|EAL85092.1 agla|OJJ79580.1 asyd|OJJ54810.1 aver|OJJ07351.1

Group10921: awen|OJJ32766.1 afla|EED53058.1 agla|OJJ78710.1 aory|EIT74762.1 asyd|OJJ60330.1 ater|EAU37693.1

Group10922: acla|EAW12890.1 afis|EAW16314.1 afum|EAL88936.1 ater|EAU33698.1 afla|EED54181.1 ater|EAU29327.1

Group10923: acla|EAW12958.1 afis|EAW17701.1 afla|EED51719.1 afum|EAL91516.1 anid|CBF90385.1 aory|EIT80750.1

Group10924: acla|EAW12998.1 agla|OJJ78993.1 ater|EAU38104.1 awen|OJJ32609.1 awen|OJJ38387.1 agla|OJJ78996.1

Group10925: awen|OJJ33442.1 anid|CBF87306.1 aory|EIT73950.1 asyd|OJJ59787.1 ater|EAU34820.1 aver|OJJ02501.1

Group10926: acla|EAW13298.1 afis|EAW17313.1 afum|EAL91148.1 asyd|OJJ58460.1 aver|OJJ01060.1 ater|EAU32991.1

Group10927: acla|EAW13303.1 afis|EAW23646.1 afla|EED52625.1 anid|CBF84294.1 awen|OJJ29668.1 azon|OJJ42658.1

Group10928: acla|EAW13454.1 awen|OJJ30236.1 afis|EAW20562.1 ater|EAU37116.1 afla|EED52342.1 aory|EIT74436.1

Group10929: acla|EAW13561.1 afis|EAW20680.1 afum|EAL84416.2 akaw|GAA92612.1 aluc|OJZ80201.1 awen|OJJ30835.1

Group10930: acla|EAW13663.1 afis|EAW20292.1 afla|EED53473.1 aory|EIT75213.1 asyd|OJJ57226.1 aver|OJI98794.1

Group10931: acla|EAW13694.1 afis|EAW16422.1 agla|OJJ81391.1 ater|EAU39441.1 atub|OJI83263.1 awen|OJJ39636.1

Group10932: acla|EAW13695.1 afis|EAW19909.1 afum|EAL84997.1 anid|CBF79755.1 asyd|OJJ58099.1 aver|OJJ04305.1

Group10933: acla|EAW13732.1 afis|EAW19847.1 afum|EAL85047.1 anid|CBF82493.1 asyd|OJJ56115.1 aver|OJJ05056.1

Group10934: acla|EAW14099.1 afis|EAW25584.1 anid|CBF90365.1 asyd|OJJ61022.1 ater|EAU35436.1 aver|OJJ00343.1

Group10935: afis|EAW16156.1 asyd|OJJ52799.1 aver|OJJ08776.1 afla|EED45362.1 aory|EIT82416.1 ater|EAU32999.1

Group10936: acla|EAW14204.1 afis|EAW19078.1 afum|EAL93806.2 anid|CBF82136.1 asyd|OJJ54317.1 aver|OJJ05271.1

Group10937: acla|EAW14205.1 afis|EAW19077.1 afum|EAL93805.2 anid|CBF82138.1 asyd|OJJ54318.1 aver|OJJ05272.1

Group10938: acla|EAW14265.1 afis|EAW19014.1 afum|EBA27501.1 agla|OJJ81766.1 awen|OJJ31648.1 azon|OJJ43169.1

Group10939: acla|EAW14279.1 afis|EAW16812.1 afum|EAL86741.1 ater|EAU32794.1 afis|EAW20666.1 afis|EAW22497.1

Group10940: acla|EAW14280.1 afla|EED46986.1 afum|EAL85151.1 aory|EIT72610.1 asyd|OJJ62552.1 aver|OJI96678.1

Group10941: acla|EAW14560.1 afis|EAW19273.1 afum|EAL93999.2 awen|OJJ37634.1 ater|EAU29491.1 agla|OJJ88400.1

Group10942: acla|EAW14640.1 afis|EAW19553.1 afla|EED49441.1 afum|EAL86956.1 aory|EIT79519.1 asyd|OJJ55492.1

Group10943: acla|EAW14641.1 afla|EED53091.1 agla|OJJ87615.1 aory|EIT74804.1 asyd|OJJ57692.1 azon|OJJ42297.1

Group10944: acla|EAW14719.1 afis|EAW24255.1 afum|EAL85923.1 agla|OJJ79618.1 awen|OJJ33045.1 azon|OJJ48474.1

Group10945: acla|EAW14742.1 afis|EAW24273.1 afla|EED48003.1 afum|EAL85941.2 aory|EIT82895.1 awen|OJJ33024.1

Group10946: acla|EAW14928.1 afis|EAW17061.1 afla|EED49820.1 aory|EIT77613.1 aory|EIT77787.1 ater|EAU36520.1

Group10947: acla|EAW14972.1 afis|EAW24497.1 afum|EAL86167.1 anid|CBF75739.1 asyd|OJJ57515.1 aver|OJJ05664.1

Group10948: anid|CBF69383.1 anid|CBF85125.1 afum|EBA27267.1 ater|EAU32996.1 asyd|OJJ57407.1 aver|OJJ05774.1

Group10949: acla|EAW15252.1 afis|EAW20225.1 afla|EED53436.1 afum|EAL85342.1 aory|EIT75219.1 awen|OJJ31965.1

Group10950: acla|EAW15258.1 afis|EAW20237.1 afla|EED45968.1 afum|EAL85355.1 anid|CBF82916.1 aory|EIT75571.1

Group10951: acla|EAW15273.1 afis|EAW20243.1 afla|EED47509.1 afum|EAL85360.1 aory|EIT74551.1 azon|OJJ51776.1

Group10952: anid|CBF80134.1 ater|EAU35874.1 afla|EED50206.1 aory|EIT77322.1 asyd|OJJ57903.1 aver|OJJ04491.1

Group10953: afis|EAW15501.1 agla|OJJ80179.1 asyd|OJJ56010.1 ater|EAU32618.1 aver|OJJ04955.1 awen|OJJ38161.1

Group10954: afis|EAW16171.1 afla|EED49648.1 aory|EIT81001.1 aver|OJJ00805.1 awen|OJJ30152.1 azon|OJJ46420.1

Group10955: afis|EAW16211.1 afum|EAL88838.1 asyd|OJJ57149.1 aver|OJJ05568.1 aver|OJJ03161.1 asyd|OJJ57618.1

Group10956: afis|EAW16252.1 anid|CBF89278.1 asyd|OJJ55340.1 ater|EAU35484.1 aver|OJJ07022.1 asyd|OJJ55185.1

Group10957: afis|EAW16254.1 ater|EAU31545.1 afis|EAW16410.1 afum|EAL89034.1 anid|CBF73415.1 awen|OJJ41052.1

Group10958: afis|EAW16405.1 afla|EED47774.1 afum|EAL89023.1 aory|EIT77129.1 awen|OJJ30220.1 azon|OJJ43942.1

Group10959: afis|EAW16792.1 afum|EAL86361.1 agla|OJJ86551.1 asyd|OJJ54826.1 aver|OJJ07334.1 awen|OJJ38000.1

Group10960: afis|EAW16793.1 afla|EED45972.1 afum|EAL86362.1 agla|OJJ79964.1 aory|EIT75672.1 awen|OJJ33624.1

Group10961: afis|EAW16856.1 afla|EED50959.1 afum|EAL86399.1 asyd|OJJ61379.1 aver|OJJ00706.1 aory|EIT76949.1

Group10962: afis|EAW16860.1 afla|EED45902.1 aory|EIT75712.1 asyd|OJJ59235.1 aver|OJJ01845.1 awen|OJJ36994.1

Group10963: afis|EAW16919.1 afla|EED56829.1 aory|EIT83113.1 asyd|OJJ62993.1 aver|OJI97975.1 afla|EED51185.1

Group10964: afis|EAW17029.1 aluc|OJZ80159.1 asyd|OJJ55864.1 aver|OJJ04805.1 awen|OJJ33827.1 asyd|OJJ59624.1

Group10965: afis|EAW17030.1 agla|OJJ79118.1 asyd|OJJ60335.1 aver|OJI99680.1 awen|OJJ36176.1 aver|OJJ03410.1

Group10966: afis|EAW17053.1 afla|EED53168.1 afum|EAL86537.1 akaw|GAA88276.1 anig|EHA28495.1 atub|OJI80950.1

Group10967: afis|EAW17203.1 afla|EED52909.1 afum|EAL86629.2 aory|EIT78608.1 asyd|OJJ64727.1 aver|OJJ01308.1

Group10968: afis|EAW17297.1 asyd|OJJ55437.1 ater|EAU29435.1 aver|OJJ06513.1 awen|OJJ36111.1 anid|CBF83929.1

Group10969: afis|EAW17762.1 afum|EAL91572.1 anid|CBF82293.1 asyd|OJJ52732.1 ater|EAU32267.1 aver|OJJ05349.1

Group10970: afis|EAW18258.1 afum|EAL87621.1 anid|CBF74968.1 asyd|OJJ64754.1 ater|EAU35673.1 aver|OJI98856.1

Group10971: afis|EAW18822.1 afum|EAL93559.1 anid|CBF84664.1 asyd|OJJ57152.1 ater|EAU37633.1 aver|OJJ03158.1

Group10972: afis|EAW19506.1 afla|EED56797.1 aory|EIT73919.1 asyd|OJJ52890.1 ater|EAU30294.1 aver|OJJ08861.1

Group10973: afis|EAW19731.1 afum|EAL86797.1 asyd|OJJ54950.1 ater|EAU30302.1 aver|OJJ07245.1 anid|CBF71453.1

Group10974: afla|EED51942.1 aory|EIT81380.1 azon|OJJ47544.1 asyd|OJJ52400.1 aver|OJJ01920.1 awen|OJJ33501.1

Group10975: afis|EAW19739.1 afla|EED56527.1 afum|EAL86790.1 agla|OJJ79781.1 aory|EIT82923.1 awen|OJJ34644.1

Group10976: afis|EAW19858.1 afla|EED56900.1 afum|EAL85037.1 aory|EIT77529.1 asyd|OJJ52512.1 aver|OJJ03479.1

Group10977: afis|EAW19869.1 afum|EAL85028.1 agla|OJJ89321.1 akaw|GAA85276.1 aluc|OJZ90635.1 aver|OJJ05207.1

Group10978: afis|EAW19905.1 afla|EED54743.1 afum|EAL85002.1 aory|EIT78929.1 ater|EAU34056.1 azon|OJJ51042.1

Group10979: afis|EAW19981.1 afum|EBA27211.1 agla|OJJ79259.1 asyd|OJJ57156.1 azon|OJJ42436.1 azon|OJJ45351.1

Group10980: afis|EAW20393.1 afla|EED54889.1 anid|CBF73690.1 aory|EIT79424.1 asyd|OJJ56652.1 aver|OJJ06073.1

Group10981: afis|EAW20699.1 afum|EAL84874.1 asyd|OJJ56274.1 atub|OJI79832.1 aver|OJJ08491.1 anid|CBF82275.1

Group10982: afis|EAW20819.1 afum|EAL84810.1 akaw|GAA82837.1 aluc|OJZ87398.1 atub|OJI81248.1 awen|OJJ33644.1

Group10983: afis|EAW21074.1 afla|EED48209.1 anid|CBF74242.1 aory|EIT72914.1 asyd|OJJ53972.1 aver|OJJ08071.1

Group10984: afla|EED49491.1 agla|OJJ81021.1 anid|CBF83099.1 aory|EIT82648.1 asyd|OJJ60676.1 aver|OJI99999.1

Group10985: afis|EAW21209.1 afum|EAL92220.1 agla|OJJ79293.1 asyd|OJJ54584.1 atub|OJI84637.1 aver|OJJ07573.1

Group10986: afis|EAW22001.1 afum|EAL93016.1 anid|CBF82668.1 asyd|OJJ61663.1 ater|EAU31120.1 aver|OJJ00984.1

Group10987: afis|EAW22099.1 afla|EED52304.1 afum|EAL91090.1 aory|EIT74339.1 ater|EAU32748.1 aver|OJJ05878.1

Group10988: afis|EAW22300.1 agla|OJJ82722.1 anid|CBF85182.1 awen|OJJ31412.1 aver|OJI99533.1 anid|CBF80147.1

Group10989: afis|EAW22746.1 agla|OJJ88065.1 aory|EIT80580.1 asyd|OJJ55934.1 aver|OJJ04883.1 awen|OJJ36947.1

Group10990: asyd|OJJ55935.1 afla|EED52577.1 agla|OJJ85232.1 aory|EIT74273.1 aver|OJJ04884.1 awen|OJJ36948.1

Group10991: akaw|GAA85275.1 aluc|OJZ80498.1 awen|OJJ31084.1 afum|EAL90263.1 asyd|OJJ62321.1 ater|EAU33523.1

Group10992: afis|EAW23618.1 afla|EED48177.1 afum|EAL87844.1 asyd|OJJ55362.1 aver|OJJ08472.1 aver|OJJ06444.1

Group10993: afis|EAW23644.1 afla|EED46050.1 aory|EIT83494.1 ater|EAU35170.1 awen|OJJ39955.1 ater|EAU30989.1

Group10994: afis|EAW23689.1 afla|EED52257.1 afum|EAL87185.1 anid|CBF76147.1 asyd|OJJ54885.1 ater|EAU34905.1

Group10995: afis|EAW24014.1 afum|EAL87444.1 agla|OJJ82998.1 asyd|OJJ53236.1 awen|OJJ35486.1 aver|OJJ03615.1

Group10996: afis|EAW24587.1 agla|OJJ82375.1 anid|CBF87337.1 asyd|OJJ52811.1 aver|OJI96481.1 aver|OJJ08789.1

Group10997: afis|EAW24677.1 asyd|OJJ57400.1 awen|OJJ40793.1 afis|EAW24707.1 afum|EAL89349.1 ater|EAU31629.1

Group10998: afis|EAW24790.1 aluc|OJZ92173.1 afla|EED47192.1 aory|EIT79308.1 awen|OJJ40376.1 azon|OJJ49227.1

Group10999: afis|EAW25607.1 afla|EED50615.1 agla|OJJ82289.1 aory|EIT78044.1 asyd|OJJ52088.1 awen|OJJ36106.1

Group11000: afis|EAW25731.1 afum|EAL84402.2 anid|CBF78390.1 asyd|OJJ62330.1 aver|OJJ04860.1 aver|OJI96436.1

Group11001: afla|EED44731.1 anid|CBF87318.1 aory|EIT77804.1 asyd|OJJ59108.1 atub|OJI79309.1 aver|OJJ01699.1

Group11002: afla|EED45037.1 aory|EIT73555.1 asyd|OJJ59695.1 ater|EAU31333.1 aver|OJJ02592.1 azon|OJJ42416.1

Group11003: afla|EED45640.1 anid|CBF78492.1 aory|EIT75641.1 asyd|OJJ56057.1 aver|OJJ04999.1 awen|OJJ33618.1

Group11004: afla|EED45653.1 anid|CBF78350.1 aory|EIT75430.1 asyd|OJJ56142.1 aver|OJJ05081.1 awen|OJJ35378.1

Group11005: afla|EED45770.1 anid|CBF71764.1 aory|EIT75526.1 asyd|OJJ61724.1 aver|OJI95822.1 awen|OJJ30372.1

Group11006: afla|EED45876.1 anid|CBF84189.1 aory|EIT75523.1 asyd|OJJ57133.1 ater|EAU39448.1 aver|OJJ03170.1

Group11007: afla|EED45896.1 anid|CBF87396.1 aory|EIT75706.1 asyd|OJJ52859.1 aver|OJJ08836.1 awen|OJJ38132.1

Group11008: asyd|OJJ65304.1 agla|OJJ89018.1 anid|CBF82272.1 ater|EAU35670.1 aver|OJI99442.1 awen|OJJ38658.1

Group11009: afla|EED46391.1 anid|CBF80108.1 aory|EIT72424.1 asyd|OJJ58327.1 atub|OJI84525.1 aver|OJJ02084.1

Group11010: afla|EED46481.1 anid|CBF83918.1 aory|EIT72434.1 asyd|OJJ55497.1 aver|OJJ06575.1 awen|OJJ36917.1

Group11011: afla|EED46519.1 akaw|GAA83305.1 aluc|OJZ83801.1 anig|EHA18949.1 aory|EIT72402.1 atub|OJI90169.1

Group11012: afla|EED47258.1 afum|EAL91081.1 anid|CBF80296.1 aory|EIT79204.1 asyd|OJJ60438.1 aver|OJJ02284.1

Group11013: afla|EED47352.1 agla|OJJ84843.1 aory|EIT82876.1 asyd|OJJ62340.1 aver|OJI96445.1 awen|OJJ33169.1

Group11014: afla|EED47650.1 akaw|GAA85926.1 aory|EIT80193.1 ater|EAU30408.1 awen|OJJ30178.1 azon|OJJ46788.1

Group11015: afla|EED48125.1 atub|OJI80333.1 afla|EED56928.1 aory|EIT77474.1 aory|EIT76262.1 awen|OJJ36397.1

Group11016: afla|EED48193.1 aluc|OJZ87032.1 anig|EHA25216.1 aory|EIT72958.1 asyd|OJJ62483.1 atub|OJI83483.1

Group11017: afla|EED48238.1 agla|OJJ78755.1 anid|CBF78629.1 aory|EIT72904.1 aver|OJJ08130.1 awen|OJJ30816.1

Group11018: afla|EED48294.1 akaw|GAA86603.1 aluc|OJZ85151.1 anig|EHA25384.1 ater|EAU37315.1 atub|OJI80425.1

Group11019: afla|EED48298.1 aory|EIT76301.1 afla|EED52299.1 anig|EHA25382.1 ater|EAU37313.1 atub|OJI80427.1

Group11020: afla|EED48322.1 asyd|OJJ56109.1 atub|OJI89796.1 aver|OJJ05052.1 awen|OJJ33527.1 azon|OJJ48590.1

Group11021: afla|EED48325.1 anid|CBF75766.1 aory|EIT76277.1 atub|OJI89793.1 awen|OJJ33528.1 azon|OJJ45976.1

Group11022: afla|EED48383.1 aory|EIT83046.1 asyd|OJJ62117.1 ater|EAU35082.1 aver|OJI96224.1 awen|OJJ31903.1

Group11023: afla|EED48415.1 agla|OJJ85774.1 anid|CBF78041.1 aory|EIT83080.1 asyd|OJJ59809.1 aver|OJJ02481.1

Group11024: afla|EED48454.1 aory|EIT83471.1 asyd|OJJ59209.1 aver|OJJ01820.1 awen|OJJ37836.1 aory|EIT83466.1

Group11025: afla|EED48504.1 aory|EIT73089.1 asyd|OJJ53410.1 aver|OJJ08341.1 azon|OJJ45927.1 anid|CBF87460.1

Group11026: aver|OJJ03959.1 anid|CBF82936.1 anig|EHA24789.1 asyd|OJJ65139.1 ater|EAU35235.1 awen|OJJ37939.1

Group11027: afla|EED48928.1 anid|CBF79039.1 aory|EIT80013.1 asyd|OJJ52411.1 ater|EAU29323.1 aver|OJJ08489.1

Group11028: afla|EED49097.1 aory|EIT82511.1 asyd|OJJ59250.1 ater|EAU33805.1 aver|OJJ01859.1 awen|OJJ29929.1

Group11029: afla|EED49378.1 aory|EIT79460.1 asyd|OJJ63498.1 ater|EAU36892.1 aver|OJI97431.1 awen|OJJ41769.1

Group11030: afla|EED49492.1 agla|OJJ81022.1 anid|CBF83101.1 aory|EIT82626.1 asyd|OJJ64761.1 aver|OJI98862.1

Group11031: afla|EED49493.1 agla|OJJ81023.1 anid|CBF83103.1 aory|EIT82653.1 asyd|OJJ64760.1 aver|OJI98861.1

Group11032: afla|EED49613.1 akaw|GAA89215.1 aluc|OJZ81614.1 aory|EIT80961.1 atub|OJI88568.1 aver|OJJ02004.1

Group11033: afla|EED50603.1 anig|EHA18745.1 ater|EAU31539.1 aver|OJI95798.1 atub|OJI86838.1 azon|OJJ42280.1

Group11034: afla|EED50843.1 anid|CBF75998.1 aory|EIT79006.1 asyd|OJJ53689.1 ater|EAU30180.1 awen|OJJ37948.1

Group11035: afla|EED51050.1 anid|CBF85593.1 aory|EIT77066.1 asyd|OJJ64191.1 aver|OJI98268.1 azon|OJJ46837.1

Group11036: afla|EED51087.1 anid|CBF83993.1 aory|EIT76986.1 agla|OJJ80680.1 azon|OJJ42245.1 agla|OJJ80682.1

Group11037: afla|EED51130.1 agla|OJJ81310.1 anid|CBF90354.1 asyd|OJJ55574.1 ater|EAU37577.1 aver|OJJ06653.1

Group11038: afla|EED51150.1 agla|OJJ88366.1 anid|CBF80151.1 aory|EIT81301.1 ater|EAU38048.1 aver|OJI99511.1

Group11039: afla|EED51495.1 akaw|GAA91746.1 aluc|OJZ92757.1 aory|EIT80437.1 atub|OJI87201.1 asyd|OJJ54974.1

Group11040: afla|EED51504.1 anid|CBF75008.1 aory|EIT80576.1 asyd|OJJ60054.1 ater|EAU31282.1 aver|OJJ02235.1

Group11041: afla|EED51555.1 anid|CBF89723.1 aory|EIT80555.1 asyd|OJJ62236.1 ater|EAU36623.1 aver|OJI96345.1

Group11042: afla|EED51557.1 anid|CBF79921.1 aory|EIT80713.1 asyd|OJJ58199.1 ater|EAU38230.1 aver|OJJ04209.1

Group11043: afla|EED51558.1 anid|CBF77716.1 aory|EIT80698.1 asyd|OJJ53065.1 ater|EAU33176.1 aver|OJJ03780.1

Group11044: afla|EED51665.1 aluc|OJZ80141.1 anid|CBF70597.1 aory|EIT80426.1 ater|EAU30913.1 anid|CBF82265.1

Group11045: afla|EED51776.1 agla|OJJ85279.1 aory|EIT81743.1 asyd|OJJ56966.1 aver|OJJ03321.1 anid|CBF73576.1

Group11046: afla|EED51898.1 akaw|GAA92702.1 aluc|OJZ88741.1 aory|EIT76175.1 atub|OJI91313.1 awen|OJJ36085.1

Group11047: afla|EED52020.1 anid|CBF88297.1 aory|EIT81381.1 ater|EAU31924.1 aver|OJJ07787.1 anid|CBF86048.1

Group11048: afla|EED52140.1 anid|CBF89346.1 aory|EIT74427.1 asyd|OJJ62504.1 ater|EAU37454.1 aver|OJI96625.1

Group11049: afla|EED52145.1 anid|CBF87484.1 asyd|OJJ52948.1 aver|OJJ08915.1 anid|CBF87483.1 aory|EIT74429.1

Group11050: afla|EED52195.1 agla|OJJ80687.1 aory|EIT74488.1 asyd|OJJ61472.1 aver|OJJ00813.1 awen|OJJ32527.1

Group11051: afla|EED52335.1 akaw|GAA86512.1 aluc|OJZ85057.1 anid|CBF70200.1 aory|EIT74469.1 atub|OJI80543.1

Group11052: afla|EED52795.1 anid|CBF88875.1 aory|EIT78179.1 asyd|OJJ54653.1 ater|EAU30966.1 aver|OJJ07508.1

Group11053: afla|EED52803.1 akaw|GAA89087.1 aluc|OJZ80504.1 anig|EHA25689.1 aory|EIT78152.1 atub|OJI88426.1

Group11054: afla|EED52978.1 akaw|GAA86389.1 anid|CBF70037.1 ater|EAU29853.1 aver|OJJ01468.1 azon|OJJ50476.1

Group11055: afla|EED53135.1 aory|EIT75248.1 asyd|OJJ57974.1 ater|EAU31427.1 aver|OJJ04427.1 azon|OJJ45208.1

Group11056: afla|EED53154.1 anid|CBF84351.1 aory|EIT75092.1 asyd|OJJ59650.1 aver|OJJ02632.1 ater|EAU34773.1

Group11057: afla|EED53327.1 aory|EIT74832.1 asyd|OJJ65383.1 ater|EAU32671.1 aver|OJI99553.1 awen|OJJ38738.1

Group11058: afla|EED54054.1 agla|OJJ82673.1 anid|CBF73933.1 aory|EIT76065.1 asyd|OJJ53673.1 ater|EAU30472.1

Group11059: afla|EED54192.1 anid|CBF76177.1 asyd|OJJ54095.1 ater|EAU33851.1 aver|OJJ08644.1 azon|OJJ45224.1

Group11060: afla|EED54706.1 anid|CBF73852.1 aory|EIT78865.1 asyd|OJJ56517.1 ater|EAU29954.1 aver|OJJ06205.1

Group11061: afla|EED55081.1 akaw|GAA84903.1 aluc|OJZ86767.1 anig|EHA21913.1 aory|EIT81657.1 atub|OJI83761.1

Group11062: afla|EED55149.1 anid|CBF83895.1 aory|EIT81662.1 ater|EAU33495.1 aver|OJJ05759.1 asyd|OJJ57419.1

Group11063: afla|EED55261.1 anid|CBF82107.1 aory|EIT83554.1 asyd|OJJ54273.1 aver|OJJ05234.1 aver|OJJ05229.1

Group11064: afla|EED55433.1 agla|OJJ82942.1 aory|EIT80832.1 awen|OJJ36301.1 awen|OJJ31586.1 asyd|OJJ55472.1

Group11065: afla|EED56595.1 agla|OJJ83770.1 aory|EIT79684.1 asyd|OJJ54325.1 aver|OJJ05279.1 awen|OJJ30241.1

Group11066: afla|EED56601.1 aver|OJI98848.1 agla|OJJ83888.1 asyd|OJJ64744.1 aory|EIT80964.1 awen|OJJ33407.1

Group11067: afla|EED56604.1 agla|OJJ81898.1 aory|EIT79637.1 asyd|OJJ65335.1 aver|OJI99481.1 awen|OJJ33513.1

Group11068: afla|EED56871.1 anid|CBF74702.1 aory|EIT77445.1 asyd|OJJ60217.1 ater|EAU31411.1 aver|OJJ04051.1

Group11069: afla|EED57053.1 agla|OJJ78600.1 aory|EIT82308.1 atub|OJI86755.1 awen|OJJ35633.1 azon|OJJ42411.1

Group11070: afla|EED57075.1 aory|EIT82287.1 asyd|OJJ61458.1 aver|OJJ00796.1 awen|OJJ41744.1 azon|OJJ46568.1

Group11071: afla|EED57078.1 agla|OJJ79341.1 anid|CBF70707.1 aory|EIT82310.1 asyd|OJJ58454.1 aver|OJJ01054.1

Group11072: afla|EED57381.1 aory|EIT80125.1 asyd|OJJ63943.1 aver|OJI96999.1 awen|OJJ35021.1 azon|OJJ47389.1

Group11073: afla|EED57513.1 aory|EIT81860.1 asyd|OJJ64585.1 aver|OJI98683.1 awen|OJJ29792.1 azon|OJJ43594.1

Group11074: afla|EED57915.1 afum|EAL86571.1 aory|EIT82347.1 asyd|OJJ62083.1 ater|EAU31850.1 awen|OJJ38259.1

Group11075: agla|OJJ78820.1 akaw|GAA88081.1 awen|OJJ33747.1 azon|OJJ42201.1 aluc|OJZ81345.1 atub|OJI91549.1

Group11076: agla|OJJ80670.1 anid|CBF80575.1 asyd|OJJ55187.1 ater|EAU37974.1 aver|OJJ07020.1 awen|OJJ31432.1

Group11077: agla|OJJ84711.1 akaw|GAA91832.1 aluc|OJZ92854.1 asyd|OJJ57293.1 atub|OJI87311.1 aver|OJJ05863.1

Group11078: agla|OJJ86472.1 anid|CBF74893.1 asyd|OJJ60104.1 ater|EAU35597.1 aver|OJJ04158.1 awen|OJJ39535.1

Group11079: agla|OJJ88688.1 anid|CBF82376.1 asyd|OJJ54195.1 ater|EAU29280.1 aver|OJJ08531.1 awen|OJJ31675.1

Group11080: akaw|GAA83149.1 aluc|OJZ79892.1 anig|EHA18390.1 ater|EAU33733.1 atub|OJI91287.1 awen|OJJ35353.1

Group11081: akaw|GAA83832.1 aluc|OJZ90886.1 anig|EHA19605.1 atub|OJI82880.1 awen|OJJ29664.1 azon|OJJ47401.1

Group11082: akaw|GAA85147.1 aluc|OJZ86491.1 asyd|OJJ55409.1 atub|OJI84037.1 aver|OJJ06488.1 awen|OJJ35933.1

Group11083: akaw|GAA88766.1 anid|CBF69550.1 asyd|OJJ64983.1 ater|EAU38122.1 aver|OJI99103.1 awen|OJJ31666.1

Group11084: akaw|GAA88968.1 aluc|OJZ89435.1 anig|EHA23152.1 asyd|OJJ54808.1 atub|OJI85358.1 aver|OJJ07353.1

Group11085: akaw|GAA90690.1 aluc|OJZ84664.1 asyd|OJJ62586.1 ater|EAU29540.1 atub|OJI89416.1 aver|OJI96705.1

Group11086: akaw|GAA92619.1 aluc|OJZ80493.1 asyd|OJJ56761.1 akaw|GAA93044.1 anig|EHA21838.1 aluc|OJZ80238.1

Group11087: anig|EHA22606.1 aluc|OJZ91579.1 aory|EIT73465.1 aver|OJJ02479.1 ater|EAU31719.1 azon|OJJ45984.1

Group11088: asyd|OJJ53494.1 aluc|OJZ89757.1 anid|CBF79128.1 atub|OJI85043.1 aver|OJJ08254.1 awen|OJJ36424.1

Group11089: anid|CBF82200.1 asyd|OJJ52182.1 aver|OJJ05963.1 awen|OJJ34428.1 anid|CBF82202.1 ater|EAU29292.1

Group11090: anid|CBF82938.1 asyd|OJJ54755.1 ater|EAU31004.1 aver|OJJ08348.1 asyd|OJJ54754.1 aver|OJJ05259.1

Group11091: abra|OJJ65464.1 abra|OJJ67272.1 abra|OJJ65463.1 afla|EED49681.1 afla|EED50600.1

Group11092: abra|OJJ65468.1 abra|OJJ65659.1 abra|OJJ65707.1 akaw|GAA92647.1 ater|EAU30113.1

Group11093: abra|OJJ65510.1 abra|OJJ65823.1 ater|EAU30110.1 agla|OJJ78479.1 akaw|GAA93290.1

Group11094: abra|OJJ67520.1 abra|OJJ70182.1 acar|OOF94082.1 acar|OOF94152.1 afum|EAL94105.1

Group11095: abra|OJJ67522.1 abra|OJJ70180.1 acar|OOF94081.1 acar|OOF94151.1 awen|OJJ34532.1

Group11096: acar|OOF90423.1 acar|OOF90427.1 anid|CBF71777.1 asyd|OJJ61717.1 aver|OJI95811.1

Group11097: acar|OOF93737.1 acar|OOF93826.1 abra|OJJ66741.1 anig|EHA27310.1 atub|OJI87565.1

Group11098: acar|OOF94048.1 acar|OOF94118.1 akaw|GAA82257.1 aluc|OJZ91887.1 atub|OJI86337.1

Group11099: acar|OOF95666.1 acar|OOF95717.1 agla|OJJ82099.1 akaw|GAA81913.1 azon|OJJ49331.1

Group11100: acar|OOF97363.1 acar|OOF97647.1 abra|OJJ72277.1 asyd|OJJ55823.1 aver|OJJ04765.1

Group11101: afis|EAW18100.1 afis|EAW20670.1 agla|OJJ83479.1 agla|OJJ83543.1 aluc|OJZ91233.1

Group11102: afis|EAW19854.1 afis|EAW24644.1 acar|OOF91482.1 acla|EAW13497.1 afum|EAL85041.1

Group11103: afum|EAL86748.1 afum|EAL86770.1 afla|EED48246.1 azon|OJJ45213.1 agla|OJJ78812.1

Group11104: afum|EAL90936.1 afum|EAL91120.1 afis|EAW15511.1 agla|OJJ87696.1 asyd|OJJ52157.1

Group11105: agla|OJJ78802.1 agla|OJJ83553.1 agla|OJJ86307.1 agla|OJJ79435.1 agla|OJJ79130.1

Group11106: agla|OJJ79528.1 agla|OJJ80559.1 agla|OJJ88962.1 acar|OOF93971.1 awen|OJJ32693.1

Group11107: agla|OJJ79590.1 agla|OJJ82596.1 agla|OJJ81185.1 afis|EAW25162.1 awen|OJJ33247.1

Group11108: agla|OJJ79775.1 awen|OJJ30128.1 awen|OJJ31420.1 asyd|OJJ55627.1 aver|OJJ06704.1

Group11109: agla|OJJ80714.1 agla|OJJ86337.1 agla|OJJ81203.1 afis|EAW16748.1 azon|OJJ50284.1

Group11110: agla|OJJ83216.1 agla|OJJ86811.1 awen|OJJ38792.1 afla|EED51649.1 aory|EIT80589.1

Group11111: agla|OJJ84034.1 agla|OJJ84149.1 agla|OJJ84040.1 agla|OJJ84035.1 agla|OJJ84033.1

Group11112: agla|OJJ87384.1 agla|OJJ87389.1 agla|OJJ87390.1 asyd|OJJ64720.1 awen|OJJ37149.1

Group11113: akaw|GAA88992.1 akaw|GAA93235.1 acar|OOF90174.1 afla|EED51669.1 aluc|OJZ80347.1

Group11114: akaw|GAA92348.1 akaw|GAA93073.1 aluc|OJZ80226.1 akaw|GAA93134.1 asyd|OJJ52326.1

Group11115: akaw|GAA92964.1 akaw|GAA93161.1 aluc|OJZ79723.1 aluc|OJZ79794.1 aluc|OJZ80328.1

Group11116: akaw|GAA93122.1 akaw|GAA93331.1 awen|OJJ29555.1 awen|OJJ29643.1 ater|EAU30106.1

Group11117: aluc|OJZ79486.1 aluc|OJZ79630.1 afla|EED50609.1 aory|EIT78131.1 asyd|OJJ53649.1

Group11118: aluc|OJZ79526.1 aluc|OJZ79769.1 aluc|OJZ79876.1 aluc|OJZ79665.1 aluc|OJZ80045.1

Group11119: aluc|OJZ79583.1 aluc|OJZ80492.1 akaw|GAA93323.1 anig|EHA17686.1 akaw|GAA93329.1

Group11120: aluc|OJZ79649.1 aluc|OJZ79994.1 atub|OJI84503.1 awen|OJJ29797.1 azon|OJJ48188.1

Group11121: aluc|OJZ79672.1 aluc|OJZ88805.1 abra|OJJ65594.1 akaw|GAA93282.1 asyd|OJJ51939.1

Group11122: aluc|OJZ79742.1 aluc|OJZ79985.1 aluc|OJZ79903.1 aluc|OJZ80180.1 aluc|OJZ80259.1

Group11123: aluc|OJZ80133.1 aluc|OJZ81119.1 abra|OJJ65481.1 akaw|GAA93068.1 akaw|GAA93334.1

Group11124: aluc|OJZ80447.1 aluc|OJZ80484.1 acla|EAW11472.1 asyd|OJJ52151.1 akaw|GAA93182.1

Group11125: anid|CBF74637.1 anid|CBF76235.1 asyd|OJJ54460.1 ater|EAU35373.1 aver|OJJ07709.1

Group11126: anid|CBF80277.1 anid|CBF89297.1 afla|EED48478.1 aory|EIT73190.1 awen|OJJ40130.1

Group11127: anig|EHA19678.1 anig|EHA23791.1 akaw|GAA83755.1 aluc|OJZ90977.1 atub|OJI82967.1

Group11128: asyd|OJJ51883.1 asyd|OJJ51885.1 asyd|OJJ51984.1 asyd|OJJ51927.1 akaw|GAA93093.1

Group11129: asyd|OJJ51922.1 asyd|OJJ52313.1 abra|OJJ67771.1 anig|EHA21860.1 ater|EAU30370.1

Group11130: asyd|OJJ51938.1 asyd|OJJ52131.1 asyd|OJJ52144.1 asyd|OJJ52241.1 acla|EAW11452.1

Group11131: asyd|OJJ51946.1 asyd|OJJ56266.1 agla|OJJ84078.1 aluc|OJZ80204.1 anid|CBF84331.1

Group11132: asyd|OJJ52079.1 asyd|OJJ54403.1 anid|CBF78256.1 ater|EAU32805.1 aver|OJJ03025.1

Group11133: asyd|OJJ52519.1 asyd|OJJ55475.1 azon|OJJ42592.1 ater|EAU33746.1 ater|EAU29634.1

Group11134: asyd|OJJ52522.1 asyd|OJJ55476.1 azon|OJJ42595.1 asyd|OJJ58317.1 aver|OJJ02093.1

Group11135: asyd|OJJ54544.1 asyd|OJJ56714.1 aver|OJJ06011.1 aver|OJJ07633.1 anid|CBF85159.1

Group11136: asyd|OJJ55441.1 asyd|OJJ58383.1 asyd|OJJ56929.1 asyd|OJJ55467.1 asyd|OJJ61692.1

Group11137: asyd|OJJ56624.1 asyd|OJJ58589.1 asyd|OJJ59716.1 afis|EAW22933.1 ater|EAU38744.1

Group11138: asyd|OJJ57919.1 asyd|OJJ61729.1 afla|EED47666.1 akaw|GAA92535.1 aory|EIT80270.1

Group11139: asyd|OJJ58691.1 asyd|OJJ65367.1 agla|OJJ88816.1 aver|OJJ01303.1 azon|OJJ50313.1

Group11140: ater|EAU29356.1 aory|EIT81756.1 acar|OOF92526.1 awen|OJJ32241.1 afla|EED51789.1

Group11141: ater|EAU29429.1 ater|EAU39348.1 anid|CBF80712.1 asyd|OJJ58389.1 aver|OJJ02029.1

Group11142: ater|EAU33399.1 ater|EAU33406.1 afis|EAW24578.1 agla|OJJ86014.1 asyd|OJJ61757.1

Group11143: ater|EAU33758.1 ater|EAU36615.1 akaw|GAA88720.1 aluc|OJZ92677.1 asyd|OJJ56928.1

Group11144: ater|EAU36874.1 ater|EAU39017.1 acla|EAW13653.1 afis|EAW19952.1 afum|EAL84963.1

Group11145: aver|OJI96461.1 aver|OJI97960.1 anid|CBF74677.1 asyd|OJJ55664.1 awen|OJJ31385.1

Group11146: aver|OJI98000.1 aver|OJJ07714.1 asyd|OJJ62967.1 afis|EAW16726.1 azon|OJJ42979.1

Group11147: awen|OJJ29963.1 awen|OJJ29965.1 awen|OJJ29964.1 afis|EAW17001.1 azon|OJJ48032.1

Group11148: awen|OJJ30772.1 awen|OJJ31709.1 agla|OJJ87853.1 azon|OJJ48233.1 azon|OJJ45039.1

Group11149: awen|OJJ30848.1 awen|OJJ36211.1 abra|OJJ77767.1 acar|OOF97503.1 anig|EHA20913.1

Group11150: awen|OJJ31095.1 abra|OJJ69639.1 afum|EAL86340.1 anid|CBF89403.1 azon|OJJ50089.1

Group11151: awen|OJJ31343.1 awen|OJJ37498.1 acla|EAW12411.1 agla|OJJ81568.1 acla|EAW13606.1

Group11152: awen|OJJ36251.1 asyd|OJJ64970.1 aver|OJI99094.1 ater|EAU33751.1 azon|OJJ45933.1

Group11153: awen|OJJ32516.1 awen|OJJ36007.1 awen|OJJ40099.1 agla|OJJ87733.1 azon|OJJ42078.1

Group11154: awen|OJJ33331.1 abra|OJJ70319.1 aluc|OJZ82588.1 anig|EHA27487.1 atub|OJI88823.1

Group11155: awen|OJJ33365.1 awen|OJJ40869.1 awen|OJJ35876.1 awen|OJJ35792.1 awen|OJJ30542.1

Group11156: awen|OJJ36468.1 awen|OJJ38291.1 agla|OJJ88684.1 ater|EAU31889.1 awen|OJJ35857.1

Group11157: awen|OJJ36943.1 awen|OJJ39647.1 abra|OJJ65478.1 ater|EAU31613.1 azon|OJJ42377.1

Group11158: azon|OJJ42212.1 azon|OJJ46790.1 afla|EED57029.1 aory|EIT82293.1 agla|OJJ79829.1

Group11159: azon|OJJ42813.1 azon|OJJ49487.1 azon|OJJ44476.1 azon|OJJ49473.1 azon|OJJ47564.1

Group11160: azon|OJJ44447.1 azon|OJJ46469.1 agla|OJJ84395.1 agla|OJJ87726.1 awen|OJJ38734.1

Group11161: azon|OJJ44853.1 azon|OJJ45564.1 akaw|GAA84667.1 aluc|OJZ87022.1 atub|OJI83496.1

Group11162: azon|OJJ45685.1 azon|OJJ46495.1 azon|OJJ48980.1 azon|OJJ46509.1 azon|OJJ51111.1

Group11163: anid|CBF71593.1 anid|CBF80250.1 anid|CBF80255.1 anid|CBF82190.1 anid|CBF89813.1

Group11164: awen|OJJ29550.1 awen|OJJ29593.1 awen|OJJ29573.1 awen|OJJ29637.1 awen|OJJ29616.1

Group11165: azon|OJJ45072.1 azon|OJJ49483.1 acla|EAW11913.1 asyd|OJJ54173.1 aver|OJJ08566.1

Group11166: agla|OJJ82005.1 agla|OJJ89325.1 agla|OJJ85454.1 agla|OJJ86193.1 agla|OJJ88741.1

Group11167: aluc|OJZ79474.1 aluc|OJZ79524.1 aluc|OJZ79717.1 aluc|OJZ79872.1 aluc|OJZ88834.1

Group11168: aluc|OJZ79511.1 aluc|OJZ79780.1 aluc|OJZ79860.1 aluc|OJZ88829.1 aluc|OJZ80243.1

Group11169: aluc|OJZ79557.1 aluc|OJZ79777.1 aluc|OJZ79864.1 aluc|OJZ80213.1 aluc|OJZ79798.1

Group11170: aluc|OJZ79567.1 aluc|OJZ79846.1 aluc|OJZ88814.1 aluc|OJZ80289.1 aluc|OJZ79716.1

Group11171: abra|OJJ65418.1 aory|AFM82510.1 abra|OJJ65582.1 akaw|BAL04872.1 azon|OJJ41972.1

Group11172: abra|OJJ65422.1 acar|OOF89753.1 akaw|BAL04879.1 aluc|OJZ79502.1 aory|AFM82520.1

Group11173: abra|OJJ65436.1 acar|OOF89789.1 afis|EAW25780.1 akaw|BAL04882.1 aory|AFM82516.1

Group11174: abra|OJJ65546.1 akaw|GAA92849.1 aluc|OJZ88791.1 asyd|OJJ55711.1 aver|OJJ06797.1

Group11175: awen|OJJ41835.1 afis|EAW17010.1 afla|EED48916.1 afum|EAL86530.1 aory|EIT81216.1

Group11176: abra|OJJ65610.1 agla|OJJ88301.1 anid|CBF83166.1 asyd|OJJ55505.1 aver|OJJ06582.1

Group11177: abra|OJJ65619.1 acar|OOF98688.1 afum|EAL87790.1 asyd|OJJ62681.1 ater|EAU37844.1

Group11178: abra|OJJ65653.1 acar|OOF97622.1 agla|OJJ88776.1 ater|EAU37667.1 awen|OJJ36082.1

Group11179: abra|OJJ65678.1 akaw|GAA84549.1 aluc|OJZ89281.1 anig|EHA23668.1 atub|OJI85518.1

Group11180: abra|OJJ65737.1 afla|EED56167.1 aory|EIT76771.1 ater|EAU31845.1 awen|OJJ36040.1

Group11181: abra|OJJ65745.1 acar|OOG00551.1 acla|EAW14212.1 afis|EAW19069.1 agla|OJJ84536.1

Group11182: abra|OJJ65789.1 acla|EAW13288.1 afis|EAW17331.1 afum|EAL91168.1 anig|EHA18727.1

Group11183: abra|OJJ65804.1 akaw|GAA92932.1 aluc|OJZ82525.1 anid|CBF85283.1 anig|EHA22489.1

Group11184: abra|OJJ65865.1 akaw|GAA92279.1 aluc|OJZ90176.1 anig|EHA23675.1 atub|OJI81603.1

Group11185: abra|OJJ65875.1 asyd|OJJ53362.1 aver|OJJ07648.1 awen|OJJ36102.1 azon|OJJ42113.1

Group11186: abra|OJJ65883.1 afis|EAW21015.1 afum|EAL92053.1 asyd|OJJ53345.1 azon|OJJ50731.1

Group11187: abra|OJJ65886.1 acar|OOF94946.1 akaw|GAA86018.1 aluc|OJZ90246.1 atub|OJI81674.1

Group11188: asyd|OJJ63946.1 aory|EIT75260.1 atub|OJI83153.1 aver|OJI96996.1 awen|OJJ39627.1

Group11189: abra|OJJ65931.1 akaw|GAA92304.1 aluc|OJZ90201.1 anig|EHA23698.1 atub|OJI81629.1

Group11190: ater|EAU37748.1 aver|OJJ04225.1 afla|EED44873.1 aory|EIT82724.1 asyd|OJJ58179.1

Group11191: acar|OOF95830.1 akaw|GAA89761.1 aluc|OJZ80622.1 anig|EHA27209.1 atub|OJI87476.1

Group11192: abra|OJJ66015.1 ater|EAU32220.1 aver|OJJ05325.1 abra|OJJ66460.1 awen|OJJ32689.1

Group11193: abra|OJJ66021.1 afla|EED56777.1 aory|EIT73924.1 asyd|OJJ61424.1 atub|OJI82845.1

Group11194: awen|OJJ34670.1 anid|CBF79112.1 afum|EAL86464.1 ater|EAU37684.1 agla|OJJ81793.1

Group11195: abra|OJJ66034.1 afla|EED45347.1 anig|EHA18396.1 ater|EAU31821.1 atub|OJI87433.1

Group11196: abra|OJJ66049.1 atub|OJI87255.1 abra|OJJ66143.1 afis|EAW16402.1 afum|EAL89024.1

Group11197: abra|OJJ66051.1 anid|CBF70059.1 anig|EHA27185.1 aver|OJJ07386.1 awen|OJJ31006.1

Group11198: abra|OJJ66057.1 akaw|GAA93127.1 aluc|OJZ81107.1 anig|EHA19262.1 atub|OJI82472.1

Group11199: afla|EED54828.1 agla|OJJ81753.1 aory|EIT79363.1 azon|OJJ46462.1 awen|OJJ33382.1

Group11200: agla|OJJ82272.1 aluc|OJZ87835.1 anid|CBF76060.1 afla|EED51101.1 aory|EIT81302.1

Group11201: abra|OJJ66102.1 akaw|GAA92669.1 aluc|OJZ90123.1 anig|EHA20947.1 atub|OJI84672.1

Group11202: abra|OJJ66121.1 afis|EAW16828.1 aluc|OJZ90140.1 anig|EHA20928.1 atub|OJI84658.1

Group11203: ater|EAU37651.1 acar|OOG00118.1 acla|EAW07060.1 azon|OJJ45217.1 afla|EED48491.1

Group11204: abra|OJJ66148.1 afla|EED53484.1 akaw|GAA88366.1 aluc|OJZ82289.1 atub|OJI88150.1

Group11205: abra|OJJ66156.1 ater|EAU33999.1 afum|EAL90467.1 afis|EAW22728.1 awen|OJJ31976.1

Group11206: akaw|GAA84885.1 aluc|OJZ86786.1 anig|EHA21896.1 awen|OJJ29810.1 atub|OJI83733.1

Group11207: abra|OJJ66198.1 acar|OOF91932.1 akaw|GAA89091.1 anig|EHA25693.1 atub|OJI88430.1

Group11208: abra|OJJ66255.1 acar|OOF90726.1 aluc|OJZ80563.1 anig|EHA25635.1 atub|OJI88368.1

Group11209: abra|OJJ66259.1 afla|EED53427.1 aluc|OJZ80566.1 anig|EHA25632.1 atub|OJI88365.1

Group11210: afis|EAW24560.1 asyd|OJJ55938.1 aver|OJJ04887.1 anid|CBF78919.1 azon|OJJ43547.1

Group11211: abra|OJJ66281.1 anig|EHA23661.1 ater|EAU38972.1 atub|OJI89942.1 azon|OJJ44258.1

Group11212: abra|OJJ66284.1 anig|EHA23664.1 atub|OJI89939.1 azon|OJJ42281.1 acar|OOF94958.1

Group11213: abra|OJJ66329.1 akaw|GAA90206.1 aluc|OJZ83139.1 anig|EHA28198.1 atub|OJI87795.1

Group11214: acar|OOF99381.1 afis|EAW17074.1 anig|EHA19928.1 asyd|OJJ59188.1 aver|OJJ01805.1

Group11215: abra|OJJ66396.1 akaw|GAA90261.1 aluc|OJZ83079.1 anig|EHA28142.1 atub|OJI87866.1

Group11216: abra|OJJ66402.1 akaw|GAA90270.1 aluc|OJZ83071.1 anig|EHA28135.1 atub|OJI87875.1

Group11217: abra|OJJ66413.1 aluc|OJZ83062.1 atub|OJI87883.1 aver|OJJ07281.1 azon|OJJ43650.1

Group11218: abra|OJJ66435.1 akaw|GAA83942.1 aluc|OJZ80824.1 anig|EHA28110.1 atub|OJI87912.1

Group11219: abra|OJJ66446.1 akaw|GAA86651.1 aluc|OJZ85209.1 anig|EHA26089.1 aver|OJJ07917.1

Group11220: abra|OJJ66452.1 anid|CBF75802.1 awen|OJJ37490.1 asyd|OJJ57536.1 aver|OJJ05643.1

Group11221: abra|OJJ66502.1 akaw|GAA91208.1 aluc|OJZ81543.1 atub|OJI88641.1 asyd|OJJ57671.1

Group11222: abra|OJJ66515.1 akaw|GAA91196.1 aluc|OJZ81530.1 anig|EHA25906.1 atub|OJI88656.1

Group11223: abra|OJJ66516.1 akaw|GAA91195.1 aluc|OJZ81529.1 anig|EHA25907.1 atub|OJI88657.1

Group11224: abra|OJJ66771.1 akaw|GAA89708.1 aluc|OJZ80676.1 anig|EHA27336.1 atub|OJI87535.1

Group11225: abra|OJJ66775.1 acar|OOF93854.1 akaw|GAA89710.1 aluc|OJZ80674.1 atub|OJI87531.1

Group11226: abra|OJJ66796.1 akaw|GAA86861.1 aluc|OJZ89985.1 anig|EHA27356.1 atub|OJI84825.1

Group11227: abra|OJJ66825.1 awen|OJJ31935.1 azon|OJJ46601.1 afum|EAL86851.1 agla|OJJ82611.1

Group11228: abra|OJJ66920.1 afis|EAW19565.1 afla|EED48985.1 agla|OJJ82090.1 aory|EIT79976.1

Group11229: abra|OJJ66945.1 akaw|GAA91475.1 aluc|OJZ85636.1 anig|EHA24896.1 atub|OJI79426.1

Group11230: abra|OJJ66960.1 akaw|GAA92201.1 aluc|OJZ85616.1 anig|EHA24910.1 atub|OJI79409.1

Group11231: abra|OJJ66973.1 afum|EAL85069.1 agla|OJJ82219.1 anig|EHA24920.1 agla|OJJ88219.1

Group11232: afis|EAW19734.1 afum|EAL86795.1 agla|OJJ89293.1 asyd|OJJ57616.1 aver|OJJ05570.1

Group11233: ater|EAU37976.1 awen|OJJ31433.1 agla|OJJ80671.1 asyd|OJJ55186.1 aver|OJJ07021.1

Group11234: afla|EED46927.1 aory|EIT72776.1 afis|EAW24754.1 awen|OJJ40022.1 aver|OJJ08132.1

Group11235: abra|OJJ67075.1 afla|EED57257.1 ater|EAU31941.1 atub|OJI89956.1 aory|EIT74064.1

Group11236: abra|OJJ67131.1 agla|OJJ80718.1 asyd|OJJ59557.1 aver|OJJ02711.1 anid|CBF81676.1

Group11237: abra|OJJ67234.1 acar|OOF98318.1 akaw|GAA91658.1 aluc|OJZ87741.1 anig|EHA20233.1

Group11238: abra|OJJ67257.1 akaw|GAA91672.1 aluc|OJZ87722.1 anig|EHA20216.1 atub|OJI80920.1

Group11239: abra|OJJ67258.1 akaw|GAA91673.1 aluc|OJZ87721.1 anig|EHA20215.1 atub|OJI80921.1

Group11240: azon|OJJ42454.1 acar|OOF94552.1 acla|EAW12195.1 afis|EAW16908.1 afum|EAL86444.1

Group11241: abra|OJJ67277.1 akaw|GAA92070.1 aluc|OJZ81157.1 asyd|OJJ52004.1 asyd|OJJ52002.1

Group11242: abra|OJJ67282.1 akaw|GAA86527.1 aluc|OJZ85075.1 anig|EHA25290.1 atub|OJI80523.1

Group11243: abra|OJJ67301.1 afla|EED57136.1 akaw|GAA86515.1 aluc|OJZ85060.1 atub|OJI80538.1

Group11244: abra|OJJ67363.1 akaw|GAA86458.1 aluc|OJZ85000.1 anig|EHA25218.1 atub|OJI80600.1

Group11245: abra|OJJ67404.1 anig|EHA25177.1 atub|OJI80636.1 akaw|GAA86423.1 aluc|OJZ84964.1

Group11246: abra|OJJ67405.1 akaw|GAA86422.1 aluc|OJZ84963.1 anig|EHA25176.1 atub|OJI80637.1

Group11247: abra|OJJ67432.1 akaw|GAA86397.1 aluc|OJZ84932.1 anig|EHA25150.1 atub|OJI80667.1

Group11248: abra|OJJ67479.1 anig|EHA25117.1 akaw|GAA86359.1 aluc|OJZ84888.1 atub|OJI80712.1

Group11249: abra|OJJ67566.1 acar|OOG00284.1 akaw|GAA88099.1 aluc|OJZ81366.1 atub|OJI91570.1

Group11250: abra|OJJ67579.1 akaw|GAA90578.1 aluc|OJZ86036.1 anig|EHA24579.1 atub|OJI79823.1

Group11251: acar|OOF91826.1 afla|EED51021.1 aory|EIT77019.1 anig|EHA26199.1 atub|OJI87261.1

Group11252: abra|OJJ67583.1 akaw|GAA85346.1 aluc|OJZ90708.1 anig|EHA24575.1 atub|OJI82135.1

Group11253: abra|OJJ67602.1 akaw|GAA89120.1 aluc|OJZ81719.1 anig|EHA25720.1 atub|OJI88463.1

Group11254: abra|OJJ67605.1 akaw|GAA89122.1 aluc|OJZ81716.1 anig|EHA25723.1 atub|OJI88466.1

Group11255: abra|OJJ67618.1 akaw|GAA89135.1 aluc|OJZ81701.1 anig|EHA25733.1 atub|OJI88480.1

Group11256: abra|OJJ67619.1 akaw|GAA89136.1 aluc|OJZ81700.1 anig|EHA25734.1 atub|OJI88481.1

Group11257: abra|OJJ67620.1 aluc|OJZ81698.1 anig|EHA25735.1 atub|OJI88482.1 akaw|GAA89138.1

Group11258: agla|OJJ79401.1 afla|EED45284.1 aory|EIT73654.1 asyd|OJJ52807.1 aver|OJJ08787.1

Group11259: abra|OJJ67641.1 aluc|OJZ81674.1 anig|EHA25766.1 atub|OJI88507.1 akaw|GAA89160.1

Group11260: abra|OJJ67644.1 akaw|GAA89163.1 aluc|OJZ81671.1 anig|EHA25771.1 atub|OJI88511.1

Group11261: abra|OJJ67646.1 akaw|GAA89164.1 aluc|OJZ81670.1 anig|EHA25774.1 atub|OJI88514.1

Group11262: abra|OJJ67651.1 akaw|GAA89167.1 aluc|OJZ81666.1 anig|EHA25777.1 atub|OJI88517.1

Group11263: abra|OJJ67657.1 akaw|GAA89173.1 aluc|OJZ81660.1 anig|EHA25783.1 atub|OJI88523.1

Group11264: abra|OJJ67666.1 akaw|GAA89182.1 aluc|OJZ81650.1 anig|EHA25793.1 atub|OJI88532.1

Group11265: abra|OJJ67672.1 akaw|GAA89188.1 aluc|OJZ81643.1 anig|EHA25800.1 atub|OJI88539.1

Group11266: abra|OJJ67697.1 acar|OOF90177.1 aluc|OJZ81623.1 anig|EHA25820.1 atub|OJI88557.1

Group11267: abra|OJJ67698.1 akaw|GAA89207.1 aluc|OJZ81622.1 anig|EHA25821.1 atub|OJI88558.1

Group11268: abra|OJJ67722.1 acar|OOF93285.1 anig|EHA25835.1 atub|OJI88572.1 ater|EAU31223.1

Group11269: abra|OJJ67732.1 agla|OJJ85827.1 atub|OJI88583.1 akaw|GAA92423.1 aluc|OJZ81599.1

Group11270: abra|OJJ67754.1 afla|EED49704.1 aory|EIT77676.1 atub|OJI84568.1 awen|OJJ36304.1

Group11271: abra|OJJ67762.1 akaw|GAA92446.1 aluc|OJZ81572.1 anig|EHA25866.1 atub|OJI88611.1

Group11272: abra|OJJ67775.1 afis|EAW20789.1 ater|EAU33383.1 atub|OJI80490.1 ater|EAU30087.1

Group11273: abra|OJJ67787.1 acar|OOF94974.1 aluc|OJZ81080.1 anig|EHA20679.1 ater|EAU30330.1

Group11274: abra|OJJ67807.1 aluc|OJZ90850.1 asyd|OJJ52899.1 atub|OJI82855.1 azon|OJJ42358.1

Group11275: abra|OJJ67822.1 akaw|GAA91459.1 aluc|OJZ82062.1 anig|EHA22191.1 atub|OJI84620.1

Group11276: abra|OJJ67841.1 acar|OOF92018.1 akaw|GAA91847.1 aluc|OJZ82051.1 atub|OJI84609.1

Group11277: abra|OJJ67867.1 acar|OOF92848.1 akaw|GAA91874.1 aluc|OJZ82025.1 atub|OJI84578.1

Group11278: abra|OJJ67899.1 akaw|GAA91903.1 aluc|OJZ81991.1 anig|EHA22265.1 atub|OJI84542.1

Group11279: azon|OJJ44460.1 ater|EAU37624.1 acla|EAW12451.1 afis|EAW16755.1 afum|EAL86327.1

Group11280: anid|CBF79879.1 awen|OJJ33768.1 asyd|OJJ58171.1 aver|OJJ04232.1 azon|OJJ42872.1

Group11281: abra|OJJ67995.1 acar|OOF92038.1 akaw|GAA90954.1 aluc|OJZ82870.1 atub|OJI89093.1

Group11282: ater|EAU36713.1 akaw|GAA92596.1 aluc|OJZ82613.1 atub|OJI88848.1 anid|CBF87736.1

Group11283: abra|OJJ68018.1 akaw|GAA90933.1 aluc|OJZ82892.1 atub|OJI89116.1 awen|OJJ32559.1

Group11284: abra|OJJ68028.1 akaw|GAA90925.1 aluc|OJZ82901.1 anig|EHA21326.1 atub|OJI89127.1

Group11285: abra|OJJ68103.1 akaw|GAA86887.1 aluc|OJZ90011.1 anig|EHA27386.1 atub|OJI84796.1

Group11286: abra|OJJ68113.1 afis|EAW19855.1 afum|EAL85040.1 asyd|OJJ54979.1 aver|OJJ03937.1

Group11287: abra|OJJ68120.1 akaw|GAA86901.1 aluc|OJZ90024.1 anig|EHA27400.1 atub|OJI84783.1

Group11288: abra|OJJ68142.1 akaw|GAA92376.1 aluc|OJZ83235.1 anig|EHA21092.1 atub|OJI87701.1

Group11289: abra|OJJ68202.1 acar|OOF96118.1 afla|EED46966.1 atub|OJI90439.1 awen|OJJ39623.1

Group11290: abra|OJJ68209.1 akaw|GAA83237.1 aluc|OJZ83736.1 anig|EHA18885.1 atub|OJI90114.1

Group11291: abra|OJJ68220.1 acar|OOF97510.1 akaw|GAA83225.1 anig|EHA18873.1 atub|OJI90101.1

Group11292: abra|OJJ68272.1 akaw|GAA83197.1 aluc|OJZ83699.1 anig|EHA18850.1 atub|OJI90070.1

Group11293: abra|OJJ68275.1 aluc|OJZ83695.1 anig|EHA18847.1 atub|OJI90067.1 azon|OJJ48131.1

Group11294: afum|EAL85008.1 aory|EIT83549.1 acla|EAW12413.1 afis|EAW18941.1 afla|EED55208.1

Group11295: abra|OJJ68311.1 acar|OOF96697.1 akaw|GAA83162.1 aluc|OJZ83662.1 atub|OJI90033.1

Group11296: abra|OJJ68317.1 akaw|GAA83156.1 aluc|OJZ83656.1 anig|EHA18808.1 atub|OJI90026.1

Group11297: abra|OJJ68324.1 acar|OOF97618.1 akaw|GAA92075.1 aluc|OJZ83649.1 atub|OJI90018.1

Group11298: abra|OJJ68347.1 acar|OOF90953.1 akaw|GAA92114.1 aluc|OJZ83611.1 atub|OJI89984.1

Group11299: abra|OJJ68350.1 akaw|GAA92118.1 aluc|OJZ83605.1 atub|OJI89977.1 awen|OJJ30193.1

Group11300: abra|OJJ68419.1 akaw|GAA92529.1 aluc|OJZ90078.1 anig|EHA20995.1 atub|OJI84728.1

Group11301: abra|OJJ68508.1 akaw|GAA92763.1 aluc|OJZ82722.1 anig|EHA21176.1 atub|OJI88956.1

Group11302: aory|EIT81282.1 afla|EED48834.1 agla|OJJ81089.1 ater|EAU34703.1 azon|OJJ45534.1

Group11303: abra|OJJ68571.1 afis|EAW25567.1 afla|EED54379.1 aory|EIT75856.1 awen|OJJ37970.1

Group11304: abra|OJJ68572.1 anid|CBF75888.1 asyd|OJJ60259.1 aver|OJJ03997.1 awen|OJJ33511.1

Group11305: abra|OJJ68587.1 akaw|GAA91919.1 aluc|OJZ81961.1 anig|EHA22286.1 atub|OJI84512.1

Group11306: abra|OJJ68589.1 acar|OOF92453.1 anid|CBF69459.1 ater|EAU35432.1 ater|EAU32764.1

Group11307: abra|OJJ68590.1 afla|EED49872.1 aory|EIT83406.1 ater|EAU35717.1 azon|OJJ50748.1

Group11308: abra|OJJ68594.1 acla|EAW11910.1 afis|EAW20746.1 afla|EED46090.1 azon|OJJ45786.1

Group11309: abra|OJJ68627.1 aluc|OJZ81928.1 anig|EHA22321.1 atub|OJI84480.1 awen|OJJ39968.1

Group11310: abra|OJJ68632.1 akaw|GAA89771.1 aluc|OJZ81924.1 anig|EHA22324.1 atub|OJI84476.1

Group11311: abra|OJJ68653.1 asyd|OJJ57278.1 atub|OJI81011.1 abra|OJJ69048.1 azon|OJJ48229.1

Group11312: abra|OJJ68811.1 acar|OOF90614.1 akaw|GAA90443.1 aluc|OJZ85784.1 atub|OJI79568.1

Group11313: ater|EAU29276.1 akaw|GAA84591.1 aluc|OJZ89336.1 atub|OJI85463.1 aver|OJJ01766.1

Group11314: afis|EAW15414.1 afum|EAL84232.2 akaw|GAA82234.1 aluc|OJZ91913.1 azon|OJJ50410.1

Group11315: abra|OJJ68860.1 acar|OOF90529.1 akaw|GAA90393.1 aluc|OJZ85838.1 atub|OJI79626.1

Group11316: abra|OJJ68883.1 akaw|GAA90373.1 aluc|OJZ85859.1 atub|OJI79645.1 aver|OJJ03405.1

Group11317: abra|OJJ68884.1 akaw|GAA90372.1 aluc|OJZ85860.1 anig|EHA28092.1 atub|OJI79646.1

Group11318: abra|OJJ68885.1 akaw|GAA90371.1 aluc|OJZ85861.1 anig|EHA28091.1 atub|OJI79647.1

Group11319: abra|OJJ68886.1 akaw|GAA90370.1 aluc|OJZ85862.1 anig|EHA28090.1 atub|OJI79648.1

Group11320: abra|OJJ68888.1 akaw|GAA90368.1 aluc|OJZ85864.1 anig|EHA28088.1 atub|OJI79650.1

Group11321: abra|OJJ68889.1 akaw|GAA90367.1 aluc|OJZ85865.1 anig|EHA28087.1 atub|OJI79651.1

Group11322: afis|EAW19830.1 afum|EAL85062.1 anid|CBF76075.1 ater|EAU32496.1 awen|OJJ36314.1

Group11323: abra|OJJ68947.1 akaw|GAA90307.1 aluc|OJZ85933.1 anig|EHA24665.1 atub|OJI79722.1

Group11324: abra|OJJ68954.1 acar|OOF91532.1 aluc|OJZ85940.1 anig|EHA24659.1 atub|OJI79728.1

Group11325: abra|OJJ69014.1 akaw|GAA90531.1 aluc|OJZ86000.1 anig|EHA24610.1 atub|OJI79787.1

Group11326: abra|OJJ69045.1 akaw|GAA90574.1 aluc|OJZ86030.1 anig|EHA24582.1 atub|OJI79819.1

Group11327: abra|OJJ69134.1 akaw|GAA86322.1 aluc|OJZ86122.1 anig|EHA24499.1 atub|OJI79918.1

Group11328: abra|OJJ69163.1 acar|OOF91142.1 aluc|OJZ86150.1 atub|OJI79942.1 azon|OJJ43763.1

Group11329: abra|OJJ69437.1 akaw|GAA88379.1 aluc|OJZ82273.1 anig|EHA28472.1 atub|OJI88166.1

Group11330: abra|OJJ69489.1 akaw|GAA88331.1 aluc|OJZ82328.1 anig|EHA28430.1 atub|OJI88110.1

Group11331: abra|OJJ69528.1 akaw|GAA88380.1 aluc|OJZ82368.1 anig|EHA28397.1 atub|OJI88074.1

Group11332: abra|OJJ69533.1 agla|OJJ86580.1 akaw|GAA83223.1 atub|OJI90099.1 aluc|OJZ83724.1

Group11333: acar|OOF95575.1 afla|EED48703.1 akaw|GAA86620.1 aluc|OJZ85175.1 aory|EIT73233.1

Group11334: abra|OJJ69569.1 acla|EAW09065.1 agla|OJJ79943.1 awen|OJJ41337.1 azon|OJJ42268.1

Group11335: abra|OJJ69605.1 acar|OOF98997.1 asyd|OJJ65118.1 aver|OJI99244.1 aver|OJI99245.1

Group11336: abra|OJJ69607.1 acar|OOF92700.1 akaw|GAA91592.1 aluc|OJZ82415.1 atub|OJI88026.1

Group11337: abra|OJJ69654.1 akaw|GAA91633.1 aluc|OJZ82460.1 anig|EHA28315.1 atub|OJI87978.1

Group11338: azon|OJJ48026.1 anid|CBF70510.1 asyd|OJJ58574.1 aver|OJJ01177.1 awen|OJJ33414.1

Group11339: abra|OJJ69695.1 akaw|GAA88815.1 aluc|OJZ82498.1 anig|EHA28274.1 atub|OJI87932.1

Group11340: abra|OJJ69729.1 akaw|GAA90182.1 aluc|OJZ83168.1 anig|EHA28233.1 atub|OJI87769.1

Group11341: abra|OJJ69730.1 akaw|GAA90183.1 aluc|OJZ83167.1 anig|EHA28232.1 atub|OJI87770.1

Group11342: abra|OJJ69737.1 acar|OOF94183.1 akaw|GAA90190.1 aluc|OJZ83159.1 atub|OJI87778.1

Group11343: abra|OJJ69806.1 akaw|GAA87853.1 aluc|OJZ85495.1 anig|EHA25028.1 atub|OJI80070.1

Group11344: acla|EAW14882.1 afis|EAW24401.1 afum|EAL86072.1 agla|OJJ79322.1 awen|OJJ30172.1

Group11345: abra|OJJ69848.1 acar|OOF93370.1 akaw|GAA87810.1 aluc|OJZ85445.1 anig|EHA24989.1

Group11346: asyd|OJJ65102.1 akaw|GAA92969.1 aver|OJI99229.1 awen|OJJ38709.1 awen|OJJ31619.1

Group11347: abra|OJJ69862.1 akaw|GAA87794.1 aluc|OJZ85429.1 atub|OJI80146.1 anig|EHA25608.1

Group11348: abra|OJJ69863.1 akaw|GAA87793.1 aluc|OJZ85428.1 anig|EHA25607.1 atub|OJI80147.1

Group11349: aory|EIT81676.1 afis|EAW15461.1 afla|EED48436.1 asyd|OJJ54362.1 aver|OJJ05315.1

Group11350: abra|OJJ69891.1 akaw|GAA87759.1 aluc|OJZ85393.1 anig|EHA25577.1 atub|OJI80178.1

Group11351: abra|OJJ70027.1 acar|OOF98709.1 akaw|GAA92697.1 aluc|OJZ90155.1 atub|OJI84644.1

Group11352: abra|OJJ70039.1 afis|EAW24622.1 afum|EAL86250.1 anig|EHA18721.1 atub|OJI79323.1

Group11353: abra|OJJ70044.1 akaw|GAA87631.1 aluc|OJZ85247.1 anig|EHA25447.1 atub|OJI80322.1

Group11354: abra|OJJ70053.1 akaw|GAA85318.1 aluc|OJZ90679.1 anig|EHA27496.1 awen|OJJ30546.1

Group11355: afis|EAW25602.1 awen|OJJ40144.1 agla|OJJ80364.1 anid|CBF82322.1 ater|EAU32209.1

Group11356: abra|OJJ70069.1 akaw|GAA87610.1 aluc|OJZ85231.1 anig|EHA25434.1 atub|OJI80349.1

Group11357: abra|OJJ70072.1 afla|EED51154.1 anid|CBF80156.1 aory|EIT81342.1 aver|OJI99515.1

Group11358: abra|OJJ70096.1 akaw|GAA87599.1 aluc|OJZ85219.1 anig|EHA25423.1 atub|OJI80364.1

Group11359: anig|EHA22568.1 afla|EED55094.1 aory|EIT81630.1 asyd|OJJ62607.1 aver|OJI96470.1

Group11360: abra|OJJ70123.1 acar|OOF98698.1 akaw|GAA86646.1 aluc|OJZ85203.1 atub|OJI80398.1

Group11361: abra|OJJ70150.1 afla|EED52386.1 akaw|GAA90557.1 aory|EIT83290.1 ater|EAU35374.1

Group11362: abra|OJJ70151.1 agla|OJJ83851.1 akaw|GAA86599.1 aluc|OJZ85146.1 atub|OJI80420.1

Group11363: abra|OJJ70164.1 akaw|GAA86611.1 aluc|OJZ85162.1 anig|EHA25360.1 atub|OJI80438.1

Group11364: abra|OJJ70169.1 acar|OOG00401.1 akaw|GAA86617.1 aluc|OJZ85170.1 atub|OJI80446.1

Group11365: abra|OJJ70175.1 acar|OOF93302.1 akaw|GAA86597.1 aluc|OJZ85144.1 atub|OJI80449.1

Group11366: abra|OJJ70186.1 akaw|GAA86589.1 aluc|OJZ85136.1 anig|EHA25368.1 atub|OJI80456.1

Group11367: abra|OJJ70194.1 akaw|GAA86581.1 aluc|OJZ85132.1 anig|EHA25351.1 atub|OJI80461.1

Group11368: abra|OJJ70204.1 akaw|GAA86573.1 aluc|OJZ85123.1 atub|OJI80470.1 azon|OJJ47991.1

Group11369: abra|OJJ70206.1 acar|OOF92573.1 akaw|GAA86571.1 aluc|OJZ85121.1 atub|OJI80472.1

Group11370: abra|OJJ70230.1 afla|EED46888.1 aluc|OJZ90860.1 anig|EHA18748.1 aory|EIT72671.1

Group11371: abra|OJJ70252.1 akaw|GAA86539.1 aluc|OJZ85087.1 atub|OJI80512.1 awen|OJJ37027.1

Group11372: abra|OJJ70278.1 akaw|GAA92871.1 aluc|OJZ82540.1 anig|EHA22508.1 atub|OJI88784.1

Group11373: abra|OJJ70291.1 acar|OOF91745.1 akaw|GAA92857.1 aluc|OJZ82556.1 atub|OJI88798.1

Group11374: abra|OJJ70292.1 acar|OOF91746.1 akaw|GAA92856.1 aluc|OJZ82557.1 atub|OJI88799.1

Group11375: abra|OJJ70299.1 akaw|GAA93022.1 aluc|OJZ82563.1 anig|EHA27511.1 atub|OJI88806.1

Group11376: abra|OJJ70301.1 akaw|GAA93021.1 aluc|OJZ82564.1 anig|EHA27510.1 atub|OJI88808.1

Group11377: abra|OJJ70314.1 akaw|GAA92570.1 aluc|OJZ82580.1 anig|EHA27491.1 atub|OJI88821.1

Group11378: akaw|GAA92764.1 aluc|OJZ82721.1 atub|OJI88955.1 acar|OOF96899.1 anig|EHA21175.1

Group11379: abra|OJJ70357.1 akaw|GAA92602.1 aluc|OJZ82619.1 anig|EHA27458.1 atub|OJI88854.1

Group11380: acar|OOF95923.1 akaw|GAA83489.1 aluc|OJZ92504.1 aory|EIT82472.1 atub|OJI86936.1

Group11381: abra|OJJ70441.1 afla|EED45987.1 aory|EIT75643.1 aver|OJI98304.1 awen|OJJ37918.1

Group11382: abra|OJJ70444.1 akaw|GAA90136.1 aluc|OJZ83211.1 anig|EHA21071.1 atub|OJI87726.1

Group11383: abra|OJJ70458.1 akaw|GAA83304.1 aluc|OJZ83799.1 anig|EHA18947.1 atub|OJI90168.1

Group11384: abra|OJJ70531.1 acar|OOF97401.1 akaw|GAA83376.1 aluc|OJZ83874.1 atub|OJI90242.1

Group11385: abra|OJJ70534.1 acar|OOF97398.1 akaw|GAA83379.1 aluc|OJZ83877.1 atub|OJI90245.1

Group11386: abra|OJJ70581.1 akaw|GAA83425.1 aluc|OJZ83926.1 anig|EHA19058.1 atub|OJI90298.1

Group11387: abra|OJJ70621.1 akaw|GAA89960.1 aluc|OJZ83967.1 anig|EHA19095.1 atub|OJI90340.1

Group11388: abra|OJJ70662.1 akaw|GAA89998.1 aluc|OJZ84012.1 anig|EHA19135.1 atub|OJI90381.1

Group11389: abra|OJJ70679.1 akaw|GAA90011.1 aluc|OJZ84027.1 anig|EHA19148.1 atub|OJI90399.1

Group11390: abra|OJJ70831.1 anig|EHA20973.1 asyd|OJJ52722.1 aver|OJJ04486.1 awen|OJJ35066.1

Group11391: abra|OJJ70885.1 acar|OOF94897.1 akaw|GAA87403.1 aluc|OJZ87637.1 atub|OJI81010.1

Group11392: abra|OJJ71071.1 acar|OOF90617.1 akaw|GAA82891.1 aluc|OJZ87460.1 anig|EHA20682.1

Group11393: abra|OJJ71073.1 akaw|GAA82889.1 aluc|OJZ87458.1 anig|EHA20680.1 atub|OJI81190.1

Group11394: acla|EAW13667.1 afis|EAW17064.1 aver|OJJ00804.1 afla|EED52444.1 aory|EIT82750.1

Group11395: abra|OJJ71107.1 akaw|GAA82862.1 aluc|OJZ87428.1 anig|EHA20650.1 atub|OJI81221.1

Group11396: afis|EAW19925.1 afum|EAL84984.2 acar|OOF93586.1 acla|EAW13681.1 azon|OJJ50108.1

Group11397: anid|CBF85314.1 asyd|OJJ64008.1 aver|OJI98095.1 afla|EED53707.1 aory|EIT75285.1

Group11398: abra|OJJ71217.1 acar|OOF94575.1 acla|EAW15346.1 anig|EHA20561.1 aver|OJJ03893.1

Group11399: abra|OJJ71224.1 akaw|GAA82765.1 aluc|OJZ87323.1 anig|EHA20558.1 atub|OJI81322.1

Group11400: akaw|GAA82745.1 anid|CBF85404.1 asyd|OJJ64063.1 aver|OJI98149.1 awen|OJJ29831.1

Group11401: abra|OJJ71269.1 acar|OOF97904.1 akaw|GAA82721.1 aluc|OJZ87275.1 atub|OJI81366.1

Group11402: afum|EAL90001.1 agla|OJJ83867.1 afis|EAW25338.1 awen|OJJ41033.1 azon|OJJ44632.1

Group11403: abra|OJJ71343.1 akaw|GAA82649.1 aluc|OJZ87198.1 anig|EHA20452.1 atub|OJI81441.1

Group11404: abra|OJJ71566.1 akaw|GAA89358.1 aluc|OJZ86303.1 anig|EHA21569.1 atub|OJI84233.1

Group11405: abra|OJJ71619.1 afis|EAW21025.1 aluc|OJZ86360.1 atub|OJI84174.1 acar|OOF98296.1

Group11406: azon|OJJ49119.1 acar|OOF98229.1 aluc|OJZ79831.1 anig|EHA26123.1 ater|EAU32273.1

Group11407: abra|OJJ71629.1 akaw|GAA89296.1 aluc|OJZ86371.1 anig|EHA21521.1 atub|OJI84163.1

Group11408: abra|OJJ71735.1 akaw|GAA85149.1 aluc|OJZ86488.1 anig|EHA22181.1 atub|OJI84041.1

Group11409: abra|OJJ71764.1 acar|OOF93297.1 agla|OJJ82501.1 asyd|OJJ65291.1 atub|OJI84034.1

Group11410: abra|OJJ71768.1 akaw|GAA85139.1 aluc|OJZ86498.1 anig|EHA22146.1 atub|OJI84028.1

Group11411: abra|OJJ71784.1 akaw|GAA85126.1 aluc|OJZ86514.1 anig|EHA22130.1 atub|OJI84010.1

Group11412: abra|OJJ71822.1 akaw|GAA85088.1 aluc|OJZ86554.1 anig|EHA22095.1 atub|OJI83968.1

Group11413: abra|OJJ71944.1 akaw|GAA84982.1 aluc|OJZ86680.1 anig|EHA21994.1 atub|OJI83847.1

Group11414: abra|OJJ71945.1 akaw|GAA84981.1 aluc|OJZ86681.1 anig|EHA21993.1 atub|OJI83846.1

Group11415: abra|OJJ71976.1 akaw|GAA84954.1 aluc|OJZ86713.1 anig|EHA21965.1 atub|OJI83816.1

Group11416: abra|OJJ71992.1 akaw|GAA84937.1 aluc|OJZ86729.1 anig|EHA21950.1 atub|OJI83799.1

Group11417: abra|OJJ72002.1 akaw|GAA84929.1 aluc|OJZ86739.1 anig|EHA21941.1 atub|OJI83789.1

Group11418: abra|OJJ72039.1 akaw|GAA84895.1 aluc|OJZ86776.1 anig|EHA21905.1 atub|OJI83745.1

Group11419: abra|OJJ72130.1 akaw|GAA84792.1 aluc|OJZ86888.1 anig|EHA21755.1 atub|OJI83631.1

Group11420: ater|EAU36011.1 abra|OJJ76598.1 agla|OJJ83625.1 afla|EED54877.1 aory|EIT79400.1

Group11421: anid|CBF83143.1 acla|EAW12872.1 aluc|OJZ89213.1 anig|EHA23345.1 atub|OJI85579.1

Group11422: abra|OJJ72212.1 acar|OOF96833.1 akaw|GAA84722.1 aluc|OJZ86964.1 atub|OJI83559.1

Group11423: anid|CBF85310.1 aory|EIT80481.1 afla|EED51662.1 asyd|OJJ64004.1 awen|OJJ33578.1

Group11424: abra|OJJ72258.1 acar|OOF96890.1 afla|EED52818.1 aluc|OJZ87009.1 atub|OJI83510.1

Group11425: abra|OJJ72279.1 akaw|GAA86031.1 aluc|OJZ90258.1 anig|EHA23753.1 atub|OJI81684.1

Group11426: abra|OJJ72307.1 akaw|GAA86058.1 aluc|OJZ90288.1 anig|EHA23778.1 atub|OJI81712.1

Group11427: abra|OJJ72309.1 aluc|OJZ90290.1 anig|EHA23780.1 asyd|OJJ65159.1 azon|OJJ42572.1

Group11428: aory|EIT81786.1 acar|OOF90668.1 afis|EAW16639.1 afum|EAL89241.1 afla|EED55658.1

Group11429: abra|OJJ72376.1 akaw|GAA86124.1 aluc|OJZ90363.1 anig|EHA23838.1 acar|OOF94842.1

Group11430: abra|OJJ72419.1 akaw|GAA86161.1 aluc|OJZ90402.1 atub|OJI81833.1 azon|OJJ45687.1

Group11431: afis|EAW17832.1 afla|EED46174.1 afum|EAL91642.1 aory|EIT76434.1 acla|EAW12668.1

Group11432: agla|OJJ87942.1 acla|EAW07125.1 afis|EAW22135.1 afum|EAL91055.1 awen|OJJ33870.1

Group11433: azon|OJJ43491.1 afla|EED57238.1 aory|EIT73982.1 asyd|OJJ53398.1 aver|OJJ02219.1

Group11434: afis|EAW17889.1 afum|EAL91698.1 acla|EAW12726.1 awen|OJJ41670.1 azon|OJJ48385.1

Group11435: abra|OJJ72607.1 acar|OOF94648.1 aory|EIT74208.1 afis|EAW21219.1 afla|EED52549.1

Group11436: awen|OJJ39812.1 acla|EAW14121.1 afla|EED46083.1 aory|EIT76421.1 awen|OJJ39863.1

Group11437: awen|OJJ30861.1 acar|OOF98794.1 akaw|GAA87803.1 aluc|OJZ85438.1 atub|OJI80130.1

Group11438: abra|OJJ72694.1 agla|OJJ79706.1 asyd|OJJ60096.1 aver|OJJ04182.1 awen|OJJ33888.1

Group11439: abra|OJJ72708.1 akaw|GAA85312.1 aluc|OJZ90673.1 anig|EHA24106.1 atub|OJI82103.1

Group11440: abra|OJJ72746.1 akaw|GAA85358.1 aluc|OJZ90720.1 anig|EHA24137.1 atub|OJI82148.1

Group11441: abra|OJJ72805.1 akaw|GAA85408.1 aluc|OJZ83303.1 anig|EHA24186.1 atub|OJI82203.1

Group11442: abra|OJJ72841.1 akaw|GAA85443.1 aluc|OJZ83348.1 anig|EHA24216.1 atub|OJI82242.1

Group11443: abra|OJJ72974.1 afla|EED54542.1 aory|EIT78763.1 ater|EAU38893.1 awen|OJJ30812.1

Group11444: abra|OJJ73012.1 akaw|GAA85591.1 aluc|OJZ83527.1 anig|EHA24350.1 atub|OJI82416.1

Group11445: abra|OJJ73078.1 acar|OOG00313.1 akaw|GAA88084.1 aluc|OJZ81348.1 atub|OJI91552.1

Group11446: abra|OJJ73112.1 afla|EED48401.1 anig|EHA24048.1 aory|EIT83068.1 atub|OJI82037.1

Group11447: abra|OJJ73114.1 akaw|GAA88055.1 aluc|OJZ81317.1 atub|OJI91520.1 anig|EHA18622.1

Group11448: abra|OJJ73117.1 akaw|GAA88051.1 aluc|OJZ81313.1 anig|EHA18617.1 atub|OJI91517.1

Group11449: abra|OJJ73123.1 akaw|GAA88042.1 aluc|OJZ81301.1 anig|EHA18610.1 atub|OJI91510.1

Group11450: awen|OJJ34469.1 akaw|GAA81896.1 aluc|OJZ92304.1 anig|EHA26669.1 atub|OJI86734.1

Group11451: abra|OJJ73139.1 akaw|GAA88033.1 aluc|OJZ81292.1 anig|EHA18591.1 atub|OJI91500.1

Group11452: ater|EAU38088.1 aver|OJJ05860.1 afla|EED53213.1 aory|EIT74847.1 asyd|OJJ57295.1

Group11453: abra|OJJ73291.1 akaw|GAA92712.1 aluc|OJZ88757.1 atub|OJI91327.1 aver|OJJ03840.1

Group11454: afis|EAW19379.1 afla|EED48339.1 acla|EAW08559.1 anid|CBF71405.1 aver|OJJ07174.1

Group11455: abra|OJJ73333.1 acar|OOF91494.1 agla|OJJ81112.1 asyd|OJJ57264.1 aver|OJJ03397.1

Group11456: abra|OJJ73367.1 akaw|GAA83117.1 aluc|OJZ88673.1 anig|EHA18355.1 atub|OJI91254.1

Group11457: abra|OJJ73394.1 aluc|OJZ85152.1 abra|OJJ73395.1 anig|EHA25379.1 atub|OJI80430.1

Group11458: abra|OJJ73397.1 aluc|OJZ88637.1 anig|EHA18333.1 atub|OJI91222.1 azon|OJJ42512.1

Group11459: afum|EAL90873.1 acla|EAW07669.1 afis|EAW22317.1 ater|EAU31005.1 aver|OJJ07473.1

Group11460: abra|OJJ73482.1 akaw|GAA87431.1 aluc|OJZ88566.1 anig|EHA18263.1 atub|OJI91152.1

Group11461: abra|OJJ73542.1 acar|OOG00717.1 akaw|GAA88609.1 aluc|OJZ88517.1 atub|OJI91102.1

Group11462: abra|OJJ73548.1 akaw|GAA88604.1 aluc|OJZ88512.1 anig|EHA18218.1 atub|OJI91097.1

Group11463: azon|OJJ43669.1 anid|CBF82169.1 asyd|OJJ54346.1 ater|EAU32687.1 aver|OJJ05298.1

Group11464: abra|OJJ73615.1 akaw|GAA88538.1 aluc|OJZ88434.1 anig|EHA18158.1 atub|OJI91031.1

Group11465: abra|OJJ73635.1 aluc|OJZ88415.1 atub|OJI91011.1 asyd|OJJ59092.1 aver|OJJ01682.1

Group11466: abra|OJJ73706.1 anig|EHA18078.1 akaw|GAA88457.1 aluc|OJZ88345.1 atub|OJI90941.1

Group11467: abra|OJJ73713.1 afis|EAW21078.1 akaw|GAA88450.1 aluc|OJZ88337.1 anig|EHA18072.1

Group11468: abra|OJJ73722.1 acar|OOG00891.1 akaw|GAA88440.1 aluc|OJZ88328.1 atub|OJI90923.1

Group11469: abra|OJJ73725.1 afla|EED53600.1 aory|EIT75231.1 aver|OJJ08871.1 azon|OJJ45997.1

Group11470: abra|OJJ73737.1 akaw|GAA88427.1 aluc|OJZ88314.1 ater|EAU39057.1 atub|OJI90909.1

Group11471: abra|OJJ73755.1 afis|EAW24620.1 afum|EAL86248.1 anig|EHA19510.1 aver|OJJ08548.1

Group11472: afla|EED56729.1 aory|EIT73953.1 asyd|OJJ64768.1 aver|OJI98868.1 ater|EAU33981.1

Group11473: abra|OJJ73811.1 akaw|GAA89430.1 aluc|OJZ88239.1 anig|EHA17986.1 atub|OJI90841.1

Group11474: abra|OJJ73853.1 afla|EED57229.1 anig|EHA21757.1 aory|EIT74043.1 azon|OJJ42633.1

Group11475: abra|OJJ73942.1 akaw|GAA89547.1 aluc|OJZ88102.1 anig|EHA17871.1 atub|OJI90710.1

Group11476: abra|OJJ73948.1 acar|OOG01076.1 akaw|GAA89553.1 aluc|OJZ88096.1 atub|OJI90704.1

Group11477: abra|OJJ73954.1 akaw|GAA89559.1 aluc|OJZ88090.1 anig|EHA17860.1 atub|OJI90698.1

Group11478: abra|OJJ73957.1 akaw|GAA85099.1 atub|OJI83978.1 agla|OJJ89635.1 aluc|OJZ86545.1

Group11479: abra|OJJ74026.1 akaw|GAA87508.1 aluc|OJZ88012.1 anig|EHA17792.1 atub|OJI90621.1

Group11480: abra|OJJ74112.1 acar|OOF90000.1 asyd|OJJ63248.1 ater|EAU30630.1 atub|OJI90534.1

Group11481: agla|OJJ86579.1 aluc|OJZ82541.1 akaw|GAA92870.1 atub|OJI88785.1 anig|EHA22509.1

Group11482: abra|OJJ74151.1 akaw|GAA89732.1 aluc|OJZ80651.1 anig|EHA27181.1 atub|OJI87506.1

Group11483: abra|OJJ74160.1 akaw|GAA89724.1 aluc|OJZ80659.1 anig|EHA27173.1 atub|OJI87515.1

Group11484: abra|OJJ74214.1 akaw|GAA86804.1 aluc|OJZ89920.1 anig|EHA27120.1 atub|OJI84885.1

Group11485: anid|CBF84306.1 asyd|OJJ56674.1 aver|OJJ06050.1 afla|EED49875.1 aory|EIT83387.1

Group11486: abra|OJJ74402.1 akaw|GAA87297.1 aluc|OJZ89721.1 anig|EHA26948.1 atub|OJI85079.1

Group11487: asyd|OJJ65130.1 afla|EED47450.1 agla|OJJ88202.1 anig|EHA25401.1 aory|EIT81897.1

Group11488: abra|OJJ74430.1 afla|EED57468.1 anig|EHA26929.1 aory|EIT80153.1 afis|EAW24539.1

Group11489: agla|OJJ81569.1 asyd|OJJ54208.1 ater|EAU38806.1 aver|OJJ01904.1 awen|OJJ30144.1

Group11490: abra|OJJ74474.1 akaw|GAA87368.1 aluc|OJZ89644.1 anig|EHA26883.1 atub|OJI85159.1

Group11491: abra|OJJ74475.1 akaw|GAA87369.1 aluc|OJZ89643.1 anig|EHA26882.1 atub|OJI85160.1

Group11492: afis|EAW22195.1 afum|EAL90996.1 afla|EED57582.1 aory|EIT82945.1 awen|OJJ34778.1

Group11493: abra|OJJ74514.1 akaw|GAA82085.1 aluc|OJZ92095.1 anig|EHA26846.1 atub|OJI86528.1

Group11494: abra|OJJ74518.1 afla|EED54605.1 akaw|GAA92800.1 aluc|OJZ88722.1 anig|EHA28205.1

Group11495: abra|OJJ74534.1 akaw|GAA82066.1 aluc|OJZ92116.1 anig|EHA26829.1 atub|OJI86549.1

Group11496: abra|OJJ74575.1 akaw|GAA82030.1 aluc|OJZ92153.1 anig|EHA26796.1 atub|OJI86581.1

Group11497: abra|OJJ74597.1 aluc|OJZ92172.1 akaw|GAA82011.1 anig|EHA26777.1 atub|OJI86604.1

Group11498: abra|OJJ74660.1 asyd|OJJ55546.1 atub|OJI86670.1 awen|OJJ30940.1 aver|OJJ06623.1

Group11499: abra|OJJ74850.1 akaw|GAA83572.1 aluc|OJZ92426.1 anig|EHA26541.1 atub|OJI86856.1

Group11500: abra|OJJ74886.1 akaw|GAA83533.1 aluc|OJZ92463.1 anig|EHA26504.1 atub|OJI86892.1

Group11501: azon|OJJ42866.1 acla|EAW11650.1 afis|EAW23805.1 afum|EBA27306.1 awen|OJJ33762.1

Group11502: abra|OJJ74951.1 afla|EED51748.1 anig|EHA26451.1 aory|EIT80488.1 atub|OJI86958.1

Group11503: abra|OJJ74973.1 anig|EHA21622.1 asyd|OJJ55445.1 aver|OJJ06518.1 awen|OJJ36163.1

Group11504: abra|OJJ75008.1 akaw|GAA88631.1 aluc|OJZ92573.1 anig|EHA26408.1 atub|OJI87009.1

Group11505: abra|OJJ75118.1 akaw|GAA88728.1 aluc|OJZ92685.1 anig|EHA26315.1 atub|OJI87113.1

Group11506: abra|OJJ75140.1 akaw|GAA88747.1 aluc|OJZ92707.1 anig|EHA26293.1 atub|OJI87133.1

Group11507: abra|OJJ75196.1 akaw|GAA91748.1 aluc|OJZ92759.1 anig|EHA26247.1 atub|OJI87203.1

Group11508: abra|OJJ75224.1 akaw|GAA91772.1 aluc|OJZ92787.1 anig|EHA26221.1 atub|OJI87230.1

Group11509: agla|OJJ80388.1 afla|EED49688.1 aory|EIT80937.1 awen|OJJ32520.1 agla|OJJ82223.1

Group11510: abra|OJJ75250.1 afla|EED47799.1 aluc|OJZ92809.1 atub|OJI87266.1 atub|OJI87264.1

Group11511: abra|OJJ75294.1 aluc|OJZ88715.1 anid|CBF79577.1 azon|OJJ42162.1 aluc|OJZ79599.1

Group11512: abra|OJJ75316.1 acar|OOF94514.1 aluc|OJZ92870.1 anig|EHA26095.1 atub|OJI87328.1

Group11513: abra|OJJ75320.1 akaw|GAA92015.1 aluc|OJZ92881.1 anig|EHA26079.1 atub|OJI87340.1

Group11514: abra|OJJ75360.1 aluc|OJZ92927.1 anig|EHA26043.1 atub|OJI87387.1 akaw|GAA92055.1

Group11515: abra|OJJ75479.1 acar|OOF90815.1 akaw|GAA84248.1 aluc|OJZ88937.1 atub|OJI85853.1

Group11516: abra|OJJ75534.1 akaw|GAA91954.1 aluc|OJZ82664.1 anig|EHA27438.1 atub|OJI88897.1

Group11517: abra|OJJ75571.1 akaw|GAA84322.1 aluc|OJZ89019.1 anig|EHA23511.1 atub|OJI85770.1

Group11518: abra|OJJ75585.1 akaw|GAA84337.1 aluc|OJZ89036.1 anig|EHA23498.1 atub|OJI85754.1

Group11519: abra|OJJ75586.1 akaw|GAA84338.1 aluc|OJZ89037.1 anig|EHA23497.1 atub|OJI85753.1

Group11520: afis|EAW19826.1 afum|EAL85066.1 acla|EAW15058.1 agla|OJJ81773.1 awen|OJJ40152.1

Group11521: abra|OJJ75597.1 akaw|GAA84348.1 aluc|OJZ89048.1 anig|EHA23488.1 atub|OJI85739.1

Group11522: abra|OJJ75600.1 akaw|GAA84351.1 aluc|OJZ89051.1 anig|EHA23487.1 atub|OJI85736.1

Group11523: abra|OJJ75608.1 akaw|GAA84358.1 aluc|OJZ89060.1 anig|EHA23478.1 atub|OJI85726.1

Group11524: abra|OJJ75750.1 akaw|GAA84483.1 aluc|OJZ89203.1 anig|EHA23355.1 atub|OJI85588.1

Group11525: azon|OJJ42173.1 afla|EED45357.1 aory|EIT82400.1 awen|OJJ35918.1 aver|OJJ07430.1

Group11526: abra|OJJ75767.1 asyd|OJJ58673.1 atub|OJI87191.1 aver|OJJ01284.1 awen|OJJ30921.1

Group11527: abra|OJJ75772.1 asyd|OJJ58677.1 atub|OJI87186.1 aver|OJJ01289.1 awen|OJJ30926.1

Group11528: abra|OJJ75837.1 afis|EAW25763.1 aluc|OJZ88051.1 anig|EHA17823.1 atub|OJI90659.1

Group11529: abra|OJJ75860.1 akaw|GAA84565.1 aluc|OJZ89298.1 anig|EHA23265.1 atub|OJI85500.1

Group11530: abra|OJJ75993.1 akaw|GAA88965.1 aluc|OJZ89438.1 anig|EHA23148.1 atub|OJI85355.1

Group11531: abra|OJJ76014.1 akaw|GAA88946.1 aluc|OJZ89460.1 atub|OJI85334.1 azon|OJJ50328.1

Group11532: abra|OJJ76162.1 akaw|GAA82109.1 aluc|OJZ92067.1 anig|EHA22994.1 atub|OJI86498.1

Group11533: abra|OJJ76168.1 akaw|GAA91444.1 aluc|OJZ84721.1 anig|EHA27994.1 atub|OJI89362.1

Group11534: abra|OJJ76277.1 akaw|GAA82209.1 aluc|OJZ91943.1 anig|EHA22902.1 atub|OJI86386.1

Group11535: abra|OJJ76289.1 acar|OOF99531.1 akaw|GAA82218.1 aluc|OJZ91933.1 atub|OJI86375.1

Group11536: abra|OJJ76329.1 afum|EAL93528.1 aluc|OJZ91884.1 asyd|OJJ58950.1 atub|OJI86335.1

Group11537: abra|OJJ76445.1 akaw|GAA82357.1 aluc|OJZ91765.1 atub|OJI86223.1 acar|OOF99688.1

Group11538: abra|OJJ76592.1 acar|OOF99395.1 akaw|GAA82496.1 aluc|OJZ91614.1 atub|OJI86067.1

Group11539: abra|OJJ76629.1 acar|OOF99369.1 akaw|GAA92828.1 aluc|OJZ91583.1 atub|OJI86032.1

Group11540: abra|OJJ76645.1 akaw|GAA92815.1 aluc|OJZ91566.1 anig|EHA22595.1 atub|OJI86016.1

Group11541: abra|OJJ76666.1 acar|OOF99532.1 afla|EED49840.1 anig|EHA21628.1 aory|EIT83401.1

Group11542: abra|OJJ76676.1 acar|OOF99333.1 akaw|GAA92516.1 aluc|OJZ91529.1 atub|OJI85983.1

Group11543: abra|OJJ76723.1 akaw|GAA92478.1 aluc|OJZ91488.1 anig|EHA22532.1 atub|OJI85943.1

Group11544: abra|OJJ76854.1 akaw|GAA87168.1 aluc|OJZ84266.1 anig|EHA27608.1 atub|OJI89817.1

Group11545: abra|OJJ76872.1 anig|EHA24716.1 acar|OOF90111.1 akaw|GAA89114.1 aluc|OJZ81726.1

Group11546: abra|OJJ76883.1 acar|OOF90100.1 anid|CBF87196.1 asyd|OJJ56897.1 awen|OJJ30941.1

Group11547: abra|OJJ76891.1 akaw|GAA87151.1 aluc|OJZ84284.1 atub|OJI89787.1 awen|OJJ36187.1

Group11548: abra|OJJ76892.1 acar|OOF92132.1 anig|EHA25830.1 ater|EAU36567.1 azon|OJJ43582.1

Group11549: awen|OJJ35548.1 afla|EED56941.1 akaw|GAA86007.1 atub|OJI81663.1 aluc|OJZ90233.1

Group11550: abra|OJJ76990.1 akaw|GAA87056.1 aluc|OJZ84390.1 atub|OJI89681.1 azon|OJJ49723.1

Group11551: abra|OJJ77017.1 afla|EED53227.1 anig|EHA20227.1 aory|EIT75128.1 aver|OJJ02005.1

Group11552: abra|OJJ77020.1 akaw|GAA87032.1 aluc|OJZ84420.1 anig|EHA27742.1 atub|OJI89653.1

Group11553: abra|OJJ77022.1 akaw|GAA87030.1 aluc|OJZ84422.1 anig|EHA27744.1 atub|OJI89651.1

Group11554: abra|OJJ77027.1 akaw|GAA87024.1 aluc|OJZ84429.1 anig|EHA27748.1 atub|OJI89645.1

Group11555: abra|OJJ77170.1 akaw|GAA90763.1 aluc|OJZ84583.1 anig|EHA27876.1 atub|OJI89497.1

Group11556: abra|OJJ77185.1 akaw|GAA90754.1 aluc|OJZ84594.1 anig|EHA27883.1 atub|OJI89485.1

Group11557: abra|OJJ77278.1 akaw|GAA90671.1 aluc|OJZ84688.1 anig|EHA27963.1 atub|OJI89394.1

Group11558: abra|OJJ77307.1 akaw|GAA87995.1 aluc|OJZ81250.1 anig|EHA18558.1 atub|OJI91447.1

Group11559: abra|OJJ77387.1 acar|OOF93256.1 akaw|GAA91374.1 aluc|OJZ84795.1 atub|OJI89287.1

Group11560: awen|OJJ35376.1 acar|OOF90041.1 akaw|GAA85851.1 aluc|OJZ91248.1 atub|OJI83245.1

Group11561: abra|OJJ77458.1 acar|OOG00737.1 acla|EAW07629.1 anig|EHA19292.1 asyd|OJJ60066.1

Group11562: abra|OJJ77465.1 acar|OOF99270.1 akaw|GAA84152.1 aluc|OJZ81073.1 atub|OJI82513.1

Group11563: abra|OJJ77500.1 akaw|GAA84121.1 aluc|OJZ81037.1 anig|EHA19329.1 atub|OJI82546.1

Group11564: abra|OJJ77551.1 akaw|GAA84072.1 aluc|OJZ80986.1 anig|EHA19374.1 atub|OJI82600.1

Group11565: abra|OJJ77552.1 akaw|GAA84071.1 aluc|OJZ80985.1 anig|EHA19375.1 atub|OJI82601.1

Group11566: abra|OJJ77606.1 acla|EAW12878.1 asyd|OJJ57436.1 ater|EAU36087.1 aver|OJJ05742.1

Group11567: abra|OJJ77641.1 akaw|GAA83993.1 aluc|OJZ80889.1 anig|EHA19449.1 atub|OJI82689.1

Group11568: abra|OJJ77646.1 aluc|OJZ80883.1 atub|OJI82694.1 awen|OJJ37814.1 acar|OOF91727.1

Group11569: abra|OJJ77738.1 akaw|GAA83891.1 aluc|OJZ90804.1 atub|OJI82793.1 awen|OJJ35813.1

Group11570: awen|OJJ34501.1 afla|EED44857.1 anid|CBF78811.1 asyd|OJJ65261.1 aver|OJI99412.1

Group11571: anid|CBF83029.1 afis|EAW25582.1 afum|EAL84292.1 asyd|OJJ56287.1 aver|OJJ08461.1

Group11572: abra|OJJ77805.1 akaw|GAA83844.1 aluc|OJZ90871.1 anig|EHA19591.1 atub|OJI82871.1

Group11573: abra|OJJ77935.1 acar|OOF98871.1 akaw|GAA83728.1 aluc|OJZ91003.1 atub|OJI82994.1

Group11574: azon|OJJ45370.1 agla|OJJ86118.1 asyd|OJJ59985.1 aver|OJJ02306.1 awen|OJJ30345.1

Group11575: abra|OJJ77942.1 afis|EAW16408.1 akaw|GAA83723.1 aluc|OJZ91010.1 atub|OJI83001.1

Group11576: abra|OJJ77958.1 akaw|GAA83725.1 aluc|OJZ91006.1 anig|EHA19711.1 atub|OJI82998.1

Group11577: abra|OJJ77961.1 acar|OOF90714.1 anid|CBF81947.1 asyd|OJJ57369.1 aver|OJJ05799.1

Group11578: abra|OJJ77979.1 akaw|GAA86001.1 aluc|OJZ91054.1 anig|EHA19757.1 atub|OJI83050.1

Group11579: abra|OJJ77981.1 akaw|GAA85999.1 aluc|OJZ91056.1 anig|EHA19759.1 atub|OJI83052.1

Group11580: abra|OJJ78031.1 akaw|GAA85954.1 aluc|OJZ91108.1 anig|EHA19806.1 atub|OJI83105.1

Group11581: abra|OJJ78058.1 agla|OJJ85719.1 akaw|GAA85934.1 aluc|OJZ91128.1 atub|OJI83139.1

Group11582: abra|OJJ78069.1 akaw|GAA85922.1 aluc|OJZ91145.1 anig|EHA19841.1 atub|OJI83158.1

Group11583: abra|OJJ78080.1 akaw|GAA85913.1 aluc|OJZ91153.1 anig|EHA19850.1 atub|OJI83168.1

Group11584: abra|OJJ78081.1 akaw|GAA85911.1 aluc|OJZ91155.1 anig|EHA19851.1 atub|OJI83170.1

Group11585: abra|OJJ78089.1 akaw|GAA85905.1 aluc|OJZ91162.1 anig|EHA19856.1 atub|OJI83177.1

Group11586: abra|OJJ78106.1 acar|OOF98759.1 aluc|OJZ91171.1 atub|OJI83185.1 awen|OJJ30277.1

Group11587: anid|CBF73972.1 asyd|OJJ62367.1 afis|EAW17323.1 afum|EAL91159.1 aver|OJI96501.1

Group11588: abra|OJJ78159.1 akaw|GAA85867.1 aluc|OJZ91212.1 anig|EHA19918.1 atub|OJI83230.1

Group11589: abra|OJJ78161.1 akaw|GAA85864.1 aluc|OJZ91215.1 anig|EHA19921.1 atub|OJI83233.1

Group11590: abra|OJJ78171.1 agla|OJJ80024.1 ater|EAU30166.1 asyd|OJJ59097.1 aver|OJJ01688.1

Group11591: abra|OJJ78176.1 akaw|GAA85857.1 aluc|OJZ91225.1 anig|EHA19931.1 atub|OJI83238.1

Group11592: abra|OJJ78184.1 acar|OOF93286.1 akaw|GAA85850.1 aluc|OJZ91249.1 atub|OJI83247.1

Group11593: abra|OJJ78197.1 acar|OOF98658.1 aluc|OJZ91268.1 anig|EHA19956.1 atub|OJI83257.1

Group11594: acla|EAW07044.1 afla|EED44975.1 aory|EIT73531.1 ater|EAU30335.1 azon|OJJ46852.1

Group11595: azon|OJJ47989.1 acar|OOG00346.1 acla|EAW13485.1 akaw|GAA88048.1 aluc|OJZ81305.1

Group11596: acar|OOF89858.1 akaw|GAA87911.1 aluc|OJZ88781.1 anig|EHA18475.1 atub|OJI91353.1

Group11597: acar|OOF89868.1 acla|EAW13237.1 afla|EED49459.1 aory|EIT79486.1 ater|EAU38425.1

Group11598: acar|OOF90158.1 aory|EIT73596.1 awen|OJJ37847.1 azon|OJJ43182.1 afla|EED45042.1

Group11599: azon|OJJ50402.1 acla|EAW07416.1 afis|EAW15632.1 asyd|OJJ60062.1 aver|OJJ02228.1

Group11600: acar|OOF90558.1 asyd|OJJ53527.1 ater|EAU37836.1 aver|OJJ08223.1 anid|CBF84497.1

Group11601: acar|OOF90580.1 afla|EED45486.1 asyd|OJJ54094.1 aver|OJJ08645.1 aory|EIT75714.1

Group11602: acar|OOF90600.1 aluc|OJZ85172.1 afla|EED48699.1 anid|CBF78730.1 aory|EIT73228.1

Group11603: acar|OOF91083.1 acla|EAW08612.1 afla|EED51225.1 akaw|GAA92245.1 aory|EIT83192.1

Group11604: acar|OOF91240.1 akaw|GAA90581.1 aluc|OJZ86039.1 anig|EHA24572.1 atub|OJI79826.1

Group11605: acar|OOF91741.1 afla|EED48871.1 anig|EHA22517.1 aory|EIT81191.1 awen|OJJ38799.1

Group11606: acar|OOF91793.1 anig|EHA25412.1 awen|OJJ32139.1 azon|OJJ43633.1 ater|EAU34523.1

Group11607: acar|OOF91815.1 afum|EAL86657.1 anid|CBF89379.1 asyd|OJJ52303.1 aver|OJJ02178.1

Group11608: acar|OOF91816.1 agla|OJJ79106.1 asyd|OJJ55392.1 aver|OJJ08265.1 awen|OJJ34588.1

Group11609: aluc|OJZ88613.1 anid|CBF87199.1 afis|EAW16201.1 afum|EAL88828.1 atub|OJI91198.1

Group11610: acar|OOF92086.1 afis|EAW25709.1 afum|EAL84382.1 afla|EED50159.1 aory|EIT76929.1

Group11611: acar|OOF92135.1 afis|EAW19863.1 ater|EAU31608.1 awen|OJJ41891.1 afum|EAL85032.1

Group11612: acar|OOF92451.1 anid|CBF80402.1 ater|EAU32762.1 aver|OJJ07408.1 ater|EAU35437.1

Group11613: acar|OOF92581.1 akaw|GAA86577.1 aluc|OJZ85128.1 anig|EHA25347.1 atub|OJI80465.1

Group11614: acar|OOF92905.1 afis|EAW16769.1 afla|EED53153.1 afum|EAL86337.1 aory|EIT75263.1

Group11615: ater|EAU29916.1 awen|OJJ32299.1 afis|EAW19915.1 afla|EED45532.1 afum|EAL84992.1

Group11616: afis|EAW23680.1 afum|EAL87123.1 asyd|OJJ54288.1 ater|EAU37627.1 aver|OJJ05242.1

Group11617: acar|OOF93145.1 agla|OJJ80311.1 aory|EIT73225.1 ater|EAU37753.1 awen|OJJ38068.1

Group11618: acar|OOF93265.1 afla|EED56025.1 afum|EAL84407.1 aory|EIT76725.1 awen|OJJ30874.1

Group11619: acar|OOF93293.1 afla|EED54454.1 agla|OJJ78525.1 aory|EIT81155.1 awen|OJJ41198.1

Group11620: acla|EAW08970.1 agla|OJJ86272.1 afis|EAW15449.1 afum|EAL84197.1 ater|EAU30839.1

Group11621: acar|OOF93346.1 akaw|GAA86108.1 aluc|OJZ90340.1 atub|OJI81767.1 ater|EAU36841.1

Group11622: acar|OOF93932.1 aluc|OJZ83621.1 anig|EHA27513.1 atub|OJI87848.1 akaw|GAA92104.1

Group11623: acar|OOF93934.1 akaw|GAA92105.1 aluc|OJZ83620.1 anig|EHA27512.1 atub|OJI87847.1

Group11624: acar|OOF94197.1 afla|EED47770.1 asyd|OJJ59945.1 ater|EAU30396.1 aver|OJJ02343.1

Group11625: asyd|OJJ65336.1 aver|OJI99482.1 afla|EED48619.1 awen|OJJ33514.1 aory|EIT79599.1

Group11626: acar|OOF94512.1 akaw|GAA92000.1 aluc|OJZ92871.1 anid|CBF88861.1 atub|OJI87329.1

Group11627: afis|EAW16981.1 afum|EAL86508.1 agla|OJJ79788.1 awen|OJJ35995.1 acla|EAW12255.1

Group11628: afis|EAW16973.1 afla|EED51778.1 aory|EIT81699.1 aver|OJJ02700.1 azon|OJJ42647.1

Group11629: akaw|GAA84852.1 aluc|OJZ86819.1 atub|OJI83698.1 agla|OJJ83637.1 aver|OJJ04869.1

Group11630: acar|OOF94677.1 anid|CBF87491.1 anig|EHA26147.1 asyd|OJJ52951.1 aver|OJJ08917.1

Group11631: azon|OJJ50289.1 acla|EAW14837.1 afis|EAW24365.1 afum|EAL86037.1 awen|OJJ32849.1

Group11632: anig|EHA28515.1 ater|EAU31655.1 afla|EED53334.1 aory|EIT75105.1 atub|OJI82827.1

Group11633: acar|OOF94960.1 aory|EIT73121.1 atub|OJI88446.1 aver|OJJ07461.1 afla|EED48595.1

Group11634: acar|OOF95008.1 acla|EAW12216.1 afis|EAW25557.1 afum|EBA27171.1 awen|OJJ31977.1

Group11635: acar|OOF95009.1 acla|EAW12219.1 afis|EAW25560.1 afum|EAL84284.1 awen|OJJ31979.1

Group11636: acar|OOF95022.1 afla|EED53083.1 aory|EIT74806.1 asyd|OJJ62857.1 aver|OJI96909.1

Group11637: ater|EAU30958.1 afla|EED46244.1 anid|CBF80415.1 aory|EIT76385.1 azon|OJJ50735.1

Group11638: afla|EED45397.1 ater|EAU33508.1 aory|EIT82396.1 asyd|OJJ54513.1 aver|OJI95770.1

Group11639: acar|OOF95599.1 akaw|GAA88039.1 aluc|OJZ81298.1 anig|EHA18607.1 atub|OJI91507.1

Group11640: acar|OOF95837.1 afla|EED49641.1 anid|CBF70130.1 aory|EIT80985.1 ater|EAU32469.1

Group11641: aory|EIT81704.1 afla|EED51881.1 afla|EED51882.1 anig|EHA22155.1 azon|OJJ42106.1

Group11642: acar|OOF96501.1 anid|CBF74630.1 asyd|OJJ57583.1 aver|OJJ05598.1 azon|OJJ51751.1

Group11643: acar|OOF96692.1 afla|EED54107.1 aory|EIT75906.1 asyd|OJJ57567.1 aver|OJJ05612.1

Group11644: acar|OOF96790.1 akaw|GAA84770.1 aluc|OJZ86913.1 anig|EHA21731.1 atub|OJI83608.1

Group11645: acar|OOF97570.1 acla|EAW09121.1 atub|OJI83131.1 aver|OJJ08167.1 awen|OJJ29722.1

Group11646: acar|OOF97637.1 akaw|GAA92099.1 aluc|OJZ83626.1 anig|EHA18779.1 atub|OJI89995.1

Group11647: acar|OOF97927.1 akaw|GAA82747.1 aluc|OJZ87303.1 anig|EHA20542.1 awen|OJJ38050.1

Group11648: azon|OJJ45429.1 aluc|OJZ85169.1 anid|CBF81952.1 anig|EHA25363.1 awen|OJJ32776.1

Group11649: acar|OOF98232.1 awen|OJJ38716.1 acar|OOF99029.1 aory|EIT72316.1 afla|EED46400.1

Group11650: acar|OOF98332.1 afla|EED47391.1 atub|OJI87441.1 aver|OJI97587.1 azon|OJJ42119.1

Group11651: acar|OOF98719.1 anig|EHA22488.1 asyd|OJJ55518.1 ater|EAU34694.1 aver|OJJ06596.1

Group11652: acar|OOF98727.1 afis|EAW21205.1 afis|EAW21206.1 afum|EAL92216.1 afum|EAL92217.1

Group11653: acar|OOF98795.1 akaw|GAA87802.1 aluc|OJZ85437.1 atub|OJI80131.1 azon|OJJ45923.1

Group11654: acar|OOF98989.1 agla|OJJ78509.1 asyd|OJJ60481.1 ater|EAU29496.1 aver|OJI99802.1

Group11655: asyd|OJJ64276.1 aver|OJI98368.1 afla|EED56303.1 aory|EIT73418.1 anid|CBF85721.1

Group11656: acar|OOF99109.1 afla|EED51168.1 anid|CBF80169.1 aory|EIT81340.1 aver|OJI99522.1

Group11657: acar|OOF99111.1 awen|OJJ30483.1 acla|EAW06491.1 afis|EAW15987.1 afum|EAL88619.1

Group11658: acar|OOF99135.1 afla|EED48438.1 agla|OJJ84624.1 asyd|OJJ61434.1 ater|EAU29760.1

Group11659: acar|OOF99286.1 akaw|GAA84163.1 aluc|OJZ81088.1 anig|EHA19282.1 atub|OJI82496.1

Group11660: acar|OOF99304.1 akaw|GAA91081.1 aluc|OJZ87897.1 anig|EHA20195.1 atub|OJI80753.1

Group11661: acar|OOF99861.1 aory|EIT74417.1 acar|OOG00208.1 azon|OJJ42204.1 afla|EED52215.1

Group11662: acar|OOG00291.1 acla|EAW12453.1 afla|EED51635.1 atub|OJI84634.1 azon|OJJ46528.1

Group11663: acar|OOG00296.1 acla|EAW15040.1 anid|CBF80514.1 asyd|OJJ54737.1 awen|OJJ36952.1

Group11664: acar|OOG00321.1 akaw|GAA89088.1 aluc|OJZ80503.1 anig|EHA25690.1 atub|OJI88427.1

Group11665: acar|OOG00408.1 afum|EAL84201.1 agla|OJJ88745.1 anig|EHA23823.1 azon|OJJ50323.1

Group11666: acar|OOG00534.1 afum|EAL86635.1 anid|CBF86043.1 asyd|OJJ54464.1 aver|OJJ07790.1

Group11667: acar|OOG00539.1 akaw|GAA92714.1 aluc|OJZ88760.1 anig|EHA18456.1 atub|OJI91330.1

Group11668: acar|OOG00636.1 afum|EAL87770.2 anig|EHA18412.1 asyd|OJJ54770.1 aver|OJI99272.1

Group11669: acar|OOG00696.1 agla|OJJ84879.1 aluc|OJZ80909.1 asyd|OJJ58133.1 aver|OJJ04272.1

Group11670: acar|OOG00736.1 afis|EAW16805.1 afla|EED49871.1 anid|CBF80485.1 aory|EIT83385.1

Group11671: acla|EAW06330.1 anig|EHA24793.1 akaw|GAA90461.1 atub|OJI79547.1 aluc|OJZ85763.1

Group11672: acla|EAW06451.1 afis|EAW15948.1 afla|EED58111.1 afum|EAL88581.1 awen|OJJ35636.1

Group11673: acla|EAW06593.1 afis|EAW16016.1 afum|EAL88645.1 awen|OJJ30448.1 azon|OJJ45440.1

Group11674: acla|EAW06760.1 afis|EAW16310.1 afum|EAL88932.1 awen|OJJ37710.1 azon|OJJ44969.1

Group11675: asyd|OJJ59634.1 afla|EED49577.1 aory|EIT82583.1 ater|EAU29693.1 aver|OJJ02647.1

Group11676: acla|EAW07111.1 afis|EAW22123.1 afla|EED46075.1 afum|EAL91064.1 aory|EIT83491.1

Group11677: acla|EAW07112.1 afla|EED49434.1 asyd|OJJ56891.1 aver|OJJ07732.1 aver|OJJ07733.1

Group11678: acla|EAW07205.1 afis|EAW22217.1 afum|EAL90977.1 agla|OJJ87827.1 awen|OJJ34803.1

Group11679: acla|EAW07212.1 afis|EAW22225.1 afum|EAL90969.1 aver|OJJ01781.1 azon|OJJ42276.1

Group11680: acla|EAW07266.1 afis|EAW22287.1 afum|EAL90894.1 awen|OJJ34860.1 agla|OJJ87762.1

Group11681: acla|EAW07770.1 agla|OJJ86197.1 agla|OJJ88487.1 afis|EAW18238.1 afum|EBA27320.1

Group11682: acla|EAW08000.1 afum|EAL88446.1 afis|EAW23035.1 awen|OJJ35706.1 agla|OJJ89442.1

Group11683: acla|EAW08471.1 anid|CBF83637.1 asyd|OJJ60784.1 aver|OJJ00117.1 awen|OJJ39842.1

Group11684: acla|EAW08548.1 afis|EAW23585.1 afum|EAL87879.1 awen|OJJ39912.1 azon|OJJ45888.1

Group11685: acla|EAW08573.1 afis|EAW23631.1 afla|EED45134.1 afum|EBA27324.1 aory|EIT73823.1

Group11686: acla|EAW08739.1 afis|EAW15697.1 afum|EAL85651.2 anig|EHA24533.1 afla|EED51356.1

Group11687: acla|EAW08945.1 afum|EAL91099.1 akaw|GAA83285.1 anig|EHA18931.1 awen|OJJ34617.1

Group11688: acla|EAW08947.1 afum|EBA27420.1 akaw|GAA83287.1 anig|EHA18932.1 awen|OJJ34619.1

Group11689: acla|EAW08948.1 afum|EAL91101.1 akaw|GAA83288.1 anig|EHA18933.1 awen|OJJ34620.1

Group11690: azon|OJJ43668.1 anid|CBF84735.1 asyd|OJJ53301.1 aver|OJJ08440.1 awen|OJJ30041.1

Group11691: acla|EAW09046.1 agla|OJJ89274.1 anid|CBF81248.1 asyd|OJJ59324.1 aver|OJJ02950.1

Group11692: acla|EAW09067.1 afis|EAW25642.1 afum|EAL84329.1 anid|CBF80275.1 awen|OJJ36095.1

Group11693: acla|EAW09081.1 afla|EED46121.1 aory|EIT73727.1 afis|EAW17182.1 aory|EIT76387.1

Group11694: acla|EAW09115.1 afis|EAW22087.1 afla|EED49612.1 afum|EAL85696.1 aory|EIT80971.1

Group11695: acla|EAW09339.1 afis|EAW21794.1 afum|EAL92819.1 awen|OJJ34078.1 azon|OJJ44263.1

Group11696: azon|OJJ51423.1 afla|EED57812.1 awen|OJJ39091.1 asyd|OJJ60990.1 aver|OJJ00310.1

Group11697: acla|EAW09827.1 afis|EAW21306.1 afum|EAL92322.1 agla|OJJ87433.1 awen|OJJ39392.1

Group11698: aluc|OJZ88612.1 anid|CBF87197.1 atub|OJI91197.1 afis|EAW16200.1 afum|EAL88827.1

Group11699: acla|EAW10017.1 agla|OJJ78661.1 aory|EIT77856.1 asyd|OJJ57247.1 aver|OJJ03047.1

Group11700: acla|EAW10115.1 anid|CBF77102.1 asyd|OJJ55094.1 aver|OJJ07114.1 awen|OJJ32552.1

Group11701: acla|EAW10400.1 afis|EAW18584.1 agla|OJJ86755.1 awen|OJJ39597.1 azon|OJJ41988.1

Group11702: acla|EAW10648.1 ater|EAU30169.1 acla|EAW11698.1 afis|EAW24243.1 afum|EAL85911.1

Group11703: acla|EAW10834.1 afis|EAW24730.1 afum|EAL89377.1 agla|OJJ85245.1 awen|OJJ37856.1

Group11704: acla|EAW11069.1 afis|EAW20805.1 afum|EAL84823.1 asyd|OJJ54168.1 aver|OJJ08571.1

Group11705: acla|EAW11093.1 afis|EAW21814.1 afum|EBA27228.1 agla|OJJ82561.1 asyd|OJJ62723.1

Group11706: afla|EED52113.1 anid|CBF80518.1 aory|EIT76897.1 asyd|OJJ65293.1 aver|OJJ01917.1

Group11707: acla|EAW11331.1 afis|EAW18164.1 afum|EAL87524.1 awen|OJJ30510.1 azon|OJJ50193.1

Group11708: acla|EAW11517.1 afis|EAW23936.1 afum|EAL87361.1 agla|OJJ79417.1 azon|OJJ51344.1

Group11709: acla|EAW11623.1 ater|EAU31391.1 asyd|OJJ58185.1 aver|OJJ04220.1 azon|OJJ48754.1

Group11710: acla|EAW11706.1 agla|OJJ86727.1 anid|CBF73384.1 ater|EAU32802.1 ater|EAU37150.1

Group11711: acla|EAW11769.1 afla|EED45360.1 aory|EIT82406.1 ater|EAU34823.1 awen|OJJ33643.1

Group11712: acla|EAW11905.1 afla|EED48384.1 agla|OJJ85502.1 aory|EIT83062.1 awen|OJJ37790.1

Group11713: acla|EAW11924.1 afis|EAW19558.1 afla|EED49684.1 afum|EAL86953.1 aory|EIT80976.1

Group11714: acla|EAW11992.1 afis|EAW19632.1 afum|EAL86890.1 awen|OJJ41876.1 azon|OJJ47075.1

Group11715: acla|EAW12277.1 afis|EAW16963.1 afla|EED45225.1 afum|EAL86490.1 awen|OJJ40084.1

Group11716: acla|EAW12348.1 afum|EAL85144.1 afis|EAW19753.1 agla|OJJ80816.1 awen|OJJ29875.1

Group11717: acla|EAW12892.1 afis|EAW20657.1 anid|CBF87746.1 asyd|OJJ52092.1 atub|OJI88906.1

Group11718: anid|CBF82320.1 afis|EAW25603.1 agla|OJJ80365.1 ater|EAU32208.1 awen|OJJ40145.1

Group11719: acla|EAW13289.1 afis|EAW17330.1 afum|EAL91167.2 ater|EAU29262.1 azon|OJJ43172.1

Group11720: acla|EAW13621.1 afis|EAW19662.1 afum|EAL86860.2 awen|OJJ30664.1 azon|OJJ44765.1

Group11721: acla|EAW13658.1 afis|EAW20398.1 afum|EAL85497.1 anid|CBF85425.1 aver|OJI99263.1

Group11722: acla|EAW13714.1 afis|EAW23648.1 afla|EED48131.1 aory|EIT76223.1 ater|EAU31787.1

Group11723: acla|EAW13842.1 anid|CBF84469.1 asyd|OJJ53536.1 ater|EAU33155.1 aver|OJJ08215.1

Group11724: acla|EAW13898.1 afis|EAW18610.1 afum|EAL93320.1 agla|OJJ81663.1 awen|OJJ32071.1

Group11725: acla|EAW14024.1 afis|EAW18730.1 afum|EAL93455.2 awen|OJJ32272.1 azon|OJJ42749.1

Group11726: acla|EAW14025.1 afis|EAW18731.1 afum|EAL93456.1 awen|OJJ32273.1 azon|OJJ42750.1

Group11727: acla|EAW14135.1 afis|EAW18848.1 afla|EED50815.1 akaw|GAA82477.1 aory|EIT78994.1

Group11728: awen|OJJ36044.1 afis|EAW17301.1 afla|EED45574.1 aory|EIT75612.1 ater|EAU36622.1

Group11729: afla|EED52864.1 anid|CBF71142.1 aory|EIT78636.1 aver|OJJ04778.1 azon|OJJ43924.1

Group11730: acla|EAW14638.1 afis|EAW23679.1 afum|EAL87122.2 ater|EAU35562.1 azon|OJJ47910.1

Group11731: acla|EAW14643.1 afla|EED48987.1 aory|EIT80004.1 asyd|OJJ63817.1 aver|OJI97121.1

Group11732: acla|EAW14718.1 afis|EAW24254.1 afum|EAL85922.1 ater|EAU31196.1 awen|OJJ33046.1

Group11733: acla|EAW14823.1 afis|EAW24352.1 afum|EAL86022.1 agla|OJJ81769.1 ater|EAU30387.1

Group11734: acla|EAW14988.1 afis|EAW24513.1 afum|EBA27261.1 agla|OJJ83395.1 aver|OJJ08783.1

Group11735: acla|EAW15032.1 afis|EAW19345.1 afla|EED52029.1 anig|EHA24138.1 aory|EIT81473.1

Group11736: acla|EAW15342.1 afis|EAW20365.1 afla|EED45742.1 afum|EAL85481.2 aory|EIT75784.1

Group11737: afis|EAW15410.1 afla|EED51877.1 afum|EAL84236.1 aory|EIT81759.1 ater|EAU37984.1

Group11738: afis|EAW15648.1 anid|CBF85168.1 asyd|OJJ60245.1 ater|EAU30251.1 aver|OJJ04016.1

Group11739: awen|OJJ29791.1 afla|EED57511.1 aory|EIT81867.1 asyd|OJJ64582.1 aver|OJI98680.1

Group11740: afis|EAW16253.1 afla|EED56873.1 ater|EAU31543.1 aory|EIT77464.1 asyd|OJJ54141.1

Group11741: afis|EAW16389.1 afum|EAL89015.1 anid|CBF84475.1 awen|OJJ32037.1 aver|OJJ08217.1

Group11742: afis|EAW16456.1 afum|EAL89053.1 asyd|OJJ52206.1 ater|EAU39537.1 aver|OJI95787.1

Group11743: afis|EAW16706.1 asyd|OJJ53363.1 aver|OJJ07649.1 awen|OJJ36104.1 azon|OJJ42114.1

Group11744: afis|EAW16711.1 afum|EAL89299.1 anid|CBF70512.1 asyd|OJJ54856.1 ater|EAU29466.1

Group11745: afum|EAL86306.2 aluc|OJZ86493.1 anig|EHA22151.1 atub|OJI84033.1 akaw|GAA85144.1

Group11746: afis|EAW16764.1 afla|EED47351.1 afum|EAL86335.1 asyd|OJJ54357.1 aver|OJJ05311.1

Group11747: afis|EAW16910.1 afum|EAL86446.1 asyd|OJJ56951.1 aver|OJJ03334.1 azon|OJJ42448.1

Group11748: afis|EAW16911.1 afum|EAL86447.1 asyd|OJJ56952.1 aver|OJJ03333.1 azon|OJJ42449.1

Group11749: afis|EAW17007.1 akaw|GAA90343.1 aluc|OJZ85889.1 awen|OJJ37866.1 azon|OJJ42655.1

Group11750: afis|EAW17060.1 afum|EAL86556.1 anid|CBF88300.1 ater|EAU31926.1 azon|OJJ43509.1

Group11751: afis|EAW17078.1 akaw|GAA92154.1 aluc|OJZ85561.1 anig|EHA24967.1 atub|OJI79344.1

Group11752: afis|EAW17141.1 afla|EED56546.1 agla|OJJ82273.1 aory|EIT79701.1 awen|OJJ35659.1

Group11753: afis|EAW17150.1 afum|EAL86594.1 anid|CBF86604.1 asyd|OJJ64824.1 aver|OJI98937.1

Group11754: afis|EAW17310.1 afla|EED49774.1 afum|EAL91145.1 aory|EIT77689.1 awen|OJJ32902.1

Group11755: afis|EAW17311.1 afum|EAL91146.1 asyd|OJJ64145.1 aver|OJI98226.1 azon|OJJ42533.1

Group11756: afla|EED47547.1 anid|CBF69509.1 aory|EIT74573.1 asyd|OJJ59216.1 aver|OJJ01826.1

Group11757: afis|EAW17422.1 afum|EAL91247.1 asyd|OJJ61568.1 ater|EAU37823.1 aver|OJJ00888.1

Group11758: afis|EAW17520.1 anid|CBF71793.1 asyd|OJJ63026.1 ater|EAU35354.1 aver|OJI97942.1

Group11759: afis|EAW17521.1 anid|CBF71792.1 asyd|OJJ63027.1 ater|EAU35353.1 aver|OJI97941.1

Group11760: afis|EAW17763.1 afum|EAL91573.1 anid|CBF82290.1 asyd|OJJ52734.1 ater|EAU32266.1

Group11761: afis|EAW17852.1 afla|EED46206.1 afum|EAL91660.1 awen|OJJ41721.1 aory|EIT76447.1

Group11762: afis|EAW18093.1 aory|EIT78055.1 aver|OJI97252.1 azon|OJJ42725.1 agla|OJJ89387.1

Group11763: afis|EAW18324.1 anig|EHA23342.1 asyd|OJJ59740.1 ater|EAU29684.1 awen|OJJ32314.1

Group11764: afis|EAW18555.1 afum|EAL93255.1 aory|EIT80620.1 ater|EAU39464.1 afla|EED51439.1

Group11765: afis|EAW19009.1 afum|EAL93743.1 akaw|GAA90359.1 aluc|OJZ85873.1 atub|OJI79659.1

Group11766: afis|EAW19140.1 afla|EED53206.1 agla|OJJ80464.1 asyd|OJJ53821.1 ater|EAU35706.1

Group11767: afis|EAW19372.1 afla|EED45374.1 aory|EIT82425.1 asyd|OJJ60288.1 aver|OJI99620.1

Group11768: afis|EAW19517.1 afla|EED45882.1 afum|EAL86980.2 aory|EIT75651.1 ater|EAU33655.1

Group11769: afis|EAW19522.1 afla|EED46522.1 aory|EIT72363.1 asyd|OJJ64208.1 akaw|GAA92090.1

Group11770: afis|EAW19728.1 afum|EAL86799.1 anid|CBF71457.1 asyd|OJJ54949.1 aver|OJJ07246.1

Group11771: afis|EAW19740.1 afla|EED56528.1 afum|EAL86789.1 aory|EIT82925.1 awen|OJJ34641.1

Group11772: afis|EAW19770.1 afum|EAL85130.1 asyd|OJJ55399.1 ater|EAU32214.1 aver|OJJ06479.1

Group11773: afis|EAW19796.1 afum|EAL85094.1 aory|EIT81735.1 ater|EAU31582.1 awen|OJJ41524.1

Group11774: afis|EAW19808.1 afum|EAL85084.1 asyd|OJJ61798.1 ater|EAU33903.1 aver|OJI95897.1

Group11775: afis|EAW19861.1 afum|EAL85034.1 anid|CBF71554.1 asyd|OJJ56281.1 ater|EAU38548.1

Group11776: afis|EAW20216.1 afla|EED55532.1 aory|EIT80814.1 asyd|OJJ56329.1 aver|OJJ06384.1

Group11777: afis|EAW20254.1 afum|EAL85371.1 anid|CBF90399.1 ater|EAU35996.1 awen|OJJ38310.1

Group11778: agla|OJJ86325.1 asyd|OJJ56104.1 ater|EAU31560.1 aver|OJJ05046.1 awen|OJJ40020.1

Group11779: afis|EAW20381.1 afla|EED46310.1 afum|EAL85494.1 aory|EIT76336.1 azon|OJJ42056.1

Group11780: afis|EAW20580.1 afla|EED53405.1 afum|EAL84484.1 aory|EIT75078.1 awen|OJJ33280.1

Group11781: afis|EAW20705.1 afum|EAL87726.1 agla|OJJ82562.1 ater|EAU39566.1 awen|OJJ31381.1

Group11782: afis|EAW20844.1 afla|EED50399.1 afum|EAL84789.1 asyd|OJJ54456.1 ater|EAU32497.1

Group11783: afis|EAW21031.1 afla|EED55087.1 afum|EAL92070.1 agla|OJJ88642.1 aver|OJJ08193.1

Group11784: afis|EAW21040.1 afum|EAL92079.1 anid|CBF69836.1 asyd|OJJ59037.1 aver|OJJ01632.1

Group11785: afis|EAW21058.1 afum|EAL92093.2 afla|EED56621.1 asyd|OJJ64179.1 ater|EAU32060.1

Group11786: afis|EAW21203.1 anid|CBF86720.1 asyd|OJJ64716.1 ater|EAU32104.1 aver|OJI98819.1

Group11787: afis|EAW21216.1 afum|EAL92227.2 aluc|OJZ79469.1 anid|CBF73942.1 azon|OJJ46547.1

Group11788: afis|EAW21805.1 afum|EAL85391.1 agla|OJJ80479.1 akaw|GAA87895.1 asyd|OJJ62695.1

Group11789: anig|EHA17701.1 asyd|OJJ52138.1 akaw|GAA93338.1 aluc|OJZ79977.1 aluc|OJZ80376.1

Group11790: afis|EAW22563.1 afla|EED53099.1 afum|EAL90632.1 aory|EIT74819.1 awen|OJJ29708.1

Group11791: afis|EAW23612.1 afum|EAL87853.1 anid|CBF74000.1 asyd|OJJ61471.1 aver|OJJ00812.1

Group11792: afis|EAW23664.1 afla|EED50427.1 agla|OJJ83622.1 anid|CBF75930.1 aory|EIT82076.1

Group11793: afis|EAW23692.1 akaw|GAA92975.1 anig|EHA18086.1 atub|OJI89952.1 awen|OJJ35047.1

Group11794: afis|EAW23920.1 afum|EBA27311.1 asyd|OJJ57137.1 aver|OJJ03166.1 azon|OJJ46493.1

Group11795: afis|EAW24470.1 afla|EED48147.1 afum|EAL86142.1 aory|EIT76229.1 ater|EAU37772.1

Group11796: afis|EAW24481.1 akaw|GAA82567.1 aluc|OJZ87100.1 anig|EHA20370.1 atub|OJI81538.1

Group11797: afis|EAW24509.1 afla|EED48172.1 afum|EAL86178.1 aory|EIT83034.1 ater|EAU32786.1

Group11798: afis|EAW24514.1 afum|EAL86180.1 aory|EIT81903.1 asyd|OJJ56876.1 ater|EAU39544.1

Group11799: afis|EAW24542.1 asyd|OJJ54767.1 aver|OJJ07400.1 afla|EED44757.1 awen|OJJ37968.1

Group11800: afis|EAW24549.1 afla|EED53107.1 afum|EAL86213.1 agla|OJJ84626.1 aory|EIT74771.1

Group11801: afis|EAW24600.1 agla|OJJ80222.1 anig|EHA25388.1 aver|OJI96490.1 awen|OJJ29703.1

Group11802: afis|EAW24618.1 afla|EED54044.1 afum|EAL86246.1 aory|EIT76068.1 ater|EAU29289.1

Group11803: afis|EAW24772.1 afum|EAL89414.2 asyd|OJJ58395.1 ater|EAU34033.1 aver|OJJ03009.1

Group11804: afis|EAW24897.1 afla|EED52558.1 afum|EAL89551.1 aory|EIT74317.1 ater|EAU37216.1

Group11805: asyd|OJJ52783.1 afla|EED52088.1 aory|EIT76916.1 aver|OJJ04877.1 awen|OJJ32286.1

Group11806: afis|EAW25262.1 afum|EBA27391.1 asyd|OJJ64042.1 aver|OJI98126.1 awen|OJJ40974.1

Group11807: afis|EAW25583.1 afum|EAL84293.1 anid|CBF83031.1 asyd|OJJ56286.1 aver|OJJ08463.1

Group11808: afis|EAW25672.1 anid|CBF70504.1 asyd|OJJ58577.1 ater|EAU33457.1 aver|OJJ01181.1

Group11809: afis|EAW25703.1 afum|EAL84376.1 ater|EAU33410.1 azon|OJJ50782.1 azon|OJJ50781.1

Group11810: afis|EAW25712.1 afla|EED55198.1 aory|EIT83573.1 ater|EAU36724.1 azon|OJJ42322.1

Group11811: afla|EED44726.1 akaw|GAA92138.1 aory|EIT77794.1 asyd|OJJ59109.1 aver|OJJ01701.1

Group11812: afla|EED44753.1 akaw|GAA93270.1 aluc|OJZ81133.1 asyd|OJJ51865.1 ater|EAU35395.1

Group11813: afla|EED44867.1 aory|EIT82720.1 atub|OJI83117.1 awen|OJJ32254.1 aory|EIT82723.1

Group11814: afla|EED44919.1 agla|OJJ79288.1 aory|EIT73519.1 asyd|OJJ60041.1 aver|OJJ02244.1

Group11815: afla|EED45069.1 agla|OJJ84403.1 aory|EIT73671.1 aory|EIT73806.1 awen|OJJ40092.1

Group11816: afla|EED45070.1 agla|OJJ83555.1 anid|CBF81787.1 asyd|OJJ57968.1 atub|OJI83167.1

Group11817: afla|EED45231.1 aory|EIT73618.1 asyd|OJJ53708.1 aver|OJJ07832.1 awen|OJJ32557.1

Group11818: afla|EED45257.1 aluc|OJZ79657.1 anid|CBF76103.1 aory|EIT73624.1 ater|EAU36306.1

Group11819: afla|EED45269.1 anid|CBF80768.1 aory|EIT73800.1 asyd|OJJ54508.1 aver|OJJ07674.1

Group11820: awen|OJJ30964.1 afla|EED55970.1 aory|EIT76658.1 asyd|OJJ56496.1 aver|OJJ06223.1

Group11821: afla|EED45434.1 anid|CBF79176.1 aory|EIT82413.1 asyd|OJJ56413.1 aver|OJJ06302.1

Group11822: afla|EED45440.1 agla|OJJ88689.1 aory|EIT82381.1 awen|OJJ33936.1 aory|EIT82432.1

Group11823: afla|EED45485.1 anig|EHA23321.1 aory|EIT75381.1 ater|EAU36549.1 awen|OJJ33563.1

Group11824: afla|EED45498.1 anid|CBF84307.1 aory|EIT75762.1 asyd|OJJ57248.1 aver|OJJ03046.1

Group11825: afla|EED45534.1 anid|CBF83282.1 aory|EIT75631.1 asyd|OJJ61443.1 aver|OJJ00779.1

Group11826: afla|EED45577.1 agla|OJJ79089.1 aory|EIT75751.1 ater|EAU36771.1 awen|OJJ36955.1

Group11827: afla|EED45593.1 aory|EIT75639.1 asyd|OJJ52188.1 aver|OJJ03864.1 asyd|OJJ52540.1

Group11828: afla|EED45631.1 aory|EIT75596.1 awen|OJJ30950.1 asyd|OJJ61500.1 aver|OJJ00837.1

Group11829: afla|EED45649.1 aory|EIT75389.1 asyd|OJJ57902.1 aver|OJJ00712.1 aver|OJJ04492.1

Group11830: afla|EED45699.1 anid|CBF89939.1 aory|EIT75576.1 asyd|OJJ62112.1 aver|OJI96220.1

Group11831: afla|EED45702.1 aory|EIT75839.1 ater|EAU31907.1 awen|OJJ29761.1 azon|OJJ50387.1

Group11832: afla|EED45778.1 aory|EIT75841.1 asyd|OJJ60500.1 aver|OJI99820.1 awen|OJJ36013.1

Group11833: afla|EED45810.1 agla|OJJ89200.1 aory|EIT75390.1 aver|OJI96467.1 awen|OJJ40811.1

Group11834: afla|EED46052.1 aory|EIT83481.1 asyd|OJJ55010.1 aver|OJI99609.1 azon|OJJ45209.1

Group11835: afla|EED46076.1 aory|EIT83474.1 asyd|OJJ56875.1 ater|EAU30975.1 aver|OJI99559.1

Group11836: afla|EED46218.1 aory|EIT76420.1 asyd|OJJ56438.1 aver|OJJ06282.1 azon|OJJ48040.1

Group11837: afla|EED46362.1 aory|EIT81177.1 asyd|OJJ59950.1 aver|OJJ02339.1 awen|OJJ40929.1

Group11838: afla|EED46385.1 agla|OJJ78982.1 aluc|OJZ85543.1 aory|EIT72373.1 aver|OJI99260.1

Group11839: afla|EED46573.1 afum|EAL86977.1 anid|CBF76642.1 aory|EIT77846.1 asyd|OJJ61202.1

Group11840: afla|EED46873.1 aory|EIT72685.1 asyd|OJJ62345.1 ater|EAU32997.1 aver|OJI96450.1

Group11841: afla|EED46882.1 aory|EIT72695.1 asyd|OJJ52080.1 ater|EAU29585.1 aver|OJJ03022.1

Group11842: afla|EED47079.1 anid|CBF89959.1 aory|EIT72483.1 asyd|OJJ54597.1 aver|OJJ07555.1

Group11843: afla|EED47257.1 aory|EIT79263.1 asyd|OJJ60435.1 atub|OJI87259.1 aver|OJJ02287.1

Group11844: afla|EED47471.1 agla|OJJ78721.1 aory|EIT79952.1 asyd|OJJ61786.1 aver|OJI95886.1

Group11845: afla|EED47519.1 aory|EIT74561.1 asyd|OJJ58132.1 aver|OJJ04273.1 awen|OJJ31322.1

Group11846: awen|OJJ31280.1 agla|OJJ85293.1 akaw|GAA87343.1 aluc|OJZ89671.1 atub|OJI85129.1

Group11847: afla|EED47576.1 aory|EIT74523.1 asyd|OJJ65246.1 ater|EAU29464.1 aver|OJI99398.1

Group11848: afla|EED47689.1 awen|OJJ34121.1 akaw|GAA88330.1 aluc|OJZ82330.1 ater|EAU34061.1

Group11849: afla|EED47695.1 agla|OJJ85098.1 aory|EIT80204.1 awen|OJJ29798.1 agla|OJJ80341.1

Group11850: ater|EAU33650.1 akaw|GAA82961.1 aluc|OJZ87542.1 atub|OJI81105.1 azon|OJJ43564.1

Group11851: afla|EED48090.1 aory|EIT76228.1 asyd|OJJ56497.1 aver|OJJ06222.1 azon|OJJ48158.1

Group11852: afla|EED48104.1 anid|CBF87341.1 aory|EIT76235.1 asyd|OJJ52802.1 aver|OJJ08779.1

Group11853: afla|EED48175.1 anid|CBF79169.1 aory|EIT83041.1 aver|OJJ07814.1 anig|EHA28264.1

Group11854: afla|EED48211.1 aory|EIT72923.1 ater|EAU33167.1 awen|OJJ41708.1 azon|OJJ42458.1

Group11855: afla|EED48244.1 anid|CBF71807.1 aory|EIT72954.1 ater|EAU32958.1 azon|OJJ46507.1

Group11856: afla|EED48280.1 afum|EAL89209.1 agla|OJJ86338.1 anid|CBF71527.1 asyd|OJJ56408.1

Group11857: afla|EED48310.1 anid|CBF79065.1 aory|EIT76293.1 asyd|OJJ56228.1 aver|OJJ05163.1

Group11858: afla|EED48311.1 anid|CBF79063.1 aory|EIT76314.1 aver|OJJ06244.1 asyd|OJJ56474.1

Group11859: afla|EED48331.1 aory|EIT76282.1 asyd|OJJ63901.1 aver|OJI97039.1 anid|CBF88681.1

Group11860: afla|EED48434.1 anid|CBF81924.1 aory|EIT81670.1 asyd|OJJ58559.1 aver|OJJ01158.1

Group11861: afla|EED48725.1 aory|EIT73267.1 asyd|OJJ56161.1 aver|OJJ05100.1 awen|OJJ39996.1

Group11862: afla|EED48861.1 aory|EIT81281.1 ater|EAU34441.1 awen|OJJ36401.1 awen|OJJ30100.1

Group11863: afla|EED49429.1 aory|EIT79459.1 akaw|GAA90294.1 aluc|OJZ83043.1 atub|OJI87907.1

Group11864: afla|EED49435.1 aluc|OJZ85524.1 anig|EHA25056.1 aory|EIT79500.1 atub|OJI80043.1

Group11865: afla|EED49576.1 aory|EIT82580.1 asyd|OJJ59635.1 ater|EAU29694.1 aver|OJJ02646.1

Group11866: afla|EED49581.1 aory|EIT82659.1 asyd|OJJ59629.1 ater|EAU29691.1 aver|OJJ02651.1

Group11867: aluc|OJZ85541.1 aory|EIT74277.1 asyd|OJJ55206.1 ater|EAU38342.1 aver|OJJ07010.1

Group11868: anid|CBF73592.1 afla|EED51112.1 aory|EIT81359.1 asyd|OJJ56219.1 aver|OJJ05151.1

Group11869: afla|EED49646.1 ater|EAU32755.1 aory|EIT81019.1 asyd|OJJ59191.1 aver|OJJ01808.1

Group11870: afla|EED49657.1 anid|CBF89998.1 aory|EIT81009.1 asyd|OJJ62079.1 aver|OJI96186.1

Group11871: afla|EED49658.1 anid|CBF84187.1 aory|EIT81006.1 asyd|OJJ57138.1 aver|OJJ03164.1

Group11872: afla|EED49689.1 agla|OJJ83406.1 anid|CBF84423.1 asyd|OJJ53546.1 aver|OJJ08207.1

Group11873: afla|EED49743.1 aory|EIT77678.1 afla|EED49744.1 anid|CBF82185.1 aory|EIT77611.1

Group11874: afla|EED49769.1 anid|CBF87979.1 aory|EIT77605.1 asyd|OJJ63491.1 aver|OJI97438.1

Group11875: afla|EED49823.1 akaw|GAA93231.1 aory|EIT77637.1 awen|OJJ33486.1 aluc|OJZ79995.1

Group11876: afla|EED50209.1 anid|CBF74144.1 aory|EIT77294.1 asyd|OJJ53918.1 aver|OJJ08016.1

Group11877: afla|EED50230.1 aory|EIT77259.1 asyd|OJJ58371.1 aver|OJJ02035.1 awen|OJJ31991.1

Group11878: afla|EED50266.1 anid|CBF84346.1 aory|EIT77293.1 asyd|OJJ55455.1 aver|OJI99494.1

Group11879: afla|EED50478.1 aory|EIT82033.1 aver|OJI96169.1 awen|OJJ37168.1 azon|OJJ50864.1

Group11880: afla|EED50960.1 aory|EIT77007.1 asyd|OJJ61376.1 aver|OJJ00703.1 awen|OJJ41772.1

Group11881: afla|EED51120.1 anid|CBF71493.1 aory|EIT81358.1 asyd|OJJ54935.1 aver|OJJ07262.1

Group11882: afla|EED51138.1 agla|OJJ88618.1 aory|EIT81357.1 asyd|OJJ54549.1 aver|OJJ07610.1

Group11883: afla|EED51152.1 anid|CBF80142.1 aory|EIT81333.1 aver|OJI99536.1 ater|EAU31007.1

Group11884: afla|EED51160.1 anid|CBF80160.1 aory|EIT81318.1 atub|OJI79835.1 aver|OJI99517.1

Group11885: afla|EED51493.1 aory|EIT80757.1 afla|EED51494.1 agla|OJJ80050.1 awen|OJJ33324.1

Group11886: afla|EED51559.1 aory|EIT80591.1 asyd|OJJ53064.1 ater|EAU33177.1 aver|OJJ03781.1

Group11887: afla|EED51638.1 aory|EIT80492.1 asyd|OJJ63610.1 aver|OJI97327.1 awen|OJJ32918.1

Group11888: afla|EED51645.1 aory|EIT80490.1 ater|EAU33075.1 azon|OJJ42229.1 aver|OJJ02052.1

Group11889: afla|EED51660.1 aory|EIT80764.1 asyd|OJJ64003.1 aver|OJI98092.1 asyd|OJJ54855.1

Group11890: afla|EED51720.1 aory|EIT80575.1 asyd|OJJ54175.1 ater|EAU30767.1 aver|OJJ08564.1

Group11891: afla|EED51726.1 akaw|GAA93164.1 aluc|OJZ83023.1 aory|EIT80422.1 aver|OJJ07427.1

Group11892: afla|EED51756.1 aory|EIT80424.1 asyd|OJJ63601.1 ater|EAU29426.1 aver|OJI97335.1

Group11893: afla|EED51833.1 akaw|GAA87143.1 aluc|OJZ84294.1 atub|OJI89775.1 aver|OJI95766.1

Group11894: afla|EED52071.1 anig|EHA27863.1 aory|EIT72307.1 asyd|OJJ65089.1 aver|OJI99215.1

Group11895: afla|EED52151.1 anid|CBF83081.1 aory|EIT74369.1 asyd|OJJ61481.1 aver|OJJ00823.1

Group11896: afla|EED52183.1 anid|CBF90370.1 aory|EIT74342.1 ater|EAU33485.1 azon|OJJ43603.1

Group11897: afla|EED52207.1 aory|EIT74361.1 asyd|OJJ58612.1 ater|EAU37642.1 aver|OJJ01218.1

Group11898: afla|EED52300.1 anig|EHA25381.1 aory|EIT74422.1 ater|EAU37312.1 atub|OJI80428.1

Group11899: afla|EED52384.1 aory|EIT83311.1 afla|EED52517.1 aory|EIT82810.1 awen|OJJ33739.1

Group11900: agla|OJJ83635.1 aluc|OJZ86817.1 atub|OJI83700.1 akaw|GAA84854.1 aver|OJJ04861.1

Group11901: afla|EED52623.1 akaw|GAA82380.1 aluc|OJZ91741.1 aory|EIT74196.1 atub|OJI86199.1

Group11902: afla|EED53052.1 aory|EIT74776.1 asyd|OJJ57577.1 anid|CBF75853.1 aver|OJJ05603.1

Group11903: afla|EED53129.1 aory|EIT75192.1 asyd|OJJ54657.1 ater|EAU33824.1 aver|OJJ07502.1

Group11904: afla|EED53199.1 agla|OJJ89089.1 aory|EIT74875.1 asyd|OJJ55201.1 awen|OJJ32919.1

Group11905: afla|EED53336.1 agla|OJJ84992.1 aory|EIT74873.1 ater|EAU30433.1 awen|OJJ31390.1

Group11906: afla|EED53409.1 asyd|OJJ54988.1 aver|OJJ07213.1 anid|CBF71416.1 ater|EAU34394.1

Group11907: afla|EED53693.1 anid|CBF83124.1 aory|EIT75336.1 asyd|OJJ64576.1 aver|OJI98673.1

Group11908: afla|EED53766.1 aory|EIT78434.1 akaw|GAA83259.1 aluc|OJZ83765.1 atub|OJI90136.1

Group11909: afla|EED54105.1 aory|EIT76089.1 asyd|OJJ62066.1 aver|OJI96172.1 anid|CBF90025.1

Group11910: afla|EED54187.1 agla|OJJ78606.1 aory|EIT76005.1 awen|OJJ34610.1 azon|OJJ43040.1

Group11911: afla|EED54311.1 agla|OJJ80604.1 aory|EIT76144.1 ater|EAU37851.1 awen|OJJ29576.1

Group11912: afla|EED54354.1 anid|CBF77242.1 aory|EIT76130.1 asyd|OJJ56916.1 aver|OJJ03376.1

Group11913: afla|EED55062.1 agla|OJJ80727.1 anid|CBF76079.1 aory|EIT81970.1 awen|OJJ33518.1

Group11914: afla|EED55105.1 aory|EIT81572.1 asyd|OJJ62957.1 aver|OJI98010.1 azon|OJJ49874.1

Group11915: afla|EED55109.1 aory|EIT81600.1 asyd|OJJ54953.1 ater|EAU33570.1 aver|OJJ07243.1

Group11916: afla|EED55111.1 anid|CBF75002.1 aory|EIT81625.1 asyd|OJJ60077.1 aver|OJJ04186.1

Group11917: afla|EED55150.1 agla|OJJ83986.1 aory|EIT81654.1 ater|EAU37414.1 awen|OJJ34907.1

Group11918: afla|EED55186.1 anid|CBF86702.1 aory|EIT83622.1 asyd|OJJ64741.1 aver|OJI98845.1

Group11919: ater|EAU34574.1 agla|OJJ79310.1 awen|OJJ29582.1 azon|OJJ50267.1 asyd|OJJ54280.1

Group11920: afla|EED55530.1 aory|EIT80818.1 asyd|OJJ54311.1 aver|OJJ05264.1 azon|OJJ43673.1

Group11921: afla|EED55971.1 aory|EIT76727.1 asyd|OJJ56495.1 aver|OJJ06224.1 awen|OJJ30966.1

Group11922: afla|EED56578.1 afum|EAL88808.1 agla|OJJ82557.1 aory|EIT79608.1 awen|OJJ30511.1

Group11923: afla|EED56614.1 anid|CBF80621.1 aory|EIT79658.1 asyd|OJJ54629.1 aver|OJJ07527.1

Group11924: afla|EED56667.1 anid|CBF77228.1 asyd|OJJ60222.1 awen|OJJ34637.1 aver|OJJ04046.1

Group11925: afla|EED56788.1 aory|EIT73914.1 ater|EAU38499.1 awen|OJJ36167.1 azon|OJJ47806.1

Group11926: afla|EED56790.1 asyd|OJJ54993.1 aver|OJJ07208.1 awen|OJJ31994.1 azon|OJJ46049.1

Group11927: afla|EED57035.1 aory|EIT82272.1 asyd|OJJ65363.1 ater|EAU32831.1 aver|OJI99537.1

Group11928: afla|EED57108.1 aluc|OJZ84843.1 aory|EIT74030.1 asyd|OJJ57405.1 awen|OJJ35020.1

Group11929: afla|EED57402.1 aory|EIT80141.1 ater|EAU29522.1 awen|OJJ36984.1 azon|OJJ43866.1

Group11930: afla|EED57512.1 aory|EIT81849.1 aver|OJI98682.1 azon|OJJ42241.1 asyd|OJJ64584.1

Group11931: afla|EED57520.1 aory|EIT81842.1 asyd|OJJ54648.1 aver|OJJ07514.1 azon|OJJ42441.1

Group11932: afum|EAL92631.1 asyd|OJJ53615.1 aver|OJJ05935.1 asyd|OJJ54764.1 aver|OJJ07403.1

Group11933: agla|OJJ79107.1 asyd|OJJ55394.1 aver|OJJ08263.1 aluc|OJZ80184.1 azon|OJJ48017.1

Group11934: agla|OJJ79256.1 anid|CBF77657.1 asyd|OJJ53116.1 ater|EAU31054.1 aver|OJJ03738.1

Group11935: atub|OJI89780.1 anid|CBF84139.1 asyd|OJJ62468.1 ater|EAU35753.1 aver|OJI96603.1

Group11936: agla|OJJ80979.1 anid|CBF78501.1 asyd|OJJ61884.1 ater|EAU32639.1 aver|OJI95982.1

Group11937: agla|OJJ81526.1 anid|CBF81072.1 asyd|OJJ56890.1 ater|EAU29637.1 awen|OJJ37975.1

Group11938: agla|OJJ82215.1 anig|EHA20957.1 ater|EAU29924.1 atub|OJI84688.1 awen|OJJ38718.1

Group11939: agla|OJJ83164.1 anid|CBF87407.1 asyd|OJJ52891.1 ater|EAU38998.1 aver|OJJ08862.1

Group11940: agla|OJJ83648.1 anid|CBF78632.1 asyd|OJJ65122.1 aver|OJI99250.1 awen|OJJ40566.1

Group11941: agla|OJJ84618.1 aluc|OJZ81023.1 aory|EIT74823.1 atub|OJI82560.1 awen|OJJ31265.1

Group11942: agla|OJJ85280.1 asyd|OJJ56964.1 aver|OJJ03322.1 anid|CBF73577.1 asyd|OJJ56965.1

Group11943: agla|OJJ85482.1 akaw|GAA92275.1 aluc|OJZ90169.1 atub|OJI81597.1 aver|OJI99606.1

Group11944: agla|OJJ88529.1 aluc|OJZ79622.1 aory|EIT81220.1 atub|OJI82655.1 anid|CBF83743.1

Group11945: akaw|GAA81958.1 aluc|OJZ92231.1 anig|EHA26726.1 asyd|OJJ54870.1 atub|OJI86661.1

Group11946: akaw|GAA83226.1 aluc|OJZ83726.1 anig|EHA18874.1 asyd|OJJ54660.1 atub|OJI90102.1

Group11947: akaw|GAA83699.1 aluc|OJZ91034.1 ater|EAU30331.1 atub|OJI83030.1 aver|OJJ05853.1

Group11948: aver|OJJ02026.1 anid|CBF86673.1 asyd|OJJ64776.1 ater|EAU30444.1 aver|OJI98876.1

Group11949: akaw|GAA83900.1 aluc|OJZ90795.1 asyd|OJJ55180.1 atub|OJI82784.1 aver|OJJ07027.1

Group11950: akaw|GAA84677.1 aluc|OJZ87012.1 asyd|OJJ53593.1 atub|OJI83507.1 aver|OJJ08155.1

Group11951: akaw|GAA85946.1 aluc|OJZ91116.1 anig|EHA19815.1 atub|OJI83120.1 azon|OJJ42804.1

Group11952: akaw|GAA89126.1 aluc|OJZ81712.1 anig|EHA25727.1 aory|EIT75894.1 atub|OJI88472.1

Group11953: akaw|GAA89307.1 aluc|OJZ86362.1 anig|EHA21529.1 atub|OJI84172.1 aver|OJJ01270.1

Group11954: akaw|GAA89766.1 aluc|OJZ80617.1 anig|EHA27215.1 atub|OJI87471.1 azon|OJJ48139.1

Group11955: akaw|GAA92120.1 aluc|OJZ83603.1 ater|EAU35151.1 azon|OJJ51594.1 atub|OJI89975.1

Group11956: akaw|GAA92141.1 anid|CBF87326.1 asyd|OJJ59112.1 atub|OJI79313.1 aver|OJJ01704.1

Group11957: akaw|GAA92558.1 aluc|OJZ90105.1 anig|EHA20965.1 atub|OJI84699.1 azon|OJJ45560.1

Group11958: anid|CBF69440.1 asyd|OJJ52394.1 ater|EAU38171.1 aver|OJJ01929.1 azon|OJJ43196.1

Group11959: anid|CBF69492.1 asyd|OJJ63452.1 ater|EAU33838.1 aver|OJI97480.1 ater|EAU31074.1

Group11960: anid|CBF74998.1 asyd|OJJ63341.1 aver|OJI97620.1 awen|OJJ37042.1 azon|OJJ43648.1

Group11961: anid|CBF79085.1 asyd|OJJ62952.1 ater|EAU38541.1 aver|OJJ01750.1 asyd|OJJ59160.1

Group11962: anid|CBF80082.1 asyd|OJJ58308.1 aver|OJJ02104.1 awen|OJJ31908.1 azon|OJJ50391.1

Group11963: anid|CBF82195.1 asyd|OJJ52185.1 ater|EAU29295.1 aver|OJJ05960.1 awen|OJJ34425.1

Group11964: anid|CBF83948.1 anig|EHA19749.1 asyd|OJJ61436.1 aver|OJJ00770.1 azon|OJJ43215.1

Group11965: anid|CBF83950.1 anig|EHA19750.1 asyd|OJJ61437.1 aver|OJJ00771.1 azon|OJJ43214.1

Group11966: anid|CBF87478.1 asyd|OJJ52943.1 ater|EAU31727.1 aver|OJJ08910.1 awen|OJJ35977.1

Group11967: anid|CBF88188.1 asyd|OJJ63632.1 aver|OJI97304.1 awen|OJJ35273.1 azon|OJJ49834.1

Group11968: anig|EHA25383.1 aory|EIT74367.1 ater|EAU37314.1 atub|OJI80426.1 aory|EIT74355.1

Group11969: asyd|OJJ63014.1 aver|OJI99848.1 asyd|OJJ65343.1 aver|OJI99489.1 azon|OJJ45198.1

Group11970: abra|OJJ65450.1 abra|OJJ65503.1 abra|OJJ65815.1 akaw|GAA93239.1

Group11971: abra|OJJ65455.1 abra|OJJ65567.1 abra|OJJ65524.1 aluc|OJZ80442.1

Group11972: abra|OJJ65499.1 abra|OJJ75563.1 afum|EAL84359.1 afis|EAW25690.1

Group11973: abra|OJJ65530.1 abra|OJJ65571.1 akaw|GAA93121.1 akaw|GAA93276.1

Group11974: abra|OJJ65663.1 abra|OJJ66171.1 acar|OOF93533.1 aluc|OJZ80020.1

Group11975: abra|OJJ67054.1 abra|OJJ72693.1 asyd|OJJ60495.1 aver|OJI99818.1

Group11976: abra|OJJ67724.1 agla|OJJ84234.1 atub|OJI89641.1 aluc|OJZ84432.1

Group11977: abra|OJJ75756.1 afla|EED54065.1 aluc|OJZ85789.1 akaw|GAA90438.1

Group11978: abra|OJJ70079.1 abra|OJJ70081.1 abra|OJJ70080.1 afla|EED45677.1

Group11979: acar|OOF89763.1 acar|OOF89772.1 acar|OOF89773.1 anig|EHA21431.1

Group11980: acar|OOF90488.1 acar|OOF90495.1 abra|OJJ65752.1 agla|OJJ87639.1

Group11981: acar|OOF90679.1 acar|OOF91584.1 asyd|OJJ54337.1 aver|OJJ05289.1

Group11982: acar|OOF93347.1 acar|OOF93422.1 asyd|OJJ53700.1 aver|OJJ07817.1

Group11983: acar|OOF94030.1 acar|OOF94100.1 abra|OJJ66423.1 anig|EHA28117.1

Group11984: acar|OOF94078.1 acar|OOF94148.1 abra|OJJ72265.1 azon|OJJ45046.1

Group11985: acar|OOF94625.1 acar|OOG00106.1 awen|OJJ32529.1 azon|OJJ44254.1

Group11986: acar|OOF95056.1 acar|OOF95123.1 afla|EED51489.1 aory|EIT80720.1

Group11987: acar|OOF95057.1 acar|OOF95124.1 acla|EAW12275.1 anid|CBF79677.1

Group11988: afis|EAW20337.1 afis|EAW23183.1 anid|CBF80591.1 anid|CBF86728.1

Group11989: afla|EED54186.1 afla|EED56602.1 aory|EIT75929.1 aory|EIT79673.1

Group11990: afum|EAL86280.1 afum|EAL87741.1 akaw|GAA93041.1 anig|EHA21840.1

Group11991: afum|EAL86726.1 afum|EAL87779.1 afla|EED53571.1 ater|EAU32014.1

Group11992: afum|EAL86747.1 afum|EAL86768.1 aluc|OJZ91229.1 anig|EHA21822.1

Group11993: agla|OJJ78414.1 agla|OJJ87181.1 awen|OJJ33558.1 azon|OJJ42434.1

Group11994: agla|OJJ78835.1 agla|OJJ88736.1 agla|OJJ78836.1 agla|OJJ78837.1

Group11995: agla|OJJ79277.1 agla|OJJ84151.1 asyd|OJJ52300.1 aluc|OJZ79966.1

Group11996: agla|OJJ79632.1 agla|OJJ86973.1 awen|OJJ35800.1 awen|OJJ39643.1

Group11997: agla|OJJ80316.1 agla|OJJ88819.1 agla|OJJ84494.1 afla|EED51747.1

Group11998: agla|OJJ80558.1 agla|OJJ88963.1 afla|EED48400.1 aory|EIT83045.1

Group11999: agla|OJJ81222.1 agla|OJJ84586.1 afis|EAW19758.1 agla|OJJ81890.1

Group12000: agla|OJJ82937.1 agla|OJJ82943.1 agla|OJJ82939.1 agla|OJJ82938.1

Group12001: agla|OJJ84643.1 agla|OJJ85058.1 abra|OJJ69548.1 afis|EAW17238.1

Group12002: agla|OJJ84783.1 agla|OJJ85242.1 asyd|OJJ53820.1 awen|OJJ40826.1

Group12003: agla|OJJ84794.1 agla|OJJ89484.1 agla|OJJ89485.1 agla|OJJ84924.1

Group12004: agla|OJJ85011.1 agla|OJJ85273.1 abra|OJJ68079.1 afis|EAW24646.1

Group12005: agla|OJJ86309.1 agla|OJJ89019.1 akaw|GAA92981.1 akaw|GAA93104.1

Group12006: agla|OJJ89335.1 agla|OJJ89338.1 agla|OJJ84372.1 agla|OJJ85303.1

Group12007: akaw|GAA87902.1 akaw|GAA93074.1 afla|EED44728.1 aory|EIT77793.1

Group12008: akaw|GAA87905.1 akaw|GAA93251.1 akaw|GAA93054.1 abra|OJJ65620.1

Group12009: akaw|GAA87907.1 akaw|GAA93051.1 asyd|OJJ52066.1 azon|OJJ42053.1

Group12010: akaw|GAA92610.1 akaw|GAA93007.1 afis|EAW20664.1 agla|OJJ87703.1

Group12011: akaw|GAA92936.1 akaw|GAA93100.1 akaw|GAA93315.1 ater|EAU36045.1

Group12012: akaw|GAA93180.1 akaw|GAA93249.1 aluc|OJZ79979.1 aluc|OJZ88816.1

Group12013: akaw|GAA93325.1 akaw|GAA93327.1 aluc|OJZ79910.1 aluc|OJZ80098.1

Group12014: aluc|OJZ79478.1 aluc|OJZ80166.1 aluc|OJZ80255.1 aluc|OJZ79698.1

Group12015: aluc|OJZ79496.1 aluc|OJZ80390.1 aluc|OJZ80500.1 aluc|OJZ79915.1

Group12016: aluc|OJZ79514.1 aluc|OJZ88821.1 akaw|GAA93339.1 asyd|OJJ51841.1

Group12017: aluc|OJZ79528.1 aluc|OJZ79826.1 acla|EAW11471.1 afla|EED47390.1

Group12018: aluc|OJZ79577.1 aluc|OJZ80088.1 aluc|OJZ80410.1 asyd|OJJ52327.1

Group12019: aluc|OJZ79616.1 aluc|OJZ80095.1 atub|OJI87461.1 akaw|GAA92888.1

Group12020: aluc|OJZ79668.1 aluc|OJZ88809.1 aluc|OJZ79755.1 aluc|OJZ80111.1

Group12021: aluc|OJZ79726.1 aluc|OJZ79935.1 aluc|OJZ80138.1 aluc|OJZ80230.1

Group12022: aluc|OJZ79778.1 aluc|OJZ80325.1 aluc|OJZ79799.1 aluc|OJZ80212.1

Group12023: aluc|OJZ79914.1 aluc|OJZ79932.1 aluc|OJZ80387.1 aluc|OJZ80132.1

Group12024: aluc|OJZ80114.1 aluc|OJZ80449.1 aluc|OJZ80482.1 asyd|OJJ52328.1

Group12025: aluc|OJZ80125.1 aluc|OJZ88795.1 anig|EHA20892.1 asyd|OJJ51844.1

Group12026: aluc|OJZ80208.1 aluc|OJZ80427.1 aluc|OJZ79856.1 abra|OJJ65562.1

Group12027: aluc|OJZ80244.1 aluc|OJZ80431.1 akaw|GAA93345.1 atub|OJI80387.1

Group12028: anid|CBF70185.1 anid|CBF78519.1 afla|EED55118.1 aory|EIT81628.1

Group12029: anid|CBF71474.1 anid|CBF75356.1 anid|CBF78886.1 agla|OJJ89171.1

Group12030: anid|CBF75354.1 anid|CBF78887.1 ater|EAU34810.1 agla|OJJ78411.1

Group12031: anid|CBF76137.1 anid|CBF89568.1 asyd|OJJ62338.1 aver|OJI96443.1

Group12032: anid|CBF76145.1 anid|CBF84592.1 anid|CBF86034.1 awen|OJJ33509.1

Group12033: anid|CBF84381.1 anid|CBF89336.1 afis|EAW17051.1 awen|OJJ33549.1

Group12034: anig|EHA28193.1 anid|CBF69487.1 aver|OJJ01853.1 asyd|OJJ59243.1

Group12035: asyd|OJJ51842.1 asyd|OJJ52214.1 asyd|OJJ52137.1 afis|EAW22018.1

Group12036: asyd|OJJ51897.1 asyd|OJJ52288.1 asyd|OJJ51994.1 asyd|OJJ52133.1

Group12037: asyd|OJJ51945.1 asyd|OJJ51999.1 asyd|OJJ52122.1 ater|EAU35389.1

Group12038: asyd|OJJ52014.1 asyd|OJJ53701.1 afis|EAW25566.1 aver|OJJ07819.1

Group12039: asyd|OJJ52030.1 asyd|OJJ53580.1 anid|CBF76225.1 aver|OJI98041.1

Group12040: asyd|OJJ52168.1 asyd|OJJ52217.1 abra|OJJ65495.1 acla|EAW10811.1

Group12041: asyd|OJJ52433.1 asyd|OJJ61678.1 asyd|OJJ59286.1 aver|OJJ04341.1

Group12042: asyd|OJJ52491.1 asyd|OJJ54423.1 asyd|OJJ54421.1 asyd|OJJ53822.1

Group12043: asyd|OJJ52934.1 asyd|OJJ62631.1 aver|OJI96746.1 aver|OJJ08901.1

Group12044: asyd|OJJ55073.1 asyd|OJJ57666.1 aver|OJJ07135.1 anid|CBF69565.1

Group12045: asyd|OJJ62610.1 asyd|OJJ63015.1 azon|OJJ42120.1 agla|OJJ85425.1

Group12046: ater|EAU31389.1 ater|EAU36655.1 asyd|OJJ54720.1 aver|OJJ07448.1

Group12047: ater|EAU31436.1 ater|EAU35967.1 afla|EED47221.1 aory|EIT79302.1

Group12048: ater|EAU34021.1 ater|EAU37626.1 asyd|OJJ57223.1 aver|OJJ03064.1

Group12049: aver|OJI96458.1 aver|OJI97961.1 awen|OJJ29735.1 ater|EAU39428.1

Group12050: aver|OJI96729.1 aver|OJJ07512.1 afla|EED45673.1 aory|EIT75609.1

Group12051: aver|OJI98278.1 aver|OJJ07749.1 abra|OJJ74635.1 asyd|OJJ54868.1

Group12052: aver|OJJ00699.1 aver|OJJ02112.1 asyd|OJJ61372.1 awen|OJJ36386.1

Group12053: aver|OJJ01960.1 aver|OJJ07459.1 acar|OOF94959.1 afla|EED53665.1

Group12054: awen|OJJ29570.1 awen|OJJ29596.1 awen|OJJ29619.1 awen|OJJ29634.1

Group12055: awen|OJJ29580.1 awen|OJJ32581.1 awen|OJJ29609.1 abra|OJJ65511.1

Group12056: awen|OJJ29821.1 awen|OJJ31315.1 agla|OJJ84599.1 azon|OJJ42606.1

Group12057: awen|OJJ29982.1 awen|OJJ30082.1 awen|OJJ29981.1 afla|EED49718.1

Group12058: awen|OJJ30179.1 awen|OJJ30184.1 awen|OJJ31476.1 acar|OOF99002.1

Group12059: awen|OJJ30183.1 awen|OJJ36369.1 ater|EAU29248.1 asyd|OJJ57921.1

Group12060: awen|OJJ30788.1 awen|OJJ31693.1 afla|EED55298.1 aory|EIT83596.1

Group12061: awen|OJJ30821.1 awen|OJJ37783.1 afum|EAL84796.1 azon|OJJ45547.1

Group12062: awen|OJJ32288.1 awen|OJJ37493.1 awen|OJJ39605.1 acar|OOF96541.1

Group12063: awen|OJJ32517.1 awen|OJJ36006.1 abra|OJJ77882.1 azon|OJJ43500.1

Group12064: awen|OJJ34405.1 awen|OJJ38782.1 afis|EAW20916.1 azon|OJJ43629.1

Group12065: awen|OJJ37436.1 awen|OJJ37812.1 agla|OJJ86965.1 aver|OJJ07576.1

Group12066: azon|OJJ42089.1 azon|OJJ45168.1 asyd|OJJ57262.1 aver|OJJ03401.1

Group12067: azon|OJJ42306.1 azon|OJJ44448.1 afla|EED47777.1 aory|EIT77154.1

Group12068: azon|OJJ42403.1 azon|OJJ50426.1 azon|OJJ43600.1 azon|OJJ42643.1

Group12069: azon|OJJ42585.1 azon|OJJ45996.1 asyd|OJJ53498.1 aver|OJJ08247.1

Group12070: azon|OJJ42781.1 azon|OJJ46443.1 afla|EED47659.1 aory|EIT80208.1

Group12071: azon|OJJ43528.1 azon|OJJ44455.1 azon|OJJ50088.1 azon|OJJ45123.1

Group12072: azon|OJJ45268.1 azon|OJJ45309.1 azon|OJJ45693.1 ater|EAU36691.1

Group12073: azon|OJJ46178.1 azon|OJJ50239.1 aory|EIT77223.1 aver|OJJ03856.1

Group12074: azon|OJJ50228.1 azon|OJJ51244.1 acar|OOF90576.1 anig|EHA21042.1

Group12075: anid|CBF69387.1 anid|CBF79009.1 anid|CBF84587.1 anid|CBF84335.1

Group12076: awen|OJJ31896.1 awen|OJJ31899.1 awen|OJJ31898.1 awen|OJJ31912.1

Group12077: agla|OJJ78542.1 agla|OJJ86873.1 agla|OJJ80124.1 agla|OJJ88958.1

Group12078: agla|OJJ79465.1 agla|OJJ87519.1 agla|OJJ82006.1 agla|OJJ89694.1

Group12079: agla|OJJ79774.1 agla|OJJ80630.1 agla|OJJ85469.1 agla|OJJ85034.1

Group12080: agla|OJJ83779.1 agla|OJJ83780.1 agla|OJJ83790.1 agla|OJJ83789.1

Group12081: agla|OJJ89404.1 agla|OJJ89405.1 agla|OJJ89406.1 agla|OJJ89407.1

Group12082: aluc|OJZ79471.1 aluc|OJZ79552.1 aluc|OJZ80341.1 aluc|OJZ79814.1

Group12083: aluc|OJZ79522.1 aluc|OJZ80415.1 aluc|OJZ79873.1 aluc|OJZ88835.1

Group12084: aluc|OJZ79527.1 aluc|OJZ79768.1 aluc|OJZ80044.1 aluc|OJZ88811.1

Group12085: aluc|OJZ79556.1 aluc|OJZ79757.1 aluc|OJZ79666.1 aluc|OJZ79893.1

Group12086: aluc|OJZ79588.1 aluc|OJZ79732.1 aluc|OJZ80062.1 aluc|OJZ79702.1

Group12087: aluc|OJZ79764.1 aluc|OJZ79912.1 aluc|OJZ80496.1 aluc|OJZ81122.1

Group12088: abra|OJJ65423.1 akaw|BAL04884.1 aory|AFM82511.1 azon|OJJ41969.1

Group12089: abra|OJJ65432.1 akaw|BAL04876.1 aluc|OJZ79481.1 aory|AFM82513.1

Group12090: akaw|GAA92350.1 anig|EHA17688.1 akaw|GAA93097.1 aluc|OJZ80224.1

Group12091: abra|OJJ65581.1 asyd|OJJ60475.1 aver|OJI99795.1 awen|OJJ30890.1

Group12092: abra|OJJ65595.1 akaw|GAA92337.1 aluc|OJZ80456.1 asyd|OJJ52119.1

Group12093: afum|EAL88797.1 afis|EAW20720.1 asyd|OJJ54640.1 aver|OJJ07517.1

Group12094: abra|OJJ65596.1 akaw|GAA92336.1 aluc|OJZ80457.1 asyd|OJJ52118.1

Group12095: abra|OJJ65601.1 agla|OJJ80066.1 asyd|OJJ52028.1 aver|OJI98896.1

Group12096: abra|OJJ65642.1 acar|OOF97539.1 asyd|OJJ60383.1 awen|OJJ32712.1

Group12097: abra|OJJ65656.1 anid|CBF87109.1 asyd|OJJ60336.1 aver|OJI99681.1

Group12098: abra|OJJ65690.1 atub|OJI83490.1 aver|OJI96727.1 awen|OJJ30912.1

Group12099: afum|EBA27227.1 acar|OOF91401.1 akaw|GAA84023.1 azon|OJJ49807.1

Group12100: abra|OJJ65713.1 agla|OJJ84755.1 akaw|GAA93053.1 awen|OJJ29561.1

Group12101: anig|EHA21376.1 aluc|OJZ82960.1 atub|OJI89192.1 awen|OJJ40824.1

Group12102: abra|OJJ65759.1 afum|EAL84942.1 ater|EAU31474.1 awen|OJJ30961.1

Group12103: abra|OJJ65805.1 aluc|OJZ82065.1 anig|EHA22487.1 atub|OJI84623.1

Group12104: abra|OJJ65866.1 aluc|OJZ90174.1 anig|EHA23673.1 atub|OJI81602.1

Group12105: abra|OJJ65893.1 akaw|GAA86012.1 aluc|OJZ90239.1 atub|OJI81668.1

Group12106: abra|OJJ65927.1 aluc|OJZ90204.1 anig|EHA23701.1 atub|OJI81633.1

Group12107: abra|OJJ65979.1 akaw|GAA90852.1 aluc|OJZ82981.1 atub|OJI89214.1

Group12108: abra|OJJ66016.1 akaw|GAA92451.1 aluc|OJZ81567.1 atub|OJI88618.1

Group12109: abra|OJJ66029.1 atub|OJI87437.1 abra|OJJ76717.1 awen|OJJ38001.1

Group12110: abra|OJJ66033.1 atub|OJI87434.1 abra|OJJ67821.1 ater|EAU31099.1

Group12111: abra|OJJ66090.1 anid|CBF78152.1 asyd|OJJ59152.1 aver|OJJ01742.1

Group12112: abra|OJJ66101.1 akaw|GAA92667.1 aluc|OJZ90121.1 atub|OJI84674.1

Group12113: azon|OJJ44973.1 anid|CBF73584.1 asyd|OJJ56214.1 aver|OJJ05147.1

Group12114: anid|CBF84391.1 afla|EED48183.1 aory|EIT72918.1 aver|OJJ08168.1

Group12115: abra|OJJ66181.1 akaw|GAA93311.1 aluc|OJZ81145.1 aluc|OJZ80118.1

Group12116: abra|OJJ66194.1 akaw|GAA89094.1 anig|EHA25696.1 atub|OJI88433.1

Group12117: abra|OJJ66224.1 acar|OOF92074.1 anig|EHA25665.1 asyd|OJJ60327.1

Group12118: abra|OJJ66225.1 aluc|OJZ80530.1 anig|EHA25664.1 atub|OJI88400.1

Group12119: abra|OJJ66267.1 aluc|OJZ85746.1 anig|EHA28104.1 atub|OJI79531.1

Group12120: awen|OJJ31447.1 akaw|GAA84548.1 aluc|OJZ89280.1 anig|EHA23284.1

Group12121: abra|OJJ66276.1 afum|EAL85123.1 anig|EHA23657.1 atub|OJI89946.1

Group12122: abra|OJJ66303.1 afla|EED47881.1 aluc|OJZ80190.1 asyd|OJJ62690.1

Group12123: abra|OJJ66337.1 aluc|OJZ83132.1 anig|EHA28192.1 atub|OJI87803.1

Group12124: abra|OJJ66426.1 akaw|GAA90290.1 aluc|OJZ83048.1 atub|OJI87900.1

Group12125: anid|CBF89343.1 ater|EAU29255.1 akaw|GAA82057.1 aluc|OJZ92123.1

Group12126: abra|OJJ66780.1 akaw|GAA89715.1 aluc|OJZ80669.1 atub|OJI87523.1

Group12127: abra|OJJ66816.1 akaw|GAA90454.1 aluc|OJZ85770.1 atub|OJI79555.1

Group12128: abra|OJJ66904.1 aluc|OJZ85679.1 anig|EHA24862.1 atub|OJI79469.1

Group12129: abra|OJJ66939.1 afla|EED57211.1 aory|EIT74028.1 atub|OJI79434.1

Group12130: acla|EAW12298.1 afis|EAW16933.1 agla|OJJ86890.1 afum|EAL86458.1

Group12131: abra|OJJ67039.1 akaw|GAA93111.1 anig|EHA25080.1 aluc|OJZ80592.1

Group12132: abra|OJJ67040.1 ater|EAU34935.1 afis|EAW17158.1 agla|OJJ88355.1

Group12133: abra|OJJ67072.1 akaw|GAA91083.1 aluc|OJZ87900.1 atub|OJI80751.1

Group12134: azon|OJJ46831.1 afla|EED55084.1 aory|EIT81591.1 ater|EAU35954.1

Group12135: abra|OJJ67098.1 akaw|GAA91063.1 aluc|OJZ87875.1 atub|OJI80774.1

Group12136: abra|OJJ67123.1 aory|EIT78609.1 atub|OJI80795.1 awen|OJJ32528.1

Group12137: acar|OOF99013.1 afla|EED53375.1 aory|EIT74863.1 awen|OJJ39794.1

Group12138: abra|OJJ67188.1 acar|OOF98375.1 anig|EHA20268.1 aory|EIT80257.1

Group12139: abra|OJJ67218.1 asyd|OJJ52203.1 ater|EAU37937.1 aver|OJI99452.1

Group12140: abra|OJJ67219.1 ater|EAU37938.1 aver|OJI99453.1 anid|CBF77069.1

Group12141: abra|OJJ67243.1 acla|EAW13444.1 ater|EAU33582.1 atub|OJI80906.1

Group12142: awen|OJJ33399.1 asyd|OJJ58504.1 ater|EAU38213.1 aver|OJJ01105.1

Group12143: abra|OJJ67416.1 anig|EHA25168.1 atub|OJI80650.1 azon|OJJ43512.1

Group12144: awen|OJJ29833.1 afla|EED46100.1 aory|EIT76358.1 afla|EED46101.1

Group12145: awen|OJJ30909.1 acla|EAW07065.1 akaw|GAA87319.1 anig|EHA19568.1

Group12146: abra|OJJ67590.1 akaw|GAA89109.1 anig|EHA25709.1 atub|OJI88450.1

Group12147: abra|OJJ67591.1 akaw|GAA89110.1 atub|OJI88451.1 aver|OJJ05785.1

Group12148: abra|OJJ67592.1 akaw|GAA89111.1 anig|EHA25710.1 atub|OJI88452.1

Group12149: abra|OJJ67594.1 akaw|GAA89113.1 anig|EHA25712.1 atub|OJI88454.1

Group12150: abra|OJJ67632.1 akaw|GAA89152.1 aluc|OJZ81684.1 atub|OJI88497.1

Group12151: abra|OJJ67650.1 aluc|OJZ81667.1 anig|EHA25776.1 atub|OJI88516.1

Group12152: abra|OJJ67658.1 akaw|GAA85195.1 aluc|OJZ90550.1 atub|OJI81972.1

Group12153: abra|OJJ67733.1 akaw|GAA92782.1 aluc|OJZ82701.1 atub|OJI88934.1

Group12154: abra|OJJ67760.1 acar|OOF94973.1 aluc|OJZ81573.1 atub|OJI88608.1

Group12155: abra|OJJ67809.1 acar|OOF94199.1 aluc|OJZ90849.1 atub|OJI80340.1

Group12156: abra|OJJ67811.1 aluc|OJZ81262.1 ater|EAU29357.1 aver|OJJ07825.1

Group12157: abra|OJJ67820.1 anig|EHA20934.1 ater|EAU31098.1 aver|OJJ07619.1

Group12158: abra|OJJ67835.1 afum|EAL87811.1 atub|OJI89150.1 awen|OJJ41604.1

Group12159: abra|OJJ67855.1 akaw|GAA91861.1 aluc|OJZ82037.1 atub|OJI84592.1

Group12160: abra|OJJ67886.1 aluc|OJZ81411.1 atub|OJI91617.1 akaw|GAA88139.1

Group12161: abra|OJJ67934.1 akaw|GAA91262.1 aluc|OJZ82806.1 atub|OJI89037.1

Group12162: anid|CBF80171.1 afla|EED51161.1 aory|EIT81309.1 aver|OJI99523.1

Group12163: anid|CBF71799.1 asyd|OJJ64765.1 aver|OJI98865.1 awen|OJJ33438.1

Group12164: agla|OJJ78806.1 anid|CBF88783.1 asyd|OJJ63994.1 aver|OJI96958.1

Group12165: abra|OJJ68051.1 akaw|GAA90904.1 aluc|OJZ82922.1 anig|EHA21345.1

Group12166: abra|OJJ68052.1 akaw|GAA90903.1 aluc|OJZ82923.1 anig|EHA21346.1

Group12167: abra|OJJ68067.1 afla|EED47513.1 aory|EIT74557.1 aver|OJJ07731.1

Group12168: abra|OJJ68137.1 akaw|GAA92380.1 aluc|OJZ83239.1 anig|EHA25842.1

Group12169: abra|OJJ68138.1 akaw|GAA92379.1 aluc|OJZ83238.1 atub|OJI87698.1

Group12170: azon|OJJ50303.1 agla|OJJ84356.1 afla|EED49719.1 aory|EIT77707.1

Group12171: abra|OJJ68150.1 agla|OJJ87675.1 asyd|OJJ52382.1 aver|OJJ01941.1

Group12172: abra|OJJ68153.1 akaw|GAA90123.1 aluc|OJZ83227.1 atub|OJI87709.1

Group12173: abra|OJJ68165.1 acar|OOF90662.1 anig|EHA18925.1 atub|OJI90155.1

Group12174: abra|OJJ68222.1 akaw|GAA83222.1 aluc|OJZ83723.1 atub|OJI90098.1

Group12175: abra|OJJ68241.1 acar|OOG00393.1 atub|OJI82859.1 awen|OJJ29876.1

Group12176: abra|OJJ68243.1 agla|OJJ79094.1 asyd|OJJ65316.1 aver|OJI96899.1

Group12177: abra|OJJ68260.1 aluc|OJZ83705.1 anig|EHA18855.1 atub|OJI90077.1

Group12178: abra|OJJ68269.1 akaw|GAA91933.1 aluc|OJZ82689.1 atub|OJI88921.1

Group12179: abra|OJJ68276.1 akaw|GAA83194.1 aluc|OJZ83694.1 atub|OJI90066.1

Group12180: abra|OJJ68279.1 akaw|GAA83193.1 aluc|OJZ83692.1 atub|OJI90064.1

Group12181: abra|OJJ68346.1 akaw|GAA92113.1 aluc|OJZ83612.1 atub|OJI89985.1

Group12182: abra|OJJ68404.1 aory|EIT73532.1 awen|OJJ38103.1 azon|OJJ42686.1

Group12183: abra|OJJ68405.1 asyd|OJJ65107.1 aver|OJI99234.1 awen|OJJ38104.1

Group12184: abra|OJJ68408.1 ater|EAU31646.1 aver|OJI96707.1 awen|OJJ39972.1

Group12185: abra|OJJ68551.1 akaw|GAA91302.1 aluc|OJZ82768.1 atub|OJI88999.1

Group12186: agla|OJJ83094.1 akaw|GAA87849.1 aluc|OJZ85489.1 aver|OJJ05536.1

Group12187: azon|OJJ50789.1 anid|CBF84430.1 asyd|OJJ53638.1 aver|OJJ01168.1

Group12188: abra|OJJ68628.1 aluc|OJZ81927.1 anig|EHA22322.1 atub|OJI84479.1

Group12189: anig|EHA28294.1 akaw|GAA88798.1 aluc|OJZ82481.1 atub|OJI87957.1

Group12190: aver|OJJ05253.1 agla|OJJ80152.1 asyd|OJJ56241.1 asyd|OJJ54307.1

Group12191: abra|OJJ68718.1 anig|EHA22406.1 aluc|OJZ81811.1 atub|OJI84383.1

Group12192: abra|OJJ68882.1 akaw|GAA90374.1 aluc|OJZ85858.1 atub|OJI79644.1

Group12193: abra|OJJ68896.1 anig|EHA24709.1 asyd|OJJ62624.1 ater|EAU33937.1

Group12194: abra|OJJ68940.1 aluc|OJZ85925.1 anig|EHA24671.1 atub|OJI79715.1

Group12195: abra|OJJ69046.1 akaw|GAA90575.1 aluc|OJZ86032.1 atub|OJI79820.1

Group12196: abra|OJJ69069.1 akaw|GAA90600.1 aluc|OJZ86058.1 atub|OJI79855.1

Group12197: abra|OJJ69295.1 aluc|OJZ82124.1 anig|EHA28608.1 atub|OJI88314.1

Group12198: afis|EAW23683.1 afum|EBA27300.1 ater|EAU32096.1 aver|OJI97352.1

Group12199: afum|EAL85009.1 acar|OOF91888.1 ater|EAU33735.1 agla|OJJ83850.1

Group12200: abra|OJJ69527.1 aluc|OJZ82367.1 anig|EHA28398.1 atub|OJI88075.1

Group12201: abra|OJJ69546.1 afis|EAW17239.1 agla|OJJ84360.1 anig|EHA19681.1

Group12202: abra|OJJ69562.1 aluc|OJZ82388.1 anig|EHA28375.1 atub|OJI88053.1

Group12203: abra|OJJ69598.1 aver|OJJ08325.1 awen|OJJ38316.1 asyd|OJJ53422.1

Group12204: abra|OJJ69604.1 asyd|OJJ52865.1 aver|OJJ08846.1 azon|OJJ46011.1

Group12205: ater|EAU36838.1 acla|EAW15300.1 afis|EAW25737.1 anid|CBF69359.1

Group12206: abra|OJJ69670.1 akaw|GAA88794.1 aluc|OJZ82477.1 anig|EHA28298.1

Group12207: abra|OJJ69671.1 akaw|GAA88795.1 aluc|OJZ82478.1 anig|EHA28297.1

Group12208: abra|OJJ69710.1 akaw|GAA90160.1 aluc|OJZ83187.1 atub|OJI87752.1

Group12209: anid|CBF80602.1 acar|OOF95026.1 aver|OJJ07474.1 asyd|OJJ54700.1

Group12210: aver|OJJ04448.1 anid|CBF80288.1 asyd|OJJ61491.1 aver|OJJ00832.1

Group12211: abra|OJJ69767.1 akaw|GAA87886.1 aluc|OJZ85531.1 atub|OJI80037.1

Group12212: asyd|OJJ56551.1 azon|OJJ51771.1 agla|OJJ85390.1 aver|OJJ06171.1

Group12213: awen|OJJ41949.1 asyd|OJJ61040.1 ater|EAU38818.1 azon|OJJ46802.1

Group12214: abra|OJJ69847.1 akaw|GAA87811.1 aluc|OJZ85446.1 atub|OJI80121.1

Group12215: abra|OJJ69853.1 akaw|GAA87805.1 aluc|OJZ85440.1 atub|OJI80128.1

Group12216: abra|OJJ69883.1 atub|OJI80167.1 akaw|GAA87775.1 aluc|OJZ85410.1

Group12217: awen|OJJ33592.1 afla|EED55228.1 aory|EIT83513.1 azon|OJJ43670.1

Group12218: afla|EED52111.1 anid|CBF80516.1 asyd|OJJ65294.1 aver|OJJ01916.1

Group12219: afum|EAL84424.1 afis|EAW20649.1 afla|EED50438.1 azon|OJJ50344.1

Group12220: agla|OJJ78862.1 aver|OJI99695.1 awen|OJJ34612.1 asyd|OJJ60357.1

Group12221: abra|OJJ70100.1 agla|OJJ88106.1 awen|OJJ31426.1 awen|OJJ37816.1

Group12222: abra|OJJ70101.1 afis|EAW19516.1 asyd|OJJ52930.1 awen|OJJ37567.1

Group12223: abra|OJJ70127.1 akaw|GAA86643.1 aluc|OJZ85200.1 atub|OJI80401.1

Group12224: abra|OJJ70133.1 anig|EHA25398.1 asyd|OJJ60983.1 atub|OJI80410.1

Group12225: abra|OJJ70134.1 anig|EHA25397.1 asyd|OJJ60981.1 atub|OJI80411.1

Group12226: abra|OJJ70184.1 anid|CBF89258.1 anig|EHA25370.1 ater|EAU30711.1

Group12227: acar|OOF92595.1 afla|EED52024.1 afum|EAL87780.1 aory|EIT81433.1

Group12228: abra|OJJ70207.1 atub|OJI88819.1 acla|EAW08957.1 agla|OJJ80659.1

Group12229: abra|OJJ70262.1 agla|OJJ83806.1 akaw|GAA86535.1 atub|OJI80516.1

Group12230: abra|OJJ70318.1 akaw|GAA92575.1 aluc|OJZ82586.1 anig|EHA27488.1

Group12231: abra|OJJ70358.1 aluc|OJZ82620.1 anig|EHA27457.1 atub|OJI88855.1

Group12232: ater|EAU36616.1 afla|EED47543.1 agla|OJJ85092.1 aory|EIT74593.1

Group12233: abra|OJJ70400.1 acar|OOF91877.1 aluc|OJZ82659.1 atub|OJI88891.1

Group12234: abra|OJJ70416.1 anig|EHA21049.1 asyd|OJJ61785.1 aver|OJI95885.1

Group12235: abra|OJJ70826.1 acar|OOG00734.1 acla|EAW09024.1 anig|EHA17997.1

Group12236: abra|OJJ70871.1 aluc|OJZ87649.1 anig|EHA20843.1 atub|OJI80994.1

Group12237: abra|OJJ70884.1 afla|EED52123.1 agla|OJJ82944.1 aory|EIT74366.1

Group12238: agla|OJJ89557.1 aluc|OJZ85807.1 atub|OJI79593.1 akaw|GAA90422.1

Group12239: abra|OJJ71033.1 akaw|GAA82921.1 aluc|OJZ87496.1 atub|OJI81152.1

Group12240: agla|OJJ80140.1 acla|EAW08981.1 afis|EAW15428.1 afum|EAL84215.1

Group12241: abra|OJJ71220.1 acar|OOF94577.1 acla|EAW15345.1 anig|EHA20564.1

Group12242: abra|OJJ71408.1 acar|OOF97763.1 aluc|OJZ87134.1 atub|OJI81508.1

Group12243: abra|OJJ71440.1 afla|EED45386.1 agla|OJJ82298.1 awen|OJJ30854.1

Group12244: afla|EED51190.1 agla|OJJ85275.1 asyd|OJJ61033.1 aver|OJJ00351.1

Group12245: abra|OJJ71644.1 acar|OOF96360.1 aluc|OJZ86387.1 atub|OJI84146.1

Group12246: abra|OJJ71706.1 ater|EAU29929.1 akaw|GAA91388.1 awen|OJJ33559.1

Group12247: abra|OJJ71729.1 akaw|GAA85164.1 aluc|OJZ86472.1 atub|OJI84057.1

Group12248: aory|EIT75143.1 acar|OOF95276.1 afla|EED53387.1 awen|OJJ32232.1

Group12249: acla|EAW10853.1 acar|OOF95927.1 asyd|OJJ65215.1 aver|OJI99365.1

Group12250: abra|OJJ71778.1 aluc|OJZ86509.1 atub|OJI84018.1 awen|OJJ33357.1

Group12251: abra|OJJ71838.1 aluc|OJZ86570.1 anig|EHA22085.1 atub|OJI83952.1

Group12252: afis|EAW21523.1 acla|EAW09616.1 afum|EAL92544.1 anid|CBF76412.1

Group12253: abra|OJJ72031.1 acar|OOF96681.1 aluc|OJZ86769.1 atub|OJI83759.1

Group12254: afum|EAL85413.1 afis|EAW20278.1 agla|OJJ85651.1 awen|OJJ40959.1

Group12255: abra|OJJ72220.1 anig|EHA21678.1 aver|OJJ02620.1 asyd|OJJ59663.1

Group12256: aory|EIT79295.1 acar|OOF93160.1 afla|EED47161.1 ater|EAU38820.1

Group12257: agla|OJJ86669.1 akaw|GAA90291.1 aluc|OJZ83047.1 atub|OJI87901.1

Group12258: abra|OJJ72266.1 akaw|GAA86020.1 aluc|OJZ90248.1 acla|EAW10035.1

Group12259: azon|OJJ42730.1 afis|EAW21275.1 afum|EAL92290.2 ater|EAU31601.1

Group12260: abra|OJJ72455.1 akaw|GAA86191.1 aluc|OJZ90440.1 atub|OJI81870.1

Group12261: anig|EHA22067.1 akaw|GAA85058.1 aluc|OJZ86593.1 atub|OJI83930.1

Group12262: abra|OJJ72677.1 afis|EAW20000.1 afum|EAL84919.1 anig|EHA24090.1

Group12263: abra|OJJ72710.1 akaw|GAA85313.1 aluc|OJZ90674.1 atub|OJI82105.1

Group12264: abra|OJJ72730.1 akaw|GAA85336.1 aluc|OJZ90699.1 atub|OJI82125.1

Group12265: abra|OJJ72778.1 aluc|OJZ83276.1 anig|EHA24164.1 atub|OJI82180.1

Group12266: abra|OJJ72929.1 akaw|GAA85511.1 aluc|OJZ83439.1 atub|OJI82330.1

Group12267: abra|OJJ72963.1 acar|OOF96145.1 aluc|OJZ83472.1 anid|CBF81007.1

Group12268: abra|OJJ72996.1 akaw|GAA85570.1 aluc|OJZ83504.1 atub|OJI82398.1

Group12269: abra|OJJ73094.1 aluc|OJZ81331.1 anig|EHA18635.1 atub|OJI91534.1

Group12270: abra|OJJ73095.1 aluc|OJZ81330.1 anig|EHA18634.1 atub|OJI91533.1

Group12271: abra|OJJ73111.1 agla|OJJ85316.1 aluc|OJZ90610.1 atub|OJI82036.1

Group12272: abra|OJJ73163.1 agla|OJJ89559.1 ater|EAU29644.1 awen|OJJ38359.1

Group12273: abra|OJJ73293.1 akaw|GAA92710.1 aluc|OJZ88754.1 atub|OJI91324.1

Group12274: abra|OJJ73295.1 acar|OOF98236.1 agla|OJJ88327.1 awen|OJJ32708.1

Group12275: abra|OJJ73306.1 afla|EED53246.1 anig|EHA21032.1 aory|EIT74893.1

Group12276: acla|EAW09266.1 afum|EAL92896.1 awen|OJJ34171.1 afis|EAW21882.1

Group12277: abra|OJJ73347.1 akaw|GAA83135.1 aluc|OJZ88692.1 atub|OJI91272.1

Group12278: abra|OJJ73410.1 akaw|GAA83081.1 aluc|OJZ88628.1 atub|OJI91212.1

Group12279: abra|OJJ73462.1 acar|OOG00663.1 afis|EAW16338.1 akaw|GAA83036.1

Group12280: abra|OJJ73473.1 akaw|GAA83028.1 aluc|OJZ88574.1 atub|OJI91160.1

Group12281: aory|EIT74414.1 afla|EED52374.1 agla|OJJ81201.1 awen|OJJ32686.1

Group12282: azon|OJJ45023.1 afla|EED56165.1 awen|OJJ36243.1 agla|OJJ85808.1

Group12283: abra|OJJ73669.1 akaw|GAA88488.1 anig|EHA18111.1 atub|OJI90978.1

Group12284: abra|OJJ73696.1 anid|CBF87426.1 asyd|OJJ52905.1 aver|OJJ08875.1

Group12285: asyd|OJJ53741.1 aver|OJJ07854.1 awen|OJJ37845.1 acar|OOF92059.1

Group12286: azon|OJJ42669.1 aory|EIT80250.1 asyd|OJJ52363.1 aver|OJJ01983.1

Group12287: afis|EAW24660.1 afum|EAL89310.1 asyd|OJJ59254.1 aver|OJJ01864.1

Group12288: abra|OJJ73878.1 akaw|GAA89490.1 aluc|OJZ88171.1 atub|OJI90769.1

Group12289: abra|OJJ73911.1 anig|EHA17900.1 aver|OJI96669.1 awen|OJJ32659.1

Group12290: abra|OJJ73913.1 akaw|GAA89519.1 aluc|OJZ88135.1 atub|OJI90738.1

Group12291: abra|OJJ73963.1 afla|EED50152.1 aluc|OJZ88081.1 atub|OJI90690.1

Group12292: agla|OJJ86140.1 anid|CBF73932.1 asyd|OJJ53675.1 aver|OJJ07791.1

Group12293: abra|OJJ74108.1 aluc|OJZ87926.1 atub|OJI90538.1 awen|OJJ38575.1

Group12294: abra|OJJ74122.1 aluc|OJZ84286.1 anig|EHA19565.1 atub|OJI89785.1

Group12295: ater|EAU30162.1 awen|OJJ37805.1 afla|EED49587.1 aory|EIT82584.1

Group12296: agla|OJJ82688.1 awen|OJJ32588.1 afis|EAW16732.1 ater|EAU33546.1

Group12297: akaw|GAA86784.1 anig|EHA27097.1 atub|OJI84910.1 aluc|OJZ89897.1

Group12298: abra|OJJ74371.1 akaw|GAA86657.1 aluc|OJZ89753.1 atub|OJI85049.1

Group12299: abra|OJJ74403.1 afla|EED57260.1 aory|EIT73978.1 aver|OJJ07367.1

Group12300: abra|OJJ74405.1 akaw|GAA87299.1 aluc|OJZ89719.1 atub|OJI85081.1

Group12301: afum|EAL86359.1 acar|OOF90166.1 afis|EAW16789.1 azon|OJJ50670.1

Group12302: afis|EAW22189.1 acla|EAW07175.1 awen|OJJ34769.1 azon|OJJ45777.1

Group12303: abra|OJJ74632.1 akaw|GAA81977.1 anig|EHA26745.1 atub|OJI86642.1

Group12304: azon|OJJ49805.1 akaw|GAA92413.1 aluc|OJZ83271.1 asyd|OJJ53099.1

Group12305: abra|OJJ74703.1 akaw|GAA81916.1 aluc|OJZ92281.1 atub|OJI86713.1

Group12306: abra|OJJ74719.1 acar|OOF95730.1 akaw|GAA81897.1 aluc|OJZ92303.1

Group12307: abra|OJJ74733.1 akaw|GAA81886.1 aluc|OJZ92315.1 atub|OJI86745.1

Group12308: abra|OJJ74799.1 asyd|OJJ60237.1 agla|OJJ86754.1 aver|OJJ04028.1

Group12309: abra|OJJ75232.1 acar|OOF94560.1 aluc|OJZ92791.1 atub|OJI87235.1

Group12310: abra|OJJ75253.1 akaw|GAA91792.1 aluc|OJZ92812.1 atub|OJI87269.1

Group12311: abra|OJJ75332.1 acar|OOF94497.1 aluc|OJZ92896.1 atub|OJI87354.1

Group12312: asyd|OJJ56170.1 afis|EAW16915.1 anid|CBF81997.1 aver|OJJ05103.1

Group12313: asyd|OJJ62927.1 agla|OJJ88008.1 aluc|OJZ89301.1 atub|OJI85497.1

Group12314: abra|OJJ75392.1 afla|EED46229.1 acla|EAW06360.1 aory|EIT76459.1

Group12315: abra|OJJ75467.1 anig|EHA23589.1 atub|OJI85869.1 aluc|OJZ88923.1

Group12316: abra|OJJ75573.1 akaw|GAA84324.1 aluc|OJZ89021.1 atub|OJI85768.1

Group12317: abra|OJJ75594.1 akaw|GAA84345.1 aluc|OJZ89046.1 atub|OJI85741.1

Group12318: abra|OJJ75598.1 akaw|GAA84349.1 aluc|OJZ89049.1 atub|OJI85738.1

Group12319: acar|OOF95347.1 afla|EED44864.1 aory|EIT82715.1 anig|EHA22132.1

Group12320: abra|OJJ75619.1 aluc|OJZ89072.1 atub|OJI85716.1 akaw|GAA84368.1

Group12321: abra|OJJ75733.1 akaw|GAA84466.1 aluc|OJZ89184.1 atub|OJI85605.1

Group12322: abra|OJJ75761.1 aluc|OJZ89212.1 anig|EHA23346.1 atub|OJI85580.1

Group12323: ater|EAU32203.1 asyd|OJJ63062.1 atub|OJI87199.1 aver|OJI97900.1

Group12324: abra|OJJ75823.1 akaw|GAA84536.1 aluc|OJZ89269.1 atub|OJI85530.1

Group12325: abra|OJJ75838.1 akaw|GAA84551.1 aluc|OJZ89283.1 atub|OJI85516.1

Group12326: abra|OJJ75849.1 acar|OOF99376.1 anig|EHA23276.1 azon|OJJ42506.1

Group12327: abra|OJJ75852.1 akaw|GAA84557.1 aluc|OJZ89289.1 atub|OJI85510.1

Group12328: abra|OJJ75918.1 aluc|OJZ89367.1 anig|EHA23216.1 atub|OJI85431.1

Group12329: abra|OJJ75937.1 akaw|GAA84638.1 aluc|OJZ89384.1 atub|OJI85414.1

Group12330: abra|OJJ75948.1 akaw|GAA84651.1 aluc|OJZ89397.1 atub|OJI85400.1

Group12331: abra|OJJ75955.1 aluc|OJZ89404.1 anig|EHA23183.1 atub|OJI85393.1

Group12332: abra|OJJ75969.1 akaw|GAA88987.1 aluc|OJZ89416.1 atub|OJI85378.1

Group12333: anig|EHA21692.1 akaw|GAA84731.1 aluc|OJZ86955.1 atub|OJI83568.1

Group12334: abra|OJJ75994.1 aluc|OJZ89439.1 anig|EHA23146.1 atub|OJI85354.1

Group12335: abra|OJJ76032.1 aluc|OJZ89484.1 anig|EHA23110.1 atub|OJI85317.1

Group12336: anid|CBF74612.1 aver|OJJ08260.1 asyd|OJJ59625.1 aver|OJJ03071.1

Group12337: aory|EIT74283.1 afla|EED52635.1 asyd|OJJ58195.1 awen|OJJ32822.1

Group12338: abra|OJJ76163.1 aluc|OJZ92066.1 anig|EHA22993.1 atub|OJI86497.1

Group12339: abra|OJJ76209.1 akaw|GAA82149.1 aluc|OJZ92019.1 atub|OJI86454.1

Group12340: abra|OJJ76223.1 aluc|OJZ92008.1 anig|EHA22947.1 atub|OJI86443.1

Group12341: abra|OJJ76241.1 acar|OOF99486.1 aluc|OJZ91990.1 atub|OJI86425.1

Group12342: abra|OJJ76304.1 akaw|GAA82236.1 aluc|OJZ91911.1 atub|OJI86361.1

Group12343: abra|OJJ76306.1 akaw|GAA82237.1 aluc|OJZ91909.1 atub|OJI86359.1

Group12344: abra|OJJ76388.1 aluc|OJZ91825.1 atub|OJI86280.1 acar|OOF99629.1

Group12345: abra|OJJ76400.1 akaw|GAA82319.1 aluc|OJZ91813.1 atub|OJI86268.1

Group12346: aory|EIT82085.1 akaw|GAA86610.1 aluc|OJZ85161.1 atub|OJI80437.1

Group12347: abra|OJJ76586.1 akaw|GAA82490.1 aluc|OJZ91620.1 atub|OJI86073.1

Group12348: abra|OJJ76589.1 akaw|GAA82493.1 aluc|OJZ91617.1 atub|OJI86070.1

Group12349: abra|OJJ76643.1 akaw|GAA92986.1 aluc|OJZ91564.1 asyd|OJJ56045.1

Group12350: abra|OJJ76644.1 akaw|GAA92987.1 aluc|OJZ91563.1 atub|OJI86014.1

Group12351: abra|OJJ76726.1 anig|EHA28101.1 asyd|OJJ55397.1 awen|OJJ30161.1

Group12352: abra|OJJ76814.1 afla|EED44775.1 akaw|GAA87207.1 atub|OJI89860.1

Group12353: abra|OJJ76873.1 akaw|GAA85847.1 aluc|OJZ91252.1 anig|EHA28100.1

Group12354: abra|OJJ76885.1 afla|EED52848.1 aory|EIT78641.1 asyd|OJJ56895.1

Group12355: abra|OJJ77009.1 akaw|GAA87039.1 aluc|OJZ84408.1 atub|OJI89663.1

Group12356: abra|OJJ77025.1 akaw|GAA87026.1 aluc|OJZ84427.1 atub|OJI89647.1

Group12357: abra|OJJ77045.1 akaw|GAA87008.1 aluc|OJZ84448.1 atub|OJI89627.1

Group12358: abra|OJJ77060.1 akaw|GAA91853.1 aluc|OJZ84467.1 atub|OJI89609.1

Group12359: abra|OJJ77061.1 akaw|GAA86992.1 aluc|OJZ84468.1 anig|EHA27779.1

Group12360: abra|OJJ77097.1 akaw|GAA86959.1 aluc|OJZ84505.1 atub|OJI89575.1

Group12361: acla|EAW09034.1 afis|EAW22734.1 afum|EAL90461.1 agla|OJJ78520.1

Group12362: abra|OJJ77292.1 anig|EHA27978.1 atub|OJI89379.1 aluc|OJZ84703.1

Group12363: abra|OJJ77393.1 acar|OOF93261.1 aluc|OJZ84801.1 azon|OJJ50266.1

Group12364: abra|OJJ77413.1 asyd|OJJ62931.1 aver|OJI98059.1 awen|OJJ41770.1

Group12365: abra|OJJ77416.1 akaw|GAA91345.1 aluc|OJZ84828.1 atub|OJI89259.1

Group12366: abra|OJJ77427.1 anid|CBF84180.1 asyd|OJJ57157.1 aver|OJJ03156.1

Group12367: aory|EIT81327.1 afla|EED51158.1 anid|CBF80154.1 aver|OJI99514.1

Group12368: awen|OJJ29986.1 agla|OJJ86405.1 asyd|OJJ54664.1 aver|OJJ02025.1

Group12369: azon|OJJ43921.1 afis|EAW25723.1 afum|EAL84396.1 asyd|OJJ57275.1

Group12370: abra|OJJ77482.1 aluc|OJZ81055.1 anig|EHA19312.1 atub|OJI82529.1

Group12371: abra|OJJ77593.1 aluc|OJZ80943.1 anig|EHA19410.1 akaw|GAA84035.1

Group12372: abra|OJJ77611.1 acar|OOF99117.1 anig|EHA19424.1 awen|OJJ30422.1

Group12373: ater|EAU37176.1 afla|EED47697.1 aory|EIT80212.1 awen|OJJ31251.1

Group12374: afis|EAW22093.1 afla|EED50926.1 aory|EIT78978.1 ater|EAU31790.1

Group12375: abra|OJJ77711.1 afla|EED45384.1 aory|EIT82436.1 atub|OJI82763.1

Group12376: abra|OJJ77726.1 aluc|OJZ88662.1 aory|EIT83094.1 aver|OJJ05763.1

Group12377: abra|OJJ77737.1 akaw|GAA83892.1 aluc|OJZ90803.1 atub|OJI82792.1

Group12378: abra|OJJ77742.1 anig|EHA18408.1 abra|OJJ78141.1 aver|OJJ06437.1

Group12379: afla|EED51780.1 agla|OJJ87025.1 asyd|OJJ54713.1 aver|OJJ07453.1

Group12380: abra|OJJ77813.1 afis|EAW16857.1 afum|EAL86400.1 aluc|OJZ88123.1

Group12381: abra|OJJ77885.1 akaw|GAA82229.1 aluc|OJZ91920.1 ater|EAU37964.1

Group12382: abra|OJJ77930.1 akaw|GAA83733.1 aluc|OJZ90998.1 atub|OJI82988.1

Group12383: abra|OJJ77933.1 akaw|GAA83730.1 aluc|OJZ91001.1 atub|OJI82991.1

Group12384: abra|OJJ77943.1 akaw|GAA83714.1 atub|OJI83011.1 aluc|OJZ91019.1

Group12385: acar|OOF96773.1 anid|CBF74011.1 asyd|OJJ53753.1 aver|OJJ07865.1

Group12386: abra|OJJ78037.1 aluc|OJZ87777.1 ater|EAU31866.1 agla|OJJ81396.1

Group12387: abra|OJJ78043.1 akaw|GAA89418.1 aluc|OJZ88255.1 atub|OJI90853.1

Group12388: abra|OJJ78047.1 anig|EHA19819.1 acla|EAW15271.1 afis|EAW20238.1

Group12389: abra|OJJ78048.1 acar|OOF95284.1 anid|CBF86030.1 anig|EHA19820.1

Group12390: anig|EHA27897.1 akaw|GAA90740.1 aluc|OJZ84608.1 atub|OJI89472.1

Group12391: abra|OJJ78092.1 akaw|GAA85906.1 aluc|OJZ91161.1 atub|OJI83176.1

Group12392: abra|OJJ78099.1 aluc|OJZ81973.1 anig|EHA26142.1 atub|OJI79695.1

Group12393: acar|OOF91491.1 akaw|GAA90345.1 aluc|OJZ85887.1 atub|OJI79674.1

Group12394: ater|EAU33250.1 afla|EED45087.1 agla|OJJ80643.1 aory|EIT73753.1

Group12395: abra|OJJ78143.1 anid|CBF73530.1 asyd|OJJ56724.1 aver|OJJ06001.1

Group12396: abra|OJJ78167.1 agla|OJJ84481.1 ater|EAU30097.1 awen|OJJ31321.1

Group12397: agla|OJJ78789.1 awen|OJJ36114.1 asyd|OJJ64786.1 aver|OJI98898.1

Group12398: abra|OJJ78204.1 akaw|GAA89022.1 aluc|OJZ80582.1 atub|OJI88348.1

Group12399: awen|OJJ36508.1 afla|EED53594.1 aory|EIT74996.1 atub|OJI83687.1

Group12400: abra|OJJ78335.1 atub|OJI83401.1 abra|OJJ78336.1 aluc|OJZ91408.1

Group12401: acar|OOF89790.1 akaw|BAL04883.1 aory|AFM82514.1 azon|OJJ41970.1

Group12402: acar|OOF90014.1 afla|EED57084.1 agla|OJJ80260.1 aory|EIT82295.1

Group12403: acar|OOF90099.1 anid|CBF84624.1 asyd|OJJ54553.1 aver|OJI96747.1

Group12404: acar|OOF90172.1 afla|EED53602.1 aory|EIT75188.1 aver|OJI98824.1

Group12405: acar|OOF90231.1 afla|EED53290.1 aory|EIT75111.1 awen|OJJ31455.1

Group12406: acar|OOF90239.1 anid|CBF80663.1 asyd|OJJ54607.1 aver|OJJ07549.1

Group12407: acar|OOF90390.1 awen|OJJ35044.1 azon|OJJ45353.1 agla|OJJ88295.1

Group12408: acar|OOF90510.1 afla|EED47874.1 agla|OJJ85526.1 azon|OJJ48718.1

Group12409: awen|OJJ34540.1 atub|OJI86046.1 akaw|GAA92837.1 anig|EHA22624.1

Group12410: acar|OOF90570.1 acla|EAW13530.1 afla|EED45167.1 aory|EIT73590.1

Group12411: aluc|OJZ82067.1 awen|OJJ32571.1 akaw|GAA91462.1 anig|EHA19360.1

Group12412: acar|OOF90626.1 akaw|GAA90434.1 aluc|OJZ85793.1 atub|OJI79579.1

Group12413: acar|OOF90642.1 akaw|GAA84624.1 aluc|OJZ89370.1 awen|OJJ32892.1

Group12414: aver|OJJ07190.1 afis|EAW19849.1 afum|EAL85044.1 agla|OJJ81562.1

Group12415: acar|OOF90911.1 agla|OJJ87940.1 aory|EIT79034.1 afla|EED50872.1

Group12416: acar|OOF90913.1 acla|EAW11057.1 asyd|OJJ56444.1 ater|EAU37670.1

Group12417: aory|EIT73656.1 afla|EED45080.1 asyd|OJJ54725.1 aver|OJJ07442.1

Group12418: agla|OJJ81919.1 asyd|OJJ52649.1 aver|OJJ04678.1 awen|OJJ38464.1

Group12419: acar|OOF91242.1 aver|OJJ05125.1 awen|OJJ32857.1 asyd|OJJ56198.1

Group12420: acar|OOF91356.1 acla|EAW12402.1 afis|EAW16850.1 afum|EAL86396.1

Group12421: acar|OOF91378.1 anid|CBF87269.1 ater|EAU29527.1 awen|OJJ38078.1

Group12422: aver|OJJ00844.1 anid|CBF83013.1 asyd|OJJ61507.1 ater|EAU30759.1

Group12423: aver|OJJ08465.1 azon|OJJ47777.1 afla|EED47903.1 aory|EIT77245.1

Group12424: acar|OOF91729.1 anid|CBF78054.1 aver|OJJ04810.1 azon|OJJ45180.1

Group12425: acar|OOF91794.1 acla|EAW15266.1 afis|EAW17057.1 afum|EAL86553.1

Group12426: afis|EAW17028.1 aluc|OJZ79598.1 asyd|OJJ58569.1 aver|OJI98840.1

Group12427: acar|OOF91817.1 agla|OJJ79125.1 asyd|OJJ55390.1 aver|OJJ08267.1

Group12428: acar|OOF91842.1 afla|EED45903.1 aory|EIT75562.1 awen|OJJ31983.1

Group12429: acar|OOF91867.1 afis|EAW19200.1 afum|EAL93931.1 asyd|OJJ58347.1

Group12430: acar|OOF91911.1 aluc|OJZ85606.1 azon|OJJ42118.1 aluc|OJZ90141.1

Group12431: acar|OOF91926.1 afis|EAW22249.1 afum|EAL90925.1 atub|OJI82858.1

Group12432: anid|CBF78495.1 asyd|OJJ61882.1 ater|EAU32641.1 aver|OJI95980.1

Group12433: acla|EAW11046.1 afis|EAW23433.1 afla|EED49629.1 aory|EIT80998.1

Group12434: acar|OOF92452.1 afla|EED53907.1 aory|EIT78482.1 ater|EAU32763.1

Group12435: acar|OOF92538.1 anid|CBF71805.1 asyd|OJJ65350.1 aver|OJI99498.1

Group12436: acla|EAW14292.1 afis|EAW18991.1 afum|EAL93727.1 awen|OJJ31613.1

Group12437: acar|OOF92542.1 asyd|OJJ65177.1 aver|OJI99330.1 awen|OJJ39690.1

Group12438: acar|OOF92793.1 asyd|OJJ57284.1 aver|OJJ05872.1 azon|OJJ50348.1

Group12439: acar|OOF92869.1 akaw|GAA86457.1 aluc|OJZ84999.1 atub|OJI80601.1

Group12440: acar|OOF93070.1 anig|EHA19733.1 atub|OJI83028.1 awen|OJJ33370.1

Group12441: acar|OOF93174.1 afla|EED47511.1 aory|EIT74594.1 ater|EAU31933.1

Group12442: acar|OOF93176.1 acla|EAW06584.1 asyd|OJJ56443.1 aver|OJJ06278.1

Group12443: acar|OOF93298.1 anid|CBF80338.1 asyd|OJJ54838.1 aver|OJJ07323.1

Group12444: afla|EED56666.1 aory|EIT79625.1 awen|OJJ34581.1 azon|OJJ42192.1

Group12445: acar|OOF93401.1 akaw|GAA87774.1 aluc|OJZ85409.1 anig|EHA25588.1

Group12446: acar|OOF93615.1 akaw|GAA92165.1 aluc|OJZ85573.1 atub|OJI79368.1

Group12447: anig|EHA25152.1 akaw|GAA86399.1 aluc|OJZ84934.1 atub|OJI80665.1

Group12448: asyd|OJJ59768.1 ater|EAU34798.1 aver|OJJ02517.1 awen|OJJ36898.1

Group12449: agla|OJJ88964.1 ater|EAU36815.1 afla|EED45989.1 aory|EIT75588.1

Group12450: acar|OOF94005.1 acla|EAW08898.1 anid|CBF73449.1 aver|OJJ08118.1

Group12451: agla|OJJ79280.1 asyd|OJJ64891.1 ater|EAU30236.1 aver|OJI99008.1

Group12452: acar|OOF94191.1 afla|EED48648.1 aory|EIT73046.1 ater|EAU35422.1

Group12453: acar|OOF94508.1 acla|EAW15351.1 afis|EAW20362.1 afum|EAL85478.2

Group12454: acar|OOF94532.1 afis|EAW17963.1 afum|EAL91773.1 azon|OJJ42388.1

Group12455: acar|OOF94921.1 agla|OJJ79305.1 aver|OJJ03053.1 awen|OJJ36160.1

Group12456: acar|OOF95007.1 acla|EAW12212.1 afis|EAW25552.1 afum|EAL84275.1

Group12457: afum|EAL85116.1 anid|CBF82795.1 asyd|OJJ55217.1 aver|OJJ06999.1

Group12458: acar|OOF95425.1 acla|EAW07351.1 ater|EAU39514.1 azon|OJJ45605.1

Group12459: acar|OOF95541.1 afis|EAW20728.1 acla|EAW15263.1 azon|OJJ45015.1

Group12460: acla|EAW12174.1 akaw|GAA84861.1 aluc|OJZ86810.1 atub|OJI83707.1

Group12461: acar|OOF95834.1 akaw|GAA86064.1 aluc|OJZ90298.1 atub|OJI81726.1

Group12462: acar|OOF95869.1 afla|EED47529.1 agla|OJJ79290.1 awen|OJJ32584.1

Group12463: awen|OJJ39220.1 agla|OJJ86772.1 afla|EED57423.1 aory|EIT80052.1

Group12464: acar|OOF95942.1 azon|OJJ42256.1 asyd|OJJ61387.1 aver|OJJ00715.1

Group12465: acar|OOF96412.1 agla|OJJ82203.1 aory|EIT81293.1 afla|EED51118.1

Group12466: acar|OOF96447.1 asyd|OJJ64726.1 ater|EAU34580.1 aver|OJI98827.1

Group12467: acar|OOF96590.1 anid|CBF74614.1 anig|EHA27723.1 awen|OJJ29532.1

Group12468: acar|OOF96638.1 afla|EED57109.1 aory|EIT74163.1 awen|OJJ29687.1

Group12469: acar|OOF96776.1 anid|CBF73993.1 asyd|OJJ53752.1 aver|OJJ07864.1

Group12470: acar|OOF96876.1 aory|EIT75481.1 awen|OJJ38802.1 azon|OJJ43573.1

Group12471: ater|EAU32315.1 acla|EAW11534.1 afis|EAW23916.1 afum|EAL87343.1

Group12472: acar|OOF97584.1 aory|EIT73196.1 azon|OJJ50755.1 afla|EED48473.1

Group12473: ater|EAU30463.1 afla|EED50987.1 anid|CBF73678.1 aory|EIT77020.1

Group12474: acar|OOF97613.1 agla|OJJ88994.1 atub|OJI80140.1 awen|OJJ37277.1

Group12475: acar|OOF97614.1 atub|OJI80141.1 aver|OJJ05134.1 awen|OJJ35442.1

Group12476: acar|OOF97625.1 afum|EAL86146.1 agla|OJJ85955.1 awen|OJJ36071.1

Group12477: acar|OOF98017.1 acla|EAW10736.1 aory|EIT75944.1 awen|OJJ36178.1

Group12478: acar|OOF98056.1 agla|OJJ88710.1 asyd|OJJ64700.1 aver|OJI98804.1

Group12479: acar|OOF98331.1 afis|EAW21475.1 afum|EAL92495.1 agla|OJJ83522.1

Group12480: anig|EHA24934.1 akaw|GAA92177.1 aluc|OJZ85587.1 atub|OJI79382.1

Group12481: afla|EED49099.1 aory|EIT82529.1 afis|EAW23421.1 afum|EAL88048.2

Group12482: acar|OOF98853.1 akaw|GAA93109.1 aluc|OJZ80594.1 anig|EHA22188.1

Group12483: acar|OOF98854.1 aluc|OJZ80596.1 anig|EHA22186.1 awen|OJJ29869.1

Group12484: acar|OOF98976.1 akaw|GAA92125.1 aluc|OJZ83597.1 atub|OJI89969.1

Group12485: acar|OOF98986.1 aory|EIT76158.1 awen|OJJ36025.1 azon|OJJ43605.1

Group12486: acar|OOF99008.1 afla|EED45162.1 aver|OJJ06556.1 azon|OJJ50000.1

Group12487: acar|OOF99051.1 akaw|GAA83947.1 atub|OJI82738.1 aluc|OJZ80828.1

Group12488: acar|OOF99184.1 aluc|OJZ80183.1 asyd|OJJ59803.1 aver|OJJ02488.1

Group12489: acar|OOF99284.1 aluc|OJZ81086.1 atub|OJI82498.1 azon|OJJ50600.1

Group12490: acar|OOF99301.1 agla|OJJ79461.1 asyd|OJJ54303.1 atub|OJI80756.1

Group12491: afum|EAL85338.1 afis|EAW20223.1 aver|OJJ08352.1 anig|EHA25399.1

Group12492: afum|EAL88831.1 awen|OJJ37829.1 afis|EAW16203.1 agla|OJJ81274.1

Group12493: acar|OOG00065.1 aluc|OJZ82066.1 ater|EAU31588.1 awen|OJJ32570.1

Group12494: acar|OOG00092.1 akaw|GAA84508.1 aluc|OJZ89237.1 anig|EHA23329.1

Group12495: acar|OOG00126.1 acla|EAW10036.1 akaw|GAA86021.1 aluc|OJZ90249.1

Group12496: afla|EED45048.1 aory|EIT73760.1 azon|OJJ42418.1 awen|OJJ32633.1

Group12497: acla|EAW09451.1 awen|OJJ33954.1 agla|OJJ79915.1 azon|OJJ48616.1

Group12498: acar|OOG00224.1 asyd|OJJ54072.1 aver|OJJ08667.1 awen|OJJ40009.1

Group12499: acar|OOG00433.1 anid|CBF69470.1 asyd|OJJ59274.1 aver|OJJ01887.1

Group12500: acar|OOG00435.1 anid|CBF69468.1 asyd|OJJ59275.1 aver|OJJ01888.1

Group12501: acar|OOG00543.1 afla|EED47887.1 aory|EIT77233.1 aver|OJJ07415.1

Group12502: acla|EAW06296.1 afis|EAW18402.1 afum|EAL93080.1 agla|OJJ84962.1

Group12503: ater|EAU31426.1 aory|EIT75187.1 asyd|OJJ57975.1 aver|OJJ04426.1

Group12504: acla|EAW06548.1 aluc|OJZ80952.1 anig|EHA19402.1 atub|OJI82631.1

Group12505: acla|EAW06579.1 afis|EAW16687.1 afla|EED51145.1 aory|EIT81337.1

Group12506: acla|EAW06681.1 afis|EAW16084.1 afum|EAL88716.1 awen|OJJ30348.1

Group12507: anig|EHA28235.1 aver|OJJ06471.1 ater|EAU36717.1 atub|OJI87768.1

Group12508: acla|EAW07098.1 afis|EAW22110.1 afum|EAL91074.1 aver|OJJ02734.1

Group12509: acla|EAW07106.1 afis|EAW22118.1 afum|EAL91068.1 awen|OJJ34701.1

Group12510: acla|EAW07142.1 awen|OJJ37872.1 agla|OJJ79600.1 anid|CBF69373.1

Group12511: acla|EAW07214.1 afis|EAW22227.1 afum|EAL90967.1 azon|OJJ42275.1

Group12512: acla|EAW07350.1 afis|EAW22380.1 afum|EAL90811.1 awen|OJJ34955.1

Group12513: acla|EAW07368.1 afis|EAW19831.1 afum|EAL85061.1 agla|OJJ87290.1

Group12514: acla|EAW08184.1 afla|EED47032.1 agla|OJJ86375.1 asyd|OJJ60176.1

Group12515: acla|EAW08499.1 afis|EAW23544.1 afum|EAL87922.1 ater|EAU35330.1

Group12516: acla|EAW08732.1 afis|EAW15706.1 afum|EAL85641.1 awen|OJJ38399.1

Group12517: acla|EAW08774.1 afum|EBA27238.1 asyd|OJJ53503.1 awen|OJJ32666.1

Group12518: agla|OJJ81399.1 asyd|OJJ53832.1 aver|OJJ07937.1 azon|OJJ45799.1

Group12519: acla|EAW08914.1 afis|EAW15480.1 agla|OJJ80159.1 azon|OJJ43151.1

Group12520: acla|EAW08946.1 afum|EAL91100.1 akaw|GAA83286.1 awen|OJJ34618.1

Group12521: acla|EAW08952.1 afis|EAW15460.1 afum|EAL85859.1 ater|EAU30837.1

Group12522: acla|EAW08962.1 afis|EAW15443.1 afum|EAL84204.1 ater|EAU35570.1

Group12523: acla|EAW08998.1 afis|EAW22069.1 agla|OJJ78772.1 aluc|OJZ79993.1

Group12524: acla|EAW09003.1 afla|EED57115.1 aory|EIT74101.1 awen|OJJ36036.1

Group12525: acla|EAW09037.1 afis|EAW22731.1 afum|EAL90464.2 agla|OJJ78523.1

Group12526: acla|EAW09351.1 afis|EAW21816.1 afum|EAL92832.1 ater|EAU36734.1

Group12527: acla|EAW09418.1 afis|EAW21718.1 awen|OJJ33991.1 azon|OJJ48562.1

Group12528: acla|EAW09512.1 afis|EAW21620.1 afum|EAL92656.1 agla|OJJ82577.1

Group12529: acla|EAW09753.1 afis|EAW21388.1 afum|EAL92403.2 anid|CBF71411.1

Group12530: acla|EAW09794.1 afis|EAW21342.1 afum|EAL92359.1 ater|EAU35966.1

Group12531: acla|EAW09868.1 agla|OJJ87500.1 awen|OJJ36572.1 azon|OJJ44316.1

Group12532: acla|EAW10015.1 afis|EAW21039.1 afum|EAL92078.1 awen|OJJ35723.1

Group12533: acla|EAW10024.1 aory|EIT81883.1 afla|EED57488.1 afum|EAL89366.2

Group12534: acla|EAW10196.1 afla|EED53877.1 aory|EIT78437.1 ater|EAU34550.1

Group12535: acla|EAW10299.1 afis|EAW25293.1 afum|EAL89956.1 awen|OJJ29696.1

Group12536: acla|EAW10406.1 afis|EAW20556.1 afum|EAL84505.1 anid|CBF73656.1

Group12537: acla|EAW10412.1 afla|EED55128.1 asyd|OJJ52933.1 aver|OJJ08899.1

Group12538: agla|OJJ85499.1 asyd|OJJ65169.1 aver|OJI99321.1 anid|CBF80667.1

Group12539: acla|EAW10839.1 afla|EED55833.1 aory|EIT76690.1 azon|OJJ43607.1

Group12540: acla|EAW10848.1 afis|EAW20282.1 afum|EAL85416.1 agla|OJJ80881.1

Group12541: acla|EAW10876.1 afis|EAW20982.1 afum|EAL84648.2 awen|OJJ34329.1

Group12542: acla|EAW11021.1 afis|EAW20835.1 afla|EED47444.1 aory|EIT81899.1

Group12543: acla|EAW11032.1 afis|EAW23432.1 afla|EED49638.1 aory|EIT81010.1

Group12544: acla|EAW11056.1 afis|EAW20820.1 afum|EAL84809.1 ater|EAU39614.1

Group12545: acla|EAW11257.1 afis|EAW18242.1 afum|EAL87606.1 ater|EAU37479.1

Group12546: acla|EAW11532.1 afis|EAW23922.1 afum|EAL87347.1 awen|OJJ35398.1

Group12547: acla|EAW11670.1 asyd|OJJ56436.1 aver|OJJ06285.1 awen|OJJ38759.1

Group12548: acla|EAW11728.1 asyd|OJJ56132.1 aver|OJJ05073.1 azon|OJJ46136.1

Group12549: acla|EAW11776.1 anid|CBF70506.1 asyd|OJJ58576.1 aver|OJJ01180.1

Group12550: afla|EED49645.1 asyd|OJJ60260.1 ater|EAU36670.1 aver|OJJ03996.1

Group12551: acla|EAW11978.1 agla|OJJ80762.1 aver|OJJ07084.1 awen|OJJ41863.1

Group12552: acla|EAW12029.1 anid|CBF81953.1 asyd|OJJ52952.1 aver|OJJ08918.1

Group12553: acla|EAW12092.1 afla|EED48247.1 ater|EAU32454.1 azon|OJJ47354.1

Group12554: acla|EAW12127.1 afis|EAW17246.1 afum|EBA27285.1 ater|EAU38592.1

Group12555: acla|EAW12128.1 afis|EAW17249.1 afum|EAL86668.1 ater|EAU38590.1

Group12556: awen|OJJ33498.1 afla|EED46280.1 agla|OJJ84142.1 aory|EIT76348.1

Group12557: acla|EAW12185.1 afis|EAW17121.1 afum|EAL86582.1 awen|OJJ34474.1

Group12558: acla|EAW12198.1 anid|CBF69436.1 asyd|OJJ52389.1 aver|OJJ01934.1

Group12559: acla|EAW12202.1 afla|EED50214.1 aory|EIT77340.1 asyd|OJJ54578.1

Group12560: acla|EAW12215.1 afis|EAW25556.1 afum|EAL84279.2 awen|OJJ31978.1

Group12561: acla|EAW12294.1 afis|EAW16925.1 afum|EAL86450.1 ater|EAU32508.1

Group12562: acla|EAW12311.1 afis|EAW24615.1 afla|EED50193.1 afum|EAL86244.1

Group12563: acla|EAW12329.1 agla|OJJ79949.1 asyd|OJJ54495.1 aver|OJJ07685.1

Group12564: acla|EAW12342.1 afis|EAW16878.1 afum|EAL86415.1 agla|OJJ78656.1

Group12565: acla|EAW12417.1 afla|EED52219.1 anid|CBF83053.1 aory|EIT74492.1

Group12566: acla|EAW12423.1 afla|EED52221.1 anid|CBF83042.1 aory|EIT74446.1

Group12567: acla|EAW12438.1 afis|EAW16750.1 afum|EAL86323.1 agla|OJJ79212.1

Group12568: acla|EAW12448.1 afla|EED48612.1 aory|EIT74314.1 aory|EIT73183.1

Group12569: acla|EAW12598.1 afum|EAL88792.1 akaw|GAA92360.1 aluc|OJZ79830.1

Group12570: acla|EAW12670.1 agla|OJJ86842.1 ater|EAU34483.1 aver|OJJ08589.1

Group12571: acla|EAW12685.1 afis|EAW17848.1 afum|EAL91657.1 awen|OJJ41725.1

Group12572: acla|EAW12780.1 anig|EHA21241.1 asyd|OJJ58201.1 azon|OJJ48322.1

Group12573: acla|EAW12905.1 afis|EAW17783.1 afla|EED51534.1 afum|EAL91591.2

Group12574: acla|EAW12960.1 aory|EIT82286.1 awen|OJJ35545.1 azon|OJJ42179.1

Group12575: acla|EAW13122.1 afis|EAW17542.1 afum|EAL91356.1 azon|OJJ51003.1

Group12576: acla|EAW13222.1 agla|OJJ85887.1 aory|EIT73584.1 afla|EED45038.1

Group12577: acla|EAW13227.1 afis|EAW17409.1 afum|EAL91243.1 ater|EAU37361.1

Group12578: acla|EAW13228.1 afis|EAW17407.1 afum|EAL91242.1 asyd|OJJ52224.1

Group12579: anig|EHA22106.1 afla|EED45487.1 aory|EIT75569.1 awen|OJJ30902.1

Group12580: acla|EAW13447.1 afla|EED46203.1 aory|EIT76386.1 ater|EAU29432.1

Group12581: acla|EAW13449.1 afis|EAW20555.1 afum|EBA27183.1 azon|OJJ43198.1

Group12582: acla|EAW13522.1 anid|CBF87074.1 asyd|OJJ61529.1 awen|OJJ34704.1

Group12583: acla|EAW13547.1 agla|OJJ88374.1 ater|EAU29445.1 afla|EED46256.1

Group12584: acla|EAW13650.1 afis|EAW19955.1 afum|EAL84959.1 ater|EAU36871.1

Group12585: acla|EAW13705.1 afis|EAW21064.1 afum|EBA27264.1 aluc|OJZ83579.1

Group12586: afla|EED56408.1 aory|EIT82999.1 anid|CBF81958.1 ater|EAU30737.1

Group12587: acla|EAW14133.1 afis|EAW18846.1 afum|EAL93577.1 agla|OJJ83523.1

Group12588: awen|OJJ38284.1 aver|OJI97629.1 anid|CBF87706.1 asyd|OJJ63333.1

Group12589: acla|EAW14281.1 afum|EAL85155.2 asyd|OJJ62554.1 aver|OJI96680.1

Group12590: acla|EAW14510.1 afis|EAW19228.1 afum|EAL93958.1 awen|OJJ37586.1

Group12591: acla|EAW14592.1 afla|EED55209.1 anig|EHA27992.1 aory|EIT83610.1

Group12592: acla|EAW14809.1 afis|EAW24338.1 awen|OJJ32880.1 azon|OJJ48978.1

Group12593: acla|EAW14840.1 afum|EBA27256.1 ater|EAU30371.1 awen|OJJ32846.1

Group12594: acla|EAW14842.1 afis|EAW24369.1 afum|EAL86041.1 ater|EAU30369.1

Group12595: acla|EAW14924.1 azon|OJJ42508.1 afla|EED52070.1 aory|EIT72305.1

Group12596: acla|EAW14926.1 afla|EED49818.1 aory|EIT77663.1 asyd|OJJ62980.1

Group12597: acla|EAW14978.1 afla|EED56850.1 aory|EIT77569.1 awen|OJJ36317.1

Group12598: acla|EAW15003.1 afis|EAW24538.1 afum|EAL86202.1 azon|OJJ48341.1

Group12599: acla|EAW15008.1 afis|EAW15437.1 afum|EAL84208.2 anid|CBF71217.1

Group12600: acla|EAW15028.1 afis|EAW24558.1 afum|EAL86221.1 awen|OJJ31238.1

Group12601: acla|EAW15029.1 afla|EED57259.1 aory|EIT74138.1 awen|OJJ36137.1

Group12602: acla|EAW15050.1 afis|EAW17412.1 agla|OJJ89306.1 anig|EHA23279.1

Group12603: acla|EAW15151.1 afis|EAW20119.1 afum|EAL85236.2 agla|OJJ88830.1

Group12604: ater|EAU32829.1 asyd|OJJ60433.1 anid|CBF87063.1 aver|OJI99755.1

Group12605: acla|EAW15259.1 afis|EAW20236.1 afum|EAL85353.1 awen|OJJ37031.1

Group12606: acla|EAW15341.1 afis|EAW20367.1 afum|EAL85483.1 azon|OJJ50763.1

Group12607: acla|EAW15359.1 afis|EAW20352.1 afum|EAL85470.1 ater|EAU32215.1

Group12608: agla|OJJ85076.1 asyd|OJJ60488.1 aver|OJI99814.1 awen|OJJ33534.1

Group12609: afis|EAW15403.1 anid|CBF80699.1 ater|EAU33084.1 ater|EAU35084.1

Group12610: afis|EAW15406.1 afla|EED51744.1 afum|EAL84240.1 aory|EIT80709.1

Group12611: afis|EAW15408.1 afla|EED47794.1 afum|EAL84239.1 aory|EIT77179.1

Group12612: afis|EAW15442.1 afum|EAL84205.1 asyd|OJJ64163.1 aver|OJI98240.1

Group12613: afis|EAW15629.1 afum|EAL85705.1 anid|CBF73658.1 asyd|OJJ54444.1

Group12614: afla|EED46061.1 aory|EIT83479.1 aver|OJI99637.1 anid|CBF87165.1

Group12615: afla|EED53178.1 aory|EIT74964.1 asyd|OJJ53365.1 aver|OJJ08382.1

Group12616: asyd|OJJ55027.1 aver|OJJ07177.1 ater|EAU36685.1 awen|OJJ41484.1

Group12617: afis|EAW16166.1 afum|EAL88801.1 aver|OJJ08858.1 azon|OJJ49589.1

Group12618: afis|EAW16179.1 aory|EIT82839.1 afla|EED47366.1 agla|OJJ79066.1

Group12619: afis|EAW16181.1 ater|EAU29303.1 aver|OJJ07307.1 ater|EAU33872.1

Group12620: afla|EED48450.1 awen|OJJ31972.1 agla|OJJ86297.1 afum|EAL86288.2

Group12621: azon|OJJ44590.1 anid|CBF88935.1 asyd|OJJ62854.1 aver|OJI96907.1

Group12622: afis|EAW16424.1 afum|EAL85137.1 anig|EHA19882.1 azon|OJJ48340.1

Group12623: aver|OJI99614.1 aluc|OJZ88481.1 asyd|OJJ52498.1 awen|OJJ38950.1

Group12624: afis|EAW16708.1 afla|EED46491.1 anid|CBF71843.1 aory|EIT72371.1

Group12625: afis|EAW16728.1 afum|EAL86303.2 afis|EAW16730.1 awen|OJJ38303.1

Group12626: afis|EAW16760.1 agla|OJJ87811.1 asyd|OJJ56901.1 awen|OJJ41527.1

Group12627: afis|EAW16819.1 afum|EBA27270.1 asyd|OJJ56741.1 aver|OJJ05986.1

Group12628: afis|EAW16859.1 asyd|OJJ62623.1 aver|OJI99301.1 awen|OJJ36991.1

Group12629: afis|EAW16899.1 afla|EED53179.1 afum|EAL86434.1 aory|EIT75198.1

Group12630: ater|EAU31586.1 anid|CBF73499.1 asyd|OJJ59268.1 aver|OJJ01880.1

Group12631: anid|CBF87043.1 asyd|OJJ60446.1 aver|OJI99764.1 awen|OJJ36367.1

Group12632: afis|EAW16971.1 afla|EED57581.1 aory|EIT82962.1 ater|EAU37145.1

Group12633: asyd|OJJ54420.1 aver|OJJ06548.1 agla|OJJ87528.1 awen|OJJ36611.1

Group12634: afis|EAW17008.1 akaw|GAA90342.1 aluc|OJZ85890.1 awen|OJJ37867.1

Group12635: afis|EAW17023.1 akaw|GAA89823.1 aluc|OJZ81867.1 anig|EHA25072.1

Group12636: afis|EAW17039.1 aluc|OJZ79601.1 aver|OJI98841.1 awen|OJJ33826.1

Group12637: atub|OJI91281.1 akaw|GAA83143.1 aluc|OJZ88700.1 anig|EHA18380.1

Group12638: afis|EAW17125.1 anid|CBF83967.1 asyd|OJJ59818.1 aver|OJJ00761.1

Group12639: afis|EAW17220.1 anid|CBF75874.1 asyd|OJJ60233.1 aver|OJJ04032.1

Group12640: atub|OJI84038.1 asyd|OJJ55410.1 aver|OJJ06489.1 awen|OJJ35934.1

Group12641: anig|EHA21859.1 akaw|GAA93038.1 aluc|OJZ82091.1 aory|EIT72971.1

Group12642: afis|EAW17526.1 afum|EAL86501.1 anid|CBF82920.1 ater|EAU39525.1

Group12643: afis|EAW17774.1 anid|CBF88655.1 asyd|OJJ63886.1 aver|OJI97053.1

Group12644: afis|EAW17798.1 afum|EAL91606.1 asyd|OJJ57628.1 ater|EAU33841.1

Group12645: afis|EAW18181.1 afum|EAL87542.1 awen|OJJ30529.1 azon|OJJ50211.1

Group12646: afis|EAW18554.1 afla|EED48113.1 afum|EAL93254.1 aory|EIT76197.1

Group12647: afis|EAW18556.1 afla|EED45068.1 afum|EAL93256.1 aory|EIT73780.1

Group12648: asyd|OJJ60429.1 aver|OJI99752.1 ater|EAU36739.1 anid|CBF87059.1

Group12649: afis|EAW19319.1 afum|EAL94036.1 ater|EAU29615.1 azon|OJJ45035.1

Group12650: afis|EAW19332.1 afum|EAL94049.1 ater|EAU29937.1 azon|OJJ44394.1

Group12651: afis|EAW19346.1 afla|EED54638.1 afum|EAL94066.1 aory|EIT78912.1

Group12652: asyd|OJJ61754.1 aver|OJI95852.1 awen|OJJ33647.1 azon|OJJ42778.1

Group12653: afis|EAW19369.1 afum|EAL94088.1 aory|EIT76838.1 afla|EED56076.1

Group12654: anid|CBF78721.1 ater|EAU29521.1 asyd|OJJ63988.1 aver|OJI96965.1

Group12655: aver|OJJ07067.1 agla|OJJ80745.1 anid|CBF71279.1 asyd|OJJ55136.1

Group12656: afis|EAW19669.1 afum|EAL86854.2 awen|OJJ31936.1 azon|OJJ46600.1

Group12657: afis|EAW19671.1 afla|EED46494.1 agla|OJJ81540.1 aory|EIT72433.1

Group12658: afis|EAW19741.1 afum|EAL86788.1 afla|EED56529.1 aory|EIT82922.1

Group12659: afis|EAW19760.1 ater|EAU36857.1 aory|EIT83283.1 afla|EED52427.1

Group12660: afis|EAW19788.1 afla|EED47534.1 afum|EAL85106.2 aory|EIT74580.1

Group12661: afis|EAW19805.1 afla|EED51065.1 afum|EAL85088.2 aory|EIT76967.1

Group12662: asyd|OJJ54160.1 aver|OJJ08584.1 anid|CBF82401.1 ater|EAU31753.1

Group12663: afis|EAW19967.1 afum|EAL84949.1 atub|OJI83199.1 awen|OJJ41818.1

Group12664: afis|EAW19969.1 afla|EED45226.1 afum|EAL84951.2 atub|OJI83198.1

Group12665: awen|OJJ32248.1 aluc|OJZ83022.1 asyd|OJJ56052.1 aver|OJJ07433.1

Group12666: afis|EAW19971.1 afum|EAL84953.1 atub|OJI83196.1 aver|OJJ07431.1

Group12667: afis|EAW20264.1 afla|EED45398.1 ater|EAU32478.1 awen|OJJ33383.1

Group12668: afis|EAW20272.1 afla|EED52438.1 afum|EAL85406.1 asyd|OJJ60800.1

Group12669: afis|EAW20287.1 afum|EAL85419.1 ater|EAU31587.1 azon|OJJ42679.1

Group12670: afis|EAW20330.1 asyd|OJJ63985.1 aver|OJI96968.1 azon|OJJ45655.1

Group12671: afis|EAW20558.1 afum|EAL84503.1 ater|EAU35417.1 aory|EIT74154.1

Group12672: afis|EAW20630.1 afum|EAL84437.1 agla|OJJ80446.1 azon|OJJ50721.1

Group12673: afis|EAW20645.1 afla|EED50442.1 afum|EAL84428.1 aory|EIT82074.1

Group12674: afis|EAW20646.1 afum|EAL84427.1 afis|EAW22096.1 afla|EED50441.1

Group12675: afis|EAW20647.1 afla|EED50440.1 afum|EAL84426.1 aory|EIT82099.1

Group12676: afis|EAW20648.1 afla|EED50439.1 afum|EAL84425.1 aory|EIT82155.1

Group12677: afis|EAW20712.1 agla|OJJ85311.1 aver|OJJ03120.1 asyd|OJJ51870.1

Group12678: afis|EAW20827.1 akaw|GAA83753.1 asyd|OJJ56855.1 ater|EAU35992.1

Group12679: afis|EAW20843.1 aory|EIT83050.1 ater|EAU37767.1 awen|OJJ33415.1

Group12680: afis|EAW21028.1 agla|OJJ89399.1 awen|OJJ36154.1 azon|OJJ50772.1

Group12681: afis|EAW21217.1 afum|EAL92229.1 ater|EAU31939.1 azon|OJJ42770.1

Group12682: afis|EAW21279.1 afum|EAL92294.1 ater|EAU31597.1 azon|OJJ42726.1

Group12683: afis|EAW22019.1 akaw|GAA93155.1 aluc|OJZ80421.1 akaw|GAA93175.1

Group12684: afis|EAW22086.1 afla|EED49609.1 afum|EAL85695.1 aory|EIT81016.1

Group12685: afis|EAW22092.1 afla|EED45137.1 aory|EIT73670.1 azon|OJJ46491.1

Group12686: anig|EHA22333.1 akaw|GAA89780.1 aluc|OJZ81911.1 atub|OJI84464.1

Group12687: afis|EAW22250.1 afla|EED44890.1 agla|OJJ89328.1 awen|OJJ37802.1

Group12688: afis|EAW22302.1 agla|OJJ82724.1 anid|CBF85186.1 awen|OJJ31410.1

Group12689: afis|EAW22305.1 agla|OJJ82732.1 anid|CBF85179.1 awen|OJJ31405.1

Group12690: afis|EAW22307.1 agla|OJJ82733.1 anid|CBF85177.1 awen|OJJ31404.1

Group12691: afis|EAW22481.1 afum|EAL90714.1 agla|OJJ81329.1 asyd|OJJ55316.1

Group12692: afla|EED48517.1 aory|EIT73055.1 asyd|OJJ64974.1 ater|EAU38745.1

Group12693: afis|EAW23632.1 agla|OJJ83716.1 anig|EHA18445.1 awen|OJJ30960.1

Group12694: afis|EAW23640.1 akaw|GAA85577.1 aluc|OJZ83513.1 ater|EAU35167.1

Group12695: afis|EAW23665.1 anid|CBF86366.1 asyd|OJJ64967.1 aver|OJI99091.1

Group12696: aluc|OJZ88354.1 akaw|GAA88465.1 anig|EHA18087.1 atub|OJI90949.1

Group12697: afis|EAW24054.1 afum|EAL91908.2 aory|EIT76118.1 ater|EAU29610.1

Group12698: afis|EAW24057.1 afum|EAL91913.1 aory|EIT80623.1 afla|EED51497.1

Group12699: afis|EAW24471.1 afum|EAL86144.1 asyd|OJJ53920.1 aver|OJJ08018.1

Group12700: afis|EAW24528.1 afum|EAL86193.1 asyd|OJJ55346.1 aver|OJJ06432.1

Group12701: afis|EAW24529.1 afum|EAL86194.1 asyd|OJJ55347.1 aver|OJJ06433.1

Group12702: afis|EAW24530.1 afum|EAL86195.1 asyd|OJJ55348.1 aver|OJJ06434.1

Group12703: afis|EAW24561.1 afum|EAL86222.1 anid|CBF83496.1 ater|EAU34205.1

Group12704: afis|EAW24626.1 anid|CBF79093.1 aver|OJJ08545.1 awen|OJJ29934.1

Group12705: afis|EAW24630.1 afla|EED50781.1 aory|EIT78119.1 ater|EAU34864.1

Group12706: afis|EAW24665.1 asyd|OJJ57399.1 afis|EAW24667.1 anid|CBF84287.1

Group12707: anid|CBF82396.1 azon|OJJ51730.1 asyd|OJJ54155.1 aver|OJJ08578.1

Group12708: afis|EAW25175.1 afum|EAL89848.1 akaw|GAA82884.1 azon|OJJ42794.1

Group12709: afis|EAW25576.1 afum|EAL84286.1 asyd|OJJ55342.1 aver|OJJ06428.1

Group12710: afis|EAW25577.1 afum|EAL84287.2 asyd|OJJ55343.1 aver|OJJ06429.1

Group12711: afis|EAW25578.1 afum|EAL84289.1 asyd|OJJ55344.1 aver|OJJ06430.1

Group12712: afis|EAW25579.1 afum|EAL84290.1 asyd|OJJ55345.1 aver|OJJ06431.1

Group12713: afis|EAW25587.1 afla|EED46133.1 afum|EAL84294.1 aory|EIT76488.1

Group12714: ater|EAU37771.1 anid|CBF82097.1 asyd|OJJ54260.1 aver|OJJ05213.1

Group12715: afis|EAW25733.1 anid|CBF69362.1 anig|EHA24047.1 ater|EAU30733.1

Group12716: afla|EED44727.1 anid|CBF87319.1 atub|OJI79310.1 aver|OJJ01700.1

Group12717: afla|EED44732.1 asyd|OJJ59105.1 atub|OJI79306.1 aver|OJJ01696.1

Group12718: afla|EED44811.1 agla|OJJ84865.1 aory|EIT77726.1 awen|OJJ33147.1

Group12719: afla|EED44963.1 aory|EIT73469.1 ater|EAU39567.1 azon|OJJ48659.1

Group12720: afla|EED45045.1 aory|EIT73660.1 asyd|OJJ56220.1 aver|OJJ05152.1

Group12721: afla|EED45098.1 aory|EIT73650.1 asyd|OJJ54179.1 aver|OJJ08561.1

Group12722: afla|EED45099.1 aory|EIT73692.1 asyd|OJJ54180.1 aver|OJJ08560.1

Group12723: afla|EED45112.1 anig|EHA26087.1 awen|OJJ31618.1 atub|OJI87333.1

Group12724: afla|EED45287.1 agla|OJJ88708.1 aory|EIT73683.1 awen|OJJ38832.1

Group12725: afla|EED45426.1 aory|EIT82451.1 ater|EAU34031.1 awen|OJJ37935.1

Group12726: afla|EED45451.1 agla|OJJ83723.1 aory|EIT82450.1 awen|OJJ34521.1

Group12727: afla|EED45507.1 anig|EHA20199.1 aory|EIT75575.1 ater|EAU30776.1

Group12728: afla|EED45817.1 aory|EIT75396.1 asyd|OJJ64188.1 aver|OJI98265.1

Group12729: afla|EED45919.1 afum|EAL94094.1 anig|EHA18741.1 aver|OJI96911.1

Group12730: afla|EED45991.1 aory|EIT75747.1 ater|EAU33128.1 awen|OJJ30976.1

Group12731: afla|EED46066.1 aory|EIT83502.1 asyd|OJJ56239.1 aory|EIT74811.1

Group12732: afla|EED46071.1 asyd|OJJ62490.1 aver|OJI95771.1 aver|OJI95940.1

Group12733: afla|EED46128.1 agla|OJJ86613.1 aory|EIT76505.1 ater|EAU38911.1

Group12734: afla|EED46224.1 aory|EIT76406.1 asyd|OJJ61419.1 aver|OJJ00754.1

Group12735: afla|EED46304.1 aory|EIT76339.1 aluc|OJZ92003.1 asyd|OJJ53539.1

Group12736: afla|EED46593.1 aory|EIT77886.1 awen|OJJ37810.1 aory|EIT77644.1

Group12737: afla|EED47256.1 aory|EIT79235.1 asyd|OJJ60436.1 aver|OJJ02286.1

Group12738: afla|EED47264.1 asyd|OJJ53566.1 ater|EAU30453.1 aver|OJJ08188.1

Group12739: afla|EED47265.1 aory|EIT79272.1 aver|OJJ07848.1 asyd|OJJ53733.1

Group12740: afla|EED47437.1 anid|CBF74960.1 asyd|OJJ52444.1 aver|OJJ03972.1

Group12741: afla|EED47492.1 asyd|OJJ52775.1 afla|EED47493.1 aory|EIT74556.1

Group12742: afla|EED47527.1 aory|EIT74558.1 ater|EAU29487.1 awen|OJJ33445.1

Group12743: afla|EED47574.1 aluc|OJZ80355.1 afla|EED50194.1 aory|EIT77390.1

Group12744: aver|OJJ07953.1 awen|OJJ36960.1 anid|CBF74035.1 asyd|OJJ53851.1

Group12745: afla|EED47758.1 aory|EIT77198.1 asyd|OJJ63034.1 aver|OJI97933.1

Group12746: afla|EED47767.1 aory|EIT77174.1 asyd|OJJ52370.1 aver|OJJ01978.1

Group12747: afla|EED47780.1 agla|OJJ83889.1 asyd|OJJ62411.1 aver|OJI96545.1

Group12748: afla|EED47864.1 agla|OJJ84437.1 aory|EIT77217.1 awen|OJJ35950.1

Group12749: afla|EED48095.1 aory|EIT76230.1 asyd|OJJ60351.1 aver|OJI99690.1

Group12750: afla|EED48148.1 agla|OJJ78447.1 aory|EIT76244.1 awen|OJJ35062.1

Group12751: afla|EED48367.1 afum|EAL94106.1 anid|CBF80285.1 aory|EIT83106.1

Group12752: afla|EED48476.1 anid|CBF89293.1 aory|EIT73248.1 ater|EAU36719.1

Group12753: afla|EED48494.1 aory|EIT73138.1 ater|EAU33398.1 aver|OJI99309.1

Group12754: afla|EED48900.1 aory|EIT81199.1 ater|EAU39620.1 awen|OJJ30204.1

Group12755: afla|EED48999.1 aluc|OJZ83983.1 aory|EIT79987.1 anig|EHA19110.1

Group12756: afla|EED49056.1 agla|OJJ84409.1 aory|EIT82484.1 asyd|OJJ56097.1

Group12757: afla|EED49072.1 aory|EIT82562.1 asyd|OJJ57956.1 aver|OJI99543.1

Group12758: afla|EED49450.1 aory|EIT79466.1 asyd|OJJ52788.1 azon|OJJ42629.1

Group12759: afla|EED49466.1 agla|OJJ82725.1 awen|OJJ33412.1 azon|OJJ46198.1

Group12760: afla|EED49541.1 aory|EIT82616.1 asyd|OJJ54938.1 aver|OJJ07258.1

Group12761: afla|EED49579.1 aory|EIT82607.1 asyd|OJJ59619.1 ater|EAU29692.1

Group12762: afla|EED49580.1 aory|EIT82572.1 asyd|OJJ59630.1 aver|OJJ02650.1

Group12763: afla|EED49672.1 anid|CBF84329.1 aory|EIT80965.1 aver|OJI96140.1

Group12764: afla|EED49698.1 asyd|OJJ56434.1 aver|OJJ06286.1 awen|OJJ29977.1

Group12765: afla|EED49745.1 agla|OJJ79839.1 anid|CBF82183.1 aory|EIT77674.1

Group12766: afla|EED49830.1 asyd|OJJ64836.1 aver|OJI98949.1 aory|EIT76349.1

Group12767: afla|EED50153.1 aory|EIT76928.1 asyd|OJJ63401.1 aver|OJI97536.1

Group12768: azon|OJJ42664.1 anid|CBF83288.1 asyd|OJJ57256.1 aver|OJJ03038.1

Group12769: afla|EED50222.1 asyd|OJJ54609.1 aver|OJJ07547.1 anid|CBF80660.1

Group12770: afla|EED50264.1 anid|CBF84349.1 aory|EIT77324.1 asyd|OJJ55453.1

Group12771: afla|EED50265.1 anid|CBF84347.1 aory|EIT77375.1 asyd|OJJ55454.1

Group12772: afla|EED50400.1 aory|EIT82118.1 atub|OJI86190.1 awen|OJJ37001.1

Group12773: afla|EED50617.1 agla|OJJ79385.1 aory|EIT78122.1 awen|OJJ39037.1

Group12774: afla|EED50777.1 agla|OJJ86278.1 aory|EIT78080.1 awen|OJJ29910.1

Group12775: afla|EED51095.1 aory|EIT81993.1 asyd|OJJ56754.1 aver|OJJ05969.1

Group12776: afla|EED51117.1 agla|OJJ82663.1 aory|EIT81364.1 aver|OJJ06746.1

Group12777: afla|EED51135.1 aory|EIT81360.1 asyd|OJJ53806.1 aver|OJJ07904.1

Group12778: afla|EED51148.1 aory|EIT81317.1 asyd|OJJ56869.1 awen|OJJ30235.1

Group12779: afla|EED51153.1 anid|CBF80182.1 aory|EIT81363.1 aver|OJI99529.1

Group12780: afla|EED51456.1 aluc|OJZ87836.1 aory|EIT80639.1 aver|OJJ08844.1

Group12781: afla|EED51462.1 aory|EIT80610.1 ater|EAU30456.1 awen|OJJ41461.1

Group12782: afla|EED51488.1 aory|EIT80469.1 asyd|OJJ54741.1 aver|OJJ07423.1

Group12783: afla|EED51664.1 aory|EIT80716.1 asyd|OJJ65172.1 aver|OJI99324.1

Group12784: afla|EED51711.1 agla|OJJ81858.1 aory|EIT80556.1 awen|OJJ35917.1

Group12785: afla|EED51783.1 aory|EIT81719.1 aver|OJJ05159.1 awen|OJJ29734.1

Group12786: afla|EED51847.1 anid|CBF80437.1 asyd|OJJ54782.1 aver|OJJ07384.1

Group12787: afla|EED52001.1 agla|OJJ81472.1 aory|EIT81458.1 awen|OJJ34615.1

Group12788: afla|EED52053.1 asyd|OJJ58713.1 aver|OJJ01326.1 anid|CBF70330.1

Group12789: afla|EED52095.1 anid|CBF82418.1 aory|EIT76927.1 awen|OJJ29694.1

Group12790: afla|EED52125.1 aluc|OJZ87029.1 aory|EIT74420.1 atub|OJI83485.1

Group12791: aory|EIT74348.1 asyd|OJJ58698.1 aver|OJJ01310.1 ater|EAU31702.1

Group12792: afla|EED52208.1 agla|OJJ79200.1 aory|EIT74458.1 awen|OJJ36202.1

Group12793: afla|EED52275.1 aory|EIT74459.1 aver|OJJ04004.1 awen|OJJ41101.1

Group12794: akaw|GAA85319.1 aluc|OJZ90680.1 anig|EHA27495.1 awen|OJJ30547.1

Group12795: afla|EED52377.1 aory|EIT82372.1 asyd|OJJ61399.1 aver|OJJ00728.1

Group12796: afla|EED52414.1 agla|OJJ88597.1 asyd|OJJ52083.1 awen|OJJ37428.1

Group12797: afla|EED52580.1 aory|EIT74291.1 ater|EAU34999.1 awen|OJJ35810.1

Group12798: afla|EED52644.1 aory|EIT74240.1 ater|EAU38850.1 afla|EED52645.1

Group12799: afla|EED52841.1 asyd|OJJ59114.1 awen|OJJ40871.1 aver|OJI99246.1

Group12800: afla|EED52961.1 akaw|GAA90411.1 aory|EIT78735.1 azon|OJJ46261.1

Group12801: afla|EED53060.1 akaw|GAA88009.1 aluc|OJZ81267.1 aory|EIT74799.1

Group12802: afla|EED53077.1 akaw|GAA83284.1 anig|EHA18930.1 aory|EIT74781.1

Group12803: afla|EED53101.1 aory|EIT74764.1 asyd|OJJ62819.1 aver|OJI96864.1

Group12804: afla|EED53141.1 aory|EIT74966.1 asyd|OJJ52378.1 aver|OJJ01969.1

Group12805: afla|EED53183.1 agla|OJJ89057.1 aory|EIT75233.1 aver|OJJ07600.1

Group12806: afla|EED53185.1 agla|OJJ82951.1 aory|EIT75081.1 awen|OJJ31589.1

Group12807: afla|EED53218.1 aory|EIT75254.1 ater|EAU32833.1 awen|OJJ31960.1

Group12808: afla|EED53267.1 aory|EIT75238.1 asyd|OJJ64360.1 aver|OJI98456.1

Group12809: afla|EED53303.1 aory|EIT74980.1 asyd|OJJ59258.1 aver|OJJ01869.1

Group12810: afla|EED53406.1 asyd|OJJ59968.1 aver|OJJ02323.1 awen|OJJ41951.1

Group12811: anid|CBF70080.1 asyd|OJJ55930.1 aver|OJJ04876.1 awen|OJJ33306.1

Group12812: afla|EED53574.1 agla|OJJ85440.1 aluc|OJZ84845.1 aory|EIT75215.1

Group12813: afla|EED53620.1 aory|EIT75225.1 afla|EED54853.1 awen|OJJ37957.1

Group12814: afla|EED53952.1 asyd|OJJ54541.1 aver|OJJ07640.1 anid|CBF79654.1

Group12815: afla|EED54102.1 agla|OJJ81289.1 aory|EIT76135.1 aver|OJJ04436.1

Group12816: afla|EED54258.1 agla|OJJ85932.1 aory|EIT76102.1 awen|OJJ37460.1

Group12817: afla|EED54341.1 aory|EIT76063.1 asyd|OJJ59957.1 aver|OJJ02332.1

Group12818: afla|EED54403.1 akaw|GAA91213.1 aluc|OJZ82858.1 aory|EIT75845.1

Group12819: afla|EED54890.1 akaw|GAA83675.1 anig|EHA24762.1 aory|EIT79402.1

Group12820: afla|EED55108.1 aory|EIT81569.1 ater|EAU30760.1 asyd|OJJ54918.1

Group12821: afla|EED55117.1 anid|CBF78521.1 aory|EIT81618.1 asyd|OJJ53434.1

Group12822: afla|EED55167.1 aory|EIT81576.1 ater|EAU32711.1 azon|OJJ45273.1

Group12823: afla|EED55191.1 asyd|OJJ64707.1 aver|OJI98809.1 awen|OJJ30080.1

Group12824: afla|EED55708.1 aory|EIT83224.1 asyd|OJJ60059.1 aver|OJJ02230.1

Group12825: afla|EED55752.1 agla|OJJ85247.1 aory|EIT76751.1 awen|OJJ39612.1

Group12826: asyd|OJJ52170.1 akaw|GAA93141.1 aluc|OJZ80034.1 aluc|OJZ79854.1

Group12827: afla|EED56533.1 aory|EIT79691.1 asyd|OJJ52844.1 aver|OJJ08821.1

Group12828: afla|EED56751.1 anid|CBF73547.1 aory|EIT73872.1 ater|EAU34582.1

Group12829: afla|EED56794.1 aory|EIT73868.1 asyd|OJJ52881.1 aver|OJJ08852.1

Group12830: afla|EED56880.1 aory|EIT77482.1 asyd|OJJ54818.1 aver|OJJ07343.1

Group12831: afla|EED56890.1 aory|EIT77486.1 ater|EAU30300.1 awen|OJJ35824.1

Group12832: afla|EED56922.1 aory|EIT77562.1 awen|OJJ36057.1 azon|OJJ45172.1

Group12833: afla|EED57043.1 akaw|GAA85824.1 aory|EIT82269.1 agla|OJJ83700.1

Group12834: afla|EED57065.1 aory|EIT73508.1 ater|EAU33400.1 azon|OJJ46513.1

Group12835: afla|EED57101.1 aory|EIT82266.1 ater|EAU33029.1 aver|OJJ03088.1

Group12836: afla|EED57122.1 aory|EIT74103.1 asyd|OJJ52875.1 awen|OJJ35395.1

Group12837: afla|EED57232.1 aory|EIT74132.1 asyd|OJJ54475.1 aver|OJJ05161.1

Group12838: afla|EED57524.1 aory|EIT81870.1 aver|OJJ08135.1 awen|OJJ36252.1

Group12839: afla|EED57525.1 aory|EIT81871.1 asyd|OJJ53613.1 aver|OJJ08136.1

Group12840: afla|EED57550.1 aory|EIT81839.1 asyd|OJJ52828.1 aver|OJJ08804.1

Group12841: afla|EED57551.1 aory|EIT81855.1 asyd|OJJ52826.1 aver|OJJ08802.1

Group12842: afla|EED57632.1 agla|OJJ81254.1 aory|EIT82966.1 awen|OJJ36467.1

Group12843: afla|EED57766.1 aory|EIT81077.1 asyd|OJJ53424.1 aver|OJJ08321.1

Group12844: afum|EAL84947.1 anid|CBF86370.1 asyd|OJJ56510.1 aver|OJJ06209.1

Group12845: afum|EAL85443.1 anid|CBF83211.1 asyd|OJJ55200.1 ater|EAU35711.1

Group12846: afum|EAL86284.1 akaw|GAA92938.1 aluc|OJZ82094.1 anig|EHA21849.1

Group12847: anid|CBF78540.1 asyd|OJJ56048.1 atub|OJI89154.1 aver|OJJ04991.1

Group12848: afum|EAL87851.1 anid|CBF73999.1 asyd|OJJ61470.1 aver|OJJ00811.1

Group12849: afum|EAL92213.1 agla|OJJ88491.1 ater|EAU36105.1 aver|OJI97250.1

Group12850: aver|OJJ07490.1 anid|CBF80543.1 asyd|OJJ54677.1 awen|OJJ31818.1

Group12851: anig|EHA18621.1 ater|EAU29574.1 asyd|OJJ64223.1 awen|OJJ31928.1

Group12852: agla|OJJ78711.1 ater|EAU37692.1 awen|OJJ32765.1 azon|OJJ46796.1

Group12853: agla|OJJ78775.1 asyd|OJJ52064.1 aver|OJJ04996.1 awen|OJJ30066.1

Group12854: agla|OJJ78790.1 asyd|OJJ65113.1 ater|EAU34531.1 aver|OJI99239.1

Group12855: agla|OJJ78861.1 akaw|GAA92188.1 aluc|OJZ85599.1 atub|OJI79394.1

Group12856: agla|OJJ79126.1 asyd|OJJ55387.1 aver|OJJ08270.1 awen|OJJ30822.1

Group12857: agla|OJJ79243.1 asyd|OJJ56595.1 aver|OJJ06130.1 awen|OJJ32773.1

Group12858: agla|OJJ79347.1 asyd|OJJ52458.1 aory|EIT77530.1 awen|OJJ31958.1

Group12859: agla|OJJ80035.1 awen|OJJ30090.1 anig|EHA24070.1 atub|OJI82062.1

Group12860: akaw|GAA88217.1 aluc|OJZ82180.1 anig|EHA28554.1 atub|OJI88257.1

Group12861: agla|OJJ80205.1 awen|OJJ36373.1 azon|OJJ42211.1 azon|OJJ45684.1

Group12862: agla|OJJ80669.1 asyd|OJJ55188.1 aver|OJJ07019.1 awen|OJJ31430.1

Group12863: agla|OJJ81535.1 anid|CBF74184.1 asyd|OJJ53944.1 aver|OJJ08041.1

Group12864: agla|OJJ81536.1 anid|CBF74182.1 asyd|OJJ53945.1 aver|OJJ08042.1

Group12865: azon|OJJ50332.1 asyd|OJJ54963.1 aver|OJJ07228.1 awen|OJJ38736.1

Group12866: agla|OJJ82209.1 asyd|OJJ60323.1 aver|OJI99671.1 awen|OJJ37959.1

Group12867: agla|OJJ82210.1 asyd|OJJ60322.1 aver|OJI99670.1 awen|OJJ37958.1

Group12868: agla|OJJ82738.1 asyd|OJJ59814.1 aver|OJJ02472.1 awen|OJJ29907.1

Group12869: agla|OJJ83650.1 anid|CBF87155.1 asyd|OJJ54413.1 aver|OJJ08133.1

Group12870: agla|OJJ83683.1 awen|OJJ35414.1 azon|OJJ49762.1 anig|EHA17981.1

Group12871: agla|OJJ84737.1 asyd|OJJ54201.1 ater|EAU36886.1 aver|OJJ08523.1

Group12872: agla|OJJ85077.1 asyd|OJJ60487.1 aver|OJI99813.1 awen|OJJ33533.1

Group12873: agla|OJJ85842.1 anid|CBF71750.1 asyd|OJJ61732.1 aver|OJI95826.1

Group12874: agla|OJJ85847.1 asyd|OJJ62352.1 atub|OJI84594.1 azon|OJJ42787.1

Group12875: agla|OJJ86250.1 anig|EHA22184.1 asyd|OJJ53782.1 aver|OJJ07888.1

Group12876: agla|OJJ86416.1 asyd|OJJ52979.1 aver|OJJ08945.1 awen|OJJ36945.1

Group12877: agla|OJJ86672.1 akaw|GAA90292.1 aluc|OJZ83046.1 atub|OJI87902.1

Group12878: agla|OJJ87087.1 asyd|OJJ53137.1 aver|OJJ03715.1 azon|OJJ44748.1

Group12879: agla|OJJ87284.1 asyd|OJJ62327.1 aver|OJI96432.1 azon|OJJ51823.1

Group12880: anid|CBF78499.1 asyd|OJJ61883.1 ater|EAU32640.1 aver|OJI95981.1

Group12881: agla|OJJ88170.1 asyd|OJJ57612.1 aver|OJJ05573.1 awen|OJJ31659.1

Group12882: agla|OJJ88376.1 anid|CBF83745.1 asyd|OJJ65110.1 aver|OJI99237.1

Group12883: akaw|GAA81959.1 aluc|OJZ92230.1 anig|EHA26727.1 atub|OJI86660.1

Group12884: akaw|GAA82113.1 aluc|OJZ92063.1 atub|OJI86495.1 anig|EHA22992.1

Group12885: akaw|GAA82568.1 aluc|OJZ87102.1 anig|EHA20371.1 atub|OJI81537.1

Group12886: akaw|GAA84296.1 aluc|OJZ88990.1 anig|EHA23536.1 atub|OJI85797.1

Group12887: akaw|GAA84297.1 aluc|OJZ88991.1 anig|EHA23535.1 atub|OJI85796.1

Group12888: akaw|GAA84513.1 aluc|OJZ89242.1 anig|EHA23325.1 atub|OJI85556.1

Group12889: akaw|GAA84858.1 aluc|OJZ86813.1 atub|OJI83704.1 awen|OJJ33631.1

Group12890: akaw|GAA84936.1 aluc|OJZ86730.1 anig|EHA21949.1 atub|OJI83798.1

Group12891: akaw|GAA85316.1 aluc|OJZ90677.1 anig|EHA21626.1 awen|OJJ30545.1

Group12892: akaw|GAA85845.1 aluc|OJZ91254.1 asyd|OJJ53340.1 aver|OJJ08405.1

Group12893: akaw|GAA85939.1 aluc|OJZ91123.1 anid|CBF76247.1 atub|OJI83128.1

Group12894: akaw|GAA85943.1 aluc|OJZ91119.1 anig|EHA19817.1 atub|OJI83123.1

Group12895: akaw|GAA85945.1 aluc|OJZ91117.1 anig|EHA19816.1 atub|OJI83121.1

Group12896: akaw|GAA85951.1 aluc|OJZ91111.1 anig|EHA19808.1 atub|OJI83108.1

Group12897: akaw|GAA86065.1 aluc|OJZ90299.1 anig|EHA23787.1 atub|OJI81724.1

Group12898: akaw|GAA86634.1 aluc|OJZ85191.1 atub|OJI80406.1 awen|OJJ39940.1

Group12899: akaw|GAA86654.1 aluc|OJZ89748.1 anig|EHA26971.1 atub|OJI85054.1

Group12900: akaw|GAA86674.1 aluc|OJZ89771.1 atub|OJI85032.1 azon|OJJ48042.1

Group12901: akaw|GAA87092.1 aluc|OJZ84346.1 atub|OJI89722.1 azon|OJJ43063.1

Group12902: akaw|GAA87509.1 aluc|OJZ88011.1 anig|EHA17791.1 atub|OJI90620.1

Group12903: akaw|GAA87609.1 aluc|OJZ85230.1 anig|EHA25433.1 atub|OJI80350.1

Group12904: akaw|GAA87812.1 aluc|OJZ85447.1 anig|EHA24990.1 atub|OJI80120.1

Group12905: akaw|GAA87848.1 aluc|OJZ85488.1 anig|EHA25025.1 atub|OJI80076.1

Group12906: akaw|GAA88045.1 aluc|OJZ81308.1 anig|EHA18616.1 atub|OJI91516.1

Group12907: akaw|GAA88435.1 aluc|OJZ88323.1 anig|EHA18059.1 atub|OJI90918.1

Group12908: akaw|GAA88799.1 aluc|OJZ82482.1 anig|EHA28293.1 atub|OJI87956.1

Group12909: akaw|GAA89216.1 aluc|OJZ81613.1 atub|OJI88569.1 aver|OJJ02003.1

Group12910: akaw|GAA89763.1 aluc|OJZ80620.1 anig|EHA27211.1 atub|OJI87474.1

Group12911: akaw|GAA89767.1 aluc|OJZ80616.1 anig|EHA27216.1 atub|OJI87470.1

Group12912: akaw|GAA90344.1 aluc|OJZ85888.1 anig|EHA24699.1 atub|OJI79675.1

Group12913: akaw|GAA90439.1 aluc|OJZ85788.1 atub|OJI79574.1 awen|OJJ35350.1

Group12914: akaw|GAA90539.1 aluc|OJZ86009.1 anig|EHA24601.1 atub|OJI79798.1

Group12915: akaw|GAA90859.1 aluc|OJZ82974.1 anig|EHA21390.1 atub|OJI89207.1

Group12916: akaw|GAA91038.1 aluc|OJZ87848.1 atub|OJI80800.1 azon|OJJ46806.1

Group12917: akaw|GAA91770.1 aluc|OJZ92784.1 anig|EHA26225.1 atub|OJI87226.1

Group12918: akaw|GAA91836.1 aluc|OJZ92858.1 anig|EHA26148.1 atub|OJI87316.1

Group12919: akaw|GAA91877.1 aluc|OJZ82022.1 anig|EHA22233.1 atub|OJI84575.1

Group12920: akaw|GAA91906.1 aluc|OJZ81988.1 atub|OJI84539.1 azon|OJJ42605.1

Group12921: akaw|GAA92050.1 aluc|OJZ92920.1 anig|EHA26049.1 atub|OJI87379.1

Group12922: akaw|GAA92102.1 aluc|OJZ83623.1 anig|EHA27515.1 atub|OJI87850.1

Group12923: akaw|GAA92148.1 aluc|OJZ85553.1 atub|OJI79352.1 anig|EHA24962.1

Group12924: asyd|OJJ52316.1 akaw|GAA92939.1 anig|EHA21843.1 azon|OJJ42036.1

Group12925: akaw|GAA92557.1 aluc|OJZ90104.1 anig|EHA20966.1 atub|OJI84700.1

Group12926: akaw|GAA92563.1 aluc|OJZ90110.1 anig|EHA20958.1 atub|OJI84689.1

Group12927: akaw|GAA92598.1 aluc|OJZ82615.1 anig|EHA27464.1 atub|OJI88850.1

Group12928: akaw|GAA92716.1 aluc|OJZ88762.1 anig|EHA18458.1 atub|OJI91332.1

Group12929: akaw|GAA92876.1 aluc|OJZ82535.1 anig|EHA22503.1 atub|OJI88779.1

Group12930: akaw|GAA93110.1 aluc|OJZ80593.1 anig|EHA22189.1 awen|OJJ29865.1

Group12931: aluc|OJZ80303.1 atub|OJI87456.1 aver|OJJ02221.1 asyd|OJJ53396.1

Group12932: aluc|OJZ83761.1 atub|OJI90134.1 azon|OJJ42427.1 aver|OJJ05049.1

Group12933: aluc|OJZ84345.1 anig|EHA27681.1 atub|OJI89723.1 awen|OJJ32234.1

Group12934: aluc|OJZ87025.1 anig|EHA21633.1 atub|OJI83493.1 azon|OJJ45794.1

Group12935: aluc|OJZ87031.1 anig|EHA25217.1 asyd|OJJ54379.1 atub|OJI83484.1

Group12936: awen|OJJ36949.1 aory|EIT74207.1 aver|OJJ04885.1 asyd|OJJ55936.1

Group12937: aluc|OJZ89699.1 ater|EAU39615.1 atub|OJI85102.1 awen|OJJ40949.1

Group12938: aluc|OJZ91141.1 asyd|OJJ55528.1 atub|OJI83151.1 aver|OJJ05144.1

Group12939: anid|CBF69434.1 asyd|OJJ52387.1 ater|EAU36795.1 aver|OJJ01936.1

Group12940: anid|CBF69494.1 asyd|OJJ59238.1 ater|EAU29440.1 aver|OJJ01848.1

Group12941: anid|CBF70703.1 asyd|OJJ58457.1 ater|EAU37970.1 aver|OJJ01057.1

Group12942: anid|CBF71430.1 asyd|OJJ52412.1 ater|EAU37827.1 aver|OJJ01946.1

Group12943: anid|CBF71499.1 asyd|OJJ54931.1 aver|OJJ07625.1 aver|OJJ07266.1

Group12944: anid|CBF71523.1 asyd|OJJ53435.1 aver|OJJ08314.1 azon|OJJ43609.1

Group12945: anid|CBF73648.1 asyd|OJJ65134.1 aver|OJI95772.1 asyd|OJJ52404.1

Group12946: anid|CBF74618.1 asyd|OJJ57622.1 ater|EAU36113.1 aver|OJJ05564.1

Group12947: anid|CBF74675.1 asyd|OJJ55665.1 aver|OJI97962.1 azon|OJJ46185.1

Group12948: anid|CBF74691.1 asyd|OJJ60211.1 aver|OJJ04057.1 awen|OJJ40117.1

Group12949: anid|CBF74971.1 asyd|OJJ60087.1 ater|EAU31429.1 aver|OJJ04174.1

Group12950: anid|CBF77110.1 asyd|OJJ62964.1 aver|OJI98003.1 azon|OJJ43190.1

Group12951: anid|CBF78148.1 asyd|OJJ65379.1 ater|EAU32100.1 aver|OJI99549.1

Group12952: anid|CBF78238.1 asyd|OJJ56040.1 ater|EAU37375.1 aver|OJJ04986.1

Group12953: anid|CBF78472.1 asyd|OJJ54762.1 ater|EAU30253.1 aver|OJJ07405.1

Group12954: anid|CBF78534.1 aory|EIT74360.1 asyd|OJJ65297.1 aver|OJI99437.1

Group12955: anid|CBF78610.1 asyd|OJJ65111.1 ater|EAU32524.1 aver|OJI99238.1

Group12956: anid|CBF78782.1 asyd|OJJ55215.1 aver|OJJ07001.1 azon|OJJ43631.1

Group12957: anid|CBF80106.1 asyd|OJJ58324.1 atub|OJI84524.1 aver|OJJ02087.1

Group12958: anid|CBF80121.1 asyd|OJJ58341.1 aver|OJJ02070.1 azon|OJJ51179.1

Group12959: anid|CBF80287.1 asyd|OJJ56857.1 ater|EAU36075.1 aver|OJJ03470.1

Group12960: anid|CBF80469.1 asyd|OJJ54772.1 aver|OJJ07394.1 ater|EAU32686.1

Group12961: anid|CBF81872.1 asyd|OJJ62653.1 ater|EAU30094.1 aver|OJI96752.1

Group12962: anid|CBF82023.1 atub|OJI80139.1 aver|OJJ05135.1 awen|OJJ37276.1

Group12963: anid|CBF82124.1 asyd|OJJ54294.1 ater|EAU29308.1 aver|OJJ05249.1

Group12964: anid|CBF82205.1 asyd|OJJ52180.1 aver|OJJ05965.1 awen|OJJ34424.1

Group12965: anid|CBF82775.1 asyd|OJJ58566.1 aver|OJJ01173.1 awen|OJJ34605.1

Group12966: anid|CBF82808.1 asyd|OJJ57639.1 ater|EAU37477.1 aver|OJJ01945.1

Group12967: anid|CBF83909.1 asyd|OJJ55420.1 ater|EAU36531.1 aver|OJJ06497.1

Group12968: anid|CBF84008.1 asyd|OJJ61381.1 ater|EAU37360.1 aver|OJJ00708.1

Group12969: anid|CBF84017.1 asyd|OJJ61378.1 aver|OJJ00705.1 awen|OJJ29825.1

Group12970: anid|CBF84158.1 asyd|OJJ57125.1 ater|EAU31584.1 aver|OJJ03179.1

Group12971: anid|CBF84267.1 asyd|OJJ57207.1 aver|OJJ03082.1 awen|OJJ41829.1

Group12972: anid|CBF84454.1 asyd|OJJ52036.1 azon|OJJ42035.1 awen|OJJ33685.1

Group12973: anid|CBF84576.1 asyd|OJJ53383.1 ater|EAU32828.1 aver|OJJ08360.1

Group12974: anid|CBF84699.1 asyd|OJJ53315.1 aver|OJJ08428.1 azon|OJJ42544.1

Group12975: anid|CBF85024.1 asyd|OJJ63109.1 aver|OJI97851.1 azon|OJJ42614.1

Group12976: anid|CBF86758.1 asyd|OJJ64710.1 aver|OJI98812.1 ater|EAU36702.1

Group12977: anid|CBF87067.1 asyd|OJJ60427.1 aver|OJI99749.1 asyd|OJJ56866.1

Group12978: anid|CBF87151.1 asyd|OJJ60306.1 aver|OJI99647.1 azon|OJJ42490.1

Group12979: anid|CBF87158.1 asyd|OJJ60307.1 aver|OJI99648.1 ater|EAU32325.1

Group12980: anid|CBF87316.1 asyd|OJJ59107.1 atub|OJI79308.1 aver|OJJ01698.1

Group12981: anid|CBF87330.1 asyd|OJJ59104.1 atub|OJI79305.1 aver|OJJ01695.1

Group12982: anid|CBF87371.1 aver|OJJ06191.1 azon|OJJ42109.1 asyd|OJJ56532.1

Group12983: anid|CBF87456.1 asyd|OJJ52925.1 ater|EAU38161.1 aver|OJJ08893.1

Group12984: anid|CBF87496.1 asyd|OJJ52967.1 aver|OJJ08933.1 awen|OJJ36410.1

Group12985: anid|CBF88657.1 asyd|OJJ63887.1 ater|EAU37502.1 aver|OJI97052.1

Group12986: anid|CBF88756.1 asyd|OJJ63975.1 ater|EAU34975.1 aver|OJI96979.1

Group12987: anid|CBF89119.1 asyd|OJJ62658.1 ater|EAU30179.1 aver|OJI96756.1

Group12988: anid|CBF89120.1 asyd|OJJ62659.1 ater|EAU30178.1 aver|OJI96757.1

Group12989: anid|CBF89129.1 asyd|OJJ62662.1 ater|EAU30175.1 aver|OJI96760.1

Group12990: anid|CBF89373.1 asyd|OJJ62936.1 ater|EAU39611.1 aver|OJI98055.1

Group12991: anig|EHA19812.1 ater|EAU29875.1 atub|OJI83112.1 awen|OJJ34678.1

Group12992: anig|EHA21835.1 asyd|OJJ55805.1 aver|OJJ04753.1 awen|OJJ29600.1

Group12993: ater|EAU33817.1 asyd|OJJ52684.1 aver|OJJ08670.1 azon|OJJ45037.1

Group12994: anig|EHA27967.1 asyd|OJJ58982.1 ater|EAU36532.1 aver|OJJ01577.1

Group12995: aory|EIT74786.1 ater|EAU31573.1 awen|OJJ33259.1 azon|OJJ50633.1

Group12996: asyd|OJJ52199.1 ater|EAU32864.1 aver|OJJ05951.1 awen|OJJ34088.1

Group12997: asyd|OJJ52544.1 aver|OJJ03860.1 awen|OJJ40970.1 azon|OJJ44556.1

Group12998: asyd|OJJ52744.1 awen|OJJ41771.1 asyd|OJJ62932.1 aver|OJI98058.1

Group12999: asyd|OJJ53630.1 atub|OJI91469.1 aver|OJJ05947.1 awen|OJJ30587.1

Group13000: asyd|OJJ53632.1 atub|OJI91471.1 aver|OJJ05949.1 awen|OJJ30583.1

Group13001: asyd|OJJ54491.1 aver|OJJ07688.1 awen|OJJ36286.1 azon|OJJ46776.1

Group13002: asyd|OJJ55095.1 atub|OJI83026.1 aver|OJJ07113.1 awen|OJJ30194.1

Group13003: asyd|OJJ55422.1 aver|OJJ06500.1 asyd|OJJ62319.1 azon|OJJ45048.1

Group13004: asyd|OJJ55697.1 aver|OJJ06782.1 awen|OJJ33233.1 azon|OJJ46571.1

Group13005: asyd|OJJ57371.1 ater|EAU37762.1 aver|OJJ05797.1 awen|OJJ35959.1

Group13006: asyd|OJJ57372.1 ater|EAU37761.1 aver|OJJ05796.1 awen|OJJ35958.1

Group13007: asyd|OJJ59106.1 aver|OJJ01697.1 asyd|OJJ63962.1 atub|OJI79307.1

Group13008: asyd|OJJ60935.1 ater|EAU30756.1 aver|OJJ00261.1 awen|OJJ29908.1

Group13009: asyd|OJJ61511.1 ater|EAU30984.1 aver|OJJ00848.1 awen|OJJ34636.1

Group13010: asyd|OJJ61753.1 ater|EAU30962.1 aver|OJI95851.1 awen|OJJ33646.1

Group13011: asyd|OJJ62359.1 atub|OJI86338.1 aver|OJI96454.1 awen|OJJ32290.1

Group13012: ater|EAU33663.1 azon|OJJ44486.1 aver|OJJ06542.1 awen|OJJ29834.1

Group13013: abra|OJJ65453.1 abra|OJJ65526.1 abra|OJJ65569.1

Group13014: abra|OJJ65473.1 abra|OJJ65558.1 abra|OJJ67124.1

Group13015: abra|OJJ65559.1 abra|OJJ65829.1 akaw|GAA93246.1

Group13016: abra|OJJ65754.1 abra|OJJ65846.1 aluc|OJZ88783.1

Group13017: abra|OJJ65827.1 abra|OJJ65828.1 akaw|GAA93245.1

Group13018: abra|OJJ66017.1 abra|OJJ66458.1 ater|EAU32217.1

Group13019: abra|OJJ76729.1 abra|OJJ77448.1 afla|EED54075.1

Group13020: acar|OOF90195.1 acar|OOF90197.1 aluc|OJZ79950.1

Group13021: acar|OOF90264.1 acar|OOG00798.1 acar|OOF93084.1

Group13022: acar|OOF90438.1 acar|OOF90445.1 asyd|OJJ55926.1

Group13023: acar|OOF90561.1 acar|OOF90562.1 acar|OOF90563.1

Group13024: acar|OOF90631.1 acar|OOF90634.1 afis|EAW25541.1

Group13025: acar|OOF94097.1 acar|OOF94167.1 agla|OJJ78969.1

Group13026: acar|OOF95058.1 acar|OOF95125.1 acla|EAW08663.1

Group13027: acar|OOF95060.1 acar|OOF95127.1 anid|CBF79679.1

Group13028: acar|OOG00250.1 acar|OOG00268.1 azon|OJJ42098.1

Group13029: afis|EAW18108.1 afis|EAW24036.1 afis|EAW25563.1

Group13030: afum|EAL90931.1 afum|EAL91126.1 acla|EAW14873.1

Group13031: agla|OJJ78408.1 agla|OJJ78919.1 agla|OJJ82792.1

Group13032: agla|OJJ78433.1 agla|OJJ85459.1 agla|OJJ89640.1

Group13033: agla|OJJ79065.1 agla|OJJ87918.1 awen|OJJ36226.1

Group13034: agla|OJJ79274.1 agla|OJJ88809.1 agla|OJJ82201.1

Group13035: agla|OJJ81537.1 agla|OJJ86488.1 agla|OJJ87265.1

Group13036: agla|OJJ83424.1 agla|OJJ89193.1 ater|EAU34811.1

Group13037: agla|OJJ83549.1 agla|OJJ89316.1 afum|EAL87730.1

Group13038: agla|OJJ83609.1 agla|OJJ83610.1 azon|OJJ47419.1

Group13039: agla|OJJ84374.1 agla|OJJ85305.1 agla|OJJ89339.1

Group13040: agla|OJJ86585.1 agla|OJJ87693.1 aver|OJJ04106.1

Group13041: agla|OJJ88218.1 asyd|OJJ63999.1 aver|OJI98088.1

Group13042: agla|OJJ88811.1 agla|OJJ89486.1 anid|CBF88644.1

Group13043: akaw|GAA92347.1 akaw|GAA93057.1 acla|EAW09608.1

Group13044: akaw|GAA92952.1 akaw|GAA93250.1 asyd|OJJ52135.1

Group13045: akaw|GAA93243.1 akaw|GAA93291.1 abra|OJJ65732.1

Group13046: aluc|OJZ79470.1 aluc|OJZ79695.1 aluc|OJZ79900.1

Group13047: aluc|OJZ79477.1 aluc|OJZ80340.1 aluc|OJZ79553.1

Group13048: aluc|OJZ79532.1 aluc|OJZ80330.1 aluc|OJZ79907.1

Group13049: aluc|OJZ79546.1 aluc|OJZ80342.1 aluc|OJZ79842.1

Group13050: aluc|OJZ79572.1 aluc|OJZ81135.1 aluc|OJZ81136.1

Group13051: aluc|OJZ79578.1 aluc|OJZ80409.1 aluc|OJZ80089.1

Group13052: aluc|OJZ79590.1 aluc|OJZ80060.1 aluc|OJZ80407.1

Group13053: aluc|OJZ79621.1 aluc|OJZ80053.1 aluc|OJZ80216.1

Group13054: aluc|OJZ79633.1 aluc|OJZ79682.1 aluc|OJZ79632.1

Group13055: aluc|OJZ79661.1 aluc|OJZ80110.1 akaw|GAA89010.1

Group13056: aluc|OJZ79662.1 aluc|OJZ88847.1 aluc|OJZ80305.1

Group13057: aluc|OJZ79688.1 aluc|OJZ80464.1 aluc|OJZ79840.1

Group13058: aluc|OJZ79690.1 aluc|OJZ80036.1 aory|EIT75739.1

Group13059: aluc|OJZ79722.1 aluc|OJZ79868.1 akaw|GAA93162.1

Group13060: aluc|OJZ79725.1 aluc|OJZ80229.1 aluc|OJZ80137.1

Group13061: aluc|OJZ79759.1 aluc|OJZ80331.1 akaw|GAA93330.1

Group13062: aluc|OJZ79784.1 aluc|OJZ80378.1 akaw|GAA93174.1

Group13063: aluc|OJZ79819.1 aluc|OJZ80413.1 akaw|GAA93221.1

Group13064: aluc|OJZ79823.1 aluc|OJZ88820.1 anig|EHA17703.1

Group13065: aluc|OJZ79850.1 aluc|OJZ80295.1 akaw|GAA93153.1

Group13066: aluc|OJZ79972.1 aluc|OJZ80476.1 aluc|OJZ88836.1

Group13067: aluc|OJZ80116.1 aluc|OJZ80451.1 akaw|GAA93356.1

Group13068: aluc|OJZ80261.1 aluc|OJZ80343.1 akaw|GAA88995.1

Group13069: aluc|OJZ80448.1 aluc|OJZ80483.1 akaw|GAA93359.1

Group13070: anid|CBF79007.1 anid|CBF79011.1 anid|CBF84589.1

Group13071: anid|CBF84377.1 anid|CBF89339.1 anig|EHA19814.1

Group13072: anid|CBF87241.1 anid|CBF87256.1 agla|OJJ82041.1

Group13073: anig|EHA21127.1 anig|EHA21136.1 acla|EAW08997.1

Group13074: anig|EHA21832.1 anig|EHA21864.1 anid|CBF84450.1

Group13075: asyd|OJJ51839.1 asyd|OJJ52329.1 aluc|OJZ79788.1

Group13076: asyd|OJJ51921.1 asyd|OJJ52312.1 aluc|OJZ79961.1

Group13077: asyd|OJJ51923.1 asyd|OJJ52311.1 acla|EAW11463.1

Group13078: asyd|OJJ51928.1 asyd|OJJ51983.1 asyd|OJJ51962.1

Group13079: asyd|OJJ51941.1 asyd|OJJ52142.1 asyd|OJJ52127.1

Group13080: asyd|OJJ51950.1 asyd|OJJ62679.1 aluc|OJZ80358.1

Group13081: asyd|OJJ51959.1 asyd|OJJ62654.1 aver|OJI96753.1

Group13082: asyd|OJJ51964.1 asyd|OJJ62581.1 aver|OJI96701.1

Group13083: asyd|OJJ51989.1 asyd|OJJ54463.1 aver|OJJ07707.1

Group13084: asyd|OJJ52018.1 asyd|OJJ52146.1 asyd|OJJ52218.1

Group13085: asyd|OJJ52024.1 asyd|OJJ52283.1 afum|EAL87731.1

Group13086: asyd|OJJ52194.1 asyd|OJJ54732.1 asyd|OJJ54368.1

Group13087: asyd|OJJ52219.1 asyd|OJJ52305.1 ater|EAU35390.1

Group13088: asyd|OJJ52266.1 asyd|OJJ52318.1 asyd|OJJ62696.1

Group13089: asyd|OJJ53280.1 asyd|OJJ56764.1 aver|OJJ03569.1

Group13090: asyd|OJJ53535.1 asyd|OJJ63010.1 aver|OJJ08216.1

Group13091: asyd|OJJ53552.1 asyd|OJJ63020.1 ater|EAU31402.1

Group13092: asyd|OJJ53793.1 asyd|OJJ57301.1 aver|OJJ07903.1

Group13093: asyd|OJJ54694.1 asyd|OJJ62715.1 awen|OJJ38794.1

Group13094: asyd|OJJ54863.1 asyd|OJJ62955.1 aver|OJI98011.1

Group13095: asyd|OJJ58372.1 asyd|OJJ62488.1 agla|OJJ80481.1

Group13096: asyd|OJJ59285.1 asyd|OJJ65369.1 aver|OJJ07161.1

Group13097: ater|EAU37462.1 afla|EED47164.1 aory|EIT79262.1

Group13098: ater|EAU29535.1 ater|EAU34637.1 anid|CBF80205.1

Group13099: ater|EAU29605.1 ater|EAU30191.1 asyd|OJJ52414.1

Group13100: ater|EAU39200.1 asyd|OJJ55519.1 aver|OJJ06597.1

Group13101: atub|OJI87528.1 atub|OJI87530.1 atub|OJI87529.1

Group13102: aver|OJI97246.1 aver|OJJ08769.1 asyd|OJJ63689.1

Group13103: aver|OJJ02492.1 aver|OJJ08766.1 asyd|OJJ59797.1

Group13104: aver|OJJ05160.1 aver|OJJ07378.1 asyd|OJJ54718.1

Group13105: aver|OJJ06703.1 aver|OJJ07808.1 asyd|OJJ55626.1

Group13106: awen|OJJ29575.1 awen|OJJ29614.1 akaw|GAA92645.1

Group13107: awen|OJJ30087.1 awen|OJJ38073.1 agla|OJJ85527.1

Group13108: awen|OJJ30778.1 awen|OJJ31703.1 agla|OJJ82180.1

Group13109: awen|OJJ30790.1 awen|OJJ31691.1 anid|CBF80735.1

Group13110: awen|OJJ30819.1 awen|OJJ38002.1 atub|OJI84643.1

Group13111: awen|OJJ31595.1 awen|OJJ40839.1 azon|OJJ49490.1

Group13112: awen|OJJ31918.1 awen|OJJ31919.1 afis|EAW20128.1

Group13113: awen|OJJ31970.1 awen|OJJ35520.1 afis|EAW19977.1

Group13114: awen|OJJ32042.1 awen|OJJ34523.1 awen|OJJ34522.1

Group13115: awen|OJJ32342.1 aver|OJI98034.1 asyd|OJJ56969.1

Group13116: awen|OJJ32514.1 awen|OJJ36009.1 abra|OJJ77883.1

Group13117: awen|OJJ35967.1 awen|OJJ35968.1 afum|EAL86713.1

Group13118: awen|OJJ36925.1 awen|OJJ36930.1 azon|OJJ43654.1

Group13119: awen|OJJ40017.1 awen|OJJ40018.1 awen|OJJ31326.1

Group13120: azon|OJJ42236.1 azon|OJJ45313.1 azon|OJJ46424.1

Group13121: azon|OJJ42670.1 azon|OJJ45164.1 azon|OJJ48054.1

Group13122: azon|OJJ43556.1 azon|OJJ45986.1 azon|OJJ46494.1

Group13123: azon|OJJ43576.1 azon|OJJ45574.1 azon|OJJ47406.1

Group13124: azon|OJJ43643.1 azon|OJJ48721.1 agla|OJJ84717.1

Group13125: azon|OJJ44501.1 azon|OJJ45177.1 acar|OOF94767.1

Group13126: azon|OJJ46560.1 azon|OJJ51073.1 awen|OJJ32916.1

Group13127: abra|OJJ65433.1 abra|OJJ65439.1 aluc|OJZ79731.1

Group13128: abra|OJJ65454.1 abra|OJJ65568.1 abra|OJJ65525.1

Group13129: abra|OJJ65629.1 abra|OJJ65662.1 abra|OJJ65730.1

Group13130: anid|CBF69925.1 anid|CBF73430.1 anid|CBF84241.1

Group13131: anid|CBF71863.1 anid|CBF76093.1 anid|CBF80239.1

Group13132: asyd|OJJ51880.1 asyd|OJJ52236.1 asyd|OJJ52277.1

Group13133: asyd|OJJ51890.1 asyd|OJJ52307.1 asyd|OJJ62687.1

Group13134: asyd|OJJ51896.1 asyd|OJJ51997.1 asyd|OJJ52308.1

Group13135: asyd|OJJ51937.1 asyd|OJJ52242.1 asyd|OJJ52132.1

Group13136: asyd|OJJ51940.1 asyd|OJJ52143.1 asyd|OJJ52128.1

Group13137: asyd|OJJ51951.1 asyd|OJJ62678.1 afum|EAL85395.1

Group13138: asyd|OJJ51958.1 asyd|OJJ52278.1 asyd|OJJ62655.1

Group13139: asyd|OJJ51986.1 asyd|OJJ62708.1 afis|EAW20717.1

Group13140: ater|EAU32912.1 ater|EAU32920.1 ater|EAU32919.1

Group13141: atub|OJI83367.1 atub|OJI83369.1 atub|OJI83368.1

Group13142: aver|OJI96127.1 aver|OJJ04447.1 ater|EAU36835.1

Group13143: awen|OJJ29530.1 awen|OJJ29608.1 awen|OJJ29531.1

Group13144: awen|OJJ29549.1 awen|OJJ29620.1 awen|OJJ29633.1

Group13145: azon|OJJ42045.1 azon|OJJ42048.1 azon|OJJ42047.1

Group13146: azon|OJJ42570.1 azon|OJJ46523.1 azon|OJJ46435.1

Group13147: acla|EAW08958.1 acla|EAW10829.1 acla|EAW14874.1

Group13148: afis|EAW16431.1 afis|EAW18465.1 afis|EAW20644.1

Group13149: agla|OJJ78941.1 agla|OJJ87858.1 agla|OJJ86639.1

Group13150: agla|OJJ79794.1 agla|OJJ83136.1 agla|OJJ87697.1

Group13151: agla|OJJ79795.1 agla|OJJ81748.1 agla|OJJ86424.1

Group13152: agla|OJJ80621.1 agla|OJJ80623.1 agla|OJJ80622.1

Group13153: agla|OJJ81012.1 agla|OJJ83432.1 aver|OJI96722.1

Group13154: agla|OJJ81343.1 agla|OJJ83434.1 agla|OJJ82253.1

Group13155: agla|OJJ82004.1 agla|OJJ86549.1 agla|OJJ83785.1

Group13156: agla|OJJ82220.1 agla|OJJ89483.1 agla|OJJ87263.1

Group13157: agla|OJJ82842.1 agla|OJJ82844.1 ater|EAU38146.1

Group13158: agla|OJJ84373.1 agla|OJJ89336.1 agla|OJJ85304.1

Group13159: agla|OJJ85904.1 agla|OJJ85907.1 agla|OJJ85905.1

Group13160: aluc|OJZ79510.1 aluc|OJZ80402.1 aluc|OJZ79862.1

Group13161: aluc|OJZ79525.1 aluc|OJZ79980.1 aluc|OJZ80369.1

Group13162: aluc|OJZ79550.1 aluc|OJZ80078.1 aluc|OJZ79744.1

Group13163: aluc|OJZ79568.1 aluc|OJZ80459.1 aluc|OJZ80288.1

Group13164: aluc|OJZ79647.1 aluc|OJZ80336.1 aluc|OJZ80043.1

Group13165: aluc|OJZ79708.1 aluc|OJZ79953.1 aluc|OJZ80152.1

Group13166: aluc|OJZ79728.1 aluc|OJZ80232.1 aluc|OJZ79938.1

Group13167: aluc|OJZ79791.1 aluc|OJZ80333.1 aluc|OJZ80048.1

Group13168: aluc|OJZ79816.1 aluc|OJZ80082.1 aluc|OJZ80251.1

Group13169: aluc|OJZ79818.1 aluc|OJZ80105.1 aluc|OJZ80084.1

Group13170: aluc|OJZ79981.1 aluc|OJZ80489.1 aluc|OJZ80169.1

Group13171: abra|OJJ65419.1 aory|EIT83501.1 asyd|OJJ52296.1

Group13172: abra|OJJ65427.1 agla|OJJ86334.1 akaw|GAA92644.1

Group13173: abra|OJJ65442.1 akaw|BAL04887.1 azon|OJJ41976.1

Group13174: anig|EHA17698.1 akaw|GAA92965.1 aluc|OJZ80329.1

Group13175: abra|OJJ65538.1 afum|EAL91912.1 asyd|OJJ57188.1

Group13176: abra|OJJ65565.1 akaw|GAA93355.1 awen|OJJ32280.1

Group13177: abra|OJJ65593.1 aluc|OJZ80057.1 akaw|GAA93258.1

Group13178: awen|OJJ32908.1 atub|OJI86040.1 asyd|OJJ54901.1

Group13179: abra|OJJ65615.1 aluc|OJZ80006.1 akaw|GAA93150.1

Group13180: akaw|GAA92882.1 abra|OJJ70029.1 atub|OJI79317.1

Group13181: abra|OJJ65641.1 acar|OOF97538.1 awen|OJJ32714.1

Group13182: abra|OJJ65646.1 acla|EAW06926.1 awen|OJJ35899.1

Group13183: abra|OJJ65667.1 aluc|OJZ81152.1 atub|OJI88769.1

Group13184: abra|OJJ65669.1 anid|CBF76111.1 atub|OJI82833.1

Group13185: abra|OJJ65674.1 aluc|OJZ79541.1 aver|OJJ03423.1

Group13186: abra|OJJ65675.1 aluc|OJZ79542.1 aver|OJJ03430.1

Group13187: abra|OJJ65677.1 afis|EAW20905.1 afum|EAL84728.2

Group13188: asyd|OJJ54177.1 aver|OJJ08562.1 azon|OJJ42582.1

Group13189: abra|OJJ65704.1 aluc|OJZ79923.1 ater|EAU35383.1

Group13190: abra|OJJ65705.1 acar|OOF90744.1 aluc|OJZ79926.1

Group13191: agla|OJJ83803.1 acar|OOF90629.1 aluc|OJZ79482.1

Group13192: abra|OJJ65757.1 afla|EED46308.1 aory|EIT76345.1

Group13193: aory|EIT78110.1 acla|EAW14267.1 afla|EED50758.1

Group13194: asyd|OJJ54316.1 ater|EAU32012.1 aver|OJJ05270.1

Group13195: acla|EAW13704.1 afis|EAW19882.1 afum|EAL85016.1

Group13196: abra|OJJ65790.1 anig|EHA18728.1 anig|EHA18729.1

Group13197: abra|OJJ65793.1 asyd|OJJ57208.1 aver|OJJ03081.1

Group13198: anid|CBF80352.1 asyd|OJJ54830.1 aver|OJJ07331.1

Group13199: abra|OJJ65798.1 asyd|OJJ54902.1 aver|OJJ02468.1

Group13200: afla|EED47497.1 aory|EIT74577.1 azon|OJJ42013.1

Group13201: abra|OJJ65811.1 anig|EHA22497.1 atub|OJI88776.1

Group13202: awen|OJJ37048.1 afla|EED53430.1 aory|EIT74995.1

Group13203: abra|OJJ65842.1 akaw|GAA92841.1 ater|EAU33404.1

Group13204: abra|OJJ65845.1 akaw|GAA92839.1 aluc|OJZ88784.1

Group13205: ater|EAU36824.1 afla|EED54136.1 aory|EIT75925.1

Group13206: anid|CBF80140.1 agla|OJJ85369.1 aver|OJJ01889.1

Group13207: acla|EAW11042.1 afla|EED49625.1 aory|EIT80945.1

Group13208: afla|EED54351.1 afis|EAW20559.1 afum|EAL84502.1

Group13209: abra|OJJ65908.1 anig|EHA21129.1 awen|OJJ33613.1

Group13210: ater|EAU29593.1 afum|EAL85115.1 aluc|OJZ81563.1

Group13211: abra|OJJ66031.1 afla|EED52375.1 atub|OJI87436.1

Group13212: abra|OJJ66032.1 atub|OJI87435.1 awen|OJJ37489.1

Group13213: abra|OJJ66035.1 atub|OJI87432.1 awen|OJJ37488.1

Group13214: abra|OJJ66037.1 ater|EAU32812.1 atub|OJI82469.1

Group13215: abra|OJJ66095.1 asyd|OJJ55458.1 aver|OJJ03133.1

Group13216: abra|OJJ66097.1 aver|OJJ03134.1 asyd|OJJ55457.1

Group13217: anid|CBF82365.1 asyd|OJJ54199.1 aver|OJJ08525.1

Group13218: acar|OOF98883.1 aver|OJI97586.1 asyd|OJJ63362.1

Group13219: aver|OJJ06719.1 aluc|OJZ89983.1 asyd|OJJ52508.1

Group13220: acla|EAW13555.1 aver|OJJ07460.1 afla|EED49761.1

Group13221: abra|OJJ66164.1 afla|EED47402.1 agla|OJJ79412.1

Group13222: abra|OJJ66174.1 asyd|OJJ61492.1 aver|OJJ00833.1

Group13223: abra|OJJ66178.1 akaw|GAA93275.1 aluc|OJZ79677.1

Group13224: abra|OJJ66184.1 acar|OOF96460.1 asyd|OJJ65284.1

Group13225: abra|OJJ66188.1 afla|EED48374.1 aory|EIT83087.1

Group13226: abra|OJJ66205.1 aluc|OJZ89829.1 atub|OJI84977.1

Group13227: afum|EAL92110.2 acla|EAW13633.1 afis|EAW21087.1

Group13228: abra|OJJ66272.1 anig|EHA23653.1 atub|OJI89950.1

Group13229: abra|OJJ66273.1 acla|EAW10348.1 afis|EAW19777.1

Group13230: abra|OJJ66274.1 anig|EHA23655.1 atub|OJI89948.1

Group13231: abra|OJJ66280.1 anig|EHA23660.1 atub|OJI89943.1

Group13232: abra|OJJ66287.1 anig|EHA23667.1 atub|OJI89936.1

Group13233: abra|OJJ66298.1 asyd|OJJ56621.1 azon|OJJ42429.1

Group13234: abra|OJJ66299.1 acla|EAW10957.1 afum|EAL87796.1

Group13235: abra|OJJ66301.1 afla|EED47877.1 aluc|OJZ80192.1

Group13236: abra|OJJ66309.1 asyd|OJJ52054.1 anig|EHA21831.1

Group13237: abra|OJJ66316.1 afla|EED52351.1 akaw|GAA93076.1

Group13238: abra|OJJ66380.1 asyd|OJJ54799.1 aver|OJJ07360.1

Group13239: abra|OJJ66381.1 asyd|OJJ54798.1 aver|OJJ07361.1

Group13240: abra|OJJ66382.1 asyd|OJJ54797.1 aver|OJJ07362.1

Group13241: abra|OJJ66432.1 acar|OOF90242.1 agla|OJJ84658.1

Group13242: abra|OJJ66436.1 akaw|GAA83941.1 aluc|OJZ80823.1

Group13243: abra|OJJ66495.1 asyd|OJJ54146.1 aver|OJI96128.1

Group13244: abra|OJJ66602.1 aluc|OJZ81444.1 atub|OJI88734.1

Group13245: abra|OJJ66663.1 aluc|OJZ80784.1 anig|EHA27245.1

Group13246: abra|OJJ66730.1 aluc|OJZ80719.1 atub|OJI87575.1

Group13247: abra|OJJ66805.1 afla|EED53420.1 atub|OJI79339.1

Group13248: abra|OJJ66814.1 afla|EED53416.1 atub|OJI79327.1

Group13249: abra|OJJ66885.1 anig|EHA24845.1 atub|OJI79487.1

Group13250: abra|OJJ66913.1 aluc|OJZ85667.1 atub|OJI79460.1

Group13251: abra|OJJ66930.1 afla|EED49766.1 awen|OJJ41938.1

Group13252: agla|OJJ89296.1 aver|OJJ08535.1 asyd|OJJ54192.1

Group13253: abra|OJJ67002.1 afis|EAW25644.1 agla|OJJ83006.1

Group13254: abra|OJJ67025.1 akaw|GAA83297.1 anig|EHA24959.1

Group13255: abra|OJJ67042.1 afla|EED52066.1 aory|EIT72304.1

Group13256: abra|OJJ67049.1 agla|OJJ82956.1 anig|EHA18594.1

Group13257: azon|OJJ42485.1 afla|EED52158.1 aory|EIT74503.1

Group13258: abra|OJJ67051.1 asyd|OJJ62602.1 awen|OJJ33861.1

Group13259: abra|OJJ67055.1 afla|EED52906.1 aory|EIT78640.1

Group13260: abra|OJJ67063.1 acar|OOF95670.1 anig|EHA18943.1

Group13261: abra|OJJ67107.1 aluc|OJZ81114.1 atub|OJI86628.1

Group13262: abra|OJJ67135.1 agla|OJJ78765.1 anig|EHA20146.1

Group13263: abra|OJJ67136.1 agla|OJJ86210.1 anig|EHA20315.1

Group13264: acla|EAW08555.1 afum|EAL87869.1 afis|EAW23597.1

Group13265: abra|OJJ67220.1 ater|EAU37939.1 aver|OJI99454.1

Group13266: abra|OJJ67221.1 afla|EED50961.1 atub|OJI81177.1

Group13267: abra|OJJ67289.1 akaw|GAA87404.1 atub|OJI81012.1

Group13268: abra|OJJ67315.1 afla|EED46372.1 aory|EIT72289.1

Group13269: abra|OJJ67355.1 anig|EHA25225.1 asyd|OJJ54611.1

Group13270: abra|OJJ67362.1 acar|OOF92460.1 aluc|OJZ85001.1

Group13271: asyd|OJJ58445.1 aver|OJJ01045.1 anid|CBF70718.1

Group13272: abra|OJJ67603.1 anig|EHA25721.1 atub|OJI88464.1

Group13273: abra|OJJ67613.1 akaw|GAA89131.1 aluc|OJZ81707.1

Group13274: acar|OOF91967.1 asyd|OJJ60369.1 aver|OJI99705.1

Group13275: anid|CBF73891.1 asyd|OJJ53661.1 aver|OJJ07769.1

Group13276: abra|OJJ67626.1 akaw|GAA89146.1 aluc|OJZ81690.1

Group13277: abra|OJJ67705.1 atub|OJI88564.1 aver|OJI99277.1

Group13278: abra|OJJ67712.1 ater|EAU33736.1 acar|OOF94899.1

Group13279: abra|OJJ67768.1 afla|EED55026.1 aory|EIT81984.1

Group13280: abra|OJJ67800.1 aluc|OJZ90856.1 afla|EED50187.1

Group13281: acar|OOF99271.1 asyd|OJJ52758.1 aver|OJJ08740.1

Group13282: abra|OJJ67834.1 afum|EAL87812.1 atub|OJI89151.1

Group13283: abra|OJJ67895.1 aluc|OJZ81995.1 atub|OJI84545.1

Group13284: abra|OJJ67946.1 aluc|OJZ82818.1 anig|EHA21258.1

Group13285: abra|OJJ67960.1 atub|OJI89063.1 aluc|OJZ82833.1

Group13286: abra|OJJ68026.1 acar|OOF92069.1 anig|EHA21323.1

Group13287: acla|EAW08928.1 afla|EED45085.1 aory|EIT73673.1

Group13288: abra|OJJ68158.1 aluc|OJZ83223.1 atub|OJI87713.1

Group13289: abra|OJJ68213.1 akaw|GAA83233.1 atub|OJI90110.1

Group13290: abra|OJJ68247.1 akaw|GAA83214.1 aluc|OJZ83716.1

Group13291: agla|OJJ80188.1 akaw|GAA89978.1 aluc|OJZ83989.1

Group13292: anid|CBF86783.1 aver|OJI98782.1 asyd|OJJ64680.1

Group13293: anid|CBF76040.1 asyd|OJJ52460.1 aver|OJJ03955.1

Group13294: anid|CBF82267.1 asyd|OJJ59955.1 aver|OJJ02334.1

Group13295: ater|EAU35301.1 asyd|OJJ58134.1 aver|OJJ04271.1

Group13296: anid|CBF84434.1 asyd|OJJ53635.1 aver|OJJ01171.1

Group13297: ater|EAU36743.1 afum|EAL88826.2 afis|EAW16195.1

Group13298: awen|OJJ36254.1 asyd|OJJ64168.1 aver|OJI98243.1

Group13299: abra|OJJ68629.1 aluc|OJZ81926.1 atub|OJI84478.1

Group13300: anid|CBF76074.1 asyd|OJJ55071.1 aver|OJJ07140.1

Group13301: aory|EIT72254.1 afla|EED46756.1 awen|OJJ41298.1

Group13302: azon|OJJ42015.1 asyd|OJJ65326.1 aver|OJI99471.1

Group13303: abra|OJJ68831.1 aluc|OJZ85809.1 atub|OJI79595.1

Group13304: afum|EAL84273.1 acla|EAW12210.1 afis|EAW25550.1

Group13305: afis|EAW17014.1 akaw|GAA89813.1 aluc|OJZ81877.1

Group13306: afum|EAL89943.1 afis|EAW25280.1 azon|OJJ50338.1

Group13307: abra|OJJ68946.1 aluc|OJZ85932.1 atub|OJI79721.1

Group13308: abra|OJJ68974.1 aluc|OJZ85963.1 atub|OJI79750.1

Group13309: aory|EIT82518.1 afla|EED49095.1 anid|CBF80558.1

Group13310: abra|OJJ69076.1 acar|OOF91215.1 asyd|OJJ55841.1

Group13311: afis|EAW19978.1 acla|EAW14481.1 afum|EAL94012.2

Group13312: abra|OJJ69338.1 aluc|OJZ82170.1 atub|OJI88267.1

Group13313: abra|OJJ69360.1 aluc|OJZ82193.1 atub|OJI88242.1

Group13314: abra|OJJ69495.1 aluc|OJZ82335.1 atub|OJI88105.1

Group13315: abra|OJJ69549.1 aluc|OJZ82379.1 atub|OJI88062.1

Group13316: abra|OJJ69586.1 aluc|OJZ82403.1 atub|OJI88038.1

Group13317: abra|OJJ69601.1 acar|OOF93268.1 awen|OJJ38319.1

Group13318: abra|OJJ69610.1 aluc|OJZ82418.1 atub|OJI88023.1

Group13319: agla|OJJ88060.1 asyd|OJJ56268.1 aver|OJJ08499.1

Group13320: atub|OJI87960.1 akaw|GAA88793.1 aluc|OJZ82476.1

Group13321: abra|OJJ69740.1 aluc|OJZ83156.1 atub|OJI87781.1

Group13322: agla|OJJ82665.1 asyd|OJJ60040.1 aver|OJJ02245.1

Group13323: abra|OJJ69781.1 afis|EAW24837.1 aver|OJJ05999.1

Group13324: asyd|OJJ59192.1 afla|EED53431.1 aory|EIT75228.1

Group13325: anid|CBF84269.1 asyd|OJJ57206.1 aver|OJJ03083.1

Group13326: aory|EIT74838.1 afis|EAW25613.1 afla|EED53589.1

Group13327: asyd|OJJ64743.1 anid|CBF86696.1 aver|OJI98847.1

Group13328: azon|OJJ42398.1 aver|OJI99434.1 asyd|OJJ65283.1

Group13329: abra|OJJ70032.1 anig|EHA18717.1 atub|OJI79321.1

Group13330: abra|OJJ70040.1 anig|EHA18722.1 atub|OJI79326.1

Group13331: abra|OJJ70091.1 acar|OOF90921.1 atub|OJI80359.1

Group13332: abra|OJJ70092.1 atub|OJI80360.1 atub|OJI88455.1

Group13333: abra|OJJ70099.1 afla|EED48387.1 aory|EIT83070.1

Group13334: aory|EIT75501.1 afla|EED45516.1 agla|OJJ79484.1

Group13335: abra|OJJ70191.1 akaw|GAA86585.1 azon|OJJ43536.1

Group13336: aver|OJJ07775.1 ater|EAU33242.1 awen|OJJ36207.1

Group13337: abra|OJJ70259.1 aluc|OJZ85085.1 anig|EHA25300.1

Group13338: anid|CBF74552.1 asyd|OJJ57684.1 aver|OJJ05490.1

Group13339: abra|OJJ70271.1 agla|OJJ82947.1 awen|OJJ34691.1

Group13340: abra|OJJ70344.1 acar|OOF91922.1 anig|EHA27467.1

Group13341: abra|OJJ70362.1 acar|OOF89853.1 ater|EAU29960.1

Group13342: abra|OJJ70393.1 akaw|GAA91966.1 anig|EHA27421.1

Group13343: abra|OJJ70405.1 acar|OOF93981.1 anig|EHA21037.1

Group13344: abra|OJJ70408.1 acar|OOF94952.1 awen|OJJ32276.1

Group13345: abra|OJJ70479.1 asyd|OJJ55444.1 aver|OJJ06517.1

Group13346: abra|OJJ70507.1 anig|EHA18994.1 atub|OJI90219.1

Group13347: abra|OJJ70545.1 aluc|OJZ83890.1 atub|OJI90257.1

Group13348: abra|OJJ70635.1 ater|EAU33583.1 awen|OJJ31594.1

Group13349: afla|EED56727.1 aory|EIT73927.1 azon|OJJ48357.1

Group13350: abra|OJJ70737.1 aluc|OJZ84082.1 atub|OJI90452.1

Group13351: abra|OJJ70771.1 aluc|OJZ84117.1 atub|OJI90490.1

Group13352: abra|OJJ70822.1 acla|EAW09020.1 anig|EHA18000.1

Group13353: abra|OJJ70825.1 acla|EAW09023.1 anig|EHA17998.1

Group13354: afum|EAL88031.1 afis|EAW23430.1 azon|OJJ45956.1

Group13355: abra|OJJ70886.1 acla|EAW10809.1 aluc|OJZ87636.1

Group13356: abra|OJJ70903.1 akaw|GAA87417.1 aluc|OJZ87621.1

Group13357: abra|OJJ70908.1 anig|EHA20814.1 atub|OJI81033.1

Group13358: abra|OJJ70909.1 aluc|OJZ87615.1 anig|EHA20812.1

Group13359: abra|OJJ71197.1 akaw|GAA82786.1 anig|EHA20582.1

Group13360: abra|OJJ71215.1 anig|EHA20559.1 aver|OJJ03896.1

Group13361: aluc|OJZ81564.1 akaw|GAA92454.1 atub|OJI88621.1

Group13362: abra|OJJ71431.1 akaw|GAA82572.1 aluc|OJZ87107.1

Group13363: abra|OJJ71437.1 asyd|OJJ55468.1 aver|OJJ06551.1

Group13364: abra|OJJ71448.1 aluc|OJZ87093.1 atub|OJI81545.1

Group13365: abra|OJJ71459.1 aluc|OJZ87084.1 azon|OJJ44805.1

Group13366: abra|OJJ71530.1 anig|EHA21600.1 atub|OJI84272.1

Group13367: abra|OJJ71569.1 aluc|OJZ86306.1 atub|OJI84230.1

Group13368: abra|OJJ71604.1 awen|OJJ38715.1 azon|OJJ42405.1

Group13369: abra|OJJ71605.1 aluc|OJZ86345.1 atub|OJI84189.1

Group13370: abra|OJJ71606.1 aluc|OJZ86346.1 anig|EHA21540.1

Group13371: abra|OJJ71640.1 aluc|OJZ86383.1 atub|OJI84150.1

Group13372: abra|OJJ71683.1 aluc|OJZ86425.1 atub|OJI84106.1

Group13373: abra|OJJ71776.1 akaw|GAA85132.1 aluc|OJZ86506.1

Group13374: abra|OJJ71785.1 aluc|OJZ86515.1 atub|OJI84009.1

Group13375: anid|CBF70301.1 aver|OJJ01341.1 asyd|OJJ58728.1

Group13376: abra|OJJ71833.1 atub|OJI83956.1 aluc|OJZ86565.1

Group13377: abra|OJJ71835.1 acar|OOF96509.1 aluc|OJZ86566.1

Group13378: afum|EAL90928.1 acar|OOF90709.1 awen|OJJ36067.1

Group13379: abra|OJJ71927.1 acar|OOF96584.1 agla|OJJ85699.1

Group13380: abra|OJJ71943.1 aluc|OJZ86679.1 atub|OJI83848.1

Group13381: awen|OJJ35354.1 afla|EED47438.1 aory|EIT81895.1

Group13382: abra|OJJ72057.1 acar|OOF96714.1 atub|OJI83725.1

Group13383: abra|OJJ72072.1 afis|EAW19914.1 ater|EAU32298.1

Group13384: abra|OJJ72104.1 akaw|GAA84818.1 aluc|OJZ86859.1

Group13385: anid|CBF73513.1 asyd|OJJ55808.1 aver|OJJ04756.1

Group13386: atub|OJI88063.1 acar|OOF90605.1 anig|EHA28386.1

Group13387: abra|OJJ72275.1 anig|EHA19912.1 awen|OJJ35392.1

Group13388: abra|OJJ72276.1 anig|EHA19913.1 awen|OJJ35393.1

Group13389: abra|OJJ72288.1 aluc|OJZ90267.1 atub|OJI81692.1

Group13390: azon|OJJ51716.1 afla|EED49861.1 aory|EIT83391.1

Group13391: azon|OJJ42108.1 asyd|OJJ56533.1 aver|OJJ06190.1

Group13392: abra|OJJ72390.1 atub|OJI81802.1 aluc|OJZ90377.1

Group13393: abra|OJJ72500.1 anig|EHA23941.1 atub|OJI81908.1

Group13394: abra|OJJ72526.1 aluc|OJZ90506.1 atub|OJI81930.1

Group13395: abra|OJJ72597.1 aluc|OJZ90578.1 atub|OJI82003.1

Group13396: abra|OJJ72612.1 anig|EHA24033.1 atub|OJI82017.1

Group13397: aver|OJJ03327.1 anid|CBF77092.1 asyd|OJJ56958.1

Group13398: awen|OJJ39033.1 aory|EIT78028.1 afla|EED50605.1

Group13399: abra|OJJ72691.1 asyd|OJJ52468.1 aver|OJJ03947.1

Group13400: abra|OJJ72711.1 anig|EHA24108.1 atub|OJI82106.1

Group13401: abra|OJJ72766.1 aluc|OJZ90741.1 atub|OJI82168.1

Group13402: abra|OJJ72812.1 aluc|OJZ83310.1 atub|OJI82210.1

Group13403: abra|OJJ72832.1 aluc|OJZ83333.1 atub|OJI82229.1

Group13404: anid|CBF85195.1 asyd|OJJ63017.1 aver|OJI97947.1

Group13405: abra|OJJ72958.1 acar|OOF96140.1 atub|OJI82360.1

Group13406: abra|OJJ73045.1 aluc|OJZ83556.1 anig|EHA24374.1

Group13407: abra|OJJ73060.1 aluc|OJZ80026.1 anig|EHA26112.1

Group13408: abra|OJJ73061.1 aluc|OJZ90872.1 anig|EHA19592.1

Group13409: abra|OJJ73064.1 aluc|OJZ81362.1 atub|OJI91566.1

Group13410: abra|OJJ73070.1 aluc|OJZ81355.1 atub|OJI91559.1

Group13411: abra|OJJ73102.1 akaw|GAA88063.1 aluc|OJZ81326.1

Group13412: abra|OJJ73129.1 anig|EHA18703.1 azon|OJJ42287.1

Group13413: abra|OJJ73165.1 asyd|OJJ58360.1 aver|OJJ02046.1

Group13414: aory|EIT72407.1 afla|EED46508.1 asyd|OJJ54747.1

Group13415: abra|OJJ73292.1 aluc|OJZ88756.1 atub|OJI91326.1

Group13416: abra|OJJ73307.1 anig|EHA21033.1 awen|OJJ35330.1

Group13417: anid|CBF71403.1 asyd|OJJ55029.1 aver|OJJ07175.1

Group13418: anid|CBF74654.1 asyd|OJJ61480.1 aver|OJJ00821.1

Group13419: afla|EED48949.1 aory|EIT79994.1 ater|EAU31698.1

Group13420: abra|OJJ73419.1 acar|OOG00621.1 akaw|GAA83072.1

Group13421: acar|OOG00676.1 afla|EED53517.1 aory|EIT74919.1

Group13422: abra|OJJ73521.1 acar|OOF90602.1 aver|OJJ04731.1

Group13423: asyd|OJJ55175.1 acar|OOF98340.1 aver|OJJ07032.1

Group13424: abra|OJJ73523.1 aory|EIT72915.1 afla|EED48257.1

Group13425: agla|OJJ81320.1 acar|OOF94217.1 azon|OJJ48031.1

Group13426: abra|OJJ73703.1 anig|EHA18081.1 awen|OJJ33597.1

Group13427: anid|CBF89961.1 aver|OJI96211.1 asyd|OJJ62105.1

Group13428: abra|OJJ73709.1 aluc|OJZ88341.1 atub|OJI90937.1

Group13429: abra|OJJ73749.1 aluc|OJZ88303.1 atub|OJI90898.1

Group13430: akaw|GAA82096.1 aluc|OJZ92083.1 atub|OJI86513.1

Group13431: abra|OJJ73893.1 aluc|OJZ88155.1 atub|OJI90755.1

Group13432: abra|OJJ74121.1 afum|EAL85398.1 aluc|OJZ84287.1

Group13433: abra|OJJ74187.1 afum|EAL86235.1 ater|EAU32781.1

Group13434: agla|OJJ79816.1 afla|EED51187.1 asyd|OJJ57220.1

Group13435: awen|OJJ37003.1 asyd|OJJ52473.1 aver|OJJ03943.1

Group13436: abra|OJJ74500.1 aluc|OJZ89618.1 atub|OJI85184.1

Group13437: abra|OJJ74516.1 asyd|OJJ54858.1 atub|OJI86530.1

Group13438: abra|OJJ74517.1 afis|EAW24049.1 asyd|OJJ54860.1

Group13439: abra|OJJ74567.1 aluc|OJZ92146.1 anig|EHA26803.1

Group13440: anid|CBF85874.1 aver|OJI98470.1 asyd|OJJ64374.1

Group13441: aver|OJI96389.1 anid|CBF89655.1 asyd|OJJ62280.1

Group13442: abra|OJJ74902.1 akaw|GAA86630.1 aluc|OJZ85185.1

Group13443: atub|OJI86906.1 akaw|GAA83520.1 aluc|OJZ92475.1

Group13444: abra|OJJ75050.1 aluc|OJZ92616.1 anig|EHA26372.1

Group13445: abra|OJJ75051.1 aluc|OJZ92617.1 anig|EHA26371.1

Group13446: asyd|OJJ55809.1 aver|OJJ04757.1 anid|CBF73511.1

Group13447: abra|OJJ75184.1 aluc|OJZ92748.1 anig|EHA26255.1

Group13448: abra|OJJ75193.1 acar|OOF92441.1 azon|OJJ43635.1

Group13449: abra|OJJ75221.1 anig|EHA26224.1 atub|OJI87227.1

Group13450: anig|EHA26223.1 atub|OJI87228.1 anid|CBF84476.1

Group13451: abra|OJJ75225.1 afla|EED48955.1 aory|EIT80016.1

Group13452: abra|OJJ75226.1 acar|OOF94563.1 anig|EHA26220.1

Group13453: abra|OJJ75259.1 aluc|OJZ92816.1 atub|OJI87274.1

Group13454: abra|OJJ75277.1 aluc|OJZ92839.1 atub|OJI87295.1

Group13455: abra|OJJ75343.1 akaw|GAA92036.1 atub|OJI87365.1

Group13456: aver|OJJ04537.1 anid|CBF79456.1 asyd|OJJ57857.1

Group13457: abra|OJJ75481.1 aluc|OJZ88941.1 atub|OJI85849.1

Group13458: abra|OJJ75542.1 akaw|GAA84298.1 aluc|OJZ88992.1

Group13459: abra|OJJ75587.1 aluc|OJZ89038.1 atub|OJI85752.1

Group13460: abra|OJJ75648.1 acar|OOG00192.1 aluc|OJZ89097.1

Group13461: abra|OJJ75685.1 aluc|OJZ89136.1 atub|OJI85649.1

Group13462: abra|OJJ75691.1 akaw|GAA84428.1 aluc|OJZ89142.1

Group13463: abra|OJJ75703.1 akaw|GAA84439.1 aluc|OJZ89154.1

Group13464: abra|OJJ75706.1 aluc|OJZ89157.1 atub|OJI85634.1

Group13465: afla|EED54451.1 aory|EIT81153.1 ater|EAU39266.1

Group13466: abra|OJJ75730.1 aluc|OJZ89182.1 atub|OJI85607.1

Group13467: ater|EAU30356.1 afla|EED47831.1 aory|EIT77106.1

Group13468: abra|OJJ75872.1 aluc|OJZ89315.1 atub|OJI85482.1

Group13469: aver|OJI99935.1 anid|CBF83343.1 asyd|OJJ60611.1

Group13470: aver|OJI98895.1 asyd|OJJ64784.1 anid|CBF83888.1

Group13471: abra|OJJ76136.1 aluc|OJZ89599.1 atub|OJI85205.1

Group13472: abra|OJJ76169.1 acar|OOF89917.1 anig|EHA27995.1

Group13473: abra|OJJ76237.1 agla|OJJ87582.1 awen|OJJ31905.1

Group13474: abra|OJJ76282.1 anig|EHA22898.1 atub|OJI86381.1

Group13475: abra|OJJ76305.1 aluc|OJZ91910.1 atub|OJI86360.1

Group13476: aver|OJJ01556.1 anid|CBF69791.1 asyd|OJJ58962.1

Group13477: aory|EIT81951.1 afla|EED55068.1 azon|OJJ50761.1

Group13478: abra|OJJ76668.1 anig|EHA22360.1 anid|CBF83130.1

Group13479: azon|OJJ49992.1 asyd|OJJ60963.1 aver|OJJ00288.1

Group13480: abra|OJJ76681.1 anig|EHA28109.1 atub|OJI87913.1

Group13481: abra|OJJ76735.1 afis|EAW21008.1 aluc|OJZ80161.1

Group13482: abra|OJJ76755.1 aluc|OJZ84158.1 anig|EHA27520.1

Group13483: abra|OJJ76879.1 asyd|OJJ60060.1 ater|EAU38554.1

Group13484: abra|OJJ76881.1 agla|OJJ83398.1 asyd|OJJ56899.1

Group13485: abra|OJJ76894.1 afla|EED57871.1 aluc|OJZ82046.1

Group13486: ater|EAU31578.1 afla|EED54079.1 aory|EIT75980.1

Group13487: abra|OJJ76963.1 acla|EAW15301.1 anid|CBF69364.1

Group13488: abra|OJJ76998.1 acar|OOF91444.1 anig|EHA27727.1

Group13489: asyd|OJJ52268.1 akaw|GAA91194.1 aluc|OJZ81528.1

Group13490: ater|EAU29539.1 afla|EED55090.1 aory|EIT81609.1

Group13491: abra|OJJ77083.1 aory|EIT82369.1 aver|OJJ08346.1

Group13492: asyd|OJJ52100.1 afla|EED49838.1 aory|EIT83390.1

Group13493: abra|OJJ77234.1 aluc|OJZ84644.1 atub|OJI89437.1

Group13494: agla|OJJ87589.1 acar|OOF93156.1 aluc|OJZ80473.1

Group13495: aver|OJI99401.1 agla|OJJ79236.1 asyd|OJJ65249.1

Group13496: abra|OJJ77436.1 acar|OOF94972.1 agla|OJJ82718.1

Group13497: abra|OJJ77462.1 asyd|OJJ56401.1 aver|OJJ06311.1

Group13498: abra|OJJ77563.1 aluc|OJZ80975.1 anig|EHA19384.1

Group13499: abra|OJJ77592.1 aluc|OJZ80944.1 atub|OJI82639.1

Group13500: abra|OJJ77696.1 aluc|OJZ90759.1 atub|OJI82746.1

Group13501: asyd|OJJ60258.1 acla|EAW15293.1 aver|OJJ03998.1

Group13502: abra|OJJ77766.1 acar|OOF97504.1 anig|EHA20912.1

Group13503: abra|OJJ77775.1 anig|EHA19551.1 atub|OJI82814.1

Group13504: afis|EAW16159.1 agla|OJJ85019.1 ater|EAU35526.1

Group13505: abra|OJJ77795.1 acar|OOF90712.1 awen|OJJ38008.1

Group13506: abra|OJJ77828.1 atub|OJI82895.1 aluc|OJZ90902.1

Group13507: abra|OJJ77872.1 afis|EAW16223.1 awen|OJJ30238.1

Group13508: agla|OJJ87728.1 awen|OJJ40103.1 azon|OJJ42373.1

Group13509: abra|OJJ77907.1 aluc|OJZ90978.1 atub|OJI82968.1

Group13510: abra|OJJ77914.1 afla|EED51611.1 aory|EIT80662.1

Group13511: abra|OJJ77915.1 afla|EED51609.1 aory|EIT80525.1

Group13512: abra|OJJ77941.1 aluc|OJZ91008.1 atub|OJI83000.1

Group13513: abra|OJJ77998.1 aluc|OJZ91074.1 atub|OJI83069.1

Group13514: abra|OJJ78071.1 anid|CBF87123.1 aver|OJJ08736.1

Group13515: abra|OJJ78090.1 aluc|OJZ91159.1 atub|OJI83174.1

Group13516: abra|OJJ78100.1 aluc|OJZ81974.1 atub|OJI79694.1

Group13517: aory|EIT76397.1 afla|EED46251.1 ater|EAU37247.1

Group13518: abra|OJJ78165.1 akaw|GAA92954.1 ater|EAU30328.1

Group13519: abra|OJJ78180.1 afis|EAW19763.1 ater|EAU36859.1

Group13520: acar|OOF89771.1 acla|EAW08168.1 anig|EHA21138.1

Group13521: acar|OOF89783.1 anid|CBF87194.1 awen|OJJ30942.1

Group13522: acar|OOF90007.1 afla|EED45511.1 aory|EIT75792.1

Group13523: acar|OOF90016.1 acla|EAW07718.1 afla|EED47873.1

Group13524: afum|EAL88059.1 afla|EED57311.1 aory|EIT74057.1

Group13525: acar|OOF90039.1 afla|EED56935.1 aory|EIT77574.1

Group13526: acar|OOF90089.1 acla|EAW15338.1 afla|EED53069.1

Group13527: acar|OOF90171.1 acla|EAW09325.1 awen|OJJ33546.1

Group13528: acar|OOF90201.1 afla|EED51998.1 aory|EIT81476.1

Group13529: acar|OOF90286.1 aluc|OJZ84547.1 atub|OJI89533.1

Group13530: acar|OOF90368.1 anig|EHA23742.1 azon|OJJ49782.1

Group13531: agla|OJJ82267.1 awen|OJJ30817.1 azon|OJJ46539.1

Group13532: acar|OOF90450.1 anid|CBF82346.1 ater|EAU33653.1

Group13533: asyd|OJJ52812.1 aver|OJJ08791.1 azon|OJJ43865.1

Group13534: acar|OOF90468.1 afla|EED53720.1 aory|EIT75294.1

Group13535: acar|OOF90536.1 acla|EAW11292.1 azon|OJJ50401.1

Group13536: aver|OJJ05145.1 anid|CBF73623.1 asyd|OJJ56212.1

Group13537: anid|CBF85208.1 aver|OJI97937.1 asyd|OJJ63031.1

Group13538: anid|CBF83801.1 aver|OJJ00214.1 asyd|OJJ60890.1

Group13539: acar|OOF90572.1 afla|EED45166.1 aory|EIT73787.1

Group13540: acar|OOF90610.1 asyd|OJJ54124.1 aver|OJJ08613.1

Group13541: acar|OOF90623.1 afla|EED49835.1 afum|EAL84843.2

Group13542: asyd|OJJ54479.1 afla|EED52150.1 aory|EIT74494.1

Group13543: agla|OJJ79092.1 aory|EIT83085.1 awen|OJJ31975.1

Group13544: acar|OOF90685.1 afla|EED55099.1 aory|EIT81646.1

Group13545: acar|OOF90903.1 anig|EHA25865.1 atub|OJI88610.1

Group13546: acla|EAW10870.1 afis|EAW20988.1 afum|EAL84641.1

Group13547: acar|OOF91760.1 afum|EAL85381.1 azon|OJJ42426.1

Group13548: agla|OJJ88057.1 asyd|OJJ56264.1 aver|OJJ08502.1

Group13549: acar|OOF91785.1 afla|EED50928.1 aory|EIT79038.1

Group13550: acar|OOF91853.1 asyd|OJJ58351.1 aver|OJJ02056.1

Group13551: acar|OOF91912.1 afla|EED48618.1 aory|EIT73257.1

Group13552: acar|OOF91915.1 afis|EAW17742.1 aver|OJJ03427.1

Group13553: acar|OOF91919.1 afla|EED52010.1 aory|EIT81397.1

Group13554: acar|OOF92527.1 afla|EED52078.1 aory|EIT76921.1

Group13555: acar|OOF92558.1 agla|OJJ87573.1 awen|OJJ40605.1

Group13556: acar|OOF92697.1 afla|EED48163.1 aory|EIT83039.1

Group13557: acar|OOF93146.1 afla|EED49522.1 aory|EIT82655.1

Group13558: acar|OOF93186.1 aluc|OJZ92894.1 anig|EHA26069.1

Group13559: acla|EAW10018.1 afis|EAW20416.1 afum|EAL89229.1

Group13560: azon|OJJ43593.1 asyd|OJJ61555.1 aver|OJJ00878.1

Group13561: acar|OOF93559.1 aver|OJI96960.1 awen|OJJ41961.1

Group13562: acar|OOF93563.1 acla|EAW15372.1 azon|OJJ49996.1

Group13563: aver|OJI95853.1 agla|OJJ79832.1 asyd|OJJ61755.1

Group13564: acar|OOF93634.1 aory|EIT73965.1 afla|EED57139.1

Group13565: acar|OOF93878.1 afis|EAW19288.1 asyd|OJJ63019.1

Group13566: acar|OOF94173.1 anig|EHA28282.1 aver|OJJ07860.1

Group13567: acar|OOF94382.1 asyd|OJJ55055.1 aver|OJJ07929.1

Group13568: acar|OOF94383.1 asyd|OJJ55054.1 aver|OJJ07155.1

Group13569: acar|OOF94488.1 aluc|OJZ92910.1 atub|OJI87370.1

Group13570: akaw|GAA89159.1 atub|OJI88506.1 aluc|OJZ81675.1

Group13571: acar|OOF94642.1 afla|EED54694.1 aory|EIT78888.1

Group13572: acar|OOF94680.1 acla|EAW11915.1 ater|EAU33994.1

Group13573: anid|CBF84386.1 asyd|OJJ59691.1 aver|OJJ02596.1

Group13574: aver|OJI96462.1 ater|EAU34576.1 asyd|OJJ55663.1

Group13575: acar|OOF95235.1 aluc|OJZ89608.1 atub|OJI85195.1

Group13576: acar|OOF95248.1 aver|OJI99580.1 awen|OJJ38747.1

Group13577: acar|OOF95249.1 aver|OJI99579.1 awen|OJJ38745.1

Group13578: acar|OOF95250.1 aver|OJI99578.1 awen|OJJ38746.1

Group13579: awen|OJJ38895.1 afla|EED48421.1 aory|EIT83063.1

Group13580: acar|OOF95613.1 afla|EED51176.1 aory|EIT81334.1

Group13581: afis|EAW20917.1 afum|EAL84717.1 awen|OJJ34404.1

Group13582: anig|EHA26755.1 atub|OJI86630.1 aluc|OJZ92202.1

Group13583: anid|CBF84438.1 asyd|OJJ53637.1 aver|OJJ01169.1

Group13584: anid|CBF84987.1 asyd|OJJ63123.1 aver|OJI97831.1

Group13585: agla|OJJ83747.1 asyd|OJJ64380.1 aver|OJI98476.1

Group13586: acar|OOF96794.1 asyd|OJJ60411.1 azon|OJJ45234.1

Group13587: acar|OOF96891.1 afla|EED52817.1 aory|EIT78210.1

Group13588: acar|OOF97088.1 agla|OJJ85080.1 aver|OJI96137.1

Group13589: aver|OJI98818.1 anid|CBF86722.1 asyd|OJJ64715.1

Group13590: acar|OOF97275.1 afis|EAW19853.1 anig|EHA18369.1

Group13591: afis|EAW23441.1 awen|OJJ39758.1 acla|EAW08398.1

Group13592: acar|OOF97567.1 awen|OJJ29727.1 azon|OJJ44461.1

Group13593: ater|EAU32638.1 afla|EED52564.1 aory|EIT74252.1

Group13594: acar|OOF97588.1 afla|EED52459.1 aory|EIT82742.1

Group13595: acar|OOF97619.1 agla|OJJ85960.1 awen|OJJ36079.1

Group13596: acar|OOF97643.1 afla|EED45610.1 aory|EIT75686.1

Group13597: acar|OOF98057.1 agla|OJJ88711.1 asyd|OJJ64699.1

Group13598: acar|OOF98084.1 akaw|GAA82903.1 atub|OJI81176.1

Group13599: acar|OOF98168.1 afla|EED55408.1 aory|EIT80856.1

Group13600: acar|OOF98352.1 akaw|GAA87700.1 azon|OJJ44814.1

Group13601: anid|CBF78625.1 asyd|OJJ65126.1 aver|OJI99254.1

Group13602: acar|OOF98696.1 aver|OJJ05894.1 azon|OJJ46431.1

Group13603: acar|OOF98899.1 afum|EAL88851.1 atub|OJI82938.1

Group13604: acar|OOF98996.1 asyd|OJJ65117.1 aver|OJI99243.1

Group13605: awen|OJJ35026.1 acla|EAW15296.1 ater|EAU37654.1

Group13606: acar|OOF99082.1 aory|EIT72397.1 afla|EED46487.1

Group13607: aver|OJI98313.1 agla|OJJ79990.1 awen|OJJ36308.1

Group13608: acar|OOF99296.1 afla|EED46492.1 aory|EIT72427.1

Group13609: asyd|OJJ64553.1 anid|CBF86213.1 aver|OJI98650.1

Group13610: afis|EAW21266.1 acla|EAW09860.1 afum|EAL92281.1

Group13611: anid|CBF83725.1 asyd|OJJ60842.1 aver|OJJ00167.1

Group13612: ater|EAU39618.1 afla|EED52447.1 aory|EIT82816.1

Group13613: acar|OOG00489.1 afis|EAW16897.1 agla|OJJ84612.1

Group13614: ater|EAU34443.1 asyd|OJJ56610.1 aver|OJJ06113.1

Group13615: anid|CBF89226.1 asyd|OJJ62517.1 aver|OJI96636.1

Group13616: acar|OOG00733.1 afis|EAW16800.1 anid|CBF80491.1

Group13617: acar|OOG00885.1 afla|EED55019.1 azon|OJJ51782.1

Group13618: acar|OOG00910.1 agla|OJJ81043.1 aver|OJI97286.1

Group13619: acla|EAW06521.1 afla|EED56607.1 aory|EIT79647.1

Group13620: ater|EAU33705.1 afla|EED53340.1 aory|EIT74976.1

Group13621: acla|EAW06585.1 agla|OJJ79821.1 ater|EAU37856.1

Group13622: acla|EAW06633.1 afis|EAW16132.1 afum|EAL88762.1

Group13623: aver|OJI98210.1 anid|CBF85520.1 asyd|OJJ64127.1

Group13624: anid|CBF85538.1 aver|OJI98221.1 asyd|OJJ64140.1

Group13625: acla|EAW06669.1 afis|EAW16096.1 afum|EAL88728.1

Group13626: acla|EAW06679.1 afis|EAW16086.1 afum|EAL88718.1

Group13627: acla|EAW06838.1 afis|EAW16458.1 afum|EAL89054.1

Group13628: acla|EAW06839.1 afum|EAL89056.2 afis|EAW16460.1

Group13629: acla|EAW06978.1 afis|EAW16592.1 afum|EAL89188.1

Group13630: acla|EAW07058.1 anid|CBF73843.1 aver|OJJ06166.1

Group13631: afum|EAL89048.1 ater|EAU36716.1 anid|CBF87741.1

Group13632: acla|EAW07087.1 afum|EAL89046.1 aver|OJJ03912.1

Group13633: acla|EAW07090.1 asyd|OJJ60593.1 aver|OJI99917.1

Group13634: aory|EIT75269.1 afla|EED53642.1 awen|OJJ30138.1

Group13635: acla|EAW07108.1 afis|EAW22120.1 afum|EAL91066.1

Group13636: acla|EAW07110.1 afis|EAW22122.1 afum|EAL91065.1

Group13637: acla|EAW07172.1 afis|EAW22187.1 afum|EAL91003.1

Group13638: acla|EAW07211.1 afis|EAW22224.1 afum|EAL90970.1

Group13639: acla|EAW07285.1 afis|EAW22318.1 afum|EAL90872.2

Group13640: anid|CBF73621.1 asyd|OJJ56213.1 aver|OJJ05146.1

Group13641: acla|EAW07487.1 afis|EAW22520.1 afum|EAL90673.1

Group13642: acla|EAW07493.1 afis|EAW22526.1 afum|EAL90668.1

Group13643: acla|EAW07587.1 afis|EAW22618.1 afum|EAL90578.1

Group13644: asyd|OJJ60310.1 aver|OJI99651.1 awen|OJJ34012.1

Group13645: acla|EAW07719.1 afis|EAW22750.1 afum|EAL90449.1

Group13646: aory|EIT81780.1 afla|EED55635.1 azon|OJJ47755.1

Group13647: acla|EAW08401.1 afis|EAW23468.1 afum|EAL87998.1

Group13648: aver|OJI96373.1 asyd|OJJ62264.1 azon|OJJ43240.1

Group13649: acla|EAW08484.1 afis|EAW23530.1 afum|EAL87936.1

Group13650: acla|EAW08518.1 afis|EAW23556.1 afum|EAL87910.1

Group13651: acla|EAW08699.1 afis|EAW15736.1 afum|EAL85612.2

Group13652: acla|EAW08766.1 afis|EAW15664.1 afum|EAL85684.1

Group13653: acla|EAW08778.1 afis|EAW15623.1 afum|EAL85711.1

Group13654: acla|EAW08974.1 afis|EAW15453.1 afum|EAL84192.1

Group13655: acla|EAW08978.1 afis|EAW15431.1 afum|EAL84212.1

Group13656: acla|EAW09015.1 afis|EAW20065.1 ater|EAU30766.1

Group13657: acla|EAW09062.1 agla|OJJ88760.1 anid|CBF78316.1

Group13658: acla|EAW09076.1 asyd|OJJ59977.1 aver|OJJ02314.1

Group13659: acla|EAW09097.1 afis|EAW25691.1 afum|EAL84361.1

Group13660: acla|EAW09113.1 afis|EAW16774.1 afum|EAL86343.2

Group13661: acla|EAW09125.1 afum|EAL94068.1 ater|EAU34936.1

Group13662: asyd|OJJ60859.1 anid|CBF83755.1 aver|OJJ00184.1

Group13663: acla|EAW09443.1 afis|EAW21693.1 afum|EAL92727.1

Group13664: acla|EAW09543.1 afis|EAW21590.1 afum|EAL92608.1

Group13665: acla|EAW09597.1 afis|EAW21540.1 afum|EAL92560.1

Group13666: ater|EAU30139.1 afla|EED48248.1 akaw|GAA92066.1

Group13667: acla|EAW09758.1 afis|EAW21383.1 afum|EAL92398.1

Group13668: acla|EAW09836.1 afis|EAW21298.1 afum|EAL92315.1

Group13669: acla|EAW09884.1 afis|EAW21242.1 afum|EAL92255.1

Group13670: acla|EAW09996.1 afis|EAW21077.1 afum|EAL92100.1

Group13671: acla|EAW10103.1 afis|EAW25484.1 afum|EAL90149.1

Group13672: acla|EAW10227.1 afis|EAW25365.1 ater|EAU30940.1

Group13673: acla|EAW10395.1 afis|EAW25182.1 afum|EAL89856.1

Group13674: acla|EAW10443.1 afis|EAW25124.1 afum|EAL89795.1

Group13675: acla|EAW10511.1 afis|EAW25056.1 afum|EAL89724.1

Group13676: acla|EAW10718.1 afis|EAW25571.1 afum|EAL89289.1

Group13677: acla|EAW10733.1 afis|EAW24842.1 afum|EAL89488.1

Group13678: acla|EAW10763.1 afis|EAW24813.1 afum|EAL89457.1

Group13679: acla|EAW10784.1 afla|EED57526.1 aory|EIT81845.1

Group13680: acla|EAW10817.1 afis|EAW24757.1 afum|EAL89399.1

Group13681: acla|EAW10828.1 afis|EAW24736.1 afum|EAL89383.2

Group13682: anid|CBF74920.1 asyd|OJJ60079.1 aver|OJJ04166.1

Group13683: acla|EAW10833.1 afis|EAW24731.1 afum|EAL89378.1

Group13684: acla|EAW10835.1 afis|EAW24728.1 afum|EBA27377.1

Group13685: acla|EAW10837.1 afis|EAW24726.1 afum|EAL89374.1

Group13686: ater|EAU35994.1 afis|EAW20252.1 afum|EAL85370.1

Group13687: acla|EAW10838.1 afla|EED55832.1 aory|EIT76669.1

Group13688: anid|CBF82831.1 aver|OJJ00904.1 asyd|OJJ61582.1

Group13689: acla|EAW11033.1 afla|EED49637.1 aory|EIT81003.1

Group13690: acla|EAW11043.1 afla|EED49626.1 aory|EIT80954.1

Group13691: acla|EAW11045.1 afla|EED49628.1 aory|EIT81000.1

Group13692: agla|OJJ86141.1 asyd|OJJ55482.1 aver|OJJ06561.1

Group13693: acla|EAW11047.1 afla|EED49630.1 aory|EIT80951.1

Group13694: asyd|OJJ55039.1 aver|OJJ07168.1 ater|EAU35915.1

Group13695: anig|EHA17707.1 akaw|GAA92950.1 aluc|OJZ81139.1

Group13696: acla|EAW11468.1 afis|EAW20727.1 afum|EBA27361.1

Group13697: acla|EAW11516.1 afis|EAW23937.1 afum|EAL87362.1

Group13698: anid|CBF77100.1 aver|OJJ03332.1 asyd|OJJ56953.1

Group13699: aluc|OJZ88122.1 asyd|OJJ61460.1 aver|OJJ00797.1

Group13700: acla|EAW11682.1 afis|EAW23699.1 afum|EAL87140.1

Group13701: acla|EAW11695.1 afis|EAW23712.1 afum|EAL87156.2

Group13702: ater|EAU37842.1 afla|EED54051.1 aory|EIT75938.1

Group13703: acla|EAW11712.1 afis|EAW23727.1 afum|EBA27301.1

Group13704: anid|CBF85113.1 asyd|OJJ63050.1 aver|OJI97913.1

Group13705: ater|EAU34966.1 asyd|OJJ61455.1 aver|OJJ00791.1

Group13706: acla|EAW12034.1 afum|EBA27231.1 afis|EAW20285.1

Group13707: acla|EAW12058.1 afis|EAW19699.1 afum|EAL86828.1

Group13708: acla|EAW12064.1 afla|EED54089.1 aory|EIT76091.1

Group13709: acla|EAW12200.1 afis|EAW25310.1 agla|OJJ89131.1

Group13710: acla|EAW12209.1 afis|EAW25549.1 afum|EAL84272.1

Group13711: acla|EAW12211.1 afis|EAW25551.1 afum|EAL84274.1

Group13712: acla|EAW12213.1 afis|EAW25553.1 afum|EAL84277.1

Group13713: acla|EAW12214.1 afis|EAW25554.1 afum|EAL84278.1

Group13714: acla|EAW12217.1 afis|EAW25558.1 afum|EAL84282.1

Group13715: acla|EAW12218.1 afis|EAW25559.1 afum|EAL84283.1

Group13716: acla|EAW12251.1 anid|CBF74952.1 awen|OJJ38332.1

Group13717: afis|EAW16957.1 aver|OJJ02654.1 anid|CBF81774.1

Group13718: acla|EAW12287.1 afis|EAW16951.1 afum|EAL86478.1

Group13719: acla|EAW12356.1 akaw|GAA92919.1 asyd|OJJ52439.1

Group13720: asyd|OJJ65081.1 anid|CBF78585.1 aver|OJI99206.1

Group13721: acla|EAW12424.1 anid|CBF83040.1 ater|EAU36465.1

Group13722: acla|EAW12445.1 aory|EIT74205.1 azon|OJJ51540.1

Group13723: acla|EAW12646.1 afis|EAW18039.1 afum|EAL91850.1

Group13724: acla|EAW12877.1 afis|EAW18833.1 afum|EAL93569.1

Group13725: acla|EAW12889.1 afis|EAW17787.1 afum|EAL91596.1

Group13726: acla|EAW12896.1 afis|EAW17771.1 afum|EAL91580.1

Group13727: acla|EAW12930.1 afis|EAW17707.1 afum|EAL91521.2

Group13728: awen|OJJ37129.1 afla|EED45709.1 aory|EIT75795.1

Group13729: aory|EIT82271.1 afla|EED57073.1 awen|OJJ35544.1

Group13730: acla|EAW13181.1 afis|EAW17459.1 afum|EAL91287.1

Group13731: acla|EAW13234.1 afis|EAW17396.1 afum|EAL91234.2

Group13732: acla|EAW13269.1 afis|EAW17350.1 afum|EAL91193.1

Group13733: acla|EAW13471.1 afum|EBA27181.1 awen|OJJ33289.1

Group13734: acla|EAW13488.1 asyd|OJJ57241.1 azon|OJJ42369.1

Group13735: afla|EED52641.1 aory|EIT74265.1 awen|OJJ30501.1

Group13736: acla|EAW13609.1 afis|EAW20001.1 afum|EAL84918.1

Group13737: acla|EAW13622.1 afis|EAW22102.1 afum|EAL91088.1

Group13738: acla|EAW13631.1 afis|EAW21089.1 afum|EAL92112.1

Group13739: acla|EAW13668.1 afum|EAL85439.1 ater|EAU37463.1

Group13740: agla|OJJ78676.1 azon|OJJ48734.1 ater|EAU30333.1

Group13741: acla|EAW13745.1 afum|EBA27295.1 aver|OJJ05558.1

Group13742: acla|EAW13755.1 afis|EAW19811.1 afum|EAL85081.1

Group13743: acla|EAW13765.1 afis|EAW19794.1 afum|EAL85096.2

Group13744: acla|EAW13859.1 afis|EAW15438.1 afum|EAL84207.1

Group13745: acla|EAW14081.1 afis|EAW18788.1 afum|EAL93522.2

Group13746: acla|EAW14095.1 afum|EBA27362.1 agla|OJJ86300.1

Group13747: acla|EAW14261.1 afis|EAW19019.1 awen|OJJ31307.1

Group13748: aory|EIT80629.1 afla|EED51732.1 anid|CBF79136.1

Group13749: acla|EAW14273.1 aluc|OJZ87300.1 atub|OJI81340.1

Group13750: acla|EAW14458.1 afis|EAW19166.1 afum|EAL93896.1

Group13751: acla|EAW14536.1 aory|EIT78882.1 afla|EED54702.1

Group13752: anid|CBF78490.1 aver|OJI99474.1 asyd|OJJ65329.1

Group13753: acla|EAW14584.1 afis|EAW19317.1 afum|EAL94034.1

Group13754: aver|OJI99654.1 asyd|OJJ60313.1 awen|OJJ34015.1

Group13755: acla|EAW14881.1 afis|EAW24400.1 afum|EAL86071.1

Group13756: acla|EAW14925.1 afla|EED49817.1 aory|EIT77657.1

Group13757: acla|EAW14951.1 afis|EAW24468.1 afum|EAL86140.1

Group13758: acla|EAW14955.1 afla|EED57791.1 aory|EIT81078.1

Group13759: acla|EAW14984.1 afis|EAW24506.1 afum|EAL86174.1

Group13760: acla|EAW14995.1 afis|EAW24521.1 afum|EAL86186.1

Group13761: acla|EAW15025.1 afum|EAL86638.1 awen|OJJ35955.1

Group13762: acla|EAW15057.1 afis|EAW19780.1 afum|EAL85109.1

Group13763: ater|EAU34054.1 afla|EED45170.1 aory|EIT73812.1

Group13764: acla|EAW15260.1 afis|EAW20235.1 afum|EAL85352.1

Group13765: acla|EAW15269.1 afis|EAW20240.1 afum|EAL85358.1

Group13766: acla|EAW15288.1 afum|EAL86978.1 afis|EAW19519.1

Group13767: acla|EAW15292.1 agla|OJJ84603.1 awen|OJJ36128.1

Group13768: acla|EAW15299.1 afis|EAW25741.1 anid|CBF69357.1

Group13769: afis|EAW15385.1 asyd|OJJ60081.1 aver|OJJ05930.1

Group13770: afis|EAW15464.1 asyd|OJJ57379.1 aver|OJJ05779.1

Group13771: aory|EIT77913.1 afla|EED46602.1 azon|OJJ46007.1

Group13772: afis|EAW15639.1 ater|EAU30734.1 azon|OJJ42124.1

Group13773: afis|EAW15649.1 asyd|OJJ53725.1 aver|OJI98280.1

Group13774: afis|EAW16068.1 afum|EAL88700.2 awen|OJJ30367.1

Group13775: afis|EAW16153.1 afla|EED45106.1 aory|EIT73674.1

Group13776: afis|EAW16154.1 afla|EED45107.1 aory|EIT73576.1

Group13777: afis|EAW16168.1 afla|EED57466.1 awen|OJJ38679.1

Group13778: afum|EAL87138.1 afla|EED48463.1 aory|EIT76273.1

Group13779: afis|EAW16172.1 afla|EED50601.1 aory|EIT78015.1

Group13780: afis|EAW16197.1 aver|OJJ08287.1 asyd|OJJ53469.1

Group13781: afis|EAW16228.1 agla|OJJ88692.1 awen|OJJ37851.1

Group13782: afis|EAW16413.1 afum|EAL89037.1 anid|CBF73424.1

Group13783: afis|EAW16426.1 asyd|OJJ61463.1 aver|OJJ05755.1

Group13784: anig|EHA19925.1 akaw|GAA92002.1 awen|OJJ32680.1

Group13785: afis|EAW16441.1 afum|EAL89042.1 azon|OJJ45297.1

Group13786: afis|EAW16442.1 afum|EAL89043.1 azon|OJJ45296.1

Group13787: afis|EAW16633.1 afla|EED53429.1 aory|EIT74831.1

Group13788: afis|EAW16635.1 afum|EAL89237.1 ater|EAU37744.1

Group13789: aver|OJJ08811.1 anid|CBF87376.1 asyd|OJJ52833.1

Group13790: afis|EAW16797.1 afla|EED46068.1 aory|EIT83498.1

Group13791: afis|EAW16896.1 afum|EAL86433.1 awen|OJJ32236.1

Group13792: afis|EAW16907.1 afla|EED51714.1 aory|EIT80505.1

Group13793: afis|EAW16924.1 afla|EED45474.1 aory|EIT75616.1

Group13794: afis|EAW16942.1 afum|EBA27274.1 anid|CBF84649.1

Group13795: afis|EAW16958.1 awen|OJJ32043.1 ater|EAU37671.1

Group13796: afis|EAW16970.1 afum|EAL86498.1 ater|EAU37853.1

Group13797: afla|EED45041.1 awen|OJJ37846.1 azon|OJJ43183.1

Group13798: afis|EAW17013.1 akaw|GAA89812.1 aluc|OJZ81878.1

Group13799: afis|EAW17021.1 akaw|GAA89820.1 aluc|OJZ81870.1

Group13800: afis|EAW17027.1 aluc|OJZ79603.1 awen|OJJ40157.1

Group13801: afis|EAW17066.1 afum|EAL90927.1 asyd|OJJ52120.1

Group13802: afis|EAW17115.1 afum|EAL86577.1 awen|OJJ35966.1

Group13803: afis|EAW17133.1 afla|EED52045.1 awen|OJJ37886.1

Group13804: afis|EAW17157.1 agla|OJJ88354.1 ater|EAU34934.1

Group13805: afis|EAW17244.1 afum|EBA27284.1 ater|EAU37685.1

Group13806: afis|EAW17254.1 asyd|OJJ59282.1 aver|OJJ01901.1

Group13807: awen|OJJ34566.1 afla|EED45153.1 aory|EIT73773.1

Group13808: afis|EAW17374.1 afum|EAL91210.1 anid|CBF88126.1

Group13809: anid|CBF71585.1 asyd|OJJ61872.1 aver|OJI95970.1

Group13810: afis|EAW17527.1 afum|EAL86502.1 ater|EAU39524.1

Group13811: afis|EAW17752.1 afla|EED48584.1 asyd|OJJ54474.1

Group13812: afis|EAW18091.1 agla|OJJ79756.1 asyd|OJJ62670.1

Group13813: afis|EAW18095.1 afla|EED56535.1 aory|EIT79697.1

Group13814: afis|EAW18133.1 afla|EED47886.1 aory|EIT77249.1

Group13815: anid|CBF76044.1 aver|OJJ03953.1 asyd|OJJ52462.1

Group13816: ater|EAU32196.1 asyd|OJJ54758.1 aver|OJJ07412.1

Group13817: ater|EAU39465.1 afla|EED46932.1 aory|EIT72641.1

Group13818: aory|EIT83066.1 afla|EED48392.1 awen|OJJ30962.1

Group13819: afis|EAW19227.1 afla|EED49597.1 awen|OJJ37949.1

Group13820: afis|EAW19255.1 afla|EED56899.1 awen|OJJ32282.1

Group13821: afis|EAW19385.1 afla|EED52380.1 aory|EIT82371.1

Group13822: afis|EAW19518.1 afum|EAL86979.1 aluc|OJZ81079.1

Group13823: afis|EAW19534.1 afum|EAL86972.1 ater|EAU36060.1

Group13824: afis|EAW19535.1 afum|EBA27296.1 awen|OJJ29902.1

Group13825: afis|EAW19559.1 afum|EAL86952.1 azon|OJJ47318.1

Group13826: afis|EAW19561.1 awen|OJJ35389.1 azon|OJJ50605.1

Group13827: akaw|GAA93094.1 afla|EED49495.1 aory|EIT82621.1

Group13828: azon|OJJ46905.1 afla|EED50996.1 aory|EIT76979.1

Group13829: anid|CBF73989.1 aver|OJI97958.1 awen|OJJ38741.1

Group13830: afis|EAW19747.1 afum|EAL91595.1 afla|EED47526.1

Group13831: afis|EAW19766.1 ater|EAU36862.1 azon|OJJ42249.1

Group13832: afis|EAW19824.1 afum|EAL85068.1 aory|EIT83367.1

Group13833: anid|CBF71501.1 asyd|OJJ54930.1 aver|OJJ07267.1

Group13834: afis|EAW19889.1 afla|EED51139.1 azon|OJJ42765.1

Group13835: afis|EAW19968.1 afum|EAL84950.2 atub|OJI83197.1

Group13836: aver|OJJ07821.1 afla|EED55889.1 asyd|OJJ52017.1

Group13837: anid|CBF87204.1 afum|EAL84869.1 ater|EAU37637.1

Group13838: afis|EAW20260.1 afum|EAL94037.1 agla|OJJ88690.1

Group13839: afis|EAW20270.1 afum|EAL85404.1 asyd|OJJ60802.1

Group13840: afis|EAW20290.1 afum|EAL85421.1 azon|OJJ50178.1

Group13841: afis|EAW20321.1 asyd|OJJ65357.1 aver|OJI99506.1

Group13842: awen|OJJ34570.1 asyd|OJJ55125.1 aver|OJJ07079.1

Group13843: afla|EED56668.1 aory|EIT79657.1 ater|EAU31814.1

Group13844: azon|OJJ46411.1 afla|EED51754.1 aory|EIT80606.1

Group13845: afis|EAW20411.1 afum|EAL89233.1 ater|EAU30196.1

Group13846: afis|EAW20673.1 agla|OJJ83423.1 anid|CBF84447.1

Group13847: afis|EAW20710.1 afum|EAL87791.1 awen|OJJ29623.1

Group13848: afis|EAW20731.1 afla|EED54426.1 aory|EIT75889.1

Group13849: afis|EAW20778.1 afla|EED50592.1 aory|EIT82087.1

Group13850: afis|EAW20786.1 afum|EAL84838.2 azon|OJJ42752.1

Group13851: afis|EAW20823.1 afum|EAL84806.1 aver|OJJ00337.1

Group13852: afis|EAW20826.1 afum|EAL84803.1 asyd|OJJ56856.1

Group13853: afis|EAW20855.1 awen|OJJ40931.1 azon|OJJ42353.1

Group13854: afis|EAW21004.1 agla|OJJ82565.1 awen|OJJ37980.1

Group13855: afis|EAW21016.1 afum|EAL92054.1 ater|EAU37795.1

Group13856: afis|EAW21027.1 afum|EAL92065.1 ater|EAU32574.1

Group13857: afis|EAW21063.1 ater|EAU31346.1 awen|OJJ30101.1

Group13858: agla|OJJ78558.1 awen|OJJ37794.1 asyd|OJJ53111.1

Group13859: afis|EAW21196.1 asyd|OJJ59259.1 aver|OJJ01871.1

Group13860: afis|EAW21200.1 awen|OJJ32935.1 ater|EAU31235.1

Group13861: afis|EAW21201.1 awen|OJJ32936.1 azon|OJJ45228.1

Group13862: afis|EAW21214.1 afum|EAL92225.1 awen|OJJ29901.1

Group13863: afis|EAW21277.1 afum|EAL92292.1 ater|EAU31599.1

Group13864: aver|OJI97601.1 agla|OJJ82388.1 asyd|OJJ53439.1

Group13865: afis|EAW21808.1 agla|OJJ78924.1 awen|OJJ35922.1

Group13866: afis|EAW22081.1 afum|EAL85690.1 aory|EIT81007.1

Group13867: anid|CBF88955.1 asyd|OJJ62807.1 aver|OJI96853.1

Group13868: asyd|OJJ62622.1 awen|OJJ36992.1 aver|OJI99300.1

Group13869: afis|EAW22747.1 akaw|GAA91088.1 aluc|OJZ87905.1

Group13870: afis|EAW23233.1 afum|EAL88249.1 ater|EAU36362.1

Group13871: afis|EAW23458.1 afum|EAL88009.1 ater|EAU39520.1

Group13872: afis|EAW23669.1 afum|EAL87114.1 asyd|OJJ53732.1

Group13873: aory|EIT79602.1 afla|EED56554.1 ater|EAU37799.1

Group13874: afis|EAW23702.1 afum|EAL87143.1 asyd|OJJ52253.1

Group13875: afis|EAW23714.1 afum|EAL87159.1 aver|OJI95857.1

Group13876: akaw|GAA92444.1 aluc|OJZ81575.1 atub|OJI88606.1

Group13877: afis|EAW23845.1 afla|EED46471.1 aory|EIT72380.1

Group13878: afis|EAW23914.1 awen|OJJ35045.1 agla|OJJ82266.1

Group13879: azon|OJJ48047.1 atub|OJI89954.1 awen|OJJ37779.1

Group13880: afis|EAW24397.1 agla|OJJ78877.1 awen|OJJ32808.1

Group13881: afla|EED45372.1 asyd|OJJ54839.1 aver|OJJ07322.1

Group13882: afis|EAW24594.1 aver|OJI96486.1 agla|OJJ80220.1

Group13883: afis|EAW24604.1 agla|OJJ82372.1 aver|OJI96495.1

Group13884: atub|OJI79830.1 afla|EED45394.1 aory|EIT82440.1

Group13885: aluc|OJZ85564.1 akaw|GAA92157.1 atub|OJI79358.1

Group13886: afis|EAW24903.1 asyd|OJJ60234.1 aver|OJJ04031.1

Group13887: afis|EAW25174.1 afum|EAL89847.1 azon|OJJ42795.1

Group13888: afis|EAW25193.1 afum|EBA27388.1 ater|EAU31574.1

Group13889: akaw|GAA92422.1 aluc|OJZ81600.1 atub|OJI88582.1

Group13890: afis|EAW25608.1 afla|EED53585.1 aory|EIT74870.1

Group13891: afis|EAW25610.1 afla|EED53587.1 aory|EIT74929.1

Group13892: afis|EAW25615.1 afla|EED53581.1 aory|EIT75131.1

Group13893: afis|EAW25662.1 asyd|OJJ53541.1 aver|OJJ08212.1

Group13894: aluc|OJZ81310.1 akaw|GAA88049.1 aver|OJJ03854.1

Group13895: afis|EAW25685.1 afum|EAL84354.1 awen|OJJ30831.1

Group13896: afis|EAW25722.1 afum|EBA27176.1 agla|OJJ78648.1

Group13897: asyd|OJJ55726.1 agla|OJJ84802.1 anid|CBF79202.1

Group13898: afla|EED44751.1 aory|EIT77783.1 azon|OJJ50358.1

Group13899: afla|EED44758.1 asyd|OJJ54766.1 aver|OJJ07401.1

Group13900: awen|OJJ34013.1 asyd|OJJ60311.1 aver|OJI99652.1

Group13901: afla|EED44925.1 agla|OJJ84469.1 aory|EIT73467.1

Group13902: afla|EED44978.1 aory|EIT73711.1 awen|OJJ40147.1

Group13903: afla|EED45028.1 aory|EIT73644.1 asyd|OJJ54865.1

Group13904: afla|EED45034.1 aory|EIT73640.1 asyd|OJJ56414.1

Group13905: afla|EED45057.1 aory|EIT73762.1 aver|OJI98828.1

Group13906: afla|EED45100.1 asyd|OJJ54181.1 aver|OJJ08559.1

Group13907: afla|EED45118.1 aory|EIT73538.1 ater|EAU39527.1

Group13908: afla|EED45152.1 aory|EIT73552.1 awen|OJJ39619.1

Group13909: afla|EED45189.1 anid|CBF78859.1 aory|EIT73774.1

Group13910: afla|EED45356.1 agla|OJJ80032.1 aory|EIT82403.1

Group13911: afla|EED45388.1 asyd|OJJ54926.1 aver|OJJ07269.1

Group13912: afla|EED45404.1 afum|EAL86631.1 aory|EIT82448.1

Group13913: afla|EED45405.1 aory|EIT82464.1 aver|OJJ03841.1

Group13914: asyd|OJJ54712.1 aver|OJJ07454.1 awen|OJJ36968.1

Group13915: afla|EED45431.1 aory|EIT82405.1 awen|OJJ35314.1

Group13916: afla|EED45535.1 aory|EIT75788.1 awen|OJJ31940.1

Group13917: afla|EED45582.1 aory|EIT75789.1 ater|EAU34196.1

Group13918: afla|EED45607.1 aory|EIT75825.1 asyd|OJJ64153.1

Group13919: afla|EED45669.1 agla|OJJ83733.1 aory|EIT75418.1

Group13920: afla|EED45680.1 aory|EIT75533.1 azon|OJJ45980.1

Group13921: afla|EED45691.1 aory|EIT75579.1 azon|OJJ42599.1

Group13922: afla|EED45695.1 anid|CBF71634.1 aory|EIT75487.1

Group13923: afla|EED45779.1 agla|OJJ87547.1 awen|OJJ29653.1

Group13924: aver|OJJ05867.1 asyd|OJJ57290.1 azon|OJJ44495.1

Group13925: anid|CBF85471.1 asyd|OJJ53605.1 aver|OJJ08143.1

Group13926: afla|EED45819.1 asyd|OJJ64187.1 aver|OJI98264.1

Group13927: afla|EED45852.1 aory|EIT75764.1 awen|OJJ29657.1

Group13928: afla|EED45853.1 aory|EIT75708.1 awen|OJJ29658.1

Group13929: afla|EED45868.1 aory|EIT75716.1 asyd|OJJ54396.1

Group13930: afla|EED45893.1 aory|EIT75699.1 awen|OJJ30143.1

Group13931: afla|EED45914.1 aory|EIT75544.1 ater|EAU31437.1

Group13932: afla|EED46145.1 anig|EHA25875.1 aory|EIT76435.1

Group13933: afla|EED46156.1 aory|EIT76418.1 awen|OJJ33423.1

Group13934: afla|EED46157.1 aory|EIT76509.1 azon|OJJ44013.1

Group13935: afla|EED46158.1 aory|EIT76399.1 agla|OJJ88969.1

Group13936: afla|EED46194.1 aory|EIT76432.1 azon|OJJ43521.1

Group13937: agla|OJJ79525.1 akaw|GAA88578.1 aluc|OJZ88482.1

Group13938: afla|EED46257.1 agla|OJJ88352.1 aory|EIT76446.1

Group13939: afla|EED46383.1 aluc|OJZ85542.1 aver|OJI99261.1

Group13940: afla|EED46390.1 aory|EIT72400.1 azon|OJJ45861.1

Group13941: afla|EED46556.1 aory|EIT77947.1 ater|EAU31401.1

Group13942: afla|EED46601.1 aory|EIT77830.1 awen|OJJ40507.1

Group13943: afla|EED46869.1 aory|EIT72688.1 anid|CBF85340.1

Group13944: afla|EED46913.1 aory|EIT72615.1 azon|OJJ42573.1

Group13945: afla|EED46916.1 agla|OJJ84152.1 aory|EIT72616.1

Group13946: afla|EED46985.1 aory|EIT72729.1 ater|EAU29557.1

Group13947: afum|EAL92633.1 asyd|OJJ53616.1 aver|OJJ05936.1

Group13948: afla|EED47105.1 agla|OJJ85115.1 aory|EIT72783.1

Group13949: afla|EED47129.1 aory|EIT79269.1 ater|EAU36462.1

Group13950: afla|EED47246.1 aory|EIT79318.1 awen|OJJ40109.1

Group13951: afla|EED47270.1 aory|EIT79271.1 awen|OJJ31415.1

Group13952: aluc|OJZ83993.1 ater|EAU38493.1 anig|EHA19117.1

Group13953: afla|EED47309.1 aory|EIT82849.1 azon|OJJ45902.1

Group13954: afla|EED47317.1 aory|EIT82831.1 asyd|OJJ54266.1

Group13955: asyd|OJJ54594.1 aver|OJJ07558.1 anid|CBF80684.1

Group13956: afla|EED47367.1 agla|OJJ88086.1 aory|EIT82852.1

Group13957: afla|EED47368.1 aory|EIT82844.1 awen|OJJ37989.1

Group13958: afla|EED47393.1 aory|EIT81933.1 ater|EAU36823.1

Group13959: afla|EED47397.1 aory|EIT81926.1 asyd|OJJ62949.1

Group13960: afla|EED47465.1 asyd|OJJ58367.1 aver|OJJ02039.1

Group13961: afla|EED47467.1 aory|EIT79951.1 awen|OJJ31953.1

Group13962: anid|CBF73902.1 asyd|OJJ63009.1 aver|OJI97949.1

Group13963: awen|OJJ35836.1 asyd|OJJ53337.1 aver|OJJ08407.1

Group13964: afla|EED47499.1 aory|EIT74533.1 aver|OJJ07503.1

Group13965: afla|EED47504.1 aory|EIT74515.1 ater|EAU34202.1

Group13966: afla|EED47525.1 aluc|OJZ89487.1 ater|EAU33899.1

Group13967: afla|EED47536.1 aory|EIT74587.1 awen|OJJ30811.1

Group13968: afla|EED47544.1 aory|EIT74568.1 ater|EAU29443.1

Group13969: afla|EED47667.1 aory|EIT80171.1 asyd|OJJ57918.1

Group13970: afla|EED47744.1 anid|CBF85568.1 aory|EIT77100.1

Group13971: afla|EED47759.1 aory|EIT77166.1 azon|OJJ46421.1

Group13972: afla|EED47871.1 agla|OJJ89267.1 aory|EIT77244.1

Group13973: afla|EED47891.1 aluc|OJZ89215.1 atub|OJI85577.1

Group13974: afla|EED48120.1 agla|OJJ87525.1 aory|EIT76208.1

Group13975: afla|EED48170.1 agla|OJJ84365.1 aory|EIT83043.1

Group13976: afla|EED48173.1 aory|EIT83038.1 azon|OJJ45859.1

Group13977: afla|EED48182.1 aory|EIT83037.1 awen|OJJ31671.1

Group13978: afla|EED48198.1 aory|EIT72911.1 ater|EAU35692.1

Group13979: afla|EED48215.1 aory|EIT72898.1 awen|OJJ30535.1

Group13980: afla|EED48276.1 aory|EIT72942.1 awen|OJJ39634.1

Group13981: afla|EED48326.1 aory|EIT76286.1 atub|OJI89792.1

Group13982: afla|EED48359.1 aory|EIT83101.1 awen|OJJ35382.1

Group13983: afla|EED48377.1 aory|EIT83090.1 awen|OJJ40870.1

Group13984: afla|EED48417.1 akaw|GAA83298.1 aory|EIT83105.1

Group13985: afla|EED48422.1 agla|OJJ84728.1 aory|EIT83071.1

Group13986: afla|EED48425.1 aory|EIT83108.1 aver|OJJ05981.1

Group13987: afla|EED48440.1 aory|EIT81674.1 ater|EAU31013.1

Group13988: afla|EED48465.1 aory|EIT76266.1 awen|OJJ31684.1

Group13989: afla|EED48479.1 anid|CBF89299.1 aory|EIT73101.1

Group13990: afla|EED48482.1 anid|CBF89305.1 aory|EIT73266.1

Group13991: afla|EED48484.1 anid|CBF89309.1 aory|EIT73170.1

Group13992: afla|EED48585.1 aory|EIT73268.1 azon|OJJ46653.1

Group13993: afla|EED48718.1 aory|EIT73184.1 awen|OJJ36199.1

Group13994: afla|EED48881.1 aory|EIT81249.1 awen|OJJ34564.1

Group13995: afla|EED48890.1 agla|OJJ85104.1 aory|EIT81255.1

Group13996: afla|EED48922.1 aory|EIT80031.1 ater|EAU29374.1

Group13997: afla|EED48933.1 agla|OJJ87939.1 awen|OJJ34700.1

Group13998: afla|EED48981.1 aory|EIT80006.1 awen|OJJ29942.1

Group13999: afla|EED48998.1 aory|EIT80030.1 azon|OJJ47608.1

Group14000: afla|EED49071.1 aory|EIT82493.1 awen|OJJ33719.1

Group14001: aver|OJJ05747.1 asyd|OJJ57433.1 awen|OJJ34682.1

Group14002: afla|EED49377.1 aver|OJJ05892.1 awen|OJJ31163.1

Group14003: afla|EED49471.1 aory|EIT79522.1 awen|OJJ35944.1

Group14004: afla|EED49563.1 aory|EIT82630.1 azon|OJJ46532.1

Group14005: awen|OJJ36196.1 asyd|OJJ60086.1 aver|OJJ04173.1

Group14006: afla|EED49686.1 aory|EIT81015.1 aver|OJJ08573.1

Group14007: afla|EED49692.1 aory|EIT77652.1 ater|EAU30297.1

Group14008: afla|EED49693.1 aory|EIT77661.1 ater|EAU30298.1

Group14009: asyd|OJJ56420.1 aver|OJJ06294.1 awen|OJJ33800.1

Group14010: ater|EAU33986.1 afla|EED55030.1 aory|EIT81959.1

Group14011: afla|EED49841.1 aory|EIT83419.1 aver|OJJ02119.1

Group14012: afla|EED49843.1 aory|EIT83422.1 aver|OJJ02121.1

Group14013: afla|EED49866.1 aory|EIT83386.1 awen|OJJ34965.1

Group14014: afla|EED49874.1 asyd|OJJ57240.1 azon|OJJ42368.1

Group14015: afla|EED49892.1 agla|OJJ86460.1 aory|EIT72466.1

Group14016: afla|EED49896.1 aory|EIT72453.1 awen|OJJ32243.1

Group14017: afla|EED49897.1 aory|EIT72469.1 awen|OJJ36267.1

Group14018: akaw|GAA89231.1 aluc|OJZ86449.1 atub|OJI84075.1

Group14019: afla|EED49995.1 akaw|GAA83022.1 aory|EIT74711.1

Group14020: afla|EED50042.1 aluc|OJZ80205.1 awen|OJJ29563.1

Group14021: afla|EED50238.1 agla|OJJ82252.1 aory|EIT77299.1

Group14022: asyd|OJJ54064.1 ater|EAU38597.1 aver|OJJ08678.1

Group14023: afla|EED50611.1 aory|EIT77999.1 aver|OJI98679.1

Group14024: afla|EED50616.1 aory|EIT78097.1 ater|EAU38695.1

Group14025: afla|EED50783.1 aory|EIT78089.1 awen|OJJ36389.1

Group14026: afla|EED50786.1 aory|EIT78092.1 asyd|OJJ61708.1

Group14027: afla|EED50929.1 aory|EIT79026.1 aver|OJJ00175.1

Group14028: anid|CBF87724.1 asyd|OJJ58694.1 aver|OJJ01306.1

Group14029: afla|EED51103.1 aory|EIT81296.1 azon|OJJ43532.1

Group14030: afla|EED51105.1 asyd|OJJ54972.1 aver|OJJ07219.1

Group14031: afla|EED51133.1 aory|EIT81325.1 aver|OJJ05879.1

Group14032: afla|EED51438.1 aory|EIT80705.1 ater|EAU31576.1

Group14033: afla|EED51445.1 aory|EIT80618.1 asyd|OJJ56235.1

Group14034: afla|EED51447.1 aory|EIT80463.1 asyd|OJJ56230.1

Group14035: afla|EED51448.1 aory|EIT80724.1 asyd|OJJ56231.1

Group14036: afla|EED51449.1 aory|EIT80727.1 asyd|OJJ56237.1

Group14037: afla|EED51452.1 aory|EIT80545.1 awen|OJJ29933.1

Group14038: afla|EED51466.1 agla|OJJ85775.1 aory|EIT80542.1

Group14039: afla|EED51485.1 aory|EIT80685.1 asyd|OJJ57966.1

Group14040: afla|EED51578.1 afum|EAL87212.1 aory|EIT80719.1

Group14041: afla|EED51610.1 agla|OJJ81284.1 aory|EIT80485.1

Group14042: afla|EED51668.1 aory|EIT80726.1 ater|EAU31404.1

Group14043: asyd|OJJ62620.1 aver|OJI99298.1 awen|OJJ36995.1

Group14044: afla|EED51708.1 aory|EIT80687.1 ater|EAU35483.1

Group14045: afla|EED51731.1 anid|CBF79141.1 ater|EAU33338.1

Group14046: afla|EED51758.1 agla|OJJ82195.1 aory|EIT80634.1

Group14047: anid|CBF76115.1 atub|OJI80381.1 azon|OJJ42566.1

Group14048: afla|EED51831.1 aory|EIT81752.1 aver|OJI95764.1

Group14049: afla|EED51838.1 aory|EIT81767.1 asyd|OJJ65402.1

Group14050: asyd|OJJ56394.1 aver|OJJ06319.1 anid|CBF73503.1

Group14051: asyd|OJJ65292.1 anid|CBF80520.1 aver|OJJ01918.1

Group14052: awen|OJJ34471.1 asyd|OJJ52871.1 aver|OJJ08937.1

Group14053: afla|EED52253.1 aory|EIT74479.1 ater|EAU31604.1

Group14054: afla|EED52260.1 anig|EHA28242.1 aory|EIT74443.1

Group14055: afla|EED52354.1 aluc|OJZ79664.1 akaw|GAA93089.1

Group14056: afla|EED52362.1 afum|EAL86495.1 aver|OJI95834.1

Group14057: afla|EED52396.1 atub|OJI88385.1 awen|OJJ35569.1

Group14058: afla|EED52404.1 aory|EIT83297.1 awen|OJJ38809.1

Group14059: afla|EED52405.1 asyd|OJJ62985.1 aver|OJI97983.1

Group14060: afla|EED52794.1 asyd|OJJ54652.1 aver|OJJ07509.1

Group14061: afla|EED52851.1 aory|EIT78662.1 atub|OJI84703.1

Group14062: afla|EED53065.1 aory|EIT74800.1 asyd|OJJ56238.1

Group14063: afla|EED53067.1 aory|EIT74807.1 asyd|OJJ56236.1

Group14064: afla|EED53151.1 aory|EIT75197.1 aver|OJJ01958.1

Group14065: akaw|GAA88050.1 aluc|OJZ81312.1 aver|OJJ03853.1

Group14066: afla|EED53189.1 asyd|OJJ54352.1 aver|OJJ05305.1

Group14067: afla|EED53266.1 aory|EIT74910.1 asyd|OJJ64359.1

Group14068: afla|EED53390.1 asyd|OJJ54121.1 aver|OJJ08616.1

Group14069: afla|EED53572.1 aory|EIT75190.1 awen|OJJ32942.1

Group14070: afla|EED53697.1 aory|EIT75282.1 ater|EAU34795.1

Group14071: anid|CBF85420.1 asyd|OJJ64074.1 aver|OJI98161.1

Group14072: afla|EED54046.1 aory|EIT75926.1 aver|OJJ06475.1

Group14073: afla|EED54049.1 aory|EIT76151.1 asyd|OJJ52553.1

Group14074: anid|CBF71465.1 asyd|OJJ54947.1 aver|OJJ07248.1

Group14075: afla|EED54069.1 aory|EIT76098.1 ater|EAU39444.1

Group14076: afla|EED54077.1 aory|EIT76056.1 ater|EAU29609.1

Group14077: afla|EED54082.1 aory|EIT75954.1 awen|OJJ37943.1

Group14078: afla|EED54133.1 agla|OJJ78605.1 aory|EIT76110.1

Group14079: afla|EED54141.1 aory|EIT76122.1 aver|OJI97560.1

Group14080: afla|EED54148.1 aory|EIT76121.1 awen|OJJ36292.1

Group14081: afla|EED54158.1 aory|EIT76107.1 awen|OJJ36153.1

Group14082: afla|EED54255.1 aory|EIT75999.1 ater|EAU32330.1

Group14083: afla|EED54352.1 aory|EIT75958.1 anig|EHA26174.1

Group14084: anid|CBF78743.1 aver|OJI95877.1 aory|EIT72648.1

Group14085: afla|EED54471.1 aory|EIT81142.1 awen|OJJ37950.1

Group14086: afla|EED54672.1 aory|EIT78906.1 ater|EAU29968.1

Group14087: afla|EED54687.1 aory|EIT78755.1 awen|OJJ31386.1

Group14088: afla|EED54737.1 aory|EIT78897.1 awen|OJJ33333.1

Group14089: asyd|OJJ53397.1 akaw|GAA90558.1 aver|OJJ02220.1

Group14090: afla|EED54926.1 aory|EIT80352.1 azon|OJJ49999.1

Group14091: afla|EED54956.1 aory|EIT80283.1 azon|OJJ50603.1

Group14092: afla|EED55036.1 aory|EIT81985.1 awen|OJJ39967.1

Group14093: afla|EED55037.1 aory|EIT81981.1 awen|OJJ39966.1

Group14094: afla|EED55040.1 aory|EIT81942.1 ater|EAU37350.1

Group14095: agla|OJJ79292.1 asyd|OJJ54794.1 aver|OJJ02015.1

Group14096: afla|EED55115.1 aory|EIT81665.1 awen|OJJ37018.1

Group14097: afla|EED55163.1 agla|OJJ84027.1 ater|EAU37668.1

Group14098: afla|EED55262.1 aory|EIT83606.1 awen|OJJ32898.1

Group14099: afla|EED55340.1 aver|OJJ05801.1 asyd|OJJ57367.1

Group14100: afla|EED55507.1 anid|CBF81667.1 aory|EIT80843.1

Group14101: afla|EED55969.1 aory|EIT76555.1 asyd|OJJ56494.1

Group14102: afla|EED56419.1 agla|OJJ82193.1 aory|EIT72880.1

Group14103: afla|EED56539.1 agla|OJJ80935.1 aory|EIT79699.1

Group14104: afla|EED56556.1 aory|EIT79616.1 ater|EAU37798.1

Group14105: afla|EED56586.1 awen|OJJ40102.1 agla|OJJ87729.1

Group14106: akaw|GAA84494.1 aluc|OJZ89217.1 atub|OJI85575.1

Group14107: afla|EED56759.1 aory|EIT73844.1 azon|OJJ50645.1

Group14108: afla|EED56779.1 aory|EIT73878.1 awen|OJJ38022.1

Group14109: afla|EED56786.1 aory|EIT73896.1 awen|OJJ36374.1

Group14110: afla|EED56799.1 aory|EIT73883.1 awen|OJJ33574.1

Group14111: afla|EED56801.1 aory|EIT73929.1 azon|OJJ44088.1

Group14112: afla|EED56816.1 akaw|GAA86081.1 atub|OJI81741.1

Group14113: afla|EED56849.1 aory|EIT77509.1 azon|OJJ46001.1

Group14114: afla|EED56907.1 aory|EIT77496.1 awen|OJJ36148.1

Group14115: akaw|GAA92512.1 aluc|OJZ91525.1 azon|OJJ45173.1

Group14116: afla|EED56952.1 aory|EIT77554.1 awen|OJJ41846.1

Group14117: afla|EED57020.1 asyd|OJJ57948.1 aver|OJJ04450.1

Group14118: aver|OJJ01355.1 anid|CBF70275.1 asyd|OJJ58745.1

Group14119: afla|EED57231.1 aory|EIT74156.1 asyd|OJJ54480.1

Group14120: afla|EED57241.1 aory|EIT74062.1 atub|OJI87884.1

Group14121: afla|EED57301.1 aory|EIT74115.1 ater|EAU30083.1

Group14122: afla|EED57483.1 aory|EIT80093.1 ater|EAU33069.1

Group14123: afla|EED57600.1 aory|EIT82927.1 ater|EAU31886.1

Group14124: afla|EED57732.1 aory|EIT79865.1 awen|OJJ32620.1

Group14125: afla|EED57734.1 aory|EIT79908.1 azon|OJJ51788.1

Group14126: afla|EED57870.1 asyd|OJJ55541.1 aver|OJJ06619.1

Group14127: afum|EAL84360.1 aver|OJI97999.1 asyd|OJJ62968.1

Group14128: afum|EAL84870.1 anid|CBF87206.1 aver|OJI99462.1

Group14129: afum|EAL85150.1 anid|CBF85132.1 aver|OJI97925.1

Group14130: afum|EAL85390.1 akaw|GAA87896.1 asyd|OJJ62698.1

Group14131: afum|EAL85430.2 agla|OJJ85933.1 ater|EAU30731.1

Group14132: afum|EAL86281.1 akaw|GAA93040.1 anig|EHA21841.1

Group14133: afum|EAL86283.1 agla|OJJ86638.1 aluc|OJZ82093.1

Group14134: afum|EAL86637.1 anid|CBF86054.1 aver|OJJ07784.1

Group14135: afum|EAL86725.1 aluc|OJZ79964.1 asyd|OJJ56759.1

Group14136: afum|EAL86769.1 aluc|OJZ79808.1 anig|EHA21842.1

Group14137: afum|EAL86774.1 agla|OJJ78482.1 awen|OJJ31337.1

Group14138: afum|EAL87145.1 aory|EIT74463.1 awen|OJJ41935.1

Group14139: afum|EAL87748.1 aluc|OJZ80160.1 awen|OJJ40154.1

Group14140: afum|EAL87797.1 awen|OJJ37931.1 azon|OJJ41994.1

Group14141: aluc|OJZ87034.1 anig|EHA25214.1 atub|OJI83481.1

Group14142: aver|OJI95816.1 anid|CBF83098.1 asyd|OJJ61721.1

Group14143: asyd|OJJ53198.1 aver|OJJ03652.1 azon|OJJ45242.1

Group14144: anig|EHA17685.1 akaw|GAA92626.1 aluc|OJZ80122.1

Group14145: aluc|OJZ86453.1 akaw|GAA89227.1 atub|OJI84079.1

Group14146: anid|CBF73495.1 asyd|OJJ60828.1 aver|OJJ00154.1

Group14147: afum|EAL92636.2 asyd|OJJ53627.1 aver|OJJ05944.1

Group14148: afum|EAL94098.1 asyd|OJJ53727.1 aver|OJI98282.1

Group14149: agla|OJJ78528.1 azon|OJJ51356.1 awen|OJJ30176.1

Group14150: agla|OJJ78531.1 asyd|OJJ65330.1 aver|OJI99475.1

Group14151: agla|OJJ78658.1 aver|OJJ03762.1 awen|OJJ36609.1

Group14152: agla|OJJ78854.1 asyd|OJJ53815.1 aver|OJJ04321.1

Group14153: asyd|OJJ62063.1 agla|OJJ87806.1 aver|OJI96162.1

Group14154: anid|CBF82173.1 asyd|OJJ54355.1 aver|OJJ05309.1

Group14155: agla|OJJ79112.1 asyd|OJJ55386.1 aver|OJJ08271.1

Group14156: awen|OJJ34017.1 asyd|OJJ60316.1 aver|OJI99657.1

Group14157: agla|OJJ79119.1 asyd|OJJ55395.1 aver|OJJ08262.1

Group14158: agla|OJJ79121.1 asyd|OJJ55380.1 awen|OJJ41413.1

Group14159: ater|EAU37988.1 agla|OJJ80065.1 awen|OJJ30906.1

Group14160: agla|OJJ79845.1 asyd|OJJ60830.1 aver|OJJ00155.1

Group14161: agla|OJJ80046.1 asyd|OJJ62577.1 aver|OJI96698.1

Group14162: agla|OJJ80193.1 asyd|OJJ58368.1 aver|OJJ02038.1

Group14163: agla|OJJ80333.1 atub|OJI81898.1 awen|OJJ32600.1

Group14164: agla|OJJ80606.1 asyd|OJJ53692.1 aver|OJJ07809.1

Group14165: agla|OJJ80632.1 aluc|OJZ80312.1 atub|OJI80385.1

Group14166: agla|OJJ80870.1 asyd|OJJ53599.1 aver|OJJ08149.1

Group14167: agla|OJJ81070.1 awen|OJJ35777.1 azon|OJJ43900.1

Group14168: anid|CBF83076.1 aver|OJJ00824.1 asyd|OJJ61482.1

Group14169: asyd|OJJ65115.1 aver|OJI99241.1 anid|CBF76239.1

Group14170: agla|OJJ81474.1 asyd|OJJ54227.1 aver|OJJ05180.1

Group14171: agla|OJJ81543.1 anid|CBF79166.1 aver|OJI96475.1

Group14172: agla|OJJ81706.1 asyd|OJJ55015.1 aver|OJJ07196.1

Group14173: agla|OJJ82007.1 awen|OJJ36165.1 aver|OJJ07896.1

Group14174: agla|OJJ82211.1 asyd|OJJ60321.1 awen|OJJ37955.1

Group14175: agla|OJJ82464.1 asyd|OJJ56947.1 aver|OJJ03338.1

Group14176: agla|OJJ82476.1 asyd|OJJ57274.1 ater|EAU30951.1

Group14177: agla|OJJ82477.1 ater|EAU30950.1 aver|OJJ03447.1

Group14178: agla|OJJ82504.1 anid|CBF81857.1 ater|EAU31863.1

Group14179: agla|OJJ82615.1 ater|EAU30080.1 awen|OJJ40131.1

Group14180: agla|OJJ82664.1 asyd|OJJ53573.1 aver|OJJ08184.1

Group14181: agla|OJJ82666.1 awen|OJJ29652.1 azon|OJJ45047.1

Group14182: agla|OJJ82760.1 asyd|OJJ52976.1 aver|OJJ08942.1

Group14183: agla|OJJ83096.1 akaw|GAA92801.1 aluc|OJZ88723.1

Group14184: agla|OJJ83480.1 aver|OJI96894.1 asyd|OJJ53795.1

Group14185: agla|OJJ83558.1 ater|EAU38736.1 awen|OJJ34150.1

Group14186: agla|OJJ84447.1 asyd|OJJ52111.1 aver|OJJ05255.1

Group14187: agla|OJJ84465.1 aory|EIT74901.1 ater|EAU31930.1

Group14188: agla|OJJ84472.1 asyd|OJJ54483.1 aver|OJJ07696.1

Group14189: agla|OJJ84569.1 akaw|GAA89015.1 aluc|OJZ79786.1

Group14190: agla|OJJ84723.1 aver|OJI99734.1 asyd|OJJ60404.1

Group14191: agla|OJJ84978.1 aver|OJJ01042.1 azon|OJJ50498.1

Group14192: agla|OJJ85295.1 asyd|OJJ65028.1 aver|OJI99147.1

Group14193: agla|OJJ85331.1 asyd|OJJ61769.1 aver|OJI95866.1

Group14194: agla|OJJ85704.1 awen|OJJ39151.1 azon|OJJ51479.1

Group14195: agla|OJJ86222.1 awen|OJJ30264.1 ater|EAU33253.1

Group14196: agla|OJJ86592.1 asyd|OJJ61475.1 aver|OJJ00816.1

Group14197: ater|EAU37272.1 aver|OJJ06012.1 asyd|OJJ56713.1

Group14198: agla|OJJ87380.1 anid|CBF86671.1 ater|EAU33420.1

Group14199: agla|OJJ87731.1 awen|OJJ40101.1 azon|OJJ42077.1

Group14200: agla|OJJ88186.1 anid|CBF82335.1 azon|OJJ41996.1

Group14201: agla|OJJ88188.1 aver|OJJ05943.1 asyd|OJJ53626.1

Group14202: agla|OJJ88230.1 awen|OJJ40915.1 azon|OJJ43558.1

Group14203: agla|OJJ88336.1 aver|OJJ05347.1 awen|OJJ30917.1

Group14204: aluc|OJZ86812.1 atub|OJI83705.1 akaw|GAA84859.1

Group14205: agla|OJJ88812.1 asyd|OJJ54515.1 aver|OJJ07669.1

Group14206: agla|OJJ88813.1 asyd|OJJ55402.1 aver|OJJ06482.1

Group14207: agla|OJJ88814.1 asyd|OJJ54516.1 aver|OJJ07667.1

Group14208: agla|OJJ88815.1 asyd|OJJ54517.1 aver|OJJ07666.1

Group14209: agla|OJJ88848.1 aluc|OJZ88659.1 aver|OJJ05765.1

Group14210: asyd|OJJ56750.1 anig|EHA21627.1 aver|OJJ05980.1

Group14211: agla|OJJ88983.1 awen|OJJ41815.1 azon|OJJ43557.1

Group14212: agla|OJJ89314.1 asyd|OJJ57225.1 aver|OJJ03062.1

Group14213: akaw|BAL04873.1 aory|AFM82512.1 azon|OJJ41982.1

Group14214: akaw|BAL04881.1 aory|AFM82508.1 azon|OJJ41987.1

Group14215: akaw|GAA82390.1 aluc|OJZ91730.1 anig|EHA22737.1

Group14216: akaw|GAA82413.1 aluc|OJZ91705.1 atub|OJI90105.1

Group14217: akaw|GAA82771.1 aluc|OJZ87330.1 atub|OJI81314.1

Group14218: akaw|GAA82772.1 aluc|OJZ87331.1 atub|OJI81313.1

Group14219: akaw|GAA82957.1 aluc|OJZ87538.1 atub|OJI81109.1

Group14220: akaw|GAA83094.1 aluc|OJZ88641.1 atub|OJI91226.1

Group14221: akaw|GAA83224.1 aluc|OJZ83725.1 atub|OJI90100.1

Group14222: akaw|GAA83895.1 aluc|OJZ90800.1 atub|OJI82789.1

Group14223: akaw|GAA83896.1 aluc|OJZ90799.1 atub|OJI82788.1

Group14224: akaw|GAA83899.1 aluc|OJZ90796.1 atub|OJI82785.1

Group14225: akaw|GAA83933.1 aluc|OJZ90762.1 atub|OJI82750.1

Group14226: akaw|GAA84052.1 aluc|OJZ80964.1 atub|OJI82622.1

Group14227: akaw|GAA84086.1 aluc|OJZ81001.1 atub|OJI82584.1

Group14228: akaw|GAA84167.1 aluc|OJZ81092.1 atub|OJI82491.1

Group14229: akaw|GAA84219.1 aluc|OJZ88897.1 atub|OJI85892.1

Group14230: akaw|GAA84274.1 aluc|OJZ88967.1 atub|OJI85823.1

Group14231: akaw|GAA84737.1 aluc|OJZ86949.1 atub|OJI83574.1

Group14232: akaw|GAA84847.1 aluc|OJZ86824.1 atub|OJI83694.1

Group14233: asyd|OJJ53606.1 anid|CBF85469.1 aver|OJJ08142.1

Group14234: akaw|GAA84850.1 aluc|OJZ86821.1 atub|OJI83696.1

Group14235: akaw|GAA84855.1 aluc|OJZ86816.1 atub|OJI83701.1

Group14236: akaw|GAA84857.1 aluc|OJZ86814.1 atub|OJI83703.1

Group14237: akaw|GAA84862.1 aluc|OJZ86809.1 atub|OJI83708.1

Group14238: akaw|GAA84966.1 aluc|OJZ86697.1 atub|OJI83828.1

Group14239: akaw|GAA85047.1 aluc|OJZ86606.1 anid|CBF80329.1

Group14240: ater|EAU37352.1 anig|EHA26088.1 atub|OJI87332.1

Group14241: akaw|GAA85277.1 aluc|OJZ90636.1 atub|OJI82064.1

Group14242: akaw|GAA85307.1 aluc|OJZ90667.1 atub|OJI82097.1

Group14243: akaw|GAA85348.1 aluc|OJZ90710.1 atub|OJI82137.1

Group14244: akaw|GAA85353.1 aluc|OJZ90715.1 atub|OJI82141.1

Group14245: akaw|GAA85437.1 aluc|OJZ83341.1 atub|OJI82236.1

Group14246: akaw|GAA85438.1 aluc|OJZ83342.1 atub|OJI82237.1

Group14247: akaw|GAA85439.1 aluc|OJZ83343.1 atub|OJI82238.1

Group14248: akaw|GAA85440.1 aluc|OJZ83344.1 atub|OJI82239.1

Group14249: akaw|GAA85520.1 atub|OJI82340.1 aluc|OJZ83448.1

Group14250: akaw|GAA85821.1 aluc|OJZ91278.1 atub|OJI83267.1

Group14251: akaw|GAA85846.1 aluc|OJZ91253.1 awen|OJJ34438.1

Group14252: akaw|GAA85944.1 aluc|OJZ91118.1 atub|OJI83122.1

Group14253: akaw|GAA85953.1 aluc|OJZ91109.1 atub|OJI83106.1

Group14254: akaw|GAA86040.1 aluc|OJZ90268.1 atub|OJI81693.1

Group14255: akaw|GAA86048.1 aluc|OJZ90277.1 atub|OJI81701.1

Group14256: akaw|GAA86107.1 aluc|OJZ90339.1 atub|OJI81766.1

Group14257: akaw|GAA86113.1 aluc|OJZ90348.1 atub|OJI81774.1

Group14258: akaw|GAA86368.1 aluc|OJZ84897.1 atub|OJI80701.1

Group14259: akaw|GAA86412.1 aluc|OJZ84952.1 anig|EHA23275.1

Group14260: akaw|GAA86413.1 aluc|OJZ84954.1 atub|OJI80646.1

Group14261: akaw|GAA86583.1 anig|EHA25353.1 atub|OJI80459.1

Group14262: akaw|GAA86598.1 aluc|OJZ85145.1 anig|EHA27469.1

Group14263: akaw|GAA86639.1 aluc|OJZ85196.1 atub|OJI80405.1

Group14264: aluc|OJZ89899.1 anig|EHA27099.1 atub|OJI84908.1

Group14265: akaw|GAA86922.1 aluc|OJZ90046.1 atub|OJI84760.1

Group14266: akaw|GAA87150.1 aluc|OJZ84285.1 atub|OJI89786.1

Group14267: akaw|GAA87301.1 aluc|OJZ89717.1 atub|OJI85083.1

Group14268: akaw|GAA87303.1 aluc|OJZ89715.1 atub|OJI85087.1

Group14269: akaw|GAA87454.1 aluc|OJZ88541.1 atub|OJI91126.1

Group14270: akaw|GAA87468.1 aluc|OJZ88528.1 atub|OJI91112.1

Group14271: akaw|GAA87630.1 atub|OJI80324.1 awen|OJJ36371.1

Group14272: akaw|GAA87709.1 aluc|OJZ85335.1 atub|OJI80236.1

Group14273: akaw|GAA87763.1 aluc|OJZ85398.1 atub|OJI80173.1

Group14274: akaw|GAA87872.1 aluc|OJZ85513.1 ater|EAU30245.1

Group14275: akaw|GAA88052.1 aluc|OJZ81314.1 anig|EHA18618.1

Group14276: akaw|GAA88104.1 aluc|OJZ81371.1 atub|OJI91576.1

Group14277: akaw|GAA88263.1 aluc|OJZ82232.1 atub|OJI88205.1

Group14278: akaw|GAA88998.1 asyd|OJJ51942.1 aluc|OJZ79920.1

Group14279: atub|OJI80388.1 akaw|GAA92354.1 aluc|OJZ80394.1

Group14280: akaw|GAA89140.1 aluc|OJZ81696.1 atub|OJI88484.1

Group14281: akaw|GAA89218.1 aluc|OJZ81611.1 atub|OJI88574.1

Group14282: akaw|GAA89226.1 aluc|OJZ86454.1 atub|OJI84080.1

Group14283: akaw|GAA89229.1 aluc|OJZ86451.1 atub|OJI84077.1

Group14284: akaw|GAA89306.1 aluc|OJZ86363.1 atub|OJI84171.1

Group14285: akaw|GAA89822.1 aluc|OJZ81868.1 anig|EHA25071.1

Group14286: ater|EAU33239.1 anid|CBF75898.1 aver|OJJ08122.1

Group14287: akaw|GAA90163.1 aluc|OJZ83184.1 anig|EHA18398.1

Group14288: akaw|GAA90164.1 aluc|OJZ83183.1 anig|EHA18397.1

Group14289: akaw|GAA90168.1 asyd|OJJ57147.1 azon|OJJ42371.1

Group14290: akaw|GAA90236.1 aluc|OJZ83104.1 atub|OJI87831.1

Group14291: akaw|GAA90304.1 aluc|OJZ83033.1 atub|OJI87915.1

Group14292: akaw|GAA90327.1 aluc|OJZ85906.1 anig|EHA24686.1

Group14293: akaw|GAA90556.1 asyd|OJJ53399.1 aver|OJJ02218.1

Group14294: akaw|GAA90598.1 aluc|OJZ86056.1 atub|OJI79853.1

Group14295: akaw|GAA90675.1 aluc|OJZ84683.1 atub|OJI89399.1

Group14296: akaw|GAA90691.1 aluc|OJZ84663.1 atub|OJI89417.1

Group14297: akaw|GAA91037.1 aluc|OJZ87847.1 atub|OJI80801.1

Group14298: akaw|GAA91078.1 aluc|OJZ87893.1 atub|OJI80759.1

Group14299: akaw|GAA91164.1 aluc|OJZ81491.1 atub|OJI88689.1

Group14300: akaw|GAA91349.1 aluc|OJZ84825.1 atub|OJI89262.1

Group14301: akaw|GAA91371.1 aluc|OJZ84798.1 atub|OJI89284.1

Group14302: akaw|GAA91401.1 aluc|OJZ84770.1 atub|OJI89311.1

Group14303: akaw|GAA91648.1 aluc|OJZ82475.1 atub|OJI87961.1

Group14304: akaw|GAA91804.1 aluc|OJZ92826.1 atub|OJI87283.1

Group14305: asyd|OJJ56129.1 anid|CBF78341.1 aver|OJJ05070.1

Group14306: akaw|GAA92271.1 aluc|OJZ90165.1 atub|OJI81593.1

Group14307: akaw|GAA92453.1 aluc|OJZ81565.1 atub|OJI88620.1

Group14308: akaw|GAA92457.1 aluc|OJZ81560.1 atub|OJI88625.1

Group14309: akaw|GAA92508.1 aluc|OJZ91521.1 atub|OJI85976.1

Group14310: akaw|GAA92536.1 aluc|OJZ90084.1 atub|OJI84722.1

Group14311: akaw|GAA92701.1 aluc|OJZ88740.1 atub|OJI91312.1

Group14312: akaw|GAA92797.1 aluc|OJZ88719.1 anig|EHA28202.1

Group14313: akaw|GAA92798.1 aluc|OJZ88720.1 anig|EHA28203.1

Group14314: akaw|GAA92827.1 aluc|OJZ91582.1 atub|OJI86031.1

Group14315: akaw|GAA92830.1 aory|EIT79180.1 ater|EAU39332.1

Group14316: akaw|GAA92852.1 aluc|OJZ80011.1 anig|EHA26119.1

Group14317: akaw|GAA93078.1 aluc|OJZ80097.1 anig|EHA17687.1

Group14318: akaw|GAA93102.1 aluc|OJZ79586.1 awen|OJJ36956.1

Group14319: akaw|GAA93203.1 aluc|OJZ79706.1 anid|CBF81148.1

Group14320: akaw|GAA93280.1 atub|OJI88197.1 aluc|OJZ79973.1

Group14321: akaw|GAA93284.1 aluc|OJZ90160.1 atub|OJI84640.1

Group14322: akaw|GAA93296.1 anig|EHA25615.1 ater|EAU32741.1

Group14323: asyd|OJJ61845.1 aver|OJI95945.1 anid|CBF71636.1

Group14324: aluc|OJZ79924.1 ater|EAU35382.1 awen|OJJ34694.1

Group14325: aluc|OJZ80147.1 asyd|OJJ64228.1 aver|OJI98321.1

Group14326: aluc|OJZ81328.1 anig|EHA18632.1 atub|OJI91531.1

Group14327: aluc|OJZ81358.1 anig|EHA18658.1 atub|OJI91562.1

Group14328: aluc|OJZ81648.1 atub|OJI88534.1 azon|OJJ43499.1

Group14329: aluc|OJZ82223.1 anig|EHA28517.1 atub|OJI88212.1

Group14330: aver|OJI97549.1 anid|CBF85247.1 asyd|OJJ63386.1

Group14331: aluc|OJZ83413.1 anig|EHA24262.1 atub|OJI82300.1

Group14332: aluc|OJZ83479.1 anig|EHA24316.1 atub|OJI82370.1

Group14333: aluc|OJZ83607.1 anig|EHA18766.1 atub|OJI89980.1

Group14334: aluc|OJZ85333.1 anig|EHA25521.1 atub|OJI80240.1

Group14335: aluc|OJZ85377.1 anig|EHA25560.1 atub|OJI80192.1

Group14336: aluc|OJZ85745.1 atub|OJI79530.1 anig|EHA28105.1

Group14337: aluc|OJZ86064.1 aver|OJJ04784.1 awen|OJJ38355.1

Group14338: aluc|OJZ86131.1 atub|OJI79927.1 anig|EHA24490.1

Group14339: aluc|OJZ87027.1 anig|EHA21631.1 atub|OJI83491.1

Group14340: aluc|OJZ87707.1 anig|EHA20889.1 atub|OJI80936.1

Group14341: aluc|OJZ88661.1 aver|OJJ05767.1 asyd|OJJ61523.1

Group14342: aluc|OJZ88841.1 aory|EIT83506.1 asyd|OJJ52035.1

Group14343: aluc|OJZ89304.1 atub|OJI85495.1 anig|EHA23262.1

Group14344: aluc|OJZ89489.1 asyd|OJJ61847.1 atub|OJI85314.1

Group14345: aluc|OJZ89584.1 anig|EHA23025.1 atub|OJI85219.1

Group14346: aluc|OJZ89609.1 anig|EHA26855.1 atub|OJI85193.1

Group14347: aluc|OJZ89909.1 anig|EHA27109.1 atub|OJI84897.1

Group14348: aluc|OJZ90839.1 ater|EAU38786.1 atub|OJI82826.1

Group14349: aluc|OJZ91209.1 anig|EHA19914.1 atub|OJI83227.1

Group14350: aluc|OJZ91244.1 anig|EHA19943.1 atub|OJI83241.1

Group14351: azon|OJJ43146.1 asyd|OJJ57172.1 awen|OJJ32420.1

Group14352: asyd|OJJ64430.1 anid|CBF85981.1 aver|OJI98531.1

Group14353: aluc|OJZ91572.1 anig|EHA22601.1 atub|OJI86023.1

Group14354: aluc|OJZ92424.1 anig|EHA26543.1 atub|OJI86854.1

Group14355: anid|CBF69423.1 asyd|OJJ57143.1 aver|OJJ04751.1

Group14356: anid|CBF69429.1 asyd|OJJ52371.1 aver|OJJ01977.1

Group14357: anid|CBF69475.1 asyd|OJJ59269.1 aver|OJJ01882.1

Group14358: anid|CBF69479.1 asyd|OJJ59271.1 aver|OJJ01884.1

Group14359: anid|CBF69496.1 asyd|OJJ59236.1 aver|OJJ01846.1

Group14360: anid|CBF69542.1 asyd|OJJ59169.1 aver|OJJ01759.1

Group14361: anid|CBF69576.1 asyd|OJJ59148.1 aver|OJJ01738.1

Group14362: anid|CBF69646.1 asyd|OJJ59079.1 aver|OJJ01672.1

Group14363: anid|CBF69926.1 asyd|OJJ59038.1 aver|OJJ01633.1

Group14364: ater|EAU34088.1 asyd|OJJ52862.1 aver|OJJ08841.1

Group14365: anid|CBF69969.1 asyd|OJJ58904.1 ater|EAU31720.1

Group14366: anid|CBF70094.1 asyd|OJJ58836.1 aver|OJJ01439.1

Group14367: anid|CBF70112.1 asyd|OJJ58821.1 aver|OJJ01425.1

Group14368: anid|CBF70219.1 asyd|OJJ52962.1 aver|OJJ08928.1

Group14369: anid|CBF70263.1 asyd|OJJ58752.1 aver|OJJ01996.1

Group14370: anid|CBF70518.1 asyd|OJJ60538.1 aver|OJI99865.1

Group14371: anid|CBF70526.1 asyd|OJJ52963.1 aver|OJJ08929.1

Group14372: anid|CBF70531.1 asyd|OJJ58555.1 aver|OJJ01153.1

Group14373: anid|CBF70614.1 asyd|OJJ58507.1 aver|OJJ01108.1

Group14374: anid|CBF70724.1 asyd|OJJ55959.1 aver|OJJ04905.1

Group14375: anid|CBF71062.1 asyd|OJJ52681.1 aver|OJJ04707.1

Group14376: anid|CBF71165.1 asyd|OJJ55846.1 aver|OJJ04789.1

Group14377: anid|CBF71247.1 asyd|OJJ55158.1 aver|OJJ07050.1

Group14378: anid|CBF71285.1 asyd|OJJ55130.1 aver|OJJ07073.1

Group14379: anid|CBF71399.1 asyd|OJJ55025.1 aver|OJJ07180.1

Group14380: anid|CBF71401.1 asyd|OJJ55026.1 aver|OJJ07178.1

Group14381: anid|CBF71449.1 asyd|OJJ54959.1 aver|OJJ07232.1

Group14382: anid|CBF71551.1 asyd|OJJ56284.1 aver|OJJ08476.1

Group14383: anid|CBF71760.1 asyd|OJJ61728.1 aver|OJI95824.1

Group14384: anid|CBF71766.1 asyd|OJJ61722.1 aver|OJI95817.1

Group14385: anid|CBF71779.1 asyd|OJJ61741.1 aver|OJI95838.1

Group14386: anid|CBF73403.1 asyd|OJJ54745.1 aver|OJJ07420.1

Group14387: anid|CBF73411.1 asyd|OJJ55942.1 aver|OJJ04891.1

Group14388: anid|CBF73509.1 asyd|OJJ55810.1 aver|OJJ04758.1

Group14389: anid|CBF73526.1 asyd|OJJ61525.1 aver|OJJ00857.1

Group14390: anid|CBF73528.1 asyd|OJJ56726.1 aver|OJJ05997.1

Group14391: anid|CBF73551.1 asyd|OJJ63991.1 aver|OJI96963.1

Group14392: anid|CBF73572.1 asyd|OJJ56689.1 aver|OJJ06034.1

Group14393: anid|CBF73588.1 asyd|OJJ56217.1 aver|OJJ05149.1

Group14394: anid|CBF73609.1 asyd|OJJ56675.1 aver|OJJ06049.1

Group14395: anid|CBF73676.1 asyd|OJJ56661.1 aver|OJJ06063.1

Group14396: anid|CBF73696.1 asyd|OJJ56651.1 aver|OJJ06074.1

Group14397: anid|CBF73737.1 asyd|OJJ56633.1 aver|OJJ06093.1

Group14398: anid|CBF73786.1 asyd|OJJ56601.1 aver|OJJ06123.1

Group14399: anid|CBF73848.1 asyd|OJJ56553.1 aver|OJJ06168.1

Group14400: anid|CBF73952.1 asyd|OJJ53711.1 aver|OJJ07837.1

Group14401: anid|CBF73954.1 asyd|OJJ53712.1 aver|OJJ07838.1

Group14402: anid|CBF73955.1 asyd|OJJ53713.1 aver|OJJ07839.1

Group14403: anid|CBF73957.1 asyd|OJJ53714.1 aver|OJJ07840.1

Group14404: anid|CBF73963.1 asyd|OJJ62372.1 aver|OJI96506.1

Group14405: anid|CBF73967.1 asyd|OJJ62370.1 aver|OJI96504.1

Group14406: anid|CBF74146.1 asyd|OJJ53919.1 aver|OJJ08017.1

Group14407: anid|CBF74401.1 aver|OJJ05401.1 asyd|OJJ57765.1

Group14408: anid|CBF74469.1 asyd|OJJ57701.1 aver|OJJ05463.1

Group14409: anid|CBF74503.1 asyd|OJJ57724.1 aver|OJJ05445.1

Group14410: anid|CBF74505.1 asyd|OJJ57725.1 aver|OJJ05444.1

Group14411: anid|CBF74507.1 asyd|OJJ57727.1 aver|OJJ05442.1

Group14412: anid|CBF74509.1 asyd|OJJ57728.1 aver|OJJ05441.1

Group14413: anid|CBF74542.1 asyd|OJJ57689.1 aver|OJJ05476.1

Group14414: anid|CBF74641.1 asyd|OJJ59265.1 aver|OJJ01875.1

Group14415: anid|CBF74722.1 asyd|OJJ60199.1 aver|OJJ04066.1

Group14416: anid|CBF74781.1 aver|OJJ04102.1 asyd|OJJ60162.1

Group14417: anid|CBF74892.1 asyd|OJJ52155.1 aver|OJJ00746.1

Group14418: anid|CBF75026.1 asyd|OJJ60075.1 aver|OJJ04187.1

Group14419: anid|CBF75028.1 asyd|OJJ60053.1 aver|OJJ02236.1

Group14420: anid|CBF75248.1 asyd|OJJ59906.1 aver|OJJ02379.1

Group14421: anid|CBF75862.1 asyd|OJJ60229.1 aver|OJJ04038.1

Group14422: anid|CBF75870.1 asyd|OJJ60242.1 aver|OJJ04024.1

Group14423: anid|CBF75894.1 asyd|OJJ60267.1 aver|OJJ03987.1

Group14424: anid|CBF75996.1 asyd|OJJ59249.1 aver|OJJ01787.1

Group14425: anid|CBF76042.1 asyd|OJJ52461.1 aver|OJJ03954.1

Group14426: anid|CBF76046.1 asyd|OJJ52463.1 aver|OJJ03952.1

Group14427: anid|CBF76048.1 asyd|OJJ52464.1 aver|OJJ03951.1

Group14428: anid|CBF76273.1 asyd|OJJ60976.1 aver|OJJ00300.1

Group14429: anid|CBF76370.1 asyd|OJJ61063.1 aver|OJJ00380.1

Group14430: anid|CBF76500.1 asyd|OJJ61129.1 aver|OJJ00450.1

Group14431: anid|CBF76729.1 asyd|OJJ61249.1 aver|OJJ00573.1

Group14432: anid|CBF76731.1 asyd|OJJ61250.1 aver|OJJ00574.1

Group14433: anid|CBF76753.1 asyd|OJJ61261.1 aver|OJJ00587.1

Group14434: anid|CBF76928.1 asyd|OJJ57068.1 aver|OJJ03230.1

Group14435: anid|CBF77020.1 asyd|OJJ57015.1 aver|OJJ03273.1

Group14436: anid|CBF77135.1 asyd|OJJ56946.1 aver|OJJ03339.1

Group14437: anid|CBF77162.1 asyd|OJJ56862.1 aver|OJJ03463.1

Group14438: anid|CBF77520.1 aver|OJJ03670.1 asyd|OJJ53181.1

Group14439: anid|CBF77992.1 asyd|OJJ55253.1 aver|OJJ06971.1

Group14440: anid|CBF78052.1 aver|OJI97562.1 azon|OJJ43663.1

Group14441: anid|CBF78269.1 asyd|OJJ56078.1 aver|OJJ05021.1

Group14442: anid|CBF78271.1 asyd|OJJ56076.1 aver|OJJ05019.1

Group14443: anid|CBF78303.1 asyd|OJJ56098.1 aver|OJJ05041.1

Group14444: anid|CBF78309.1 asyd|OJJ56092.1 aver|OJJ05035.1

Group14445: anid|CBF78329.1 asyd|OJJ56084.1 aver|OJJ05027.1

Group14446: anid|CBF78335.1 asyd|OJJ56126.1 aver|OJJ05068.1

Group14447: anid|CBF78425.1 asyd|OJJ58319.1 aver|OJJ02092.1

Group14448: anid|CBF78457.1 asyd|OJJ64189.1 aver|OJI98266.1

Group14449: anid|CBF78577.1 asyd|OJJ65071.1 aver|OJI99196.1

Group14450: anid|CBF78579.1 asyd|OJJ65072.1 aver|OJI99197.1

Group14451: anid|CBF78844.1 asyd|OJJ65276.1 aver|OJI99428.1

Group14452: anid|CBF79020.1 asyd|OJJ57252.1 aver|OJJ03042.1

Group14453: anid|CBF79099.1 asyd|OJJ55028.1 aver|OJJ07176.1

Group14454: anid|CBF79180.1 aver|OJJ04747.1 asyd|OJJ52718.1

Group14455: anid|CBF79425.1 asyd|OJJ57838.1 aver|OJJ04552.1

Group14456: anid|CBF79568.1 aver|OJJ04475.1 asyd|OJJ57916.1

Group14457: anid|CBF79589.1 asyd|OJJ57980.1 aver|OJJ04420.1

Group14458: anid|CBF79624.1 asyd|OJJ58005.1 aver|OJJ04396.1

Group14459: anid|CBF79744.1 asyd|OJJ58087.1 aver|OJJ04322.1

Group14460: anid|CBF79749.1 asyd|OJJ58097.1 aver|OJJ04307.1

Group14461: anid|CBF79917.1 asyd|OJJ58192.1 aver|OJJ04213.1

Group14462: anid|CBF79937.1 asyd|OJJ58209.1 aver|OJJ02213.1

Group14463: anid|CBF80069.1 asyd|OJJ58295.1 aver|OJJ02129.1

Group14464: anid|CBF80115.1 asyd|OJJ58330.1 aver|OJJ02081.1

Group14465: anid|CBF80119.1 asyd|OJJ58340.1 aver|OJJ02071.1

Group14466: anid|CBF80432.1 asyd|OJJ54778.1 aver|OJJ07387.1

Group14467: anid|CBF80445.1 asyd|OJJ54795.1 aver|OJJ07370.1

Group14468: anid|CBF80448.1 asyd|OJJ54756.1 aver|OJJ07414.1

Group14469: anid|CBF80506.1 asyd|OJJ54735.1 aver|OJJ07436.1

Group14470: anid|CBF80531.1 asyd|OJJ55618.1 aver|OJJ06693.1

Group14471: anid|CBF80580.1 asyd|OJJ56927.1 aver|OJJ03367.1

Group14472: anid|CBF80716.1 asyd|OJJ58387.1 aver|OJJ02031.1

Group14473: anid|CBF80718.1 aver|OJJ02032.1 asyd|OJJ58386.1

Group14474: anid|CBF80925.1 asyd|OJJ56318.1 aver|OJJ06396.1

Group14475: anid|CBF80984.1 asyd|OJJ53565.1 aver|OJJ08189.1

Group14476: anid|CBF81046.1 asyd|OJJ56439.1 aver|OJJ06281.1

Group14477: anid|CBF81475.1 asyd|OJJ59449.1 aver|OJJ02823.1

Group14478: anid|CBF81477.1 asyd|OJJ59450.1 aver|OJJ02822.1

Group14479: anid|CBF81778.1 asyd|OJJ54866.1 aver|OJI96473.1

Group14480: anid|CBF81781.1 asyd|OJJ58364.1 aver|OJJ02042.1

Group14481: anid|CBF81895.1 asyd|OJJ59681.1 aver|OJJ02607.1

Group14482: anid|CBF81929.1 asyd|OJJ57420.1 aver|OJJ05758.1

Group14483: anid|CBF81931.1 asyd|OJJ57417.1 aver|OJJ05760.1

Group14484: awen|OJJ35532.1 asyd|OJJ54281.1 aver|OJJ05233.1

Group14485: anid|CBF82112.1 asyd|OJJ54287.1 aver|OJJ05241.1

Group14486: anid|CBF82115.1 asyd|OJJ54290.1 aver|OJJ05245.1

Group14487: anid|CBF82167.1 asyd|OJJ54345.1 aver|OJJ05297.1

Group14488: anid|CBF82171.1 asyd|OJJ54354.1 aver|OJJ05308.1

Group14489: anid|CBF82292.1 asyd|OJJ52733.1 aver|OJJ05348.1

Group14490: anid|CBF82306.1 asyd|OJJ61693.1 aver|OJJ08127.1

Group14491: anid|CBF82372.1 aory|EIT75863.1 azon|OJJ42562.1

Group14492: anid|CBF82440.1 asyd|OJJ54126.1 aver|OJJ08610.1

Group14493: anid|CBF82604.1 asyd|OJJ54027.1 aver|OJJ08713.1

Group14494: anid|CBF82777.1 asyd|OJJ53737.1 aver|OJJ07852.1

Group14495: anid|CBF82883.1 asyd|OJJ55064.1 aver|OJJ07147.1

Group14496: anid|CBF83007.1 asyd|OJJ56437.1 aver|OJJ06284.1

Group14497: anid|CBF83070.1 asyd|OJJ61487.1 aver|OJJ00829.1

Group14498: anid|CBF83078.1 asyd|OJJ52798.1 aver|OJJ08775.1

Group14499: anid|CBF83356.1 asyd|OJJ60617.1 aver|OJI99941.1

Group14500: anid|CBF83386.1 asyd|OJJ60633.1 aver|OJI99956.1

Group14501: anid|CBF83704.1 asyd|OJJ52494.1 aver|OJJ06530.1

Group14502: anid|CBF83831.1 asyd|OJJ60908.1 aver|OJJ00235.1

Group14503: anid|CBF83917.1 asyd|OJJ55493.1 aver|OJJ06569.1

Group14504: anid|CBF83942.1 asyd|OJJ61447.1 aver|OJJ00783.1

Group14505: anid|CBF83984.1 asyd|OJJ57947.1 aver|OJJ04451.1

Group14506: anid|CBF84006.1 aver|OJI97563.1 azon|OJJ42261.1

Group14507: anid|CBF84229.1 asyd|OJJ57180.1 aver|OJJ03110.1

Group14508: anid|CBF84236.1 asyd|OJJ57187.1 aver|OJJ03102.1

Group14509: anid|CBF84379.1 asyd|OJJ62965.1 aver|OJI98002.1

Group14510: anid|CBF84398.1 asyd|OJJ53579.1 aver|OJJ08179.1

Group14511: anid|CBF84406.1 asyd|OJJ53569.1 aver|OJJ08186.1

Group14512: anid|CBF84417.1 asyd|OJJ53558.1 aver|OJJ08196.1

Group14513: anid|CBF84428.1 asyd|OJJ53639.1 aver|OJJ01167.1

Group14514: anid|CBF84484.1 asyd|OJJ64713.1 aver|OJI98816.1

Group14515: anid|CBF84535.1 asyd|OJJ55561.1 aver|OJJ06642.1

Group14516: aver|OJJ08390.1 asyd|OJJ53353.1 awen|OJJ36233.1

Group14517: anid|CBF84843.1 asyd|OJJ63205.1 aver|OJI97750.1

Group14518: anid|CBF85117.1 asyd|OJJ63048.1 aver|OJI97915.1

Group14519: anid|CBF85140.1 asyd|OJJ63036.1 aver|OJI97931.1

Group14520: anid|CBF85146.1 asyd|OJJ63003.1 aver|OJI97957.1

Group14521: anid|CBF85161.1 asyd|OJJ54545.1 aver|OJJ07632.1

Group14522: anid|CBF85226.1 asyd|OJJ57608.1 aver|OJJ05576.1

Group14523: anid|CBF85286.1 asyd|OJJ56752.1 aver|OJJ05972.1

Group14524: anid|CBF85288.1 asyd|OJJ56753.1 aver|OJJ05970.1

Group14525: anid|CBF85424.1 asyd|OJJ64076.1 aver|OJI98163.1

Group14526: anid|CBF85774.1 asyd|OJJ64309.1 aver|OJI98402.1

Group14527: anid|CBF86059.1 asyd|OJJ54469.1 aver|OJJ07780.1

Group14528: anid|CBF86346.1 asyd|OJJ64977.1 aver|OJI99099.1

Group14529: anid|CBF86377.1 asyd|OJJ64963.1 aver|OJI99086.1

Group14530: anid|CBF86407.1 asyd|OJJ64946.1 aver|OJI99068.1

Group14531: anid|CBF86615.1 asyd|OJJ64816.1 aver|OJI98929.1

Group14532: anid|CBF86645.1 asyd|OJJ64797.1 aver|OJI98911.1

Group14533: anid|CBF86749.1 asyd|OJJ64706.1 aver|OJI98808.1

Group14534: anid|CBF86768.1 asyd|OJJ64689.1 aver|OJI98792.1

Group14535: anid|CBF86873.1 asyd|OJJ64629.1 aver|OJI98727.1

Group14536: anid|CBF86921.1 asyd|OJJ55624.1 aver|OJJ06699.1

Group14537: anid|CBF87048.1 asyd|OJJ60443.1 aver|OJI99761.1

Group14538: anid|CBF87065.1 asyd|OJJ53736.1 aver|OJJ07851.1

Group14539: anid|CBF87086.1 asyd|OJJ60412.1 aver|OJI99739.1

Group14540: anid|CBF87118.1 asyd|OJJ60358.1 aver|OJI99696.1

Group14541: anid|CBF87135.1 asyd|OJJ60371.1 aver|OJI99706.1

Group14542: anid|CBF87167.1 asyd|OJJ60294.1 aver|OJI99632.1

Group14543: anid|CBF87183.1 asyd|OJJ55021.1 aver|OJI96404.1

Group14544: awen|OJJ36059.1 asyd|OJJ62966.1 aver|OJI98001.1

Group14545: anid|CBF87444.1 asyd|OJJ63360.1 aver|OJI97589.1

Group14546: anid|CBF87446.1 asyd|OJJ63361.1 aver|OJI97588.1

Group14547: anid|CBF87449.1 asyd|OJJ58570.1 aver|OJJ01174.1

Group14548: anid|CBF87518.1 asyd|OJJ52981.1 aver|OJJ08950.1

Group14549: anid|CBF87520.1 asyd|OJJ52982.1 aver|OJJ08951.1

Group14550: anid|CBF87722.1 asyd|OJJ63340.1 aver|OJI97621.1

Group14551: anid|CBF87751.1 asyd|OJJ56865.1 aver|OJJ06180.1

Group14552: anid|CBF88207.1 aver|OJI97290.1 asyd|OJJ63644.1

Group14553: anid|CBF88271.1 asyd|OJJ63681.1 aver|OJI97258.1

Group14554: anid|CBF88397.1 aver|OJI97194.1 asyd|OJJ63746.1

Group14555: anid|CBF88461.1 asyd|OJJ63775.1 aver|OJI97162.1

Group14556: anid|CBF88562.1 asyd|OJJ63834.1 aver|OJI97103.1

Group14557: anid|CBF88678.1 asyd|OJJ63899.1 aver|OJI97041.1

Group14558: anid|CBF88772.1 asyd|OJJ63984.1 aver|OJI96969.1

Group14559: anid|CBF88776.1 asyd|OJJ63472.1 aver|OJI97459.1

Group14560: anid|CBF88778.1 asyd|OJJ63473.1 aver|OJI97457.1

Group14561: anid|CBF88843.1 asyd|OJJ62868.1 aver|OJI96921.1

Group14562: anid|CBF88871.1 asyd|OJJ64581.1 aver|OJI98678.1

Group14563: anid|CBF89117.1 asyd|OJJ62657.1 aver|OJI96755.1

Group14564: anid|CBF89122.1 asyd|OJJ62660.1 aver|OJI96758.1

Group14565: anid|CBF89131.1 asyd|OJJ62663.1 aver|OJI96761.1

Group14566: anid|CBF89135.1 asyd|OJJ62651.1 aver|OJI96751.1

Group14567: anid|CBF89173.1 aver|OJI96666.1 asyd|OJJ62540.1

Group14568: anid|CBF89175.1 asyd|OJJ62539.1 aver|OJI96665.1

Group14569: anid|CBF89375.1 asyd|OJJ62472.1 aver|OJI96607.1

Group14570: anid|CBF89410.1 asyd|OJJ62456.1 aver|OJI96591.1

Group14571: anid|CBF89557.1 asyd|OJJ62342.1 aver|OJI96446.1

Group14572: anid|CBF89565.1 asyd|OJJ62346.1 aver|OJI96451.1

Group14573: anid|CBF89566.1 asyd|OJJ62339.1 aver|OJI96444.1

Group14574: anid|CBF89570.1 asyd|OJJ65271.1 aver|OJI99423.1

Group14575: anid|CBF89604.1 asyd|OJJ62308.1 aver|OJI96416.1

Group14576: anid|CBF89835.1 asyd|OJJ62179.1 aver|OJI96288.1

Group14577: anid|CBF90092.1 asyd|OJJ62035.1 aver|OJI96125.1

Group14578: anid|CBF90311.1 asyd|OJJ61914.1 aver|OJI96007.1

Group14579: anid|CBF90378.1 asyd|OJJ62125.1 aver|OJI96233.1

Group14580: anig|EHA22529.1 ater|EAU33342.1 aver|OJJ01797.1

Group14581: anig|EHA23285.1 ater|EAU29434.1 azon|OJJ42274.1

Group14582: anig|EHA23408.1 asyd|OJJ60337.1 aver|OJI99682.1

Group14583: anig|EHA27750.1 atub|OJI89643.1 aver|OJJ08328.1

Group14584: aory|EIT75673.1 ater|EAU32771.1 aver|OJI97598.1

Group14585: aory|EIT77107.1 asyd|OJJ59778.1 aver|OJJ02508.1

Group14586: aory|EIT82307.1 asyd|OJJ57951.1 aver|OJJ04446.1

Group14587: asyd|OJJ52205.1 ater|EAU36867.1 aver|OJI95788.1

Group14588: asyd|OJJ52351.1 ater|EAU33822.1 aver|OJJ07897.1

Group14589: asyd|OJJ52373.1 ater|EAU38987.1 aver|OJJ01976.1

Group14590: asyd|OJJ52474.1 aver|OJJ03942.1 awen|OJJ37004.1

Group14591: asyd|OJJ52479.1 aver|OJJ05893.1 awen|OJJ31164.1

Group14592: asyd|OJJ52761.1 aver|OJJ08743.1 awen|OJJ40744.1

Group14593: asyd|OJJ52774.1 ater|EAU38646.1 aver|OJJ08757.1

Group14594: asyd|OJJ52814.1 aver|OJJ08793.1 awen|OJJ40868.1

Group14595: asyd|OJJ52848.1 aver|OJJ08826.1 awen|OJJ37375.1

Group14596: asyd|OJJ53447.1 aver|OJJ08305.1 awen|OJJ41339.1

Group14597: asyd|OJJ53619.1 atub|OJI91473.1 awen|OJJ30586.1

Group14598: asyd|OJJ53744.1 ater|EAU30917.1 aver|OJJ07858.1

Group14599: asyd|OJJ53771.1 ater|EAU30267.1 aver|OJJ07879.1

Group14600: asyd|OJJ53814.1 ater|EAU31387.1 aver|OJJ07915.1

Group14601: asyd|OJJ54096.1 ater|EAU36840.1 aver|OJJ08643.1

Group14602: asyd|OJJ54187.1 ater|EAU35237.1 aver|OJJ08555.1

Group14603: asyd|OJJ54327.1 ater|EAU29306.1 aver|OJJ05281.1

Group14604: asyd|OJJ54391.1 ater|EAU32810.1 aver|OJI99555.1

Group14605: asyd|OJJ54507.1 ater|EAU33037.1 aver|OJJ07675.1

Group14606: asyd|OJJ54650.1 ater|EAU30281.1 aver|OJJ07511.1

Group14607: asyd|OJJ54651.1 ater|EAU30280.1 aver|OJJ07510.1

Group14608: asyd|OJJ54666.1 aver|OJJ02023.1 awen|OJJ34681.1

Group14609: asyd|OJJ54669.1 aver|OJJ02020.1 azon|OJJ50312.1

Group14610: asyd|OJJ54825.1 ater|EAU34239.1 aver|OJJ07335.1

Group14611: asyd|OJJ54850.1 ater|EAU35599.1 aver|OJJ07316.1

Group14612: asyd|OJJ54923.1 aver|OJJ07272.1 azon|OJJ45632.1

Group14613: asyd|OJJ55408.1 aver|OJJ06487.1 awen|OJJ35932.1

Group14614: asyd|OJJ56041.1 ater|EAU37376.1 aver|OJJ04987.1

Group14615: asyd|OJJ56059.1 aver|OJJ05001.1 awen|OJJ33508.1

Group14616: asyd|OJJ56124.1 aver|OJJ05066.1 awen|OJJ35913.1

Group14617: asyd|OJJ56140.1 aver|OJJ05079.1 awen|OJJ38744.1

Group14618: asyd|OJJ56160.1 aver|OJJ05099.1 awen|OJJ39997.1

Group14619: asyd|OJJ56207.1 ater|EAU36042.1 aver|OJJ05138.1

Group14620: asyd|OJJ56503.1 ater|EAU31639.1 aver|OJJ06216.1

Group14621: asyd|OJJ56976.1 aver|OJJ03312.1 awen|OJJ33540.1

Group14622: asyd|OJJ57086.1 aver|OJJ03215.1 awen|OJJ36210.1

Group14623: asyd|OJJ57127.1 ater|EAU36117.1 aver|OJJ03178.1

Group14624: asyd|OJJ57283.1 aver|OJJ05873.1 azon|OJJ46800.1

Group14625: asyd|OJJ57408.1 ater|EAU34547.1 aver|OJJ05773.1

Group14626: asyd|OJJ58044.1 ater|EAU38803.1 azon|OJJ50745.1

Group14627: asyd|OJJ58197.1 ater|EAU38231.1 aver|OJJ04210.1

Group14628: asyd|OJJ58353.1 aver|OJJ02054.1 awen|OJJ36814.1

Group14629: asyd|OJJ58354.1 aver|OJJ02053.1 azon|OJJ42228.1

Group14630: asyd|OJJ58556.1 aver|OJJ01154.1 azon|OJJ42243.1

Group14631: asyd|OJJ58560.1 ater|EAU38542.1 aver|OJJ01159.1

Group14632: asyd|OJJ59257.1 aver|OJJ01868.1 awen|OJJ36073.1

Group14633: asyd|OJJ59278.1 aver|OJJ01893.1 awen|OJJ40926.1

Group14634: asyd|OJJ59579.1 aver|OJJ02690.1 azon|OJJ43186.1

Group14635: asyd|OJJ60281.1 aver|OJI99610.1 awen|OJJ39505.1

Group14636: asyd|OJJ60308.1 aver|OJI99649.1 awen|OJJ34010.1

Group14637: asyd|OJJ60309.1 aver|OJI99650.1 awen|OJJ34011.1

Group14638: asyd|OJJ60312.1 aver|OJI99653.1 awen|OJJ34014.1

Group14639: asyd|OJJ60315.1 aver|OJI99656.1 awen|OJJ34016.1

Group14640: asyd|OJJ60477.1 ater|EAU39530.1 aver|OJI99797.1

Group14641: asyd|OJJ60499.1 aver|OJI99821.1 awen|OJJ33596.1

Group14642: asyd|OJJ60515.1 aver|OJI99841.1 azon|OJJ45783.1

Group14643: asyd|OJJ61039.1 ater|EAU38819.1 azon|OJJ46801.1

Group14644: asyd|OJJ61400.1 aver|OJJ00729.1 awen|OJJ33520.1

Group14645: asyd|OJJ61409.1 aver|OJJ00739.1 azon|OJJ45992.1

Group14646: asyd|OJJ61695.1 ater|EAU39523.1 aver|OJI95779.1

Group14647: asyd|OJJ62167.1 aver|OJI96278.1 awen|OJJ40170.1

Group14648: asyd|OJJ62329.1 ater|EAU32507.1 aver|OJI96435.1

Group14649: asyd|OJJ62457.1 ater|EAU33527.1 aver|OJI96592.1

Group14650: asyd|OJJ62466.1 aver|OJI96601.1 awen|OJJ34590.1

Group14651: asyd|OJJ62496.1 ater|EAU36884.1 aver|OJJ07650.1

Group14652: asyd|OJJ62621.1 aver|OJI99299.1 awen|OJJ36993.1

Group14653: asyd|OJJ62929.1 aver|OJI98061.1 awen|OJJ41768.1

Group14654: asyd|OJJ63061.1 ater|EAU30423.1 aver|OJI97902.1

Group14655: asyd|OJJ63173.1 aver|OJI97781.1 ater|EAU38713.1

Group14656: asyd|OJJ63495.1 ater|EAU31462.1 aver|OJI97434.1

Group14657: asyd|OJJ63987.1 ater|EAU32301.1 aver|OJI96966.1

Group14658: asyd|OJJ64183.1 ater|EAU39599.1 aver|OJI98260.1

Group14659: asyd|OJJ64320.1 ater|EAU32177.1 aver|OJI98413.1

Group14660: asyd|OJJ64790.1 aver|OJI98902.1 awen|OJJ41509.1

Group14661: asyd|OJJ64795.1 aver|OJI98910.1 awen|OJJ36234.1

Group14662: asyd|OJJ64971.1 ater|EAU33961.1 aver|OJI99095.1

Group14663: asyd|OJJ65243.1 aver|OJI99395.1 awen|OJJ38300.1

Group14664: asyd|OJJ65288.1 aver|OJJ08735.1 awen|OJJ31971.1

Group14665: asyd|OJJ65299.1 aver|OJI99439.1 azon|OJJ51833.1

Group14666: asyd|OJJ65365.1 ater|EAU35430.1 aver|OJJ01304.1

Group14667: ater|EAU31954.1 atub|OJI88933.1 aver|OJJ03916.1

Group14668: abra|OJJ65465.1 abra|OJJ65474.1

Group14669: abra|OJJ65494.1 abra|OJJ65649.1

Group14670: abra|OJJ65549.1 abra|OJJ65555.1

Group14671: abra|OJJ65650.1 abra|OJJ75560.1

Group14672: abra|OJJ65652.1 abra|OJJ75561.1

Group14673: abra|OJJ67053.1 abra|OJJ68239.1

Group14674: acar|OOF89765.1 acar|OOF89774.1

Group14675: acar|OOF90106.1 acar|OOF98053.1

Group14676: acar|OOF90422.1 acar|OOF90426.1

Group14677: acar|OOF90439.1 acar|OOF90446.1

Group14678: acar|OOF90440.1 acar|OOF90447.1

Group14679: acar|OOF90473.1 acar|OOF90480.1

Group14680: acar|OOF90474.1 acar|OOF90481.1

Group14681: acar|OOF90475.1 acar|OOF90482.1

Group14682: acar|OOF90476.1 acar|OOF90483.1

Group14683: acar|OOF90477.1 acar|OOF90484.1

Group14684: acar|OOF90478.1 acar|OOF90485.1

Group14685: acar|OOF90494.1 acar|OOF90503.1

Group14686: acar|OOF91720.1 acar|OOF91724.1

Group14687: acar|OOF91722.1 acar|OOF91726.1

Group14688: acar|OOF94045.1 acar|OOF94115.1

Group14689: acar|OOF94072.1 acar|OOF94142.1

Group14690: acar|OOF94080.1 acar|OOF94150.1

Group14691: acar|OOF94096.1 acar|OOF94166.1

Group14692: acar|OOF95044.1 acar|OOF95111.1

Group14693: acar|OOF95052.1 acar|OOF95119.1

Group14694: acar|OOF95053.1 acar|OOF95120.1

Group14695: acar|OOF95059.1 acar|OOF95126.1

Group14696: acar|OOF95622.1 acar|OOF95674.1

Group14697: acar|OOG00253.1 acar|OOG00271.1

Group14698: acla|EAW12036.1 acla|EAW12879.1

Group14699: afla|EED45097.1 afla|EED45491.1

Group14700: afla|EED49210.1 afla|EED56618.1

Group14701: agla|OJJ78406.1 agla|OJJ78764.1

Group14702: agla|OJJ78457.1 agla|OJJ83498.1

Group14703: agla|OJJ78465.1 agla|OJJ80594.1

Group14704: agla|OJJ78481.1 agla|OJJ79796.1

Group14705: agla|OJJ78540.1 agla|OJJ78728.1

Group14706: agla|OJJ79010.1 agla|OJJ80225.1

Group14707: agla|OJJ79091.1 agla|OJJ81300.1

Group14708: agla|OJJ79759.1 agla|OJJ88429.1

Group14709: agla|OJJ79780.1 agla|OJJ85457.1

Group14710: agla|OJJ79844.1 agla|OJJ85998.1

Group14711: agla|OJJ80514.1 agla|OJJ85398.1

Group14712: agla|OJJ80605.1 agla|OJJ82356.1

Group14713: agla|OJJ80624.1 agla|OJJ80625.1

Group14714: agla|OJJ81291.1 agla|OJJ89319.1

Group14715: agla|OJJ87859.1 agla|OJJ78921.1

Group14716: agla|OJJ82050.1 agla|OJJ82055.1

Group14717: agla|OJJ82066.1 agla|OJJ82954.1

Group14718: agla|OJJ82254.1 agla|OJJ83433.1

Group14719: agla|OJJ82847.1 agla|OJJ88204.1

Group14720: agla|OJJ82922.1 agla|OJJ83800.1

Group14721: agla|OJJ84077.1 agla|OJJ87264.1

Group14722: agla|OJJ84907.1 agla|OJJ83287.1

Group14723: agla|OJJ85486.1 agla|OJJ85488.1

Group14724: agla|OJJ86336.1 agla|OJJ87980.1

Group14725: agla|OJJ86550.1 agla|OJJ87009.1

Group14726: agla|OJJ86648.1 agla|OJJ89311.1

Group14727: agla|OJJ86729.1 agla|OJJ88110.1

Group14728: agla|OJJ87097.1 agla|OJJ87099.1

Group14729: agla|OJJ87637.1 agla|OJJ88095.1

Group14730: akaw|GAA92068.1 akaw|GAA93092.1

Group14731: akaw|GAA92349.1 akaw|GAA93135.1

Group14732: akaw|GAA92368.1 akaw|GAA93353.1

Group14733: akaw|GAA92611.1 akaw|GAA93006.1

Group14734: akaw|GAA93028.1 akaw|GAA93067.1

Group14735: akaw|GAA93184.1 akaw|GAA93265.1

Group14736: akaw|GAA93294.1 akaw|GAA93312.1

Group14737: aluc|OJZ79483.1 aluc|OJZ79694.1

Group14738: aluc|OJZ79495.1 aluc|OJZ80499.1

Group14739: aluc|OJZ79533.1 aluc|OJZ79761.1

Group14740: aluc|OJZ79566.1 aluc|OJZ79848.1

Group14741: aluc|OJZ79573.1 aluc|OJZ81137.1

Group14742: aluc|OJZ79610.1 aluc|OJZ80080.1

Group14743: aluc|OJZ79635.1 aluc|OJZ80075.1

Group14744: aluc|OJZ79671.1 aluc|OJZ79751.1

Group14745: aluc|OJZ79707.1 aluc|OJZ80153.1

Group14746: aluc|OJZ79718.1 aluc|OJZ79871.1

Group14747: aluc|OJZ79727.1 aluc|OJZ80231.1

Group14748: aluc|OJZ79745.1 aluc|OJZ79983.1

Group14749: aluc|OJZ79763.1 aluc|OJZ79911.1

Group14750: aluc|OJZ79775.1 aluc|OJZ80136.1

Group14751: aluc|OJZ79820.1 aluc|OJZ80412.1

Group14752: aluc|OJZ79929.1 aluc|OJZ80389.1

Group14753: aluc|OJZ79931.1 aluc|OJZ80388.1

Group14754: aluc|OJZ79965.1 aluc|OJZ80475.1

Group14755: aluc|OJZ79982.1 aluc|OJZ80079.1

Group14756: aluc|OJZ79990.1 aluc|OJZ91545.1

Group14757: aluc|OJZ80021.1 aluc|OJZ80176.1

Group14758: aluc|OJZ80041.1 aluc|OJZ80337.1

Group14759: aluc|OJZ80054.1 aluc|OJZ80215.1

Group14760: aluc|OJZ80157.1 aluc|OJZ80319.1

Group14761: aluc|OJZ80245.1 aluc|OJZ80432.1

Group14762: anid|CBF69531.1 anid|CBF86063.1

Group14763: anid|CBF71837.1 anid|CBF82958.1

Group14764: anid|CBF80694.1 anid|CBF84322.1

Group14765: anid|CBF81146.1 anid|CBF89248.1

Group14766: anid|CBF82215.1 anid|CBF82216.1

Group14767: anid|CBF82240.1 anid|CBF87228.1

Group14768: anid|CBF86379.1 anid|CBF89320.1

Group14769: aory|EIT78835.1 afla|EED54654.1

Group14770: asyd|OJJ51850.1 asyd|OJJ52246.1

Group14771: asyd|OJJ51873.1 asyd|OJJ51981.1

Group14772: asyd|OJJ51874.1 asyd|OJJ51980.1

Group14773: asyd|OJJ51884.1 asyd|OJJ52234.1

Group14774: asyd|OJJ51926.1 asyd|OJJ52309.1

Group14775: asyd|OJJ51953.1 asyd|OJJ62676.1

Group14776: asyd|OJJ51969.1 asyd|OJJ51991.1

Group14777: asyd|OJJ52023.1 asyd|OJJ52284.1

Group14778: asyd|OJJ52116.1 asyd|OJJ52171.1

Group14779: asyd|OJJ52195.1 asyd|OJJ54370.1

Group14780: asyd|OJJ52238.1 asyd|OJJ52267.1

Group14781: asyd|OJJ52251.1 asyd|OJJ52258.1

Group14782: asyd|OJJ52295.1 asyd|OJJ62685.1

Group14783: asyd|OJJ52776.1 asyd|OJJ52777.1

Group14784: asyd|OJJ52809.1 asyd|OJJ65405.1

Group14785: asyd|OJJ53799.1 asyd|OJJ54445.1

Group14786: asyd|OJJ54140.1 asyd|OJJ55189.1

Group14787: ater|EAU29592.1 ater|EAU30415.1

Group14788: ater|EAU29979.1 ater|EAU32471.1

Group14789: ater|EAU31049.1 ater|EAU34695.1

Group14790: ater|EAU31653.1 ater|EAU39310.1

Group14791: ater|EAU31728.1 ater|EAU39616.1

Group14792: ater|EAU31744.1 ater|EAU33403.1

Group14793: ater|EAU33564.1 ater|EAU37222.1

Group14794: ater|EAU34493.1 ater|EAU37114.1

Group14795: aver|OJI95753.1 aver|OJJ06438.1

Group14796: aver|OJI96735.1 aver|OJJ03399.1

Group14797: aver|OJI97923.1 aver|OJI97924.1

Group14798: aver|OJI98315.1 aver|OJJ07713.1

Group14799: aver|OJI98892.1 aver|OJJ05269.1

Group14800: aver|OJI99302.1 aver|OJJ07469.1

Group14801: aver|OJI99629.1 aver|OJJ07369.1

Group14802: aver|OJJ00340.1 aver|OJJ05776.1

Group14803: aver|OJJ03365.1 aver|OJJ07296.1

Group14804: aver|OJJ03398.1 aver|OJJ08768.1

Group14805: aver|OJJ03444.1 aver|OJJ03886.1

Group14806: aver|OJJ07615.1 aver|OJJ07623.1

Group14807: awen|OJJ29556.1 awen|OJJ29644.1

Group14808: awen|OJJ29557.1 awen|OJJ29646.1

Group14809: awen|OJJ29606.1 awen|OJJ32582.1

Group14810: awen|OJJ29710.1 awen|OJJ29794.1

Group14811: awen|OJJ30791.1 awen|OJJ31690.1

Group14812: awen|OJJ30793.1 awen|OJJ31688.1

Group14813: awen|OJJ30795.1 awen|OJJ31686.1

Group14814: awen|OJJ31284.1 awen|OJJ37716.1

Group14815: awen|OJJ31944.1 awen|OJJ31945.1

Group14816: awen|OJJ32039.1 awen|OJJ32999.1

Group14817: awen|OJJ32565.1 awen|OJJ37837.1

Group14818: awen|OJJ32783.1 awen|OJJ33782.1

Group14819: awen|OJJ32940.1 awen|OJJ37849.1

Group14820: awen|OJJ33282.1 awen|OJJ33283.1

Group14821: awen|OJJ33400.1 awen|OJJ36228.1

Group14822: awen|OJJ33581.1 awen|OJJ35019.1

Group14823: awen|OJJ33694.1 awen|OJJ38676.1

Group14824: awen|OJJ34110.1 awen|OJJ34111.1

Group14825: awen|OJJ35406.1 awen|OJJ40654.1

Group14826: awen|OJJ37974.1 awen|OJJ37977.1

Group14827: azon|OJJ42072.1 azon|OJJ48733.1

Group14828: azon|OJJ42273.1 azon|OJJ42642.1

Group14829: azon|OJJ42400.1 azon|OJJ51826.1

Group14830: azon|OJJ42439.1 azon|OJJ46535.1

Group14831: azon|OJJ42453.1 azon|OJJ46972.1

Group14832: azon|OJJ42465.1 azon|OJJ45246.1

Group14833: azon|OJJ42469.1 azon|OJJ45911.1

Group14834: azon|OJJ42565.1 azon|OJJ42571.1

Group14835: azon|OJJ44500.1 azon|OJJ45797.1

Group14836: azon|OJJ45075.1 azon|OJJ50103.1

Group14837: azon|OJJ45269.1 azon|OJJ45308.1

Group14838: azon|OJJ45900.1 azon|OJJ51057.1

Group14839: azon|OJJ46559.1 azon|OJJ50752.1

Group14840: azon|OJJ47255.1 azon|OJJ48052.1

Group14841: azon|OJJ47256.1 azon|OJJ47344.1

Group14842: azon|OJJ47341.1 azon|OJJ48413.1

Group14843: azon|OJJ47807.1 azon|OJJ51804.1

Group14844: azon|OJJ50234.1 azon|OJJ51283.1

Group14845: abra|OJJ65445.1 abra|OJJ65480.1

Group14846: abra|OJJ65447.1 abra|OJJ65563.1

Group14847: abra|OJJ65448.1 abra|OJJ65564.1

Group14848: abra|OJJ65467.1 abra|OJJ65660.1

Group14849: abra|OJJ65469.1 abra|OJJ65706.1

Group14850: abra|OJJ65504.1 abra|OJJ65816.1

Group14851: abra|OJJ65505.1 abra|OJJ65817.1

Group14852: abra|OJJ65523.1 abra|OJJ65566.1

Group14853: abra|OJJ65543.1 abra|OJJ67280.1

Group14854: abra|OJJ65544.1 abra|OJJ68291.1

Group14855: abra|OJJ65618.1 abra|OJJ65716.1

Group14856: abra|OJJ65664.1 abra|OJJ66172.1

Group14857: abra|OJJ66290.1 abra|OJJ69757.1

Group14858: anid|CBF70348.1 anid|CBF73871.1

Group14859: anid|CBF74009.1 anid|CBF79031.1

Group14860: anid|CBF75900.1 anid|CBF75902.1

Group14861: anid|CBF80274.1 anid|CBF81843.1

Group14862: anid|CBF80692.1 anid|CBF84326.1

Group14863: anid|CBF81151.1 anid|CBF86888.1

Group14864: anid|CBF82313.1 anid|CBF84375.1

Group14865: anid|CBF82962.1 anid|CBF89290.1

Group14866: anid|CBF84591.1 anid|CBF86032.1

Group14867: acar|OOF89762.1 acar|OOF89775.1

Group14868: asyd|OJJ51846.1 asyd|OJJ52141.1

Group14869: asyd|OJJ51853.1 asyd|OJJ51915.1

Group14870: asyd|OJJ51855.1 asyd|OJJ54877.1

Group14871: asyd|OJJ51881.1 asyd|OJJ52154.1

Group14872: asyd|OJJ51886.1 asyd|OJJ52169.1

Group14873: asyd|OJJ51899.1 asyd|OJJ52150.1

Group14874: asyd|OJJ51924.1 asyd|OJJ52310.1

Group14875: asyd|OJJ51933.1 asyd|OJJ52322.1

Group14876: asyd|OJJ51934.1 asyd|OJJ52323.1

Group14877: asyd|OJJ51935.1 asyd|OJJ52324.1

Group14878: asyd|OJJ51936.1 asyd|OJJ52325.1

Group14879: asyd|OJJ51952.1 asyd|OJJ62677.1

Group14880: asyd|OJJ51954.1 asyd|OJJ62675.1

Group14881: asyd|OJJ51985.1 asyd|OJJ52304.1

Group14882: asyd|OJJ51996.1 asyd|OJJ52314.1

Group14883: asyd|OJJ52020.1 asyd|OJJ52274.1

Group14884: asyd|OJJ52022.1 asyd|OJJ52285.1

Group14885: asyd|OJJ52048.1 asyd|OJJ65411.1

Group14886: asyd|OJJ52049.1 asyd|OJJ62684.1

Group14887: asyd|OJJ52147.1 asyd|OJJ52317.1

Group14888: asyd|OJJ52153.1 asyd|OJJ52280.1

Group14889: asyd|OJJ52164.1 asyd|OJJ62673.1

Group14890: asyd|OJJ52405.1 asyd|OJJ57627.1

Group14891: asyd|OJJ53068.1 asyd|OJJ56149.1

Group14892: asyd|OJJ53279.1 asyd|OJJ56763.1

Group14893: asyd|OJJ53281.1 asyd|OJJ56765.1

Group14894: asyd|OJJ53364.1 asyd|OJJ63018.1

Group14895: asyd|OJJ53642.1 asyd|OJJ58062.1

Group14896: asyd|OJJ54645.1 asyd|OJJ54646.1

Group14897: asyd|OJJ54695.1 asyd|OJJ54696.1

Group14898: asyd|OJJ55603.1 asyd|OJJ55606.1

Group14899: asyd|OJJ55604.1 asyd|OJJ55607.1

Group14900: acar|OOF90179.1 acar|OOF95343.1

Group14901: acar|OOF90489.1 acar|OOF90496.1

Group14902: acar|OOF90490.1 acar|OOF90497.1

Group14903: asyd|OJJ61553.1 asyd|OJJ61554.1

Group14904: acar|OOF91719.1 acar|OOF91723.1

Group14905: ater|EAU32502.1 ater|EAU35993.1

Group14906: ater|EAU32796.1 ater|EAU34346.1

Group14907: ater|EAU33996.1 ater|EAU33997.1

Group14908: ater|EAU36312.1 ater|EAU37068.1

Group14909: atub|OJI85695.1 atub|OJI85696.1

Group14910: aver|OJI96218.1 aver|OJJ02089.1

Group14911: acar|OOF94068.1 acar|OOF94138.1

Group14912: acar|OOF94083.1 acar|OOF94153.1

Group14913: acar|OOF94495.1 acar|OOF95399.1

Group14914: acar|OOF94692.1 acar|OOF94693.1

Group14915: awen|OJJ29533.1 awen|OJJ29641.1

Group14916: awen|OJJ29543.1 awen|OJJ29648.1

Group14917: awen|OJJ29545.1 awen|OJJ32591.1

Group14918: awen|OJJ29554.1 awen|OJJ29642.1

Group14919: awen|OJJ29579.1 awen|OJJ29604.1

Group14920: awen|OJJ29591.1 awen|OJJ29639.1

Group14921: awen|OJJ29743.1 awen|OJJ37019.1

Group14922: awen|OJJ30129.1 awen|OJJ34493.1

Group14923: awen|OJJ30748.1 awen|OJJ31733.1

Group14924: awen|OJJ30749.1 awen|OJJ31732.1

Group14925: awen|OJJ30751.1 awen|OJJ31730.1

Group14926: awen|OJJ30754.1 awen|OJJ31727.1

Group14927: awen|OJJ30761.1 awen|OJJ31720.1

Group14928: awen|OJJ30770.1 awen|OJJ31711.1

Group14929: awen|OJJ30842.1 awen|OJJ32917.1

Group14930: awen|OJJ31325.1 awen|OJJ35387.1

Group14931: awen|OJJ31419.1 awen|OJJ40216.1

Group14932: awen|OJJ31425.1 awen|OJJ35805.1

Group14933: awen|OJJ31900.1 awen|OJJ38092.1

Group14934: awen|OJJ31901.1 awen|OJJ31915.1

Group14935: awen|OJJ31957.1 awen|OJJ36420.1

Group14936: awen|OJJ32038.1 awen|OJJ37770.1

Group14937: awen|OJJ32247.1 awen|OJJ37028.1

Group14938: acar|OOF95041.1 acar|OOF95108.1

Group14939: awen|OJJ35795.1 awen|OJJ38065.1

Group14940: awen|OJJ35868.1 awen|OJJ38662.1

Group14941: awen|OJJ36478.1 awen|OJJ36479.1

Group14942: awen|OJJ37813.1 awen|OJJ40904.1

Group14943: awen|OJJ37820.1 awen|OJJ37822.1

Group14944: awen|OJJ37937.1 awen|OJJ37938.1

Group14945: awen|OJJ39718.1 awen|OJJ39720.1

Group14946: azon|OJJ42017.1 azon|OJJ45014.1

Group14947: azon|OJJ42188.1 azon|OJJ48025.1

Group14948: azon|OJJ42277.1 azon|OJJ46531.1

Group14949: azon|OJJ42342.1 azon|OJJ46520.1

Group14950: azon|OJJ42464.1 azon|OJJ45245.1

Group14951: azon|OJJ42466.1 azon|OJJ45247.1

Group14952: azon|OJJ42516.1 azon|OJJ42578.1

Group14953: azon|OJJ45025.1 azon|OJJ50324.1

Group14954: azon|OJJ45185.1 azon|OJJ50400.1

Group14955: azon|OJJ46141.1 azon|OJJ46142.1

Group14956: azon|OJJ46422.1 azon|OJJ51802.1

Group14957: azon|OJJ47563.1 azon|OJJ47811.1

Group14958: acar|OOF97689.1 acar|OOF97739.1

Group14959: acla|EAW08289.1 acla|EAW12997.1

Group14960: acla|EAW11777.1 acla|EAW15026.1

Group14961: afis|EAW15485.1 afis|EAW19658.1

Group14962: afis|EAW16170.1 afis|EAW18840.1

Group14963: afis|EAW17171.1 afis|EAW19371.1

Group14964: afla|EED45528.1 afla|EED45619.1

Group14965: afum|EAL86285.2 afum|EAL86738.1

Group14966: abra|OJJ71213.1 abra|OJJ75255.1

Group14967: agla|OJJ78454.1 agla|OJJ87959.1

Group14968: agla|OJJ78455.1 agla|OJJ81298.1

Group14969: agla|OJJ78464.1 agla|OJJ79291.1

Group14970: agla|OJJ85036.1 agla|OJJ78643.1

Group14971: agla|OJJ78556.1 agla|OJJ88574.1

Group14972: agla|OJJ78729.1 agla|OJJ80326.1

Group14973: agla|OJJ78771.1 agla|OJJ82972.1

Group14974: agla|OJJ79080.1 agla|OJJ86872.1

Group14975: agla|OJJ79279.1 agla|OJJ88489.1

Group14976: agla|OJJ79365.1 agla|OJJ85429.1

Group14977: agla|OJJ79383.1 agla|OJJ79384.1

Group14978: agla|OJJ79468.1 agla|OJJ79470.1

Group14979: agla|OJJ79872.1 agla|OJJ85343.1

Group14980: agla|OJJ79908.1 agla|OJJ80988.1

Group14981: agla|OJJ80062.1 agla|OJJ83548.1

Group14982: agla|OJJ80313.1 agla|OJJ88794.1

Group14983: agla|OJJ80317.1 agla|OJJ83606.1

Group14984: agla|OJJ80627.1 agla|OJJ80628.1

Group14985: agla|OJJ80686.1 agla|OJJ83810.1

Group14986: agla|OJJ81336.1 agla|OJJ88494.1

Group14987: agla|OJJ81481.1 agla|OJJ81825.1

Group14988: agla|OJJ82687.1 agla|OJJ86252.1

Group14989: agla|OJJ83431.1 agla|OJJ88334.1

Group14990: agla|OJJ83484.1 agla|OJJ88341.1

Group14991: agla|OJJ83551.1 agla|OJJ89315.1

Group14992: agla|OJJ83782.1 agla|OJJ86854.1

Group14993: agla|OJJ83784.1 agla|OJJ84565.1

Group14994: agla|OJJ83786.1 agla|OJJ86853.1

Group14995: agla|OJJ84757.1 agla|OJJ85318.1

Group14996: agla|OJJ84887.1 agla|OJJ88098.1

Group14997: agla|OJJ84908.1 agla|OJJ88820.1

Group14998: agla|OJJ84923.1 agla|OJJ86841.1

Group14999: agla|OJJ85443.1 agla|OJJ88023.1

Group15000: agla|OJJ85959.1 agla|OJJ87960.1

Group15001: agla|OJJ86651.1 agla|OJJ87958.1

Group15002: agla|OJJ87026.1 agla|OJJ89634.1

Group15003: agla|OJJ87394.1 agla|OJJ88697.1

Group15004: agla|OJJ87765.1 agla|OJJ88707.1

Group15005: agla|OJJ87851.1 agla|OJJ87852.1

Group15006: agla|OJJ87981.1 agla|OJJ88089.1

Group15007: agla|OJJ88094.1 agla|OJJ88096.1

Group15008: akaw|GAA92660.1 akaw|GAA93117.1

Group15009: aluc|OJZ79475.1 aluc|OJZ79908.1

Group15010: aluc|OJZ79494.1 aluc|OJZ80223.1

Group15011: aluc|OJZ79497.1 aluc|OJZ80391.1

Group15012: aluc|OJZ79515.1 aluc|OJZ79519.1

Group15013: aluc|OJZ79523.1 aluc|OJZ79770.1

Group15014: aluc|OJZ79535.1 aluc|OJZ80009.1

Group15015: aluc|OJZ79548.1 aluc|OJZ79812.1

Group15016: aluc|OJZ79555.1 aluc|OJZ79767.1

Group15017: aluc|OJZ79585.1 aluc|OJZ80490.1

Group15018: aluc|OJZ79589.1 aluc|OJZ80061.1

Group15019: aluc|OJZ79592.1 aluc|OJZ80058.1

Group15020: aluc|OJZ79620.1 aluc|OJZ80502.1

Group15021: aluc|OJZ79639.1 aluc|OJZ88798.1

Group15022: aluc|OJZ79640.1 aluc|OJZ80348.1

Group15023: aluc|OJZ79641.1 aluc|OJZ79997.1

Group15024: aluc|OJZ79719.1 aluc|OJZ79772.1

Group15025: aluc|OJZ79730.1 aluc|OJZ80233.1

Group15026: aluc|OJZ79741.1 aluc|OJZ79986.1

Group15027: aluc|OJZ79758.1 aluc|OJZ80332.1

Group15028: aluc|OJZ79776.1 aluc|OJZ88832.1

Group15029: aluc|OJZ79804.1 aluc|OJZ80037.1

Group15030: aluc|OJZ79809.1 aluc|OJZ79956.1

Group15031: aluc|OJZ79815.1 aluc|OJZ80081.1

Group15032: aluc|OJZ79817.1 aluc|OJZ80083.1

Group15033: aluc|OJZ79833.1 aluc|OJZ80220.1

Group15034: aluc|OJZ79882.1 aluc|OJZ91290.1

Group15035: aluc|OJZ79943.1 aluc|OJZ79970.1

Group15036: aluc|OJZ80051.1 aluc|OJZ80372.1

Group15037: aluc|OJZ80096.1 aluc|OJZ80130.1

Group15038: aluc|OJZ80103.1 aluc|OJZ80368.1

Group15039: aluc|OJZ80104.1 aluc|OJZ81117.1

Group15040: aluc|OJZ80126.1 aluc|OJZ88794.1

Group15041: aluc|OJZ80148.1 aluc|OJZ80264.1

Group15042: aluc|OJZ80150.1 aluc|OJZ80349.1

Group15043: aluc|OJZ80156.1 aluc|OJZ80320.1

Group15044: aluc|OJZ80158.1 aluc|OJZ80318.1

Group15045: aluc|OJZ80165.1 aluc|OJZ80256.1

Group15046: aluc|OJZ80297.1 aluc|OJZ88822.1

Group15047: aluc|OJZ80380.1 aluc|OJZ88826.1

Group15048: aluc|OJZ80450.1 aluc|OJZ81141.1

Group15049: abra|OJJ65415.1 azon|OJJ41986.1

Group15050: abra|OJJ65428.1 acar|OOF89752.1

Group15051: abra|OJJ65443.1 atub|OJI87361.1

Group15052: abra|OJJ65459.1 aluc|OJZ85536.1

Group15053: abra|OJJ65479.1 akaw|GAA92641.1

Group15054: abra|OJJ65507.1 acar|OOF93531.1

Group15055: abra|OJJ65512.1 atub|OJI82850.1

Group15056: abra|OJJ65513.1 atub|OJI82849.1

Group15057: abra|OJJ65514.1 atub|OJI82844.1

Group15058: abra|OJJ65516.1 atub|OJI82842.1

Group15059: abra|OJJ65518.1 atub|OJI82841.1

Group15060: abra|OJJ65519.1 atub|OJI82840.1

Group15061: abra|OJJ65520.1 akaw|GAA93118.1

Group15062: abra|OJJ65535.1 akaw|GAA93332.1

Group15063: abra|OJJ65537.1 afla|EED53063.1

Group15064: azon|OJJ43209.1 awen|OJJ33443.1

Group15065: abra|OJJ65587.1 afum|EAL91137.2

Group15066: akaw|GAA92356.1 aluc|OJZ80396.1

Group15067: abra|OJJ65611.1 afla|EED54386.1

Group15068: abra|OJJ65614.1 afla|EED47876.1

Group15069: aluc|OJZ80005.1 akaw|GAA93151.1

Group15070: abra|OJJ65627.1 agla|OJJ78557.1

Group15071: abra|OJJ65630.1 afla|EED52038.1

Group15072: abra|OJJ65631.1 afla|EED52039.1

Group15073: abra|OJJ65638.1 ater|EAU35407.1

Group15074: abra|OJJ65643.1 awen|OJJ36081.1

Group15075: abra|OJJ65668.1 aluc|OJZ81151.1

Group15076: abra|OJJ65681.1 anig|EHA25308.1

Group15077: aory|EIT72290.1 afla|EED46378.1

Group15078: abra|OJJ65692.1 aver|OJJ08176.1

Group15079: abra|OJJ65693.1 aory|EIT73820.1

Group15080: abra|OJJ65703.1 azon|OJJ46175.1

Group15081: abra|OJJ65715.1 aluc|OJZ80206.1

Group15082: abra|OJJ65736.1 afum|EAL86728.1

Group15083: azon|OJJ41997.1 agla|OJJ79521.1

Group15084: abra|OJJ65749.1 aluc|OJZ79838.1

Group15085: abra|OJJ65755.1 ater|EAU35418.1

Group15086: abra|OJJ65761.1 ater|EAU33983.1

Group15087: abra|OJJ65783.1 aluc|OJZ81113.1

Group15088: abra|OJJ65786.1 aluc|OJZ81111.1

Group15089: abra|OJJ65838.1 ater|EAU31694.1

Group15090: abra|OJJ65840.1 awen|OJJ29610.1

Group15091: abra|OJJ65878.1 aver|OJJ03995.1

Group15092: abra|OJJ65935.1 atub|OJI81625.1

Group15093: abra|OJJ65960.1 anig|EHA21413.1

Group15094: abra|OJJ65978.1 atub|OJI89215.1

Group15095: abra|OJJ66007.1 anig|EHA21369.1

Group15096: abra|OJJ66024.1 azon|OJJ45088.1

Group15097: abra|OJJ66050.1 atub|OJI87254.1

Group15098: abra|OJJ66150.1 afla|EED54060.1

Group15099: abra|OJJ66151.1 afla|EED54061.1

Group15100: abra|OJJ66159.1 ater|EAU39622.1

Group15101: abra|OJJ66192.1 acar|OOF91938.1

Group15102: abra|OJJ66226.1 acar|OOF92073.1

Group15103: afum|EAL85124.1 afis|EAW19778.1

Group15104: abra|OJJ66291.1 aver|OJJ02331.1

Group15105: abra|OJJ66302.1 aluc|OJZ80191.1

Group15106: abra|OJJ66304.1 azon|OJJ42058.1

Group15107: abra|OJJ66305.1 aluc|OJZ91227.1

Group15108: abra|OJJ66311.1 aluc|OJZ80188.1

Group15109: abra|OJJ66318.1 afla|EED47883.1

Group15110: abra|OJJ66321.1 afla|EED47880.1

Group15111: abra|OJJ66327.1 afis|EAW17211.1

Group15112: abra|OJJ66334.1 atub|OJI87800.1

Group15113: abra|OJJ66408.1 acla|EAW15052.1

Group15114: abra|OJJ66416.1 atub|OJI87889.1

Group15115: ater|EAU29256.1 akaw|GAA82058.1

Group15116: afla|EED47387.1 aory|EIT82828.1

Group15117: ater|EAU31817.1 asyd|OJJ54867.1

Group15118: abra|OJJ66466.1 aver|OJI96977.1

Group15119: agla|OJJ81843.1 ater|EAU36163.1

Group15120: abra|OJJ66497.1 asyd|OJJ54147.1

Group15121: abra|OJJ66517.1 aluc|OJZ81527.1

Group15122: abra|OJJ66582.1 akaw|GAA91136.1

Group15123: abra|OJJ66584.1 anig|EHA25967.1

Group15124: abra|OJJ66673.1 anig|EHA27255.1

Group15125: abra|OJJ66790.1 aluc|OJZ89977.1

Group15126: abra|OJJ66804.1 azon|OJJ41968.1

Group15127: abra|OJJ66806.1 atub|OJI79337.1

Group15128: abra|OJJ66809.1 atub|OJI79333.1

Group15129: abra|OJJ66812.1 atub|OJI79330.1

Group15130: abra|OJJ66813.1 atub|OJI79329.1

Group15131: abra|OJJ66821.1 anig|EHA24787.1

Group15132: abra|OJJ66914.1 atub|OJI79459.1

Group15133: abra|OJJ67000.1 aluc|OJZ85572.1

Group15134: abra|OJJ67003.1 atub|OJI79366.1

Group15135: abra|OJJ67010.1 anig|EHA25580.1

Group15136: abra|OJJ67017.1 acar|OOF93605.1

Group15137: abra|OJJ67038.1 akaw|GAA93112.1

Group15138: afla|EED52069.1 aory|EIT72302.1

Group15139: abra|OJJ67045.1 anig|EHA18598.1

Group15140: abra|OJJ67047.1 anig|EHA18596.1

Group15141: ater|EAU31768.1 azon|OJJ42008.1

Group15142: abra|OJJ67226.1 aluc|OJZ87748.1

Group15143: abra|OJJ67241.1 anig|EHA20225.1

Group15144: abra|OJJ67254.1 anig|EHA20218.1

Group15145: abra|OJJ67297.1 atub|OJI79589.1

Group15146: abra|OJJ67304.1 afla|EED45664.1

Group15147: aver|OJJ03935.1 asyd|OJJ52477.1

Group15148: asyd|OJJ61365.1 aver|OJJ00692.1

Group15149: abra|OJJ67678.1 afis|EAW25667.1

Group15150: aory|EIT80657.1 afla|EED51441.1

Group15151: aory|EIT75589.1 afla|EED45855.1

Group15152: abra|OJJ67707.1 atub|OJI88566.1

Group15153: abra|OJJ67711.1 afis|EAW20749.1

Group15154: abra|OJJ67717.1 aver|OJJ07194.1

Group15155: abra|OJJ67721.1 anig|EHA25834.1

Group15156: asyd|OJJ65406.1 aver|OJJ05896.1

Group15157: abra|OJJ67782.1 awen|OJJ36194.1

Group15158: abra|OJJ67783.1 agla|OJJ78614.1

Group15159: abra|OJJ67784.1 agla|OJJ78613.1

Group15160: abra|OJJ67804.1 aluc|OJZ90853.1

Group15161: abra|OJJ67825.1 anig|EHA22194.1

Group15162: abra|OJJ67827.1 anig|EHA22197.1

Group15163: abra|OJJ67828.1 anig|EHA22198.1

Group15164: agla|OJJ79083.1 awen|OJJ41028.1

Group15165: abra|OJJ67831.1 anig|EHA22201.1

Group15166: anid|CBF88293.1 ater|EAU31922.1

Group15167: abra|OJJ67970.1 acar|OOF95032.1

Group15168: abra|OJJ67994.1 aluc|OJZ82868.1

Group15169: abra|OJJ68024.1 atub|OJI89123.1

Group15170: afla|EED49506.1 aory|EIT82599.1

Group15171: abra|OJJ68151.1 aver|OJJ07754.1

Group15172: abra|OJJ68175.1 acla|EAW14646.1

Group15173: abra|OJJ68231.1 acar|OOF95815.1

Group15174: abra|OJJ68367.1 asyd|OJJ52876.1

Group15175: afis|EAW21273.1 afum|EAL92288.1

Group15176: abra|OJJ68504.1 acar|OOF90926.1

Group15177: abra|OJJ68540.1 atub|OJI88989.1

Group15178: aory|EIT82773.1 afla|EED52527.1

Group15179: abra|OJJ68573.1 awen|OJJ32253.1

Group15180: abra|OJJ68580.1 atub|OJI91462.1

Group15181: akaw|GAA91964.1 aluc|OJZ82653.1

Group15182: abra|OJJ68630.1 anig|EHA22323.1

Group15183: abra|OJJ68631.1 aluc|OJZ81925.1

Group15184: abra|OJJ68658.1 agla|OJJ83715.1

Group15185: abra|OJJ68667.1 acar|OOF97782.1

Group15186: abra|OJJ68712.1 atub|OJI84388.1

Group15187: abra|OJJ68816.1 atub|OJI79577.1

Group15188: abra|OJJ68872.1 atub|OJI79637.1

Group15189: abra|OJJ68907.1 anig|EHA21438.1

Group15190: abra|OJJ68936.1 aluc|OJZ85921.1

Group15191: abra|OJJ68939.1 atub|OJI79714.1

Group15192: afla|EED50398.1 aory|EIT82071.1

Group15193: abra|OJJ68964.1 aluc|OJZ85951.1

Group15194: abra|OJJ68981.1 atub|OJI79755.1

Group15195: abra|OJJ69148.1 anig|EHA24487.1

Group15196: aory|EIT79487.1 afla|EED49464.1

Group15197: abra|OJJ69213.1 asyd|OJJ52248.1

Group15198: abra|OJJ69260.1 atub|OJI80032.1

Group15199: asyd|OJJ53388.1 aver|OJJ08355.1

Group15200: asyd|OJJ53888.1 aver|OJJ07986.1

Group15201: abra|OJJ69464.1 aluc|OJZ82307.1

Group15202: abra|OJJ69472.1 anig|EHA28443.1

Group15203: abra|OJJ69487.1 atub|OJI88112.1

Group15204: abra|OJJ69547.1 anig|EHA19682.1

Group15205: asyd|OJJ64577.1 aver|OJI98674.1

Group15206: abra|OJJ69589.1 aluc|OJZ82407.1

Group15207: abra|OJJ69591.1 anig|EHA28358.1

Group15208: afla|EED47671.1 aory|EIT80190.1

Group15209: abra|OJJ69622.1 aluc|OJZ82520.1

Group15210: abra|OJJ69756.1 anig|EHA21152.1

Group15211: abra|OJJ69762.1 anig|EHA28305.1

Group15212: aver|OJJ08201.1 asyd|OJJ53553.1

Group15213: abra|OJJ70028.1 anig|EHA28108.1

Group15214: abra|OJJ70052.1 anig|EHA21624.1

Group15215: abra|OJJ70054.1 anid|CBF82503.1

Group15216: aory|EIT82305.1 afla|EED57087.1

Group15217: abra|OJJ70112.1 anig|EHA26093.1

Group15218: abra|OJJ70113.1 anig|EHA26092.1

Group15219: afum|EAL85603.1 afis|EAW15745.1

Group15220: abra|OJJ70125.1 acar|OOF98335.1

Group15221: abra|OJJ70145.1 akaw|GAA90565.1

Group15222: abra|OJJ70157.1 aluc|OJZ85154.1

Group15223: abra|OJJ70163.1 anig|EHA25801.1

Group15224: awen|OJJ40746.1 aver|OJJ08744.1

Group15225: abra|OJJ70192.1 akaw|GAA86584.1

Group15226: aver|OJI95876.1 asyd|OJJ61774.1

Group15227: abra|OJJ70246.1 acar|OOF92552.1

Group15228: abra|OJJ70258.1 agla|OJJ78611.1

Group15229: akaw|GAA92006.1 ater|EAU33099.1

Group15230: asyd|OJJ60045.1 aver|OJJ02242.1

Group15231: afla|EED48314.1 anig|EHA27468.1

Group15232: abra|OJJ70359.1 anig|EHA27456.1

Group15233: abra|OJJ70361.1 aory|EIT73560.1

Group15234: abra|OJJ70642.1 aluc|OJZ83992.1

Group15235: asyd|OJJ64580.1 aver|OJI98677.1

Group15236: abra|OJJ70723.1 atub|OJI90441.1

Group15237: abra|OJJ70740.1 atub|OJI90455.1

Group15238: abra|OJJ70823.1 acla|EAW09021.1

Group15239: abra|OJJ70889.1 anig|EHA20831.1

Group15240: abra|OJJ70896.1 aluc|OJZ87627.1

Group15241: abra|OJJ70927.1 aluc|OJZ87600.1

Group15242: abra|OJJ70937.1 aluc|OJZ87591.1

Group15243: abra|OJJ71018.1 atub|OJI81136.1

Group15244: abra|OJJ71074.1 aluc|OJZ87456.1

Group15245: abra|OJJ71093.1 anig|EHA25713.1

Group15246: abra|OJJ71137.1 atub|OJI86689.1

Group15247: abra|OJJ71222.1 anig|EHA20566.1

Group15248: asyd|OJJ52506.1 aver|OJJ06718.1

Group15249: abra|OJJ71240.1 aluc|OJZ87305.1

Group15250: afla|EED53806.1 aory|EIT78431.1

Group15251: abra|OJJ71391.1 aluc|OJZ87149.1

Group15252: abra|OJJ71392.1 anig|EHA20408.1

Group15253: abra|OJJ71436.1 atub|OJI81536.1

Group15254: abra|OJJ71659.1 atub|OJI84131.1

Group15255: abra|OJJ71696.1 azon|OJJ42070.1

Group15256: abra|OJJ71730.1 awen|OJJ31956.1

Group15257: abra|OJJ71750.1 acar|OOF93993.1

Group15258: asyd|OJJ60380.1 aver|OJI99709.1

Group15259: abra|OJJ71781.1 anig|EHA22133.1

Group15260: afum|EAL87904.1 afis|EAW23564.1

Group15261: afla|EED51273.1 aory|EIT81556.1

Group15262: abra|OJJ71834.1 atub|OJI83955.1

Group15263: abra|OJJ71912.1 aluc|OJZ86649.1

Group15264: azon|OJJ42210.1 acar|OOF99438.1

Group15265: abra|OJJ71950.1 atub|OJI83842.1

Group15266: abra|OJJ71999.1 akaw|GAA84930.1

Group15267: abra|OJJ72000.1 anig|EHA21942.1

Group15268: abra|OJJ72029.1 aluc|OJZ86766.1

Group15269: abra|OJJ72077.1 aluc|OJZ86831.1

Group15270: akaw|GAA84839.1 anig|EHA21800.1

Group15271: aver|OJI99827.1 asyd|OJJ60503.1

Group15272: abra|OJJ72138.1 anig|EHA21747.1

Group15273: aver|OJJ05769.1 asyd|OJJ57412.1

Group15274: abra|OJJ72195.1 atub|OJI83576.1

Group15275: abra|OJJ72199.1 anig|EHA24125.1

Group15276: abra|OJJ72320.1 anig|EHA23793.1

Group15277: abra|OJJ72331.1 ater|EAU30183.1

Group15278: abra|OJJ72365.1 aluc|OJZ90352.1

Group15279: abra|OJJ72449.1 aluc|OJZ90433.1

Group15280: abra|OJJ72522.1 agla|OJJ78796.1

Group15281: afis|EAW21066.1 afum|EAL86269.1

Group15282: afla|EED49868.1 aory|EIT83418.1

Group15283: abra|OJJ72646.1 atub|OJI82051.1

Group15284: abra|OJJ72652.1 anig|EHA24067.1

Group15285: abra|OJJ72667.1 atub|OJI82076.1

Group15286: abra|OJJ72742.1 anig|EHA24134.1

Group15287: abra|OJJ72785.1 akaw|GAA85392.1

Group15288: abra|OJJ72839.1 aluc|OJZ83339.1

Group15289: abra|OJJ72877.1 aluc|OJZ83388.1

Group15290: abra|OJJ72881.1 atub|OJI82278.1

Group15291: abra|OJJ72884.1 atub|OJI82279.1

Group15292: abra|OJJ73055.1 aluc|OJZ83566.1

Group15293: abra|OJJ73096.1 afla|EED55097.1

Group15294: abra|OJJ73101.1 atub|OJI91528.1

Group15295: abra|OJJ73131.1 anig|EHA18705.1

Group15296: aory|EIT73695.1 afla|EED45126.1

Group15297: abra|OJJ73213.1 atub|OJI91411.1

Group15298: abra|OJJ73296.1 acar|OOF98237.1

Group15299: anid|CBF88298.1 ater|EAU31925.1

Group15300: abra|OJJ73358.1 akaw|GAA83126.1

Group15301: asyd|OJJ52822.1 aver|OJJ08798.1

Group15302: abra|OJJ73418.1 akaw|GAA83073.1

Group15303: abra|OJJ73421.1 acar|OOG00624.1

Group15304: abra|OJJ73525.1 acar|OOF90212.1

Group15305: abra|OJJ73572.1 anig|EHA18197.1

Group15306: abra|OJJ73625.1 akaw|GAA88528.1

Group15307: asyd|OJJ59214.1 aver|OJJ01824.1

Group15308: abra|OJJ73760.1 anig|EHA18033.1

Group15309: abra|OJJ73918.1 anig|EHA17894.1

Group15310: abra|OJJ73979.1 atub|OJI90674.1

Group15311: abra|OJJ73999.1 aluc|OJZ88044.1

Group15312: abra|OJJ74124.1 afla|EED52214.1

Group15313: abra|OJJ74128.1 anig|EHA27205.1

Group15314: abra|OJJ74181.1 azon|OJJ47414.1

Group15315: abra|OJJ74263.1 atub|OJI84938.1

Group15316: abra|OJJ74274.1 aluc|OJZ89859.1

Group15317: abra|OJJ74412.1 atub|OJI85088.1

Group15318: abra|OJJ74427.1 aluc|OJZ89693.1

Group15319: abra|OJJ74448.1 aver|OJI98050.1

Group15320: abra|OJJ74476.1 acar|OOF95270.1

Group15321: afla|EED47163.1 aory|EIT79316.1

Group15322: aver|OJJ06705.1 asyd|OJJ55628.1

Group15323: abra|OJJ74737.1 anig|EHA26652.1

Group15324: abra|OJJ74813.1 afla|EED56019.1

Group15325: asyd|OJJ65385.1 anid|CBF82911.1

Group15326: abra|OJJ74970.1 aver|OJJ02992.1

Group15327: abra|OJJ75018.1 aluc|OJZ92584.1

Group15328: abra|OJJ75028.1 aluc|OJZ92595.1

Group15329: abra|OJJ75041.1 anig|EHA26379.1

Group15330: abra|OJJ75049.1 atub|OJI87050.1

Group15331: abra|OJJ75167.1 atub|OJI87162.1

Group15332: abra|OJJ75264.1 aluc|OJZ92822.1

Group15333: abra|OJJ75278.1 anig|EHA23286.1

Group15334: abra|OJJ75298.1 aver|OJI95773.1

Group15335: aver|OJI98032.1 asyd|OJJ62951.1

Group15336: abra|OJJ75470.1 atub|OJI85864.1

Group15337: abra|OJJ75509.1 acla|EAW06520.1

Group15338: asyd|OJJ64192.1 aver|OJI98269.1

Group15339: abra|OJJ75601.1 acar|OOF98769.1

Group15340: abra|OJJ75640.1 aluc|OJZ89089.1

Group15341: abra|OJJ75661.1 atub|OJI85672.1

Group15342: abra|OJJ75678.1 aluc|OJZ89128.1

Group15343: abra|OJJ75683.1 ater|EAU32978.1

Group15344: afla|EED46499.1 aory|EIT72317.1

Group15345: abra|OJJ75839.1 acar|OOF96619.1

Group15346: abra|OJJ75870.1 aluc|OJZ89313.1

Group15347: abra|OJJ75923.1 anig|EHA23211.1

Group15348: abra|OJJ75927.1 anid|CBF83333.1

Group15349: afla|EED55816.1 aory|EIT76764.1

Group15350: abra|OJJ76155.1 aluc|OJZ92073.1

Group15351: abra|OJJ76185.1 aluc|OJZ92044.1

Group15352: abra|OJJ76206.1 aluc|OJZ92022.1

Group15353: abra|OJJ76221.1 anig|EHA22949.1

Group15354: abra|OJJ76232.1 anig|EHA18751.1

Group15355: abra|OJJ76264.1 akaw|GAA82196.1

Group15356: abra|OJJ76281.1 aluc|OJZ91939.1

Group15357: abra|OJJ76407.1 atub|OJI86259.1

Group15358: abra|OJJ76488.1 aluc|OJZ91717.1

Group15359: abra|OJJ76562.1 atub|OJI86098.1

Group15360: abra|OJJ76696.1 agla|OJJ81055.1

Group15361: abra|OJJ76711.1 acar|OOF92525.1

Group15362: abra|OJJ76713.1 aver|OJJ01955.1

Group15363: asyd|OJJ53478.1 aver|OJJ08279.1

Group15364: abra|OJJ76742.1 azon|OJJ45657.1

Group15365: abra|OJJ76747.1 azon|OJJ42802.1

Group15366: abra|OJJ76748.1 acar|OOF89842.1

Group15367: abra|OJJ76886.1 anig|EHA26976.1

Group15368: abra|OJJ76890.1 afis|EAW16701.1

Group15369: abra|OJJ76925.1 aver|OJI98300.1

Group15370: anid|CBF82270.1 afum|EAL94104.1

Group15371: abra|OJJ77255.1 anig|EHA22320.1

Group15372: abra|OJJ77308.1 anig|EHA28071.1

Group15373: abra|OJJ77337.1 aluc|OJZ84745.1

Group15374: abra|OJJ77357.1 anig|EHA28036.1

Group15375: abra|OJJ77445.1 ater|EAU35413.1

Group15376: abra|OJJ77449.1 aver|OJJ04308.1

Group15377: afla|EED53270.1 aory|EIT75001.1

Group15378: abra|OJJ77741.1 anig|EHA18409.1

Group15379: abra|OJJ77745.1 anig|EHA20900.1

Group15380: abra|OJJ77753.1 agla|OJJ83618.1

Group15381: abra|OJJ77755.1 aluc|OJZ90812.1

Group15382: abra|OJJ77764.1 anig|EHA20910.1

Group15383: abra|OJJ77791.1 acla|EAW12266.1

Group15384: abra|OJJ77800.1 anig|EHA19584.1

Group15385: afum|EAL87846.1 afis|EAW23616.1

Group15386: abra|OJJ77850.1 acar|OOF99855.1

Group15387: abra|OJJ77886.1 aluc|OJZ91919.1

Group15388: abra|OJJ77891.1 aluc|OJZ90962.1

Group15389: aver|OJI98085.1 asyd|OJJ62904.1

Group15390: abra|OJJ77923.1 atub|OJI81810.1

Group15391: abra|OJJ77999.1 acar|OOF98823.1

Group15392: abra|OJJ78022.1 aluc|OJZ91097.1

Group15393: abra|OJJ78067.1 aver|OJI97461.1

Group15394: abra|OJJ78133.1 anig|EHA19892.1

Group15395: abra|OJJ78140.1 aluc|OJZ82074.1

Group15396: abra|OJJ78148.1 ater|EAU33812.1

Group15397: abra|OJJ78170.1 ater|EAU30167.1

Group15398: abra|OJJ78172.1 ater|EAU30165.1

Group15399: abra|OJJ78173.1 agla|OJJ84480.1

Group15400: afla|EED46351.1 aory|EIT81214.1

Group15401: abra|OJJ78338.1 aluc|OJZ91410.1

Group15402: acar|OOF89769.1 afis|EAW21001.1

Group15403: acar|OOF89778.1 azon|OJJ47002.1

Group15404: acar|OOF89779.1 afla|EED45938.1

Group15405: acar|OOF89787.1 anig|EHA21132.1

Group15406: asyd|OJJ62550.1 aver|OJI96675.1

Group15407: acar|OOF89879.1 afla|EED44756.1

Group15408: aver|OJI98235.1 asyd|OJJ64158.1

Group15409: asyd|OJJ58088.1 aver|OJJ04313.1

Group15410: acar|OOF90010.1 ater|EAU35968.1

Group15411: afla|EED45506.1 aory|EIT75735.1

Group15412: aory|EIT74425.1 afla|EED52288.1

Group15413: acar|OOF90120.1 aory|EIT76247.1

Group15414: acar|OOF90240.1 anig|EHA19270.1

Group15415: acar|OOF90350.1 azon|OJJ48146.1

Group15416: acar|OOF90353.1 azon|OJJ49967.1

Group15417: acar|OOF90386.1 asyd|OJJ58091.1

Group15418: aver|OJJ00867.1 asyd|OJJ61540.1

Group15419: acar|OOF90581.1 ater|EAU30980.1

Group15420: afla|EED48258.1 aory|EIT72901.1

Group15421: asyd|OJJ56445.1 aver|OJJ06277.1

Group15422: asyd|OJJ54366.1 aver|OJJ05319.1

Group15423: acar|OOF90696.1 aver|OJJ07890.1

Group15424: asyd|OJJ63397.1 aver|OJI97539.1

Group15425: acar|OOF90879.1 anig|EHA28165.1

Group15426: afla|EED46578.1 aory|EIT77897.1

Group15427: acar|OOF90905.1 atub|OJI88612.1

Group15428: acar|OOF90923.1 ater|EAU36146.1

Group15429: atub|OJI88950.1 aluc|OJZ82716.1

Group15430: acar|OOF91228.1 afla|EED50198.1

Group15431: acar|OOF91349.1 afum|EAL91904.2

Group15432: afla|EED50444.1 aory|EIT82168.1

Group15433: acar|OOF91455.1 afla|EED54129.1

Group15434: aver|OJJ05593.1 asyd|OJJ57590.1

Group15435: aver|OJI99666.1 asyd|OJJ52516.1

Group15436: acar|OOF91829.1 afla|EED57242.1

Group15437: afla|EED48360.1 aory|EIT83089.1

Group15438: akaw|GAA89132.1 aluc|OJZ81704.1

Group15439: aory|EIT83052.1 afla|EED48361.1

Group15440: asyd|OJJ56634.1 aver|OJJ06092.1

Group15441: acar|OOF92007.1 awen|OJJ36186.1

Group15442: aory|EIT83215.1 afla|EED55711.1

Group15443: acar|OOF92400.1 aluc|OJZ84945.1

Group15444: asyd|OJJ65374.1 aver|OJI99541.1

Group15445: awen|OJJ33635.1 aver|OJI95802.1

Group15446: acar|OOF92529.1 akaw|GAA88767.1

Group15447: acar|OOF92550.1 agla|OJJ82743.1

Group15448: acar|OOF92583.1 asyd|OJJ59221.1

Group15449: acar|OOF92585.1 azon|OJJ44382.1

Group15450: acar|OOF92588.1 azon|OJJ45971.1

Group15451: acar|OOF92589.1 azon|OJJ45972.1

Group15452: acar|OOF92591.1 azon|OJJ45973.1

Group15453: acar|OOF92656.1 anid|CBF82954.1

Group15454: acar|OOF92666.1 afis|EAW19289.1

Group15455: aory|EIT74892.1 afla|EED53130.1

Group15456: acar|OOF92823.1 aver|OJJ05975.1

Group15457: acar|OOF92943.1 azon|OJJ42244.1

Group15458: acar|OOF93023.1 awen|OJJ29706.1

Group15459: acar|OOF93316.1 ater|EAU31546.1

Group15460: afis|EAW16773.1 anid|CBF75884.1

Group15461: acar|OOF93323.1 ater|EAU31652.1

Group15462: agla|OJJ82678.1 awen|OJJ40229.1

Group15463: asyd|OJJ61771.1 aver|OJI95873.1

Group15464: acar|OOF93374.1 aluc|OJZ87910.1

Group15465: aver|OJJ01830.1 acla|EAW11969.1

Group15466: acar|OOF93425.1 atub|OJI80196.1

Group15467: acar|OOF93558.1 azon|OJJ50777.1

Group15468: afla|EED56033.1 aory|EIT76821.1

Group15469: aory|EIT74853.1 afla|EED53123.1

Group15470: acar|OOF93921.1 azon|OJJ46794.1

Group15471: acar|OOF93944.1 anig|EHA21026.1

Group15472: acar|OOF93978.1 anig|EHA19268.1

Group15473: aver|OJJ08182.1 asyd|OJJ53575.1

Group15474: acar|OOF94017.1 azon|OJJ50606.1

Group15475: acar|OOF94207.1 azon|OJJ43894.1

Group15476: afum|EAL92223.1 afis|EAW16436.1

Group15477: acar|OOF94482.1 azon|OJJ42800.1

Group15478: acar|OOF94706.1 aluc|OJZ90534.1

Group15479: acla|EAW11911.1 ater|EAU33993.1

Group15480: acar|OOF94811.1 ater|EAU38811.1

Group15481: aver|OJJ06614.1 asyd|OJJ55536.1

Group15482: acar|OOF94870.1 ater|EAU32909.1

Group15483: acar|OOF94911.1 aluc|OJZ79927.1

Group15484: asyd|OJJ57514.1 aver|OJJ05665.1

Group15485: asyd|OJJ59919.1 aver|OJJ02367.1

Group15486: acar|OOF94977.1 awen|OJJ40090.1

Group15487: asyd|OJJ58859.1 aver|OJJ01460.1

Group15488: acar|OOF94979.1 atub|OJI83024.1

Group15489: acar|OOF95018.1 awen|OJJ34375.1

Group15490: acar|OOF95030.1 azon|OJJ42314.1

Group15491: acar|OOF95197.1 agla|OJJ84713.1

Group15492: acar|OOF95245.1 awen|OJJ38748.1

Group15493: acar|OOF95282.1 atub|OJI85148.1

Group15494: aver|OJJ07589.1 asyd|OJJ54565.1

Group15495: acar|OOF95336.1 azon|OJJ42567.1

Group15496: aory|EIT79480.1 afla|EED49470.1

Group15497: acar|OOF95595.1 anig|EHA24753.1

Group15498: acar|OOF95804.1 anig|EHA26572.1

Group15499: acar|OOF95832.1 agla|OJJ79166.1

Group15500: acar|OOF96404.1 akaw|GAA89239.1

Group15501: acar|OOF96413.1 aory|EIT81338.1

Group15502: acar|OOF96443.1 awen|OJJ34651.1

Group15503: anid|CBF73447.1 aver|OJJ08117.1

Group15504: acar|OOF97140.1 anig|EHA23741.1

Group15505: acar|OOF97517.1 awen|OJJ38800.1

Group15506: acar|OOF97568.1 acla|EAW12162.1

Group15507: akaw|GAA85697.1 aluc|OJZ91406.1

Group15508: acar|OOF97582.1 aver|OJJ01908.1

Group15509: acar|OOF97586.1 awen|OJJ31355.1

Group15510: acar|OOF97604.1 akaw|GAA92086.1

Group15511: acar|OOF97623.1 agla|OJJ89302.1

Group15512: acar|OOF97645.1 awen|OJJ37239.1

Group15513: acar|OOF97841.1 akaw|GAA84347.1

Group15514: acar|OOF97995.1 aory|EIT74596.1

Group15515: atub|OJI84462.1 aluc|OJZ81909.1

Group15516: acar|OOF98129.1 acla|EAW13630.1

Group15517: acar|OOF98182.1 afum|EAL86714.1

Group15518: agla|OJJ80661.1 awen|OJJ35940.1

Group15519: aory|EIT73714.1 afla|EED45047.1

Group15520: acar|OOF98456.1 azon|OJJ50966.1

Group15521: acar|OOF98462.1 afla|EED45452.1

Group15522: acar|OOF98568.1 aluc|OJZ91374.1

Group15523: acar|OOF98684.1 awen|OJJ32681.1

Group15524: acar|OOF98692.1 azon|OJJ46426.1

Group15525: acar|OOF98693.1 azon|OJJ46427.1

Group15526: acar|OOF98695.1 azon|OJJ46429.1

Group15527: acar|OOF98701.1 atub|OJI79833.1

Group15528: acar|OOF98725.1 aver|OJJ07746.1

Group15529: acar|OOF98797.1 agla|OJJ83928.1

Group15530: acar|OOF98897.1 azon|OJJ42503.1

Group15531: aver|OJJ05334.1 asyd|OJJ54398.1

Group15532: aory|EIT83044.1 afla|EED48365.1

Group15533: aver|OJJ08426.1 asyd|OJJ53317.1

Group15534: acar|OOF99160.1 acla|EAW12191.1

Group15535: acar|OOF99180.1 atub|OJI82592.1

Group15536: acar|OOF99489.1 afla|EED50915.1

Group15537: acar|OOF99540.1 awen|OJJ29880.1

Group15538: afla|EED57864.1 aory|EIT81087.1

Group15539: acar|OOF99894.1 awen|OJJ36412.1

Group15540: afum|EAL84209.1 afis|EAW15436.1

Group15541: afis|EAW19521.1 aory|EIT72421.1

Group15542: acar|OOG00124.1 aver|OJJ07152.1

Group15543: acar|OOG00293.1 atub|OJI84638.1

Group15544: acar|OOG00397.1 asyd|OJJ58095.1

Group15545: acar|OOG00398.1 asyd|OJJ58092.1

Group15546: asyd|OJJ54220.1 aver|OJJ05172.1

Group15547: aver|OJJ08916.1 asyd|OJJ52950.1

Group15548: acar|OOG00613.1 azon|OJJ43531.1

Group15549: acar|OOG00614.1 azon|OJJ47351.1

Group15550: acar|OOG01135.1 atub|OJI90636.1

Group15551: acla|EAW06549.1 azon|OJJ45408.1

Group15552: acla|EAW06580.1 awen|OJJ34550.1

Group15553: acla|EAW06614.1 afis|EAW16151.1

Group15554: acla|EAW06733.1 ater|EAU37615.1

Group15555: asyd|OJJ59229.1 aver|OJJ01839.1

Group15556: acla|EAW07068.1 aver|OJJ07755.1

Group15557: afla|EED53644.1 aory|EIT75348.1

Group15558: acla|EAW07099.1 afis|EAW22114.1

Group15559: acla|EAW07101.1 afis|EAW22116.1

Group15560: afla|EED45536.1 aory|EIT75510.1

Group15561: acla|EAW07119.1 afis|EAW17373.1

Group15562: acla|EAW07209.1 afis|EAW22221.1

Group15563: awen|OJJ38305.1 asyd|OJJ57343.1

Group15564: acla|EAW07226.1 afis|EAW22240.1

Group15565: acla|EAW07615.1 awen|OJJ35279.1

Group15566: acla|EAW07627.1 asyd|OJJ60064.1

Group15567: acla|EAW08257.1 afla|EED55681.1

Group15568: afis|EAW19752.1 afum|EAL85145.1

Group15569: afla|EED55271.1 aory|EIT83516.1

Group15570: aory|EIT78685.1 afla|EED52931.1

Group15571: acla|EAW08927.1 azon|OJJ48235.1

Group15572: acla|EAW08941.1 afis|EAW15458.1

Group15573: acla|EAW08943.1 azon|OJJ47384.1

Group15574: agla|OJJ79122.1 asyd|OJJ55381.1

Group15575: acla|EAW09017.1 ater|EAU30726.1

Group15576: acla|EAW09053.1 aory|EIT81629.1

Group15577: afla|EED48938.1 aory|EIT80040.1

Group15578: acla|EAW09116.1 asyd|OJJ54394.1

Group15579: asyd|OJJ53855.1 aver|OJJ07955.1

Group15580: acla|EAW09584.1 afum|EBA27464.1

Group15581: acla|EAW09622.1 awen|OJJ37330.1

Group15582: acla|EAW09695.1 awen|OJJ39254.1

Group15583: acla|EAW10032.1 atub|OJI79836.1

Group15584: acla|EAW10041.1 ater|EAU38848.1

Group15585: afis|EAW25399.1 afum|EAL90058.1

Group15586: acla|EAW10256.1 agla|OJJ83865.1

Group15587: afis|EAW25285.1 afum|EAL89948.1

Group15588: afla|EED55838.1 aory|EIT76731.1

Group15589: acla|EAW10832.1 anid|CBF78968.1

Group15590: asyd|OJJ56902.1 aver|OJJ03391.1

Group15591: acla|EAW10858.1 awen|OJJ29625.1

Group15592: asyd|OJJ56062.1 aver|OJJ05004.1

Group15593: acla|EAW11053.1 aory|EIT81014.1

Group15594: acla|EAW11071.1 agla|OJJ85146.1

Group15595: acla|EAW11110.1 anid|CBF75394.1

Group15596: asyd|OJJ59611.1 aver|OJJ02658.1

Group15597: aver|OJJ02688.1 asyd|OJJ59581.1

Group15598: aluc|OJZ79802.1 akaw|GAA89001.1

Group15599: acla|EAW11457.1 aluc|OJZ83375.1

Group15600: acla|EAW11469.1 afis|EAW19375.1

Group15601: akaw|GAA92658.1 aluc|OJZ79613.1

Group15602: awen|OJJ30139.1 agla|OJJ79603.1

Group15603: acla|EAW11703.1 afis|EAW25204.1

Group15604: acla|EAW11705.1 ater|EAU32801.1

Group15605: acla|EAW11916.1 ater|EAU33995.1

Group15606: afis|EAW19557.1 afum|EAL86954.1

Group15607: acla|EAW12165.1 awen|OJJ37889.1

Group15608: acla|EAW12181.1 afis|EAW16678.1

Group15609: acla|EAW12197.1 afis|EAW25311.1

Group15610: acla|EAW12290.1 azon|OJJ43174.1

Group15611: acla|EAW12328.1 afis|EAW17183.1

Group15612: acla|EAW12373.1 awen|OJJ33598.1

Group15613: acla|EAW12394.1 afis|EAW16842.1

Group15614: acla|EAW12415.1 anid|CBF83057.1

Group15615: acla|EAW12418.1 anid|CBF83051.1

Group15616: acla|EAW12419.1 anid|CBF83049.1

Group15617: acla|EAW12422.1 anid|CBF83044.1

Group15618: asyd|OJJ53555.1 aver|OJJ08200.1

Group15619: acla|EAW12907.1 asyd|OJJ60508.1

Group15620: aver|OJJ00748.1 awen|OJJ38294.1

Group15621: aluc|OJZ84202.1 atub|OJI89882.1

Group15622: acla|EAW13436.1 ater|EAU29243.1

Group15623: afis|EAW20850.1 afum|EBA27205.1

Group15624: aver|OJJ01788.1 awen|OJJ35126.1

Group15625: acla|EAW13448.1 ater|EAU29433.1

Group15626: acla|EAW13516.1 ater|EAU29415.1

Group15627: acla|EAW13519.1 ater|EAU29411.1

Group15628: acla|EAW13535.1 anid|CBF69540.1

Group15629: acla|EAW13537.1 anid|CBF69538.1

Group15630: acla|EAW13539.1 azon|OJJ45983.1

Group15631: aory|EIT74516.1 afla|EED47582.1

Group15632: aory|EIT74514.1 afla|EED47540.1

Group15633: acla|EAW13623.1 atub|OJI83192.1

Group15634: acla|EAW13636.1 aory|EIT83099.1

Group15635: afla|EED48983.1 aory|EIT80026.1

Group15636: acla|EAW13686.1 afis|EAW19920.1

Group15637: aver|OJJ07855.1 asyd|OJJ53742.1

Group15638: acla|EAW13806.1 awen|OJJ30706.1

Group15639: acla|EAW14026.1 afum|EAL93457.1

Group15640: acla|EAW14253.1 afla|EED56617.1

Group15641: acla|EAW14285.1 afum|EAL86336.1

Group15642: acla|EAW14312.1 afum|EAL93709.1

Group15643: acla|EAW14515.1 afla|EED47655.1

Group15644: aver|OJJ03142.1 asyd|OJJ57174.1

Group15645: acla|EAW14587.1 afis|EAW19318.1

Group15646: acla|EAW14621.1 ater|EAU32945.1

Group15647: aver|OJJ00030.1 asyd|OJJ60705.1

Group15648: anid|CBF80180.1 aver|OJI99528.1

Group15649: anig|EHA19881.1 afis|EAW16423.1

Group15650: acla|EAW15018.1 afis|EAW24550.1

Group15651: acla|EAW15021.1 awen|OJJ29754.1

Group15652: acla|EAW15059.1 afum|EAL85110.1

Group15653: acla|EAW15061.1 afum|EAL85112.1

Group15654: afis|EAW16192.1 afum|EAL88823.1

Group15655: acla|EAW15268.1 afis|EAW20241.1

Group15656: aver|OJI99626.1 anid|CBF82284.1

Group15657: acla|EAW15321.1 afis|EAW20322.1

Group15658: acla|EAW15340.1 afis|EAW20369.1

Group15659: acla|EAW15369.1 ater|EAU38090.1

Group15660: afis|EAW15384.1 aver|OJJ05931.1

Group15661: afis|EAW15386.1 aver|OJJ05929.1

Group15662: afis|EAW15391.1 aver|OJJ05928.1

Group15663: afis|EAW15435.1 afum|EBA27169.1

Group15664: afis|EAW15439.1 afum|EBA27168.1

Group15665: afis|EAW15507.1 afum|EBA27421.1

Group15666: aver|OJJ03125.1 asyd|OJJ52046.1

Group15667: afis|EAW15551.1 asyd|OJJ65145.1

Group15668: afis|EAW15624.1 afum|EAL85710.1

Group15669: afis|EAW15635.1 ater|EAU35924.1

Group15670: afis|EAW15986.1 afum|EAL88618.1

Group15671: afis|EAW16158.1 ater|EAU35525.1

Group15672: afis|EAW16183.1 afum|EAL88814.1

Group15673: afis|EAW16212.1 agla|OJJ79505.1

Group15674: afis|EAW16255.1 ater|EAU31544.1

Group15675: afis|EAW16303.1 afum|EAL88925.1

Group15676: afis|EAW16349.1 afum|EAL88974.1

Group15677: asyd|OJJ57380.1 aver|OJJ05778.1

Group15678: afis|EAW16404.1 agla|OJJ78759.1

Group15679: afis|EAW16438.1 afum|EAL89041.1

Group15680: afis|EAW16440.1 ater|EAU32056.1

Group15681: afis|EAW16452.1 anid|CBF74895.1

Group15682: afis|EAW16692.1 aver|OJI99220.1

Group15683: afis|EAW16712.1 afum|EAL89300.1

Group15684: afis|EAW16719.1 anid|CBF75716.1

Group15685: afis|EAW16722.1 afum|EAL86296.2

Group15686: afis|EAW16729.1 afum|EAL86304.2

Group15687: afis|EAW16731.1 afum|EAL86307.1

Group15688: afis|EAW16756.1 afum|EAL86328.1

Group15689: afis|EAW16757.1 afum|EAL86329.1

Group15690: afis|EAW16766.1 asyd|OJJ64182.1

Group15691: afis|EAW16796.1 afum|EAL86365.1

Group15692: afis|EAW16799.1 afla|EED46871.1

Group15693: afis|EAW16814.1 asyd|OJJ62350.1

Group15694: afis|EAW16818.1 afum|EAL86740.1

Group15695: afis|EAW16830.1 afum|EAL86378.1

Group15696: afis|EAW16868.1 afum|EAL86407.1

Group15697: afis|EAW16877.1 afum|EAL86414.1

Group15698: afis|EAW16898.1 agla|OJJ84611.1

Group15699: afis|EAW16929.1 afum|EAL86454.1

Group15700: afis|EAW16935.1 afum|EAL86460.1

Group15701: asyd|OJJ52545.1 aver|OJJ03859.1

Group15702: aver|OJJ03765.1 asyd|OJJ53082.1

Group15703: afis|EAW17002.1 asyd|OJJ60318.1

Group15704: afis|EAW17006.1 awen|OJJ37865.1

Group15705: afla|EED54038.1 aory|EIT74599.1

Group15706: afis|EAW17070.1 azon|OJJ46013.1

Group15707: awen|OJJ31608.1 ater|EAU37817.1

Group15708: asyd|OJJ53080.1 aver|OJJ03767.1

Group15709: afis|EAW17087.1 afum|EAL86564.2

Group15710: afis|EAW17100.1 aver|OJJ08552.1

Group15711: afis|EAW17140.1 awen|OJJ34743.1

Group15712: afis|EAW17143.1 akaw|GAA92902.1

Group15713: afis|EAW17208.1 asyd|OJJ57945.1

Group15714: afis|EAW17209.1 agla|OJJ79438.1

Group15715: afis|EAW17215.1 afum|EAL86641.1

Group15716: afis|EAW17233.1 aver|OJI98012.1

Group15717: afis|EAW17248.1 ater|EAU38591.1

Group15718: afis|EAW17303.1 afum|EAL91112.2

Group15719: afis|EAW17358.1 ater|EAU37994.1

Group15720: afis|EAW17434.1 afum|EAL91261.1

Group15721: afis|EAW17485.1 afum|EBA27326.1

Group15722: afis|EAW17773.1 afum|EAL91582.1

Group15723: afis|EAW18131.1 afum|EAL87490.1

Group15724: afis|EAW18153.1 afum|EAL87512.1

Group15725: afis|EAW18256.1 afum|EAL87619.1

Group15726: afis|EAW18289.1 afum|EBA27322.1

Group15727: afis|EAW18316.1 afum|EAL87683.1

Group15728: afla|EED52830.1 aory|EIT78185.1

Group15729: afla|EED52789.1 aory|EIT78211.1

Group15730: afis|EAW18831.1 afum|EAL93565.2

Group15731: afis|EAW19017.1 afum|EBA27503.1

Group15732: afis|EAW19043.1 afum|EAL93772.1

Group15733: afis|EAW19112.1 afum|EAL93843.2

Group15734: afis|EAW19256.1 azon|OJJ44577.1

Group15735: afis|EAW19298.1 afum|EAL94017.1

Group15736: afis|EAW19300.1 afum|EAL94019.1

Group15737: afis|EAW19301.1 afum|EAL94020.1

Group15738: afis|EAW19306.1 afum|EAL94024.1

Group15739: afis|EAW19307.1 afum|EAL94025.1

Group15740: afis|EAW19365.1 afum|EAL94084.2

Group15741: afis|EAW19366.1 afum|EAL94085.1

Group15742: afis|EAW19485.1 afum|EAL87010.1

Group15743: afis|EAW19542.1 afum|EAL86966.1

Group15744: afis|EAW19596.1 afum|EAL86922.1

Group15745: afis|EAW19667.1 afum|EAL86856.1

Group15746: afis|EAW19678.1 aver|OJJ08345.1

Group15747: afis|EAW19685.1 afum|EAL86839.1

Group15748: afis|EAW19702.1 afum|EAL86824.1

Group15749: afis|EAW19755.1 afum|EAL85142.1

Group15750: afis|EAW19756.1 afum|EAL85141.1

Group15751: afis|EAW19757.1 afum|EAL85140.1

Group15752: afis|EAW19762.1 ater|EAU36858.1

Group15753: afis|EAW19765.1 ater|EAU36861.1

Group15754: afis|EAW19767.1 ater|EAU36863.1

Group15755: afis|EAW19768.1 ater|EAU36864.1

Group15756: aory|EIT80917.1 afla|EED55488.1

Group15757: afis|EAW19774.1 afum|EAL85125.1

Group15758: afis|EAW19793.1 afum|EAL85097.1

Group15759: afis|EAW19821.1 ater|EAU29480.1

Group15760: afis|EAW19828.1 azon|OJJ42656.1

Group15761: afis|EAW19840.1 anig|EHA26574.1

Group15762: agla|OJJ79742.1 awen|OJJ33426.1

Group15763: afis|EAW19859.1 afum|EAL85036.1

Group15764: afis|EAW19860.1 afum|EAL85035.1

Group15765: afis|EAW19898.1 ater|EAU29942.1

Group15766: afis|EAW19902.1 ater|EAU34043.1

Group15767: afis|EAW19910.1 afum|EAL84996.2

Group15768: afis|EAW19916.1 afum|EAL84991.1

Group15769: aver|OJJ05136.1 atub|OJI80142.1

Group15770: afis|EAW19919.1 afum|EAL84989.2

Group15771: afis|EAW19970.1 afum|EAL84952.1

Group15772: afis|EAW19986.1 afum|EAL84929.1

Group15773: afis|EAW20277.1 afum|EAL85411.1

Group15774: afis|EAW20332.1 akaw|GAA88765.1

Group15775: afis|EAW20373.1 afum|EAL85487.1

Group15776: afis|EAW20410.1 afum|EAL89234.2

Group15777: afis|EAW20413.1 afum|EBA27372.1

Group15778: afis|EAW20491.1 afum|EAL84563.1

Group15779: afis|EAW20576.1 afum|EAL84489.1

Group15780: afis|EAW20639.1 afum|EAL84430.1

Group15781: afis|EAW20658.1 asyd|OJJ52093.1

Group15782: afis|EAW20663.1 azon|OJJ42049.1

Group15783: afis|EAW20672.1 anid|CBF84449.1

Group15784: afis|EAW20679.1 afum|EAL84418.2

Group15785: afis|EAW20683.1 anid|CBF87355.1

Group15786: afis|EAW20708.1 azon|OJJ42239.1

Group15787: afis|EAW20716.1 asyd|OJJ52042.1

Group15788: afis|EAW20753.1 afum|EAL84864.1

Group15789: afis|EAW20759.1 afum|EAL84861.1

Group15790: afis|EAW20760.1 afum|EAL84859.1

Group15791: asyd|OJJ60271.1 aver|OJJ03979.1

Group15792: afis|EAW20795.1 azon|OJJ46413.1

Group15793: afis|EAW20804.1 afum|EAL84824.1

Group15794: afis|EAW20817.1 afum|EBA27206.1

Group15795: afis|EAW20828.1 awen|OJJ41727.1

Group15796: afis|EAW20851.1 afum|EBA27204.1

Group15797: afis|EAW20857.1 awen|OJJ40933.1

Group15798: afis|EAW20889.1 afum|EAL84744.1

Group15799: afis|EAW21000.1 afla|EED44720.1

Group15800: afla|EED48316.1 aory|EIT76291.1

Group15801: afis|EAW21017.1 afum|EAL92055.1

Group15802: afis|EAW21042.1 ater|EAU31433.1

Group15803: afis|EAW21053.1 afum|EAL92089.1

Group15804: afis|EAW21059.1 afla|EED48133.1

Group15805: afis|EAW21071.1 ater|EAU31984.1

Group15806: afis|EAW21130.1 afum|EAL92148.1

Group15807: asyd|OJJ59446.1 aver|OJJ02826.1

Group15808: afis|EAW21189.1 afla|EED54139.1

Group15809: afis|EAW21199.1 awen|OJJ32934.1

Group15810: afis|EAW21204.1 ater|EAU32103.1

Group15811: afis|EAW21207.1 afum|EAL92218.1

Group15812: afis|EAW21270.1 afum|EBA27458.1

Group15813: afis|EAW21304.1 awen|OJJ39394.1

Group15814: afis|EAW21322.1 afum|EAL92339.1

Group15815: aory|EIT76643.1 afla|EED55809.1

Group15816: afis|EAW21444.1 afum|EAL92462.2

Group15817: afis|EAW21544.1 afum|EAL92564.1

Group15818: afis|EAW21630.1 afum|EAL92664.1

Group15819: afis|EAW21686.1 afum|EAL92720.1

Group15820: afis|EAW21748.1 afum|EBA27472.1

Group15821: asyd|OJJ63023.1 aver|OJI97945.1

Group15822: afis|EAW21839.1 afum|EBA27476.1

Group15823: asyd|OJJ61235.1 aver|OJJ00558.1

Group15824: afis|EAW21845.1 afum|EAL92857.2

Group15825: asyd|OJJ64549.1 aver|OJI98645.1

Group15826: afis|EAW22053.1 awen|OJJ31621.1

Group15827: afis|EAW22094.1 afum|EAL87821.1

Group15828: anid|CBF87084.1 aver|OJJ07534.1

Group15829: afis|EAW22100.1 afum|EAL91089.1

Group15830: afis|EAW22154.1 afum|EAL91036.1

Group15831: afis|EAW22477.1 afum|EAL90715.1

Group15832: afis|EAW22566.1 awen|OJJ36325.1

Group15833: anid|CBF80824.1 asyd|OJJ59908.1

Group15834: agla|OJJ79472.1 awen|OJJ38894.1

Group15835: afis|EAW22673.1 afum|EAL90519.1

Group15836: afis|EAW22710.1 agla|OJJ78681.1

Group15837: afla|EED46247.1 aory|EIT76483.1

Group15838: afis|EAW23081.1 afum|EAL88398.1

Group15839: afis|EAW23121.1 afum|EAL88359.1

Group15840: afis|EAW23531.1 afum|EAL87935.1

Group15841: afis|EAW23593.1 awen|OJJ34756.1

Group15842: awen|OJJ40950.1 agla|OJJ87027.1

Group15843: afis|EAW23615.1 afum|EAL87847.2

Group15844: afis|EAW23637.1 afum|EAL86632.1

Group15845: afis|EAW23653.1 ater|EAU32808.1

Group15846: afis|EAW23693.1 afum|EAL87136.1

Group15847: afis|EAW23694.1 ater|EAU30657.1

Group15848: afis|EAW23812.1 afum|EAL87247.1

Group15849: afis|EAW24020.1 afum|EAL87450.1

Group15850: afis|EAW24030.1 afum|EAL87461.1

Group15851: afis|EAW24241.1 afum|EAL85909.2

Group15852: afis|EAW24355.1 afum|EBA27253.1

Group15853: afis|EAW24407.1 ater|EAU29649.1

Group15854: aory|EIT77209.1 anid|CBF80266.1

Group15855: afla|EED55093.1 aory|EIT81574.1

Group15856: afis|EAW24534.1 ater|EAU29282.1

Group15857: afis|EAW24541.1 afum|EBA27262.1

Group15858: afis|EAW24545.1 afum|EAL86209.1

Group15859: afis|EAW24590.1 aver|OJI96484.1

Group15860: afis|EAW24592.1 aver|OJI96485.1

Group15861: afis|EAW24601.1 aver|OJI96491.1

Group15862: afis|EAW24605.1 agla|OJJ82373.1

Group15863: afis|EAW24613.1 afum|EAL86243.1

Group15864: afis|EAW24616.1 afum|EAL86245.1

Group15865: afis|EAW24643.1 anid|CBF82188.1

Group15866: afis|EAW24645.1 afum|EAL90944.1

Group15867: afis|EAW24648.1 afum|EAL86749.1

Group15868: afis|EAW24652.1 afum|EAL89318.1

Group15869: afis|EAW24653.1 afum|EAL89317.1

Group15870: afis|EAW24654.1 afum|EAL89316.1

Group15871: afis|EAW24655.1 afum|EAL89315.1

Group15872: afis|EAW24656.1 afum|EAL89314.2

Group15873: afis|EAW24657.1 afum|EAL89313.1

Group15874: afis|EAW24658.1 afum|EAL89312.1

Group15875: afis|EAW24659.1 afum|EAL89311.1

Group15876: afis|EAW24676.1 asyd|OJJ57384.1

Group15877: afis|EAW24680.1 asyd|OJJ57386.1

Group15878: afis|EAW24681.1 asyd|OJJ57387.1

Group15879: afis|EAW24729.1 afum|EBA27378.1

Group15880: afis|EAW25009.1 afla|EED55429.1

Group15881: afis|EAW25189.1 agla|OJJ79841.1

Group15882: afis|EAW25239.1 awen|OJJ35620.1

Group15883: afis|EAW25466.1 afum|EAL90129.2

Group15884: afis|EAW25564.1 asyd|OJJ57957.1

Group15885: afis|EAW25595.1 afum|EAL84301.1

Group15886: afis|EAW25598.1 afum|EAL84304.1

Group15887: aory|EIT75745.1 afla|EED45529.1

Group15888: afis|EAW25606.1 aver|OJJ07494.1

Group15889: afis|EAW25612.1 afla|EED53578.1

Group15890: afis|EAW25660.1 afum|EAL84334.1

Group15891: afis|EAW25684.1 awen|OJJ33833.1

Group15892: aver|OJJ03421.1 afum|EAL86563.2

Group15893: afis|EAW25718.1 afum|EAL84393.2

Group15894: afis|EAW25719.1 afum|EAL84394.2

Group15895: afla|EED44729.1 aory|EIT77802.1

Group15896: afla|EED44750.1 aory|EIT77761.1

Group15897: afla|EED44764.1 aory|EIT77782.1

Group15898: afla|EED44768.1 aory|EIT77764.1

Group15899: afla|EED44821.1 aory|EIT77714.1

Group15900: afla|EED44868.1 aory|EIT82725.1

Group15901: afla|EED44869.1 aory|EIT82721.1

Group15902: afla|EED44870.1 aory|EIT82726.1

Group15903: afla|EED44874.1 aory|EIT83253.1

Group15904: afla|EED44882.1 aory|EIT83258.1

Group15905: afla|EED44887.1 aory|EIT83243.1

Group15906: afla|EED44888.1 aory|EIT83242.1

Group15907: afla|EED44900.1 aory|EIT83239.1

Group15908: afla|EED44902.1 aory|EIT73501.1

Group15909: afla|EED44904.1 aory|EIT73473.1

Group15910: afla|EED44912.1 aory|EIT73484.1

Group15911: afla|EED44913.1 aory|EIT73495.1

Group15912: afla|EED44917.1 aory|EIT73474.1

Group15913: afla|EED44927.1 aory|EIT73523.1

Group15914: afla|EED44933.1 aory|EIT73513.1

Group15915: afla|EED44934.1 aory|EIT73504.1

Group15916: afla|EED44937.1 ater|EAU29359.1

Group15917: afla|EED44953.1 aory|EIT73483.1

Group15918: afla|EED44957.1 aory|EIT73471.1

Group15919: afla|EED44968.1 aory|EIT73502.1

Group15920: afla|EED44995.1 aory|EIT73578.1

Group15921: afla|EED45006.1 aory|EIT73822.1

Group15922: afla|EED45007.1 aory|EIT73602.1

Group15923: afla|EED45050.1 aory|EIT73799.1

Group15924: afla|EED45051.1 aory|EIT73581.1

Group15925: afla|EED45053.1 aory|EIT73785.1

Group15926: afla|EED45056.1 aory|EIT73580.1

Group15927: afla|EED45058.1 ater|EAU37132.1

Group15928: afla|EED45061.1 aory|EIT73603.1

Group15929: afla|EED45086.1 aory|EIT73585.1

Group15930: afla|EED45089.1 aory|EIT73598.1

Group15931: afla|EED45091.1 aory|EIT73789.1

Group15932: afla|EED45092.1 aory|EIT73550.1

Group15933: afla|EED45103.1 aory|EIT73777.1

Group15934: afla|EED45115.1 aory|EIT73763.1

Group15935: afla|EED45125.1 azon|OJJ45672.1

Group15936: afla|EED45135.1 aory|EIT73599.1

Group15937: afla|EED45139.1 aory|EIT73748.1

Group15938: afla|EED45142.1 aory|EIT73655.1

Group15939: afla|EED45143.1 aory|EIT73592.1

Group15940: afla|EED45157.1 aory|EIT73623.1

Group15941: afla|EED45158.1 aory|EIT73819.1

Group15942: afla|EED45165.1 aory|EIT73527.1

Group15943: afla|EED45181.1 aory|EIT73690.1

Group15944: afla|EED45187.1 aory|EIT73627.1

Group15945: afla|EED45190.1 aory|EIT73571.1

Group15946: afla|EED45198.1 aory|EIT73718.1

Group15947: afla|EED45200.1 aory|EIT73788.1

Group15948: afla|EED45203.1 aory|EIT73810.1

Group15949: afla|EED45208.1 aluc|OJZ89967.1

Group15950: afla|EED45209.1 aory|EIT73634.1

Group15951: afla|EED45219.1 aory|EIT73587.1

Group15952: afla|EED45220.1 aory|EIT73684.1

Group15953: afla|EED45222.1 aory|EIT73784.1

Group15954: afla|EED45230.1 ater|EAU30187.1

Group15955: afla|EED45254.1 aory|EIT73693.1

Group15956: afla|EED45255.1 aory|EIT73535.1

Group15957: afla|EED45283.1 aory|EIT73557.1

Group15958: afla|EED45301.1 aory|EIT73704.1

Group15959: afla|EED45365.1 aory|EIT82430.1

Group15960: afla|EED45367.1 aory|EIT82461.1

Group15961: afla|EED45385.1 aory|EIT82454.1

Group15962: afla|EED45391.1 aory|EIT82419.1

Group15963: afla|EED45416.1 aory|EIT82390.1

Group15964: afla|EED45422.1 aory|EIT82417.1

Group15965: afla|EED45425.1 aory|EIT82453.1

Group15966: afla|EED45432.1 aory|EIT82422.1

Group15967: afla|EED45441.1 aory|EIT82397.1

Group15968: asyd|OJJ61484.1 aver|OJJ00826.1

Group15969: afla|EED45456.1 aory|EIT82407.1

Group15970: afla|EED45463.1 aory|EIT82402.1

Group15971: afla|EED45468.1 aory|EIT75640.1

Group15972: afla|EED45476.1 aory|EIT75404.1

Group15973: afla|EED45480.1 aory|EIT75530.1

Group15974: afla|EED45484.1 aory|EIT75722.1

Group15975: afla|EED45495.1 aory|EIT75570.1

Group15976: afla|EED45508.1 aory|EIT75495.1

Group15977: afla|EED45509.1 aory|EIT75516.1

Group15978: afla|EED45510.1 aory|EIT75653.1

Group15979: afla|EED45522.1 aory|EIT75812.1

Group15980: afla|EED45526.1 aory|EIT75401.1

Group15981: afla|EED45546.1 aory|EIT75548.1

Group15982: afla|EED45580.1 aory|EIT75467.1

Group15983: afla|EED45591.1 aory|EIT75635.1

Group15984: afla|EED45603.1 aory|EIT75402.1

Group15985: afla|EED45612.1 azon|OJJ47860.1

Group15986: afla|EED45613.1 aory|EIT75458.1

Group15987: afla|EED45614.1 aory|EIT75414.1

Group15988: afla|EED45623.1 aory|EIT75445.1

Group15989: afla|EED45626.1 aory|EIT75729.1

Group15990: afla|EED45627.1 ater|EAU32690.1

Group15991: afla|EED45630.1 asyd|OJJ56234.1

Group15992: afla|EED45637.1 aory|EIT75478.1

Group15993: afla|EED45650.1 aory|EIT75494.1

Group15994: afla|EED45651.1 aory|EIT75652.1

Group15995: afla|EED45660.1 aory|EIT75660.1

Group15996: afla|EED45671.1 aory|EIT75785.1

Group15997: afla|EED45686.1 aory|EIT75624.1

Group15998: afla|EED45687.1 aory|EIT75738.1

Group15999: afla|EED45689.1 aory|EIT75691.1

Group16000: afla|EED45705.1 aory|EIT75453.1

Group16001: afla|EED45749.1 aory|EIT75472.1

Group16002: afla|EED45761.1 aory|EIT75568.1

Group16003: azon|OJJ43180.1 awen|OJJ32632.1

Group16004: afla|EED45801.1 aory|EIT75497.1

Group16005: afla|EED45814.1 aory|EIT75529.1

Group16006: afla|EED45824.1 aory|EIT75633.1

Group16007: afla|EED45833.1 aory|EIT75505.1

Group16008: afla|EED45856.1 aory|EIT75490.1

Group16009: afla|EED45859.1 aory|EIT75665.1

Group16010: afla|EED45873.1 aory|EIT75410.1

Group16011: afla|EED45898.1 aory|EIT75506.1

Group16012: aver|OJJ08365.1 asyd|OJJ53378.1

Group16013: afla|EED45956.1 aory|EIT75695.1

Group16014: afla|EED45966.1 aory|EIT75806.1

Group16015: afla|EED45978.1 aory|EIT75437.1

Group16016: afla|EED45979.1 aory|EIT75807.1

Group16017: afla|EED45980.1 aory|EIT75834.1

Group16018: afla|EED45995.1 aory|EIT75687.1

Group16019: afla|EED46011.1 aory|EIT75573.1

Group16020: afla|EED46056.1 aory|EIT83493.1

Group16021: afla|EED46062.1 aory|EIT83496.1

Group16022: afla|EED46064.1 aory|EIT83487.1

Group16023: afla|EED46065.1 aory|EIT83480.1

Group16024: afla|EED46074.1 aory|EIT83483.1

Group16025: afla|EED46092.1 aory|EIT76502.1

Group16026: afla|EED46135.1 aory|EIT76361.1

Group16027: afla|EED46152.1 aory|EIT76467.1

Group16028: afla|EED46189.1 aory|EIT76374.1

Group16029: afla|EED46198.1 aory|EIT76454.1

Group16030: afla|EED46222.1 aory|EIT76363.1

Group16031: afla|EED46228.1 aory|EIT76359.1

Group16032: afla|EED46233.1 aory|EIT76382.1

Group16033: afla|EED46249.1 aory|EIT76370.1

Group16034: afla|EED46250.1 aory|EIT76484.1

Group16035: afla|EED46255.1 aory|EIT76360.1

Group16036: afla|EED46281.1 aory|EIT76338.1

Group16037: afla|EED46292.1 aory|EIT76337.1

Group16038: afla|EED46299.1 aory|EIT76340.1

Group16039: afla|EED46303.1 aory|EIT76330.1

Group16040: afla|EED46306.1 atub|OJI80332.1

Group16041: afla|EED46319.1 aory|EIT81234.1

Group16042: afla|EED46343.1 aory|EIT81262.1

Group16043: afla|EED46349.1 aory|EIT81190.1

Group16044: afla|EED46357.1 aory|EIT81175.1

Group16045: afla|EED46358.1 aory|EIT81259.1

Group16046: afla|EED46375.1 aory|EIT72288.1

Group16047: afla|EED46380.1 aory|EIT72293.1

Group16048: afla|EED46384.1 aory|EIT72372.1

Group16049: afla|EED46444.1 anid|CBF87453.1

Group16050: afla|EED46445.1 aory|EIT72423.1

Group16051: afla|EED46500.1 aory|EIT72367.1

Group16052: afla|EED46501.1 aory|EIT72408.1

Group16053: afla|EED46503.1 aory|EIT72422.1

Group16054: afla|EED46504.1 aory|EIT72417.1

Group16055: afla|EED46505.1 aory|EIT72413.1

Group16056: afla|EED46517.1 aory|EIT72445.1

Group16057: afla|EED46521.1 aory|EIT72348.1

Group16058: afla|EED46530.1 aory|EIT72429.1

Group16059: afla|EED46531.1 aory|EIT72315.1

Group16060: afla|EED46536.1 aory|EIT72323.1

Group16061: afla|EED46552.1 agla|OJJ81457.1

Group16062: afla|EED46579.1 aory|EIT77957.1

Group16063: afla|EED46580.1 aory|EIT77892.1

Group16064: afla|EED46581.1 aory|EIT77861.1

Group16065: akaw|GAA86637.1 aluc|OJZ85194.1

Group16066: afla|EED46592.1 aory|EIT77836.1

Group16067: afla|EED46596.1 aory|EIT77922.1

Group16068: afla|EED46613.1 aory|EIT77841.1

Group16069: afla|EED46647.1 aory|EIT77944.1

Group16070: afla|EED46743.1 aory|EIT77860.1

Group16071: afla|EED46786.1 aory|EIT72753.1

Group16072: afla|EED46817.1 aory|EIT72534.1

Group16073: afla|EED46840.1 aory|EIT72555.1

Group16074: afla|EED46841.1 aory|EIT72612.1

Group16075: afla|EED46842.1 aory|EIT72657.1

Group16076: afla|EED46843.1 aory|EIT72734.1

Group16077: afla|EED46853.1 aory|EIT72773.1

Group16078: afla|EED46875.1 aory|EIT72549.1

Group16079: afla|EED46880.1 aory|EIT72717.1

Group16080: afla|EED46881.1 aory|EIT72751.1

Group16081: afla|EED46893.1 aory|EIT72591.1

Group16082: afla|EED46895.1 aory|EIT72676.1

Group16083: afla|EED46896.1 aory|EIT72719.1

Group16084: afla|EED46908.1 aory|EIT72722.1

Group16085: ater|EAU31347.1 agla|OJJ80843.1

Group16086: aver|OJJ03000.1 asyd|OJJ56209.1

Group16087: afla|EED46928.1 aory|EIT72558.1

Group16088: afla|EED46929.1 aory|EIT72702.1

Group16089: afla|EED46950.1 aory|EIT72731.1

Group16090: afla|EED46963.1 anid|CBF78261.1

Group16091: afla|EED46987.1 aory|EIT72561.1

Group16092: afla|EED47034.1 aory|EIT72556.1

Group16093: afla|EED47065.1 aory|EIT72502.1

Group16094: afla|EED47074.1 aory|EIT72475.1

Group16095: afla|EED47133.1 aory|EIT79253.1

Group16096: afla|EED47220.1 aory|EIT79176.1

Group16097: afla|EED47228.1 aory|EIT79194.1

Group16098: afla|EED47235.1 aory|EIT79305.1

Group16099: afla|EED47241.1 aory|EIT79208.1

Group16100: afla|EED47245.1 aory|EIT79257.1

Group16101: afla|EED47247.1 aory|EIT79236.1

Group16102: afla|EED47250.1 aory|EIT79285.1

Group16103: afla|EED47262.1 aory|EIT79317.1

Group16104: afla|EED47267.1 aory|EIT79281.1

Group16105: afla|EED47275.1 aory|EIT79301.1

Group16106: afla|EED47280.1 aory|EIT79213.1

Group16107: afla|EED47287.1 aory|EIT79224.1

Group16108: afla|EED47290.1 aory|EIT79278.1

Group16109: afla|EED47297.1 aory|EIT82856.1

Group16110: afla|EED47302.1 aory|EIT82865.1

Group16111: afla|EED47303.1 aory|EIT82892.1

Group16112: afla|EED47363.1 aory|EIT82847.1

Group16113: afla|EED47364.1 aory|EIT82829.1

Group16114: afla|EED47385.1 aory|EIT82836.1

Group16115: afla|EED47386.1 aory|EIT82835.1

Group16116: afla|EED47398.1 aory|EIT81935.1

Group16117: afla|EED47399.1 aory|EIT81931.1

Group16118: afla|EED47401.1 aory|EIT83381.1

Group16119: afla|EED47416.1 aory|EIT81910.1

Group16120: afla|EED47417.1 aory|EIT81915.1

Group16121: afla|EED47419.1 aory|EIT81908.1

Group16122: afla|EED47422.1 aory|EIT81923.1

Group16123: afla|EED47453.1 aory|EIT79968.1

Group16124: afla|EED47455.1 aory|EIT79954.1

Group16125: afla|EED47475.1 aory|EIT79949.1

Group16126: afla|EED47489.1 aory|EIT79971.1

Group16127: asyd|OJJ64154.1 aver|OJI98233.1

Group16128: afla|EED47518.1 aory|EIT74550.1

Group16129: afla|EED47535.1 aory|EIT74559.1

Group16130: afla|EED47563.1 aory|EIT74582.1

Group16131: afla|EED47564.1 agla|OJJ86149.1

Group16132: afla|EED47571.1 aluc|OJZ80352.1

Group16133: afla|EED47572.1 aluc|OJZ80353.1

Group16134: afla|EED47585.1 aory|EIT74528.1

Group16135: afla|EED47622.1 aory|EIT80268.1

Group16136: afla|EED47648.1 aory|EIT80216.1

Group16137: afla|EED47662.1 aory|EIT80254.1

Group16138: afla|EED47677.1 aory|EIT80275.1

Group16139: afla|EED47680.1 aory|EIT80203.1

Group16140: afla|EED47693.1 aory|EIT80184.1

Group16141: afla|EED47721.1 aory|EIT77145.1

Group16142: afla|EED47727.1 aory|EIT77199.1

Group16143: afla|EED47741.1 aory|EIT77162.1

Group16144: afla|EED47743.1 aory|EIT77157.1

Group16145: afla|EED47756.1 aory|EIT77156.1

Group16146: afla|EED47768.1 aory|EIT77188.1

Group16147: afla|EED47787.1 aory|EIT77183.1

Group16148: afla|EED47801.1 aory|EIT77138.1

Group16149: afla|EED47806.1 aory|EIT77190.1

Group16150: afla|EED47812.1 aory|EIT77151.1

Group16151: afla|EED47841.1 aory|EIT77213.1

Group16152: afla|EED47843.1 aory|EIT77239.1

Group16153: afla|EED47861.1 aory|EIT77235.1

Group16154: afla|EED47866.1 aory|EIT77250.1

Group16155: afla|EED47882.1 asyd|OJJ62689.1

Group16156: afla|EED47895.1 aory|EIT77211.1

Group16157: afla|EED47901.1 ater|EAU30658.1

Group16158: afla|EED47925.1 aory|EIT79809.1

Group16159: afla|EED47934.1 aory|EIT79860.1

Group16160: afla|EED47959.1 aory|EIT79814.1

Group16161: afla|EED47962.1 aory|EIT79837.1

Group16162: afla|EED48078.1 aory|EIT82478.1

Group16163: afla|EED48102.1 anid|CBF84010.1

Group16164: afla|EED48111.1 aory|EIT76224.1

Group16165: afla|EED48114.1 aory|EIT76231.1

Group16166: afla|EED48122.1 aory|EIT76202.1

Group16167: afla|EED48123.1 aory|EIT76251.1

Group16168: afla|EED48124.1 aory|EIT76189.1

Group16169: afla|EED48134.1 aory|EIT76256.1

Group16170: afla|EED48138.1 aory|EIT76211.1

Group16171: afla|EED48139.1 aory|EIT76236.1

Group16172: aver|OJI96141.1 asyd|OJJ53760.1

Group16173: afla|EED48152.1 aory|EIT76199.1

Group16174: afla|EED48157.1 aory|EIT83033.1

Group16175: afla|EED48162.1 aory|EIT83040.1

Group16176: afla|EED48212.1 aory|EIT72924.1

Group16177: afla|EED48213.1 aory|EIT72895.1

Group16178: afla|EED48218.1 aory|EIT72930.1

Group16179: afla|EED48223.1 aory|EIT72937.1

Group16180: afla|EED48224.1 aory|EIT72941.1

Group16181: afla|EED48225.1 aory|EIT72953.1

Group16182: afla|EED48237.1 aory|EIT72900.1

Group16183: afla|EED48255.1 aory|EIT72892.1

Group16184: afla|EED48282.1 aory|EIT72913.1

Group16185: afla|EED48283.1 aory|EIT72943.1

Group16186: afla|EED48297.1 aory|EIT76313.1

Group16187: afla|EED48307.1 aory|EIT76279.1

Group16188: afla|EED48312.1 aory|EIT76298.1

Group16189: afla|EED48323.1 atub|OJI89795.1

Group16190: afla|EED48333.1 aory|EIT76290.1

Group16191: afla|EED48335.1 aory|EIT76316.1

Group16192: afla|EED48348.1 aory|EIT76305.1

Group16193: afla|EED48356.1 aory|EIT72299.1

Group16194: afla|EED48358.1 aory|EIT83100.1

Group16195: afla|EED48368.1 aory|EIT83069.1

Group16196: afla|EED48369.1 aory|EIT83054.1

Group16197: afla|EED48371.1 aory|EIT83058.1

Group16198: afla|EED48375.1 aory|EIT83075.1

Group16199: afla|EED48376.1 aory|EIT83072.1

Group16200: afla|EED48380.1 aory|EIT83081.1

Group16201: afla|EED48398.1 awen|OJJ32511.1

Group16202: afla|EED48404.1 aory|EIT83103.1

Group16203: afla|EED48426.1 aory|EIT83102.1

Group16204: afla|EED48445.1 aory|EIT81668.1

Group16205: afla|EED48446.1 aory|EIT83467.1

Group16206: afla|EED48451.1 aory|EIT83469.1

Group16207: afla|EED48452.1 aory|EIT83473.1

Group16208: afla|EED48453.1 aory|EIT83470.1

Group16209: afla|EED48458.1 aory|EIT76275.1

Group16210: afla|EED48459.1 aory|EIT76267.1

Group16211: afla|EED48460.1 aory|EIT76271.1

Group16212: afla|EED48461.1 aory|EIT76274.1

Group16213: afla|EED48462.1 aory|EIT76268.1

Group16214: afla|EED48470.1 aory|EIT73062.1

Group16215: afla|EED48488.1 aory|EIT73044.1

Group16216: afla|EED48505.1 aory|EIT73262.1

Group16217: afla|EED48518.1 aory|EIT73229.1

Group16218: afla|EED48520.1 aory|EIT73080.1

Group16219: afla|EED48532.1 aory|EIT73237.1

Group16220: afla|EED48606.1 aory|EIT73081.1

Group16221: afla|EED48607.1 aory|EIT73065.1

Group16222: afla|EED48611.1 aory|EIT73091.1

Group16223: afla|EED48614.1 aory|EIT73104.1

Group16224: afla|EED48620.1 aory|EIT73039.1

Group16225: afla|EED48621.1 aory|EIT73162.1

Group16226: afla|EED48646.1 ater|EAU37621.1

Group16227: afla|EED48649.1 aory|EIT73132.1

Group16228: afla|EED48671.1 aory|EIT73133.1

Group16229: afla|EED48693.1 ater|EAU33235.1

Group16230: afla|EED48734.1 aory|EIT73040.1

Group16231: afla|EED48764.1 aory|EIT73025.1

Group16232: afla|EED48804.1 aory|EIT73026.1

Group16233: afla|EED48883.1 aory|EIT81258.1

Group16234: afla|EED48884.1 aory|EIT81218.1

Group16235: awen|OJJ39625.1 agla|OJJ78435.1

Group16236: afla|EED48891.1 aory|EIT81223.1

Group16237: afla|EED48898.1 aory|EIT81231.1

Group16238: afla|EED48904.1 aory|EIT81164.1

Group16239: afla|EED48921.1 aory|EIT80046.1

Group16240: afla|EED48932.1 aory|EIT79984.1

Group16241: afla|EED48934.1 aory|EIT80009.1

Group16242: afla|EED48940.1 ater|EAU29925.1

Group16243: afla|EED48941.1 ater|EAU29926.1

Group16244: afla|EED48954.1 aory|EIT80000.1

Group16245: afla|EED48973.1 aory|EIT79998.1

Group16246: afla|EED48975.1 aory|EIT79978.1

Group16247: afla|EED48982.1 aory|EIT80036.1

Group16248: afla|EED49031.1 aory|EIT82556.1

Group16249: afla|EED49070.1 aory|EIT82501.1

Group16250: agla|OJJ89202.1 awen|OJJ40105.1

Group16251: afla|EED49136.1 aory|EIT78587.1

Group16252: afla|EED49139.1 awen|OJJ35365.1

Group16253: afla|EED49146.1 aory|EIT78524.1

Group16254: afla|EED49160.1 aory|EIT78577.1

Group16255: afla|EED49168.1 aory|EIT78521.1

Group16256: afla|EED49172.1 aory|EIT78549.1

Group16257: afla|EED49185.1 aory|EIT78533.1

Group16258: afla|EED49270.1 aory|EIT82206.1

Group16259: afla|EED49285.1 aory|EIT82209.1

Group16260: afla|EED49292.1 aory|EIT82225.1

Group16261: afla|EED49346.1 aory|EIT79514.1

Group16262: afla|EED49351.1 aory|EIT79555.1

Group16263: afla|EED49395.1 aory|EIT79532.1

Group16264: afla|EED49415.1 aory|EIT79589.1

Group16265: afla|EED49417.1 aory|EIT79501.1

Group16266: afla|EED49420.1 aory|EIT79476.1

Group16267: afla|EED49426.1 aory|EIT79518.1

Group16268: afla|EED49430.1 aory|EIT79547.1

Group16269: afla|EED49431.1 aory|EIT79481.1

Group16270: afla|EED49442.1 aory|EIT79566.1

Group16271: afla|EED49446.1 aory|EIT79541.1

Group16272: afla|EED49447.1 aory|EIT79478.1

Group16273: afla|EED49451.1 aory|EIT79550.1

Group16274: afla|EED49456.1 aory|EIT79533.1

Group16275: afla|EED49472.1 aory|EIT79588.1

Group16276: afla|EED49475.1 aory|EIT79577.1

Group16277: afla|EED49479.1 aory|EIT79539.1

Group16278: afla|EED49484.1 aory|EIT79557.1

Group16279: afla|EED49527.1 aory|EIT82627.1

Group16280: afla|EED49531.1 aory|EIT82596.1

Group16281: afla|EED49538.1 aory|EIT82636.1

Group16282: afla|EED49542.1 aory|EIT82631.1

Group16283: afla|EED49564.1 aory|EIT82632.1

Group16284: afla|EED49565.1 aory|EIT82622.1

Group16285: afla|EED49571.1 aory|EIT82646.1

Group16286: afla|EED49578.1 aory|EIT82608.1

Group16287: afla|EED49600.1 ater|EAU35384.1

Group16288: afla|EED49605.1 aory|EIT80986.1

Group16289: afla|EED49620.1 aory|EIT80978.1

Group16290: afla|EED49644.1 aory|EIT81021.1

Group16291: afla|EED49647.1 aory|EIT80980.1

Group16292: afla|EED49655.1 aory|EIT80946.1

Group16293: afla|EED49661.1 aory|EIT80936.1

Group16294: afla|EED49663.1 aory|EIT80957.1

Group16295: afla|EED49664.1 aory|EIT81020.1

Group16296: afla|EED49669.1 aory|EIT80952.1

Group16297: afla|EED49673.1 aory|EIT80988.1

Group16298: afla|EED49696.1 aory|EIT77640.1

Group16299: afla|EED49717.1 aory|EIT77618.1

Group16300: afla|EED49720.1 aory|EIT77645.1

Group16301: afla|EED49726.1 aory|EIT77680.1

Group16302: afla|EED49728.1 agla|OJJ82708.1

Group16303: afla|EED49730.1 aory|EIT77630.1

Group16304: afla|EED49738.1 aory|EIT77701.1

Group16305: afla|EED49748.1 aory|EIT77639.1

Group16306: afla|EED49760.1 aory|EIT77673.1

Group16307: afla|EED49765.1 aory|EIT77672.1

Group16308: anig|EHA18923.1 ater|EAU39470.1

Group16309: afla|EED49775.1 aory|EIT77636.1

Group16310: afla|EED49783.1 aory|EIT77685.1

Group16311: afla|EED49790.1 asyd|OJJ56233.1

Group16312: afla|EED49800.1 aory|EIT77627.1

Group16313: afla|EED49813.1 aory|EIT77669.1

Group16314: afla|EED49829.1 aory|EIT77656.1

Group16315: afla|EED49834.1 aory|EIT83393.1

Group16316: afla|EED49836.1 aory|EIT83388.1

Group16317: afla|EED49845.1 aver|OJJ02117.1

Group16318: afla|EED49847.1 awen|OJJ36077.1

Group16319: afla|EED49849.1 awen|OJJ36075.1

Group16320: afla|EED49856.1 aory|EIT78828.1

Group16321: afla|EED49860.1 aory|EIT83416.1

Group16322: afla|EED49865.1 aory|EIT83420.1

Group16323: afla|EED49869.1 aory|EIT83394.1

Group16324: afla|EED49900.1 aory|EIT72457.1

Group16325: afla|EED49919.1 aory|EIT72273.1

Group16326: afla|EED49928.1 aory|EIT72266.1

Group16327: afla|EED49943.1 aory|EIT83443.1

Group16328: afla|EED49948.1 aory|EIT83430.1

Group16329: afla|EED49970.1 aory|EIT83453.1

Group16330: afla|EED49979.1 aory|EIT74644.1

Group16331: afla|EED49981.1 aory|EIT74651.1

Group16332: afla|EED50014.1 aory|EIT74624.1

Group16333: afla|EED50039.1 aory|EIT74715.1

Group16334: afla|EED50047.1 aory|EIT74680.1

Group16335: afla|EED50053.1 aory|EIT74706.1

Group16336: afla|EED50067.1 aory|EIT74668.1

Group16337: afla|EED50083.1 aory|EIT74672.1

Group16338: afla|EED50101.1 aory|EIT74678.1

Group16339: afla|EED50107.1 aory|EIT74709.1

Group16340: afla|EED50127.1 aory|EIT74612.1

Group16341: afla|EED50131.1 aory|EIT74730.1

Group16342: afla|EED50138.1 aory|EIT74694.1

Group16343: afla|EED50139.1 aory|EIT74729.1

Group16344: afla|EED50140.1 aory|EIT74641.1

Group16345: afla|EED50161.1 aory|EIT76915.1

Group16346: afla|EED50177.1 aory|EIT72284.1

Group16347: afla|EED50185.1 aory|EIT77378.1

Group16348: afla|EED50186.1 aory|EIT77386.1

Group16349: afla|EED50213.1 aory|EIT77274.1

Group16350: aver|OJI97120.1 asyd|OJJ63818.1

Group16351: afla|EED50217.1 aory|EIT77319.1

Group16352: afla|EED50233.1 aory|EIT77285.1

Group16353: afla|EED50234.1 aory|EIT77346.1

Group16354: afla|EED50241.1 aory|EIT77354.1

Group16355: afla|EED50267.1 aory|EIT77288.1

Group16356: afla|EED50290.1 aory|EIT77359.1

Group16357: afla|EED50416.1 anid|CBF78083.1

Group16358: afla|EED50422.1 aory|EIT82052.1

Group16359: afla|EED50428.1 aory|EIT82012.1

Group16360: afla|EED50443.1 aory|EIT82006.1

Group16361: afla|EED50459.1 aory|EIT82126.1

Group16362: afla|EED50473.1 aory|EIT82005.1

Group16363: afla|EED50515.1 aory|EIT82157.1

Group16364: afla|EED50519.1 aory|EIT82089.1

Group16365: afla|EED50526.1 aory|EIT82073.1

Group16366: afla|EED50559.1 aory|EIT82004.1

Group16367: afla|EED50572.1 aory|EIT82161.1

Group16368: afla|EED50580.1 aory|EIT82114.1

Group16369: afla|EED50598.1 aory|EIT78091.1

Group16370: afla|EED50606.1 aory|EIT78118.1

Group16371: afla|EED50610.1 aory|EIT78113.1

Group16372: afla|EED50659.1 aory|EIT78034.1

Group16373: afla|EED50697.1 aory|EIT78098.1

Group16374: afla|EED50704.1 aory|EIT77997.1

Group16375: afla|EED50751.1 aory|EIT78120.1

Group16376: afla|EED50757.1 aory|EIT78063.1

Group16377: afla|EED50760.1 aory|EIT77979.1

Group16378: afla|EED50784.1 aory|EIT78018.1

Group16379: afla|EED50789.1 aory|EIT77984.1

Group16380: afla|EED50810.1 aory|EIT79063.1

Group16381: afla|EED50811.1 aory|EIT79000.1

Group16382: afla|EED50812.1 aory|EIT79039.1

Group16383: afla|EED50842.1 aory|EIT79003.1

Group16384: afla|EED50849.1 aory|EIT79037.1

Group16385: afla|EED50850.1 aory|EIT79055.1

Group16386: afla|EED50856.1 aory|EIT79020.1

Group16387: afla|EED50857.1 aory|EIT79052.1

Group16388: afla|EED50859.1 aory|EIT79017.1

Group16389: afla|EED50930.1 aory|EIT78996.1

Group16390: afla|EED50931.1 aory|EIT79019.1

Group16391: afla|EED50934.1 aory|EIT79002.1

Group16392: afla|EED50945.1 aory|EIT77008.1

Group16393: afla|EED50951.1 aory|EIT77039.1

Group16394: afla|EED50958.1 aory|EIT77029.1

Group16395: afla|EED50963.1 aory|EIT77004.1

Group16396: afla|EED50968.1 aory|EIT76957.1

Group16397: afla|EED50994.1 aory|EIT76976.1

Group16398: afla|EED51000.1 aory|EIT76969.1

Group16399: afla|EED51015.1 aory|EIT77011.1

Group16400: afla|EED51016.1 aory|EIT76984.1

Group16401: afla|EED51017.1 aory|EIT77033.1

Group16402: afla|EED51020.1 aory|EIT76994.1

Group16403: afla|EED51025.1 aory|EIT76974.1

Group16404: afla|EED51041.1 aory|EIT76965.1

Group16405: afla|EED51044.1 aory|EIT76940.1

Group16406: afla|EED51046.1 aory|EIT76989.1

Group16407: afla|EED51048.1 aory|EIT76991.1

Group16408: afla|EED51053.1 aory|EIT77003.1

Group16409: afla|EED51060.1 aory|EIT77035.1

Group16410: afla|EED51061.1 aory|EIT76945.1

Group16411: afla|EED51067.1 aory|EIT76977.1

Group16412: afla|EED51069.1 aory|EIT76944.1

Group16413: afla|EED51088.1 aory|EIT81995.1

Group16414: afla|EED51104.1 aory|EIT81304.1

Group16415: afla|EED51108.1 aory|EIT81313.1

Group16416: afla|EED51111.1 aory|EIT81336.1

Group16417: afla|EED51122.1 aory|EIT81343.1

Group16418: afla|EED51136.1 aory|EIT81306.1

Group16419: afla|EED51143.1 awen|OJJ38780.1

Group16420: afla|EED51149.1 aory|EIT81298.1

Group16421: afla|EED51155.1 aory|EIT81332.1

Group16422: afla|EED51164.1 aver|OJI99532.1

Group16423: afla|EED51179.1 aory|EIT81328.1

Group16424: afla|EED51181.1 aory|EIT81351.1

Group16425: afla|EED51184.1 aory|EIT81345.1

Group16426: afla|EED51188.1 aver|OJJ07635.1

Group16427: afla|EED51267.1 aory|EIT81519.1

Group16428: afla|EED51304.1 aory|EIT81559.1

Group16429: afla|EED51352.1 aory|EIT80461.1

Group16430: afla|EED51355.1 aory|EIT80608.1

Group16431: afla|EED51375.1 aory|EIT80504.1

Group16432: afla|EED51377.1 aory|EIT80593.1

Group16433: afla|EED51378.1 aory|EIT80484.1

Group16434: afla|EED51382.1 aory|EIT80712.1

Group16435: afla|EED51383.1 aory|EIT80548.1

Group16436: afla|EED51435.1 aory|EIT80475.1

Group16437: afla|EED51455.1 aory|EIT80454.1

Group16438: afla|EED51482.1 aory|EIT80696.1

Group16439: afla|EED51490.1 aory|EIT80615.1

Group16440: afla|EED51491.1 aory|EIT80648.1

Group16441: afla|EED51492.1 aory|EIT80518.1

Group16442: afla|EED51518.1 aory|EIT80611.1

Group16443: afla|EED51529.1 aory|EIT80536.1

Group16444: afla|EED51549.1 aory|EIT80429.1

Group16445: afla|EED51554.1 aory|EIT80602.1

Group16446: afla|EED51566.1 aory|EIT80440.1

Group16447: afla|EED51587.1 aory|EIT80590.1

Group16448: afla|EED51588.1 aory|EIT80681.1

Group16449: afla|EED51589.1 aluc|OJZ92919.1

Group16450: afla|EED51601.1 aory|EIT80732.1

Group16451: afla|EED51604.1 aory|EIT80423.1

Group16452: afla|EED51628.1 aory|EIT80586.1

Group16453: afla|EED51630.1 aluc|OJZ87907.1

Group16454: afla|EED51646.1 aory|EIT80701.1

Group16455: afla|EED51648.1 aory|EIT80676.1

Group16456: afla|EED51661.1 aory|EIT80568.1

Group16457: afla|EED51666.1 aory|EIT80513.1

Group16458: afla|EED51676.1 aory|EIT80595.1

Group16459: afla|EED51698.1 anid|CBF83162.1

Group16460: afla|EED51707.1 aory|EIT80447.1

Group16461: afla|EED51709.1 aory|EIT80672.1

Group16462: afla|EED51721.1 aory|EIT80669.1

Group16463: afla|EED51723.1 aory|EIT80613.1

Group16464: afla|EED51724.1 aory|EIT80677.1

Group16465: afla|EED51725.1 aory|EIT80546.1

Group16466: afla|EED51728.1 aory|EIT80747.1

Group16467: afla|EED51733.1 anid|CBF79139.1

Group16468: afla|EED51746.1 aory|EIT80691.1

Group16469: afla|EED51753.1 aory|EIT80759.1

Group16470: afla|EED51769.1 aory|EIT80018.1

Group16471: afla|EED51772.1 aory|EIT81737.1

Group16472: afla|EED51777.1 aory|EIT81774.1

Group16473: afla|EED51819.1 aory|EIT81718.1

Group16474: afla|EED51823.1 aory|EIT81729.1

Group16475: afla|EED51828.1 aory|EIT81714.1

Group16476: afla|EED51834.1 aory|EIT81727.1

Group16477: afla|EED51842.1 aory|EIT81739.1

Group16478: afla|EED51858.1 aory|EIT81745.1

Group16479: afla|EED51859.1 aory|EIT81702.1

Group16480: afla|EED51933.1 aory|EIT81477.1

Group16481: afla|EED51941.1 aory|EIT81464.1

Group16482: afla|EED51968.1 aory|EIT81405.1

Group16483: afla|EED51972.1 aory|EIT81455.1

Group16484: afla|EED51992.1 aory|EIT81456.1

Group16485: afla|EED51993.1 aory|EIT81444.1

Group16486: afla|EED51994.1 aory|EIT81410.1

Group16487: afla|EED52000.1 aory|EIT81470.1

Group16488: afla|EED52003.1 aory|EIT81393.1

Group16489: afla|EED52011.1 aory|EIT81437.1

Group16490: afla|EED52012.1 aory|EIT81449.1

Group16491: afla|EED52014.1 aory|EIT81378.1

Group16492: afla|EED52018.1 aory|EIT81428.1

Group16493: asyd|OJJ55491.1 anid|CBF83145.1

Group16494: afla|EED52037.1 agla|OJJ78909.1

Group16495: afla|EED52043.1 aory|EIT81411.1

Group16496: afla|EED52052.1 aory|EIT81479.1

Group16497: afla|EED52056.1 aory|EIT81384.1

Group16498: afla|EED52062.1 aory|EIT81386.1

Group16499: afla|EED52072.1 aory|EIT72303.1

Group16500: afla|EED52074.1 aory|EIT76935.1

Group16501: afla|EED52075.1 afum|EAL86754.2

Group16502: afla|EED52076.1 aory|EIT76925.1

Group16503: afla|EED52080.1 aory|EIT76937.1

Group16504: afla|EED52092.1 aory|EIT76906.1

Group16505: afla|EED52102.1 aory|EIT76920.1

Group16506: afla|EED52106.1 aory|EIT76939.1

Group16507: afla|EED52148.1 awen|OJJ32932.1

Group16508: afla|EED52155.1 aory|EIT74335.1

Group16509: afla|EED52160.1 azon|OJJ42481.1

Group16510: afla|EED52166.1 azon|OJJ50339.1

Group16511: afla|EED52169.1 aory|EIT74467.1

Group16512: afla|EED52170.1 aory|EIT74462.1

Group16513: afla|EED52174.1 aory|EIT74330.1

Group16514: afla|EED52189.1 aory|EIT74457.1

Group16515: afla|EED52193.1 aory|EIT74511.1

Group16516: afla|EED52217.1 aory|EIT74468.1

Group16517: afla|EED52223.1 aory|EIT74364.1

Group16518: afla|EED52226.1 aory|EIT74484.1

Group16519: afla|EED52235.1 aory|EIT74508.1

Group16520: afla|EED52240.1 aory|EIT74444.1

Group16521: afla|EED52249.1 aory|EIT74341.1

Group16522: afla|EED52250.1 aory|EIT74431.1

Group16523: afla|EED52256.1 aory|EIT74398.1

Group16524: afla|EED52258.1 awen|OJJ33778.1

Group16525: afla|EED52272.1 aory|EIT74380.1

Group16526: afla|EED52280.1 aory|EIT74509.1

Group16527: afla|EED52283.1 aory|EIT74470.1

Group16528: afla|EED52286.1 aory|EIT74363.1

Group16529: afla|EED52295.1 aory|EIT74437.1

Group16530: afla|EED52307.1 aory|EIT74421.1

Group16531: afla|EED52308.1 aory|EIT74321.1

Group16532: afla|EED52309.1 aory|EIT74426.1

Group16533: afla|EED52316.1 aory|EIT74329.1

Group16534: afla|EED52327.1 aory|EIT74455.1

Group16535: afla|EED52333.1 aory|EIT74389.1

Group16536: afla|EED52334.1 aory|EIT74512.1

Group16537: afla|EED52338.1 aory|EIT74388.1

Group16538: afla|EED52340.1 aory|EIT74439.1

Group16539: asyd|OJJ61886.1 aver|OJI95984.1

Group16540: afla|EED52369.1 aory|EIT82367.1

Group16541: afla|EED52378.1 aory|EIT82370.1

Group16542: afla|EED52407.1 ater|EAU37126.1

Group16543: afla|EED52411.1 aory|EIT83279.1

Group16544: afla|EED52416.1 aory|EIT83317.1

Group16545: afla|EED52419.1 aory|EIT83319.1

Group16546: atub|OJI88613.1 anig|EHA25867.1

Group16547: afla|EED52460.1 aory|EIT82802.1

Group16548: afla|EED52512.1 aory|EIT82782.1

Group16549: afla|EED52516.1 aory|EIT82818.1

Group16550: afla|EED52518.1 aory|EIT82806.1

Group16551: afla|EED52523.1 aory|EIT82747.1

Group16552: afla|EED52532.1 aory|EIT74182.1

Group16553: afla|EED52533.1 aory|EIT74258.1

Group16554: afla|EED52539.1 awen|OJJ38790.1

Group16555: afla|EED52542.1 awen|OJJ38791.1

Group16556: afla|EED52557.1 aory|EIT74223.1

Group16557: afla|EED52560.1 azon|OJJ43114.1

Group16558: afla|EED52579.1 aory|EIT74281.1

Group16559: afla|EED52581.1 aory|EIT74217.1

Group16560: afla|EED52652.1 atub|OJI82035.1

Group16561: afla|EED52677.1 aory|EIT74220.1

Group16562: afla|EED52722.1 aory|EIT78155.1

Group16563: afla|EED52796.1 aory|EIT78242.1

Group16564: afla|EED52805.1 aory|EIT78213.1

Group16565: afla|EED52826.1 aory|EIT78238.1

Group16566: afla|EED52835.1 aory|EIT78603.1

Group16567: afla|EED52837.1 aory|EIT78667.1

Group16568: afla|EED52839.1 aory|EIT78670.1

Group16569: afla|EED52840.1 aory|EIT78610.1

Group16570: afla|EED52842.1 aory|EIT78647.1

Group16571: afla|EED52846.1 aory|EIT78665.1

Group16572: afla|EED52847.1 aory|EIT78605.1

Group16573: afla|EED52850.1 aory|EIT78649.1

Group16574: afla|EED52859.1 aory|EIT78657.1

Group16575: afla|EED52869.1 afum|EAL93229.1

Group16576: afla|EED52885.1 aory|EIT78677.1

Group16577: afla|EED52886.1 aory|EIT78589.1

Group16578: afla|EED52901.1 aory|EIT78626.1

Group16579: afla|EED52964.1 aory|EIT78720.1

Group16580: afla|EED53007.1 aory|EIT77420.1

Group16581: afla|EED53046.1 azon|OJJ49872.1

Group16582: afla|EED53070.1 aory|EIT74815.1

Group16583: afla|EED53076.1 aory|EIT74769.1

Group16584: aver|OJJ01855.1 asyd|OJJ59245.1

Group16585: afla|EED53080.1 aory|EIT74778.1

Group16586: afla|EED53087.1 aory|EIT74766.1

Group16587: afla|EED53138.1 aory|EIT74862.1

Group16588: afla|EED53149.1 aory|EIT75019.1

Group16589: afla|EED53165.1 aory|EIT75096.1

Group16590: afla|EED53186.1 aory|EIT75227.1

Group16591: afla|EED53195.1 aory|EIT75020.1

Group16592: afla|EED53204.1 aory|EIT75181.1

Group16593: afla|EED53223.1 aory|EIT74936.1

Group16594: afla|EED53229.1 aory|EIT75218.1

Group16595: afla|EED53238.1 aory|EIT75258.1

Group16596: afla|EED53245.1 aory|EIT74975.1

Group16597: afla|EED53247.1 aory|EIT75014.1

Group16598: afla|EED53278.1 aory|EIT74915.1

Group16599: afla|EED53291.1 aory|EIT75033.1

Group16600: afla|EED53314.1 aory|EIT74856.1

Group16601: afla|EED53320.1 aory|EIT75170.1

Group16602: afla|EED53329.1 aory|EIT75097.1

Group16603: afla|EED53335.1 aory|EIT75139.1

Group16604: afla|EED53365.1 aory|EIT74941.1

Group16605: afla|EED53366.1 aory|EIT74858.1

Group16606: afla|EED53367.1 aory|EIT75159.1

Group16607: afla|EED53397.1 aory|EIT74835.1

Group16608: afla|EED53408.1 aory|EIT75114.1

Group16609: afla|EED53413.1 aory|EIT75222.1

Group16610: aver|OJJ08577.1 asyd|OJJ54163.1

Group16611: afla|EED53433.1 aory|EIT75157.1

Group16612: afla|EED53448.1 aory|EIT74932.1

Group16613: afla|EED53453.1 aory|EIT75203.1

Group16614: afla|EED53454.1 aory|EIT75264.1

Group16615: afla|EED53456.1 aory|EIT74959.1

Group16616: afla|EED53457.1 aory|EIT74912.1

Group16617: afla|EED53459.1 aory|EIT75255.1

Group16618: afla|EED53474.1 aory|EIT74827.1

Group16619: afla|EED53477.1 aory|EIT74937.1

Group16620: afla|EED53486.1 aory|EIT75040.1

Group16621: afla|EED53492.1 aory|EIT75257.1

Group16622: afla|EED53495.1 aory|EIT74973.1

Group16623: afla|EED53498.1 azon|OJJ44464.1

Group16624: afla|EED53499.1 aory|EIT74979.1

Group16625: afla|EED53504.1 aory|EIT75088.1

Group16626: afla|EED53551.1 aory|EIT75137.1

Group16627: afla|EED53579.1 aory|EIT74839.1

Group16628: afla|EED53580.1 aory|EIT75006.1

Group16629: afla|EED53593.1 aory|EIT74946.1

Group16630: afla|EED53601.1 aory|EIT74826.1

Group16631: afla|EED53611.1 aory|EIT75206.1

Group16632: afla|EED53619.1 aory|EIT74864.1

Group16633: afla|EED53630.1 aory|EIT75136.1

Group16634: afla|EED53634.1 aory|EIT75208.1

Group16635: afla|EED53638.1 aory|EIT75183.1

Group16636: afla|EED53655.1 aory|EIT75280.1

Group16637: afla|EED53656.1 aory|EIT75276.1

Group16638: afla|EED53695.1 aory|EIT75312.1

Group16639: aver|OJI95755.1 anid|CBF76153.1

Group16640: afla|EED53710.1 aory|EIT75274.1

Group16641: afla|EED53717.1 aory|EIT75283.1

Group16642: afla|EED53751.1 aory|EIT78411.1

Group16643: afla|EED53760.1 aory|EIT78436.1

Group16644: afla|EED53787.1 atub|OJI86834.1

Group16645: afla|EED53801.1 aory|EIT78373.1

Group16646: afla|EED53837.1 aory|EIT78398.1

Group16647: afla|EED53908.1 aory|EIT78496.1

Group16648: afla|EED53918.1 aory|EIT78258.1

Group16649: afla|EED53982.1 aory|EIT78278.1

Group16650: afla|EED53990.1 aory|EIT78332.1

Group16651: afla|EED54020.1 aory|EIT78337.1

Group16652: afla|EED54025.1 aory|EIT78266.1

Group16653: afla|EED54045.1 aver|OJJ06476.1

Group16654: afla|EED54048.1 aory|EIT76086.1

Group16655: afla|EED54052.1 aory|EIT75895.1

Group16656: afla|EED54056.1 aory|EIT76108.1

Group16657: afla|EED54067.1 aory|EIT76064.1

Group16658: afla|EED54081.1 aory|EIT76055.1

Group16659: afla|EED54090.1 aory|EIT76136.1

Group16660: afla|EED54096.1 aory|EIT76016.1

Group16661: afla|EED54099.1 aory|EIT76127.1

Group16662: afla|EED54101.1 aory|EIT76012.1

Group16663: asyd|OJJ60827.1 aver|OJJ00153.1

Group16664: afla|EED54106.1 aory|EIT75948.1

Group16665: afla|EED54113.1 aory|EIT76049.1

Group16666: afla|EED54132.1 aory|EIT75946.1

Group16667: afla|EED54147.1 aory|EIT76129.1

Group16668: afla|EED54150.1 aory|EIT75994.1

Group16669: afla|EED54152.1 atub|OJI91291.1

Group16670: afla|EED54157.1 aory|EIT75962.1

Group16671: afla|EED54165.1 aory|EIT76077.1

Group16672: afla|EED54173.1 aory|EIT75960.1

Group16673: afla|EED54174.1 aory|EIT76117.1

Group16674: afla|EED54190.1 ater|EAU33850.1

Group16675: aver|OJI97593.1 asyd|OJJ52103.1

Group16676: afla|EED54236.1 aory|EIT76097.1

Group16677: afla|EED54250.1 aory|EIT75995.1

Group16678: afla|EED54312.1 aory|EIT75936.1

Group16679: afla|EED54323.1 aory|EIT75913.1

Group16680: afla|EED54335.1 aory|EIT75972.1

Group16681: afla|EED54342.1 aory|EIT75981.1

Group16682: afla|EED54346.1 aory|EIT76163.1

Group16683: afla|EED54359.1 anid|CBF80458.1

Group16684: afla|EED54360.1 aory|EIT75959.1

Group16685: afla|EED54368.1 aory|EIT76160.1

Group16686: afla|EED54382.1 aory|EIT75851.1

Group16687: afla|EED54383.1 aory|EIT75864.1

Group16688: afla|EED54393.1 aory|EIT75874.1

Group16689: afla|EED54395.1 aory|EIT75868.1

Group16690: afla|EED54402.1 aory|EIT75865.1

Group16691: atub|OJI90163.1 aluc|OJZ79712.1

Group16692: afla|EED54406.1 aory|EIT75850.1

Group16693: afla|EED54419.1 aory|EIT75867.1

Group16694: afla|EED54475.1 aory|EIT81128.1

Group16695: afla|EED54497.1 aory|EIT78764.1

Group16696: afla|EED54541.1 aory|EIT78883.1

Group16697: afla|EED54551.1 aory|EIT78940.1

Group16698: afla|EED54613.1 agla|OJJ87531.1

Group16699: afla|EED54619.1 aory|EIT78953.1

Group16700: afla|EED54623.1 aory|EIT78938.1

Group16701: afla|EED54657.1 aory|EIT78842.1

Group16702: afla|EED54659.1 aory|EIT78928.1

Group16703: afla|EED54660.1 aory|EIT78804.1

Group16704: afla|EED54667.1 aory|EIT78909.1

Group16705: afla|EED54684.1 aory|EIT78878.1

Group16706: afla|EED54690.1 aory|EIT78847.1

Group16707: afla|EED54733.1 aory|EIT78778.1

Group16708: afla|EED54745.1 aory|EIT78855.1

Group16709: afla|EED54747.1 aory|EIT78793.1

Group16710: afla|EED54759.1 aory|EIT79403.1

Group16711: afla|EED54772.1 aory|EIT79397.1

Group16712: afla|EED54801.1 aory|EIT79344.1

Group16713: afla|EED54814.1 aory|EIT79375.1

Group16714: afla|EED54830.1 aory|EIT79371.1

Group16715: afla|EED54866.1 aory|EIT79414.1

Group16716: afla|EED54876.1 aory|EIT79354.1

Group16717: afla|EED54891.1 aory|EIT79452.1

Group16718: afla|EED54895.1 aory|EIT79418.1

Group16719: afla|EED54903.1 aory|EIT80360.1

Group16720: afla|EED54910.1 aory|EIT80364.1

Group16721: afla|EED54914.1 aory|EIT80342.1

Group16722: afla|EED54925.1 aory|EIT80311.1

Group16723: afla|EED55008.1 aory|EIT80348.1

Group16724: afla|EED55022.1 aory|EIT81980.1

Group16725: afla|EED55029.1 aory|EIT81955.1

Group16726: afla|EED55038.1 aory|EIT81978.1

Group16727: afla|EED55041.1 aory|EIT81975.1

Group16728: afla|EED55043.1 aory|EIT81982.1

Group16729: afla|EED55066.1 aory|EIT81963.1

Group16730: afla|EED55069.1 aory|EIT81990.1

Group16731: afla|EED55104.1 aory|EIT81632.1

Group16732: afla|EED55114.1 aory|EIT81626.1

Group16733: afla|EED55124.1 aory|EIT81615.1

Group16734: afla|EED55136.1 aory|EIT81599.1

Group16735: afla|EED55139.1 aory|EIT81651.1

Group16736: afla|EED55142.1 aory|EIT81652.1

Group16737: afla|EED55151.1 aory|EIT81623.1

Group16738: afla|EED55169.1 aory|EIT81655.1

Group16739: afla|EED55173.1 aory|EIT81638.1

Group16740: afla|EED55195.1 aory|EIT83609.1

Group16741: afla|EED55196.1 aory|EIT83567.1

Group16742: afla|EED55227.1 aory|EIT83623.1

Group16743: afla|EED55234.1 aory|EIT83541.1

Group16744: afla|EED55255.1 aory|EIT83558.1

Group16745: afla|EED55260.1 aory|EIT83624.1

Group16746: afla|EED55284.1 aory|EIT83602.1

Group16747: afla|EED55295.1 aory|EIT83607.1

Group16748: afla|EED55296.1 aory|EIT83540.1

Group16749: afla|EED55299.1 aory|EIT83508.1

Group16750: afla|EED55307.1 agla|OJJ89029.1

Group16751: afla|EED55346.1 aory|EIT80905.1

Group16752: afla|EED55365.1 aory|EIT80930.1

Group16753: afla|EED55494.1 aory|EIT80907.1

Group16754: afla|EED55603.1 aory|EIT82698.1

Group16755: afla|EED55670.1 aory|EIT77098.1

Group16756: afla|EED55785.1 aory|EIT76539.1

Group16757: afla|EED55787.1 aory|EIT76738.1

Group16758: afla|EED55813.1 aory|EIT76575.1

Group16759: afla|EED55852.1 aory|EIT76760.1

Group16760: afla|EED55872.1 aory|EIT76613.1

Group16761: asyd|OJJ60516.1 aver|OJI99842.1

Group16762: afla|EED55905.1 aory|EIT76845.1

Group16763: afla|EED55942.1 aory|EIT76639.1

Group16764: afla|EED55968.1 aory|EIT76618.1

Group16765: afla|EED55972.1 aory|EIT76808.1

Group16766: afla|EED55974.1 aory|EIT76695.1

Group16767: afla|EED55977.1 aory|EIT76653.1

Group16768: afla|EED55995.1 aory|EIT76611.1

Group16769: afla|EED56026.1 aory|EIT76675.1

Group16770: afla|EED56030.1 aory|EIT76812.1

Group16771: afla|EED56032.1 aory|EIT76786.1

Group16772: afla|EED56075.1 aory|EIT76684.1

Group16773: afla|EED56084.1 aory|EIT76565.1

Group16774: afla|EED56124.1 aory|EIT76772.1

Group16775: afla|EED56168.1 aory|EIT76878.1

Group16776: afla|EED56197.1 aory|EIT73392.1

Group16777: afla|EED56202.1 aory|EIT73349.1

Group16778: afla|EED56209.1 aory|EIT73435.1

Group16779: afla|EED56305.1 aory|EIT73385.1

Group16780: afla|EED56316.1 aory|EIT73419.1

Group16781: afla|EED56317.1 aory|EIT73452.1

Group16782: afla|EED56324.1 aory|EIT73453.1

Group16783: afla|EED56334.1 aory|EIT73416.1

Group16784: afla|EED56343.1 aory|EIT73451.1

Group16785: afla|EED56356.1 aory|EIT73334.1

Group16786: afla|EED56358.1 aory|EIT73294.1

Group16787: afla|EED56381.1 aory|EIT73364.1

Group16788: afla|EED56382.1 aory|EIT73384.1

Group16789: afla|EED56409.1 aory|EIT83007.1

Group16790: afla|EED56417.1 aory|EIT72884.1

Group16791: afla|EED56428.1 ater|EAU33598.1

Group16792: afla|EED56448.1 aory|EIT72858.1

Group16793: afla|EED56462.1 aory|EIT72806.1

Group16794: afla|EED56532.1 aory|EIT79692.1

Group16795: afla|EED56537.1 aory|EIT79690.1

Group16796: afla|EED56543.1 aory|EIT79696.1

Group16797: afla|EED56547.1 aory|EIT79689.1

Group16798: afla|EED56552.1 ater|EAU37801.1

Group16799: afla|EED56565.1 aory|EIT79598.1

Group16800: afla|EED56587.1 aory|EIT79611.1

Group16801: afla|EED56594.1 aory|EIT79609.1

Group16802: afla|EED56606.1 aory|EIT79592.1

Group16803: afla|EED56608.1 aory|EIT79622.1

Group16804: afla|EED56611.1 aory|EIT79683.1

Group16805: afla|EED56615.1 aory|EIT79643.1

Group16806: afla|EED56628.1 aory|EIT79664.1

Group16807: afla|EED56629.1 aory|EIT79593.1

Group16808: afla|EED56642.1 aory|EIT79661.1

Group16809: afla|EED56717.1 aory|EIT73832.1

Group16810: afla|EED56740.1 aory|EIT73909.1

Group16811: afla|EED56753.1 aory|EIT73925.1

Group16812: afla|EED56754.1 aory|EIT73913.1

Group16813: afla|EED56756.1 aory|EIT73852.1

Group16814: afla|EED56763.1 aory|EIT73933.1

Group16815: afla|EED56770.1 aory|EIT73839.1

Group16816: afla|EED56772.1 aory|EIT73932.1

Group16817: afla|EED56774.1 aory|EIT73886.1

Group16818: afla|EED56780.1 aory|EIT73922.1

Group16819: afla|EED56784.1 aory|EIT73859.1

Group16820: afla|EED56793.1 aory|EIT73855.1

Group16821: afla|EED56807.1 aory|EIT73936.1

Group16822: asyd|OJJ52739.1 aver|OJJ02996.1

Group16823: afla|EED56826.1 aory|EIT73890.1

Group16824: afla|EED56838.1 aory|EIT82823.1

Group16825: afla|EED56844.1 aory|EIT77527.1

Group16826: afla|EED56854.1 aory|EIT77581.1

Group16827: afla|EED56863.1 aory|EIT77579.1

Group16828: afla|EED56866.1 aory|EIT77485.1

Group16829: afla|EED56867.1 aory|EIT77588.1

Group16830: afla|EED56887.1 aory|EIT77495.1

Group16831: afla|EED56892.1 agla|OJJ80498.1

Group16832: afla|EED56894.1 aory|EIT77453.1

Group16833: afla|EED56901.1 aory|EIT77452.1

Group16834: afla|EED56923.1 aory|EIT77539.1

Group16835: afla|EED56924.1 azon|OJJ45174.1

Group16836: afla|EED56931.1 aory|EIT77456.1

Group16837: afla|EED56942.1 aory|EIT77572.1

Group16838: afla|EED57037.1 aory|EIT82278.1

Group16839: afla|EED57041.1 aory|EIT82296.1

Group16840: afla|EED57058.1 aory|EIT82259.1

Group16841: afla|EED57081.1 aory|EIT82281.1

Group16842: afla|EED57094.1 aory|EIT82300.1

Group16843: afla|EED57114.1 aory|EIT74124.1

Group16844: afla|EED57116.1 aory|EIT74004.1

Group16845: afla|EED57123.1 aory|EIT74023.1

Group16846: afla|EED57161.1 aory|EIT74072.1

Group16847: afla|EED57208.1 aory|EIT73966.1

Group16848: afla|EED57213.1 aory|EIT74044.1

Group16849: afla|EED57226.1 aory|EIT74142.1

Group16850: afla|EED57239.1 aory|EIT74063.1

Group16851: afla|EED57244.1 aory|EIT74131.1

Group16852: afla|EED57258.1 aory|EIT74100.1

Group16853: afla|EED57264.1 aory|EIT74070.1

Group16854: afla|EED57267.1 aory|EIT73962.1

Group16855: afla|EED57268.1 aory|EIT74068.1

Group16856: afla|EED57272.1 aory|EIT73983.1

Group16857: afla|EED57300.1 aory|EIT74061.1

Group16858: afla|EED57305.1 aory|EIT73996.1

Group16859: afla|EED57332.1 aory|EIT73986.1

Group16860: afla|EED57343.1 aory|EIT80157.1

Group16861: afla|EED57360.1 aory|EIT80119.1

Group16862: afla|EED57414.1 aory|EIT80154.1

Group16863: afla|EED57455.1 aory|EIT80145.1

Group16864: afla|EED57471.1 aory|EIT80144.1

Group16865: afla|EED57482.1 aory|EIT80087.1

Group16866: afla|EED57486.1 aory|EIT81878.1

Group16867: afla|EED57490.1 aory|EIT81879.1

Group16868: afla|EED57492.1 aory|EIT81881.1

Group16869: afla|EED57514.1 aluc|OJZ80818.1

Group16870: afla|EED57540.1 aory|EIT81859.1

Group16871: afla|EED57564.1 aory|EIT81856.1

Group16872: afla|EED57569.1 aory|EIT82928.1

Group16873: afla|EED57570.1 aory|EIT82980.1

Group16874: afla|EED57573.1 aory|EIT82982.1

Group16875: afla|EED57580.1 aory|EIT82979.1

Group16876: afla|EED57617.1 aory|EIT82994.1

Group16877: afla|EED57618.1 aory|EIT82933.1

Group16878: afla|EED57642.1 aory|EIT79879.1

Group16879: afla|EED57675.1 aory|EIT79932.1

Group16880: afla|EED57676.1 aory|EIT79883.1

Group16881: afla|EED57711.1 aory|EIT79912.1

Group16882: afla|EED57801.1 aory|EIT81049.1

Group16883: afla|EED57825.1 aory|EIT81114.1

Group16884: afla|EED57856.1 aory|EIT81121.1

Group16885: afla|EED57863.1 awen|OJJ37046.1

Group16886: afla|EED57869.1 aory|EIT81033.1

Group16887: afla|EED57874.1 aory|EIT83268.1

Group16888: afla|EED57882.1 aory|EIT83267.1

Group16889: afla|EED57888.1 aory|EIT83274.1

Group16890: afla|EED57889.1 aory|EIT82738.1

Group16891: afla|EED57908.1 aory|EIT82354.1

Group16892: afla|EED57911.1 aory|EIT82321.1

Group16893: afla|EED57917.1 aory|EIT82337.1

Group16894: afla|EED57939.1 aory|EIT82361.1

Group16895: afla|EED58000.1 aory|EIT79807.1

Group16896: afla|EED58005.1 aory|EIT79733.1

Group16897: afla|EED58012.1 aory|EIT79792.1

Group16898: afla|EED58035.1 aory|EIT79740.1

Group16899: afla|EED58079.1 aory|EIT79124.1

Group16900: afla|EED58086.1 aory|EIT79108.1

Group16901: afla|EED58159.1 aory|EIT79106.1

Group16902: afum|EAL84781.1 aver|OJJ00745.1

Group16903: afum|EAL84868.1 anid|CBF87203.1

Group16904: afum|EAL84981.1 anig|EHA23978.1

Group16905: afum|EAL85103.1 ater|EAU36159.1

Group16906: afum|EAL85126.1 aory|EIT72581.1

Group16907: afum|EAL85340.1 aluc|OJZ80252.1

Group16908: afum|EAL85380.1 aluc|OJZ79828.1

Group16909: afum|EAL85388.1 akaw|GAA87897.1

Group16910: agla|OJJ83629.1 awen|OJJ31324.1

Group16911: afum|EAL85422.1 azon|OJJ50177.1

Group16912: afum|EAL86277.1 ater|EAU32476.1

Group16913: afum|EAL86402.1 anig|EHA24468.1

Group16914: afum|EAL86442.1 aory|EIT73553.1

Group16915: afum|EAL86536.1 akaw|GAA88275.1

Group16916: afum|EAL86544.1 ater|EAU36563.1

Group16917: afum|EAL86721.1 aory|EIT80420.1

Group16918: afum|EAL86745.1 anig|EHA21857.1

Group16919: afum|EAL87133.1 anig|EHA23472.1

Group16920: asyd|OJJ62933.1 aver|OJI98057.1

Group16921: afum|EAL87783.1 anig|EHA21824.1

Group16922: afum|EAL87805.1 akaw|GAA83754.1

Group16923: afum|EAL88866.1 agla|OJJ85627.1

Group16924: afum|EAL89008.2 aver|OJJ08172.1

Group16925: afum|EAL89014.2 aluc|OJZ90828.1

Group16926: asyd|OJJ65079.1 aver|OJI99204.1

Group16927: afum|EAL91123.1 anid|CBF73643.1

Group16928: afum|EAL91129.1 agla|OJJ83276.1

Group16929: afum|EAL91130.1 aluc|OJZ80471.1

Group16930: azon|OJJ51754.1 aver|OJJ08137.1

Group16931: afum|EAL91910.1 ater|EAU29608.1

Group16932: afum|EAL92628.1 asyd|OJJ60378.1

Group16933: afum|EAL92630.1 aver|OJJ05865.1

Group16934: anig|EHA25785.1 awen|OJJ34652.1

Group16935: afum|EAL94099.1 ater|EAU32776.1

Group16936: agla|OJJ78462.1 awen|OJJ30562.1

Group16937: agla|OJJ78483.1 awen|OJJ31338.1

Group16938: agla|OJJ78563.1 azon|OJJ42288.1

Group16939: agla|OJJ78649.1 awen|OJJ34647.1

Group16940: aver|OJI97953.1 asyd|OJJ63007.1

Group16941: agla|OJJ78784.1 awen|OJJ38720.1

Group16942: agla|OJJ78808.1 ater|EAU36741.1

Group16943: agla|OJJ78811.1 asyd|OJJ61464.1

Group16944: agla|OJJ78823.1 atub|OJI91290.1

Group16945: asyd|OJJ55466.1 aver|OJJ04320.1

Group16946: agla|OJJ79168.1 atub|OJI81723.1

Group16947: agla|OJJ79255.1 ater|EAU31055.1

Group16948: agla|OJJ79268.1 awen|OJJ36379.1

Group16949: agla|OJJ79273.1 awen|OJJ29898.1

Group16950: agla|OJJ79304.1 asyd|OJJ61694.1

Group16951: agla|OJJ79355.1 azon|OJJ41992.1

Group16952: agla|OJJ79395.1 awen|OJJ31596.1

Group16953: agla|OJJ79474.1 awen|OJJ41515.1

Group16954: agla|OJJ79764.1 awen|OJJ30388.1

Group16955: agla|OJJ79867.1 anig|EHA18423.1

Group16956: agla|OJJ79970.1 awen|OJJ40935.1

Group16957: aluc|OJZ90633.1 atub|OJI82060.1

Group16958: agla|OJJ80047.1 awen|OJJ39631.1

Group16959: aver|OJJ07233.1 ater|EAU36848.1

Group16960: agla|OJJ80211.1 asyd|OJJ54600.1

Group16961: agla|OJJ80324.1 awen|OJJ32723.1

Group16962: agla|OJJ80392.1 awen|OJJ29731.1

Group16963: agla|OJJ80516.1 ater|EAU33706.1

Group16964: agla|OJJ80551.1 ater|EAU33818.1

Group16965: agla|OJJ80676.1 aory|EIT82796.1

Group16966: agla|OJJ80681.1 aory|EIT77045.1

Group16967: agla|OJJ80807.1 awen|OJJ37979.1

Group16968: agla|OJJ80832.1 aver|OJJ08250.1

Group16969: agla|OJJ80839.1 asyd|OJJ62682.1

Group16970: agla|OJJ81000.1 asyd|OJJ53485.1

Group16971: agla|OJJ81001.1 asyd|OJJ53432.1

Group16972: agla|OJJ81473.1 azon|OJJ42205.1

Group16973: agla|OJJ81559.1 aver|OJJ07189.1

Group16974: agla|OJJ81563.1 aver|OJI98884.1

Group16975: agla|OJJ81741.1 aory|EIT80044.1

Group16976: agla|OJJ81751.1 awen|OJJ37976.1

Group16977: agla|OJJ81929.1 asyd|OJJ60095.1

Group16978: agla|OJJ82214.1 awen|OJJ31963.1

Group16979: agla|OJJ82329.1 ater|EAU32947.1

Group16980: agla|OJJ82378.1 atub|OJI87465.1

Group16981: agla|OJJ82458.1 anig|EHA24122.1

Group16982: agla|OJJ82524.1 awen|OJJ36131.1

Group16983: agla|OJJ82593.1 atub|OJI83010.1

Group16984: agla|OJJ82599.1 asyd|OJJ56418.1

Group16985: agla|OJJ82721.1 awen|OJJ31413.1

Group16986: agla|OJJ82726.1 awen|OJJ31409.1

Group16987: agla|OJJ82734.1 awen|OJJ31403.1

Group16988: agla|OJJ82825.1 azon|OJJ45349.1

Group16989: agla|OJJ83024.1 azon|OJJ48352.1

Group16990: agla|OJJ83077.1 awen|OJJ36193.1

Group16991: agla|OJJ83080.1 azon|OJJ42083.1

Group16992: agla|OJJ83429.1 aver|OJI96720.1

Group16993: agla|OJJ83430.1 aver|OJI96721.1

Group16994: agla|OJJ83547.1 aory|EIT72973.1

Group16995: agla|OJJ83626.1 ater|EAU29764.1

Group16996: agla|OJJ83631.1 azon|OJJ43580.1

Group16997: agla|OJJ83645.1 ater|EAU32793.1

Group16998: agla|OJJ83649.1 awen|OJJ40565.1

Group16999: aver|OJJ04762.1 asyd|OJJ55818.1

Group17000: agla|OJJ83708.1 awen|OJJ32719.1

Group17001: agla|OJJ83734.1 anid|CBF87185.1

Group17002: agla|OJJ83898.1 ater|EAU37775.1

Group17003: agla|OJJ83934.1 ater|EAU34916.1

Group17004: agla|OJJ83942.1 anig|EHA22258.1

Group17005: agla|OJJ84017.1 ater|EAU30204.1

Group17006: agla|OJJ84265.1 awen|OJJ35916.1

Group17007: agla|OJJ84357.1 anid|CBF76139.1

Group17008: agla|OJJ84466.1 awen|OJJ38772.1

Group17009: agla|OJJ84474.1 awen|OJJ32544.1

Group17010: agla|OJJ84475.1 awen|OJJ32545.1

Group17011: asyd|OJJ52391.1 aver|OJJ01932.1

Group17012: agla|OJJ84753.1 awen|OJJ37025.1

Group17013: agla|OJJ85026.1 awen|OJJ31416.1

Group17014: asyd|OJJ64578.1 aver|OJI98675.1

Group17015: agla|OJJ85120.1 aluc|OJZ80067.1

Group17016: aver|OJJ01266.1 asyd|OJJ58658.1

Group17017: agla|OJJ85333.1 awen|OJJ31554.1

Group17018: agla|OJJ85395.1 ater|EAU31593.1

Group17019: agla|OJJ85430.1 awen|OJJ30931.1

Group17020: agla|OJJ85433.1 awen|OJJ29698.1

Group17021: agla|OJJ85452.1 asyd|OJJ52561.1

Group17022: agla|OJJ85456.1 aluc|OJZ80472.1

Group17023: agla|OJJ85641.1 azon|OJJ42208.1

Group17024: agla|OJJ85642.1 azon|OJJ42206.1

Group17025: agla|OJJ85920.1 awen|OJJ37065.1

Group17026: agla|OJJ86108.1 ater|EAU30188.1

Group17027: agla|OJJ86351.1 aver|OJJ07137.1

Group17028: agla|OJJ86397.1 awen|OJJ39468.1

Group17029: agla|OJJ86417.1 aory|EIT72727.1

Group17030: agla|OJJ86477.1 awen|OJJ39542.1

Group17031: agla|OJJ86505.1 azon|OJJ46434.1

Group17032: agla|OJJ86570.1 awen|OJJ38740.1

Group17033: agla|OJJ86663.1 awen|OJJ33830.1

Group17034: agla|OJJ86804.1 awen|OJJ38958.1

Group17035: agla|OJJ86865.1 awen|OJJ38037.1

Group17036: agla|OJJ86883.1 awen|OJJ33366.1

Group17037: agla|OJJ86985.1 anid|CBF80063.1

Group17038: agla|OJJ87308.1 awen|OJJ41239.1

Group17039: aver|OJJ01914.1 asyd|OJJ56226.1

Group17040: agla|OJJ87732.1 awen|OJJ40100.1

Group17041: agla|OJJ87746.1 awen|OJJ34884.1

Group17042: agla|OJJ87925.1 azon|OJJ46449.1

Group17043: agla|OJJ88091.1 atub|OJI80497.1

Group17044: asyd|OJJ58702.1 aver|OJJ01314.1

Group17045: asyd|OJJ57605.1 aver|OJJ05578.1

Group17046: agla|OJJ88604.1 ater|EAU29842.1

Group17047: agla|OJJ88744.1 azon|OJJ50322.1

Group17048: agla|OJJ88992.1 awen|OJJ40801.1

Group17049: agla|OJJ89072.1 anig|EHA19731.1

Group17050: agla|OJJ89197.1 asyd|OJJ52441.1

Group17051: agla|OJJ89219.1 awen|OJJ30972.1

Group17052: agla|OJJ89307.1 ater|EAU31833.1

Group17053: asyd|OJJ60961.1 aver|OJJ00286.1

Group17054: akaw|BAL04875.1 aluc|OJZ79480.1

Group17055: akaw|BAL04877.1 aory|AFM82521.1

Group17056: akaw|BAL04880.1 azon|OJJ41981.1

Group17057: akaw|BAL04885.1 azon|OJJ41975.1

Group17058: akaw|GAA82072.1 aluc|OJZ92110.1

Group17059: akaw|GAA82111.1 aluc|OJZ92065.1

Group17060: akaw|GAA82141.1 aluc|OJZ92030.1

Group17061: akaw|GAA82415.1 atub|OJI86162.1

Group17062: akaw|GAA82583.1 aluc|OJZ87120.1

Group17063: akaw|GAA82587.1 aluc|OJZ87126.1

Group17064: akaw|GAA82791.1 aluc|OJZ87350.1

Group17065: akaw|GAA82792.1 aluc|OJZ87351.1

Group17066: akaw|GAA82881.1 aluc|OJZ87449.1

Group17067: akaw|GAA82937.1 aluc|OJZ87514.1

Group17068: akaw|GAA83048.1 atub|OJI91179.1

Group17069: aver|OJJ03449.1 asyd|OJJ62612.1

Group17070: akaw|GAA83093.1 aluc|OJZ88640.1

Group17071: akaw|GAA83527.1 atub|OJI86898.1

Group17072: akaw|GAA83592.1 aluc|OJZ92405.1

Group17073: akaw|GAA83607.1 atub|OJI86816.1

Group17074: akaw|GAA83676.1 anig|EHA24763.1

Group17075: akaw|GAA83680.1 anig|EHA25617.1

Group17076: akaw|GAA83825.1 aluc|OJZ90893.1

Group17077: akaw|GAA83964.1 aluc|OJZ80853.1

Group17078: akaw|GAA83967.1 aluc|OJZ80856.1

Group17079: akaw|GAA83968.1 aluc|OJZ80860.1

Group17080: akaw|GAA84022.1 aluc|OJZ80926.1

Group17081: akaw|GAA84224.1 aluc|OJZ88904.1

Group17082: akaw|GAA84272.1 aluc|OJZ88965.1

Group17083: akaw|GAA84441.1 atub|OJI85633.1

Group17084: akaw|GAA84482.1 aluc|OJZ89202.1

Group17085: akaw|GAA84590.1 aluc|OJZ89335.1

Group17086: akaw|GAA84678.1 aluc|OJZ87011.1

Group17087: akaw|GAA84730.1 aluc|OJZ86956.1

Group17088: akaw|GAA84848.1 aluc|OJZ86823.1

Group17089: akaw|GAA84902.1 anig|EHA21912.1

Group17090: akaw|GAA85223.1 atub|OJI82001.1

Group17091: akaw|GAA85297.1 aluc|OJZ90657.1

Group17092: akaw|GAA85335.1 aluc|OJZ90698.1

Group17093: akaw|GAA85436.1 aluc|OJZ83340.1

Group17094: akaw|GAA85709.1 aluc|OJZ91394.1

Group17095: akaw|GAA85756.1 aluc|OJZ91344.1

Group17096: akaw|GAA85809.1 atub|OJI83280.1

Group17097: akaw|GAA85842.1 aluc|OJZ91257.1

Group17098: akaw|GAA85844.1 aluc|OJZ91255.1

Group17099: akaw|GAA85848.1 aluc|OJZ91251.1

Group17100: akaw|GAA86082.1 atub|OJI81742.1

Group17101: asyd|OJJ62465.1 aver|OJI96600.1

Group17102: akaw|GAA86258.1 aluc|OJZ90516.1

Group17103: akaw|GAA86460.1 atub|OJI80596.1

Group17104: akaw|GAA86614.1 aluc|OJZ85166.1

Group17105: akaw|GAA86619.1 aluc|OJZ85173.1

Group17106: akaw|GAA86636.1 aluc|OJZ85193.1

Group17107: akaw|GAA86991.1 aluc|OJZ84469.1

Group17108: akaw|GAA87108.1 aluc|OJZ84330.1

Group17109: akaw|GAA87398.1 aluc|OJZ89614.1

Group17110: akaw|GAA87494.1 aluc|OJZ88028.1

Group17111: akaw|GAA87615.1 aluc|OJZ85236.1

Group17112: akaw|GAA87708.1 aluc|OJZ85331.1

Group17113: akaw|GAA87743.1 atub|OJI80194.1

Group17114: akaw|GAA87782.1 anig|EHA25596.1

Group17115: akaw|GAA87797.1 aluc|OJZ85432.1

Group17116: akaw|GAA88008.1 atub|OJI91466.1

Group17117: akaw|GAA88137.1 aluc|OJZ81408.1

Group17118: aluc|OJZ82372.1 atub|OJI88070.1

Group17119: akaw|GAA88449.1 aluc|OJZ88336.1

Group17120: akaw|GAA88462.1 asyd|OJJ54534.1

Group17121: akaw|GAA88579.1 aluc|OJZ88483.1

Group17122: akaw|GAA88580.1 aluc|OJZ88484.1

Group17123: akaw|GAA88582.1 aluc|OJZ88486.1

Group17124: akaw|GAA89000.1 aluc|OJZ80236.1

Group17125: akaw|GAA89004.1 aluc|OJZ88833.1

Group17126: akaw|GAA89009.1 aluc|OJZ79999.1

Group17127: akaw|GAA89020.1 aluc|OJZ80584.1

Group17128: akaw|GAA89137.1 aluc|OJZ81699.1

Group17129: akaw|GAA89214.1 aluc|OJZ81615.1

Group17130: akaw|GAA89320.1 aluc|OJZ86350.1

Group17131: akaw|GAA89353.1 atub|OJI84227.1

Group17132: akaw|GAA89383.1 aluc|OJZ86276.1

Group17133: akaw|GAA89469.1 aluc|OJZ88195.1

Group17134: akaw|GAA89524.1 aluc|OJZ88128.1

Group17135: akaw|GAA89768.1 aluc|OJZ80615.1

Group17136: akaw|GAA89769.1 aluc|OJZ80614.1

Group17137: akaw|GAA89770.1 aluc|OJZ80613.1

Group17138: akaw|GAA89821.1 aluc|OJZ81869.1

Group17139: akaw|GAA89874.1 aluc|OJZ81805.1

Group17140: akaw|GAA89979.1 atub|OJI90361.1

Group17141: akaw|GAA90140.1 aluc|OJZ83207.1

Group17142: akaw|GAA90180.1 aluc|OJZ83170.1

Group17143: akaw|GAA90181.1 aluc|OJZ83169.1

Group17144: akaw|GAA90263.1 atub|OJI87868.1

Group17145: akaw|GAA90296.1 aluc|OJZ83041.1

Group17146: akaw|GAA90302.1 aluc|OJZ83035.1

Group17147: akaw|GAA90313.1 aluc|OJZ85926.1

Group17148: akaw|GAA90338.1 aluc|OJZ85894.1

Group17149: akaw|GAA90429.1 aluc|OJZ85800.1

Group17150: akaw|GAA90689.1 aluc|OJZ84665.1

Group17151: aory|EIT79695.1 aver|OJJ07425.1

Group17152: akaw|GAA90891.1 aluc|OJZ82937.1

Group17153: akaw|GAA90892.1 aluc|OJZ82938.1

Group17154: akaw|GAA90901.1 aluc|OJZ82925.1

Group17155: akaw|GAA91087.1 aluc|OJZ87904.1

Group17156: akaw|GAA91091.1 aluc|OJZ87911.1

Group17157: akaw|GAA91211.1 aluc|OJZ82860.1

Group17158: akaw|GAA91214.1 aluc|OJZ82857.1

Group17159: akaw|GAA91361.1 aluc|OJZ84810.1

Group17160: akaw|GAA91450.1 aluc|OJZ84714.1

Group17161: akaw|GAA91843.1 atub|OJI84613.1

Group17162: akaw|GAA91981.1 aluc|OJZ82632.1

Group17163: akaw|GAA92047.1 aluc|OJZ92918.1

Group17164: akaw|GAA92048.1 aory|EIT80479.1

Group17165: akaw|GAA92135.1 aluc|OJZ83588.1

Group17166: akaw|GAA92173.1 atub|OJI79378.1

Group17167: akaw|GAA92339.1 aluc|OJZ80455.1

Group17168: akaw|GAA92343.1 aluc|OJZ79595.1

Group17169: akaw|GAA92355.1 aluc|OJZ80395.1

Group17170: akaw|GAA92363.1 aluc|OJZ79881.1

Group17171: akaw|GAA92365.1 asyd|OJJ52301.1

Group17172: akaw|GAA92369.1 aluc|OJZ79660.1

Group17173: akaw|GAA92387.1 aluc|OJZ83247.1

Group17174: akaw|GAA92395.1 aluc|OJZ83255.1

Group17175: akaw|GAA92482.1 aluc|OJZ91494.1

Group17176: akaw|GAA92483.1 aluc|OJZ91495.1

Group17177: akaw|GAA92493.1 aluc|OJZ91505.1

Group17178: akaw|GAA92559.1 atub|OJI84698.1

Group17179: akaw|GAA92572.1 aluc|OJZ82583.1

Group17180: akaw|GAA92607.1 aluc|OJZ80064.1

Group17181: akaw|GAA92618.1 aluc|OJZ79584.1

Group17182: akaw|GAA92620.1 aluc|OJZ79534.1

Group17183: akaw|GAA92622.1 aluc|OJZ80497.1

Group17184: akaw|GAA92623.1 aluc|OJZ80438.1

Group17185: akaw|GAA92624.1 aluc|OJZ79837.1

Group17186: akaw|GAA92632.1 aluc|OJZ79472.1

Group17187: akaw|GAA92635.1 aluc|OJZ79739.1

Group17188: akaw|GAA92637.1 ater|EAU34939.1

Group17189: akaw|GAA92659.1 azon|OJJ42651.1

Group17190: akaw|GAA92699.1 aluc|OJZ88738.1

Group17191: akaw|GAA92717.1 atub|OJI91334.1

Group17192: akaw|GAA92813.1 aluc|OJZ88736.1

Group17193: akaw|GAA92843.1 ater|EAU35394.1

Group17194: akaw|GAA92851.1 atub|OJI87464.1

Group17195: akaw|GAA92854.1 awen|OJJ30191.1

Group17196: akaw|GAA92931.1 aluc|OJZ82524.1

Group17197: akaw|GAA92949.1 aluc|OJZ88806.1

Group17198: akaw|GAA92966.1 aluc|OJZ79870.1

Group17199: akaw|GAA92976.1 atub|OJI89951.1

Group17200: akaw|GAA93027.1 aluc|OJZ80440.1

Group17201: akaw|GAA93058.1 aluc|OJZ80491.1

Group17202: akaw|GAA93061.1 aluc|OJZ80495.1

Group17203: akaw|GAA93066.1 aluc|OJZ80435.1

Group17204: akaw|GAA93081.1 aluc|OJZ80227.1

Group17205: akaw|GAA93084.1 aluc|OJZ80481.1

Group17206: aluc|OJZ80237.1 asyd|OJJ56760.1

Group17207: akaw|GAA93133.1 aluc|OJZ80250.1

Group17208: akaw|GAA93136.1 aluc|OJZ80225.1

Group17209: akaw|GAA93138.1 aluc|OJZ80221.1

Group17210: akaw|GAA93139.1 aluc|OJZ79834.1

Group17211: akaw|GAA93152.1 aluc|OJZ80002.1

Group17212: akaw|GAA93163.1 aluc|OJZ80436.1

Group17213: akaw|GAA93166.1 aluc|OJZ83025.1

Group17214: akaw|GAA93167.1 aluc|OJZ83026.1

Group17215: akaw|GAA93169.1 aluc|OJZ83028.1

Group17216: akaw|GAA93202.1 aluc|OJZ79705.1

Group17217: akaw|GAA93210.1 aluc|OJZ79489.1

Group17218: akaw|GAA93225.1 anig|EHA21869.1

Group17219: akaw|GAA93236.1 aluc|OJZ80346.1

Group17220: akaw|GAA93254.1 aluc|OJZ79904.1

Group17221: akaw|GAA93260.1 aluc|OJZ80403.1

Group17222: akaw|GAA93264.1 aluc|OJZ79570.1

Group17223: akaw|GAA93266.1 aluc|OJZ80063.1

Group17224: akaw|GAA93273.1 aluc|OJZ79674.1

Group17225: akaw|GAA93281.1 aluc|OJZ79576.1

Group17226: akaw|GAA93287.1 aluc|OJZ90157.1

Group17227: akaw|GAA93297.1 anig|EHA25614.1

Group17228: akaw|GAA93298.1 anig|EHA25613.1

Group17229: akaw|GAA93303.1 aluc|OJZ81120.1

Group17230: akaw|GAA93304.1 aluc|OJZ80444.1

Group17231: akaw|GAA93305.1 aluc|OJZ80446.1

Group17232: akaw|GAA93313.1 aluc|OJZ80123.1

Group17233: akaw|GAA93314.1 aluc|OJZ80124.1

Group17234: akaw|GAA93322.1 aluc|OJZ79531.1

Group17235: akaw|GAA93336.1 aluc|OJZ79562.1

Group17236: akaw|GAA93337.1 aluc|OJZ79563.1

Group17237: akaw|GAA93347.1 aluc|OJZ79737.1

Group17238: akaw|GAA93348.1 aluc|OJZ79736.1

Group17239: aluc|OJZ79536.1 asyd|OJJ52063.1

Group17240: aluc|OJZ79537.1 atub|OJI80384.1

Group17241: aluc|OJZ79827.1 anig|EHA26122.1

Group17242: aluc|OJZ79888.1 atub|OJI80540.1

Group17243: aluc|OJZ79918.1 asyd|OJJ51943.1

Group17244: aluc|OJZ79971.1 anig|EHA20893.1

Group17245: aluc|OJZ80146.1 atub|OJI85103.1

Group17246: aluc|OJZ80461.1 atub|OJI87466.1

Group17247: aluc|OJZ80462.1 atub|OJI87467.1

Group17248: aluc|OJZ80590.1 anig|EHA25074.1

Group17249: aluc|OJZ80799.1 anig|EHA27235.1

Group17250: aluc|OJZ80840.1 atub|OJI82727.1

Group17251: aluc|OJZ80907.1 atub|OJI82671.1

Group17252: aluc|OJZ80910.1 atub|OJI82669.1

Group17253: aluc|OJZ81012.1 atub|OJI82572.1

Group17254: aluc|OJZ81085.1 atub|OJI82499.1

Group17255: aluc|OJZ81187.1 atub|OJI91383.1

Group17256: aluc|OJZ81223.1 atub|OJI91421.1

Group17257: aver|OJJ05989.1 asyd|OJJ56734.1

Group17258: anid|CBF79121.1 atub|OJI85465.1

Group17259: aluc|OJZ81652.1 anid|CBF83094.1

Group17260: aluc|OJZ81706.1 atub|OJI88477.1

Group17261: aluc|OJZ81781.1 atub|OJI84354.1

Group17262: aluc|OJZ81790.1 anig|EHA22425.1

Group17263: aluc|OJZ81792.1 anig|EHA22423.1

Group17264: aluc|OJZ81827.1 atub|OJI84397.1

Group17265: aluc|OJZ81914.1 atub|OJI84466.1

Group17266: aluc|OJZ81933.1 anig|EHA22315.1

Group17267: aluc|OJZ82100.1 atub|OJI88337.1

Group17268: aluc|OJZ82192.1 atub|OJI88243.1

Group17269: aluc|OJZ82212.1 anig|EHA28528.1

Group17270: aluc|OJZ82275.1 atub|OJI88164.1

Group17271: aluc|OJZ82325.1 atub|OJI88115.1

Group17272: aluc|OJZ82356.1 atub|OJI88084.1

Group17273: aluc|OJZ82452.1 atub|OJI87988.1

Group17274: aluc|OJZ82511.1 anig|EHA20907.1

Group17275: asyd|OJJ61545.1 aver|OJJ00872.1

Group17276: aluc|OJZ82587.1 atub|OJI88822.1

Group17277: aluc|OJZ82595.1 atub|OJI88830.1

Group17278: aluc|OJZ82600.1 atub|OJI88836.1

Group17279: aluc|OJZ82639.1 anig|EHA27431.1

Group17280: aluc|OJZ82650.1 atub|OJI88882.1

Group17281: aluc|OJZ82669.1 atub|OJI88904.1

Group17282: aluc|OJZ82733.1 atub|OJI88966.1

Group17283: aluc|OJZ82761.1 atub|OJI88994.1

Group17284: aluc|OJZ82855.1 atub|OJI89085.1

Group17285: aluc|OJZ82933.1 atub|OJI89162.1

Group17286: aluc|OJZ83020.1 anid|CBF81988.1

Group17287: aluc|OJZ83032.1 atub|OJI87916.1

Group17288: aluc|OJZ83049.1 atub|OJI87899.1

Group17289: aluc|OJZ83050.1 atub|OJI87898.1

Group17290: aluc|OJZ83143.1 anig|EHA23288.1

Group17291: aluc|OJZ83209.1 atub|OJI87728.1

Group17292: aluc|OJZ83437.1 atub|OJI82328.1

Group17293: aluc|OJZ83507.1 atub|OJI82401.1

Group17294: aluc|OJZ83529.1 atub|OJI82419.1

Group17295: aluc|OJZ83547.1 atub|OJI82438.1

Group17296: aluc|OJZ83563.1 atub|OJI82453.1

Group17297: aluc|OJZ83578.1 atub|OJI89953.1

Group17298: aluc|OJZ83586.1 atub|OJI89959.1

Group17299: aluc|OJZ83636.1 atub|OJI90005.1

Group17300: aluc|OJZ83679.1 atub|OJI90050.1

Group17301: aluc|OJZ83792.1 atub|OJI90160.1

Group17302: aluc|OJZ83973.1 atub|OJI90346.1

Group17303: aluc|OJZ83990.1 anig|EHA19115.1

Group17304: aluc|OJZ84015.1 atub|OJI90386.1

Group17305: aluc|OJZ84087.1 atub|OJI90459.1

Group17306: aluc|OJZ84096.1 atub|OJI90469.1

Group17307: asyd|OJJ55756.1 aver|OJJ06841.1

Group17308: aluc|OJZ84310.1 atub|OJI89758.1

Group17309: aluc|OJZ84544.1 atub|OJI89536.1

Group17310: aluc|OJZ84614.1 atub|OJI89467.1

Group17311: aluc|OJZ84639.1 atub|OJI89441.1

Group17312: aluc|OJZ84670.1 atub|OJI89410.1

Group17313: aluc|OJZ84696.1 atub|OJI89386.1

Group17314: aluc|OJZ84710.1 atub|OJI89373.1

Group17315: aluc|OJZ84743.1 atub|OJI89338.1

Group17316: aluc|OJZ84879.1 anig|EHA25108.1

Group17317: aluc|OJZ84906.1 atub|OJI80691.1

Group17318: aluc|OJZ84909.1 atub|OJI80689.1

Group17319: aluc|OJZ85241.1 atub|OJI80334.1

Group17320: aluc|OJZ85334.1 atub|OJI80239.1

Group17321: aluc|OJZ85360.1 atub|OJI80211.1

Group17322: aluc|OJZ85454.1 atub|OJI80110.1

Group17323: aluc|OJZ85486.1 atub|OJI80078.1

Group17324: aluc|OJZ85493.1 atub|OJI80072.1

Group17325: aver|OJJ05941.1 asyd|OJJ53624.1

Group17326: aluc|OJZ85557.1 atub|OJI79348.1

Group17327: aluc|OJZ85638.1 atub|OJI79429.1

Group17328: aluc|OJZ85669.1 atub|OJI79461.1

Group17329: aluc|OJZ85697.1 atub|OJI79486.1

Group17330: aluc|OJZ85700.1 atub|OJI79490.1

Group17331: aluc|OJZ85710.1 atub|OJI79501.1

Group17332: aluc|OJZ85749.1 atub|OJI79533.1

Group17333: aluc|OJZ85923.1 atub|OJI79713.1

Group17334: aluc|OJZ85936.1 atub|OJI79725.1

Group17335: aluc|OJZ85987.1 atub|OJI79776.1

Group17336: aluc|OJZ86077.1 atub|OJI79874.1

Group17337: aluc|OJZ86084.1 atub|OJI79880.1

Group17338: aluc|OJZ86264.1 atub|OJI84279.1

Group17339: aluc|OJZ86332.1 atub|OJI84203.1

Group17340: aluc|OJZ86341.1 atub|OJI84195.1

Group17341: aluc|OJZ86388.1 atub|OJI84145.1

Group17342: aluc|OJZ86389.1 anig|EHA21508.1

Group17343: aluc|OJZ86433.1 atub|OJI84099.1

Group17344: aluc|OJZ86434.1 atub|OJI84097.1

Group17345: aluc|OJZ86458.1 atub|OJI84069.1

Group17346: aluc|OJZ86540.1 atub|OJI83983.1

Group17347: aluc|OJZ86560.1 atub|OJI83961.1

Group17348: aluc|OJZ86599.1 atub|OJI83924.1

Group17349: aluc|OJZ86737.1 atub|OJI83791.1

Group17350: aluc|OJZ86997.1 atub|OJI83522.1

Group17351: aluc|OJZ87068.1 atub|OJI81568.1

Group17352: aluc|OJZ87098.1 atub|OJI81540.1

Group17353: aluc|OJZ87242.1 atub|OJI81400.1

Group17354: aluc|OJZ87345.1 atub|OJI81299.1

Group17355: aluc|OJZ87375.1 atub|OJI81271.1

Group17356: aluc|OJZ87414.1 atub|OJI81234.1

Group17357: aluc|OJZ87415.1 atub|OJI81233.1

Group17358: aluc|OJZ87416.1 atub|OJI81232.1

Group17359: aluc|OJZ87494.1 atub|OJI81155.1

Group17360: aluc|OJZ87566.1 anig|EHA20768.1

Group17361: aluc|OJZ87616.1 anig|EHA20813.1

Group17362: aluc|OJZ87633.1 atub|OJI81016.1

Group17363: aluc|OJZ87783.1 atub|OJI80864.1

Group17364: aluc|OJZ87819.1 atub|OJI80825.1

Group17365: aluc|OJZ87822.1 atub|OJI80822.1

Group17366: aluc|OJZ87837.1 anid|CBF76058.1

Group17367: aluc|OJZ87916.1 anig|EHA17711.1

Group17368: aluc|OJZ87928.1 atub|OJI90540.1

Group17369: aluc|OJZ88082.1 atub|OJI90691.1

Group17370: aluc|OJZ88146.1 atub|OJI90750.1

Group17371: aluc|OJZ88147.1 atub|OJI90751.1

Group17372: aluc|OJZ88148.1 anig|EHA17909.1

Group17373: aluc|OJZ88208.1 atub|OJI90809.1

Group17374: aluc|OJZ88266.1 atub|OJI90866.1

Group17375: aluc|OJZ88268.1 atub|OJI90868.1

Group17376: aluc|OJZ88293.1 atub|OJI90889.1

Group17377: aluc|OJZ88306.1 atub|OJI90901.1

Group17378: aluc|OJZ88492.1 atub|OJI91079.1

Group17379: aluc|OJZ88547.1 atub|OJI91132.1

Group17380: aluc|OJZ88658.1 aver|OJJ05764.1

Group17381: aluc|OJZ88660.1 aver|OJJ05766.1

Group17382: aluc|OJZ88711.1 atub|OJI91298.1

Group17383: aluc|OJZ88725.1 atub|OJI91302.1

Group17384: aluc|OJZ88887.1 atub|OJI85902.1

Group17385: aluc|OJZ88910.1 atub|OJI85881.1

Group17386: aluc|OJZ88924.1 atub|OJI85868.1

Group17387: aluc|OJZ88938.1 atub|OJI85852.1

Group17388: aluc|OJZ88940.1 atub|OJI85850.1

Group17389: aluc|OJZ88944.1 atub|OJI85845.1

Group17390: aluc|OJZ89026.1 anig|EHA23506.1

Group17391: aluc|OJZ89027.1 atub|OJI85764.1

Group17392: aluc|OJZ89057.1 atub|OJI85729.1

Group17393: aluc|OJZ89101.1 atub|OJI85680.1

Group17394: aluc|OJZ89190.1 atub|OJI85600.1

Group17395: aluc|OJZ89220.1 atub|OJI85572.1

Group17396: aluc|OJZ89314.1 atub|OJI85483.1

Group17397: aluc|OJZ89462.1 anig|EHA23126.1

Group17398: aluc|OJZ89491.1 atub|OJI85312.1

Group17399: aluc|OJZ89582.1 atub|OJI85221.1

Group17400: aluc|OJZ89585.1 atub|OJI85217.1

Group17401: aluc|OJZ89826.1 anig|EHA27032.1

Group17402: aluc|OJZ89948.1 anig|EHA27149.1

Group17403: aluc|OJZ89975.1 anig|EHA27349.1

Group17404: aluc|OJZ90354.1 atub|OJI81781.1

Group17405: aluc|OJZ90409.1 atub|OJI81841.1

Group17406: aluc|OJZ90434.1 atub|OJI81865.1

Group17407: aluc|OJZ90482.1 atub|OJI81907.1

Group17408: aluc|OJZ90486.1 anig|EHA23944.1

Group17409: aluc|OJZ90530.1 atub|OJI81952.1

Group17410: aluc|OJZ90742.1 atub|OJI82169.1

Group17411: aluc|OJZ90760.1 atub|OJI82747.1

Group17412: aluc|OJZ90844.1 atub|OJI82831.1

Group17413: aluc|OJZ90898.1 atub|OJI82890.1

Group17414: aluc|OJZ91075.1 atub|OJI83070.1

Group17415: aluc|OJZ91134.1 atub|OJI83144.1

Group17416: aluc|OJZ91140.1 aver|OJJ05143.1

Group17417: anid|CBF80174.1 aver|OJI99525.1

Group17418: aluc|OJZ91320.1 atub|OJI83313.1

Group17419: aluc|OJZ91426.1 atub|OJI83419.1

Group17420: aluc|OJZ91437.1 atub|OJI83431.1

Group17421: aluc|OJZ91439.1 atub|OJI83433.1

Group17422: aluc|OJZ91441.1 anig|EHA20105.1

Group17423: aluc|OJZ91511.1 atub|OJI85963.1

Group17424: aluc|OJZ91546.1 atub|OJI85999.1

Group17425: aluc|OJZ91554.1 anig|EHA22589.1

Group17426: aluc|OJZ91558.1 atub|OJI86010.1

Group17427: aluc|OJZ91601.1 atub|OJI86053.1

Group17428: aluc|OJZ91637.1 atub|OJI86090.1

Group17429: aluc|OJZ91665.1 anig|EHA22685.1

Group17430: aluc|OJZ91668.1 atub|OJI86123.1

Group17431: aluc|OJZ91681.1 atub|OJI86134.1

Group17432: aluc|OJZ91682.1 atub|OJI86135.1

Group17433: aluc|OJZ91757.1 ater|EAU38397.1

Group17434: aluc|OJZ91766.1 anig|EHA22766.1

Group17435: aluc|OJZ91772.1 anig|EHA22772.1

Group17436: aluc|OJZ91979.1 ater|EAU30952.1

Group17437: aluc|OJZ91980.1 anig|EHA20390.1

Group17438: aluc|OJZ91996.1 atub|OJI86431.1

Group17439: aluc|OJZ92101.1 atub|OJI86535.1

Group17440: aluc|OJZ92194.1 atub|OJI87955.1

Group17441: aluc|OJZ92215.1 atub|OJI86646.1

Group17442: aluc|OJZ92228.1 atub|OJI86658.1

Group17443: aluc|OJZ92300.1 atub|OJI86729.1

Group17444: aluc|OJZ92302.1 atub|OJI86732.1

Group17445: aluc|OJZ92390.1 atub|OJI86820.1

Group17446: aluc|OJZ92532.1 anig|EHA26442.1

Group17447: aluc|OJZ92533.1 atub|OJI86969.1

Group17448: aluc|OJZ92554.1 atub|OJI86990.1

Group17449: aluc|OJZ92623.1 anig|EHA26366.1

Group17450: aluc|OJZ92624.1 anig|EHA26365.1

Group17451: aluc|OJZ92697.1 atub|OJI87125.1

Group17452: aluc|OJZ92726.1 anig|EHA26275.1

Group17453: aluc|OJZ92739.1 atub|OJI87164.1

Group17454: aluc|OJZ92753.1 atub|OJI87178.1

Group17455: aluc|OJZ92794.1 atub|OJI87237.1

Group17456: aluc|OJZ92821.1 anig|EHA26187.1

Group17457: aluc|OJZ92836.1 asyd|OJJ53763.1

Group17458: aluc|OJZ92885.1 atub|OJI87344.1

Group17459: anid|CBF69445.1 aver|OJJ06465.1

Group17460: aver|OJJ06643.1 asyd|OJJ55562.1

Group17461: anid|CBF70352.1 aver|OJJ06047.1

Group17462: anid|CBF70804.1 awen|OJJ36116.1

Group17463: azon|OJJ42432.1 aver|OJI95858.1

Group17464: anid|CBF71203.1 asyd|OJJ55036.1

Group17465: anid|CBF71207.1 aver|OJI96130.1

Group17466: anid|CBF71438.1 asyd|OJJ56672.1

Group17467: anid|CBF71480.1 aver|OJJ01276.1

Group17468: anid|CBF73396.1 aver|OJJ03014.1

Group17469: anid|CBF73399.1 asyd|OJJ52427.1

Group17470: anid|CBF73435.1 aver|OJJ08110.1

Group17471: anid|CBF73440.1 aver|OJJ08113.1

Group17472: anid|CBF73451.1 aver|OJJ08119.1

Group17473: anid|CBF73469.1 aver|OJJ08121.1

Group17474: anid|CBF73596.1 azon|OJJ44481.1

Group17475: aver|OJJ08771.1 asyd|OJJ52793.1

Group17476: anid|CBF73654.1 aver|OJI99659.1

Group17477: anid|CBF74660.1 asyd|OJJ54389.1

Group17478: anid|CBF75167.1 asyd|OJJ59958.1

Group17479: anid|CBF75411.1 asyd|OJJ56173.1

Group17480: anid|CBF75892.1 aver|OJJ08873.1

Group17481: anid|CBF76078.1 ater|EAU33662.1

Group17482: asyd|OJJ54531.1 aver|OJJ07657.1

Group17483: anid|CBF77678.1 aver|OJJ04867.1

Group17484: asyd|OJJ55288.1 aver|OJJ06940.1

Group17485: anid|CBF78129.1 asyd|OJJ55909.1

Group17486: anid|CBF78403.1 aver|OJI99308.1

Group17487: anid|CBF78542.1 azon|OJJ44497.1

Group17488: anid|CBF78716.1 anig|EHA26173.1

Group17489: anid|CBF78942.1 aver|OJJ01865.1

Group17490: anid|CBF78966.1 aver|OJJ06747.1

Group17491: anid|CBF79122.1 atub|OJI85464.1

Group17492: anid|CBF79240.1 awen|OJJ36954.1

Group17493: anid|CBF79296.1 aver|OJJ06859.1

Group17494: anid|CBF79919.1 asyd|OJJ58198.1

Group17495: anid|CBF80173.1 aver|OJI99524.1

Group17496: anid|CBF80355.1 asyd|OJJ54827.1

Group17497: anid|CBF80421.1 ater|EAU29528.1

Group17498: aver|OJJ08828.1 asyd|OJJ52850.1

Group17499: anid|CBF81163.1 anig|EHA21126.1

Group17500: anid|CBF81768.1 asyd|OJJ56691.1

Group17501: anid|CBF81993.1 asyd|OJJ52780.1

Group17502: anid|CBF82041.1 aver|OJJ05175.1

Group17503: anid|CBF82299.1 asyd|OJJ52728.1

Group17504: anid|CBF82475.1 aver|OJJ08635.1

Group17505: anid|CBF83022.1 asyd|OJJ61032.1

Group17506: anid|CBF83035.1 asyd|OJJ53749.1

Group17507: anid|CBF83093.1 atub|OJI88530.1

Group17508: anid|CBF83261.1 asyd|OJJ60582.1

Group17509: anid|CBF83739.1 awen|OJJ39941.1

Group17510: aver|OJJ02108.1 asyd|OJJ58304.1

Group17511: anid|CBF84493.1 anig|EHA19601.1

Group17512: anid|CBF84557.1 asyd|OJJ62587.1

Group17513: anid|CBF84611.1 ater|EAU36755.1

Group17514: anid|CBF85199.1 ater|EAU32058.1

Group17515: anid|CBF85290.1 anig|EHA18732.1

Group17516: anid|CBF85323.1 azon|OJJ42069.1

Group17517: anid|CBF86658.1 asyd|OJJ64787.1

Group17518: anid|CBF86718.1 aver|OJJ01995.1

Group17519: anid|CBF86734.1 aver|OJJ03078.1

Group17520: aver|OJJ08358.1 asyd|OJJ53384.1

Group17521: anid|CBF87251.1 ater|EAU29531.1

Group17522: anid|CBF87359.1 awen|OJJ39789.1

Group17523: anid|CBF87362.1 awen|OJJ39788.1

Group17524: anid|CBF87364.1 awen|OJJ39787.1

Group17525: asyd|OJJ52945.1 aver|OJJ08912.1

Group17526: anid|CBF87572.1 asyd|OJJ63245.1

Group17527: anid|CBF87729.1 aver|OJJ08124.1

Group17528: anig|EHA22620.1 asyd|OJJ53393.1

Group17529: anid|CBF88304.1 ater|EAU31928.1

Group17530: anid|CBF88820.1 atub|OJI85023.1

Group17531: anid|CBF89267.1 aver|OJJ06189.1

Group17532: anid|CBF89292.1 ater|EAU33883.1

Group17533: anid|CBF89408.1 awen|OJJ37801.1

Group17534: anig|EHA17689.1 aory|EIT76245.1

Group17535: anig|EHA17690.1 aver|OJJ07504.1

Group17536: anig|EHA17972.1 atub|OJI90825.1

Group17537: anig|EHA18411.1 awen|OJJ41830.1

Group17538: anig|EHA18473.1 atub|OJI91351.1

Group17539: anig|EHA18791.1 atub|OJI90008.1

Group17540: anig|EHA18927.1 atub|OJI90158.1

Group17541: anig|EHA19666.1 atub|OJI82953.1

Group17542: anig|EHA20156.1 awen|OJJ33216.1

Group17543: anig|EHA20660.1 azon|OJJ45830.1

Group17544: anig|EHA20956.1 atub|OJI84687.1

Group17545: anig|EHA21514.1 atub|OJI84154.1

Group17546: anig|EHA21666.1 ater|EAU37750.1

Group17547: anig|EHA21667.1 atub|OJI87921.1

Group17548: anig|EHA21756.1 ater|EAU30221.1

Group17549: anig|EHA21938.1 atub|OJI83786.1

Group17550: anig|EHA22022.1 atub|OJI83879.1

Group17551: anig|EHA22139.1 atub|OJI84021.1

Group17552: anig|EHA22429.1 atub|OJI84360.1

Group17553: anig|EHA22551.1 asyd|OJJ61683.1

Group17554: anig|EHA22998.1 atub|OJI86501.1

Group17555: anig|EHA23038.1 atub|OJI85235.1

Group17556: anig|EHA23473.1 ater|EAU36242.1

Group17557: anig|EHA23493.1 atub|OJI85746.1

Group17558: anig|EHA23654.1 atub|OJI89949.1

Group17559: anig|EHA23993.1 asyd|OJJ56936.1

Group17560: anig|EHA24193.1 atub|OJI82211.1

Group17561: anig|EHA24245.1 atub|OJI82276.1

Group17562: anig|EHA24778.1 atub|OJI79561.1

Group17563: anig|EHA24783.1 azon|OJJ51811.1

Group17564: anig|EHA24863.1 atub|OJI79468.1

Group17565: ater|EAU36850.1 aver|OJJ07234.1

Group17566: anig|EHA25153.1 atub|OJI80664.1

Group17567: anig|EHA25829.1 ater|EAU30173.1

Group17568: anig|EHA25862.1 atub|OJI88604.1

Group17569: anig|EHA26152.1 azon|OJJ45940.1

Group17570: anig|EHA26571.1 ater|EAU31075.1

Group17571: anig|EHA26576.1 ater|EAU30921.1

Group17572: anig|EHA26577.1 ater|EAU38785.1

Group17573: anig|EHA26783.1 atub|OJI86597.1

Group17574: anig|EHA26854.1 atub|OJI85194.1

Group17575: anig|EHA26974.1 atub|OJI85050.1

Group17576: anig|EHA27361.1 atub|OJI84818.1

Group17577: anig|EHA27465.1 atub|OJI88849.1

Group17578: anig|EHA27584.1 atub|OJI89847.1

Group17579: anig|EHA28206.1 awen|OJJ32250.1

Group17580: asyd|OJJ62668.1 aver|OJI96766.1

Group17581: anig|EHA28359.1 atub|OJI88033.1

Group17582: aory|EIT72738.1 ater|EAU30545.1

Group17583: aory|EIT74344.1 awen|OJJ39516.1

Group17584: aory|EIT75337.1 asyd|OJJ60852.1

Group17585: aory|EIT77273.1 aver|OJJ06755.1

Group17586: aory|EIT77983.1 ater|EAU36271.1

Group17587: asyd|OJJ56701.1 aver|OJJ06024.1

Group17588: aory|EIT80754.1 awen|OJJ41599.1

Group17589: aory|EIT80933.1 atub|OJI88345.1

Group17590: aory|EIT81297.1 ater|EAU31887.1

Group17591: aory|EIT81834.1 ater|EAU36137.1

Group17592: asyd|OJJ53427.1 aver|OJJ08318.1

Group17593: aory|EIT83114.1 asyd|OJJ62486.1

Group17594: asyd|OJJ52076.1 aver|OJJ03027.1

Group17595: asyd|OJJ52078.1 aver|OJJ03026.1

Group17596: asyd|OJJ52089.1 aver|OJJ03478.1

Group17597: asyd|OJJ52105.1 aver|OJI97595.1

Group17598: asyd|OJJ52148.1 ater|EAU30127.1

Group17599: asyd|OJJ52191.1 aver|OJJ05955.1

Group17600: asyd|OJJ52345.1 aver|OJJ06549.1

Group17601: asyd|OJJ52347.1 aver|OJI99660.1

Group17602: asyd|OJJ52348.1 aver|OJJ05332.1

Group17603: asyd|OJJ52352.1 aver|OJJ01994.1

Group17604: asyd|OJJ52356.1 aver|OJJ01993.1

Group17605: asyd|OJJ52357.1 aver|OJJ01992.1

Group17606: asyd|OJJ52379.1 aver|OJJ01968.1

Group17607: asyd|OJJ52380.1 aver|OJJ01967.1

Group17608: asyd|OJJ52381.1 aver|OJJ01966.1

Group17609: asyd|OJJ52383.1 aver|OJJ01940.1

Group17610: asyd|OJJ52388.1 aver|OJJ01935.1

Group17611: asyd|OJJ52392.1 aver|OJJ01931.1

Group17612: asyd|OJJ52393.1 aver|OJJ01930.1

Group17613: asyd|OJJ52406.1 aver|OJI98316.1

Group17614: asyd|OJJ52413.1 aver|OJJ00342.1

Group17615: asyd|OJJ52415.1 aver|OJJ08922.1

Group17616: asyd|OJJ52428.1 aver|OJJ05543.1

Group17617: asyd|OJJ52430.1 aver|OJJ05541.1

Group17618: asyd|OJJ52445.1 aver|OJJ03971.1

Group17619: asyd|OJJ52454.1 aver|OJJ03961.1

Group17620: asyd|OJJ52472.1 aver|OJJ03944.1

Group17621: asyd|OJJ52478.1 aver|OJJ03934.1

Group17622: asyd|OJJ52483.1 aver|OJJ03924.1

Group17623: asyd|OJJ52484.1 aver|OJJ03923.1

Group17624: asyd|OJJ52488.1 aver|OJJ03915.1

Group17625: asyd|OJJ52490.1 aver|OJJ03914.1

Group17626: asyd|OJJ52495.1 aver|OJJ06531.1

Group17627: asyd|OJJ52496.1 aver|OJJ03888.1

Group17628: asyd|OJJ52497.1 aver|OJI99615.1

Group17629: asyd|OJJ52499.1 aver|OJJ03884.1

Group17630: asyd|OJJ52515.1 aver|OJI99667.1

Group17631: asyd|OJJ52528.1 aver|OJJ03872.1

Group17632: asyd|OJJ52541.1 aver|OJJ03863.1

Group17633: asyd|OJJ52542.1 aver|OJJ03862.1

Group17634: asyd|OJJ52550.1 aver|OJJ03850.1

Group17635: asyd|OJJ52552.1 aver|OJJ03848.1

Group17636: asyd|OJJ52719.1 ater|EAU38047.1

Group17637: asyd|OJJ52743.1 aver|OJJ02999.1

Group17638: asyd|OJJ52750.1 aver|OJJ04009.1

Group17639: asyd|OJJ52760.1 aver|OJJ08742.1

Group17640: asyd|OJJ52767.1 aver|OJJ08750.1

Group17641: asyd|OJJ52772.1 aver|OJJ08755.1

Group17642: asyd|OJJ52773.1 aver|OJJ08756.1

Group17643: asyd|OJJ52782.1 aver|OJJ08760.1

Group17644: asyd|OJJ52790.1 aver|OJJ08765.1

Group17645: asyd|OJJ52795.1 aver|OJJ08773.1

Group17646: asyd|OJJ52806.1 aver|OJJ08786.1

Group17647: asyd|OJJ52813.1 aver|OJJ08792.1

Group17648: asyd|OJJ52816.1 aver|OJJ08795.1

Group17649: asyd|OJJ52832.1 aver|OJJ08807.1

Group17650: asyd|OJJ52846.1 aver|OJJ08824.1

Group17651: asyd|OJJ52852.1 aver|OJJ08830.1

Group17652: asyd|OJJ52853.1 aver|OJJ08831.1

Group17653: asyd|OJJ52863.1 aver|OJJ08842.1

Group17654: asyd|OJJ52870.1 aver|OJI98021.1

Group17655: asyd|OJJ52884.1 aver|OJJ08855.1

Group17656: asyd|OJJ52885.1 aver|OJJ08856.1

Group17657: asyd|OJJ52886.1 aver|OJJ08857.1

Group17658: asyd|OJJ52894.1 aver|OJJ08865.1

Group17659: asyd|OJJ52900.1 aver|OJJ08869.1

Group17660: asyd|OJJ52908.1 aver|OJJ08877.1

Group17661: asyd|OJJ52909.1 aver|OJJ08878.1

Group17662: asyd|OJJ52910.1 aver|OJJ08879.1

Group17663: asyd|OJJ52911.1 aver|OJJ08880.1

Group17664: asyd|OJJ52917.1 aver|OJJ08887.1

Group17665: asyd|OJJ52922.1 aver|OJJ08891.1

Group17666: asyd|OJJ52923.1 aver|OJJ08892.1

Group17667: asyd|OJJ52947.1 aver|OJJ08914.1

Group17668: asyd|OJJ52955.1 aver|OJJ08921.1

Group17669: asyd|OJJ52957.1 aver|OJJ08923.1

Group17670: asyd|OJJ52971.1 aver|OJJ08938.1

Group17671: asyd|OJJ52972.1 aver|OJJ08939.1

Group17672: asyd|OJJ52974.1 aver|OJJ08940.1

Group17673: asyd|OJJ53040.1 aver|OJJ03804.1

Group17674: asyd|OJJ53069.1 aver|OJJ03776.1

Group17675: asyd|OJJ53113.1 aver|OJJ03742.1

Group17676: asyd|OJJ53346.1 aver|OJJ08399.1

Group17677: asyd|OJJ53351.1 aver|OJJ08393.1

Group17678: asyd|OJJ53354.1 aver|OJJ08389.1

Group17679: asyd|OJJ53355.1 aver|OJJ08386.1

Group17680: asyd|OJJ53359.1 aver|OJJ07645.1

Group17681: asyd|OJJ53377.1 aver|OJJ08366.1

Group17682: asyd|OJJ53381.1 aver|OJJ08362.1

Group17683: asyd|OJJ53412.1 aver|OJJ08339.1

Group17684: asyd|OJJ53413.1 aver|OJJ08338.1

Group17685: asyd|OJJ53415.1 aver|OJJ08336.1

Group17686: asyd|OJJ53416.1 aver|OJJ08335.1

Group17687: asyd|OJJ53417.1 aver|OJJ08334.1

Group17688: asyd|OJJ53418.1 aver|OJJ08333.1

Group17689: asyd|OJJ53420.1 aver|OJJ08331.1

Group17690: asyd|OJJ53421.1 aver|OJJ08330.1

Group17691: asyd|OJJ53425.1 aver|OJJ08320.1

Group17692: asyd|OJJ53430.1 aver|OJJ08315.1

Group17693: asyd|OJJ53440.1 aver|OJJ08311.1

Group17694: asyd|OJJ53442.1 aver|OJJ08309.1

Group17695: asyd|OJJ53450.1 aver|OJJ08302.1

Group17696: asyd|OJJ53460.1 aver|OJJ08295.1

Group17697: asyd|OJJ53489.1 aver|OJJ03353.1

Group17698: asyd|OJJ53491.1 aver|OJJ03351.1

Group17699: asyd|OJJ53495.1 aver|OJJ08253.1

Group17700: asyd|OJJ53497.1 aver|OJJ08248.1

Group17701: asyd|OJJ53500.1 aver|OJJ08245.1

Group17702: asyd|OJJ53504.1 aver|OJJ08242.1

Group17703: asyd|OJJ53506.1 aver|OJJ08240.1

Group17704: asyd|OJJ53520.1 aver|OJJ08230.1

Group17705: asyd|OJJ53521.1 aver|OJJ08229.1

Group17706: asyd|OJJ53522.1 aver|OJJ08228.1

Group17707: asyd|OJJ53524.1 aver|OJJ08226.1

Group17708: asyd|OJJ53533.1 awen|OJJ33416.1

Group17709: asyd|OJJ53550.1 aver|OJJ06170.1

Group17710: asyd|OJJ53554.1 aver|OJJ08202.1

Group17711: asyd|OJJ53556.1 aver|OJJ08199.1

Group17712: asyd|OJJ53568.1 aver|OJJ08187.1

Group17713: asyd|OJJ53595.1 aver|OJJ08153.1

Group17714: asyd|OJJ53600.1 aver|OJJ08148.1

Group17715: asyd|OJJ53607.1 aver|OJJ08141.1

Group17716: asyd|OJJ53608.1 aver|OJJ08140.1

Group17717: asyd|OJJ53620.1 awen|OJJ34583.1

Group17718: asyd|OJJ53621.1 aver|OJJ05938.1

Group17719: asyd|OJJ53629.1 aver|OJJ05946.1

Group17720: asyd|OJJ53686.1 aver|OJJ07804.1

Group17721: asyd|OJJ53709.1 aver|OJJ07835.1

Group17722: asyd|OJJ53710.1 aver|OJJ07836.1

Group17723: asyd|OJJ53726.1 aver|OJI98281.1

Group17724: asyd|OJJ53728.1 aver|OJI98283.1

Group17725: asyd|OJJ53746.1 aver|OJJ07861.1

Group17726: asyd|OJJ53755.1 aver|OJJ07866.1

Group17727: asyd|OJJ53756.1 aver|OJJ07867.1

Group17728: asyd|OJJ53757.1 aver|OJJ07868.1

Group17729: asyd|OJJ53766.1 aver|OJJ07874.1

Group17730: asyd|OJJ53767.1 aver|OJJ07875.1

Group17731: asyd|OJJ53768.1 aver|OJJ07876.1

Group17732: asyd|OJJ53792.1 aver|OJJ03441.1

Group17733: asyd|OJJ53797.1 aver|OJI96897.1

Group17734: asyd|OJJ53807.1 aver|OJJ07907.1

Group17735: asyd|OJJ53818.1 aver|OJJ07533.1

Group17736: asyd|OJJ53826.1 aver|OJJ07927.1

Group17737: asyd|OJJ53827.1 aver|OJJ07931.1

Group17738: asyd|OJJ53828.1 aver|OJJ07932.1

Group17739: asyd|OJJ53829.1 aver|OJJ07933.1

Group17740: asyd|OJJ53837.1 aver|OJJ07940.1

Group17741: asyd|OJJ53854.1 aver|OJJ07957.1

Group17742: asyd|OJJ53929.1 aver|OJJ08028.1

Group17743: asyd|OJJ53934.1 aver|OJJ08032.1

Group17744: asyd|OJJ53943.1 aver|OJJ08040.1

Group17745: awen|OJJ37049.1 ater|EAU39531.1

Group17746: asyd|OJJ54091.1 azon|OJJ46447.1

Group17747: asyd|OJJ54093.1 aver|OJJ08646.1

Group17748: asyd|OJJ54123.1 aver|OJJ08614.1

Group17749: asyd|OJJ54131.1 aver|OJJ08604.1

Group17750: asyd|OJJ54132.1 aver|OJJ08603.1

Group17751: asyd|OJJ54170.1 aver|OJJ08569.1

Group17752: asyd|OJJ54184.1 aver|OJJ08556.1

Group17753: asyd|OJJ54190.1 aver|OJJ08549.1

Group17754: asyd|OJJ54202.1 aver|OJJ08522.1

Group17755: asyd|OJJ54212.1 aver|OJJ08519.1

Group17756: asyd|OJJ54216.1 aver|OJJ08515.1

Group17757: asyd|OJJ54217.1 aver|OJJ08514.1

Group17758: asyd|OJJ54223.1 aver|OJJ05176.1

Group17759: asyd|OJJ54228.1 aver|OJJ05181.1

Group17760: asyd|OJJ54309.1 aver|OJJ05257.1

Group17761: asyd|OJJ54319.1 aver|OJJ05273.1

Group17762: asyd|OJJ54338.1 aver|OJJ05290.1

Group17763: asyd|OJJ54341.1 aver|OJJ05293.1

Group17764: asyd|OJJ54342.1 aver|OJJ05294.1

Group17765: asyd|OJJ54347.1 aver|OJJ05299.1

Group17766: asyd|OJJ54349.1 aver|OJJ05301.1

Group17767: asyd|OJJ54356.1 aver|OJJ05310.1

Group17768: asyd|OJJ54363.1 aver|OJJ05316.1

Group17769: asyd|OJJ54364.1 aver|OJJ05317.1

Group17770: asyd|OJJ54365.1 aver|OJJ05318.1

Group17771: asyd|OJJ54371.1 aver|OJJ05326.1

Group17772: asyd|OJJ54373.1 aver|OJJ05327.1

Group17773: asyd|OJJ54374.1 aver|OJJ05328.1

Group17774: asyd|OJJ54414.1 aver|OJJ05557.1

Group17775: asyd|OJJ54416.1 aver|OJJ05556.1

Group17776: asyd|OJJ54417.1 aver|OJJ05555.1

Group17777: asyd|OJJ54431.1 ater|EAU31020.1

Group17778: asyd|OJJ54432.1 awen|OJJ39736.1

Group17779: asyd|OJJ54437.1 aver|OJJ04033.1

Group17780: asyd|OJJ54452.1 azon|OJJ50686.1

Group17781: asyd|OJJ54453.1 aver|OJJ07702.1

Group17782: asyd|OJJ54454.1 aver|OJJ07701.1

Group17783: asyd|OJJ54484.1 aver|OJJ07695.1

Group17784: asyd|OJJ54489.1 aver|OJJ07690.1

Group17785: asyd|OJJ54498.1 aver|OJJ07682.1

Group17786: asyd|OJJ54505.1 aver|OJJ07677.1

Group17787: asyd|OJJ54514.1 aver|OJJ07670.1

Group17788: asyd|OJJ54519.1 aver|OJJ07665.1

Group17789: asyd|OJJ54526.1 aver|OJJ05306.1

Group17790: asyd|OJJ54559.1 aver|OJJ07604.1

Group17791: asyd|OJJ54561.1 aver|OJJ07595.1

Group17792: ater|EAU37339.1 aver|OJI99597.1

Group17793: asyd|OJJ54575.1 aver|OJJ07579.1

Group17794: asyd|OJJ54593.1 aver|OJJ07559.1

Group17795: asyd|OJJ54595.1 aver|OJJ07557.1

Group17796: asyd|OJJ54604.1 aver|OJJ07551.1

Group17797: asyd|OJJ54605.1 aver|OJJ07550.1

Group17798: asyd|OJJ54617.1 aver|OJJ07542.1

Group17799: asyd|OJJ54623.1 aver|OJJ07535.1

Group17800: asyd|OJJ54625.1 aver|OJJ07531.1

Group17801: asyd|OJJ54636.1 aver|OJJ07520.1

Group17802: asyd|OJJ54637.1 aver|OJJ07519.1

Group17803: asyd|OJJ54679.1 aver|OJJ07488.1

Group17804: asyd|OJJ54709.1 aver|OJJ07457.1

Group17805: asyd|OJJ54717.1 aver|OJJ07450.1

Group17806: asyd|OJJ54721.1 aver|OJJ07447.1

Group17807: asyd|OJJ54723.1 aver|OJJ07443.1

Group17808: asyd|OJJ54746.1 aver|OJJ07419.1

Group17809: asyd|OJJ54763.1 aver|OJJ07404.1

Group17810: asyd|OJJ54777.1 aver|OJJ07388.1

Group17811: asyd|OJJ54803.1 aver|OJI95781.1

Group17812: asyd|OJJ54817.1 aver|OJJ07344.1

Group17813: asyd|OJJ54820.1 aver|OJJ07341.1

Group17814: asyd|OJJ54843.1 aver|OJJ07319.1

Group17815: asyd|OJJ54846.1 aver|OJJ07636.1

Group17816: asyd|OJJ54873.1 aver|OJJ03889.1

Group17817: asyd|OJJ54874.1 aver|OJJ03890.1

Group17818: asyd|OJJ54875.1 aver|OJJ03891.1

Group17819: asyd|OJJ54882.1 ater|EAU37340.1

Group17820: asyd|OJJ54906.1 aver|OJJ07295.1

Group17821: asyd|OJJ54933.1 aver|OJJ07265.1

Group17822: asyd|OJJ54964.1 aver|OJJ07227.1

Group17823: asyd|OJJ54965.1 aver|OJJ07226.1

Group17824: asyd|OJJ54966.1 aver|OJJ07225.1

Group17825: asyd|OJJ54968.1 aver|OJJ07223.1

Group17826: asyd|OJJ54969.1 aver|OJJ07222.1

Group17827: asyd|OJJ54990.1 aver|OJJ07211.1

Group17828: asyd|OJJ55005.1 aver|OJJ07202.1

Group17829: asyd|OJJ55044.1 aver|OJJ07163.1

Group17830: asyd|OJJ55053.1 aver|OJJ07156.1

Group17831: asyd|OJJ55056.1 aver|OJJ07154.1

Group17832: asyd|OJJ55058.1 aver|OJJ07607.1

Group17833: asyd|OJJ55059.1 aver|OJJ07153.1

Group17834: asyd|OJJ55061.1 aver|OJJ07150.1

Group17835: asyd|OJJ55063.1 aver|OJJ07148.1

Group17836: asyd|OJJ55070.1 aver|OJJ07141.1

Group17837: asyd|OJJ55162.1 aver|OJJ07045.1

Group17838: asyd|OJJ55177.1 aver|OJJ07030.1

Group17839: asyd|OJJ55195.1 aver|OJJ07018.1

Group17840: asyd|OJJ55211.1 aver|OJJ07006.1

Group17841: asyd|OJJ55273.1 aver|OJJ06954.1

Group17842: asyd|OJJ55274.1 aver|OJJ06953.1

Group17843: asyd|OJJ55341.1 aver|OJI98025.1

Group17844: asyd|OJJ55352.1 aver|OJJ06441.1

Group17845: asyd|OJJ55356.1 atub|OJI80374.1

Group17846: asyd|OJJ55357.1 aver|OJJ01942.1

Group17847: asyd|OJJ55400.1 aver|OJJ06480.1

Group17848: asyd|OJJ55401.1 aver|OJJ06481.1

Group17849: asyd|OJJ55403.1 aver|OJJ06483.1

Group17850: asyd|OJJ55404.1 aver|OJJ06484.1

Group17851: asyd|OJJ55407.1 aver|OJJ03918.1

Group17852: asyd|OJJ55416.1 aver|OJJ06493.1

Group17853: asyd|OJJ55421.1 aver|OJJ06498.1

Group17854: asyd|OJJ55443.1 aver|OJJ06516.1

Group17855: asyd|OJJ55461.1 aver|OJJ06527.1

Group17856: asyd|OJJ55473.1 aver|OJJ08258.1

Group17857: asyd|OJJ55483.1 aver|OJJ06562.1

Group17858: asyd|OJJ55502.1 aver|OJJ06580.1

Group17859: asyd|OJJ55534.1 aver|OJJ06612.1

Group17860: asyd|OJJ55557.1 aver|OJJ06638.1

Group17861: asyd|OJJ55563.1 aver|OJJ06644.1

Group17862: asyd|OJJ55571.1 aver|OJJ06650.1

Group17863: asyd|OJJ55605.1 aver|OJJ06682.1

Group17864: asyd|OJJ55613.1 aver|OJJ06688.1

Group17865: asyd|OJJ55619.1 aver|OJJ06694.1

Group17866: asyd|OJJ55620.1 aver|OJJ06695.1

Group17867: asyd|OJJ55621.1 aver|OJJ06696.1

Group17868: asyd|OJJ55636.1 aver|OJJ06715.1

Group17869: asyd|OJJ55661.1 aver|OJJ06744.1

Group17870: asyd|OJJ55662.1 aver|OJJ06745.1

Group17871: asyd|OJJ55677.1 aver|OJJ06760.1

Group17872: asyd|OJJ55712.1 aver|OJJ06798.1

Group17873: asyd|OJJ55806.1 aver|OJJ04754.1

Group17874: asyd|OJJ55812.1 aver|OJJ04760.1

Group17875: asyd|OJJ55848.1 aver|OJJ04791.1

Group17876: asyd|OJJ55865.1 aver|OJJ04808.1

Group17877: asyd|OJJ55884.1 aver|OJJ04829.1

Group17878: asyd|OJJ55897.1 aver|OJJ04842.1

Group17879: asyd|OJJ55917.1 aver|OJJ04857.1

Group17880: asyd|OJJ55927.1 aver|OJI99310.1

Group17881: asyd|OJJ55931.1 aver|OJJ04880.1

Group17882: asyd|OJJ55932.1 aver|OJJ04881.1

Group17883: asyd|OJJ55953.1 aver|OJJ04899.1

Group17884: asyd|OJJ55954.1 aver|OJJ04900.1

Group17885: asyd|OJJ56038.1 aver|OJJ04984.1

Group17886: asyd|OJJ56042.1 aver|OJJ04988.1

Group17887: asyd|OJJ56053.1 aver|OJJ04994.1

Group17888: asyd|OJJ56061.1 aver|OJJ05003.1

Group17889: asyd|OJJ56068.1 aver|OJJ05011.1

Group17890: asyd|OJJ56077.1 aver|OJJ05020.1

Group17891: asyd|OJJ56080.1 aver|OJJ05023.1

Group17892: asyd|OJJ56088.1 aver|OJJ05031.1

Group17893: asyd|OJJ56091.1 aver|OJJ05034.1

Group17894: asyd|OJJ56099.1 aver|OJJ05042.1

Group17895: asyd|OJJ56108.1 aver|OJJ05051.1

Group17896: asyd|OJJ56112.1 aver|OJJ05055.1

Group17897: asyd|OJJ56114.1 aver|OJJ07446.1

Group17898: asyd|OJJ56134.1 aver|OJJ05074.1

Group17899: asyd|OJJ56135.1 aver|OJJ05075.1

Group17900: asyd|OJJ56136.1 aver|OJJ05076.1

Group17901: asyd|OJJ56137.1 aver|OJJ05077.1

Group17902: asyd|OJJ56150.1 aver|OJJ05089.1

Group17903: asyd|OJJ56151.1 aver|OJJ05090.1

Group17904: asyd|OJJ56171.1 aver|OJJ05104.1

Group17905: asyd|OJJ56172.1 aver|OJJ05105.1

Group17906: asyd|OJJ56174.1 aver|OJJ05106.1

Group17907: asyd|OJJ56176.1 aver|OJJ05107.1

Group17908: asyd|OJJ56177.1 aver|OJJ05109.1

Group17909: asyd|OJJ56182.1 aver|OJJ07199.1

Group17910: asyd|OJJ56183.1 aver|OJJ05111.1

Group17911: asyd|OJJ56184.1 aver|OJJ05113.1

Group17912: asyd|OJJ56188.1 aver|OJJ05116.1

Group17913: asyd|OJJ56201.1 aver|OJJ05128.1

Group17914: asyd|OJJ56206.1 aver|OJJ05137.1

Group17915: asyd|OJJ56210.1 aver|OJJ05139.1

Group17916: asyd|OJJ56218.1 aver|OJJ05150.1

Group17917: asyd|OJJ56221.1 aver|OJJ05153.1

Group17918: asyd|OJJ56227.1 azon|OJJ51831.1

Group17919: asyd|OJJ56245.1 aver|OJJ05168.1

Group17920: asyd|OJJ56247.1 aver|OJJ08513.1

Group17921: asyd|OJJ56248.1 aver|OJJ08512.1

Group17922: asyd|OJJ56250.1 aver|OJJ08510.1

Group17923: asyd|OJJ56251.1 aver|OJJ08509.1

Group17924: asyd|OJJ56259.1 aver|OJJ08505.1

Group17925: asyd|OJJ56260.1 aver|OJJ08504.1

Group17926: asyd|OJJ56270.1 aver|OJJ08495.1

Group17927: asyd|OJJ56390.1 aver|OJJ06323.1

Group17928: asyd|OJJ56410.1 awen|OJJ34673.1

Group17929: asyd|OJJ56412.1 aver|OJJ06303.1

Group17930: asyd|OJJ56419.1 aver|OJJ06295.1

Group17931: asyd|OJJ56424.1 aver|OJJ06293.1

Group17932: asyd|OJJ56433.1 aver|OJJ06287.1

Group17933: asyd|OJJ56471.1 aver|OJJ06248.1

Group17934: asyd|OJJ56472.1 aver|OJJ06247.1

Group17935: asyd|OJJ56511.1 aver|OJJ06208.1

Group17936: asyd|OJJ56513.1 aver|OJJ06207.1

Group17937: asyd|OJJ56527.1 aver|OJJ06196.1

Group17938: asyd|OJJ56539.1 aver|OJJ06183.1

Group17939: asyd|OJJ56541.1 aver|OJJ06181.1

Group17940: asyd|OJJ56547.1 aver|OJJ06175.1

Group17941: asyd|OJJ56629.1 aver|OJJ06097.1

Group17942: asyd|OJJ56637.1 aver|OJJ06089.1

Group17943: asyd|OJJ56668.1 aver|OJJ06055.1

Group17944: asyd|OJJ56683.1 aver|OJJ06040.1

Group17945: asyd|OJJ56703.1 aver|OJJ06022.1

Group17946: asyd|OJJ56705.1 aver|OJJ06020.1

Group17947: asyd|OJJ56706.1 aver|OJJ06019.1

Group17948: asyd|OJJ56711.1 aver|OJJ06014.1

Group17949: asyd|OJJ56727.1 aver|OJJ05993.1

Group17950: asyd|OJJ56728.1 aver|OJJ05992.1

Group17951: asyd|OJJ56733.1 aver|OJJ05990.1

Group17952: asyd|OJJ56743.1 aver|OJJ08349.1

Group17953: asyd|OJJ56755.1 aver|OJJ05968.1

Group17954: asyd|OJJ56800.1 aver|OJJ03534.1

Group17955: asyd|OJJ56846.1 aver|OJJ04250.1

Group17956: asyd|OJJ56850.1 aver|OJJ03464.1

Group17957: asyd|OJJ56872.1 aver|OJI96732.1

Group17958: asyd|OJJ56887.1 awen|OJJ30429.1

Group17959: asyd|OJJ56894.1 aver|OJJ07734.1

Group17960: asyd|OJJ56905.1 aver|OJJ03388.1

Group17961: asyd|OJJ56918.1 aver|OJJ03374.1

Group17962: asyd|OJJ56919.1 aver|OJJ03373.1

Group17963: asyd|OJJ56924.1 aver|OJJ03370.1

Group17964: asyd|OJJ56925.1 aver|OJJ03369.1

Group17965: asyd|OJJ56926.1 aver|OJJ03368.1

Group17966: asyd|OJJ56935.1 aver|OJJ03349.1

Group17967: asyd|OJJ56957.1 aver|OJJ03328.1

Group17968: asyd|OJJ57003.1 aver|OJJ03284.1

Group17969: asyd|OJJ57007.1 aver|OJJ03280.1

Group17970: asyd|OJJ57117.1 aver|OJJ03183.1

Group17971: asyd|OJJ57123.1 aver|OJJ03181.1

Group17972: asyd|OJJ57135.1 aver|OJJ03168.1

Group17973: asyd|OJJ57150.1 aver|OJJ03159.1

Group17974: asyd|OJJ57153.1 aver|OJJ07869.1

Group17975: asyd|OJJ57173.1 aver|OJJ03143.1

Group17976: asyd|OJJ57186.1 aver|OJJ03104.1

Group17977: asyd|OJJ57192.1 aver|OJJ03099.1

Group17978: asyd|OJJ57217.1 aver|OJJ07885.1

Group17979: asyd|OJJ57222.1 aver|OJJ03065.1

Group17980: asyd|OJJ57231.1 aver|OJJ03058.1

Group17981: asyd|OJJ57235.1 aver|OJJ03050.1

Group17982: asyd|OJJ57239.1 azon|OJJ42367.1

Group17983: asyd|OJJ57243.1 azon|OJJ42363.1

Group17984: asyd|OJJ57250.1 aver|OJJ03044.1

Group17985: asyd|OJJ57253.1 aver|OJJ03041.1

Group17986: asyd|OJJ57254.1 aver|OJJ03040.1

Group17987: asyd|OJJ57265.1 aver|OJJ03396.1

Group17988: asyd|OJJ57267.1 aver|OJJ03394.1

Group17989: asyd|OJJ57269.1 aver|OJI96734.1

Group17990: asyd|OJJ57270.1 aver|OJI96733.1

Group17991: asyd|OJJ57279.1 aver|OJJ05877.1

Group17992: asyd|OJJ57286.1 aver|OJJ05870.1

Group17993: asyd|OJJ57289.1 aver|OJJ05868.1

Group17994: asyd|OJJ57291.1 aver|OJJ05866.1

Group17995: asyd|OJJ57296.1 aver|OJJ05854.1

Group17996: asyd|OJJ57307.1 aver|OJI99611.1

Group17997: asyd|OJJ57353.1 aver|OJJ05812.1

Group17998: asyd|OJJ57376.1 aver|OJJ05792.1

Group17999: asyd|OJJ57377.1 aver|OJJ05781.1

Group18000: asyd|OJJ57382.1 aver|OJJ05775.1

Group18001: asyd|OJJ57421.1 aver|OJJ05757.1

Group18002: asyd|OJJ57422.1 aver|OJJ05756.1

Group18003: asyd|OJJ57423.1 aver|OJJ05754.1

Group18004: asyd|OJJ57434.1 aver|OJJ05744.1

Group18005: asyd|OJJ57440.1 aver|OJJ05738.1

Group18006: asyd|OJJ57566.1 aver|OJJ05613.1

Group18007: asyd|OJJ57570.1 aver|OJJ05609.1

Group18008: asyd|OJJ57573.1 aver|OJJ05606.1

Group18009: asyd|OJJ57575.1 aver|OJJ05605.1

Group18010: asyd|OJJ57576.1 aver|OJJ05604.1

Group18011: asyd|OJJ57578.1 aver|OJJ05602.1

Group18012: asyd|OJJ57591.1 aver|OJJ05591.1

Group18013: asyd|OJJ57593.1 aver|OJJ05590.1

Group18014: asyd|OJJ57595.1 aver|OJJ05588.1

Group18015: asyd|OJJ57597.1 aver|OJJ05586.1

Group18016: asyd|OJJ57598.1 aver|OJJ05584.1

Group18017: asyd|OJJ57604.1 aver|OJJ05579.1

Group18018: asyd|OJJ57617.1 aver|OJJ05569.1

Group18019: asyd|OJJ57619.1 aver|OJJ05567.1

Group18020: asyd|OJJ57630.1 awen|OJJ33428.1

Group18021: asyd|OJJ57631.1 awen|OJJ33429.1

Group18022: asyd|OJJ57636.1 aver|OJJ01943.1

Group18023: asyd|OJJ57641.1 aver|OJJ05497.1

Group18024: asyd|OJJ57643.1 aver|OJJ05500.1

Group18025: asyd|OJJ57674.1 aver|OJJ05480.1

Group18026: asyd|OJJ57682.1 aver|OJJ05488.1

Group18027: asyd|OJJ57711.1 aver|OJJ05454.1

Group18028: asyd|OJJ57738.1 aver|OJJ05429.1

Group18029: asyd|OJJ57909.1 aver|OJJ04483.1

Group18030: asyd|OJJ57941.1 aver|OJJ04455.1

Group18031: asyd|OJJ57976.1 aver|OJJ04425.1

Group18032: asyd|OJJ58040.1 ater|EAU38802.1

Group18033: asyd|OJJ58042.1 aver|OJJ04359.1

Group18034: asyd|OJJ58043.1 ater|EAU38804.1

Group18035: asyd|OJJ58109.1 aver|OJJ04295.1

Group18036: asyd|OJJ58144.1 aver|OJJ04259.1

Group18037: asyd|OJJ58163.1 aver|OJJ04239.1

Group18038: asyd|OJJ58196.1 aver|OJJ04211.1

Group18039: asyd|OJJ58223.1 aver|OJJ02199.1

Group18040: asyd|OJJ58236.1 aver|OJJ02189.1

Group18041: asyd|OJJ58253.1 aver|OJJ02170.1

Group18042: asyd|OJJ58311.1 aver|OJJ02100.1

Group18043: asyd|OJJ58312.1 aver|OJJ02099.1

Group18044: asyd|OJJ58333.1 aver|OJJ02077.1

Group18045: asyd|OJJ58335.1 aver|OJJ02075.1

Group18046: asyd|OJJ58336.1 aver|OJJ02074.1

Group18047: asyd|OJJ58359.1 aver|OJJ02048.1

Group18048: asyd|OJJ58363.1 aver|OJJ02043.1

Group18049: asyd|OJJ58366.1 aver|OJJ02040.1

Group18050: asyd|OJJ58447.1 aver|OJJ01049.1

Group18051: asyd|OJJ58509.1 aver|OJJ01110.1

Group18052: asyd|OJJ58554.1 aver|OJJ01152.1

Group18053: asyd|OJJ58557.1 aver|OJJ01156.1

Group18054: asyd|OJJ58572.1 aver|OJJ01175.1

Group18055: asyd|OJJ58573.1 aver|OJJ01176.1

Group18056: asyd|OJJ58591.1 aver|OJJ01193.1

Group18057: asyd|OJJ58602.1 aver|OJJ01204.1

Group18058: asyd|OJJ58663.1 aver|OJJ01269.1

Group18059: asyd|OJJ58684.1 aver|OJJ01296.1

Group18060: asyd|OJJ58688.1 aver|OJJ01300.1

Group18061: asyd|OJJ58693.1 aver|OJJ01305.1

Group18062: asyd|OJJ58701.1 aver|OJJ01313.1

Group18063: asyd|OJJ58711.1 aver|OJJ01324.1

Group18064: asyd|OJJ58856.1 aver|OJJ01457.1

Group18065: asyd|OJJ58940.1 aver|OJJ01536.1

Group18066: asyd|OJJ59083.1 aver|OJJ01676.1

Group18067: asyd|OJJ59086.1 aver|OJJ01678.1

Group18068: asyd|OJJ59095.1 aver|OJJ01685.1

Group18069: asyd|OJJ59102.1 aver|OJJ01693.1

Group18070: asyd|OJJ59103.1 aver|OJJ01694.1

Group18071: asyd|OJJ59113.1 aver|OJJ01705.1

Group18072: asyd|OJJ59116.1 aver|OJJ01706.1

Group18073: asyd|OJJ59153.1 aver|OJJ01743.1

Group18074: asyd|OJJ59179.1 aver|OJJ01799.1

Group18075: asyd|OJJ59195.1 aver|OJJ01811.1

Group18076: asyd|OJJ59212.1 aver|OJJ01822.1

Group18077: asyd|OJJ59213.1 aver|OJJ01823.1

Group18078: asyd|OJJ59225.1 aver|OJJ01835.1

Group18079: asyd|OJJ59256.1 aver|OJJ01867.1

Group18080: asyd|OJJ59283.1 aver|OJJ01902.1

Group18081: asyd|OJJ59284.1 aver|OJJ01903.1

Group18082: asyd|OJJ59436.1 aver|OJJ02835.1

Group18083: asyd|OJJ59482.1 aver|OJJ02792.1

Group18084: asyd|OJJ59509.1 aver|OJJ02765.1

Group18085: asyd|OJJ59574.1 aver|OJJ02694.1

Group18086: asyd|OJJ59575.1 aver|OJJ02693.1

Group18087: asyd|OJJ59618.1 aver|OJJ02652.1

Group18088: asyd|OJJ59623.1 aver|OJJ01913.1

Group18089: asyd|OJJ59805.1 aver|OJJ02485.1

Group18090: asyd|OJJ59806.1 aver|OJJ02484.1

Group18091: asyd|OJJ59807.1 aver|OJJ02482.1

Group18092: asyd|OJJ59811.1 aver|OJJ02477.1

Group18093: asyd|OJJ59812.1 aver|OJJ02476.1

Group18094: asyd|OJJ59827.1 aver|OJJ02460.1

Group18095: asyd|OJJ59902.1 aver|OJJ02384.1

Group18096: asyd|OJJ59903.1 aver|OJJ02383.1

Group18097: asyd|OJJ59948.1 aver|OJJ02341.1

Group18098: asyd|OJJ59954.1 aver|OJJ02335.1

Group18099: asyd|OJJ59963.1 aver|OJJ02327.1

Group18100: asyd|OJJ59964.1 aver|OJJ02326.1

Group18101: asyd|OJJ59965.1 aver|OJJ02325.1

Group18102: asyd|OJJ59987.1 aver|OJJ02304.1

Group18103: asyd|OJJ60000.1 aver|OJJ02291.1

Group18104: asyd|OJJ60044.1 aver|OJJ02243.1

Group18105: asyd|OJJ60046.1 aver|OJJ02241.1

Group18106: asyd|OJJ60071.1 aver|OJJ04190.1

Group18107: asyd|OJJ60078.1 aver|OJJ04165.1

Group18108: asyd|OJJ60092.1 aver|OJJ04179.1

Group18109: asyd|OJJ60198.1 aver|OJJ04067.1

Group18110: asyd|OJJ60204.1 atub|OJI90227.1

Group18111: asyd|OJJ60208.1 aver|OJJ07742.1

Group18112: asyd|OJJ60228.1 aver|OJJ04039.1

Group18113: asyd|OJJ60240.1 aver|OJJ04026.1

Group18114: asyd|OJJ60248.1 aver|OJJ08134.1

Group18115: asyd|OJJ60283.1 aver|OJI99613.1

Group18116: asyd|OJJ60286.1 aver|OJI99618.1

Group18117: asyd|OJJ60287.1 aver|OJI99619.1

Group18118: asyd|OJJ60295.1 aver|OJI99633.1

Group18119: asyd|OJJ60339.1 azon|OJJ47734.1

Group18120: asyd|OJJ60344.1 aver|OJI99685.1

Group18121: asyd|OJJ60347.1 aver|OJI99689.1

Group18122: asyd|OJJ60348.1 aver|OJI99645.1

Group18123: asyd|OJJ60353.1 aver|OJI99692.1

Group18124: asyd|OJJ60356.1 aver|OJI99694.1

Group18125: asyd|OJJ60374.1 ater|EAU33978.1

Group18126: asyd|OJJ60384.1 aver|OJI99712.1

Group18127: asyd|OJJ60385.1 aver|OJI99714.1

Group18128: asyd|OJJ60393.1 aver|OJI99722.1

Group18129: asyd|OJJ60394.1 aver|OJI99723.1

Group18130: asyd|OJJ60400.1 aver|OJI99729.1

Group18131: asyd|OJJ60401.1 aver|OJI99730.1

Group18132: asyd|OJJ60406.1 aver|OJI99736.1

Group18133: asyd|OJJ60408.1 aver|OJI99737.1

Group18134: asyd|OJJ60409.1 aver|OJI99738.1

Group18135: asyd|OJJ60417.1 aver|OJI99741.1

Group18136: asyd|OJJ60419.1 aver|OJI99743.1

Group18137: asyd|OJJ60422.1 aver|OJI99745.1

Group18138: asyd|OJJ60423.1 aver|OJI99747.1

Group18139: asyd|OJJ60431.1 aver|OJI99754.1

Group18140: asyd|OJJ60437.1 aver|OJJ02285.1

Group18141: asyd|OJJ60479.1 aver|OJI99799.1

Group18142: asyd|OJJ60482.1 aver|OJI99807.1

Group18143: asyd|OJJ60483.1 aver|OJI99808.1

Group18144: asyd|OJJ60484.1 aver|OJI99809.1

Group18145: asyd|OJJ60501.1 aver|OJI99819.1

Group18146: asyd|OJJ60505.1 aver|OJI99830.1

Group18147: asyd|OJJ60514.1 aver|OJI99840.1

Group18148: asyd|OJJ60566.1 aver|OJI99892.1

Group18149: asyd|OJJ60585.1 aver|OJI99908.1

Group18150: asyd|OJJ60604.1 aver|OJI99930.1

Group18151: asyd|OJJ60623.1 aver|OJI99947.1

Group18152: asyd|OJJ60667.1 aver|OJI99990.1

Group18153: asyd|OJJ60680.1 aver|OJJ00005.1

Group18154: asyd|OJJ60726.1 azon|OJJ48654.1

Group18155: asyd|OJJ60774.1 aver|OJJ00108.1

Group18156: asyd|OJJ60775.1 aver|OJJ00109.1

Group18157: asyd|OJJ60866.1 aver|OJJ00192.1

Group18158: asyd|OJJ60907.1 aver|OJJ00234.1

Group18159: asyd|OJJ60910.1 aver|OJJ00237.1

Group18160: asyd|OJJ60928.1 aver|OJJ00254.1

Group18161: asyd|OJJ60945.1 aver|OJJ00271.1

Group18162: asyd|OJJ60946.1 aver|OJJ00273.1

Group18163: asyd|OJJ60950.1 aver|OJJ00277.1

Group18164: asyd|OJJ60951.1 aver|OJJ00278.1

Group18165: asyd|OJJ60986.1 aver|OJJ00305.1

Group18166: asyd|OJJ61005.1 aver|OJJ00323.1

Group18167: asyd|OJJ61021.1 azon|OJJ45275.1

Group18168: asyd|OJJ61041.1 aver|OJJ00357.1

Group18169: asyd|OJJ61086.1 aver|OJJ00401.1

Group18170: asyd|OJJ61143.1 awen|OJJ39232.1

Group18171: asyd|OJJ61163.1 aver|OJJ00487.1

Group18172: asyd|OJJ61192.1 aver|OJJ00516.1

Group18173: asyd|OJJ61215.1 aver|OJJ00539.1

Group18174: asyd|OJJ61331.1 aver|OJJ00656.1

Group18175: asyd|OJJ61377.1 aver|OJJ00704.1

Group18176: asyd|OJJ61390.1 aver|OJJ00719.1

Group18177: asyd|OJJ61394.1 aver|OJJ00723.1

Group18178: asyd|OJJ61402.1 aver|OJJ00730.1

Group18179: asyd|OJJ61418.1 aver|OJJ00753.1

Group18180: asyd|OJJ61429.1 aver|OJJ00765.1

Group18181: asyd|OJJ61444.1 aver|OJJ00780.1

Group18182: asyd|OJJ61446.1 aver|OJJ00782.1

Group18183: asyd|OJJ61449.1 aver|OJJ00784.1

Group18184: asyd|OJJ61450.1 aver|OJJ00785.1

Group18185: asyd|OJJ61461.1 aver|OJJ00800.1

Group18186: asyd|OJJ61462.1 aver|OJJ00807.1

Group18187: asyd|OJJ61468.1 aver|OJJ00808.1

Group18188: asyd|OJJ61478.1 aver|OJJ00819.1

Group18189: asyd|OJJ61479.1 aver|OJJ00820.1

Group18190: asyd|OJJ61483.1 aver|OJJ00825.1

Group18191: asyd|OJJ61485.1 aver|OJJ00827.1

Group18192: asyd|OJJ61493.1 awen|OJJ41827.1

Group18193: asyd|OJJ61497.1 aver|OJJ00834.1

Group18194: asyd|OJJ61501.1 aver|OJJ00839.1

Group18195: asyd|OJJ61504.1 aver|OJJ00841.1

Group18196: asyd|OJJ61512.1 aver|OJJ00849.1

Group18197: asyd|OJJ61513.1 aver|OJJ00850.1

Group18198: asyd|OJJ61514.1 aver|OJJ00851.1

Group18199: asyd|OJJ61533.1 aver|OJJ00861.1

Group18200: awen|OJJ30110.1 ater|EAU31339.1

Group18201: asyd|OJJ61541.1 aver|OJJ00868.1

Group18202: asyd|OJJ61542.1 aver|OJJ00869.1

Group18203: asyd|OJJ61543.1 aver|OJJ00870.1

Group18204: asyd|OJJ61544.1 aver|OJJ00871.1

Group18205: asyd|OJJ61552.1 aver|OJJ00877.1

Group18206: asyd|OJJ61556.1 aver|OJJ00879.1

Group18207: asyd|OJJ61557.1 aver|OJJ00880.1

Group18208: asyd|OJJ61558.1 aver|OJJ00881.1

Group18209: asyd|OJJ61559.1 aver|OJJ00882.1

Group18210: asyd|OJJ61560.1 aver|OJJ00883.1

Group18211: asyd|OJJ61585.1 aver|OJJ00907.1

Group18212: asyd|OJJ61615.1 aver|OJJ00938.1

Group18213: asyd|OJJ61703.1 aver|OJI95792.1

Group18214: asyd|OJJ61707.1 aver|OJI95804.1

Group18215: asyd|OJJ61709.1 aver|OJI95805.1

Group18216: asyd|OJJ61740.1 aver|OJI95837.1

Group18217: asyd|OJJ61742.1 aver|OJI95839.1

Group18218: asyd|OJJ61768.1 aver|OJI95865.1

Group18219: asyd|OJJ61773.1 aver|OJI95875.1

Group18220: asyd|OJJ61846.1 aver|OJI95946.1

Group18221: asyd|OJJ61888.1 aver|OJI95985.1

Group18222: asyd|OJJ61902.1 aver|OJI95995.1

Group18223: asyd|OJJ62009.1 aver|OJI96101.1

Group18224: asyd|OJJ62096.1 aver|OJI96202.1

Group18225: asyd|OJJ62130.1 aver|OJI96239.1

Group18226: asyd|OJJ62142.1 aver|OJI96252.1

Group18227: asyd|OJJ62242.1 aver|OJI96352.1

Group18228: asyd|OJJ62267.1 aver|OJI96376.1

Group18229: asyd|OJJ62320.1 azon|OJJ45049.1

Group18230: asyd|OJJ62328.1 aver|OJI96434.1

Group18231: asyd|OJJ62416.1 aver|OJI96551.1

Group18232: asyd|OJJ62461.1 aver|OJI96596.1

Group18233: asyd|OJJ62478.1 aver|OJI96615.1

Group18234: asyd|OJJ62479.1 aver|OJI96616.1

Group18235: asyd|OJJ62548.1 aver|OJI96676.1

Group18236: asyd|OJJ62556.1 aver|OJI96681.1

Group18237: asyd|OJJ62558.1 aver|OJI96683.1

Group18238: asyd|OJJ62578.1 aver|OJI96699.1

Group18239: asyd|OJJ62617.1 aver|OJJ03452.1

Group18240: asyd|OJJ62618.1 aver|OJJ03453.1

Group18241: asyd|OJJ62656.1 aver|OJI96754.1

Group18242: asyd|OJJ62664.1 aver|OJI96762.1

Group18243: asyd|OJJ62686.1 aver|OJJ03115.1

Group18244: asyd|OJJ62719.1 awen|OJJ32592.1

Group18245: asyd|OJJ62779.1 aver|OJI96824.1

Group18246: asyd|OJJ62805.1 aver|OJI96851.1

Group18247: asyd|OJJ62900.1 aver|OJI96954.1

Group18248: asyd|OJJ62914.1 aver|OJI98075.1

Group18249: asyd|OJJ62926.1 aver|OJI98063.1

Group18250: asyd|OJJ62930.1 aver|OJI98060.1

Group18251: asyd|OJJ62938.1 aver|OJI98053.1

Group18252: asyd|OJJ62942.1 aver|OJI98048.1

Group18253: asyd|OJJ62959.1 aver|OJI98008.1

Group18254: asyd|OJJ62977.1 aver|OJI97992.1

Group18255: asyd|OJJ62983.1 aver|OJI97986.1

Group18256: asyd|OJJ63001.1 aver|OJI97965.1

Group18257: asyd|OJJ63005.1 aver|OJI97955.1

Group18258: asyd|OJJ63006.1 aver|OJI97954.1

Group18259: asyd|OJJ63011.1 ater|EAU33154.1

Group18260: asyd|OJJ63042.1 aver|OJI97926.1

Group18261: asyd|OJJ63132.1 aver|OJI97822.1

Group18262: asyd|OJJ63334.1 aver|OJI97628.1

Group18263: asyd|OJJ63351.1 aver|OJI97610.1

Group18264: asyd|OJJ63352.1 aver|OJI97609.1

Group18265: asyd|OJJ63357.1 aver|OJI97604.1

Group18266: asyd|OJJ63358.1 aver|OJI97597.1

Group18267: asyd|OJJ63415.1 aver|OJI97523.1

Group18268: asyd|OJJ63470.1 aver|OJI97462.1

Group18269: asyd|OJJ63510.1 aver|OJI97420.1

Group18270: asyd|OJJ63526.1 aver|OJI97405.1

Group18271: asyd|OJJ63527.1 aver|OJI97404.1

Group18272: asyd|OJJ63646.1 aver|OJI97288.1

Group18273: asyd|OJJ63688.1 aver|OJI97247.1

Group18274: asyd|OJJ63778.1 aver|OJI97159.1

Group18275: asyd|OJJ63815.1 aver|OJI97123.1

Group18276: asyd|OJJ63937.1 aver|OJI97005.1

Group18277: asyd|OJJ63948.1 aver|OJI96994.1

Group18278: asyd|OJJ63949.1 aver|OJI96992.1

Group18279: asyd|OJJ63971.1 ater|EAU31414.1

Group18280: asyd|OJJ64024.1 aver|OJI98110.1

Group18281: asyd|OJJ64029.1 aver|OJI98115.1

Group18282: asyd|OJJ64129.1 aver|OJI98212.1

Group18283: asyd|OJJ64138.1 aver|OJI98220.1

Group18284: asyd|OJJ64159.1 aver|OJI98236.1

Group18285: asyd|OJJ64161.1 aver|OJI98238.1

Group18286: asyd|OJJ64169.1 aver|OJI98244.1

Group18287: asyd|OJJ64170.1 aver|OJI98245.1

Group18288: asyd|OJJ64173.1 aver|OJI98248.1

Group18289: asyd|OJJ64184.1 aver|OJI98261.1

Group18290: asyd|OJJ64185.1 aver|OJI98262.1

Group18291: asyd|OJJ64217.1 azon|OJJ50408.1

Group18292: asyd|OJJ64220.1 aver|OJI98303.1

Group18293: asyd|OJJ64224.1 ater|EAU29575.1

Group18294: asyd|OJJ64234.1 aver|OJI98326.1

Group18295: asyd|OJJ64236.1 aver|OJI98329.1

Group18296: asyd|OJJ64245.1 aver|OJI98336.1

Group18297: asyd|OJJ64287.1 aver|OJI98379.1

Group18298: asyd|OJJ64292.1 aver|OJI98385.1

Group18299: asyd|OJJ64315.1 aver|OJI98408.1

Group18300: asyd|OJJ64321.1 aver|OJI98414.1

Group18301: asyd|OJJ64323.1 aver|OJI98416.1

Group18302: asyd|OJJ64324.1 aver|OJI98417.1

Group18303: asyd|OJJ64329.1 aver|OJI98422.1

Group18304: asyd|OJJ64338.1 aver|OJI98431.1

Group18305: asyd|OJJ64438.1 aver|OJI98539.1

Group18306: asyd|OJJ64568.1 aver|OJI98665.1

Group18307: asyd|OJJ64573.1 aver|OJI98670.1

Group18308: asyd|OJJ64579.1 aver|OJI98676.1

Group18309: asyd|OJJ64648.1 aver|OJI98748.1

Group18310: asyd|OJJ64679.1 aver|OJI98781.1

Group18311: asyd|OJJ64681.1 aver|OJI98783.1

Group18312: asyd|OJJ64711.1 aver|OJI98813.1

Group18313: asyd|OJJ64728.1 aver|OJI98832.1

Group18314: asyd|OJJ64746.1 aver|OJI98849.1

Group18315: asyd|OJJ64756.1 aver|OJI98858.1

Group18316: asyd|OJJ64762.1 aver|OJI98863.1

Group18317: asyd|OJJ64764.1 aver|OJI98864.1

Group18318: asyd|OJJ64769.1 aver|OJI98869.1

Group18319: asyd|OJJ64789.1 aver|OJI98901.1

Group18320: asyd|OJJ64804.1 aver|OJI98918.1

Group18321: asyd|OJJ64805.1 aver|OJI98919.1

Group18322: asyd|OJJ64823.1 aver|OJI98936.1

Group18323: asyd|OJJ64845.1 aver|OJI98956.1

Group18324: asyd|OJJ64964.1 aver|OJI99087.1

Group18325: asyd|OJJ64965.1 aver|OJI99088.1

Group18326: asyd|OJJ65066.1 aver|OJI99190.1

Group18327: asyd|OJJ65077.1 aver|OJI99202.1

Group18328: asyd|OJJ65087.1 aver|OJI99213.1

Group18329: asyd|OJJ65125.1 aver|OJI99253.1

Group18330: asyd|OJJ65142.1 awen|OJJ30555.1

Group18331: asyd|OJJ65161.1 aver|OJI99312.1

Group18332: asyd|OJJ65168.1 aver|OJI99320.1

Group18333: asyd|OJJ65173.1 aver|OJI99326.1

Group18334: asyd|OJJ65174.1 aver|OJI99327.1

Group18335: asyd|OJJ65175.1 aver|OJI99328.1

Group18336: asyd|OJJ65176.1 aver|OJI99329.1

Group18337: asyd|OJJ65180.1 aver|OJI99333.1

Group18338: asyd|OJJ65248.1 aver|OJI99400.1

Group18339: asyd|OJJ65250.1 aver|OJI99402.1

Group18340: asyd|OJJ65251.1 aver|OJI99403.1

Group18341: asyd|OJJ65253.1 aver|OJI99404.1

Group18342: asyd|OJJ65254.1 aver|OJI99405.1

Group18343: asyd|OJJ65255.1 aver|OJI99406.1

Group18344: asyd|OJJ65256.1 aver|OJI99407.1

Group18345: asyd|OJJ65266.1 aver|OJI99418.1

Group18346: asyd|OJJ65277.1 awen|OJJ37825.1

Group18347: asyd|OJJ65278.1 ater|EAU37989.1

Group18348: asyd|OJJ65280.1 aver|OJI99430.1

Group18349: asyd|OJJ65281.1 aver|OJI99432.1

Group18350: asyd|OJJ65282.1 aver|OJI99433.1

Group18351: asyd|OJJ65300.1 aver|OJI99440.1

Group18352: asyd|OJJ65319.1 aver|OJI99464.1

Group18353: asyd|OJJ65321.1 aver|OJI99466.1

Group18354: asyd|OJJ65328.1 aver|OJJ07721.1

Group18355: asyd|OJJ65338.1 aver|OJI99485.1

Group18356: asyd|OJJ65339.1 aver|OJI99486.1

Group18357: asyd|OJJ65344.1 aver|OJI99491.1

Group18358: asyd|OJJ65345.1 aver|OJI99492.1

Group18359: asyd|OJJ65346.1 aver|OJI99493.1

Group18360: asyd|OJJ65348.1 aver|OJI99495.1

Group18361: asyd|OJJ65352.1 aver|OJI99500.1

Group18362: asyd|OJJ65368.1 aver|OJJ07158.1

Group18363: asyd|OJJ65371.1 aver|OJI99538.1

Group18364: asyd|OJJ65372.1 aver|OJI99539.1

Group18365: asyd|OJJ65380.1 aver|OJI99550.1

Group18366: asyd|OJJ65396.1 aver|OJI99568.1

Group18367: asyd|OJJ65397.1 aver|OJI99569.1

Group18368: ater|EAU29231.1 aver|OJJ05341.1

Group18369: ater|EAU29234.1 aver|OJJ05345.1

Group18370: ater|EAU29235.1 aver|OJJ05346.1

Group18371: ater|EAU29236.1 aver|OJJ05336.1

Group18372: ater|EAU29273.1 aver|OJJ01770.1

Group18373: ater|EAU29274.1 aver|OJJ01768.1

Group18374: ater|EAU29275.1 aver|OJJ01767.1

Group18375: ater|EAU29277.1 aver|OJJ01764.1

Group18376: ater|EAU29278.1 aver|OJJ01769.1

Group18377: ater|EAU29311.1 awen|OJJ33834.1

Group18378: ater|EAU29572.1 awen|OJJ35316.1

Group18379: ater|EAU29964.1 awen|OJJ35823.1

Group18380: ater|EAU30217.1 aver|OJJ07306.1

Group18381: ater|EAU30323.1 azon|OJJ43176.1

Group18382: ater|EAU31067.1 azon|OJJ51655.1

Group18383: ater|EAU31202.1 awen|OJJ34630.1

Group18384: ater|EAU31340.1 awen|OJJ30108.1

Group18385: ater|EAU31341.1 awen|OJJ30107.1

Group18386: ater|EAU31343.1 awen|OJJ30104.1

Group18387: ater|EAU31408.1 awen|OJJ35957.1

Group18388: ater|EAU31460.1 aver|OJJ03431.1

Group18389: ater|EAU31500.1 azon|OJJ43624.1

Group18390: ater|EAU31767.1 azon|OJJ42006.1

Group18391: ater|EAU31844.1 aver|OJJ08411.1

Group18392: ater|EAU31942.1 awen|OJJ39688.1

Group18393: ater|EAU32082.1 aver|OJJ04405.1

Group18394: ater|EAU32156.1 awen|OJJ40036.1

Group18395: ater|EAU32226.1 aver|OJJ05223.1

Group18396: ater|EAU32514.1 aver|OJI96493.1

Group18397: ater|EAU32643.1 aver|OJI98830.1

Group18398: ater|EAU32645.1 awen|OJJ33503.1

Group18399: ater|EAU32816.1 awen|OJJ32541.1

Group18400: ater|EAU33085.1 azon|OJJ43687.1

Group18401: ater|EAU33127.1 awen|OJJ30977.1

Group18402: ater|EAU33622.1 aver|OJJ06435.1

Group18403: ater|EAU33623.1 aver|OJJ06439.1

Group18404: aver|OJJ01876.1 awen|OJJ37972.1

Group18405: ater|EAU33847.1 azon|OJJ47376.1

Group18406: ater|EAU33867.1 azon|OJJ43524.1

Group18407: ater|EAU34046.1 awen|OJJ40110.1

Group18408: ater|EAU35705.1 aver|OJJ07434.1

Group18409: ater|EAU35749.1 aver|OJJ07760.1

Group18410: ater|EAU35856.1 awen|OJJ29672.1

Group18411: ater|EAU36048.1 awen|OJJ30956.1

Group18412: ater|EAU36049.1 awen|OJJ30955.1

Group18413: ater|EAU36073.1 awen|OJJ35067.1

Group18414: ater|EAU36632.1 aver|OJJ07139.1

Group18415: ater|EAU36708.1 aver|OJJ06458.1

Group18416: ater|EAU37130.1 azon|OJJ42059.1

Group18417: ater|EAU37404.1 aver|OJJ00327.1

Group18418: ater|EAU37682.1 azon|OJJ42271.1

Group18419: ater|EAU38632.1 azon|OJJ46583.1

Group18420: ater|EAU39020.1 awen|OJJ35319.1

Group18421: ater|EAU39067.1 azon|OJJ43881.1

Group18422: ater|EAU39607.1 awen|OJJ34657.1

Group18423: ater|EAU39621.1 awen|OJJ30205.1

Group18424: atub|OJI79324.1 awen|OJJ37861.1

Group18425: atub|OJI80138.1 aver|OJJ05133.1

Group18426: atub|OJI80488.1 awen|OJJ30846.1

Group18427: atub|OJI81832.1 awen|OJJ32217.1

Group18428: atub|OJI82286.1 awen|OJJ37002.1

Group18429: atub|OJI83113.1 awen|OJJ33409.1

Group18430: atub|OJI85941.1 azon|OJJ42295.1

Group18431: awen|OJJ35371.1 aver|OJJ08813.1

Group18432: atub|OJI91292.1 awen|OJJ40849.1

Group18433: aver|OJI95801.1 awen|OJJ33636.1

Group18434: aver|OJI95803.1 awen|OJJ33638.1

Group18435: awen|OJJ41693.1 azon|OJJ43939.1

Group18436: aver|OJJ00456.1 awen|OJJ39222.1

Group18437: aver|OJJ00747.1 awen|OJJ38295.1

Group18438: aver|OJJ01155.1 azon|OJJ42242.1

Group18439: aver|OJJ02009.1 azon|OJJ47857.1

Group18440: aver|OJJ02011.1 azon|OJJ47855.1

Group18441: aver|OJJ03475.1 azon|OJJ45210.1

Group18442: aver|OJJ03855.1 awen|OJJ37016.1

Group18443: aver|OJJ03887.1 awen|OJJ32656.1

Group18444: aver|OJJ03905.1 azon|OJJ44713.1

Group18445: aver|OJJ04012.1 azon|OJJ42213.1

Group18446: aver|OJJ06257.1 awen|OJJ41171.1

Group18447: aver|OJJ06609.1 azon|OJJ42588.1

Group18448: aver|OJJ08606.1 azon|OJJ48023.1

Group18449: aver|OJJ08745.1 awen|OJJ40748.1

Group18450: aver|OJJ08814.1 awen|OJJ35372.1

Group18451: awen|OJJ30852.1 azon|OJJ43035.1

Group18452: awen|OJJ30860.1 azon|OJJ50318.1

Group18453: awen|OJJ31336.1 azon|OJJ47386.1

Group18454: awen|OJJ31678.1 azon|OJJ45087.1

Group18455: awen|OJJ31681.1 azon|OJJ42499.1

Group18456: awen|OJJ31937.1 azon|OJJ46599.1

Group18457: awen|OJJ32697.1 azon|OJJ51791.1

Group18458: awen|OJJ32805.1 azon|OJJ49027.1

Group18459: awen|OJJ33693.1 azon|OJJ46548.1

Group18460: awen|OJJ34034.1 azon|OJJ48206.1

Group18461: awen|OJJ34125.1 azon|OJJ44223.1

Group18462: awen|OJJ34491.1 azon|OJJ46008.1

Group18463: awen|OJJ34631.1 azon|OJJ43116.1

Group18464: awen|OJJ35355.1 azon|OJJ43597.1

Group18465: awen|OJJ35832.1 azon|OJJ45773.1

Group18466: awen|OJJ36291.1 azon|OJJ50428.1

Group18467: awen|OJJ37017.1 azon|OJJ42676.1

Group18468: awen|OJJ37464.1 azon|OJJ51142.1

Group18469: awen|OJJ38086.1 azon|OJJ46847.1

Group18470: awen|OJJ38304.1 azon|OJJ45932.1

Group18471: awen|OJJ38829.1 azon|OJJ49625.1

Group18472: awen|OJJ39499.1 azon|OJJ47701.1

Group18473: awen|OJJ40027.1 azon|OJJ42011.1

Group18474: awen|OJJ41079.1 azon|OJJ44661.1

**Additional data file 2:** Core genome of Aspergillus. The file consists of all the clusters containing at least one protein from all the analysed 18 species. Abbreviations - abra : *Aspergillus brasiliensis*, acar : *A. carbonarius*, acla : *A. clavatus*, afis : *A. fischeri*, afla : *A. flavus*, afum : *A. fumigatus*, agla : *A. glaucus*, akaw : *A. kawachii*, alue : *A. luchuensis*, anid : *A. nidulans*, anig : *A. niger*, aora : *A. oryzae*, asyd : *A. sydowii*, ater : *A. terreus*, atub : *A. tubingenesis*, aver : *A. versicolor*, awen : *A. wentii*, azon : *A. zonatus*.

Group1: abra|OJJ66207.1 abra|OJJ67944.1 abra|OJJ69734.1 abra|OJJ73921.1 abra|OJJ75305.1 abra|OJJ75840.1 abra|OJJ73699.1 abra|OJJ69537.1 abra|OJJ78094.1 acar|OOF90653.1 acar|OOF94187.1 acar|OOG00776.1 acar|OOG01052.1 acar|OOF92566.1 acar|OOF93189.1 acar|OOF97512.1 acar|OOF98777.1 acla|EAW07798.1 acla|EAW08899.1 acla|EAW11681.1 akaw|GAA89528.1 akaw|GAA90187.1 akaw|GAA91252.1 aluc|OJZ80511.1 aluc|OJZ82816.1 aluc|OJZ88125.1 aluc|OJZ89282.1 aluc|OJZ83162.1 anid|CBF73441.1 anid|CBF90374.1 anid|CBF87872.1 anig|EHA17891.1 anig|EHA21256.1 anig|EHA21434.1 anig|EHA25711.1 anig|EHA28226.1 anig|EHA18088.1 anig|EHA19858.1 anig|EHA25622.1 anig|EHA25682.1 anig|EHA23283.1 aory|EIT74664.1 aory|EIT83097.1 aory|EIT80579.1 aory|EIT81467.1 aory|EIT83286.1 asyd|OJJ56148.1 asyd|OJJ63431.1 atub|OJI87775.1 atub|OJI89047.1 atub|OJI90732.1 atub|OJI88419.1 aver|OJI97506.1 aver|OJI99587.1 aver|OJJ03355.1 aver|OJJ05088.1 aver|OJJ08114.1 aver|OJJ03487.1 awen|OJJ29812.1 awen|OJJ35864.1 awen|OJJ38889.1 azon|OJJ42489.1 azon|OJJ44494.1 azon|OJJ46205.1 azon|OJJ47369.1 azon|OJJ45854.1 afis|EAW22835.1 afla|EED50110.1 afla|EED51745.1 afum|EAL90367.2 agla|OJJ78560.1 akaw|GAA89080.1 aory|EIT81736.1 ater|EAU35730.1 abra|OJJ67593.1 akaw|GAA89112.1 atub|OJI88453.1 acar|OOF91540.1 abra|OJJ70223.1 afla|EED52040.1 afum|EAL91343.1 akaw|GAA85904.1 aluc|OJZ91163.1 anid|CBF80212.1 atub|OJI83179.1 acar|OOF96620.1 akaw|GAA84550.1 atub|OJI85517.1 afla|EED52433.1 acla|EAW10758.1 afis|EAW16932.1 afla|EED48405.1 afla|EED51820.1 afla|EED51822.1 afla|EED52154.1 aory|EIT74440.1 anid|CBF73982.1 ater|EAU33387.1

Group2: abra|OJJ67588.1 abra|OJJ73922.1 abra|OJJ75304.1 acar|OOF99263.1 acar|OOG01053.1 acar|OOF99291.1 acar|OOF99559.1 acla|EAW07086.1 acla|EAW08665.1 acla|EAW08900.1 acla|EAW11679.1 acla|EAW15286.1 acla|EAW07799.1 afis|EAW16180.1 afis|EAW16446.1 afis|EAW20684.1 afla|EED44993.1 afla|EED47804.1 afla|EED49850.1 afla|EED50111.1 afla|EED52159.1 afum|EAL89049.1 afum|EAL91342.1 akaw|GAA82422.1 akaw|GAA89107.1 akaw|GAA89529.1 anid|CBF73453.1 anid|CBF87069.1 anid|CBF76036.1 anid|CBF87869.1 anid|CBF90372.1 anig|EHA17890.1 anig|EHA25707.1 anig|EHA20896.1 anig|EHA21433.1 anig|EHA24938.1 aory|EIT73739.1 aory|EIT74412.1 aory|EIT74615.1 aory|EIT80956.1 asyd|OJJ56147.1 asyd|OJJ63429.1 ater|EAU29302.1 ater|EAU30156.1 ater|EAU37759.1 ater|EAU30201.1 atub|OJI86150.1 atub|OJI88448.1 atub|OJI88518.1 atub|OJI90731.1 aver|OJI97508.1 aver|OJI99595.1 aver|OJJ03909.1 aver|OJJ05087.1 aver|OJI99581.1 aver|OJI99596.1 aver|OJJ07364.1 aver|OJJ08120.1 aver|OJJ08464.1 aver|OJI99625.1 aver|OJJ07311.1 aver|OJJ08781.1 awen|OJJ36076.1 awen|OJJ38362.1 awen|OJJ32929.1 awen|OJJ38888.1 azon|OJJ42030.1 azon|OJJ42483.1 azon|OJJ46579.1 azon|OJJ48580.1 afis|EAW22836.1 afum|EAL90366.1 agla|OJJ80563.1 aluc|OJZ85583.1 aluc|OJZ88124.1 asyd|OJJ61536.1 ater|EAU38874.1 awen|OJJ32292.1 abra|OJJ67652.1 afis|EAW16453.1 afis|EAW25605.1 agla|OJJ79232.1 akaw|GAA89168.1 aluc|OJZ81665.1 anig|EHA25778.1 ater|EAU33874.1 aver|OJJ07366.1 azon|OJJ50363.1 acla|EAW14636.1 ater|EAU36802.1 awen|OJJ34576.1 afum|EAL88812.1 aver|OJI99592.1 asyd|OJJ52456.1 aver|OJJ03957.1 awen|OJJ34650.1

Group3: abra|OJJ67584.1 abra|OJJ67975.1 abra|OJJ75778.1 abra|OJJ77754.1 acar|OOF92816.1 acar|OOF96723.1 acla|EAW08895.1 acla|EAW11760.1 acla|EAW12170.1 acla|EAW12336.1 acla|EAW14856.1 afla|EED47354.1 afla|EED52019.1 afla|EED47890.1 afla|EED48583.1 agla|OJJ79933.1 agla|OJJ80111.1 agla|OJJ83619.1 agla|OJJ86349.1 akaw|GAA84491.1 akaw|GAA85345.1 akaw|GAA87318.1 akaw|GAA91228.1 akaw|GAA92525.1 akaw|GAA90346.1 akaw|GAA85925.1 aluc|OJZ82842.1 aluc|OJZ85886.1 aluc|OJZ89214.1 aluc|OJZ90707.1 aluc|OJZ91539.1 aluc|OJZ91139.1 anid|CBF71467.1 anid|CBF83055.1 anid|CBF86052.1 anid|CBF87072.1 anid|CBF88289.1 anig|EHA19567.1 anig|EHA20193.1 anig|EHA21280.1 anig|EHA22576.1 anig|EHA24574.1 anig|EHA24700.1 anig|EHA28244.1 aory|EIT73118.1 aory|EIT76300.1 aory|EIT77964.1 aory|EIT81441.1 aory|EIT82867.1 ater|EAU29417.1 ater|EAU31921.1 ater|EAU32663.1 ater|EAU36720.1 atub|OJI79672.1 atub|OJI83150.1 atub|OJI85993.1 atub|OJI91114.1 atub|OJI82134.1 atub|OJI85578.1 atub|OJI89072.1 atub|OJI82802.1 aver|OJI96730.1 aver|OJJ05906.1 awen|OJJ29758.1 awen|OJJ34705.1 awen|OJJ40111.1 awen|OJJ29917.1 awen|OJJ30910.1 awen|OJJ38044.1 awen|OJJ39589.1 awen|OJJ29944.1 afis|EAW23793.1 afum|EAL87227.2 asyd|OJJ62861.1 aver|OJI96914.1 asyd|OJJ61528.1 abra|OJJ70374.1 acar|OOF91895.1 afis|EAW16886.1 afum|EAL86424.2 ater|EAU30225.1 azon|OJJ48577.1 acar|OOF93599.1 azon|OJJ49991.1 afla|EED46560.1 akaw|GAA87467.1 aluc|OJZ88529.1

Group4: abra|OJJ67662.1 abra|OJJ77815.1 abra|OJJ78181.1 abra|OJJ74373.1 abra|OJJ78098.1 abra|OJJ76619.1 acar|OOF91828.1 acar|OOF91847.1 acar|OOF92934.1 acar|OOF94993.1 acar|OOF97632.1 acar|OOF95055.1 acar|OOF95122.1 acla|EAW07064.1 afis|EAW19759.1 afis|EAW23638.1 akaw|GAA83602.1 akaw|GAA84851.1 akaw|GAA92094.1 akaw|GAA83831.1 akaw|GAA85928.1 akaw|GAA89177.1 akaw|GAA86655.1 akaw|GAA85575.1 akaw|GAA85859.1 aluc|OJZ81656.1 aluc|OJZ83631.1 aluc|OJZ90887.1 aluc|OJZ91221.1 aluc|OJZ81972.1 aluc|OJZ86820.1 aluc|OJZ89749.1 aluc|OJZ92392.1 aluc|OJZ83511.1 aluc|OJZ89332.1 aluc|OJZ91137.1 anid|CBF80211.1 anid|CBF82304.1 anig|EHA19289.1 anig|EHA24757.1 anig|EHA19606.1 anig|EHA25789.1 anig|EHA26583.1 aory|EIT74370.1 aory|EIT76972.1 ater|EAU35165.1 ater|EAU36856.1 atub|OJI79585.1 atub|OJI79696.1 atub|OJI83697.1 atub|OJI85053.1 atub|OJI86822.1 atub|OJI82881.1 atub|OJI88527.1 atub|OJI90000.1 atub|OJI83148.1 aver|OJJ04870.1 aver|OJJ06306.1

aver|OJJ07785.1 awen|OJJ29662.1 awen|OJJ29874.1 awen|OJJ33672.1 agla|OJJ83634.1 asyd|OJJ52727.1 aver|OJJ02001.1 azon|OJJ47400.1 afla|EED51023.1 abra|OJJ74811.1 acar|OOF95802.1 afla|EED52130.1 afum|EAL85129.1 atub|OJI85468.1 abra|OJJ77455.1 afla|EED49595.1 anig|EHA19288.1 asyd|OJJ56406.1 ater|EAU32666.1 ater|EAU38928.1 ater|EAU39559.1 anid|CBF79115.1 afis|EAW19771.1 akaw|GAA84588.1 akaw|GAA84589.1 aluc|OJZ89334.1 atub|OJI82404.1

Group5: abra|OJJ67238.1 abra|OJJ68732.1 acar|OOF92227.1 acar|OOF98313.1 acla|EAW12167.1 acla|EAW12516.1 afis|EAW15603.1 afis|EAW17072.1 afis|EAW19893.1 afis|EAW21005.1 afis|EAW24131.1 afis|EAW24651.1 afis|EAW22083.1 afis|EAW24734.1 afla|EED46089.1 afla|EED46801.1 afla|EED48098.1 afla|EED52234.1 afla|EED48355.1 afla|EED49615.1 afla|EED55083.1 afla|EED56403.1 afum|EAL85692.1 afum|EAL89381.1 afum|EAL91987.1 afum|EAL85730.1 afum|EAL86560.1 agla|OJJ79085.1 agla|OJJ82566.1 agla|OJJ86784.1 agla|OJJ87207.1 akaw|GAA89881.1 akaw|GAA91663.1 aluc|OJZ81798.1 aluc|OJZ87737.1 anid|CBF75787.1 anid|CBF86562.1 anid|CBF86663.1 anig|EHA20229.1 anig|EHA22418.1 aory|EIT72725.1 aory|EIT74466.1 aory|EIT76413.1 aory|EIT80974.1 aory|EIT81644.1 aory|EIT83015.1 asyd|OJJ59270.1 asyd|OJJ64851.1 asyd|OJJ60947.1 asyd|OJJ64782.1 asyd|OJJ65051.1 ater|EAU30561.1 ater|EAU31344.1 ater|EAU38103.1 atub|OJI80903.1 atub|OJI84369.1 aver|OJI98893.1 aver|OJI99174.1 aver|OJJ00274.1 aver|OJJ08481.1 aver|OJI98963.1 aver|OJJ01883.1 aver|OJJ07136.1 awen|OJJ30103.1 awen|OJJ40222.1 awen|OJJ41358.1 awen|OJJ31256.1 awen|OJJ37981.1 azon|OJJ46971.1 azon|OJJ48822.1 abra|OJJ73660.1 acar|OOG00838.1 akaw|GAA88498.1 aluc|OJZ88393.1 anig|EHA18121.1 aory|EIT72296.1 atub|OJI90988.1 awen|OJJ38335.1 anid|CBF74724.1 aluc|OJZ80164.1 aory|EIT76217.1 ater|EAU32520.1

Group7: abra|OJJ65820.1 abra|OJJ66803.1 acar|OOF90643.1 acar|OOF94182.1 afis|EAW16457.1 afis|EAW21767.1 afla|EED55145.1 afla|EED55917.1 akaw|GAA92417.1 akaw|GAA93242.1 anid|CBF75176.1 anid|CBF84668.1 anid|CBF85153.1 anid|CBF88209.1 anid|CBF75912.1 anid|CBF80593.1 anid|CBF80303.1 anid|CBF80770.1 anig|EHA18419.1 anig|EHA24965.1 anig|EHA21352.1 aory|EIT76564.1 aory|EIT81636.1 aory|EIT81823.1 asyd|OJJ52880.1 asyd|OJJ54446.1 asyd|OJJ60235.1 asyd|OJJ61610.1 ater|EAU32131.1 ater|EAU35090.1 ater|EAU37273.1 aver|OJJ04030.1 aver|OJJ08401.1 aver|OJJ08851.1 aver|OJJ07396.1 azon|OJJ42892.1 azon|OJJ45918.1 azon|OJJ48221.1 azon|OJJ48680.1 azon|OJJ49289.1 agla|OJJ83550.1 anig|EHA26118.1 awen|OJJ41193.1 azon|OJJ42522.1 abra|OJJ67032.1 acla|EAW08788.1 afis|EAW19897.1 afis|EAW23754.1 agla|OJJ82646.1 akaw|GAA92151.1 aluc|OJZ85558.1 ater|EAU30865.1 atub|OJI79347.1 abra|OJJ76132.1 acar|OOF99791.1 acla|EAW09686.1 afis|EAW21453.1 afla|EED55883.1 afum|EAL92472.1 agla|OJJ85599.1 akaw|GAA88835.1 aluc|OJZ89593.1 anid|CBF76549.1 anig|EHA23018.1 aory|EIT76608.1 asyd|OJJ61156.1 atub|OJI85209.1 aver|OJJ00478.1 awen|OJJ39247.1 azon|OJJ51608.1 abra|OJJ77926.1 acar|OOF89902.1 afum|EAL87190.2 akaw|GAA83738.1 aluc|OJZ90994.1 atub|OJI82984.1 aver|OJJ07910.1 aver|OJJ00933.1

Group8: abra|OJJ66275.1 abra|OJJ77982.1 abra|OJJ70821.1 abra|OJJ71091.1 abra|OJJ75537.1 abra|OJJ78258.1 acar|OOF97571.1 acar|OOF98050.1 acar|OOG00735.1 acar|OOF98604.1 acla|EAW07624.1 acla|EAW08950.1 acla|EAW09019.1 acla|EAW09117.1 acla|EAW13520.1 acla|EAW15062.1 afla|EED45116.1 afla|EED47649.1 afla|EED49862.1 afla|EED51183.1 akaw|GAA82870.1 akaw|GAA91957.1 akaw|GAA85998.1 aluc|OJZ82661.1 aluc|OJZ84344.1 aluc|OJZ87437.1 aluc|OJZ91057.1 aluc|OJZ91332.1 anig|EHA18001.1 anig|EHA18935.1 anig|EHA20663.1 anig|EHA27435.1 anig|EHA27680.1 anig|EHA19760.1 anig|EHA23147.1 anig|EHA23656.1 anig|EHA20014.1 aory|EIT73707.1 aory|EIT80187.1 aory|EIT83384.1 asyd|OJJ57246.1 asyd|OJJ60063.1 asyd|OJJ63687.1 atub|OJI81211.1 atub|OJI83130.1 atub|OJI83325.1 atub|OJI88893.1 atub|OJI89724.1 atub|OJI83053.1 atub|OJI89947.1 aver|OJI99011.1 aver|OJJ08178.1 awen|OJJ29725.1 awen|OJJ32235.1 awen|OJJ34622.1 afla|EED49876.1 afum|EAL85113.2 akaw|GAA87094.1 aory|EIT83412.1 ater|EAU38971.1 afis|EAW16804.1 agla|OJJ85462.1 anid|CBF80487.1 ater|EAU29410.1 azon|OJJ42361.1 acla|EAW10019.1 afis|EAW20415.1 afum|EAL89230.2 anid|CBF85555.1 ater|EAU29808.1 azon|OJJ43495.1 aver|OJI97248.1 afum|EAL91103.2 akaw|GAA83290.1 akaw|GAA83291.1 afla|EED45117.1 aver|OJJ08105.1

Group9: abra|OJJ68001.1 abra|OJJ69377.1 abra|OJJ77432.1 abra|OJJ78055.1 acar|OOF92044.1 acar|OOF94347.1 afis|EAW19338.1 afis|EAW24682.1 afis|EAW20700.1 afis|EAW24697.1 afis|EAW25724.1 afla|EED45393.1 afla|EED47975.1 afla|EED51175.1 afla|EED57518.1 afum|EAL84397.1 afum|EAL84875.1 afum|EAL89339.1 afum|EAL94057.1 agla|OJJ83163.1 agla|OJJ85718.1 agla|OJJ88362.1 akaw|GAA85937.1 akaw|GAA88581.1 akaw|GAA88246.1 akaw|GAA90949.1 aluc|OJZ82213.1 aluc|OJZ82875.1 aluc|OJZ88485.1 aluc|OJZ91125.1 anid|CBF70387.1 anid|CBF79143.1 anid|CBF90097.1 anid|CBF74114.1 anid|CBF80184.1 anig|EHA21301.1 anig|EHA28527.1 aory|EIT75060.1 aory|EIT80443.1 aory|EIT79828.1 aory|EIT81356.1 aory|EIT81872.1 aory|EIT82404.1 asyd|OJJ53903.1 asyd|OJJ56275.1 asyd|OJJ57401.1 asyd|OJJ57276.1 asyd|OJJ58638.1 asyd|OJJ62031.1 ater|EAU31624.1 ater|EAU37396.1 ater|EAU31762.1 ater|EAU38791.1 atub|OJI79831.1 atub|OJI83136.1 atub|OJI88222.1 atub|OJI89099.1 aver|OJI96122.1 aver|OJJ01245.1 aver|OJI99530.1 aver|OJJ08000.1 aver|OJJ08490.1 awen|OJJ32995.1 awen|OJJ37839.1 awen|OJJ40791.1 azon|OJJ48191.1 azon|OJJ48520.1 acla|EAW14609.1 azon|OJJ43922.1 acla|EAW13612.1 afla|EED53479.1 anig|EHA20150.1 anig|EHA19826.1 afla|EED51738.1 afla|EED51739.1 ater|EAU33967.1

Group10: abra|OJJ73857.1 abra|OJJ76179.1 acar|OOF94216.1 acar|OOF99761.1 acar|OOG01005.1 acla|EAW07747.1 acla|EAW09647.1 afis|EAW17299.1 afis|EAW22784.1 afis|EAW20842.1 afis|EAW21493.1 afla|EED48870.1 afla|EED50046.1 afum|EAL84790.1 afum|EAL92512.1 afum|EAL90415.1 agla|OJJ83596.1 agla|OJJ85655.1 akaw|GAA82124.1 akaw|GAA89472.1 akaw|GAA92487.1 aluc|OJZ80154.1 aluc|OJZ88191.1 aluc|OJZ92051.1 aluc|OJZ91499.1 anid|CBF76107.1 anid|CBF87969.1 anid|CBF76478.1 anid|CBF80461.1 anid|CBF83927.1 anig|EHA17943.1 anig|EHA22980.1 anig|EHA22539.1 aory|EIT74611.1 aory|EIT76730.1

aory|EIT80155.1 aory|EIT81260.1 asyd|OJJ53335.1 asyd|OJJ61118.1 asyd|OJJ55439.1 asyd|OJJ60355.1 asyd|OJJ63488.1 ater|EAU29437.1 ater|EAU35045.1 ater|EAU38925.1 ater|EAU29951.1 atub|OJI80378.1 atub|OJI90790.1 atub|OJI85952.1 atub|OJI86482.1 aver|OJI97441.1 aver|OJJ06514.1 aver|OJI99693.1 aver|OJJ08409.1 aver|OJJ00436.1 awen|OJJ36113.1 awen|OJJ38951.1 awen|OJJ39202.1 awen|OJJ40168.1 azon|OJJ49724.1 azon|OJJ51661.1 abra|OJJ70289.1 acar|OOF91742.1 akaw|GAA92859.1 aluc|OJZ82554.1 anig|EHA22518.1 atub|OJI88795.1 awen|OJJ38795.1 afla|EED57355.1 afla|EED55928.1

Group11: abra|OJJ66566.1 abra|OJJ74913.1 acar|OOF91007.1 acar|OOF95904.1 acla|EAW08458.1 acla|EAW10929.1 afis|EAW20932.1 afis|EAW23503.1 afla|EED50286.1 afla|EED57404.1 afum|EAL84702.1 afum|EAL87958.1 agla|OJJ83956.1 agla|OJJ86749.1 akaw|GAA83509.1 akaw|GAA91151.1 aluc|OJZ81479.1 aluc|OJZ92486.1 anid|CBF82508.1 anid|CBF89725.1 anig|EHA18724.1 anig|EHA18871.1 anig|EHA25948.1 anig|EHA26487.1 aory|EIT77296.1 aory|EIT80109.1 asyd|OJJ54080.1 asyd|OJJ62235.1 ater|EAU35292.1 ater|EAU38577.1 atub|OJI86917.1 atub|OJI88701.1 aver|OJI96344.1 aver|OJJ08658.1 awen|OJJ34386.1 awen|OJJ39829.1 azon|OJJ47952.1 azon|OJJ48118.1 acar|OOF91006.1 acla|EAW10930.1 afis|EAW20931.1 afla|EED50285.1 afum|EAL84703.2 agla|OJJ83957.1 akaw|GAA91152.1 anid|CBF82510.1 anig|EHA25947.1 aory|EIT77323.1 asyd|OJJ54081.1 ater|EAU38576.1 aver|OJJ08657.1 awen|OJJ34387.1 azon|OJJ48119.1 abra|OJJ68224.1 acar|OOF95905.1 acla|EAW08457.1 afis|EAW23498.1 afla|EED57403.1 afum|EAL87965.1 akaw|GAA83219.1 aluc|OJZ83721.1 anig|EHA18870.1 aory|EIT80136.1 ater|EAU35291.1 atub|OJI90095.1 awen|OJJ39827.1 anig|EHA26486.1 atub|OJI86918.1 agla|OJJ83079.1

Group12: abra|OJJ67365.1 abra|OJJ70373.1 abra|OJJ72695.1 abra|OJJ77105.1 abra|OJJ78039.1 abra|OJJ77002.1 acar|OOF90082.1 acar|OOF91896.1 acar|OOF92990.1 acar|OOF91448.1 acla|EAW08775.1 acla|EAW12337.1 afis|EAW16885.1 afis|EAW20732.1 afis|EAW25626.1 afla|EED47350.1 afla|EED50231.1 afla|EED54427.1 afum|EAL84314.2 afum|EAL86423.2 agla|OJJ78903.1 agla|OJJ79932.1 agla|OJJ85297.1 akaw|GAA86951.1 akaw|GAA91977.1 aluc|OJZ80600.1 aluc|OJZ84401.1 aluc|OJZ82637.1 aluc|OJZ84514.1 anid|CBF80656.1 anid|CBF82934.1 anid|CBF83105.1 anig|EHA27442.1 anig|EHA27812.1 aory|EIT75862.1 aory|EIT77345.1 aory|EIT81172.1 aory|EIT81924.1 aory|EIT82860.1 asyd|OJJ53449.1 asyd|OJJ54612.1 asyd|OJJ61456.1 ater|EAU30228.1 ater|EAU38239.1 atub|OJI88868.1 atub|OJI89567.1 atub|OJI89671.1 aver|OJJ00792.1 aver|OJJ05324.1 aver|OJJ07546.1 aver|OJJ08303.1 awen|OJJ29598.1 awen|OJJ29759.1 awen|OJJ31482.1 awen|OJJ34518.1 awen|OJJ36047.1 azon|OJJ42524.1 afla|EED55182.1 akaw|GAA87046.1 anig|EHA27731.1 aory|EIT81588.1 asyd|OJJ55521.1 ater|EAU32705.1 aver|OJJ06599.1 azon|OJJ50398.1 afla|EED48862.1 ater|EAU32222.1

Group13: abra|OJJ74143.1 abra|OJJ77919.1 acar|OOF95048.1 acar|OOF95115.1 acar|OOF95586.1 acar|OOF98862.1 acar|OOF99364.1 afla|EED47372.1 afla|EED48594.1 akaw|GAA83744.1 akaw|GAA89741.1 akaw|GAA92822.1 akaw|GAA92977.1 aluc|OJZ80240.1 aluc|OJZ91576.1 aluc|OJZ80643.1 aluc|OJZ90987.1 anid|CBF75107.1 anid|CBF76688.1 anid|CBF85354.1 anid|CBF86773.1 anig|EHA19692.1 anig|EHA27189.1 aory|EIT73093.1 aory|EIT82880.1 asyd|OJJ59996.1 asyd|OJJ61226.1 asyd|OJJ64036.1 asyd|OJJ64686.1 ater|EAU32382.1 ater|EAU33556.1 atub|OJI82978.1 atub|OJI87498.1 aver|OJI98121.1 aver|OJJ00549.1 aver|OJJ02295.1 aver|OJI98789.1 abra|OJJ69652.1 acar|OOF98861.1 acla|EAW06691.1 afis|EAW16074.1 afla|EED48593.1 afum|EAL88706.2 agla|OJJ86094.1 akaw|GAA91631.1 aluc|OJZ82458.1 anid|CBF75109.1 anig|EHA28317.1 aory|EIT73047.1 ater|EAU33557.1 atub|OJI87980.1 awen|OJJ30359.1 azon|OJJ45380.1 acla|EAW06690.1 afis|EAW16075.1 afum|EAL88707.1 awen|OJJ30358.1 abra|OJJ76634.1 afla|EED53145.1 agla|OJJ82986.1 anig|EHA22603.1 aory|EIT75089.1 ater|EAU37809.1 atub|OJI86025.1 awen|OJJ31622.1 azon|OJJ47320.1

Group14: abra|OJJ65763.1 abra|OJJ65764.1 abra|OJJ68458.1 abra|OJJ70187.1 acar|OOF91603.1 acar|OOF93288.1 acar|OOF99308.1 acar|OOF99313.1 afis|EAW16734.1 afis|EAW19114.1 afla|EED45012.1 afla|EED57380.1 afla|EED57383.1 afum|EAL86261.1 afum|EAL86308.1 afum|EAL86264.1 afum|EAL93846.1 akaw|GAA83845.1 akaw|GAA86588.1 akaw|GAA92381.1 aluc|OJZ83240.1 aluc|OJZ85135.1 aluc|OJZ90870.1 anid|CBF74944.1 anid|CBF83824.1 anid|CBF84265.1 anig|EHA19590.1 anig|EHA25367.1 anig|EHA21095.1 anig|EHA22863.1 aory|EIT73582.1 aory|EIT75916.1 aory|EIT80067.1 aory|EIT80069.1 asyd|OJJ57214.1 asyd|OJJ58070.1 asyd|OJJ63941.1 asyd|OJJ60903.1 asyd|OJJ63609.1 asyd|OJJ63938.1 atub|OJI80457.1 atub|OJI82868.1 atub|OJI87697.1 aver|OJI97001.1 aver|OJJ00229.1 aver|OJJ03075.1 aver|OJJ04337.1 aver|OJJ07009.1 awen|OJJ30374.1 awen|OJJ30814.1 awen|OJJ32530.1 acla|EAW14406.1 agla|OJJ89198.1 azon|OJJ47391.1 aver|OJI97004.1 aver|OJI97328.1 awen|OJJ40814.1 azon|OJJ50090.1 ater|EAU37377.1 azon|OJJ42904.1 ater|EAU38814.1 asyd|OJJ55207.1

Group15: abra|OJJ69361.1 abra|OJJ71787.1 abra|OJJ75622.1 acar|OOF94364.1 acar|OOF96466.1 acar|OOG00218.1 acla|EAW08453.1 acla|EAW09357.1 acla|EAW14751.1 afis|EAW21780.1 afis|EAW23502.1 afis|EAW24282.1 afla|EED46347.1 afla|EED47994.1 afum|EAL85951.1 afum|EAL87959.1 afum|EAL92809.1 agla|OJJ80637.1 agla|OJJ84344.1 akaw|GAA84369.1 akaw|GAA85124.1 akaw|GAA88230.1 aluc|OJZ82194.1 aluc|OJZ86517.1 aluc|OJZ89073.1 anig|EHA22129.1 anig|EHA23464.1 anig|EHA28543.1 aory|EIT79847.1 aory|EIT81244.1 asyd|OJJ53917.1 asyd|OJJ56885.1 asyd|OJJ61185.1 asyd|OJJ62326.1 ater|EAU31230.1 ater|EAU38460.1 atub|OJI84007.1 atub|OJI85713.1 atub|OJI88241.1 aver|OJI96431.1 aver|OJJ00509.1 aver|OJJ08015.1 awen|OJJ33015.1 awen|OJJ34069.1 awen|OJJ39823.1 azon|OJJ47947.1 azon|OJJ48162.1 azon|OJJ48502.1 afla|EED57398.1 anid|CBF76607.1 agla|OJJ86745.1 abra|OJJ74918.1 acar|OOF95909.1 akaw|GAA83505.1 aluc|OJZ92489.1 anid|CBF89735.1 anig|EHA26483.1 aory|EIT80098.1 asyd|OJJ62232.1 ater|EAU35287.1 atub|OJI86922.1 aver|OJI96341.1

Group16: abra|OJJ69532.1 abra|OJJ73245.1 abra|OJJ75746.1 abra|OJJ71793.1 acar|OOF90431.1 acar|OOF92663.1 acar|OOG00484.1 acar|OOF96473.1 acla|EAW06919.1 acla|EAW08051.1 afis|EAW16536.1 afis|EAW23087.1 afla|EED48561.1 afla|EED48728.1 afla|EED56133.1 afum|EAL88392.1 afum|EAL89133.1 agla|OJJ82786.1 agla|OJJ89597.1 akaw|GAA84479.1 akaw|GAA85117.1 akaw|GAA87935.1 akaw|GAA88383.1 aluc|OJZ81182.1 aluc|OJZ82371.1 aluc|OJZ86523.1 aluc|OJZ89199.1 anid|CBF75152.1 anid|CBF81741.1 anid|CBF82143.1 anig|EHA18500.1 anig|EHA23358.1 anig|EHA28393.1 anig|EHA22123.1 aory|EIT73145.1 aory|EIT73234.1 aory|EIT76572.1 aory|EIT79614.1 asyd|OJJ54322.1 asyd|OJJ59597.1 asyd|OJJ59975.1 asyd|OJJ56163.1 atub|OJI83999.1 atub|OJI91379.1 atub|OJI85591.1 aver|OJJ02316.1 aver|OJJ05102.1 aver|OJJ05276.1 aver|OJJ02671.1 awen|OJJ30245.1 awen|OJJ39992.1 awen|OJJ40286.1 ater|EAU36211.1 ater|EAU36599.1 atub|OJI88071.1 awen|OJJ36697.1 azon|OJJ47500.1 azon|OJJ49147.1 afla|EED56590.1 agla|OJJ79942.1 anig|EHA23359.1

Group17: abra|OJJ67548.1 abra|OJJ67738.1 abra|OJJ69732.1 abra|OJJ75389.1 abra|OJJ67871.1 acar|OOF90253.1 acar|OOF90937.1 acar|OOF94189.1 acar|OOF94221.1 acar|OOF94457.1 acla|EAW11677.1 acla|EAW12103.1 afla|EED45500.1 afla|EED48836.1 afla|EED51171.1 afla|EED52156.1 akaw|GAA88112.1 akaw|GAA90185.1 akaw|GAA92429.1 akaw|GAA92752.1 akaw|GAA91880.1 aluc|OJZ81382.1 aluc|OJZ81593.1 aluc|OJZ83164.1 aluc|OJZ92960.1 aluc|OJZ82019.1 anid|CBF73431.1 anid|CBF87554.1 anig|EHA18677.1 anig|EHA25848.1 anig|EHA26015.1 anig|EHA28228.1 anig|EHA22237.1 aory|EIT74393.1 aory|EIT75554.1 aory|EIT81271.1 aory|EIT82420.1 aory|EIT81347.1 ater|EAU30799.1 ater|EAU34705.1 atub|OJI84570.1 atub|OJI87418.1 atub|OJI87773.1 atub|OJI88590.1 atub|OJI91588.1 aver|OJJ08108.1 aver|OJJ08968.1 azon|OJJ42487.1 azon|OJJ45532.1 afis|EAW17283.1 afum|EAL86701.1 agla|OJJ81086.1 anid|CBF82821.1 asyd|OJJ52999.1 asyd|OJJ61577.1 aver|OJJ00899.1 awen|OJJ35752.1 afla|EED45454.1 anid|CBF80162.1 aver|OJI99518.1

Group18: abra|OJJ67797.1 abra|OJJ73515.1 abra|OJJ68679.1 abra|OJJ70328.1 abra|OJJ76402.1 acar|OOF91783.1 acar|OOF92181.1 acar|OOG00703.1 acla|EAW07449.1 acla|EAW12568.1 afis|EAW18727.1 afis|EAW22482.1 afis|EAW24082.1 afla|EED45270.1 afla|EED51391.1 afum|EAL90712.1 afum|EAL91937.1 agla|OJJ79953.1 agla|OJJ82286.1 agla|OJJ87548.1 akaw|GAA82322.1 akaw|GAA89832.1 akaw|GAA92583.1 akaw|GAA87464.1 aluc|OJZ81855.1 aluc|OJZ82597.1 aluc|OJZ88532.1 aluc|OJZ91810.1 anig|EHA18233.1 anig|EHA22369.1 anig|EHA27480.1 anig|EHA22803.1 aory|EIT73742.1 aory|EIT78822.1 aory|EIT80707.1 asyd|OJJ53887.1 asyd|OJJ62829.1 ater|EAU30215.1 ater|EAU30505.1 ater|EAU39231.1 ater|EAU38177.1 atub|OJI84423.1 atub|OJI88833.1 atub|OJI91117.1 atub|OJI86266.1 aver|OJI96872.1 aver|OJJ07985.1 awen|OJJ29718.1 awen|OJJ32268.1 awen|OJJ35101.1 acla|EAW14021.1 aluc|OJZ90857.1 anid|CBF88889.1 atub|OJI82853.1 azon|OJJ50007.1 afla|EED50684.1 afla|EED54508.1 acar|OOF99644.1 afum|EAL93452.1 aory|EIT78049.1

Group19: abra|OJJ67556.1 abra|OJJ67739.1 abra|OJJ69721.1 abra|OJJ75388.1 abra|OJJ67870.1 acar|OOF90043.1 acar|OOF90938.1 acar|OOF94195.1 acar|OOF94224.1 acar|OOF94458.1 acla|EAW11676.1 acla|EAW12104.1 afla|EED45513.1 afla|EED48837.1 afla|EED51172.1 afla|EED52163.1 akaw|GAA88108.1 akaw|GAA90174.1 akaw|GAA92430.1 akaw|GAA92751.1 akaw|GAA91879.1 aluc|OJZ81378.1 aluc|OJZ81592.1 aluc|OJZ92959.1 aluc|OJZ82020.1 aluc|OJZ83176.1 anid|CBF73444.1 anid|CBF82822.1 anid|CBF87553.1 anig|EHA18673.1 anig|EHA25849.1 anig|EHA26016.1 anig|EHA28250.1 anig|EHA22236.1 aory|EIT74418.1 aory|EIT75385.1 aory|EIT81285.1 aory|EIT82421.1 aory|EIT81323.1 asyd|OJJ52998.1 asyd|OJJ61578.1 atub|OJI84572.1 atub|OJI87417.1 atub|OJI87761.1 atub|OJI91584.1 atub|OJI88591.1 aver|OJJ00900.1 aver|OJJ08115.1 aver|OJJ08967.1 azon|OJJ42478.1 azon|OJJ45531.1 afis|EAW17282.1 afum|EAL86700.1 agla|OJJ81085.1 ater|EAU30800.1 ater|EAU34706.1 awen|OJJ35753.1 anid|CBF80164.1 aver|OJI99519.1

Group20: abra|OJJ67608.1 abra|OJJ70431.1 acar|OOF89840.1 acar|OOF91955.1 acla|EAW11023.1 acla|EAW14999.1 afis|EAW18722.1 afis|EAW19401.1 afis|EAW20837.1 afis|EAW24523.1 afla|EED45470.1 afla|EED46489.1 afla|EED50679.1 afum|EAL84797.1 afum|EAL86188.1 afum|EAL93446.1 agla|OJJ79209.1 agla|OJJ83714.1 akaw|GAA89124.1 akaw|GAA90149.1 aluc|OJZ81714.1 aluc|OJZ83199.1 anid|CBF74338.1 anid|CBF82877.1 aory|EIT72354.1 aory|EIT75736.1 asyd|OJJ57954.1 asyd|OJJ59255.1 asyd|OJJ61569.1 ater|EAU30785.1 ater|EAU36144.1 ater|EAU36801.1 atub|OJI87739.1 atub|OJI88468.1 aver|OJJ00889.1 aver|OJJ01866.1 awen|OJJ32698.1 awen|OJJ34547.1 azon|OJJ42767.1 azon|OJJ43003.1 anig|EHA25725.1 agla|OJJ78774.1 anig|EHA21062.1 abra|OJJ76408.1 acar|OOF99650.1 acla|EAW14020.1 agla|OJJ80462.1 akaw|GAA82327.1 aluc|OJZ91804.1 anid|CBF70690.1 anig|EHA22798.1 asyd|OJJ58469.1 ater|EAU38182.1 atub|OJI86258.1 aver|OJJ01070.1 awen|OJJ32266.1 azon|OJJ46779.1 azon|OJJ51803.1 asyd|OJJ51967.1

Group21: abra|OJJ67735.1 abra|OJJ68593.1 abra|OJJ73309.1 abra|OJJ77765.1 acar|OOF90932.1 acar|OOF97505.1 acla|EAW12049.1 acla|EAW12177.1 afis|EAW17018.1 afis|EAW19983.1 afla|EED45460.1 afla|EED45667.1 afla|EED48486.1 afla|EED52023.1 afum|EAL84933.1 afum|EAL86540.1 akaw|GAA89817.1 akaw|GAA92425.1 aluc|OJZ81597.1 aluc|OJZ81873.1 anid|CBF69451.1 anid|CBF80428.1 anid|CBF82812.1 anid|CBF83139.1 anid|CBF88295.1 anid|CBF89312.1 anid|CBF86046.1 anig|EHA20911.1 anig|EHA25844.1 anig|EHA28237.1 aory|EIT73076.1 aory|EIT75517.1 aory|EIT81451.1 aory|EIT82410.1 aory|EIT79328.1 asyd|OJJ61573.1 asyd|OJJ62373.1 ater|EAU29529.1 ater|EAU30794.1 ater|EAU31923.1 ater|EAU35431.1 ater|EAU36706.1 atub|OJI80342.1 atub|OJI87755.1 aver|OJI96507.1 aver|OJJ00893.1 aver|OJJ07788.1 aver|OJJ06454.1 agla|OJJ79464.1 atub|OJI88585.1 awen|OJJ38079.1 azon|OJJ46787.1 azon|OJJ47411.1 akaw|GAA90162.1 aluc|OJZ83185.1 abra|OJJ70061.1 anig|EHA21035.1 acar|OOF91380.1 anig|EHA28257.1

Group23: abra|OJJ70376.1 abra|OJJ76944.1 acar|OOF91394.1 acar|OOF91893.1 acar|OOF94281.1 acla|EAW09924.1 acla|EAW12333.1 acla|EAW12657.1 acla|EAW13468.1 afis|EAW16889.1 afis|EAW18023.1 afis|EAW20581.1 afis|EAW21172.1 afla|EED47357.1 afla|EED52279.1 afla|EED54755.1 afum|EAL84483.1 afum|EAL86427.2 afum|EAL92192.1 afum|EAL91835.1 agla|OJJ79937.1 agla|OJJ80007.1 agla|OJJ88693.1 akaw|GAA87096.1 akaw|GAA88352.1 akaw|GAA91976.1 aluc|OJZ82303.1 aluc|OJZ84342.1 aluc|OJZ82638.1 aory|EIT79353.1 aory|EIT82885.1 asyd|OJJ54585.1 asyd|OJJ55452.1 ater|EAU35839.1 ater|EAU37103.1 atub|OJI88134.1 atub|OJI89726.1 atub|OJI88871.1 aver|OJJ06525.1 aver|OJJ07569.1 awen|OJJ29755.1 awen|OJJ33285.1 awen|OJJ36510.1 azon|OJJ42518.1 azon|OJJ46073.1 azon|OJJ50660.1 abra|OJJ69460.1 anid|CBF83321.1 anig|EHA28452.1 asyd|OJJ60544.1 aver|OJI99870.1 azon|OJJ44385.1 anid|CBF76070.1 anid|CBF80669.1 anig|EHA27432.1 ater|EAU29931.1 ater|EAU30227.1 anig|EHA27678.1 aory|EIT74323.1

Group24: abra|OJJ69144.1 abra|OJJ70352.1 abra|OJJ77612.1 acar|OOF91158.1 acar|OOF99116.1 acla|EAW06570.1 acla|EAW08667.1 acla|EAW13589.1 afis|EAW15744.1 afis|EAW16026.1 afis|EAW20014.1 afum|EAL84904.1 afum|EAL85604.1 afum|EAL88655.1 agla|OJJ81906.1 agla|OJJ86054.1 agla|OJJ87984.1 akaw|GAA84019.1 akaw|GAA86313.1 aluc|OJZ80922.1 aluc|OJZ86130.1 anid|CBF71034.1 anid|CBF77968.1 anid|CBF84719.1 anig|EHA19425.1 anig|EHA24491.1 aory|EIT75778.1 aory|EIT81521.1 aory|EIT83013.1 asyd|OJJ52664.1 asyd|OJJ53328.1 asyd|OJJ55266.1 asyd|OJJ60058.1 ater|EAU29366.1 ater|EAU30124.1 ater|EAU32401.1 atub|OJI79926.1 atub|OJI82658.1 aver|OJJ02231.1 aver|OJJ04691.1 aver|OJJ06960.1 aver|OJJ08416.1 awen|OJJ30028.1 awen|OJJ38444.1 azon|OJJ43750.1 azon|OJJ45430.1 azon|OJJ48427.1 ater|EAU33615.1 awen|OJJ30431.1 awen|OJJ41731.1 afla|EED46021.1 afla|EED56407.1 anid|CBF75022.1 acla|EAW12683.1 afis|EAW17846.1 afum|EAL91655.1 ater|EAU33048.1 afla|EED51306.1

Group25: abra|OJJ70961.1 abra|OJJ76031.1 acar|OOF98169.1 acar|OOF99897.1 acla|EAW09101.1 acla|EAW09782.1 acla|EAW10522.1 afis|EAW21356.1 afis|EAW25044.1 afis|EAW25699.1 afis|EAW25729.1 afla|EED45847.1 afla|EED55407.1 afla|EED55771.1 afum|EAL84400.2 afum|EAL89712.1 afum|EAL92371.2 agla|OJJ78663.1 agla|OJJ87377.1 agla|OJJ89150.1 agla|OJJ88275.1 akaw|GAA82986.1 akaw|GAA91201.1 aluc|OJZ81536.1 aluc|OJZ87571.1 aluc|OJZ89483.1 anid|CBF76249.1 anid|CBF85912.1 anig|EHA20773.1 anig|EHA23111.1 anig|EHA25900.1 aory|EIT75461.1 aory|EIT80822.1 aory|EIT75500.1 aory|EIT76538.1 asyd|OJJ59042.1 asyd|OJJ60969.1 asyd|OJJ64398.1 ater|EAU35202.1 ater|EAU35794.1 ater|EAU36445.1 ater|EAU37742.1 atub|OJI81078.1 atub|OJI85318.1 aver|OJI98494.1 aver|OJJ00294.1 aver|OJJ01637.1 awen|OJJ37057.1 awen|OJJ39342.1 awen|OJJ40655.1 azon|OJJ51688.1 ater|EAU36998.1 afla|EED54858.1 anid|CBF69709.1 atub|OJI88650.1 aory|EIT79380.1

Group26: abra|OJJ71513.1 abra|OJJ73019.1 abra|OJJ77809.1 acar|OOF96188.1 acar|OOF96228.1 acar|OOF96266.1 acar|OOF98958.1 acla|EAW07995.1 acla|EAW11308.1 acla|EAW15070.1 afis|EAW18195.1 afis|EAW23030.1 afis|EAW20042.1 afla|EED45337.1 afla|EED54986.1 afla|EED56290.1 afum|EAL85162.1 afum|EAL88452.1 afum|EAL87556.1 agla|OJJ85896.1 agla|OJJ89448.1 agla|OJJ88920.1 akaw|GAA83840.1 akaw|GAA85596.1 akaw|GAA89403.1 aluc|OJZ83533.1 aluc|OJZ86251.1 aluc|OJZ90876.1 anid|CBF80914.1 anid|CBF86102.1 anig|EHA19596.1 anig|EHA24355.1 anig|EHA21614.1 aory|EIT73286.1 aory|EIT80288.1 asyd|OJJ56326.1 asyd|OJJ64481.1 ater|EAU29627.1 ater|EAU33801.1 atub|OJI82424.1 atub|OJI82873.1 atub|OJI84291.1 aver|OJI98582.1 aver|OJJ06387.1 awen|OJJ30557.1 awen|OJJ35716.1 azon|OJJ44174.1 azon|OJJ45510.1 azon|OJJ46319.1 anid|CBF84784.1 anid|CBF84786.1 aory|EIT73629.1 asyd|OJJ63232.1 ater|EAU38649.1 aver|OJI97724.1 awen|OJJ31798.1

Group27: abra|OJJ67095.1 abra|OJJ67686.1 abra|OJJ69631.1 abra|OJJ76432.1 acar|OOF92679.1 acar|OOF94074.1 acar|OOF94144.1 acar|OOF99676.1 acla|EAW10021.1 acla|EAW12377.1 acla|EAW13992.1 afis|EAW16823.1 afis|EAW18700.1 afis|EAW22316.1 afla|EED50648.1 afla|EED55075.1 afum|EAL86374.1 afum|EAL90874.1 afum|EAL93422.1 agla|OJJ79102.1 agla|OJJ82361.1 akaw|GAA82347.1 akaw|GAA89198.1 akaw|GAA91066.1 akaw|GAA91611.1 aluc|OJZ81632.1 aluc|OJZ82438.1 aluc|OJZ87880.1 aluc|OJZ91780.1 anid|CBF70637.1 anid|CBF82989.1 anig|EHA20183.1 anig|EHA25811.1 anig|EHA28335.1 aory|EIT78077.1 aory|EIT81953.1 aory|EIT82431.1 asyd|OJJ52074.1 asyd|OJJ54406.1 asyd|OJJ58493.1 asyd|OJJ64734.1 ater|EAU29368.1 ater|EAU35424.1 ater|EAU38206.1 atub|OJI80771.1 atub|OJI86236.1 atub|OJI88003.1 atub|OJI88548.1 aver|OJI98838.1 aver|OJJ01096.1 aver|OJJ03029.1 awen|OJJ32183.1 awen|OJJ40928.1 azon|OJJ46764.1 anig|EHA22777.1

Group28: abra|OJJ68117.1 abra|OJJ77078.1 abra|OJJ68815.1 acar|OOF90616.1 acar|OOF92960.1 acar|OOF93911.1 afis|EAW17741.1 afis|EAW17974.1 afla|EED45883.1 afla|EED55321.1 afum|EAL91555.1 afum|EAL91783.1 agla|OJJ81756.1 agla|OJJ85465.1 agla|OJJ89268.1 akaw|GAA86898.1 akaw|GAA86975.1 akaw|GAA90440.1 aluc|OJZ84488.1 aluc|OJZ85787.1 aluc|OJZ88708.1 aluc|OJZ90021.1 anid|CBF71442.1 anid|CBF84572.1 anig|EHA24771.1 anig|EHA27397.1 anig|EHA27792.1 aory|EIT75581.1 aory|EIT76204.1 asyd|OJJ53389.1 asyd|OJJ54954.1 ater|EAU30960.1 ater|EAU35114.1 ater|EAU36891.1 atub|OJI79571.1 atub|OJI84786.1 atub|OJI89592.1 atub|OJI91296.1 aver|OJI99661.1 aver|OJJ03907.1 aver|OJJ07242.1 aver|OJJ08354.1 awen|OJJ32944.1 awen|OJJ39601.1 azon|OJJ42107.1 azon|OJJ42757.1 azon|OJJ42786.1 azon|OJJ44596.1 azon|OJJ47560.1 acla|EAW11105.1 afla|EED52131.1 acla|EAW12809.1 ater|EAU36942.1 awen|OJJ32177.1 aory|EIT74465.1

Group30: abra|OJJ68096.1 abra|OJJ70248.1 abra|OJJ73110.1 abra|OJJ74007.1 acar|OOF92551.1 acar|OOF93899.1 acar|OOG00337.1 acar|OOG01129.1 afla|EED44907.1 afla|EED44932.1 afla|EED56562.1 agla|OJJ78703.1 agla|OJJ88418.1 akaw|GAA86542.1 akaw|GAA86880.1 akaw|GAA88056.1 akaw|GAA92925.1 akaw|GAA87488.1 aluc|OJZ81318.1 aluc|OJZ82516.1 aluc|OJZ85090.1 aluc|OJZ88037.1 aluc|OJZ90004.1

anid|CBF74975.1 anid|CBF78035.1 anid|CBF81473.1 anig|EHA17810.1 anig|EHA18624.1 anig|EHA25311.1 anig|EHA27379.1 anig|EHA20903.1 aory|EIT73500.1 aory|EIT73507.1 asyd|OJJ55224.1 asyd|OJJ59448.1 asyd|OJJ60089.1 asyd|OJJ61384.1 asyd|OJJ57439.1 ater|EAU39548.1 ater|EAU39587.1 atub|OJI80509.1 atub|OJI84804.1 atub|OJI91521.1 atub|OJI90644.1 aver|OJJ00711.1 aver|OJJ04176.1 aver|OJJ05739.1 aver|OJJ02824.1 awen|OJJ33639.1 awen|OJJ40047.1 acla|EAW07039.1 afis|EAW16665.1 afum|EAL89269.1 azon|OJJ50123.1

Group31: abra|OJJ69067.1 abra|OJJ70519.1 abra|OJJ77965.1 acar|OOF91229.1 acar|OOF97409.1 acar|OOF98850.1 acla|EAW08156.1 acla|EAW08763.1 afis|EAW15666.1 afis|EAW16089.1 afis|EAW23194.1 afla|EED46889.1 afla|EED47060.1 afla|EED48625.1 afum|EAL85682.1 afum|EAL88289.1 afum|EAL88721.1 agla|OJJ81804.1 agla|OJJ86130.1 agla|OJJ86345.1 akaw|GAA83366.1 akaw|GAA83698.1 akaw|GAA90597.1 aluc|OJZ83864.1 aluc|OJZ91035.1 aluc|OJZ86055.1 anid|CBF71178.1 anid|CBF74704.1 anid|CBF75129.1 anig|EHA19005.1 anig|EHA19734.1 anig|EHA24556.1 aory|EIT72496.1 aory|EIT73097.1 asyd|OJJ55863.1 asyd|OJJ60218.1 asyd|OJJ59982.1 ater|EAU32505.1 ater|EAU33161.1 ater|EAU36321.1 atub|OJI79852.1 atub|OJI83031.1 atub|OJI90232.1 aver|OJJ02309.1 aver|OJJ04050.1 aver|OJJ04804.1 awen|OJJ30341.1 awen|OJJ38341.1 awen|OJJ36585.1 azon|OJJ42517.1 azon|OJJ43041.1 aory|EIT72708.1 acla|EAW06676.1 azon|OJJ47622.1

Group33: abra|OJJ66203.1 abra|OJJ68990.1 abra|OJJ74346.1 acar|OOF91561.1 acar|OOF91930.1 acar|OOF95383.1 acla|EAW07400.1 acla|EAW14202.1 afis|EAW19080.1 afis|EAW22426.1 afla|EED50168.1 afla|EED52504.1 afum|EAL91073.1 afum|EAL90765.1 afum|EAL93809.1 agla|OJJ84526.1 agla|OJJ87597.1 agla|OJJ84718.1 akaw|GAA86684.1 akaw|GAA89082.1 aluc|OJZ80508.1 aluc|OJZ85975.1 aluc|OJZ89784.1 anid|CBF76870.1 anid|CBF79672.1 anid|CBF88817.1 anig|EHA24631.1 anig|EHA26999.1 anig|EHA25684.1 aory|EIT76001.1 aory|EIT82781.1 asyd|OJJ52200.1 asyd|OJJ58034.1 asyd|OJJ61321.1 asyd|OJJ62887.1 ater|EAU32126.1 ater|EAU32863.1 ater|EAU32522.1 ater|EAU39281.1 atub|OJI85021.1 atub|OJI88421.1 aver|OJI96939.1 aver|OJJ00646.1 aver|OJJ04365.1 aver|OJJ05950.1 aver|OJJ07745.1 awen|OJJ32472.1 awen|OJJ35005.1 awen|OJJ34090.1 azon|OJJ45647.1 azon|OJJ46232.1 akaw|GAA90510.1 atub|OJI79765.1

Group34: abra|OJJ71614.1 abra|OJJ77069.1 acar|OOF92952.1 acar|OOF96330.1 acla|EAW13507.1 acla|EAW15156.1 afis|EAW20123.1 afis|EAW21377.1 afis|EAW20124.1 afis|EAW20605.1 afla|EED47726.1 afla|EED57252.1 afum|EAL84455.1 afum|EAL85242.1 agla|OJJ88636.1 agla|OJJ88823.1 akaw|GAA86984.1 akaw|GAA89313.1 aluc|OJZ84477.1 aluc|OJZ86355.1 anid|CBF84947.1 anid|CBF86944.1 anig|EHA21535.1 anig|EHA27785.1 aory|EIT73980.1 aory|EIT77148.1 asyd|OJJ55590.1 asyd|OJJ63148.1 ater|EAU35261.1 ater|EAU38740.1 atub|OJI84179.1 atub|OJI89602.1 aver|OJI97806.1 aver|OJJ06669.1 awen|OJJ31893.1 awen|OJJ33448.1 azon|OJJ46243.1 azon|OJJ50705.1 abra|OJJ71611.1 acar|OOF96327.1 acla|EAW15155.1 afla|EED57253.1 afum|EAL85241.2 agla|OJJ88828.1 akaw|GAA89316.1 anid|CBF84945.1 aory|EIT74161.1 asyd|OJJ63149.1 atub|OJI84182.1 aver|OJI97805.1 awen|OJJ31892.1 azon|OJJ42421.1 azon|OJJ46244.1

Group35: afis|EAW16487.1 afis|EAW19960.1 afla|EED48721.1 afla|EED49671.1 afum|EAL84955.1 afum|EAL89081.1 agla|OJJ79324.1 agla|OJJ82857.1 agla|OJJ88164.1 aory|EIT73140.1 aory|EIT80950.1 aory|EIT75645.1 aory|EIT79202.1 ater|EAU35270.1 ater|EAU36551.1 abra|OJJ68065.1 acla|EAW06870.1 afla|EED45908.1 akaw|GAA90894.1 aluc|OJZ82935.1 anid|CBF75512.1 anid|CBF84598.1 anig|EHA21355.1 anig|EHA22760.1 asyd|OJJ53379.1 atub|OJI89164.1 aver|OJJ02547.1 awen|OJJ29993.1 azon|OJJ49197.1 abra|OJJ71770.1 akaw|GAA85137.1 aluc|OJZ86500.1 anig|EHA22144.1 atub|OJI84026.1 azon|OJJ48613.1 abra|OJJ78207.1 acar|OOF98647.1 acla|EAW11241.1 afis|EAW18262.1 afum|EAL87626.1 akaw|GAA85819.1 aluc|OJZ91280.1 anid|CBF79531.1 anig|EHA19965.1 asyd|OJJ57895.1 ater|EAU33270.1 atub|OJI83270.1 aver|OJJ04500.1 awen|OJJ30636.1 afla|EED47227.1 asyd|OJJ59741.1 awen|OJJ40346.1 aver|OJJ08364.1

Group37: abra|OJJ72940.1 abra|OJJ77077.1 acar|OOF92763.1 acar|OOF92959.1 acar|OOF97146.1 afis|EAW18980.1 afis|EAW21099.1 afis|EAW24628.1 afla|EED53187.1 afla|EED54887.1 agla|OJJ80287.1 agla|OJJ82940.1 akaw|GAA86976.1 akaw|GAA91285.1 aluc|OJZ82784.1 aluc|OJZ84487.1 anig|EHA18922.1 anig|EHA21226.1 aory|EIT75171.1 aory|EIT79411.1 asyd|OJJ58358.1 asyd|OJJ60157.1 ater|EAU34862.1 ater|EAU37837.1 atub|OJI89016.1 atub|OJI89593.1 aver|OJJ02049.1 aver|OJJ04107.1 awen|OJJ31588.1 awen|OJJ37322.1 azon|OJJ47304.1 azon|OJJ47677.1 abra|OJJ70575.1 acla|EAW08374.1 afis|EAW23403.1 afla|EED47010.1 afum|EAL88067.1 agla|OJJ86391.1 akaw|GAA83419.1 aluc|OJZ83919.1 anid|CBF74791.1 anig|EHA19052.1 aory|EIT72601.1 ater|EAU36372.1 atub|OJI90290.1 awen|OJJ39474.1 acla|EAW14304.1 afum|EAL93716.2 anid|CBF69653.1 asyd|OJJ59075.1 aver|OJJ01668.1 anig|EHA27791.1

Group38: abra|OJJ65887.1 abra|OJJ73728.1 abra|OJJ73840.1 abra|OJJ75062.1 acar|OOF94948.1 acar|OOF97272.1 acar|OOG00894.1 acar|OOG00991.1 acla|EAW07735.1 acla|EAW09321.1 afis|EAW21824.1 afis|EAW24409.1 afla|EED45566.1 afla|EED50029.1 afum|EAL86078.1 afum|EAL92840.1 agla|OJJ83601.1 agla|OJJ85455.1 agla|OJJ89020.1 akaw|GAA86017.1 akaw|GAA88434.1 akaw|GAA88674.1 akaw|GAA89458.1 aluc|OJZ88209.1 aluc|OJZ88322.1 aluc|OJZ90245.1 aluc|OJZ92628.1 anid|CBF84638.1 anid|CBF87998.1 anig|EHA17956.1 anig|EHA18058.1 anig|EHA26362.1 aory|EIT74733.1 aory|EIT75750.1 asyd|OJJ53366.1 asyd|OJJ63506.1 ater|EAU38301.1 ater|EAU38950.1 atub|OJI81673.1 atub|OJI87058.1 atub|OJI90810.1 atub|OJI90917.1 aver|OJI97424.1 aver|OJJ08381.1 awen|OJJ32729.1 awen|OJJ37942.1 awen|OJJ38966.1 awen|OJJ41210.1 azon|OJJ46022.1 azon|OJJ48676.1 aory|EIT76918.1

Group39: abra|OJJ66342.1 abra|OJJ71162.1 abra|OJJ75572.1 acar|OOF89825.1 acar|OOF94094.1 acar|OOF94165.1 acar|OOF97998.1 acla|EAW09994.1 acla|EAW10337.1 afis|EAW21096.1 afis|EAW25252.1

afla|EED51678.1 afla|EED53719.1 afla|EED54679.1 afum|EAL86170.1 afum|EAL89914.1 afum|EAL92119.1 agla|OJJ80114.1 agla|OJJ89225.1 akaw|GAA82817.1 akaw|GAA84323.1 akaw|GAA90216.1 aluc|OJZ83127.1 aluc|OJZ87378.1 aluc|OJZ89020.1 anid|CBF80117.1 anid|CBF85345.1 anig|EHA20610.1 anig|EHA23510.1 anig|EHA28188.1 aory|EIT75350.1 aory|EIT78900.1 aory|EIT80684.1 asyd|OJJ54726.1 asyd|OJJ64027.1 atub|OJI81268.1 atub|OJI85769.1 atub|OJI87806.1 aver|OJI98113.1 aver|OJJ07441.1 awen|OJJ36352.1 awen|OJJ40966.1 awen|OJJ38784.1 ater|EAU34396.1 azon|OJJ44550.1 abra|OJJ67575.1 acar|OOF90092.1 akaw|GAA89104.1 anig|EHA25706.1 atub|OJI88443.1 afis|EAW24500.1

Group40: abra|OJJ73400.1 abra|OJJ73868.1 acar|OOG00598.1 acar|OOG01011.1 acla|EAW07007.1 acla|EAW07756.1 afis|EAW16623.1 afis|EAW19494.1 afis|EAW22793.1 afla|EED44972.1 afla|EED48978.1 afla|EED50061.1 afla|EED56714.1 afum|EAL87000.1 afum|EAL89214.1 afum|EAL90406.1 agla|OJJ83569.1 agla|OJJ85298.1 akaw|GAA83092.1 akaw|GAA89474.1 akaw|GAA89692.1 aluc|OJZ80692.1 aluc|OJZ88189.1 aluc|OJZ88639.1 anid|CBF81984.1 anid|CBF87954.1 anig|EHA17940.1 anig|EHA18328.1 anig|EHA27323.1 aory|EIT73515.1 aory|EIT73856.1 aory|EIT74647.1 aory|EIT79983.1 asyd|OJJ56154.1 asyd|OJJ59777.1 asyd|OJJ63479.1 ater|EAU34806.1 ater|EAU36732.1 ater|EAU38918.1 atub|OJI87552.1 atub|OJI90788.1 atub|OJI91225.1 aver|OJI97451.1 aver|OJJ02509.1 aver|OJJ05094.1 awen|OJJ36912.1 awen|OJJ38940.1 azon|OJJ45543.1 azon|OJJ49714.1 anid|CBF75442.1

Group41: abra|OJJ65606.1 abra|OJJ68131.1 abra|OJJ68372.1 abra|OJJ69834.1 acar|OOF90085.1 acar|OOF93362.1 acar|OOG00535.1 afla|EED45615.1 afla|EED45682.1 afla|EED54532.1 agla|OJJ80470.1 agla|OJJ80842.1 agla|OJJ83418.1 akaw|GAA86911.1 akaw|GAA87826.1 aluc|OJZ85462.1 aluc|OJZ90034.1 anig|EHA25004.1 anig|EHA25073.1 anig|EHA27410.1 aory|EIT75511.1 aory|EIT75540.1 aory|EIT78869.1 aory|EIT81830.1 asyd|OJJ52349.1 asyd|OJJ56735.1 asyd|OJJ60489.1 atub|OJI80101.1 atub|OJI84772.1 aver|OJI99815.1 aver|OJJ05882.1 aver|OJJ05988.1 awen|OJJ32535.1 awen|OJJ33535.1 awen|OJJ34624.1 awen|OJJ35129.1 awen|OJJ40106.1 acla|EAW07472.1 afis|EAW17054.1 afum|EAL86551.2 akaw|GAA83834.1 aluc|OJZ90883.1 anid|CBF86106.1 ater|EAU30279.1 atub|OJI82467.1 anig|EHA19603.1 atub|OJI82878.1 azon|OJJ47403.1 afla|EED57508.1

Group42: abra|OJJ65781.1 abra|OJJ69519.1 abra|OJJ69805.1 acar|OOF93570.1 acar|OOF94236.1 acla|EAW09965.1 acla|EAW15041.1 afis|EAW17149.1 afis|EAW21123.1 afis|EAW24564.1 afla|EED50957.1 afla|EED51796.1 afla|EED54117.1 afla|EED57909.1 afum|EAL86225.1 afum|EAL92146.1 agla|OJJ78698.1 agla|OJJ79809.1 agla|OJJ80088.1 akaw|GAA87854.1 akaw|GAA88307.1 akaw|GAA90882.1 aluc|OJZ82358.1 aluc|OJZ82950.1 aluc|OJZ85496.1 anid|CBF77670.1 anid|CBF87020.1 anig|EHA21430.1 anig|EHA25029.1 anig|EHA28405.1 aory|EIT76140.1 aory|EIT77042.1 aory|EIT81685.1 aory|EIT82334.1 asyd|OJJ53106.1 asyd|OJJ60460.1 ater|EAU32188.1 ater|EAU36777.1 atub|OJI80069.1 atub|OJI88083.1 atub|OJI89182.1 aver|OJI99780.1 aver|OJJ03746.1 awen|OJJ36453.1 awen|OJJ37008.1 awen|OJJ40041.1 azon|OJJ44424.1 azon|OJJ50127.1 aver|OJI99293.1

Group43: abra|OJJ67343.1 abra|OJJ74161.1 abra|OJJ74904.1 acar|OOF92471.1 acar|OOF95566.1 acar|OOF95902.1 afis|EAW18497.1 afis|EAW23515.1 afis|EAW24769.1 afum|EAL89410.1 afum|EAL93187.1 agla|OJJ83565.1 agla|OJJ85096.1 agla|OJJ86763.1 agla|OJJ88214.1 akaw|GAA83518.1 akaw|GAA86474.1 akaw|GAA89723.1 aluc|OJZ80660.1 aluc|OJZ85015.1 aluc|OJZ92477.1 anid|CBF70261.1 anid|CBF89705.1 anig|EHA25236.1 anig|EHA26493.1 anig|EHA27172.1 aory|EIT77595.1 aory|EIT78639.1 aory|EIT80122.1 asyd|OJJ58753.1 asyd|OJJ62252.1 asyd|OJJ62467.1 atub|OJI80582.1 atub|OJI86908.1 atub|OJI87516.1 aver|OJI96362.1 aver|OJJ01361.1 awen|OJJ31051.1 awen|OJJ37928.1 awen|OJJ39840.1 acla|EAW08468.1 afla|EED49731.1 afum|EAL87948.1 anid|CBF89382.1 ater|EAU35304.1 azon|OJJ43252.1 ater|EAU34034.1 aver|OJI96602.1 afla|EED57415.1

Group44: abra|OJJ68123.1 abra|OJJ69603.1 abra|OJJ72218.1 acar|OOF92701.1 acar|OOF93917.1 acar|OOF96841.1 acla|EAW12866.1 acla|EAW14975.1 afis|EAW17815.1 afis|EAW24499.1 afla|EED45599.1 afla|EED51849.1 afla|EED54703.1 afum|EAL86169.1 afum|EAL91624.1 agla|OJJ78971.1 agla|OJJ85550.1 agla|OJJ81686.1 akaw|GAA84717.1 akaw|GAA86904.1 akaw|GAA91591.1 aluc|OJZ82414.1 aluc|OJZ86970.1 aluc|OJZ90027.1 anig|EHA21680.1 anig|EHA27403.1 anig|EHA28353.1 aory|EIT75721.1 aory|EIT78771.1 aory|EIT81733.1 asyd|OJJ54895.1 asyd|OJJ58303.1 asyd|OJJ56516.1 asyd|OJJ56725.1 atub|OJI83551.1 atub|OJI84780.1 atub|OJI88027.1 aver|OJJ02109.1 aver|OJJ05998.1 aver|OJJ06206.1 awen|OJJ32626.1 awen|OJJ36932.1 anid|CBF73850.1 ater|EAU30287.1 azon|OJJ50345.1 anid|CBF80089.1 ater|EAU31562.1 awen|OJJ33657.1 ater|EAU29952.1

Group45: abra|OJJ68536.1 abra|OJJ72481.1 acar|OOF90150.1 acar|OOF94753.1 acla|EAW10385.1 acla|EAW12826.1 afis|EAW17874.1 afis|EAW17991.1 afis|EAW25211.1 afla|EED45114.1 afla|EED51912.1 afla|EED46241.1 afum|EAL89872.1 afum|EAL91801.1 afum|EAL91682.1 agla|OJJ80251.1 agla|OJJ83074.1 akaw|GAA86216.1 akaw|GAA91315.1 aluc|OJZ82750.1 aluc|OJZ90468.1 anid|CBF79986.1 anid|CBF89933.1 anig|EHA21199.1 anig|EHA23927.1 aory|EIT73554.1 aory|EIT76181.1 aory|EIT76362.1 ater|EAU31453.1 ater|EAU37217.1 ater|EAU31502.1 ater|EAU33093.1 atub|OJI81892.1 atub|OJI88986.1 aver|OJI96221.1 aver|OJJ02185.1 awen|OJJ41004.1 awen|OJJ41694.1 azon|OJJ45116.1 azon|OJJ46814.1 azon|OJJ48270.1 azon|OJJ48393.1 acla|EAW12711.1 asyd|OJJ58240.1 asyd|OJJ62114.1 awen|OJJ41555.1 anid|CBF69461.1 aver|OJJ01892.1 asyd|OJJ59277.1

Group46: abra|OJJ72575.1 abra|OJJ77969.1 acar|OOF94684.1 acar|OOF98846.1 akaw|GAA82423.1 akaw|GAA87621.1 akaw|GAA83693.1 akaw|GAA85201.1 aluc|OJZ90556.1 aluc|OJZ91039.1 anig|EHA19738.1

anig|EHA24003.1 asyd|OJJ54419.1 asyd|OJJ59096.1 atub|OJI81979.1 atub|OJI83035.1 acla|EAW06671.1 afis|EAW16094.1 afla|EED48630.1 afla|EED56583.1 afum|EAL88726.1 agla|OJJ88578.1 anid|CBF85458.1 aory|EIT73102.1 aory|EIT79674.1 asyd|OJJ58704.1 asyd|OJJ64100.1 ater|EAU30075.1 ater|EAU33170.1 aver|OJI98183.1 aver|OJJ01316.1 awen|OJJ30336.1 awen|OJJ40588.1 azon|OJJ42310.1 azon|OJJ45364.1 abra|OJJ76510.1 acar|OOF99461.1 afla|EED54610.1 agla|OJJ87529.1 aluc|OJZ91695.1 anig|EHA22708.1 aory|EIT78921.1 ater|EAU38073.1 atub|OJI86149.1 aver|OJJ01687.1 aver|OJJ06547.1 awen|OJJ36612.1 agla|OJJ86127.1 anid|CBF70342.1

Group47: acar|OOF91492.1 acar|OOF98333.1 acar|OOF92665.1 acar|OOF93877.1 afis|EAW15510.1 afis|EAW17137.1 afis|EAW16415.1 afis|EAW16568.1 afis|EAW17138.1 afis|EAW25618.1 afum|EAL89039.1 afum|EAL89165.1 anid|CBF80635.1 anid|CBF89254.1 ater|EAU31707.1 ater|EAU36836.1 abra|OJJ74339.1 acla|EAW07389.1 afis|EAW22415.1 afla|EED52493.1 afum|EAL90777.1 agla|OJJ87679.1 akaw|GAA86691.1 aluc|OJZ89792.1 anid|CBF88800.1 anig|EHA27005.1 aory|EIT82748.1 asyd|OJJ62896.1 ater|EAU39292.1 atub|OJI85014.1 aver|OJI96948.1 awen|OJJ34994.1 azon|OJJ45637.1 agla|OJJ82753.1 anig|EHA22499.1 atub|OJI84627.1 afis|EAW19287.1 acla|EAW06953.1 awen|OJJ40251.1 azon|OJJ49111.1 acar|OOG00127.1 akaw|GAA86019.1 aluc|OJZ90247.1 asyd|OJJ61228.1 aver|OJJ00551.1 anid|CBF76692.1 akaw|GAA91465.1 aluc|OJZ82070.1 afum|EAL84308.2

Group48: abra|OJJ66388.1 abra|OJJ69074.1 acla|EAW08755.1 acla|EAW11763.1 afis|EAW15677.1 afis|EAW23796.1 afis|EAW25170.1 afla|EED46854.1 afla|EED51603.1 afla|EED53628.1 afum|EAL85670.1 afum|EAL87230.1 afum|EAL89843.1 akaw|GAA90254.1 akaw|GAA90605.1 aluc|OJZ83086.1 aluc|OJZ86063.1 anid|CBF71157.1 anid|CBF79858.1 anid|CBF85251.1 aory|EIT72587.1 aory|EIT74962.1 aory|EIT80702.1 asyd|OJJ51987.1 asyd|OJJ54461.1 asyd|OJJ54551.1 asyd|OJJ55842.1 asyd|OJJ58159.1 asyd|OJJ59172.1 asyd|OJJ62925.1 asyd|OJJ64763.1 ater|EAU31334.1 ater|EAU32484.1 ater|EAU34334.1 atub|OJI79860.1 atub|OJI87858.1 aver|OJI98064.1 aver|OJJ04243.1 aver|OJJ04785.1 aver|OJJ07708.1 awen|OJJ30141.1 awen|OJJ33753.1 awen|OJJ38354.1 azon|OJJ42861.1 azon|OJJ43857.1 acar|OOF91216.1 agla|OJJ81826.1 anig|EHA24550.1

Group49: abra|OJJ66054.1 abra|OJJ67991.1 abra|OJJ68633.1 acar|OOF92033.1 acar|OOF92137.1 acla|EAW07055.1 acla|EAW12619.1 afis|EAW16409.1 afis|EAW16681.1 afis|EAW18065.1 afla|EED45379.1 afla|EED47440.1 afla|EED51440.1 afum|EAL84867.1 afum|EAL89033.1 afum|EAL89285.1 afum|EAL91877.1 akaw|GAA89772.1 akaw|GAA90957.1 aluc|OJZ81922.1 aluc|OJZ82865.1 anid|CBF73413.1 anid|CBF86422.1 anid|CBF87201.1 anig|EHA21295.1 anig|EHA22325.1 aory|EIT80529.1 aory|EIT81893.1 aory|EIT82380.1 asyd|OJJ59980.1 asyd|OJJ64938.1 ater|EAU30466.1 ater|EAU35871.1 atub|OJI84474.1 atub|OJI89090.1 aver|OJI99059.1 aver|OJJ02311.1 awen|OJJ36169.1 awen|OJJ41452.1 azon|OJJ47178.1 azon|OJJ48910.1 agla|OJJ87352.1 anid|CBF75131.1 aver|OJJ01785.1 asyd|OJJ59979.1 anid|CBF75132.1 aver|OJJ02312.1

Group50: abra|OJJ69511.1 abra|OJJ74314.1 acar|OOF94242.1 acar|OOF95413.1 acla|EAW07365.1 acla|EAW09962.1 afis|EAW21133.1 afis|EAW22392.1 afla|EED46140.1 afla|EED56055.1 afum|EAL90799.1 afum|EAL92151.1 agla|OJJ80083.1 agla|OJJ87654.1 akaw|GAA86711.1 akaw|GAA88313.1 aluc|OJZ82349.1 aluc|OJZ89812.1 anid|CBF87003.1 anid|CBF88741.1 anig|EHA27022.1 anig|EHA28411.1 aory|EIT76474.1 aory|EIT76698.1 asyd|OJJ60468.1 asyd|OJJ63953.1 ater|EAU36765.1 ater|EAU39326.1 atub|OJI84993.1 atub|OJI88091.1 aver|OJI96988.1 aver|OJI99788.1 awen|OJJ34969.1 awen|OJJ36463.1 acar|OOF94243.1 acla|EAW09961.1 afis|EAW21134.1 afum|EAL92153.1 agla|OJJ80077.1 anid|CBF87002.1 aory|EIT76417.1 ater|EAU36766.1 awen|OJJ36462.1 azon|OJJ44415.1 azon|OJJ44422.1 azon|OJJ45618.1 afla|EED46139.1

Group55: abra|OJJ66953.1 abra|OJJ70025.1 acar|OOF93535.1 acar|OOF93655.1 agla|OJJ79210.1 agla|OJJ85149.1 akaw|GAA87635.1 akaw|GAA92208.1 aluc|OJZ85251.1 aluc|OJZ85625.1 anig|EHA24903.1 anig|EHA25451.1 aory|EIT80438.1 aory|EIT82112.1 asyd|OJJ55882.1 asyd|OJJ65094.1 ater|EAU32531.1 ater|EAU37243.1 atub|OJI79418.1 atub|OJI80318.1 aver|OJI99218.1 aver|OJJ04827.1 awen|OJJ30173.1 awen|OJJ38273.1 awen|OJJ41598.1 acla|EAW08800.1 afis|EAW15597.1 afla|EED50419.1 afum|EBA27239.1 anid|CBF78088.1 azon|OJJ43909.1 asyd|OJJ55868.1 acar|OOF93534.1 acla|EAW08799.1 afis|EAW15598.1 afum|EAL85734.1 agla|OJJ88612.1 akaw|GAA87636.1 anid|CBF78056.1 anig|EHA25452.1 aory|EIT82031.1 ater|EAU32530.1 atub|OJI80317.1 aver|OJJ04811.1 azon|OJJ43905.1

Group56: agla|OJJ81571.1 agla|OJJ85255.1 asyd|OJJ54206.1 asyd|OJJ59064.1 asyd|OJJ54427.1 ater|EAU31423.1 ater|EAU38101.1 aver|OJJ01658.1 aver|OJJ01906.1 awen|OJJ30145.1 awen|OJJ31520.1 abra|OJJ76240.1 acar|OOF99485.1 acla|EAW14198.1 afis|EAW18923.1 afla|EED50910.1 afum|EAL93655.1 agla|OJJ85250.1 akaw|GAA82174.1 aluc|OJZ91991.1 anid|CBF69674.1 anig|EHA22937.1 aory|EIT79045.1 asyd|OJJ59065.1 ater|EAU38098.1 atub|OJI86426.1 awen|OJJ31518.1 azon|OJJ47234.1 abra|OJJ76243.1 acar|OOF99488.1 acla|EAW06837.1 afis|EAW18926.1 afla|EED50916.1 afum|EAL93657.1 akaw|GAA82176.1 aluc|OJZ91988.1 anid|CBF69676.1 anig|EHA22935.1 aory|EIT78980.1 atub|OJI86423.1 azon|OJJ47236.1 afis|EAW16385.1 afla|EED48287.1 afum|EAL89013.2 aory|EIT72910.1

Group57: abra|OJJ69646.1 abra|OJJ77015.1 abra|OJJ77136.1 acar|OOF90285.1 acar|OOF91423.1 acar|OOF91458.1 acla|EAW09001.1 acla|EAW10860.1 acla|EAW12947.1 acla|EAW13503.1 afis|EAW17722.1 afis|EAW20602.1 afis|EAW20998.1 afum|EAL84459.1 afum|EAL91537.1 agla|OJJ78915.1 agla|OJJ79354.1 agla|OJJ88647.1 akaw|GAA87035.1 akaw|GAA90793.1 akaw|GAA91624.1 aluc|OJZ82450.1 aluc|OJZ84416.1 aluc|OJZ84546.1 anig|EHA27739.1 anig|EHA27843.1 anig|EHA28324.1 aory|EIT75439.1 aory|EIT77113.1 asyd|OJJ52539.1 asyd|OJJ55599.1 atub|OJI87989.1 atub|OJI89534.1 atub|OJI89658.1 aver|OJJ03865.1

aver|OJJ06679.1 awen|OJJ33394.1 awen|OJJ35790.1 afla|EED45724.1 afla|EED47731.1 anid|CBF86932.1 ater|EAU35247.1 azon|OJJ43330.1 agla|OJJ84075.1

Group58: acar|OOF92599.1 acar|OOF92620.1 abra|OJJ72332.1 acar|OOF94882.1 acla|EAW11990.1 afis|EAW19630.1 afla|EED56982.1 afum|EAL86892.1 agla|OJJ80775.1 akaw|GAA86084.1 aluc|OJZ90316.1 anid|CBF71323.1 anig|EHA23803.1 aory|EIT77586.1 asyd|OJJ55109.1 ater|EAU32896.1 atub|OJI81744.1 aver|OJJ07097.1 awen|OJJ41874.1 azon|OJJ47072.1 abra|OJJ72333.1 acar|OOF94880.1 acla|EAW11989.1 afis|EAW19629.1 afla|EED56981.1 afum|EAL86893.1 akaw|GAA86085.1 anid|CBF71321.1 aory|EIT77447.1 ater|EAU32897.1 azon|OJJ47071.1 abra|OJJ72335.1 acla|EAW11987.1 afis|EAW19627.1 afla|EED56979.1 afum|EAL86894.1 agla|OJJ80773.1 akaw|GAA86087.1 anid|CBF71319.1 aory|EIT77480.1 asyd|OJJ55111.1 ater|EAU32899.1 aver|OJJ07095.1 azon|OJJ47069.1

Group59: acar|OOF92600.1 acar|OOF92621.1 abra|OJJ76130.1 acar|OOF99793.1 acla|EAW09689.1 afis|EAW21450.1 afla|EED55879.1 afum|EAL92469.1 agla|OJJ85598.1 akaw|GAA88837.1 aluc|OJZ89591.1 anid|CBF76555.1 anig|EHA23020.1 aory|EIT76712.1 asyd|OJJ61157.1 ater|EAU35092.1 atub|OJI85211.1 aver|OJJ00479.1 awen|OJJ39248.1 azon|OJJ51607.1 abra|OJJ76131.1 acar|OOF99792.1 acla|EAW09687.1 afis|EAW21452.1 afla|EED55881.1 afum|EAL92471.1 akaw|GAA88836.1 aluc|OJZ89592.1 anid|CBF76551.1 anig|EHA23019.1 aory|EIT76739.1 ater|EAU35091.1 atub|OJI85210.1 abra|OJJ77110.1 acla|EAW09688.1 afis|EAW21451.1 afla|EED55880.1 afum|EAL92470.1 akaw|GAA90818.1 aluc|OJZ84519.1 anid|CBF76553.1 anig|EHA27817.1 aory|EIT76614.1 atub|OJI89561.1

Group61: abra|OJJ65992.1 abra|OJJ66994.1 abra|OJJ68819.1 acar|OOF90628.1 acar|OOF92880.1 afis|EAW20003.1 afis|EAW20740.1 afla|EED46008.1 afla|EED47482.1 afla|EED49530.1 agla|OJJ87844.1 agla|OJJ87962.1 akaw|GAA90432.1 akaw|GAA90866.1 akaw|GAA92170.1 aluc|OJZ82967.1 aluc|OJZ85578.1 aluc|OJZ85795.1 anig|EHA18425.1 anig|EHA21383.1 anig|EHA24761.1 anig|EHA24941.1 aory|EIT75392.1 aory|EIT79965.1 aory|EIT82595.1 asyd|OJJ52756.1 asyd|OJJ53312.1 asyd|OJJ54834.1 atub|OJI79374.1 atub|OJI79581.1 atub|OJI89200.1 atub|OJI91621.1 aver|OJJ07327.1 aver|OJJ08431.1 aver|OJJ08738.1 azon|OJJ43639.1 azon|OJJ50209.1 acla|EAW13602.1 afum|EAL84916.1 anid|CBF80344.1 anid|CBF84693.1 ater|EAU37349.1 awen|OJJ30010.1

Group62: abra|OJJ68708.1 abra|OJJ69824.1 acar|OOF92210.1 acar|OOF93339.1 acla|EAW08971.1 acla|EAW12537.1 afis|EAW15450.1 afis|EAW24112.1 afla|EED46821.1 afla|EED56823.1 akaw|GAA87836.1 akaw|GAA89862.1 aluc|OJZ81821.1 aluc|OJZ85472.1 anig|EHA22396.1 anig|EHA25013.1 atub|OJI80091.1 atub|OJI84392.1 awen|OJJ41377.1 awen|OJJ41767.1 abra|OJJ68707.1 acar|OOF92209.1 acla|EAW12538.1 afis|EAW24111.1 afum|EAL91965.1 agla|OJJ87188.1 akaw|GAA89861.1 aluc|OJZ81823.1 anid|CBF86525.1 anig|EHA22395.1 aory|EIT72582.1 asyd|OJJ64873.1 ater|EAU30539.1 atub|OJI84393.1 aver|OJI98986.1 azon|OJJ48844.1 afum|EAL91966.1 agla|OJJ87189.1 anid|CBF86527.1 aory|EIT72529.1 asyd|OJJ64872.1 ater|EAU30540.1 aver|OJI98984.1

Group64: agla|OJJ84606.1 agla|OJJ86279.1 anid|CBF78809.1 anid|CBF85739.1 asyd|OJJ64288.1 asyd|OJJ65260.1 aver|OJI98381.1 aver|OJI99411.1 awen|OJJ30086.1 awen|OJJ30833.1 awen|OJJ38668.1 azon|OJJ44554.1 azon|OJJ44649.1 abra|OJJ67129.1 acar|OOF98424.1 afis|EAW19329.1 afla|EED47551.1 afum|EAL94046.1 akaw|GAA91039.1 aluc|OJZ87849.1 anig|EHA20152.1 aory|EIT74544.1 ater|EAU29453.1 atub|OJI80799.1 abra|OJJ67137.1 acar|OOF98421.1 acla|EAW14610.1 afis|EAW19339.1 afla|EED47549.1 afum|EAL94058.1 akaw|GAA91032.1 aluc|OJZ87840.1 anid|CBF78807.1 anig|EHA20314.1 aory|EIT74530.1 asyd|OJJ65259.1 ater|EAU29452.1 atub|OJI80806.1 aver|OJI99410.1 agla|OJJ83890.1 awen|OJJ41067.1 agla|OJJ89228.1 awen|OJJ40968.1

Group65: abra|OJJ65919.1 abra|OJJ71178.1 acar|OOF94968.1 acar|OOF97988.1 acla|EAW10328.1 acla|EAW13616.1 afis|EAW19973.1 afis|EAW25266.1 afla|EED50456.1 afla|EED53730.1 afum|EAL84937.1 afum|EAL89929.1 agla|OJJ80057.1 agla|OJJ89242.1 akaw|GAA82807.1 akaw|GAA92313.1 aluc|OJZ87365.1 aluc|OJZ90211.1 anid|CBF84421.1 anid|CBF85372.1 anig|EHA20600.1 anig|EHA23709.1 aory|EIT75304.1 aory|EIT82028.1 asyd|OJJ53545.1 asyd|OJJ57927.1 asyd|OJJ64047.1 ater|EAU31473.1 ater|EAU34407.1 atub|OJI81279.1 atub|OJI81642.1 aver|OJI97591.1 aver|OJI98132.1 aver|OJI98886.1 aver|OJJ08208.1 awen|OJJ36011.1 awen|OJJ40979.1 azon|OJJ42397.1 azon|OJJ44566.1 azon|OJJ45171.1 azon|OJJ50333.1 asyd|OJJ54388.1

Group66: abra|OJJ66757.1 abra|OJJ68421.1 abra|OJJ70242.1 acar|OOF92554.1 acar|OOF93838.1 afis|EAW19495.1 afis|EAW21208.1 afum|EAL86999.1 afum|EAL92219.1 agla|OJJ78585.1 agla|OJJ81224.1 agla|OJJ86343.1 agla|OJJ87022.1 akaw|GAA86545.1 akaw|GAA89691.1 akaw|GAA92527.1 aluc|OJZ80693.1 aluc|OJZ85093.1 aluc|OJZ90076.1 anig|EHA20997.1 anig|EHA27322.1 anig|EHA25314.1 aory|EIT73935.1 aory|EIT77370.1 asyd|OJJ59201.1 asyd|OJJ59780.1 ater|EAU34261.1 ater|EAU34807.1 atub|OJI80506.1 atub|OJI87553.1 atub|OJI84632.1 atub|OJI84730.1 aver|OJJ01814.1 aver|OJJ02507.1 awen|OJJ36122.1 awen|OJJ36911.1 azon|OJJ42905.1 azon|OJJ45981.1 acla|EAW11883.1 afla|EED50220.1 afla|EED56713.1 anid|CBF75444.1

Group67: abra|OJJ67357.1 abra|OJJ69620.1 abra|OJJ69925.1 acar|OOF92464.1 acar|OOF92686.1 acar|OOF93436.1 acla|EAW06385.1 acla|EAW08905.1 afis|EAW15491.1 afis|EAW18491.1 afla|EED49005.1 afla|EED52913.1 afum|EAL85832.1 afum|EAL93181.1 agla|OJJ80168.1 agla|OJJ85086.1 akaw|GAA86464.1 akaw|GAA87732.1 akaw|GAA91604.1 aluc|OJZ82429.1 aluc|OJZ85004.1 aluc|OJZ85363.1 anig|EHA25223.1

anig|EHA25546.1 anig|EHA28342.1 aory|EIT78593.1 aory|EIT82534.1 asyd|OJJ53723.1 asyd|OJJ56020.1 asyd|OJJ58801.1 ater|EAU29917.1 ater|EAU32628.1 atub|OJI80207.1 atub|OJI80594.1 atub|OJI88012.1 awen|OJJ31044.1 awen|OJJ38147.1 azon|OJJ43140.1 azon|OJJ50555.1 anid|CBF70161.1 aver|OJJ01405.1 aver|OJJ04967.1

Group68: abra|OJJ69659.1 abra|OJJ71032.1 acar|OOF92597.1 acar|OOF92618.1 acar|OOF92644.1 acar|OOF98104.1 aluc|OJZ82466.1 aluc|OJZ87497.1 anig|EHA20713.1 anig|EHA28309.1 atub|OJI81151.1 atub|OJI87972.1 acla|EAW08698.1 acla|EAW12766.1 afis|EAW15737.1 afis|EAW17928.1 afla|EED49112.1 afla|EED51315.1 afum|EAL85611.1 afum|EAL91739.1 agla|OJJ81894.1 agla|OJJ83005.1 akaw|GAA91638.1 akaw|GAA91639.1 aluc|OJZ82465.1 anid|CBF71049.1 anid|CBF80811.1 anig|EHA28310.1 aory|EIT78548.1 aory|EIT81553.1 asyd|OJJ52673.1 asyd|OJJ54450.1 ater|EAU32405.1 ater|EAU33543.1 atub|OJI87973.1 aver|OJJ04699.1 aver|OJJ07704.1 awen|OJJ38433.1 awen|OJJ41618.1 azon|OJJ43810.1 azon|OJJ48339.1 akaw|GAA82922.1

Group69: abra|OJJ70622.1 abra|OJJ71755.1 abra|OJJ72061.1 abra|OJJ76523.1 acar|OOF96715.1 acar|OOF96808.1 acar|OOF99444.1 acla|EAW13671.1 acla|EAW14172.1 afis|EAW15618.1 afis|EAW18886.1 afla|EED50879.1 afla|EED53904.1 afla|EED57853.1 agla|OJJ80215.1 agla|OJJ80629.1 akaw|GAA82436.1 akaw|GAA84874.1 aluc|OJZ84834.1 aluc|OJZ86797.1 aluc|OJZ91685.1 anig|EHA21886.1 anig|EHA22697.1 anig|EHA25744.1 aory|EIT78491.1 aory|EIT78970.1 aory|EIT81115.1 ater|EAU31982.1 ater|EAU32969.1 ater|EAU38058.1 atub|OJI83721.1 atub|OJI86137.1 aver|OJJ01614.1 aver|OJJ02222.1 awen|OJJ32444.1 awen|OJJ38463.1 azon|OJJ47202.1 azon|OJJ47982.1 afum|EAL93619.1 anid|CBF69896.1 asyd|OJJ59001.1

Group71: abra|OJJ65673.1 abra|OJJ68238.1 abra|OJJ66038.1 abra|OJJ68515.1 abra|OJJ74212.1 abra|OJJ74240.1 acar|OOF92683.1 acar|OOF95490.1 acar|OOF95508.1 afla|EED47448.1 afla|EED57720.1 akaw|GAA86780.1 akaw|GAA86806.1 akaw|GAA91332.1 aluc|OJZ79540.1 aluc|OJZ89922.1 aluc|OJZ82729.1 aluc|OJZ89892.1 anig|EHA21182.1 anig|EHA27091.1 anig|EHA27122.1 aory|EIT79930.1 aory|EIT81920.1 atub|OJI84883.1 atub|OJI84915.1 atub|OJI88962.1 aver|OJI97104.1 aver|OJJ03419.1 acla|EAW07265.1 afum|EAL90895.1 ater|EAU34298.1 agla|OJJ87719.1 anid|CBF88605.1 awen|OJJ34900.1 azon|OJJ50067.1 afis|EAW22286.1 anid|CBF88560.1 asyd|OJJ63833.1 ater|EAU34323.1 asyd|OJJ63857.1

Group72: abra|OJJ66697.1 abra|OJJ68381.1 acar|OOF93705.1 acar|OOF93791.1 acar|OOF93987.1 acla|EAW11832.1 acla|EAW12357.1 afis|EAW16780.1 afis|EAW19443.1 afla|EED56190.1 afla|EED56651.1 afum|EAL86351.1 afum|EAL87052.1 agla|OJJ79770.1 agla|OJJ81157.1 akaw|GAA89641.1 akaw|GAA92564.1 aluc|OJZ80748.1 aluc|OJZ90112.1 anid|CBF80588.1 anid|CBF81398.1 anig|EHA20960.1 anig|EHA27274.1 aory|EIT73439.1 aory|EIT79687.1 asyd|OJJ54689.1 asyd|OJJ59403.1 ater|EAU30272.1 ater|EAU34745.1 atub|OJI84692.1 atub|OJI87197.1 atub|OJI87605.1 aver|OJI99850.1 aver|OJJ02870.1 aver|OJJ07480.1 aver|OJJ07987.1 awen|OJJ29847.1 awen|OJJ36844.1 azon|OJJ42956.1 azon|OJJ50350.1

Group73: abra|OJJ68502.1 abra|OJJ73653.1 acar|OOF98748.1 acar|OOG00831.1 acla|EAW07563.1 acla|EAW12139.1 acla|EAW15367.1 afis|EAW17197.1 afis|EAW17227.1 afis|EAW22597.1 afla|EED45179.1 afla|EED51197.1 afla|EED51520.1 afla|EED54722.1 afum|EAL86625.1 afum|EAL86649.1 afum|EAL90601.1 agla|OJJ80961.1 agla|OJJ86657.1 akaw|GAA88505.1 akaw|GAA92769.1 aluc|OJZ82715.1 aluc|OJZ88400.1 anid|CBF69884.1 anid|CBF89553.1 anig|EHA18128.1 anig|EHA21170.1 aory|EIT73539.1 aory|EIT78945.1 aory|EIT80690.1 asyd|OJJ59011.1 asyd|OJJ65377.1 ater|EAU31751.1 ater|EAU39106.1 atub|OJI88949.1 atub|OJI90995.1 aver|OJI99545.1 aver|OJJ01604.1 awen|OJJ35225.1 azon|OJJ51827.1

Group74: abra|OJJ66378.1 abra|OJJ68574.1 abra|OJJ75363.1 acar|OOF89872.1 acar|OOF92870.1 akaw|GAA90247.1 akaw|GAA91914.1 aluc|OJZ81966.1 aluc|OJZ83093.1 atub|OJI84517.1 atub|OJI87392.1 atub|OJI87843.1 acla|EAW08967.1 agla|OJJ89584.1 anid|CBF87061.1 anig|EHA22281.1 anig|EHA28155.1 asyd|OJJ60430.1 ater|EAU36814.1 aver|OJI99753.1 afla|EED48897.1 afum|EAL93812.1 akaw|GAA92059.1 aluc|OJZ92932.1 azon|OJJ46231.1 afis|EAW19083.1 agla|OJJ81069.1 anig|EHA26039.1 aory|EIT81242.1 ater|EAU36156.1 awen|OJJ35776.1 azon|OJJ45936.1 acla|EAW14375.1 afla|EED54315.1 aory|EIT76109.1 ater|EAU32130.1 awen|OJJ32478.1 afla|EED46053.1 aory|EIT83489.1

Group75: abra|OJJ66645.1 abra|OJJ77622.1 acar|OOF93665.1 acar|OOF93749.1 acar|OOF99113.1 afis|EAW19398.1 afis|EAW25441.1 afla|EED46395.1 afla|EED56341.1 afum|EAL87096.1 afum|EAL90099.1 agla|OJJ81109.1 agla|OJJ84218.1 akaw|GAA84009.1 akaw|GAA89598.1 aluc|OJZ80804.1 aluc|OJZ80911.1 anid|CBF74274.1 anid|CBF85786.1 anig|EHA19435.1 anig|EHA27230.1 aory|EIT72345.1 aory|EIT73378.1 aory|EIT83124.1 ater|EAU33678.1 ater|EAU36124.1 atub|OJI82668.1 atub|OJI87652.1 acla|EAW10153.1 acla|EAW11788.1 afla|EED50354.1 asyd|OJJ51908.1 asyd|OJJ64317.1 aver|OJJ05362.1 awen|OJJ36793.1 azon|OJJ43006.1 azon|OJJ43662.1 aver|OJI98410.1 awen|OJJ30031.1

Group77: abra|OJJ68971.1 abra|OJJ74920.1 acar|OOF91548.1 acar|OOF95911.1 acla|EAW08451.1 acla|EAW14216.1 afis|EAW19065.1 afis|EAW23496.1 afla|EED54431.1 afla|EED57396.1 afum|EAL87967.1 afum|EAL93794.1 agla|OJJ83427.1 agla|OJJ84542.1 agla|OJJ86743.1 akaw|GAA83503.1 akaw|GAA90498.1 aluc|OJZ85958.1 aluc|OJZ92491.1 anid|CBF79648.1 anid|CBF89739.1 anig|EHA24643.1 anig|EHA26481.1 aory|EIT75884.1 aory|EIT80127.1 asyd|OJJ52407.1 asyd|OJJ58022.1 asyd|OJJ62230.1 ater|EAU32108.1 ater|EAU35285.1 atub|OJI79746.1 atub|OJI86924.1 aver|OJI96339.1 aver|OJJ04377.1 awen|OJJ31497.1 awen|OJJ39821.1 azon|OJJ46209.1 azon|OJJ47945.1 anid|CBF76133.1

Group79: abra|OJJ69282.1 abra|OJJ71129.1 acar|OOF90163.1 acar|OOF98023.1 acar|OOF94440.1 acla|EAW10369.1 acla|EAW14671.1 afis|EAW24202.1 afis|EAW25223.1 afla|EED48070.1 afla|EED53683.1 afum|EAL85872.1 afum|EAL89884.1 agla|OJJ79190.1 agla|OJJ79675.1 akaw|GAA82845.1 akaw|GAA88154.1 aluc|OJZ82110.1 aluc|OJZ87406.1 anid|CBF74328.1 anid|CBF85271.1 anig|EHA20636.1 anig|EHA28620.1 aory|EIT75297.1 aory|EIT83329.1 asyd|OJJ53993.1 asyd|OJJ62910.1 ater|EAU31153.1 ater|EAU34382.1 atub|OJI81240.1 atub|OJI88326.1 aver|OJI98079.1 aver|OJJ08091.1 awen|OJJ33096.1 awen|OJJ40889.1 azon|OJJ42425.1 azon|OJJ44518.1 azon|OJJ44851.1

Group81: abra|OJJ72988.1 abra|OJJ77279.1 acar|OOF93153.1 acar|OOF96164.1 acla|EAW08023.1 acla|EAW13278.1 afis|EAW17339.1 afis|EAW23057.1 afum|EAL88423.2 afum|EAL91183.1 akaw|GAA85562.1 akaw|GAA90670.1 aluc|OJZ83496.1 aluc|OJZ84689.1 anig|EHA24331.1 anig|EHA27964.1 asyd|OJJ54336.1 asyd|OJJ56391.1 ater|EAU33849.1 ater|EAU37607.1 atub|OJI82390.1 atub|OJI89393.1 aver|OJJ05288.1 aver|OJJ06322.1 awen|OJJ35679.1 awen|OJJ40816.1 afla|EED51042.1 afla|EED54960.1 agla|OJJ79345.1 agla|OJJ89418.1 anid|CBF81037.1 aory|EIT80299.1 azon|OJJ44147.1 azon|OJJ50224.1 anid|CBF84215.1 aory|EIT77041.1 asyd|OJJ57175.1 aver|OJJ03139.1

Group82: abra|OJJ73798.1 abra|OJJ76038.1 acar|OOF99891.1 acar|OOG00949.1 acla|EAW07105.1 acla|EAW07691.1 afla|EED45540.1 afla|EED49992.1 agla|OJJ86348.1 agla|OJJ87936.1 akaw|GAA88927.1 akaw|GAA89416.1 aluc|OJZ88257.1 aluc|OJZ89493.1 anid|CBF71497.1 anid|CBF88069.1 anig|EHA18002.1 anig|EHA23106.1 aory|EIT74740.1 aory|EIT75450.1 asyd|OJJ54934.1 asyd|OJJ60320.1 asyd|OJJ63552.1 ater|EAU32047.1 ater|EAU38995.1 atub|OJI85309.1 atub|OJI90856.1 aver|OJI97382.1 aver|OJI99669.1 aver|OJJ07264.1 awen|OJJ34497.1 awen|OJJ34697.1 awen|OJJ35343.1 azon|OJJ45277.1 azon|OJJ49485.1 azon|OJJ49775.1 afis|EAW22706.1 afum|EAL90486.1

Group83: abra|OJJ74861.1 abra|OJJ78108.1 aluc|OJZ91173.1 aluc|OJZ92439.1 anig|EHA19866.1 anig|EHA26528.1 atub|OJI83187.1 atub|OJI86868.1 acar|OOF95861.1 agla|OJJ86823.1 akaw|GAA83559.1 akaw|GAA85898.1 awen|OJJ39889.1 azon|OJJ43220.1 acar|OOF98757.1 acla|EAW06608.1 afis|EAW16215.1 afla|EED48709.1 afum|EAL88842.1 agla|OJJ86220.1 anid|CBF79542.1 aory|EIT73181.1 asyd|OJJ57900.1 ater|EAU33259.1 aver|OJJ04494.1 awen|OJJ30257.1 azon|OJJ43953.1 abra|OJJ78109.1 acla|EAW06607.1 afis|EAW16216.1 afla|EED48710.1 afum|EAL88843.1 akaw|GAA85897.1 anid|CBF79540.1 aory|EIT73231.1 ater|EAU33260.1 awen|OJJ30256.1 azon|OJJ43954.1

Group85: abra|OJJ69585.1 acar|OOF91569.1 acla|EAW14381.1 acla|EAW14383.1 acla|EAW14384.1 afis|EAW19087.1 afis|EAW19089.1 afis|EAW19090.1 afla|EED54319.1 afla|EED54321.1 afla|EED54322.1 afum|EAL93817.1 afum|EAL93819.1 afum|EAL93820.1 agla|OJJ84514.1 agla|OJJ84515.1 agla|OJJ84517.1 akaw|GAA91580.1 akaw|GAA91581.1 akaw|GAA91583.1 aluc|OJZ82402.1 anid|CBF79686.1 anid|CBF79690.1 anid|CBF79692.1 anig|EHA28363.1 aory|EIT76048.1 aory|EIT76070.1 aory|EIT76088.1 asyd|OJJ58048.1 ater|EAU32135.1 ater|EAU32138.1 ater|EAU32139.1 atub|OJI88039.1 aver|OJJ04355.1 awen|OJJ32484.1 azon|OJJ46222.1 azon|OJJ46228.1 azon|OJJ46230.1

Group87: abra|OJJ73172.1 abra|OJJ75139.1 acar|OOF97197.1 acar|OOG00411.1 acla|EAW06855.1 acla|EAW08318.1 afis|EAW16472.1 afis|EAW23353.1 afla|EED47207.1 afla|EED55602.1 afum|EAL88123.1 afum|EAL89066.1 agla|OJJ82874.1 agla|OJJ86473.1 akaw|GAA87999.1 akaw|GAA88746.1 aluc|OJZ81255.1 aluc|OJZ92706.1 anid|CBF74897.1 anid|CBF81437.1 anig|EHA18563.1 anig|EHA26294.1 aory|EIT79222.1 aory|EIT82665.1 asyd|OJJ59426.1 asyd|OJJ60103.1 ater|EAU35596.1 ater|EAU36570.1 atub|OJI87132.1 atub|OJI91451.1 aver|OJJ02845.1 aver|OJJ04159.1 awen|OJJ39536.1 awen|OJJ40364.1 azon|OJJ47731.1 azon|OJJ49213.1 anid|CBF74899.1

Group89: aory|EIT73009.1 aory|EIT82639.1 abra|OJJ75430.1 acar|OOF90776.1 acla|EAW13800.1 acla|EAW13801.1 acla|EAW13802.1 afis|EAW18330.1 afis|EAW18331.1 afis|EAW18332.1 afla|EED48794.1 afla|EED48795.1 afla|EED49574.1 afum|EAL87695.1 afum|EAL87696.1 afum|EAL87697.1 agla|OJJ88054.1 akaw|GAA84206.1 akaw|GAA84207.1 aluc|OJZ88883.1 aluc|OJZ88884.1 anid|CBF79398.1 anid|CBF79400.1 anid|CBF79402.1 anig|EHA23623.1 aory|EIT72994.1 aory|EIT73029.1 asyd|OJJ57827.1 ater|EAU33343.1 ater|EAU33344.1 atub|OJI85907.1 atub|OJI85908.1 atub|OJI85909.1 aver|OJJ04565.1 awen|OJJ30712.1 azon|OJJ43305.1 afla|EED48796.1

Group90: aory|EIT73779.1 aory|EIT77467.1 awen|OJJ29827.1 awen|OJJ36939.1 abra|OJJ65853.1 acar|OOF95014.1 acla|EAW11610.1 afis|EAW23843.1 afla|EED56987.1 afum|EAL93778.1 agla|OJJ78690.1 akaw|GAA92291.1 aluc|OJZ90188.1 anig|EHA23687.1 ater|EAU31468.1 atub|OJI81615.1 aver|OJI99116.1 awen|OJJ29704.1 azon|OJJ46986.1 abra|OJJ77924.1 acar|OOF90880.1 afla|EED45127.1 akaw|GAA83740.1 aluc|OJZ90991.1 anid|CBF86319.1 anig|EHA19699.1 asyd|OJJ64996.1 atub|OJI82982.1 acar|OOG00327.1 asyd|OJJ55713.1 atub|OJI87196.1 aver|OJJ06799.1 afum|EAL87274.1 agla|OJJ86442.1 asyd|OJJ62873.1 ater|EAU31778.1 aver|OJI96926.1

Group91: abra|OJJ67337.1 acar|OOF92476.1 acla|EAW06397.1 afis|EAW18501.1 afla|EED52900.1 afum|EAL93192.1 agla|OJJ85126.1 akaw|GAA86479.1 aluc|OJZ85020.1 anid|CBF75182.1 anig|EHA25241.1 aory|EIT78666.1 asyd|OJJ59951.1 ater|EAU30070.1 atub|OJI80577.1 aver|OJJ02338.1 awen|OJJ31055.1 azon|OJJ50565.1 abra|OJJ69524.1 acar|OOF94230.1 acla|EAW11627.1 afis|EAW23829.1 afla|EED54410.1 afum|EAL87261.1 agla|OJJ82483.1 akaw|GAA88301.1 aluc|OJZ82364.1 anid|CBF75186.1 anig|EHA28401.1

aory|EIT75866.1 ater|EAU31384.1 atub|OJI88078.1 awen|OJJ38733.1 azon|OJJ50221.1 afla|EED51976.1 aory|EIT81396.1 agla|OJJ78561.1

Group94: abra|OJJ68844.1 abra|OJJ73357.1 acar|OOF90545.1 acar|OOG00565.1 acla|EAW06977.1 acla|EAW15277.1 afis|EAW16591.1 afis|EAW20248.1 afla|EED45005.1 afla|EED57097.1 afum|EAL85366.1 afum|EAL89187.1 akaw|GAA83127.1 akaw|GAA90404.1 aluc|OJZ85824.1 aluc|OJZ88683.1 anid|CBF78712.1 anid|CBF81894.1 anig|EHA18363.1 anig|EHA24737.1 aory|EIT73755.1 aory|EIT82298.1 asyd|OJJ59680.1 asyd|OJJ63029.1 ater|EAU32039.1 ater|EAU36671.1 atub|OJI79611.1 atub|OJI91263.1 aver|OJI97939.1 aver|OJJ02608.1 awen|OJJ31246.1 awen|OJJ40224.1 azon|OJJ48439.1 azon|OJJ49082.1 agla|OJJ82668.1 azon|OJJ45914.1

Group95: abra|OJJ69553.1 abra|OJJ70394.1 aory|EIT78629.1 aory|EIT82274.1 acar|OOF91880.1 afla|EED57090.1 agla|OJJ82606.1 akaw|GAA91965.1 aluc|OJZ82652.1 anid|CBF75146.1 anig|EHA27420.1 asyd|OJJ59972.1 ater|EAU30264.1 atub|OJI88884.1 aver|OJJ02319.1 awen|OJJ41493.1 abra|OJJ71088.1 acar|OOF98047.1 acla|EAW06408.1 afis|EAW18512.1 afla|EED52888.1 afum|EAL93207.1 agla|OJJ85018.1 akaw|GAA82873.1 aluc|OJZ87440.1 anid|CBF70237.1 anig|EHA20665.1 asyd|OJJ58768.1 ater|EAU29455.1 atub|OJI81208.1 aver|OJJ01377.1 awen|OJJ31066.1 azon|OJJ46021.1 acar|OOF98419.1 acar|OOF99514.1 azon|OJJ42344.1

Group96: abra|OJJ71749.1 abra|OJJ72735.1 acar|OOF90204.1 acar|OOF95966.1 afla|EED52022.1 afla|EED52611.1 afum|EAL85108.1 afum|EAL87310.1 agla|OJJ78437.1 agla|OJJ82556.1 akaw|GAA85163.1 akaw|GAA85341.1 aluc|OJZ86473.1 aluc|OJZ90703.1 anig|EHA22158.1 anig|EHA24129.1 aory|EIT73811.1 aory|EIT74178.1 aory|EIT81402.1 asyd|OJJ54106.1 asyd|OJJ56840.1 aver|OJJ03498.1 aver|OJJ08629.1 awen|OJJ31677.1 awen|OJJ33864.1 afis|EAW23882.1 agla|OJJ80999.1 anid|CBF77189.1 ater|EAU33464.1 atub|OJI82130.1 atub|OJI84056.1 azon|OJJ42521.1 azon|OJJ48689.1 acla|EAW11571.1 afis|EAW19784.1 afla|EED45206.1

Group97: abra|OJJ71850.1 abra|OJJ73479.1 acar|OOF96520.1 acar|OOG00674.1 acla|EAW06803.1 acla|EAW13882.1 afis|EAW16352.1 afis|EAW18594.1 afla|EED52753.1 afla|EED53519.1 afum|EAL88977.1 afum|EAL93302.1 agla|OJJ81646.1 agla|OJJ88248.1 akaw|GAA85067.1 akaw|GAA87429.1 aluc|OJZ86584.1 aluc|OJZ88568.1 anid|CBF70389.1 anid|CBF89223.1 anig|EHA18265.1 anig|EHA22076.1 aory|EIT75113.1 aory|EIT78157.1 asyd|OJJ58637.1 asyd|OJJ62519.1 ater|EAU34109.1 ater|EAU34888.1 atub|OJI83939.1 atub|OJI91154.1 aver|OJI96638.1 aver|OJJ01243.1 awen|OJJ31157.1 awen|OJJ37755.1 azon|OJJ45016.1 azon|OJJ46632.1

Group98: abra|OJJ72143.1 abra|OJJ77114.1 acar|OOF93003.1 acar|OOF96781.1 acla|EAW12957.1 acla|EAW14937.1 afis|EAW17702.1 afis|EAW24453.1 afum|EAL86122.1 afum|EAL91517.1 agla|OJJ79467.1 agla|OJJ80300.1 akaw|GAA84780.1 akaw|GAA90815.1 aluc|OJZ84523.1 aluc|OJZ86900.1 anid|CBF73820.1 anid|CBF89927.1 anig|EHA21742.1 anig|EHA27821.1 aory|EIT75637.1 aory|EIT83612.1 asyd|OJJ56569.1 asyd|OJJ62120.1 atub|OJI83619.1 atub|OJI89556.1 aver|OJI96228.1 aver|OJJ06153.1 awen|OJJ32744.1 awen|OJJ37133.1 azon|OJJ45294.1 azon|OJJ50831.1 afla|EED55221.1 ater|EAU29992.1 ater|EAU31909.1 afla|EED45698.1

Group99: abra|OJJ72622.1 abra|OJJ76815.1 acar|OOF91310.1 acar|OOF94641.1 acla|EAW11602.1 acla|EAW13375.1 afis|EAW20489.1 afis|EAW23856.1 afum|EAL84566.1 afum|EAL87284.1 agla|OJJ82510.1 agla|OJJ84827.1 akaw|GAA85243.1 akaw|GAA87206.1 aluc|OJZ84225.1 aluc|OJZ90600.1 anid|CBF77244.1 anid|CBF79220.1 anig|EHA24038.1 anig|EHA27574.1 aory|EIT74289.1 aory|EIT77786.1 asyd|OJJ55735.1 asyd|OJJ56812.1 ater|EAU29567.1 ater|EAU33499.1 atub|OJI82024.1 atub|OJI89859.1 aver|OJJ03523.1 aver|OJJ06821.1 awen|OJJ33189.1 awen|OJJ33817.1 azon|OJJ46391.1 azon|OJJ48719.1 afla|EED52576.1 afla|EED44774.1

Group100: abra|OJJ73827.1 abra|OJJ74268.1 acar|OOF95463.1 acar|OOG00979.1 acla|EAW07312.1 acla|EAW07722.1 afis|EAW22334.1 afis|EAW22754.1 afla|EED50015.1 afla|EED57748.1 afum|EAL90445.1 afum|EAL90858.1 agla|OJJ83671.1 agla|OJJ87714.1 akaw|GAA86755.1 akaw|GAA89445.1 aluc|OJZ88222.1 aluc|OJZ89865.1 anid|CBF88019.1 anid|CBF88645.1 anig|EHA17969.1 anig|EHA27065.1 aory|EIT74682.1 aory|EIT81059.1 asyd|OJJ63522.1 asyd|OJJ63881.1 ater|EAU38966.1 ater|EAU39471.1 atub|OJI84943.1 atub|OJI90822.1 aver|OJI97058.1 aver|OJI97408.1 awen|OJJ34912.1 awen|OJJ35426.1 azon|OJJ45572.1 azon|OJJ49750.1

Group101: abra|OJJ76503.1 abra|OJJ76504.1 acar|OOF99462.1 acar|OOF99465.1 afis|EAW18906.1 afis|EAW18907.1 afla|EED50840.1 afla|EED50841.1 afum|EAL93638.1 afum|EAL93639.1 agla|OJJ81511.1 agla|OJJ81512.1 akaw|GAA82419.1 akaw|GAA82420.1 aluc|OJZ91697.1 aluc|OJZ91699.1 anid|CBF69713.1 anid|CBF69715.1 anig|EHA22712.1 anig|EHA22713.1 aory|EIT78986.1 aory|EIT79030.1 asyd|OJJ59043.1 asyd|OJJ59044.1 ater|EAU38075.1 ater|EAU38076.1 atub|OJI86152.1 atub|OJI86154.1 aver|OJJ01638.1 aver|OJJ01639.1 awen|OJJ32463.1 awen|OJJ32464.1 azon|OJJ47220.1 azon|OJJ47221.1 acla|EAW14191.1 acla|EAW14192.1

Group103: afla|EED46026.1 afla|EED49987.1 akaw|GAA89413.1 akaw|GAA90839.1 aluc|OJZ82994.1 aluc|OJZ88260.1 aory|EIT74653.1 aory|EIT75776.1 atub|OJI89228.1 atub|OJI90859.1 awen|OJJ30039.1 awen|OJJ35338.1 abra|OJJ65964.1 acar|OOF92910.1 acla|EAW13583.1 afis|EAW20018.1 afis|EAW22701.1 afum|EAL84899.2 afum|EAL90491.1 agla|OJJ87998.1 anid|CBF84731.1 anig|EHA21408.1 asyd|OJJ53303.1

ater|EAU29378.1 ater|EAU39001.1 aver|OJJ08438.1 azon|OJJ43685.1 abra|OJJ73794.1 acar|OOG00953.1 acla|EAW07686.1 agla|OJJ79008.1 anid|CBF88080.1 anig|EHA18005.1 asyd|OJJ63557.1 aver|OJI97375.1 azon|OJJ49780.1

Group105: abra|OJJ76497.1 acar|OOF99470.1 acla|EAW14363.1 afis|EAW18913.1 afla|EED50836.1 afum|EAL93644.1 agla|OJJ81517.1 akaw|GAA82414.1 aluc|OJZ91703.1 anid|CBF69695.1 anig|EHA22717.1 aory|EIT79010.1 asyd|OJJ59053.1 ater|EAU38080.1 atub|OJI86163.1 aver|OJJ01647.1 awen|OJJ31506.1 azon|OJJ41989.1 azon|OJJ47222.1 abra|OJJ76500.1 acar|OOF99467.1 acla|EAW14196.1 afis|EAW18912.1 afla|EED50837.1 afum|EAL93643.1 akaw|GAA82416.1 aluc|OJZ91702.1 anid|CBF69697.1 anig|EHA22716.1 aory|EIT79064.1 asyd|OJJ59051.1 ater|EAU38079.1 atub|OJI86161.1 awen|OJJ31505.1 azon|OJJ47223.1 azon|OJJ41991.1

Group106: abra|OJJ72950.1 abra|OJJ74416.1 acla|EAW06831.1 acla|EAW11196.1 afis|EAW15926.1 afis|EAW16381.1 afis|EAW19938.1 afum|EAL88562.1 afum|EAL89006.1 akaw|GAA85531.1 akaw|GAA87307.1 aluc|OJZ83460.1 aluc|OJZ89710.1 anid|CBF81021.1 anid|CBF89371.1 atub|OJI82352.1 atub|OJI85092.1 aver|OJI96612.1 aver|OJJ06329.1 awen|OJJ35654.1 awen|OJJ37914.1 acar|OOF96132.1 afla|EED54935.1 agla|OJJ89379.1 anig|EHA24304.1 aory|EIT80345.1 asyd|OJJ56383.1 ater|EAU35529.1 azon|OJJ48008.1 acar|OOF95327.1 afla|EED49933.1 anig|EHA26941.1 aory|EIT83444.1 asyd|OJJ62476.1 ater|EAU34071.1

Group107: abra|OJJ68266.1 acar|OOF97546.1 acla|EAW06826.1 afis|EAW16377.1 afla|EED53460.1 afum|EAL89002.1 agla|OJJ78595.1 akaw|GAA91930.1 aluc|OJZ82692.1 anid|CBF74547.1 anig|EHA27369.1 aory|EIT75115.1 asyd|OJJ57687.1 ater|EAU34075.1 atub|OJI88924.1 aver|OJJ05493.1 awen|OJJ37911.1 azon|OJJ45090.1 abra|OJJ68268.1 acar|OOF97548.1 acla|EAW07026.1 afis|EAW16652.1 afla|EED44948.1 afum|EAL89256.1 agla|OJJ78685.1 akaw|GAA91932.1 aluc|OJZ82690.1 anid|CBF74545.1 anig|EHA27367.1 aory|EIT73496.1 ater|EAU39576.1 atub|OJI88922.1 aver|OJJ05494.1 awen|OJJ40062.1 azon|OJJ45565.1

Group108: abra|OJJ69218.1 acar|OOF91100.1 acla|EAW08627.1 afis|EAW15804.1 afla|EED51241.1 afum|EAL85546.1 agla|OJJ81971.1 akaw|GAA92227.1 aluc|OJZ86200.1 anid|CBF70912.1 anig|EHA24423.1 aory|EIT83194.1 asyd|OJJ52602.1 ater|EAU33444.1 atub|OJI79993.1 aver|OJJ04628.1 awen|OJJ38514.1 azon|OJJ43738.1 abra|OJJ69219.1 acar|OOF91099.1 acla|EAW08626.1 afis|EAW15805.1 afla|EED51240.1 afum|EAL85545.1 akaw|GAA92228.1 aluc|OJZ86201.1 anid|CBF70910.1 anig|EHA24422.1 aory|EIT83177.1 asyd|OJJ52601.1 ater|EAU33443.1 atub|OJI79994.1 aver|OJJ04627.1 awen|OJJ38516.1 azon|OJJ43737.1

Group109: abra|OJJ73774.1 acar|OOG00931.1 acla|EAW07658.1 afis|EAW22676.1 afla|EED49957.1 afum|EAL90516.1 agla|OJJ79041.1 akaw|GAA88402.1 aluc|OJZ88280.1 anid|CBF88133.1 anig|EHA18025.1 aory|EIT83452.1 asyd|OJJ63590.1 ater|EAU39033.1 atub|OJI90880.1 aver|OJI97343.1 awen|OJJ35305.1 azon|OJJ49811.1 abra|OJJ73775.1 acar|OOG00932.1 acla|EAW07660.1 afis|EAW22677.1 afla|EED49958.1 afum|EAL90515.1 agla|OJJ79040.1 akaw|GAA88401.1 aluc|OJZ88279.1 anid|CBF88131.1 anig|EHA18024.1 aory|EIT83440.1 asyd|OJJ63589.1 ater|EAU39032.1 atub|OJI90879.1 aver|OJI97344.1 azon|OJJ49810.1

Group110: abra|OJJ74284.1 acar|OOF89817.1 acla|EAW07328.1 afis|EAW22360.1 afla|EED57768.1 afum|EAL90831.1 agla|OJJ87609.1 akaw|GAA86740.1 aluc|OJZ89849.1 anid|CBF88679.1 anig|EHA27050.1 aory|EIT81097.1 asyd|OJJ63900.1 ater|EAU39494.1 atub|OJI84957.1 aver|OJI97040.1 awen|OJJ34934.1 azon|OJJ45587.1 abra|OJJ74285.1 acar|OOF95446.1 acla|EAW07329.1 afis|EAW22361.1 afla|EED57769.1 afum|EAL90830.1 agla|OJJ87610.1 akaw|GAA86739.1 aluc|OJZ89848.1 anid|CBF88686.1 anig|EHA27049.1 aory|EIT81110.1 asyd|OJJ63902.1 ater|EAU39495.1 atub|OJI84958.1 aver|OJI97038.1 awen|OJJ34935.1

Group111: abra|OJJ74383.1 acar|OOF95354.1 acla|EAW07424.1 afis|EAW22450.1 afla|EED54465.1 afum|EAL90741.1 agla|OJJ87572.1 akaw|GAA87284.1 aluc|OJZ89735.1 anid|CBF88925.1 anig|EHA26961.1 aory|EIT81125.1 asyd|OJJ62853.1 ater|EAU39257.1 atub|OJI85066.1 aver|OJI96906.1 awen|OJJ35075.1 azon|OJJ50032.1 abra|OJJ74384.1 acar|OOF95353.1 acla|EAW07423.1 afis|EAW22449.1 afla|EED54467.1 afum|EAL90742.1 akaw|GAA87285.1 aluc|OJZ89734.1 anid|CBF88927.1 anig|EHA26960.1 aory|EIT81127.1 asyd|OJJ62852.1 ater|EAU39255.1 atub|OJI85067.1 aver|OJI96905.1 awen|OJJ35076.1 azon|OJJ50031.1

Group112: abra|OJJ75811.1 acar|OOG00066.1 acla|EAW14862.1 afis|EAW24390.1 afla|EED47820.1 afum|EAL86062.1 agla|OJJ78870.1 akaw|GAA84525.1 aluc|OJZ89258.1 anid|CBF73709.1 anig|EHA23310.1 aory|EIT77118.1 asyd|OJJ56644.1 ater|EAU30344.1 atub|OJI85541.1 aver|OJJ06082.1 awen|OJJ32814.1 azon|OJJ49018.1 abra|OJJ75812.1 acar|OOG00063.1 acla|EAW14861.1 afis|EAW24389.1 afum|EAL86061.1 agla|OJJ83285.1 akaw|GAA84526.1 aluc|OJZ89259.1 anid|CBF73707.1 anig|EHA23309.1 aory|EIT77172.1 asyd|OJJ56645.1 ater|EAU30345.1 atub|OJI85540.1 aver|OJJ06081.1 awen|OJJ32815.1 azon|OJJ49017.1

Group113: abra|OJJ77674.1 acar|OOF99062.1 acla|EAW06583.1 afis|EAW16009.1 afla|EED56386.1 afum|EAL88639.1 agla|OJJ86034.1 akaw|GAA83958.1 aluc|OJZ80845.1 anid|CBF74571.1 anig|EHA19478.1 aory|EIT83011.1 asyd|OJJ57651.1 ater|EAU33634.1 atub|OJI82722.1 aver|OJJ05511.1 awen|OJJ30456.1 azon|OJJ45447.1 abra|OJJ77675.1 acar|OOF99061.1 acla|EAW06587.1 afis|EAW16010.1 afla|EED56387.1 afum|EAL88640.1 agla|OJJ86035.1 akaw|GAA83957.1 aluc|OJZ80844.1 anid|CBF74569.1 anig|EHA19479.1 aory|EIT82998.1 ater|EAU33633.1 atub|OJI82723.1 aver|OJJ05510.1 awen|OJJ30455.1 azon|OJJ45446.1

Group114: abra|OJJ65981.1 abra|OJJ73719.1 acar|OOF92892.1 acar|OOG00881.1 acla|EAW07608.1 acla|EAW13600.1 afis|EAW20005.1 afis|EAW22638.1 afla|EED46007.1 afla|EED49889.1 afum|EAL84913.1 afum|EAL90554.2 akaw|GAA88444.1 akaw|GAA90854.1 aluc|OJZ82979.1 aluc|OJZ88332.1 anid|CBF84697.1

anid|CBF88205.1 anig|EHA18067.1 anig|EHA21394.1 asyd|OJJ53314.1 asyd|OJJ63639.1 atub|OJI89212.1 atub|OJI90927.1 aver|OJI97297.1 aver|OJJ08429.1 awen|OJJ30019.1 awen|OJJ35261.1 agla|OJJ80980.1 aory|EIT72460.1 aory|EIT75522.1 ater|EAU29347.1 ater|EAU39068.1 azon|OJJ44553.1

Group115: abra|OJJ66667.1 abra|OJJ72896.1 acar|OOF93682.1 acar|OOF93767.1 acar|OOF96087.1 acla|EAW11355.1 acla|EAW11808.1 afis|EAW18139.1 afis|EAW19418.1 afla|EED50375.1 afla|EED58065.1 afum|EAL87076.1 afum|EAL87496.1 agla|OJJ81131.1 agla|OJJ83822.1 akaw|GAA85483.1 akaw|GAA89616.1 aluc|OJZ80780.1 aluc|OJZ83405.1 anig|EHA24256.1 anig|EHA27249.1 aory|EIT79750.1 aory|EIT83148.1 atub|OJI82294.1 atub|OJI87632.1 awen|OJJ35581.1 awen|OJJ36816.1 azon|OJJ42986.1 azon|OJJ46941.1 anid|CBF74365.1 asyd|OJJ57784.1 ater|EAU34724.1 aver|OJJ05380.1 ater|EAU35462.1

Group116: abra|OJJ69228.1 abra|OJJ71754.1 asyd|OJJ52597.1 asyd|OJJ53400.1 aver|OJJ04201.1 aver|OJJ04624.1 abra|OJJ69224.1 acar|OOF91096.1 acla|EAW08622.1 afis|EAW15810.1 afla|EED51237.1 afum|EAL85541.1 agla|OJJ81976.1 akaw|GAA92232.1 aluc|OJZ86204.1 anid|CBF70905.1 anig|EHA24418.1 aory|EIT83188.1 ater|EAU33440.1 atub|OJI79998.1 awen|OJJ38521.1 azon|OJJ43734.1 acar|OOF91092.1 acla|EAW08620.1 afis|EAW15813.1 afla|EED51233.1 afum|EAL85538.1 akaw|GAA92236.1 aluc|OJZ86208.1 anid|CBF70903.1 aory|EIT83195.1 ater|EAU33435.1 atub|OJI80002.1 awen|OJJ38526.1

Group117: abra|OJJ69289.1 abra|OJJ70324.1 acar|OOF91780.1 acar|OOF94433.1 acla|EAW12318.1 acla|EAW14678.1 afis|EAW16903.1 afis|EAW24209.1 afla|EED45264.1 afla|EED48062.1 afum|EAL85879.1 afum|EAL86439.1 akaw|GAA88161.1 akaw|GAA92580.1 aluc|OJZ82118.1 aluc|OJZ82593.1 anig|EHA27482.1 anig|EHA28613.1 asyd|OJJ53986.1 asyd|OJJ54375.1 ater|EAU30213.1 ater|EAU31159.1 atub|OJI88319.1 atub|OJI88828.1 aver|OJJ05329.1 aver|OJJ08084.1 awen|OJJ29716.1 awen|OJJ33087.1 azon|OJJ42612.1 azon|OJJ44859.1 agla|OJJ79662.1 anid|CBF74267.1 aory|EIT83358.1 aory|EIT73698.1

Group118: abra|OJJ69846.1 abra|OJJ72321.1 acar|OOF94891.1 acar|OOF97566.1 afis|EAW19582.1 afis|EAW19642.1 afla|EED47805.1 afla|EED56993.1 agla|OJJ79220.1 agla|OJJ80784.1 akaw|GAA86072.1 akaw|GAA87813.1 aluc|OJZ85448.1 aluc|OJZ90307.1 anig|EHA23794.1 anig|EHA24991.1 aory|EIT77130.1 aory|EIT77504.1 atub|OJI80119.1 atub|OJI81731.1 aver|OJJ05597.1 aver|OJJ07110.1 acla|EAW12002.1 afum|EAL86880.1 anid|CBF71346.1 asyd|OJJ55096.1 ater|EAU32886.1 awen|OJJ41887.1 azon|OJJ42708.1 azon|OJJ46844.1 acla|EAW07815.1 anid|CBF74628.1 asyd|OJJ57585.1 ater|EAU30349.1

Group119: abra|OJJ70036.1 abra|OJJ72033.1 acla|EAW08504.1 acla|EAW09509.1 afis|EAW17096.1 afis|EAW21624.1 afla|EED48527.1 afla|EED57027.1 afla|EED57851.1 akaw|GAA84899.1 akaw|GAA88365.1 aluc|OJZ82290.1 aluc|OJZ86772.1 anig|EHA18733.1 anig|EHA21909.1 anig|EHA26569.1 aory|EIT72294.1 aory|EIT73160.1 aory|EIT81085.1 ater|EAU38312.1 ater|EAU38711.1 atub|OJI83756.1 atub|OJI88149.1 aver|OJJ03867.1 aver|OJJ08171.1 acar|OOF96691.1 afum|EAL92660.1 agla|OJJ79695.1 anid|CBF71556.1 asyd|OJJ52536.1 awen|OJJ33874.1 azon|OJJ48675.1 afla|EED46381.1 afla|EED46382.1

Group121: abra|OJJ74623.1 abra|OJJ74837.1 acar|OOF95085.1 acar|OOF95151.1 acar|OOF95841.1 acla|EAW10761.1 acla|EAW14613.1 afis|EAW19343.1 afis|EAW24815.1 afla|EED47168.1 afla|EED50136.1 afla|EED56891.1 afum|EAL89459.1 afum|EAL94062.1 akaw|GAA81986.1 akaw|GAA83583.1 aluc|OJZ92204.1 aluc|OJZ92414.1 anig|EHA26551.1 anig|EHA26754.1 aory|EIT74736.1 aory|EIT79324.1 aory|EIT77583.1 ater|EAU36492.1 ater|EAU39073.1 atub|OJI86632.1 atub|OJI86844.1 azon|OJJ48665.1 azon|OJJ49253.1 agla|OJJ82911.1 anid|CBF85881.1 asyd|OJJ64378.1 aver|OJI98474.1 awen|OJJ40401.1

Group122: agla|OJJ84376.1 agla|OJJ85709.1 asyd|OJJ57959.1 asyd|OJJ61061.1 abra|OJJ71937.1 acar|OOF96594.1 acar|OOF96595.1 acla|EAW09592.1 acla|EAW09593.1 afis|EAW21545.1 afis|EAW21546.1 afla|EED55996.1 afla|EED55997.1 afum|EAL92565.1 afum|EAL92566.1 agla|OJJ85708.1 akaw|GAA84987.1 aluc|OJZ86674.1 anid|CBF76366.1 anid|CBF76368.1 anig|EHA21999.1 anig|EHA22000.1 aory|EIT76685.1 aory|EIT76776.1 asyd|OJJ61062.1 ater|EAU34993.1 ater|EAU34994.1 atub|OJI83854.1 aver|OJJ00378.1 aver|OJJ00379.1 awen|OJJ39144.1 awen|OJJ39146.1 azon|OJJ51476.1 azon|OJJ51477.1

Group123: abra|OJJ67331.1 acar|OOF92482.1 acla|EAW06403.1 afis|EAW18507.1 afla|EED52893.1 afum|EAL93201.1 agla|OJJ85114.1 akaw|GAA86485.1 aluc|OJZ85026.1 anid|CBF70247.1 anig|EHA25247.1 aory|EIT78591.1 asyd|OJJ58761.1 ater|EAU30064.1 atub|OJI80571.1 aver|OJJ01371.1 awen|OJJ31060.1 azon|OJJ50571.1 abra|OJJ67332.1 acar|OOF92481.1 acla|EAW06402.1 afis|EAW18506.1 afla|EED52895.1 afum|EAL93200.1 akaw|GAA86484.1 aluc|OJZ85025.1 anig|EHA25246.1 aory|EIT78664.1 asyd|OJJ58760.1 ater|EAU30065.1 atub|OJI80572.1 aver|OJJ01368.1 awen|OJJ31059.1 azon|OJJ50569.1

Group124: abra|OJJ68518.1 acar|OOF92724.1 acla|EAW12845.1 afis|EAW18008.1 afla|EED51887.1 afum|EAL91820.1 agla|OJJ80274.1 akaw|GAA91330.1 aluc|OJZ82732.1 anid|CBF80034.1 anig|EHA21184.1 aory|EIT81755.1 asyd|OJJ58272.1 ater|EAU31524.1 atub|OJI88965.1 aver|OJJ02152.1 awen|OJJ41532.1 azon|OJJ48252.1 atub|OJI79785.1 abra|OJJ69012.1 acar|OOF92723.1 acla|EAW12846.1 afis|EAW18009.1 afla|EED51886.1 afum|EAL91821.1 agla|OJJ80275.1 akaw|GAA90529.1 aluc|OJZ85998.1 anid|CBF80035.1 anig|EHA24612.1 aory|EIT81715.1 ater|EAU31525.1 awen|OJJ41531.1 azon|OJJ48251.1

Group125: abra|OJJ69958.1 acar|OOF93468.1 acla|EAW08829.1 afis|EAW15568.1 afla|EED49078.1 afum|EAL85765.1 agla|OJJ85182.1 akaw|GAA87702.1 aluc|OJZ85324.1 anid|CBF78110.1 anig|EHA25514.1 aory|EIT82568.1 ater|EAU32562.1 atub|OJI80248.1 abra|OJJ69959.1 acla|EAW08833.1 afis|EAW15567.1 afla|EED49076.1 afla|EED49077.1 afum|EAL85766.1 agla|OJJ85184.1 akaw|GAA87701.1 anid|CBF78112.1

anid|CBF78114.1 aory|EIT82498.1 aory|EIT82525.1 asyd|OJJ55899.1 ater|EAU32566.1 atub|OJI80249.1 aver|OJJ04844.1 awen|OJJ38227.1 azon|OJJ43068.1 acla|EAW08832.1 azon|OJJ47853.1

Group126: abra|OJJ71382.1 acar|OOF97787.1 acla|EAW10157.1 afis|EAW25437.1 afla|EED53925.1 afum|EAL90095.1 agla|OJJ87082.1 akaw|GAA82612.1 aluc|OJZ87158.1 anid|CBF77596.1 anig|EHA20417.1 aory|EIT78347.1 asyd|OJJ53142.1 ater|EAU34609.1 atub|OJI81482.1 aver|OJJ03710.1 awen|OJJ41165.1 azon|OJJ44743.1 abra|OJJ71383.1 acar|OOF97786.1 acla|EAW10158.1 afis|EAW25436.1 afla|EED53924.1 afum|EAL90094.1 agla|OJJ87081.1 akaw|GAA82611.1 aluc|OJZ87157.1 anid|CBF77594.1 anig|EHA20416.1 aory|EIT78262.1 asyd|OJJ53143.1 ater|EAU34608.1 atub|OJI81483.1 aver|OJJ03709.1

Group127: abra|OJJ72469.1 acar|OOF94768.1 acla|EAW12702.1 afis|EAW17865.1 afla|EED46230.1 afum|EAL91673.1 agla|OJJ83099.1 akaw|GAA86207.1 aluc|OJZ90456.1 anid|CBF76910.1 anig|EHA23918.1 aory|EIT76409.1 asyd|OJJ57089.1 ater|EAU33080.1 atub|OJI81883.1 aver|OJJ03211.1 awen|OJJ41706.1 azon|OJJ48402.1 abra|OJJ72470.1 acar|OOF94765.1 acla|EAW12703.1 afis|EAW17866.1 afla|EED46231.1 afum|EAL91674.1 agla|OJJ83086.1 akaw|GAA86208.1 aluc|OJZ90457.1 anid|CBF76912.1 anig|EHA23919.1 aory|EIT76527.1 ater|EAU33081.1 atub|OJI81884.1 awen|OJJ41705.1 azon|OJJ48401.1

Group128: abra|OJJ72932.1 acar|OOF96112.1 acla|EAW06458.1 afis|EAW15954.1 afla|EED58104.1 afum|EAL88587.1 agla|OJJ89359.1 akaw|GAA85514.1 aluc|OJZ83442.1 anid|CBF81063.1 anig|EHA24288.1 aory|EIT79164.1 asyd|OJJ56456.1 ater|EAU35503.1 atub|OJI82333.1 aver|OJJ06266.1 awen|OJJ35622.1 azon|OJJ46902.1 abra|OJJ72934.1 acar|OOF96115.1 acla|EAW06457.1 afis|EAW15953.1 afla|EED58105.1 afum|EAL88586.1 akaw|GAA85515.1 aluc|OJZ83443.1 anid|CBF81061.1 anig|EHA24289.1 aory|EIT79130.1 asyd|OJJ56455.1 ater|EAU35504.1 atub|OJI82334.1 aver|OJJ06267.1 awen|OJJ35623.1

Group129: abra|OJJ76465.1 acar|OOF99703.1 acla|EAW13966.1 afis|EAW18675.1 afla|EED50624.1 afum|EAL93395.2 agla|OJJ82401.1 aluc|OJZ91745.1 anid|CBF70588.1 anig|EHA22748.1 aory|EIT78145.1 asyd|OJJ58523.1 ater|EAU38235.1 atub|OJI86203.1 aver|OJJ01124.1 awen|OJJ32147.1 azon|OJJ46731.1 abra|OJJ76466.1 acar|OOF99704.1 acla|EAW13965.1 afis|EAW18674.1 afla|EED50625.1 afum|EAL93394.1 akaw|GAA82377.1 aluc|OJZ91744.1 anid|CBF70586.1 anig|EHA22747.1 aory|EIT77989.1 asyd|OJJ58524.1 ater|EAU38236.1 atub|OJI86202.1 aver|OJJ01125.1 awen|OJJ32146.1 azon|OJJ46730.1

Group130: abra|OJJ66702.1 abra|OJJ76322.1 acar|OOF93706.1 acar|OOF93792.1 afla|EED45227.1 afla|EED56652.1 anid|CBF78509.1 anid|CBF81402.1 anig|EHA22872.1 anig|EHA27277.1 aory|EIT73751.1 aory|EIT79654.1 asyd|OJJ59404.1 asyd|OJJ65313.1 ater|EAU29738.1 ater|EAU34746.1 atub|OJI86345.1 atub|OJI87602.1 acla|EAW11833.1 afis|EAW19444.1 afum|EAL87051.1 agla|OJJ81158.1 akaw|GAA89644.1 aluc|OJZ80745.1 aver|OJJ02869.1 awen|OJJ36845.1 azon|OJJ42319.1 azon|OJJ42955.1 acar|OOF97765.1 akaw|GAA82251.1 aluc|OJZ91893.1 aver|OJI99451.1 awen|OJJ40922.1

Group131: abra|OJJ68714.1 abra|OJJ76939.1 acar|OOF91389.1 acar|OOF92214.1 afum|EAL84486.2 afum|EAL91972.2 aluc|OJZ81814.1 aluc|OJZ84338.1 anig|EHA22401.1 anig|EHA27672.1 atub|OJI84386.1 atub|OJI89730.1 awen|OJJ33278.1 awen|OJJ41371.1 azon|OJJ48840.1 azon|OJJ50652.1 acla|EAW12532.1 acla|EAW13466.1 afis|EAW20578.1 afis|EAW24117.1 afla|EED46815.1 afla|EED52313.1 agla|OJJ88717.1 akaw|GAA87100.1 akaw|GAA89867.1 anid|CBF86537.1 aory|EIT74384.1 asyd|OJJ64866.1 ater|EAU37106.1 aver|OJI98979.1 anid|CBF78940.1 asyd|OJJ55646.1 aver|OJJ06731.1

Group132: abra|OJJ69799.1 abra|OJJ75720.1 acar|OOF90260.1 acar|OOF93578.1 afis|EAW21759.1 afis|EAW24741.1 afum|EAL89387.1 afum|EAL92789.1 akaw|GAA84456.1 akaw|GAA87860.1 aluc|OJZ85501.1 aluc|OJZ89172.1 anid|CBF80400.1 anid|CBF83678.1 anig|EHA23383.1 anig|EHA25034.1 aory|EIT75373.1 aory|EIT79769.1 asyd|OJJ54761.1 asyd|OJJ55920.1 asyd|OJJ60823.1 atub|OJI80064.1 atub|OJI85618.1 aver|OJJ00149.1 aver|OJJ04873.1 aver|OJJ07409.1 azon|OJJ45021.1 azon|OJJ48200.1 acla|EAW09376.1 afla|EED45538.1 agla|OJJ84391.1 ater|EAU38431.1 awen|OJJ34043.1

Group133: abra|OJJ71453.1 abra|OJJ72857.1 acar|OOF96054.1 acar|OOF97693.1 acar|OOF97744.1 acar|OOF98763.1 afis|EAW22749.1 afis|EAW24033.1 afla|EED45033.1 afla|EED49120.1 afum|EAL87464.1 afum|EAL90450.1 akaw|GAA82558.1 akaw|GAA85456.1 aluc|OJZ83364.1 aluc|OJZ87090.1 anig|EHA20361.1 anig|EHA24229.1 aory|EIT73568.1 aory|EIT78580.1 asyd|OJJ59869.1 asyd|OJJ65349.1 ater|EAU29414.1 ater|EAU29711.1 atub|OJI81548.1 atub|OJI82255.1 aver|OJI99497.1 aver|OJJ02418.1 acla|EAW11389.1 agla|OJJ86999.1 anid|CBF75320.1 awen|OJJ35505.1 azon|OJJ46979.1

Group134: abra|OJJ72895.1 abra|OJJ73247.1 acar|OOF96086.1 acar|OOG00487.1 acla|EAW06921.1 acla|EAW11356.1 afis|EAW16538.1 afis|EAW18138.1 afla|EED56131.1 afla|EED58064.1 afum|EAL87495.1 afum|EAL89135.1 agla|OJJ82784.1 agla|OJJ83821.1 akaw|GAA85482.1 akaw|GAA87933.1 aluc|OJZ81180.1 aluc|OJZ83404.1 anig|EHA18498.1 anig|EHA24255.1 aory|EIT76761.1 aory|EIT79757.1 ater|EAU35461.1 ater|EAU36601.1 atub|OJI82293.1 atub|OJI91377.1 awen|OJJ35580.1 awen|OJJ40283.1 azon|OJJ46943.1 azon|OJJ49145.1 anid|CBF81745.1 asyd|OJJ59599.1 aver|OJJ02669.1

Group135: anid|CBF74543.1 anid|CBF80800.1 asyd|OJJ54487.1 asyd|OJJ57688.1 aver|OJJ05495.1 aver|OJJ07692.1 awen|OJJ30771.1 awen|OJJ31710.1 abra|OJJ67440.1 acar|OOF92387.1 acar|OOF92388.1 acla|EAW06324.1 afis|EAW18438.1 afla|EED52977.1 afum|EAL93120.1 agla|OJJ85004.1 akaw|GAA86390.1 aluc|OJZ84924.1 anid|CBF70039.1 anig|EHA25142.1 anig|EHA25143.1 aory|EIT78691.1 aory|EIT78713.1

asyd|OJJ58866.1 ater|EAU29862.1 atub|OJI80675.1 aver|OJJ01467.1 azon|OJJ50475.1 acla|EAW06323.1 afis|EAW18437.1 afum|EAL93119.1 ater|EAU29854.1 afla|EED52975.1

Group136: aory|EIT76219.1 abra|OJJ66160.1 acla|EAW07091.1 ater|EAU30184.1 abra|OJJ76943.1 acar|OOF91392.1 acla|EAW14645.1 afis|EAW23759.1 afla|EED45469.1 afum|EAL87193.2 agla|OJJ78794.1 akaw|GAA87097.1 aluc|OJZ84341.1 anid|CBF86208.1 anig|EHA27675.1 anig|EHA27676.1 aory|EIT75611.1 asyd|OJJ64548.1 atub|OJI89727.1 aver|OJI98644.1 awen|OJJ40873.1 afis|EAW21192.1 anid|CBF78488.1 asyd|OJJ58352.1 ater|EAU36866.1 aver|OJJ02055.1 awen|OJJ31298.1 afis|EAW22111.1 afla|EED48116.1 aver|OJJ07112.1 azon|OJJ50585.1 awen|OJJ29841.1 awen|OJJ30060.1

Group137: abra|OJJ68073.1 acar|OOF92659.1 acla|EAW14013.1 afis|EAW18719.1 afla|EED50674.1 afum|EAL93443.2 agla|OJJ82339.1 akaw|GAA92398.1 aluc|OJZ83259.1 anid|CBF70677.1 anig|EHA21113.1 aory|EIT78139.1 asyd|OJJ58472.1 ater|EAU38185.1 atub|OJI87680.1 aver|OJJ01073.1 awen|OJJ32224.1 azon|OJJ46775.1 abra|OJJ68477.1 acar|OOF92676.1 acla|EAW07215.1 afis|EAW22228.1 afla|EED57619.1 afum|EAL90966.1 agla|OJJ87821.1 anid|CBF88467.1 anig|EHA21112.1 aory|EIT82940.1 asyd|OJJ63779.1 ater|EAU34233.1 aver|OJI97158.1 awen|OJJ34810.1 azon|OJJ45740.1

Group138: abra|OJJ71306.1 acar|OOF97864.1 acla|EAW10234.1 afis|EAW25358.1 afla|EED53840.1 afum|EAL90020.1 agla|OJJ83908.1 akaw|GAA82685.1 aluc|OJZ87238.1 anid|CBF77463.1 anig|EHA20484.1 aory|EIT78472.1 asyd|OJJ53218.1 ater|EAU34506.1 atub|OJI81404.1 aver|OJJ03634.1 awen|OJJ41084.1 azon|OJJ44664.1 abra|OJJ71307.1 acla|EAW10233.1 afis|EAW25359.1 afla|EED53841.1 afum|EAL90021.1 agla|OJJ83909.1 akaw|GAA82684.1 aluc|OJZ87237.1 anid|CBF77461.1 aory|EIT78435.1 asyd|OJJ53217.1 ater|EAU34507.1 atub|OJI81405.1 awen|OJJ41085.1 azon|OJJ44665.1

Group139: abra|OJJ71587.1 acar|OOF96317.1 acla|EAW15133.1 afis|EAW20105.1 afla|EED53369.1 afum|EAL85222.1 agla|OJJ88850.1 akaw|GAA89336.1 aluc|OJZ86327.1 anid|CBF84906.1 anig|EHA21551.1 aory|EIT75191.1 asyd|OJJ63169.1 ater|EAU38724.1 atub|OJI84207.1 aver|OJI97785.1 awen|OJJ31876.1 azon|OJJ46259.1 abra|OJJ71588.1 acla|EAW15132.1 afis|EAW20104.1 afum|EAL85221.1 agla|OJJ88849.1 akaw|GAA89335.1 aluc|OJZ86328.1 anid|CBF84908.1 anig|EHA21550.1 aory|EIT75224.1 asyd|OJJ63168.1 atub|OJI84206.1 aver|OJI97786.1 awen|OJJ31877.1 azon|OJJ46257.1

Group140: abra|OJJ72666.1 acar|OOF94609.1 acla|EAW11529.1 afis|EAW23925.1 afla|EED50596.1 afum|EAL87350.1 agla|OJJ79403.1 akaw|GAA85287.1 aluc|OJZ90645.1 anid|CBF77250.1 anig|EHA24080.1 aory|EIT82072.1 asyd|OJJ56809.1 ater|EAU32310.1 atub|OJI82075.1 aver|OJJ03525.1 awen|OJJ39029.1 azon|OJJ51358.1 abra|OJJ72668.1 acla|EAW11530.1 afis|EAW23924.1 afla|EED50597.1 afum|EAL87349.1 agla|OJJ79402.1 akaw|GAA85288.1 aluc|OJZ90646.1 anid|CBF77248.1 anig|EHA24081.1 aory|EIT78140.1 asyd|OJJ56810.1 ater|EAU32312.1 awen|OJJ39031.1 azon|OJJ51359.1

Group141: abra|OJJ74273.1 acar|OOF95457.1 acla|EAW07318.1 afis|EAW22350.1 afla|EED57754.1 afum|EAL90842.1 agla|OJJ87709.1 akaw|GAA86750.1 aluc|OJZ89860.1 anid|CBF88653.1 anig|EHA27060.1 aory|EIT81054.1 ater|EAU39489.1 aver|OJI97054.1 awen|OJJ34904.1 azon|OJJ45578.1 abra|OJJ74279.1 acar|OOF95452.1 acla|EAW07323.1 afis|EAW22355.1 afla|EED57761.1 afum|EAL90836.1 agla|OJJ87608.1 akaw|GAA86745.1 aluc|OJZ89854.1 anid|CBF88666.1 anig|EHA27055.1 aory|EIT81031.1 asyd|OJJ63892.1 atub|OJI84952.1 aver|OJI97047.1 awen|OJJ34929.1 azon|OJJ45582.1

Group142: abra|OJJ68218.1 abra|OJJ69789.1 acar|OOF93590.1 acar|OOF97507.1 agla|OJJ78902.1 agla|OJJ86249.1 akaw|GAA83228.1 akaw|GAA87869.1 aluc|OJZ83728.1 aluc|OJZ85510.1 anid|CBF84393.1 anid|CBF86693.1 atub|OJI80056.1 atub|OJI90104.1 acla|EAW09054.1 acla|EAW12123.1 afis|EAW25629.1 afla|EED55183.1 afum|EAL84317.1 anig|EHA18876.1 aory|EIT80665.1 aory|EIT81643.1 asyd|OJJ55507.1 ater|EAU32706.1 aver|OJJ06584.1 aver|OJJ08175.1 awen|OJJ36048.1 azon|OJJ43184.1 afla|EED51722.1 anig|EHA25042.1 asyd|OJJ53581.1 anid|CBF83169.1

Group143: abra|OJJ68305.1 abra|OJJ68976.1 afla|EED55146.1 afla|EED57528.1 agla|OJJ80112.1 agla|OJJ84537.1 akaw|GAA83169.1 akaw|GAA90501.1 aluc|OJZ83668.1 aluc|OJZ85964.1 aory|EIT81602.1 aory|EIT81847.1 asyd|OJJ59831.1 asyd|OJJ65307.1 atub|OJI79752.1 atub|OJI90039.1 awen|OJJ31501.1 awen|OJJ36340.1 azon|OJJ46215.1 azon|OJJ48729.1 acar|OOF91551.1 acar|OOF96683.1 acla|EAW14213.1 afis|EAW19068.1 afum|EAL93797.1 anid|CBF78527.1 anig|EHA18823.1 anig|EHA24640.1 aver|OJI99445.1 afla|EED54434.1 aory|EIT75875.1 ater|EAU32117.1

Group145: abra|OJJ70543.1 abra|OJJ73773.1 acar|OOF94608.1 acar|OOG00929.1 afis|EAW15411.1 afis|EAW22675.1 afum|EAL84235.1 afum|EAL90517.1 akaw|GAA83389.1 akaw|GAA88403.1 aluc|OJZ83888.1 aluc|OJZ88281.1 anid|CBF74491.1 anid|CBF88137.1 anig|EHA18026.1 anig|EHA19025.1 aory|EIT72732.1 aory|EIT83438.1 asyd|OJJ57716.1 asyd|OJJ63598.1 ater|EAU36301.1 ater|EAU39034.1 atub|OJI90255.1 atub|OJI90881.1 azon|OJJ45068.1 azon|OJJ49812.1 acla|EAW07657.1 afla|EED49956.1 agla|OJJ79042.1 aver|OJI97338.1 aver|OJJ05451.1 awen|OJJ35304.1

Group147: abra|OJJ73071.1 abra|OJJ77012.1 acla|EAW09005.1 acla|EAW15322.1 afis|EAW20323.1 afis|EAW25545.1 akaw|GAA87037.1 akaw|GAA88090.1 aluc|OJZ81354.1 aluc|OJZ84412.1 asyd|OJJ54270.1 asyd|OJJ55537.1 ater|EAU32181.1 ater|EAU39595.1 atub|OJI89661.1 atub|OJI91558.1 aver|OJJ05226.1 aver|OJJ06615.1 acar|OOG00304.1 afla|EED44875.1 afum|EAL85448.1 agla|OJJ78910.1 anid|CBF82101.1

anid|CBF83180.1 anig|EHA18655.1 aory|EIT83261.1 awen|OJJ36038.1 azon|OJJ51343.1 afla|EED54130.1 afum|EAL84263.1 anig|EHA27738.1 aory|EIT76050.1

Group149: acar|OOF94510.1 acar|OOG00617.1 agla|OJJ78704.1 agla|OJJ82589.1 agla|OJJ88673.1 awen|OJJ33431.1 awen|OJJ40048.1 abra|OJJ73366.1 acar|OOG00573.1 acla|EAW12876.1 afis|EAW17800.1 afum|EAL91608.1 akaw|GAA83118.1 aluc|OJZ88674.1 anid|CBF87487.1 anig|EHA18356.1 asyd|OJJ52958.1 ater|EAU31398.1 atub|OJI91255.1 aver|OJJ08926.1 abra|OJJ73415.1 afla|EED44896.1 akaw|GAA83076.1 aluc|OJZ88622.1 anig|EHA18315.1 aory|EIT83248.1 asyd|OJJ62646.1 atub|OJI91207.1 aver|OJJ07782.1 azon|OJJ50122.1 afis|EAW20384.1 asyd|OJJ54467.1

Group150: acar|OOF95065.1 acar|OOF95131.1 acar|OOF95066.1 acar|OOF95132.1 abra|OJJ74648.1 acla|EAW10738.1 acla|EAW10739.1 afis|EAW24835.1 afis|EAW24836.1 afla|EED47139.1 afla|EED47140.1 afum|EAL89480.1 afum|EAL89481.2 agla|OJJ82028.1 akaw|GAA81962.1 akaw|GAA81963.1 aluc|OJZ92226.1 anid|CBF74584.1 anid|CBF74586.1 anig|EHA26730.1 aory|EIT79207.1 aory|EIT79226.1 asyd|OJJ57656.1 ater|EAU36469.1 ater|EAU36470.1 atub|OJI86656.1 aver|OJJ05516.1 aver|OJJ05517.1 awen|OJJ40422.1 awen|OJJ40423.1 azon|OJJ49280.1 azon|OJJ49281.1

Group151: acar|OOF95215.1 acar|OOG00549.1 afla|EED45224.1 afla|EED52255.1 afla|EED57542.1 afum|EAL86491.1 afum|EAL91022.1 aory|EIT73609.1 aory|EIT74331.1 aory|EIT81831.1 asyd|OJJ52204.1 asyd|OJJ54774.1 asyd|OJJ64753.1 ater|EAU30189.1 ater|EAU32767.1 ater|EAU34175.1 aver|OJI98855.1 aver|OJJ07392.1 aver|OJJ07305.1 abra|OJJ74541.1 acla|EAW07155.1 acla|EAW12276.1 afis|EAW16964.1 agla|OJJ87885.1 akaw|GAA82061.1 aluc|OJZ92121.1 anid|CBF86686.1 anig|EHA26824.1 atub|OJI86554.1 awen|OJJ34744.1 azon|OJJ42779.1 anid|CBF80465.1

Group153: asyd|OJJ60702.1 asyd|OJJ64821.1 aver|OJI98934.1 aver|OJJ00028.1 abra|OJJ72028.1 acar|OOF96677.1 afis|EAW21619.1 afla|EED57848.1 akaw|GAA84904.1 aluc|OJZ86765.1 anid|CBF83493.1 anig|EHA21914.1 aory|EIT81067.1 ater|EAU38309.1 atub|OJI83762.1 awen|OJJ37096.1 azon|OJJ48684.1 abra|OJJ77154.1 acar|OOF93040.1 acla|EAW12917.1 afis|EAW17740.1 afla|EED45741.1 afum|EAL91554.1 agla|OJJ81711.1 akaw|GAA90776.1 aluc|OJZ84568.1 anid|CBF86610.1 anig|EHA27862.1 aory|EIT75552.1 ater|EAU31867.1 atub|OJI89514.1 azon|OJJ50798.1

Group154: atub|OJI81651.1 atub|OJI86062.1 abra|OJJ65911.1 acar|OOF94961.1 acla|EAW12054.1 afis|EAW19697.1 afla|EED57032.1 afum|EAL86830.1 agla|OJJ80819.1 akaw|GAA92321.1 aluc|OJZ90219.1 anid|CBF71392.1 anig|EHA23717.1 aory|EIT82283.1 asyd|OJJ55041.1 ater|EAU33691.1 aver|OJI98387.1 awen|OJJ41939.1 azon|OJJ50749.1 abra|OJJ76597.1 akaw|GAA82501.1 aluc|OJZ91610.1 anig|EHA22639.1 acla|EAW11329.1 afis|EAW18166.1 afla|EED56329.1 afum|EAL87526.1 agla|OJJ85952.1 anid|CBF85749.1 aory|EIT73387.1 asyd|OJJ64295.1 aver|OJJ07167.1

Group155: abra|OJJ66622.1 acar|OOF91045.1 acla|EAW10875.1 afis|EAW20983.1 afla|EED50338.1 afum|EAL84647.2 agla|OJJ84060.1 akaw|GAA91103.1 aluc|OJZ81427.1 anid|CBF82618.1 anig|EHA25995.1 aory|EIT77260.1 asyd|OJJ54020.1 ater|EAU38633.1 atub|OJI88753.1 aver|OJJ08720.1 awen|OJJ34328.1 azon|OJJ48073.1 afla|EED50340.1 akaw|GAA91102.1 aluc|OJZ81426.1 acar|OOF91046.1 acla|EAW10874.1 afis|EAW20984.1 afla|EED50339.1 afum|EAL84646.1 agla|OJJ84063.1 anid|CBF82622.1 aory|EIT77350.1 atub|OJI88754.1 awen|OJJ34327.1 azon|OJJ48069.1

Group156: abra|OJJ68143.1 acar|OOF97481.1 acla|EAW13069.1 afis|EAW17595.1 afla|EED49225.1 afum|EAL91408.1 agla|OJJ81333.1 akaw|GAA92375.1 aluc|OJZ83234.1 anid|CBF90206.1 anig|EHA21091.1 aory|EIT83153.1 asyd|OJJ61974.1 ater|EAU37033.1 atub|OJI87702.1 aver|OJI96067.1 awen|OJJ37251.1 azon|OJJ50946.1 abra|OJJ68185.1 acla|EAW13066.1 afis|EAW17598.1 afla|EED49223.1 afum|EAL91411.1 agla|OJJ81335.1 akaw|GAA83261.1 aluc|OJZ83767.1 anid|CBF90203.1 asyd|OJJ61976.1 ater|EAU37035.1 atub|OJI90138.1 aver|OJI96069.1 awen|OJJ37248.1

Group157: abra|OJJ70602.1 acar|OOF97117.1 acla|EAW08180.1 afis|EAW23215.1 afla|EED47037.1 afum|EAL88269.1 akaw|GAA89942.1 aluc|OJZ83947.1 anig|EHA19076.1 aory|EIT72544.1 asyd|OJJ60180.1 ater|EAU36345.1 atub|OJI90318.1 awen|OJJ39449.1 abra|OJJ70604.1 acar|OOF97116.1 acla|EAW08178.1 afis|EAW23213.1 afum|EBA27340.1 agla|OJJ78731.1 akaw|GAA89944.1 aluc|OJZ83949.1 anid|CBF74742.1 anig|EHA19078.1 aory|EIT72666.1 asyd|OJJ60182.1 ater|EAU36343.1 atub|OJI90320.1 aver|OJJ04082.1 awen|OJJ39447.1 azon|OJJ47649.1 anid|CBF74745.1

Group158: abra|OJJ71268.1 acar|OOF97905.1 acar|OOF97907.1 acla|EAW10269.1 acla|EAW10270.1 afis|EAW25324.1 afis|EAW25325.1 afla|EED53799.1 afla|EED53800.1 afum|EAL89985.1 afum|EAL89986.1 agla|OJJ83862.1 agla|OJJ83863.1 akaw|GAA82722.1 aluc|OJZ87276.1 anid|CBF75258.1 anid|CBF75260.1 anig|EHA20517.1 anig|EHA20518.1 aory|EIT78367.1 aory|EIT78397.1 asyd|OJJ59900.1 asyd|OJJ59901.1 ater|EAU34470.1 atub|OJI81365.1 aver|OJJ02385.1 aver|OJJ02386.1 awen|OJJ41038.1 awen|OJJ41039.1 azon|OJJ44627.1 ater|EAU34472.1 ater|EAU34471.1

Group159: abra|OJJ72382.1 acar|OOF94837.1 acla|EAW11953.1 acla|EAW11954.1 afis|EAW19590.1 afis|EAW19591.1 afla|EED56945.1 afum|EAL86927.1 afum|EAL86928.1 agla|OJJ80730.1 agla|OJJ80732.1 akaw|GAA86129.1 akaw|GAA86131.1 aluc|OJZ90369.1 anid|CBF71264.1 anid|CBF71266.1 anig|EHA23843.1 anig|EHA23844.1 aory|EIT77519.1 aory|EIT77568.1 asyd|OJJ55144.1 asyd|OJJ55146.1 ater|EAU32934.1

ater|EAU32936.1 atub|OJI81795.1 aver|OJJ07061.1 aver|OJJ07063.1 awen|OJJ41805.1 awen|OJJ41808.1 azon|OJJ47024.1 azon|OJJ47026.1 acar|OOF94836.1

Group160: abra|OJJ72953.1 acar|OOF96134.1 acla|EAW11198.1 afis|EAW15928.1 afla|EED54933.1 afum|EAL88564.1 agla|OJJ89377.1 akaw|GAA85534.1 aluc|OJZ83463.1 anid|CBF81024.1 anig|EHA24306.1 aory|EIT80331.1 asyd|OJJ56385.1 ater|EAU35524.1 atub|OJI82355.1 aver|OJJ06327.1 awen|OJJ35652.1 azon|OJJ46886.1 abra|OJJ72954.1 acar|OOF96135.1 acla|EAW11199.1 afis|EAW15929.1 afla|EED54932.1 afum|EAL88565.1 akaw|GAA85535.1 anid|CBF81026.1 anig|EHA24307.1 aory|EIT80358.1 asyd|OJJ56386.1 ater|EAU35523.1 awen|OJJ35651.1 azon|OJJ46887.1

Group161: abra|OJJ74876.1 acar|OOF95878.1 acar|OOF95879.1 acla|EAW08494.1 acla|EAW08495.1 afis|EAW23540.1 afis|EAW23541.1 afla|EED57438.1 afla|EED57439.1 afum|EAL87925.1 afum|EAL87926.1 agla|OJJ86795.1 akaw|GAA83544.1 akaw|GAA83545.1 aluc|OJZ92453.1 anid|CBF89661.1 anid|CBF89663.1 anig|EHA26514.1 anig|EHA26515.1 aory|EIT80116.1 aory|EIT80129.1 asyd|OJJ62276.1 asyd|OJJ62277.1 ater|EAU35326.1 ater|EAU35327.1 atub|OJI86881.1 atub|OJI86882.1 aver|OJI96385.1 aver|OJI96386.1 awen|OJJ39871.1 awen|OJJ39872.1 azon|OJJ43230.1

Group162: abra|OJJ75341.1 acar|OOF89856.1 acla|EAW09853.1 afis|EAW21282.1 afla|EED54833.1 afum|EAL92299.1 akaw|GAA92034.1 aluc|OJZ92904.1 anid|CBF83359.1 anig|EHA26059.1 aory|EIT79370.1 ater|EAU35911.1 atub|OJI87363.1 awen|OJJ39417.1 abra|OJJ75940.1 acar|OOF99970.1 acla|EAW09854.1 afis|EAW21281.1 afla|EED54832.1 afum|EAL92298.1 agla|OJJ87478.1 akaw|GAA84642.1 aluc|OJZ89387.1 anid|CBF83358.1 anig|EHA23197.1 aory|EIT79445.1 asyd|OJJ60618.1 ater|EAU35910.1 atub|OJI85410.1 aver|OJI99942.1 awen|OJJ39418.1 azon|OJJ44300.1

Group163: abra|OJJ75791.1 acar|OOG00087.1 acar|OOG00097.1 acla|EAW14890.1 acla|EAW14891.1 afis|EAW24412.1 afis|EAW24413.1 afla|EED47763.1 afla|EED47764.1 afum|EAL86081.1 afum|EAL86082.1 agla|OJJ79215.1 agla|OJJ79216.1 akaw|GAA84506.1 akaw|GAA84510.1 aluc|OJZ89235.1 aluc|OJZ89239.1 anid|CBF73743.1 anid|CBF73745.1 anig|EHA23328.1 aory|EIT77117.1 aory|EIT77194.1 asyd|OJJ56623.1 ater|EAU30316.1 ater|EAU30317.1 atub|OJI85559.1 aver|OJJ06102.1 aver|OJJ06103.1 awen|OJJ32796.1 awen|OJJ32798.1 azon|OJJ49036.1 azon|OJJ49037.1

Group164: abra|OJJ76816.1 acar|OOF91311.1 acla|EAW13376.1 afis|EAW20490.1 afla|EED44773.1 afum|EAL84564.1 agla|OJJ84837.1 akaw|GAA87205.1 aluc|OJZ84226.1 anig|EHA27575.1 aory|EIT77777.1 asyd|OJJ55730.1 atub|OJI89858.1 aver|OJJ06816.1 awen|OJJ33190.1 azon|OJJ46392.1 abra|OJJ76822.1 acar|OOF91317.1 acla|EAW13382.1 afis|EAW20496.1 afla|EED44772.1 afum|EAL84558.1 agla|OJJ84806.1 akaw|GAA87200.1 aluc|OJZ84233.1 anid|CBF79209.1 anig|EHA27580.1 aory|EIT77769.1 ater|EAU37190.1 atub|OJI89852.1 awen|OJJ33196.1 azon|OJJ46400.1

Group165: abra|OJJ78214.1 acla|EAW11236.1 afis|EAW18269.1 afla|EED48735.1 afum|EAL87634.1 agla|OJJ88158.1 akaw|GAA85813.1 aluc|OJZ91286.1 anid|CBF79514.1 anig|EHA19971.1 aory|EIT73143.1 asyd|OJJ57887.1 ater|EAU33276.1 atub|OJI83276.1 aver|OJJ04507.1 awen|OJJ30643.1 azon|OJJ47972.1 abra|OJJ78215.1 acar|OOF98641.1 acla|EAW11235.1 afis|EAW18270.1 afla|EED48736.1 afum|EAL87635.1 agla|OJJ88157.1 akaw|GAA85812.1 aluc|OJZ91287.1 anid|CBF79512.1 anig|EHA19972.1 aory|EIT73147.1 asyd|OJJ57886.1 atub|OJI83277.1 aver|OJJ04508.1

Group168: abra|OJJ66675.1 abra|OJJ75504.1 acar|OOF90835.1 acar|OOF93687.1 acar|OOF93772.1 afis|EAW16470.1 afis|EAW19425.1 afum|EAL87069.1 afum|EAL89064.1 akaw|GAA84268.1 akaw|GAA89623.1 aluc|OJZ80773.1 aluc|OJZ88960.1 anig|EHA23560.1 anig|EHA27257.1 atub|OJI85829.1 atub|OJI87625.1 acla|EAW06851.1 afla|EED50383.1 agla|OJJ81137.1 anid|CBF74378.1 aory|EIT83131.1 asyd|OJJ57778.1 ater|EAU34729.1 aver|OJJ05387.1 awen|OJJ36823.1 azon|OJJ42980.1 afla|EED49104.1 asyd|OJJ57938.1 aver|OJJ04456.1 aory|EIT82490.1

Group169: abra|OJJ67685.1 abra|OJJ69630.1 acar|OOF92680.1 acar|OOF94075.1 acar|OOF94145.1 acar|OOF96015.1 afum|EAL86373.1 afum|EAL87420.1 akaw|GAA89197.1 akaw|GAA91610.1 aluc|OJZ81633.1 aluc|OJZ82435.1 anig|EHA25810.1 anig|EHA28336.1 atub|OJI88006.1 atub|OJI88547.1 afla|EED55077.1 aory|EIT81950.1 acla|EAW12376.1 afis|EAW16822.1 agla|OJJ86924.1 anid|CBF82987.1 asyd|OJJ64732.1 ater|EAU35425.1 aver|OJI98836.1 awen|OJJ35458.1 azon|OJJ51285.1 anid|CBF77369.1 asyd|OJJ53260.1 aver|OJJ03591.1 acla|EAW11432.1

Group170: acar|OOF91973.1 acar|OOF94079.1 acar|OOF94149.1 acar|OOF99292.1 afla|EED45949.1 afla|EED47803.1 aluc|OJZ79501.1 aluc|OJZ79636.1 aluc|OJZ81691.1 aluc|OJZ80076.1 aory|EIT75701.1 aory|EIT77131.1 asyd|OJJ52720.1 asyd|OJJ61396.1 asyd|OJJ61537.1 ater|EAU29987.1 ater|EAU30155.1 aver|OJJ00725.1 aver|OJJ04752.1 awen|OJJ32293.1 awen|OJJ35892.1 abra|OJJ67625.1 acla|EAW15287.1 afis|EAW20776.1 afum|EAL84844.1 agla|OJJ85083.1 akaw|GAA89145.1 anid|CBF87439.1 anig|EHA25753.1 atub|OJI88489.1 azon|OJJ42146.1

Group171: acar|OOF92319.1 acar|OOF92361.1 abra|OJJ67475.1 acla|EAW06297.1 afis|EAW18404.1 afla|EED53004.1 afum|EAL93084.1 agla|OJJ84961.1 akaw|GAA86362.1 aluc|OJZ84889.1 anid|CBF70745.1 anig|EHA25118.1 aory|EIT77437.1 asyd|OJJ58438.1 ater|EAU29821.1 atub|OJI80708.1 aver|OJJ01038.1 awen|OJJ31746.1 abra|OJJ67478.1 acla|EAW06294.1 afis|EAW18400.1 afla|EED53006.1 afum|EAL93078.1

agla|OJJ84971.1 akaw|GAA86360.1 anid|CBF70749.1 aory|EIT77424.1 ater|EAU29816.1 atub|OJI80711.1 awen|OJJ31749.1 azon|OJJ50474.1

Group172: acar|OOF98484.1 acar|OOF98497.1 abra|OJJ78390.1 acar|OOF98498.1 acla|EAW09142.1 acla|EAW09143.1 afis|EAW22004.1 afis|EAW22005.1 afla|EED46729.1 afum|EAL93019.1 afum|EAL93020.1 agla|OJJ84093.1 agla|OJJ84094.1 akaw|GAA85641.1 akaw|GAA85642.1 aluc|OJZ91469.1 anid|CBF82660.1 anid|CBF82662.1 anig|EHA20131.1 aory|EIT77865.1 aory|EIT77967.1 asyd|OJJ61666.1 asyd|OJJ61667.1 ater|EAU31123.1 ater|EAU31125.1 atub|OJI83462.1 aver|OJJ00987.1 awen|OJJ34299.1 awen|OJJ34300.1 azon|OJJ43338.1 azon|OJJ43339.1

Group174: afla|EED46122.1 afla|EED53117.1 afla|EED48931.1 afla|EED51100.1 akaw|GAA86153.1 akaw|GAA92990.1 aory|EIT74792.1 aory|EIT79985.1 aory|EIT81307.1 atub|OJI81824.1 atub|OJI86011.1 abra|OJJ72410.1 acar|OOF94819.1 acla|EAW14266.1 afis|EAW19012.1 afum|EAL93746.1 agla|OJJ81770.1 aluc|OJZ90394.1 anig|EHA23863.1 aory|EIT76367.1 ater|EAU37790.1 awen|OJJ31644.1 abra|OJJ76649.1 acar|OOF99352.1 aluc|OJZ91559.1 anid|CBF86747.1 anig|EHA22591.1 asyd|OJJ64703.1 aver|OJI98807.1 azon|OJJ47337.1 afis|EAW16798.1

Group176: ater|EAU33722.1 ater|EAU38738.1 abra|OJJ71609.1 acar|OOF96326.1 acla|EAW15150.1 acla|EAW15152.1 afis|EAW20118.1 afis|EAW20120.1 afla|EED53353.1 afla|EED57256.1 afum|EAL85235.1 afum|EBA27224.1 agla|OJJ88826.1 agla|OJJ88831.1 akaw|GAA89318.1 akaw|GAA89319.1 aluc|OJZ86351.1 aluc|OJZ86353.1 anid|CBF84937.1 anid|CBF84939.1 anig|EHA21536.1 aory|EIT74846.1 asyd|OJJ63156.1 ater|EAU38737.1 atub|OJI84184.1 aver|OJI97800.1 aver|OJI97801.1 awen|OJJ31888.1 azon|OJJ46246.1 aory|EIT74143.1 asyd|OJJ63154.1

Group177: awen|OJJ33301.1 awen|OJJ37090.1 abra|OJJ76956.1 acar|OOF91397.1 acla|EAW13475.1 afla|EED52244.1 agla|OJJ88687.1 akaw|GAA87087.1 aluc|OJZ84351.1 anid|CBF78932.1 anig|EHA27688.1 aory|EIT74404.1 asyd|OJJ62348.1 ater|EAU37096.1 atub|OJI89715.1 azon|OJJ50794.1 abra|OJJ77159.1 acar|OOF93045.1 acla|EAW12913.1 afis|EAW17747.1 afla|EED45747.1 afum|EAL91559.1 agla|OJJ81707.1 akaw|GAA90772.1 aluc|OJZ84572.1 anid|CBF86617.1 aory|EIT75770.1 asyd|OJJ64815.1 ater|EAU31860.1 atub|OJI89510.1 aver|OJI98928.1

Group178: abra|OJJ65971.1 acar|OOF92902.1 acla|EAW13592.1 afis|EAW20012.1 afla|EED46018.1 afum|EAL84906.1 akaw|GAA90846.1 aluc|OJZ82987.1 anid|CBF84721.1 anig|EHA21402.1 aory|EIT75680.1 asyd|OJJ53329.1 ater|EAU29354.1 atub|OJI89221.1 aver|OJJ08415.1 awen|OJJ30025.1 abra|OJJ65972.1 acar|OOF92900.1 acla|EAW13593.1 afis|EAW20011.1 afla|EED46017.1 afum|EAL84907.1 agla|OJJ87976.1 anid|CBF83963.1 anig|EHA21401.1 aory|EIT75560.1 asyd|OJJ61425.1 ater|EAU29353.1 aver|OJJ00759.1 awen|OJJ30024.1 azon|OJJ43649.1

Group179: abra|OJJ66528.1 acar|OOF90981.1 acla|EAW10962.1 afis|EAW20897.1 afla|EED50247.1 afum|EAL84735.1 agla|OJJ84007.1 akaw|GAA91185.1 aluc|OJZ81518.1 anid|CBF82460.1 anig|EHA25916.1 aory|EIT77290.1 ater|EAU38531.1 atub|OJI88667.1 awen|OJJ34450.1 azon|OJJ51738.1 abra|OJJ66529.1 acar|OOF90980.1 acla|EAW10961.1 afis|EAW20898.1 afla|EED50248.1 afum|EAL84734.2 agla|OJJ84008.1 akaw|GAA91184.1 aluc|OJZ81519.1 anid|CBF82458.1 aory|EIT77356.1 asyd|OJJ54115.1 atub|OJI88668.1 aver|OJJ08620.1 azon|OJJ51737.1

Group180: abra|OJJ66543.1 acar|OOF90995.1 afla|EED50933.1 agla|OJJ78717.1 akaw|GAA91168.1 aluc|OJZ81496.1 anid|CBF71517.1 anig|EHA25934.1 aory|EIT79048.1 asyd|OJJ54917.1 ater|EAU29874.1 atub|OJI88685.1 aver|OJJ07286.1 awen|OJJ30168.1 abra|OJJ67418.1 acar|OOF92403.1 acla|EAW06346.1 afis|EAW18457.1 afla|EED52951.1 afum|EAL93142.1 agla|OJJ85040.1 akaw|GAA86410.1 aluc|OJZ84950.1 anid|CBF70055.1 anig|EHA25165.1 aory|EIT78701.1 asyd|OJJ58858.1 atub|OJI80652.1 aver|OJJ01459.1 awen|OJJ31009.1 azon|OJJ50523.1

Group181: abra|OJJ67409.1 acar|OOF92406.1 acla|EAW06349.1 afis|EAW18461.1 afla|EED52947.1 afum|EAL93147.1 agla|OJJ85045.1 akaw|GAA86418.1 aluc|OJZ84959.1 anid|CBF70089.1 anig|EHA25173.1 aory|EIT78700.1 asyd|OJJ58841.1 ater|EAU29879.1 atub|OJI80641.1 aver|OJJ01443.1 awen|OJJ31013.1 azon|OJJ50526.1 abra|OJJ67410.1 acar|OOF92404.1 afis|EAW18460.1 afla|EED52948.1 afum|EAL93146.1 akaw|GAA86417.1 aluc|OJZ84958.1 anid|CBF70087.1 anig|EHA25172.1 aory|EIT78733.1 ater|EAU29878.1 atub|OJI80642.1 awen|OJJ31012.1

Group182: abra|OJJ67954.1 acar|OOF92799.1 acla|EAW11644.1 afis|EAW23813.1 afis|EAW23814.1 afla|EED51991.1 afla|EED52002.1 afum|EAL87248.1 agla|OJJ82453.1 akaw|GAA91242.1 akaw|GAA91243.1 aluc|OJZ82827.1 anid|CBF79881.1 anig|EHA21264.1 anig|EHA21265.1 aory|EIT81469.1 aory|EIT81487.1 asyd|OJJ58172.1 ater|EAU31366.1 atub|OJI89057.1 aver|OJJ04231.1 awen|OJJ33769.1 azon|OJJ46009.1 anid|CBF79883.1 asyd|OJJ58173.1 acla|EAW11643.1 afum|EAL87249.1 agla|OJJ82454.1 ater|EAU31369.1 awen|OJJ33770.1 azon|OJJ42873.1

Group183: abra|OJJ68614.1 acar|OOF92124.1 acar|OOF92125.1 acla|EAW12605.1 acla|EAW12606.1 afis|EAW18078.1 afis|EAW18079.1 afla|EED51472.1 afla|EED51473.1 afum|EAL91890.1 afum|EAL91891.1 agla|OJJ87372.1 akaw|GAA92907.1 akaw|GAA92908.1 aluc|OJZ81940.1 aluc|OJZ81941.1 anid|CBF86395.1 anid|CBF86397.1 anig|EHA22308.1 aory|EIT80638.1 aory|EIT80738.1 asyd|OJJ64952.1 asyd|OJJ64954.1

ater|EAU30443.1 ater|EAU30445.1 atub|OJI84493.1 atub|OJI84494.1 aver|OJI99074.1 aver|OJI99076.1 awen|OJJ41467.1 azon|OJJ48924.1

Group186: abra|OJJ74487.1 acar|OOF95262.1 acla|EAW07208.1 afis|EAW22220.1 afla|EED57613.1 afum|EAL90974.1 agla|OJJ87824.1 akaw|GAA87381.1 aluc|OJZ89631.1 anig|EHA26874.1 ater|EAU34230.1 atub|OJI85171.1 aver|OJI97165.1 awen|OJJ34805.1 abra|OJJ74488.1 acar|OOF95261.1 acla|EAW07207.1 afis|EAW22219.1 afla|EED57612.1 afum|EAL90975.1 agla|OJJ87825.1 akaw|GAA87382.1 aluc|OJZ89630.1 anid|CBF88457.1 anig|EHA26873.1 aory|EIT82983.1 asyd|OJJ63772.1 ater|EAU34229.1 atub|OJI85172.1 aver|OJI97166.1 azon|OJJ45743.1

Group188: abra|OJJ74864.1 acar|OOF95864.1 acar|OOF95866.1 acla|EAW08515.1 acla|EAW08517.1 afis|EAW23553.1 afis|EAW23555.1 afla|EED57448.1 afla|EED57450.1 afum|EAL87911.1 afum|EAL87913.1 agla|OJJ86813.1 agla|OJJ86815.1 akaw|GAA83554.1 akaw|GAA83556.1 aluc|OJZ92442.1 anid|CBF89641.1 anid|CBF89643.1 anig|EHA26525.1 aory|EIT80074.1 aory|EIT80092.1 asyd|OJJ62288.1 asyd|OJJ62290.1 ater|EAU35336.1 ater|EAU35338.1 atub|OJI86871.1 aver|OJI96396.1 aver|OJI96398.1 awen|OJJ39884.1 awen|OJJ39886.1 azon|OJJ43221.1

Group189: abra|OJJ75158.1 acar|OOF97180.1 acar|OOF97181.1 acla|EAW08333.1 acla|EAW08334.1 afis|EAW23368.1 afis|EAW23369.1 afla|EED55588.1 afla|EED55589.1 afum|EAL88104.2 afum|EAL88105.1 agla|OJJ86455.1 agla|OJJ86456.1 akaw|GAA88764.1 akaw|GAA88769.1 aluc|OJZ92728.1 anid|CBF74819.1 anid|CBF74821.1 anig|EHA26273.1 aory|EIT82662.1 aory|EIT82702.1 asyd|OJJ60141.1 asyd|OJJ60142.1 ater|EAU35619.1 ater|EAU35620.1 atub|OJI87152.1 aver|OJJ04121.1 awen|OJJ39518.1 awen|OJJ39519.1 azon|OJJ47713.1 azon|OJJ47715.1

Group190: abra|OJJ75500.1 acar|OOF90833.1 acla|EAW09294.1 afis|EAW21853.1 afla|EED46457.1 afum|EAL92866.1 agla|OJJ84283.1 akaw|GAA84266.1 aluc|OJZ88959.1 anid|CBF76739.1 anig|EHA23561.1 aory|EIT72412.1 asyd|OJJ61254.1 ater|EAU30852.1 atub|OJI85830.1 aver|OJJ00577.1 awen|OJJ34137.1 azon|OJJ44207.1 abra|OJJ75502.1 acar|OOF90834.1 acla|EAW09295.1 afis|EAW21852.1 afla|EED46458.1 afum|EAL92865.1 akaw|GAA84267.1 anid|CBF76737.1 aory|EIT72344.1 asyd|OJJ61253.1 ater|EAU30851.1 awen|OJJ34135.1 azon|OJJ44209.1

Group191: abra|OJJ76457.1 acar|OOF99697.1 acla|EAW13972.1 acla|EAW13973.1 afis|EAW18681.1 afis|EAW18682.1 afla|EED50628.1 afum|EAL93402.1 afum|EAL93403.1 agla|OJJ82393.1 agla|OJJ82394.1 akaw|GAA82369.1 akaw|GAA82370.1 aluc|OJZ91752.1 aluc|OJZ91753.1 anid|CBF70601.1 anid|CBF70603.1 anig|EHA22755.1 anig|EHA22756.1 aory|EIT78067.1 asyd|OJJ58516.1 asyd|OJJ58517.1 ater|EAU38228.1 atub|OJI86210.1 atub|OJI86211.1 aver|OJJ01117.1 aver|OJJ01118.1 awen|OJJ32153.1 azon|OJJ46737.1 afla|EED50629.1 aory|EIT78047.1

Group192: abra|OJJ77384.1 acla|EAW09036.1 afla|EED56897.1 akaw|GAA91377.1 aluc|OJZ84792.1 anig|EHA28060.1 aory|EIT77450.1 asyd|OJJ57351.1 ater|EAU38783.1 atub|OJI89290.1 aver|OJJ05814.1 awen|OJJ35827.1 abra|OJJ78137.1 acar|OOF99382.1 acla|EAW14634.1 afis|EAW22732.1 afla|EED55765.1 afum|EAL90463.1 agla|OJJ78522.1 aluc|OJZ82078.1 anid|CBF78393.1 ater|EAU30739.1 awen|OJJ32040.1 azon|OJJ47350.1 acar|OOF96813.1 anig|EHA25049.1 aluc|OJZ82079.1 atub|OJI88447.1 akaw|GAA87875.1 aluc|OJZ85516.1 atub|OJI80050.1

Group195: abra|OJJ68217.1 abra|OJJ69796.1 acar|OOF92535.1 acar|OOF97501.1 akaw|GAA83229.1 akaw|GAA87863.1 aluc|OJZ83729.1 aluc|OJZ85504.1 anig|EHA18877.1 anig|EHA25036.1 aory|EIT80435.1 aory|EIT81864.1 atub|OJI80061.1 atub|OJI90106.1 awen|OJJ31685.1 awen|OJJ32019.1 azon|OJJ42231.1 azon|OJJ46071.1 acla|EAW15175.1 afis|EAW20145.1 afla|EED51644.1 afum|EAL85260.1 agla|OJJ89015.1 anid|CBF81813.1 asyd|OJJ59648.1 ater|EAU39446.1 aver|OJJ02634.1 asyd|OJJ58357.1 ater|EAU33077.1 aver|OJJ02050.1

Group196: abra|OJJ68439.1 abra|OJJ71007.1 acar|OOF93955.1 acar|OOF98135.1 afis|EAW19801.1 afis|EAW25026.1 agla|OJJ79158.1 agla|OJJ87950.1 akaw|GAA82946.1 akaw|GAA86933.1 aluc|OJZ87524.1 aluc|OJZ90058.1 anid|CBF81549.1 anid|CBF82379.1 anig|EHA20734.1 anig|EHA21013.1 asyd|OJJ54138.1 asyd|OJJ59494.1 ater|EAU30748.1 ater|EAU35761.1 atub|OJI81124.1 atub|OJI84749.1 aver|OJJ02780.1 aver|OJJ08598.1 acla|EAW10559.1 afla|EED55448.1 afum|EAL89669.1 aory|EIT80786.1 awen|OJJ40613.1 azon|OJJ49462.1

Group197: abra|OJJ69676.1 abra|OJJ72100.1 acar|OOF90056.1 acar|OOF96745.1 aluc|OJZ82484.1 aluc|OJZ86855.1 anig|EHA21784.1 anig|EHA28290.1 asyd|OJJ60799.1 asyd|OJJ62562.1 atub|OJI83664.1 atub|OJI87948.1 awen|OJJ31984.1 awen|OJJ34027.1 afis|EAW21744.1 afum|EAL92777.1 agla|OJJ81779.1 agla|OJJ84428.1 akaw|GAA84821.1 akaw|GAA88801.1 ater|EAU32512.1 aver|OJI96686.1 azon|OJJ42257.1 acla|EAW09389.1 afla|EED58033.1 anid|CBF83660.1 aory|EIT79754.1 ater|EAU38444.1 aver|OJJ00132.1 acar|OOF95939.1

Group198: abra|OJJ72836.1 abra|OJJ73712.1 agla|OJJ78798.1 agla|OJJ86951.1 akaw|GAA85433.1 akaw|GAA88451.1 aluc|OJZ83336.1 aluc|OJZ88338.1 anig|EHA18073.1 anig|EHA24212.1 aory|EIT75956.1 aory|EIT78575.1 asyd|OJJ53233.1 asyd|OJJ54785.1 ater|EAU29729.1 ater|EAU32826.1 atub|OJI82233.1 atub|OJI90934.1 aver|OJJ03618.1 aver|OJJ07380.1 awen|OJJ35489.1 awen|OJJ38098.1 acar|OOF96038.1

acla|EAW11404.1 afis|EAW24017.1 afla|EED49142.1 afum|EAL87447.1 anid|CBF77418.1 azon|OJJ47000.1 afla|EED54085.1

Group200: acar|OOF98095.1 acar|OOF99166.1 acla|EAW06529.1 acla|EAW10590.1 afis|EAW16066.1 afis|EAW24978.1 afla|EED48587.1 afla|EED55481.1 afum|EAL88698.1 afum|EAL89638.1 anid|CBF75098.1 anid|CBF81606.1 asyd|OJJ59522.1 asyd|OJJ60007.1 ater|EAU33563.1 ater|EAU35727.1 aver|OJJ02278.1 aver|OJJ02750.1 awen|OJJ30369.1 awen|OJJ40575.1 azon|OJJ45388.1 azon|OJJ49435.1 abra|OJJ71044.1 agla|OJJ79139.1 akaw|GAA82912.1 aluc|OJZ87482.1 anig|EHA20704.1 aory|EIT73215.1 aory|EIT80785.1 atub|OJI81166.1

Group206: abra|OJJ66942.1 acar|OOF91706.1 acar|OOF91708.1 acla|EAW14500.1 acla|EAW14501.1 afis|EAW19214.1 afis|EAW19215.1 afla|EED47685.1 afla|EED47686.1 afum|EAL93947.1 afum|EAL93948.1 agla|OJJ88482.1 akaw|GAA91478.1 akaw|GAA91479.1 aluc|OJZ85640.1 anid|CBF78591.1 anig|EHA24894.1 aory|EIT80213.1 aory|EIT80244.1 asyd|OJJ65083.1 ater|EAU37253.1 atub|OJI79431.1 aver|OJI99209.1 awen|OJJ37572.1 azon|OJJ51231.1 anid|CBF78589.1 ater|EAU37254.1 aver|OJI99208.1 awen|OJJ37571.1 asyd|OJJ65084.1

Group207: abra|OJJ67140.1 acar|OOF98417.1 acla|EAW14599.1 afis|EAW19328.1 afum|EAL94045.1 agla|OJJ86277.1 akaw|GAA91029.1 aluc|OJZ87832.1 anid|CBF83940.1 anig|EHA20311.1 aory|EIT74592.1 asyd|OJJ53609.1 ater|EAU29456.1 atub|OJI80809.1 aver|OJJ08139.1 awen|OJJ30097.1 azon|OJJ44450.1 abra|OJJ67993.1 acar|OOF92035.1 acla|EAW13634.1 afis|EAW23673.1 afla|EED47394.1 afum|EAL87116.1 akaw|GAA90955.1 aluc|OJZ82867.1 anig|EHA21297.1 aory|EIT81925.1 atub|OJI89092.1 azon|OJJ45993.1 afla|EED47553.1

Group208: abra|OJJ70660.1 acar|OOF97061.1 acla|EAW15046.1 afis|EAW23152.1 afla|EED48515.1 afum|EAL86237.1 agla|OJJ89650.1 akaw|GAA89996.1 aluc|OJZ84010.1 anid|CBF74423.1 anig|EHA19132.1 aory|EIT73250.1 asyd|OJJ57754.1 ater|EAU36259.1 atub|OJI90379.1 aver|OJJ05413.1 awen|OJJ36648.1 azon|OJJ47567.1 acla|EAW08098.1 afum|EAL88329.1 agla|OJJ89649.1 anig|EHA19133.1 azon|OJJ47566.1 acar|OOF97062.1 afis|EAW24607.1 anid|CBF87133.1 aory|EIT73278.1 asyd|OJJ60373.1 ater|EAU36260.1 aver|OJI99708.1

Group209: abra|OJJ71126.1 acar|OOF98025.1 acar|OOF98026.1 acla|EAW10372.1 acla|EAW10373.1 afis|EAW25219.1 afis|EAW25220.1 afla|EED53686.1 afla|EED53687.1 afum|EAL89880.1 afum|EAL89881.1 agla|OJJ79187.1 akaw|GAA82848.1 akaw|GAA82849.1 aluc|OJZ87409.1 anid|CBF85301.1 anig|EHA20639.1 aory|EIT75287.1 aory|EIT75345.1 asyd|OJJ62915.1 asyd|OJJ62916.1 ater|EAU34378.1 ater|EAU34379.1 atub|OJI81236.1 atub|OJI81237.1 aver|OJI98074.1 awen|OJJ40885.1 awen|OJJ40886.1 azon|OJJ44513.1 azon|OJJ44514.1

Group211: abra|OJJ74957.1 acar|OOF97356.1 acla|EAW08414.1 afis|EAW23455.1 afla|EED57346.1 afum|EAL88011.1 agla|OJJ86695.1 akaw|GAA83465.1 akaw|GAA83466.1 aluc|OJZ92529.1 anid|CBF89806.1 anig|EHA26445.1 anig|EHA26446.1 aory|EIT80091.1 asyd|OJJ62193.1 ater|EAU36921.1 atub|OJI86965.1 atub|OJI86966.1 aver|OJI96302.1 awen|OJJ39774.1 azon|OJJ47901.1 acla|EAW08415.1 afis|EAW23454.1 afla|EED57349.1 afum|EAL88012.1 agla|OJJ86696.1 anid|CBF89804.1 ater|EAU36920.1 aver|OJI96305.1 awen|OJJ39775.1

Group212: abra|OJJ75967.1 acar|OOF99952.1 acla|EAW09831.1 afis|EAW21302.1 afla|EED57262.1 afum|EAL92319.1 agla|OJJ87435.1 akaw|GAA88989.1 aluc|OJZ89414.1 anid|CBF83395.1 anig|EHA23173.1 aory|EIT74086.1 asyd|OJJ60638.1 ater|EAU35935.1 atub|OJI85381.1 aver|OJI99960.1 awen|OJJ39396.1 azon|OJJ44280.1 acla|EAW09826.1 afis|EAW21307.1 afum|EAL92323.1 anid|CBF83397.1 asyd|OJJ60645.1 ater|EAU35937.1 aver|OJI99963.1 afis|EAW21308.1 afum|EAL92324.1 anid|CBF83399.1 asyd|OJJ60644.1 ater|EAU35938.1

Group213: abra|OJJ76386.1 acar|OOF99627.1 acla|EAW14037.1 acla|EAW14038.1 afis|EAW18743.1 afis|EAW18744.1 afla|EED50699.1 afla|EED50700.1 afum|EAL93469.1 afum|EAL93470.1 agla|OJJ82309.1 akaw|GAA82305.1 akaw|GAA82306.1 aluc|OJZ91827.1 aluc|OJZ91828.1 anid|CBF69994.1 anid|CBF69996.1 anig|EHA22816.1 aory|EIT78103.1 aory|EIT78138.1 asyd|OJJ58888.1 asyd|OJJ58889.1 ater|EAU37870.1 ater|EAU37873.1 atub|OJI86282.1 atub|OJI86283.1 aver|OJJ01489.1 awen|OJJ32308.1 awen|OJJ32309.1 azon|OJJ42718.1

Group214: abra|OJJ65770.1 abra|OJJ69062.1 acar|OOF91221.1 acar|OOG00531.1 afla|EED46866.1 afla|EED56884.1 akaw|GAA90592.1 akaw|GAA90890.1 aluc|OJZ82939.1 aluc|OJZ86050.1 anig|EHA21359.1 anig|EHA24561.1 aory|EIT72522.1 aory|EIT77460.1 atub|OJI79847.1 atub|OJI89171.1 aver|OJJ03011.1 aver|OJJ07939.1 afis|EAW15670.1 afum|EAL85677.1 anid|CBF81578.1 asyd|OJJ53834.1 ater|EAU32494.1 awen|OJJ38346.1 acla|EAW11671.1 agla|OJJ81808.1 agla|OJJ85313.1 awen|OJJ38760.1 azon|OJJ42640.1

Group216: abra|OJJ67846.1 abra|OJJ69869.1 acar|OOF90337.1 acar|OOF92827.1 acla|EAW11753.1 acla|EAW15317.1 akaw|GAA87787.1 akaw|GAA91854.1 aluc|OJZ82045.1 aluc|OJZ85422.1 anig|EHA22212.1 anig|EHA25601.1 asyd|OJJ52765.1 asyd|OJJ58155.1 ater|EAU31327.1 ater|EAU32679.1 aver|OJJ04248.1 aver|OJJ08748.1 awen|OJJ31347.1 awen|OJJ33731.1 afis|EAW23786.1 afla|EED51595.1 afum|EAL87220.2 agla|OJJ78939.1 anid|CBF79847.1 atub|OJI84604.1 azon|OJJ42847.1 atub|OJI80153.1 aory|EIT80419.1

Group217: abra|OJJ69627.1 abra|OJJ76818.1 acar|OOF91313.1 acar|OOF92684.1 acla|EAW08738.1 acla|EAW13378.1 agla|OJJ84834.1 agla|OJJ86329.1 akaw|GAA87204.1 akaw|GAA91607.1 anig|EHA27577.1 anig|EHA28339.1 atub|OJI88009.1 atub|OJI89856.1 awen|OJJ33192.1 awen|OJJ38390.1 azon|OJJ43848.1 azon|OJJ46477.1 afis|EAW15698.1 afla|EED51354.1 afum|EAL85650.1 aluc|OJZ82432.1 anid|CBF78282.1 aory|EIT80486.1 asyd|OJJ56072.1 ater|EAU39440.1 aver|OJJ05015.1 aluc|OJZ84228.1 asyd|OJJ52533.1

Group218: abra|OJJ69629.1 abra|OJJ74245.1 acar|OOF92681.1 acar|OOF95485.1 afla|EED55078.1 afla|EED57736.1 aluc|OJZ82434.1 aluc|OJZ89887.1 asyd|OJJ63867.1 asyd|OJJ64731.1 acla|EAW07299.1 agla|OJJ87688.1 akaw|GAA91609.1 anig|EHA28337.1 aory|EIT81976.1 atub|OJI84920.1 atub|OJI88007.1 aver|OJI98835.1 awen|OJJ34926.1 afis|EAW22346.1 afum|EAL90846.1 akaw|GAA86775.1 anid|CBF82985.1 anig|EHA27086.1 ater|EAU39484.1 aver|OJI97072.1 azon|OJJ50069.1 ater|EAU35426.1 aory|EIT79895.1

Group221: acar|OOF92608.1 acar|OOF92629.1 abra|OJJ67199.1 acar|OOF98365.1 acla|EAW14525.1 acla|EAW14526.1 afis|EAW19243.1 afis|EAW19244.1 afla|EED47633.1 afla|EED47634.1 afum|EAL93972.1 afum|EAL93973.1 agla|OJJ88454.1 akaw|GAA90979.1 akaw|GAA90980.1 aluc|OJZ87775.1 anid|CBF78652.1 anig|EHA20259.1 aory|EIT80236.1 asyd|OJJ65195.1 ater|EAU29500.1 atub|OJI80870.1 aver|OJI99344.1 awen|OJJ37599.1 azon|OJJ51254.1 agla|OJJ88453.1 anid|CBF78654.1 azon|OJJ51255.1 ater|EAU29499.1

Group222: afis|EAW21841.1 afis|EAW25659.1 agla|OJJ84294.1 agla|OJJ87901.1 aory|EIT72356.1 aory|EIT79167.1 asyd|OJJ52337.1 asyd|OJJ61239.1 asyd|OJJ54547.1 aver|OJI96725.1 aver|OJI96959.1 aver|OJJ07630.1 aver|OJJ00562.1 abra|OJJ68267.1 acar|OOF97547.1 acla|EAW09304.1 afla|EED46529.1 afum|EAL92853.1 akaw|GAA91931.1 aluc|OJZ82691.1 anid|CBF76711.1 anig|EHA27368.1 ater|EAU38518.1 atub|OJI88923.1 awen|OJJ34124.1 azon|OJJ44222.1 aluc|OJZ80200.1 afla|EED47266.1 anid|CBF85164.1

Group223: afla|EED48230.1 afla|EED58028.1 anid|CBF83667.1 anid|CBF86363.1 aory|EIT72902.1 aory|EIT79778.1 asyd|OJJ60808.1 asyd|OJJ64969.1 aver|OJI99093.1 aver|OJJ00136.1 awen|OJJ30134.1 awen|OJJ35830.1 abra|OJJ72106.1 acar|OOF96749.1 acla|EAW09385.1 afis|EAW21750.1 afum|EAL92781.2 agla|OJJ84424.1 akaw|GAA84816.1 aluc|OJZ86861.1 anig|EHA21779.1 ater|EAU38440.1 atub|OJI83659.1 awen|OJJ34031.1 azon|OJJ48204.1 anid|CBF83471.1 ater|EAU34242.1 aver|OJJ00014.1 asyd|OJJ60689.1

Group224: azon|OJJ42769.1 azon|OJJ50372.1 abra|OJJ71617.1 acar|OOF96333.1 afla|EED57118.1 agla|OJJ80138.1 akaw|GAA89310.1 aluc|OJZ86358.1 anig|EHA21532.1 aory|EIT73985.1 atub|OJI84176.1 awen|OJJ40249.1 abra|OJJ77100.1 acar|OOF92984.1 acla|EAW12282.1 afis|EAW16959.1 afla|EED45239.1 afum|EAL86486.1 akaw|GAA86956.1 aluc|OJZ84508.1 anid|CBF80512.1 anig|EHA27807.1 aory|EIT73744.1 asyd|OJJ54736.1 ater|EAU30185.1 atub|OJI89572.1 aver|OJJ07435.1 agla|OJJ85349.1 awen|OJJ30386.1

Group225: azon|OJJ46793.1 azon|OJJ51796.1 abra|OJJ77127.1 acar|OOF93013.1 acla|EAW12902.1 afis|EAW16407.1 afla|EED51674.1 agla|OJJ81607.1 akaw|GAA90801.1 aluc|OJZ84537.1 anid|CBF82038.1 anig|EHA27835.1 aory|EIT79575.1 asyd|OJJ59200.1 ater|EAU36805.1 atub|OJI89543.1 aver|OJJ01812.1 awen|OJJ36343.1 acar|OOF93300.1 afla|EED49754.1 aory|EIT81671.1 asyd|OJJ54221.1 acar|OOF99299.1 aory|EIT77631.1 afla|EED48439.1 awen|OJJ38768.1 aory|EIT80498.1 afum|EAL91133.1 aver|OJJ05173.1

Group226: abra|OJJ67152.1 acar|OOF90071.1 acar|OOF98406.1 acla|EAW14572.1 acla|EAW14573.1 afis|EAW19295.1 afis|EAW19296.1 afla|EED47590.1 afla|EED47591.1 afum|EAL94014.1 afum|EAL94015.1 agla|OJJ88385.1 akaw|GAA91020.1 akaw|GAA91021.1 aluc|OJZ87823.1 aluc|OJZ87824.1 anid|CBF78768.1 anid|CBF78770.1 anig|EHA20303.1 aory|EIT74518.1 aory|EIT74526.1 asyd|OJJ65238.1 ater|EAU29476.1 ater|EAU29477.1 atub|OJI80820.1 atub|OJI80821.1 aver|OJI99388.1 awen|OJJ37646.1 azon|OJJ44911.1

Group228: abra|OJJ68668.1 acar|OOF92168.1 acla|EAW12580.1 afis|EAW24068.1 afla|EED51407.1 afum|EAL91924.1 agla|OJJ87316.1 aluc|OJZ81882.1 anid|CBF86457.1 anig|EHA22358.1 aory|EIT80674.1 asyd|OJJ64915.1 ater|EAU30490.1 atub|OJI84435.1 aver|OJI99035.1 awen|OJJ41422.1 azon|OJJ48887.1 acla|EAW12579.1 afis|EAW24069.1 afla|EED51406.1 afum|EAL91925.1 agla|OJJ87315.1 akaw|GAA89809.1 aluc|OJZ81881.1 anid|CBF86458.1 aory|EIT80695.1 asyd|OJJ64914.1 awen|OJJ41421.1 azon|OJJ48886.1

Group229: abra|OJJ68715.1 acar|OOF92215.1 acar|OOF92216.1 acla|EAW12530.1 acla|EAW12531.1 afis|EAW24118.1 afis|EAW24119.1 afla|EED46814.1 afum|EAL91973.1 afum|EAL91974.1 agla|OJJ87193.1 akaw|GAA89868.1 aluc|OJZ81813.1 anid|CBF86539.1 anid|CBF86540.1 anig|EHA22402.1 anig|EHA22403.1 aory|EIT72603.1 aory|EIT72683.1 asyd|OJJ64863.1 asyd|OJJ64865.1 ater|EAU30546.1 ater|EAU30547.1 atub|OJI84385.1 aver|OJI98974.1 aver|OJI98978.1 awen|OJJ41370.1 azon|OJJ48839.1 afla|EED46813.1

Group230: abra|OJJ68757.1 acar|OOF92245.1 acla|EAW12494.1 afis|EAW24151.1 afla|EED46780.1 afum|EAL92006.1 akaw|GAA89901.1 aluc|OJZ81770.1 anid|CBF77913.1 anig|EHA22446.1 aory|EIT72705.1 asyd|OJJ55298.1 ater|EAU30583.1 atub|OJI84342.1 aver|OJJ06928.1 awen|OJJ41327.1 abra|OJJ68758.1 acla|EAW12493.1 afis|EAW24152.1 afla|EED46779.1 afum|EAL92007.1 agla|OJJ89543.1 akaw|GAA89902.1 aluc|OJZ81769.1 anid|CBF77911.1 aory|EIT72741.1 atub|OJI84341.1 awen|OJJ41326.1 azon|OJJ48800.1

Group231: abra|OJJ70392.1 acar|OOF90061.1 acla|EAW12387.1 acla|EAW12388.1 afis|EAW16834.1 afis|EAW16835.1 afla|EED47328.1 afum|EAL86381.1 afum|EAL86382.2 agla|OJJ81545.1 agla|OJJ89426.1 akaw|GAA91967.1 aluc|OJZ82649.1 anid|CBF74687.1 anid|CBF74689.1 anig|EHA27422.1 anig|EHA27423.1 aory|EIT82824.1 asyd|OJJ60210.1 ater|EAU30239.1 atub|OJI88881.1 aver|OJJ04058.1 awen|OJJ35685.1 azon|OJJ44153.1 acar|OOF90060.1 afla|EED47327.1 aory|EIT82846.1 ater|EAU30242.1 acar|OOF91882.1

Group232: abra|OJJ70540.1 acar|OOF97393.1 acla|EAW08138.1 acla|EAW08139.1 afis|EAW23174.1 afis|EAW23175.1 afla|EED46944.1 afla|EED46945.1 afum|EAL88308.1 afum|EAL88309.1 agla|OJJ89686.1 agla|OJJ89687.1 akaw|GAA83385.1 akaw|GAA83386.1 aluc|OJZ83885.1 anid|CBF74496.1 anid|CBF74498.1 anig|EHA19022.1 aory|EIT72667.1 aory|EIT72762.1 asyd|OJJ57719.1 asyd|OJJ57720.1 ater|EAU36303.1 ater|EAU36304.1 atub|OJI90252.1 aver|OJJ05448.1 awen|OJJ36603.1 azon|OJJ47603.1 azon|OJJ47604.1

Group233: abra|OJJ73887.1 acar|OOG01025.1 acar|OOG01026.1 acla|EAW07769.1 acla|EAW07771.1 afis|EAW22805.1 afis|EAW22807.1 afla|EED50075.1 afum|EAL90394.1 afum|EBA27407.1 agla|OJJ83582.1 akaw|GAA89498.1 akaw|GAA89499.1 aluc|OJZ88162.1 anid|CBF87924.1 anig|EHA17916.1 anig|EHA17917.1 aory|EIT74704.1 asyd|OJJ63459.1 asyd|OJJ63460.1 ater|EAU38903.1 ater|EAU38904.1 atub|OJI90761.1 aver|OJI97472.1 aver|OJI97473.1 awen|OJJ38923.1 azon|OJJ49700.1 aory|EIT74626.1 afla|EED50076.1

Group234: abra|OJJ74769.1 acar|OOF95760.1 acla|EAW10642.1 afis|EAW24924.1 afla|EED55549.1 afum|EAL89579.1 agla|OJJ82153.1 akaw|GAA83643.1 aluc|OJZ92352.1 anid|CBF75699.1 anig|EHA26622.1 aory|EIT82671.1 asyd|OJJ57496.1 atub|OJI86780.1 aver|OJJ05684.1 awen|OJJ40521.1 azon|OJJ49380.1 abra|OJJ74770.1 acla|EAW10641.1 afis|EAW24925.1 afla|EED55548.1 afum|EAL89580.1 akaw|GAA83642.1 anid|CBF75701.1 aory|EIT82683.1 ater|EAU35655.1 atub|OJI86781.1 aver|OJJ05683.1 azon|OJJ49381.1

Group235: abra|OJJ74915.1 acar|OOF95906.1 acar|OOF95907.1 acla|EAW08455.1 acla|EAW08456.1 afis|EAW23499.1 afis|EAW23500.1 afla|EED57400.1 afla|EED57401.1 afum|EAL87962.2 afum|EAL87963.1 agla|OJJ86747.1 agla|OJJ86748.1 akaw|GAA83507.1 akaw|GAA83508.1 aluc|OJZ92487.1 anid|CBF89730.1 anid|CBF89731.1 anig|EHA26485.1 aory|EIT80151.1 aory|EIT80164.1 asyd|OJJ62234.1 ater|EAU35289.1 ater|EAU35290.1 atub|OJI86920.1 aver|OJI96343.1 awen|OJJ39825.1 azon|OJJ47949.1 azon|OJJ47950.1

Group236: abra|OJJ75075.1 acar|OOF97258.1 acar|OOF97259.1 acla|EAW08256.1 afis|EAW23290.1 afla|EED55682.1 afum|EAL88189.1 agla|OJJ86548.1 akaw|GAA88687.1 akaw|GAA88688.1 aluc|OJZ92641.1 anid|CBF89509.1 anig|EHA26351.1 aory|EIT77074.1 asyd|OJJ62397.1 ater|EAU37583.1 atub|OJI87072.1 aver|OJI96531.1 awen|OJJ39667.1 azon|OJJ47797.1 aory|EIT77075.1 acla|EAW08258.1 afis|EAW23291.1 afla|EED55680.1 afum|EAL88188.1 anid|CBF89508.1 asyd|OJJ62398.1 ater|EAU37582.1 azon|OJJ47796.1

Group238: abra|OJJ75492.1 acar|OOF90824.1 acar|OOF90825.1 acla|EAW11224.1 acla|EAW11225.1 afis|EAW18278.1 afis|EAW18279.1 afla|EED48745.1 afla|EED48746.1 afum|EAL87644.1 afum|EAL87645.1 agla|OJJ88147.1 agla|OJJ88148.1 akaw|GAA84257.1 akaw|GAA84258.1 aluc|OJZ88951.1 anid|CBF79492.1 anid|CBF79494.1 anig|EHA23569.1 anig|EHA23570.1 aory|EIT73072.1 aory|EIT73086.1 asyd|OJJ57877.1 ater|EAU33284.1 ater|EAU33285.1 atub|OJI85838.1 aver|OJJ04517.1 awen|OJJ30651.1 azon|OJJ47964.1

Group239: abra|OJJ75717.1 acar|OOG00139.1 acar|OOG00141.1 acla|EAW09392.1 acla|EAW09394.1 afis|EAW21739.1 afis|EAW21741.1 afla|EED58014.1 afla|EED58016.1 afum|EAL92773.1 afum|EAL92774.1 agla|OJJ84432.1 agla|OJJ84449.1 akaw|GAA84452.1 akaw|GAA84453.1 aluc|OJZ89169.1 anid|CBF83696.1 anid|CBF83699.1 anig|EHA23386.1 aory|EIT79752.1 aory|EIT79775.1 asyd|OJJ60825.1 ater|EAU38420.1 ater|EAU38429.1 atub|OJI85621.1 aver|OJJ00151.1 awen|OJJ34021.1 awen|OJJ34023.1 azon|OJJ48535.1

Group240: abra|OJJ77333.1 acar|OOF93207.1 acla|EAW13139.1 afis|EAW17511.1 afla|EED49310.1 afum|EAL91331.1 agla|OJJ81447.1 akaw|GAA91425.1 aluc|OJZ84741.1 anid|CBF71568.1 anig|EHA28015.1 aory|EIT82226.1 asyd|OJJ61881.1 ater|EAU36952.1 atub|OJI89340.1 aver|OJI95979.1 awen|OJJ37333.1 azon|OJJ51021.1 afla|EED49307.1 afum|EAL91330.1 anid|CBF71531.1 asyd|OJJ61893.1 aver|OJI95989.1 azon|OJJ51020.1 acla|EAW13140.1 afis|EAW17512.1 anig|EHA28014.1 aory|EIT82236.1 awen|OJJ37334.1

Group243: abra|OJJ66383.1 abra|OJJ71952.1 acar|OOF94059.1 acar|OOF94129.1 acar|OOF96603.1 afla|EED51643.1 afla|EED53089.1 akaw|GAA84976.1 akaw|GAA90249.1 aluc|OJZ83091.1 aluc|OJZ86686.1 aory|EIT74787.1 aory|EIT80741.1 atub|OJI83839.1 atub|OJI87845.1 azon|OJJ48234.1 azon|OJJ51069.1 acla|EAW12654.1 afis|EAW18031.1 afum|EAL91842.1 agla|OJJ85401.1 anid|CBF76050.1 anig|EHA21987.1 asyd|OJJ52485.1 ater|EAU36811.1 aver|OJJ03921.1 awen|OJJ41506.1 ater|EAU35537.1

Group245: abra|OJJ68302.1 abra|OJJ74425.1 acar|OOF90436.1 acar|OOF90443.1 acar|OOF96685.1 afis|EAW24469.1 afis|EAW25516.1 aluc|OJZ80145.1 aluc|OJZ83671.1 aluc|OJZ89698.1 atub|OJI85105.1 atub|OJI90042.1 awen|OJJ34908.1 awen|OJJ41259.1 acla|EAW14952.1 afla|EED55056.1 afum|EAL86141.1 afum|EAL90181.1 agla|OJJ87287.1 akaw|GAA83172.1 anid|CBF77786.1 anig|EHA18826.1 aory|EIT81972.1 asyd|OJJ56269.1 ater|EAU33362.1 aver|OJJ08496.1 azon|OJJ42850.1 asyd|OJJ54439.1

Group247: abra|OJJ71231.1 abra|OJJ72448.1 acar|OOF94788.1 acar|OOF97944.1 afla|EED46190.1 afla|EED56273.1 akaw|GAA82757.1 akaw|GAA86186.1 aluc|OJZ87316.1 aluc|OJZ90432.1 ater|EAU33047.1 ater|EAU33789.1 atub|OJI81329.1 atub|OJI81864.1 awen|OJJ31638.1 awen|OJJ41732.1 acla|EAW12682.1 afis|EAW17845.1 afum|EAL91654.1 agla|OJJ81772.1 agla|OJJ83123.1 anig|EHA20550.1 asyd|OJJ55268.1 aver|OJJ06959.1 azon|OJJ48428.1 anid|CBF77966.1 anig|EHA23901.1 aory|EIT76458.1

Group248: acar|OOF95098.1 acar|OOF95164.1 acar|OOF96577.1 afis|EAW21529.1 afis|EAW24614.1 agla|OJJ80793.1 agla|OJJ81846.1 agla|OJJ85693.1 agla|OJJ84478.1 aory|EIT76721.1 aory|EIT79947.1 asyd|OJJ52471.1 asyd|OJJ61079.1 ater|EAU31519.1 ater|EAU35012.1 abra|OJJ71920.1 acla|EAW09610.1 afla|EED47452.1 afla|EED55978.1 afum|EAL92550.1 akaw|GAA85002.1 aluc|OJZ86657.1 anid|CBF76400.1 anig|EHA22014.1 atub|OJI83870.1 aver|OJJ00395.1 awen|OJJ39164.1 azon|OJJ51493.1

Group250: afis|EAW18309.1 afis|EAW25696.1 awen|OJJ30683.1 awen|OJJ32214.1 abra|OJJ75441.1 acar|OOF90786.1 acla|EAW13822.1 afla|EED53243.1 afum|EAL87675.1 agla|OJJ88078.1 akaw|GAA84216.1 aluc|OJZ88894.1 anid|CBF79436.1 anig|EHA23612.1 aory|EIT72989.1 asyd|OJJ54980.1 asyd|OJJ57846.1 ater|EAU33319.1 atub|OJI85895.1 aver|OJJ04318.1 aver|OJJ04546.1 awen|OJJ32538.1 azon|OJJ43103.1 afla|EED50918.1 aory|EIT74822.1 aory|EIT78997.1 asyd|OJJ52725.1 ater|EAU32817.1

Group252: agla|OJJ82640.1 agla|OJJ85276.1 asyd|OJJ57347.1 asyd|OJJ65114.1 aver|OJI99240.1 aver|OJJ05818.1 aver|OJJ00352.1 aver|OJJ05340.1 awen|OJJ30078.1 awen|OJJ32593.1 abra|OJJ78114.1 acar|OOF98754.1 acla|EAW07003.1 afis|EAW16620.1 afla|EED48645.1 afum|EAL89217.1 agla|OJJ84764.1 akaw|GAA85893.1 aluc|OJZ91179.1 anid|CBF81971.1 anig|EHA19870.1 aory|EIT73182.1 ater|EAU36705.1 atub|OJI83191.1 azon|OJJ42184.1 aory|EIT73798.1 asyd|OJJ61034.1 ater|EAU37620.1

Group256: abra|OJJ67322.1 acar|OOF92491.1 acar|OOF92492.1 acla|EAW06413.1 acla|EAW06414.1 afis|EAW18517.1 afis|EAW18518.1 afla|EED52880.1 afla|EED52881.1 afum|EAL93212.1 afum|EAL93213.1 agla|OJJ85130.1 akaw|GAA86495.1 akaw|GAA86496.1 aluc|OJZ85037.1 aluc|OJZ85038.1 anid|CBF70228.1 anid|CBF70230.1 anig|EHA25256.1 aory|EIT78607.1 aory|EIT78620.1 asyd|OJJ58772.1 ater|EAU30052.1 ater|EAU30053.1 atub|OJI80559.1 aver|OJJ01381.1 awen|OJJ31070.1 azon|OJJ50580.1

Group257: abra|OJJ67358.1 acar|OOF92463.1 acla|EAW06384.1 afis|EAW18490.1 afla|EED52916.1 afum|EAL93180.1 agla|OJJ85085.1 akaw|GAA86463.1 aluc|OJZ85003.1 anid|CBF70157.1 anig|EHA25222.1 aory|EIT78645.1 asyd|OJJ58803.1 ater|EAU29914.1 atub|OJI80595.1 aver|OJJ01407.1 awen|OJJ31043.1 azon|OJJ50554.1 abra|OJJ67360.1 acla|EAW06383.1 afis|EAW18489.1 afla|EED52917.1 afum|EAL93179.1 akaw|GAA86461.1 anid|CBF70155.1 anig|EHA25220.1 aory|EIT78598.1 asyd|OJJ58804.1

Group259: abra|OJJ68687.1 acar|OOF92189.1 acar|OOF92190.1 acla|EAW12558.1 acla|EAW12560.1 afis|EAW24090.1 afis|EAW24092.1 afla|EED46847.1 afla|EED46849.1 afum|EAL91945.1 afum|EAL91947.2 agla|OJJ87160.1 agla|OJJ87166.1 akaw|GAA89840.1 akaw|GAA89842.1 aluc|OJZ81846.1 anid|CBF86494.1 anig|EHA22376.1 aory|EIT72742.1 asyd|OJJ64893.1 ater|EAU30514.1 ater|EAU30516.1 atub|OJI84415.1 aver|OJI99014.1 awen|OJJ41397.1 awen|OJJ41399.1 azon|OJJ48865.1 aory|EIT72593.1

Group260: abra|OJJ68761.1 acar|OOF92248.1 acla|EAW12490.1 afis|EAW24155.1 afla|EED46777.1 afum|EAL92010.1 akaw|GAA89905.1 aluc|OJZ81765.1 anid|CBF77904.1 anig|EHA22449.1 asyd|OJJ55301.1 atub|OJI84337.1 abra|OJJ68762.1 acla|EAW12489.1 afis|EAW24156.1 afum|EAL92011.1 agla|OJJ89525.1 akaw|GAA89906.1 aluc|OJZ81764.1 anid|CBF77902.1 anig|EHA22450.1 aory|EIT72652.1 asyd|OJJ55302.1 ater|EAU30587.1 atub|OJI84336.1 aver|OJJ06924.1 awen|OJJ41322.1 azon|OJJ48795.1

Group261: abra|OJJ69707.1 acar|OOF94204.1 acla|EAW14911.1 afla|EED55248.1 agla|OJJ79247.1 akaw|GAA90156.1 aluc|OJZ83191.1 anid|CBF73771.1 anig|EHA28261.1 aory|EIT77697.1 asyd|OJJ56592.1 atub|OJI87748.1 awen|OJJ35897.1 azon|OJJ50407.1 abra|OJJ72111.1 acar|OOF90025.1 afis|EAW24432.1 afum|EAL86100.1 akaw|GAA84811.1 aluc|OJZ86866.1 anig|EHA21775.1 ater|EAU30013.1 atub|OJI83654.1 aver|OJJ06133.1 awen|OJJ32770.1 aory|EIT83627.1 azon|OJJ44000.1 afla|EED49732.1

Group263: abra|OJJ70208.1 acar|OOF90365.1 acla|EAW13177.1 afis|EAW22620.1 afla|EED51827.1 afum|EAL87187.2 agla|OJJ85385.1 akaw|GAA86568.1 aluc|OJZ85119.1 anid|CBF84539.1 anig|EHA25339.1 aory|EIT81723.1 asyd|OJJ53483.1 ater|EAU32773.1 atub|OJI80474.1 aver|OJJ08276.1 awen|OJJ33335.1 azon|OJJ51798.1 acla|EAW11725.1 afis|EAW23750.1 asyd|OJJ64104.1 aver|OJI98187.1 azon|OJJ51793.1 afum|EAL90576.1 afla|EED53104.1 anid|CBF85467.1 agla|OJJ87544.1 aory|EIT74796.1

Group264: anig|EHA20613.1 atub|OJI81264.1 abra|OJJ71157.1 acar|OOF98002.1 acla|EAW10341.1 afis|EAW25250.1 afla|EED53714.1 afum|EAL89912.1 agla|OJJ89220.1 akaw|GAA82821.1 aluc|OJZ87382.1 anid|CBF85305.1 aory|EIT74933.1 asyd|OJJ63998.1 ater|EAU34356.1 aver|OJI98087.1 awen|OJJ40962.1 azon|OJJ44543.1 acla|EAW10342.1 afis|EAW25247.1 afla|EED53713.1 afum|EAL89909.1 agla|OJJ89214.1 akaw|GAA82822.1 aluc|OJZ87383.1 anid|CBF85306.1 awen|OJJ40961.1 azon|OJJ44529.1

Group265: abra|OJJ71667.1 acar|OOF96377.1 acar|OOF96378.1 acla|EAW15205.1 acla|EAW15206.1 afis|EAW20170.1 afis|EAW20171.1 afla|EED57177.1 afla|EED57179.1 afum|EAL85288.1 afum|EAL85289.1 agla|OJJ88759.1 akaw|GAA89264.1 akaw|GAA89265.1 aluc|OJZ86409.1 anid|CBF85026.1 anid|CBF85028.1 anig|EHA21490.1 aory|EIT73988.1 aory|EIT74058.1 asyd|OJJ63103.1 ater|EAU35205.1 ater|EAU35206.1 atub|OJI84122.1 aver|OJI97858.1 aver|OJI97859.1 awen|OJJ32067.1 azon|OJJ46099.1

Group267: abra|OJJ75148.1 acar|OOF97189.1 acar|OOF97190.1 acla|EAW08321.1 acla|EAW08322.1 afis|EAW23356.1 afis|EAW23357.1 afla|EED55606.1 afla|EED55607.1 afum|EAL88119.2 afum|EAL88120.2 agla|OJJ86469.1 akaw|GAA88753.1 akaw|GAA88754.1 aluc|OJZ92713.1 aluc|OJZ92714.1 anid|CBF74884.1 anig|EHA26286.1 aory|EIT81807.1 asyd|OJJ60109.1 ater|EAU35607.1 ater|EAU35608.1 atub|OJI87139.1 atub|OJI87140.1 aver|OJJ04153.1 awen|OJJ39530.1 azon|OJJ47726.1 azon|OJJ47728.1

Group269: abra|OJJ76491.1 acar|OOF99722.1 acar|OOF99723.1 acla|EAW13941.1 acla|EAW13944.1 afis|EAW18652.1 afis|EAW18653.1 afla|EED52680.1 afla|EED52681.1 afum|EAL93368.1 afum|EAL93369.1 agla|OJJ82422.1 akaw|GAA82406.1 akaw|GAA82407.1 aluc|OJZ91713.1 aluc|OJZ91714.1 anid|CBF70539.1 anid|CBF70540.1 anig|EHA22724.1 aory|EIT74312.1 aory|EIT74318.1 asyd|OJJ58549.1 ater|EAU38259.1 ater|EAU38260.1 atub|OJI86172.1 aver|OJJ01147.1 awen|OJJ32120.1 azon|OJJ46706.1

Group270: abra|OJJ76517.1 acar|OOF99456.1 acla|EAW14184.1 acla|EAW14185.1 afis|EAW18899.1 afis|EAW18900.1 afla|EED50848.1 afla|EED50853.1 afum|EAL93631.1 afum|EAL93632.1 agla|OJJ81505.1 akaw|GAA82429.1 akaw|GAA82430.1 aluc|OJZ91689.1 anid|CBF69917.1 anid|CBF69919.1 anig|EHA22702.1 aory|EIT78983.1 aory|EIT79046.1 asyd|OJJ58989.1 ater|EAU38067.1 ater|EAU38068.1 atub|OJI86142.1 aver|OJJ01626.1 aver|OJJ01627.1 awen|OJJ32457.1 azon|OJJ47214.1 azon|OJJ47215.1

Group271: abra|OJJ76807.1 acar|OOF91299.1 acla|EAW13366.1 afis|EAW20480.1 afla|EED44787.1 afum|EAL84574.1 agla|OJJ84832.1 akaw|GAA87216.1 aluc|OJZ84214.1 anid|CBF79227.1 anig|EHA27567.1 aory|EIT77778.1 asyd|OJJ55739.1 ater|EAU29559.1 atub|OJI89869.1 aver|OJJ06825.1 awen|OJJ33179.1 azon|OJJ46382.1 afla|EED44788.1 aory|EIT77780.1 awen|OJJ33175.1 azon|OJJ46828.1 acar|OOF91301.1 acla|EAW13367.1 afis|EAW20481.1 afum|EAL84573.1 akaw|GAA87215.1 aluc|OJZ84215.1

Group272: abra|OJJ77199.1 acar|OOF93082.1 acar|OOF93087.1 acla|EAW13188.1 acla|EAW13189.1 afis|EAW17452.1 afis|EAW17453.1 afla|EED49369.1 afla|EED49370.1 afum|EAL91280.1 afum|EAL91281.2 agla|OJJ80350.1 agla|OJJ80360.1 akaw|GAA90742.1 aluc|OJZ84606.1 anid|CBF71675.1 anid|CBF71677.1 anig|EHA27895.1 anig|EHA27896.1 aory|EIT79469.1 aory|EIT79540.1 asyd|OJJ61825.1 ater|EAU33932.1 ater|EAU33933.1 atub|OJI89473.1 aver|OJI95925.1 awen|OJJ37394.1 azon|OJJ51067.1

Group273: abra|OJJ77369.1 acar|OOF93239.1 acla|EAW13106.1 acla|EAW13107.1 afis|EAW17558.1 afla|EED49266.1 afla|EED49267.1 afum|EAL91372.1 agla|OJJ81410.1 akaw|GAA91392.1 akaw|GAA91393.1 aluc|OJZ84778.1 aluc|OJZ84779.1 anid|CBF90279.1 anig|EHA28046.1 aory|EIT82182.1 aory|EIT82245.1 asyd|OJJ61916.1 ater|EAU36989.1 atub|OJI89303.1 aver|OJI96009.1 awen|OJJ37296.1 azon|OJJ50989.1 afis|EAW17557.1 afum|EAL91371.1 agla|OJJ81411.1 anid|CBF90281.1 asyd|OJJ61917.1

Group276: acar|OOF92615.1 acar|OOF92636.1 abra|OJJ69082.1 acar|OOF91209.1 acla|EAW08752.1 afis|EAW15684.1 afla|EED51370.1 afum|EAL85664.1 agla|OJJ81833.1 akaw|GAA90611.1 akaw|GAA90612.1 aluc|OJZ86070.1 anid|CBF71148.1 anig|EHA24544.1 aory|EIT80661.1 asyd|OJJ55837.1 ater|EAU32473.1 atub|OJI79866.1 atub|OJI79867.1 aver|OJJ04780.1 awen|OJJ38376.1 azon|OJJ43860.1 acla|EAW08751.1 afis|EAW15685.1 afla|EED51369.1 afum|EAL85663.1 anid|CBF71146.1

Group277: acar|OOF93141.1 acar|OOF96894.1 asyd|OJJ55471.1 asyd|OJJ61030.1 asyd|OJJ57185.1 abra|OJJ72261.1 acla|EAW13265.1 afis|EAW16849.1 afla|EED49789.1 afum|EAL86395.1 agla|OJJ88181.1 akaw|GAA84669.1 aluc|OJZ87020.1 anid|CBF84231.1 anig|EHA21635.1 aory|EIT77692.1 ater|EAU33865.1 atub|OJI83498.1 aver|OJJ00350.1 aver|OJJ03105.1 awen|OJJ36910.1 azon|OJJ50296.1 acla|EAW12401.1 afla|EED51180.1 aory|EIT81355.1 anid|CBF89260.1 azon|OJJ50917.1

Group278: acar|OOF94414.1 acar|OOF97591.1 afis|EAW24229.1 afis|EAW24933.1 afla|EED46551.1 afla|EED48042.1 afum|EAL85898.1 afum|EAL89589.1 aory|EIT77904.1 aory|EIT83324.1 asyd|OJJ53967.1 asyd|OJJ64678.1 ater|EAU29958.1 ater|EAU31178.1 aver|OJI98780.1 aver|OJJ08063.1 anig|EHA28592.1 abra|OJJ69311.1 acla|EAW14698.1 agla|OJJ79642.1 akaw|GAA88180.1 aluc|OJZ82140.1 anid|CBF74228.1 anig|EHA24084.1 atub|OJI88297.1 awen|OJJ33068.1 azon|OJJ48455.1

Group280: afis|EAW18048.1 afis|EAW24488.1 afum|EAL86157.1 afum|EAL86730.1 afum|EAL91860.1 abra|OJJ68602.1 acar|OOF92112.1 acla|EAW12637.1 afla|EED51498.1 agla|OJJ85419.1 akaw|GAA91926.1 aluc|OJZ81954.1 anid|CBF79800.1 anig|EHA22293.1 aory|EIT80704.1 asyd|OJJ58129.1 ater|EAU31275.1 atub|OJI84505.1 aver|OJJ04276.1 awen|OJJ32709.1 awen|OJJ41486.1 azon|OJJ48216.1 afla|EED48171.1 agla|OJJ83081.1 aory|EIT83032.1 ater|EAU38954.1 aver|OJI98885.1

Group282: afla|EED51568.1 afla|EED51716.1 afla|EED56540.1 agla|OJJ78952.1 agla|OJJ80278.1 aory|EIT79700.1 aory|EIT80654.1 aory|EIT80755.1 asyd|OJJ57607.1 asyd|OJJ58114.1 aver|OJJ04291.1 aver|OJJ05577.1 awen|OJJ33707.1 awen|OJJ34440.1 abra|OJJ67863.1 acar|OOF92843.1 acla|EAW11735.1 afis|EAW23768.1 afum|EAL87202.1 agla|OJJ85386.1 akaw|GAA91870.1 aluc|OJZ82029.1 anid|CBF79783.1 anig|EHA22227.1 ater|EAU31312.1 atub|OJI84583.1 azon|OJJ42833.1

Group284: afum|EAL86514.1 afum|EAL91321.1 asyd|OJJ52759.1 asyd|OJJ54478.1 asyd|OJJ61874.1 azon|OJJ42558.1 azon|OJJ51475.1 azon|OJJ51028.1 abra|OJJ74834.1 acar|OOF95824.1 acla|EAW13149.1 afis|EAW17498.1 afla|EED49318.1 agla|OJJ81459.1 akaw|GAA83586.1 aluc|OJZ92411.1 anid|CBF71582.1 anig|EHA26554.1 aory|EIT82205.1 ater|EAU34444.1 ater|EAU36943.1 atub|OJI86841.1 aver|OJI95972.1 awen|OJJ37343.1 acla|EAW12237.1 afis|EAW16990.1 aver|OJJ08741.1

Group285: agla|OJJ79575.1 agla|OJJ86866.1 agla|OJJ88176.1 agla|OJJ89239.1 asyd|OJJ62473.1 asyd|OJJ64049.1 ater|EAU34049.1 ater|EAU34409.1 aver|OJI96608.1 aver|OJI98135.1 awen|OJJ38034.1 awen|OJJ40680.1 awen|OJJ40981.1 abra|OJJ74157.1 acar|OOF95570.1 acla|EAW12279.1 afis|EAW16961.1 afla|EED49740.1 afum|EAL86487.2 akaw|GAA89727.1 aluc|OJZ80656.1 anid|CBF89363.1 anig|EHA27176.1 aory|EIT75292.1 atub|OJI87511.1 azon|OJJ44568.1 anid|CBF85376.1

Group289: azon|OJJ42747.1 azon|OJJ51180.1 abra|OJJ76401.1 acar|OOF99642.1 acla|EAW14023.1 afis|EAW18729.1 afla|EED50685.1 afum|EAL93454.1 agla|OJJ82287.1 akaw|GAA82321.1 aluc|OJZ91811.1 anid|CBF70016.1 anig|EHA22804.1 aory|EIT78004.1 asyd|OJJ58877.1 ater|EAU38175.1 atub|OJI86267.1 aver|OJJ01479.1 awen|OJJ32270.1 akaw|GAA82320.1 aluc|OJZ91812.1 acar|OOF99643.1 acla|EAW14022.1 afis|EAW18728.1 afum|EAL93453.1 anid|CBF70018.1 ater|EAU38176.1

Group290: abra|OJJ65930.1 acar|OOF94991.1 acla|EAW12067.1 afis|EAW19710.1 afla|EED57049.1 afum|EAL86817.2 agla|OJJ82841.1 akaw|GAA92305.1 akaw|GAA92306.1 aluc|OJZ90202.1 anid|CBF84278.1 anig|EHA23699.1 aory|EIT82284.1 asyd|OJJ57230.1 ater|EAU32834.1 atub|OJI81631.1 aver|OJJ03059.1 awen|OJJ32205.1 azon|OJJ42677.1 afla|EED57054.1 acar|OOF94992.1 acla|EAW12066.1 afis|EAW19709.1 afum|EAL86818.1 aory|EIT82258.1 ater|EAU32836.1 azon|OJJ42678.1

Group291: abra|OJJ66585.1 acar|OOF91022.1 acla|EAW10906.1 acla|EAW10907.1 afis|EAW20951.1 afis|EAW20952.1 afla|EED50307.1 afla|EED50308.1 afum|EAL84682.1 afum|EAL84683.2 agla|OJJ83929.1 akaw|GAA91132.1 akaw|GAA91134.1 aluc|OJZ81461.1 anid|CBF82552.1 anid|CBF82554.1 anig|EHA25968.1 aory|EIT77335.1 aory|EIT77337.1 asyd|OJJ54058.1 ater|EAU38602.1 ater|EAU38603.1 atub|OJI88718.1 aver|OJJ08683.1 awen|OJJ34364.1 awen|OJJ34365.1 azon|OJJ48100.1

Group293: abra|OJJ67205.1 acar|OOF98357.1 acar|OOF98358.1 acla|EAW14519.1 afis|EAW19237.1 afla|EED47640.1 afum|EAL93965.2 agla|OJJ88462.1 akaw|GAA90972.1 akaw|GAA90973.1 aluc|OJZ87769.1 anid|CBF78640.1 anig|EHA20253.1 aory|EIT80182.1 asyd|OJJ65189.1 ater|EAU37226.1 atub|OJI80876.1 aver|OJI99338.1 awen|OJJ37593.1 azon|OJJ51248.1 acla|EAW14518.1 afis|EAW19236.1 afla|EED47646.1 afum|EAL93964.1 anid|CBF78638.1 aory|EIT80263.1 ater|EAU37227.1

Group295: abra|OJJ68390.1 afla|EED49558.1 agla|OJJ87147.1 aory|EIT82645.1 atub|OJI84701.1 abra|OJJ77648.1 acar|OOF99088.1 acla|EAW12091.1 afis|EAW19985.1 afum|EBA27209.1 akaw|GAA83987.1 aluc|OJZ80881.1 anid|CBF87267.1 anig|EHA19455.1 asyd|OJJ62371.1 ater|EAU33659.1 atub|OJI82696.1 aver|OJI98437.1 awen|OJJ38077.1 azon|OJJ42589.1 anid|CBF73965.1 aver|OJI96505.1 afla|EED52196.1 aory|EIT74391.1 ater|EAU29811.1 aluc|OJZ83787.1 anid|CBF85559.1

Group296: abra|OJJ69188.1 acar|OOF91124.1 acla|EAW08657.1 afis|EAW15774.1 afla|EED51274.1 afum|EAL85575.1 agla|OJJ81937.1 akaw|GAA86276.1 akaw|GAA86277.1 aluc|OJZ86174.1 anid|CBF70974.1 anig|EHA24450.1 anig|EHA24451.1 aory|EIT81499.1 asyd|OJJ52632.1 ater|EAU32367.1 atub|OJI79966.1 aver|OJJ04660.1 awen|OJJ38481.1 azon|OJJ43782.1 acla|EAW08658.1 afis|EAW15773.1 afla|EED49465.1 anid|CBF70972.1 ater|EAU32368.1 azon|OJJ43781.1 aory|EIT79571.1

Group297: abra|OJJ69236.1 acar|OOF91082.1 acar|OOF91084.1 acla|EAW08611.1 acla|EAW08613.1 afis|EAW15821.1 afis|EAW15822.1 afla|EED51224.1 afla|EED51226.1 afum|EAL85529.1 afum|EAL85530.1 agla|OJJ81985.1 akaw|GAA92244.1 akaw|GAA92246.1 aluc|OJZ86216.1 anid|CBF70886.1 anid|CBF70887.1 anig|EHA24410.1 aory|EIT83181.1 aory|EIT83189.1 asyd|OJJ52589.1 ater|EAU33426.1 ater|EAU33427.1 atub|OJI80011.1 aver|OJJ04616.1 awen|OJJ38535.1 azon|OJJ43727.1

Group298: abra|OJJ70462.1 acar|OOF97462.1 acar|OOF97465.1 acla|EAW13051.1 acla|EAW13053.1 afis|EAW17613.1 afis|EAW17616.1 afla|EED49204.1 afla|EED49207.1 afum|EAL91426.1 afum|EAL91429.1 agla|OJJ81324.1 akaw|GAA83308.1 akaw|GAA83311.1 aluc|OJZ83805.1 anid|CBF90169.1 anid|CBF90172.1 anig|EHA18952.1 aory|EIT78583.1 aory|EIT83172.1 asyd|OJJ61991.1 ater|EAU37048.1 ater|EAU37050.1 atub|OJI90172.1 aver|OJI96083.1 awen|OJJ37229.1 azon|OJJ50932.1

Group299: abra|OJJ71130.1 acar|OOF98022.1 acla|EAW10367.1 acla|EAW10368.1 afis|EAW25224.1 afis|EAW25225.1 afla|EED53669.1 afla|EED53682.1 afum|EAL89885.1 afum|EAL89886.1 akaw|GAA82844.1 aluc|OJZ87405.1 anid|CBF85264.1 anid|CBF85265.1 anig|EHA20634.1 anig|EHA20635.1 aory|EIT75307.1 aory|EIT75353.1 ater|EAU34383.1 ater|EAU34384.1 atub|OJI81241.1 aver|OJI98082.1 aver|OJI98083.1 agla|OJJ79183.1 asyd|OJJ62906.1 awen|OJJ40890.1 azon|OJJ44519.1

Group301: abra|OJJ72938.1 acar|OOF96120.1 acar|OOF96121.1 acla|EAW06452.1 acla|EAW06453.1 afis|EAW15949.1 afla|EED58109.1 afla|EED58110.1 afum|EAL88582.1 agla|OJJ89365.1 akaw|GAA85519.1 aluc|OJZ83447.1 anid|CBF81052.1 anig|EHA24293.1 anig|EHA24294.1 aory|EIT79139.1 aory|EIT79141.1 asyd|OJJ56450.1 ater|EAU35509.1 atub|OJI82338.1 atub|OJI82339.1 aver|OJJ06272.1 awen|OJJ35627.1 azon|OJJ46898.1 anid|CBF81054.1 ater|EAU35508.1 aver|OJJ06271.1

Group302: abra|OJJ73010.1 acar|OOF96182.1 acar|OOF96183.1 acla|EAW08002.1 acla|EAW08003.1 afis|EAW23037.1 afis|EAW23038.1 afla|EED54980.1 afum|EAL88443.1 afum|EAL88444.1 agla|OJJ89440.1 akaw|GAA85588.1 akaw|GAA85589.1 aluc|OJZ83525.1 anid|CBF80929.1 anid|CBF80931.1 anig|EHA24348.1 aory|EIT80318.1 aory|EIT80383.1 asyd|OJJ56316.1 ater|EAU29618.1 ater|EAU29621.1 atub|OJI82414.1 aver|OJJ06398.1 awen|OJJ35704.1 azon|OJJ44166.1 afla|EED54979.1

Group303: abra|OJJ73242.1 acar|OOG00479.1 acar|OOG00481.1 acla|EAW06915.1 acla|EAW06916.1 afis|EAW16532.1 afis|EAW16533.1 afla|EED56136.1 afla|EED56140.1 afum|EAL89129.1 afum|EAL89130.2 agla|OJJ82788.1 akaw|GAA87938.1 akaw|GAA87939.1 aluc|OJZ81186.1 anid|CBF81734.1 anid|CBF81736.1 anig|EHA18503.1 aory|EIT76580.1 aory|EIT76630.1 asyd|OJJ59594.1 ater|EAU36595.1 ater|EAU36596.1 atub|OJI91382.1 aver|OJJ02674.1 awen|OJJ40289.1 azon|OJJ49150.1

Group304: abra|OJJ73503.1 acar|OOG00688.1 acla|EAW07435.1 acla|EAW07436.1 afis|EAW22461.1 afis|EAW22462.1 afla|EED54482.1 afla|EED54483.1 afum|EAL90729.1 afum|EAL90730.1 agla|OJJ87561.1 akaw|GAA87450.1 akaw|GAA87451.1 aluc|OJZ88545.1 anid|CBF88911.1 anid|CBF88912.1 anig|EHA18245.1 aory|EIT81150.1 asyd|OJJ62841.1 ater|EAU39244.1 atub|OJI91130.1 aver|OJI96884.1 aver|OJI96885.1 awen|OJJ35087.1 awen|OJJ35088.1 azon|OJJ50020.1 aory|EIT81157.1

Group306: abra|OJJ76286.1 acar|OOF99527.1 acla|EAW14330.1 afis|EAW18958.1 afum|EAL93691.1 akaw|GAA82215.1 aory|EIT76999.1 asyd|OJJ59136.1 awen|OJJ31557.1 azon|OJJ47274.1 abra|OJJ76287.1 acar|OOF99530.1 acla|EAW14329.1 afis|EAW18959.1 afla|EED51005.1 afum|EAL93692.1 agla|OJJ85336.1 akaw|GAA82216.1 aluc|OJZ91935.1 anid|CBF69598.1 anig|EHA22895.1 aory|EIT76941.1 ater|EAU38141.1 atub|OJI86377.1 aver|OJJ01725.1 awen|OJJ31558.1 azon|OJJ47275.1

Group307: abra|OJJ77576.1 acar|OOF99151.1 acla|EAW06541.1 acla|EAW06542.1 afis|EAW16053.1 afis|EAW16054.1 afla|EED56444.1 afla|EED56445.1 afum|EAL88685.1 afum|EAL88686.1 agla|OJJ86087.1 akaw|GAA84049.1 akaw|GAA84050.1 aluc|OJZ80961.1 anid|CBF75070.1 anid|CBF75072.1 anig|EHA19396.1 aory|EIT72818.1 asyd|OJJ60021.1 ater|EAU33580.1 ater|EAU33584.1 atub|OJI82625.1 aver|OJJ02264.1 awen|OJJ30385.1 awen|OJJ30389.1 azon|OJJ45401.1 aory|EIT72866.1

Group308: abra|OJJ77839.1 acar|OOF98932.1 acla|EAW11284.1 afis|EAW18216.1 afla|EED56254.1 afum|EAL87582.1 akaw|GAA83808.1 aory|EIT73366.1 ater|EAU33774.1 azon|OJJ50174.1 abra|OJJ77840.1 acar|OOF98931.1 acla|EAW11283.1 afis|EAW18217.1 afla|EED56253.1 afum|EAL87583.1 agla|OJJ85865.1 akaw|GAA83807.1 aluc|OJZ90915.1 anid|CBF86146.1 anig|EHA19627.1 aory|EIT73438.1 asyd|OJJ64511.1 atub|OJI82907.1 aver|OJI98609.1 awen|OJJ30595.1 azon|OJJ50175.1

Group309: abra|OJJ65742.1 abra|OJJ77048.1 acar|OOF91730.1 acar|OOF92937.1 aluc|OJZ84452.1 aluc|OJZ86322.1 atub|OJI84212.1 atub|OJI89624.1 acla|EAW13550.1 afis|EAW20627.1 afla|EED49777.1 afum|EAL84441.1 agla|OJJ86946.1 akaw|GAA87005.1 anid|CBF83228.1 anig|EHA27767.1 aory|EIT77606.1 asyd|OJJ55581.1 ater|EAU32164.1 aver|OJJ06660.1 awen|OJJ33462.1 azon|OJJ50718.1 afla|EED52520.1 anig|EHA19838.1 aory|EIT82787.1 awen|OJJ33741.1

Group313: abra|OJJ71059.1 abra|OJJ73085.1 akaw|GAA82900.1 akaw|GAA88075.1 anig|EHA18644.1 anig|EHA20691.1 ater|EAU29720.1 ater|EAU35713.1 atub|OJI81179.1 atub|OJI91543.1 aver|OJJ02730.1 aver|OJJ06544.1 awen|OJJ34513.1 awen|OJJ40561.1 acar|OOF98083.1 acla|EAW10600.1 afis|EAW24968.1 afum|EAL89627.1 agla|OJJ82234.1 aluc|OJZ87469.1 anid|CBF81643.1 aory|EIT80808.1 asyd|OJJ59538.1 azon|OJJ49422.1 afla|EED57716.1 aory|EIT79920.1

Group316: acla|EAW06814.1 acla|EAW11026.1 afis|EAW16363.1 afis|EAW20840.1 afum|EAL84793.1 afum|EAL88988.1 agla|OJJ78575.1 agla|OJJ80753.1 aver|OJI96276.1 aver|OJI96627.1 awen|OJJ37768.1 awen|OJJ40172.1 abra|OJJ73491.1 acar|OOG00684.1 afla|EED53503.1 agla|OJJ79965.1 akaw|GAA87440.1 aluc|OJZ88557.1 anid|CBF89245.1 anig|EHA18255.1 aory|EIT74859.1 asyd|OJJ62507.1 ater|EAU34089.1 atub|OJI91142.1 azon|OJJ45066.1 asyd|OJJ62165.1

Group317: acla|EAW11040.1 acla|EAW11477.1 afla|EED47466.1 afla|EED49623.1 afla|EED50532.1 agla|OJJ82839.1 agla|OJJ86906.1 aory|EIT79957.1 aory|EIT81005.1 aory|EIT82111.1 awen|OJJ31954.1 awen|OJJ35441.1 azon|OJJ45954.1 azon|OJJ51301.1 abra|OJJ72795.1 acar|OOF89880.1 afis|EAW23976.1 afum|EAL87406.1 akaw|GAA85398.1 aluc|OJZ83292.1 anid|CBF77345.1 anig|EHA24176.1 asyd|OJJ53272.1 ater|EAU29776.1 atub|OJI82193.1 aver|OJJ03577.1

Group318: afla|EED44930.1 afla|EED51540.1 ater|EAU31301.1 ater|EAU38559.1 awen|OJJ33684.1 awen|OJJ36221.1 abra|OJJ67873.1 acar|OOF92853.1 acla|EAW11724.1 afis|EAW23749.1 afum|EAL87183.1 agla|OJJ78962.1 agla|OJJ89691.1 akaw|GAA91882.1 aluc|OJZ82015.1 anid|CBF79817.1 anig|EHA22241.1 aory|EIT80515.1 asyd|OJJ58137.1 atub|OJI84564.1 aver|OJJ04268.1 azon|OJJ42827.1 aver|OJJ03949.1 afis|EAW24762.1 afum|EAL89404.1 asyd|OJJ52466.1

Group319: afla|EED50195.1 afla|EED57633.1 anid|CBF78576.1 anid|CBF84581.1 anid|CBF88483.1 aory|EIT77388.1 aory|EIT82958.1 asyd|OJJ59798.1 asyd|OJJ63787.1 asyd|OJJ65070.1 aver|OJI97151.1 aver|OJI99195.1 aver|OJJ02491.1 abra|OJJ74472.1 acar|OOF95272.1 acla|EAW07225.1 afis|EAW22239.1 afum|EAL90953.1 agla|OJJ87812.1 akaw|GAA87366.1 aluc|OJZ89647.1 anig|EHA26885.1 ater|EAU34246.1 atub|OJI85157.1 awen|OJJ34817.1 azon|OJJ45732.1

Group323: abra|OJJ66489.1 acar|OOF90965.1 acar|OOF90966.1 acla|EAW10979.1 acla|EAW10980.1 afis|EAW20879.1 afis|EAW20880.1 afla|EED45565.1 afum|EAL84752.1 afum|EAL84753.1 agla|OJJ83989.1 akaw|GAA92472.1 akaw|GAA92473.1 aluc|OJZ81546.1 anid|CBF82862.1 anid|CBF82864.1 anig|EHA25889.1 aory|EIT75388.1 aory|EIT75432.1 asyd|OJJ61598.1 ater|EAU30811.1 atub|OJI88638.1 aver|OJJ00922.1 awen|OJJ34480.1 azon|OJJ51757.1 afla|EED45564.1

Group324: abra|OJJ67172.1 acar|OOF98391.1 acla|EAW14551.1 afis|EAW19268.1 afla|EED47606.1 afum|EAL93995.1 agla|OJJ88427.1 akaw|GAA91002.1 aluc|OJZ87803.1 anid|CBF78699.1 anig|EHA20284.1 aory|EIT80198.1 asyd|OJJ65220.1 ater|EAU29549.1 atub|OJI80844.1 aver|OJI99369.1 awen|OJJ37629.1 azon|OJJ47017.1 afla|EED47607.1 anid|CBF78697.1 aory|EIT80260.1 acar|OOF98392.1 acla|EAW14552.1 afis|EAW19269.1 afum|EAL93996.1 anig|EHA20285.1

Group326: abra|OJJ68235.1 acar|OOF93581.1 acla|EAW10392.1 afis|EAW25199.1 afum|EBA27389.1 agla|OJJ85249.1 anid|CBF85213.1 asyd|OJJ54665.1 ater|EAU34341.1 atub|OJI83115.1 aver|OJI97964.1 awen|OJJ33408.1 abra|OJJ71100.1 afla|EED53646.1 akaw|GAA82868.1 aluc|OJZ87435.1 anig|EHA20657.1 aory|EIT75355.1 atub|OJI81213.1 azon|OJJ47678.1 aver|OJJ02024.1 awen|OJJ31517.1 asyd|OJJ63002.1 agla|OJJ86390.1 awen|OJJ39475.1 agla|OJJ89313.1

Group327: abra|OJJ68548.1 acar|OOF92744.1 acla|EAW12819.1 acla|EAW12820.1 afis|EAW17984.1 afis|EAW17985.1 afla|EED51920.1 afla|EED51921.1 afum|EAL91793.1 afum|EAL91794.1 agla|OJJ80241.1 akaw|GAA91305.1 akaw|GAA91306.1 aluc|OJZ82765.1 anid|CBF79988.1 anid|CBF79990.1 anig|EHA21208.1 aory|EIT81392.1 aory|EIT81447.1 asyd|OJJ58239.1 ater|EAU31494.1 ater|EAU31495.1 atub|OJI88997.1 aver|OJJ02186.1 awen|OJJ41561.1 azon|OJJ48276.1

Group328: abra|OJJ68713.1 acar|OOF92213.1 acla|EAW11027.1 afis|EAW20841.1 afla|EED46816.1 afum|EAL84792.1 agla|OJJ80754.1 akaw|GAA89866.1 aluc|OJZ81815.1 anid|CBF86535.1 anig|EHA22400.1 aory|EIT72698.1 asyd|OJJ62166.1 ater|EAU30544.1 atub|OJI84387.1 aver|OJI96277.1 awen|OJJ40171.1 azon|OJJ50392.1 afis|EAW24116.1 anid|CBF84526.1 acla|EAW12533.1 agla|OJJ88735.1 afum|EAL91971.1 awen|OJJ41372.1 aver|OJI98980.1 asyd|OJJ64867.1

Group330: abra|OJJ70396.1 acar|OOG01144.1 acla|EAW11244.1 afis|EAW19549.1 afla|EED49895.1 afum|EAL89219.1 agla|OJJ88525.1 akaw|GAA91962.1 aluc|OJZ82655.1 anig|EHA24571.1 asyd|OJJ65105.1 ater|EAU33680.1 atub|OJI88886.1 aver|OJI99232.1 awen|OJJ31981.1 azon|OJJ47790.1 afis|EAW18105.1 akaw|GAA92884.1 aory|EIT77372.1 afis|EAW16617.1 afum|EAL86960.1 anig|EHA27418.1 afla|EED50268.1 anid|CBF84502.1 asyd|OJJ53512.1 aver|OJI96187.1

Group331: abra|OJJ71980.1 acar|OOF96631.1 acla|EAW09559.1 acla|EAW09561.1 afis|EAW21576.1 afis|EAW21578.1 afla|EED57795.1 afum|EAL92595.1 afum|EAL92596.1 agla|OJJ85761.1 akaw|GAA84949.1 akaw|GAA84950.1 aluc|OJZ86717.1 anid|CBF76292.1 anid|CBF76294.1 anig|EHA21962.1 aory|EIT81026.1 aory|EIT81117.1 asyd|OJJ61004.1 ater|EAU34955.1 ater|EAU34956.1 atub|OJI83812.1 aver|OJJ00322.1 awen|OJJ39103.1 azon|OJJ51431.1 afla|EED57794.1

Group332: abra|OJJ75736.1 acar|OOF89930.1 acla|EAW14905.1 afis|EAW24426.1 afla|EED55252.1 afum|EAL86094.1 agla|OJJ79240.1 akaw|GAA84469.1 aluc|OJZ89188.1 anid|CBF73780.1 anig|EHA23369.1 aory|EIT83550.1 asyd|OJJ56598.1 ater|EAU30018.1 atub|OJI85602.1 aver|OJJ06127.1 awen|OJJ32778.1 azon|OJJ44006.1 acla|EAW14906.1 afis|EAW24427.1 afla|EED55251.1 afum|EAL86095.1 aluc|OJZ89187.1 anig|EHA23370.1 ater|EAU30019.1 awen|OJJ32779.1

Group333: abra|OJJ75938.1 acar|OOF99972.1 acar|OOF99973.1 acla|EAW09855.1 acla|EAW09856.1 afis|EAW21271.1 afis|EAW21280.1 afla|EED54831.1 afum|EAL92286.1 afum|EAL92296.1 agla|OJJ87477.1 akaw|GAA84639.1 akaw|GAA84640.1 aluc|OJZ89385.1 anid|CBF83352.1 anid|CBF83354.1 anig|EHA23199.1 aory|EIT79381.1 aory|EIT79404.1 asyd|OJJ60616.1 ater|EAU35907.1 ater|EAU35908.1 atub|OJI85413.1 aver|OJI99940.1 awen|OJJ39421.1 azon|OJJ44303.1

Group334: abra|OJJ76077.1 acar|OOF99848.1 acar|OOF99849.1 acla|EAW09743.1 acla|EAW09744.1 afis|EAW21398.1 afis|EAW21399.1 afla|EED55817.1 afla|EED55818.1 afum|EAL92412.1 afum|EAL92413.1 agla|OJJ85524.1 akaw|GAA88889.1 akaw|GAA88890.1 aluc|OJZ89533.1 anid|CBF83833.1 anig|EHA23070.1 aory|EIT76532.1 aory|EIT76875.1 asyd|OJJ60909.1 ater|EAU35159.1 atub|OJI85272.1 aver|OJJ00236.1 awen|OJJ39304.1 azon|OJJ51548.1 ater|EAU35160.1

Group335: abra|OJJ78321.1 acar|OOF98553.1 acla|EAW09204.1 acla|EAW09205.1 afis|EAW21942.1 afis|EAW21943.1 afla|EED46659.1 afla|EED46660.1 afum|EAL92956.1 afum|EAL92957.1 agla|OJJ84181.1 akaw|GAA85707.1 akaw|GAA85708.1 aluc|OJZ91396.1 anid|CBF84053.1 anid|CBF84055.1 anig|EHA20073.1 aory|EIT77914.1 aory|EIT77936.1 asyd|OJJ61344.1 ater|EAU30947.1 ater|EAU30991.1 atub|OJI83388.1 aver|OJJ00669.1 awen|OJJ34234.1 azon|OJJ43402.1

Group337: abra|OJJ66243.1 abra|OJJ72838.1 afla|EED45789.1 afla|EED49140.1 aluc|OJZ80548.1 aluc|OJZ83338.1 anig|EHA24214.1 anig|EHA25648.1 aory|EIT75668.1 aory|EIT78573.1 atub|OJI82235.1 atub|OJI88383.1 acar|OOF96040.1 acla|EAW11403.1 afis|EAW24018.1 afum|EAL87448.1 akaw|GAA85435.1 akaw|GAA89048.1 anid|CBF87091.1 asyd|OJJ55451.1 ater|EAU29728.1 awen|OJJ35491.1 azon|OJJ46999.1 aver|OJJ06524.1 agla|OJJ78562.1

Group340: abra|OJJ69049.1 abra|OJJ77527.1 akaw|GAA84093.1 akaw|GAA90580.1 aluc|OJZ81008.1 aluc|OJZ86038.1 anig|EHA19352.1 anig|EHA24573.1 asyd|OJJ51977.1 asyd|OJJ54328.1 atub|OJI79825.1 atub|OJI82576.1 acar|OOF99199.1 acla|EAW06702.1 afis|EAW16247.1 afum|EAL88875.1 agla|OJJ86256.1 anid|CBF82150.1 aory|EIT75005.1 ater|EAU37457.1 aver|OJJ05282.1 awen|OJJ30209.1 azon|OJJ43933.1 afla|EED53308.1 asyd|OJJ57961.1

Group342: abra|OJJ70211.1 abra|OJJ76495.1 acar|OOF92571.1 acar|OOF99727.1 akaw|GAA82411.1 akaw|GAA86565.1 aluc|OJZ85115.1 aluc|OJZ91709.1 azon|OJJ46702.1 azon|OJJ50101.1 acla|EAW13937.1 afis|EAW18648.1 afla|EED52691.1 afum|EAL93364.1 agla|OJJ82426.1 anid|CBF70533.1 anig|EHA22720.1 anig|EHA25335.1 aory|EIT74236.1 asyd|OJJ58553.1 atub|OJI80479.1 atub|OJI86168.1 aver|OJJ01151.1 awen|OJJ32116.1 ater|EAU38263.1

Group343: abra|OJJ74685.1 abra|OJJ77443.1 acar|OOF95646.1 acar|OOF95697.1 asyd|OJJ52766.1 asyd|OJJ57547.1 asyd|OJJ56548.1 aver|OJJ05631.1 aver|OJJ06174.1 aver|OJJ08749.1 awen|OJJ30371.1 awen|OJJ40449.1 acla|EAW10711.1 afis|EAW24859.1 afla|EED47099.1 afum|EAL89507.1 agla|OJJ82077.1 akaw|GAA81929.1 aluc|OJZ92262.1 anid|CBF75821.1 anig|EHA26697.1 aory|EIT72503.1 ater|EAU36441.1 atub|OJI86696.1 azon|OJJ49305.1

Group344: abra|OJJ75156.1 abra|OJJ77714.1 acar|OOF97183.1 acar|OOF99026.1 akaw|GAA83918.1 akaw|GAA88762.1 aluc|OJZ90779.1 aluc|OJZ92725.1 anig|EHA19512.1 anig|EHA26276.1 ater|EAU35617.1 ater|EAU38821.1 atub|OJI82766.1 atub|OJI87150.1 acla|EAW08331.1 afis|EAW23366.1 afla|EED55592.1 afum|EAL88107.1 agla|OJJ86458.1 anid|CBF74825.1 aory|EIT82696.1 asyd|OJJ60139.1 aver|OJJ04123.1 awen|OJJ39521.1 azon|OJJ47717.1

Group345: acar|OOF90695.1 acar|OOF91261.1 afla|EED44834.1 afla|EED46120.1 agla|OJJ84890.1 agla|OJJ88358.1 anid|CBF79319.1 anid|CBF85127.1 asyd|OJJ55782.1 asyd|OJJ63353.1 aver|OJI97608.1 aver|OJJ06871.1 aver|OJJ07891.1 abra|OJJ76767.1 acla|EAW13321.1 afis|EAW20436.1 afum|EAL84616.1 akaw|GAA87259.1 aluc|OJZ84170.1 anig|EHA27531.1 aory|EIT77716.1 ater|EAU37722.1 atub|OJI89914.1 awen|OJJ33127.1 azon|OJJ46343.1

Group347: acla|EAW06619.1 acla|EAW13553.1 awen|OJJ30278.1 awen|OJJ33599.1 azon|OJJ42199.1 azon|OJJ50720.1 abra|OJJ77044.1 acar|OOF90272.1 afis|EAW16146.1 afis|EAW20629.1 afla|EED49763.1 afum|EAL84438.1 afum|EAL88775.1 agla|OJJ80445.1 agla|OJJ86199.1 akaw|GAA87009.1 aluc|OJZ84447.1 anid|CBF84471.1 anig|EHA27764.1 aory|EIT77658.1 asyd|OJJ53537.1 ater|EAU32165.1 atub|OJI89628.1 aver|OJJ08214.1 azon|OJJ43971.1

Group348: acla|EAW07744.1 acla|EAW10138.1 afis|EAW20748.1 afis|EAW22781.1 afla|EED47130.1 afla|EED50040.1 afum|EAL84866.1 afum|EAL90418.1 anid|CBF71564.1 anid|CBF87981.1 ater|EAU33992.1 ater|EAU38933.1 awen|OJJ36890.1 awen|OJJ38956.1 abra|OJJ73849.1 acar|OOG00999.1 agla|OJJ83643.1 akaw|GAA89466.1 aluc|OJZ88200.1 anig|EHA17948.1 aory|EIT79245.1 asyd|OJJ63496.1 atub|OJI90798.1 aver|OJI97433.1 azon|OJJ49728.1

Group349: afis|EAW15424.1 afis|EAW16124.1 afum|EAL84220.1 afum|EAL88754.1 aory|EIT73054.1 aory|EIT80955.1 abra|OJJ69804.1 acar|OOF93572.1 acla|EAW06641.1 afla|EED48664.1 agla|OJJ86166.1 akaw|GAA87855.1 aluc|OJZ85497.1 anid|CBF85532.1 anig|EHA25030.1 asyd|OJJ64135.1 ater|EAU37174.1 atub|OJI80068.1 aver|OJI98218.1 awen|OJJ30306.1 azon|OJJ48681.1 acar|OOF91913.1 afla|EED47503.1 aory|EIT74578.1 ater|EAU33207.1

Group350: afis|EAW20695.1 afis|EAW25202.1 afum|EAL86271.1 afum|EAL89865.1 anid|CBF80363.1 anid|CBF85220.1 asyd|OJJ54819.1 asyd|OJJ60277.1 asyd|OJJ62998.1 ater|EAU34342.1 ater|EAU35428.1 aver|OJI97968.1 aver|OJJ07342.1 abra|OJJ71102.1 acar|OOF98041.1 acla|EAW10389.1 afla|EED53649.1 agla|OJJ88973.1 akaw|GAA82866.1 aluc|OJZ87433.1 anig|EHA20655.1 aory|EIT75301.1 atub|OJI81215.1 awen|OJJ40837.1 azon|OJJ46857.1

Group354: afla|EED45547.1 afla|EED48860.1 afum|EAL84770.1 afum|EAL86315.1 aory|EIT75539.1 aory|EIT81273.1 awen|OJJ29954.1 awen|OJJ34505.1 abra|OJJ67746.1 acar|OOF90945.1 acla|EAW12433.1 afis|EAW20865.1 agla|OJJ83914.1 akaw|GAA92437.1 aluc|OJZ81583.1 anid|CBF80387.1 anig|EHA25856.1 asyd|OJJ61074.1 ater|EAU30808.1 atub|OJI88598.1 aver|OJJ00391.1 azon|OJJ51781.1 acla|EAW10996.1 agla|OJJ79837.1 afis|EAW16740.1

Group355: afla|EED45554.1 afla|EED57298.1 aory|EIT74121.1 aory|EIT75629.1 asyd|OJJ56283.1 asyd|OJJ61592.1 asyd|OJJ61824.1 ater|EAU30081.1 ater|EAU30816.1 aver|OJI95924.1 aver|OJJ00914.1 aver|OJJ08478.1 abra|OJJ67752.1 acar|OOF90956.1 acla|EAW10988.1 afis|EAW20872.1 afum|EAL84762.1 agla|OJJ83979.1 akaw|GAA92442.1 aluc|OJZ81577.1 anid|CBF82847.1 anig|EHA25861.1 atub|OJI88603.1 awen|OJJ34489.1 azon|OJJ51766.1

Group357: afum|EAL90938.1 afum|EAL91118.1 abra|OJJ70518.1 acar|OOF97412.1 acla|EAW08157.1 afis|EAW23195.1 afla|EED57873.1 afum|EAL88288.2 agla|OJJ86332.1 akaw|GAA83365.1 aluc|OJZ83863.1 anid|CBF74706.1 anig|EHA19003.1 aory|EIT83278.1 asyd|OJJ60219.1 ater|EAU36323.1 atub|OJI90231.1 aver|OJJ04049.1 awen|OJJ39425.1 azon|OJJ47626.1 afis|EAW20299.1 agla|OJJ84725.1 asyd|OJJ62722.1 aver|OJJ01792.1 azon|OJJ49994.1

Group359: agla|OJJ83698.1 agla|OJJ89439.1 akaw|GAA85587.1 akaw|GAA85825.1 aluc|OJZ83524.1 aluc|OJZ91276.1 anig|EHA19962.1 anig|EHA24347.1 asyd|OJJ56315.1 asyd|OJJ58149.1 atub|OJI82413.1 atub|OJI83265.1 abra|OJJ73009.1 acar|OOF96181.1 acla|EAW08004.1 afis|EAW23039.1 afla|EED45399.1 afum|EAL88442.1 anid|CBF80933.1 aory|EIT82471.1 ater|EAU29619.1 aver|OJJ06399.1 awen|OJJ35703.1 azon|OJJ44165.1 aver|OJJ04255.1

Group360: aluc|OJZ83601.1 aluc|OJZ88595.1 anig|EHA18289.1 anig|EHA18760.1 asyd|OJJ56054.1 asyd|OJJ61370.1 atub|OJI89973.1 atub|OJI91180.1 aver|OJJ00697.1 aver|OJJ04995.1 awen|OJJ35022.1 awen|OJJ37732.1 abra|OJJ73445.1 acar|OOG00650.1 acla|EAW06781.1 afis|EAW16331.1 afla|EED53547.1 afum|EAL88956.1 agla|OJJ88261.1 akaw|GAA83049.1 anid|CBF84027.1 aory|EIT75029.1 ater|EAU34127.1 azon|OJJ45215.1 akaw|GAA92121.1

Group367: abra|OJJ67359.1 acar|OOF93335.1 acla|EAW10444.1 afla|EED55318.1 agla|OJJ89047.1 akaw|GAA86462.1 anid|CBF87506.1 anig|EHA25221.1 aory|EIT76259.1 asyd|OJJ58362.1 ater|EAU39395.1 aver|OJJ02044.1 awen|OJJ40742.1 abra|OJJ70853.1 acar|OOF98263.1 afis|EAW25123.1 afum|EAL89794.1 akaw|GAA91720.1 aluc|OJZ87668.1 anid|CBF85688.1 anig|EHA20859.1 asyd|OJJ64259.1 atub|OJI80975.1 aver|OJI98351.1 azon|OJJ44113.1

Group372: abra|OJJ70648.1 acar|OOF97073.1 acla|EAW08111.1 acla|EAW08112.1 afis|EAW23139.1 afis|EAW23140.1 afla|EED48502.1 afum|EAL88341.1 afum|EAL88342.1 agla|OJJ89661.1 akaw|GAA89984.1 aluc|OJZ83998.1 anid|CBF74444.1 anid|CBF74446.1 anig|EHA19121.1 aory|EIT73212.1 aory|EIT73270.1 asyd|OJJ57742.1 ater|EAU36277.1 atub|OJI90367.1 aver|OJJ05425.1 awen|OJJ36636.1 azon|OJJ47580.1 afla|EED48503.1 ater|EAU36278.1

Group373: abra|OJJ70697.1 acar|OOF90045.1 acla|EAW08067.1 afis|EAW23103.1 afla|EED48544.1 afum|EAL88376.1 agla|OJJ89639.1 akaw|GAA90028.1 akaw|GAA90029.1 aluc|OJZ84044.1 anid|CBF81363.1 anig|EHA19166.1 aory|EIT73061.1 asyd|OJJ59383.1 ater|EAU36232.1 atub|OJI90416.1 aver|OJJ02890.1 awen|OJJ36677.1 azon|OJJ47521.1 afla|EED48545.1 acla|EAW08068.1 afis|EAW23104.1 afum|EAL88375.1 anid|CBF81365.1 aory|EIT73082.1

Group376: abra|OJJ72657.1 acar|OOF94617.1 acla|EAW11521.1 afis|EAW23933.1 afla|EED50585.1 afum|EAL87357.1 agla|OJJ79413.1 akaw|GAA85279.1 akaw|GAA85280.1 aluc|OJZ90638.1 anid|CBF77273.1 anig|EHA24073.1 aory|EIT82169.1 asyd|OJJ56797.1 ater|EAU32304.1 atub|OJI82066.1 aver|OJJ03537.1 awen|OJJ39022.1 azon|OJJ51347.1 acla|EAW11520.1 afum|EAL87358.1 aory|EIT82139.1 afis|EAW23932.1 afla|EED50586.1 azon|OJJ51348.1

Group378: abra|OJJ74406.1 acar|OOF95334.1 acla|EAW06822.1 afis|EAW16185.1 afla|EED53465.1 afum|EAL88816.2 agla|OJJ78591.1 akaw|GAA87300.1 aluc|OJZ89718.1 anid|CBF89935.1 anig|EHA26946.1 aory|EIT74868.1 asyd|OJJ62121.1 ater|EAU34079.1 atub|OJI85082.1 aver|OJI96229.1 awen|OJJ37907.1 azon|OJJ45085.1 afis|EAW16370.1 acla|EAW11051.1 afla|EED49634.1 aory|EIT80960.1 azon|OJJ45955.1 ater|EAU36747.1 azon|OJJ45949.1

Group379: abra|OJJ76313.1 acar|OOF99561.1 acar|OOF99562.1 acla|EAW14106.1 afis|EAW18806.1 afis|EAW18807.1 afla|EED50770.1 afum|EAL93542.1 afum|EAL93543.1 agla|OJJ80536.1 akaw|GAA82241.1 aluc|OJZ91904.1 anid|CBF69790.1 anig|EHA22880.1 aory|EIT78083.1 asyd|OJJ58961.1 ater|EAU37958.1 atub|OJI86354.1 aver|OJJ01555.1 awen|OJJ32382.1 azon|OJJ47143.1 agla|OJJ80538.1 acla|EAW14105.1 anid|CBF69788.1 awen|OJJ32381.1

Group380: abra|OJJ76772.1 acar|OOF91266.1 acla|EAW13327.1 afis|EAW20442.1 afla|EED44828.1 afum|EAL84610.2 agla|OJJ84883.1 akaw|GAA87253.1 akaw|GAA87254.1 aluc|OJZ84176.1 anid|CBF79308.1 anig|EHA27535.1 aory|EIT77723.1 asyd|OJJ55777.1 ater|EAU37717.1 atub|OJI89908.1 atub|OJI89909.1 aver|OJJ06866.1 awen|OJJ33132.1 azon|OJJ46347.1 acla|EAW13326.1 afis|EAW20441.1 afla|EED44829.1 afum|EAL84611.1 anid|CBF79310.1

Group381: abra|OJJ77345.1 acar|OOF93215.1 acla|EAW13129.1 acla|EAW13130.1 afis|EAW17533.1 afis|EAW17534.1 afla|EED49295.1 afla|EED49296.1 afum|EAL91347.1 afum|EAL91348.1 agla|OJJ81437.1 akaw|GAA91415.1 akaw|GAA91416.1 aluc|OJZ84756.1 anid|CBF90328.1 anid|CBF90329.1 anig|EHA28025.1 aory|EIT82179.1 asyd|OJJ61906.1 ater|EAU36962.1 ater|EAU36963.1 atub|OJI89328.1 aver|OJI95999.1 awen|OJJ37319.1 azon|OJJ51011.1

Group384: abra|OJJ66370.1 abra|OJJ77693.1 acar|OOF98983.1 acar|OOF99046.1 akaw|GAA83937.1 akaw|GAA90239.1 atub|OJI82743.1 atub|OJI87834.1 acla|EAW06572.1 afis|EAW16024.1 afla|EED56405.1 afum|EAL88653.1 agla|OJJ81572.1 aluc|OJZ83101.1 anid|CBF75012.1 anig|EHA19493.1 anig|EHA28166.1 aory|EIT83000.1 asyd|OJJ60056.1 ater|EAU33617.1 aver|OJJ02233.1 awen|OJJ30433.1 azon|OJJ45432.1 aluc|OJZ80819.1

Group385: abra|OJJ67097.1 abra|OJJ77658.1 aluc|OJZ80870.1 aluc|OJZ87876.1 anig|EHA19465.1 anig|EHA20181.1 atub|OJI80773.1 atub|OJI82707.1 acar|OOF98457.1 acar|OOF99078.1 acla|EAW06500.1 afis|EAW15996.1 afla|EED56370.1 afum|EAL88628.2 agla|OJJ86011.1 akaw|GAA83977.1 akaw|GAA91064.1 anid|CBF81691.1 aory|EIT73327.1 asyd|OJJ59562.1 ater|EAU33651.1 aver|OJJ02706.1 awen|OJJ30473.1 azon|OJJ45465.1

Group387: abra|OJJ68864.1 abra|OJJ75043.1 acar|OOF90525.1 acar|OOF97290.1 akaw|GAA88660.1 akaw|GAA90389.1 aluc|OJZ85842.1 aluc|OJZ92609.1 atub|OJI79630.1 atub|OJI87043.1 awen|OJJ31662.1 awen|OJJ39696.1 acla|EAW08232.1 afis|EAW23266.1 afla|EED55747.1 afum|EAL88213.1 agla|OJJ86586.1 anid|CBF84130.1 anig|EHA24722.1 aory|EIT83228.1 asyd|OJJ57114.1 ater|EAU37532.1 aver|OJJ03186.1 azon|OJJ47828.1

Group389: abra|OJJ70579.1 abra|OJJ75835.1 akaw|GAA83423.1 akaw|GAA84545.1 aluc|OJZ83923.1 aluc|OJZ89278.1 anig|EHA19056.1 anig|EHA23291.1 atub|OJI85521.1 atub|OJI90295.1 awen|OJJ32048.1 awen|OJJ39470.1 acar|OOF97142.1 acla|EAW08201.1 afis|EAW23235.1 afla|EED47014.1 afum|EAL88247.1 agla|OJJ86395.1 anid|CBF75150.1 aory|EIT72630.1 asyd|OJJ59974.1 ater|EAU36368.1 aver|OJJ02317.1 azon|OJJ47672.1

Group391: acar|OOF90694.1 acar|OOF95969.1 afla|EED46110.1 afla|EED52607.1 agla|OJJ82549.1 agla|OJJ84416.1 awen|OJJ31372.1 awen|OJJ33859.1 abra|OJJ72741.1 acla|EAW11574.1 afis|EAW23880.1 afum|EAL87308.1 akaw|GAA85352.1 aluc|OJZ90714.1 anid|CBF77193.1 anig|EHA24133.1 aory|EIT74176.1 asyd|OJJ56836.1 ater|EAU33467.1 atub|OJI82140.1 aver|OJJ03501.1 azon|OJJ48693.1 asyd|OJJ58816.1 aory|EIT76373.1

Group392: acar|OOF92653.1 acar|OOG00348.1 abra|OJJ69030.1 afla|EED56357.1 akaw|GAA90546.1 aluc|OJZ86016.1 anig|EHA24595.1 aory|EIT73420.1 ater|EAU31400.1 atub|OJI79805.1 awen|OJJ34779.1 abra|OJJ73125.1 acla|EAW07183.1 afis|EAW22196.1 afum|EAL90995.1 agla|OJJ87849.1 akaw|GAA88040.1 aluc|OJZ81299.1 anid|CBF88408.1 anig|EHA18608.1 asyd|OJJ63753.1 atub|OJI91508.1 aver|OJI97188.1 azon|OJJ45760.1

Group394: acar|OOF95062.1 acar|OOF95129.1 afla|EED47135.1 afla|EED49651.1 anid|CBF74606.1 anid|CBF84541.1 asyd|OJJ53476.1 asyd|OJJ57659.1 aver|OJJ05520.1 aver|OJJ08280.1 abra|OJJ74650.1 acla|EAW10735.1 afis|EAW24839.1 afum|EAL89485.1 agla|OJJ82030.1 akaw|GAA81960.1 aluc|OJZ92229.1 anig|EHA26728.1 aory|EIT79283.1 ater|EAU36464.1 atub|OJI86659.1 awen|OJJ40425.1 azon|OJJ49283.1 ater|EAU29279.1

Group403: aory|EIT74183.1 aory|EIT82897.1 ater|EAU31229.1 ater|EAU37935.1 aver|OJJ03325.1 aver|OJJ08019.1 abra|OJJ69359.1 acar|OOF94365.1 acla|EAW14750.1 afis|EAW24281.1 afla|EED47995.1 afum|EAL85950.1 agla|OJJ80635.1 akaw|GAA88229.1 aluc|OJZ82191.1 anid|CBF74148.1 anig|EHA28544.1 asyd|OJJ53921.1 asyd|OJJ56960.1 atub|OJI88244.1 awen|OJJ33016.1 azon|OJJ48501.1 acar|OOF90185.1 afla|EED52650.1

Group406: abra|OJJ66593.1 acar|OOF91028.1 acla|EAW10898.1 afis|EAW20960.1 afum|EAL84673.1 agla|OJJ83918.1 akaw|GAA91124.1 aluc|OJZ81454.1 anid|CBF82567.1 anig|EHA25974.1 aory|EIT77369.1 asyd|OJJ54050.1 ater|EAU38610.1 atub|OJI88727.1 aver|OJJ08692.1 awen|OJJ34357.1 azon|OJJ48093.1 acla|EAW10899.1 afis|EAW20959.1 afla|EED50315.1 afum|EAL84674.1 akaw|GAA91125.1 aluc|OJZ81455.1 aory|EIT77298.1

Group409: abra|OJJ66987.1 acar|OOF93625.1 acla|EAW08983.1 afis|EAW24689.1 afla|EED48347.1 afum|EAL89330.1 agla|OJJ83266.1 akaw|GAA92176.1 aluc|OJZ85586.1 anid|CBF73671.1 anig|EHA24935.1 aory|EIT76285.1 asyd|OJJ56664.1 ater|EAU31416.1 atub|OJI79381.1 aver|OJJ06059.1 awen|OJJ32833.1 azon|OJJ49002.1 acla|EAW14852.1 afis|EAW24379.1 afla|EED47832.1 afum|EAL86051.1 aory|EIT77109.1 ater|EAU30357.1

Group410: abra|OJJ67104.1 acar|OOF95836.1 afla|EED47889.1 aluc|OJZ82016.1 anig|EHA22240.1 aory|EIT77218.1 atub|OJI84565.1 abra|OJJ75997.1 acar|OOF99928.1 acla|EAW09808.1 afis|EAW21328.1 afla|EED57286.1 afum|EAL92346.2 agla|OJJ87411.1 akaw|GAA88962.1 aluc|OJZ89443.1 anid|CBF83430.1 anig|EHA23143.1 aory|EIT74039.1 asyd|OJJ60660.1 ater|EAU35953.1 aver|OJI99983.1 awen|OJJ39370.1 azon|OJJ51710.1

Group412: afis|EAW15422.1 afum|EAL84222.1 abra|OJJ74863.1 acar|OOF95863.1 acla|EAW08764.1 afla|EED57451.1 agla|OJJ81802.1 akaw|GAA83557.1 aluc|OJZ92441.1 anid|CBF89639.1 anig|EHA26526.1 aory|EIT80148.1 asyd|OJJ62291.1 ater|EAU32509.1 atub|OJI86870.1 aver|OJI96399.1 awen|OJJ39887.1 azon|OJJ50235.1 afum|EAL85683.1 afis|EAW15665.1 afla|EED46890.1 awen|OJJ38340.1 acla|EAW08989.1 aory|EIT72560.1

Group414: abra|OJJ68074.1 acar|OOF96032.1 acla|EAW11411.1 afis|EAW24010.1 afla|EED49151.1 afum|EAL87440.1 agla|OJJ86942.1 akaw|GAA92399.1 aluc|OJZ83260.1 anid|CBF77404.1 anig|EHA21114.1 aory|EIT79051.1 asyd|OJJ53240.1 ater|EAU29741.1 atub|OJI87679.1 aver|OJJ03610.1 awen|OJJ35482.1 azon|OJJ51266.1 aory|EIT78571.1 afla|EED50884.1 ater|EAU38050.1 asyd|OJJ59006.1 aver|OJJ01610.1 awen|OJJ37601.1

Group415: abra|OJJ68223.1 acla|EAW07100.1 afis|EAW20575.1 afum|EAL84490.1 akaw|GAA83221.1 aluc|OJZ83722.1 anig|EHA18872.1 atub|OJI90097.1 azon|OJJ50650.1 abra|OJJ76992.1 acar|OOF91437.1 afla|EED52310.1 agla|OJJ88718.1 akaw|GAA87054.1 aluc|OJZ84392.1 anid|CBF78937.1 anig|EHA27721.1 aory|EIT74487.1 asyd|OJJ55645.1 ater|EAU37109.1 atub|OJI89679.1 aver|OJJ06730.1 awen|OJJ33275.1 acla|EAW13463.1

Group418: abra|OJJ70549.1 afla|EED53198.1 agla|OJJ89090.1 akaw|GAA83393.1 aluc|OJZ83893.1 asyd|OJJ55203.1 atub|OJI90260.1 aver|OJJ07013.1 abra|OJJ70550.1 acar|OOF97171.1 acla|EAW08341.1 afis|EAW23376.1 afla|EED55581.1 afum|EAL88095.1 agla|OJJ86447.1 anid|CBF74843.1 anig|EHA19029.1 asyd|OJJ60130.1 ater|EAU35630.1 atub|OJI90261.1 aver|OJJ04131.1 awen|OJJ39507.1 azon|OJJ47706.1 aory|EIT74938.1

Group421: abra|OJJ71639.1 acar|OOF96356.1 acla|EAW15180.1 afis|EAW20150.1 afla|EED57200.1 afum|EAL85264.1 agla|OJJ88784.1 akaw|GAA89286.1 akaw|GAA89287.1 aluc|OJZ86382.1 anid|CBF84986.1 anig|EHA21512.1 aory|EIT73993.1 asyd|OJJ63125.1 ater|EAU38765.1 atub|OJI84151.1 aver|OJI97830.1 awen|OJJ32023.1 azon|OJJ46067.1 acla|EAW15179.1 afis|EAW20149.1 anid|CBF84984.1 aory|EIT74054.1 afum|EAL85265.1

Group422: abra|OJJ71666.1 acar|OOF96376.1 acla|EAW15203.1 acla|EAW15204.1 afis|EAW20168.1 afis|EAW20169.1 afla|EED57180.1 afum|EAL85286.1 afum|EAL85287.1 agla|OJJ88761.1 akaw|GAA89266.1 akaw|GAA89267.1 aluc|OJZ86408.1 anid|CBF85015.1 anid|CBF85017.1 anig|EHA21491.1 aory|EIT74089.1 asyd|OJJ63104.1 ater|EAU38800.1 atub|OJI84123.1 aver|OJI97857.1 awen|OJJ32065.1 awen|OJJ32066.1 azon|OJJ46098.1

Group426: abra|OJJ74271.1 acar|OOF95459.1 acla|EAW07315.1 afis|EAW22331.1 afla|EED57752.1 afum|EAL90861.1 agla|OJJ87711.1 akaw|GAA86752.1 aluc|OJZ89862.1 anid|CBF88649.1 anig|EHA27062.1 aory|EIT81123.1 asyd|OJJ63884.1 ater|EAU39460.1 atub|OJI84946.1 aver|OJI97056.1 awen|OJJ34909.1 azon|OJJ45576.1 abra|OJJ74564.1 acar|OOF95196.1 akaw|GAA82037.1 aluc|OJZ92145.1 anig|EHA26804.1 atub|OJI86574.1

Group427: abra|OJJ74337.1 acar|OOF95390.1 acla|EAW07388.1 afis|EAW22412.1 afla|EED52492.1 afum|EAL90780.1 agla|OJJ80127.1 akaw|GAA86692.1 aluc|OJZ89793.1 anid|CBF88798.1 anig|EHA27006.1 aory|EIT82746.1 asyd|OJJ64791.1 ater|EAU39293.1 atub|OJI85013.1 aver|OJI98903.1 awen|OJJ34993.1 azon|OJJ45636.1 afis|EAW15417.1 afla|EED50613.1 afum|EAL84228.1 aory|EIT78052.1 ater|EAU37467.1 awen|OJJ32510.1

Group429: abra|OJJ77163.1 acar|OOF93049.1 acla|EAW11721.1 afis|EAW23742.1 afla|EED45436.1 afum|EAL87179.1 agla|OJJ81702.1 akaw|GAA90769.1 aluc|OJZ84576.1 anid|CBF86622.1 anig|EHA27869.1 aory|EIT82382.1 asyd|OJJ65088.1 ater|EAU37973.1 atub|OJI89506.1 aver|OJI99214.1 awen|OJJ37083.1 azon|OJJ42895.1 anid|CBF87116.1 awen|OJJ35821.1 ater|EAU31855.1 asyd|OJJ64812.1 aver|OJI98925.1 anid|CBF71823.1

Group430: abra|OJJ65741.1 abra|OJJ71669.1 asyd|OJJ61538.1 asyd|OJJ63100.1 aver|OJI97862.1 aver|OJI98822.1 aver|OJJ00865.1 acar|OOF96381.1 acla|EAW15209.1 afis|EAW20174.1 afla|EED57174.1 afum|EAL85292.1 agla|OJJ88756.1 akaw|GAA89262.1 aluc|OJZ86412.1 anid|CBF85033.1 anig|EHA21487.1 aory|EIT74095.1 ater|EAU35210.1 atub|OJI84119.1 awen|OJJ41764.1 azon|OJJ46103.1 afla|EED54878.1

Group434: abra|OJJ67714.1 abra|OJJ68753.1 akaw|GAA89213.1 akaw|GAA89897.1 aluc|OJZ81616.1 aluc|OJZ81774.1 anig|EHA22442.1 anig|EHA25827.1 atub|OJI84347.1 atub|OJI88567.1 acar|OOF92242.1 acla|EAW12498.1 afis|EAW24147.1 afla|EED46784.1 afum|EAL92002.1 agla|OJJ87227.1 anid|CBF77921.1 aory|EIT72760.1 asyd|OJJ55295.1 ater|EAU30579.1 aver|OJJ06931.1 awen|OJJ41332.1 azon|OJJ48802.1

Group435: abra|OJJ70999.1 abra|OJJ75834.1 acar|OOF98141.1 acla|EAW10556.1 afis|EAW25023.1 afla|EED55445.1 afum|EAL89672.1 agla|OJJ89191.1 akaw|GAA82949.1 aluc|OJZ87528.1 anid|CBF81543.1 anig|EHA20738.1 aory|EIT80817.1 asyd|OJJ59491.1 ater|EAU35764.1 atub|OJI81120.1 aver|OJJ02783.1 awen|OJJ40617.1 azon|OJJ49465.1 akaw|GAA84544.1 aluc|OJZ89277.1 anig|EHA23292.1 atub|OJI85522.1

Group436: abra|OJJ73227.1 abra|OJJ77180.1 acar|OOF91798.1 acar|OOG00465.1 agla|OJJ82802.1 agla|OJJ84567.1 asyd|OJJ52864.1 asyd|OJJ59702.1 acla|EAW06901.1 afis|EAW16518.1 afla|EED56153.1 afum|EAL89113.1 akaw|GAA87953.1 aluc|OJZ81201.1 anid|CBF75551.1 anig|EHA18516.1 aory|EIT76710.1 ater|EAU36584.1 atub|OJI91396.1 aver|OJJ02585.1 aver|OJJ08843.1 awen|OJJ40305.1 azon|OJJ49166.1

Group438: abra|OJJ75609.1 abra|OJJ76068.1 akaw|GAA84359.1 akaw|GAA88899.1 aluc|OJZ89061.1 aluc|OJZ89522.1 anig|EHA23079.1 anig|EHA23477.1 atub|OJI85281.1 atub|OJI85725.1 acar|OOG00229.1 acla|EAW09371.1 afis|EAW21764.1 afla|EED46335.1 afum|EAL92794.1 agla|OJJ84387.1 anid|CBF83753.1 aory|EIT81195.1 asyd|OJJ60858.1 ater|EAU38451.1 aver|OJJ00182.1 awen|OJJ34047.1 azon|OJJ48184.1

Group441: acar|OOF94062.1 acar|OOF94132.1 abra|OJJ66364.1 acla|EAW08975.1 afis|EAW15454.1 afla|EED54752.1 afum|EAL84191.1 agla|OJJ80556.1 akaw|GAA90237.1 aluc|OJZ83103.1 anid|CBF80456.1 anig|EHA28167.1 aory|EIT78886.1 asyd|OJJ54768.1 ater|EAU32265.1 atub|OJI87832.1 aver|OJJ07399.1 awen|OJJ36431.1 azon|OJJ43598.1 atub|OJI80330.1 afla|EED48876.1 aory|EIT81202.1 awen|OJJ31672.1

Group442: acla|EAW10867.1 acla|EAW11076.1 afla|EED50345.1 afla|EED55539.1 afum|EAL84638.1 afum|EAL87765.1 aory|EIT77302.1 aory|EIT80884.1 asyd|OJJ54018.1 asyd|OJJ56874.1 abra|OJJ66629.1 acar|OOF91053.1 afis|EAW20991.1 agla|OJJ84069.1 akaw|GAA91096.1 aluc|OJZ81419.1 anid|CBF82616.1 anig|EHA26002.1 ater|EAU38640.1 atub|OJI88762.1 aver|OJJ08722.1 awen|OJJ34319.1 azon|OJJ48065.1

Group446: afla|EED44969.1 afla|EED46569.1 afla|EED51147.1 aory|EIT73475.1 aory|EIT77864.1 aory|EIT81352.1 asyd|OJJ56870.1 asyd|OJJ61208.1 abra|OJJ75570.1 acar|OOF90888.1 acla|EAW09352.1 afis|EAW21817.1 afum|EAL92833.1 agla|OJJ84325.1 akaw|GAA84321.1 aluc|OJZ89018.1 anid|CBF76652.1 anig|EHA23512.1 ater|EAU38486.1 atub|OJI85771.1 aver|OJJ00532.1 awen|OJJ34091.1 azon|OJJ44255.1

Group447: afla|EED47292.1 afla|EED54844.1 agla|OJJ87461.1 agla|OJJ89110.1 aory|EIT79178.1 aory|EIT79336.1 asyd|OJJ60317.1 asyd|OJJ60627.1 asyd|OJJ62712.1 abra|OJJ75953.1 acar|OOF99959.1 acla|EAW09843.1 afis|EAW21292.1 afum|EAL92309.1 akaw|GAA84656.1 aluc|OJZ89402.1 anid|CBF83376.1 anig|EHA23185.1 ater|EAU35925.1 atub|OJI85395.1 aver|OJI99951.1 awen|OJJ39407.1 azon|OJJ44290.1

Group448: afla|EED48892.1 afla|EED55381.1 aory|EIT80798.1 aory|EIT81174.1 asyd|OJJ56534.1 asyd|OJJ64421.1 aver|OJI98520.1 aver|OJJ06188.1 abra|OJJ70928.1 acar|OOF98198.1 acla|EAW10494.1 afis|EAW25073.1 afum|EAL89742.1 agla|OJJ89119.1 akaw|GAA83014.1 aluc|OJZ87599.1 anid|CBF85962.1 anig|EHA20799.1 ater|EAU39552.1 atub|OJI81048.1 awen|OJJ40684.1 azon|OJJ44061.1 awen|OJJ34526.1

Group449: afla|EED54572.1 afla|EED56566.1 asyd|OJJ53770.1 asyd|OJJ62761.1 ater|EAU30266.1 ater|EAU39167.1 aver|OJI96804.1 aver|OJJ07878.1 abra|OJJ73591.1 acar|OOG00775.1 acla|EAW07510.1

afis|EAW22541.1 afum|EAL90653.1 agla|OJJ80900.1 akaw|GAA88561.1 aluc|OJZ88459.1 anid|CBF89038.1 anig|EHA18179.1 aory|EIT78798.1 aory|EIT79646.1 atub|OJI91057.1 awen|OJJ35170.1 azon|OJJ49937.1

Group452: akaw|GAA83965.1 akaw|GAA90739.1 aluc|OJZ80854.1 aluc|OJZ84609.1 abra|OJJ77200.1 acar|OOF93088.1 acla|EAW13191.1 afis|EAW17450.1 afla|EED49372.1 afum|EAL91278.2 agla|OJJ80362.1 akaw|GAA90738.1 aluc|OJZ84610.1 anid|CBF71683.1 anig|EHA27899.1 aory|EIT79587.1 asyd|OJJ61822.1 ater|EAU33930.1 atub|OJI89471.1 aver|OJI95922.1 awen|OJJ37396.1 azon|OJJ51068.1 anig|EHA27898.1

Group455: aory|EIT75378.1 aory|EIT78539.1 awen|OJJ34056.1 awen|OJJ35494.1 azon|OJJ46995.1 azon|OJJ50589.1 abra|OJJ72842.1 acar|OOF96043.1 acla|EAW11400.1 afis|EAW24021.1 afla|EED45752.1 afla|EED49134.1 afla|EED49135.1 afum|EAL87451.1 agla|OJJ86954.1 akaw|GAA85444.1 aluc|OJZ83350.1 anid|CBF77424.1 anig|EHA24217.1 asyd|OJJ53230.1 ater|EAU29725.1 atub|OJI82243.1 aver|OJJ03622.1

Group456: asyd|OJJ51978.1 asyd|OJJ52250.1 asyd|OJJ56074.1 abra|OJJ76969.1 acar|OOF91414.1 acla|EAW13273.1 acla|EAW13494.1 afis|EAW20594.1 afla|EED47642.1 afum|EAL84467.1 agla|OJJ88662.1 akaw|GAA87075.1 aluc|OJZ84365.1 anid|CBF78279.1 anig|EHA27700.1 aory|EIT80239.1 ater|EAU35240.1 atub|OJI89703.1 aver|OJJ05017.1 awen|OJJ33318.1 azon|OJJ50690.1 afis|EAW17345.1 afum|EAL91189.2

Group461: ater|EAU29737.1 ater|EAU36828.1 abra|OJJ68505.1 acar|OOF90180.1 acar|OOF91789.1 acla|EAW11100.1 afis|EAW20763.1 afla|EED53200.1 afum|EAL84856.1 agla|OJJ84006.1 akaw|GAA92768.1 aluc|OJZ82717.1 anid|CBF80354.1 anig|EHA21171.1 aory|EIT75122.1 asyd|OJJ54828.1 atub|OJI88951.1 aver|OJJ07333.1 awen|OJJ29787.1 azon|OJJ42797.1 asyd|OJJ58563.1 aver|OJJ01162.1 azon|OJJ42773.1

Group467: abra|OJJ66772.1 acar|OOF93853.1 acla|EAW08992.1 afis|EAW15416.1 afla|EED48724.1 afum|EAL84229.2 agla|OJJ79941.1 anid|CBF81992.1 anig|EHA27337.1 aory|EIT73206.1 asyd|OJJ56162.1 ater|EAU32712.1 aver|OJJ05101.1 awen|OJJ39993.1 azon|OJJ42917.1 abra|OJJ75745.1 akaw|GAA84478.1 aluc|OJZ89198.1 anig|EHA23360.1 atub|OJI85592.1 afla|EED55174.1 aory|EIT81585.1 awen|OJJ36897.1

Group471: abra|OJJ67925.1 acar|OOF92776.1 afis|EAW17948.1 afum|EAL91758.1 agla|OJJ89340.1 akaw|GAA91272.1 aluc|OJZ82797.1 anid|CBF80065.1 anig|EHA21238.1 ater|EAU31422.1 atub|OJI89028.1 aver|OJJ02091.1 awen|OJJ41595.1 azon|OJJ48310.1 abra|OJJ74125.1 acla|EAW14963.1 afla|EED48217.1 akaw|GAA89760.1 aluc|OJZ80623.1 anig|EHA27208.1 aory|EIT72959.1 asyd|OJJ58320.1 awen|OJJ31633.1

Group473: awen|OJJ41504.1 abra|OJJ71797.1 acar|OOF96477.1 acla|EAW13728.1 afis|EAW18033.1 afum|EAL91844.1 agla|OJJ85408.1 akaw|GAA85113.1 aluc|OJZ86527.1 anid|CBF81681.1 anig|EHA22119.1 aory|EIT81720.1 asyd|OJJ53625.1 asyd|OJJ58292.1 atub|OJI83995.1 aver|OJJ05942.1 awen|OJJ31297.1 azon|OJJ50337.1 acar|OOF92709.1 afla|EED51862.1 anid|CBF80076.1 ater|EAU31550.1 aver|OJJ02132.1

Group476: abra|OJJ68512.1 akaw|GAA91334.1 aluc|OJZ82727.1 anig|EHA21180.1 atub|OJI88960.1 abra|OJJ69632.1 acar|OOF92677.1 acla|EAW10562.1 afis|EAW25029.1 afla|EED55452.1 afum|EAL89666.1 agla|OJJ79155.1 akaw|GAA91612.1 aluc|OJZ82439.1 anid|CBF81553.1 anig|EHA28334.1 aory|EIT80876.1 asyd|OJJ59496.1 ater|EAU35758.1 atub|OJI88002.1 aver|OJJ02778.1 awen|OJJ40610.1 azon|OJJ49459.1

Group478: abra|OJJ72302.1 acar|OOF94922.1 acla|EAW10039.1 afis|EAW24476.1 afla|EED56937.1 afum|EAL86149.1 agla|OJJ86412.1 akaw|GAA86053.1 aluc|OJZ90283.1 anid|CBF78363.1 anig|EHA23773.1 aory|EIT77524.1 asyd|OJJ57323.1 ater|EAU29927.1 atub|OJI81707.1 aver|OJJ05837.1 awen|OJJ40788.1 acla|EAW12030.1 afis|EAW24671.1 afum|EAL89324.1 anig|EHA28157.1 azon|OJJ48190.1 awen|OJJ34571.1

Group480: abra|OJJ73797.1 acar|OOG00952.1 acla|EAW07688.1 afis|EAW22703.1 afla|EED49989.1 afum|EAL90489.1 agla|OJJ79006.1 akaw|GAA89415.1 aluc|OJZ88258.1 anid|CBF88074.1 anig|EHA18003.1 aory|EIT73485.1 asyd|OJJ63555.1 ater|EAU38999.1 atub|OJI90857.1 aver|OJI97379.1 awen|OJJ35340.1 azon|OJJ49778.1 aory|EIT74604.1 afis|EAW21081.1 afum|EAL92102.1 agla|OJJ79284.1 afla|EED44920.1

Group481: abra|OJJ74536.1 acar|OOF96722.1 afla|EED46724.1 akaw|GAA82065.1 aluc|OJZ92117.1 anig|EHA26827.1 aory|EIT77932.1 asyd|OJJ61662.1 ater|EAU31119.1 atub|OJI86550.1 abra|OJJ78387.1 acar|OOF98501.1 acla|EAW09146.1 afis|EAW22000.1 afum|EAL93015.1 agla|OJJ84097.1 akaw|GAA85645.1 aluc|OJZ91466.1 anid|CBF82670.1 atub|OJI83459.1 aver|OJJ00983.1 awen|OJJ34296.1 azon|OJJ43342.1

Group484: abra|OJJ76940.1 acar|OOF91390.1 acla|EAW09287.1 afis|EAW21860.1 afla|EED45974.1 afum|EAL92873.1 agla|OJJ79533.1 akaw|GAA87099.1 aluc|OJZ84339.1 anid|CBF89674.1 anig|EHA27673.1 aory|EIT75604.1 asyd|OJJ62270.1 ater|EAU29774.1 atub|OJI89729.1 aver|OJI96379.1 awen|OJJ34144.1 azon|OJJ50245.1 afla|EED46450.1 agla|OJJ84275.1 aory|EIT72411.1 ater|EAU30859.1 azon|OJJ43482.1

Group486: abra|OJJ78343.1 acar|OOF98537.1 acla|EAW13688.1 afis|EAW19912.1 afla|EED47242.1 afum|EAL84995.1 agla|OJJ84607.1 akaw|GAA85690.1 aluc|OJZ91415.1 anid|CBF87843.1 anig|EHA20088.1 aory|EIT79169.1 asyd|OJJ61361.1 ater|EAU31064.1 atub|OJI83410.1 aver|OJJ00688.1 awen|OJJ33622.1 azon|OJJ43388.1 afla|EED46677.1 anid|CBF84039.1 aory|EIT77968.1 awen|OJJ34250.1 agla|OJJ88201.1

Group489: abra|OJJ68479.1 abra|OJJ69702.1 akaw|GAA88821.1 akaw|GAA92402.1 aluc|OJZ82505.1 aluc|OJZ83263.1 atub|OJI87674.1 atub|OJI87926.1 acar|OOF92670.1 acla|EAW15184.1 afis|EAW20154.1 afla|EED57195.1 afum|EAL85269.1 agla|OJJ88775.1 anid|CBF84996.1 anig|EHA28265.1 aory|EIT74166.1 asyd|OJJ63119.1 ater|EAU38769.1 aver|OJI97839.1 awen|OJJ32030.1 azon|OJJ46082.1

Group490: abra|OJJ69764.1 abra|OJJ76049.1 acla|EAW08772.1 acla|EAW09780.1 aluc|OJZ85534.1 aluc|OJZ89505.1 ater|EAU29246.1 ater|EAU35201.1 acar|OOF99880.1 afis|EAW21358.1 afla|EED55773.1 afum|EAL92373.1 agla|OJJ85477.1 akaw|GAA88916.1 anid|CBF76263.1 anig|EHA23095.1 aory|EIT76750.1 asyd|OJJ60971.1 atub|OJI85298.1 aver|OJJ00296.1 awen|OJJ39340.1 azon|OJJ51686.1

Group492: acar|OOF93735.1 acar|OOF93823.1 abra|OJJ66740.1 acla|EAW07107.1 afis|EAW19478.1 afla|EED56686.1 afum|EAL87018.1 agla|OJJ78581.1 akaw|GAA89678.1 aluc|OJZ80708.1 anid|CBF69805.1 anig|EHA27309.1 aory|EIT73937.1 asyd|OJJ58969.1 ater|EAU30765.1 atub|OJI87566.1 aver|OJJ01564.1 awen|OJJ40135.1 azon|OJJ45831.1 afis|EAW22119.1 agla|OJJ81206.1 afum|EAL91067.1

Group496: acla|EAW12269.1 acla|EAW15163.1 agla|OJJ78609.1 agla|OJJ88803.1 ater|EAU31506.1 ater|EAU38748.1 aver|OJI97813.1 aver|OJI98043.1 abra|OJJ71622.1 acar|OOF96339.1 afis|EAW20132.1 afla|EED57220.1 afum|EAL85248.1 akaw|GAA89303.1 aluc|OJZ86365.1 anid|CBF84960.1 anig|EHA21527.1 aory|EIT74164.1 asyd|OJJ63141.1 atub|OJI84169.1 awen|OJJ31998.1 azon|OJJ50773.1

Group500: afis|EAW18163.1 afis|EAW21088.1 afum|EAL87523.1 afum|EAL92111.1 abra|OJJ77694.1 acar|OOF99045.1 acla|EAW11332.1 afla|EED56330.1 agla|OJJ85962.1 akaw|GAA83936.1 aluc|OJZ90757.1 anid|CBF85753.1 anig|EHA19494.1 aory|EIT73367.1 asyd|OJJ64297.1 ater|EAU33690.1 atub|OJI82744.1 aver|OJI98390.1 awen|OJJ30508.1 azon|OJJ50317.1 acla|EAW10845.1 acla|EAW13632.1

Group501: afla|EED45701.1 afla|EED55177.1 aory|EIT75733.1 aory|EIT81570.1 ater|EAU31908.1 ater|EAU32716.1 azon|OJJ50388.1 azon|OJJ50758.1 abra|OJJ68251.1 acar|OOF97530.1 acla|EAW09063.1 afis|EAW25635.1 afum|EAL84323.1 agla|OJJ78897.1 akaw|GAA83210.1 aluc|OJZ83712.1 anid|CBF83153.1 anig|EHA18863.1 asyd|OJJ55515.1 atub|OJI90087.1 aver|OJJ06592.1 awen|OJJ36092.1

Group502: afla|EED49422.1 afla|EED56716.1 afla|EED53576.1 aory|EIT73869.1 aory|EIT74843.1 aory|EIT79474.1 abra|OJJ66759.1 acar|OOF93842.1 acla|EAW11885.1 afis|EAW19497.1 afum|EAL86996.1 agla|OJJ81661.1 akaw|GAA89696.1 aluc|OJZ80689.1 anid|CBF75434.1 anig|EHA27326.1 asyd|OJJ59785.1 ater|EAU34809.1 atub|OJI87548.1 aver|OJJ02503.1 awen|OJJ36914.1 azon|OJJ42897.1

Group504: afla|EED52027.1 afla|EED55465.1 aory|EIT80878.1 aory|EIT81484.1 awen|OJJ35115.1 awen|OJJ40599.1 abra|OJJ77085.1 acar|OOF92969.1 acla|EAW10573.1 afis|EAW24994.1 afum|EAL89655.1 agla|OJJ82261.1 akaw|GAA86970.1 aluc|OJZ84494.1 anid|CBF81588.1 anig|EHA27797.1 asyd|OJJ59513.1 ater|EAU35745.1 atub|OJI89586.1 aver|OJJ02760.1 azon|OJJ49449.1 azon|OJJ49978.1

Group509: agla|OJJ79128.1 agla|OJJ84540.1 asyd|OJJ53810.1 asyd|OJJ58024.1 aver|OJJ04376.1 aver|OJJ07911.1 awen|OJJ31498.1 awen|OJJ36278.1 abra|OJJ68972.1 acar|OOF91549.1 acla|EAW14215.1 afis|EAW19066.1 afla|EED54432.1 afum|EAL93795.1 akaw|GAA90499.1 aluc|OJZ85961.1 anid|CBF79650.1 anig|EHA24642.1 aory|EIT75890.1 ater|EAU32114.1 atub|OJI79748.1 azon|OJJ46211.1

Group510: agla|OJJ81570.1 agla|OJJ84260.1 asyd|OJJ54207.1 asyd|OJJ61270.1 awen|OJJ30146.1 awen|OJJ34161.1 abra|OJJ78235.1 acar|OOF98625.1 acla|EAW09275.1 afis|EAW21872.1 afla|EED46436.1 afum|EAL92886.1 akaw|GAA85790.1 aluc|OJZ91309.1 anid|CBF76772.1 anig|EHA19993.1 aory|EIT72312.1 ater|EAU30870.1 atub|OJI83302.1 aver|OJJ00599.1 aver|OJJ01907.1 azon|OJJ43468.1

Group511: agla|OJJ82328.1 agla|OJJ89180.1 ater|EAU35772.1 ater|EAU39458.1 atub|OJI81101.1 atub|OJI87185.1 awen|OJJ30919.1 awen|OJJ40626.1 abra|OJJ70986.1 acar|OOF98145.1 acla|EAW10546.1 afis|EAW25014.1 afla|EED55435.1 afum|EAL89682.1 akaw|GAA82963.1 aluc|OJZ87544.1 anid|CBF81525.1 anig|EHA20748.1 aory|EIT80926.1 asyd|OJJ59480.1 aver|OJJ02794.1 azon|OJJ49057.1

Group512: agla|OJJ83608.1 agla|OJJ87442.1 anig|EHA21812.1 anig|EHA23178.1 ater|EAU35930.1 ater|EAU39415.1 awen|OJJ36034.1 awen|OJJ39401.1 abra|OJJ75961.1 acar|OOF99954.1 acla|EAW09837.1 afis|EAW21297.1 afla|EED54848.1 afum|EAL92314.1 akaw|GAA84661.1 aluc|OJZ89409.1 anid|CBF83385.1 aory|EIT79420.1 asyd|OJJ60632.1 atub|OJI85387.1 aver|OJI99955.1 azon|OJJ44285.1

Group513: akaw|GAA83114.1 akaw|GAA92362.1 aluc|OJZ79905.1 aluc|OJZ81127.1 aluc|OJZ88669.1 aluc|OJZ88850.1 abra|OJJ73371.1 acar|OOG00577.1 acla|EAW06992.1 afis|EAW16606.1 afla|EED45025.1 afum|EAL89203.1 agla|OJJ82647.1 anid|CBF81913.1 anig|EHA18352.1 aory|EIT73821.1 asyd|OJJ59694.1 ater|EAU36688.1 atub|OJI91251.1 aver|OJJ02593.1 awen|OJJ40200.1 azon|OJJ49067.1

Group515: asyd|OJJ54189.1 asyd|OJJ55024.1 ater|EAU32854.1 ater|EAU36887.1 aver|OJJ07182.1 aver|OJJ08550.1 abra|OJJ69557.1 acar|OOF91593.1 acla|EAW12881.1 afis|EAW17796.1 afla|EED57033.1 afum|EAL91604.2 agla|OJJ89112.1 akaw|GAA91566.1 aluc|OJZ82384.1 anid|CBF71395.1 anig|EHA28379.1 aory|EIT82249.1 atub|OJI88056.1 awen|OJJ41608.1 azon|OJJ48327.1 agla|OJJ84736.1

Group517: asyd|OJJ55069.1 asyd|OJJ55250.1 ater|EAU31464.1 ater|EAU32149.1 aver|OJJ06973.1 aver|OJJ07142.1 abra|OJJ77233.1 acar|OOF93115.1 acla|EAW12666.1 afis|EAW17830.1 afla|EED46172.1 afum|EAL91640.1 agla|OJJ80651.1 akaw|GAA90709.1 aluc|OJZ84643.1 anid|CBF77996.1 anig|EHA27929.1 aory|EIT74935.1 atub|OJI89438.1 awen|OJJ41759.1 azon|OJJ48451.1 afla|EED53219.1

Group526: acla|EAW15302.1 anid|CBF84723.1 abra|OJJ76196.1 acar|OOF99748.1 afla|EED52713.1 agla|OJJ81243.1 akaw|GAA82138.1 aluc|OJZ92033.1 anig|EHA22966.1 aory|EIT78196.1 ater|EAU38283.1

atub|OJI86466.1 awen|OJJ32095.1 azon|OJJ46673.1 afis|EAW20227.1 afum|EAL85344.1 aver|OJJ08433.1 afis|EAW25742.1 afla|EED46022.1 aory|EIT75659.1 asyd|OJJ53309.1 awen|OJJ30032.1

Group539: abra|OJJ69330.1 acar|OOF94397.1 acla|EAW14710.1 afis|EAW24245.1 afla|EED48029.1 afum|EAL85913.2 agla|OJJ79629.1 akaw|GAA88201.1 aluc|OJZ82162.1 anid|CBF86924.1 anig|EHA28570.1 aory|EIT83325.1 asyd|OJJ55608.1 ater|EAU32738.1 atub|OJI88275.1 aver|OJJ06683.1 awen|OJJ38017.1 azon|OJJ51787.1 asyd|OJJ53731.1 afis|EAW23670.1 afum|EAL87115.1 agla|OJJ81340.1

Group542: abra|OJJ70674.1 acar|OOF97053.1 acla|EAW08090.1 afis|EAW23125.1 afla|EED48525.1 afum|EAL88355.1 agla|OJJ87945.1 akaw|GAA90006.1 aluc|OJZ84022.1 anid|CBF74410.1 anig|EHA19143.1 aory|EIT73109.1 asyd|OJJ57761.1 ater|EAU36253.1 atub|OJI90393.1 aver|OJJ05405.1 awen|OJJ36655.1 azon|OJJ47551.1 awen|OJJ33435.1 afla|EED47672.1 aory|EIT80214.1 ater|EAU37239.1

Group543: abra|OJJ71147.1 acar|OOF98008.1 acla|EAW10357.1 afis|EAW25237.1 afla|EED53704.1 afum|EAL89898.1 agla|OJJ89205.1 akaw|GAA82830.1 aluc|OJZ87391.1 anid|CBF85312.1 anig|EHA20620.1 aory|EIT75340.1 asyd|OJJ64005.1 ater|EAU34361.1 atub|OJI81256.1 aver|OJI98093.1 awen|OJJ40948.1 azon|OJJ44541.1 afis|EAW25236.1 afum|EAL89897.1 awen|OJJ40953.1 azon|OJJ44540.1

Group544: abra|OJJ72457.1 acar|OOF94781.1 acla|EAW12689.1 afis|EAW17853.1 afla|EED46207.1 afum|EAL91661.1 agla|OJJ83117.1 akaw|GAA86193.1 aluc|OJZ90443.1 anid|CBF77956.1 anig|EHA23907.1 aory|EIT76431.1 asyd|OJJ61702.1 ater|EAU33055.1 atub|OJI81872.1 aver|OJI95791.1 awen|OJJ41720.1 azon|OJJ48418.1 afla|EED45133.1 aory|EIT73615.1 awen|OJJ30412.1 agla|OJJ86835.1

Group545: abra|OJJ73099.1 acar|OOG00332.1 acla|EAW07049.1 afis|EAW16675.1 afla|EED49659.1 afum|EBA27373.1 agla|OJJ78688.1 akaw|GAA88064.1 aluc|OJZ81327.1 anid|CBF89880.1 anig|EHA18631.1 aory|EIT80967.1 asyd|OJJ54564.1 ater|EAU33734.1 atub|OJI91530.1 aver|OJJ07590.1 awen|OJJ40031.1 azon|OJJ50640.1 acla|EAW08376.1 afis|EAW23405.1 afum|EAL88065.2 ater|EAU37518.1

Group547: abra|OJJ76094.1 acar|OOF99830.1 acar|OOF99831.1 acla|EAW09725.1 afis|EAW21416.1 afla|EED55842.1 afum|EAL92431.1 agla|OJJ85546.1 akaw|GAA88872.1 akaw|GAA88873.1 aluc|OJZ89550.1 anid|CBF83794.1 anig|EHA23056.1 aory|EIT76615.1 asyd|OJJ60885.1 ater|EAU35132.1 atub|OJI85255.1 atub|OJI85256.1 aver|OJJ00209.1 awen|OJJ39286.1 azon|OJJ51568.1 aory|EIT76662.1

Group548: abra|OJJ76170.1 acar|OOF99769.1 acla|EAW13698.1 afis|EAW21484.1 afla|EED55916.1 afum|EAL92504.1 agla|OJJ85638.1 akaw|GAA82116.1 aluc|OJZ92060.1 anid|CBF76494.1 anig|EHA22989.1 aory|EIT76674.1 asyd|OJJ61126.1 ater|EAU35054.1 atub|OJI86492.1 aver|OJJ00447.1 awen|OJJ39213.1 azon|OJJ51641.1 acla|EAW09655.1 afla|EED49764.1 aory|EIT77609.1 awen|OJJ29776.1

Group550: abra|OJJ68047.1 abra|OJJ70802.1 acar|OOF96908.1 acar|OOF96943.1 aory|EIT72863.1 aory|EIT81579.1 acla|EAW11122.1 afis|EAW15851.1 afla|EED56522.1 afum|EAL88487.1 agla|OJJ87257.1 akaw|GAA90118.1 aluc|OJZ84148.1 anid|CBF81176.1 anig|EHA21341.1 asyd|OJJ59292.1 ater|EAU31666.1 atub|OJI90520.1 aver|OJJ02983.1 awen|OJJ36776.1 azon|OJJ47422.1

Group552: abra|OJJ72113.1 abra|OJJ72712.1 anig|EHA21773.1 anig|EHA24109.1 atub|OJI82107.1 atub|OJI83652.1 acar|OOF96753.1 acla|EAW14914.1 afis|EAW24435.1 afla|EED55246.1 afum|EAL86103.1 agla|OJJ79250.1 akaw|GAA84809.1 aluc|OJZ86868.1 anid|CBF73765.1 aory|EIT83588.1 asyd|OJJ56590.1 ater|EAU30011.1 aver|OJJ06135.1 awen|OJJ32764.1 azon|OJJ43998.1

Group554: acar|OOF91686.1 acar|OOF91797.1 ater|EAU30834.1 ater|EAU37279.1 awen|OJJ37554.1 awen|OJJ37864.1 abra|OJJ66921.1 acla|EAW14487.1 afis|EAW19196.1 afla|EED47712.1 afum|EAL93928.1 agla|OJJ88511.1 akaw|GAA91498.1 aluc|OJZ85659.1 anid|CBF78550.1 anig|EHA24874.1 aory|EIT80197.1 asyd|OJJ65057.1 atub|OJI79452.1 aver|OJI99182.1 azon|OJJ51215.1

Group555: acar|OOF93691.1 acar|OOF93775.1 abra|OJJ66681.1 acla|EAW11818.1 afis|EAW19429.1 afla|EED56633.1 afum|EAL87065.1 agla|OJJ81141.1 akaw|GAA89626.1 aluc|OJZ80768.1 anid|CBF74384.1 anig|EHA27260.1 aory|EIT79639.1 asyd|OJJ57774.1 ater|EAU34732.1 atub|OJI87621.1 awen|OJJ36828.1 azon|OJJ42976.1 aver|OJJ05392.1 aver|OJJ05391.1 awen|OJJ36827.1

Group559: acla|EAW09341.1 acla|EAW11685.1 afis|EAW21796.1 afis|EAW23700.1 afum|EAL87141.1 afum|EAL92821.1 abra|OJJ75576.1 acar|OOF90894.1 afla|EED46599.1 agla|OJJ84330.1 akaw|GAA84328.1 aluc|OJZ89025.1 anid|CBF76627.1 anig|EHA23507.1 aory|EIT77837.1 asyd|OJJ61195.1 ater|EAU38478.1 atub|OJI85765.1 aver|OJJ00519.1 awen|OJJ34081.1 azon|OJJ44259.1

Group560: afis|EAW16155.1 afis|EAW23200.1 afum|EAL88027.1 afum|EAL88283.1 ater|EAU36328.1 ater|EAU36668.1 abra|OJJ70513.1 acar|OOF97416.1 acla|EAW08162.1 afla|EED47054.1 agla|OJJ78748.1 akaw|GAA83360.1 aluc|OJZ83858.1 anid|CBF74719.1 anig|EHA18999.1 aory|EIT72491.1 asyd|OJJ60202.1 atub|OJI90225.1 aver|OJJ04064.1 awen|OJJ39430.1 azon|OJJ47634.1

Group561: afis|EAW17104.1 afis|EAW25308.1 agla|OJJ88961.1 agla|OJJ89287.1 abra|OJJ71251.1 acar|OOF97919.1 acla|EAW10284.1 afum|EAL89971.1 akaw|GAA82736.1 aluc|OJZ87291.1 anid|CBF75291.1 anig|EHA20532.1 aory|EIT78474.1 asyd|OJJ59887.1 ater|EAU34456.1 atub|OJI81350.1 aver|OJJ02399.1 awen|OJJ41022.1 azon|OJJ44609.1 asyd|OJJ65171.1 afla|EED53785.1

Group563: afis|EAW22109.1 afis|EAW22582.1 afum|EAL90617.1 afum|EAL91075.1 abra|OJJ73636.1 acar|OOG00816.1 acla|EAW07548.1 afla|EED54627.1 agla|OJJ80942.1 akaw|GAA88519.1 aluc|OJZ88414.1 anid|CBF88345.1 anig|EHA18141.1 aory|EIT78908.1 asyd|OJJ63718.1 ater|EAU39122.1 atub|OJI91010.1 aver|OJI97222.1 awen|OJJ35210.1 azon|OJJ49899.1 anid|CBF89563.1

Group566: afla|EED45981.1 afla|EED55382.1 aory|EIT75435.1 aory|EIT80896.1 ater|EAU33990.1 ater|EAU39553.1 abra|OJJ70929.1 acar|OOF98196.1 acla|EAW10495.1 afis|EAW25072.1 afum|EAL89741.1 agla|OJJ89122.1 akaw|GAA83013.1 aluc|OJZ87598.1 anid|CBF85960.1 anig|EHA20798.1 asyd|OJJ64420.1 atub|OJI81049.1 aver|OJI98519.1 awen|OJJ40683.1 azon|OJJ44060.1

Group567: afla|EED48911.1 afla|EED51260.1 aory|EIT81176.1 aory|EIT81506.1 awen|OJJ29671.1 awen|OJJ38491.1 abra|OJJ69199.1 acar|OOF91116.1 acla|EAW08647.1 afis|EAW15784.1 afum|EAL85566.1 agla|OJJ81947.1 akaw|GAA92212.1 aluc|OJZ86184.1 anid|CBF70950.1 anig|EHA24440.1 asyd|OJJ52622.1 ater|EAU32357.1 atub|OJI79976.1 aver|OJJ04649.1 azon|OJJ43792.1

Group568: afla|EED51201.1 afla|EED53566.1 asyd|OJJ62648.1 asyd|OJJ65376.1 aver|OJI96748.1 aver|OJI99544.1 abra|OJJ73167.1 acar|OOG00407.1 acla|EAW06285.1 afis|EAW22140.1 afum|EAL91050.1 agla|OJJ87935.1 akaw|GAA88004.1 aluc|OJZ81260.1 anid|CBF89140.1 anig|EHA18567.1 aory|EIT75205.1 ater|EAU34142.1 atub|OJI91456.1 awen|OJJ34714.1 azon|OJJ50465.1

Group570: agla|OJJ78645.1 agla|OJJ81233.1 agla|OJJ83403.1 abra|OJJ71870.1 acar|OOF96539.1 acla|EAW13900.1 afis|EAW18612.1 afla|EED52733.1 afum|EAL93322.1 akaw|GAA85045.1 aluc|OJZ86609.1 anid|CBF70422.1 anig|EHA22057.1 aory|EIT78158.1 asyd|OJJ58619.1 ater|EAU34909.1 atub|OJI83917.1 aver|OJJ01226.1 awen|OJJ32073.1 azon|OJJ46654.1 agla|OJJ81232.1

Group573: akaw|GAA82012.1 akaw|GAA83300.1 anig|EHA23471.1 anig|EHA26778.1 azon|OJJ46821.1 azon|OJJ49225.1 abra|OJJ74596.1 acar|OOF95173.1 acla|EAW10785.1 afis|EAW24793.1 afla|EED47193.1 afum|EAL89435.1 agla|OJJ82887.1 aluc|OJZ92171.1 anid|CBF81462.1 aory|EIT79250.1 asyd|OJJ59441.1 ater|EAU36515.1 atub|OJI86603.1 aver|OJJ02832.1 awen|OJJ40374.1

Group574: aluc|OJZ79634.1 aluc|OJZ79637.1 aluc|OJZ79681.1 aluc|OJZ90921.1 abra|OJJ77846.1 acar|OOF98926.1 acla|EAW11277.1 afis|EAW18223.1 afla|EED56246.1 afum|EAL87589.2 agla|OJJ85859.1 akaw|GAA83801.1 anid|CBF86151.1 anig|EHA19633.1 aory|EIT73368.1 asyd|OJJ64517.1 ater|EAU33768.1 atub|OJI82913.1 aver|OJI98615.1 awen|OJJ30601.1 azon|OJJ50167.1

Group576: aluc|OJZ80315.1 aluc|OJZ86416.1 anid|CBF76119.1 anid|CBF85040.1 atub|OJI80382.1 atub|OJI84115.1 abra|OJJ71674.1 acar|OOF96385.1 acla|EAW15213.1 afis|EAW20178.1 afla|EED57170.1 afum|EAL85296.1 agla|OJJ84688.1 akaw|GAA89258.1 anig|EHA21483.1 aory|EIT74051.1 asyd|OJJ63096.1 ater|EAU35214.1 aver|OJI97866.1 awen|OJJ31177.1 azon|OJJ46108.1

Group580: anid|CBF71344.1 anid|CBF71770.1 asyd|OJJ55097.1 asyd|OJJ55529.1 aver|OJJ07109.1 aver|OJJ07313.1 abra|OJJ72319.1 acar|OOF94892.1 acla|EAW12001.1 afis|EAW19641.1 afla|EED56992.1 afum|EAL86881.1 agla|OJJ80783.1 akaw|GAA86071.1 aluc|OJZ90306.1 anig|EHA23792.1 aory|EIT77578.1 ater|EAU32888.1 atub|OJI81730.1 awen|OJJ41886.1 azon|OJJ50727.1

Group584: asyd|OJJ53054.1 asyd|OJJ62331.1 aver|OJI96437.1 aver|OJJ03790.1 abra|OJJ67212.1 acar|OOF98347.1 acla|EAW08554.1 afis|EAW23596.1 afla|EED55272.1 afum|EAL87871.1 agla|OJJ86856.1 akaw|GAA90965.1 aluc|OJZ87761.1 anid|CBF89583.1 anig|EHA20246.1 aory|EIT83575.1 ater|EAU35356.1 atub|OJI80883.1 awen|OJJ39918.1 azon|OJJ45892.1 anid|CBF77738.1

Group585: asyd|OJJ53773.1 asyd|OJJ64196.1 aver|OJI98285.1 aver|OJJ01986.1 abra|OJJ71111.1 acar|OOF98036.1 acla|EAW10381.1 afis|EAW25212.1 afla|EED53666.1 afum|EAL89873.1 agla|OJJ88949.1 akaw|GAA82858.1 aluc|OJZ87423.1 anid|CBF85604.1 anig|EHA19933.1 anig|EHA20646.1 aory|EIT75278.1 ater|EAU34369.1 atub|OJI81225.1 awen|OJJ40874.1 azon|OJJ46813.1

Group592: abra|OJJ65854.1 acar|OOF95015.1 acla|EAW13528.1 afis|EAW20729.1 afla|EED48906.1 afum|EAL84879.1 agla|OJJ84616.1 akaw|GAA92290.1 aluc|OJZ90187.1 anid|CBF85584.1 anig|EHA23686.1 aory|EIT81194.1 asyd|OJJ64174.1 atub|OJI81614.1 aver|OJJ06463.1 awen|OJJ29911.1 azon|OJJ42620.1 afla|EED55134.1 aory|EIT81587.1 ater|EAU32788.1 aver|OJI98249.1

Group594: abra|OJJ66784.1 afla|EED47234.1 anid|CBF87107.1 aory|EIT79229.1 abra|OJJ75747.1 acar|OOG00125.1 acla|EAW09092.1 afla|EED52418.1 agla|OJJ82218.1 akaw|GAA84480.1 aluc|OJZ89200.1 anig|EHA23357.1 aory|EIT83316.1 asyd|OJJ56738.1 ater|EAU31807.1 atub|OJI85590.1 aver|OJJ05987.1 awen|OJJ41471.1 azon|OJJ44445.1 afum|EAL84355.1 afis|EAW25686.1

Group600: abra|OJJ69337.1 acar|OOF94389.1 acla|EAW06601.1 afis|EAW16003.1 afla|EED48032.1 afum|EAL88635.2 agla|OJJ79631.1 akaw|GAA88207.1 aluc|OJZ82169.1 anid|CBF85846.1 anig|EHA28563.1 aory|EIT83338.1 asyd|OJJ53958.1 ater|EAU31185.1 atub|OJI88268.1 aver|OJJ08054.1 awen|OJJ33053.1 azon|OJJ48464.1 asyd|OJJ64357.1 ater|EAU33640.1 aver|OJI98453.1

Group605: abra|OJJ70312.1 acar|OOF93014.1 acla|EAW13439.1 afis|EAW20553.1 afla|EED45906.1 afum|EAL84508.1 agla|OJJ80823.1 akaw|GAA93011.1 aluc|OJZ82574.1 anid|CBF78981.1 anig|EHA27499.1

aory|EIT75600.1 asyd|OJJ55678.1 ater|EAU37124.1 atub|OJI88818.1 aver|OJJ06761.1 awen|OJJ41921.1 azon|OJJ42685.1 anig|EHA23882.1 afla|EED52345.1 aory|EIT74410.1

Group606: abra|OJJ70650.1 acar|OOF97072.1 acla|EAW08109.1 afis|EAW23142.1 afla|EED47570.1 afum|EAL88339.1 agla|OJJ89659.1 akaw|GAA89986.1 aluc|OJZ84000.1 anid|CBF74440.1 anig|EHA19122.1 aory|EIT73130.1 asyd|OJJ57744.1 ater|EAU36275.1 atub|OJI90369.1 aver|OJJ05423.1 awen|OJJ36638.1 azon|OJJ47578.1 afla|EED44702.1 aluc|OJZ80351.1 aory|EIT79586.1

Group607: abra|OJJ70731.1 acar|OOF97007.1 acla|EAW11182.1 afis|EAW15913.1 afla|EED57767.1 afum|EAL88548.2 agla|OJJ89577.1 akaw|GAA90059.1 aluc|OJZ84076.1 anid|CBF81305.1 anig|EHA19195.1 aory|EIT81027.1 asyd|OJJ59352.1 ater|EAU36192.1 atub|OJI90448.1 aver|OJJ02922.1 awen|OJJ36717.1 azon|OJJ47476.1 acla|EAW10824.1 afum|EAL89393.1 awen|OJJ36142.1

Group610: abra|OJJ71163.1 acar|OOF97997.1 acla|EAW14151.1 afis|EAW18864.1 afla|EED53721.1 afum|EAL93596.1 agla|OJJ85239.1 akaw|GAA82816.1 aluc|OJZ87377.1 anid|CBF85347.1 anig|EHA20609.1 aory|EIT75273.1 asyd|OJJ64028.1 ater|EAU34397.1 atub|OJI81269.1 aver|OJI98114.1 awen|OJJ32721.1 azon|OJJ50355.1 afla|EED45395.1 aory|EIT82387.1 awen|OJJ39952.1

Group612: abra|OJJ71741.1 acar|OOF96432.1 acla|EAW06445.1 afis|EAW18550.1 afla|EED52825.1 afum|EAL93250.1 agla|OJJ81594.1 akaw|GAA85155.1 aluc|OJZ86482.1 anid|CBF70273.1 anig|EHA22165.1 aory|EIT78166.1 asyd|OJJ58746.1 ater|EAU33222.1 atub|OJI84048.1 aver|OJJ01356.1 awen|OJJ31108.1 azon|OJJ46554.1 ater|EAU31802.1 afla|EED56761.1 aory|EIT73941.1

Group613: abra|OJJ72042.1 acar|OOF96701.1 acla|EAW09503.1 afis|EAW21633.1 afla|EED57861.1 afum|EAL92666.2 agla|OJJ79697.1 akaw|GAA84891.1 aluc|OJZ86780.1 anid|CBF83510.1 anig|EHA21902.1 aory|EIT81075.1 asyd|OJJ60709.1 ater|EAU38319.1 atub|OJI83740.1 aver|OJJ00035.1 awen|OJJ33882.1 azon|OJJ48672.1 afum|EAL91062.1 acla|EAW07114.1 afis|EAW22126.1

Group617: abra|OJJ72859.1 acar|OOF96056.1 acla|EAW11387.1 afis|EAW18109.1 afla|EED49118.1 afum|EAL87466.1 agla|OJJ86997.1 akaw|GAA85458.1 aluc|OJZ83366.1 anid|CBF75326.1 anig|EHA24231.1 aory|EIT78544.1 asyd|OJJ59867.1 ater|EAU29709.1 atub|OJI82257.1 aver|OJJ02421.1 awen|OJJ35507.1 azon|OJJ46977.1 afla|EED55499.1 agla|OJJ87955.1 aory|EIT80816.1

Group619: abra|OJJ73104.1 acar|OOG00342.1 acla|EAW12431.1 afis|EAW16738.1 afla|EED44914.1 afum|EAL86313.2 agla|OJJ82713.1 akaw|GAA88061.1 aluc|OJZ81323.1 anid|CBF82088.1 anig|EHA18629.1 aory|EIT73472.1 asyd|OJJ54256.1 ater|EAU39589.1 atub|OJI91526.1 aver|OJJ05209.1 awen|OJJ39338.1 anid|CBF75878.1 asyd|OJJ60230.1 aver|OJJ04036.1 azon|OJJ51650.1

Group623: abra|OJJ74191.1 acar|OOF95530.1 acla|EAW07243.1 afis|EAW22264.1 afla|EED57656.1 afum|EAL90916.1 agla|OJJ87791.1 akaw|GAA86826.1 aluc|OJZ89944.1 anid|CBF88515.1 anig|EHA27143.1 aory|EIT80760.1 asyd|OJJ63805.1 ater|EAU34274.1 atub|OJI84862.1 aver|OJI97133.1 awen|OJJ34837.1 azon|OJJ45710.1 afla|EED51484.1 asyd|OJJ58154.1 aver|OJJ04249.1

Group624: abra|OJJ74556.1 acar|OOF95199.1 acla|EAW12292.1 afis|EAW16945.1 afla|EED46082.1 afum|EAL86472.2 agla|OJJ89555.1 akaw|GAA82040.1 aluc|OJZ92139.1 anid|CBF88435.1 anig|EHA26810.1 aory|EIT76401.1 asyd|OJJ63762.1 ater|EAU35227.1 atub|OJI86571.1 aver|OJI97178.1 awen|OJJ36332.1 azon|OJJ46487.1 acla|EAW11680.1 afis|EAW23697.1 anid|CBF76117.1

Group628: abra|OJJ76531.1 acar|OOF99442.1 acla|EAW12929.1 afis|EAW17727.1 afla|EED48895.1 afum|EAL91542.1 agla|OJJ87872.1 akaw|GAA82443.1 aluc|OJZ91673.1 anid|CBF90286.1 anig|EHA22691.1 aory|EIT81184.1 asyd|OJJ61920.1 ater|EAU36831.1 atub|OJI86128.1 aver|OJI96012.1 awen|OJJ39163.1 azon|OJJ44435.1 acar|OOF93188.1 acla|EAW12526.1 awen|OJJ40114.1

Group630: abra|OJJ76842.1 acar|OOF89970.1 acla|EAW13405.1 afis|EAW20516.1 afla|EED52466.1 afum|EAL84541.1 agla|OJJ84774.1 akaw|GAA87180.1 aluc|OJZ84253.1 anid|CBF79059.1 anig|EHA27598.1 aory|EIT82819.1 asyd|OJJ55708.1 ater|EAU37185.1 atub|OJI89830.1 aver|OJJ06794.1 awen|OJJ33222.1 azon|OJJ45121.1 aluc|OJZ87023.1 anig|EHA21634.1 atub|OJI83495.1

Group631: abra|OJJ77007.1 acar|OOF91453.1 acla|EAW15047.1 afis|EAW24611.1 afla|EED47377.1 afum|EAL86241.1 agla|OJJ78906.1 akaw|GAA87041.1 aluc|OJZ84406.1 anid|CBF83177.1 anig|EHA27736.1 aory|EIT75955.1 asyd|OJJ55523.1 ater|EAU32183.1 atub|OJI89666.1 aver|OJJ06601.1 awen|OJJ36042.1 azon|OJJ45129.1 acla|EAW15048.1 aory|EIT82833.1 afla|EED54123.1

Group632: abra|OJJ77033.1 acar|OOF91472.1 acla|EAW07271.1 afis|EAW22292.1 afla|EED57692.1 afum|EAL90889.1 agla|OJJ87756.1 akaw|GAA87018.1 aluc|OJZ84436.1 anid|CBF88573.1 anig|EHA27755.1 aory|EIT79900.1 asyd|OJJ63839.1 ater|EAU34305.1 atub|OJI89637.1 aver|OJI97098.1 awen|OJJ34866.1 azon|OJJ45662.1 afla|EED51864.1 aory|EIT81771.1 awen|OJJ30800.1

Group633: abra|OJJ77037.1 acar|OOF90079.1 acla|EAW13558.1 afis|EAW20634.1 afla|EED49758.1 afum|EAL84434.1 agla|OJJ86283.1 akaw|GAA87014.1 aluc|OJZ84440.1 anid|CBF83213.1 anig|EHA27759.1 aory|EIT77698.1 asyd|OJJ65203.1 ater|EAU32169.1 atub|OJI89633.1 aver|OJJ06646.1 awen|OJJ33605.1 azon|OJJ50726.1 asyd|OJJ55565.1 azon|OJJ42610.1 aver|OJI99352.1

Group635: abra|OJJ77191.1 acar|OOF93077.1 acla|EAW13182.1 afis|EAW17458.1 afla|EED49364.1 afum|EAL91286.1 agla|OJJ80049.1 akaw|GAA90749.1 aluc|OJZ84599.1 anid|CBF71666.1 anig|EHA27889.1 aory|EIT79494.1 asyd|OJJ59263.1 ater|EAU33943.1 atub|OJI89480.1 aver|OJJ01874.1 awen|OJJ37389.1 azon|OJJ51062.1 agla|OJJ80359.1 asyd|OJJ61831.1 aver|OJI95930.1

Group636: abra|OJJ77325.1 acar|OOF93201.1 acla|EAW13147.1 afis|EAW17500.1 afla|EED45479.1 afum|EAL91323.1 agla|OJJ81453.1 akaw|GAA91431.1 aluc|OJZ84735.1 anid|CBF71578.1 anig|EHA28008.1 aory|EIT82181.1 asyd|OJJ61876.1 ater|EAU36946.1 atub|OJI89346.1 aver|OJI95974.1 awen|OJJ37341.1 azon|OJJ51026.1 agla|OJJ78829.1 aory|EIT75365.1 asyd|OJJ62947.1

Group637: abra|OJJ77475.1 acar|OOF99250.1 acla|EAW06757.1 afis|EAW16307.1 afla|EED53242.1 afum|EAL88929.1 agla|OJJ88269.1 akaw|GAA84143.1 aluc|OJZ81063.1 anid|CBF69567.1 anig|EHA19305.1 aory|EIT75142.1 asyd|OJJ59151.1 ater|EAU29335.1 atub|OJI82522.1 aver|OJJ08866.1 awen|OJJ34880.1 azon|OJJ44967.1 aver|OJJ01741.1 awen|OJJ36323.1 asyd|OJJ52895.1

Group638: abra|OJJ78236.1 acar|OOF98624.1 acla|EAW09274.1 afis|EAW21874.1 afla|EED46435.1 afum|EAL92888.1 agla|OJJ84259.1 akaw|GAA85789.1 aluc|OJZ91310.1 anid|CBF76774.1 anig|EHA19994.1 aory|EIT72385.1 asyd|OJJ61272.1 ater|EAU30873.1 atub|OJI83303.1 aver|OJJ00600.1 awen|OJJ34162.1 azon|OJJ43467.1 afla|EED46434.1 awen|OJJ34163.1 asyd|OJJ61271.1

Group644: abra|OJJ74604.1 abra|OJJ77441.1 acar|OOF95100.1 acar|OOF95166.1 acla|EAW10775.1 afis|EAW24801.1 afla|EED47186.1 afum|EAL89442.1 agla|OJJ82895.1 akaw|GAA82004.1 aluc|OJZ92183.1 anid|CBF81510.1 anig|EHA26770.1 aory|EIT79225.1 asyd|OJJ59469.1 ater|EAU36506.1 atub|OJI86611.1 aver|OJJ02805.1 awen|OJJ40384.1 azon|OJJ49233.1

Group646: acar|OOF90425.1 acar|OOF90429.1 acar|OOF93386.1 abra|OJJ69872.1 acla|EAW08934.1 afis|EAW15465.1 afla|EED46918.1 afum|EAL85857.1 agla|OJJ78859.1 akaw|GAA87784.1 aluc|OJZ85419.1 anid|CBF71773.1 anig|EHA25598.1 aory|EIT72659.1 asyd|OJJ61719.1 ater|EAU32668.1 atub|OJI80156.1 aver|OJI95813.1 awen|OJJ38110.1 azon|OJJ43167.1

Group648: acar|OOF90442.1 acar|OOF90448.1 abra|OJJ78364.1 acar|OOF98523.1 acla|EAW09171.1 afis|EAW21975.1 afla|EED46698.1 afum|EAL92990.1 agla|OJJ84127.1 akaw|GAA85669.1 aluc|OJZ91440.1 anid|CBF82718.1 anig|EHA20104.1 aory|EIT77954.1 asyd|OJJ61636.1 ater|EAU31093.1 atub|OJI83434.1 aver|OJJ00959.1 awen|OJJ34268.1 azon|OJJ43367.1

Group649: acar|OOF90479.1 acar|OOF90486.1 acar|OOF96185.1 abra|OJJ73013.1 acla|EAW07999.1 afis|EAW23034.1 afla|EED54982.1 afum|EAL88447.1 agla|OJJ89443.1 akaw|GAA85592.1 aluc|OJZ83528.1 anid|CBF80924.1 anig|EHA24351.1 aory|EIT80284.1 asyd|OJJ56320.1 ater|EAU29623.1 atub|OJI82417.1 aver|OJJ06394.1 awen|OJJ35708.1 azon|OJJ44168.1

Group652: acar|OOF92603.1 acar|OOF92624.1 asyd|OJJ51990.1 asyd|OJJ56630.1 abra|OJJ68460.1 acla|EAW14885.1 afis|EAW24404.1 afla|EED47809.1 afum|EAL86075.1 agla|OJJ89341.1 akaw|GAA92382.1 aluc|OJZ83241.1 anid|CBF73735.1 anig|EHA21096.1 aory|EIT77126.1 ater|EAU30327.1 atub|OJI87696.1 aver|OJJ06096.1 awen|OJJ32804.1 azon|OJJ49024.1

Group653: acar|OOF92614.1 acar|OOF92635.1 awen|OJJ30756.1 awen|OJJ31725.1 abra|OJJ67454.1 acla|EAW06311.1 afis|EAW18420.1 afla|EED52988.1 afum|EAL93104.1 agla|OJJ84988.1 akaw|GAA86377.1 aluc|OJZ84911.1 anid|CBF70700.1 anig|EHA25132.1 aory|EIT78732.1 asyd|OJJ58461.1 ater|EAU29838.1 atub|OJI80687.1 aver|OJJ01061.1 azon|OJJ50488.1

Group655: acar|OOF93670.1 acar|OOF93754.1 abra|OJJ66651.1 acla|EAW11794.1 afis|EAW19404.1 afla|EED50359.1 afum|EAL87091.1 agla|OJJ81118.1 akaw|GAA89603.1 aluc|OJZ80795.1 anid|CBF74343.1 anig|EHA27236.1 aory|EIT83145.1 asyd|OJJ51900.1 asyd|OJJ57797.1 ater|EAU36141.1 atub|OJI87646.1 aver|OJJ05367.1 awen|OJJ36800.1 azon|OJJ42998.1

Group656: acar|OOF93714.1 acar|OOF93801.1 agla|OJJ81170.1 agla|OJJ82654.1 abra|OJJ66712.1 acla|EAW11843.1 afis|EAW19454.1 afla|EED56662.1 afum|EAL87042.1 akaw|GAA89654.1 aluc|OJZ80735.1 anid|CBF81419.1 anig|EHA27286.1 aory|EIT79629.1 asyd|OJJ59414.1 ater|EAU34757.1 atub|OJI87592.1 aver|OJJ02859.1 awen|OJJ36855.1 azon|OJJ42945.1

Group659: acar|OOF95099.1 acar|OOF95165.1 asyd|OJJ59458.1 asyd|OJJ62484.1 abra|OJJ74605.1 acla|EAW10774.1 afis|EAW24802.1 afla|EED47184.1 afum|EAL89443.1 agla|OJJ82896.1 akaw|GAA82003.1 aluc|OJZ92184.1 anid|CBF81490.1 anig|EHA26769.1 aory|EIT79286.1 ater|EAU36505.1 atub|OJI86612.1 aver|OJJ02815.1 awen|OJJ40385.1 azon|OJJ49235.1

Group661: acar|OOF95637.1 acar|OOF95688.1 aver|OJJ05622.1 aver|OJJ07803.1 abra|OJJ74674.1 acla|EAW10723.1 afis|EAW24850.1 afla|EED47121.1 afum|EAL89496.1 agla|OJJ82064.1 akaw|GAA81939.1 aluc|OJZ92251.1 anid|CBF75837.1 anig|EHA26705.1 aory|EIT79282.1 asyd|OJJ57556.1 ater|EAU36453.1 atub|OJI86684.1 awen|OJJ40435.1 azon|OJJ49296.1

Group662: acar|OOF96241.1 acar|OOF96278.1 afum|EAL85174.2 afum|EAL87772.2 abra|OJJ71525.1 acla|EAW15082.1 afis|EAW20054.1 afla|EED45326.1 agla|OJJ88906.1 akaw|GAA89392.1 aluc|OJZ86265.1

anid|CBF84810.1 anig|EHA21603.1 aory|EIT73801.1 asyd|OJJ63220.1 ater|EAU38663.1 atub|OJI84278.1 aver|OJI97735.1 awen|OJJ31812.1 azon|OJJ46310.1

Group663: acar|OOF96244.1 acar|OOF96281.1 afum|EAL85177.1 afum|EAL87761.1 abra|OJJ71529.1 acla|EAW15085.1 afis|EAW20057.1 afla|EED45323.1 agla|OJJ88903.1 akaw|GAA89389.1 aluc|OJZ86269.1 anid|CBF84842.1 anig|EHA21601.1 aory|EIT73791.1 asyd|OJJ63216.1 ater|EAU38666.1 atub|OJI84273.1 aver|OJI97739.1 awen|OJJ31816.1 azon|OJJ46307.1

Group664: acar|OOF96246.1 acar|OOF96283.1 afum|EAL85179.1 afum|EAL87759.1 abra|OJJ71533.1 acla|EAW15087.1 afis|EAW20059.1 afla|EED45321.1 agla|OJJ88901.1 akaw|GAA89387.1 aluc|OJZ86271.1 anid|CBF84838.1 anig|EHA21598.1 aory|EIT73620.1 asyd|OJJ63214.1 ater|EAU38668.1 atub|OJI84270.1 aver|OJI97741.1 awen|OJJ31819.1 azon|OJJ46305.1

Group665: acar|OOF96247.1 acar|OOF96284.1 afum|EAL85180.1 afum|EAL87758.1 abra|OJJ71534.1 acla|EAW15088.1 afis|EAW20060.1 afla|EED45319.1 agla|OJJ88899.1 akaw|GAA89386.1 aluc|OJZ86272.1 anid|CBF84836.1 anig|EHA21597.1 aory|EIT73579.1 asyd|OJJ63213.1 ater|EAU38669.1 atub|OJI84268.1 aver|OJI97742.1 awen|OJJ31820.1 azon|OJJ46304.1

Group666: acar|OOF97654.1 acar|OOF97701.1 abra|OJJ71503.1 acla|EAW10045.1 afis|EAW25539.1 afla|EED44716.1 afum|EAL90202.1 agla|OJJ87266.1 akaw|GAA82517.1 aluc|OJZ87044.1 anid|CBF77824.1 anig|EHA20317.1 aory|EIT83461.1 asyd|OJJ53006.1 ater|EAU33380.1 atub|OJI81591.1 aver|OJJ03837.1 awen|OJJ41281.1 azon|OJJ49499.1 afla|EED44715.1

Group671: acla|EAW10425.1 acla|EAW11189.1 afum|EAL88556.2 afum|EAL89817.2 abra|OJJ70818.1 acar|OOF98284.1 afis|EAW25144.1 afla|EED55301.1 agla|OJJ89017.1 akaw|GAA91697.1 aluc|OJZ87695.1 anid|CBF85681.1 anig|EHA20883.1 aory|EIT83608.1 asyd|OJJ64255.1 ater|EAU39416.1 atub|OJI80947.1 aver|OJI98346.1 awen|OJJ40769.1 azon|OJJ46879.1

Group672: afis|EAW16434.1 afis|EAW20510.1 agla|OJJ84787.1 agla|OJJ85600.1 abra|OJJ76837.1 acar|OOF91329.1 acla|EAW13399.1 afla|EED52451.1 afum|EAL84546.1 akaw|GAA87186.1 aluc|OJZ84248.1 anid|CBF79187.1 anig|EHA27593.1 aory|EIT82791.1 asyd|OJJ55716.1 ater|EAU37204.1 atub|OJI89835.1 aver|OJJ06802.1 awen|OJJ33211.1 azon|OJJ46484.1

Group673: afis|EAW17153.1 afis|EAW21838.1 afum|EAL87745.2 afum|EBA27475.1 abra|OJJ75520.1 acar|OOF90847.1 acla|EAW09307.1 afla|EED46537.1 agla|OJJ84300.1 akaw|GAA84284.1 aluc|OJZ88977.1 anid|CBF76703.1 anig|EHA23548.1 aory|EIT72350.1 asyd|OJJ61234.1 ater|EAU38513.1 atub|OJI85812.1 aver|OJJ00557.1 awen|OJJ34114.1 azon|OJJ44226.1

Group674: afis|EAW17990.1 afis|EAW20711.1 afum|EAL91127.1 afum|EAL91800.1 abra|OJJ68535.1 acar|OOF92737.1 acla|EAW12825.1 afla|EED51913.1 agla|OJJ80250.1 akaw|GAA91316.1 aluc|OJZ82748.1 anid|CBF79983.1 anig|EHA21198.1 aory|EIT81369.1 asyd|OJJ58245.1 ater|EAU31501.1 atub|OJI88983.1 aver|OJJ02181.1 awen|OJJ41556.1 azon|OJJ48271.1

Group677: afla|EED45834.1 afla|EED51243.1 aory|EIT75702.1 aory|EIT83187.1 abra|OJJ69216.1 acar|OOF91102.1 acla|EAW08629.1 afis|EAW15802.1 afum|EAL85548.1 agla|OJJ81969.1 akaw|GAA92225.1 aluc|OJZ86198.1 anid|CBF70916.1 anig|EHA24425.1 asyd|OJJ52604.1 ater|EAU33446.1 atub|OJI79990.1 aver|OJJ04630.1 awen|OJJ38512.1 azon|OJJ43740.1

Group678: afla|EED46513.1 afla|EED55226.1 aory|EIT72395.1 aory|EIT83617.1 abra|OJJ72126.1 acar|OOF96766.1 acla|EAW14932.1 afis|EAW24447.1 afum|EAL86116.1 agla|OJJ80293.1 akaw|GAA84797.1 aluc|OJZ86883.1 anid|CBF73809.1 anig|EHA21761.1 asyd|OJJ56577.1 ater|EAU29999.1 atub|OJI83636.1 aver|OJJ06146.1 awen|OJJ32750.1 azon|OJJ43988.1

Group681: agla|OJJ78415.1 agla|OJJ84293.1 abra|OJJ69363.1 acar|OOF94362.1 acla|EAW14753.1 afis|EAW24284.1 afla|EED47992.1 afum|EAL85953.1 akaw|GAA88232.1 aluc|OJZ82196.1 anid|CBF74135.1 anig|EHA28541.1 aory|EIT79812.1 asyd|OJJ53915.1 ater|EAU31232.1 atub|OJI88239.1 aver|OJJ08013.1 awen|OJJ33012.1 azon|OJJ48503.1 agla|OJJ83146.1

Group684: agla|OJJ79930.1 agla|OJJ82748.1 awen|OJJ29762.1 awen|OJJ33576.1 abra|OJJ70370.1 acar|OOF91900.1 acla|EAW12338.1 afis|EAW16884.1 afla|EED47358.1 afum|EAL86420.1 akaw|GAA91980.1 aluc|OJZ82634.1 anid|CBF80682.1 anig|EHA27445.1 aory|EIT82887.1 asyd|OJJ54592.1 ater|EAU30230.1 atub|OJI88865.1 aver|OJJ07560.1 azon|OJJ50386.1

Group688: aory|EIT72414.1 aory|EIT75970.1 abra|OJJ66829.1 acar|OOF91607.1 acla|EAW14412.1 afis|EAW19120.1 afla|EED46512.1 afla|EED54294.1 afum|EAL93852.1 agla|OJJ80435.1 akaw|GAA90466.1 aluc|OJZ85758.1 anid|CBF75403.1 anig|EHA24798.1 asyd|OJJ59823.1 ater|EAU37383.1 atub|OJI79543.1 aver|OJJ02464.1 awen|OJJ37451.1 azon|OJJ51128.1

Group690: asyd|OJJ52010.1 asyd|OJJ52511.1 asyd|OJJ58584.1 abra|OJJ76189.1 acar|OOF99753.1 acla|EAW13913.1 afis|EAW18624.1 afum|EAL93335.1 agla|OJJ81236.1 akaw|GAA82133.1 aluc|OJZ92039.1 anid|CBF70490.1 anig|EHA22972.1 aory|EIT78164.1 ater|EAU34924.1 atub|OJI86473.1 aver|OJJ01188.1 awen|OJJ32087.1 azon|OJJ46666.1 afla|EED52723.1

Group691: asyd|OJJ52365.1 asyd|OJJ62740.1 aver|OJI96783.1 aver|OJJ01982.1 abra|OJJ74531.1 acar|OOF95221.1 acla|EAW07161.1 afis|EAW22176.1 afla|EED57555.1 afum|EAL91013.1 agla|OJJ87878.1 akaw|GAA82069.1 aluc|OJZ92113.1 anid|CBF89075.1 anig|EHA26832.1 aory|EIT81844.1 ater|EAU34181.1 atub|OJI86546.1 awen|OJJ34753.1 azon|OJJ45807.1

Group692: asyd|OJJ52442.1 asyd|OJJ59129.1 ater|EAU33481.1 ater|EAU38150.1 abra|OJJ76293.1 acar|OOF99538.1 acla|EAW14323.1 afis|EAW18965.1 afla|EED51014.1 afum|EAL93697.2 agla|OJJ85347.1 akaw|GAA82221.1 aluc|OJZ91929.1 anid|CBF69609.1 anig|EHA22891.1 aory|EIT76953.1 atub|OJI86372.1 aver|OJJ01717.1 awen|OJJ31565.1 azon|OJJ46798.1

Group693: asyd|OJJ52683.1 asyd|OJJ58527.1 aver|OJJ01128.1 aver|OJJ08669.1 abra|OJJ76471.1 acar|OOF99708.1 acla|EAW13962.1 afis|EAW18671.1 afla|EED50620.1 afum|EAL93390.1 agla|OJJ82404.1 akaw|GAA82383.1 aluc|OJZ91738.1 anid|CBF70581.1 anig|EHA22744.1 aory|EIT78086.1 ater|EAU38240.1 atub|OJI86196.1 awen|OJJ32143.1 azon|OJJ46724.1

Group694: asyd|OJJ52787.1 asyd|OJJ58807.1 aver|OJJ01410.1 aver|OJJ08762.1 abra|OJJ67374.1 acar|OOF92437.1 acla|EAW06381.1 afis|EAW18487.1 afla|EED52921.1 afum|EAL93176.1 agla|OJJ85082.1 akaw|GAA86448.1 aluc|OJZ84991.1 anid|CBF70149.1 anig|EHA25203.1 aory|EIT78597.1 ater|EAU29912.1 atub|OJI80609.1 awen|OJJ31041.1 azon|OJJ48423.1

Group695: asyd|OJJ54188.1 asyd|OJJ58628.1 aver|OJJ01234.1 aver|OJJ08551.1 abra|OJJ71860.1 acar|OOF96532.1 acla|EAW13891.1 afis|EAW18603.1 afla|EED52744.1 afum|EAL93311.1 agla|OJJ81655.1 akaw|GAA85055.1 aluc|OJZ86597.1 anid|CBF70407.1 anig|EHA22065.1 aory|EIT78216.1 ater|EAU34898.1 atub|OJI83926.1 awen|OJJ31169.1 azon|OJJ46646.1

Group697: asyd|OJJ55918.1 asyd|OJJ60822.1 aver|OJJ00148.1 aver|OJJ04871.1 abra|OJJ75721.1 acar|OOF90261.1 acla|EAW09377.1 afis|EAW21758.1 afla|EED58019.1 afum|EAL92788.2 agla|OJJ84392.1 akaw|GAA84457.1 aluc|OJZ89173.1 anid|CBF83680.1 anig|EHA23382.1 aory|EIT79759.1 ater|EAU38432.1 atub|OJI85617.1 awen|OJJ34042.1 azon|OJJ48199.1

Group698: asyd|OJJ57403.1 asyd|OJJ62790.1 aver|OJI96837.1 aver|OJJ07752.1 abra|OJJ73561.1 acar|OOG00743.1 acla|EAW07478.1 afis|EAW22510.1 afla|EED54538.1 afum|EAL90683.1 agla|OJJ80868.1 akaw|GAA88593.1 aluc|OJZ88500.1 anid|CBF88979.1 anig|EHA18209.1 aory|EIT78737.1 ater|EAU39204.1 atub|OJI91087.1 awen|OJJ35136.1 azon|OJJ49963.1

Group699: asyd|OJJ61696.1 asyd|OJJ63657.1 aver|OJI95780.1 aver|OJI97277.1 abra|OJJ73686.1 acar|OOG00858.1 acla|EAW07601.1 afis|EAW22632.1 afla|EED49882.1 afum|EAL90561.1 agla|OJJ80987.1 akaw|GAA88473.1 aluc|OJZ88363.1 anid|CBF88232.1 anig|EHA18095.1 aory|EIT72462.1 ater|EAU39076.1 atub|OJI90959.1 awen|OJJ35254.1 azon|OJJ49852.1

Group701: ater|EAU29422.1 ater|EAU29976.1 awen|OJJ32730.1 awen|OJJ40859.1 abra|OJJ72200.1 acar|OOF96820.1 acla|EAW14947.1 afis|EAW24462.1 afla|EED55204.1 afum|EAL86133.1 agla|OJJ80320.1 akaw|GAA84736.1 aluc|OJZ86950.1 anid|CBF73838.1 anig|EHA21697.1 aory|EIT83625.1 asyd|OJJ56557.1 atub|OJI83573.1 aver|OJJ06163.1 azon|OJJ45283.1

Group702: aver|OJI99622.1 aver|OJJ08782.1 abra|OJJ69941.1 acar|OOF93451.1 acla|EAW08881.1 afis|EAW15514.1 afla|EED49021.1 agla|OJJ80185.1 akaw|GAA87718.1 aluc|OJZ85345.1 anid|CBF87265.1 anig|EHA25533.1 asyd|OJJ60386.1 ater|EAU32610.1 atub|OJI80225.1 aver|OJI99715.1 awen|OJJ38168.1 azon|OJJ43128.1 afum|EAL85817.1 aory|EIT82522.1

Group703: aver|OJJ00302.1 aver|OJJ08608.1 abra|OJJ68423.1 acar|OOF93930.1 acla|EAW11595.1 afis|EAW23861.1 afla|EED51802.1 afum|EAL87289.1 agla|OJJ87920.1 akaw|GAA86949.1 aluc|OJZ90074.1 anid|CBF82425.1 anig|EHA20999.1 aory|EIT81707.1 asyd|OJJ54128.1 asyd|OJJ60984.1 ater|EAU30753.1 atub|OJI84732.1 awen|OJJ36149.1 azon|OJJ43896.1

Group705: azon|OJJ42149.1 azon|OJJ43551.1 abra|OJJ67976.1 acar|OOF92817.1 acla|EAW11759.1 afis|EAW23792.1 afla|EED51600.1 afum|EAL87226.1 agla|OJJ78821.1 akaw|GAA91227.1 aluc|OJZ82843.1 anid|CBF79854.1 anig|EHA21281.1 aory|EIT80489.1 asyd|OJJ58158.1 ater|EAU31332.1 atub|OJI89073.1 aver|OJJ04245.1 awen|OJJ33749.1 awen|OJJ34607.1

Group716: abra|OJJ66628.1 acar|OOF91051.1 acla|EAW10868.1 afis|EAW20990.1 afla|EED46119.1 afum|EAL84639.1 agla|OJJ84068.1 akaw|GAA91097.1 aluc|OJZ81420.1 anid|CBF82620.1 anig|EHA26001.1 aory|EIT77263.1 asyd|OJJ54019.1 ater|EAU38639.1 atub|OJI88761.1 aver|OJJ08721.1 awen|OJJ34320.1 azon|OJJ48067.1 aory|EIT76479.1 afla|EED50344.1

Group719: abra|OJJ66927.1 acar|OOF91695.1 acla|EAW14492.1 afis|EAW19206.1 afla|EED47706.1 afum|EAL93936.1 agla|OJJ88505.1 akaw|GAA91490.1 aluc|OJZ85652.1 anid|CBF78561.1 anig|EHA24881.1 aory|EIT80267.1 asyd|OJJ65063.1 ater|EAU37266.1 atub|OJI79443.1 aver|OJI99187.1 awen|OJJ37559.1 azon|OJJ51221.1 ater|EAU31748.1 awen|OJJ37930.1

Group720: abra|OJJ66958.1 acar|OOF93650.1 acla|EAW14962.1 afis|EAW24485.1 afla|EED45449.1 afum|EAL86154.1 agla|OJJ88956.1 akaw|GAA92203.1 aluc|OJZ85618.1 anid|CBF82826.1 anig|EHA24908.1

aory|EIT82465.1 asyd|OJJ55929.1 ater|EAU37956.1 atub|OJI79411.1 aver|OJJ04879.1 awen|OJJ34538.1 azon|OJJ51794.1 agla|OJJ83743.1 aver|OJJ00902.1

Group731: abra|OJJ69242.1 acar|OOF91077.1 acla|EAW08606.1 afis|EAW15827.1 afla|EED51218.1 afum|EAL85524.1 agla|OJJ81990.1 akaw|GAA92251.1 aluc|OJZ86222.1 anid|CBF70872.1 anig|EHA24405.1 aory|EIT80412.1 asyd|OJJ52582.1 ater|EAU33421.1 atub|OJI80016.1 aver|OJJ04610.1 awen|OJJ38541.1 azon|OJJ43722.1 asyd|OJJ54750.1 aver|OJJ07417.1

Group740: abra|OJJ70960.1 acar|OOF98170.1 acla|EAW10521.1 afis|EAW25045.1 afla|EED48351.1 afum|EAL89713.1 agla|OJJ89149.1 akaw|GAA82987.1 aluc|OJZ87572.1 anid|CBF85914.1 anig|EHA20774.1 aory|EIT80893.1 asyd|OJJ64399.1 ater|EAU35795.1 atub|OJI81077.1 aver|OJI98495.1 awen|OJJ40656.1 azon|OJJ44034.1 aory|EIT72298.1 afla|EED55406.1

Group741: abra|OJJ71146.1 acar|OOF98009.1 acar|OOF98010.1 acla|EAW10360.1 afis|EAW25233.1 afla|EED53679.1 afum|EAL89894.1 agla|OJJ79174.1 akaw|GAA82831.1 aluc|OJZ87392.1 anid|CBF85327.1 anig|EHA20621.1 anig|EHA20622.1 aory|EIT75303.1 asyd|OJJ64017.1 ater|EAU34392.1 atub|OJI81254.1 aver|OJI98103.1 awen|OJJ40899.1 azon|OJJ44526.1

Group743: abra|OJJ71853.1 acar|OOF96523.1 afla|EED52750.1 afum|EAL93305.1 agla|OJJ81650.1 akaw|GAA85064.1 aluc|OJZ86587.1 anid|CBF70395.1 anig|EHA22073.1 aory|EIT78183.1 asyd|OJJ58634.1 atub|OJI83936.1 aver|OJJ01240.1 awen|OJJ31160.1 acla|EAW13885.1 afis|EAW18597.1 ater|EAU34891.1 azon|OJJ46636.1 asyd|OJJ54226.1 aver|OJJ05179.1

Group745: abra|OJJ72207.1 acar|OOF96828.1 acla|EAW14969.1 afis|EAW24492.1 afla|EED54697.1 afum|EAL86161.1 agla|OJJ80329.1 akaw|GAA84727.1 aluc|OJZ86959.1 anid|CBF71478.1 anig|EHA21689.1 aory|EIT78783.1 asyd|OJJ54943.1 ater|EAU29944.1 atub|OJI83564.1 aver|OJJ07252.1 awen|OJJ33962.1 azon|OJJ42613.1 azon|OJJ48606.1 awen|OJJ32703.1

Group746: abra|OJJ72535.1 acar|OOF94717.1 acla|EAW12742.1 afis|EAW17905.1 afla|EED46297.1 afum|EAL91714.1 agla|OJJ83037.1 akaw|GAA86257.1 aluc|OJZ90515.1 anid|CBF76985.1 anig|EHA23970.1 aory|EIT76354.1 asyd|OJJ57035.1 ater|EAU33123.1 atub|OJI81939.1 aver|OJJ03257.1 awen|OJJ41645.1 azon|OJJ48367.1 aluc|OJZ90514.1 aory|EIT76342.1

Group747: abra|OJJ72848.1 acar|OOF96049.1 acla|EAW11395.1 afis|EAW24026.1 afla|EED49128.1 afum|EAL87457.1 agla|OJJ86959.1 akaw|GAA85449.1 aluc|OJZ83357.1 anid|CBF84482.1 anig|EHA24222.1 aory|EIT78568.1 asyd|OJJ53530.1 ater|EAU29718.1 atub|OJI82248.1 aver|OJJ08220.1 awen|OJJ35499.1 azon|OJJ43175.1 acla|EAW12378.1 awen|OJJ40008.1

Group748: abra|OJJ73145.1 acar|OOG00357.1 acla|EAW07021.1 afis|EAW16647.1 afla|EED45442.1 afum|EAL89251.1 agla|OJJ82595.1 akaw|GAA88027.1 aluc|OJZ81286.1 anid|CBF82058.1 anig|EHA18585.1 aory|EIT82414.1 asyd|OJJ54238.1 ater|EAU39572.1 atub|OJI91493.1 aver|OJJ05190.1 awen|OJJ40067.1 azon|OJJ45557.1 aver|OJJ01178.1 afla|EED50972.1

Group749: abra|OJJ73181.1 acar|OOG00417.1 afla|EED47213.1 afum|EAL89070.1 akaw|GAA87994.1 aluc|OJZ81249.1 anig|EHA18557.1 ater|EAU36561.1 awen|OJJ40360.1 azon|OJJ49208.1 acla|EAW06859.1 afis|EAW16476.1 agla|OJJ82869.1 anid|CBF75495.1 aory|EIT79277.1 asyd|OJJ59750.1 atub|OJI91445.1 aver|OJJ02537.1 acar|OOG00418.1 atub|OJI91444.1

Group750: abra|OJJ73280.1 acar|OOG00516.1 acla|EAW06955.1 afis|EAW16570.1 afla|EED56094.1 afum|EAL89167.1 agla|OJJ82745.1 akaw|GAA92725.1 aluc|OJZ88769.1 anid|CBF81824.1 anig|EHA18465.1 aory|EIT76687.1 asyd|OJJ59656.1 ater|EAU36638.1 atub|OJI91342.1 aver|OJJ02626.1 awen|OJJ40248.1 azon|OJJ49110.1 afum|EAL87777.1 aory|EIT72969.1

Group751: abra|OJJ73388.1 acar|OOG00590.1 acla|EAW07000.1 afis|EAW16615.1 afla|EED44990.1 afum|EAL89221.1 agla|OJJ82629.1 akaw|GAA83101.1 aluc|OJZ88649.1 anid|CBF82980.1 anig|EHA18340.1 aory|EIT73600.1 asyd|OJJ53668.1 ater|EAU36700.1 atub|OJI91236.1 aver|OJJ07774.1 awen|OJJ40187.1 azon|OJJ49474.1 anig|EHA18404.1 ater|EAU37639.1

Group752: abra|OJJ73407.1 acar|OOG00604.1 acla|EAW07012.1 afis|EAW16632.1 afla|EED48324.1 afum|EAL89228.1 agla|OJJ82617.1 akaw|GAA83084.1 aluc|OJZ88631.1 anid|CBF82011.1 anig|EHA18321.1 aory|EIT73498.1 asyd|OJJ56190.1 ater|EAU36752.1 atub|OJI91215.1 aver|OJJ05118.1 awen|OJJ40077.1 azon|OJJ43490.1 aory|EIT76284.1 afla|EED48295.1

Group753: abra|OJJ73423.1 acar|OOG00627.1 acla|EAW07036.1 afis|EAW16662.1 afla|EED44895.1 afum|EAL89266.1 agla|OJJ78707.1 akaw|GAA83069.1 aluc|OJZ88620.1 anid|CBF84550.1 anig|EHA18311.1 aory|EIT83238.1 asyd|OJJ53473.1 ater|EAU31782.1 atub|OJI91205.1 aver|OJJ08283.1 awen|OJJ40051.1 azon|OJJ47485.1 afla|EED56569.1 aory|EIT79633.1

Group754: abra|OJJ73685.1 acar|OOG00855.1 acar|OOG00856.1 acla|EAW07600.1 afis|EAW22631.1 afla|EED49880.1 afla|EED49881.1 afum|EBA27410.1 agla|OJJ80989.1 akaw|GAA88474.1 aluc|OJZ88364.1 anid|CBF88234.1 anig|EHA18096.1 aory|EIT72461.1 ater|EAU39077.1 atub|OJI90961.1 aver|OJI97276.1 awen|OJJ35252.1 azon|OJJ49854.1 asyd|OJJ63658.1

Group755: abra|OJJ73793.1 acar|OOG00954.1 acla|EAW07685.1 afis|EAW22700.1 afla|EED49986.1 afum|EAL90492.1 agla|OJJ79009.1 akaw|GAA89412.1 aluc|OJZ88261.1 anid|CBF88081.1 anig|EHA18006.1 aory|EIT74617.1 asyd|OJJ63558.1 ater|EAU39002.1 atub|OJI90860.1 aver|OJI97374.1 awen|OJJ35337.1 azon|OJJ49781.1 ater|EAU30260.1 awen|OJJ38024.1

Group756: abra|OJJ73803.1 acar|OOG00958.1 acla|EAW07675.1 afis|EAW22690.1 afla|EED49983.1 afum|EAL90502.1 agla|OJJ79015.1 akaw|GAA89422.1 aluc|OJZ88250.1 anid|CBF88089.1 anig|EHA17993.1 aory|EIT74622.1 asyd|OJJ63564.1 ater|EAU39009.1 atub|OJI90849.1 aver|OJI97369.1 awen|OJJ33510.1 azon|OJJ49784.1 acar|OOG00352.1 afla|EED46078.1

Group757: abra|OJJ73866.1 acar|OOF97590.1 acla|EAW06976.1 afis|EAW16590.1 afla|EED46107.1 afum|EAL89185.1 agla|OJJ83704.1 akaw|GAA89476.1 aluc|OJZ88187.1 anid|CBF76004.1 anig|EHA17938.1 aory|EIT76398.1 asyd|OJJ57426.1 ater|EAU29326.1 atub|OJI90786.1 aver|OJJ03990.1 awen|OJJ40225.1 azon|OJJ49083.1 anid|CBF75976.1 asyd|OJJ60263.1

Group758: abra|OJJ74117.1 acar|OOG01222.1 acla|EAW07959.1 afis|EAW22994.1 afla|EED58187.1 afum|EAL90205.1 agla|OJJ83290.1 akaw|GAA87590.1 aluc|OJZ87915.1 anid|CBF87568.1 anig|EHA17710.1 aory|EIT79090.1 asyd|OJJ63243.1 ater|EAU30624.1 atub|OJI90527.1 aver|OJI97713.1 awen|OJJ38567.1 azon|OJJ50434.1 anig|EHA17709.1 afum|EAL90206.1

Group760: abra|OJJ74757.1 acar|OOF95750.1 acla|EAW10654.1 afis|EAW24913.1 afla|EED46998.1 afum|EAL89567.1 agla|OJJ82924.1 akaw|GAA83654.1 aluc|OJZ92341.1 anid|CBF75675.1 anig|EHA26633.1 aory|EIT72680.1 asyd|OJJ57485.1 ater|EAU36385.1 atub|OJI86767.1 aver|OJJ05695.1 awen|OJJ40509.1 azon|OJJ49368.1 afla|EED48195.1 aory|EIT72956.1

Group761: abra|OJJ74761.1 acar|OOF95754.1 acla|EAW10650.1 afis|EAW24917.1 afla|EED47002.1 afum|EAL89572.2 agla|OJJ82145.1 akaw|GAA83650.1 aluc|OJZ92345.1 anid|CBF75683.1 anig|EHA26629.1 aory|EIT72569.1 asyd|OJJ57489.1 ater|EAU36381.1 atub|OJI86771.1 aver|OJJ05691.1 awen|OJJ40513.1 azon|OJJ49372.1 anid|CBF75685.1 atub|OJI86772.1

Group762: abra|OJJ74833.1 acar|OOF95823.1 acla|EAW11247.1 afis|EAW18253.1 afla|EED56199.1 afum|EAL87616.1 agla|OJJ88191.1 akaw|GAA83587.1 aluc|OJZ92410.1 anid|CBF86203.1 anig|EHA26555.1 aory|EIT73440.1 asyd|OJJ58331.1 ater|EAU33725.1 atub|OJI86840.1 aver|OJJ02080.1 awen|OJJ30627.1 azon|OJJ47986.1 aver|OJI98640.1 asyd|OJJ64544.1

Group764: abra|OJJ74964.1 acar|OOF97350.1 acla|EAW08410.1 afis|EAW23459.1 afla|EED57342.1 afum|EAL88007.2 agla|OJJ86690.1 akaw|GAA83462.1 aluc|OJZ92534.1 anid|CBF89818.1 anig|EHA26441.1 aory|EIT80102.1 asyd|OJJ62188.1 ater|EAU37472.1 atub|OJI86970.1 aver|OJI96297.1 awen|OJJ39771.1 azon|OJJ47905.1 afla|EED57341.1 aory|EIT80055.1

Group766: abra|OJJ75618.1 acar|OOG00217.1 acla|EAW09356.1 afis|EAW21782.1 afla|EED46350.1 afum|EAL92810.1 agla|OJJ84341.1 akaw|GAA84367.1 aluc|OJZ89071.1 anid|CBF76611.1 anig|EHA23465.1 aory|EIT81207.1 asyd|OJJ61187.1 ater|EAU38459.1 atub|OJI85717.1 aver|OJJ00510.1 awen|OJJ34070.1 azon|OJJ48159.1 afum|EAL85431.1 anid|CBF76609.1

Group769: abra|OJJ76070.1 acar|OOF99859.1 acla|EAW09749.1 afis|EAW21393.1 afla|EED55810.1 afum|EAL92407.1 agla|OJJ85517.1 akaw|GAA88897.1 aluc|OJZ89524.1 anid|CBF71432.1 anig|EHA23077.1 aory|EIT76869.1 asyd|OJJ61713.1 ater|EAU35164.1 atub|OJI85279.1 aver|OJJ07221.1 awen|OJJ39311.1 azon|OJJ51541.1 asyd|OJJ54970.1 ater|EAU32747.1

Group771: abra|OJJ76117.1 acar|OOF99808.1 acla|EAW09703.1 afis|EAW21437.1 afla|EED45636.1 afum|EAL92452.2 agla|OJJ85581.1 akaw|GAA88850.1 aluc|OJZ89574.1 anid|CBF76585.1 anig|EHA23033.1 aory|EIT76558.1 asyd|OJJ61172.1 ater|EAU35108.1 atub|OJI85230.1 aver|OJJ00497.1 awen|OJJ39261.1 azon|OJJ51593.1 aory|EIT75602.1 afla|EED55865.1

Group772: abra|OJJ76529.1 acar|OOF99449.1 acla|EAW14176.1 afis|EAW18891.1 afla|EED50873.1 afum|EBA27495.1 agla|OJJ81497.1 akaw|GAA82440.1 akaw|GAA82441.1 aluc|OJZ91675.1 aluc|OJZ91676.1 anid|CBF69904.1 anig|EHA22693.1 aory|EIT79015.1 asyd|OJJ58997.1 ater|EAU38063.1 atub|OJI86130.1 aver|OJJ01619.1 awen|OJJ32450.1 azon|OJJ47208.1

Group773: abra|OJJ77055.1 acar|OOF92945.1 acla|EAW13514.1 afis|EAW19907.1 afla|EED54205.1 afum|EAL84999.1 agla|OJJ88625.1 akaw|GAA86997.1 aluc|OJZ84460.1 anid|CBF86958.1 anig|EHA27775.1 aory|EIT76071.1 asyd|OJJ55597.1 ater|EAU35255.1 atub|OJI89615.1 aver|OJJ06677.1 awen|OJJ33456.1 azon|OJJ50712.1 afis|EAW20612.1 acla|EAW13697.1

Group775: abra|OJJ77859.1 acar|OOF98914.1 acla|EAW11267.1 afis|EAW18231.1 afla|EED52136.1 afum|EAL87599.1 agla|OJJ85846.1 akaw|GAA83790.1 aluc|OJZ90933.1 anid|CBF86176.1 anig|EHA19643.1 aory|EIT73335.1 asyd|OJJ64527.1 ater|EAU33760.1 atub|OJI82925.1 aver|OJI98625.1 awen|OJJ30611.1 azon|OJJ48005.1 azon|OJJ42776.1 awen|OJJ36209.1

Group776: abra|OJJ77974.1 acar|OOF98840.1 acla|EAW06664.1 afis|EAW16101.1 afla|EED48636.1 afum|EAL88733.1 agla|OJJ86121.1 akaw|GAA83686.1 aluc|OJZ91047.1 anid|CBF85446.1 anig|EHA19744.1

aory|EIT73060.1 asyd|OJJ64094.1 ater|EAU33175.1 aver|OJI98176.1 awen|OJJ30331.1 azon|OJJ45359.1 anid|CBF85444.1 asyd|OJJ64092.1 atub|OJI83043.1

Group778: abra|OJJ78259.1 acar|OOF98601.1 acla|EAW09251.1 afis|EAW21896.1 afla|EED46412.1 afum|EBA27479.1 agla|OJJ84236.1 akaw|GAA85766.1 aluc|OJZ91333.1 anig|EHA20015.1 aory|EIT72351.1 asyd|OJJ61292.1 ater|EAU30891.1 atub|OJI83326.1 aver|OJJ00618.1 awen|OJJ34185.1 azon|OJJ43448.1 aluc|OJZ91334.1 akaw|GAA85767.1 anid|CBF76812.1

Group783: abra|OJJ68371.1 abra|OJJ72091.1 acar|OOF96736.1 acla|EAW09472.1 afis|EAW21661.1 afla|EED57928.1 afum|EAL92696.1 agla|OJJ79735.1 akaw|GAA84829.1 aluc|OJZ86846.1 anid|CBF83566.1 anig|EHA21792.1 aory|EIT82318.1 asyd|OJJ60745.1 ater|EAU38349.1 atub|OJI83673.1 aver|OJJ00077.1 awen|OJJ33927.1 azon|OJJ48635.1

Group784: acar|OOF90452.1 acar|OOF90459.1 abra|OJJ71882.1 acla|EAW13911.1 afis|EAW18623.1 afla|EED52721.1 afum|EAL93333.1 agla|OJJ81237.1 akaw|GAA85034.1 aluc|OJZ86620.1 anid|CBF70493.1 anig|EHA22046.1 aory|EIT78225.1 asyd|OJJ58583.1 ater|EAU34923.1 atub|OJI83905.1 aver|OJJ01187.1 awen|OJJ32086.1 azon|OJJ46665.1

Group785: acar|OOF90454.1 acar|OOF90462.1 abra|OJJ71884.1 acla|EAW09637.1 afis|EAW21503.1 afla|EED55939.1 afum|EAL92522.1 agla|OJJ85666.1 akaw|GAA85032.1 aluc|OJZ86622.1 anid|CBF76453.1 anig|EHA22044.1 aory|EIT76619.1 asyd|OJJ61107.1 ater|EAU35035.1 atub|OJI83903.1 aver|OJJ00424.1 awen|OJJ39192.1 azon|OJJ51508.1

Group786: acar|OOF90455.1 acar|OOF90461.1 abra|OJJ71885.1 acla|EAW09636.1 afis|EAW21504.1 afla|EED55940.1 afum|EAL92523.1 agla|OJJ85667.1 akaw|GAA85031.1 aluc|OJZ86623.1 anid|CBF76451.1 anig|EHA22043.1 aory|EIT76886.1 asyd|OJJ61106.1 ater|EAU35034.1 atub|OJI83902.1 aver|OJJ00423.1 awen|OJJ39191.1 azon|OJJ51507.1

Group787: acar|OOF90456.1 acar|OOF90463.1 abra|OJJ71886.1 acla|EAW09635.1 afis|EAW21505.1 afla|EED55941.1 afum|EAL92524.1 agla|OJJ85668.1 akaw|GAA85030.1 aluc|OJZ86624.1 anid|CBF76449.1 anig|EHA22042.1 aory|EIT76597.1 asyd|OJJ61105.1 ater|EAU35033.1 atub|OJI83901.1 aver|OJJ00422.1 awen|OJJ39190.1 azon|OJJ51506.1

Group790: acar|OOF92286.1 acar|OOF92327.1 abra|OJJ67516.1 acla|EAW06255.1 afis|EAW18362.1 afla|EED53043.1 afum|EAL93033.1 agla|OJJ84926.1 akaw|GAA86324.1 aluc|OJZ84848.1 anid|CBF70825.1 anig|EHA25082.1 aory|EIT77408.1 asyd|OJJ58399.1 ater|EAU32230.1 atub|OJI80748.1 aver|OJJ01000.1 awen|OJJ31788.1 azon|OJJ50438.1

Group791: acar|OOF92287.1 acar|OOF92328.1 abra|OJJ67515.1 acla|EAW06254.1 afis|EAW18361.1 afla|EED53044.1 afum|EAL93032.2 agla|OJJ84925.1 akaw|GAA86325.1 aluc|OJZ84849.1 anid|CBF70827.1 anig|EHA25083.1 aory|EIT77438.1 asyd|OJJ58398.1 ater|EAU32229.1 atub|OJI80747.1 aver|OJJ00999.1 awen|OJJ31789.1 azon|OJJ42001.1

Group792: acar|OOF92288.1 acar|OOF92329.1 abra|OJJ67514.1 acla|EAW06260.1 afis|EAW18367.1 afla|EED53038.1 afum|EAL93041.1 agla|OJJ84931.1 akaw|GAA86326.1 aluc|OJZ84850.1 anid|CBF70815.1 anig|EHA25084.1 aory|EIT77399.1 asyd|OJJ58404.1 ater|EAU32236.1 atub|OJI80746.1 aver|OJJ01005.1 awen|OJJ31783.1 azon|OJJ50443.1

Group793: acar|OOF92289.1 acar|OOF92330.1 abra|OJJ67513.1 acla|EAW06259.1 afis|EAW18366.1 afla|EED53039.1 afum|EAL93040.1 agla|OJJ84930.1 akaw|GAA86327.1 aluc|OJZ84851.1 anid|CBF70817.1 anig|EHA25085.1 aory|EIT77396.1 asyd|OJJ58403.1 ater|EAU32235.1 atub|OJI80745.1 aver|OJJ01004.1 awen|OJJ31784.1 azon|OJJ50442.1

Group794: acar|OOF92291.1 acar|OOF92332.1 abra|OJJ67510.1 afis|EAW18368.1 afla|EED53037.1 afum|EAL93043.1 agla|OJJ84933.1 akaw|GAA86329.1 aluc|OJZ84854.1 anid|CBF70814.1 anig|EHA25087.1 aory|EIT77401.1 asyd|OJJ58405.1 ater|EAU32238.1 atub|OJI80742.1 aver|OJJ01006.1 awen|OJJ31781.1 azon|OJJ43320.1 acla|EAW06262.1

Group795: acar|OOF92293.1 acar|OOF92334.1 abra|OJJ67507.1 acla|EAW06264.1 afis|EAW18370.1 afla|EED53036.1 afum|EAL93045.1 agla|OJJ84935.1 akaw|GAA86331.1 aluc|OJZ84856.1 anid|CBF70808.1 anig|EHA25089.1 aory|EIT77405.1 asyd|OJJ58408.1 ater|EAU32240.1 atub|OJI80740.1 aver|OJJ01010.1 awen|OJJ31779.1 azon|OJJ50445.1

Group796: acar|OOF92297.1 acar|OOF92338.1 abra|OJJ67503.1 acla|EAW06267.1 afis|EAW18373.1 afla|EED53033.1 afum|EAL93048.1 agla|OJJ84937.1 akaw|GAA86335.1 aluc|OJZ84860.1 anid|CBF70800.1 anig|EHA25093.1 aory|EIT77403.1 asyd|OJJ58411.1 ater|EAU32243.1 atub|OJI80736.1 aver|OJJ01013.1 awen|OJJ31777.1 azon|OJJ50448.1

Group797: acar|OOF92298.1 acar|OOF92339.1 abra|OJJ67502.1 acla|EAW06268.1 afis|EAW18374.1 afla|EED53032.1 afum|EAL93049.1 agla|OJJ84938.1 akaw|GAA86336.1 aluc|OJZ84861.1 anid|CBF70798.1 anig|EHA25094.1 aory|EIT77421.1 asyd|OJJ58412.1 ater|EAU32244.1 atub|OJI80735.1 aver|OJJ01014.1 awen|OJJ31766.1 azon|OJJ50449.1

Group798: acar|OOF92299.1 acar|OOF92340.1 abra|OJJ67501.1 acla|EAW06269.1 afis|EAW18375.1 afla|EED53031.1 afum|EAL93050.1 agla|OJJ84939.1 akaw|GAA86337.1 aluc|OJZ84862.1 anid|CBF70797.1 anig|EHA25095.1 aory|EIT77415.1 asyd|OJJ58413.1 ater|EAU32245.1 atub|OJI80734.1 aver|OJJ01015.1 awen|OJJ31767.1 azon|OJJ50450.1

Group799: acar|OOF92301.1 acar|OOF92342.1 abra|OJJ67499.1 acla|EAW06271.1 afis|EAW18377.1 afla|EED53029.1 afum|EAL93052.1 agla|OJJ84941.1 akaw|GAA86339.1 aluc|OJZ84864.1 anid|CBF70793.1 anig|EHA25096.1 aory|EIT77426.1 asyd|OJJ58415.1 ater|EAU32247.1 atub|OJI80732.1 aver|OJJ01017.1 awen|OJJ31769.1 azon|OJJ50452.1

Group800: acar|OOF92303.1 acar|OOF92343.1 abra|OJJ67498.1 acla|EAW06272.1 afis|EAW18378.1 afla|EED53028.1 afum|EAL93053.1 agla|OJJ84942.1 akaw|GAA86340.1 aluc|OJZ84865.1 anid|CBF70791.1 anig|EHA25097.1 aory|EIT77397.1 asyd|OJJ58416.1 ater|EAU32248.1 atub|OJI80731.1 aver|OJJ01018.1 awen|OJJ31770.1 azon|OJJ50453.1

Group801: acar|OOF92304.1 acar|OOF92345.1 abra|OJJ67496.1 acla|EAW06274.1 afis|EAW18380.1 afla|EED53026.1 afum|EAL93055.1 agla|OJJ84944.1 akaw|GAA86342.1 aluc|OJZ84867.1 anid|CBF70787.1 anig|EHA25099.1 aory|EIT77406.1 asyd|OJJ58418.1 ater|EAU32249.1 atub|OJI80729.1 aver|OJJ01020.1 awen|OJJ31772.1 azon|OJJ50455.1

Group802: acar|OOF92305.1 acar|OOF92346.1 abra|OJJ67495.1 acla|EAW06275.1 afis|EAW18381.1 afla|EED53025.1 afum|EAL93056.1 agla|OJJ84945.1 akaw|GAA86343.1 aluc|OJZ84869.1 anid|CBF70785.1 anig|EHA25100.1 aory|EIT77407.1 asyd|OJJ58419.1 ater|EAU32250.1 atub|OJI80728.1 aver|OJJ01021.1 awen|OJJ31773.1 azon|OJJ50456.1

Group803: acar|OOF92306.1 acar|OOF92347.1 abra|OJJ67494.1 acla|EAW06276.1 afis|EAW18382.1 afla|EED53024.1 afum|EAL93057.1 agla|OJJ84946.1 akaw|GAA86344.1 aluc|OJZ84870.1 anid|CBF70783.1 anig|EHA25101.1 aory|EIT77418.1 asyd|OJJ58420.1 ater|EAU32251.1 atub|OJI80727.1 aver|OJJ01022.1 awen|OJJ31774.1 azon|OJJ50457.1

Group804: acar|OOF92307.1 acar|OOF92348.1 abra|OJJ67492.1 acla|EAW06278.1 afis|EAW18384.1 afla|EED53022.1 afum|EAL93059.1 agla|OJJ84948.1 akaw|GAA86346.1 aluc|OJZ84872.1 anid|CBF70781.1 anig|EHA25103.1 aory|EIT77395.1 asyd|OJJ58422.1 ater|EAU32253.1 atub|OJI80726.1 aver|OJJ01023.1 awen|OJJ31775.1 azon|OJJ50458.1

Group805: acar|OOF92308.1 acar|OOF92349.1 abra|OJJ67491.1 acla|EAW06279.1 afis|EAW18385.1 afla|EED53021.1 afum|EAL93060.1 agla|OJJ84949.1 akaw|GAA86347.1 aluc|OJZ84874.1 anid|CBF70779.1 anig|EHA25104.1 aory|EIT77414.1 asyd|OJJ58423.1 ater|EAU32254.1 atub|OJI80725.1 aver|OJJ01024.1 awen|OJJ31776.1 azon|OJJ45926.1

Group806: acar|OOF92310.1 acar|OOF92351.1 abra|OJJ67490.1 acla|EAW06282.1 afis|EAW18388.1 afla|EED53019.1 afum|EAL93063.1 agla|OJJ84951.1 akaw|GAA86348.1 aluc|OJZ84875.1 anid|CBF70772.1 anig|EHA25106.1 aory|EIT77433.1 asyd|OJJ58427.1 ater|EAU32261.1 atub|OJI80723.1 aver|OJJ01028.1 awen|OJJ31763.1 azon|OJJ50461.1

Group807: acar|OOF92311.1 acar|OOF92352.1 abra|OJJ67489.1 acla|EAW06284.1 afis|EAW18390.1 afla|EED53017.1 afum|EAL93065.1 agla|OJJ84953.1 akaw|GAA86350.1 aluc|OJZ84878.1 anid|CBF70776.1 anig|EHA25107.1 aory|EIT77439.1 asyd|OJJ58425.1 ater|EAU32263.1 atub|OJI80721.1 aver|OJJ01026.1 awen|OJJ31761.1 azon|OJJ50464.1

Group808: acar|OOF92312.1 acar|OOF92353.1 abra|OJJ67488.1 acla|EAW06286.1 afis|EAW18391.1 afla|EED53016.1 afum|EAL93066.1 agla|OJJ84955.1 akaw|GAA86351.1 aluc|OJZ84880.1 anid|CBF70777.1 anig|EHA25110.1 aory|EIT77398.1 asyd|OJJ58424.1 ater|EAU32278.1 atub|OJI80719.1 aver|OJJ01025.1 awen|OJJ31758.1 azon|OJJ50466.1

Group809: acar|OOF92314.1 acar|OOF92355.1 abra|OJJ67485.1 acla|EAW06289.1 afis|EAW18394.1 afla|EED53013.1 afum|EAL93070.1 agla|OJJ84966.1 akaw|GAA86354.1 aluc|OJZ84883.1 anid|CBF70762.1 anig|EHA25112.1 aory|EIT77419.1 asyd|OJJ58430.1 ater|EAU32282.1 atub|OJI80717.1 aver|OJJ01032.1 awen|OJJ31756.1 azon|OJJ50472.1

Group810: acar|OOF92317.1 acar|OOF92359.1 abra|OJJ67482.1 acla|EAW06291.1 afis|EAW18397.1 afla|EED53009.1 afum|EAL93074.1 agla|OJJ84968.1 akaw|GAA86357.1 aluc|OJZ84886.1 anid|CBF70756.1 anig|EHA25115.1 aory|EIT77428.1 asyd|OJJ58434.1 ater|EAU32285.1 atub|OJI80714.1 aver|OJJ01035.1 awen|OJJ31754.1 azon|OJJ50470.1

Group812: acar|OOF92320.1 acar|OOF92362.1 abra|OJJ67474.1 acla|EAW06298.1 afis|EAW18405.1 afla|EED53003.1 afum|EAL93085.1 agla|OJJ84956.1 akaw|GAA86363.1 aluc|OJZ84891.1 anid|CBF69754.1 anig|EHA25120.1 aory|EIT77409.1 asyd|OJJ58942.1 ater|EAU29822.1 atub|OJI80707.1 aver|OJJ01538.1 awen|OJJ31745.1 azon|OJJ50505.1

Group813: acar|OOF92596.1 acar|OOF92617.1 abra|OJJ68969.1 acla|EAW14218.1 afis|EAW19063.1 afla|EED54429.1 afum|EAL93792.1 agla|OJJ84544.1 akaw|GAA90496.1 aluc|OJZ85956.1 anid|CBF79645.1

anig|EHA24645.1 aory|EIT75848.1 asyd|OJJ58020.1 ater|EAU32106.1 atub|OJI79744.1 aver|OJJ04379.1 awen|OJJ31494.1 azon|OJJ46207.1

Group814: acar|OOF92598.1 acar|OOF92619.1 abra|OJJ68487.1 acla|EAW06807.1 afis|EAW16356.1 afla|EED53514.1 afum|EAL88981.1 agla|OJJ88237.1 akaw|GAA92784.1 aluc|OJZ82699.1 anid|CBF89230.1 anig|EHA21155.1 aory|EIT74940.1 asyd|OJJ62515.1 ater|EAU34103.1 atub|OJI88931.1 aver|OJI96634.1 awen|OJJ37760.1 azon|OJJ45058.1

Group815: acar|OOF92601.1 acar|OOF92622.1 abra|OJJ77122.1 acla|EAW12935.1 afis|EAW17711.1 afla|EED45713.1 afum|EAL91526.1 agla|OJJ81737.1 akaw|GAA90806.1 aluc|OJZ84532.1 anid|CBF76922.1 anig|EHA27830.1 aory|EIT75682.1 asyd|OJJ57071.1 ater|EAU31890.1 atub|OJI89548.1 aver|OJJ03227.1 awen|OJJ37125.1 azon|OJJ50824.1

Group816: acar|OOF92602.1 acar|OOF92623.1 abra|OJJ75110.1 acla|EAW08316.1 afis|EAW23324.1 afla|EED55642.1 afum|EAL88155.1 agla|OJJ86519.1 akaw|GAA88719.1 aluc|OJZ92676.1 anid|CBF89451.1 anig|EHA26322.1 aory|EIT81779.1 asyd|OJJ62432.1 ater|EAU35572.1 atub|OJI87104.1 aver|OJI96568.1 awen|OJJ39570.1 azon|OJJ47763.1

Group817: acar|OOF92604.1 acar|OOF92625.1 abra|OJJ78212.1 acla|EAW11239.1 afis|EAW18266.1 afla|EED48732.1 afum|EAL87631.1 agla|OJJ88160.1 akaw|GAA85815.1 aluc|OJZ91284.1 anid|CBF79520.1 anig|EHA19969.1 aory|EIT73261.1 asyd|OJJ57890.1 ater|EAU33273.1 atub|OJI83274.1 aver|OJJ04504.1 awen|OJJ30640.1 azon|OJJ47975.1

Group818: acar|OOF92606.1 acar|OOF92627.1 abra|OJJ72009.1 acla|EAW09531.1 afis|EAW21601.1 afla|EED57828.1 afum|EAL92620.1 agla|OJJ85796.1 akaw|GAA84924.1 aluc|OJZ86745.1 anid|CBF83460.1 anig|EHA21934.1 aory|EIT81025.1 asyd|OJJ60683.1 ater|EAU38291.1 atub|OJI83782.1 aver|OJJ00008.1 awen|OJJ39081.1 azon|OJJ51413.1

Group819: acar|OOF92607.1 acar|OOF92628.1 abra|OJJ71278.1 acla|EAW10260.1 afis|EAW25333.1 afla|EED53813.1 afum|EAL89996.1 agla|OJJ83874.1 akaw|GAA82710.1 aluc|OJZ87265.1 anid|CBF75238.1 anig|EHA20507.1 aory|EIT78421.1 asyd|OJJ59912.1 ater|EAU34479.1 atub|OJI81377.1 aver|OJJ02374.1 awen|OJJ41027.1 azon|OJJ44637.1

Group820: acar|OOF92609.1 acar|OOF92630.1 abra|OJJ74612.1 acla|EAW10770.1 afis|EAW24806.1 afla|EED47177.1 afum|EAL89448.1 agla|OJJ82901.1 akaw|GAA81996.1 aluc|OJZ92192.1 anid|CBF85894.1 anig|EHA26762.1 aory|EIT79234.1 asyd|OJJ64389.1 ater|EAU36500.1 atub|OJI86621.1 aver|OJI98483.1 awen|OJJ40390.1 azon|OJJ49240.1

Group821: acar|OOF92610.1 acar|OOF92631.1 abra|OJJ71181.1 acla|EAW10323.1 afis|EAW25269.1 afla|EED53734.1 afum|EAL89932.1 agla|OJJ89237.1 akaw|GAA82804.1 aluc|OJZ87362.1 anid|CBF85380.1 anig|EHA20597.1 aory|EIT75356.1 asyd|OJJ64050.1 ater|EAU34412.1 atub|OJI81283.1 aver|OJI98136.1 awen|OJJ40983.1 azon|OJJ44569.1

Group822: acar|OOF92611.1 acar|OOF92632.1 abra|OJJ72794.1 acla|EAW11478.1 afis|EAW23975.1 afla|EED50533.1 afum|EAL87405.1 agla|OJJ86905.1 akaw|GAA85397.1 aluc|OJZ83290.1 anid|CBF77343.1 anig|EHA24175.1 aory|EIT82080.1 asyd|OJJ53273.1 ater|EAU29777.1 atub|OJI82192.1 aver|OJJ03576.1 awen|OJJ35440.1 azon|OJJ51303.1

Group823: acar|OOF92612.1 acar|OOF92633.1 abra|OJJ66525.1 acla|EAW10966.1 afis|EAW20894.1 afla|EED50243.1 afum|EAL84739.1 agla|OJJ84001.1 akaw|GAA91188.1 aluc|OJZ81515.1 anid|CBF82444.1 anig|EHA25913.1 aory|EIT77371.1 asyd|OJJ54119.1 ater|EAU38528.1 atub|OJI88664.1 aver|OJJ08617.1 awen|OJJ34454.1 azon|OJJ51741.1

Group824: acar|OOF92616.1 acar|OOF92637.1 abra|OJJ76860.1 acla|EAW13421.1 afis|EAW20532.1 afla|EED52482.1 afum|EAL84525.1 agla|OJJ84699.1 akaw|GAA87162.1 aluc|OJZ84272.1 anid|CBF79000.1 anig|EHA27614.1 aory|EIT82763.1 asyd|OJJ55688.1 ater|EAU37152.1 atub|OJI89811.1 aver|OJJ06772.1 awen|OJJ33239.1 azon|OJJ50618.1

Group825: acar|OOF93657.1 acar|OOF93741.1 abra|OJJ66636.1 acla|EAW11779.1 afis|EAW19389.1 afla|EED50347.1 afum|EAL87106.1 agla|OJJ81100.1 akaw|GAA89589.1 aluc|OJZ80813.1 anid|CBF74291.1 anig|EHA27221.1 aory|EIT81936.1 asyd|OJJ51912.1 ater|EAU36052.1 atub|OJI87662.1 aver|OJJ05353.1 awen|OJJ36782.1 azon|OJJ43016.1

Group826: acar|OOF93659.1 acar|OOF93743.1 abra|OJJ66639.1 acla|EAW11782.1 afis|EAW19392.1 afla|EED50350.1 afum|EAL87102.1 agla|OJJ81103.1 akaw|GAA89592.1 aluc|OJZ80810.1 anid|CBF74285.1 anig|EHA27224.1 aory|EIT81937.1 asyd|OJJ51914.1 ater|EAU36055.1 atub|OJI87658.1 aver|OJJ05356.1 awen|OJJ36785.1 azon|OJJ43013.1

Group827: acar|OOF93660.1 acar|OOF93744.1 abra|OJJ66640.1 acla|EAW11783.1 afis|EAW19393.1 afla|EED50351.1 agla|OJJ81104.1 akaw|GAA89593.1 aluc|OJZ80809.1 anig|EHA27225.1 aory|EIT81941.1 asyd|OJJ51863.1 ater|EAU36056.1 atub|OJI87657.1 aver|OJJ05357.1 awen|OJJ36787.1 azon|OJJ43012.1 afum|EAL87101.1 anid|CBF74283.1

Group828: acar|OOF93663.1 acar|OOF93747.1 abra|OJJ66643.1 acla|EAW11786.1 afis|EAW19396.1 afla|EED50352.1 afum|EAL87098.1 agla|OJJ81107.1 akaw|GAA89596.1 aluc|OJZ80806.1 anid|CBF74278.1 anig|EHA27228.1 aory|EIT83119.1 asyd|OJJ51859.1 ater|EAU36122.1 atub|OJI87654.1 aver|OJJ05360.1 awen|OJJ36791.1 azon|OJJ43008.1

Group829: acar|OOF93667.1 acar|OOF93751.1 abra|OJJ66647.1 acla|EAW11790.1 afis|EAW19400.1 afla|EED50355.1 afum|EAL87094.1 agla|OJJ81111.1 akaw|GAA89600.1 aluc|OJZ80802.1 anid|CBF74336.1 anig|EHA27232.1 aory|EIT83130.1 asyd|OJJ51906.1 ater|EAU36126.1 atub|OJI87650.1 aver|OJJ05363.1 awen|OJJ36795.1 azon|OJJ43004.1

Group830: acar|OOF93669.1 acar|OOF93753.1 abra|OJJ66649.1 acla|EAW11792.1 afis|EAW19403.1 afla|EED50358.1 afum|EAL87092.1 agla|OJJ81117.1 akaw|GAA89602.1 aluc|OJZ80800.1 anid|CBF74298.1 anig|EHA27234.1 aory|EIT83146.1 asyd|OJJ51902.1 ater|EAU36143.1 atub|OJI87648.1 aver|OJJ05365.1 awen|OJJ36798.1 azon|OJJ43001.1

Group831: acar|OOF93671.1 acar|OOF93755.1 abra|OJJ66653.1 acla|EAW11795.1 afis|EAW19405.1 afla|EED50361.1 afum|EAL87090.1 agla|OJJ81119.1 akaw|GAA89604.1 aluc|OJZ80794.1 anid|CBF74345.1 anig|EHA27237.1 aory|EIT83126.1 asyd|OJJ57796.1 ater|EAU36140.1 atub|OJI87645.1 aver|OJJ05368.1 awen|OJJ36801.1 azon|OJJ42997.1

Group832: acar|OOF93672.1 acar|OOF93756.1 abra|OJJ66655.1 acla|EAW11796.1 afis|EAW19406.1 afla|EED50362.1 afum|EAL87088.1 agla|OJJ81120.1 akaw|GAA89605.1 aluc|OJZ80793.1 anid|CBF74347.1 anig|EHA27238.1 aory|EIT83144.1 asyd|OJJ57795.1 ater|EAU36139.1 atub|OJI87644.1 aver|OJJ05369.1 awen|OJJ36802.1 azon|OJJ42996.1

Group833: acar|OOF93673.1 acar|OOF93757.1 abra|OJJ66656.1 acla|EAW11797.1 afis|EAW19407.1 afla|EED50363.1 afum|EAL87087.1 agla|OJJ81121.1 akaw|GAA89606.1 aluc|OJZ80792.1 anid|CBF74349.1 anig|EHA27239.1 aory|EIT83134.1 asyd|OJJ57794.1 ater|EAU36138.1 atub|OJI87643.1 aver|OJJ05370.1 awen|OJJ36803.1 azon|OJJ42995.1

Group834: acar|OOF93674.1 acar|OOF93758.1 abra|OJJ66657.1 acla|EAW11798.1 afis|EAW19408.1 afla|EED50364.1 afum|EAL87086.1 agla|OJJ81122.1 akaw|GAA89607.1 aluc|OJZ80791.1 anid|CBF74356.1 anig|EHA27240.1 aory|EIT83115.1 asyd|OJJ57793.1 ater|EAU34715.1 atub|OJI87642.1 aver|OJJ05371.1 awen|OJJ36804.1 azon|OJJ42994.1

Group835: acar|OOF93676.1 acar|OOF93761.1 abra|OJJ66660.1 acla|EAW11802.1 afis|EAW19412.1 afla|EED50367.1 afum|EAL87082.1 agla|OJJ81126.1 akaw|GAA89610.1 aluc|OJZ80788.1 anid|CBF74358.1 anig|EHA27242.1 aory|EIT83138.1 asyd|OJJ57789.1 ater|EAU34718.1 atub|OJI87639.1 aver|OJJ05375.1 awen|OJJ36808.1 azon|OJJ42992.1

Group836: acar|OOF93677.1 acar|OOF93762.1 abra|OJJ66661.1 acla|EAW11803.1 afis|EAW19413.1 afla|EED50368.1 afum|EAL87081.1 agla|OJJ81127.1 akaw|GAA89611.1 aluc|OJZ80787.1 anid|CBF74360.1 anig|EHA27243.1 aory|EIT83132.1 asyd|OJJ57788.1 ater|EAU34719.1 atub|OJI87638.1 aver|OJJ05376.1 awen|OJJ36809.1 azon|OJJ42991.1

Group837: acar|OOF93678.1 acar|OOF93763.1 abra|OJJ66662.1 acla|EAW11804.1 afis|EAW19414.1 afla|EED50369.1 afum|EAL87080.2 agla|OJJ81128.1 akaw|GAA89612.1 aluc|OJZ80786.1 anid|CBF74362.1 anig|EHA27244.1 aory|EIT83123.1 asyd|OJJ57787.1 ater|EAU34720.1 atub|OJI87636.1 aver|OJJ05377.1 awen|OJJ36810.1 azon|OJJ42990.1

Group838: acar|OOF93679.1 acar|OOF93765.1 abra|OJJ66664.1 acla|EAW11805.1 afis|EAW19415.1 afla|EED50371.1 afum|EAL87079.1 agla|OJJ81129.1 akaw|GAA89613.1 aluc|OJZ80783.1 anid|CBF74364.1 anig|EHA27246.1 aory|EIT83116.1 asyd|OJJ57785.1 ater|EAU34721.1 atub|OJI87635.1 aver|OJJ05379.1 awen|OJJ36812.1 azon|OJJ42989.1

Group839: acar|OOF93681.1 acar|OOF93764.1 abra|OJJ66666.1 acla|EAW11807.1 afis|EAW19417.1 afla|EED50373.1 afum|EAL87077.1 agla|OJJ81130.1 akaw|GAA89615.1 aluc|OJZ80781.1 anid|CBF86348.1 anig|EHA27248.1 aory|EIT83140.1 asyd|OJJ64978.1 ater|EAU34723.1 atub|OJI87633.1 aver|OJI99100.1 awen|OJJ36815.1 azon|OJJ42987.1

Group840: acar|OOF93685.1 acar|OOF93770.1 abra|OJJ66671.1 acla|EAW11812.1 afis|EAW19422.1 afla|EED50379.1 afum|EAL87072.1 agla|OJJ81134.1 akaw|GAA89620.1 aluc|OJZ80776.1 anid|CBF74373.1 anig|EHA27253.1 aory|EIT83120.1 asyd|OJJ57781.1 ater|EAU34726.1 atub|OJI87628.1 aver|OJJ05384.1 awen|OJJ36820.1 azon|OJJ42982.1

Group841: acar|OOF93686.1 acar|OOF93771.1 abra|OJJ66672.1 acla|EAW11813.1 afis|EAW19423.1 afla|EED50380.1 afum|EAL87071.1 agla|OJJ81135.1 akaw|GAA89621.1 aluc|OJZ80775.1 anid|CBF74375.1 anig|EHA27254.1 aory|EIT83122.1 asyd|OJJ57780.1 ater|EAU34727.1 atub|OJI87627.1 aver|OJJ05385.1 awen|OJJ36821.1 azon|OJJ42981.1

Group842: acar|OOF93690.1 acar|OOF93776.1 abra|OJJ66680.1 acla|EAW11817.1 afis|EAW19428.1 afla|EED56632.1 afum|EAL87066.1 agla|OJJ81140.1 akaw|GAA89625.1 aluc|OJZ80769.1 anid|CBF74382.1

anig|EHA27259.1 aory|EIT79645.1 asyd|OJJ57775.1 ater|EAU34731.1 atub|OJI87622.1 aver|OJJ05390.1 awen|OJJ36826.1 azon|OJJ42977.1

Group843: acar|OOF93692.1 acar|OOF93778.1 abra|OJJ66683.1 acla|EAW11820.1 afis|EAW19431.1 afla|EED56635.1 afum|EAL87063.1 agla|OJJ81143.1 akaw|GAA89628.1 aluc|OJZ80766.1 anid|CBF81374.1 anig|EHA27262.1 aory|EIT79669.1 asyd|OJJ59388.1 ater|EAU34734.1 atub|OJI87619.1 aver|OJJ02885.1 awen|OJJ36832.1 azon|OJJ42974.1

Group844: acar|OOF93693.1 acar|OOF93777.1 abra|OJJ66682.1 acla|EAW11819.1 afis|EAW19430.1 afla|EED56634.1 afum|EAL87064.1 agla|OJJ81142.1 akaw|GAA89627.1 aluc|OJZ80767.1 anid|CBF74386.1 anig|EHA27261.1 aory|EIT79632.1 asyd|OJJ57773.1 ater|EAU34733.1 atub|OJI87620.1 aver|OJJ05393.1 awen|OJJ36829.1 azon|OJJ42975.1

Group845: acar|OOF93696.1 acar|OOF93781.1 abra|OJJ66687.1 acla|EAW11822.1 afis|EAW19433.1 afla|EED56638.1 afum|EAL87062.1 agla|OJJ81145.1 akaw|GAA89631.1 aluc|OJZ80762.1 anid|CBF81380.1 anig|EHA27265.1 aory|EIT79671.1 asyd|OJJ59391.1 ater|EAU34735.1 atub|OJI87616.1 aver|OJJ02881.1 awen|OJJ36833.1 azon|OJJ42972.1

Group846: acar|OOF93699.1 acar|OOF93784.1 abra|OJJ66690.1 acla|EAW11825.1 afis|EAW19436.1 afla|EED56641.1 afum|EAL87059.1 agla|OJJ81148.1 akaw|GAA89634.1 aluc|OJZ80759.1 anid|CBF81383.1 anig|EHA27268.1 aory|EIT79678.1 asyd|OJJ59395.1 ater|EAU34738.1 atub|OJI87613.1 aver|OJJ02877.1 awen|OJJ36834.1 azon|OJJ42969.1

Group847: acar|OOF93701.1 acar|OOF93787.1 abra|OJJ66693.1 acla|EAW11828.1 afis|EAW19440.1 afla|EED56647.1 afum|EAL87056.1 agla|OJJ81155.1 akaw|GAA89637.1 aluc|OJZ80752.1 anid|CBF79760.1 anig|EHA27271.1 aory|EIT79660.1 asyd|OJJ58101.1 ater|EAU34741.1 atub|OJI87609.1 aver|OJJ04303.1 awen|OJJ36840.1 azon|OJJ42961.1

Group848: acar|OOF93703.1 acar|OOF93790.1 abra|OJJ66695.1 acla|EAW11830.1 afis|EAW19442.1 afla|EED56649.1 afum|EAL87054.1 agla|OJJ81153.1 akaw|GAA89639.1 aluc|OJZ80750.1 anid|CBF81395.1 anig|EHA27272.1 aory|EIT79630.1 asyd|OJJ59401.1 ater|EAU34743.1 atub|OJI87607.1 aver|OJJ02872.1 awen|OJJ36842.1 azon|OJJ42958.1

Group849: acar|OOF93707.1 acar|OOF93793.1 abra|OJJ66703.1 acla|EAW11834.1 afis|EAW19445.1 afla|EED56653.1 afum|EAL87050.1 agla|OJJ81160.1 akaw|GAA89645.1 aluc|OJZ80744.1 anid|CBF81404.1 anig|EHA27278.1 aory|EIT79624.1 asyd|OJJ59405.1 ater|EAU34747.1 atub|OJI87601.1 aver|OJJ02868.1 awen|OJJ36846.1 azon|OJJ42954.1

Group850: acar|OOF93708.1 acar|OOF93794.1 abra|OJJ66704.1 acla|EAW11835.1 afis|EAW19446.1 afla|EED56654.1 afum|EAL87049.1 agla|OJJ81161.1 akaw|GAA89646.1 aluc|OJZ80743.1 anid|CBF81405.1 anig|EHA27279.1 aory|EIT79662.1 asyd|OJJ59406.1 ater|EAU34748.1 atub|OJI87600.1 aver|OJJ02867.1 awen|OJJ36847.1 azon|OJJ42953.1

Group851: acar|OOF93709.1 acar|OOF93795.1 abra|OJJ66706.1 acla|EAW11837.1 afis|EAW19448.1 afla|EED56656.1 afum|EBA27299.1 agla|OJJ81163.1 akaw|GAA89648.1 aluc|OJZ80741.1 anid|CBF81409.1 anig|EHA27281.1 aory|EIT79680.1 asyd|OJJ59408.1 ater|EAU34750.1 atub|OJI87598.1 aver|OJJ02865.1 awen|OJJ36849.1 azon|OJJ42951.1

Group852: acar|OOF93710.1 acar|OOF93796.1 abra|OJJ66708.1 acla|EAW11839.1 afis|EAW19450.1 afla|EED56658.1 afum|EAL87046.1 agla|OJJ81165.1 akaw|GAA89650.1 aluc|OJZ80739.1 anid|CBF81412.1 anig|EHA27283.1 aory|EIT79655.1 asyd|OJJ59410.1 ater|EAU34752.1 atub|OJI87596.1 aver|OJJ02863.1 awen|OJJ36851.1 azon|OJJ42949.1

Group853: acar|OOF93711.1 acar|OOF93797.1 abra|OJJ66709.1 acla|EAW11840.1 afis|EAW19451.1 afla|EED56659.1 afum|EAL87045.1 agla|OJJ81166.1 akaw|GAA89651.1 aluc|OJZ80738.1 anid|CBF81414.1 anig|EHA27284.1 aory|EIT79675.1 asyd|OJJ59411.1 ater|EAU34753.1 atub|OJI87595.1 aver|OJJ02862.1 awen|OJJ36852.1 azon|OJJ42948.1

Group854: acar|OOF93715.1 acar|OOF93802.1 abra|OJJ66713.1 acla|EAW11844.1 afis|EAW19455.1 afla|EED56663.1 afum|EAL87041.1 agla|OJJ81171.1 akaw|GAA89655.1 aluc|OJZ80734.1 anid|CBF81421.1 anig|EHA27287.1 aory|EIT79656.1 asyd|OJJ59415.1 ater|EAU34758.1 atub|OJI87591.1 aver|OJJ02858.1 awen|OJJ36856.1 azon|OJJ42944.1

Group855: acar|OOF93716.1 acar|OOF93803.1 abra|OJJ66714.1 acla|EAW11845.1 afis|EAW19456.1 afla|EED56664.1 afum|EAL87040.1 agla|OJJ81173.1 akaw|GAA89656.1 aluc|OJZ80733.1 anid|CBF81423.1 anig|EHA27288.1 aory|EIT79663.1 asyd|OJJ59416.1 ater|EAU34759.1 atub|OJI87590.1 aver|OJJ02857.1 awen|OJJ36858.1 azon|OJJ42943.1

Group856: acar|OOF93717.1 acar|OOF93804.1 abra|OJJ66715.1 acla|EAW11846.1 afis|EAW19457.1 afla|EED56665.1 afum|EAL87039.1 agla|OJJ81174.1 akaw|GAA89657.1 aluc|OJZ80732.1 anid|CBF81425.1 anig|EHA27289.1 aory|EIT79668.1 asyd|OJJ59417.1 ater|EAU34760.1 atub|OJI87589.1 aver|OJJ02856.1 awen|OJJ36859.1 azon|OJJ42942.1

Group857: acar|OOF93718.1 acar|OOF93805.1 abra|OJJ66716.1 acla|EAW11847.1 afis|EAW19458.1 afla|EED56669.1 afum|EAL87038.1 agla|OJJ81175.1 akaw|GAA89658.1 aluc|OJZ80731.1 anid|CBF81427.1 anig|EHA27290.1 aory|EIT79631.1 asyd|OJJ59418.1 ater|EAU34761.1 atub|OJI87588.1 aver|OJJ02851.1 awen|OJJ36860.1 azon|OJJ42941.1

Group858: acar|OOF93721.1 acar|OOF93808.1 abra|OJJ66719.1 acla|EAW11850.1 afis|EAW19461.1 afla|EED56672.1 afum|EAL87035.1 agla|OJJ81178.1 akaw|GAA89661.1 aluc|OJZ80728.1 anid|CBF74529.1 anig|EHA27293.1 aory|EIT82995.1 asyd|OJJ57737.1 ater|EAU34764.1 atub|OJI87584.1 aver|OJJ05430.1 awen|OJJ36863.1 azon|OJJ42938.1

Group859: acar|OOF93723.1 acar|OOF93810.1 abra|OJJ66721.1 acla|EAW11852.1 afis|EAW19463.1 afla|EED56674.1 afum|EAL87033.1 agla|OJJ81180.1 akaw|GAA89663.1 aluc|OJZ80726.1 anid|CBF75491.1 anig|EHA27294.1 aory|EIT73836.1 asyd|OJJ59753.1 ater|EAU34766.1 atub|OJI87582.1 aver|OJJ02535.1 awen|OJJ36865.1 azon|OJJ42936.1

Group860: acar|OOF93724.1 acar|OOF93811.1 abra|OJJ66722.1 acla|EAW11853.1 afis|EAW19464.1 afla|EED56675.1 afum|EAL87032.1 agla|OJJ81181.1 akaw|GAA89664.1 aluc|OJZ80725.1 anid|CBF75490.1 anig|EHA27295.1 aory|EIT73882.1 asyd|OJJ59754.1 ater|EAU34768.1 atub|OJI87581.1 aver|OJJ02534.1 awen|OJJ36866.1 azon|OJJ42935.1

Group861: acar|OOF93726.1 acar|OOF93814.1 abra|OJJ66726.1 acla|EAW11856.1 afis|EAW19467.1 afla|EED56677.1 afum|EAL87029.1 agla|OJJ81187.1 akaw|GAA89667.1 aluc|OJZ80722.1 anid|CBF75486.1 anig|EHA27298.1 aory|EIT73931.1 asyd|OJJ59756.1 ater|EAU34777.1 atub|OJI87578.1 aver|OJJ02532.1 awen|OJJ36872.1 azon|OJJ42932.1

Group862: acar|OOF93727.1 acar|OOF93815.1 abra|OJJ66727.1 acla|EAW11857.1 afis|EAW19468.1 afla|EED56678.1 afum|EAL87028.1 agla|OJJ81188.1 akaw|GAA89668.1 aluc|OJZ80721.1 anid|CBF75484.1 anig|EHA27299.1 aory|EIT73958.1 asyd|OJJ59757.1 ater|EAU34778.1 atub|OJI87577.1 aver|OJJ02531.1 awen|OJJ36873.1 azon|OJJ42931.1

Group863: acar|OOF93728.1 acar|OOF93816.1 abra|OJJ66728.1 acla|EAW11858.1 afis|EAW19469.1 afla|EED56679.1 afum|EAL87027.1 agla|OJJ81190.1 akaw|GAA89669.1 aluc|OJZ80720.1 anid|CBF75482.1 anig|EHA27300.1 aory|EIT73843.1 asyd|OJJ59758.1 ater|EAU34779.1 atub|OJI87576.1 aver|OJJ02530.1 awen|OJJ36874.1 azon|OJJ42930.1

Group864: acar|OOF93729.1 acar|OOF93817.1 abra|OJJ66731.1 acla|EAW11859.1 afis|EAW19470.1 afla|EED56680.1 afum|EAL87026.1 agla|OJJ81191.1 akaw|GAA89670.1 aluc|OJZ80718.1 anid|CBF75480.1 anig|EHA27301.1 aory|EIT73854.1 asyd|OJJ59759.1 ater|EAU34780.1 atub|OJI87574.1 aver|OJJ02529.1 awen|OJJ36875.1 azon|OJJ42929.1

Group865: acar|OOF93731.1 acar|OOF93819.1 abra|OJJ66733.1 acla|EAW11861.1 afis|EAW19472.1 afla|EED56682.1 afum|EAL87024.1 agla|OJJ81193.1 akaw|GAA89672.1 aluc|OJZ80716.1 anid|CBF75474.1 anig|EHA27303.1 aory|EIT73961.1 asyd|OJJ59761.1 ater|EAU34782.1 atub|OJI87572.1 aver|OJJ02527.1 awen|OJJ36877.1 azon|OJJ42927.1

Group866: acar|OOF93734.1 acar|OOF93822.1 abra|OJJ66736.1 acla|EAW11864.1 afis|EAW19475.1 afla|EED56683.1 afum|EAL87021.1 agla|OJJ81198.1 akaw|GAA89675.1 aluc|OJZ80711.1 anid|CBF75470.1 anig|EHA27306.1 aory|EIT73904.1 asyd|OJJ59764.1 ater|EAU34787.1 atub|OJI87569.1 aver|OJJ02524.1 awen|OJJ36880.1 azon|OJJ42925.1

Group867: acar|OOF93736.1 acar|OOF93825.1 abra|OJJ66739.1 acla|EAW11866.1 afis|EAW19477.1 afla|EED56685.1 afum|EAL87019.1 agla|OJJ81207.1 akaw|GAA89677.1 aluc|OJZ80709.1 anid|CBF71519.1 anig|EHA27308.1 aory|EIT73831.1 asyd|OJJ54916.1 ater|EAU34789.1 atub|OJI87567.1 aver|OJJ07287.1 awen|OJJ36882.1 azon|OJJ42923.1

Group868: acar|OOF93739.1 acar|OOF93829.1 abra|OJJ66743.1 acla|EAW11872.1 afis|EAW19483.1 afla|EED56701.1 afum|EAL87012.1 agla|OJJ81208.1 akaw|GAA89681.1 aluc|OJZ80704.1 anid|CBF75464.1 anig|EHA27313.1 aory|EIT73864.1 asyd|OJJ59767.1 ater|EAU34796.1 atub|OJI87562.1 aver|OJJ02520.1 awen|OJJ36895.1 azon|OJJ42918.1

Group870: acar|OOF94064.1 acar|OOF94134.1 abra|OJJ66360.1 acla|EAW10009.1 afis|EAW21047.1 afla|EED51623.1 afum|EAL92083.1 agla|OJJ82365.1 akaw|GAA90233.1 aluc|OJZ83108.1 anid|CBF87099.1 anig|EHA28171.1 aory|EIT80605.1 asyd|OJJ60391.1 ater|EAU36819.1 atub|OJI87826.1 aver|OJI99720.1 awen|OJJ32175.1 azon|OJJ46754.1

Group871: acar|OOF94087.1 acar|OOF94157.1 abra|OJJ66351.1 acla|EAW06467.1 afis|EAW15962.1 afla|EED58095.1 afum|EAL88595.1 agla|OJJ89350.1 akaw|GAA90224.1 aluc|OJZ83117.1 anid|CBF81081.1 anig|EHA28180.1 aory|EIT79147.1 asyd|OJJ56464.1 ater|EAU35492.1 atub|OJI87817.1 aver|OJJ06256.1 awen|OJJ35612.1 azon|OJJ46912.1

Group872: acar|OOF94090.1 acar|OOF94160.1 abra|OJJ66347.1 acla|EAW09988.1 afis|EAW21105.1 afla|EED51687.1 afum|EAL92126.1 agla|OJJ80110.1 akaw|GAA90221.1 aluc|OJZ83120.1 anid|CBF87045.1

anig|EHA28184.1 aory|EIT80607.1 asyd|OJJ60445.1 ater|EAU36794.1 atub|OJI87813.1 aver|OJI99763.1 awen|OJJ36366.1 azon|OJJ43506.1

Group874: acar|OOF95064.1 acar|OOF95130.1 abra|OJJ74649.1 acla|EAW10737.1 afis|EAW24838.1 afla|EED47138.1 afum|EAL89484.1 agla|OJJ82029.1 akaw|GAA81961.1 aluc|OJZ92227.1 anid|CBF74603.1 anig|EHA26729.1 aory|EIT79201.1 asyd|OJJ57657.1 ater|EAU36468.1 atub|OJI86657.1 aver|OJJ05518.1 awen|OJJ40424.1 azon|OJJ49282.1

Group875: acar|OOF95067.1 acar|OOF95133.1 abra|OJJ74647.1 acla|EAW10740.1 afis|EAW24834.1 afla|EED47141.1 afum|EAL89479.1 agla|OJJ82027.1 akaw|GAA81964.1 aluc|OJZ92225.1 anid|CBF74582.1 anig|EHA26731.1 aory|EIT79244.1 asyd|OJJ57655.1 ater|EAU36471.1 atub|OJI86655.1 aver|OJJ05515.1 awen|OJJ40421.1 azon|OJJ49279.1

Group876: acar|OOF95068.1 acar|OOF95134.1 abra|OJJ74646.1 acla|EAW10741.1 afis|EAW24833.1 afla|EED47142.1 afum|EAL89478.1 agla|OJJ82026.1 akaw|GAA81965.1 aluc|OJZ92224.1 anid|CBF74580.1 anig|EHA26732.1 aory|EIT79261.1 asyd|OJJ57654.1 ater|EAU36472.1 atub|OJI86654.1 aver|OJJ05514.1 awen|OJJ40420.1 azon|OJJ49278.1

Group877: acar|OOF95069.1 acar|OOF95135.1 abra|OJJ68490.1 acla|EAW10742.1 afis|EAW24832.1 afla|EED47143.1 afum|EAL89477.1 agla|OJJ82025.1 akaw|GAA92780.1 aluc|OJZ82703.1 anid|CBF74578.1 anig|EHA21158.1 aory|EIT79304.1 asyd|OJJ57653.1 ater|EAU36473.1 atub|OJI88936.1 aver|OJJ05513.1 awen|OJJ40419.1 azon|OJJ49277.1

Group878: acar|OOF95070.1 acar|OOF95136.1 abra|OJJ74645.1 acla|EAW10743.1 afis|EAW24831.1 afla|EED47144.1 afum|EAL89476.1 agla|OJJ82019.1 akaw|GAA81966.1 aluc|OJZ92223.1 anid|CBF74576.1 anig|EHA26733.1 aory|EIT79280.1 asyd|OJJ57652.1 ater|EAU36474.1 atub|OJI86653.1 aver|OJJ05512.1 awen|OJJ40418.1 azon|OJJ49276.1

Group879: acar|OOF95076.1 acar|OOF95140.1 abra|OJJ74640.1 acla|EAW10748.1 afis|EAW24826.1 afla|EED47149.1 afum|EAL89470.1 agla|OJJ82015.1 akaw|GAA81971.1 aluc|OJZ92219.1 anid|CBF85858.1 anig|EHA26739.1 aory|EIT79249.1 asyd|OJJ64367.1 ater|EAU36479.1 atub|OJI86650.1 aver|OJI98462.1 awen|OJJ40412.1 azon|OJJ49262.1

Group880: acar|OOF95080.1 acar|OOF95145.1 abra|OJJ74631.1 acla|EAW10753.1 afis|EAW24822.1 afla|EED47153.1 afum|EAL89466.1 agla|OJJ82920.1 akaw|GAA81978.1 aluc|OJZ92212.1 anid|CBF85851.1 anig|EHA26746.1 aory|EIT79200.1 asyd|OJJ64363.1 ater|EAU36483.1 atub|OJI86641.1 aver|OJI98458.1 awen|OJJ40408.1 azon|OJJ49257.1

Group882: acar|OOF95084.1 acar|OOF95149.1 abra|OJJ74624.1 acla|EAW10760.1 afis|EAW24816.1 afla|EED47167.1 afum|EAL89460.2 agla|OJJ82910.1 akaw|GAA81984.1 aluc|OJZ92206.1 anid|CBF85879.1 anig|EHA26752.1 aory|EIT79187.1 asyd|OJJ64377.1 ater|EAU39452.1 atub|OJI86635.1 aver|OJI98473.1 awen|OJJ40402.1 azon|OJJ43613.1

Group883: acar|OOF95086.1 acar|OOF95152.1 abra|OJJ74618.1 acla|EAW10764.1 afis|EAW24812.1 afla|EED47171.1 afum|EAL89455.1 agla|OJJ82907.1 akaw|GAA81988.1 aluc|OJZ92201.1 anid|CBF85885.1 anig|EHA26756.1 aory|EIT79186.1 asyd|OJJ64381.1 ater|EAU36495.1 atub|OJI86629.1 aver|OJI98477.1 awen|OJJ40396.1 azon|OJJ49249.1

Group884: acar|OOF95087.1 acar|OOF95153.1 abra|OJJ74617.1 acla|EAW10765.1 afis|EAW24811.1 afla|EED47172.1 afum|EAL89454.1 agla|OJJ82906.1 akaw|GAA81989.1 aluc|OJZ92200.1 anid|CBF85886.1 anig|EHA26757.1 aory|EIT79291.1 asyd|OJJ64382.1 ater|EAU36496.1 atub|OJI86627.1 aver|OJI98478.1 awen|OJJ40395.1 azon|OJJ49248.1

Group885: acar|OOF95089.1 acar|OOF95155.1 abra|OJJ74615.1 acla|EAW10767.1 afis|EAW24809.1 afla|EED47174.1 afum|EAL89452.1 agla|OJJ82904.1 akaw|GAA81991.1 aluc|OJZ92198.1 anid|CBF85888.1 anig|EHA26759.1 aory|EIT79300.1 asyd|OJJ64384.1 ater|EAU36497.1 atub|OJI86625.1 aver|OJI98480.1 awen|OJJ40393.1 azon|OJJ49246.1

Group886: acar|OOF95090.1 acar|OOF95156.1 abra|OJJ74614.1 acla|EAW10768.1 afis|EAW24808.1 afla|EED47175.1 afum|EAL89450.2 agla|OJJ82903.1 akaw|GAA81992.1 aluc|OJZ92197.1 anid|CBF85890.1 anig|EHA26760.1 aory|EIT79254.1 asyd|OJJ64385.1 ater|EAU36498.1 atub|OJI86624.1 aver|OJI98481.1 awen|OJJ40392.1 azon|OJJ49245.1

Group887: acar|OOF95091.1 acar|OOF95157.1 abra|OJJ74613.1 acla|EAW10769.1 afis|EAW24807.1 afla|EED47176.1 afum|EAL89449.1 agla|OJJ82902.1 akaw|GAA81995.1 aluc|OJZ92193.1 anid|CBF85892.1 anig|EHA26761.1 aory|EIT79197.1 asyd|OJJ64387.1 ater|EAU36499.1 atub|OJI86622.1 aver|OJI98482.1 awen|OJJ40391.1 azon|OJJ49241.1

Group888: acar|OOF95092.1 acar|OOF95158.1 abra|OJJ74611.1 acla|EAW10771.1 afis|EAW24805.1 afla|EED47180.1 afum|EAL89447.1 agla|OJJ82900.1 akaw|GAA81997.1 aluc|OJZ92191.1 anid|CBF81492.1 anig|EHA26763.1 aory|EIT79217.1 asyd|OJJ59459.1 ater|EAU36502.1 atub|OJI86620.1 aver|OJJ02814.1 awen|OJJ40389.1 azon|OJJ49239.1

Group889: acar|OOF95093.1 acar|OOF95159.1 abra|OJJ74610.1 acla|EAW10772.1 afis|EAW24804.1 afla|EED47181.1 afum|EAL89446.1 agla|OJJ82899.1 akaw|GAA81998.1 aluc|OJZ92190.1 anid|CBF81493.1 anig|EHA26764.1 aory|EIT79264.1 asyd|OJJ59460.1 ater|EAU36503.1 atub|OJI86619.1 aver|OJJ02813.1 awen|OJJ40388.1 azon|OJJ49238.1

Group890: acar|OOF95095.1 acar|OOF95162.1 abra|OJJ74608.1 acla|EAW10773.1 afis|EAW24803.1 afla|EED47183.1 afum|EAL89444.1 agla|OJJ82897.1 akaw|GAA82000.1 aluc|OJZ92188.1 anid|CBF81495.1 anig|EHA26765.1 aory|EIT79220.1 asyd|OJJ59463.1 ater|EAU36504.1 atub|OJI86617.1 aver|OJJ02811.1 azon|OJJ49236.1 awen|OJJ40386.1

Group891: acar|OOF95101.1 acar|OOF95167.1 abra|OJJ74603.1 acla|EAW10776.1 afis|EAW24798.1 afla|EED47189.1 afum|EAL89440.1 agla|OJJ82894.1 akaw|GAA82005.1 aluc|OJZ92182.1 anid|CBF81508.1 anig|EHA26771.1 aory|EIT79258.1 asyd|OJJ59468.1 ater|EAU36507.1 atub|OJI86610.1 aver|OJJ02806.1 awen|OJJ40383.1 azon|OJJ49232.1

Group893: acar|OOF95104.1 acar|OOF95170.1 abra|OJJ74600.1 acla|EAW10779.1 afis|EAW24795.1 afla|EED47187.1 afum|EAL89437.1 agla|OJJ82891.1 akaw|GAA82008.1 aluc|OJZ92179.1 anid|CBF81503.1 anig|EHA26774.1 aory|EIT79230.1 asyd|OJJ59466.1 ater|EAU36510.1 atub|OJI86607.1 aver|OJJ02808.1 awen|OJJ40380.1 azon|OJJ49230.1

Group894: acar|OOF95106.1 acar|OOF95172.1 abra|OJJ74598.1 acla|EAW10781.1 afis|EAW24788.1 afla|EED47191.1 afum|EAL89427.1 agla|OJJ82890.1 akaw|GAA82010.1 aluc|OJZ92176.1 anid|CBF81466.1 anig|EHA26776.1 aory|EIT79298.1 asyd|OJJ59443.1 ater|EAU36513.1 atub|OJI86605.1 aver|OJJ02830.1 awen|OJJ40378.1 azon|OJJ49229.1

Group895: acar|OOF95629.1 acar|OOF95680.1 abra|OJJ74664.1 acla|EAW10731.1 afis|EAW24844.1 afla|EED47127.1 afum|EAL89490.1 agla|OJJ82044.1 akaw|GAA81947.1 aluc|OJZ92241.1 anid|CBF75849.1 anig|EHA26713.1 aory|EIT79274.1 asyd|OJJ57571.1 ater|EAU36460.1 atub|OJI86675.1 aver|OJJ05608.1 awen|OJJ40429.1 azon|OJJ49290.1

Group896: acar|OOF95630.1 acar|OOF95681.1 abra|OJJ74665.1 acla|EAW10730.1 afis|EAW24845.1 afla|EED47126.1 afum|EAL89491.1 agla|OJJ82045.1 akaw|GAA81946.1 aluc|OJZ92242.1 anid|CBF75847.1 anig|EHA26712.1 aory|EIT79168.1 asyd|OJJ57562.1 ater|EAU36459.1 atub|OJI86676.1 aver|OJJ05616.1 awen|OJJ40430.1 azon|OJJ49291.1

Group897: acar|OOF95631.1 acar|OOF95683.1 abra|OJJ74666.1 acla|EAW10729.1 afis|EAW24846.1 afla|EED47125.1 afum|EAL89492.1 agla|OJJ82058.1 akaw|GAA81945.1 aluc|OJZ92244.1 anid|CBF75845.1 anig|EHA26711.1 aory|EIT79177.1 asyd|OJJ57561.1 ater|EAU36457.1 atub|OJI86678.1 aver|OJJ05617.1 awen|OJJ40431.1 azon|OJJ49292.1

Group898: acar|OOF95632.1 acar|OOF95684.1 abra|OJJ74671.1 acla|EAW10726.1 afis|EAW24849.1 afla|EED47122.1 afum|EAL89495.1 agla|OJJ82061.1 akaw|GAA81942.1 aluc|OJZ92248.1 anid|CBF75839.1 anig|EHA26708.1 aory|EIT79246.1 asyd|OJJ57557.1 ater|EAU36454.1 atub|OJI86681.1 aver|OJJ05621.1 awen|OJJ40434.1 azon|OJJ49295.1

Group899: acar|OOF95633.1 acar|OOF95685.1 abra|OJJ74668.1 acla|EAW10728.1 afis|EAW24847.1 afla|EED47124.1 afum|EAL89493.1 agla|OJJ82059.1 akaw|GAA81944.1 aluc|OJZ92246.1 anid|CBF75843.1 anig|EHA26710.1 aory|EIT79296.1 asyd|OJJ57560.1 ater|EAU36456.1 atub|OJI86679.1 aver|OJJ05618.1 awen|OJJ40432.1 azon|OJJ49293.1

Group900: acar|OOF95638.1 acar|OOF95689.1 abra|OJJ74675.1 acla|EAW10722.1 afis|EAW24851.1 afla|EED47120.1 afum|EAL89498.1 agla|OJJ82065.1 akaw|GAA81938.1 aluc|OJZ92252.1 anid|CBF75835.1 anig|EHA26704.1 aory|EIT79290.1 asyd|OJJ57555.1 ater|EAU36452.1 atub|OJI86685.1 aver|OJJ05623.1 awen|OJJ40436.1 azon|OJJ49297.1

Group901: acar|OOF95639.1 acar|OOF95690.1 abra|OJJ74677.1 acla|EAW10720.1 afis|EAW24853.1 afla|EED47118.1 afum|EAL89500.1 agla|OJJ82067.1 akaw|GAA81937.1 aluc|OJZ92254.1 anid|CBF75831.1 anig|EHA26703.1 aory|EIT79297.1 asyd|OJJ57553.1 ater|EAU36449.1 atub|OJI86687.1 aver|OJJ05625.1 awen|OJJ40438.1 azon|OJJ49299.1

Group902: acar|OOF95640.1 acar|OOF95691.1 abra|OJJ74678.1 acla|EAW10719.1 afis|EAW24854.1 afla|EED47117.1 afum|EAL89501.1 agla|OJJ82068.1 akaw|GAA81936.1 aluc|OJZ92255.1 anid|CBF90310.1 anig|EHA26702.1 aory|EIT79215.1 asyd|OJJ61915.1 ater|EAU36448.1 atub|OJI86688.1 aver|OJI96008.1 awen|OJJ40439.1 azon|OJJ46990.1

Group903: acar|OOF95641.1 acar|OOF95692.1 abra|OJJ74679.1 acla|EAW10716.1 afis|EAW24855.1 afla|EED47094.1 afum|EAL89503.2 agla|OJJ82069.1 akaw|GAA81935.1 aluc|OJZ92256.1 anid|CBF75829.1 anig|EHA26701.1 aory|EIT72488.1 asyd|OJJ57552.1 ater|EAU36447.1 atub|OJI86690.1 aver|OJJ05626.1 awen|OJJ40443.1 azon|OJJ49301.1

Group904: acar|OOF95643.1 acar|OOF95694.1 abra|OJJ74681.1 acla|EAW10714.1 afis|EAW24856.1 afla|EED47096.1 afum|EAL89504.1 agla|OJJ82071.1 akaw|GAA81933.1 aluc|OJZ92258.1 anid|CBF75827.1

anig|EHA26700.1 aory|EIT72489.1 asyd|OJJ57550.1 ater|EAU36446.1 atub|OJI86692.1 aver|OJJ05628.1 awen|OJJ40445.1 azon|OJJ49302.1

Group905: acar|OOF95644.1 acar|OOF95695.1 abra|OJJ74683.1 acla|EAW10713.1 afis|EAW24857.1 afla|EED47097.1 afum|EAL89505.2 agla|OJJ82072.1 akaw|GAA81931.1 aluc|OJZ92260.1 anid|CBF75825.1 anig|EHA26699.1 aory|EIT72494.1 asyd|OJJ57549.1 ater|EAU36443.1 atub|OJI86694.1 aver|OJJ05629.1 awen|OJJ40446.1 azon|OJJ49303.1

Group906: acar|OOF95645.1 acar|OOF95696.1 abra|OJJ74684.1 acla|EAW10712.1 afis|EAW24858.1 afla|EED47098.1 afum|EAL89506.2 agla|OJJ82073.1 akaw|GAA81930.1 aluc|OJZ92261.1 anid|CBF75823.1 anig|EHA26698.1 aory|EIT72492.1 asyd|OJJ57548.1 ater|EAU36442.1 atub|OJI86695.1 aver|OJJ05630.1 awen|OJJ40447.1 azon|OJJ49304.1

Group907: acar|OOF95647.1 acar|OOF95698.1 abra|OJJ74686.1 acla|EAW10710.1 afis|EAW24860.1 afla|EED47100.1 afum|EAL89509.1 agla|OJJ82078.1 akaw|GAA81928.1 aluc|OJZ92263.1 anid|CBF75819.1 anig|EHA26696.1 aory|EIT72779.1 asyd|OJJ57546.1 ater|EAU36440.1 atub|OJI86697.1 aver|OJJ05632.1 awen|OJJ40450.1 azon|OJJ49306.1

Group908: acar|OOF95648.1 acar|OOF95699.1 abra|OJJ74687.1 acla|EAW10709.1 afis|EAW24861.1 afla|EED47101.1 afum|EAL89510.1 agla|OJJ82079.1 akaw|GAA81927.1 aluc|OJZ92264.1 anid|CBF75817.1 anig|EHA26695.1 aory|EIT72784.1 asyd|OJJ57545.1 ater|EAU36439.1 atub|OJI86698.1 aver|OJJ05633.1 awen|OJJ40451.1 azon|OJJ49307.1

Group909: acar|OOF95649.1 acar|OOF95700.1 abra|OJJ74688.1 acla|EAW10708.1 afis|EAW24862.1 afla|EED47102.1 afum|EAL89511.1 agla|OJJ82080.1 akaw|GAA81926.1 aluc|OJZ92265.1 anid|CBF75815.1 anig|EHA26694.1 aory|EIT72778.1 asyd|OJJ57544.1 ater|EAU36438.1 atub|OJI86699.1 aver|OJJ05634.1 awen|OJJ40452.1 azon|OJJ49308.1

Group910: acar|OOF95650.1 acar|OOF95701.1 abra|OJJ74689.1 acla|EAW10707.1 afis|EAW24863.1 afla|EED47103.1 afum|EAL89513.1 agla|OJJ82081.1 akaw|GAA81925.1 aluc|OJZ92266.1 anid|CBF75813.1 anig|EHA26693.1 aory|EIT72782.1 asyd|OJJ57543.1 ater|EAU36437.1 atub|OJI86700.1 aver|OJJ05635.1 awen|OJJ40453.1 azon|OJJ49309.1

Group911: acar|OOF95651.1 acar|OOF95702.1 abra|OJJ74690.1 acla|EAW10706.1 afis|EAW24864.1 afla|EED47104.1 afum|EAL89514.1 agla|OJJ82082.1 akaw|GAA81924.1 aluc|OJZ92267.1 anid|CBF75811.1 anig|EHA26692.1 aory|EIT72780.1 asyd|OJJ57542.1 ater|EAU36436.1 atub|OJI86701.1 aver|OJJ05636.1 awen|OJJ40454.1 azon|OJJ49310.1

Group912: acar|OOF95653.1 acar|OOF95704.1 abra|OJJ74693.1 acla|EAW10705.1 afis|EAW24865.1 afla|EED47106.1 afum|EAL89515.1 agla|OJJ82083.1 akaw|GAA81923.1 aluc|OJZ92270.1 anid|CBF75809.1 anig|EHA26691.1 aory|EIT72781.1 asyd|OJJ57541.1 ater|EAU36435.1 atub|OJI86704.1 aver|OJJ05637.1 awen|OJJ40455.1 azon|OJJ49311.1

Group913: acar|OOF95660.1 acar|OOF95711.1 abra|OJJ68465.1 acla|EAW10698.1 afis|EAW24872.1 afla|EED47112.1 afum|EAL89522.1 agla|OJJ82093.1 akaw|GAA92385.1 aluc|OJZ83244.1 anid|CBF75794.1 anig|EHA21099.1 aory|EIT79170.1 asyd|OJJ57530.1 ater|EAU36429.1 atub|OJI87692.1 aver|OJJ05648.1 awen|OJJ40463.1 azon|OJJ49323.1

Group914: acar|OOF95661.1 acar|OOF95712.1 abra|OJJ68466.1 acla|EAW10697.1 afis|EAW24873.1 afla|EED47113.1 afum|EAL89523.1 agla|OJJ82094.1 akaw|GAA92386.1 aluc|OJZ83245.1 anid|CBF75792.1 anig|EHA21100.1 aory|EIT79294.1 asyd|OJJ57531.1 ater|EAU36428.1 atub|OJI87691.1 aver|OJJ05647.1 awen|OJJ40464.1 azon|OJJ49324.1

Group915: acar|OOF95662.1 acar|OOF95713.1 abra|OJJ68467.1 acla|EAW10696.1 afis|EAW24874.1 afla|EED47114.1 afum|EAL89524.1 agla|OJJ82095.1 akaw|GAA92388.1 aluc|OJZ83248.1 anid|CBF75790.1 anig|EHA21101.1 aory|EIT79175.1 asyd|OJJ57532.1 ater|EAU36427.1 atub|OJI87689.1 aver|OJJ05646.1 awen|OJJ40465.1 azon|OJJ49325.1

Group916: acar|OOF95663.1 acar|OOF95714.1 abra|OJJ74702.1 acla|EAW10695.1 afis|EAW24875.1 afla|EED47115.1 afum|EAL89525.1 agla|OJJ82096.1 akaw|GAA81917.1 aluc|OJZ92279.1 anid|CBF75789.1 anig|EHA26685.1 aory|EIT79273.1 asyd|OJJ57533.1 ater|EAU36426.1 atub|OJI86712.1 aver|OJJ05645.1 awen|OJJ40466.1 azon|OJJ49328.1

Group917: acar|OOF95665.1 acar|OOF95716.1 abra|OJJ74704.1 acla|EAW10693.1 afis|EAW24877.1 afla|EED47089.1 afum|EAL89527.1 agla|OJJ82098.1 akaw|GAA81914.1 aluc|OJZ92284.1 anid|CBF75758.1 anig|EHA26684.1 aory|EIT72484.1 asyd|OJJ57527.1 ater|EAU36424.1 atub|OJI86715.1 aver|OJJ05651.1 awen|OJJ40467.1 azon|OJJ49330.1

Group918: acar|OOF96224.1 acar|OOF96262.1 abra|OJJ71508.1 acla|EAW15066.1 afis|EAW20038.1 afla|EED45342.1 afum|EAL85159.1 agla|OJJ88925.1 akaw|GAA89405.1 aluc|OJZ86247.1 anid|CBF84777.1 anig|EHA21617.1 aory|EIT73643.1 asyd|OJJ63237.1 ater|EAU38653.1 atub|OJI84295.1 aver|OJI97719.1 awen|OJJ31793.1 azon|OJJ43703.1

Group919: acar|OOF96229.1 acar|OOF96267.1 abra|OJJ71515.1 acla|EAW15072.1 afis|EAW20044.1 afla|EED45336.1 afum|EAL85164.2 agla|OJJ88918.1 akaw|GAA89401.1 aluc|OJZ86253.1 anid|CBF84789.1 anig|EHA21612.1 aory|EIT73570.1 asyd|OJJ63230.1 ater|EAU38648.1 atub|OJI84290.1 aver|OJI97726.1 awen|OJJ31800.1 azon|OJJ46321.1

Group920: acar|OOF96233.1 acar|OOF96270.1 abra|OJJ71517.1 acla|EAW15075.1 afis|EAW20046.1 afla|EED45334.1 afum|EAL85167.1 agla|OJJ88914.1 akaw|GAA89400.1 aluc|OJZ86256.1 anid|CBF84795.1 anig|EHA21610.1 aory|EIT73786.1 asyd|OJJ63227.1 ater|EAU38656.1 atub|OJI84287.1 aver|OJI97728.1 awen|OJJ31804.1 azon|OJJ46318.1

Group921: acar|OOF96234.1 acar|OOF96271.1 abra|OJJ71518.1 acla|EAW15076.1 afis|EAW20047.1 afla|EED45333.1 afum|EAL85168.2 agla|OJJ88913.1 akaw|GAA89399.1 aluc|OJZ86257.1 anid|CBF84797.1 anig|EHA21609.1 aory|EIT73741.1 asyd|OJJ63226.1 ater|EAU38657.1 atub|OJI84286.1 aver|OJI97729.1 awen|OJJ31805.1 azon|OJJ46317.1

Group922: acar|OOF96235.1 acar|OOF96272.1 abra|OJJ71519.1 acla|EAW15077.1 afis|EAW20048.1 afla|EED45332.1 afum|EAL85169.1 agla|OJJ88912.1 akaw|GAA89398.1 aluc|OJZ86258.1 anid|CBF84799.1 anig|EHA21608.1 aory|EIT73661.1 asyd|OJJ63225.1 ater|EAU38658.1 atub|OJI84285.1 aver|OJI97730.1 awen|OJJ31806.1 azon|OJJ46316.1

Group923: acar|OOF96236.1 acar|OOF96273.1 abra|OJJ71520.1 acla|EAW15078.1 afis|EAW20049.1 afum|EAL85170.2 agla|OJJ88911.1 akaw|GAA89397.1 aluc|OJZ86259.1 anid|CBF84801.1 anig|EHA21607.1 aory|EIT73827.1 asyd|OJJ63224.1 ater|EAU38659.1 atub|OJI84284.1 aver|OJI97731.1 awen|OJJ31807.1 azon|OJJ46315.1 afla|EED45331.1

Group924: acar|OOF96239.1 acar|OOF96276.1 abra|OJJ71523.1 acla|EAW15080.1 afis|EAW20052.1 afla|EED45328.1 afum|EAL85172.1 agla|OJJ88908.1 akaw|GAA89394.1 aluc|OJZ86262.1 anid|CBF84806.1 anig|EHA21605.1 aory|EIT73685.1 asyd|OJJ63222.1 ater|EAU38661.1 atub|OJI84281.1 aver|OJI97733.1 awen|OJJ31810.1 azon|OJJ46312.1

Group925: acar|OOF96240.1 acar|OOF96277.1 abra|OJJ71524.1 acla|EAW15081.1 afis|EAW20053.1 afla|EED45327.1 afum|EAL85173.2 agla|OJJ88907.1 akaw|GAA89393.1 aluc|OJZ86263.1 anid|CBF84808.1 anig|EHA21604.1 aory|EIT73613.1 asyd|OJJ63221.1 ater|EAU38662.1 atub|OJI84280.1 aver|OJI97734.1 awen|OJJ31811.1 azon|OJJ46311.1

Group926: acar|OOF96242.1 acar|OOF96279.1 abra|OJJ71527.1 acla|EAW15083.1 afis|EAW20055.1 afla|EED45325.1 afum|EAL85175.2 agla|OJJ88905.1 akaw|GAA89391.1 aluc|OJZ86266.1 anid|CBF84813.1 anig|EHA21602.1 aory|EIT73716.1 asyd|OJJ63218.1 ater|EAU38664.1 atub|OJI84275.1 aver|OJI97737.1 awen|OJJ31814.1 azon|OJJ46309.1

Group928: acar|OOF96245.1 acar|OOF96282.1 abra|OJJ71531.1 acla|EAW15086.1 afis|EAW20058.1 afla|EED45322.1 afum|EAL85178.1 agla|OJJ88902.1 akaw|GAA89388.1 aluc|OJZ86270.1 anid|CBF84840.1 anig|EHA21599.1 aory|EIT73696.1 asyd|OJJ63215.1 ater|EAU38667.1 atub|OJI84271.1 aver|OJI97740.1 awen|OJJ31817.1 azon|OJJ46306.1

Group930: acar|OOF96249.1 acar|OOF96286.1 abra|OJJ71537.1 acla|EAW15090.1 afis|EAW20062.1 afla|EED45317.1 afum|EAL85182.2 agla|OJJ88897.1 akaw|GAA89384.1 aluc|OJZ86274.1 anid|CBF84832.1 anig|EHA21595.1 aory|EIT73715.1 asyd|OJJ63211.1 ater|EAU38671.1 atub|OJI84266.1 aver|OJI97744.1 awen|OJJ31822.1 azon|OJJ46303.1

Group931: acar|OOF96250.1 acar|OOF96287.1 abra|OJJ71540.1 acla|EAW15095.1 afis|EAW20064.1 afla|EED45313.1 afum|EAL85184.1 agla|OJJ88893.1 akaw|GAA89381.1 aluc|OJZ86278.1 anid|CBF84828.1 anig|EHA21593.1 aory|EIT73604.1 asyd|OJJ63209.1 ater|EAU38673.1 atub|OJI84261.1 aver|OJI97746.1 awen|OJJ31825.1 azon|OJJ46300.1

Group932: acar|OOF96251.1 acar|OOF96290.1 abra|OJJ71543.1 acla|EAW15097.1 afis|EAW20069.1 afla|EED45310.1 afum|EAL85187.1 agla|OJJ88891.1 akaw|GAA89379.1 aluc|OJZ86281.1 anid|CBF84823.1 anig|EHA21590.1 aory|EIT73765.1 asyd|OJJ63206.1 ater|EAU38675.1 atub|OJI84258.1 aver|OJI97749.1 awen|OJJ31828.1 azon|OJJ46296.1

Group933: acar|OOF96252.1 acar|OOF96291.1 abra|OJJ71545.1 acla|EAW15099.1 afis|EAW20071.1 afla|EED45308.1 afum|EAL85189.1 agla|OJJ88889.1 akaw|GAA89377.1 aluc|OJZ86283.1 anid|CBF84845.1 anig|EHA21588.1 aory|EIT73815.1 asyd|OJJ63204.1 ater|EAU38676.1 atub|OJI84255.1 aver|OJI97751.1 awen|OJJ31830.1 azon|OJJ46294.1

Group934: acar|OOF96254.1 acar|OOF96293.1 abra|OJJ71549.1 acla|EAW15103.1 afis|EAW20075.1 afla|EED45304.1 afum|EAL85193.2 agla|OJJ88885.1 akaw|GAA89373.1 aluc|OJZ86287.1 anid|CBF84853.1 anig|EHA21584.1 aory|EIT73551.1 asyd|OJJ63200.1 ater|EAU38683.1 atub|OJI84251.1 aver|OJI97755.1 awen|OJJ31834.1 azon|OJJ46290.1

Group935: acar|OOF96255.1 acar|OOF96294.1 abra|OJJ71550.1 acla|EAW15104.1 afis|EAW20076.1 afla|EED45302.1 afum|EAL85194.1 agla|OJJ88884.1 akaw|GAA89372.1 aluc|OJZ86288.1 anid|CBF84855.1

anig|EHA21583.1 aory|EIT73645.1 asyd|OJJ63199.1 ater|EAU38684.1 atub|OJI84250.1 aver|OJI97756.1 awen|OJJ31835.1 azon|OJJ46289.1

Group936: acar|OOF96256.1 acar|OOF96295.1 abra|OJJ71551.1 acla|EAW15105.1 afis|EAW20077.1 afla|EED45298.1 afum|EAL85195.1 agla|OJJ88883.1 akaw|GAA89371.1 aluc|OJZ86289.1 anid|CBF84867.1 anig|EHA21582.1 aory|EIT73681.1 asyd|OJJ63198.1 ater|EAU38685.1 atub|OJI84249.1 aver|OJI97757.1 awen|OJJ31836.1 azon|OJJ46288.1

Group937: acar|OOF96257.1 acar|OOF96296.1 abra|OJJ71554.1 acla|EAW15108.1 afis|EAW20079.1 afla|EED45296.1 afum|EAL85198.1 agla|OJJ88880.1 akaw|GAA89368.1 aluc|OJZ86292.1 anid|CBF84858.1 anig|EHA21579.1 aory|EIT73808.1 asyd|OJJ63195.1 ater|EAU38687.1 atub|OJI84246.1 aver|OJI97760.1 awen|OJJ31839.1 azon|OJJ46284.1

Group938: acar|OOF96906.1 acar|OOF96942.1 abra|OJJ70803.1 acla|EAW11121.1 afis|EAW15850.1 afla|EED56523.1 afum|EAL88486.1 agla|OJJ87258.1 akaw|GAA90119.1 aluc|OJZ84149.1 anid|CBF81174.1 anig|EHA19253.1 aory|EIT72791.1 asyd|OJJ59291.1 ater|EAU31665.1 atub|OJI90521.1 aver|OJJ02984.1 awen|OJJ36777.1 azon|OJJ47421.1

Group939: acar|OOF96913.1 acar|OOF96948.1 abra|OJJ70797.1 acla|EAW11126.1 afis|EAW15855.1 afla|EED56517.1 afum|EAL88491.1 agla|OJJ87252.1 akaw|GAA90113.1 aluc|OJZ84142.1 anid|CBF81185.1 anig|EHA19250.1 aory|EIT72794.1 asyd|OJJ59297.1 ater|EAU31670.1 atub|OJI90516.1 aver|OJJ02978.1 awen|OJJ36772.1 azon|OJJ47426.1

Group940: acar|OOF96914.1 acar|OOF96949.1 abra|OJJ70796.1 acla|EAW11127.1 afis|EAW15856.1 afla|EED56516.1 afum|EAL88492.1 agla|OJJ87251.1 akaw|GAA90112.1 aluc|OJZ84141.1 anid|CBF81187.1 anig|EHA19249.1 aory|EIT72847.1 asyd|OJJ59298.1 ater|EAU31671.1 atub|OJI90515.1 aver|OJJ02977.1 awen|OJJ36771.1 azon|OJJ47427.1

Group941: acar|OOF96916.1 acar|OOF96951.1 abra|OJJ70794.1 acla|EAW11129.1 afis|EAW15858.1 afla|EED56514.1 afum|EAL88494.1 agla|OJJ87249.1 akaw|GAA90110.1 aluc|OJZ84139.1 anid|CBF81191.1 anig|EHA19247.1 aory|EIT72852.1 asyd|OJJ59300.1 ater|EAU31673.1 atub|OJI90513.1 aver|OJJ02975.1 awen|OJJ36769.1 azon|OJJ47429.1

Group942: acar|OOF96917.1 acar|OOF96952.1 abra|OJJ70793.1 acla|EAW11130.1 afis|EAW15861.1 afla|EED56513.1 afum|EAL88496.1 agla|OJJ87248.1 akaw|GAA90109.1 aluc|OJZ84138.1 anid|CBF81193.1 anig|EHA19246.1 aory|EIT72867.1 asyd|OJJ59301.1 ater|EAU31674.1 atub|OJI90512.1 aver|OJJ02974.1 awen|OJJ36768.1 azon|OJJ47430.1

Group943: acar|OOF96918.1 acar|OOF96953.1 abra|OJJ70792.1 acla|EAW11131.1 afis|EAW15862.1 afla|EED56512.1 afum|EAL88497.1 agla|OJJ87247.1 akaw|GAA90108.1 aluc|OJZ84137.1 anid|CBF81194.1 anig|EHA19245.1 aory|EIT72817.1 asyd|OJJ59302.1 ater|EAU31675.1 atub|OJI90511.1 aver|OJJ02973.1 awen|OJJ36767.1 azon|OJJ48009.1

Group944: acar|OOF96919.1 acar|OOF96954.1 abra|OJJ70791.1 acla|EAW11132.1 afis|EAW15863.1 afla|EED56511.1 afum|EAL88498.1 agla|OJJ87246.1 akaw|GAA90107.1 aluc|OJZ84136.1 anid|CBF81196.1 anig|EHA19244.1 aory|EIT72814.1 asyd|OJJ59303.1 ater|EAU31676.1 atub|OJI90510.1 aver|OJJ02972.1 awen|OJJ36766.1 azon|OJJ47431.1

Group945: acar|OOF96921.1 acar|OOF96956.1 abra|OJJ70789.1 acla|EAW11133.1 afis|EAW15865.1 afla|EED56510.1 afum|EAL88499.1 agla|OJJ87244.1 akaw|GAA90105.1 aluc|OJZ84134.1 anid|CBF81200.1 anig|EHA19242.1 aory|EIT72849.1 asyd|OJJ59304.1 ater|EAU31678.1 atub|OJI90508.1 aver|OJJ02971.1 awen|OJJ36765.1 azon|OJJ47433.1

Group946: acar|OOF96924.1 acar|OOF96959.1 abra|OJJ70786.1 acla|EAW11135.1 afis|EAW15867.1 afla|EED56508.1 afum|EAL88501.1 agla|OJJ87241.1 akaw|GAA90103.1 aluc|OJZ84131.1 anid|CBF81205.1 anig|EHA19239.1 aory|EIT72789.1 asyd|OJJ59307.1 ater|EAU31680.1 atub|OJI90505.1 aver|OJJ02968.1 awen|OJJ36762.1 azon|OJJ47436.1

Group947: acar|OOF96927.1 acar|OOF96962.1 abra|OJJ70781.1 acla|EAW11138.1 afis|EAW15869.1 afla|EED56505.1 afum|EAL88503.1 agla|OJJ87238.1 akaw|GAA90100.1 aluc|OJZ84128.1 anid|CBF81211.1 anig|EHA19236.1 aory|EIT72793.1 asyd|OJJ59310.1 ater|EAU31686.1 atub|OJI90502.1 aver|OJJ02965.1 awen|OJJ36760.1 azon|OJJ47439.1

Group948: acar|OOF96928.1 acar|OOF96963.1 abra|OJJ70780.1 acla|EAW11139.1 afis|EAW15870.1 afla|EED56504.1 afum|EAL88504.1 agla|OJJ87237.1 akaw|GAA90099.1 aluc|OJZ84127.1 anid|CBF81213.1 anig|EHA19235.1 aory|EIT72861.1 asyd|OJJ59311.1 ater|EAU31687.1 atub|OJI90501.1 aver|OJJ02964.1 awen|OJJ36759.1 azon|OJJ47440.1

Group949: acar|OOF96930.1 acar|OOF96965.1 abra|OJJ70778.1 acla|EAW11141.1 afis|EAW15872.1 afla|EED56503.1 afum|EAL88506.1 agla|OJJ87235.1 akaw|GAA90097.1 aluc|OJZ84125.1 anid|CBF81219.1 anig|EHA19233.1 aory|EIT72819.1 asyd|OJJ59312.1 ater|EAU31689.1 atub|OJI90499.1 aver|OJJ02963.1 awen|OJJ36758.1 azon|OJJ47442.1

Group951: acar|OOF96932.1 acar|OOF96967.1 abra|OJJ70776.1 acla|EAW11143.1 afis|EAW15874.1 afla|EED56501.1 afum|EAL88508.1 agla|OJJ87234.1 akaw|GAA90095.1 aluc|OJZ84123.1 anid|CBF81222.1 anig|EHA19231.1 aory|EIT72827.1 asyd|OJJ59314.1 ater|EAU31691.1 atub|OJI90495.1 aver|OJJ02961.1 awen|OJJ36756.1 azon|OJJ47444.1

Group952: acar|OOF97657.1 acar|OOF97704.1 abra|OJJ71500.1 acla|EAW10048.1 afis|EAW25536.1 afla|EED44712.1 afum|EAL90200.1 agla|OJJ87267.1 akaw|GAA82520.1 aluc|OJZ87047.1 anid|CBF77819.1 anig|EHA20320.1 aory|EIT83462.1 asyd|OJJ53009.1 ater|EAU33377.1 atub|OJI81588.1 aver|OJJ03834.1 awen|OJJ41278.1 azon|OJJ49501.1

Group953: acar|OOF97658.1 acar|OOF97705.1 abra|OJJ71499.1 acla|EAW10050.1 afis|EAW25534.1 afla|EED44709.1 afum|EAL90198.1 agla|OJJ87269.1 akaw|GAA82521.1 aluc|OJZ87048.1 anid|CBF77817.1 anig|EHA20321.1 aory|EIT83458.1 asyd|OJJ53010.1 ater|EAU33376.1 atub|OJI81587.1 aver|OJJ03833.1 awen|OJJ41276.1 azon|OJJ49508.1

Group954: acar|OOF97660.1 acar|OOF97707.1 abra|OJJ71497.1 acla|EAW10052.1 afis|EAW25532.1 afla|EED54033.1 afum|EAL90196.2 agla|OJJ87270.1 akaw|GAA82523.1 aluc|OJZ87050.1 anid|CBF77813.1 anig|EHA20323.1 aory|EIT78299.1 asyd|OJJ53012.1 ater|EAU33374.1 atub|OJI81585.1 aver|OJJ03831.1 awen|OJJ41274.1 azon|OJJ49506.1

Group955: acar|OOF97661.1 acar|OOF97708.1 abra|OJJ71496.1 acla|EAW10053.1 afis|EAW25531.1 afla|EED54032.1 afum|EBA27398.1 agla|OJJ87271.1 akaw|GAA82524.1 aluc|OJZ87051.1 anid|CBF77811.1 anig|EHA20324.1 aory|EIT78333.1 asyd|OJJ53013.1 ater|EAU33373.1 atub|OJI81584.1 aver|OJJ03830.1 awen|OJJ41273.1 azon|OJJ49505.1

Group956: acar|OOF97663.1 acar|OOF97710.1 abra|OJJ71494.1 acla|EAW10055.1 afis|EAW25529.1 afla|EED54030.1 afum|EAL90194.1 agla|OJJ87273.1 akaw|GAA82526.1 aluc|OJZ87053.1 anid|CBF77821.1 anig|EHA20326.1 aory|EIT78259.1 asyd|OJJ53015.1 ater|EAU33372.1 atub|OJI81582.1 aver|OJJ03828.1 awen|OJJ41271.1 azon|OJJ49509.1

Group957: acar|OOF97664.1 acar|OOF97711.1 abra|OJJ71493.1 acla|EAW10057.1 afis|EAW25527.1 afla|EED54028.1 afum|EAL90192.1 agla|OJJ87275.1 akaw|GAA82527.1 aluc|OJZ87054.1 anid|CBF77805.1 anig|EHA20327.1 aory|EIT78292.1 asyd|OJJ53017.1 ater|EAU33370.1 atub|OJI81581.1 aver|OJJ03826.1 awen|OJJ41269.1 azon|OJJ49511.1

Group958: acar|OOF97666.1 acar|OOF97713.1 abra|OJJ71491.1 acla|EAW10059.1 afis|EAW25525.1 afla|EED55229.1 afum|EAL90190.2 agla|OJJ81067.1 akaw|GAA82529.1 aluc|OJZ87056.1 anid|CBF77802.1 anig|EHA20329.1 aory|EIT83576.1 asyd|OJJ53019.1 ater|EAU32314.1 atub|OJI81579.1 aver|OJJ03824.1 awen|OJJ40768.1 azon|OJJ44041.1

Group959: acar|OOF97669.1 acar|OOF97715.1 abra|OJJ71489.1 acla|EAW10061.1 afis|EAW25523.1 afla|EED54026.1 afum|EAL90188.2 agla|OJJ87278.1 akaw|GAA82531.1 aluc|OJZ87058.1 anid|CBF77800.1 anig|EHA20331.1 aory|EIT78257.1 asyd|OJJ53021.1 ater|EAU33368.1 atub|OJI81577.1 aver|OJJ03822.1 awen|OJJ41267.1 azon|OJJ44832.1

Group960: acar|OOF97671.1 acar|OOF97718.1 abra|OJJ71486.1 acla|EAW10064.1 afis|EAW25521.1 afla|EED54023.1 afum|EAL90186.2 agla|OJJ87281.1 akaw|GAA82534.1 aluc|OJZ87061.1 anid|CBF77796.1 anig|EHA20334.1 aory|EIT78303.1 asyd|OJJ53024.1 ater|EAU33367.1 atub|OJI81574.1 aver|OJJ03819.1 awen|OJJ41264.1 azon|OJJ44829.1

Group961: acar|OOF97672.1 acar|OOF97719.1 abra|OJJ71485.1 acla|EAW10065.1 afis|EAW25520.1 afla|EED54022.1 afum|EAL90185.1 agla|OJJ87282.1 akaw|GAA82535.1 aluc|OJZ87062.1 anid|CBF77794.1 anig|EHA20335.1 aory|EIT78334.1 asyd|OJJ53025.1 ater|EAU33366.1 atub|OJI81573.1 aver|OJJ03818.1 awen|OJJ41263.1 azon|OJJ44828.1

Group962: acar|OOF97673.1 acar|OOF97720.1 abra|OJJ71481.1 acla|EAW10068.1 afis|EAW25517.1 afla|EED54017.1 afum|EAL90182.1 agla|OJJ87286.1 akaw|GAA82538.1 aluc|OJZ87066.1 anid|CBF77788.1 anig|EHA20338.1 aory|EIT78273.1 asyd|OJJ53028.1 ater|EAU33363.1 atub|OJI81570.1 aver|OJJ03816.1 awen|OJJ41260.1 azon|OJJ44825.1

Group963: acar|OOF97674.1 acar|OOF97721.1 abra|OJJ71478.1 acla|EAW10071.1 afis|EAW25515.1 afla|EED54016.1 afum|EAL90180.1 agla|OJJ87288.1 akaw|GAA82539.1 aluc|OJZ87067.1 anid|CBF77785.1 anig|EHA20339.1 aory|EIT78267.1 asyd|OJJ53029.1 ater|EAU33361.1 atub|OJI81569.1 aver|OJJ03815.1 awen|OJJ41258.1 azon|OJJ44824.1

Group964: acar|OOF97675.1 acar|OOF97723.1 abra|OJJ71476.1 acla|EAW10073.1 afis|EAW25513.1 afla|EED54014.1 afum|EAL90178.1 agla|OJJ87291.1 akaw|GAA82540.1 aluc|OJZ87070.1 anid|CBF77781.1 anig|EHA20340.1 aory|EIT78254.1 asyd|OJJ53031.1 ater|EAU33359.1 atub|OJI81567.1 aver|OJJ03813.1 awen|OJJ41256.1 azon|OJJ44821.1

Group965: acar|OOF97678.1 acar|OOF97726.1 abra|OJJ71474.1 acla|EAW10076.1 afis|EAW25510.1 afla|EED54011.1 afum|EAL90176.2 agla|OJJ87295.1 akaw|GAA82542.1 aluc|OJZ87072.1 anid|CBF77775.1

anig|EHA20342.1 aory|EIT78253.1 asyd|OJJ53034.1 ater|EAU33357.1 atub|OJI81565.1 aver|OJJ03810.1 awen|OJJ41252.1 azon|OJJ44819.1

Group966: acar|OOF97679.1 acar|OOF97727.1 abra|OJJ71472.1 acla|EAW10078.1 afis|EAW25508.1 afla|EED54008.1 afum|EAL90174.1 agla|OJJ87297.1 akaw|GAA82544.1 aluc|OJZ87074.1 anid|CBF77772.1 anig|EHA20343.1 aory|EIT78280.1 asyd|OJJ53036.1 ater|EAU33355.1 atub|OJI81563.1 aver|OJJ03808.1 awen|OJJ41250.1 azon|OJJ44817.1

Group967: acar|OOF97680.1 acar|OOF97729.1 abra|OJJ71470.1 acla|EAW10081.1 afis|EAW25506.1 afla|EED54006.1 afum|EAL90172.1 agla|OJJ87298.1 akaw|GAA82545.1 aluc|OJZ87075.1 anid|CBF77768.1 anig|EHA20345.1 aory|EIT78338.1 asyd|OJJ53037.1 ater|EAU33354.1 atub|OJI81561.1 aver|OJJ03807.1 awen|OJJ41249.1 azon|OJJ44816.1

Group968: acar|OOF97681.1 acar|OOF97730.1 abra|OJJ71469.1 acla|EAW10082.1 afis|EAW25505.1 afla|EED54005.1 afum|EAL90171.1 agla|OJJ87299.1 akaw|GAA82546.1 aluc|OJZ87076.1 anid|CBF77766.1 anig|EHA20346.1 aory|EIT78265.1 asyd|OJJ53038.1 ater|EAU33353.1 atub|OJI81560.1 aver|OJJ03806.1 awen|OJJ41248.1 azon|OJJ44813.1

Group969: acar|OOF97682.1 acar|OOF97731.1 abra|OJJ71468.1 acla|EAW10083.1 afis|EAW25504.1 afla|EED54003.1 afum|EAL90170.1 agla|OJJ87300.1 akaw|GAA82547.1 aluc|OJZ87077.1 anid|CBF77764.1 anig|EHA20347.1 aory|EIT78261.1 asyd|OJJ53039.1 ater|EAU33352.1 atub|OJI81559.1 aver|OJJ03805.1 awen|OJJ41247.1 azon|OJJ44812.1

Group970: acar|OOF97684.1 acar|OOF97734.1 abra|OJJ71464.1 acla|EAW10086.1 afis|EAW25501.1 afla|EED54000.1 afum|EAL90167.1 agla|OJJ87303.1 akaw|GAA82550.1 aluc|OJZ87080.1 anid|CBF77756.1 anig|EHA20351.1 aory|EIT78290.1 asyd|OJJ53043.1 ater|EAU34671.1 atub|OJI81557.1 aver|OJJ03801.1 awen|OJJ41243.1 azon|OJJ44809.1

Group971: acar|OOF97685.1 acar|OOF97735.1 abra|OJJ71465.1 acla|EAW10085.1 afis|EAW25502.1 afla|EED54001.1 afum|EAL90168.1 agla|OJJ87302.1 akaw|GAA82549.1 aluc|OJZ87079.1 anid|CBF77758.1 anig|EHA20350.1 aory|EIT78293.1 asyd|OJJ53042.1 ater|EAU34672.1 atub|OJI81558.1 aver|OJJ03802.1 awen|OJJ41244.1 azon|OJJ44810.1

Group972: acar|OOF97686.1 acar|OOF97736.1 abra|OJJ71463.1 acla|EAW10088.1 afis|EAW25499.1 afla|EED53999.1 afum|EAL90165.1 agla|OJJ87305.1 akaw|GAA82551.1 aluc|OJZ87081.1 anid|CBF77754.1 anig|EHA20352.1 aory|EIT78297.1 asyd|OJJ53044.1 ater|EAU34670.1 atub|OJI81556.1 aver|OJJ03800.1 awen|OJJ41241.1 azon|OJJ44807.1

Group973: acar|OOF97690.1 acar|OOF97740.1 abra|OJJ71458.1 acla|EAW10091.1 afis|EAW25496.1 afla|EED53996.1 afum|EAL90162.1 agla|OJJ87309.1 akaw|GAA82553.1 aluc|OJZ87085.1 anid|CBF77749.1 anig|EHA20355.1 aory|EIT78288.1 asyd|OJJ53047.1 ater|EAU34667.1 atub|OJI81553.1 aver|OJJ03798.1 awen|OJJ41238.1 azon|OJJ44804.1

Group974: acar|OOF97691.1 acar|OOF97741.1 abra|OJJ71456.1 acla|EAW10093.1 afis|EAW25494.1 afla|EED53994.1 afum|EAL90160.1 agla|OJJ87311.1 akaw|GAA82555.1 aluc|OJZ87087.1 anid|CBF77745.1 anig|EHA20357.1 aory|EIT78318.1 asyd|OJJ53049.1 ater|EAU34665.1 atub|OJI81551.1 aver|OJJ03796.1 awen|OJJ41236.1 azon|OJJ44802.1

Group976: acar|OOF97694.1 acar|OOF97743.1 abra|OJJ71454.1 acla|EAW10095.1 afis|EAW25492.1 afla|EED53992.1 afum|EAL90157.1 agla|OJJ87140.1 akaw|GAA82557.1 aluc|OJZ87089.1 anid|CBF77743.1 anig|EHA20360.1 aory|EIT78276.1 asyd|OJJ53051.1 ater|EAU34663.1 atub|OJI81549.1 aver|OJJ03794.1 awen|OJJ41234.1 azon|OJJ44799.1

Group977: acar|OOF97695.1 acar|OOF97745.1 abra|OJJ71451.1 acla|EAW10096.1 afis|EAW25491.1 afla|EED53991.1 afum|EAL90156.1 agla|OJJ87139.1 akaw|GAA82559.1 aluc|OJZ87091.1 anid|CBF77741.1 anig|EHA20362.1 aory|EIT78277.1 asyd|OJJ53052.1 ater|EAU34662.1 atub|OJI81547.1 aver|OJJ03792.1 awen|OJJ41233.1 azon|OJJ44798.1

Group978: acar|OOF97697.1 acar|OOF97747.1 abra|OJJ71447.1 acla|EAW10099.1 afis|EAW25488.1 afla|EED53986.1 afum|EAL90153.1 agla|OJJ87136.1 akaw|GAA82562.1 aluc|OJZ87094.1 anid|CBF77724.1 anig|EHA20365.1 aory|EIT78270.1 asyd|OJJ53061.1 ater|EAU34659.1 atub|OJI81544.1 aver|OJJ03783.1 awen|OJJ41229.1 azon|OJJ44796.1

Group979: acar|OOF97698.1 acar|OOF97748.1 abra|OJJ71446.1 acla|EAW10100.1 afis|EAW25487.1 afla|EED53985.1 afum|EAL90152.1 agla|OJJ87135.1 akaw|GAA82563.1 aluc|OJZ87095.1 anid|CBF77722.1 anig|EHA20366.1 aory|EIT78345.1 asyd|OJJ53062.1 ater|EAU34658.1 atub|OJI81543.1 aver|OJJ03782.1 awen|OJJ41228.1 azon|OJJ44795.1

Group980: acar|OOF97699.1 acar|OOF97749.1 abra|OJJ71444.1 acla|EAW10102.1 afis|EAW25485.1 afla|EED53983.1 afum|EAL90150.1 agla|OJJ87133.1 akaw|GAA82565.1 aluc|OJZ87097.1 anid|CBF77713.1 anig|EHA20368.1 aory|EIT78331.1 asyd|OJJ53067.1 ater|EAU34656.1 atub|OJI81541.1 aver|OJJ03778.1 awen|OJJ41226.1 azon|OJJ44781.1

Group981: acar|OOF98471.1 acar|OOF98473.1 abra|OJJ78401.1 acla|EAW09131.1 afis|EAW22015.1 afla|EED46740.1 afum|EAL93030.1 agla|OJJ84082.1 akaw|GAA85630.1 aluc|OJZ91480.1 anid|CBF82625.1 anig|EHA20142.1 aory|EIT77878.1 asyd|OJJ54012.1 ater|EAU31136.1 atub|OJI83473.1 aver|OJJ08730.1 awen|OJJ34311.1 azon|OJJ43327.1

Group982: acar|OOF98472.1 acar|OOF98486.1 abra|OJJ78402.1 acla|EAW09130.1 afis|EAW22016.1 afla|EED46741.1 afum|EAL93031.1 agla|OJJ84081.1 akaw|GAA85629.1 aluc|OJZ91481.1 anid|CBF82627.1 anig|EHA20143.1 aory|EIT77969.1 asyd|OJJ54011.1 ater|EAU31137.1 atub|OJI83474.1 aver|OJJ08731.1 awen|OJJ34312.1 azon|OJJ43328.1

Group983: acar|OOF98475.1 acar|OOF98487.1 abra|OJJ78399.1 acla|EAW09133.1 afis|EAW22013.1 afla|EED46738.1 afum|EAL93028.1 agla|OJJ84084.1 akaw|GAA85632.1 aluc|OJZ91478.1 anid|CBF82644.1 anig|EHA20140.1 aory|EIT77900.1 asyd|OJJ61676.1 ater|EAU31135.1 atub|OJI83471.1 aver|OJJ00997.1 awen|OJJ34309.1 azon|OJJ48062.1

Group984: acar|OOF98476.1 acar|OOF98489.1 abra|OJJ78398.1 acla|EAW09134.1 afis|EAW22012.1 afla|EED46737.1 afum|EAL93027.1 agla|OJJ84085.1 akaw|GAA85633.1 aluc|OJZ91477.1 anid|CBF82646.1 anig|EHA20139.1 aory|EIT77930.1 asyd|OJJ61675.1 ater|EAU31134.1 atub|OJI83470.1 aver|OJJ00996.1 awen|OJJ34308.1 azon|OJJ48061.1

Group985: acar|OOF98478.1 acar|OOF98491.1 abra|OJJ78396.1 acla|EAW09136.1 afis|EAW22011.1 afla|EED46735.1 afum|EAL93026.1 agla|OJJ84087.1 akaw|GAA85635.1 aluc|OJZ91475.1 anid|CBF82649.1 anig|EHA20137.1 aory|EIT77821.1 asyd|OJJ61673.1 ater|EAU31132.1 atub|OJI83468.1 aver|OJJ00994.1 awen|OJJ34306.1 azon|OJJ43333.1

Group986: acar|OOF98481.1 acar|OOF98494.1 abra|OJJ78393.1 acla|EAW09139.1 afis|EAW22008.1 afla|EED46732.1 afum|EAL93023.1 agla|OJJ84090.1 akaw|GAA85638.1 aluc|OJZ91472.1 anid|CBF82655.1 anig|EHA20134.1 aory|EIT77839.1 asyd|OJJ61670.1 ater|EAU31130.1 atub|OJI83465.1 aver|OJJ00991.1 awen|OJJ34303.1 azon|OJJ43335.1

Group987: acar|OOF98482.1 acar|OOF98495.1 abra|OJJ78391.1 acla|EAW09141.1 afis|EAW22006.1 afla|EED46730.1 afum|EAL93021.2 agla|OJJ84092.1 akaw|GAA85640.1 aluc|OJZ91470.1 anid|CBF82659.1 anig|EHA20132.1 aory|EIT77908.1 asyd|OJJ61668.1 ater|EAU31128.1 atub|OJI83463.1 aver|OJJ00989.1 awen|OJJ34301.1 azon|OJJ43337.1

Group988: acar|OOF98483.1 acar|OOF98496.1 abra|OJJ78392.1 acla|EAW09140.1 afis|EAW22007.1 afla|EED46731.1 afum|EAL93022.1 agla|OJJ84091.1 akaw|GAA85639.1 aluc|OJZ91471.1 anid|CBF82657.1 anig|EHA20133.1 aory|EIT77818.1 asyd|OJJ61669.1 ater|EAU31129.1 atub|OJI83464.1 aver|OJJ00990.1 awen|OJJ34302.1 azon|OJJ43336.1

Group989: acar|OOF98531.1 acar|OOF98532.1 abra|OJJ78355.1 acla|EAW09180.1 afis|EAW21966.1 afla|EED46687.1 afum|EAL92980.1 agla|OJJ84137.1 akaw|GAA85679.1 aluc|OJZ91430.1 anid|CBF82734.1 anig|EHA20099.1 aory|EIT77938.1 asyd|OJJ61626.1 ater|EAU31086.1 atub|OJI83423.1 aver|OJJ00951.1 awen|OJJ34260.1 azon|OJJ43377.1

Group991: afis|EAW15984.1 afis|EAW22037.1 abra|OJJ77639.1 acar|OOF99107.1 acla|EAW06489.1 afla|EED56354.1 afum|EAL88616.1 agla|OJJ85987.1 akaw|GAA83995.1 aluc|OJZ80891.1 anid|CBF85810.1 anig|EHA19447.1 aory|EIT73316.1 asyd|OJJ64340.1 ater|EAU33668.1 atub|OJI82687.1 aver|OJI98433.1 awen|OJJ30486.1 azon|OJJ45477.1

Group992: afis|EAW16169.1 afis|EAW20696.1 abra|OJJ78079.1 acar|OOF98785.1 acla|EAW06612.1 afla|EED48696.1 afum|EAL88803.1 agla|OJJ86214.1 akaw|GAA85915.1 aluc|OJZ91151.1 anid|CBF79561.1 anig|EHA19849.1 aory|EIT73253.1 asyd|OJJ57924.1 ater|EAU33245.1 atub|OJI83165.1 aver|OJJ04468.1 awen|OJJ30269.1 azon|OJJ43965.1

Group994: afis|EAW16703.1 afis|EAW22511.1 abra|OJJ73562.1 acar|OOG00744.1 acla|EAW07479.1 afla|EED54539.1 afum|EAL90682.1 agla|OJJ80876.1 akaw|GAA88592.1 aluc|OJZ88499.1 anid|CBF88981.1 anig|EHA18208.1 aory|EIT78923.1 asyd|OJJ62789.1 ater|EAU39198.1 atub|OJI91086.1 aver|OJI96836.1 awen|OJJ35137.1 azon|OJJ49962.1

Group995: afis|EAW17302.1 afis|EAW21312.1 abra|OJJ75975.1 acar|OOF99944.1 acla|EAW09822.1 afla|EED57269.1 afum|EAL92328.1 agla|OJJ87430.1 akaw|GAA88981.1 aluc|OJZ89422.1 anid|CBF83405.1 anig|EHA23165.1 aory|EIT74144.1 asyd|OJJ60646.1 ater|EAU35941.1 atub|OJI85371.1 aver|OJI99964.1 awen|OJJ39388.1 azon|OJJ44276.1

Group996: afis|EAW20681.1 afis|EAW24898.1 abra|OJJ74730.1 acar|OOF89795.1 acla|EAW10669.1 afla|EED46977.1 afum|EAL89552.2 agla|OJJ82121.1 akaw|GAA81887.1 aluc|OJZ92314.1 anid|CBF75653.1 anig|EHA26655.1 aory|EIT72604.1 asyd|OJJ57466.1 ater|EAU36401.1 atub|OJI86743.1 aver|OJJ05715.1 awen|OJJ40490.1 azon|OJJ49353.1

Group997: afis|EAW21461.1 afis|EAW22039.1 abra|OJJ76153.1 acar|OOF89976.1 acla|EAW09678.1 afla|EED55893.1 afum|EAL92480.1 agla|OJJ85609.1 akaw|GAA82102.1 aluc|OJZ92076.1 anid|CBF76534.1

anig|EHA23002.1 aory|EIT76550.1 asyd|OJJ61148.1 ater|EAU35074.1 atub|OJI86506.1 aver|OJJ00470.1 awen|OJJ39238.1 azon|OJJ51616.1

Group1000: afla|EED47569.1 afla|EED49784.1 abra|OJJ77270.1 acar|OOF93140.1 acla|EAW13268.1 afis|EAW17349.1 afum|EAL91192.2 agla|OJJ88227.1 akaw|GAA90678.1 aluc|OJZ84677.1 anid|CBF84233.1 anig|EHA27955.1 aory|EIT77705.1 asyd|OJJ57184.1 ater|EAU33864.1 atub|OJI89403.1 aver|OJJ03106.1 awen|OJJ34563.1 azon|OJJ50311.1

Group1003: agla|OJJ78416.1 agla|OJJ80642.1 abra|OJJ72427.1 acar|OOF94806.1 acla|EAW12660.1 afis|EAW17824.1 afla|EED46164.1 afum|EAL91634.1 akaw|GAA86168.1 aluc|OJZ90411.1 anid|CBF78007.1 anig|EHA23877.1 aory|EIT76471.1 asyd|OJJ55244.1 ater|EAU33025.1 atub|OJI81842.1 aver|OJJ06980.1 awen|OJJ41754.1 azon|OJJ48444.1

Group1004: agla|OJJ78484.1 agla|OJJ87510.1 abra|OJJ75909.1 acar|OOF99994.1 acla|EAW09880.1 afis|EAW21246.1 afla|EED54806.1 afum|EAL92259.1 akaw|GAA84615.1 aluc|OJZ89359.1 anid|CBF83238.1 anig|EHA23222.1 aory|EIT79446.1 asyd|OJJ60599.1 ater|EAU35886.1 atub|OJI85439.1 aver|OJI99922.1 awen|OJJ36559.1 azon|OJJ44332.1

Group1005: agla|OJJ78485.1 agla|OJJ87511.1 abra|OJJ75908.1 acar|OOF99995.1 acla|EAW09881.1 afis|EAW21245.1 afla|EED54805.1 afum|EAL92258.1 akaw|GAA84614.1 aluc|OJZ89358.1 anid|CBF83236.1 anig|EHA23223.1 aory|EIT79346.1 asyd|OJJ60600.1 ater|EAU35885.1 atub|OJI85440.1 aver|OJI99923.1 awen|OJJ36558.1 azon|OJJ44333.1

Group1006: agla|OJJ78486.1 agla|OJJ87512.1 abra|OJJ75907.1 acar|OOF99996.1 acla|EAW09882.1 afis|EAW21244.1 afla|EED54804.1 afum|EAL92257.1 akaw|GAA84613.1 aluc|OJZ89357.1 anid|CBF83253.1 anig|EHA23224.1 aory|EIT79338.1 asyd|OJJ60588.1 ater|EAU35884.1 atub|OJI85441.1 aver|OJI99911.1 awen|OJJ36555.1 azon|OJJ44334.1

Group1009: agla|OJJ82043.1 agla|OJJ82886.1 abra|OJJ74586.1 acar|OOF95180.1 acla|EAW10792.1 afis|EAW24784.1 afla|EED47201.1 afum|EAL89423.1 akaw|GAA82019.1 aluc|OJZ92164.1 anid|CBF81451.1 anig|EHA26785.1 aory|EIT79216.1 asyd|OJJ59433.1 ater|EAU36525.1 atub|OJI86593.1 aver|OJJ02838.1 awen|OJJ40369.1 azon|OJJ49218.1

Group1010: agla|OJJ82106.1 agla|OJJ85287.1 abra|OJJ74712.1 acar|OOF95721.1 acla|EAW10686.1 afis|EAW24884.1 afla|EED47082.1 afum|EAL89534.1 akaw|GAA81905.1 aluc|OJZ92293.1 anid|CBF75745.1 anig|EHA26676.1 aory|EIT72490.1 asyd|OJJ57520.1 ater|EAU36417.1 atub|OJI86723.1 aver|OJJ05658.1 awen|OJJ40474.1 azon|OJJ49338.1

Group1011: agla|OJJ82204.1 agla|OJJ88613.1 abra|OJJ66850.1 acar|OOF91626.1 acla|EAW14428.1 afis|EAW19135.1 afla|EED54279.1 afum|EAL93867.1 akaw|GAA91556.1 aluc|OJZ85734.1 anid|CBF86332.1 anig|EHA24816.1 aory|EIT76149.1 asyd|OJJ64990.1 ater|EAU37406.1 atub|OJI79519.1 aver|OJI99110.1 awen|OJJ37471.1 azon|OJJ51151.1

Group1012: agla|OJJ87407.1 agla|OJJ87526.1 abra|OJJ76001.1 acar|OOF99924.1 acla|EAW09804.1 afis|EAW21332.1 afla|EED57290.1 afum|EAL92350.1 akaw|GAA88959.1 aluc|OJZ89446.1 anid|CBF83435.1 anig|EHA23140.1 aory|EIT74092.1 asyd|OJJ60664.1 ater|EAU35958.1 atub|OJI85347.1 aver|OJI99987.1 awen|OJJ39366.1 azon|OJJ51705.1

Group1013: akaw|GAA87623.1 akaw|GAA91524.1 abra|OJJ66888.1 acar|OOF91659.1 acla|EAW14460.1 afis|EAW19168.1 afla|EED54229.1 afum|EAL93898.1 agla|OJJ88556.1 aluc|OJZ85694.1 anid|CBF86275.1 anig|EHA24847.1 aory|EIT75898.1 asyd|OJJ65023.1 ater|EAU37305.1 atub|OJI79484.1 aver|OJI99142.1 awen|OJJ37521.1 azon|OJJ51187.1

Group1014: akaw|GAA90564.1 akaw|GAA91190.1 abra|OJJ66522.1 acar|OOF90976.1 acla|EAW10968.1 afis|EAW20892.1 afla|EED45587.1 afum|EAL84741.1 agla|OJJ84000.1 aluc|OJZ81513.1 anid|CBF82751.1 anig|EHA25911.1 aory|EIT75801.1 asyd|OJJ61613.1 ater|EAU30831.1 atub|OJI88662.1 aver|OJJ00936.1 awen|OJJ34455.1 azon|OJJ51742.1

Group1017: anid|CBF75896.1 anid|CBF87037.1 abra|OJJ68334.1 acar|OOF97603.1 acla|EAW09978.1 afis|EAW21111.1 afla|EED51763.1 afum|EAL92134.1 agla|OJJ80102.1 akaw|GAA92087.1 aluc|OJZ83637.1 anig|EHA18789.1 aory|EIT79989.1 asyd|OJJ60451.1 ater|EAU36787.1 atub|OJI90006.1 aver|OJI99769.1 awen|OJJ36440.1 azon|OJJ44433.1

Group1018: anid|CBF76123.1 anid|CBF81569.1 abra|OJJ71017.1 acar|OOF98123.1 acla|EAW10567.1 afis|EAW25033.1 afla|EED55458.1 afum|EAL89661.1 agla|OJJ82245.1 akaw|GAA82938.1 aluc|OJZ87515.1 anig|EHA20726.1 aory|EIT80849.1 asyd|OJJ59506.1 ater|EAU35752.1 atub|OJI81135.1 aver|OJJ02768.1 awen|OJJ40606.1 azon|OJJ49455.1

Group1023: aory|EIT76145.1 aory|EIT78526.1 abra|OJJ72823.1 acar|OOF96028.1 acla|EAW11415.1 afis|EAW24006.1 afla|EED49155.1 afum|EAL87436.1 agla|OJJ86937.1 akaw|GAA85424.1 aluc|OJZ83325.1 anid|CBF77396.1 anig|EHA24203.1 asyd|OJJ53244.1 ater|EAU29745.1 atub|OJI82222.1 aver|OJJ03606.1 awen|OJJ35478.1 azon|OJJ51270.1

Group1024: aory|EIT76394.1 aory|EIT77963.1 abra|OJJ78294.1 acar|OOF98573.1 acla|EAW09222.1 afis|EAW21926.1 afla|EED46618.1 afum|EAL92939.1 agla|OJJ84196.1 akaw|GAA85733.1 aluc|OJZ91368.1 anid|CBF76880.1 anig|EHA20046.1 asyd|OJJ61326.1 ater|EAU30932.1 atub|OJI83361.1 aver|OJJ00651.1 awen|OJJ34216.1 azon|OJJ43418.1

Group1026: asyd|OJJ51877.1 asyd|OJJ54109.1 abra|OJJ66556.1 acar|OOF91001.1 acla|EAW10938.1 afis|EAW20923.1 afla|EED50277.1 afum|EAL84711.1 agla|OJJ83966.1 akaw|GAA91159.1 aluc|OJZ81487.1 anid|CBF82451.1 anig|EHA25940.1 aory|EIT77347.1 ater|EAU38570.1 atub|OJI88693.1 aver|OJJ08625.1 awen|OJJ34397.1 azon|OJJ48133.1

Group1027: asyd|OJJ51975.1 asyd|OJJ60994.1 abra|OJJ71988.1 acar|OOF96641.1 acla|EAW09550.1 afis|EAW21586.1 afla|EED57807.1 afum|EAL92604.1 agla|OJJ85779.1 akaw|GAA84941.1 aluc|OJZ86725.1 anid|CBF76287.1 anig|EHA21954.1 aory|EIT81038.1 ater|EAU34945.1 atub|OJI83803.1 aver|OJJ00314.1 awen|OJJ39095.1 azon|OJJ51428.1

Group1028: asyd|OJJ52256.1 asyd|OJJ61135.1 abra|OJJ76154.1 acar|OOF99776.1 acla|EAW09663.1 afis|EAW21474.1 afla|EED46116.1 afum|EAL92493.1 agla|OJJ85624.1 akaw|GAA82103.1 aluc|OJZ92074.1 anid|CBF76511.1 anig|EHA23001.1 aory|EIT76380.1 ater|EAU35059.1 atub|OJI86505.1 aver|OJJ00457.1 awen|OJJ39223.1 azon|OJJ51633.1

Group1029: asyd|OJJ52749.1 asyd|OJJ64102.1 abra|OJJ77967.1 acar|OOF98848.1 acla|EAW06673.1 afis|EAW16092.1 afla|EED48628.1 afum|EAL88724.1 agla|OJJ86129.1 akaw|GAA83695.1 aluc|OJZ91037.1 anid|CBF85462.1 anig|EHA19736.1 aory|EIT73247.1 ater|EAU33164.1 atub|OJI83033.1 aver|OJI98185.1 awen|OJJ30338.1 azon|OJJ45366.1

Group1031: asyd|OJJ53278.1 asyd|OJJ56762.1 abra|OJJ72786.1 acar|OOF96003.1 acla|EAW11483.1 afis|EAW23970.1 afla|EED50538.1 afum|EAL87400.1 agla|OJJ86899.1 akaw|GAA85393.1 aluc|OJZ83284.1 anid|CBF77331.1 anig|EHA24170.1 aory|EIT82142.1 ater|EAU29782.1 atub|OJI82187.1 aver|OJJ03570.1 awen|OJJ38973.1 azon|OJJ51308.1

Group1032: asyd|OJJ53282.1 asyd|OJJ56766.1 abra|OJJ72783.1 acar|OOF96001.1 acla|EAW11484.1 afis|EAW23969.1 afla|EED50540.1 afum|EAL87399.1 agla|OJJ86898.1 akaw|GAA85391.1 aluc|OJZ83282.1 anid|CBF77330.1 anig|EHA24169.1 aory|EIT82164.1 ater|EAU29783.1 atub|OJI82186.1 aver|OJJ03568.1 awen|OJJ38974.1 azon|OJJ51309.1

Group1033: asyd|OJJ53391.1 asyd|OJJ55131.1 abra|OJJ72369.1 acar|OOF94851.1 acla|EAW11966.1 afis|EAW19602.1 afla|EED56958.1 afum|EAL86916.1 agla|OJJ80744.1 akaw|GAA86119.1 aluc|OJZ90357.1 anid|CBF71283.1 anig|EHA23833.1 aory|EIT77537.1 ater|EAU32923.1 atub|OJI81785.1 aver|OJJ07072.1 awen|OJJ41838.1 azon|OJJ47049.1

Group1035: asyd|OJJ53802.1 asyd|OJJ61125.1 abra|OJJ76171.1 acar|OOF99768.1 acla|EAW09654.1 afis|EAW21485.1 afla|EED55918.1 afum|EAL92505.1 agla|OJJ85644.1 akaw|GAA82117.1 aluc|OJZ92059.1 anid|CBF76492.1 anig|EHA22988.1 aory|EIT76579.1 ater|EAU35053.1 atub|OJI86491.1 aver|OJJ00446.1 awen|OJJ39212.1 azon|OJJ51642.1

Group1038: asyd|OJJ57870.1 asyd|OJJ61020.1 abra|OJJ75464.1 acar|OOF90276.1 acla|EAW13831.1 afis|EAW18300.1 afla|EED48769.1 afum|EAL87666.1 agla|OJJ88123.1 akaw|GAA84237.1 aluc|OJZ88920.1 anid|CBF79478.1 anig|EHA23592.1 aory|EIT73032.1 ater|EAU33308.1 atub|OJI85872.1 aver|OJJ04524.1 awen|OJJ30675.1 azon|OJJ43275.1

Group1040: ater|EAU30138.1 ater|EAU38860.1 abra|OJJ73936.1 acar|OOG01067.1 acla|EAW07813.1 afis|EAW22849.1 afla|EED50124.1 afum|EAL90351.1 agla|OJJ83475.1 akaw|GAA89542.1 aluc|OJZ88108.1 anid|CBF87853.1 anig|EHA17877.1 aory|EIT74737.1 asyd|OJJ63417.1 atub|OJI90715.1 aver|OJI97521.1 awen|OJJ38872.1 azon|OJJ49662.1

Group1041: ater|EAU30572.1 ater|EAU36084.1 abra|OJJ68744.1 acar|OOF90309.1 acla|EAW14290.1 afis|EAW18993.1 afla|EED46791.1 afum|EAL93728.1 agla|OJJ82981.1 akaw|GAA89891.1 aluc|OJZ81782.1 anid|CBF77932.1 anig|EHA22433.1 aory|EIT72766.1 asyd|OJJ52417.1 atub|OJI84356.1 aver|OJJ03362.1 awen|OJJ31615.1 azon|OJJ47315.1

Group1043: atub|OJI80331.1 atub|OJI90561.1 abra|OJJ74085.1 acar|OOG01195.1 acla|EAW07929.1 afis|EAW22967.1 afla|EED54404.1 afum|EAL90230.1 agla|OJJ83320.1 akaw|GAA87565.1 aluc|OJZ87950.1 anid|CBF87610.1 anig|EHA17736.1 aory|EIT75873.1 asyd|OJJ63271.1 ater|EAU30655.1 aver|OJI97686.1 awen|OJJ38596.1 azon|OJJ47331.1

Group1045: aver|OJI96684.1 aver|OJJ05918.1 abra|OJJ73439.1 acar|OOG00645.1 acla|EAW06773.1 afis|EAW16324.1 afla|EED53558.1 afum|EAL88947.1 agla|OJJ88286.1 akaw|GAA83054.1 aluc|OJZ88601.1 anid|CBF89160.1 anig|EHA18295.1 aory|EIT74902.1 asyd|OJJ62559.1 ater|EAU34133.1 atub|OJI91186.1 awen|OJJ37724.1 azon|OJJ44983.1

Group1046: aver|OJI98887.1 aver|OJJ06757.1 abra|OJJ77001.1 acar|OOF91446.1 acla|EAW13450.1 afis|EAW20561.1 afla|EED52343.1 afum|EAL84500.1 agla|OJJ88729.1 akaw|GAA87047.1 aluc|OJZ84400.1

anid|CBF78975.1 anig|EHA27730.1 aory|EIT74333.1 asyd|OJJ55674.1 ater|EAU37119.1 atub|OJI89672.1 awen|OJJ33268.1 azon|OJJ50642.1

Group1047: aver|OJI98940.1 aver|OJI99461.1 abra|OJJ69903.1 acar|OOF93421.1 acla|EAW08920.1 afis|EAW15474.1 afla|EED46925.1 afum|EAL85847.1 agla|OJJ89052.1 akaw|GAA87749.1 aluc|OJZ85382.1 anid|CBF86601.1 anig|EHA25565.1 aory|EIT72600.1 asyd|OJJ64827.1 ater|EAU32654.1 atub|OJI80188.1 awen|OJJ36395.1 azon|OJJ44109.1

Group1049: aver|OJJ01895.1 aver|OJJ04283.1 abra|OJJ68598.1 acar|OOF92108.1 acla|EAW12641.1 afis|EAW18044.1 afla|EED51502.1 afum|EAL91855.2 agla|OJJ85415.1 akaw|GAA91922.1 aluc|OJZ81958.1 anid|CBF79781.1 anig|EHA22289.1 aory|EIT80736.1 asyd|OJJ58122.1 ater|EAU31279.1 atub|OJI84509.1 awen|OJJ41490.1 azon|OJJ48220.1

Group1050: aver|OJJ05206.1 aver|OJJ06610.1 aluc|OJZ88621.1 anig|EHA18312.1 atub|OJI91206.1 abra|OJJ73422.1 acar|OOG00625.1 acla|EAW07037.1 afis|EAW16663.1 afla|EED44906.1 afum|EAL89267.1 agla|OJJ78706.1 akaw|GAA83070.1 anid|CBF82078.1 aory|EIT73489.1 asyd|OJJ54252.1 ater|EAU39585.1 awen|OJJ40050.1 azon|OJJ50117.1

Group1052: awen|OJJ29714.1 awen|OJJ31500.1 abra|OJJ72682.1 acar|OOF95931.1 acla|EAW12316.1 afis|EAW16904.1 afla|EED45262.1 afum|EAL86440.1 agla|OJJ84538.1 akaw|GAA85300.1 aluc|OJZ90660.1 anid|CBF80774.1 anig|EHA24095.1 aory|EIT73651.1 asyd|OJJ54502.1 ater|EAU30211.1 atub|OJI82089.1 aver|OJJ07679.1 azon|OJJ46214.1

Group1053: awen|OJJ30059.1 awen|OJJ32960.1 abra|OJJ69424.1 acar|OOF94312.1 acla|EAW14803.1 afis|EAW24334.1 afla|EED47930.1 afum|EAL86004.1 agla|OJJ83203.1 akaw|GAA88290.1 aluc|OJZ82260.1 anig|EHA28482.1 aory|EIT79819.1 ater|EAU30414.1 atub|OJI88178.1 anid|CBF74037.1 aver|OJJ07954.1 azon|OJJ48958.1 asyd|OJJ53852.1

Group1054: awen|OJJ30747.1 awen|OJJ31734.1 abra|OJJ67459.1 acar|OOF92371.1 acla|EAW06307.1 afis|EAW18416.1 afla|EED52993.1 afum|EAL93098.1 agla|OJJ84982.1 akaw|GAA86374.1 aluc|OJZ84905.1 anid|CBF70716.1 anig|EHA25129.1 aory|EIT77436.1 asyd|OJJ58446.1 ater|EAU29831.1 atub|OJI80693.1 aver|OJJ01046.1 azon|OJJ50492.1

Group1055: awen|OJJ30760.1 awen|OJJ31721.1 abra|OJJ67449.1 acar|OOF90341.1 acla|EAW06315.1 afis|EAW18429.1 afla|EED52984.1 afum|EAL93109.1 agla|OJJ84994.1 akaw|GAA86381.1 aluc|OJZ84915.1 anid|CBF70683.1 anig|EHA25136.1 aory|EIT78705.1 asyd|OJJ58466.1 ater|EAU29845.1 atub|OJI80683.1 aver|OJJ01065.1 azon|OJJ50482.1

Group1056: awen|OJJ30764.1 awen|OJJ31717.1 abra|OJJ67446.1 acar|OOF90343.1 acla|EAW06317.1 afis|EAW18431.1 afla|EED52982.1 afum|EAL93111.2 agla|OJJ84997.1 akaw|GAA86383.1 aluc|OJZ84917.1 anid|CBF70025.1 anig|EHA25137.1 aory|EIT78725.1 asyd|OJJ58872.1 ater|EAU29847.1 atub|OJI80681.1 aver|OJJ01475.1 azon|OJJ50481.1

Group1057: awen|OJJ30768.1 awen|OJJ31713.1 abra|OJJ67442.1 acar|OOF92386.1 acla|EAW06321.1 afis|EAW18435.1 afla|EED52980.1 afum|EAL93116.1 agla|OJJ85002.1 akaw|GAA86387.1 aluc|OJZ84922.1 anid|CBF70033.1 anig|EHA25140.1 aory|EIT78731.1 asyd|OJJ58868.1 ater|EAU29851.1 atub|OJI80677.1 aver|OJJ01470.1 azon|OJJ50478.1

Group1060: awen|OJJ32955.1 awen|OJJ34548.1 abra|OJJ69452.1 acar|OOF94289.1 acla|EAW14816.1 afis|EAW24346.1 afla|EED47917.1 afum|EAL86016.1 agla|OJJ83207.1 akaw|GAA88367.1 aluc|OJZ82288.1 anid|CBF74019.1 anig|EHA28460.1 aory|EIT83370.1 asyd|OJJ53841.1 ater|EAU30400.1 atub|OJI88151.1 aver|OJJ07944.1 azon|OJJ48966.1

Group1061: awen|OJJ35632.1 awen|OJJ41410.1 abra|OJJ68673.1 acar|OOF92174.1 acla|EAW12575.1 afis|EAW24073.1 afla|EED51400.1 afum|EAL91929.1 agla|OJJ87142.1 akaw|GAA89826.1 aluc|OJZ81862.1 anid|CBF86467.1 anig|EHA22364.1 aory|EIT80717.1 asyd|OJJ64908.1 ater|EAU30495.1 atub|OJI84429.1 aver|OJI99030.1 azon|OJJ48882.1

Group1062: awen|OJJ35956.1 awen|OJJ39245.1 abra|OJJ76134.1 acar|OOF99789.1 acla|EAW09684.1 afis|EAW21455.1 afla|EED55885.1 afum|EAL92474.1 agla|OJJ85603.1 akaw|GAA88833.1 aluc|OJZ89596.1 anid|CBF76546.1 anig|EHA23016.1 aory|EIT76856.1 asyd|OJJ61154.1 ater|EAU35080.1 atub|OJI85207.1 aver|OJJ00476.1 azon|OJJ51610.1

Group1063: awen|OJJ35989.1 awen|OJJ38023.1 abra|OJJ75270.1 acar|OOF94533.1 acla|EAW12238.1 afis|EAW16989.1 afla|EED48968.1 afum|EAL86513.1 agla|OJJ79827.1 akaw|GAA91808.1 aluc|OJZ92830.1 anid|CBF87435.1 anig|EHA26179.1 aory|EIT79996.1 asyd|OJJ52914.1 ater|EAU31738.1 atub|OJI87287.1 aver|OJJ08884.1 azon|OJJ50274.1

Group1064: awen|OJJ36336.1 awen|OJJ40727.1 abra|OJJ70865.1 acar|OOF98251.1 acla|EAW10457.1 afis|EAW25110.1 afla|EED55332.1 afum|EAL89781.1 agla|OJJ89071.1 akaw|GAA91732.1 aluc|OJZ87654.1 anid|CBF86071.1 anig|EHA20848.1 aory|EIT76213.1 asyd|OJJ64460.1 ater|EAU39380.1 atub|OJI80988.1 aver|OJI98563.1 azon|OJJ44099.1

Group1065: awen|OJJ38099.1 awen|OJJ40555.1 abra|OJJ74809.1 acar|OOF95801.1 acla|EAW10605.1 afis|EAW24962.1 afla|EED55500.1 afum|EAL89620.1 agla|OJJ82197.1 akaw|GAA83603.1 aluc|OJZ92391.1 anid|CBF81656.1 anig|EHA26584.1 aory|EIT80890.1 asyd|OJJ59547.1 ater|EAU35693.1 atub|OJI86821.1 aver|OJJ02721.1 azon|OJJ49417.1

Group1066: awen|OJJ40158.1 awen|OJJ41199.1 abra|OJJ71411.1 acar|OOF97761.1 acla|EAW10126.1 afis|EAW25462.1 afla|EED53951.1 afum|EAL90125.1 agla|OJJ87114.1 akaw|GAA82589.1 aluc|OJZ87129.1 anid|CBF77649.1 anig|EHA20391.1 aory|EIT78320.1 asyd|OJJ53118.1 ater|EAU34631.1 atub|OJI81511.1 aver|OJJ03736.1 azon|OJJ44770.1

Group1068: azon|OJJ43179.1 azon|OJJ47539.1 abra|OJJ70678.1 acar|OOF97049.1 acla|EAW08086.1 afis|EAW23122.1 afla|EED48531.1 afum|EAL88358.1 agla|OJJ89617.1 akaw|GAA90010.1 aluc|OJZ84026.1 anid|CBF74402.1 anig|EHA19147.1 aory|EIT73068.1 asyd|OJJ57764.1 ater|EAU36250.1 atub|OJI90398.1 aver|OJJ05402.1 awen|OJJ36659.1

Group1069: azon|OJJ44484.1 azon|OJJ48556.1 abra|OJJ75695.1 acar|OOG00157.1 acla|EAW09413.1 afis|EAW21723.1 afla|EED57992.1 afum|EAL92756.1 agla|OJJ79861.1 akaw|GAA84431.1 aluc|OJZ89146.1 anid|CBF83716.1 anig|EHA23402.1 aory|EIT79772.1 asyd|OJJ60837.1 ater|EAU38407.1 atub|OJI85642.1 aver|OJJ00161.1 awen|OJJ33996.1

Group1070: azon|OJJ45306.1 azon|OJJ46721.1 abra|OJJ76474.1 acar|OOF99711.1 acla|EAW13959.1 afis|EAW18668.1 afla|EED52660.1 afum|EAL93386.1 agla|OJJ82407.1 akaw|GAA82386.1 aluc|OJZ91735.1 anid|CBF70575.1 anig|EHA22741.1 aory|EIT74271.1 asyd|OJJ58530.1 ater|EAU38243.1 atub|OJI86193.1 aver|OJJ01132.1 awen|OJJ32140.1

Group1072: azon|OJJ45841.1 azon|OJJ45942.1 abra|OJJ69362.1 acar|OOF94363.1 acla|EAW14752.1 afis|EAW24283.1 afla|EED47993.1 afum|EAL85952.1 agla|OJJ83145.1 akaw|GAA88231.1 aluc|OJZ82195.1 anid|CBF74136.1 anig|EHA28542.1 aory|EIT79842.1 asyd|OJJ53916.1 ater|EAU31231.1 atub|OJI88240.1 aver|OJJ08014.1 awen|OJJ33013.1

Group1073: azon|OJJ50179.1 azon|OJJ50395.1 abra|OJJ68426.1 acar|OOF93927.1 acla|EAW11928.1 afis|EAW19562.1 afla|EED50583.1 afum|EAL86950.1 agla|OJJ81713.1 akaw|GAA86946.1 aluc|OJZ90071.1 anid|CBF89380.1 anig|EHA21002.1 aory|EIT82053.1 asyd|OJJ62464.1 ater|EAU31050.1 atub|OJI84735.1 aver|OJI96599.1 awen|OJJ35963.1

Group1077: abra|OJJ65944.1 acar|OOF92929.1 acla|EAW13567.1 afis|EAW20033.1 afla|EED46044.1 afum|EAL84883.1 agla|OJJ88021.1 akaw|GAA90822.1 aluc|OJZ83013.1 anid|CBF84765.1 anig|EHA21428.1 aory|EIT75479.1 asyd|OJJ53284.1 ater|EAU29399.1 atub|OJI89248.1 aver|OJJ08457.1 awen|OJJ30056.1 azon|OJJ46327.1 afla|EED46043.1

Group1083: abra|OJJ66594.1 acar|OOF91029.1 acla|EAW10897.1 afis|EAW20961.1 afla|EED50316.1 afum|EAL84672.1 agla|OJJ83917.1 akaw|GAA91123.1 aluc|OJZ81453.1 anid|CBF82569.1 anig|EHA25975.1 aory|EIT77366.1 asyd|OJJ54049.1 ater|EAU38611.1 atub|OJI88728.1 aver|OJJ08694.1 awen|OJJ34356.1 azon|OJJ48092.1 aory|EIT77330.1

Group1084: abra|OJJ66648.1 acar|OOF93752.1 acla|EAW11791.1 afis|EAW19402.1 afla|EED50356.1 afum|EAL87093.1 agla|OJJ81115.1 akaw|GAA89601.1 aluc|OJZ80801.1 anid|CBF74340.1 anig|EHA27233.1 aory|EIT83133.1 asyd|OJJ51904.1 ater|EAU36127.1 atub|OJI87649.1 aver|OJJ05364.1 awen|OJJ36797.1 azon|OJJ43002.1 acar|OOF93668.1

Group1085: abra|OJJ66851.1 acar|OOF91627.1 acla|EAW14429.1 afis|EAW19136.1 afla|EED54277.1 afum|EAL93868.1 agla|OJJ88615.1 akaw|GAA91555.1 aluc|OJZ85733.1 anid|CBF86327.1 anig|EHA24817.1 aory|EIT75982.1 asyd|OJJ64992.1 ater|EAU37407.1 atub|OJI79518.1 aver|OJI99111.1 awen|OJJ37473.1 azon|OJJ51152.1 anid|CBF86329.1

Group1086: abra|OJJ66857.1 acar|OOF91634.1 acla|EAW14436.1 afis|EAW19144.1 afla|EED54269.1 afum|EAL93875.1 agla|OJJ88587.1 akaw|GAA91548.1 aluc|OJZ85726.1 anid|CBF86314.1 anig|EHA24822.1 aory|EIT75975.1 asyd|OJJ64999.1 ater|EAU37416.1 atub|OJI79512.1 aver|OJI99119.1 awen|OJJ37483.1 azon|OJJ51160.1 aory|EIT76087.1

Group1087: abra|OJJ66865.1 acar|OOF91641.1 acla|EAW14441.1 afis|EAW19149.1 afla|EED54264.1 afum|EAL93880.1 agla|OJJ88575.1 akaw|GAA91543.1 aluc|OJZ85717.1 anid|CBF86306.1 anig|EHA24827.1 aory|EIT76024.1 asyd|OJJ65005.1 ater|EAU37422.1 atub|OJI79507.1 aver|OJI99124.1 awen|OJJ37502.1 azon|OJJ51167.1 afla|EED54263.1

Group1088: abra|OJJ67116.1 acar|OOF98436.1 acla|EAW13746.1 afis|EAW19825.1 afla|EED56208.1 afum|EAL85067.1 agla|OJJ83144.1 akaw|GAA91049.1 aluc|OJZ87859.1 anid|CBF83120.1 anig|EHA20165.1 aory|EIT73304.1 asyd|OJJ61473.1 ater|EAU31280.1 atub|OJI80788.1 aver|OJJ00814.1 awen|OJJ36528.1 azon|OJJ47562.1 agla|OJJ79980.1

Group1093: abra|OJJ67523.1 awen|OJJ30978.1 abra|OJJ69776.1 acar|OOF93595.1 acla|EAW11058.1 afis|EAW20815.1 afla|EED53102.1 afum|EBA27207.1 agla|OJJ81282.1 akaw|GAA87877.1 aluc|OJZ85518.1

anid|CBF86687.1 anig|EHA25051.1 aory|EIT74763.1 asyd|OJJ64752.1 ater|EAU38934.1 atub|OJI80048.1 aver|OJI98853.1 azon|OJJ45137.1

Group1098: abra|OJJ67796.1 agla|OJJ84477.1 abra|OJJ75071.1 acar|OOF97263.1 acla|EAW08252.1 afis|EAW23286.1 afla|EED55688.1 afum|EAL88193.1 akaw|GAA88683.1 aluc|OJZ92637.1 anid|CBF89516.1 anig|EHA26354.1 aory|EIT77084.1 asyd|OJJ62394.1 ater|EAU37586.1 atub|OJI87068.1 aver|OJI96527.1 awen|OJJ39670.1 azon|OJJ47800.1

Group1099: aory|EIT77379.1 abra|OJJ71274.1 acar|OOF97900.1 acla|EAW10254.1 afis|EAW25337.1 afla|EED53807.1 afum|EAL90000.1 agla|OJJ83868.1 akaw|GAA82716.1 aluc|OJZ87270.1 anid|CBF75246.1 anig|EHA20511.1 asyd|OJJ59907.1 ater|EAU34477.1 atub|OJI81372.1 aver|OJJ02378.1 awen|OJJ41034.1 azon|OJJ44631.1 aory|EIT78377.1

Group1104: abra|OJJ68219.1 acar|OOF97509.1 acla|EAW09056.1 afis|EAW25630.1 afla|EED55187.1 afum|EAL84318.1 agla|OJJ78900.1 akaw|GAA83227.1 aluc|OJZ83727.1 anid|CBF83164.1 anig|EHA18875.1 aory|EIT83560.1 asyd|OJJ55506.1 ater|EAU32707.1 atub|OJI90103.1 aver|OJJ06581.1 awen|OJJ36050.1 azon|OJJ50396.1 acla|EAW09057.1

Group1106: abra|OJJ68422.1 acar|OOF97621.1 acla|EAW10999.1 afis|EAW20862.1 afla|EED45444.1 afum|EAL84773.1 agla|OJJ83749.1 akaw|GAA92526.1 aluc|OJZ90075.1 anid|CBF71444.1 anig|EHA20998.1 aory|EIT82447.1 asyd|OJJ60710.1 ater|EAU30805.1 atub|OJI84731.1 aver|OJJ00036.1 awen|OJJ34508.1 azon|OJJ51786.1 agla|OJJ89303.1

Group1108: abra|OJJ68521.1 acar|OOF92726.1 acla|EAW12843.1 afis|EAW18006.1 afla|EED51889.1 afum|EAL91818.1 agla|OJJ80269.1 akaw|GAA91328.1 aluc|OJZ82735.1 anid|CBF80030.1 anig|EHA21186.1 aory|EIT82919.1 asyd|OJJ58269.1 ater|EAU31522.1 atub|OJI88969.1 aver|OJJ02154.1 awen|OJJ41536.1 azon|OJJ48254.1 agla|OJJ80268.1

Group1109: abra|OJJ68529.1 acar|OOF92733.1 acla|EAW12835.1 afis|EAW18000.1 afla|EED51896.1 afum|EAL91811.1 agla|OJJ80263.1 akaw|GAA91321.1 aluc|OJZ82742.1 anid|CBF80021.1 anig|EHA21193.1 aory|EIT76177.1 asyd|OJJ58263.1 ater|EAU31512.1 atub|OJI88977.1 aver|OJJ02161.1 awen|OJJ41542.1 azon|OJJ48260.1 aory|EIT76174.1

Group1110: abra|OJJ68552.1 acar|OOF92747.1 acla|EAW12815.1 afis|EAW17980.1 afla|EED51925.1 afum|EAL91789.1 agla|OJJ80237.1 akaw|GAA91301.1 aluc|OJZ82769.1 anid|CBF79979.1 anig|EHA21211.1 aory|EIT81371.1 asyd|OJJ58235.1 ater|EAU31490.1 atub|OJI89000.1 aver|OJJ02190.1 awen|OJJ41565.1 azon|OJJ48280.1 akaw|GAA92005.1

Group1111: abra|OJJ68565.1 acar|OOF92758.1 acla|EAW12801.1 afis|EAW17967.1 afla|EED51939.1 afum|EAL91776.1 agla|OJJ80213.1 akaw|GAA91289.1 aluc|OJZ82780.1 anid|CBF79959.1 anig|EHA21223.1 aory|EIT81465.1 asyd|OJJ58221.1 ater|EAU31477.1 atub|OJI89013.1 aver|OJJ02201.1 awen|OJJ41577.1 azon|OJJ48290.1 akaw|GAA91288.1

Group1112: abra|OJJ68634.1 acar|OOF92138.1 acla|EAW12618.1 afis|EAW18066.1 afla|EED51450.1 afum|EBA27443.1 agla|OJJ87353.1 akaw|GAA89773.1 aluc|OJZ81921.1 anid|CBF86420.1 anig|EHA22326.1 anig|EHA22327.1 aory|EIT80631.1 asyd|OJJ64939.1 ater|EAU30462.1 atub|OJI84473.1 aver|OJI99061.1 awen|OJJ41453.1 azon|OJJ48911.1

Group1113: abra|OJJ68759.1 acar|OOF92246.1 acla|EAW12492.1 afis|EAW24153.1 afla|EED46778.1 afum|EAL92008.1 agla|OJJ89544.1 akaw|GAA89903.1 aluc|OJZ81767.1 anid|CBF77909.1 anig|EHA22447.1 aory|EIT72606.1 asyd|OJJ55299.1 ater|EAU30585.1 atub|OJI84340.1 aver|OJJ06926.1 awen|OJJ41324.1 azon|OJJ48798.1 ater|EAU30584.1

Group1114: abra|OJJ68992.1 acar|OOF92704.1 acla|EAW12644.1 afis|EAW18041.1 afla|EED51855.1 afla|EED51856.1 afum|EAL91852.1 agla|OJJ85413.1 akaw|GAA90512.1 aluc|OJZ85977.1 anid|CBF80101.1 anig|EHA24629.1 aory|EIT81757.1 asyd|OJJ58297.1 ater|EAU31556.1 atub|OJI79767.1 aver|OJJ02127.1 awen|OJJ41496.1 azon|OJJ48224.1

Group1115: abra|OJJ69111.1 acar|OOF91183.1 acla|EAW08721.1 afis|EAW15726.1 afla|EED51338.1 afum|EAL85621.1 agla|OJJ81865.1 akaw|GAA90636.1 aluc|OJZ86100.1 anid|CBF71089.1 anig|EHA24519.1 aory|EIT81534.1 asyd|OJJ52700.1 ater|EAU32431.1 atub|OJI79895.1 aver|OJJ04728.1 awen|OJJ38410.1 azon|OJJ43832.1 afla|EED51337.1

Group1116: abra|OJJ69231.1 acar|OOF91089.1 acla|EAW08618.1 afis|EAW15816.1 afla|EED51231.1 afum|EAL85535.1 agla|OJJ81980.1 akaw|GAA92239.1 aluc|OJZ86211.1 anid|CBF70897.1 anig|EHA24415.1 aory|EIT83178.1 asyd|OJJ52594.1 ater|EAU33432.1 atub|OJI80005.1 aver|OJJ04621.1 awen|OJJ38530.1 azon|OJJ43732.1 atub|OJI80006.1

Group1117: abra|OJJ69301.1 acar|OOF94424.1 acla|EAW14687.1 afis|EAW24218.1 afla|EED48053.1 afum|EAL85887.1 agla|OJJ79652.1 akaw|GAA88170.1 aluc|OJZ82129.1 anid|CBF74250.1 anig|EHA28602.1 aory|EIT83328.1 asyd|OJJ53976.1 ater|EAU31168.1 atub|OJI88307.1 aver|OJJ08075.1 awen|OJJ33078.1 azon|OJJ44868.1 aory|EIT83336.1

Group1118: abra|OJJ69328.1 acar|OOF94399.1 acla|EAW14712.1 afis|EAW24247.1 afla|EED48026.1 afum|EAL85915.1 agla|OJJ79626.1 akaw|GAA88199.1 aluc|OJZ82160.1 anid|CBF74202.1 anig|EHA28572.1 aory|EIT83332.1 asyd|OJJ53955.1 ater|EAU31188.1 atub|OJI88277.1 aver|OJJ08051.1 awen|OJJ33059.1 azon|OJJ48467.1 afla|EED48027.1

Group1119: abra|OJJ69488.1 acar|OOF94260.1 acla|EAW09905.1 afis|EAW21154.1 afla|EED54658.1 afum|EAL92172.1 agla|OJJ80039.1 akaw|GAA88332.1 aluc|OJZ82327.1 anid|CBF86989.1 anig|EHA28431.1 aory|EIT78787.1 asyd|OJJ60510.1 ater|EAU39556.1 atub|OJI88111.1 aver|OJI99836.1 awen|OJJ36482.1 azon|OJJ44363.1 anig|EHA21818.1

Group1120: abra|OJJ69514.1 acar|OOF94239.1 acla|EAW09954.1 afis|EAW21142.1 afla|EED46141.1 afum|EAL92161.1 agla|OJJ80069.1 akaw|GAA88310.1 aluc|OJZ82352.1 anid|CBF87008.1 anig|EHA28408.1 aory|EIT76511.1 asyd|OJJ60471.1 ater|EAU36763.1 atub|OJI88088.1 aver|OJI99791.1 awen|OJJ36466.1 azon|OJJ44412.1 azon|OJJ42893.1

Group1121: abra|OJJ69523.1 acar|OOF94231.1 acla|EAW09971.1 afis|EAW21117.1 afla|EED51790.1 afum|EAL92141.1 agla|OJJ80094.1 akaw|GAA88302.1 aluc|OJZ82363.1 anid|CBF87028.1 anig|EHA28402.1 aory|EIT81753.1 asyd|OJJ60456.1 ater|EAU36782.1 atub|OJI88079.1 aver|OJJ06754.1 awen|OJJ36447.1 azon|OJJ44428.1 aver|OJI99774.1

Group1122: abra|OJJ69615.1 acar|OOF92691.1 acla|EAW12075.1 afis|EAW19716.1 afla|EED53109.1 afum|EAL86811.1 agla|OJJ80862.1 akaw|GAA91599.1 aluc|OJZ82423.1 anid|CBF80080.1 anig|EHA28346.1 aory|EIT74768.1 asyd|OJJ58309.1 ater|EAU37777.1 atub|OJI88017.1 aver|OJJ02103.1 awen|OJJ35106.1 azon|OJJ45858.1 awen|OJJ34696.1

Group1123: abra|OJJ69668.1 acar|OOF95768.1 acla|EAW10632.1 afis|EAW24935.1 afla|EED55537.1 afum|EAL87763.1 agla|OJJ82162.1 akaw|GAA91646.1 aluc|OJZ82473.1 anid|CBF75718.1 anig|EHA28300.1 aory|EIT80819.1 asyd|OJJ57503.1 ater|EAU35664.1 atub|OJI87964.1 aver|OJJ05675.1 awen|OJJ40530.1 azon|OJJ49390.1 afum|EAL89592.1

Group1131: abra|OJJ70466.1 acar|OOF97458.1 acla|EAW13048.1 afis|EAW17619.1 afla|EED49199.1 afla|EED49200.1 afum|EAL91432.1 agla|OJJ81306.1 akaw|GAA83315.1 aluc|OJZ83810.1 anid|CBF90163.1 anig|EHA18955.1 aory|EIT78513.1 asyd|OJJ61994.1 ater|EAU37053.1 atub|OJI90177.1 aver|OJI96086.1 awen|OJJ37225.1 azon|OJJ50928.1

Group1132: abra|OJJ70467.1 acar|OOF97457.1 acla|EAW13047.1 afis|EAW17620.1 afla|EED49197.1 afla|EED49198.1 afum|EAL91433.2 agla|OJJ81305.1 akaw|GAA83316.1 aluc|OJZ83811.1 anid|CBF90161.1 anig|EHA18956.1 aory|EIT78563.1 asyd|OJJ61995.1 ater|EAU37054.1 atub|OJI90178.1 aver|OJI96087.1 awen|OJJ37224.1 azon|OJJ50927.1

Group1133: abra|OJJ70612.1 acar|OOF97110.1 acla|EAW13015.1 afis|EAW17652.1 afla|EED50506.1 afum|EAL91467.1 agla|OJJ79552.1 akaw|GAA89951.1 aluc|OJZ83958.1 anid|CBF90075.1 anig|EHA19086.1 aory|EIT82132.1 asyd|OJJ62041.1 ater|EAU37088.1 atub|OJI90330.1 aver|OJI96147.1 awen|OJJ37193.1 azon|OJJ50892.1 aver|OJJ07605.1

Group1134: abra|OJJ70647.1 acar|OOF97074.1 acla|EAW08113.1 afis|EAW23138.1 afla|EED48501.1 afum|EAL88343.1 agla|OJJ89662.1 akaw|GAA89983.1 aluc|OJZ83997.1 anid|CBF74448.1 anig|EHA19120.1 aory|EIT73103.1 asyd|OJJ57741.1 ater|EAU36279.1 atub|OJI90366.1 aver|OJJ05426.1 awen|OJJ36635.1 azon|OJJ47581.1 awen|OJJ36634.1

Group1135: abra|OJJ70656.1 acar|OOF97066.1 acla|EAW08103.1 afis|EAW23148.1 afla|EED48511.1 afum|EAL88333.1 agla|OJJ89654.1 akaw|GAA89992.1 aluc|OJZ84006.1 anid|CBF74430.1 anig|EHA19128.1 aory|EIT73155.1 asyd|OJJ57750.1 ater|EAU32975.1 atub|OJI90375.1 aver|OJJ05417.1 awen|OJJ36643.1 azon|OJJ47571.1 acar|OOG00309.1

Group1136: abra|OJJ70745.1 acar|OOF96999.1 acla|EAW11175.1 afis|EAW15906.1 afla|EED56463.1 afum|EAL88541.1 agla|OJJ89566.1 akaw|GAA90066.1 aluc|OJZ84090.1 anid|CBF81270.1 anig|EHA19203.1 aory|EIT72857.1 asyd|OJJ59333.1 ater|EAU36183.1 atub|OJI90461.1 aver|OJJ02941.1 awen|OJJ36725.1 azon|OJJ47484.1 ater|EAU30153.1

Group1137: abra|OJJ70747.1 acar|OOF96998.1 acla|EAW11174.1 afis|EAW15905.1 afla|EED56464.1 afum|EAL88540.1 agla|OJJ89565.1 akaw|GAA90067.1 aluc|OJZ84091.1 anid|CBF81274.1 anig|EHA19204.1 aory|EIT72825.1 asyd|OJJ59334.1 ater|EAU36182.1 atub|OJI90462.1 aver|OJJ02940.1 awen|OJJ36726.1 azon|OJJ47471.1 aory|EIT72838.1

Group1138: abra|OJJ70748.1 acar|OOF96997.1 acla|EAW11173.1 afis|EAW15904.1 afla|EED56466.1 afum|EAL88539.1 agla|OJJ89564.1 akaw|GAA90068.1 aluc|OJZ84092.1 anid|CBF81276.1 anig|EHA19205.1 aory|EIT72390.1 asyd|OJJ59335.1 ater|EAU36181.1 atub|OJI90463.1 aver|OJJ02939.1 awen|OJJ36727.1 azon|OJJ47470.1 afla|EED46475.1

Group1140: abra|OJJ70840.1 acar|OOF98274.1 acla|EAW13281.1 afis|EAW17336.1 afla|EED46105.1 afum|EAL91180.1 agla|OJJ82057.1 akaw|GAA91708.1 aluc|OJZ87681.1 anid|CBF85633.1 anig|EHA20871.1

aory|EIT76521.1 asyd|OJJ56056.1 ater|EAU33888.1 atub|OJI80962.1 aver|OJJ04998.1 awen|OJJ33506.1 azon|OJJ50300.1 azon|OJJ51817.1

Group1141: abra|OJJ70845.1 acar|OOF98270.1 acla|EAW10436.1 afis|EAW25131.1 afla|EED55314.1 afum|EAL89802.1 agla|OJJ89035.1 akaw|GAA91713.1 aluc|OJZ87676.1 anid|CBF86816.1 anig|EHA20866.1 aory|EIT83519.1 asyd|OJJ64657.1 ater|EAU39403.1 atub|OJI80967.1 aver|OJI98760.1 awen|OJJ40756.1 azon|OJJ44121.1 aver|OJJ02113.1

Group1142: abra|OJJ70882.1 acar|OOF98239.1 acla|EAW10470.1 afis|EAW25097.1 afla|EED55351.1 afum|EAL89768.1 agla|OJJ85476.1 akaw|GAA91744.1 aluc|OJZ87640.1 anid|CBF86010.1 anig|EHA20834.1 aory|EIT80916.1 asyd|OJJ64447.1 ater|EAU39355.1 atub|OJI81005.1 aver|OJI98550.1 awen|OJJ40711.1 azon|OJJ44087.1 ater|EAU29372.1

Group1143: abra|OJJ70985.1 acar|OOF98146.1 acla|EAW10545.1 afis|EAW25013.1 afla|EED55432.1 afum|EAL89683.1 agla|OJJ89177.1 akaw|GAA82964.1 aluc|OJZ87545.1 aluc|OJZ87546.1 anid|CBF81523.1 anig|EHA20749.1 aory|EIT80770.1 asyd|OJJ59478.1 ater|EAU35773.1 atub|OJI81100.1 aver|OJJ02795.1 awen|OJJ40627.1 azon|OJJ49056.1

Group1144: abra|OJJ71009.1 acar|OOF98133.1 acla|EAW10561.1 afis|EAW25028.1 afla|EED55451.1 afum|EAL89667.1 agla|OJJ79156.1 agla|OJJ82560.1 akaw|GAA82944.1 aluc|OJZ87522.1 anid|CBF81555.1 anig|EHA20732.1 aory|EIT80765.1 asyd|OJJ59497.1 ater|EAU35759.1 atub|OJI81127.1 aver|OJJ02777.1 awen|OJJ40611.1 azon|OJJ49460.1

Group1145: abra|OJJ71029.1 acar|OOF98110.1 acla|EAW10582.1 afis|EAW24987.1 afla|EED55471.1 afum|EAL89647.1 agla|OJJ82275.1 akaw|GAA82926.1 aluc|OJZ87501.1 anid|CBF81601.1 anig|EHA20716.1 aory|EIT80792.1 asyd|OJJ59520.1 ater|EAU35738.1 atub|OJI81148.1 aver|OJJ02753.1 awen|OJJ40583.1 azon|OJJ41984.1 azon|OJJ49442.1

Group1146: abra|OJJ71043.1 acar|OOF98096.1 acla|EAW10591.1 afis|EAW24977.1 afla|EED55482.1 afum|EAL89637.1 agla|OJJ79140.1 akaw|GAA82913.1 aluc|OJZ87485.1 anid|CBF81627.1 anig|EHA20706.1 aory|EIT80881.1 asyd|OJJ59529.1 ater|EAU35726.1 atub|OJI81163.1 aver|OJJ02741.1 awen|OJJ40574.1 azon|OJJ49436.1 afla|EED55483.1

Group1147: abra|OJJ71134.1 acar|OOF98018.1 acla|EAW10363.1 afis|EAW25229.1 afla|EED53673.1 afum|EAL89890.2 agla|OJJ79179.1 akaw|GAA82840.1 aluc|OJZ87401.1 anid|CBF85338.1 anig|EHA20630.1 aory|EIT75324.1 asyd|OJJ64023.1 ater|EAU34387.1 atub|OJI81245.1 aver|OJI98109.1 awen|OJJ40894.1 azon|OJJ44523.1 aory|EIT75321.1

Group1148: abra|OJJ71194.1 acar|OOF97972.1 acla|EAW10310.1 afis|EAW25283.1 afla|EED53749.1 afum|EAL89946.1 agla|OJJ89257.1 akaw|GAA82789.1 aluc|OJZ87348.1 anid|CBF85422.1 anig|EHA20584.1 aory|EIT78394.1 asyd|OJJ64075.1 ater|EAU34425.1 atub|OJI81295.1 aver|OJI98162.1 awen|OJJ40996.1 azon|OJJ44586.1 asyd|OJJ60350.1

Group1149: abra|OJJ71328.1 acar|OOF97846.1 acla|EAW10211.1 afis|EAW25383.1 afla|EED53860.1 afum|EAL90041.1 agla|OJJ83775.1 akaw|GAA82664.1 aluc|OJZ87214.1 anid|CBF77488.1 anig|EHA20465.1 aory|EIT78406.1 asyd|OJJ53195.1 ater|EAU34529.1 atub|OJI81425.1 aver|OJJ03655.1 awen|OJJ41109.1 azon|OJJ44687.1 aory|EIT78415.1

Group1150: abra|OJJ71401.1 acar|OOF97769.1 acla|EAW10133.1 afis|EAW25454.1 afla|EED53942.1 afla|EED53943.1 afum|EAL90113.1 agla|OJJ87107.1 akaw|GAA82596.1 aluc|OJZ87141.1 anid|CBF77635.1 anig|EHA20399.1 aory|EIT78263.1 asyd|OJJ53125.1 ater|EAU34624.1 atub|OJI81501.1 aver|OJJ03729.1 awen|OJJ41186.1 azon|OJJ44762.1

Group1152: abra|OJJ71994.1 acar|OOF96648.1 acla|EAW09538.1 afis|EAW21594.1 afla|EED57816.1 afum|EAL92612.1 agla|OJJ85787.1 akaw|GAA84934.1 aluc|OJZ86732.1 anid|CBF83446.1 anig|EHA21947.1 aory|EIT81030.1 asyd|OJJ60671.1 ater|EAU34930.1 atub|OJI83796.1 aver|OJI99994.1 awen|OJJ39087.1 azon|OJJ51418.1 ater|EAU32744.1

Group1153: abra|OJJ72045.1 acar|OOF96703.1 acla|EAW09501.1 afis|EAW21635.1 afla|EED48487.1 afum|EAL92669.1 agla|OJJ79700.1 akaw|GAA84889.1 aluc|OJZ86782.1 anid|CBF83498.1 anig|EHA21900.1 aory|EIT83266.1 asyd|OJJ60712.1 ater|EAU38321.1 atub|OJI83738.1 aver|OJJ00045.1 awen|OJJ33884.1 azon|OJJ48669.1 aory|EIT73220.1

Group1156: abra|OJJ72423.1 acar|OOF90372.1 acla|EAW12860.1 afis|EAW17809.1 afla|EED46159.1 afum|EAL91617.1 agla|OJJ80653.1 akaw|GAA86165.1 aluc|OJZ90406.1 anid|CBF78020.1 anig|EHA23874.1 aory|EIT76396.1 asyd|OJJ55235.1 ater|EAU33015.1 atub|OJI81837.1 aver|OJJ06988.1 awen|OJJ41763.1 azon|OJJ44903.1 afla|EED46160.1

Group1157: abra|OJJ72434.1 acar|OOF94801.1 acla|EAW12667.1 afis|EAW17831.1 afla|EED46173.1 afum|EAL91641.1 agla|OJJ80640.1 akaw|GAA86174.1 aluc|OJZ90417.1 anid|CBF77994.1 anig|EHA23885.1 aory|EIT76438.1 asyd|OJJ55252.1 ater|EAU33034.1 atub|OJI81850.1 aver|OJI99643.1 awen|OJJ41750.1 azon|OJJ48442.1 aver|OJJ06972.1

Group1159: abra|OJJ72549.1 acar|OOF94703.1 acla|EAW12754.1 afis|EAW17917.1 afla|EED46320.1 afum|EBA27439.1 agla|OJJ83026.1 akaw|GAA85181.1 aluc|OJZ90537.1 anid|CBF76959.1 anig|EHA23985.1 aory|EIT81197.1 atub|OJI81958.1 awen|OJJ41628.1 asyd|OJJ57052.1 aver|OJJ03244.1 azon|OJJ48353.1 ater|EAU33139.1 asyd|OJJ57051.1

Group1161: abra|OJJ72629.1 acar|OOF94635.1 acla|EAW11593.1 afis|EAW23864.1 afla|EED52589.1 afum|EAL87292.1 agla|OJJ82532.1 akaw|GAA85249.1 aluc|OJZ90606.1 anid|CBF77222.1 anig|EHA24044.1 aory|EIT74309.1 asyd|OJJ56820.1 ater|EAU33487.1 atub|OJI82030.1 aver|OJJ03515.1 awen|OJJ33839.1 azon|OJJ48708.1 awen|OJJ33432.1

Group1162: abra|OJJ72716.1 acar|OOF95949.1 acla|EAW11552.1 afis|EAW23901.1 afla|EED52636.1 afum|EAL87328.1 agla|OJJ79367.1 akaw|GAA85324.1 aluc|OJZ90685.1 anid|CBF77143.1 anig|EHA24113.1 aory|EIT74168.1 asyd|OJJ56912.1 ater|EAU32337.1 atub|OJI82111.1 aver|OJJ03380.1 awen|OJJ39048.1 azon|OJJ51378.1 afis|EAW25771.1

Group1163: abra|OJJ72751.1 acar|OOF95974.1 acla|EAW11510.1 afis|EAW23943.1 afla|EED50573.1 afum|EAL87368.1 agla|OJJ79431.1 akaw|GAA85363.1 aluc|OJZ90725.1 anid|CBF77256.1 anig|EHA24143.1 aory|EIT82138.1 asyd|OJJ56806.1 ater|EAU36036.1 atub|OJI82153.1 aver|OJJ03528.1 awen|OJJ39012.1 azon|OJJ51337.1 awen|OJJ40567.1

Group1166: abra|OJJ72907.1 acar|OOF96090.1 acla|EAW11346.1 afis|EAW18148.1 afla|EED58076.1 afum|EAL87506.1 agla|OJJ83833.1 akaw|GAA85492.1 aluc|OJZ83417.1 anid|CBF81114.1 anig|EHA24266.1 aory|EIT79127.1 ater|EAU35470.1 atub|OJI82304.1 aver|OJJ06234.1 awen|OJJ35591.1 azon|OJJ46913.1 asyd|OJJ56485.1 asyd|OJJ57077.1

Group1167: abra|OJJ73268.1 acar|OOG00508.1 acla|EAW06945.1 afis|EAW16559.1 afla|EED56102.1 afum|EAL89157.1 agla|OJJ82763.1 akaw|GAA87914.1 aluc|OJZ81161.1 anid|CBF81806.1 anig|EHA18478.1 aory|EIT76895.1 asyd|OJJ59644.1 ater|EAU36627.1 atub|OJI91356.1 aver|OJJ02638.1 awen|OJJ40257.1 azon|OJJ49121.1 acar|OOF91712.1

Group1168: abra|OJJ73284.1 acar|OOG00520.1 acla|EAW06959.1 afis|EAW16574.1 afla|EED56090.1 afum|EAL89171.2 agla|OJJ82706.1 akaw|GAA92721.1 aluc|OJZ88765.1 anid|CBF81830.1 anig|EHA18461.1 aory|EIT76622.1 asyd|OJJ59660.1 ater|EAU36642.1 atub|OJI91338.1 aver|OJJ02622.1 awen|OJJ40244.1 azon|OJJ49105.1 asyd|OJJ52085.1

Group1169: abra|OJJ73346.1 acar|OOG00556.1 acla|EAW06973.1 afis|EAW16587.1 afla|EED56078.1 afum|EAL89182.1 agla|OJJ82679.1 akaw|GAA83136.1 aluc|OJZ88693.1 anid|CBF81890.1 anig|EHA18373.1 aory|EIT76830.1 asyd|OJJ59676.1 ater|EAU36664.1 ater|EAU39117.1 atub|OJI91273.1 aver|OJJ02610.1 awen|OJJ40228.1 azon|OJJ49093.1

Group1170: abra|OJJ73352.1 acar|OOG00563.1 acla|EAW12784.1 afis|EAW19851.1 afla|EED52646.1 afum|EAL85042.1 agla|OJJ81301.1 akaw|GAA83131.1 aluc|OJZ88687.1 anid|CBF73979.1 anig|EHA18367.1 aory|EIT74299.1 asyd|OJJ54702.1 ater|EAU32479.1 atub|OJI91267.1 aver|OJJ07471.1 awen|OJJ32014.1 azon|OJJ46075.1 afis|EAW17946.1

Group1171: abra|OJJ73409.1 acar|OOG00606.1 acla|EAW07013.1 afis|EAW16640.1 afla|EED44962.1 afum|EAL89242.1 agla|OJJ82610.1 akaw|GAA83082.1 aluc|OJZ88629.1 anid|CBF82018.1 anig|EHA18319.1 aory|EIT73512.1 asyd|OJJ56191.1 ater|EAU36751.1 atub|OJI91213.1 aver|OJJ05120.1 awen|OJJ40074.1 azon|OJJ45550.1 anid|CBF76269.1

Group1172: abra|OJJ73438.1 acar|OOG00644.1 acla|EAW06774.1 afis|EAW16325.1 afla|EED53556.1 afum|EAL88949.1 agla|OJJ88288.1 akaw|GAA83055.1 aluc|OJZ88602.1 anid|CBF89162.1 anig|EHA18296.1 aory|EIT75182.1 asyd|OJJ62547.1 ater|EAU34132.1 atub|OJI91187.1 aver|OJI96673.1 awen|OJJ37725.1 azon|OJJ44985.1 afla|EED53557.1

Group1175: abra|OJJ73649.1 acar|OOG00830.1 acla|EAW07562.1 afis|EAW22596.1 afla|EED54646.1 afum|EAL90602.1 agla|OJJ80960.1 akaw|GAA88506.1 aluc|OJZ88401.1 anid|CBF88373.1 anig|EHA18129.1 aory|EIT78830.1 asyd|OJJ63733.1 ater|EAU39107.1 atub|OJI90997.1 aver|OJI97208.1 awen|OJJ35224.1 azon|OJJ49885.1 afla|EED54647.1

Group1177: abra|OJJ73743.1 acar|OOG00906.1 acla|EAW07637.1 afis|EAW22657.1 afla|EED49925.1 afum|EAL90535.1 agla|OJJ81040.1 akaw|GAA88422.1 aluc|OJZ88309.1 anid|CBF80225.1 anig|EHA18046.1 aory|EIT72274.1 asyd|OJJ54399.1 ater|EAU39050.1 atub|OJI90904.1 aver|OJJ05335.1 awen|OJJ35266.1 azon|OJJ49829.1 ater|EAU30036.1

Group1179: abra|OJJ73822.1 acar|OOG00974.1 acla|EAW07711.1 afis|EAW22727.1 afla|EED50013.1 afum|EAL90468.2 agla|OJJ83675.1 akaw|GAA89441.1 aluc|OJZ88226.1 anid|CBF88026.1 anig|EHA17974.1 aory|EIT74607.1 asyd|OJJ63528.1 ater|EAU38970.1 atub|OJI90827.1 aver|OJI97403.1 awen|OJJ35422.1 azon|OJJ49754.1 aory|EIT74669.1

Group1180: abra|OJJ73906.1 acar|OOG01039.1 acla|EAW07784.1 afis|EAW22820.1 afla|EED50095.1 afum|EBA27405.1 agla|OJJ83518.1 akaw|GAA89513.1 aluc|OJZ88141.1 anid|CBF87897.1 anig|EHA17904.1

aory|EIT74756.1 asyd|OJJ63443.1 ater|EAU38890.1 atub|OJI90744.1 aver|OJI97493.1 awen|OJJ38907.1 azon|OJJ49688.1 afla|EED50094.1

Group1182: abra|OJJ73974.1 acar|OOG01101.1 acla|EAW07845.1 afis|EAW22881.1 afla|EED50179.1 afum|EAL90318.1 agla|OJJ83449.1 akaw|GAA89577.1 aluc|OJZ88070.1 anid|CBF87807.1 anig|EHA17843.1 aory|EIT77377.1 asyd|OJJ63387.1 ater|EAU38845.1 atub|OJI90679.1 aver|OJI97548.1 awen|OJJ38841.1 azon|OJJ49638.1 agla|OJJ83448.1

Group1183: abra|OJJ73990.1 acar|OOG01116.1 acla|EAW07861.1 afis|EAW22896.1 afla|EED54867.1 afum|EAL90302.1 agla|OJJ83416.1 akaw|GAA87473.1 aluc|OJZ88055.1 anid|CBF87777.1 anig|EHA17827.1 aory|EIT79357.1 asyd|OJJ63370.1 atub|OJI90663.1 aver|OJI97576.1 awen|OJJ38822.1 azon|OJJ49621.1 ater|EAU35980.1 awen|OJJ38821.1

Group1185: abra|OJJ74469.1 acar|OOF95278.1 acla|EAW07228.1 afis|EAW22242.1 afla|EED57635.1 afum|EAL90951.1 agla|OJJ87809.1 akaw|GAA87362.1 aluc|OJZ89651.1 anid|CBF88489.1 anig|EHA26889.1 aory|EIT82991.1 asyd|OJJ63790.1 ater|EAU34249.1 atub|OJI85153.1 aver|OJI97147.1 awen|OJJ34820.1 azon|OJJ45730.1 aver|OJJ06299.1

Group1187: abra|OJJ74764.1 acar|OOF95757.1 acla|EAW10646.1 afis|EAW24920.1 afla|EED55553.1 afum|EAL89575.1 agla|OJJ82149.1 akaw|GAA83647.1 aluc|OJZ92348.1 anig|EHA26626.1 aory|EIT82708.1 asyd|OJJ57492.1 asyd|OJJ62481.1 ater|EAU35652.1 atub|OJI86775.1 aver|OJJ05688.1 awen|OJJ40517.1 azon|OJJ49376.1 anid|CBF75691.1

Group1188: abra|OJJ74777.1 acar|OOF95766.1 acla|EAW10634.1 afis|EAW24932.1 afla|EED55541.1 afum|EAL89588.1 agla|OJJ82160.1 akaw|GAA83635.1 aluc|OJZ92358.1 anid|CBF75714.1 anig|EHA26615.1 aory|EIT82707.1 asyd|OJJ57501.1 ater|EAU35662.1 atub|OJI86788.1 aver|OJJ05677.1 awen|OJJ40528.1 azon|OJJ49389.1 afum|EAL87767.1

Group1190: abra|OJJ74880.1 acar|OOF95883.1 acla|EAW08490.1 afis|EAW23536.1 afla|EED57434.1 afum|EAL87930.1 agla|OJJ86791.1 akaw|GAA83540.1 aluc|OJZ92457.1 anid|CBF89670.1 anig|EHA26510.1 aory|EIT80126.1 asyd|OJJ62272.1 ater|EAU35322.1 atub|OJI86886.1 aver|OJI96381.1 awen|OJJ39866.1 azon|OJJ43234.1 afla|EED57433.1

Group1191: abra|OJJ74903.1 acar|OOF95901.1 acla|EAW08469.1 afis|EAW23516.1 afla|EED57417.1 afum|EAL87947.2 agla|OJJ86764.1 akaw|GAA83519.1 aluc|OJZ92476.1 anid|CBF89704.1 anig|EHA26494.1 aory|EIT80064.1 asyd|OJJ62253.1 ater|EAU35305.1 atub|OJI86907.1 aver|OJI96363.1 awen|OJJ39841.1 azon|OJJ43251.1 aory|EIT80158.1

Group1192: abra|OJJ74942.1 acar|OOF97373.1 acla|EAW08428.1 afis|EAW23472.1 afla|EED57368.1 afum|EAL87994.1 agla|OJJ86711.1 akaw|GAA83479.1 aluc|OJZ92516.1 anid|CBF89779.1 anig|EHA26459.1 aory|EIT80072.1 asyd|OJJ62207.1 ater|EAU36908.1 atub|OJI86949.1 aver|OJI96319.1 awen|OJJ39798.1 azon|OJJ47919.1 awen|OJJ40002.1

Group1193: abra|OJJ75019.1 acar|OOF89964.1 akaw|GAA88640.1 aluc|OJZ92586.1 anid|CBF84112.1 anig|EHA26399.1 atub|OJI87018.1 abra|OJJ75020.1 acla|EAW08212.1 afis|EAW23246.1 afla|EED55727.1 afum|EAL88235.1 agla|OJJ86614.1 aory|EIT83220.1 asyd|OJJ57103.1 ater|EAU37528.1 aver|OJJ03196.1 awen|OJJ39724.1 azon|OJJ47850.1

Group1194: abra|OJJ75054.1 acar|OOF97283.1 acla|EAW08239.1 afis|EAW23273.1 afla|EED51546.1 afum|EAL88206.1 agla|OJJ86575.1 akaw|GAA88667.1 aluc|OJZ92619.1 anid|CBF89542.1 anig|EHA26369.1 aory|EIT77094.1 asyd|OJJ62380.1 ater|EAU37558.1 atub|OJI87053.1 aver|OJI96514.1 awen|OJJ39684.1 azon|OJJ47821.1 aory|EIT80554.1

Group1195: abra|OJJ75063.1 acar|OOF97271.1 acla|EAW08246.1 afis|EAW23280.1 afla|EED55696.1 afum|EAL88199.1 agla|OJJ86567.1 akaw|GAA88675.1 aluc|OJZ92629.1 anid|CBF89529.1 anig|EHA26361.1 aory|EIT77085.1 asyd|OJJ62387.1 ater|EAU37578.1 atub|OJI87059.1 aver|OJI96520.1 awen|OJJ39677.1 azon|OJJ43147.1 awen|OJJ31335.1

Group1199: abra|OJJ75173.1 acar|OOF94605.1 acla|EAW13003.1 afis|EAW17665.1 afla|EED50492.1 afum|EAL91479.1 agla|OJJ79538.1 akaw|GAA88780.1 aluc|OJZ92741.1 anid|CBF90049.1 anig|EHA26262.1 aory|EIT82090.1 asyd|OJJ62054.1 ater|EAU31988.1 atub|OJI87166.1 aver|OJI96160.1 awen|OJJ37176.1 azon|OJJ50876.1 asyd|OJJ61465.1

Group1202: abra|OJJ75302.1 acar|OOF92541.1 acla|EAW09084.1 afis|EAW25675.1 afla|EED47643.1 afum|EAL84345.1 agla|OJJ86330.1 akaw|GAA91835.1 aluc|OJZ92857.1 anid|CBF78280.1 anig|EHA26149.1 aory|EIT76311.1 asyd|OJJ52225.1 ater|EAU36081.1 atub|OJI87314.1 aver|OJJ05016.1 awen|OJJ35856.1 azon|OJJ42600.1 asyd|OJJ56073.1

Group1203: abra|OJJ75308.1 acar|OOF94521.1 acla|EAW12232.1 afis|EAW16996.1 afla|EED48959.1 afum|EAL86520.1 agla|OJJ79823.1 akaw|GAA91992.1 aluc|OJZ92862.1 anid|CBF87466.1 anig|EHA26103.1 aory|EIT80011.1 asyd|OJJ52936.1 ater|EAU31732.1 atub|OJI87320.1 aver|OJJ08903.1 awen|OJJ35985.1 azon|OJJ47407.1 azon|OJJ50280.1

Group1204: abra|OJJ75752.1 acar|OOG00116.1 acla|EAW09398.1 afis|EAW16786.1 afla|EED47412.1 afum|EAL86356.1 agla|OJJ78549.1 akaw|GAA84485.1 aluc|OJZ89206.1 anid|CBF84608.1 anig|EHA23353.1 aory|EIT81898.1 asyd|OJJ60434.1 ater|EAU37659.1 atub|OJI85586.1 aver|OJI99756.1 awen|OJJ40856.1 azon|OJJ42408.1 ater|EAU30953.1

Group1206: abra|OJJ75996.1 acar|OOF99929.1 acla|EAW09809.1 afis|EAW21327.1 afla|EED57285.1 afum|EAL92345.1 agla|OJJ87412.1 akaw|GAA88963.1 aluc|OJZ89442.1 anid|CBF83428.1 anig|EHA23144.1 aory|EIT74010.1 asyd|OJJ60659.1 ater|EAU35952.1 atub|OJI85351.1 aver|OJI99982.1 awen|OJJ39371.1 azon|OJJ51711.1 afla|EED57284.1

Group1208: abra|OJJ76593.1 acar|OOF99394.1 acla|EAW14308.1 afis|EAW18977.1 afla|EED53193.1 afum|EAL93713.2 agla|OJJ82935.1 akaw|GAA82497.1 aluc|OJZ91613.1 anid|CBF69648.1 anig|EHA22642.1 aory|EIT74974.1 asyd|OJJ59078.1 ater|EAU38168.1 atub|OJI86066.1 aver|OJJ01671.1 awen|OJJ31583.1 azon|OJJ47298.1 afla|EED53194.1

Group1211: abra|OJJ76859.1 acar|OOF91347.1 acla|EAW13420.1 afis|EAW20531.1 afla|EED52481.1 afum|EAL84526.1 agla|OJJ84700.1 akaw|GAA87163.1 aluc|OJZ84271.1 anid|CBF79002.1 anig|EHA27613.1 aory|EIT82813.1 asyd|OJJ55689.1 ater|EAU32506.1 atub|OJI89812.1 aver|OJJ06773.1 awen|OJJ33238.1 azon|OJJ50610.1 afis|EAW15383.1

Group1212: abra|OJJ76952.1 acar|OOF91399.1 acla|EAW13478.1 afis|EAW20585.1 afla|EED52259.1 afum|EAL84480.1 agla|OJJ80829.1 akaw|GAA87090.1 aluc|OJZ84348.1 anid|CBF78926.1 anig|EHA27685.1 aory|EIT76185.1 asyd|OJJ55629.1 ater|EAU37094.1 atub|OJI89720.1 aver|OJJ06706.1 awen|OJJ33298.1 azon|OJJ50675.1 afis|EAW16394.1

Group1219: abra|OJJ77364.1 acar|OOF93233.1 acla|EAW13112.1 afis|EAW17552.1 afla|EED49273.1 afum|EAL91366.1 agla|OJJ81416.1 akaw|GAA91398.1 aluc|OJZ84773.1 anid|CBF90298.1 anig|EHA28041.1 aory|EIT82235.1 asyd|OJJ61925.1 ater|EAU36982.1 atub|OJI89308.1 aver|OJI96017.1 awen|OJJ37301.1 azon|OJJ50994.1 anid|CBF87295.1

Group1220: abra|OJJ77453.1 acar|OOF99273.1 acla|EAW12649.1 afis|EAW18036.1 afla|EED54071.1 afum|EAL91847.1 agla|OJJ78571.1 akaw|GAA84155.1 aluc|OJZ81076.1 anid|CBF81938.1 anig|EHA19286.1 aory|EIT76067.1 asyd|OJJ57409.1 ater|EAU29318.1 ater|EAU31609.1 atub|OJI82509.1 aver|OJJ05772.1 awen|OJJ34693.1 azon|OJJ48230.1

Group1224: abra|OJJ77564.1 acar|OOF99168.1 acla|EAW06526.1 afis|EAW16070.1 afla|EED48589.1 afum|EAL88702.1 agla|OJJ86098.1 akaw|GAA84062.1 aluc|OJZ80974.1 anid|CBF75103.1 anig|EHA19385.1 aory|EIT83270.1 asyd|OJJ60005.1 ater|EAU33561.1 atub|OJI82611.1 aver|OJJ02281.1 awen|OJJ30365.1 azon|OJJ45384.1 afla|EED57884.1

Group1226: abra|OJJ77599.1 acar|OOF99127.1 acla|EAW06563.1 afis|EAW16033.1 afla|EED56415.1 afla|EED56416.1 afum|EAL88664.1 agla|OJJ86061.1 akaw|GAA84029.1 aluc|OJZ80934.1 anid|CBF75034.1 anig|EHA19416.1 aory|EIT72886.1 asyd|OJJ60050.1 ater|EAU33608.1 atub|OJI82646.1 aver|OJJ02239.1 awen|OJJ30414.1 azon|OJJ45422.1

Group1230: abra|OJJ77818.1 acar|OOF98950.1 acla|EAW11305.1 afis|EAW18192.1 afla|EED56282.1 afum|EAL87560.1 agla|OJJ85885.1 akaw|GAA83828.1 aluc|OJZ90890.1 anid|CBF86109.1 anig|EHA19609.1 aory|EIT73431.1 asyd|OJJ64488.1 ater|EAU33797.1 atub|OJI82884.1 aver|OJI98589.1 awen|OJJ30564.1 azon|OJJ45507.1 aory|EIT73332.1

Group1231: abra|OJJ77831.1 acar|OOF98939.1 acla|EAW11295.1 afis|EAW18207.1 afla|EED56264.1 afum|EAL87572.1 agla|OJJ85875.1 akaw|GAA83816.1 aluc|OJZ90905.1 anid|CBF86133.1 anig|EHA19620.1 aory|EIT73347.1 asyd|OJJ64503.1 ater|EAU33782.1 atub|OJI82898.1 aver|OJI98602.1 awen|OJJ30579.1 azon|OJJ45497.1 afla|EED56263.1

Group1233: abra|OJJ77901.1 acar|OOF98882.1 acla|EAW06511.1 afis|EAW16229.1 afla|EED53352.1 afum|EAL88852.1 agla|OJJ86237.1 akaw|GAA83761.1 aluc|OJZ90970.1 anid|CBF86221.1 anig|EHA19672.1 aory|EIT75062.1 asyd|OJJ64562.1 ater|EAU33715.1 atub|OJI82961.1 aver|OJI98659.1 awen|OJJ30233.1 azon|OJJ43950.1 anid|CBF86220.1

Group1235: abra|OJJ78155.1 acar|OOF90381.1 acla|EAW11243.1 afis|EAW18260.1 afla|EED48719.1 afum|EAL87623.1 agla|OJJ88167.1 akaw|GAA85871.1 aluc|OJZ91207.1 anid|CBF79523.1 anig|EHA19911.1 aory|EIT73142.1 asyd|OJJ57893.1 ater|EAU33268.1 atub|OJI83223.1 aver|OJJ04502.1 awen|OJJ30632.1 awen|OJJ30633.1 azon|OJJ47984.1

Group1236: abra|OJJ78254.1 acar|OOF98608.1 acla|EAW09255.1 afis|EAW21892.1 afla|EED46416.1 afum|EAL92906.1 agla|OJJ84240.1 akaw|GAA85772.1 aluc|OJZ91328.1 anid|CBF76804.1 anig|EHA20010.1 aory|EIT72374.1 asyd|OJJ61288.1 atub|OJI83321.1 aver|OJJ00614.1 awen|OJJ34181.1 azon|OJJ43451.1 acar|OOF99280.1 ater|EAU36657.1

Group1237: abra|OJJ78362.1 acar|OOF98525.1 acla|EAW09173.1 afis|EAW21973.1 afla|EED46696.1 afum|EAL92988.1 agla|OJJ84129.1 akaw|GAA85671.1 aluc|OJZ91436.1 anid|CBF82714.1 anig|EHA20102.1

aory|EIT77832.1 asyd|OJJ61638.1 ater|EAU31091.1 atub|OJI83430.1 aver|OJJ00961.1 awen|OJJ34266.1 azon|OJJ43370.1 aory|EIT77819.1

Group1328: anig|EHA20833.1 afum|EAL89767.1 abra|OJJ70883.1 acar|OOF98234.1 acla|EAW10471.1 afis|EAW25096.1 afla|EED55352.1 agla|OJJ89081.1 akaw|GAA87402.1 aluc|OJZ87638.1 anid|CBF86008.1 aory|EIT80853.1 asyd|OJJ64446.1 ater|EAU39354.1 atub|OJI81009.1 aver|OJI98549.1 awen|OJJ40710.1 azon|OJJ44086.1

Group1340: asyd|OJJ60628.1 abra|OJJ75954.1 acar|OOF99958.1 acla|EAW09842.1 afis|EAW21293.1 afla|EED54845.1 afum|EAL92310.1 agla|OJJ87460.1 akaw|GAA84657.1 aluc|OJZ89403.1 anid|CBF83378.1 anig|EHA23184.1 aory|EIT79413.1 ater|EAU35926.1 atub|OJI85394.1 aver|OJI99952.1 awen|OJJ39406.1 azon|OJJ44289.1

Group1365: abra|OJJ65896.1 acar|OOF94956.1 acla|EAW11696.1 afis|EAW23713.1 afla|EED44871.1 afum|EAL87157.1 agla|OJJ81717.1 akaw|GAA86009.1 aluc|OJZ90235.1 anid|CBF83004.1 anig|EHA23735.1 aory|EIT82717.1 asyd|OJJ61502.1 ater|EAU31796.1 atub|OJI81665.1 aver|OJJ00840.1 awen|OJJ37102.1 azon|OJJ50803.1

Group1366: abra|OJJ65913.1 acar|OOF94964.1 acla|EAW12019.1 afis|EAW19660.1 afla|EED57030.1 afum|EAL86863.1 agla|OJJ80822.1 akaw|GAA92319.1 aluc|OJZ90217.1 anid|CBF71390.1 anig|EHA23715.1 aory|EIT82277.1 asyd|OJJ55042.1 ater|EAU32852.1 atub|OJI81649.1 aver|OJJ07166.1 awen|OJJ41922.1 azon|OJJ42684.1

Group1367: abra|OJJ65945.1 acar|OOF92928.1 acla|EAW13569.1 afis|EAW20032.1 afla|EED46042.1 afum|EAL84884.1 agla|OJJ88019.1 akaw|GAA90823.1 aluc|OJZ83012.1 anid|CBF84764.1 anig|EHA21427.1 aory|EIT75566.1 asyd|OJJ53285.1 ater|EAU29398.1 atub|OJI89247.1 aver|OJJ08456.1 awen|OJJ30055.1 azon|OJJ43701.1

Group1368: abra|OJJ65946.1 acar|OOF92925.1 acla|EAW13570.1 afis|EAW20031.1 afla|EED46041.1 afum|EAL84885.1 agla|OJJ88018.1 akaw|GAA90824.1 aluc|OJZ83011.1 anid|CBF84760.1 anig|EHA21426.1 aory|EIT75583.1 asyd|OJJ53286.1 ater|EAU29395.1 atub|OJI89246.1 aver|OJJ08455.1 awen|OJJ30054.1 azon|OJJ43702.1

Group1369: abra|OJJ65948.1 acar|OOF92924.1 acla|EAW13571.1 afis|EAW20030.1 afla|EED46039.1 afum|EAL84887.1 agla|OJJ88016.1 akaw|GAA90826.1 aluc|OJZ83009.1 anid|CBF84756.1 anig|EHA21424.1 aory|EIT75781.1 asyd|OJJ53288.1 ater|EAU29393.1 atub|OJI89244.1 aver|OJJ08453.1 awen|OJJ30052.1 azon|OJJ43700.1

Group1370: abra|OJJ65949.1 acar|OOF92923.1 acla|EAW13572.1 afis|EAW20029.1 afla|EED46038.1 afum|EAL84888.1 agla|OJJ88015.1 akaw|GAA90827.1 aluc|OJZ83008.1 anid|CBF84754.1 anig|EHA21423.1 aory|EIT75542.1 asyd|OJJ53289.1 ater|EAU29392.1 atub|OJI89243.1 aver|OJJ08452.1 awen|OJJ30051.1 azon|OJJ43698.1

Group1371: abra|OJJ65951.1 acar|OOF92921.1 acla|EAW13573.1 afis|EAW20028.1 afla|EED46037.1 afum|EAL84889.1 agla|OJJ88000.1 akaw|GAA90828.1 aluc|OJZ83006.1 anid|CBF84753.1 anig|EHA21422.1 aory|EIT75829.1 asyd|OJJ53290.1 ater|EAU29391.1 atub|OJI89241.1 aver|OJJ08450.1 awen|OJJ30042.1 azon|OJJ43688.1

Group1372: abra|OJJ65952.1 acar|OOF92920.1 acla|EAW13574.1 afis|EAW20027.1 afla|EED46036.1 afum|EAL84890.1 agla|OJJ88001.1 akaw|GAA90829.1 aluc|OJZ83005.1 anid|CBF84751.1 anig|EHA21421.1 aory|EIT75375.1 asyd|OJJ53291.1 ater|EAU29390.1 atub|OJI89240.1 aver|OJJ08449.1 awen|OJJ30043.1 azon|OJJ43689.1

Group1373: abra|OJJ65954.1 acar|OOF92918.1 acla|EAW13575.1 afis|EAW20026.1 afla|EED46034.1 afum|EAL84891.1 agla|OJJ88004.1 akaw|GAA90831.1 aluc|OJZ83003.1 anid|CBF84749.1 anig|EHA21419.1 aory|EIT75468.1 asyd|OJJ53293.1 ater|EAU29388.1 atub|OJI89237.1 aver|OJJ08447.1 awen|OJJ30045.1 azon|OJJ43691.1

Group1374: abra|OJJ65956.1 acar|OOF92916.1 acla|EAW13577.1 afis|EAW20024.1 afla|EED46032.1 afum|EAL84893.1 agla|OJJ88006.1 akaw|GAA90833.1 aluc|OJZ83001.1 anid|CBF84745.1 anig|EHA21417.1 aory|EIT75584.1 asyd|OJJ53295.1 ater|EAU29386.1 atub|OJI89235.1 aver|OJJ08445.1 awen|OJJ30046.1 azon|OJJ43693.1

Group1375: abra|OJJ65957.1 acar|OOF92915.1 acla|EAW13578.1 afis|EAW20023.1 afla|EED46031.1 afum|EAL84894.1 agla|OJJ88007.1 akaw|GAA90834.1 aluc|OJZ83000.1 anid|CBF84743.1 anig|EHA21416.1 aory|EIT75718.1 asyd|OJJ53296.1 ater|EAU29385.1 atub|OJI89234.1 aver|OJJ08444.1 awen|OJJ30047.1 azon|OJJ43694.1

Group1376: abra|OJJ65958.1 acar|OOF92914.1 acla|EAW13579.1 afis|EAW20022.1 afla|EED46030.1 afum|EAL84895.1 agla|OJJ88011.1 akaw|GAA90835.1 aluc|OJZ82999.1 anid|CBF84741.1 anig|EHA21415.1 aory|EIT75399.1 asyd|OJJ53297.1 ater|EAU29384.1 atub|OJI89233.1 aver|OJJ08443.1 awen|OJJ30048.1 azon|OJJ43695.1

Group1377: abra|OJJ65959.1 acar|OOF92913.1 acla|EAW13580.1 afis|EAW20021.1 afla|EED46029.1 afum|EAL84896.2 agla|OJJ88012.1 akaw|GAA90836.1 aluc|OJZ82998.1 anid|CBF84739.1 anig|EHA21414.1 aory|EIT75790.1 asyd|OJJ53298.1 ater|EAU29383.1 atub|OJI89232.1 aver|OJJ08442.1 awen|OJJ30049.1 azon|OJJ43696.1

Group1378: abra|OJJ65963.1 acar|OOF92911.1 acla|EAW13582.1 afis|EAW20019.1 afla|EED46027.1 afum|EAL84898.1 agla|OJJ87997.1 akaw|GAA90838.1 aluc|OJZ82995.1 anid|CBF84733.1 anig|EHA21410.1 aory|EIT75391.1 asyd|OJJ53302.1 ater|EAU29380.1 atub|OJI89229.1 aver|OJJ08439.1 awen|OJJ30040.1 azon|OJJ43684.1

Group1379: abra|OJJ65969.1 acar|OOF92904.1 acla|EAW13590.1 afis|EAW20013.1 afla|EED46020.1 afum|EAL84905.1 agla|OJJ87978.1 akaw|GAA90844.1 aluc|OJZ82989.1 anid|CBF84715.1 anig|EHA21404.1 aory|EIT75372.1 asyd|OJJ53326.1 ater|EAU29355.1 atub|OJI89223.1 aver|OJJ08417.1 awen|OJJ30027.1 azon|OJJ43661.1

Group1380: abra|OJJ65976.1 acar|OOF92896.1 acla|EAW13597.1 afis|EAW20007.1 afla|EED46013.1 afum|EAL84911.1 agla|OJJ87972.1 akaw|GAA90850.1 aluc|OJZ82983.1 anid|CBF84706.1 anig|EHA21397.1 aory|EIT75456.1 asyd|OJJ53321.1 ater|EAU29349.1 atub|OJI89217.1 aver|OJJ08422.1 awen|OJJ30020.1 azon|OJJ49472.1

Group1385: abra|OJJ66483.1 acar|OOF90960.1 acla|EAW10986.1 afis|EAW20874.1 afla|EED45556.1 afum|EAL84760.1 agla|OJJ83981.1 akaw|GAA92468.1 aluc|OJZ81551.1 anid|CBF82851.1 anig|EHA25883.1 aory|EIT75688.1 asyd|OJJ61594.1 ater|EAU30815.1 atub|OJI88634.1 aver|OJJ00916.1 awen|OJJ34486.1 azon|OJJ51763.1

Group1386: akaw|GAA92469.1 aluc|OJZ81550.1 atub|OJI88635.1 abra|OJJ66485.1 acar|OOF90962.1 acla|EAW10984.1 afis|EAW20876.1 afla|EED45558.1 afum|EAL84757.1 agla|OJJ83983.1 anid|CBF82855.1 anig|EHA25885.1 aory|EIT75434.1 asyd|OJJ61595.1 ater|EAU30813.1 aver|OJJ00918.1 awen|OJJ34484.1 azon|OJJ51761.1

Group1387: abra|OJJ66488.1 acar|OOF90964.1 acla|EAW10981.1 afis|EAW20878.1 afla|EED45563.1 afum|EAL84754.1 agla|OJJ83988.1 akaw|GAA92471.1 aluc|OJZ81547.1 anid|CBF82860.1 anig|EHA25888.1 aory|EIT75724.1 asyd|OJJ61597.1 ater|EAU38706.1 atub|OJI88637.1 aver|OJJ00921.1 awen|OJJ34481.1 azon|OJJ51758.1

Group1388: abra|OJJ66491.1 acar|OOF90967.1 acla|EAW10978.1 afis|EAW20881.1 afla|EED45568.1 afum|EAL84751.1 agla|OJJ83990.1 akaw|GAA91209.1 aluc|OJZ81544.1 anid|CBF82866.1 anig|EHA25891.1 aory|EIT75376.1 asyd|OJJ61599.1 ater|EAU30823.1 atub|OJI88640.1 aver|OJJ00923.1 awen|OJJ34479.1 azon|OJJ51756.1

Group1389: abra|OJJ66504.1 acar|OOF90968.1 acla|EAW10977.1 afis|EAW20882.1 afla|EED45567.1 afum|EAL84750.1 agla|OJJ83991.1 akaw|GAA91206.1 aluc|OJZ81541.1 anid|CBF82764.1 anig|EHA25895.1 aory|EIT75654.1 asyd|OJJ61604.1 ater|EAU30824.1 atub|OJI88645.1 aver|OJJ00927.1 awen|OJJ34478.1 azon|OJJ51755.1

Group1390: abra|OJJ66506.1 acar|OOF90970.1 acla|EAW10975.1 afis|EAW20885.1 afla|EED45570.1 afum|EAL84748.1 agla|OJJ83992.1 akaw|GAA91204.1 aluc|OJZ81539.1 anid|CBF82758.1 anig|EHA25897.1 aory|EIT75723.1 asyd|OJJ61607.1 ater|EAU30825.1 atub|OJI88647.1 aver|OJJ00930.1 awen|OJJ34462.1 azon|OJJ51752.1

Group1391: abra|OJJ66507.1 acar|OOF90971.1 acla|EAW10974.1 afis|EAW20886.1 afla|EED45571.1 afum|EAL84747.1 agla|OJJ83993.1 akaw|GAA91203.1 aluc|OJZ81538.1 anid|CBF82760.1 anig|EHA25898.1 aory|EIT75837.1 asyd|OJJ61606.1 ater|EAU30826.1 atub|OJI88648.1 aver|OJJ00929.1 awen|OJJ34461.1 azon|OJJ51750.1

Group1392: abra|OJJ66508.1 acar|OOF90972.1 acla|EAW10973.1 afis|EAW20887.1 afla|EED45572.1 afum|EAL84746.1 agla|OJJ83994.1 akaw|GAA91202.1 aluc|OJZ81537.1 anid|CBF82762.1 anig|EHA25899.1 aory|EIT75742.1 asyd|OJJ61605.1 ater|EAU30827.1 atub|OJI88649.1 aver|OJJ00928.1 awen|OJJ34460.1 azon|OJJ51749.1

Group1393: abra|OJJ66518.1 acar|OOF90973.1 acla|EAW10971.1 afis|EAW20888.1 afla|EED45576.1 afum|EAL84745.1 agla|OJJ83996.1 akaw|GAA91193.1 aluc|OJZ81526.1 anid|CBF82768.1 anig|EHA25908.1 aory|EIT75514.1 asyd|OJJ61601.1 ater|EAU30828.1 atub|OJI88659.1 aver|OJJ00925.1 awen|OJJ34459.1 azon|OJJ51748.1

Group1394: abra|OJJ66532.1 acar|OOF90983.1 acla|EAW10960.1 afis|EAW20899.1 afla|EED50249.1 afum|EAL84733.1 agla|OJJ84012.1 akaw|GAA91183.1 aluc|OJZ81520.1 anid|CBF82455.1 anig|EHA25917.1 aory|EIT77292.1 asyd|OJJ54114.1 ater|EAU38532.1 atub|OJI88669.1 aver|OJJ08621.1 awen|OJJ34449.1 azon|OJJ51735.1

Group1395: abra|OJJ66534.1 acar|OOF90986.1 acla|EAW10958.1 afis|EAW20901.1 afla|EED50251.1 afum|EAL84730.1 agla|OJJ84015.1 akaw|GAA91181.1 aluc|OJZ81522.1 anid|CBF82453.1 anig|EHA25920.1

aory|EIT77336.1 asyd|OJJ54112.1 ater|EAU38534.1 atub|OJI88671.1 aver|OJJ08623.1 awen|OJJ34447.1 azon|OJJ51733.1

Group1396: abra|OJJ66536.1 acar|OOF90987.1 acla|EAW10955.1 afis|EAW20906.1 afla|EED50254.1 afum|EAL84727.1 agla|OJJ84022.1 akaw|GAA91179.1 aluc|OJZ81524.1 anid|CBF82467.1 anig|EHA25922.1 aory|EIT77306.1 asyd|OJJ54098.1 ater|EAU38536.1 atub|OJI88673.1 aver|OJJ08640.1 awen|OJJ34445.1 azon|OJJ51728.1

Group1397: abra|OJJ66538.1 acar|OOF90989.1 acla|EAW10953.1 afis|EAW20908.1 afla|EED50256.1 afum|EAL84725.1 agla|OJJ84024.1 akaw|GAA91177.1 aluc|OJZ81509.1 anid|CBF82471.1 anig|EHA25924.1 aory|EIT77316.1 asyd|OJJ54100.1 ater|EAU38538.1 atub|OJI88675.1 aver|OJJ08638.1 awen|OJJ34443.1 azon|OJJ51720.1

Group1398: abra|OJJ66542.1 acar|OOF90994.1 acla|EAW09002.1 afis|EAW25543.1 afla|EED50261.1 afum|EAL84261.1 agla|OJJ84031.1 akaw|GAA91167.1 aluc|OJZ81495.1 anid|CBF80805.1 anig|EHA25935.1 aory|EIT77304.1 asyd|OJJ54485.1 ater|EAU38562.1 atub|OJI88686.1 aver|OJJ07694.1 awen|OJJ34423.1 azon|OJJ49761.1

Group1399: abra|OJJ66544.1 acar|OOF90996.1 acla|EAW10945.1 afis|EAW20913.1 afla|EED50263.1 afum|EAL84721.1 agla|OJJ79574.1 akaw|GAA91169.1 aluc|OJZ81497.1 anid|CBF84410.1 anig|EHA25933.1 aory|EIT77352.1 asyd|OJJ53562.1 ater|EAU38564.1 atub|OJI88684.1 aver|OJJ08190.1 awen|OJJ34421.1 azon|OJJ48150.1

Group1401: abra|OJJ66558.1 acar|OOF90013.1 acla|EAW10937.1 afis|EAW20924.1 afla|EED50278.1 afum|EAL84710.1 agla|OJJ83965.1 akaw|GAA91158.1 aluc|OJZ81486.1 anid|CBF82449.1 anig|EHA25941.1 aory|EIT77317.1 asyd|OJJ54111.1 ater|EAU38571.1 atub|OJI88694.1 aver|OJJ08624.1 awen|OJJ34396.1 azon|OJJ48132.1

Group1402: abra|OJJ66562.1 acar|OOF91003.1 acla|EAW10934.1 afis|EAW20927.1 afla|EED50281.1 afum|EAL84707.1 agla|OJJ83961.1 akaw|GAA91155.1 aluc|OJZ81483.1 anid|CBF82497.1 anig|EHA25944.1 aory|EIT77267.1 asyd|OJJ54088.1 ater|EAU38573.1 atub|OJI88698.1 aver|OJJ08650.1 awen|OJJ34392.1 azon|OJJ48123.1

Group1403: abra|OJJ66563.1 acar|OOF91004.1 acla|EAW10933.1 afis|EAW20928.1 afla|EED50282.1 afum|EAL84706.1 agla|OJJ83960.1 akaw|GAA91154.1 aluc|OJZ81482.1 anid|CBF82515.1 anig|EHA25945.1 aory|EIT77363.1 asyd|OJJ54084.1 ater|EAU38574.1 atub|OJI88699.1 aver|OJJ08654.1 awen|OJJ34390.1 azon|OJJ48122.1

Group1404: abra|OJJ66564.1 acar|OOF91005.1 acla|EAW10932.1 afis|EAW20929.1 afla|EED50283.1 afum|EAL84705.1 agla|OJJ83959.1 akaw|GAA91153.1 aluc|OJZ81481.1 anid|CBF82513.1 anig|EHA25946.1 aory|EIT77279.1 asyd|OJJ54083.1 ater|EAU38575.1 atub|OJI88700.1 aver|OJJ08655.1 awen|OJJ34389.1 azon|OJJ48121.1

Group1405: abra|OJJ66568.1 acar|OOF91009.1 acla|EAW10927.1 afis|EAW20402.1 afla|EED50288.1 afum|EAL85501.1 agla|OJJ83952.1 akaw|GAA91149.1 aluc|OJZ81476.1 anid|CBF82504.1 anig|EHA25950.1 aory|EIT77254.1 asyd|OJJ54078.1 ater|EAU38579.1 atub|OJI88703.1 aver|OJJ08660.1 awen|OJJ34384.1 azon|OJJ48115.1

Group1406: abra|OJJ66570.1 acar|OOF91010.1 acla|EAW10924.1 afis|EAW20934.1 afla|EED50291.1 afum|EAL84700.1 agla|OJJ83951.1 akaw|GAA91148.1 aluc|OJZ81474.1 anid|CBF82532.1 anig|EHA25951.1 aory|EIT77301.1 asyd|OJJ54076.1 ater|EAU38580.1 atub|OJI88704.1 aver|OJJ08663.1 awen|OJJ34382.1 azon|OJJ48114.1

Group1407: abra|OJJ66573.1 acar|OOF91014.1 acla|EAW10917.1 afis|EAW20941.1 afla|EED50296.1 afum|EAL84693.1 agla|OJJ83946.1 akaw|GAA91145.1 aluc|OJZ81471.1 anid|CBF82534.1 anig|EHA25956.1 aory|EIT77257.1 asyd|OJJ54069.1 ater|EAU38586.1 atub|OJI88707.1 aver|OJJ08673.1 awen|OJJ34376.1 azon|OJJ50768.1

Group1408: abra|OJJ66576.1 acar|OOF91017.1 acla|EAW10914.1 afis|EAW20944.1 afla|EED50298.1 afum|EAL84690.2 agla|OJJ83943.1 akaw|GAA91142.1 aluc|OJZ81468.1 anid|CBF82539.1 anig|EHA25959.1 aory|EIT77376.1 asyd|OJJ54066.1 ater|EAU38588.1 atub|OJI88710.1 aver|OJJ08676.1 awen|OJJ34372.1 azon|OJJ48108.1

Group1409: abra|OJJ66577.1 acar|OOF91018.1 acla|EAW10913.1 afis|EAW20945.1 afla|EED50299.1 afum|EAL84689.2 agla|OJJ83941.1 akaw|GAA91141.1 aluc|OJZ81467.1 anid|CBF82541.1 anig|EHA25960.1 aory|EIT77303.1 asyd|OJJ54065.1 ater|EAU38596.1 atub|OJI88711.1 aver|OJJ08677.1 awen|OJJ34371.1 azon|OJJ48107.1

Group1410: abra|OJJ66579.1 acar|OOF91019.1 acla|EAW10911.1 afis|EAW20947.1 afla|EED50301.1 afum|EAL84687.1 agla|OJJ83939.1 akaw|GAA91139.1 aluc|OJZ81465.1 anid|CBF82545.1 anig|EHA25962.1 aory|EIT77341.1 asyd|OJJ54063.1 ater|EAU38598.1 atub|OJI88713.1 aver|OJJ08679.1 awen|OJJ34369.1 azon|OJJ48105.1

Group1411: abra|OJJ66580.1 acar|OOF91020.1 acla|EAW10910.1 afis|EAW20948.1 afla|EED50302.1 afum|EAL84686.1 agla|OJJ83938.1 akaw|GAA91138.1 aluc|OJZ81464.1 anid|CBF82547.1 anig|EHA25963.1 aory|EIT77278.1 asyd|OJJ54062.1 ater|EAU38599.1 atub|OJI88714.1 aver|OJJ08680.1 awen|OJJ34368.1 azon|OJJ48104.1

Group1412: abra|OJJ66581.1 acar|OOF89819.1 acla|EAW10909.1 afis|EAW20949.1 afla|EED50303.1 afum|EAL84685.1 agla|OJJ83936.1 akaw|GAA91137.1 aluc|OJZ81463.1 anid|CBF82549.1 anig|EHA25964.1 aory|EIT77276.1 asyd|OJJ54061.1 ater|EAU38600.1 atub|OJI88715.1 aver|OJJ08681.1 awen|OJJ34367.1 azon|OJJ48103.1

Group1413: abra|OJJ66583.1 acar|OOF91021.1 acla|EAW10908.1 afis|EAW20950.1 afla|EED50305.1 afum|EAL84684.1 agla|OJJ83930.1 akaw|GAA91135.1 aluc|OJZ81462.1 anid|CBF82550.1 anig|EHA25966.1 aory|EIT77342.1 asyd|OJJ54060.1 ater|EAU38601.1 atub|OJI88716.1 aver|OJJ08682.1 awen|OJJ34366.1 azon|OJJ48101.1

Group1414: abra|OJJ66587.1 acar|OOF91024.1 acla|EAW10904.1 afis|EAW20954.1 afla|EED50310.1 afum|EAL84680.2 agla|OJJ83925.1 akaw|GAA91130.1 aluc|OJZ81459.1 anid|CBF82558.1 anig|EHA25970.1 aory|EIT77289.1 asyd|OJJ54056.1 ater|EAU38605.1 atub|OJI88720.1 aver|OJJ08685.1 awen|OJJ34362.1 azon|OJJ48098.1

Group1415: abra|OJJ66588.1 acar|OOF91025.1 acla|EAW10903.1 afis|EAW20955.1 afla|EED50311.1 afum|EAL84679.1 agla|OJJ83924.1 akaw|GAA91129.1 aluc|OJZ81458.1 anid|CBF82560.1 anig|EHA25971.1 aory|EIT77266.1 asyd|OJJ54054.1 ater|EAU38606.1 atub|OJI88721.1 aver|OJJ08687.1 awen|OJJ34361.1 azon|OJJ48097.1

Group1416: abra|OJJ66589.1 acar|OOF91026.1 acla|EAW10902.1 afis|EAW20956.1 afla|EED50312.1 afum|EAL84677.1 agla|OJJ83922.1 akaw|GAA91128.1 aluc|OJZ81457.1 anid|CBF82562.1 anig|EHA25972.1 aory|EIT77295.1 asyd|OJJ54052.1 ater|EAU38607.1 atub|OJI88722.1 aver|OJJ08688.1 awen|OJJ34360.1 azon|OJJ48096.1

Group1417: abra|OJJ66590.1 acar|OOF91027.1 acla|EAW10901.1 afis|EAW20957.1 afla|EED50313.1 afum|EAL84676.1 agla|OJJ83920.1 akaw|GAA91127.1 aluc|OJZ81456.1 anid|CBF82564.1 anig|EHA25973.1 aory|EIT77362.1 asyd|OJJ54051.1 ater|EAU38608.1 atub|OJI88723.1 aver|OJJ08689.1 awen|OJJ34359.1 azon|OJJ48095.1

Group1418: abra|OJJ66596.1 acar|OOF91030.1 acla|EAW10896.1 afis|EAW20962.1 afla|EED50317.1 afum|EAL84671.1 agla|OJJ83916.1 akaw|GAA91122.1 aluc|OJZ81450.1 anid|CBF82571.1 anig|EHA25976.1 aory|EIT77307.1 asyd|OJJ54048.1 ater|EAU38612.1 atub|OJI88729.1 aver|OJJ08695.1 awen|OJJ34354.1 azon|OJJ48091.1

Group1419: abra|OJJ66598.1 acar|OOF91031.1 acla|EAW10895.1 afis|EAW20963.1 afla|EED50318.1 afum|EAL84670.1 agla|OJJ83915.1 akaw|GAA91121.1 aluc|OJZ81449.1 anid|CBF82572.1 anig|EHA25977.1 aory|EIT77348.1 asyd|OJJ54047.1 ater|EAU38613.1 atub|OJI88730.1 aver|OJJ08696.1 awen|OJJ34352.1 azon|OJJ48089.1

Group1420: abra|OJJ66600.1 acar|OOF91033.1 acla|EAW10892.1 afis|EAW20966.1 afla|EED50320.1 afum|EAL84667.1 agla|OJJ84043.1 akaw|GAA91119.1 aluc|OJZ81446.1 anid|CBF82578.1 anig|EHA25979.1 aory|EIT77255.1 asyd|OJJ54042.1 ater|EAU38616.1 atub|OJI88732.1 aver|OJJ08699.1 awen|OJJ34349.1 azon|OJJ48086.1

Group1421: abra|OJJ66601.1 acar|OOF91034.1 acla|EAW10891.1 afis|EAW20967.1 afla|EED50321.1 afum|EAL84666.1 agla|OJJ84044.1 akaw|GAA91118.1 aluc|OJZ81445.1 anid|CBF82580.1 anig|EHA25980.1 aory|EIT77332.1 asyd|OJJ54041.1 ater|EAU38617.1 atub|OJI88733.1 aver|OJJ08700.1 awen|OJJ34348.1 azon|OJJ48085.1

Group1422: abra|OJJ66604.1 acar|OOF90114.1 acla|EAW10889.1 afis|EAW20969.1 afla|EED50323.1 afum|EAL84665.1 agla|OJJ84046.1 akaw|GAA91116.1 aluc|OJZ81442.1 anid|CBF82584.1 anig|EHA25982.1 aory|EIT77349.1 asyd|OJJ54039.1 ater|EAU38619.1 atub|OJI88736.1 aver|OJJ08701.1 awen|OJJ34346.1 azon|OJJ48084.1

Group1423: abra|OJJ66606.1 acar|OOF90112.1 acla|EAW10887.1 afis|EAW20971.1 afla|EED50325.1 afum|EAL84663.1 agla|OJJ84048.1 akaw|GAA91114.1 aluc|OJZ81440.1 anid|CBF82585.1 anig|EHA25984.1 aory|EIT77291.1 asyd|OJJ54037.1 ater|EAU38620.1 atub|OJI88738.1 aver|OJJ08703.1 awen|OJJ34344.1 azon|OJJ48083.1

Group1424: abra|OJJ66610.1 acar|OOF91035.1 acla|EAW10886.1 afis|EAW20972.1 afla|EED50326.1 afum|EAL84661.1 agla|OJJ84050.1 akaw|GAA91113.1 aluc|OJZ81439.1 anid|CBF82587.1 anig|EHA25985.1 aory|EIT77361.1 asyd|OJJ54036.1 ater|EAU38622.1 atub|OJI88742.1 aver|OJJ08704.1 awen|OJJ34341.1 azon|OJJ48081.1

Group1425: abra|OJJ66612.1 acar|OOF91036.1 acla|EAW10884.1 afis|EAW20974.1 afla|EED50329.1 afum|EAL84659.1 agla|OJJ84052.1 akaw|GAA91111.1 aluc|OJZ81437.1 anid|CBF82593.1 anig|EHA25987.1

aory|EIT77331.1 asyd|OJJ54033.1 ater|EAU38624.1 atub|OJI88744.1 aver|OJJ08707.1 awen|OJJ34339.1 azon|OJJ48079.1

Group1426: abra|OJJ66613.1 acar|OOF91038.1 acla|EAW10883.1 afis|EAW20975.1 afla|EED50330.1 afum|EAL84658.1 agla|OJJ84053.1 akaw|GAA91110.1 aluc|OJZ81436.1 anid|CBF82595.1 anig|EHA25988.1 aory|EIT77270.1 asyd|OJJ54032.1 ater|EAU38625.1 atub|OJI88745.1 aver|OJJ08708.1 awen|OJJ34337.1 azon|OJJ48078.1

Group1427: abra|OJJ66617.1 acar|OOF91040.1 acla|EAW10880.1 afis|EAW20978.1 afla|EED50333.1 afum|EAL84655.1 agla|OJJ84055.1 akaw|GAA91108.1 aluc|OJZ81432.1 anid|CBF82598.1 anig|EHA25990.1 aory|EIT77374.1 asyd|OJJ54030.1 ater|EAU38627.1 atub|OJI88748.1 aver|OJJ08710.1 awen|OJJ34334.1 azon|OJJ48076.1

Group1428: abra|OJJ66619.1 acar|OOF91042.1 acla|EAW10878.1 afis|EAW20980.1 afla|EED50335.1 afum|EAL84651.2 agla|OJJ84057.1 akaw|GAA91106.1 aluc|OJZ81430.1 anid|CBF71772.1 anig|EHA25992.1 aory|EIT77286.1 asyd|OJJ61720.1 ater|EAU38628.1 atub|OJI88750.1 aver|OJI95814.1 awen|OJJ34332.1 azon|OJJ48074.1

Group1429: abra|OJJ66624.1 acar|OOF91048.1 acla|EAW10872.1 afis|EAW20986.1 afla|EED50341.1 afum|EAL84644.1 agla|OJJ84065.1 akaw|GAA91101.1 aluc|OJZ81425.1 anid|CBF82611.1 anig|EHA25997.1 aory|EIT77327.1 asyd|OJJ54022.1 ater|EAU38635.1 atub|OJI88756.1 aver|OJJ08718.1 awen|OJJ34325.1 azon|OJJ48071.1

Group1430: abra|OJJ66627.1 acar|OOF91052.1 acla|EAW10869.1 afis|EAW20989.1 afla|EED50343.1 afum|EAL84640.2 agla|OJJ84067.1 akaw|GAA91098.1 aluc|OJZ81421.1 anid|CBF82605.1 anig|EHA26000.1 aory|EIT77281.1 asyd|OJJ54026.1 ater|EAU38638.1 atub|OJI88760.1 aver|OJJ08714.1 awen|OJJ34321.1 azon|OJJ48066.1

Group1431: abra|OJJ66749.1 acar|OOF93832.1 acla|EAW11875.1 afis|EAW19486.1 afla|EED56704.1 afum|EAL87008.1 agla|OJJ81211.1 akaw|GAA89685.1 aluc|OJZ80700.1 anid|CBF75460.1 anig|EHA27316.1 aory|EIT73946.1 asyd|OJJ59769.1 ater|EAU34799.1 atub|OJI87559.1 aver|OJJ02516.1 awen|OJJ36902.1 azon|OJJ42915.1

Group1432: abra|OJJ66751.1 acar|OOF93833.1 acla|EAW11876.1 afis|EAW19487.1 afla|EED56705.1 afum|EAL87007.1 agla|OJJ81213.1 akaw|GAA89686.1 aluc|OJZ80699.1 anid|CBF75458.1 anig|EHA27317.1 aory|EIT73903.1 asyd|OJJ59771.1 ater|EAU34800.1 atub|OJI87558.1 aver|OJJ02515.1 awen|OJJ36903.1 azon|OJJ42913.1

Group1433: abra|OJJ66752.1 acar|OOF93834.1 acla|EAW11877.1 afis|EAW19488.1 afla|EED56706.1 afum|EAL87006.1 agla|OJJ81214.1 akaw|GAA89687.1 aluc|OJZ80698.1 anid|CBF75456.1 anig|EHA27318.1 aory|EIT73834.1 asyd|OJJ59772.1 ater|EAU34801.1 atub|OJI87557.1 aver|OJJ02514.1 awen|OJJ36904.1 azon|OJJ42912.1

Group1434: abra|OJJ66753.1 acar|OOF93835.1 acla|EAW11880.1 afis|EAW19491.1 afla|EED56710.1 afum|EAL87003.1 agla|OJJ81220.1 akaw|GAA89688.1 aluc|OJZ80697.1 anid|CBF75452.1 anig|EHA27319.1 aory|EIT73847.1 asyd|OJJ59774.1 ater|EAU34803.1 atub|OJI87556.1 aver|OJJ02512.1 awen|OJJ36907.1 azon|OJJ42909.1

Group1435: abra|OJJ66755.1 acar|OOF93836.1 acla|EAW11881.1 afis|EAW19492.1 afla|EED56711.1 afum|EAL87002.1 agla|OJJ81221.1 akaw|GAA89689.1 aluc|OJZ80696.1 anid|CBF75450.1 anig|EHA27320.1 aory|EIT73907.1 asyd|OJJ59775.1 ater|EAU34804.1 atub|OJI87555.1 aver|OJJ02511.1 awen|OJJ36908.1 azon|OJJ42908.1

Group1437: abra|OJJ66819.1 acar|OOF91602.1 acla|EAW14405.1 afis|EAW19113.1 afla|EED54301.1 afum|EAL93845.1 agla|OJJ84489.1 akaw|GAA90457.1 aluc|OJZ85767.1 anid|CBF79734.1 anig|EHA24786.1 aory|EIT76106.1 asyd|OJJ58069.1 ater|EAU37372.1 atub|OJI79552.1 aver|OJJ04338.1 awen|OJJ32523.1 azon|OJJ50791.1

Group1438: abra|OJJ66828.1 acar|OOF91606.1 acla|EAW14411.1 afis|EAW19119.1 afla|EED54295.1 afum|EAL93851.1 agla|OJJ80434.1 akaw|GAA90464.1 aluc|OJZ85759.1 anid|CBF79724.1 anig|EHA24797.1 aory|EIT76143.1 asyd|OJJ58075.1 ater|EAU37382.1 atub|OJI79544.1 aver|OJJ04332.1 awen|OJJ37450.1 azon|OJJ51127.1

Group1439: abra|OJJ66830.1 acar|OOF91608.1 acla|EAW14413.1 afis|EAW19121.1 afla|EED54293.1 afum|EAL93853.1 agla|OJJ80436.1 akaw|GAA90467.1 aluc|OJZ85757.1 anid|CBF79722.1 anig|EHA24799.1 aory|EIT76028.1 asyd|OJJ58077.1 ater|EAU37384.1 atub|OJI79542.1 aver|OJJ04330.1 awen|OJJ37452.1 azon|OJJ51129.1

Group1440: abra|OJJ66832.1 acar|OOF91609.1 acla|EAW14414.1 afis|EAW19122.1 afla|EED54292.1 afum|EAL93854.1 agla|OJJ80437.1 akaw|GAA90468.1 aluc|OJZ85756.1 anid|CBF79720.1 anig|EHA24800.1 aory|EIT76096.1 asyd|OJJ58078.1 ater|EAU37385.1 atub|OJI79541.1 aver|OJJ04329.1 awen|OJJ37453.1 azon|OJJ51130.1

Group1441: abra|OJJ66833.1 acar|OOF91610.1 acla|EAW14415.1 afis|EAW19123.1 afla|EED54291.1 afum|EAL93855.1 agla|OJJ80438.1 akaw|GAA90469.1 aluc|OJZ85755.1 anid|CBF79718.1 anig|EHA24801.1 aory|EIT76111.1 asyd|OJJ58079.1 ater|EAU37386.1 atub|OJI79540.1 aver|OJJ04328.1 awen|OJJ37454.1 azon|OJJ51131.1

Group1442: abra|OJJ66834.1 acar|OOF91611.1 acla|EAW14416.1 afis|EAW19124.1 afla|EED54290.1 afum|EAL93856.1 agla|OJJ80439.1 akaw|GAA90470.1 aluc|OJZ85754.1 anid|CBF79717.1 anig|EHA24802.1 aory|EIT76047.1 asyd|OJJ58080.1 ater|EAU37387.1 atub|OJI79538.1 aver|OJJ04327.1 awen|OJJ37455.1 azon|OJJ51134.1

Group1443: abra|OJJ66835.1 acar|OOF91613.1 acla|EAW14419.1 afis|EAW19126.1 afla|EED54288.1 afum|EAL93858.1 agla|OJJ80441.1 akaw|GAA90472.1 aluc|OJZ85752.1 anid|CBF79737.1 anig|EHA24804.1 aory|EIT75920.1 asyd|OJJ58083.1 ater|EAU37393.1 atub|OJI79535.1 aver|OJJ04325.1 awen|OJJ37457.1 azon|OJJ51137.1

Group1444: abra|OJJ66836.1 acar|OOF91614.1 acla|EAW14420.1 afis|EAW19127.1 afla|EED54287.1 afum|EAL93859.1 agla|OJJ88596.1 akaw|GAA90473.1 aluc|OJZ85751.1 anid|CBF79739.1 anig|EHA24805.1 aory|EIT75974.1 asyd|OJJ58084.1 ater|EAU37394.1 atub|OJI79534.1 aver|OJJ04324.1 awen|OJJ37459.1 azon|OJJ51138.1

Group1446: abra|OJJ66843.1 acar|OOF91620.1 acla|EAW14417.1 afis|EAW19130.1 afla|EED54284.1 afum|EAL93862.1 agla|OJJ88602.1 akaw|GAA90479.1 aluc|OJZ85740.1 anid|CBF79758.1 anig|EHA24811.1 aory|EIT76041.1 asyd|OJJ58100.1 ater|EAU37398.1 atub|OJI79525.1 aver|OJJ04304.1 awen|OJJ37465.1 azon|OJJ51143.1

Group1447: abra|OJJ66846.1 acar|OOF91624.1 acla|EAW14425.1 afis|EAW19132.1 afla|EED54281.1 afum|EAL93864.1 agla|OJJ88607.1 akaw|GAA91559.1 aluc|OJZ85737.1 anid|CBF86338.1 anig|EHA24813.1 aory|EIT76060.1 asyd|OJJ64988.1 ater|EAU37405.1 atub|OJI79522.1 aver|OJI99108.1 awen|OJJ37468.1 azon|OJJ51146.1

Group1448: abra|OJJ66847.1 acar|OOF91625.1 acla|EAW14426.1 afis|EAW19133.1 afla|EED54280.1 afum|EAL93865.1 agla|OJJ88608.1 akaw|GAA91558.1 aluc|OJZ85736.1 anid|CBF86336.1 anig|EHA24814.1 aory|EIT76085.1 asyd|OJJ64989.1 ater|EAU37401.1 atub|OJI79521.1 aver|OJI99109.1 awen|OJJ37469.1 azon|OJJ51147.1

Group1449: abra|OJJ66853.1 acar|OOF91629.1 acla|EAW14431.1 afis|EAW19138.1 afla|EED54275.1 afum|EAL93870.1 agla|OJJ88617.1 akaw|GAA91553.1 aluc|OJZ85731.1 anid|CBF86324.1 anig|EHA24818.1 aory|EIT75921.1 asyd|OJJ64993.1 ater|EAU37408.1 atub|OJI79516.1 aver|OJI99112.1 awen|OJJ37475.1 azon|OJJ51155.1

Group1450: abra|OJJ66854.1 acar|OOF91630.1 acla|EAW14432.1 afis|EAW19139.1 afla|EED54274.1 afum|EAL93871.2 agla|OJJ88591.1 akaw|GAA91552.1 aluc|OJZ85730.1 anid|CBF86322.1 anig|EHA24819.1 aory|EIT75977.1 asyd|OJJ64994.1 ater|EAU37410.1 atub|OJI79515.1 aver|OJI99113.1 awen|OJJ37476.1 azon|OJJ51157.1

Group1451: abra|OJJ66855.1 acar|OOF91632.1 acla|EAW14433.1 afis|EAW19141.1 afla|EED54272.1 afum|EAL93872.1 agla|OJJ88589.1 akaw|GAA91551.1 aluc|OJZ85728.1 anid|CBF86321.1 anig|EHA24820.1 aory|EIT76058.1 asyd|OJJ64995.1 ater|EAU37412.1 atub|OJI79514.1 aver|OJI99115.1 awen|OJJ37481.1 azon|OJJ51158.1

Group1452: abra|OJJ66856.1 acar|OOF91633.1 acla|EAW14434.1 afis|EAW19142.1 afla|EED54271.1 afum|EAL93873.1 agla|OJJ88588.1 akaw|GAA91550.1 aluc|OJZ85727.1 anid|CBF86317.1 anig|EHA24821.1 aory|EIT76153.1 asyd|OJJ64997.1 ater|EAU37413.1 atub|OJI79513.1 aver|OJI99117.1 awen|OJJ37482.1 azon|OJJ51159.1

Group1453: abra|OJJ66858.1 acar|OOF91636.1 acla|EAW14437.1 afis|EAW19145.1 afla|EED54268.1 afum|EAL93876.1 agla|OJJ88586.1 akaw|GAA91547.1 aluc|OJZ85725.1 anid|CBF86313.1 anig|EHA24823.1 aory|EIT76154.1 asyd|OJJ65000.1 ater|EAU37417.1 atub|OJI79511.1 aver|OJI99120.1 awen|OJJ37484.1 azon|OJJ51161.1

Group1454: abra|OJJ66861.1 acar|OOF91638.1 acla|EAW14438.1 afis|EAW19146.1 afla|EED54267.1 afum|EAL93877.1 agla|OJJ88584.1 akaw|GAA91546.1 aluc|OJZ85721.1 anid|CBF86311.1 anig|EHA24824.1 aory|EIT76119.1 asyd|OJJ65001.1 ater|EAU37420.1 atub|OJI79510.1 aver|OJI99121.1 awen|OJJ37485.1 azon|OJJ51163.1

Group1455: abra|OJJ66864.1 acar|OOF91640.1 acla|EAW14440.1 afis|EAW19148.1 afla|EED54265.1 afum|EAL93879.1 agla|OJJ88576.1 akaw|GAA91544.1 aluc|OJZ85719.1 anid|CBF86308.1 anig|EHA24826.1 aory|EIT75979.1 asyd|OJJ65004.1 ater|EAU37421.1 atub|OJI79508.1 aver|OJI99123.1 awen|OJJ37501.1 azon|OJJ51166.1

Group1456: abra|OJJ66866.1 acar|OOF91643.1 acla|EAW14442.1 afis|EAW19150.1 afla|EED54257.1 afum|EAL93881.1 agla|OJJ88572.1 akaw|GAA91542.1 aluc|OJZ85716.1 anid|CBF86305.1 anig|EHA24828.1

aory|EIT76051.1 asyd|OJJ65006.1 ater|EAU37424.1 atub|OJI79506.1 aver|OJI99125.1 awen|OJJ37503.1 azon|OJJ51168.1

Group1457: abra|OJJ66869.1 acar|OOF91646.1 acla|EAW14445.1 afis|EAW19153.1 afla|EED54252.1 afum|EAL93884.1 agla|OJJ88569.1 akaw|GAA91539.1 aluc|OJZ85712.1 anid|CBF86299.1 anig|EHA24831.1 aory|EIT76023.1 asyd|OJJ65009.1 ater|EAU37426.1 atub|OJI79503.1 aver|OJI99128.1 awen|OJJ37506.1 azon|OJJ51172.1

Group1458: abra|OJJ66871.1 acar|OOF91647.1 acla|EAW14446.1 afis|EAW19154.1 afla|EED54251.1 afum|EAL93885.1 agla|OJJ88568.1 akaw|GAA91538.1 aluc|OJZ85711.1 anid|CBF86297.1 anig|EHA24832.1 aory|EIT75939.1 asyd|OJJ65010.1 ater|EAU37427.1 atub|OJI79502.1 aver|OJI99129.1 awen|OJJ37507.1 azon|OJJ51173.1

Group1459: abra|OJJ66873.1 acar|OOF91649.1 acla|EAW14448.1 afis|EAW19156.1 afla|EED54246.1 afum|EAL93887.1 agla|OJJ88566.1 akaw|GAA91536.1 aluc|OJZ85708.1 anid|CBF86294.1 anig|EHA24834.1 aory|EIT75988.1 asyd|OJJ65012.1 ater|EAU37429.1 atub|OJI79499.1 aver|OJI99131.1 awen|OJJ37509.1 azon|OJJ51174.1

Group1460: abra|OJJ66874.1 acar|OOF91650.1 acla|EAW14449.1 afis|EAW19157.1 afla|EED54245.1 afum|EBA27505.1 agla|OJJ88565.1 akaw|GAA91535.1 aluc|OJZ85707.1 anid|CBF86292.1 anig|EHA24835.1 aory|EIT75937.1 asyd|OJJ65013.1 ater|EAU37430.1 atub|OJI79498.1 aver|OJI99132.1 awen|OJJ37510.1 azon|OJJ51175.1

Group1461: abra|OJJ66875.1 acar|OOF91651.1 acla|EAW14450.1 afis|EAW19158.1 afla|EED54244.1 afum|EBA27506.1 agla|OJJ88564.1 akaw|GAA91534.1 aluc|OJZ85706.1 anid|CBF86291.1 anig|EHA24836.1 aory|EIT76133.1 asyd|OJJ65014.1 ater|EAU37431.1 atub|OJI79497.1 aver|OJI99133.1 awen|OJJ37511.1 azon|OJJ51176.1

Group1462: abra|OJJ66879.1 acar|OOF91654.1 acla|EAW14454.1 afis|EAW19162.1 afla|EED54240.1 afum|EAL93892.2 agla|OJJ88562.1 akaw|GAA91530.1 aluc|OJZ85703.1 anid|CBF86285.1 anig|EHA24840.1 aory|EIT76157.1 asyd|OJJ65017.1 ater|EAU37311.1 atub|OJI79493.1 aver|OJI99135.1 awen|OJJ37515.1 azon|OJJ51178.1

Group1463: abra|OJJ66880.1 acar|OOF91655.1 acla|EAW14455.1 afis|EAW19163.1 afla|EED54239.1 afum|EAL93893.1 agla|OJJ88561.1 akaw|GAA91529.1 aluc|OJZ85702.1 anid|CBF86282.1 anig|EHA24841.1 aory|EIT76014.1 asyd|OJJ65018.1 ater|EAU37310.1 atub|OJI79492.1 aver|OJI99136.1 awen|OJJ37516.1 azon|OJJ51181.1

Group1464: abra|OJJ66883.1 acar|OOF91657.1 acla|EAW14457.1 afis|EAW19165.1 afla|EED54232.1 afum|EAL93895.1 agla|OJJ88559.1 akaw|GAA91527.1 aluc|OJZ85699.1 anid|CBF86279.1 anig|EHA24843.1 aory|EIT75967.1 asyd|OJJ65020.1 ater|EAU37308.1 atub|OJI79489.1 aver|OJI99138.1 awen|OJJ37518.1 azon|OJJ51185.1

Group1465: abra|OJJ66887.1 acar|OOF91660.1 acla|EAW14459.1 afis|EAW19167.1 afla|EED54230.1 afum|EAL93897.1 agla|OJJ88557.1 akaw|GAA91525.1 aluc|OJZ85696.1 anid|CBF86276.1 anig|EHA24846.1 aory|EIT75971.1 asyd|OJJ65022.1 ater|EAU37306.1 atub|OJI79485.1 aver|OJI99141.1 awen|OJJ37520.1 azon|OJJ51186.1

Group1466: abra|OJJ66889.1 acar|OOF91661.1 acla|EAW14461.1 afis|EAW19169.1 afla|EED54228.1 afum|EAL93899.1 agla|OJJ88555.1 akaw|GAA91523.1 aluc|OJZ85693.1 anid|CBF86273.1 anig|EHA24848.1 aory|EIT76026.1 asyd|OJJ65024.1 ater|EAU37304.1 atub|OJI79483.1 aver|OJI99143.1 awen|OJJ37522.1 azon|OJJ51188.1

Group1467: abra|OJJ66890.1 acar|OOF91662.1 acla|EAW14462.1 afis|EAW19170.1 afla|EED54226.1 afum|EAL93900.1 agla|OJJ88554.1 akaw|GAA91522.1 aluc|OJZ85692.1 anid|CBF86271.1 anig|EHA24849.1 aory|EIT75952.1 asyd|OJJ65025.1 ater|EAU37303.1 atub|OJI79482.1 aver|OJI99144.1 awen|OJJ37523.1 azon|OJJ51189.1

Group1468: abra|OJJ66891.1 acar|OOF91663.1 acla|EAW14463.1 afis|EAW19171.1 afla|EED54224.1 afum|EAL93901.1 agla|OJJ88553.1 akaw|GAA91521.1 aluc|OJZ85691.1 anid|CBF86270.1 anig|EHA24850.1 aory|EIT75965.1 asyd|OJJ65026.1 ater|EAU37302.1 atub|OJI79481.1 aver|OJI99145.1 awen|OJJ37524.1 azon|OJJ51190.1

Group1469: abra|OJJ66893.1 acar|OOF91664.1 acla|EAW14465.1 afis|EAW19173.1 afla|EED54222.1 afum|EAL93903.1 agla|OJJ88551.1 akaw|GAA91519.1 aluc|OJZ85689.1 anid|CBF86266.1 anig|EHA24852.1 aory|EIT76009.1 asyd|OJJ65029.1 ater|EAU37300.1 atub|OJI79479.1 aver|OJI99148.1 awen|OJJ37526.1 azon|OJJ51193.1

Group1470: abra|OJJ66894.1 acar|OOF91665.1 acla|EAW14466.1 afis|EAW19174.1 afla|EED54221.1 afum|EAL93904.1 agla|OJJ88550.1 akaw|GAA91518.1 aluc|OJZ85688.1 anid|CBF86265.1 anig|EHA24853.1 aory|EIT76027.1 asyd|OJJ65030.1 ater|EAU37299.1 atub|OJI79478.1 aver|OJI99149.1 awen|OJJ37527.1 azon|OJJ51194.1

Group1471: abra|OJJ66895.1 acar|OOF91666.1 acla|EAW14467.1 afis|EAW19175.1 afla|EED54220.1 afum|EAL93905.1 agla|OJJ88548.1 akaw|GAA91517.1 aluc|OJZ85687.1 anid|CBF86263.1 anig|EHA24854.1 aory|EIT75931.1 asyd|OJJ65031.1 ater|EAU37298.1 atub|OJI79477.1 aver|OJI99150.1 awen|OJJ37528.1 azon|OJJ51195.1

Group1472: abra|OJJ66896.1 acar|OOF91667.1 acla|EAW14468.1 afis|EAW19176.1 afla|EED54219.1 afum|EAL93906.1 agla|OJJ88547.1 akaw|GAA91516.1 aluc|OJZ85686.1 anid|CBF86262.1 anig|EHA24855.1 aory|EIT76150.1 asyd|OJJ65032.1 ater|EAU37297.1 atub|OJI79476.1 aver|OJI99151.1 awen|OJJ37530.1 azon|OJJ51196.1

Group1473: abra|OJJ66897.1 acar|OOF91668.1 acla|EAW14469.1 afis|EAW19177.1 afla|EED54218.1 afum|EAL93907.1 agla|OJJ88546.1 akaw|GAA91515.1 aluc|OJZ85685.1 anid|CBF86260.1 anig|EHA24856.1 aory|EIT75968.1 asyd|OJJ65033.1 ater|EAU37296.1 atub|OJI79475.1 aver|OJI99152.1 awen|OJJ37531.1 azon|OJJ51197.1

Group1474: abra|OJJ66899.1 acar|OOF91671.1 acla|EAW14470.1 afis|EAW19179.1 afla|EED54217.1 afum|EAL93909.1 agla|OJJ88544.1 akaw|GAA91513.1 aluc|OJZ85683.1 anid|CBF86259.1 anig|EHA24858.1 aory|EIT76147.1 asyd|OJJ65035.1 ater|EAU37294.1 atub|OJI79473.1 aver|OJI99154.1 awen|OJJ37533.1 azon|OJJ51199.1

Group1475: abra|OJJ66900.1 acar|OOF91672.1 acla|EAW14471.1 afis|EAW19180.1 afla|EED54216.1 afum|EAL93910.1 agla|OJJ88543.1 akaw|GAA91512.1 aluc|OJZ85682.1 anid|CBF86257.1 anig|EHA24859.1 aory|EIT76034.1 asyd|OJJ65036.1 ater|EAU37293.1 atub|OJI79472.1 aver|OJI99155.1 awen|OJJ37534.1 azon|OJJ51200.1

Group1476: abra|OJJ66903.1 acar|OOF91673.1 acla|EAW14473.1 afis|EAW19182.1 afla|EED54213.1 afum|EAL93912.1 agla|OJJ88540.1 akaw|GAA91510.1 aluc|OJZ85680.1 anid|CBF86253.1 anig|EHA24861.1 aory|EIT76125.1 asyd|OJJ65038.1 ater|EAU37291.1 atub|OJI79470.1 aver|OJI99157.1 awen|OJJ37536.1 azon|OJJ51202.1

Group1477: abra|OJJ66905.1 acar|OOF91675.1 acla|EAW14474.1 afis|EAW19183.1 afla|EED54212.1 afum|EAL93913.1 agla|OJJ88539.1 akaw|GAA91509.1 aluc|OJZ85676.1 anid|CBF86252.1 anig|EHA24864.1 aory|EIT75985.1 asyd|OJJ65041.1 ater|EAU37290.1 atub|OJI79467.1 aver|OJI99159.1 awen|OJJ37538.1 azon|OJJ51203.1

Group1478: abra|OJJ66906.1 acar|OOF91676.1 acla|EAW14475.1 afis|EAW19184.1 afla|EED54211.1 afum|EAL93915.1 akaw|GAA91508.1 aluc|OJZ85675.1 anid|CBF86250.1 anig|EHA24865.1 aory|EIT76099.1 asyd|OJJ65042.1 ater|EAU37289.1 atub|OJI79466.1 aver|OJI99160.1 awen|OJJ37539.1 azon|OJJ51204.1 agla|OJJ88538.1

Group1479: abra|OJJ66907.1 acar|OOF91677.1 acla|EAW14476.1 afla|EED54210.1 afum|EAL93916.1 agla|OJJ88537.1 akaw|GAA91507.1 aluc|OJZ85673.1 anid|CBF86248.1 anig|EHA24866.1 aory|EIT76017.1 asyd|OJJ65043.1 ater|EAU37287.1 atub|OJI79465.1 aver|OJI99162.1 awen|OJJ37542.1 afis|EAW19185.1 azon|OJJ51206.1

Group1480: abra|OJJ66909.1 acar|OOF91679.1 acla|EAW14478.1 afis|EAW19187.1 afla|EED47724.1 afum|EAL93918.1 agla|OJJ88534.1 akaw|GAA91505.1 aluc|OJZ85671.1 anid|CBF86246.1 anig|EHA24868.1 aory|EIT77160.1 asyd|OJJ65044.1 ater|EAU37286.1 atub|OJI79463.1 aver|OJI99165.1 awen|OJJ37545.1 azon|OJJ51210.1

Group1481: abra|OJJ66911.1 acar|OOF91680.1 acla|EAW14480.1 afis|EAW19189.1 afla|EED47722.1 afum|EAL93921.1 agla|OJJ88532.1 akaw|GAA91504.1 aluc|OJZ85670.1 anid|CBF86243.1 anig|EHA24869.1 aory|EIT77116.1 asyd|OJJ65046.1 ater|EAU37285.1 atub|OJI79462.1 aver|OJI99167.1 awen|OJJ37549.1 azon|OJJ51211.1

Group1482: abra|OJJ66915.1 acar|OOF91682.1 acla|EAW14482.1 afis|EAW19191.1 afla|EED47718.1 afum|EAL93922.1 agla|OJJ88518.1 akaw|GAA91503.1 aluc|OJZ85665.1 anid|CBF86239.1 anig|EHA24870.1 aory|EIT80246.1 asyd|OJJ65049.1 ater|EAU37284.1 atub|OJI79458.1 aver|OJI99171.1 awen|OJJ37551.1 azon|OJJ51212.1

Group1483: abra|OJJ66918.1 acar|OOF91685.1 acla|EAW14485.1 afis|EAW19194.1 afla|EED47714.1 afum|EAL93925.2 agla|OJJ88515.1 akaw|GAA91500.1 aluc|OJZ85661.1 anid|CBF78544.1 anig|EHA24872.1 aory|EIT80273.1 asyd|OJJ65054.1 ater|EAU37281.1 atub|OJI79455.1 aver|OJI99177.1 awen|OJJ37552.1 azon|OJJ51213.1

Group1484: abra|OJJ66919.1 acar|OOF91687.1 acla|EAW14486.1 afis|EAW19195.1 afla|EED47713.1 afum|EAL93927.2 agla|OJJ88514.1 akaw|GAA91499.1 aluc|OJZ85660.1 anid|CBF78546.1 anig|EHA24873.1 aory|EIT80279.1 asyd|OJJ65055.1 ater|EAU37280.1 atub|OJI79454.1 aver|OJI99178.1 awen|OJJ37553.1 azon|OJJ51214.1

Group1485: abra|OJJ66923.1 acar|OOF91690.1 acla|EAW14489.1 afis|EAW19203.1 afla|EED47709.1 afum|EAL93933.1 agla|OJJ88509.1 akaw|GAA91496.1 aluc|OJZ85657.1 anid|CBF78554.1 anig|EHA24876.1

aory|EIT80207.1 asyd|OJJ65059.1 ater|EAU37277.1 atub|OJI79449.1 aver|OJI99184.1 awen|OJJ37556.1 azon|OJJ51219.1

Group1486: abra|OJJ66924.1 acar|OOF91692.1 acla|EAW14490.1 afis|EAW19204.1 afla|EED47708.1 afum|EAL93934.1 agla|OJJ88508.1 akaw|GAA91493.1 aluc|OJZ85654.1 anid|CBF78558.1 anig|EHA24878.1 aory|EIT80227.1 asyd|OJJ65061.1 ater|EAU37268.1 atub|OJI79446.1 aver|OJI99185.1 awen|OJJ37557.1 azon|OJJ51220.1

Group1487: abra|OJJ66928.1 acar|OOF91696.1 acla|EAW14493.1 afis|EAW19207.1 afla|EED47705.1 afum|EAL93937.1 agla|OJJ88504.1 akaw|GAA91489.1 aluc|OJZ85651.1 anid|CBF78563.1 anig|EHA24882.1 aory|EIT80245.1 asyd|OJJ65064.1 ater|EAU37265.1 atub|OJI79442.1 aver|OJI99188.1 awen|OJJ37560.1 azon|OJJ51222.1

Group1488: abra|OJJ66931.1 acar|OOF91698.1 acla|EAW14494.1 afis|EAW19208.1 afla|EED47703.1 afum|EAL93938.2 agla|OJJ88501.1 akaw|GAA91487.1 aluc|OJZ85649.1 anid|CBF78566.1 anig|EHA24884.1 aory|EIT80217.1 asyd|OJJ65067.1 ater|EAU37260.1 atub|OJI79440.1 aver|OJI99191.1 awen|OJJ37562.1 azon|OJJ51225.1

Group1489: abra|OJJ66934.1 acar|OOF91701.1 acla|EAW14496.1 afis|EAW19210.1 afla|EED47702.1 afum|EAL93940.1 agla|OJJ88490.1 akaw|GAA91485.1 aluc|OJZ85647.1 anid|CBF78570.1 anig|EHA24887.1 aory|EIT80269.1 asyd|OJJ65075.1 ater|EAU37258.1 atub|OJI79438.1 aver|OJI99200.1 awen|OJJ37564.1 azon|OJJ51227.1

Group1490: afum|EAL92928.1 abra|OJJ78283.1 acar|OOF98585.1 acla|EAW09232.1 afis|EAW21916.1 afla|EED46633.1 agla|OJJ84207.1 akaw|GAA85744.1 aluc|OJZ91357.1 anig|EHA20035.1 aory|EIT77912.1 asyd|OJJ61312.1 ater|EAU30918.1 atub|OJI83350.1 aver|OJJ00637.1 awen|OJJ34206.1 azon|OJJ43428.1 anid|CBF76852.1

Group1492: abra|OJJ66938.1 acar|OOF91704.1 acla|EAW14497.1 afis|EAW19211.1 afla|EED47699.1 afum|EAL93941.1 agla|OJJ88486.1 akaw|GAA91482.1 aluc|OJZ85644.1 anid|CBF78583.1 anig|EHA24891.1 aory|EIT80265.1 asyd|OJJ65078.1 ater|EAU37256.1 atub|OJI79435.1 aver|OJI99203.1 awen|OJJ37565.1 azon|OJJ51228.1

Group1493: abra|OJJ66940.1 acar|OOF91705.1 acla|EAW14498.1 afis|EAW19212.1 afla|EED47698.1 afum|EBA27508.1 agla|OJJ88485.1 akaw|GAA91481.1 aluc|OJZ85642.1 anid|CBF78581.1 anig|EHA24892.1 aory|EIT80243.1 asyd|OJJ65076.1 ater|EAU37255.1 atub|OJI79433.1 aver|OJI99201.1 awen|OJJ37568.1 azon|OJJ51229.1

Group1494: abra|OJJ66948.1 acar|OOF91714.1 acla|EAW14503.1 afis|EAW19217.1 afla|EED47682.1 afum|EAL93950.1 agla|OJJ88478.1 akaw|GAA91472.1 aluc|OJZ85632.1 anid|CBF78594.1 anig|EHA24899.1 aory|EIT80231.1 asyd|OJJ65091.1 ater|EAU37251.1 atub|OJI79422.1 aver|OJI99216.1 awen|OJJ37574.1 azon|OJJ51234.1

Group1495: abra|OJJ66952.1 acar|OOF93656.1 acla|EAW14505.1 afis|EAW19218.1 afla|EED47681.1 afum|EAL93951.2 akaw|GAA91468.1 aluc|OJZ85626.1 anid|CBF78597.1 anig|EHA24902.1 aory|EIT80228.1 asyd|OJJ65093.1 ater|EAU37250.1 atub|OJI79419.1 aver|OJI99217.1 azon|OJJ51235.1 agla|OJJ88477.1 awen|OJJ37576.1

Group1496: abra|OJJ66955.1 acar|OOF93653.1 acla|EAW14506.1 afis|EAW19219.1 afla|EED47670.1 afum|EAL93952.1 agla|OJJ88476.1 akaw|GAA92206.1 aluc|OJZ85621.1 anid|CBF78603.1 anig|EHA24905.1 aory|EIT80271.1 asyd|OJJ65097.1 ater|EAU37238.1 atub|OJI79414.1 aver|OJI99223.1 awen|OJJ37577.1 azon|OJJ51236.1

Group1497: abra|OJJ66956.1 acar|OOF93652.1 acla|EAW14507.1 afis|EAW19220.1 afla|EED47669.1 afum|EAL93953.1 agla|OJJ88464.1 akaw|GAA92205.1 aluc|OJZ85620.1 anid|CBF78605.1 anig|EHA24906.1 aory|EIT80172.1 asyd|OJJ65098.1 ater|EAU37237.1 atub|OJI79413.1 aver|OJI99224.1 awen|OJJ37583.1 azon|OJJ51237.1

Group1498: abra|OJJ66962.1 acar|OOF93645.1 acla|EAW14512.1 afis|EAW19230.1 afla|EED47658.1 afum|EAL93959.1 agla|OJJ88473.1 akaw|GAA92199.1 aluc|OJZ85614.1 anid|CBF78614.1 anig|EHA24912.1 aory|EIT80215.1 asyd|OJJ65178.1 ater|EAU37233.1 atub|OJI79407.1 aver|OJI99331.1 awen|OJJ37588.1 azon|OJJ51242.1

Group1499: abra|OJJ66980.1 acar|OOF93631.1 acla|EAW11770.1 afis|EAW23678.1 afla|EED52197.1 afum|EAL87121.1 agla|OJJ78538.1 akaw|GAA92184.1 aluc|OJZ85594.1 anid|CBF89572.1 anig|EHA24928.1 aory|EIT74397.1 asyd|OJJ62335.1 ater|EAU35375.1 atub|OJI79390.1 aver|OJI96441.1 awen|OJJ35575.1 azon|OJJ43197.1

Group1501: abra|OJJ66992.1 acar|OOF93622.1 acla|EAW15343.1 afis|EAW20364.1 afla|EED51949.1 afum|EAL85480.2 agla|OJJ81790.1 akaw|GAA92172.1 aluc|OJZ85580.1 anid|CBF69463.1 anig|EHA24939.1 aory|EIT81480.1 asyd|OJJ59276.1 ater|EAU36758.1 atub|OJI79376.1 aver|OJJ01891.1 awen|OJJ32524.1 azon|OJJ50243.1

Group1503: abra|OJJ67118.1 acar|OOF98434.1 acla|EAW11710.1 afis|EAW23729.1 afla|EED47523.1 afum|EAL87167.1 agla|OJJ88177.1 akaw|GAA91047.1 aluc|OJZ87857.1 anid|CBF69394.1 anig|EHA20163.1 aory|EIT74567.1 asyd|OJJ52918.1 ater|EAU30755.1 atub|OJI80790.1 aver|OJJ08888.1 awen|OJJ32546.1 azon|OJJ51795.1

Group1504: abra|OJJ67150.1 acar|OOF98408.1 acla|EAW14575.1 afis|EAW19299.1 afla|EED47588.1 afum|EAL94018.1 agla|OJJ88383.1 akaw|GAA91023.1 aluc|OJZ87826.1 anid|CBF78774.1 anig|EHA20305.1 aory|EIT74538.1 asyd|OJJ65240.1 ater|EAU29474.1 atub|OJI80817.1 aver|OJI99390.1 awen|OJJ37648.1 azon|OJJ44913.1

Group1505: abra|OJJ67151.1 acar|OOF98407.1 acla|EAW14574.1 afis|EAW19297.1 afla|EED47589.1 afum|EAL94016.1 agla|OJJ88384.1 akaw|GAA91022.1 aluc|OJZ87825.1 anid|CBF78772.1 anig|EHA20304.1 aory|EIT74535.1 asyd|OJJ65239.1 ater|EAU29475.1 atub|OJI80819.1 aver|OJI99389.1 awen|OJJ37647.1 azon|OJJ44912.1

Group1506: abra|OJJ67153.1 acar|OOF98405.1 acla|EAW14571.1 afis|EAW19294.1 afla|EED47592.1 afum|EAL94013.1 agla|OJJ88386.1 akaw|GAA91019.1 aluc|OJZ87821.1 anid|CBF78767.1 anig|EHA20302.1 aory|EIT74534.1 asyd|OJJ65237.1 ater|EAU29478.1 atub|OJI80823.1 aver|OJI99386.1 awen|OJJ37645.1 azon|OJJ44910.1

Group1507: abra|OJJ67154.1 acar|OOF98404.1 acla|EAW14570.1 afis|EAW19283.1 afla|EED47593.1 afum|EAL94009.1 agla|OJJ88395.1 akaw|GAA91018.1 aluc|OJZ87820.1 anid|CBF78763.1 anig|EHA20301.1 aory|EIT80189.1 asyd|OJJ65235.1 ater|EAU29479.1 atub|OJI80824.1 aver|OJI99384.1 awen|OJJ37644.1 azon|OJJ44909.1

Group1508: abra|OJJ67157.1 acar|OOF98402.1 acla|EAW14568.1 afis|EAW19281.1 afla|EED47596.1 afum|EAL94007.1 agla|OJJ88393.1 akaw|GAA91015.1 aluc|OJZ87816.1 anid|CBF78760.1 anig|EHA20297.1 aory|EIT80256.1 asyd|OJJ65233.1 ater|EAU29482.1 atub|OJI80828.1 aver|OJI99382.1 awen|OJJ37642.1 azon|OJJ44906.1

Group1509: abra|OJJ67159.1 acar|OOF98400.1 acla|EAW14567.1 afis|EAW19280.1 afla|EED47598.1 afum|EAL94006.1 agla|OJJ88391.1 akaw|GAA91013.1 aluc|OJZ87814.1 anid|CBF78756.1 anig|EHA20295.1 aory|EIT80259.1 asyd|OJJ65231.1 ater|EAU29483.1 atub|OJI80830.1 aver|OJI99380.1 awen|OJJ37640.1 azon|OJJ44904.1

Group1510: abra|OJJ67163.1 acar|OOF98397.1 acla|EAW14565.1 afis|EAW19278.1 afla|EED47600.1 afum|EAL94004.1 agla|OJJ88389.1 akaw|GAA91011.1 aluc|OJZ87812.1 anid|CBF78752.1 anig|EHA20294.1 aory|EIT80253.1 asyd|OJJ65229.1 ater|EAU29485.1 atub|OJI80832.1 aver|OJI99378.1 awen|OJJ37638.1 azon|OJJ44882.1

Group1511: abra|OJJ67164.1 acar|OOF98398.1 acla|EAW14564.1 afis|EAW19277.1 afla|EED47601.1 afum|EAL94003.1 agla|OJJ88396.1 akaw|GAA91010.1 aluc|OJZ87811.1 anid|CBF78750.1 anig|EHA20293.1 aory|EIT80205.1 asyd|OJJ65228.1 ater|EAU29486.1 atub|OJI80833.1 aver|OJI99377.1 awen|OJJ37637.1 azon|OJJ44883.1

Group1512: abra|OJJ67173.1 acar|OOF98390.1 acla|EAW14550.1 afis|EAW19267.1 afla|EED47608.1 afum|EAL93994.1 agla|OJJ88426.1 akaw|GAA91001.1 aluc|OJZ87802.1 anid|CBF78695.1 anig|EHA20283.1 aory|EIT80277.1 asyd|OJJ65218.1 ater|EAU29548.1 atub|OJI80845.1 aver|OJI99368.1 awen|OJJ37628.1 azon|OJJ47016.1

Group1513: abra|OJJ67175.1 acar|OOF98388.1 acla|EAW14548.1 afis|EAW19265.1 afla|EED47609.1 afum|EAL93992.1 agla|OJJ88424.1 akaw|GAA90999.1 aluc|OJZ87800.1 anid|CBF78691.1 anig|EHA20281.1 aory|EIT80181.1 asyd|OJJ65216.1 ater|EAU29546.1 atub|OJI80847.1 aver|OJI99366.1 awen|OJJ37626.1 azon|OJJ47015.1

Group1514: abra|OJJ67176.1 acar|OOF98386.1 acla|EAW14547.1 afis|EAW19264.1 afla|EED47610.1 afum|EAL93991.1 agla|OJJ88423.1 akaw|GAA90998.1 aluc|OJZ87799.1 anid|CBF78688.1 anig|EHA20280.1 aory|EIT80241.1 asyd|OJJ65214.1 ater|EAU29545.1 atub|OJI80848.1 aver|OJI99364.1 awen|OJJ37625.1 azon|OJJ47014.1

Group1516: abra|OJJ67179.1 acar|OOF98384.1 acla|EAW14544.1 afis|EAW19262.1 afla|EED47613.1 afum|EAL93989.1 agla|OJJ88420.1 akaw|GAA90997.1 aluc|OJZ87796.1 anid|CBF78682.1 anig|EHA20277.1 aory|EIT80281.1 asyd|OJJ65211.1 ater|EAU29542.1 atub|OJI80851.1 aver|OJI99361.1 awen|OJJ37622.1 azon|OJJ47011.1

Group1517: abra|OJJ67180.1 acar|OOF98382.1 acla|EAW14543.1 afis|EAW19261.1 afla|EED47614.1 afum|EAL93988.1 agla|OJJ88433.1 akaw|GAA90996.1 aluc|OJZ87795.1 anid|CBF78680.1 anig|EHA20276.1 aory|EIT80274.1 asyd|OJJ65210.1 ater|EAU29514.1 atub|OJI80852.1 aver|OJI99360.1 awen|OJJ37621.1 azon|OJJ47010.1

Group1518: abra|OJJ67182.1 acar|OOF98380.1 acla|EAW14541.1 afis|EAW19259.1 afla|EED47616.1 afum|EAL93986.1 agla|OJJ88436.1 akaw|GAA90994.1 aluc|OJZ87793.1 anid|CBF78678.1 anig|EHA20274.1

aory|EIT80183.1 asyd|OJJ65209.1 ater|EAU29516.1 atub|OJI80854.1 aver|OJI99358.1 awen|OJJ37620.1 azon|OJJ47009.1

Group1519: abra|OJJ67183.1 acar|OOF98379.1 acla|EAW14540.1 afis|EAW19258.1 afla|EED47618.1 afum|EAL93985.1 agla|OJJ88437.1 akaw|GAA90993.1 aluc|OJZ87792.1 anid|CBF78676.1 anig|EHA20273.1 aory|EIT80276.1 asyd|OJJ65208.1 ater|EAU29517.1 atub|OJI80855.1 aver|OJI99357.1 awen|OJJ37619.1 azon|OJJ47008.1

Group1520: abra|OJJ67187.1 acar|OOF98377.1 acla|EAW14537.1 afis|EAW19253.1 afla|EED47621.1 afum|EAL93982.1 agla|OJJ88444.1 akaw|GAA90990.1 aluc|OJZ87788.1 anid|CBF78671.1 anig|EHA20270.1 aory|EIT80226.1 asyd|OJJ65205.1 ater|EAU29510.1 atub|OJI80859.1 aver|OJI99354.1 awen|OJJ37616.1 azon|OJJ47005.1

Group1521: abra|OJJ67189.1 acar|OOF98374.1 acla|EAW14534.1 afis|EAW19252.1 afla|EED47626.1 afum|EAL93981.1 agla|OJJ88445.1 akaw|GAA90988.1 aluc|OJZ87786.1 anid|CBF78669.1 anig|EHA20266.1 aory|EIT80202.1 asyd|OJJ65202.1 ater|EAU29506.1 atub|OJI80861.1 aver|OJI99351.1 awen|OJJ37615.1 azon|OJJ47004.1

Group1522: abra|OJJ67190.1 acar|OOF98373.1 acla|EAW14533.1 afis|EAW19251.1 afla|EED47627.1 afum|EAL93980.1 agla|OJJ88446.1 akaw|GAA90987.1 aluc|OJZ87785.1 anid|CBF78667.1 anig|EHA20265.1 aory|EIT80224.1 asyd|OJJ65201.1 ater|EAU29505.1 atub|OJI80862.1 aver|OJI99350.1 awen|OJJ37614.1 azon|OJJ51262.1

Group1523: abra|OJJ67192.1 acar|OOF98372.1 acla|EAW14532.1 afis|EAW19250.1 afla|EED47628.1 afum|EAL93979.1 agla|OJJ88447.1 akaw|GAA90986.1 aluc|OJZ87784.1 anid|CBF78663.1 anig|EHA20264.1 aory|EIT80188.1 asyd|OJJ65199.1 ater|EAU29504.1 atub|OJI80863.1 aver|OJI99348.1 awen|OJJ37612.1 azon|OJJ51261.1

Group1524: abra|OJJ67193.1 acar|OOF98371.1 acla|EAW14531.1 afis|EAW19249.1 afla|EED47629.1 afum|EAL93978.1 agla|OJJ88448.1 akaw|GAA90985.1 aluc|OJZ87782.1 anid|CBF78665.1 anig|EHA20263.1 aory|EIT80199.1 asyd|OJJ65200.1 ater|EAU29503.1 atub|OJI80865.1 aver|OJI99349.1 awen|OJJ37611.1 azon|OJJ51260.1

Group1525: abra|OJJ67195.1 acar|OOF98369.1 acla|EAW14529.1 afis|EAW19247.1 afla|EED47631.1 afum|EAL93976.1 agla|OJJ88450.1 akaw|GAA90983.1 aluc|OJZ87780.1 anid|CBF78658.1 anig|EHA20262.1 aory|EIT80177.1 asyd|OJJ65197.1 ater|EAU29501.1 atub|OJI80867.1 aver|OJI99346.1 awen|OJJ37604.1 azon|OJJ51257.1

Group1526: abra|OJJ67200.1 acar|OOF98364.1 acla|EAW14524.1 afis|EAW19242.1 afla|EED47635.1 afum|EAL93971.2 agla|OJJ88455.1 akaw|GAA90978.1 aluc|OJZ87774.1 anid|CBF78650.1 anig|EHA20258.1 aory|EIT80223.1 asyd|OJJ65194.1 ater|EAU29498.1 atub|OJI80871.1 aver|OJI99343.1 awen|OJJ37598.1 azon|OJJ51253.1

Group1527: abra|OJJ67201.1 acar|OOF98362.1 acla|EAW14523.1 afis|EAW19241.1 afla|EED47636.1 afum|EAL93970.2 agla|OJJ88456.1 akaw|GAA90977.1 aluc|OJZ87773.1 anid|CBF78648.1 anig|EHA20257.1 aory|EIT80218.1 asyd|OJJ65193.1 ater|EAU29497.1 atub|OJI80872.1 aver|OJI99342.1 awen|OJJ37597.1 azon|OJJ51252.1

Group1528: abra|OJJ67202.1 acar|OOF98361.1 acla|EAW14522.1 afis|EAW19240.1 afla|EED47637.1 afum|EAL93969.1 agla|OJJ88458.1 akaw|GAA90976.1 aluc|OJZ87772.1 anid|CBF78646.1 anig|EHA20256.1 aory|EIT80185.1 asyd|OJJ65192.1 ater|EAU37223.1 atub|OJI80873.1 aver|OJI99341.1 awen|OJJ37596.1 azon|OJJ51251.1

Group1529: abra|OJJ67203.1 acar|OOF98360.1 acla|EAW14521.1 afis|EAW19239.1 afla|EED47638.1 afum|EBA27509.1 agla|OJJ88459.1 akaw|GAA90975.1 aluc|OJZ87771.1 anid|CBF78644.1 anig|EHA20255.1 aory|EIT80222.1 asyd|OJJ65191.1 ater|EAU37224.1 atub|OJI80874.1 aver|OJI99340.1 awen|OJJ37595.1 azon|OJJ51250.1

Group1530: abra|OJJ67204.1 acar|OOF98359.1 acla|EAW14520.1 afis|EAW19238.1 afla|EED47639.1 afum|EAL93966.1 agla|OJJ88461.1 akaw|GAA90974.1 aluc|OJZ87770.1 anid|CBF78642.1 anig|EHA20254.1 aory|EIT80175.1 asyd|OJJ65190.1 ater|EAU37225.1 atub|OJI80875.1 aver|OJI99339.1 awen|OJJ37594.1 azon|OJJ51249.1

Group1531: abra|OJJ67213.1 acar|OOF98346.1 acla|EAW08553.1 afis|EAW23595.1 afla|EED47505.1 afum|EAL87872.1 agla|OJJ86855.1 akaw|GAA90964.1 aluc|OJZ87760.1 anid|CBF89585.1 anig|EHA20245.1 aory|EIT74581.1 asyd|OJJ62318.1 ater|EAU35355.1 atub|OJI80884.1 aver|OJI96426.1 awen|OJJ39917.1 azon|OJJ45891.1

Group1532: abra|OJJ67215.1 acar|OOF98343.1 acla|EAW10167.1 afis|EAW25427.1 afla|EED53915.1 afum|EAL90086.1 agla|OJJ87071.1 akaw|GAA90962.1 aluc|OJZ87757.1 anid|CBF77551.1 anig|EHA20243.1 aory|EIT78350.1 asyd|OJJ53152.1 ater|EAU34597.1 atub|OJI80886.1 aver|OJJ03699.1 awen|OJJ41156.1 azon|OJJ44734.1

Group1533: abra|OJJ67225.1 acar|OOF98326.1 acla|EAW10408.1 afis|EAW25160.1 afla|EED53622.1 afum|EAL89835.1 agla|OJJ88637.1 akaw|GAA91652.1 aluc|OJZ87749.1 anid|CBF80654.1 anig|EHA20237.1 aory|EIT75132.1 asyd|OJJ54613.1 ater|EAU34333.1 atub|OJI80894.1 aver|OJJ07545.1 awen|OJJ33447.1 azon|OJJ50704.1

Group1534: abra|OJJ67239.1 acar|OOF89889.1 acla|EAW13993.1 afis|EAW18701.1 afla|EED50650.1 afum|EAL93424.1 agla|OJJ82360.1 akaw|GAA91664.1 aluc|OJZ87736.1 anid|CBF70640.1 anig|EHA20228.1 aory|EIT78150.1 asyd|OJJ58491.1 ater|EAU38205.1 atub|OJI80904.1 aver|OJJ01094.1 awen|OJJ32184.1 azon|OJJ48705.1

Group1535: abra|OJJ67298.1 acar|OOF92514.1 acla|EAW15241.1 afis|EAW20208.1 afla|EED57135.1 afum|EAL85325.1 agla|OJJ84656.1 akaw|GAA86516.1 aluc|OJZ85061.1 anid|CBF85098.1 anig|EHA25276.1 aory|EIT74134.1 asyd|OJJ63059.1 ater|EAU32018.1 atub|OJI80536.1 aver|OJI97904.1 awen|OJJ31212.1 azon|OJJ46148.1

Group1536: abra|OJJ67306.1 acar|OOF92507.1 acla|EAW06427.1 afis|EAW18531.1 afla|EED52870.1 afum|EAL93228.1 agla|OJJ85144.1 akaw|GAA86509.1 aluc|OJZ85052.1 anid|CBF70204.1 anig|EHA25269.1 aory|EIT78588.1 asyd|OJJ58784.1 ater|EAU30039.1 atub|OJI80546.1 aver|OJJ01392.1 awen|OJJ31089.1 azon|OJJ50594.1

Group1537: abra|OJJ67308.1 acar|OOF92504.1 acla|EAW06424.1 afis|EAW18528.1 afla|EED52873.1 afum|EAL93225.1 agla|OJJ85141.1 akaw|GAA86507.1 aluc|OJZ85050.1 anid|CBF70208.1 anig|EHA25267.1 aory|EIT78611.1 asyd|OJJ58781.1 ater|EAU30041.1 atub|OJI80548.1 aver|OJJ01389.1 awen|OJJ31086.1 azon|OJJ50591.1

Group1538: abra|OJJ67310.1 acar|OOF92502.1 acla|EAW06422.1 afis|EAW18526.1 afla|EED52875.1 afum|EAL93221.1 agla|OJJ85140.1 akaw|GAA86505.1 aluc|OJZ85048.1 anid|CBF70211.1 anig|EHA25265.1 aory|EIT78615.1 asyd|OJJ58779.1 ater|EAU30043.1 atub|OJI80550.1 aver|OJJ01388.1 awen|OJJ31080.1 azon|OJJ50588.1

Group1539: abra|OJJ67311.1 acar|OOF92501.1 acla|EAW06421.1 afis|EAW18525.1 afla|EED52876.1 afum|EAL93220.1 agla|OJJ85139.1 akaw|GAA86504.1 aluc|OJZ85047.1 anid|CBF70213.1 anig|EHA25264.1 aory|EIT78600.1 asyd|OJJ58778.1 ater|EAU30044.1 atub|OJI80551.1 aver|OJJ01387.1 awen|OJJ31079.1 azon|OJJ50587.1

Group1540: abra|OJJ67319.1 acar|OOF92496.1 acla|EAW06417.1 afis|EAW18521.1 afla|EED52877.1 afum|EAL93216.1 agla|OJJ85133.1 akaw|GAA86499.1 aluc|OJZ85041.1 anid|CBF70223.1 anig|EHA25259.1 aory|EIT78623.1 asyd|OJJ58774.1 ater|EAU30049.1 atub|OJI80556.1 aver|OJJ01383.1 awen|OJJ31073.1 azon|OJJ49993.1

Group1541: abra|OJJ67321.1 acar|OOF92493.1 acla|EAW06415.1 afis|EAW18519.1 afla|EED52879.1 afum|EAL93214.1 agla|OJJ85131.1 akaw|GAA86497.1 aluc|OJZ85039.1 anid|CBF70226.1 anig|EHA25257.1 aory|EIT78673.1 asyd|OJJ58773.1 ater|EAU30051.1 atub|OJI80558.1 aver|OJJ01382.1 awen|OJJ31071.1 azon|OJJ50581.1

Group1542: abra|OJJ67323.1 acar|OOF92490.1 acla|EAW06412.1 afis|EAW18516.1 afla|EED52882.1 afum|EAL93211.1 agla|OJJ85129.1 akaw|GAA86493.1 aluc|OJZ85035.1 anid|CBF70232.1 anig|EHA25255.1 aory|EIT78656.1 asyd|OJJ58771.1 ater|EAU30054.1 atub|OJI80562.1 aver|OJJ01380.1 awen|OJJ31592.1 azon|OJJ50579.1

Group1543: abra|OJJ67325.1 acar|OOF92488.1 acla|EAW06410.1 afis|EAW18514.1 afla|EED52884.1 afum|EAL93209.1 agla|OJJ85107.1 akaw|GAA86491.1 aluc|OJZ85033.1 anid|CBF70234.1 anig|EHA25253.1 aory|EIT78594.1 asyd|OJJ58770.1 ater|EAU30055.1 atub|OJI80564.1 aver|OJJ01379.1 awen|OJJ31068.1 azon|OJJ50577.1

Group1544: abra|OJJ67326.1 acar|OOF92487.1 acla|EAW06409.1 afis|EAW18513.1 afla|EED52887.1 afum|EAL93208.1 agla|OJJ85108.1 akaw|GAA86490.1 aluc|OJZ85032.1 anid|CBF70236.1 anig|EHA25252.1 aory|EIT78631.1 asyd|OJJ58769.1 ater|EAU30057.1 atub|OJI80565.1 aver|OJJ01378.1 awen|OJJ31067.1 azon|OJJ50576.1

Group1545: abra|OJJ67327.1 acar|OOF92486.1 acla|EAW06407.1 afis|EAW18511.1 afla|EED52889.1 afum|EAL93206.1 agla|OJJ85109.1 akaw|GAA86489.1 aluc|OJZ85030.1 anid|CBF70239.1 anig|EHA25251.1 aory|EIT78676.1 asyd|OJJ58767.1 ater|EAU30058.1 atub|OJI80567.1 aver|OJJ01376.1 awen|OJJ31065.1 azon|OJJ50575.1

Group1546: abra|OJJ67328.1 acar|OOF92485.1 acla|EAW06406.1 afis|EAW18510.1 afla|EED52890.1 afum|EAL93205.1 agla|OJJ85110.1 akaw|GAA86488.1 aluc|OJZ85029.1 anid|CBF70241.1 anig|EHA25250.1 aory|EIT78606.1 asyd|OJJ58766.1 ater|EAU30059.1 atub|OJI80568.1 aver|OJJ01375.1 awen|OJJ31064.1 azon|OJJ50574.1

Group1547: abra|OJJ67329.1 acar|OOF92483.1 acla|EAW06405.1 afis|EAW18509.1 afla|EED52891.1 afum|EAL93204.1 agla|OJJ85111.1 akaw|GAA86487.1 aluc|OJZ85028.1 anid|CBF70243.1 anig|EHA25249.1

aory|EIT78599.1 asyd|OJJ58765.1 ater|EAU30060.1 atub|OJI80569.1 aver|OJJ01374.1 awen|OJJ31063.1 azon|OJJ50573.1

Group1548: abra|OJJ67330.1 acar|OOF92484.1 acla|EAW06404.1 afis|EAW18508.1 afla|EED52894.1 afum|EAL93203.1 agla|OJJ85112.1 akaw|GAA86486.1 aluc|OJZ85027.1 anid|CBF70245.1 anig|EHA25248.1 aory|EIT78590.1 asyd|OJJ58762.1 ater|EAU30063.1 atub|OJI80570.1 aver|OJJ01372.1 awen|OJJ31061.1 azon|OJJ50572.1

Group1549: abra|OJJ67334.1 acar|OOF92479.1 acla|EAW06400.1 afis|EAW18504.1 afla|EED52897.1 afum|EAL93197.1 agla|OJJ85119.1 akaw|GAA86482.1 aluc|OJZ85023.1 anid|CBF70249.1 anig|EHA25244.1 aory|EIT78613.1 asyd|OJJ58759.1 ater|EAU30067.1 atub|OJI80574.1 aver|OJJ01367.1 awen|OJJ31058.1 azon|OJJ50567.1

Group1550: abra|OJJ67335.1 acar|OOF92478.1 acla|EAW06399.1 afis|EAW18503.1 afla|EED52898.1 afum|EAL93196.2 agla|OJJ85118.1 akaw|GAA86481.1 aluc|OJZ85022.1 anid|CBF70251.1 anig|EHA25243.1 aory|EIT78654.1 asyd|OJJ58758.1 ater|EAU30068.1 atub|OJI80575.1 aver|OJJ01366.1 awen|OJJ31057.1 azon|OJJ50568.1

Group1551: abra|OJJ67336.1 acar|OOF92477.1 acla|EAW06398.1 afis|EAW18502.1 afla|EED52899.1 afum|EAL93195.1 agla|OJJ85123.1 akaw|GAA86480.1 aluc|OJZ85021.1 anid|CBF70253.1 anig|EHA25242.1 aory|EIT78660.1 asyd|OJJ58757.1 ater|EAU30069.1 atub|OJI80576.1 aver|OJJ01365.1 awen|OJJ31056.1 azon|OJJ50566.1

Group1552: abra|OJJ67339.1 acar|OOF92474.1 acla|EAW06395.1 afis|EAW18499.1 afla|EED52902.1 afum|EAL93190.1 agla|OJJ85128.1 akaw|GAA86477.1 aluc|OJZ85018.1 anid|CBF70257.1 anig|EHA25239.1 aory|EIT78634.1 asyd|OJJ58755.1 ater|EAU30071.1 atub|OJI80579.1 aver|OJJ01363.1 awen|OJJ31053.1 azon|OJJ50563.1

Group1553: abra|OJJ67340.1 acar|OOF92473.1 acla|EAW06394.1 afis|EAW18498.1 afla|EED52903.1 afum|EAL93188.1 agla|OJJ85105.1 akaw|GAA86476.1 aluc|OJZ85017.1 anid|CBF70259.1 anig|EHA25238.1 aory|EIT78622.1 asyd|OJJ58754.1 ater|EAU30072.1 atub|OJI80580.1 aver|OJJ01362.1 awen|OJJ31052.1 azon|OJJ50564.1

Group1554: abra|OJJ67346.1 acar|OOF92469.1 acla|EAW06390.1 afis|EAW18496.1 afla|EED52907.1 afum|EAL93186.1 agla|OJJ85091.1 akaw|GAA86472.1 aluc|OJZ85013.1 anid|CBF70170.1 anig|EHA25234.1 aory|EIT78612.1 asyd|OJJ58795.1 ater|EAU30073.1 atub|OJI80584.1 aver|OJJ01400.1 awen|OJJ31049.1 azon|OJJ50560.1

Group1555: abra|OJJ67347.1 acar|OOF92468.1 acla|EAW06389.1 afis|EAW18495.1 afla|EED52908.1 afum|EAL93185.1 agla|OJJ85090.1 akaw|GAA86471.1 aluc|OJZ85012.1 anid|CBF70168.1 anig|EHA25233.1 aory|EIT78663.1 asyd|OJJ58797.1 ater|EAU29921.1 atub|OJI80585.1 aver|OJJ01401.1 awen|OJJ31048.1 azon|OJJ50559.1

Group1556: abra|OJJ67350.1 acar|OOF92466.1 acla|EAW06386.1 afis|EAW18492.1 afla|EED52912.1 afum|EAL93182.1 agla|OJJ85087.1 akaw|GAA86468.1 aluc|OJZ85009.1 anid|CBF70162.1 anig|EHA25230.1 aory|EIT78650.1 asyd|OJJ58800.1 ater|EAU29918.1 atub|OJI80588.1 aver|OJJ01404.1 awen|OJJ31045.1 azon|OJJ50556.1

Group1557: abra|OJJ67375.1 acar|OOF92436.1 acla|EAW06380.1 afis|EAW18486.1 afla|EED52922.1 afum|EAL93175.1 agla|OJJ85081.1 akaw|GAA86447.1 aluc|OJZ84990.1 anid|CBF70147.1 anig|EHA25202.1 aory|EIT78638.1 asyd|OJJ58808.1 ater|EAU29911.1 atub|OJI80610.1 aver|OJJ01411.1 awen|OJJ31033.1 azon|OJJ50552.1

Group1558: abra|OJJ67377.1 acar|OOF92435.1 acla|EAW06379.1 afis|EAW18485.1 afla|EED52923.1 afum|EBA27486.1 agla|OJJ85074.1 akaw|GAA86446.1 aluc|OJZ84988.1 anid|CBF70145.1 anig|EHA25201.1 aory|EIT78668.1 asyd|OJJ58809.1 ater|EAU29910.1 atub|OJI80611.1 aver|OJJ01412.1 awen|OJJ31035.1 azon|OJJ50550.1

Group1559: abra|OJJ67380.1 acar|OOF92432.1 acla|EAW06377.1 afis|EAW18483.1 afla|EED52925.1 afum|EAL93172.1 agla|OJJ85072.1 akaw|GAA86444.1 aluc|OJZ84986.1 anid|CBF86341.1 anig|EHA25199.1 aory|EIT78658.1 asyd|OJJ64985.1 ater|EAU29909.1 atub|OJI80614.1 aver|OJI99105.1 awen|OJJ31037.1 azon|OJJ50548.1

Group1560: abra|OJJ67381.1 acar|OOF92433.1 acla|EAW06376.1 afis|EAW18482.1 afla|EED52926.1 afum|EAL93171.1 agla|OJJ85063.1 akaw|GAA86443.1 aluc|OJZ84985.1 anid|CBF86343.1 anig|EHA25198.1 aory|EIT78671.1 asyd|OJJ64984.1 ater|EAU29908.1 atub|OJI80615.1 aver|OJI99104.1 awen|OJJ31038.1 azon|OJJ50547.1

Group1561: abra|OJJ67385.1 acar|OOF92428.1 acla|EAW06374.1 afis|EAW18480.1 afla|EED52929.1 afum|EAL93169.1 agla|OJJ85066.1 akaw|GAA86439.1 aluc|OJZ84981.1 anid|CBF70139.1 anig|EHA25194.1 aory|EIT78702.1 asyd|OJJ58812.1 ater|EAU29896.1 atub|OJI80620.1 aver|OJJ01415.1 awen|OJJ31040.1 azon|OJJ50545.1

Group1562: abra|OJJ67387.1 acar|OOF92427.1 acla|EAW06372.1 afis|EAW18478.1 afla|EED52932.1 afum|EAL93166.1 agla|OJJ85068.1 akaw|GAA86437.1 aluc|OJZ84980.1 anid|CBF70136.1 anig|EHA25192.1 aory|EIT78730.1 asyd|OJJ58814.1 ater|EAU29895.1 atub|OJI80622.1 aver|OJJ01417.1 awen|OJJ31031.1 azon|OJJ50544.1

Group1563: abra|OJJ67393.1 acar|OOF92423.1 acla|EAW06370.1 afis|EAW18476.1 afla|EED52934.1 afum|EAL93163.1 agla|OJJ85071.1 akaw|GAA86434.1 aluc|OJZ84976.1 anid|CBF70132.1 anig|EHA25188.1 aory|EIT78693.1 asyd|OJJ58817.1 ater|EAU29893.1 atub|OJI80625.1 aver|OJJ01421.1 awen|OJJ31029.1 azon|OJJ50542.1

Group1564: abra|OJJ67394.1 acar|OOF92422.1 acla|EAW06369.1 afis|EAW18475.1 afla|EED52935.1 afum|EAL93162.1 agla|OJJ85062.1 akaw|GAA86433.1 aluc|OJZ84975.1 anid|CBF70116.1 anig|EHA25187.1 aory|EIT78723.1 asyd|OJJ58819.1 ater|EAU29892.1 atub|OJI80626.1 aver|OJJ01423.1 awen|OJJ31028.1 azon|OJJ50541.1

Group1565: abra|OJJ67395.1 acar|OOF92421.1 acla|EAW06368.1 afis|EAW18474.1 afla|EED52936.1 afum|EAL93161.1 agla|OJJ85061.1 akaw|GAA86432.1 aluc|OJZ84974.1 anid|CBF70108.1 anig|EHA25186.1 aory|EIT78688.1 asyd|OJJ58823.1 ater|EAU29891.1 atub|OJI80627.1 aver|OJJ01427.1 awen|OJJ31026.1 azon|OJJ50540.1

Group1566: abra|OJJ67396.1 acar|OOF92419.1 acla|EAW06364.1 afis|EAW18473.1 afla|EED52937.1 afum|EAL93160.2 agla|OJJ85060.1 akaw|GAA86431.1 aluc|OJZ84973.1 anid|CBF70107.1 anig|EHA25185.1 aory|EIT78722.1 asyd|OJJ58827.1 ater|EAU29890.1 atub|OJI80628.1 aver|OJJ01430.1 awen|OJJ31025.1 azon|OJJ50539.1

Group1567: abra|OJJ67398.1 acar|OOF92418.1 acla|EAW06362.1 afis|EAW18471.1 afla|EED52939.1 afum|EAL93158.1 agla|OJJ85056.1 akaw|GAA86429.1 aluc|OJZ84971.1 anid|CBF70105.1 anig|EHA25183.1 aory|EIT78698.1 asyd|OJJ58829.1 ater|EAU29888.1 atub|OJI80630.1 aver|OJJ01432.1 awen|OJJ31023.1 azon|OJJ50537.1

Group1568: abra|OJJ67399.1 acar|OOF92417.1 acla|EAW06359.1 afis|EAW18470.1 afla|EED52940.1 afum|EAL93156.1 agla|OJJ85055.1 akaw|GAA86428.1 aluc|OJZ84969.1 anid|CBF70103.1 anig|EHA25182.1 aory|EIT78679.1 asyd|OJJ58830.1 ater|EAU29887.1 atub|OJI80631.1 aver|OJJ01433.1 awen|OJJ31022.1 azon|OJJ50536.1

Group1569: abra|OJJ67400.1 acar|OOF92416.1 acla|EAW06358.1 afis|EAW18469.1 afla|EED52941.1 afum|EAL93155.1 agla|OJJ85054.1 akaw|GAA86427.1 aluc|OJZ84968.1 anid|CBF70101.1 anig|EHA25181.1 aory|EIT78712.1 asyd|OJJ58831.1 ater|EAU29886.1 atub|OJI80632.1 aver|OJJ01434.1 awen|OJJ31021.1 azon|OJJ50535.1

Group1570: abra|OJJ67401.1 acar|OOF92415.1 acla|EAW06357.1 afis|EAW18468.1 afla|EED52942.1 afum|EAL93154.1 agla|OJJ85053.1 akaw|GAA86426.1 aluc|OJZ84967.1 anid|CBF70100.1 anig|EHA25180.1 aory|EIT78694.1 asyd|OJJ58832.1 ater|EAU29885.1 atub|OJI80633.1 aver|OJJ01435.1 awen|OJJ31020.1 azon|OJJ50534.1

Group1571: abra|OJJ67403.1 acar|OOF92413.1 acla|EAW06355.1 afis|EAW18466.1 afla|EED52944.1 afum|EAL93152.1 agla|OJJ85051.1 akaw|GAA86424.1 aluc|OJZ84965.1 anid|CBF70096.1 anig|EHA25178.1 aory|EIT78707.1 asyd|OJJ58834.1 ater|EAU29883.1 atub|OJI80635.1 aver|OJJ01437.1 awen|OJJ31018.1 azon|OJJ50532.1

Group1572: abra|OJJ67411.1 acar|OOF89994.1 acla|EAW06348.1 afis|EAW18459.1 afla|EED52949.1 afum|EAL93145.1 agla|OJJ85042.1 akaw|GAA86416.1 aluc|OJZ84957.1 anid|CBF70085.1 anig|EHA25171.1 aory|EIT78721.1 asyd|OJJ58842.1 ater|EAU29877.1 atub|OJI80643.1 aver|OJJ01444.1 awen|OJJ31011.1 azon|OJJ50525.1

Group1573: abra|OJJ67412.1 acar|OOF89992.1 acla|EAW06347.1 afis|EAW18458.1 afla|EED52950.1 afum|EAL93143.1 agla|OJJ85041.1 akaw|GAA86415.1 aluc|OJZ84956.1 anid|CBF70084.1 anig|EHA25170.1 aory|EIT78692.1 asyd|OJJ58843.1 ater|EAU29876.1 atub|OJI80644.1 aver|OJJ01445.1 awen|OJJ31010.1 azon|OJJ50524.1

Group1574: abra|OJJ67419.1 acar|OOF92402.1 acla|EAW06345.1 afis|EAW18456.1 afla|EED52952.1 afum|EAL93141.1 agla|OJJ85039.1 akaw|GAA86409.1 aluc|OJZ84949.1 anid|CBF70053.1 anig|EHA25164.1 aory|EIT78736.1 asyd|OJJ58857.1 ater|EAU29873.1 atub|OJI80653.1 aver|OJJ01458.1 awen|OJJ31008.1 azon|OJJ50522.1

Group1575: abra|OJJ67421.1 acar|OOF92401.1 acla|EAW06344.1 afis|EAW18455.1 afla|EED52953.1 afum|EAL93140.1 agla|OJJ85038.1 akaw|GAA86408.1 aluc|OJZ84947.1 anid|CBF70057.1 anig|EHA25163.1 aory|EIT78696.1 asyd|OJJ58855.1 ater|EAU29872.1 atub|OJI80654.1 aver|OJJ01456.1 awen|OJJ31007.1 azon|OJJ50521.1

Group1576: abra|OJJ67423.1 acar|OOF89873.1 acla|EAW06341.1 afis|EAW18453.1 afla|EED52955.1 afum|EAL93138.1 agla|OJJ85032.1 akaw|GAA86406.1 aluc|OJZ84944.1 anid|CBF70061.1 anig|EHA25161.1

aory|EIT78714.1 asyd|OJJ58853.1 ater|EAU29870.1 atub|OJI80656.1 aver|OJJ01454.1 awen|OJJ31005.1 azon|OJJ50519.1

Group1577: abra|OJJ67424.1 acar|OOF89874.1 acla|EAW06340.1 afis|EAW18452.1 afla|EED52956.1 afum|EAL93137.1 agla|OJJ85030.1 akaw|GAA86405.1 aluc|OJZ84943.1 anid|CBF70062.1 anig|EHA25160.1 aory|EIT78680.1 asyd|OJJ58852.1 ater|EAU29869.1 atub|OJI80657.1 aver|OJJ01453.1 awen|OJJ31004.1 azon|OJJ50517.1

Group1578: abra|OJJ67426.1 acar|OOF92399.1 acla|EAW06338.1 afis|EAW18450.1 afla|EED52958.1 afum|EAL93135.1 agla|OJJ85028.1 akaw|GAA86404.1 aluc|OJZ84941.1 anid|CBF70064.1 anig|EHA25158.1 aory|EIT78699.1 asyd|OJJ58850.1 ater|EAU29868.1 atub|OJI80659.1 aver|OJJ01451.1 awen|OJJ31002.1 azon|OJJ50516.1

Group1579: abra|OJJ67429.1 acar|OOF92397.1 acla|EAW06335.1 afis|EAW18447.1 afla|EED52963.1 afum|EAL93132.1 agla|OJJ85024.1 akaw|GAA86401.1 aluc|OJZ84937.1 anid|CBF70070.1 anig|EHA25155.1 aory|EIT78728.1 asyd|OJJ58847.1 ater|EAU29866.1 atub|OJI80662.1 aver|OJJ01448.1 awen|OJJ30999.1 azon|OJJ50513.1

Group1581: abra|OJJ67434.1 acar|OOF92394.1 acla|EAW06332.1 afis|EAW18445.1 afla|EED52967.1 afum|EAL93130.1 agla|OJJ85021.1 akaw|GAA86396.1 aluc|OJZ84930.1 anid|CBF70041.1 anig|EHA25149.1 aory|EIT78729.1 asyd|OJJ58865.1 ater|EAU29864.1 atub|OJI80669.1 aver|OJJ01466.1 awen|OJJ30993.1 azon|OJJ50510.1

Group1582: abra|OJJ67435.1 acar|OOF92393.1 acla|EAW06331.1 afis|EAW18444.1 afla|EED52968.1 afum|EAL93128.1 agla|OJJ85020.1 akaw|GAA86395.1 aluc|OJZ84929.1 anid|CBF70043.1 anig|EHA25148.1 aory|EIT78686.1 asyd|OJJ58864.1 ater|EAU29863.1 atub|OJI80670.1 aver|OJJ01465.1 awen|OJJ30992.1 azon|OJJ50511.1

Group1583: abra|OJJ67437.1 acar|OOF92391.1 acla|EAW06327.1 afis|EAW18441.1 afla|EED52972.1 afum|EAL93123.1 agla|OJJ85006.1 akaw|GAA86393.1 aluc|OJZ84927.1 anid|CBF70049.1 anig|EHA25146.1 aory|EIT78690.1 asyd|OJJ58861.1 ater|EAU29860.1 atub|OJI80672.1 aver|OJJ01462.1 awen|OJJ30989.1 azon|OJJ50507.1

Group1584: abra|OJJ67438.1 acar|OOF92389.1 acla|EAW06328.1 afis|EAW18442.1 afla|EED52971.1 afum|EAL93125.1 agla|OJJ85008.1 akaw|GAA86392.1 aluc|OJZ84926.1 anid|CBF70047.1 anig|EHA25145.1 aory|EIT78682.1 asyd|OJJ58862.1 ater|EAU29859.1 atub|OJI80673.1 aver|OJJ01463.1 awen|OJJ30990.1 azon|OJJ50508.1

Group1585: abra|OJJ67439.1 acar|OOF92390.1 acla|EAW06329.1 afis|EAW18443.1 afla|EED52970.1 afum|EAL93126.1 agla|OJJ85009.1 akaw|GAA86391.1 aluc|OJZ84925.1 anid|CBF70045.1 anig|EHA25144.1 aory|EIT78727.1 asyd|OJJ58863.1 ater|EAU29858.1 atub|OJI80674.1 aver|OJJ01464.1 awen|OJJ30991.1 azon|OJJ50509.1

Group1587: abra|OJJ67471.1 acar|OOF89932.1 acla|EAW06300.1 afis|EAW18407.1 afla|EED52997.1 afum|EAL93087.1 agla|OJJ84959.1 akaw|GAA86365.1 aluc|OJZ84895.1 anid|CBF70741.1 anig|EHA25122.1 aory|EIT77423.1 asyd|OJJ58439.1 ater|EAU29825.1 atub|OJI80705.1 aver|OJJ01039.1 awen|OJJ31743.1 azon|OJJ50501.1

Group1589: abra|OJJ67742.1 acar|OOF90949.1 acla|EAW10991.1 afis|EAW20869.1 afla|EED45539.1 afum|EAL84765.1 agla|OJJ83977.1 akaw|GAA92433.1 aluc|OJZ81588.1 anid|CBF82842.1 anig|EHA25852.1 aory|EIT75440.1 asyd|OJJ61589.1 ater|EAU30819.1 atub|OJI88594.1 aver|OJJ00911.1 awen|OJJ34495.1 azon|OJJ51769.1

Group1590: abra|OJJ67743.1 acar|OOF90948.1 acla|EAW10992.1 afis|EAW20868.1 afla|EED45543.1 afum|EAL84766.2 agla|OJJ83976.1 akaw|GAA92434.1 aluc|OJZ81586.1 anid|CBF82840.1 anig|EHA25853.1 aory|EIT75711.1 asyd|OJJ61588.1 ater|EAU30820.1 atub|OJI88595.1 aver|OJJ00910.1 awen|OJJ34498.1 azon|OJJ51773.1

Group1591: abra|OJJ67744.1 acar|OOF90947.1 acla|EAW10993.1 afis|EAW20867.1 afla|EED45544.1 afum|EAL84767.1 agla|OJJ83975.1 akaw|GAA92435.1 aluc|OJZ81585.1 anid|CBF82838.1 anig|EHA25854.1 aory|EIT75422.1 asyd|OJJ61587.1 ater|EAU30821.1 atub|OJI88596.1 aver|OJJ00909.1 awen|OJJ34499.1 azon|OJJ51774.1

Group1592: abra|OJJ67747.1 acar|OOF90944.1 acla|EAW10997.1 afis|EAW20864.1 afla|EED45548.1 afum|EAL84771.1 agla|OJJ83751.1 akaw|GAA92438.1 aluc|OJZ81582.1 anid|CBF82835.1 anig|EHA25857.1 aory|EIT75692.1 asyd|OJJ61584.1 ater|EAU30807.1 atub|OJI88599.1 aver|OJJ00906.1 awen|OJJ34506.1 azon|OJJ51784.1

Group1593: abra|OJJ67748.1 acar|OOF90943.1 acla|EAW10998.1 afis|EAW20863.1 afla|EED45549.1 afum|EAL84772.1 agla|OJJ83750.1 akaw|GAA92439.1 aluc|OJZ81581.1 anid|CBF82833.1 anig|EHA25858.1 aory|EIT75610.1 asyd|OJJ61583.1 ater|EAU30806.1 atub|OJI88600.1 aver|OJJ00905.1 awen|OJJ34507.1 azon|OJJ51785.1

Group1594: abra|OJJ67749.1 acar|OOF90950.1 acla|EAW10990.1 afis|EAW20870.1 afla|EED45552.1 afum|EAL84764.1 agla|OJJ83978.1 akaw|GAA92440.1 aluc|OJZ81580.1 anid|CBF82844.1 anig|EHA25859.1 aory|EIT75541.1 asyd|OJJ61590.1 ater|EAU30818.1 atub|OJI88601.1 aver|OJJ00912.1 awen|OJJ34494.1 azon|OJJ51768.1

Group1596: abra|OJJ67847.1 acar|OOF92828.1 acla|EAW11752.1 afis|EAW23785.1 afla|EED51594.1 afum|EAL87219.2 agla|OJJ78938.1 akaw|GAA91855.1 aluc|OJZ82044.1 anid|CBF79845.1 anig|EHA22213.1 aory|EIT80570.1 asyd|OJJ58153.1 ater|EAU31326.1 atub|OJI84603.1 aver|OJJ04251.1 awen|OJJ33730.1 azon|OJJ42852.1

Group1597: abra|OJJ67848.1 acar|OOF92829.1 acla|EAW11751.1 afis|EAW23784.1 afla|EED51593.1 afum|EAL87218.2 agla|OJJ78937.1 akaw|GAA91856.1 aluc|OJZ82043.1 anid|CBF79841.1 anig|EHA22214.1 aory|EIT80650.1 asyd|OJJ58152.1 ater|EAU31325.1 atub|OJI84602.1 aver|OJJ04252.1 awen|OJJ33729.1 azon|OJJ42853.1

Group1598: abra|OJJ67851.1 acar|OOF92832.1 acla|EAW11748.1 afis|EAW23781.1 afla|EED51585.1 afum|EAL87215.1 agla|OJJ78933.1 akaw|GAA91859.1 aluc|OJZ82040.1 anid|CBF79839.1 anig|EHA22216.1 aory|EIT80522.1 asyd|OJJ58148.1 ater|EAU31322.1 atub|OJI84599.1 aver|OJJ04256.1 awen|OJJ33724.1 azon|OJJ42856.1

Group1599: abra|OJJ67859.1 acar|OOF92840.1 acla|EAW11739.1 afis|EAW23772.1 afla|EED51573.1 afum|EAL87206.1 agla|OJJ78947.1 akaw|GAA91866.1 aluc|OJZ82033.1 anid|CBF79823.1 anig|EHA22223.1 aory|EIT80715.1 asyd|OJJ58141.1 ater|EAU31316.1 atub|OJI84588.1 aver|OJJ04264.1 awen|OJJ33703.1 azon|OJJ42837.1

Group1600: abra|OJJ67861.1 acar|OOF92839.1 acla|EAW11737.1 afis|EAW23770.1 afla|EED51571.1 afum|EAL87204.1 agla|OJJ78949.1 anid|CBF79821.1 anig|EHA22225.1 aory|EIT80633.1 asyd|OJJ58138.1 ater|EAU31314.1 aver|OJJ04267.1 awen|OJJ33705.1 azon|OJJ42836.1 akaw|GAA91868.1 aluc|OJZ82031.1 atub|OJI84586.1

Group1601: abra|OJJ67865.1 acar|OOF92845.1 acla|EAW11734.1 afis|EAW23766.1 afla|EED51552.1 afum|EAL87199.1 agla|OJJ78956.1 akaw|GAA91872.1 aluc|OJZ82027.1 anid|CBF79770.1 anig|EHA22229.1 aory|EIT80651.1 asyd|OJJ58112.1 ater|EAU31310.1 atub|OJI84581.1 aver|OJJ04293.1 awen|OJJ33692.1 azon|OJJ42832.1

Group1602: abra|OJJ67866.1 acar|OOF92846.1 acla|EAW11733.1 afis|EAW23765.1 afla|EED51550.1 afum|EAL87198.1 agla|OJJ78957.1 akaw|GAA91873.1 aluc|OJZ82026.1 anid|CBF79772.1 anig|EHA22230.1 aory|EIT80718.1 asyd|OJJ58110.1 ater|EAU31311.1 atub|OJI84579.1 aver|OJJ04294.1 awen|OJJ33689.1 azon|OJJ42830.1

Group1603: abra|OJJ67881.1 acar|OOF92865.1 acla|EAW11722.1 afis|EAW23744.1 afla|EED51536.1 afum|EAL87180.1 agla|OJJ78963.1 akaw|GAA91888.1 aluc|OJZ82009.1 anid|CBF81714.1 anig|EHA22248.1 aory|EIT80594.1 asyd|OJJ59582.1 ater|EAU31299.1 atub|OJI84558.1 aver|OJJ02687.1 awen|OJJ33679.1 azon|OJJ42820.1

Group1604: abra|OJJ67913.1 acar|OOF92766.1 acla|EAW12796.1 afis|EAW17959.1 afla|EED51948.1 afum|EAL91770.1 agla|OJJ80207.1 akaw|GAA91283.1 aluc|OJZ82786.1 anid|CBF79950.1 anig|EHA21227.1 aory|EIT81390.1 asyd|OJJ58216.1 ater|EAU31448.1 atub|OJI89018.1 aver|OJJ02206.1 awen|OJJ41583.1 azon|OJJ48295.1

Group1605: abra|OJJ67915.1 acar|OOF92767.1 acla|EAW12794.1 afis|EAW17957.1 afla|EED51950.1 afum|EAL91768.1 agla|OJJ80204.1 akaw|GAA91281.1 aluc|OJZ82788.1 anid|CBF79947.1 anig|EHA21229.1 aory|EIT81406.1 asyd|OJJ58214.1 ater|EAU31445.1 atub|OJI89020.1 aver|OJJ02208.1 awen|OJJ41585.1 azon|OJJ48297.1

Group1606: abra|OJJ67917.1 acar|OOF92769.1 acla|EAW12792.1 afis|EAW17955.1 afla|EED51952.1 afum|EAL91766.1 agla|OJJ80202.1 akaw|GAA91279.1 aluc|OJZ82790.1 anid|CBF79943.1 anig|EHA21231.1 aory|EIT81414.1 asyd|OJJ58212.1 ater|EAU31443.1 atub|OJI89021.1 aver|OJJ02210.1 awen|OJJ41587.1 azon|OJJ48302.1

Group1607: abra|OJJ67918.1 acar|OOF92770.1 acla|EAW12791.1 afis|EAW17951.1 afla|EED51953.1 afum|EAL91761.1 agla|OJJ80201.1 akaw|GAA91278.1 aluc|OJZ82791.1 anid|CBF79941.1 anig|EHA21232.1 aory|EIT81400.1 asyd|OJJ58211.1 ater|EAU31442.1 atub|OJI89022.1 aver|OJJ02211.1 awen|OJJ41588.1 azon|OJJ48303.1

Group1608: abra|OJJ67919.1 acar|OOF92771.1 acla|EAW12790.1 afis|EAW17952.1 afla|EED51954.1 afum|EAL91762.1 agla|OJJ80200.1 akaw|GAA91277.1 aluc|OJZ82792.1 anid|CBF79939.1 anig|EHA21233.1 aory|EIT81379.1 asyd|OJJ58210.1 ater|EAU31441.1 atub|OJI89023.1 aver|OJJ02212.1 awen|OJJ41589.1 azon|OJJ48304.1

Group1609: abra|OJJ67920.1 acar|OOF92772.1 acla|EAW12789.1 afis|EAW17953.1 afla|EED51955.1 afum|EAL91763.1 agla|OJJ80199.1 akaw|GAA91276.1 aluc|OJZ82793.1 anid|CBF79934.1 anig|EHA21234.1

aory|EIT81461.1 asyd|OJJ58207.1 ater|EAU31440.1 atub|OJI89024.1 aver|OJJ02215.1 awen|OJJ41590.1 azon|OJJ48305.1

Group1610: abra|OJJ67922.1 acar|OOF92773.1 acla|EAW12788.1 afis|EAW17954.1 afla|EED51956.1 afum|EAL91765.1 agla|OJJ80198.1 akaw|GAA91275.1 aluc|OJZ82794.1 anid|CBF79932.1 anig|EHA21235.1 aory|EIT81403.1 asyd|OJJ58206.1 ater|EAU31439.1 atub|OJI89025.1 aver|OJJ02216.1 awen|OJJ41591.1 azon|OJJ48306.1

Group1611: abra|OJJ67923.1 acar|OOF92774.1 acla|EAW12787.1 afis|EAW17950.1 afla|EED51957.1 afum|EAL91760.1 agla|OJJ80197.1 akaw|GAA91274.1 aluc|OJZ82795.1 anid|CBF79931.1 anig|EHA21236.1 aory|EIT81374.1 asyd|OJJ58205.1 ater|EAU31438.1 atub|OJI89026.1 aver|OJJ02217.1 awen|OJJ41592.1 azon|OJJ48308.1

Group1612: abra|OJJ67926.1 acar|OOF92775.1 acla|EAW12785.1 afis|EAW17947.1 afla|EED51959.1 afum|EAL91757.1 agla|OJJ80195.1 akaw|GAA91271.1 aluc|OJZ82798.1 anid|CBF79929.1 anig|EHA21239.1 aory|EIT81382.1 asyd|OJJ58204.1 ater|EAU31431.1 atub|OJI89029.1 aver|OJJ04202.1 awen|OJJ41596.1 azon|OJJ48312.1

Group1613: abra|OJJ67928.1 acar|OOF92777.1 acla|EAW12781.1 afis|EAW17943.1 afla|EED51961.1 afum|EAL91754.1 agla|OJJ82989.1 akaw|GAA91269.1 aluc|OJZ82800.1 anid|CBF79925.1 anig|EHA21240.1 aory|EIT81385.1 asyd|OJJ58202.1 ater|EAU31396.1 atub|OJI89031.1 aver|OJJ04207.1 awen|OJJ41600.1 azon|OJJ48323.1

Group1614: abra|OJJ67929.1 acar|OOF92779.1 acla|EAW11622.1 afis|EAW23836.1 afla|EED51962.1 afum|EAL87267.1 agla|OJJ82498.1 akaw|GAA91268.1 aluc|OJZ82801.1 anid|CBF79923.1 anig|EHA21242.1 aory|EIT81370.1 asyd|OJJ58200.1 ater|EAU31395.1 atub|OJI89032.1 aver|OJJ04208.1 awen|OJJ33798.1 azon|OJJ48745.1

Group1615: abra|OJJ67936.1 acar|OOF92785.1 acla|EAW11625.1 afis|EAW23831.1 afla|EED51974.1 afum|EAL87263.1 agla|OJJ82481.1 akaw|GAA91260.1 aluc|OJZ82808.1 anid|CBF79904.1 anig|EHA21248.1 aory|EIT81407.1 asyd|OJJ58187.1 ater|EAU31385.1 atub|OJI89039.1 aver|OJJ04218.1 awen|OJJ33789.1 azon|OJJ48757.1

Group1616: abra|OJJ67938.1 acar|OOF92787.1 acla|EAW11628.1 afis|EAW23828.1 afla|EED51977.1 afum|EAL87260.1 agla|OJJ82484.1 akaw|GAA91258.1 aluc|OJZ82810.1 anid|CBF79908.1 anig|EHA21249.1 aory|EIT81387.1 asyd|OJJ58189.1 ater|EAU31383.1 atub|OJI89040.1 aver|OJJ04216.1 awen|OJJ33787.1 azon|OJJ48759.1

Group1617: abra|OJJ67940.1 acar|OOF92789.1 acla|EAW11630.1 afis|EAW23826.1 afla|EED51980.1 afum|EAL87258.1 agla|OJJ82486.1 akaw|GAA91256.1 aluc|OJZ82812.1 anid|CBF79911.1 anig|EHA21252.1 aory|EIT81452.1 asyd|OJJ58190.1 ater|EAU31381.1 atub|OJI89043.1 aver|OJJ04215.1 awen|OJJ33784.1 azon|OJJ42886.1

Group1618: abra|OJJ67942.1 acar|OOF92791.1 acla|EAW11636.1 afis|EAW23821.1 afla|EED51983.1 afum|EAL87256.1 agla|OJJ82468.1 akaw|GAA91254.1 aluc|OJZ82814.1 anid|CBF79894.1 anig|EHA21254.1 aory|EIT81391.1 asyd|OJJ58182.1 ater|EAU31373.1 atub|OJI89045.1 aver|OJJ04224.1 awen|OJJ33777.1 azon|OJJ42883.1

Group1619: abra|OJJ67951.1 acar|OOF92798.1 acla|EAW11641.1 afis|EAW23816.1 afla|EED51989.1 afum|EAL87251.1 agla|OJJ82462.1 akaw|GAA91246.1 aluc|OJZ82824.1 anid|CBF79887.1 anig|EHA21262.1 aory|EIT81439.1 asyd|OJJ58175.1 ater|EAU31371.1 atub|OJI89053.1 aver|OJJ04229.1 awen|OJJ33773.1 azon|OJJ42877.1

Group1620: abra|OJJ67957.1 acar|OOF92802.1 acla|EAW11646.1 afis|EAW23810.1 afla|EED52005.1 afum|EAL87244.1 agla|OJJ82450.1 akaw|GAA91239.1 aluc|OJZ82830.1 anid|CBF79876.1 anig|EHA21268.1 aory|EIT81434.1 asyd|OJJ58169.1 ater|EAU31363.1 atub|OJI89060.1 aver|OJJ04234.1 awen|OJJ33766.1 azon|OJJ42870.1

Group1621: abra|OJJ67958.1 acar|OOF92803.1 acla|EAW11647.1 afis|EAW23809.1 afla|EED52006.1 afum|EAL87243.1 agla|OJJ82449.1 akaw|GAA91238.1 aluc|OJZ82831.1 anid|CBF79874.1 anig|EHA21269.1 aory|EIT81404.1 asyd|OJJ58168.1 ater|EAU31362.1 atub|OJI89061.1 aver|OJJ04235.1 awen|OJJ33765.1 azon|OJJ42868.1

Group1622: abra|OJJ67977.1 acar|OOF92819.1 acla|EAW11758.1 afis|EAW23791.1 afla|EED51599.1 afum|EAL87225.1 agla|OJJ78819.1 akaw|GAA91226.1 aluc|OJZ82844.1 anid|CBF79853.1 anig|EHA21282.1 aory|EIT80507.1 asyd|OJJ58157.1 ater|EAU31330.1 atub|OJI89074.1 aver|OJJ04246.1 awen|OJJ33746.1 azon|OJJ42843.1

Group1623: abra|OJJ67978.1 acar|OOF92818.1 acla|EAW11757.1 afis|EAW23790.1 afla|EED51598.1 afum|EAL87224.1 agla|OJJ78817.1 akaw|GAA91225.1 aluc|OJZ82845.1 anid|CBF79851.1 anig|EHA21283.1 aory|EIT80526.1 asyd|OJJ58156.1 ater|EAU31329.1 atub|OJI89075.1 aver|OJJ04247.1 awen|OJJ33745.1 azon|OJJ42844.1

Group1624: abra|OJJ68015.1 acar|OOF92057.1 acla|EAW08795.1 afis|EAW15605.1 afla|EED55861.1 afum|EAL85728.1 agla|OJJ79064.1 akaw|GAA90936.1 aluc|OJZ82888.1 anid|CBF69625.1 anig|EHA21314.1 aory|EIT76549.1 asyd|OJJ59120.1 ater|EAU37248.1 atub|OJI89113.1 aver|OJJ01709.1 awen|OJJ32398.1 azon|OJJ43874.1

Group1626: abra|OJJ68060.1 acar|OOF92081.1 acla|EAW13655.1 afis|EAW19935.1 afla|EED45904.1 afum|EAL84975.2 agla|OJJ79327.1 akaw|GAA90896.1 aluc|OJZ82932.1 anid|CBF84602.1 anig|EHA21353.1 aory|EIT75822.1 asyd|OJJ53375.1 ater|EAU36898.1 atub|OJI89161.1 aver|OJJ08369.1 awen|OJJ29978.1 azon|OJJ50215.1

Group1627: abra|OJJ68075.1 acar|OOF94470.1 acla|EAW12117.1 afis|EAW17265.1 afla|EED48857.1 afum|EAL86682.1 agla|OJJ85378.1 akaw|GAA92736.1 aluc|OJZ92944.1 anid|CBF87528.1 anig|EHA26028.1 aory|EIT81286.1 asyd|OJJ52984.1 ater|EAU36128.1 atub|OJI87403.1 aver|OJJ08953.1 awen|OJJ35769.1 azon|OJJ50248.1

Group1628: abra|OJJ68089.1 acar|OOF93893.1 acla|EAW15283.1 afis|EAW17127.1 afla|EED48286.1 afum|EAL86587.1 agla|OJJ78597.1 akaw|GAA86874.1 aluc|OJZ89998.1 anid|CBF80813.1 anig|EHA27373.1 aory|EIT72919.1 asyd|OJJ54451.1 ater|EAU34073.1 atub|OJI84810.1 aver|OJJ07703.1 awen|OJJ29885.1 azon|OJJ50148.1

Group1629: abra|OJJ68125.1 acar|OOF93919.1 acla|EAW12869.1 afis|EAW17817.1 afla|EED45601.1 afum|EAL91627.1 agla|OJJ81689.1 akaw|GAA86906.1 aluc|OJZ90029.1 anid|CBF80748.1 anig|EHA27405.1 aory|EIT75765.1 asyd|OJJ54539.1 ater|EAU30285.1 atub|OJI84778.1 aver|OJJ07642.1 awen|OJJ36934.1 azon|OJJ42822.1

Group1631: abra|OJJ68139.1 acar|OOF98356.1 acla|EAW14517.1 afis|EAW19235.1 afla|EED47647.1 afum|EAL93963.1 agla|OJJ88463.1 akaw|GAA92378.1 aluc|OJZ83237.1 anid|CBF78634.1 anig|EHA21094.1 aory|EIT80221.1 asyd|OJJ65123.1 ater|EAU37228.1 atub|OJI87699.1 aver|OJI99251.1 awen|OJJ37592.1 azon|OJJ51245.1

Group1632: abra|OJJ68145.1 acar|OOF89988.1 acla|EAW07390.1 afis|EAW22416.1 afla|EED52494.1 afum|EAL90776.1 agla|OJJ87680.1 akaw|GAA92373.1 aluc|OJZ83231.1 anid|CBF88053.1 anig|EHA21089.1 aory|EIT82805.1 asyd|OJJ63540.1 ater|EAU39291.1 atub|OJI87705.1 aver|OJI97393.1 awen|OJJ34995.1 azon|OJJ45638.1

Group1634: abra|OJJ68168.1 acar|OOF97466.1 acla|EAW13054.1 afis|EAW17612.1 afla|EED49203.1 afum|EAL91425.1 agla|OJJ81325.1 akaw|GAA83275.1 aluc|OJZ83781.1 anid|CBF90174.1 anig|EHA18921.1 aory|EIT78579.1 asyd|OJJ61990.1 ater|EAU37047.1 atub|OJI90151.1 aver|OJI96082.1 awen|OJJ37230.1 azon|OJJ50933.1

Group1635: abra|OJJ68169.1 acar|OOF97467.1 acla|EAW13055.1 afis|EAW17611.1 afla|EED49202.1 afum|EAL91424.1 agla|OJJ81326.1 akaw|GAA83274.1 aluc|OJZ83780.1 anid|CBF90175.1 anig|EHA18920.1 aory|EIT78562.1 asyd|OJJ61989.1 ater|EAU37046.1 atub|OJI90150.1 aver|OJI96081.1 awen|OJJ37231.1 azon|OJJ50934.1

Group1636: abra|OJJ68171.1 acar|OOF97468.1 acla|EAW13056.1 afis|EAW17610.1 afla|EED49201.1 afum|EAL91423.1 agla|OJJ81331.1 akaw|GAA83273.1 aluc|OJZ83779.1 anid|CBF90177.1 anig|EHA18919.1 aory|EIT78514.1 asyd|OJJ61988.1 ater|EAU37045.1 atub|OJI90149.1 aver|OJI96080.1 awen|OJJ37232.1 azon|OJJ50935.1

Group1637: abra|OJJ68172.1 acar|OOF97470.1 acla|EAW13057.1 afis|EAW17609.1 afla|EED49213.1 afum|EAL91422.1 agla|OJJ81356.1 akaw|GAA83272.1 aluc|OJZ83778.1 anid|CBF90179.1 anig|EHA18918.1 aory|EIT83168.1 asyd|OJJ61987.1 ater|EAU37044.1 atub|OJI90148.1 aver|OJI96079.1 awen|OJJ37233.1 azon|OJJ50936.1

Group1638: abra|OJJ68173.1 acar|OOF97471.1 acla|EAW13058.1 afis|EAW17608.1 afla|EED49214.1 afum|EAL91421.1 agla|OJJ81355.1 akaw|GAA83271.1 aluc|OJZ83777.1 anid|CBF90181.1 anig|EHA18917.1 aory|EIT83162.1 asyd|OJJ61986.1 ater|EAU37043.1 atub|OJI90147.1 aver|OJI96078.1 awen|OJJ37234.1 azon|OJJ50937.1

Group1639: abra|OJJ68176.1 acar|OOF97473.1 acla|EAW13060.1 afis|EAW17604.1 afla|EED49217.1 afum|EAL91417.1 agla|OJJ81352.1 akaw|GAA83268.1 aluc|OJZ83774.1 anid|CBF90186.1 anig|EHA18914.1 aory|EIT83159.1 asyd|OJJ61983.1 ater|EAU37041.1 atub|OJI90145.1 aver|OJI96075.1 awen|OJJ37237.1 azon|OJJ50939.1

Group1640: abra|OJJ68177.1 acar|OOF97474.1 acla|EAW13061.1 afis|EAW17603.1 afla|EED49218.1 afum|EAL91416.1 agla|OJJ81351.1 akaw|GAA83267.1 aluc|OJZ83773.1 anid|CBF90188.1 anig|EHA18913.1 aory|EIT83167.1 asyd|OJJ61982.1 ater|EAU37040.1 atub|OJI90144.1 aver|OJI96074.1 awen|OJJ37238.1 azon|OJJ50940.1

Group1641: abra|OJJ68181.1 acar|OOF97477.1 acla|EAW13063.1 afis|EAW17601.1 afla|EED49220.1 afum|EBA27434.1 agla|OJJ81348.1 akaw|GAA83265.1 aluc|OJZ83771.1 anid|CBF90190.1 anig|EHA18911.1

aory|EIT83161.1 asyd|OJJ61981.1 ater|EAU37038.1 atub|OJI90142.1 aver|OJI96073.1 awen|OJJ37244.1 azon|OJJ50942.1

Group1642: abra|OJJ68182.1 acar|OOF97476.1 acla|EAW13064.1 afis|EAW17600.1 afla|EED49221.1 afum|EAL91414.1 agla|OJJ81347.1 akaw|GAA83264.1 aluc|OJZ83770.1 anid|CBF90192.1 anig|EHA18910.1 aory|EIT83165.1 asyd|OJJ61980.1 ater|EAU37037.1 atub|OJI90141.1 aver|OJI96072.1 awen|OJJ37245.1 azon|OJJ50943.1

Group1643: abra|OJJ68184.1 acar|OOF97479.1 acla|EAW13065.1 afis|EAW17599.1 afla|EED49222.1 afum|EAL91412.2 agla|OJJ81345.1 akaw|GAA83262.1 aluc|OJZ83768.1 anig|EHA18908.1 aory|EIT83170.1 asyd|OJJ61978.1 ater|EAU37036.1 atub|OJI90139.1 aver|OJI96070.1 awen|OJJ37247.1 azon|OJJ50945.1 anid|CBF90194.1

Group1644: abra|OJJ68190.1 acar|OOF97486.1 acla|EAW13072.1 afis|EAW17592.1 afla|EED49230.1 afum|EAL91405.1 agla|OJJ81357.1 akaw|GAA83255.1 aluc|OJZ83756.1 anid|CBF90212.1 anig|EHA18903.1 aory|EIT83166.1 asyd|OJJ61969.1 ater|EAU37029.1 atub|OJI90131.1 aver|OJI96062.1 awen|OJJ37254.1 azon|OJJ50949.1

Group1645: abra|OJJ68193.1 acar|OOF97489.1 acla|EAW13076.1 afis|EAW17588.1 afla|EED49234.1 afum|EAL91401.1 agla|OJJ81364.1 akaw|GAA83252.1 aluc|OJZ83753.1 anid|CBF90220.1 anig|EHA18900.1 aory|EIT83171.1 asyd|OJJ61966.1 ater|EAU37023.1 atub|OJI90128.1 aver|OJI96058.1 awen|OJJ37258.1 azon|OJJ50953.1

Group1646: abra|OJJ68194.1 acar|OOF97490.1 acla|EAW13077.1 afis|EAW17587.1 afla|EED49235.1 afum|EAL91400.1 agla|OJJ81365.1 akaw|GAA83251.1 aluc|OJZ83752.1 anid|CBF90222.1 anig|EHA18899.1 aory|EIT83154.1 asyd|OJJ61965.1 ater|EAU37022.1 atub|OJI90127.1 aver|OJI96057.1 awen|OJJ37259.1 azon|OJJ50954.1

Group1647: abra|OJJ68196.1 acar|OOF97491.1 acla|EAW13079.1 afis|EAW17585.1 afla|EED49237.1 afum|EAL91398.2 agla|OJJ81368.1 akaw|GAA83249.1 aluc|OJZ83750.1 anid|CBF90225.1 anig|EHA18897.1 aory|EIT83169.1 asyd|OJJ61963.1 ater|EAU37020.1 atub|OJI90125.1 aver|OJI96055.1 awen|OJJ37261.1 azon|OJJ50956.1

Group1648: abra|OJJ68197.1 acar|OOF89886.1 acla|EAW13080.1 afis|EAW17584.1 afla|EED49239.1 afum|EAL91397.1 agla|OJJ81369.1 akaw|GAA83248.1 aluc|OJZ83749.1 anid|CBF90227.1 anig|EHA18896.1 aory|EIT82222.1 asyd|OJJ61962.1 ater|EAU37019.1 atub|OJI90124.1 aver|OJI96054.1 awen|OJJ37262.1 azon|OJJ50957.1

Group1649: abra|OJJ68199.1 acar|OOF90119.1 acla|EAW13081.1 afis|EAW17583.1 afla|EED49240.1 afum|EAL91396.1 agla|OJJ81370.1 akaw|GAA83247.1 aluc|OJZ83748.1 anid|CBF90229.1 anig|EHA18895.1 aory|EIT82239.1 asyd|OJJ61961.1 ater|EAU37018.1 atub|OJI90123.1 aver|OJI96053.1 awen|OJJ37265.1 azon|OJJ50959.1

Group1650: abra|OJJ68200.1 acar|OOF90117.1 acla|EAW13083.1 afis|EAW17581.1 afla|EED49241.1 agla|OJJ81372.1 akaw|GAA83245.1 aluc|OJZ83747.1 anid|CBF90232.1 anig|EHA18893.1 aory|EIT82216.1 asyd|OJJ61959.1 ater|EAU37016.1 atub|OJI90122.1 aver|OJI96051.1 awen|OJJ37267.1 azon|OJJ50960.1 afum|EAL91394.1

Group1651: abra|OJJ68201.1 acar|OOF90116.1 acla|EAW13084.1 afis|EAW17580.1 afla|EED49242.1 afum|EAL91393.1 agla|OJJ81379.1 akaw|GAA83244.1 aluc|OJZ83746.1 anid|CBF90234.1 anig|EHA18892.1 aory|EIT82232.1 asyd|OJJ61958.1 ater|EAU37015.1 atub|OJI90121.1 aver|OJI96050.1 awen|OJJ37268.1 azon|OJJ50961.1

Group1652: abra|OJJ68203.1 acar|OOF90234.1 acla|EAW13086.1 afis|EAW17579.1 afla|EED49243.1 afum|EAL91392.1 agla|OJJ81378.1 akaw|GAA83243.1 aluc|OJZ83744.1 anid|CBF90238.1 anig|EHA18891.1 aory|EIT82211.1 asyd|OJJ61957.1 ater|EAU37014.1 atub|OJI90120.1 aver|OJI96049.1 awen|OJJ37269.1 azon|OJJ50962.1

Group1653: abra|OJJ68204.1 acar|OOF90236.1 acla|EAW13087.1 afis|EAW17577.1 afla|EED49244.1 afum|EAL91390.1 agla|OJJ81377.1 akaw|GAA83242.1 aluc|OJZ83741.1 anid|CBF90245.1 anig|EHA18890.1 aory|EIT82198.1 asyd|OJJ61951.1 ater|EAU37013.1 atub|OJI90119.1 aver|OJI96042.1 awen|OJJ37270.1 azon|OJJ50969.1

Group1654: abra|OJJ68205.1 acar|OOF90235.1 acla|EAW13088.1 afis|EAW17576.1 afla|EED49245.1 afum|EAL91389.1 agla|OJJ81381.1 akaw|GAA83241.1 aluc|OJZ83740.1 anid|CBF90247.1 anig|EHA18889.1 aory|EIT82237.1 asyd|OJJ61950.1 ater|EAU37012.1 atub|OJI90118.1 aver|OJI96041.1 awen|OJJ37271.1 azon|OJJ50971.1

Group1655: abra|OJJ68207.1 acar|OOF97493.1 acla|EAW13090.1 afis|EAW17574.1 afla|EED49247.1 afum|EAL91387.1 agla|OJJ81384.1 akaw|GAA83239.1 aluc|OJZ83738.1 anid|CBF90250.1 anig|EHA18887.1 aory|EIT82189.1 asyd|OJJ61948.1 ater|EAU37010.1 atub|OJI90116.1 aver|OJI96040.1 awen|OJJ37273.1 azon|OJJ50973.1

Group1656: abra|OJJ68211.1 acar|OOF97495.1 acla|EAW12973.1 afis|EAW17691.1 afla|EED50453.1 afum|EAL91505.1 agla|OJJ79488.1 akaw|GAA83235.1 aluc|OJZ83734.1 anid|CBF89980.1 anig|EHA18883.1 aory|EIT82051.1 asyd|OJJ62090.1 ater|EAU31953.1 atub|OJI90112.1 aver|OJI96196.1 awen|OJJ37145.1 azon|OJJ50845.1

Group1657: abra|OJJ68212.1 acar|OOF97496.1 acla|EAW12972.1 afis|EAW17692.1 afla|EED50452.1 afum|EAL91506.1 agla|OJJ79487.1 akaw|GAA83234.1 aluc|OJZ83733.1 anid|CBF89978.1 anig|EHA18882.1 aory|EIT82145.1 asyd|OJJ62091.1 ater|EAU31958.1 atub|OJI90111.1 aver|OJI96197.1 awen|OJJ37144.1 azon|OJJ50844.1

Group1658: abra|OJJ68249.1 acar|OOF97528.1 acla|EAW09061.1 afis|EAW25634.1 afla|EED55178.1 afum|EAL84322.1 agla|OJJ78898.1 akaw|GAA83212.1 aluc|OJZ83714.1 anid|CBF83155.1 anig|EHA18865.1 aory|EIT81648.1 asyd|OJJ55514.1 ater|EAU32715.1 atub|OJI90089.1 aver|OJJ06591.1 awen|OJJ36053.1 azon|OJJ50759.1

Group1659: abra|OJJ68257.1 acar|OOF97534.1 acla|EAW09064.1 afis|EAW25636.1 afla|EED55176.1 afum|EAL84324.1 agla|OJJ78896.1 akaw|GAA83205.1 aluc|OJZ83707.1 anid|CBF83151.1 anig|EHA18858.1 aory|EIT81641.1 asyd|OJJ55516.1 ater|EAU32717.1 atub|OJI90079.1 aver|OJJ06593.1 awen|OJJ36093.1 azon|OJJ50756.1

Group1660: abra|OJJ68259.1 acar|OOF97536.1 acla|EAW14109.1 afis|EAW18811.1 afla|EED50761.1 afum|EAL93547.1 agla|OJJ80544.1 akaw|GAA92415.1 aluc|OJZ83273.1 anid|CBF69795.1 anig|EHA18856.1 aory|EIT78026.1 asyd|OJJ58964.1 ater|EAU37966.1 atub|OJI87668.1 aver|OJJ01558.1 awen|OJJ32387.1 azon|OJJ47147.1

Group1662: abra|OJJ68281.1 acar|OOF97555.1 acla|EAW09093.1 afis|EAW25687.1 afla|EED55157.1 afum|EAL84356.1 agla|OJJ78886.1 akaw|GAA83191.1 aluc|OJZ83690.1 anid|CBF83933.1 anig|EHA18844.1 aory|EIT81633.1 asyd|OJJ55435.1 ater|EAU32721.1 atub|OJI90062.1 aver|OJJ06512.1 awen|OJJ36120.1 azon|OJJ50742.1

Group1663: abra|OJJ68301.1 acar|OOF96686.1 acla|EAW12631.1 afis|EAW18053.1 afla|EED56195.1 afum|EAL91864.1 agla|OJJ87336.1 akaw|GAA83173.1 aluc|OJZ83672.1 anid|CBF83350.1 anig|EHA18827.1 aory|EIT73338.1 asyd|OJJ60615.1 ater|EAU34830.1 atub|OJI90043.1 aver|OJI99939.1 awen|OJJ41435.1 azon|OJJ48899.1

Group1666: abra|OJJ68449.1 acar|OOF93946.1 acla|EAW11772.1 afis|EAW23677.1 afla|EED54146.1 afum|EAL87120.1 agla|OJJ78443.1 akaw|GAA86924.1 aluc|OJZ90048.1 anid|CBF75094.1 anig|EHA21022.1 aory|EIT75945.1 asyd|OJJ60010.1 ater|EAU29439.1 atub|OJI84758.1 aver|OJJ02275.1 awen|OJJ38063.1 azon|OJJ50775.1

Group1667: abra|OJJ68459.1 acar|OOF92643.1 acla|EAW08639.1 afis|EAW15792.1 afla|EED51253.1 afum|EAL85558.1 agla|OJJ81955.1 akaw|GAA92421.1 aluc|OJZ80815.1 anid|CBF70968.1 anig|EHA21125.1 aory|EIT81555.1 asyd|OJJ52614.1 ater|EAU32350.1 atub|OJI87664.1 aver|OJJ04641.1 awen|OJJ38500.1 azon|OJJ43801.1

Group1668: abra|OJJ68464.1 acar|OOF94169.1 acla|EAW10056.1 afis|EAW25528.1 afla|EED54029.1 afum|EAL90193.2 agla|OJJ87274.1 akaw|GAA92384.1 aluc|OJZ83243.1 anid|CBF77807.1 anig|EHA21098.1 aory|EIT78344.1 asyd|OJJ53016.1 ater|EAU33371.1 atub|OJI87693.1 aver|OJJ03827.1 awen|OJJ41270.1 azon|OJJ49510.1

Group1669: abra|OJJ68468.1 acar|OOF89893.1 acla|EAW10429.1 afis|EAW25140.1 afla|EED55305.1 afum|EAL89812.1 agla|OJJ89027.1 akaw|GAA92389.1 aluc|OJZ83249.1 anid|CBF86095.1 anig|EHA21102.1 aory|EIT83530.1 asyd|OJJ64474.1 ater|EAU39409.1 atub|OJI87688.1 aver|OJI98578.1 awen|OJJ40764.1 azon|OJJ44129.1

Group1670: abra|OJJ68475.1 acar|OOF92975.1 acla|EAW06477.1 afis|EAW15972.1 afla|EED58084.1 afum|EAL88604.1 agla|OJJ83843.1 akaw|GAA92396.1 aluc|OJZ83256.1 anid|CBF81100.1 anig|EHA21110.1 aory|EIT79134.1 asyd|OJJ56477.1 ater|EAU35476.1 atub|OJI87682.1 aver|OJJ06241.1 awen|OJJ35600.1 azon|OJJ46920.1

Group1671: abra|OJJ68476.1 acar|OOF99927.1 acla|EAW09807.1 afis|EAW21329.1 afla|EED57287.1 afum|EAL92347.1 agla|OJJ87410.1 akaw|GAA92397.1 aluc|OJZ83258.1 anid|CBF83431.1 anig|EHA21111.1 aory|EIT74125.1 asyd|OJJ60661.1 ater|EAU35955.1 atub|OJI87681.1 aver|OJI99984.1 awen|OJJ39369.1 azon|OJJ51709.1

Group1672: abra|OJJ68478.1 acar|OOF96519.1 acla|EAW13881.1 afis|EAW18593.1 afla|EED52754.1 afum|EAL93301.1 agla|OJJ81645.1 akaw|GAA92401.1 aluc|OJZ83262.1 anid|CBF70381.1 anig|EHA21115.1 aory|EIT78198.1 asyd|OJJ58642.1 ater|EAU34887.1 atub|OJI87675.1 aver|OJJ01248.1 awen|OJJ31156.1 azon|OJJ46631.1

Group1673: abra|OJJ68485.1 acar|OOG00353.1 acla|EAW07017.1 afis|EAW16643.1 afla|EED44966.1 afum|EAL89246.1 agla|OJJ82604.1 akaw|GAA92786.1 aluc|OJZ82697.1 anid|CBF82045.1 anig|EHA21146.1

aory|EIT73490.1 asyd|OJJ54225.1 ater|EAU39568.1 atub|OJI88929.1 aver|OJJ05178.1 awen|OJJ40071.1 azon|OJJ45553.1

Group1674: abra|OJJ68486.1 acar|OOF95442.1 acla|EAW07333.1 afis|EAW22365.1 afla|EED57773.1 afum|EBA27416.1 agla|OJJ87616.1 akaw|GAA92785.1 aluc|OJZ82698.1 anid|CBF88690.1 anig|EHA21153.1 aory|EIT81113.1 asyd|OJJ63908.1 ater|EAU39499.1 atub|OJI88930.1 aver|OJI97033.1 awen|OJJ34939.1 azon|OJJ45591.1

Group1675: abra|OJJ68497.1 acar|OOF99362.1 acla|EAW14277.1 afis|EAW18997.1 afla|EED53139.1 afum|EAL93732.1 agla|OJJ80191.1 akaw|GAA92773.1 aluc|OJZ82710.1 anid|CBF86779.1 anig|EHA21165.1 aory|EIT74918.1 asyd|OJJ64682.1 ater|EAU37806.1 atub|OJI88943.1 aver|OJI98784.1 awen|OJJ31625.1 azon|OJJ47325.1

Group1676: abra|OJJ68499.1 acar|OOF93648.1 acla|EAW14509.1 afis|EAW19222.1 afla|EED47663.1 afum|EAL93954.1 agla|OJJ88466.1 akaw|GAA92772.1 aluc|OJZ82712.1 anid|CBF78608.1 anig|EHA21167.1 aory|EIT80278.1 asyd|OJJ65103.1 ater|EAU37236.1 atub|OJI88945.1 aver|OJI99230.1 awen|OJJ37585.1 azon|OJJ51239.1

Group1677: abra|OJJ68500.1 acar|OOF92674.1 acla|EAW08059.1 afis|EAW23095.1 afla|EED48553.1 afum|EAL88383.1 agla|OJJ89607.1 akaw|GAA92771.1 aluc|OJZ82713.1 anid|CBF75180.1 anig|EHA21168.1 aory|EIT73191.1 asyd|OJJ59952.1 ater|EAU36223.1 atub|OJI88946.1 aver|OJJ02337.1 awen|OJJ36686.1 azon|OJJ47510.1

Group1678: abra|OJJ68501.1 acar|OOF89894.1 acla|EAW11963.1 afis|EAW19600.1 afla|EED56956.1 afum|EAL86918.1 agla|OJJ80742.1 akaw|GAA92770.1 aluc|OJZ82714.1 anid|CBF71291.1 anig|EHA21169.1 aory|EIT77492.1 asyd|OJJ55128.1 ater|EAU32925.1 atub|OJI88947.1 aver|OJJ07076.1 awen|OJJ41841.1 azon|OJJ47038.1

Group1679: abra|OJJ68507.1 acar|OOF92954.1 acla|EAW07233.1 afis|EAW22254.1 afla|EED57640.1 afum|EAL90924.1 agla|OJJ87802.1 akaw|GAA92766.1 aluc|OJZ82719.1 anid|CBF88500.1 anig|EHA21173.1 aory|EIT82931.1 asyd|OJJ63795.1 ater|EAU34255.1 atub|OJI88953.1 aver|OJI97142.1 awen|OJJ34826.1 azon|OJJ45723.1

Group1680: abra|OJJ68510.1 acar|OOF92672.1 acla|EAW07690.1 afis|EAW22705.1 afla|EED49991.1 afum|EAL90487.1 agla|OJJ79004.1 akaw|GAA92761.1 aluc|OJZ82724.1 anid|CBF88071.1 anig|EHA21178.1 aory|EIT74613.1 asyd|OJJ63553.1 ater|EAU38996.1 atub|OJI88958.1 aver|OJI97381.1 awen|OJJ35342.1 azon|OJJ49776.1

Group1681: abra|OJJ68511.1 acar|OOF92678.1 acla|EAW14940.1 afis|EAW24455.1 afla|EED55219.1 afum|EAL86124.1 agla|OJJ80302.1 akaw|GAA91335.1 aluc|OJZ82725.1 anid|CBF75392.1 anig|EHA21179.1 aory|EIT83615.1 asyd|OJJ59834.1 ater|EAU29438.1 atub|OJI88959.1 aver|OJJ02454.1 awen|OJJ32742.1 azon|OJJ45292.1

Group1682: abra|OJJ68517.1 acar|OOF99154.1 acla|EAW06536.1 afis|EAW16059.1 afla|EED48573.1 afum|EAL88692.1 agla|OJJ86106.1 akaw|GAA91331.1 aluc|OJZ82731.1 anid|CBF75082.1 anig|EHA21183.1 aory|EIT73064.1 asyd|OJJ60019.1 ater|EAU33575.1 atub|OJI88964.1 aver|OJJ02266.1 awen|OJJ30380.1 azon|OJJ45396.1

Group1683: abra|OJJ68524.1 acar|OOF92729.1 acla|EAW12839.1 afis|EAW18004.1 afla|EED51892.1 afum|EAL91815.2 agla|OJJ80272.1 akaw|GAA91325.1 aluc|OJZ82738.1 anid|CBF80032.1 anig|EHA21189.1 aory|EIT76182.1 asyd|OJJ58271.1 ater|EAU31516.1 atub|OJI88973.1 aver|OJJ02153.1 awen|OJJ41538.1 azon|OJJ48256.1

Group1684: abra|OJJ68526.1 acar|OOF92730.1 acla|EAW12838.1 afis|EAW18003.1 afla|EED51893.1 afum|EAL91814.2 agla|OJJ80266.1 akaw|GAA91324.1 aluc|OJZ82739.1 anid|CBF80026.1 anig|EHA21190.1 aory|EIT76172.1 asyd|OJJ58266.1 ater|EAU31515.1 atub|OJI88974.1 aver|OJJ02157.1 awen|OJJ41539.1 azon|OJJ48257.1

Group1685: abra|OJJ68527.1 acar|OOF92731.1 acla|EAW12837.1 afis|EAW18002.1 afla|EED51894.1 afum|EAL91813.1 agla|OJJ80265.1 akaw|GAA91323.1 aluc|OJZ82740.1 anid|CBF80025.1 anig|EHA21191.1 aory|EIT76169.1 asyd|OJJ58265.1 ater|EAU31514.1 atub|OJI88975.1 aver|OJJ02158.1 awen|OJJ41540.1 azon|OJJ48258.1

Group1686: abra|OJJ68528.1 acar|OOF92732.1 acla|EAW12836.1 afis|EAW18001.1 afla|EED51895.1 afum|EAL91812.1 agla|OJJ80264.1 akaw|GAA91322.1 aluc|OJZ82741.1 anid|CBF80023.1 anig|EHA21192.1 aory|EIT76176.1 asyd|OJJ58264.1 ater|EAU31513.1 atub|OJI88976.1 aver|OJJ02160.1 awen|OJJ41541.1 azon|OJJ48259.1

Group1687: abra|OJJ68531.1 acar|OOF92734.1 acla|EAW12834.1 afis|EAW17999.1 afla|EED51899.1 afum|EAL91810.1 agla|OJJ80262.1 akaw|GAA91320.1 aluc|OJZ82743.1 anid|CBF80019.1 anig|EHA21194.1 aory|EIT76173.1 asyd|OJJ58262.1 ater|EAU31511.1 atub|OJI88978.1 aver|OJJ02162.1 awen|OJJ41543.1 azon|OJJ48261.1

Group1688: abra|OJJ68532.1 acar|OOF92735.1 acla|EAW12833.1 afis|EAW17998.1 afla|EED51900.1 afum|EAL91809.2 agla|OJJ80261.1 akaw|GAA91319.1 aluc|OJZ82744.1 anid|CBF80017.1 anig|EHA21195.1 aory|EIT76171.1 asyd|OJJ58261.1 ater|EAU31510.1 atub|OJI88979.1 aver|OJJ02163.1 awen|OJJ41544.1 azon|OJJ48262.1

Group1689: abra|OJJ68533.1 acar|OOF92736.1 afis|EAW17997.1 afla|EED51903.1 afum|EAL91807.2 akaw|GAA91318.1 aluc|OJZ82746.1 anig|EHA21196.1 aory|EIT76183.1 asyd|OJJ58260.1 atub|OJI88980.1 aver|OJJ02164.1 awen|OJJ41545.1 acla|EAW12832.1 agla|OJJ80257.1 ater|EAU31509.1 azon|OJJ48263.1 anid|CBF80015.1

Group1690: abra|OJJ68537.1 acar|OOF90151.1 acla|EAW12827.1 afis|EAW17992.1 afla|EED51911.1 afum|EAL91802.1 agla|OJJ80253.1 akaw|GAA91314.1 aluc|OJZ82751.1 anid|CBF80004.1 anig|EHA21200.1 aory|EIT76179.1 asyd|OJJ58250.1 ater|EAU31503.1 atub|OJI88987.1 aver|OJJ02174.1 awen|OJJ41554.1 azon|OJJ48269.1

Group1691: abra|OJJ68538.1 acar|OOF90152.1 acla|EAW12828.1 afis|EAW17993.1 afla|EED51908.1 afum|EAL91803.1 agla|OJJ80254.1 akaw|GAA91313.1 aluc|OJZ82752.1 anid|CBF80005.1 anig|EHA21201.1 aory|EIT76170.1 asyd|OJJ58251.1 ater|EAU31504.1 atub|OJI88988.1 aver|OJJ02173.1 awen|OJJ41553.1 azon|OJJ48268.1

Group1692: abra|OJJ68541.1 acar|OOF92739.1 acla|EAW12829.1 afis|EAW17994.1 afla|EED51907.1 afum|EAL91804.1 agla|OJJ80255.1 akaw|GAA91312.1 aluc|OJZ82753.1 anid|CBF80009.1 anig|EHA21202.1 aory|EIT76184.1 asyd|OJJ58256.1 ater|EAU31505.1 atub|OJI88990.1 aver|OJJ02167.1 awen|OJJ41551.1 azon|OJJ48267.1

Group1693: abra|OJJ68542.1 acar|OOF92740.1 acla|EAW12830.1 afis|EAW17995.1 afla|EED51906.1 afum|EAL91805.1 agla|OJJ80259.1 akaw|GAA91311.1 aluc|OJZ82755.1 anid|CBF80011.1 anig|EHA21203.1 aory|EIT76180.1 asyd|OJJ58257.1 ater|EAU31507.1 atub|OJI88991.1 aver|OJJ02166.1 awen|OJJ41550.1 azon|OJJ48266.1

Group1694: abra|OJJ68544.1 acar|OOF92742.1 acla|EAW12823.1 afis|EAW17988.1 afla|EED51915.1 afum|EAL91797.1 agla|OJJ80248.1 akaw|GAA91309.1 aluc|OJZ82759.1 anid|CBF79996.1 anig|EHA21205.1 aory|EIT81365.1 asyd|OJJ58247.1 ater|EAU31498.1 atub|OJI88993.1 aver|OJJ02179.1 awen|OJJ41558.1 azon|OJJ48273.1

Group1695: abra|OJJ68546.1 acar|OOF92743.1 acla|EAW12822.1 afis|EAW17987.1 afla|EED51916.1 afum|EAL91796.1 agla|OJJ80242.1 akaw|GAA91308.1 aluc|OJZ82762.1 anid|CBF80000.1 anig|EHA21206.1 aory|EIT81367.1 asyd|OJJ58248.1 ater|EAU31497.1 atub|OJI88995.1 aver|OJJ02176.1 awen|OJJ41559.1 azon|OJJ48274.1

Group1696: abra|OJJ68549.1 acar|OOF92745.1 acla|EAW12818.1 afis|EAW17983.1 afla|EED51922.1 afum|EAL91792.1 agla|OJJ80240.1 akaw|GAA91304.1 aluc|OJZ82766.1 anid|CBF79992.1 anig|EHA21209.1 aory|EIT81388.1 asyd|OJJ58238.1 ater|EAU31493.1 atub|OJI88998.1 aver|OJJ02187.1 awen|OJJ41562.1 azon|OJJ48277.1

Group1697: abra|OJJ68554.1 acar|OOF92749.1 acla|EAW12813.1 afis|EAW17978.1 afla|EED51927.1 afum|EAL91787.1 agla|OJJ80235.1 akaw|GAA91299.1 aluc|OJZ82771.1 anid|CBF79976.1 anig|EHA21213.1 aory|EIT81454.1 asyd|OJJ58232.1 ater|EAU31489.1 atub|OJI89002.1 aver|OJJ02192.1 awen|OJJ41567.1 azon|OJJ44836.1

Group1698: abra|OJJ68555.1 acar|OOF92750.1 acla|EAW12812.1 afis|EAW17977.1 afla|EED51928.1 afum|EAL91786.1 agla|OJJ80234.1 akaw|GAA91298.1 aluc|OJZ82772.1 anid|CBF79975.1 anig|EHA21214.1 aory|EIT81416.1 asyd|OJJ58231.1 ater|EAU31488.1 atub|OJI89003.1 aver|OJJ02193.1 awen|OJJ41568.1 azon|OJJ44837.1

Group1699: abra|OJJ68556.1 acar|OOF92751.1 acla|EAW12811.1 afis|EAW17976.1 afla|EED51929.1 afum|EAL91785.1 agla|OJJ80233.1 akaw|GAA91297.1 aluc|OJZ82773.1 anid|CBF79973.1 anig|EHA21215.1 aory|EIT81423.1 asyd|OJJ58230.1 ater|EAU31487.1 atub|OJI89004.1 aver|OJJ02194.1 awen|OJJ41569.1 azon|OJJ44838.1

Group1700: abra|OJJ68557.1 acar|OOF92752.1 acla|EAW12810.1 afis|EAW17975.1 afla|EED51930.1 afum|EAL91784.1 agla|OJJ80232.1 akaw|GAA91296.1 aluc|OJZ82774.1 anid|CBF79972.1 anig|EHA21216.1 aory|EIT81446.1 asyd|OJJ58229.1 ater|EAU31486.1 atub|OJI89005.1 aver|OJJ02195.1 awen|OJJ41570.1 azon|OJJ48283.1

Group1701: abra|OJJ68559.1 acar|OOF92754.1 acla|EAW12806.1 afis|EAW17972.1 afla|EED51932.1 afum|EAL91781.1 agla|OJJ80230.1 akaw|GAA91294.1 aluc|OJZ82775.1 anid|CBF79968.1 anig|EHA21218.1 aory|EIT81462.1 asyd|OJJ58227.1 ater|EAU31482.1 atub|OJI89007.1 aver|OJJ02197.1 awen|OJJ41572.1 azon|OJJ48285.1

Group1702: abra|OJJ68561.1 acar|OOF92755.1 acla|EAW12804.1 afis|EAW17970.1 afla|EED51934.1 afum|EAL91779.1 agla|OJJ80228.1 akaw|GAA91292.1 aluc|OJZ82777.1 anid|CBF79964.1 anig|EHA21220.1

aory|EIT81427.1 asyd|OJJ58224.1 ater|EAU31480.1 atub|OJI89009.1 aver|OJJ02198.1 awen|OJJ41574.1 azon|OJJ48287.1

Group1703: abra|OJJ68564.1 acar|OOF92757.1 acla|EAW12802.1 afis|EAW17968.1 afla|EED51937.1 afum|EAL91777.1 agla|OJJ80226.1 akaw|GAA91290.1 aluc|OJZ82779.1 anid|CBF79961.1 anig|EHA21222.1 aory|EIT81398.1 asyd|OJJ58222.1 ater|EAU31478.1 atub|OJI89012.1 aver|OJJ02200.1 awen|OJJ41576.1 azon|OJJ48289.1

Group1704: abra|OJJ68567.1 acar|OOF92761.1 acla|EAW12798.1 afis|EAW17964.1 afla|EED51946.1 afum|EAL91774.1 agla|OJJ80209.1 akaw|GAA91286.1 aluc|OJZ82782.1 anid|CBF79954.1 anig|EHA21225.1 aory|EIT81431.1 asyd|OJJ58218.1 ater|EAU31450.1 atub|OJI89015.1 aver|OJJ02204.1 awen|OJJ41580.1 azon|OJJ48293.1

Group1705: abra|OJJ68599.1 acar|OOF92109.1 acla|EAW12640.1 afis|EAW18045.1 afla|EED51501.1 afum|EAL91857.1 agla|OJJ85416.1 akaw|GAA91923.1 aluc|OJZ81957.1 anid|CBF79779.1 anig|EHA22290.1 aory|EIT80534.1 asyd|OJJ58121.1 ater|EAU31278.1 atub|OJI84508.1 aver|OJJ04284.1 awen|OJJ41489.1 azon|OJJ48219.1

Group1706: abra|OJJ68601.1 acar|OOF92111.1 acla|EAW12638.1 afis|EAW18047.1 afla|EED51499.1 afum|EAL91859.1 agla|OJJ85418.1 akaw|GAA91925.1 aluc|OJZ81955.1 anid|CBF79776.1 anig|EHA22292.1 aory|EIT80455.1 asyd|OJJ58119.1 ater|EAU31276.1 atub|OJI84506.1 aver|OJJ04286.1 awen|OJJ41487.1 azon|OJJ48217.1

Group1707: abra|OJJ68605.1 acar|OOF92114.1 acla|EAW12588.1 afis|EAW18087.1 afla|EED51483.1 afum|EAL91899.1 agla|OJJ85421.1 akaw|GAA92916.1 aluc|OJZ81952.1 anid|CBF79764.1 anig|EHA22301.1 aory|EIT80442.1 asyd|OJJ58103.1 ater|EAU30435.1 atub|OJI84502.1 aver|OJJ04301.1 awen|OJJ41480.1 azon|OJJ48212.1

Group1708: abra|OJJ68606.1 acar|OOF92117.1 acla|EAW12587.1 afis|EAW18086.1 afla|EED51480.1 afum|EAL91898.1 agla|OJJ85422.1 akaw|GAA92915.1 aluc|OJZ81948.1 anid|CBF79762.1 anig|EHA22302.1 aory|EIT80466.1 asyd|OJJ58102.1 ater|EAU30436.1 atub|OJI84501.1 aver|OJJ04302.1 awen|OJJ41479.1 azon|OJJ48211.1

Group1710: abra|OJJ68608.1 acar|OOF92119.1 acla|EAW12594.1 afis|EAW18084.1 afla|EED51478.1 afum|EAL91896.1 agla|OJJ85424.1 akaw|GAA92913.1 aluc|OJZ81946.1 anid|CBF86386.1 anig|EHA22303.1 aory|EIT80678.1 asyd|OJJ64959.1 ater|EAU30438.1 atub|OJI84499.1 aver|OJI99081.1 awen|OJJ41477.1 azon|OJJ48209.1

Group1711: abra|OJJ68610.1 acar|OOF92120.1 acla|EAW12601.1 afis|EAW18083.1 afla|EED51477.1 afum|EAL91895.1 agla|OJJ85446.1 akaw|GAA92912.1 aluc|OJZ81945.1 anid|CBF86388.1 anig|EHA22304.1 aory|EIT80541.1 asyd|OJJ64958.1 ater|EAU30439.1 atub|OJI84498.1 aver|OJI99080.1 awen|OJJ41472.1 azon|OJJ48931.1

Group1712: abra|OJJ68611.1 acar|OOF92121.1 acla|EAW12602.1 afis|EAW18082.1 afla|EED51476.1 afum|EAL91894.1 agla|OJJ85447.1 akaw|GAA92911.1 aluc|OJZ81944.1 anid|CBF86390.1 anig|EHA22305.1 aory|EIT80460.1 asyd|OJJ64957.1 ater|EAU30440.1 atub|OJI84497.1 aver|OJI99079.1 awen|OJJ41470.1 azon|OJJ48927.1

Group1713: abra|OJJ68612.1 acar|OOF92122.1 acla|EAW12603.1 afis|EAW18081.1 afla|EED51475.1 afum|EAL91893.1 agla|OJJ87374.1 akaw|GAA92910.1 aluc|OJZ81943.1 anid|CBF86391.1 anig|EHA22306.1 aory|EIT80723.1 asyd|OJJ64956.1 ater|EAU30441.1 atub|OJI84496.1 aver|OJI99078.1 awen|OJJ41469.1 azon|OJJ48926.1

Group1714: abra|OJJ68613.1 acar|OOF92123.1 acla|EAW12604.1 afis|EAW18080.1 afla|EED51474.1 afum|EAL91892.1 agla|OJJ87373.1 akaw|GAA92909.1 aluc|OJZ81942.1 anid|CBF86393.1 anig|EHA22307.1 aory|EIT80697.1 asyd|OJJ64955.1 ater|EAU30442.1 atub|OJI84495.1 aver|OJI99077.1 awen|OJJ41468.1 azon|OJJ48925.1

Group1715: abra|OJJ68615.1 acar|OOF92126.1 acla|EAW12607.1 afis|EAW18077.1 afla|EED51471.1 afum|EAL91889.1 agla|OJJ87370.1 akaw|GAA92906.1 aluc|OJZ81939.1 anid|CBF86404.1 anig|EHA22309.1 aory|EIT80562.1 asyd|OJJ64951.1 ater|EAU30446.1 atub|OJI84492.1 aver|OJI99073.1 awen|OJJ41466.1 azon|OJJ48923.1

Group1716: abra|OJJ68617.1 acar|OOF92128.1 acla|EAW12609.1 afis|EAW18075.1 afla|EED51467.1 afum|EAL91887.1 agla|OJJ87367.1 akaw|GAA92904.1 aluc|OJZ81937.1 anid|CBF86400.1 anig|EHA22311.1 aory|EIT80468.1 asyd|OJJ64949.1 ater|EAU30452.1 atub|OJI84490.1 aver|OJI99071.1 awen|OJJ41464.1 azon|OJJ48919.1

Group1717: abra|OJJ68618.1 acar|OOF92129.1 acla|EAW12610.1 afis|EAW18074.1 afla|EED51464.1 afum|EAL91886.1 agla|OJJ87366.1 akaw|GAA92903.1 aluc|OJZ81936.1 anid|CBF86398.1 anig|EHA22312.1 aory|EIT80711.1 asyd|OJJ64948.1 ater|EAU30454.1 atub|OJI84489.1 aver|OJI99070.1 awen|OJJ41463.1 azon|OJJ48918.1

Group1718: abra|OJJ68619.1 acar|OOF92130.1 acla|EAW12611.1 afis|EAW18073.1 afla|EED51463.1 afum|EAL91885.1 agla|OJJ87365.1 akaw|GAA92899.1 aluc|OJZ81935.1 anid|CBF86406.1 anig|EHA22313.1 aory|EIT80619.1 asyd|OJJ64947.1 ater|EAU30455.1 atub|OJI84488.1 aver|OJI99069.1 awen|OJJ41462.1 azon|OJJ48917.1

Group1719: abra|OJJ68620.1 acar|OOF92131.1 acla|EAW13703.1 afis|EAW19885.1 afla|EED57031.1 afum|EAL85012.1 agla|OJJ89079.1 akaw|GAA92898.1 aluc|OJZ81934.1 anid|CBF82756.1 anig|EHA22314.1 aory|EIT82313.1 asyd|OJJ61608.1 ater|EAU33697.1 atub|OJI84487.1 aver|OJJ00931.1 awen|OJJ36996.1 azon|OJJ42189.1

Group1720: abra|OJJ68623.1 acar|OOF92133.1 acla|EAW12612.1 afis|EAW18072.1 afla|EED51461.1 afum|EAL91884.1 agla|OJJ87364.1 akaw|GAA92897.1 aluc|OJZ81932.1 anid|CBF86411.1 anig|EHA22316.1 aory|EIT80603.1 asyd|OJJ64944.1 ater|EAU30457.1 atub|OJI84485.1 aver|OJI99066.1 awen|OJJ41459.1 azon|OJJ48916.1

Group1721: abra|OJJ68626.1 acar|OOF92136.1 acla|EAW12613.1 afis|EAW18071.1 afla|EED51460.1 afum|EAL91883.1 agla|OJJ87363.1 akaw|GAA92895.1 aluc|OJZ81929.1 anid|CBF86413.1 anig|EHA22318.1 aory|EIT80647.1 asyd|OJJ64943.1 ater|EAU30458.1 atub|OJI84481.1 aver|OJI99065.1 awen|OJJ41458.1 azon|OJJ48915.1

Group1722: abra|OJJ68636.1 acar|OOF92140.1 acla|EAW12615.1 afis|EAW18069.1 afla|EED51458.1 afum|EAL91881.2 agla|OJJ87356.1 akaw|GAA89775.1 aluc|OJZ81919.1 anid|CBF86416.1 anig|EHA22328.1 aory|EIT80491.1 asyd|OJJ64941.1 ater|EAU30460.1 atub|OJI84471.1 aver|OJI99063.1 awen|OJJ41456.1 azon|OJJ48913.1

Group1723: abra|OJJ68637.1 acar|OOF92141.1 acla|EAW12614.1 afis|EAW18070.1 afla|EED51459.1 afum|EAL91882.1 agla|OJJ87357.1 akaw|GAA89776.1 aluc|OJZ81918.1 anid|CBF86415.1 anig|EHA22329.1 aory|EIT80742.1 asyd|OJJ64942.1 ater|EAU30459.1 atub|OJI84470.1 aver|OJI99064.1 awen|OJJ41457.1 azon|OJJ48914.1

Group1724: abra|OJJ68642.1 acar|OOF92146.1 acla|EAW12620.1 afis|EAW18064.1 afla|EED51433.1 afum|EAL91876.1 akaw|GAA89782.1 aluc|OJZ81910.1 anid|CBF86424.1 anig|EHA22334.1 aory|EIT80728.1 asyd|OJJ64937.1 ater|EAU30467.1 atub|OJI84463.1 aver|OJI99058.1 awen|OJJ41451.1 azon|OJJ48909.1 agla|OJJ87350.1

Group1725: abra|OJJ68643.1 acar|OOF92145.1 acla|EAW12621.1 afis|EAW18063.1 afla|EED51432.1 afum|EAL91875.1 agla|OJJ87349.1 akaw|GAA89783.1 aluc|OJZ81908.1 anid|CBF86426.1 anig|EHA22335.1 aory|EIT80621.1 asyd|OJJ64936.1 ater|EAU30468.1 atub|OJI84461.1 aver|OJI99057.1 awen|OJJ41449.1 azon|OJJ48908.1

Group1726: abra|OJJ68647.1 acar|OOF92149.1 acla|EAW12623.1 afis|EAW18061.1 afla|EED51429.1 afum|EAL91872.1 agla|OJJ87325.1 akaw|GAA89787.1 aluc|OJZ81904.1 anid|CBF86430.1 anig|EHA22339.1 aory|EIT80637.1 asyd|OJJ64934.1 ater|EAU30473.1 atub|OJI84457.1 aver|OJI99054.1 awen|OJJ41445.1 azon|OJJ48907.1

Group1727: abra|OJJ68648.1 acar|OOF92150.1 acla|EAW12624.1 afis|EAW18060.1 afla|EED51428.1 afum|EAL91871.1 agla|OJJ87326.1 akaw|GAA89788.1 aluc|OJZ81903.1 anid|CBF86432.1 anig|EHA22340.1 aory|EIT80552.1 asyd|OJJ64933.1 ater|EAU30474.1 atub|OJI84456.1 aver|OJI99053.1 awen|OJJ41444.1 azon|OJJ48905.1

Group1728: abra|OJJ68649.1 acar|OOF92151.1 acla|EAW12625.1 afis|EAW18059.1 afla|EED51427.1 afum|EAL91870.1 agla|OJJ87327.1 akaw|GAA89789.1 aluc|OJZ81902.1 anid|CBF86434.1 anig|EHA22341.1 aory|EIT80495.1 asyd|OJJ64932.1 ater|EAU30475.1 atub|OJI84455.1 aver|OJI99051.1 awen|OJJ41442.1 azon|OJJ48904.1

Group1729: abra|OJJ68650.1 acar|OOF92152.1 acla|EAW12626.1 afis|EAW18058.1 afla|EED51426.1 afum|EBA27442.1 agla|OJJ87328.1 akaw|GAA89790.1 aluc|OJZ81901.1 anid|CBF86435.1 anig|EHA22342.1 aory|EIT80761.1 asyd|OJJ64931.1 ater|EAU30476.1 atub|OJI84454.1 aver|OJI99050.1 awen|OJJ41441.1 azon|OJJ48903.1

Group1730: abra|OJJ68652.1 acar|OOF92153.1 acla|EAW12628.1 afis|EAW18056.1 afla|EED51423.1 afum|EAL91867.1 agla|OJJ87330.1 akaw|GAA89792.1 aluc|OJZ81899.1 anid|CBF86443.1 anig|EHA22344.1 aory|EIT80509.1 asyd|OJJ64928.1 ater|EAU30478.1 atub|OJI84452.1 aver|OJI99047.1 awen|OJJ41439.1 azon|OJJ48901.1

Group1731: abra|OJJ68655.1 acar|OOF92155.1 acla|EAW12630.1 afis|EAW18054.1 afla|EED51421.1 afum|EAL91865.1 agla|OJJ87335.1 akaw|GAA89794.1 aluc|OJZ81897.1 anid|CBF86438.1 anig|EHA22345.1 aory|EIT80746.1 asyd|OJJ64927.1 ater|EAU30479.1 atub|OJI84450.1 aver|OJI99045.1 awen|OJJ41436.1 azon|OJJ48900.1

Group1732: abra|OJJ68656.1 acar|OOF92156.1 acla|EAW12632.1 afis|EAW18052.1 afla|EED51418.1 afum|EAL91863.1 agla|OJJ87339.1 akaw|GAA89795.1 aluc|OJZ81896.1 anid|CBF86439.1 anig|EHA22346.1

aory|EIT80553.1 asyd|OJJ64925.1 ater|EAU30480.1 atub|OJI84449.1 aver|OJI99044.1 awen|OJJ41433.1 azon|OJJ48898.1

Group1733: abra|OJJ68657.1 acar|OOF92157.1 acla|EAW12633.1 afis|EAW18051.1 afla|EED51417.1 afum|EAL91862.1 agla|OJJ87340.1 akaw|GAA89796.1 aluc|OJZ81895.1 anid|CBF86440.1 anig|EHA22347.1 aory|EIT80630.1 asyd|OJJ64924.1 ater|EAU30481.1 atub|OJI84448.1 aver|OJI99043.1 awen|OJJ41432.1 azon|OJJ48897.1

Group1734: abra|OJJ68659.1 acar|OOF92159.1 acla|EAW12590.1 afis|EAW24059.1 afla|EED51416.1 afum|EAL91914.1 agla|OJJ87341.1 akaw|GAA89797.1 aluc|OJZ81894.1 anid|CBF86444.1 anig|EHA22348.1 aory|EIT80652.1 asyd|OJJ64923.1 ater|EAU30482.1 atub|OJI84447.1 aver|OJI99042.1 awen|OJJ41431.1 azon|OJJ48896.1

Group1735: abra|OJJ68660.1 acar|OOF92158.1 acla|EAW12591.1 afis|EAW24060.1 afla|EED51415.1 afum|EAL91915.1 agla|OJJ87342.1 akaw|GAA89798.1 aluc|OJZ81893.1 anid|CBF86446.1 anig|EHA22349.1 aory|EIT80731.1 asyd|OJJ64922.1 ater|EAU30483.1 atub|OJI84446.1 aver|OJI99041.1 awen|OJJ41430.1 azon|OJJ48895.1

Group1736: abra|OJJ68662.1 acar|OOF92162.1 acla|EAW12586.1 afis|EAW24062.1 afla|EED51412.1 afum|EAL91917.1 agla|OJJ87344.1 akaw|GAA89802.1 aluc|OJZ81889.1 anid|CBF86449.1 anig|EHA22352.1 aory|EIT80533.1 asyd|OJJ64920.1 ater|EAU30484.1 atub|OJI84442.1 aver|OJI99040.1 awen|OJJ41428.1 azon|OJJ48893.1

Group1737: abra|OJJ68664.1 acar|OOF92165.1 acla|EAW12585.1 afis|EAW24063.1 afla|EED51411.1 afum|EAL91918.1 agla|OJJ87345.1 akaw|GAA89805.1 aluc|OJZ81886.1 anid|CBF86451.1 anig|EHA22354.1 aory|EIT80561.1 asyd|OJJ64919.1 ater|EAU30485.1 atub|OJI84439.1 aver|OJI99039.1 awen|OJJ41426.1 azon|OJJ48891.1

Group1738: abra|OJJ68665.1 acar|OOF92167.1 acla|EAW12584.1 afis|EAW24064.1 afla|EED51410.1 afum|EAL91919.1 agla|OJJ87346.1 akaw|GAA89806.1 aluc|OJZ81885.1 anid|CBF86452.1 anig|EHA22355.1 aory|EIT80617.1 asyd|OJJ64918.1 ater|EAU30486.1 atub|OJI84438.1 aver|OJI99038.1 awen|OJJ41425.1 azon|OJJ48890.1

Group1739: abra|OJJ68666.1 acar|OOF92166.1 acla|EAW12583.1 afis|EAW24065.1 afla|EED51409.1 afum|EAL91920.1 agla|OJJ87347.1 akaw|GAA89807.1 aluc|OJZ81884.1 anid|CBF86454.1 anig|EHA22356.1 aory|EIT80471.1 asyd|OJJ64917.1 ater|EAU30487.1 atub|OJI84437.1 aver|OJI99037.1 awen|OJJ41424.1 azon|OJJ48889.1

Group1740: abra|OJJ68670.1 acar|OOF92170.1 acla|EAW12577.1 afis|EAW24071.1 afla|EED51404.1 afum|EAL91927.1 agla|OJJ87312.1 akaw|GAA89811.1 aluc|OJZ81879.1 anid|CBF86462.1 anig|EHA22359.1 aory|EIT80762.1 asyd|OJJ64912.1 ater|EAU30492.1 atub|OJI84433.1 aver|OJI99033.1 awen|OJJ41419.1 azon|OJJ48884.1

Group1741: abra|OJJ68677.1 acar|OOF92179.1 acla|EAW12570.1 afis|EAW24080.1 afla|EED51393.1 afum|EAL91935.1 agla|OJJ87151.1 akaw|GAA89830.1 aluc|OJZ81857.1 anid|CBF86478.1 anig|EHA22367.1 aory|EIT80640.1 asyd|OJJ64902.1 ater|EAU30501.1 atub|OJI84425.1 aver|OJI99024.1 awen|OJJ41418.1 azon|OJJ48875.1

Group1742: abra|OJJ68678.1 acar|OOF92180.1 acla|EAW12569.1 afis|EAW24081.1 afla|EED51392.1 afum|EAL91936.1 agla|OJJ87152.1 akaw|GAA89831.1 aluc|OJZ81856.1 anid|CBF86480.1 anig|EHA22368.1 aory|EIT80524.1 asyd|OJJ64901.1 ater|EAU30504.1 atub|OJI84424.1 aver|OJI99022.1 awen|OJJ41407.1 azon|OJJ48873.1

Group1743: abra|OJJ68681.1 acar|OOF92183.1 acla|EAW12566.1 afis|EAW24084.1 afla|EED51389.1 afum|EAL91939.1 agla|OJJ87154.1 akaw|GAA89834.1 aluc|OJZ81852.1 anid|CBF86484.1 anig|EHA22370.1 aory|EIT80477.1 asyd|OJJ64899.1 ater|EAU30507.1 atub|OJI84421.1 aver|OJI99020.1 awen|OJJ41405.1 azon|OJJ48871.1

Group1744: abra|OJJ68682.1 acar|OOF92184.1 acla|EAW12565.1 afis|EAW24085.1 afla|EED51388.1 afum|EAL91940.1 agla|OJJ87155.1 akaw|GAA89835.1 aluc|OJZ81851.1 anid|CBF86486.1 anig|EHA22371.1 aory|EIT80535.1 asyd|OJJ64898.1 ater|EAU30508.1 atub|OJI84420.1 aver|OJI99019.1 awen|OJJ41404.1 azon|OJJ48870.1

Group1745: abra|OJJ68684.1 acar|OOF92186.1 acla|EAW12563.1 afis|EAW24087.1 afla|EED51386.1 afum|EAL91942.1 agla|OJJ87157.1 akaw|GAA89837.1 aluc|OJZ81849.1 anid|CBF86489.1 anig|EHA22373.1 aory|EIT80476.1 asyd|OJJ64896.1 ater|EAU30511.1 atub|OJI84418.1 aver|OJI99017.1 awen|OJJ41402.1 azon|OJJ48868.1

Group1746: abra|OJJ68685.1 acar|OOF92187.1 acla|EAW12562.1 afis|EAW24088.1 afla|EED46851.1 afum|EAL91943.1 agla|OJJ87158.1 akaw|GAA89838.1 aluc|OJZ81848.1 anid|CBF86491.1 anig|EHA22374.1 aory|EIT72584.1 asyd|OJJ64895.1 ater|EAU30512.1 atub|OJI84417.1 aver|OJI99016.1 awen|OJJ41401.1 azon|OJJ48867.1

Group1747: abra|OJJ68686.1 acar|OOF92188.1 acla|EAW12561.1 afis|EAW24089.1 afla|EED46850.1 afum|EAL91944.1 agla|OJJ87159.1 akaw|GAA89839.1 aluc|OJZ81847.1 anid|CBF86493.1 anig|EHA22375.1 aory|EIT72520.1 asyd|OJJ64894.1 ater|EAU30513.1 atub|OJI84416.1 aver|OJI99015.1 awen|OJJ41400.1 azon|OJJ48866.1

Group1748: abra|OJJ68689.1 acar|OOF92193.1 acla|EAW12555.1 afis|EAW24096.1 afla|EED46838.1 afum|EAL91950.1 agla|OJJ87169.1 akaw|GAA89844.1 aluc|OJZ81843.1 anid|CBF86498.1 anig|EHA22379.1 aory|EIT72590.1 asyd|OJJ64889.1 ater|EAU30523.1 atub|OJI84411.1 aver|OJI99003.1 awen|OJJ41394.1 azon|OJJ48862.1

Group1749: abra|OJJ68690.1 acar|OOF92194.1 acla|EAW12554.1 afis|EAW24097.1 afla|EED46837.1 afum|EAL91951.2 agla|OJJ87171.1 akaw|GAA89845.1 aluc|OJZ81842.1 anid|CBF86500.1 anig|EHA22380.1 aory|EIT72693.1 asyd|OJJ64888.1 ater|EAU30524.1 atub|OJI84410.1 aver|OJI99002.1 awen|OJJ41392.1 azon|OJJ48861.1

Group1750: abra|OJJ68691.1 acar|OOF92195.1 acla|EAW12553.1 afis|EAW24098.1 afla|EED46836.1 afum|EAL91952.1 agla|OJJ87172.1 akaw|GAA89846.1 aluc|OJZ81841.1 anid|CBF86501.1 anig|EHA22381.1 aory|EIT72579.1 asyd|OJJ64887.1 ater|EAU30525.1 atub|OJI84409.1 aver|OJI99001.1 awen|OJJ41391.1 azon|OJJ48860.1

Group1751: abra|OJJ68692.1 acar|OOF92196.1 acla|EAW12552.1 afis|EAW24099.1 afla|EED46835.1 afum|EAL91953.2 agla|OJJ87173.1 akaw|GAA89847.1 aluc|OJZ81840.1 anid|CBF86503.1 anig|EHA22382.1 aory|EIT72533.1 asyd|OJJ64886.1 ater|EAU30526.1 atub|OJI84408.1 aver|OJI99000.1 awen|OJJ41390.1 azon|OJJ48859.1

Group1752: abra|OJJ68693.1 acar|OOF92197.1 acla|EAW12551.1 afis|EAW24100.1 afla|EED46834.1 afum|EAL91954.1 agla|OJJ87174.1 akaw|GAA89848.1 aluc|OJZ81838.1 anid|CBF86505.1 anig|EHA22383.1 aory|EIT72706.1 asyd|OJJ64885.1 ater|EAU30527.1 atub|OJI84407.1 aver|OJI98999.1 awen|OJJ41389.1 azon|OJJ48858.1

Group1753: abra|OJJ68695.1 acar|OOF92199.1 acla|EAW12549.1 afis|EAW24101.1 afla|EED46832.1 afum|EAL91955.1 agla|OJJ87176.1 akaw|GAA89850.1 aluc|OJZ81836.1 anid|CBF86507.1 anig|EHA22385.1 aory|EIT72575.1 asyd|OJJ64883.1 ater|EAU30529.1 atub|OJI84405.1 aver|OJI98997.1 awen|OJJ41387.1 azon|OJJ48856.1

Group1754: abra|OJJ68696.1 acar|OOF92200.1 acla|EAW12548.1 afis|EAW24102.1 afla|EED46831.1 afum|EAL91956.1 agla|OJJ87177.1 akaw|GAA89851.1 aluc|OJZ81835.1 anid|CBF86509.1 anig|EHA22386.1 aory|EIT72546.1 asyd|OJJ64882.1 ater|EAU30530.1 atub|OJI84404.1 aver|OJI98996.1 awen|OJJ41386.1 azon|OJJ48855.1

Group1755: abra|OJJ68697.1 acar|OOF92201.1 acla|EAW12547.1 afis|EAW24103.1 afla|EED46830.1 afum|EAL91957.1 agla|OJJ87178.1 akaw|GAA89852.1 aluc|OJZ81834.1 anid|CBF86511.1 anig|EHA22387.1 aory|EIT72771.1 asyd|OJJ64881.1 ater|EAU30531.1 atub|OJI84403.1 aver|OJI98995.1 awen|OJJ41385.1 azon|OJJ48854.1

Group1756: abra|OJJ68698.1 acar|OOF92202.1 acla|EAW12546.1 afis|EAW24104.1 afla|EED46829.1 afum|EAL91958.1 agla|OJJ87179.1 akaw|GAA89853.1 aluc|OJZ81833.1 anid|CBF86513.1 anig|EHA22388.1 aory|EIT72709.1 asyd|OJJ64880.1 ater|EAU30532.1 atub|OJI84402.1 aver|OJI98994.1 awen|OJJ41384.1 azon|OJJ48853.1

Group1757: abra|OJJ68703.1 acar|OOF92205.1 acla|EAW12542.1 afis|EAW24107.1 afla|EED46826.1 afum|EAL91961.1 agla|OJJ87182.1 akaw|GAA89857.1 aluc|OJZ81828.1 anid|CBF86518.1 anig|EHA22391.1 aory|EIT72757.1 asyd|OJJ64877.1 ater|EAU30535.1 atub|OJI84398.1 aver|OJI98990.1 awen|OJJ41381.1 azon|OJJ48850.1

Group1758: abra|OJJ68704.1 acar|OOF92206.1 acla|EAW12541.1 afis|EAW24108.1 afla|EED46825.1 afum|EAL91962.1 agla|OJJ87183.1 akaw|GAA89858.1 aluc|OJZ81826.1 anid|CBF86520.1 anig|EHA22392.1 aory|EIT72547.1 asyd|OJJ64876.1 ater|EAU30536.1 atub|OJI84396.1 aver|OJI98989.1 awen|OJJ41380.1 azon|OJJ48848.1

Group1759: abra|OJJ68706.1 acar|OOF92208.1 acla|EAW12539.1 afis|EAW24110.1 afla|EED46822.1 afum|EAL91964.1 agla|OJJ87185.1 akaw|GAA89860.1 aluc|OJZ81824.1 anid|CBF86523.1 anig|EHA22394.1 aory|EIT72674.1 asyd|OJJ64874.1 ater|EAU30538.1 atub|OJI84394.1 aver|OJI98987.1 awen|OJJ41378.1 azon|OJJ48846.1

Group1760: abra|OJJ68709.1 acar|OOF92211.1 acla|EAW12536.1 afis|EAW24113.1 afla|EED46820.1 afum|EAL91967.1 agla|OJJ87190.1 akaw|GAA89863.1 aluc|OJZ81818.1 anid|CBF86529.1 anig|EHA22397.1 aory|EIT72599.1 asyd|OJJ64871.1 ater|EAU30541.1 atub|OJI84391.1 aver|OJI98983.1 awen|OJJ41375.1 azon|OJJ48843.1

Group1761: abra|OJJ68710.1 acar|OOF92212.1 acla|EAW12535.1 afis|EAW24114.1 afla|EED46819.1 afum|EAL91968.1 agla|OJJ87191.1 akaw|GAA89864.1 aluc|OJZ81817.1 anid|CBF86531.1 anig|EHA22398.1

aory|EIT72684.1 asyd|OJJ64870.1 ater|EAU30542.1 atub|OJI84390.1 aver|OJI98982.1 awen|OJJ41374.1 azon|OJJ48842.1

Group1762: abra|OJJ68717.1 acar|OOF92217.1 acla|EAW12529.1 afis|EAW24120.1 afla|EED46812.1 afum|EAL91975.1 agla|OJJ87194.1 akaw|GAA89869.1 aluc|OJZ81812.1 anid|CBF86542.1 anig|EHA22405.1 aory|EIT72532.1 asyd|OJJ64862.1 ater|EAU30548.1 atub|OJI84384.1 aver|OJI98973.1 awen|OJJ41369.1 azon|OJJ48836.1

Group1763: abra|OJJ68719.1 acar|OOF92218.1 acla|EAW12528.1 afis|EAW24121.1 afla|EED46810.1 afum|EAL91977.1 agla|OJJ87196.1 akaw|GAA89870.1 aluc|OJZ81810.1 anid|CBF86546.1 anig|EHA22407.1 aory|EIT72617.1 asyd|OJJ64860.1 ater|EAU30549.1 atub|OJI84382.1 aver|OJI98972.1 awen|OJJ41368.1 azon|OJJ48835.1

Group1764: abra|OJJ68722.1 acar|OOF92221.1 acla|EAW12524.1 afis|EAW24124.1 afla|EED46809.1 afum|EAL91980.1 agla|OJJ87199.1 akaw|GAA89872.1 aluc|OJZ81807.1 anid|CBF86551.1 anig|EHA22410.1 aory|EIT72759.1 asyd|OJJ64857.1 ater|EAU30551.1 atub|OJI84379.1 aver|OJI98969.1 awen|OJJ41365.1 azon|OJJ48832.1

Group1765: abra|OJJ68723.1 acar|OOF92222.1 acla|EAW12523.1 afis|EAW24125.1 afla|EED46808.1 afum|EAL91981.1 agla|OJJ87200.1 akaw|GAA89873.1 aluc|OJZ81806.1 anid|CBF86553.1 anig|EHA22411.1 aory|EIT72530.1 asyd|OJJ64856.1 ater|EAU30552.1 atub|OJI84378.1 aver|OJI98968.1 awen|OJJ41364.1 azon|OJJ48831.1

Group1766: abra|OJJ68725.1 acar|OOF92224.1 acla|EAW12521.1 afis|EAW24126.1 afla|EED46806.1 afum|EAL91982.1 agla|OJJ87202.1 akaw|GAA89876.1 aluc|OJZ81803.1 anid|CBF86555.1 anig|EHA22413.1 aory|EIT72636.1 asyd|OJJ64855.1 ater|EAU30553.1 atub|OJI84375.1 aver|OJI98967.1 awen|OJJ41363.1 azon|OJJ48829.1

Group1767: abra|OJJ68728.1 acar|OOF92225.1 acla|EAW12520.1 afis|EAW24127.1 afla|EED46805.1 afum|EAL91983.1 agla|OJJ87203.1 akaw|GAA89877.1 aluc|OJZ81802.1 anid|CBF86557.1 anig|EHA22414.1 aory|EIT72714.1 asyd|OJJ64854.1 ater|EAU30557.1 atub|OJI84374.1 aver|OJI98966.1 awen|OJJ41362.1 azon|OJJ48827.1

Group1768: abra|OJJ68729.1 acar|OOF92226.1 acla|EAW12519.1 afis|EAW24128.1 afla|EED46804.1 afum|EAL91984.1 agla|OJJ87204.1 akaw|GAA89878.1 aluc|OJZ81801.1 anid|CBF86559.1 anig|EHA22415.1 aory|EIT72675.1 asyd|OJJ64853.1 ater|EAU30558.1 atub|OJI84372.1 aver|OJI98965.1 awen|OJJ41361.1 azon|OJJ48825.1

Group1769: abra|OJJ68730.1 acar|OOF92228.1 acla|EAW12518.1 afis|EAW24129.1 afla|EED46803.1 afum|EAL91985.1 agla|OJJ87205.1 akaw|GAA89879.1 aluc|OJZ81800.1 anid|CBF86560.1 anig|EHA22416.1 aory|EIT72669.1 asyd|OJJ64852.1 ater|EAU30559.1 atub|OJI84371.1 aver|OJI98964.1 awen|OJJ41360.1 azon|OJJ48824.1

Group1770: abra|OJJ68734.1 acar|OOF92230.1 acla|EAW12513.1 afis|EAW24133.1 afla|EED46800.1 afum|EAL91988.1 agla|OJJ87210.1 akaw|GAA89882.1 aluc|OJZ81796.1 anid|CBF86566.1 anig|EHA22420.1 aory|EIT72682.1 asyd|OJJ64849.1 ater|EAU30562.1 atub|OJI84367.1 aver|OJI98961.1 awen|OJJ41356.1 azon|OJJ48819.1

Group1771: abra|OJJ68735.1 acar|OOF92231.1 acla|EAW12512.1 afis|EAW24134.1 afla|EED46799.1 afum|EAL91989.2 agla|OJJ87211.1 akaw|GAA89883.1 aluc|OJZ81794.1 anid|CBF86568.1 anig|EHA22421.1 aory|EIT72755.1 asyd|OJJ64848.1 ater|EAU30563.1 atub|OJI84366.1 aver|OJI98960.1 awen|OJJ41355.1 azon|OJJ48818.1

Group1772: abra|OJJ68736.1 acar|OOF92232.1 acla|EAW12511.1 afis|EAW24135.1 afla|EED46798.1 afum|EAL91990.1 agla|OJJ87212.1 akaw|GAA89884.1 aluc|OJZ81793.1 anid|CBF86570.1 anig|EHA22422.1 aory|EIT72542.1 asyd|OJJ64847.1 ater|EAU30564.1 atub|OJI84365.1 aver|OJI98959.1 awen|OJJ41354.1 azon|OJJ48817.1

Group1773: abra|OJJ68738.1 acar|OOF92234.1 afis|EAW24137.1 afla|EED46797.1 afum|EAL91992.1 agla|OJJ87215.1 akaw|GAA89886.1 aluc|OJZ81789.1 anid|CBF77943.1 anig|EHA22426.1 aory|EIT72631.1 asyd|OJJ55282.1 ater|EAU30566.1 atub|OJI84363.1 aver|OJJ06946.1 awen|OJJ41352.1 azon|OJJ48815.1 acla|EAW12509.1

Group1774: abra|OJJ68739.1 acar|OOF92235.1 acla|EAW12508.1 afis|EAW24138.1 afla|EED46796.1 afum|EAL91993.1 agla|OJJ87216.1 akaw|GAA89887.1 aluc|OJZ81788.1 anid|CBF77941.1 anig|EHA22427.1 aory|EIT72772.1 asyd|OJJ55283.1 ater|EAU30567.1 atub|OJI84362.1 aver|OJJ06945.1 awen|OJJ41351.1 azon|OJJ48814.1

Group1775: abra|OJJ68742.1 acar|OOF90307.1 acla|EAW12506.1 afis|EAW24141.1 afla|EED46793.1 afum|EAL91996.1 agla|OJJ87220.1 akaw|GAA89889.1 aluc|OJZ81784.1 anid|CBF77936.1 anig|EHA22431.1 aory|EIT72574.1 asyd|OJJ55286.1 ater|EAU30570.1 atub|OJI84358.1 aver|OJJ06942.1 awen|OJJ41347.1 azon|OJJ48811.1

Group1776: abra|OJJ68743.1 acar|OOF90308.1 acla|EAW12505.1 afis|EAW24142.1 afla|EED46792.1 afum|EAL91997.1 agla|OJJ87221.1 akaw|GAA89890.1 aluc|OJZ81783.1 anid|CBF77934.1 anig|EHA22432.1 aory|EIT72571.1 asyd|OJJ55287.1 ater|EAU30571.1 atub|OJI84357.1 aver|OJJ06941.1 awen|OJJ41346.1 azon|OJJ48810.1

Group1777: abra|OJJ68748.1 acar|OOF92236.1 acla|EAW12503.1 afis|EAW24143.1 afla|EED46789.1 afum|EAL91998.1 agla|OJJ87222.1 akaw|GAA89892.1 aluc|OJZ81779.1 anid|CBF77930.1 anig|EHA22436.1 aory|EIT72633.1 asyd|OJJ55290.1 ater|EAU30574.1 atub|OJI84353.1 aver|OJJ06939.1 awen|OJJ41345.1 azon|OJJ48809.1

Group1778: abra|OJJ68749.1 acar|OOF92237.1 acla|EAW12502.1 afis|EAW24144.1 afla|EED46788.1 afum|EAL91999.1 agla|OJJ87223.1 akaw|GAA89893.1 aluc|OJZ81778.1 anid|CBF77928.1 anig|EHA22438.1 aory|EIT72562.1 asyd|OJJ55291.1 ater|EAU30575.1 atub|OJI84351.1 aver|OJJ06936.1 awen|OJJ41336.1 azon|OJJ48806.1

Group1779: abra|OJJ68750.1 acar|OOF92238.1 acla|EAW12501.1 afis|EAW24145.1 afla|EED46787.1 afum|EAL92000.1 agla|OJJ87224.1 akaw|GAA89894.1 aluc|OJZ81777.1 anid|CBF77926.1 anig|EHA22439.1 aory|EIT72718.1 asyd|OJJ55292.1 ater|EAU30576.1 atub|OJI84350.1 aver|OJJ06935.1 awen|OJJ41335.1 azon|OJJ48804.1

Group1780: abra|OJJ68752.1 acar|OOF92239.1 acla|EAW12499.1 afis|EAW24146.1 afla|EED46783.1 afum|EAL92001.1 agla|OJJ87226.1 akaw|GAA89896.1 aluc|OJZ81775.1 anid|CBF77923.1 anig|EHA22441.1 aory|EIT72696.1 asyd|OJJ55294.1 ater|EAU30578.1 atub|OJI84348.1 aver|OJJ06932.1 awen|OJJ41333.1 azon|OJJ48803.1

Group1781: abra|OJJ68756.1 acar|OOF92244.1 acla|EAW12495.1 afis|EAW24150.1 afla|EED46781.1 afum|EAL92005.1 agla|OJJ89542.1 akaw|GAA89900.1 aluc|OJZ81771.1 anid|CBF77915.1 anig|EHA22445.1 aory|EIT72518.1 asyd|OJJ55297.1 ater|EAU30582.1 atub|OJI84344.1 aver|OJJ06929.1 awen|OJJ41328.1 azon|OJJ48801.1

Group1782: abra|OJJ68763.1 acar|OOF92249.1 acla|EAW12488.1 afis|EAW24157.1 afla|EED46776.1 afum|EAL92012.2 agla|OJJ89524.1 akaw|GAA89907.1 aluc|OJZ81762.1 anid|CBF77900.1 anig|EHA22451.1 aory|EIT72774.1 asyd|OJJ55303.1 ater|EAU30588.1 atub|OJI84334.1 aver|OJJ06923.1 awen|OJJ41321.1 azon|OJJ48794.1

Group1783: abra|OJJ68766.1 acar|OOF92252.1 acla|EAW12485.1 afis|EAW24160.1 afla|EED46773.1 afum|EAL92016.1 agla|OJJ89521.1 akaw|GAA89910.1 aluc|OJZ81759.1 anid|CBF77894.1 anig|EHA22453.1 aory|EIT72644.1 asyd|OJJ55306.1 ater|EAU30591.1 atub|OJI84331.1 aver|OJJ06920.1 awen|OJJ41318.1 azon|OJJ48791.1

Group1784: abra|OJJ68768.1 acar|OOF92253.1 acla|EAW12484.1 afis|EAW24161.1 afla|EED46772.1 afum|EAL92017.1 agla|OJJ89519.1 akaw|GAA89911.1 aluc|OJZ81758.1 anid|CBF77892.1 anig|EHA22454.1 aory|EIT72707.1 asyd|OJJ55307.1 ater|EAU30592.1 atub|OJI84329.1 aver|OJJ06919.1 awen|OJJ41316.1 azon|OJJ48790.1

Group1785: abra|OJJ68769.1 acar|OOF92254.1 acla|EAW12483.1 afis|EAW24162.1 afla|EED46771.1 afum|EAL92018.1 agla|OJJ89518.1 akaw|GAA89912.1 aluc|OJZ81757.1 anid|CBF77890.1 anig|EHA22455.1 aory|EIT72701.1 asyd|OJJ55308.1 ater|EAU30593.1 atub|OJI84328.1 aver|OJJ06918.1 awen|OJJ41315.1 azon|OJJ48789.1

Group1786: abra|OJJ68770.1 acar|OOF92255.1 acla|EAW12482.1 afis|EAW24163.1 afla|EED46770.1 afum|EAL92020.1 agla|OJJ89517.1 akaw|GAA89913.1 aluc|OJZ81756.1 anid|CBF77888.1 anig|EHA22456.1 aory|EIT72763.1 asyd|OJJ55309.1 ater|EAU30594.1 atub|OJI84327.1 aver|OJJ06917.1 awen|OJJ41314.1 azon|OJJ48788.1

Group1787: abra|OJJ68785.1 acar|OOF92269.1 acla|EAW12469.1 afis|EAW24172.1 afla|EED46757.1 afum|EAL92034.1 agla|OJJ89503.1 akaw|GAA89927.1 aluc|OJZ81743.1 anid|CBF77861.1 anig|EHA22469.1 aory|EIT72255.1 asyd|OJJ55323.1 ater|EAU30607.1 atub|OJI84314.1 aver|OJJ06905.1 awen|OJJ41300.1 azon|OJJ48774.1

Group1788: abra|OJJ68788.1 acar|OOF92272.1 acla|EAW12466.1 afis|EAW24175.1 afla|EED46754.1 afum|EAL92037.1 agla|OJJ89499.1 akaw|GAA89929.1 aluc|OJZ81740.1 anid|CBF77855.1 anig|EHA22472.1 aory|EIT72259.1 asyd|OJJ55326.1 ater|EAU30610.1 atub|OJI84311.1 aver|OJJ06902.1 awen|OJJ41295.1 azon|OJJ48771.1

Group1789: abra|OJJ68790.1 acar|OOF92274.1 acla|EAW12463.1 afis|EAW24178.1 afla|EED46751.1 afum|EAL92040.1 agla|OJJ89496.1 akaw|GAA89932.1 aluc|OJZ81736.1 anid|CBF77849.1 anig|EHA22475.1 aory|EIT72252.1 asyd|OJJ55329.1 ater|EAU30613.1 atub|OJI84308.1 aver|OJJ06899.1 awen|OJJ41291.1 azon|OJJ48765.1

Group1790: abra|OJJ68793.1 acar|OOF92278.1 acla|EAW12460.1 afis|EAW24182.1 afla|EED46749.1 afum|EAL92042.1 agla|OJJ89493.1 akaw|GAA89933.1 aluc|OJZ81734.1 anid|CBF77844.1 anig|EHA22478.1

aory|EIT72258.1 asyd|OJJ55332.1 ater|EAU30615.1 atub|OJI84305.1 aver|OJJ06896.1 awen|OJJ41288.1 azon|OJJ48762.1

Group1791: abra|OJJ68794.1 acar|OOF92277.1 acla|EAW12459.1 afis|EAW24183.1 afla|EED46748.1 afum|EAL92043.1 agla|OJJ89492.1 akaw|GAA89934.1 aluc|OJZ81733.1 anid|CBF77842.1 anig|EHA22479.1 aory|EIT72260.1 asyd|OJJ55333.1 ater|EAU30616.1 atub|OJI84304.1 aver|OJJ06895.1 awen|OJJ41287.1 azon|OJJ48769.1

Group1792: abra|OJJ68797.1 acar|OOF92281.1 acla|EAW14657.1 afis|EAW24186.1 afla|EED46746.1 afum|EAL92046.1 agla|OJJ89489.1 akaw|GAA89936.1 aluc|OJZ81730.1 anid|CBF77836.1 anig|EHA22482.1 aory|EIT72257.1 asyd|OJJ55336.1 ater|EAU30619.1 atub|OJI84301.1 aver|OJJ06891.1 awen|OJJ41284.1 azon|OJJ43704.1

Group1794: abra|OJJ68855.1 acar|OOF90533.1 acla|EAW15256.1 afis|EAW20230.1 afla|EED57124.1 afum|EAL85347.2 agla|OJJ84632.1 akaw|GAA90398.1 aluc|OJZ85833.1 anid|CBF85122.1 anig|EHA24730.1 aory|EIT74034.1 asyd|OJJ63044.1 ater|EAU32032.1 atub|OJI79621.1 aver|OJI97919.1 awen|OJJ31233.1 azon|OJJ46156.1

Group1795: abra|OJJ68856.1 acar|OOF90532.1 afla|EED57125.1 afum|EAL85346.1 agla|OJJ84633.1 akaw|GAA90397.1 aluc|OJZ85834.1 anid|CBF85120.1 anig|EHA24729.1 aory|EIT74077.1 ater|EAU32031.1 atub|OJI79622.1 aver|OJI97918.1 awen|OJJ31232.1 azon|OJJ46155.1 acla|EAW15255.1 afis|EAW20229.1 asyd|OJJ63045.1

Group1796: abra|OJJ68859.1 acar|OOF90530.1 acla|EAW15250.1 afis|EAW20218.1 afla|EED57129.1 afum|EAL85333.1 agla|OJJ84638.1 akaw|GAA90394.1 aluc|OJZ85837.1 anid|CBF85115.1 anig|EHA24726.1 aory|EIT74090.1 asyd|OJJ63049.1 ater|EAU32027.1 atub|OJI79625.1 aver|OJI97914.1 awen|OJJ31228.1 azon|OJJ46151.1

Group1797: abra|OJJ68871.1 acar|OOF89965.1 acla|EAW15248.1 afis|EAW20214.1 afla|EED57130.1 afum|EAL85331.1 agla|OJJ84640.1 akaw|GAA90383.1 aluc|OJZ85848.1 anid|CBF85111.1 anig|EHA28098.1 aory|EIT74018.1 asyd|OJJ63051.1 ater|EAU32024.1 atub|OJI79636.1 aver|OJI97912.1 awen|OJJ31220.1 azon|OJJ46137.1

Group1798: abra|OJJ68913.1 acar|OOF91500.1 acla|EAW15323.1 afis|EAW20327.1 afla|EED57093.1 afum|EAL85453.1 agla|OJJ86344.1 akaw|GAA90336.1 aluc|OJZ85896.1 anid|CBF87781.1 anig|EHA24696.1 aory|EIT82282.1 asyd|OJJ63373.1 ater|EAU32055.1 atub|OJI79681.1 aver|OJI97573.1 awen|OJJ39915.1 azon|OJJ45889.1

Group1799: abra|OJJ68918.1 acar|OOF91506.1 acla|EAW14259.1 afis|EAW19021.1 afla|EED56623.1 afum|EAL93752.1 agla|OJJ84614.1 akaw|GAA90331.1 aluc|OJZ85901.1 anid|CBF85579.1 anig|EHA24691.1 aory|EIT79672.1 asyd|OJJ64177.1 ater|EAU32063.1 atub|OJI79686.1 aver|OJI98253.1 awen|OJJ31311.1 azon|OJJ46160.1

Group1800: abra|OJJ68919.1 acar|OOF91507.1 acla|EAW14260.1 afis|EAW19020.1 afla|EED56622.1 afum|EAL93751.1 agla|OJJ84615.1 akaw|GAA90330.1 aluc|OJZ85902.1 anid|CBF85577.1 anig|EHA24690.1 aory|EIT79600.1 asyd|OJJ64178.1 ater|EAU32062.1 atub|OJI79687.1 aver|OJI98254.1 awen|OJJ31310.1 azon|OJJ46161.1

Group1801: abra|OJJ68926.1 acar|OOF91512.1 acla|EAW14248.1 afis|EAW19032.1 afla|EED50385.1 afum|EAL93761.1 agla|OJJ84590.1 akaw|GAA90322.1 aluc|OJZ85911.1 anid|CBF79596.1 anig|EHA24681.1 aory|EIT83141.1 asyd|OJJ57985.1 ater|EAU32071.1 atub|OJI79701.1 aver|OJJ04415.1 awen|OJJ31461.1 azon|OJJ46163.1

Group1803: abra|OJJ68932.1 acar|OOF91515.1 acla|EAW14247.1 afis|EAW19033.1 afla|EED50387.1 afum|EAL93762.2 agla|OJJ84588.1 akaw|GAA90317.1 aluc|OJZ85916.1 anid|CBF79600.1 anig|EHA24675.1 aory|EIT83129.1 asyd|OJJ57990.1 ater|EAU32073.1 atub|OJI79707.1 aver|OJJ04412.1 awen|OJJ31463.1 azon|OJJ46164.1

Group1804: abra|OJJ68933.1 acar|OOF91516.1 acla|EAW14246.1 afla|EED50389.1 agla|OJJ84587.1 akaw|GAA90316.1 aluc|OJZ85917.1 anig|EHA24674.1 aory|EIT83117.1 asyd|OJJ57991.1 ater|EAU32074.1 atub|OJI79710.1 aver|OJJ04411.1 awen|OJJ31464.1 azon|OJJ46166.1 afum|EAL93763.1 anid|CBF79601.1 afis|EAW19034.1

Group1805: abra|OJJ68937.1 acar|OOF91518.1 acla|EAW14243.1 afis|EAW19037.1 afla|EED50391.1 afum|EAL93766.1 agla|OJJ84584.1 akaw|GAA90314.1 aluc|OJZ85922.1 anid|CBF79605.1 anig|EHA24672.1 aory|EIT82128.1 asyd|OJJ57993.1 ater|EAU32076.1 atub|OJI79712.1 aver|OJJ04409.1 awen|OJJ31466.1 azon|OJJ46169.1

Group1806: abra|OJJ68941.1 acar|OOF91521.1 acla|EAW14242.1 afis|EAW19038.1 afla|EED50393.1 afum|EAL93767.1 agla|OJJ84582.1 akaw|GAA90312.1 aluc|OJZ85927.1 anid|CBF79609.1 anig|EHA24670.1 aory|EIT82042.1 asyd|OJJ57994.1 ater|EAU32077.1 atub|OJI79716.1 aver|OJJ04408.1 awen|OJJ31468.1 azon|OJJ46172.1

Group1807: abra|OJJ68943.1 acar|OOF91522.1 acla|EAW14241.1 afis|EAW19039.1 afla|EED50394.1 afum|EAL93768.1 agla|OJJ84573.1 akaw|GAA90310.1 aluc|OJZ85929.1 anid|CBF79610.1 anig|EHA24669.1 aory|EIT82167.1 asyd|OJJ57995.1 ater|EAU32078.1 atub|OJI79717.1 aver|OJJ04407.1 awen|OJJ31469.1 azon|OJJ46173.1

Group1808: abra|OJJ68944.1 acar|OOF91524.1 acla|EAW14239.1 afis|EAW19041.1 afla|EED50396.1 afum|EAL93770.1 agla|OJJ84564.1 akaw|GAA90309.1 aluc|OJZ85930.1 anid|CBF79612.1 anig|EHA24667.1 aory|EIT82062.1 asyd|OJJ57999.1 ater|EAU32080.1 atub|OJI79719.1 aver|OJJ04403.1 awen|OJJ31470.1 azon|OJJ46179.1

Group1809: abra|OJJ68945.1 acar|OOF91525.1 acla|EAW14238.1 afis|EAW19042.1 afla|EED50397.1 afum|EAL93771.1 agla|OJJ84563.1 akaw|GAA90308.1 aluc|OJZ85931.1 anid|CBF79613.1 anig|EHA24666.1 aory|EIT82121.1 asyd|OJJ57998.1 ater|EAU32081.1 atub|OJI79720.1 aver|OJJ04404.1 awen|OJJ31471.1 azon|OJJ46181.1

Group1810: abra|OJJ68948.1 acar|OOF91526.1 acla|EAW14236.1 afis|EAW19044.1 afla|EED50401.1 afum|EAL93773.1 agla|OJJ84561.1 akaw|GAA90306.1 aluc|OJZ85934.1 anig|EHA24664.1 aory|EIT82075.1 ater|EAU32083.1 atub|OJI79723.1 azon|OJJ46184.1 anid|CBF79617.1 asyd|OJJ58000.1 aver|OJJ04402.1 awen|OJJ31473.1

Group1811: abra|OJJ68949.1 acar|OOF91527.1 acla|EAW14235.1 afis|EAW19045.1 afla|EED50402.1 afum|EAL93774.1 agla|OJJ84560.1 akaw|GAA90305.1 aluc|OJZ85935.1 anid|CBF79619.1 anig|EHA24663.1 aory|EIT82000.1 asyd|OJJ58001.1 ater|EAU32084.1 atub|OJI79724.1 aver|OJJ04400.1 awen|OJJ31474.1 azon|OJJ46188.1

Group1812: abra|OJJ68952.1 acar|OOF91530.1 acla|EAW14234.1 afis|EAW19046.1 afla|EED50405.1 afum|EAL93775.1 agla|OJJ84559.1 akaw|GAA90481.1 aluc|OJZ85938.1 anid|CBF79621.1 anig|EHA24661.1 aory|EIT82119.1 asyd|OJJ58003.1 ater|EAU32085.1 atub|OJI79726.1 aver|OJJ04398.1 awen|OJJ31475.1 azon|OJJ46189.1

Group1813: abra|OJJ68953.1 acar|OOF91531.1 acla|EAW14233.1 afis|EAW19047.1 afla|EED50406.1 afum|EAL93776.1 agla|OJJ84558.1 akaw|GAA90482.1 aluc|OJZ85939.1 anid|CBF79622.1 anig|EHA24660.1 aory|EIT82158.1 asyd|OJJ58004.1 ater|EAU32086.1 atub|OJI79727.1 aver|OJJ04397.1 awen|OJJ31477.1 azon|OJJ46190.1

Group1814: abra|OJJ68956.1 acar|OOF91535.1 acla|EAW14229.1 afis|EAW19052.1 afla|EED54409.1 afum|EAL93781.1 agla|OJJ84555.1 akaw|GAA90484.1 aluc|OJZ85942.1 anid|CBF79627.1 anig|EHA24657.1 aory|EIT75857.1 asyd|OJJ58009.1 ater|EAU32089.1 atub|OJI79730.1 aver|OJJ04391.1 awen|OJJ31483.1 azon|OJJ46193.1

Group1815: abra|OJJ68957.1 acar|OOF91534.1 acla|EAW14228.1 afis|EAW19053.1 afla|EED54411.1 afum|EAL93782.1 agla|OJJ84554.1 akaw|GAA90485.1 aluc|OJZ85943.1 anid|CBF79628.1 anig|EHA24656.1 aory|EIT75871.1 asyd|OJJ58010.1 ater|EAU32090.1 atub|OJI79731.1 aver|OJJ04390.1 awen|OJJ31484.1 azon|OJJ46194.1

Group1816: abra|OJJ68959.1 acar|OOF91537.1 acla|EAW14226.1 afis|EAW19055.1 afla|EED54413.1 afum|EAL93784.1 agla|OJJ84552.1 akaw|GAA90487.1 aluc|OJZ85945.1 anid|CBF79630.1 anig|EHA24654.1 aory|EIT75877.1 asyd|OJJ58012.1 ater|EAU32091.1 atub|OJI79733.1 aver|OJJ04388.1 awen|OJJ31486.1 azon|OJJ46196.1

Group1817: abra|OJJ68960.1 acar|OOF91538.1 acla|EAW14225.1 afis|EAW19056.1 afla|EED54414.1 afum|EAL93785.1 agla|OJJ84551.1 akaw|GAA90488.1 aluc|OJZ85946.1 anid|CBF79632.1 anig|EHA24653.1 aory|EIT75852.1 asyd|OJJ58013.1 ater|EAU32092.1 atub|OJI79734.1 aver|OJJ04387.1 awen|OJJ31487.1 azon|OJJ46197.1

Group1818: abra|OJJ68961.1 acar|OOF91539.1 acla|EAW14224.1 afis|EAW19057.1 afla|EED54415.1 afum|EAL93786.1 agla|OJJ84550.1 akaw|GAA90489.1 aluc|OJZ85947.1 anid|CBF79634.1 anig|EHA24652.1 aory|EIT75893.1 asyd|OJJ58014.1 ater|EAU32093.1 atub|OJI79735.1 aver|OJJ04386.1 awen|OJJ31488.1 azon|OJJ46199.1

Group1819: abra|OJJ68963.1 acar|OOF91541.1 acla|EAW14223.1 afis|EAW19058.1 afla|EED54418.1 afum|EAL93787.1 agla|OJJ84549.1 akaw|GAA90491.1 aluc|OJZ85950.1 anid|CBF79638.1 anig|EHA24650.1 aory|EIT75892.1 asyd|OJJ58015.1 ater|EAU32094.1 atub|OJI79738.1 aver|OJJ04385.1 awen|OJJ31489.1 azon|OJJ46201.1

Group1820: abra|OJJ68965.1 acar|OOF91542.1 acla|EAW14222.1 afis|EAW19059.1 afla|EED54421.1 afum|EAL93788.1 agla|OJJ84548.1 akaw|GAA90492.1 aluc|OJZ85952.1 anid|CBF79639.1 anig|EHA24649.1 aory|EIT75888.1 asyd|OJJ58016.1 ater|EAU32095.1 atub|OJI79740.1 aver|OJJ04384.1 awen|OJJ31490.1 azon|OJJ46202.1

Group1821: abra|OJJ68966.1 acar|OOF91544.1 acla|EAW14221.1 afis|EAW19060.1 afla|EED54422.1 afum|EAL93789.1 agla|OJJ84547.1 akaw|GAA90493.1 aluc|OJZ85953.1 anid|CBF79640.1 anig|EHA24648.1

aory|EIT75846.1 asyd|OJJ58017.1 ater|EAU32098.1 atub|OJI79741.1 aver|OJJ04382.1 awen|OJJ31491.1 azon|OJJ46203.1

Group1822: abra|OJJ68967.1 acar|OOF91545.1 acla|EAW14220.1 afis|EAW19061.1 afla|EED54423.1 afum|EAL93790.1 agla|OJJ84546.1 akaw|GAA90494.1 aluc|OJZ85954.1 anid|CBF79641.1 anig|EHA24647.1 aory|EIT75876.1 asyd|OJJ58018.1 ater|EAU32099.1 atub|OJI79742.1 aver|OJJ04381.1 awen|OJJ31492.1 azon|OJJ46204.1

Group1823: abra|OJJ68968.1 acar|OOF91546.1 acla|EAW14219.1 afis|EAW19062.1 afla|EED54428.1 afum|EAL93791.1 agla|OJJ84545.1 akaw|GAA90495.1 aluc|OJZ85955.1 anid|CBF79643.1 anig|EHA24646.1 aory|EIT75878.1 asyd|OJJ58019.1 ater|EAU32105.1 atub|OJI79743.1 aver|OJJ04380.1 awen|OJJ31493.1 azon|OJJ46206.1

Group1824: abra|OJJ68970.1 acar|OOF91547.1 acla|EAW14217.1 afis|EAW19064.1 afla|EED54430.1 afum|EAL93793.2 agla|OJJ84543.1 akaw|GAA90497.1 aluc|OJZ85957.1 anid|CBF79646.1 anig|EHA24644.1 aory|EIT75886.1 asyd|OJJ58021.1 ater|EAU32107.1 atub|OJI79745.1 aver|OJJ04378.1 awen|OJJ31496.1 azon|OJJ46208.1

Group1825: abra|OJJ68973.1 acar|OOF91550.1 acla|EAW14214.1 afis|EAW19067.1 afla|EED54433.1 afum|EAL93796.1 agla|OJJ84539.1 akaw|GAA90500.1 aluc|OJZ85962.1 anid|CBF79652.1 anig|EHA24641.1 aory|EIT75842.1 asyd|OJJ58025.1 ater|EAU32115.1 atub|OJI79749.1 aver|OJJ04374.1 awen|OJJ31499.1 azon|OJJ46213.1

Group1826: abra|OJJ68978.1 acar|OOF91553.1 acla|EAW14210.1 afis|EAW19071.1 afla|EED54436.1 afum|EAL93799.1 agla|OJJ84534.1 akaw|GAA90503.1 aluc|OJZ85966.1 anid|CBF79663.1 anig|EHA24638.1 aory|EIT75872.1 asyd|OJJ58028.1 ater|EAU32119.1 atub|OJI79754.1 aver|OJJ04371.1 awen|OJJ31503.1 azon|OJJ46218.1

Group1827: abra|OJJ68982.1 acar|OOF91554.1 acla|EAW14209.1 afis|EAW19072.1 afla|EED54437.1 afum|EAL93800.1 agla|OJJ84532.1 akaw|GAA90504.1 aluc|OJZ85968.1 anid|CBF79661.1 anig|EHA24637.1 aory|EIT75860.1 asyd|OJJ58029.1 ater|EAU32120.1 atub|OJI79756.1 aver|OJJ04370.1 awen|OJJ32466.1 azon|OJJ46219.1

Group1828: abra|OJJ68984.1 acar|OOF91555.1 acla|EAW14207.1 afis|EAW19074.1 afla|EED54439.1 afum|EAL93802.1 agla|OJJ84529.1 akaw|GAA90506.1 aluc|OJZ85970.1 anid|CBF79667.1 anig|EHA24635.1 aory|EIT75870.1 asyd|OJJ58031.1 ater|EAU32122.1 atub|OJI79758.1 aver|OJJ04368.1 awen|OJJ32468.1 azon|OJJ46238.1

Group1829: abra|OJJ68991.1 acar|OOF92703.1 acla|EAW12643.1 afis|EAW18042.1 afla|EED51854.1 afum|EAL91853.1 agla|OJJ85414.1 akaw|GAA90511.1 aluc|OJZ85976.1 anid|CBF80097.1 anig|EHA24630.1 aory|EIT81766.1 asyd|OJJ58299.1 ater|EAU31557.1 atub|OJI79766.1 aver|OJJ02125.1 awen|OJJ41492.1 azon|OJJ48223.1

Group1830: abra|OJJ68994.1 acar|OOF92705.1 acla|EAW12645.1 afis|EAW18040.1 afla|EED51857.1 afum|EAL91851.1 agla|OJJ85412.1 akaw|GAA90513.1 aluc|OJZ85978.1 anid|CBF80103.1 anig|EHA24628.1 aory|EIT81744.1 asyd|OJJ58296.1 ater|EAU31555.1 atub|OJI79768.1 aver|OJJ02128.1 awen|OJJ41497.1 azon|OJJ48225.1

Group1831: abra|OJJ68996.1 acar|OOF92707.1 acla|EAW12647.1 afis|EAW18038.1 afla|EED51860.1 afum|EAL91849.1 agla|OJJ85410.1 akaw|GAA90515.1 aluc|OJZ85980.1 anid|CBF80073.1 anig|EHA24626.1 aory|EIT81689.1 asyd|OJJ58294.1 ater|EAU31552.1 atub|OJI79770.1 aver|OJJ02130.1 awen|OJJ41499.1 azon|OJJ48227.1

Group1832: abra|OJJ68997.1 acar|OOF92708.1 acla|EAW12648.1 afis|EAW18037.1 afla|EED51861.1 afum|EAL91848.1 agla|OJJ85409.1 akaw|GAA90516.1 aluc|OJZ85981.1 anid|CBF80074.1 anig|EHA24625.1 aory|EIT81730.1 asyd|OJJ58293.1 ater|EAU31551.1 atub|OJI79771.1 aver|OJJ02131.1 awen|OJJ41500.1 azon|OJJ48228.1

Group1833: abra|OJJ69004.1 acar|OOF92717.1 acla|EAW12855.1 afis|EAW18018.1 afla|EED51873.1 afum|EAL91830.1 agla|OJJ85372.1 akaw|GAA90522.1 aluc|OJZ85990.1 anid|CBF80052.1 anig|EHA24620.1 aory|EIT81725.1 asyd|OJJ58280.1 ater|EAU31534.1 atub|OJI79779.1 aver|OJJ02143.1 awen|OJJ41521.1 azon|OJJ48243.1

Group1834: abra|OJJ69005.1 acar|OOF92718.1 acla|EAW12854.1 afla|EED51874.1 afum|EAL91829.1 akaw|GAA90523.1 aluc|OJZ85991.1 anig|EHA24619.1 aory|EIT81686.1 ater|EAU31533.1 atub|OJI79780.1 afis|EAW18017.1 asyd|OJJ58279.1 aver|OJJ02144.1 awen|OJJ41522.1 agla|OJJ85371.1 anid|CBF80050.1 azon|OJJ48244.1

Group1835: abra|OJJ69008.1 acar|OOF92719.1 acla|EAW12850.1 afis|EAW18013.1 afla|EED51883.1 afum|EAL91825.1 agla|OJJ80286.1 akaw|GAA90525.1 aluc|OJZ85994.1 anid|CBF80040.1 anig|EHA24616.1 aory|EIT81726.1 asyd|OJJ58275.1 ater|EAU31529.1 atub|OJI79782.1 aver|OJJ02149.1 awen|OJJ41528.1 azon|OJJ48248.1

Group1836: abra|OJJ69010.1 acar|OOF92721.1 acla|EAW12848.1 afis|EAW18011.1 afla|EED51885.1 afum|EAL91823.1 agla|OJJ80283.1 akaw|GAA90527.1 aluc|OJZ85996.1 anid|CBF80037.1 anig|EHA24614.1 aory|EIT81760.1 asyd|OJJ58273.1 ater|EAU31527.1 atub|OJI79784.1 aver|OJJ02151.1 awen|OJJ41530.1 azon|OJJ48250.1

Group1837: azon|OJJ51098.1 abra|OJJ74349.1 acar|OOF95381.1 acla|EAW07403.1 afis|EAW22429.1 afla|EED52507.1 afum|EAL90762.1 agla|OJJ87594.1 akaw|GAA86681.1 aluc|OJZ89781.1 anid|CBF88824.1 anig|EHA26996.1 aory|EIT82792.1 asyd|OJJ62884.1 ater|EAU39278.1 atub|OJI85025.1 aver|OJI96936.1 awen|OJJ35008.1

Group1838: abra|OJJ69024.1 acar|OOF92985.1 acla|EAW06466.1 afis|EAW15961.1 afla|EED58096.1 afum|EAL88594.1 agla|OJJ89351.1 akaw|GAA90541.1 aluc|OJZ86011.1 anid|CBF81079.1 anig|EHA24600.1 aory|EIT79116.1 asyd|OJJ56463.1 ater|EAU35493.1 atub|OJI79800.1 aver|OJJ06258.1 awen|OJJ35613.1 azon|OJJ46911.1

Group1839: abra|OJJ69026.1 acar|OOF92660.1 acla|EAW12556.1 afis|EAW24095.1 afla|EED46839.1 afum|EAL91949.1 agla|OJJ87168.1 akaw|GAA92787.1 aluc|OJZ82696.1 anid|CBF75138.1 anig|EHA21145.1 aory|EIT72775.1 asyd|OJJ59978.1 ater|EAU30520.1 atub|OJI88928.1 aver|OJJ02313.1 awen|OJJ41395.1 azon|OJJ48863.1

Group1840: abra|OJJ69029.1 acar|OOF92658.1 acla|EAW10072.1 afis|EAW25514.1 afla|EED54015.1 afum|EAL90179.1 agla|OJJ87289.1 akaw|GAA90545.1 aluc|OJZ86015.1 anid|CBF77783.1 anig|EHA24596.1 aory|EIT78264.1 asyd|OJJ53030.1 ater|EAU33360.1 atub|OJI79804.1 aver|OJJ03814.1 awen|OJJ41257.1 azon|OJJ44822.1

Group1841: abra|OJJ69034.1 acar|OOF89828.1 acla|EAW08636.1 afis|EAW15795.1 afla|EED51250.1 afum|EAL85555.1 agla|OJJ81958.1 akaw|GAA90549.1 aluc|OJZ86019.1 anid|CBF70929.1 anig|EHA24592.1 aory|EIT81525.1 asyd|OJJ52611.1 ater|EAU32347.1 atub|OJI79808.1 aver|OJJ04638.1 awen|OJJ38503.1 azon|OJJ43748.1

Group1842: abra|OJJ69036.1 acar|OOF92649.1 acla|EAW12687.1 afis|EAW17850.1 afla|EED46204.1 afum|EAL91659.1 agla|OJJ83119.1 akaw|GAA90551.1 aluc|OJZ86021.1 anid|CBF77960.1 anig|EHA24590.1 aory|EIT76404.1 asyd|OJJ55272.1 ater|EAU33052.1 atub|OJI79810.1 aver|OJJ06955.1 awen|OJJ41723.1 azon|OJJ48421.1

Group1843: abra|OJJ69037.1 acar|OOF92647.1 acla|EAW14104.1 afis|EAW18805.1 afla|EED50771.1 afum|EAL93541.1 agla|OJJ80535.1 akaw|GAA90552.1 aluc|OJZ86022.1 anid|CBF88044.1 anig|EHA24589.1 aory|EIT78003.1 asyd|OJJ63545.1 ater|EAU37943.1 atub|OJI79811.1 aver|OJI97389.1 awen|OJJ32380.1 azon|OJJ47142.1

Group1844: abra|OJJ69038.1 acar|OOF92646.1 acla|EAW06768.1 afis|EAW16318.1 afla|EED53563.1 afum|EAL88942.1 agla|OJJ88279.1 akaw|GAA90553.1 aluc|OJZ86023.1 anid|CBF89147.1 anig|EHA24588.1 aory|EIT74896.1 asyd|OJJ62568.1 ater|EAU34139.1 atub|OJI79812.1 aver|OJI96692.1 awen|OJJ37718.1 azon|OJJ44975.1

Group1845: abra|OJJ69039.1 acar|OOF92645.1 acla|EAW11330.1 afis|EAW18165.1 afla|EED53435.1 afum|EAL87525.2 agla|OJJ85953.1 akaw|GAA90554.1 aluc|OJZ86024.1 anid|CBF85751.1 anig|EHA24587.1 aory|EIT74836.1 asyd|OJJ64296.1 ater|EAU36013.1 atub|OJI79813.1 aver|OJI98389.1 awen|OJJ30512.1 azon|OJJ50291.1

Group1846: abra|OJJ69057.1 acar|OOF91226.1 acla|EAW08757.1 afis|EAW15674.1 afla|EED46856.1 afum|EAL85673.1 agla|OJJ81821.1 akaw|GAA90587.1 aluc|OJZ86045.1 anid|CBF71161.1 anig|EHA24566.1 aory|EIT72754.1 asyd|OJJ55844.1 ater|EAU32486.1 atub|OJI79842.1 aver|OJJ04787.1 awen|OJJ38352.1 azon|OJJ42790.1

Group1847: abra|OJJ69058.1 acar|OOF91225.1 acla|EAW08756.1 afis|EAW15673.1 afla|EED46857.1 afum|EAL85675.1 agla|OJJ81820.1 akaw|GAA90588.1 aluc|OJZ86046.1 anid|CBF71163.1 anig|EHA24565.1 aory|EIT72609.1 asyd|OJJ55845.1 ater|EAU32487.1 atub|OJI79843.1 aver|OJJ04788.1 awen|OJJ38351.1 azon|OJJ42791.1

Group1848: abra|OJJ69064.1 acar|OOF91219.1 acla|EAW08760.1 afis|EAW15668.1 afla|EED46876.1 afum|EAL85680.1 agla|OJJ81806.1 akaw|GAA90594.1 aluc|OJZ86052.1 anid|CBF71174.1 anig|EHA24559.1 aory|EIT72756.1 asyd|OJJ55853.1 ater|EAU32503.1 atub|OJI79849.1 aver|OJJ04796.1 awen|OJJ38344.1 azon|OJJ43043.1

Group1849: abra|OJJ69065.1 acar|OOF91218.1 acla|EAW08759.1 afis|EAW15667.1 afla|EED46877.1 afum|EAL85681.1 agla|OJJ81805.1 akaw|GAA90595.1 aluc|OJZ86053.1 anid|CBF71176.1 anig|EHA24558.1 aory|EIT72761.1 asyd|OJJ55854.1 ater|EAU32504.1 atub|OJI79850.1 aver|OJJ04797.1 awen|OJJ38343.1 azon|OJJ43042.1

Group1850: abra|OJJ69071.1 acar|OOF91231.1 acla|EAW08767.1 afis|EAW15663.1 afla|EED46886.1 afum|EAL85685.1 agla|OJJ81785.1 akaw|GAA90602.1 aluc|OJZ86060.1 anid|CBF71180.1 anig|EHA24553.1

aory|EIT72594.1 asyd|OJJ55862.1 ater|EAU32510.1 atub|OJI79857.1 aver|OJJ04803.1 awen|OJJ38339.1 azon|OJJ43024.1

Group1852: abra|OJJ69080.1 acar|OOF91211.1 acla|EAW08754.1 afis|EAW15682.1 afla|EED51372.1 afum|EAL85666.1 agla|OJJ81831.1 akaw|GAA90609.1 aluc|OJZ86068.1 anid|CBF71152.1 anig|EHA24546.1 aory|EIT80462.1 asyd|OJJ55839.1 ater|EAU32475.1 atub|OJI79864.1 aver|OJJ04782.1 awen|OJJ38374.1 azon|OJJ43858.1

Group1853: abra|OJJ69081.1 acar|OOF91210.1 acla|EAW08753.1 afis|EAW15683.1 afla|EED51371.1 afum|EAL85665.1 agla|OJJ81832.1 akaw|GAA90610.1 aluc|OJZ86069.1 anid|CBF71150.1 anig|EHA24545.1 aory|EIT80441.1 asyd|OJJ55838.1 ater|EAU32474.1 atub|OJI79865.1 aver|OJJ04781.1 awen|OJJ38375.1 azon|OJJ43859.1

Group1854: abra|OJJ69084.1 acar|OOF91208.1 acla|EAW08750.1 afis|EAW15686.1 afla|EED51368.1 afum|EAL85662.1 agla|OJJ81834.1 akaw|GAA90613.1 aluc|OJZ86071.1 anid|CBF71144.1 anig|EHA24543.1 aory|EIT80616.1 asyd|OJJ55836.1 ater|EAU32467.1 atub|OJI79868.1 aver|OJJ04779.1 awen|OJJ38377.1 azon|OJJ43861.1

Group1855: abra|OJJ69089.1 acar|OOF91204.1 acla|EAW08743.1 afis|EAW15693.1 afla|EED51360.1 afum|EAL85655.1 agla|OJJ81840.1 akaw|GAA90618.1 aluc|OJZ86078.1 anid|CBF71129.1 anig|EHA24538.1 aory|EIT80527.1 asyd|OJJ55829.1 ater|EAU32462.1 atub|OJI79875.1 aver|OJJ04771.1 awen|OJJ38384.1 azon|OJJ43853.1

Group1856: abra|OJJ69090.1 acar|OOF91203.1 acla|EAW08742.1 afis|EAW15694.1 afla|EED51359.1 afum|EAL85654.1 agla|OJJ81841.1 akaw|GAA90619.1 aluc|OJZ86079.1 anid|CBF71127.1 anig|EHA24537.1 aory|EIT80431.1 asyd|OJJ55828.1 ater|EAU32461.1 atub|OJI79876.1 aver|OJJ04770.1 awen|OJJ38385.1 azon|OJJ43852.1

Group1857: abra|OJJ69091.1 acar|OOF91202.1 acla|EAW08741.1 afis|EAW15695.1 afla|EED51358.1 afum|EAL85653.1 agla|OJJ81842.1 akaw|GAA90620.1 aluc|OJZ86080.1 anid|CBF71119.1 anig|EHA24535.1 aory|EIT80729.1 asyd|OJJ55827.1 ater|EAU32460.1 atub|OJI79877.1 aver|OJJ04769.1 awen|OJJ38386.1 azon|OJJ43850.1

Group1858: abra|OJJ69093.1 acar|OOF91201.1 acla|EAW08740.1 afis|EAW15696.1 afla|EED51357.1 afum|EAL85652.2 agla|OJJ81850.1 akaw|GAA90621.1 aluc|OJZ86081.1 anid|CBF71121.1 anig|EHA24534.1 aory|EIT80641.1 asyd|OJJ55826.1 ater|EAU32455.1 atub|OJI79878.1 aver|OJJ04768.1 awen|OJJ38389.1 azon|OJJ43849.1

Group1859: abra|OJJ69094.1 acar|OOF91200.1 acla|EAW08737.1 afis|EAW15699.1 afla|EED51353.1 afum|EAL85649.1 agla|OJJ81849.1 akaw|GAA90622.1 aluc|OJZ86083.1 anid|CBF71123.1 anig|EHA24532.1 aory|EIT80448.1 asyd|OJJ55825.1 ater|EAU32452.1 atub|OJI79879.1 aver|OJJ04767.1 awen|OJJ38391.1 azon|OJJ43847.1

Group1860: abra|OJJ69099.1 acar|OOF91197.1 acla|EAW08734.1 afis|EAW15702.1 afla|EED51349.1 afum|EAL85645.1 agla|OJJ81844.1 akaw|GAA90625.1 aluc|OJZ86088.1 anid|CBF71108.1 anig|EHA24529.1 aory|EIT80559.1 asyd|OJJ52712.1 ater|EAU32442.1 atub|OJI79884.1 aver|OJJ04741.1 awen|OJJ38396.1 azon|OJJ43845.1

Group1861: abra|OJJ69100.1 acar|OOF91196.1 acla|EAW08733.1 afis|EAW15707.1 afla|EED51348.1 afum|EAL85640.1 agla|OJJ81851.1 akaw|GAA90626.1 aluc|OJZ86089.1 anid|CBF71106.1 anig|EHA24528.1 aory|EIT80499.1 asyd|OJJ52711.1 atub|OJI79885.1 aver|OJJ04740.1 awen|OJJ38397.1 ater|EAU32441.1 azon|OJJ43844.1

Group1862: abra|OJJ69104.1 acar|OOF91194.1 acla|EAW08728.1 afis|EAW15703.1 afla|EED51345.1 afum|EAL85644.1 agla|OJJ81855.1 akaw|GAA90629.1 aluc|OJZ86092.1 anid|CBF71100.1 anig|EHA24525.1 aory|EIT80418.1 asyd|OJJ52707.1 ater|EAU32438.1 atub|OJI79888.1 aver|OJJ04736.1 awen|OJJ38402.1 azon|OJJ43841.1

Group1863: abra|OJJ69108.1 acar|OOF91181.1 acla|EAW08724.1 afis|EAW15732.1 afla|EED51341.1 afum|EAL85615.2 agla|OJJ81861.1 akaw|GAA90633.1 aluc|OJZ86096.1 anid|CBF71094.1 anig|EHA24522.1 aory|EIT81523.1 asyd|OJJ52704.1 ater|EAU32434.1 atub|OJI79892.1 aver|OJJ04733.1 awen|OJJ38406.1 azon|OJJ43836.1

Group1864: abra|OJJ69110.1 acar|OOF91184.1 acla|EAW08722.1 afis|EAW15730.1 afla|EED51339.1 afum|EAL85617.1 agla|OJJ81864.1 akaw|GAA90635.1 aluc|OJZ86098.1 anid|CBF71090.1 anig|EHA24520.1 aory|EIT81546.1 asyd|OJJ52701.1 ater|EAU32432.1 atub|OJI79894.1 aver|OJJ04729.1 awen|OJJ38409.1 azon|OJJ43833.1

Group1865: abra|OJJ69113.1 acar|OOF91185.1 acla|EAW08719.1 afis|EAW15728.1 afla|EED51335.1 afum|EAL85619.1 agla|OJJ81867.1 akaw|GAA90638.1 aluc|OJZ86102.1 anid|CBF71085.1 anig|EHA24517.1 aory|EIT81524.1 asyd|OJJ52698.1 ater|EAU32430.1 atub|OJI79897.1 aver|OJJ04726.1 awen|OJJ38412.1 azon|OJJ43830.1

Group1866: abra|OJJ69116.1 acar|OOF91188.1 acla|EAW08716.1 afis|EAW15724.1 afla|EED51332.1 afum|EAL85623.1 agla|OJJ81870.1 akaw|GAA90641.1 aluc|OJZ86105.1 anid|CBF71079.1 anig|EHA24515.1 aory|EIT81503.1 asyd|OJJ52695.1 ater|EAU32427.1 atub|OJI79900.1 aver|OJJ04723.1 awen|OJJ38415.1 azon|OJJ43827.1

Group1867: abra|OJJ69119.1 acar|OOF91191.1 acla|EAW08713.1 afis|EAW15721.1 afla|EED51329.1 afum|EAL85626.1 agla|OJJ81873.1 akaw|GAA90644.1 aluc|OJZ86108.1 anid|CBF71075.1 anig|EHA24513.1 aory|EIT81515.1 asyd|OJJ52692.1 ater|EAU32424.1 atub|OJI79903.1 aver|OJJ04720.1 awen|OJJ38418.1 azon|OJJ43824.1

Group1868: abra|OJJ69120.1 acar|OOF91192.1 acla|EAW08712.1 afis|EAW15720.1 afla|EED51328.1 afum|EAL85628.1 agla|OJJ81874.1 akaw|GAA90645.1 aluc|OJZ86109.1 anid|CBF71073.1 anig|EHA24512.1 aory|EIT81554.1 asyd|OJJ52691.1 ater|EAU32423.1 atub|OJI79904.1 aver|OJJ04719.1 awen|OJJ38419.1 azon|OJJ43823.1

Group1869: abra|OJJ69121.1 acar|OOF91193.1 acla|EAW08711.1 afis|EAW15719.1 afla|EED51327.1 afum|EAL85629.2 agla|OJJ81875.1 akaw|GAA90646.1 aluc|OJZ86110.1 anid|CBF71071.1 anig|EHA24511.1 aory|EIT81517.1 asyd|OJJ52690.1 ater|EAU32420.1 atub|OJI79905.1 aver|OJJ04718.1 awen|OJJ38420.1 azon|OJJ43822.1

Group1870: abra|OJJ69122.1 acar|OOF91178.1 acla|EAW08710.1 afis|EAW15718.1 afla|EED51326.1 afum|EAL85630.2 agla|OJJ81877.1 akaw|GAA90647.1 aluc|OJZ86111.1 anid|CBF71070.1 anig|EHA24510.1 aory|EIT81526.1 asyd|OJJ52689.1 ater|EAU32421.1 atub|OJI79906.1 aver|OJJ04717.1 awen|OJJ38421.1 azon|OJJ43821.1

Group1871: abra|OJJ69123.1 acar|OOF91177.1 acla|EAW08709.1 afis|EAW15717.1 afla|EED51325.1 afum|EAL85631.1 agla|OJJ81878.1 akaw|GAA90648.1 aluc|OJZ86112.1 anid|CBF71068.1 anig|EHA24509.1 aory|EIT81505.1 asyd|OJJ52688.1 ater|EAU32422.1 atub|OJI79907.1 aver|OJJ04716.1 awen|OJJ38422.1 azon|OJJ43820.1

Group1872: abra|OJJ69125.1 acar|OOF91175.1 acla|EAW08707.1 afis|EAW15715.1 afla|EED51323.1 afum|EAL85633.1 agla|OJJ81886.1 akaw|GAA90650.1 aluc|OJZ86114.1 anid|CBF71064.1 anig|EHA24507.1 aory|EIT81539.1 asyd|OJJ52685.1 ater|EAU32419.1 atub|OJI79909.1 aver|OJJ04713.1 awen|OJJ38425.1 azon|OJJ43818.1

Group1873: abra|OJJ69126.1 acar|OOF91174.1 acla|EAW08706.1 afis|EAW15714.1 afla|EED51322.1 afum|EAL85634.1 agla|OJJ81887.1 akaw|GAA90651.1 aluc|OJZ86115.1 anid|CBF71058.1 anig|EHA24506.1 aory|EIT81529.1 asyd|OJJ52680.1 ater|EAU32418.1 atub|OJI79910.1 aver|OJJ04706.1 awen|OJJ38426.1 azon|OJJ43817.1

Group1874: abra|OJJ69127.1 acar|OOF91173.1 acla|EAW08705.1 afis|EAW15713.1 afla|EED51321.1 afum|EAL85635.1 agla|OJJ81888.1 akaw|GAA90652.1 aluc|OJZ86116.1 anid|CBF71060.1 anig|EHA24505.1 aory|EIT81530.1 asyd|OJJ52679.1 ater|EAU32417.1 atub|OJI79911.1 aver|OJJ04705.1 awen|OJJ38427.1 azon|OJJ43816.1

Group1875: abra|OJJ69128.1 acar|OOF91172.1 acla|EAW08704.1 afis|EAW15712.1 afla|EED51320.1 afum|EAL85636.1 agla|OJJ81889.1 akaw|GAA90653.1 aluc|OJZ86117.1 anid|CBF71057.1 anig|EHA24504.1 aory|EIT81550.1 asyd|OJJ52678.1 ater|EAU32416.1 atub|OJI79912.1 aver|OJJ04704.1 awen|OJJ38428.1 azon|OJJ43815.1

Group1876: abra|OJJ69131.1 acar|OOF91169.1 acla|EAW08701.1 afis|EAW15709.1 afla|EED51317.1 afum|EAL85638.1 agla|OJJ81892.1 akaw|GAA90656.1 aluc|OJZ86120.1 anid|CBF71053.1 anig|EHA24501.1 aory|EIT81516.1 asyd|OJJ52675.1 ater|EAU32414.1 atub|OJI79916.1 aver|OJJ04701.1 awen|OJJ38431.1 azon|OJJ43812.1

Group1877: abra|OJJ69132.1 acar|OOF91168.1 acla|EAW08700.1 afis|EAW15708.1 afla|EED51316.1 afum|EAL85639.1 agla|OJJ81893.1 akaw|GAA90657.1 aluc|OJZ86121.1 anid|CBF71051.1 anig|EHA24500.1 aory|EIT81557.1 asyd|OJJ52674.1 ater|EAU32413.1 atub|OJI79917.1 aver|OJJ04700.1 awen|OJJ38432.1 azon|OJJ43811.1

Group1878: abra|OJJ69136.1 acar|OOF91164.1 acla|EAW08697.1 afis|EAW15738.1 afla|EED51314.1 afum|EAL85610.1 agla|OJJ81895.1 akaw|GAA86320.1 aluc|OJZ86124.1 anid|CBF71047.1 anig|EHA24497.1 aory|EIT81566.1 asyd|OJJ52672.1 ater|EAU32406.1 atub|OJI79920.1 aver|OJJ04698.1 awen|OJJ38434.1 azon|OJJ43808.1

Group1879: abra|OJJ69138.1 acar|OOF91163.1 acla|EAW08696.1 afis|EAW15739.1 afla|EED51313.1 afum|EAL85609.1 agla|OJJ81900.1 akaw|GAA86319.1 aluc|OJZ86125.1 anid|CBF71045.1 anig|EHA24496.1 aory|EIT81532.1 asyd|OJJ52671.1 ater|EAU32408.1 atub|OJI79921.1 aver|OJJ04697.1 awen|OJJ38438.1 azon|OJJ43807.1

Group1880: abra|OJJ69143.1 acar|OOF91159.1 acla|EAW08666.1 afis|EAW15743.1 afla|EED51309.1 afum|EAL85605.1 agla|OJJ81904.1 akaw|GAA86314.1 aluc|OJZ86129.1 anid|CBF71038.1 anig|EHA24492.1

aory|EIT81535.1 asyd|OJJ52666.1 ater|EAU32402.1 atub|OJI79925.1 aver|OJJ04693.1 awen|OJJ38443.1 azon|OJJ43749.1

Group1881: abra|OJJ69147.1 acar|OOF91156.1 acla|EAW08669.1 afis|EAW15757.1 afla|EED51302.1 afum|EAL85591.1 agla|OJJ81905.1 akaw|GAA86311.1 aluc|OJZ86133.1 anid|CBF71025.1 anig|EHA24488.1 aory|EIT81545.1 asyd|OJJ52660.1 ater|EAU32400.1 atub|OJI79929.1 aver|OJJ04688.1 awen|OJJ38445.1 azon|OJJ43751.1

Group1882: abra|OJJ69154.1 acar|OOF91149.1 acla|EAW08671.1 afis|EAW15753.1 afla|EED51298.1 afum|EAL85596.1 agla|OJJ81909.1 akaw|GAA86308.1 aluc|OJZ86141.1 anid|CBF71023.1 anig|EHA24483.1 aory|EIT81538.1 asyd|OJJ52659.1 ater|EAU32397.1 atub|OJI79933.1 aver|OJJ04687.1 awen|OJJ38453.1 azon|OJJ43753.1

Group1883: abra|OJJ69155.1 acar|OOF91148.1 acla|EAW08672.1 afis|EAW15752.1 afla|EED51297.1 afum|EAL85597.1 agla|OJJ81910.1 akaw|GAA86307.1 aluc|OJZ86142.1 anid|CBF71021.1 anig|EHA24482.1 aory|EIT81540.1 asyd|OJJ52658.1 ater|EAU32396.1 atub|OJI79934.1 aver|OJJ04686.1 awen|OJJ38454.1 azon|OJJ43754.1

Group1884: abra|OJJ69156.1 acar|OOF91147.1 acla|EAW08673.1 afis|EAW15751.1 afla|EED51296.1 afum|EAL85598.1 agla|OJJ81911.1 akaw|GAA86306.1 aluc|OJZ86143.1 anid|CBF71019.1 anig|EHA24481.1 aory|EIT81520.1 asyd|OJJ52657.1 ater|EAU32395.1 atub|OJI79935.1 aver|OJJ04685.1 awen|OJJ38455.1 azon|OJJ43755.1

Group1885: abra|OJJ69157.1 acar|OOF91146.1 acla|EAW08674.1 afis|EAW15750.1 afum|EAL85599.1 agla|OJJ81913.1 akaw|GAA86305.1 aluc|OJZ86144.1 anid|CBF71017.1 anig|EHA24480.1 aory|EIT81492.1 asyd|OJJ52656.1 ater|EAU32394.1 atub|OJI79936.1 aver|OJJ04684.1 awen|OJJ38456.1 azon|OJJ43756.1 afla|EED51295.1

Group1886: abra|OJJ69158.1 acar|OOF91145.1 acla|EAW08675.1 afis|EAW15749.1 afla|EED51294.1 afum|EAL85600.1 agla|OJJ81914.1 akaw|GAA86304.1 aluc|OJZ86145.1 anid|CBF71015.1 anig|EHA24479.1 aory|EIT81541.1 asyd|OJJ52655.1 ater|EAU32393.1 atub|OJI79937.1 aver|OJJ04683.1 awen|OJJ38457.1 azon|OJJ43757.1

Group1887: abra|OJJ69159.1 acar|OOF91144.1 acla|EAW08676.1 afis|EAW15748.1 afla|EED51293.1 afum|EAL85601.1 agla|OJJ81915.1 akaw|GAA86303.1 aluc|OJZ86146.1 anid|CBF71013.1 anig|EHA24478.1 aory|EIT81547.1 asyd|OJJ52654.1 ater|EAU32392.1 atub|OJI79938.1 aver|OJJ04682.1 awen|OJJ38458.1 azon|OJJ43759.1

Group1888: abra|OJJ69161.1 acar|OOF91141.1 acla|EAW08678.1 afis|EAW15746.1 afla|EED51291.1 afum|EAL85602.1 agla|OJJ81917.1 akaw|GAA86301.1 aluc|OJZ86148.1 anid|CBF71009.1 anig|EHA24476.1 aory|EIT81488.1 asyd|OJJ52651.1 ater|EAU32390.1 atub|OJI79940.1 aver|OJJ04680.1 awen|OJJ38460.1 azon|OJJ43761.1

Group1889: abra|OJJ69162.1 acar|OOF91140.1 acla|EAW08679.1 afis|EAW15754.1 afla|EED51290.1 afum|EAL85595.1 agla|OJJ81918.1 akaw|GAA86300.1 aluc|OJZ86149.1 anid|CBF71007.1 anig|EHA24475.1 aory|EIT81493.1 asyd|OJJ52650.1 ater|EAU32389.1 atub|OJI79941.1 aver|OJJ04679.1 awen|OJJ38462.1 azon|OJJ43762.1

Group1890: abra|OJJ69164.1 acar|OOF91139.1 acla|EAW08680.1 afis|EAW15758.1 afla|EED51289.1 afum|EAL85590.1 agla|OJJ81920.1 akaw|GAA86299.1 aluc|OJZ86151.1 anid|CBF71006.1 anig|EHA24474.1 aory|EIT81496.1 asyd|OJJ52648.1 ater|EAU32388.1 atub|OJI79943.1 aver|OJJ04677.1 awen|OJJ38466.1 azon|OJJ43764.1

Group1891: abra|OJJ69165.1 acar|OOF91138.1 acla|EAW08681.1 afis|EAW15759.1 afla|EED51288.1 afum|EAL85589.1 agla|OJJ81921.1 akaw|GAA86298.1 aluc|OJZ86152.1 anid|CBF71004.1 anig|EHA24473.1 aory|EIT81511.1 asyd|OJJ52647.1 ater|EAU32387.1 atub|OJI79944.1 aver|OJJ04676.1 awen|OJJ38467.1 azon|OJJ43765.1

Group1892: abra|OJJ69166.1 acar|OOF91137.1 acla|EAW08682.1 afis|EAW15760.1 afla|EED51287.1 afum|EAL85588.1 agla|OJJ81922.1 akaw|GAA86297.1 aluc|OJZ86153.1 anid|CBF71002.1 anig|EHA24472.1 aory|EIT81510.1 asyd|OJJ52646.1 ater|EAU32386.1 atub|OJI79945.1 aver|OJJ04675.1 awen|OJJ38468.1 azon|OJJ43766.1

Group1893: abra|OJJ69167.1 acar|OOF91136.1 acla|EAW08683.1 afis|EAW15761.1 afla|EED51286.1 afum|EAL85587.1 agla|OJJ81923.1 akaw|GAA86296.1 aluc|OJZ86154.1 anid|CBF71000.1 anig|EHA24471.1 aory|EIT81497.1 asyd|OJJ52645.1 ater|EAU32385.1 atub|OJI79946.1 aver|OJJ04674.1 awen|OJJ38469.1 azon|OJJ43767.1

Group1894: abra|OJJ69168.1 acar|OOF91135.1 acla|EAW08684.1 afis|EAW15762.1 afla|EED51285.1 afum|EAL85586.1 agla|OJJ81924.1 akaw|GAA86295.1 aluc|OJZ86155.1 anid|CBF70998.1 anig|EHA24470.1 aory|EIT81522.1 asyd|OJJ52644.1 ater|EAU32384.1 atub|OJI79947.1 aver|OJJ04673.1 awen|OJJ38470.1 azon|OJJ43769.1

Group1895: abra|OJJ69177.1 acar|OOF91129.1 acla|EAW08686.1 afis|EAW15763.1 afla|EED51284.1 afum|EAL85585.1 agla|OJJ81925.1 akaw|GAA86287.1 aluc|OJZ86163.1 anid|CBF70996.1 anig|EHA24461.1 aory|EIT81489.1 asyd|OJJ52643.1 ater|EAU32378.1 atub|OJI79955.1 aver|OJJ04672.1 awen|OJJ38471.1 azon|OJJ43770.1

Group1896: abra|OJJ69178.1 acar|OOF90317.1 acla|EAW08687.1 afis|EAW15764.1 afla|EED51283.1 afum|EAL85584.1 agla|OJJ81927.1 akaw|GAA86286.1 aluc|OJZ86164.1 anid|CBF70994.1 anig|EHA24460.1 aory|EIT81513.1 asyd|OJJ52642.1 ater|EAU32377.1 atub|OJI79956.1 aver|OJJ04670.1 awen|OJJ38472.1 azon|OJJ43772.1

Group1897: abra|OJJ69179.1 acar|OOF90318.1 acla|EAW08688.1 afis|EAW15765.1 afla|EED51282.1 afum|EAL85583.1 agla|OJJ81928.1 akaw|GAA86285.1 aluc|OJZ86165.1 anid|CBF70993.1 anig|EHA24459.1 aory|EIT81558.1 asyd|OJJ52641.1 ater|EAU32376.1 atub|OJI79957.1 aver|OJJ04669.1 awen|OJJ38473.1 azon|OJJ43773.1

Group1898: abra|OJJ69180.1 acar|OOF90319.1 acla|EAW08689.1 afis|EAW15766.1 afla|EED51281.1 afum|EAL85582.1 agla|OJJ81930.1 akaw|GAA86284.1 aluc|OJZ86166.1 anig|EHA24458.1 aory|EIT81544.1 asyd|OJJ52640.1 ater|EAU32375.1 atub|OJI79958.1 awen|OJJ38474.1 anid|CBF70991.1 aver|OJJ04668.1 azon|OJJ43774.1

Group1899: abra|OJJ69181.1 acar|OOF90321.1 acla|EAW08690.1 afis|EAW15767.1 afla|EED51280.1 afum|EAL85581.1 agla|OJJ81931.1 akaw|GAA86283.1 aluc|OJZ86167.1 anid|CBF70989.1 anig|EHA24457.1 aory|EIT81536.1 asyd|OJJ52639.1 ater|EAU32374.1 atub|OJI79959.1 aver|OJJ04667.1 awen|OJJ38475.1 azon|OJJ43775.1

Group1900: abra|OJJ69183.1 acar|OOF90320.1 acla|EAW08691.1 afis|EAW15768.1 afla|EED51279.1 afum|EAL85580.1 agla|OJJ81932.1 akaw|GAA86282.1 aluc|OJZ86168.1 anid|CBF70987.1 anig|EHA24456.1 aory|EIT81531.1 asyd|OJJ52638.1 ater|EAU32373.1 atub|OJI79960.1 aver|OJJ04666.1 awen|OJJ38476.1 azon|OJJ43776.1

Group1901: abra|OJJ69184.1 acar|OOF91128.1 acla|EAW08662.1 afis|EAW15769.1 afla|EED51278.1 afum|EAL85579.1 agla|OJJ81933.1 akaw|GAA86281.1 aluc|OJZ86170.1 anid|CBF70983.1 anig|EHA24455.1 aory|EIT81490.1 asyd|OJJ52636.1 ater|EAU32372.1 atub|OJI79962.1 aver|OJJ04664.1 awen|OJJ38477.1 azon|OJJ43777.1

Group1902: abra|OJJ69186.1 acar|OOF91126.1 acla|EAW08660.1 afis|EAW15771.1 afla|EED51276.1 afum|EAL85577.1 agla|OJJ81935.1 akaw|GAA86279.1 aluc|OJZ86172.1 anid|CBF70979.1 anig|EHA24453.1 aory|EIT81507.1 asyd|OJJ52634.1 ater|EAU32370.1 atub|OJI79964.1 aver|OJJ04662.1 awen|OJJ38479.1 azon|OJJ43779.1

Group1903: abra|OJJ69187.1 acar|OOF91125.1 acla|EAW08659.1 afis|EAW15772.1 afla|EED51275.1 afum|EAL85576.1 agla|OJJ81936.1 akaw|GAA86278.1 aluc|OJZ86173.1 anid|CBF70977.1 anig|EHA24452.1 aory|EIT81560.1 asyd|OJJ52633.1 ater|EAU32369.1 atub|OJI79965.1 aver|OJJ04661.1 awen|OJJ38480.1 azon|OJJ43780.1

Group1904: abra|OJJ69189.1 acar|OOF91122.1 acla|EAW08656.1 afis|EAW15775.1 afla|EED51271.1 afum|EAL85574.1 agla|OJJ81938.1 akaw|GAA86275.1 aluc|OJZ86175.1 anid|CBF70976.1 anig|EHA24449.1 aory|EIT81533.1 asyd|OJJ52631.1 ater|EAU32366.1 atub|OJI79967.1 aver|OJJ04659.1 awen|OJJ38482.1 azon|OJJ43783.1

Group1905: abra|OJJ69191.1 acar|OOF91120.1 acla|EAW08654.1 afis|EAW15777.1 afla|EED51268.1 afum|EAL85572.1 agla|OJJ81940.1 akaw|GAA86273.1 aluc|OJZ86177.1 anid|CBF70933.1 anig|EHA24447.1 aory|EIT81563.1 asyd|OJJ52629.1 ater|EAU32364.1 atub|OJI79969.1 aver|OJJ04657.1 awen|OJJ38484.1 azon|OJJ43785.1

Group1906: abra|OJJ69192.1 acar|OOF91121.1 acla|EAW08653.1 afis|EAW15778.1 afla|EED51266.1 afum|EAL85571.1 agla|OJJ81941.1 akaw|GAA86272.1 aluc|OJZ86178.1 anid|CBF70935.1 anig|EHA24446.1 aory|EIT81565.1 asyd|OJJ52628.1 ater|EAU32363.1 atub|OJI79970.1 aver|OJJ04656.1 awen|OJJ38485.1 azon|OJJ43786.1

Group1907: abra|OJJ69193.1 acar|OOF91119.1 acla|EAW08652.1 afis|EAW15779.1 afla|EED51265.1 afum|EAL85570.1 agla|OJJ81942.1 akaw|GAA86271.1 aluc|OJZ86179.1 anid|CBF70937.1 anig|EHA24445.1 aory|EIT81542.1 asyd|OJJ52627.1 ater|EAU32362.1 atub|OJI79971.1 aver|OJJ04655.1 awen|OJJ38486.1 azon|OJJ43787.1

Group1908: abra|OJJ69194.1 acar|OOF91118.1 acla|EAW08651.1 afis|EAW15780.1 afla|EED51264.1 afum|EBA27235.1 agla|OJJ81943.1 akaw|GAA86270.1 aluc|OJZ86180.1 anid|CBF70939.1 anig|EHA24444.1 aory|EIT81528.1 asyd|OJJ52626.1 ater|EAU32361.1 atub|OJI79972.1 aver|OJJ04654.1 awen|OJJ38487.1 azon|OJJ43788.1

Group1909: abra|OJJ69196.1 acar|OOF90027.1 acla|EAW08650.1 afis|EAW15781.1 afla|EED51263.1 afum|EAL85569.1 agla|OJJ81944.1 akaw|GAA92209.1 aluc|OJZ86181.1 anid|CBF70941.1 anig|EHA24443.1

aory|EIT81567.1 asyd|OJJ52625.1 ater|EAU32360.1 atub|OJI79973.1 aver|OJJ04652.1 awen|OJJ38488.1 azon|OJJ43789.1

Group1910: abra|OJJ69197.1 acar|OOF90026.1 acla|EAW08649.1 afis|EAW15782.1 afla|EED51262.1 afum|EAL85568.1 agla|OJJ81945.1 akaw|GAA92210.1 aluc|OJZ86182.1 anid|CBF70945.1 anig|EHA24442.1 aory|EIT81562.1 asyd|OJJ52624.1 ater|EAU32359.1 atub|OJI79974.1 aver|OJJ04651.1 awen|OJJ38489.1 azon|OJJ43790.1

Group1911: abra|OJJ69198.1 acar|OOF91117.1 acla|EAW08648.1 afis|EAW15783.1 afla|EED51261.1 afum|EAL85567.1 agla|OJJ81946.1 akaw|GAA92211.1 aluc|OJZ86183.1 anid|CBF70946.1 anig|EHA24441.1 aory|EIT81552.1 asyd|OJJ52623.1 ater|EAU32358.1 atub|OJI79975.1 aver|OJJ04650.1 awen|OJJ38490.1 azon|OJJ43791.1

Group1912: abra|OJJ69200.1 acar|OOF91115.1 acla|EAW08646.1 afis|EAW15785.1 afla|EED51259.1 afum|EAL85565.1 agla|OJJ81948.1 akaw|GAA92213.1 aluc|OJZ86185.1 anid|CBF70954.1 anig|EHA24439.1 aory|EIT81568.1 asyd|OJJ52621.1 ater|EAU32356.1 atub|OJI79977.1 aver|OJJ04648.1 awen|OJJ38492.1 azon|OJJ43794.1

Group1913: abra|OJJ69204.1 acar|OOF91112.1 acla|EAW08643.1 afis|EAW15788.1 afla|EED51257.1 afum|EAL85562.1 agla|OJJ81951.1 akaw|GAA92216.1 aluc|OJZ86188.1 anid|CBF70960.1 anig|EHA24435.1 aory|EIT81501.1 asyd|OJJ52618.1 ater|EAU32353.1 atub|OJI79980.1 aver|OJJ04645.1 awen|OJJ38496.1 azon|OJJ43797.1

Group1914: abra|OJJ69205.1 acar|OOF91109.1 acla|EAW08642.1 afis|EAW15789.1 afla|EED51256.1 afum|EAL85561.1 agla|OJJ81952.1 akaw|GAA92217.1 aluc|OJZ86189.1 anid|CBF70962.1 anig|EHA24434.1 aory|EIT81537.1 asyd|OJJ52617.1 ater|EAU32352.1 atub|OJI79981.1 aver|OJJ04644.1 awen|OJJ38497.1 azon|OJJ43798.1

Group1915: abra|OJJ69206.1 acar|OOF91111.1 acla|EAW08641.1 afis|EAW15790.1 afla|EED51255.1 afum|EAL85560.1 agla|OJJ81953.1 akaw|GAA92218.1 aluc|OJZ86190.1 anid|CBF70964.1 anig|EHA24433.1 aory|EIT81514.1 asyd|OJJ52616.1 ater|EAU32351.1 atub|OJI79982.1 aver|OJJ04643.1 awen|OJJ38498.1 azon|OJJ43799.1

Group1916: abra|OJJ69208.1 acar|OOF91108.1 acla|EAW08638.1 afis|EAW15793.1 afla|EED51252.1 afum|EAL85557.1 agla|OJJ81956.1 akaw|GAA92219.1 aluc|OJZ86191.1 anid|CBF70970.1 anig|EHA24431.1 aory|EIT81504.1 asyd|OJJ52613.1 ater|EAU32349.1 atub|OJI79983.1 aver|OJJ04640.1 awen|OJJ38501.1 azon|OJJ43802.1

Group1917: abra|OJJ69209.1 acar|OOF91107.1 acla|EAW08637.1 afis|EAW15794.1 afla|EED51251.1 afum|EAL85556.1 agla|OJJ81957.1 akaw|GAA92220.1 aluc|OJZ86192.1 anid|CBF70931.1 anig|EHA24430.1 aory|EIT81500.1 asyd|OJJ52612.1 ater|EAU32348.1 atub|OJI79984.1 aver|OJJ04639.1 awen|OJJ38502.1 azon|OJJ43803.1

Group1918: abra|OJJ69210.1 acar|OOF91106.1 acla|EAW08635.1 afis|EAW15796.1 afla|EED51249.1 afum|EAL85554.1 agla|OJJ81960.1 akaw|GAA92221.1 aluc|OJZ86193.1 anid|CBF70927.1 anig|EHA24429.1 aory|EIT81508.1 asyd|OJJ52610.1 ater|EAU32346.1 atub|OJI79985.1 aver|OJJ04637.1 awen|OJJ38504.1 azon|OJJ43747.1

Group1919: abra|OJJ69211.1 acar|OOF91105.1 acla|EAW08634.1 afis|EAW15797.1 afla|EED51248.1 afum|EAL85553.2 agla|OJJ81961.1 akaw|GAA92222.1 aluc|OJZ86194.1 anid|CBF70925.1 anig|EHA24428.1 aory|EIT81561.1 asyd|OJJ52609.1 ater|EAU32345.1 atub|OJI79986.1 aver|OJJ04636.1 awen|OJJ38505.1 azon|OJJ43746.1

Group1920: abra|OJJ69212.1 acar|OOF91104.1 acla|EAW08633.1 afis|EAW15798.1 afla|EED51247.1 afum|EAL85552.1 agla|OJJ81962.1 akaw|GAA92223.1 aluc|OJZ86195.1 anid|CBF70923.1 anig|EHA24427.1 aory|EIT81502.1 asyd|OJJ52608.1 ater|EAU32344.1 atub|OJI79988.1 aver|OJJ04635.1 awen|OJJ38506.1 azon|OJJ43745.1

Group1921: abra|OJJ69215.1 acar|OOF91103.1 acla|EAW08630.1 afis|EAW15801.1 afla|EED51244.1 afum|EAL85549.2 agla|OJJ81968.1 akaw|GAA92224.1 aluc|OJZ86196.1 anid|CBF70918.1 anig|EHA24426.1 aory|EIT83186.1 asyd|OJJ52605.1 ater|EAU33447.1 atub|OJI79989.1 aver|OJJ04632.1 awen|OJJ38510.1 azon|OJJ43742.1

Group1922: abra|OJJ69217.1 acar|OOF91101.1 acla|EAW08628.1 afis|EAW15803.1 afla|EED51242.1 afum|EAL85547.1 agla|OJJ81970.1 akaw|GAA92226.1 aluc|OJZ86199.1 anid|CBF70914.1 anig|EHA24424.1 aory|EIT83176.1 asyd|OJJ52603.1 ater|EAU33445.1 atub|OJI79991.1 aver|OJJ04629.1 awen|OJJ38513.1 azon|OJJ43739.1

Group1923: abra|OJJ69222.1 acar|OOF91098.1 acla|EAW12739.1 afis|EAW17902.1 afla|EED51238.1 afum|EAL91710.1 agla|OJJ81973.1 akaw|GAA92230.1 aluc|OJZ86202.1 anid|CBF70908.1 anig|EHA24420.1 aory|EIT83180.1 asyd|OJJ52599.1 ater|EAU33438.1 atub|OJI79996.1 aver|OJJ04626.1 awen|OJJ38518.1 azon|OJJ43736.1

Group1924: abra|OJJ69230.1 acar|OOF91090.1 acla|EAW08619.1 afis|EAW15815.1 afla|EED51232.1 afum|EAL85536.1 agla|OJJ81979.1 akaw|GAA92238.1 aluc|OJZ86210.1 anid|CBF70899.1 anig|EHA24416.1 aory|EIT83190.1 asyd|OJJ52595.1 ater|EAU33433.1 atub|OJI80004.1 aver|OJJ04622.1 awen|OJJ38528.1 azon|OJJ43733.1

Group1925: abra|OJJ69232.1 acar|OOF91088.1 acla|EAW08617.1 afis|EAW15817.1 afla|EED51230.1 afum|EAL85534.1 agla|OJJ81981.1 akaw|GAA92240.1 aluc|OJZ86212.1 anid|CBF70895.1 anig|EHA24414.1 aory|EIT83179.1 asyd|OJJ52593.1 ater|EAU33431.1 atub|OJI80007.1 aver|OJJ04620.1 awen|OJJ38531.1 azon|OJJ43731.1

Group1926: abra|OJJ69234.1 acar|OOF91086.1 acla|EAW08615.1 afis|EAW15819.1 afla|EED51228.1 afum|EAL85532.1 agla|OJJ81983.1 akaw|GAA92242.1 aluc|OJZ86214.1 anid|CBF70891.1 anig|EHA24412.1 aory|EIT83174.1 asyd|OJJ52591.1 ater|EAU33429.1 atub|OJI80009.1 aver|OJJ04618.1 awen|OJJ38533.1 azon|OJJ43729.1

Group1927: abra|OJJ69235.1 acar|OOF91085.1 acla|EAW08614.1 afis|EAW15820.1 afla|EED51227.1 afum|EAL85531.1 agla|OJJ81984.1 akaw|GAA92243.1 aluc|OJZ86215.1 anid|CBF70889.1 anig|EHA24411.1 aory|EIT83196.1 asyd|OJJ52590.1 ater|EAU33428.1 atub|OJI80010.1 aver|OJJ04617.1 awen|OJJ38534.1 azon|OJJ43728.1

Group1928: abra|OJJ69237.1 acar|OOF91081.1 acla|EAW08610.1 afis|EAW15823.1 afla|EED51223.1 afum|EAL85528.1 agla|OJJ81986.1 akaw|GAA92247.1 aluc|OJZ86217.1 anid|CBF70884.1 anig|EHA24409.1 aory|EIT83191.1 asyd|OJJ52588.1 ater|EAU33425.1 atub|OJI80012.1 aver|OJJ04615.1 awen|OJJ38536.1 azon|OJJ43726.1

Group1929: abra|OJJ69238.1 acar|OOF91080.1 acla|EAW08609.1 afis|EAW15824.1 afla|EED51221.1 afum|EAL85527.1 agla|OJJ81987.1 akaw|GAA92248.1 aluc|OJZ86218.1 anid|CBF70878.1 anig|EHA24408.1 aory|EIT80400.1 asyd|OJJ52586.1 ater|EAU33424.1 atub|OJI80013.1 aver|OJJ04613.1 awen|OJJ38538.1 azon|OJJ43725.1

Group1930: abra|OJJ69239.1 acar|OOF91079.1 acla|EAW08608.1 afis|EAW15825.1 afla|EED51220.1 afum|EAL85526.1 agla|OJJ81988.1 akaw|GAA92249.1 aluc|OJZ86220.1 anid|CBF70876.1 anig|EHA24407.1 aory|EIT80394.1 asyd|OJJ52584.1 ater|EAU33423.1 atub|OJI80014.1 aver|OJJ04612.1 awen|OJJ38539.1 azon|OJJ43724.1

Group1931: abra|OJJ69240.1 acar|OOF91078.1 acla|EAW08607.1 afis|EAW15826.1 afla|EED51219.1 afum|EAL85525.1 agla|OJJ81989.1 akaw|GAA92250.1 aluc|OJZ86221.1 anid|CBF70874.1 anig|EHA24406.1 aory|EIT80397.1 asyd|OJJ52583.1 ater|EAU33422.1 atub|OJI80015.1 aver|OJJ04611.1 awen|OJJ38540.1 azon|OJJ43723.1

Group1932: abra|OJJ69244.1 acar|OOF91076.1 acla|EAW08605.1 afis|EAW15828.1 afla|EED51217.1 afum|EAL85523.1 agla|OJJ81992.1 akaw|GAA92252.1 aluc|OJZ86225.1 anid|CBF70869.1 anig|EHA24403.1 aory|EIT80398.1 asyd|OJJ52580.1 ater|EAU33419.1 atub|OJI80018.1 aver|OJJ04608.1 awen|OJJ38542.1 azon|OJJ43721.1

Group1934: abra|OJJ69248.1 acar|OOF91073.1 acla|EAW08601.1 afis|EAW15832.1 afla|EED51214.1 afum|EAL85520.1 agla|OJJ81996.1 akaw|GAA92255.1 aluc|OJZ86229.1 anid|CBF70861.1 anig|EHA24400.1 aory|EIT80409.1 asyd|OJJ52576.1 ater|EAU33416.1 atub|OJI80022.1 aver|OJJ04605.1 awen|OJJ38545.1 azon|OJJ43717.1

Group1935: abra|OJJ69249.1 acar|OOF91072.1 acla|EAW08600.1 afis|EAW15833.1 afla|EED51213.1 afum|EAL85519.1 agla|OJJ81997.1 akaw|GAA92256.1 aluc|OJZ86230.1 anid|CBF70859.1 anig|EHA24399.1 aory|EIT80405.1 asyd|OJJ52575.1 ater|EAU33415.1 atub|OJI80023.1 aver|OJJ04604.1 awen|OJJ38546.1 azon|OJJ43716.1

Group1936: abra|OJJ69250.1 acar|OOF91071.1 acla|EAW08599.1 afis|EAW15834.1 afla|EED51212.1 afum|EAL85518.1 agla|OJJ81998.1 akaw|GAA92257.1 aluc|OJZ86231.1 anid|CBF70858.1 anig|EHA24398.1 aory|EIT80395.1 asyd|OJJ52574.1 ater|EAU33414.1 atub|OJI80024.1 aver|OJJ04603.1 awen|OJJ38547.1 azon|OJJ43715.1

Group1937: abra|OJJ69252.1 acar|OOF91069.1 acla|EAW08597.1 afis|EAW15836.1 afla|EED51210.1 afum|EAL85515.1 agla|OJJ82000.1 akaw|GAA92259.1 aluc|OJZ86232.1 anid|CBF70854.1 anig|EHA24397.1 aory|EIT80401.1 asyd|OJJ52573.1 ater|EAU33412.1 atub|OJI80026.1 aver|OJJ04602.1 awen|OJJ38550.1 azon|OJJ43713.1

Group1938: atub|OJI80028.1 abra|OJJ69256.1 acar|OOF91065.1 acla|EAW08593.1 afis|EAW15841.1 afla|EED51208.1 afum|EAL85510.1 agla|OJJ84917.1 akaw|GAA92262.1 aluc|OJZ86236.1 anid|CBF70845.1 anig|EHA24392.1 aory|EIT80413.1 asyd|OJJ52568.1 ater|EAU33394.1 aver|OJJ04597.1 awen|OJJ38555.1 azon|OJJ43711.1

Group1939: abra|OJJ69257.1 acar|OOF91064.1 acla|EAW08592.1 afis|EAW15842.1 afla|EED51207.1 afum|EAL85509.1 agla|OJJ84916.1 akaw|GAA92263.1 aluc|OJZ86237.1 anid|CBF70843.1 anig|EHA24391.1

aory|EIT80396.1 asyd|OJJ52567.1 ater|EAU33393.1 atub|OJI80029.1 aver|OJJ04596.1 awen|OJJ38556.1 azon|OJJ43710.1

Group1941: abra|OJJ69262.1 acar|OOF91060.1 acla|EAW08589.1 afis|EAW15846.1 afla|EED51204.1 afum|EAL85505.1 agla|OJJ84913.1 akaw|GAA92267.1 aluc|OJZ86241.1 anid|CBF70835.1 anig|EHA24388.1 aory|EIT80414.1 asyd|OJJ52564.1 ater|EAU33390.1 atub|OJI80033.1 aver|OJJ04592.1 awen|OJJ38561.1 azon|OJJ48761.1

Group1942: abra|OJJ69263.1 acar|OOF91059.1 acla|EAW08588.1 afis|EAW15847.1 afla|EED51203.1 afum|EAL85504.1 agla|OJJ84911.1 akaw|GAA92268.1 aluc|OJZ86242.1 anid|CBF70833.1 anig|EHA24387.1 aory|EIT80410.1 asyd|OJJ52563.1 ater|EAU33389.1 atub|OJI80034.1 aver|OJJ04591.1 awen|OJJ38562.1 azon|OJJ43615.1

Group1943: abra|OJJ69268.1 acar|OOF89861.1 acla|EAW14653.1 afis|EAW24188.1 afla|EED48081.1 afum|EAL85861.1 agla|OJJ79691.1 akaw|GAA88143.1 aluc|OJZ82097.1 anid|CBF74306.1 anig|EHA28632.1 aory|EIT82476.1 asyd|OJJ54008.1 ater|EAU31140.1 atub|OJI88340.1 aver|OJJ08103.1 awen|OJJ33111.1 azon|OJJ41999.1

Group1944: abra|OJJ69275.1 acar|OOF94447.1 acla|EAW14663.1 afis|EAW24194.1 afla|EED48077.1 afum|EAL85866.1 agla|OJJ79684.1 akaw|GAA88148.1 aluc|OJZ82102.1 anid|CBF74313.1 anig|EHA28627.1 aory|EIT82481.1 asyd|OJJ54002.1 ater|EAU31145.1 atub|OJI88333.1 aver|OJJ08099.1 awen|OJJ33105.1 azon|OJJ44844.1

Group1945: abra|OJJ69276.1 acar|OOF94446.1 acla|EAW14664.1 afis|EAW24195.1 afla|EED48076.1 afum|EAL85867.1 agla|OJJ79683.1 akaw|GAA88149.1 aluc|OJZ82103.1 anid|CBF74315.1 anig|EHA28626.1 aory|EIT82479.1 asyd|OJJ54001.1 ater|EAU31146.1 atub|OJI88332.1 aver|OJJ08098.1 awen|OJJ33104.1 azon|OJJ44845.1

Group1946: abra|OJJ69277.1 acar|OOF94445.1 acla|EAW14665.1 afis|EAW24196.1 afla|EED48075.1 afum|EAL85868.1 agla|OJJ79682.1 akaw|GAA88150.1 aluc|OJZ82104.1 anid|CBF74317.1 anig|EHA28625.1 aory|EIT82482.1 asyd|OJJ54000.1 ater|EAU31147.1 atub|OJI88331.1 aver|OJJ08097.1 awen|OJJ33103.1 azon|OJJ44846.1

Group1947: abra|OJJ69278.1 acar|OOF94444.1 acla|EAW14666.1 afis|EAW24197.1 afla|EED48074.1 afum|EAL85869.1 agla|OJJ79681.1 akaw|GAA88151.1 aluc|OJZ82105.1 anid|CBF74319.1 anig|EHA28624.1 aory|EIT82477.1 asyd|OJJ53999.1 ater|EAU31148.1 atub|OJI88330.1 aver|OJJ08096.1 awen|OJJ33102.1 azon|OJJ44847.1

Group1948: abra|OJJ69279.1 acar|OOF94443.1 acla|EAW14668.1 afis|EAW24199.1 afla|EED48073.1 afum|EAL85871.1 agla|OJJ79678.1 akaw|GAA88152.1 aluc|OJZ82106.1 anid|CBF74323.1 anig|EHA28623.1 aory|EIT83321.1 asyd|OJJ53996.1 ater|EAU31150.1 atub|OJI88329.1 aver|OJJ08094.1 awen|OJJ33099.1 azon|OJJ44848.1

Group1949: abra|OJJ69281.1 acar|OOF94441.1 acla|EAW14670.1 afis|EAW24201.1 afla|EED48071.1 afum|EBA27248.1 agla|OJJ79676.1 akaw|GAA88153.1 aluc|OJZ82109.1 anid|CBF74326.1 anig|EHA28621.1 aory|EIT83320.1 asyd|OJJ53994.1 ater|EAU31152.1 atub|OJI88327.1 aver|OJJ08092.1 awen|OJJ33097.1 azon|OJJ44850.1

Group1950: abra|OJJ69283.1 acar|OOF94439.1 acla|EAW14672.1 afis|EAW24203.1 afla|EED48069.1 afum|EAL85873.2 agla|OJJ79674.1 akaw|GAA88155.1 aluc|OJZ82112.1 anid|CBF74330.1 anig|EHA28619.1 aory|EIT83340.1 asyd|OJJ53992.1 ater|EAU31154.1 atub|OJI88325.1 aver|OJJ08090.1 awen|OJJ33095.1 azon|OJJ44852.1

Group1951: abra|OJJ69284.1 acar|OOF94438.1 acla|EAW14673.1 afis|EAW24204.1 afla|EED48067.1 afum|EAL85874.2 agla|OJJ79673.1 akaw|GAA88156.1 aluc|OJZ82113.1 anid|CBF74332.1 anig|EHA28618.1 aory|EIT83335.1 asyd|OJJ53991.1 ater|EAU31155.1 atub|OJI88324.1 aver|OJJ08089.1 awen|OJJ33094.1 azon|OJJ44854.1

Group1952: abra|OJJ69285.1 acar|OOF94436.1 acla|EAW14674.1 afis|EAW24205.1 afla|EED48066.1 afum|EAL85875.1 agla|OJJ79672.1 akaw|GAA88157.1 aluc|OJZ82114.1 anid|CBF74272.1 anig|EHA28617.1 aory|EIT83356.1 asyd|OJJ53990.1 ater|EAU31156.1 atub|OJI88323.1 aver|OJJ08088.1 awen|OJJ33093.1 azon|OJJ44855.1

Group1953: abra|OJJ69286.1 acar|OOF94437.1 acla|EAW14675.1 afis|EAW24206.1 afla|EED48065.1 afum|EAL85876.1 agla|OJJ79671.1 akaw|GAA88158.1 aluc|OJZ82115.1 anid|CBF74271.1 anig|EHA28616.1 aory|EIT83359.1 asyd|OJJ53989.1 ater|EAU31157.1 atub|OJI88322.1 aver|OJJ08087.1 awen|OJJ33092.1 azon|OJJ44856.1

Group1954: abra|OJJ69288.1 acar|OOF94434.1 acla|EAW14677.1 afis|EAW24208.1 afla|EED48063.1 afum|EAL85878.1 agla|OJJ79669.1 akaw|GAA88160.1 aluc|OJZ82117.1 anid|CBF74269.1 anig|EHA28614.1 aory|EIT83351.1 asyd|OJJ53987.1 ater|EAU31158.1 atub|OJI88320.1 aver|OJJ08085.1 awen|OJJ33089.1 azon|OJJ44858.1

Group1955: abra|OJJ69292.1 acar|OOF94431.1 acla|EAW14680.1 afis|EAW24211.1 afla|EED48060.1 afum|EAL85880.1 agla|OJJ79654.1 akaw|GAA88163.1 aluc|OJZ82121.1 anid|CBF74263.1 anig|EHA28611.1 aory|EIT83350.1 asyd|OJJ53983.1 ater|EAU31161.1 atub|OJI88317.1 aver|OJJ08082.1 awen|OJJ33085.1 azon|OJJ44861.1

Group1956: abra|OJJ69293.1 acar|OOF94430.1 acla|EAW14681.1 afis|EAW24212.1 afla|EED48059.1 afum|EAL85881.1 agla|OJJ79663.1 akaw|GAA88164.1 aluc|OJZ82122.1 anid|CBF74261.1 anig|EHA28610.1 aory|EIT83323.1 asyd|OJJ53982.1 ater|EAU31162.1 atub|OJI88316.1 aver|OJJ08081.1 awen|OJJ33084.1 azon|OJJ44862.1

Group1957: abra|OJJ69294.1 acar|OOF94429.1 acla|EAW14682.1 afis|EAW24213.1 afla|EED48058.1 afum|EAL85882.1 agla|OJJ79665.1 akaw|GAA88165.1 aluc|OJZ82123.1 anid|CBF74260.1 anig|EHA28609.1 aory|EIT83345.1 asyd|OJJ53981.1 ater|EAU31163.1 atub|OJI88315.1 aver|OJJ08080.1 awen|OJJ33083.1 azon|OJJ44863.1

Group1958: abra|OJJ69296.1 acar|OOF94428.1 acla|EAW14683.1 afis|EAW24214.1 afla|EED48057.1 afum|EAL85883.1 agla|OJJ79666.1 akaw|GAA88166.1 aluc|OJZ82125.1 anid|CBF74258.1 anig|EHA28607.1 aory|EIT83357.1 asyd|OJJ53980.1 ater|EAU31164.1 atub|OJI88313.1 aver|OJJ08079.1 awen|OJJ33082.1 azon|OJJ44864.1

Group1959: abra|OJJ69297.1 acar|OOF94427.1 acla|EAW14684.1 afis|EAW24215.1 afla|EED48056.1 afum|EAL85884.1 agla|OJJ79667.1 akaw|GAA88167.1 aluc|OJZ82126.1 anid|CBF74256.1 anig|EHA28606.1 aory|EIT83352.1 asyd|OJJ53979.1 ater|EAU31165.1 atub|OJI88312.1 aver|OJJ08078.1 awen|OJJ33081.1 azon|OJJ44865.1

Group1960: abra|OJJ69298.1 acar|OOF94425.1 acla|EAW14685.1 afis|EAW24216.1 afla|EED48055.1 afum|EAL85885.1 agla|OJJ79668.1 akaw|GAA88168.1 aluc|OJZ82127.1 anid|CBF74254.1 anig|EHA28605.1 aory|EIT83339.1 asyd|OJJ53978.1 ater|EAU31166.1 atub|OJI88311.1 aver|OJJ08077.1 awen|OJJ33080.1 azon|OJJ44866.1

Group1961: abra|OJJ69299.1 acar|OOF94426.1 acla|EAW14686.1 afis|EAW24217.1 afla|EED48054.1 afum|EAL85886.1 agla|OJJ79653.1 akaw|GAA88169.1 aluc|OJZ82128.1 anid|CBF74252.1 anig|EHA28603.1 aory|EIT83360.1 asyd|OJJ53977.1 ater|EAU31167.1 atub|OJI88310.1 aver|OJJ08076.1 awen|OJJ33079.1 azon|OJJ44867.1

Group1962: abra|OJJ69302.1 acar|OOF94423.1 acla|EAW14688.1 afis|EAW24219.1 afla|EED48052.1 afum|EAL85888.1 agla|OJJ79651.1 akaw|GAA88171.1 aluc|OJZ82130.1 anid|CBF74248.1 anig|EHA28601.1 aory|EIT83354.1 asyd|OJJ53975.1 ater|EAU31169.1 atub|OJI88306.1 aver|OJJ08074.1 awen|OJJ33077.1 azon|OJJ44869.1

Group1963: abra|OJJ69305.1 acar|OOF94420.1 acla|EAW14691.1 afis|EAW24222.1 afla|EED48050.1 afum|EAL85891.2 agla|OJJ79648.1 akaw|GAA88174.1 aluc|OJZ82134.1 anid|CBF74239.1 anig|EHA28597.1 aory|EIT83349.1 asyd|OJJ53971.1 ater|EAU31172.1 atub|OJI88303.1 aver|OJJ08070.1 awen|OJJ33074.1 azon|OJJ44873.1

Group1964: abra|OJJ69307.1 acar|OOF94419.1 acla|EAW14694.1 afis|EAW24225.1 afla|EED48046.1 afum|EAL85893.1 agla|OJJ79646.1 akaw|GAA88176.1 aluc|OJZ82136.1 anid|CBF74234.1 anig|EHA28595.1 aory|EIT83343.1 asyd|OJJ53969.1 ater|EAU31174.1 atub|OJI88301.1 aver|OJJ08067.1 awen|OJJ33071.1 azon|OJJ44875.1

Group1965: abra|OJJ69308.1 acar|OOF94418.1 acla|EAW14695.1 afis|EAW24226.1 afla|EED48045.1 afum|EAL85894.1 agla|OJJ79645.1 akaw|GAA88177.1 aluc|OJZ82137.1 anid|CBF74232.1 anig|EHA28594.1 aory|EIT83341.1 asyd|OJJ53968.1 ater|EAU31175.1 atub|OJI88300.1 aver|OJJ08066.1 awen|OJJ33070.1 azon|OJJ44877.1

Group1966: abra|OJJ69312.1 acar|OOF94415.1 acla|EAW14738.1 afis|EAW24269.1 afla|EED48008.1 afum|EAL85937.1 agla|OJJ79596.1 akaw|GAA88181.1 aluc|OJZ82141.1 anid|CBF74167.1 anig|EHA28591.1 aory|EIT82915.1 asyd|OJJ53936.1 ater|EAU31214.1 atub|OJI88296.1 aver|OJJ08034.1 awen|OJJ33031.1 azon|OJJ48488.1

Group1967: abra|OJJ69313.1 acar|OOF94413.1 acla|EAW14737.1 afis|EAW24268.1 afla|EED48009.1 afum|EAL85936.1 agla|OJJ79597.1 akaw|GAA88182.1 aluc|OJZ82142.1 anid|CBF74169.1 anig|EHA28589.1 aory|EIT82903.1 asyd|OJJ53937.1 ater|EAU31213.1 atub|OJI88295.1 aver|OJJ08035.1 awen|OJJ33032.1 azon|OJJ48487.1

Group1968: abra|OJJ69314.1 acar|OOF94412.1 acla|EAW14736.1 afis|EAW24267.1 afla|EED48010.1 afum|EAL85935.1 agla|OJJ79598.1 akaw|GAA88183.1 aluc|OJZ82143.1 anid|CBF74171.1 anig|EHA28588.1 aory|EIT82909.1 asyd|OJJ53938.1 ater|EAU31212.1 atub|OJI88294.1 aver|OJJ08036.1 awen|OJJ33033.1 azon|OJJ48486.1

Group1969: abra|OJJ69315.1 acar|OOF90302.1 acla|EAW14734.1 afis|EAW24266.1 afla|EED48011.1 afum|EAL85934.1 agla|OJJ79606.1 akaw|GAA88184.1 aluc|OJZ82144.1 anid|CBF74173.1 anig|EHA28586.1

aory|EIT82916.1 asyd|OJJ53940.1 ater|EAU31210.1 atub|OJI88292.1 aver|OJJ08037.1 awen|OJJ33034.1 azon|OJJ48485.1

Group1970: abra|OJJ69316.1 acar|OOF90303.1 acla|EAW14732.1 afis|EAW24265.1 afla|EED48012.1 afum|EAL85933.1 agla|OJJ79607.1 akaw|GAA88185.1 aluc|OJZ82145.1 anid|CBF74175.1 anig|EHA28585.1 aory|EIT82905.1 asyd|OJJ53941.1 ater|EAU31209.1 atub|OJI88291.1 aver|OJJ08038.1 awen|OJJ33035.1 azon|OJJ48484.1

Group1971: abra|OJJ69319.1 acar|OOF94411.1 acla|EAW14726.1 afis|EAW24262.1 akaw|GAA88188.1 aluc|OJZ82148.1 ater|EAU31207.1 atub|OJI88287.1 aver|OJJ08043.1 awen|OJJ33039.1 azon|OJJ48479.1 afum|EAL85930.1 agla|OJJ79610.1 anid|CBF74186.1 anig|EHA28582.1 aory|EIT82904.1 asyd|OJJ53946.1 afla|EED48015.1

Group1972: abra|OJJ69329.1 acar|OOF94398.1 acla|EAW14711.1 afis|EAW24246.1 afla|EED48028.1 afum|EAL85914.1 agla|OJJ79627.1 akaw|GAA88200.1 aluc|OJZ82161.1 anid|CBF74210.1 anig|EHA28571.1 aory|EIT83355.1 asyd|OJJ53956.1 ater|EAU31187.1 atub|OJI88276.1 aver|OJJ08052.1 awen|OJJ33058.1 azon|OJJ48466.1

Group1973: abra|OJJ69332.1 acar|OOF94395.1 acla|EAW14708.1 afis|EAW24239.1 afla|EED48031.1 afum|EAL85907.1 agla|OJJ79630.1 akaw|GAA88203.1 aluc|OJZ82164.1 anid|CBF74213.1 anig|EHA28568.1 aory|EIT83322.1 asyd|OJJ53957.1 ater|EAU31186.1 atub|OJI88273.1 aver|OJJ08053.1 awen|OJJ33054.1 azon|OJJ48465.1

Group1974: anig|EHA28560.1 abra|OJJ69342.1 acar|OOF94385.1 acla|EAW14704.1 afis|EAW24235.1 afla|EED48036.1 afum|EAL85904.1 agla|OJJ79636.1 akaw|GAA88211.1 aluc|OJZ82174.1 anid|CBF74219.1 aory|EIT83346.1 asyd|OJJ53961.1 ater|EAU31183.1 atub|OJI88263.1 aver|OJJ08057.1 awen|OJJ33063.1 azon|OJJ48461.1

Group1975: abra|OJJ69343.1 acar|OOF94386.1 acla|EAW14703.1 afis|EAW24234.1 afla|EED48037.1 afum|EAL85903.1 agla|OJJ79637.1 akaw|GAA88212.1 aluc|OJZ82175.1 anid|CBF74221.1 anig|EHA28559.1 aory|EIT83326.1 asyd|OJJ53962.1 ater|EAU31182.1 atub|OJI88262.1 aver|OJJ08058.1 awen|OJJ33064.1 azon|OJJ48460.1

Group1976: abra|OJJ69345.1 acar|OOF94379.1 acla|EAW14701.1 afis|EAW24232.1 afla|EED48039.1 afum|EAL85901.2 agla|OJJ79639.1 akaw|GAA88214.1 aluc|OJZ82177.1 anid|CBF74223.1 anig|EHA28557.1 aory|EIT83353.1 asyd|OJJ53964.1 ater|EAU31180.1 atub|OJI88260.1 aver|OJJ08060.1 awen|OJJ33065.1 azon|OJJ48458.1

Group1977: abra|OJJ69346.1 acar|OOF94378.1 acla|EAW14700.1 afis|EAW24231.1 afla|EED48040.1 afum|EAL85900.1 agla|OJJ79640.1 akaw|GAA88215.1 aluc|OJZ82178.1 anid|CBF74224.1 anig|EHA28556.1 aory|EIT83342.1 asyd|OJJ53965.1 ater|EAU31179.1 atub|OJI88259.1 aver|OJJ08061.1 awen|OJJ33066.1 azon|OJJ48457.1

Group1978: abra|OJJ69349.1 acar|OOF94376.1 acla|EAW14739.1 afis|EAW24270.1 afla|EED48007.1 afum|EAL85938.1 agla|OJJ79595.1 akaw|GAA88219.1 aluc|OJZ82181.1 anid|CBF74165.1 anig|EHA28553.1 aory|EIT82896.1 asyd|OJJ53935.1 ater|EAU31215.1 atub|OJI88254.1 aver|OJJ08033.1 awen|OJJ33030.1 azon|OJJ48489.1

Group1979: abra|OJJ69351.1 acar|OOF94375.1 acla|EAW14740.1 afis|EAW24271.1 afla|EED48006.1 afum|EAL85939.1 agla|OJJ79594.1 akaw|GAA88220.1 aluc|OJZ82183.1 anid|CBF74163.1 anig|EHA28552.1 aory|EIT82918.1 asyd|OJJ53933.1 ater|EAU31216.1 atub|OJI88253.1 aver|OJJ08031.1 awen|OJJ33028.1 azon|OJJ48490.1

Group1980: abra|OJJ69353.1 acar|OOF94372.1 acla|EAW14743.1 afis|EAW24274.1 afla|EED48002.1 afum|EAL85942.1 agla|OJJ79587.1 akaw|GAA88222.1 aluc|OJZ82186.1 anid|CBF74158.1 anig|EHA28550.1 aory|EIT82912.1 asyd|OJJ53927.1 ater|EAU31221.1 atub|OJI88251.1 aver|OJJ08026.1 awen|OJJ33023.1 azon|OJJ48494.1

Group1981: abra|OJJ69355.1 acar|OOF94370.1 acla|EAW14745.1 afis|EAW24276.1 afla|EED48000.1 afum|EAL85945.1 agla|OJJ79585.1 akaw|GAA88224.1 aluc|OJZ82187.1 anid|CBF74156.1 anig|EHA28548.1 aory|EIT82907.1 asyd|OJJ53925.1 ater|EAU31219.1 atub|OJI88249.1 aver|OJJ08024.1 awen|OJJ33021.1 azon|OJJ48496.1

Group1982: abra|OJJ69357.1 acar|OOF94367.1 acla|EAW14748.1 afis|EAW24279.1 afla|EED47997.1 afum|EAL85948.1 agla|OJJ80633.1 akaw|GAA88227.1 aluc|OJZ82189.1 anid|CBF74151.1 anig|EHA28546.1 aory|EIT82902.1 asyd|OJJ53923.1 ater|EAU31228.1 atub|OJI88246.1 aver|OJJ08021.1 awen|OJJ33018.1 azon|OJJ48499.1

Group1983: abra|OJJ69366.1 acar|OOF94359.1 acla|EAW14756.1 afis|EAW24287.1 afla|EED47989.1 afum|EAL85956.1 agla|OJJ83152.1 akaw|GAA88235.1 aluc|OJZ82200.1 anid|CBF74129.1 anig|EHA28539.1 aory|EIT79810.1 asyd|OJJ53912.1 ater|EAU31238.1 atub|OJI88236.1 aver|OJJ08010.1 awen|OJJ33009.1 azon|OJJ48506.1

Group1984: abra|OJJ69367.1 acar|OOF94358.1 acla|EAW14757.1 afis|EAW24288.1 afla|EED47988.1 afum|EAL85957.1 agla|OJJ83153.1 akaw|GAA88236.1 aluc|OJZ82201.1 anid|CBF74127.1 anig|EHA28538.1 aory|EIT79844.1 asyd|OJJ53911.1 ater|EAU31239.1 atub|OJI88235.1 aver|OJJ08009.1 awen|OJJ33008.1 azon|OJJ48507.1

Group1985: abra|OJJ69369.1 acar|OOF89837.1 acla|EAW14759.1 afis|EAW24290.1 afla|EED47986.1 afum|EAL85958.1 agla|OJJ83155.1 akaw|GAA88238.1 aluc|OJZ82203.1 anid|CBF74125.1 anig|EHA28536.1 aory|EIT79849.1 asyd|OJJ53909.1 ater|EAU31240.1 atub|OJI88233.1 aver|OJJ08007.1 awen|OJJ33006.1 azon|OJJ48509.1

Group1986: abra|OJJ69370.1 acar|OOF94357.1 acla|EAW14760.1 afis|EAW24291.1 afla|EED47985.1 afum|EAL85959.1 agla|OJJ83156.1 akaw|GAA88239.1 aluc|OJZ82204.1 anid|CBF74123.1 anig|EHA28535.1 aory|EIT79854.1 asyd|OJJ53908.1 ater|EAU31241.1 atub|OJI88232.1 aver|OJJ08006.1 awen|OJJ33005.1 azon|OJJ48510.1

Group1987: abra|OJJ69371.1 acar|OOF94355.1 acla|EAW14761.1 afis|EAW24292.1 afla|EED47984.1 afum|EAL85960.1 agla|OJJ83157.1 akaw|GAA88240.1 aluc|OJZ82205.1 anid|CBF74121.1 anig|EHA28534.1 aory|EIT79808.1 asyd|OJJ53907.1 ater|EAU31242.1 atub|OJI88231.1 aver|OJJ08005.1 awen|OJJ33004.1 azon|OJJ48511.1

Group1988: abra|OJJ69373.1 acar|OOF94354.1 acla|EAW14763.1 afis|EAW24294.1 afla|EED47982.1 afum|EAL85962.1 agla|OJJ83159.1 akaw|GAA88242.1 aluc|OJZ82208.1 anid|CBF74118.1 anig|EHA28532.1 aory|EIT79845.1 asyd|OJJ53905.1 ater|EAU31244.1 atub|OJI88228.1 aver|OJJ08003.1 awen|OJJ33001.1 azon|OJJ48513.1

Group1989: abra|OJJ69375.1 acar|OOF94351.1 acla|EAW14765.1 afis|EAW24296.1 afla|EED47980.1 afum|EAL85964.1 agla|OJJ83161.1 akaw|GAA88244.1 aluc|OJZ82210.1 anid|CBF75402.1 anig|EHA28530.1 aory|EIT79827.1 asyd|OJJ59822.1 ater|EAU31246.1 atub|OJI88226.1 aver|OJJ02465.1 awen|OJJ32998.1 azon|OJJ48514.1

Group1990: abra|OJJ69376.1 acar|OOF94349.1 acla|EAW14766.1 afis|EAW24297.1 afla|EED47979.1 afum|EAL85965.1 agla|OJJ83162.1 akaw|GAA88245.1 aluc|OJZ82211.1 anid|CBF74116.1 anig|EHA28529.1 aory|EIT79833.1 asyd|OJJ53904.1 ater|EAU31248.1 atub|OJI88225.1 aver|OJJ08001.1 awen|OJJ32997.1 azon|OJJ48515.1

Group1991: abra|OJJ69381.1 acar|OOF94344.1 acla|EAW14768.1 afis|EAW24299.1 afla|EED47973.1 afum|EAL85967.1 agla|OJJ83167.1 akaw|GAA88249.1 aluc|OJZ82216.1 anid|CBF74108.1 anig|EHA28524.1 aory|EIT79831.1 asyd|OJJ53901.1 ater|EAU31250.1 atub|OJI88219.1 aver|OJJ07998.1 awen|OJJ32993.1 azon|OJJ48523.1

Group1992: abra|OJJ69383.1 acar|OOF94343.1 acla|EAW14770.1 afis|EAW24301.1 afla|EED47971.1 afum|EAL85969.1 agla|OJJ83169.1 akaw|GAA88251.1 aluc|OJZ82217.1 anid|CBF74104.1 anig|EHA28523.1 aory|EIT79840.1 asyd|OJJ53899.1 ater|EAU31252.1 atub|OJI88218.1 aver|OJJ07996.1 awen|OJJ32991.1 azon|OJJ48525.1

Group1993: abra|OJJ69384.1 acar|OOF94342.1 acla|EAW14771.1 afis|EAW24302.1 afla|EED47970.1 afum|EAL85970.1 agla|OJJ83170.1 akaw|GAA88252.1 aluc|OJZ82218.1 anid|CBF74102.1 anig|EHA28522.1 aory|EIT79832.1 asyd|OJJ53898.1 ater|EAU31253.1 atub|OJI88217.1 aver|OJJ07995.1 awen|OJJ32990.1 azon|OJJ45840.1

Group1994: abra|OJJ69385.1 acar|OOF94341.1 acla|EAW14772.1 afis|EAW24303.1 afla|EED47969.1 afum|EAL85971.1 agla|OJJ83171.1 akaw|GAA88253.1 aluc|OJZ82219.1 anid|CBF74100.1 anig|EHA28521.1 aory|EIT79817.1 asyd|OJJ53897.1 ater|EAU31256.1 atub|OJI88216.1 aver|OJJ07994.1 awen|OJJ32989.1 azon|OJJ48526.1

Group1995: abra|OJJ69386.1 acar|OOF94340.1 acla|EAW14773.1 afis|EAW24304.1 afla|EED47968.1 afum|EAL85972.1 agla|OJJ83172.1 akaw|GAA88254.1 aluc|OJZ82220.1 anid|CBF79548.1 anig|EHA28520.1 aory|EIT79824.1 asyd|OJJ62162.1 ater|EAU31257.1 atub|OJI88215.1 aver|OJI96272.1 awen|OJJ32988.1 azon|OJJ48527.1

Group1996: abra|OJJ69387.1 acar|OOF94339.1 acla|EAW14774.1 afis|EAW24305.1 afla|EED47965.1 afum|EAL85973.1 agla|OJJ83173.1 akaw|GAA88255.1 aluc|OJZ82221.1 anid|CBF74098.1 anig|EHA28519.1 aory|EIT79856.1 asyd|OJJ53896.1 ater|EAU31258.1 atub|OJI88214.1 aver|OJJ07993.1 awen|OJJ32987.1 azon|OJJ48932.1

Group1997: abra|OJJ69398.1 acar|OOF94332.1 acla|EAW14780.1 afis|EAW24311.1 afla|EED47956.1 afum|EAL85979.1 agla|OJJ83179.1 akaw|GAA88264.1 aluc|OJZ82234.1 anid|CBF74081.1 anig|EHA28507.1 aory|EIT79851.1 asyd|OJJ53882.1 ater|EAU31264.1 atub|OJI88204.1 aver|OJJ07980.1 awen|OJJ32981.1 azon|OJJ48936.1

Group1998: abra|OJJ69400.1 acar|OOF94330.1 acla|EAW14782.1 afis|EAW24313.1 afla|EED47954.1 afum|EAL85981.1 agla|OJJ83181.1 akaw|GAA88266.1 aluc|OJZ82236.1 anid|CBF74079.1 anig|EHA28505.1

aory|EIT79862.1 asyd|OJJ53880.1 ater|EAU31266.1 atub|OJI88202.1 aver|OJJ07978.1 awen|OJJ32979.1 azon|OJJ48938.1

Group1999: abra|OJJ69402.1 acar|OOF94329.1 acla|EAW14784.1 afis|EAW24315.1 afla|EED47952.1 afum|EAL85983.1 agla|OJJ83183.1 akaw|GAA88267.1 aluc|OJZ82238.1 anid|CBF74078.1 anig|EHA28503.1 aory|EIT79841.1 asyd|OJJ53878.1 ater|EAU31267.1 atub|OJI88200.1 aver|OJJ07976.1 awen|OJJ32977.1 azon|OJJ48940.1

Group2000: abra|OJJ69404.1 acar|OOF94327.1 acla|EAW14786.1 afis|EAW24317.1 afla|EED47950.1 afum|EAL85985.1 agla|OJJ83185.1 akaw|GAA88269.1 aluc|OJZ82240.1 anid|CBF74074.1 anig|EHA28501.1 aory|EIT79859.1 asyd|OJJ53876.1 ater|EAU31270.1 atub|OJI88198.1 aver|OJJ07974.1 awen|OJJ32975.1 azon|OJJ48942.1

Group2001: abra|OJJ69405.1 acar|OOF94326.1 acla|EAW14787.1 afis|EAW24318.1 afla|EED47949.1 afum|EAL85987.1 agla|OJJ83186.1 akaw|GAA88270.1 aluc|OJZ82242.1 anid|CBF74073.1 anig|EHA28500.1 aory|EIT79816.1 asyd|OJJ53875.1 ater|EAU31271.1 atub|OJI88196.1 aver|OJJ07973.1 awen|OJJ32974.1 azon|OJJ48943.1

Group2002: abra|OJJ69410.1 acar|OOF94323.1 afla|EED47946.1 afum|EAL85990.1 akaw|GAA88277.1 aluc|OJZ82247.1 anid|CBF74069.1 anig|EHA28494.1 aory|EIT79850.1 asyd|OJJ53873.1 ater|EAU31273.1 atub|OJI88191.1 aver|OJJ07971.1 azon|OJJ48945.1 acla|EAW14789.1 afis|EAW24320.1 agla|OJJ83189.1 awen|OJJ32972.1

Group2003: abra|OJJ69411.1 acar|OOF94322.1 acla|EAW14790.1 afis|EAW24321.1 afla|EED47945.1 afum|EAL85991.1 agla|OJJ83190.1 akaw|GAA88278.1 aluc|OJZ82248.1 anid|CBF74068.1 anig|EHA28493.1 aory|EIT79836.1 asyd|OJJ53872.1 ater|EAU31274.1 atub|OJI88190.1 aver|OJJ07970.1 awen|OJJ32971.1 azon|OJJ48946.1

Group2004: abra|OJJ69412.1 acar|OOF94321.1 acla|EAW14791.1 afis|EAW24322.1 afla|EED47944.1 afum|EAL85992.1 agla|OJJ83191.1 akaw|GAA88279.1 aluc|OJZ82249.1 anid|CBF74066.1 anig|EHA28492.1 aory|EIT79820.1 asyd|OJJ53871.1 ater|EAU30430.1 atub|OJI88189.1 aver|OJJ07969.1 awen|OJJ32970.1 azon|OJJ48947.1

Group2006: abra|OJJ69414.1 acar|OOF94318.1 acla|EAW14795.1 afis|EAW24326.1 afla|EED47940.1 afum|EAL85996.1 agla|OJJ83193.1 akaw|GAA88282.1 aluc|OJZ82251.1 anid|CBF74056.1 anig|EHA28490.1 aory|EIT79834.1 asyd|OJJ53868.1 ater|EAU30427.1 atub|OJI88186.1 aver|OJJ07966.1 awen|OJJ32968.1 azon|OJJ48949.1

Group2007: abra|OJJ69415.1 acar|OOF94317.1 acla|EAW14796.1 afis|EAW24327.1 afla|EED47939.1 afum|EAL85997.1 agla|OJJ83194.1 akaw|GAA88283.1 aluc|OJZ82252.1 anid|CBF74054.1 anig|EHA28489.1 aory|EIT79852.1 asyd|OJJ53867.1 ater|EAU30426.1 atub|OJI88185.1 aver|OJJ07965.1 awen|OJJ32967.1 azon|OJJ48950.1

Group2008: abra|OJJ69417.1 acar|OOF94315.1 acla|EAW14798.1 afis|EAW24329.1 afla|EED47937.1 afum|EAL85999.1 agla|OJJ83196.1 akaw|GAA88285.1 aluc|OJZ82254.1 anid|CBF74052.1 anig|EHA28487.1 aory|EIT79861.1 asyd|OJJ53865.1 atub|OJI88183.1 aver|OJJ07963.1 awen|OJJ32965.1 azon|OJJ48953.1 ater|EAU30424.1

Group2009: abra|OJJ69419.1 acar|OOF90227.1 acla|EAW14799.1 afis|EAW24330.1 afla|EED47935.1 afum|EAL86000.1 agla|OJJ83197.1 akaw|GAA88286.1 aluc|OJZ82255.1 anid|CBF74050.1 anig|EHA28486.1 aory|EIT79863.1 asyd|OJJ53863.1 ater|EAU30422.1 atub|OJI88182.1 aver|OJJ07962.1 awen|OJJ32964.1 azon|OJJ48954.1

Group2010: abra|OJJ69420.1 acar|OOF90228.1 acla|EAW14800.1 afis|EAW24331.1 afla|EED47933.1 afum|EAL86001.1 agla|OJJ83198.1 akaw|GAA88287.1 aluc|OJZ82256.1 anid|CBF74048.1 anig|EHA28485.1 aory|EIT79848.1 asyd|OJJ53862.1 ater|EAU30418.1 atub|OJI88181.1 aver|OJJ07961.1 awen|OJJ32963.1 azon|OJJ48955.1

Group2011: abra|OJJ69421.1 acar|OOF90226.1 acla|EAW14801.1 afis|EAW24332.1 afla|EED47932.1 afum|EAL86002.1 agla|OJJ83199.1 akaw|GAA88288.1 aluc|OJZ82258.1 anid|CBF74045.1 anig|EHA28484.1 aory|EIT79839.1 asyd|OJJ53857.1 ater|EAU30417.1 atub|OJI88180.1 aver|OJJ07958.1 awen|OJJ32962.1 azon|OJJ48960.1

Group2012: abra|OJJ69423.1 acar|OOF94313.1 acla|EAW14802.1 afis|EAW24333.1 afla|EED47931.1 afum|EAL86003.1 agla|OJJ83202.1 akaw|GAA88289.1 aluc|OJZ82259.1 anid|CBF74039.1 anig|EHA28483.1 aory|EIT79843.1 asyd|OJJ53859.1 ater|EAU30416.1 atub|OJI88179.1 aver|OJJ07959.1 awen|OJJ32961.1 azon|OJJ48959.1

Group2013: abra|OJJ69446.1 acar|OOF94294.1 acla|EAW14811.1 afis|EAW24341.1 afla|EED47922.1 afum|EAL86012.1 agla|OJJ83224.1 akaw|GAA88372.1 aluc|OJZ82282.1 anid|CBF74027.1 anig|EHA28465.1 aory|EIT83372.1 asyd|OJJ53846.1 ater|EAU30406.1 atub|OJI88157.1 aver|OJJ07948.1 awen|OJJ32950.1 azon|OJJ48970.1

Group2014: abra|OJJ69448.1 acar|OOF94293.1 acla|EAW14812.1 afis|EAW24342.1 afla|EED47920.1 afum|EAL86013.1 agla|OJJ83223.1 akaw|GAA88371.1 aluc|OJZ82283.1 anid|CBF74025.1 anig|EHA28464.1 aory|EIT83362.1 asyd|OJJ53845.1 ater|EAU30404.1 atub|OJI88155.1 aver|OJJ07947.1 awen|OJJ32951.1 azon|OJJ48969.1

Group2015: abra|OJJ69449.1 acar|OOF94292.1 acla|EAW14813.1 afis|EAW24343.1 afla|EED47919.1 afum|EAL86014.1 agla|OJJ83210.1 akaw|GAA88370.1 aluc|OJZ82285.1 anid|CBF74028.1 anig|EHA28463.1 aory|EIT83364.1 asyd|OJJ53847.1 ater|EAU30403.1 atub|OJI88154.1 aver|OJJ07949.1 awen|OJJ32952.1 azon|OJJ48968.1

Group2016: abra|OJJ69453.1 acar|OOF94288.1 acla|EAW14817.1 afis|EAW24347.1 afla|EED47916.1 afum|EAL86017.1 agla|OJJ83206.1 akaw|GAA88359.1 aluc|OJZ82296.1 anid|CBF74017.1 anig|EHA28459.1 aory|EIT83378.1 asyd|OJJ53840.1 ater|EAU30399.1 atub|OJI88143.1 aver|OJJ07943.1 awen|OJJ32956.1 azon|OJJ48964.1

Group2017: abra|OJJ69456.1 acar|OOF94285.1 acla|EAW14821.1 afis|EAW24349.1 afla|EED47914.1 afum|EAL86019.2 agla|OJJ81762.1 akaw|GAA88356.1 aluc|OJZ82299.1 anid|CBF76155.1 anig|EHA28456.1 aory|EIT83376.1 asyd|OJJ52360.1 ater|EAU30395.1 atub|OJI88140.1 aver|OJJ01988.1 awen|OJJ32875.1 azon|OJJ48976.1

Group2018: abra|OJJ69457.1 acar|OOF94284.1 acla|EAW14820.1 afis|EAW24350.1 afla|EED47913.1 afum|EAL86020.2 agla|OJJ81761.1 akaw|GAA88355.1 aluc|OJZ82300.1 anid|CBF76157.1 anig|EHA28455.1 aory|EIT83375.1 asyd|OJJ52359.1 ater|EAU30394.1 atub|OJI88139.1 aver|OJJ01990.1 awen|OJJ32874.1 azon|OJJ48975.1

Group2019: abra|OJJ69458.1 acar|OOF94283.1 acla|EAW14819.1 afis|EAW24351.1 afla|EED47912.1 afum|EAL86021.1 agla|OJJ81760.1 akaw|GAA88354.1 aluc|OJZ82301.1 anid|CBF76159.1 anig|EHA28454.1 aory|EIT83371.1 asyd|OJJ52358.1 ater|EAU30393.1 atub|OJI88137.1 aver|OJJ01991.1 awen|OJJ32873.1 azon|OJJ48974.1

Group2020: abra|OJJ69459.1 acar|OOF94282.1 acla|EAW09925.1 afis|EAW21173.1 afla|EED54758.1 afum|EAL92193.1 agla|OJJ80005.1 akaw|GAA88353.1 aluc|OJZ82302.1 anid|CBF83317.1 anig|EHA28453.1 aory|EIT79382.1 asyd|OJJ60559.1 ater|EAU35840.1 atub|OJI88135.1 aver|OJI99885.1 awen|OJJ36511.1 azon|OJJ44387.1

Group2021: abra|OJJ69463.1 acar|OOF94278.1 acla|EAW09922.1 afis|EAW21170.1 afla|EED54735.1 afum|EAL92189.2 agla|OJJ80010.1 akaw|GAA88349.1 aluc|OJZ82306.1 anid|CBF83232.1 anig|EHA28450.1 aory|EIT78745.1 asyd|OJJ60541.1 ater|EAU35837.1 atub|OJI88132.1 aver|OJI99868.1 awen|OJJ36507.1 azon|OJJ44379.1

Group2022: abra|OJJ69467.1 acar|OOF94275.1 acla|EAW09920.1 afis|EAW21168.1 afla|EED54732.1 afum|EAL92187.1 agla|OJJ80012.1 akaw|GAA88346.1 aluc|OJZ82311.1 anid|CBF86960.1 anig|EHA28447.1 aory|EIT78891.1 asyd|OJJ60539.1 ater|EAU35835.1 atub|OJI88129.1 aver|OJI99866.1 awen|OJJ36505.1 azon|OJJ44377.1

Group2023: abra|OJJ69468.1 acar|OOF94274.1 acla|EAW09919.1 afis|EAW21167.1 afla|EED54731.1 afum|EAL92186.1 agla|OJJ80013.1 akaw|GAA88345.1 aluc|OJZ82312.1 anid|CBF86961.1 anig|EHA28446.1 aory|EIT78775.1 asyd|OJJ60537.1 ater|EAU35834.1 atub|OJI88128.1 aver|OJI99864.1 awen|OJJ36504.1 azon|OJJ44376.1

Group2024: abra|OJJ69469.1 acar|OOF94273.1 acla|EAW09918.1 afis|EAW21166.1 afla|EED54730.1 afum|EAL92185.1 agla|OJJ80015.1 akaw|GAA88344.1 aluc|OJZ82313.1 anid|CBF86963.1 anig|EHA28445.1 aory|EIT78843.1 asyd|OJJ60536.1 ater|EAU35833.1 atub|OJI88127.1 aver|OJI99863.1 awen|OJJ36503.1 azon|OJJ44375.1

Group2025: abra|OJJ69471.1 acar|OOF94272.1 acla|EAW09917.1 afis|EAW21165.1 afla|EED54729.1 afum|EAL92184.1 agla|OJJ80016.1 akaw|GAA88343.1 aluc|OJZ82314.1 anid|CBF86964.1 anig|EHA28444.1 aory|EIT78934.1 asyd|OJJ60535.1 ater|EAU35832.1 atub|OJI88126.1 aver|OJI99862.1 awen|OJJ36500.1 azon|OJJ44374.1

Group2026: abra|OJJ69473.1 acar|OOF94270.1 acla|EAW09915.1 afis|EAW21163.1 afla|EED54726.1 afum|EAL92182.1 agla|OJJ80020.1 akaw|GAA88342.1 aluc|OJZ82316.1 anid|CBF86968.1 anig|EHA28442.1 aory|EIT78844.1 asyd|OJJ60532.1 ater|EAU35829.1 atub|OJI88125.1 aver|OJI99859.1 awen|OJJ36497.1 azon|OJJ44372.1

Group2027: abra|OJJ69474.1 acar|OOF94268.1 acla|EAW09914.1 afis|EAW21162.1 afla|EED54725.1 afum|EAL92181.1 agla|OJJ80021.1 akaw|GAA88341.1 aluc|OJZ82317.1 anid|CBF86970.1 anig|EHA28441.1 aory|EIT78833.1 asyd|OJJ60531.1 ater|EAU35828.1 atub|OJI88124.1 aver|OJI99858.1 awen|OJJ36496.1 azon|OJJ44371.1

Group2028: abra|OJJ69475.1 acar|OOF94269.1 acla|EAW09913.1 afis|EAW21161.1 afla|EED54724.1 afum|EAL92180.1 agla|OJJ80022.1 akaw|GAA88340.1 aluc|OJZ82318.1 anid|CBF86972.1 anig|EHA28440.1

aory|EIT78951.1 asyd|OJJ60530.1 ater|EAU35827.1 atub|OJI88123.1 aver|OJI99857.1 awen|OJJ36495.1 azon|OJJ44370.1

Group2029: abra|OJJ69477.1 acar|OOF94265.1 acla|EAW09911.1 afis|EAW21160.1 afla|EED54649.1 afum|EAL92178.1 agla|OJJ80028.1 akaw|GAA88338.1 aluc|OJZ82320.1 anid|CBF86977.1 anig|EHA28438.1 aory|EIT78896.1 asyd|OJJ60527.1 ater|EAU35822.1 atub|OJI88120.1 aver|OJI99855.1 awen|OJJ36492.1 azon|OJJ44365.1

Group2030: abra|OJJ69481.1 acar|OOF94263.1 acla|EAW09908.1 akaw|GAA88335.1 aluc|OJZ82323.1 anig|EHA28435.1 atub|OJI88117.1 awen|OJJ36488.1 agla|OJJ80036.1 afis|EAW21157.1 afum|EAL92175.1 afla|EED54653.1 aory|EIT78820.1 ater|EAU35819.1 anid|CBF86982.1 asyd|OJJ60523.1 aver|OJI99847.1 azon|OJJ44359.1

Group2031: abra|OJJ69483.1 acar|OOF94262.1 acla|EAW09907.1 afis|EAW21156.1 afla|EED54652.1 afum|EAL92174.1 agla|OJJ80037.1 akaw|GAA88334.1 aluc|OJZ82324.1 anid|CBF86984.1 anig|EHA28434.1 aory|EIT78871.1 asyd|OJJ60518.1 ater|EAU35818.1 atub|OJI88116.1 aver|OJI99845.1 awen|OJJ36487.1 azon|OJJ44360.1

Group2032: abra|OJJ69485.1 acar|OOF94261.1 acla|EAW09906.1 afis|EAW21155.1 afla|EED54661.1 afum|EAL92173.1 agla|OJJ80038.1 akaw|GAA88333.1 aluc|OJZ82326.1 anid|CBF86986.1 anig|EHA28432.1 aory|EIT78925.1 asyd|OJJ60520.1 ater|EAU35817.1 atub|OJI88114.1 aver|OJI99846.1 awen|OJJ36486.1 azon|OJJ44362.1

Group2034: abra|OJJ69496.1 acar|OOF94254.1 acla|EAW09950.1 afis|EAW21146.1 afla|EED46146.1 afum|EAL92165.1 agla|OJJ80070.1 akaw|GAA88325.1 aluc|OJZ82336.1 anid|CBF87010.1 anig|EHA28424.1 aory|EIT76496.1 asyd|OJJ60472.1 ater|EAU36761.1 atub|OJI88104.1 aver|OJI99792.1 awen|OJJ36469.1 azon|OJJ44354.1

Group2035: abra|OJJ69497.1 acar|OOF94256.1 acla|EAW09951.1 afis|EAW21145.1 afla|EED46147.1 afum|EAL92164.1 agla|OJJ80071.1 akaw|GAA88324.1 aluc|OJZ82337.1 anid|CBF86995.1 anig|EHA28423.1 aory|EIT76462.1 asyd|OJJ60473.1 ater|EAU36757.1 atub|OJI88103.1 aver|OJI99793.1 awen|OJJ36470.1 azon|OJJ44355.1

Group2036: abra|OJJ69498.1 acar|OOF94257.1 acla|EAW09952.1 afis|EAW21144.1 afla|EED46148.1 afum|EAL92163.1 agla|OJJ80072.1 akaw|GAA88323.1 aluc|OJZ82338.1 anid|CBF86997.1 anig|EHA28422.1 aory|EIT76514.1 asyd|OJJ60474.1 ater|EAU36756.1 atub|OJI88102.1 aver|OJI99794.1 awen|OJJ36471.1 azon|OJJ44356.1

Group2037: abra|OJJ69505.1 acar|OOF94247.1 acla|EAW09957.1 afis|EAW21136.1 afla|EED51811.1 afum|EAL92155.1 agla|OJJ80076.1 akaw|GAA88318.1 aluc|OJZ82344.1 anid|CBF87016.1 anig|EHA28416.1 aory|EIT81758.1 asyd|OJJ60463.1 ater|EAU36774.1 atub|OJI88096.1 aver|OJI99783.1 awen|OJJ36456.1 azon|OJJ44420.1

Group2038: abra|OJJ69506.1 acar|OOF94246.1 acla|EAW09958.1 afis|EAW21137.1 afla|EED51812.1 afum|EAL92156.1 agla|OJJ80081.1 akaw|GAA88317.1 aluc|OJZ82345.1 anid|CBF87014.1 anig|EHA28415.1 aory|EIT81705.1 asyd|OJJ60464.1 ater|EAU36773.1 atub|OJI88095.1 aver|OJI99784.1 awen|OJJ36457.1 azon|OJJ44418.1

Group2039: abra|OJJ69508.1 acar|OOF94245.1 acla|EAW09959.1 afis|EAW21138.1 afla|EED51813.1 afum|EAL92157.2 agla|OJJ80080.1 akaw|GAA88316.1 aluc|OJZ82346.1 anid|CBF87012.1 anig|EHA28414.1 aory|EIT81694.1 asyd|OJJ60465.1 ater|EAU36772.1 atub|OJI88094.1 aver|OJI99785.1 awen|OJJ36458.1 azon|OJJ44419.1

Group2040: abra|OJJ69510.1 acar|OOF94244.1 acla|EAW09960.1 afis|EAW21135.1 afla|EED46138.1 afum|EAL92154.1 agla|OJJ80078.1 akaw|GAA88314.1 aluc|OJZ82348.1 anid|CBF87000.1 anig|EHA28412.1 aory|EIT76494.1 asyd|OJJ60467.1 ater|EAU36767.1 atub|OJI88092.1 aver|OJI99787.1 awen|OJJ36461.1 azon|OJJ44416.1

Group2041: abra|OJJ69513.1 acar|OOF94240.1 acla|EAW09955.1 afis|EAW21141.1 afla|EED46142.1 afum|EAL92160.1 agla|OJJ80068.1 akaw|GAA88311.1 aluc|OJZ82351.1 anid|CBF87007.1 anig|EHA28409.1 aory|EIT76415.1 asyd|OJJ60470.1 ater|EAU36762.1 atub|OJI88089.1 aver|OJI99790.1 awen|OJJ36465.1 azon|OJJ44413.1

Group2042: abra|OJJ69516.1 acar|OOF94238.1 acla|EAW09964.1 afis|EAW21124.1 afla|EED51799.1 afum|EAL92147.1 agla|OJJ80086.1 akaw|GAA88308.1 aluc|OJZ82354.1 anid|CBF87018.1 anig|EHA28406.1 aory|EIT81724.1 asyd|OJJ60461.1 ater|EAU36776.1 atub|OJI88086.1 aver|OJI99781.1 awen|OJJ36454.1 azon|OJJ44423.1

Group2043: abra|OJJ69520.1 acar|OOF94237.1 acla|EAW09966.1 afis|EAW21122.1 afla|EED51795.1 afum|EAL92145.1 agla|OJJ80089.1 akaw|GAA88306.1 aluc|OJZ82359.1 anid|CBF87021.1 anig|EHA28404.1 aory|EIT81713.1 asyd|OJJ60459.1 ater|EAU36778.1 atub|OJI88082.1 aver|OJI99779.1 awen|OJJ36452.1 azon|OJJ44425.1

Group2044: abra|OJJ69522.1 acar|OOF94233.1 acla|EAW09968.1 afis|EAW21120.1 afla|EED51793.1 afum|EBA27448.1 agla|OJJ80091.1 akaw|GAA88305.1 aluc|OJZ82362.1 anid|CBF87025.1 anig|EHA28403.1 aory|EIT81754.1 asyd|OJJ60457.1 ater|EAU36779.1 atub|OJI88080.1 aver|OJI99777.1 awen|OJJ36450.1 azon|OJJ44427.1

Group2045: abra|OJJ69525.1 acar|OOF94229.1 acla|EAW09972.1 afis|EAW21116.1 afla|EED51767.1 afum|EAL92139.1 agla|OJJ80095.1 akaw|GAA88300.1 aluc|OJZ82365.1 anid|CBF87030.1 anig|EHA28400.1 aory|EIT79993.1 asyd|OJJ60455.1 ater|EAU36783.1 atub|OJI88077.1 aver|OJI99773.1 awen|OJJ36446.1 azon|OJJ44429.1

Group2047: abra|OJJ69535.1 acar|OOF95965.1 acla|EAW11570.1 afis|EAW23883.1 afla|EED52612.1 afum|EAL87311.1 agla|OJJ82570.1 akaw|GAA88385.1 aluc|OJZ82373.1 anid|CBF77187.1 anig|EHA28391.1 aory|EIT74250.1 asyd|OJJ56841.1 ater|EAU33463.1 atub|OJI88069.1 aver|OJJ03497.1 awen|OJJ33865.1 azon|OJJ48688.1

Group2048: abra|OJJ69536.1 acar|OOF91604.1 acla|EAW14409.1 afis|EAW19117.1 afla|EED54297.1 afum|EAL93849.1 agla|OJJ80432.1 akaw|GAA91927.1 aluc|OJZ82695.1 anid|CBF79728.1 anig|EHA21143.1 aory|EIT75966.1 asyd|OJJ58073.1 ater|EAU37379.1 atub|OJI88927.1 aver|OJJ04334.1 awen|OJJ37447.1 azon|OJJ51125.1

Group2049: abra|OJJ69575.1 acar|OOF91580.1 acla|EAW14394.1 afis|EAW19102.1 afla|EED54308.1 afum|EAL93834.1 agla|OJJ84500.1 akaw|GAA91572.1 aluc|OJZ82392.1 anid|CBF79696.1 anig|EHA28372.1 aory|EIT75942.1 asyd|OJJ58050.1 ater|EAU32151.1 atub|OJI88049.1 aver|OJJ04353.1 awen|OJJ32495.1 azon|OJJ46034.1

Group2050: abra|OJJ69580.1 acar|OOF91573.1 acla|EAW14388.1 afis|EAW19095.1 afla|EED54328.1 afum|EAL93826.1 agla|OJJ84506.1 akaw|GAA91576.1 aluc|OJZ82398.1 anid|CBF79705.1 anig|EHA28367.1 aory|EIT75940.1 asyd|OJJ58055.1 ater|EAU32143.1 atub|OJI88044.1 aver|OJJ04348.1 awen|OJJ32488.1 azon|OJJ46043.1

Group2052: abra|OJJ69584.1 acar|OOF91570.1 acla|EAW14385.1 afis|EAW19091.1 afla|EED54326.1 afum|EAL93821.1 agla|OJJ84512.1 akaw|GAA91579.1 aluc|OJZ82401.1 anid|CBF79711.1 anig|EHA28364.1 aory|EIT76092.1 asyd|OJJ58058.1 ater|EAU32140.1 atub|OJI88040.1 aver|OJJ04345.1 awen|OJJ32485.1 azon|OJJ46046.1

Group2053: abra|OJJ69587.1 acar|OOF91568.1 acla|EAW14380.1 afis|EAW19086.1 afla|EED54318.1 afum|EAL93816.1 agla|OJJ84518.1 akaw|GAA91584.1 aluc|OJZ82404.1 anid|CBF79684.1 anig|EHA28362.1 aory|EIT76159.1 asyd|OJJ58047.1 ater|EAU32134.1 atub|OJI88037.1 aver|OJJ04356.1 awen|OJJ32483.1 azon|OJJ46227.1

Group2054: abra|OJJ69588.1 acar|OOF91567.1 acla|EAW14379.1 afis|EAW19085.1 afla|EED54317.1 afum|EAL93815.1 agla|OJJ84521.1 akaw|GAA91585.1 aluc|OJZ82406.1 anid|CBF79683.1 anig|EHA28361.1 aory|EIT75897.1 asyd|OJJ58046.1 ater|EAU32133.1 atub|OJI88036.1 aver|OJJ04357.1 awen|OJJ32482.1 azon|OJJ46226.1

Group2055: abra|OJJ69590.1 acar|OOF91565.1 acla|EAW14378.1 afis|EAW19084.1 afla|EED54316.1 afum|EAL93814.1 agla|OJJ84522.1 akaw|GAA91586.1 aluc|OJZ82408.1 anid|CBF79676.1 anig|EHA28360.1 aory|EIT75902.1 asyd|OJJ58038.1 ater|EAU32132.1 atub|OJI88034.1 aver|OJJ04362.1 awen|OJJ32481.1 azon|OJJ46224.1

Group2057: abra|OJJ69602.1 acar|OOF92702.1 acla|EAW10013.1 afis|EAW21041.1 afla|EED51373.1 afum|EAL92080.1 agla|OJJ84213.1 akaw|GAA91590.1 aluc|OJZ82413.1 anid|CBF80091.1 anig|EHA28354.1 aory|EIT80745.1 asyd|OJJ58302.1 ater|EAU31725.1 atub|OJI88028.1 aver|OJJ02110.1 awen|OJJ32696.1 azon|OJJ50700.1

Group2058: abra|OJJ69616.1 acar|OOF92690.1 acla|EAW12076.1 afis|EAW19717.1 afla|EED53108.1 afum|EAL86810.1 agla|OJJ80861.1 akaw|GAA91600.1 aluc|OJZ82424.1 anid|CBF80078.1 anig|EHA28345.1 aory|EIT74760.1 asyd|OJJ58310.1 ater|EAU37776.1 atub|OJI88016.1 aver|OJJ02101.1 awen|OJJ35107.1 azon|OJJ49987.1

Group2059: abra|OJJ69628.1 afla|EED55079.1 agla|OJJ86032.1 aluc|OJZ82433.1 anid|CBF82983.1 anig|EHA28338.1 asyd|OJJ64730.1 ater|EAU30171.1 atub|OJI88008.1 aver|OJI98834.1 awen|OJJ30457.1 azon|OJJ45448.1 acar|OOF92682.1 afis|EAW16821.1 afum|EAL86371.1 acla|EAW12375.1 aory|EIT81573.1 akaw|GAA91608.1

Group2061: abra|OJJ69637.1 acar|OOF92675.1 acla|EAW12571.1 afis|EAW24079.1 afla|EED51394.1 afum|EAL91934.1 agla|OJJ87150.1 akaw|GAA91617.1 aluc|OJZ82443.1 anid|CBF86476.1 anig|EHA28330.1 aory|EIT80502.1 asyd|OJJ64903.1 ater|EAU30500.1 atub|OJI87997.1 aver|OJI99025.1 awen|OJJ41417.1 azon|OJJ48876.1

Group2062: azon|OJJ44974.1 abra|OJJ73161.1 acar|OOG00404.1 acla|EAW06767.1 afis|EAW16317.1 afla|EED53564.1 afum|EAL88940.1 agla|OJJ88278.1 akaw|GAA88011.1 aluc|OJZ81269.1 anid|CBF89145.1

anig|EHA18569.1 aory|EIT75175.1 asyd|OJJ62569.1 ater|EAU34140.1 atub|OJI91476.1 aver|OJI96693.1 awen|OJJ37717.1

Group2063: abra|OJJ69655.1 acar|OOG00750.1 acla|EAW07485.1 afis|EAW22518.1 afla|EED54547.1 afum|EAL90675.1 agla|OJJ80878.1 akaw|GAA91634.1 aluc|OJZ82461.1 anid|CBF88995.1 anig|EHA28314.1 aory|EIT78864.1 asyd|OJJ62783.1 ater|EAU39191.1 atub|OJI87977.1 aver|OJI96828.1 awen|OJJ35145.1 azon|OJJ49955.1

Group2064: abra|OJJ69657.1 acar|OOF95821.1 acla|EAW11286.1 afis|EAW18213.1 afla|EED56258.1 afum|EAL87579.1 agla|OJJ85869.1 akaw|GAA91636.1 aluc|OJZ82463.1 anid|CBF75160.1 anig|EHA28312.1 aory|EIT73422.1 asyd|OJJ59962.1 ater|EAU33776.1 atub|OJI87975.1 aver|OJJ02328.1 awen|OJJ30591.1 azon|OJJ50185.1

Group2065: abra|OJJ69663.1 acar|OOF98113.1 acla|EAW10577.1 afis|EAW24990.1 afla|EED55469.1 afum|EAL89650.1 agla|OJJ82264.1 akaw|GAA91641.1 aluc|OJZ82468.1 anid|CBF81596.1 anig|EHA28307.1 aory|EIT80882.1 asyd|OJJ59517.1 ater|EAU35741.1 atub|OJI87970.1 aver|OJJ02756.1 awen|OJJ40595.1 azon|OJJ49446.1

Group2066: abra|OJJ69664.1 acar|OOG00819.1 acla|EAW07551.1 afis|EAW22585.1 afla|EED54630.1 afum|EAL90614.1 agla|OJJ80945.1 akaw|GAA91642.1 aluc|OJZ82469.1 anid|CBF88352.1 anig|EHA28306.1 aory|EIT78765.1 asyd|OJJ63721.1 ater|EAU39114.1 atub|OJI87968.1 aver|OJI97219.1 awen|OJJ35213.1 azon|OJJ49896.1

Group2067: abra|OJJ69665.1 acar|OOG00018.1 acla|EAW09937.1 afis|EAW21182.1 afla|EED54777.1 afum|EAL92203.1 agla|OJJ79988.1 akaw|GAA91643.1 aluc|OJZ82470.1 anid|CBF83310.1 anig|EHA28303.1 aory|EIT79439.1 asyd|OJJ60549.1 ater|EAU35848.1 atub|OJI87967.1 aver|OJI99875.1 awen|OJJ36521.1 azon|OJJ44397.1

Group2068: abra|OJJ69666.1 acar|OOG00019.1 acla|EAW09936.1 afis|EAW21181.1 afla|EED54778.1 afum|EAL92202.1 agla|OJJ79998.1 akaw|GAA91644.1 aluc|OJZ82471.1 anid|CBF83308.1 anig|EHA28302.1 aory|EIT79454.1 asyd|OJJ60550.1 ater|EAU35847.1 atub|OJI87966.1 aver|OJI99876.1 awen|OJJ36520.1 azon|OJJ44396.1

Group2069: abra|OJJ69667.1 acar|OOF97488.1 acla|EAW13075.1 afis|EAW17589.1 afla|EED49233.1 afum|EAL91402.1 agla|OJJ81360.1 akaw|GAA91645.1 aluc|OJZ82472.1 anid|CBF90218.1 anig|EHA28301.1 aory|EIT83160.1 asyd|OJJ61967.1 ater|EAU37024.1 atub|OJI87965.1 aver|OJI96059.1 awen|OJJ37257.1 azon|OJJ50952.1

Group2070: abra|OJJ69669.1 acar|OOF95822.1 acla|EAW06663.1 afis|EAW16102.1 afla|EED48637.1 afum|EAL88734.1 agla|OJJ86131.1 akaw|GAA91647.1 aluc|OJZ82474.1 anid|CBF85442.1 anig|EHA28299.1 aory|EIT73069.1 asyd|OJJ64091.1 ater|EAU33179.1 atub|OJI87963.1 aver|OJI98175.1 awen|OJJ30330.1 azon|OJJ45358.1

Group2071: abra|OJJ69677.1 acar|OOF92983.1 acla|EAW14808.1 afis|EAW24337.1 afla|EED47927.1 afum|EAL86008.1 agla|OJJ83205.1 akaw|GAA88802.1 aluc|OJZ82485.1 anid|CBF74030.1 anig|EHA28289.1 aory|EIT79853.1 asyd|OJJ53849.1 ater|EAU30411.1 atub|OJI87947.1 aver|OJJ07950.1 awen|OJJ32957.1 azon|OJJ48963.1

Group2072: abra|OJJ69682.1 acar|OOF93221.1 acla|EAW13124.1 afis|EAW17540.1 afla|EED49287.1 afum|EAL91354.1 agla|OJJ81429.1 akaw|GAA92416.1 aluc|OJZ83274.1 anid|CBF90315.1 anig|EHA28284.1 aory|EIT82215.1 asyd|OJJ61912.1 ater|EAU36970.1 atub|OJI87666.1 aver|OJI96005.1 awen|OJJ37313.1 azon|OJJ51005.1

Group2073: abra|OJJ69687.1 acar|OOF92648.1 acla|EAW10319.1 afis|EAW25273.1 afla|EED53738.1 afum|EAL89936.1 agla|OJJ89247.1 akaw|GAA88811.1 aluc|OJZ82494.1 anid|CBF85387.1 anig|EHA28278.1 aory|EIT75306.1 asyd|OJJ64055.1 ater|EAU34416.1 atub|OJI87938.1 aver|OJI98140.1 awen|OJJ40987.1 azon|OJJ44574.1

Group2074: abra|OJJ69698.1 acar|OOF90521.1 acla|EAW15242.1 afis|EAW20209.1 afla|EED57134.1 afum|EAL85326.1 agla|OJJ84655.1 akaw|GAA88816.1 aluc|OJZ82500.1 anid|CBF85100.1 anig|EHA28270.1 aory|EIT74032.1 asyd|OJJ63057.1 ater|EAU32019.1 atub|OJI87931.1 aver|OJI97905.1 awen|OJJ31213.1 azon|OJJ46146.1

Group2075: abra|OJJ69699.1 acar|OOF95410.1 acla|EAW07370.1 afis|EAW22395.1 afla|EED56059.1 afum|EAL90796.1 agla|OJJ87658.1 akaw|GAA88818.1 aluc|OJZ82502.1 anid|CBF88747.1 anig|EHA28269.1 aory|EIT76693.1 asyd|OJJ63956.1 ater|EAU39322.1 atub|OJI87929.1 aver|OJI96985.1 awen|OJJ34973.1 azon|OJJ45622.1

Group2076: abra|OJJ69701.1 acar|OOF93222.1 acla|EAW13123.1 afis|EAW17541.1 afla|EED49286.1 afum|EAL91355.1 agla|OJJ81428.1 akaw|GAA88820.1 aluc|OJZ82504.1 anid|CBF90313.1 anig|EHA28266.1 aory|EIT82202.1 asyd|OJJ61913.1 ater|EAU36971.1 atub|OJI87927.1 aver|OJI96006.1 awen|OJJ37312.1 azon|OJJ51004.1

Group2077: abra|OJJ69703.1 acar|OOF98863.1 acla|EAW06689.1 afis|EAW16076.1 afla|EED48596.1 afum|EAL88708.1 agla|OJJ86093.1 akaw|GAA88822.1 aluc|OJZ82506.1 anid|CBF75115.1 anig|EHA28263.1 aory|EIT73056.1 asyd|OJJ59995.1 ater|EAU33555.1 atub|OJI87925.1 aver|OJJ02296.1 awen|OJJ30357.1 azon|OJJ45379.1

Group2078: abra|OJJ69718.1 acar|OOF94196.1 acla|EAW09976.1 afis|EAW21113.1 afla|EED51764.1 afum|EAL92136.1 agla|OJJ80098.1 akaw|GAA90171.1 aluc|OJZ83179.1 anid|CBF87036.1 anig|EHA28253.1 aory|EIT80019.1 asyd|OJJ60452.1 ater|EAU36786.1 atub|OJI87758.1 aver|OJI99770.1 awen|OJJ36441.1 azon|OJJ44432.1

Group2079: abra|OJJ69746.1 acar|OOF94176.1 acla|EAW09985.1 afis|EAW21108.1 afla|EED51696.1 afum|EAL92129.1 agla|OJJ80107.1 akaw|GAA90198.1 aluc|OJZ83150.1 anid|CBF87041.1 anig|EHA28216.1 aory|EIT80588.1 asyd|OJJ60448.1 ater|EAU36792.1 atub|OJI87787.1 aver|OJI99766.1 awen|OJJ36382.1 azon|OJJ51534.1

Group2080: aver|OJJ05453.1 abra|OJJ70544.1 acar|OOF94607.1 acla|EAW08135.1 afis|EAW23171.1 afla|EED46949.1 afum|EAL88312.2 agla|OJJ89683.1 akaw|GAA83390.1 aluc|OJZ83889.1 anid|CBF74485.1 anig|EHA19026.1 aory|EIT72627.1 asyd|OJJ57712.1 ater|EAU36300.1 atub|OJI90256.1 awen|OJJ36606.1 azon|OJJ47601.1

Group2082: abra|OJJ69822.1 acar|OOF93341.1 acla|EAW08969.1 afis|EAW15448.1 afla|EED56825.1 afum|EAL84198.2 agla|OJJ86271.1 akaw|GAA87838.1 aluc|OJZ85474.1 anid|CBF78305.1 anig|EHA25015.1 aory|EIT73942.1 asyd|OJJ56090.1 ater|EAU30838.1 atub|OJI80089.1 aver|OJJ05033.1 awen|OJJ31223.1 azon|OJJ45071.1

Group2085: abra|OJJ69898.1 acar|OOF93417.1 acla|EAW08922.1 afis|EAW15471.1 afla|EED46922.1 afum|EAL85850.1 agla|OJJ80151.1 akaw|GAA87752.1 aluc|OJZ85385.1 anid|CBF78234.1 anig|EHA25569.1 aory|EIT72668.1 asyd|OJJ56036.1 ater|EAU32657.1 atub|OJI80185.1 aver|OJJ04982.1 awen|OJJ38121.1 azon|OJJ48022.1

Group2086: abra|OJJ69901.1 acar|OOF93419.1 acla|EAW08921.1 afis|EAW15473.1 afla|EED46923.1 afum|EAL85848.1 akaw|GAA87750.1 aluc|OJZ85383.1 anid|CBF78228.1 anig|EHA25567.1 aory|EIT72551.1 asyd|OJJ56034.1 ater|EAU32656.1 atub|OJI80187.1 aver|OJJ04980.1 awen|OJJ38126.1 azon|OJJ43160.1 agla|OJJ80154.1

Group2087: abra|OJJ69906.1 acar|OOF93423.1 acla|EAW08918.1 afis|EAW15476.1 afla|EED46939.1 afum|EAL85845.1 agla|OJJ80155.1 akaw|GAA87747.1 aluc|OJZ85380.1 anid|CBF78226.1 anig|EHA25563.1 aory|EIT72564.1 asyd|OJJ56033.1 ater|EAU32651.1 atub|OJI80190.1 aver|OJJ04979.1 awen|OJJ38133.1 azon|OJJ43159.1

Group2088: abra|OJJ69914.1 acar|OOF93426.1 acla|EAW08915.1 afis|EAW15479.1 afla|EED46934.1 afum|EAL85842.1 agla|OJJ80158.1 akaw|GAA87742.1 aluc|OJZ85374.1 anid|CBF78221.1 anig|EHA25558.1 aory|EIT72728.1 asyd|OJJ56028.1 ater|EAU32648.1 atub|OJI80195.1 aver|OJJ04975.1 awen|OJJ38138.1 azon|OJJ43152.1

Group2090: abra|OJJ69940.1 acar|OOF93449.1 acla|EAW08883.1 afis|EAW15506.1 afla|EED49019.1 afum|EAL85818.1 agla|OJJ80183.1 akaw|GAA87719.1 aluc|OJZ85347.1 anid|CBF84473.1 anig|EHA25534.1 aory|EIT82494.1 asyd|OJJ53540.1 ater|EAU32612.1 atub|OJI80223.1 aver|OJJ08213.1 awen|OJJ38166.1 azon|OJJ45800.1

Group2092: abra|OJJ69953.1 acar|OOF93463.1 acla|EAW08869.1 afis|EAW15525.1 afla|EED49036.1 afum|EAL85803.1 agla|OJJ85221.1 akaw|GAA87706.1 aluc|OJZ85329.1 anid|CBF78217.1 anig|EHA25518.1 aory|EIT82537.1 asyd|OJJ55991.1 ater|EAU32598.1 atub|OJI80243.1 aver|OJJ04939.1 awen|OJJ38189.1 azon|OJJ43106.1

Group2093: abra|OJJ69955.1 acar|OOF93465.1 acla|EAW08867.1 afis|EAW15527.1 afla|EED49038.1 afum|EAL85801.1 agla|OJJ85219.1 akaw|GAA87704.1 aluc|OJZ85327.1 anid|CBF78212.1 anig|EHA25517.1 aory|EIT82564.1 asyd|OJJ55989.1 ater|EAU32597.1 atub|OJI80245.1 aver|OJJ04937.1 awen|OJJ38191.1 azon|OJJ43104.1

Group2094: abra|OJJ69962.1 acar|OOF93470.1 acla|EAW08834.1 afis|EAW15566.1 afla|EED49075.1 afum|EAL85767.2 agla|OJJ85185.1 akaw|GAA87697.1 aluc|OJZ85321.1 anid|CBF78116.1 anig|EHA25511.1 aory|EIT82536.1 asyd|OJJ55900.1 ater|EAU32567.1 atub|OJI80252.1 aver|OJJ04845.1 awen|OJJ38226.1 azon|OJJ43069.1

Group2095: abra|OJJ69963.1 acar|OOF93471.1 acla|EAW08835.1 afis|EAW15565.1 afla|EED49074.1 agla|OJJ85186.1 akaw|GAA87696.1 aluc|OJZ85320.1 anid|CBF78118.1 anig|EHA25510.1 aory|EIT82524.1 asyd|OJJ55901.1 ater|EAU32568.1 atub|OJI80253.1 aver|OJJ04846.1 awen|OJJ38225.1 azon|OJJ43070.1 afum|EAL85768.1

Group2096: abra|OJJ69971.1 acar|OOF93475.1 acla|EAW08842.1 afis|EAW15559.1 afla|EED49063.1 afum|EAL85773.1 agla|OJJ85195.1 akaw|GAA87690.1 aluc|OJZ85311.1 anid|CBF78159.1 anig|EHA25506.1

aory|EIT82560.1 asyd|OJJ55960.1 ater|EAU32575.1 atub|OJI80260.1 aver|OJJ04906.1 awen|OJJ38217.1 azon|OJJ43077.1

Group2097: abra|OJJ69974.1 acar|OOF93479.1 acla|EAW08844.1 afis|EAW15556.1 afla|EED49061.1 afum|EAL85776.1 agla|OJJ85198.1 akaw|GAA87687.1 aluc|OJZ85307.1 anid|CBF78163.1 anig|EHA25503.1 aory|EIT82517.1 asyd|OJJ55964.1 ater|EAU32577.1 atub|OJI80264.1 aver|OJJ04910.1 awen|OJJ38214.1 azon|OJJ43081.1

Group2098: abra|OJJ69975.1 acar|OOF93481.1 acla|EAW08846.1 afis|EAW15554.1 afla|EED49057.1 afum|EAL85779.1 agla|OJJ85201.1 akaw|GAA87686.1 aluc|OJZ85306.1 anid|CBF78167.1 anig|EHA25502.1 aory|EIT82497.1 asyd|OJJ55968.1 ater|EAU32580.1 atub|OJI80265.1 aver|OJJ04915.1 awen|OJJ38210.1 azon|OJJ43084.1

Group2099: abra|OJJ69978.1 acar|OOF93484.1 acla|EAW08847.1 afis|EAW15549.1 afla|EED49055.1 afum|EAL85783.1 agla|OJJ85202.1 akaw|GAA87683.1 aluc|OJZ85302.1 anid|CBF78165.1 anig|EHA25499.1 aory|EIT82558.1 asyd|OJJ55969.1 ater|EAU32583.1 atub|OJI80268.1 aver|OJJ04916.1 awen|OJJ38209.1 azon|OJJ43086.1

Group2100: abra|OJJ69979.1 acar|OOF93483.1 acla|EAW08849.1 afis|EAW15547.1 afla|EED49054.1 afum|EAL85785.1 agla|OJJ85205.1 akaw|GAA87682.1 aluc|OJZ85301.1 anid|CBF78176.1 anig|EHA25498.1 aory|EIT82504.1 asyd|OJJ55970.1 ater|EAU32582.1 atub|OJI80269.1 aver|OJJ04917.1 awen|OJJ38208.1 azon|OJJ43087.1

Group2101: abra|OJJ69982.1 acar|OOF93486.1 acla|EAW08852.1 afis|EAW15544.1 afla|EED49051.1 afum|EAL85787.1 agla|OJJ85207.1 akaw|GAA87680.1 aluc|OJZ85298.1 anid|CBF78182.1 anig|EHA25496.1 aory|EIT82527.1 asyd|OJJ55973.1 ater|EAU32584.1 atub|OJI80272.1 aver|OJJ04921.1 awen|OJJ38205.1 azon|OJJ43090.1

Group2102: abra|OJJ69983.1 acar|OOF93487.1 acla|EAW08853.1 afis|EAW15543.1 afla|EED49050.1 afum|EAL85788.1 agla|OJJ85208.1 akaw|GAA87679.1 aluc|OJZ85297.1 anid|CBF78183.1 anig|EHA25495.1 aory|EIT82528.1 asyd|OJJ55974.1 ater|EAU32585.1 atub|OJI80273.1 aver|OJJ04922.1 awen|OJJ38204.1 azon|OJJ43091.1

Group2103: abra|OJJ69984.1 acar|OOF93489.1 acla|EAW08855.1 afis|EAW15541.1 afla|EED49049.1 afum|EAL85789.1 agla|OJJ85209.1 akaw|GAA87677.1 aluc|OJZ85295.1 anid|CBF78187.1 anig|EHA25493.1 aory|EIT82548.1 asyd|OJJ55976.1 ater|EAU32586.1 atub|OJI80275.1 aver|OJJ04924.1 awen|OJJ38202.1 azon|OJJ43093.1

Group2104: abra|OJJ69985.1 acar|OOF93490.1 acla|EAW08856.1 afis|EAW15540.1 afla|EED49048.1 afum|EAL85790.1 agla|OJJ85210.1 akaw|GAA87676.1 aluc|OJZ85294.1 anid|CBF78189.1 anig|EHA25492.1 aory|EIT82530.1 asyd|OJJ55977.1 ater|EAU32587.1 atub|OJI80276.1 aver|OJJ04925.1 awen|OJJ38201.1 azon|OJJ43094.1

Group2105: abra|OJJ69987.1 acar|OOF93491.1 acla|EAW08858.1 afis|EAW15538.1 afla|EED49046.1 afum|EAL85792.1 agla|OJJ85211.1 akaw|GAA87674.1 aluc|OJZ85292.1 anid|CBF78193.1 anig|EHA25490.1 aory|EIT82570.1 asyd|OJJ55979.1 ater|EAU32589.1 atub|OJI80278.1 aver|OJJ04927.1 awen|OJJ38200.1 azon|OJJ43095.1

Group2106: abra|OJJ69989.1 acar|OOF93494.1 acla|EAW08861.1 afis|EAW15533.1 afla|EED49044.1 afum|EAL85795.1 agla|OJJ85214.1 akaw|GAA87672.1 aluc|OJZ85290.1 anid|CBF78199.1 anig|EHA25488.1 aory|EIT82552.1 asyd|OJJ55983.1 ater|EAU32592.1 atub|OJI80280.1 aver|OJJ04930.1 awen|OJJ38197.1 azon|OJJ43096.1

Group2107: abra|OJJ69990.1 acar|OOF93496.1 acla|EAW08863.1 afis|EAW15531.1 afla|EED49042.1 afum|EAL85797.1 agla|OJJ85216.1 akaw|GAA87671.1 aluc|OJZ85289.1 anid|CBF78202.1 anig|EHA25486.1 aory|EIT82550.1 asyd|OJJ55985.1 ater|EAU32594.1 atub|OJI80281.1 aver|OJJ04932.1 awen|OJJ38195.1 azon|OJJ43099.1

Group2108: abra|OJJ69991.1 acar|OOF93497.1 acla|EAW08864.1 afis|EAW15530.1 afla|EED49041.1 afum|EAL85798.1 agla|OJJ85217.1 akaw|GAA87670.1 aluc|OJZ85288.1 anid|CBF78204.1 anig|EHA25485.1 aory|EIT82567.1 asyd|OJJ55986.1 ater|EAU32595.1 atub|OJI80282.1 aver|OJJ04933.1 awen|OJJ38194.1 azon|OJJ43100.1

Group2109: abra|OJJ69995.1 acar|OOF93502.1 acla|EAW08824.1 afis|EAW15573.1 afla|EED49082.1 afum|EAL85761.1 agla|OJJ85179.1 akaw|GAA87666.1 aluc|OJZ85284.1 anid|CBF78105.1 anig|EHA25481.1 aory|EIT82549.1 asyd|OJJ55895.1 ater|EAU32558.1 atub|OJI80286.1 aver|OJJ04840.1 awen|OJJ38232.1 azon|OJJ43064.1

Group2110: abra|OJJ70001.1 acar|OOF93507.1 acla|EAW08818.1 afis|EAW15579.1 afla|EED49093.1 afum|EAL85755.1 agla|OJJ85173.1 akaw|GAA87660.1 aluc|OJZ85278.1 anid|CBF78100.1 anig|EHA25475.1 aory|EIT82531.1 asyd|OJJ55889.1 ater|EAU32551.1 atub|OJI80292.1 aver|OJJ04834.1 awen|OJJ38241.1 azon|OJJ43058.1

Group2111: abra|OJJ70003.1 acar|OOF93509.1 acla|EAW08816.1 afis|EAW15581.1 afla|EED49090.1 afum|EAL85753.2 agla|OJJ85170.1 akaw|GAA87657.1 aluc|OJZ85276.1 anid|CBF78096.1 anig|EHA25473.1 aory|EIT82539.1 asyd|OJJ55886.1 ater|EAU32549.1 atub|OJI80294.1 aver|OJJ04831.1 awen|OJJ38243.1 azon|OJJ43055.1

Group2112: abra|OJJ70008.1 acar|OOF93514.1 acla|EAW08815.1 afis|EAW15582.1 afla|EED49088.1 afum|EAL85752.1 agla|OJJ85169.1 akaw|GAA87654.1 aluc|OJZ85271.1 anid|CBF78090.1 anig|EHA25470.1 aory|EIT82507.1 asyd|OJJ55883.1 ater|EAU32547.1 atub|OJI80300.1 aver|OJJ04828.1 awen|OJJ38246.1 azon|OJJ43052.1

Group2114: abra|OJJ70013.1 acar|OOF93519.1 acla|EAW08811.1 afis|EAW15586.1 afla|EED54443.1 afum|EAL85746.1 agla|OJJ85166.1 akaw|GAA87649.1 aluc|OJZ85264.1 anid|CBF78063.1 anig|EHA25464.1 aory|EIT81154.1 asyd|OJJ55870.1 ater|EAU32543.1 atub|OJI80305.1 aver|OJJ04815.1 awen|OJJ38252.1 azon|OJJ43032.1

Group2115: abra|OJJ70014.1 acar|OOF93520.1 acla|EAW08809.1 afis|EAW15587.1 afla|EED50410.1 afum|EAL85745.1 agla|OJJ85161.1 akaw|GAA87648.1 aluc|OJZ85263.1 anid|CBF78069.1 anig|EHA25463.1 aory|EIT82131.1 asyd|OJJ55872.1 ater|EAU32541.1 atub|OJI80306.1 aver|OJJ04817.1 awen|OJJ38254.1 azon|OJJ43033.1

Group2116: abra|OJJ70015.1 acar|OOF93521.1 acla|EAW08808.1 afis|EAW15588.1 afla|EED50411.1 afum|EAL85744.1 agla|OJJ85160.1 akaw|GAA87646.1 aluc|OJZ85261.1 anid|CBF78071.1 anig|EHA25462.1 aory|EIT82015.1 asyd|OJJ55873.1 ater|EAU32539.1 atub|OJI80307.1 aver|OJJ04818.1 awen|OJJ38255.1 azon|OJJ43038.1

Group2117: abra|OJJ70016.1 acar|OOF93522.1 acla|EAW08807.1 afis|EAW15589.1 afla|EED50412.1 afum|EAL85743.1 agla|OJJ85159.1 akaw|GAA87645.1 aluc|OJZ85260.1 anid|CBF78072.1 anig|EHA25461.1 aory|EIT82097.1 asyd|OJJ55874.1 ater|EAU32538.1 atub|OJI80308.1 aver|OJJ04819.1 awen|OJJ38262.1 azon|OJJ43023.1

Group2118: abra|OJJ70018.1 acar|OOF93524.1 acla|EAW08805.1 afis|EAW15591.1 afla|EED50414.1 afum|EAL85741.1 agla|OJJ85157.1 akaw|GAA87643.1 aluc|OJZ85258.1 anid|CBF78076.1 anig|EHA25459.1 aory|EIT82095.1 asyd|OJJ55876.1 ater|EAU32537.1 atub|OJI80310.1 aver|OJJ04821.1 awen|OJJ38264.1 azon|OJJ43021.1

Group2119: abra|OJJ70026.1 acar|OOF93536.1 acla|EAW08802.1 afis|EAW15595.1 afla|EED50418.1 afum|EAL85736.2 agla|OJJ85151.1 akaw|GAA87634.1 aluc|OJZ85250.1 anid|CBF78087.1 anig|EHA25450.1 aory|EIT82070.1 asyd|OJJ55881.1 ater|EAU32532.1 atub|OJI80319.1 aver|OJJ04826.1 awen|OJJ38271.1 azon|OJJ43910.1

Group2120: abra|OJJ70042.1 acar|OOF93537.1 acla|EAW08803.1 afis|EAW15594.1 afla|EED50417.1 afum|EAL85737.2 agla|OJJ85153.1 akaw|GAA87633.1 aluc|OJZ85249.1 anid|CBF78085.1 anig|EHA25449.1 aory|EIT82162.1 asyd|OJJ55880.1 ater|EAU32533.1 atub|OJI80320.1 aver|OJJ04825.1 awen|OJJ38270.1 azon|OJJ43912.1

Group2123: abra|OJJ70159.1 acar|OOF93314.1 acla|EAW06525.1 afis|EAW16241.1 afla|EED57104.1 afum|EAL88863.1 agla|OJJ88977.1 akaw|GAA86606.1 aluc|OJZ85156.1 anid|CBF78253.1 anig|EHA25372.1 aory|EIT73977.1 asyd|OJJ56050.1 ater|EAU34336.1 atub|OJI80433.1 aver|OJJ04993.1 awen|OJJ36019.1 azon|OJJ42535.1

Group2125: abra|OJJ70321.1 acar|OOF91776.1 acla|EAW12313.1 afis|EAW16939.1 afla|EED45249.1 afum|EAL86466.1 agla|OJJ79973.1 akaw|GAA92577.1 aluc|OJZ82590.1 anid|CBF75409.1 anig|EHA27485.1 aory|EIT73687.1 asyd|OJJ59815.1 ater|EAU30208.1 atub|OJI88825.1 aver|OJJ02473.1 awen|OJJ29713.1 azon|OJJ45106.1

Group2127: abra|OJJ70367.1 acar|OOF91904.1 acla|EAW12332.1 afis|EAW16890.1 afla|EED53384.1 afum|EAL86428.2 agla|OJJ79944.1 akaw|GAA91984.1 aluc|OJZ82629.1 anid|CBF80798.1 anig|EHA27448.1 aory|EIT75200.1 asyd|OJJ54488.1 ater|EAU30224.1 atub|OJI88862.1 aver|OJJ07691.1 awen|OJJ29748.1 azon|OJJ50203.1

Group2128: abra|OJJ70389.1 acar|OOF90063.1 acla|EAW12390.1 afis|EAW16837.1 afla|EED47319.1 afum|EAL86385.1 agla|OJJ83707.1 akaw|GAA91969.1 aluc|OJZ82647.1 anid|CBF80639.1 anig|EHA27425.1 aory|EIT82834.1 asyd|OJJ54621.1 ater|EAU30244.1 atub|OJI88879.1 aver|OJJ07537.1 awen|OJJ33871.1 azon|OJJ43960.1

Group2129: abra|OJJ70449.1 acar|OOF91831.1 acla|EAW09124.1 afis|EAW24708.1 afla|EED52336.1 afum|EAL89350.1 agla|OJJ82814.1 akaw|GAA90132.1 aluc|OJZ83215.1 anid|CBF83955.1 anig|EHA21075.1 aory|EIT74337.1 asyd|OJJ61431.1 ater|EAU37947.1 atub|OJI87721.1 aver|OJJ00766.1 awen|OJJ39949.1 azon|OJJ51122.1

Group2130: abra|OJJ70463.1 acar|OOF97461.1 acla|EAW13050.1 afis|EAW17617.1 afla|EED49208.1 afum|EAL91430.1 agla|OJJ81323.1 akaw|GAA83312.1 aluc|OJZ83806.1 anid|CBF90167.1 anig|EHA18953.1

aory|EIT83163.1 asyd|OJJ61992.1 ater|EAU37051.1 atub|OJI90173.1 aver|OJI96084.1 awen|OJJ37228.1 azon|OJJ50931.1

Group2131: abra|OJJ70464.1 acar|OOF97459.1 acla|EAW13049.1 afis|EAW17618.1 afla|EED49209.1 afum|EAL91431.1 agla|OJJ81322.1 akaw|GAA83313.1 aluc|OJZ83807.1 anid|CBF90165.1 anig|EHA18954.1 aory|EIT83149.1 asyd|OJJ61993.1 ater|EAU37052.1 atub|OJI90174.1 aver|OJI96085.1 awen|OJJ37227.1 azon|OJJ50930.1

Group2132: abra|OJJ70468.1 acar|OOF97456.1 acla|EAW13046.1 afis|EAW17621.1 afla|EED49196.1 afum|EAL91434.1 agla|OJJ81304.1 akaw|GAA83317.1 aluc|OJZ83812.1 anid|CBF90159.1 anig|EHA18957.1 aory|EIT78509.1 asyd|OJJ61996.1 ater|EAU37055.1 atub|OJI90179.1 aver|OJI96088.1 awen|OJJ37223.1 azon|OJJ50926.1

Group2133: abra|OJJ70469.1 acar|OOF97455.1 acla|EAW13045.1 afis|EAW17622.1 afla|EED49195.1 afum|EAL91435.1 agla|OJJ81303.1 akaw|GAA83318.1 aluc|OJZ83813.1 anid|CBF90157.1 anig|EHA18958.1 aory|EIT78576.1 asyd|OJJ61997.1 ater|EAU37056.1 atub|OJI90180.1 aver|OJI96089.1 awen|OJJ37222.1 azon|OJJ50925.1

Group2134: abra|OJJ70470.1 acar|OOF97453.1 acla|EAW13044.1 afis|EAW17623.1 afla|EED49194.1 afum|EAL91437.1 agla|OJJ81302.1 akaw|GAA83319.1 aluc|OJZ83814.1 anid|CBF90156.1 anig|EHA18959.1 aory|EIT78552.1 asyd|OJJ61998.1 ater|EAU37057.1 atub|OJI90181.1 aver|OJI96090.1 awen|OJJ37221.1 azon|OJJ50924.1

Group2135: abra|OJJ70473.1 acar|OOF97452.1 acla|EAW13041.1 afis|EAW17626.1 afla|EED49192.1 afum|EAL91440.1 agla|OJJ81276.1 akaw|GAA83322.1 aluc|OJZ83817.1 anid|CBF90150.1 anig|EHA18962.1 aory|EIT78505.1 asyd|OJJ62001.1 ater|EAU37060.1 atub|OJI90184.1 aver|OJI96093.1 awen|OJJ37218.1 azon|OJJ50921.1

Group2136: abra|OJJ70475.1 acar|OOF97450.1 acla|EAW13039.1 afis|EAW17628.1 afla|EED49190.1 afum|EAL91442.1 agla|OJJ81275.1 akaw|GAA83324.1 aluc|OJZ83819.1 anid|CBF90147.1 anig|EHA18964.1 aory|EIT78550.1 asyd|OJJ62002.1 ater|EAU37062.1 atub|OJI90186.1 aver|OJI96094.1 awen|OJJ37216.1 azon|OJJ50919.1

Group2137: abra|OJJ70476.1 acar|OOF97449.1 acla|EAW13038.1 afis|EAW17629.1 afla|EED49189.1 afum|EAL91443.1 agla|OJJ81272.1 akaw|GAA83325.1 aluc|OJZ83820.1 anid|CBF90141.1 anig|EHA18965.1 aory|EIT78529.1 asyd|OJJ62006.1 ater|EAU37063.1 atub|OJI90187.1 aver|OJI96098.1 awen|OJJ37215.1 azon|OJJ50916.1

Group2138: abra|OJJ70477.1 acar|OOF97448.1 acla|EAW13037.1 afis|EAW17630.1 afla|EED49188.1 afum|EAL91445.1 agla|OJJ81271.1 akaw|GAA83326.1 aluc|OJZ83822.1 anid|CBF90143.1 anig|EHA18966.1 aory|EIT78536.1 asyd|OJJ62005.1 ater|EAU37065.1 atub|OJI90189.1 aver|OJI96096.1 awen|OJJ37214.1 azon|OJJ50914.1

Group2139: abra|OJJ70478.1 acar|OOF97447.1 acla|EAW13036.1 afis|EAW17631.1 afla|EED49187.1 afum|EAL91446.1 agla|OJJ81270.1 akaw|GAA83327.1 aluc|OJZ83823.1 anid|CBF90145.1 anig|EHA18967.1 aory|EIT78534.1 asyd|OJJ62004.1 ater|EAU37066.1 atub|OJI90190.1 aver|OJI96095.1 awen|OJJ37213.1 azon|OJJ50913.1

Group2140: abra|OJJ70481.1 acar|OOF97444.1 acla|EAW13035.1 afis|EAW17632.1 afla|EED49186.1 afum|EAL91447.1 agla|OJJ81269.1 akaw|GAA83329.1 aluc|OJZ83825.1 anid|CBF90139.1 anig|EHA18969.1 aory|EIT78523.1 asyd|OJJ62011.1 ater|EAU37067.1 atub|OJI90192.1 aver|OJI96103.1 awen|OJJ37212.1 azon|OJJ50912.1

Group2141: abra|OJJ70482.1 acar|OOF97443.1 acla|EAW13034.1 afis|EAW17633.1 afla|EED50531.1 afum|EAL91448.1 agla|OJJ81266.1 akaw|GAA83330.1 aluc|OJZ83826.1 anid|CBF90137.1 anig|EHA18970.1 aory|EIT82093.1 asyd|OJJ62012.1 ater|EAU37069.1 atub|OJI90193.1 aver|OJI96104.1 awen|OJJ37211.1 azon|OJJ50911.1

Group2142: abra|OJJ70483.1 acar|OOF97442.1 acla|EAW13033.1 afis|EAW17634.1 afla|EED50530.1 afum|EAL91449.1 agla|OJJ81265.1 akaw|GAA83331.1 aluc|OJZ83827.1 anid|CBF90135.1 anig|EHA18971.1 aory|EIT82030.1 asyd|OJJ62013.1 ater|EAU37070.1 atub|OJI90194.1 aver|OJI96105.1 awen|OJJ37210.1 azon|OJJ50910.1

Group2143: abra|OJJ70484.1 acar|OOF97441.1 acla|EAW13031.1 afis|EAW17636.1 afla|EED50528.1 afum|EAL91451.2 agla|OJJ81263.1 akaw|GAA83333.1 aluc|OJZ83829.1 anid|CBF90132.1 anig|EHA18972.1 aory|EIT82038.1 asyd|OJJ62015.1 ater|EAU37072.1 atub|OJI90196.1 aver|OJI96106.1 awen|OJJ37208.1 azon|OJJ50908.1

Group2144: abra|OJJ70488.1 acar|OOF97438.1 acla|EAW13028.1 afis|EAW17639.1 afla|EED50524.1 afum|EAL91454.1 agla|OJJ79567.1 akaw|GAA83336.1 aluc|OJZ83832.1 anid|CBF90125.1 anig|EHA18975.1 aory|EIT82049.1 asyd|OJJ62018.1 ater|EAU37075.1 atub|OJI90199.1 aver|OJI96109.1 awen|OJJ37206.1 azon|OJJ50905.1

Group2145: abra|OJJ70489.1 acar|OOF97437.1 acla|EAW13027.1 afis|EAW17640.1 afla|EED50523.1 afum|EAL91455.1 agla|OJJ79568.1 akaw|GAA83337.1 aluc|OJZ83833.1 anig|EHA18976.1 aory|EIT82102.1 asyd|OJJ62019.1 ater|EAU37076.1 atub|OJI90200.1 aver|OJI96110.1 awen|OJJ37205.1 azon|OJJ50904.1 anid|CBF90123.1

Group2146: abra|OJJ70490.1 acar|OOF97436.1 acla|EAW13026.1 afis|EAW17641.1 afla|EED50522.1 afum|EAL91456.1 agla|OJJ79569.1 akaw|GAA83338.1 aluc|OJZ83834.1 anid|CBF90121.1 anig|EHA18977.1 aory|EIT82016.1 asyd|OJJ62020.1 ater|EAU37077.1 atub|OJI90201.1 aver|OJI96111.1 awen|OJJ37204.1 azon|OJJ50903.1

Group2147: abra|OJJ70492.1 acar|OOF97435.1 acla|EAW13025.1 afis|EAW17642.1 afla|EED50518.1 afum|EAL91457.1 agla|OJJ79562.1 akaw|GAA83340.1 aluc|OJZ83836.1 anid|CBF90119.1 anig|EHA18979.1 aory|EIT82059.1 asyd|OJJ62021.1 ater|EAU37078.1 atub|OJI90203.1 aver|OJI96112.1 awen|OJJ37203.1 azon|OJJ50902.1

Group2148: abra|OJJ70493.1 acar|OOF97432.1 acla|EAW13024.1 afis|EAW17643.1 afla|EED50517.1 afum|EAL91458.1 agla|OJJ79561.1 akaw|GAA83341.1 aluc|OJZ83837.1 anid|CBF90118.1 anig|EHA18980.1 aory|EIT82149.1 asyd|OJJ62022.1 ater|EAU37079.1 atub|OJI90204.1 aver|OJI96113.1 awen|OJJ37202.1 azon|OJJ50901.1

Group2149: abra|OJJ70495.1 acar|OOF97431.1 acla|EAW13022.1 afis|EAW17645.1 afla|EED50512.1 afum|EAL91460.1 agla|OJJ79559.1 akaw|GAA83343.1 aluc|OJZ83840.1 anid|CBF90114.1 anig|EHA18982.1 aory|EIT82064.1 asyd|OJJ62024.1 ater|EAU37081.1 atub|OJI90206.1 aver|OJI96115.1 awen|OJJ37200.1 azon|OJJ50899.1

Group2150: abra|OJJ70498.1 acar|OOF97428.1 acla|EAW08176.1 afis|EAW23212.1 afla|EED47039.1 afum|EAL88272.1 agla|OJJ78733.1 akaw|GAA83346.1 aluc|OJZ83843.1 anid|CBF74740.1 anig|EHA18985.1 aory|EIT72595.1 asyd|OJJ60184.1 ater|EAU36340.1 atub|OJI90209.1 aver|OJJ04080.1 awen|OJJ39445.1 azon|OJJ47647.1

Group2151: abra|OJJ70499.1 acar|OOF97427.1 acla|EAW08175.1 afis|EAW23211.1 afla|EED47040.1 afum|EAL88273.1 agla|OJJ78734.1 akaw|GAA83347.1 aluc|OJZ83844.1 anid|CBF74738.1 anig|EHA18986.1 aory|EIT72653.1 asyd|OJJ60185.1 ater|EAU36339.1 atub|OJI90210.1 aver|OJJ04079.1 awen|OJJ39444.1 azon|OJJ47646.1

Group2152: abra|OJJ70500.1 acar|OOF97425.1 acla|EAW08174.1 afis|EAW23210.1 afla|EED47041.1 afum|EAL88274.1 agla|OJJ78735.1 akaw|GAA83348.1 aluc|OJZ83845.1 anid|CBF74736.1 anig|EHA18987.1 aory|EIT72592.1 asyd|OJJ60186.1 ater|EAU36338.1 atub|OJI90211.1 aver|OJJ04078.1 awen|OJJ39443.1 azon|OJJ47645.1

Group2153: abra|OJJ70504.1 acar|OOF97421.1 acla|EAW08170.1 afis|EAW23206.1 afla|EED47047.1 afum|EAL88277.1 agla|OJJ78739.1 akaw|GAA83353.1 aluc|OJZ83849.1 anid|CBF74734.1 anig|EHA18991.1 aory|EIT72747.1 asyd|OJJ60189.1 ater|EAU36334.1 atub|OJI90215.1 aver|OJJ04075.1 awen|OJJ39438.1 azon|OJJ47641.1

Group2154: abra|OJJ70509.1 acar|OOF97420.1 acla|EAW08166.1 afis|EAW23204.1 afla|EED47050.1 afum|EAL88279.1 agla|OJJ78743.1 akaw|GAA83356.1 aluc|OJZ83854.1 anid|CBF74730.1 anig|EHA18995.1 aory|EIT72509.1 asyd|OJJ60194.1 ater|EAU36332.1 atub|OJI90221.1 aver|OJJ04071.1 awen|OJJ39434.1 azon|OJJ47638.1

Group2155: abra|OJJ70510.1 acar|OOF97419.1 acla|EAW08165.1 afis|EAW23203.1 afla|EED47051.1 afum|EAL88280.1 agla|OJJ78745.1 akaw|GAA83357.1 aluc|OJZ83855.1 anid|CBF74728.1 anig|EHA18996.1 aory|EIT72511.1 asyd|OJJ60195.1 ater|EAU36331.1 atub|OJI90222.1 aver|OJJ04070.1 awen|OJJ39433.1 azon|OJJ47637.1

Group2156: abra|OJJ70512.1 acar|OOF97417.1 acla|EAW08163.1 afis|EAW23201.1 afla|EED47053.1 afum|EAL88282.1 agla|OJJ78747.1 akaw|GAA83359.1 aluc|OJZ83857.1 anid|CBF74721.1 anig|EHA18998.1 aory|EIT72498.1 asyd|OJJ60201.1 ater|EAU36329.1 atub|OJI90224.1 aver|OJJ04065.1 awen|OJJ39431.1 azon|OJJ47635.1

Group2157: abra|OJJ70514.1 acar|OOF97415.1 acla|EAW08161.1 afis|EAW23199.1 afla|EED47055.1 afum|EAL88284.1 agla|OJJ78749.1 akaw|GAA83361.1 aluc|OJZ83859.1 anid|CBF74717.1 anig|EHA19000.1 aory|EIT72508.1 asyd|OJJ60203.1 ater|EAU36327.1 atub|OJI90226.1 aver|OJJ04063.1 awen|OJJ39429.1 azon|OJJ47633.1

Group2158: abra|OJJ70515.1 acar|OOF97414.1 acla|EAW08160.1 afis|EAW23198.1 afla|EED47056.1 afum|EAL88285.1 agla|OJJ86331.1 akaw|GAA83362.1 aluc|OJZ83860.1 anid|CBF74715.1 anig|EHA19001.1 aory|EIT72476.1 asyd|OJJ60205.1 ater|EAU36325.1 atub|OJI90228.1 aver|OJJ04062.1 awen|OJJ39428.1 azon|OJJ47632.1

Group2159: abra|OJJ70521.1 acar|OOF97407.1 acla|EAW08154.1 afis|EAW23185.1 afla|EED47062.1 afum|EAL88298.1 agla|OJJ86360.1 akaw|GAA83368.1 aluc|OJZ83866.1 anid|CBF74698.1 anig|EHA19007.1

aory|EIT72505.1 asyd|OJJ60215.1 ater|EAU36320.1 atub|OJI90234.1 aver|OJJ04053.1 awen|OJJ36587.1 azon|OJJ47619.1

Group2160: abra|OJJ70522.1 acar|OOF97406.1 acla|EAW08153.1 afis|EAW23186.1 afla|EED47064.1 afum|EAL88297.1 agla|OJJ86361.1 akaw|GAA83369.1 aluc|OJZ83867.1 anid|CBF74696.1 anig|EHA19008.1 aory|EIT72495.1 asyd|OJJ60214.1 ater|EAU36319.1 atub|OJI90235.1 aver|OJJ04054.1 awen|OJJ36588.1 azon|OJJ47618.1

Group2161: abra|OJJ70529.1 acar|OOF97402.1 acla|EAW08142.1 afis|EAW23178.1 afla|EED49000.1 afum|EAL88305.1 agla|OJJ89690.1 akaw|GAA83374.1 aluc|OJZ83872.1 anid|CBF74512.1 anig|EHA19013.1 aory|EIT80028.1 asyd|OJJ57730.1 ater|EAU36311.1 atub|OJI90240.1 aver|OJJ05439.1 awen|OJJ36601.1 azon|OJJ47607.1

Group2162: abra|OJJ70532.1 acar|OOF97400.1 acla|EAW08145.1 afis|EAW23181.1 afla|EED47073.1 afum|EAL88302.1 akaw|GAA83377.1 aluc|OJZ83875.1 anid|CBF74516.1 anig|EHA19015.1 aory|EIT72487.1 asyd|OJJ57732.1 ater|EAU36314.1 atub|OJI90243.1 aver|OJJ05437.1 awen|OJJ36597.1 azon|OJJ47611.1 agla|OJJ86370.1

Group2163: abra|OJJ70536.1 acar|OOF97397.1 acla|EAW08148.1 afis|EAW23191.1 afla|EED47070.1 afum|EAL88292.1 agla|OJJ86367.1 akaw|GAA83381.1 aluc|OJZ83880.1 anid|CBF74521.1 anig|EHA19018.1 aory|EIT72473.1 asyd|OJJ57733.1 ater|EAU36315.1 atub|OJI90247.1 aver|OJJ05434.1 awen|OJJ36593.1 azon|OJJ47612.1

Group2164: abra|OJJ70537.1 acar|OOF97396.1 acla|EAW08149.1 afis|EAW23190.1 afla|EED47069.1 afum|EAL88293.1 agla|OJJ86365.1 akaw|GAA83382.1 aluc|OJZ83881.1 anid|CBF74523.1 anig|EHA19019.1 aory|EIT72486.1 asyd|OJJ57734.1 ater|EAU36309.1 atub|OJI90249.1 aver|OJJ05433.1 awen|OJJ36592.1 azon|OJJ47614.1

Group2165: abra|OJJ70539.1 acar|OOF97394.1 acla|EAW08151.1 afis|EAW23188.1 afla|EED47067.1 afum|EAL88295.1 agla|OJJ86363.1 akaw|GAA83384.1 aluc|OJZ83884.1 anid|CBF74527.1 anig|EHA19021.1 aory|EIT72500.1 asyd|OJJ57736.1 ater|EAU36310.1 atub|OJI90251.1 aver|OJJ05431.1 awen|OJJ36590.1 azon|OJJ47616.1

Group2166: abra|OJJ70541.1 acar|OOF97392.1 acla|EAW08137.1 afis|EAW23173.1 afla|EED46946.1 afum|EAL88310.1 agla|OJJ89685.1 akaw|GAA83387.1 aluc|OJZ83886.1 anid|CBF74495.1 anig|EHA19023.1 aory|EIT72576.1 asyd|OJJ57718.1 ater|EAU36302.1 atub|OJI90253.1 aver|OJJ05449.1 awen|OJJ36604.1 azon|OJJ47602.1

Group2167: abra|OJJ70552.1 acar|OOF97168.1 acla|EAW08344.1 afis|EAW23379.1 afla|EED55578.1 afum|EAL88092.1 agla|OJJ86444.1 akaw|GAA83396.1 aluc|OJZ83895.1 anid|CBF74849.1 anig|EHA19031.1 aory|EIT82685.1 asyd|OJJ60128.1 ater|EAU35633.1 atub|OJI90263.1 aver|OJJ04133.1 awen|OJJ39503.1 azon|OJJ47704.1

Group2168: abra|OJJ70554.1 acar|OOF97165.1 acla|EAW08350.1 afis|EAW23383.1 afla|EED55571.1 afum|EAL88088.1 agla|OJJ86437.1 akaw|GAA83399.1 aluc|OJZ83897.1 anid|CBF74854.1 anig|EHA19034.1 aory|EIT82660.1 asyd|OJJ60126.1 ater|EAU35636.1 atub|OJI90267.1 aver|OJJ04136.1 awen|OJJ39498.1 azon|OJJ47700.1

Group2169: abra|OJJ70555.1 acar|OOF97164.1 acla|EAW08353.1 afis|EAW23384.1 afla|EED55572.1 afum|EAL88087.1 agla|OJJ86436.1 akaw|GAA83400.1 aluc|OJZ83898.1 anid|CBF74856.1 anig|EHA19035.1 aory|EIT82663.1 asyd|OJJ60125.1 ater|EAU35637.1 atub|OJI90268.1 aver|OJJ04137.1 awen|OJJ39497.1 azon|OJJ47699.1

Group2170: abra|OJJ70556.1 acar|OOF97163.1 acla|EAW08354.1 afis|EAW23385.1 afla|EED55573.1 afum|EAL88086.2 agla|OJJ86435.1 akaw|GAA83401.1 aluc|OJZ83899.1 anid|CBF74875.1 anig|EHA19036.1 aory|EIT82690.1 asyd|OJJ60114.1 ater|EAU35638.1 atub|OJI90269.1 aver|OJJ04148.1 awen|OJJ39496.1 azon|OJJ47698.1

Group2171: abra|OJJ70557.1 acar|OOF97162.1 acla|EAW08355.1 afis|EAW23386.1 afla|EED55574.1 afum|EAL88085.1 agla|OJJ86432.1 akaw|GAA83402.1 aluc|OJZ83900.1 anid|CBF74873.1 anig|EHA19037.1 aory|EIT82673.1 asyd|OJJ60115.1 ater|EAU35639.1 atub|OJI90270.1 aver|OJJ04147.1 awen|OJJ39495.1 azon|OJJ47696.1

Group2172: abra|OJJ70559.1 acar|OOF97161.1 acla|EAW08357.1 afis|EAW23388.1 afla|EED55576.1 afum|EAL88083.1 agla|OJJ86430.1 akaw|GAA83404.1 aluc|OJZ83903.1 anid|CBF74870.1 anig|EHA19038.1 aory|EIT82706.1 asyd|OJJ60117.1 ater|EAU35640.1 atub|OJI90272.1 aver|OJJ04145.1 awen|OJJ39493.1 azon|OJJ47694.1

Group2173: abra|OJJ70560.1 acar|OOF97160.1 acla|EAW08358.1 afis|EAW23389.1 afla|EED55568.1 afum|EAL88082.1 agla|OJJ86428.1 akaw|GAA83405.1 aluc|OJZ83904.1 anid|CBF74868.1 anig|EHA19039.1 aory|EIT82664.1 asyd|OJJ60118.1 ater|EAU35642.1 atub|OJI90273.1 aver|OJJ04144.1 awen|OJJ39492.1 azon|OJJ47693.1

Group2174: abra|OJJ70561.1 acar|OOF97158.1 acla|EAW08360.1 afis|EAW23391.1 afla|EED55566.1 afum|EAL88080.1 agla|OJJ86426.1 akaw|GAA83406.1 aluc|OJZ83905.1 anid|CBF74864.1 anig|EHA19040.1 aory|EIT82703.1 asyd|OJJ60121.1 ater|EAU35644.1 atub|OJI90276.1 aver|OJJ04141.1 awen|OJJ39488.1 azon|OJJ47691.1

Group2175: abra|OJJ70564.1 acar|OOF97156.1 acla|EAW08362.1 afis|EAW23393.1 afla|EED55564.1 afum|EAL88077.1 agla|OJJ86420.1 akaw|GAA83408.1 aluc|OJZ83907.1 anid|CBF74860.1 anig|EHA19042.1 aory|EIT82692.1 asyd|OJJ60123.1 ater|EAU35646.1 atub|OJI90278.1 aver|OJJ04139.1 awen|OJJ39486.1 azon|OJJ47689.1

Group2176: abra|OJJ70567.1 acar|OOF97155.1 acla|EAW08366.1 afis|EAW23396.1 afla|EED55560.1 afum|EAL88074.1 agla|OJJ86410.1 akaw|GAA83411.1 aluc|OJZ83910.1 anid|CBF74807.1 anig|EHA19044.1 aory|EIT82680.1 asyd|OJJ60148.1 ater|EAU35649.1 atub|OJI90281.1 aver|OJJ04115.1 awen|OJJ39483.1 azon|OJJ48236.1

Group2177: abra|OJJ70568.1 acar|OOF97154.1 acla|EAW08367.1 afis|EAW23397.1 afla|EED55559.1 afum|EAL88073.1 agla|OJJ86409.1 akaw|GAA83412.1 aluc|OJZ83911.1 anid|CBF74805.1 anig|EHA19045.1 aory|EIT82705.1 asyd|OJJ60149.1 ater|EAU35650.1 atub|OJI90282.1 aver|OJJ04114.1 awen|OJJ39482.1 azon|OJJ47684.1

Group2178: abra|OJJ70569.1 acar|OOF97152.1 acla|EAW08371.1 afis|EAW23398.1 afla|EED55554.1 afum|EAL88072.1 agla|OJJ86403.1 akaw|GAA83413.1 aluc|OJZ83913.1 anid|CBF74803.1 anig|EHA19046.1 aory|EIT82712.1 asyd|OJJ60150.1 ater|EAU35651.1 atub|OJI90283.1 aver|OJJ04113.1 awen|OJJ39481.1 azon|OJJ47683.1

Group2179: abra|OJJ70570.1 acar|OOF97151.1 acla|EAW08370.1 afis|EAW23399.1 afla|EED47005.1 afum|EAL88071.1 agla|OJJ86402.1 akaw|GAA83414.1 aluc|OJZ83914.1 anid|CBF74801.1 anig|EHA19047.1 aory|EIT72537.1 asyd|OJJ60152.1 ater|EAU36378.1 atub|OJI90284.1 aver|OJJ04112.1 awen|OJJ39480.1 azon|OJJ47682.1

Group2180: abra|OJJ70571.1 acar|OOF97150.1 acla|EAW08369.1 afis|EAW23400.1 afla|EED47006.1 afum|EAL88070.1 agla|OJJ86401.1 akaw|GAA83415.1 aluc|OJZ83915.1 anid|CBF73601.1 anig|EHA19048.1 aory|EIT72638.1 asyd|OJJ56680.1 ater|EAU36377.1 atub|OJI90285.1 aver|OJJ06043.1 awen|OJJ39479.1 azon|OJJ47681.1

Group2181: abra|OJJ70574.1 acar|OOF97147.1 acla|EAW08373.1 afis|EAW23402.1 afla|EED47009.1 afum|EAL88068.1 agla|OJJ86389.1 akaw|GAA83418.1 aluc|OJZ83918.1 anid|CBF74795.1 anig|EHA19051.1 aory|EIT72635.1 asyd|OJJ60155.1 ater|EAU36373.1 atub|OJI90289.1 aver|OJJ04109.1 awen|OJJ39476.1 azon|OJJ47679.1

Group2182: abra|OJJ70576.1 acar|OOF97145.1 acla|EAW08375.1 afis|EAW23404.1 afla|EED47011.1 afum|EAL88066.1 agla|OJJ86392.1 akaw|GAA83420.1 aluc|OJZ83920.1 anid|CBF74793.1 anig|EHA19053.1 aory|EIT72673.1 asyd|OJJ60156.1 ater|EAU36371.1 atub|OJI90292.1 aver|OJJ04108.1 awen|OJJ39473.1 azon|OJJ47675.1

Group2183: abra|OJJ70577.1 acar|OOF97144.1 acla|EAW08203.1 afis|EAW23237.1 afla|EED47012.1 afum|EAL88245.1 agla|OJJ86393.1 akaw|GAA83421.1 aluc|OJZ83921.1 anid|CBF74787.1 anig|EHA19054.1 aory|EIT72690.1 asyd|OJJ60159.1 ater|EAU36370.1 atub|OJI90293.1 aver|OJJ04105.1 awen|OJJ39472.1 azon|OJJ47674.1

Group2184: abra|OJJ70578.1 acar|OOF97143.1 acla|EAW08202.1 afis|EAW23236.1 afla|EED47013.1 afum|EAL88246.1 agla|OJJ86394.1 akaw|GAA83422.1 aluc|OJZ83922.1 anid|CBF74785.1 anig|EHA19055.1 aory|EIT72768.1 asyd|OJJ60160.1 ater|EAU36369.1 atub|OJI90294.1 aver|OJJ04104.1 awen|OJJ39471.1 azon|OJJ47673.1

Group2185: abra|OJJ70580.1 acar|OOF97138.1 acla|EAW08200.1 afis|EAW23234.1 afla|EED47015.1 afum|EAL88248.1 agla|OJJ86396.1 akaw|GAA83424.1 aluc|OJZ83925.1 anid|CBF74783.1 anig|EHA19057.1 aory|EIT72553.1 asyd|OJJ60161.1 ater|EAU36367.1 atub|OJI90297.1 aver|OJJ04103.1 awen|OJJ39469.1 azon|OJJ47669.1

Group2186: abra|OJJ70582.1 acar|OOF97137.1 acla|EAW08198.1 afis|EAW23232.1 afla|EED47016.1 afum|EAL88250.1 agla|OJJ86398.1 akaw|GAA83426.1 aluc|OJZ83927.1 anid|CBF74779.1 anig|EHA19059.1 aory|EIT72651.1 asyd|OJJ60163.1 ater|EAU36361.1 atub|OJI90299.1 aver|OJJ04101.1 awen|OJJ39467.1 azon|OJJ47667.1

Group2187: abra|OJJ70584.1 acar|OOF97136.1 acla|EAW08197.1 afis|EAW23231.1 afla|EED47019.1 afum|EAL88252.1 agla|OJJ86399.1 akaw|GAA83427.1 aluc|OJZ83928.1 anid|CBF74777.1 anig|EHA19060.1 aory|EIT72752.1 asyd|OJJ60164.1 ater|EAU36360.1 atub|OJI90300.1 aver|OJJ04100.1 awen|OJJ39466.1 azon|OJJ47666.1

Group2188: abra|OJJ70585.1 acar|OOF97135.1 acla|EAW08196.1 afis|EAW23230.1 afla|EED47020.1 afum|EAL88253.1 agla|OJJ86400.1 akaw|GAA83428.1 aluc|OJZ83929.1 anid|CBF74775.1 anig|EHA19061.1

aory|EIT72543.1 asyd|OJJ60165.1 ater|EAU36359.1 atub|OJI90301.1 aver|OJJ04099.1 awen|OJJ39465.1 azon|OJJ47665.1

Group2189: abra|OJJ70589.1 acar|OOF97128.1 acla|EAW08192.1 afis|EAW23226.1 afla|EED47024.1 afum|EAL88257.1 agla|OJJ86383.1 akaw|GAA83433.1 aluc|OJZ83934.1 anid|CBF74767.1 anig|EHA19065.1 aory|EIT72765.1 asyd|OJJ60169.1 ater|EAU36356.1 atub|OJI90306.1 aver|OJJ04095.1 awen|OJJ39461.1 azon|OJJ47660.1

Group2190: abra|OJJ70592.1 acar|OOF97126.1 acla|EAW08188.1 afis|EAW23222.1 afla|EED47028.1 afum|EAL88262.1 agla|OJJ86379.1 akaw|GAA83437.1 aluc|OJZ83938.1 anid|CBF74759.1 anig|EHA19068.1 aory|EIT72692.1 asyd|OJJ60173.1 ater|EAU36353.1 atub|OJI90311.1 aver|OJJ04091.1 awen|OJJ39458.1 azon|OJJ47656.1

Group2191: abra|OJJ70594.1 acar|OOF97124.1 acla|EAW08186.1 afis|EAW23220.1 afla|EED47030.1 afum|EAL88264.1 agla|OJJ86377.1 akaw|GAA83439.1 aluc|OJZ83940.1 anid|CBF74757.1 anig|EHA19070.1 aory|EIT72550.1 asyd|OJJ60174.1 ater|EAU36351.1 atub|OJI90313.1 aver|OJJ04089.1 awen|OJJ39456.1 azon|OJJ47654.1

Group2192: abra|OJJ70595.1 acar|OOF97123.1 acla|EAW08185.1 afis|EAW23219.1 afla|EED47031.1 afum|EAL88265.1 agla|OJJ86376.1 akaw|GAA83440.1 aluc|OJZ83941.1 anid|CBF74755.1 anig|EHA19071.1 aory|EIT72573.1 asyd|OJJ60175.1 ater|EAU36350.1 atub|OJI90314.1 aver|OJJ04088.1 awen|OJJ39455.1 azon|OJJ47653.1

Group2193: abra|OJJ70600.1 acar|OOF97120.1 acla|EAW08182.1 afis|EAW23217.1 afla|EED47035.1 afum|EAL88267.1 agla|OJJ86373.1 akaw|GAA89940.1 aluc|OJZ83945.1 anid|CBF74749.1 anig|EHA19074.1 aory|EIT72655.1 asyd|OJJ60178.1 ater|EAU36347.1 atub|OJI90316.1 aver|OJJ04084.1 awen|OJJ39451.1 azon|OJJ47651.1

Group2194: abra|OJJ70606.1 acar|OOF97115.1 acla|EAW13020.1 afis|EAW17647.1 afla|EED50511.1 afum|EAL91462.1 agla|OJJ79557.1 akaw|GAA89946.1 aluc|OJZ83952.1 anid|CBF90111.1 anig|EHA19080.1 aory|EIT82166.1 asyd|OJJ62026.1 ater|EAU37083.1 atub|OJI90323.1 aver|OJI96117.1 awen|OJJ37198.1 azon|OJJ50897.1

Group2195: abra|OJJ70607.1 acar|OOF97114.1 acla|EAW13019.1 afis|EAW17648.1 afla|EED50510.1 afum|EAL91464.1 agla|OJJ79556.1 akaw|GAA89947.1 aluc|OJZ83954.1 anid|CBF90087.1 anig|EHA19081.1 aory|EIT82125.1 asyd|OJJ62036.1 ater|EAU37084.1 atub|OJI90324.1 aver|OJI96142.1 awen|OJJ37197.1 azon|OJJ50896.1

Group2196: abra|OJJ70608.1 acar|OOF97113.1 acla|EAW13018.1 afis|EAW17649.1 afla|EED50509.1 afum|EAL91465.1 agla|OJJ79555.1 akaw|GAA89948.1 aluc|OJZ83955.1 anid|CBF90085.1 anig|EHA19082.1 aory|EIT82043.1 asyd|OJJ62037.1 ater|EAU37085.1 atub|OJI90325.1 aver|OJI96143.1 awen|OJJ37196.1 azon|OJJ50895.1

Group2197: abra|OJJ70611.1 acar|OOF97111.1 acla|EAW13016.1 afis|EAW17651.1 afla|EED50508.1 afum|EAL91466.1 agla|OJJ79553.1 akaw|GAA89950.1 aluc|OJZ83957.1 anid|CBF90077.1 anig|EHA19084.1 aory|EIT82060.1 asyd|OJJ62040.1 ater|EAU37087.1 atub|OJI90327.1 aver|OJI96146.1 awen|OJJ37194.1 azon|OJJ50893.1

Group2198: abra|OJJ70613.1 acar|OOF97109.1 acla|EAW13014.1 afis|EAW17653.1 afla|EED50505.1 afum|EAL91468.1 agla|OJJ79551.1 akaw|GAA89952.1 aluc|OJZ83959.1 anid|CBF90074.1 anig|EHA19087.1 aory|EIT82010.1 asyd|OJJ62042.1 ater|EAU37089.1 atub|OJI90331.1 aver|OJI96148.1 awen|OJJ37192.1 azon|OJJ50891.1

Group2199: abra|OJJ70616.1 acar|OOF97105.1 acla|EAW13011.1 afis|EAW17656.1 afla|EED50502.1 afum|EAL91471.1 agla|OJJ79549.1 akaw|GAA89955.1 aluc|OJZ83962.1 anid|CBF90066.1 anig|EHA19090.1 aory|EIT82103.1 asyd|OJJ62045.1 ater|EAU31997.1 atub|OJI90334.1 aver|OJI96151.1 awen|OJJ37187.1 azon|OJJ50889.1

Group2200: abra|OJJ70617.1 acar|OOF97104.1 acla|EAW13010.1 afis|EAW17657.1 afla|EED50501.1 afum|EAL91472.1 agla|OJJ79548.1 akaw|GAA89956.1 aluc|OJZ83963.1 anid|CBF90065.1 anig|EHA19091.1 aory|EIT82023.1 asyd|OJJ62046.1 ater|EAU31996.1 atub|OJI90335.1 aver|OJI96152.1 awen|OJJ37186.1 azon|OJJ50888.1

Group2201: abra|OJJ70618.1 acar|OOF97103.1 acla|EAW13009.1 afis|EAW17658.1 afla|EED50500.1 afum|EAL91473.1 agla|OJJ79547.1 akaw|GAA89957.1 aluc|OJZ83964.1 anid|CBF90063.1 anig|EHA19092.1 aory|EIT82147.1 asyd|OJJ62048.1 ater|EAU31994.1 atub|OJI90337.1 aver|OJI96153.1 awen|OJJ37185.1 azon|OJJ50886.1

Group2202: abra|OJJ70619.1 acar|OOF97102.1 acla|EAW13008.1 afis|EAW17659.1 afla|EED50499.1 afum|EAL91474.1 agla|OJJ79546.1 akaw|GAA89958.1 aluc|OJZ83965.1 anid|CBF90061.1 anig|EHA19093.1 aory|EIT82055.1 asyd|OJJ62049.1 ater|EAU31993.1 atub|OJI90338.1 aver|OJI96154.1 awen|OJJ37184.1 azon|OJJ50885.1

Group2203: abra|OJJ70620.1 acar|OOF97101.1 acla|EAW13007.1 afis|EAW17660.1 afla|EED50497.1 afum|EAL91475.1 agla|OJJ79529.1 akaw|GAA89959.1 aluc|OJZ83966.1 anid|CBF90059.1 anig|EHA19094.1 aory|EIT82172.1 asyd|OJJ62050.1 ater|EAU31992.1 atub|OJI90339.1 aver|OJI96155.1 awen|OJJ37182.1 azon|OJJ50884.1

Group2204: abra|OJJ70625.1 acar|OOF97099.1 acla|EAW13005.1 afis|EAW17663.1 afla|EED50495.1 afum|EAL91477.2 agla|OJJ79532.1 akaw|GAA89963.1 aluc|OJZ83970.1 anid|CBF90056.1 anig|EHA19099.1 aory|EIT82160.1 asyd|OJJ62052.1 ater|EAU31990.1 atub|OJI90343.1 aver|OJI96158.1 awen|OJJ37179.1 azon|OJJ50881.1

Group2205: abra|OJJ70626.1 acar|OOF97098.1 acla|EAW08131.1 afis|EAW23168.1 afla|EED46953.1 afum|EAL88316.1 agla|OJJ89679.1 akaw|GAA89964.1 aluc|OJZ83971.1 anid|CBF74478.1 anig|EHA19100.1 aory|EIT72624.1 asyd|OJJ57707.1 ater|EAU36297.1 atub|OJI90344.1 aver|OJJ05457.1 awen|OJJ36615.1 azon|OJJ47598.1

Group2206: abra|OJJ70627.1 acar|OOF97097.1 acla|EAW08130.1 afis|EAW23167.1 afla|EED46954.1 afum|EAL88317.2 agla|OJJ89678.1 akaw|GAA89965.1 aluc|OJZ83972.1 anid|CBF74476.1 anig|EHA19101.1 aory|EIT72656.1 asyd|OJJ57706.1 ater|EAU36296.1 atub|OJI90345.1 aver|OJJ05458.1 awen|OJJ36616.1 azon|OJJ41979.1

Group2207: abra|OJJ70628.1 acar|OOF97096.1 acla|EAW08129.1 afis|EAW23166.1 afla|EED46955.1 afum|EAL88318.1 agla|OJJ89677.1 akaw|GAA89966.1 aluc|OJZ83974.1 anid|CBF74475.1 anig|EHA19102.1 aory|EIT72623.1 asyd|OJJ57705.1 ater|EAU36295.1 atub|OJI90347.1 aver|OJJ05459.1 awen|OJJ36617.1 azon|OJJ47597.1

Group2208: abra|OJJ70629.1 acar|OOF97095.1 acla|EAW08128.1 afis|EAW23165.1 afla|EED46957.1 afum|EBA27343.1 agla|OJJ89676.1 akaw|GAA89967.1 aluc|OJZ83976.1 anid|CBF74473.1 anig|EHA19103.1 aory|EIT72540.1 asyd|OJJ57704.1 ater|EAU36294.1 atub|OJI90349.1 aver|OJJ05460.1 awen|OJJ36618.1 azon|OJJ47595.1

Group2209: abra|OJJ70630.1 acar|OOF97093.1 acla|EAW08126.1 afis|EAW23163.1 afla|EED46958.1 afum|EAL88320.1 agla|OJJ89674.1 akaw|GAA89969.1 aluc|OJZ83978.1 anid|CBF74467.1 anig|EHA19105.1 aory|EIT72637.1 asyd|OJJ57700.1 ater|EAU36293.1 atub|OJI90351.1 aver|OJJ05464.1 awen|OJJ36620.1 azon|OJJ47593.1

Group2210: abra|OJJ70639.1 acar|OOF97080.1 acla|EAW08118.1 afis|EAW23133.1 afla|EED48995.1 afum|EAL88348.1 agla|OJJ89666.1 akaw|GAA89975.1 aluc|OJZ83986.1 anid|CBF74456.1 anig|EHA19112.1 aory|EIT80032.1 asyd|OJJ57677.1 ater|EAU36286.1 atub|OJI90358.1 aver|OJJ05483.1 awen|OJJ36627.1 azon|OJJ47586.1

Group2211: abra|OJJ70640.1 acar|OOF97081.1 acla|EAW08117.1 afis|EAW23134.1 afla|EED48497.1 afum|EAL88347.1 agla|OJJ89665.1 akaw|GAA89976.1 aluc|OJZ83987.1 anid|CBF74455.1 anig|EHA19113.1 aory|EIT73203.1 asyd|OJJ57676.1 ater|EAU36282.1 atub|OJI90359.1 aver|OJJ05482.1 awen|OJJ36630.1 azon|OJJ47584.1

Group2212: abra|OJJ70646.1 acar|OOF97075.1 acla|EAW08114.1 afis|EAW23137.1 afla|EED48500.1 afum|EAL88344.1 agla|OJJ89663.1 akaw|GAA89982.1 aluc|OJZ83996.1 anid|CBF74450.1 anig|EHA19119.1 aory|EIT73193.1 asyd|OJJ57740.1 ater|EAU36280.1 atub|OJI90365.1 aver|OJJ05427.1 awen|OJJ36633.1 azon|OJJ47582.1

Group2213: abra|OJJ70653.1 acar|OOF97070.1 acla|EAW08106.1 afis|EAW23145.1 afla|EED48508.1 afum|EAL88336.1 agla|OJJ89657.1 akaw|GAA89989.1 aluc|OJZ84003.1 anid|CBF74436.1 anig|EHA19125.1 aory|EIT73083.1 asyd|OJJ57747.1 ater|EAU36266.1 atub|OJI90372.1 aver|OJJ05420.1 awen|OJJ36640.1 azon|OJJ47574.1

Group2214: abra|OJJ70654.1 acar|OOF97068.1 acla|EAW08105.1 afis|EAW23146.1 afla|EED48509.1 afum|EAL88335.1 agla|OJJ89656.1 akaw|GAA89990.1 aluc|OJZ84004.1 anid|CBF74434.1 anig|EHA19126.1 aory|EIT73165.1 asyd|OJJ57748.1 ater|EAU36265.1 atub|OJI90373.1 aver|OJJ05419.1 awen|OJJ36641.1 azon|OJJ47573.1

Group2215: abra|OJJ70657.1 acar|OOF97065.1 acla|EAW08102.1 afis|EAW23149.1 afla|EED48512.1 afum|EAL88332.1 agla|OJJ89653.1 akaw|GAA89993.1 aluc|OJZ84007.1 anid|CBF74428.1 anig|EHA19129.1 aory|EIT73157.1 asyd|OJJ57751.1 ater|EAU36263.1 atub|OJI90376.1 aver|OJJ05416.1 awen|OJJ36645.1 azon|OJJ47570.1

Group2216: abra|OJJ70659.1 acar|OOF97063.1 acla|EAW08100.1 afis|EAW23151.1 afla|EED48514.1 afum|EAL88330.2 agla|OJJ89651.1 akaw|GAA89995.1 aluc|OJZ84009.1 anid|CBF74424.1 anig|EHA19131.1 aory|EIT73110.1 asyd|OJJ57753.1 ater|EAU36261.1 atub|OJI90378.1 aver|OJJ05414.1 awen|OJJ36647.1 azon|OJJ47568.1

Group2217: abra|OJJ70661.1 acar|OOF97060.1 acla|EAW08097.1 afis|EAW23153.1 afla|EED48516.1 afum|EAL88328.1 agla|OJJ89648.1 akaw|GAA89997.1 aluc|OJZ84011.1 anid|CBF74421.1 anig|EHA19134.1

aory|EIT73108.1 asyd|OJJ57755.1 ater|EAU36258.1 atub|OJI90380.1 aver|OJJ05412.1 awen|OJJ36649.1 azon|OJJ47553.1

Group2218: abra|OJJ70666.1 acar|OOF97058.1 acla|EAW08096.1 afis|EAW23154.1 afla|EED48519.1 afum|EAL88327.2 agla|OJJ89646.1 akaw|GAA90001.1 aluc|OJZ84016.1 anid|CBF74419.1 anig|EHA19138.1 aory|EIT73174.1 asyd|OJJ57756.1 ater|EAU36257.1 atub|OJI90387.1 aver|OJJ05411.1 awen|OJJ36650.1 azon|OJJ47545.1

Group2219: abra|OJJ70669.1 acar|OOF97057.1 acla|EAW08094.1 afis|EAW23129.1 afla|EED48521.1 afum|EAL88351.1 agla|OJJ89645.1 akaw|GAA90002.1 aluc|OJZ84017.1 anid|CBF74417.1 anig|EHA19139.1 aory|EIT73127.1 asyd|OJJ57757.1 ater|EAU36256.1 atub|OJI90389.1 aver|OJJ05410.1 awen|OJJ36651.1 azon|OJJ47547.1

Group2220: abra|OJJ70671.1 acar|OOF97055.1 acla|EAW08093.1 afis|EAW23128.1 afla|EED48522.1 afum|EAL88352.1 agla|OJJ89644.1 akaw|GAA90003.1 aluc|OJZ84019.1 anid|CBF74413.1 anig|EHA19140.1 aory|EIT73195.1 asyd|OJJ57758.1 ater|EAU36255.1 atub|OJI90390.1 aver|OJJ05408.1 awen|OJJ36652.1 azon|OJJ47548.1

Group2221: abra|OJJ70672.1 acar|OOF97054.1 acla|EAW08092.1 afis|EAW23127.1 afla|EED48523.1 afum|EAL88353.1 agla|OJJ89643.1 akaw|GAA90004.1 aluc|OJZ84020.1 anid|CBF74412.1 anig|EHA19141.1 aory|EIT73139.1 asyd|OJJ57759.1 ater|EAU36254.1 atub|OJI90391.1 aver|OJJ05407.1 awen|OJJ36653.1 azon|OJJ47549.1

Group2222: abra|OJJ70675.1 acar|OOF97052.1 acla|EAW08089.1 afis|EAW23124.1 afla|EED48528.1 afum|EAL88356.1 agla|OJJ89641.1 akaw|GAA90007.1 aluc|OJZ84023.1 anid|CBF74406.1 anig|EHA19144.1 aory|EIT73244.1 asyd|OJJ57762.1 ater|EAU36252.1 atub|OJI90395.1 aver|OJJ05404.1 awen|OJJ36656.1 azon|OJJ47552.1

Group2223: abra|OJJ70677.1 acar|OOF97050.1 acla|EAW08087.1 afis|EAW23123.1 afla|EED48530.1 afum|EAL88357.1 agla|OJJ89616.1 akaw|GAA90009.1 aluc|OJZ84025.1 anid|CBF74404.1 anig|EHA19146.1 aory|EIT73116.1 asyd|OJJ57763.1 ater|EAU36251.1 atub|OJI90397.1 aver|OJJ05403.1 awen|OJJ36658.1 azon|OJJ47540.1

Group2224: abra|OJJ70680.1 acar|OOF97047.1 acla|EAW08084.1 afis|EAW23120.1 afla|EED48533.1 afum|EAL88360.1 agla|OJJ89619.1 akaw|GAA90012.1 aluc|OJZ84028.1 anid|CBF74398.1 anig|EHA19149.1 aory|EIT73210.1 asyd|OJJ57766.1 ater|EAU36249.1 atub|OJI90400.1 aver|OJJ05400.1 awen|OJJ36661.1 azon|OJJ47537.1

Group2225: abra|OJJ70681.1 acar|OOF97046.1 acla|EAW08083.1 afis|EAW23119.1 afla|EED48534.1 afum|EAL88361.1 agla|OJJ89621.1 akaw|GAA90013.1 aluc|OJZ84029.1 anid|CBF74396.1 anig|EHA19150.1 aory|EIT73258.1 asyd|OJJ57767.1 ater|EAU36248.1 atub|OJI90401.1 aver|OJJ05399.1 awen|OJJ36662.1 azon|OJJ47536.1

Group2226: abra|OJJ70684.1 acar|OOF97043.1 acla|EAW08078.1 afis|EAW23117.1 afla|EED48535.1 afum|EAL88363.1 agla|OJJ89624.1 akaw|GAA90016.1 aluc|OJZ84032.1 anid|CBF74393.1 anig|EHA19153.1 aory|EIT73171.1 asyd|OJJ57769.1 ater|EAU36245.1 atub|OJI90404.1 aver|OJJ05397.1 awen|OJJ36665.1 azon|OJJ47534.1

Group2227: abra|OJJ70685.1 acar|OOF97042.1 acla|EAW08077.1 afis|EAW23116.1 afla|EED48536.1 afum|EAL88364.1 agla|OJJ89625.1 akaw|GAA90017.1 aluc|OJZ84033.1 anid|CBF74391.1 anig|EHA19154.1 aory|EIT73186.1 asyd|OJJ57770.1 ater|EAU36244.1 atub|OJI90405.1 aver|OJJ05396.1 awen|OJJ36666.1 azon|OJJ47533.1

Group2229: abra|OJJ70689.1 acar|OOF97038.1 acla|EAW08076.1 afis|EAW23111.1 afla|EED48537.1 afum|EAL88368.1 agla|OJJ89630.1 akaw|GAA90021.1 aluc|OJZ84037.1 anid|CBF74389.1 anig|EHA19158.1 aory|EIT73070.1 asyd|OJJ57771.1 ater|EAU36239.1 atub|OJI90409.1 aver|OJJ05395.1 awen|OJJ36670.1 azon|OJJ47529.1

Group2230: abra|OJJ70694.1 acar|OOF97035.1 acla|EAW08071.1 afis|EAW23107.1 afla|EED48541.1 afum|EAL88372.1 agla|OJJ89636.1 akaw|GAA90025.1 aluc|OJZ84041.1 anid|CBF81370.1 anig|EHA19163.1 aory|EIT73129.1 asyd|OJJ59386.1 ater|EAU36235.1 atub|OJI90413.1 aver|OJJ02887.1 awen|OJJ36674.1 azon|OJJ47524.1

Group2231: abra|OJJ70696.1 acar|OOF97033.1 acla|EAW08069.1 afis|EAW23105.1 afla|EED48543.1 afum|EAL88374.1 agla|OJJ89638.1 akaw|GAA90027.1 aluc|OJZ84043.1 anid|CBF81367.1 anig|EHA19165.1 aory|EIT73169.1 asyd|OJJ59384.1 ater|EAU36233.1 atub|OJI90415.1 aver|OJJ02889.1 awen|OJJ36676.1 azon|OJJ47522.1

Group2232: abra|OJJ70700.1 acar|OOF90048.1 acla|EAW08065.1 afis|EAW23101.1 afla|EED48549.1 afum|EAL88377.1 agla|OJJ89613.1 akaw|GAA90031.1 aluc|OJZ84046.1 anid|CBF81359.1 anig|EHA19167.1 aory|EIT73156.1 asyd|OJJ59381.1 ater|EAU36227.1 atub|OJI90418.1 aver|OJJ02892.1 awen|OJJ36680.1 azon|OJJ47517.1

Group2233: abra|OJJ70701.1 acar|OOF89978.1 acla|EAW08064.1 afis|EAW23100.1 afla|EED48548.1 afum|EAL88378.2 agla|OJJ89612.1 akaw|GAA90032.1 aluc|OJZ84047.1 anig|EHA19168.1 aory|EIT73049.1 asyd|OJJ59380.1 ater|EAU36228.1 atub|OJI90419.1 aver|OJJ02893.1 awen|OJJ36681.1 azon|OJJ47516.1 anid|CBF81358.1

Group2234: abra|OJJ70702.1 acar|OOF89980.1 acla|EAW08063.1 afis|EAW23099.1 afla|EED48547.1 afum|EAL88379.1 agla|OJJ89611.1 akaw|GAA90033.1 aluc|OJZ84049.1 anid|CBF81356.1 anig|EHA19169.1 aory|EIT73063.1 asyd|OJJ59379.1 ater|EAU36230.1 atub|OJI90420.1 aver|OJJ02894.1 awen|OJJ36682.1 azon|OJJ47515.1

Group2235: abra|OJJ70704.1 acar|OOF97032.1 acla|EAW08061.1 afis|EAW23097.1 afla|EED48551.1 afum|EAL88381.2 agla|OJJ89609.1 akaw|GAA90035.1 aluc|OJZ84051.1 anid|CBF81352.1 anig|EHA19171.1 aory|EIT73218.1 asyd|OJJ59377.1 ater|EAU36225.1 atub|OJI90422.1 aver|OJJ02896.1 awen|OJJ36684.1 azon|OJJ47512.1

Group2236: abra|OJJ70705.1 acar|OOF97031.1 acla|EAW08060.1 afis|EAW23096.1 afla|EED48552.1 afum|EAL88382.2 agla|OJJ89608.1 akaw|GAA90036.1 aluc|OJZ84052.1 anid|CBF81350.1 anig|EHA19172.1 aory|EIT73272.1 asyd|OJJ59376.1 ater|EAU36224.1 atub|OJI90423.1 aver|OJJ02897.1 awen|OJJ36685.1 azon|OJJ47511.1

Group2238: abra|OJJ70708.1 acar|OOF97028.1 acla|EAW08056.1 afis|EAW23092.1 afla|EED48556.1 afum|EBA27345.1 agla|OJJ89604.1 akaw|GAA90039.1 aluc|OJZ84055.1 anid|CBF81348.1 anig|EHA19176.1 aory|EIT73227.1 asyd|OJJ59374.1 ater|EAU36215.1 atub|OJI90426.1 aver|OJJ02900.1 awen|OJJ36690.1 azon|OJJ47506.1

Group2239: abra|OJJ70710.1 acar|OOF97026.1 acla|EAW08054.1 afis|EAW23090.1 afla|EED48558.1 afum|EAL88389.1 agla|OJJ89602.1 akaw|GAA90041.1 aluc|OJZ84057.1 anid|CBF81345.1 anig|EHA19177.1 aory|EIT73059.1 asyd|OJJ59372.1 ater|EAU36213.1 atub|OJI90428.1 aver|OJJ02902.1 awen|OJJ36692.1 azon|OJJ47504.1

Group2240: abra|OJJ70711.1 acar|OOF97025.1 acla|EAW08053.1 afis|EAW23089.1 afla|EED48559.1 afum|EAL88390.1 agla|OJJ89601.1 akaw|GAA90042.1 aluc|OJZ84058.1 anid|CBF81343.1 anig|EHA19178.1 aory|EIT73173.1 asyd|OJJ59371.1 ater|EAU36212.1 atub|OJI90429.1 aver|OJJ02903.1 awen|OJJ36694.1 azon|OJJ47503.1

Group2241: abra|OJJ70714.1 acar|OOF97022.1 acla|EAW08049.1 afis|EAW23085.1 afla|EED48563.1 afum|EAL88394.1 agla|OJJ89595.1 akaw|GAA90045.1 aluc|OJZ84061.1 anid|CBF81339.1 anig|EHA19181.1 aory|EIT73232.1 asyd|OJJ59369.1 ater|EAU36209.1 atub|OJI90432.1 aver|OJJ02905.1 awen|OJJ36699.1 azon|OJJ47498.1

Group2242: abra|OJJ70715.1 acar|OOF97021.1 acla|EAW08048.1 afis|EAW23084.1 afla|EED48564.1 afum|EAL88395.1 agla|OJJ89594.1 akaw|GAA90046.1 aluc|OJZ84062.1 anid|CBF81337.1 anig|EHA19182.1 aory|EIT73208.1 asyd|OJJ59368.1 ater|EAU36208.1 atub|OJI90433.1 aver|OJJ02906.1 awen|OJJ36700.1 azon|OJJ47497.1

Group2243: abra|OJJ70716.1 acar|OOF97020.1 acla|EAW08047.1 afis|EAW23083.1 afla|EED48565.1 afum|EAL88396.1 agla|OJJ89592.1 akaw|GAA90047.1 aluc|OJZ84063.1 anid|CBF81335.1 anig|EHA19183.1 aory|EIT73176.1 asyd|OJJ59367.1 ater|EAU36207.1 atub|OJI90434.1 aver|OJJ02907.1 awen|OJJ36701.1 azon|OJJ47496.1

Group2244: abra|OJJ70717.1 acar|OOF97019.1 acla|EAW08046.1 afis|EAW23082.1 afla|EED48566.1 afum|EAL88397.1 agla|OJJ89591.1 akaw|GAA90048.1 aluc|OJZ84064.1 anid|CBF81333.1 anig|EHA19184.1 aory|EIT73236.1 asyd|OJJ59366.1 ater|EAU36206.1 atub|OJI90435.1 aver|OJJ02908.1 awen|OJJ36702.1 azon|OJJ47495.1

Group2245: abra|OJJ70718.1 acar|OOF97018.1 acla|EAW08044.1 afis|EAW23080.1 afla|EED48567.1 afum|EAL88399.1 agla|OJJ89588.1 akaw|GAA90049.1 aluc|OJZ84065.1 anid|CBF81330.1 anig|EHA19185.1 aory|EIT73202.1 asyd|OJJ59364.1 ater|EAU36205.1 atub|OJI90436.1 aver|OJJ02910.1 awen|OJJ36704.1 azon|OJJ47494.1

Group2246: abra|OJJ70719.1 acar|OOF97017.1 acla|EAW08043.1 afis|EAW23079.1 afla|EED48568.1 afum|EAL88400.1 agla|OJJ89587.1 akaw|GAA90050.1 aluc|OJZ84066.1 anid|CBF81328.1 anig|EHA19186.1 aory|EIT73095.1 asyd|OJJ59363.1 ater|EAU36204.1 atub|OJI90437.1 aver|OJJ02911.1 awen|OJJ36705.1 azon|OJJ47493.1

Group2247: abra|OJJ70720.1 acar|OOF97016.1 acla|EAW08042.1 afis|EAW23078.1 afla|EED48569.1 afum|EAL88401.1 agla|OJJ89586.1 akaw|GAA90051.1 aluc|OJZ84067.1 anid|CBF81326.1 anig|EHA19187.1 aory|EIT73149.1 asyd|OJJ59362.1 ater|EAU36202.1 atub|OJI90438.1 aver|OJJ02912.1 awen|OJJ36706.1 azon|OJJ47489.1

Group2249: abra|OJJ70726.1 acar|OOF97013.1 acla|EAW08040.1 afis|EAW23075.1 afla|EED56447.1 afum|EAL88404.1 agla|OJJ89582.1 akaw|GAA90054.1 aluc|OJZ84071.1 anid|CBF81318.1 anig|EHA19190.1

aory|EIT72820.1 asyd|OJJ59359.1 ater|EAU36199.1 atub|OJI90443.1 aver|OJJ02915.1 awen|OJJ36709.1 azon|OJJ47492.1

Group2250: abra|OJJ70727.1 acar|OOF97012.1 acla|EAW08039.1 afis|EAW23074.1 afla|EED56449.1 afum|EAL88405.1 agla|OJJ89581.1 akaw|GAA90055.1 aluc|OJZ84072.1 anid|CBF81316.1 anig|EHA19191.1 aory|EIT72790.1 asyd|OJJ59358.1 ater|EAU36198.1 atub|OJI90444.1 aver|OJJ02916.1 awen|OJJ36710.1 azon|OJJ47487.1

Group2251: abra|OJJ70728.1 acar|OOF97011.1 acla|EAW08038.1 afis|EAW23073.1 afla|EED56450.1 afum|EAL88406.1 agla|OJJ89580.1 akaw|GAA90056.1 aluc|OJZ84073.1 anid|CBF81314.1 anig|EHA19192.1 aory|EIT72843.1 asyd|OJJ59357.1 ater|EAU36197.1 atub|OJI90445.1 aver|OJJ02917.1 awen|OJJ36711.1 azon|OJJ47488.1

Group2252: abra|OJJ70729.1 acar|OOF97009.1 acla|EAW11184.1 afis|EAW15915.1 afla|EED56455.1 afum|EAL88550.2 agla|OJJ89579.1 akaw|GAA90057.1 aluc|OJZ84074.1 anid|CBF81309.1 anig|EHA19193.1 aory|EIT72851.1 asyd|OJJ59354.1 ater|EAU36193.1 atub|OJI90446.1 aver|OJJ02920.1 awen|OJJ36715.1 azon|OJJ47474.1

Group2253: abra|OJJ70733.1 acar|OOF97005.1 acla|EAW11180.1 afis|EAW15911.1 afla|EED56457.1 afum|EAL88546.2 agla|OJJ89575.1 akaw|GAA90061.1 aluc|OJZ84078.1 anid|CBF81301.1 anig|EHA19197.1 aory|EIT72816.1 asyd|OJJ59350.1 ater|EAU36191.1 atub|OJI90450.1 aver|OJJ02924.1 awen|OJJ36719.1 azon|OJJ47478.1

Group2254: abra|OJJ70735.1 acar|OOF97004.1 acla|EAW11179.1 afis|EAW15910.1 afla|EED56458.1 afum|EAL88545.1 agla|OJJ89574.1 akaw|GAA90062.1 aluc|OJZ84080.1 anid|CBF81299.1 anig|EHA19198.1 aory|EIT72821.1 asyd|OJJ59349.1 ater|EAU36189.1 atub|OJI90451.1 aver|OJJ02925.1 awen|OJJ36720.1 azon|OJJ47479.1

Group2255: abra|OJJ70738.1 acar|OOF97003.1 acla|EAW11178.1 afis|EAW15909.1 afla|EED56459.1 afum|EAL88544.1 agla|OJJ89573.1 akaw|GAA90063.1 aluc|OJZ84083.1 anid|CBF81298.1 anig|EHA19199.1 aory|EIT72803.1 asyd|OJJ59348.1 ater|EAU36188.1 atub|OJI90453.1 aver|OJJ02926.1 awen|OJJ36721.1 azon|OJJ47480.1

Group2256: abra|OJJ70741.1 acar|OOF97002.1 acla|EAW11176.1 afis|EAW15907.1 afla|EED56461.1 afum|EAL88542.1 agla|OJJ78421.1 akaw|GAA90065.1 aluc|OJZ84086.1 anid|CBF81263.1 anig|EHA19202.1 aory|EIT72874.1 asyd|OJJ59332.1 ater|EAU36186.1 atub|OJI90457.1 aver|OJJ02942.1 awen|OJJ36723.1 azon|OJJ47482.1

Group2257: abra|OJJ70751.1 acar|OOF96994.1 acla|EAW11170.1 afis|EAW15901.1 afla|EED56470.1 afum|EAL88536.1 agla|OJJ89561.1 akaw|GAA90071.1 aluc|OJZ84095.1 anid|CBF81281.1 anig|EHA19208.1 aory|EIT72864.1 asyd|OJJ59338.1 ater|EAU36180.1 atub|OJI90468.1 aver|OJJ02936.1 awen|OJJ36730.1 azon|OJJ47467.1

Group2258: abra|OJJ70752.1 acar|OOF96992.1 acla|EAW11169.1 afis|EAW15900.1 afla|EED56471.1 afum|EAL88535.1 agla|OJJ89560.1 akaw|GAA90072.1 aluc|OJZ84097.1 anid|CBF81283.1 anig|EHA19209.1 aory|EIT72802.1 asyd|OJJ59340.1 ater|EAU36179.1 atub|OJI90470.1 aver|OJJ02935.1 awen|OJJ36731.1 azon|OJJ47466.1

Group2259: abra|OJJ70753.1 acar|OOF96991.1 acla|EAW11168.1 afis|EAW15899.1 afla|EED56472.1 afum|EAL88534.1 agla|OJJ89553.1 akaw|GAA90073.1 aluc|OJZ84098.1 anid|CBF81285.1 anig|EHA19210.1 aory|EIT72807.1 asyd|OJJ59341.1 ater|EAU36178.1 atub|OJI90471.1 aver|OJJ02934.1 awen|OJJ36732.1 azon|OJJ47465.1

Group2260: abra|OJJ70754.1 acar|OOF96990.1 acla|EAW11167.1 afis|EAW15898.1 afla|EED56473.1 afum|EAL88533.1 agla|OJJ89552.1 akaw|GAA90074.1 aluc|OJZ84099.1 anid|CBF81287.1 anig|EHA19211.1 aory|EIT72796.1 asyd|OJJ59342.1 ater|EAU36177.1 atub|OJI90472.1 aver|OJJ02933.1 awen|OJJ36733.1 azon|OJJ47464.1

Group2261: abra|OJJ70755.1 acar|OOF96989.1 acla|EAW11166.1 afis|EAW15897.1 afla|EED56474.1 afum|EAL88532.1 agla|OJJ89551.1 akaw|GAA90075.1 aluc|OJZ84100.1 anid|CBF81288.1 anig|EHA19212.1 aory|EIT72834.1 asyd|OJJ59343.1 ater|EAU36176.1 atub|OJI90473.1 aver|OJJ02931.1 awen|OJJ36734.1 azon|OJJ47463.1

Group2262: abra|OJJ70756.1 acar|OOF96988.1 acla|EAW11165.1 afis|EAW15896.1 afla|EED56475.1 afum|EAL88531.1 agla|OJJ89550.1 akaw|GAA90076.1 aluc|OJZ84101.1 anid|CBF81290.1 anig|EHA19213.1 aory|EIT72841.1 asyd|OJJ59344.1 ater|EAU36175.1 atub|OJI90474.1 aver|OJJ02930.1 awen|OJJ36735.1 azon|OJJ47462.1

Group2263: abra|OJJ70758.1 acar|OOF96986.1 acla|EAW11163.1 afis|EAW15894.1 afla|EED56477.1 afum|EAL88529.1 agla|OJJ89548.1 akaw|GAA90078.1 aluc|OJZ84103.1 anid|CBF81294.1 anig|EHA19215.1 aory|EIT72822.1 asyd|OJJ59346.1 ater|EAU36173.1 atub|OJI90476.1 aver|OJJ02928.1 awen|OJJ36737.1 azon|OJJ47460.1

Group2264: abra|OJJ70760.1 acar|OOF96984.1 acla|EAW11161.1 afis|EAW15892.1 afla|EED56479.1 afum|EAL88527.1 agla|OJJ89530.1 akaw|GAA90080.1 aluc|OJZ84105.1 anid|CBF81259.1 anig|EHA19217.1 aory|EIT72795.1 asyd|OJJ59330.1 ater|EAU31714.1 atub|OJI90479.1 aver|OJJ02944.1 awen|OJJ36739.1 azon|OJJ47459.1

Group2265: abra|OJJ70762.1 acar|OOF96982.1 acla|EAW11159.1 afis|EAW15890.1 afla|EED56482.1 afum|EAL88525.2 agla|OJJ89532.1 akaw|GAA90082.1 aluc|OJZ84107.1 anid|CBF81256.1 anig|EHA19219.1 aory|EIT72811.1 asyd|OJJ59328.1 ater|EAU31712.1 atub|OJI90481.1 aver|OJJ02946.1 awen|OJJ36740.1 azon|OJJ47457.1

Group2266: abra|OJJ70766.1 acar|OOF96979.1 acla|EAW11156.1 afis|EAW15887.1 afla|EED56485.1 afum|EAL88521.2 agla|OJJ89534.1 akaw|GAA90086.1 aluc|OJZ84112.1 anid|CBF81250.1 anig|EHA19222.1 aory|EIT72797.1 asyd|OJJ59325.1 ater|EAU31709.1 atub|OJI90485.1 aver|OJJ02949.1 awen|OJJ36743.1 azon|OJJ47454.1

Group2267: abra|OJJ70767.1 acar|OOF96977.1 acla|EAW11154.1 afis|EAW15885.1 afla|EED56487.1 afum|EAL88519.1 agla|OJJ89536.1 akaw|GAA90087.1 aluc|OJZ84113.1 anid|CBF81243.1 anig|EHA19223.1 aory|EIT72824.1 asyd|OJJ59321.1 ater|EAU31705.1 atub|OJI90486.1 aver|OJJ02952.1 awen|OJJ36746.1 azon|OJJ47453.1

Group2268: abra|OJJ70768.1 acar|OOF96978.1 acla|EAW11153.1 afis|EAW15884.1 afla|EED56488.1 afum|EAL88518.1 agla|OJJ89537.1 akaw|GAA90088.1 aluc|OJZ84114.1 anid|CBF81241.1 anig|EHA19224.1 aory|EIT72873.1 asyd|OJJ59320.1 ater|EAU31704.1 atub|OJI90487.1 aver|OJJ02953.1 awen|OJJ36747.1 azon|OJJ47452.1

Group2269: abra|OJJ70770.1 acar|OOF96976.1 acla|EAW11151.1 afis|EAW15882.1 afla|EED56489.1 afum|EAL88516.1 agla|OJJ89539.1 akaw|GAA90090.1 aluc|OJZ84116.1 anid|CBF81237.1 anig|EHA19226.1 aory|EIT72859.1 asyd|OJJ59319.1 ater|EAU31703.1 atub|OJI90489.1 aver|OJJ02954.1 awen|OJJ36749.1 azon|OJJ47450.1

Group2270: abra|OJJ70774.1 acar|OOF96936.1 acla|EAW11147.1 afis|EAW15878.1 afla|EED56497.1 afum|EAL88512.1 agla|OJJ87230.1 akaw|GAA90094.1 aluc|OJZ84121.1 anid|CBF81230.1 anig|EHA19229.1 aory|EIT72868.1 asyd|OJJ59316.1 ater|EAU31699.1 atub|OJI90493.1 aver|OJJ02957.1 awen|OJJ36752.1 azon|OJJ47447.1

Group2271: abra|OJJ70795.1 acar|OOF96950.1 acla|EAW11128.1 afis|EAW15857.1 afla|EED56515.1 afum|EAL88493.1 agla|OJJ87250.1 akaw|GAA90111.1 aluc|OJZ84140.1 anid|CBF81189.1 anig|EHA19248.1 aory|EIT72805.1 asyd|OJJ59299.1 ater|EAU31672.1 atub|OJI90514.1 aver|OJJ02976.1 awen|OJJ36770.1 azon|OJJ47428.1

Group2272: abra|OJJ70809.1 acar|OOF98293.1 acla|EAW10418.1 afis|EAW25151.1 afla|EED55282.1 afum|EAL89826.2 agla|OJJ89004.1 akaw|GAA91689.1 aluc|OJZ87705.1 anid|CBF81138.1 anig|EHA20888.1 aory|EIT83553.1 asyd|OJJ56507.1 ater|EAU39430.1 atub|OJI80938.1 aver|OJJ06212.1 awen|OJJ40780.1 azon|OJJ46872.1

Group2273: abra|OJJ70815.1 acar|OOF98285.1 acla|EAW10424.1 afis|EAW25145.1 afla|EED55294.1 afum|EAL89818.1 agla|OJJ89008.1 akaw|GAA91695.1 aluc|OJZ87698.1 anid|CBF85679.1 anig|EHA20885.1 aory|EIT83533.1 asyd|OJJ64254.1 ater|EAU39423.1 atub|OJI80945.1 aver|OJI98344.1 awen|OJJ40770.1 azon|OJJ46878.1

Group2274: abra|OJJ70833.1 acar|OOF98281.1 acla|EAW10427.1 afis|EAW25142.1 afla|EED55303.1 afum|EAL89814.1 agla|OJJ89025.1 akaw|GAA91701.1 aluc|OJZ87689.1 anid|CBF85686.1 anig|EHA20878.1 aory|EIT83517.1 asyd|OJJ64258.1 ater|EAU39411.1 atub|OJI80951.1 aver|OJI98350.1 awen|OJJ40766.1 azon|OJJ44131.1

Group2275: abra|OJJ70834.1 acar|OOF98280.1 acla|EAW10428.1 afis|EAW25141.1 afla|EED55304.1 afum|EAL89813.1 agla|OJJ89026.1 akaw|GAA91702.1 aluc|OJZ87688.1 anid|CBF86093.1 anig|EHA20877.1 aory|EIT83571.1 asyd|OJJ64473.1 ater|EAU39410.1 atub|OJI80952.1 aver|OJI98577.1 awen|OJJ40765.1 azon|OJJ44130.1

Group2276: abra|OJJ70837.1 acar|OOF98277.1 acla|EAW10432.1 afis|EAW25137.1 afla|EED55309.1 afum|EAL89808.2 agla|OJJ89030.1 akaw|GAA91705.1 aluc|OJZ87684.1 anid|CBF85710.1 anig|EHA20874.1 aory|EIT83613.1 asyd|OJJ64270.1 ater|EAU39407.1 atub|OJI80959.1 aver|OJI98362.1 awen|OJJ40761.1 azon|OJJ44125.1

Group2277: abra|OJJ70839.1 acar|OOF98275.1 acla|EAW10433.1 afis|EAW25135.1 afla|EED55311.1 afum|EAL89806.2 agla|OJJ89032.1 akaw|GAA91707.1 aluc|OJZ87682.1 anid|CBF85706.1 anig|EHA20872.1 aory|EIT83546.1 asyd|OJJ64268.1 ater|EAU39406.1 atub|OJI80961.1 aver|OJI98360.1 awen|OJJ40759.1 azon|OJJ44123.1

Group2278: abra|OJJ70843.1 acar|OOF98272.1 acla|EAW10434.1 afis|EAW25134.1 afla|EED55312.1 afum|EAL89805.1 agla|OJJ89033.1 akaw|GAA91711.1 aluc|OJZ87678.1 anid|CBF85705.1 anig|EHA20868.1

aory|EIT83614.1 asyd|OJJ64267.1 ater|EAU39405.1 atub|OJI80965.1 aver|OJI98359.1 awen|OJJ40758.1 azon|OJJ44122.1

Group2279: abra|OJJ70848.1 acar|OOF98266.1 acla|EAW10439.1 afis|EAW25128.1 afla|EED55316.1 afum|EAL89799.1 agla|OJJ89042.1 akaw|GAA91716.1 aluc|OJZ87672.1 anid|CBF85697.1 anig|EHA20863.1 aory|EIT83568.1 asyd|OJJ64263.1 ater|EAU39401.1 atub|OJI80970.1 aver|OJI98355.1 awen|OJJ40753.1 azon|OJJ44117.1

Group2280: abra|OJJ70851.1 acar|OOF98265.1 acla|EAW10441.1 afis|EAW25126.1 afla|EED55317.1 afum|EAL89797.1 agla|OJJ89044.1 akaw|GAA91718.1 aluc|OJZ87670.1 anid|CBF85694.1 anig|EHA20861.1 aory|EIT83542.1 asyd|OJJ64261.1 ater|EAU39399.1 atub|OJI80972.1 aver|OJI98353.1 awen|OJJ40750.1 azon|OJJ44115.1

Group2281: abra|OJJ70854.1 acar|OOF98262.1 acla|EAW10447.1 afis|EAW25120.1 afla|EED55322.1 afum|EAL89791.1 agla|OJJ89053.1 akaw|GAA91721.1 aluc|OJZ87667.1 anid|CBF86087.1 anig|EHA20858.1 aory|EIT76263.1 asyd|OJJ64469.1 ater|EAU39392.1 atub|OJI80976.1 aver|OJI98573.1 awen|OJJ40739.1 azon|OJJ44108.1

Group2282: abra|OJJ70855.1 acar|OOF98261.1 acla|EAW10446.1 afis|EAW25121.1 afla|EED55320.1 afum|EAL89792.1 agla|OJJ89049.1 akaw|GAA91722.1 aluc|OJZ87665.1 anid|CBF86089.1 anig|EHA20857.1 aory|EIT76239.1 asyd|OJJ64471.1 ater|EAU39393.1 atub|OJI80978.1 aver|OJI98575.1 awen|OJJ40740.1 azon|OJJ44110.1

Group2283: abra|OJJ70856.1 acar|OOF98260.1 acla|EAW10445.1 afis|EAW25122.1 afla|EED55319.1 afum|EAL89793.1 agla|OJJ89048.1 akaw|GAA91723.1 aluc|OJZ87664.1 anid|CBF86091.1 anig|EHA20856.1 aory|EIT76220.1 asyd|OJJ64472.1 ater|EAU39394.1 atub|OJI80979.1 aver|OJI98576.1 awen|OJJ40741.1 azon|OJJ44112.1

Group2284: abra|OJJ70857.1 acar|OOF98259.1 acla|EAW10449.1 afis|EAW25118.1 afla|EED55324.1 afum|EAL89789.1 agla|OJJ89059.1 akaw|GAA91724.1 aluc|OJZ87663.1 anid|CBF86086.1 anig|EHA20855.1 aory|EIT76226.1 asyd|OJJ64468.1 ater|EAU39386.1 atub|OJI80980.1 aver|OJI98572.1 awen|OJJ40737.1 azon|OJJ44107.1

Group2285: abra|OJJ70858.1 acar|OOF98258.1 acla|EAW10450.1 afis|EAW25117.1 afla|EED55325.1 afum|EAL89788.1 agla|OJJ89063.1 akaw|GAA91725.1 aluc|OJZ87662.1 anid|CBF86084.1 anig|EHA20854.1 aory|EIT76188.1 asyd|OJJ64467.1 ater|EAU39387.1 atub|OJI80981.1 aver|OJI98571.1 awen|OJJ40734.1 azon|OJJ44106.1

Group2286: abra|OJJ70859.1 acar|OOF98257.1 acla|EAW10451.1 afis|EAW25116.1 afla|EED55326.1 afum|EAL89787.1 agla|OJJ89064.1 akaw|GAA91726.1 aluc|OJZ87661.1 anid|CBF86082.1 anig|EHA20853.1 aory|EIT76238.1 asyd|OJJ64466.1 ater|EAU39388.1 atub|OJI80982.1 aver|OJI98570.1 awen|OJJ40733.1 azon|OJJ44105.1

Group2287: abra|OJJ70860.1 acar|OOF98256.1 acla|EAW10452.1 afis|EAW25115.1 afla|EED55327.1 afum|EAL89786.1 agla|OJJ89065.1 akaw|GAA91727.1 aluc|OJZ87660.1 anid|CBF86080.1 anig|EHA20852.1 aory|EIT76212.1 asyd|OJJ64465.1 ater|EAU39389.1 atub|OJI80983.1 aver|OJI98569.1 awen|OJJ40732.1 azon|OJJ44104.1

Group2288: abra|OJJ70861.1 acar|OOF98255.1 acla|EAW10453.1 afis|EAW25114.1 afla|EED55328.1 afum|EAL89785.1 agla|OJJ89066.1 akaw|GAA91728.1 aluc|OJZ87659.1 anid|CBF86079.1 anig|EHA20851.1 aory|EIT76255.1 asyd|OJJ64464.1 ater|EAU39390.1 atub|OJI80984.1 aver|OJI98568.1 awen|OJJ40731.1 azon|OJJ44103.1

Group2289: abra|OJJ70862.1 acar|OOF98254.1 acla|EAW10454.1 afis|EAW25113.1 afla|EED55329.1 afum|EAL89784.1 agla|OJJ89067.1 akaw|GAA91729.1 aluc|OJZ87658.1 anid|CBF86077.1 anig|EHA20850.1 aory|EIT76257.1 asyd|OJJ64463.1 ater|EAU39383.1 atub|OJI80985.1 aver|OJI98567.1 awen|OJJ40730.1 azon|OJJ44102.1

Group2290: abra|OJJ70864.1 acar|OOF98252.1 acla|EAW10456.1 afis|EAW25111.1 afla|EED55331.1 afum|EAL89782.2 agla|OJJ89070.1 akaw|GAA91731.1 aluc|OJZ87656.1 anid|CBF86073.1 anig|EHA20849.1 aory|EIT76253.1 asyd|OJJ64461.1 ater|EAU39381.1 atub|OJI80987.1 aver|OJI98564.1 awen|OJJ40728.1 azon|OJJ44100.1

Group2291: abra|OJJ70878.1 acar|OOF98242.1 acla|EAW10467.1 afis|EAW25100.1 afla|EED55345.1 afum|EAL89771.1 agla|OJJ85468.1 akaw|GAA91741.1 aluc|OJZ87644.1 anid|CBF86015.1 anig|EHA20837.1 aory|EIT80879.1 asyd|OJJ64450.1 ater|EAU39362.1 atub|OJI81001.1 aver|OJI98553.1 awen|OJJ40714.1 azon|OJJ44091.1

Group2292: abra|OJJ70880.1 acar|OOF98241.1 acla|EAW10468.1 afis|EAW25099.1 afla|EED55347.1 afum|EAL89770.1 agla|OJJ85472.1 akaw|GAA91742.1 aluc|OJZ87643.1 anid|CBF86013.1 anig|EHA20836.1 aory|EIT80895.1 asyd|OJJ64449.1 ater|EAU39361.1 atub|OJI81002.1 aver|OJI98552.1 awen|OJJ40713.1 azon|OJJ44090.1

Group2293: abra|OJJ70890.1 acar|OOF98227.1 acla|EAW10472.1 afis|EAW25095.1 afla|EED55353.1 afum|EAL89766.2 agla|OJJ89082.1 akaw|GAA87406.1 aluc|OJZ87634.1 anid|CBF86004.1 anig|EHA20829.1 aory|EIT80918.1 asyd|OJJ64445.1 ater|EAU39353.1 atub|OJI81015.1 aver|OJI98548.1 awen|OJJ40709.1 azon|OJJ44085.1

Group2294: abra|OJJ70892.1 acar|OOF90217.1 acla|EAW10474.1 afis|EAW25093.1 afla|EED55354.1 afum|EAL89764.1 agla|OJJ89084.1 akaw|GAA87408.1 aluc|OJZ87631.1 anid|CBF86001.1 anig|EHA20827.1 aory|EIT80886.1 asyd|OJJ64443.1 ater|EAU39351.1 atub|OJI81018.1 aver|OJI98545.1 awen|OJJ40707.1 azon|OJJ44083.1

Group2295: abra|OJJ70895.1 acar|OOF90219.1 acla|EAW10482.1 afis|EAW25085.1 afla|EED55362.1 afum|EAL89754.1 agla|OJJ89097.1 akaw|GAA87411.1 aluc|OJZ87628.1 anid|CBF85988.1 anig|EHA20824.1 aory|EIT80873.1 asyd|OJJ64434.1 ater|EAU39341.1 atub|OJI81022.1 aver|OJI98535.1 awen|OJJ40697.1 azon|OJJ44075.1

Group2296: abra|OJJ70897.1 acar|OOF90220.1 acla|EAW10481.1 afis|EAW25086.1 afla|EED55361.1 afum|EAL89755.1 agla|OJJ89096.1 akaw|GAA87412.1 aluc|OJZ87626.1 anid|CBF85990.1 anig|EHA20823.1 aory|EIT80880.1 asyd|OJJ64435.1 ater|EAU39342.1 atub|OJI81023.1 aver|OJI98536.1 awen|OJJ40698.1 azon|OJJ44076.1

Group2297: abra|OJJ70898.1 acar|OOF98228.1 acla|EAW10480.1 afis|EAW25087.1 afla|EED55360.1 afum|EAL89756.1 agla|OJJ89095.1 akaw|GAA87413.1 aluc|OJZ87625.1 anid|CBF85992.1 anig|EHA20822.1 aory|EIT80799.1 asyd|OJJ64436.1 ater|EAU39343.1 atub|OJI81024.1 aver|OJI98537.1 awen|OJJ40700.1 azon|OJJ44077.1

Group2298: abra|OJJ70899.1 acar|OOF98221.1 acla|EAW10479.1 afis|EAW25088.1 afla|EED55359.1 afum|EAL89757.1 agla|OJJ89094.1 akaw|GAA87414.1 aluc|OJZ87624.1 anid|CBF85994.1 anig|EHA20821.1 aory|EIT80835.1 asyd|OJJ64437.1 ater|EAU39344.1 atub|OJI81025.1 aver|OJI98538.1 awen|OJJ40701.1 azon|OJJ44078.1

Group2299: abra|OJJ70901.1 acar|OOF98218.1 acla|EAW10477.1 afis|EAW25090.1 afla|EED55357.1 afum|EAL89760.2 agla|OJJ89087.1 akaw|GAA87416.1 aluc|OJZ87622.1 anid|CBF85997.1 anig|EHA20819.1 aory|EIT80906.1 asyd|OJJ64440.1 ater|EAU39345.1 atub|OJI81027.1 aver|OJI98542.1 awen|OJJ40704.1 azon|OJJ44080.1

Group2300: abra|OJJ70906.1 acar|OOF98214.1 acla|EAW10484.1 afis|EAW25083.1 afla|EED55367.1 afum|EAL89752.1 agla|OJJ89102.1 akaw|GAA87420.1 aluc|OJZ87618.1 anid|CBF85985.1 anig|EHA20816.1 aory|EIT80861.1 asyd|OJJ64432.1 ater|EAU39339.1 atub|OJI81031.1 aver|OJI98533.1 awen|OJJ40695.1 azon|OJJ44072.1

Group2301: abra|OJJ70911.1 acar|OOF98211.1 acla|EAW10486.1 afis|EAW25081.1 afla|EED55369.1 afum|EAL89750.1 agla|OJJ89105.1 akaw|GAA87422.1 aluc|OJZ87614.1 anid|CBF85977.1 anig|EHA20811.1 aory|EIT80845.1 asyd|OJJ64429.1 ater|EAU39519.1 atub|OJI81035.1 aver|OJI98529.1 awen|OJJ40693.1 azon|OJJ44069.1

Group2302: abra|OJJ70912.1 acar|OOF98210.1 acla|EAW10487.1 afis|EAW25080.1 afla|EED55370.1 afum|EAL89749.1 agla|OJJ89106.1 akaw|GAA87423.1 aluc|OJZ87613.1 anid|CBF85975.1 anig|EHA20810.1 aory|EIT80866.1 asyd|OJJ64428.1 ater|EAU39539.1 atub|OJI81036.1 aver|OJI98528.1 awen|OJJ40692.1 azon|OJJ44068.1

Group2303: abra|OJJ70913.1 acar|OOF89971.1 acla|EAW10488.1 afis|EAW25079.1 afla|EED55371.1 afum|EAL89748.1 agla|OJJ89107.1 akaw|GAA87424.1 aluc|OJZ87612.1 anid|CBF85973.1 anig|EHA20809.1 aory|EIT80872.1 asyd|OJJ64427.1 ater|EAU39540.1 atub|OJI81037.1 aver|OJI98527.1 awen|OJJ40691.1 azon|OJJ44067.1

Group2304: abra|OJJ70915.1 acar|OOF98209.1 acla|EAW10489.1 afis|EAW25078.1 afla|EED55375.1 afum|EAL89747.1 agla|OJJ89115.1 akaw|GAA87425.1 aluc|OJZ87610.1 anid|CBF85970.1 anig|EHA20808.1 aory|EIT80925.1 asyd|OJJ64426.1 ater|EAU39545.1 atub|OJI81038.1 aver|OJI98526.1 awen|OJJ40689.1 azon|OJJ44066.1

Group2305: abra|OJJ70916.1 acar|OOF98208.1 acla|EAW10490.1 afis|EAW25077.1 afla|EED55376.1 afum|EAL89746.1 agla|OJJ89116.1 akaw|GAA87426.1 aluc|OJZ87609.1 anid|CBF85968.1 anig|EHA20807.1 aory|EIT80812.1 asyd|OJJ64425.1 ater|EAU39546.1 atub|OJI81039.1 aver|OJI98524.1 awen|OJJ40688.1 azon|OJJ44065.1

Group2306: abra|OJJ70918.1 acar|OOF98206.1 acla|EAW10492.1 afis|EAW25075.1 afla|EED55378.1 afum|EAL89744.1 agla|OJJ89117.1 akaw|GAA87428.1 aluc|OJZ87608.1 anid|CBF85964.1 anig|EHA20806.1 aory|EIT80830.1 asyd|OJJ64423.1 ater|EAU39550.1 atub|OJI81040.1 aver|OJI98522.1 awen|OJJ40686.1 azon|OJJ44063.1

Group2307: abra|OJJ70931.1 acar|OOF98195.1 acla|EAW10497.1 afis|EAW25070.1 afla|EED55384.1 afum|EAL89739.1 agla|OJJ89124.1 akaw|GAA83011.1 aluc|OJZ87596.1 anid|CBF85958.1 anig|EHA20796.1

aory|EIT80805.1 asyd|OJJ64418.1 ater|EAU39555.1 atub|OJI81051.1 aver|OJI98517.1 awen|OJJ40681.1 azon|OJJ44058.1

Group2308: abra|OJJ70933.1 acar|OOF89985.1 acla|EAW10499.1 afis|EAW25068.1 afla|EED55386.1 afum|EAL89737.2 agla|OJJ89126.1 akaw|GAA83009.1 aluc|OJZ87594.1 anid|CBF85954.1 anig|EHA20794.1 aory|EIT80870.1 asyd|OJJ64416.1 ater|EAU35815.1 atub|OJI81053.1 aver|OJI98515.1 awen|OJJ40675.1 azon|OJJ44056.1

Group2309: abra|OJJ70940.1 acar|OOF98192.1 acla|EAW10505.1 afis|EAW25062.1 afla|EED55391.1 afum|EAL89731.2 agla|OJJ89133.1 akaw|GAA83004.1 aluc|OJZ87588.1 anid|CBF85943.1 anig|EHA20789.1 aory|EIT80778.1 asyd|OJJ64412.1 ater|EAU35810.1 atub|OJI81060.1 aver|OJI98511.1 awen|OJJ40669.1 azon|OJJ44050.1

Group2310: abra|OJJ70941.1 acar|OOF98191.1 acla|EAW10506.1 afis|EAW25061.1 afla|EED55392.1 afum|EAL89729.1 agla|OJJ89134.1 akaw|GAA83003.1 aluc|OJZ87587.1 anid|CBF85941.1 anig|EHA20788.1 aory|EIT80784.1 asyd|OJJ64411.1 ater|EAU35809.1 atub|OJI81061.1 aver|OJI98510.1 awen|OJJ40668.1 azon|OJJ44049.1

Group2311: abra|OJJ70942.1 acar|OOF98190.1 acla|EAW10507.1 afis|EAW25060.1 afla|EED55393.1 afum|EAL89728.1 agla|OJJ89135.1 akaw|GAA83002.1 aluc|OJZ87586.1 anid|CBF85940.1 anig|EHA20787.1 aory|EIT80892.1 asyd|OJJ64410.1 ater|EAU35808.1 atub|OJI81062.1 aver|OJI98509.1 awen|OJJ40667.1 azon|OJJ44048.1

Group2312: abra|OJJ70944.1 acar|OOF98187.1 acla|EAW10512.1 afis|EAW25055.1 afla|EED55396.1 afum|EAL89723.2 agla|OJJ89137.1 akaw|GAA82999.1 aluc|OJZ87584.1 anid|CBF85933.1 anig|EHA20785.1 aory|EIT80802.1 asyd|OJJ64408.1 ater|EAU35805.1 atub|OJI81064.1 aver|OJI98506.1 awen|OJJ40665.1 azon|OJJ44044.1

Group2313: abra|OJJ70945.1 acar|OOF98186.1 acla|EAW10513.1 afis|EAW25054.1 afla|EED55397.1 afum|EAL89722.1 agla|OJJ89138.1 akaw|GAA82998.1 aluc|OJZ87583.1 anid|CBF85931.1 anig|EHA20784.1 aory|EIT80901.1 asyd|OJJ64407.1 ater|EAU35804.1 atub|OJI81065.1 aver|OJI98505.1 awen|OJJ40664.1 azon|OJJ44045.1

Group2314: abra|OJJ70946.1 acar|OOF98185.1 acla|EAW10514.1 afis|EAW25053.1 afla|EED55398.1 afum|EAL89721.1 agla|OJJ89139.1 akaw|GAA82997.1 aluc|OJZ87582.1 anid|CBF85929.1 anig|EHA20783.1 aory|EIT80773.1 asyd|OJJ64406.1 ater|EAU35803.1 atub|OJI81066.1 aver|OJI98504.1 awen|OJJ40663.1 azon|OJJ44043.1

Group2315: abra|OJJ70954.1 acar|OOF98176.1 acla|EAW10515.1 afis|EAW25051.1 afla|EED55400.1 afum|EAL89719.1 agla|OJJ89141.1 akaw|GAA82993.1 aluc|OJZ87578.1 anid|CBF85925.1 anig|EHA20780.1 aory|EIT80833.1 asyd|OJJ64405.1 ater|EAU35801.1 atub|OJI81071.1 aver|OJI98501.1 awen|OJJ40661.1 azon|OJJ44040.1

Group2316: abra|OJJ70955.1 acar|OOF98175.1 acla|EAW10516.1 afis|EAW25050.1 afla|EED55401.1 afum|EAL89718.1 agla|OJJ89142.1 akaw|GAA82992.1 aluc|OJZ87577.1 anid|CBF85923.1 anig|EHA20779.1 aory|EIT80841.1 asyd|OJJ64404.1 ater|EAU35800.1 atub|OJI81072.1 aver|OJI98500.1 awen|OJJ40660.1 azon|OJJ44039.1

Group2317: abra|OJJ70957.1 acar|OOF98173.1 acla|EAW10518.1 afis|EAW25048.1 afla|EED55403.1 afum|EAL89716.1 agla|OJJ89144.1 akaw|GAA82990.1 aluc|OJZ87575.1 anid|CBF85920.1 anig|EHA20777.1 aory|EIT80783.1 asyd|OJJ64402.1 ater|EAU35798.1 atub|OJI81074.1 aver|OJI98498.1 awen|OJJ40659.1 azon|OJJ44037.1

Group2318: abra|OJJ70958.1 acar|OOF98172.1 acla|EAW10519.1 afis|EAW25047.1 afla|EED55404.1 afum|EAL89715.1 agla|OJJ89145.1 akaw|GAA82989.1 aluc|OJZ87574.1 anid|CBF85918.1 anig|EHA20776.1 aory|EIT80839.1 asyd|OJJ64401.1 ater|EAU35797.1 atub|OJI81075.1 aver|OJI98497.1 awen|OJJ40658.1 azon|OJJ44036.1

Group2319: abra|OJJ70959.1 acar|OOF98171.1 acla|EAW10520.1 afis|EAW25046.1 afla|EED55405.1 afum|EAL89714.1 agla|OJJ89148.1 akaw|GAA82988.1 aluc|OJZ87573.1 anid|CBF85916.1 anig|EHA20775.1 aory|EIT80804.1 asyd|OJJ64400.1 ater|EAU35796.1 atub|OJI81076.1 aver|OJI98496.1 awen|OJJ40657.1 azon|OJJ44035.1

Group2320: abra|OJJ70963.1 acar|OOF98167.1 acla|EAW10524.1 afis|EAW25042.1 afla|EED55410.1 afum|EAL89710.1 agla|OJJ89152.1 akaw|GAA82984.1 aluc|OJZ87569.1 anid|CBF85908.1 anig|EHA20771.1 aory|EIT80900.1 asyd|OJJ64396.1 ater|EAU35792.1 atub|OJI81080.1 aver|OJI98492.1 awen|OJJ40647.1 azon|OJJ44032.1

Group2321: abra|OJJ70964.1 acar|OOF98166.1 acla|EAW10525.1 afis|EAW25041.1 afla|EED55411.1 afum|EAL89709.1 agla|OJJ89153.1 akaw|GAA82983.1 aluc|OJZ87568.1 anid|CBF85907.1 anig|EHA20770.1 aory|EIT80795.1 asyd|OJJ64395.1 ater|EAU35791.1 atub|OJI81081.1 aver|OJI98490.1 awen|OJJ40646.1 azon|OJJ44031.1

Group2322: abra|OJJ70968.1 acar|OOF98162.1 acla|EAW10528.1 afis|EAW25038.1 afla|EED55415.1 afum|EAL89706.1 agla|OJJ89155.1 akaw|GAA82980.1 aluc|OJZ87564.1 anid|CBF85901.1 anig|EHA20766.1 aory|EIT80889.1 asyd|OJJ64392.1 ater|EAU35788.1 atub|OJI81083.1 aver|OJI98487.1 awen|OJJ40644.1 azon|OJJ44028.1

Group2323: abra|OJJ70971.1 acar|OOF98159.1 acla|EAW10531.1 afis|EAW24998.1 afla|EED55418.1 afum|EAL89703.1 agla|OJJ89158.1 akaw|GAA82977.1 aluc|OJZ87561.1 anid|CBF85898.1 anig|EHA20763.1 aory|EIT80823.1 asyd|OJJ64390.1 ater|EAU35786.1 atub|OJI81086.1 aver|OJI98485.1 awen|OJJ40641.1 azon|OJJ44026.1

Group2324: abra|OJJ70973.1 acar|OOF98158.1 acla|EAW10532.1 afis|EAW24999.1 afla|EED55419.1 afum|EAL89699.1 agla|OJJ89161.1 akaw|GAA82976.1 aluc|OJZ87559.1 anid|CBF81499.1 anig|EHA20762.1 aory|EIT80834.1 asyd|OJJ59465.1 ater|EAU35785.1 atub|OJI81088.1 aver|OJJ02809.1 awen|OJJ40640.1 azon|OJJ44024.1

Group2325: abra|OJJ70976.1 acar|OOF98154.1 acla|EAW10535.1 afis|EAW25002.1 afla|EED55422.1 afum|EAL89696.1 agla|OJJ89165.1 akaw|GAA82973.1 aluc|OJZ87556.1 anid|CBF81486.1 anig|EHA20759.1 aory|EIT80883.1 ater|EAU35783.1 atub|OJI81091.1 awen|OJJ40637.1 azon|OJJ49046.1 asyd|OJJ59457.1 aver|OJJ02816.1

Group2326: abra|OJJ70977.1 acar|OOF98155.1 acla|EAW10536.1 afis|EAW25003.1 afla|EED55423.1 afum|EAL89695.1 agla|OJJ89166.1 akaw|GAA82972.1 aluc|OJZ87555.1 anid|CBF81484.1 anig|EHA20758.1 aory|EIT80840.1 asyd|OJJ59456.1 ater|EAU35782.1 atub|OJI81092.1 aver|OJJ02817.1 awen|OJJ40636.1 azon|OJJ49047.1

Group2327: abra|OJJ70979.1 acar|OOF98153.1 acla|EAW10538.1 afis|EAW25005.1 afla|EED55425.1 afum|EAL89693.1 agla|OJJ89168.1 akaw|GAA82970.1 aluc|OJZ87553.1 anid|CBF81481.1 anig|EHA20756.1 aory|EIT80871.1 asyd|OJJ59454.1 ater|EAU35780.1 atub|OJI81093.1 aver|OJJ02819.1 awen|OJJ40634.1 azon|OJJ49049.1

Group2328: abra|OJJ70980.1 acar|OOF98151.1 acla|EAW10539.1 afis|EAW25006.1 afla|EED55426.1 afum|EAL89691.2 agla|OJJ89169.1 akaw|GAA82969.1 aluc|OJZ87551.1 anid|CBF81512.1 anig|EHA20754.1 aory|EIT80772.1 asyd|OJJ59470.1 ater|EAU35779.1 atub|OJI81094.1 aver|OJJ02804.1 awen|OJJ40633.1 azon|OJJ49050.1

Group2329: abra|OJJ70981.1 acar|OOF98150.1 acla|EAW10540.1 afis|EAW25007.1 afla|EED55427.1 afum|EAL89690.1 agla|OJJ89170.1 akaw|GAA82968.1 aluc|OJZ87550.1 anid|CBF81514.1 anig|EHA20753.1 aory|EIT80826.1 asyd|OJJ59472.1 ater|EAU35778.1 atub|OJI81095.1 aver|OJJ02802.1 awen|OJJ40632.1 azon|OJJ49051.1

Group2330: abra|OJJ70982.1 acar|OOF98148.1 acla|EAW10541.1 afis|EAW25008.1 afla|EED55428.1 afum|EBA27387.1 agla|OJJ89173.1 akaw|GAA82967.1 aluc|OJZ87549.1 anid|CBF81516.1 anig|EHA20752.1 aory|EIT80794.1 asyd|OJJ59473.1 ater|EAU35777.1 atub|OJI81097.1 aver|OJJ02801.1 awen|OJJ40631.1 azon|OJJ49052.1

Group2331: abra|OJJ70983.1 acar|OOF98149.1 acla|EAW10542.1 afis|EAW25010.1 afla|EED55430.1 afum|EAL89686.1 agla|OJJ89174.1 akaw|GAA82966.1 aluc|OJZ87548.1 anid|CBF81518.1 anig|EHA20751.1 aory|EIT80803.1 asyd|OJJ59474.1 ater|EAU35776.1 atub|OJI81098.1 aver|OJJ02799.1 awen|OJJ40630.1 azon|OJJ49053.1

Group2332: abra|OJJ70984.1 acar|OOF98147.1 acla|EAW10543.1 afis|EAW25011.1 afla|EED55431.1 afum|EAL89685.1 agla|OJJ89175.1 akaw|GAA82965.1 aluc|OJZ87547.1 anid|CBF81519.1 anig|EHA20750.1 aory|EIT80824.1 asyd|OJJ59475.1 ater|EAU35775.1 atub|OJI81099.1 aver|OJJ02798.1 awen|OJJ40629.1 azon|OJJ49054.1

Group2333: abra|OJJ70989.1 acar|OOF98144.1 acla|EAW10547.1 afis|EAW25015.1 afla|EED55437.1 afum|EAL89681.1 agla|OJJ89181.1 akaw|GAA82962.1 aluc|OJZ87543.1 anid|CBF81527.1 anig|EHA20747.1 aory|EIT80927.1 asyd|OJJ59481.1 ater|EAU35771.1 atub|OJI81104.1 aver|OJJ02793.1 awen|OJJ40625.1 azon|OJJ49058.1

Group2334: abra|OJJ70991.1 acar|OOF98143.1 acla|EAW10548.1 afis|EAW25016.1 afla|EED55439.1 afum|EAL89680.1 agla|OJJ89182.1 akaw|GAA82960.1 aluc|OJZ87541.1 anid|CBF81529.1 anig|EHA20746.1 aory|EIT80844.1 asyd|OJJ59483.1 ater|EAU35770.1 atub|OJI81106.1 aver|OJJ02791.1 awen|OJJ40624.1 azon|OJJ49059.1

Group2335: abra|OJJ70996.1 acar|OOF90034.1 acla|EAW10552.1 afis|EAW25019.1 afla|EED55442.1 afum|EAL89676.1 agla|OJJ89187.1 akaw|GAA82952.1 aluc|OJZ87532.1 anid|CBF81538.1 anig|EHA20741.1 aory|EIT80858.1 asyd|OJJ59488.1 ater|EAU35767.1 atub|OJI81117.1 aver|OJJ02786.1 awen|OJJ40621.1 azon|OJJ49469.1

Group2336: abra|OJJ70998.1 acar|OOF98142.1 acla|EAW10555.1 afis|EAW25022.1 afla|EED55444.1 afum|EAL89673.1 agla|OJJ89190.1 akaw|GAA82950.1 aluc|OJZ87529.1 anid|CBF81541.1 anig|EHA20739.1

aory|EIT80864.1 asyd|OJJ59490.1 ater|EAU35765.1 atub|OJI81119.1 aver|OJJ02784.1 awen|OJJ40618.1 azon|OJJ49466.1

Group2337: abra|OJJ71001.1 acar|OOF98139.1 acla|EAW10558.1 afis|EAW25025.1 afla|EED55447.1 afum|EAL89670.1 agla|OJJ79159.1 akaw|GAA82947.1 aluc|OJZ87525.1 anid|CBF81547.1 anig|EHA20736.1 aory|EIT80809.1 asyd|OJJ59493.1 ater|EAU35762.1 atub|OJI81123.1 aver|OJJ02781.1 awen|OJJ40614.1 azon|OJJ49463.1

Group2338: abra|OJJ71008.1 acar|OOF98134.1 acla|EAW10560.1 afis|EAW25027.1 afla|EED55449.1 afum|EAL89668.1 agla|OJJ79157.1 akaw|GAA82945.1 aluc|OJZ87523.1 anid|CBF81551.1 anig|EHA20733.1 aory|EIT80929.1 asyd|OJJ59495.1 ater|EAU35760.1 atub|OJI81125.1 aver|OJJ02779.1 awen|OJJ40612.1 azon|OJJ49461.1

Group2339: abra|OJJ71013.1 acar|OOF98126.1 acla|EAW10564.1 afis|EAW25031.1 afla|EED55454.1 afum|EAL89664.1 agla|OJJ82242.1 akaw|GAA82941.1 aluc|OJZ87518.1 anid|CBF81564.1 anig|EHA20729.1 aory|EIT80774.1 asyd|OJJ59502.1 ater|EAU35756.1 atub|OJI81131.1 aver|OJJ02772.1 awen|OJJ40608.1 azon|OJJ49458.1

Group2340: abra|OJJ71016.1 acar|OOF98124.1 acla|EAW10566.1 afis|EAW25032.1 afla|EED55457.1 afum|EAL89662.1 agla|OJJ82244.1 akaw|GAA82939.1 aluc|OJZ87516.1 anid|CBF81567.1 anig|EHA20727.1 aory|EIT80779.1 asyd|OJJ59504.1 ater|EAU35755.1 atub|OJI81134.1 aver|OJJ02770.1 awen|OJJ40607.1 azon|OJJ49456.1

Group2341: abra|OJJ71019.1 acar|OOF98122.1 acla|EAW10568.1 afis|EAW25034.1 afla|EED55459.1 afum|EAL89660.1 agla|OJJ82255.1 akaw|GAA82936.1 aluc|OJZ87513.1 anid|CBF81574.1 anig|EHA20725.1 aory|EIT80788.1 asyd|OJJ59507.1 ater|EAU35751.1 atub|OJI81138.1 aver|OJJ02767.1 awen|OJJ40604.1 azon|OJJ49454.1

Group2342: abra|OJJ71021.1 acar|OOF98119.1 acla|EAW10570.1 afis|EAW25036.1 afla|EED55462.1 afum|EAL89658.1 agla|OJJ82257.1 akaw|GAA82934.1 aluc|OJZ87511.1 anid|CBF81583.1 anig|EHA20723.1 aory|EIT80855.1 asyd|OJJ59510.1 ater|EAU35747.1 atub|OJI81140.1 aver|OJJ02764.1 awen|OJJ40602.1 azon|OJJ49452.1

Group2343: abra|OJJ71022.1 acar|OOF98117.1 acla|EAW10571.1 afis|EAW25037.1 afla|EED55463.1 afum|EAL89657.1 agla|OJJ82258.1 akaw|GAA82933.1 aluc|OJZ87510.1 anid|CBF81585.1 anig|EHA20722.1 aory|EIT80922.1 asyd|OJJ59511.1 ater|EAU38720.1 atub|OJI81141.1 aver|OJJ02762.1 awen|OJJ40601.1 azon|OJJ49451.1

Group2344: abra|OJJ71023.1 acar|OOF98118.1 acla|EAW10572.1 afis|EAW24995.1 afla|EED55464.1 afum|EAL89656.1 agla|OJJ82259.1 akaw|GAA82932.1 aluc|OJZ87509.1 anid|CBF81586.1 anig|EHA20721.1 aory|EIT80875.1 asyd|OJJ59512.1 ater|EAU35746.1 atub|OJI81142.1 aver|OJJ02761.1 awen|OJJ40600.1 azon|OJJ49450.1

Group2345: abra|OJJ71028.1 acar|OOF98111.1 acla|EAW10579.1 afis|EAW24988.1 afla|EED55470.1 afum|EAL89648.1 agla|OJJ82274.1 akaw|GAA82927.1 aluc|OJZ87502.1 anid|CBF81599.1 anig|EHA20717.1 aory|EIT80811.1 asyd|OJJ59519.1 ater|EAU35739.1 atub|OJI81147.1 aver|OJJ02754.1 awen|OJJ40593.1 azon|OJJ49443.1

Group2346: abra|OJJ71030.1 acar|OOF98109.1 acla|EAW10583.1 afis|EAW24986.1 afla|EED55472.1 afum|EAL89646.1 agla|OJJ82276.1 akaw|GAA82925.1 aluc|OJZ87498.1 anid|CBF81603.1 anig|EHA20715.1 aory|EIT80838.1 asyd|OJJ59521.1 ater|EAU35737.1 atub|OJI81149.1 aver|OJJ02752.1 awen|OJJ40582.1 azon|OJJ49441.1

Group2347: abra|OJJ71036.1 acar|OOF98101.1 acla|EAW10586.1 afis|EAW24982.1 afla|EED55477.1 afum|EAL89642.1 agla|OJJ82279.1 akaw|GAA82919.1 aluc|OJZ87491.1 anid|CBF81614.1 anig|EHA20711.1 aory|EIT80831.1 asyd|OJJ59526.1 ater|EAU35734.1 atub|OJI81157.1 aver|OJJ02746.1 awen|OJJ40579.1 azon|OJJ49430.1

Group2348: abra|OJJ71038.1 acar|OOF98100.1 acla|EAW10587.1 afis|EAW24981.1 afla|EED55478.1 afum|EAL89641.1 agla|OJJ79136.1 akaw|GAA82918.1 aluc|OJZ87490.1 anid|CBF81612.1 anig|EHA20710.1 aory|EIT80854.1 asyd|OJJ59525.1 ater|EAU35733.1 atub|OJI81158.1 aver|OJJ02747.1 awen|OJJ40578.1 azon|OJJ49431.1

Group2349: abra|OJJ71040.1 acar|OOF98098.1 acla|EAW10589.1 afis|EAW24979.1 afla|EED55480.1 afum|EAL89639.1 agla|OJJ79138.1 akaw|GAA82916.1 aluc|OJZ87488.1 anid|CBF81608.1 anig|EHA20708.1 aory|EIT80915.1 asyd|OJJ59523.1 ater|EAU35729.1 atub|OJI81160.1 aver|OJJ02749.1 awen|OJJ40576.1 azon|OJJ49434.1

Group2350: abra|OJJ71042.1 acar|OOF98097.1 acla|EAW10592.1 afis|EAW24976.1 afla|EED55484.1 afum|EAL89636.1 agla|OJJ79141.1 akaw|GAA82914.1 aluc|OJZ87486.1 anid|CBF81629.1 anig|EHA20707.1 aory|EIT80868.1 asyd|OJJ59530.1 ater|EAU35725.1 atub|OJI81162.1 aver|OJJ02740.1 awen|OJJ40573.1 azon|OJJ49438.1

Group2351: abra|OJJ71045.1 acar|OOF98094.1 acla|EAW10593.1 afis|EAW24975.1 afla|EED55485.1 afum|EAL89635.1 agla|OJJ82241.1 akaw|GAA82911.1 aluc|OJZ87481.1 anid|CBF81630.1 anig|EHA20703.1 aory|EIT80859.1 asyd|OJJ59531.1 ater|EAU35723.1 atub|OJI81167.1 aver|OJJ02739.1 awen|OJJ40572.1 azon|OJJ49429.1

Group2352: abra|OJJ71047.1 acar|OOF98092.1 acla|EAW10595.1 afis|EAW24973.1 afla|EED55486.1 afum|EAL89633.1 agla|OJJ82239.1 akaw|GAA82909.1 aluc|OJZ87479.1 anid|CBF81634.1 anig|EHA20701.1 aory|EIT80885.1 asyd|OJJ59533.1 ater|EAU35721.1 atub|OJI81169.1 aver|OJJ02737.1 awen|OJJ40570.1 azon|OJJ49427.1

Group2353: abra|OJJ71048.1 acar|OOF98091.1 acla|EAW10596.1 afis|EAW24972.1 afla|EED55487.1 afum|EBA27384.1 agla|OJJ82238.1 akaw|GAA82908.1 aluc|OJZ87478.1 anid|CBF81636.1 anig|EHA20700.1 aory|EIT80911.1 asyd|OJJ59534.1 ater|EAU35720.1 atub|OJI81170.1 aver|OJJ02736.1 awen|OJJ40569.1 azon|OJJ49426.1

Group2354: abra|OJJ71049.1 acar|OOF98090.1 acla|EAW10597.1 afis|EAW24971.1 afla|EED55490.1 afum|EAL89630.2 agla|OJJ82237.1 akaw|GAA82907.1 aluc|OJZ87477.1 anid|CBF81638.1 anig|EHA20699.1 aory|EIT80796.1 asyd|OJJ59535.1 ater|EAU35719.1 atub|OJI81171.1 aver|OJJ02735.1 awen|OJJ40568.1 azon|OJJ49425.1

Group2355: abra|OJJ71052.1 acar|OOF98087.1 acla|EAW10603.1 afis|EAW24965.1 afla|EED55495.1 afum|EAL89623.1 agla|OJJ82224.1 akaw|GAA82905.1 aluc|OJZ87475.1 anid|CBF81647.1 anig|EHA20695.1 aory|EIT80847.1 asyd|OJJ59541.1 ater|EAU35709.1 atub|OJI81173.1 aver|OJJ02727.1 awen|OJJ40558.1 azon|OJJ49419.1

Group2357: abra|OJJ71061.1 acar|OOF98082.1 acla|EAW10598.1 afis|EAW24970.1 afla|EED55491.1 afum|EAL89629.1 agla|OJJ82236.1 akaw|GAA82898.1 aluc|OJZ87467.1 anid|CBF81640.1 anig|EHA20689.1 aory|EIT80891.1 asyd|OJJ59536.1 ater|EAU35715.1 atub|OJI81181.1 aver|OJJ02733.1 awen|OJJ40564.1 azon|OJJ49424.1

Group2358: abra|OJJ71075.1 acar|OOF98069.1 acla|EAW10604.1 afis|EAW24964.1 afla|EED55497.1 afum|EAL89622.1 agla|OJJ82226.1 akaw|GAA82887.1 aluc|OJZ87455.1 anid|CBF81649.1 anig|EHA20676.1 aory|EIT80869.1 asyd|OJJ59542.1 ater|EAU35707.1 atub|OJI81193.1 aver|OJJ02726.1 awen|OJJ40556.1 azon|OJJ49418.1

Group2359: abra|OJJ71103.1 acar|OOF90221.1 acla|EAW10388.1 afis|EAW25205.1 afla|EED53664.1 afum|EAL89866.2 agla|OJJ88951.1 akaw|GAA82865.1 aluc|OJZ87432.1 anid|CBF85221.1 anig|EHA20653.1 aory|EIT75288.1 asyd|OJJ62997.1 ater|EAU34349.1 atub|OJI81216.1 aver|OJI97970.1 awen|OJJ40840.1 azon|OJJ46822.1

Group2360: abra|OJJ71104.1 acar|OOF90222.1 acla|EAW10387.1 afis|EAW25206.1 afla|EED53662.1 afum|EAL89867.1 agla|OJJ88952.1 akaw|GAA82864.1 aluc|OJZ87431.1 anid|CBF85223.1 anig|EHA20652.1 aory|EIT75326.1 asyd|OJJ62996.1 ater|EAU34348.1 atub|OJI81217.1 aver|OJI97971.1 awen|OJJ40841.1 azon|OJJ46823.1

Group2361: abra|OJJ71105.1 acar|OOF90223.1 acla|EAW10386.1 afis|EAW25207.1 afla|EED53660.1 afum|EAL89868.1 agla|OJJ79791.1 akaw|GAA82863.1 aluc|OJZ87430.1 anid|CBF85225.1 anig|EHA20651.1 aory|EIT75316.1 asyd|OJJ62995.1 ater|EAU34347.1 atub|OJI81218.1 aver|OJI97972.1 awen|OJJ35846.1 azon|OJJ45132.1

Group2362: abra|OJJ71112.1 acar|OOF98038.1 acla|EAW10380.1 afis|EAW25213.1 afla|EED53667.1 afum|EAL89874.1 agla|OJJ88948.1 akaw|GAA82857.1 aluc|OJZ87422.1 anid|CBF85606.1 anig|EHA20645.1 aory|EIT75347.1 asyd|OJJ64197.1 ater|EAU34370.1 atub|OJI81226.1 aver|OJI98286.1 awen|OJJ40875.1 azon|OJJ46812.1

Group2363: abra|OJJ71117.1 acar|OOF98033.1 acla|EAW10378.1 afis|EAW25215.1 afla|EED53692.1 afum|EAL89876.1 agla|OJJ88944.1 akaw|GAA82853.1 aluc|OJZ87418.1 anid|CBF85254.1 anig|EHA20642.1 aory|EIT75323.1 asyd|OJJ62921.1 ater|EAU34372.1 atub|OJI81230.1 aver|OJI98068.1 awen|OJJ40878.1 azon|OJJ44506.1

Group2364: abra|OJJ71119.1 acar|OOF98031.1 acla|EAW10376.1 afis|EAW25217.1 afla|EED53690.1 afum|EAL89878.1 agla|OJJ79184.1 akaw|GAA82851.1 aluc|OJZ87417.1 anid|CBF85273.1 anig|EHA20641.1 aory|EIT75328.1 asyd|OJJ62920.1 ater|EAU34374.1 atub|OJI81231.1 aver|OJI98070.1 awen|OJJ40880.1 azon|OJJ44508.1

Group2365: abra|OJJ71125.1 acar|OOF98028.1 acla|EAW10374.1 afis|EAW25218.1 afla|EED53688.1 afum|EAL89879.1 agla|OJJ79186.1 akaw|GAA82850.1 aluc|OJZ87410.1 anid|CBF85299.1 anig|EHA20640.1 aory|EIT75354.1 asyd|OJJ62917.1 ater|EAU34377.1 atub|OJI81235.1 aver|OJI98073.1 awen|OJJ40883.1 azon|OJJ44512.1

Group2366: abra|OJJ71128.1 acar|OOF98024.1 acla|EAW10370.1 afis|EAW25222.1 afla|EED53684.1 afum|EAL89883.1 agla|OJJ79189.1 akaw|GAA82846.1 aluc|OJZ87407.1 anid|CBF85258.1 anig|EHA20637.1

aory|EIT75281.1 asyd|OJJ62911.1 ater|EAU34381.1 atub|OJI81239.1 aver|OJI98078.1 awen|OJJ40888.1 azon|OJJ44517.1

Group2367: abra|OJJ71132.1 acar|OOF98020.1 acla|EAW10365.1 afis|EAW25227.1 afla|EED53671.1 afum|EAL89888.1 agla|OJJ79181.1 akaw|GAA82842.1 aluc|OJZ87403.1 anid|CBF85269.1 anig|EHA20632.1 aory|EIT75290.1 asyd|OJJ62908.1 ater|EAU34385.1 atub|OJI81243.1 aver|OJI98080.1 awen|OJJ40892.1 azon|OJJ44521.1

Group2368: abra|OJJ71133.1 acar|OOF98019.1 acla|EAW10364.1 afis|EAW25228.1 afla|EED53672.1 afum|EAL89889.1 agla|OJJ79180.1 akaw|GAA82841.1 aluc|OJZ87402.1 anid|CBF85262.1 anig|EHA20631.1 aory|EIT75302.1 asyd|OJJ62905.1 ater|EAU34386.1 atub|OJI81244.1 aver|OJI98084.1 awen|OJJ40893.1 azon|OJJ44522.1

Group2369: abra|OJJ71142.1 acar|OOF98014.1 acla|EAW10359.1 afis|EAW25234.1 afla|EED53675.1 afum|EAL89895.1 agla|OJJ89203.1 akaw|GAA82835.1 aluc|OJZ87396.1 anid|CBF85334.1 anig|EHA20626.1 aory|EIT75320.1 asyd|OJJ64021.1 ater|EAU34389.1 atub|OJI81250.1 aver|OJI98107.1 awen|OJJ40946.1 azon|OJJ44527.1

Group2370: abra|OJJ71143.1 acar|OOF98013.1 acla|EAW10358.1 afis|EAW25235.1 afla|EED53676.1 afum|EAL89896.2 agla|OJJ89204.1 akaw|GAA82834.1 aluc|OJZ87395.1 anid|CBF85332.1 anig|EHA20625.1 aory|EIT75309.1 asyd|OJJ64020.1 ater|EAU34390.1 atub|OJI81251.1 aver|OJI98106.1 awen|OJJ40947.1 azon|OJJ44528.1

Group2371: abra|OJJ71145.1 acar|OOF98011.1 acla|EAW10361.1 afis|EAW25232.1 afla|EED53678.1 afum|EAL89893.1 agla|OJJ79175.1 akaw|GAA82832.1 aluc|OJZ87393.1 anid|CBF85329.1 anig|EHA20623.1 aory|EIT75284.1 asyd|OJJ64018.1 ater|EAU34391.1 atub|OJI81253.1 aver|OJI98104.1 awen|OJJ40898.1 azon|OJJ44525.1

Group2372: abra|OJJ71150.1 acar|OOF98007.1 acla|EAW10352.1 afis|EAW25242.1 afla|EED53702.1 afum|EAL89904.1 agla|OJJ89208.1 akaw|GAA82827.1 aluc|OJZ87388.1 anid|CBF85319.1 anig|EHA20617.1 aory|EIT75296.1 asyd|OJJ64011.1 ater|EAU34367.1 atub|OJI81259.1 aver|OJI98099.1 awen|OJJ40955.1 azon|OJJ44538.1

Group2373: abra|OJJ71151.1 acar|OOF98006.1 acla|EAW10346.1 afis|EAW25243.1 afla|EED53700.1 afum|EAL89905.1 agla|OJJ89209.1 akaw|GAA82826.1 aluc|OJZ87387.1 anid|CBF85321.1 anig|EHA20616.1 aory|EIT75325.1 asyd|OJJ64012.1 ater|EAU34360.1 atub|OJI81260.1 aver|OJI98100.1 awen|OJJ40956.1 azon|OJJ44537.1

Group2374: abra|OJJ71158.1 acar|OOF98000.1 acla|EAW10339.1 afis|EAW25248.1 afla|EED53716.1 afum|EAL89910.1 agla|OJJ89222.1 akaw|GAA82820.1 aluc|OJZ87381.1 anid|CBF85341.1 anig|EHA20612.1 aory|EIT75359.1 asyd|OJJ64025.1 ater|EAU34355.1 atub|OJI81265.1 aver|OJI98111.1 awen|OJJ40964.1 azon|OJJ44545.1

Group2375: abra|OJJ71167.1 acar|OOF97996.1 acla|EAW10336.1 afis|EAW25253.1 afla|EED53722.1 afum|EAL89916.1 agla|OJJ89227.1 akaw|GAA82814.1 aluc|OJZ87374.1 anid|CBF85349.1 anig|EHA20607.1 aory|EIT75275.1 asyd|OJJ64033.1 ater|EAU34398.1 atub|OJI81272.1 aver|OJI98117.1 awen|OJJ40967.1 azon|OJJ44551.1

Group2376: abra|OJJ71169.1 acar|OOF97994.1 acla|EAW10333.1 afis|EAW25259.1 afla|EED53724.1 afum|EAL89921.1 agla|OJJ89230.1 akaw|GAA82812.1 aluc|OJZ87372.1 anid|CBF85360.1 anig|EHA20605.1 aory|EIT75308.1 asyd|OJJ64039.1 ater|EAU34400.1 atub|OJI81274.1 aver|OJI98123.1 awen|OJJ40972.1 azon|OJJ44558.1

Group2377: abra|OJJ71170.1 acar|OOF97993.1 acla|EAW10332.1 afla|EED53725.1 agla|OJJ89231.1 akaw|GAA82811.1 aluc|OJZ87371.1 anid|CBF85362.1 anig|EHA20604.1 aory|EIT75319.1 asyd|OJJ64040.1 ater|EAU34401.1 atub|OJI81275.1 aver|OJI98124.1 awen|OJJ40973.1 afis|EAW25260.1 afum|EAL89922.1 azon|OJJ44559.1

Group2378: abra|OJJ71173.1 acar|OOF97992.1 acla|EAW10331.1 afis|EAW25263.1 afla|EED53727.1 afum|EAL89926.2 agla|OJJ89233.1 akaw|GAA82810.1 aluc|OJZ87369.1 anid|CBF85365.1 anig|EHA20603.1 aory|EIT75277.1 asyd|OJJ64043.1 ater|EAU34402.1 atub|OJI81276.1 aver|OJI98127.1 awen|OJJ40975.1 azon|OJJ47355.1

Group2379: abra|OJJ71174.1 acar|OOF97991.1 acla|EAW10330.1 afis|EAW25264.1 afla|EED53728.1 afum|EAL89927.1 agla|OJJ89234.1 akaw|GAA82809.1 aluc|OJZ87368.1 anid|CBF85367.1 anig|EHA20602.1 aory|EIT75295.1 asyd|OJJ64045.1 ater|EAU34403.1 atub|OJI81277.1 aver|OJI98128.1 awen|OJJ40976.1 azon|OJJ44563.1

Group2380: abra|OJJ71179.1 acar|OOF97987.1 acla|EAW10327.1 afis|EAW25267.1 afla|EED53731.1 afum|EAL89930.1 agla|OJJ89241.1 akaw|GAA82806.1 aluc|OJZ87364.1 anid|CBF85374.1 anig|EHA20599.1 aory|EIT75331.1 asyd|OJJ64048.1 ater|EAU34408.1 atub|OJI81280.1 aver|OJI98133.1 awen|OJJ40980.1 azon|OJJ44567.1

Group2381: abra|OJJ71182.1 acar|OOF97984.1 acla|EAW10322.1 afis|EAW25270.1 afla|EED53735.1 afum|EAL89933.1 agla|OJJ89243.1 akaw|GAA82803.1 aluc|OJZ87361.1 anid|CBF85382.1 anig|EHA20596.1 aory|EIT75338.1 asyd|OJJ64051.1 ater|EAU34413.1 atub|OJI81284.1 aver|OJI98137.1 awen|OJJ40984.1 azon|OJJ44570.1

Group2382: abra|OJJ71183.1 acar|OOF97983.1 acla|EAW10321.1 afis|EAW25271.1 afla|EED53736.1 afum|EAL89934.1 agla|OJJ89244.1 akaw|GAA82802.1 aluc|OJZ87360.1 anid|CBF85384.1 anig|EHA20595.1 aory|EIT75335.1 asyd|OJJ64053.1 ater|EAU34414.1 atub|OJI81285.1 aver|OJI98138.1 awen|OJJ40985.1 azon|OJJ44572.1

Group2383: abra|OJJ71184.1 acar|OOF97982.1 acla|EAW10320.1 afis|EAW25272.1 afla|EED53737.1 afum|EAL89935.1 agla|OJJ89245.1 akaw|GAA82801.1 aluc|OJZ87359.1 anid|CBF85385.1 anig|EHA20594.1 aory|EIT75311.1 asyd|OJJ64054.1 ater|EAU34415.1 atub|OJI81286.1 aver|OJI98139.1 awen|OJJ40986.1 azon|OJJ44573.1

Group2384: abra|OJJ71185.1 acar|OOF97981.1 acla|EAW10318.1 afis|EAW25274.1 afla|EED53739.1 afum|EAL89937.1 agla|OJJ89248.1 akaw|GAA82800.1 aluc|OJZ87358.1 anid|CBF85389.1 anig|EHA20593.1 aory|EIT75291.1 asyd|OJJ64056.1 ater|EAU34417.1 atub|OJI81287.1 aver|OJI98141.1 awen|OJJ40988.1 azon|OJJ44575.1

Group2385: abra|OJJ71187.1 acar|OOF97980.1 acla|EAW10316.1 afis|EAW25276.1 afla|EED53742.1 afum|EAL89939.1 agla|OJJ89250.1 akaw|GAA82798.1 aluc|OJZ87357.1 anid|CBF85392.1 anig|EHA20591.1 aory|EIT75317.1 asyd|OJJ64058.1 ater|EAU34419.1 atub|OJI81288.1 aver|OJI98143.1 awen|OJJ40990.1 azon|OJJ44578.1

Group2386: abra|OJJ71188.1 acar|OOF97979.1 acla|EAW10315.1 afis|EAW25277.1 afla|EED53744.1 afum|EAL89940.1 agla|OJJ89251.1 akaw|GAA82797.1 aluc|OJZ87356.1 anid|CBF85394.1 anig|EHA20590.1 aory|EIT75351.1 asyd|OJJ64059.1 ater|EAU34420.1 atub|OJI81289.1 aver|OJI98145.1 awen|OJJ40991.1 azon|OJJ44580.1

Group2387: abra|OJJ71189.1 acar|OOF97978.1 acla|EAW10314.1 afis|EAW25278.1 afla|EED53745.1 afum|EAL89941.1 agla|OJJ89252.1 akaw|GAA82796.1 aluc|OJZ87355.1 anid|CBF85396.1 anig|EHA20589.1 aory|EIT75313.1 asyd|OJJ64060.1 ater|EAU34421.1 atub|OJI81290.1 aver|OJI98146.1 awen|OJJ40992.1 azon|OJJ44581.1

Group2388: abra|OJJ71192.1 acar|OOF97975.1 acla|EAW10312.1 afis|EAW25281.1 afla|EED53747.1 afum|EAL89944.1 agla|OJJ89255.1 akaw|GAA82793.1 aluc|OJZ87352.1 anid|CBF85418.1 anig|EHA20586.1 aory|EIT75318.1 asyd|OJJ64073.1 ater|EAU34422.1 atub|OJI81293.1 aver|OJI98160.1 awen|OJJ40994.1 azon|OJJ44583.1

Group2389: abra|OJJ71198.1 acar|OOF97970.1 acla|EAW10305.1 afis|EAW25287.1 afla|EED53752.1 afum|EAL89950.1 agla|OJJ89260.1 akaw|GAA82785.1 aluc|OJZ87344.1 anid|CBF85429.1 anig|EHA20581.1 asyd|OJJ64083.1 ater|EAU34428.1 atub|OJI81300.1 aver|OJI98168.1 awen|OJJ40999.1 azon|OJJ44591.1 aory|EIT78420.1

Group2390: abra|OJJ71203.1 acar|OOF97968.1 acla|EAW10304.1 afis|EAW25288.1 afla|EED53753.1 afum|EAL89951.1 agla|OJJ89261.1 akaw|GAA82781.1 aluc|OJZ87340.1 anid|CBF85431.1 anig|EHA20577.1 aory|EIT78400.1 asyd|OJJ64084.1 ater|EAU34429.1 atub|OJI81304.1 aver|OJI98169.1 awen|OJJ41000.1 azon|OJJ44594.1

Group2391: abra|OJJ71204.1 acar|OOF97967.1 acla|EAW10303.1 afis|EAW25289.1 afla|EED53754.1 afum|EAL89952.1 agla|OJJ89262.1 akaw|GAA82780.1 aluc|OJZ87339.1 anid|CBF85433.1 anig|EHA20576.1 aory|EIT78451.1 asyd|OJJ64085.1 ater|EAU34430.1 atub|OJI81305.1 aver|OJI98170.1 awen|OJJ41001.1 azon|OJJ44593.1

Group2392: abra|OJJ71205.1 acar|OOF97965.1 acla|EAW10302.1 afis|EAW25290.1 afla|EED53755.1 afum|EAL89953.1 agla|OJJ89263.1 akaw|GAA82779.1 aluc|OJZ87338.1 anid|CBF85435.1 anig|EHA20575.1 aory|EIT78386.1 asyd|OJJ64086.1 ater|EAU34431.1 atub|OJI81306.1 aver|OJI98171.1 awen|OJJ41002.1 azon|OJJ44592.1

Group2393: abra|OJJ71206.1 acar|OOF97966.1 acla|EAW10301.1 afis|EAW25291.1 afla|EED53756.1 afum|EAL89954.1 agla|OJJ89265.1 akaw|GAA82778.1 aluc|OJZ87337.1 anid|CBF85413.1 anig|EHA20574.1 aory|EIT78379.1 asyd|OJJ64069.1 ater|EAU34432.1 atub|OJI81307.1 aver|OJI98156.1 awen|OJJ41003.1 azon|OJJ44595.1

Group2394: abra|OJJ71210.1 acar|OOF97956.1 acla|EAW10300.1 afis|EAW25292.1 afla|EED53757.1 afum|EAL89955.1 agla|OJJ89269.1 akaw|GAA82774.1 aluc|OJZ87333.1 anid|CBF85411.1 anig|EHA20569.1 aory|EIT78389.1 asyd|OJJ64068.1 ater|EAU34433.1 atub|OJI81311.1 aver|OJI98155.1 awen|OJJ41006.1 azon|OJJ44602.1

Group2395: abra|OJJ71211.1 acar|OOF97955.1 acla|EAW10298.1 afis|EAW25294.1 afla|EED53759.1 afum|EAL89957.1 agla|OJJ89270.1 akaw|GAA82773.1 aluc|OJZ87332.1 anid|CBF85409.1 anig|EHA20568.1

aory|EIT78363.1 asyd|OJJ64067.1 ater|EAU34434.1 atub|OJI81312.1 aver|OJI98154.1 awen|OJJ41008.1 azon|OJJ44601.1

Group2396: abra|OJJ71227.1 acar|OOF97952.1 acla|EAW10295.1 afis|EAW25297.1 afla|EED53763.1 afum|EAL89960.1 agla|OJJ89273.1 akaw|GAA82762.1 aluc|OJZ87320.1 anid|CBF85405.1 anig|EHA20554.1 aory|EIT78455.1 asyd|OJJ64064.1 ater|EAU34438.1 atub|OJI81325.1 aver|OJI98151.1 awen|OJJ41011.1 azon|OJJ44598.1

Group2397: abra|OJJ71241.1 acar|OOF97926.1 acla|EAW09108.1 afis|EAW25717.1 afla|EED53769.1 afum|EAL84392.2 agla|OJJ85442.1 akaw|GAA82746.1 aluc|OJZ87301.1 anid|CBF77088.1 anig|EHA20541.1 aory|EIT78470.1 asyd|OJJ56975.1 ater|EAU34440.1 atub|OJI81339.1 aver|OJJ03313.1 awen|OJJ33601.1 azon|OJJ44472.1

Group2398: abra|OJJ71244.1 acar|OOF97924.1 acla|EAW10291.1 afis|EAW25300.1 afla|EED53774.1 afum|EAL89963.2 agla|OJJ89280.1 akaw|GAA82743.1 aluc|OJZ87298.1 anid|CBF75193.1 anig|EHA20539.1 aory|EIT78388.1 asyd|OJJ59938.1 ater|EAU34448.1 atub|OJI81342.1 aver|OJJ02349.1 awen|OJJ41015.1 azon|OJJ44603.1

Group2399: abra|OJJ71245.1 acar|OOF97923.1 acla|EAW10290.1 afis|EAW25301.1 afla|EED53776.1 afum|EAL89964.1 agla|OJJ89281.1 akaw|GAA82742.1 aluc|OJZ87297.1 anid|CBF75195.1 anig|EHA20538.1 aory|EIT78454.1 asyd|OJJ59937.1 ater|EAU34449.1 atub|OJI81343.1 aver|OJJ02350.1 awen|OJJ41016.1 azon|OJJ42625.1

Group2400: abra|OJJ71248.1 acar|OOF97920.1 acla|EAW10287.1 afis|EAW25305.1 afla|EED53783.1 afum|EAL89968.2 agla|OJJ89284.1 akaw|GAA82739.1 aluc|OJZ87294.1 anid|CBF75296.1 anig|EHA20535.1 aory|EIT78359.1 asyd|OJJ59884.1 ater|EAU34453.1 atub|OJI81346.1 aver|OJJ02402.1 awen|OJJ41019.1 azon|OJJ44606.1

Group2401: abra|OJJ71255.1 acar|OOF97916.1 acla|EAW10281.1 afis|EAW25313.1 afla|EED53788.1 afum|EAL89974.1 agla|OJJ83848.1 akaw|GAA82733.1 aluc|OJZ87288.1 anid|CBF75287.1 anig|EHA20529.1 aory|EIT78401.1 asyd|OJJ59889.1 ater|EAU34458.1 atub|OJI81353.1 aver|OJJ02397.1 awen|OJJ41053.1 azon|OJJ44614.1

Group2402: abra|OJJ71256.1 acar|OOF97915.1 acla|EAW10280.1 afis|EAW25314.1 afla|EED53789.1 afum|EAL89975.1 agla|OJJ83849.1 akaw|GAA82732.1 aluc|OJZ87287.1 anid|CBF75285.1 anig|EHA20528.1 aory|EIT78413.1 asyd|OJJ59890.1 ater|EAU34459.1 atub|OJI81354.1 aver|OJJ02396.1 awen|OJJ41050.1 azon|OJJ44612.1

Group2403: abra|OJJ71258.1 acar|OOF97914.1 acla|EAW10278.1 afis|EAW25316.1 afla|EED53791.1 afum|EAL89977.2 agla|OJJ83854.1 akaw|GAA82731.1 aluc|OJZ87285.1 anid|CBF75283.1 anig|EHA20527.1 aory|EIT78447.1 asyd|OJJ59892.1 ater|EAU34460.1 atub|OJI81356.1 aver|OJJ02394.1 awen|OJJ41047.1 azon|OJJ44616.1

Group2404: abra|OJJ71259.1 acar|OOF97913.1 acla|EAW10277.1 afis|EAW25317.1 afla|EED53792.1 afum|EAL89978.2 agla|OJJ83855.1 akaw|GAA82730.1 aluc|OJZ87284.1 anid|CBF75281.1 anig|EHA20526.1 aory|EIT78371.1 asyd|OJJ59893.1 ater|EAU34464.1 atub|OJI81357.1 aver|OJJ02393.1 awen|OJJ41046.1 azon|OJJ44617.1

Group2405: abra|OJJ71262.1 acar|OOF97912.1 acla|EAW10275.1 afis|EAW25319.1 afla|EED53794.1 afum|EAL89980.2 agla|OJJ83857.1 akaw|GAA82727.1 aluc|OJZ87281.1 anid|CBF75269.1 anig|EHA20523.1 aory|EIT78456.1 asyd|OJJ59895.1 ater|EAU34465.1 atub|OJI81360.1 aver|OJJ02391.1 awen|OJJ41044.1 azon|OJJ44619.1

Group2406: abra|OJJ71263.1 acar|OOF97911.1 acla|EAW10274.1 afis|EAW25320.1 afla|EED53795.1 afum|EAL89981.1 agla|OJJ83858.1 akaw|GAA82726.1 aluc|OJZ87280.1 anid|CBF75267.1 anig|EHA20522.1 aory|EIT78395.1 asyd|OJJ59896.1 ater|EAU34466.1 atub|OJI81361.1 aver|OJJ02390.1 awen|OJJ41043.1 azon|OJJ44620.1

Group2407: abra|OJJ71264.1 acar|OOF97910.1 acla|EAW10273.1 afis|EAW25321.1 afla|EED53796.1 afum|EAL89982.1 agla|OJJ83859.1 akaw|GAA82725.1 aluc|OJZ87279.1 anid|CBF75265.1 anig|EHA20521.1 aory|EIT78365.1 asyd|OJJ59897.1 ater|EAU34467.1 atub|OJI81362.1 aver|OJJ02389.1 awen|OJJ41042.1 azon|OJJ44621.1

Group2408: abra|OJJ71265.1 acar|OOF97909.1 acla|EAW10272.1 afis|EAW25322.1 afla|EED53797.1 afum|EAL89983.1 agla|OJJ83860.1 akaw|GAA82724.1 aluc|OJZ87278.1 anid|CBF75263.1 anig|EHA20520.1 aory|EIT78429.1 asyd|OJJ59898.1 ater|EAU34468.1 atub|OJI81363.1 aver|OJJ02388.1 awen|OJJ41041.1 azon|OJJ44624.1

Group2409: abra|OJJ71267.1 acar|OOF97908.1 acla|EAW10271.1 afis|EAW25323.1 afla|EED53798.1 afum|EAL89984.1 agla|OJJ83861.1 akaw|GAA82723.1 aluc|OJZ87277.1 anid|CBF75261.1 anig|EHA20519.1 aory|EIT78438.1 asyd|OJJ59899.1 ater|EAU34469.1 atub|OJI81364.1 aver|OJJ02387.1 awen|OJJ41040.1 azon|OJJ44626.1

Group2410: abra|OJJ71270.1 acar|OOF97902.1 acla|EAW10268.1 afis|EAW25326.1 afla|EED53803.1 afum|EAL89987.1 agla|OJJ83870.1 akaw|GAA82720.1 aluc|OJZ87274.1 anid|CBF75254.1 anig|EHA20515.1 aory|EIT78467.1 asyd|OJJ59904.1 ater|EAU34473.1 atub|OJI81368.1 aver|OJJ02381.1 awen|OJJ41031.1 azon|OJJ44634.1

Group2411: abra|OJJ71271.1 acar|OOF97903.1 acla|EAW10267.1 afis|EAW25327.1 afla|EED53804.1 afum|EAL89988.1 agla|OJJ83869.1 akaw|GAA82719.1 aluc|OJZ87273.1 anid|CBF75252.1 anig|EHA20514.1 aory|EIT78360.1 asyd|OJJ59905.1 ater|EAU34474.1 atub|OJI81369.1 aver|OJJ02380.1 awen|OJJ41032.1 azon|OJJ44633.1

Group2413: abra|OJJ71277.1 acar|OOF97896.1 acla|EAW10258.1 afis|EAW25335.1 afla|EED53811.1 afum|EAL89998.1 agla|OJJ83872.1 akaw|GAA82712.1 aluc|OJZ87266.1 anid|CBF75240.1 anig|EHA20508.1 aory|EIT78414.1 asyd|OJJ59911.1 ater|EAU34478.1 atub|OJI81376.1 aver|OJJ02375.1 awen|OJJ41029.1 azon|OJJ44636.1

Group2414: abra|OJJ71279.1 acar|OOF97892.1 acla|EAW10261.1 afis|EAW25332.1 afla|EED53814.1 afum|EAL89994.1 agla|OJJ83875.1 akaw|GAA82709.1 aluc|OJZ87264.1 anid|CBF75236.1 anig|EHA20506.1 aory|EIT78424.1 asyd|OJJ59913.1 ater|EAU34480.1 atub|OJI81378.1 aver|OJJ02373.1 awen|OJJ41026.1 azon|OJJ44638.1

Group2415: abra|OJJ71280.1 acar|OOF97893.1 acla|EAW10262.1 afis|EAW25331.1 afla|EED53815.1 afum|EAL89993.1 agla|OJJ83877.1 akaw|GAA82708.1 aluc|OJZ87263.1 anid|CBF75234.1 anig|EHA20505.1 aory|EIT78448.1 asyd|OJJ59914.1 ater|EAU34481.1 atub|OJI81379.1 aver|OJJ02372.1 awen|OJJ41025.1 azon|OJJ44639.1

Group2416: abra|OJJ71286.1 acar|OOF97886.1 acla|EAW10251.1 afis|EAW25341.1 afla|EED53822.1 afum|EBA27392.1 agla|OJJ83884.1 akaw|GAA82703.1 aluc|OJZ87257.1 anid|CBF75213.1 anig|EHA20502.1 aory|EIT78427.1 asyd|OJJ59928.1 ater|EAU34487.1 atub|OJI81385.1 aver|OJJ02358.1 awen|OJJ41062.1 azon|OJJ44645.1

Group2417: abra|OJJ71287.1 acar|OOF97884.1 acla|EAW10250.1 afis|EAW25342.1 afla|EED53823.1 afum|EAL90005.2 agla|OJJ83885.1 akaw|GAA82702.1 aluc|OJZ87256.1 anid|CBF75211.1 anig|EHA20501.1 aory|EIT78440.1 asyd|OJJ59927.1 ater|EAU34488.1 atub|OJI81386.1 aver|OJJ02359.1 awen|OJJ41063.1 azon|OJJ44646.1

Group2418: abra|OJJ71288.1 acar|OOF97885.1 acla|EAW10249.1 afis|EAW25343.1 afla|EED53824.1 afum|EAL90006.1 agla|OJJ83886.1 akaw|GAA82701.1 aluc|OJZ87255.1 anid|CBF75209.1 anig|EHA20500.1 aory|EIT78410.1 asyd|OJJ59926.1 ater|EAU34489.1 atub|OJI81387.1 aver|OJJ02360.1 awen|OJJ41065.1 azon|OJJ44647.1

Group2419: abra|OJJ71289.1 acar|OOF97883.1 acla|EAW10248.1 afis|EAW25344.1 afla|EED53825.1 afum|EAL90007.1 agla|OJJ83887.1 akaw|GAA82700.1 aluc|OJZ87254.1 anid|CBF75207.1 anig|EHA20499.1 aory|EIT78419.1 asyd|OJJ59925.1 ater|EAU34490.1 atub|OJI81388.1 aver|OJJ02361.1 awen|OJJ41066.1 azon|OJJ44648.1

Group2420: abra|OJJ71290.1 acar|OOF97882.1 acla|EAW10247.1 afis|EAW25345.1 afla|EED53826.1 afum|EAL90008.2 agla|OJJ83891.1 akaw|GAA82699.1 aluc|OJZ87253.1 anid|CBF75205.1 anig|EHA20498.1 aory|EIT78441.1 asyd|OJJ59924.1 ater|EAU34491.1 atub|OJI81389.1 aver|OJJ02362.1 awen|OJJ41068.1 azon|OJJ44650.1

Group2421: abra|OJJ71291.1 acar|OOF97881.1 acla|EAW10246.1 afis|EAW25346.1 afla|EED53827.1 afum|EAL90009.1 agla|OJJ83892.1 akaw|GAA82698.1 aluc|OJZ87252.1 anid|CBF75203.1 anig|EHA20497.1 aory|EIT78374.1 asyd|OJJ59923.1 ater|EAU34492.1 atub|OJI81390.1 aver|OJJ02363.1 awen|OJJ41069.1 azon|OJJ44651.1

Group2422: abra|OJJ71293.1 acar|OOF97877.1 acla|EAW10245.1 afis|EAW25347.1 afla|EED53828.1 afum|EAL90010.2 agla|OJJ83893.1 akaw|GAA82696.1 aluc|OJZ87250.1 anid|CBF77426.1 anig|EHA20495.1 aory|EIT78392.1 asyd|OJJ53229.1 ater|EAU34495.1 atub|OJI81392.1 aver|OJJ03623.1 awen|OJJ41070.1 azon|OJJ44652.1

Group2423: abra|OJJ71294.1 acar|OOF97876.1 acla|EAW10244.1 afis|EAW25348.1 afla|EED53829.1 afum|EAL90011.1 agla|OJJ83894.1 akaw|GAA82695.1 aluc|OJZ87249.1 anid|CBF77428.1 anig|EHA20494.1 aory|EIT78450.1 asyd|OJJ53228.1 ater|EAU34496.1 atub|OJI81393.1 aver|OJJ03624.1 awen|OJJ41071.1 azon|OJJ44653.1

Group2424: abra|OJJ71295.1 acar|OOF97875.1 acla|EAW10243.1 afis|EAW25349.1 afla|EED53830.1 afum|EAL90012.1 agla|OJJ83895.1 akaw|GAA82694.1 aluc|OJZ87248.1 anid|CBF77430.1 anig|EHA20493.1 asyd|OJJ53227.1 ater|EAU34497.1 atub|OJI81394.1 aver|OJJ03625.1 awen|OJJ41072.1 azon|OJJ44654.1 aory|EIT78356.1

Group2425: abra|OJJ71296.1 acar|OOF97874.1 acla|EAW10242.1 afis|EAW25350.1 afla|EED53831.1 afum|EAL90013.1 agla|OJJ83896.1 akaw|GAA82693.1 aluc|OJZ87247.1 anid|CBF77432.1 anig|EHA20492.1

aory|EIT78425.1 asyd|OJJ53226.1 ater|EAU34498.1 atub|OJI81395.1 aver|OJJ03626.1 awen|OJJ41073.1 azon|OJJ44655.1

Group2426: abra|OJJ71297.1 acar|OOF97873.1 acla|EAW10241.1 afis|EAW25351.1 afla|EED53832.1 afum|EAL90014.1 agla|OJJ83897.1 akaw|GAA82692.1 aluc|OJZ87246.1 anid|CBF77434.1 anig|EHA20491.1 aory|EIT78366.1 asyd|OJJ53225.1 ater|EAU34499.1 atub|OJI81396.1 aver|OJJ03627.1 awen|OJJ41074.1 azon|OJJ44656.1

Group2427: abra|OJJ71300.1 acar|OOF97870.1 acla|EAW10238.1 afis|EAW25354.1 afla|EED53835.1 afum|EAL90016.1 agla|OJJ83902.1 akaw|GAA82689.1 aluc|OJZ87243.1 anid|CBF77471.1 anig|EHA20488.1 aory|EIT78370.1 asyd|OJJ53222.1 ater|EAU34502.1 atub|OJI81399.1 aver|OJJ03630.1 awen|OJJ41077.1 azon|OJJ44659.1

Group2428: abra|OJJ71302.1 acar|OOF97869.1 acla|EAW10237.1 afis|EAW25355.1 afla|EED53836.1 afum|EAL90017.1 agla|OJJ83903.1 akaw|GAA82688.1 aluc|OJZ87241.1 anid|CBF77469.1 anig|EHA20487.1 aory|EIT78369.1 asyd|OJJ53221.1 ater|EAU34503.1 atub|OJI81401.1 aver|OJJ03631.1 awen|OJJ41078.1 azon|OJJ44660.1

Group2429: abra|OJJ71303.1 acar|OOF97868.1 acla|EAW10236.1 afis|EAW25356.1 afla|EED53838.1 afum|EAL90018.1 agla|OJJ83905.1 akaw|GAA82687.1 aluc|OJZ87240.1 anid|CBF77467.1 anig|EHA20486.1 aory|EIT78449.1 asyd|OJJ53220.1 ater|EAU34504.1 atub|OJI81402.1 aver|OJJ03632.1 awen|OJJ41081.1 azon|OJJ44662.1

Group2430: abra|OJJ71304.1 acar|OOF97867.1 acla|EAW10235.1 afis|EAW25357.1 afla|EED53839.1 afum|EAL90019.1 agla|OJJ83906.1 akaw|GAA82686.1 aluc|OJZ87239.1 anid|CBF77465.1 anig|EHA20485.1 aory|EIT78442.1 asyd|OJJ53219.1 ater|EAU34505.1 atub|OJI81403.1 aver|OJJ03633.1 awen|OJJ41082.1 azon|OJJ44663.1

Group2431: abra|OJJ71308.1 acar|OOF97863.1 acla|EAW10232.1 afis|EAW25360.1 afum|EAL90022.1 agla|OJJ83910.1 akaw|GAA82683.1 aluc|OJZ87236.1 anid|CBF77459.1 anig|EHA20483.1 aory|EIT78368.1 asyd|OJJ53216.1 ater|EAU34508.1 atub|OJI81406.1 aver|OJJ03635.1 awen|OJJ41086.1 azon|OJJ44666.1 afla|EED53842.1

Group2432: abra|OJJ71310.1 acar|OOF97862.1 acla|EAW10230.1 afis|EAW25362.1 afla|EED53844.1 afum|EAL90024.1 agla|OJJ83912.1 akaw|GAA82681.1 aluc|OJZ87234.1 anid|CBF77455.1 anig|EHA20481.1 aory|EIT78471.1 asyd|OJJ53214.1 ater|EAU34509.1 atub|OJI81408.1 aver|OJJ03637.1 awen|OJJ41088.1 azon|OJJ44668.1

Group2433: abra|OJJ71311.1 acar|OOF97861.1 acla|EAW10229.1 afis|EAW25363.1 afla|EED53845.1 afum|EAL90025.1 agla|OJJ83913.1 akaw|GAA82680.1 aluc|OJZ87233.1 anid|CBF77453.1 anig|EHA20480.1 aory|EIT78383.1 asyd|OJJ53213.1 ater|EAU34510.1 atub|OJI81409.1 aver|OJJ03638.1 awen|OJJ41089.1 azon|OJJ44669.1

Group2434: abra|OJJ71312.1 acar|OOF97860.1 acla|EAW10228.1 afis|EAW25364.1 afla|EED53846.1 afum|EAL90026.1 agla|OJJ83752.1 akaw|GAA82679.1 aluc|OJZ87232.1 anid|CBF77452.1 anig|EHA20479.1 aory|EIT78459.1 asyd|OJJ53211.1 ater|EAU34511.1 atub|OJI81410.1 aver|OJJ03639.1 awen|OJJ41091.1 azon|OJJ44670.1

Group2435: abra|OJJ71315.1 acar|OOF97857.1 acla|EAW10224.1 afis|EAW25368.1 afla|EED53849.1 afum|EAL90028.2 agla|OJJ83755.1 akaw|GAA82676.1 aluc|OJZ87229.1 anid|CBF77446.1 anig|EHA20476.1 aory|EIT78461.1 asyd|OJJ53208.1 ater|EAU34513.1 atub|OJI81413.1 aver|OJJ03642.1 awen|OJJ41094.1 azon|OJJ44673.1

Group2436: abra|OJJ71316.1 acar|OOF97856.1 acla|EAW10223.1 afis|EAW25369.1 afla|EED53850.1 afum|EAL90029.2 agla|OJJ83756.1 akaw|GAA82675.1 aluc|OJZ87228.1 anid|CBF77444.1 anig|EHA20475.1 aory|EIT78384.1 asyd|OJJ53207.1 ater|EAU34514.1 atub|OJI81414.1 aver|OJJ03643.1 awen|OJJ41095.1 azon|OJJ44674.1

Group2437: abra|OJJ71317.1 acar|OOF97855.1 acla|EAW10222.1 afis|EAW25370.1 afla|EED53851.1 afum|EAL90030.1 agla|OJJ83757.1 akaw|GAA82674.1 aluc|OJZ87227.1 anid|CBF77442.1 anig|EHA20474.1 aory|EIT78393.1 asyd|OJJ53206.1 ater|EAU34515.1 atub|OJI81415.1 aver|OJJ03644.1 awen|OJJ41096.1 azon|OJJ44675.1

Group2438: abra|OJJ71320.1 acar|OOF97854.1 acla|EAW10219.1 afis|EAW25373.1 afla|EED53854.1 afum|EAL90033.1 agla|OJJ83760.1 akaw|GAA82671.1 aluc|OJZ87224.1 anid|CBF77475.1 anig|EHA20471.1 aory|EIT78376.1 asyd|OJJ53203.1 ater|EAU34518.1 atub|OJI81418.1 aver|OJJ03647.1 awen|OJJ41099.1 azon|OJJ44678.1

Group2439: abra|OJJ71321.1 acar|OOF97852.1 acla|EAW10218.1 afis|EAW25375.1 afla|EED53855.1 afum|EAL90034.1 agla|OJJ83761.1 akaw|GAA82670.1 aluc|OJZ87223.1 anid|CBF77477.1 anig|EHA20470.1 aory|EIT78390.1 asyd|OJJ53202.1 ater|EAU34519.1 atub|OJI81419.1 aver|OJJ03648.1 awen|OJJ41100.1 azon|OJJ44679.1

Group2440: abra|OJJ71322.1 acar|OOF97853.1 acla|EAW10217.1 afis|EAW25376.1 afla|EED53856.1 afum|EAL90035.1 agla|OJJ83765.1 akaw|GAA82669.1 aluc|OJZ87222.1 anid|CBF77479.1 anig|EHA20469.1 aory|EIT78453.1 asyd|OJJ53201.1 ater|EAU34520.1 atub|OJI81420.1 aver|OJJ03649.1 awen|OJJ41103.1 azon|OJJ44680.1

Group2441: abra|OJJ71324.1 acar|OOF97850.1 acla|EAW10215.1 afis|EAW25378.1 afla|EED53858.1 afum|EAL90037.1 agla|OJJ83767.1 akaw|GAA82667.1 aluc|OJZ87220.1 anid|CBF77483.1 anig|EHA20468.1 aory|EIT78426.1 asyd|OJJ53199.1 ater|EAU34522.1 atub|OJI81421.1 aver|OJJ03651.1 awen|OJJ41105.1 azon|OJJ44682.1

Group2442: abra|OJJ71325.1 acar|OOF97849.1 acla|EAW10214.1 afis|EAW25379.1 afla|EED53859.1 afum|EAL90038.1 agla|OJJ83772.1 akaw|GAA82666.1 aluc|OJZ87219.1 anid|CBF77485.1 anig|EHA20467.1 aory|EIT78378.1 asyd|OJJ53197.1 ater|EAU34527.1 atub|OJI81422.1 aver|OJJ03653.1 awen|OJJ41106.1 azon|OJJ44683.1

Group2444: abra|OJJ71333.1 acar|OOF97842.1 acla|EAW10206.1 afis|EAW25389.1 afla|EED53866.1 afum|EAL90048.2 agla|OJJ83792.1 akaw|GAA82659.1 aluc|OJZ87209.1 anid|CBF77500.1 anig|EHA20461.1 aory|EIT78380.1 asyd|OJJ53190.1 ater|EAU34537.1 atub|OJI81430.1 aver|OJJ03660.1 awen|OJJ41116.1 azon|OJJ44692.1

Group2445: abra|OJJ71334.1 acar|OOF97838.1 acla|EAW10205.1 afis|EAW25390.1 afla|EED53867.1 afum|EAL90049.1 agla|OJJ87028.1 akaw|GAA82658.1 aluc|OJZ87208.1 anid|CBF77502.1 anig|EHA20459.1 aory|EIT78439.1 asyd|OJJ53189.1 ater|EAU34538.1 atub|OJI81431.1 aver|OJJ03661.1 awen|OJJ41117.1 azon|OJJ44693.1

Group2446: abra|OJJ71337.1 acar|OOF97835.1 acla|EAW10202.1 afis|EAW25393.1 afla|EED53870.1 afum|EAL90052.1 agla|OJJ87031.1 akaw|GAA82655.1 aluc|OJZ87205.1 anid|CBF77508.1 anig|EHA20458.1 aory|EIT78391.1 asyd|OJJ53186.1 ater|EAU34541.1 atub|OJI81435.1 aver|OJJ03665.1 awen|OJJ41119.1 azon|OJJ43212.1

Group2447: abra|OJJ71339.1 acar|OOF97833.1 acla|EAW10200.1 afis|EAW25395.1 afla|EED53872.1 afum|EAL90054.1 agla|OJJ87033.1 akaw|GAA82653.1 aluc|OJZ87203.1 anid|CBF77512.1 anig|EHA20456.1 aory|EIT78468.1 asyd|OJJ53185.1 ater|EAU34543.1 atub|OJI81437.1 aver|OJJ03666.1 awen|OJJ41121.1 azon|OJJ44696.1

Group2448: abra|OJJ71340.1 acar|OOF97832.1 acla|EAW10199.1 afis|EAW25396.1 afla|EED53873.1 afum|EAL90055.1 agla|OJJ87034.1 akaw|GAA82652.1 aluc|OJZ87202.1 anid|CBF77514.1 anig|EHA20455.1 aory|EIT78362.1 asyd|OJJ53184.1 ater|EAU34544.1 atub|OJI81438.1 aver|OJJ03667.1 awen|OJJ41122.1 azon|OJJ44697.1

Group2449: abra|OJJ71341.1 acar|OOF97831.1 acla|EAW10198.1 afis|EAW25397.1 afla|EED53874.1 afum|EAL90056.1 agla|OJJ87035.1 akaw|GAA82651.1 aluc|OJZ87201.1 anid|CBF77516.1 anig|EHA20454.1 aory|EIT78399.1 asyd|OJJ53183.1 ater|EAU34545.1 atub|OJI81439.1 aver|OJJ03668.1 awen|OJJ41123.1 azon|OJJ44698.1

Group2450: abra|OJJ71342.1 acar|OOF97830.1 acla|EAW10197.1 afis|EAW25398.1 afla|EED53875.1 afum|EAL90057.1 agla|OJJ87036.1 akaw|GAA82650.1 aluc|OJZ87200.1 anid|CBF77518.1 anig|EHA20453.1 aory|EIT78445.1 asyd|OJJ53182.1 ater|EAU34549.1 atub|OJI81440.1 aver|OJJ03669.1 awen|OJJ41124.1 azon|OJJ44699.1

Group2451: abra|OJJ71345.1 acar|OOF97827.1 acla|EAW10194.1 afis|EAW25401.1 afla|EED53879.1 afum|EAL90060.1 agla|OJJ87042.1 akaw|GAA82647.1 aluc|OJZ87196.1 anid|CBF77524.1 anig|EHA20451.1 aory|EIT78402.1 asyd|OJJ53180.1 ater|EAU34552.1 atub|OJI81443.1 aver|OJJ03671.1 awen|OJJ41128.1 azon|OJJ44702.1

Group2452: abra|OJJ71346.1 acar|OOF97824.1 acla|EAW10192.1 afis|EAW25403.1 afla|EED53882.1 afum|EAL90062.1 agla|OJJ87044.1 akaw|GAA82646.1 aluc|OJZ87193.1 anid|CBF77530.1 anig|EHA20450.1 aory|EIT78396.1 asyd|OJJ53177.1 ater|EAU34554.1 atub|OJI81445.1 aver|OJJ03675.1 awen|OJJ41130.1 azon|OJJ44704.1

Group2453: abra|OJJ71347.1 acar|OOF97823.1 acla|EAW10191.1 afis|EAW25404.1 afla|EED53883.1 afum|EAL90063.1 agla|OJJ87045.1 akaw|GAA82645.1 aluc|OJZ87192.1 anid|CBF77532.1 anig|EHA20449.1 aory|EIT78372.1 asyd|OJJ53176.1 ater|EAU34555.1 atub|OJI81446.1 aver|OJJ03676.1 awen|OJJ41131.1 azon|OJJ44705.1

Group2454: abra|OJJ71349.1 acar|OOF97821.1 acla|EAW10189.1 afis|EAW25406.1 afla|EED53885.1 afum|EAL90065.1 agla|OJJ87047.1 akaw|GAA82643.1 aluc|OJZ87190.1 anid|CBF77536.1 anig|EHA20447.1 aory|EIT78483.1 asyd|OJJ53174.1 ater|EAU34557.1 atub|OJI81448.1 aver|OJJ03678.1 awen|OJJ41133.1 azon|OJJ44707.1

Group2455: abra|OJJ71351.1 acar|OOF97818.1 acla|EAW10187.1 afis|EAW25408.1 afla|EED53887.1 afum|EAL90067.1 agla|OJJ87049.1 akaw|GAA82641.1 aluc|OJZ87188.1 anid|CBF77540.1 anig|EHA20445.1

aory|EIT78477.1 asyd|OJJ53172.1 ater|EAU34559.1 atub|OJI81450.1 aver|OJJ03680.1 awen|OJJ41135.1 azon|OJJ44709.1

Group2456: abra|OJJ71352.1 acar|OOF97819.1 acla|EAW10186.1 afis|EAW25409.1 afla|EED53888.1 afum|EAL90068.1 agla|OJJ87050.1 akaw|GAA82640.1 aluc|OJZ87187.1 anid|CBF77588.1 anig|EHA20444.1 aory|EIT78490.1 asyd|OJJ53171.1 ater|EAU34560.1 atub|OJI81451.1 aver|OJJ03681.1 awen|OJJ41136.1 azon|OJJ44715.1

Group2457: abra|OJJ71353.1 acar|OOF97817.1 acla|EAW10183.1 afis|EAW25411.1 afla|EED53890.1 afum|EAL90070.1 agla|OJJ87052.1 akaw|GAA82639.1 aluc|OJZ87186.1 anid|CBF77584.1 anig|EHA20443.1 aory|EIT78494.1 asyd|OJJ53169.1 ater|EAU34562.1 atub|OJI81452.1 aver|OJJ03683.1 awen|OJJ41138.1 azon|OJJ44717.1

Group2458: abra|OJJ71354.1 acar|OOF97816.1 acla|EAW10182.1 afis|EAW25412.1 afla|EED53891.1 afum|EAL90071.1 agla|OJJ87054.1 akaw|GAA82638.1 aluc|OJZ87185.1 anid|CBF77582.1 anig|EHA20442.1 aory|EIT78487.1 asyd|OJJ53168.1 ater|EAU34563.1 atub|OJI81453.1 aver|OJJ03684.1 awen|OJJ41139.1 azon|OJJ44719.1

Group2459: abra|OJJ71355.1 acar|OOF97815.1 acla|EAW10181.1 afis|EAW25413.1 afla|EED53892.1 afum|EAL90072.1 agla|OJJ87055.1 akaw|GAA82637.1 aluc|OJZ87184.1 anid|CBF77580.1 anig|EHA20441.1 aory|EIT78502.1 asyd|OJJ53167.1 ater|EAU34564.1 atub|OJI81455.1 aver|OJJ03685.1 awen|OJJ41140.1 azon|OJJ44720.1

Group2460: abra|OJJ71357.1 acar|OOF97813.1 acla|EAW10179.1 afis|EAW25415.1 afla|EED53894.1 afum|EAL90074.1 agla|OJJ87057.1 akaw|GAA82635.1 aluc|OJZ87182.1 anid|CBF77576.1 anig|EHA20440.1 aory|EIT78485.1 asyd|OJJ53165.1 ater|EAU34566.1 atub|OJI81457.1 aver|OJJ03687.1 awen|OJJ41142.1 azon|OJJ44722.1

Group2461: abra|OJJ71358.1 acar|OOF97812.1 acla|EAW10178.1 afis|EAW25416.1 afla|EED53895.1 afum|EAL90075.1 agla|OJJ87058.1 akaw|GAA82634.1 aluc|OJZ87181.1 anid|CBF77574.1 anig|EHA20439.1 aory|EIT78501.1 asyd|OJJ53164.1 ater|EAU34567.1 atub|OJI81458.1 aver|OJJ03688.1 awen|OJJ41144.1 azon|OJJ44723.1

Group2462: abra|OJJ71359.1 acar|OOF97811.1 acla|EAW10177.1 afis|EAW25417.1 afla|EED53896.1 afum|EAL90076.1 agla|OJJ87059.1 akaw|GAA82633.1 aluc|OJZ87180.1 anid|CBF77573.1 anig|EHA20438.1 aory|EIT78493.1 asyd|OJJ53163.1 ater|EAU34568.1 atub|OJI81459.1 aver|OJJ03689.1 awen|OJJ41145.1 azon|OJJ44724.1

Group2463: abra|OJJ71360.1 acar|OOF97810.1 acla|EAW10176.1 afis|EAW25418.1 afla|EED53897.1 afum|EAL90077.1 agla|OJJ87060.1 akaw|GAA82632.1 aluc|OJZ87179.1 anid|CBF77571.1 anig|EHA20437.1 aory|EIT78486.1 asyd|OJJ53162.1 ater|EAU34569.1 atub|OJI81460.1 aver|OJJ03690.1 awen|OJJ41146.1 azon|OJJ44725.1

Group2464: abra|OJJ71361.1 acar|OOF97809.1 acla|EAW10175.1 afis|EAW25419.1 afla|EED53898.1 afum|EAL90078.1 agla|OJJ87061.1 akaw|GAA82631.1 aluc|OJZ87178.1 anid|CBF77569.1 anig|EHA20436.1 aory|EIT78503.1 asyd|OJJ53161.1 ater|EAU34570.1 atub|OJI81461.1 aver|OJJ03691.1 awen|OJJ41147.1 azon|OJJ44726.1

Group2465: abra|OJJ71363.1 acar|OOF97808.1 acla|EAW10174.1 afis|EAW25420.1 afla|EED53899.1 afum|EAL90079.1 agla|OJJ87064.1 akaw|GAA82630.1 aluc|OJZ87177.1 anid|CBF77567.1 anig|EHA20435.1 aory|EIT78497.1 asyd|OJJ53160.1 ater|EAU34571.1 atub|OJI81462.1 aver|OJJ03692.1 awen|OJJ41148.1 azon|OJJ44727.1

Group2466: abra|OJJ71365.1 acar|OOF97807.1 acla|EAW10173.1 afis|EAW25421.1 afla|EED53900.1 afum|EAL90080.2 agla|OJJ87063.1 akaw|GAA82629.1 aluc|OJZ87175.1 anid|CBF77563.1 anig|EHA20433.1 aory|EIT78488.1 asyd|OJJ53158.1 ater|EAU34572.1 atub|OJI81463.1 aver|OJJ03693.1 awen|OJJ41149.1 azon|OJJ44728.1

Group2467: abra|OJJ71366.1 acar|OOF97806.1 acla|EAW10172.1 afis|EAW25422.1 afla|EED53901.1 afum|EAL90081.1 agla|OJJ87062.1 akaw|GAA82628.1 aluc|OJZ87174.1 anid|CBF77561.1 anig|EHA20432.1 aory|EIT78499.1 asyd|OJJ53157.1 ater|EAU34573.1 atub|OJI81464.1 aver|OJJ03694.1 awen|OJJ41150.1 azon|OJJ44729.1

Group2468: abra|OJJ71367.1 acar|OOF97805.1 acla|EAW10171.1 afis|EAW25423.1 afla|EED53902.1 afum|EAL90082.1 agla|OJJ87065.1 akaw|GAA82627.1 aluc|OJZ87173.1 anid|CBF77559.1 anig|EHA20431.1 aory|EIT78495.1 asyd|OJJ53156.1 ater|EAU34577.1 atub|OJI81465.1 aver|OJJ03695.1 awen|OJJ41151.1 azon|OJJ44730.1

Group2469: abra|OJJ71368.1 acar|OOF97804.1 acla|EAW10170.1 afis|EAW25424.1 afla|EED53903.1 afum|EAL90083.1 agla|OJJ87066.1 akaw|GAA82626.1 aluc|OJZ87172.1 anid|CBF77557.1 anig|EHA20430.1 aory|EIT78480.1 asyd|OJJ53155.1 ater|EAU34578.1 atub|OJI81466.1 aver|OJJ03696.1 awen|OJJ41152.1 azon|OJJ44731.1

Group2470: abra|OJJ71370.1 acar|OOF97798.1 acla|EAW10168.1 afis|EAW25426.1 afla|EED53914.1 afum|EAL90085.1 agla|OJJ87070.1 akaw|GAA82624.1 aluc|OJZ87170.1 anid|CBF77553.1 anig|EHA20428.1 aory|EIT78484.1 asyd|OJJ53153.1 ater|EAU34591.1 atub|OJI81469.1 aver|OJJ03698.1 awen|OJJ41155.1 azon|OJJ44733.1

Group2471: abra|OJJ71371.1 acar|OOF97797.1 acla|EAW10166.1 afis|EAW25428.1 afla|EED53916.1 afum|EAL90087.1 agla|OJJ87073.1 akaw|GAA82623.1 aluc|OJZ87169.1 anid|CBF77549.1 anig|EHA20427.1 aory|EIT78314.1 asyd|OJJ53151.1 ater|EAU34598.1 atub|OJI81470.1 aver|OJJ03700.1 awen|OJJ41157.1 azon|OJJ44735.1

Group2472: abra|OJJ71372.1 acar|OOF97796.1 acla|EAW10165.1 afis|EAW25429.1 afla|EED53917.1 afum|EAL90088.1 agla|OJJ87074.1 akaw|GAA82622.1 aluc|OJZ87168.1 anid|CBF77547.1 anig|EHA20426.1 aory|EIT78324.1 asyd|OJJ53150.1 ater|EAU34600.1 atub|OJI81471.1 aver|OJJ03701.1 awen|OJJ41158.1 azon|OJJ44736.1

Group2473: abra|OJJ71374.1 acar|OOF97794.1 acla|EAW10163.1 afis|EAW25431.1 afla|EED53919.1 afum|EAL90089.1 agla|OJJ87076.1 akaw|GAA82620.1 aluc|OJZ87166.1 anid|CBF77543.1 anig|EHA20425.1 aory|EIT78269.1 asyd|OJJ53148.1 ater|EAU34603.1 atub|OJI81473.1 aver|OJJ03703.1 awen|OJJ41160.1 azon|OJJ44738.1

Group2474: abra|OJJ71376.1 acar|OOF97792.1 acla|EAW10148.1 afis|EAW25445.1 afla|EED53931.1 afum|EAL90103.1 agla|OJJ87089.1 akaw|GAA82618.1 aluc|OJZ87164.1 anid|CBF77610.1 anig|EHA20423.1 aory|EIT78328.1 asyd|OJJ53135.1 ater|EAU34615.1 atub|OJI81475.1 aver|OJJ03717.1 awen|OJJ41174.1 azon|OJJ44750.1

Group2475: abra|OJJ71378.1 acar|OOF97791.1 acla|EAW10146.1 afis|EAW25447.1 afla|EED53929.1 afum|EAL90105.1 agla|OJJ87086.1 akaw|GAA82616.1 aluc|OJZ87162.1 anid|CBF77606.1 anig|EHA20421.1 aory|EIT78352.1 asyd|OJJ53138.1 ater|EAU34613.1 atub|OJI81477.1 aver|OJJ03714.1 awen|OJJ41169.1 azon|OJJ44747.1

Group2476: abra|OJJ71379.1 acar|OOF97790.1 acla|EAW10154.1 afis|EAW25440.1 afla|EED53928.1 afum|EAL90098.1 agla|OJJ87085.1 akaw|GAA82615.1 aluc|OJZ87161.1 anid|CBF77602.1 anig|EHA20420.1 aory|EIT78271.1 asyd|OJJ53139.1 ater|EAU34612.1 atub|OJI81478.1 aver|OJJ03713.1 awen|OJJ41168.1 azon|OJJ44746.1

Group2477: abra|OJJ71380.1 acar|OOF97789.1 acla|EAW10155.1 afis|EAW25439.1 afla|EED53927.1 afum|EAL90097.1 agla|OJJ87084.1 akaw|GAA82614.1 aluc|OJZ87160.1 anid|CBF77600.1 anig|EHA20419.1 aory|EIT78298.1 asyd|OJJ53140.1 ater|EAU34611.1 atub|OJI81480.1 aver|OJJ03712.1 awen|OJJ41167.1 azon|OJJ44745.1

Group2478: abra|OJJ71384.1 acar|OOF97785.1 acla|EAW10159.1 afis|EAW25435.1 afla|EED53923.1 afum|EAL90093.1 agla|OJJ87080.1 akaw|GAA82610.1 aluc|OJZ87156.1 anid|CBF77592.1 anig|EHA20415.1 aory|EIT78272.1 asyd|OJJ53144.1 ater|EAU34607.1 atub|OJI81484.1 aver|OJJ03708.1 awen|OJJ41164.1 azon|OJJ44742.1

Group2479: abra|OJJ71386.1 acar|OOF97783.1 acla|EAW10161.1 afis|EAW25433.1 afla|EED53921.1 afum|EAL90091.1 agla|OJJ87078.1 akaw|GAA82608.1 aluc|OJZ87154.1 anid|CBF77590.1 anig|EHA20413.1 aory|EIT78355.1 asyd|OJJ53146.1 ater|EAU34605.1 atub|OJI81486.1 aver|OJJ03706.1 awen|OJJ41162.1 azon|OJJ44740.1

Group2480: abra|OJJ71389.1 acar|OOF97779.1 acla|EAW10150.1 afis|EAW25443.1 afla|EED53933.1 afum|EAL90101.1 agla|OJJ87092.1 akaw|GAA82605.1 aluc|OJZ87151.1 anid|CBF77614.1 anig|EHA20410.1 aory|EIT78302.1 asyd|OJJ53133.1 ater|EAU34616.1 atub|OJI81489.1 aver|OJJ03719.1 awen|OJJ41176.1 azon|OJJ44752.1

Group2481: abra|OJJ71390.1 acar|OOF97778.1 acla|EAW10151.1 afis|EAW25442.1 afla|EED53934.1 afum|EAL90100.1 agla|OJJ87094.1 akaw|GAA82604.1 aluc|OJZ87150.1 anid|CBF77616.1 anig|EHA20409.1 aory|EIT78339.1 asyd|OJJ53132.1 ater|EAU34617.1 atub|OJI81490.1 aver|OJJ03720.1 awen|OJJ41177.1 azon|OJJ44753.1

Group2482: abra|OJJ71393.1 acar|OOF97775.1 acla|EAW10145.1 afis|EAW25448.1 afla|EED53935.1 afum|EAL90106.1 agla|OJJ87095.1 akaw|GAA82603.1 aluc|OJZ87148.1 anid|CBF77624.1 anig|EHA20407.1 aory|EIT78256.1 asyd|OJJ53131.1 ater|EAU34618.1 atub|OJI81492.1 aver|OJJ03723.1 awen|OJJ41179.1 azon|OJJ44755.1

Group2483: abra|OJJ71395.1 acar|OOF97774.1 acla|EAW10144.1 afis|EAW25449.1 afla|EED53936.1 afum|EAL90108.1 agla|OJJ87101.1 akaw|GAA82602.1 aluc|OJZ87147.1 anid|CBF77626.1 anig|EHA20406.1 aory|EIT78279.1 asyd|OJJ53130.1 ater|EAU34619.1 atub|OJI81494.1 aver|OJJ03724.1 awen|OJJ41180.1 azon|OJJ44756.1

Group2484: abra|OJJ71397.1 acar|OOF97772.1 acla|EAW10142.1 afis|EAW25451.1 afla|EED53938.1 afum|EAL90110.1 agla|OJJ87103.1 akaw|GAA82600.1 aluc|OJZ87145.1 anid|CBF77629.1 anig|EHA20404.1

aory|EIT78274.1 asyd|OJJ53128.1 ater|EAU34621.1 atub|OJI81496.1 aver|OJJ03726.1 awen|OJJ41182.1 azon|OJJ44758.1

Group2485: abra|OJJ71399.1 acar|OOF97771.1 acla|EAW10135.1 afis|EAW25452.1 afla|EED53940.1 afum|EAL90111.1 agla|OJJ87105.1 akaw|GAA82598.1 aluc|OJZ87143.1 anid|CBF77056.1 anig|EHA20403.1 aory|EIT78282.1 asyd|OJJ56993.1 ater|EAU34622.1 atub|OJI81498.1 aver|OJJ03294.1 awen|OJJ41184.1 azon|OJJ44760.1

Group2486: abra|OJJ71400.1 acar|OOF97770.1 acla|EAW10134.1 afis|EAW25453.1 afla|EED53941.1 afum|EAL90112.1 agla|OJJ87106.1 akaw|GAA82597.1 aluc|OJZ87142.1 anid|CBF77633.1 anig|EHA20402.1 aory|EIT78319.1 asyd|OJJ53126.1 ater|EAU34623.1 atub|OJI81500.1 aver|OJJ03728.1 awen|OJJ41185.1 azon|OJJ44761.1

Group2487: abra|OJJ71402.1 acar|OOF97768.1 acla|EAW10132.1 afis|EAW25455.1 afla|EED53944.1 afum|EAL90114.1 agla|OJJ87108.1 akaw|GAA82595.1 aluc|OJZ87139.1 anid|CBF77637.1 anig|EHA20398.1 aory|EIT78284.1 asyd|OJJ53124.1 ater|EAU34625.1 atub|OJI81502.1 aver|OJJ03730.1 awen|OJJ41187.1 azon|OJJ44763.1

Group2488: abra|OJJ71404.1 acar|OOF97767.1 acla|EAW10131.1 afis|EAW25456.1 afla|EED53945.1 afum|EAL90115.2 agla|OJJ87109.1 akaw|GAA82594.1 aluc|OJZ87138.1 anid|CBF77639.1 anig|EHA20397.1 aory|EIT78325.1 asyd|OJJ53123.1 ater|EAU34626.1 atub|OJI81503.1 aver|OJJ03731.1 awen|OJJ41188.1 azon|OJJ44764.1

Group2489: abra|OJJ71405.1 acar|OOF97766.1 acla|EAW10130.1 afis|EAW25457.1 afla|EED53946.1 afum|EAL90117.1 agla|OJJ87110.1 akaw|GAA82593.1 aluc|OJZ87137.1 anid|CBF77641.1 anig|EHA20396.1 aory|EIT78306.1 asyd|OJJ53122.1 ater|EAU34627.1 atub|OJI81504.1 aver|OJJ03732.1 awen|OJJ41189.1 azon|OJJ44767.1

Group2490: abra|OJJ71407.1 acar|OOF97764.1 acla|EAW10128.1 afis|EAW25459.1 afla|EED53948.1 afum|EAL90119.1 agla|OJJ87112.1 akaw|GAA82592.1 aluc|OJZ87135.1 anid|CBF77645.1 anig|EHA20394.1 aory|EIT78349.1 asyd|OJJ53120.1 ater|EAU34629.1 atub|OJI81507.1 aver|OJJ03734.1 awen|OJJ41191.1 azon|OJJ44768.1

Group2492: abra|OJJ71413.1 acar|OOF90159.1 acla|EAW10125.1 afis|EAW25463.1 afla|EED53955.1 afum|EAL90126.1 agla|OJJ87115.1 akaw|GAA82588.1 aluc|OJZ87128.1 anid|CBF77653.1 anig|EHA20389.1 aory|EIT78291.1 asyd|OJJ53114.1 ater|EAU34632.1 atub|OJI81512.1 aver|OJJ03741.1 awen|OJJ41200.1 azon|OJJ44771.1

Group2493: abra|OJJ71416.1 acar|OOF90160.1 acla|EAW10123.1 afis|EAW25464.1 afla|EED53958.1 afum|EAL90127.1 agla|OJJ87116.1 akaw|GAA82586.1 aluc|OJZ87124.1 anid|CBF77655.1 anig|EHA20387.1 aory|EIT78307.1 asyd|OJJ53115.1 ater|EAU34634.1 atub|OJI81516.1 aver|OJJ03740.1 awen|OJJ41201.1 azon|OJJ44774.1

Group2494: abra|OJJ71418.1 acar|OOF97758.1 acla|EAW10122.1 afis|EAW25465.1 afla|EED53960.1 afum|EAL90128.2 agla|OJJ87117.1 akaw|GAA82585.1 aluc|OJZ87122.1 anid|CBF77651.1 anig|EHA20386.1 aory|EIT78354.1 asyd|OJJ53117.1 ater|EAU34635.1 atub|OJI81519.1 aver|OJJ03737.1 awen|OJJ41205.1 azon|OJJ44775.1

Group2495: abra|OJJ71420.1 acar|OOF97757.1 acla|EAW10120.1 afis|EAW25467.1 afla|EED53963.1 afum|EAL90131.1 agla|OJJ87118.1 akaw|GAA82582.1 aluc|OJZ87118.1 anid|CBF77666.1 anig|EHA20384.1 aory|EIT78330.1 asyd|OJJ53102.1 ater|EAU34636.1 atub|OJI81522.1 aver|OJJ03751.1 awen|OJJ41206.1 azon|OJJ44777.1

Group2496: abra|OJJ71421.1 acar|OOF97756.1 acla|EAW10111.1 afis|EAW25476.1 afla|EED53964.1 afum|EAL90139.1 agla|OJJ87120.1 akaw|GAA82581.1 aluc|OJZ87117.1 anid|CBF77674.1 anig|EHA20383.1 aory|EIT78326.1 asyd|OJJ53091.1 ater|EAU34640.1 atub|OJI81523.1 aver|OJJ03755.1 awen|OJJ41207.1 azon|OJJ44779.1

Group2497: abra|OJJ71423.1 acar|OOF97755.1 acla|EAW10116.1 afis|EAW25471.1 afla|EED53968.1 afum|EAL90135.1 agla|OJJ87122.1 akaw|GAA82579.1 aluc|OJZ87115.1 anid|CBF77682.1 anig|EHA20381.1 aory|EIT78300.1 asyd|OJJ53088.1 ater|EAU34644.1 atub|OJI81525.1 aver|OJJ03758.1 awen|OJJ41212.1 azon|OJJ44793.1

Group2498: abra|OJJ71424.1 acar|OOF97754.1 acla|EAW10119.1 afis|EAW25470.1 afla|EED53969.1 afum|EAL90134.1 agla|OJJ87123.1 akaw|GAA82578.1 aluc|OJZ87114.1 anid|CBF77684.1 anig|EHA20380.1 aory|EIT78285.1 asyd|OJJ53087.1 ater|EAU34646.1 atub|OJI81526.1 aver|OJJ03759.1 awen|OJJ41213.1 azon|OJJ44792.1

Group2499: abra|OJJ71425.1 acar|OOF97753.1 acla|EAW10118.1 afis|EAW25469.1 afla|EED53970.1 afum|EAL90133.1 agla|OJJ87124.1 akaw|GAA82577.1 aluc|OJZ87113.1 anid|CBF77686.1 anig|EHA20379.1 aory|EIT78275.1 asyd|OJJ53086.1 ater|EAU34647.1 atub|OJI81527.1 aver|OJJ03760.1 awen|OJJ41214.1 azon|OJJ44791.1

Group2500: abra|OJJ71426.1 acar|OOF97752.1 acla|EAW10117.1 afis|EAW25468.1 afla|EED53971.1 afum|EAL90132.1 agla|OJJ87125.1 akaw|GAA82576.1 aluc|OJZ87112.1 anid|CBF77688.1 anig|EHA20378.1 aory|EIT78308.1 asyd|OJJ53085.1 ater|EAU34648.1 atub|OJI81528.1 aver|OJJ03761.1 awen|OJJ41215.1 azon|OJJ44790.1

Group2501: abra|OJJ71428.1 acar|OOF97750.1 acla|EAW10110.1 afis|EAW25477.1 afla|EED53972.1 afum|EAL90140.1 agla|OJJ87126.1 akaw|GAA82575.1 aluc|OJZ87110.1 anid|CBF77709.1 anig|EHA20377.1 aory|EIT78305.1 asyd|OJJ53077.1 ater|EAU34649.1 atub|OJI81529.1 aver|OJJ03771.1 awen|OJJ41216.1 azon|OJJ44789.1

Group2502: abra|OJJ71432.1 acar|OOF90401.1 acla|EAW10106.1 afis|EAW25481.1 afla|EED53979.1 afum|EAL90146.1 agla|OJJ87129.1 akaw|GAA82571.1 aluc|OJZ87106.1 anid|CBF77701.1 anig|EHA20374.1 aory|EIT78311.1 asyd|OJJ53072.1 ater|EAU34652.1 atub|OJI81532.1 aver|OJJ03773.1 awen|OJJ41222.1 azon|OJJ44785.1

Group2503: abra|OJJ71435.1 acar|OOF90399.1 acla|EAW10104.1 afis|EAW25483.1 afla|EED53981.1 afum|EAL90148.1 agla|OJJ87131.1 akaw|GAA82569.1 aluc|OJZ87103.1 anid|CBF77698.1 anig|EHA20372.1 aory|EIT78335.1 asyd|OJJ53070.1 ater|EAU34654.1 atub|OJI81535.1 aver|OJJ03775.1 awen|OJJ41224.1 azon|OJJ44783.1

Group2504: abra|OJJ71507.1 acar|OOF96261.1 acla|EAW15065.1 afis|EAW20037.1 afla|EED45343.1 afum|EAL85158.1 agla|OJJ88926.1 akaw|GAA89406.1 aluc|OJZ86246.1 anid|CBF84775.1 anig|EHA21618.1 aory|EIT73818.1 asyd|OJJ63238.1 ater|EAU38647.1 atub|OJI84296.1 aver|OJI97718.1 awen|OJJ31792.1 azon|OJJ46326.1

Group2505: abra|OJJ71542.1 acar|OOF96289.1 acla|EAW15096.1 afis|EAW20068.1 afla|EED45311.1 afum|EAL85186.1 agla|OJJ88892.1 akaw|GAA89380.1 aluc|OJZ86280.1 anid|CBF84825.1 anig|EHA21591.1 aory|EIT73564.1 asyd|OJJ63207.1 ater|EAU38674.1 atub|OJI84259.1 aver|OJI97748.1 awen|OJJ31827.1 azon|OJJ46297.1

Group2506: abra|OJJ71556.1 acar|OOF90418.1 acla|EAW15109.1 afis|EAW20080.1 afla|EED45295.1 afum|EAL85199.1 agla|OJJ88879.1 akaw|GAA89367.1 aluc|OJZ86293.1 anid|CBF84857.1 anig|EHA21578.1 aory|EIT73533.1 asyd|OJJ63194.1 ater|EAU38688.1 atub|OJI84243.1 aver|OJI97761.1 awen|OJJ31840.1 azon|OJJ46283.1

Group2507: abra|OJJ71560.1 acar|OOF90420.1 acla|EAW15112.1 afis|EAW20083.1 afla|EED45291.1 afum|EAL85202.1 agla|OJJ88875.1 akaw|GAA89364.1 aluc|OJZ86297.1 anid|CBF84871.1 anig|EHA21575.1 aory|EIT73626.1 asyd|OJJ63191.1 ater|EAU38691.1 atub|OJI84240.1 aver|OJI97765.1 awen|OJJ31843.1 azon|OJJ46280.1

Group2508: abra|OJJ71561.1 acar|OOF90421.1 acla|EAW15113.1 afis|EAW20084.1 afla|EED45290.1 afum|EAL85203.1 agla|OJJ88874.1 akaw|GAA89363.1 aluc|OJZ86298.1 anid|CBF84873.1 anig|EHA21574.1 aory|EIT73721.1 asyd|OJJ63190.1 ater|EAU38692.1 atub|OJI84239.1 aver|OJI97766.1 awen|OJJ31844.1 azon|OJJ46279.1

Group2509: abra|OJJ71562.1 acar|OOF96298.1 acla|EAW15114.1 afis|EAW20085.1 afla|EED45289.1 afum|EAL85204.1 agla|OJJ88873.1 akaw|GAA89362.1 aluc|OJZ86299.1 anid|CBF84875.1 anig|EHA21573.1 aory|EIT73546.1 asyd|OJJ63189.1 ater|EAU38693.1 atub|OJI84237.1 aver|OJI97767.1 awen|OJJ31845.1 azon|OJJ46278.1

Group2510: abra|OJJ71563.1 acar|OOF96299.1 acla|EAW15115.1 afis|EAW20086.1 afla|EED45288.1 afum|EAL85205.1 agla|OJJ88872.1 akaw|GAA89361.1 aluc|OJZ86300.1 anid|CBF84877.1 anig|EHA21572.1 aory|EIT73633.1 asyd|OJJ63188.1 ater|EAU38694.1 atub|OJI84236.1 aver|OJI97768.1 awen|OJJ31846.1 azon|OJJ46277.1

Group2511: abra|OJJ71568.1 acar|OOF96302.1 acla|EAW15118.1 afis|EAW20088.1 afla|EED45282.1 afum|EAL85207.1 agla|OJJ88868.1 akaw|GAA89356.1 aluc|OJZ86305.1 anid|CBF84880.1 anig|EHA21567.1 aory|EIT73561.1 asyd|OJJ63185.1 ater|EAU38697.1 atub|OJI84231.1 aver|OJI97770.1 awen|OJJ31849.1 azon|OJJ46274.1

Group2512: abra|OJJ71570.1 acar|OOF96303.1 acla|EAW15119.1 afis|EAW20089.1 afla|EED45281.1 afum|EAL85208.1 agla|OJJ88867.1 akaw|GAA89355.1 aluc|OJZ86307.1 anid|CBF84882.1 anig|EHA21566.1 aory|EIT73628.1 asyd|OJJ63184.1 ater|EAU38698.1 atub|OJI84229.1 aver|OJI97771.1 awen|OJJ31851.1 azon|OJJ46273.1

Group2513: abra|OJJ71573.1 acar|OOF96305.1 acla|EAW15122.1 afis|EAW20092.1 afla|EED45278.1 afum|EAL85211.1 agla|OJJ88862.1 akaw|GAA89351.1 aluc|OJZ86310.1 anid|CBF84897.1 anig|EHA21563.1 aory|EIT73632.1 asyd|OJJ63175.1 ater|EAU38701.1 atub|OJI84225.1 aver|OJI97779.1 awen|OJJ31855.1 azon|OJJ46270.1

Group2514: abra|OJJ71577.1 acar|OOF96306.1 acla|EAW15124.1 afis|EAW20094.1 afla|EED45276.1 afum|EAL85212.1 agla|OJJ88860.1 akaw|GAA89347.1 aluc|OJZ86314.1 anid|CBF84893.1 anig|EHA21561.1

aory|EIT73817.1 asyd|OJJ63177.1 ater|EAU38704.1 atub|OJI84221.1 aver|OJI97777.1 awen|OJJ31857.1 azon|OJJ46268.1

Group2515: abra|OJJ71580.1 acar|OOF96310.1 acla|EAW15127.1 afis|EAW20097.1 afla|EED45273.1 afum|EAL85215.1 agla|OJJ88857.1 akaw|GAA89344.1 aluc|OJZ86317.1 anid|CBF84888.1 anig|EHA21558.1 aory|EIT73668.1 asyd|OJJ63180.1 ater|EAU38705.1 atub|OJI84218.1 aver|OJI97774.1 awen|OJJ31860.1 azon|OJJ46265.1

Group2516: abra|OJJ71584.1 acar|OOF96314.1 acla|EAW15129.1 afis|EAW20101.1 afla|EED53372.1 afum|EAL85218.1 agla|OJJ88854.1 akaw|GAA89339.1 aluc|OJZ86324.1 anid|CBF84901.1 anig|EHA21554.1 aory|EIT74990.1 asyd|OJJ63172.1 ater|EAU38721.1 atub|OJI84210.1 aver|OJI97782.1 awen|OJJ31871.1 azon|OJJ46263.1

Group2517: abra|OJJ71585.1 acar|OOF96315.1 acla|EAW15130.1 afis|EAW20102.1 afla|EED53371.1 afum|EAL85219.1 agla|OJJ88853.1 akaw|GAA89338.1 aluc|OJZ86325.1 anid|CBF84903.1 anig|EHA21553.1 aory|EIT75070.1 asyd|OJJ63171.1 ater|EAU38722.1 atub|OJI84209.1 aver|OJI97783.1 awen|OJJ31872.1 azon|OJJ46262.1

Group2518: abra|OJJ71586.1 acar|OOF96316.1 acla|EAW15131.1 afis|EAW20103.1 afla|EED53370.1 afum|EAL85220.1 agla|OJJ88851.1 akaw|GAA89337.1 aluc|OJZ86326.1 anid|CBF84905.1 anig|EHA21552.1 aory|EIT75110.1 asyd|OJJ63170.1 ater|EAU38723.1 atub|OJI84208.1 aver|OJI97784.1 awen|OJJ31875.1 azon|OJJ46260.1

Group2519: abra|OJJ71593.1 acar|OOF96321.1 acla|EAW15138.1 afis|EAW20110.1 afla|EED53362.1 afum|EAL85227.1 agla|OJJ88842.1 akaw|GAA89330.1 aluc|OJZ86335.1 anid|CBF84919.1 anig|EHA21546.1 aory|EIT74851.1 asyd|OJJ63164.1 ater|EAU38728.1 atub|OJI84200.1 aver|OJI97791.1 awen|OJJ31881.1 azon|OJJ46253.1

Group2520: abra|OJJ71594.1 acar|OOF96322.1 acla|EAW15139.1 afis|EAW20111.1 afla|EED53361.1 afum|EAL85228.2 agla|OJJ88841.1 akaw|GAA89329.1 aluc|OJZ86336.1 anid|CBF84921.1 anig|EHA21545.1 aory|EIT75176.1 asyd|OJJ63163.1 ater|EAU38729.1 atub|OJI84199.1 aver|OJI97792.1 awen|OJJ31882.1 azon|OJJ46252.1

Group2521: abra|OJJ71595.1 acar|OOF96323.1 acla|EAW15140.1 afis|EAW20112.1 afla|EED53360.1 afum|EAL85229.1 agla|OJJ88840.1 akaw|GAA89328.1 aluc|OJZ86337.1 anid|CBF84923.1 anig|EHA21544.1 aory|EIT75201.1 asyd|OJJ63162.1 ater|EAU38730.1 atub|OJI84198.1 aver|OJI97793.1 awen|OJJ31883.1 azon|OJJ46251.1

Group2522: abra|OJJ71596.1 acar|OOF96324.1 acla|EAW15141.1 afis|EAW20113.1 afla|EED53357.1 afum|EAL85230.1 agla|OJJ82514.1 akaw|GAA89327.1 aluc|OJZ86338.1 anid|CBF84924.1 anig|EHA21543.1 aory|EIT75098.1 asyd|OJJ63161.1 ater|EAU38732.1 atub|OJI84197.1 aver|OJI97794.1 azon|OJJ46250.1 awen|OJJ31884.1

Group2523: abra|OJJ71615.1 acar|OOF96331.1 acla|EAW15157.1 afis|EAW20125.1 afla|EED57251.1 afum|EAL85243.1 agla|OJJ88822.1 akaw|GAA89312.1 aluc|OJZ86356.1 anid|CBF84949.1 anig|EHA21534.1 aory|EIT74081.1 asyd|OJJ63147.1 ater|EAU38741.1 atub|OJI84178.1 aver|OJI97807.1 awen|OJJ31894.1 azon|OJJ46242.1

Group2524: abra|OJJ71621.1 acar|OOF96338.1 acla|EAW15161.1 afis|EAW20130.1 afla|EED57222.1 afum|EAL85246.1 agla|OJJ88805.1 akaw|GAA89305.1 aluc|OJZ86364.1 anid|CBF84956.1 anig|EHA21528.1 aory|EIT74137.1 asyd|OJJ63143.1 ater|EAU38747.1 atub|OJI84170.1 aver|OJI97811.1 awen|OJJ31996.1 azon|OJJ46051.1

Group2525: abra|OJJ71623.1 acar|OOF96340.1 acla|EAW15164.1 afis|EAW20133.1 afla|EED57219.1 afum|EAL85249.1 agla|OJJ88802.1 akaw|GAA89302.1 aluc|OJZ86366.1 anid|CBF84962.1 anig|EHA21526.1 aory|EIT74020.1 asyd|OJJ63140.1 ater|EAU38749.1 atub|OJI84168.1 aver|OJI97814.1 awen|OJJ32001.1 azon|OJJ46054.1

Group2526: abra|OJJ71624.1 acar|OOF96341.1 acla|EAW15165.1 afis|EAW20134.1 afla|EED57218.1 afum|EAL85250.1 agla|OJJ88801.1 akaw|GAA89301.1 aluc|OJZ86367.1 anid|CBF84964.1 anig|EHA21525.1 aory|EIT74013.1 asyd|OJJ63139.1 ater|EAU38750.1 atub|OJI84167.1 aver|OJI97815.1 awen|OJJ32002.1 azon|OJJ46055.1

Group2527: abra|OJJ71626.1 acar|OOF96343.1 acla|EAW15166.1 afis|EAW20135.1 afla|EED57217.1 afum|EBA27225.1 agla|OJJ88799.1 akaw|GAA89300.1 aluc|OJZ86368.1 anid|CBF84965.1 anig|EHA21524.1 aory|EIT73997.1 asyd|OJJ63138.1 ater|EAU38758.1 atub|OJI84166.1 aver|OJI97816.1 awen|OJJ32003.1 azon|OJJ46056.1

Group2528: abra|OJJ71627.1 acar|OOF96345.1 acla|EAW15168.1 afis|EAW20137.1 afla|EED57212.1 afum|EAL85252.1 agla|OJJ88798.1 akaw|GAA89298.1 aluc|OJZ86369.1 anid|CBF84968.1 anig|EHA21523.1 aory|EIT74094.1 asyd|OJJ63136.1 ater|EAU38756.1 atub|OJI84165.1 aver|OJI97818.1 awen|OJJ32006.1 azon|OJJ46058.1

Group2529: abra|OJJ71628.1 acar|OOF96346.1 acla|EAW15169.1 afis|EAW20138.1 afla|EED57210.1 afum|EAL85253.1 agla|OJJ88797.1 akaw|GAA89297.1 aluc|OJZ86370.1 anid|CBF84970.1 anig|EHA21522.1 aory|EIT74150.1 asyd|OJJ63135.1 ater|EAU38755.1 atub|OJI84164.1 aver|OJI97819.1 awen|OJJ32007.1 azon|OJJ46063.1

Group2530: abra|OJJ71630.1 acar|OOF96347.1 acla|EAW15172.1 afis|EAW20141.1 afla|EED57209.1 afum|EAL85256.1 agla|OJJ88791.1 akaw|GAA89295.1 aluc|OJZ86372.1 anid|CBF84973.1 anig|EHA21520.1 aory|EIT74038.1 asyd|OJJ63133.1 ater|EAU38753.1 atub|OJI84162.1 aver|OJI97821.1 awen|OJJ32010.1 azon|OJJ46079.1

Group2531: abra|OJJ71633.1 acar|OOF96350.1 acla|EAW15174.1 afis|EAW20143.1 afla|EED57205.1 afum|EAL85258.1 agla|OJJ88789.1 akaw|GAA89293.1 aluc|OJZ86376.1 anid|CBF84978.1 anig|EHA21518.1 aory|EIT74107.1 asyd|OJJ63130.1 atub|OJI84158.1 aver|OJI97824.1 awen|OJJ32013.1 azon|OJJ46076.1 ater|EAU38760.1

Group2532: abra|OJJ71637.1 acar|OOF96354.1 acla|EAW15177.1 afis|EAW20147.1 afla|EED57202.1 afum|EAL85262.1 agla|OJJ88786.1 akaw|GAA89289.1 aluc|OJZ86380.1 anid|CBF84980.1 anig|EHA21513.1 aory|EIT73995.1 asyd|OJJ63127.1 ater|EAU38763.1 atub|OJI84153.1 aver|OJI97828.1 awen|OJJ32021.1 azon|OJJ46069.1

Group2533: abra|OJJ71643.1 acar|OOF96359.1 acla|EAW15183.1 afis|EAW20153.1 afla|EED57197.1 afum|EAL85268.1 agla|OJJ88781.1 akaw|GAA89283.1 aluc|OJZ86386.1 anid|CBF84993.1 anig|EHA21509.1 aory|EIT74014.1 asyd|OJJ63121.1 ater|EAU38767.1 atub|OJI84147.1 aver|OJI97837.1 awen|OJJ32026.1 azon|OJJ46080.1

Group2534: abra|OJJ71649.1 acar|OOF96363.1 acla|EAW15186.1 afis|EAW20156.1 afla|EED57194.1 afum|EAL85271.1 agla|OJJ88774.1 akaw|GAA89280.1 aluc|OJZ86393.1 anid|CBF84998.1 anig|EHA21505.1 aory|EIT74147.1 asyd|OJJ63118.1 ater|EAU38770.1 atub|OJI84140.1 aver|OJI97841.1 awen|OJJ32031.1 azon|OJJ46083.1

Group2535: abra|OJJ71655.1 acar|OOF96369.1 acla|EAW15192.1 afis|EAW20161.1 afla|EED57187.1 afum|EAL85279.1 agla|OJJ88769.1 akaw|GAA89274.1 aluc|OJZ86401.1 anid|CBF85009.1 anig|EHA21499.1 aory|EIT74084.1 asyd|OJJ63112.1 ater|EAU38778.1 atub|OJI84134.1 aver|OJI97847.1 awen|OJJ32056.1 azon|OJJ46090.1

Group2536: abra|OJJ71656.1 acar|OOF96370.1 acla|EAW15194.1 afis|EAW20163.1 afla|EED57186.1 afum|EAL85281.1 agla|OJJ88767.1 akaw|GAA89273.1 aluc|OJZ86402.1 anid|CBF85011.1 anig|EHA21498.1 aory|EIT74088.1 asyd|OJJ63111.1 ater|EAU38794.1 atub|OJI84133.1 aver|OJI97849.1 awen|OJJ32058.1 azon|OJJ46092.1

Group2537: abra|OJJ71663.1 acar|OOF96373.1 acla|EAW15199.1 afis|EAW20165.1 afla|EED57183.1 afum|EAL85283.1 agla|OJJ88764.1 akaw|GAA89270.1 aluc|OJZ86405.1 anid|CBF85022.1 anig|EHA21494.1 aory|EIT74158.1 asyd|OJJ63107.1 ater|EAU38797.1 atub|OJI84126.1 aver|OJI97854.1 awen|OJJ32062.1 azon|OJJ46095.1

Group2538: abra|OJJ71665.1 acar|OOF96375.1 acla|EAW15202.1 afis|EAW20167.1 afla|EED57181.1 afum|EAL85285.1 agla|OJJ88762.1 akaw|GAA89268.1 aluc|OJZ86407.1 anid|CBF85019.1 anig|EHA21492.1 aory|EIT74007.1 asyd|OJJ63105.1 ater|EAU38799.1 atub|OJI84124.1 aver|OJI97856.1 awen|OJJ32064.1 azon|OJJ46097.1

Group2539: abra|OJJ71668.1 acar|OOF96380.1 acla|EAW15208.1 afis|EAW20173.1 afla|EED57175.1 afum|EAL85291.1 agla|OJJ88757.1 akaw|GAA89263.1 aluc|OJZ86411.1 anid|CBF85031.1 anig|EHA21488.1 aory|EIT74021.1 asyd|OJJ63101.1 ater|EAU35209.1 atub|OJI84120.1 aver|OJI97861.1 awen|OJJ32068.1 azon|OJJ46101.1

Group2540: abra|OJJ71670.1 acar|OOF96382.1 acla|EAW15210.1 afis|EAW20175.1 afla|EED57173.1 afum|EAL85293.1 agla|OJJ84691.1 akaw|GAA89261.1 aluc|OJZ86413.1 anid|CBF85035.1 anig|EHA21486.1 aory|EIT73987.1 asyd|OJJ63099.1 ater|EAU35211.1 atub|OJI84118.1 aver|OJI97863.1 awen|OJJ31174.1 azon|OJJ46104.1

Group2541: abra|OJJ71671.1 acar|OOF96383.1 acla|EAW15211.1 afis|EAW20176.1 afla|EED57172.1 afum|EAL85294.1 agla|OJJ84690.1 akaw|GAA89260.1 aluc|OJZ86414.1 anid|CBF85037.1 anig|EHA21485.1 aory|EIT73964.1 asyd|OJJ63098.1 ater|EAU35212.1 atub|OJI84117.1 aver|OJI97864.1 awen|OJJ31175.1 azon|OJJ46105.1

Group2542: abra|OJJ71672.1 acar|OOF96384.1 acla|EAW15212.1 afis|EAW20177.1 afla|EED57171.1 afum|EAL85295.1 agla|OJJ84689.1 akaw|GAA89259.1 aluc|OJZ86415.1 anid|CBF85039.1 anig|EHA21484.1 aory|EIT74066.1 asyd|OJJ63097.1 ater|EAU35213.1 atub|OJI84116.1 aver|OJI97865.1 awen|OJJ31176.1 azon|OJJ46106.1

Group2543: abra|OJJ71675.1 acar|OOF96386.1 acla|EAW15214.1 afis|EAW20179.1 afla|EED57169.1 afum|EAL85297.1 agla|OJJ84687.1 akaw|GAA89257.1 aluc|OJZ86417.1 anid|CBF85042.1 anig|EHA21482.1

aory|EIT74151.1 asyd|OJJ63095.1 ater|EAU35215.1 atub|OJI84114.1 aver|OJI97867.1 awen|OJJ31178.1 azon|OJJ46109.1

Group2544: abra|OJJ71676.1 acar|OOF96387.1 acla|EAW15215.1 afis|EAW20180.1 afla|EED57168.1 afum|EAL85298.1 agla|OJJ84686.1 akaw|GAA89256.1 aluc|OJZ86418.1 anid|CBF85044.1 anig|EHA21481.1 aory|EIT73991.1 asyd|OJJ63094.1 ater|EAU35216.1 atub|OJI84113.1 aver|OJI97868.1 awen|OJJ31179.1 azon|OJJ46110.1

Group2545: abra|OJJ71677.1 acar|OOF96388.1 acla|EAW15216.1 afis|EAW20181.1 afla|EED57167.1 afum|EAL85299.1 agla|OJJ84685.1 akaw|GAA89255.1 aluc|OJZ86419.1 anid|CBF85046.1 anig|EHA21480.1 aory|EIT74127.1 asyd|OJJ63093.1 ater|EAU35217.1 atub|OJI84112.1 aver|OJI97869.1 awen|OJJ31180.1 azon|OJJ46111.1

Group2546: abra|OJJ71678.1 acar|OOF96389.1 acla|EAW15217.1 afis|EAW20182.1 afla|EED57166.1 afum|EAL85300.1 agla|OJJ84684.1 akaw|GAA89254.1 aluc|OJZ86420.1 anid|CBF85048.1 anig|EHA21479.1 aory|EIT74003.1 asyd|OJJ63092.1 ater|EAU35218.1 atub|OJI84111.1 aver|OJI97870.1 awen|OJJ31181.1 azon|OJJ46112.1

Group2547: abra|OJJ71679.1 acar|OOF96390.1 acla|EAW15218.1 afis|EAW20183.1 afla|EED57165.1 afum|EAL85301.1 agla|OJJ84683.1 akaw|GAA89253.1 aluc|OJZ86421.1 anid|CBF85050.1 anig|EHA21478.1 aory|EIT74002.1 asyd|OJJ63091.1 ater|EAU35219.1 atub|OJI84110.1 aver|OJI97871.1 awen|OJJ31182.1 azon|OJJ46113.1

Group2548: abra|OJJ71682.1 acar|OOF96393.1 acla|EAW15221.1 afis|EAW20186.1 afla|EED57164.1 afum|EAL85305.1 agla|OJJ84680.1 akaw|GAA89250.1 aluc|OJZ86424.1 anid|CBF85054.1 anig|EHA21474.1 aory|EIT74048.1 asyd|OJJ63088.1 ater|EAU35222.1 atub|OJI84107.1 aver|OJI97874.1 awen|OJJ31186.1 azon|OJJ46116.1

Group2549: abra|OJJ71685.1 acar|OOF96395.1 acla|EAW15223.1 afis|EAW20188.1 afla|EED57158.1 afum|EAL85307.1 agla|OJJ84678.1 akaw|GAA89248.1 aluc|OJZ86427.1 anid|CBF85058.1 anig|EHA21472.1 aory|EIT73974.1 asyd|OJJ63086.1 ater|EAU35223.1 atub|OJI84104.1 aver|OJI97876.1 awen|OJJ31188.1 azon|OJJ46118.1

Group2550: abra|OJJ71686.1 acar|OOF96396.1 acla|EAW15224.1 afis|EAW20189.1 afla|EED57157.1 afum|EAL85308.1 agla|OJJ84677.1 akaw|GAA89247.1 aluc|OJZ86428.1 anid|CBF85060.1 anig|EHA21471.1 aory|EIT74135.1 asyd|OJJ63085.1 ater|EAU35224.1 atub|OJI84103.1 aver|OJI97877.1 awen|OJJ31189.1 azon|OJJ46119.1

Group2551: abra|OJJ71687.1 acar|OOF96397.1 acla|EAW15225.1 afis|EAW20190.1 afla|EED57156.1 afum|EAL85309.1 agla|OJJ84676.1 akaw|GAA89246.1 aluc|OJZ86429.1 anid|CBF85062.1 anig|EHA21470.1 aory|EIT74033.1 asyd|OJJ63084.1 ater|EAU35225.1 atub|OJI84102.1 aver|OJI97879.1 awen|OJJ31190.1 azon|OJJ46124.1

Group2552: abra|OJJ71689.1 acar|OOF96398.1 acla|EAW15226.1 afis|EAW20191.1 afla|EED57155.1 afum|EAL85310.1 agla|OJJ84675.1 akaw|GAA89245.1 aluc|OJZ86431.1 anid|CBF85064.1 anig|EHA21469.1 aory|EIT74046.1 asyd|OJJ63083.1 ater|EAU35226.1 atub|OJI84100.1 aver|OJI97881.1 awen|OJJ31196.1 azon|OJJ46120.1

Group2553: abra|OJJ71691.1 acar|OOF96399.1 acla|EAW15228.1 afis|EAW20192.1 afla|EED57152.1 afum|EAL85312.2 agla|OJJ84673.1 akaw|GAA89244.1 aluc|OJZ86435.1 anid|CBF85065.1 anig|EHA21468.1 aory|EIT74165.1 asyd|OJJ63082.1 ater|EAU32000.1 atub|OJI84095.1 aver|OJI97882.1 awen|OJJ31197.1 azon|OJJ46125.1

Group2554: abra|OJJ71692.1 acar|OOF96400.1 acla|EAW15229.1 afis|EAW20193.1 afla|EED57151.1 afum|EAL85313.1 agla|OJJ84672.1 akaw|GAA89243.1 aluc|OJZ86436.1 anid|CBF85067.1 anig|EHA21467.1 aory|EIT74001.1 asyd|OJJ63081.1 ater|EAU32001.1 atub|OJI84094.1 aver|OJI97883.1 awen|OJJ31198.1 azon|OJJ46126.1

Group2555: abra|OJJ71693.1 acar|OOF96402.1 acla|EAW15230.1 afis|EAW20194.1 afla|EED57150.1 afum|EAL85314.1 agla|OJJ84671.1 akaw|GAA89242.1 aluc|OJZ86437.1 anid|CBF85069.1 anig|EHA21466.1 aory|EIT74019.1 asyd|OJJ63080.1 ater|EAU32002.1 atub|OJI84093.1 aver|OJI97884.1 awen|OJJ31199.1 azon|OJJ46127.1

Group2556: abra|OJJ71697.1 acar|OOF96405.1 acla|EAW15233.1 afis|EAW20197.1 afla|EED57147.1 afum|EAL85317.1 agla|OJJ84668.1 akaw|GAA89238.1 aluc|OJZ86440.1 anid|CBF85078.1 anig|EHA21461.1 aory|EIT74146.1 asyd|OJJ63076.1 ater|EAU32005.1 atub|OJI84088.1 aver|OJI97888.1 awen|OJJ31203.1 azon|OJJ46128.1

Group2557: abra|OJJ71699.1 acar|OOF96407.1 acla|EAW15235.1 afis|EAW20199.1 afla|EED57145.1 afum|EAL85319.1 agla|OJJ84666.1 akaw|GAA89236.1 aluc|OJZ86442.1 anid|CBF85082.1 anig|EHA21459.1 aory|EIT74005.1 asyd|OJJ63073.1 ater|EAU32007.1 atub|OJI84086.1 aver|OJI97890.1 awen|OJJ31205.1 azon|OJJ41971.1

Group2558: abra|OJJ71700.1 acar|OOF96408.1 acla|EAW15236.1 afis|EAW20200.1 afla|EED57144.1 afum|EAL85320.1 agla|OJJ84665.1 akaw|GAA89235.1 aluc|OJZ86443.1 anid|CBF85084.1 anig|EHA21458.1 aory|EIT74097.1 asyd|OJJ63072.1 ater|EAU32008.1 atub|OJI84085.1 aver|OJI97891.1 awen|OJJ31206.1 azon|OJJ46130.1

Group2559: abra|OJJ71701.1 acar|OOF96409.1 acla|EAW15237.1 afis|EAW20201.1 afla|EED57143.1 afum|EAL85321.1 agla|OJJ84663.1 akaw|GAA89234.1 aluc|OJZ86444.1 anid|CBF85085.1 anig|EHA21457.1 aory|EIT73998.1 asyd|OJJ63070.1 ater|EAU32009.1 atub|OJI84084.1 aver|OJI97892.1 awen|OJJ31207.1 azon|OJJ46131.1

Group2560: abra|OJJ71705.1 acar|OOF96415.1 afis|EAW20206.1 afla|EED57137.1 afum|EAL85324.1 agla|OJJ84661.1 akaw|GAA89232.1 aluc|OJZ86447.1 anig|EHA21454.1 aory|EIT74126.1 asyd|OJJ63064.1 ater|EAU32017.1 atub|OJI84081.1 aver|OJI97897.1 awen|OJJ31210.1 acla|EAW15239.1 anid|CBF85093.1 azon|OJJ46134.1

Group2561: abra|OJJ71714.1 acar|OOF96419.1 acla|EAW06442.1 afis|EAW18547.1 afla|EED52833.1 afum|EAL93247.1 agla|OJJ81584.1 akaw|GAA85175.1 aluc|OJZ86459.1 anid|CBF70177.1 anig|EHA21449.1 aory|EIT78161.1 asyd|OJJ58793.1 ater|EAU30022.1 atub|OJI84068.1 aver|OJJ01397.1 awen|OJJ31099.1 azon|OJJ46565.1

Group2562: abra|OJJ71715.1 acar|OOF96420.1 acla|EAW06441.1 afis|EAW18546.1 afla|EED52832.1 afum|EAL93246.1 agla|OJJ81587.1 akaw|GAA85174.1 aluc|OJZ86461.1 anid|CBF70265.1 anig|EHA21448.1 aory|EIT78207.1 asyd|OJJ58750.1 ater|EAU34832.1 atub|OJI84067.1 aver|OJJ01360.1 awen|OJJ31100.1 azon|OJJ46562.1

Group2563: abra|OJJ71716.1 acar|OOF96421.1 acla|EAW06440.1 afis|EAW18545.1 afla|EED52831.1 afum|EAL93245.1 agla|OJJ81589.1 akaw|GAA85173.1 aluc|OJZ86462.1 anid|CBF70267.1 anig|EHA21447.1 aory|EIT78194.1 asyd|OJJ58749.1 ater|EAU34833.1 atub|OJI84066.1 aver|OJJ01359.1 awen|OJJ31101.1 azon|OJJ46561.1

Group2564: abra|OJJ71719.1 acar|OOF96423.1 acla|EAW06439.1 afis|EAW18543.1 afla|EED52829.1 afum|EAL93243.1 agla|OJJ81591.1 akaw|GAA85171.1 aluc|OJZ86465.1 anid|CBF70269.1 anig|EHA21445.1 aory|EIT78205.1 asyd|OJJ58748.1 ater|EAU34834.1 atub|OJI84064.1 aver|OJJ01358.1 awen|OJJ31103.1 azon|OJJ46558.1

Group2565: abra|OJJ71720.1 acar|OOF96425.1 acla|EAW06444.1 afis|EAW18549.1 afla|EED52828.1 afum|EAL93249.1 agla|OJJ81592.1 akaw|GAA85170.1 aluc|OJZ86466.1 anid|CBF70271.1 anig|EHA21444.1 aory|EIT78197.1 asyd|OJJ58747.1 ater|EAU34836.1 atub|OJI84063.1 aver|OJJ01357.1 awen|OJJ31105.1 azon|OJJ46557.1

Group2566: abra|OJJ71742.1 acar|OOF96433.1 acla|EAW06446.1 afis|EAW18551.1 afla|EED52823.1 afum|EAL93251.1 agla|OJJ81605.1 akaw|GAA85156.1 aluc|OJZ86481.1 anid|CBF70277.1 anig|EHA22164.1 aory|EIT78237.1 asyd|OJJ58744.1 ater|EAU34839.1 atub|OJI84049.1 aver|OJJ01354.1 awen|OJJ31111.1 azon|OJJ46587.1

Group2567: abra|OJJ71745.1 acar|OOF96436.1 acla|EAW06449.1 afis|EAW18553.1 afla|EED52822.1 afum|EAL93253.1 agla|OJJ81603.1 akaw|GAA85159.1 aluc|OJZ86478.1 anid|CBF70280.1 anig|EHA22162.1 aory|EIT78230.1 asyd|OJJ58741.1 ater|EAU34841.1 atub|OJI84052.1 aver|OJJ01351.1 awen|OJJ31114.1 azon|OJJ46590.1

Group2568: abra|OJJ71803.1 acar|OOF96483.1 acla|EAW13844.1 afis|EAW18558.1 afla|EED52808.1 afum|EAL93258.1 agla|OJJ81601.1 akaw|GAA85107.1 aluc|OJZ86533.1 anid|CBF70286.1 anig|EHA22113.1 aory|EIT78195.1 asyd|OJJ58737.1 ater|EAU34847.1 atub|OJI83988.1 aver|OJJ01348.1 awen|OJJ31116.1 azon|OJJ46596.1

Group2569: abra|OJJ71806.1 acar|OOF96484.1 acla|EAW13845.1 afis|EAW18559.1 afla|EED52807.1 afum|EAL93259.1 agla|OJJ81600.1 akaw|GAA85106.1 aluc|OJZ86536.1 anid|CBF70288.1 anig|EHA22112.1 aory|EIT78169.1 asyd|OJJ58735.1 ater|EAU34848.1 atub|OJI83987.1 aver|OJJ01347.1 awen|OJJ31117.1 azon|OJJ46595.1

Group2570: abra|OJJ71809.1 acar|OOF96487.1 acla|EAW13848.1 afis|EAW18562.1 afla|EED52792.1 afum|EAL93263.1 agla|OJJ81609.1 akaw|GAA85103.1 aluc|OJZ86539.1 anid|CBF70294.1 anig|EHA22110.1 aory|EIT78177.1 asyd|OJJ58732.1 ater|EAU34850.1 atub|OJI83984.1 aver|OJJ01344.1 awen|OJJ31120.1 azon|OJJ46597.1

Group2571: abra|OJJ71810.1 acar|OOF96488.1 acla|EAW13849.1 afis|EAW18563.1 afla|EED52791.1 afum|EAL93265.2 agla|OJJ81612.1 akaw|GAA85102.1 aluc|OJZ86541.1 anid|CBF70298.1 anig|EHA22109.1 aory|EIT78239.1 asyd|OJJ58730.1 ater|EAU34851.1 atub|OJI83982.1 aver|OJJ01343.1 awen|OJJ31121.1 azon|OJJ46602.1

Group2572: abra|OJJ71811.1 acar|OOF96489.1 acla|EAW13850.1 afis|EAW18564.1 afla|EED52790.1 afum|EAL93266.1 agla|OJJ81613.1 akaw|GAA85101.1 aluc|OJZ86542.1 anid|CBF70300.1 anig|EHA22108.1

aory|EIT78209.1 asyd|OJJ58729.1 ater|EAU34852.1 atub|OJI83981.1 aver|OJJ01342.1 awen|OJJ31122.1 azon|OJJ46603.1

Group2573: abra|OJJ71813.1 acar|OOF96491.1 acla|EAW13853.1 afis|EAW18567.1 afla|EED52786.1 afum|EAL93269.2 agla|OJJ81616.1 akaw|GAA85097.1 aluc|OJZ86546.1 anid|CBF70305.1 anig|EHA22104.1 aory|EIT78200.1 asyd|OJJ58726.1 ater|EAU34854.1 atub|OJI83977.1 aver|OJJ01339.1 awen|OJJ31124.1 azon|OJJ46605.1

Group2574: abra|OJJ71816.1 acar|OOF96493.1 acla|EAW13856.1 afis|EAW18570.1 afla|EED52781.1 afum|EAL93273.1 agla|OJJ81621.1 akaw|GAA85094.1 aluc|OJZ86548.1 anid|CBF70311.1 anig|EHA22101.1 aory|EIT78224.1 asyd|OJJ58723.1 ater|EAU34855.1 atub|OJI83974.1 aver|OJJ01336.1 awen|OJJ31127.1 azon|OJJ46608.1

Group2575: abra|OJJ71817.1 acar|OOF96494.1 acla|EAW13857.1 afis|EAW18571.1 afla|EED52780.1 afum|EAL93274.1 agla|OJJ81622.1 akaw|GAA85093.1 aluc|OJZ86549.1 anid|CBF70313.1 anig|EHA22100.1 aory|EIT78222.1 asyd|OJJ58722.1 ater|EAU34856.1 atub|OJI83973.1 aver|OJJ01335.1 awen|OJJ31128.1 azon|OJJ46609.1

Group2576: abra|OJJ71818.1 acar|OOF96495.1 acla|EAW13860.1 afis|EAW18572.1 afla|EED52776.1 afum|EAL93275.1 agla|OJJ81623.1 akaw|GAA85092.1 aluc|OJZ86550.1 anid|CBF70315.1 anig|EHA22099.1 aory|EIT78170.1 asyd|OJJ58721.1 ater|EAU34857.1 atub|OJI83972.1 aver|OJJ01334.1 awen|OJJ31129.1 azon|OJJ46610.1

Group2577: abra|OJJ71819.1 acar|OOF96496.1 acla|EAW13861.1 afis|EAW18573.1 afla|EED52775.1 afum|EAL93276.1 agla|OJJ81624.1 akaw|GAA85091.1 aluc|OJZ86551.1 anid|CBF70317.1 anig|EHA22098.1 aory|EIT78219.1 asyd|OJJ58720.1 ater|EAU34858.1 atub|OJI83971.1 aver|OJJ01333.1 awen|OJJ31130.1 azon|OJJ46611.1

Group2578: abra|OJJ71820.1 acar|OOF96497.1 acla|EAW13862.1 afis|EAW18574.1 afla|EED52774.1 afum|EAL93277.1 agla|OJJ81625.1 akaw|GAA85090.1 aluc|OJZ86552.1 anid|CBF70319.1 anig|EHA22097.1 aory|EIT78236.1 asyd|OJJ58719.1 ater|EAU34869.1 atub|OJI83970.1 aver|OJJ01332.1 awen|OJJ31131.1 azon|OJJ46612.1

Group2579: abra|OJJ71821.1 acar|OOF96498.1 acla|EAW13863.1 afis|EAW18575.1 afla|EED52773.1 afum|EAL93278.1 agla|OJJ81626.1 akaw|GAA85089.1 aluc|OJZ86553.1 anid|CBF70320.1 anig|EHA22096.1 aory|EIT78159.1 asyd|OJJ58718.1 ater|EAU34868.1 atub|OJI83969.1 aver|OJJ01331.1 awen|OJJ31132.1 azon|OJJ46613.1

Group2580: abra|OJJ71823.1 acar|OOF96499.1 acla|EAW13864.1 afis|EAW18576.1 afla|EED52772.1 afum|EAL93279.1 agla|OJJ81627.1 akaw|GAA85087.1 aluc|OJZ86555.1 anid|CBF70322.1 anig|EHA22094.1 aory|EIT78160.1 asyd|OJJ58717.1 ater|EAU34867.1 atub|OJI83967.1 aver|OJJ01330.1 awen|OJJ31133.1 azon|OJJ46614.1

Group2581: abra|OJJ71824.1 acar|OOF96500.1 acla|EAW13865.1 afis|EAW18577.1 afla|EED52771.1 afum|EAL93280.1 agla|OJJ81628.1 akaw|GAA85086.1 aluc|OJZ86556.1 anid|CBF70324.1 anig|EHA22093.1 aory|EIT78235.1 asyd|OJJ58716.1 ater|EAU34866.1 atub|OJI83966.1 aver|OJJ01329.1 awen|OJJ31134.1 azon|OJJ46615.1

Group2582: abra|OJJ71829.1 acar|OOF96505.1 acla|EAW13866.1 afis|EAW18578.1 afla|EED52770.1 afum|EAL93281.1 agla|OJJ81629.1 akaw|GAA85082.1 aluc|OJZ86561.1 anid|CBF70354.1 anig|EHA22089.1 aory|EIT78184.1 asyd|OJJ58657.1 ater|EAU34861.1 atub|OJI83960.1 aver|OJJ01265.1 awen|OJJ31135.1 azon|OJJ46616.1

Group2583: abra|OJJ71837.1 acar|OOF90146.1 acla|EAW13868.1 afis|EAW18580.1 afla|EED52766.1 afum|EAL93284.1 agla|OJJ81631.1 akaw|GAA85078.1 aluc|OJZ86568.1 anid|CBF70360.1 anig|EHA22086.1 aory|EIT78154.1 asyd|OJJ58653.1 ater|EAU34873.1 atub|OJI83953.1 aver|OJJ01262.1 awen|OJJ31143.1 azon|OJJ46619.1

Group2584: abra|OJJ71840.1 acar|OOF90149.1 acla|EAW13871.1 afis|EAW18582.1 afla|EED52762.1 afum|EAL93286.1 agla|OJJ81634.1 akaw|GAA85077.1 aluc|OJZ86573.1 anid|CBF70362.1 anig|EHA22084.1 aory|EIT78249.1 asyd|OJJ58652.1 ater|EAU34876.1 atub|OJI83950.1 aver|OJJ01258.1 awen|OJJ31146.1 azon|OJJ46621.1

Group2585: abra|OJJ71842.1 acar|OOF96511.1 acla|EAW13873.1 afis|EAW18585.1 afla|EED52761.1 afum|EAL93289.1 agla|OJJ81636.1 akaw|GAA85075.1 aluc|OJZ86575.1 anid|CBF70366.1 anig|EHA22082.1 aory|EIT78178.1 asyd|OJJ58650.1 ater|EAU34878.1 atub|OJI83948.1 aver|OJJ01256.1 awen|OJJ31148.1 azon|OJJ46623.1

Group2586: abra|OJJ71843.1 acar|OOF96512.1 acla|EAW13874.1 afis|EAW18586.1 afla|EED52760.1 afum|EAL93290.1 agla|OJJ81637.1 akaw|GAA85074.1 aluc|OJZ86576.1 anid|CBF70370.1 anig|EHA22081.1 aory|EIT78214.1 asyd|OJJ58649.1 ater|EAU34879.1 atub|OJI83946.1 aver|OJJ01255.1 awen|OJJ31149.1 azon|OJJ46624.1

Group2587: abra|OJJ71844.1 acar|OOF96513.1 acla|EAW13875.1 afis|EAW18587.1 afla|EED52759.1 afum|EAL93294.2 agla|OJJ81638.1 akaw|GAA85073.1 aluc|OJZ86577.1 anid|CBF70372.1 anig|EHA22080.1 aory|EIT78199.1 asyd|OJJ58648.1 ater|EAU34880.1 atub|OJI83945.1 aver|OJJ01254.1 awen|OJJ31150.1 azon|OJJ46625.1

Group2588: abra|OJJ71845.1 acar|OOF96514.1 acla|EAW13876.1 afis|EAW18588.1 afla|EED52758.1 afum|EAL93295.2 agla|OJJ81640.1 akaw|GAA85072.1 aluc|OJZ86578.1 anid|CBF70374.1 anig|EHA22079.1 aory|EIT78218.1 asyd|OJJ58647.1 ater|EAU34881.1 atub|OJI83944.1 aver|OJJ01253.1 awen|OJJ31151.1 azon|OJJ46626.1

Group2589: abra|OJJ71846.1 acar|OOF96515.1 acla|EAW13877.1 afis|EAW18589.1 afla|EED52757.1 afum|EAL93296.2 agla|OJJ81641.1 akaw|GAA85071.1 aluc|OJZ86579.1 anid|CBF70375.1 anig|EHA22078.1 aory|EIT78231.1 asyd|OJJ58646.1 ater|EAU34882.1 atub|OJI83943.1 aver|OJJ01252.1 awen|OJJ31152.1 azon|OJJ46627.1

Group2590: abra|OJJ71851.1 acar|OOF96521.1 acla|EAW13883.1 afis|EAW18595.1 afla|EED52752.1 afum|EAL93303.1 agla|OJJ81647.1 akaw|GAA85066.1 aluc|OJZ86585.1 anid|CBF70391.1 anig|EHA22075.1 aory|EIT78172.1 asyd|OJJ58636.1 ater|EAU34889.1 atub|OJI83938.1 aver|OJJ01242.1 awen|OJJ31158.1 azon|OJJ46633.1

Group2591: abra|OJJ71854.1 acar|OOF96524.1 acla|EAW13886.1 afis|EAW18598.1 afla|EED52749.1 afum|EAL93306.1 agla|OJJ81651.1 akaw|GAA85063.1 aluc|OJZ86588.1 anid|CBF70397.1 anig|EHA22072.1 aory|EIT78201.1 asyd|OJJ58633.1 ater|EAU34892.1 atub|OJI83935.1 aver|OJJ01239.1 awen|OJJ31161.1 azon|OJJ46637.1

Group2592: abra|OJJ71855.1 acar|OOF96525.1 acla|EAW13887.1 afis|EAW18599.1 afla|EED52748.1 afum|EAL93307.1 agla|OJJ81652.1 akaw|GAA85062.1 aluc|OJZ86589.1 anid|CBF70399.1 anig|EHA22071.1 aory|EIT78171.1 asyd|OJJ58632.1 ater|EAU34893.1 atub|OJI83934.1 aver|OJJ01238.1 awen|OJJ31162.1 azon|OJJ46638.1

Group2593: abra|OJJ71857.1 acar|OOF96527.1 acla|EAW13889.1 afis|EAW18601.1 afla|EED52746.1 afum|EAL93309.1 agla|OJJ81653.1 akaw|GAA85060.1 aluc|OJZ86591.1 anid|CBF70403.1 anig|EHA22069.1 aory|EIT78188.1 asyd|OJJ58630.1 ater|EAU34895.1 atub|OJI83932.1 aver|OJJ01236.1 awen|OJJ31166.1 azon|OJJ46640.1

Group2594: abra|OJJ71858.1 acar|OOF96528.1 acla|EAW13890.1 afis|EAW18602.1 afla|EED52745.1 afum|EAL93310.1 agla|OJJ81654.1 akaw|GAA85059.1 aluc|OJZ86592.1 anid|CBF70405.1 anig|EHA22068.1 aory|EIT78217.1 asyd|OJJ58629.1 ater|EAU34896.1 atub|OJI83931.1 aver|OJJ01235.1 awen|OJJ31167.1 azon|OJJ46641.1

Group2595: abra|OJJ71861.1 acar|OOF96533.1 acla|EAW13892.1 afis|EAW18604.1 afla|EED52743.1 afum|EAL93312.1 agla|OJJ81656.1 akaw|GAA85054.1 aluc|OJZ86598.1 anid|CBF70409.1 anig|EHA22064.1 aory|EIT78243.1 asyd|OJJ58627.1 ater|EAU34899.1 atub|OJI83925.1 aver|OJJ01233.1 awen|OJJ31170.1 azon|OJJ46647.1

Group2596: abra|OJJ71864.1 acar|OOF96534.1 acla|EAW13894.1 afis|EAW18606.1 afla|EED52740.1 afum|EBA27488.1 agla|OJJ81660.1 akaw|GAA85052.1 aluc|OJZ86601.1 anig|EHA22062.1 aory|EIT78163.1 ater|EAU34901.1 atub|OJI83922.1 awen|OJJ31172.1 azon|OJJ46649.1 anid|CBF70412.1 asyd|OJJ58625.1 aver|OJJ01231.1

Group2597: abra|OJJ71865.1 acar|OOF96535.1 acla|EAW13895.1 afis|EAW18607.1 afla|EED52739.1 afum|EAL93317.1 agla|OJJ81666.1 akaw|GAA85051.1 aluc|OJZ86602.1 anid|CBF70414.1 anig|EHA22061.1 aory|EIT78220.1 asyd|OJJ58624.1 ater|EAU34902.1 atub|OJI83921.1 aver|OJJ01230.1 awen|OJJ31173.1 azon|OJJ46650.1

Group2598: abra|OJJ71866.1 acar|OOF96536.1 acla|EAW13896.1 afis|EAW18608.1 afla|EED52738.1 afum|EAL93318.1 agla|OJJ81665.1 akaw|GAA85049.1 aluc|OJZ86603.1 anid|CBF70416.1 anig|EHA22060.1 aory|EIT78223.1 asyd|OJJ58623.1 ater|EAU34903.1 atub|OJI83920.1 aver|OJJ01229.1 awen|OJJ32069.1 azon|OJJ46651.1

Group2599: abra|OJJ71869.1 acar|OOF96540.1 acla|EAW13899.1 afis|EAW18611.1 afla|EED52736.1 afum|EAL93321.1 agla|OJJ81234.1 akaw|GAA85046.1 aluc|OJZ86608.1 anid|CBF70420.1 anig|EHA22058.1 aory|EIT78190.1 asyd|OJJ58620.1 ater|EAU34908.1 atub|OJI83918.1 aver|OJJ01227.1 awen|OJJ32072.1 azon|OJJ46652.1

Group2600: abra|OJJ71873.1 acar|OOF96544.1 acla|EAW13903.1 afis|EAW18615.1 afla|EED48366.1 afum|EAL93325.1 agla|OJJ81229.1 akaw|GAA85042.1 aluc|OJZ86612.1 anid|CBF70428.1 anig|EHA22054.1 aory|EIT83110.1 asyd|OJJ58616.1 ater|EAU34912.1 atub|OJI83914.1 aver|OJJ01223.1 awen|OJJ32076.1 azon|OJJ46656.1

Group2601: abra|OJJ71874.1 acar|OOF96545.1 acla|EAW13904.1 afis|EAW18616.1 afla|EED52730.1 afum|EAL93326.1 agla|OJJ81228.1 akaw|GAA85041.1 aluc|OJZ86613.1 anid|CBF70426.1 anig|EHA22053.1

aory|EIT78192.1 asyd|OJJ58617.1 ater|EAU34913.1 atub|OJI83913.1 aver|OJJ01224.1 awen|OJJ32077.1 azon|OJJ46657.1

Group2603: abra|OJJ71878.1 acar|OOF96546.1 acla|EAW13907.1 afis|EAW18619.1 afla|EED52726.1 afum|EAL93329.1 agla|OJJ81241.1 akaw|GAA85038.1 aluc|OJZ86616.1 anid|CBF70501.1 anig|EHA22050.1 aory|EIT78153.1 asyd|OJJ58579.1 ater|EAU34919.1 atub|OJI83910.1 aver|OJJ01183.1 awen|OJJ32081.1 azon|OJJ46660.1

Group2604: abra|OJJ71879.1 acar|OOF96547.1 acla|EAW13908.1 afis|EAW18620.1 afla|EED52725.1 afum|EAL93330.1 agla|OJJ81240.1 akaw|GAA85037.1 aluc|OJZ86617.1 anid|CBF70499.1 anig|EHA22049.1 aory|EIT78189.1 asyd|OJJ58580.1 ater|EAU34920.1 atub|OJI83909.1 aver|OJJ01184.1 awen|OJJ32082.1 azon|OJJ46661.1

Group2605: abra|OJJ71880.1 acar|OOF96548.1 acla|EAW13909.1 afis|EAW18621.1 afla|EED52724.1 afum|EAL93331.1 agla|OJJ81239.1 akaw|GAA85036.1 aluc|OJZ86618.1 anid|CBF70497.1 anig|EHA22048.1 aory|EIT78181.1 asyd|OJJ58581.1 ater|EAU34921.1 atub|OJI83908.1 aver|OJJ01185.1 awen|OJJ32084.1 azon|OJJ46663.1

Group2606: abra|OJJ71889.1 acar|OOF96551.1 acla|EAW09632.1 afis|EAW21508.1 afla|EED55945.1 afum|EAL92528.1 agla|OJJ85671.1 akaw|GAA85028.1 aluc|OJZ86626.1 anid|CBF76443.1 anig|EHA22039.1 aory|EIT76729.1 asyd|OJJ61102.1 ater|EAU35030.1 atub|OJI83899.1 aver|OJJ00419.1 awen|OJJ39187.1 azon|OJJ51504.1

Group2607: abra|OJJ71891.1 acar|OOF96552.1 acla|EAW09631.1 afis|EAW21509.1 afla|EED55947.1 afum|EAL92529.1 agla|OJJ85673.1 akaw|GAA85027.1 aluc|OJZ86630.1 anid|CBF76441.1 anig|EHA22038.1 aory|EIT76718.1 asyd|OJJ61101.1 ater|EAU35029.1 atub|OJI83898.1 aver|OJJ00416.1 awen|OJJ39185.1 azon|OJJ51503.1

Group2608: abra|OJJ71895.1 acar|OOF96556.1 acla|EAW09627.1 afis|EAW21513.1 afla|EED55951.1 afum|EAL92533.1 agla|OJJ85677.1 akaw|GAA85023.1 aluc|OJZ86634.1 anid|CBF76433.1 anig|EHA22035.1 aory|EIT76705.1 asyd|OJJ61096.1 ater|EAU35027.1 atub|OJI83894.1 aver|OJJ00411.1 awen|OJJ39181.1 azon|OJJ51512.1

Group2609: abra|OJJ71896.1 acar|OOF96557.1 acla|EAW09626.1 afis|EAW21514.1 afla|EED55952.1 afum|EAL92534.1 agla|OJJ85678.1 akaw|GAA85022.1 aluc|OJZ86635.1 anid|CBF76431.1 anig|EHA22034.1 aory|EIT76554.1 asyd|OJJ61095.1 ater|EAU35026.1 atub|OJI83893.1 aver|OJJ00410.1 awen|OJJ39180.1 azon|OJJ51513.1

Group2610: abra|OJJ71898.1 acar|OOF96559.1 acla|EAW09624.1 afis|EAW21516.1 afla|EED55954.1 afum|EAL92536.1 agla|OJJ85680.1 akaw|GAA85020.1 aluc|OJZ86637.1 anid|CBF76427.1 anig|EHA22032.1 aory|EIT76837.1 asyd|OJJ61093.1 ater|EAU35024.1 atub|OJI83891.1 aver|OJJ00408.1 awen|OJJ39178.1 azon|OJJ51516.1

Group2611: abra|OJJ71902.1 acar|OOF96560.1 acla|EAW09621.1 afis|EAW21518.1 afla|EED55956.1 afum|EAL92539.1 agla|OJJ85681.1 akaw|GAA85018.1 aluc|OJZ86639.1 anid|CBF76423.1 anig|EHA22030.1 aory|EIT76726.1 asyd|OJJ61091.1 ater|EAU35022.1 atub|OJI83888.1 aver|OJJ00406.1 awen|OJJ39175.1 azon|OJJ51515.1

Group2612: abra|OJJ71903.1 acar|OOF96561.1 acla|EAW09620.1 afis|EAW21519.1 afla|EED55957.1 afum|EAL92540.1 agla|OJJ85682.1 akaw|GAA85017.1 aluc|OJZ86640.1 anid|CBF76419.1 anig|EHA22029.1 aory|EIT76756.1 asyd|OJJ61090.1 ater|EAU35021.1 atub|OJI83887.1 aver|OJJ00405.1 awen|OJJ39174.1 azon|OJJ51514.1

Group2613: abra|OJJ71905.1 acar|OOF96562.1 acla|EAW09618.1 afis|EAW21521.1 afla|EED55959.1 afum|EAL92542.1 agla|OJJ85684.1 akaw|GAA85015.1 aluc|OJZ86642.1 anid|CBF76416.1 anig|EHA22027.1 aory|EIT76533.1 asyd|OJJ61088.1 ater|EAU35020.1 atub|OJI83885.1 aver|OJJ00403.1 awen|OJJ39172.1 azon|OJJ51501.1

Group2614: abra|OJJ71908.1 acar|OOF96568.1 acla|EAW09615.1 afis|EAW21524.1 afla|EED55962.1 afum|EAL92545.1 agla|OJJ85687.1 akaw|GAA85011.1 aluc|OJZ86646.1 anid|CBF76410.1 anig|EHA22024.1 aory|EIT76688.1 asyd|OJJ61085.1 ater|EAU35017.1 atub|OJI83881.1 aver|OJJ00400.1 awen|OJJ39169.1 azon|OJJ51498.1

Group2615: abra|OJJ71917.1 acar|OOF96574.1 acla|EAW09613.1 afis|EAW21526.1 afla|EED55965.1 afum|EAL92547.1 agla|OJJ85689.1 akaw|GAA85005.1 aluc|OJZ86654.1 anid|CBF76406.1 anig|EHA22017.1 aory|EIT76778.1 asyd|OJJ61082.1 ater|EAU35015.1 atub|OJI83873.1 aver|OJJ00398.1 awen|OJJ39167.1 azon|OJJ51496.1

Group2616: abra|OJJ71918.1 acar|OOF96575.1 acla|EAW09612.1 afis|EAW21527.1 afla|EED55966.1 afum|EAL92548.1 agla|OJJ85690.1 akaw|GAA85004.1 aluc|OJZ86655.1 anid|CBF76404.1 anig|EHA22016.1 aory|EIT76686.1 asyd|OJJ61081.1 ater|EAU35014.1 atub|OJI83872.1 aver|OJJ00397.1 awen|OJJ39166.1 azon|OJJ51495.1

Group2617: abra|OJJ71921.1 acar|OOF96578.1 acla|EAW09607.1 afis|EAW21530.1 afla|EED55979.1 afum|EAL92551.1 agla|OJJ85694.1 akaw|GAA85001.1 aluc|OJZ86658.1 anid|CBF76398.1 anig|EHA22013.1 aory|EIT76728.1 asyd|OJJ61078.1 ater|EAU35009.1 atub|OJI83869.1 aver|OJJ00394.1 awen|OJJ39162.1 azon|OJJ51492.1

Group2618: abra|OJJ71922.1 acar|OOF96579.1 acla|EAW09606.1 afis|EAW21531.1 afla|EED55980.1 afum|EAL92552.1 agla|OJJ85695.1 akaw|GAA85000.1 aluc|OJZ86659.1 anid|CBF76396.1 anig|EHA22012.1 aory|EIT76765.1 asyd|OJJ61077.1 ater|EAU35008.1 atub|OJI83868.1 aver|OJJ00393.1 awen|OJJ39161.1 azon|OJJ51491.1

Group2619: abra|OJJ71923.1 acar|OOF96580.1 acla|EAW09605.1 afis|EAW21532.1 afla|EED55981.1 afum|EAL92553.1 agla|OJJ85696.1 akaw|GAA84999.1 aluc|OJZ86660.1 anid|CBF76394.1 anig|EHA22011.1 aory|EIT76691.1 asyd|OJJ61076.1 ater|EAU35007.1 atub|OJI83867.1 aver|OJJ00392.1 awen|OJJ39160.1 azon|OJJ51490.1

Group2620: abra|OJJ71924.1 acar|OOF96581.1 acla|EAW09604.1 afis|EAW21533.1 afla|EED55982.1 afum|EAL92554.1 agla|OJJ85697.1 akaw|GAA84998.1 aluc|OJZ86661.1 anid|CBF76392.1 anig|EHA22010.1 aory|EIT76892.1 asyd|OJJ61073.1 ater|EAU35006.1 atub|OJI83864.1 aver|OJJ00390.1 awen|OJJ39159.1 azon|OJJ51489.1

Group2621: abra|OJJ71928.1 acar|OOF96585.1 afis|EAW21536.1 afla|EED55986.1 afum|EAL92556.2 agla|OJJ85700.1 anid|CBF76386.1 aory|EIT76707.1 aver|OJJ00386.1 acla|EAW09601.1 akaw|GAA84995.1 aluc|OJZ86665.1 anig|EHA22008.1 asyd|OJJ61070.1 ater|EAU35003.1 atub|OJI83861.1 awen|OJJ39155.1 azon|OJJ51486.1

Group2622: abra|OJJ71929.1 acar|OOF96586.1 acla|EAW09600.1 afis|EAW21537.1 afla|EED55989.1 afum|EAL92557.1 agla|OJJ85701.1 akaw|GAA84994.1 aluc|OJZ86666.1 anid|CBF76382.1 anig|EHA22007.1 aory|EIT76583.1 asyd|OJJ61068.1 ater|EAU35002.1 atub|OJI83860.1 aver|OJJ00384.1 awen|OJJ39154.1 azon|OJJ51485.1

Group2623: abra|OJJ71931.1 acar|OOF96588.1 acla|EAW09598.1 afis|EAW21539.1 afla|EED55991.1 afum|EAL92559.2 agla|OJJ85703.1 akaw|GAA84992.1 aluc|OJZ86668.1 anid|CBF76378.1 anig|EHA22005.1 aory|EIT76546.1 asyd|OJJ61066.1 ater|EAU35001.1 atub|OJI83859.1 aver|OJJ00382.1 awen|OJJ39152.1 azon|OJJ51478.1

Group2624: abra|OJJ71935.1 acar|OOF96591.1 acla|EAW09595.1 afis|EAW21542.1 afla|EED55993.1 afum|EAL92562.1 agla|OJJ85705.1 akaw|GAA84989.1 aluc|OJZ86672.1 anid|CBF76374.1 anig|EHA22002.1 aory|EIT76774.1 asyd|OJJ61064.1 ater|EAU34997.1 atub|OJI83856.1 aver|OJJ00381.1 awen|OJJ39148.1 azon|OJJ51482.1

Group2625: abra|OJJ71938.1 acar|OOF96596.1 acla|EAW09591.1 afis|EAW21547.1 afla|EED55999.1 afum|EAL92567.1 agla|OJJ85710.1 akaw|GAA84986.1 aluc|OJZ86675.1 anid|CBF76364.1 anig|EHA21998.1 aory|EIT76832.1 asyd|OJJ61060.1 ater|EAU34992.1 atub|OJI83853.1 aver|OJJ00377.1 awen|OJJ39143.1 azon|OJJ51464.1

Group2626: abra|OJJ71940.1 acar|OOF96598.1 acla|EAW09590.1 afis|EAW21548.1 afla|EED56001.1 afum|EAL92568.1 agla|OJJ85726.1 akaw|GAA84985.1 aluc|OJZ86676.1 anid|CBF76362.1 anig|EHA21997.1 aory|EIT76581.1 asyd|OJJ61059.1 ater|EAU34991.1 atub|OJI83852.1 aver|OJJ00376.1 awen|OJJ39142.1 azon|OJJ51465.1

Group2627: abra|OJJ71942.1 acar|OOF96599.1 acla|EAW09588.1 afis|EAW21550.1 afla|EED56002.1 afum|EAL92570.1 agla|OJJ85728.1 akaw|GAA84983.1 aluc|OJZ86678.1 anid|CBF76358.1 anig|EHA21995.1 aory|EIT76540.1 asyd|OJJ61057.1 ater|EAU34989.1 atub|OJI83849.1 aver|OJJ00374.1 awen|OJJ39140.1 azon|OJJ51467.1

Group2628: abra|OJJ71946.1 acar|OOF96600.1 acla|EAW09587.1 afis|EAW21551.1 afla|EED56003.1 afum|EAL92571.1 agla|OJJ85729.1 akaw|GAA84980.1 aluc|OJZ86682.1 anid|CBF76356.1 anig|EHA21992.1 aory|EIT76839.1 asyd|OJJ61056.1 ater|EAU34988.1 atub|OJI83845.1 aver|OJJ00373.1 awen|OJJ39139.1 azon|OJJ51470.1

Group2629: abra|OJJ71951.1 acar|OOF96602.1 acla|EAW09583.1 afis|EAW21554.1 afla|EED56006.1 afum|EAL92574.1 agla|OJJ85731.1 akaw|GAA84977.1 aluc|OJZ86685.1 anid|CBF76350.1 anig|EHA21988.1 aory|EIT76802.1 asyd|OJJ61053.1 ater|EAU34980.1 atub|OJI83841.1 aver|OJJ00370.1 awen|OJJ39136.1 azon|OJJ51474.1

Group2630: abra|OJJ71956.1 acar|OOF96615.1 acla|EAW09579.1 afis|EAW21558.1 afla|EED56010.1 afum|EAL92577.1 agla|OJJ85736.1 akaw|GAA84971.1 aluc|OJZ86691.1 anid|CBF76344.1 anig|EHA21982.1 aory|EIT76530.1 asyd|OJJ61049.1 ater|EAU34976.1 atub|OJI83833.1 aver|OJJ00365.1 awen|OJJ39132.1 azon|OJJ51460.1

Group2631: abra|OJJ71957.1 acar|OOF96613.1 acla|EAW09578.1 afis|EAW21559.1 afla|EED56011.1 afum|EAL92578.1 agla|OJJ85737.1 akaw|GAA84970.1 aluc|OJZ86692.1 anid|CBF76342.1 anig|EHA21981.1

aory|EIT76831.1 asyd|OJJ61048.1 ater|EAU34974.1 atub|OJI83832.1 aver|OJJ00364.1 awen|OJJ39131.1 azon|OJJ51459.1

Group2632: abra|OJJ71959.1 acar|OOF96611.1 acla|EAW09577.1 afis|EAW21560.1 afla|EED56012.1 afum|EAL92579.1 agla|OJJ85739.1 akaw|GAA84969.1 aluc|OJZ86693.1 anid|CBF76340.1 anig|EHA21980.1 aory|EIT76670.1 asyd|OJJ61046.1 atub|OJI83831.1 aver|OJJ00363.1 awen|OJJ39129.1 azon|OJJ51458.1 ater|EAU34973.1

Group2633: abra|OJJ71960.1 acar|OOF96610.1 acla|EAW09576.1 afis|EAW21561.1 afla|EED56014.1 afum|EAL92580.1 agla|OJJ85740.1 akaw|GAA84968.1 aluc|OJZ86694.1 anid|CBF76338.1 anig|EHA21979.1 aory|EIT76536.1 asyd|OJJ61045.1 ater|EAU34972.1 atub|OJI83830.1 aver|OJJ00362.1 awen|OJJ39128.1 azon|OJJ51456.1

Group2634: abra|OJJ71961.1 acar|OOF96609.1 acla|EAW09575.1 afis|EAW21562.1 afla|EED56015.1 afum|EAL92581.1 agla|OJJ85741.1 akaw|GAA84967.1 aluc|OJZ86696.1 anid|CBF76336.1 anig|EHA21978.1 aory|EIT76848.1 asyd|OJJ61044.1 ater|EAU34971.1 atub|OJI83829.1 aver|OJJ00361.1 awen|OJJ39127.1 azon|OJJ51455.1

Group2635: abra|OJJ71966.1 acar|OOF89890.1 acla|EAW09571.1 afis|EAW21565.1 afum|EAL92584.1 agla|OJJ85750.1 akaw|GAA84962.1 aluc|OJZ86702.1 anid|CBF76330.1 anig|EHA21974.1 aory|EIT76870.1 asyd|OJJ61038.1 ater|EAU34967.1 atub|OJI83824.1 aver|OJJ00356.1 awen|OJJ39120.1 azon|OJJ51450.1 afla|EED56022.1

Group2636: abra|OJJ71969.1 acar|OOF96604.1 acla|EAW09570.1 afis|EAW21566.1 afla|EED56031.1 afum|EAL92585.1 agla|OJJ85751.1 akaw|GAA84961.1 aluc|OJZ86704.1 anid|CBF76328.1 anig|EHA21972.1 aory|EIT76642.1 asyd|OJJ61016.1 ater|EAU34965.1 atub|OJI83823.1 aver|OJJ00336.1 awen|OJJ39115.1 azon|OJJ51449.1

Group2637: abra|OJJ71970.1 acar|OOF96621.1 acla|EAW09569.1 afis|EAW21567.1 afla|EED56034.1 afum|EAL92586.1 agla|OJJ85752.1 akaw|GAA84960.1 aluc|OJZ86705.1 anid|CBF76326.1 anig|EHA21971.1 aory|EIT76629.1 asyd|OJJ61015.1 ater|EAU34964.1 atub|OJI83822.1 aver|OJJ00334.1 awen|OJJ39113.1 azon|OJJ51448.1

Group2638: abra|OJJ71971.1 acar|OOF96622.1 acla|EAW09568.1 afis|EAW21568.1 afla|EED56035.1 afum|EAL92587.1 agla|OJJ85753.1 akaw|GAA84959.1 aluc|OJZ86706.1 anid|CBF76324.1 anig|EHA21970.1 aory|EIT76766.1 asyd|OJJ61014.1 ater|EAU34963.1 atub|OJI83821.1 aver|OJJ00333.1 awen|OJJ39112.1 azon|OJJ51447.1

Group2639: abra|OJJ71972.1 acar|OOF96623.1 acla|EAW09567.1 afis|EAW21569.1 afla|EED56036.1 afum|EBA27467.1 agla|OJJ85754.1 akaw|GAA84958.1 aluc|OJZ86707.1 anid|CBF76322.1 anig|EHA21969.1 aory|EIT76617.1 asyd|OJJ61013.1 ater|EAU34962.1 atub|OJI83820.1 aver|OJJ00332.1 awen|OJJ39111.1 azon|OJJ51446.1

Group2640: abra|OJJ71973.1 acar|OOF96625.1 acla|EAW09565.1 afis|EAW21571.1 afla|EED56037.1 afum|EAL92589.1 agla|OJJ85771.1 akaw|GAA84957.1 aluc|OJZ86708.1 anid|CBF76318.1 anig|EHA21968.1 aory|EIT76745.1 asyd|OJJ61011.1 ater|EAU34961.1 atub|OJI83819.1 aver|OJJ00330.1 awen|OJJ39110.1 azon|OJJ51445.1

Group2641: abra|OJJ71974.1 acar|OOF96624.1 acla|EAW09564.1 afis|EAW21572.1 afla|EED56038.1 afum|EAL92590.1 agla|OJJ85768.1 akaw|GAA84956.1 aluc|OJZ86709.1 anid|CBF76316.1 anig|EHA21967.1 aory|EIT76889.1 asyd|OJJ61010.1 ater|EAU34960.1 atub|OJI83818.1 aver|OJJ00329.1 awen|OJJ39109.1 azon|OJJ51444.1

Group2642: abra|OJJ71975.1 acar|OOF96626.1 acla|EAW09563.1 afis|EAW21573.1 afla|EED57792.1 afum|EAL92591.2 agla|OJJ85766.1 akaw|GAA84955.1 aluc|OJZ86710.1 anid|CBF76314.1 anig|EHA21966.1 aory|EIT81122.1 asyd|OJJ61009.1 ater|EAU34959.1 atub|OJI83817.1 aver|OJJ00328.1 awen|OJJ39108.1 azon|OJJ51443.1

Group2643: abra|OJJ71978.1 acar|OOF96628.1 acla|EAW09562.1 afis|EAW21575.1 afla|EED57793.1 afum|EAL92593.1 agla|OJJ85764.1 akaw|GAA84952.1 aluc|OJZ86715.1 anid|CBF76306.1 anig|EHA21963.1 aory|EIT81108.1 asyd|OJJ61006.1 ater|EAU34957.1 atub|OJI83814.1 aver|OJJ00324.1 awen|OJJ39105.1 azon|OJJ51438.1

Group2644: abra|OJJ71983.1 acar|OOF96634.1 acla|EAW09556.1 afis|EAW21581.1 afla|EED57799.1 afum|EAL92599.2 agla|OJJ85757.1 akaw|GAA84946.1 aluc|OJZ86720.1 anid|CBF76300.1 anig|EHA21959.1 aory|EIT81064.1 asyd|OJJ61001.1 ater|EAU34953.1 atub|OJI83809.1 aver|OJJ00319.1 awen|OJJ39100.1 azon|OJJ51434.1

Group2645: abra|OJJ71987.1 acar|OOF96642.1 acla|EAW09551.1 afis|EAW21585.1 afla|EED57806.1 afum|EAL92603.2 agla|OJJ85777.1 akaw|GAA84942.1 aluc|OJZ86724.1 anid|CBF76289.1 anig|EHA21955.1 aory|EIT81058.1 asyd|OJJ60995.1 ater|EAU34946.1 atub|OJI83804.1 aver|OJJ00315.1 awen|OJJ39096.1 azon|OJJ51429.1

Group2646: abra|OJJ71990.1 acar|OOF96644.1 acla|EAW09545.1 afis|EAW21588.1 afla|EED57809.1 afum|EAL92606.1 agla|OJJ85781.1 akaw|GAA84939.1 aluc|OJZ86727.1 anid|CBF76285.1 anig|EHA21952.1 aory|EIT81040.1 asyd|OJJ60992.1 ater|EAU34943.1 atub|OJI83801.1 aver|OJJ00312.1 awen|OJJ39093.1 azon|OJJ51426.1

Group2647: abra|OJJ71991.1 acar|OOF96645.1 acla|EAW09544.1 afis|EAW21589.1 afla|EED57811.1 afum|EAL92607.1 agla|OJJ85782.1 akaw|GAA84938.1 aluc|OJZ86728.1 anid|CBF76283.1 anig|EHA21951.1 aory|EIT81035.1 asyd|OJJ60991.1 ater|EAU34942.1 atub|OJI83800.1 aver|OJJ00311.1 awen|OJJ39092.1 azon|OJJ51425.1

Group2648: abra|OJJ71993.1 acar|OOF96647.1 acla|EAW09542.1 afis|EAW21591.1 afla|EED57813.1 afum|EAL92609.1 agla|OJJ85784.1 akaw|GAA84935.1 aluc|OJZ86731.1 anid|CBF83440.1 anig|EHA21948.1 aory|EIT81099.1 asyd|OJJ60668.1 ater|EAU34940.1 atub|OJI83797.1 aver|OJI99991.1 awen|OJJ39090.1 azon|OJJ51421.1

Group2649: abra|OJJ71995.1 acar|OOF96649.1 acla|EAW09540.1 afis|EAW21593.1 afla|EED57815.1 afum|EAL92611.1 agla|OJJ85788.1 akaw|GAA84933.1 aluc|OJZ86733.1 anid|CBF83444.1 anig|EHA21946.1 aory|EIT81101.1 asyd|OJJ60670.1 ater|EAU34931.1 atub|OJI83795.1 aver|OJI99993.1 awen|OJJ39088.1 azon|OJJ51419.1

Group2650: abra|OJJ71997.1 acar|OOF96651.1 acla|EAW09541.1 afis|EAW21592.1 afla|EED57814.1 afum|EAL92610.1 agla|OJJ85789.1 akaw|GAA84932.1 aluc|OJZ86735.1 anid|CBF83442.1 anig|EHA21944.1 aory|EIT81111.1 asyd|OJJ60669.1 ater|EAU34932.1 atub|OJI83793.1 aver|OJI99992.1 awen|OJJ39089.1 azon|OJJ51420.1

Group2651: abra|OJJ71998.1 acar|OOF96653.1 acla|EAW09536.1 afis|EAW21596.1 afla|EED57819.1 afum|EAL92614.1 agla|OJJ85790.1 akaw|GAA84931.1 aluc|OJZ86736.1 anid|CBF83453.1 anig|EHA21943.1 aory|EIT81091.1 asyd|OJJ60672.1 ater|EAU34928.1 atub|OJI83792.1 aver|OJI99995.1 awen|OJJ39086.1 azon|OJJ51417.1

Group2652: abra|OJJ72007.1 acar|OOF96657.1 acla|EAW09534.1 afis|EAW21599.1 afla|EED57824.1 afum|EAL92618.1 agla|OJJ85794.1 akaw|GAA84926.1 aluc|OJZ86743.1 anid|CBF83457.1 anig|EHA21936.1 aory|EIT81107.1 asyd|OJJ60679.1 ater|EAU38289.1 atub|OJI83784.1 aver|OJJ00004.1 awen|OJJ39083.1 azon|OJJ51415.1

Group2654: abra|OJJ72010.1 acar|OOF96658.1 acla|EAW09529.1 afis|EAW21603.1 afla|EED57829.1 afum|EAL92622.1 agla|OJJ85797.1 akaw|GAA84923.1 aluc|OJZ86746.1 anid|CBF83462.1 anig|EHA21933.1 aory|EIT81061.1 asyd|OJJ60684.1 ater|EAU38292.1 atub|OJI83781.1 aver|OJJ00009.1 awen|OJJ39080.1 azon|OJJ51412.1

Group2655: abra|OJJ72014.1 acar|OOF96663.1 acla|EAW09526.1 afis|EAW21606.1 afla|EED57833.1 afum|EAL92625.1 agla|OJJ85801.1 akaw|GAA84919.1 aluc|OJZ86750.1 anid|CBF83469.1 anig|EHA21929.1 aory|EIT81024.1 asyd|OJJ60688.1 ater|EAU38295.1 atub|OJI83777.1 aver|OJJ00013.1 awen|OJJ39076.1 azon|OJJ51408.1

Group2656: abra|OJJ72015.1 acar|OOF96664.1 acla|EAW09525.1 afis|EAW21607.1 afla|EED57834.1 afum|EAL92626.1 agla|OJJ85802.1 akaw|GAA84918.1 aluc|OJZ86751.1 anid|CBF83473.1 anig|EHA21928.1 aory|EIT81105.1 asyd|OJJ60690.1 ater|EAU38296.1 atub|OJI83775.1 aver|OJJ00015.1 awen|OJJ39075.1 azon|OJJ51407.1

Group2657: abra|OJJ72016.1 acar|OOF96666.1 acla|EAW09524.1 afis|EAW21608.1 afla|EED57837.1 afum|EAL92642.1 agla|OJJ85803.1 akaw|GAA84917.1 aluc|OJZ86752.1 anid|CBF83475.1 anig|EHA21927.1 aory|EIT81080.1 asyd|OJJ60691.1 ater|EAU38297.1 atub|OJI83774.1 aver|OJJ00016.1 awen|OJJ39074.1 azon|OJJ51406.1

Group2658: abra|OJJ72017.1 acar|OOF96665.1 acla|EAW09523.1 afis|EAW21609.1 afla|EED57838.1 afum|EAL92643.1 agla|OJJ85804.1 akaw|GAA84916.1 aluc|OJZ86753.1 anid|CBF83477.1 anig|EHA21926.1 aory|EIT81100.1 asyd|OJJ60692.1 ater|EAU38298.1 atub|OJI83773.1 aver|OJJ00017.1 awen|OJJ39073.1 azon|OJJ51405.1

Group2659: abra|OJJ72018.1 acar|OOF96667.1 acla|EAW09521.1 afis|EAW21611.1 afla|EED57840.1 afum|EAL92645.1 agla|OJJ78616.1 akaw|GAA84914.1 aluc|OJZ86755.1 anid|CBF83480.1 anig|EHA21924.1 aory|EIT81096.1 asyd|OJJ60694.1 ater|EAU38302.1 atub|OJI83772.1 aver|OJJ00020.1 awen|OJJ39071.1 azon|OJJ51402.1

Group2660: abra|OJJ72019.1 acar|OOF96669.1 acla|EAW09520.1 afis|EAW21612.1 afla|EED57841.1 afum|EAL92646.1 agla|OJJ78617.1 akaw|GAA84913.1 aluc|OJZ86756.1 anid|CBF83482.1 anig|EHA21923.1 aory|EIT81062.1 asyd|OJJ60695.1 ater|EAU38303.1 atub|OJI83771.1 aver|OJJ00021.1 awen|OJJ39070.1 azon|OJJ51401.1

Group2661: abra|OJJ72020.1 acar|OOF96668.1 acla|EAW09515.1 afis|EAW21617.1 afla|EED57842.1 afum|EAL92652.1 agla|OJJ78622.1 akaw|GAA84912.1 aluc|OJZ86757.1 anid|CBF83484.1 anig|EHA21922.1

aory|EIT81032.1 asyd|OJJ60696.1 ater|EAU38304.1 atub|OJI83770.1 aver|OJJ00022.1 awen|OJJ39069.1 azon|OJJ51400.1

Group2662: abra|OJJ72022.1 acar|OOF96671.1 acla|EAW09517.1 afis|EAW21615.1 afla|EED57844.1 afum|EAL92649.1 agla|OJJ78624.1 akaw|GAA84910.1 aluc|OJZ86759.1 anid|CBF83485.1 anig|EHA21920.1 aory|EIT81023.1 asyd|OJJ60698.1 ater|EAU38305.1 atub|OJI83768.1 aver|OJJ00024.1 awen|OJJ39067.1 azon|OJJ51398.1

Group2663: abra|OJJ72024.1 acar|OOF96673.1 acla|EAW09519.1 afis|EAW21613.1 afla|EED57846.1 afum|EAL92647.1 agla|OJJ78626.1 akaw|GAA84908.1 aluc|OJZ86761.1 anid|CBF83489.1 anig|EHA21918.1 aory|EIT81029.1 asyd|OJJ60700.1 ater|EAU38307.1 atub|OJI83766.1 aver|OJJ00026.1 awen|OJJ39065.1 azon|OJJ51396.1

Group2664: abra|OJJ72032.1 acar|OOF96682.1 acla|EAW09511.1 afis|EAW21621.1 afla|EED57850.1 afum|EAL92657.2 agla|OJJ82578.1 akaw|GAA84900.1 aluc|OJZ86770.1 anid|CBF83495.1 anig|EHA21910.1 aory|EIT81046.1 asyd|OJJ60704.1 ater|EAU38310.1 atub|OJI83757.1 aver|OJJ00029.1 awen|OJJ33873.1 azon|OJJ48677.1

Group2665: abra|OJJ72044.1 acar|OOF96702.1 acla|EAW09502.1 afis|EAW21634.1 afla|EED57876.1 afum|EAL92667.1 agla|OJJ79699.1 akaw|GAA84890.1 aluc|OJZ86781.1 anid|CBF83500.1 anig|EHA21901.1 aory|EIT83264.1 asyd|OJJ60711.1 ater|EAU38320.1 atub|OJI83739.1 aver|OJJ00038.1 awen|OJJ33883.1 azon|OJJ48670.1

Group2666: abra|OJJ72046.1 acar|OOF96704.1 acla|EAW09500.1 afis|EAW21636.1 afla|EED57879.1 afum|EAL92670.1 agla|OJJ79701.1 akaw|GAA84888.1 aluc|OJZ86783.1 anid|CBF83517.1 anig|EHA21899.1 aory|EIT83275.1 asyd|OJJ60714.1 ater|EAU38322.1 atub|OJI83737.1 aver|OJJ00047.1 awen|OJJ33885.1 azon|OJJ48668.1

Group2667: abra|OJJ72048.1 acar|OOF96706.1 acla|EAW09498.1 afis|EAW21638.1 afla|EED57881.1 afum|EAL92672.2 agla|OJJ79703.1 akaw|GAA84886.1 aluc|OJZ86785.1 anid|CBF83521.1 anig|EHA21897.1 aory|EIT83269.1 asyd|OJJ60716.1 ater|EAU38323.1 atub|OJI83735.1 aver|OJJ00049.1 awen|OJJ33887.1 azon|OJJ48666.1

Group2668: abra|OJJ72052.1 acar|OOF96709.1 acla|EAW09495.1 afis|EAW21641.1 afla|EED57902.1 afum|EAL92675.1 agla|OJJ79708.1 akaw|GAA84882.1 aluc|OJZ86789.1 anid|CBF83526.1 anig|EHA21894.1 aory|EIT82320.1 asyd|OJJ60718.1 ater|EAU38326.1 atub|OJI83730.1 aver|OJJ00051.1 awen|OJJ33898.1 azon|OJJ48662.1

Group2669: abra|OJJ72054.1 acar|OOF96711.1 acla|EAW09493.1 afis|EAW21643.1 afla|EED57900.1 afum|EAL92677.1 agla|OJJ79710.1 akaw|GAA84880.1 aluc|OJZ86791.1 anid|CBF83528.1 anig|EHA21892.1 aory|EIT82343.1 asyd|OJJ60720.1 ater|EAU38328.1 atub|OJI83728.1 aver|OJJ00053.1 awen|OJJ33900.1 azon|OJJ48660.1

Group2670: abra|OJJ72056.1 acar|OOF96713.1 acla|EAW09492.1 afis|EAW21644.1 afla|EED57899.1 afum|EAL92678.1 agla|OJJ79711.1 akaw|GAA84878.1 aluc|OJZ86793.1 anid|CBF83530.1 anig|EHA21890.1 aory|EIT82331.1 asyd|OJJ60721.1 ater|EAU38329.1 atub|OJI83726.1 aver|OJJ00054.1 awen|OJJ33901.1 azon|OJJ48658.1

Group2671: abra|OJJ72062.1 acar|OOF96716.1 acla|EAW09489.1 afis|EAW21647.1 afla|EED57896.1 afum|EAL92681.1 agla|OJJ79718.1 akaw|GAA84873.1 aluc|OJZ86798.1 anid|CBF83537.1 anig|EHA21885.1 aory|EIT82735.1 asyd|OJJ60725.1 ater|EAU38332.1 atub|OJI83720.1 aver|OJJ00058.1 awen|OJJ33907.1 azon|OJJ48655.1

Group2672: abra|OJJ72063.1 acar|OOF96717.1 acla|EAW09479.1 afis|EAW21657.1 afla|EED57895.1 afum|EAL92692.1 agla|OJJ79719.1 akaw|GAA84872.1 aluc|OJZ86799.1 anid|CBF83539.1 anig|EHA21884.1 aory|EIT82741.1 asyd|OJJ60727.1 ater|EAU38333.1 atub|OJI83719.1 aver|OJJ00059.1 awen|OJJ33908.1 azon|OJJ48653.1

Group2673: abra|OJJ72064.1 acar|OOF96718.1 acla|EAW09478.1 afis|EAW21658.1 afla|EED57894.1 afum|EAL92693.1 agla|OJJ79720.1 akaw|GAA84871.1 aluc|OJZ86800.1 anid|CBF83541.1 anig|EHA21883.1 aory|EIT82736.1 asyd|OJJ60728.1 ater|EAU38334.1 atub|OJI83718.1 aver|OJJ00060.1 awen|OJJ33909.1 azon|OJJ48652.1

Group2674: abra|OJJ72065.1 acar|OOF96719.1 acla|EAW09477.1 afis|EAW21659.1 afla|EED57893.1 afum|EAL92694.1 agla|OJJ79721.1 akaw|GAA84870.1 aluc|OJZ86801.1 anid|CBF83542.1 anig|EHA21882.1 aory|EIT82740.1 asyd|OJJ60729.1 ater|EAU38335.1 atub|OJI83717.1 aver|OJJ00061.1 awen|OJJ33910.1 azon|OJJ48651.1

Group2675: abra|OJJ72068.1 acar|OOF96724.1 acla|EAW09483.1 afis|EAW21653.1 afla|EED57886.1 afum|EAL92688.1 agla|OJJ79724.1 akaw|GAA84867.1 aluc|OJZ86803.1 anid|CBF83550.1 anig|EHA21879.1 aory|EIT83265.1 asyd|OJJ60733.1 ater|EAU38339.1 atub|OJI83715.1 aver|OJJ00065.1 awen|OJJ33914.1 azon|OJJ48647.1

Group2676: abra|OJJ72075.1 acar|OOF96727.1 acla|EAW09485.1 afis|EAW21651.1 afla|EED57918.1 afum|EAL92685.2 agla|OJJ79727.1 akaw|GAA84842.1 aluc|OJZ86829.1 anid|CBF83553.1 anig|EHA21806.1 aory|EIT82364.1 asyd|OJJ60735.1 ater|EAU38343.1 atub|OJI83689.1 aver|OJJ00067.1 awen|OJJ33917.1 azon|OJJ48643.1

Group2677: abra|OJJ72085.1 acar|OOF96730.1 acla|EAW09488.1 afis|EAW21648.1 afla|EED57923.1 afum|EAL92682.2 agla|OJJ79730.1 akaw|GAA84835.1 aluc|OJZ86839.1 anid|CBF83559.1 anig|EHA21798.1 aory|EIT82355.1 asyd|OJJ60738.1 ater|EAU38345.1 atub|OJI83679.1 aver|OJJ00070.1 awen|OJJ33919.1 azon|OJJ48645.1

Group2678: abra|OJJ72086.1 acar|OOF96732.1 acla|EAW09467.1 afis|EAW21669.1 afla|EED57933.1 afum|EAL92704.1 agla|OJJ79740.1 akaw|GAA84834.1 aluc|OJZ86840.1 anid|CBF83582.1 anig|EHA21797.1 aory|EIT82344.1 asyd|OJJ60781.1 ater|EAU38353.1 atub|OJI83678.1 aver|OJJ00114.1 awen|OJJ33932.1 azon|OJJ48640.1

Group2679: abra|OJJ72087.1 acar|OOF96731.1 acla|EAW09468.1 afis|EAW21668.1 afla|EED57932.1 afum|EAL92703.1 agla|OJJ79739.1 akaw|GAA84833.1 aluc|OJZ86841.1 anid|CBF83580.1 anig|EHA21796.1 aory|EIT82317.1 asyd|OJJ60749.1 ater|EAU38352.1 atub|OJI83677.1 aver|OJJ00081.1 awen|OJJ33931.1 azon|OJJ48639.1

Group2680: abra|OJJ72089.1 acar|OOF96735.1 acla|EAW09470.1 afis|EAW21666.1 afla|EED57930.1 afum|EAL92701.1 agla|OJJ79737.1 akaw|GAA84831.1 aluc|OJZ86843.1 anid|CBF83569.1 anig|EHA21794.1 aory|EIT82323.1 asyd|OJJ60747.1 ater|EAU38351.1 atub|OJI83675.1 aver|OJJ00079.1 awen|OJJ33930.1 azon|OJJ48637.1

Group2681: abra|OJJ72090.1 acar|OOF96734.1 acla|EAW09471.1 afis|EAW21665.1 afla|EED57929.1 afum|EAL92700.1 agla|OJJ79736.1 akaw|GAA84830.1 aluc|OJZ86844.1 anid|CBF83568.1 anig|EHA21793.1 aory|EIT82319.1 asyd|OJJ60746.1 ater|EAU38350.1 atub|OJI83674.1 aver|OJJ00078.1 awen|OJJ33929.1 azon|OJJ48636.1

Group2682: abra|OJJ72092.1 acar|OOF96737.1 acla|EAW09473.1 afis|EAW21662.1 afla|EED57927.1 afum|EAL92697.1 agla|OJJ79734.1 akaw|GAA84828.1 aluc|OJZ86847.1 anid|CBF83564.1 anig|EHA21791.1 aory|EIT82336.1 asyd|OJJ60744.1 ater|EAU38348.1 atub|OJI83672.1 aver|OJJ00076.1 awen|OJJ33926.1 azon|OJJ48634.1

Group2683: abra|OJJ72094.1 acar|OOF96739.1 acla|EAW09475.1 afis|EAW21664.1 afla|EED57925.1 afum|EAL92699.1 agla|OJJ79732.1 akaw|GAA84826.1 aluc|OJZ86849.1 anid|CBF83562.1 anig|EHA21789.1 aory|EIT82328.1 asyd|OJJ60742.1 ater|EAU38347.1 atub|OJI83670.1 aver|OJJ00074.1 awen|OJJ33924.1 azon|OJJ48632.1

Group2684: abra|OJJ72095.1 acar|OOF96740.1 acla|EAW09476.1 afis|EAW21660.1 afla|EED57924.1 afum|EAL92695.1 agla|OJJ79731.1 akaw|GAA84825.1 aluc|OJZ86850.1 anid|CBF83560.1 anig|EHA21788.1 aory|EIT82356.1 asyd|OJJ60741.1 ater|EAU38346.1 atub|OJI83669.1 aver|OJJ00073.1 awen|OJJ33923.1 azon|OJJ48631.1

Group2685: abra|OJJ72096.1 acar|OOF96741.1 acla|EAW09391.1 afis|EAW21742.1 afla|EED58036.1 afum|EAL92775.1 agla|OJJ84431.1 akaw|GAA84824.1 aluc|OJZ86851.1 anid|CBF83748.1 anig|EHA21787.1 aory|EIT79774.1 asyd|OJJ60855.1 ater|EAU38446.1 atub|OJI83668.1 aver|OJJ00179.1 awen|OJJ34024.1 azon|OJJ48534.1

Group2686: abra|OJJ72099.1 acar|OOF96744.1 acla|EAW09390.1 afis|EAW21743.1 afla|EED58034.1 afum|EAL92776.1 agla|OJJ84427.1 akaw|GAA84822.1 aluc|OJZ86854.1 anid|CBF83738.1 anig|EHA21785.1 aory|EIT79762.1 asyd|OJJ60853.1 ater|EAU38445.1 atub|OJI83665.1 aver|OJJ00177.1 awen|OJJ34026.1 azon|OJJ48530.1

Group2687: abra|OJJ72101.1 acar|OOF96746.1 acla|EAW09388.1 afis|EAW21746.1 afla|EED58032.1 afum|EAL92778.1 agla|OJJ84430.1 akaw|GAA84820.1 aluc|OJZ86856.1 anid|CBF83662.1 anig|EHA21783.1 aory|EIT79789.1 asyd|OJJ60804.1 ater|EAU38443.1 atub|OJI83663.1 aver|OJJ00133.1 awen|OJJ34028.1 azon|OJJ48201.1

Group2688: abra|OJJ72105.1 acar|OOF96748.1 acla|EAW09386.1 afis|EAW21749.1 afla|EED58029.1 afum|EAL92780.2 agla|OJJ84425.1 akaw|GAA84817.1 aluc|OJZ86860.1 anid|CBF83665.1 anig|EHA21780.1 aory|EIT79704.1 asyd|OJJ60806.1 ater|EAU38441.1 atub|OJI83660.1 aver|OJJ00135.1 awen|OJJ34030.1 azon|OJJ48203.1

Group2689: abra|OJJ72110.1 acar|OOF90022.1 acla|EAW14910.1 afis|EAW24431.1 afla|EED55244.1 afum|EAL86099.1 agla|OJJ79245.1 akaw|GAA84812.1 aluc|OJZ86865.1 anid|CBF73773.1 anig|EHA21776.1 aory|EIT83529.1 asyd|OJJ56593.1 ater|EAU30014.1 atub|OJI83655.1 aver|OJJ06132.1 awen|OJJ32771.1 azon|OJJ44001.1

Group2690: abra|OJJ72112.1 acar|OOF96752.1 acla|EAW14912.1 afis|EAW24433.1 afla|EED55247.1 afum|EAL86101.1 agla|OJJ79248.1 akaw|GAA84810.1 aluc|OJZ86867.1 anid|CBF73767.1 anig|EHA21774.1

aory|EIT83512.1 asyd|OJJ56591.1 ater|EAU30012.1 atub|OJI83653.1 aver|OJJ06134.1 awen|OJJ32768.1 azon|OJJ43999.1

Group2691: abra|OJJ72115.1 acar|OOF96755.1 acla|EAW14916.1 afis|EAW24438.1 afla|EED55241.1 afum|EAL86106.1 agla|OJJ78777.1 akaw|GAA84807.1 aluc|OJZ86870.1 anid|CBF73762.1 anig|EHA21771.1 aory|EIT83515.1 asyd|OJJ56588.1 ater|EAU30009.1 atub|OJI83649.1 aver|OJJ06137.1 awen|OJJ32761.1 azon|OJJ43996.1

Group2692: abra|OJJ72116.1 acar|OOF96756.1 acla|EAW14917.1 afis|EAW24439.1 afla|EED55239.1 afum|EAL86107.1 agla|OJJ78778.1 akaw|GAA84806.1 aluc|OJZ86871.1 anid|CBF73760.1 anig|EHA21770.1 aory|EIT83598.1 asyd|OJJ56587.1 ater|EAU30008.1 atub|OJI83648.1 aver|OJJ06138.1 awen|OJJ32760.1 azon|OJJ43995.1

Group2693: abra|OJJ72117.1 acar|OOF96758.1 acla|EAW14918.1 afis|EAW24440.1 afla|EED55238.1 afum|EAL86108.1 agla|OJJ78779.1 akaw|GAA84805.1 aluc|OJZ86872.1 anid|CBF73758.1 anig|EHA21769.1 aory|EIT83628.1 asyd|OJJ56586.1 ater|EAU30007.1 atub|OJI83647.1 aver|OJJ06139.1 awen|OJJ32759.1 azon|OJJ43994.1

Group2694: abra|OJJ72118.1 acar|OOF96759.1 acla|EAW14919.1 afis|EAW24441.1 afla|EED55237.1 afum|EAL86109.1 agla|OJJ78780.1 akaw|GAA84804.1 aluc|OJZ86873.1 anid|CBF73756.1 anig|EHA21768.1 aory|EIT83564.1 asyd|OJJ56585.1 ater|EAU30006.1 atub|OJI83645.1 aver|OJJ06140.1 awen|OJJ32758.1 azon|OJJ43993.1

Group2695: abra|OJJ72119.1 acar|OOF96760.1 acla|EAW14920.1 afis|EAW24442.1 afla|EED55236.1 afum|EAL86110.1 agla|OJJ78783.1 akaw|GAA84803.1 aluc|OJZ86875.1 anid|CBF73754.1 anig|EHA21767.1 aory|EIT83587.1 asyd|OJJ56584.1 ater|EAU30005.1 atub|OJI83643.1 aver|OJJ06141.1 awen|OJJ32757.1 azon|OJJ43991.1

Group2696: abra|OJJ72123.1 acar|OOF96763.1 acla|EAW14923.1 afis|EAW24445.1 afla|EED55232.1 afum|EAL86114.1 agla|OJJ80290.1 akaw|GAA84800.1 aluc|OJZ86880.1 anid|CBF73749.1 anig|EHA21764.1 aory|EIT83556.1 asyd|OJJ56580.1 ater|EAU30002.1 atub|OJI83639.1 aver|OJJ06144.1 awen|OJJ32754.1 azon|OJJ43989.1

Group2697: abra|OJJ72124.1 acar|OOF96764.1 acla|EAW14930.1 afis|EAW24446.1 afla|EED55231.1 afum|EAL86115.1 agla|OJJ80291.1 akaw|GAA84799.1 aluc|OJZ86881.1 anid|CBF73806.1 anig|EHA21763.1 aory|EIT83582.1 asyd|OJJ56579.1 ater|EAU30001.1 atub|OJI83638.1 aver|OJJ06145.1 awen|OJJ32753.1 azon|OJJ43992.1

Group2699: abra|OJJ72139.1 acar|OOF96777.1 acla|EAW14933.1 afis|EAW24448.1 afla|EED55225.1 afum|EAL86117.1 agla|OJJ80294.1 akaw|GAA84783.1 aluc|OJZ86897.1 anid|CBF73811.1 anig|EHA21746.1 aory|EIT83551.1 asyd|OJJ56576.1 ater|EAU29998.1 atub|OJI83622.1 aver|OJJ06147.1 awen|OJJ32749.1 azon|OJJ45300.1

Group2700: abra|OJJ72140.1 acar|OOF96778.1 acla|EAW14934.1 afis|EAW24449.1 afla|EED55224.1 afum|EAL86118.1 agla|OJJ80295.1 akaw|GAA84784.1 aluc|OJZ86896.1 anid|CBF73813.1 anig|EHA21745.1 aory|EIT83509.1 asyd|OJJ56575.1 ater|EAU29997.1 atub|OJI83623.1 aver|OJJ06148.1 awen|OJJ32748.1 azon|OJJ45301.1

Group2701: abra|OJJ72141.1 acar|OOF96779.1 acla|EAW14935.1 afis|EAW24450.1 afla|EED55223.1 afum|EAL86119.1 agla|OJJ80296.1 akaw|GAA84782.1 aluc|OJZ86898.1 anid|CBF73815.1 anig|EHA21744.1 aory|EIT83569.1 asyd|OJJ56574.1 ater|EAU29996.1 atub|OJI83621.1 aver|OJJ06149.1 awen|OJJ32747.1 azon|OJJ45299.1

Group2702: abra|OJJ72145.1 acar|OOF96782.1 acla|EAW14939.1 afis|EAW24454.1 afla|EED55220.1 afum|EAL86123.1 agla|OJJ80301.1 akaw|GAA84779.1 aluc|OJZ86902.1 anid|CBF73822.1 anig|EHA21741.1 aory|EIT83594.1 asyd|OJJ56567.1 ater|EAU29991.1 atub|OJI83618.1 aver|OJJ06154.1 awen|OJJ32743.1 azon|OJJ45293.1

Group2703: abra|OJJ72146.1 acar|OOF96783.1 acla|EAW14941.1 afis|EAW24456.1 afla|EED55218.1 afum|EAL86126.1 agla|OJJ80303.1 akaw|GAA84778.1 aluc|OJZ86903.1 anid|CBF73824.1 anig|EHA21740.1 aory|EIT83584.1 asyd|OJJ56566.1 ater|EAU29988.1 atub|OJI83617.1 aver|OJJ06155.1 awen|OJJ32741.1 azon|OJJ45291.1

Group2704: abra|OJJ72147.1 acar|OOF96784.1 acla|EAW14942.1 afis|EAW24457.1 afla|EED55217.1 afum|EAL86127.1 agla|OJJ80304.1 akaw|GAA84777.1 aluc|OJZ86904.1 anid|CBF73826.1 anig|EHA21739.1 aory|EIT83603.1 asyd|OJJ56564.1 ater|EAU29985.1 atub|OJI83616.1 aver|OJJ06157.1 awen|OJJ32740.1 azon|OJJ45290.1

Group2705: abra|OJJ72148.1 acar|OOF96785.1 acla|EAW14943.1 afis|EAW24458.1 afla|EED55216.1 afum|EAL86128.1 agla|OJJ80305.1 akaw|GAA84776.1 aluc|OJZ86905.1 anid|CBF73828.1 anig|EHA21738.1 aory|EIT83601.1 asyd|OJJ56563.1 ater|EAU29984.1 atub|OJI83615.1 aver|OJJ06158.1 awen|OJJ32739.1 azon|OJJ45289.1

Group2706: abra|OJJ72149.1 acar|OOF96786.1 acla|EAW14944.1 afis|EAW24459.1 afla|EED55215.1 afum|EAL86129.1 agla|OJJ80306.1 akaw|GAA84775.1 aluc|OJZ86906.1 anid|CBF73830.1 anig|EHA21737.1 aory|EIT83574.1 asyd|OJJ56562.1 ater|EAU29983.1 atub|OJI83614.1 aver|OJJ06159.1 awen|OJJ32738.1 azon|OJJ45288.1

Group2707: abra|OJJ72150.1 acar|OOF96787.1 acla|EAW14945.1 afis|EAW24460.1 afla|EED55214.1 afum|EAL86130.1 agla|OJJ80308.1 akaw|GAA84774.1 aluc|OJZ86909.1 anid|CBF73832.1 anig|EHA21736.1 aory|EIT83521.1 asyd|OJJ56561.1 ater|EAU29982.1 atub|OJI83613.1 aver|OJJ06160.1 awen|OJJ32736.1 azon|OJJ45287.1

Group2708: abra|OJJ72209.1 acar|OOF96830.1 acla|EAW14967.1 afis|EAW24490.1 afla|EED54675.1 afum|EAL86159.1 agla|OJJ81567.1 akaw|GAA84725.1 aluc|OJZ86961.1 anid|CBF89709.1 anig|EHA21687.1 aory|EIT78852.1 asyd|OJJ62250.1 ater|EAU29966.1 atub|OJI83562.1 aver|OJI96360.1 awen|OJJ32706.1 azon|OJJ45248.1

Group2711: abra|OJJ72290.1 acar|OOF94933.1 acla|EAW12012.1 afis|EAW19652.1 afla|EED57003.1 afum|EAL86870.1 agla|OJJ80798.1 akaw|GAA86042.1 aluc|OJZ90271.1 anid|CBF71361.1 anig|EHA23763.1 aory|EIT77490.1 asyd|OJJ55084.1 ater|EAU32876.1 atub|OJI81695.1 aver|OJJ07123.1 awen|OJJ41903.1 azon|OJJ42697.1

Group2712: abra|OJJ72296.1 acar|OOF94928.1 acla|EAW12007.1 afis|EAW19647.1 afla|EED56998.1 afum|EAL86875.1 agla|OJJ80789.1 akaw|GAA86047.1 aluc|OJZ90276.1 anid|CBF71354.1 anig|EHA23768.1 aory|EIT77515.1 asyd|OJJ55089.1 ater|EAU32881.1 atub|OJI81700.1 aver|OJJ07118.1 awen|OJJ41898.1 azon|OJJ42702.1

Group2713: abra|OJJ72298.1 acar|OOF94926.1 acla|EAW12006.1 afis|EAW19646.1 afla|EED56997.1 afum|EAL86876.1 agla|OJJ80788.1 akaw|GAA86049.1 aluc|OJZ90278.1 anid|CBF71352.1 anig|EHA23769.1 aory|EIT77535.1 asyd|OJJ55091.1 ater|EAU32883.1 atub|OJI81703.1 aver|OJJ07117.1 awen|OJJ41897.1 azon|OJJ42703.1

Group2714: abra|OJJ72299.1 acar|OOF94924.1 acla|EAW12005.1 afis|EAW19645.1 afla|EED56996.1 afum|EAL86877.1 agla|OJJ80787.1 akaw|GAA86050.1 aluc|OJZ90280.1 anid|CBF71350.1 anig|EHA23770.1 aory|EIT77477.1 asyd|OJJ55092.1 ater|EAU32884.1 atub|OJI81704.1 aver|OJJ07116.1 awen|OJJ41896.1 azon|OJJ42704.1

Group2715: abra|OJJ72318.1 acar|OOF94893.1 acla|EAW12000.1 afis|EAW19640.1 afla|EED56991.1 afum|EAL86882.1 agla|OJJ80782.1 akaw|GAA86067.1 aluc|OJZ90301.1 anid|CBF71343.1 anig|EHA23789.1 aory|EIT77491.1 asyd|OJJ55098.1 ater|EAU32889.1 atub|OJI81727.1 aver|OJJ07108.1 awen|OJJ41885.1 azon|OJJ42710.1

Group2716: abra|OJJ72324.1 acar|OOF89799.1 acla|EAW11999.1 afis|EAW19639.1 afla|EED56990.1 afum|EAL86883.1 agla|OJJ80781.1 akaw|GAA86074.1 aluc|OJZ90309.1 anid|CBF71341.1 anig|EHA23796.1 aory|EIT77478.1 asyd|OJJ55099.1 ater|EAU32890.1 atub|OJI81734.1 aver|OJJ07107.1 awen|OJJ41884.1 azon|OJJ42711.1

Group2717: abra|OJJ72327.1 acar|OOF94887.1 acla|EAW11996.1 afis|EAW19636.1 afla|EED56988.1 afum|EAL86886.1 agla|OJJ80778.1 akaw|GAA86077.1 aluc|OJZ90312.1 anid|CBF71335.1 anig|EHA23799.1 aory|EIT77538.1 asyd|OJJ55103.1 ater|EAU32892.1 atub|OJI81737.1 aver|OJJ07104.1 awen|OJJ41881.1 azon|OJJ47079.1

Group2718: abra|OJJ72328.1 acar|OOF94886.1 acla|EAW11995.1 afis|EAW19635.1 afla|EED56986.1 afum|EAL86887.1 agla|OJJ80777.1 akaw|GAA86078.1 aluc|OJZ90313.1 anid|CBF71333.1 anig|EHA23800.1 aory|EIT77532.1 asyd|OJJ55104.1 atub|OJI81738.1 aver|OJJ07103.1 awen|OJJ41880.1 azon|OJJ47078.1 ater|EAU32893.1

Group2719: abra|OJJ72338.1 acar|OOF94878.1 acla|EAW11985.1 afis|EAW19625.1 afla|EED56977.1 afum|EBA27293.1 agla|OJJ80768.1 akaw|GAA86089.1 aluc|OJZ90318.1 anid|CBF71315.1 anig|EHA23804.1 aory|EIT77446.1 asyd|OJJ55113.1 ater|EAU32903.1 atub|OJI81747.1 aver|OJJ07093.1 awen|OJJ41872.1 azon|OJJ47067.1

Group2720: abra|OJJ72339.1 acar|OOF94877.1 acla|EAW11984.1 afis|EAW19624.1 afla|EED56976.1 afum|EAL86898.1 agla|OJJ80767.1 akaw|GAA86090.1 aluc|OJZ90319.1 anid|CBF71313.1 anig|EHA23805.1 aory|EIT77523.1 asyd|OJJ55114.1 ater|EAU32902.1 atub|OJI81748.1 aver|OJJ07092.1 awen|OJJ41871.1 azon|OJJ47066.1

Group2721: abra|OJJ72343.1 acar|OOF94874.1 acla|EAW11983.1 afis|EAW19621.1 afla|EED56973.1 afum|EAL86900.1 agla|OJJ80766.1 akaw|GAA86092.1 aluc|OJZ90322.1 anid|CBF71293.1 anig|EHA23808.1 aory|EIT77575.1 asyd|OJJ55117.1 ater|EAU32905.1 atub|OJI81749.1 aver|OJJ07089.1 awen|OJJ41868.1 azon|OJJ47061.1

Group2722: abra|OJJ72344.1 acar|OOF94873.1 acla|EAW11982.1 afis|EAW19620.1 afla|EED56972.1 afum|EAL86902.1 agla|OJJ80765.1 akaw|GAA86093.1 aluc|OJZ90323.1 anid|CBF71296.1 anig|EHA23809.1

aory|EIT77533.1 asyd|OJJ55118.1 ater|EAU32906.1 atub|OJI81750.1 aver|OJJ07088.1 awen|OJJ41867.1 azon|OJJ47060.1

Group2723: abra|OJJ72345.1 acar|OOF94872.1 acla|EAW11981.1 afis|EAW19619.1 afla|EED56967.1 afum|EAL86903.1 agla|OJJ80764.1 akaw|GAA86094.1 aluc|OJZ90324.1 anid|CBF71298.1 anig|EHA23810.1 aory|EIT77589.1 asyd|OJJ55119.1 ater|EAU32907.1 atub|OJI81751.1 aver|OJJ07087.1 awen|OJJ41866.1 azon|OJJ47059.1

Group2724: abra|OJJ72346.1 acar|OOF94871.1 acla|EAW11980.1 afis|EAW19618.1 afla|EED56968.1 afum|EAL86904.1 agla|OJJ80763.1 akaw|GAA86095.1 aluc|OJZ90325.1 anid|CBF71300.1 anig|EHA23811.1 aory|EIT77479.1 asyd|OJJ55120.1 ater|EAU32908.1 atub|OJI81752.1 aver|OJJ07086.1 awen|OJJ41865.1 azon|OJJ47058.1

Group2725: abra|OJJ72349.1 acar|OOF94869.1 acla|EAW11977.1 afis|EAW19609.1 afla|EED56969.1 afum|EAL86913.1 agla|OJJ80761.1 akaw|GAA86096.1 aluc|OJZ90328.1 anid|CBF71302.1 anig|EHA23812.1 aory|EIT77540.1 asyd|OJJ55121.1 ater|EAU32910.1 atub|OJI81755.1 aver|OJJ07083.1 awen|OJJ41862.1 azon|OJJ47055.1

Group2726: abra|OJJ72351.1 acar|OOF94867.1 acla|EAW11975.1 afis|EAW19611.1 afla|EED56966.1 afum|EAL86911.1 agla|OJJ80759.1 akaw|GAA86098.1 aluc|OJZ90330.1 anid|CBF71306.1 anig|EHA23814.1 aory|EIT77494.1 asyd|OJJ55123.1 ater|EAU32921.1 atub|OJI81757.1 aver|OJJ07080.1 awen|OJJ41859.1 azon|OJJ47053.1

Group2727: abra|OJJ72352.1 acar|OOF94866.1 acla|EAW11973.1 afis|EAW19613.1 afla|EED56961.1 afum|EAL86909.1 agla|OJJ80758.1 akaw|GAA86099.1 aluc|OJZ90331.1 anid|CBF71308.1 anig|EHA23815.1 aory|EIT77512.1 asyd|OJJ55126.1 ater|EAU32914.1 atub|OJI81758.1 aver|OJJ07078.1 awen|OJJ41857.1 azon|OJJ47051.1

Group2728: abra|OJJ72353.1 acar|OOF94865.1 acla|EAW11972.1 afis|EAW19614.1 afla|EED56962.1 afum|EAL86908.1 agla|OJJ80757.1 akaw|GAA86100.1 aluc|OJZ90332.1 anid|CBF71310.1 anig|EHA23816.1 aory|EIT77559.1 asyd|OJJ55127.1 ater|EAU32915.1 atub|OJI81759.1 aver|OJJ07077.1 awen|OJJ41856.1 azon|OJJ47050.1

Group2729: abra|OJJ72355.1 acar|OOF94863.1 acla|EAW11961.1 afis|EAW19598.1 afla|EED56954.1 afum|EAL86920.1 agla|OJJ80740.1 akaw|GAA86102.1 aluc|OJZ90334.1 anid|CBF71287.1 anig|EHA23818.1 aory|EIT77544.1 asyd|OJJ55129.1 ater|EAU32927.1 atub|OJI81761.1 aver|OJJ07074.1 awen|OJJ41844.1 azon|OJJ44888.1

Group2730: abra|OJJ72368.1 acar|OOF94849.1 acla|EAW11964.1 afis|EAW19601.1 afla|EED56957.1 afum|EAL86917.1 agla|OJJ80743.1 akaw|GAA86118.1 aluc|OJZ90356.1 anid|CBF71281.1 anig|EHA23834.1 aory|EIT77555.1 asyd|OJJ55133.1 ater|EAU32924.1 atub|OJI81783.1 aver|OJJ07071.1 awen|OJJ41839.1 azon|OJJ47044.1

Group2732: abra|OJJ72373.1 acar|OOF94855.1 acla|EAW11967.1 afis|EAW19617.1 afla|EED56965.1 afum|EAL86905.1 agla|OJJ80746.1 akaw|GAA86123.1 aluc|OJZ90362.1 anid|CBF71277.1 anig|EHA23829.1 aory|EIT77547.1 asyd|OJJ55139.1 ater|EAU32917.1 atub|OJI81789.1 aver|OJJ07066.1 awen|OJJ41850.1 azon|OJJ47046.1

Group2733: abra|OJJ72377.1 acar|OOF94841.1 acla|EAW11958.1 afis|EAW19595.1 afla|EED56951.1 afum|EAL86923.1 agla|OJJ80736.1 akaw|GAA86125.1 aluc|OJZ90364.1 anid|CBF71262.1 anig|EHA23839.1 aory|EIT77501.1 asyd|OJJ55150.1 ater|EAU32929.1 atub|OJI81791.1 aver|OJJ07057.1 awen|OJJ41849.1 azon|OJJ47020.1

Group2734: abra|OJJ72379.1 acar|OOF94840.1 acla|EAW11957.1 afis|EAW19594.1 afla|EED56950.1 afum|EAL86924.1 agla|OJJ80735.1 akaw|GAA86127.1 aluc|OJZ90366.1 anid|CBF71275.1 anig|EHA23841.1 aory|EIT77449.1 asyd|OJJ55149.1 ater|EAU32930.1 atub|OJI81793.1 aver|OJJ07058.1 awen|OJJ41811.1 azon|OJJ47021.1

Group2735: abra|OJJ72381.1 acar|OOF94838.1 acla|EAW11955.1 afis|EAW19592.1 afla|EED56948.1 afum|EAL86926.1 agla|OJJ80733.1 akaw|GAA86128.1 aluc|OJZ90368.1 anid|CBF71268.1 anig|EHA23842.1 aory|EIT77465.1 asyd|OJJ55147.1 ater|EAU32933.1 atub|OJI81794.1 aver|OJJ07060.1 awen|OJJ41809.1 azon|OJJ47023.1

Group2736: abra|OJJ72388.1 acar|OOF94835.1 acla|EAW11949.1 afis|EAW19586.1 afla|EED56914.1 afum|EAL86932.1 agla|OJJ80723.1 akaw|GAA86136.1 aluc|OJZ90374.1 anid|CBF71254.1 anig|EHA23848.1 aory|EIT77498.1 asyd|OJJ55153.1 ater|EAU32962.1 atub|OJI81800.1 aver|OJJ07054.1 awen|OJJ41800.1 azon|OJJ47031.1

Group2737: abra|OJJ72389.1 acar|OOF94834.1 acla|EAW11948.1 afis|EAW19585.1 afla|EED56913.1 afum|EAL86933.1 agla|OJJ80722.1 akaw|GAA86137.1 aluc|OJZ90376.1 anid|CBF71253.1 anig|EHA23849.1 aory|EIT77525.1 asyd|OJJ55155.1 ater|EAU32963.1 atub|OJI81801.1 aver|OJJ07053.1 awen|OJJ41799.1 azon|OJJ47032.1

Group2738: abra|OJJ72391.1 acar|OOF94833.1 acla|EAW11947.1 afis|EAW19584.1 afla|EED56912.1 afum|EAL86934.1 agla|OJJ80712.1 akaw|GAA86138.1 aluc|OJZ90378.1 anid|CBF71251.1 anig|EHA23850.1 aory|EIT77463.1 asyd|OJJ55156.1 ater|EAU32967.1 atub|OJI81803.1 aver|OJJ07052.1 awen|OJJ41798.1 azon|OJJ47033.1

Group2739: abra|OJJ72394.1 acar|OOF94832.1 acla|EAW11946.1 afis|EAW19583.1 afla|EED56911.1 afum|EAL86935.2 agla|OJJ80711.1 akaw|GAA86139.1 aluc|OJZ90379.1 anid|CBF71249.1 anig|EHA23851.1 aory|EIT77567.1 asyd|OJJ55157.1 ater|EAU32968.1 atub|OJI81804.1 aver|OJJ07051.1 awen|OJJ41797.1 azon|OJJ47034.1

Group2740: abra|OJJ72395.1 acar|OOF94831.1 acla|EAW11945.1 afis|EAW19581.1 afla|EED56910.1 afum|EAL86937.1 agla|OJJ80710.1 akaw|GAA86140.1 aluc|OJZ90380.1 anid|CBF71245.1 anig|EHA23852.1 aory|EIT77560.1 asyd|OJJ55159.1 ater|EAU32972.1 atub|OJI81806.1 aver|OJJ07049.1 awen|OJJ41795.1 azon|OJJ47036.1

Group2741: abra|OJJ72396.1 acar|OOF94830.1 acla|EAW11944.1 afis|EAW19580.1 afla|EED56909.1 afum|EAL86938.1 agla|OJJ80709.1 akaw|GAA86141.1 aluc|OJZ90381.1 anid|CBF71243.1 anig|EHA23853.1 aory|EIT77483.1 asyd|OJJ55160.1 ater|EAU32973.1 atub|OJI81807.1 aver|OJJ07047.1 awen|OJJ41794.1 azon|OJJ47037.1

Group2742: abra|OJJ72397.1 acar|OOF94829.1 acla|EAW11943.1 afis|EAW19579.1 afla|EED56804.1 afum|EAL86939.1 agla|OJJ80708.1 akaw|GAA86142.1 aluc|OJZ90382.1 anid|CBF71239.1 anig|EHA23854.1 aory|EIT73874.1 asyd|OJJ55163.1 ater|EAU32976.1 atub|OJI81808.1 aver|OJJ07044.1 awen|OJJ41792.1 azon|OJJ47041.1

Group2743: abra|OJJ72399.1 acar|OOF94828.1 acla|EAW11941.1 afis|EAW19577.1 afla|EED56805.1 afum|EAL86941.2 agla|OJJ80706.1 akaw|GAA86144.1 aluc|OJZ90384.1 anid|CBF71236.1 anig|EHA23856.1 aory|EIT73858.1 asyd|OJJ55165.1 ater|EAU33003.1 atub|OJI81811.1 aver|OJJ07042.1 awen|OJJ41790.1 azon|OJJ47039.1

Group2744: abra|OJJ72401.1 acar|OOF94827.1 acla|EAW11940.1 afis|EAW19575.1 afla|EED56808.1 afum|EAL86942.1 agla|OJJ80704.1 akaw|GAA86145.1 aluc|OJZ90386.1 anid|CBF71234.1 anig|EHA23857.1 aory|EIT73879.1 asyd|OJJ55166.1 ater|EAU33002.1 atub|OJI81814.1 aver|OJJ07040.1 awen|OJJ41788.1 azon|OJJ44890.1

Group2745: abra|OJJ72402.1 acar|OOF94826.1 acla|EAW11939.1 afis|EAW19574.1 afla|EED56809.1 afum|EAL86943.1 agla|OJJ80702.1 akaw|GAA86146.1 aluc|OJZ90387.1 anid|CBF71232.1 anig|EHA23858.1 aory|EIT73889.1 asyd|OJJ55167.1 ater|EAU33001.1 atub|OJI81815.1 aver|OJJ07039.1 awen|OJJ41787.1 azon|OJJ44891.1

Group2746: abra|OJJ72404.1 acar|OOF94824.1 acla|EAW11937.1 afis|EAW19572.1 afla|EED56811.1 afum|EAL86945.1 agla|OJJ80700.1 akaw|GAA86148.1 aluc|OJZ90389.1 anid|CBF71228.1 anig|EHA23859.1 aory|EIT73949.1 asyd|OJJ55170.1 ater|EAU32995.1 atub|OJI81818.1 aver|OJJ07037.1 awen|OJJ41785.1 azon|OJJ44893.1

Group2747: abra|OJJ72405.1 acar|OOF94823.1 acla|EAW11936.1 afis|EAW19571.1 afla|EED56812.1 afum|EAL86946.1 agla|OJJ80699.1 akaw|GAA86149.1 aluc|OJZ90390.1 anid|CBF71226.1 anig|EHA23860.1 aory|EIT73876.1 asyd|OJJ55171.1 ater|EAU32994.1 atub|OJI81819.1 aver|OJJ07036.1 awen|OJJ41784.1 azon|OJJ44894.1

Group2748: abra|OJJ72408.1 acar|OOF94821.1 acla|EAW11933.1 afis|EAW19569.1 afla|EED56814.1 afum|EAL86947.1 agla|OJJ80697.1 akaw|GAA86151.1 aluc|OJZ90392.1 anid|CBF78033.1 anig|EHA23862.1 aory|EIT73916.1 asyd|OJJ55227.1 ater|EAU32992.1 atub|OJI81821.1 aver|OJJ06994.1 awen|OJJ41782.1 azon|OJJ44896.1

Group2749: abra|OJJ72412.1 acar|OOF94817.1 acla|EAW11929.1 afis|EAW19563.1 afla|EED56817.1 afum|EAL86949.1 agla|OJJ80692.1 akaw|GAA86155.1 aluc|OJZ90396.1 anid|CBF78024.1 anig|EHA23866.1 aory|EIT73850.1 asyd|OJJ55233.1 ater|EAU32986.1 atub|OJI81827.1 aver|OJJ06990.1 awen|OJJ41778.1 azon|OJJ44899.1

Group2750: abra|OJJ72413.1 acar|OOF94816.1 acla|EAW12857.1 afis|EAW17821.1 afla|EED56818.1 afum|EAL91631.1 agla|OJJ80690.1 akaw|GAA86156.1 aluc|OJZ90397.1 anid|CBF78026.1 anig|EHA23867.1 aory|EIT73888.1 asyd|OJJ55232.1 ater|EAU32985.1 atub|OJI81828.1 aver|OJJ06991.1 awen|OJJ41777.1 azon|OJJ44900.1

Group2751: abra|OJJ72414.1 acar|OOF94815.1 acla|EAW12858.1 afis|EAW17820.1 afla|EED56819.1 afum|EAL91630.1 agla|OJJ80689.1 akaw|GAA86157.1 aluc|OJZ90398.1 anid|CBF78027.1 anig|EHA23868.1 aory|EIT73947.1 asyd|OJJ55231.1 ater|EAU32984.1 atub|OJI81829.1 aver|OJJ06992.1 awen|OJJ41776.1 azon|OJJ44901.1

Group2752: abra|OJJ72415.1 acar|OOF94814.1 acla|EAW12859.1 afis|EAW17808.1 afla|EED56820.1 afum|EAL91616.1 agla|OJJ80688.1 akaw|GAA86158.1 aluc|OJZ90399.1 anid|CBF78029.1 anig|EHA23869.1

aory|EIT73877.1 asyd|OJJ55230.1 ater|EAU32982.1 atub|OJI81830.1 aver|OJJ06993.1 awen|OJJ41775.1 azon|OJJ44902.1

Group2754: abra|OJJ72421.1 acar|OOF90370.1 acla|EAW12862.1 afis|EAW17811.1 afla|EED46162.1 afum|EAL91619.1 agla|OJJ80655.1 akaw|GAA86163.1 aluc|OJZ90404.1 anid|CBF78013.1 anig|EHA23872.1 aory|EIT76383.1 asyd|OJJ55240.1 ater|EAU33017.1 atub|OJI81835.1 aver|OJJ06983.1 awen|OJJ41761.1 azon|OJJ44878.1

Group2755: abra|OJJ72422.1 acar|OOF90371.1 acla|EAW12861.1 afis|EAW17810.1 afla|EED46161.1 afum|EAL91618.1 agla|OJJ80654.1 akaw|GAA86164.1 aluc|OJZ90405.1 anid|CBF78014.1 anig|EHA23873.1 aory|EIT76437.1 asyd|OJJ55239.1 ater|EAU33016.1 atub|OJI81836.1 aver|OJJ06984.1 awen|OJJ41762.1 azon|OJJ44879.1

Group2757: abra|OJJ72425.1 acar|OOF94810.1 acla|EAW12659.1 afis|EAW17823.1 afla|EED46166.1 afum|EAL91633.1 agla|OJJ80641.1 akaw|GAA86167.1 aluc|OJZ90408.1 anid|CBF78009.1 anig|EHA23876.1 aory|EIT76477.1 asyd|OJJ55242.1 ater|EAU33024.1 atub|OJI81840.1 aver|OJJ06982.1 awen|OJJ41751.1 azon|OJJ48443.1

Group2758: abra|OJJ72428.1 acar|OOF94804.1 acla|EAW12661.1 afis|EAW17825.1 afla|EED46163.1 afum|EAL91635.1 agla|OJJ80646.1 akaw|GAA86169.1 aluc|OJZ90412.1 anid|CBF78005.1 anig|EHA23878.1 aory|EIT76524.1 asyd|OJJ55245.1 ater|EAU33026.1 atub|OJI81843.1 aver|OJJ06979.1 awen|OJJ41755.1 azon|OJJ48445.1

Group2759: abra|OJJ72429.1 acar|OOF94805.1 acla|EAW12662.1 afis|EAW17826.1 afla|EED46168.1 afum|EAL91636.1 agla|OJJ80647.1 akaw|GAA86170.1 aluc|OJZ90413.1 anid|CBF78003.1 anig|EHA23879.1 aory|EIT76408.1 asyd|OJJ55246.1 ater|EAU33027.1 atub|OJI81845.1 aver|OJJ06978.1 awen|OJJ41756.1 azon|OJJ48446.1

Group2760: abra|OJJ72430.1 acar|OOF94803.1 acla|EAW12663.1 afis|EAW17827.1 afla|EED46169.1 afum|EAL91637.1 agla|OJJ80648.1 akaw|GAA86171.1 aluc|OJZ90414.1 anid|CBF78001.1 anig|EHA23880.1 aory|EIT76516.1 asyd|OJJ55247.1 ater|EAU33031.1 atub|OJI81846.1 aver|OJJ06977.1 awen|OJJ41757.1 azon|OJJ48447.1

Group2761: abra|OJJ72439.1 acar|OOF94798.1 acla|EAW12672.1 afis|EAW17835.1 afla|EED46180.1 afum|EAL91645.1 agla|OJJ83131.1 akaw|GAA86178.1 aluc|OJZ90423.1 anid|CBF77987.1 anig|EHA23892.1 aory|EIT76377.1 asyd|OJJ55256.1 ater|EAU33039.1 atub|OJI81855.1 aver|OJJ06968.1 awen|OJJ41741.1 azon|OJJ48437.1

Group2762: abra|OJJ72440.1 acar|OOF94797.1 acla|EAW12673.1 afis|EAW17836.1 afla|EED46181.1 afum|EAL91646.1 agla|OJJ83130.1 akaw|GAA86179.1 aluc|OJZ90424.1 anid|CBF77985.1 anig|EHA23893.1 aory|EIT76515.1 asyd|OJJ55257.1 ater|EAU33040.1 atub|OJI81856.1 aver|OJJ06967.1 awen|OJJ41740.1 azon|OJJ48436.1

Group2763: abra|OJJ72443.1 acar|OOF94793.1 acla|EAW12676.1 afis|EAW17839.1 afla|EED46184.1 afum|EAL91649.1 agla|OJJ83128.1 akaw|GAA86182.1 aluc|OJZ90427.1 anid|CBF77979.1 anig|EHA23897.1 aory|EIT76493.1 asyd|OJJ55260.1 ater|EAU33042.1 atub|OJI81859.1 aver|OJJ06965.1 awen|OJJ41737.1 azon|OJJ48434.1

Group2764: abra|OJJ72445.1 acar|OOF94791.1 acla|EAW12678.1 afis|EAW17841.1 afla|EED46186.1 afum|EAL91650.1 agla|OJJ83126.1 akaw|GAA86183.1 aluc|OJZ90429.1 anid|CBF77975.1 anig|EHA23898.1 aory|EIT76513.1 asyd|OJJ55262.1 ater|EAU33043.1 atub|OJI81861.1 aver|OJJ06963.1 awen|OJJ41735.1 azon|OJJ48432.1

Group2765: abra|OJJ72446.1 acar|OOF94790.1 acla|EAW12679.1 afis|EAW17842.1 afla|EED46187.1 afum|EAL91651.1 agla|OJJ83125.1 akaw|GAA86184.1 aluc|OJZ90430.1 anid|CBF77973.1 anig|EHA23899.1 aory|EIT76407.1 asyd|OJJ55263.1 ater|EAU33044.1 atub|OJI81862.1 aver|OJJ06962.1 awen|OJJ41734.1 azon|OJJ48431.1

Group2766: abra|OJJ72450.1 acar|OOF94786.1 acla|EAW12684.1 afis|EAW17847.1 afla|EED46201.1 afum|EAL91656.1 agla|OJJ83122.1 akaw|GAA86187.1 aluc|OJZ90435.1 anid|CBF77964.1 anig|EHA23902.1 aory|EIT76452.1 asyd|OJJ55269.1 ater|EAU33049.1 atub|OJI81866.1 aver|OJJ06958.1 awen|OJJ41726.1 azon|OJJ48425.1

Group2767: abra|OJJ72458.1 acar|OOF94779.1 acla|EAW12690.1 afis|EAW17854.1 afla|EED46208.1 afum|EAL91662.1 agla|OJJ83116.1 akaw|GAA86194.1 aluc|OJZ90445.1 anid|CBF77955.1 anig|EHA23908.1 aory|EIT76378.1 asyd|OJJ55275.1 ater|EAU33056.1 atub|OJI81873.1 aver|OJJ06952.1 awen|OJJ41719.1 azon|OJJ48416.1

Group2768: abra|OJJ72459.1 acar|OOF94780.1 acla|EAW12691.1 afis|EAW17855.1 afla|EED46209.1 afum|EAL91663.1 agla|OJJ83115.1 akaw|GAA86195.1 aluc|OJZ90446.1 anid|CBF77953.1 anig|EHA23909.1 aory|EIT76395.1 asyd|OJJ55276.1 ater|EAU33057.1 atub|OJI81874.1 aver|OJJ06951.1 awen|OJJ41718.1 azon|OJJ48415.1

Group2769: abra|OJJ72460.1 acar|OOF94778.1 acla|EAW12692.1 afis|EAW17856.1 afla|EED46210.1 afum|EAL91664.1 agla|OJJ83113.1 akaw|GAA86196.1 aluc|OJZ90447.1 anid|CBF77951.1 anig|EHA23910.1 aory|EIT76442.1 asyd|OJJ55277.1 ater|EAU33058.1 atub|OJI81875.1 aver|OJJ06950.1 awen|OJJ41716.1 azon|OJJ48411.1

Group2770: abra|OJJ72461.1 acar|OOF94777.1 acla|EAW12693.1 afis|EAW17857.1 afla|EED46212.1 afum|EAL91665.1 agla|OJJ83112.1 akaw|GAA86197.1 aluc|OJZ90448.1 anid|CBF77949.1 anig|EHA23911.1 aory|EIT76364.1 asyd|OJJ55278.1 ater|EAU33059.1 atub|OJI81876.1 aver|OJJ06949.1 awen|OJJ41715.1 azon|OJJ48410.1

Group2771: abra|OJJ72462.1 acar|OOF94775.1 acla|EAW12694.1 afis|EAW17858.1 afla|EED46213.1 afum|EAL91666.1 agla|OJJ83111.1 akaw|GAA86198.1 aluc|OJZ90449.1 anid|CBF77947.1 anig|EHA23912.1 aory|EIT76487.1 asyd|OJJ55279.1 ater|EAU33064.1 atub|OJI81877.1 aver|OJJ06948.1 awen|OJJ41714.1 azon|OJJ48409.1

Group2772: abra|OJJ72467.1 acar|OOF94773.1 acla|EAW12699.1 afis|EAW17862.1 afla|EED46225.1 afum|EAL91670.2 agla|OJJ83101.1 akaw|GAA86203.1 aluc|OJZ90454.1 anid|CBF88078.1 anig|EHA23916.1 aory|EIT76473.1 asyd|OJJ57087.1 ater|EAU33070.1 atub|OJI81881.1 aver|OJJ03214.1 awen|OJJ41709.1 azon|OJJ48404.1

Group2773: abra|OJJ72468.1 acar|OOF94772.1 acla|EAW12700.1 afis|EAW17863.1 afla|EED46226.1 afum|EAL91671.1 agla|OJJ83100.1 akaw|GAA86204.1 aluc|OJZ90455.1 anid|CBF84085.1 anig|EHA23917.1 aory|EIT76453.1 asyd|OJJ57088.1 ater|EAU33078.1 atub|OJI81882.1 aver|OJJ03213.1 awen|OJJ41707.1 azon|OJJ48403.1

Group2774: abra|OJJ72471.1 acar|OOF94764.1 acla|EAW12704.1 afis|EAW17867.1 afla|EED46236.1 afum|EAL91675.1 agla|OJJ83087.1 akaw|GAA86209.1 aluc|OJZ90458.1 anid|CBF89893.1 anig|EHA23920.1 aory|EIT76466.1 asyd|OJJ62139.1 ater|EAU33082.1 atub|OJI81885.1 aver|OJI96249.1 awen|OJJ41704.1 azon|OJJ48400.1

Group2775: abra|OJJ72476.1 acar|OOF94760.1 acla|EAW12707.1 afis|EAW17870.1 afla|EED46238.1 afum|EAL91678.1 agla|OJJ83090.1 akaw|GAA86212.1 aluc|OJZ90462.1 anid|CBF89899.1 anig|EHA23923.1 aory|EIT76429.1 asyd|OJJ62136.1 ater|EAU33088.1 atub|OJI81888.1 aver|OJI96246.1 awen|OJJ41700.1 azon|OJJ48397.1

Group2776: abra|OJJ72478.1 acar|OOF94756.1 acla|EAW12709.1 afis|EAW17872.1 afla|EED46239.1 afum|EAL91680.1 agla|OJJ83092.1 akaw|GAA86214.1 aluc|OJZ90465.1 anid|CBF89903.1 anig|EHA23925.1 aory|EIT76379.1 asyd|OJJ62134.1 ater|EAU33090.1 atub|OJI81890.1 aver|OJI96244.1 awen|OJJ41698.1 azon|OJJ48395.1

Group2778: abra|OJJ72488.1 acar|OOF94747.1 acla|EAW12719.1 afis|EAW17882.1 afla|EED46269.1 afum|EAL91690.1 agla|OJJ83065.1 akaw|GAA86223.1 aluc|OJZ90475.1 anig|EHA23934.1 aory|EIT76427.1 asyd|OJJ57008.1 ater|EAU33102.1 atub|OJI81900.1 aver|OJJ03279.1 awen|OJJ41684.1 azon|OJJ48392.1 anid|CBF77035.1

Group2779: abra|OJJ72489.1 acar|OOF92642.1 acla|EAW12720.1 afis|EAW17883.1 afla|EED46270.1 afum|EAL91691.1 agla|OJJ83064.1 akaw|GAA86224.1 aluc|OJZ90476.1 anid|CBF77033.1 anig|EHA23935.1 aory|EIT76482.1 asyd|OJJ57009.1 ater|EAU33103.1 atub|OJI81901.1 aver|OJJ03278.1 awen|OJJ41682.1 azon|OJJ48391.1

Group2780: abra|OJJ72493.1 acar|OOF94745.1 acla|EAW12722.1 afis|EAW17885.1 afla|EED46272.1 afum|EAL91693.1 agla|OJJ83062.1 akaw|GAA86227.1 aluc|OJZ90479.1 anid|CBF77029.1 anig|EHA23938.1 aory|EIT76321.1 asyd|OJJ57011.1 ater|EAU33105.1 atub|OJI81904.1 aver|OJJ03276.1 awen|OJJ41674.1 azon|OJJ48389.1

Group2781: abra|OJJ72494.1 acar|OOF94744.1 acla|EAW12723.1 afis|EAW17886.1 afla|EED46273.1 afum|EAL91694.1 agla|OJJ83061.1 akaw|GAA86228.1 aluc|OJZ90480.1 anid|CBF77027.1 anig|EHA23939.1 aory|EIT76334.1 asyd|OJJ57012.1 ater|EAU33106.1 atub|OJI81905.1 aver|OJJ03275.1 awen|OJJ41673.1 azon|OJJ48388.1

Group2782: abra|OJJ72497.1 acar|OOF94742.1 acla|EAW12725.1 afis|EAW17888.1 afla|EED46275.1 afum|EAL91697.1 agla|OJJ83059.1 akaw|GAA86229.1 aluc|OJZ90481.1 anid|CBF77026.1 anig|EHA23940.1 aory|EIT76333.1 asyd|OJJ57013.1 ater|EAU33107.1 atub|OJI81906.1 aver|OJJ03274.1 awen|OJJ41671.1 azon|OJJ48386.1

Group2783: abra|OJJ72503.1 acar|OOF94738.1 acla|EAW12727.1 afis|EAW17890.1 afla|EED46282.1 afum|EAL91699.1 agla|OJJ83055.1 akaw|GAA86231.1 aluc|OJZ90485.1 anid|CBF77016.1 anig|EHA23943.1 aory|EIT76352.1 asyd|OJJ57017.1 ater|EAU33110.1 atub|OJI81910.1 aver|OJJ03272.1 awen|OJJ41667.1 azon|OJJ48384.1

Group2784: abra|OJJ72507.1 acar|OOF94735.1 acla|EAW12730.1 afis|EAW17893.1 afla|EED46286.1 afum|EAL91702.1 agla|OJJ83049.1 akaw|GAA86235.1 aluc|OJZ90490.1 anid|CBF77006.1 anig|EHA23948.1

aory|EIT76341.1 asyd|OJJ57021.1 ater|EAU33114.1 atub|OJI81914.1 aver|OJJ03268.1 awen|OJJ41659.1 azon|OJJ48379.1

Group2785: abra|OJJ72509.1 acar|OOF94733.1 acla|EAW12732.1 afis|EAW17895.1 afla|EED46267.1 afum|EAL91703.1 agla|OJJ83047.1 akaw|GAA86237.1 aluc|OJZ90492.1 anid|CBF77004.1 anig|EHA23950.1 aory|EIT76414.1 asyd|OJJ57023.1 ater|EAU33115.1 atub|OJI81916.1 aver|OJJ03266.1 awen|OJJ41657.1 azon|OJJ48377.1

Group2786: abra|OJJ72511.1 acar|OOF94732.1 acla|EAW12733.1 afis|EAW17896.1 afla|EED46266.1 afum|EAL91704.1 agla|OJJ83046.1 akaw|GAA86238.1 aluc|OJZ90493.1 anid|CBF77002.1 anig|EHA23951.1 aory|EIT76411.1 asyd|OJJ57025.1 ater|EAU33116.1 atub|OJI81917.1 aver|OJJ03265.1 awen|OJJ41655.1 azon|OJJ48375.1

Group2787: abra|OJJ72513.1 acar|OOF94730.1 acla|EAW12734.1 afis|EAW17897.1 afla|EED46265.1 afum|EAL91705.1 agla|OJJ83045.1 akaw|GAA86240.1 aluc|OJZ90495.1 anid|CBF76998.1 anig|EHA23954.1 aory|EIT76381.1 asyd|OJJ57027.1 ater|EAU33117.1 atub|OJI81920.1 aver|OJJ03263.1 awen|OJJ41654.1 azon|OJJ48374.1

Group2788: abra|OJJ72514.1 acar|OOF94729.1 acla|EAW12735.1 afis|EAW17898.1 afla|EED46264.1 afum|EAL91706.1 agla|OJJ83044.1 akaw|GAA86241.1 aluc|OJZ90496.1 anid|CBF76996.1 anig|EHA23955.1 aory|EIT76461.1 asyd|OJJ57028.1 ater|EAU33118.1 atub|OJI81921.1 aver|OJJ03262.1 awen|OJJ41653.1 azon|OJJ48373.1

Group2789: abra|OJJ72515.1 acar|OOF94728.1 afis|EAW17899.1 afla|EED46263.1 agla|OJJ83043.1 akaw|GAA86242.1 aluc|OJZ90497.1 anid|CBF76994.1 anig|EHA23956.1 aory|EIT76422.1 asyd|OJJ57029.1 atub|OJI81922.1 aver|OJJ03261.1 awen|OJJ41652.1 azon|OJJ48372.1 acla|EAW12736.1 afum|EAL91707.1 ater|EAU33119.1

Group2790: abra|OJJ72516.1 acar|OOF94727.1 acla|EAW12737.1 afis|EAW17900.1 afla|EED46262.1 afum|EAL91708.1 agla|OJJ83042.1 akaw|GAA86243.1 aluc|OJZ90498.1 anid|CBF76992.1 anig|EHA23957.1 aory|EIT76428.1 asyd|OJJ57030.1 ater|EAU33120.1 atub|OJI81923.1 aver|OJJ03260.1 awen|OJJ41651.1 azon|OJJ48371.1

Group2791: abra|OJJ72518.1 acar|OOF94726.1 acla|EAW12738.1 afis|EAW17901.1 afla|EED46261.1 afum|EAL91709.1 agla|OJJ83041.1 akaw|GAA86244.1 aluc|OJZ90499.1 anid|CBF76990.1 anig|EHA23958.1 aory|EIT76470.1 asyd|OJJ57032.1 ater|EAU33121.1 atub|OJI81924.1 aver|OJJ03259.1 awen|OJJ41648.1 azon|OJJ48370.1

Group2792: abra|OJJ72538.1 acar|OOF89947.1 acla|EAW12744.1 afis|EAW17907.1 afla|EED46301.1 afum|EAL91716.2 agla|OJJ83035.1 akaw|GAA86262.1 aluc|OJZ90520.1 anid|CBF76981.1 anig|EHA23974.1 aory|EIT76350.1 asyd|OJJ57037.1 ater|EAU33125.1 atub|OJI81944.1 aver|OJJ03255.1 awen|OJJ41643.1 azon|OJJ48364.1

Group2793: abra|OJJ72539.1 acar|OOF89942.1 acla|EAW12745.1 afis|EAW17908.1 afla|EED46302.1 afum|EAL91717.2 agla|OJJ83034.1 akaw|GAA86263.1 aluc|OJZ90522.1 anid|CBF76979.1 anig|EHA23975.1 aory|EIT76327.1 asyd|OJJ57038.1 ater|EAU33126.1 atub|OJI81945.1 aver|OJJ03253.1 awen|OJJ41641.1 azon|OJJ48363.1

Group2794: abra|OJJ72543.1 acar|OOF94712.1 acla|EAW12747.1 afis|EAW17911.1 afla|EED46313.1 afum|EAL91720.1 agla|OJJ83032.1 akaw|GAA86267.1 aluc|OJZ90527.1 anid|CBF76967.1 anig|EHA23979.1 aory|EIT76323.1 asyd|OJJ57041.1 ater|EAU33131.1 atub|OJI81949.1 aver|OJJ03252.1 awen|OJJ41638.1 azon|OJJ48360.1

Group2795: abra|OJJ72544.1 acar|OOF94711.1 acla|EAW12748.1 afis|EAW17912.1 afla|EED46314.1 afum|EAL91721.1 agla|OJJ83031.1 akaw|GAA86268.1 aluc|OJZ90528.1 anid|CBF76969.1 anig|EHA23980.1 aory|EIT76335.1 asyd|OJJ57042.1 ater|EAU33132.1 atub|OJI81950.1 aver|OJJ03251.1 awen|OJJ41637.1 azon|OJJ48359.1

Group2796: abra|OJJ72545.1 acar|OOF94710.1 acla|EAW12749.1 afis|EAW17913.1 afla|EED46315.1 afum|EAL91722.1 agla|OJJ83030.1 akaw|GAA86269.1 aluc|OJZ90529.1 anid|CBF76971.1 anig|EHA23981.1 aory|EIT76319.1 asyd|OJJ57043.1 ater|EAU33133.1 atub|OJI81951.1 aver|OJJ03250.1 awen|OJJ41636.1 azon|OJJ48356.1

Group2797: abra|OJJ72551.1 acar|OOF94702.1 acla|EAW12756.1 afis|EAW17919.1 afla|EED46322.1 afum|EAL91730.1 agla|OJJ83023.1 akaw|GAA85183.1 aluc|OJZ90538.1 anid|CBF76955.1 anig|EHA23986.1 aory|EIT81188.1 asyd|OJJ57054.1 ater|EAU33554.1 atub|OJI81959.1 aver|OJJ03243.1 awen|OJJ41627.1 azon|OJJ48350.1

Group2798: abra|OJJ72554.1 acar|OOF94700.1 acla|EAW12759.1 afis|EAW17921.1 afla|EED46324.1 afum|EAL91732.1 agla|OJJ83020.1 akaw|GAA85186.1 aluc|OJZ90541.1 anid|CBF76951.1 anig|EHA23989.1 aory|EIT81201.1 asyd|OJJ57057.1 ater|EAU33550.1 atub|OJI81963.1 aver|OJJ03241.1 awen|OJJ41624.1 azon|OJJ48347.1

Group2799: abra|OJJ72555.1 acar|OOF94698.1 acla|EAW12760.1 afis|EAW17922.1 afla|EED46325.1 afum|EAL91733.1 agla|OJJ83019.1 akaw|GAA85187.1 aluc|OJZ90542.1 anid|CBF76949.1 anig|EHA23990.1 aory|EIT81261.1 asyd|OJJ57058.1 ater|EAU33549.1 atub|OJI81964.1 aver|OJJ03240.1 awen|OJJ41623.1 azon|OJJ48346.1

Group2801: abra|OJJ72564.1 acar|OOF94690.1 acla|EAW12764.1 afis|EAW17926.1 afla|EED46328.1 afum|EAL91737.1 agla|OJJ83013.1 akaw|GAA92409.1 aluc|OJZ83268.1 anid|CBF76942.1 anig|EHA21121.1 aory|EIT81217.1 asyd|OJJ57062.1 ater|EAU33545.1 atub|OJI87669.1 aver|OJJ03236.1 awen|OJJ41620.1 azon|OJJ48343.1

Group2802: abra|OJJ72565.1 acar|OOF94689.1 acla|EAW12765.1 afis|EAW17927.1 afla|EED58037.1 afum|EAL91738.1 agla|OJJ83007.1 akaw|GAA92406.1 aluc|OJZ83267.1 anid|CBF76940.1 anig|EHA21120.1 aory|EIT79801.1 asyd|OJJ57063.1 ater|EAU33544.1 atub|OJI87670.1 aver|OJJ03235.1 awen|OJJ41619.1 azon|OJJ48342.1

Group2803: abra|OJJ72567.1 acar|OOF94687.1 acla|EAW12767.1 afis|EAW17930.1 afla|EED49111.1 afum|EAL91741.1 agla|OJJ83003.1 akaw|GAA85194.1 aluc|OJZ90549.1 anid|CBF76936.1 anig|EHA23997.1 aory|EIT78530.1 asyd|OJJ57064.1 ater|EAU33540.1 atub|OJI81971.1 aver|OJJ03234.1 awen|OJJ41616.1 azon|OJJ48337.1

Group2805: abra|OJJ72581.1 acar|OOF94671.1 acla|EAW12768.1 afis|EAW17931.1 afla|EED49110.1 afum|EAL91742.1 agla|OJJ83002.1 akaw|GAA85207.1 aluc|OJZ90562.1 anid|CBF76934.1 anig|EHA24008.1 aory|EIT78507.1 asyd|OJJ57065.1 ater|EAU33539.1 atub|OJI81985.1 aver|OJJ03233.1 awen|OJJ41615.1 azon|OJJ48336.1

Group2806: abra|OJJ72582.1 acar|OOF94670.1 acla|EAW12769.1 afis|EAW17932.1 afla|EED49109.1 afum|EAL91743.1 agla|OJJ83001.1 akaw|GAA85208.1 aluc|OJZ90563.1 anid|CBF76932.1 anig|EHA24009.1 aory|EIT78546.1 asyd|OJJ57066.1 ater|EAU33538.1 atub|OJI81986.1 aver|OJJ03232.1 awen|OJJ41614.1 azon|OJJ48335.1

Group2807: abra|OJJ72583.1 acar|OOF94669.1 acla|EAW12770.1 afis|EAW17933.1 afla|EED49108.1 afum|EAL91744.1 agla|OJJ83000.1 akaw|GAA85209.1 aluc|OJZ90564.1 anid|CBF76930.1 anig|EHA24010.1 aory|EIT78564.1 asyd|OJJ57067.1 ater|EAU33537.1 atub|OJI81987.1 aver|OJJ03231.1 awen|OJJ41613.1 azon|OJJ48333.1

Group2808: abra|OJJ72585.1 acar|OOF94667.1 acla|EAW12771.1 afis|EAW17934.1 afla|EED49107.1 afum|EAL91745.1 agla|OJJ82999.1 akaw|GAA85211.1 aluc|OJZ90566.1 anid|CBF77039.1 anig|EHA24012.1 aory|EIT78519.1 asyd|OJJ57002.1 ater|EAU33536.1 atub|OJI81989.1 aver|OJJ03285.1 awen|OJJ41612.1 azon|OJJ48331.1

Group2809: abra|OJJ72588.1 acar|OOF94662.1 acla|EAW12774.1 afis|EAW17937.1 afla|EED49105.1 afum|EAL91748.1 agla|OJJ82995.1 akaw|GAA85214.1 aluc|OJZ90569.1 anid|CBF77044.1 anig|EHA24014.1 aory|EIT78518.1 asyd|OJJ56999.1 ater|EAU33533.1 atub|OJI81992.1 aver|OJJ03288.1 awen|OJJ41609.1 azon|OJJ48328.1

Group2810: abra|OJJ72589.1 acar|OOF94661.1 acla|EAW12775.1 afis|EAW17938.1 afla|EED49103.1 afum|EAL91749.1 agla|OJJ82994.1 akaw|GAA85215.1 aluc|OJZ90570.1 anid|CBF77046.1 anig|EHA24015.1 aory|EIT82547.1 asyd|OJJ56998.1 ater|EAU33530.1 atub|OJI81993.1 aver|OJJ03289.1 awen|OJJ41607.1 azon|OJJ48325.1

Group2811: abra|OJJ72592.1 acar|OOF94658.1 acla|EAW12778.1 afis|EAW17941.1 afla|EED52574.1 afum|EAL91752.1 agla|OJJ82991.1 akaw|GAA85219.1 aluc|OJZ90573.1 anid|CBF77049.1 anig|EHA24018.1 aory|EIT74256.1 asyd|OJJ56996.1 ater|EAU33524.1 atub|OJI81997.1 aver|OJJ03291.1 awen|OJJ41602.1 azon|OJJ48320.1

Group2812: abra|OJJ72596.1 acar|OOF94657.1 acla|EAW11620.1 afla|EED52571.1 akaw|GAA85224.1 aluc|OJZ90577.1 anig|EHA24021.1 aory|EIT74280.1 ater|EAU33521.1 atub|OJI82002.1 azon|OJJ48744.1 afis|EAW23838.1 afum|EAL87269.1 asyd|OJJ56992.1 aver|OJJ03295.1 anid|CBF77058.1 agla|OJJ82497.1 awen|OJJ33801.1

Group2813: abra|OJJ72598.1 acar|OOF94656.1 acla|EAW11619.1 afis|EAW23853.1 afla|EED52570.1 afum|EAL87281.1 agla|OJJ82496.1 akaw|GAA85225.1 aluc|OJZ90579.1 anid|CBF77060.1 anig|EHA24022.1 aory|EIT74300.1 asyd|OJJ56991.1 ater|EAU33520.1 atub|OJI82004.1 aver|OJJ03296.1 awen|OJJ33802.1 azon|OJJ48743.1

Group2814: abra|OJJ72599.1 acar|OOF94655.1 acla|EAW11618.1 afis|EAW23852.1 afla|EED52569.1 afum|EAL87280.1 agla|OJJ82495.1 akaw|GAA85226.1 aluc|OJZ90580.1 anid|CBF77062.1 anig|EHA24023.1 aory|EIT74211.1 asyd|OJJ56990.1 ater|EAU33519.1 atub|OJI82005.1 aver|OJJ03297.1 awen|OJJ33803.1 azon|OJJ48742.1

Group2815: abra|OJJ72600.1 acar|OOF94653.1 acla|EAW11617.1 afis|EAW23851.1 afla|EED52568.1 afum|EAL87279.1 agla|OJJ82494.1 akaw|GAA85228.1 aluc|OJZ90583.1 anid|CBF77063.1 anig|EHA24024.1

aory|EIT74296.1 asyd|OJJ56989.1 ater|EAU33518.1 atub|OJI82006.1 aver|OJJ03298.1 awen|OJJ33804.1 azon|OJJ48740.1

Group2816: abra|OJJ72603.1 acar|OOF94650.1 acla|EAW11611.1 afis|EAW23844.1 afla|EED52553.1 afum|EAL87275.1 agla|OJJ82490.1 akaw|GAA85231.1 aluc|OJZ90586.1 anid|CBF77071.1 anig|EHA24026.1 aory|EIT74313.1 asyd|OJJ56986.1 ater|EAU33511.1 atub|OJI82009.1 aver|OJJ03301.1 awen|OJJ33807.1 azon|OJJ48737.1

Group2817: abra|OJJ72605.1 acar|OOF94649.1 afis|EAW23848.1 afla|EED52565.1 agla|OJJ82491.1 akaw|GAA85232.1 aluc|OJZ90588.1 anig|EHA24027.1 aory|EIT74169.1 atub|OJI82011.1 aver|OJJ03300.1 acla|EAW11614.1 afum|EAL87276.2 anid|CBF77067.1 asyd|OJJ56987.1 ater|EAU33515.1 awen|OJJ33806.1 azon|OJJ48727.1

Group2818: abra|OJJ72625.1 acar|OOF94639.1 acla|EAW11599.1 afis|EAW23858.1 afla|EED52583.1 afum|EAL87286.1 agla|OJJ82525.1 akaw|GAA85245.1 aluc|OJZ90602.1 anid|CBF77236.1 anig|EHA24039.1 aory|EIT74247.1 asyd|OJJ56814.1 ater|EAU33493.1 atub|OJI82026.1 aver|OJJ03521.1 awen|OJJ33819.1 azon|OJJ48716.1

Group2819: abra|OJJ72626.1 acar|OOF94638.1 acla|EAW11598.1 afis|EAW23859.1 afla|EED52584.1 afum|EAL87287.1 agla|OJJ82526.1 akaw|GAA85246.1 aluc|OJZ90603.1 anid|CBF77234.1 anig|EHA24040.1 aory|EIT74284.1 asyd|OJJ56815.1 ater|EAU33492.1 atub|OJI82027.1 aver|OJJ03520.1 awen|OJJ33820.1 azon|OJJ48715.1

Group2820: abra|OJJ72627.1 acar|OOF94637.1 acla|EAW11596.1 afis|EAW23860.1 afla|EED52585.1 afum|EAL87288.1 agla|OJJ82527.1 akaw|GAA85247.1 aluc|OJZ90604.1 anid|CBF77232.1 anig|EHA24041.1 aory|EIT74206.1 asyd|OJJ56816.1 ater|EAU33490.1 atub|OJI82028.1 awen|OJJ33821.1 azon|OJJ48714.1 aver|OJJ03519.1

Group2822: abra|OJJ72640.1 acar|OOF94631.1 acla|EAW11584.1 afis|EAW23869.1 afla|EED52597.1 afum|EAL87297.1 agla|OJJ82538.1 akaw|GAA85261.1 aluc|OJZ90619.1 anid|CBF77217.1 anig|EHA24056.1 aory|EIT74235.1 asyd|OJJ56823.1 ater|EAU33478.1 atub|OJI82045.1 aver|OJJ03512.1 awen|OJJ33845.1 azon|OJJ48702.1

Group2823: abra|OJJ72641.1 acar|OOF94630.1 acla|EAW11583.1 afis|EAW23870.1 afla|EED52598.1 afum|EAL87298.1 agla|OJJ82539.1 akaw|GAA85262.1 aluc|OJZ90620.1 anid|CBF77219.1 anig|EHA24057.1 aory|EIT74294.1 asyd|OJJ56822.1 ater|EAU33479.1 atub|OJI82046.1 aver|OJJ03513.1 awen|OJJ33846.1 azon|OJJ48701.1

Group2824: abra|OJJ72642.1 acar|OOF94629.1 acla|EAW11582.1 afis|EAW23871.1 afla|EED52599.1 afum|EAL87299.1 agla|OJJ82540.1 akaw|GAA85263.1 aluc|OJZ90621.1 anid|CBF77220.1 anig|EHA24058.1 aory|EIT74246.1 asyd|OJJ56821.1 ater|EAU33475.1 atub|OJI82047.1 aver|OJJ03514.1 awen|OJJ33847.1 azon|OJJ48700.1

Group2825: abra|OJJ72647.1 acar|OOF94623.1 acla|EAW11581.1 afis|EAW23872.1 afla|EED52600.1 afum|EAL87300.1 agla|OJJ82542.1 akaw|GAA85267.1 aluc|OJZ90625.1 anid|CBF84521.1 anig|EHA24062.1 aory|EIT74293.1 asyd|OJJ60278.1 ater|EAU33474.1 atub|OJI82052.1 aver|OJJ03978.1 awen|OJJ33848.1 azon|OJJ48699.1

Group2826: abra|OJJ72648.1 acar|OOF94622.1 acla|EAW11580.1 afis|EAW23874.1 afla|EED52602.1 afum|EAL87302.1 agla|OJJ82544.1 akaw|GAA85269.1 aluc|OJZ90627.1 anid|CBF77209.1 anig|EHA24063.1 aory|EIT74307.1 asyd|OJJ56829.1 ater|EAU33472.1 atub|OJI82053.1 aver|OJJ03507.1 awen|OJJ33850.1 azon|OJJ48698.1

Group2827: abra|OJJ72653.1 acar|OOF94620.1 afis|EAW23877.1 afla|EED52605.1 afum|EAL87305.1 agla|OJJ82547.1 akaw|GAA85272.1 aluc|OJZ90631.1 anid|CBF77201.1 anig|EHA24068.1 aory|EIT74181.1 asyd|OJJ56833.1 ater|EAU33469.1 atub|OJI82057.1 aver|OJJ03504.1 awen|OJJ33854.1 azon|OJJ48695.1 acla|EAW11577.1

Group2828: abra|OJJ72654.1 acar|OOF94619.1 acla|EAW11576.1 afis|EAW23878.1 afla|EED52606.1 afum|EAL87306.1 agla|OJJ82548.1 akaw|GAA85273.1 aluc|OJZ90632.1 anid|CBF77199.1 anig|EHA24069.1 aory|EIT74260.1 asyd|OJJ56834.1 ater|EAU33468.1 atub|OJI82059.1 aver|OJJ03503.1 awen|OJJ33858.1 azon|OJJ48694.1

Group2829: abra|OJJ72658.1 acar|OOF94615.1 acla|EAW11522.1 afis|EAW23931.1 afla|EED50587.1 afum|EAL87356.1 agla|OJJ79411.1 akaw|GAA85281.1 aluc|OJZ90639.1 anid|CBF77275.1 anig|EHA24074.1 aory|EIT82143.1 asyd|OJJ56796.1 ater|EAU32296.1 atub|OJI82067.1 aver|OJJ03538.1 awen|OJJ39023.1 azon|OJJ51349.1

Group2830: abra|OJJ72659.1 acar|OOF94614.1 acla|EAW11523.1 afis|EAW23930.1 afla|EED50588.1 afum|EAL87355.1 agla|OJJ79407.1 akaw|GAA85282.1 aluc|OJZ90640.1 anid|CBF77277.1 anig|EHA24075.1 aory|EIT82106.1 asyd|OJJ56795.1 ater|EAU32306.1 atub|OJI82068.1 aver|OJJ03539.1 awen|OJJ39024.1 azon|OJJ51350.1

Group2831: abra|OJJ72661.1 acar|OOF94612.1 acla|EAW11526.1 afis|EAW23928.1 afla|EED50590.1 afum|EAL87353.1 agla|OJJ79406.1 akaw|GAA85284.1 aluc|OJZ90642.1 anid|CBF77281.1 anig|EHA24077.1 aory|EIT82136.1 asyd|OJJ56793.1 ater|EAU32307.1 atub|OJI82070.1 aver|OJJ03541.1 awen|OJJ39026.1 azon|OJJ51352.1

Group2832: abra|OJJ72662.1 acar|OOF94611.1 acla|EAW11527.1 afis|EAW23927.1 afla|EED50591.1 afum|EAL87352.1 agla|OJJ79405.1 akaw|GAA85285.1 aluc|OJZ90643.1 anid|CBF77254.1 anig|EHA24078.1 aory|EIT82117.1 asyd|OJJ56807.1 ater|EAU32308.1 atub|OJI82071.1 aver|OJJ03527.1 awen|OJJ39027.1 azon|OJJ51353.1

Group2833: abra|OJJ72663.1 acar|OOF94610.1 acla|EAW11528.1 afis|EAW23926.1 afla|EED50594.1 afum|EAL87351.1 agla|OJJ79404.1 akaw|GAA85286.1 aluc|OJZ90644.1 anid|CBF77252.1 anig|EHA24079.1 aory|EIT82151.1 asyd|OJJ56808.1 ater|EAU32309.1 atub|OJI82072.1 aver|OJJ03526.1 awen|OJJ39028.1 azon|OJJ51354.1

Group2834: abra|OJJ72669.1 acar|OOF97389.1 acla|EAW11531.1 afis|EAW23923.1 afla|EED50599.1 afum|EAL87348.1 agla|OJJ79397.1 akaw|GAA85289.1 aluc|OJZ90647.1 anid|CBF77090.1 anig|EHA24082.1 aory|EIT78106.1 asyd|OJJ56959.1 ater|EAU32313.1 atub|OJI82077.1 aver|OJJ03326.1 awen|OJJ39032.1 azon|OJJ51360.1

Group2835: abra|OJJ72670.1 acar|OOF97388.1 acla|EAW11537.1 afis|EAW23919.1 afla|EED50608.1 afum|EAL87345.1 agla|OJJ79386.1 akaw|GAA85290.1 aluc|OJZ90648.1 anid|CBF77096.1 anig|EHA24083.1 aory|EIT78069.1 asyd|OJJ56955.1 ater|EAU32318.1 atub|OJI82078.1 aver|OJJ03330.1 awen|OJJ39036.1 azon|OJJ51364.1

Group2836: abra|OJJ72675.1 acar|OOF97380.1 acla|EAW11538.1 afis|EAW23913.1 afla|EED50618.1 afum|EAL87342.1 agla|OJJ79382.1 akaw|GAA85294.1 aluc|OJZ90653.1 anid|CBF77098.1 anig|EHA24088.1 aory|EIT78036.1 asyd|OJJ56954.1 ater|EAU32319.1 atub|OJI82082.1 aver|OJJ03331.1 awen|OJJ39038.1 azon|OJJ51365.1

Group2837: abra|OJJ72683.1 acar|OOF95932.1 acla|EAW13159.1 afis|EAW17487.1 afla|EED49329.1 afum|EAL91310.1 agla|OJJ81475.1 akaw|GAA85301.1 aluc|OJZ90661.1 anid|CBF71613.1 anig|EHA24096.1 aory|EIT79552.1 asyd|OJJ61862.1 ater|EAU36932.1 atub|OJI82090.1 aver|OJI95960.1 awen|OJJ37355.1 azon|OJJ51040.1

Group2838: abra|OJJ72684.1 acar|OOF95933.1 acla|EAW11540.1 afis|EAW23911.1 afla|EED52657.1 afum|EAL87340.1 agla|OJJ79380.1 akaw|GAA85302.1 aluc|OJZ90662.1 anid|CBF77104.1 anig|EHA24097.1 aory|EIT74175.1 asyd|OJJ56950.1 ater|EAU32321.1 atub|OJI82091.1 aver|OJJ03335.1 awen|OJJ39040.1 azon|OJJ51367.1

Group2839: abra|OJJ72685.1 acar|OOF95934.1 acla|EAW11541.1 afis|EAW23910.1 afla|EED52655.1 afum|EAL87339.1 agla|OJJ79379.1 akaw|GAA85303.1 aluc|OJZ90663.1 anid|CBF77106.1 anig|EHA24098.1 aory|EIT74315.1 asyd|OJJ56949.1 ater|EAU32322.1 atub|OJI82092.1 aver|OJJ03336.1 awen|OJJ39041.1 azon|OJJ51368.1

Group2840: abra|OJJ72687.1 acar|OOF95936.1 acla|EAW11542.1 afis|EAW23909.1 afla|EED52654.1 afum|EAL87337.1 agla|OJJ79378.1 akaw|GAA85305.1 aluc|OJZ90665.1 anid|CBF77108.1 anig|EHA24100.1 aory|EIT74231.1 asyd|OJJ56948.1 ater|EAU32324.1 atub|OJI82094.1 aver|OJJ03337.1 awen|OJJ39042.1 azon|OJJ51370.1

Group2841: abra|OJJ72706.1 acar|OOF95943.1 acla|EAW11546.1 afis|EAW23907.1 afla|EED52642.1 afum|EAL87335.1 agla|OJJ79369.1 akaw|GAA85310.1 aluc|OJZ90671.1 anid|CBF77127.1 anig|EHA24104.1 aory|EIT74245.1 asyd|OJJ56941.1 ater|EAU32331.1 atub|OJI82101.1 aver|OJJ03344.1 awen|OJJ39043.1 azon|OJJ51372.1

Group2842: abra|OJJ72707.1 acar|OOF95944.1 acla|EAW11548.1 afis|EAW23905.1 afla|EED52640.1 afum|EAL87333.1 agla|OJJ79370.1 akaw|GAA85311.1 aluc|OJZ90672.1 anid|CBF77125.1 anig|EHA24105.1 aory|EIT74243.1 asyd|OJJ56940.1 ater|EAU32332.1 atub|OJI82102.1 aver|OJJ03345.1 awen|OJJ39044.1 azon|OJJ51373.1

Group2843: abra|OJJ72714.1 acar|OOF95946.1 acla|EAW11550.1 afis|EAW23903.1 afla|EED52638.1 afum|EAL87331.1 agla|OJJ79372.1 akaw|GAA85322.1 aluc|OJZ90683.1 anid|CBF77139.1 anig|EHA24111.1 aory|EIT74295.1 asyd|OJJ56914.1 ater|EAU32335.1 atub|OJI82109.1 aver|OJJ03378.1 awen|OJJ39046.1 azon|OJJ51375.1

Group2844: abra|OJJ72715.1 acar|OOF95947.1 acla|EAW11551.1 afis|EAW23902.1 afla|EED52637.1 afum|EAL87329.1 agla|OJJ79373.1 akaw|GAA85323.1 aluc|OJZ90684.1 anid|CBF77141.1 anig|EHA24112.1 aory|EIT74272.1 asyd|OJJ56913.1 ater|EAU32336.1 atub|OJI82110.1 aver|OJJ03379.1 awen|OJJ39047.1 azon|OJJ51376.1

Group2845: abra|OJJ72719.1 acar|OOF95952.1 acla|EAW11555.1 afis|EAW23898.1 afla|EED52632.1 afum|EAL87325.1 agla|OJJ78640.1 akaw|GAA85326.1 aluc|OJZ90688.1 anid|CBF77148.1 anig|EHA24116.1

aory|EIT74173.1 asyd|OJJ56910.1 ater|EAU32339.1 atub|OJI82113.1 aver|OJJ03383.1 awen|OJJ39050.1 azon|OJJ51382.1

Group2846: abra|OJJ72721.1 acar|OOF95954.1 acla|EAW11557.1 afis|EAW23896.1 afla|EED52630.1 afum|EAL87324.1 agla|OJJ78637.1 akaw|GAA85327.1 aluc|OJZ90690.1 anid|CBF77152.1 anig|EHA24117.1 aory|EIT74255.1 asyd|OJJ56908.1 ater|EAU32340.1 atub|OJI82115.1 aver|OJJ03385.1 awen|OJJ39052.1 azon|OJJ51384.1

Group2847: abra|OJJ72722.1 acar|OOF95955.1 acla|EAW11558.1 afis|EAW23895.1 afla|EED52629.1 afum|EAL87323.1 agla|OJJ78636.1 akaw|GAA85328.1 aluc|OJZ90691.1 anid|CBF77154.1 anig|EHA24118.1 aory|EIT74200.1 asyd|OJJ56907.1 ater|EAU32341.1 atub|OJI82116.1 aver|OJJ03386.1 awen|OJJ39053.1 azon|OJJ51385.1

Group2848: abra|OJJ72725.1 acar|OOF95958.1 acla|EAW11562.1 afis|EAW23891.1 afla|EED52621.1 afum|EAL87319.1 agla|OJJ78631.1 akaw|GAA85331.1 aluc|OJZ90694.1 anid|CBF77171.1 anig|EHA24119.1 aory|EIT74302.1 asyd|OJJ56851.1 ater|EAU33451.1 atub|OJI82119.1 aver|OJJ03466.1 awen|OJJ39058.1 azon|OJJ51389.1

Group2849: abra|OJJ72727.1 acar|OOF95960.1 acla|EAW11564.1 afis|EAW23889.1 afla|EED52619.1 afum|EBA27309.1 agla|OJJ78629.1 akaw|GAA85333.1 aluc|OJZ90696.1 anid|CBF77175.1 anig|EHA24121.1 aory|EIT74266.1 asyd|OJJ56848.1 ater|EAU33454.1 atub|OJI82122.1 aver|OJJ03491.1 awen|OJJ39061.1 azon|OJJ51391.1

Group2850: abra|OJJ72729.1 acar|OOF95961.1 acla|EAW11565.1 afis|EAW23888.1 afla|EED52617.1 afum|EAL87315.1 agla|OJJ78628.1 akaw|GAA85334.1 aluc|OJZ90697.1 anid|CBF77177.1 anig|EHA24123.1 aory|EIT74304.1 asyd|OJJ56847.1 ater|EAU33456.1 atub|OJI82123.1 aver|OJJ03492.1 awen|OJJ39063.1 azon|OJJ51392.1

Group2851: abra|OJJ72733.1 acar|OOF95963.1 acla|EAW11568.1 afis|EAW23885.1 afla|EED52614.1 afum|EBA27307.1 agla|OJJ82572.1 akaw|GAA85339.1 aluc|OJZ90701.1 anid|CBF77183.1 anig|EHA24127.1 aory|EIT74261.1 asyd|OJJ56843.1 ater|EAU33461.1 atub|OJI82128.1 aver|OJJ03495.1 awen|OJJ33867.1 azon|OJJ48687.1

Group2852: abra|OJJ72736.1 acar|OOF95967.1 acla|EAW11572.1 afis|EAW23881.1 afla|EED52610.1 afum|EAL87309.1 agla|OJJ82558.1 akaw|GAA85342.1 aluc|OJZ90704.1 anid|CBF77191.1 anig|EHA24130.1 aory|EIT74269.1 asyd|OJJ56839.1 ater|EAU33465.1 atub|OJI82131.1 aver|OJJ03499.1 awen|OJJ33863.1 azon|OJJ48690.1

Group2853: abra|OJJ72748.1 acar|OOF95971.1 acla|EAW11515.1 afis|EAW23938.1 afla|EED50579.1 afum|EAL87363.1 agla|OJJ79426.1 akaw|GAA85360.1 aluc|OJZ90722.1 anid|CBF77265.1 anig|EHA24140.1 aory|EIT82037.1 asyd|OJJ56801.1 ater|EAU32295.1 atub|OJI82150.1 aver|OJJ03533.1 awen|OJJ39017.1 azon|OJJ51342.1

Group2854: abra|OJJ72749.1 acar|OOF95972.1 acla|EAW11514.1 afis|EAW23939.1 afla|EED50578.1 afum|EAL87364.1 agla|OJJ79427.1 akaw|GAA85361.1 aluc|OJZ90723.1 anid|CBF77263.1 anig|EHA24141.1 aory|EIT82108.1 asyd|OJJ56802.1 ater|EAU32294.1 atub|OJI82151.1 aver|OJJ03532.1 awen|OJJ39016.1 azon|OJJ51341.1

Group2855: abra|OJJ72750.1 acar|OOF95973.1 acla|EAW11513.1 afis|EAW23940.1 afla|EED50577.1 afum|EAL87365.1 agla|OJJ79428.1 akaw|GAA85362.1 aluc|OJZ90724.1 anid|CBF77262.1 anig|EHA24142.1 aory|EIT82017.1 asyd|OJJ56803.1 ater|EAU32293.1 atub|OJI82152.1 aver|OJJ03531.1 awen|OJJ39015.1 azon|OJJ51340.1

Group2856: abra|OJJ72755.1 acar|OOF95976.1 acla|EAW11512.1 afis|EAW23941.1 afla|EED50576.1 afum|EAL87366.1 agla|OJJ79429.1 akaw|GAA85365.1 aluc|OJZ90728.1 anid|CBF77260.1 anig|EHA24145.1 aory|EIT82066.1 asyd|OJJ56804.1 ater|EAU32292.1 atub|OJI82155.1 aver|OJJ03530.1 awen|OJJ39014.1 azon|OJJ51339.1

Group2857: abra|OJJ72756.1 acar|OOF95977.1 acla|EAW11508.1 afis|EAW23945.1 afla|EED50571.1 afum|EAL87370.1 agla|OJJ79436.1 akaw|GAA85366.1 aluc|OJZ90729.1 anid|CBF77290.1 anig|EHA24146.1 aory|EIT82165.1 asyd|OJJ56784.1 ater|EAU32289.1 atub|OJI82156.1 aver|OJJ03549.1 awen|OJJ39009.1 azon|OJJ51335.1

Group2858: abra|OJJ72761.1 acar|OOF95980.1 acla|EAW11503.1 afis|EAW23949.1 afla|EED50567.1 afum|EAL87375.1 agla|OJJ79442.1 akaw|GAA85371.1 aluc|OJZ90735.1 anid|CBF77283.1 anig|EHA24151.1 aory|EIT82002.1 asyd|OJJ56792.1 ater|EAU29803.1 atub|OJI82163.1 aver|OJJ03542.1 awen|OJJ39000.1 azon|OJJ51331.1

Group2859: abra|OJJ72763.1 acar|OOF95983.1 acla|EAW11501.1 afis|EAW23951.1 afla|EED50565.1 afum|EAL87379.1 agla|OJJ79444.1 akaw|GAA85373.1 aluc|OJZ90738.1 anid|CBF77296.1 anig|EHA24153.1 aory|EIT82176.1 asyd|OJJ56782.1 ater|EAU29801.1 atub|OJI82165.1 aver|OJJ03551.1 awen|OJJ38997.1 azon|OJJ51329.1

Group2860: abra|OJJ72764.1 acar|OOF95984.1 acla|EAW11500.1 afis|EAW23952.1 afla|EED50564.1 afum|EAL87380.1 agla|OJJ79445.1 akaw|GAA85374.1 aluc|OJZ90739.1 anid|CBF77298.1 anig|EHA24154.1 aory|EIT82174.1 asyd|OJJ56781.1 ater|EAU29800.1 atub|OJI82166.1 aver|OJJ03552.1 awen|OJJ38996.1 azon|OJJ51328.1

Group2861: abra|OJJ72765.1 acar|OOF95985.1 acla|EAW11499.1 afis|EAW23953.1 afla|EED50562.1 afum|EAL87381.1 agla|OJJ79446.1 akaw|GAA85375.1 aluc|OJZ90740.1 anid|CBF77300.1 anig|EHA24155.1 aory|EIT82008.1 asyd|OJJ56780.1 ater|EAU29799.1 atub|OJI82167.1 aver|OJJ03553.1 awen|OJJ38995.1 azon|OJJ51327.1

Group2862: abra|OJJ72767.1 acar|OOF95986.1 acla|EAW11498.1 afis|EAW23954.1 afla|EED50561.1 afum|EAL87382.1 agla|OJJ79447.1 akaw|GAA85376.1 aluc|OJZ90743.1 anid|CBF77302.1 anig|EHA24156.1 aory|EIT82091.1 asyd|OJJ56779.1 ater|EAU29798.1 atub|OJI82170.1 aver|OJJ03554.1 awen|OJJ38994.1 azon|OJJ51326.1

Group2863: abra|OJJ72768.1 acar|OOF95987.1 acla|EAW11497.1 afis|EAW23955.1 afla|EED50557.1 afum|EAL87383.1 agla|OJJ86878.1 akaw|GAA85377.1 aluc|OJZ90745.1 anid|CBF77304.1 anig|EHA24157.1 aory|EIT82133.1 asyd|OJJ56778.1 ater|EAU29797.1 atub|OJI82171.1 aver|OJJ03555.1 awen|OJJ38993.1 azon|OJJ51324.1

Group2864: abra|OJJ72769.1 acar|OOF95988.1 acla|EAW11496.1 afis|EAW23956.1 afla|EED50556.1 afum|EAL87384.1 agla|OJJ86879.1 akaw|GAA85378.1 aluc|OJZ90746.1 anid|CBF77306.1 anig|EHA24158.1 aory|EIT82152.1 asyd|OJJ56777.1 ater|EAU29796.1 atub|OJI82172.1 aver|OJJ03556.1 awen|OJJ38991.1 azon|OJJ51323.1

Group2865: abra|OJJ72770.1 acar|OOF95989.1 acla|EAW11495.1 afis|EAW23957.1 afla|EED50555.1 afum|EAL87385.1 agla|OJJ86880.1 akaw|GAA85379.1 aluc|OJZ90747.1 anid|CBF77308.1 anig|EHA24159.1 aory|EIT82065.1 asyd|OJJ56776.1 ater|EAU29795.1 atub|OJI82173.1 aver|OJJ03557.1 awen|OJJ38990.1 azon|OJJ51322.1

Group2866: abra|OJJ72773.1 acar|OOF95992.1 acla|EAW11493.1 afis|EAW23960.1 afla|EED50552.1 afum|EAL87389.1 agla|OJJ86885.1 akaw|GAA85382.1 aluc|OJZ90750.1 anid|CBF77314.1 anig|EHA24161.1 aory|EIT82096.1 asyd|OJJ56773.1 ater|EAU29792.1 atub|OJI82176.1 aver|OJJ03560.1 awen|OJJ38987.1 azon|OJJ51319.1

Group2867: abra|OJJ72775.1 acar|OOF95993.1 acla|EAW11491.1 afis|EAW23962.1 afla|EED50548.1 afum|EAL87391.2 agla|OJJ86887.1 akaw|GAA85384.1 anig|EHA24162.1 aory|EIT82083.1 ater|EAU29790.1 atub|OJI82177.1 awen|OJJ38985.1 azon|OJJ51317.1 anid|CBF83986.1 asyd|OJJ61403.1 aver|OJJ00731.1 aluc|OJZ90752.1

Group2868: abra|OJJ72776.1 acar|OOF95994.1 acla|EAW11490.1 afis|EAW23963.1 afla|EED50547.1 afum|EAL87393.1 agla|OJJ86888.1 akaw|GAA85385.1 aluc|OJZ90754.1 anid|CBF77316.1 anig|EHA24163.1 aory|EIT82035.1 asyd|OJJ56772.1 ater|EAU29789.1 atub|OJI82178.1 aver|OJJ03561.1 awen|OJJ38984.1 azon|OJJ51316.1

Group2869: abra|OJJ72779.1 acar|OOF95996.1 acla|EAW11488.1 afis|EAW23965.1 afla|EED50545.1 afum|EAL87395.1 agla|OJJ86894.1 akaw|GAA85387.1 aluc|OJZ83277.1 anid|CBF77322.1 anig|EHA24165.1 aory|EIT82007.1 asyd|OJJ56770.1 ater|EAU29787.1 atub|OJI82181.1 aver|OJJ03563.1 awen|OJJ38978.1 azon|OJJ51314.1

Group2870: abra|OJJ72780.1 acar|OOF95999.1 acla|EAW11487.1 afis|EAW23966.1 afla|EED50544.1 afum|EAL87396.1 agla|OJJ86895.1 akaw|GAA85388.1 aluc|OJZ83279.1 anid|CBF77324.1 anig|EHA24166.1 aory|EIT82088.1 asyd|OJJ56769.1 ater|EAU29786.1 atub|OJI82183.1 aver|OJJ03565.1 awen|OJJ38977.1 azon|OJJ51313.1

Group2871: abra|OJJ72781.1 acar|OOF96000.1 acla|EAW11486.1 afis|EAW23967.1 afla|EED50542.1 afum|EAL87397.1 agla|OJJ86896.1 akaw|GAA85389.1 aluc|OJZ83280.1 anid|CBF77326.1 anig|EHA24167.1 aory|EIT82086.1 asyd|OJJ56768.1 ater|EAU29785.1 atub|OJI82184.1 aver|OJJ03566.1 awen|OJJ38976.1 azon|OJJ51312.1

Group2872: abra|OJJ72782.1 acar|OOF96002.1 acla|EAW11485.1 afis|EAW23968.1 afla|EED50541.1 afum|EAL87398.2 agla|OJJ86897.1 akaw|GAA85390.1 aluc|OJZ83281.1 anid|CBF77328.1 anig|EHA24168.1 aory|EIT82024.1 asyd|OJJ56767.1 ater|EAU29784.1 atub|OJI82185.1 aver|OJJ03567.1 awen|OJJ38975.1 azon|OJJ51310.1

Group2873: abra|OJJ72792.1 acar|OOF90123.1 acla|EAW11480.1 afis|EAW23973.1 afla|EED50535.1 afum|EAL87403.1 agla|OJJ86902.1 akaw|GAA85395.1 aluc|OJZ83288.1 anid|CBF77339.1 anig|EHA24173.1 aory|EIT82084.1 asyd|OJJ53275.1 ater|EAU29779.1 atub|OJI82190.1 aver|OJJ03574.1 awen|OJJ35438.1 azon|OJJ51305.1

Group2874: abra|OJJ72797.1 acar|OOF96005.1 acla|EAW11475.1 afis|EAW23978.1 afla|EED49183.1 afum|EAL87408.1 agla|OJJ86908.1 akaw|GAA85400.1 aluc|OJZ83294.1 anid|CBF77349.1 anig|EHA24178.1

aory|EIT78581.1 asyd|OJJ53271.1 ater|EAU29772.1 atub|OJI82195.1 aver|OJJ03579.1 awen|OJJ35446.1 azon|OJJ51300.1

Group2875: abra|OJJ72798.1 acar|OOF96006.1 acla|EAW11474.1 afis|EAW23979.1 afla|EED49182.1 afum|EAL87409.1 agla|OJJ86910.1 akaw|GAA85401.1 aluc|OJZ83295.1 anid|CBF77351.1 anig|EHA24179.1 aory|EIT78531.1 asyd|OJJ53270.1 ater|EAU29771.1 atub|OJI82196.1 aver|OJJ03580.1 awen|OJJ35447.1 azon|OJJ51299.1

Group2876: abra|OJJ72800.1 acar|OOF96008.1 acla|EAW11444.1 afis|EAW23981.1 afla|EED49178.1 afum|EAL87411.1 agla|OJJ86914.1 akaw|GAA85403.1 aluc|OJZ83298.1 anid|CBF77356.1 anig|EHA24181.1 aory|EIT78527.1 asyd|OJJ53268.1 ater|EAU29770.1 atub|OJI82198.1 aver|OJJ03583.1 awen|OJJ35449.1 azon|OJJ51297.1

Group2877: abra|OJJ72801.1 acar|OOF96009.1 acla|EAW11443.1 afis|EAW23982.1 afla|EED49177.1 afum|EAL87412.1 agla|OJJ86916.1 akaw|GAA85404.1 aluc|OJZ83299.1 anid|CBF77358.1 anig|EHA24182.1 aory|EIT78537.1 asyd|OJJ53267.1 ater|EAU29769.1 atub|OJI82199.1 aver|OJJ03584.1 awen|OJJ35450.1 azon|OJJ51293.1

Group2878: abra|OJJ72803.1 acar|OOF96011.1 acla|EAW11441.1 afis|EAW23984.1 afla|EED49175.1 afum|EAL87413.1 agla|OJJ86918.1 akaw|GAA85406.1 aluc|OJZ83301.1 anid|CBF77362.1 anig|EHA24183.1 aory|EIT78558.1 asyd|OJJ53265.1 ater|EAU29767.1 atub|OJI82201.1 aver|OJJ03586.1 awen|OJJ35452.1 azon|OJJ51291.1

Group2879: abra|OJJ72804.1 acar|OOF96012.1 acla|EAW11440.1 afis|EAW23985.1 afla|EED49173.1 afum|EAL87414.1 agla|OJJ86919.1 akaw|GAA85407.1 aluc|OJZ83302.1 anid|CBF77364.1 anig|EHA24184.1 aory|EIT78557.1 asyd|OJJ53264.1 ater|EAU29766.1 atub|OJI82202.1 aver|OJJ03587.1 awen|OJJ35453.1 azon|OJJ51290.1

Group2880: abra|OJJ72808.1 acar|OOF96014.1 acla|EAW11433.1 afis|EAW23989.1 afla|EED49169.1 afum|EAL87419.1 agla|OJJ86923.1 akaw|GAA85411.1 aluc|OJZ83306.1 anid|CBF77367.1 anig|EHA24189.1 aory|EIT78567.1 asyd|OJJ53261.1 ater|EAU29761.1 atub|OJI82206.1 aver|OJJ03590.1 awen|OJJ35457.1 azon|OJJ51286.1

Group2881: abra|OJJ72810.1 acar|OOF96017.1 acla|EAW11430.1 afis|EAW23993.1 afla|EED49167.1 afum|EAL87423.1 agla|OJJ86926.1 akaw|GAA85413.1 aluc|OJZ83308.1 anid|CBF77373.1 anig|EHA24191.1 aory|EIT78512.1 asyd|OJJ53258.1 ater|EAU29758.1 atub|OJI82208.1 aver|OJJ03593.1 awen|OJJ35462.1 azon|OJJ51281.1

Group2882: abra|OJJ72811.1 acar|OOF96018.1 acla|EAW11429.1 afis|EAW23994.1 afla|EED49166.1 afum|EAL87424.1 agla|OJJ86927.1 akaw|GAA85414.1 aluc|OJZ83309.1 anid|CBF77375.1 anig|EHA24192.1 aory|EIT78551.1 asyd|OJJ53257.1 ater|EAU29757.1 atub|OJI82209.1 aver|OJJ03594.1 awen|OJJ35463.1 azon|OJJ51280.1

Group2883: abra|OJJ72814.1 acar|OOF96020.1 acla|EAW11428.1 afis|EAW23995.1 afla|EED49165.1 afum|EAL87425.1 agla|OJJ86928.1 akaw|GAA85415.1 aluc|OJZ83312.1 anid|CBF77378.1 anig|EHA24194.1 aory|EIT78566.1 asyd|OJJ53255.1 ater|EAU29756.1 atub|OJI82212.1 aver|OJJ03597.1 awen|OJJ35465.1 azon|OJJ51278.1

Group2884: abra|OJJ72815.1 acar|OOF96021.1 acla|EAW11427.1 afis|EAW23996.1 afla|EED49164.1 afum|EAL87426.1 agla|OJJ86929.1 akaw|GAA85416.1 aluc|OJZ83314.1 anid|CBF77379.1 anig|EHA24195.1 aory|EIT78572.1 asyd|OJJ53254.1 ater|EAU29755.1 atub|OJI82213.1 aver|OJJ03599.1 awen|OJJ35467.1 azon|OJJ51277.1

Group2885: abra|OJJ72818.1 acar|OOF96024.1 acla|EAW11422.1 afis|EAW23999.1 afla|EED49162.1 afum|EAL87429.1 agla|OJJ86933.1 akaw|GAA85419.1 aluc|OJZ83320.1 anid|CBF77385.1 anig|EHA24198.1 aory|EIT78542.1 asyd|OJJ53250.1 ater|EAU29750.1 atub|OJI82217.1 awen|OJJ35470.1 azon|OJJ51274.1 aver|OJJ03602.1

Group2886: abra|OJJ72820.1 acar|OOF96025.1 acla|EAW11419.1 afis|EAW24003.1 afla|EED49158.1 afum|EAL87433.1 agla|OJJ86935.1 akaw|GAA85421.1 aluc|OJZ83322.1 anid|CBF77390.1 anig|EHA24200.1 aory|EIT78547.1 asyd|OJJ53247.1 ater|EAU29748.1 atub|OJI82219.1 aver|OJJ03604.1 awen|OJJ35474.1 azon|OJJ51272.1

Group2887: abra|OJJ72822.1 acar|OOF96027.1 acla|EAW11417.1 afis|EAW24005.1 afla|EED49156.1 afum|EAL87435.1 agla|OJJ86936.1 akaw|GAA85423.1 aluc|OJZ83324.1 anid|CBF77394.1 anig|EHA24202.1 aory|EIT78532.1 asyd|OJJ53245.1 ater|EAU29746.1 atub|OJI82221.1 aver|OJJ03605.1 awen|OJJ35477.1 azon|OJJ51271.1

Group2888: abra|OJJ72824.1 acar|OOF96029.1 acla|EAW11414.1 afis|EAW24007.1 afla|EED49154.1 afum|EAL87437.1 agla|OJJ86938.1 akaw|GAA85425.1 aluc|OJZ83326.1 anid|CBF77398.1 anig|EHA24204.1 aory|EIT78570.1 asyd|OJJ53243.1 ater|EAU29744.1 atub|OJI82223.1 aver|OJJ03607.1 awen|OJJ35479.1 azon|OJJ51269.1

Group2889: abra|OJJ72825.1 acar|OOF96030.1 acla|EAW11413.1 afis|EAW24008.1 afla|EED49153.1 afum|EAL87438.1 agla|OJJ86939.1 akaw|GAA85426.1 aluc|OJZ83327.1 anid|CBF77400.1 anig|EHA24205.1 aory|EIT78506.1 asyd|OJJ53242.1 ater|EAU29743.1 atub|OJI82224.1 aver|OJJ03608.1 awen|OJJ35480.1 azon|OJJ51268.1

Group2890: abra|OJJ72826.1 acar|OOF96031.1 acla|EAW11412.1 afis|EAW24009.1 afla|EED49152.1 afum|EAL87439.1 agla|OJJ86940.1 akaw|GAA85427.1 aluc|OJZ83328.1 anid|CBF77402.1 anig|EHA24206.1 aory|EIT78555.1 asyd|OJJ53241.1 ater|EAU29742.1 atub|OJI82225.1 aver|OJJ03609.1 awen|OJJ35481.1 azon|OJJ51267.1

Group2891: abra|OJJ72827.1 acar|OOF96033.1 acla|EAW11410.1 afis|EAW24011.1 afla|EED49150.1 afum|EAL87441.1 agla|OJJ86943.1 akaw|GAA85428.1 aluc|OJZ83330.1 anid|CBF77406.1 anig|EHA24207.1 aory|EIT78559.1 asyd|OJJ53239.1 ater|EAU29740.1 atub|OJI82226.1 aver|OJJ03611.1 awen|OJJ35483.1 azon|OJJ51265.1

Group2892: abra|OJJ72831.1 acar|OOF96035.1 acla|EAW11408.1 afis|EAW24013.1 afla|EED49147.1 afum|EAL87443.1 agla|OJJ86945.1 akaw|GAA85430.1 aluc|OJZ83332.1 anid|CBF77410.1 anig|EHA24209.1 aory|EIT78522.1 asyd|OJJ53237.1 ater|EAU29734.1 atub|OJI82228.1 aver|OJJ03614.1 awen|OJJ35485.1 azon|OJJ51263.1

Group2893: abra|OJJ72833.1 acar|OOF96036.1 acla|EAW11406.1 afis|EAW24015.1 afla|EED49144.1 afum|EAL87445.1 agla|OJJ86949.1 akaw|GAA85431.1 aluc|OJZ83334.1 anid|CBF77414.1 anig|EHA24210.1 aory|EIT78508.1 asyd|OJJ53235.1 ater|EAU29731.1 atub|OJI82230.1 aver|OJJ03616.1 awen|OJJ35487.1 azon|OJJ47003.1

Group2894: abra|OJJ72835.1 acar|OOF96037.1 acla|EAW11405.1 afis|EAW24016.1 afla|EED49143.1 afum|EAL87446.1 agla|OJJ86950.1 akaw|GAA85432.1 aluc|OJZ83335.1 anid|CBF77416.1 anig|EHA24211.1 aory|EIT78565.1 asyd|OJJ53234.1 ater|EAU29730.1 atub|OJI82232.1 aver|OJJ03617.1 awen|OJJ35488.1 azon|OJJ47001.1

Group2895: abra|OJJ72837.1 acar|OOF96039.1 acla|EAW06923.1 afis|EAW16540.1 afla|EED53743.1 afum|EAL89137.1 agla|OJJ85576.1 akaw|GAA85434.1 aluc|OJZ83337.1 anid|CBF69989.1 anig|EHA24213.1 aory|EIT75293.1 asyd|OJJ58892.1 ater|EAU39054.1 atub|OJI82234.1 aver|OJJ01493.1 awen|OJJ40280.1 azon|OJJ49143.1

Group2896: abra|OJJ72840.1 acar|OOF96042.1 acla|EAW11402.1 afis|EAW24019.1 afla|EED49137.1 afum|EAL87449.1 agla|OJJ86952.1 akaw|GAA85442.1 aluc|OJZ83347.1 anid|CBF77420.1 anig|EHA24215.1 aory|EIT78560.1 asyd|OJJ53232.1 ater|EAU29727.1 atub|OJI82241.1 aver|OJJ03619.1 awen|OJJ35492.1 azon|OJJ46998.1

Group2897: abra|OJJ72844.1 acar|OOF96045.1 acla|EAW11399.1 afis|EAW24022.1 afla|EED49132.1 afum|EAL87453.1 agla|OJJ86955.1 akaw|GAA85445.1 aluc|OJZ83352.1 anid|CBF75300.1 anig|EHA24218.1 aory|EIT78561.1 asyd|OJJ59881.1 ater|EAU29724.1 atub|OJI82244.1 aver|OJJ02405.1 awen|OJJ35495.1 azon|OJJ46994.1

Group2898: abra|OJJ72847.1 acar|OOF96048.1 acla|EAW11396.1 afis|EAW24025.1 afla|EED49129.1 afum|EAL87456.1 agla|OJJ86958.1 akaw|GAA85448.1 aluc|OJZ83355.1 anid|CBF75305.1 anig|EHA24221.1 aory|EIT78574.1 asyd|OJJ59876.1 ater|EAU29719.1 atub|OJI82247.1 aver|OJJ02410.1 awen|OJJ35498.1 azon|OJJ46991.1

Group2899: abra|OJJ72850.1 acar|OOF96051.1 acla|EAW11393.1 afis|EAW24028.1 afla|EED49126.1 afum|EAL87459.1 agla|OJJ86961.1 akaw|GAA85451.1 aluc|OJZ83359.1 anid|CBF75310.1 anig|EHA24224.1 aory|EIT78543.1 asyd|OJJ59874.1 ater|EAU29716.1 atub|OJI82250.1 aver|OJJ02413.1 awen|OJJ35501.1 azon|OJJ46988.1

Group2900: abra|OJJ72853.1 acar|OOF96052.1 acla|EAW11391.1 afis|EAW24031.1 afla|EED49123.1 afum|EAL87462.1 agla|OJJ87001.1 akaw|GAA85454.1 aluc|OJZ83362.1 anid|CBF75316.1 anig|EHA24227.1 aory|EIT78584.1 asyd|OJJ59871.1 ater|EAU29713.1 atub|OJI82253.1 aver|OJJ02416.1 awen|OJJ35503.1 azon|OJJ46983.1

Group2901: abra|OJJ72854.1 acar|OOF96053.1 acla|EAW11390.1 afis|EAW24032.1 afla|EED49121.1 afum|EAL87463.1 agla|OJJ87000.1 akaw|GAA85455.1 aluc|OJZ83363.1 anid|CBF75318.1 anig|EHA24228.1 aory|EIT78515.1 asyd|OJJ59870.1 ater|EAU29712.1 atub|OJI82254.1 aver|OJJ02417.1 awen|OJJ35504.1 azon|OJJ46981.1

Group2902: abra|OJJ72858.1 acar|OOF96055.1 acla|EAW11388.1 afis|EAW24034.1 afla|EED49119.1 afum|EAL87465.1 agla|OJJ86998.1 akaw|GAA85457.1 aluc|OJZ83365.1 anid|CBF75324.1 anig|EHA24230.1 aory|EIT78541.1 asyd|OJJ59868.1 ater|EAU29710.1 atub|OJI82256.1 aver|OJJ02419.1 awen|OJJ35506.1 azon|OJJ46978.1

Group2903: abra|OJJ72860.1 acar|OOF96057.1 acla|EAW11386.1 afis|EAW18110.1 afla|EED49117.1 afum|EAL87467.1 agla|OJJ86996.1 akaw|GAA85459.1 aluc|OJZ83367.1 anid|CBF75328.1 anig|EHA24232.1

aory|EIT78582.1 asyd|OJJ59865.1 ater|EAU29708.1 atub|OJI82259.1 aver|OJJ02422.1 awen|OJJ35508.1 azon|OJJ47630.1

Group2904: abra|OJJ72861.1 acar|OOF96058.1 acla|EAW11385.1 afis|EAW18111.1 afla|EED49114.1 afum|EAL87468.1 agla|OJJ86995.1 akaw|GAA85460.1 aluc|OJZ83368.1 anid|CBF75330.1 anig|EHA24233.1 aory|EIT78525.1 asyd|OJJ59864.1 ater|EAU29707.1 atub|OJI82260.1 aver|OJJ02423.1 awen|OJJ35509.1 azon|OJJ46976.1

Group2905: abra|OJJ72862.1 acar|OOF96060.1 acla|EAW11384.1 afis|EAW18112.1 afla|EED49113.1 afum|EAL87469.1 agla|OJJ86994.1 akaw|GAA85461.1 aluc|OJZ83369.1 anid|CBF75332.1 anig|EHA24234.1 aory|EIT78586.1 asyd|OJJ59863.1 ater|EAU29706.1 atub|OJI82261.1 aver|OJJ02424.1 awen|OJJ35510.1 azon|OJJ46975.1

Group2906: abra|OJJ72863.1 acar|OOF96062.1 acla|EAW11379.1 afis|EAW18116.1 afla|EED58041.1 afum|EAL87473.1 agla|OJJ86980.1 akaw|GAA85462.1 aluc|OJZ83376.1 anid|CBF75340.1 anig|EHA24235.1 aory|EIT79770.1 asyd|OJJ59859.1 ater|EAU29701.1 atub|OJI82262.1 aver|OJJ02428.1 awen|OJJ35515.1 azon|OJJ46970.1

Group2907: abra|OJJ72865.1 acar|OOF96064.1 acla|EAW11382.1 afis|EAW18114.1 afla|EED58039.1 afum|EAL87471.1 agla|OJJ86982.1 akaw|GAA85464.1 aluc|OJZ83378.1 anid|CBF75336.1 anig|EHA24237.1 aory|EIT79795.1 asyd|OJJ59861.1 ater|EAU29703.1 atub|OJI82264.1 aver|OJJ02426.1 awen|OJJ35512.1 azon|OJJ46973.1

Group2908: abra|OJJ72866.1 acar|OOF96065.1 acla|EAW11383.1 afis|EAW18113.1 afla|EED58038.1 afum|EAL87470.1 agla|OJJ86991.1 akaw|GAA85465.1 aluc|OJZ83379.1 anid|CBF75334.1 anig|EHA24238.1 aory|EIT79794.1 asyd|OJJ59862.1 ater|EAU29705.1 atub|OJI82265.1 aver|OJJ02425.1 awen|OJJ35511.1 azon|OJJ46974.1

Group2909: abra|OJJ72869.1 acar|OOF96068.1 acla|EAW11376.1 afis|EAW18119.1 afla|EED58044.1 afum|EAL87477.1 agla|OJJ87011.1 akaw|GAA85468.1 aluc|OJZ83382.1 anid|CBF75346.1 anig|EHA24241.1 aory|EIT79722.1 asyd|OJJ59854.1 ater|EAU29698.1 atub|OJI82268.1 aver|OJJ02431.1 awen|OJJ35518.1 azon|OJJ46966.1

Group2910: abra|OJJ72871.1 acar|OOF96069.1 acla|EAW11375.1 afis|EAW18120.1 afla|EED58045.1 afum|EAL87478.1 agla|OJJ87014.1 akaw|GAA85469.1 aluc|OJZ83385.1 anid|CBF75348.1 anig|EHA24243.1 aory|EIT79798.1 asyd|OJJ59853.1 ater|EAU29697.1 atub|OJI82271.1 aver|OJJ02432.1 awen|OJJ35521.1 azon|OJJ46965.1

Group2911: abra|OJJ72874.1 acar|OOF96071.1 acla|EAW11374.1 afis|EAW18121.1 afla|EED58046.1 afum|EAL87479.1 agla|OJJ87015.1 akaw|GAA85470.1 aluc|OJZ83386.1 anid|CBF75352.1 anig|EHA24244.1 aory|EIT79741.1 asyd|OJJ59852.1 ater|EAU29695.1 atub|OJI82272.1 aver|OJJ02435.1 awen|OJJ35523.1 azon|OJJ46963.1

Group2912: abra|OJJ72880.1 acar|OOF96073.1 acla|EAW11366.1 afis|EAW18129.1 afla|EED58053.1 afum|EAL87486.1 agla|OJJ83795.1 akaw|GAA85472.1 aluc|OJZ83391.1 anid|CBF75366.1 anig|EHA24246.1 aory|EIT79707.1 asyd|OJJ59848.1 ater|EAU35448.1 atub|OJI82277.1 aver|OJJ02440.1 awen|OJJ35557.1 azon|OJJ46953.1

Group2913: abra|OJJ72886.1 acar|OOF96076.1 acla|EAW11364.1 afis|EAW18130.1 afla|EED58054.1 afum|EAL87487.1 agla|OJJ83802.1 akaw|GAA85473.1 aluc|OJZ83394.1 anid|CBF75370.1 anig|EHA24247.1 aory|EIT79721.1 asyd|OJJ59847.1 ater|EAU35452.1 atub|OJI82280.1 aver|OJJ02441.1 awen|OJJ35561.1 azon|OJJ46951.1

Group2914: abra|OJJ72887.1 acar|OOF96077.1 acla|EAW11363.1 afis|EAW18132.1 afla|EED58056.1 afum|EAL87489.1 agla|OJJ83814.1 akaw|GAA85474.1 aluc|OJZ83395.1 anid|CBF75372.1 anig|EHA24248.1 aory|EIT79771.1 asyd|OJJ59846.1 ater|EAU35454.1 atub|OJI82281.1 aver|OJJ02442.1 awen|OJJ35571.1 azon|OJJ46949.1

Group2915: abra|OJJ72890.1 acar|OOF96080.1 acla|EAW11360.1 afis|EAW18134.1 afla|EED58059.1 afum|EAL87491.1 agla|OJJ83818.1 akaw|GAA85477.1 aluc|OJZ83398.1 anid|CBF75380.1 anig|EHA24251.1 aory|EIT79745.1 asyd|OJJ59843.1 ater|EAU35457.1 atub|OJI82284.1 aver|OJJ02446.1 awen|OJJ35576.1 azon|OJJ46946.1

Group2916: abra|OJJ72892.1 acar|OOF96083.1 acla|EAW11358.1 afis|EAW18136.1 afla|EED58061.1 afum|EAL87493.1 agla|OJJ83819.1 akaw|GAA85479.1 aluc|OJZ83400.1 anid|CBF75376.1 anig|EHA24252.1 aory|EIT79756.1 asyd|OJJ59840.1 ater|EAU35459.1 atub|OJI82288.1 aver|OJJ02449.1 awen|OJJ35578.1 azon|OJJ46945.1

Group2917: abra|OJJ72894.1 acar|OOF96085.1 acla|EAW11357.1 afis|EAW18137.1 afla|EED58062.1 afum|EAL87494.1 agla|OJJ83820.1 akaw|GAA85481.1 aluc|OJZ83403.1 anid|CBF75384.1 anig|EHA24254.1 aory|EIT79803.1 asyd|OJJ59838.1 ater|EAU35460.1 atub|OJI82291.1 aver|OJJ02450.1 awen|OJJ35579.1 azon|OJJ46944.1

Group2918: abra|OJJ72898.1 acar|OOF96088.1 acla|EAW11353.1 afis|EAW18141.1 afla|EED58067.1 afum|EAL87498.1 agla|OJJ83826.1 akaw|GAA85485.1 aluc|OJZ83407.1 anid|CBF81129.1 anig|EHA24258.1 aory|EIT79748.1 asyd|OJJ56491.1 ater|EAU35464.1 atub|OJI82296.1 aver|OJJ06227.1 awen|OJJ35583.1 azon|OJJ46939.1

Group2919: abra|OJJ72904.1 acar|OOF89820.1 acla|EAW11350.1 afis|EAW18145.1 afla|EED58072.1 afum|EAL87502.1 agla|OJJ83828.1 akaw|GAA85489.1 aluc|OJZ83414.1 anid|CBF81120.1 anig|EHA24263.1 aory|EIT79155.1 asyd|OJJ56488.1 ater|EAU35468.1 atub|OJI82301.1 aver|OJJ06231.1 awen|OJJ35588.1 azon|OJJ46937.1

Group2920: abra|OJJ72906.1 acar|OOF96089.1 acla|EAW11348.1 afis|EAW18147.1 afla|EED58075.1 afum|EAL87505.1 agla|OJJ83832.1 akaw|GAA85491.1 aluc|OJZ83416.1 anid|CBF81116.1 anig|EHA24265.1 aory|EIT79152.1 asyd|OJJ56486.1 ater|EAU35446.1 atub|OJI82303.1 aver|OJJ06233.1 awen|OJJ35590.1 azon|OJJ46935.1

Group2921: abra|OJJ72909.1 acar|OOF96092.1 acla|EAW11344.1 afis|EAW18150.1 afla|EED58078.1 afum|EAL87508.1 agla|OJJ83835.1 akaw|GAA85494.1 aluc|OJZ83419.1 anid|CBF81110.1 anig|EHA24268.1 aory|EIT79111.1 asyd|OJJ56483.1 ater|EAU35472.1 atub|OJI82306.1 aver|OJJ06236.1 awen|OJJ35593.1 azon|OJJ46915.1

Group2922: abra|OJJ72910.1 acar|OOF96093.1 acla|EAW11343.1 afis|EAW18151.1 afla|EED58080.1 afum|EAL87509.1 agla|OJJ83836.1 akaw|GAA85495.1 aluc|OJZ83421.1 anid|CBF81108.1 anig|EHA24269.1 aory|EIT79142.1 asyd|OJJ56481.1 ater|EAU35473.1 atub|OJI82307.1 aver|OJJ06238.1 awen|OJJ35594.1 azon|OJJ46916.1

Group2923: abra|OJJ72912.1 acar|OOF96094.1 acla|EAW06479.1 afis|EAW15974.1 afla|EED58082.1 afum|EAL88606.1 agla|OJJ83838.1 akaw|GAA85496.1 aluc|OJZ83422.1 anid|CBF81105.1 anig|EHA24270.1 aory|EIT79133.1 asyd|OJJ56479.1 ater|EAU35474.1 atub|OJI82308.1 aver|OJJ06239.1 awen|OJJ35597.1 azon|OJJ46918.1

Group2924: abra|OJJ72913.1 acar|OOF96095.1 acla|EAW06478.1 afis|EAW15973.1 afla|EED58083.1 afum|EAL88605.1 agla|OJJ83842.1 akaw|GAA85497.1 aluc|OJZ83423.1 anid|CBF81101.1 anig|EHA24271.1 aory|EIT79117.1 asyd|OJJ56478.1 ater|EAU35475.1 atub|OJI82309.1 aver|OJJ06240.1 awen|OJJ35599.1 azon|OJJ46919.1

Group2925: abra|OJJ72914.1 acar|OOF96096.1 acla|EAW06476.1 afis|EAW15971.1 afla|EED58085.1 afum|EAL88603.1 agla|OJJ83844.1 akaw|GAA85498.1 aluc|OJZ83424.1 anid|CBF81098.1 anig|EHA24272.1 aory|EIT79112.1 asyd|OJJ56476.1 ater|EAU35477.1 atub|OJI82310.1 aver|OJJ06242.1 awen|OJJ35601.1 azon|OJJ46921.1

Group2926: abra|OJJ72917.1 acar|OOF96099.1 acla|EAW06474.1 afis|EAW15968.1 afla|EED58089.1 afum|EAL88601.1 agla|OJJ89290.1 akaw|GAA85500.1 aluc|OJZ83427.1 anid|CBF81094.1 anig|EHA24279.1 aory|EIT79146.1 asyd|OJJ56470.1 ater|EAU35480.1 atub|OJI82315.1 aver|OJJ06249.1 awen|OJJ35605.1 azon|OJJ46925.1

Group2927: abra|OJJ72920.1 acar|OOF96102.1 acla|EAW06471.1 afis|EAW15965.1 afla|EED58091.1 afum|EAL88598.1 agla|OJJ89346.1 akaw|GAA85503.1 aluc|OJZ83430.1 anid|CBF81089.1 anig|EHA24281.1 aory|EIT79110.1 asyd|OJJ56467.1 ater|EAU35487.1 atub|OJI82318.1 aver|OJJ06252.1 awen|OJJ35608.1 azon|OJJ46928.1

Group2928: abra|OJJ72922.1 acar|OOF96103.1 acla|EAW06469.1 afis|EAW15964.1 afla|EED58093.1 afum|EAL88597.1 agla|OJJ89348.1 akaw|GAA85504.1 aluc|OJZ83432.1 anid|CBF81085.1 anig|EHA24282.1 aory|EIT79125.1 asyd|OJJ56465.1 ater|EAU35488.1 atub|OJI82321.1 aver|OJJ06254.1 awen|OJJ35610.1 azon|OJJ46930.1

Group2929: abra|OJJ72930.1 acar|OOF96110.1 acla|EAW06460.1 afis|EAW15956.1 afla|EED58102.1 afum|EAL88589.1 agla|OJJ89357.1 akaw|GAA85512.1 aluc|OJZ83440.1 anid|CBF81067.1 anig|EHA24286.1 aory|EIT79135.1 asyd|OJJ56458.1 ater|EAU35501.1 atub|OJI82331.1 aver|OJJ06264.1 awen|OJJ35618.1 azon|OJJ46904.1

Group2930: abra|OJJ72931.1 acar|OOF96111.1 acla|EAW06459.1 afis|EAW15955.1 afla|EED58103.1 afum|EAL88588.1 agla|OJJ89358.1 akaw|GAA85513.1 aluc|OJZ83441.1 anid|CBF81065.1 anig|EHA24287.1 aory|EIT79109.1 asyd|OJJ56457.1 ater|EAU35502.1 atub|OJI82332.1 aver|OJJ06265.1 awen|OJJ35619.1 azon|OJJ46903.1

Group2931: abra|OJJ72935.1 acar|OOF96116.1 acla|EAW06456.1 afis|EAW15952.1 afla|EED58106.1 afum|EAL88585.1 agla|OJJ89362.1 akaw|GAA85516.1 aluc|OJZ83444.1 anid|CBF81059.1 anig|EHA24290.1 aory|EIT79114.1 asyd|OJJ56454.1 ater|EAU35505.1 atub|OJI82335.1 aver|OJJ06268.1 awen|OJJ35624.1 azon|OJJ46901.1

Group2932: abra|OJJ72936.1 acar|OOF96117.1 acla|EAW06455.1 afis|EAW15951.1 afla|EED58107.1 afum|EAL88584.1 agla|OJJ89363.1 akaw|GAA85517.1 aluc|OJZ83445.1 anid|CBF81057.1 anig|EHA24291.1

aory|EIT79166.1 asyd|OJJ56453.1 ater|EAU35506.1 atub|OJI82336.1 aver|OJJ06269.1 awen|OJJ35625.1 azon|OJJ46900.1

Group2933: abra|OJJ72937.1 acar|OOF96119.1 acla|EAW06454.1 afis|EAW15950.1 afla|EED58108.1 afum|EAL88583.1 agla|OJJ89364.1 akaw|GAA85518.1 aluc|OJZ83446.1 anid|CBF81056.1 anig|EHA24292.1 aory|EIT79143.1 asyd|OJJ56451.1 ater|EAU35507.1 atub|OJI82337.1 aver|OJJ06270.1 awen|OJJ35626.1 azon|OJJ46899.1

Group2934: abra|OJJ72942.1 acar|OOF96123.1 acla|EAW11209.1 afis|EAW15942.1 afla|EED58116.1 afum|EAL88576.1 agla|OJJ89368.1 akaw|GAA85524.1 aluc|OJZ83452.1 anid|CBF80959.1 anig|EHA24298.1 aory|EIT79144.1 asyd|OJJ56346.1 ater|EAU35514.1 atub|OJI82344.1 aver|OJJ06367.1 awen|OJJ35640.1 azon|OJJ46895.1

Group2935: abra|OJJ72945.1 acar|OOF96126.1 acla|EAW11205.1 afis|EAW15935.1 afla|EED58120.1 afum|EAL88571.1 agla|OJJ89371.1 akaw|GAA85527.1 aluc|OJZ83455.1 anid|CBF80966.1 anig|EHA24301.1 aory|EIT79156.1 asyd|OJJ56349.1 ater|EAU35517.1 atub|OJI82347.1 aver|OJJ06364.1 awen|OJJ35644.1 azon|OJJ46892.1

Group2936: abra|OJJ72949.1 acar|OOF96130.1 acla|EAW11195.1 afis|EAW15925.1 afla|EED54936.1 afum|EAL88561.1 agla|OJJ89380.1 akaw|GAA85530.1 aluc|OJZ83459.1 anid|CBF81019.1 anig|EHA24303.1 aory|EIT80380.1 asyd|OJJ56382.1 ater|EAU35530.1 atub|OJI82351.1 aver|OJJ06331.1 awen|OJJ35655.1 azon|OJJ46884.1

Group2937: abra|OJJ72956.1 acar|OOF96137.1 acla|EAW11201.1 afis|EAW15931.1 afla|EED54930.1 afum|EAL88567.1 agla|OJJ89375.1 akaw|GAA85537.1 aluc|OJZ83465.1 anid|CBF81029.1 anig|EHA24309.1 aory|EIT80323.1 asyd|OJJ56388.1 ater|EAU35521.1 atub|OJI82358.1 aver|OJJ06325.1 awen|OJJ35648.1 azon|OJJ46889.1

Group2938: abra|OJJ72960.1 acar|OOF96142.1 acla|EAW11192.1 afis|EAW15922.1 afla|EED54938.1 afum|EAL88559.1 agla|OJJ89386.1 akaw|GAA85540.1 aluc|OJZ83468.1 anid|CBF81013.1 anig|EHA24312.1 aory|EIT80338.1 asyd|OJJ56379.1 ater|EAU35533.1 atub|OJI82362.1 aver|OJJ06334.1 awen|OJJ35658.1 azon|OJJ46881.1

Group2939: abra|OJJ72967.1 acar|OOF96149.1 acla|EAW11187.1 afis|EAW15918.1 afla|EED54943.1 afum|EAL88554.1 agla|OJJ89393.1 akaw|GAA85546.1 aluc|OJZ83478.1 anid|CBF81002.1 anig|EHA24315.1 aory|EIT80355.1 asyd|OJJ56373.1 ater|EAU35538.1 atub|OJI82368.1 aver|OJJ06340.1 awen|OJJ35661.1 azon|OJJ44132.1

Group2940: abra|OJJ72971.1 acar|OOF96150.1 acla|EAW08037.1 afis|EAW23072.1 afla|EED54944.1 afum|EAL88407.1 agla|OJJ89394.1 akaw|GAA85547.1 aluc|OJZ83482.1 anid|CBF80992.1 anig|EHA24317.1 aory|EIT80389.1 asyd|OJJ56368.1 ater|EAU35539.1 atub|OJI82372.1 aver|OJJ06346.1 awen|OJJ35663.1 azon|OJJ44133.1

Group2941: abra|OJJ72972.1 acar|OOF96151.1 acla|EAW08036.1 afis|EAW23071.1 afla|EED54945.1 afum|EAL88408.1 agla|OJJ89395.1 akaw|GAA85548.1 aluc|OJZ83483.1 anid|CBF80994.1 anig|EHA24318.1 aory|EIT80359.1 asyd|OJJ56369.1 ater|EAU35540.1 atub|OJI82373.1 aver|OJJ06345.1 awen|OJJ35664.1 azon|OJJ44134.1

Group2942: abra|OJJ72973.1 acar|OOF96152.1 acla|EAW08035.1 afis|EAW23070.1 afla|EED54946.1 afum|EAL88409.1 agla|OJJ89396.1 akaw|GAA85549.1 aluc|OJZ83484.1 anid|CBF80996.1 anig|EHA24319.1 aory|EIT80382.1 asyd|OJJ56370.1 ater|EAU37593.1 atub|OJI82374.1 aver|OJJ06344.1 awen|OJJ35665.1 azon|OJJ44135.1

Group2943: abra|OJJ72975.1 acar|OOF96153.1 acla|EAW08034.1 afis|EAW23069.1 afla|EED54947.1 afum|EAL88410.1 agla|OJJ89397.1 akaw|GAA85550.1 aluc|OJZ83485.1 anid|CBF80998.1 anig|EHA24320.1 aory|EIT80368.1 asyd|OJJ56371.1 ater|EAU37596.1 atub|OJI82376.1 aver|OJJ06343.1 awen|OJJ35666.1 azon|OJJ44137.1

Group2944: abra|OJJ72976.1 acar|OOF96154.1 acla|EAW08033.1 afis|EAW23068.1 afla|EED54948.1 afum|EAL88411.1 agla|OJJ89398.1 akaw|GAA85551.1 aluc|OJZ83486.1 anid|CBF81000.1 anig|EHA24321.1 aory|EIT80295.1 asyd|OJJ56372.1 ater|EAU37597.1 atub|OJI82377.1 aver|OJJ06342.1 awen|OJJ35667.1 azon|OJJ44138.1

Group2945: abra|OJJ72979.1 acar|OOF96158.1 acla|EAW08031.1 afis|EAW23065.1 afla|EED54952.1 afum|EAL88414.1 agla|OJJ89409.1 akaw|GAA85554.1 aluc|OJZ83489.1 anid|CBF80981.1 anig|EHA24324.1 aory|EIT80321.1 asyd|OJJ56358.1 ater|EAU37601.1 atub|OJI82381.1 aver|OJJ06356.1 awen|OJJ35670.1 azon|OJJ44140.1

Group2946: abra|OJJ72980.1 acar|OOF96156.1 acla|EAW08030.1 afis|EAW23064.1 afla|EED54953.1 afum|EAL88415.1 agla|OJJ89410.1 akaw|GAA85555.1 aluc|OJZ83490.1 anid|CBF80979.1 anig|EHA24325.1 aory|EIT80361.1 asyd|OJJ56357.1 ater|EAU37602.1 atub|OJI82382.1 aver|OJJ06357.1 awen|OJJ35671.1 azon|OJJ44141.1

Group2947: abra|OJJ72982.1 acar|OOF96159.1 acla|EAW08028.1 afis|EAW23062.1 afla|EED54955.1 afum|EAL88417.1 agla|OJJ89412.1 akaw|GAA85557.1 aluc|OJZ83492.1 anid|CBF80975.1 anig|EHA24327.1 aory|EIT80363.1 asyd|OJJ56355.1 ater|EAU37603.1 atub|OJI82386.1 aver|OJJ06359.1 awen|OJJ35673.1 azon|OJJ44142.1

Group2948: abra|OJJ72983.1 acar|OOF96160.1 acla|EAW08027.1 afis|EAW23061.1 afla|EED54957.1 afum|EAL88418.1 agla|OJJ89414.1 akaw|GAA85558.1 aluc|OJZ83493.1 anid|CBF80973.1 anig|EHA24328.1 aory|EIT80353.1 asyd|OJJ56354.1 ater|EAU37604.1 atub|OJI82387.1 aver|OJJ06360.1 awen|OJJ35676.1 azon|OJJ44143.1

Group2949: abra|OJJ72984.1 acar|OOF96161.1 acla|EAW08026.1 afis|EAW23060.1 afla|EED54958.1 afum|EAL88419.1 agla|OJJ89415.1 akaw|GAA85559.1 aluc|OJZ83494.1 anid|CBF80972.1 anig|EHA24329.1 aory|EIT80387.1 asyd|OJJ56353.1 ater|EAU37605.1 atub|OJI82388.1 aver|OJJ06361.1 awen|OJJ35677.1 azon|OJJ44144.1

Group2950: abra|OJJ72985.1 acar|OOF96162.1 acla|EAW08025.1 afis|EAW23059.1 afla|EED54959.1 afum|EAL88420.1 agla|OJJ89417.1 akaw|GAA85560.1 aluc|OJZ83495.1 anid|CBF80970.1 anig|EHA24330.1 aory|EIT80388.1 asyd|OJJ56352.1 ater|EAU37606.1 atub|OJI82389.1 aver|OJJ06362.1 awen|OJJ35678.1 azon|OJJ44146.1

Group2951: abra|OJJ72989.1 acar|OOF96165.1 acla|EAW08022.1 afis|EAW23056.1 afla|EED54961.1 afum|EAL88424.1 agla|OJJ89419.1 akaw|GAA85563.1 aluc|OJZ83497.1 anid|CBF81039.1 anig|EHA24332.1 aory|EIT80362.1 asyd|OJJ56392.1 ater|EAU37608.1 atub|OJI82391.1 aver|OJJ06321.1 awen|OJJ35680.1 azon|OJJ44148.1

Group2952: abra|OJJ72990.1 acar|OOF96166.1 acla|EAW08021.1 afis|EAW23055.1 afla|EED54962.1 afum|EAL88425.1 agla|OJJ89420.1 akaw|GAA85564.1 aluc|OJZ83498.1 anid|CBF81041.1 anig|EHA24333.1 aory|EIT80303.1 asyd|OJJ56393.1 ater|EAU37609.1 atub|OJI82392.1 aver|OJJ06320.1 awen|OJJ35681.1 azon|OJJ44149.1

Group2953: abra|OJJ72991.1 acar|OOF96167.1 acla|EAW08020.1 afis|EAW23054.1 afla|EED54964.1 afum|EAL88426.1 agla|OJJ89422.1 akaw|GAA85565.1 aluc|OJZ83499.1 anid|CBF81043.1 anig|EHA24334.1 aory|EIT80378.1 asyd|OJJ56441.1 ater|EAU37610.1 atub|OJI82393.1 aver|OJJ06280.1 awen|OJJ35682.1 azon|OJJ44150.1

Group2954: abra|OJJ72997.1 acar|OOF96173.1 acla|EAW08016.1 afis|EAW23050.1 afla|EED54968.1 afum|EAL88430.1 agla|OJJ89429.1 akaw|GAA85571.1 aluc|OJZ83505.1 anid|CBF80953.1 anig|EHA24338.1 aory|EIT80340.1 asyd|OJJ56343.1 ater|EAU29602.1 atub|OJI82399.1 aver|OJJ06370.1 awen|OJJ35689.1 azon|OJJ44154.1

Group2955: abra|OJJ72998.1 acar|OOF89938.1 acla|EAW08015.1 afis|EAW23049.1 afla|EED54969.1 afum|EAL88431.1 agla|OJJ89430.1 akaw|GAA85572.1 aluc|OJZ83506.1 anid|CBF80951.1 anig|EHA24339.1 aory|EIT80346.1 asyd|OJJ56342.1 ater|EAU29601.1 atub|OJI82400.1 aver|OJJ06371.1 awen|OJJ35690.1 azon|OJJ44155.1

Group2956: abra|OJJ72999.1 acar|OOF89939.1 acla|EAW08014.1 afis|EAW23048.1 afla|EED54970.1 afum|EAL88432.1 agla|OJJ89431.1 akaw|GAA85573.1 aluc|OJZ83509.1 anid|CBF80949.1 anig|EHA24340.1 aory|EIT80379.1 asyd|OJJ56341.1 ater|EAU29600.1 atub|OJI82402.1 aver|OJJ06372.1 awen|OJJ35691.1 azon|OJJ44157.1

Group2957: abra|OJJ73002.1 acar|OOF96175.1 acla|EAW08011.1 afis|EAW23046.1 afla|EED54973.1 afum|EAL88435.1 agla|OJJ89432.1 akaw|GAA85580.1 aluc|OJZ83516.1 anid|CBF80946.1 anig|EHA24341.1 aory|EIT80343.1 asyd|OJJ56339.1 ater|EAU29614.1 atub|OJI82407.1 aver|OJJ06374.1 awen|OJJ35694.1 azon|OJJ44158.1

Group2958: abra|OJJ73003.1 acar|OOF96176.1 acla|EAW08010.1 afis|EAW23045.1 afla|EED54974.1 afum|EAL88436.1 agla|OJJ89433.1 akaw|GAA85581.1 aluc|OJZ83517.1 anid|CBF80944.1 anig|EHA24342.1 aory|EIT80339.1 asyd|OJJ56338.1 ater|EAU29613.1 atub|OJI82408.1 aver|OJJ06375.1 awen|OJJ35695.1 azon|OJJ44159.1

Group2959: abra|OJJ73006.1 acar|OOF96178.1 acla|EAW08007.1 afis|EAW23042.1 afla|EED54976.1 afum|EAL88439.1 agla|OJJ89436.1 akaw|GAA85584.1 aluc|OJZ83521.1 anid|CBF80938.1 anig|EHA24344.1 aory|EIT80316.1 asyd|OJJ56312.1 ater|EAU29617.1 atub|OJI82410.1 aver|OJJ06402.1 awen|OJJ35700.1 azon|OJJ44162.1

Group2960: abra|OJJ73008.1 acar|OOF96180.1 acla|EAW08005.1 afis|EAW23040.1 afla|EED54978.1 afum|EAL88441.1 agla|OJJ89438.1 akaw|GAA85586.1 aluc|OJZ83523.1 anid|CBF80935.1 anig|EHA24346.1 aory|EIT80392.1 asyd|OJJ56314.1 ater|EAU29603.1 atub|OJI82412.1 aver|OJJ06400.1 awen|OJJ35702.1 azon|OJJ44164.1

Group2961: abra|OJJ73016.1 acar|OOF96186.1 acla|EAW07997.1 afis|EAW23032.1 afla|EED54984.1 afum|EAL88450.1 agla|OJJ89445.1 akaw|GAA85594.1 aluc|OJZ83531.1 anid|CBF80910.1 anig|EHA24353.1

aory|EIT80327.1 asyd|OJJ56328.1 ater|EAU29625.1 atub|OJI82421.1 aver|OJJ06385.1 awen|OJJ35713.1 azon|OJJ44172.1

Group2962: abra|OJJ73022.1 acar|OOF96190.1 acla|EAW07993.1 afis|EAW23028.1 afla|EED54987.1 afum|EAL88453.2 agla|OJJ89451.1 akaw|GAA85597.1 aluc|OJZ83535.1 anid|CBF80920.1 anig|EHA24356.1 aory|EIT80371.1 asyd|OJJ56322.1 ater|EAU29630.1 atub|OJI82426.1 aver|OJJ06391.1 awen|OJJ35717.1 azon|OJJ44176.1

Group2963: abra|OJJ73023.1 acar|OOF96191.1 acla|EAW07992.1 afis|EAW23027.1 afla|EED54988.1 afum|EAL88455.1 agla|OJJ89452.1 akaw|GAA85598.1 aluc|OJZ83536.1 anid|CBF80918.1 anig|EHA24357.1 aory|EIT80332.1 asyd|OJJ56323.1 ater|EAU29629.1 atub|OJI82427.1 aver|OJJ06390.1 awen|OJJ35718.1 azon|OJJ44177.1

Group2964: abra|OJJ73025.1 acar|OOF96192.1 acla|EAW07991.1 afis|EAW23026.1 afla|EED54989.1 afum|EAL88456.1 agla|OJJ89453.1 akaw|GAA85599.1 aluc|OJZ83537.1 anid|CBF80916.1 anig|EHA24358.1 aory|EIT80337.1 asyd|OJJ56324.1 ater|EAU29628.1 atub|OJI82428.1 aver|OJJ06389.1 awen|OJJ35719.1 azon|OJJ44178.1

Group2965: abra|OJJ73029.1 acar|OOF96195.1 acla|EAW07988.1 afis|EAW23023.1 afla|EED54993.1 afum|EAL88459.1 agla|OJJ89456.1 akaw|GAA85602.1 aluc|OJZ83540.1 anid|CBF80905.1 anig|EHA24360.1 aory|EIT80347.1 asyd|OJJ56332.1 ater|EAU29651.1 atub|OJI82432.1 aver|OJJ06382.1 awen|OJJ30720.1 azon|OJJ44181.1

Group2966: abra|OJJ73030.1 acar|OOF96196.1 acla|EAW07987.1 afis|EAW23022.1 afla|EED54994.1 afum|EAL88460.1 agla|OJJ89457.1 akaw|GAA85603.1 aluc|OJZ83541.1 anid|CBF80903.1 anig|EHA24361.1 aory|EIT80374.1 asyd|OJJ56333.1 ater|EAU29253.1 atub|OJI82433.1 aver|OJJ06381.1 awen|OJJ30721.1 azon|OJJ44182.1

Group2967: abra|OJJ73031.1 acar|OOF96197.1 acla|EAW07986.1 afis|EAW23021.1 afla|EED54995.1 afum|EAL88461.1 agla|OJJ89458.1 akaw|GAA85604.1 aluc|OJZ83542.1 anid|CBF80901.1 anig|EHA24362.1 aory|EIT80385.1 asyd|OJJ56334.1 ater|EAU29652.1 atub|OJI82434.1 aver|OJJ06380.1 awen|OJJ30722.1 azon|OJJ44183.1

Group2968: abra|OJJ73032.1 acar|OOF96198.1 acla|EAW07985.1 afis|EAW23020.1 afla|EED54996.1 afum|EAL88462.1 agla|OJJ89459.1 akaw|GAA85605.1 aluc|OJZ83543.1 anid|CBF80900.1 anig|EHA24363.1 aory|EIT80298.1 asyd|OJJ56335.1 ater|EAU29653.1 atub|OJI82435.1 aver|OJJ06379.1 awen|OJJ30723.1 azon|OJJ44184.1

Group2969: abra|OJJ73037.1 acar|OOF96200.1 acla|EAW07982.1 afis|EAW23017.1 afla|EED54999.1 afum|EAL88464.1 agla|OJJ89463.1 akaw|GAA85608.1 aluc|OJZ83548.1 anig|EHA24366.1 aory|EIT80350.1 asyd|OJJ56309.1 ater|EAU29657.1 atub|OJI82439.1 aver|OJJ06405.1 awen|OJJ29535.1 azon|OJJ44187.1 anid|CBF80894.1

Group2970: abra|OJJ73038.1 acar|OOF96201.1 acla|EAW07981.1 afis|EAW23016.1 afla|EED55000.1 afum|EAL88465.1 agla|OJJ89464.1 akaw|GAA85609.1 aluc|OJZ83549.1 anid|CBF80896.1 anig|EHA24367.1 aory|EIT80294.1 asyd|OJJ56310.1 ater|EAU29658.1 atub|OJI82440.1 aver|OJJ06404.1 awen|OJJ29534.1 azon|OJJ44188.1

Group2971: abra|OJJ73039.1 acar|OOF96204.1 acla|EAW07980.1 afis|EAW23015.1 afla|EED55001.1 afum|EAL88466.1 agla|OJJ89465.1 akaw|GAA85610.1 aluc|OJZ83550.1 anid|CBF80890.1 anig|EHA24368.1 aory|EIT80296.1 asyd|OJJ56307.1 ater|EAU29659.1 atub|OJI82441.1 aver|OJJ06408.1 awen|OJJ30728.1 azon|OJJ44189.1

Group2972: abra|OJJ73040.1 acar|OOF96203.1 acla|EAW07979.1 afis|EAW23014.1 afla|EED55002.1 afum|EAL88467.2 agla|OJJ89466.1 akaw|GAA85611.1 aluc|OJZ83551.1 anid|CBF80888.1 anig|EHA24369.1 aory|EIT80286.1 asyd|OJJ56306.1 ater|EAU29660.1 atub|OJI82442.1 aver|OJJ06409.1 awen|OJJ30729.1 azon|OJJ44190.1

Group2973: abra|OJJ73041.1 acar|OOF96202.1 acla|EAW07978.1 afis|EAW23013.1 afla|EED55003.1 afum|EAL88468.1 agla|OJJ89467.1 akaw|GAA85612.1 aluc|OJZ83552.1 anid|CBF80886.1 anig|EHA24370.1 aory|EIT80391.1 asyd|OJJ56305.1 ater|EAU29661.1 atub|OJI82443.1 aver|OJJ06410.1 awen|OJJ30730.1 azon|OJJ44191.1

Group2974: abra|OJJ73042.1 acar|OOF96205.1 acla|EAW07977.1 afis|EAW23012.1 afla|EED55004.1 afum|EAL88469.1 agla|OJJ89468.1 akaw|GAA85613.1 aluc|OJZ83553.1 anid|CBF80885.1 anig|EHA24371.1 aory|EIT80335.1 asyd|OJJ56304.1 ater|EAU29662.1 atub|OJI82444.1 aver|OJJ06411.1 awen|OJJ30731.1 azon|OJJ44192.1

Group2975: abra|OJJ73046.1 acar|OOF96208.1 acla|EAW07973.1 afis|EAW23008.1 afla|EED55007.1 afum|EAL88472.1 agla|OJJ89472.1 akaw|GAA85617.1 aluc|OJZ83557.1 anid|CBF80879.1 anig|EHA24375.1 aory|EIT80310.1 asyd|OJJ56302.1 ater|EAU29665.1 atub|OJI82447.1 aver|OJJ06415.1 awen|OJJ30733.1 azon|OJJ44196.1

Group2976: abra|OJJ73048.1 acar|OOF96211.1 acla|EAW07971.1 afis|EAW23006.1 afla|EED55009.1 afum|EAL88474.1 agla|OJJ89473.1 akaw|GAA85619.1 aluc|OJZ83559.1 anid|CBF80875.1 anig|EHA24377.1 aory|EIT80349.1 asyd|OJJ56300.1 ater|EAU29667.1 atub|OJI82449.1 aver|OJJ06417.1 awen|OJJ30735.1 azon|OJJ44197.1

Group2977: abra|OJJ73049.1 acar|OOF96212.1 acla|EAW07970.1 afis|EAW23005.1 afla|EED55010.1 afum|EAL88475.1 agla|OJJ89474.1 akaw|GAA85620.1 aluc|OJZ83560.1 anid|CBF80873.1 anig|EHA24378.1 aory|EIT80326.1 asyd|OJJ56299.1 ater|EAU29668.1 atub|OJI82450.1 aver|OJJ06418.1 awen|OJJ30736.1 azon|OJJ44198.1

Group2978: abra|OJJ73050.1 acar|OOF96213.1 acla|EAW07969.1 afis|EAW23004.1 afla|EED55011.1 afum|EAL88476.2 agla|OJJ89475.1 akaw|GAA85621.1 aluc|OJZ83561.1 anid|CBF80871.1 anig|EHA24379.1 aory|EIT80312.1 asyd|OJJ56298.1 ater|EAU29669.1 atub|OJI82451.1 aver|OJJ06419.1 awen|OJJ30737.1 azon|OJJ44199.1

Group2979: abra|OJJ73051.1 acar|OOF96214.1 acla|EAW07968.1 afis|EAW23003.1 afla|EED55012.1 afum|EAL88477.1 agla|OJJ89476.1 akaw|GAA85622.1 aluc|OJZ83562.1 anid|CBF80869.1 anig|EHA24380.1 aory|EIT80369.1 asyd|OJJ56297.1 ater|EAU29670.1 atub|OJI82452.1 aver|OJJ06420.1 awen|OJJ30738.1 azon|OJJ44200.1

Group2980: abra|OJJ73052.1 acar|OOF96215.1 acla|EAW07967.1 afis|EAW23002.1 afla|EED55013.1 afum|EAL88478.1 agla|OJJ89477.1 akaw|GAA85623.1 aluc|OJZ83564.1 anid|CBF80868.1 anig|EHA24381.1 aory|EIT80329.1 asyd|OJJ56296.1 ater|EAU29671.1 atub|OJI82454.1 aver|OJJ06421.1 awen|OJJ30739.1 azon|OJJ44201.1

Group2981: abra|OJJ73056.1 acar|OOF96217.1 acla|EAW07965.1 afis|EAW23000.1 afla|EED55015.1 afum|EAL88480.1 agla|OJJ89479.1 akaw|GAA85624.1 aluc|OJZ83567.1 anid|CBF80864.1 anig|EHA24383.1 aory|EIT80314.1 asyd|OJJ56294.1 ater|EAU29673.1 atub|OJI82456.1 aver|OJJ06423.1 awen|OJJ30741.1 azon|OJJ44204.1

Group2982: abra|OJJ73058.1 acar|OOF96218.1 acla|EAW07963.1 afis|EAW22998.1 afla|EED55017.1 afum|EAL88482.1 agla|OJJ89481.1 akaw|GAA85626.1 aluc|OJZ83569.1 anid|CBF80860.1 anig|EHA24385.1 aory|EIT80381.1 asyd|OJJ56292.1 ater|EAU29674.1 atub|OJI82458.1 aver|OJJ06425.1 awen|OJJ30743.1 azon|OJJ44202.1

Group2983: abra|OJJ73059.1 acar|OOF96220.1 acla|EAW07962.1 afis|EAW22997.1 afla|EED55018.1 afum|EAL88483.1 agla|OJJ89482.1 akaw|GAA85627.1 aluc|OJZ83570.1 anid|CBF80856.1 anig|EHA24386.1 aory|EIT80291.1 asyd|OJJ56291.1 ater|EAU29675.1 atub|OJI82459.1 aver|OJJ06426.1 awen|OJJ30744.1 azon|OJJ44205.1

Group2985: abra|OJJ73142.1 acar|OOG00354.1 acla|EAW07018.1 afis|EAW16644.1 afla|EED44956.1 afum|EAL89248.1 agla|OJJ82603.1 akaw|GAA88030.1 aluc|OJZ81289.1 anid|CBF82053.1 anig|EHA18588.1 aory|EIT73503.1 asyd|OJJ54235.1 ater|EAU39569.1 atub|OJI91497.1 aver|OJJ05187.1 awen|OJJ40070.1 azon|OJJ45554.1

Group2986: abra|OJJ73143.1 acar|OOG00355.1 acla|EAW07019.1 afis|EAW16645.1 afla|EED44955.1 afum|EAL89249.1 agla|OJJ82602.1 akaw|GAA88029.1 aluc|OJZ81288.1 anid|CBF82055.1 anig|EHA18587.1 aory|EIT73491.1 asyd|OJJ54236.1 ater|EAU39570.1 atub|OJI91496.1 aver|OJJ05188.1 awen|OJJ40069.1 azon|OJJ45555.1

Group2987: abra|OJJ73147.1 acar|OOG00358.1 acla|EAW07023.1 afis|EAW16649.1 afla|EED44951.1 afum|EAL89253.1 agla|OJJ78458.1 akaw|GAA88025.1 aluc|OJZ81284.1 anid|CBF82061.1 anig|EHA18583.1 aory|EIT73464.1 asyd|OJJ54240.1 ater|EAU39573.1 atub|OJI91491.1 aver|OJJ05192.1 awen|OJJ40065.1 azon|OJJ45561.1

Group2988: abra|OJJ73148.1 acar|OOG00360.1 acla|EAW07024.1 afis|EAW16650.1 afla|EED44950.1 afum|EAL89254.1 agla|OJJ78459.1 akaw|GAA88024.1 aluc|OJZ81282.1 anid|CBF82063.1 anig|EHA18582.1 aory|EIT73478.1 asyd|OJJ54241.1 ater|EAU39574.1 atub|OJI91490.1 aver|OJJ05193.1 awen|OJJ40064.1 azon|OJJ45562.1

Group2989: abra|OJJ73149.1 acar|OOG00361.1 acla|EAW07025.1 afis|EAW16651.1 afla|EED44949.1 afum|EAL89255.1 agla|OJJ78460.1 akaw|GAA88023.1 aluc|OJZ81281.1 anid|CBF82064.1 anig|EHA18581.1 aory|EIT73505.1 asyd|OJJ54242.1 ater|EAU39575.1 atub|OJI91489.1 aver|OJJ05194.1 awen|OJJ40063.1 azon|OJJ45563.1

Group2990: abra|OJJ73151.1 acar|OOG00363.1 acla|EAW07028.1 afis|EAW16654.1 afla|EED44946.1 afum|EAL89258.1 agla|OJJ78687.1 akaw|GAA88021.1 aluc|OJZ81279.1 anid|CBF82067.1 anig|EHA18579.1 aory|EIT73521.1 asyd|OJJ54244.1 ater|EAU39578.1 atub|OJI91487.1 aver|OJJ05196.1 awen|OJJ40060.1 azon|OJJ45567.1

Group2991: abra|OJJ73156.1 acar|OOG00371.1 acla|EAW07030.1 afis|EAW16656.1 afla|EED44941.1 afum|EAL89260.1 agla|OJJ78716.1 akaw|GAA88016.1 aluc|OJZ81274.1 anid|CBF82070.1 anig|EHA18574.1

aory|EIT73470.1 asyd|OJJ54246.1 ater|EAU39579.1 atub|OJI91481.1 aver|OJJ05199.1 awen|OJJ40058.1 azon|OJJ50081.1

Group2992: abra|OJJ73158.1 acar|OOG00369.1 acla|EAW07032.1 afis|EAW16658.1 afla|EED44944.1 afum|EAL89262.1 agla|OJJ78714.1 akaw|GAA88014.1 aluc|OJZ81272.1 anid|CBF82072.1 anig|EHA18572.1 aory|EIT73479.1 asyd|OJJ54248.1 ater|EAU39581.1 atub|OJI91479.1 aver|OJJ05202.1 awen|OJJ40056.1 azon|OJJ50105.1

Group2993: abra|OJJ73159.1 acar|OOG00368.1 acla|EAW07033.1 afis|EAW16659.1 afla|EED44936.1 afum|EAL89263.1 agla|OJJ78713.1 akaw|GAA88013.1 aluc|OJZ81271.1 anid|CBF82073.1 anig|EHA18571.1 aory|EIT73463.1 asyd|OJJ54249.1 ater|EAU39582.1 atub|OJI91478.1 aver|OJJ05203.1 awen|OJJ40055.1 azon|OJJ50106.1

Group2994: abra|OJJ73160.1 acar|OOG00405.1 acla|EAW06769.1 afis|EAW16319.1 afla|EED53562.1 afum|EAL88943.1 agla|OJJ88280.1 akaw|GAA88012.1 aluc|OJZ81270.1 anid|CBF89151.1 anig|EHA18570.1 aory|EIT74869.1 asyd|OJJ62566.1 ater|EAU34138.1 atub|OJI91477.1 aver|OJI96690.1 awen|OJJ37720.1 azon|OJJ44978.1

Group2995: abra|OJJ73168.1 acar|OOG00409.1 acla|EAW07130.1 afis|EAW22141.1 afla|EED53567.1 afum|EAL91049.1 agla|OJJ87934.1 akaw|GAA88003.1 aluc|OJZ81259.1 anid|CBF89137.1 anig|EHA18566.1 aory|EIT75049.1 asyd|OJJ62649.1 ater|EAU34143.1 atub|OJI91455.1 aver|OJI96749.1 awen|OJJ34715.1 azon|OJJ45847.1

Group2996: abra|OJJ73170.1 acar|OOG00410.1 acla|EAW10796.1 afis|EAW24780.1 afla|EED47205.1 afum|EAL89419.1 agla|OJJ82876.1 akaw|GAA88001.1 aluc|OJZ81257.1 anid|CBF81439.1 anig|EHA18564.1 aory|EIT79182.1 asyd|OJJ59428.1 ater|EAU36528.1 atub|OJI91453.1 aver|OJJ02843.1 awen|OJJ40366.1 azon|OJJ49215.1

Group2997: abra|OJJ73173.1 acar|OOG00413.1 acla|EAW06856.1 afis|EAW16473.1 afla|EED47208.1 afum|EAL89067.1 agla|OJJ82873.1 akaw|GAA87998.1 aluc|OJZ81254.1 anid|CBF81435.1 anig|EHA18562.1 aory|EIT79309.1 asyd|OJJ59424.1 ater|EAU36569.1 atub|OJI91450.1 aver|OJJ02846.1 awen|OJJ40363.1 azon|OJJ49211.1

Group2998: abra|OJJ73176.1 acar|OOG00416.1 acla|EAW06858.1 afis|EAW16475.1 afla|EED47210.1 afum|EAL89069.1 agla|OJJ82871.1 akaw|GAA87996.1 aluc|OJZ81251.1 anid|CBF81431.1 anig|EHA18560.1 aory|EIT79330.1 asyd|OJJ59422.1 ater|EAU36562.1 atub|OJI91448.1 aver|OJJ02848.1 awen|OJJ40361.1 azon|OJJ49209.1

Group2999: abra|OJJ73183.1 acar|OOG00420.1 acla|EAW06861.1 afis|EAW16478.1 afla|EED47215.1 afum|EAL89072.1 agla|OJJ82867.1 akaw|GAA87992.1 aluc|OJZ81246.1 anid|CBF75501.1 anig|EHA18555.1 aory|EIT79241.1 asyd|OJJ59747.1 ater|EAU36560.1 atub|OJI91442.1 aver|OJJ02541.1 awen|OJJ40357.1 azon|OJJ49206.1

Group3000: abra|OJJ73184.1 acar|OOG00421.1 acla|EAW06862.1 afis|EAW16479.1 afla|EED47216.1 afum|EAL89073.1 agla|OJJ82866.1 akaw|GAA87991.1 aluc|OJZ81245.1 anid|CBF75499.1 anig|EHA18554.1 aory|EIT79319.1 asyd|OJJ59748.1 ater|EAU36559.1 atub|OJI91441.1 aver|OJJ02540.1 awen|OJJ40356.1 azon|OJJ49205.1

Group3001: abra|OJJ73185.1 acar|OOG00422.1 acla|EAW06863.1 afis|EAW16480.1 afla|EED47217.1 afum|EAL89074.1 agla|OJJ82865.1 akaw|GAA87990.1 aluc|OJZ81244.1 anid|CBF75503.1 anig|EHA18553.1 aory|EIT79172.1 asyd|OJJ59746.1 ater|EAU36558.1 atub|OJI91440.1 aver|OJJ02542.1 awen|OJJ40354.1 azon|OJJ49204.1

Group3002: abra|OJJ73189.1 acar|OOG00425.1 acla|EAW06865.1 afis|EAW16482.1 afla|EED47222.1 afum|EAL89076.1 agla|OJJ82862.1 akaw|GAA87988.1 aluc|OJZ81241.1 anid|CBF75506.1 anig|EHA18551.1 aory|EIT79303.1 asyd|OJJ59744.1 ater|EAU36556.1 atub|OJI91438.1 aver|OJJ02544.1 awen|OJJ40350.1 azon|OJJ49202.1

Group3003: abra|OJJ73190.1 acar|OOG00426.1 acla|EAW06866.1 afis|EAW16483.1 afla|EED47223.1 afum|EAL89077.1 agla|OJJ82861.1 akaw|GAA87987.1 aluc|OJZ81240.1 anid|CBF75508.1 anig|EHA18550.1 aory|EIT79190.1 asyd|OJJ59743.1 ater|EAU36555.1 atub|OJI91437.1 aver|OJJ02545.1 awen|OJJ40348.1 azon|OJJ49201.1

Group3004: abra|OJJ73191.1 acar|OOG00428.1 acla|EAW06867.1 afis|EAW16484.1 afla|EED47224.1 afum|EAL89078.1 agla|OJJ82860.1 akaw|GAA87986.1 aluc|OJZ81239.1 anid|CBF75520.1 anig|EHA18549.1 aory|EIT79323.1 asyd|OJJ59736.1 ater|EAU36554.1 atub|OJI91436.1 aver|OJJ02551.1 awen|OJJ40343.1 azon|OJJ49200.1

Group3005: abra|OJJ73193.1 acar|OOG00429.1 acla|EAW06869.1 afis|EAW16486.1 afla|EED47226.1 afum|EAL89080.1 agla|OJJ82858.1 akaw|GAA87984.1 aluc|OJZ81236.1 anid|CBF75516.1 anig|EHA18547.1 aory|EIT79231.1 asyd|OJJ59738.1 ater|EAU36552.1 atub|OJI91434.1 aver|OJJ02549.1 awen|OJJ40345.1 azon|OJJ49198.1

Group3006: abra|OJJ73194.1 acar|OOG00431.1 acla|EAW06871.1 afis|EAW16488.1 afla|EED47229.1 afum|EAL89082.1 agla|OJJ82852.1 akaw|GAA87983.1 aluc|OJZ81235.1 anid|CBF75510.1 anig|EHA18546.1 aory|EIT79266.1 asyd|OJJ59742.1 ater|EAU36548.1 atub|OJI91432.1 aver|OJJ02546.1 awen|OJJ40342.1 azon|OJJ49196.1

Group3007: abra|OJJ73196.1 acar|OOG00434.1 acla|EAW06873.1 afis|EAW16490.1 afla|EED47231.1 afum|EAL89084.1 agla|OJJ82829.1 akaw|GAA87981.1 aluc|OJZ81233.1 anid|CBF75522.1 anig|EHA18544.1 aory|EIT79209.1 asyd|OJJ59735.1 ater|EAU36547.1 atub|OJI91430.1 aver|OJJ02552.1 awen|OJJ40339.1 azon|OJJ49193.1

Group3008: abra|OJJ73199.1 acar|OOG00438.1 acla|EAW06876.1 afis|EAW16493.1 afla|EED47237.1 afum|EAL89087.1 agla|OJJ82832.1 akaw|GAA87978.1 aluc|OJZ81230.1 anid|CBF75527.1 anig|EHA18541.1 aory|EIT79313.1 asyd|OJJ59732.1 ater|EAU36545.1 atub|OJI91427.1 aver|OJJ02555.1 awen|OJJ40336.1 azon|OJJ49190.1

Group3009: abra|OJJ73202.1 acar|OOG00441.1 acla|EAW06879.1 afis|EAW16496.1 afla|EED47251.1 afum|EAL89090.1 agla|OJJ82835.1 akaw|GAA87975.1 aluc|OJZ81227.1 anid|CBF75606.1 anig|EHA18538.1 aory|EIT79275.1 asyd|OJJ59727.1 ater|EAU36543.1 atub|OJI91424.1 aver|OJJ02559.1 awen|OJJ40333.1 azon|OJJ49187.1

Group3010: abra|OJJ73203.1 acar|OOG00442.1 acla|EAW06880.1 afis|EAW16497.1 afla|EED56182.1 afum|EBA27367.1 agla|OJJ82836.1 akaw|GAA87974.1 aluc|OJZ81224.1 anid|CBF75605.1 anig|EHA18537.1 aory|EIT73365.1 asyd|OJJ59726.1 ater|EAU36542.1 atub|OJI91422.1 aver|OJJ02560.1 awen|OJJ40332.1 azon|OJJ49186.1

Group3011: abra|OJJ73204.1 acar|OOG00443.1 acla|EAW06881.1 afis|EAW16498.1 afla|EED56181.1 afum|EAL89093.1 agla|OJJ82838.1 akaw|GAA87973.1 aluc|OJZ81222.1 anid|CBF75604.1 anig|EHA18536.1 aory|EIT73379.1 asyd|OJJ59725.1 ater|EAU36541.1 atub|OJI91420.1 aver|OJJ02561.1 awen|OJJ40331.1 azon|OJJ49185.1

Group3012: abra|OJJ73205.1 acar|OOG00444.1 acla|EAW06882.1 afis|EAW16499.1 afla|EED56180.1 afum|EAL89094.1 agla|OJJ82826.1 akaw|GAA87972.1 aluc|OJZ81221.1 anid|CBF75602.1 anig|EHA18535.1 aory|EIT76650.1 asyd|OJJ59724.1 ater|EAU36540.1 atub|OJI91419.1 aver|OJJ02562.1 awen|OJJ40330.1 azon|OJJ49184.1

Group3013: abra|OJJ73206.1 acar|OOG00445.1 acla|EAW06883.1 afis|EAW16500.1 afla|EED56179.1 afum|EAL89095.1 agla|OJJ82827.1 akaw|GAA87971.1 aluc|OJZ81220.1 anid|CBF75600.1 anig|EHA18534.1 aory|EIT76763.1 asyd|OJJ59723.1 ater|EAU36539.1 atub|OJI91418.1 aver|OJJ02563.1 awen|OJJ40328.1 azon|OJJ49183.1

Group3014: abra|OJJ73207.1 acar|OOG00446.1 acla|EAW06884.1 afis|EAW16501.1 afla|EED56178.1 afum|EAL89096.1 agla|OJJ82828.1 akaw|GAA87970.1 aluc|OJZ81219.1 anid|CBF75598.1 anig|EHA18533.1 aory|EIT76626.1 asyd|OJJ59722.1 ater|EAU36538.1 atub|OJI91417.1 aver|OJJ02564.1 awen|OJJ40327.1 azon|OJJ49182.1

Group3015: abra|OJJ73209.1 acar|OOG00447.1 acla|EAW06885.1 afis|EAW16502.1 afla|EED56177.1 afum|EAL89097.1 agla|OJJ82815.1 akaw|GAA87969.1 aluc|OJZ81218.1 anid|CBF75589.1 anig|EHA18532.1 aory|EIT76868.1 asyd|OJJ59721.1 ater|EAU36537.1 atub|OJI91416.1 aver|OJJ02565.1 awen|OJJ40326.1 azon|OJJ49181.1

Group3016: abra|OJJ73210.1 acar|OOG00449.1 acla|EAW06886.1 afis|EAW16503.1 afla|EED56175.1 afum|EAL89098.1 agla|OJJ82816.1 akaw|GAA87968.1 aluc|OJZ81217.1 anid|CBF75591.1 anig|EHA18531.1 aory|EIT76792.1 asyd|OJJ59720.1 ater|EAU36536.1 atub|OJI91415.1 aver|OJJ02566.1 awen|OJJ40324.1 azon|OJJ49180.1

Group3017: abra|OJJ73212.1 acar|OOG00448.1 acla|EAW06888.1 afis|EAW16505.1 afla|EED56173.1 afum|EAL89100.1 agla|OJJ82818.1 akaw|GAA87966.1 aluc|OJZ81215.1 anid|CBF75594.1 anig|EHA18529.1 aory|EIT76612.1 asyd|OJJ59718.1 ater|EAU36535.1 atub|OJI91412.1 aver|OJJ02568.1 awen|OJJ40321.1 azon|OJJ49178.1

Group3018: abra|OJJ73216.1 acar|OOG00453.1 acla|EAW06891.1 afis|EAW16508.1 afla|EED56170.1 afum|EAL89103.1 agla|OJJ82822.1 akaw|GAA87963.1 aluc|OJZ81212.1 anid|CBF75578.1 anig|EHA18526.1 aory|EIT76719.1 asyd|OJJ59714.1 ater|EAU36572.1 atub|OJI91407.1 aver|OJJ02572.1 awen|OJJ40317.1 azon|OJJ47186.1

Group3019: abra|OJJ73218.1 acar|OOG00455.1 acla|EAW06892.1 afis|EAW16509.1 afla|EED56164.1 afum|EAL89104.2 agla|OJJ82812.1 akaw|GAA87961.1 aluc|OJZ81210.1 anid|CBF75572.1 anig|EHA18524.1 aory|EIT76824.1 asyd|OJJ59711.1 ater|EAU36575.1 atub|OJI91405.1 aver|OJJ02575.1 awen|OJJ40315.1 azon|OJJ49175.1

Group3020: abra|OJJ73219.1 acar|OOG00457.1 acla|EAW06894.1 afis|EAW16511.1 afla|EED56162.1 afum|EAL89106.1 agla|OJJ82811.1 akaw|GAA87959.1 aluc|OJZ81208.1 anid|CBF75568.1 anig|EHA18523.1

aory|EIT76692.1 asyd|OJJ59709.1 ater|EAU36577.1 atub|OJI91404.1 aver|OJJ02576.1 awen|OJJ40313.1 azon|OJJ49173.1

Group3021: abra|OJJ73220.1 acar|OOG00458.1 acla|EAW06895.1 afis|EAW16512.1 afla|EED56161.1 afum|EAL89107.2 agla|OJJ82810.1 akaw|GAA87958.1 aluc|OJZ81207.1 anid|CBF75566.1 anig|EHA18522.1 aory|EIT76654.1 asyd|OJJ59708.1 ater|EAU36578.1 atub|OJI91403.1 aver|OJJ02577.1 awen|OJJ40312.1 azon|OJJ49172.1

Group3022: abra|OJJ73222.1 acar|OOG00460.1 acla|EAW06897.1 afis|EAW16514.1 afla|EED56159.1 afum|EAL89109.1 agla|OJJ82808.1 akaw|GAA87956.1 aluc|OJZ81205.1 anid|CBF75562.1 anig|EHA18520.1 aory|EIT76696.1 asyd|OJJ59706.1 ater|EAU36580.1 atub|OJI91401.1 aver|OJJ02579.1 awen|OJJ40309.1 azon|OJJ49170.1

Group3023: abra|OJJ73223.1 acar|OOG00461.1 acla|EAW06898.1 afis|EAW16515.1 afla|EED56158.1 afum|EAL89110.2 agla|OJJ82807.1 akaw|GAA87955.1 aluc|OJZ81204.1 anid|CBF75560.1 anig|EHA18519.1 aory|EIT76553.1 asyd|OJJ59705.1 ater|EAU36581.1 atub|OJI91400.1 aver|OJJ02580.1 awen|OJJ40308.1 azon|OJJ49169.1

Group3024: abra|OJJ73224.1 acar|OOG00462.1 acla|EAW06899.1 afis|EAW16516.1 afla|EED56157.1 afum|EAL89111.2 agla|OJJ82806.1 akaw|GAA87954.1 aluc|OJZ81203.1 anid|CBF75558.1 anig|EHA18518.1 aory|EIT76604.1 asyd|OJJ59704.1 ater|EAU36582.1 atub|OJI91398.1 aver|OJJ02582.1 awen|OJJ40307.1 azon|OJJ49168.1

Group3025: abra|OJJ73228.1 acar|OOG00466.1 acla|EAW06902.1 afis|EAW16519.1 afla|EED56152.1 afum|EAL89114.1 agla|OJJ82801.1 akaw|GAA87952.1 aluc|OJZ81200.1 anid|CBF75555.1 anig|EHA18515.1 aory|EIT76840.1 asyd|OJJ57431.1 ater|EAU36585.1 atub|OJI91395.1 aver|OJJ05749.1 awen|OJJ40303.1 azon|OJJ49165.1

Group3026: abra|OJJ73229.1 acar|OOG00468.1 acla|EAW06904.1 afis|EAW16521.1 afla|EED56150.1 afum|EAL89116.1 agla|OJJ82800.1 akaw|GAA87951.1 aluc|OJZ81199.1 anid|CBF75547.1 anig|EHA18514.1 aory|EIT76779.1 asyd|OJJ57441.1 ater|EAU36586.1 atub|OJI91394.1 aver|OJJ05737.1 awen|OJJ40302.1 azon|OJJ49164.1

Group3027: abra|OJJ73230.1 acar|OOG00469.1 acla|EAW06905.1 afis|EAW16522.1 afla|EED56149.1 afum|EAL89117.1 agla|OJJ82799.1 akaw|GAA87950.1 aluc|OJZ81198.1 anid|CBF75545.1 anig|EHA18513.1 aory|EIT76528.1 asyd|OJJ57442.1 ater|EAU36587.1 atub|OJI91393.1 aver|OJJ05736.1 awen|OJJ40301.1 azon|OJJ49163.1

Group3028: abra|OJJ73231.1 acar|OOG00470.1 acla|EAW06906.1 afis|EAW16523.1 afla|EED56148.1 afum|EAL89118.1 agla|OJJ82798.1 akaw|GAA87949.1 aluc|OJZ81197.1 anid|CBF75543.1 anig|EHA18512.1 aory|EIT76854.1 asyd|OJJ57443.1 ater|EAU36588.1 atub|OJI91392.1 aver|OJJ05735.1 awen|OJJ40300.1 azon|OJJ49162.1

Group3029: abra|OJJ73232.1 acar|OOG00471.1 acla|EAW06907.1 afis|EAW16524.1 afla|EED56147.1 afum|EAL89119.1 agla|OJJ82797.1 akaw|GAA87948.1 aluc|OJZ81196.1 anid|CBF75541.1 anig|EHA18511.1 aory|EIT76769.1 asyd|OJJ57446.1 ater|EAU36589.1 atub|OJI91391.1 aver|OJJ05734.1 awen|OJJ40299.1 azon|OJJ49161.1

Group3030: abra|OJJ73234.1 acar|OOG00472.1 acla|EAW06909.1 afis|EAW16526.1 afla|EED56145.1 afum|EAL89122.1 agla|OJJ82796.1 akaw|GAA87946.1 aluc|OJZ81194.1 anid|CBF75537.1 anig|EHA18509.1 aory|EIT76537.1 asyd|OJJ57447.1 ater|EAU36590.1 atub|OJI91389.1 aver|OJJ05732.1 awen|OJJ40297.1 azon|OJJ49159.1

Group3031: abra|OJJ73236.1 acar|OOG00473.1 acla|EAW06910.1 afis|EAW16527.1 afla|EED56144.1 afum|EAL89123.2 agla|OJJ82795.1 akaw|GAA87945.1 aluc|OJZ81193.1 anid|CBF75535.1 anig|EHA18508.1 aory|EIT76806.1 asyd|OJJ57448.1 ater|EAU36591.1 atub|OJI91388.1 aver|OJJ05731.1 awen|OJJ40296.1 azon|OJJ49158.1

Group3032: abra|OJJ73237.1 acar|OOG00474.1 acla|EAW06911.1 afis|EAW16528.1 afla|EED56143.1 afum|EAL89124.1 agla|OJJ82793.1 akaw|GAA87944.1 aluc|OJZ81191.1 anid|CBF75533.1 anig|EHA18507.1 aory|EIT76880.1 asyd|OJJ57449.1 ater|EAU36592.1 atub|OJI91387.1 aver|OJJ05730.1 awen|OJJ40295.1 azon|OJJ49157.1

Group3033: abra|OJJ73239.1 acar|OOG00476.1 acla|EAW06912.1 afis|EAW16529.1 afla|EED56141.1 afum|EAL89125.1 agla|OJJ82791.1 akaw|GAA87942.1 aluc|OJZ81189.1 anid|CBF75529.1 anig|EHA18505.1 aory|EIT76699.1 asyd|OJJ57450.1 ater|EAU36593.1 atub|OJI91385.1 aver|OJJ05728.1 awen|OJJ40293.1 azon|OJJ49154.1

Group3034: abra|OJJ73240.1 acar|OOG00477.1 acla|EAW06913.1 afis|EAW16530.1 afla|EED56138.1 afum|EAL89127.1 agla|OJJ82790.1 akaw|GAA87941.1 aluc|OJZ81188.1 anid|CBF81731.1 anig|EHA18504.1 aory|EIT76798.1 asyd|OJJ59592.1 ater|EAU36594.1 atub|OJI91384.1 aver|OJJ02677.1 awen|OJJ40291.1 azon|OJJ49153.1

Group3035: abra|OJJ73246.1 acar|OOG00485.1 acla|EAW06920.1 afis|EAW16537.1 afla|EED56132.1 afum|EAL89134.1 agla|OJJ82785.1 akaw|GAA87934.1 aluc|OJZ81181.1 anid|CBF81743.1 anig|EHA18499.1 aory|EIT76834.1 asyd|OJJ59598.1 ater|EAU36600.1 atub|OJI91378.1 aver|OJJ02670.1 awen|OJJ40285.1 azon|OJJ49146.1

Group3036: abra|OJJ73248.1 acar|OOG00488.1 acla|EAW06922.1 afis|EAW16539.1 afla|EED56130.1 afum|EAL89136.1 agla|OJJ82783.1 akaw|GAA87932.1 aluc|OJZ81179.1 anid|CBF81746.1 anig|EHA18497.1 aory|EIT76822.1 asyd|OJJ59600.1 ater|EAU36602.1 atub|OJI91375.1 aver|OJJ02668.1 awen|OJJ40282.1 azon|OJJ49144.1

Group3037: abra|OJJ73249.1 acar|OOG00491.1 acla|EAW06924.1 afis|EAW16541.1 afla|EED56127.1 afum|EAL89138.1 agla|OJJ82782.1 akaw|GAA87931.1 aluc|OJZ81178.1 anid|CBF81748.1 anig|EHA18496.1 aory|EIT76785.1 asyd|OJJ59601.1 ater|EAU36603.1 atub|OJI91374.1 aver|OJJ02667.1 awen|OJJ40279.1 azon|OJJ49142.1

Group3038: abra|OJJ73251.1 acar|OOG00495.1 acla|EAW06929.1 afis|EAW16544.1 afla|EED56121.1 afum|EAL89142.2 agla|OJJ82779.1 akaw|GAA87929.1 aluc|OJZ81176.1 anid|CBF81754.1 anig|EHA18494.1 aory|EIT76628.1 asyd|OJJ59604.1 ater|EAU36607.1 atub|OJI91372.1 aver|OJJ02665.1 awen|OJJ40276.1 azon|OJJ49139.1

Group3039: abra|OJJ73252.1 acar|OOG00494.1 acla|EAW06930.1 afis|EAW16545.1 afla|EED56120.1 afum|EAL89143.1 agla|OJJ82778.1 akaw|GAA87928.1 aluc|OJZ81175.1 anid|CBF81755.1 anig|EHA18493.1 aory|EIT76556.1 asyd|OJJ59605.1 ater|EAU36608.1 atub|OJI91371.1 aver|OJJ02664.1 awen|OJJ40275.1 azon|OJJ49138.1

Group3040: abra|OJJ73253.1 acar|OOG00496.1 acla|EAW06931.1 afis|EAW16546.1 afum|EAL89144.1 agla|OJJ82777.1 akaw|GAA87927.1 aluc|OJZ81174.1 anid|CBF81757.1 anig|EHA18492.1 aory|EIT76862.1 asyd|OJJ59606.1 ater|EAU36609.1 atub|OJI91370.1 aver|OJJ02663.1 awen|OJJ40271.1 azon|OJJ49137.1 afla|EED56118.1

Group3041: abra|OJJ73256.1 acar|OOG00499.1 acla|EAW06933.1 afis|EAW16548.1 afla|EED56116.1 afum|EAL89146.1 agla|OJJ82775.1 akaw|GAA87925.1 aluc|OJZ81172.1 anid|CBF81761.1 anig|EHA18490.1 aory|EIT76624.1 asyd|OJJ59607.1 ater|EAU36611.1 atub|OJI91367.1 aver|OJJ02662.1 awen|OJJ40269.1 azon|OJJ49133.1

Group3042: abra|OJJ73257.1 acar|OOG00498.1 acla|EAW06934.1 afis|EAW16549.1 afla|EED56115.1 afum|EAL89147.1 agla|OJJ82774.1 akaw|GAA87924.1 aluc|OJZ81171.1 anid|CBF81763.1 anig|EHA18489.1 aory|EIT76873.1 asyd|OJJ59608.1 ater|EAU36612.1 atub|OJI91366.1 aver|OJJ02661.1 awen|OJJ40268.1 azon|OJJ49132.1

Group3043: abra|OJJ73258.1 acar|OOG00500.1 acla|EAW06935.1 afis|EAW16550.1 afla|EED56112.1 afum|EAL89148.1 agla|OJJ82773.1 akaw|GAA87923.1 aluc|OJZ81170.1 anid|CBF81765.1 anig|EHA18488.1 aory|EIT76657.1 ater|EAU36613.1 atub|OJI91365.1 awen|OJJ40267.1 azon|OJJ49131.1 asyd|OJJ59609.1 aver|OJJ02660.1

Group3044: abra|OJJ73259.1 acar|OOG00501.1 acla|EAW06936.1 afis|EAW16551.1 afla|EED56111.1 afum|EAL89149.1 agla|OJJ82772.1 akaw|GAA87922.1 aluc|OJZ81169.1 anid|CBF81766.1 anig|EHA18487.1 aory|EIT76585.1 asyd|OJJ59610.1 ater|EAU36617.1 atub|OJI91364.1 aver|OJJ02659.1 awen|OJJ40266.1 azon|OJJ49130.1

Group3045: abra|OJJ73262.1 acar|OOG00502.1 acla|EAW06939.1 afis|EAW16554.1 afla|EED56108.1 afum|EAL89152.1 agla|OJJ82769.1 akaw|GAA87920.1 aluc|OJZ81167.1 anid|CBF81795.1 anig|EHA18484.1 aory|EIT76711.1 asyd|OJJ59637.1 ater|EAU36619.1 atub|OJI91362.1 aver|OJJ02644.1 awen|OJJ40264.1 azon|OJJ49127.1

Group3046: abra|OJJ73263.1 acar|OOG00503.1 acla|EAW06940.1 afis|EAW16555.1 afla|EED56107.1 afum|EAL89153.1 agla|OJJ82768.1 akaw|GAA87919.1 aluc|OJZ81166.1 anid|CBF81796.1 anig|EHA18483.1 aory|EIT76566.1 asyd|OJJ59638.1 ater|EAU36620.1 atub|OJI91361.1 aver|OJJ02643.1 awen|OJJ40263.1 azon|OJJ49126.1

Group3047: abra|OJJ73264.1 acar|OOG00504.1 acla|EAW06941.1 afis|EAW16556.1 afla|EED56106.1 afum|EAL89154.1 agla|OJJ82767.1 akaw|GAA87918.1 aluc|OJZ81165.1 anid|CBF81798.1 anig|EHA18482.1 aory|EIT76736.1 asyd|OJJ59639.1 ater|EAU36621.1 atub|OJI91360.1 aver|OJJ02642.1 awen|OJJ40262.1 azon|OJJ49125.1

Group3048: abra|OJJ73265.1 acar|OOG00505.1 acla|EAW06942.1 afis|EAW16557.1 afla|EED56105.1 afum|EAL89155.1 agla|OJJ82766.1 akaw|GAA87917.1 aluc|OJZ81164.1 anid|CBF81802.1 anig|EHA18481.1 aory|EIT76677.1 asyd|OJJ59642.1 ater|EAU36624.1 atub|OJI91359.1 aver|OJJ02640.1 awen|OJJ40260.1 azon|OJJ49122.1

Group3049: abra|OJJ73266.1 acar|OOG00506.1 acla|EAW06943.1 afis|EAW16558.1 afla|EED56104.1 afum|EAL89156.1 agla|OJJ82765.1 akaw|GAA87916.1 aluc|OJZ81163.1 anid|CBF81800.1 anig|EHA18480.1

aory|EIT76573.1 asyd|OJJ59640.1 ater|EAU36625.1 atub|OJI91358.1 aver|OJJ02641.1 awen|OJJ40259.1 azon|OJJ49123.1

Group3050: abra|OJJ73269.1 acar|OOG00509.1 acla|EAW06946.1 afis|EAW16561.1 afla|EED56101.1 afum|EAL89159.1 agla|OJJ82762.1 akaw|GAA87913.1 aluc|OJZ81160.1 anid|CBF81808.1 anig|EHA18477.1 aory|EIT76636.1 asyd|OJJ59645.1 ater|EAU36628.1 atub|OJI91355.1 aver|OJJ02637.1 awen|OJJ40256.1 azon|OJJ49120.1

Group3051: abra|OJJ73275.1 acar|OOG00514.1 acla|EAW06950.1 afis|EAW16565.1 afla|EED56098.1 afum|EAL89162.1 agla|OJJ82756.1 akaw|GAA92730.1 aluc|OJZ88774.1 anid|CBF81817.1 anig|EHA18468.1 aory|EIT76722.1 asyd|OJJ59651.1 ater|EAU36634.1 atub|OJI91345.1 aver|OJJ02631.1 awen|OJJ40253.1 azon|OJJ49115.1

Group3052: abra|OJJ73278.1 acar|OOG00515.1 acla|EAW06954.1 afis|EAW16569.1 afla|EED56095.1 afum|EAL89166.1 agla|OJJ82751.1 akaw|GAA92727.1 aluc|OJZ88771.1 anid|CBF81822.1 anig|EHA18467.1 aory|EIT82359.1 asyd|OJJ59654.1 ater|EAU36637.1 atub|OJI91344.1 aver|OJJ02628.1 awen|OJJ40250.1 azon|OJJ41980.1

Group3053: abra|OJJ73281.1 acar|OOG00517.1 acla|EAW06956.1 afis|EAW16571.1 afla|EED56093.1 afum|EAL89168.1 agla|OJJ82703.1 akaw|GAA92724.1 aluc|OJZ88768.1 anid|CBF81826.1 anig|EHA18464.1 aory|EIT76754.1 asyd|OJJ59657.1 ater|EAU36639.1 atub|OJI91341.1 aver|OJJ02625.1 awen|OJJ40247.1 azon|OJJ49108.1

Group3054: abra|OJJ73283.1 acar|OOG00519.1 acla|EAW06958.1 afis|EAW16573.1 afla|EED56091.1 afum|EAL89170.2 agla|OJJ82705.1 akaw|GAA92722.1 aluc|OJZ88766.1 anid|CBF81828.1 anig|EHA18462.1 aory|EIT76851.1 asyd|OJJ59659.1 ater|EAU36641.1 atub|OJI91339.1 aver|OJJ02623.1 awen|OJJ40245.1 azon|OJJ49106.1

Group3055: abra|OJJ73287.1 acar|OOG00522.1 acla|EAW06962.1 afis|EAW16577.1 afla|EED56087.1 afum|EAL89173.1 agla|OJJ82700.1 akaw|GAA92718.1 aluc|OJZ88763.1 anid|CBF81876.1 anig|EHA18459.1 aory|EIT76884.1 asyd|OJJ59665.1 ater|EAU36645.1 atub|OJI91335.1 aver|OJJ02618.1 awen|OJJ40240.1 azon|OJJ49103.1

Group3057: abra|OJJ73338.1 acar|OOG00523.1 acla|EAW06963.1 afis|EAW16578.1 afla|EED56086.1 afum|EAL89174.1 agla|OJJ82699.1 akaw|GAA83146.1 aluc|OJZ88703.1 anid|CBF81878.1 anig|EHA18387.1 aory|EIT76746.1 asyd|OJJ59666.1 ater|EAU36646.1 atub|OJI91284.1 aver|OJJ02617.1 awen|OJJ40239.1 azon|OJJ49102.1

Group3058: abra|OJJ73339.1 acar|OOG00524.1 acla|EAW06964.1 afis|EAW16579.1 afla|EED56085.1 afum|EAL89175.1 agla|OJJ82698.1 akaw|GAA83145.1 aluc|OJZ88702.1 anid|CBF81880.1 anig|EHA18386.1 aory|EIT76894.1 asyd|OJJ59667.1 ater|EAU36647.1 atub|OJI91283.1 aver|OJJ02616.1 awen|OJJ40238.1 azon|OJJ49101.1

Group3059: abra|OJJ73340.1 acar|OOG00527.1 acla|EAW06965.1 afis|EAW16580.1 afla|EED56082.1 afum|EAL89176.1 agla|OJJ82695.1 akaw|GAA83142.1 aluc|OJZ88699.1 anid|CBF81883.1 anig|EHA18379.1 aory|EIT76744.1 asyd|OJJ59669.1 ater|EAU36649.1 atub|OJI91280.1 aver|OJJ02614.1 awen|OJJ40234.1 azon|OJJ49100.1

Group3060: abra|OJJ73343.1 acar|OOG00553.1 acla|EAW06970.1 afis|EAW16584.1 afla|EED56081.1 afum|EAL89179.1 agla|OJJ82694.1 akaw|GAA83139.1 aluc|OJZ88696.1 anid|CBF81885.1 anig|EHA18376.1 aory|EIT76749.1 asyd|OJJ59671.1 ater|EAU36661.1 atub|OJI91277.1 aver|OJJ02613.1 awen|OJJ40232.1 azon|OJJ49096.1

Group3061: abra|OJJ73344.1 acar|OOG00554.1 acla|EAW06971.1 afis|EAW16585.1 afla|EED56080.1 afum|EAL89180.1 agla|OJJ82693.1 akaw|GAA83138.1 aluc|OJZ88695.1 anid|CBF81887.1 anig|EHA18375.1 aory|EIT76713.1 asyd|OJJ59672.1 ater|EAU36662.1 atub|OJI91275.1 aver|OJJ02612.1 awen|OJJ40231.1 azon|OJJ49095.1

Group3062: abra|OJJ73345.1 acar|OOG00555.1 acla|EAW06972.1 afis|EAW16586.1 afla|EED56079.1 afum|EAL89181.2 agla|OJJ82692.1 akaw|GAA83137.1 aluc|OJZ88694.1 anid|CBF81889.1 anig|EHA18374.1 aory|EIT76590.1 asyd|OJJ59673.1 ater|EAU36663.1 atub|OJI91274.1 aver|OJJ02611.1 awen|OJJ40230.1 azon|OJJ49094.1

Group3063: abra|OJJ73359.1 acar|OOG00567.1 acla|EAW06979.1 afis|EAW16593.1 afla|EED45008.1 afum|EAL89189.1 agla|OJJ82660.1 akaw|GAA83125.1 aluc|OJZ88681.1 anid|CBF81897.1 anig|EHA18362.1 aory|EIT73639.1 asyd|OJJ59682.1 ater|EAU36673.1 atub|OJI91262.1 aver|OJJ02606.1 awen|OJJ40214.1 azon|OJJ49080.1

Group3064: abra|OJJ73361.1 acar|OOG00568.1 acla|EAW06981.1 afis|EAW16595.1 afla|EED45014.1 afum|EAL89191.1 agla|OJJ82659.1 akaw|GAA83123.1 aluc|OJZ88679.1 anid|CBF81901.1 anig|EHA18360.1 aory|EIT73637.1 asyd|OJJ59685.1 ater|EAU36674.1 atub|OJI91260.1 aver|OJJ02602.1 awen|OJJ40213.1 azon|OJJ49079.1

Group3065: abra|OJJ73362.1 acar|OOG00569.1 acla|EAW06982.1 afis|EAW16596.1 afla|EED45015.1 afum|EAL89192.1 agla|OJJ82658.1 akaw|GAA83122.1 aluc|OJZ88678.1 anid|CBF81902.1 anig|EHA18359.1 aory|EIT73689.1 asyd|OJJ59686.1 ater|EAU36675.1 atub|OJI91259.1 aver|OJJ02601.1 awen|OJJ40212.1 azon|OJJ49078.1

Group3066: abra|OJJ73365.1 acar|OOG00572.1 acla|EAW06985.1 afis|EAW16599.1 afla|EED45018.1 afum|EAL89195.1 agla|OJJ82655.1 akaw|GAA83119.1 aluc|OJZ88675.1 anid|CBF81905.1 anig|EHA18357.1 aory|EIT73617.1 asyd|OJJ59688.1 ater|EAU36680.1 atub|OJI91256.1 aver|OJJ02599.1 awen|OJJ40209.1 azon|OJJ49075.1

Group3067: abra|OJJ73368.1 acar|OOG00574.1 acla|EAW06988.1 afis|EAW16602.1 afla|EED45021.1 afum|EAL89198.1 agla|OJJ82650.1 akaw|GAA83116.1 aluc|OJZ88672.1 anid|CBF81909.1 anig|EHA18354.1 aory|EIT73783.1 asyd|OJJ59690.1 ater|EAU36683.1 atub|OJI91253.1 aver|OJJ02597.1 awen|OJJ40204.1 azon|OJJ49071.1

Group3068: abra|OJJ73378.1 acar|OOF90413.1 acla|EAW06994.1 afis|EAW16609.1 afla|EED45027.1 afum|EAL89205.1 agla|OJJ82638.1 akaw|GAA83111.1 aluc|OJZ88665.1 anid|CBF81965.1 anig|EHA18350.1 aory|EIT73642.1 asyd|OJJ57355.1 ater|EAU36693.1 atub|OJI91247.1 aver|OJJ05810.1 awen|OJJ40198.1 azon|OJJ49069.1

Group3069: abra|OJJ73382.1 acar|OOG00581.1 acla|EAW06997.1 afis|EAW16611.1 afla|EED45000.1 afum|EAL89207.1 agla|OJJ82634.1 akaw|GAA83107.1 aluc|OJZ88656.1 anid|CBF81963.1 anig|EHA18347.1 aory|EIT73611.1 asyd|OJJ57358.1 ater|EAU36696.1 atub|OJI91242.1 aver|OJJ05808.1 awen|OJJ40193.1 azon|OJJ49061.1

Group3070: abra|OJJ73383.1 acar|OOG00582.1 acla|EAW06998.1 afis|EAW16613.1 afla|EED44999.1 afum|EAL89223.1 agla|OJJ82633.1 akaw|GAA83106.1 aluc|OJZ88654.1 anid|CBF81961.1 anig|EHA18346.1 aory|EIT73720.1 asyd|OJJ57359.1 ater|EAU36697.1 atub|OJI91241.1 aver|OJJ05807.1 awen|OJJ40192.1 azon|OJJ49060.1

Group3071: abra|OJJ73402.1 acar|OOG00600.1 acla|EAW07005.1 afis|EAW16622.1 afla|EED44970.1 afum|EAL89215.1 agla|OJJ82622.1 akaw|GAA83088.1 aluc|OJZ88635.1 anid|CBF81987.1 anig|EHA18325.1 aory|EIT73499.1 asyd|OJJ56156.1 ater|EAU36736.1 atub|OJI91219.1 aver|OJJ05096.1 awen|OJJ40086.1 azon|OJJ49496.1

Group3072: abra|OJJ73424.1 acar|OOG00628.1 acla|EAW07034.1 afis|EAW16660.1 afla|EED44893.1 afum|EAL89264.1 agla|OJJ78709.1 akaw|GAA83068.1 aluc|OJZ88619.1 anid|CBF82075.1 anig|EHA18310.1 aory|EIT83240.1 asyd|OJJ54250.1 ater|EAU39583.1 atub|OJI91204.1 aver|OJJ05204.1 awen|OJJ40053.1 azon|OJJ50109.1

Group3073: abra|OJJ73425.1 acar|OOG00629.1 acla|EAW07035.1 afis|EAW16661.1 afla|EED44894.1 afum|EAL89265.1 agla|OJJ78708.1 akaw|GAA83067.1 aluc|OJZ88618.1 anid|CBF82076.1 anig|EHA18309.1 aory|EIT83251.1 asyd|OJJ54251.1 ater|EAU39584.1 atub|OJI91203.1 aver|OJJ05205.1 awen|OJJ40052.1 azon|OJJ50116.1

Group3074: abra|OJJ73432.1 acar|OOG00639.1 acla|EAW06771.1 afis|EAW16322.1 afla|EED53560.1 afum|EAL88945.1 agla|OJJ88284.1 akaw|GAA83060.1 aluc|OJZ88609.1 anid|CBF89157.1 anig|EHA18302.1 aory|EIT75216.1 asyd|OJJ62563.1 ater|EAU34135.1 atub|OJI91194.1 aver|OJI96687.1 awen|OJJ37722.1 azon|OJJ44981.1

Group3075: abra|OJJ73437.1 acar|OOG00643.1 acla|EAW06775.1 afis|EAW16326.1 afla|EED53555.1 afum|EAL88950.1 agla|OJJ88266.1 akaw|GAA83056.1 aluc|OJZ88604.1 anid|CBF89164.1 anig|EHA18297.1 aory|EIT75038.1 asyd|OJJ62546.1 ater|EAU34131.1 atub|OJI91188.1 aver|OJI96672.1 awen|OJJ37726.1 azon|OJJ44986.1

Group3076: abra|OJJ73440.1 acar|OOG00646.1 acla|EAW06772.1 afis|EAW16323.1 afla|EED53559.1 afum|EAL88946.1 agla|OJJ88285.1 akaw|GAA83053.1 aluc|OJZ88600.1 anid|CBF89158.1 anig|EHA18294.1 aory|EIT74917.1 asyd|OJJ62560.1 ater|EAU34134.1 atub|OJI91185.1 aver|OJI96685.1 awen|OJJ37723.1 azon|OJJ44982.1

Group3077: abra|OJJ73442.1 acar|OOG00648.1 acla|EAW06777.1 afis|EAW16328.1 afla|EED53553.1 afum|EAL88952.1 agla|OJJ88264.1 akaw|GAA83052.1 aluc|OJZ88598.1 anid|CBF89168.1 anig|EHA18292.1 aory|EIT74949.1 asyd|OJJ62543.1 ater|EAU34129.1 atub|OJI91183.1 aver|OJI96670.1 awen|OJJ37729.1 azon|OJJ44988.1

Group3078: abra|OJJ73451.1 acar|OOG00654.1 acla|EAW06782.1 afis|EAW16332.1 afla|EED53545.1 afum|EAL88957.1 agla|OJJ88260.1 akaw|GAA83045.1 aluc|OJZ88590.1 anid|CBF89183.1 anig|EHA18285.1 aory|EIT75091.1 asyd|OJJ62537.1 ater|EAU34126.1 atub|OJI91176.1 aver|OJI96663.1 awen|OJJ37733.1 azon|OJJ44990.1

Group3079: abra|OJJ73456.1 acar|OOG00656.1 acla|EAW06783.1 afis|EAW16333.1 afla|EED53543.1 afum|EAL88958.1 agla|OJJ88259.1 akaw|GAA83042.1 aluc|OJZ88587.1 anid|CBF89188.1 anig|EHA18281.1

aory|EIT75083.1 asyd|OJJ62536.1 ater|EAU34125.1 atub|OJI91173.1 aver|OJI96661.1 awen|OJJ37734.1 azon|OJJ44993.1

Group3080: abra|OJJ73458.1 acar|OOG00659.1 acla|EAW06785.1 afis|EAW16335.1 afla|EED53541.1 afum|EAL88960.1 agla|OJJ88251.1 akaw|GAA83040.1 aluc|OJZ88585.1 anid|CBF89192.1 anig|EHA18280.1 aory|EIT74882.1 asyd|OJJ62534.1 ater|EAU34123.1 atub|OJI91171.1 aver|OJI96659.1 awen|OJJ37737.1 azon|OJJ44997.1

Group3081: abra|OJJ73460.1 acar|OOG00661.1 acla|EAW06787.1 afis|EAW16336.1 afla|EED53539.1 afum|EAL88962.1 agla|OJJ88254.1 akaw|GAA83038.1 aluc|OJZ88583.1 anid|CBF89196.1 anig|EHA18278.1 aory|EIT75013.1 asyd|OJJ62532.1 ater|EAU34121.1 atub|OJI91169.1 aver|OJI96657.1 awen|OJJ37739.1 azon|OJJ45000.1

Group3082: abra|OJJ73463.1 acar|OOG00664.1 acla|EAW06789.1 afis|EAW16339.1 afla|EED53536.1 afum|EAL88964.1 agla|OJJ88256.1 akaw|GAA83035.1 aluc|OJZ88581.1 anid|CBF89199.1 anig|EHA18277.1 aory|EIT74845.1 asyd|OJJ62530.1 ater|EAU34119.1 atub|OJI91167.1 aver|OJI96655.1 awen|OJJ37741.1 azon|OJJ45002.1

Group3083: abra|OJJ73464.1 acar|OOG00665.1 acla|EAW06790.1 afis|EAW16340.1 afla|EED53535.1 afum|EAL88965.1 agla|OJJ88257.1 akaw|GAA83034.1 aluc|OJZ88580.1 anid|CBF89201.1 anig|EHA18276.1 aory|EIT75162.1 asyd|OJJ62529.1 ater|EAU34117.1 atub|OJI91166.1 aver|OJI96654.1 awen|OJJ37742.1 azon|OJJ45003.1

Group3084: abra|OJJ73465.1 acar|OOG00666.1 acla|EAW06792.1 afis|EAW16341.1 afla|EED53534.1 afum|EAL88966.1 agla|OJJ88247.1 akaw|GAA83033.1 aluc|OJZ88579.1 anid|CBF89203.1 anig|EHA18275.1 aory|EIT75067.1 asyd|OJJ62528.1 ater|EAU34116.1 atub|OJI91165.1 aver|OJI96651.1 awen|OJJ37743.1 azon|OJJ45004.1

Group3085: abra|OJJ73466.1 acar|OOG00667.1 acla|EAW06793.1 afis|EAW16342.1 afla|EED53533.1 afum|EAL88967.1 agla|OJJ88246.1 akaw|GAA83032.1 aluc|OJZ88578.1 anid|CBF89205.1 anig|EHA18274.1 aory|EIT75133.1 asyd|OJJ62527.1 ater|EAU34115.1 atub|OJI91164.1 aver|OJI96650.1 awen|OJJ37744.1 azon|OJJ45005.1

Group3086: abra|OJJ73467.1 acar|OOG00668.1 acla|EAW06794.1 afis|EAW16343.1 afla|EED53532.1 afum|EAL88968.1 agla|OJJ88245.1 akaw|GAA83031.1 aluc|OJZ88577.1 anid|CBF89206.1 anig|EHA18273.1 aory|EIT74961.1 asyd|OJJ62526.1 ater|EAU34114.1 atub|OJI91163.1 aver|OJI96649.1 awen|OJJ37745.1 azon|OJJ45006.1

Group3087: abra|OJJ73470.1 acar|OOG00669.1 acla|EAW06795.1 afis|EAW16344.1 afla|EED53526.1 afum|EAL88969.1 agla|OJJ88244.1 akaw|GAA83030.1 aluc|OJZ88576.1 anid|CBF89217.1 anig|EHA18272.1 aory|EIT74890.1 asyd|OJJ62524.1 ater|EAU34113.1 atub|OJI91162.1 aver|OJI96645.1 awen|OJJ37747.1 azon|OJJ45008.1

Group3088: abra|OJJ73474.1 acar|OOG00670.1 acla|EAW06799.1 afis|EAW16348.1 afla|EED53523.1 afum|EAL88973.1 agla|OJJ88239.1 akaw|GAA83027.1 aluc|OJZ88573.1 anid|CBF89210.1 anig|EHA18271.1 aory|EIT75044.1 asyd|OJJ62522.1 ater|EAU34111.1 atub|OJI91159.1 aver|OJI96641.1 awen|OJJ37751.1 azon|OJJ45010.1

Group3089: abra|OJJ73476.1 acar|OOG00673.1 acla|EAW06802.1 afis|EAW16351.1 afla|EED53521.1 afum|EAL88976.1 agla|OJJ88249.1 akaw|GAA83025.1 aluc|OJZ88571.1 anid|CBF89221.1 anig|EHA18269.1 aory|EIT75100.1 asyd|OJJ62520.1 ater|EAU34110.1 atub|OJI91157.1 aver|OJI96639.1 awen|OJJ37754.1 azon|OJJ45013.1

Group3090: abra|OJJ73484.1 acar|OOG00678.1 acla|EAW06808.1 afis|EAW16357.1 afla|EED53513.1 afum|EAL88982.1 agla|OJJ88236.1 akaw|GAA87433.1 aluc|OJZ88564.1 anid|CBF89232.1 anig|EHA18261.1 aory|EIT75027.1 asyd|OJJ62514.1 ater|EAU34102.1 atub|OJI91150.1 aver|OJI96633.1 awen|OJJ37761.1 azon|OJJ45061.1

Group3091: abra|OJJ73485.1 acar|OOG00680.1 acla|EAW06809.1 afis|EAW16358.1 afla|EED53512.1 afum|EAL88983.1 agla|OJJ88235.1 akaw|GAA87434.1 aluc|OJZ88563.1 anid|CBF89234.1 anig|EHA18260.1 aory|EIT75048.1 asyd|OJJ62513.1 ater|EAU34101.1 atub|OJI91149.1 aver|OJI96632.1 awen|OJJ37762.1 azon|OJJ45062.1

Group3092: abra|OJJ73486.1 acar|OOG00679.1 acla|EAW06810.1 afis|EAW16359.1 afla|EED53510.1 afum|EAL88984.1 agla|OJJ88233.1 akaw|GAA87435.1 aluc|OJZ88562.1 anid|CBF89236.1 anig|EHA18259.1 aory|EIT75112.1 asyd|OJJ62512.1 ater|EAU34100.1 atub|OJI91147.1 aver|OJI96631.1 awen|OJJ37763.1 azon|OJJ45063.1

Group3093: abra|OJJ73487.1 acar|OOG00681.1 acla|EAW06811.1 afis|EAW16360.1 afla|EED53508.1 afum|EAL88985.1 agla|OJJ88232.1 akaw|GAA87436.1 aluc|OJZ88561.1 anid|CBF89237.1 anig|EHA18258.1 aory|EIT75230.1 asyd|OJJ62510.1 ater|EAU34092.1 atub|OJI91146.1 aver|OJI96630.1 awen|OJJ37764.1 azon|OJJ45064.1

Group3094: abra|OJJ73488.1 acar|OOG00682.1 acla|EAW06812.1 afis|EAW16361.1 afla|EED53507.1 afum|EAL88986.1 agla|OJJ88231.1 akaw|GAA87437.1 aluc|OJZ88560.1 anid|CBF89239.1 anig|EHA18257.1 aory|EIT75024.1 asyd|OJJ62509.1 ater|EAU34091.1 atub|OJI91145.1 aver|OJI96629.1 awen|OJJ37765.1 azon|OJJ45065.1

Group3095: abra|OJJ73496.1 acar|OOG00694.1 acla|EAW07430.1 afis|EAW22456.1 afla|EED54488.1 afum|EAL90735.1 agla|OJJ87568.1 akaw|GAA87445.1 aluc|OJZ88552.1 anid|CBF88921.1 anig|EHA18250.1 aory|EIT78863.1 asyd|OJJ62846.1 ater|EAU39249.1 atub|OJI91137.1 aver|OJI96890.1 awen|OJJ35081.1 azon|OJJ50026.1

Group3096: abra|OJJ73497.1 acar|OOG00693.1 acla|EAW07431.1 afis|EAW22457.1 afla|EED54487.1 afum|EAL90734.1 agla|OJJ87567.1 akaw|GAA87446.1 aluc|OJZ88551.1 anid|CBF88920.1 anig|EHA18249.1 aory|EIT78747.1 asyd|OJJ62845.1 ater|EAU39248.1 atub|OJI91136.1 aver|OJI96889.1 awen|OJJ35082.1 azon|OJJ50025.1

Group3097: abra|OJJ73507.1 acar|OOG00697.1 acla|EAW07440.1 afis|EAW22467.1 afla|EED54499.1 afum|EAL90724.1 agla|OJJ87557.1 akaw|GAA87455.1 aluc|OJZ88540.1 anid|CBF88903.1 anig|EHA18241.1 aory|EIT78913.1 asyd|OJJ62837.1 ater|EAU39241.1 atub|OJI91125.1 aver|OJI96880.1 awen|OJJ35093.1 azon|OJJ50016.1

Group3098: abra|OJJ73508.1 acar|OOG00698.1 acla|EAW07441.1 afis|EAW22468.1 afla|EED54500.1 afum|EAL90723.1 agla|OJJ87556.1 akaw|GAA87456.1 aluc|OJZ88539.1 anid|CBF88902.1 anig|EHA18240.1 aory|EIT78813.1 asyd|OJJ62836.1 ater|EAU39240.1 atub|OJI91124.1 aver|OJI96879.1 awen|OJJ35094.1 azon|OJJ50015.1

Group3099: abra|OJJ73510.1 acar|OOG00699.1 acla|EAW07443.1 afis|EAW22470.1 afla|EED54502.1 afum|EAL90721.1 agla|OJJ87554.1 akaw|GAA87458.1 aluc|OJZ88537.1 anid|CBF88900.1 anig|EHA18238.1 aory|EIT78910.1 asyd|OJJ62834.1 ater|EAU39238.1 atub|OJI91122.1 aver|OJI96877.1 awen|OJJ35096.1 azon|OJJ50013.1

Group3100: abra|OJJ73512.1 acar|OOG00701.1 acla|EAW07445.1 afis|EAW22472.1 afla|EED54504.1 afum|EAL90719.1 agla|OJJ87552.1 akaw|GAA87460.1 aluc|OJZ88535.1 anid|CBF88896.1 anig|EHA18236.1 aory|EIT78949.1 asyd|OJJ62832.1 ater|EAU39236.1 atub|OJI91120.1 aver|OJI96875.1 awen|OJJ35098.1 azon|OJJ50011.1

Group3101: abra|OJJ73513.1 acar|OOG00700.1 acla|EAW07447.1 afis|EAW22474.1 afla|EED54506.1 afum|EAL90717.1 agla|OJJ87550.1 akaw|GAA87462.1 aluc|OJZ88534.1 anid|CBF88892.1 anig|EHA18235.1 aory|EIT78850.1 asyd|OJJ62831.1 ater|EAU39235.1 atub|OJI91119.1 aver|OJI96874.1 awen|OJJ35099.1 azon|OJJ50010.1

Group3102: abra|OJJ73516.1 acar|OOG00706.1 acla|EAW07451.1 afis|EAW22483.1 afla|EED54509.1 afum|EAL90711.1 agla|OJJ87543.1 akaw|GAA87465.1 aluc|OJZ88531.1 anid|CBF88887.1 anig|EHA18232.1 aory|EIT78939.1 asyd|OJJ62826.1 ater|EAU39232.1 atub|OJI91116.1 aver|OJI96871.1 awen|OJJ35102.1 azon|OJJ50006.1

Group3103: abra|OJJ73529.1 acar|OOG00710.1 acla|EAW07454.1 afis|EAW22486.1 afla|EED54512.1 afum|EAL90707.1 agla|OJJ80863.1 akaw|GAA88617.1 aluc|OJZ88526.1 anid|CBF88942.1 anig|EHA18228.1 aory|EIT78750.1 asyd|OJJ62815.1 ater|EAU39228.1 atub|OJI91111.1 aver|OJI96859.1 awen|OJJ35105.1 azon|OJJ49989.1

Group3104: abra|OJJ73534.1 acar|OOG00712.1 acla|EAW07456.1 afis|EAW22487.1 afla|EED54516.1 afum|EAL90706.1 agla|OJJ80860.1 akaw|GAA88616.1 aluc|OJZ88525.1 anid|CBF88944.1 anig|EHA18227.1 aory|EIT78867.1 asyd|OJJ62814.1 ater|EAU39227.1 atub|OJI91110.1 aver|OJI96858.1 awen|OJJ35108.1 azon|OJJ49986.1

Group3105: abra|OJJ73537.1 acar|OOG00713.1 acla|EAW07459.1 afis|EAW22490.1 afla|EED54519.1 afum|EAL90703.1 agla|OJJ80856.1 akaw|GAA88613.1 aluc|OJZ88522.1 anid|CBF88950.1 anig|EHA18225.1 aory|EIT78823.1 asyd|OJJ62811.1 ater|EAU39225.1 atub|OJI91107.1 aver|OJI96857.1 awen|OJJ35110.1 azon|OJJ49983.1

Group3106: abra|OJJ73539.1 acar|OOG00715.1 acla|EAW07460.1 afis|EAW22491.1 afla|EED54520.1 afum|EAL90702.1 agla|OJJ80854.1 akaw|GAA88612.1 aluc|OJZ88520.1 anid|CBF88951.1 anig|EHA18224.1 aory|EIT78935.1 asyd|OJJ62809.1 ater|EAU39224.1 atub|OJI91105.1 aver|OJI96855.1 awen|OJJ35112.1 azon|OJJ49981.1

Group3107: abra|OJJ73541.1 acar|OOG00716.1 acla|EAW07461.1 afis|EAW22492.1 afla|EED54521.1 afum|EAL90701.1 agla|OJJ80853.1 akaw|GAA88610.1 aluc|OJZ88518.1 anid|CBF88953.1 anig|EHA18223.1 aory|EIT78920.1 asyd|OJJ62808.1 ater|EAU39223.1 atub|OJI91103.1 aver|OJI96854.1 awen|OJJ35113.1 azon|OJJ49980.1

Group3108: abra|OJJ73547.1 acar|OOG00721.1 acla|EAW07466.1 afis|EAW22499.1 afla|EED54526.1 afum|EAL90694.1 agla|OJJ80847.1 akaw|GAA88605.1 aluc|OJZ88513.1 anid|CBF88960.1 anig|EHA18219.1

aory|EIT78800.1 asyd|OJJ62804.1 ater|EAU39219.1 atub|OJI91098.1 aver|OJI96850.1 awen|OJJ35122.1 azon|OJJ49976.1

Group3109: abra|OJJ73549.1 acar|OOG00723.1 acla|EAW07468.1 afis|EAW22501.1 afla|EED54529.1 afum|EAL90692.1 agla|OJJ80851.1 akaw|GAA88603.1 aluc|OJZ88511.1 anid|CBF88962.1 anig|EHA18217.1 aory|EIT78773.1 asyd|OJJ62799.1 ater|EAU39217.1 atub|OJI91096.1 aver|OJI96846.1 awen|OJJ35124.1 azon|OJJ49974.1

Group3110: abra|OJJ73550.1 acar|OOG00724.1 acla|EAW07469.1 afis|EAW22502.1 afla|EED54530.1 afum|EAL90691.1 agla|OJJ80852.1 akaw|GAA88602.1 aluc|OJZ88510.1 anid|CBF88965.1 anig|EHA18216.1 aory|EIT78738.1 asyd|OJJ62798.1 ater|EAU39216.1 atub|OJI91095.1 aver|OJI96845.1 awen|OJJ35125.1 azon|OJJ49973.1

Group3111: abra|OJJ73555.1 acar|OOG00726.1 acla|EAW07473.1 afis|EAW22505.1 afla|EED54533.1 afum|EAL90688.1 agla|OJJ87536.1 akaw|GAA88598.1 aluc|OJZ88506.1 anid|CBF88970.1 anig|EHA18213.1 aory|EIT78812.1 asyd|OJJ62795.1 ater|EAU39210.1 atub|OJI91092.1 aver|OJI96842.1 awen|OJJ35130.1 azon|OJJ49969.1

Group3112: abra|OJJ73556.1 acar|OOG00728.1 acla|EAW07474.1 afis|EAW22506.1 afla|EED54534.1 afum|EAL90687.1 agla|OJJ87537.1 akaw|GAA88597.1 aluc|OJZ88505.1 anid|CBF88972.1 anig|EHA18212.1 aory|EIT78875.1 asyd|OJJ62794.1 ater|EAU39209.1 atub|OJI91091.1 aver|OJI96841.1 awen|OJJ35131.1 azon|OJJ49968.1

Group3113: abra|OJJ73560.1 acar|OOG00731.1 acla|EAW07477.1 afis|EAW22509.1 afla|EED54537.1 afum|EAL90684.1 agla|OJJ80867.1 akaw|GAA88594.1 aluc|OJZ88502.1 anid|CBF88977.1 anig|EHA18210.1 aory|EIT78879.1 asyd|OJJ62791.1 ater|EAU39205.1 atub|OJI91088.1 aver|OJI96838.1 awen|OJJ35134.1 azon|OJJ49964.1

Group3114: abra|OJJ73563.1 acar|OOG00746.1 acla|EAW07480.1 afis|EAW22512.1 afla|EED54540.1 afum|EAL90681.1 agla|OJJ80875.1 akaw|GAA88591.1 aluc|OJZ88498.1 anid|CBF88983.1 anig|EHA18207.1 aory|EIT78924.1 asyd|OJJ62788.1 ater|EAU39197.1 atub|OJI91085.1 aver|OJI96835.1 awen|OJJ35138.1 azon|OJJ49961.1

Group3115: abra|OJJ73566.1 acar|OOG00748.1 acla|EAW07482.1 afis|EAW22514.1 afla|EED54544.1 afum|EAL90679.1 agla|OJJ80873.1 akaw|GAA88589.1 aluc|OJZ88496.1 anid|CBF88984.1 anig|EHA18205.1 aory|EIT78876.1 asyd|OJJ62786.1 ater|EAU39195.1 atub|OJI91083.1 aver|OJI96833.1 awen|OJJ35140.1 azon|OJJ49959.1

Group3116: abra|OJJ73567.1 acar|OOG00745.1 acla|EAW07483.1 afis|EAW22515.1 afla|EED54545.1 afum|EAL90678.1 agla|OJJ80872.1 akaw|GAA88588.1 aluc|OJZ88495.1 anid|CBF88986.1 anig|EHA18204.1 aory|EIT78884.1 asyd|OJJ62785.1 ater|EAU39194.1 atub|OJI91082.1 aver|OJI96831.1 awen|OJJ35141.1 azon|OJJ49958.1

Group3117: abra|OJJ73568.1 acar|OOG00749.1 acla|EAW07484.1 afis|EAW22516.1 afla|EED54546.1 afum|EAL90677.1 agla|OJJ80877.1 akaw|GAA88587.1 aluc|OJZ88494.1 anid|CBF88988.1 anig|EHA18203.1 aory|EIT78756.1 asyd|OJJ62784.1 ater|EAU39193.1 atub|OJI91081.1 aver|OJI96830.1 awen|OJJ35142.1 azon|OJJ49957.1

Group3118: abra|OJJ73573.1 acar|OOG00754.1 acla|EAW07490.1 afis|EAW22523.1 afla|EED54548.1 afum|EAL90671.1 agla|OJJ80884.1 akaw|GAA88577.1 aluc|OJZ88480.1 anid|CBF89007.1 anig|EHA18196.1 aory|EIT78743.1 asyd|OJJ62776.1 ater|EAU39187.1 atub|OJI91074.1 aver|OJI96821.1 awen|OJJ35150.1 azon|OJJ49951.1

Group3119: abra|OJJ73574.1 acar|OOG00755.1 acla|EAW07491.1 afis|EAW22524.1 afla|EED54549.1 afum|EAL90670.1 agla|OJJ80885.1 akaw|GAA88576.1 aluc|OJZ88479.1 anid|CBF89009.1 anig|EHA18195.1 aory|EIT78837.1 asyd|OJJ62775.1 ater|EAU39186.1 atub|OJI91073.1 aver|OJI96820.1 awen|OJJ35151.1 azon|OJJ49950.1

Group3120: abra|OJJ73575.1 acar|OOG00756.1 acla|EAW07492.1 afis|EAW22525.1 afla|EED54553.1 afum|EAL90669.1 agla|OJJ80886.1 akaw|GAA88575.1 aluc|OJZ88477.1 anid|CBF89011.1 anig|EHA18194.1 aory|EIT78752.1 asyd|OJJ62774.1 ater|EAU39185.1 atub|OJI91072.1 aver|OJI96819.1 awen|OJJ35152.1 azon|OJJ49949.1

Group3121: abra|OJJ73578.1 acar|OOG00759.1 acla|EAW07496.1 afis|EAW22529.1 afla|EED54557.1 afum|EAL90665.1 agla|OJJ80890.1 akaw|GAA88573.1 aluc|OJZ88474.1 anid|CBF89014.1 anig|EHA18191.1 aory|EIT78796.1 asyd|OJJ62772.1 ater|EAU39182.1 atub|OJI91069.1 aver|OJI96817.1 awen|OJJ35156.1 azon|OJJ49945.1

Group3122: abra|OJJ73580.1 acar|OOG00761.1 acla|EAW07498.1 afis|EAW22531.1 afla|EED54559.1 afum|EAL90663.1 agla|OJJ80892.1 akaw|GAA88571.1 aluc|OJZ88472.1 anid|CBF89018.1 anig|EHA18189.1 aory|EIT78759.1 asyd|OJJ62770.1 ater|EAU39178.1 atub|OJI91067.1 aver|OJI96815.1 awen|OJJ35158.1 azon|OJJ49944.1

Group3123: abra|OJJ73581.1 acar|OOG00762.1 acla|EAW07499.1 afis|EAW22532.1 afla|EED54560.1 afum|EAL90662.1 agla|OJJ80893.1 akaw|GAA88570.1 aluc|OJZ88471.1 anid|CBF89020.1 anig|EHA18188.1 aory|EIT78827.1 asyd|OJJ62769.1 ater|EAU39177.1 atub|OJI91066.1 aver|OJI96814.1 awen|OJJ35159.1 azon|OJJ49943.1

Group3124: abra|OJJ73582.1 acar|OOG00763.1 acla|EAW07500.1 afis|EAW22533.1 afla|EED54561.1 afum|EAL90661.1 agla|OJJ80894.1 akaw|GAA88569.1 aluc|OJZ88470.1 anid|CBF89023.1 anig|EHA18187.1 aory|EIT78937.1 asyd|OJJ62768.1 ater|EAU39176.1 atub|OJI91065.1 aver|OJI96813.1 awen|OJJ35160.1 azon|OJJ49942.1

Group3125: abra|OJJ73583.1 acar|OOG00764.1 acla|EAW07501.1 afis|EAW22534.1 afla|EED54562.1 afum|EAL90660.1 agla|OJJ80895.1 akaw|GAA88568.1 aluc|OJZ88469.1 anid|CBF89025.1 anig|EHA18186.1 aory|EIT78784.1 asyd|OJJ62767.1 ater|EAU39175.1 atub|OJI91064.1 aver|OJI96812.1 awen|OJJ35161.1 azon|OJJ49941.1

Group3126: abra|OJJ73585.1 acar|OOG00766.1 acla|EAW07503.1 afis|EAW22536.1 afla|EED54564.1 afum|EAL90658.1 agla|OJJ80897.1 akaw|GAA88566.1 aluc|OJZ88467.1 anid|CBF89029.1 anig|EHA18184.1 aory|EIT78903.1 asyd|OJJ62765.1 ater|EAU39173.1 atub|OJI91062.1 aver|OJI96810.1 awen|OJJ35163.1 azon|OJJ49940.1

Group3127: abra|OJJ73586.1 acar|OOG00767.1 acla|EAW07504.1 afis|EAW22537.1 afla|EED54565.1 afum|EAL90657.1 agla|OJJ80898.1 akaw|GAA88565.1 aluc|OJZ88466.1 anid|CBF89031.1 anig|EHA18183.1 aory|EIT78873.1 asyd|OJJ62764.1 ater|EAU39172.1 atub|OJI91061.1 aver|OJI96809.1 awen|OJJ35165.1 azon|OJJ49939.1

Group3128: abra|OJJ73589.1 acar|OOG00771.1 acla|EAW07507.1 afis|EAW22539.1 afla|EED54568.1 afum|EAL90655.1 agla|OJJ80899.1 akaw|GAA88563.1 aluc|OJZ88461.1 anid|CBF89036.1 anig|EHA18181.1 aory|EIT78782.1 asyd|OJJ62762.1 ater|EAU39170.1 atub|OJI91059.1 aver|OJI96805.1 awen|OJJ35168.1 azon|OJJ49938.1

Group3129: abra|OJJ73592.1 acar|OOG00777.1 acla|EAW07511.1 afis|EAW22542.1 afla|EED54573.1 afum|EAL90652.1 agla|OJJ80904.1 akaw|GAA88560.1 aluc|OJZ88458.1 anid|CBF89040.1 anig|EHA18178.1 aory|EIT78776.1 asyd|OJJ62760.1 ater|EAU39166.1 atub|OJI91056.1 aver|OJI96803.1 awen|OJJ35171.1 azon|OJJ49936.1

Group3131: abra|OJJ73598.1 acar|OOG00784.1 acla|EAW07516.1 afis|EAW22546.1 afla|EED54578.1 afum|EAL90649.1 agla|OJJ80908.1 akaw|GAA88554.1 aluc|OJZ88452.1 anid|CBF89049.1 anig|EHA18173.1 aory|EIT78955.1 asyd|OJJ62755.1 ater|EAU39159.1 atub|OJI91050.1 aver|OJI96797.1 awen|OJJ35177.1 azon|OJJ49931.1

Group3132: abra|OJJ73599.1 acar|OOG00785.1 acla|EAW07517.1 afis|EAW22547.1 afla|EED54579.1 afum|EAL90648.1 agla|OJJ80909.1 akaw|GAA88553.1 aluc|OJZ88451.1 anid|CBF89051.1 anig|EHA18172.1 aory|EIT78753.1 asyd|OJJ62754.1 ater|EAU39158.1 atub|OJI91049.1 aver|OJI96796.1 awen|OJJ35178.1 azon|OJJ49930.1

Group3133: abra|OJJ73601.1 acar|OOG00786.1 acla|EAW07519.1 afis|EAW22549.1 afla|EED54582.1 afum|EAL90646.1 agla|OJJ80911.1 akaw|GAA88551.1 aluc|OJZ88447.1 anid|CBF89057.1 anig|EHA18170.1 aory|EIT78932.1 asyd|OJJ62752.1 ater|EAU39155.1 atub|OJI91046.1 aver|OJI96794.1 awen|OJJ35180.1 azon|OJJ49927.1

Group3134: abra|OJJ73606.1 acar|OOG00790.1 acla|EAW07523.1 afis|EAW22553.1 afla|EED54585.1 afum|EAL90642.1 agla|OJJ80915.1 akaw|GAA88547.1 aluc|OJZ88443.1 anid|CBF89062.1 anig|EHA18167.1 aory|EIT78786.1 asyd|OJJ62748.1 ater|EAU39151.1 atub|OJI91041.1 aver|OJI96788.1 awen|OJJ35184.1 azon|OJJ49924.1

Group3135: abra|OJJ73607.1 acar|OOG00791.1 acla|EAW07524.1 afis|EAW22554.1 afla|EED54586.1 afum|EAL90641.1 agla|OJJ80916.1 akaw|GAA88546.1 aluc|OJZ88442.1 anid|CBF89063.1 anig|EHA18166.1 aory|EIT78927.1 asyd|OJJ62747.1 ater|EAU39150.1 atub|OJI91040.1 aver|OJI96787.1 awen|OJJ35185.1 azon|OJJ49923.1

Group3136: abra|OJJ73609.1 acar|OOG00793.1 acla|EAW07526.1 afis|EAW22556.1 afla|EED54587.1 afum|EAL90639.1 agla|OJJ80917.1 akaw|GAA88544.1 aluc|OJZ88440.1 anid|CBF89065.1 anig|EHA18164.1 aory|EIT78856.1 asyd|OJJ62745.1 ater|EAU39148.1 atub|OJI91038.1 aver|OJI96785.1 awen|OJJ35187.1 azon|OJJ49921.1

Group3137: abra|OJJ73611.1 acar|OOG00794.1 acla|EAW07528.1 afis|EAW22558.1 afla|EED54589.1 afum|EAL90637.1 agla|OJJ80918.1 akaw|GAA88542.1 aluc|OJZ88439.1 anid|CBF89071.1 anig|EHA18162.1 aory|EIT78846.1 asyd|OJJ62743.1 ater|EAU39146.1 atub|OJI91036.1 aver|OJI96784.1 awen|OJJ35188.1 azon|OJJ49918.1

Group3138: abra|OJJ73612.1 acar|OOG00795.1 acla|EAW07529.1 afis|EAW22559.1 afla|EED54590.1 afum|EAL90636.1 agla|OJJ80919.1 akaw|GAA88541.1 aluc|OJZ88437.1 anid|CBF88377.1 anig|EHA18161.1

aory|EIT78952.1 asyd|OJJ63734.1 ater|EAU39145.1 atub|OJI91034.1 aver|OJI97206.1 awen|OJJ35189.1 azon|OJJ49917.1

Group3139: abra|OJJ73616.1 acar|OOG00800.1 acla|EAW07531.1 afis|EAW22562.1 afla|EED54592.1 afum|EAL90633.1 agla|OJJ80921.1 akaw|GAA88537.1 aluc|OJZ88433.1 anid|CBF88314.1 anig|EHA18157.1 aory|EIT78915.1 asyd|OJJ63700.1 ater|EAU39142.1 atub|OJI91030.1 aver|OJI97239.1 awen|OJJ35191.1 azon|OJJ49915.1

Group3140: abra|OJJ73617.1 acar|OOG00799.1 acla|EAW07532.1 afis|EAW22561.1 afla|EED54596.1 afum|EAL90634.1 agla|OJJ80922.1 akaw|GAA88536.1 aluc|OJZ88432.1 anid|CBF88316.1 anig|EHA18156.1 aory|EIT78857.1 asyd|OJJ63701.1 ater|EAU39141.1 atub|OJI91029.1 aver|OJI97238.1 awen|OJJ35192.1 azon|OJJ49914.1

Group3141: abra|OJJ73618.1 acar|OOG00801.1 acla|EAW07533.1 afis|EAW22567.1 afla|EED54597.1 afum|EAL90631.1 agla|OJJ80923.1 akaw|GAA88535.1 aluc|OJZ88431.1 anid|CBF88318.1 anig|EHA18155.1 aory|EIT78761.1 asyd|OJJ63702.1 ater|EAU39140.1 atub|OJI91028.1 aver|OJI97237.1 awen|OJJ35193.1 azon|OJJ49913.1

Group3142: abra|OJJ73619.1 acar|OOG00802.1 acla|EAW07534.1 afis|EAW22568.1 afla|EED54598.1 afum|EAL90630.1 agla|OJJ80924.1 akaw|GAA88534.1 aluc|OJZ88430.1 anid|CBF88320.1 anig|EHA18154.1 aory|EIT78762.1 asyd|OJJ63703.1 ater|EAU39139.1 atub|OJI91027.1 aver|OJI97236.1 awen|OJJ35194.1 azon|OJJ49912.1

Group3143: abra|OJJ73621.1 acar|OOG00805.1 acla|EAW07536.1 afis|EAW22570.1 afla|EED54600.1 afum|EAL90628.1 agla|OJJ80926.1 akaw|GAA88532.1 aluc|OJZ88427.1 anid|CBF88323.1 anig|EHA18152.1 aory|EIT78841.1 asyd|OJJ63705.1 ater|EAU39137.1 atub|OJI91025.1 aver|OJI97234.1 awen|OJJ35196.1 azon|OJJ49911.1

Group3144: abra|OJJ73622.1 acar|OOG00804.1 acla|EAW07537.1 afis|EAW22571.1 afla|EED54601.1 afum|EAL90627.1 agla|OJJ80928.1 akaw|GAA88531.1 aluc|OJZ88426.1 anid|CBF88325.1 anig|EHA18151.1 aory|EIT78890.1 asyd|OJJ63706.1 ater|EAU39136.1 atub|OJI91024.1 aver|OJI97233.1 awen|OJJ35198.1 azon|OJJ49910.1

Group3145: abra|OJJ73624.1 acar|OOG00807.1 acla|EAW07539.1 afis|EAW22573.1 afla|EED54602.1 afum|EAL90625.1 agla|OJJ80930.1 akaw|GAA88529.1 aluc|OJZ88424.1 anid|CBF88328.1 anig|EHA18149.1 aory|EIT78905.1 asyd|OJJ63708.1 ater|EAU39134.1 atub|OJI91020.1 aver|OJI97230.1 awen|OJJ35200.1 azon|OJJ49907.1

Group3146: abra|OJJ73627.1 acar|OOG00809.1 acla|EAW07541.1 afis|EAW22575.1 afla|EED54604.1 afum|EAL90623.1 agla|OJJ80931.1 akaw|GAA88526.1 aluc|OJZ88422.1 anid|CBF88332.1 anig|EHA18147.1 aory|EIT78895.1 asyd|OJJ63711.1 ater|EAU39129.1 atub|OJI91018.1 aver|OJI97228.1 awen|OJJ35203.1 azon|OJJ49905.1

Group3147: abra|OJJ73632.1 acar|OOG00814.1 acla|EAW07546.1 afis|EAW22580.1 afla|EED54625.1 afum|EAL90619.2 agla|OJJ80940.1 akaw|GAA88521.1 aluc|OJZ88417.1 anid|CBF88341.1 anig|EHA18143.1 aory|EIT78840.1 asyd|OJJ63716.1 ater|EAU39124.1 atub|OJI91013.1 aver|OJI97224.1 awen|OJJ35208.1 azon|OJJ49901.1

Group3148: abra|OJJ73633.1 acar|OOG00815.1 acla|EAW07547.1 afis|EAW22581.1 afla|EED54626.1 afum|EAL90618.1 agla|OJJ80941.1 akaw|GAA88520.1 aluc|OJZ88416.1 anid|CBF88343.1 anig|EHA18142.1 aory|EIT78811.1 asyd|OJJ63717.1 ater|EAU39123.1 atub|OJI91012.1 aver|OJI97223.1 awen|OJJ35209.1 azon|OJJ49900.1

Group3149: abra|OJJ73637.1 acar|OOG00817.1 acla|EAW07549.1 afis|EAW22583.1 afla|EED54628.1 afum|EAL90616.1 agla|OJJ80943.1 akaw|GAA88518.1 aluc|OJZ88413.1 anid|CBF88349.1 anig|EHA18140.1 aory|EIT78801.1 asyd|OJJ63719.1 ater|EAU39121.1 atub|OJI91009.1 aver|OJI97221.1 awen|OJJ35211.1 azon|OJJ49898.1

Group3150: abra|OJJ73638.1 acar|OOG00818.1 acla|EAW07550.1 afis|EAW22584.1 afla|EED54629.1 afum|EAL90615.1 agla|OJJ80944.1 akaw|GAA88517.1 aluc|OJZ88412.1 anid|CBF88350.1 anig|EHA18139.1 aory|EIT78791.1 asyd|OJJ63720.1 ater|EAU39120.1 atub|OJI91007.1 aver|OJI97220.1 awen|OJJ35212.1 azon|OJJ49897.1

Group3151: abra|OJJ73640.1 acar|OOG00820.1 acla|EAW07553.1 afis|EAW22587.1 afla|EED54633.1 afum|EAL90612.1 agla|OJJ80947.1 akaw|GAA88515.1 aluc|OJZ88410.1 anid|CBF88356.1 anig|EHA18137.1 aory|EIT78894.1 asyd|OJJ63723.1 ater|EAU39113.1 atub|OJI91006.1 aver|OJI97217.1 awen|OJJ35215.1 azon|OJJ49894.1

Group3152: abra|OJJ73641.1 acar|OOG00822.1 acla|EAW07554.1 afis|EAW22588.1 afla|EED54634.1 afum|EAL90611.1 agla|OJJ80948.1 akaw|GAA88514.1 aluc|OJZ88409.1 anid|CBF88358.1 anig|EHA18136.1 aory|EIT78862.1 asyd|OJJ63724.1 ater|EAU39112.1 atub|OJI91005.1 aver|OJI97216.1 awen|OJJ35216.1 azon|OJJ49893.1

Group3153: abra|OJJ73643.1 acar|OOG00824.1 acla|EAW07556.1 afis|EAW22590.1 afla|EED54636.1 afum|EAL90609.1 agla|OJJ80950.1 akaw|GAA88512.1 aluc|OJZ88407.1 anid|CBF88360.1 anig|EHA18134.1 aory|EIT78889.1 asyd|OJJ63726.1 ater|EAU39119.1 atub|OJI91003.1 aver|OJI97214.1 awen|OJJ35218.1 azon|OJJ49890.1

Group3154: abra|OJJ73644.1 acar|OOG00825.1 acla|EAW07557.1 afis|EAW22591.1 afla|EED54641.1 afum|EAL90608.1 agla|OJJ80951.1 akaw|GAA88511.1 aluc|OJZ88406.1 anid|CBF88364.1 anig|EHA18133.1 aory|EIT78790.1 asyd|OJJ63727.1 ater|EAU39111.1 atub|OJI91002.1 aver|OJI97213.1 awen|OJJ35219.1 azon|OJJ49889.1

Group3155: abra|OJJ73645.1 acar|OOG00826.1 acla|EAW07558.1 afis|EAW22592.1 afla|EED54642.1 afum|EAL90607.1 agla|OJJ80952.1 akaw|GAA88510.1 aluc|OJZ88405.1 anid|CBF88365.1 anig|EHA18132.1 aory|EIT78880.1 asyd|OJJ63728.1 ater|EAU39110.1 atub|OJI91001.1 aver|OJI97212.1 awen|OJJ35220.1 azon|OJJ49888.1

Group3156: abra|OJJ73647.1 acar|OOG00829.1 acla|EAW07560.1 afis|EAW22594.1 afla|EED54644.1 afum|EAL90605.1 agla|OJJ80954.1 akaw|GAA88508.1 aluc|OJZ88403.1 anid|CBF88369.1 anig|EHA18131.1 aory|EIT78768.1 asyd|OJJ63730.1 ater|EAU39109.1 atub|OJI90999.1 aver|OJI97210.1 awen|OJJ35222.1 azon|OJJ49886.1

Group3157: abra|OJJ73654.1 acar|OOG00832.1 acla|EAW07564.1 afis|EAW22598.1 afla|EED54720.1 afum|EAL90600.1 agla|OJJ80966.1 akaw|GAA88504.1 aluc|OJZ88399.1 anid|CBF88312.1 anig|EHA18127.1 aory|EIT78821.1 asyd|OJJ63698.1 ater|EAU39105.1 atub|OJI90994.1 aver|OJI97241.1 awen|OJJ35226.1 azon|OJJ49884.1

Group3158: abra|OJJ73655.1 acar|OOG00833.1 acla|EAW07565.1 afis|EAW22599.1 afla|EED54719.1 afum|EAL90599.1 agla|OJJ80967.1 akaw|GAA88503.1 aluc|OJZ88398.1 anid|CBF88310.1 anig|EHA18126.1 aory|EIT78916.1 asyd|OJJ63697.1 ater|EAU39104.1 atub|OJI90993.1 aver|OJI97242.1 awen|OJJ35228.1 azon|OJJ49883.1

Group3159: abra|OJJ73656.1 acar|OOG00834.1 acla|EAW07566.1 afis|EAW22600.1 afla|EED54718.1 afum|EAL90598.1 agla|OJJ80968.1 akaw|GAA88502.1 aluc|OJZ88397.1 anid|CBF88308.1 anig|EHA18125.1 aory|EIT78770.1 asyd|OJJ63696.1 ater|EAU39103.1 atub|OJI90992.1 aver|OJI97243.1 awen|OJJ35229.1 azon|OJJ49882.1

Group3160: abra|OJJ73657.1 acar|OOG00835.1 acla|EAW07567.1 afis|EAW22601.1 afla|EED54717.1 afum|EAL90597.1 agla|OJJ80971.1 akaw|GAA88501.1 aluc|OJZ88396.1 anid|CBF88256.1 anig|EHA18124.1 aory|EIT78942.1 asyd|OJJ63672.1 ater|EAU39102.1 atub|OJI90991.1 aver|OJI97266.1 awen|OJJ35230.1 azon|OJJ49881.1

Group3161: abra|OJJ73658.1 acar|OOG00836.1 acla|EAW07568.1 afis|EAW22602.1 afla|EED54716.1 afum|EAL90596.1 agla|OJJ80972.1 akaw|GAA88500.1 aluc|OJZ88395.1 anid|CBF88257.1 anig|EHA18123.1 aory|EIT78742.1 asyd|OJJ63673.1 ater|EAU39101.1 atub|OJI90990.1 aver|OJI97265.1 awen|OJJ35231.1 azon|OJJ49880.1

Group3162: abra|OJJ73665.1 acar|OOG00843.1 acla|EAW07572.1 afis|EAW22606.1 afla|EED49514.1 afum|EAL90592.1 agla|OJJ80975.1 akaw|GAA88491.1 aluc|OJZ88387.1 anid|CBF88263.1 anig|EHA18114.1 aory|EIT82633.1 asyd|OJJ63677.1 ater|EAU39097.1 atub|OJI90982.1 aver|OJI97262.1 awen|OJJ35234.1 azon|OJJ49877.1

Group3163: abra|OJJ73667.1 acar|OOG00844.1 acla|EAW07573.1 afis|EAW22607.1 afla|EED49805.1 afum|EAL90591.1 agla|OJJ80977.1 akaw|GAA88490.1 aluc|OJZ88385.1 anid|CBF88265.1 anig|EHA18113.1 aory|EIT77683.1 asyd|OJJ63678.1 ater|EAU39096.1 atub|OJI90980.1 aver|OJI97261.1 awen|OJJ35235.1 azon|OJJ49876.1

Group3164: abra|OJJ73668.1 acar|OOG00845.1 acla|EAW07575.1 afis|EAW22609.1 afla|EED49807.1 afum|EAL90589.1 agla|OJJ81009.1 akaw|GAA88489.1 aluc|OJZ88384.1 anid|CBF88269.1 anig|EHA18112.1 aory|EIT77641.1 asyd|OJJ63680.1 ater|EAU39095.1 atub|OJI90979.1 aver|OJI97259.1 awen|OJJ35237.1 azon|OJJ49870.1

Group3165: abra|OJJ73671.1 acar|OOG00846.1 acla|EAW07583.1 afis|EAW22611.1 afla|EED49809.1 afum|EAL90587.1 agla|OJJ81005.1 akaw|GAA88486.1 aluc|OJZ88382.1 anid|CBF88281.1 anig|EHA18109.1 aory|EIT77633.1 asyd|OJJ63684.1 ater|EAU39092.1 atub|OJI90976.1 aver|OJI97255.1 awen|OJJ35239.1 azon|OJJ49868.1

Group3166: abra|OJJ73672.1 acar|OOG00847.1 acla|EAW07582.1 afis|EAW22612.1 afla|EED49810.1 afum|EAL90586.1 agla|OJJ81004.1 akaw|GAA88485.1 aluc|OJZ88381.1 anid|CBF88283.1 anig|EHA18108.1 aory|EIT77693.1 asyd|OJJ63685.1 ater|EAU39091.1 atub|OJI90975.1 aver|OJI97254.1 awen|OJJ35240.1 azon|OJJ49867.1

Group3167: abra|OJJ73673.1 acar|OOG00848.1 acla|EAW07581.1 afis|EAW22613.1 afla|EED49811.1 afum|EAL90585.1 agla|OJJ81003.1 akaw|GAA88484.1 aluc|OJZ88379.1 anid|CBF88287.1 anig|EHA18106.1

aory|EIT77666.1 asyd|OJJ63686.1 ater|EAU39090.1 atub|OJI90973.1 aver|OJI97253.1 awen|OJJ35241.1 azon|OJJ49866.1

Group3168: abra|OJJ73681.1 acar|OOG00853.1 acla|EAW07596.1 afis|EAW22627.1 afla|EED49827.1 afum|EAL90569.1 agla|OJJ80994.1 akaw|GAA88478.1 aluc|OJZ88368.1 anid|CBF88241.1 anig|EHA18101.1 aory|EIT77681.1 asyd|OJJ63662.1 ater|EAU39082.1 atub|OJI90966.1 aver|OJI97272.1 awen|OJJ35248.1 azon|OJJ49858.1

Group3169: abra|OJJ73684.1 acar|OOG00854.1 acla|EAW07599.1 afis|EAW22630.1 afla|EED49879.1 afum|EAL90565.1 agla|OJJ80990.1 akaw|GAA88475.1 aluc|OJZ88365.1 anid|CBF88236.1 anig|EHA18097.1 aory|EIT72464.1 asyd|OJJ63659.1 ater|EAU39078.1 atub|OJI90963.1 aver|OJI97275.1 awen|OJJ35251.1 azon|OJJ49855.1

Group3170: abra|OJJ73688.1 acar|OOG00860.1 acla|EAW07603.1 afis|EAW22634.1 afla|EED49884.1 afum|EAL90559.1 agla|OJJ80985.1 akaw|GAA88471.1 aluc|OJZ88361.1 anid|CBF88229.1 anig|EHA18093.1 aory|EIT72459.1 asyd|OJJ63655.1 ater|EAU39074.1 atub|OJI90957.1 aver|OJI97279.1 awen|OJJ35256.1 azon|OJJ49851.1

Group3171: abra|OJJ73689.1 acar|OOG00861.1 acla|EAW07604.1 afis|EAW22635.1 afum|EAL90558.2 akaw|GAA88470.1 aluc|OJZ88360.1 anid|CBF88227.1 anig|EHA18092.1 asyd|OJJ63654.1 ater|EAU39071.1 atub|OJI90956.1 aver|OJI97280.1 awen|OJJ35257.1 azon|OJJ49850.1 afla|EED49885.1 aory|EIT72467.1 agla|OJJ80984.1

Group3172: abra|OJJ73692.1 acar|OOG00863.1 acla|EAW07607.1 afis|EAW22637.1 afla|EED49888.1 afum|EAL90555.1 agla|OJJ80981.1 akaw|GAA88468.1 aluc|OJZ88357.1 anid|CBF88221.1 anig|EHA18090.1 aory|EIT72465.1 asyd|OJJ63652.1 ater|EAU39069.1 atub|OJI90952.1 aver|OJI97282.1 awen|OJJ35260.1 azon|OJJ49847.1

Group3173: abra|OJJ73724.1 acar|OOG00893.1 acla|EAW07613.1 afis|EAW22643.1 afla|EED49912.1 afum|EAL90549.2 agla|OJJ81025.1 akaw|GAA88438.1 aluc|OJZ88326.1 anid|CBF88175.1 anig|EHA18062.1 aory|EIT72277.1 asyd|OJJ63625.1 ater|EAU39064.1 atub|OJI90921.1 aver|OJI97311.1 awen|OJJ35281.1 azon|OJJ49845.1

Group3174: abra|OJJ73731.1 acar|OOG00897.1 acla|EAW07616.1 afis|EAW22645.1 afla|EED49914.1 afum|EAL90547.1 agla|OJJ81028.1 akaw|GAA88432.1 aluc|OJZ88319.1 anid|CBF88179.1 anig|EHA18055.1 aory|EIT72268.1 asyd|OJJ63627.1 ater|EAU39062.1 atub|OJI90914.1 aver|OJI97309.1 awen|OJJ35278.1 azon|OJJ49841.1

Group3175: abra|OJJ73733.1 acar|OOG00898.1 acla|EAW07617.1 afis|EAW22646.1 afla|EED49915.1 afum|EAL90546.1 agla|OJJ81029.1 akaw|GAA88431.1 aluc|OJZ88318.1 anid|CBF88180.1 anig|EHA18054.1 aory|EIT72267.1 asyd|OJJ63628.1 ater|EAU39061.1 atub|OJI90913.1 aver|OJI97308.1 awen|OJJ35277.1 azon|OJJ49840.1

Group3176: abra|OJJ73734.1 acar|OOG00899.1 acla|EAW07618.1 afis|EAW22647.1 afla|EED49916.1 afum|EAL90545.1 agla|OJJ81030.1 akaw|GAA88430.1 aluc|OJZ88317.1 anid|CBF88182.1 anig|EHA18053.1 aory|EIT72271.1 asyd|OJJ63629.1 ater|EAU39060.1 atub|OJI90912.1 aver|OJI97307.1 awen|OJJ35276.1 azon|OJJ49839.1

Group3177: abra|OJJ73735.1 acar|OOG00900.1 acla|EAW07619.1 afis|EAW22648.1 afla|EED49917.1 afum|EAL90544.1 agla|OJJ81031.1 akaw|GAA88429.1 aluc|OJZ88316.1 anid|CBF88184.1 anig|EHA18052.1 aory|EIT72269.1 asyd|OJJ63630.1 ater|EAU39059.1 atub|OJI90911.1 aver|OJI97306.1 awen|OJJ35275.1 azon|OJJ49832.1

Group3178: abra|OJJ73736.1 acar|OOG00901.1 acla|EAW07620.1 afis|EAW22649.1 afla|EED49918.1 afum|EAL90543.1 agla|OJJ81032.1 akaw|GAA88428.1 aluc|OJZ88315.1 anid|CBF88186.1 anig|EHA18051.1 aory|EIT72278.1 asyd|OJJ63631.1 ater|EAU39058.1 atub|OJI90910.1 aver|OJI97305.1 awen|OJJ35274.1 azon|OJJ49833.1

Group3179: abra|OJJ73739.1 acar|OOG00902.1 acla|EAW07633.1 afis|EAW22653.1 afla|EED49921.1 afum|EAL90539.1 agla|OJJ81036.1 akaw|GAA88425.1 aluc|OJZ88312.1 anid|CBF88197.1 anig|EHA18049.1 aory|EIT72265.1 asyd|OJJ63634.1 ater|EAU39053.1 atub|OJI90907.1 aver|OJI97302.1 awen|OJJ35270.1 azon|OJJ49837.1

Group3180: abra|OJJ73741.1 acar|OOG00904.1 acla|EAW07635.1 afis|EAW22655.1 afla|EED49923.1 afum|EAL90537.1 agla|OJJ81038.1 akaw|GAA88424.1 aluc|OJZ88311.1 anid|CBF88201.1 anig|EHA18048.1 aory|EIT72272.1 asyd|OJJ63637.1 ater|EAU39051.1 atub|OJI90906.1 aver|OJI97299.1 awen|OJJ35268.1 azon|OJJ49831.1

Group3181: abra|OJJ73744.1 acar|OOG00908.1 acla|EAW07640.1 afis|EAW22660.1 afla|EED49930.1 afum|EAL90532.1 agla|OJJ81041.1 akaw|GAA88421.1 aluc|OJZ88308.1 anid|CBF88218.1 anig|EHA18045.1 aory|EIT72279.1 asyd|OJJ63649.1 ater|EAU39048.1 atub|OJI90903.1 aver|OJI97284.1 awen|OJJ35265.1 azon|OJJ49828.1

Group3182: abra|OJJ73745.1 acar|OOG00909.1 acla|EAW07639.1 afis|EAW22659.1 afla|EED49931.1 afum|EAL90533.1 agla|OJJ81042.1 akaw|GAA88420.1 aluc|OJZ88307.1 anid|CBF88215.1 anig|EHA18044.1 aory|EIT72275.1 asyd|OJJ63648.1 ater|EAU39049.1 atub|OJI90902.1 aver|OJI97285.1 awen|OJJ35283.1 azon|OJJ49827.1

Group3183: abra|OJJ73751.1 acar|OOG00914.1 acla|EAW07642.1 afis|EAW22662.1 afla|EED49935.1 afum|EAL90530.1 agla|OJJ79062.1 akaw|GAA88416.1 aluc|OJZ88301.1 anid|CBF88168.1 anig|EHA18040.1 aory|EIT83423.1 asyd|OJJ63622.1 ater|EAU39046.1 atub|OJI90896.1 aver|OJI97315.1 awen|OJJ35287.1 azon|OJJ49826.1

Group3184: abra|OJJ73752.1 acar|OOG00915.1 acla|EAW07643.1 afis|EAW22663.1 afla|EED49937.1 afum|EAL90529.1 akaw|GAA88415.1 aluc|OJZ88300.1 anid|CBF88166.1 anig|EHA18039.1 aory|EIT83433.1 asyd|OJJ63621.1 ater|EAU39045.1 atub|OJI90894.1 aver|OJI97316.1 awen|OJJ35288.1 azon|OJJ49825.1 agla|OJJ79061.1

Group3185: abra|OJJ73753.1 acar|OOG00916.1 acla|EAW07644.1 afis|EAW22664.1 afla|EED49938.1 afum|EAL90528.1 agla|OJJ79060.1 akaw|GAA88414.1 aluc|OJZ88298.1 anid|CBF88164.1 anig|EHA18038.1 aory|EIT83434.1 asyd|OJJ63620.1 ater|EAU39044.1 atub|OJI90893.1 aver|OJI97318.1 awen|OJJ35290.1 azon|OJJ49824.1

Group3186: abra|OJJ73756.1 acar|OOG00918.1 acla|EAW07646.1 afis|EAW22666.1 afla|EED49940.1 afum|EAL90526.1 agla|OJJ79058.1 akaw|GAA88412.1 aluc|OJZ88296.1 anid|CBF88161.1 anig|EHA18036.1 aory|EIT83451.1 asyd|OJJ63619.1 ater|EAU39042.1 atub|OJI90892.1 aver|OJI97320.1 awen|OJJ35292.1 azon|OJJ49822.1

Group3187: abra|OJJ73757.1 acar|OOG00919.1 acla|EAW07647.1 afis|EAW22667.1 afla|EED49941.1 afum|EAL90525.1 agla|OJJ79057.1 akaw|GAA88411.1 aluc|OJZ88295.1 anid|CBF88159.1 anig|EHA18035.1 aory|EIT83456.1 asyd|OJJ63618.1 ater|EAU39041.1 atub|OJI90891.1 aver|OJI97321.1 awen|OJJ35293.1 azon|OJJ49821.1

Group3188: abra|OJJ73758.1 acar|OOG00920.1 acla|EAW07648.1 afis|EAW22668.1 afla|EED49942.1 afum|EAL90524.1 agla|OJJ79056.1 akaw|GAA88410.1 aluc|OJZ88294.1 anid|CBF88157.1 anig|EHA18034.1 aory|EIT83429.1 asyd|OJJ63617.1 ater|EAU39040.1 atub|OJI90890.1 aver|OJI97322.1 awen|OJJ35294.1 azon|OJJ49820.1

Group3189: abra|OJJ73766.1 acar|OOG00925.1 acla|EAW07652.1 afis|EAW22670.1 afla|EED49947.1 afum|EAL90522.1 agla|OJJ79049.1 akaw|GAA88408.1 aluc|OJZ88290.1 anid|CBF88155.1 anig|EHA18031.1 aory|EIT83455.1 asyd|OJJ63608.1 ater|EAU39038.1 atub|OJI90887.1 aver|OJI97329.1 awen|OJJ35298.1 azon|OJJ49817.1

Group3190: abra|OJJ73768.1 acar|OOG00927.1 acla|EAW07653.1 afis|EAW22671.1 afla|EED49949.1 afum|EAL90521.1 agla|OJJ79048.1 akaw|GAA88407.1 aluc|OJZ88289.1 anid|CBF88151.1 anig|EHA18030.1 aory|EIT83435.1 asyd|OJJ63607.1 ater|EAU39037.1 atub|OJI90886.1 aver|OJI97330.1 awen|OJJ35299.1 azon|OJJ49816.1

Group3191: abra|OJJ73769.1 acar|OOG00926.1 acla|EAW07654.1 afis|EAW22672.1 afla|EED49951.1 afum|EAL90520.1 agla|OJJ79046.1 akaw|GAA88406.1 aluc|OJZ88288.1 anid|CBF88145.1 anig|EHA18029.1 aory|EIT83454.1 asyd|OJJ63604.1 ater|EAU39036.1 atub|OJI90885.1 aver|OJI97332.1 awen|OJJ35300.1 azon|OJJ49815.1

Group3192: abra|OJJ73776.1 acar|OOG00933.1 acla|EAW07661.1 afis|EAW22678.1 afla|EED49959.1 afum|EAL90514.1 agla|OJJ79038.1 akaw|GAA88400.1 aluc|OJZ88278.1 anid|CBF88130.1 anig|EHA18023.1 aory|EIT83426.1 asyd|OJJ63588.1 ater|EAU39031.1 atub|OJI90878.1 aver|OJI97345.1 awen|OJJ35306.1 azon|OJJ49809.1

Group3193: abra|OJJ73778.1 acar|OOG00934.1 acla|EAW07662.1 afis|EAW22679.1 afla|EED49960.1 afum|EAL90513.1 agla|OJJ79037.1 akaw|GAA88399.1 aluc|OJZ88277.1 anid|CBF88128.1 anig|EHA18022.1 aory|EIT83445.1 asyd|OJJ63587.1 ater|EAU39030.1 atub|OJI90877.1 aver|OJI97347.1 awen|OJJ35307.1 azon|OJJ49806.1

Group3194: abra|OJJ73781.1 acar|OOG00937.1 acla|EAW07666.1 afis|EAW22683.1 afla|EED49967.1 afum|EAL90509.1 agla|OJJ79035.1 akaw|GAA88396.1 aluc|OJZ88274.1 anid|CBF88119.1 anig|EHA18019.1 aory|EIT83437.1 asyd|OJJ63583.1 ater|EAU39026.1 atub|OJI90874.1 aver|OJI97351.1 awen|OJJ35317.1 azon|OJJ49799.1

Group3195: abra|OJJ73783.1 acar|OOG00939.1 acla|EAW07668.1 afis|EAW22685.1 afla|EED49969.1 afum|EAL90508.1 agla|OJJ79032.1 akaw|GAA88394.1 aluc|OJZ88272.1 anid|CBF88111.1 anig|EHA18017.1 aory|EIT83432.1 asyd|OJJ63580.1 ater|EAU39019.1 atub|OJI90872.1 aver|OJI97356.1 awen|OJJ35318.1 azon|OJJ49798.1

Group3196: abra|OJJ73784.1 acar|OOG00940.1 acla|EAW07671.1 afis|EAW22686.1 afla|EED49971.1 afum|EAL90507.1 agla|OJJ79031.1 akaw|GAA88393.1 aluc|OJZ88271.1 anid|CBF88109.1 anig|EHA18016.1

aory|EIT74696.1 asyd|OJJ63578.1 ater|EAU39021.1 atub|OJI90871.1 aver|OJI97357.1 awen|OJJ35320.1 azon|OJJ49796.1

Group3197: abra|OJJ73785.1 acar|OOG00941.1 acla|EAW07672.1 afis|EAW22687.1 afla|EED49972.1 afum|EAL90506.1 agla|OJJ79029.1 akaw|GAA88392.1 aluc|OJZ88270.1 anid|CBF88107.1 anig|EHA18014.1 aory|EIT74698.1 asyd|OJJ63577.1 ater|EAU39022.1 atub|OJI90870.1 aver|OJI97359.1 awen|OJJ35321.1 azon|OJJ49794.1

Group3198: abra|OJJ73786.1 acar|OOG00942.1 acla|EAW07673.1 afis|EAW22688.1 afla|EED49973.1 afum|EAL90505.1 agla|OJJ79028.1 akaw|GAA88391.1 aluc|OJZ88269.1 anid|CBF88106.1 anig|EHA18013.1 aory|EIT74642.1 asyd|OJJ63576.1 ater|EAU39023.1 atub|OJI90869.1 aver|OJI97360.1 awen|OJJ35322.1 azon|OJJ49793.1

Group3199: abra|OJJ73787.1 acar|OOF90345.1 acla|EAW07674.1 afis|EAW22689.1 afla|EED49974.1 afum|EAL90504.1 agla|OJJ79024.1 akaw|GAA88390.1 aluc|OJZ88267.1 anid|CBF88104.1 anig|EHA18012.1 aory|EIT74677.1 asyd|OJJ63575.1 ater|EAU39024.1 atub|OJI90867.1 aver|OJI97361.1 awen|OJJ35323.1 azon|OJJ49792.1

Group3200: abra|OJJ73791.1 acar|OOF90347.1 acla|EAW07680.1 afis|EAW22695.1 afla|EED49977.1 afum|EAL90498.1 agla|OJJ79021.1 akaw|GAA89410.1 aluc|OJZ88263.1 anid|CBF88098.1 anig|EHA18009.1 aory|EIT74603.1 asyd|OJJ63570.1 ater|EAU39015.1 atub|OJI90862.1 aver|OJI97364.1 awen|OJJ35325.1 azon|OJJ49789.1

Group3201: abra|OJJ73801.1 acar|OOG00957.1 acla|EAW07683.1 afis|EAW22698.1 afla|EED49984.1 afum|EAL90494.1 agla|OJJ79012.1 akaw|GAA89421.1 aluc|OJZ88252.1 anid|CBF88087.1 anig|EHA17994.1 aory|EIT74667.1 asyd|OJJ63563.1 ater|EAU39008.1 atub|OJI90850.1 aver|OJI97370.1 awen|OJJ35334.1 azon|OJJ49783.1

Group3202: abra|OJJ73804.1 acar|OOG00959.1 acla|EAW07676.1 afis|EAW22691.1 afla|EED49982.1 afum|EAL90501.1 agla|OJJ79016.1 akaw|GAA89423.1 aluc|OJZ88249.1 anid|CBF88090.1 anig|EHA17992.1 aory|EIT74721.1 asyd|OJJ63565.1 ater|EAU39010.1 atub|OJI90848.1 aver|OJI97368.1 awen|OJJ35333.1 azon|OJJ49785.1

Group3203: abra|OJJ73805.1 acar|OOG00960.1 acla|EAW07677.1 afis|EAW22692.1 afla|EED49980.1 afum|EBA27409.1 agla|OJJ79018.1 akaw|GAA89425.1 aluc|OJZ88247.1 anid|CBF88095.1 anig|EHA17991.1 aory|EIT74734.1 asyd|OJJ63566.1 ater|EAU39012.1 atub|OJI90847.1 aver|OJI97367.1 awen|OJJ35332.1 azon|OJJ49786.1

Group3204: abra|OJJ73806.1 acar|OOG00961.1 acla|EAW07694.1 afis|EAW22709.1 afla|EED49996.1 afum|EAL90483.1 agla|OJJ79001.1 akaw|GAA89426.1 aluc|OJZ88246.1 anid|CBF88064.1 anig|EHA17990.1 aory|EIT74673.1 asyd|OJJ63549.1 ater|EAU38992.1 atub|OJI90846.1 aver|OJI97385.1 awen|OJJ35346.1 azon|OJJ49771.1

Group3205: abra|OJJ73807.1 acar|OOG00963.1 acla|EAW07695.1 afis|EAW22712.1 afla|EED49997.1 afum|EAL90482.1 agla|OJJ79000.1 akaw|GAA89427.1 aluc|OJZ88243.1 anid|CBF88062.1 anig|EHA17989.1 aory|EIT74610.1 asyd|OJJ63548.1 ater|EAU38993.1 atub|OJI90845.1 aver|OJI97386.1 awen|OJJ35347.1 azon|OJJ49770.1

Group3206: abra|OJJ73816.1 acar|OOG00970.1 acla|EAW07703.1 afis|EAW22721.1 afla|EED50006.1 afum|EAL90474.1 agla|OJJ83682.1 akaw|GAA89435.1 aluc|OJZ88234.1 anid|CBF88036.1 anig|EHA17980.1 aory|EIT74720.1 asyd|OJJ63535.1 ater|EAU38983.1 atub|OJI90836.1 aver|OJI97397.1 awen|OJJ35415.1 azon|OJJ49760.1

Group3207: abra|OJJ73821.1 acar|OOG00973.1 acla|EAW07710.1 afis|EAW22726.1 afla|EED50012.1 afum|EAL90469.1 agla|OJJ83676.1 akaw|GAA89440.1 aluc|OJZ88227.1 anid|CBF88028.1 anig|EHA17975.1 aory|EIT74707.1 asyd|OJJ63529.1 ater|EAU38978.1 atub|OJI90828.1 aver|OJI97402.1 awen|OJJ35421.1 azon|OJJ49755.1

Group3208: abra|OJJ73830.1 acar|OOG00981.1 acla|EAW07725.1 afis|EAW22757.1 afla|EED50018.1 afum|EAL90442.2 agla|OJJ83666.1 akaw|GAA89448.1 aluc|OJZ88219.1 anid|CBF88015.1 anig|EHA17966.1 aory|EIT74656.1 asyd|OJJ63519.1 ater|EAU38962.1 atub|OJI90819.1 aver|OJI97411.1 awen|OJJ35429.1 azon|OJJ49747.1

Group3210: abra|OJJ73832.1 acar|OOG00983.1 acla|EAW07727.1 afis|EAW22759.1 afla|EED50020.1 afum|EAL90440.1 agla|OJJ83661.1 akaw|GAA89450.1 aluc|OJZ88217.1 anid|CBF88013.1 anig|EHA17964.1 aory|EIT74627.1 asyd|OJJ63517.1 ater|EAU38960.1 atub|OJI90817.1 aver|OJI97413.1 awen|OJJ35431.1 azon|OJJ49745.1

Group3211: abra|OJJ73833.1 acar|OOG00984.1 acla|EAW07728.1 afis|EAW22760.1 afla|EED50021.1 afum|EAL90439.1 agla|OJJ83660.1 akaw|GAA89451.1 aluc|OJZ88216.1 anid|CBF88011.1 anig|EHA17963.1 aory|EIT74690.1 asyd|OJJ63516.1 ater|EAU38959.1 atub|OJI90816.1 aver|OJI97414.1 awen|OJJ35433.1 azon|OJJ49744.1

Group3212: abra|OJJ73834.1 acar|OOG00985.1 acla|EAW07729.1 afis|EAW22761.1 afla|EED50022.1 afum|EAL90438.1 agla|OJJ83659.1 akaw|GAA89452.1 aluc|OJZ88215.1 anid|CBF88009.1 anig|EHA17962.1 aory|EIT74638.1 asyd|OJJ63515.1 ater|EAU38958.1 atub|OJI90815.1 aver|OJI97415.1 awen|OJJ35434.1 azon|OJJ49743.1

Group3213: abra|OJJ73836.1 acar|OOG00987.1 acla|EAW07731.1 afis|EAW22763.1 afla|EED50025.1 afum|EAL90436.2 agla|OJJ83657.1 akaw|GAA89454.1 aluc|OJZ88213.1 anid|CBF88005.1 anig|EHA17961.1 aory|EIT74679.1 asyd|OJJ63513.1 ater|EAU38956.1 atub|OJI90813.1 aver|OJI97417.1 awen|OJJ35436.1 azon|OJJ49741.1

Group3214: abra|OJJ73837.1 acar|OOG00988.1 acla|EAW07732.1 afis|EAW22764.1 afla|EED50026.1 afum|EAL90435.1 agla|OJJ83656.1 akaw|GAA89455.1 aluc|OJZ88212.1 anid|CBF88003.1 anig|EHA17960.1 aory|EIT74726.1 asyd|OJJ63509.1 ater|EAU38953.1 atub|OJI90812.1 aver|OJI97421.1 awen|OJJ38969.1 azon|OJJ49740.1

Group3215: abra|OJJ73838.1 acar|OOG00989.1 acla|EAW07733.1 afis|EAW22765.1 afla|EED50027.1 afum|EAL90434.1 agla|OJJ83655.1 akaw|GAA89456.1 aluc|OJZ88211.1 anid|CBF88001.1 anig|EHA17958.1 aory|EIT74637.1 asyd|OJJ63508.1 ater|EAU38952.1 atub|OJI90811.1 aver|OJI97422.1 awen|OJJ38968.1 azon|OJJ49738.1

Group3216: abra|OJJ73841.1 acar|OOG00992.1 acla|EAW07736.1 afis|EAW22772.1 afla|EED50031.1 afum|EAL90426.1 agla|OJJ83602.1 akaw|GAA89459.1 aluc|OJZ88207.1 anid|CBF87996.1 anig|EHA17955.1 aory|EIT74633.1 asyd|OJJ63505.1 ater|EAU38941.1 atub|OJI90808.1 aver|OJI97425.1 awen|OJJ38965.1 azon|OJJ49736.1

Group3217: abra|OJJ73842.1 acar|OOG00993.1 acla|EAW07737.1 afis|EAW22773.1 afla|EED50032.1 afum|EAL90425.1 agla|OJJ83603.1 akaw|GAA89460.1 aluc|OJZ88206.1 anid|CBF87994.1 anig|EHA17954.1 aory|EIT74732.1 asyd|OJJ63504.1 ater|EAU38940.1 atub|OJI90807.1 aver|OJI97426.1 awen|OJJ38964.1 azon|OJJ49735.1

Group3218: abra|OJJ73843.1 acar|OOG00994.1 acla|EAW07738.1 afis|EAW22774.1 afla|EED50033.1 afum|EAL90424.1 agla|OJJ83604.1 akaw|GAA89461.1 aluc|OJZ88205.1 anid|CBF87992.1 anig|EHA17953.1 aory|EIT74665.1 asyd|OJJ63503.1 ater|EAU38939.1 atub|OJI90806.1 aver|OJI97427.1 awen|OJJ38963.1 azon|OJJ49734.1

Group3219: abra|OJJ73844.1 acar|OOG00995.1 acla|EAW07739.1 afis|EAW22775.1 afla|EED50034.1 afum|EAL90423.1 agla|OJJ83605.1 akaw|GAA89462.1 aluc|OJZ88204.1 anid|CBF87991.1 anig|EHA17952.1 aory|EIT74686.1 asyd|OJJ63502.1 ater|EAU38938.1 atub|OJI90805.1 aver|OJI97428.1 awen|OJJ38962.1 azon|OJJ49733.1

Group3220: abra|OJJ73846.1 acar|OOG00997.1 acla|EAW07741.1 afis|EAW22777.1 afla|EED50037.1 afum|EAL90421.1 agla|OJJ83628.1 akaw|GAA89464.1 aluc|OJZ88202.1 anid|CBF87987.1 anig|EHA17950.1 aory|EIT74713.1 asyd|OJJ63500.1 ater|EAU38937.1 atub|OJI90802.1 aver|OJI97430.1 awen|OJJ38960.1 azon|OJJ49731.1

Group3221: abra|OJJ73848.1 acar|OOG00998.1 acla|EAW07742.1 afis|EAW22778.1 afla|EED50038.1 afum|EAL90420.1 agla|OJJ83632.1 akaw|GAA89465.1 aluc|OJZ88201.1 anid|CBF87983.1 anig|EHA17949.1 aory|EIT74616.1 asyd|OJJ63497.1 ater|EAU38936.1 atub|OJI90799.1 aver|OJI97432.1 awen|OJJ38959.1 azon|OJJ49730.1

Group3222: abra|OJJ73850.1 acar|OOG01000.1 acla|EAW07743.1 afis|EAW22780.1 afla|EED50041.1 afum|EAL90419.1 agla|OJJ83633.1 akaw|GAA89467.1 aluc|OJZ88199.1 anid|CBF87970.1 anig|EHA17947.1 aory|EIT74714.1 asyd|OJJ63494.1 ater|EAU38932.1 atub|OJI90797.1 aver|OJI97435.1 awen|OJJ38957.1 azon|OJJ49729.1

Group3223: abra|OJJ73851.1 acar|OOG01001.1 acla|EAW07745.1 afis|EAW22782.1 afla|EED50044.1 afum|EAL90417.1 agla|OJJ83644.1 akaw|GAA89468.1 aluc|OJZ88197.1 anid|CBF87972.1 anig|EHA17946.1 aory|EIT74643.1 asyd|OJJ63493.1 ater|EAU38930.1 atub|OJI90796.1 aver|OJI97436.1 awen|OJJ38955.1 azon|OJJ49727.1

Group3224: abra|OJJ73854.1 acar|OOG01002.1 acla|EAW07746.1 afis|EAW22783.1 afla|EED50045.1 afum|EAL90416.1 agla|OJJ83653.1 akaw|GAA89470.1 aluc|OJZ88194.1 anid|CBF87974.1 anig|EHA17945.1 aory|EIT74701.1 asyd|OJJ63492.1 ater|EAU38926.1 atub|OJI90793.1 aver|OJI97437.1 awen|OJJ38954.1 azon|OJJ49726.1

Group3225: abra|OJJ73861.1 acar|OOG01007.1 acla|EAW07751.1 afis|EAW22788.1 afla|EED50052.1 afum|EAL90411.1 agla|OJJ83588.1 akaw|GAA89481.1 aluc|OJZ88182.1 anid|CBF87963.1 anig|EHA17934.1 aory|EIT74670.1 asyd|OJJ63484.1 ater|EAU38922.1 atub|OJI90781.1 aver|OJI97445.1 awen|OJJ38946.1 azon|OJJ49718.1

Group3226: abra|OJJ73864.1 acar|OOF89914.1 acla|EAW07754.1 afis|EAW22791.1 afla|EED50058.1 afum|EAL90408.1 agla|OJJ83567.1 akaw|GAA89478.1 aluc|OJZ88185.1 anid|CBF87957.1 anig|EHA17936.1

aory|EIT74695.1 asyd|OJJ63481.1 ater|EAU38920.1 atub|OJI90784.1 aver|OJI97449.1 awen|OJJ38943.1 azon|OJJ49716.1

Group3227: abra|OJJ73867.1 acar|OOF89912.1 acla|EAW07755.1 afis|EAW22792.1 afla|EED50059.1 afum|EAL90407.2 agla|OJJ83568.1 akaw|GAA89475.1 aluc|OJZ88188.1 anid|CBF87955.1 anig|EHA17939.1 aory|EIT74728.1 asyd|OJJ63480.1 ater|EAU38919.1 atub|OJI90787.1 aver|OJI97450.1 awen|OJJ38941.1 azon|OJJ49715.1

Group3228: abra|OJJ73869.1 acar|OOG01012.1 acla|EAW07757.1 afis|EAW22794.1 afla|EED50062.1 afum|EAL90405.1 agla|OJJ83570.1 akaw|GAA89473.1 aluc|OJZ88190.1 anid|CBF87952.1 anig|EHA17941.1 aory|EIT74606.1 asyd|OJJ63478.1 ater|EAU38917.1 atub|OJI90789.1 aver|OJI97452.1 awen|OJJ38939.1 azon|OJJ49713.1

Group3229: abra|OJJ73874.1 acar|OOG01014.1 acla|EAW07759.1 afis|EAW22795.1 afla|EED50063.1 afum|EAL90404.1 agla|OJJ83571.1 akaw|GAA89486.1 aluc|OJZ88176.1 anid|CBF87950.1 anig|EHA17928.1 aory|EIT74699.1 asyd|OJJ63477.1 ater|EAU38915.1 atub|OJI90774.1 aver|OJI97453.1 awen|OJJ38938.1 azon|OJJ49712.1

Group3230: abra|OJJ73876.1 acar|OOG01016.1 acla|EAW07761.1 afis|EAW22797.1 afla|EED50066.1 afum|EAL90402.1 agla|OJJ83574.1 akaw|GAA89488.1 aluc|OJZ88173.1 anid|CBF87946.1 anig|EHA17926.1 aory|EIT74703.1 asyd|OJJ63475.1 ater|EAU38913.1 atub|OJI90771.1 aver|OJI97455.1 awen|OJJ38932.1 azon|OJJ49708.1

Group3231: abra|OJJ73881.1 acar|OOG01020.1 acla|EAW07763.1 afis|EAW22799.1 afla|EED50069.1 afum|EAL90400.1 agla|OJJ83576.1 akaw|GAA89492.1 aluc|OJZ88169.1 anid|CBF87936.1 anig|EHA17923.1 aory|EIT74640.1 asyd|OJJ63466.1 ater|EAU38910.1 atub|OJI90767.1 aver|OJI97466.1 awen|OJJ38930.1 azon|OJJ49706.1

Group3232: abra|OJJ73882.1 acar|OOG01021.1 acla|EAW07764.1 afis|EAW22800.1 afla|EED50070.1 afum|EAL90399.1 agla|OJJ83577.1 akaw|GAA89493.1 aluc|OJZ88168.1 anid|CBF87934.1 anig|EHA17922.1 aory|EIT74735.1 asyd|OJJ63465.1 ater|EAU38909.1 atub|OJI90766.1 aver|OJI97467.1 awen|OJJ38929.1 azon|OJJ49705.1

Group3233: abra|OJJ73883.1 acar|OOG01022.1 acla|EAW07765.1 afis|EAW22801.1 afla|EED50071.1 afum|EAL90398.1 agla|OJJ83578.1 akaw|GAA89494.1 aluc|OJZ88167.1 anid|CBF87932.1 anig|EHA17921.1 aory|EIT74619.1 asyd|OJJ63464.1 ater|EAU38908.1 atub|OJI90765.1 aver|OJI97468.1 awen|OJJ38928.1 azon|OJJ49704.1

Group3234: abra|OJJ73884.1 acar|OOG01023.1 acla|EAW07766.1 afis|EAW22802.1 afla|EED50072.1 afum|EAL90397.1 agla|OJJ83579.1 akaw|GAA89495.1 aluc|OJZ88166.1 anid|CBF87930.1 anig|EHA17920.1 aory|EIT74719.1 asyd|OJJ63463.1 ater|EAU38907.1 atub|OJI90764.1 aver|OJI97469.1 awen|OJJ38927.1 azon|OJJ49703.1

Group3235: abra|OJJ73886.1 acar|OOG01024.1 acla|EAW07768.1 afis|EAW22804.1 afla|EED50074.1 afum|EAL90395.1 agla|OJJ83581.1 akaw|GAA89497.1 aluc|OJZ88164.1 anid|CBF87926.1 anig|EHA17918.1 aory|EIT74660.1 asyd|OJJ63461.1 ater|EAU38905.1 atub|OJI90762.1 aver|OJI97471.1 awen|OJJ38924.1 azon|OJJ49701.1

Group3236: abra|OJJ73889.1 acar|OOG01027.1 acla|EAW07773.1 afis|EAW22809.1 afla|EED50078.1 afum|EAL90392.1 agla|OJJ83584.1 akaw|GAA89501.1 aluc|OJZ88160.1 anid|CBF87904.1 anig|EHA17915.1 aory|EIT74748.1 asyd|OJJ63457.1 ater|EAU38902.1 atub|OJI90759.1 aver|OJI97475.1 awen|OJJ38921.1 azon|OJJ49698.1

Group3237: abra|OJJ73890.1 acar|OOG01030.1 acla|EAW07774.1 afis|EAW22810.1 afla|EED50079.1 afum|EAL90391.1 agla|OJJ83585.1 akaw|GAA89502.1 aluc|OJZ88159.1 anid|CBF87906.1 anig|EHA17914.1 aory|EIT74744.1 asyd|OJJ63456.1 ater|EAU38901.1 atub|OJI90758.1 aver|OJI97476.1 awen|OJJ38920.1 azon|OJJ49697.1

Group3238: abra|OJJ73891.1 acar|OOG01029.1 acla|EAW07775.1 afis|EAW22811.1 afla|EED50080.1 afum|EAL90390.1 agla|OJJ83566.1 akaw|GAA89503.1 aluc|OJZ88158.1 anid|CBF87908.1 anig|EHA17913.1 aory|EIT74652.1 asyd|OJJ63455.1 ater|EAU38900.1 atub|OJI90757.1 aver|OJI97477.1 awen|OJJ38915.1 azon|OJJ49696.1

Group3239: abra|OJJ73897.1 acar|OOG01034.1 acla|EAW07778.1 afis|EAW22815.1 afla|EED50088.1 afum|EAL90386.1 agla|OJJ83527.1 akaw|GAA89507.1 aluc|OJZ88149.1 anid|CBF87915.1 anig|EHA17910.1 aory|EIT74671.1 asyd|OJJ63448.1 ater|EAU38896.1 atub|OJI90752.1 aver|OJI97486.1 awen|OJJ38913.1 azon|OJJ49693.1

Group3240: abra|OJJ73901.1 acar|OOG01036.1 acla|EAW07780.1 afis|EAW22816.1 afla|EED50089.1 afum|EAL90385.1 agla|OJJ83524.1 akaw|GAA89508.1 aluc|OJZ88145.1 anid|CBF87918.1 anig|EHA17908.1 aory|EIT74605.1 asyd|OJJ63447.1 ater|EAU38895.1 atub|OJI90749.1 aver|OJI97489.1 awen|OJJ38911.1 azon|OJJ49692.1

Group3241: abra|OJJ73902.1 acar|OOG01037.1 acla|EAW07781.1 afis|EAW22817.1 afla|EED50090.1 afum|EAL90384.1 agla|OJJ83521.1 akaw|GAA89509.1 aluc|OJZ88144.1 anid|CBF87920.1 anig|EHA17907.1 aory|EIT74746.1 asyd|OJJ63446.1 ater|EAU38894.1 atub|OJI90748.1 aver|OJI97490.1 awen|OJJ38910.1 azon|OJJ49691.1

Group3242: abra|OJJ73904.1 acar|OOG01038.1 acla|EAW07783.1 afis|EAW22819.1 afla|EED50092.1 afum|EAL90382.1 agla|OJJ83519.1 akaw|GAA89511.1 aluc|OJZ88142.1 anid|CBF87899.1 anig|EHA17905.1 aory|EIT74751.1 asyd|OJJ63444.1 ater|EAU38891.1 atub|OJI90746.1 aver|OJI97492.1 awen|OJJ38908.1 azon|OJJ49689.1

Group3243: abra|OJJ73908.1 acar|OOG01041.1 acla|EAW07786.1 afis|EAW22822.1 afla|EED50097.1 afum|EAL90378.2 agla|OJJ83516.1 akaw|GAA89515.1 aluc|OJZ88139.1 anid|CBF87894.1 anig|EHA17903.1 aory|EIT74681.1 asyd|OJJ63441.1 ater|EAU38888.1 atub|OJI90742.1 aver|OJI97495.1 awen|OJJ38905.1 azon|OJJ49686.1

Group3244: abra|OJJ73909.1 acar|OOG01042.1 acla|EAW07787.1 afis|EAW22823.1 afla|EED50098.1 afum|EAL90377.1 agla|OJJ83515.1 akaw|GAA89516.1 aluc|OJZ88138.1 anid|CBF87892.1 anig|EHA17902.1 aory|EIT74745.1 asyd|OJJ63440.1 ater|EAU38887.1 atub|OJI90741.1 aver|OJI97496.1 awen|OJJ38904.1 azon|OJJ49685.1

Group3245: abra|OJJ73910.1 acar|OOG01043.1 acla|EAW07788.1 afis|EAW22824.1 afla|EED50099.1 afum|EAL90376.1 agla|OJJ83514.1 akaw|GAA89517.1 aluc|OJZ88137.1 anid|CBF87890.1 anig|EHA17901.1 aory|EIT74620.1 asyd|OJJ63439.1 ater|EAU38886.1 atub|OJI90740.1 aver|OJI97497.1 awen|OJJ38903.1 azon|OJJ49684.1

Group3246: abra|OJJ73912.1 acar|OOG01044.1 acla|EAW07789.1 afis|EAW22825.1 afla|EED50100.1 afum|EAL90375.2 agla|OJJ83513.1 akaw|GAA89518.1 aluc|OJZ88136.1 anid|CBF87888.1 anig|EHA17899.1 aory|EIT74685.1 asyd|OJJ63438.1 ater|EAU38885.1 atub|OJI90739.1 aver|OJI97499.1 awen|OJJ38902.1 azon|OJJ49683.1

Group3247: abra|OJJ73915.1 acar|OOG01047.1 acla|EAW07791.1 afis|EAW22829.1 afla|EED50104.1 afum|EAL90373.2 agla|OJJ83510.1 akaw|GAA89521.1 aluc|OJZ88133.1 anid|CBF87886.1 anig|EHA17897.1 aory|EIT74692.1 asyd|OJJ63437.1 ater|EAU38882.1 atub|OJI90737.1 aver|OJI97500.1 awen|OJJ38899.1 azon|OJJ49680.1

Group3248: abra|OJJ73916.1 acar|OOG01048.1 acla|EAW07792.1 afis|EAW22830.1 afla|EED50105.1 afum|EAL90372.2 agla|OJJ83509.1 akaw|GAA89522.1 aluc|OJZ88131.1 anid|CBF87883.1 anig|EHA17896.1 aory|EIT74639.1 asyd|OJJ63435.1 ater|EAU38881.1 atub|OJI90736.1 aver|OJI97502.1 awen|OJJ38898.1 azon|OJJ49679.1

Group3249: abra|OJJ73920.1 acar|OOG01051.1 acla|EAW07797.1 afis|EAW22834.1 afla|EED50109.1 afum|EAL90368.1 agla|OJJ83505.1 akaw|GAA89527.1 aluc|OJZ88126.1 anid|CBF87876.1 anig|EHA17892.1 aory|EIT74609.1 asyd|OJJ63432.1 ater|EAU38875.1 atub|OJI90733.1 aver|OJI97505.1 awen|OJJ38890.1 azon|OJJ49676.1

Group3250: abra|OJJ73923.1 acar|OOG01056.1 acla|EAW07800.1 afis|EAW22837.1 afla|EED50112.1 afum|EAL90365.2 agla|OJJ83504.1 akaw|GAA89530.1 aluc|OJZ88120.1 anid|CBF87867.1 anig|EHA17889.1 aory|EIT74684.1 asyd|OJJ63428.1 ater|EAU38873.1 atub|OJI90730.1 aver|OJI97509.1 awen|OJJ38887.1 azon|OJJ49675.1

Group3251: abra|OJJ73924.1 acar|OOG01057.1 acla|EAW07801.1 afis|EAW22838.1 afla|EED50113.1 afum|EAL90364.2 agla|OJJ83503.1 akaw|GAA89531.1 aluc|OJZ88119.1 anid|CBF87865.1 anig|EHA17888.1 aory|EIT74621.1 asyd|OJJ63427.1 ater|EAU38872.1 atub|OJI90729.1 aver|OJI97510.1 awen|OJJ38885.1 azon|OJJ49674.1

Group3252: abra|OJJ73926.1 acar|OOG01058.1 acla|EAW07803.1 afis|EAW22840.1 afla|EED50115.1 afum|EAL90363.1 agla|OJJ83501.1 akaw|GAA89533.1 aluc|OJZ88117.1 anid|CBF87861.1 anig|EHA17887.1 aory|EIT74702.1 asyd|OJJ63425.1 ater|EAU38870.1 atub|OJI90727.1 aver|OJI97512.1 awen|OJJ38883.1 azon|OJJ49672.1

Group3253: abra|OJJ73927.1 acar|OOG01060.1 acla|EAW07804.1 afis|EAW22841.1 afla|EED50116.1 afum|EAL90362.2 agla|OJJ83500.1 akaw|GAA89534.1 aluc|OJZ88116.1 anid|CBF87860.1 anig|EHA17886.1 aory|EIT74608.1 asyd|OJJ63424.1 ater|EAU38869.1 atub|OJI90725.1 aver|OJI97513.1 awen|OJJ38882.1 azon|OJJ49671.1

Group3254: abra|OJJ73928.1 acar|OOG01061.1 acla|EAW07805.1 afis|EAW22842.1 afla|EED50117.1 afum|EAL90361.1 agla|OJJ83496.1 akaw|GAA89535.1 aluc|OJZ88115.1 anid|CBF87858.1 anig|EHA17885.1 aory|EIT74650.1 asyd|OJJ63423.1 ater|EAU38868.1 atub|OJI90724.1 aver|OJI97514.1 awen|OJJ38881.1 azon|OJJ49670.1

Group3255: abra|OJJ73930.1 acar|OOG01062.1 acla|EAW07808.1 afis|EAW22845.1 afla|EED50120.1 afum|EAL90357.1 agla|OJJ83492.1 akaw|GAA89538.1 aluc|OJZ88112.1 anid|CBF87846.1 anig|EHA17882.1

aory|EIT74689.1 asyd|OJJ63421.1 ater|EAU38865.1 atub|OJI90720.1 aver|OJI97517.1 awen|OJJ38878.1 azon|OJJ49667.1

Group3256: abra|OJJ73931.1 acar|OOG01063.1 acla|EAW07809.1 afis|EAW22846.1 afla|EED50121.1 afum|EAL90356.1 agla|OJJ83485.1 akaw|GAA89539.1 aluc|OJZ88111.1 anid|CBF87848.1 anig|EHA17881.1 aory|EIT74731.1 asyd|OJJ63420.1 ater|EAU38864.1 atub|OJI90719.1 aver|OJI97518.1 awen|OJJ38877.1 azon|OJJ49666.1

Group3257: abra|OJJ73938.1 acar|OOG01069.1 afis|EAW22850.1 afla|EED50125.1 afum|EAL90350.1 agla|OJJ83473.1 akaw|GAA89543.1 aluc|OJZ88106.1 anid|CBF87855.1 anig|EHA17875.1 aory|EIT74705.1 asyd|OJJ63416.1 ater|EAU38859.1 atub|OJI90714.1 aver|OJI97522.1 awen|OJJ38871.1 azon|OJJ49663.1 acla|EAW07814.1

Group3258: abra|OJJ73939.1 acar|OOG01068.1 acla|EAW07816.1 afis|EAW22851.1 afla|EED50126.1 afum|EAL90349.1 agla|OJJ83471.1 akaw|GAA89544.1 aluc|OJZ88105.1 anid|CBF87841.1 anig|EHA17874.1 aory|EIT74661.1 asyd|OJJ63414.1 ater|EAU38858.1 atub|OJI90713.1 aver|OJI97524.1 awen|OJJ38867.1 azon|OJJ49661.1

Group3259: abra|OJJ73944.1 acar|OOG01072.1 acla|EAW07817.1 afis|EAW22852.1 afla|EED50132.1 afum|EAL90348.1 agla|OJJ83469.1 akaw|GAA89549.1 aluc|OJZ88100.1 anid|CBF87839.1 anig|EHA17869.1 aory|EIT74662.1 asyd|OJJ63413.1 ater|EAU38857.1 atub|OJI90708.1 aver|OJI97525.1 awen|OJJ38866.1 azon|OJJ49660.1

Group3260: abra|OJJ73945.1 acar|OOG01073.1 acla|EAW07818.1 afis|EAW22853.1 afla|EED50133.1 afum|EAL90347.1 agla|OJJ83468.1 akaw|GAA89550.1 aluc|OJZ88099.1 anid|CBF87838.1 anig|EHA17868.1 aory|EIT74757.1 asyd|OJJ63412.1 ater|EAU38856.1 atub|OJI90707.1 aver|OJI97526.1 awen|OJJ38865.1 azon|OJJ49659.1

Group3261: abra|OJJ73946.1 acar|OOG01074.1 acla|EAW07819.1 afis|EAW22854.1 afla|EED50134.1 afum|EAL90346.1 agla|OJJ83467.1 akaw|GAA89551.1 aluc|OJZ88098.1 anid|CBF87836.1 anig|EHA17867.1 aory|EIT74630.1 asyd|OJJ63410.1 atub|OJI90706.1 aver|OJI97527.1 awen|OJJ38864.1 azon|OJJ49658.1 ater|EAU38807.1

Group3262: abra|OJJ73949.1 acar|OOG01077.1 acla|EAW07823.1 afis|EAW22859.1 afla|EED50137.1 afum|EAL90341.1 agla|OJJ83465.1 akaw|GAA89554.1 aluc|OJZ88095.1 anid|CBF87833.1 anig|EHA17865.1 aory|EIT74739.1 asyd|OJJ63408.1 ater|EAU38809.1 atub|OJI90703.1 aver|OJI97529.1 awen|OJJ38861.1 azon|OJJ49655.1

Group3263: abra|OJJ73952.1 acar|OOG01080.1 acla|EAW07826.1 afis|EAW22862.1 afla|EED50146.1 afum|EAL90338.1 agla|OJJ83462.1 akaw|GAA89557.1 aluc|OJZ88092.1 anid|CBF87832.1 anig|EHA17862.1 aory|EIT74749.1 asyd|OJJ63407.1 ater|EAU38816.1 atub|OJI90700.1 aver|OJI97530.1 awen|OJJ38858.1 azon|OJJ49656.1

Group3264: abra|OJJ73959.1 acar|OOG01086.1 acla|EAW07828.1 afis|EAW22864.1 afla|EED50148.1 afum|EAL90336.1 agla|OJJ83460.1 akaw|GAA89562.1 aluc|OJZ88086.1 anid|CBF87829.1 anig|EHA17857.1 aory|EIT74755.1 asyd|OJJ63405.1 ater|EAU38824.1 atub|OJI90695.1 aver|OJI97531.1 awen|OJJ38856.1 azon|OJJ49654.1

Group3265: abra|OJJ73961.1 acar|OOG01087.1 acla|EAW07830.1 afis|EAW22866.1 afla|EED50150.1 afum|EAL90333.1 agla|OJJ83458.1 akaw|GAA89564.1 aluc|OJZ88084.1 anid|CBF87827.1 anig|EHA17855.1 aory|EIT74659.1 asyd|OJJ63403.1 ater|EAU38827.1 atub|OJI90693.1 aver|OJI97533.1 awen|OJJ38854.1 azon|OJJ49652.1

Group3266: abra|OJJ73962.1 acar|OOG01089.1 acla|EAW07831.1 afis|EAW22867.1 afla|EED50151.1 afum|EAL90332.1 agla|OJJ83457.1 akaw|GAA89565.1 aluc|OJZ88083.1 anid|CBF87826.1 anig|EHA17854.1 aory|EIT74722.1 asyd|OJJ63402.1 ater|EAU38828.1 atub|OJI90692.1 aver|OJI97534.1 awen|OJJ38853.1 azon|OJJ49651.1

Group3267: abra|OJJ73964.1 acar|OOG01090.1 acla|EAW07834.1 afis|EAW22871.1 afla|EED50154.1 afum|EAL90327.1 agla|OJJ83452.1 akaw|GAA89567.1 aluc|OJZ88080.1 anid|CBF87822.1 anig|EHA17852.1 aory|EIT76938.1 asyd|OJJ63400.1 ater|EAU38835.1 atub|OJI90689.1 aver|OJI97537.1 awen|OJJ38851.1 azon|OJJ49647.1

Group3268: abra|OJJ73967.1 acar|OOG01093.1 acla|EAW07837.1 afis|EAW22874.1 afla|EED50157.1 afum|EAL90324.1 agla|OJJ83443.1 akaw|GAA89570.1 aluc|OJZ88077.1 anid|CBF87816.1 anig|EHA17849.1 aory|EIT76898.1 asyd|OJJ63395.1 ater|EAU38838.1 atub|OJI90686.1 aver|OJI97541.1 awen|OJJ38848.1 azon|OJJ49644.1

Group3269: abra|OJJ73971.1 acar|OOG01097.1 acla|EAW07842.1 afis|EAW22878.1 afla|EED50183.1 afum|EAL90321.1 agla|OJJ83446.1 akaw|GAA89574.1 aluc|OJZ88073.1 anid|CBF87811.1 anig|EHA17846.1 aory|EIT77380.1 asyd|OJJ63390.1 ater|EAU38842.1 atub|OJI90682.1 aver|OJI97545.1 awen|OJJ38844.1 azon|OJJ49641.1

Group3270: abra|OJJ73976.1 acar|OOG01103.1 acla|EAW07847.1 afis|EAW22883.1 afla|EED50176.1 afum|EAL90316.1 agla|OJJ83441.1 akaw|GAA89579.1 aluc|OJZ88068.1 anid|CBF87803.1 anig|EHA17841.1 aory|EIT72282.1 asyd|OJJ63384.1 ater|EAU38846.1 atub|OJI90677.1 aver|OJI97551.1 awen|OJJ38839.1 azon|OJJ49636.1

Group3271: abra|OJJ73978.1 acar|OOG01104.1 acla|EAW07849.1 afis|EAW22885.1 afla|EED50174.1 afum|EAL90313.1 agla|OJJ83438.1 akaw|GAA89581.1 aluc|OJZ88066.1 anid|CBF87801.1 anig|EHA17839.1 aory|EIT72285.1 asyd|OJJ63382.1 ater|EAU35978.1 atub|OJI90675.1 aver|OJI97553.1 awen|OJJ38837.1 azon|OJJ49634.1

Group3272: abra|OJJ73980.1 acar|OOG01107.1 acla|EAW07851.1 afis|EAW22887.1 afla|EED50172.1 afum|EAL90311.1 agla|OJJ83436.1 akaw|GAA89583.1 aluc|OJZ88064.1 anid|CBF87798.1 anig|EHA17837.1 aory|EIT72283.1 asyd|OJJ63380.1 ater|EAU35985.1 atub|OJI90673.1 aver|OJI97555.1 awen|OJJ38835.1 azon|OJJ49632.1

Group3273: abra|OJJ73982.1 acar|OOG01109.1 acla|EAW07854.1 afis|EAW22890.1 afla|EED54859.1 afum|EAL90308.1 agla|OJJ83420.1 akaw|GAA89586.1 aluc|OJZ88062.1 anid|CBF87791.1 anig|EHA17835.1 aory|EIT79448.1 asyd|OJJ63377.1 ater|EAU35982.1 atub|OJI90671.1 aver|OJI97569.1 awen|OJJ38831.1 azon|OJJ49630.1

Group3274: abra|OJJ73983.1 acar|OOG01110.1 acla|EAW07855.1 afis|EAW22891.1 afla|EED54860.1 afum|EAL90307.1 agla|OJJ83407.1 akaw|GAA89587.1 aluc|OJZ88061.1 anid|CBF87790.1 anig|EHA17834.1 aory|EIT79392.1 asyd|OJJ63376.1 ater|EAU36000.1 atub|OJI90670.1 aver|OJI97570.1 awen|OJJ38830.1 azon|OJJ49629.1

Group3275: abra|OJJ73986.1 acar|OOG01111.1 acla|EAW07857.1 afis|EAW22893.1 afla|EED54862.1 afum|EAL90305.2 agla|OJJ83410.1 akaw|GAA87469.1 aluc|OJZ88059.1 anid|CBF87783.1 anig|EHA17831.1 aory|EIT79335.1 asyd|OJJ63374.1 ater|EAU35990.1 atub|OJI90667.1 aver|OJI97572.1 awen|OJJ38827.1 azon|OJJ49627.1

Group3276: abra|OJJ73989.1 acar|OOG01115.1 acla|EAW07860.1 afis|EAW22895.1 afla|EED54864.1 afum|EAL90303.1 agla|OJJ83415.1 akaw|GAA87472.1 aluc|OJZ88056.1 anid|CBF74977.1 anig|EHA17828.1 aory|EIT79412.1 asyd|OJJ60090.1 ater|EAU35987.1 atub|OJI90664.1 aver|OJJ04177.1 awen|OJJ38823.1 azon|OJJ49622.1

Group3277: abra|OJJ73991.1 acar|OOG01117.1 acla|EAW07862.1 afis|EAW22897.1 afla|EED54868.1 afum|EAL90301.1 agla|OJJ83417.1 akaw|GAA87474.1 aluc|OJZ88054.1 anid|CBF87775.1 anig|EHA17826.1 aory|EIT79359.1 asyd|OJJ63369.1 ater|EAU35981.1 atub|OJI90662.1 aver|OJI97577.1 awen|OJJ38820.1 azon|OJJ49620.1

Group3279: abra|OJJ73993.1 acar|OOG01118.1 acla|EAW07864.1 afis|EAW22899.1 afla|EED54870.1 afum|EAL90298.1 agla|OJJ83392.1 akaw|GAA87476.1 aluc|OJZ88052.1 anid|CBF87771.1 anig|EHA17824.1 aory|EIT79377.1 asyd|OJJ63367.1 ater|EAU36002.1 atub|OJI90660.1 aver|OJI97579.1 awen|OJJ38731.1 azon|OJJ49618.1

Group3280: abra|OJJ74000.1 acar|OOG01121.1 acla|EAW07866.1 afis|EAW22901.1 afla|EED54872.1 afum|EAL90296.1 agla|OJJ83389.1 akaw|GAA87482.1 aluc|OJZ88043.1 anid|CBF87767.1 anig|EHA17816.1 aory|EIT79383.1 asyd|OJJ63365.1 ater|EAU36006.1 atub|OJI90650.1 aver|OJI97582.1 awen|OJJ38729.1 azon|OJJ49615.1

Group3281: abra|OJJ74002.1 acar|OOG01122.1 acla|EAW07867.1 afis|EAW22902.1 afla|EED54873.1 afum|EAL90295.1 agla|OJJ83388.1 akaw|GAA87483.1 aluc|OJZ88042.1 anid|CBF87766.1 anig|EHA17815.1 aory|EIT79350.1 asyd|OJJ63364.1 ater|EAU36007.1 atub|OJI90649.1 aver|OJI97583.1 awen|OJJ38728.1 azon|OJJ49614.1

Group3282: abra|OJJ74003.1 acar|OOG01124.1 acla|EAW07868.1 afis|EAW22903.1 afla|EED54874.1 afum|EAL90294.1 agla|OJJ83387.1 akaw|GAA87484.1 aluc|OJZ88041.1 anid|CBF87764.1 anig|EHA17814.1 aory|EIT79433.1 asyd|OJJ63363.1 ater|EAU36010.1 atub|OJI90648.1 aver|OJI97584.1 awen|OJJ38727.1 azon|OJJ49613.1

Group3283: abra|OJJ74016.1 acar|OOG01137.1 acla|EAW07870.1 afis|EAW22905.1 afla|EED54897.1 afum|EAL90292.1 agla|OJJ83384.1 akaw|GAA87498.1 aluc|OJZ88024.1 anid|CBF87713.1 anig|EHA17801.1 aory|EIT80309.1 asyd|OJJ63338.1 ater|EAU36034.1 atub|OJI90632.1 aver|OJI97623.1 awen|OJJ38725.1 azon|OJJ49609.1

Group3284: abra|OJJ74017.1 acar|OOG01138.1 acla|EAW07871.1 afis|EAW22906.1 afla|EED54898.1 afum|EAL90291.1 agla|OJJ83383.1 akaw|GAA87499.1 aluc|OJZ88022.1 anid|CBF87712.1 anig|EHA17800.1 aory|EIT80293.1 asyd|OJJ63337.1 ater|EAU36033.1 atub|OJI90631.1 aver|OJI97625.1 awen|OJJ38724.1 azon|OJJ49608.1

Group3285: abra|OJJ74020.1 acar|OOG01139.1 acla|EAW07874.1 afis|EAW22909.1 afla|EED54901.1 afum|EAL90288.1 agla|OJJ83380.1 akaw|GAA87502.1 aluc|OJZ88019.1 anid|CBF87705.1 anig|EHA17798.1

aory|EIT80370.1 asyd|OJJ63332.1 ater|EAU36030.1 atub|OJI90628.1 aver|OJI97630.1 awen|OJJ38656.1 azon|OJJ49605.1

Group3286: abra|OJJ74021.1 acar|OOG01140.1 acla|EAW07875.1 afis|EAW22910.1 afla|EED54902.1 afum|EAL90287.1 agla|OJJ83379.1 akaw|GAA87503.1 aluc|OJZ88018.1 anid|CBF87703.1 anig|EHA17797.1 aory|EIT80365.1 asyd|OJJ63331.1 ater|EAU36029.1 atub|OJI90627.1 aver|OJI97631.1 awen|OJJ38655.1 azon|OJJ49604.1

Group3287: abra|OJJ74022.1 acar|OOG01141.1 acla|EAW07876.1 afis|EAW22911.1 afla|EED54904.1 afum|EAL90286.1 agla|OJJ83378.1 akaw|GAA87505.1 aluc|OJZ88015.1 anid|CBF87694.1 anig|EHA17796.1 aory|EIT80313.1 asyd|OJJ63326.1 ater|EAU36028.1 atub|OJI90625.1 aver|OJI97636.1 awen|OJJ38654.1 azon|OJJ49603.1

Group3288: abra|OJJ74023.1 acar|OOG01142.1 acla|EAW07877.1 afis|EAW22912.1 afla|EED54905.1 afum|EAL90285.1 agla|OJJ83377.1 akaw|GAA87504.1 aluc|OJZ88017.1 anid|CBF87695.1 anig|EHA17795.1 aory|EIT80317.1 asyd|OJJ63327.1 ater|EAU36027.1 atub|OJI90626.1 aver|OJI97635.1 awen|OJJ38653.1 azon|OJJ49602.1

Group3289: abra|OJJ74024.1 acar|OOG01143.1 acla|EAW07878.1 afis|EAW22913.1 afla|EED54906.1 afum|EAL90284.1 agla|OJJ83376.1 akaw|GAA87506.1 aluc|OJZ88014.1 anid|CBF87699.1 anig|EHA17794.1 aory|EIT80324.1 asyd|OJJ63329.1 ater|EAU36026.1 atub|OJI90623.1 aver|OJI97633.1 awen|OJJ38652.1 azon|OJJ49601.1

Group3290: abra|OJJ74027.1 acar|OOG01146.1 acla|EAW07879.1 afis|EAW22914.1 afla|EED54907.1 afum|EAL90283.1 agla|OJJ83375.1 akaw|GAA87510.1 aluc|OJZ88010.1 anid|CBF87697.1 anig|EHA17790.1 aory|EIT80315.1 asyd|OJJ63328.1 ater|EAU30708.1 atub|OJI90619.1 aver|OJI97634.1 awen|OJJ38651.1 azon|OJJ49599.1

Group3291: abra|OJJ74029.1 acar|OOG01147.1 acla|EAW07880.1 afis|EAW22915.1 afla|EED54911.1 afum|EAL90282.1 agla|OJJ83373.1 akaw|GAA87512.1 aluc|OJZ88008.1 anid|CBF87690.1 anig|EHA17788.1 aory|EIT80322.1 asyd|OJJ63325.1 ater|EAU30707.1 atub|OJI90617.1 aver|OJI97637.1 awen|OJJ38649.1 azon|OJJ49595.1

Group3292: abra|OJJ74032.1 acar|OOG01151.1 acla|EAW07883.1 afis|EAW22918.1 afla|EED54915.1 afum|EAL90279.1 agla|OJJ83371.1 akaw|GAA87516.1 aluc|OJZ88005.1 anid|CBF87687.1 anig|EHA17786.1 aory|EIT80289.1 asyd|OJJ63320.1 ater|EAU30705.1 atub|OJI90613.1 aver|OJI97639.1 awen|OJJ38646.1 azon|OJJ49588.1

Group3293: abra|OJJ74033.1 acar|OOG01152.1 acla|EAW07884.1 afis|EAW22919.1 afla|EED54916.1 afum|EAL90278.1 agla|OJJ83369.1 akaw|GAA87517.1 aluc|OJZ88004.1 anid|CBF87685.1 anig|EHA17785.1 aory|EIT80336.1 asyd|OJJ63319.1 ater|EAU30704.1 atub|OJI90612.1 aver|OJI97640.1 awen|OJJ38645.1 azon|OJJ49587.1

Group3294: abra|OJJ74035.1 acla|EAW07885.1 afis|EAW22920.1 afla|EED54917.1 afum|EAL90277.1 agla|OJJ83368.1 akaw|GAA87518.1 anid|CBF87683.1 aory|EIT80285.1 asyd|OJJ63318.1 ater|EAU30703.1 aver|OJI97641.1 awen|OJJ38644.1 azon|OJJ49586.1 acar|OOG01153.1 aluc|OJZ88003.1 anig|EHA17784.1 atub|OJI90611.1

Group3295: abra|OJJ74036.1 acar|OOG01155.1 acla|EAW07886.1 afis|EAW22921.1 afla|EED54918.1 afum|EAL90276.1 agla|OJJ83367.1 akaw|GAA87519.1 aluc|OJZ88000.1 anid|CBF87681.1 anig|EHA17782.1 aory|EIT80300.1 asyd|OJJ63317.1 ater|EAU30702.1 atub|OJI90609.1 aver|OJI97642.1 awen|OJJ38643.1 azon|OJJ49585.1

Group3296: abra|OJJ74037.1 acar|OOG01156.1 acla|EAW07887.1 afis|EAW22922.1 afla|EED54919.1 afum|EAL90275.1 agla|OJJ83366.1 akaw|GAA87520.1 aluc|OJZ87999.1 anid|CBF87680.1 anig|EHA17781.1 aory|EIT80390.1 asyd|OJJ63316.1 ater|EAU30701.1 atub|OJI90608.1 aver|OJI97643.1 awen|OJJ38642.1 azon|OJJ49584.1

Group3297: abra|OJJ74038.1 acar|OOG01157.1 acla|EAW07888.1 afis|EAW22923.1 afla|EED54920.1 afum|EAL90274.1 agla|OJJ83365.1 akaw|GAA87521.1 aluc|OJZ87998.1 anid|CBF87678.1 anig|EHA17780.1 aory|EIT80297.1 asyd|OJJ63315.1 ater|EAU30700.1 atub|OJI90607.1 aver|OJI97644.1 awen|OJJ38641.1 azon|OJJ49583.1

Group3298: abra|OJJ74039.1 acar|OOG01158.1 acla|EAW07889.1 afis|EAW22924.1 afla|EED54921.1 afum|EAL90273.1 agla|OJJ83364.1 akaw|GAA87522.1 aluc|OJZ87997.1 anid|CBF87676.1 anig|EHA17779.1 aory|EIT80292.1 asyd|OJJ63314.1 ater|EAU30699.1 atub|OJI90606.1 aver|OJI97645.1 awen|OJJ38640.1 azon|OJJ49582.1

Group3299: abra|OJJ74040.1 acar|OOG01159.1 acla|EAW07890.1 afis|EAW22925.1 afla|EED54922.1 afum|EAL90272.1 agla|OJJ83363.1 akaw|GAA87523.1 aluc|OJZ87996.1 anid|CBF87674.1 anig|EHA17778.1 aory|EIT80341.1 asyd|OJJ63313.1 ater|EAU30698.1 atub|OJI90605.1 aver|OJI97646.1 awen|OJJ38639.1 azon|OJJ49581.1

Group3300: abra|OJJ74043.1 acar|OOG01161.1 acla|EAW07892.1 afis|EAW22927.1 afla|EED54924.1 afum|EAL90270.1 agla|OJJ83361.1 akaw|GAA87525.1 aluc|OJZ87994.1 anid|CBF87671.1 anig|EHA17776.1 aory|EIT80287.1 asyd|OJJ63311.1 ater|EAU30696.1 atub|OJI90603.1 aver|OJI97648.1 awen|OJJ38637.1 azon|OJJ49579.1

Group3301: abra|OJJ74045.1 acar|OOG01164.1 acla|EAW07895.1 afis|EAW22930.1 afla|EED58123.1 afum|EAL90267.1 agla|OJJ83358.1 akaw|GAA87528.1 aluc|OJZ87990.1 anid|CBF87667.1 anig|EHA17773.1 aory|EIT79163.1 asyd|OJJ63308.1 ater|EAU30694.1 atub|OJI90600.1 aver|OJI97651.1 awen|OJJ38634.1 azon|OJJ49576.1

Group3302: abra|OJJ74048.1 acar|OOF89921.1 acla|EAW07897.1 afis|EAW22932.1 afla|EED58126.1 afum|EAL90264.1 agla|OJJ83355.1 akaw|GAA87530.1 aluc|OJZ87987.1 anid|CBF87662.1 anig|EHA17770.1 aory|EIT79157.1 asyd|OJJ63305.1 ater|EAU30692.1 atub|OJI90597.1 aver|OJI97654.1 awen|OJJ38631.1 azon|OJJ49573.1

Group3303: abra|OJJ74049.1 acar|OOF89922.1 acla|EAW07898.1 afis|EAW22934.1 afla|EED58127.1 afum|EAL90262.1 agla|OJJ83354.1 akaw|GAA87531.1 aluc|OJZ87986.1 anid|CBF87661.1 anig|EHA17769.1 aory|EIT79153.1 asyd|OJJ63304.1 ater|EAU30691.1 atub|OJI90596.1 aver|OJI97655.1 awen|OJJ38630.1 azon|OJJ49572.1

Group3304: abra|OJJ74050.1 acar|OOF89923.1 acla|EAW07899.1 afis|EAW22935.1 afla|EED58128.1 afum|EAL90261.1 agla|OJJ83353.1 akaw|GAA87532.1 aluc|OJZ87985.1 anid|CBF87659.1 anig|EHA17768.1 aory|EIT79137.1 asyd|OJJ63303.1 ater|EAU30690.1 atub|OJI90595.1 aver|OJI97656.1 awen|OJJ38629.1 azon|OJJ49571.1

Group3305: abra|OJJ74053.1 acar|OOG01166.1 acla|EAW07901.1 afis|EAW22938.1 afla|EED58131.1 afum|EAL90258.1 agla|OJJ83351.1 akaw|GAA87535.1 aluc|OJZ87982.1 anid|CBF87654.1 anig|EHA17765.1 aory|EIT79120.1 asyd|OJJ63299.1 ater|EAU30687.1 atub|OJI90592.1 aver|OJI97659.1 awen|OJJ38626.1 azon|OJJ49568.1

Group3306: abra|OJJ74054.1 acar|OOG01167.1 acla|EAW07902.1 afis|EAW22939.1 afla|EED58133.1 afum|EAL90257.1 agla|OJJ83349.1 akaw|GAA87536.1 aluc|OJZ87981.1 anid|CBF87652.1 anig|EHA17764.1 aory|EIT79138.1 asyd|OJJ63298.1 ater|EAU30686.1 atub|OJI90591.1 aver|OJI97660.1 awen|OJJ38625.1 azon|OJJ49567.1

Group3307: abra|OJJ74056.1 acar|OOG01169.1 acla|EAW07903.1 afis|EAW22940.1 afla|EED58154.1 afum|EAL90256.1 agla|OJJ83348.1 akaw|GAA87538.1 aluc|OJZ87979.1 anid|CBF87650.1 anig|EHA17762.1 aory|EIT79084.1 asyd|OJJ63297.1 ater|EAU30684.1 atub|OJI90589.1 aver|OJI97661.1 awen|OJJ38624.1 azon|OJJ49565.1

Group3308: abra|OJJ74060.1 acar|OOG01172.1 acla|EAW07905.1 afis|EAW22943.1 afla|EED58153.1 afum|EAL90254.1 agla|OJJ83345.1 akaw|GAA87541.1 aluc|OJZ87976.1 anid|CBF87646.1 anig|EHA17760.1 aory|EIT79073.1 asyd|OJJ63295.1 ater|EAU30681.1 atub|OJI90586.1 aver|OJI97663.1 awen|OJJ38621.1 azon|OJJ49562.1

Group3309: abra|OJJ74061.1 acar|OOG01173.1 acla|EAW07906.1 afis|EAW22944.1 afla|EED58151.1 afum|EAL90253.1 agla|OJJ83344.1 akaw|GAA87542.1 aluc|OJZ87975.1 anid|CBF87644.1 anig|EHA17759.1 aory|EIT79080.1 asyd|OJJ63294.1 ater|EAU30680.1 atub|OJI90585.1 aver|OJI97664.1 awen|OJJ38620.1 azon|OJJ49561.1

Group3310: abra|OJJ74062.1 acar|OOG01175.1 acla|EAW07907.1 afis|EAW22945.1 afla|EED58150.1 afum|EAL90252.1 agla|OJJ83343.1 akaw|GAA87543.1 aluc|OJZ87974.1 anid|CBF87642.1 anig|EHA17758.1 aory|EIT79097.1 asyd|OJJ63293.1 ater|EAU30679.1 atub|OJI90584.1 aver|OJI97665.1 awen|OJJ38619.1 azon|OJJ49560.1

Group3311: abra|OJJ74063.1 acar|OOG01176.1 acla|EAW07908.1 afis|EAW22946.1 afla|EED58149.1 afum|EAL90251.1 agla|OJJ83342.1 akaw|GAA87544.1 aluc|OJZ87973.1 anid|CBF87640.1 anig|EHA17757.1 aory|EIT79094.1 asyd|OJJ63292.1 ater|EAU30678.1 atub|OJI90583.1 aver|OJI97666.1 awen|OJJ38618.1 azon|OJJ49559.1

Group3312: abra|OJJ74064.1 acar|OOG01174.1 acla|EAW07909.1 afis|EAW22947.1 afla|EED58148.1 afum|EAL90250.1 agla|OJJ83341.1 akaw|GAA87545.1 aluc|OJZ87972.1 anid|CBF87639.1 anig|EHA17756.1 aory|EIT79089.1 asyd|OJJ63291.1 ater|EAU30677.1 atub|OJI90582.1 aver|OJI97667.1 awen|OJJ38617.1 azon|OJJ49558.1

Group3313: abra|OJJ74065.1 acar|OOG01177.1 acla|EAW07910.1 afis|EAW22948.1 afla|EED58147.1 afum|EAL90249.1 agla|OJJ83339.1 akaw|GAA87546.1 aluc|OJZ87971.1 anid|CBF87637.1 anig|EHA17755.1 aory|EIT79088.1 asyd|OJJ63290.1 ater|EAU30676.1 atub|OJI90581.1 aver|OJI97668.1 awen|OJJ38616.1 azon|OJJ49557.1

Group3314: abra|OJJ74066.1 acar|OOG01179.1 acla|EAW07911.1 afis|EAW22949.1 afla|EED58146.1 afum|EAL90248.1 agla|OJJ83338.1 akaw|GAA87547.1 aluc|OJZ87970.1 anid|CBF87636.1 anig|EHA17754.1

aory|EIT79070.1 asyd|OJJ63289.1 ater|EAU30675.1 atub|OJI90580.1 aver|OJI97669.1 awen|OJJ38615.1 azon|OJJ49556.1

Group3315: abra|OJJ74067.1 acar|OOG01178.1 acla|EAW07912.1 afis|EAW22950.1 afla|EED58145.1 afum|EAL90247.1 agla|OJJ83337.1 akaw|GAA87548.1 aluc|OJZ87969.1 anid|CBF87634.1 anig|EHA17753.1 aory|EIT79067.1 asyd|OJJ63288.1 ater|EAU30674.1 atub|OJI90579.1 aver|OJI97670.1 awen|OJJ38614.1 azon|OJJ49555.1

Group3316: abra|OJJ74068.1 acar|OOG01180.1 acla|EAW07913.1 afis|EAW22951.1 afla|EED58144.1 afum|EAL90246.1 agla|OJJ83336.1 akaw|GAA87549.1 aluc|OJZ87968.1 anid|CBF87632.1 anig|EHA17752.1 aory|EIT79092.1 asyd|OJJ63287.1 ater|EAU30673.1 atub|OJI90578.1 aver|OJI97671.1 awen|OJJ38613.1 azon|OJJ49554.1

Group3317: abra|OJJ74069.1 acar|OOG01181.1 acla|EAW07914.1 afis|EAW22952.1 afla|EED58143.1 afum|EAL90245.1 agla|OJJ83335.1 akaw|GAA87550.1 aluc|OJZ87967.1 anid|CBF87630.1 anig|EHA17751.1 aory|EIT79082.1 asyd|OJJ63286.1 ater|EAU30672.1 atub|OJI90577.1 aver|OJI97672.1 awen|OJJ38612.1 azon|OJJ49553.1

Group3318: abra|OJJ74070.1 acar|OOG01182.1 acla|EAW07915.1 afis|EAW22953.1 afla|EED58142.1 afum|EAL90244.1 agla|OJJ83334.1 akaw|GAA87551.1 aluc|OJZ87966.1 anid|CBF87629.1 anig|EHA17750.1 aory|EIT79065.1 asyd|OJJ63285.1 ater|EAU30671.1 atub|OJI90576.1 aver|OJI97673.1 awen|OJJ38611.1 azon|OJJ49552.1

Group3319: abra|OJJ74071.1 acar|OOG01183.1 acla|EAW07916.1 afis|EAW22954.1 afla|EED58141.1 afum|EAL90243.1 agla|OJJ83333.1 akaw|GAA87552.1 aluc|OJZ87965.1 anid|CBF87627.1 anig|EHA17749.1 aory|EIT79103.1 asyd|OJJ63284.1 ater|EAU30670.1 atub|OJI90575.1 aver|OJI97674.1 awen|OJJ38610.1 azon|OJJ49551.1

Group3320: abra|OJJ74072.1 acar|OOG01184.1 acla|EAW07917.1 afis|EAW22955.1 afla|EED58140.1 afum|EAL90242.1 agla|OJJ83332.1 akaw|GAA87553.1 aluc|OJZ87964.1 anid|CBF87625.1 anig|EHA17748.1 aory|EIT79149.1 asyd|OJJ63283.1 ater|EAU30669.1 atub|OJI90574.1 aver|OJI97675.1 awen|OJJ38608.1 azon|OJJ49550.1

Group3321: abra|OJJ74075.1 acar|OOG01187.1 acla|EAW07920.1 afis|EAW22958.1 afla|EED58138.1 afum|EAL90239.1 agla|OJJ83330.1 akaw|GAA87556.1 aluc|OJZ87961.1 anid|CBF87624.1 anig|EHA17745.1 aory|EIT79160.1 asyd|OJJ63281.1 ater|EAU30667.1 atub|OJI90571.1 aver|OJI97677.1 awen|OJJ38605.1 azon|OJJ49547.1

Group3322: abra|OJJ74076.1 acar|OOG01188.1 acla|EAW07921.1 afis|EAW22959.1 afla|EED58137.1 afum|EAL90238.1 agla|OJJ83329.1 akaw|GAA87557.1 aluc|OJZ87960.1 anid|CBF87622.1 anig|EHA17744.1 aory|EIT79140.1 asyd|OJJ63280.1 ater|EAU30666.1 atub|OJI90570.1 aver|OJI97678.1 awen|OJJ38604.1 azon|OJJ49546.1

Group3323: abra|OJJ74078.1 acar|OOG01190.1 acla|EAW07923.1 afis|EAW22961.1 afla|EED58135.1 afum|EAL90236.1 agla|OJJ83327.1 akaw|GAA87559.1 aluc|OJZ87957.1 anid|CBF87621.1 anig|EHA17742.1 aory|EIT79121.1 asyd|OJJ63278.1 ater|EAU30664.1 atub|OJI90568.1 aver|OJI97680.1 awen|OJJ38602.1 azon|OJJ49544.1

Group3324: abra|OJJ74079.1 acar|OOG01191.1 acla|EAW07924.1 afis|EAW22962.1 afla|EED58155.1 afum|EAL90235.1 agla|OJJ83326.1 akaw|GAA87560.1 aluc|OJZ87956.1 anid|CBF87619.1 anig|EHA17741.1 aory|EIT79083.1 asyd|OJJ63277.1 ater|EAU30663.1 atub|OJI90567.1 aver|OJI97681.1 awen|OJJ38601.1 azon|OJJ49543.1

Group3325: abra|OJJ74080.1 acar|OOG01192.1 acla|EAW07925.1 afis|EAW22963.1 afla|EED58156.1 afum|EAL90234.1 agla|OJJ83324.1 akaw|GAA87561.1 aluc|OJZ87955.1 anid|CBF87617.1 anig|EHA17740.1 aory|EIT79101.1 asyd|OJJ63276.1 ater|EAU30662.1 atub|OJI90566.1 aver|OJI97682.1 awen|OJJ38600.1 azon|OJJ49542.1

Group3326: abra|OJJ74081.1 acar|OOG01193.1 acla|EAW07926.1 afis|EAW22964.1 afla|EED58157.1 afum|EAL90233.1 agla|OJJ83323.1 akaw|GAA87562.1 aluc|OJZ87954.1 anid|CBF87615.1 anig|EHA17739.1 aory|EIT79095.1 asyd|OJJ63275.1 ater|EAU30661.1 atub|OJI90565.1 aver|OJI97683.1 awen|OJJ38599.1 azon|OJJ49541.1

Group3327: abra|OJJ74082.1 acar|OOG01194.1 acla|EAW07927.1 afis|EAW22965.1 afla|EED58158.1 afum|EAL90232.1 agla|OJJ83322.1 akaw|GAA87563.1 aluc|OJZ87953.1 anid|CBF87613.1 anig|EHA17738.1 aory|EIT79104.1 asyd|OJJ63274.1 ater|EAU30653.1 atub|OJI90564.1 aver|OJI97684.1 awen|OJJ38598.1 azon|OJJ49540.1

Group3328: abra|OJJ74084.1 acar|OOG01196.1 acla|EAW07928.1 afis|EAW22966.1 afla|EED58160.1 afum|EAL90231.1 agla|OJJ83321.1 akaw|GAA87564.1 aluc|OJZ87951.1 anid|CBF87612.1 anig|EHA17737.1 aory|EIT79091.1 asyd|OJJ63272.1 ater|EAU30654.1 atub|OJI90563.1 aver|OJI97685.1 awen|OJJ38597.1 azon|OJJ49538.1

Group3329: abra|OJJ74086.1 acar|OOG01197.1 acla|EAW07930.1 afis|EAW22968.1 afla|EED58161.1 afum|EAL90229.1 agla|OJJ83319.1 akaw|GAA87566.1 aluc|OJZ87949.1 anid|CBF87609.1 anig|EHA17735.1 aory|EIT79100.1 asyd|OJJ63270.1 ater|EAU30659.1 atub|OJI90560.1 aver|OJI97687.1 awen|OJJ38595.1 azon|OJJ49537.1

Group3330: abra|OJJ74089.1 acar|OOG01198.1 acla|EAW07932.1 afis|EAW22970.1 afla|EED58163.1 afum|EAL90227.2 agla|OJJ83317.1 akaw|GAA87568.1 aluc|OJZ87947.1 anid|CBF74548.1 anig|EHA17733.1 aory|EIT79078.1 asyd|OJJ57686.1 ater|EAU30652.1 atub|OJI90558.1 aver|OJJ05492.1 awen|OJJ38593.1 azon|OJJ49536.1

Group3331: abra|OJJ74090.1 acar|OOG01199.1 acla|EAW07933.1 afis|EAW22971.1 afla|EED58164.1 afum|EAL90226.1 agla|OJJ83316.1 akaw|GAA87569.1 aluc|OJZ87946.1 anid|CBF87607.1 anig|EHA17732.1 aory|EIT79076.1 asyd|OJJ63269.1 ater|EAU30651.1 atub|OJI90557.1 aver|OJI97688.1 awen|OJJ38591.1 azon|OJJ49535.1

Group3332: abra|OJJ74091.1 acar|OOG01200.1 acla|EAW07934.1 afis|EAW22972.1 afla|EED58166.1 afum|EAL90225.1 agla|OJJ83315.1 akaw|GAA87570.1 aluc|OJZ87945.1 anid|CBF87606.1 anig|EHA17731.1 aory|EIT79075.1 asyd|OJJ63268.1 ater|EAU30650.1 atub|OJI90556.1 aver|OJI97689.1 awen|OJJ38590.1 azon|OJJ49533.1

Group3333: abra|OJJ74093.1 acar|OOG01202.1 acla|EAW07936.1 afis|EAW22974.1 afla|EED58168.1 afum|EAL90223.1 agla|OJJ83313.1 akaw|GAA87572.1 aluc|OJZ87943.1 anid|CBF87602.1 anig|EHA17730.1 aory|EIT79071.1 asyd|OJJ63266.1 ater|EAU30647.1 atub|OJI90554.1 aver|OJI97691.1 awen|OJJ38589.1 azon|OJJ49532.1

Group3334: abra|OJJ74094.1 acar|OOG01204.1 acla|EAW07937.1 afis|EAW22975.1 afla|EED58169.1 afum|EAL90222.1 agla|OJJ83312.1 akaw|GAA87573.1 aluc|OJZ87942.1 anid|CBF87600.1 anig|EHA17729.1 aory|EIT79096.1 asyd|OJJ63265.1 ater|EAU30646.1 atub|OJI90553.1 aver|OJI97692.1 awen|OJJ38588.1 azon|OJJ49531.1

Group3335: abra|OJJ74095.1 acar|OOG01203.1 acla|EAW07938.1 afis|EAW22976.1 afla|EED58170.1 afum|EAL90221.1 agla|OJJ83311.1 akaw|GAA87574.1 aluc|OJZ87941.1 anid|CBF87598.1 anig|EHA17728.1 aory|EIT79085.1 asyd|OJJ63264.1 ater|EAU30645.1 atub|OJI90552.1 aver|OJI97693.1 awen|OJJ38587.1 azon|OJJ49530.1

Group3336: abra|OJJ74096.1 acar|OOG01205.1 acla|EAW07940.1 afis|EAW22978.1 afla|EED58171.1 afum|EAL90219.1 agla|OJJ83310.1 akaw|GAA87575.1 aluc|OJZ87940.1 anid|CBF87596.1 anig|EHA17727.1 aory|EIT79081.1 asyd|OJJ63262.1 ater|EAU30644.1 atub|OJI90551.1 aver|OJI97695.1 awen|OJJ38586.1 azon|OJJ49528.1

Group3337: abra|OJJ74100.1 acar|OOG01209.1 acla|EAW07943.1 afis|EAW22981.1 afla|EED58174.1 afum|EAL90216.1 agla|OJJ83306.1 akaw|GAA87578.1 aluc|OJZ87936.1 anid|CBF87589.1 anig|EHA17724.1 aory|EIT79069.1 asyd|OJJ63258.1 ater|EAU30640.1 atub|OJI90548.1 aver|OJI97699.1 awen|OJJ38582.1 azon|OJJ49524.1

Group3338: abra|OJJ74102.1 acar|OOG01211.1 acla|EAW07945.1 afis|EAW22983.1 afla|EED58176.1 afum|EAL90215.1 agla|OJJ83304.1 akaw|GAA87580.1 aluc|OJZ87934.1 anid|CBF87597.1 anig|EHA17723.1 aory|EIT79086.1 asyd|OJJ63263.1 ater|EAU30638.1 atub|OJI90546.1 aver|OJI97694.1 awen|OJJ38580.1 azon|OJJ49522.1

Group3339: abra|OJJ74103.1 acar|OOG01213.1 acla|EAW07946.1 afis|EAW22985.1 afla|EED58177.1 afum|EAL90214.1 agla|OJJ83302.1 akaw|GAA87581.1 aluc|OJZ87933.1 anid|CBF87584.1 anig|EHA17721.1 aory|EIT79072.1 asyd|OJJ63255.1 ater|EAU30637.1 atub|OJI90545.1 aver|OJI97702.1 awen|OJJ38579.1 azon|OJJ49520.1

Group3340: abra|OJJ74109.1 acar|OOG01217.1 acla|EAW07953.1 afis|EAW22990.1 afla|EED58182.1 afum|EAL90210.1 agla|OJJ83296.1 akaw|GAA87585.1 aluc|OJZ87925.1 anid|CBF87579.1 anig|EHA17717.1 aory|EIT79105.1 asyd|OJJ63251.1 ater|EAU30632.1 atub|OJI90537.1 aver|OJI97707.1 awen|OJJ38574.1 azon|OJJ49515.1

Group3341: abra|OJJ74116.1 acar|OOG01220.1 acla|EAW07958.1 afis|EAW22993.1 afla|EED58186.1 afum|EAL90208.1 agla|OJJ83291.1 akaw|GAA87589.1 aluc|OJZ87918.1 anid|CBF87570.1 anig|EHA17712.1 aory|EIT79074.1 asyd|OJJ63244.1 ater|EAU30627.1 atub|OJI90529.1 aver|OJI97712.1 awen|OJJ38568.1 azon|OJJ44834.1

Group3342: abra|OJJ74118.1 acar|OOG01223.1 acla|EAW07960.1 afis|EAW22995.1 afla|EED58188.1 afum|EAL90204.1 agla|OJJ83289.1 akaw|GAA87591.1 aluc|OJZ87914.1 anid|CBF87567.1 anig|EHA17708.1 aory|EIT79079.1 asyd|OJJ63242.1 ater|EAU30623.1 atub|OJI90526.1 aver|OJI97714.1 awen|OJJ38566.1 azon|OJJ50435.1

Group3345: abra|OJJ74188.1 acar|OOF95533.1 acla|EAW07238.1 afis|EAW22261.1 afla|EED57649.1 afum|EAL90919.1 agla|OJJ87796.1 akaw|GAA86829.1 aluc|OJZ89947.1 anid|CBF88508.1 anig|EHA27148.1

aory|EIT79898.1 asyd|OJJ63801.1 ater|EAU34268.1 atub|OJI84859.1 aver|OJI97138.1 awen|OJJ34833.1 azon|OJJ45713.1

Group3346: abra|OJJ74189.1 acar|OOF95532.1 acla|EAW07241.1 afis|EAW22262.1 afla|EED57651.1 afum|EAL90918.1 agla|OJJ87794.1 akaw|GAA86828.1 aluc|OJZ89946.1 anid|CBF88511.1 anig|EHA27145.1 aory|EIT79914.1 asyd|OJJ63803.1 ater|EAU34269.1 atub|OJI84860.1 aver|OJI97135.1 awen|OJJ34835.1 azon|OJJ45712.1

Group3347: abra|OJJ74190.1 acar|OOF95531.1 acla|EAW07242.1 afis|EAW22263.1 afla|EED57655.1 afum|EAL90917.1 agla|OJJ87792.1 akaw|GAA86827.1 aluc|OJZ89945.1 anid|CBF88513.1 anig|EHA27144.1 aory|EIT79925.1 asyd|OJJ63804.1 ater|EAU34273.1 atub|OJI84861.1 aver|OJI97134.1 awen|OJJ34836.1 azon|OJJ45711.1

Group3348: abra|OJJ74193.1 acar|OOF95529.1 acla|EAW07245.1 afis|EAW22266.1 afla|EED57658.1 afum|EAL90914.1 agla|OJJ87789.1 akaw|GAA86824.1 aluc|OJZ89942.1 anid|CBF88519.1 anig|EHA27141.1 aory|EIT79939.1 asyd|OJJ63807.1 ater|EAU34277.1 atub|OJI84864.1 aver|OJI97131.1 awen|OJJ34838.1 azon|OJJ45708.1

Group3349: abra|OJJ74194.1 acar|OOF95527.1 acla|EAW07247.1 afis|EAW22268.1 afla|EED57660.1 afum|EAL90912.1 agla|OJJ87787.1 akaw|GAA86823.1 aluc|OJZ89941.1 anid|CBF88523.1 anig|EHA27140.1 aory|EIT79938.1 asyd|OJJ63808.1 ater|EAU34279.1 atub|OJI84865.1 aver|OJI97129.1 awen|OJJ34841.1 azon|OJJ45706.1

Group3350: abra|OJJ74195.1 acar|OOF95526.1 acla|EAW07248.1 afis|EAW22269.1 afla|EED57661.1 afum|EAL90911.1 agla|OJJ87786.1 akaw|GAA86822.1 aluc|OJZ89939.1 anid|CBF88527.1 anig|EHA27139.1 aory|EIT79909.1 asyd|OJJ63810.1 ater|EAU34280.1 atub|OJI84866.1 aver|OJI97127.1 awen|OJJ34842.1 azon|OJJ45705.1

Group3351: abra|OJJ74196.1 acar|OOF95524.1 acla|EAW07249.1 afis|EAW22270.1 afla|EED57662.1 afum|EAL90910.1 agla|OJJ87785.1 akaw|GAA86821.1 aluc|OJZ89938.1 anid|CBF88528.1 anig|EHA27138.1 aory|EIT79867.1 asyd|OJJ63811.1 ater|EAU34281.1 atub|OJI84868.1 aver|OJI97126.1 awen|OJJ34843.1 azon|OJJ45704.1

Group3352: abra|OJJ74197.1 acar|OOF95523.1 acla|EAW07250.1 afis|EAW22271.1 afla|EED57663.1 afum|EAL90909.1 agla|OJJ87784.1 akaw|GAA86820.1 aluc|OJZ89937.1 anid|CBF88530.1 anig|EHA27137.1 aory|EIT79903.1 asyd|OJJ63812.1 ater|EAU34282.1 atub|OJI84869.1 aver|OJI97125.1 awen|OJJ34844.1 azon|OJJ45703.1

Group3353: abra|OJJ74198.1 acar|OOF95521.1 acla|EAW07252.1 afis|EAW22273.1 afla|EED57665.1 afum|EAL90908.1 agla|OJJ87781.1 akaw|GAA86819.1 aluc|OJZ89936.1 anid|CBF88536.1 anig|EHA27136.1 aory|EIT79943.1 asyd|OJJ63820.1 ater|EAU34283.1 atub|OJI84870.1 aver|OJI97118.1 awen|OJJ34846.1 azon|OJJ45702.1

Group3354: abra|OJJ74199.1 acar|OOF95520.1 acla|EAW07253.1 afis|EAW22274.1 afla|EED57666.1 afum|EAL90907.1 agla|OJJ87780.1 akaw|GAA86818.1 aluc|OJZ89935.1 anid|CBF88537.1 anig|EHA27135.1 aory|EIT79937.1 asyd|OJJ63821.1 ater|EAU34284.1 atub|OJI84871.1 aver|OJI97117.1 awen|OJJ34847.1 azon|OJJ45700.1

Group3355: abra|OJJ74201.1 acar|OOF95519.1 acla|EAW07254.1 afis|EAW22275.1 afla|EED57667.1 afum|EAL90906.1 agla|OJJ87779.1 akaw|GAA86817.1 aluc|OJZ89934.1 anid|CBF88539.1 anig|EHA27134.1 aory|EIT79875.1 asyd|OJJ63822.1 ater|EAU34285.1 atub|OJI84872.1 aver|OJI97116.1 awen|OJJ34848.1 azon|OJJ45699.1

Group3356: abra|OJJ74202.1 acar|OOF95518.1 afla|EED57668.1 afum|EAL90905.1 agla|OJJ87777.1 akaw|GAA86816.1 aluc|OJZ89933.1 anid|CBF88541.1 anig|EHA27133.1 aory|EIT79884.1 ater|EAU34286.1 atub|OJI84873.1 aver|OJI97115.1 azon|OJJ45698.1 acla|EAW07255.1 afis|EAW22276.1 asyd|OJJ63823.1 awen|OJJ34849.1

Group3357: abra|OJJ74203.1 acar|OOF95517.1 acla|EAW07256.1 afis|EAW22277.1 afla|EED57669.1 afum|EAL90904.1 agla|OJJ87776.1 akaw|GAA86815.1 aluc|OJZ89932.1 anid|CBF88543.1 anig|EHA27132.1 aory|EIT79926.1 asyd|OJJ63824.1 ater|EAU34287.1 atub|OJI84874.1 aver|OJI97114.1 awen|OJJ34850.1 azon|OJJ45697.1

Group3358: abra|OJJ74204.1 acar|OOF95516.1 acla|EAW07257.1 afis|EAW22278.1 afla|EED57671.1 afum|EAL90903.1 agla|OJJ87775.1 akaw|GAA86814.1 aluc|OJZ89931.1 anid|CBF88545.1 anig|EHA27131.1 aory|EIT79927.1 asyd|OJJ63825.1 ater|EAU34288.1 atub|OJI84875.1 aver|OJI97113.1 awen|OJJ34851.1 azon|OJJ45696.1

Group3359: abra|OJJ74206.1 acar|OOF95514.1 acla|EAW07259.1 afis|EAW22280.1 afla|EED57677.1 afum|EAL90901.1 agla|OJJ87773.1 akaw|GAA86812.1 aluc|OJZ89929.1 anid|CBF88548.1 anig|EHA27129.1 aory|EIT79874.1 asyd|OJJ63827.1 ater|EAU34290.1 atub|OJI84877.1 aver|OJI97111.1 awen|OJJ34853.1 azon|OJJ45677.1

Group3360: abra|OJJ74207.1 acar|OOF95513.1 acla|EAW07260.1 afis|EAW22281.1 afla|EED57678.1 afum|EAL90900.1 agla|OJJ87768.1 akaw|GAA86811.1 aluc|OJZ89928.1 anid|CBF88550.1 anig|EHA27128.1 aory|EIT79890.1 asyd|OJJ63828.1 ater|EAU34291.1 atub|OJI84878.1 aver|OJI97110.1 awen|OJJ34854.1 azon|OJJ45676.1

Group3361: abra|OJJ74208.1 acar|OOF95512.1 acla|EAW07261.1 afis|EAW22282.1 afla|EED57679.1 afum|EAL90899.1 agla|OJJ87767.1 akaw|GAA86810.1 aluc|OJZ89926.1 anid|CBF88552.1 anig|EHA27127.1 aory|EIT79907.1 asyd|OJJ63829.1 ater|EAU34292.1 atub|OJI84879.1 aver|OJI97109.1 awen|OJJ34855.1 azon|OJJ45674.1

Group3362: abra|OJJ74209.1 acar|OOF95511.1 acla|EAW07262.1 afis|EAW22283.1 afla|EED57680.1 afum|EAL90898.1 agla|OJJ87766.1 akaw|GAA86809.1 aluc|OJZ89925.1 anid|CBF88554.1 anig|EHA27126.1 aory|EIT79921.1 asyd|OJJ63830.1 ater|EAU34293.1 atub|OJI84880.1 aver|OJI97108.1 awen|OJJ34856.1 azon|OJJ45673.1

Group3363: abra|OJJ74210.1 acar|OOF95510.1 acla|EAW07263.1 afis|EAW22284.1 afla|EED57682.1 afum|EAL90897.1 agla|OJJ87764.1 akaw|GAA86808.1 aluc|OJZ89924.1 anid|CBF88556.1 anig|EHA27124.1 aory|EIT79901.1 asyd|OJJ63831.1 ater|EAU34296.1 atub|OJI84881.1 aver|OJI97107.1 awen|OJJ34857.1 azon|OJJ45671.1

Group3364: abra|OJJ74211.1 acar|OOF95509.1 acla|EAW07264.1 afis|EAW22285.1 afla|EED57685.1 afum|EAL90896.1 agla|OJJ87763.1 akaw|GAA86807.1 aluc|OJZ89923.1 anid|CBF88558.1 anig|EHA27123.1 aory|EIT79887.1 asyd|OJJ63832.1 ater|EAU34297.1 atub|OJI84882.1 aver|OJI97106.1 awen|OJJ34858.1 azon|OJJ45670.1

Group3365: abra|OJJ74218.1 acar|OOF95503.1 acla|EAW07273.1 afis|EAW22294.1 afla|EED57690.1 afum|EAL90887.1 agla|OJJ87758.1 akaw|GAA86800.1 aluc|OJZ89916.1 anid|CBF88569.1 anig|EHA27116.1 aory|EIT79904.1 asyd|OJJ63837.1 ater|EAU34303.1 atub|OJI84889.1 aver|OJI97100.1 awen|OJJ34864.1 azon|OJJ45665.1

Group3366: abra|OJJ74220.1 acar|OOF90433.1 acla|EAW07284.1 afis|EAW22315.1 afla|EED57695.1 afum|EAL90875.1 agla|OJJ87735.1 akaw|GAA86798.1 aluc|OJZ89914.1 anid|CBF88577.1 anig|EHA27114.1 aory|EIT79936.1 asyd|OJJ63841.1 ater|EAU34308.1 atub|OJI84891.1 aver|OJI97096.1 awen|OJJ34868.1 azon|OJJ50045.1

Group3367: abra|OJJ74223.1 acar|OOF90434.1 acla|EAW07282.1 afis|EAW22313.1 afla|EED57697.1 afum|EAL90877.1 agla|OJJ87737.1 akaw|GAA86796.1 aluc|OJZ89911.1 anid|CBF88580.1 anig|EHA27111.1 aory|EIT79934.1 asyd|OJJ63843.1 ater|EAU34310.1 atub|OJI84895.1 aver|OJI97094.1 awen|OJJ34872.1 azon|OJJ50047.1

Group3368: abra|OJJ74224.1 acar|OOF90435.1 acla|EAW07281.1 afis|EAW22312.1 afla|EED57698.1 afum|EAL90878.1 agla|OJJ87738.1 akaw|GAA86795.1 aluc|OJZ89910.1 anid|CBF88582.1 anig|EHA27110.1 aory|EIT79916.1 asyd|OJJ63844.1 ater|EAU34311.1 atub|OJI84896.1 aver|OJI97093.1 awen|OJJ34873.1 azon|OJJ50048.1

Group3369: abra|OJJ74225.1 acar|OOF95502.1 acla|EAW07280.1 afis|EAW22311.1 afla|EED57701.1 afum|EAL90879.1 agla|OJJ87739.1 akaw|GAA86794.1 aluc|OJZ89908.1 anid|CBF88584.1 anig|EHA27108.1 aory|EIT79931.1 asyd|OJJ63846.1 ater|EAU34313.1 atub|OJI84898.1 aver|OJI97090.1 awen|OJJ34876.1 azon|OJJ50049.1

Group3370: abra|OJJ74228.1 acar|OOF95498.1 acla|EAW07276.1 afis|EAW22297.1 afla|EED57705.1 afum|EAL90884.1 agla|OJJ87748.1 akaw|GAA86791.1 aluc|OJZ89905.1 anid|CBF88592.1 anig|EHA27105.1 aory|EIT79881.1 asyd|OJJ63850.1 ater|EAU34317.1 atub|OJI84901.1 aver|OJI97087.1 awen|OJJ34881.1 azon|OJJ50050.1

Group3371: abra|OJJ74231.1 acar|OOF95495.1 acla|EAW07286.1 afis|EAW22319.1 afla|EED57710.1 afum|EAL90871.1 agla|OJJ87745.1 akaw|GAA86788.1 aluc|OJZ89902.1 anid|CBF88599.1 anig|EHA27102.1 aory|EIT79923.1 asyd|OJJ63854.1 ater|EAU34320.1 atub|OJI84905.1 aver|OJI97084.1 awen|OJJ34885.1 azon|OJJ50053.1

Group3372: abra|OJJ74237.1 acar|OOF95492.1 acla|EAW07288.1 afis|EAW22321.1 afla|EED57718.1 afum|EAL90870.1 agla|OJJ87749.1 akaw|GAA86783.1 aluc|OJZ89895.1 anid|CBF88602.1 anig|EHA27094.1 aory|EIT79929.1 asyd|OJJ63855.1 ater|EAU34321.1 atub|OJI84912.1 aver|OJI97083.1 awen|OJJ34888.1 azon|OJJ50055.1

Group3373: abra|OJJ74242.1 acar|OOF95488.1 acla|EAW07291.1 afis|EAW22325.1 afla|EED57723.1 afum|EAL90867.1 agla|OJJ87721.1 akaw|GAA86778.1 aluc|OJZ89890.1 anid|CBF88609.1 anig|EHA27089.1 aory|EIT79878.1 asyd|OJJ63859.1 ater|EAU34324.1 atub|OJI84917.1 aver|OJI97080.1 awen|OJJ34896.1 azon|OJJ50065.1

Group3374: abra|OJJ74246.1 acar|OOF95484.1 acla|EAW07298.1 afis|EAW22347.1 afla|EED57735.1 afum|EAL90845.1 agla|OJJ87687.1 akaw|GAA86774.1 aluc|OJZ89886.1 anid|CBF88623.1 anig|EHA27085.1

aory|EIT79913.1 asyd|OJJ63868.1 ater|EAU39483.1 atub|OJI84921.1 aver|OJI97071.1 awen|OJJ34928.1 azon|OJJ50070.1

Group3375: abra|OJJ74249.1 acar|OOF95479.1 acla|EAW07293.1 afis|EAW22327.1 afla|EED57726.1 afum|EAL90865.1 agla|OJJ87723.1 akaw|GAA86772.1 aluc|OJZ89883.1 anid|CBF88613.1 anig|EHA27082.1 aory|EIT79866.1 asyd|OJJ63861.1 ater|EAU34326.1 atub|OJI84924.1 aver|OJI97078.1 awen|OJJ34894.1 azon|OJJ50063.1

Group3376: abra|OJJ74250.1 acar|OOF95480.1 acla|EAW07294.1 afis|EAW22328.1 afla|EED57727.1 afum|EAL90864.1 agla|OJJ87724.1 akaw|GAA86771.1 aluc|OJZ89882.1 anid|CBF88615.1 anig|EHA27081.1 aory|EIT79906.1 asyd|OJJ63862.1 ater|EAU34327.1 atub|OJI84925.1 aver|OJI97077.1 awen|OJJ34893.1 azon|OJJ50062.1

Group3377: abra|OJJ74252.1 acar|OOF95477.1 acla|EAW07296.1 afis|EAW22330.1 afla|EED57729.1 afum|EAL90862.1 agla|OJJ87752.1 akaw|GAA86769.1 aluc|OJZ89880.1 anid|CBF88619.1 anig|EHA27079.1 aory|EIT79894.1 asyd|OJJ63864.1 ater|EAU34331.1 atub|OJI84927.1 aver|OJI97075.1 awen|OJJ34891.1 azon|OJJ50060.1

Group3378: abra|OJJ74256.1 acar|OOF95472.1 acla|EAW07302.1 afis|EAW22343.1 afla|EED57738.1 afum|EAL90848.1 agla|OJJ87691.1 akaw|GAA86765.1 aluc|OJZ89876.1 anid|CBF88627.1 anig|EHA27075.1 aory|EIT79915.1 asyd|OJJ63870.1 ater|EAU39481.1 atub|OJI84931.1 aver|OJI97069.1 awen|OJJ34922.1 azon|OJJ50072.1

Group3379: abra|OJJ74257.1 acar|OOF95471.1 acla|EAW07303.1 afis|EAW22342.1 afla|EED57739.1 afum|EAL90849.1 agla|OJJ87692.1 akaw|GAA86764.1 aluc|OJZ89875.1 anid|CBF88628.1 anig|EHA27074.1 aory|EIT79892.1 asyd|OJJ63871.1 ater|EAU39480.1 atub|OJI84932.1 aver|OJI97068.1 awen|OJJ34921.1 azon|OJJ50073.1

Group3380: abra|OJJ74260.1 acar|OOF89900.1 acla|EAW07305.1 afis|EAW22340.1 afla|EED57741.1 afum|EAL90852.1 agla|OJJ87698.1 akaw|GAA86761.1 aluc|OJZ89871.1 anid|CBF88632.1 anig|EHA27071.1 aory|EIT79893.1 asyd|OJJ63874.1 ater|EAU39478.1 atub|OJI84936.1 aver|OJI97064.1 awen|OJJ34919.1 azon|OJJ50075.1

Group3381: abra|OJJ74261.1 acar|OOF89901.1 acla|EAW07306.1 afis|EAW22339.1 afla|EED57742.1 afum|EAL90853.1 agla|OJJ87699.1 akaw|GAA86760.1 aluc|OJZ89870.1 anid|CBF88634.1 anig|EHA27070.1 aory|EIT79922.1 asyd|OJJ63875.1 ater|EAU39477.1 atub|OJI84937.1 aver|OJI97063.1 awen|OJJ34918.1 azon|OJJ50076.1

Group3382: abra|OJJ74264.1 acar|OOF95466.1 acla|EAW07308.1 afis|EAW22338.1 afla|EED57743.1 afum|EAL90854.1 agla|OJJ87700.1 akaw|GAA86759.1 aluc|OJZ89869.1 anid|CBF88636.1 anig|EHA27069.1 aory|EIT81083.1 asyd|OJJ63877.1 ater|EAU39475.1 atub|OJI84939.1 aver|OJI97062.1 awen|OJJ34916.1 azon|OJJ50079.1

Group3383: abra|OJJ74265.1 acar|OOF95465.1 acla|EAW07309.1 afis|EAW22337.1 afla|EED57744.1 afum|EAL90855.1 agla|OJJ87701.1 akaw|GAA86758.1 aluc|OJZ89868.1 anid|CBF88638.1 anig|EHA27068.1 aory|EIT81047.1 asyd|OJJ63878.1 ater|EAU39474.1 atub|OJI84940.1 aver|OJI97061.1 awen|OJJ34915.1 azon|OJJ45569.1

Group3384: abra|OJJ74266.1 acar|OOF95464.1 acla|EAW07310.1 afis|EAW22336.1 afla|EED57746.1 afum|EAL90856.1 agla|OJJ87702.1 akaw|GAA86757.1 aluc|OJZ89867.1 anid|CBF88640.1 anig|EHA27067.1 aory|EIT81092.1 asyd|OJJ63879.1 ater|EAU39473.1 atub|OJI84941.1 aver|OJI97060.1 awen|OJJ34914.1 azon|OJJ45570.1

Group3385: abra|OJJ74267.1 acar|OOF89814.1 acla|EAW07311.1 afis|EAW22335.1 afla|EED57747.1 afum|EAL90857.2 agla|OJJ87715.1 akaw|GAA86756.1 aluc|OJZ89866.1 anid|CBF88641.1 anig|EHA27066.1 aory|EIT81022.1 asyd|OJJ63880.1 ater|EAU39472.1 atub|OJI84942.1 aver|OJI97059.1 awen|OJJ34913.1 azon|OJJ45571.1

Group3386: abra|OJJ74270.1 acar|OOF95460.1 acla|EAW07314.1 afis|EAW22332.1 afla|EED57751.1 afum|EAL90860.1 agla|OJJ87712.1 akaw|GAA86753.1 aluc|OJZ89863.1 anid|CBF88647.1 anig|EHA27063.1 aory|EIT81116.1 asyd|OJJ63882.1 ater|EAU39461.1 atub|OJI84945.1 aver|OJI97057.1 awen|OJJ34910.1 azon|OJJ45575.1

Group3387: abra|OJJ74272.1 acar|OOF95458.1 acla|EAW07317.1 afis|EAW22349.1 afla|EED57753.1 afum|EAL90843.1 agla|OJJ87710.1 akaw|GAA86751.1 aluc|OJZ89861.1 anid|CBF88651.1 anig|EHA27061.1 aory|EIT81051.1 asyd|OJJ63885.1 ater|EAU39459.1 atub|OJI84947.1 aver|OJI97055.1 awen|OJJ34905.1 azon|OJJ45577.1

Group3388: abra|OJJ74276.1 acar|OOF95455.1 acla|EAW07320.1 afis|EAW22352.1 afla|EED57756.1 afum|EAL90840.1 agla|OJJ87707.1 akaw|GAA86748.1 aluc|OJZ89857.1 anid|CBF88661.1 anig|EHA27058.1 aory|EIT81039.1 asyd|OJJ63889.1 ater|EAU39488.1 atub|OJI84949.1 aver|OJI97050.1 awen|OJJ34903.1 azon|OJJ45579.1

Group3389: abra|OJJ74277.1 acar|OOF95454.1 acla|EAW07321.1 afis|EAW22353.1 afla|EED57759.1 afum|EAL90839.1 agla|OJJ87716.1 akaw|GAA86747.1 aluc|OJZ89856.1 anid|CBF88662.1 anig|EHA27057.1 aory|EIT81042.1 asyd|OJJ63890.1 ater|EAU39487.1 atub|OJI84950.1 aver|OJI97049.1 awen|OJJ34902.1 azon|OJJ45580.1

Group3390: abra|OJJ74278.1 acar|OOF95453.1 acla|EAW07322.1 afis|EAW22354.1 afla|EED57760.1 afum|EAL90838.1 agla|OJJ87717.1 akaw|GAA86746.1 aluc|OJZ89855.1 anid|CBF88664.1 anig|EHA27056.1 aory|EIT81028.1 asyd|OJJ63891.1 ater|EAU39486.1 atub|OJI84951.1 aver|OJI97048.1 awen|OJJ34901.1 azon|OJJ45581.1

Group3391: abra|OJJ74280.1 acar|OOF95451.1 acla|EAW07324.1 afis|EAW22356.1 afla|EED57762.1 afum|EAL90835.1 agla|OJJ87598.1 akaw|GAA86744.1 aluc|OJZ89853.1 anid|CBF88668.1 anig|EHA27054.1 aory|EIT81074.1 asyd|OJJ63893.1 ater|EAU39490.1 atub|OJI84953.1 aver|OJI97046.1 awen|OJJ34930.1 azon|OJJ45583.1

Group3392: abra|OJJ74281.1 acar|OOF95450.1 acla|EAW07325.1 afis|EAW22357.1 afla|EED57763.1 afum|EAL90834.1 agla|OJJ87682.1 akaw|GAA86743.1 aluc|OJZ89852.1 anid|CBF88672.1 anig|EHA27053.1 aory|EIT81043.1 asyd|OJJ63895.1 ater|EAU39491.1 atub|OJI84954.1 aver|OJI97044.1 awen|OJJ34931.1 azon|OJJ45584.1

Group3393: abra|OJJ74282.1 acar|OOF95449.1 acla|EAW07326.1 afis|EAW22358.1 afla|EED57764.1 afum|EAL90833.1 agla|OJJ87683.1 akaw|GAA86742.1 aluc|OJZ89851.1 anid|CBF88674.1 anig|EHA27052.1 aory|EIT81053.1 asyd|OJJ63896.1 ater|EAU39492.1 atub|OJI84955.1 aver|OJI97043.1 awen|OJJ34932.1 azon|OJJ45585.1

Group3394: abra|OJJ74283.1 acar|OOF95447.1 acla|EAW07327.1 afis|EAW22359.1 afla|EED57765.1 afum|EAL90832.1 agla|OJJ87684.1 akaw|GAA86741.1 aluc|OJZ89850.1 anid|CBF88676.1 anig|EHA27051.1 aory|EIT81093.1 asyd|OJJ63897.1 ater|EAU39493.1 atub|OJI84956.1 aver|OJI97042.1 awen|OJJ34933.1 azon|OJJ45586.1

Group3395: abra|OJJ74290.1 acar|OOF95441.1 acla|EAW07334.1 afis|EAW22366.1 afla|EED57774.1 afum|EBA27415.1 agla|OJJ87624.1 akaw|GAA86734.1 aluc|OJZ89844.1 anid|CBF88692.1 anig|EHA27045.1 aory|EIT81109.1 asyd|OJJ63910.1 ater|EAU39501.1 atub|OJI84964.1 aver|OJI97031.1 awen|OJJ34940.1 azon|OJJ45592.1

Group3396: abra|OJJ74291.1 acar|OOF95440.1 acla|EAW07335.1 afis|EAW22367.1 afla|EED57775.1 afum|EAL90825.1 agla|OJJ87622.1 akaw|GAA86733.1 aluc|OJZ89843.1 anid|CBF88694.1 anig|EHA27044.1 aory|EIT81063.1 asyd|OJJ63911.1 ater|EAU39502.1 atub|OJI84965.1 aver|OJI97030.1 awen|OJJ34941.1 azon|OJJ45593.1

Group3397: abra|OJJ74292.1 acar|OOF95439.1 acla|EAW07336.1 afis|EAW22368.1 afla|EED57776.1 afum|EAL90824.1 akaw|GAA86732.1 aluc|OJZ89842.1 anid|CBF88695.1 anig|EHA27043.1 aory|EIT81045.1 asyd|OJJ63913.1 ater|EAU39503.1 atub|OJI84966.1 aver|OJI97029.1 awen|OJJ34942.1 azon|OJJ45594.1 agla|OJJ87621.1

Group3398: abra|OJJ74293.1 acar|OOF95438.1 acla|EAW07337.1 afis|EAW22369.1 afla|EED57777.1 afum|EAL90823.1 agla|OJJ87620.1 akaw|GAA86731.1 aluc|OJZ89841.1 anid|CBF88697.1 anig|EHA27042.1 aory|EIT81034.1 asyd|OJJ63914.1 ater|EAU39504.1 atub|OJI84967.1 aver|OJI97028.1 awen|OJJ34944.1 azon|OJJ45595.1

Group3399: abra|OJJ74294.1 acar|OOF95437.1 acla|EAW07339.1 afis|EAW22370.1 afla|EED57778.1 afum|EAL90822.2 agla|OJJ87619.1 akaw|GAA86730.1 aluc|OJZ89840.1 anid|CBF88699.1 anig|EHA27041.1 aory|EIT81037.1 asyd|OJJ63915.1 ater|EAU39505.1 atub|OJI84968.1 aver|OJI97027.1 awen|OJJ34945.1 azon|OJJ45596.1

Group3400: abra|OJJ74295.1 acar|OOF95436.1 acla|EAW07340.1 afis|EAW22372.1 afla|EED57779.1 afum|EAL90820.2 agla|OJJ87617.1 akaw|GAA86729.1 aluc|OJZ89839.1 anid|CBF88703.1 anig|EHA27040.1 aory|EIT81082.1 asyd|OJJ63919.1 ater|EAU39506.1 atub|OJI84969.1 aver|OJI97024.1 awen|OJJ34947.1 azon|OJJ45597.1

Group3401: abra|OJJ74296.1 acar|OOF95435.1 acla|EAW07341.1 afis|EAW22373.1 afla|EED57780.1 afum|EAL90819.1 agla|OJJ87626.1 akaw|GAA86728.1 aluc|OJZ89838.1 anid|CBF88705.1 anig|EHA27039.1 aory|EIT81056.1 asyd|OJJ63920.1 ater|EAU39507.1 atub|OJI84970.1 aver|OJI97023.1 awen|OJJ34948.1 azon|OJJ45598.1

Group3402: abra|OJJ74297.1 acar|OOF95434.1 acla|EAW07342.1 afis|EAW22374.1 afla|EED57781.1 afum|EAL90818.1 agla|OJJ87627.1 akaw|GAA86727.1 aluc|OJZ89837.1 anid|CBF88706.1 anig|EHA27038.1 aory|EIT81052.1 asyd|OJJ63923.1 ater|EAU39508.1 atub|OJI84971.1 aver|OJI97022.1 awen|OJJ34949.1 azon|OJJ45599.1

Group3403: abra|OJJ74298.1 acar|OOF95433.1 acla|EAW07343.1 afis|EAW22375.1 afum|EAL90817.1 agla|OJJ87628.1 akaw|GAA86726.1 aluc|OJZ89835.1 anid|CBF88710.1 anig|EHA27037.1 aory|EIT81071.1

asyd|OJJ63924.1 ater|EAU39509.1 atub|OJI84972.1 aver|OJI97020.1 awen|OJJ34950.1 azon|OJJ45600.1 afla|EED57783.1

Group3404: abra|OJJ74299.1 acar|OOF95431.1 acla|EAW07344.1 afis|EAW22376.1 afla|EED57784.1 afum|EAL90816.1 agla|OJJ87629.1 akaw|GAA86725.1 aluc|OJZ89834.1 anid|CBF88712.1 anig|EHA27036.1 aory|EIT81070.1 asyd|OJJ63925.1 ater|EAU39510.1 atub|OJI84973.1 aver|OJI97019.1 awen|OJJ34951.1 azon|OJJ45601.1

Group3405: abra|OJJ74301.1 acar|OOF95429.1 acla|EAW07346.1 afis|EAW22378.1 afla|EED57786.1 afum|EAL90814.1 agla|OJJ87631.1 akaw|GAA86724.1 aluc|OJZ89832.1 anid|CBF88714.1 anig|EHA27034.1 aory|EIT81069.1 asyd|OJJ63926.1 ater|EAU39512.1 atub|OJI84974.1 aver|OJI97018.1 awen|OJJ34952.1 azon|OJJ45602.1

Group3406: abra|OJJ74302.1 acar|OOF95428.1 acla|EAW07349.1 afis|EAW22379.1 afla|EED57787.1 afum|EAL90812.1 agla|OJJ87635.1 akaw|GAA86723.1 aluc|OJZ89830.1 anid|CBF88716.1 anig|EHA27033.1 aory|EIT81084.1 asyd|OJJ63927.1 ater|EAU39513.1 atub|OJI84975.1 aver|OJI97017.1 awen|OJJ34953.1 azon|OJJ45603.1

Group3407: abra|OJJ74304.1 acar|OOF95424.1 acla|EAW07352.1 afis|EAW22381.1 afla|EED57790.1 afum|EAL90810.1 agla|OJJ87640.1 akaw|GAA86720.1 aluc|OJZ89824.1 anid|CBF88718.1 anig|EHA27031.1 aory|EIT81089.1 asyd|OJJ63928.1 ater|EAU39515.1 atub|OJI84982.1 aver|OJI97016.1 awen|OJJ34956.1 azon|OJJ45608.1

Group3408: abra|OJJ74305.1 acar|OOF95423.1 acla|EAW07353.1 afis|EAW22382.1 afla|EED56040.1 afum|EAL90809.1 agla|OJJ87641.1 akaw|GAA86719.1 aluc|OJZ89822.1 anid|CBF88721.1 anig|EHA27030.1 aory|EIT76582.1 asyd|OJJ63929.1 ater|EAU39516.1 atub|OJI84983.1 aver|OJI97014.1 awen|OJJ34957.1 azon|OJJ45609.1

Group3409: abra|OJJ74306.1 acar|OOF95422.1 acla|EAW07354.1 afis|EAW22383.1 afla|EED56041.1 afum|EAL90808.1 agla|OJJ87642.1 akaw|GAA86718.1 aluc|OJZ89821.1 anid|CBF88723.1 anig|EHA27029.1 aory|EIT76563.1 asyd|OJJ63930.1 ater|EAU39517.1 atub|OJI84984.1 aver|OJI97013.1 awen|OJJ34958.1 azon|OJJ45610.1

Group3410: abra|OJJ74307.1 acar|OOF95420.1 acla|EAW07355.1 afis|EAW22384.1 afla|EED56044.1 afum|EAL90806.1 agla|OJJ87644.1 akaw|GAA86717.1 aluc|OJZ89820.1 anid|CBF88725.1 anig|EHA27028.1 aory|EIT76594.1 asyd|OJJ63931.1 ater|EAU39337.1 atub|OJI84985.1 aver|OJI97012.1 awen|OJJ34959.1 azon|OJJ45611.1

Group3411: abra|OJJ74310.1 acar|OOF95419.1 acla|EAW07357.1 afis|EAW22386.1 afla|EED56046.1 afum|EAL90804.1 agla|OJJ87647.1 akaw|GAA86714.1 aluc|OJZ89816.1 anid|CBF88730.1 anig|EHA27025.1 aory|EIT76610.1 asyd|OJJ63934.1 ater|EAU39335.1 atub|OJI84989.1 aver|OJI97008.1 awen|OJJ34961.1 azon|OJJ45613.1

Group3412: abra|OJJ74311.1 acar|OOF95417.1 acla|EAW07358.1 afis|EAW22387.1 afla|EED56047.1 afum|EAL90803.1 agla|OJJ87648.1 akaw|GAA86713.1 aluc|OJZ89815.1 anid|CBF88732.1 anig|EHA27024.1 aory|EIT76591.1 asyd|OJJ63935.1 ater|EAU39334.1 atub|OJI84990.1 aver|OJI97007.1 awen|OJJ34962.1 azon|OJJ45614.1

Group3413: abra|OJJ74315.1 acar|OOF95412.1 acla|EAW07366.1 afis|EAW22393.1 afla|EED56056.1 afum|EAL90798.1 agla|OJJ87655.1 akaw|GAA86710.1 aluc|OJZ89811.1 anid|CBF88743.1 anig|EHA27021.1 aory|EIT76567.1 asyd|OJJ63954.1 ater|EAU39325.1 atub|OJI84994.1 aver|OJI96987.1 awen|OJJ34970.1 azon|OJJ45619.1

Group3414: abra|OJJ74316.1 acar|OOF95411.1 acla|EAW07367.1 afis|EAW22394.1 afla|EED56057.1 afum|EAL90797.1 agla|OJJ87657.1 akaw|GAA86709.1 aluc|OJZ89810.1 anid|CBF88745.1 anig|EHA27020.1 aory|EIT76865.1 asyd|OJJ63955.1 ater|EAU39324.1 atub|OJI84995.1 aver|OJI96986.1 awen|OJJ34971.1 azon|OJJ45620.1

Group3415: abra|OJJ74320.1 acar|OOF95409.1 acla|EAW07372.1 afis|EAW22397.1 afla|EED56061.1 afum|EAL90794.1 agla|OJJ87660.1 akaw|GAA86706.1 aluc|OJZ89808.1 anid|CBF88750.1 anig|EHA27018.1 aory|EIT76888.1 asyd|OJJ63958.1 ater|EAU39320.1 atub|OJI84998.1 aver|OJI96983.1 awen|OJJ34975.1 azon|OJJ45624.1

Group3416: abra|OJJ74325.1 acar|OOF95406.1 acla|EAW07377.1 afis|EAW22402.1 afla|EED56066.1 afum|EAL90789.1 agla|OJJ87665.1 akaw|GAA86701.1 aluc|OJZ89803.1 anid|CBF88765.1 anig|EHA27016.1 aory|EIT76819.1 asyd|OJJ63981.1 ater|EAU39306.1 atub|OJI85003.1 aver|OJI96972.1 awen|OJJ34981.1 azon|OJJ45629.1

Group3417: abra|OJJ74326.1 acar|OOF95405.1 acla|EAW07378.1 afis|EAW22403.1 afla|EED56067.1 afum|EAL90788.1 agla|OJJ87666.1 akaw|GAA86700.1 aluc|OJZ89802.1 anid|CBF88763.1 anig|EHA27015.1 aory|EIT76544.1 asyd|OJJ63982.1 ater|EAU39305.1 atub|OJI85004.1 aver|OJI96971.1 awen|OJJ34982.1 azon|OJJ45631.1

Group3418: abra|OJJ74329.1 acar|OOF95402.1 acla|EAW07379.1 afis|EAW22404.1 afla|EED56068.1 afum|EAL90787.1 agla|OJJ87667.1 akaw|GAA86699.1 aluc|OJZ89801.1 anid|CBF88785.1 anig|EHA27014.1 aory|EIT76781.1 asyd|OJJ62903.1 ater|EAU39304.1 atub|OJI85005.1 aver|OJI96957.1 awen|OJJ34983.1 azon|OJJ45633.1

Group3419: abra|OJJ74336.1 acar|OOF95391.1 acla|EAW07386.1 afis|EAW22411.1 afla|EED52491.1 afum|EAL90781.1 agla|OJJ87678.1 akaw|GAA86693.1 aluc|OJZ89794.1 anid|CBF88796.1 anig|EHA27007.1 aory|EIT82814.1 asyd|OJJ62897.1 ater|EAU39294.1 atub|OJI85012.1 aver|OJI96950.1 awen|OJJ34992.1 azon|OJJ45635.1

Group3420: abra|OJJ74340.1 acar|OOF89991.1 acla|EAW07391.1 afis|EAW22417.1 afla|EED52495.1 afum|EAL90775.1 agla|OJJ87599.1 akaw|GAA86690.1 aluc|OJZ89791.1 anid|CBF88802.1 anig|EHA27004.1 aory|EIT82790.1 asyd|OJJ62895.1 ater|EAU39289.1 atub|OJI85015.1 aver|OJI96947.1 awen|OJJ34996.1 azon|OJJ45639.1

Group3421: abra|OJJ74341.1 acar|OOF95389.1 acla|EAW07392.1 afis|EAW22418.1 afla|EED52496.1 afum|EAL90774.1 agla|OJJ87600.1 akaw|GAA86689.1 aluc|OJZ89790.1 anid|CBF88804.1 anig|EHA27003.1 aory|EIT82800.1 asyd|OJJ62894.1 ater|EAU39288.1 atub|OJI85016.1 aver|OJI96946.1 awen|OJJ34997.1 azon|OJJ45646.1

Group3422: abra|OJJ74343.1 acar|OOF95387.1 acla|EAW07397.1 afis|EAW22423.1 afla|EED52501.1 afum|EAL90768.2 agla|OJJ87605.1 akaw|GAA86687.1 aluc|OJZ89788.1 anid|CBF88813.1 anig|EHA27002.1 aory|EIT82762.1 asyd|OJJ62889.1 ater|EAU39284.1 atub|OJI85018.1 aver|OJI96941.1 awen|OJJ35002.1 azon|OJJ45642.1

Group3423: abra|OJJ74344.1 acar|OOF95386.1 acla|EAW07398.1 afis|EAW22424.1 afla|EED52502.1 afum|EAL90767.1 agla|OJJ87606.1 akaw|GAA86686.1 aluc|OJZ89787.1 anid|CBF75162.1 anig|EHA27001.1 aory|EIT82765.1 asyd|OJJ59961.1 ater|EAU39283.1 atub|OJI85019.1 aver|OJJ02329.1 awen|OJJ35003.1 azon|OJJ45641.1

Group3424: abra|OJJ74347.1 acar|OOF95382.1 acla|EAW07401.1 afis|EAW22427.1 afla|EED52505.1 afum|EAL90764.1 agla|OJJ87596.1 akaw|GAA86683.1 aluc|OJZ89783.1 anid|CBF88819.1 anig|EHA26998.1 aory|EIT82749.1 asyd|OJJ62886.1 ater|EAU39280.1 atub|OJI85022.1 aver|OJI96938.1 awen|OJJ35006.1 azon|OJJ45648.1

Group3425: abra|OJJ74348.1 acar|OOF95380.1 acla|EAW07402.1 afis|EAW22428.1 afla|EED52506.1 afum|EAL90763.1 agla|OJJ87595.1 akaw|GAA86682.1 aluc|OJZ89782.1 anid|CBF88822.1 anig|EHA26997.1 aory|EIT82757.1 asyd|OJJ62885.1 ater|EAU39279.1 atub|OJI85024.1 aver|OJI96937.1 awen|OJJ35007.1 azon|OJJ45649.1

Group3426: abra|OJJ74352.1 acar|OOF95378.1 acla|EAW07405.1 afis|EAW22431.1 afla|EED52510.1 afum|EAL90760.1 agla|OJJ87592.1 akaw|GAA86679.1 aluc|OJZ89776.1 anid|CBF88828.1 anig|EHA26994.1 aory|EIT82794.1 asyd|OJJ62882.1 ater|EAU39276.1 atub|OJI85027.1 aver|OJI96934.1 awen|OJJ35010.1 azon|OJJ45651.1

Group3427: abra|OJJ74353.1 acar|OOF95377.1 acla|EAW07406.1 afis|EAW22432.1 afla|EED52513.1 afum|EAL90759.1 agla|OJJ87590.1 akaw|GAA86678.1 aluc|OJZ89775.1 anid|CBF88832.1 anig|EHA26993.1 aory|EIT82776.1 asyd|OJJ62880.1 ater|EAU39275.1 atub|OJI85028.1 aver|OJI96932.1 awen|OJJ35011.1 azon|OJJ45652.1

Group3428: abra|OJJ74355.1 acar|OOF95375.1 acla|EAW07409.1 afis|EAW22435.1 afla|EED52515.1 afum|EAL90756.1 agla|OJJ87587.1 akaw|GAA86676.1 aluc|OJZ89773.1 anid|CBF88835.1 anig|EHA26991.1 aory|EIT82785.1 asyd|OJJ62878.1 ater|EAU39273.1 atub|OJI85030.1 aver|OJI96930.1 awen|OJJ35013.1 azon|OJJ45654.1

Group3429: abra|OJJ74359.1 acar|OOF95370.1 acla|EAW07410.1 afis|EAW22436.1 afla|EED54446.1 afum|EAL90755.1 agla|OJJ87586.1 akaw|GAA86672.1 aluc|OJZ89769.1 anid|CBF88882.1 anig|EHA26989.1 asyd|OJJ62877.1 ater|EAU39272.1 atub|OJI85034.1 aver|OJI96929.1 awen|OJJ35014.1 azon|OJJ45656.1 aory|EIT81137.1

Group3430: abra|OJJ74361.1 acar|OOF95368.1 acla|EAW07413.1 afis|EAW22439.1 afla|EED54449.1 afum|EAL90752.1 agla|OJJ87584.1 akaw|GAA86669.1 aluc|OJZ89766.1 anid|CBF88862.1 anig|EHA26986.1 aory|EIT81135.1 asyd|OJJ62874.1 ater|EAU39269.1 atub|OJI85036.1 aver|OJI96927.1 awen|OJJ35016.1 azon|OJJ45661.1

Group3431: abra|OJJ74364.1 acar|OOF95366.1 acla|EAW07414.1 afis|EAW22441.1 afla|EED54457.1 afum|EAL90750.1 agla|OJJ87583.1 akaw|GAA86666.1 aluc|OJZ89763.1 anid|CBF88845.1 anig|EHA26983.1 aory|EIT81148.1 asyd|OJJ62867.1 ater|EAU39263.1 atub|OJI85039.1 aver|OJI96920.1 awen|OJJ35018.1 azon|OJJ50043.1

Group3432: abra|OJJ74386.1 acar|OOF95352.1 acla|EAW07427.1 afis|EAW22453.1 afla|EED54469.1 afum|EAL90738.1 agla|OJJ87570.1 akaw|GAA87287.1 aluc|OJZ89732.1 anid|CBF88931.1 anig|EHA26958.1

aory|EIT81158.1 asyd|OJJ62850.1 ater|EAU39253.1 atub|OJI85069.1 aver|OJI96903.1 awen|OJJ35078.1 azon|OJJ50029.1

Group3433: abra|OJJ74395.1 acar|OOF95341.1 acla|EAW06819.1 afis|EAW16367.1 afla|EED53469.1 afum|EAL88992.2 agla|OJJ78588.1 akaw|GAA87293.1 aluc|OJZ89725.1 anid|CBF89350.1 anig|EHA26953.1 aory|EIT75003.1 asyd|OJJ62502.1 ater|EAU34081.1 atub|OJI85075.1 aver|OJI96623.1 awen|OJJ37772.1 azon|OJJ45083.1

Group3434: abra|OJJ74414.1 acar|OOF95329.1 acla|EAW06828.1 afis|EAW16378.1 afla|EED53458.1 afum|EAL89003.1 agla|OJJ78596.1 akaw|GAA87305.1 aluc|OJZ89712.1 anid|CBF88042.1 anig|EHA26943.1 aory|EIT75071.1 asyd|OJJ63543.1 ater|EAU34074.1 atub|OJI85090.1 aver|OJI97391.1 awen|OJJ37912.1 azon|OJJ45091.1

Group3435: abra|OJJ74417.1 acar|OOF95326.1 acla|EAW06832.1 afis|EAW16382.1 afla|EED53452.1 afum|EAL89007.1 agla|OJJ88205.1 akaw|GAA87308.1 aluc|OJZ89709.1 anid|CBF89369.1 anig|EHA26940.1 aory|EIT75129.1 asyd|OJJ62475.1 ater|EAU34070.1 atub|OJI85093.1 aver|OJI96611.1 awen|OJJ37915.1 azon|OJJ46177.1

Group3436: abra|OJJ74438.1 acar|OOF95308.1 acla|EAW12299.1 afis|EAW23841.1 afla|EED49145.1 afum|EAL87272.1 agla|OJJ79053.1 akaw|GAA87331.1 aluc|OJZ89683.1 anid|CBF89626.1 anig|EHA26919.1 aory|EIT78517.1 asyd|OJJ62298.1 ater|EAU29595.1 atub|OJI85117.1 aver|OJI96406.1 awen|OJJ35935.1 azon|OJJ49112.1

Group3437: abra|OJJ74456.1 acar|OOF95294.1 acla|EAW07237.1 afis|EAW22260.1 afla|EED57647.1 afum|EAL90920.1 agla|OJJ87795.1 akaw|GAA87350.1 aluc|OJZ89664.1 anid|CBF88509.1 anig|EHA26901.1 aory|EIT79886.1 asyd|OJJ63802.1 ater|EAU34263.1 atub|OJI85136.1 aver|OJI97136.1 awen|OJJ34832.1 azon|OJJ45714.1

Group3438: abra|OJJ74457.1 acar|OOF95293.1 acla|EAW07236.1 afis|EAW22259.1 afla|EED57646.1 afum|EAL90921.1 agla|OJJ87799.1 akaw|GAA87351.1 aluc|OJZ89663.1 anid|CBF88506.1 anig|EHA26900.1 aory|EIT79868.1 asyd|OJJ63799.1 ater|EAU34262.1 atub|OJI85137.1 aver|OJI97139.1 awen|OJJ34831.1 azon|OJJ45715.1

Group3439: abra|OJJ74463.1 acar|OOF95288.1 acla|EAW07235.1 afis|EAW22256.1 afla|EED57643.1 afum|EAL90922.1 agla|OJJ87800.1 akaw|GAA87355.1 aluc|OJZ89659.1 anid|CBF88504.1 anig|EHA26896.1 aory|EIT79873.1 asyd|OJJ63798.1 ater|EAU34257.1 atub|OJI85141.1 aver|OJI97140.1 awen|OJJ34828.1 azon|OJJ45720.1

Group3440: abra|OJJ74464.1 acar|OOF95286.1 acla|EAW07234.1 afis|EAW22255.1 afla|EED57641.1 afum|EAL90923.1 agla|OJJ87801.1 akaw|GAA87356.1 aluc|OJZ89658.1 anid|CBF88502.1 anig|EHA26895.1 aory|EIT82944.1 asyd|OJJ63796.1 ater|EAU34256.1 atub|OJI85143.1 aver|OJI97141.1 awen|OJJ34827.1 azon|OJJ45722.1

Group3441: abra|OJJ74466.1 acar|OOF95281.1 acla|EAW07231.1 afis|EAW22245.1 afla|EED57638.1 afum|EAL90948.1 agla|OJJ87805.1 akaw|GAA87359.1 aluc|OJZ89654.1 anid|CBF88496.1 anig|EHA26892.1 aory|EIT82948.1 asyd|OJJ63793.1 ater|EAU34253.1 atub|OJI85150.1 aver|OJI97144.1 awen|OJJ34823.1 azon|OJJ45727.1

Group3442: abra|OJJ74467.1 acar|OOF95280.1 acla|EAW07230.1 afis|EAW22244.1 afla|EED57637.1 afum|EAL90949.1 agla|OJJ87807.1 akaw|GAA87360.1 aluc|OJZ89653.1 anid|CBF88493.1 anig|EHA26891.1 aory|EIT82977.1 asyd|OJJ63792.1 ater|EAU34252.1 atub|OJI85151.1 aver|OJI97145.1 awen|OJJ34822.1 azon|OJJ45728.1

Group3443: abra|OJJ74468.1 acar|OOF95279.1 acla|EAW07229.1 afis|EAW22243.1 afla|EED57636.1 afum|EAL90950.1 agla|OJJ87808.1 akaw|GAA87361.1 aluc|OJZ89652.1 anid|CBF88491.1 anig|EHA26890.1 aory|EIT82978.1 asyd|OJJ63791.1 ater|EAU34251.1 atub|OJI85152.1 aver|OJI97146.1 awen|OJJ34821.1 azon|OJJ45729.1

Group3444: abra|OJJ74473.1 acar|OOF95271.1 acla|EAW07224.1 afis|EAW22238.1 afla|EED57627.1 afum|EAL90955.1 agla|OJJ87813.1 akaw|GAA87367.1 aluc|OJZ89645.1 anid|CBF88481.1 anig|EHA26884.1 aory|EIT82957.1 asyd|OJJ63785.1 ater|EAU34245.1 atub|OJI85158.1 aver|OJI97152.1 awen|OJJ34816.1 azon|OJJ45733.1

Group3445: abra|OJJ74477.1 acar|OOF95269.1 acla|EAW07223.1 afis|EAW22237.1 afla|EED57626.1 afum|EAL90956.1 agla|OJJ87814.1 akaw|GAA87370.1 aluc|OJZ89642.1 anid|CBF88479.1 anig|EHA26881.1 aory|EIT82989.1 asyd|OJJ63784.1 ater|EAU34244.1 atub|OJI85161.1 aver|OJI97153.1 awen|OJJ34815.1 azon|OJJ45734.1

Group3446: abra|OJJ74478.1 acar|OOF95268.1 acla|EAW07222.1 afis|EAW22236.1 afla|EED57625.1 afum|EAL90957.1 agla|OJJ87815.1 akaw|GAA87371.1 aluc|OJZ89641.1 anid|CBF88477.1 anig|EHA26880.1 aory|EIT82990.1 asyd|OJJ63783.1 ater|EAU34243.1 atub|OJI85162.1 aver|OJI97154.1 awen|OJJ34814.1 azon|OJJ45735.1

Group3447: abra|OJJ74481.1 acar|OOF95267.1 acla|EAW07220.1 afis|EAW22234.1 afla|EED57623.1 afum|EAL90959.1 agla|OJJ87817.1 akaw|GAA87374.1 aluc|OJZ89638.1 anid|CBF88474.1 anig|EHA26879.1 aory|EIT82951.1 asyd|OJJ63782.1 ater|EAU34237.1 atub|OJI85165.1 aver|OJI97155.1 awen|OJJ34813.1 azon|OJJ45736.1

Group3448: abra|OJJ74484.1 acar|OOF95266.1 acla|EAW07217.1 afis|EAW22231.1 afla|EED57620.1 afum|EAL90962.1 agla|OJJ87820.1 akaw|GAA87377.1 aluc|OJZ89635.1 anid|CBF88469.1 anig|EHA26877.1 aory|EIT82992.1 asyd|OJJ63780.1 ater|EAU34234.1 atub|OJI85168.1 aver|OJI97157.1 awen|OJJ34811.1 azon|OJJ45738.1

Group3449: abra|OJJ74489.1 acar|OOF95260.1 acla|EAW07206.1 afis|EAW22218.1 afla|EED57611.1 afum|EAL90976.1 agla|OJJ87826.1 akaw|GAA87383.1 aluc|OJZ89629.1 anid|CBF88456.1 anig|EHA26872.1 aory|EIT82939.1 asyd|OJJ63771.1 ater|EAU34228.1 atub|OJI85173.1 aver|OJI97167.1 awen|OJJ34804.1 azon|OJJ45749.1

Group3450: abra|OJJ74490.1 acar|OOF95259.1 acla|EAW07204.1 afis|EAW22216.1 afla|EED57610.1 afum|EAL90978.1 agla|OJJ87830.1 akaw|GAA87384.1 aluc|OJZ89628.1 anid|CBF88454.1 anig|EHA26871.1 aory|EIT82985.1 asyd|OJJ63770.1 ater|EAU34224.1 atub|OJI85174.1 aver|OJI97168.1 awen|OJJ34801.1 azon|OJJ45748.1

Group3451: abra|OJJ74491.1 acar|OOF95258.1 acla|EAW07203.1 afis|EAW22215.1 afla|EED57609.1 afum|EAL90979.1 agla|OJJ87829.1 akaw|GAA87385.1 aluc|OJZ89627.1 anid|CBF88452.1 anig|EHA26870.1 aory|EIT82971.1 asyd|OJJ63769.1 ater|EAU34223.1 atub|OJI85175.1 aver|OJI97169.1 awen|OJJ34800.1 azon|OJJ45747.1

Group3452: abra|OJJ74493.1 acar|OOF95256.1 acla|EAW07201.1 afis|EAW22213.1 afla|EED57607.1 afum|EAL90981.2 agla|OJJ87831.1 akaw|GAA87386.1 aluc|OJZ89625.1 anid|CBF88448.1 anig|EHA26869.1 aory|EIT82952.1 asyd|OJJ63768.1 ater|EAU34222.1 atub|OJI85177.1 aver|OJI97170.1 awen|OJJ34798.1 azon|OJJ45745.1

Group3453: abra|OJJ74494.1 acar|OOF95255.1 acla|EAW07200.1 afis|EAW22212.1 afla|EED57606.1 afum|EAL90982.1 agla|OJJ87832.1 akaw|GAA87387.1 aluc|OJZ89624.1 anid|CBF88446.1 anig|EHA26868.1 aory|EIT82953.1 asyd|OJJ63767.1 ater|EAU34221.1 atub|OJI85178.1 aver|OJI97171.1 awen|OJJ34797.1 azon|OJJ45744.1

Group3454: abra|OJJ74496.1 acar|OOF95254.1 acla|EAW07197.1 afis|EAW22209.1 afla|EED57603.1 afum|EAL90985.1 agla|OJJ87835.1 akaw|GAA87390.1 aluc|OJZ89622.1 anig|EHA26866.1 asyd|OJJ63765.1 ater|EAU34219.1 atub|OJI85180.1 aver|OJI97174.1 awen|OJJ34794.1 anid|CBF88442.1 aory|EIT82943.1 azon|OJJ45750.1

Group3455: abra|OJJ74497.1 acar|OOF95253.1 acla|EAW07195.1 afis|EAW22207.1 afla|EED57602.1 afum|EAL90986.2 agla|OJJ87837.1 akaw|GAA87392.1 aluc|OJZ89621.1 anid|CBF88439.1 anig|EHA26865.1 aory|EIT82955.1 asyd|OJJ63764.1 ater|EAU34218.1 atub|OJI85181.1 aver|OJI97176.1 awen|OJJ34793.1 azon|OJJ45753.1

Group3456: abra|OJJ74498.1 acar|OOF95252.1 acla|EAW13756.1 afis|EAW19810.1 afla|EED57601.1 afum|EAL85082.1 agla|OJJ87838.1 akaw|GAA87393.1 aluc|OJZ89620.1 anid|CBF88437.1 anig|EHA26864.1 aory|EIT82959.1 asyd|OJJ63763.1 ater|EAU34217.1 atub|OJI85182.1 aver|OJI97177.1 awen|OJJ34792.1 azon|OJJ45754.1

Group3457: abra|OJJ74503.1 acar|OOF95241.1 acla|EAW07189.1 afis|EAW22202.1 afla|EED57593.1 afum|EAL90990.1 agla|OJJ87842.1 akaw|GAA87397.1 aluc|OJZ89615.1 anid|CBF88430.1 anig|EHA26860.1 aory|EIT82963.1 asyd|OJJ63760.1 ater|EAU34212.1 atub|OJI85188.1 aver|OJI97180.1 awen|OJJ34787.1 azon|OJJ45771.1

Group3458: abra|OJJ74506.1 acar|OOF89839.1 acla|EAW07192.1 afis|EAW22205.1 afla|EED57588.1 afum|EAL90988.1 agla|OJJ87845.1 akaw|GAA88823.1 aluc|OJZ89611.1 anid|CBF88417.1 anig|EHA26856.1 aory|EIT82968.1 asyd|OJJ63756.1 ater|EAU34209.1 atub|OJI85192.1 aver|OJI97184.1 awen|OJJ34786.1 azon|OJJ45766.1

Group3459: abra|OJJ74508.1 acar|OOF95234.1 acla|EAW07184.1 afis|EAW22197.1 afla|EED57584.1 afum|EAL90994.1 agla|OJJ87848.1 akaw|GAA82091.1 aluc|OJZ92089.1 anid|CBF88409.1 anig|EHA26852.1 aory|EIT82932.1 asyd|OJJ63754.1 ater|EAU34207.1 atub|OJI86519.1 aver|OJI97187.1 awen|OJJ34780.1 azon|OJJ45761.1

Group3460: abra|OJJ74513.1 acar|OOF95230.1 acla|EAW07177.1 afis|EAW22191.1 afla|EED57575.1 afum|EAL91000.1 agla|OJJ87856.1 akaw|GAA82086.1 aluc|OJZ92094.1 anid|CBF88400.1 anig|EHA26847.1 aory|EIT82946.1 asyd|OJJ63748.1 ater|EAU34199.1 atub|OJI86527.1 aver|OJI97192.1 awen|OJJ34771.1 azon|OJJ45775.1

Group3461: abra|OJJ74519.1 acar|OOF95229.1 acla|EAW07174.1 afis|EAW22188.1 afla|EED57572.1 afum|EAL91002.1 agla|OJJ87861.1 akaw|GAA82081.1 aluc|OJZ92098.1 anid|CBF88395.1 anig|EHA26844.1

aory|EIT82984.1 asyd|OJJ63745.1 ater|EAU34195.1 atub|OJI86531.1 aver|OJI97195.1 awen|OJJ34768.1 azon|OJJ45778.1

Group3462: abra|OJJ74523.1 acar|OOF95226.1 acla|EAW07170.1 afis|EAW22185.1 afla|EED57563.1 afum|EAL91004.1 agla|OJJ87870.1 akaw|GAA82078.1 aluc|OJZ92102.1 anid|CBF88391.1 anig|EHA26841.1 aory|EIT81827.1 asyd|OJJ63742.1 ater|EAU34193.1 atub|OJI86536.1 aver|OJI97198.1 awen|OJJ34765.1 azon|OJJ45780.1

Group3463: abra|OJJ74524.1 acar|OOF95225.1 acla|EAW07169.1 afis|EAW22184.1 afla|EED57562.1 afum|EAL91005.1 agla|OJJ87869.1 akaw|GAA82077.1 aluc|OJZ92104.1 anid|CBF75419.1 anig|EHA26840.1 aory|EIT81846.1 asyd|OJJ59792.1 ater|EAU34188.1 atub|OJI86538.1 aver|OJJ02495.1 awen|OJJ34764.1 azon|OJJ45781.1

Group3464: abra|OJJ74527.1 acar|OOF95223.1 acla|EAW07164.1 afis|EAW22179.1 afla|EED57558.1 afum|EAL91010.1 agla|OJJ87876.1 akaw|GAA82074.1 aluc|OJZ92108.1 anid|CBF88384.1 anig|EHA26837.1 aory|EIT81836.1 asyd|OJJ63738.1 ater|EAU34184.1 atub|OJI86542.1 aver|OJI97202.1 awen|OJJ34760.1 azon|OJJ45791.1

Group3465: abra|OJJ74532.1 acar|OOF95220.1 acla|EAW07160.1 afis|EAW22175.1 afla|EED57554.1 afum|EAL91014.1 agla|OJJ87879.1 akaw|GAA82068.1 aluc|OJZ92114.1 anid|CBF89077.1 anig|EHA26831.1 aory|EIT81832.1 asyd|OJJ62739.1 ater|EAU34180.1 atub|OJI86547.1 aver|OJI96782.1 awen|OJJ39689.1 azon|OJJ45808.1

Group3466: abra|OJJ74533.1 acar|OOF95219.1 acla|EAW07159.1 afis|EAW22174.1 afla|EED57553.1 afum|EAL91015.1 agla|OJJ87880.1 akaw|GAA82067.1 aluc|OJZ92115.1 anid|CBF89079.1 anig|EHA26830.1 aory|EIT81853.1 asyd|OJJ62738.1 ater|EAU34179.1 atub|OJI86548.1 aver|OJI96781.1 awen|OJJ34749.1 azon|OJJ45810.1

Group3467: abra|OJJ74546.1 acar|OOF95210.1 acla|EAW07152.1 afis|EAW22166.1 afla|EED57536.1 afum|EAL91025.1 agla|OJJ87891.1 akaw|GAA82052.1 aluc|OJZ92128.1 anid|CBF89087.1 anig|EHA26820.1 aory|EIT81873.1 asyd|OJJ62734.1 ater|EAU34173.1 atub|OJI86560.1 aver|OJI96777.1 awen|OJJ34740.1 azon|OJJ45819.1

Group3468: abra|OJJ74548.1 acar|OOF95209.1 acla|EAW07150.1 afis|EAW22164.1 afla|EED57535.1 afum|EAL91027.1 agla|OJJ87893.1 akaw|GAA82050.1 aluc|OJZ92130.1 anid|CBF89090.1 anig|EHA26818.1 aory|EIT81851.1 asyd|OJJ62732.1 ater|EAU34171.1 atub|OJI86562.1 aver|OJI96775.1 awen|OJJ34738.1 azon|OJJ45821.1

Group3469: abra|OJJ74549.1 acar|OOF95208.1 acla|EAW07149.1 afis|EAW22163.1 afla|EED57534.1 afum|EAL91028.1 agla|OJJ87894.1 akaw|GAA82049.1 aluc|OJZ92131.1 anid|CBF89094.1 anig|EHA26817.1 aory|EIT81858.1 asyd|OJJ62731.1 ater|EAU34166.1 atub|OJI86563.1 aver|OJI96774.1 awen|OJJ34737.1 azon|OJJ45822.1

Group3470: abra|OJJ74553.1 acar|OOF95202.1 acla|EAW07145.1 afis|EAW22159.1 afla|EED57531.1 afum|EAL91031.1 agla|OJJ87905.1 akaw|GAA82044.1 aluc|OJZ92136.1 anid|CBF89101.1 anig|EHA26813.1 aory|EIT81829.1 asyd|OJJ62727.1 ater|EAU34162.1 atub|OJI86568.1 aver|OJI96770.1 awen|OJJ34733.1 azon|OJJ45826.1

Group3471: abra|OJJ74569.1 acar|OOF95192.1 acla|EAW07140.1 afis|EAW22153.1 afla|EED53614.1 afum|EAL91037.1 agla|OJJ87909.1 akaw|GAA82036.1 aluc|OJZ92147.1 anid|CBF89105.1 anig|EHA26802.1 aory|EIT75057.1 asyd|OJJ62725.1 ater|EAU34157.1 atub|OJI86575.1 aver|OJI96768.1 awen|OJJ34729.1 azon|OJJ45834.1

Group3472: abra|OJJ74571.1 acar|OOF95191.1 acla|EAW07138.1 afis|EAW22151.1 afla|EED53613.1 afum|EAL91039.1 agla|OJJ87911.1 akaw|GAA82034.1 aluc|OJZ92149.1 anid|CBF89108.1 anig|EHA26800.1 aory|EIT75195.1 asyd|OJJ62667.1 ater|EAU34156.1 atub|OJI86577.1 aver|OJI96765.1 awen|OJJ34727.1 azon|OJJ45835.1

Group3473: abra|OJJ74573.1 acar|OOF95189.1 acla|EAW07136.1 afis|EAW22149.1 afla|EED53610.1 afum|EAL91041.1 agla|OJJ87914.1 akaw|GAA82032.1 aluc|OJZ92151.1 anid|CBF89112.1 anig|EHA26798.1 aory|EIT74840.1 asyd|OJJ62665.1 ater|EAU34155.1 atub|OJI86579.1 aver|OJI96763.1 awen|OJJ34725.1 azon|OJJ45839.1

Group3474: abra|OJJ74577.1 acar|OOF95187.1 acla|EAW07133.1 afis|EAW22146.1 afla|EED45848.1 afum|EAL91044.1 agla|OJJ87922.1 akaw|GAA82028.1 aluc|OJZ92155.1 anid|CBF78365.1 anig|EHA26794.1 aory|EIT75625.1 asyd|OJJ57324.1 ater|EAU34149.1 atub|OJI86583.1 aver|OJJ05836.1 awen|OJJ34721.1 azon|OJJ46642.1

Group3475: abra|OJJ74578.1 acar|OOF95186.1 acla|EAW07132.1 afis|EAW22145.1 afla|EED45849.1 afum|EAL91045.1 agla|OJJ87923.1 akaw|GAA82027.1 aluc|OJZ92156.1 anid|CBF78367.1 anig|EHA26793.1 aory|EIT75756.1 asyd|OJJ57325.1 ater|EAU34148.1 atub|OJI86584.1 aver|OJJ05835.1 awen|OJJ34720.1 azon|OJJ46643.1

Group3476: abra|OJJ74581.1 acar|OOF95183.1 acla|EAW07131.1 afis|EAW22142.1 afla|EED53568.1 afum|EAL91048.2 agla|OJJ87932.1 akaw|GAA82024.1 aluc|OJZ92159.1 anid|CBF89555.1 anig|EHA26790.1 aory|EIT75148.1 asyd|OJJ62347.1 ater|EAU31752.1 atub|OJI86587.1 aver|OJI96453.1 awen|OJJ34717.1 azon|OJJ45846.1

Group3477: abra|OJJ74584.1 acar|OOF95182.1 acla|EAW10794.1 afis|EAW24782.1 afla|EED47203.1 afum|EAL89421.1 agla|OJJ82878.1 akaw|GAA82021.1 aluc|OJZ92162.1 anid|CBF81444.1 anig|EHA26787.1 aory|EIT79171.1 asyd|OJJ59430.1 ater|EAU36526.1 atub|OJI86590.1 aver|OJJ02841.1 awen|OJJ40368.1 azon|OJJ49217.1

Group3479: abra|OJJ74588.1 acar|OOF95178.1 acla|EAW10790.1 afis|EAW24786.1 afla|EED47199.1 afum|EAL89425.1 agla|OJJ82884.1 akaw|GAA82017.1 aluc|OJZ92165.1 anid|CBF81455.1 anig|EHA26784.1 aory|EIT79195.1 asyd|OJJ59435.1 ater|EAU36523.1 atub|OJI86595.1 aver|OJJ02836.1 awen|OJJ40370.1 azon|OJJ49220.1

Group3480: abra|OJJ74589.1 acar|OOF95177.1 acla|EAW10789.1 afis|EAW24787.1 afla|EED47197.1 afum|EAL89426.1 agla|OJJ82883.1 akaw|GAA82016.1 aluc|OJZ92167.1 anid|CBF81457.1 anig|EHA26782.1 aory|EIT79260.1 asyd|OJJ59437.1 ater|EAU36522.1 atub|OJI86598.1 aver|OJJ02834.1 awen|OJJ40371.1 azon|OJJ49221.1

Group3481: abra|OJJ74594.1 acar|OOF95175.1 acla|EAW10788.1 afis|EAW24791.1 afla|EED47195.1 afum|EAL89433.1 agla|OJJ82881.1 akaw|GAA82014.1 aluc|OJZ92169.1 anid|CBF81460.1 anig|EHA26780.1 aory|EIT79181.1 asyd|OJJ59439.1 ater|EAU36521.1 atub|OJI86601.1 aver|OJJ02833.1 awen|OJJ40372.1 azon|OJJ49223.1

Group3482: abra|OJJ74595.1 acar|OOF95174.1 acla|EAW10786.1 afis|EAW24792.1 afla|EED47194.1 afum|EAL89434.1 agla|OJJ82879.1 akaw|GAA82013.1 aluc|OJZ92170.1 anid|CBF81464.1 anig|EHA26779.1 aory|EIT79314.1 asyd|OJJ59442.1 ater|EAU36516.1 atub|OJI86602.1 aver|OJJ02831.1 awen|OJJ40373.1 azon|OJJ49224.1

Group3484: abra|OJJ74705.1 acar|OOF89903.1 acla|EAW10692.1 afis|EAW24878.1 afla|EED47088.1 afum|EAL89528.1 agla|OJJ82100.1 akaw|GAA81912.1 aluc|OJZ92285.1 anid|CBF75756.1 anig|EHA26683.1 aory|EIT72477.1 asyd|OJJ57526.1 ater|EAU36423.1 atub|OJI86716.1 aver|OJJ05652.1 awen|OJJ40468.1 azon|OJJ49332.1

Group3485: abra|OJJ74707.1 acar|OOF89905.1 acla|EAW10691.1 afis|EAW24879.1 afla|EED47087.1 afum|EAL89529.1 agla|OJJ82101.1 akaw|GAA81910.1 aluc|OJZ92288.1 anid|CBF75754.1 anig|EHA26681.1 aory|EIT72507.1 asyd|OJJ57525.1 ater|EAU36422.1 atub|OJI86718.1 aver|OJJ05653.1 awen|OJJ40469.1 azon|OJJ49333.1

Group3486: abra|OJJ74708.1 acar|OOF95718.1 acla|EAW10690.1 afis|EAW24880.1 afla|EED47086.1 afum|EAL89530.1 agla|OJJ82102.1 akaw|GAA81909.1 aluc|OJZ92289.1 anid|CBF75752.1 anig|EHA26680.1 aory|EIT72478.1 asyd|OJJ57524.1 ater|EAU36421.1 atub|OJI86719.1 aver|OJJ05654.1 awen|OJJ40470.1 azon|OJJ49334.1

Group3487: abra|OJJ74709.1 acar|OOF95719.1 acla|EAW10689.1 afis|EAW24881.1 afla|EED47085.1 afum|EAL89531.1 agla|OJJ82103.1 akaw|GAA81908.1 aluc|OJZ92290.1 anid|CBF75750.1 anig|EHA26679.1 aory|EIT72499.1 asyd|OJJ57523.1 ater|EAU36420.1 atub|OJI86720.1 aver|OJJ05655.1 awen|OJJ40471.1 azon|OJJ49335.1

Group3488: abra|OJJ74711.1 acar|OOF95722.1 acla|EAW10687.1 afis|EAW24883.1 afla|EED47083.1 afum|EAL89533.1 agla|OJJ82105.1 akaw|GAA81906.1 aluc|OJZ92292.1 anid|CBF75747.1 anig|EHA26677.1 aory|EIT72512.1 asyd|OJJ57521.1 ater|EAU36418.1 atub|OJI86722.1 aver|OJJ05657.1 awen|OJJ40473.1 azon|OJJ49337.1

Group3489: abra|OJJ74714.1 acar|OOF95723.1 acla|EAW10685.1 afis|EAW24885.1 afla|EED47081.1 afum|EAL89535.2 agla|OJJ82108.1 akaw|GAA81904.1 aluc|OJZ92294.1 anid|CBF75617.1 anig|EHA26675.1 aory|EIT72482.1 asyd|OJJ57452.1 ater|EAU36416.1 atub|OJI86724.1 aver|OJJ05726.1 awen|OJJ40475.1 azon|OJJ49339.1

Group3490: abra|OJJ74715.1 acar|OOF95724.1 acla|EAW10684.1 afis|EAW24886.1 afla|EED47080.1 afum|EAL89537.1 agla|OJJ82109.1 akaw|GAA81903.1 aluc|OJZ92295.1 anid|CBF75619.1 anig|EHA26674.1 aory|EIT72515.1 asyd|OJJ57453.1 ater|EAU36415.1 atub|OJI86725.1 aver|OJJ05725.1 awen|OJJ40476.1 azon|OJJ49340.1

Group3491: abra|OJJ74717.1 acar|OOF95728.1 acla|EAW10681.1 afis|EAW24889.1 afla|EED46965.1 afum|EAL89540.1 agla|OJJ82112.1 akaw|GAA81899.1 aluc|OJZ92299.1 anid|CBF75625.1 anig|EHA26672.1 aory|EIT72622.1 asyd|OJJ57456.1 ater|EAU36412.1 atub|OJI86728.1 aver|OJJ05723.1 awen|OJJ40479.1 azon|OJJ49343.1

Group3492: abra|OJJ74722.1 acar|OOF95731.1 acla|EAW10675.1 afis|EAW24891.1 afla|EED46970.1 afum|EAL89544.1 agla|OJJ82115.1 akaw|GAA81894.1 aluc|OJZ92306.1 anid|CBF75630.1 anig|EHA26662.1

aory|EIT72703.1 asyd|OJJ57459.1 ater|EAU36407.1 atub|OJI86736.1 aver|OJJ05721.1 awen|OJJ40484.1 azon|OJJ49346.1

Group3493: abra|OJJ74723.1 acar|OOF95732.1 acla|EAW10674.1 afis|EAW24892.1 afla|EED46971.1 afum|EAL89545.1 agla|OJJ82116.1 akaw|GAA81893.1 aluc|OJZ92307.1 anid|CBF75632.1 anig|EHA26661.1 aory|EIT72525.1 asyd|OJJ57460.1 ater|EAU36406.1 atub|OJI86737.1 aver|OJJ05720.1 awen|OJJ40485.1 azon|OJJ49347.1

Group3494: abra|OJJ74724.1 acar|OOF95733.1 acla|EAW10673.1 afis|EAW24893.1 afla|EED46972.1 afum|EAL89546.1 agla|OJJ82117.1 akaw|GAA81892.1 aluc|OJZ92308.1 anid|CBF75634.1 anig|EHA26660.1 aory|EIT72626.1 asyd|OJJ57461.1 ater|EAU36405.1 atub|OJI86738.1 aver|OJJ05719.1 awen|OJJ40486.1 azon|OJJ49348.1

Group3495: abra|OJJ74727.1 acar|OOF90030.1 acla|EAW10671.1 afis|EAW24895.1 afla|EED46975.1 afum|EAL89549.1 agla|OJJ82119.1 akaw|GAA81890.1 aluc|OJZ92311.1 anid|CBF75657.1 anig|EHA26658.1 aory|EIT72598.1 asyd|OJJ57463.1 ater|EAU36403.1 atub|OJI86740.1 aver|OJJ05717.1 awen|OJJ40488.1 azon|OJJ49351.1

Group3496: abra|OJJ74735.1 acar|OOF95734.1 acla|EAW10667.1 afis|EAW24899.1 afla|EED46978.1 afum|EAL89553.1 agla|OJJ82124.1 akaw|GAA83674.1 aluc|OJZ92317.1 anid|CBF75649.1 anig|EHA26654.1 aory|EIT72632.1 asyd|OJJ57470.1 ater|EAU36400.1 atub|OJI86746.1 aver|OJJ05712.1 awen|OJJ40493.1 azon|OJJ49355.1

Group3497: abra|OJJ74736.1 acar|OOF95735.1 acla|EAW10666.1 afis|EAW24900.1 afla|EED46980.1 afum|EAL89554.1 agla|OJJ82125.1 akaw|GAA83673.1 aluc|OJZ92318.1 anid|CBF75648.1 anig|EHA26653.1 aory|EIT72726.1 asyd|OJJ57472.1 ater|EAU36399.1 atub|OJI86747.1 aver|OJJ05711.1 awen|OJJ40494.1 azon|OJJ49356.1

Group3498: abra|OJJ74738.1 acar|OOF95736.1 acla|EAW10665.1 afis|EAW24901.1 afla|EED46981.1 afum|EAL89555.1 agla|OJJ82126.1 akaw|GAA83672.1 aluc|OJZ92319.1 anid|CBF75646.1 anig|EHA26651.1 aory|EIT72528.1 asyd|OJJ57473.1 ater|EAU36398.1 atub|OJI86748.1 aver|OJJ05710.1 awen|OJJ40495.1 azon|OJJ49357.1

Group3499: abra|OJJ74739.1 acar|OOF95737.1 acla|EAW10664.1 afis|EAW24902.1 afla|EED46982.1 afum|EAL89556.1 agla|OJJ82127.1 akaw|GAA83671.1 aluc|OJZ92320.1 anid|CBF75644.1 anig|EHA26650.1 aory|EIT72548.1 asyd|OJJ57474.1 ater|EAU36397.1 atub|OJI86749.1 aver|OJJ05707.1 awen|OJJ40496.1 azon|OJJ49358.1

Group3500: abra|OJJ74749.1 acar|OOF95742.1 acla|EAW10661.1 afis|EAW24906.1 afla|EED46989.1 afum|EAL89559.1 agla|OJJ82130.1 akaw|GAA83662.1 aluc|OJZ92330.1 anid|CBF75661.1 anig|EHA26641.1 aory|EIT72764.1 asyd|OJJ57478.1 ater|EAU36392.1 atub|OJI86759.1 aver|OJJ05703.1 awen|OJJ40499.1 azon|OJJ49361.1

Group3501: abra|OJJ74750.1 acar|OOF95743.1 acla|EAW10660.1 afis|EAW24907.1 afla|EED46991.1 afum|EAL89560.1 agla|OJJ82131.1 akaw|GAA83661.1 aluc|OJZ92331.1 anid|CBF75665.1 anig|EHA26640.1 aory|EIT72700.1 asyd|OJJ57479.1 ater|EAU36391.1 atub|OJI86760.1 aver|OJJ05702.1 awen|OJJ40500.1 azon|OJJ49362.1

Group3502: abra|OJJ74751.1 acar|OOF95744.1 acla|EAW10659.1 afis|EAW24908.1 afla|EED46992.1 afum|EAL89561.1 agla|OJJ82132.1 akaw|GAA83660.1 aluc|OJZ92332.1 anid|CBF75667.1 anig|EHA26639.1 aory|EIT72681.1 asyd|OJJ57480.1 ater|EAU36390.1 atub|OJI86761.1 aver|OJJ05701.1 awen|OJJ40501.1 azon|OJJ49363.1

Group3503: abra|OJJ74752.1 acar|OOF95745.1 acla|EAW10658.1 afis|EAW24909.1 afla|EED46994.1 afum|EAL89562.1 agla|OJJ82133.1 akaw|GAA83659.1 aluc|OJZ92333.1 anid|CBF75669.1 anig|EHA26638.1 aory|EIT72691.1 asyd|OJJ57481.1 ater|EAU36389.1 atub|OJI86762.1 aver|OJJ05700.1 awen|OJJ40503.1 azon|OJJ49364.1

Group3504: abra|OJJ74754.1 acar|OOF95747.1 acla|EAW10656.1 afis|EAW24911.1 afla|EED46996.1 afum|EAL89565.1 agla|OJJ82135.1 akaw|GAA83657.1 aluc|OJZ92335.1 anid|CBF75671.1 anig|EHA26636.1 aory|EIT72517.1 asyd|OJJ57483.1 ater|EAU36387.1 atub|OJI86764.1 aver|OJJ05698.1 awen|OJJ40506.1 azon|OJJ49366.1

Group3505: abra|OJJ74756.1 acar|OOF95749.1 acla|EAW10655.1 afis|EAW24912.1 afla|EED46997.1 afum|EAL89566.2 agla|OJJ82141.1 akaw|GAA83655.1 aluc|OJZ92338.1 anid|CBF75673.1 anig|EHA26634.1 aory|EIT72758.1 asyd|OJJ57484.1 ater|EAU36386.1 atub|OJI86766.1 aver|OJJ05697.1 awen|OJJ40508.1 azon|OJJ49367.1

Group3506: abra|OJJ74758.1 acar|OOF95751.1 acla|EAW10653.1 afis|EAW24914.1 afla|EED46999.1 afum|EAL89568.1 agla|OJJ82142.1 akaw|GAA83653.1 aluc|OJZ92342.1 anid|CBF75677.1 anig|EHA26632.1 aory|EIT72541.1 asyd|OJJ57486.1 ater|EAU36384.1 atub|OJI86768.1 aver|OJJ05694.1 awen|OJJ40510.1 azon|OJJ49369.1

Group3507: abra|OJJ74759.1 acar|OOF95752.1 acla|EAW10652.1 afis|EAW24915.1 afla|EED47000.1 afum|EAL89569.1 agla|OJJ82143.1 akaw|GAA83652.1 aluc|OJZ92343.1 anid|CBF75679.1 anig|EHA26631.1 aory|EIT72710.1 asyd|OJJ57487.1 ater|EAU36383.1 atub|OJI86769.1 aver|OJJ05693.1 awen|OJJ40511.1 azon|OJJ49370.1

Group3508: abra|OJJ74760.1 acar|OOF95753.1 acla|EAW10651.1 afis|EAW24916.1 afla|EED47001.1 afum|EAL89571.1 agla|OJJ82144.1 akaw|GAA83651.1 aluc|OJZ92344.1 anid|CBF75681.1 anig|EHA26630.1 aory|EIT72621.1 asyd|OJJ57488.1 ater|EAU36382.1 atub|OJI86770.1 aver|OJJ05692.1 awen|OJJ40512.1 azon|OJJ49371.1

Group3509: abra|OJJ74762.1 acar|OOF95755.1 acla|EAW10649.1 afis|EAW24918.1 afla|EED47003.1 afum|EAL89573.1 agla|OJJ82147.1 akaw|GAA83649.1 aluc|OJZ92346.1 anid|CBF75687.1 anig|EHA26628.1 aory|EIT72583.1 asyd|OJJ57490.1 ater|EAU36380.1 atub|OJI86773.1 aver|OJJ05690.1 awen|OJJ40515.1 azon|OJJ49374.1

Group3510: abra|OJJ74763.1 acar|OOF95756.1 acla|EAW10647.1 afis|EAW24919.1 afla|EED47004.1 afum|EAL89574.1 agla|OJJ82148.1 akaw|GAA83648.1 aluc|OJZ92347.1 anid|CBF75689.1 anig|EHA26627.1 aory|EIT72654.1 asyd|OJJ57491.1 ater|EAU36379.1 atub|OJI86774.1 aver|OJJ05689.1 awen|OJJ40516.1 azon|OJJ49375.1

Group3511: abra|OJJ74766.1 acar|OOF95758.1 acla|EAW10645.1 afis|EAW24921.1 afla|EED55552.1 afum|EAL89576.1 agla|OJJ82150.1 akaw|GAA83646.1 aluc|OJZ92349.1 anid|CBF75693.1 anig|EHA26625.1 aory|EIT82689.1 asyd|OJJ57493.1 ater|EAU35653.1 atub|OJI86777.1 aver|OJJ05687.1 awen|OJJ40518.1 azon|OJJ49377.1

Group3512: abra|OJJ74767.1 acar|OOF95759.1 acla|EAW10644.1 afis|EAW24922.1 afla|EED55551.1 afum|EAL89577.2 agla|OJJ82151.1 akaw|GAA83645.1 aluc|OJZ92350.1 anid|CBF75695.1 anig|EHA26624.1 aory|EIT82687.1 asyd|OJJ57494.1 ater|EAU35654.1 atub|OJI86778.1 aver|OJJ05686.1 awen|OJJ40519.1 azon|OJJ49378.1

Group3513: abra|OJJ74772.1 acar|OOF95762.1 acla|EAW10639.1 afis|EAW24927.1 afla|EED55546.1 afum|EAL89583.1 agla|OJJ82155.1 akaw|GAA83640.1 aluc|OJZ92353.1 anid|CBF75704.1 anig|EHA26621.1 aory|EIT82681.1 asyd|OJJ57497.1 ater|EAU35657.1 atub|OJI86783.1 aver|OJJ05681.1 awen|OJJ40523.1 azon|OJJ49383.1

Group3514: abra|OJJ74774.1 acar|OOF95763.1 acla|EAW10637.1 afis|EAW24929.1 afla|EED55544.1 afum|EAL89585.1 agla|OJJ82157.1 akaw|GAA83638.1 aluc|OJZ92355.1 anid|CBF75708.1 anig|EHA26619.1 aory|EIT82710.1 asyd|OJJ57499.1 ater|EAU35659.1 atub|OJI86785.1 aver|OJJ05679.1 awen|OJJ40525.1 azon|OJJ49385.1

Group3515: abra|OJJ74775.1 acar|OOF95764.1 acla|EAW10636.1 afis|EAW24930.1 afla|EED55543.1 afum|EAL89586.1 agla|OJJ82158.1 akaw|GAA83637.1 aluc|OJZ92356.1 anid|CBF75710.1 anig|EHA26618.1 aory|EIT82674.1 asyd|OJJ57500.1 ater|EAU35660.1 atub|OJI86786.1 aver|OJJ05678.1 awen|OJJ40526.1 azon|OJJ49386.1

Group3516: abra|OJJ74776.1 acar|OOF95765.1 acla|EAW10635.1 afis|EAW24931.1 afla|EED55542.1 afum|EAL89587.1 agla|OJJ82159.1 akaw|GAA83636.1 aluc|OJZ92357.1 anid|CBF75712.1 anig|EHA26617.1 aory|EIT82695.1 asyd|OJJ57502.1 ater|EAU35661.1 atub|OJI86787.1 aver|OJJ05676.1 awen|OJJ40527.1 azon|OJJ49387.1

Group3517: abra|OJJ74779.1 acar|OOF95769.1 acla|EAW10631.1 afis|EAW24936.1 afla|EED55536.1 afum|EAL89593.1 agla|OJJ82163.1 akaw|GAA83633.1 aluc|OJZ92360.1 anid|CBF75723.1 anig|EHA26613.1 aory|EIT80806.1 asyd|OJJ57506.1 ater|EAU35665.1 atub|OJI86790.1 aver|OJJ05672.1 awen|OJJ40531.1 azon|OJJ49391.1

Group3518: abra|OJJ74780.1 acar|OOF95771.1 acla|EAW10630.1 afis|EAW24937.1 afla|EED55535.1 afum|EAL89594.1 agla|OJJ82164.1 akaw|GAA83632.1 aluc|OJZ92361.1 anid|CBF75725.1 anig|EHA26612.1 aory|EIT80813.1 asyd|OJJ57507.1 ater|EAU35666.1 atub|OJI86791.1 aver|OJJ05671.1 awen|OJJ40532.1 azon|OJJ49392.1

Group3519: abra|OJJ74781.1 acar|OOF95772.1 acla|EAW10629.1 afis|EAW24938.1 afla|EED55534.1 afum|EAL89595.1 agla|OJJ82165.1 akaw|GAA83631.1 aluc|OJZ92362.1 anid|CBF75727.1 anig|EHA26611.1 aory|EIT80903.1 asyd|OJJ57508.1 ater|EAU35667.1 atub|OJI86792.1 aver|OJJ05670.1 awen|OJJ40533.1 azon|OJJ49393.1

Group3520: abra|OJJ74784.1 acar|OOF95774.1 acla|EAW10627.1 afis|EAW24940.1 afla|EED55525.1 afum|EAL89597.1 agla|OJJ82167.1 akaw|GAA83629.1 aluc|OJZ92364.1 anid|CBF75735.1 anig|EHA26609.1 aory|EIT80775.1 asyd|OJJ57512.1 ater|EAU35669.1 atub|OJI86794.1 aver|OJJ05667.1 awen|OJJ40536.1 azon|OJJ49395.1

Group3521: abra|OJJ74786.1 acar|OOF95776.1 acla|EAW10625.1 afis|EAW24942.1 afla|EED55523.1 afum|EAL89599.1 agla|OJJ82169.1 akaw|GAA83627.1 aluc|OJZ92366.1 anid|CBF75741.1 anig|EHA26607.1

aory|EIT80928.1 asyd|OJJ57517.1 ater|EAU35674.1 atub|OJI86796.1 aver|OJJ05662.1 awen|OJJ40537.1 azon|OJJ49397.1

Group3522: abra|OJJ74788.1 acar|OOF95777.1 acla|EAW10623.1 afis|EAW24944.1 afla|EED55521.1 afum|EAL89601.1 agla|OJJ82171.1 akaw|GAA83625.1 aluc|OJZ92368.1 anid|CBF75743.1 anig|EHA26605.1 aory|EIT80857.1 asyd|OJJ57519.1 ater|EAU35675.1 atub|OJI86798.1 aver|OJJ05659.1 awen|OJJ40539.1 azon|OJJ49399.1

Group3523: abra|OJJ74789.1 acar|OOF95779.1 acla|EAW10622.1 afis|EAW24945.1 afla|EED55520.1 afum|EAL89602.1 agla|OJJ82173.1 akaw|GAA83624.1 aluc|OJZ92369.1 anid|CBF75615.1 anig|EHA26604.1 aory|EIT80768.1 asyd|OJJ57451.1 ater|EAU35676.1 atub|OJI86799.1 aver|OJJ05727.1 awen|OJJ40540.1 azon|OJJ49400.1

Group3524: abra|OJJ74790.1 acar|OOF95780.1 acla|EAW10621.1 afis|EAW24946.1 afla|EED55519.1 afum|EAL89603.1 agla|OJJ82174.1 akaw|GAA83623.1 aluc|OJZ92370.1 anid|CBF81729.1 anig|EHA26603.1 aory|EIT80898.1 asyd|OJJ59591.1 ater|EAU35678.1 atub|OJI86800.1 aver|OJJ02678.1 awen|OJJ40541.1 azon|OJJ49401.1

Group3525: abra|OJJ74791.1 acar|OOF95781.1 acla|EAW10620.1 afis|EAW24947.1 afla|EED55518.1 afum|EAL89604.1 agla|OJJ82175.1 akaw|GAA83622.1 aluc|OJZ92371.1 anid|CBF81728.1 anig|EHA26602.1 aory|EIT80815.1 asyd|OJJ59590.1 ater|EAU35679.1 atub|OJI86801.1 aver|OJJ02679.1 awen|OJJ40542.1 azon|OJJ49402.1

Group3526: abra|OJJ74792.1 acar|OOF95782.1 acla|EAW10619.1 afis|EAW24948.1 afla|EED55517.1 afum|EAL89605.1 agla|OJJ82176.1 akaw|GAA83621.1 aluc|OJZ92372.1 anid|CBF81726.1 anig|EHA26601.1 aory|EIT80913.1 asyd|OJJ59589.1 ater|EAU35680.1 atub|OJI86802.1 aver|OJJ02680.1 awen|OJJ40543.1 azon|OJJ49403.1

Group3527: abra|OJJ74797.1 acar|OOF95789.1 acla|EAW10609.1 afis|EAW24958.1 afla|EED55505.1 afum|EAL89616.1 agla|OJJ82189.1 akaw|GAA83615.1 aluc|OJZ92378.1 anid|CBF81663.1 anig|EHA26595.1 aory|EIT80867.1 asyd|OJJ59551.1 ater|EAU35697.1 atub|OJI86808.1 aver|OJJ02717.1 awen|OJJ40552.1 azon|OJJ49411.1

Group3528: abra|OJJ74802.1 acar|OOF95795.1 acla|EAW10614.1 afis|EAW24953.1 afla|EED55511.1 afum|EAL89610.1 agla|OJJ82184.1 akaw|GAA83611.1 aluc|OJZ92383.1 anid|CBF81717.1 anig|EHA26592.1 aory|EIT80894.1 asyd|OJJ59584.1 ater|EAU35685.1 atub|OJI86812.1 aver|OJJ02686.1 awen|OJJ40548.1 azon|OJJ49408.1

Group3529: abra|OJJ74804.1 acar|OOF95796.1 acla|EAW10616.1 afis|EAW24951.1 afla|EED55513.1 afum|EAL89608.1 agla|OJJ82187.1 akaw|GAA83609.1 aluc|OJZ92385.1 anid|CBF81721.1 anig|EHA26590.1 aory|EIT80836.1 asyd|OJJ59586.1 atub|OJI86814.1 aver|OJJ02683.1 awen|OJJ40546.1 azon|OJJ49406.1 ater|EAU35683.1

Group3530: abra|OJJ74805.1 acar|OOF95797.1 acla|EAW10617.1 afis|EAW24950.1 afla|EED55514.1 afum|EAL89607.1 agla|OJJ82179.1 akaw|GAA83608.1 aluc|OJZ92386.1 anid|CBF81722.1 anig|EHA26589.1 aory|EIT80921.1 asyd|OJJ59587.1 ater|EAU35682.1 atub|OJI86815.1 aver|OJJ02682.1 awen|OJJ40545.1 azon|OJJ49405.1

Group3531: abra|OJJ74806.1 acar|OOF95798.1 acla|EAW10618.1 afis|EAW24949.1 afla|EED55516.1 afum|EAL89606.1 agla|OJJ82177.1 akaw|GAA83606.1 aluc|OJZ92387.1 anid|CBF81724.1 anig|EHA26588.1 aory|EIT80820.1 asyd|OJJ59588.1 ater|EAU35681.1 atub|OJI86817.1 aver|OJJ02681.1 awen|OJJ40544.1 azon|OJJ49404.1

Group3532: abra|OJJ74808.1 acar|OOF95800.1 acla|EAW10606.1 afis|EAW24961.1 afla|EED55501.1 afum|EAL89619.1 agla|OJJ82196.1 akaw|GAA83604.1 aluc|OJZ92389.1 anid|CBF81658.1 anig|EHA26585.1 aory|EIT80766.1 asyd|OJJ59548.1 ater|EAU35694.1 atub|OJI86819.1 aver|OJJ02720.1 awen|OJJ40554.1 azon|OJJ49416.1

Group3533: abra|OJJ74830.1 acar|OOF95817.1 acla|EAW11373.1 afis|EAW18122.1 afla|EED58048.1 afum|EAL87480.1 agla|OJJ87016.1 akaw|GAA83590.1 aluc|OJZ92407.1 anid|CBF75358.1 anig|EHA26558.1 aory|EIT79720.1 asyd|OJJ59851.1 ater|EAU29689.1 atub|OJI86835.1 aver|OJJ02436.1 awen|OJJ35524.1 azon|OJJ46961.1

Group3534: abra|OJJ74831.1 acar|OOF95819.1 acla|EAW11372.1 afis|EAW18123.1 afla|EED58049.1 afum|EAL87481.1 agla|OJJ87017.1 akaw|GAA83589.1 aluc|OJZ92408.1 anid|CBF80348.1 anig|EHA26557.1 aory|EIT79773.1 asyd|OJJ54832.1 ater|EAU29688.1 atub|OJI86836.1 aver|OJJ07329.1 awen|OJJ35525.1 azon|OJJ46959.1

Group3535: abra|OJJ74840.1 acar|OOF95843.1 acla|EAW09369.1 afis|EAW21766.1 afla|EED46332.1 afum|EAL92796.1 agla|OJJ84385.1 akaw|GAA83581.1 aluc|OJZ92416.1 anid|CBF83756.1 anig|EHA26550.1 aory|EIT81192.1 asyd|OJJ60860.1 ater|EAU38449.1 atub|OJI86846.1 aver|OJJ00185.1 awen|OJJ34049.1 azon|OJJ48182.1

Group3536: abra|OJJ74841.1 acar|OOF94168.1 acla|EAW08544.1 afis|EAW23581.1 afla|EED57489.1 afum|EAL87884.1 agla|OJJ86848.1 akaw|GAA83580.1 aluc|OJZ92417.1 anid|CBF89912.1 anig|EHA26549.1 aory|EIT81889.1 asyd|OJJ62129.1 ater|EAU35371.1 atub|OJI86847.1 aver|OJI96238.1 awen|OJJ39909.1 azon|OJJ45885.1

Group3537: abra|OJJ74847.1 acar|OOF95848.1 acla|EAW08541.1 afis|EAW23578.1 afla|EED57477.1 afum|EAL87887.1 agla|OJJ86843.1 akaw|GAA83575.1 aluc|OJZ92422.1 anid|CBF89598.1 anig|EHA26545.1 aory|EIT80110.1 asyd|OJJ62311.1 ater|EAU35366.1 atub|OJI86852.1 aver|OJI96419.1 awen|OJJ39906.1 azon|OJJ45883.1

Group3538: abra|OJJ74848.1 acar|OOF95849.1 acla|EAW08540.1 afis|EAW23577.1 afla|EED57476.1 afum|EAL87888.1 agla|OJJ86838.1 akaw|GAA83574.1 aluc|OJZ92423.1 anid|CBF89600.1 anig|EHA26544.1 aory|EIT80147.1 asyd|OJJ62310.1 ater|EAU35367.1 atub|OJI86853.1 aver|OJI96418.1 awen|OJJ39905.1 azon|OJJ45882.1

Group3539: abra|OJJ74852.1 acar|OOF95851.1 acla|EAW08535.1 afis|EAW23573.1 afla|EED50962.1 afum|EAL87892.2 agla|OJJ86833.1 akaw|GAA83570.1 aluc|OJZ92428.1 anid|CBF89610.1 anig|EHA26539.1 aory|EIT77015.1 asyd|OJJ62305.1 ater|EAU35343.1 atub|OJI86858.1 aver|OJI96413.1 awen|OJJ39901.1 azon|OJJ42316.1

Group3540: abra|OJJ74854.1 acar|OOF95854.1 acla|EAW08531.1 afis|EAW23570.1 afla|EED57463.1 afum|EAL87897.2 agla|OJJ86831.1 akaw|GAA83568.1 aluc|OJZ92430.1 anid|CBF89617.1 anig|EHA26536.1 aory|EIT80101.1 asyd|OJJ62302.1 ater|EAU35346.1 atub|OJI86860.1 aver|OJI96411.1 awen|OJJ39899.1 azon|OJJ45876.1

Group3541: abra|OJJ74856.1 acar|OOF95857.1 acla|EAW08528.1 afis|EAW23568.1 afla|EED57459.1 afum|EAL87899.1 agla|OJJ86829.1 akaw|GAA83566.1 aluc|OJZ92432.1 anid|CBF89621.1 anig|EHA26534.1 aory|EIT80063.1 asyd|OJJ62300.1 ater|EAU35349.1 atub|OJI86862.1 aver|OJI96408.1 awen|OJJ39896.1 azon|OJJ45874.1

Group3542: abra|OJJ74859.1 acar|OOF95859.1 acla|EAW08521.1 afis|EAW23559.1 afla|EED57454.1 afum|EAL87907.1 agla|OJJ86825.1 akaw|GAA83561.1 aluc|OJZ92437.1 anid|CBF89634.1 anig|EHA26531.1 aory|EIT80133.1 asyd|OJJ62295.1 ater|EAU35342.1 atub|OJI86866.1 aver|OJI96402.1 awen|OJJ39893.1 azon|OJJ43217.1

Group3543: abra|OJJ74860.1 acar|OOF95860.1 acla|EAW08520.1 afis|EAW23558.1 afla|EED57453.1 afum|EAL87908.1 agla|OJJ86826.1 akaw|GAA83560.1 aluc|OJZ92438.1 anid|CBF89636.1 anig|EHA26530.1 aory|EIT80071.1 asyd|OJJ62294.1 ater|EAU35341.1 atub|OJI86867.1 aver|OJI96401.1 awen|OJJ39891.1 azon|OJJ43218.1

Group3544: abra|OJJ74862.1 acar|OOF95862.1 acla|EAW08519.1 afis|EAW23557.1 afla|EED57452.1 afum|EAL87909.1 agla|OJJ86819.1 akaw|GAA83558.1 aluc|OJZ92440.1 anid|CBF89638.1 anig|EHA26527.1 aory|EIT80146.1 asyd|OJJ62292.1 ater|EAU35340.1 atub|OJI86869.1 aver|OJI96400.1 awen|OJJ39888.1 azon|OJJ43219.1

Group3545: abra|OJJ74867.1 acar|OOF95868.1 acla|EAW08513.1 afis|EAW23551.1 afla|EED57446.1 afum|EAL87915.1 agla|OJJ86807.1 akaw|GAA83553.1 aluc|OJZ92445.1 anid|CBF89646.1 anig|EHA26523.1 aory|EIT80111.1 asyd|OJJ62286.1 ater|EAU35335.1 atub|OJI86873.1 aver|OJI96394.1 awen|OJJ39882.1 azon|OJJ43223.1

Group3546: abra|OJJ74869.1 acar|OOF95870.1 acla|EAW08511.1 afis|EAW23549.1 afla|EED57444.1 afum|EAL87917.2 agla|OJJ86805.1 akaw|GAA83551.1 aluc|OJZ92447.1 anid|CBF89650.1 anig|EHA26521.1 aory|EIT80076.1 asyd|OJJ62284.1 ater|EAU35334.1 atub|OJI86875.1 aver|OJI96392.1 awen|OJJ39880.1 azon|OJJ43224.1

Group3547: abra|OJJ74872.1 acar|OOF95873.1 acla|EAW08508.1 afis|EAW23546.1 afla|EED57441.1 afum|EAL87920.1 agla|OJJ86800.1 akaw|GAA83548.1 aluc|OJZ92450.1 anid|CBF89654.1 anig|EHA26518.1 aory|EIT80061.1 asyd|OJJ62282.1 ater|EAU35331.1 atub|OJI86878.1 aver|OJI96390.1 awen|OJJ39877.1 azon|OJJ43226.1

Group3548: abra|OJJ74874.1 acar|OOF95875.1 acla|EAW08498.1 afis|EAW23543.1 afla|EED57440.1 afum|EAL87923.2 agla|OJJ86797.1 akaw|GAA83546.1 aluc|OJZ92452.1 anid|CBF89657.1 anig|EHA26516.1 aory|EIT80096.1 asyd|OJJ62279.1 ater|EAU35329.1 atub|OJI86880.1 aver|OJI96388.1 awen|OJJ39875.1 azon|OJJ43227.1

Group3549: abra|OJJ74877.1 acar|OOF95880.1 acla|EAW08493.1 afis|EAW23539.1 afla|EED57437.1 afum|EAL87927.1 agla|OJJ86794.1 akaw|GAA83543.1 aluc|OJZ92454.1 anid|CBF89665.1 anig|EHA26513.1 aory|EIT80117.1 asyd|OJJ62275.1 ater|EAU35325.1 atub|OJI86883.1 aver|OJI96384.1 awen|OJJ39870.1 azon|OJJ43231.1

Group3550: abra|OJJ74879.1 acar|OOF95882.1 acla|EAW08491.1 afis|EAW23537.1 afla|EED57435.1 afum|EAL87929.1 agla|OJJ86792.1 akaw|GAA83541.1 aluc|OJZ92456.1 anid|CBF89668.1 anig|EHA26511.1

aory|EIT80166.1 asyd|OJJ62273.1 ater|EAU35323.1 atub|OJI86885.1 aver|OJI96382.1 awen|OJJ39867.1 azon|OJJ43233.1

Group3551: abra|OJJ74881.1 acar|OOF95884.1 acla|EAW08489.1 afis|EAW23535.1 afla|EED57432.1 afum|EAL87931.1 agla|OJJ86790.1 akaw|GAA83539.1 aluc|OJZ92458.1 anid|CBF89672.1 anig|EHA26509.1 aory|EIT80162.1 asyd|OJJ62271.1 ater|EAU35321.1 atub|OJI86887.1 aver|OJI96380.1 awen|OJJ39865.1 azon|OJJ43235.1

Group3552: abra|OJJ74882.1 acar|OOF95886.1 acla|EAW08487.1 afis|EAW23533.1 afla|EED57430.1 afum|EAL87933.1 agla|OJJ86782.1 akaw|GAA83537.1 aluc|OJZ92459.1 anid|CBF89678.1 anig|EHA26508.1 aory|EIT80077.1 asyd|OJJ62269.1 ater|EAU35319.1 atub|OJI86888.1 aver|OJI96378.1 awen|OJJ39862.1 azon|OJJ43236.1

Group3553: abra|OJJ74884.1 akaw|GAA83535.1 aluc|OJZ92461.1 anig|EHA26506.1 atub|OJI86890.1 acar|OOF95889.1 acla|EAW08483.1 afis|EAW23529.1 afla|EED57428.1 afum|EAL87937.1 agla|OJJ86778.1 anid|CBF89683.1 asyd|OJJ62266.1 ater|EAU35317.1 aver|OJI96375.1 awen|OJJ39858.1 azon|OJJ43239.1 aory|EIT80152.1

Group3554: abra|OJJ74888.1 acar|OOF95893.1 acla|EAW08479.1 afis|EAW23525.1 afla|EED57424.1 afum|EAL87940.1 agla|OJJ86774.1 akaw|GAA83531.1 aluc|OJZ92465.1 anid|CBF89691.1 anig|EHA26502.1 aory|EIT80080.1 asyd|OJJ62262.1 ater|EAU35313.1 atub|OJI86894.1 aver|OJI96371.1 awen|OJJ39854.1 azon|OJJ43245.1

Group3555: abra|OJJ74889.1 acar|OOF95892.1 acla|EAW08478.1 afis|EAW23524.1 afla|EED50993.1 afum|EAL87941.1 agla|OJJ86773.1 akaw|GAA83530.1 aluc|OJZ92466.1 anid|CBF76066.1 anig|EHA26501.1 aory|EIT76998.1 asyd|OJJ60396.1 ater|EAU31002.1 atub|OJI86895.1 aver|OJI99725.1 awen|OJJ39853.1 azon|OJJ50688.1

Group3556: abra|OJJ74897.1 acar|OOF95897.1 acla|EAW08475.1 afis|EAW23521.1 afum|EAL87943.1 agla|OJJ86768.1 akaw|GAA83524.1 aluc|OJZ92471.1 anig|EHA26497.1 ater|EAU35309.1 atub|OJI86901.1 awen|OJJ39846.1 afla|EED57421.1 aory|EIT80149.1 aver|OJI96367.1 azon|OJJ43248.1 anid|CBF89696.1 asyd|OJJ62258.1

Group3557: abra|OJJ74901.1 acar|OOF95900.1 acla|EAW08472.1 afis|EAW23518.1 afla|EED57418.1 afum|EAL87946.1 agla|OJJ86765.1 akaw|GAA83521.1 aluc|OJZ92474.1 anid|CBF89702.1 anig|EHA26495.1 aory|EIT80062.1 asyd|OJJ62254.1 atub|OJI86905.1 aver|OJI96364.1 awen|OJJ39843.1 azon|OJJ43250.1 ater|EAU35306.1

Group3558: abra|OJJ74905.1 acar|OOF90301.1 acla|EAW08467.1 afis|EAW23514.1 afla|EED57413.1 afum|EAL87949.1 agla|OJJ86762.1 akaw|GAA83517.1 aluc|OJZ92478.1 anig|EHA26492.1 aory|EIT80167.1 asyd|OJJ59973.1 ater|EAU35303.1 atub|OJI86909.1 aver|OJJ02318.1 awen|OJJ39839.1 azon|OJJ43253.1 anid|CBF75148.1

Group3559: abra|OJJ74907.1 acar|OOF90299.1 acla|EAW08465.1 afis|EAW23510.1 afla|EED57410.1 afum|EAL87951.1 agla|OJJ86760.1 akaw|GAA83515.1 aluc|OJZ92480.1 anid|CBF89718.1 anig|EHA26490.1 aory|EIT80124.1 asyd|OJJ62245.1 ater|EAU35298.1 atub|OJI86911.1 aver|OJI96355.1 awen|OJJ39836.1 azon|OJJ43256.1

Group3560: abra|OJJ74909.1 acar|OOF90294.1 acla|EAW08463.1 afis|EAW23508.1 afla|EED57408.1 afum|EAL87953.1 agla|OJJ86758.1 akaw|GAA83513.1 aluc|OJZ92482.1 anid|CBF89714.1 anig|EHA26489.1 aory|EIT80104.1 asyd|OJJ62247.1 ater|EAU35296.1 atub|OJI86913.1 aver|OJI96357.1 awen|OJJ39834.1 azon|OJJ43258.1

Group3561: abra|OJJ74910.1 acar|OOF90297.1 acla|EAW08462.1 afis|EAW23507.1 afla|EED57407.1 afum|EAL87954.1 agla|OJJ86757.1 akaw|GAA83512.1 aluc|OJZ92483.1 anid|CBF89712.1 anig|EHA26488.1 aory|EIT80089.1 asyd|OJJ62248.1 ater|EAU35295.1 atub|OJI86914.1 aver|OJI96358.1 awen|OJJ39833.1 azon|OJJ43259.1

Group3562: abra|OJJ74916.1 acar|OOF95908.1 acla|EAW08454.1 afis|EAW23501.1 afla|EED57399.1 afum|EAL87961.1 agla|OJJ86746.1 akaw|GAA83506.1 aluc|OJZ92488.1 anid|CBF89733.1 anig|EHA26484.1 aory|EIT80085.1 asyd|OJJ62233.1 ater|EAU35288.1 atub|OJI86921.1 aver|OJI96342.1 awen|OJJ39824.1 azon|OJJ47948.1

Group3563: abra|OJJ74919.1 acar|OOF95910.1 afis|EAW23497.1 afla|EED57397.1 afum|EAL87966.1 agla|OJJ86744.1 akaw|GAA83504.1 aluc|OJZ92490.1 anid|CBF89737.1 anig|EHA26482.1 aory|EIT80105.1 asyd|OJJ62231.1 ater|EAU35286.1 atub|OJI86923.1 aver|OJI96340.1 awen|OJJ39822.1 azon|OJJ47946.1 acla|EAW08452.1

Group3564: abra|OJJ74922.1 acar|OOF95912.1 acla|EAW08449.1 afis|EAW23494.1 afla|EED57394.1 afum|EAL87969.1 agla|OJJ86741.1 akaw|GAA83501.1 aluc|OJZ92493.1 anid|CBF89742.1 anig|EHA26479.1 aory|EIT80159.1 asyd|OJJ62228.1 ater|EAU35283.1 atub|OJI86926.1 aver|OJI96337.1 awen|OJJ39819.1 azon|OJJ47943.1

Group3565: abra|OJJ74923.1 acar|OOF95914.1 acla|EAW08448.1 afis|EAW23493.1 afla|EED57393.1 afum|EAL87970.1 agla|OJJ86740.1 akaw|GAA83500.1 aluc|OJZ92494.1 anid|CBF89744.1 anig|EHA26478.1 aory|EIT80132.1 asyd|OJJ62227.1 ater|EAU35282.1 atub|OJI86927.1 aver|OJI96336.1 awen|OJJ39818.1 azon|OJJ47941.1

Group3566: abra|OJJ74924.1 acar|OOF95915.1 acla|EAW08447.1 afis|EAW23492.1 afla|EED57391.1 afum|EAL87971.1 agla|OJJ86737.1 akaw|GAA83499.1 aluc|OJZ92495.1 anid|CBF89746.1 anig|EHA26477.1 aory|EIT80165.1 asyd|OJJ62225.1 ater|EAU35279.1 atub|OJI86928.1 aver|OJI96334.1 awen|OJJ39816.1 azon|OJJ47939.1

Group3567: abra|OJJ74925.1 acar|OOF95916.1 acla|EAW08446.1 afis|EAW23491.1 afla|EED57390.1 afum|EAL87972.1 agla|OJJ86736.1 akaw|GAA83498.1 aluc|OJZ92496.1 anid|CBF89747.1 anig|EHA26476.1 aory|EIT80058.1 asyd|OJJ62224.1 ater|EAU35278.1 atub|OJI86929.1 aver|OJI96333.1 awen|OJJ39815.1 azon|OJJ47938.1

Group3568: abra|OJJ74928.1 acar|OOF95917.1 acla|EAW08442.1 afis|EAW23487.1 afla|EED57386.1 afum|EAL87976.1 agla|OJJ86732.1 akaw|GAA83494.1 aluc|OJZ92499.1 anid|CBF89755.1 anig|EHA26473.1 aory|EIT80123.1 asyd|OJJ62220.1 ater|EAU35274.1 atub|OJI86932.1 aver|OJI96330.1 awen|OJJ39810.1 azon|OJJ47934.1

Group3569: abra|OJJ74930.1 acar|OOF95920.1 acla|EAW08439.1 afis|EAW23484.1 afla|EED57379.1 afum|EAL87979.1 agla|OJJ86720.1 akaw|GAA83491.1 aluc|OJZ92502.1 anid|CBF89760.1 anig|EHA26471.1 aory|EIT80056.1 asyd|OJJ62217.1 ater|EAU35271.1 atub|OJI86934.1 aver|OJI96328.1 awen|OJJ39808.1 azon|OJJ47929.1

Group3570: abra|OJJ74931.1 acar|OOF95921.1 acla|EAW08438.1 afis|EAW23483.1 afla|EED57378.1 afum|EAL87980.1 agla|OJJ86719.1 akaw|GAA83490.1 aluc|OJZ92503.1 anid|CBF89762.1 anig|EHA26470.1 aory|EIT80160.1 asyd|OJJ62216.1 ater|EAU36899.1 atub|OJI86935.1 aver|OJI96327.1 awen|OJJ39807.1 azon|OJJ47928.1

Group3571: abra|OJJ74932.1 acar|OOF95922.1 acla|EAW08437.1 afis|EAW23482.1 afla|EED57377.1 afum|EAL87982.1 agla|OJJ86718.1 akaw|GAA83488.1 aluc|OJZ92505.1 anid|CBF89764.1 anig|EHA26468.1 aory|EIT80168.1 asyd|OJJ62215.1 ater|EAU36900.1 atub|OJI86937.1 aver|OJI96326.1 awen|OJJ39806.1 azon|OJJ47927.1

Group3572: abra|OJJ74935.1 acar|OOF97377.1 acla|EAW08436.1 afis|EAW23481.1 afla|EED57376.1 afum|EAL87983.1 agla|OJJ86717.1 akaw|GAA83485.1 aluc|OJZ92508.1 anid|CBF89766.1 anig|EHA26465.1 aory|EIT80075.1 asyd|OJJ62214.1 ater|EAU36901.1 atub|OJI86941.1 aver|OJI96325.1 awen|OJJ39805.1 azon|OJJ47926.1

Group3573: abra|OJJ74938.1 acar|OOF97376.1 acla|EAW08433.1 afis|EAW23476.1 afla|EED57372.1 afum|EAL87989.2 agla|OJJ86715.1 akaw|GAA83482.1 aluc|OJZ92513.1 anid|CBF89771.1 anig|EHA26462.1 aory|EIT80114.1 asyd|OJJ62212.1 ater|EAU36904.1 atub|OJI86944.1 aver|OJI96323.1 awen|OJJ39802.1 azon|OJJ47923.1

Group3574: abra|OJJ74939.1 acar|OOF97375.1 acla|EAW08431.1 afis|EAW23474.1 afla|EED57370.1 afum|EAL87991.2 agla|OJJ86713.1 akaw|GAA83481.1 aluc|OJZ92514.1 anid|CBF89775.1 anig|EHA26461.1 aory|EIT80156.1 asyd|OJJ62209.1 ater|EAU36906.1 atub|OJI86945.1 aver|OJI96321.1 awen|OJJ39800.1 azon|OJJ47922.1

Group3575: abra|OJJ74941.1 acar|OOF97374.1 acla|EAW08429.1 afis|EAW23473.1 afla|EED57369.1 afum|EAL87993.1 agla|OJJ86712.1 akaw|GAA83480.1 aluc|OJZ92515.1 anid|CBF89777.1 anig|EHA26460.1 aory|EIT80066.1 asyd|OJJ62208.1 ater|EAU36907.1 atub|OJI86948.1 aver|OJI96320.1 awen|OJJ39799.1 azon|OJJ47920.1

Group3576: abra|OJJ74943.1 acar|OOF97372.1 acla|EAW08427.1 afis|EAW23471.1 afla|EED57367.1 afum|EAL87995.1 agla|OJJ86710.1 akaw|GAA83478.1 aluc|OJZ92517.1 anid|CBF89782.1 anig|EHA26458.1 aory|EIT80170.1 asyd|OJJ62205.1 ater|EAU36909.1 atub|OJI86950.1 aver|OJI96317.1 awen|OJJ39796.1 azon|OJJ47918.1

Group3577: abra|OJJ74945.1 acar|OOF97371.1 acla|EAW08426.1 afis|EAW23442.1 afla|EED57366.1 afum|EAL88021.1 agla|OJJ86709.1 akaw|GAA83477.1 aluc|OJZ92518.1 anid|CBF89784.1 anig|EHA26457.1 aory|EIT80053.1 asyd|OJJ62204.1 ater|EAU36910.1 atub|OJI86952.1 aver|OJI96316.1 awen|OJJ39790.1 azon|OJJ47917.1

Group3578: abra|OJJ74946.1 acar|OOF97370.1 acla|EAW08425.1 afis|EAW23444.1 afla|EED57365.1 afum|EAL88020.1 agla|OJJ86707.1 akaw|GAA83476.1 aluc|OJZ92519.1 anid|CBF89786.1 anig|EHA26456.1 aory|EIT80115.1 asyd|OJJ62203.1 ater|EAU36911.1 atub|OJI86953.1 aver|OJI96315.1 awen|OJJ39786.1 azon|OJJ47916.1

Group3579: abra|OJJ74947.1 acar|OOF97369.1 acla|EAW08424.1 afis|EAW23445.1 afla|EED57364.1 afum|EAL88019.1 agla|OJJ86706.1 akaw|GAA83475.1 aluc|OJZ92520.1 anid|CBF89788.1 anig|EHA26455.1

aory|EIT80057.1 asyd|OJJ62202.1 ater|EAU36912.1 atub|OJI86954.1 aver|OJI96314.1 awen|OJJ39785.1 azon|OJJ47915.1

Group3580: abra|OJJ74949.1 acar|OOF97367.1 acla|EAW08422.1 afis|EAW23447.1 afla|EED57362.1 afum|EAL88018.1 agla|OJJ86702.1 akaw|GAA83473.1 aluc|OJZ92522.1 anid|CBF89791.1 anig|EHA26453.1 aory|EIT80100.1 asyd|OJJ62200.1 ater|EAU36913.1 atub|OJI86956.1 aver|OJI96312.1 awen|OJJ39782.1 azon|OJJ47913.1

Group3581: abra|OJJ74952.1 acar|OOF97362.1 acla|EAW08421.1 afis|EAW23448.1 afla|EED57359.1 afum|EAL88017.1 agla|OJJ86701.1 akaw|GAA83471.1 aluc|OJZ92524.1 anid|CBF89793.1 anig|EHA26450.1 aory|EIT80142.1 asyd|OJJ62199.1 ater|EAU36914.1 atub|OJI86959.1 aver|OJI96311.1 awen|OJJ39781.1 azon|OJJ47912.1

Group3582: abra|OJJ74953.1 acar|OOF97360.1 acla|EAW08420.1 afis|EAW23449.1 afla|EED57358.1 afum|EBA27332.1 agla|OJJ86700.1 akaw|GAA83470.1 aluc|OJZ92525.1 anid|CBF89795.1 anig|EHA26449.1 aory|EIT80082.1 asyd|OJJ62198.1 ater|EAU36916.1 atub|OJI86961.1 awen|OJJ39780.1 azon|OJJ47911.1 aver|OJI96310.1

Group3583: abra|OJJ74954.1 acar|OOF97359.1 acla|EAW08418.1 afis|EAW23451.1 afla|EED57352.1 afum|EAL88015.1 agla|OJJ86699.1 akaw|GAA83469.1 aluc|OJZ92526.1 anid|CBF89799.1 anig|EHA26448.1 aory|EIT80134.1 asyd|OJJ62196.1 ater|EAU36917.1 atub|OJI86962.1 aver|OJI96308.1 awen|OJJ39778.1 azon|OJJ47908.1

Group3584: abra|OJJ74956.1 acar|OOF97357.1 acla|EAW08416.1 afis|EAW23453.1 afla|EED57350.1 afum|EAL88013.1 agla|OJJ86697.1 akaw|GAA83467.1 aluc|OJZ92528.1 anid|CBF89802.1 anig|EHA26447.1 aory|EIT80106.1 asyd|OJJ62194.1 ater|EAU36919.1 atub|OJI86964.1 aver|OJI96306.1 awen|OJJ39776.1 azon|OJJ47906.1

Group3585: abra|OJJ74958.1 acar|OOF97355.1 acla|EAW08413.1 afis|EAW23456.1 afla|EED57345.1 afum|EBA27330.1 akaw|GAA83464.1 aluc|OJZ92530.1 anid|CBF89808.1 anig|EHA26444.1 aory|EIT80113.1 asyd|OJJ62191.1 ater|EAU36922.1 atub|OJI86967.1 aver|OJI96301.1 awen|OJJ39773.1 azon|OJJ47902.1 agla|OJJ86693.1

Group3586: abra|OJJ74959.1 acar|OOF97354.1 acla|EAW08412.1 afis|EAW23457.1 afla|EED57344.1 afum|EBA27329.1 agla|OJJ86692.1 akaw|GAA83463.1 aluc|OJZ92531.1 anid|CBF89810.1 anig|EHA26443.1 aory|EIT80054.1 asyd|OJJ62190.1 ater|EAU36923.1 atub|OJI86968.1 aver|OJI96300.1 awen|OJJ39772.1 azon|OJJ47903.1

Group3587: abra|OJJ74965.1 acar|OOF97349.1 acla|EAW08409.1 afis|EAW23460.1 afla|EED57340.1 afum|EAL88006.1 agla|OJJ86689.1 akaw|GAA83461.1 aluc|OJZ92535.1 anid|CBF89820.1 anig|EHA26440.1 aory|EIT74102.1 asyd|OJJ62187.1 ater|EAU37473.1 atub|OJI86971.1 aver|OJI96296.1 awen|OJJ39770.1 azon|OJJ47900.1

Group3588: abra|OJJ74966.1 acar|OOF97348.1 acla|EAW08408.1 afis|EAW23461.1 afla|EED57339.1 afum|EAL88005.1 agla|OJJ86688.1 akaw|GAA83460.1 aluc|OJZ92536.1 anid|CBF89850.1 anig|EHA26439.1 aory|EIT74042.1 asyd|OJJ62186.1 ater|EAU37474.1 atub|OJI86972.1 aver|OJI96295.1 awen|OJJ39769.1 azon|OJJ47899.1

Group3589: abra|OJJ74967.1 acar|OOF97346.1 acla|EAW08407.1 afis|EAW23462.1 afla|EED57338.1 afum|EAL88004.1 agla|OJJ86687.1 akaw|GAA83459.1 aluc|OJZ92537.1 anid|CBF89848.1 anig|EHA26438.1 aory|EIT74136.1 asyd|OJJ62185.1 ater|EAU37475.1 atub|OJI86973.1 aver|OJI96294.1 awen|OJJ39768.1 azon|OJJ47898.1

Group3590: abra|OJJ74969.1 acar|OOF97345.1 acla|EAW08405.1 afis|EAW23464.1 afla|EED57335.1 afum|EAL88002.1 agla|OJJ86685.1 akaw|GAA83457.1 aluc|OJZ92539.1 anid|CBF89843.1 anig|EHA26437.1 aory|EIT74036.1 asyd|OJJ62183.1 ater|EAU37480.1 atub|OJI86975.1 aver|OJI96292.1 awen|OJJ39766.1 azon|OJJ47896.1

Group3591: abra|OJJ74975.1 acar|OOF97344.1 acla|EAW08404.1 afis|EAW23465.1 afla|EED57334.1 afum|EAL88001.1 agla|OJJ86684.1 akaw|GAA83456.1 aluc|OJZ92540.1 anid|CBF89841.1 anig|EHA26436.1 aory|EIT73981.1 asyd|OJJ62182.1 ater|EAU37481.1 atub|OJI86976.1 aver|OJI96291.1 awen|OJJ39765.1 azon|OJJ47895.1

Group3592: abra|OJJ74976.1 acar|OOF97343.1 acla|EAW08403.1 afis|EAW23466.1 afla|EED57333.1 afum|EAL88000.1 agla|OJJ86683.1 akaw|GAA83455.1 aluc|OJZ92541.1 anid|CBF89839.1 anig|EHA26435.1 aory|EIT74149.1 asyd|OJJ62181.1 ater|EAU37482.1 atub|OJI86977.1 aver|OJI96290.1 awen|OJJ39764.1 azon|OJJ47894.1

Group3593: abra|OJJ74980.1 acar|OOF97340.1 acla|EAW08400.1 afis|EAW23469.1 afla|EED57331.1 afum|EAL87997.1 agla|OJJ86680.1 akaw|GAA83452.1 aluc|OJZ92544.1 anid|CBF89833.1 anig|EHA26432.1 aory|EIT74122.1 asyd|OJJ62178.1 ater|EAU37485.1 atub|OJI86980.1 aver|OJI96287.1 awen|OJJ39760.1 azon|OJJ47889.1

Group3594: abra|OJJ74981.1 acar|OOF97339.1 acla|EAW08399.1 afis|EAW23470.1 afla|EED57330.1 afum|EAL87996.1 agla|OJJ86679.1 akaw|GAA83451.1 aluc|OJZ92545.1 anid|CBF89831.1 anig|EHA26431.1 aory|EIT74162.1 asyd|OJJ62177.1 ater|EAU37486.1 atub|OJI86981.1 aver|OJI96286.1 awen|OJJ39759.1 azon|OJJ47888.1

Group3596: abra|OJJ74988.1 acar|OOF97333.1 acla|EAW08394.1 afis|EAW23437.1 afla|EED57327.1 afum|EAL88025.1 agla|OJJ86674.1 akaw|GAA83446.1 aluc|OJZ92550.1 anid|CBF89852.1 anig|EHA26427.1 aory|EIT74016.1 asyd|OJJ62176.1 ater|EAU37489.1 atub|OJI86987.1 aver|OJI96285.1 awen|OJJ39754.1 azon|OJJ47881.1

Group3597: abra|OJJ74991.1 acar|OOF97329.1 acla|EAW08392.1 afis|EAW23423.1 afla|EED57322.1 afum|EAL88033.1 agla|OJJ86673.1 akaw|GAA83444.1 aluc|OJZ92555.1 anid|CBF89824.1 anig|EHA26424.1 aory|EIT74106.1 asyd|OJJ62171.1 ater|EAU37494.1 atub|OJI86991.1 aver|OJI96283.1 awen|OJJ39753.1 azon|OJJ47880.1

Group3598: abra|OJJ74993.1 acar|OOF97326.1 acla|EAW08390.1 afis|EAW23419.1 afla|EED57321.1 afum|EAL88050.1 agla|OJJ86644.1 akaw|GAA83442.1 aluc|OJZ92559.1 anid|CBF89854.1 anig|EHA26421.1 aory|EIT74073.1 asyd|OJJ62161.1 ater|EAU37503.1 atub|OJI86995.1 aver|OJI96271.1 awen|OJJ39752.1 azon|OJJ47878.1

Group3599: abra|OJJ74996.1 acar|OOF97324.1 acla|EAW08388.1 afis|EAW23417.1 afla|EED57318.1 afum|EAL88052.2 agla|OJJ86634.1 akaw|GAA88620.1 aluc|OJZ92562.1 anid|CBF89861.1 anig|EHA26419.1 aory|EIT73973.1 asyd|OJJ62158.1 ater|EAU37505.1 atub|OJI86998.1 aver|OJI96268.1 awen|OJJ39750.1 azon|OJJ47876.1

Group3600: abra|OJJ74997.1 acar|OOF97323.1 acla|EAW08387.1 afis|EAW23416.1 afla|EED57317.1 afum|EAL88053.2 agla|OJJ86633.1 akaw|GAA88621.1 aluc|OJZ92563.1 anid|CBF89862.1 anig|EHA26418.1 aory|EIT74065.1 asyd|OJJ62157.1 ater|EAU37506.1 atub|OJI86999.1 aver|OJI96267.1 awen|OJJ39749.1 azon|OJJ47875.1

Group3601: abra|OJJ74999.1 acar|OOF97322.1 acla|EAW08386.1 afis|EAW23415.1 afla|EED57316.1 afum|EAL88054.1 agla|OJJ86632.1 akaw|GAA88622.1 aluc|OJZ92564.1 anid|CBF89864.1 anig|EHA26417.1 aory|EIT73976.1 asyd|OJJ62156.1 ater|EAU37507.1 atub|OJI87000.1 aver|OJI96266.1 awen|OJJ39748.1 azon|OJJ47874.1

Group3602: abra|OJJ75000.1 acar|OOF97321.1 acla|EAW08385.1 afis|EAW23414.1 afla|EED57315.1 afum|EAL88055.1 agla|OJJ86631.1 akaw|GAA88623.1 aluc|OJZ92565.1 anid|CBF89865.1 anig|EHA26416.1 aory|EIT74071.1 asyd|OJJ62155.1 ater|EAU37508.1 atub|OJI87001.1 aver|OJI96265.1 awen|OJJ39747.1 azon|OJJ47873.1

Group3603: abra|OJJ75002.1 acar|OOF90037.1 acla|EAW08383.1 afis|EAW23412.1 afla|EED57313.1 afum|EAL88057.1 agla|OJJ86629.1 akaw|GAA88625.1 aluc|OJZ92567.1 anid|CBF89869.1 anig|EHA26414.1 aory|EIT74133.1 asyd|OJJ62153.1 ater|EAU37510.1 atub|OJI87003.1 aver|OJI96263.1 awen|OJJ39745.1 azon|OJJ47871.1

Group3604: abra|OJJ75003.1 acar|OOF90036.1 acla|EAW08382.1 afis|EAW23411.1 afla|EED57312.1 afum|EAL88058.1 agla|OJJ86628.1 akaw|GAA88626.1 aluc|OJZ92568.1 anid|CBF89871.1 anig|EHA26413.1 aory|EIT74029.1 asyd|OJJ62152.1 ater|EAU37511.1 atub|OJI87004.1 aver|OJI96262.1 awen|OJJ39744.1 azon|OJJ47870.1

Group3606: abra|OJJ75005.1 acar|OOF97320.1 acla|EAW08380.1 afis|EAW23409.1 afla|EED57309.1 afum|EAL88061.2 agla|OJJ86625.1 akaw|GAA88628.1 aluc|OJZ92570.1 anid|CBF89873.1 anig|EHA26411.1 aory|EIT74087.1 asyd|OJJ62150.1 ater|EAU37515.1 atub|OJI87006.1 aver|OJI96260.1 awen|OJJ39742.1 azon|OJJ47868.1

Group3607: abra|OJJ75006.1 acar|OOF97319.1 acla|EAW08379.1 afis|EAW23408.1 afla|EED57308.1 afum|EAL88062.1 agla|OJJ86624.1 akaw|GAA88629.1 aluc|OJZ92571.1 anid|CBF89874.1 anig|EHA26410.1 aory|EIT74117.1 asyd|OJJ62149.1 ater|EAU37512.1 atub|OJI87007.1 aver|OJI96259.1 awen|OJJ39735.1 azon|OJJ47867.1

Group3608: abra|OJJ75007.1 acar|OOF97318.1 acla|EAW08378.1 afis|EAW23407.1 afla|EED57307.1 afum|EAL88063.2 agla|OJJ86623.1 akaw|GAA88630.1 aluc|OJZ92572.1 anid|CBF89876.1 anig|EHA26409.1 aory|EIT74022.1 asyd|OJJ62148.1 ater|EAU37513.1 atub|OJI87008.1 aver|OJI96258.1 awen|OJJ39734.1 azon|OJJ47866.1

Group3609: abra|OJJ75009.1 acar|OOF97317.1 acla|EAW08204.1 afis|EAW23238.1 afla|EED55718.1 afum|EAL88244.1 agla|OJJ86622.1 akaw|GAA88632.1 aluc|OJZ92574.1 anid|CBF89882.1 anig|EHA26407.1 aory|EIT83207.1 asyd|OJJ62146.1 ater|EAU37519.1 atub|OJI87010.1 aver|OJI96256.1 awen|OJJ39732.1 azon|OJJ47864.1

Group3610: abra|OJJ75010.1 acar|OOF97316.1 acla|EAW08205.1 afis|EAW23239.1 afla|EED55719.1 afum|EAL88243.1 agla|OJJ86620.1 akaw|GAA88633.1 aluc|OJZ92575.1 anid|CBF89884.1 anig|EHA26406.1

aory|EIT83209.1 asyd|OJJ62145.1 ater|EAU37520.1 atub|OJI87011.1 aver|OJI96255.1 awen|OJJ39731.1 azon|OJJ47863.1

Group3611: abra|OJJ75011.1 acar|OOF97315.1 acla|EAW08206.1 afis|EAW23240.1 afla|EED55720.1 afum|EAL88242.1 agla|OJJ86619.1 akaw|GAA88634.1 aluc|OJZ92576.1 anid|CBF89885.1 anig|EHA26405.1 aory|EIT83217.1 asyd|OJJ62144.1 ater|EAU37521.1 atub|OJI87012.1 aver|OJI96254.1 awen|OJJ39730.1 azon|OJJ47862.1

Group3612: abra|OJJ75012.1 acar|OOF97314.1 acla|EAW08207.1 afis|EAW23241.1 afla|EED55722.1 afum|EAL88241.1 agla|OJJ86618.1 akaw|GAA88635.1 aluc|OJZ92577.1 anid|CBF89887.1 anig|EHA26404.1 aory|EIT83231.1 asyd|OJJ62143.1 ater|EAU37522.1 atub|OJI87013.1 aver|OJI96253.1 awen|OJJ39729.1 azon|OJJ47861.1

Group3613: abra|OJJ75014.1 acar|OOF97312.1 acla|EAW08209.1 afis|EAW23243.1 afla|EED55724.1 afum|EAL88239.1 agla|OJJ86617.1 akaw|GAA88637.1 aluc|OJZ92579.1 anid|CBF89891.1 anig|EHA26402.1 aory|EIT83225.1 asyd|OJJ62140.1 ater|EAU37524.1 atub|OJI87015.1 aver|OJI96250.1 awen|OJJ39727.1 azon|OJJ47858.1

Group3614: abra|OJJ75022.1 acar|OOF89975.1 acla|EAW08214.1 afis|EAW23248.1 afla|EED55729.1 afum|EAL88233.1 agla|OJJ86609.1 akaw|GAA88642.1 aluc|OJZ92589.1 anid|CBF84108.1 anig|EHA26397.1 aory|EIT83221.1 asyd|OJJ57101.1 ater|EAU37530.1 atub|OJI87021.1 aver|OJJ03198.1 awen|OJJ39716.1 azon|OJJ47847.1

Group3615: abra|OJJ75025.1 acar|OOF97307.1 acla|EAW08217.1 afis|EAW23251.1 afla|EED55732.1 afum|EAL88230.1 agla|OJJ86603.1 akaw|GAA88645.1 aluc|OJZ92592.1 anid|CBF84103.1 anig|EHA26394.1 aory|EIT83210.1 asyd|OJJ57097.1 ater|EAU37549.1 atub|OJI87025.1 aver|OJJ03202.1 awen|OJJ39713.1 azon|OJJ47844.1

Group3616: abra|OJJ75026.1 acar|OOF97306.1 acla|EAW08218.1 afis|EAW23252.1 afla|EED55733.1 afum|EAL88229.1 agla|OJJ86602.1 akaw|GAA88646.1 aluc|OJZ92593.1 anid|CBF84101.1 anig|EHA26393.1 aory|EIT83216.1 asyd|OJJ57096.1 ater|EAU37548.1 atub|OJI87027.1 aver|OJJ03203.1 awen|OJJ39712.1 azon|OJJ47843.1

Group3617: abra|OJJ75027.1 acar|OOF97305.1 acla|EAW08219.1 afis|EAW23253.1 afla|EED55734.1 afum|EAL88228.1 agla|OJJ86601.1 akaw|GAA88647.1 aluc|OJZ92594.1 anid|CBF84100.1 anig|EHA26392.1 aory|EIT83226.1 asyd|OJJ57095.1 ater|EAU37547.1 atub|OJI87028.1 aver|OJJ03204.1 awen|OJJ39711.1 azon|OJJ47842.1

Group3618: abra|OJJ75030.1 acar|OOF97303.1 acla|EAW08221.1 afis|EAW23255.1 afla|EED55736.1 afum|EAL88226.1 agla|OJJ86599.1 akaw|GAA88649.1 aluc|OJZ92597.1 anid|CBF84096.1 anig|EHA26390.1 aory|EIT83204.1 asyd|OJJ57093.1 ater|EAU37544.1 atub|OJI87031.1 aver|OJJ03206.1 awen|OJJ39709.1 azon|OJJ47840.1

Group3619: abra|OJJ75031.1 acar|OOF97302.1 acla|EAW08222.1 afis|EAW23256.1 afla|EED55737.1 afum|EAL88225.1 agla|OJJ86598.1 akaw|GAA88650.1 aluc|OJZ92598.1 anid|CBF84092.1 anig|EHA26389.1 aory|EIT83229.1 asyd|OJJ57092.1 ater|EAU37543.1 atub|OJI87032.1 aver|OJJ03208.1 awen|OJJ39708.1 azon|OJJ47839.1

Group3620: abra|OJJ75032.1 acar|OOF97301.1 acla|EAW08223.1 afis|EAW23257.1 afla|EED55738.1 afum|EAL88224.1 agla|OJJ86597.1 akaw|GAA88651.1 aluc|OJZ92599.1 anid|CBF84094.1 anig|EHA26388.1 aory|EIT83202.1 asyd|OJJ57091.1 ater|EAU37542.1 atub|OJI87034.1 aver|OJJ03209.1 awen|OJJ39707.1 azon|OJJ47838.1

Group3621: abra|OJJ75033.1 acar|OOF97300.1 acla|EAW08224.1 afis|EAW23258.1 afla|EED55739.1 afum|EAL88223.1 agla|OJJ86596.1 akaw|GAA88652.1 aluc|OJZ92600.1 anid|CBF84116.1 anig|EHA26387.1 aory|EIT83200.1 asyd|OJJ57105.1 ater|EAU37541.1 atub|OJI87035.1 aver|OJJ03194.1 awen|OJJ39706.1 azon|OJJ47837.1

Group3622: abra|OJJ75034.1 acar|OOF97299.1 acla|EAW08225.1 afis|EAW23259.1 afla|EED55740.1 afum|EAL88222.1 agla|OJJ86595.1 akaw|GAA88653.1 aluc|OJZ92602.1 anid|CBF84117.1 anig|EHA26386.1 aory|EIT83218.1 asyd|OJJ57106.1 ater|EAU37540.1 atub|OJI87036.1 aver|OJJ03193.1 awen|OJJ39705.1 azon|OJJ47835.1

Group3623: abra|OJJ75035.1 acar|OOF97298.1 acla|EAW08226.1 afis|EAW23260.1 afla|EED55741.1 afum|EAL88221.1 agla|OJJ86594.1 akaw|GAA88654.1 aluc|OJZ92603.1 anid|CBF84119.1 anig|EHA26385.1 aory|EIT83223.1 asyd|OJJ57107.1 ater|EAU37539.1 atub|OJI87037.1 aver|OJJ03192.1 awen|OJJ39704.1 azon|OJJ47834.1

Group3624: abra|OJJ75036.1 acar|OOF97297.1 acla|EAW08231.1 afis|EAW23265.1 afla|EED55746.1 afum|EAL88215.1 agla|OJJ86587.1 akaw|GAA88655.1 aluc|OJZ92604.1 anid|CBF84128.1 anig|EHA26384.1 aory|EIT83198.1 asyd|OJJ57112.1 ater|EAU37536.1 atub|OJI87038.1 aver|OJJ03187.1 awen|OJJ39699.1 azon|OJJ47829.1

Group3625: abra|OJJ75038.1 acar|OOF97294.1 acla|EAW08229.1 afis|EAW23263.1 afla|EED55744.1 afum|EAL88217.1 agla|OJJ86589.1 akaw|GAA88657.1 aluc|OJZ92606.1 anid|CBF84124.1 anig|EHA26382.1 aory|EIT83214.1 asyd|OJJ57110.1 ater|EAU37534.1 atub|OJI87040.1 aver|OJJ03189.1 awen|OJJ39701.1 azon|OJJ47831.1

Group3626: abra|OJJ75039.1 acar|OOF97293.1 acla|EAW08227.1 afis|EAW23261.1 afla|EED55742.1 afum|EAL88219.1 agla|OJJ86591.1 akaw|GAA88658.1 aluc|OJZ92607.1 anid|CBF84123.1 anig|EHA26381.1 aory|EIT83212.1 asyd|OJJ57109.1 ater|EAU37538.1 atub|OJI87041.1 aver|OJJ03190.1 awen|OJJ39703.1 azon|OJJ47833.1

Group3627: abra|OJJ75040.1 acar|OOF97292.1 acla|EAW08228.1 afis|EAW23262.1 afla|EED55743.1 afum|EAL88218.1 agla|OJJ86590.1 akaw|GAA88659.1 aluc|OJZ92608.1 anid|CBF84121.1 anig|EHA26380.1 aory|EIT83233.1 asyd|OJJ57108.1 ater|EAU37537.1 atub|OJI87042.1 aver|OJJ03191.1 awen|OJJ39702.1 azon|OJJ47832.1

Group3628: abra|OJJ75046.1 acar|OOF89831.1 acla|EAW08235.1 afis|EAW23269.1 afla|EED55710.1 afum|EAL88210.2 agla|OJJ86583.1 akaw|GAA88663.1 aluc|OJZ92612.1 anid|CBF89550.1 anig|EHA26375.1 aory|EIT83205.1 asyd|OJJ62375.1 ater|EAU37553.1 atub|OJI87046.1 aver|OJI96509.1 awen|OJJ39692.1 azon|OJJ47825.1

Group3629: abra|OJJ75047.1 acar|OOF97288.1 acla|EAW08236.1 afis|EAW23270.1 afla|EED55709.1 afum|EAL88209.1 agla|OJJ86582.1 akaw|GAA88664.1 aluc|OJZ92613.1 anid|CBF89548.1 anig|EHA26374.1 aory|EIT83232.1 asyd|OJJ62376.1 ater|EAU37554.1 atub|OJI87047.1 aver|OJI96510.1 awen|OJJ39691.1 azon|OJJ47824.1

Group3630: abra|OJJ75048.1 acar|OOF97286.1 acla|EAW08237.1 afis|EAW23271.1 afla|EED55706.1 afum|EAL88208.1 agla|OJJ86578.1 akaw|GAA88665.1 aluc|OJZ92615.1 anid|CBF89546.1 anig|EHA26373.1 aory|EIT83206.1 asyd|OJJ62377.1 ater|EAU37556.1 atub|OJI87049.1 aver|OJI96511.1 awen|OJJ39687.1 azon|OJJ47823.1

Group3631: abra|OJJ75052.1 acar|OOF97284.1 acla|EAW08238.1 afis|EAW23272.1 afla|EED55705.1 afum|EAL88207.1 agla|OJJ86577.1 akaw|GAA88666.1 aluc|OJZ92618.1 anid|CBF89544.1 anig|EHA26370.1 aory|EIT77089.1 asyd|OJJ62379.1 ater|EAU37557.1 atub|OJI87052.1 aver|OJI96513.1 awen|OJJ39685.1 azon|OJJ47822.1

Group3632: abra|OJJ75056.1 acar|OOF97281.1 acla|EAW08241.1 afis|EAW23275.1 afla|EED55703.1 afum|EAL88204.1 agla|OJJ86573.1 akaw|GAA88669.1 aluc|OJZ92622.1 anid|CBF89539.1 anig|EHA26367.1 aory|EIT77072.1 asyd|OJJ62382.1 ater|EAU37560.1 atub|OJI87055.1 aver|OJI96516.1 awen|OJJ39682.1 azon|OJJ47819.1

Group3633: abra|OJJ75057.1 acar|OOF97279.1 acla|EAW08242.1 afis|EAW23276.1 afla|EED55701.1 afum|EAL88203.1 agla|OJJ86571.1 akaw|GAA88670.1 aluc|OJZ92626.1 anid|CBF89533.1 anig|EHA26364.1 aory|EIT77071.1 asyd|OJJ62384.1 ater|EAU37561.1 atub|OJI87056.1 aver|OJI96518.1 awen|OJJ39680.1 azon|OJJ47818.1

Group3634: abra|OJJ75064.1 acar|OOF97270.1 acla|EAW08245.1 afis|EAW23279.1 afla|EED55695.1 afum|EAL88200.1 agla|OJJ86566.1 akaw|GAA88676.1 aluc|OJZ92630.1 anid|CBF89527.1 anig|EHA26360.1 aory|EIT77073.1 asyd|OJJ62388.1 ater|EAU37592.1 atub|OJI87061.1 aver|OJI96521.1 awen|OJJ39676.1 azon|OJJ47816.1

Group3635: abra|OJJ75066.1 acar|OOF97268.1 acla|EAW08247.1 afis|EAW23281.1 afla|EED55694.1 afum|EAL88198.2 agla|OJJ86565.1 akaw|GAA88678.1 aluc|OJZ92632.1 anid|CBF89525.1 anig|EHA26359.1 aory|EIT77087.1 asyd|OJJ62389.1 ater|EAU37591.1 atub|OJI87063.1 aver|OJI96522.1 awen|OJJ39675.1 azon|OJJ47805.1

Group3636: abra|OJJ75069.1 acar|OOF97264.1 acla|EAW08250.1 afis|EAW23284.1 afla|EED55691.1 afum|EAL88195.1 agla|OJJ86562.1 akaw|GAA88681.1 aluc|OJZ92635.1 anid|CBF89520.1 anig|EHA26356.1 aory|EIT77082.1 asyd|OJJ62392.1 ater|EAU37588.1 atub|OJI87066.1 aver|OJI96525.1 awen|OJJ39672.1 azon|OJJ47802.1

Group3637: abra|OJJ75070.1 acar|OOF97265.1 acla|EAW08251.1 afis|EAW23285.1 afla|EED55690.1 afum|EAL88194.1 agla|OJJ86561.1 akaw|GAA88682.1 aluc|OJZ92636.1 anid|CBF89518.1 anig|EHA26355.1 aory|EIT77093.1 asyd|OJJ62393.1 ater|EAU37587.1 atub|OJI87067.1 aver|OJI96526.1 awen|OJJ39671.1 azon|OJJ47801.1

Group3638: abra|OJJ75077.1 acar|OOF97256.1 acla|EAW08287.1 afis|EAW23320.1 afla|EED55678.1 afum|EAL88156.2 agla|OJJ86545.1 akaw|GAA88690.1 aluc|OJZ92643.1 anid|CBF89504.1 anig|EHA26349.1 aory|EIT77070.1 asyd|OJJ62400.1 ater|EAU37581.1 atub|OJI87074.1 aver|OJI96533.1 awen|OJJ39665.1 azon|OJJ47793.1

Group3639: abra|OJJ75080.1 acar|OOF97253.1 acla|EAW08284.1 afis|EAW23317.1 afla|EED55676.1 afum|EBA27338.1 agla|OJJ86542.1 akaw|GAA88693.1 aluc|OJZ92646.1 anid|CBF89487.1 anig|EHA26346.1

aory|EIT77097.1 asyd|OJJ62404.1 ater|EAU35544.1 atub|OJI87077.1 aver|OJI96537.1 awen|OJJ39662.1 azon|OJJ47791.1

Group3640: abra|OJJ75081.1 acar|OOF97252.1 acla|EAW08283.1 afis|EAW23316.1 afla|EED55675.1 afum|EAL88161.2 agla|OJJ86541.1 akaw|GAA88694.1 aluc|OJZ92647.1 anid|CBF89489.1 anig|EHA26345.1 aory|EIT77095.1 asyd|OJJ62405.1 ater|EAU35545.1 atub|OJI87078.1 aver|OJI96538.1 awen|OJJ39661.1 azon|OJJ47789.1

Group3641: abra|OJJ75082.1 acar|OOF97251.1 acla|EAW08282.1 afis|EAW23315.1 afla|EED55674.1 afum|EAL88162.1 agla|OJJ86540.1 akaw|GAA88695.1 aluc|OJZ92648.1 anid|CBF89491.1 anig|EHA26344.1 aory|EIT77078.1 asyd|OJJ62406.1 ater|EAU35546.1 atub|OJI87079.1 aver|OJI96539.1 awen|OJJ39660.1 azon|OJJ47788.1

Group3642: abra|OJJ75084.1 acar|OOF97249.1 acla|EAW08280.1 afis|EAW23313.1 afla|EED55672.1 afum|EAL88164.1 agla|OJJ86538.1 akaw|GAA88697.1 aluc|OJZ92650.1 anid|CBF89495.1 anig|EHA26342.1 aory|EIT77083.1 asyd|OJJ62408.1 ater|EAU35548.1 atub|OJI87081.1 aver|OJI96541.1 awen|OJJ39658.1 azon|OJJ47786.1

Group3643: abra|OJJ75087.1 acar|OOF97246.1 acla|EAW08277.1 afis|EAW23310.1 afla|EED55665.1 afum|EAL88167.1 akaw|GAA88699.1 aluc|OJZ92653.1 anid|CBF89485.1 anig|EHA26341.1 asyd|OJJ62412.1 ater|EAU35551.1 atub|OJI87084.1 aver|OJI96546.1 awen|OJJ39655.1 azon|OJJ47783.1 agla|OJJ86535.1 aory|EIT81799.1

Group3644: abra|OJJ75089.1 acar|OOF97244.1 acla|EAW08275.1 afis|EAW23308.1 afla|EED55663.1 afum|EAL88169.1 agla|OJJ86533.1 akaw|GAA88701.1 aluc|OJZ92655.1 anid|CBF89482.1 anig|EHA26339.1 aory|EIT81804.1 asyd|OJJ62413.1 ater|EAU35553.1 atub|OJI87086.1 aver|OJI96548.1 awen|OJJ39653.1 azon|OJJ47781.1

Group3645: abra|OJJ75090.1 acar|OOF97243.1 acla|EAW08274.1 afis|EAW23307.1 afla|EED55662.1 afum|EAL88170.1 agla|OJJ86532.1 akaw|GAA88702.1 aluc|OJZ92656.1 anid|CBF89480.1 anig|EHA26338.1 aory|EIT81820.1 asyd|OJJ62414.1 ater|EAU35554.1 atub|OJI87087.1 aver|OJI96549.1 awen|OJJ39652.1 azon|OJJ47780.1

Group3646: abra|OJJ75091.1 acar|OOF97241.1 acla|EAW08273.1 afis|EAW23306.1 afla|EED55661.1 afum|EAL88171.1 agla|OJJ86531.1 akaw|GAA88703.1 aluc|OJZ92657.1 anid|CBF89478.1 anig|EHA26337.1 aory|EIT81795.1 asyd|OJJ62415.1 ater|EAU35569.1 atub|OJI87088.1 aver|OJI96550.1 awen|OJJ39590.1 azon|OJJ47779.1

Group3647: abra|OJJ75093.1 acar|OOF97239.1 acla|EAW08271.1 afis|EAW23304.1 afla|EED55659.1 afum|EAL88173.1 agla|OJJ86529.1 akaw|GAA88705.1 anid|CBF89474.1 anig|EHA26335.1 aory|EIT81791.1 asyd|OJJ62418.1 ater|EAU35568.1 atub|OJI87090.1 aver|OJI96553.1 awen|OJJ39586.1 azon|OJJ47766.1 aluc|OJZ92659.1

Group3648: abra|OJJ75094.1 acar|OOF97238.1 acla|EAW08270.1 afis|EAW23303.1 afla|EED55656.1 afum|EAL88175.1 agla|OJJ86528.1 akaw|GAA88706.1 aluc|OJZ92660.1 anid|CBF89472.1 anig|EHA26334.1 aory|EIT81787.1 asyd|OJJ62419.1 ater|EAU35567.1 atub|OJI87091.1 aver|OJI96555.1 awen|OJJ39585.1 azon|OJJ47767.1

Group3649: abra|OJJ75096.1 acar|OOF97236.1 acla|EAW08268.1 afis|EAW23301.1 afla|EED55653.1 afum|EAL88178.1 agla|OJJ86507.1 akaw|GAA88708.1 aluc|OJZ92662.1 anid|CBF89468.1 anig|EHA26332.1 aory|EIT81800.1 asyd|OJJ62421.1 ater|EAU35565.1 atub|OJI87093.1 aver|OJI96556.1 awen|OJJ39583.1 azon|OJJ47769.1

Group3650: abra|OJJ75102.1 acar|OOF97232.1 acla|EAW08266.1 afis|EAW23299.1 afla|EED55649.1 afum|EAL88180.1 agla|OJJ86511.1 akaw|GAA88711.1 aluc|OJZ92666.1 anid|CBF89465.1 anig|EHA26329.1 aory|EIT81776.1 asyd|OJJ62424.1 ater|EAU35561.1 atub|OJI87097.1 aver|OJI96561.1 awen|OJJ39581.1 azon|OJJ47771.1

Group3651: abra|OJJ75103.1 acar|OOF97231.1 acla|EAW08265.1 afis|EAW23298.1 afla|EED55648.1 afum|EAL88181.2 agla|OJJ86513.1 akaw|GAA88712.1 aluc|OJZ92668.1 anid|CBF89463.1 anig|EHA26328.1 aory|EIT81789.1 asyd|OJJ62425.1 ater|EAU35559.1 atub|OJI87098.1 aver|OJI96562.1 awen|OJJ39579.1 azon|OJJ47772.1

Group3652: abra|OJJ75106.1 acar|OOF97228.1 acla|EAW08262.1 afis|EAW23295.1 afla|EED55645.1 afum|EAL88183.1 agla|OJJ86516.1 akaw|GAA88715.1 aluc|OJZ92671.1 anid|CBF89459.1 anig|EHA26326.1 aory|EIT81783.1 asyd|OJJ62427.1 ater|EAU35558.1 atub|OJI87100.1 aver|OJI96564.1 awen|OJJ39576.1 azon|OJJ47775.1

Group3653: abra|OJJ75107.1 acar|OOF97227.1 acla|EAW08261.1 afis|EAW23294.1 afla|EED55644.1 afum|EAL88184.1 agla|OJJ86517.1 akaw|GAA88716.1 aluc|OJZ92672.1 anid|CBF89457.1 anig|EHA26325.1 aory|EIT81790.1 asyd|OJJ62428.1 ater|EAU35557.1 atub|OJI87101.1 aver|OJI96565.1 awen|OJJ39575.1 azon|OJJ47776.1

Group3654: abra|OJJ75109.1 acar|OOF97223.1 acla|EAW08315.1 afis|EAW23325.1 afla|EED55641.1 afum|EAL88154.1 agla|OJJ86520.1 akaw|GAA88718.1 aluc|OJZ92675.1 anid|CBF89453.1 anig|EHA26323.1 aory|EIT81818.1 asyd|OJJ62431.1 ater|EAU35573.1 atub|OJI87103.1 aver|OJI96567.1 awen|OJJ39569.1 azon|OJJ47762.1

Group3655: abra|OJJ75114.1 acar|OOF97217.1 acla|EAW08309.1 afis|EAW23330.1 afla|EED55637.1 afum|EAL88149.2 agla|OJJ86525.1 akaw|GAA88724.1 aluc|OJZ92681.1 anid|CBF89441.1 anig|EHA26318.1 aory|EIT81798.1 asyd|OJJ62439.1 ater|EAU35577.1 atub|OJI87109.1 aver|OJI96575.1 awen|OJJ39564.1 azon|OJJ47757.1

Group3656: abra|OJJ75115.1 acar|OOF97216.1 acla|EAW08308.1 afis|EAW23331.1 afla|EED55636.1 afum|EAL88148.1 agla|OJJ86503.1 akaw|GAA88725.1 aluc|OJZ92682.1 anid|CBF89439.1 anig|EHA26317.1 aory|EIT81819.1 asyd|OJJ62440.1 ater|EAU35578.1 atub|OJI87110.1 aver|OJI96576.1 awen|OJJ39563.1 azon|OJJ47756.1

Group3657: abra|OJJ75120.1 acar|OOF97213.1 acla|EAW08306.1 afis|EAW23332.1 afla|EED55634.1 afum|EAL88146.2 agla|OJJ86500.1 akaw|GAA88729.1 aluc|OJZ92687.1 anid|CBF89437.1 anig|EHA26313.1 aory|EIT81801.1 asyd|OJJ62442.1 ater|EAU35580.1 atub|OJI87115.1 aver|OJI96578.1 awen|OJJ39561.1 azon|OJJ47754.1

Group3658: abra|OJJ75122.1 acar|OOF97212.1 acla|EAW08305.1 afis|EAW23334.1 afla|EED55632.1 afum|EAL88144.1 agla|OJJ86499.1 akaw|GAA88730.1 aluc|OJZ92688.1 anid|CBF89435.1 anig|EHA26312.1 aory|EIT81824.1 asyd|OJJ62443.1 ater|EAU35581.1 atub|OJI87116.1 aver|OJI96579.1 awen|OJJ39560.1 azon|OJJ47753.1

Group3659: abra|OJJ75124.1 acar|OOF97210.1 acla|EAW08304.1 afis|EAW23336.1 afla|EED55630.1 afum|EAL88142.1 agla|OJJ86494.1 akaw|GAA88732.1 aluc|OJZ92690.1 anid|CBF89432.1 anig|EHA26310.1 aory|EIT81797.1 asyd|OJJ62444.1 ater|EAU35582.1 atub|OJI87118.1 aver|OJI96581.1 awen|OJJ39558.1 azon|OJJ47749.1

Group3660: abra|OJJ75126.1 acar|OOF97209.1 acla|EAW08303.1 afis|EAW23337.1 afla|EED55629.1 afum|EAL88141.1 agla|OJJ86493.1 akaw|GAA88733.1 aluc|OJZ92691.1 anid|CBF89430.1 anig|EHA26309.1 aory|EIT81788.1 asyd|OJJ62445.1 ater|EAU35583.1 atub|OJI87119.1 aver|OJI96582.1 awen|OJJ39557.1 azon|OJJ47750.1

Group3661: abra|OJJ75129.1 acar|OOF97207.1 acla|EAW08300.1 afis|EAW23340.1 afla|EED55626.1 afum|EAL88138.1 agla|OJJ86490.1 akaw|GAA88736.1 aluc|OJZ92694.1 anid|CBF89425.1 anig|EHA26306.1 aory|EIT81810.1 asyd|OJJ62448.1 ater|EAU35585.1 atub|OJI87122.1 aver|OJI96585.1 awen|OJJ39554.1 azon|OJJ47748.1

Group3662: abra|OJJ75130.1 acar|OOF97205.1 acla|EAW08298.1 afis|EAW23342.1 afla|EED55624.1 afum|EAL88136.2 agla|OJJ86486.1 akaw|GAA88737.1 aluc|OJZ92695.1 anid|CBF89422.1 anig|EHA26304.1 aory|EIT81811.1 asyd|OJJ62451.1 ater|EAU35586.1 atub|OJI87123.1 aver|OJI96586.1 awen|OJJ39552.1 azon|OJJ47747.1

Group3663: abra|OJJ75131.1 acar|OOF97204.1 acla|EAW08297.1 afis|EAW23343.1 afla|EED55623.1 afum|EAL88135.1 agla|OJJ86485.1 akaw|GAA88738.1 aluc|OJZ92696.1 anid|CBF89420.1 anig|EHA26302.1 aory|EIT81805.1 asyd|OJJ62452.1 ater|EAU35587.1 atub|OJI87124.1 aver|OJI96587.1 awen|OJJ39550.1 azon|OJJ47746.1

Group3664: abra|OJJ75133.1 acar|OOF97203.1 acla|EAW08294.1 afis|EAW23345.1 afla|EED55621.1 afum|EAL88132.1 agla|OJJ86482.1 akaw|GAA88740.1 aluc|OJZ92699.1 anid|CBF89412.1 anig|EHA26300.1 aory|EIT81825.1 asyd|OJJ62455.1 ater|EAU35589.1 atub|OJI87127.1 aver|OJI96590.1 awen|OJJ39547.1 azon|OJJ47744.1

Group3665: abra|OJJ75134.1 acar|OOF97202.1 acla|EAW08293.1 afis|EAW23346.1 afla|EED55620.1 afum|EAL88130.1 agla|OJJ86481.1 akaw|GAA88741.1 aluc|OJZ92700.1 anid|CBF74918.1 anig|EHA26299.1 aory|EIT81822.1 asyd|OJJ60098.1 ater|EAU35590.1 atub|OJI87128.1 aver|OJJ04164.1 awen|OJJ39545.1 azon|OJJ47743.1

Group3666: abra|OJJ75136.1 acar|OOF97201.1 acla|EAW08292.1 afis|EAW23347.1 afla|EED55619.1 afum|EAL88129.1 agla|OJJ86480.1 akaw|GAA88742.1 aluc|OJZ92702.1 anid|CBF74916.1 anig|EHA26298.1 aory|EIT81794.1 asyd|OJJ60099.1 ater|EAU35591.1 atub|OJI87129.1 aver|OJJ04163.1 awen|OJJ39544.1 azon|OJJ47742.1

Group3667: abra|OJJ75146.1 acar|OOF97192.1 acla|EAW08319.1 afis|EAW23354.1 afla|EED55604.1 afum|EAL88122.2 agla|OJJ86471.1 akaw|GAA88751.1 aluc|OJZ92711.1 anid|CBF74886.1 anig|EHA26288.1 aory|EIT82711.1 asyd|OJJ60107.1 ater|EAU35606.1 atub|OJI87137.1 aver|OJJ04155.1 awen|OJJ39532.1 azon|OJJ47729.1

Group3668: abra|OJJ75149.1 acar|OOF97188.1 acla|EAW08323.1 afis|EAW23358.1 afla|EED55608.1 afum|EAL88118.1 agla|OJJ86468.1 akaw|GAA88755.1 aluc|OJZ92715.1 anid|CBF74882.1 anig|EHA26285.1

aory|EIT81782.1 asyd|OJJ60110.1 ater|EAU35609.1 atub|OJI87141.1 aver|OJJ04152.1 awen|OJJ39529.1 azon|OJJ47725.1

Group3669: abra|OJJ75150.1 acar|OOF97187.1 acla|EAW08324.1 afis|EAW23359.1 afla|EED55610.1 afum|EBA27336.1 agla|OJJ86467.1 akaw|GAA88756.1 aluc|OJZ92716.1 anid|CBF74880.1 anig|EHA26283.1 aory|EIT81806.1 asyd|OJJ60111.1 ater|EAU35610.1 atub|OJI87142.1 aver|OJJ04151.1 awen|OJJ39528.1 azon|OJJ47724.1

Group3670: abra|OJJ75152.1 acar|OOF97185.1 acla|EAW08327.1 afis|EAW23362.1 afla|EED55596.1 afum|EAL88112.1 agla|OJJ86464.1 akaw|GAA88757.1 aluc|OJZ92720.1 anid|CBF74837.1 anig|EHA26280.1 aory|EIT82684.1 asyd|OJJ60134.1 ater|EAU35613.1 atub|OJI87145.1 aver|OJJ04128.1 awen|OJJ39525.1 azon|OJJ47721.1

Group3671: abra|OJJ75153.1 acar|OOF97184.1 acla|EAW08328.1 afis|EAW23363.1 afla|EED55595.1 afum|EBA27334.1 agla|OJJ86463.1 akaw|GAA88758.1 aluc|OJZ92721.1 anid|CBF74835.1 anig|EHA26279.1 aory|EIT82682.1 asyd|OJJ60135.1 ater|EAU35614.1 atub|OJI87146.1 aver|OJJ04127.1 awen|OJJ39524.1 azon|OJJ47720.1

Group3672: abra|OJJ75157.1 acar|OOF97182.1 acla|EAW08332.1 afis|EAW23367.1 afla|EED55590.1 afum|EAL88106.1 agla|OJJ86457.1 akaw|GAA88763.1 aluc|OJZ92727.1 anid|CBF74823.1 anig|EHA26274.1 aory|EIT82667.1 asyd|OJJ60140.1 ater|EAU35618.1 atub|OJI87151.1 aver|OJJ04122.1 awen|OJJ39520.1 azon|OJJ47716.1

Group3673: abra|OJJ75159.1 acar|OOF97179.1 acla|EAW08335.1 afis|EAW23370.1 afla|EED55587.1 afum|EAL88103.2 agla|OJJ86453.1 akaw|GAA88770.1 aluc|OJZ92729.1 anid|CBF74817.1 anig|EHA26272.1 aory|EIT82666.1 asyd|OJJ60143.1 ater|EAU35621.1 atub|OJI87153.1 aver|OJJ04120.1 awen|OJJ39513.1 azon|OJJ47712.1

Group3674: abra|OJJ75160.1 acar|OOF97178.1 acla|EAW08336.1 afis|EAW23371.1 afla|EED55586.1 afum|EAL88101.1 agla|OJJ86452.1 akaw|GAA88771.1 aluc|OJZ92730.1 anid|CBF74815.1 anig|EHA26271.1 aory|EIT82693.1 asyd|OJJ60144.1 ater|EAU35622.1 atub|OJI87154.1 aver|OJJ04119.1 awen|OJJ39512.1 azon|OJJ47711.1

Group3675: abra|OJJ75162.1 acar|OOF97176.1 acla|EAW08338.1 afis|EAW23373.1 afla|EED55585.1 afum|EAL88099.1 agla|OJJ86450.1 akaw|GAA88773.1 aluc|OJZ92732.1 anid|CBF74811.1 anig|EHA26269.1 aory|EIT82709.1 asyd|OJJ60146.1 ater|EAU35624.1 atub|OJI87156.1 aver|OJJ04117.1 awen|OJJ39510.1 azon|OJJ47709.1

Group3676: abra|OJJ75165.1 acar|OOF97173.1 acla|EAW08339.1 afis|EAW23374.1 afla|EED55584.1 afum|EAL88097.1 agla|OJJ86449.1 akaw|GAA88776.1 aluc|OJZ92736.1 anid|CBF74839.1 anig|EHA26266.1 aory|EIT82678.1 asyd|OJJ60133.1 ater|EAU35628.1 atub|OJI87160.1 aver|OJJ04129.1 awen|OJJ39509.1 azon|OJJ47708.1

Group3677: abra|OJJ75166.1 acar|OOF97172.1 acla|EAW08340.1 afis|EAW23375.1 afla|EED55583.1 afum|EAL88096.1 agla|OJJ86448.1 akaw|GAA88777.1 aluc|OJZ92737.1 anid|CBF74841.1 anig|EHA26265.1 aory|EIT82676.1 asyd|OJJ60132.1 ater|EAU35629.1 atub|OJI87161.1 aver|OJJ04130.1 awen|OJJ39508.1 azon|OJJ47707.1

Group3678: abra|OJJ75168.1 acar|OOF94606.1 acla|EAW08132.1 afis|EAW23169.1 afla|EED46952.1 afum|EAL88315.1 agla|OJJ89680.1 akaw|GAA88778.1 aluc|OJZ92738.1 anid|CBF74480.1 anig|EHA26264.1 aory|EIT72535.1 asyd|OJJ57708.1 ater|EAU36298.1 atub|OJI87163.1 aver|OJJ05456.1 awen|OJJ36614.1 azon|OJJ47599.1

Group3679: abra|OJJ75172.1 acar|OOF94604.1 acla|EAW13004.1 afis|EAW17664.1 afla|EED50493.1 afum|EAL91478.1 agla|OJJ79537.1 akaw|GAA88779.1 aluc|OJZ92740.1 anid|CBF90052.1 anig|EHA26263.1 aory|EIT82027.1 asyd|OJJ62053.1 ater|EAU31989.1 atub|OJI87165.1 aver|OJI96159.1 awen|OJJ37177.1 azon|OJJ50877.1

Group3680: abra|OJJ75177.1 acar|OOF94602.1 acla|EAW13001.1 afis|EAW17667.1 afla|EED50488.1 afum|EAL91481.1 agla|OJJ79539.1 akaw|GAA88782.1 aluc|OJZ92743.1 anid|CBF90043.1 anig|EHA26260.1 aory|EIT82124.1 asyd|OJJ62058.1 ater|EAU31980.1 atub|OJI87168.1 aver|OJI96164.1 awen|OJJ37174.1 azon|OJJ50873.1

Group3681: abra|OJJ75178.1 acar|OOF94601.1 acla|EAW13000.1 afis|EAW17668.1 afla|EED50487.1 afum|EAL91482.1 agla|OJJ79540.1 akaw|GAA88783.1 aluc|OJZ92744.1 anid|CBF90041.1 anig|EHA26259.1 aory|EIT82171.1 asyd|OJJ62059.1 ater|EAU31979.1 atub|OJI87169.1 aver|OJI96165.1 awen|OJJ37173.1 azon|OJJ50872.1

Group3682: abra|OJJ75179.1 acar|OOF94600.1 acla|EAW12999.1 afis|EAW17669.1 afla|EED50486.1 afum|EAL91483.1 agla|OJJ79541.1 akaw|GAA88784.1 aluc|OJZ92745.1 anid|CBF90039.1 anig|EHA26258.1 aory|EIT82129.1 asyd|OJJ62060.1 ater|EAU31978.1 atub|OJI87170.1 aver|OJI96166.1 awen|OJJ37172.1 azon|OJJ50871.1

Group3683: abra|OJJ75182.1 acar|OOF94598.1 acla|EAW12995.1 afis|EAW17671.1 afla|EED50482.1 afum|EAL91485.1 agla|OJJ79543.1 akaw|GAA88786.1 aluc|OJZ92746.1 anid|CBF90036.1 anig|EHA26257.1 aory|EIT82081.1 asyd|OJJ62061.1 ater|EAU31976.1 atub|OJI87173.1 aver|OJI96167.1 awen|OJJ37171.1 azon|OJJ50867.1

Group3684: abra|OJJ75186.1 acar|OOF94596.1 acla|EAW12991.1 afis|EAW17674.1 afla|EED50477.1 afum|EAL91487.1 agla|OJJ79515.1 akaw|GAA88788.1 aluc|OJZ92749.1 anid|CBF90029.1 anig|EHA26254.1 aory|EIT82056.1 asyd|OJJ62064.1 ater|EAU31972.1 atub|OJI87175.1 aver|OJI96170.1 awen|OJJ37167.1 azon|OJJ50863.1

Group3685: abra|OJJ75189.1 acar|OOF94594.1 acla|EAW12990.1 afis|EAW17675.1 afla|EED50476.1 afum|EAL91488.1 agla|OJJ79514.1 akaw|GAA88790.1 aluc|OJZ92752.1 anid|CBF90024.1 anig|EHA26252.1 aory|EIT82044.1 asyd|OJJ62067.1 ater|EAU31971.1 atub|OJI87177.1 aver|OJI96173.1 awen|OJJ37165.1 azon|OJJ50862.1

Group3686: abra|OJJ75190.1 acar|OOF94593.1 acla|EAW12989.1 afis|EAW17676.1 afla|EED50475.1 afum|EAL91489.1 agla|OJJ79500.1 akaw|GAA88791.1 aluc|OJZ92754.1 anid|CBF90022.1 anig|EHA26251.1 aory|EIT82153.1 asyd|OJJ62068.1 ater|EAU31970.1 atub|OJI87179.1 aver|OJI96174.1 awen|OJJ37164.1 azon|OJJ50860.1

Group3687: abra|OJJ75198.1 acar|OOF94588.1 acla|EAW12983.1 afis|EAW17681.1 afla|EED50470.1 afum|EAL91495.1 agla|OJJ79499.1 akaw|GAA91750.1 aluc|OJZ92761.1 anid|CBF90009.1 anig|EHA26245.1 aory|EIT82018.1 asyd|OJJ62073.1 ater|EAU31944.1 atub|OJI87205.1 aver|OJI96180.1 awen|OJJ37158.1 azon|OJJ50856.1

Group3688: abra|OJJ75199.1 acar|OOF94589.1 acla|EAW12982.1 afis|EAW17682.1 afla|EED50469.1 afum|EAL91496.1 agla|OJJ79498.1 akaw|GAA91751.1 aluc|OJZ92762.1 anid|CBF90007.1 anig|EHA26244.1 aory|EIT82107.1 asyd|OJJ62074.1 ater|EAU31945.1 atub|OJI87206.1 aver|OJI96181.1 awen|OJJ37157.1 azon|OJJ50855.1

Group3689: abra|OJJ75201.1 acar|OOF94587.1 acla|EAW12981.1 afis|EAW17683.1 afla|EED50468.1 afum|EAL91497.1 agla|OJJ79496.1 akaw|GAA91752.1 aluc|OJZ92763.1 anid|CBF90005.1 anig|EHA26243.1 aory|EIT82022.1 asyd|OJJ62075.1 ater|EAU31946.1 atub|OJI87207.1 aver|OJI96182.1 awen|OJJ37156.1 azon|OJJ50854.1

Group3690: abra|OJJ75203.1 acar|OOF94585.1 acla|EAW12979.1 afis|EAW17685.1 afla|EED50466.1 afum|EAL91499.1 agla|OJJ79494.1 akaw|GAA91754.1 aluc|OJZ92766.1 anid|CBF90002.1 anig|EHA26241.1 aory|EIT82013.1 asyd|OJJ62078.1 ater|EAU31948.1 atub|OJI87210.1 aver|OJI96185.1 awen|OJJ37154.1 azon|OJJ50852.1

Group3692: abra|OJJ75207.1 acar|OOF94582.1 acla|EAW12976.1 afis|EAW17688.1 afla|EED50463.1 afum|EAL91502.1 agla|OJJ79491.1 akaw|GAA91757.1 aluc|OJZ92769.1 anid|CBF89985.1 anig|EHA26238.1 aory|EIT82159.1 asyd|OJJ62086.1 ater|EAU31951.1 atub|OJI87213.1 aver|OJI96193.1 awen|OJJ37151.1 azon|OJJ50848.1

Group3693: abra|OJJ75208.1 acar|OOF94581.1 acla|EAW12975.1 afis|EAW17689.1 afla|EED50455.1 afum|EAL91503.1 agla|OJJ79490.1 akaw|GAA91758.1 aluc|OJZ92770.1 anid|CBF89984.1 anig|EHA26237.1 aory|EIT82021.1 asyd|OJJ62087.1 ater|EAU31952.1 atub|OJI87214.1 aver|OJI96194.1 awen|OJJ37147.1 azon|OJJ50847.1

Group3694: abra|OJJ75212.1 acar|OOF94572.1 acla|EAW13091.1 afis|EAW17572.1 afla|EED49249.1 afum|EAL91386.1 agla|OJJ81386.1 akaw|GAA91762.1 aluc|OJZ92776.1 anid|CBF90252.1 anig|EHA26234.1 aory|EIT82180.1 asyd|OJJ61946.1 ater|EAU37009.1 atub|OJI87218.1 aver|OJI96038.1 awen|OJJ37275.1 azon|OJJ50975.1

Group3695: abra|OJJ75213.1 acar|OOF94571.1 acla|EAW13092.1 afis|EAW17571.1 afla|EED49250.1 afum|EBA27432.1 agla|OJJ81387.1 akaw|GAA91763.1 aluc|OJZ92777.1 anid|CBF90254.1 anig|EHA26233.1 aory|EIT82196.1 asyd|OJJ61945.1 ater|EAU37008.1 atub|OJI87219.1 aver|OJI96037.1 awen|OJJ37278.1 azon|OJJ50976.1

Group3696: abra|OJJ75214.1 acar|OOF94570.1 acla|EAW13093.1 afis|EAW17570.1 afla|EED49251.1 afum|EAL91383.1 agla|OJJ81390.1 akaw|GAA91764.1 aluc|OJZ92778.1 anid|CBF90256.1 anig|EHA26232.1 aory|EIT82218.1 asyd|OJJ61944.1 ater|EAU37007.1 atub|OJI87220.1 aver|OJI96036.1 awen|OJJ37281.1 azon|OJJ50977.1

Group3697: abra|OJJ75215.1 acar|OOF94569.1 acla|EAW13094.1 afis|EAW17569.1 afla|EED49252.1 afum|EAL91382.1 agla|OJJ81395.1 akaw|GAA91765.1 aluc|OJZ92779.1 anid|CBF90258.1 anig|EHA26231.1 aory|EIT82201.1 asyd|OJJ61943.1 ater|EAU37006.1 atub|OJI87221.1 awen|OJJ37282.1 azon|OJJ50978.1 aver|OJI96035.1

Group3698: abra|OJJ75227.1 acar|OOF94562.1 acla|EAW12254.1 afis|EAW16982.1 afla|EED48301.1 afum|EAL86509.1 agla|OJJ79789.1 akaw|GAA91773.1 aluc|OJZ92788.1 anid|CBF86682.1 anig|EHA26219.1

aory|EIT76292.1 asyd|OJJ64757.1 ater|EAU31794.1 atub|OJI87231.1 aver|OJI98859.1 awen|OJJ35994.1 azon|OJJ50263.1

Group3699: abra|OJJ75272.1 acar|OOF94531.1 acla|EAW12236.1 afis|EAW16991.1 afla|EED48964.1 afum|EAL86515.1 agla|OJJ81058.1 akaw|GAA91810.1 aluc|OJZ92832.1 anid|CBF87436.1 anig|EHA26177.1 asyd|OJJ59098.1 ater|EAU31737.1 atub|OJI87289.1 aver|OJJ01689.1 awen|OJJ35979.1 azon|OJJ50286.1 aory|EIT80025.1

Group3700: abra|OJJ75274.1 acar|OOF94529.1 acla|EAW12235.1 afis|EAW16993.1 afla|EED48956.1 afum|EAL86517.1 agla|OJJ79819.1 akaw|GAA91812.1 aluc|OJZ92834.1 anid|CBF87472.1 anig|EHA26175.1 aory|EIT80048.1 asyd|OJJ52939.1 ater|EAU31735.1 atub|OJI87291.1 aver|OJJ08906.1 awen|OJJ35982.1 azon|OJJ50283.1

Group3702: abra|OJJ75284.1 acar|OOF94523.1 acla|EAW12234.1 afis|EAW16994.1 afla|EED48957.1 afum|EAL86518.2 agla|OJJ79820.1 akaw|GAA91822.1 aluc|OJZ92845.1 anid|CBF87470.1 anig|EHA26165.1 aory|EIT79980.1 asyd|OJJ52938.1 ater|EAU31734.1 atub|OJI87301.1 aver|OJJ08905.1 awen|OJJ35983.1 azon|OJJ50282.1

Group3703: abra|OJJ75307.1 acar|OOF94522.1 acla|EAW12233.1 afis|EAW16995.1 afla|EED48958.1 afum|EAL86519.2 agla|OJJ79822.1 akaw|GAA91991.1 aluc|OJZ92861.1 anid|CBF87468.1 anig|EHA26104.1 aory|EIT79977.1 asyd|OJJ52937.1 ater|EAU31733.1 atub|OJI87319.1 aver|OJJ08904.1 awen|OJJ35984.1 azon|OJJ50281.1

Group3704: abra|OJJ75309.1 acar|OOF94520.1 acla|EAW12231.1 afis|EAW16997.1 afla|EED48962.1 afum|EAL86522.2 agla|OJJ79824.1 akaw|GAA91993.1 aluc|OJZ92863.1 anid|CBF87465.1 anig|EHA26102.1 aory|EIT79990.1 asyd|OJJ52935.1 ater|EAU31731.1 atub|OJI87321.1 aver|OJJ08902.1 awen|OJJ35986.1 azon|OJJ42560.1

Group3705: abra|OJJ75310.1 acar|OOF94519.1 acla|EAW12230.1 afis|EAW16998.1 afla|EED48952.1 afum|EAL86523.1 agla|OJJ79825.1 akaw|GAA91994.1 aluc|OJZ92864.1 anid|CBF87476.1 anig|EHA26101.1 aory|EIT79981.1 asyd|OJJ52941.1 ater|EAU31729.1 atub|OJI87322.1 aver|OJJ08908.1 awen|OJJ35987.1 azon|OJJ50276.1

Group3706: abra|OJJ75314.1 acar|OOF94515.1 acla|EAW10816.1 afis|EAW24758.1 afla|EED56728.1 afum|EAL89400.1 agla|OJJ78831.1 akaw|GAA91998.1 aluc|OJZ92868.1 anid|CBF89399.1 anig|EHA26097.1 aory|EIT73833.1 asyd|OJJ62458.1 ater|EAU34827.1 atub|OJI87326.1 aver|OJI96593.1 awen|OJJ33340.1 azon|OJJ42602.1

Group3708: abra|OJJ75342.1 acar|OOF89833.1 afla|EED48926.1 agla|OJJ81063.1 akaw|GAA92035.1 aluc|OJZ92905.1 anid|CBF87500.1 anig|EHA26058.1 atub|OJI87364.1 aver|OJJ08935.1 awen|OJJ35965.1 azon|OJJ50308.1 acla|EAW12182.1 afis|EAW17116.1 afum|EAL86578.1 asyd|OJJ52969.1 ater|EAU36171.1 aory|EIT80020.1

Group3710: abra|OJJ75381.1 acar|OOF94466.1 acla|EAW12111.1 afis|EAW17274.1 afla|EED48845.1 afum|EAL86692.1 agla|OJJ81078.1 akaw|GAA92743.1 aluc|OJZ92951.1 anid|CBF87538.1 anig|EHA26024.1 aory|EIT81278.1 asyd|OJJ52990.1 ater|EAU34713.1 atub|OJI87410.1 aver|OJJ08959.1 awen|OJJ35760.1 azon|OJJ45524.1

Group3711: abra|OJJ75383.1 acar|OOF94464.1 acla|EAW12109.1 afis|EAW17276.1 afla|EED48843.1 afum|EAL86694.1 agla|OJJ81080.1 akaw|GAA92745.1 aluc|OJZ92953.1 anid|CBF87542.1 anig|EHA26022.1 aory|EIT81280.1 asyd|OJJ52992.1 ater|EAU34711.1 atub|OJI87412.1 aver|OJJ08961.1 awen|OJJ35758.1 azon|OJJ45526.1

Group3712: abra|OJJ75384.1 acar|OOF94463.1 acla|EAW12108.1 afis|EAW17277.1 afla|EED48842.1 afum|EAL86695.1 agla|OJJ81081.1 akaw|GAA92746.1 aluc|OJZ92954.1 anid|CBF87544.1 anig|EHA26021.1 aory|EIT81283.1 asyd|OJJ52993.1 ater|EAU34710.1 atub|OJI87413.1 aver|OJJ08962.1 awen|OJJ35757.1 azon|OJJ45527.1

Group3713: abra|OJJ75385.1 acar|OOF94461.1 acla|EAW12107.1 afis|EAW17278.1 afla|EED48841.1 afum|EAL86696.1 agla|OJJ81082.1 akaw|GAA92747.1 aluc|OJZ92955.1 anid|CBF87545.1 anig|EHA26020.1 aory|EIT81279.1 asyd|OJJ52994.1 ater|EAU34709.1 atub|OJI87414.1 aver|OJJ08963.1 awen|OJJ35756.1 azon|OJJ45528.1

Group3714: abra|OJJ75391.1 acar|OOF94456.1 acla|EAW12101.1 afis|EAW17287.1 afla|EED48833.1 afum|EAL86705.1 agla|OJJ81090.1 akaw|GAA92754.1 aluc|OJZ92962.1 anid|CBF87556.1 anig|EHA26013.1 aory|EIT81270.1 asyd|OJJ53001.1 ater|EAU34702.1 atub|OJI87420.1 aver|OJJ08970.1 awen|OJJ35750.1 azon|OJJ45536.1

Group3715: abra|OJJ75394.1 acar|OOF94455.1 acla|EAW12099.1 afis|EAW17285.1 afla|EED48831.1 afum|EAL86703.1 agla|OJJ81092.1 akaw|GAA92756.1 aluc|OJZ92965.1 anid|CBF87558.1 anig|EHA26010.1 aory|EIT81288.1 asyd|OJJ53003.1 ater|EAU34700.1 atub|OJI87423.1 aver|OJJ08971.1 awen|OJJ35749.1 azon|OJJ45542.1

Group3716: abra|OJJ75396.1 acar|OOF94453.1 acla|EAW12097.1 afis|EAW17289.1 afla|EED48829.1 afum|EAL86709.1 agla|OJJ81094.1 akaw|GAA92758.1 aluc|OJZ92968.1 anid|CBF81166.1 anig|EHA26008.1 aory|EIT81272.1 asyd|OJJ59287.1 ater|EAU34699.1 atub|OJI87425.1 aver|OJJ02988.1 awen|OJJ35747.1 azon|OJJ45540.1

Group3717: abra|OJJ75397.1 acar|OOF94451.1 acla|EAW12096.1 afis|EAW17290.1 afla|EED48828.1 afum|EAL86710.1 agla|OJJ81095.1 akaw|GAA92759.1 aluc|OJZ92969.1 anid|CBF74295.1 anig|EHA26007.1 aory|EIT81266.1 asyd|OJJ51909.1 ater|EAU34698.1 atub|OJI87426.1 aver|OJJ05350.1 awen|OJJ35746.1 azon|OJJ45539.1

Group3718: abra|OJJ75399.1 acar|OOF90746.1 acla|EAW13772.1 afis|EAW18360.1 afla|EED48826.1 afum|EAL87724.1 agla|OJJ88025.1 akaw|GAA84178.1 aluc|OJZ88851.1 anid|CBF79349.1 anig|EHA23652.1 aory|EIT72979.1 asyd|OJJ57798.1 ater|EAU34692.1 atub|OJI85939.1 aver|OJJ04589.1 awen|OJJ35744.1 azon|OJJ49497.1

Group3719: abra|OJJ75400.1 acar|OOF90747.1 acla|EAW13773.1 afis|EAW18359.1 afla|EED48825.1 afum|EAL87723.1 agla|OJJ88026.1 akaw|GAA84179.1 aluc|OJZ88852.1 anid|CBF79351.1 anig|EHA23651.1 aory|EIT73008.1 asyd|OJJ57799.1 ater|EAU34691.1 atub|OJI85938.1 aver|OJJ04588.1 awen|OJJ35743.1 azon|OJJ42023.1

Group3721: abra|OJJ75402.1 acar|OOF90749.1 acla|EAW13774.1 afis|EAW18358.1 afla|EED48823.1 afum|EAL87722.1 agla|OJJ88027.1 akaw|GAA84181.1 aluc|OJZ88854.1 anid|CBF79353.1 anig|EHA23649.1 aory|EIT72976.1 asyd|OJJ57800.1 ater|EAU34690.1 atub|OJI85936.1 aver|OJJ04587.1 awen|OJJ35742.1 azon|OJJ43326.1

Group3722: abra|OJJ75404.1 acar|OOF90751.1 acla|EAW13776.1 afis|EAW18356.1 afum|EAL87719.1 agla|OJJ88029.1 akaw|GAA84182.1 aluc|OJZ88856.1 anid|CBF79356.1 anig|EHA23647.1 aory|EIT72985.1 asyd|OJJ57802.1 ater|EAU34689.1 atub|OJI85934.1 aver|OJJ04585.1 awen|OJJ35740.1 azon|OJJ43323.1 afla|EED48821.1

Group3723: abra|OJJ75408.1 acar|OOF90754.1 acla|EAW13781.1 afis|EAW18351.1 afla|EED48816.1 afum|EAL87714.1 agla|OJJ88033.1 akaw|GAA84184.1 aluc|OJZ88859.1 anid|CBF79365.1 anig|EHA23644.1 aory|EIT72998.1 asyd|OJJ57807.1 ater|EAU34685.1 atub|OJI85931.1 aver|OJJ04581.1 awen|OJJ35735.1 azon|OJJ43321.1

Group3724: abra|OJJ75409.1 acar|OOF90755.1 acla|EAW13780.1 afis|EAW18352.1 afla|EED48817.1 afum|EAL87715.1 agla|OJJ88032.1 akaw|GAA84185.1 aluc|OJZ88860.1 anid|CBF79364.1 anig|EHA23643.1 aory|EIT73011.1 asyd|OJJ57806.1 ater|EAU34686.1 atub|OJI85930.1 aver|OJJ04582.1 awen|OJJ35736.1 azon|OJJ43322.1

Group3725: abra|OJJ75410.1 acar|OOF90756.1 acla|EAW13782.1 afis|EAW18350.1 afla|EED48815.1 afum|EAL87713.1 agla|OJJ88034.1 akaw|GAA84186.1 aluc|OJZ88861.1 anid|CBF79367.1 anig|EHA23642.1 aory|EIT73016.1 asyd|OJJ57808.1 ater|EAU34684.1 atub|OJI85929.1 aver|OJJ04580.1 awen|OJJ35734.1 azon|OJJ43319.1

Group3726: abra|OJJ75413.1 acar|OOF90757.1 acla|EAW13785.1 afis|EAW18347.1 afla|EED48812.1 afum|EAL87711.1 agla|OJJ88037.1 akaw|GAA84188.1 aluc|OJZ88864.1 anid|CBF79386.1 anig|EHA23639.1 aory|EIT73027.1 asyd|OJJ57811.1 ater|EAU34682.1 atub|OJI85926.1 aver|OJJ04577.1 awen|OJJ35731.1 azon|OJJ43318.1

Group3727: abra|OJJ75414.1 acar|OOF90760.1 acla|EAW13786.1 afis|EAW18346.1 afla|EED48811.1 afum|EAL87710.1 agla|OJJ88038.1 akaw|GAA84189.1 aluc|OJZ88865.1 anid|CBF79384.1 anig|EHA23638.1 aory|EIT73037.1 asyd|OJJ57812.1 ater|EAU34681.1 atub|OJI85925.1 aver|OJJ04576.1 awen|OJJ35730.1 azon|OJJ43317.1

Group3728: abra|OJJ75415.1 acar|OOF90759.1 acla|EAW13787.1 afis|EAW18345.1 afla|EED48810.1 afum|EAL87709.1 agla|OJJ88039.1 akaw|GAA84190.1 aluc|OJZ88866.1 anid|CBF79382.1 anig|EHA23637.1 aory|EIT72992.1 asyd|OJJ57813.1 ater|EAU34680.1 atub|OJI85924.1 aver|OJJ04575.1 awen|OJJ35729.1 azon|OJJ43316.1

Group3729: abra|OJJ75417.1 acar|OOF90762.1 acla|EAW13789.1 afis|EAW18343.1 afla|EED48808.1 afum|EAL87706.1 agla|OJJ88041.1 akaw|GAA84192.1 aluc|OJZ88868.1 anid|CBF79378.1 anig|EHA23635.1 aory|EIT73022.1 asyd|OJJ57815.1 ater|EAU34678.1 atub|OJI85922.1 aver|OJJ04573.1 awen|OJJ35727.1 azon|OJJ43315.1

Group3730: abra|OJJ75421.1 acar|OOF90765.1 acla|EAW13793.1 afis|EAW18339.1 afla|EED48803.1 afum|EAL87702.1 agla|OJJ88045.1 akaw|GAA84196.1 aluc|OJZ88873.1 anid|CBF79387.1 anig|EHA23632.1 aory|EIT73019.1 asyd|OJJ57819.1 ater|EAU34674.1 atub|OJI85918.1 aver|OJJ04572.1 awen|OJJ35722.1 azon|OJJ43312.1

Group3731: abra|OJJ75425.1 acar|OOF90770.1 acla|EAW13795.1 afis|EAW18337.1 afla|EED48801.1 afum|EAL87701.1 agla|OJJ88047.1 akaw|GAA84200.1 aluc|OJZ88877.1 anid|CBF79391.1 anig|EHA23628.1

aory|EIT73024.1 asyd|OJJ57821.1 ater|EAU33350.1 atub|OJI85914.1 aver|OJJ04570.1 awen|OJJ35720.1 azon|OJJ43310.1

Group3732: abra|OJJ75426.1 acar|OOF90771.1 acla|EAW13796.1 afis|EAW18336.1 afla|EED48800.1 afum|EAL87700.1 agla|OJJ88049.1 akaw|GAA84201.1 aluc|OJZ88878.1 anid|CBF79395.1 anig|EHA23627.1 aory|EIT73005.1 asyd|OJJ57825.1 ater|EAU33349.1 atub|OJI85913.1 aver|OJJ04567.1 awen|OJJ30717.1 azon|OJJ43309.1

Group3733: abra|OJJ75427.1 acar|OOF90773.1 acla|EAW13798.1 afis|EAW18334.1 afla|EED48798.1 afum|EAL87699.1 agla|OJJ88051.1 akaw|GAA84203.1 aluc|OJZ88880.1 anid|CBF79393.1 anig|EHA23626.1 aory|EIT73035.1 asyd|OJJ57822.1 ater|EAU33347.1 atub|OJI85912.1 aver|OJJ04569.1 awen|OJJ30715.1 azon|OJJ43307.1

Group3734: abra|OJJ75428.1 acar|OOF90774.1 acla|EAW11898.1 afis|EAW19508.1 afla|EED56888.1 afum|EAL86986.1 agla|OJJ88052.1 akaw|GAA84204.1 aluc|OJZ88881.1 anid|CBF82976.1 anig|EHA23625.1 aory|EIT77571.1 asyd|OJJ61561.1 ater|EAU33346.1 atub|OJI85911.1 aver|OJJ00884.1 awen|OJJ30714.1 azon|OJJ45886.1

Group3735: abra|OJJ75429.1 acar|OOF90775.1 acla|EAW13799.1 afis|EAW18333.1 afla|EED48797.1 afum|EAL87698.1 agla|OJJ88053.1 akaw|GAA84205.1 aluc|OJZ88882.1 anid|CBF79397.1 anig|EHA23624.1 aory|EIT73004.1 asyd|OJJ57826.1 ater|EAU33345.1 atub|OJI85910.1 aver|OJJ04566.1 awen|OJJ30713.1 azon|OJJ43306.1

Group3736: abra|OJJ75432.1 acar|OOF90777.1 acla|EAW13804.1 afis|EAW18327.1 afla|EED48793.1 afum|EAL87693.1 agla|OJJ88056.1 akaw|GAA84208.1 aluc|OJZ88885.1 anid|CBF79406.1 anig|EHA23621.1 aory|EIT72996.1 asyd|OJJ57829.1 ater|EAU33336.1 atub|OJI85905.1 aver|OJJ04563.1 awen|OJJ30711.1 azon|OJJ43303.1

Group3737: abra|OJJ75433.1 acar|OOF90778.1 acla|EAW13805.1 afis|EAW18325.1 afla|EED48792.1 afum|EAL87691.1 agla|OJJ88066.1 akaw|GAA84209.1 aluc|OJZ88886.1 anid|CBF79408.1 anig|EHA23620.1 aory|EIT72982.1 asyd|OJJ57830.1 ater|EAU33334.1 atub|OJI85904.1 aver|OJJ04562.1 awen|OJJ30707.1 azon|OJJ43302.1

Group3738: abra|OJJ75435.1 acar|OOF90779.1 acla|EAW13827.1 afis|EAW18304.1 afla|EED48773.1 afum|EAL87670.1 agla|OJJ88119.1 akaw|GAA84210.1 aluc|OJZ88888.1 anid|CBF79447.1 anig|EHA23619.1 aory|EIT73020.1 asyd|OJJ57851.1 ater|EAU33311.1 atub|OJI85901.1 aver|OJJ04541.1 awen|OJJ30678.1 azon|OJJ43279.1

Group3739: abra|OJJ75436.1 acar|OOF90780.1 acla|EAW13826.1 afis|EAW18305.1 afla|EED48774.1 afum|EAL87671.1 agla|OJJ88118.1 akaw|GAA84211.1 aluc|OJZ88889.1 anid|CBF79445.1 anig|EHA23618.1 aory|EIT73030.1 asyd|OJJ57850.1 ater|EAU33312.1 atub|OJI85900.1 aver|OJJ04542.1 awen|OJJ30679.1 azon|OJJ43280.1

Group3740: abra|OJJ75438.1 acar|OOF90783.1 acla|EAW12400.1 afis|EAW16848.1 afla|EED51031.1 afum|EAL86394.1 agla|OJJ81486.1 akaw|GAA84213.1 aluc|OJZ88891.1 anid|CBF81652.1 anig|EHA23615.1 aory|EIT77036.1 asyd|OJJ59545.1 ater|EAU30241.1 atub|OJI85898.1 aver|OJJ02722.1 awen|OJJ30081.1 azon|OJJ44766.1

Group3741: abra|OJJ75442.1 acar|OOF90787.1 acla|EAW13821.1 afis|EAW18310.1 afla|EED48779.1 afum|EAL87676.1 agla|OJJ88079.1 akaw|GAA84217.1 aluc|OJZ88895.1 anid|CBF79434.1 anig|EHA23611.1 aory|EIT73023.1 asyd|OJJ57844.1 ater|EAU33320.1 atub|OJI85894.1 aver|OJJ04548.1 awen|OJJ30685.1 azon|OJJ43284.1

Group3742: abra|OJJ75443.1 acar|OOF90788.1 acla|EAW13820.1 afis|EAW18311.1 afla|EED48780.1 afum|EAL87677.1 agla|OJJ88080.1 akaw|GAA84218.1 aluc|OJZ88896.1 anid|CBF79432.1 anig|EHA23610.1 aory|EIT72999.1 asyd|OJJ57845.1 ater|EAU33321.1 atub|OJI85893.1 aver|OJJ04547.1 awen|OJJ30686.1 azon|OJJ43285.1

Group3743: abra|OJJ75445.1 acar|OOF90789.1 acla|EAW13818.1 afis|EAW18312.1 afla|EED48781.1 afum|EAL87679.1 agla|OJJ88081.1 akaw|GAA84220.1 aluc|OJZ88900.1 anid|CBF79430.1 anig|EHA23608.1 aory|EIT73002.1 asyd|OJJ57841.1 ater|EAU33322.1 atub|OJI85891.1 aver|OJJ04549.1 awen|OJJ30688.1 azon|OJJ43286.1

Group3744: abra|OJJ75446.1 acar|OOF90790.1 acla|EAW13817.1 afis|EAW18313.1 afla|EED48782.1 afum|EAL87680.1 agla|OJJ88082.1 akaw|GAA84221.1 aluc|OJZ88901.1 anid|CBF79428.1 anig|EHA23607.1 aory|EIT73015.1 asyd|OJJ57840.1 ater|EAU33323.1 atub|OJI85890.1 aver|OJJ04550.1 awen|OJJ30689.1 azon|OJJ43287.1

Group3745: abra|OJJ75447.1 acar|OOF90791.1 acla|EAW13816.1 afis|EAW18314.1 afla|EED48783.1 afum|EAL87681.1 agla|OJJ88083.1 akaw|GAA84222.1 aluc|OJZ88902.1 anid|CBF79427.1 anig|EHA23606.1 aory|EIT73014.1 asyd|OJJ57839.1 ater|EAU33324.1 atub|OJI85889.1 aver|OJJ04551.1 awen|OJJ30690.1 azon|OJJ43288.1

Group3746: abra|OJJ75448.1 acar|OOF90792.1 acla|EAW13815.1 afis|EAW18315.1 afla|EED48784.1 afum|EAL87682.1 agla|OJJ88084.1 akaw|GAA84223.1 aluc|OJZ88903.1 anid|CBF79423.1 anig|EHA23605.1 aory|EIT72980.1 asyd|OJJ57837.1 ater|EAU33325.1 atub|OJI85888.1 aver|OJJ04553.1 awen|OJJ30691.1 azon|OJJ43289.1

Group3747: abra|OJJ75450.1 acar|OOF90794.1 acla|EAW13813.1 afis|EAW18317.1 afla|EED48785.1 afum|EAL87684.1 agla|OJJ88101.1 akaw|GAA84225.1 aluc|OJZ88905.1 anid|CBF79421.1 anig|EHA23603.1 aory|EIT73000.1 asyd|OJJ57836.1 ater|EAU33326.1 atub|OJI85886.1 aver|OJJ04554.1 awen|OJJ30699.1 azon|OJJ43292.1

Group3748: abra|OJJ75452.1 acar|OOF90796.1 acla|EAW13811.1 afis|EAW18319.1 afla|EED48787.1 afum|EAL87686.1 agla|OJJ88073.1 akaw|GAA84227.1 aluc|OJZ88907.1 anid|CBF87726.1 anig|EHA23601.1 aory|EIT72981.1 asyd|OJJ63342.1 ater|EAU33329.1 atub|OJI85884.1 aver|OJI97619.1 awen|OJJ30701.1 azon|OJJ50771.1

Group3749: abra|OJJ75454.1 acar|OOF90798.1 acla|EAW13809.1 afis|EAW18321.1 afla|EED48789.1 afum|EAL87688.1 agla|OJJ88070.1 akaw|GAA84229.1 aluc|OJZ88909.1 anid|CBF79416.1 anig|EHA23599.1 aory|EIT73021.1 asyd|OJJ57833.1 ater|EAU33331.1 atub|OJI85882.1 aver|OJJ04557.1 awen|OJJ30703.1 azon|OJJ43296.1

Group3750: abra|OJJ75457.1 acar|OOF90802.1 acla|EAW13808.1 afis|EAW18322.1 afla|EED48790.1 afum|EAL87689.2 agla|OJJ88069.1 akaw|GAA84232.1 aluc|OJZ88913.1 anid|CBF79414.1 anig|EHA23597.1 aory|EIT73033.1 asyd|OJJ57832.1 ater|EAU33332.1 atub|OJI85878.1 aver|OJJ04558.1 awen|OJJ30704.1 azon|OJJ43297.1

Group3751: abra|OJJ75460.1 acar|OOF90279.1 acla|EAW13828.1 afis|EAW18303.1 afla|EED48772.1 afum|EAL87669.1 agla|OJJ88120.1 akaw|GAA84234.1 aluc|OJZ88917.1 anid|CBF79449.1 anig|EHA23595.1 aory|EIT72974.1 asyd|OJJ57852.1 ater|EAU33310.1 atub|OJI85875.1 aver|OJJ04540.1 awen|OJJ30677.1 azon|OJJ43278.1

Group3752: abra|OJJ75463.1 acar|OOF90277.1 acla|EAW13830.1 afis|EAW18301.1 afla|EED48770.1 afum|EAL87667.2 agla|OJJ88122.1 akaw|GAA84236.1 aluc|OJZ88919.1 anid|CBF79480.1 anig|EHA23593.1 aory|EIT72978.1 asyd|OJJ57871.1 ater|EAU33309.1 atub|OJI85873.1 aver|OJJ04523.1 awen|OJJ30676.1 azon|OJJ43276.1

Group3753: abra|OJJ75465.1 acar|OOF90275.1 acla|EAW13832.1 afis|EAW18299.1 afla|EED48768.1 afum|EAL87665.1 agla|OJJ88124.1 akaw|GAA84238.1 aluc|OJZ88921.1 anid|CBF79476.1 anig|EHA23591.1 aory|EIT73017.1 asyd|OJJ57869.1 ater|EAU33307.1 atub|OJI85871.1 aver|OJJ04525.1 awen|OJJ30673.1 azon|OJJ43274.1

Group3754: abra|OJJ75466.1 acar|OOF90803.1 acla|EAW13833.1 afis|EAW18298.1 afla|EED48767.1 afum|EAL87664.2 agla|OJJ88125.1 akaw|GAA84239.1 aluc|OJZ88922.1 anid|CBF79472.1 anig|EHA23590.1 aory|EIT73036.1 asyd|OJJ57867.1 ater|EAU33306.1 atub|OJI85870.1 aver|OJJ04526.1 awen|OJJ30672.1 azon|OJJ43272.1

Group3755: abra|OJJ75468.1 acar|OOF90805.1 acla|EAW13834.1 afis|EAW18297.1 afla|EED48763.1 afum|EAL87663.1 agla|OJJ88126.1 akaw|GAA84240.1 aluc|OJZ88925.1 anid|CBF79470.1 anig|EHA23588.1 aory|EIT73013.1 asyd|OJJ57865.1 ater|EAU33305.1 atub|OJI85867.1 aver|OJJ04528.1 awen|OJJ30671.1 azon|OJJ43271.1

Group3756: abra|OJJ75472.1 acar|OOF90809.1 acla|EAW13837.1 afis|EAW18294.1 afla|EED48761.1 afum|EAL87660.1 agla|OJJ88129.1 akaw|GAA84243.1 aluc|OJZ88931.1 anid|CBF79465.1 anig|EHA23585.1 aory|EIT73038.1 asyd|OJJ57861.1 ater|EAU33303.1 atub|OJI85861.1 aver|OJJ04531.1 awen|OJJ30669.1 azon|OJJ43268.1

Group3757: abra|OJJ75473.1 acar|OOF90810.1 acla|EAW13838.1 afis|EAW18293.1 afla|EED48760.1 afum|EAL87659.1 agla|OJJ88132.1 akaw|GAA84244.1 aluc|OJZ88932.1 anid|CBF79463.1 anig|EHA23584.1 aory|EIT72991.1 asyd|OJJ57860.1 ater|EAU33302.1 atub|OJI85860.1 aver|OJJ04532.1 awen|OJJ30668.1 azon|OJJ43267.1

Group3758: abra|OJJ75474.1 acar|OOF90811.1 acla|EAW13839.1 afis|EAW18292.1 afla|EED48759.1 afum|EAL87658.1 agla|OJJ88133.1 akaw|GAA84245.1 aluc|OJZ88933.1 anid|CBF79461.1 anig|EHA23583.1 aory|EIT72984.1 asyd|OJJ57859.1 ater|EAU33301.1 atub|OJI85859.1 aver|OJJ04534.1 awen|OJJ30667.1 azon|OJJ43265.1

Group3759: abra|OJJ75476.1 acar|OOF90812.1 acla|EAW13841.1 afis|EAW18291.1 afla|EED48758.1 afum|EAL87656.1 agla|OJJ88134.1 akaw|GAA84246.1 aluc|OJZ88935.1 anid|CBF79459.1 anig|EHA23582.1 aory|EIT73034.1 asyd|OJJ57858.1 ater|EAU33299.1 atub|OJI85857.1 aver|OJJ04535.1 awen|OJJ30666.1 azon|OJJ43263.1

Group3760: abra|OJJ75480.1 acar|OOF90816.1 acla|EAW11214.1 afis|EAW18288.1 afla|EED48756.1 afum|EAL87654.2 agla|OJJ88136.1 akaw|GAA84249.1 aluc|OJZ88939.1 anid|CBF79454.1 anig|EHA23579.1

aory|EIT73128.1 asyd|OJJ57856.1 ater|EAU33296.1 atub|OJI85851.1 aver|OJJ04538.1 awen|OJJ30661.1 azon|OJJ47954.1

Group3761: abra|OJJ75486.1 acar|OOF90818.1 acla|EAW11217.1 afis|EAW18286.1 afla|EED48753.1 afum|EAL87652.1 agla|OJJ88138.1 akaw|GAA84252.1 aluc|OJZ88945.1 anid|CBF79450.1 anig|EHA23576.1 aory|EIT73238.1 asyd|OJJ57853.1 ater|EAU33292.1 atub|OJI85844.1 aver|OJJ04539.1 awen|OJJ30658.1 azon|OJJ47956.1

Group3762: abra|OJJ75487.1 acar|OOF90820.1 acla|EAW11218.1 afis|EAW18285.1 afla|EED48752.1 afum|EAL87651.1 agla|OJJ88139.1 akaw|GAA84253.1 aluc|OJZ88946.1 anid|CBF79482.1 anig|EHA23575.1 aory|EIT73219.1 asyd|OJJ57872.1 ater|EAU33291.1 atub|OJI85843.1 aver|OJJ04522.1 awen|OJJ30657.1 azon|OJJ47957.1

Group3763: abra|OJJ75488.1 acar|OOF90819.1 acla|EAW11219.1 afis|EAW18284.1 afla|EED48751.1 afum|EAL87650.1 agla|OJJ88140.1 akaw|GAA84254.1 aluc|OJZ88947.1 anid|CBF79483.1 anig|EHA23574.1 aory|EIT73141.1 asyd|OJJ57873.1 ater|EAU33290.1 atub|OJI85842.1 aver|OJJ04521.1 awen|OJJ30655.1 azon|OJJ47959.1

Group3764: abra|OJJ75491.1 acar|OOF90823.1 acla|EAW11223.1 afis|EAW18280.1 afla|EED48747.1 afum|EAL87646.1 agla|OJJ88146.1 akaw|GAA84256.1 aluc|OJZ88950.1 anid|CBF79491.1 anig|EHA23571.1 aory|EIT73260.1 asyd|OJJ57876.1 ater|EAU33286.1 atub|OJI85839.1 aver|OJJ04518.1 awen|OJJ30652.1 azon|OJJ47963.1

Group3765: abra|OJJ75493.1 acar|OOF90826.1 acla|EAW11226.1 afis|EAW18277.1 afla|EED48744.1 afum|EAL87643.1 agla|OJJ88149.1 akaw|GAA84259.1 aluc|OJZ88952.1 anid|CBF79496.1 anig|EHA23568.1 aory|EIT73150.1 asyd|OJJ57878.1 ater|EAU33283.1 atub|OJI85837.1 aver|OJJ04516.1 awen|OJJ30650.1 azon|OJJ47965.1

Group3766: abra|OJJ75494.1 acar|OOF90827.1 acla|EAW09288.1 afis|EAW21859.1 afla|EED46451.1 afum|EAL92872.1 agla|OJJ84276.1 akaw|GAA84260.1 aluc|OJZ88953.1 anid|CBF76751.1 anig|EHA23567.1 aory|EIT72375.1 asyd|OJJ61260.1 ater|EAU30858.1 atub|OJI85836.1 aver|OJJ00583.1 awen|OJJ34143.1 azon|OJJ43483.1

Group3767: abra|OJJ75495.1 acar|OOF90828.1 acla|EAW09289.1 afis|EAW21858.1 afla|EED46452.1 afum|EAL92871.1 agla|OJJ84277.1 akaw|GAA84261.1 aluc|OJZ88954.1 anid|CBF76749.1 anig|EHA23566.1 aory|EIT72324.1 asyd|OJJ61259.1 ater|EAU30857.1 atub|OJI85835.1 aver|OJJ00582.1 awen|OJJ34142.1 azon|OJJ43484.1

Group3768: abra|OJJ75496.1 acar|OOF90829.1 acla|EAW09290.1 afis|EAW21857.1 afla|EED46453.1 afum|EAL92870.1 agla|OJJ84278.1 akaw|GAA84262.1 aluc|OJZ88955.1 anid|CBF76747.1 anig|EHA23565.1 aory|EIT72382.1 asyd|OJJ61258.1 ater|EAU30856.1 atub|OJI85834.1 aver|OJJ00581.1 awen|OJJ34141.1 azon|OJJ43485.1

Group3769: abra|OJJ75497.1 acar|OOF90830.1 acla|EAW09291.1 afis|EAW21856.1 afla|EED46454.1 afum|EAL92869.1 agla|OJJ84279.1 akaw|GAA84263.1 aluc|OJZ88956.1 anid|CBF76745.1 aory|EIT72446.1 asyd|OJJ61257.1 ater|EAU30855.1 atub|OJI85833.1 aver|OJJ00580.1 awen|OJJ34140.1 azon|OJJ43486.1 anig|EHA23564.1

Group3770: abra|OJJ75506.1 acar|OOF89956.1 acla|EAW09296.1 afis|EAW21851.1 afla|EED46459.1 afum|EAL92864.1 agla|OJJ84284.1 akaw|GAA84269.1 aluc|OJZ88962.1 anid|CBF76735.1 anig|EHA23559.1 aory|EIT72419.1 asyd|OJJ61252.1 ater|EAU30850.1 atub|OJI85828.1 aver|OJJ00576.1 awen|OJJ34134.1 azon|OJJ44210.1

Group3771: abra|OJJ75507.1 acar|OOF89955.1 acla|EAW09297.1 afis|EAW21850.1 afla|EED46460.1 afum|EAL92863.2 agla|OJJ84285.1 akaw|GAA84270.1 aluc|OJZ88963.1 anid|CBF76733.1 anig|EHA23558.1 aory|EIT72388.1 asyd|OJJ61251.1 ater|EAU30849.1 atub|OJI85827.1 aver|OJJ00575.1 awen|OJJ34133.1 azon|OJJ46409.1

Group3772: abra|OJJ75511.1 acar|OOF90837.1 acla|EAW09299.1 afis|EAW21848.1 afum|EAL92861.2 agla|OJJ84287.1 akaw|GAA84275.1 aluc|OJZ88968.1 anid|CBF76727.1 anig|EHA23555.1 asyd|OJJ61248.1 ater|EAU30847.1 atub|OJI85822.1 aver|OJJ00572.1 awen|OJJ34131.1 azon|OJJ44212.1 afla|EED46464.1 aory|EIT72335.1

Group3773: abra|OJJ75512.1 acar|OOF90839.1 acla|EAW09300.1 afis|EAW21847.1 afla|EED46468.1 afum|EAL92860.1 agla|OJJ84288.1 akaw|GAA84276.1 aluc|OJZ88969.1 anid|CBF76725.1 anig|EHA23554.1 aory|EIT72394.1 asyd|OJJ61246.1 ater|EAU30846.1 atub|OJI85821.1 aver|OJJ00570.1 awen|OJJ34130.1 azon|OJJ44213.1

Group3774: abra|OJJ75518.1 acar|OOF90845.1 acla|EAW09302.1 afis|EAW21844.1 afla|EED46526.1 afum|EAL92856.1 agla|OJJ84292.1 akaw|GAA84282.1 aluc|OJZ88975.1 anid|CBF76717.1 anig|EHA23550.1 aory|EIT72426.1 asyd|OJJ61242.1 ater|EAU38521.1 atub|OJI85814.1 aver|OJJ00565.1 awen|OJJ34128.1 azon|OJJ44215.1

Group3775: abra|OJJ75521.1 acar|OOF90848.1 acla|EAW09308.1 afis|EAW21837.1 afla|EED46538.1 afum|EAL92851.1 agla|OJJ84305.1 akaw|GAA84285.1 aluc|OJZ88978.1 anid|CBF76701.1 anig|EHA23547.1 aory|EIT72362.1 asyd|OJJ61233.1 ater|EAU38512.1 atub|OJI85811.1 aver|OJJ00556.1 awen|OJJ34113.1 azon|OJJ44227.1

Group3776: abra|OJJ75523.1 acar|OOF90850.1 acla|EAW09310.1 afis|EAW21835.1 afla|EED46540.1 afum|EAL92850.1 agla|OJJ84309.1 akaw|GAA84286.1 aluc|OJZ88980.1 anid|CBF76699.1 anig|EHA23546.1 aory|EIT72364.1 asyd|OJJ61232.1 ater|EAU38510.1 atub|OJI85809.1 aver|OJJ00555.1 awen|OJJ34109.1 azon|OJJ44232.1

Group3777: abra|OJJ75527.1 acar|OOF90854.1 acla|EAW09313.1 afis|EAW21832.1 afla|EED46544.1 afum|EAL92847.1 agla|OJJ84312.1 akaw|GAA84289.1 aluc|OJZ88983.1 anid|CBF76694.1 anig|EHA23543.1 aory|EIT77942.1 asyd|OJJ61229.1 ater|EAU38508.1 atub|OJI85806.1 aver|OJJ00552.1 awen|OJJ34105.1 azon|OJJ44236.1

Group3778: abra|OJJ75528.1 acar|OOF90855.1 acla|EAW09314.1 afis|EAW21831.1 afla|EED46545.1 afum|EAL92846.1 agla|OJJ84313.1 akaw|GAA84290.1 aluc|OJZ88984.1 anid|CBF76680.1 anig|EHA23542.1 aory|EIT77881.1 asyd|OJJ61223.1 ater|EAU38507.1 atub|OJI85804.1 aver|OJJ00546.1 awen|OJJ34104.1 azon|OJJ44238.1

Group3779: abra|OJJ75539.1 acar|OOF90858.1 acla|EAW09315.1 afis|EAW21830.1 afla|EED46546.1 afum|EAL92845.1 agla|OJJ84314.1 akaw|GAA84293.1 aluc|OJZ88987.1 anid|CBF76676.1 anig|EHA23539.1 aory|EIT77971.1 asyd|OJJ61221.1 ater|EAU38506.1 atub|OJI85801.1 aver|OJJ00545.1 awen|OJJ34103.1 azon|OJJ44239.1

Group3780: abra|OJJ75544.1 acar|OOF90864.1 acla|EAW09316.1 afis|EAW21829.1 afla|EED46547.1 afum|EAL92844.1 agla|OJJ84315.1 akaw|GAA84300.1 aluc|OJZ88994.1 anid|CBF76674.1 anig|EHA23533.1 aory|EIT77834.1 asyd|OJJ61220.1 ater|EAU38505.1 atub|OJI85794.1 aver|OJJ00544.1 awen|OJJ34102.1 azon|OJJ44242.1

Group3781: abra|OJJ75545.1 acar|OOF90866.1 acla|EAW09317.1 afis|EAW21828.1 afla|EED46548.1 afum|EAL92843.1 agla|OJJ84316.1 akaw|GAA84301.1 aluc|OJZ88996.1 anid|CBF76670.1 anig|EHA23532.1 aory|EIT77919.1 asyd|OJJ61219.1 ater|EAU38504.1 atub|OJI85793.1 aver|OJJ00543.1 awen|OJJ34101.1 azon|OJJ44243.1

Group3782: abra|OJJ75548.1 acar|OOF90868.1 acla|EAW09319.1 afis|EAW21826.1 afla|EED46553.1 afum|EAL92841.1 agla|OJJ84318.1 akaw|GAA84303.1 aluc|OJZ88998.1 anid|CBF76666.1 anig|EHA23530.1 aory|EIT77928.1 asyd|OJJ61217.1 ater|EAU38502.1 atub|OJI85791.1 aver|OJJ00541.1 awen|OJJ34099.1 azon|OJJ44245.1

Group3783: abra|OJJ75552.1 acar|OOF90870.1 acla|EAW09324.1 afis|EAW21821.1 afla|EED46558.1 afum|EAL92837.1 agla|OJJ84321.1 akaw|GAA84307.1 aluc|OJZ89002.1 anid|CBF76660.1 anig|EHA23526.1 aory|EIT77966.1 asyd|OJJ61212.1 ater|EAU38495.1 atub|OJI85787.1 aver|OJJ00537.1 awen|OJJ34095.1 azon|OJJ44249.1

Group3784: abra|OJJ75553.1 acar|OOF90872.1 acla|EAW09355.1 afis|EAW21820.1 afla|EED46559.1 afum|EAL92836.1 agla|OJJ84322.1 akaw|GAA84308.1 aluc|OJZ89003.1 anid|CBF76658.1 anig|EHA23525.1 aory|EIT77877.1 asyd|OJJ61211.1 ater|EAU38494.1 atub|OJI85786.1 aver|OJJ00535.1 awen|OJJ34094.1 azon|OJJ44252.1

Group3785: abra|OJJ75554.1 acar|OOF90873.1 acla|EAW09354.1 afis|EAW21819.1 afla|EED46564.1 afum|EAL92835.1 agla|OJJ84327.1 akaw|GAA84309.1 aluc|OJZ89004.1 anid|CBF76654.1 anig|EHA23524.1 aory|EIT77873.1 asyd|OJJ61210.1 ater|EAU38490.1 atub|OJI85785.1 aver|OJJ00534.1 awen|OJJ34093.1 azon|OJJ44253.1

Group3786: abra|OJJ75574.1 acar|OOF90890.1 acla|EAW09346.1 afis|EAW21799.1 afla|EED46590.1 afum|EAL92824.2 agla|OJJ84324.1 akaw|GAA84325.1 aluc|OJZ89022.1 anid|CBF76633.1 anig|EHA23509.1 aory|EIT77934.1 asyd|OJJ61197.1 ater|EAU38481.1 atub|OJI85767.1 aver|OJJ00521.1 awen|OJJ34086.1 azon|OJJ44256.1

Group3787: abra|OJJ75575.1 acar|OOF90893.1 acla|EAW09343.1 afis|EAW21797.1 afla|EED46598.1 afum|EAL92822.2 agla|OJJ84331.1 akaw|GAA84327.1 aluc|OJZ89024.1 anid|CBF76629.1 anig|EHA23508.1 aory|EIT77937.1 asyd|OJJ61196.1 ater|EAU38479.1 atub|OJI85766.1 aver|OJJ00520.1 awen|OJJ34082.1 azon|OJJ44260.1

Group3789: abra|OJJ75579.1 acar|OOF90896.1 acla|EAW09340.1 afis|EAW21795.1 afla|EED46600.1 afum|EAL92820.1 agla|OJJ84333.1 akaw|GAA84331.1 aluc|OJZ89030.1 anid|CBF76625.1 anig|EHA23504.1 aory|EIT77931.1 asyd|OJJ61194.1 ater|EAU38477.1 atub|OJI85761.1 aver|OJJ00518.1 azon|OJJ44262.1 awen|OJJ34079.1

Group3790: abra|OJJ75581.1 acar|OOF90899.1 acla|EAW09337.1 afis|EAW21791.1 afla|EED46604.1 afum|EAL92816.1 agla|OJJ88519.1 akaw|GAA84333.1 aluc|OJZ89032.1 anid|CBF76623.1 anig|EHA23502.1

aory|EIT77970.1 asyd|OJJ61193.1 ater|EAU38475.1 atub|OJI85758.1 aver|OJJ00517.1 awen|OJJ37963.1 azon|OJJ46824.1

Group3791: abra|OJJ75592.1 acar|OOG00236.1 acla|EAW09330.1 afis|EAW21786.1 afla|EED46610.1 afum|EAL92813.1 agla|OJJ84337.1 akaw|GAA84344.1 aluc|OJZ89044.1 anid|CBF76617.1 anig|EHA23491.1 aory|EIT77835.1 asyd|OJJ61189.1 ater|EAU38469.1 atub|OJI85743.1 aver|OJJ00513.1 awen|OJJ34072.1 azon|OJJ48155.1

Group3792: abra|OJJ75602.1 acar|OOG00235.1 acla|EAW09327.1 afis|EAW21784.1 afla|EED46611.1 afum|EAL92811.1 agla|OJJ84339.1 akaw|GAA84352.1 aluc|OJZ89052.1 anid|CBF76613.1 anig|EHA23486.1 aory|EIT77805.1 asyd|OJJ61188.1 ater|EAU38467.1 atub|OJI85735.1 aver|OJJ00511.1 awen|OJJ34071.1 azon|OJJ48157.1

Group3793: abra|OJJ75605.1 acar|OOG00232.1 acla|EAW09367.1 afis|EAW21770.1 afla|EED46330.1 afum|EAL92799.1 agla|OJJ84378.1 akaw|GAA84356.1 aluc|OJZ89058.1 anid|CBF83758.1 anig|EHA23484.1 aory|EIT81243.1 asyd|OJJ60861.1 ater|EAU38448.1 atub|OJI85728.1 aver|OJJ00187.1 awen|OJJ34057.1 azon|OJJ48175.1

Group3794: abra|OJJ75610.1 acar|OOG00228.1 acla|EAW09372.1 afis|EAW21763.1 afla|EED46336.1 afum|EAL92793.1 agla|OJJ84388.1 akaw|GAA84360.1 aluc|OJZ89062.1 anid|CBF83751.1 anig|EHA23476.1 aory|EIT81185.1 asyd|OJJ60857.1 ater|EAU38452.1 atub|OJI85724.1 aver|OJJ00181.1 awen|OJJ34046.1 azon|OJJ48185.1

Group3795: abra|OJJ75611.1 acar|OOG00227.1 acla|EAW09373.1 afis|EAW21762.1 afla|EED46337.1 afum|EAL92792.1 agla|OJJ84389.1 akaw|GAA84361.1 aluc|OJZ89063.1 anid|CBF83749.1 anig|EHA23475.1 aory|EIT81205.1 asyd|OJJ60856.1 ater|EAU38453.1 atub|OJI85723.1 aver|OJJ00180.1 awen|OJJ34045.1 azon|OJJ48186.1

Group3796: abra|OJJ75615.1 acar|OOG00215.1 acla|EAW09363.1 afis|EAW21774.1 afla|EED46340.1 afum|EAL92803.1 agla|OJJ84350.1 akaw|GAA84364.1 aluc|OJZ89068.1 anid|CBF76597.1 anig|EHA23468.1 aory|EIT81181.1 asyd|OJJ61178.1 ater|EAU38464.1 atub|OJI85720.1 aver|OJJ00504.1 awen|OJJ34062.1 azon|OJJ48170.1

Group3797: abra|OJJ75625.1 acar|OOG00211.1 acla|EAW09360.1 afis|EAW21777.1 afla|EED46344.1 afum|EAL92806.1 agla|OJJ84342.1 akaw|GAA84372.1 aluc|OJZ89076.1 anid|CBF76603.1 anig|EHA23461.1 aory|EIT81165.1 asyd|OJJ61182.1 ater|EAU38462.1 atub|OJI85709.1 aver|OJJ00507.1 awen|OJJ34066.1 azon|OJJ48165.1

Group3798: abra|OJJ75628.1 acar|OOG00210.1 acla|EAW09465.1 afis|EAW21671.1 afla|EED57935.1 afum|EAL92706.1 agla|OJJ79745.1 akaw|GAA84375.1 aluc|OJZ89079.1 anid|CBF83585.1 anig|EHA23458.1 aory|EIT82333.1 asyd|OJJ60779.1 ater|EAU38355.1 atub|OJI85706.1 aver|OJJ00112.1 awen|OJJ33935.1 azon|OJJ48642.1

Group3799: abra|OJJ75630.1 acar|OOG00205.1 acla|EAW09463.1 afis|EAW21674.1 afla|EED57936.1 afum|EAL92708.1 agla|OJJ79747.1 akaw|GAA84377.1 aluc|OJZ89081.1 anid|CBF83589.1 anig|EHA23456.1 aory|EIT82341.1 asyd|OJJ60776.1 ater|EAU38357.1 atub|OJI85703.1 aver|OJJ00110.1 awen|OJJ33940.1 azon|OJJ48629.1

Group3800: abra|OJJ75633.1 acar|OOG00202.1 acla|EAW09461.1 agla|OJJ79927.1 akaw|GAA84380.1 aluc|OJZ89084.1 anig|EHA23453.1 asyd|OJJ60773.1 ater|EAU38359.1 atub|OJI85700.1 aver|OJJ00107.1 awen|OJJ33943.1 afla|EED57941.1 anid|CBF83591.1 afis|EAW21676.1 afum|EAL92710.1 aory|EIT82327.1 azon|OJJ48627.1

Group3801: abra|OJJ75634.1 acar|OOG00201.1 acla|EAW09460.1 afis|EAW21677.1 afla|EED57942.1 afum|EAL92711.1 agla|OJJ79926.1 akaw|GAA84381.1 aluc|OJZ89085.1 anid|CBF83593.1 anig|EHA23452.1 aory|EIT82349.1 asyd|OJJ60772.1 ater|EAU38360.1 atub|OJI85699.1 aver|OJJ00106.1 awen|OJJ33944.1 azon|OJJ48626.1

Group3802: abra|OJJ75636.1 acar|OOG00200.1 acla|EAW09459.1 afis|EAW21678.1 afla|EED57943.1 afum|EAL92712.1 agla|OJJ79925.1 akaw|GAA84382.1 aluc|OJZ89086.1 anid|CBF83595.1 anig|EHA23451.1 aory|EIT82316.1 asyd|OJJ60771.1 ater|EAU38361.1 atub|OJI85698.1 aver|OJJ00105.1 awen|OJJ33945.1 azon|OJJ48625.1

Group3803: abra|OJJ75637.1 acar|OOG00199.1 acla|EAW09458.1 afis|EAW21679.1 afla|EED57944.1 afum|EAL92713.1 agla|OJJ79924.1 akaw|GAA84383.1 aluc|OJZ89087.1 anid|CBF83597.1 anig|EHA23450.1 aory|EIT82325.1 asyd|OJJ60770.1 ater|EAU38362.1 atub|OJI85697.1 aver|OJJ00104.1 awen|OJJ33946.1 azon|OJJ48624.1

Group3804: abra|OJJ75639.1 acar|OOG00198.1 acla|EAW09457.1 afis|EAW21680.1 afla|EED57945.1 afum|EAL92714.1 agla|OJJ79923.1 akaw|GAA84384.1 aluc|OJZ89088.1 anid|CBF83598.1 anig|EHA23449.1 aory|EIT82335.1 asyd|OJJ60769.1 ater|EAU38363.1 atub|OJI85694.1 aver|OJJ00103.1 awen|OJJ33947.1 azon|OJJ48623.1

Group3805: abra|OJJ75642.1 acar|OOG00196.1 acla|EAW09456.1 afis|EAW21681.1 afla|EED57946.1 afum|EAL92715.1 agla|OJJ79921.1 akaw|GAA84386.1 aluc|OJZ89091.1 anid|CBF83600.1 anig|EHA23447.1 aory|EIT82326.1 asyd|OJJ60768.1 ater|EAU38364.1 atub|OJI85692.1 aver|OJJ00102.1 awen|OJJ33948.1 azon|OJJ48622.1

Group3806: abra|OJJ75644.1 acar|OOG00195.1 acla|EAW09454.1 afis|EAW21683.1 afla|EED57950.1 afum|EAL92717.1 agla|OJJ79918.1 akaw|GAA84389.1 aluc|OJZ89093.1 anid|CBF83605.1 anig|EHA23444.1 aory|EIT82322.1 asyd|OJJ60766.1 ater|EAU38367.1 atub|OJI85688.1 aver|OJJ00100.1 awen|OJJ33950.1 azon|OJJ48620.1

Group3807: abra|OJJ75645.1 acar|OOG00194.1 acla|EAW09453.1 afis|EAW21684.1 afla|EED57951.1 afum|EAL92718.2 agla|OJJ79917.1 akaw|GAA84390.1 aluc|OJZ89094.1 anid|CBF83607.1 anig|EHA23443.1 aory|EIT82348.1 asyd|OJJ60765.1 ater|EAU38368.1 atub|OJI85687.1 aver|OJJ00099.1 awen|OJJ33951.1 azon|OJJ48619.1

Group3808: abra|OJJ75650.1 acar|OOG00190.1 acla|EAW09452.1 afis|EAW21685.1 afla|EED57956.1 afum|EAL92719.1 agla|OJJ79914.1 akaw|GAA84393.1 aluc|OJZ89098.1 anid|CBF83611.1 anig|EHA23440.1 aory|EIT79731.1 asyd|OJJ60762.1 ater|EAU38371.1 atub|OJI85683.1 aver|OJJ00096.1 awen|OJJ33955.1 azon|OJJ48615.1

Group3809: abra|OJJ75651.1 acar|OOG00189.1 acla|EAW09449.1 afis|EAW21687.1 afla|EED57957.1 afum|EAL92721.2 agla|OJJ79913.1 akaw|GAA84394.1 aluc|OJZ89099.1 anid|CBF83612.1 anig|EHA23439.1 aory|EIT79765.1 asyd|OJJ60761.1 ater|EAU38372.1 atub|OJI85682.1 aver|OJJ00095.1 awen|OJJ33956.1 azon|OJJ48612.1

Group3810: abra|OJJ75652.1 acar|OOG00188.1 acla|EAW09448.1 afis|EAW21688.1 afla|EED57958.1 afum|EAL92722.1 agla|OJJ79912.1 akaw|GAA84395.1 aluc|OJZ89100.1 anid|CBF83614.1 anig|EHA23438.1 aory|EIT79734.1 asyd|OJJ60760.1 ater|EAU38373.1 atub|OJI85681.1 aver|OJJ00094.1 awen|OJJ33957.1 azon|OJJ48611.1

Group3811: abra|OJJ75654.1 acar|OOG00187.1 acla|EAW09446.1 afis|EAW21690.1 afla|EED57960.1 afum|EAL92724.1 agla|OJJ79910.1 akaw|GAA84397.1 aluc|OJZ89103.1 anid|CBF83617.1 anig|EHA23436.1 aory|EIT79781.1 asyd|OJJ60758.1 ater|EAU38375.1 atub|OJI85678.1 aver|OJJ00092.1 awen|OJJ33959.1 azon|OJJ48609.1

Group3812: abra|OJJ75656.1 acar|OOG00186.1 acla|EAW09444.1 afis|EAW21692.1 afla|EED57962.1 afum|EAL92726.1 agla|OJJ79907.1 akaw|GAA84399.1 aluc|OJZ89106.1 anid|CBF83621.1 anig|EHA23434.1 aory|EIT79782.1 asyd|OJJ60756.1 ater|EAU38377.1 atub|OJI85676.1 aver|OJJ00090.1 awen|OJJ33961.1 azon|OJJ48607.1

Group3813: abra|OJJ75657.1 acar|OOG00185.1 acla|EAW09442.1 afis|EAW21694.1 afla|EED57963.1 afum|EAL92728.1 agla|OJJ79905.1 akaw|GAA84400.1 aluc|OJZ89107.1 anid|CBF83623.1 anig|EHA23433.1 aory|EIT79708.1 asyd|OJJ60755.1 ater|EAU38379.1 atub|OJI85675.1 aver|OJJ00089.1 awen|OJJ33963.1 azon|OJJ48605.1

Group3814: abra|OJJ75658.1 acar|OOG00184.1 acla|EAW09441.1 afis|EAW21695.1 afla|EED57964.1 afum|EAL92729.1 agla|OJJ79904.1 akaw|GAA84401.1 aluc|OJZ89109.1 anid|CBF83624.1 anig|EHA23432.1 aory|EIT79788.1 asyd|OJJ60754.1 ater|EAU38380.1 atub|OJI85674.1 aver|OJJ00088.1 awen|OJJ33964.1 azon|OJJ48603.1

Group3815: abra|OJJ75662.1 acar|OOG00181.1 acla|EAW09439.1 afis|EAW21697.1 afla|EED57966.1 afum|EAL92731.1 agla|OJJ79901.1 akaw|GAA84403.1 aluc|OJZ89113.1 anid|CBF83628.1 anig|EHA23430.1 aory|EIT79705.1 asyd|OJJ60752.1 ater|EAU38382.1 atub|OJI85671.1 aver|OJJ00085.1 awen|OJJ33966.1 azon|OJJ48600.1

Group3816: abra|OJJ75664.1 acar|OOG00180.1 acla|EAW09437.1 afis|EAW21699.1 afla|EED57968.1 afum|EAL92733.1 agla|OJJ79899.1 akaw|GAA84405.1 aluc|OJZ89115.1 anid|CBF83632.1 anig|EHA23428.1 aory|EIT79768.1 asyd|OJJ60750.1 ater|EAU38384.1 atub|OJI85669.1 aver|OJJ00083.1 awen|OJJ33968.1 azon|OJJ48598.1

Group3817: abra|OJJ75665.1 acar|OOG00179.1 acla|EAW09436.1 afis|EAW21700.1 afla|EED57969.1 afum|EAL92734.1 agla|OJJ79898.1 akaw|GAA84406.1 aluc|OJZ89116.1 anid|CBF83633.1 anig|EHA23427.1 aory|EIT79802.1 asyd|OJJ60782.1 ater|EAU38385.1 atub|OJI85668.1 aver|OJJ00115.1 awen|OJJ33970.1 azon|OJJ48589.1

Group3818: abra|OJJ75667.1 acar|OOG00177.1 acla|EAW09434.1 afis|EAW21702.1 afla|EED57971.1 afum|EAL92736.1 agla|OJJ79896.1 akaw|GAA84408.1 aluc|OJZ89118.1 anid|CBF83639.1 anig|EHA23425.1 aory|EIT79763.1 asyd|OJJ60785.1 ater|EAU38387.1 atub|OJI85666.1 aver|OJJ00118.1 awen|OJJ33972.1 azon|OJJ48587.1

Group3819: abra|OJJ75668.1 acar|OOG00176.1 acla|EAW09433.1 afis|EAW21703.1 afla|EED57972.1 afum|EAL92737.1 agla|OJJ84485.1 akaw|GAA84409.1 aluc|OJZ89119.1 anid|CBF83649.1 anig|EHA23424.1

aory|EIT79804.1 asyd|OJJ60792.1 ater|EAU38388.1 atub|OJI85665.1 aver|OJJ00125.1 awen|OJJ33973.1 azon|OJJ48586.1

Group3820: abra|OJJ75670.1 acar|OOG00175.1 acla|EAW09432.1 afis|EAW21704.1 afla|EED57973.1 afum|EAL92738.1 agla|OJJ79888.1 akaw|GAA84410.1 aluc|OJZ89120.1 anid|CBF83648.1 anig|EHA23423.1 aory|EIT79806.1 asyd|OJJ60791.1 ater|EAU38389.1 atub|OJI85664.1 aver|OJJ00124.1 awen|OJJ33975.1 azon|OJJ48581.1

Group3821: abra|OJJ75672.1 acar|OOG00173.1 acla|EAW09425.1 afis|EAW21711.1 afla|EED57975.1 afum|EAL92746.1 agla|OJJ79890.1 akaw|GAA84413.1 aluc|OJZ89122.1 anid|CBF83646.1 anig|EHA23421.1 aory|EIT79737.1 asyd|OJJ60789.1 ater|EAU38391.1 atub|OJI85662.1 aver|OJJ00122.1 awen|OJJ33977.1 azon|OJJ48583.1

Group3822: abra|OJJ75673.1 acar|OOG00172.1 acla|EAW09426.1 afis|EAW21710.1 afla|EED57976.1 afum|EAL92745.1 agla|OJJ79891.1 akaw|GAA84414.1 aluc|OJZ89123.1 anid|CBF83644.1 anig|EHA23420.1 aory|EIT79753.1 asyd|OJJ60788.1 ater|EAU38392.1 atub|OJI85661.1 aver|OJJ00121.1 awen|OJJ33980.1 azon|OJJ48575.1

Group3823: abra|OJJ75674.1 acar|OOG00171.1 acla|EAW09427.1 afis|EAW21709.1 afla|EED57977.1 afum|EAL92744.1 agla|OJJ79892.1 akaw|GAA84415.1 aluc|OJZ89124.1 anid|CBF83642.1 anig|EHA23419.1 aory|EIT79730.1 asyd|OJJ60787.1 ater|EAU38393.1 atub|OJI85660.1 aver|OJJ00120.1 awen|OJJ33981.1 azon|OJJ48574.1

Group3824: abra|OJJ75675.1 acar|OOG00170.1 acla|EAW09428.1 afis|EAW21708.1 afla|EED57978.1 afum|EAL92743.1 agla|OJJ79893.1 akaw|GAA84416.1 aluc|OJZ89125.1 anid|CBF83641.1 anig|EHA23418.1 aory|EIT79718.1 asyd|OJJ60786.1 ater|EAU38394.1 atub|OJI85659.1 aver|OJJ00119.1 awen|OJJ33982.1 azon|OJJ48573.1

Group3825: abra|OJJ75677.1 acar|OOG00169.1 acla|EAW09423.1 afis|EAW21712.1 afla|EED57980.1 afum|EAL92747.1 agla|OJJ79895.1 akaw|GAA84418.1 aluc|OJZ89127.1 anid|CBF83653.1 anig|EHA23416.1 aory|EIT79727.1 asyd|OJJ60794.1 ater|EAU38396.1 atub|OJI85657.1 aver|OJJ00127.1 awen|OJJ33984.1 azon|OJJ48570.1

Group3826: abra|OJJ75679.1 acar|OOG00168.1 acla|EAW09424.1 afis|EAW21713.1 afla|EED57981.1 afum|EAL92748.1 agla|OJJ79886.1 akaw|GAA84419.1 aluc|OJZ89129.1 anid|CBF83655.1 anig|EHA23415.1 aory|EIT79767.1 asyd|OJJ60795.1 ater|EAU38398.1 atub|OJI85656.1 aver|OJJ00128.1 awen|OJJ33985.1 azon|OJJ48568.1

Group3827: abra|OJJ75680.1 acar|OOG00166.1 acla|EAW09422.1 afis|EAW21714.1 afla|EED57984.1 afum|EAL92749.1 agla|OJJ79885.1 akaw|GAA84420.1 aluc|OJZ89130.1 anid|CBF83657.1 anig|EHA23414.1 aory|EIT79791.1 asyd|OJJ60796.1 ater|EAU38399.1 atub|OJI85655.1 aver|OJJ00129.1 awen|OJJ33986.1 azon|OJJ48567.1

Group3829: abra|OJJ75686.1 acar|OOG00164.1 acla|EAW09419.1 afis|EAW21717.1 afla|EED57987.1 afum|EAL92751.1 agla|OJJ79880.1 akaw|GAA84424.1 aluc|OJZ89137.1 anid|CBF83708.1 anig|EHA23410.1 aory|EIT79780.1 asyd|OJJ60831.1 ater|EAU38402.1 atub|OJI85648.1 aver|OJJ00156.1 awen|OJJ33990.1 azon|OJJ48563.1

Group3830: abra|OJJ75687.1 acar|OOG00163.1 acla|EAW09417.1 afis|EAW21719.1 afla|EED57990.1 afum|EAL92752.1 agla|OJJ79865.1 akaw|GAA84425.1 aluc|OJZ89138.1 anid|CBF83710.1 anig|EHA23407.1 aory|EIT79712.1 asyd|OJJ60833.1 ater|EAU38403.1 atub|OJI85647.1 aver|OJJ00157.1 awen|OJJ33992.1 azon|OJJ48561.1

Group3831: abra|OJJ75688.1 acar|OOG00162.1 acla|EAW09416.1 afis|EAW21720.1 afla|EED57989.1 afum|EAL92753.1 agla|OJJ79864.1 akaw|GAA84426.1 aluc|OJZ89139.1 anid|CBF83711.1 anig|EHA23406.1 aory|EIT79747.1 asyd|OJJ60834.1 ater|EAU38404.1 atub|OJI85646.1 aver|OJJ00158.1 awen|OJJ33993.1 azon|OJJ48560.1

Group3832: abra|OJJ75689.1 acar|OOG00161.1 acla|EAW09415.1 afis|EAW21721.1 afla|EED57988.1 afum|EAL92754.1 agla|OJJ79863.1 akaw|GAA84427.1 aluc|OJZ89140.1 anid|CBF83713.1 anig|EHA23405.1 aory|EIT79743.1 asyd|OJJ60835.1 ater|EAU38405.1 atub|OJI85645.1 aver|OJJ00159.1 awen|OJJ33994.1 azon|OJJ48559.1

Group3833: abra|OJJ75692.1 acar|OOG00160.1 acla|EAW09414.1 afis|EAW21722.1 afla|EED57991.1 afum|EAL92755.1 agla|OJJ79862.1 akaw|GAA84429.1 aluc|OJZ89143.1 anid|CBF83715.1 anig|EHA23404.1 aory|EIT79797.1 asyd|OJJ60836.1 ater|EAU38406.1 atub|OJI85644.1 aver|OJJ00160.1 awen|OJJ33995.1 azon|OJJ48558.1

Group3834: abra|OJJ75696.1 acar|OOG00158.1 acla|EAW09412.1 afis|EAW21724.1 afla|EED57993.1 afum|EAL92757.1 agla|OJJ79860.1 akaw|GAA84432.1 aluc|OJZ89147.1 anid|CBF83718.1 anig|EHA23401.1 aory|EIT79706.1 asyd|OJJ60838.1 ater|EAU38408.1 atub|OJI85641.1 aver|OJJ00162.1 awen|OJJ33997.1 azon|OJJ48555.1

Group3835: abra|OJJ75698.1 acar|OOG00155.1 acla|EAW09410.1 afis|EAW21726.1 afla|EED57995.1 afum|EAL92759.1 agla|OJJ79858.1 akaw|GAA84434.1 aluc|OJZ89149.1 anid|CBF83722.1 anig|EHA23399.1 aory|EIT79805.1 asyd|OJJ60840.1 ater|EAU38410.1 atub|OJI85639.1 aver|OJJ00164.1 awen|OJJ33999.1 azon|OJJ48553.1

Group3836: abra|OJJ75701.1 acar|OOG00153.1 acla|EAW09407.1 afis|EAW21729.1 afla|EED57998.1 afum|EAL92762.1 agla|OJJ79855.1 akaw|GAA84437.1 aluc|OJZ89152.1 anid|CBF83727.1 anig|EHA23397.1 aory|EIT79787.1 asyd|OJJ60843.1 ater|EAU38413.1 atub|OJI85636.1 aver|OJJ00168.1 awen|OJJ34002.1 azon|OJJ48550.1

Group3837: abra|OJJ75707.1 acar|OOG00151.1 acla|EAW09404.1 afis|EAW21732.1 afla|EED58001.1 afum|EAL92765.1 agla|OJJ79853.1 akaw|GAA84442.1 aluc|OJZ89158.1 anid|CBF83732.1 anig|EHA23395.1 aory|EIT79716.1 asyd|OJJ60846.1 ater|EAU38415.1 atub|OJI85632.1 aver|OJJ00171.1 awen|OJJ34005.1 azon|OJJ48546.1

Group3838: abra|OJJ75709.1 acar|OOG00149.1 acla|EAW09403.1 afis|EAW21733.1 afla|EED58002.1 afum|EAL92766.1 agla|OJJ79850.1 akaw|GAA84443.1 aluc|OJZ89160.1 anid|CBF83734.1 anig|EHA23394.1 aory|EIT79783.1 asyd|OJJ60848.1 ater|EAU38416.1 atub|OJI85630.1 aver|OJJ00174.1 awen|OJJ34006.1 azon|OJJ48545.1

Group3839: abra|OJJ75710.1 acar|OOG00150.1 acla|EAW09402.1 afis|EAW21734.1 afla|EED58006.1 afum|EAL92767.1 agla|OJJ79849.1 akaw|GAA84444.1 aluc|OJZ89161.1 anid|CBF83736.1 anig|EHA23393.1 aory|EIT79710.1 asyd|OJJ60850.1 ater|EAU36017.1 atub|OJI85629.1 aver|OJJ00172.1 awen|OJJ34007.1 azon|OJJ48544.1

Group3840: abra|OJJ75716.1 acar|OOG00143.1 acla|EAW09397.1 afis|EAW21737.1 afla|EED58011.1 afum|EAL92771.1 agla|OJJ84451.1 akaw|GAA84450.1 aluc|OJZ89168.1 anid|CBF83703.1 anig|EHA23387.1 aory|EIT79723.1 asyd|OJJ60826.1 ater|EAU38419.1 atub|OJI85623.1 aver|OJJ00152.1 awen|OJJ34019.1 azon|OJJ48539.1

Group3841: abra|OJJ75719.1 acar|OOG00137.1 acla|EAW09375.1 afis|EAW21760.1 afla|EED58018.1 afum|EAL92790.2 agla|OJJ84390.1 akaw|GAA84455.1 aluc|OJZ89171.1 anid|CBF83695.1 anig|EHA23384.1 aory|EIT79793.1 asyd|OJJ60824.1 ater|EAU38430.1 atub|OJI85619.1 aver|OJJ00150.1 awen|OJJ34044.1 azon|OJJ48187.1

Group3842: abra|OJJ75722.1 acar|OOG00136.1 acla|EAW09378.1 afis|EAW21757.1 afla|EED58020.1 afum|EAL92787.1 agla|OJJ84393.1 akaw|GAA84458.1 aluc|OJZ89174.1 anid|CBF83682.1 anig|EHA23381.1 aory|EIT79724.1 asyd|OJJ60821.1 ater|EAU38433.1 atub|OJI85616.1 aver|OJJ00147.1 awen|OJJ34041.1 azon|OJJ48198.1

Group3843: abra|OJJ75723.1 acar|OOG00135.1 acla|EAW09379.1 afis|EAW21756.1 afla|EED58022.1 afum|EAL92786.1 agla|OJJ84394.1 akaw|GAA84459.1 aluc|OJZ89175.1 anid|CBF83684.1 anig|EHA23380.1 asyd|OJJ60820.1 ater|EAU38434.1 atub|OJI85615.1 aver|OJJ00146.1 awen|OJJ34040.1 azon|OJJ48197.1 aory|EIT79736.1

Group3844: abra|OJJ75724.1 acar|OOG00134.1 acla|EAW09380.1 afis|EAW21755.1 afla|EED58023.1 afum|EAL92785.1 agla|OJJ84398.1 akaw|GAA84460.1 aluc|OJZ89176.1 anid|CBF83685.1 anig|EHA23379.1 aory|EIT79744.1 asyd|OJJ60819.1 ater|EAU38435.1 atub|OJI85613.1 aver|OJJ00145.1 awen|OJJ34039.1 azon|OJJ48196.1

Group3845: abra|OJJ75726.1 acar|OOG00132.1 acla|EAW09383.1 afis|EAW21752.1 afla|EED58026.1 afum|EAL92782.1 agla|OJJ84401.1 akaw|GAA84462.1 aluc|OJZ89178.1 anid|CBF83693.1 anig|EHA23377.1 aory|EIT79749.1 asyd|OJJ60816.1 ater|EAU38438.1 atub|OJI85611.1 aver|OJJ00142.1 awen|OJJ34035.1 azon|OJJ48207.1

Group3847: abra|OJJ75732.1 acar|OOF89927.1 acla|EAW14909.1 afis|EAW24430.1 afla|EED55243.1 afum|EAL86098.1 agla|OJJ79244.1 akaw|GAA84465.1 aluc|OJZ89183.1 anid|CBF73775.1 anig|EHA23373.1 aory|EIT83557.1 asyd|OJJ56594.1 ater|EAU30015.1 atub|OJI85606.1 aver|OJJ06131.1 awen|OJJ32772.1 azon|OJJ44003.1

Group3848: abra|OJJ75734.1 acar|OOF89928.1 acla|EAW14908.1 afis|EAW24429.1 afla|EED55249.1 afum|EAL86097.1 agla|OJJ79242.1 akaw|GAA84467.1 aluc|OJZ89185.1 anid|CBF73777.1 anig|EHA23372.1 aory|EIT83561.1 asyd|OJJ56596.1 ater|EAU30016.1 atub|OJI85604.1 aver|OJJ06129.1 awen|OJJ32774.1 azon|OJJ44004.1

Group3849: abra|OJJ75735.1 acar|OOF89929.1 acla|EAW14907.1 afis|EAW24428.1 afla|EED55250.1 afum|EAL86096.1 agla|OJJ79241.1 akaw|GAA84468.1 aluc|OJZ89186.1 anid|CBF73778.1 anig|EHA23371.1 aory|EIT83631.1 asyd|OJJ56597.1 ater|EAU30017.1 atub|OJI85603.1 aver|OJJ06128.1 awen|OJJ32775.1 azon|OJJ44005.1

Group3850: abra|OJJ75737.1 acar|OOF89931.1 acla|EAW14904.1 afis|EAW24425.1 afla|EED55254.1 afum|EAL86093.1 agla|OJJ79239.1 akaw|GAA84470.1 aluc|OJZ89189.1 anid|CBF73782.1 anig|EHA23368.1

aory|EIT83565.1 asyd|OJJ56599.1 ater|EAU30020.1 atub|OJI85601.1 aver|OJJ06126.1 awen|OJJ32781.1 azon|OJJ44007.1

Group3851: abra|OJJ75738.1 acar|OOG00131.1 acla|EAW14903.1 afis|EAW24424.1 afla|EED55258.1 afum|EAL86092.1 agla|OJJ79238.1 akaw|GAA84471.1 aluc|OJZ89191.1 anid|CBF73784.1 anig|EHA23367.1 aory|EIT83539.1 asyd|OJJ56600.1 ater|EAU30021.1 atub|OJI85599.1 aver|OJJ06125.1 awen|OJJ32780.1 azon|OJJ44009.1

Group3852: abra|OJJ75784.1 acar|OOG00104.1 acla|EAW14898.1 afis|EAW24420.1 afla|EED47748.1 afum|EAL86089.1 agla|OJJ79224.1 akaw|GAA84500.1 aluc|OJZ89227.1 anid|CBF73792.1 anig|EHA23335.1 aory|EIT77177.1 asyd|OJJ56604.1 ater|EAU30307.1 atub|OJI85565.1 aver|OJJ06119.1 awen|OJJ32789.1 azon|OJJ44022.1

Group3853: abra|OJJ75785.1 acar|OOG00103.1 acla|EAW14897.1 afis|EAW24419.1 afla|EED47749.1 afum|EAL86088.1 agla|OJJ79226.1 akaw|GAA84501.1 aluc|OJZ89229.1 anid|CBF73795.1 anig|EHA23334.1 aory|EIT77122.1 asyd|OJJ56605.1 ater|EAU30308.1 atub|OJI85564.1 aver|OJJ06118.1 awen|OJJ32790.1 azon|OJJ44021.1

Group3854: abra|OJJ75786.1 acar|OOG00102.1 acla|EAW14896.1 afis|EAW24418.1 afla|EED47750.1 afum|EAL86087.1 agla|OJJ79227.1 akaw|GAA84502.1 aluc|OJZ89230.1 anid|CBF73797.1 anig|EHA23333.1 aory|EIT77133.1 asyd|OJJ56607.1 ater|EAU30309.1 atub|OJI85563.1 aver|OJJ06117.1 awen|OJJ32791.1 azon|OJJ44020.1

Group3855: abra|OJJ75787.1 acar|OOG00101.1 acla|EAW14895.1 afis|EAW24417.1 afla|EED47751.1 afum|EAL86086.1 agla|OJJ79228.1 akaw|GAA84503.1 aluc|OJZ89231.1 anid|CBF73799.1 anig|EHA23332.1 aory|EIT77182.1 asyd|OJJ56608.1 ater|EAU30310.1 atub|OJI85562.1 aver|OJJ06116.1 awen|OJJ32792.1 azon|OJJ44019.1

Group3856: abra|OJJ75788.1 acar|OOG00099.1 acla|EAW14893.1 afis|EAW24415.1 afla|EED47766.1 afum|EAL86084.1 agla|OJJ79218.1 akaw|GAA84504.1 aluc|OJZ89232.1 anid|CBF73803.1 anig|EHA23331.1 aory|EIT77193.1 asyd|OJJ56615.1 ater|EAU30314.1 atub|OJI85561.1 aver|OJJ06109.1 awen|OJJ32794.1 azon|OJJ49041.1

Group3857: abra|OJJ75789.1 acar|OOG00098.1 acla|EAW14892.1 afis|EAW24414.1 afla|EED47765.1 afum|EAL86083.1 agla|OJJ79217.1 akaw|GAA84505.1 aluc|OJZ89233.1 anid|CBF73747.1 anig|EHA23330.1 aory|EIT77149.1 asyd|OJJ56622.1 ater|EAU30315.1 atub|OJI85560.1 aver|OJJ06104.1 awen|OJJ32795.1 azon|OJJ49038.1

Group3858: abra|OJJ75792.1 acar|OOG00086.1 acla|EAW14889.1 afis|EAW24411.1 afum|EAL86080.1 agla|OJJ79214.1 akaw|GAA84511.1 aluc|OJZ89240.1 anid|CBF73741.1 anig|EHA23327.1 aory|EIT77168.1 asyd|OJJ56625.1 ater|EAU30318.1 atub|OJI85558.1 aver|OJJ06101.1 awen|OJJ32799.1 azon|OJJ49033.1 afla|EED47762.1

Group3859: abra|OJJ75796.1 acar|OOG00085.1 acla|EAW14888.1 afis|EAW24410.1 afla|EED47761.1 afum|EAL86079.1 agla|OJJ79213.1 akaw|GAA84514.1 aluc|OJZ89243.1 anid|CBF73739.1 anig|EHA23322.1 aory|EIT77110.1 asyd|OJJ56626.1 ater|EAU30319.1 atub|OJI85552.1 aver|OJJ06100.1 awen|OJJ32800.1 azon|OJJ49032.1

Group3860: abra|OJJ75804.1 acar|OOG00071.1 acla|EAW14879.1 afis|EAW24396.1 afla|EED47814.1 afum|EAL86068.1 agla|OJJ78876.1 akaw|GAA84520.1 aluc|OJZ89252.1 anid|CBF73724.1 anig|EHA23315.1 aory|EIT77191.1 asyd|OJJ56638.1 ater|EAU30339.1 atub|OJI85546.1 aver|OJJ06088.1 awen|OJJ32809.1 azon|OJJ49030.1

Group3861: abra|OJJ75805.1 acar|OOG00070.1 acla|EAW14878.1 afis|EAW24395.1 afla|EED47815.1 afum|EAL86067.1 agla|OJJ78875.1 akaw|GAA84521.1 aluc|OJZ89253.1 anid|CBF73720.1 anig|EHA23314.1 aory|EIT77139.1 asyd|OJJ56639.1 ater|EAU30340.1 atub|OJI85545.1 aver|OJJ06087.1 awen|OJJ32810.1 azon|OJJ49031.1

Group3862: abra|OJJ75808.1 acar|OOG00068.1 acla|EAW14864.1 afis|EAW24392.1 afla|EED47818.1 afum|EAL86064.1 agla|OJJ78872.1 akaw|GAA84523.1 aluc|OJZ89256.1 anid|CBF73713.1 anig|EHA23312.1 aory|EIT77112.1 asyd|OJJ56642.1 ater|EAU30342.1 atub|OJI85543.1 aver|OJJ06084.1 awen|OJJ32812.1 azon|OJJ49020.1

Group3863: abra|OJJ75810.1 acar|OOG00067.1 acla|EAW14863.1 afis|EAW24391.1 afla|EED47819.1 afum|EAL86063.1 agla|OJJ78871.1 akaw|GAA84524.1 aluc|OJZ89257.1 anid|CBF73711.1 anig|EHA23311.1 aory|EIT77158.1 asyd|OJJ56643.1 ater|EAU30343.1 atub|OJI85542.1 aver|OJJ06083.1 awen|OJJ32813.1 azon|OJJ49019.1

Group3864: abra|OJJ75813.1 acar|OOG00062.1 acla|EAW14860.1 afis|EAW24388.1 afla|EED47821.1 afum|EAL86060.1 agla|OJJ83284.1 akaw|GAA84527.1 aluc|OJZ89260.1 anid|CBF73705.1 anig|EHA23308.1 aory|EIT77124.1 asyd|OJJ56646.1 ater|EAU30346.1 atub|OJI85539.1 aver|OJJ06080.1 awen|OJJ32817.1 azon|OJJ49015.1

Group3865: abra|OJJ75814.1 acar|OOG00061.1 acla|EAW14859.1 afis|EAW24387.1 afla|EED47822.1 afum|EAL86059.1 agla|OJJ83283.1 akaw|GAA84528.1 aluc|OJZ89261.1 anid|CBF73703.1 anig|EHA23307.1 aory|EIT77142.1 asyd|OJJ56647.1 ater|EAU30347.1 atub|OJI85538.1 aver|OJJ06079.1 awen|OJJ32818.1 azon|OJJ49014.1

Group3866: abra|OJJ75817.1 acar|OOG00058.1 acla|EAW14858.1 afis|EAW24385.1 afla|EED47824.1 afum|EAL86057.1 agla|OJJ83281.1 akaw|GAA84531.1 aluc|OJZ89264.1 anid|CBF73697.1 anig|EHA23304.1 aory|EIT77137.1 asyd|OJJ56650.1 ater|EAU30350.1 atub|OJI85535.1 aver|OJJ06075.1 awen|OJJ32825.1 azon|OJJ49011.1

Group3867: abra|OJJ75821.1 acar|OOG00055.1 acla|EAW14854.1 afis|EAW24381.1 afla|EED47829.1 afum|EAL86053.1 agla|OJJ83269.1 akaw|GAA84534.1 aluc|OJZ89267.1 anid|CBF73667.1 anig|EHA23301.1 aory|EIT77192.1 asyd|OJJ56666.1 ater|EAU30354.1 atub|OJI85532.1 aver|OJJ06057.1 awen|OJJ32831.1 azon|OJJ49005.1

Group3868: abra|OJJ75822.1 acar|OOG00054.1 acla|EAW14853.1 afis|EAW24380.1 afla|EED47830.1 afum|EAL86052.1 agla|OJJ83268.1 akaw|GAA84535.1 aluc|OJZ89268.1 anid|CBF73669.1 anig|EHA23300.1 aory|EIT77144.1 asyd|OJJ56665.1 ater|EAU30355.1 atub|OJI85531.1 aver|OJJ06058.1 awen|OJJ32832.1 azon|OJJ49004.1

Group3869: abra|OJJ75828.1 acar|OOG00052.1 acla|EAW14850.1 afis|EAW24377.1 afla|EED47834.1 afum|EAL86049.1 agla|OJJ83263.1 akaw|GAA84538.1 aluc|OJZ89271.1 anid|CBF73674.1 anig|EHA23298.1 aory|EIT77135.1 asyd|OJJ56662.1 ater|EAU30359.1 atub|OJI85528.1 aver|OJJ06062.1 awen|OJJ32835.1 azon|OJJ49000.1

Group3870: abra|OJJ75829.1 acar|OOG00051.1 acla|EAW14849.1 afis|EAW24376.1 afla|EED47835.1 afum|EAL86048.1 agla|OJJ83262.1 akaw|GAA84539.1 aluc|OJZ89272.1 anid|CBF73680.1 anig|EHA23297.1 aory|EIT77202.1 asyd|OJJ56659.1 ater|EAU30360.1 atub|OJI85527.1 aver|OJJ06066.1 awen|OJJ32836.1 azon|OJJ48998.1

Group3871: abra|OJJ75831.1 acar|OOG00049.1 acla|EAW14847.1 afis|EAW24374.1 afla|EED47837.1 afum|EAL86046.1 agla|OJJ83260.1 akaw|GAA84541.1 aluc|OJZ89274.1 anid|CBF73682.1 anig|EHA23295.1 aory|EIT77185.1 asyd|OJJ56657.1 ater|EAU30362.1 atub|OJI85525.1 aver|OJJ06068.1 awen|OJJ32838.1 azon|OJJ48996.1

Group3872: abra|OJJ75832.1 acar|OOG00048.1 acla|EAW14846.1 afis|EAW24373.1 afla|EED47838.1 afum|EAL86045.1 agla|OJJ83259.1 akaw|GAA84542.1 aluc|OJZ89275.1 anid|CBF73684.1 anig|EHA23294.1 aory|EIT77136.1 asyd|OJJ56656.1 ater|EAU30363.1 atub|OJI85524.1 aver|OJJ06069.1 awen|OJJ32839.1 azon|OJJ48995.1

Group3873: abra|OJJ75843.1 acar|OOG00045.1 acla|EAW14844.1 afis|EAW24371.1 afla|EED47846.1 afum|EAL86043.1 agla|OJJ83256.1 akaw|GAA84553.1 aluc|OJZ89285.1 anid|CBF73688.1 anig|EHA23281.1 aory|EIT77207.1 asyd|OJJ56654.1 ater|EAU30368.1 atub|OJI85514.1 aver|OJJ06071.1 awen|OJJ32844.1 azon|OJJ46816.1

Group3874: abra|OJJ75851.1 acar|OOG00042.1 acla|EAW14835.1 afis|EAW24364.1 afla|EED47849.1 afum|EBA27254.1 agla|OJJ83249.1 akaw|GAA84556.1 aluc|OJZ89288.1 anid|CBF73935.1 anig|EHA23273.1 aory|EIT77237.1 asyd|OJJ53672.1 ater|EAU30373.1 atub|OJI85511.1 aver|OJJ07779.1 awen|OJJ32869.1 azon|OJJ48991.1

Group3875: abra|OJJ75854.1 acar|OOG00041.1 acla|EAW14833.1 afis|EAW24362.1 afla|EED47851.1 afum|EAL86032.1 agla|OJJ83247.1 akaw|GAA84559.1 aluc|OJZ89291.1 anid|CBF73928.1 anig|EHA23271.1 aory|EIT77246.1 asyd|OJJ53677.1 ater|EAU30376.1 atub|OJI85508.1 aver|OJJ07793.1 awen|OJJ32865.1 azon|OJJ48989.1

Group3876: abra|OJJ75855.1 acar|OOG00040.1 acla|EAW14832.1 afis|EAW24361.1 afla|EED47852.1 afum|EAL86031.1 agla|OJJ83246.1 akaw|GAA84560.1 aluc|OJZ89293.1 anid|CBF73926.1 anig|EHA23270.1 aory|EIT77210.1 asyd|OJJ53678.1 ater|EAU30377.1 atub|OJI85507.1 aver|OJJ07794.1 awen|OJJ32864.1 azon|OJJ48988.1

Group3877: abra|OJJ75856.1 acar|OOG00038.1 acla|EAW14831.1 afis|EAW24360.1 afla|EED47853.1 afum|EAL86030.1 agla|OJJ83245.1 akaw|GAA84561.1 aluc|OJZ89294.1 anid|CBF73924.1 anig|EHA23269.1 aory|EIT77236.1 asyd|OJJ53679.1 ater|EAU30378.1 atub|OJI85506.1 aver|OJJ07795.1 awen|OJJ32863.1 azon|OJJ48987.1

Group3878: abra|OJJ75858.1 acar|OOG00037.1 acla|EAW14829.1 afis|EAW24358.1 afla|EED47856.1 afum|EAL86028.1 agla|OJJ83242.1 akaw|GAA84563.1 aluc|OJZ89296.1 anid|CBF73921.1 anig|EHA23267.1 aory|EIT77221.1 asyd|OJJ53681.1 ater|EAU30381.1 atub|OJI85503.1 aver|OJJ07797.1 awen|OJJ32861.1 azon|OJJ48985.1

Group3879: abra|OJJ75864.1 acar|OOG00032.1 acla|EAW14824.1 afis|EAW24353.1 afla|EED47910.1 afum|EAL86024.2 agla|OJJ83233.1 akaw|GAA84570.1 aluc|OJZ89306.1 anid|CBF73944.1 anig|EHA23260.1

aory|EIT83380.1 asyd|OJJ53698.1 ater|EAU30386.1 atub|OJI85493.1 aver|OJJ07816.1 awen|OJJ32872.1 azon|OJJ48973.1

Group3880: abra|OJJ75865.1 acar|OOG00029.1 acla|EAW09926.1 afis|EAW21174.1 afla|EED54761.1 afum|EAL92194.1 agla|OJJ80004.1 akaw|GAA84571.1 aluc|OJZ89307.1 anid|CBF83297.1 anig|EHA23259.1 aory|EIT79426.1 asyd|OJJ60558.1 ater|EAU35841.1 atub|OJI85491.1 aver|OJI99884.1 awen|OJJ36513.1 azon|OJJ44389.1

Group3881: abra|OJJ75867.1 acar|OOG00027.1 acla|EAW09928.1 afis|EAW21176.1 afla|EED54763.1 afum|EAL92196.1 agla|OJJ80001.1 akaw|GAA84573.1 aluc|OJZ89309.1 anid|CBF83306.1 anig|EHA23257.1 aory|EIT79399.1 asyd|OJJ60555.1 ater|EAU35843.1 atub|OJI85490.1 aver|OJI99881.1 awen|OJJ36515.1 azon|OJJ44391.1

Group3882: abra|OJJ75873.1 acar|OOG00022.1 acla|EAW09932.1 afis|EAW21179.1 afla|EED54769.1 afum|EAL92199.1 agla|OJJ79993.1 akaw|GAA84576.1 aluc|OJZ89317.1 anid|CBF83303.1 anig|EHA23254.1 aory|EIT79449.1 asyd|OJJ60552.1 ater|EAU35845.1 atub|OJI85480.1 aver|OJI99878.1 awen|OJJ36518.1 azon|OJJ44393.1

Group3883: abra|OJJ75875.1 acar|OOG00020.1 acla|EAW09935.1 afis|EAW21180.1 afla|EED54785.1 afum|EAL92201.1 agla|OJJ79997.1 akaw|GAA84577.1 aluc|OJZ89319.1 anid|CBF83301.1 anig|EHA23253.1 aory|EIT79362.1 asyd|OJJ60551.1 ater|EAU35846.1 atub|OJI85479.1 aver|OJI99877.1 awen|OJJ36519.1 azon|OJJ44395.1

Group3884: abra|OJJ75876.1 acar|OOG00017.1 acla|EAW09938.1 afis|EAW21183.1 afla|EED54776.1 afum|EAL92204.1 agla|OJJ79987.1 akaw|GAA84578.1 aluc|OJZ89320.1 anid|CBF83312.1 anig|EHA23252.1 aory|EIT79349.1 asyd|OJJ60548.1 ater|EAU35849.1 atub|OJI85478.1 aver|OJI99874.1 awen|OJJ36522.1 azon|OJJ44398.1

Group3885: abra|OJJ75879.1 acar|OOG00014.1 acla|EAW09940.1 afis|EAW21185.1 afla|EED54774.1 afum|EAL92207.1 agla|OJJ79982.1 akaw|GAA84582.1 aluc|OJZ89325.1 anid|CBF83315.1 anig|EHA23249.1 aory|EIT79429.1 asyd|OJJ60547.1 ater|EAU35850.1 atub|OJI85474.1 aver|OJI99873.1 awen|OJJ36526.1 azon|OJJ44399.1

Group3886: abra|OJJ75880.1 acar|OOG00013.1 acla|EAW09942.1 afis|EAW21187.1 afla|EED54773.1 afum|EAL92209.1 agla|OJJ79979.1 akaw|GAA84583.1 aluc|OJZ89326.1 anid|CBF83319.1 anig|EHA23248.1 aory|EIT79437.1 asyd|OJJ60546.1 ater|EAU35851.1 atub|OJI85473.1 aver|OJI99872.1 awen|OJJ36529.1 azon|OJJ44401.1

Group3887: abra|OJJ75884.1 acar|OOF90248.1 acla|EAW09944.1 afis|EAW21222.1 afla|EED54784.1 afum|EBA27454.1 agla|OJJ78510.1 akaw|GAA84592.1 aluc|OJZ89337.1 anid|CBF83295.1 anig|EHA23244.1 aory|EIT79434.1 asyd|OJJ60560.1 ater|EAU35852.1 atub|OJI85462.1 aver|OJI99886.1 awen|OJJ36530.1 azon|OJJ44404.1

Group3888: abra|OJJ75886.1 acar|OOF90247.1 acla|EAW09945.1 afis|EAW21223.1 afla|EED54783.1 afum|EAL92236.1 agla|OJJ78507.1 akaw|GAA84593.1 aluc|OJZ89338.1 anid|CBF83293.1 anig|EHA23243.1 aory|EIT79443.1 asyd|OJJ60561.1 ater|EAU35853.1 atub|OJI85461.1 aver|OJI99887.1 awen|OJJ36531.1 azon|OJJ44405.1

Group3889: abra|OJJ75888.1 acar|OOF90245.1 acla|EAW09946.1 afis|EAW21224.1 afla|EED54782.1 afum|EAL92237.1 agla|OJJ78506.1 akaw|GAA84594.1 aluc|OJZ89340.1 anid|CBF83291.1 anig|EHA23242.1 aory|EIT79409.1 asyd|OJJ60562.1 ater|EAU35854.1 atub|OJI85459.1 aver|OJI99888.1 awen|OJJ36532.1 azon|OJJ44406.1

Group3890: abra|OJJ75889.1 acar|OOF90244.1 acla|EAW09948.1 afis|EAW21226.1 afla|EED54779.1 afum|EBA27456.1 agla|OJJ78504.1 akaw|GAA84596.1 aluc|OJZ89341.1 anid|CBF83289.1 anig|EHA23241.1 aory|EIT79431.1 asyd|OJJ60563.1 ater|EAU35857.1 atub|OJI85458.1 aver|OJI99890.1 awen|OJJ36534.1 azon|OJJ44408.1

Group3891: abra|OJJ75890.1 acar|OOG00012.1 acla|EAW09898.1 afis|EAW21228.1 afla|EED54786.1 afum|EAL92240.1 agla|OJJ78502.1 akaw|GAA84597.1 aluc|OJZ89342.1 anid|CBF83280.1 anig|EHA23240.1 aory|EIT79332.1 asyd|OJJ60568.1 ater|EAU35859.1 atub|OJI85457.1 aver|OJI99893.1 awen|OJJ36537.1 azon|OJJ44351.1

Group3892: abra|OJJ75891.1 acar|OOG00011.1 acla|EAW09897.1 afis|EAW21229.1 afla|EED54787.1 afum|EAL92241.2 agla|OJJ78501.1 akaw|GAA84598.1 aluc|OJZ89343.1 anid|CBF83278.1 anig|EHA23239.1 aory|EIT79422.1 asyd|OJJ60569.1 ater|EAU35860.1 atub|OJI85456.1 aver|OJI99894.1 awen|OJJ36538.1 azon|OJJ44349.1

Group3893: abra|OJJ75892.1 acar|OOG00010.1 acla|EAW09896.1 afis|EAW21230.1 afla|EED54788.1 afum|EAL92242.1 agla|OJJ78492.1 akaw|GAA84599.1 aluc|OJZ89344.1 anid|CBF83277.1 anig|EHA23238.1 aory|EIT79453.1 asyd|OJJ60570.1 ater|EAU35863.1 atub|OJI85455.1 aver|OJI99895.1 awen|OJJ36539.1 azon|OJJ44348.1

Group3894: abra|OJJ75897.1 acar|OOG00007.1 acla|EAW09894.1 afis|EAW21232.1 afla|EED54793.1 afum|EAL92244.1 agla|OJJ78494.1 akaw|GAA84603.1 aluc|OJZ89347.1 anid|CBF83273.1 anig|EHA23234.1 aory|EIT79342.1 asyd|OJJ60573.1 ater|EAU35865.1 atub|OJI85451.1 aver|OJI99898.1 awen|OJJ36542.1 azon|OJJ44346.1

Group3895: abra|OJJ75898.1 acar|OOG00006.1 acla|EAW09893.1 afis|EAW21233.1 afla|EED54794.1 afum|EAL92245.1 agla|OJJ78495.1 akaw|GAA84604.1 aluc|OJZ89348.1 anid|CBF83271.1 anig|EHA23233.1 aory|EIT79361.1 asyd|OJJ60574.1 ater|EAU35866.1 atub|OJI85450.1 aver|OJI99899.1 awen|OJJ36543.1 azon|OJJ44345.1

Group3896: abra|OJJ75899.1 acar|OOG00005.1 acla|EAW09892.1 afis|EAW21234.1 afla|EED54795.1 afum|EAL92246.1 agla|OJJ78496.1 akaw|GAA84605.1 aluc|OJZ89349.1 anid|CBF83270.1 anig|EHA23232.1 aory|EIT79442.1 asyd|OJJ60575.1 ater|EAU35867.1 atub|OJI85449.1 aver|OJI99900.1 awen|OJJ36544.1 azon|OJJ44344.1

Group3897: abra|OJJ75902.1 acar|OOG00002.1 acla|EAW09888.1 afis|EAW21238.1 afla|EED54798.1 afum|EAL92250.1 agla|OJJ78500.1 akaw|GAA84608.1 aluc|OJZ89352.1 anid|CBF83266.1 anig|EHA23229.1 aory|EIT79394.1 asyd|OJJ60579.1 ater|EAU35879.1 atub|OJI85446.1 aver|OJI99904.1 awen|OJJ36549.1 azon|OJJ44340.1

Group3898: abra|OJJ75903.1 acar|OOG00001.1 acla|EAW09887.1 afis|EAW21239.1 afla|EED54799.1 afum|EAL92252.1 agla|OJJ78491.1 akaw|GAA84609.1 aluc|OJZ89353.1 anid|CBF83264.1 anig|EHA23228.1 aory|EIT79376.1 asyd|OJJ60580.1 ater|EAU35880.1 atub|OJI85445.1 aver|OJI99905.1 awen|OJJ36550.1 azon|OJJ44339.1

Group3899: abra|OJJ75904.1 acar|OOF99999.1 acla|EAW09886.1 afis|EAW21240.1 afla|EED54800.1 afum|EAL92253.1 agla|OJJ78490.1 akaw|GAA84610.1 aluc|OJZ89354.1 anid|CBF83262.1 anig|EHA23227.1 aory|EIT79341.1 asyd|OJJ60581.1 ater|EAU35881.1 atub|OJI85444.1 aver|OJI99906.1 awen|OJJ36551.1 azon|OJJ44338.1

Group3900: abra|OJJ75905.1 acar|OOF99998.1 acla|EAW09885.1 afis|EAW21241.1 afla|EED54802.1 afum|EAL92254.1 agla|OJJ78489.1 akaw|GAA84611.1 aluc|OJZ89355.1 anid|CBF83259.1 anig|EHA23226.1 aory|EIT79395.1 asyd|OJJ60584.1 ater|EAU35882.1 atub|OJI85443.1 aver|OJI99907.1 awen|OJJ36552.1 azon|OJJ44337.1

Group3901: abra|OJJ75906.1 acar|OOF99997.1 acla|EAW09883.1 afis|EAW21243.1 afla|EED54803.1 afum|EAL92256.1 agla|OJJ78487.1 akaw|GAA84612.1 aluc|OJZ89356.1 anid|CBF83255.1 anig|EHA23225.1 aory|EIT79348.1 asyd|OJJ60587.1 ater|EAU35883.1 atub|OJI85442.1 aver|OJI99910.1 awen|OJJ36554.1 azon|OJJ44335.1

Group3902: abra|OJJ75910.1 acar|OOF99993.1 acla|EAW09879.1 afis|EAW21247.1 afla|EED54807.1 afum|EAL92260.1 agla|OJJ87509.1 akaw|GAA84616.1 aluc|OJZ89360.1 anid|CBF83240.1 anig|EHA23221.1 aory|EIT79440.1 asyd|OJJ60598.1 ater|EAU35887.1 atub|OJI85438.1 aver|OJI99921.1 awen|OJJ36560.1 azon|OJJ44331.1

Group3903: abra|OJJ75911.1 acar|OOF99992.1 acla|EAW09878.1 afis|EAW21248.1 afla|EED54808.1 afum|EAL92261.1 agla|OJJ87508.1 akaw|GAA84617.1 aluc|OJZ89361.1 anid|CBF83242.1 anig|EHA23220.1 aory|EIT79438.1 asyd|OJJ60597.1 ater|EAU35888.1 atub|OJI85437.1 aver|OJI99920.1 awen|OJJ36561.1 azon|OJJ44327.1

Group3904: abra|OJJ75914.1 acar|OOF99989.1 acla|EAW09876.1 afis|EAW21250.1 afla|EED54810.1 afum|EAL92263.1 agla|OJJ87501.1 akaw|GAA84619.1 aluc|OJZ89363.1 anid|CBF83245.1 anig|EHA23219.1 aory|EIT79398.1 asyd|OJJ60594.1 ater|EAU35890.1 atub|OJI85435.1 aver|OJI99918.1 awen|OJJ36564.1 azon|OJJ44324.1

Group3905: abra|OJJ75916.1 acar|OOF90029.1 acla|EAW09874.1 afis|EAW21252.1 afla|EED54812.1 afum|EAL92266.1 agla|OJJ87495.1 akaw|GAA84621.1 aluc|OJZ89365.1 anid|CBF83247.1 anig|EHA23217.1 aory|EIT79364.1 asyd|OJJ60592.1 ater|EAU35892.1 atub|OJI85433.1 aver|OJI99916.1 awen|OJJ36567.1 azon|OJJ44321.1

Group3906: abra|OJJ75919.1 acar|OOF90028.1 acla|EAW09873.1 afis|EAW21253.1 afla|EED54815.1 afum|EAL92268.1 agla|OJJ87497.1 akaw|GAA84622.1 aluc|OJZ89368.1 anid|CBF83249.1 anig|EHA23215.1 aory|EIT79356.1 asyd|OJJ60590.1 ater|EAU35893.1 atub|OJI85430.1 aver|OJI99913.1 awen|OJJ36568.1 azon|OJJ44319.1

Group3907: abra|OJJ75920.1 acar|OOF99987.1 acla|EAW09872.1 afis|EAW21254.1 afla|EED54816.1 afum|EAL92269.1 agla|OJJ87498.1 akaw|GAA84623.1 aluc|OJZ89369.1 anid|CBF83251.1 anig|EHA23214.1 aory|EIT79384.1 asyd|OJJ60589.1 ater|EAU35894.1 atub|OJI85429.1 aver|OJI99912.1 awen|OJJ36569.1 azon|OJJ44318.1

Group3908: abra|OJJ75922.1 acar|OOF99986.1 acla|EAW09866.1 afis|EAW21259.1 afla|EED54818.1 afum|EAL92274.1 agla|OJJ87491.1 akaw|GAA84626.1 aluc|OJZ89372.1 anid|CBF83325.1 anig|EHA23212.1

aory|EIT79360.1 asyd|OJJ60601.1 ater|EAU35896.1 atub|OJI85426.1 aver|OJI99924.1 awen|OJJ36574.1 azon|OJJ44315.1

Group3909: abra|OJJ75924.1 acar|OOF99984.1 acla|EAW09865.1 afis|EAW21260.1 afla|EED54819.1 afum|EAL92275.1 agla|OJJ87490.1 akaw|GAA84627.1 aluc|OJZ89373.1 anid|CBF83329.1 anig|EHA23210.1 aory|EIT79435.1 asyd|OJJ60605.1 ater|EAU35897.1 atub|OJI85425.1 aver|OJI99929.1 awen|OJJ36575.1 azon|OJJ44314.1

Group3910: abra|OJJ75931.1 acar|OOF99979.1 acla|EAW09862.1 afis|EAW21264.1 afla|EED54823.1 afum|EAL92279.1 agla|OJJ87471.1 akaw|GAA84632.1 aluc|OJZ89378.1 anid|CBF83340.1 anig|EHA23205.1 aory|EIT79396.1 asyd|OJJ60609.1 ater|EAU35901.1 atub|OJI85420.1 aver|OJI99933.1 awen|OJJ36581.1 azon|OJJ44309.1

Group3911: abra|OJJ75932.1 acar|OOF99978.1 acla|EAW09861.1 afis|EAW21265.1 afla|EED54824.1 afum|EAL92280.1 agla|OJJ87472.1 akaw|GAA84633.1 aluc|OJZ89379.1 anid|CBF83342.1 anig|EHA23204.1 aory|EIT79447.1 asyd|OJJ60610.1 ater|EAU35902.1 atub|OJI85419.1 aver|OJI99934.1 awen|OJJ36582.1 azon|OJJ44308.1

Group3912: abra|OJJ75934.1 acar|OOF99976.1 acla|EAW09859.1 afis|EAW21267.1 afla|EED54826.1 afum|EAL92282.1 agla|OJJ87474.1 akaw|GAA84635.1 aluc|OJZ89381.1 anid|CBF83345.1 anig|EHA23202.1 aory|EIT79337.1 asyd|OJJ60612.1 ater|EAU35903.1 atub|OJI85417.1 aver|OJI99936.1 awen|OJJ36584.1 azon|OJJ47371.1

Group3913: abra|OJJ75935.1 acar|OOF99975.1 acla|EAW09858.1 afis|EAW21268.1 afla|EED54827.1 afum|EAL92283.1 agla|OJJ87475.1 akaw|GAA84636.1 aluc|OJZ89382.1 anid|CBF83347.1 anig|EHA23201.1 aory|EIT79343.1 asyd|OJJ60613.1 ater|EAU35904.1 atub|OJI85416.1 aver|OJI99937.1 awen|OJJ39424.1 azon|OJJ44306.1

Group3914: abra|OJJ75936.1 acar|OOF99974.1 acla|EAW09857.1 afis|EAW21269.1 afla|EED54829.1 afum|EBA27457.1 agla|OJJ87476.1 akaw|GAA84637.1 aluc|OJZ89383.1 anid|CBF83349.1 anig|EHA23200.1 aory|EIT79419.1 asyd|OJJ60614.1 ater|EAU35905.1 atub|OJI85415.1 aver|OJI99938.1 awen|OJJ39423.1 azon|OJJ44305.1

Group3915: abra|OJJ75942.1 acar|OOF99968.1 acla|EAW09851.1 afis|EAW21284.1 afla|EED54835.1 afum|EAL92301.1 agla|OJJ87479.1 akaw|GAA84644.1 aluc|OJZ89389.1 anid|CBF83363.1 anig|EHA23195.1 aory|EIT79451.1 asyd|OJJ60620.1 ater|EAU35913.1 atub|OJI85408.1 aver|OJI99944.1 awen|OJJ39415.1 azon|OJJ44298.1

Group3916: abra|OJJ75944.1 acar|OOF99967.1 acla|EAW09849.1 afis|EAW21286.1 afla|EED54837.1 afum|EAL92303.1 agla|OJJ87468.1 akaw|GAA84646.1 aluc|OJZ89391.1 anid|CBF83367.1 anig|EHA23193.1 aory|EIT79347.1 asyd|OJJ60622.1 ater|EAU35918.1 atub|OJI85406.1 aver|OJI99946.1 awen|OJJ39413.1 azon|OJJ44296.1

Group3917: abra|OJJ75949.1 acar|OOF99964.1 acla|EAW09847.1 afis|EAW21288.1 afla|EED54839.1 afum|EAL92305.1 agla|OJJ87466.1 akaw|GAA84652.1 aluc|OJZ89398.1 anid|CBF83371.1 anig|EHA23189.1 aory|EIT79388.1 asyd|OJJ60624.1 ater|EAU35920.1 atub|OJI85399.1 aver|OJI99948.1 awen|OJJ39411.1 azon|OJJ44294.1

Group3918: abra|OJJ75950.1 acar|OOF99962.1 acla|EAW09846.1 afis|EAW21289.1 afla|EED54840.1 afum|EAL92306.1 agla|OJJ87464.1 akaw|GAA84653.1 aluc|OJZ89399.1 anid|CBF83372.1 anig|EHA23188.1 aory|EIT79351.1 asyd|OJJ60625.1 ater|EAU35921.1 atub|OJI85398.1 aver|OJI99949.1 awen|OJJ39410.1 azon|OJJ44293.1

Group3919: abra|OJJ75951.1 acar|OOF99961.1 acla|EAW09845.1 afis|EAW21290.1 afla|EED54842.1 afum|EAL92307.1 agla|OJJ87463.1 akaw|GAA84654.1 aluc|OJZ89400.1 anid|CBF85245.1 anig|EHA23187.1 aory|EIT79441.1 asyd|OJJ62975.1 ater|EAU35922.1 atub|OJI85397.1 aver|OJI97994.1 awen|OJJ35914.1 azon|OJJ44292.1

Group3920: abra|OJJ75952.1 acar|OOF99960.1 acla|EAW09844.1 afis|EAW21291.1 afla|EED54843.1 afum|EAL92308.1 agla|OJJ87462.1 akaw|GAA84655.1 aluc|OJZ89401.1 anid|CBF83374.1 anig|EHA23186.1 aory|EIT79331.1 asyd|OJJ60626.1 ater|EAU35923.1 atub|OJI85396.1 aver|OJI99950.1 awen|OJJ39408.1 azon|OJJ44291.1

Group3921: abra|OJJ75958.1 acar|OOF99956.1 acla|EAW09840.1 afis|EAW21295.1 afla|EED54846.1 afum|EAL92312.1 agla|OJJ87444.1 akaw|GAA84659.1 aluc|OJZ89406.1 anid|CBF83381.1 anig|EHA23181.1 aory|EIT79417.1 asyd|OJJ60629.1 ater|EAU35928.1 atub|OJI85391.1 aver|OJI99953.1 awen|OJJ39404.1 azon|OJJ44288.1

Group3922: abra|OJJ75959.1 acar|OOF99955.1 acla|EAW09839.1 afis|EAW21296.1 afla|EED54847.1 afum|EAL92313.1 agla|OJJ87443.1 akaw|GAA84660.1 aluc|OJZ89407.1 anid|CBF83383.1 anig|EHA23180.1 aory|EIT79405.1 asyd|OJJ60630.1 ater|EAU35929.1 atub|OJI85388.1 aver|OJI99954.1 awen|OJJ39403.1 azon|OJJ44287.1

Group3923: abra|OJJ75963.1 acar|OOF90049.1 acla|EAW09834.1 afis|EAW21299.1 afla|EED54851.1 afum|EAL92316.1 agla|OJJ87439.1 akaw|GAA84663.1 aluc|OJZ89411.1 anid|CBF83388.1 anig|EHA23176.1 aory|EIT79436.1 asyd|OJJ60634.1 ater|EAU35932.1 atub|OJI85385.1 aver|OJI99957.1 awen|OJJ39399.1 azon|OJJ44283.1

Group3924: abra|OJJ75965.1 acar|OOF90050.1 acla|EAW09833.1 afis|EAW21300.1 afla|EED54852.1 afum|EAL92317.1 agla|OJJ87438.1 akaw|GAA84664.1 aluc|OJZ89412.1 anid|CBF83392.1 anig|EHA23175.1 aory|EIT79390.1 asyd|OJJ60635.1 ater|EAU35933.1 atub|OJI85384.1 aver|OJI99958.1 awen|OJJ39398.1 azon|OJJ44282.1

Group3925: abra|OJJ75966.1 acar|OOF90052.1 acla|EAW09832.1 afis|EAW21301.1 afla|EED57261.1 afum|EAL92318.1 agla|OJJ87436.1 akaw|GAA84665.1 aluc|OJZ89413.1 anid|CBF83394.1 anig|EHA23174.1 aory|EIT73984.1 asyd|OJJ60636.1 ater|EAU35934.1 atub|OJI85382.1 aver|OJI99959.1 awen|OJJ39397.1 azon|OJJ44281.1

Group3926: abra|OJJ75973.1 acar|OOF99946.1 acla|EAW09823.1 afis|EAW21310.1 afla|EED57266.1 afum|EAL92326.1 agla|OJJ87432.1 akaw|GAA88983.1 aluc|OJZ89420.1 anid|CBF83404.1 anig|EHA23167.1 aory|EIT74110.1 asyd|OJJ60640.1 ater|EAU35940.1 atub|OJI85373.1 aver|OJI99961.1 awen|OJJ39390.1 azon|OJJ44278.1

Group3927: abra|OJJ75977.1 acar|OOF99943.1 acla|EAW09821.1 afis|EAW21313.1 afla|EED57270.1 afum|EAL92330.1 agla|OJJ87429.1 akaw|GAA88980.1 aluc|OJZ89423.1 anid|CBF83407.1 anig|EHA23164.1 aory|EIT74037.1 asyd|OJJ60647.1 ater|EAU35942.1 atub|OJI85370.1 aver|OJI99965.1 awen|OJJ39386.1 azon|OJJ44275.1

Group3928: abra|OJJ75978.1 acar|OOF99941.1 acla|EAW09820.1 afis|EAW21314.1 afla|EED57271.1 afum|EAL92331.1 agla|OJJ87426.1 akaw|GAA88979.1 aluc|OJZ89424.1 anid|CBF83408.1 anig|EHA23163.1 aory|EIT74076.1 asyd|OJJ60648.1 ater|EAU35943.1 atub|OJI85369.1 aver|OJI99968.1 awen|OJJ39383.1 azon|OJJ44273.1

Group3929: abra|OJJ75980.1 acar|OOF99939.1 acla|EAW09819.1 afis|EAW21315.1 afla|EED57273.1 afum|EAL92332.1 agla|OJJ87424.1 akaw|GAA88977.1 aluc|OJZ89426.1 anid|CBF83412.1 anig|EHA23160.1 aory|EIT74008.1 asyd|OJJ60650.1 ater|EAU35944.1 atub|OJI85367.1 aver|OJI99970.1 awen|OJJ39381.1 azon|OJJ44271.1

Group3930: abra|OJJ75982.1 acar|OOF99937.1 acla|EAW09817.1 afis|EAW21317.1 afla|EED57275.1 afum|EAL92334.1 agla|OJJ87422.1 akaw|GAA88975.1 aluc|OJZ89428.1 anid|CBF83416.1 anig|EHA23159.1 aory|EIT74079.1 asyd|OJJ60652.1 ater|EAU35945.1 atub|OJI85365.1 aver|OJI99972.1 awen|OJJ39379.1 azon|OJJ44269.1

Group3931: abra|OJJ75983.1 acar|OOF99936.1 acla|EAW09816.1 afis|EAW21318.1 afla|EED57276.1 afum|EAL92335.1 agla|OJJ87421.1 akaw|GAA88974.1 aluc|OJZ89429.1 anid|CBF83417.1 anig|EHA23158.1 aory|EIT74114.1 asyd|OJJ60653.1 ater|EAU35946.1 atub|OJI85364.1 aver|OJI99973.1 awen|OJJ39378.1 azon|OJJ44268.1

Group3932: abra|OJJ75984.1 acar|OOF99935.1 acla|EAW09815.1 afis|EAW21319.1 afla|EED57277.1 afum|EAL92336.1 agla|OJJ87420.1 akaw|GAA88973.1 aluc|OJZ89430.1 anid|CBF83419.1 anig|EHA23157.1 aory|EIT74055.1 asyd|OJJ60654.1 ater|EAU35947.1 atub|OJI85363.1 aver|OJI99974.1 awen|OJJ39377.1 azon|OJJ44267.1

Group3933: abra|OJJ75987.1 acar|OOF99934.1 acla|EAW09813.1 afis|EAW21321.1 afla|EED57279.1 afum|EAL92338.1 agla|OJJ87416.1 akaw|GAA88971.1 aluc|OJZ89432.1 anid|CBF83421.1 anig|EHA23155.1 aory|EIT74031.1 asyd|OJJ60655.1 ater|EAU35948.1 atub|OJI85361.1 aver|OJI99976.1 awen|OJJ39375.1 azon|OJJ44265.1

Group3934: abra|OJJ75988.1 acar|OOF99933.1 acla|EAW09812.1 afis|EAW21323.1 afla|EED57280.1 afum|EAL92340.1 agla|OJJ87415.1 akaw|GAA88970.1 aluc|OJZ89433.1 anid|CBF83423.1 anig|EHA23154.1 aory|EIT74009.1 asyd|OJJ60656.1 ater|EAU35949.1 atub|OJI85360.1 aver|OJI99977.1 awen|OJJ39374.1 azon|OJJ44264.1

Group3935: abra|OJJ75995.1 acar|OOF99930.1 acla|EAW09810.1 afis|EAW21325.1 afla|EED57283.1 afum|EAL92343.1 agla|OJJ87413.1 akaw|GAA88964.1 aluc|OJZ89441.1 anid|CBF83426.1 anig|EHA23145.1 aory|EIT74153.1 asyd|OJJ60658.1 ater|EAU35951.1 atub|OJI85352.1 aver|OJI99981.1 awen|OJJ39372.1 azon|OJJ51712.1

Group3936: abra|OJJ76003.1 acar|OOF99921.1 acla|EAW09802.1 afis|EAW21334.1 afla|EED57292.1 afum|EAL92352.1 agla|OJJ87399.1 akaw|GAA88957.1 aluc|OJZ89448.1 anid|CBF76279.1 anig|EHA23138.1 aory|EIT74027.1 asyd|OJJ60988.1 ater|EAU35959.1 atub|OJI85345.1 aver|OJJ00308.1 awen|OJJ39364.1 azon|OJJ51703.1

Group3937: abra|OJJ76004.1 acar|OOF99922.1 acla|EAW09801.1 afis|EAW21335.1 afla|EED57293.1 afum|EAL92353.1 agla|OJJ87398.1 akaw|GAA88956.1 aluc|OJZ89449.1 anid|CBF76277.1 anig|EHA23137.1

aory|EIT73971.1 asyd|OJJ60987.1 ater|EAU35960.1 atub|OJI85344.1 aver|OJJ00307.1 awen|OJJ39363.1 azon|OJJ51702.1

Group3938: abra|OJJ76005.1 acar|OOF99920.1 acla|EAW09800.1 afis|EAW21336.1 afla|EED57294.1 afum|EAL92354.1 agla|OJJ82541.1 akaw|GAA88955.1 aluc|OJZ89451.1 anid|CBF83881.1 anig|EHA23136.1 aory|EIT74083.1 asyd|OJJ60940.1 ater|EAU35961.1 atub|OJI85343.1 aver|OJJ00266.1 awen|OJJ39362.1 azon|OJJ51701.1

Group3939: abra|OJJ76007.1 acar|OOF99919.1 acla|EAW09798.1 afis|EAW21338.1 afla|EED57296.1 afum|EAL92356.1 agla|OJJ87396.1 akaw|GAA88953.1 aluc|OJZ89453.1 anid|CBF83884.1 anig|EHA23134.1 aory|EIT74155.1 asyd|OJJ60942.1 ater|EAU35963.1 atub|OJI85341.1 aver|OJJ00268.1 awen|OJJ39360.1 azon|OJJ51699.1

Group3940: abra|OJJ76008.1 acar|OOF99917.1 acla|EAW09797.1 afis|EAW21339.1 afla|EED55754.1 afum|EBA27459.1 agla|OJJ87395.1 akaw|GAA88952.1 aluc|OJZ89454.1 anid|CBF83886.1 anig|EHA23133.1 aory|EIT76828.1 asyd|OJJ60943.1 ater|EAU35964.1 atub|OJI85340.1 aver|OJJ00269.1 awen|OJJ39359.1 azon|OJJ51698.1

Group3941: abra|OJJ76010.1 acar|OOF99914.1 acla|EAW09795.1 afis|EAW21341.1 afla|EED55756.1 afum|EAL92358.1 agla|OJJ87391.1 akaw|GAA88950.1 aluc|OJZ89456.1 anid|CBF76183.1 anig|EHA23131.1 aory|EIT76602.1 asyd|OJJ60962.1 ater|EAU35965.1 atub|OJI85338.1 aver|OJJ00287.1 awen|OJJ39356.1 azon|OJJ51697.1

Group3942: abra|OJJ76023.1 acar|OOF99901.1 acla|EAW09785.1 afis|EAW21353.1 afla|EED55768.1 afum|EAL92368.1 agla|OJJ87381.1 akaw|GAA88934.1 aluc|OJZ89473.1 anid|CBF76255.1 anig|EHA23118.1 aory|EIT76752.1 asyd|OJJ60965.1 ater|EAU35974.1 atub|OJI85325.1 aver|OJJ00290.1 awen|OJJ39347.1 azon|OJJ51691.1

Group3943: abra|OJJ76025.1 acar|OOF99902.1 acla|EAW09784.1 afis|EAW21354.1 afla|EED55769.1 afum|EAL92369.1 agla|OJJ87375.1 akaw|GAA88933.1 aluc|OJZ89475.1 anid|CBF76253.1 anig|EHA23117.1 aory|EIT76709.1 asyd|OJJ60966.1 ater|EAU35204.1 atub|OJI85324.1 aver|OJJ00291.1 awen|OJJ39346.1 azon|OJJ51690.1

Group3944: abra|OJJ76035.1 acar|OOF99893.1 acla|EAW09768.1 afis|EAW21370.1 afla|EED55788.1 afum|EAL92387.1 agla|OJJ85492.1 akaw|GAA88929.1 aluc|OJZ89490.1 anid|CBF83869.1 anig|EHA23108.1 aory|EIT76876.1 asyd|OJJ60933.1 ater|EAU35186.1 atub|OJI85313.1 aver|OJJ00259.1 awen|OJJ39327.1 azon|OJJ51672.1

Group3945: abra|OJJ76037.1 acar|OOF99892.1 acla|EAW09769.1 afis|EAW21369.1 afla|EED55786.1 afum|EAL92386.1 agla|OJJ85493.1 akaw|GAA88928.1 aluc|OJZ89492.1 anid|CBF83867.1 anig|EHA23107.1 aory|EIT76715.1 asyd|OJJ60932.1 ater|EAU35187.1 atub|OJI85311.1 aver|OJJ00258.1 awen|OJJ39328.1 azon|OJJ51673.1

Group3946: abra|OJJ76039.1 acar|OOF99890.1 acla|EAW09770.1 afis|EAW21368.1 afla|EED55782.1 afum|EAL92385.2 agla|OJJ85494.1 akaw|GAA88926.1 aluc|OJZ89495.1 anid|CBF83874.1 anig|EHA23105.1 aory|EIT76864.1 asyd|OJJ60937.1 ater|EAU35189.1 atub|OJI85308.1 aver|OJJ00263.1 awen|OJJ39329.1 azon|OJJ51674.1

Group3947: abra|OJJ76040.1 acar|OOF99889.1 acla|EAW09771.1 afis|EAW21367.1 afla|EED55781.1 afum|EAL92384.1 agla|OJJ85495.1 akaw|GAA88925.1 aluc|OJZ89496.1 anid|CBF83872.1 anig|EHA23104.1 aory|EIT76541.1 asyd|OJJ60936.1 ater|EAU35190.1 atub|OJI85307.1 aver|OJJ00262.1 awen|OJJ39330.1 azon|OJJ51675.1

Group3948: abra|OJJ76044.1 acar|OOF99886.1 acla|EAW09774.1 afis|EAW21364.1 afla|EED55778.1 afum|EAL92380.1 agla|OJJ85485.1 akaw|GAA88921.1 aluc|OJZ89500.1 anid|CBF83875.1 anig|EHA23101.1 aory|EIT76770.1 asyd|OJJ60938.1 ater|EAU35196.1 atub|OJI85303.1 aver|OJJ00264.1 awen|OJJ39334.1 azon|OJJ51679.1

Group3949: abra|OJJ76046.1 acar|OOF99884.1 acla|EAW09777.1 afis|EAW21361.1 afla|EED55775.1 afum|EAL92377.1 agla|OJJ85480.1 akaw|GAA88919.1 aluc|OJZ89502.1 anid|CBF76267.1 anig|EHA23098.1 aory|EIT76605.1 asyd|OJJ60973.1 ater|EAU35199.1 atub|OJI85301.1 aver|OJJ00298.1 awen|OJJ39337.1 azon|OJJ51682.1

Group3950: abra|OJJ76048.1 acar|OOF99882.1 acla|EAW09779.1 afis|EAW21359.1 afla|EED55774.1 afum|EAL92375.1 agla|OJJ85478.1 akaw|GAA88917.1 aluc|OJZ89504.1 anid|CBF76265.1 anig|EHA23096.1 aory|EIT76598.1 asyd|OJJ60972.1 ater|EAU35200.1 atub|OJI85299.1 aver|OJJ00297.1 awen|OJJ39339.1 azon|OJJ51684.1

Group3951: abra|OJJ76056.1 acar|OOF99874.1 acla|EAW09762.1 afis|EAW21375.1 afla|EED55793.1 afum|EAL92392.1 agla|OJJ85508.1 akaw|GAA88910.1 aluc|OJZ89511.1 anid|CBF83860.1 anig|EHA23088.1 aory|EIT76584.1 asyd|OJJ60926.1 ater|EAU35182.1 atub|OJI85292.1 aver|OJJ00252.1 awen|OJJ39322.1 azon|OJJ51667.1

Group3952: abra|OJJ76057.1 acar|OOF99873.1 acla|EAW09763.1 afis|EAW21374.1 afla|EED55792.1 afum|EAL92391.1 agla|OJJ85500.1 akaw|GAA88909.1 aluc|OJZ89512.1 anid|CBF83862.1 anig|EHA23087.1 aory|EIT76633.1 asyd|OJJ60927.1 ater|EAU35183.1 atub|OJI85291.1 aver|OJJ00253.1 awen|OJJ39323.1 azon|OJJ51668.1

Group3953: abra|OJJ76058.1 acar|OOF99872.1 acla|EAW09760.1 afis|EAW21381.1 afla|EED55795.1 afum|EAL92396.1 agla|OJJ85505.1 akaw|GAA88908.1 aluc|OJZ89513.1 anid|CBF83856.1 anig|EHA23086.1 aory|EIT76717.1 asyd|OJJ60924.1 ater|EAU35180.1 atub|OJI85290.1 aver|OJJ00250.1 awen|OJJ39320.1 azon|OJJ51665.1

Group3954: abra|OJJ76059.1 acar|OOF99871.1 acla|EAW09759.1 afis|EAW21382.1 afla|EED55796.1 afum|EAL92397.1 agla|OJJ85504.1 akaw|GAA88907.1 aluc|OJZ89514.1 anid|CBF83854.1 anig|EHA23085.1 aory|EIT76703.1 asyd|OJJ60923.1 ater|EAU35179.1 atub|OJI85289.1 aver|OJJ00249.1 awen|OJJ39319.1 azon|OJJ51664.1

Group3955: abra|OJJ76061.1 acar|OOF99870.1 acla|EAW09757.1 afis|EAW21384.1 afla|EED55798.1 afum|EAL92399.1 agla|OJJ85509.1 akaw|GAA88905.1 aluc|OJZ89516.1 anid|CBF83847.1 anig|EHA23083.1 aory|EIT76813.1 asyd|OJJ60919.1 ater|EAU35178.1 atub|OJI85287.1 aver|OJJ00245.1 awen|OJJ39317.1 azon|OJJ51530.1

Group3956: abra|OJJ76062.1 acar|OOF99869.1 acla|EAW09756.1 afis|EAW21385.1 afla|EED55799.1 afum|EAL92400.1 agla|OJJ85510.1 akaw|GAA88904.1 aluc|OJZ89517.1 anid|CBF83845.1 anig|EHA23082.1 aory|EIT76797.1 asyd|OJJ60918.1 ater|EAU35171.1 atub|OJI85286.1 aver|OJJ00244.1 awen|OJJ39316.1 azon|OJJ51531.1

Group3957: abra|OJJ76064.1 acar|OOF99865.1 acla|EAW09754.1 afis|EAW21387.1 afla|EED55801.1 afum|EAL92402.1 agla|OJJ85512.1 akaw|GAA88902.1 aluc|OJZ89519.1 anid|CBF83842.1 anig|EHA23081.1 aory|EIT76678.1 asyd|OJJ60916.1 ater|EAU35173.1 atub|OJI85284.1 aver|OJJ00243.1 awen|OJJ39314.1 azon|OJJ51532.1

Group3958: abra|OJJ76069.1 acar|OOF99862.1 acla|EAW09751.1 afis|EAW21391.1 afla|EED55805.1 afum|EAL92405.1 agla|OJJ85515.1 akaw|GAA88898.1 aluc|OJZ89523.1 anid|CBF83838.1 anig|EHA23078.1 aory|EIT76825.1 asyd|OJJ60913.1 ater|EAU35176.1 atub|OJI85280.1 aver|OJJ00240.1 awen|OJJ39312.1 azon|OJJ51538.1

Group3959: abra|OJJ76071.1 acar|OOF99857.1 acla|EAW09748.1 afis|EAW21394.1 afla|EED55811.1 afum|EAL92408.1 agla|OJJ85518.1 akaw|GAA88896.1 aluc|OJZ89525.1 anid|CBF83826.1 anig|EHA23076.1 aory|EIT76796.1 asyd|OJJ60904.1 ater|EAU35162.1 atub|OJI85278.1 aver|OJJ00230.1 awen|OJJ39310.1 azon|OJJ51542.1

Group3960: abra|OJJ76079.1 acar|OOF99846.1 acla|EAW09742.1 afis|EAW21400.1 afla|EED55821.1 afum|EAL92414.1 agla|OJJ85525.1 akaw|GAA88887.1 aluc|OJZ89536.1 anid|CBF83829.1 anig|EHA23068.1 aory|EIT76664.1 asyd|OJJ60906.1 ater|EAU35150.1 atub|OJI85270.1 aver|OJJ00232.1 awen|OJJ39303.1 azon|OJJ51549.1

Group3961: abra|OJJ76080.1 acar|OOF99845.1 acla|EAW09741.1 afis|EAW21401.1 afla|EED55822.1 afum|EAL92415.1 agla|OJJ85528.1 akaw|GAA88886.1 aluc|OJZ89538.1 anid|CBF83820.1 anig|EHA23067.1 aory|EIT76768.1 asyd|OJJ60901.1 ater|EAU35149.1 atub|OJI85269.1 aver|OJJ00227.1 awen|OJJ39302.1 azon|OJJ51550.1

Group3962: abra|OJJ76082.1 acar|OOF99844.1 acla|EAW09739.1 afis|EAW21402.1 afla|EED55824.1 afum|EAL92416.1 agla|OJJ85530.1 akaw|GAA88884.1 aluc|OJZ89540.1 anid|CBF83819.1 anig|EHA23066.1 aory|EIT76826.1 asyd|OJJ60900.1 ater|EAU29588.1 atub|OJI85267.1 aver|OJJ00225.1 awen|OJJ30539.1 azon|OJJ51552.1

Group3963: abra|OJJ76083.1 acar|OOF99843.1 acla|EAW09738.1 afis|EAW21403.1 afla|EED55825.1 afum|EAL92417.1 agla|OJJ85531.1 akaw|GAA88883.1 aluc|OJZ89541.1 anid|CBF83817.1 anig|EHA23065.1 aory|EIT76589.1 asyd|OJJ60899.1 ater|EAU35147.1 atub|OJI85266.1 aver|OJJ00224.1 awen|OJJ39300.1 azon|OJJ51553.1

Group3964: abra|OJJ76084.1 acar|OOF99842.1 acla|EAW09737.1 afis|EAW21404.1 afla|EED55826.1 afum|EAL92418.1 agla|OJJ85533.1 akaw|GAA88882.1 aluc|OJZ89542.1 anid|CBF83815.1 anig|EHA23064.1 aory|EIT76656.1 asyd|OJJ60898.1 ater|EAU35146.1 atub|OJI85265.1 aver|OJJ00223.1 awen|OJJ39299.1 azon|OJJ51554.1

Group3965: abra|OJJ76088.1 acar|OOF99837.1 acla|EAW09734.1 afis|EAW21407.1 afla|EED55829.1 afum|EAL92421.1 agla|OJJ85537.1 akaw|GAA88879.1 aluc|OJZ89545.1 anid|CBF83810.1 anig|EHA23061.1 aory|EIT76811.1 asyd|OJJ60895.1 ater|EAU35141.1 atub|OJI85261.1 aver|OJJ00220.1 awen|OJJ39294.1 azon|OJJ51560.1

Group3966: abra|OJJ76090.1 acar|OOF99835.1 acla|EAW09731.1 afla|EED55831.1 agla|OJJ85541.1 akaw|GAA88877.1 aluc|OJZ89547.1 anig|EHA23059.1 aory|EIT76755.1 atub|OJI85259.1 awen|OJJ39291.1

azon|OJJ51563.1 afis|EAW21410.1 afum|EAL92424.1 anid|CBF83805.1 asyd|OJJ60892.1 ater|EAU35136.1 aver|OJJ00216.1

Group3967: abra|OJJ76092.1 acar|OOF99833.1 acla|EAW09730.1 afis|EAW21411.1 afla|EED55834.1 afum|EAL92425.1 agla|OJJ85542.1 akaw|GAA88875.1 aluc|OJZ89548.1 anid|CBF83803.1 anig|EHA23058.1 aory|EIT76568.1 asyd|OJJ60891.1 ater|EAU35135.1 atub|OJI85258.1 aver|OJJ00215.1 awen|OJJ39290.1 azon|OJJ51564.1

Group3968: abra|OJJ76095.1 acar|OOF99829.1 acla|EAW09723.1 afis|EAW21418.1 afla|EED55843.1 afum|EAL92433.1 agla|OJJ85551.1 akaw|GAA88871.1 aluc|OJZ89551.1 anid|CBF83792.1 anig|EHA23055.1 aory|EIT76877.1 asyd|OJJ60881.1 ater|EAU35131.1 atub|OJI85254.1 aver|OJJ00207.1 awen|OJJ39285.1 azon|OJJ51569.1

Group3969: abra|OJJ76096.1 acar|OOF99828.1 acla|EAW09722.1 afis|EAW21419.1 afla|EED55844.1 afum|EAL92434.1 agla|OJJ85553.1 akaw|GAA88870.1 aluc|OJZ89552.1 anid|CBF83790.1 anig|EHA23054.1 aory|EIT76586.1 asyd|OJJ60880.1 ater|EAU35130.1 atub|OJI85253.1 aver|OJJ00206.1 awen|OJJ39284.1 azon|OJJ51570.1

Group3970: abra|OJJ76098.1 acar|OOF99827.1 acla|EAW09720.1 afis|EAW21421.1 afla|EED55846.1 afum|EAL92436.1 agla|OJJ85557.1 akaw|GAA88868.1 aluc|OJZ89554.1 anid|CBF83787.1 anig|EHA23052.1 aory|EIT76632.1 asyd|OJJ60878.1 ater|EAU35128.1 atub|OJI85251.1 aver|OJJ00203.1 awen|OJJ39280.1 azon|OJJ51572.1

Group3971: abra|OJJ76099.1 acar|OOF99825.1 acla|EAW09719.1 afis|EAW21422.1 afla|EED55847.1 afum|EAL92437.1 agla|OJJ85558.1 akaw|GAA88867.1 aluc|OJZ89555.1 anid|CBF83785.1 anig|EHA23051.1 aory|EIT76561.1 asyd|OJJ60877.1 ater|EAU35127.1 atub|OJI85249.1 aver|OJJ00202.1 awen|OJJ39279.1 azon|OJJ51573.1

Group3972: abra|OJJ76100.1 acar|OOF99824.1 acla|EAW09718.1 afis|EAW21423.1 afla|EED55848.1 afum|EAL92438.1 agla|OJJ85559.1 akaw|GAA88866.1 aluc|OJZ89556.1 anid|CBF83783.1 anig|EHA23050.1 aory|EIT76818.1 asyd|OJJ60876.1 ater|EAU35126.1 atub|OJI85248.1 aver|OJJ00201.1 awen|OJJ39278.1 azon|OJJ51574.1

Group3973: abra|OJJ76102.1 acar|OOF99823.1 acla|EAW09716.1 afis|EAW21425.1 afla|EED55850.1 afum|EAL92440.1 agla|OJJ85562.1 akaw|GAA88864.1 aluc|OJZ89558.1 anid|CBF83781.1 anig|EHA23048.1 aory|EIT76846.1 asyd|OJJ60874.1 ater|EAU35124.1 atub|OJI85246.1 aver|OJJ00199.1 awen|OJJ39276.1 azon|OJJ51576.1

Group3974: abra|OJJ76103.1 acar|OOF99822.1 acla|EAW09715.1 afis|EAW21426.1 afla|EED55853.1 afum|EAL92441.1 agla|OJJ85563.1 akaw|GAA88863.1 aluc|OJZ89559.1 anid|CBF83780.1 anig|EHA23047.1 aory|EIT76704.1 asyd|OJJ60873.1 ater|EAU35123.1 atub|OJI85245.1 aver|OJJ00198.1 awen|OJJ39275.1 azon|OJJ51577.1

Group3975: abra|OJJ76104.1 acar|OOF99821.1 acla|EAW09714.1 afis|EAW21427.1 afla|EED55854.1 afum|EAL92442.1 agla|OJJ85564.1 akaw|GAA88862.1 aluc|OJZ89560.1 anid|CBF83778.1 anig|EHA23046.1 aory|EIT76600.1 asyd|OJJ60871.1 ater|EAU35121.1 atub|OJI85244.1 aver|OJJ00197.1 awen|OJJ39274.1 azon|OJJ51579.1

Group3976: abra|OJJ76107.1 acar|OOF99819.1 acla|EAW09713.1 afis|EAW21428.1 afla|EED55855.1 afum|EAL92443.1 agla|OJJ85566.1 akaw|GAA88859.1 aluc|OJZ89563.1 anid|CBF83776.1 anig|EHA23043.1 aory|EIT76883.1 asyd|OJJ60870.1 ater|EAU35120.1 atub|OJI85241.1 aver|OJJ00195.1 awen|OJJ39273.1 azon|OJJ51582.1

Group3977: abra|OJJ76108.1 acar|OOF99818.1 acla|EAW09712.1 afis|EAW21429.1 afla|EED55856.1 afum|EAL92444.1 agla|OJJ85567.1 akaw|GAA88858.1 aluc|OJZ89564.1 anid|CBF83774.1 anig|EHA23042.1 aory|EIT76794.1 asyd|OJJ60869.1 ater|EAU35119.1 atub|OJI85240.1 aver|OJJ00194.1 awen|OJJ39271.1 azon|OJJ51583.1

Group3978: abra|OJJ76109.1 acar|OOF99817.1 acla|EAW09711.1 afis|EAW21430.1 afla|EED55857.1 afum|EAL92445.1 agla|OJJ85568.1 akaw|GAA88857.1 aluc|OJZ89565.1 anid|CBF83772.1 anig|EHA23041.1 aory|EIT76635.1 asyd|OJJ60868.1 ater|EAU35118.1 atub|OJI85239.1 aver|OJJ00193.1 awen|OJJ39270.1 azon|OJJ51584.1

Group3979: abra|OJJ76111.1 acar|OOF99815.1 acla|EAW09709.1 afis|EAW21431.1 afla|EED55859.1 afum|EAL92446.2 agla|OJJ85572.1 akaw|GAA88856.1 aluc|OJZ89567.1 anid|CBF83769.1 anig|EHA23040.1 aory|EIT76616.1 asyd|OJJ60865.1 ater|EAU35117.1 atub|OJI85237.1 aver|OJJ00191.1 awen|OJJ39269.1 azon|OJJ51587.1

Group3980: abra|OJJ76112.1 acar|OOF99813.1 acla|EAW09708.1 afis|EAW21432.1 afla|EED55860.1 afum|EAL92447.1 agla|OJJ85573.1 akaw|GAA88855.1 aluc|OJZ89568.1 anid|CBF83767.1 anig|EHA23039.1 aory|EIT76829.1 asyd|OJJ60864.1 ater|EAU35116.1 atub|OJI85236.1 aver|OJJ00190.1 awen|OJJ39267.1 azon|OJJ51588.1

Group3981: abra|OJJ76113.1 acar|OOF99812.1 acla|EAW09707.1 afis|EAW21433.1 afla|EED55862.1 afum|EAL92448.2 agla|OJJ85574.1 akaw|GAA88854.1 aluc|OJZ89569.1 anid|CBF83765.1 anig|EHA23037.1 aory|EIT76807.1 asyd|OJJ60863.1 ater|EAU35111.1 atub|OJI85234.1 aver|OJJ00189.1 awen|OJJ39266.1 azon|OJJ51589.1

Group3982: abra|OJJ76114.1 acar|OOF99811.1 acla|EAW09706.1 afis|EAW21434.1 afla|EED55863.1 afum|EAL92449.1 agla|OJJ85575.1 akaw|GAA88853.1 aluc|OJZ89571.1 anid|CBF76591.1 anig|EHA23036.1 aory|EIT76762.1 asyd|OJJ61175.1 ater|EAU35110.1 atub|OJI85233.1 aver|OJJ00501.1 awen|OJJ39265.1 azon|OJJ51590.1

Group3983: abra|OJJ76116.1 acar|OOF99810.1 acla|EAW09704.1 afis|EAW21436.1 afla|EED55864.1 afum|EAL92451.1 agla|OJJ85579.1 akaw|GAA88851.1 aluc|OJZ89573.1 anid|CBF76587.1 anig|EHA23034.1 aory|EIT76708.1 asyd|OJJ61173.1 ater|EAU35109.1 atub|OJI85231.1 aver|OJJ00499.1 awen|OJJ39263.1 azon|OJJ51592.1

Group3984: abra|OJJ76118.1 acar|OOF99807.1 acla|EAW09701.1 afis|EAW21438.1 afla|EED55866.1 afum|EAL92453.2 agla|OJJ85583.1 akaw|GAA88849.1 aluc|OJZ89575.1 anid|CBF76583.1 anig|EHA23032.1 aory|EIT76853.1 asyd|OJJ61171.1 ater|EAU35107.1 atub|OJI85229.1 aver|OJJ00496.1 awen|OJJ39260.1 azon|OJJ51595.1

Group3985: abra|OJJ76120.1 acar|OOF99805.1 acla|EAW09699.1 afis|EAW21440.1 afla|EED55868.1 afum|EAL92456.1 agla|OJJ85587.1 akaw|GAA88847.1 aluc|OJZ89577.1 anid|CBF76579.1 anig|EHA23031.1 aory|EIT76773.1 asyd|OJJ61169.1 ater|EAU35105.1 atub|OJI85227.1 aver|OJJ00494.1 awen|OJJ39258.1 azon|OJJ51598.1

Group3986: abra|OJJ76121.1 acar|OOF99804.1 acla|EAW09698.1 afis|EAW21441.1 afla|EED55869.1 afum|EAL92458.2 agla|OJJ85588.1 akaw|GAA88846.1 aluc|OJZ89578.1 anid|CBF76577.1 anig|EHA23030.1 aory|EIT76534.1 asyd|OJJ61168.1 ater|EAU35104.1 atub|OJI85226.1 aver|OJJ00493.1 awen|OJJ39257.1 azon|OJJ51599.1

Group3987: abra|OJJ76122.1 acar|OOF99803.1 acla|EAW09697.1 afis|EAW21442.1 afla|EED55870.1 afum|EAL92460.1 agla|OJJ85589.1 akaw|GAA88845.1 aluc|OJZ89579.1 anid|CBF76575.1 anig|EHA23029.1 aory|EIT76841.1 asyd|OJJ61167.1 ater|EAU35103.1 atub|OJI85225.1 aver|OJJ00492.1 awen|OJJ39256.1 azon|OJJ51600.1

Group3988: abra|OJJ76123.1 acar|OOF99802.1 acla|EAW09696.1 afis|EAW21443.1 afla|EED55871.1 afum|EAL92461.2 agla|OJJ85590.1 akaw|GAA88844.1 aluc|OJZ89580.1 anid|CBF76573.1 anig|EHA23028.1 aory|EIT76804.1 asyd|OJJ61166.1 ater|EAU35102.1 atub|OJI85224.1 aver|OJJ00491.1 awen|OJJ39255.1 azon|OJJ51601.1

Group3989: abra|OJJ76125.1 acar|OOF99800.1 acla|EAW09694.1 afis|EAW21445.1 afla|EED55873.1 afum|EAL92464.2 agla|OJJ85593.1 akaw|GAA88842.1 aluc|OJZ89583.1 anid|CBF76569.1 anig|EHA23026.1 aory|EIT76681.1 asyd|OJJ61164.1 ater|EAU35099.1 atub|OJI85220.1 aver|OJJ00488.1 awen|OJJ39253.1 azon|OJJ51602.1

Group3990: abra|OJJ76126.1 acar|OOF99798.1 acla|EAW09693.1 afis|EAW21446.1 afla|EED55874.1 afum|EAL92465.2 agla|OJJ85594.1 akaw|GAA88841.1 aluc|OJZ89587.1 anid|CBF76565.1 anig|EHA23024.1 aory|EIT76673.1 asyd|OJJ61162.1 ater|EAU35097.1 atub|OJI85216.1 aver|OJJ00486.1 awen|OJJ39252.1 azon|OJJ51603.1

Group3991: abra|OJJ76127.1 acar|OOF99796.1 acla|EAW09692.1 afis|EAW21447.1 afla|EED55876.1 afum|EAL92466.1 agla|OJJ85595.1 akaw|GAA88840.1 aluc|OJZ89588.1 anid|CBF76563.1 anig|EHA23023.1 aory|EIT76593.1 asyd|OJJ61161.1 ater|EAU35096.1 atub|OJI85214.1 aver|OJJ00483.1 awen|OJJ39251.1 azon|OJJ51604.1

Group3992: abra|OJJ76128.1 acar|OOF99795.1 acla|EAW09691.1 afis|EAW21448.1 afla|EED55877.1 afum|EAL92467.1 agla|OJJ85596.1 akaw|GAA88839.1 aluc|OJZ89589.1 anid|CBF76559.1 anig|EHA23022.1 aory|EIT76881.1 asyd|OJJ61159.1 ater|EAU35095.1 atub|OJI85213.1 aver|OJJ00481.1 awen|OJJ39250.1 azon|OJJ51605.1

Group3993: abra|OJJ76129.1 acar|OOF99794.1 acla|EAW09690.1 afis|EAW21449.1 afla|EED55878.1 afum|EAL92468.1 agla|OJJ85597.1 akaw|GAA88838.1 aluc|OJZ89590.1 anid|CBF76557.1 anig|EHA23021.1 aory|EIT76638.1 asyd|OJJ61158.1 ater|EAU35094.1 atub|OJI85212.1 aver|OJJ00480.1 awen|OJJ39249.1 azon|OJJ51606.1

Group3994: abra|OJJ76133.1 acar|OOF99790.1 acla|EAW09685.1 afis|EAW21454.1 afla|EED55884.1 afum|EAL92473.1 agla|OJJ85602.1 akaw|GAA88834.1 aluc|OJZ89594.1 anid|CBF76548.1 anig|EHA23017.1 aory|EIT76723.1 asyd|OJJ61155.1 ater|EAU35081.1 atub|OJI85208.1 aver|OJJ00477.1 awen|OJJ39246.1 azon|OJJ51609.1

Group3995: abra|OJJ76138.1 acar|OOF99786.1 acla|EAW09681.1 afis|EAW21458.1 afla|EED55888.1 afum|EAL92477.1 agla|OJJ85605.1 akaw|GAA88830.1 aluc|OJZ89601.1 anid|CBF76540.1 anig|EHA23014.1

aory|EIT76702.1 asyd|OJJ61151.1 ater|EAU35077.1 atub|OJI85203.1 aver|OJJ00473.1 awen|OJJ39242.1 azon|OJJ51613.1

Group3996: abra|OJJ76140.1 acar|OOF99785.1 acla|EAW09679.1 afis|EAW21460.1 afla|EED55891.1 afum|EAL92479.1 agla|OJJ85608.1 akaw|GAA88828.1 aluc|OJZ89603.1 anid|CBF76536.1 anig|EHA23012.1 aory|EIT76836.1 asyd|OJJ61149.1 ater|EAU35075.1 atub|OJI85201.1 aver|OJJ00471.1 awen|OJJ39239.1 azon|OJJ51615.1

Group3997: abra|OJJ76141.1 acar|OOF99784.1 acla|EAW09664.1 afis|EAW21473.1 afla|EED55908.1 afum|EAL92492.1 agla|OJJ85622.1 akaw|GAA88826.1 aluc|OJZ89605.1 anid|CBF76513.1 anig|EHA23011.1 aory|EIT76607.1 asyd|OJJ61136.1 ater|EAU35060.1 atub|OJI85198.1 aver|OJJ00458.1 awen|OJJ39224.1 azon|OJJ51630.1

Group3998: abra|OJJ76142.1 acar|OOF99783.1 acla|EAW09665.1 afis|EAW21472.1 afla|EED55902.1 afum|EAL92491.1 agla|OJJ85621.1 akaw|GAA88825.1 aluc|OJZ89606.1 anid|CBF76515.1 anig|EHA23010.1 aory|EIT76799.1 asyd|OJJ61137.1 ater|EAU35061.1 atub|OJI85197.1 aver|OJJ00459.1 awen|OJJ39225.1 azon|OJJ51629.1

Group3999: abra|OJJ76143.1 acar|OOF99782.1 acla|EAW09666.1 afis|EAW21471.1 afla|EED55904.1 afum|EAL92490.1 agla|OJJ85620.1 akaw|GAA88824.1 aluc|OJZ89607.1 anid|CBF76517.1 anig|EHA23009.1 aory|EIT76648.1 asyd|OJJ61138.1 ater|EAU35062.1 atub|OJI85196.1 aver|OJJ00460.1 awen|OJJ39227.1 azon|OJJ51628.1

Group4000: abra|OJJ76145.1 acar|OOF99781.1 acla|EAW09668.1 afis|EAW21469.1 afla|EED55907.1 afum|EAL92488.1 agla|OJJ85618.1 akaw|GAA82094.1 aluc|OJZ92086.1 anid|CBF76521.1 anig|EHA23008.1 aory|EIT76871.1 asyd|OJJ61140.1 ater|EAU35064.1 atub|OJI86515.1 aver|OJJ00462.1 awen|OJJ39229.1 azon|OJJ51626.1

Group4001: abra|OJJ76146.1 acar|OOF99780.1 acla|EAW09671.1 afis|EAW21467.1 afla|EED55900.1 afum|EAL92487.1 agla|OJJ85617.1 akaw|GAA82095.1 aluc|OJZ92084.1 anid|CBF76523.1 anig|EHA23007.1 aory|EIT76694.1 asyd|OJJ61141.1 ater|EAU35066.1 atub|OJI86514.1 aver|OJJ00463.1 awen|OJJ39230.1 azon|OJJ51625.1

Group4002: abra|OJJ76150.1 acar|OOF99777.1 acla|EAW09675.1 afis|EAW21464.1 afla|EED55896.1 afum|EAL92484.1 agla|OJJ85612.1 akaw|GAA82099.1 aluc|OJZ92079.1 anid|CBF76529.1 anig|EHA23005.1 aory|EIT76592.1 asyd|OJJ61145.1 ater|EAU35071.1 atub|OJI86509.1 aver|OJJ00467.1 awen|OJJ39235.1 azon|OJJ51620.1

Group4003: abra|OJJ76151.1 acar|OOF89898.1 acla|EAW09676.1 afis|EAW21463.1 afla|EED55895.1 afum|EAL92483.1 agla|OJJ85611.1 akaw|GAA82100.1 aluc|OJZ92078.1 anid|CBF76530.1 anig|EHA23004.1 aory|EIT76817.1 asyd|OJJ61146.1 ater|EAU35072.1 atub|OJI86508.1 aver|OJJ00468.1 awen|OJJ39236.1 azon|OJJ51619.1

Group4004: abra|OJJ76152.1 acar|OOF89899.1 acla|EAW09677.1 afis|EAW21462.1 afla|EED55894.1 afum|EAL92481.1 agla|OJJ85610.1 akaw|GAA82101.1 aluc|OJZ92077.1 anid|CBF76532.1 anig|EHA23003.1 aory|EIT76700.1 asyd|OJJ61147.1 ater|EAU35073.1 atub|OJI86507.1 aver|OJJ00469.1 awen|OJJ39237.1 azon|OJJ51618.1

Group4005: abra|OJJ76157.1 acar|OOF99775.1 acla|EAW09661.1 afis|EAW21478.1 afla|EED55910.1 afum|EAL92498.1 agla|OJJ85630.1 akaw|GAA82105.1 aluc|OJZ92071.1 anid|CBF76508.1 anig|EHA23000.1 aory|EIT76734.1 asyd|OJJ61133.1 ater|EAU35058.1 atub|OJI86503.1 aver|OJJ00454.1 awen|OJJ39221.1 azon|OJJ51635.1

Group4006: abra|OJJ76160.1 acar|OOF99772.1 acla|EAW09658.1 afis|EAW21481.1 afla|EED55913.1 afum|EAL92501.1 agla|OJJ85635.1 akaw|GAA82107.1 aluc|OJZ92069.1 anid|CBF76502.1 anig|EHA22997.1 aory|EIT76621.1 asyd|OJJ61130.1 ater|EAU35056.1 atub|OJI86500.1 aver|OJJ00451.1 awen|OJJ39216.1 azon|OJJ51638.1

Group4007: abra|OJJ76166.1 acar|OOF99771.1 acla|EAW09657.1 afis|EAW21482.1 afla|EED55914.1 afum|EAL92502.1 agla|OJJ85636.1 akaw|GAA82114.1 aluc|OJZ92062.1 anid|CBF76498.1 anig|EHA22991.1 aory|EIT76649.1 asyd|OJJ61128.1 ater|EAU35055.1 atub|OJI86494.1 aver|OJJ00449.1 awen|OJJ39215.1 azon|OJJ51640.1

Group4008: abra|OJJ76172.1 acar|OOF99767.1 acla|EAW09653.1 afis|EAW21486.1 afla|EED55919.1 afum|EAL92506.1 agla|OJJ85645.1 akaw|GAA82118.1 aluc|OJZ92058.1 anid|CBF76490.1 anig|EHA22987.1 aory|EIT76891.1 asyd|OJJ61124.1 ater|EAU35051.1 atub|OJI86490.1 aver|OJJ00445.1 awen|OJJ39211.1 azon|OJJ51643.1

Group4009: abra|OJJ76174.1 acar|OOF99766.1 acla|EAW09652.1 afis|EAW21488.1 afla|EED55923.1 afum|EAL92507.1 agla|OJJ85648.1 akaw|GAA82119.1 aluc|OJZ92056.1 anid|CBF76488.1 anig|EHA22985.1 aory|EIT76645.1 asyd|OJJ61123.1 ater|EAU35050.1 atub|OJI86487.1 aver|OJJ00441.1 awen|OJJ39207.1 azon|OJJ51645.1

Group4010: abra|OJJ76175.1 acar|OOF99765.1 acla|EAW09651.1 afis|EAW21489.1 afla|EED55924.1 afum|EAL92508.1 agla|OJJ85649.1 akaw|GAA82120.1 aluc|OJZ92055.1 anid|CBF76486.1 anig|EHA22984.1 aory|EIT76531.1 asyd|OJJ61122.1 ater|EAU35049.1 atub|OJI86486.1 aver|OJJ00440.1 awen|OJJ39206.1 azon|OJJ51646.1

Group4011: abra|OJJ76176.1 acar|OOF99764.1 acla|EAW09650.1 afis|EAW21490.1 afla|EED55925.1 afum|EAL92509.1 agla|OJJ85652.1 akaw|GAA82121.1 aluc|OJZ92054.1 anid|CBF76484.1 anig|EHA22983.1 aory|EIT76596.1 asyd|OJJ61121.1 ater|EAU35048.1 atub|OJI86485.1 aver|OJJ00439.1 awen|OJJ39205.1 azon|OJJ51647.1

Group4012: abra|OJJ76177.1 acar|OOF99762.1 acla|EAW09649.1 afis|EAW21491.1 afla|EED55926.1 afum|EAL92510.1 agla|OJJ85653.1 akaw|GAA82122.1 aluc|OJZ92053.1 anid|CBF76482.1 anig|EHA22982.1 aory|EIT76879.1 asyd|OJJ61120.1 ater|EAU35047.1 atub|OJI86484.1 aver|OJJ00438.1 awen|OJJ39204.1 azon|OJJ51648.1

Group4013: abra|OJJ76178.1 acar|OOF99763.1 acla|EAW09648.1 afis|EAW21492.1 afla|EED55927.1 afum|EAL92511.1 agla|OJJ85654.1 akaw|GAA82123.1 aluc|OJZ92052.1 anid|CBF76480.1 anig|EHA22981.1 aory|EIT76815.1 asyd|OJJ61119.1 ater|EAU35046.1 atub|OJI86483.1 aver|OJJ00437.1 awen|OJJ39203.1 azon|OJJ51649.1

Group4014: abra|OJJ76180.1 acar|OOF99760.1 acla|EAW09646.1 afis|EAW21494.1 afla|EED55929.1 afum|EAL92513.1 agla|OJJ85656.1 akaw|GAA82125.1 aluc|OJZ92049.1 anid|CBF76476.1 anig|EHA22979.1 aory|EIT76689.1 asyd|OJJ61117.1 ater|EAU35044.1 atub|OJI86481.1 aver|OJJ00435.1 awen|OJJ39201.1 azon|OJJ51526.1

Group4015: abra|OJJ76181.1 acar|OOF99759.1 acla|EAW09645.1 afis|EAW21495.1 afla|EED55930.1 afum|EAL92514.1 agla|OJJ85657.1 akaw|GAA82126.1 aluc|OJZ92048.1 anid|CBF76474.1 anig|EHA22978.1 aory|EIT76659.1 asyd|OJJ61116.1 ater|EAU35043.1 atub|OJI86480.1 aver|OJJ00434.1 awen|OJJ39200.1 azon|OJJ51525.1

Group4016: abra|OJJ76182.1 acar|OOF99758.1 acla|EAW09644.1 afis|EAW21496.1 afla|EED55931.1 afum|EAL92515.1 agla|OJJ85658.1 akaw|GAA82127.1 aluc|OJZ92047.1 anid|CBF76472.1 anig|EHA22977.1 aory|EIT76569.1 asyd|OJJ61115.1 ater|EAU35042.1 atub|OJI86479.1 aver|OJJ00433.1 awen|OJJ39199.1 azon|OJJ51524.1

Group4017: abra|OJJ76183.1 acar|OOF99757.1 acla|EAW09643.1 afis|EAW21497.1 afla|EED55933.1 afum|EAL92516.1 agla|OJJ85660.1 akaw|GAA82128.1 aluc|OJZ92046.1 anid|CBF76468.1 anig|EHA22976.1 aory|EIT76844.1 asyd|OJJ61114.1 ater|EAU35041.1 atub|OJI86478.1 aver|OJJ00430.1 awen|OJJ39198.1 azon|OJJ51523.1

Group4018: abra|OJJ76187.1 acar|OOF99755.1 acla|EAW09640.1 afis|EAW21500.1 afla|EED55936.1 afum|EAL92519.1 agla|OJJ85663.1 akaw|GAA82131.1 aluc|OJZ92042.1 anid|CBF76459.1 anig|EHA22974.1 aory|EIT76547.1 asyd|OJJ61111.1 ater|EAU35038.1 atub|OJI86475.1 aver|OJJ00427.1 awen|OJJ39195.1 azon|OJJ51522.1

Group4019: abra|OJJ76188.1 acar|OOF99754.1 acla|EAW09639.1 afis|EAW21501.1 afla|EED55937.1 afum|EAL92520.1 agla|OJJ85664.1 akaw|GAA82132.1 aluc|OJZ92041.1 anid|CBF76457.1 anig|EHA22973.1 aory|EIT76679.1 asyd|OJJ61110.1 ater|EAU35037.1 atub|OJI86474.1 aver|OJJ00426.1 awen|OJJ39194.1 azon|OJJ51519.1

Group4020: abra|OJJ76191.1 acar|OOF99751.1 acla|EAW13914.1 afis|EAW18625.1 afla|EED52717.1 afum|EAL93337.1 agla|OJJ81242.1 akaw|GAA82134.1 aluc|OJZ92038.1 anid|CBF70486.1 anig|EHA22970.1 aory|EIT78208.1 asyd|OJJ58585.1 ater|EAU34925.1 atub|OJI86472.1 aver|OJJ01189.1 awen|OJJ32090.1 azon|OJJ46667.1

Group4021: abra|OJJ76192.1 acar|OOF99752.1 acla|EAW13915.1 afis|EAW18626.1 afla|EED52716.1 afum|EAL93338.1 agla|OJJ81246.1 akaw|GAA82135.1 aluc|OJZ92037.1 anid|CBF70484.1 anig|EHA22969.1 aory|EIT78180.1 asyd|OJJ58586.1 ater|EAU34926.1 atub|OJI86470.1 aver|OJJ01190.1 awen|OJJ32092.1 azon|OJJ46669.1

Group4022: abra|OJJ76193.1 acar|OOF99750.1 acla|EAW13916.1 afis|EAW18627.1 afla|EED52715.1 afum|EAL93339.1 agla|OJJ81245.1 akaw|GAA82136.1 aluc|OJZ92036.1 anid|CBF70478.1 anig|EHA22968.1 aory|EIT78215.1 asyd|OJJ58587.1 ater|EAU38285.1 atub|OJI86469.1 aver|OJJ01191.1 awen|OJJ32093.1 azon|OJJ46670.1

Group4023: abra|OJJ76194.1 acar|OOF99749.1 acla|EAW13917.1 afis|EAW18628.1 afla|EED52714.1 afum|EAL93340.1 agla|OJJ81244.1 akaw|GAA82137.1 aluc|OJZ92035.1 anid|CBF70480.1 anig|EHA22967.1 aory|EIT78226.1 asyd|OJJ58588.1 ater|EAU38284.1 atub|OJI86468.1 aver|OJJ01192.1 awen|OJJ32094.1 azon|OJJ46671.1

Group4024: abra|OJJ76197.1 acar|OOF99747.1 acla|EAW13918.1 afis|EAW18629.1 afla|EED52712.1 afum|EAL93341.1 agla|OJJ81251.1 akaw|GAA82139.1 aluc|OJZ92032.1 anid|CBF70482.1 anig|EHA22965.1

aory|EIT78174.1 asyd|OJJ58592.1 ater|EAU38282.1 atub|OJI86465.1 aver|OJJ01194.1 awen|OJJ32096.1 azon|OJJ46675.1

Group4025: abra|OJJ76200.1 acar|OOF99744.1 acla|EAW13921.1 afis|EAW18632.1 afla|EED52709.1 afum|EAL93345.1 agla|OJJ82445.1 akaw|GAA82143.1 aluc|OJZ92028.1 anid|CBF70472.1 anig|EHA22962.1 aory|EIT74199.1 asyd|OJJ58595.1 ater|EAU38279.1 atub|OJI86462.1 aver|OJJ01197.1 awen|OJJ32099.1 azon|OJJ46681.1

Group4026: abra|OJJ76201.1 acar|OOF99743.1 acla|EAW13922.1 afis|EAW18633.1 afla|EED52708.1 afum|EAL93346.1 agla|OJJ82444.1 akaw|GAA82144.1 aluc|OJZ92027.1 anid|CBF70470.1 anig|EHA22961.1 aory|EIT74198.1 asyd|OJJ58596.1 ater|EAU38278.1 atub|OJI86461.1 aver|OJJ01198.1 awen|OJJ32100.1 azon|OJJ46682.1

Group4027: abra|OJJ76202.1 acar|OOF99742.1 acla|EAW13923.1 afis|EAW18634.1 afla|EED52707.1 afum|EAL93347.1 agla|OJJ82443.1 akaw|GAA82145.1 aluc|OJZ92026.1 anid|CBF70468.1 anig|EHA22960.1 aory|EIT74275.1 asyd|OJJ58597.1 ater|EAU38277.1 atub|OJI86460.1 aver|OJJ01199.1 awen|OJJ32101.1 azon|OJJ46683.1

Group4028: abra|OJJ76204.1 acar|OOF99741.1 acla|EAW13925.1 afis|EAW18636.1 afla|EED52705.1 afum|EAL93349.1 akaw|GAA82147.1 aluc|OJZ92024.1 anid|CBF70461.1 anig|EHA22958.1 aory|EIT74316.1 asyd|OJJ58600.1 atub|OJI86458.1 aver|OJJ01202.1 awen|OJJ32103.1 azon|OJJ46685.1 agla|OJJ82439.1 ater|EAU38275.1

Group4029: abra|OJJ76210.1 acar|OOF99739.1 acla|EAW13927.1 afis|EAW18638.1 afla|EED52702.1 afum|EAL93351.1 agla|OJJ82436.1 akaw|GAA82150.1 aluc|OJZ92018.1 anid|CBF70453.1 anig|EHA22956.1 aory|EIT74209.1 asyd|OJJ58604.1 ater|EAU38273.1 atub|OJI86453.1 aver|OJJ01207.1 awen|OJJ32106.1 azon|OJJ46687.1

Group4030: abra|OJJ76213.1 acar|OOF99737.1 acla|EAW13929.1 afis|EAW18640.1 afla|EED52700.1 afum|EAL93354.2 agla|OJJ82434.1 akaw|GAA82152.1 aluc|OJZ92016.1 anid|CBF70449.1 anig|EHA22955.1 aory|EIT74286.1 asyd|OJJ58606.1 ater|EAU38271.1 atub|OJI86451.1 aver|OJJ01210.1 awen|OJJ32108.1 azon|OJJ46689.1

Group4031: abra|OJJ76214.1 acar|OOF99736.1 acla|EAW13930.1 afis|EAW18641.1 afla|EED52699.1 afum|EAL93355.1 agla|OJJ82433.1 akaw|GAA82153.1 aluc|OJZ92015.1 anid|CBF70447.1 anig|EHA22954.1 aory|EIT74167.1 asyd|OJJ58607.1 ater|EAU38270.1 atub|OJI86450.1 aver|OJJ01211.1 awen|OJJ32109.1 azon|OJJ46690.1

Group4032: abra|OJJ76215.1 acar|OOF99735.1 acla|EAW13931.1 afis|EAW18642.1 afla|EED52698.1 afum|EAL93356.1 agla|OJJ82432.1 akaw|GAA82154.1 aluc|OJZ92014.1 anid|CBF70445.1 anig|EHA22953.1 aory|EIT74249.1 asyd|OJJ58608.1 ater|EAU38269.1 atub|OJI86449.1 aver|OJJ01212.1 awen|OJJ32110.1 azon|OJJ46691.1

Group4033: abra|OJJ76220.1 acar|OOF99731.1 acla|EAW13935.1 afis|EAW18646.1 afla|EED52696.1 afum|EAL93361.1 agla|OJJ82429.1 akaw|GAA82158.1 aluc|OJZ92010.1 anid|CBF70437.1 anig|EHA22950.1 aory|EIT74305.1 asyd|OJJ58611.1 ater|EAU38265.1 atub|OJI86445.1 aver|OJJ01217.1 awen|OJJ32114.1 azon|OJJ46695.1

Group4034: abra|OJJ76225.1 acar|OOF99472.1 acla|EAW14365.1 afis|EAW18915.1 afla|EED50899.1 afum|EBA27498.1 agla|OJJ81529.1 akaw|GAA82161.1 aluc|OJZ92006.1 anid|CBF69691.1 anig|EHA22946.1 aory|EIT78987.1 asyd|OJJ59055.1 ater|EAU38082.1 atub|OJI86441.1 aver|OJJ01649.1 awen|OJJ31508.1 azon|OJJ47227.1

Group4035: abra|OJJ76226.1 acar|OOF99473.1 acla|EAW14366.1 afis|EAW18916.1 afla|EED50900.1 afum|EAL93647.1 agla|OJJ81530.1 akaw|GAA82162.1 aluc|OJZ92005.1 anid|CBF69689.1 anig|EHA22945.1 aory|EIT79047.1 asyd|OJJ59056.1 ater|EAU38083.1 atub|OJI86440.1 aver|OJJ01650.1 awen|OJJ31509.1 azon|OJJ47228.1

Group4036: abra|OJJ76227.1 acar|OOF99474.1 acla|EAW14367.1 afis|EAW18917.1 afla|EED50901.1 afum|EAL93648.1 agla|OJJ81531.1 akaw|GAA82164.1 aluc|OJZ92002.1 anid|CBF69688.1 anig|EHA22943.1 aory|EIT79014.1 asyd|OJJ59057.1 ater|EAU38084.1 atub|OJI86438.1 aver|OJJ01651.1 awen|OJJ31510.1 azon|OJJ47229.1

Group4037: abra|OJJ76228.1 acar|OOF99475.1 acla|EAW14368.1 afis|EAW18918.1 afla|EED50902.1 afum|EAL93649.2 agla|OJJ81532.1 akaw|GAA82165.1 aluc|OJZ92001.1 anid|CBF69686.1 anig|EHA22942.1 aory|EIT78973.1 asyd|OJJ59058.1 ater|EAU38085.1 atub|OJI86437.1 aver|OJJ01652.1 awen|OJJ31511.1 azon|OJJ47230.1

Group4038: abra|OJJ76242.1 acar|OOF99487.1 acla|EAW14197.1 afis|EAW18925.1 afla|EED50914.1 afum|EBA27500.1 agla|OJJ85252.1 akaw|GAA82175.1 aluc|OJZ91989.1 anid|CBF81679.1 anig|EHA22936.1 aory|EIT79011.1 asyd|OJJ59560.1 ater|EAU38100.1 atub|OJI86424.1 aver|OJJ02708.1 awen|OJJ31519.1 azon|OJJ47235.1

Group4039: abra|OJJ76245.1 acar|OOF99491.1 acla|EAW14362.1 afis|EAW18927.1 afla|EED50920.1 afum|EAL93658.1 agla|OJJ85256.1 akaw|GAA82178.1 aluc|OJZ91986.1 anid|CBF69671.1 anig|EHA22932.1 aory|EIT78966.1 asyd|OJJ59063.1 ater|EAU38106.1 atub|OJI86421.1 aver|OJJ01657.1 awen|OJJ31521.1 azon|OJJ47237.1

Group4040: abra|OJJ76246.1 acar|OOF99492.1 acla|EAW14361.1 afis|EAW18928.1 afla|EED50921.1 afum|EAL93659.1 agla|OJJ85257.1 akaw|GAA82179.1 aluc|OJZ91985.1 anid|CBF69669.1 anig|EHA22931.1 aory|EIT79013.1 asyd|OJJ59062.1 ater|EAU38107.1 atub|OJI86420.1 aver|OJJ01656.1 awen|OJJ31522.1 azon|OJJ47238.1

Group4041: abra|OJJ76248.1 acar|OOF89844.1 acla|EAW14359.1 afis|EAW18931.1 afla|EED50936.1 afum|EAL93663.1 agla|OJJ85260.1 akaw|GAA82181.1 aluc|OJZ91982.1 anid|CBF86796.1 anig|EHA22929.1 aory|EIT78972.1 asyd|OJJ64670.1 ater|EAU38110.1 atub|OJI86418.1 aver|OJI98772.1 awen|OJJ31525.1 azon|OJJ47240.1

Group4042: abra|OJJ76250.1 acar|OOF99495.1 acla|EAW14357.1 afis|EAW18933.1 afla|EED50938.1 afum|EAL93666.2 agla|OJJ85262.1 akaw|GAA82183.1 aluc|OJZ91977.1 anid|CBF86800.1 anig|EHA22927.1 aory|EIT77014.1 asyd|OJJ64668.1 ater|EAU38112.1 atub|OJI86416.1 aver|OJI98770.1 awen|OJJ31527.1 azon|OJJ47242.1

Group4043: abra|OJJ76251.1 acar|OOF99496.1 acla|EAW14356.1 afis|EAW18934.1 afla|EED50939.1 afum|EAL93667.2 agla|OJJ85263.1 akaw|GAA82184.1 aluc|OJZ91976.1 anid|CBF86802.1 anig|EHA22926.1 aory|EIT77060.1 asyd|OJJ64667.1 ater|EAU38113.1 atub|OJI86415.1 aver|OJI98769.1 awen|OJJ31528.1 azon|OJJ47243.1

Group4044: abra|OJJ76252.1 acar|OOF99497.1 acla|EAW14355.1 afis|EAW18935.1 afla|EED50940.1 afum|EAL93668.2 agla|OJJ85264.1 akaw|GAA82185.1 aluc|OJZ91975.1 anid|CBF86804.1 anig|EHA22925.1 aory|EIT76996.1 asyd|OJJ64666.1 ater|EAU38114.1 atub|OJI86414.1 aver|OJI98768.1 awen|OJJ31529.1 azon|OJJ47244.1

Group4045: abra|OJJ76253.1 acar|OOF99498.1 acla|EAW14354.1 afis|EAW18936.1 afla|EED50941.1 afum|EAL93669.2 agla|OJJ85265.1 akaw|GAA82186.1 aluc|OJZ91974.1 anid|CBF86805.1 anig|EHA22924.1 aory|EIT77063.1 asyd|OJJ64665.1 ater|EAU38115.1 atub|OJI86413.1 aver|OJI98767.1 awen|OJJ31530.1 azon|OJJ47245.1

Group4046: abra|OJJ76254.1 acar|OOF99499.1 acla|EAW14353.1 afis|EAW18937.1 afla|EED50942.1 afum|EAL93670.1 agla|OJJ85270.1 akaw|GAA82188.1 aluc|OJZ91972.1 anid|CBF86807.1 anig|EHA22918.1 aory|EIT77016.1 asyd|OJJ64664.1 ater|EAU38116.1 atub|OJI86411.1 aver|OJI98766.1 awen|OJJ31532.1 azon|OJJ47246.1

Group4047: abra|OJJ76256.1 acar|OOF99501.1 acla|EAW14350.1 afis|EAW18940.1 afla|EED50966.1 afum|EAL93673.1 agla|OJJ85271.1 akaw|GAA82190.1 aluc|OJZ91970.1 anid|CBF86811.1 anig|EHA22920.1 aory|EIT76988.1 asyd|OJJ64661.1 ater|EAU38119.1 atub|OJI86409.1 aver|OJI98763.1 awen|OJJ31534.1 azon|OJJ47250.1

Group4048: abra|OJJ76259.1 acar|OOF99504.1 acla|EAW14351.1 afis|EAW18939.1 afla|EED50969.1 afum|EAL93672.1 agla|OJJ85268.1 akaw|GAA82192.1 aluc|OJZ91964.1 anid|CBF86815.1 anig|EHA22923.1 aory|EIT77055.1 asyd|OJJ64658.1 ater|EAU38120.1 atub|OJI86407.1 aver|OJI98761.1 awen|OJJ31535.1 azon|OJJ47249.1

Group4049: abra|OJJ76262.1 acar|OOF99506.1 acla|EAW14346.1 afis|EAW18944.1 afla|EED50978.1 afum|EAL93677.1 agla|OJJ85278.1 akaw|GAA82194.1 aluc|OJZ91960.1 anid|CBF86822.1 anig|EHA22916.1 aory|EIT77009.1 asyd|OJJ64655.1 ater|EAU38124.1 atub|OJI86404.1 aver|OJI98757.1 awen|OJJ31540.1 azon|OJJ47257.1

Group4050: abra|OJJ76266.1 afla|EED50981.1 afum|EAL93681.2 anid|CBF86829.1 aory|EIT76942.1 asyd|OJJ64651.1 atub|OJI86400.1 aver|OJI98752.1 acar|OOF99509.1 acla|EAW14343.1 afis|EAW18947.1 agla|OJJ85308.1 akaw|GAA82198.1 aluc|OJZ91954.1 anig|EHA22913.1 ater|EAU38127.1 awen|OJJ31543.1 azon|OJJ47260.1

Group4051: abra|OJJ76267.1 acar|OOF99511.1 acla|EAW14341.1 afis|EAW18949.1 afla|EED50985.1 afum|EAL93683.1 agla|OJJ85320.1 akaw|GAA82200.1 aluc|OJZ91953.1 anid|CBF86833.1 anig|EHA22912.1 aory|EIT77049.1 asyd|OJJ64649.1 ater|EAU38135.1 atub|OJI86398.1 aver|OJI98749.1 awen|OJJ31547.1 azon|OJJ47263.1

Group4052: abra|OJJ76269.1 acar|OOF99517.1 acla|EAW12851.1 afis|EAW18014.1 afla|EED51880.1 afum|EAL91826.1 agla|OJJ85368.1 akaw|GAA82202.1 aluc|OJZ91951.1 anid|CBF80042.1 anig|EHA22910.1 aory|EIT81710.1 asyd|OJJ58276.1 ater|EAU31530.1 atub|OJI86396.1 aver|OJJ02148.1 awen|OJJ41526.1 azon|OJJ48247.1

Group4053: abra|OJJ76288.1 acar|OOF99529.1 acla|EAW14328.1 afis|EAW18960.1 afla|EED51006.1 afum|EAL93693.1 agla|OJJ85342.1 akaw|GAA82217.1 aluc|OJZ91934.1 anid|CBF69600.1 anig|EHA22894.1

aory|EIT77056.1 asyd|OJJ59135.1 ater|EAU38143.1 atub|OJI86376.1 aver|OJJ01724.1 awen|OJJ31559.1 azon|OJJ47276.1

Group4054: abra|OJJ76290.1 acar|OOF99535.1 acla|EAW14325.1 afis|EAW18963.1 afla|EED51010.1 afum|EAL93695.1 agla|OJJ85344.1 akaw|GAA82219.1 aluc|OJZ91932.1 anid|CBF69605.1 anig|EHA22893.1 aory|EIT76961.1 asyd|OJJ59132.1 ater|EAU38148.1 atub|OJI86374.1 aver|OJJ01721.1 awen|OJJ31562.1 azon|OJJ47283.1

Group4055: abra|OJJ76292.1 acar|OOF99537.1 acla|EAW14324.1 afis|EAW18964.1 afla|EED51011.1 afum|EAL93696.1 agla|OJJ85346.1 akaw|GAA82220.1 aluc|OJZ91930.1 anid|CBF69607.1 anig|EHA22892.1 aory|EIT77048.1 asyd|OJJ59130.1 atub|OJI86373.1 aver|OJJ01720.1 awen|OJJ31564.1 azon|OJJ47282.1 ater|EAU38149.1

Group4056: abra|OJJ76296.1 acar|OOF99542.1 acla|EAW14322.1 afis|EAW18966.1 afla|EED51032.1 afum|EAL93698.2 agla|OJJ85352.1 akaw|GAA82223.1 aluc|OJZ91926.1 anid|CBF69611.1 anig|EHA22889.1 aory|EIT76971.1 asyd|OJJ59128.1 ater|EAU38151.1 atub|OJI86370.1 aver|OJJ01716.1 awen|OJJ31566.1 azon|OJJ47284.1

Group4057: abra|OJJ76297.1 acar|OOF99545.1 acla|EAW14321.1 afis|EAW18967.1 afla|EED51033.1 afum|EAL93699.1 agla|OJJ85353.1 akaw|GAA82224.1 aluc|OJZ91925.1 anid|CBF69613.1 anig|EHA22888.1 aory|EIT76951.1 asyd|OJJ59127.1 ater|EAU38153.1 atub|OJI86369.1 aver|OJJ01715.1 awen|OJJ31567.1 azon|OJJ47285.1

Group4058: abra|OJJ76299.1 acar|OOF99546.1 acla|EAW14319.1 afis|EAW18969.1 afla|EED51035.1 afum|EAL93702.1 agla|OJJ85355.1 akaw|GAA82226.1 aluc|OJZ91923.1 anid|CBF69617.1 anig|EHA22887.1 aory|EIT77006.1 asyd|OJJ59124.1 ater|EAU38155.1 atub|OJI86367.1 aver|OJJ01713.1 awen|OJJ31569.1 azon|OJJ47287.1

Group4059: abra|OJJ76300.1 acar|OOF99544.1 acla|EAW14318.1 afis|EAW18970.1 afla|EED51036.1 afum|EAL93703.1 agla|OJJ85356.1 akaw|GAA82227.1 aluc|OJZ91922.1 anid|CBF69619.1 anig|EHA22886.1 aory|EIT77012.1 asyd|OJJ59123.1 ater|EAU38156.1 atub|OJI86366.1 aver|OJJ01712.1 awen|OJJ31570.1 azon|OJJ47288.1

Group4060: abra|OJJ76301.1 acar|OOF99547.1 acla|EAW14317.1 afis|EAW18971.1 afla|EED51037.1 afum|EAL93704.1 agla|OJJ85357.1 akaw|GAA82228.1 aluc|OJZ91921.1 anid|CBF69621.1 anig|EHA22885.1 aory|EIT77040.1 asyd|OJJ59122.1 ater|EAU38157.1 atub|OJI86365.1 aver|OJJ01711.1 awen|OJJ31571.1 azon|OJJ47289.1

Group4061: abra|OJJ76302.1 acar|OOF99548.1 acla|EAW14316.1 afis|EAW18972.1 afla|EED51038.1 afum|EAL93705.1 agla|OJJ85360.1 akaw|GAA82232.1 aluc|OJZ91916.1 anid|CBF69627.1 anig|EHA22884.1 aory|EIT77044.1 asyd|OJJ59119.1 ater|EAU38158.1 atub|OJI86364.1 aver|OJJ01708.1 awen|OJJ31572.1 azon|OJJ47291.1

Group4062: abra|OJJ76303.1 acar|OOF99549.1 acla|EAW14315.1 afis|EAW18973.1 afla|EED51039.1 afum|EAL93706.1 agla|OJJ85361.1 akaw|GAA82233.1 aluc|OJZ91914.1 anid|CBF69629.1 anig|EHA22883.1 aory|EIT77038.1 asyd|OJJ59117.1 ater|EAU38159.1 atub|OJI86363.1 aver|OJJ01707.1 awen|OJJ31573.1 azon|OJJ47290.1

Group4063: abra|OJJ76307.1 acar|OOF99552.1 acla|EAW14108.1 afis|EAW18810.1 afla|EED50762.1 afum|EAL93546.1 agla|OJJ80543.1 akaw|GAA82238.1 aluc|OJZ91908.1 anid|CBF69793.1 anig|EHA22882.1 aory|EIT78035.1 asyd|OJJ58963.1 ater|EAU37962.1 atub|OJI86358.1 aver|OJJ01557.1 awen|OJJ32385.1 azon|OJJ47146.1

Group4064: abra|OJJ76314.1 acar|OOF99566.1 acla|EAW14101.1 afis|EAW18802.1 afla|EED50774.1 afum|EAL93538.1 agla|OJJ80532.1 akaw|GAA82245.1 aluc|OJZ91900.1 anid|CBF69784.1 anig|EHA22879.1 aory|EIT77990.1 asyd|OJJ58960.1 ater|EAU37940.1 atub|OJI86353.1 aver|OJJ01554.1 awen|OJJ32376.1 azon|OJJ47139.1

Group4065: abra|OJJ76323.1 acar|OOF99575.1 acla|EAW14092.1 afis|EAW18798.1 afla|EED50791.1 afum|EAL93534.1 agla|OJJ80527.1 akaw|GAA82252.1 aluc|OJZ91892.1 anid|CBF69777.1 anig|EHA22871.1 aory|EIT78043.1 asyd|OJJ58956.1 ater|EAU37930.1 atub|OJI86344.1 aver|OJJ01550.1 awen|OJJ32372.1 azon|OJJ47135.1

Group4066: abra|OJJ76325.1 acar|OOF99577.1 acla|EAW14090.1 afis|EAW18796.1 afla|EED50793.1 afum|EAL93532.1 agla|OJJ80525.1 akaw|GAA82254.1 aluc|OJZ91890.1 anid|CBF69773.1 anig|EHA22870.1 aory|EIT78128.1 asyd|OJJ58954.1 ater|EAU37927.1 atub|OJI86342.1 aver|OJJ01548.1 awen|OJJ32370.1 azon|OJJ47133.1

Group4067: abra|OJJ76326.1 acar|OOF99578.1 afis|EAW18795.1 afla|EED50794.1 afum|EAL93531.1 agla|OJJ80524.1 akaw|GAA82255.1 aluc|OJZ91889.1 anid|CBF69771.1 anig|EHA22869.1 aory|EIT78000.1 asyd|OJJ58953.1 ater|EAU37926.1 atub|OJI86341.1 aver|OJJ01547.1 awen|OJJ32369.1 azon|OJJ47132.1 acla|EAW14089.1

Group4068: abra|OJJ76327.1 acar|OOF99580.1 acla|EAW14088.1 afis|EAW18794.1 afla|EED50795.1 afum|EAL93530.1 agla|OJJ80523.1 akaw|GAA82256.1 aluc|OJZ91888.1 anid|CBF69769.1 anig|EHA22868.1 aory|EIT78079.1 asyd|OJJ58952.1 ater|EAU37925.1 atub|OJI86340.1 aver|OJJ01546.1 awen|OJJ32367.1 azon|OJJ47131.1

Group4069: abra|OJJ76332.1 acar|OOF99584.1 acla|EAW14083.1 afis|EAW18790.1 afla|EED50748.1 afum|EAL93525.1 agla|OJJ80519.1 akaw|GAA82260.1 aluc|OJZ91881.1 anid|CBF69761.1 anig|EHA22859.1 aory|EIT78056.1 asyd|OJJ58947.1 ater|EAU37921.1 atub|OJI86332.1 aver|OJJ01542.1 awen|OJJ32363.1 azon|OJJ47126.1

Group4070: abra|OJJ76334.1 acar|OOF99587.1 acla|EAW14079.1 afis|EAW18786.1 afla|EED50745.1 afum|EAL93519.1 agla|OJJ80509.1 akaw|GAA82262.1 aluc|OJZ91878.1 anid|CBF69752.1 anig|EHA22856.1 aory|EIT78009.1 asyd|OJJ58941.1 ater|EAU37918.1 atub|OJI86329.1 aver|OJJ01537.1 awen|OJJ32360.1 azon|OJJ47123.1

Group4071: abra|OJJ76336.1 acar|OOF99588.1 acla|EAW14078.1 afis|EAW18785.1 afla|EED50744.1 afum|EAL93517.2 agla|OJJ80507.1 akaw|GAA82264.1 aluc|OJZ91876.1 anid|CBF69748.1 anig|EHA22855.1 aory|EIT78137.1 asyd|OJJ58938.1 ater|EAU37917.1 atub|OJI86327.1 aver|OJJ01534.1 awen|OJJ32358.1 azon|OJJ47121.1

Group4072: abra|OJJ76337.1 acar|OOF99589.1 acla|EAW14077.1 afis|EAW18784.1 afla|EED50743.1 afum|EAL93516.2 agla|OJJ80506.1 akaw|GAA82265.1 aluc|OJZ91875.1 anid|CBF69746.1 anig|EHA22854.1 aory|EIT78027.1 asyd|OJJ58937.1 ater|EAU37916.1 atub|OJI86326.1 aver|OJJ01533.1 awen|OJJ32357.1 azon|OJJ47120.1

Group4073: abra|OJJ76338.1 acar|OOF99590.1 acla|EAW14076.1 afis|EAW18783.1 afla|EED50742.1 afum|EAL93515.1 agla|OJJ80504.1 akaw|GAA82266.1 aluc|OJZ91874.1 anid|CBF69744.1 anig|EHA22853.1 aory|EIT78012.1 asyd|OJJ58936.1 ater|EAU37915.1 atub|OJI86325.1 aver|OJJ01532.1 awen|OJJ32356.1 azon|OJJ47119.1

Group4074: abra|OJJ76340.1 acar|OOF99591.1 acla|EAW14075.1 afis|EAW18782.1 afla|EED50741.1 afum|EAL93514.1 agla|OJJ80502.1 akaw|GAA82267.1 aluc|OJZ91873.1 anid|CBF69742.1 anig|EHA22852.1 aory|EIT78041.1 asyd|OJJ58935.1 ater|EAU37914.1 atub|OJI86324.1 aver|OJJ01531.1 awen|OJJ32354.1 azon|OJJ47118.1

Group4075: abra|OJJ76342.1 acar|OOF99593.1 acla|EAW14073.1 afis|EAW18780.1 afla|EED50739.1 afum|EAL93512.1 agla|OJJ80500.1 akaw|GAA82269.1 aluc|OJZ91871.1 anid|CBF69738.1 anig|EHA22850.1 aory|EIT77981.1 asyd|OJJ58933.1 ater|EAU37912.1 atub|OJI86322.1 aver|OJJ01529.1 awen|OJJ32352.1 azon|OJJ47116.1

Group4076: abra|OJJ76344.1 acar|OOF99595.1 acla|EAW14071.1 afis|EAW18778.1 afla|EED50737.1 afum|EAL93510.1 agla|OJJ80495.1 akaw|GAA82271.1 aluc|OJZ91869.1 anid|CBF69734.1 anig|EHA22849.1 aory|EIT78123.1 asyd|OJJ58931.1 ater|EAU37910.1 atub|OJI86320.1 aver|OJJ01527.1 awen|OJJ32349.1 azon|OJJ47114.1

Group4077: abra|OJJ76345.1 acar|OOF99596.1 acla|EAW14070.1 afis|EAW18777.1 afla|EED50736.1 afum|EAL93509.1 agla|OJJ80494.1 akaw|GAA82272.1 aluc|OJZ91868.1 anid|CBF69732.1 anig|EHA22848.1 aory|EIT78071.1 asyd|OJJ58930.1 ater|EAU37909.1 atub|OJI86319.1 aver|OJJ01526.1 awen|OJJ32348.1 azon|OJJ47113.1

Group4078: abra|OJJ76346.1 acar|OOF99597.1 acla|EAW14069.1 afis|EAW18776.1 afla|EED50735.1 afum|EAL93508.1 agla|OJJ80493.1 akaw|GAA82273.1 aluc|OJZ91867.1 anid|CBF69730.1 anig|EHA22847.1 aory|EIT78130.1 asyd|OJJ58929.1 ater|EAU37908.1 atub|OJI86318.1 aver|OJJ01525.1 awen|OJJ32347.1 azon|OJJ47112.1

Group4079: abra|OJJ76347.1 acar|OOF99598.1 acla|EAW14068.1 afis|EAW18775.1 afla|EED50734.1 afum|EAL93507.1 agla|OJJ80491.1 akaw|GAA82274.1 aluc|OJZ91866.1 anid|CBF69728.1 anig|EHA22846.1 aory|EIT78021.1 asyd|OJJ58928.1 ater|EAU37906.1 atub|OJI86317.1 aver|OJJ01524.1 awen|OJJ32346.1 azon|OJJ47111.1

Group4080: abra|OJJ76349.1 acar|OOF99600.1 acla|EAW14066.1 afis|EAW18773.1 afla|EED50732.1 afum|EAL93505.1 agla|OJJ80489.1 akaw|GAA82276.1 aluc|OJZ91864.1 anid|CBF69724.1 anig|EHA22844.1 aory|EIT78029.1 asyd|OJJ58927.1 ater|EAU37904.1 atub|OJI86315.1 aver|OJJ01523.1 awen|OJJ32344.1 azon|OJJ47110.1

Group4081: abra|OJJ76351.1 acar|OOF99603.1 acla|EAW14064.1 afis|EAW18771.1 afla|EED50730.1 afum|EAL93502.1 agla|OJJ80467.1 akaw|GAA82278.1 aluc|OJZ91862.1 anid|CBF69934.1 anig|EHA22842.1 aory|EIT78074.1 asyd|OJJ58922.1 ater|EAU37902.1 atub|OJI86314.1 aver|OJJ01519.1 awen|OJJ32340.1 azon|OJJ47106.1

Group4082: abra|OJJ76353.1 acar|OOF99604.1 acla|EAW14062.1 afis|EAW18769.1 afla|EED50728.1 afum|EAL93500.2 agla|OJJ82285.1 akaw|GAA82279.1 aluc|OJZ91860.1 anid|CBF69938.1 anig|EHA22840.1

aory|EIT78133.1 asyd|OJJ58920.1 ater|EAU37900.1 atub|OJI86311.1 aver|OJJ01517.1 awen|OJJ32338.1 azon|OJJ47103.1

Group4083: abra|OJJ76354.1 acar|OOF99605.1 acla|EAW14061.1 afis|EAW18768.1 afla|EED50727.1 afum|EAL93499.1 agla|OJJ82284.1 akaw|GAA82280.1 aluc|OJZ91859.1 anid|CBF69940.1 anig|EHA22839.1 aory|EIT78038.1 asyd|OJJ58919.1 ater|EAU37899.1 atub|OJI86310.1 aver|OJJ01516.1 awen|OJJ32337.1 azon|OJJ47102.1

Group4084: abra|OJJ76356.1 acar|OOF99607.1 acla|EAW14059.1 afis|EAW18765.1 afla|EED50725.1 afum|EAL93497.1 agla|OJJ80456.1 akaw|GAA82282.1 aluc|OJZ91857.1 anid|CBF69944.1 anig|EHA22837.1 aory|EIT78144.1 asyd|OJJ58917.1 ater|EAU37895.1 atub|OJI86308.1 aver|OJJ01514.1 awen|OJJ32335.1 azon|OJJ47101.1

Group4085: abra|OJJ76358.1 acar|OOF99609.1 acla|EAW14058.1 afis|EAW18764.1 afla|EED50724.1 afum|EAL93496.2 agla|OJJ80457.1 akaw|GAA82284.1 aluc|OJZ91855.1 anid|CBF69945.1 anig|EHA22836.1 aory|EIT78031.1 asyd|OJJ58915.1 ater|EAU37894.1 atub|OJI86306.1 aver|OJJ01512.1 awen|OJJ32333.1 azon|OJJ47099.1

Group4086: abra|OJJ76359.1 acar|OOF99610.1 acla|EAW14057.1 afis|EAW18763.1 afla|EED50723.1 afum|EAL93495.1 agla|OJJ80458.1 akaw|GAA82285.1 aluc|OJZ91854.1 anid|CBF69947.1 anig|EHA22835.1 aory|EIT78108.1 asyd|OJJ58914.1 ater|EAU37893.1 atub|OJI86305.1 aver|OJJ01511.1 awen|OJJ32332.1 azon|OJJ47098.1

Group4087: abra|OJJ76362.1 acar|OOF99613.1 acla|EAW14054.1 afis|EAW18760.1 afla|EED50720.1 afum|EAL93491.1 agla|OJJ82333.1 akaw|GAA82288.1 aluc|OJZ91850.1 anid|CBF69953.1 anig|EHA22832.1 aory|EIT78011.1 asyd|OJJ58911.1 ater|EAU37891.1 atub|OJI86302.1 aver|OJJ01508.1 awen|OJJ32330.1 azon|OJJ47095.1

Group4088: abra|OJJ76363.1 acar|OOF99614.1 acla|EAW14053.1 afis|EAW18759.1 afla|EED50719.1 afum|EAL93489.1 agla|OJJ82332.1 akaw|GAA82289.1 aluc|OJZ91849.1 anid|CBF69955.1 anig|EHA22831.1 aory|EIT78126.1 asyd|OJJ58910.1 ater|EAU37928.1 atub|OJI86301.1 aver|OJJ01507.1 awen|OJJ32329.1 azon|OJJ47094.1

Group4089: abra|OJJ76364.1 acar|OOF99615.1 acla|EAW14052.1 afis|EAW18758.1 afla|EED50718.1 afum|EAL93488.2 agla|OJJ82331.1 akaw|GAA82290.1 aluc|OJZ91848.1 anid|CBF69957.1 anig|EHA22830.1 aory|EIT78109.1 asyd|OJJ58909.1 ater|EAU37890.1 atub|OJI86300.1 aver|OJJ01506.1 awen|OJJ32328.1 azon|OJJ47093.1

Group4090: abra|OJJ76366.1 acar|OOF89987.1 acla|EAW14050.1 afis|EAW18756.1 afla|EED50716.1 afum|EAL93484.1 agla|OJJ82324.1 akaw|GAA82292.1 aluc|OJZ91846.1 anid|CBF69963.1 anig|EHA22828.1 aory|EIT78115.1 asyd|OJJ58907.1 ater|EAU37887.1 atub|OJI86298.1 aver|OJJ01504.1 awen|OJJ32325.1 azon|OJJ47092.1

Group4091: abra|OJJ76368.1 acar|OOF99617.1 acla|EAW14048.1 afis|EAW18754.1 afla|EED50714.1 afum|EAL93481.1 agla|OJJ82322.1 akaw|GAA82294.1 aluc|OJZ91844.1 anid|CBF69971.1 anig|EHA22827.1 aory|EIT77980.1 asyd|OJJ58901.1 ater|EAU37885.1 atub|OJI86296.1 aver|OJJ01501.1 awen|OJJ32323.1 azon|OJJ47090.1

Group4092: abra|OJJ76370.1 acar|OOF99619.1 acla|EAW14047.1 afis|EAW18753.1 afla|EED50712.1 afum|EAL93480.1 agla|OJJ82321.1 akaw|GAA82296.1 aluc|OJZ91842.1 anid|CBF69973.1 anig|EHA22825.1 aory|EIT78076.1 asyd|OJJ58900.1 ater|EAU37884.1 atub|OJI86294.1 aver|OJJ01500.1 awen|OJJ32322.1 azon|OJJ47089.1

Group4093: abra|OJJ76378.1 acar|OOF99623.1 acla|EAW14044.1 afis|EAW18750.1 afla|EED50709.1 afum|EAL93477.1 agla|OJJ82317.1 akaw|GAA82299.1 aluc|OJZ91836.1 anid|CBF69979.1 anig|EHA22823.1 aory|EIT78032.1 asyd|OJJ58896.1 ater|EAU37879.1 atub|OJI86290.1 aver|OJJ01497.1 awen|OJJ32317.1 azon|OJJ47085.1

Group4094: abra|OJJ76379.1 acar|OOF99624.1 acla|EAW14043.1 afis|EAW18749.1 afla|EED50708.1 afum|EAL93475.1 agla|OJJ82316.1 akaw|GAA82300.1 aluc|OJZ91835.1 anid|CBF69983.1 anig|EHA22822.1 aory|EIT77996.1 asyd|OJJ58895.1 ater|EAU37878.1 atub|OJI86289.1 aver|OJJ01496.1 awen|OJJ32316.1 azon|OJJ47084.1

Group4095: abra|OJJ76381.1 acar|OOF89925.1 acla|EAW14041.1 afis|EAW18747.1 afla|EED50706.1 afum|EAL93473.1 agla|OJJ82314.1 akaw|GAA82302.1 aluc|OJZ91833.1 anid|CBF69987.1 anig|EHA22820.1 aory|EIT78116.1 asyd|OJJ58893.1 ater|EAU37876.1 atub|OJI86287.1 aver|OJJ01494.1 awen|OJJ32313.1 azon|OJJ47082.1

Group4096: abra|OJJ76382.1 acar|OOF89924.1 acla|EAW14040.1 afis|EAW18746.1 afla|EED50705.1 afum|EAL93472.1 agla|OJJ82313.1 akaw|GAA82303.1 aluc|OJZ91832.1 anid|CBF69992.1 anig|EHA22819.1 aory|EIT78094.1 asyd|OJJ58891.1 ater|EAU37875.1 atub|OJI86286.1 aver|OJJ01492.1 awen|OJJ32312.1 azon|OJJ42712.1

Group4097: abra|OJJ76387.1 acar|OOF99628.1 acla|EAW14036.1 afis|EAW18742.1 afla|EED50698.1 afum|EAL93468.1 agla|OJJ82308.1 akaw|GAA82307.1 aluc|OJZ91826.1 anid|CBF69998.1 anig|EHA22815.1 aory|EIT78061.1 asyd|OJJ58887.1 ater|EAU37869.1 atub|OJI86281.1 aver|OJJ01488.1 awen|OJJ32307.1 azon|OJJ42719.1

Group4098: abra|OJJ76389.1 acar|OOF99630.1 acla|EAW14035.1 afis|EAW18741.1 afla|EED50696.1 afum|EAL93467.1 agla|OJJ82307.1 akaw|GAA82308.1 aluc|OJZ91824.1 anid|CBF70000.1 anig|EHA22814.1 aory|EIT78051.1 asyd|OJJ58886.1 ater|EAU37864.1 atub|OJI86279.1 aver|OJJ01487.1 awen|OJJ32306.1 azon|OJJ42720.1

Group4099: abra|OJJ76390.1 acar|OOF99631.1 acla|EAW14034.1 afis|EAW18740.1 afla|EED50695.1 afum|EAL93466.1 agla|OJJ82306.1 akaw|GAA82309.1 aluc|OJZ91823.1 anid|CBF70002.1 anig|EHA22813.1 aory|EIT78073.1 asyd|OJJ58884.1 ater|EAU37863.1 atub|OJI86278.1 aver|OJJ01486.1 awen|OJJ32305.1 azon|OJJ42721.1

Group4100: abra|OJJ76392.1 acar|OOF99632.1 acla|EAW14032.1 afis|EAW18738.1 afla|EED50693.1 afum|EAL93464.1 agla|OJJ82304.1 akaw|GAA82311.1 aluc|OJZ91821.1 anid|CBF70006.1 anig|EHA22811.1 aory|EIT78112.1 asyd|OJJ58882.1 ater|EAU37861.1 atub|OJI86276.1 aver|OJJ01484.1 awen|OJJ32302.1 azon|OJJ42723.1

Group4101: abra|OJJ76393.1 acar|OOF99633.1 acla|EAW14031.1 afis|EAW18737.1 afla|EED50692.1 afum|EAL93462.1 agla|OJJ82303.1 akaw|GAA82312.1 aluc|OJZ91820.1 anid|CBF70008.1 anig|EHA22810.1 aory|EIT78068.1 asyd|OJJ58881.1 ater|EAU37860.1 atub|OJI86275.1 aver|OJJ01483.1 awen|OJJ32301.1 azon|OJJ42724.1

Group4102: abra|OJJ76395.1 acar|OOF99635.1 acla|EAW14029.1 afis|EAW18735.1 afla|EED50690.1 afum|EAL93460.1 agla|OJJ82301.1 akaw|GAA82314.1 aluc|OJZ91818.1 anid|CBF70012.1 anig|EHA22808.1 aory|EIT78132.1 asyd|OJJ58879.1 ater|EAU37858.1 atub|OJI86273.1 aver|OJJ01481.1 awen|OJJ32298.1 azon|OJJ42734.1

Group4103: abra|OJJ76396.1 acar|OOF99637.1 acla|EAW14027.1 afis|EAW18733.1 afla|EED50688.1 afum|EAL93458.1 agla|OJJ82299.1 akaw|GAA82316.1 aluc|OJZ91816.1 anid|CBF70014.1 anig|EHA22807.1 aory|EIT78099.1 asyd|OJJ58878.1 ater|EAU37848.1 atub|OJI86272.1 aver|OJJ01480.1 awen|OJJ32296.1 azon|OJJ42736.1

Group4104: abra|OJJ76404.1 acar|OOF99647.1 acla|EAW14017.1 afis|EAW18725.1 afla|EED50682.1 afum|EAL93450.1 agla|OJJ80475.1 akaw|GAA82324.1 aluc|OJZ91808.1 anid|CBF70021.1 anig|EHA22801.1 aory|EIT77978.1 asyd|OJJ58874.1 ater|EAU38179.1 atub|OJI86263.1 aver|OJJ01477.1 awen|OJJ32229.1 azon|OJJ46783.1

Group4105: abra|OJJ76406.1 acar|OOF99649.1 acla|EAW14019.1 afis|EAW18723.1 afla|EED50680.1 afum|EAL93448.1 agla|OJJ80477.1 akaw|GAA82326.1 aluc|OJZ91806.1 anid|CBF70688.1 anig|EHA22799.1 aory|EIT78149.1 asyd|OJJ58468.1 ater|EAU38180.1 atub|OJI86261.1 aver|OJJ01067.1 awen|OJJ32231.1 azon|OJJ46781.1

Group4106: abra|OJJ76409.1 acar|OOF99651.1 acla|EAW14015.1 afis|EAW18721.1 afla|EED50678.1 afum|EAL93445.1 agla|OJJ82334.1 akaw|GAA82328.1 aluc|OJZ91802.1 anid|CBF70681.1 anig|EHA22797.1 aory|EIT78078.1 asyd|OJJ58470.1 ater|EAU38183.1 atub|OJI86257.1 aver|OJJ01071.1 awen|OJJ32227.1 azon|OJJ46778.1

Group4107: abra|OJJ76411.1 acar|OOF99652.1 acla|EAW14014.1 afis|EAW18720.1 afla|EED50677.1 afum|EAL93444.1 agla|OJJ82335.1 akaw|GAA82329.1 aluc|OJZ91801.1 anid|CBF70679.1 anig|EHA22796.1 aory|EIT78134.1 asyd|OJJ58471.1 ater|EAU38184.1 atub|OJI86256.1 aver|OJJ01072.1 awen|OJJ32226.1 azon|OJJ46777.1

Group4108: abra|OJJ76417.1 acar|OOF99659.1 acla|EAW14009.1 afis|EAW18715.1 afla|EED50669.1 afum|EAL93440.1 agla|OJJ82344.1 akaw|GAA82332.1 aluc|OJZ91795.1 anid|CBF70670.1 anig|EHA22792.1 aory|EIT78070.1 asyd|OJJ58476.1 ater|EAU38192.1 atub|OJI86251.1 aver|OJJ01079.1 awen|OJJ32202.1 azon|OJJ46771.1

Group4109: abra|OJJ76418.1 acar|OOF99660.1 acla|EAW14008.1 afis|EAW18714.1 afla|EED50667.1 afum|EAL93439.1 agla|OJJ82345.1 akaw|GAA82333.1 aluc|OJZ91794.1 anid|CBF70668.1 anig|EHA22791.1 aory|EIT78037.1 asyd|OJJ58477.1 ater|EAU38193.1 atub|OJI86250.1 aver|OJJ01080.1 awen|OJJ32201.1 azon|OJJ46770.1

Group4110: abra|OJJ76419.1 acar|OOF99661.1 acla|EAW14007.1 afis|EAW18713.1 afla|EED50666.1 afum|EAL93438.1 agla|OJJ82346.1 akaw|GAA82334.1 aluc|OJZ91793.1 anid|CBF70666.1 anig|EHA22790.1 aory|EIT78088.1 asyd|OJJ58478.1 ater|EAU38194.1 atub|OJI86249.1 aver|OJJ01081.1 awen|OJJ32200.1 azon|OJJ46769.1

Group4111: abra|OJJ76420.1 acar|OOF99662.1 acla|EAW14006.1 afis|EAW18712.1 afla|EED50665.1 afum|EAL93437.1 agla|OJJ82347.1 akaw|GAA82335.1 aluc|OJZ91792.1 anid|CBF70665.1 anig|EHA22789.1

aory|EIT77998.1 asyd|OJJ58480.1 ater|EAU38195.1 atub|OJI86248.1 aver|OJJ01082.1 awen|OJJ32199.1 azon|OJJ46768.1

Group4112: abra|OJJ76422.1 acar|OOF99665.1 acla|EAW14003.1 afis|EAW18711.1 afla|EED50662.1 afum|EAL93434.1 agla|OJJ82348.1 akaw|GAA82337.1 aluc|OJZ91790.1 anid|CBF70661.1 anig|EHA22787.1 asyd|OJJ58481.1 ater|EAU38197.1 atub|OJI86246.1 aver|OJJ01083.1 awen|OJJ32197.1 azon|OJJ46766.1 aory|EIT77986.1

Group4113: abra|OJJ76423.1 acar|OOF99667.1 acla|EAW14002.1 afis|EAW18710.1 afla|EED50660.1 afum|EAL93433.1 agla|OJJ82349.1 akaw|GAA82338.1 aluc|OJZ91789.1 anid|CBF70657.1 anig|EHA22786.1 aory|EIT78084.1 asyd|OJJ58482.1 ater|EAU38198.1 atub|OJI86245.1 aver|OJJ01085.1 awen|OJJ32196.1 azon|OJJ46765.1

Group4114: abra|OJJ76429.1 acar|OOF99673.1 acla|EAW13996.1 afis|EAW18704.1 afla|EED50653.1 afum|EAL93427.2 agla|OJJ82354.1 akaw|GAA82344.1 aluc|OJZ91783.1 anid|CBF70646.1 anig|EHA22780.1 aory|EIT78100.1 asyd|OJJ58488.1 ater|EAU38202.1 atub|OJI86239.1 aver|OJJ01091.1 awen|OJJ32187.1 azon|OJJ46760.1

Group4115: abra|OJJ76430.1 acar|OOF99674.1 acla|EAW13995.1 afis|EAW18703.1 afla|EED50652.1 afum|EAL93426.1 agla|OJJ82355.1 akaw|GAA82345.1 aluc|OJZ91782.1 anid|CBF70644.1 anig|EHA22779.1 aory|EIT78020.1 asyd|OJJ58489.1 ater|EAU38203.1 atub|OJI86238.1 aver|OJJ01092.1 awen|OJJ32186.1 azon|OJJ46761.1

Group4116: abra|OJJ76431.1 acar|OOF99675.1 acla|EAW13994.1 afis|EAW18702.1 afla|EED50651.1 afum|EAL93425.1 agla|OJJ82359.1 akaw|GAA82346.1 aluc|OJZ91781.1 anid|CBF70642.1 anig|EHA22778.1 aory|EIT78030.1 asyd|OJJ58490.1 ater|EAU38204.1 atub|OJI86237.1 aver|OJJ01093.1 awen|OJJ32185.1 azon|OJJ46762.1

Group4117: abra|OJJ76433.1 acar|OOF99677.1 acla|EAW13991.1 afis|EAW18699.1 afla|EED50647.1 afum|EAL93421.1 agla|OJJ82362.1 akaw|GAA82348.1 aluc|OJZ91779.1 anid|CBF70635.1 anig|EHA22776.1 aory|EIT78093.1 asyd|OJJ58494.1 ater|EAU38207.1 atub|OJI86235.1 aver|OJJ01097.1 awen|OJJ32181.1 azon|OJJ46756.1

Group4118: abra|OJJ76434.1 acar|OOF99679.1 acla|EAW13989.1 afis|EAW18697.1 afla|EED50645.1 afum|EAL93419.1 agla|OJJ82371.1 akaw|GAA82349.1 aluc|OJZ91778.1 anid|CBF70631.1 anig|EHA22775.1 aory|EIT78142.1 asyd|OJJ58496.1 ater|EAU38209.1 atub|OJI86234.1 aver|OJJ01099.1 awen|OJJ32168.1 azon|OJJ46752.1

Group4119: abra|OJJ76440.1 acar|OOF99685.1 acla|EAW13984.1 afis|EAW18693.1 afla|EED50642.1 afum|EAL93414.1 agla|OJJ82368.1 akaw|GAA82354.1 aluc|OJZ91769.1 anid|CBF70618.1 anig|EHA22769.1 aory|EIT78025.1 asyd|OJJ58503.1 ater|EAU38215.1 atub|OJI86227.1 aver|OJJ01104.1 awen|OJJ32164.1 azon|OJJ46745.1

Group4120: abra|OJJ76441.1 acar|OOF99686.1 acla|EAW13983.1 afis|EAW18692.1 afla|EED50641.1 afum|EAL93413.2 agla|OJJ82366.1 akaw|GAA82355.1 aluc|OJZ91768.1 anid|CBF70620.1 anig|EHA22768.1 aory|EIT78095.1 asyd|OJJ58502.1 ater|EAU38216.1 atub|OJI86226.1 aver|OJJ01103.1 awen|OJJ32163.1 azon|OJJ46746.1

Group4121: abra|OJJ76446.1 acar|OOF99689.1 acla|EAW13982.1 afis|EAW18691.1 afla|EED50638.1 afum|EAL93412.1 agla|OJJ82381.1 akaw|GAA82358.1 aluc|OJZ91764.1 anid|CBF70624.1 anig|EHA22765.1 aory|EIT78085.1 asyd|OJJ58499.1 ater|EAU38217.1 atub|OJI86222.1 aver|OJJ01102.1 awen|OJJ32162.1 azon|OJJ46748.1

Group4122: abra|OJJ76448.1 acar|OOF99690.1 acla|EAW13980.1 afis|EAW18689.1 afla|EED50637.1 afum|EAL93410.1 agla|OJJ82383.1 akaw|GAA82360.1 aluc|OJZ91762.1 anid|CBF70616.1 anig|EHA22763.1 aory|EIT78124.1 asyd|OJJ58506.1 ater|EAU38219.1 atub|OJI86220.1 aver|OJJ01107.1 awen|OJJ32160.1 azon|OJJ46744.1

Group4123: abra|OJJ76449.1 acar|OOF99692.1 acla|EAW13979.1 afis|EAW18688.1 afla|EED50636.1 afum|EAL93409.1 agla|OJJ82384.1 akaw|GAA82361.1 aluc|OJZ91761.1 anid|CBF70612.1 anig|EHA22762.1 aory|EIT78146.1 asyd|OJJ58510.1 ater|EAU38220.1 atub|OJI86219.1 aver|OJJ01111.1 awen|OJJ32159.1 azon|OJJ46743.1

Group4124: abra|OJJ76455.1 acar|OOF99696.1 acla|EAW13975.1 afis|EAW18684.1 afla|EED50622.1 afum|EAL93405.1 agla|OJJ82387.1 akaw|GAA82367.1 aluc|OJZ91755.1 anid|CBF70607.1 anig|EHA22758.1 aory|EIT78141.1 asyd|OJJ58514.1 ater|EAU38226.1 atub|OJI86213.1 aver|OJJ01115.1 awen|OJJ32155.1 azon|OJJ46739.1

Group4125: abra|OJJ76456.1 acar|OOF99695.1 acla|EAW13974.1 afis|EAW18683.1 afla|EED50627.1 afum|EAL93404.1 agla|OJJ82392.1 akaw|GAA82368.1 aluc|OJZ91754.1 anid|CBF70605.1 anig|EHA22757.1 aory|EIT77985.1 asyd|OJJ58515.1 ater|EAU38227.1 atub|OJI86212.1 aver|OJJ01116.1 awen|OJJ32154.1 azon|OJJ46738.1

Group4126: abra|OJJ76460.1 acar|OOF99699.1 acla|EAW13970.1 afis|EAW18679.1 afla|EED50631.1 afum|EAL93399.1 agla|OJJ82396.1 akaw|GAA82373.1 aluc|OJZ91749.1 anid|CBF70596.1 anig|EHA22752.1 aory|EIT78016.1 asyd|OJJ58519.1 ater|EAU38232.1 atub|OJI86207.1 aver|OJJ01120.1 awen|OJJ32151.1 azon|OJJ46735.1

Group4127: abra|OJJ76464.1 acar|OOF99702.1 acla|EAW13967.1 afis|EAW18676.1 afla|EED50634.1 afum|EAL93396.1 agla|OJJ82400.1 akaw|GAA82376.1 aluc|OJZ91746.1 anid|CBF70590.1 anig|EHA22749.1 aory|EIT78033.1 asyd|OJJ58522.1 ater|EAU38234.1 atub|OJI86204.1 aver|OJJ01123.1 awen|OJJ32148.1 azon|OJJ46732.1

Group4128: abra|OJJ76467.1 acar|OOF99705.1 acla|EAW13964.1 afis|EAW18673.1 afla|EED50626.1 afum|EAL93393.2 agla|OJJ82402.1 akaw|GAA82378.1 aluc|OJZ91743.1 anid|CBF70584.1 anig|EHA22746.1 aory|EIT78060.1 asyd|OJJ58525.1 ater|EAU38237.1 atub|OJI86201.1 aver|OJJ01126.1 awen|OJJ32145.1 azon|OJJ46729.1

Group4129: abra|OJJ76468.1 acar|OOF99706.1 acla|EAW13963.1 afis|EAW18672.1 afla|EED50621.1 afum|EAL93391.1 agla|OJJ82403.1 akaw|GAA82379.1 aluc|OJZ91742.1 anid|CBF70583.1 anig|EHA22745.1 aory|EIT78045.1 asyd|OJJ58526.1 ater|EAU38238.1 atub|OJI86200.1 aver|OJJ01127.1 awen|OJJ32144.1 azon|OJJ46728.1

Group4130: abra|OJJ76472.1 acar|OOF99709.1 acla|EAW13961.1 afis|EAW18670.1 afla|EED52658.1 afum|EAL93388.1 agla|OJJ82405.1 akaw|GAA82384.1 aluc|OJZ91737.1 anid|CBF70579.1 anig|EHA22743.1 aory|EIT74274.1 asyd|OJJ58528.1 ater|EAU38241.1 atub|OJI86195.1 aver|OJJ01130.1 awen|OJJ32142.1 azon|OJJ46723.1

Group4131: abra|OJJ76479.1 acar|OOF90192.1 acla|EAW13956.1 afis|EAW18666.1 afla|EED52663.1 afum|EAL93384.1 agla|OJJ82411.1 akaw|GAA82392.1 aluc|OJZ91728.1 anid|CBF70569.1 anig|EHA22735.1 aory|EIT74303.1 asyd|OJJ58533.1 ater|EAU38246.1 atub|OJI86186.1 aver|OJJ01135.1 awen|OJJ32134.1 azon|OJJ46717.1

Group4132: acar|OOF90191.1 aluc|OJZ91727.1 anig|EHA22734.1 atub|OJI86185.1 abra|OJJ76481.1 acla|EAW13954.1 afis|EAW18664.1 afla|EED52665.1 afum|EAL93382.1 agla|OJJ82412.1 akaw|GAA82394.1 anid|CBF70567.1 aory|EIT74310.1 asyd|OJJ58535.1 ater|EAU38248.1 aver|OJJ01136.1 awen|OJJ32132.1 azon|OJJ46715.1

Group4133: abra|OJJ76482.1 acar|OOF90190.1 acla|EAW13953.1 afis|EAW18663.1 afla|EED52666.1 afum|EBA27491.1 agla|OJJ82413.1 akaw|GAA82395.1 aluc|OJZ91726.1 anid|CBF70565.1 anig|EHA22733.1 aory|EIT74219.1 asyd|OJJ58536.1 ater|EAU38249.1 atub|OJI86184.1 aver|OJJ01137.1 awen|OJJ32131.1 azon|OJJ46714.1

Group4134: abra|OJJ76483.1 acar|OOF99714.1 acla|EAW13951.1 afis|EAW18661.1 afla|EED52669.1 afum|EAL93380.1 agla|OJJ82416.1 akaw|GAA82397.1 aluc|OJZ91725.1 anid|CBF70557.1 anig|EHA22732.1 aory|EIT74184.1 asyd|OJJ58538.1 ater|EAU38251.1 atub|OJI86182.1 aver|OJJ01138.1 awen|OJJ32128.1 azon|OJJ46711.1

Group4135: abra|OJJ76484.1 acar|OOF99715.1 acla|EAW13950.1 afis|EAW18659.1 afla|EED52670.1 afum|EAL93377.1 agla|OJJ82417.1 akaw|GAA82398.1 aluc|OJZ91724.1 anid|CBF70556.1 anig|EHA22731.1 aory|EIT74290.1 asyd|OJJ58539.1 ater|EAU38252.1 atub|OJI86181.1 aver|OJJ01139.1 awen|OJJ32127.1 azon|OJJ46710.1

Group4136: abra|OJJ76485.1 acar|OOF99718.1 acla|EAW13948.1 afis|EAW18658.1 afla|EED52672.1 afum|EAL93375.1 agla|OJJ82418.1 akaw|GAA82401.1 aluc|OJZ91721.1 anid|CBF70552.1 anig|EHA22728.1 aory|EIT74194.1 asyd|OJJ58540.1 ater|EAU38254.1 atub|OJI86178.1 aver|OJJ01141.1 awen|OJJ32125.1 azon|OJJ46709.1

Group4137: abra|OJJ76487.1 acar|OOF99720.1 acla|EAW13946.1 afis|EAW18656.1 afla|EED52674.1 afum|EAL93372.1 agla|OJJ82419.1 akaw|GAA82403.1 aluc|OJZ91718.1 anid|CBF70546.1 anig|EHA22726.1 aory|EIT74203.1 asyd|OJJ58543.1 ater|EAU38256.1 atub|OJI86175.1 aver|OJJ01143.1 awen|OJJ32123.1 azon|OJJ46708.1

Group4138: abra|OJJ76501.1 acar|OOF99466.1 acla|EAW14195.1 afis|EAW18911.1 afla|EED50838.1 afum|EAL93642.1 agla|OJJ81515.1 akaw|GAA82417.1 aluc|OJZ91701.1 anid|CBF69699.1 anig|EHA22715.1 aory|EIT79018.1 asyd|OJJ59050.1 ater|EAU38078.1 atub|OJI86160.1 aver|OJJ01646.1 awen|OJJ31504.1 azon|OJJ47224.1

Group4139: abra|OJJ76513.1 acar|OOF99459.1 acla|EAW14187.1 afis|EAW18902.1 afla|EED50846.1 afum|EAL93634.1 agla|OJJ81507.1 akaw|GAA82426.1 aluc|OJZ91692.1 anid|CBF69923.1 anig|EHA22705.1 aory|EIT79043.1 asyd|OJJ59039.1 ater|EAU38072.1 atub|OJI86145.1 aver|OJJ01634.1 awen|OJJ32459.1 azon|OJJ47217.1

Group4140: abra|OJJ76516.1 acar|OOF99457.1 acla|EAW14186.1 afis|EAW18901.1 afla|EED50847.1 afum|EAL93633.1 agla|OJJ81506.1 akaw|GAA82428.1 aluc|OJZ91690.1 anid|CBF69921.1 anig|EHA22703.1

aory|EIT79032.1 asyd|OJJ58987.1 ater|EAU38069.1 atub|OJI86143.1 aver|OJJ01629.1 awen|OJJ32458.1 azon|OJJ47216.1

Group4141: abra|OJJ76520.1 acar|OOF99453.1 acla|EAW14181.1 afis|EAW18896.1 afla|EED50867.1 afum|EAL93628.1 agla|OJJ81502.1 akaw|GAA82433.1 aluc|OJZ91688.1 anid|CBF69913.1 anig|EHA22700.1 aory|EIT79004.1 asyd|OJJ58992.1 ater|EAU38066.1 atub|OJI86139.1 aver|OJJ01623.1 awen|OJJ32454.1 azon|OJJ47211.1

Group4142: abra|OJJ76524.1 acar|OOF99445.1 acla|EAW14173.1 afis|EAW18887.1 afla|EED50878.1 afum|EAL93620.1 agla|OJJ81493.1 akaw|GAA82437.1 aluc|OJZ91684.1 anid|CBF69898.1 anig|EHA22696.1 aory|EIT79044.1 asyd|OJJ59000.1 ater|EAU38059.1 atub|OJI86136.1 aver|OJJ01615.1 awen|OJJ32445.1 azon|OJJ47203.1

Group4143: abra|OJJ76528.1 acar|OOF99447.1 acla|EAW14175.1 afis|EAW18889.1 afla|EED50874.1 afum|EAL93622.1 agla|OJJ81495.1 akaw|GAA82439.1 aluc|OJZ91677.1 anid|CBF69902.1 anig|EHA22694.1 aory|EIT78988.1 asyd|OJJ58998.1 ater|EAU38061.1 atub|OJI86132.1 aver|OJJ01617.1 awen|OJJ32448.1 azon|OJJ47206.1

Group4144: abra|OJJ76530.1 acar|OOF99443.1 acla|EAW14171.1 afis|EAW18885.1 afla|EED50880.1 afum|EAL93618.1 agla|OJJ80620.1 akaw|GAA82442.1 aluc|OJZ91674.1 anid|CBF69894.1 anig|EHA22692.1 aory|EIT78957.1 asyd|OJJ59002.1 ater|EAU38057.1 atub|OJI86129.1 aver|OJJ01613.1 awen|OJJ32443.1 azon|OJJ47201.1

Group4145: abra|OJJ76534.1 acar|OOF99440.1 acla|EAW14169.1 afis|EAW18883.1 afla|EED50882.1 afum|EAL93616.1 agla|OJJ80618.1 akaw|GAA82445.1 aluc|OJZ91671.1 anid|CBF69892.1 anig|EHA22689.1 aory|EIT79022.1 asyd|OJJ59005.1 ater|EAU38055.1 atub|OJI86126.1 aver|OJJ01611.1 awen|OJJ32440.1 azon|OJJ47199.1

Group4146: abra|OJJ76535.1 acar|OOF99439.1 acla|EAW14168.1 afis|EAW18882.1 afla|EED50883.1 afum|EAL93614.1 agla|OJJ80617.1 akaw|GAA82446.1 aluc|OJZ91670.1 anid|CBF70466.1 anig|EHA22688.1 aory|EIT78979.1 asyd|OJJ58598.1 ater|EAU38054.1 atub|OJI86125.1 aver|OJJ01200.1 awen|OJJ32441.1 azon|OJJ47198.1

Group4147: abra|OJJ76536.1 acar|OOF99432.1 acla|EAW14162.1 afis|EAW18875.1 afla|EED50892.1 afum|EAL93607.1 agla|OJJ80612.1 akaw|GAA82447.1 aluc|OJZ91669.1 anid|CBF69869.1 anig|EHA22687.1 aory|EIT79035.1 asyd|OJJ59019.1 ater|EAU38040.1 atub|OJI86124.1 aver|OJJ01597.1 awen|OJJ32434.1 azon|OJJ47192.1

Group4148: abra|OJJ76537.1 acar|OOF99433.1 acla|EAW14163.1 afis|EAW18876.1 afla|EED50891.1 afum|EAL93608.1 agla|OJJ80613.1 akaw|GAA82448.1 aluc|OJZ91667.1 anid|CBF69873.1 anig|EHA22686.1 aory|EIT79049.1 asyd|OJJ59018.1 ater|EAU38041.1 atub|OJI86122.1 aver|OJJ01598.1 awen|OJJ32435.1 azon|OJJ47191.1

Group4149: abra|OJJ76540.1 acar|OOF99435.1 acla|EAW14165.1 afis|EAW18878.1 afla|EED50888.1 afum|EAL93610.1 agla|OJJ80615.1 akaw|GAA82450.1 aluc|OJZ91663.1 anid|CBF69877.1 anig|EHA22683.1 aory|EIT79008.1 asyd|OJJ59015.1 ater|EAU38043.1 atub|OJI86120.1 aver|OJJ01600.1 awen|OJJ32437.1 azon|OJJ47196.1

Group4150: abra|OJJ76542.1 acar|OOF99437.1 acla|EAW12334.1 afis|EAW16888.1 afla|EED49676.1 afum|EAL86426.1 agla|OJJ79936.1 akaw|GAA82451.1 aluc|OJZ91661.1 anid|CBF84569.1 anig|EHA22681.1 aory|EIT80968.1 asyd|OJJ53441.1 ater|EAU39143.1 atub|OJI86117.1 aver|OJJ08310.1 awen|OJJ29756.1 azon|OJJ43187.1

Group4151: abra|OJJ76543.1 acar|OOF99431.1 acla|EAW14161.1 afis|EAW18874.1 afla|EED50893.1 afum|EAL93606.1 agla|OJJ80611.1 akaw|GAA82452.1 aluc|OJZ91660.1 anid|CBF69867.1 anig|EHA22680.1 aory|EIT79007.1 asyd|OJJ59020.1 ater|EAU38039.1 atub|OJI86116.1 aver|OJJ01596.1 awen|OJJ32433.1 azon|OJJ47193.1

Group4152: abra|OJJ76545.1 acar|OOF99429.1 acla|EAW14159.1 afis|EAW18872.1 afla|EED50895.1 afum|EAL93604.1 agla|OJJ80609.1 akaw|GAA82454.1 aluc|OJZ91658.1 anid|CBF69863.1 anig|EHA22678.1 aory|EIT79057.1 asyd|OJJ59023.1 ater|EAU38036.1 atub|OJI86114.1 aver|OJJ01594.1 awen|OJJ32431.1 azon|OJJ47188.1

Group4153: abra|OJJ76546.1 acar|OOF99428.1 acla|EAW14158.1 afis|EAW18871.1 afla|EED50831.1 afum|EAL93603.1 agla|OJJ80595.1 akaw|GAA82455.1 aluc|OJZ91657.1 anid|CBF69852.1 anig|EHA22677.1 aory|EIT79042.1 asyd|OJJ59028.1 ater|EAU38035.1 atub|OJI86112.1 aver|OJJ01588.1 awen|OJJ32430.1 azon|OJJ47185.1

Group4154: abra|OJJ76547.1 acar|OOF99427.1 acla|EAW14157.1 afis|EAW18870.1 afla|EED50832.1 afum|EAL93602.1 agla|OJJ80596.1 akaw|GAA82456.1 aluc|OJZ91656.1 anid|CBF69854.1 anig|EHA22676.1 aory|EIT79054.1 asyd|OJJ59027.1 ater|EAU38034.1 atub|OJI86111.1 aver|OJJ01589.1 awen|OJJ32429.1 azon|OJJ47184.1

Group4155: abra|OJJ76549.1 acar|OOF99426.1 acla|EAW14156.1 afis|EAW18869.1 afla|EED50833.1 afum|EAL93601.1 agla|OJJ80600.1 akaw|GAA82457.1 aluc|OJZ91654.1 anid|CBF69856.1 anig|EHA22675.1 aory|EIT78961.1 asyd|OJJ59026.1 ater|EAU38033.1 atub|OJI86108.1 aver|OJJ01591.1 awen|OJJ32428.1 azon|OJJ47183.1

Group4156: abra|OJJ76550.1 acar|OOF89946.1 acla|EAW14155.1 afis|EAW18868.1 afla|EED50834.1 afum|EAL93600.1 agla|OJJ80601.1 akaw|GAA82458.1 aluc|OJZ91653.1 anid|CBF69857.1 anig|EHA22674.1 aory|EIT78982.1 asyd|OJJ59025.1 ater|EAU38032.1 atub|OJI86107.1 aver|OJJ01592.1 awen|OJJ32427.1 azon|OJJ47182.1

Group4157: abra|OJJ76551.1 acar|OOF89945.1 acla|EAW14153.1 afis|EAW18866.1 afla|EED50897.1 afum|EAL93598.1 agla|OJJ80603.1 akaw|GAA82459.1 aluc|OJZ91652.1 anid|CBF69861.1 anig|EHA22673.1 aory|EIT79016.1 asyd|OJJ59024.1 ater|EAU38030.1 atub|OJI86106.1 aver|OJJ01593.1 awen|OJJ32425.1 azon|OJJ47180.1

Group4158: abra|OJJ76554.1 acar|OOF99424.1 acla|EAW14150.1 afis|EAW18863.1 afla|EED50830.1 afum|EAL93595.1 agla|OJJ80586.1 akaw|GAA82461.1 aluc|OJZ91650.1 anid|CBF69848.1 anig|EHA22671.1 aory|EIT79053.1 asyd|OJJ59030.1 ater|EAU38025.1 atub|OJI86104.1 aver|OJJ01586.1 awen|OJJ32419.1 azon|OJJ47176.1

Group4159: abra|OJJ76558.1 acar|OOF99421.1 acla|EAW14147.1 afis|EAW18860.1 afla|EED50827.1 afum|EAL93592.1 agla|OJJ80583.1 akaw|GAA82464.1 aluc|OJZ91647.1 anid|CBF69842.1 anig|EHA22668.1 aory|EIT78975.1 asyd|OJJ59033.1 ater|EAU38023.1 atub|OJI86101.1 aver|OJJ01583.1 awen|OJJ32416.1 azon|OJJ47173.1

Group4160: abra|OJJ76563.1 acar|OOF99419.1 acla|EAW14144.1 afis|EAW18857.1 afla|EED50824.1 afum|EAL93589.2 agla|OJJ80579.1 akaw|GAA82467.1 aluc|OJZ91643.1 anid|CBF69834.1 anig|EHA22665.1 aory|EIT78963.1 asyd|OJJ58985.1 ater|EAU38020.1 atub|OJI86097.1 aver|OJJ01580.1 awen|OJJ32413.1 azon|OJJ47170.1

Group4161: abra|OJJ76565.1 acar|OOF99418.1 acla|EAW14143.1 afis|EAW18856.1 afla|EED50823.1 afum|EAL93587.1 akaw|GAA82469.1 aluc|OJZ91641.1 anid|CBF69831.1 anig|EHA22664.1 aory|EIT79024.1 asyd|OJJ58983.1 ater|EAU38019.1 atub|OJI86095.1 aver|OJJ01578.1 awen|OJJ32411.1 azon|OJJ47168.1 agla|OJJ80577.1

Group4162: abra|OJJ76566.1 acar|OOF99355.1 acla|EAW14142.1 afis|EAW18855.1 afla|EED50822.1 afum|EAL93586.1 agla|OJJ80574.1 akaw|GAA82470.1 aluc|OJZ91640.1 anid|CBF69829.1 anig|EHA22663.1 aory|EIT78993.1 asyd|OJJ58981.1 ater|EAU38013.1 atub|OJI86094.1 aver|OJJ01576.1 awen|OJJ32410.1 azon|OJJ47167.1

Group4163: abra|OJJ76567.1 acar|OOF99417.1 acla|EAW14141.1 afis|EAW18854.1 afla|EED50821.1 afum|EAL93585.1 agla|OJJ80573.1 akaw|GAA82471.1 aluc|OJZ91639.1 anid|CBF69827.1 anig|EHA22662.1 aory|EIT78968.1 asyd|OJJ58980.1 ater|EAU38012.1 atub|OJI86093.1 aver|OJJ01575.1 awen|OJJ32409.1 azon|OJJ47166.1

Group4164: abra|OJJ76569.1 acar|OOF99416.1 acla|EAW14139.1 afis|EAW18852.1 afla|EED50819.1 afum|EAL93583.1 agla|OJJ80571.1 akaw|GAA82473.1 aluc|OJZ91636.1 anid|CBF89917.1 anig|EHA22660.1 aory|EIT79009.1 asyd|OJJ62126.1 ater|EAU38005.1 atub|OJI86089.1 aver|OJI96234.1 awen|OJJ32406.1 azon|OJJ47164.1

Group4165: abra|OJJ76570.1 acar|OOF99415.1 acla|EAW14138.1 afis|EAW18851.1 afla|EED50818.1 afum|EAL93582.1 agla|OJJ80570.1 akaw|GAA82474.1 aluc|OJZ91635.1 anid|CBF69823.1 anig|EHA22659.1 aory|EIT78969.1 asyd|OJJ58978.1 ater|EAU38004.1 atub|OJI86088.1 aver|OJJ01573.1 awen|OJJ32405.1 azon|OJJ47163.1

Group4166: abra|OJJ76571.1 acar|OOF99414.1 acla|EAW14137.1 afis|EAW18850.1 afla|EED50817.1 afum|EAL93581.1 agla|OJJ80569.1 akaw|GAA82475.1 aluc|OJZ91634.1 anid|CBF69821.1 anig|EHA22658.1 aory|EIT79031.1 asyd|OJJ58977.1 ater|EAU38003.1 atub|OJI86087.1 aver|OJJ01572.1 awen|OJJ32404.1 azon|OJJ47162.1

Group4167: abra|OJJ76572.1 acar|OOF99413.1 acla|EAW14136.1 afis|EAW18849.1 afla|EED50816.1 afum|EAL93580.1 agla|OJJ80568.1 akaw|GAA82476.1 aluc|OJZ91633.1 anid|CBF69819.1 anig|EHA22657.1 aory|EIT78989.1 asyd|OJJ58976.1 ater|EAU38002.1 atub|OJI86086.1 aver|OJJ01571.1 awen|OJJ32403.1 azon|OJJ47161.1

Group4168: abra|OJJ76573.1 acar|OOF99412.1 acla|EAW14134.1 afis|EAW18847.1 afla|EED50814.1 afum|EAL93579.1 agla|OJJ80567.1 akaw|GAA82478.1 aluc|OJZ91632.1 anid|CBF69817.1 anig|EHA22656.1 aory|EIT79029.1 asyd|OJJ58975.1 ater|EAU38001.1 atub|OJI86085.1 aver|OJJ01570.1 awen|OJJ32402.1 azon|OJJ47160.1

Group4169: abra|OJJ76576.1 acar|OOF99408.1 acla|EAW14117.1 afis|EAW18819.1 afla|EED50808.1 afum|EAL93555.1 agla|OJJ80554.1 akaw|GAA82481.1 aluc|OJZ91629.1 anid|CBF69809.1 anig|EHA22653.1

aory|EIT78964.1 asyd|OJJ58971.1 ater|EAU37998.1 atub|OJI86082.1 aver|OJJ01566.1 awen|OJJ32395.1 azon|OJJ47157.1

Group4170: abra|OJJ76577.1 acar|OOF99407.1 acla|EAW14116.1 afis|EAW18818.1 afla|EED50809.1 afum|EAL93554.1 agla|OJJ80552.1 akaw|GAA82482.1 aluc|OJZ91628.1 anid|CBF69811.1 anig|EHA22652.1 aory|EIT78995.1 asyd|OJJ58972.1 ater|EAU37997.1 atub|OJI86081.1 aver|OJJ01567.1 awen|OJJ32394.1 azon|OJJ47156.1

Group4171: abra|OJJ76579.1 acar|OOF99405.1 acla|EAW14114.1 afis|EAW18816.1 afla|EED50804.1 afum|EAL93552.1 agla|OJJ80549.1 akaw|GAA82484.1 aluc|OJZ91626.1 anid|CBF69815.1 anig|EHA22650.1 aory|EIT78104.1 asyd|OJJ58974.1 ater|EAU37991.1 atub|OJI86079.1 aver|OJJ01569.1 awen|OJJ32392.1 azon|OJJ47153.1

Group4172: abra|OJJ76581.1 acar|OOF99404.1 acla|EAW14113.1 afis|EAW18815.1 afla|EED50802.1 afum|EAL93551.1 agla|OJJ80548.1 akaw|GAA82485.1 aluc|OJZ91624.1 anid|CBF69803.1 anig|EHA22649.1 aory|EIT77988.1 asyd|OJJ58968.1 ater|EAU37982.1 atub|OJI86077.1 aver|OJJ01563.1 awen|OJJ32391.1 azon|OJJ47152.1

Group4173: abra|OJJ76582.1 acar|OOF99403.1 acla|EAW14112.1 afis|EAW18814.1 afla|EED50800.1 afum|EAL93550.2 agla|OJJ80547.1 akaw|GAA82486.1 aluc|OJZ91623.1 anid|CBF69801.1 anig|EHA22648.1 aory|EIT78081.1 asyd|OJJ58967.1 ater|EAU37981.1 atub|OJI86076.1 aver|OJJ01562.1 awen|OJJ32390.1 azon|OJJ47151.1

Group4174: abra|OJJ76588.1 acar|OOF99398.1 acla|EAW14313.1 afis|EAW18974.1 afla|EED53230.1 afum|EAL93708.1 agla|OJJ82930.1 akaw|GAA82492.1 aluc|OJZ91618.1 anid|CBF69639.1 anig|EHA22645.1 aory|EIT74874.1 asyd|OJJ59084.1 ater|EAU38166.1 atub|OJI86071.1 aver|OJJ01677.1 awen|OJJ31576.1 azon|OJJ47293.1

Group4175: abra|OJJ76590.1 acar|OOF99397.1 acla|EAW14311.1 afis|EAW18975.1 afla|EED53222.1 afum|EAL93710.1 agla|OJJ82932.1 akaw|GAA82494.1 aluc|OJZ91616.1 anid|CBF69642.1 anig|EHA22644.1 aory|EIT74886.1 asyd|OJJ59081.1 ater|EAU38164.1 atub|OJI86069.1 aver|OJJ01674.1 awen|OJJ31580.1 azon|OJJ47295.1

Group4176: abra|OJJ76591.1 acar|OOF99396.1 acla|EAW14310.1 afis|EAW18976.1 afla|EED53221.1 afum|EAL93711.2 agla|OJJ82933.1 akaw|GAA82495.1 aluc|OJZ91615.1 anid|CBF69644.1 anig|EHA22643.1 aory|EIT74842.1 asyd|OJJ59080.1 ater|EAU38163.1 atub|OJI86068.1 aver|OJJ01673.1 awen|OJJ31581.1 azon|OJJ47296.1

Group4177: abra|OJJ76594.1 acar|OOF99393.1 acla|EAW14307.1 afis|EAW18978.1 afla|EED53192.1 afum|EAL93714.1 agla|OJJ82936.1 akaw|GAA82498.1 aluc|OJZ91612.1 anid|CBF69649.1 anig|EHA22641.1 aory|EIT75152.1 asyd|OJJ59077.1 ater|EAU38169.1 atub|OJI86065.1 aver|OJJ01670.1 awen|OJJ31584.1 azon|OJJ47299.1

Group4178: abra|OJJ76601.1 acar|OOF99390.1 acla|EAW14305.1 afis|EAW18979.1 afla|EED53188.1 afum|EAL93715.1 agla|OJJ82941.1 akaw|GAA82504.1 aluc|OJZ91607.1 anid|CBF69652.1 anig|EHA22636.1 aory|EIT74907.1 asyd|OJJ59076.1 ater|EAU37838.1 atub|OJI86059.1 aver|OJJ01669.1 awen|OJJ31587.1 azon|OJJ47303.1

Group4179: abra|OJJ76605.1 acar|OOF99386.1 acla|EAW14301.1 afis|EAW18983.1 afla|EED53161.1 afum|EAL93719.1 agla|OJJ82958.1 akaw|GAA82509.1 aluc|OJZ91603.1 anid|CBF69659.1 anig|EHA22632.1 aory|EIT74944.1 asyd|OJJ59072.1 ater|EAU37822.1 atub|OJI86055.1 aver|OJJ01665.1 awen|OJJ31600.1 azon|OJJ47307.1

Group4180: abra|OJJ76606.1 acar|OOF99384.1 acla|EAW14300.1 afis|EAW18984.1 afla|EED53162.1 afum|EAL93720.1 agla|OJJ82959.1 akaw|GAA82510.1 aluc|OJZ91602.1 anid|CBF69661.1 anig|EHA22631.1 aory|EIT74985.1 asyd|OJJ59071.1 ater|EAU37821.1 atub|OJI86054.1 aver|OJJ01664.1 awen|OJJ31604.1 azon|OJJ47308.1

Group4181: abra|OJJ76609.1 acar|OOF99383.1 acla|EAW14297.1 afis|EAW18986.1 afla|EED53164.1 afum|EAL93722.1 agla|OJJ82964.1 akaw|GAA82512.1 aluc|OJZ91599.1 anid|CBF69665.1 anig|EHA22629.1 aory|EIT75120.1 asyd|OJJ59069.1 ater|EAU37819.1 atub|OJI86050.1 aver|OJJ01662.1 awen|OJJ31606.1 azon|OJJ47310.1

Group4182: abra|OJJ76624.1 acar|OOF99374.1 acla|EAW14295.1 afis|EAW18988.1 afla|EED53173.1 afum|EAL93724.1 agla|OJJ82975.1 akaw|GAA92833.1 aluc|OJZ91588.1 anid|CBF86791.1 anig|EHA22612.1 aory|EIT75118.1 asyd|OJJ64674.1 ater|EAU37816.1 atub|OJI86037.1 aver|OJI98775.1 awen|OJJ31609.1 azon|OJJ47312.1

Group4183: abra|OJJ76631.1 acar|OOF99366.1 acla|EAW08624.1 afis|EAW15808.1 afla|EED53158.1 afum|EAL85543.1 agla|OJJ81974.1 akaw|GAA92825.1 aluc|OJZ91580.1 anid|CBF86787.1 anig|EHA22607.1 aory|EIT75169.1 asyd|OJJ64676.1 ater|EAU37812.1 atub|OJI86029.1 aver|OJI98777.1 awen|OJJ38519.1 azon|OJJ50132.1

Group4187: abra|OJJ76753.1 acar|OOF91249.1 acla|EAW13307.1 afis|EAW20421.1 afla|EED44850.1 afum|EAL84629.1 agla|OJJ84904.1 akaw|GAA87271.1 aluc|OJZ84156.1 anid|CBF79342.1 anig|EHA27518.1 aory|EIT77742.1 asyd|OJJ55796.1 ater|EAU37735.1 atub|OJI89928.1 aver|OJJ06885.1 awen|OJJ33114.1 azon|OJJ46330.1

Group4188: abra|OJJ76756.1 acar|OOF91251.1 acla|EAW13310.1 afis|EAW20424.1 afla|EED44846.1 afum|EAL84624.2 agla|OJJ84901.1 akaw|GAA87269.1 aluc|OJZ84159.1 anid|CBF79340.1 anig|EHA27521.1 aory|EIT77736.1 asyd|OJJ55792.1 ater|EAU37733.1 atub|OJI89925.1 aver|OJJ06882.1 awen|OJJ33117.1 azon|OJJ46333.1

Group4189: abra|OJJ76758.1 acar|OOF91253.1 acla|EAW13312.1 afis|EAW20426.1 afla|EED44842.1 afum|EAL84623.1 agla|OJJ84899.1 akaw|GAA87268.1 aluc|OJZ84161.1 anid|CBF79338.1 anig|EHA27522.1 aory|EIT77746.1 asyd|OJJ55790.1 ater|EAU37731.1 atub|OJI89923.1 aver|OJJ06880.1 awen|OJJ33119.1 azon|OJJ46335.1

Group4190: abra|OJJ76760.1 acar|OOF91255.1 acla|EAW13314.1 afis|EAW20428.1 afla|EED44840.1 afum|EAL84622.1 agla|OJJ84897.1 akaw|GAA87266.1 aluc|OJZ84163.1 anid|CBF79334.1 anig|EHA27524.1 aory|EIT77728.1 asyd|OJJ55788.1 ater|EAU37729.1 atub|OJI89921.1 aver|OJJ06878.1 awen|OJJ33121.1 azon|OJJ46337.1

Group4191: abra|OJJ76761.1 acar|OOF91256.1 acla|EAW13315.1 afis|EAW20429.1 afla|EED44839.1 afum|EAL84621.1 agla|OJJ84896.1 akaw|GAA87265.1 aluc|OJZ84164.1 anid|CBF79332.1 anig|EHA27525.1 aory|EIT77744.1 asyd|OJJ55787.1 ater|EAU37728.1 atub|OJI89920.1 aver|OJJ06877.1 awen|OJJ33122.1 azon|OJJ46338.1

Group4192: abra|OJJ76762.1 acar|OOF91257.1 acla|EAW13316.1 afis|EAW20430.1 afla|EED44838.1 afum|EAL84620.1 agla|OJJ84895.1 akaw|GAA87264.1 aluc|OJZ84165.1 anid|CBF79330.1 anig|EHA27526.1 aory|EIT77755.1 asyd|OJJ55786.1 ater|EAU37727.1 atub|OJI89919.1 aver|OJJ06876.1 awen|OJJ33123.1 azon|OJJ46339.1

Group4193: abra|OJJ76764.1 acar|OOF91259.1 acla|EAW13318.1 afis|EAW20432.1 afla|EED44836.1 afum|EAL84619.1 agla|OJJ84893.1 akaw|GAA87262.1 aluc|OJZ84168.1 anid|CBF79325.1 anig|EHA27528.1 aory|EIT77720.1 asyd|OJJ55784.1 ater|EAU37725.1 atub|OJI89916.1 aver|OJJ06874.1 awen|OJJ33125.1 azon|OJJ46341.1

Group4194: abra|OJJ76766.1 acar|OOF91260.1 acla|EAW13320.1 afis|EAW20435.1 afla|EED44835.1 afum|EAL84617.1 agla|OJJ84891.1 akaw|GAA87260.1 aluc|OJZ84169.1 anid|CBF79321.1 anig|EHA27530.1 aory|EIT77727.1 asyd|OJJ55783.1 ater|EAU37723.1 atub|OJI89915.1 aver|OJJ06872.1 awen|OJJ33126.1 azon|OJJ46342.1

Group4195: abra|OJJ76768.1 acar|OOF91262.1 acla|EAW13322.1 afis|EAW20437.1 afla|EED44833.1 afum|EAL84615.1 agla|OJJ84886.1 akaw|GAA87258.1 aluc|OJZ84171.1 anid|CBF79317.1 anig|EHA27532.1 aory|EIT77715.1 asyd|OJJ55781.1 ater|EAU37721.1 atub|OJI89913.1 aver|OJJ06870.1 awen|OJJ33128.1 azon|OJJ46344.1

Group4196: abra|OJJ76769.1 acar|OOF91263.1 acla|EAW13323.1 afis|EAW20438.1 afla|EED44832.1 afum|EAL84614.1 agla|OJJ84885.1 akaw|GAA87257.1 aluc|OJZ84172.1 anid|CBF79312.1 anig|EHA27533.1 aory|EIT77757.1 asyd|OJJ55780.1 ater|EAU37720.1 atub|OJI89912.1 aver|OJJ06869.1 awen|OJJ33129.1 azon|OJJ46345.1

Group4197: abra|OJJ76774.1 acar|OOF91269.1 acla|EAW13331.1 afis|EAW20446.1 afla|EED44824.1 afum|EAL84607.1 agla|OJJ84875.1 akaw|GAA87250.1 aluc|OJZ84178.1 anid|CBF79292.1 anig|EHA27537.1 aory|EIT77756.1 asyd|OJJ55775.1 ater|EAU37713.1 atub|OJI89905.1 aver|OJJ06864.1 awen|OJJ33136.1 azon|OJJ46349.1

Group4198: abra|OJJ76776.1 acar|OOF91271.1 acla|EAW13335.1 afis|EAW20450.1 afla|EED44818.1 afum|EAL84603.1 agla|OJJ84872.1 akaw|GAA87248.1 aluc|OJZ84180.1 anid|CBF79297.1 anig|EHA27540.1 aory|EIT77719.1 asyd|OJJ55773.1 ater|EAU37709.1 atub|OJI89903.1 aver|OJJ06860.1 awen|OJJ33140.1 azon|OJJ46351.1

Group4199: abra|OJJ76779.1 acar|OOF91275.1 acla|EAW13338.1 afis|EAW20453.1 afla|EED44815.1 afum|EAL84600.1 agla|OJJ84869.1 akaw|GAA87245.1 aluc|OJZ84183.1 anid|CBF79284.1 anig|EHA27543.1 aory|EIT77717.1 asyd|OJJ55770.1 ater|EAU37706.1 atub|OJI89900.1 aver|OJJ06856.1 awen|OJJ33143.1 azon|OJJ46354.1

Group4200: abra|OJJ76780.1 acar|OOF91276.1 acla|EAW13339.1 afis|EAW20454.1 afla|EED44814.1 afum|EAL84599.1 agla|OJJ84868.1 akaw|GAA87244.1 aluc|OJZ84184.1 anid|CBF79286.1 anig|EHA27544.1 aory|EIT77725.1 asyd|OJJ55769.1 ater|EAU37705.1 atub|OJI89899.1 aver|OJJ06855.1 awen|OJJ33144.1 azon|OJJ46355.1

Group4201: abra|OJJ76784.1 acar|OOF91279.1 acla|EAW13342.1 afis|EAW20457.1 afla|EED44810.1 afum|EAL84595.1 agla|OJJ84862.1 akaw|GAA87241.1 aluc|OJZ84187.1 anid|CBF79283.1 anig|EHA27547.1

aory|EIT77730.1 asyd|OJJ55766.1 ater|EAU37703.1 atub|OJI89896.1 aver|OJJ06852.1 awen|OJJ33149.1 azon|OJJ46359.1

Group4202: abra|OJJ76787.1 acar|OOF91281.1 acla|EAW13345.1 afis|EAW20460.1 afla|EED44808.1 afum|EAL84593.1 agla|OJJ84858.1 akaw|GAA87238.1 aluc|OJZ84189.1 anid|CBF79277.1 anig|EHA27549.1 aory|EIT77754.1 asyd|OJJ55763.1 ater|EAU37700.1 atub|OJI89893.1 aver|OJJ06849.1 awen|OJJ33153.1 azon|OJJ46362.1

Group4203: abra|OJJ76790.1 acar|OOF91284.1 acla|EAW13348.1 afis|EAW20463.1 afla|EED44805.1 afum|EAL84590.1 agla|OJJ84854.1 akaw|GAA87235.1 aluc|OJZ84193.1 anid|CBF79267.1 anig|EHA27552.1 aory|EIT77734.1 asyd|OJJ55760.1 ater|EAU37738.1 atub|OJI89890.1 aver|OJJ06845.1 awen|OJJ33157.1 azon|OJJ46366.1

Group4204: abra|OJJ76792.1 acar|OOF91283.1 acla|EAW13351.1 afis|EAW20465.1 afla|EED44802.1 afum|EAL84588.1 agla|OJJ84852.1 akaw|GAA87233.1 aluc|OJZ84196.1 anid|CBF79269.1 anig|EHA27553.1 aory|EIT77749.1 asyd|OJJ55759.1 ater|EAU37698.1 atub|OJI89888.1 aver|OJJ06844.1 awen|OJJ33159.1 azon|OJJ46367.1

Group4205: abra|OJJ76793.1 acar|OOF91286.1 acla|EAW13352.1 afis|EAW20466.1 afla|EED44801.1 afum|EAL84586.2 agla|OJJ84851.1 akaw|GAA87232.1 aluc|OJZ84197.1 anid|CBF79271.1 anig|EHA27554.1 aory|EIT77739.1 asyd|OJJ55758.1 ater|EAU37697.1 atub|OJI89887.1 aver|OJJ06843.1 awen|OJJ33160.1 azon|OJJ46368.1

Group4206: abra|OJJ76796.1 acar|OOF91289.1 acla|EAW13355.1 afis|EAW20469.1 afla|EED44799.1 afum|EAL84583.1 agla|OJJ84849.1 akaw|GAA87227.1 aluc|OJZ84203.1 anid|CBF79257.1 anig|EHA27557.1 aory|EIT77745.1 asyd|OJJ55748.1 ater|EAU37694.1 atub|OJI89881.1 aver|OJJ06835.1 awen|OJJ33163.1 azon|OJJ46370.1

Group4207: abra|OJJ76798.1 acar|OOF91291.1 acla|EAW13357.1 afis|EAW20471.1 afla|EED44797.1 afum|EAL84582.1 agla|OJJ84847.1 akaw|GAA87225.1 aluc|OJZ84205.1 anid|CBF79258.1 anig|EHA27558.1 aory|EIT77743.1 asyd|OJJ55750.1 ater|EAU37691.1 atub|OJI89879.1 aver|OJJ06837.1 awen|OJJ33165.1 azon|OJJ46372.1

Group4208: abra|OJJ76799.1 acar|OOF91292.1 acla|EAW13358.1 afis|EAW20472.1 afla|EED44796.1 afum|EAL84580.2 agla|OJJ84846.1 akaw|GAA87224.1 aluc|OJZ84206.1 anid|CBF79260.1 anig|EHA27559.1 aory|EIT77751.1 asyd|OJJ55751.1 ater|EAU37690.1 atub|OJI89878.1 aver|OJJ06838.1 awen|OJJ33166.1 azon|OJJ46373.1

Group4209: abra|OJJ76800.1 acar|OOF91294.1 acla|EAW13359.1 afis|EAW20473.1 afla|EED44794.1 afum|EAL84579.1 agla|OJJ84845.1 akaw|GAA87223.1 aluc|OJZ84207.1 anid|CBF79262.1 anig|EHA27561.1 aory|EIT77790.1 asyd|OJJ55754.1 ater|EAU37689.1 atub|OJI89877.1 aver|OJJ06839.1 awen|OJJ33167.1 azon|OJJ46375.1

Group4210: abra|OJJ76802.1 acar|OOF91296.1 acla|EAW13361.1 afis|EAW20475.1 afla|EED44793.1 afum|EAL84578.1 agla|OJJ84842.1 akaw|GAA87221.1 aluc|OJZ84209.1 anid|CBF79255.1 anig|EHA27563.1 aory|EIT77775.1 asyd|OJJ55747.1 ater|EAU37687.1 atub|OJI89874.1 aver|OJJ06834.1 awen|OJJ33170.1 azon|OJJ46377.1

Group4211: abra|OJJ76806.1 acar|OOF91300.1 acla|EAW13365.1 afis|EAW20479.1 afla|EED44789.1 afum|EAL84575.1 agla|OJJ84831.1 akaw|GAA87217.1 aluc|OJZ84213.1 anid|CBF79213.1 anig|EHA27566.1 aory|EIT77792.1 asyd|OJJ55738.1 ater|EAU29558.1 atub|OJI89870.1 aver|OJJ06824.1 awen|OJJ33174.1 azon|OJJ46381.1

Group4212: abra|OJJ76809.1 acar|OOF91305.1 acla|EAW13370.1 afis|EAW20484.1 afla|EED44784.1 afum|EAL84571.1 agla|OJJ84813.1 akaw|GAA87212.1 aluc|OJZ84218.1 anid|CBF79235.1 anig|EHA27569.1 aory|EIT77759.1 asyd|OJJ55742.1 ater|EAU29562.1 atub|OJI89866.1 aver|OJJ06829.1 awen|OJJ33185.1 azon|OJJ46384.1

Group4213: abra|OJJ76810.1 acar|OOF91306.1 acla|EAW13371.1 afis|EAW20485.1 afla|EED44782.1 afum|EAL84570.1 agla|OJJ84822.1 akaw|GAA87211.1 aluc|OJZ84221.1 anid|CBF79237.1 anig|EHA27570.1 aory|EIT77768.1 asyd|OJJ55743.1 ater|EAU29563.1 atub|OJI89865.1 aver|OJJ06830.1 awen|OJJ33186.1 azon|OJJ46388.1

Group4214: abra|OJJ76811.1 acar|OOF91307.1 acla|EAW13372.1 afis|EAW20486.1 afla|EED44781.1 afum|EAL84569.1 agla|OJJ84825.1 akaw|GAA87210.1 aluc|OJZ84222.1 anid|CBF79215.1 anig|EHA27571.1 aory|EIT77774.1 asyd|OJJ55737.1 ater|EAU29564.1 atub|OJI89864.1 aver|OJJ06823.1 awen|OJJ33187.1 azon|OJJ46389.1

Group4215: abra|OJJ76813.1 acar|OOF91309.1 acla|EAW13374.1 afis|EAW20488.1 afla|EED44776.1 afum|EAL84567.1 agla|OJJ84826.1 akaw|GAA87208.1 aluc|OJZ84224.1 anid|CBF79219.1 anig|EHA27573.1 aory|EIT77760.1 asyd|OJJ55736.1 ater|EAU29566.1 atub|OJI89861.1 aver|OJJ06822.1 awen|OJJ33188.1 azon|OJJ46390.1

Group4216: abra|OJJ76819.1 acar|OOF91315.1 acla|EAW13379.1 afis|EAW20493.1 afla|EED44737.1 afum|EAL84561.1 agla|OJJ84833.1 akaw|GAA87203.1 aluc|OJZ84229.1 anid|CBF79226.1 anig|EHA27578.1 aory|EIT77797.1 asyd|OJJ55732.1 ater|EAU29569.1 atub|OJI89855.1 aver|OJJ06818.1 awen|OJJ33193.1 azon|OJJ46403.1

Group4217: abra|OJJ76821.1 acar|OOF91316.1 acla|EAW13381.1 afis|EAW20495.1 afla|EED44771.1 afum|EAL84559.1 agla|OJJ84807.1 akaw|GAA87201.1 aluc|OJZ84232.1 anid|CBF79222.1 anig|EHA27579.1 aory|EIT77762.1 asyd|OJJ55734.1 ater|EAU37189.1 atub|OJI89853.1 aver|OJJ06820.1 awen|OJJ33195.1 azon|OJJ46401.1

Group4218: abra|OJJ76823.1 acar|OOF91319.1 acla|EAW13383.1 afis|EAW20497.1 afla|EED44739.1 afum|EAL84557.1 agla|OJJ84805.1 akaw|GAA87199.1 aluc|OJZ84234.1 anid|CBF79208.1 anig|EHA27581.1 aory|EIT77796.1 asyd|OJJ55729.1 ater|EAU37191.1 atub|OJI89851.1 aver|OJJ06815.1 awen|OJJ33197.1 azon|OJJ46399.1

Group4219: abra|OJJ76827.1 acar|OOF91321.1 acla|EAW13387.1 afis|EAW20501.1 afla|EED44744.1 afum|EAL84554.1 agla|OJJ84797.1 akaw|GAA87195.1 aluc|OJZ84239.1 anid|CBF79200.1 anig|EHA27586.1 aory|EIT77773.1 asyd|OJJ55725.1 ater|EAU37195.1 atub|OJI89844.1 aver|OJJ06811.1 awen|OJJ33202.1 azon|OJJ46396.1

Group4220: abra|OJJ76830.1 acar|OOF91324.1 acla|EAW13392.1 afis|EAW20504.1 afla|EED44747.1 afum|EAL84552.1 agla|OJJ84800.1 akaw|GAA87192.1 aluc|OJZ84242.1 anid|CBF79195.1 anig|EHA27587.1 aory|EIT77784.1 asyd|OJJ55722.1 ater|EAU37199.1 atub|OJI89841.1 aver|OJJ06808.1 awen|OJJ33205.1 azon|OJJ46478.1

Group4222: abra|OJJ76834.1 acar|OOF91327.1 acla|EAW13396.1 afis|EAW20507.1 afla|EED52448.1 afum|EAL84549.1 agla|OJJ84784.1 akaw|GAA87189.1 aluc|OJZ84245.1 anid|CBF79193.1 anig|EHA27590.1 aory|EIT82772.1 asyd|OJJ55719.1 ater|EAU37201.1 atub|OJI89838.1 aver|OJJ06805.1 awen|OJJ33208.1 azon|OJJ46481.1

Group4223: abra|OJJ76835.1 acar|OOF91328.1 acla|EAW13397.1 afis|EAW20508.1 afla|EED52449.1 afum|EAL84548.1 agla|OJJ84785.1 akaw|GAA87188.1 aluc|OJZ84246.1 anid|CBF79190.1 anig|EHA27591.1 aory|EIT82778.1 asyd|OJJ55718.1 ater|EAU37202.1 atub|OJI89837.1 aver|OJJ06804.1 awen|OJJ33209.1 azon|OJJ46482.1

Group4224: abra|OJJ76836.1 acar|OOF90072.1 acla|EAW13398.1 afis|EAW20509.1 afla|EED52450.1 afum|EAL84547.1 agla|OJJ84786.1 akaw|GAA87187.1 aluc|OJZ84247.1 anid|CBF79189.1 anig|EHA27592.1 aory|EIT82759.1 asyd|OJJ55717.1 ater|EAU37203.1 atub|OJI89836.1 aver|OJJ06803.1 awen|OJJ33210.1 azon|OJJ46483.1

Group4225: abra|OJJ76838.1 acar|OOF91330.1 acla|EAW13400.1 afis|EAW20511.1 afla|EED52452.1 afum|EAL84545.2 agla|OJJ84788.1 akaw|GAA87185.1 aluc|OJZ84249.1 anid|CBF79186.1 anig|EHA27594.1 aory|EIT82799.1 asyd|OJJ55715.1 ater|EAU37205.1 atub|OJI89834.1 aver|OJJ06801.1 awen|OJJ33213.1 azon|OJJ46485.1

Group4226: abra|OJJ76839.1 acar|OOF91331.1 acla|EAW13401.1 afis|EAW20512.1 afla|EED52453.1 afum|EBA27192.1 agla|OJJ84789.1 akaw|GAA87184.1 aluc|OJZ84250.1 anid|CBF79184.1 anig|EHA27595.1 aory|EIT82753.1 asyd|OJJ55714.1 ater|EAU37208.1 atub|OJI89833.1 aver|OJJ06800.1 awen|OJJ33214.1 azon|OJJ46486.1

Group4227: abra|OJJ76840.1 acar|OOF91333.1 acla|EAW13403.1 afis|EAW20514.1 afla|EED52468.1 afum|EAL84543.2 agla|OJJ84776.1 akaw|GAA87182.1 aluc|OJZ84251.1 anid|CBF79055.1 anig|EHA27596.1 aory|EIT82784.1 asyd|OJJ55710.1 ater|EAU37187.1 atub|OJI89832.1 aver|OJJ06796.1 awen|OJJ33220.1 azon|OJJ46488.1

Group4228: abra|OJJ76843.1 acar|OOF89969.1 acla|EAW13406.1 afis|EAW20517.1 afla|EED52458.1 afum|EAL84540.1 agla|OJJ84769.1 akaw|GAA87179.1 aluc|OJZ84254.1 anid|CBF88058.1 anig|EHA27599.1 aory|EIT82793.1 asyd|OJJ63541.1 ater|EAU37184.1 atub|OJI89829.1 aver|OJI97392.1 awen|OJJ33223.1 azon|OJJ46553.1

Group4229: abra|OJJ76844.1 acar|OOF89968.1 acla|EAW13407.1 afis|EAW20518.1 afla|EED52457.1 afum|EAL84539.1 agla|OJJ84768.1 akaw|GAA87177.1 aluc|OJZ84256.1 anid|CBF79061.1 anig|EHA27600.1 aory|EIT82769.1 asyd|OJJ55707.1 ater|EAU37183.1 atub|OJI89827.1 aver|OJJ06793.1 awen|OJJ33224.1 azon|OJJ46581.1

Group4230: abra|OJJ76845.1 acar|OOF91335.1 acla|EAW13408.1 afis|EAW20519.1 afla|EED52455.1 afum|EAL84538.1 agla|OJJ84767.1 akaw|GAA87176.1 aluc|OJZ84257.1 anid|CBF79067.1 anig|EHA27601.1 aory|EIT82798.1 asyd|OJJ55706.1 ater|EAU37182.1 atub|OJI89826.1 aver|OJJ06792.1 awen|OJJ33225.1 azon|OJJ46580.1

Group4231: abra|OJJ76847.1 acar|OOF91337.1 acla|EAW13410.1 afis|EAW20521.1 afla|EED52470.1 afum|EAL84536.1 agla|OJJ84765.1 akaw|GAA87174.1 aluc|OJZ84259.1 anid|CBF79070.1 anig|EHA27602.1

aory|EIT82770.1 asyd|OJJ55704.1 ater|EAU37167.1 atub|OJI89824.1 aver|OJJ06790.1 awen|OJJ33227.1 azon|OJJ46577.1

Group4232: abra|OJJ76849.1 acar|OOF91338.1 acla|EAW13412.1 afis|EAW20523.1 afla|EED52472.1 afum|EAL84534.1 agla|OJJ84740.1 akaw|GAA87172.1 aluc|OJZ84261.1 anid|CBF79074.1 anig|EHA27604.1 aory|EIT82821.1 asyd|OJJ55702.1 ater|EAU37165.1 atub|OJI89822.1 aver|OJJ06787.1 awen|OJJ33229.1 azon|OJJ46575.1

Group4233: abra|OJJ76851.1 acar|OOF91339.1 acla|EAW13413.1 afis|EAW20524.1 afla|EED52473.1 afum|EAL84533.1 agla|OJJ84742.1 akaw|GAA87171.1 aluc|OJZ84262.1 anid|CBF79076.1 anig|EHA27605.1 aory|EIT82758.1 asyd|OJJ55701.1 ater|EAU37164.1 atub|OJI89821.1 aver|OJJ06786.1 awen|OJJ33230.1 azon|OJJ46574.1

Group4234: abra|OJJ76852.1 acar|OOF91340.1 acla|EAW13414.1 afis|EAW20525.1 afla|EED52474.1 afum|EAL84532.1 agla|OJJ84743.1 akaw|GAA87170.1 aluc|OJZ84264.1 anid|CBF79078.1 anig|EHA27606.1 aory|EIT82817.1 asyd|OJJ55699.1 ater|EAU37163.1 atub|OJI89819.1 aver|OJJ06784.1 awen|OJJ33231.1 azon|OJJ46573.1

Group4235: abra|OJJ76855.1 acar|OOF91341.1 acla|EAW13416.1 afis|EAW20527.1 afla|EED52476.1 afum|EAL84530.1 agla|OJJ84745.1 akaw|GAA87167.1 aluc|OJZ84267.1 anid|CBF79081.1 anig|EHA27609.1 aory|EIT82771.1 asyd|OJJ55696.1 ater|EAU37161.1 atub|OJI89816.1 aver|OJJ06781.1 awen|OJJ33234.1 azon|OJJ46570.1

Group4236: abra|OJJ76857.1 acar|OOF91343.1 acla|EAW13418.1 afis|EAW20529.1 afla|EED52478.1 afum|EAL84528.1 agla|OJJ84706.1 akaw|GAA87165.1 aluc|OJZ84269.1 anid|CBF79006.1 anig|EHA27611.1 aory|EIT82803.1 asyd|OJJ55691.1 ater|EAU37159.1 atub|OJI89814.1 aver|OJJ06776.1 awen|OJJ33236.1 azon|OJJ50608.1

Group4237: abra|OJJ76903.1 acar|OOF91351.1 acla|EAW13425.1 afis|EAW20536.1 afla|EED52442.1 afum|EAL84522.1 agla|OJJ84695.1 akaw|GAA87135.1 aluc|OJZ84302.1 anid|CBF78995.1 anig|EHA27638.1 aory|EIT82797.1 asyd|OJJ55685.1 ater|EAU37143.1 atub|OJI89766.1 aver|OJJ06769.1 awen|OJJ33243.1 azon|OJJ50622.1

Group4238: abra|OJJ76916.1 acar|OOF91363.1 acla|EAW13428.1 afis|EAW20544.1 afla|EED52424.1 afum|EAL84514.1 agla|OJJ88753.1 akaw|GAA87122.1 aluc|OJZ84316.1 anid|CBF78988.1 anig|EHA27651.1 aory|EIT83288.1 asyd|OJJ55682.1 ater|EAU37137.1 atub|OJI89752.1 aver|OJJ06766.1 awen|OJJ33252.1 azon|OJJ50638.1

Group4239: abra|OJJ76917.1 acar|OOF91364.1 acla|EAW13429.1 afis|EAW20545.1 afla|EED52423.1 afum|EBA27188.1 agla|OJJ88752.1 akaw|GAA87121.1 aluc|OJZ84317.1 anid|CBF78986.1 anig|EHA27652.1 aory|EIT83281.1 asyd|OJJ55681.1 ater|EAU37136.1 atub|OJI89751.1 aver|OJJ06765.1 awen|OJJ33253.1 azon|OJJ50637.1

Group4240: abra|OJJ76918.1 acar|OOF91365.1 acla|EAW13430.1 afis|EAW20546.1 afla|EED52422.1 afum|EAL84513.1 agla|OJJ88751.1 akaw|GAA87120.1 aluc|OJZ84318.1 anid|CBF78985.1 anig|EHA27653.1 aory|EIT83315.1 asyd|OJJ55680.1 ater|EAU37135.1 atub|OJI89750.1 aver|OJJ06763.1 awen|OJJ33254.1 azon|OJJ50636.1

Group4241: abra|OJJ76919.1 acar|OOF91366.1 acla|EAW13431.1 afis|EAW20547.1 afla|EED52421.1 afum|EAL84512.1 agla|OJJ88750.1 akaw|GAA87119.1 aluc|OJZ84319.1 anid|CBF78983.1 anig|EHA27654.1 aory|EIT83284.1 asyd|OJJ55679.1 ater|EAU37134.1 atub|OJI89749.1 aver|OJJ06762.1 awen|OJJ33255.1 azon|OJJ50635.1

Group4244: abra|OJJ76950.1 acar|OOF91396.1 acla|EAW13470.1 afis|EAW20582.1 afla|EED52277.1 afum|EAL84482.1 agla|OJJ88700.1 akaw|GAA87091.1 aluc|OJZ84347.1 anid|CBF78946.1 anig|EHA27683.1 aory|EIT74415.1 asyd|OJJ55651.1 ater|EAU37100.1 atub|OJI89721.1 aver|OJJ06736.1 awen|OJJ33287.1 azon|OJJ50663.1

Group4245: abra|OJJ76954.1 acar|OOF91398.1 acla|EAW13476.1 afis|EAW20583.1 afla|EED52245.1 afum|EBA27180.1 agla|OJJ88686.1 akaw|GAA87088.1 aluc|OJZ84350.1 anid|CBF78930.1 anig|EHA27687.1 aory|EIT74502.1 asyd|OJJ55642.1 ater|EAU37097.1 atub|OJI89717.1 aver|OJJ06725.1 awen|OJJ33300.1 azon|OJJ50671.1

Group4246: abra|OJJ76970.1 acar|OOF91415.1 acla|EAW13495.1 afis|EAW20595.1 afla|EED52229.1 afum|EAL84466.1 agla|OJJ88661.1 akaw|GAA87074.1 aluc|OJZ84366.1 anid|CBF78911.1 anig|EHA27701.1 aory|EIT74324.1 asyd|OJJ55615.1 ater|EAU35241.1 atub|OJI89702.1 aver|OJJ06690.1 awen|OJJ33385.1 azon|OJJ50691.1

Group4247: abra|OJJ76971.1 acar|OOF91416.1 acla|EAW13496.1 afis|EAW20596.1 afla|EED52228.1 afum|EAL84465.1 agla|OJJ88660.1 akaw|GAA87073.1 aluc|OJZ84367.1 anid|CBF78909.1 anig|EHA27702.1 aory|EIT74434.1 asyd|OJJ55614.1 ater|EAU35242.1 atub|OJI89701.1 aver|OJJ06689.1 awen|OJJ33388.1 azon|OJJ50692.1

Group4248: abra|OJJ76972.1 acar|OOF91417.1 acla|EAW13498.1 afis|EAW20597.1 afla|EED52213.1 afum|EAL84464.1 agla|OJJ88659.1 akaw|GAA87072.1 aluc|OJZ84368.1 anid|CBF78908.1 anig|EHA27703.1 aory|EIT74430.1 asyd|OJJ55612.1 ater|EAU35243.1 atub|OJI89700.1 aver|OJJ06687.1 awen|OJJ33389.1 azon|OJJ50693.1

Group4249: abra|OJJ76976.1 acar|OOF91419.1 acla|EAW13502.1 afis|EAW20601.1 afla|EED47729.1 afum|EAL84460.1 agla|OJJ88650.1 akaw|GAA87069.1 aluc|OJZ84372.1 anid|CBF75136.1 anig|EHA27707.1 aory|EIT77171.1 asyd|OJJ59976.1 ater|EAU35246.1 atub|OJI89695.1 aver|OJJ02315.1 awen|OJJ33393.1 azon|OJJ50697.1

Group4250: abra|OJJ76977.1 acar|OOF91422.1 acla|EAW13504.1 afis|EAW20603.1 afla|EED47732.1 afum|EAL84458.1 agla|OJJ88646.1 akaw|GAA87068.1 aluc|OJZ84374.1 anid|CBF86930.1 anig|EHA27708.1 aory|EIT77153.1 asyd|OJJ55600.1 ater|EAU35248.1 atub|OJI89694.1 aver|OJJ06680.1 awen|OJJ33395.1 azon|OJJ50698.1

Group4251: abra|OJJ76978.1 acar|OOF91424.1 acla|EAW13505.1 afis|EAW20604.1 afla|EED47733.1 afum|EAL84457.1 agla|OJJ88645.1 akaw|GAA87067.1 aluc|OJZ84376.1 anid|CBF86928.1 anig|EHA27709.1 aory|EIT77125.1 asyd|OJJ55601.1 ater|EAU35250.1 atub|OJI89693.1 aver|OJJ06681.1 awen|OJJ33396.1 azon|OJJ50394.1

Group4252: abra|OJJ76981.1 acar|OOF91427.1 acla|EAW12964.1 afis|EAW17699.1 afla|EED45692.1 afum|EAL91514.1 agla|OJJ79475.1 akaw|GAA87064.1 aluc|OJZ84381.1 anid|CBF89963.1 anig|EHA27712.1 aory|EIT75518.1 asyd|OJJ62104.1 ater|EAU31915.1 atub|OJI89690.1 aver|OJI96210.1 awen|OJJ37134.1 azon|OJJ50834.1

Group4253: abra|OJJ76984.1 acar|OOF91429.1 acla|EAW12966.1 afis|EAW17697.1 afla|EED50445.1 afum|EAL91511.1 agla|OJJ79478.1 akaw|GAA87062.1 aluc|OJZ84383.1 anid|CBF89965.1 anig|EHA27714.1 aory|EIT82014.1 asyd|OJJ62100.1 ater|EAU31918.1 atub|OJI89688.1 aver|OJI96206.1 awen|OJJ37137.1 azon|OJJ50836.1

Group4254: abra|OJJ76985.1 acar|OOF91430.1 acla|EAW12967.1 afis|EAW17696.1 afla|EED50446.1 afum|EAL91510.1 agla|OJJ79479.1 akaw|GAA87061.1 aluc|OJZ84384.1 anid|CBF89966.1 anig|EHA27715.1 aory|EIT82098.1 asyd|OJJ62099.1 ater|EAU31919.1 atub|OJI89687.1 aver|OJI96205.1 awen|OJJ37139.1 azon|OJJ50839.1

Group4255: abra|OJJ76986.1 acar|OOF92639.1 acla|EAW12968.1 afis|EAW17695.1 afla|EED50447.1 afum|EAL91509.1 agla|OJJ79483.1 akaw|GAA87060.1 aluc|OJZ84385.1 anid|CBF89968.1 anig|EHA27716.1 aory|EIT82034.1 asyd|OJJ62098.1 ater|EAU31920.1 atub|OJI89686.1 aver|OJI96204.1 awen|OJJ37140.1 azon|OJJ50840.1

Group4256: abra|OJJ76987.1 acar|OOF91431.1 acla|EAW12969.1 afis|EAW17694.1 afla|EED50448.1 afum|EAL91508.1 agla|OJJ79485.1 akaw|GAA87059.1 aluc|OJZ84386.1 anid|CBF89970.1 anig|EHA27717.1 aory|EIT82134.1 asyd|OJJ62097.1 ater|EAU31961.1 atub|OJI89685.1 aver|OJI96203.1 awen|OJJ37141.1 azon|OJJ50841.1

Group4257: abra|OJJ76993.1 acar|OOF91438.1 acla|EAW13455.1 afis|EAW20563.1 afum|EAL84498.1 agla|OJJ88728.1 akaw|GAA87053.1 aluc|OJZ84393.1 anid|CBF78959.1 anig|EHA27722.1 aory|EIT74501.1 asyd|OJJ55658.1 ater|EAU37112.1 atub|OJI89678.1 aver|OJJ06742.1 awen|OJJ33269.1 azon|OJJ50641.1 afla|EED52341.1

Group4258: abra|OJJ76996.1 acar|OOF91443.1 acla|EAW13460.1 afis|EAW20566.1 afla|EED52324.1 afum|EAL84495.1 agla|OJJ88727.1 akaw|GAA87050.1 aluc|OJZ84396.1 anid|CBF78935.1 anig|EHA27726.1 aory|EIT74477.1 asyd|OJJ55644.1 ater|EAU37111.1 atub|OJI89676.1 aver|OJJ06728.1 awen|OJJ33271.1 azon|OJJ50646.1

Group4259: abra|OJJ77016.1 acar|OOF95265.1 acla|EAW07210.1 afis|EAW22222.1 afla|EED57616.1 afum|EAL90972.1 agla|OJJ87822.1 akaw|GAA87034.1 aluc|OJZ84418.1 anid|CBF88465.1 anig|EHA27740.1 aory|EIT82988.1 asyd|OJJ63777.1 ater|EAU34232.1 atub|OJI89656.1 aver|OJI97160.1 awen|OJJ34809.1 azon|OJJ45741.1

Group4260: abra|OJJ77024.1 acar|OOF91461.1 acla|EAW12057.1 afis|EAW19698.1 afla|EED54134.1 afum|EAL86829.1 agla|OJJ80810.1 akaw|GAA87028.1 aluc|OJZ84424.1 anid|CBF81131.1 anig|EHA27746.1 aory|EIT76042.1 asyd|OJJ56500.1 ater|EAU32171.1 atub|OJI89649.1 aver|OJJ06219.1 awen|OJJ41934.1 azon|OJJ48232.1

Group4261: abra|OJJ77043.1 acar|OOF90273.1 acla|EAW13554.1 afis|EAW20631.1 afla|EED49762.1 afum|EAL84436.1 agla|OJJ80447.1 akaw|GAA87010.1 aluc|OJZ84445.1 anid|CBF83219.1 anig|EHA27763.1 aory|EIT77598.1 asyd|OJJ55568.1 ater|EAU32166.1 atub|OJI89629.1 aver|OJJ06647.1 awen|OJJ33600.1 azon|OJJ50722.1

Group4262: abra|OJJ77049.1 acar|OOF92938.1 acla|EAW13549.1 afis|EAW20628.1 afla|EED49778.1 afum|EAL84442.1 agla|OJJ88595.1 akaw|GAA87004.1 aluc|OJZ84453.1 anid|CBF83230.1 anig|EHA27768.1

aory|EIT77696.1 asyd|OJJ55582.1 ater|EAU32163.1 atub|OJI89623.1 aver|OJJ06661.1 awen|OJJ33461.1 azon|OJJ50717.1

Group4263: abra|OJJ77050.1 acar|OOF92939.1 acla|EAW13544.1 afis|EAW20616.1 afla|EED54198.1 afum|EAL84445.1 agla|OJJ88594.1 akaw|GAA87003.1 aluc|OJZ84454.1 anid|CBF86935.1 anig|EHA27769.1 aory|EIT75987.1 asyd|OJJ55584.1 ater|EAU35262.1 atub|OJI89621.1 aver|OJJ06663.1 awen|OJJ33460.1 azon|OJJ50716.1

Group4264: abra|OJJ77064.1 acar|OOF92948.1 acla|EAW13511.1 afis|EAW20609.1 afla|EED54207.1 afum|EAL84451.1 agla|OJJ88630.1 akaw|GAA86988.1 aluc|OJZ84472.1 anid|CBF86951.1 anig|EHA27781.1 aory|EIT76020.1 asyd|OJJ55594.1 ater|EAU35257.1 atub|OJI89606.1 aver|OJJ06673.1 awen|OJJ33452.1 azon|OJJ50709.1

Group4265: abra|OJJ77068.1 acar|OOF92951.1 acla|EAW13508.1 afis|EAW20606.1 afla|EED47725.1 afum|EAL84454.1 agla|OJJ88635.1 akaw|GAA86985.1 aluc|OJZ84476.1 anid|CBF86946.1 anig|EHA27784.1 aory|EIT77120.1 asyd|OJJ55591.1 ater|EAU35260.1 atub|OJI89603.1 aver|OJJ06670.1 awen|OJJ33449.1 azon|OJJ50706.1

Group4266: abra|OJJ77076.1 acar|OOF92958.1 acla|EAW11204.1 afis|EAW15934.1 afla|EED54927.1 afum|EAL88570.1 agla|OJJ89373.1 akaw|GAA86977.1 aluc|OJZ84486.1 anid|CBF86351.1 anig|EHA27790.1 aory|EIT80330.1 asyd|OJJ64980.1 ater|EAU35519.1 atub|OJI89594.1 aver|OJI99102.1 awen|OJJ35645.1 azon|OJJ46891.1

Group4267: abra|OJJ77079.1 acar|OOF92961.1 acla|EAW12041.1 afis|EAW19690.1 afla|EED57010.1 afum|EAL86836.1 agla|OJJ80804.1 akaw|GAA86974.1 aluc|OJZ84489.1 anid|CBF75140.1 anig|EHA27793.1 aory|EIT82303.1 asyd|OJJ59969.1 atub|OJI89591.1 aver|OJJ02322.1 awen|OJJ41917.1 azon|OJJ42689.1 ater|EAU32856.1

Group4268: abra|OJJ77090.1 acar|OOF92974.1 acla|EAW08632.1 afis|EAW15799.1 afla|EED51246.1 afum|EAL85551.1 agla|OJJ81963.1 akaw|GAA86965.1 aluc|OJZ84499.1 anid|CBF70921.1 anig|EHA27799.1 aory|EIT83184.1 asyd|OJJ52607.1 ater|EAU32343.1 atub|OJI89581.1 aver|OJJ04634.1 awen|OJJ38507.1 azon|OJJ43744.1

Group4269: abra|OJJ77091.1 acar|OOF92976.1 acla|EAW10011.1 afis|EAW21045.1 afla|EED51625.1 afum|EAL92081.1 agla|OJJ82363.1 akaw|GAA86964.1 aluc|OJZ84500.1 anid|CBF87097.1 anig|EHA27800.1 aory|EIT80549.1 asyd|OJJ60392.1 ater|EAU36817.1 atub|OJI89580.1 aver|OJI99721.1 awen|OJJ32179.1 azon|OJJ46755.1

Group4270: abra|OJJ77094.1 acar|OOF92977.1 acla|EAW10918.1 afis|EAW20940.1 afla|EED50295.1 afum|EAL84694.1 agla|OJJ83947.1 akaw|GAA86961.1 aluc|OJZ84503.1 anid|CBF82521.1 anig|EHA27803.1 aory|EIT77252.1 asyd|OJJ54070.1 ater|EAU38585.1 atub|OJI89577.1 aver|OJJ08672.1 awen|OJJ34377.1 azon|OJJ48111.1

Group4271: abra|OJJ77095.1 acar|OOF92978.1 acla|EAW10919.1 afis|EAW20939.1 afla|EED50294.1 afum|EAL84695.1 agla|OJJ83948.1 akaw|GAA86960.1 aluc|OJZ84504.1 anid|CBF82523.1 anig|EHA27804.1 aory|EIT77268.1 asyd|OJJ54071.1 ater|EAU38584.1 atub|OJI89576.1 aver|OJJ08671.1 awen|OJJ34378.1 azon|OJJ48112.1

Group4272: abra|OJJ77106.1 acar|OOF92991.1 acla|EAW11608.1 afis|EAW23840.1 afla|EED52551.1 afum|EAL87271.1 agla|OJJ82489.1 akaw|GAA86950.1 aluc|OJZ84515.1 anid|CBF77077.1 anig|EHA27813.1 aory|EIT74221.1 asyd|OJJ56983.1 ater|EAU33507.1 atub|OJI89566.1 aver|OJJ03304.1 awen|OJJ33810.1 azon|OJJ48724.1

Group4273: abra|OJJ77111.1 acar|OOF93001.1 acla|EAW07270.1 afis|EAW22291.1 afla|EED57693.1 afum|EAL90890.1 agla|OJJ87755.1 akaw|GAA90817.1 aluc|OJZ84520.1 anid|CBF88575.1 anig|EHA27818.1 aory|EIT79896.1 asyd|OJJ63840.1 ater|EAU34306.1 atub|OJI89560.1 aver|OJI97097.1 awen|OJJ34867.1 azon|OJJ50044.1

Group4274: abra|OJJ77116.1 acar|OOF93005.1 acla|EAW12952.1 afis|EAW17704.1 afla|EED45696.1 afum|EAL91519.2 agla|OJJ81740.1 akaw|GAA90813.1 aluc|OJZ84526.1 anid|CBF89929.1 anig|EHA27823.1 aory|EIT75367.1 asyd|OJJ62119.1 ater|EAU31912.1 atub|OJI89554.1 aver|OJI96226.1 awen|OJJ37131.1 azon|OJJ50830.1

Group4275: abra|OJJ77120.1 acar|OOF93007.1 acla|EAW12932.1 afis|EAW17709.1 afla|EED45711.1 afum|EAL91524.2 agla|OJJ81738.1 akaw|GAA90809.1 aluc|OJZ84530.1 anid|CBF89906.1 anig|EHA27827.1 aory|EIT75650.1 asyd|OJJ62132.1 ater|EAU31903.1 atub|OJI89550.1 aver|OJI96241.1 awen|OJJ37127.1 azon|OJJ50826.1

Group4276: abra|OJJ77123.1 acar|OOF93009.1 acla|EAW12936.1 afis|EAW17712.1 afla|EED45714.1 afum|EAL91527.1 agla|OJJ81736.1 akaw|GAA90805.1 aluc|OJZ84533.1 anid|CBF76920.1 anig|EHA27831.1 aory|EIT75661.1 asyd|OJJ57072.1 ater|EAU31891.1 atub|OJI89547.1 aver|OJJ03226.1 awen|OJJ37124.1 azon|OJJ50823.1

Group4277: abra|OJJ77125.1 acar|OOF93011.1 acla|EAW12938.1 afis|EAW17714.1 afla|EED45716.1 afum|EAL91529.1 agla|OJJ81734.1 akaw|GAA90803.1 aluc|OJZ84535.1 anid|CBF76916.1 anig|EHA27833.1 aory|EIT75429.1 asyd|OJJ57079.1 ater|EAU31892.1 atub|OJI89545.1 aver|OJJ03224.1 awen|OJJ37122.1 azon|OJJ50821.1

Group4278: abra|OJJ77126.1 acar|OOF93012.1 acla|EAW12939.1 afis|EAW17715.1 afla|EED45717.1 afum|EAL91530.1 agla|OJJ81733.1 akaw|GAA90802.1 aluc|OJZ84536.1 anid|CBF76914.1 anig|EHA27834.1 aory|EIT75816.1 asyd|OJJ57080.1 ater|EAU31893.1 atub|OJI89544.1 aver|OJJ03223.1 awen|OJJ37121.1 azon|OJJ50820.1

Group4279: abra|OJJ77130.1 acar|OOF93018.1 acla|EAW12941.1 afis|EAW17717.1 afla|EED45719.1 afum|EAL91532.1 agla|OJJ81731.1 akaw|GAA90798.1 aluc|OJZ84540.1 anid|CBF84088.1 anig|EHA27838.1 aory|EIT75412.1 asyd|OJJ57082.1 ater|EAU31895.1 atub|OJI89540.1 aver|OJJ03220.1 awen|OJJ37119.1 azon|OJJ50818.1

Group4280: abra|OJJ77131.1 acar|OOF93019.1 acla|EAW12942.1 afis|EAW17718.1 afla|EED45720.1 afum|EAL91533.1 agla|OJJ81728.1 akaw|GAA90797.1 aluc|OJZ84541.1 anid|CBF76908.1 anig|EHA27839.1 aory|EIT75617.1 asyd|OJJ57083.1 ater|EAU31896.1 atub|OJI89539.1 aver|OJJ03219.1 awen|OJJ37118.1 azon|OJJ50817.1

Group4281: abra|OJJ77133.1 acar|OOF93021.1 acla|EAW12944.1 afis|EAW17720.1 afla|EED45722.1 afum|EAL91535.2 agla|OJJ81730.1 akaw|GAA90795.1 aluc|OJZ84543.1 anid|CBF76904.1 anig|EHA27841.1 aory|EIT75737.1 asyd|OJJ57085.1 ater|EAU31898.1 atub|OJI89537.1 aver|OJJ03217.1 awen|OJJ37116.1 azon|OJJ50815.1

Group4282: abra|OJJ77138.1 acar|OOF90283.1 acla|EAW12948.1 afis|EAW17723.1 afla|EED45725.1 afum|EAL91538.1 agla|OJJ81727.1 akaw|GAA90791.1 aluc|OJZ84549.1 anid|CBF86584.1 anig|EHA27845.1 aory|EIT75656.1 asyd|OJJ64839.1 ater|EAU31901.1 atub|OJI89531.1 aver|OJI98950.1 awen|OJJ37115.1 azon|OJJ50813.1

Group4283: abra|OJJ77139.1 acar|OOF90282.1 acla|EAW12950.1 afis|EAW17725.1 afla|EED45726.1 afum|EAL91540.1 agla|OJJ81726.1 akaw|GAA90790.1 aluc|OJZ84552.1 anid|CBF86583.1 anig|EHA27846.1 aory|EIT75496.1 asyd|OJJ64840.1 ater|EAU31881.1 atub|OJI89530.1 aver|OJI98951.1 awen|OJJ37113.1 azon|OJJ50812.1

Group4284: abra|OJJ77141.1 acar|OOF93027.1 acla|EAW12927.1 afis|EAW17728.1 afla|EED45728.1 afum|EAL91543.1 agla|OJJ81723.1 akaw|GAA90788.1 aluc|OJZ84555.1 anid|CBF86577.1 anig|EHA27849.1 aory|EIT75836.1 asyd|OJJ64842.1 ater|EAU31879.1 atub|OJI89527.1 aver|OJI98953.1 awen|OJJ37110.1 azon|OJJ50809.1

Group4285: abra|OJJ77142.1 acar|OOF93028.1 acla|EAW12926.1 afis|EAW17729.1 afla|EED45729.1 afum|EAL91544.1 agla|OJJ81722.1 akaw|GAA90787.1 aluc|OJZ84556.1 anid|CBF86575.1 anig|EHA27850.1 aory|EIT75504.1 asyd|OJJ64843.1 ater|EAU31878.1 atub|OJI89526.1 aver|OJI98954.1 awen|OJJ37109.1 azon|OJJ50808.1

Group4286: abra|OJJ77143.1 acar|OOF93029.1 acla|EAW12925.1 afis|EAW17730.1 afla|EED45730.1 afum|EAL91545.1 agla|OJJ81721.1 akaw|GAA90786.1 aluc|OJZ84558.1 anid|CBF86573.1 anig|EHA27851.1 aory|EIT75486.1 asyd|OJJ64844.1 ater|EAU31877.1 atub|OJI89525.1 aver|OJI98955.1 awen|OJJ37108.1 azon|OJJ50807.1

Group4287: abra|OJJ77144.1 acar|OOF93030.1 acla|EAW12924.1 afis|EAW17731.1 afla|EED45731.1 afum|EAL91546.1 agla|OJJ81720.1 akaw|GAA90785.1 aluc|OJZ84559.1 anid|CBF86588.1 anig|EHA27852.1 aory|EIT75623.1 asyd|OJJ64838.1 ater|EAU31876.1 atub|OJI89524.1 aver|OJI98948.1 awen|OJJ37107.1 azon|OJJ50806.1

Group4288: abra|OJJ77145.1 acar|OOF93031.1 acla|EAW12923.1 afis|EAW17732.1 afla|EED45732.1 afum|EAL91547.1 agla|OJJ81719.1 akaw|GAA90784.1 aluc|OJZ84560.1 anid|CBF86586.1 anig|EHA27853.1 aory|EIT75469.1 asyd|OJJ64837.1 ater|EAU31875.1 atub|OJI89523.1 aver|OJI98947.1 awen|OJJ37105.1 azon|OJJ50805.1

Group4289: abra|OJJ77146.1 acar|OOF93032.1 acla|EAW12922.1 afis|EAW17733.1 afla|EED45734.1 afum|EAL91548.1 agla|OJJ81718.1 akaw|GAA90783.1 aluc|OJZ84561.1 anid|CBF86597.1 anig|EHA27855.1 aory|EIT75719.1 asyd|OJJ64833.1 ater|EAU31874.1 atub|OJI89521.1 aver|OJI98946.1 awen|OJJ37104.1 azon|OJJ50804.1

Group4290: abra|OJJ77148.1 acar|OOF93034.1 acla|EAW12921.1 afis|EAW17734.1 afla|EED45735.1 afum|EAL91549.1 agla|OJJ81716.1 akaw|GAA90782.1 aluc|OJZ84562.1 anid|CBF86595.1 anig|EHA27856.1 aory|EIT75759.1 asyd|OJJ64832.1 ater|EAU31873.1 atub|OJI89520.1 aver|OJI98945.1 azon|OJJ50802.1 awen|OJJ37101.1

Group4291: abra|OJJ77149.1 acar|OOF93035.1 acla|EAW12920.1 afis|EAW17735.1 afla|EED45736.1 afum|EAL91550.1 agla|OJJ81715.1 akaw|GAA90781.1 aluc|OJZ84563.1 anid|CBF86593.1 anig|EHA27857.1

aory|EIT75423.1 asyd|OJJ64831.1 ater|EAU31872.1 atub|OJI89519.1 aver|OJI98944.1 awen|OJJ37100.1 azon|OJJ50801.1

Group4292: abra|OJJ77157.1 acar|OOF93044.1 acla|EAW12915.1 afis|EAW17745.1 afla|EED45745.1 afum|EAL91557.1 agla|OJJ81708.1 akaw|GAA90773.1 aluc|OJZ84571.1 anid|CBF86614.1 anig|EHA27866.1 aory|EIT75582.1 asyd|OJJ64818.1 ater|EAU31862.1 atub|OJI89511.1 aver|OJI98931.1 awen|OJJ37092.1 azon|OJJ50796.1

Group4293: abra|OJJ77164.1 acar|OOF93050.1 acla|EAW12910.1 afis|EAW17754.1 afla|EED45758.1 afum|EAL91567.1 agla|OJJ81699.1 akaw|GAA90768.1 aluc|OJZ84577.1 anid|CBF86624.1 anig|EHA27870.1 aory|EIT75538.1 asyd|OJJ64811.1 ater|EAU31854.1 atub|OJI89504.1 aver|OJI98924.1 awen|OJJ37082.1 azon|OJJ50787.1

Group4294: abra|OJJ77165.1 acar|OOF93054.1 acla|EAW12901.1 afis|EAW17764.1 afla|EED45766.1 afum|EAL91574.1 agla|OJJ81698.1 akaw|GAA90767.1 aluc|OJZ84578.1 anid|CBF86628.1 anig|EHA27871.1 aory|EIT75559.1 asyd|OJJ64809.1 ater|EAU31853.1 atub|OJI89503.1 aver|OJI98922.1 awen|OJJ37081.1 azon|OJJ50784.1

Group4295: abra|OJJ77166.1 acar|OOF93055.1 acla|EAW12900.1 afis|EAW17765.1 afla|EED45767.1 afum|EAL91575.1 agla|OJJ81697.1 akaw|GAA90766.1 aluc|OJZ84579.1 anid|CBF86630.1 anig|EHA27872.1 aory|EIT75709.1 asyd|OJJ64808.1 ater|EAU31852.1 atub|OJI89502.1 aver|OJI98921.1 awen|OJJ37080.1 azon|OJJ50783.1

Group4296: abra|OJJ77176.1 acar|OOF93064.1 acla|EAW13174.1 afis|EAW17464.1 afla|EED49356.1 afum|EAL91293.1 agla|OJJ80340.1 akaw|GAA90759.1 aluc|OJZ84587.1 anid|CBF71650.1 anig|EHA27880.1 aory|EIT79568.1 asyd|OJJ61840.1 ater|EAU33949.1 atub|OJI89491.1 aver|OJI95939.1 awen|OJJ37381.1 azon|OJJ51055.1

Group4297: abra|OJJ77177.1 acar|OOF93065.1 acla|EAW13175.1 afis|EAW17463.1 afla|EED49357.1 afum|EAL91292.1 agla|OJJ80342.1 akaw|GAA90758.1 aluc|OJZ84589.1 anid|CBF71651.1 anig|EHA27881.1 aory|EIT79484.1 asyd|OJJ61839.1 ater|EAU33948.1 atub|OJI89490.1 aver|OJI95938.1 awen|OJJ37382.1 azon|OJJ51056.1

Group4298: abra|OJJ77186.1 acar|OOF93074.1 acla|EAW13176.1 afis|EAW17462.1 afla|EED49360.1 afum|EAL91291.1 agla|OJJ80343.1 akaw|GAA90753.1 aluc|OJZ84595.1 anid|CBF71661.1 anig|EHA27884.1 aory|EIT79468.1 asyd|OJJ61834.1 ater|EAU33947.1 atub|OJI89484.1 aver|OJI95933.1 awen|OJJ37384.1 azon|OJJ51059.1

Group4299: abra|OJJ77188.1 acar|OOF92641.1 acla|EAW13178.1 afis|EAW17461.1 afla|EED49361.1 afum|EAL91289.1 agla|OJJ80344.1 akaw|GAA90752.1 aluc|OJZ84596.1 anid|CBF71663.1 anig|EHA27885.1 aory|EIT79490.1 asyd|OJJ61833.1 ater|EAU33946.1 atub|OJI89483.1 aver|OJI95932.1 awen|OJJ37385.1 azon|OJJ51060.1

Group4300: abra|OJJ77190.1 acar|OOF93076.1 acla|EAW13179.1 afis|EAW17460.1 afla|EED49362.1 afum|EAL91288.1 agla|OJJ80345.1 akaw|GAA90750.1 aluc|OJZ84598.1 anid|CBF71665.1 anig|EHA27887.1 aory|EIT79506.1 asyd|OJJ61832.1 ater|EAU33945.1 atub|OJI89481.1 aver|OJI95931.1 awen|OJJ37386.1 azon|OJJ51061.1

Group4301: abra|OJJ77193.1 acar|OOF93078.1 acla|EAW13183.1 afis|EAW17457.1 afla|EED49365.1 afum|EAL91285.1 agla|OJJ80354.1 akaw|GAA90748.1 aluc|OJZ84600.1 anid|CBF71668.1 anig|EHA27890.1 aory|EIT79576.1 asyd|OJJ61829.1 ater|EAU33944.1 atub|OJI89479.1 aver|OJI95929.1 awen|OJJ37390.1 azon|OJJ51063.1

Group4302: abra|OJJ77194.1 acar|OOF93079.1 acla|EAW13184.1 afis|EAW17456.1 afla|EED49366.1 afum|EAL91284.1 agla|OJJ80353.1 akaw|GAA90747.1 aluc|OJZ84601.1 anid|CBF71670.1 anig|EHA27891.1 aory|EIT79508.1 asyd|OJJ61828.1 ater|EAU33936.1 atub|OJI89478.1 aver|OJI95928.1 awen|OJJ37391.1 azon|OJJ51064.1

Group4303: abra|OJJ77195.1 acar|OOF93080.1 acla|EAW13186.1 afis|EAW17455.1 afla|EED49367.1 afum|EAL91283.1 agla|OJJ80352.1 akaw|GAA90746.1 aluc|OJZ84602.1 anid|CBF71671.1 anig|EHA27892.1 aory|EIT79573.1 asyd|OJJ61827.1 ater|EAU33935.1 atub|OJI89477.1 aver|OJI95927.1 awen|OJJ37392.1 azon|OJJ51065.1

Group4304: abra|OJJ77198.1 acar|OOF93081.1 acla|EAW13187.1 afis|EAW17454.1 afla|EED49368.1 afum|EAL91282.1 agla|OJJ80351.1 akaw|GAA90743.1 aluc|OJZ84605.1 anid|CBF71673.1 anig|EHA27894.1 aory|EIT79544.1 asyd|OJJ61826.1 ater|EAU33934.1 atub|OJI89474.1 aver|OJI95926.1 awen|OJJ37393.1 azon|OJJ51066.1

Group4305: abra|OJJ77201.1 acar|OOF93089.1 acla|EAW13192.1 afis|EAW17449.1 afla|EED49373.1 afum|EBA27429.1 akaw|GAA90737.1 aluc|OJZ84611.1 anid|CBF71685.1 anig|EHA27900.1 aory|EIT79520.1 asyd|OJJ61821.1 ater|EAU33929.1 atub|OJI89470.1 aver|OJI95921.1 awen|OJJ37397.1 azon|OJJ51070.1 agla|OJJ80368.1

Group4306: abra|OJJ77202.1 acar|OOF93090.1 acla|EAW13193.1 afis|EAW17448.1 afla|EED49374.1 afum|EAL91275.2 agla|OJJ80369.1 akaw|GAA90736.1 aluc|OJZ84612.1 anid|CBF71687.1 anig|EHA27901.1 aory|EIT79525.1 asyd|OJJ61820.1 ater|EAU33928.1 atub|OJI89469.1 aver|OJI95920.1 awen|OJJ37398.1 azon|OJJ51071.1

Group4307: abra|OJJ77204.1 acar|OOF93092.1 acla|EAW13196.1 akaw|GAA90734.1 aluc|OJZ84615.1 anid|CBF71688.1 anig|EHA27903.1 aory|EIT79511.1 asyd|OJJ61819.1 ater|EAU33925.1 atub|OJI89466.1 aver|OJI95919.1 awen|OJJ37400.1 afis|EAW17446.1 afla|EED49381.1 afum|EAL91272.1 agla|OJJ80371.1 azon|OJJ51074.1

Group4308: abra|OJJ77205.1 acar|OOF93094.1 acla|EAW13197.1 afis|EAW17445.1 afla|EED49382.1 afum|EAL91271.1 agla|OJJ80372.1 akaw|GAA90733.1 aluc|OJZ84616.1 anid|CBF71690.1 anig|EHA27904.1 aory|EIT79527.1 asyd|OJJ61818.1 ater|EAU33924.1 atub|OJI89465.1 aver|OJI95918.1 awen|OJJ37401.1 azon|OJJ51075.1

Group4309: abra|OJJ77206.1 acar|OOF93093.1 acla|EAW13198.1 afis|EAW17444.1 afla|EED49383.1 afum|EAL91270.1 agla|OJJ80373.1 akaw|GAA90732.1 aluc|OJZ84617.1 anid|CBF71692.1 anig|EHA27905.1 aory|EIT79499.1 asyd|OJJ61817.1 ater|EAU33923.1 atub|OJI89464.1 aver|OJI95917.1 awen|OJJ37402.1 azon|OJJ51076.1

Group4310: abra|OJJ77207.1 acar|OOF93095.1 acla|EAW13199.1 afis|EAW17443.1 afla|EED49384.1 afum|EAL91269.1 agla|OJJ80374.1 akaw|GAA90731.1 aluc|OJZ84618.1 anid|CBF71694.1 anig|EHA27906.1 aory|EIT79562.1 asyd|OJJ61816.1 ater|EAU33922.1 atub|OJI89463.1 aver|OJI95916.1 awen|OJJ37403.1 azon|OJJ51077.1

Group4311: abra|OJJ77209.1 acar|OOF93096.1 acla|EAW13201.1 afis|EAW17441.1 afla|EED49386.1 afum|EAL91267.1 agla|OJJ80376.1 akaw|GAA90729.1 aluc|OJZ84620.1 anid|CBF71698.1 anig|EHA27908.1 aory|EIT79558.1 asyd|OJJ61814.1 ater|EAU33920.1 atub|OJI89461.1 aver|OJI95913.1 awen|OJJ37405.1 azon|OJJ51079.1

Group4312: abra|OJJ77210.1 acar|OOF93098.1 acla|EAW13202.1 afis|EAW17440.1 afla|EED49387.1 afum|EAL91266.1 agla|OJJ80377.1 akaw|GAA90728.1 aluc|OJZ84621.1 anid|CBF71700.1 anig|EHA27909.1 aory|EIT79563.1 asyd|OJJ61813.1 ater|EAU33919.1 atub|OJI89460.1 aver|OJI95912.1 awen|OJJ37406.1 azon|OJJ51080.1

Group4313: abra|OJJ77211.1 acar|OOF93097.1 acla|EAW13203.1 afis|EAW17439.1 afla|EED49388.1 afum|EAL91265.1 agla|OJJ80378.1 akaw|GAA90727.1 aluc|OJZ84622.1 anid|CBF71702.1 anig|EHA27910.1 aory|EIT79503.1 asyd|OJJ61812.1 ater|EAU33918.1 atub|OJI89459.1 aver|OJI95911.1 awen|OJJ37407.1 azon|OJJ51081.1

Group4315: abra|OJJ77214.1 acar|OOF93101.1 acla|EAW13206.1 afis|EAW17436.1 afla|EED49392.1 afum|EAL91263.1 agla|OJJ80381.1 akaw|GAA90724.1 aluc|OJZ84625.1 anid|CBF71706.1 anig|EHA27912.1 aory|EIT79513.1 asyd|OJJ61809.1 ater|EAU33915.1 atub|OJI89456.1 aver|OJI95908.1 awen|OJJ37410.1 azon|OJJ51084.1

Group4316: abra|OJJ77215.1 acar|OOF93102.1 acla|EAW13207.1 afis|EAW17435.1 afla|EED49409.1 afum|EAL91262.1 agla|OJJ80383.1 akaw|GAA90723.1 aluc|OJZ84626.1 anid|CBF71708.1 anig|EHA27913.1 aory|EIT79458.1 asyd|OJJ61808.1 ater|EAU33914.1 atub|OJI89455.1 aver|OJI95907.1 awen|OJJ37411.1 azon|OJJ51085.1

Group4317: abra|OJJ77216.1 acar|OOF93103.1 acla|EAW13209.1 afis|EAW17432.1 afla|EED49408.1 afum|EAL91260.1 agla|OJJ80384.1 akaw|GAA90722.1 aluc|OJZ84627.1 anid|CBF71710.1 anig|EHA27914.1 aory|EIT79545.1 asyd|OJJ61807.1 ater|EAU33913.1 atub|OJI89452.1 aver|OJI95906.1 awen|OJJ37412.1 azon|OJJ51086.1

Group4318: abra|OJJ77220.1 acar|OOF93105.1 acla|EAW13210.1 afis|EAW17431.1 afla|EED49407.1 afum|EAL91259.1 agla|OJJ80387.1 akaw|GAA90719.1 aluc|OJZ84630.1 anid|CBF71712.1 anig|EHA27918.1 aory|EIT79560.1 asyd|OJJ61806.1 ater|EAU33912.1 atub|OJI89449.1 aver|OJI95905.1 awen|OJJ37413.1 azon|OJJ51087.1

Group4319: abra|OJJ77223.1 acar|OOF93108.1 acla|EAW13214.1 afis|EAW17428.1 afla|EED49402.1 afum|EAL91256.1 agla|OJJ80396.1 akaw|GAA90716.1 aluc|OJZ84633.1 anid|CBF71716.1 anig|EHA27921.1 aory|EIT79534.1 asyd|OJJ61804.1 ater|EAU33909.1 atub|OJI89446.1 aver|OJI95903.1 awen|OJJ37415.1 azon|OJJ51089.1

Group4320: abra|OJJ77225.1 acar|OOF93110.1 acla|EAW13216.1 afis|EAW17426.1 afla|EED49400.1 afum|EAL91254.1 agla|OJJ80398.1 akaw|GAA90714.1 aluc|OJZ84635.1 anid|CBF71720.1 anig|EHA27923.1 aory|EIT79524.1 asyd|OJJ61802.1 ater|EAU33908.1 atub|OJI89444.1 aver|OJI95901.1 awen|OJJ37417.1 azon|OJJ51091.1

Group4321: abra|OJJ77226.1 acar|OOF93111.1 acla|EAW13217.1 afis|EAW17425.1 afla|EED49399.1 afum|EAL91253.1 agla|OJJ80399.1 akaw|GAA90713.1 aluc|OJZ84636.1 anid|CBF71722.1 anig|EHA27924.1

aory|EIT79475.1 asyd|OJJ61801.1 ater|EAU33907.1 atub|OJI89443.1 aver|OJI95900.1 awen|OJJ37418.1 azon|OJJ51092.1

Group4322: abra|OJJ77230.1 acar|OOF93113.1 acla|EAW13219.1 afis|EAW17423.1 afla|EED49397.1 afum|EAL91251.1 agla|OJJ80401.1 akaw|GAA90711.1 aluc|OJZ84640.1 anid|CBF71729.1 anig|EHA27926.1 aory|EIT79564.1 asyd|OJJ61799.1 ater|EAU33905.1 atub|OJI89440.1 aver|OJI95898.1 awen|OJJ37421.1 azon|OJJ51094.1

Group4323: abra|OJJ77238.1 acar|OOF93119.1 acla|EAW13245.1 afis|EAW17391.1 afla|EED49477.1 afum|EAL91228.1 agla|OJJ80406.1 akaw|GAA90705.1 aluc|OJZ84648.1 anid|CBF71739.1 anig|EHA27932.1 aory|EIT79509.1 asyd|OJJ61792.1 ater|EAU33896.1 atub|OJI89432.1 aver|OJI95892.1 awen|OJJ37424.1 azon|OJJ51104.1

Group4324: abra|OJJ77240.1 acar|OOF93121.1 acla|EAW13247.1 afis|EAW17389.1 afla|EED49481.1 afum|EAL91225.1 agla|OJJ80408.1 akaw|GAA90703.1 aluc|OJZ84650.1 anid|CBF71743.1 anig|EHA27933.1 aory|EIT79553.1 asyd|OJJ61790.1 ater|EAU33894.1 atub|OJI89430.1 aver|OJI95890.1 awen|OJJ37426.1 azon|OJJ51102.1

Group4326: abra|OJJ77244.1 acar|OOF93125.1 acla|EAW13248.1 afis|EAW17388.1 afla|EED49482.1 afum|EAL91224.1 agla|OJJ80409.1 akaw|GAA90699.1 aluc|OJZ84654.1 anid|CBF71787.1 anig|EHA27937.1 aory|EIT79464.1 asyd|OJJ61746.1 ater|EAU33893.1 atub|OJI89426.1 aver|OJI95843.1 awen|OJJ37427.1 azon|OJJ51101.1

Group4327: abra|OJJ77247.1 acar|OOF90006.1 acla|EAW13252.1 afis|EAW17384.1 afla|EED49501.1 afum|EAL91219.1 agla|OJJ80419.1 akaw|GAA90695.1 aluc|OJZ84658.1 anid|CBF84196.1 anig|EHA27940.1 aory|EIT82601.1 asyd|OJJ57160.1 ater|EAU33890.1 atub|OJI89422.1 aver|OJJ03153.1 awen|OJJ33371.1 azon|OJJ51109.1

Group4328: abra|OJJ77250.1 acar|OOF90005.1 acla|EAW13253.1 afis|EAW17383.1 afla|EED49504.1 afum|EAL91218.1 agla|OJJ80420.1 akaw|GAA90693.1 aluc|OJZ84661.1 anid|CBF84198.1 anig|EHA27943.1 aory|EIT82610.1 asyd|OJJ57161.1 ater|EAU33880.1 atub|OJI89419.1 aver|OJJ03152.1 awen|OJJ37433.1 azon|OJJ51112.1

Group4329: abra|OJJ77260.1 acar|OOF93128.1 afla|EED49508.1 akaw|GAA90687.1 aluc|OJZ84667.1 anig|EHA27946.1 aory|EIT82587.1 asyd|OJJ57166.1 ater|EAU33878.1 atub|OJI89413.1 awen|OJJ37439.1 azon|OJJ51115.1 acla|EAW13256.1 afis|EAW17380.1 afum|EAL91215.1 agla|OJJ80422.1 anid|CBF84207.1 aver|OJJ03148.1

Group4330: abra|OJJ77261.1 acar|OOF93130.1 acla|EAW13257.1 afis|EAW17379.1 afla|EED49509.1 afum|EAL91214.1 agla|OJJ80423.1 akaw|GAA90686.1 aluc|OJZ84668.1 anid|CBF84209.1 anig|EHA27947.1 aory|EIT82656.1 asyd|OJJ57167.1 ater|EAU33877.1 atub|OJI89412.1 aver|OJJ03147.1 awen|OJJ37441.1 azon|OJJ51117.1

Group4331: abra|OJJ77262.1 acar|OOF93132.1 acla|EAW13258.1 afis|EAW17378.1 afla|EED49510.1 afum|EAL91213.1 agla|OJJ80424.1 akaw|GAA90685.1 aluc|OJZ84669.1 anid|CBF84210.1 anig|EHA27948.1 aory|EIT82657.1 asyd|OJJ57168.1 ater|EAU33876.1 atub|OJI89411.1 aver|OJJ03146.1 awen|OJJ37442.1 azon|OJJ51120.1

Group4332: abra|OJJ77263.1 acar|OOF93131.1 acla|EAW13259.1 afis|EAW17377.1 afla|EED49511.1 afum|EAL91212.1 agla|OJJ80425.1 akaw|GAA90684.1 aluc|OJZ84671.1 anid|CBF84212.1 anig|EHA27949.1 aory|EIT82654.1 asyd|OJJ57169.1 ater|EAU33875.1 atub|OJI89409.1 aver|OJJ03145.1 awen|OJJ37443.1 azon|OJJ51121.1

Group4333: abra|OJJ77275.1 acar|OOF93150.1 acla|EAW13275.1 afis|EAW17342.1 afla|EED49781.1 afum|EBA27424.1 agla|OJJ79342.1 akaw|GAA90674.1 aluc|OJZ84684.1 anid|CBF84217.1 anig|EHA27960.1 aory|EIT77695.1 asyd|OJJ57179.1 ater|EAU33855.1 atub|OJI89397.1 aver|OJJ03135.1 awen|OJJ29675.1 azon|OJJ45994.1

Group4334: abra|OJJ77289.1 acar|OOF93167.1 acla|EAW13173.1 afis|EAW17465.1 afla|EED49355.1 afum|EAL91294.1 agla|OJJ80339.1 akaw|GAA90660.1 aluc|OJZ84700.1 anid|CBF71646.1 anig|EHA27975.1 aory|EIT79489.1 asyd|OJJ61841.1 ater|EAU33950.1 atub|OJI89382.1 aver|OJI95941.1 awen|OJJ37380.1 azon|OJJ51054.1

Group4335: abra|OJJ77290.1 acar|OOF93168.1 acla|EAW13172.1 afis|EAW17466.1 afla|EED49354.1 afum|EAL91295.1 agla|OJJ80338.1 akaw|GAA90659.1 aluc|OJZ84701.1 anid|CBF71638.1 anig|EHA27976.1 aory|EIT79580.1 asyd|OJJ61844.1 ater|EAU33951.1 atub|OJI89381.1 aver|OJI95944.1 awen|OJJ37379.1 azon|OJJ51053.1

Group4336: abra|OJJ77291.1 acar|OOF93169.1 acla|EAW13171.1 afis|EAW17468.1 afla|EED49353.1 afum|EAL91297.1 agla|OJJ80336.1 akaw|GAA90658.1 aluc|OJZ84702.1 anid|CBF71640.1 anig|EHA27977.1 aory|EIT79590.1 asyd|OJJ61843.1 ater|EAU33952.1 atub|OJI89380.1 aver|OJI95943.1 awen|OJJ37378.1 azon|OJJ51052.1

Group4337: abra|OJJ77295.1 acar|OOF93172.1 acla|EAW13169.1 afis|EAW17470.1 afla|EED49348.1 afum|EAL91299.1 agla|OJJ78440.1 akaw|GAA91456.1 aluc|OJZ84707.1 anid|CBF71630.1 anig|EHA27981.1 aory|EIT79461.1 asyd|OJJ61849.1 ater|EAU33954.1 atub|OJI89376.1 aver|OJI95948.1 awen|OJJ37372.1 azon|OJJ51050.1

Group4338: abra|OJJ77297.1 acar|OOF93177.1 acla|EAW13167.1 afis|EAW17472.1 afla|EED49343.1 akaw|GAA91454.1 aluc|OJZ84709.1 anid|CBF71626.1 anig|EHA27983.1 asyd|OJJ61851.1 ater|EAU33956.1 atub|OJI89374.1 aver|OJI95950.1 afum|EAL91301.1 agla|OJJ81490.1 aory|EIT79471.1 awen|OJJ37370.1 azon|OJJ51049.1

Group4339: abra|OJJ77300.1 acar|OOF93179.1 acla|EAW13165.1 afis|EAW17474.1 afla|EED49340.1 afum|EAL91303.1 agla|OJJ81488.1 akaw|GAA91452.1 aluc|OJZ84712.1 anid|CBF71622.1 anig|EHA27985.1 aory|EIT79516.1 asyd|OJJ61854.1 ater|EAU33958.1 atub|OJI89371.1 aver|OJI95953.1 awen|OJJ37368.1 azon|OJJ51047.1

Group4340: abra|OJJ77301.1 acar|OOF93180.1 acla|EAW13164.1 afis|EAW17475.1 afla|EED49339.1 afum|EAL91304.1 agla|OJJ81487.1 akaw|GAA91451.1 aluc|OJZ84713.1 anid|CBF71620.1 anig|EHA27986.1 aory|EIT79565.1 asyd|OJJ61855.1 ater|EAU33959.1 atub|OJI89370.1 aver|OJI95954.1 awen|OJJ37367.1 azon|OJJ51046.1

Group4341: abra|OJJ77303.1 acar|OOF93183.1 acla|EAW13163.1 afis|EAW17477.1 afla|EED49337.1 afum|EAL91306.1 agla|OJJ81480.1 akaw|GAA91448.1 aluc|OJZ84716.1 anid|CBF71603.1 anig|EHA27988.1 aory|EIT79497.1 asyd|OJJ61857.1 ater|EAU33693.1 atub|OJI89367.1 aver|OJI95956.1 awen|OJJ37365.1 azon|OJJ51045.1

Group4342: abra|OJJ77304.1 acar|OOF93182.1 acla|EAW13162.1 afis|EAW17478.1 afla|EED49336.1 afum|EAL91307.1 agla|OJJ81479.1 akaw|GAA91447.1 aluc|OJZ84717.1 anid|CBF71605.1 anig|EHA27989.1 aory|EIT79535.1 asyd|OJJ61858.1 ater|EAU33694.1 atub|OJI89366.1 aver|OJI95957.1 awen|OJJ37364.1 azon|OJJ51044.1

Group4343: abra|OJJ77305.1 acar|OOF93184.1 acla|EAW13161.1 afis|EAW17479.1 afla|EED49335.1 afum|EAL91308.1 agla|OJJ81478.1 akaw|GAA91446.1 aluc|OJZ84718.1 anid|CBF71607.1 anig|EHA27990.1 aory|EIT79462.1 asyd|OJJ61859.1 ater|EAU33695.1 atub|OJI89365.1 aver|OJI95958.1 awen|OJJ37362.1 azon|OJJ51043.1

Group4344: abra|OJJ77306.1 acar|OOF93185.1 acla|EAW13160.1 afis|EAW17481.1 afla|EED49334.1 afum|EAL91309.1 agla|OJJ81477.1 akaw|GAA91445.1 aluc|OJZ84720.1 anid|CBF71609.1 anig|EHA27991.1 aory|EIT79526.1 asyd|OJJ61860.1 ater|EAU36927.1 atub|OJI89363.1 aver|OJI95959.1 awen|OJJ37357.1 azon|OJJ51041.1

Group4345: abra|OJJ77315.1 acar|OOF93191.1 acla|EAW13157.1 afis|EAW17489.1 afla|EED49327.1 afum|EAL91311.1 agla|OJJ81467.1 akaw|GAA91440.1 aluc|OJZ84725.1 anid|CBF71617.1 anig|EHA27999.1 aory|EIT79549.1 asyd|OJJ61864.1 ater|EAU36934.1 atub|OJI89357.1 aver|OJI95962.1 awen|OJJ37352.1 azon|OJJ51038.1

Group4346: abra|OJJ77318.1 acar|OOF93194.1 acla|EAW13154.1 afis|EAW17492.1 afla|EED49324.1 afum|EAL91314.1 agla|OJJ81464.1 akaw|GAA91438.1 aluc|OJZ84728.1 anid|CBF71599.1 anig|EHA28001.1 aory|EIT79543.1 asyd|OJJ61866.1 ater|EAU36936.1 atub|OJI89354.1 aver|OJI95964.1 awen|OJJ37349.1 azon|OJJ51035.1

Group4347: abra|OJJ77319.1 acar|OOF93195.1 acla|EAW13153.1 afis|EAW17493.1 afla|EED49323.1 afum|EAL91315.1 agla|OJJ81463.1 akaw|GAA91437.1 aluc|OJZ84729.1 anid|CBF71595.1 anig|EHA28002.1 aory|EIT79479.1 asyd|OJJ61868.1 ater|EAU36937.1 atub|OJI89353.1 aver|OJI95966.1 awen|OJJ37348.1 azon|OJJ51034.1

Group4348: abra|OJJ77320.1 acar|OOF93196.1 acla|EAW13152.1 afis|EAW17494.1 afla|EED49322.1 afum|EAL91316.1 agla|OJJ81460.1 akaw|GAA91436.1 aluc|OJZ84730.1 anid|CBF71589.1 anig|EHA28003.1 aory|EIT79493.1 asyd|OJJ61870.1 ater|EAU36938.1 atub|OJI89352.1 aver|OJI95968.1 awen|OJJ37347.1 azon|OJJ51033.1

Group4349: abra|OJJ77326.1 acar|OOF93202.1 acla|EAW13146.1 afis|EAW17518.1 afla|EED49315.1 afum|EAL91336.1 agla|OJJ81452.1 akaw|GAA91430.1 aluc|OJZ84736.1 anid|CBF71576.1 anig|EHA28009.1 aory|EIT82190.1 asyd|OJJ61877.1 ater|EAU36948.1 atub|OJI89345.1 aver|OJI95975.1 awen|OJJ37339.1 azon|OJJ51025.1

Group4350: abra|OJJ77328.1 acar|OOF93203.1 acla|EAW13145.1 afis|EAW17517.1 afla|EED49314.1 afum|EAL91335.1 agla|OJJ81451.1 akaw|GAA91429.1 aluc|OJZ84737.1 anid|CBF71574.1 anig|EHA28010.1 aory|EIT82219.1 asyd|OJJ61878.1 ater|EAU36949.1 atub|OJI89344.1 aver|OJI95976.1 awen|OJJ37338.1 azon|OJJ51024.1

Group4351: abra|OJJ77329.1 acar|OOF93204.1 acla|EAW13144.1 afis|EAW17516.1 afla|EED49313.1 afum|EAL91334.1 agla|OJJ81450.1 akaw|GAA91428.1 aluc|OJZ84738.1 anid|CBF71572.1 anig|EHA28011.1

aory|EIT82242.1 asyd|OJJ61879.1 ater|EAU36950.1 atub|OJI89343.1 aver|OJI95977.1 awen|OJJ37337.1 azon|OJJ51023.1

Group4352: abra|OJJ77334.1 acar|OOF89885.1 acla|EAW13138.1 afis|EAW17507.1 afla|EED49308.1 afum|EAL91329.1 agla|OJJ81445.1 akaw|GAA91424.1 aluc|OJZ84742.1 anid|CBF71533.1 anig|EHA28016.1 aory|EIT82210.1 asyd|OJJ61892.1 ater|EAU36953.1 atub|OJI89339.1 aver|OJI95988.1 awen|OJJ37332.1 azon|OJJ51019.1

Group4353: abra|OJJ77338.1 acar|OOF93209.1 acla|EAW13137.1 afis|EAW17506.1 afla|EED49306.1 afum|EAL91328.2 agla|OJJ81444.1 akaw|GAA91423.1 aluc|OJZ84747.1 anid|CBF71535.1 anig|EHA28017.1 aory|EIT82204.1 asyd|OJJ61889.1 ater|EAU36955.1 atub|OJI89335.1 aver|OJI95986.1 awen|OJJ37329.1 azon|OJJ51018.1

Group4354: abra|OJJ77339.1 acar|OOF93210.1 acla|EAW13136.1 afis|EAW17505.1 afla|EED49305.1 afum|EAL91327.1 agla|OJJ81443.1 akaw|GAA91422.1 aluc|OJZ84748.1 anid|CBF71537.1 anig|EHA28018.1 aory|EIT82207.1 asyd|OJJ61894.1 ater|EAU36956.1 atub|OJI89334.1 aver|OJI95990.1 awen|OJJ37328.1 azon|OJJ51017.1

Group4355: abra|OJJ77341.1 acar|OOF93211.1 acla|EAW13134.1 afis|EAW17503.1 afla|EED49301.1 afum|EBA27430.1 agla|OJJ81441.1 akaw|GAA91420.1 aluc|OJZ84750.1 anid|CBF71541.1 anig|EHA28020.1 aory|EIT82193.1 asyd|OJJ61896.1 ater|EAU36958.1 atub|OJI89332.1 aver|OJI95992.1 awen|OJJ37326.1 azon|OJJ51015.1

Group4356: abra|OJJ77342.1 acar|OOF93212.1 acla|EAW13133.1 afis|EAW17502.1 afla|EED49300.1 afum|EAL91325.1 agla|OJJ81440.1 akaw|GAA91419.1 aluc|OJZ84751.1 anid|CBF71543.1 anig|EHA28021.1 aory|EIT82195.1 asyd|OJJ61897.1 ater|EAU36959.1 atub|OJI89331.1 aver|OJI95993.1 awen|OJJ37325.1 azon|OJJ51014.1

Group4357: abra|OJJ77343.1 acar|OOF93213.1 acla|EAW13132.1 afis|EAW17501.1 afla|EED49299.1 afum|EAL91324.1 agla|OJJ81439.1 akaw|GAA91418.1 aluc|OJZ84753.1 anid|CBF85589.1 anig|EHA28022.1 aory|EIT82233.1 asyd|OJJ64171.1 ater|EAU36960.1 atub|OJI89330.1 aver|OJI98246.1 awen|OJJ37324.1 azon|OJJ51013.1

Group4358: abra|OJJ77344.1 acar|OOF93214.1 acla|EAW13131.1 afis|EAW17532.1 afla|EED49298.1 afum|EAL91346.1 agla|OJJ81438.1 akaw|GAA91417.1 aluc|OJZ84754.1 anid|CBF90335.1 anig|EHA28023.1 aory|EIT82183.1 asyd|OJJ61898.1 ater|EAU36961.1 atub|OJI89329.1 aver|OJI95994.1 awen|OJJ37320.1 azon|OJJ51012.1

Group4359: abra|OJJ77346.1 acar|OOF93216.1 acla|EAW13128.1 afis|EAW17535.1 afla|EED49294.1 afum|EAL91349.1 agla|OJJ81436.1 akaw|GAA91414.1 aluc|OJZ84757.1 anid|CBF90326.1 anig|EHA28026.1 aory|EIT82178.1 asyd|OJJ61907.1 ater|EAU36964.1 atub|OJI89327.1 aver|OJI96000.1 awen|OJJ37318.1 azon|OJJ51010.1

Group4360: abra|OJJ77347.1 acar|OOF93217.1 acla|EAW13127.1 afis|EAW17536.1 afla|EED49293.1 afum|EAL91350.1 agla|OJJ81433.1 akaw|GAA91413.1 aluc|OJZ84758.1 anid|CBF90324.1 anig|EHA28027.1 aory|EIT82228.1 asyd|OJJ61908.1 ater|EAU36965.1 atub|OJI89326.1 aver|OJI96001.1 awen|OJJ37317.1 azon|OJJ51009.1

Group4361: abra|OJJ77348.1 acar|OOF93218.1 acla|EAW13126.1 afis|EAW17537.1 afla|EED49290.1 afum|EAL91351.1 agla|OJJ81432.1 akaw|GAA91412.1 aluc|OJZ84759.1 anid|CBF90322.1 anig|EHA28028.1 aory|EIT82241.1 asyd|OJJ61909.1 ater|EAU36967.1 atub|OJI89325.1 aver|OJI96002.1 awen|OJJ37315.1 azon|OJJ51007.1

Group4362: abra|OJJ77350.1 acar|OOF93220.1 acla|EAW13125.1 afis|EAW17539.1 afla|EED49288.1 afum|EAL91353.1 agla|OJJ81430.1 akaw|GAA91411.1 aluc|OJZ84761.1 anid|CBF90317.1 anig|EHA28030.1 aory|EIT82231.1 asyd|OJJ61911.1 ater|EAU36969.1 atub|OJI89323.1 aver|OJI96004.1 awen|OJJ37314.1 azon|OJJ51006.1

Group4363: abra|OJJ77353.1 acar|OOF93223.1 acla|EAW13121.1 afis|EAW17543.1 afla|EED49284.1 afum|EAL91357.1 agla|OJJ81426.1 akaw|GAA91408.1 aluc|OJZ84764.1 anid|CBF90308.1 anig|EHA28033.1 aory|EIT82220.1 asyd|OJJ61932.1 ater|EAU36973.1 atub|OJI89320.1 aver|OJI96024.1 awen|OJJ37310.1 azon|OJJ51002.1

Group4364: abra|OJJ77355.1 acar|OOF93225.1 acla|EAW13119.1 afis|EAW17545.1 afla|EED49282.1 afum|EAL91359.1 agla|OJJ81424.1 akaw|GAA91406.1 aluc|OJZ84765.1 anid|CBF90305.1 anig|EHA28034.1 aory|EIT82244.1 asyd|OJJ61930.1 ater|EAU36975.1 atub|OJI89319.1 aver|OJI96022.1 awen|OJJ37309.1 azon|OJJ51001.1

Group4365: abra|OJJ77356.1 acar|OOF93226.1 acla|EAW13118.1 afis|EAW17546.1 afla|EED49281.1 afum|EAL91360.1 agla|OJJ81423.1 akaw|GAA91405.1 aluc|OJZ84766.1 anid|CBF90303.1 anig|EHA28035.1 aory|EIT82243.1 asyd|OJJ61929.1 ater|EAU36976.1 atub|OJI89318.1 aver|OJI96021.1 awen|OJJ37308.1 azon|OJJ51000.1

Group4366: abra|OJJ77359.1 acar|OOF93228.1 acla|EAW13117.1 afis|EAW17547.1 afla|EED49278.1 afum|EAL91361.1 agla|OJJ81421.1 akaw|GAA91404.1 aluc|OJZ84767.1 anid|CBF90290.1 anig|EHA28037.1 aory|EIT82197.1 asyd|OJJ61921.1 ater|EAU36977.1 atub|OJI89315.1 aver|OJI96013.1 awen|OJJ37307.1 azon|OJJ50999.1

Group4367: abra|OJJ77361.1 acar|OOF93230.1 acla|EAW13115.1 afis|EAW17549.1 afla|EED49276.1 afum|EAL91363.1 agla|OJJ81419.1 akaw|GAA91402.1 aluc|OJZ84769.1 anid|CBF90294.1 anig|EHA28038.1 aory|EIT82240.1 asyd|OJJ61923.1 ater|EAU36985.1 atub|OJI89312.1 aver|OJI96015.1 awen|OJJ37305.1 azon|OJJ50997.1

Group4368: abra|OJJ77362.1 acar|OOF93231.1 acla|EAW13114.1 afis|EAW17550.1 afla|EED49275.1 afum|EAL91364.1 agla|OJJ81418.1 akaw|GAA91400.1 aluc|OJZ84771.1 anid|CBF90296.1 anig|EHA28039.1 aory|EIT82229.1 asyd|OJJ61924.1 ater|EAU36984.1 atub|OJI89310.1 aver|OJI96016.1 awen|OJJ37303.1 azon|OJJ50996.1

Group4369: abra|OJJ77365.1 acar|OOF93235.1 acla|EAW13111.1 afis|EAW17553.1 afla|EED49272.1 afum|EAL91367.1 agla|OJJ81415.1 akaw|GAA91397.1 aluc|OJZ84774.1 anid|CBF90299.1 anig|EHA28042.1 aory|EIT82208.1 asyd|OJJ61926.1 ater|EAU36981.1 atub|OJI89307.1 aver|OJI96018.1 awen|OJJ37300.1 azon|OJJ50993.1

Group4370: abra|OJJ77366.1 acar|OOF93236.1 acla|EAW13110.1 afis|EAW17554.1 afla|EED49271.1 afum|EAL91368.1 agla|OJJ81414.1 akaw|GAA91396.1 aluc|OJZ84775.1 anid|CBF90301.1 anig|EHA28043.1 aory|EIT82221.1 asyd|OJJ61927.1 ater|EAU36980.1 atub|OJI89306.1 aver|OJI96019.1 awen|OJJ37299.1 azon|OJJ50992.1

Group4371: abra|OJJ77368.1 acar|OOF93238.1 acla|EAW13108.1 afis|EAW17556.1 afla|EED49268.1 afum|EAL91370.1 agla|OJJ81412.1 akaw|GAA91394.1 aluc|OJZ84777.1 anid|CBF90282.1 anig|EHA28045.1 aory|EIT82203.1 asyd|OJJ61918.1 ater|EAU36988.1 atub|OJI89304.1 aver|OJI96010.1 awen|OJJ37297.1 azon|OJJ50990.1

Group4372: abra|OJJ77370.1 acar|OOF93241.1 acla|EAW13105.1 afis|EAW17559.1 afla|EED49265.1 afum|EAL91373.1 agla|OJJ81409.1 akaw|GAA91391.1 aluc|OJZ84780.1 anid|CBF90277.1 anig|EHA28047.1 aory|EIT82185.1 asyd|OJJ61933.1 ater|EAU36990.1 atub|OJI89302.1 aver|OJI96025.1 awen|OJJ37295.1 azon|OJJ50988.1

Group4373: abra|OJJ77371.1 acar|OOF93240.1 acla|EAW13104.1 afis|EAW17560.1 afla|EED49264.1 afum|EAL91374.1 agla|OJJ81408.1 akaw|GAA91390.1 aluc|OJZ84781.1 anid|CBF90275.1 anig|EHA28048.1 aory|EIT82187.1 asyd|OJJ61934.1 ater|EAU36991.1 atub|OJI89301.1 aver|OJI96026.1 awen|OJJ37294.1 azon|OJJ50987.1

Group4374: abra|OJJ77372.1 acar|OOF93242.1 acla|EAW13103.1 afis|EAW17561.1 afla|EED49263.1 afum|EAL91375.1 agla|OJJ81407.1 akaw|GAA91389.1 aluc|OJZ84782.1 anid|CBF90273.1 anig|EHA28049.1 aory|EIT82191.1 asyd|OJJ61935.1 ater|EAU36992.1 atub|OJI89300.1 aver|OJI96027.1 awen|OJJ37293.1 azon|OJJ50986.1

Group4375: abra|OJJ77373.1 acar|OOF93244.1 acla|EAW13102.1 afis|EAW17562.1 afla|EED49259.1 afum|EAL91376.1 agla|OJJ81406.1 akaw|GAA91387.1 aluc|OJZ84783.1 anid|CBF90271.1 anig|EHA28050.1 aory|EIT82238.1 asyd|OJJ61936.1 ater|EAU37000.1 atub|OJI89299.1 aver|OJI96028.1 awen|OJJ37292.1 azon|OJJ50985.1

Group4376: abra|OJJ77374.1 acar|OOF93243.1 acla|EAW13101.1 afis|EAW17563.1 afla|EED49258.1 afum|EAL91377.1 agla|OJJ81405.1 akaw|GAA91386.1 aluc|OJZ84784.1 anid|CBF90269.1 anig|EHA28051.1 aory|EIT82186.1 asyd|OJJ61937.1 ater|EAU37001.1 atub|OJI89298.1 aver|OJI96029.1 awen|OJJ37291.1 azon|OJJ50984.1

Group4377: abra|OJJ77375.1 acar|OOF93245.1 acla|EAW13100.1 afis|EAW17564.1 afla|EED49257.1 afum|EAL91378.1 agla|OJJ81402.1 akaw|GAA91385.1 aluc|OJZ84785.1 anid|CBF90267.1 anig|EHA28052.1 aory|EIT82246.1 asyd|OJJ61938.1 ater|EAU37002.1 atub|OJI89297.1 aver|OJI96030.1 awen|OJJ37290.1 azon|OJJ50983.1

Group4378: abra|OJJ77376.1 acar|OOF93246.1 acla|EAW13098.1 afis|EAW17565.1 afla|EED49256.1 afum|EAL91379.1 agla|OJJ81401.1 akaw|GAA91384.1 aluc|OJZ84786.1 anid|CBF90266.1 anig|EHA28053.1 aory|EIT82227.1 asyd|OJJ61939.1 ater|EAU37003.1 atub|OJI89296.1 aver|OJI96031.1 awen|OJJ37288.1 azon|OJJ50982.1

Group4379: abra|OJJ77378.1 acar|OOF93247.1 acla|EAW13096.1 afis|EAW17567.1 afla|EED49254.1 afum|EAL91381.1 agla|OJJ81398.1 akaw|GAA91382.1 aluc|OJZ84787.1 anid|CBF90262.1 anig|EHA28055.1 aory|EIT82184.1 asyd|OJJ61941.1 ater|EAU37004.1 atub|OJI89295.1 aver|OJI96033.1 awen|OJJ37286.1 azon|OJJ50980.1

Group4380: abra|OJJ77476.1 acar|OOF99249.1 acla|EAW06738.1 afis|EAW16288.1 afla|EED53265.1 afum|EAL88910.1 agla|OJJ88310.1 akaw|GAA84142.1 aluc|OJZ81062.1 anid|CBF86860.1 anig|EHA19306.1

aory|EIT75107.1 asyd|OJJ64634.1 ater|EAU37333.1 atub|OJI82523.1 aver|OJI98733.1 awen|OJJ37692.1 azon|OJJ44952.1

Group4381: abra|OJJ77477.1 acar|OOF99248.1 acla|EAW06739.1 afis|EAW16289.1 afla|EED53264.1 afum|EAL88911.1 agla|OJJ88311.1 akaw|GAA84141.1 aluc|OJZ81061.1 anid|CBF86858.1 anig|EHA19307.1 aory|EIT74825.1 asyd|OJJ64635.1 ater|EAU37334.1 atub|OJI82524.1 aver|OJI98734.1 awen|OJJ37691.1 azon|OJJ44951.1

Group4382: abra|OJJ77478.1 acar|OOF99247.1 acla|EAW06740.1 afis|EAW16290.1 afla|EED53263.1 afum|EAL88912.1 agla|OJJ88312.1 akaw|GAA84140.1 aluc|OJZ81060.1 anid|CBF86856.1 anig|EHA19308.1 aory|EIT75015.1 asyd|OJJ64636.1 ater|EAU37335.1 atub|OJI82525.1 aver|OJI98735.1 awen|OJJ37690.1 azon|OJJ44950.1

Group4383: abra|OJJ77479.1 acar|OOF99246.1 acla|EAW06741.1 afis|EAW16291.1 afla|EED53262.1 afum|EAL88913.1 agla|OJJ88313.1 akaw|GAA84139.1 aluc|OJZ81059.1 anid|CBF86854.1 anig|EHA19309.1 aory|EIT74947.1 asyd|OJJ64637.1 ater|EAU37336.1 atub|OJI82526.1 aver|OJI98736.1 awen|OJJ37689.1 azon|OJJ44949.1

Group4384: abra|OJJ77485.1 acar|OOF99240.1 acla|EAW06746.1 afis|EAW16296.1 afla|EED53258.1 afum|EAL88918.1 agla|OJJ88303.1 akaw|GAA84133.1 aluc|OJZ81051.1 anid|CBF86847.1 anig|EHA19316.1 aory|EIT75226.1 asyd|OJJ64642.1 ater|EAU29340.1 atub|OJI82533.1 aver|OJI98742.1 awen|OJJ37695.1 azon|OJJ44956.1

Group4385: abra|OJJ77489.1 acar|OOF99236.1 acla|EAW06751.1 afis|EAW16301.1 afla|EED53253.1 afum|EAL88923.1 agla|OJJ88293.1 akaw|GAA84130.1 aluc|OJZ81047.1 anid|CBF86841.1 anig|EHA19320.1 aory|EIT74998.1 asyd|OJJ64646.1 ater|EAU37346.1 atub|OJI82537.1 aver|OJI98746.1 awen|OJJ37700.1 azon|OJJ44961.1

Group4386: abra|OJJ77490.1 acar|OOF99235.1 acla|EAW06752.1 afis|EAW16302.1 afla|EED53252.1 afum|EAL88924.1 agla|OJJ88292.1 akaw|GAA84129.1 aluc|OJZ81046.1 anid|CBF86839.1 anig|EHA19321.1 aory|EIT75063.1 asyd|OJJ64647.1 ater|EAU37345.1 atub|OJI82538.1 aver|OJI98747.1 awen|OJJ37701.1 azon|OJJ44962.1

Group4387: abra|OJJ77493.1 acar|OOF99233.1 acla|EAW06754.1 afis|EAW16304.1 afla|EED53250.1 afum|EAL88926.1 agla|OJJ88290.1 akaw|GAA84127.1 aluc|OJZ81044.1 anid|CBF69582.1 anig|EHA19323.1 aory|EIT75043.1 asyd|OJJ59144.1 ater|EAU29332.1 atub|OJI82540.1 aver|OJJ01734.1 awen|OJJ37704.1 azon|OJJ44964.1

Group4388: abra|OJJ77494.1 acar|OOF99232.1 acla|EAW06755.1 afis|EAW16305.1 afla|EED53249.1 afum|EAL88927.1 agla|OJJ88267.1 akaw|GAA84126.1 aluc|OJZ81043.1 anid|CBF69583.1 anig|EHA19324.1 aory|EIT75153.1 asyd|OJJ59145.1 ater|EAU29333.1 atub|OJI82541.1 aver|OJJ01735.1 awen|OJJ37705.1 azon|OJJ44965.1

Group4390: abra|OJJ77499.1 acar|OOF99227.1 acla|EAW06736.1 afis|EAW16287.1 afla|EED53269.1 afum|EAL88909.1 agla|OJJ88309.1 akaw|GAA84122.1 aluc|OJZ81039.1 anid|CBF86861.1 anig|EHA19328.1 aory|EIT74855.1 asyd|OJJ64633.1 ater|EAU37332.1 atub|OJI82545.1 aver|OJI98732.1 awen|OJJ37682.1 azon|OJJ44954.1

Group4391: abra|OJJ77502.1 acar|OOF99225.1 acla|EAW06734.1 afis|EAW16285.1 afla|EED53271.1 afum|EAL88906.1 agla|OJJ88306.1 akaw|GAA84119.1 aluc|OJZ81035.1 anid|CBF86869.1 anig|EHA19331.1 aory|EIT75236.1 asyd|OJJ64631.1 ater|EAU37329.1 atub|OJI82548.1 aver|OJI98729.1 awen|OJJ37679.1 azon|OJJ44945.1

Group4392: abra|OJJ77503.1 acar|OOF99224.1 acla|EAW06732.1 afis|EAW16284.1 afla|EED53272.1 afum|EAL88905.1 agla|OJJ88305.1 akaw|GAA84118.1 aluc|OJZ81034.1 anid|CBF86871.1 anig|EHA19332.1 aory|EIT75050.1 asyd|OJJ64630.1 ater|EAU37328.1 atub|OJI82549.1 aver|OJI98728.1 awen|OJJ37677.1 azon|OJJ44942.1

Group4393: abra|OJJ77504.1 acar|OOF99222.1 acla|EAW06730.1 afis|EAW16283.1 afla|EED53273.1 afum|EAL88904.1 agla|OJJ88318.1 akaw|GAA84116.1 aluc|OJZ81032.1 anid|CBF86876.1 anig|EHA19333.1 aory|EIT75123.1 asyd|OJJ64627.1 ater|EAU37327.1 atub|OJI82551.1 aver|OJI98725.1 awen|OJJ37675.1 azon|OJJ44940.1

Group4394: abra|OJJ77507.1 acar|OOF99219.1 acla|EAW06727.1 afis|EAW16280.1 afla|EED53276.1 afum|EAL88901.1 agla|OJJ88324.1 akaw|GAA84113.1 aluc|OJZ81029.1 anid|CBF86880.1 anig|EHA19336.1 aory|EIT74958.1 asyd|OJJ64626.1 ater|EAU37325.1 atub|OJI82554.1 aver|OJI98722.1 awen|OJJ37672.1 azon|OJJ44937.1

Group4395: abra|OJJ77508.1 acar|OOF99218.1 acla|EAW06726.1 afis|EAW16279.1 afla|EED53277.1 afum|EAL88900.1 agla|OJJ88323.1 akaw|GAA84112.1 aluc|OJZ81028.1 anid|CBF86882.1 anig|EHA19337.1 aory|EIT74848.1 asyd|OJJ64625.1 ater|EAU37324.1 atub|OJI82555.1 aver|OJI98721.1 awen|OJJ37671.1 azon|OJJ44936.1

Group4396: abra|OJJ77509.1 acar|OOF99217.1 acla|EAW06725.1 afis|EAW16278.1 afla|EED53279.1 afum|EAL88899.1 agla|OJJ88321.1 akaw|GAA84111.1 aluc|OJZ81027.1 anid|CBF86883.1 anig|EHA19338.1 aory|EIT74824.1 asyd|OJJ64624.1 ater|EAU37323.1 atub|OJI82556.1 aver|OJI98720.1 awen|OJJ37670.1 azon|OJJ44935.1

Group4397: abra|OJJ77510.1 acar|OOF99216.1 acla|EAW06724.1 afis|EAW16277.1 afla|EED53280.1 afum|EAL88898.1 agla|OJJ88328.1 akaw|GAA84110.1 aluc|OJZ81026.1 anid|CBF86885.1 anig|EHA19339.1 aory|EIT75207.1 asyd|OJJ64623.1 ater|EAU37322.1 atub|OJI82557.1 aver|OJI98719.1 awen|OJJ37669.1 azon|OJJ44934.1

Group4398: abra|OJJ77511.1 acar|OOF99214.1 acla|EAW06723.1 afis|EAW16276.1 afla|EED53281.1 afum|EAL88897.1 agla|OJJ88329.1 akaw|GAA84109.1 aluc|OJZ81025.1 anid|CBF86892.1 anig|EHA18669.1 aory|EIT75064.1 asyd|OJJ64622.1 ater|EAU37321.1 atub|OJI82558.1 aver|OJI98718.1 awen|OJJ37668.1 azon|OJJ44933.1

Group4399: abra|OJJ77513.1 acar|OOF99211.1 acla|EAW06722.1 afis|EAW16265.1 afla|EED53285.1 afum|EAL88886.1 agla|OJJ88330.1 akaw|GAA84107.1 aluc|OJZ81022.1 anid|CBF86894.1 anig|EHA19341.1 aory|EIT74876.1 asyd|OJJ64621.1 ater|EAU37438.1 atub|OJI82561.1 aver|OJI98717.1 awen|OJJ37667.1 azon|OJJ44930.1

Group4400: abra|OJJ77514.1 acar|OOF99210.1 acla|EAW06721.1 afis|EAW16266.1 afla|EED53284.1 afum|EAL88887.1 agla|OJJ88331.1 akaw|GAA84106.1 aluc|OJZ81021.1 anid|CBF86895.1 anig|EHA19342.1 aory|EIT75173.1 asyd|OJJ64620.1 ater|EAU37439.1 atub|OJI82562.1 aver|OJI98716.1 awen|OJJ37666.1 azon|OJJ44931.1

Group4401: abra|OJJ77516.1 acar|OOF99207.1 acla|EAW06720.1 afis|EAW16267.1 afla|EED53282.1 afum|EAL88888.1 agla|OJJ88333.1 akaw|GAA84104.1 aluc|OJZ81019.1 anid|CBF86897.1 anig|EHA19343.1 aory|EIT75090.1 asyd|OJJ64618.1 ater|EAU37440.1 atub|OJI82564.1 aver|OJI98714.1 awen|OJJ37664.1 azon|OJJ44923.1

Group4402: abra|OJJ77517.1 acar|OOF99208.1 acla|EAW06719.1 afis|EAW16268.1 afla|EED53288.1 afum|EAL88889.1 agla|OJJ88337.1 akaw|GAA84103.1 aluc|OJZ81018.1 anid|CBF86899.1 anig|EHA19344.1 aory|EIT74865.1 asyd|OJJ64617.1 ater|EAU37443.1 atub|OJI82565.1 aver|OJI98713.1 awen|OJJ37663.1 azon|OJJ44924.1

Group4403: abra|OJJ77520.1 acar|OOF99203.1 acla|EAW06715.1 afla|EED53293.1 afum|EAL88893.1 akaw|GAA84100.1 aluc|OJZ81016.1 anid|CBF86904.1 anig|EHA19346.1 aory|EIT74861.1 asyd|OJJ64613.1 atub|OJI82568.1 aver|OJI98709.1 awen|OJJ37656.1 azon|OJJ44929.1 afis|EAW16272.1 agla|OJJ88342.1 ater|EAU37447.1

Group4404: abra|OJJ77522.1 acar|OOF99202.1 acla|EAW06717.1 afis|EAW16270.1 afla|EED53295.1 afum|EAL88891.1 agla|OJJ88339.1 akaw|GAA84098.1 aluc|OJZ81014.1 anid|CBF86903.1 anig|EHA19347.1 aory|EIT74914.1 asyd|OJJ64615.1 ater|EAU37445.1 atub|OJI82570.1 aver|OJI98711.1 awen|OJJ37660.1 azon|OJJ44927.1

Group4405: abra|OJJ77524.1 acar|OOF99200.1 acla|EAW06718.1 afis|EAW16269.1 afla|EED53289.1 afum|EAL88890.1 agla|OJJ88338.1 akaw|GAA84095.1 aluc|OJZ81011.1 anid|CBF86901.1 anig|EHA19350.1 aory|EIT74922.1 asyd|OJJ64616.1 ater|EAU37444.1 atub|OJI82573.1 aver|OJI98712.1 awen|OJJ37661.1 azon|OJJ44926.1

Group4406: abra|OJJ77526.1 acar|OOF99198.1 acla|EAW06712.1 afis|EAW16275.1 afla|EED53297.1 afum|EAL88896.1 agla|OJJ88345.1 akaw|GAA84094.1 aluc|OJZ81009.1 anid|CBF86909.1 anig|EHA19351.1 aory|EIT75079.1 asyd|OJJ64610.1 ater|EAU37450.1 atub|OJI82575.1 aver|OJI98706.1 awen|OJJ37658.1 azon|OJJ44920.1

Group4407: abra|OJJ77535.1 acar|OOF99189.1 acla|EAW06704.1 afis|EAW16249.1 afla|EED53313.1 afum|EAL88877.1 agla|OJJ86260.1 akaw|GAA84085.1 aluc|OJZ81000.1 anid|CBF78891.1 anig|EHA19359.1 aory|EIT74965.1 asyd|OJJ64597.1 ater|EAU37466.1 atub|OJI82585.1 aver|OJI98694.1 awen|OJJ30206.1 azon|OJJ43931.1

Group4408: abra|OJJ77554.1 acar|OOF99173.1 acla|EAW06523.1 afis|EAW16239.1 afla|EED53338.1 afum|EAL88860.1 agla|OJJ86246.1 akaw|GAA84069.1 aluc|OJZ80983.1 anid|CBF78863.1 anig|EHA19377.1 aory|EIT75140.1 asyd|OJJ64574.1 ater|EAU33704.1 atub|OJI82603.1 aver|OJI98671.1 awen|OJJ30218.1 azon|OJJ43940.1

Group4410: abra|OJJ77561.1 acar|OOF99169.1 acla|EAW06694.1 afis|EAW16072.1 afla|EED48591.1 afum|EAL88704.1 agla|OJJ86096.1 akaw|GAA84064.1 aluc|OJZ80977.1 anid|CBF75113.1 anig|EHA19382.1 aory|EIT73146.1 asyd|OJJ59998.1 ater|EAU33559.1 atub|OJI82608.1 aver|OJJ02293.1 awen|OJJ30362.1 azon|OJJ45381.1

Group4411: abra|OJJ77566.1 acar|OOF99164.1 acla|EAW06531.1 afis|EAW16064.1 afla|EED48578.1 afum|EAL88696.1 agla|OJJ86101.1 akaw|GAA84060.1 aluc|OJZ80972.1 anid|CBF75092.1 anig|EHA19387.1

aory|EIT73042.1 asyd|OJJ60011.1 ater|EAU33571.1 atub|OJI82613.1 aver|OJJ02274.1 awen|OJJ30375.1 azon|OJJ45391.1

Group4412: abra|OJJ77567.1 acar|OOF99162.1 acla|EAW06532.1 afis|EAW16063.1 afla|EED48577.1 afum|EAL88695.1 agla|OJJ86102.1 akaw|GAA84059.1 aluc|OJZ80971.1 anid|CBF75090.1 anig|EHA19388.1 aory|EIT73179.1 asyd|OJJ60012.1 ater|EAU33572.1 atub|OJI82614.1 aver|OJJ02273.1 awen|OJJ30376.1 azon|OJJ45392.1

Group4413: abra|OJJ77568.1 acar|OOF99163.1 acla|EAW06533.1 afis|EAW16062.1 afla|EED48576.1 afum|EAL88694.1 agla|OJJ86103.1 akaw|GAA84058.1 aluc|OJZ80970.1 anid|CBF75088.1 anig|EHA19389.1 aory|EIT73163.1 asyd|OJJ60013.1 ater|EAU33573.1 atub|OJI82615.1 aver|OJJ02272.1 awen|OJJ30377.1 azon|OJJ45393.1

Group4414: abra|OJJ77572.1 acar|OOF99157.1 acla|EAW06539.1 afis|EAW16056.1 afla|EED56442.1 afum|EAL88688.1 agla|OJJ86089.1 akaw|GAA84054.1 aluc|OJZ80966.1 anid|CBF75076.1 anig|EHA19392.1 aory|EIT72786.1 asyd|OJJ60016.1 ater|EAU33578.1 atub|OJI82620.1 aver|OJJ02269.1 awen|OJJ30383.1 azon|OJJ45400.1

Group4415: abra|OJJ77573.1 acar|OOF99156.1 acla|EAW06538.1 afis|EAW16057.1 afla|EED56441.1 afum|EAL88689.1 agla|OJJ86090.1 akaw|GAA84053.1 aluc|OJZ80965.1 anid|CBF75078.1 anig|EHA19393.1 aory|EIT72787.1 asyd|OJJ60017.1 ater|EAU33577.1 atub|OJI82621.1 aver|OJJ02268.1 awen|OJJ30382.1 azon|OJJ45399.1

Group4416: abra|OJJ77574.1 acar|OOF99155.1 acla|EAW06537.1 afis|EAW16058.1 afla|EED48572.1 afum|EAL88690.2 agla|OJJ86107.1 akaw|GAA84051.1 aluc|OJZ80963.1 anid|CBF75080.1 anig|EHA19394.1 aory|EIT73249.1 asyd|OJJ60018.1 ater|EAU33576.1 atub|OJI82623.1 aver|OJJ02267.1 awen|OJJ30381.1 azon|OJJ45398.1

Group4417: abra|OJJ77577.1 acar|OOF99150.1 acla|EAW06543.1 afis|EAW16052.1 afla|EED56446.1 afum|EAL88684.1 agla|OJJ86086.1 akaw|GAA84048.1 aluc|OJZ80959.1 anid|CBF75068.1 anig|EHA19397.1 aory|EIT72800.1 asyd|OJJ60022.1 ater|EAU33581.1 atub|OJI82626.1 aver|OJJ02262.1 awen|OJJ30390.1 azon|OJJ45402.1

Group4418: abra|OJJ77579.1 acar|OOF99148.1 acla|EAW06545.1 afis|EAW16049.1 afla|EED56438.1 afum|EAL88682.1 agla|OJJ86083.1 akaw|GAA84046.1 aluc|OJZ80956.1 anid|CBF75064.1 anig|EHA19399.1 aory|EIT72808.1 asyd|OJJ60024.1 ater|EAU33587.1 atub|OJI82628.1 aver|OJJ02260.1 awen|OJJ30394.1 azon|OJJ45404.1

Group4419: abra|OJJ77580.1 acar|OOF99147.1 acla|EAW06546.1 afis|EAW16048.1 afla|EED56437.1 afum|EAL88681.1 agla|OJJ86082.1 akaw|GAA84045.1 aluc|OJZ80955.1 anid|CBF75062.1 anig|EHA19400.1 aory|EIT72865.1 asyd|OJJ60025.1 ater|EAU33588.1 atub|OJI82629.1 aver|OJJ02259.1 awen|OJJ30396.1 azon|OJJ45405.1

Group4420: abra|OJJ77581.1 acar|OOF99146.1 acla|EAW06547.1 afis|EAW16047.1 afla|EED56436.1 afum|EAL88680.1 agla|OJJ86081.1 akaw|GAA84044.1 aluc|OJZ80954.1 anid|CBF75060.1 anig|EHA19401.1 aory|EIT72831.1 asyd|OJJ60026.1 ater|EAU33590.1 atub|OJI82630.1 aver|OJJ02258.1 awen|OJJ30398.1 azon|OJJ45406.1

Group4421: abra|OJJ77585.1 acar|OOF92640.1 acla|EAW06551.1 afis|EAW16044.1 afla|EED56433.1 afum|EAL88676.1 agla|OJJ86077.1 akaw|GAA84041.1 aluc|OJZ80950.1 anid|CBF75054.1 anig|EHA19404.1 aory|EIT72801.1 asyd|OJJ60029.1 ater|EAU33593.1 atub|OJI82633.1 aver|OJJ02255.1 awen|OJJ30402.1 azon|OJJ45410.1

Group4422: abra|OJJ77586.1 acar|OOF99144.1 acla|EAW06552.1 afis|EAW16043.1 afla|EED56432.1 afum|EAL88675.1 agla|OJJ86076.1 akaw|GAA84040.1 aluc|OJZ80949.1 anid|CBF75052.1 anig|EHA19405.1 aory|EIT72860.1 asyd|OJJ60030.1 ater|EAU33594.1 atub|OJI82634.1 aver|OJJ02254.1 awen|OJJ30403.1 azon|OJJ45411.1

Group4423: abra|OJJ77588.1 acar|OOF99143.1 acla|EAW06554.1 afis|EAW16041.1 afla|EED56430.1 afum|EAL88673.1 agla|OJJ86074.1 akaw|GAA84038.1 aluc|OJZ80947.1 anid|CBF75050.1 anig|EHA19407.1 aory|EIT72854.1 asyd|OJJ60032.1 ater|EAU33596.1 atub|OJI82636.1 aver|OJJ02252.1 awen|OJJ30405.1 azon|OJJ45413.1

Group4424: abra|OJJ77590.1 acar|OOF99141.1 acla|EAW06556.1 afis|EAW16039.1 afla|EED56427.1 afum|EAL88671.1 agla|OJJ86072.1 akaw|GAA84036.1 aluc|OJZ80945.1 anid|CBF75046.1 anig|EHA19409.1 aory|EIT72877.1 asyd|OJJ60034.1 ater|EAU33599.1 atub|OJI82638.1 aver|OJJ02250.1 awen|OJJ30407.1 azon|OJJ45415.1

Group4425: abra|OJJ77598.1 acar|OOF99128.1 acla|EAW06562.1 afis|EAW16034.1 afla|EED56420.1 afum|EAL88665.1 agla|OJJ86066.1 akaw|GAA84030.1 aluc|OJZ80935.1 anid|CBF75036.1 anig|EHA19415.1 aory|EIT72883.1 asyd|OJJ60049.1 ater|EAU33606.1 atub|OJI82645.1 aver|OJJ02240.1 awen|OJJ30413.1 azon|OJJ45421.1

Group4426: abra|OJJ77601.1 acar|OOF99126.1 acla|EAW06565.1 afis|EAW16031.1 afla|EED56413.1 afum|EAL88662.1 agla|OJJ86059.1 akaw|GAA84027.1 aluc|OJZ80932.1 anid|CBF75030.1 anig|EHA19418.1 aory|EIT72887.1 asyd|OJJ60052.1 ater|EAU33610.1 atub|OJI82648.1 aver|OJJ02237.1 awen|OJJ30417.1 azon|OJJ45424.1

Group4427: abra|OJJ77602.1 acar|OOF99124.1 acla|EAW06567.1 afis|EAW16029.1 afla|EED56412.1 afum|EAL88660.1 agla|OJJ86057.1 akaw|GAA84026.1 aluc|OJZ80931.1 anid|CBF75016.1 anig|EHA19419.1 aory|EIT72882.1 asyd|OJJ60070.1 ater|EAU33611.1 atub|OJI82649.1 aver|OJJ04192.1 awen|OJJ30419.1 azon|OJJ45426.1

Group4428: abra|OJJ77603.1 acar|OOF99123.1 acla|EAW06568.1 afis|EAW16028.1 afla|EED56411.1 afum|EAL88658.1 agla|OJJ86056.1 akaw|GAA84025.1 aluc|OJZ80930.1 anid|CBF75018.1 anig|EHA19420.1 aory|EIT72881.1 asyd|OJJ60069.1 ater|EAU33612.1 atub|OJI82650.1 aver|OJJ02226.1 awen|OJJ30420.1 azon|OJJ45427.1

Group4429: abra|OJJ77605.1 acar|OOF99122.1 acla|EAW06569.1 afis|EAW16027.1 afla|EED56410.1 afum|EAL88657.1 agla|OJJ86055.1 akaw|GAA84024.1 aluc|OJZ80928.1 anid|CBF75020.1 anig|EHA19421.1 aory|EIT83014.1 asyd|OJJ60068.1 ater|EAU33613.1 atub|OJI82651.1 aver|OJJ02227.1 awen|OJJ30421.1 azon|OJJ45428.1

Group4430: abra|OJJ77613.1 acar|OOF99115.1 acla|EAW06571.1 afis|EAW16025.1 afla|EED56406.1 afum|EAL88654.1 agla|OJJ86053.1 akaw|GAA84018.1 aluc|OJZ80921.1 anid|CBF75014.1 anig|EHA19426.1 aory|EIT83006.1 asyd|OJJ60057.1 ater|EAU33616.1 atub|OJI82659.1 aver|OJJ02232.1 awen|OJJ30432.1 azon|OJJ45431.1

Group4431: abra|OJJ77614.1 acar|OOF90324.1 acla|EAW11334.1 afis|EAW18161.1 afla|EED56332.1 afum|EAL87521.1 agla|OJJ85965.1 akaw|GAA84017.1 aluc|OJZ80920.1 anid|CBF85769.1 anig|EHA19427.1 aory|EIT73455.1 asyd|OJJ64306.1 ater|EAU33688.1 atub|OJI82660.1 aver|OJI98399.1 awen|OJJ30506.1 azon|OJJ45495.1

Group4432: abra|OJJ77615.1 acar|OOF90323.1 acla|EAW11335.1 afis|EAW18160.1 afla|EED56333.1 afum|EBA27314.1 agla|OJJ85966.1 akaw|GAA84016.1 aluc|OJZ80919.1 anid|CBF85772.1 anig|EHA19428.1 aory|EIT73306.1 asyd|OJJ64308.1 ater|EAU33687.1 atub|OJI82661.1 aver|OJI98400.1 awen|OJJ30505.1 azon|OJJ45494.1

Group4433: abra|OJJ77616.1 acar|OOF90325.1 acla|EAW11336.1 afis|EAW18159.1 afla|EED56335.1 afum|EAL87518.1 agla|OJJ85968.1 akaw|GAA84015.1 aluc|OJZ80918.1 anid|CBF85776.1 anig|EHA19429.1 aory|EIT73326.1 asyd|OJJ64310.1 ater|EAU33686.1 atub|OJI82662.1 aver|OJI98403.1 awen|OJJ30504.1 azon|OJJ45493.1

Group4434: abra|OJJ77617.1 acar|OOF90326.1 acla|EAW11337.1 afis|EAW18158.1 afla|EED56336.1 afum|EAL87517.1 agla|OJJ85969.1 akaw|GAA84014.1 aluc|OJZ80917.1 anid|CBF85777.1 anig|EHA19430.1 aory|EIT73425.1 asyd|OJJ64311.1 ater|EAU33685.1 atub|OJI82663.1 aver|OJI98404.1 awen|OJJ30503.1 azon|OJJ45492.1

Group4435: abra|OJJ77618.1 acar|OOF90327.1 acla|EAW11338.1 afis|EAW18157.1 afla|EED56337.1 afum|EAL87516.1 agla|OJJ85970.1 akaw|GAA84013.1 aluc|OJZ80916.1 anid|CBF85779.1 anig|EHA19431.1 aory|EIT73376.1 asyd|OJJ64312.1 ater|EAU33684.1 atub|OJI82664.1 aver|OJI98405.1 awen|OJJ30502.1 azon|OJJ45491.1

Group4436: abra|OJJ77619.1 acar|OOF90328.1 acla|EAW11339.1 afis|EAW18156.1 afla|EED56338.1 afum|EAL87515.1 agla|OJJ85971.1 akaw|GAA84012.1 aluc|OJZ80915.1 anid|CBF85781.1 anig|EHA19432.1 aory|EIT73330.1 asyd|OJJ64313.1 ater|EAU33683.1 atub|OJI82665.1 aver|OJI98406.1 awen|OJJ30500.1 azon|OJJ45490.1

Group4437: abra|OJJ77620.1 acar|OOF90329.1 acla|EAW11340.1 afis|EAW18155.1 afla|EED56339.1 afum|EAL87514.1 agla|OJJ85972.1 akaw|GAA84011.1 aluc|OJZ80914.1 anid|CBF85783.1 anig|EHA19433.1 aory|EIT73446.1 asyd|OJJ64314.1 ater|EAU33682.1 atub|OJI82666.1 aver|OJI98407.1 awen|OJJ30499.1 azon|OJJ45489.1

Group4438: abra|OJJ77621.1 acar|OOF99114.1 acla|EAW11341.1 afis|EAW18154.1 afla|EED56340.1 afum|EAL87513.1 agla|OJJ85973.1 akaw|GAA84010.1 aluc|OJZ80913.1 anid|CBF85784.1 anig|EHA19434.1 aory|EIT73328.1 asyd|OJJ64316.1 ater|EAU33681.1 atub|OJI82667.1 aver|OJI98409.1 awen|OJJ30498.1 azon|OJJ45488.1

Group4439: abra|OJJ77626.1 acar|OOF99112.1 acla|EAW11342.1 afis|EAW18152.1 afla|EED56342.1 afum|EAL87511.1 agla|OJJ85975.1 akaw|GAA84007.1 aluc|OJZ80905.1 anid|CBF85790.1 anig|EHA19436.1 aory|EIT73287.1 asyd|OJJ64325.1 ater|EAU33677.1 atub|OJI82672.1 aver|OJI98419.1 awen|OJJ30495.1 azon|OJJ45487.1

Group4440: abra|OJJ77629.1 acar|OOF99097.1 acla|EAW06480.1 afis|EAW15975.1 afla|EED56345.1 afum|EAL88607.1 agla|OJJ85980.1 akaw|GAA84005.1 aluc|OJZ80901.1 anid|CBF85795.1 anig|EHA19438.1

aory|EIT73353.1 asyd|OJJ64330.1 ater|EAU33676.1 atub|OJI82678.1 aver|OJI98423.1 awen|OJJ30494.1 azon|OJJ45485.1

Group4441: abra|OJJ77630.1 acar|OOF99098.1 acla|EAW06481.1 afis|EAW15976.1 afla|EED56346.1 afum|EAL88608.1 agla|OJJ85981.1 akaw|GAA84004.1 aluc|OJZ80900.1 anid|CBF85797.1 anig|EHA19439.1 aory|EIT73314.1 asyd|OJJ64331.1 ater|EAU33675.1 atub|OJI82679.1 aver|OJI98424.1 awen|OJJ30493.1 azon|OJJ45484.1

Group4442: abra|OJJ77631.1 acar|OOF99099.1 acla|EAW06482.1 afis|EAW15977.1 afla|EED56347.1 afum|EAL88609.1 agla|OJJ85982.1 akaw|GAA84003.1 aluc|OJZ80898.1 anid|CBF85799.1 anig|EHA19440.1 aory|EIT73397.1 asyd|OJJ64332.1 ater|EAU33674.1 atub|OJI82680.1 aver|OJI98425.1 awen|OJJ30492.1 azon|OJJ45483.1

Group4443: abra|OJJ77632.1 acar|OOF99100.1 acla|EAW06484.1 afis|EAW15979.1 afla|EED56349.1 afum|EAL88611.1 agla|OJJ85984.1 akaw|GAA84001.1 aluc|OJZ80897.1 anid|CBF85803.1 anig|EHA19442.1 aory|EIT73329.1 asyd|OJJ64334.1 ater|EAU33672.1 atub|OJI82681.1 aver|OJI98427.1 awen|OJJ30491.1 azon|OJJ45482.1

Group4444: abra|OJJ77637.1 acar|OOF99105.1 acla|EAW06487.1 afis|EAW15982.1 afla|EED56352.1 afum|EAL88614.1 agla|OJJ85985.1 akaw|GAA83997.1 aluc|OJZ80893.1 anid|CBF85806.1 anig|EHA19445.1 aory|EIT73461.1 asyd|OJJ64337.1 ater|EAU33670.1 atub|OJI82685.1 aver|OJI98430.1 awen|OJJ30488.1 azon|OJJ45479.1

Group4445: abra|OJJ77638.1 acar|OOF99106.1 acla|EAW06488.1 afis|EAW15983.1 afla|EED56353.1 afum|EAL88615.1 agla|OJJ85986.1 akaw|GAA83996.1 aluc|OJZ80892.1 anid|CBF85808.1 anig|EHA19446.1 aory|EIT73319.1 asyd|OJJ64339.1 ater|EAU33669.1 atub|OJI82686.1 aver|OJI98432.1 awen|OJJ30487.1 azon|OJJ45478.1

Group4446: abra|OJJ77640.1 acar|OOF99108.1 acla|EAW06490.1 afis|EAW15985.1 afla|EED56355.1 afum|EAL88617.1 agla|OJJ85988.1 akaw|GAA83994.1 aluc|OJZ80890.1 anid|CBF85812.1 anig|EHA19448.1 aory|EIT73389.1 asyd|OJJ64341.1 ater|EAU33667.1 atub|OJI82688.1 aver|OJI98434.1 awen|OJJ30485.1 azon|OJJ45476.1

Group4447: abra|OJJ77642.1 acar|OOF99094.1 acla|EAW06492.1 afis|EAW15988.1 afla|EED56360.1 afum|EAL88620.1 agla|OJJ85990.1 akaw|GAA83992.1 aluc|OJZ80888.1 anid|CBF85815.1 anig|EHA19450.1 aory|EIT73300.1 asyd|OJJ64342.1 ater|EAU33661.1 atub|OJI82690.1 aver|OJI98435.1 awen|OJJ30482.1 azon|OJJ45473.1

Group4449: abra|OJJ77652.1 acar|OOF99084.1 acla|EAW06494.1 afis|EAW15990.1 afla|EED56362.1 afum|EAL88622.1 agla|OJJ85992.1 akaw|GAA83983.1 aluc|OJZ80877.1 anid|CBF85819.1 anig|EHA19459.1 aory|EIT73339.1 asyd|OJJ64343.1 ater|EAU33658.1 atub|OJI82700.1 aver|OJI98440.1 awen|OJJ30480.1 azon|OJJ45471.1

Group4450: abra|OJJ77654.1 acar|OOF99083.1 acla|EAW06496.1 afis|EAW15992.1 afla|EED56364.1 afum|EAL88624.2 agla|OJJ86003.1 akaw|GAA83981.1 aluc|OJZ80875.1 anid|CBF85823.1 anig|EHA19460.1 aory|EIT73404.1 asyd|OJJ64344.1 ater|EAU33657.1 atub|OJI82702.1 aver|OJI98442.1 awen|OJJ30478.1 azon|OJJ45470.1

Group4451: abra|OJJ77655.1 acar|OOF99081.1 acla|EAW06497.1 afis|EAW15993.1 afla|EED56365.1 afum|EAL88625.1 agla|OJJ86004.1 akaw|GAA83980.1 aluc|OJZ80874.1 anid|CBF85824.1 anig|EHA19462.1 aory|EIT73307.1 asyd|OJJ64345.1 ater|EAU33656.1 atub|OJI82704.1 aver|OJI98443.1 awen|OJJ30477.1 azon|OJJ45469.1

Group4452: abra|OJJ77656.1 acar|OOF99080.1 acla|EAW06498.1 afis|EAW15994.1 afla|EED56367.1 afum|EAL88626.1 agla|OJJ86008.1 akaw|GAA83979.1 aluc|OJZ80872.1 anid|CBF85828.1 anig|EHA19463.1 asyd|OJJ64348.1 ater|EAU33654.1 atub|OJI82705.1 aver|OJI98445.1 awen|OJJ30475.1 azon|OJJ45467.1 aory|EIT73315.1

Group4453: abra|OJJ77657.1 acar|OOF99079.1 acla|EAW06499.1 afis|EAW15995.1 afla|EED56369.1 afum|EAL88627.1 agla|OJJ86009.1 akaw|GAA83978.1 aluc|OJZ80871.1 anid|CBF85830.1 anig|EHA19464.1 aory|EIT73436.1 asyd|OJJ64349.1 ater|EAU33652.1 atub|OJI82706.1 aver|OJI98446.1 awen|OJJ30474.1 azon|OJJ45466.1

Group4454: abra|OJJ77659.1 acar|OOF99076.1 acla|EAW06501.1 afis|EAW15997.1 afla|EED56371.1 afum|EAL88629.1 agla|OJJ86017.1 akaw|GAA83976.1 aluc|OJZ80869.1 anid|CBF85832.1 anig|EHA19466.1 aory|EIT73442.1 asyd|OJJ64350.1 ater|EAU33649.1 atub|OJI82708.1 aver|OJI98447.1 awen|OJJ30472.1 azon|OJJ45464.1

Group4455: abra|OJJ77660.1 acar|OOF99075.1 acla|EAW06504.1 afis|EAW15998.1 afla|EED56372.1 afum|EAL88630.1 agla|OJJ86013.1 akaw|GAA83975.1 aluc|OJZ80868.1 anid|CBF85834.1 anig|EHA19467.1 aory|EIT73360.1 asyd|OJJ64351.1 ater|EAU33648.1 atub|OJI82709.1 aver|OJI98448.1 awen|OJJ30471.1 azon|OJJ45463.1

Group4456: abra|OJJ77661.1 acar|OOF99073.1 acla|EAW06598.1 afis|EAW16006.1 afla|EED56380.1 afum|EAL88638.2 agla|OJJ86025.1 akaw|GAA83974.1 aluc|OJZ80867.1 anid|CBF85868.1 anig|EHA19468.1 aory|EIT73448.1 asyd|OJJ64371.1 ater|EAU33637.1 atub|OJI82710.1 aver|OJI98467.1 awen|OJJ30461.1 azon|OJJ45451.1

Group4457: abra|OJJ77665.1 acar|OOF99070.1 acla|EAW06507.1 afis|EAW16002.1 afla|EED56376.1 afum|EAL88634.1 agla|OJJ86024.1 akaw|GAA83970.1 aluc|OJZ80863.1 anid|CBF85844.1 anig|EHA19472.1 aory|EIT73423.1 asyd|OJJ64356.1 ater|EAU33643.1 atub|OJI82714.1 aver|OJI98452.1 awen|OJJ30466.1 azon|OJJ45457.1

Group4458: abra|OJJ77667.1 acar|OOF99068.1 acla|EAW06506.1 afis|EAW16000.1 afla|EED56375.1 afum|EAL88633.1 agla|OJJ86020.1 akaw|GAA83962.1 aluc|OJZ80850.1 anid|CBF85840.1 anig|EHA19474.1 aory|EIT73310.1 asyd|OJJ64354.1 ater|EAU33645.1 atub|OJI82718.1 aver|OJI98450.1 awen|OJJ30469.1 azon|OJJ45461.1

Group4459: abra|OJJ77671.1 acar|OOF99066.1 acla|EAW06505.1 afis|EAW15999.1 afla|EED56374.1 afum|EAL88631.1 agla|OJJ86019.1 akaw|GAA83961.1 aluc|OJZ80848.1 anid|CBF85835.1 anig|EHA19475.1 aory|EIT73377.1 asyd|OJJ64352.1 ater|EAU33647.1 atub|OJI82719.1 aver|OJI98449.1 awen|OJJ30470.1 azon|OJJ45462.1

Group4460: abra|OJJ77680.1 acar|OOF99060.1 acla|EAW06591.1 afis|EAW16014.1 afla|EED56391.1 afum|EAL88643.1 agla|OJJ86040.1 akaw|GAA83954.1 aluc|OJZ80838.1 anid|CBF74562.1 anig|EHA19482.1 aory|EIT83009.1 asyd|OJJ57647.1 ater|EAU33630.1 atub|OJI82730.1 aver|OJJ05504.1 awen|OJJ30450.1 azon|OJJ45442.1

Group4461: abra|OJJ77681.1 acar|OOF99058.1 acla|EAW06592.1 afis|EAW16015.1 afla|EED56392.1 afum|EAL88644.1 agla|OJJ86041.1 akaw|GAA83952.1 aluc|OJZ80836.1 anid|CBF74560.1 anig|EHA19483.1 aory|EIT83002.1 asyd|OJJ57646.1 ater|EAU33629.1 atub|OJI82732.1 aver|OJJ05503.1 awen|OJJ30449.1 azon|OJJ45441.1

Group4462: abra|OJJ77682.1 acar|OOF99059.1 acla|EAW06594.1 afis|EAW16017.1 afla|EED56393.1 afum|EAL88646.1 agla|OJJ86042.1 akaw|GAA83951.1 aluc|OJZ80835.1 anid|CBF74558.1 anig|EHA19484.1 aory|EIT83003.1 asyd|OJJ57645.1 ater|EAU33628.1 atub|OJI82733.1 aver|OJJ05502.1 awen|OJJ30447.1 azon|OJJ45439.1

Group4463: abra|OJJ77683.1 acar|OOF99057.1 acla|EAW06595.1 afis|EAW16018.1 afla|EED56394.1 afum|EAL88647.1 agla|OJJ86045.1 akaw|GAA83950.1 aluc|OJZ80834.1 anid|CBF74555.1 anig|EHA19485.1 aory|EIT83010.1 asyd|OJJ57644.1 ater|EAU33627.1 atub|OJI82734.1 aver|OJJ05501.1 awen|OJJ30446.1 azon|OJJ45438.1

Group4464: abra|OJJ77684.1 acar|OOF99056.1 acla|EAW06596.1 afis|EAW16019.1 afla|EED56396.1 afum|EAL88648.1 agla|OJJ86046.1 akaw|GAA83949.1 aluc|OJZ80833.1 anid|CBF74553.1 anig|EHA19486.1 aory|EIT83017.1 asyd|OJJ57642.1 ater|EAU33626.1 atub|OJI82735.1 aver|OJJ05499.1 awen|OJJ30445.1 azon|OJJ45437.1

Group4465: abra|OJJ77690.1 acar|OOF99049.1 acla|EAW06575.1 afis|EAW16021.1 afla|EED56399.1 afum|EAL88650.1 agla|OJJ86048.1 akaw|GAA83944.1 aluc|OJZ80825.1 anid|CBF74538.1 anig|EHA19490.1 aory|EIT83016.1 asyd|OJJ57695.1 ater|EAU33619.1 atub|OJI82740.1 aver|OJJ05471.1 awen|OJJ30443.1 azon|OJJ45435.1

Group4466: abra|OJJ77691.1 acar|OOF99048.1 acla|EAW06574.1 afis|EAW16022.1 afla|EED56401.1 afum|EAL88651.1 agla|OJJ86051.1 akaw|GAA83939.1 aluc|OJZ80821.1 anid|CBF74540.1 anig|EHA19491.1 aory|EIT83005.1 asyd|OJJ57694.1 ater|EAU33618.1 atub|OJI82741.1 aver|OJJ05472.1 awen|OJJ30439.1 azon|OJJ45434.1

Group4467: abra|OJJ77695.1 acar|OOF99044.1 acla|EAW11326.1 afis|EAW18169.1 afla|EED56325.1 afum|EBA27315.1 agla|OJJ85948.1 akaw|GAA83935.1 aluc|OJZ90758.1 anid|CBF85744.1 anig|EHA19495.1 aory|EIT73371.1 asyd|OJJ64290.1 ater|EAU33835.1 atub|OJI82745.1 aver|OJI98383.1 awen|OJJ30516.1 azon|OJJ50197.1

Group4468: abra|OJJ77697.1 acar|OOF99043.1 acla|EAW11327.1 afis|EAW18168.1 afla|EED56326.1 afum|EAL87529.2 agla|OJJ85949.1 akaw|GAA83934.1 aluc|OJZ90761.1 anid|CBF85746.1 anig|EHA19496.1 aory|EIT73447.1 asyd|OJJ64291.1 ater|EAU33836.1 atub|OJI82748.1 aver|OJI98384.1 awen|OJJ30514.1 azon|OJJ50195.1

Group4469: abra|OJJ77699.1 acar|OOF99040.1 acla|EAW11324.1 afis|EAW18171.1 afla|EED56322.1 afum|EAL87533.1 agla|OJJ85935.1 akaw|GAA83931.1 aluc|OJZ90764.1 anid|CBF85742.1 anig|EHA19498.1 aory|EIT73405.1 asyd|OJJ64289.1 ater|EAU33833.1 atub|OJI82752.1 aver|OJI98382.1 awen|OJJ30518.1 azon|OJJ50199.1

Group4470: abra|OJJ77704.1 acar|OOF99036.1 acla|EAW11315.1 afis|EAW18183.1 afla|EED56304.1 afum|EAL87545.1 agla|OJJ85903.1 akaw|GAA83926.1 aluc|OJZ90769.1 anid|CBF85723.1 anig|EHA19504.1

aory|EIT73331.1 asyd|OJJ64277.1 ater|EAU33819.1 atub|OJI82757.1 aver|OJI98369.1 awen|OJJ30531.1 azon|OJJ50212.1

Group4471: abra|OJJ77708.1 acar|OOF99032.1 acla|EAW11317.1 afis|EAW18179.1 afla|EED56307.1 afum|EAL87540.1 agla|OJJ85910.1 akaw|GAA83922.1 aluc|OJZ90773.1 anid|CBF85729.1 anig|EHA19507.1 aory|EIT73380.1 asyd|OJJ64280.1 ater|EAU33821.1 atub|OJI82760.1 aver|OJI98372.1 awen|OJJ30527.1 azon|OJJ50208.1

Group4472: abra|OJJ77709.1 acar|OOF99031.1 acla|EAW11318.1 afis|EAW18178.1 afla|EED56306.1 afum|EAL87539.1 agla|OJJ85912.1 akaw|GAA83921.1 aluc|OJZ90775.1 anid|CBF85732.1 anig|EHA19508.1 aory|EIT73415.1 asyd|OJJ64283.1 ater|EAU33827.1 atub|OJI82761.1 aver|OJI98375.1 awen|OJJ30526.1 azon|OJJ50207.1

Group4473: abra|OJJ77710.1 acar|OOF99030.1 acla|EAW11319.1 afis|EAW18177.1 afla|EED56312.1 afum|EAL87538.1 agla|OJJ85913.1 akaw|GAA83920.1 aluc|OJZ90776.1 anid|CBF85733.1 anig|EHA19509.1 aory|EIT73345.1 asyd|OJJ64284.1 ater|EAU33828.1 atub|OJI82762.1 aver|OJI98376.1 awen|OJJ30525.1 azon|OJJ50206.1

Group4474: abra|OJJ77717.1 acar|OOF99022.1 acla|EAW11320.1 afis|EAW18176.1 afla|EED56313.1 afum|EAL87537.1 agla|OJJ85914.1 akaw|GAA83915.1 aluc|OJZ90782.1 anid|CBF85735.1 anig|EHA19515.1 aory|EIT73299.1 asyd|OJJ64285.1 ater|EAU33829.1 atub|OJI82769.1 aver|OJI98377.1 awen|OJJ30524.1 azon|OJJ50205.1

Group4475: abra|OJJ77718.1 acar|OOF99019.1 acla|EAW11321.1 afis|EAW18175.1 afla|EED56314.1 afum|EAL87536.1 agla|OJJ85915.1 akaw|GAA83914.1 aluc|OJZ90783.1 anid|CBF85737.1 anig|EHA19516.1 aory|EIT73381.1 asyd|OJJ64286.1 ater|EAU33830.1 atub|OJI82770.1 aver|OJI98378.1 awen|OJJ30523.1 azon|OJJ50204.1

Group4476: abra|OJJ77810.1 acar|OOF98957.1 acla|EAW11307.1 afis|EAW18194.1 afla|EED56284.1 afum|EBA27318.1 agla|OJJ85895.1 akaw|GAA83839.1 aluc|OJZ90877.1 anid|CBF86104.1 anig|EHA19597.1 aory|EIT73433.1 asyd|OJJ64482.1 ater|EAU33799.1 atub|OJI82874.1 aver|OJI98583.1 awen|OJJ30558.1 azon|OJJ45509.1

Group4477: abra|OJJ77811.1 acar|OOF98956.1 acla|EAW11306.1 afis|EAW18193.1 afla|EED56283.1 afum|EAL87559.1 agla|OJJ85893.1 akaw|GAA83838.1 aluc|OJZ90878.1 anid|CBF86108.1 anig|EHA19599.1 aory|EIT73325.1 asyd|OJJ64487.1 ater|EAU33798.1 atub|OJI82875.1 aver|OJI98588.1 awen|OJJ30559.1 azon|OJJ45508.1

Group4478: abra|OJJ77819.1 acar|OOF98949.1 acla|EAW11304.1 afis|EAW18191.1 afla|EED56281.1 afum|EAL87562.2 agla|OJJ85884.1 akaw|GAA83827.1 aluc|OJZ90891.1 anid|CBF86111.1 anig|EHA19610.1 aory|EIT73396.1 asyd|OJJ64489.1 ater|EAU33796.1 atub|OJI82885.1 aver|OJI98590.1 awen|OJJ30565.1 azon|OJJ45506.1

Group4479: abra|OJJ77821.1 acar|OOF98945.1 acla|EAW11303.1 afis|EAW18199.1 afla|EED56279.1 afum|EAL87564.1 agla|OJJ85883.1 akaw|GAA83824.1 aluc|OJZ90894.1 anid|CBF86115.1 anig|EHA19613.1 aory|EIT73401.1 asyd|OJJ64491.1 ater|EAU33795.1 atub|OJI82887.1 aver|OJI98593.1 awen|OJJ30567.1 azon|OJJ45505.1

Group4480: abra|OJJ77824.1 acar|OOF98944.1 acla|EAW11301.1 afis|EAW18201.1 afla|EED56277.1 afum|EAL87566.1 agla|OJJ85881.1 akaw|GAA83822.1 aluc|OJZ90897.1 anid|CBF86122.1 anig|EHA19615.1 aory|EIT73449.1 asyd|OJJ64496.1 ater|EAU33793.1 atub|OJI82889.1 aver|OJI98598.1 awen|OJJ30569.1 azon|OJJ45503.1

Group4481: abra|OJJ77826.1 acar|OOF98942.1 acla|EAW11299.1 afis|EAW18203.1 afla|EED56269.1 afum|EAL87568.1 agla|OJJ85879.1 akaw|GAA83820.1 aluc|OJZ90900.1 anid|CBF86119.1 anig|EHA19616.1 aory|EIT73413.1 asyd|OJJ64494.1 ater|EAU33786.1 atub|OJI82893.1 aver|OJI98596.1 awen|OJJ30571.1 azon|OJJ45501.1

Group4482: abra|OJJ77827.1 acar|OOF98941.1 acla|EAW11298.1 afis|EAW18204.1 afla|EED56268.1 afum|EAL87569.1 agla|OJJ85878.1 akaw|GAA83819.1 aluc|OJZ90901.1 anid|CBF86124.1 anig|EHA19617.1 aory|EIT73341.1 asyd|OJJ64497.1 ater|EAU33785.1 atub|OJI82894.1 aver|OJI98599.1 awen|OJJ30572.1 azon|OJJ45500.1

Group4483: abra|OJJ77830.1 acar|OOF98940.1 acla|EAW11296.1 afis|EAW18206.1 afla|EED56265.1 afum|EAL87571.1 agla|OJJ85876.1 akaw|GAA83817.1 aluc|OJZ90904.1 anid|CBF86132.1 anig|EHA19619.1 aory|EIT73429.1 asyd|OJJ64502.1 ater|EAU33783.1 atub|OJI82897.1 aver|OJI98601.1 awen|OJJ30578.1 azon|OJJ45498.1

Group4484: abra|OJJ77834.1 acar|OOF98937.1 acla|EAW11289.1 afis|EAW18210.1 afla|EED56260.1 afum|EAL87576.2 agla|OJJ85872.1 akaw|GAA83813.1 aluc|OJZ90908.1 anid|CBF86139.1 anig|EHA19623.1 aory|EIT73444.1 asyd|OJJ64506.1 ater|EAU33778.1 atub|OJI82901.1 aver|OJI98605.1 awen|OJJ30588.1 azon|OJJ50189.1

Group4485: abra|OJJ77841.1 acar|OOF98930.1 acla|EAW11282.1 afis|EAW18218.1 afla|EED56252.1 afum|EAL87584.1 agla|OJJ85864.1 akaw|GAA83806.1 aluc|OJZ90916.1 anid|CBF86148.1 anig|EHA19628.1 aory|EIT73348.1 asyd|OJJ64512.1 ater|EAU33773.1 atub|OJI82908.1 aver|OJI98610.1 awen|OJJ30596.1 azon|OJJ50172.1

Group4486: abra|OJJ77844.1 acar|OOF98928.1 acla|EAW11279.1 afis|EAW18221.1 afla|EED56249.1 afum|EAL87587.1 agla|OJJ85861.1 akaw|GAA83803.1 aluc|OJZ90919.1 anid|CBF86155.1 anig|EHA19631.1 aory|EIT73362.1 asyd|OJJ64515.1 ater|EAU33770.1 atub|OJI82911.1 aver|OJI98613.1 awen|OJJ30599.1 azon|OJJ50169.1

Group4487: abra|OJJ77845.1 acar|OOF98927.1 acla|EAW11278.1 afis|EAW18222.1 afla|EED56247.1 afum|EAL87588.1 agla|OJJ85860.1 akaw|GAA83802.1 aluc|OJZ90920.1 anid|CBF86153.1 anig|EHA19632.1 aory|EIT73414.1 asyd|OJJ64516.1 ater|EAU33769.1 atub|OJI82912.1 aver|OJI98614.1 awen|OJJ30600.1 azon|OJJ50168.1

Group4488: abra|OJJ77847.1 acar|OOF98925.1 acla|EAW11276.1 afis|EAW18224.1 afla|EED56245.1 afum|EAL87590.1 agla|OJJ85858.1 akaw|GAA83800.1 aluc|OJZ90922.1 anid|CBF86159.1 anig|EHA19634.1 aory|EIT73297.1 asyd|OJJ64518.1 ater|EAU33767.1 atub|OJI82914.1 aver|OJI98616.1 awen|OJJ30602.1 azon|OJJ50166.1

Group4489: abra|OJJ77851.1 acar|OOF98922.1 acla|EAW11275.1 afis|EAW18225.1 afla|EED56244.1 afum|EAL87591.1 agla|OJJ85857.1 akaw|GAA83797.1 aluc|OJZ90925.1 anid|CBF86161.1 anig|EHA19637.1 aory|EIT73369.1 asyd|OJJ64519.1 ater|EAU33766.1 atub|OJI82917.1 aver|OJI98617.1 awen|OJJ30603.1 azon|OJJ50165.1

Group4490: abra|OJJ77852.1 acar|OOF98921.1 acla|EAW11274.1 afis|EAW18226.1 afla|EED56243.1 afum|EAL87593.2 agla|OJJ85856.1 akaw|GAA83796.1 aluc|OJZ90926.1 anid|CBF86162.1 anig|EHA19638.1 aory|EIT73293.1 asyd|OJJ64520.1 ater|EAU33765.1 atub|OJI82918.1 aver|OJI98618.1 awen|OJJ30604.1 azon|OJJ50164.1

Group4491: abra|OJJ77853.1 acar|OOF98920.1 acla|EAW11273.1 afis|EAW18227.1 afla|EED56242.1 afum|EAL87594.1 agla|OJJ85855.1 akaw|GAA83795.1 aluc|OJZ90927.1 anid|CBF86164.1 anig|EHA19639.1 aory|EIT73313.1 asyd|OJJ64521.1 ater|EAU33764.1 atub|OJI82919.1 aver|OJI98619.1 awen|OJJ30605.1 azon|OJJ50163.1

Group4492: abra|OJJ77854.1 acar|OOF98919.1 acla|EAW11272.1 afis|EAW18228.1 afla|EED56241.1 afum|EAL87595.1 agla|OJJ85854.1 akaw|GAA83794.1 aluc|OJZ90928.1 anid|CBF86166.1 anig|EHA19640.1 aory|EIT73375.1 asyd|OJJ64522.1 ater|EAU33763.1 atub|OJI82920.1 aver|OJI98620.1 awen|OJJ30606.1 azon|OJJ50162.1

Group4493: abra|OJJ77857.1 acar|OOF98917.1 acla|EAW11269.1 afis|EAW18230.1 afla|EED56238.1 afum|EAL87597.1 agla|OJJ85844.1 akaw|GAA83792.1 aluc|OJZ90931.1 anid|CBF86172.1 anig|EHA19642.1 aory|EIT73450.1 asyd|OJJ64525.1 ater|EAU33761.1 atub|OJI82923.1 aver|OJI98623.1 awen|OJJ30609.1 azon|OJJ50159.1

Group4494: abra|OJJ77862.1 acar|OOF98911.1 acla|EAW11264.1 afis|EAW18234.1 afla|EED56232.1 afum|EBA27319.1 agla|OJJ85850.1 akaw|GAA83787.1 aluc|OJZ90937.1 anid|CBF86182.1 anig|EHA19645.1 aory|EIT73318.1 asyd|OJJ64530.1 ater|EAU33752.1 atub|OJI82928.1 aver|OJI98628.1 awen|OJJ30614.1 azon|OJJ48003.1

Group4495: abra|OJJ77864.1 acar|OOF98909.1 acla|EAW11262.1 afis|EAW18236.1 afla|EED56230.1 afum|EAL87603.1 agla|OJJ85838.1 akaw|GAA83785.1 aluc|OJZ90939.1 anid|CBF86185.1 anig|EHA19647.1 aory|EIT73458.1 asyd|OJJ64532.1 ater|EAU33749.1 atub|OJI82930.1 aver|OJI98630.1 awen|OJJ30616.1 azon|OJJ48001.1

Group4496: abra|OJJ77865.1 acar|OOF98907.1 acla|EAW11261.1 afis|EAW18237.1 afla|EED56229.1 afum|EAL87604.2 agla|OJJ85837.1 akaw|GAA83784.1 aluc|OJZ90940.1 anid|CBF86187.1 anig|EHA19648.1 aory|EIT73322.1 asyd|OJJ64534.1 ater|EAU33748.1 atub|OJI82931.1 aver|OJI98631.1 awen|OJJ30617.1 azon|OJJ48000.1

Group4497: abra|OJJ77866.1 acar|OOF98906.1 acla|EAW11259.1 afis|EAW18239.1 afla|EED56227.1 afum|EBA27321.1 agla|OJJ85831.1 akaw|GAA83782.1 aluc|OJZ90942.1 anid|CBF86191.1 anig|EHA19650.1 aory|EIT73427.1 asyd|OJJ64536.1 ater|EAU33744.1 atub|OJI82933.1 aver|OJI98633.1 awen|OJJ30619.1 azon|OJJ47998.1

Group4498: abra|OJJ77876.1 acar|OOF98895.1 acla|EAW06509.1 afis|EAW16222.1 afla|EED50188.1 afum|EAL88850.1 agla|OJJ86234.1 akaw|GAA83776.1 aluc|OJZ90951.1 anid|CBF86216.1 anig|EHA19657.1 aory|EIT77385.1 asyd|OJJ64558.1 ater|EAU33719.1 atub|OJI82942.1 aver|OJI98655.1 awen|OJJ30248.1 azon|OJJ43951.1

Group4499: abra|OJJ77878.1 acar|OOF98894.1 acla|EAW06602.1 afis|EAW16221.1 afla|EED50189.1 afum|EAL88849.1 agla|OJJ86233.1 akaw|GAA83775.1 aluc|OJZ90952.1 anid|CBF86215.1 anig|EHA19658.1

aory|EIT77381.1 asyd|OJJ64557.1 ater|EAU33720.1 atub|OJI82943.1 aver|OJI98654.1 awen|OJJ30250.1 azon|OJJ43952.1

Group4501: abra|OJJ77890.1 acar|OOF98890.1 acla|EAW11245.1 afis|EAW18255.1 afla|EED56196.1 afum|EAL87618.1 agla|OJJ88189.1 akaw|GAA83768.1 aluc|OJZ90960.1 anid|CBF86206.1 anig|EHA19665.1 aory|EIT73437.1 asyd|OJJ64546.1 ater|EAU33727.1 atub|OJI82951.1 aver|OJI98642.1 awen|OJJ30629.1 azon|OJJ46567.1

Group4502: abra|OJJ77893.1 acar|OOF98889.1 acla|EAW11246.1 afis|EAW18254.1 afla|EED56198.1 afum|EAL87617.1 agla|OJJ88190.1 akaw|GAA83767.1 aluc|OJZ90963.1 anid|CBF86205.1 anig|EHA19667.1 aory|EIT73303.1 asyd|OJJ64545.1 ater|EAU33726.1 atub|OJI82954.1 aver|OJI98641.1 awen|OJJ30628.1 azon|OJJ47987.1

Group4503: abra|OJJ77897.1 acar|OOF98887.1 acla|EAW11248.1 afis|EAW18249.1 afla|EED56215.1 afum|EAL87614.1 agla|OJJ85821.1 akaw|GAA83765.1 aluc|OJZ90967.1 anid|CBF86200.1 anig|EHA19669.1 aory|EIT73361.1 asyd|OJJ64542.1 ater|EAU33731.1 atub|OJI82957.1 aver|OJI98638.1 awen|OJJ30625.1 azon|OJJ47993.1

Group4504: abra|OJJ77898.1 acar|OOF98886.1 acla|EAW11249.1 afis|EAW18248.1 afla|EED56216.1 afum|EAL87613.1 agla|OJJ85822.1 akaw|GAA83764.1 aluc|OJZ90968.1 anid|CBF86198.1 anig|EHA19670.1 aory|EIT73459.1 asyd|OJJ64541.1 ater|EAU33732.1 atub|OJI82958.1 aver|OJI98637.1 awen|OJJ30624.1 azon|OJJ47994.1

Group4505: abra|OJJ77902.1 acar|OOF98881.1 acla|EAW06512.1 afis|EAW16230.1 afla|EED53351.1 afum|EAL88853.1 agla|OJJ86245.1 akaw|GAA83760.1 aluc|OJZ90971.1 anid|CBF86223.1 anig|EHA19673.1 aory|EIT75232.1 asyd|OJJ64563.1 ater|EAU33714.1 atub|OJI82962.1 aver|OJI98660.1 awen|OJJ30232.1 azon|OJJ43949.1

Group4506: abra|OJJ77903.1 acar|OOF98879.1 acla|EAW06513.1 afis|EAW16231.1 afla|EED53350.1 afum|EAL88854.1 agla|OJJ86244.1 akaw|GAA83759.1 aluc|OJZ90972.1 anid|CBF86224.1 anig|EHA19674.1 aory|EIT74926.1 asyd|OJJ64564.1 ater|EAU33713.1 atub|OJI82963.1 aver|OJI98661.1 awen|OJJ30231.1 azon|OJJ43948.1

Group4507: abra|OJJ77910.1 acar|OOF98875.1 acla|EAW06518.1 afis|EAW16236.1 afla|EED53345.1 afum|EAL88857.1 agla|OJJ86239.1 akaw|GAA83749.1 aluc|OJZ90982.1 anid|CBF86232.1 anig|EHA19687.1 aory|EIT74884.1 asyd|OJJ64566.1 ater|EAU33708.1 atub|OJI82973.1 aver|OJI98663.1 awen|OJJ30224.1 azon|OJJ43944.1

Group4508: anig|EHA19693.1 abra|OJJ77921.1 acar|OOF98864.1 acla|EAW06687.1 afis|EAW16078.1 afla|EED48598.1 afum|EAL88710.1 agla|OJJ86111.1 akaw|GAA83742.1 aluc|OJZ90989.1 anid|CBF75119.1 aory|EIT73211.1 asyd|OJJ59993.1 ater|EAU33144.1 atub|OJI82980.1 aver|OJJ02298.1 awen|OJJ30355.1 azon|OJJ45377.1

Group4509: abra|OJJ77936.1 acar|OOF98872.1 acla|EAW06678.1 afis|EAW16087.1 afla|EED48608.1 afum|EAL88719.1 agla|OJJ86119.1 akaw|GAA83727.1 aluc|OJZ91004.1 anid|CBF75125.1 anig|EHA19709.1 aory|EIT73213.1 asyd|OJJ59984.1 ater|EAU33157.1 atub|OJI82995.1 aver|OJJ02307.1 awen|OJJ30344.1 azon|OJJ45369.1

Group4510: abra|OJJ77937.1 acar|OOF98873.1 acla|EAW06677.1 afis|EAW16088.1 afla|EED48609.1 afum|EAL88720.1 agla|OJJ86120.1 akaw|GAA83726.1 aluc|OJZ91005.1 anid|CBF75127.1 anig|EHA19710.1 aory|EIT73207.1 asyd|OJJ59983.1 ater|EAU33159.1 atub|OJI82996.1 aver|OJJ02308.1 awen|OJJ30343.1 azon|OJJ45368.1

Group4511: abra|OJJ77968.1 acar|OOF98847.1 acla|EAW06672.1 afis|EAW16093.1 afla|EED48629.1 afum|EAL88725.1 akaw|GAA83694.1 aluc|OJZ91038.1 anid|CBF85460.1 anig|EHA19737.1 aory|EIT73172.1 asyd|OJJ64101.1 ater|EAU33165.1 atub|OJI83034.1 aver|OJI98184.1 awen|OJJ30337.1 azon|OJJ45367.1 agla|OJJ86128.1

Group4512: abra|OJJ77971.1 acar|OOF98843.1 acla|EAW06670.1 afis|EAW16095.1 afla|EED48631.1 afum|EAL88727.1 agla|OJJ86126.1 akaw|GAA83691.1 aluc|OJZ91041.1 anid|CBF85455.1 anig|EHA19740.1 aory|EIT73094.1 asyd|OJJ64098.1 ater|EAU33171.1 atub|OJI83037.1 aver|OJI98181.1 awen|OJJ30335.1 azon|OJJ45363.1

Group4513: abra|OJJ77972.1 acar|OOF98842.1 acla|EAW06667.1 afis|EAW16098.1 afla|EED48633.1 afum|EAL88730.1 agla|OJJ86124.1 akaw|GAA83689.1 aluc|OJZ91043.1 anid|CBF85451.1 anig|EHA19742.1 aory|EIT73279.1 asyd|OJJ64096.1 ater|EAU33173.1 atub|OJI83039.1 aver|OJI98179.1 awen|OJJ30333.1 azon|OJJ45362.1

Group4514: abra|OJJ77973.1 acar|OOF98841.1 acla|EAW06665.1 afis|EAW16100.1 afla|EED48635.1 afum|EAL88732.1 agla|OJJ86122.1 anid|CBF85447.1 anig|EHA19743.1 aory|EIT73131.1 asyd|OJJ64095.1 ater|EAU33174.1 aver|OJI98177.1 awen|OJJ30332.1 azon|OJJ45360.1 akaw|GAA83687.1 aluc|OJZ91045.1 atub|OJI83041.1

Group4515: abra|OJJ77988.1 acar|OOF98835.1 acla|EAW06661.1 afis|EAW16104.1 afla|EED48640.1 afum|EAL88736.1 agla|OJJ86133.1 akaw|GAA85992.1 aluc|OJZ91063.1 anid|CBF85480.1 anig|EHA19767.1 aory|EIT73283.1 asyd|OJJ64108.1 ater|EAU33183.1 atub|OJI83059.1 aver|OJI98190.1 awen|OJJ30328.1 azon|OJJ45345.1

Group4516: abra|OJJ77989.1 acar|OOF98834.1 acla|EAW06660.1 afis|EAW16105.1 afla|EED48641.1 afum|EAL88737.1 agla|OJJ86134.1 akaw|GAA85991.1 aluc|OJZ91064.1 anid|CBF85482.1 anig|EHA19768.1 aory|EIT73197.1 asyd|OJJ64109.1 ater|EAU33184.1 atub|OJI83060.1 aver|OJI98191.1 awen|OJJ30327.1 azon|OJJ45344.1

Group4517: abra|OJJ77990.1 acar|OOF98833.1 acla|EAW06659.1 afis|EAW16106.1 afla|EED48642.1 afum|EAL88738.1 agla|OJJ86135.1 akaw|GAA85990.1 aluc|OJZ91066.1 anid|CBF85484.1 anig|EHA19769.1 aory|EIT73122.1 asyd|OJJ64110.1 ater|EAU33185.1 atub|OJI83061.1 aver|OJI98192.1 awen|OJJ30325.1 azon|OJJ45343.1

Group4518: abra|OJJ77991.1 acar|OOF98831.1 acla|EAW06658.1 afis|EAW16107.1 afla|EED48643.1 afum|EAL88739.2 agla|OJJ86136.1 akaw|GAA85989.1 aluc|OJZ91067.1 anid|CBF85486.1 anig|EHA19770.1 aory|EIT73085.1 asyd|OJJ64111.1 ater|EAU33190.1 atub|OJI83062.1 aver|OJI98193.1 awen|OJJ30324.1 azon|OJJ45342.1

Group4519: abra|OJJ77992.1 acar|OOF98830.1 acla|EAW06657.1 afis|EAW16108.1 afla|EED48644.1 afum|EAL88740.1 agla|OJJ86137.1 akaw|GAA85988.1 aluc|OJZ91068.1 anid|CBF85487.1 anig|EHA19771.1 aory|EIT73264.1 asyd|OJJ64112.1 ater|EAU33191.1 atub|OJI83063.1 aver|OJI98194.1 awen|OJJ30323.1 azon|OJJ45341.1

Group4520: abra|OJJ77993.1 acar|OOF98829.1 acla|EAW06655.1 afis|EAW16110.1 afla|EED48650.1 afum|EAL88741.1 agla|OJJ86145.1 akaw|GAA85987.1 aluc|OJZ91069.1 anid|CBF85492.1 anig|EHA19772.1 aory|EIT73074.1 asyd|OJJ64113.1 ater|EAU33193.1 atub|OJI83064.1 aver|OJI98196.1 awen|OJJ30320.1 azon|OJJ45339.1

Group4521: abra|OJJ77994.1 acar|OOF98828.1 acla|EAW06654.1 afis|EAW16111.1 afla|EED48651.1 afum|EAL88742.1 agla|OJJ86146.1 akaw|GAA85986.1 aluc|OJZ91070.1 anid|CBF85494.1 anig|EHA19773.1 aory|EIT73265.1 asyd|OJJ64114.1 ater|EAU33194.1 atub|OJI83065.1 aver|OJI98197.1 awen|OJJ30319.1 azon|OJJ45338.1

Group4522: abra|OJJ77995.1 acar|OOF98827.1 acla|EAW06653.1 afis|EAW16112.1 afla|EED48652.1 afum|EAL88743.1 agla|OJJ86151.1 akaw|GAA85985.1 aluc|OJZ91071.1 anid|CBF85496.1 anig|EHA19774.1 aory|EIT73201.1 asyd|OJJ64115.1 ater|EAU33195.1 atub|OJI83066.1 aver|OJI98198.1 awen|OJJ30318.1 azon|OJJ45337.1

Group4523: abra|OJJ78001.1 acar|OOF98821.1 acla|EAW06649.1 afis|EAW16116.1 afla|EED48657.1 afum|EAL88747.1 agla|OJJ86155.1 akaw|GAA85982.1 aluc|OJZ91077.1 anid|CBF85505.1 anig|EHA19778.1 aory|EIT73053.1 asyd|OJJ64119.1 ater|EAU33199.1 atub|OJI83072.1 aver|OJI98202.1 awen|OJJ30314.1 azon|OJJ45331.1

Group4524: abra|OJJ78004.1 acar|OOF98819.1 acla|EAW06646.1 afis|EAW16119.1 afla|EED48660.1 afum|EAL88750.1 agla|OJJ86160.1 akaw|GAA85979.1 aluc|OJZ91080.1 anid|CBF85543.1 anig|EHA19781.1 aory|EIT73098.1 asyd|OJJ64143.1 ater|EAU33202.1 atub|OJI83075.1 aver|OJI98224.1 awen|OJJ30311.1 azon|OJJ45328.1

Group4525: abra|OJJ78005.1 acar|OOF98818.1 acla|EAW06645.1 afis|EAW16120.1 afla|EED48661.1 afum|EAL88751.1 agla|OJJ86161.1 akaw|GAA85978.1 aluc|OJZ91081.1 anid|CBF85541.1 anig|EHA19782.1 aory|EIT73067.1 asyd|OJJ64142.1 ater|EAU33203.1 atub|OJI83077.1 aver|OJI98223.1 awen|OJJ30310.1 azon|OJJ45327.1

Group4526: abra|OJJ78008.1 acar|OOF98815.1 acla|EAW06642.1 afis|EAW16123.1 afla|EED48663.1 afum|EAL88753.1 agla|OJJ86164.1 akaw|GAA85975.1 aluc|OJZ91084.1 anid|CBF85534.1 anig|EHA19785.1 aory|EIT73192.1 asyd|OJJ64136.1 ater|EAU33206.1 atub|OJI83080.1 aver|OJI98219.1 awen|OJJ30307.1 azon|OJJ45310.1

Group4527: abra|OJJ78010.1 acar|OOF98813.1 acla|EAW06639.1 afis|EAW16126.1 afla|EED48666.1 afum|EAL88756.1 agla|OJJ86168.1 akaw|GAA85973.1 aluc|OJZ91086.1 anid|CBF85529.1 anig|EHA19787.1 aory|EIT73204.1 asyd|OJJ64133.1 ater|EAU33209.1 atub|OJI83082.1 aver|OJI98216.1 awen|OJJ30303.1 azon|OJJ45304.1

Group4528: abra|OJJ78011.1 acar|OOF98812.1 acla|EAW06638.1 afis|EAW16127.1 afla|EED48667.1 afum|EAL88757.1 agla|OJJ86169.1 akaw|GAA85972.1 aluc|OJZ91087.1 anid|CBF85527.1 anig|EHA19788.1 aory|EIT73187.1 asyd|OJJ64132.1 ater|EAU33210.1 atub|OJI83083.1 aver|OJI98215.1 awen|OJJ30302.1 azon|OJJ45303.1

Group4529: abra|OJJ78012.1 acar|OOF98811.1 acla|EAW06637.1 afis|EAW16128.1 afla|EED48668.1 afum|EAL88758.1 agla|OJJ86174.1 akaw|GAA85971.1 aluc|OJZ91088.1 anid|CBF85525.1 anig|EHA19789.1

aory|EIT73230.1 asyd|OJJ64131.1 ater|EAU33211.1 atub|OJI83085.1 aver|OJI98214.1 awen|OJJ30301.1 azon|OJJ45302.1

Group4530: abra|OJJ78013.1 acar|OOF98810.1 acla|EAW06635.1 afis|EAW16130.1 afla|EED48670.1 afum|EAL88760.1 agla|OJJ86175.1 akaw|GAA85970.1 aluc|OJZ91089.1 anid|CBF85523.1 anig|EHA19790.1 aory|EIT73154.1 asyd|OJJ64130.1 ater|EAU33212.1 atub|OJI83086.1 aver|OJI98213.1 awen|OJJ30299.1 azon|OJJ43987.1

Group4531: abra|OJJ78014.1 acar|OOF98809.1 acla|EAW06634.1 afis|EAW16131.1 afla|EED48669.1 afum|EAL88761.1 agla|OJJ86176.1 akaw|GAA85969.1 aluc|OJZ91090.1 anid|CBF85521.1 anig|EHA19791.1 aory|EIT73222.1 asyd|OJJ64128.1 ater|EAU33213.1 atub|OJI83087.1 aver|OJI98211.1 awen|OJJ30298.1 azon|OJJ43986.1

Group4532: abra|OJJ78018.1 acar|OOF98805.1 acla|EAW06629.1 afis|EAW16136.1 afla|EED48676.1 afum|EAL88766.2 agla|OJJ86181.1 akaw|GAA85965.1 aluc|OJZ91094.1 anid|CBF85512.1 anig|EHA19795.1 aory|EIT73051.1 asyd|OJJ64123.1 ater|EAU33218.1 atub|OJI83091.1 aver|OJI98206.1 awen|OJJ30292.1 azon|OJJ43981.1

Group4533: abra|OJJ78021.1 acar|OOF98803.1 acla|EAW06627.1 afis|EAW16138.1 afla|EED48678.1 afum|EAL88767.1 agla|OJJ86183.1 akaw|GAA85964.1 aluc|OJZ91096.1 anid|CBF85545.1 anig|EHA19797.1 aory|EIT73188.1 asyd|OJJ64144.1 ater|EAU33219.1 atub|OJI83094.1 aver|OJI98225.1 awen|OJJ30290.1 azon|OJJ43979.1

Group4534: abra|OJJ78023.1 acar|OOF98802.1 acla|EAW06626.1 afis|EAW16139.1 afla|EED48679.1 afum|EAL88768.1 agla|OJJ86185.1 akaw|GAA85963.1 aluc|OJZ91098.1 anid|CBF85546.1 anig|EHA19798.1 aory|EIT73152.1 asyd|OJJ64148.1 ater|EAU33220.1 atub|OJI83095.1 aver|OJI98228.1 awen|OJJ30289.1 azon|OJJ43978.1

Group4535: abra|OJJ78024.1 acar|OOF98801.1 acla|EAW06625.1 afis|EAW16140.1 afla|EED48681.1 afum|EAL88769.2 agla|OJJ86189.1 akaw|GAA85962.1 aluc|OJZ91099.1 anid|CBF85548.1 anig|EHA19799.1 aory|EIT73077.1 asyd|OJJ64149.1 ater|EAU33225.1 atub|OJI83096.1 aver|OJI98229.1 awen|OJJ30288.1 azon|OJJ43977.1

Group4536: abra|OJJ78026.1 acar|OOF98799.1 acla|EAW06623.1 afis|EAW16142.1 afla|EED48683.1 afum|EAL88771.1 agla|OJJ86190.1 akaw|GAA85960.1 aluc|OJZ91101.1 anid|CBF85551.1 anig|EHA19801.1 aory|EIT73175.1 asyd|OJJ64151.1 ater|EAU33227.1 atub|OJI83098.1 aver|OJI98231.1 awen|OJJ30286.1 azon|OJJ43975.1

Group4537: abra|OJJ78027.1 acar|OOF98796.1 acla|EAW06622.1 afis|EAW16143.1 afla|EED48684.1 afum|EAL88772.1 agla|OJJ86191.1 akaw|GAA85959.1 aluc|OJZ91102.1 anid|CBF79592.1 anig|EHA19802.1 aory|EIT73194.1 asyd|OJJ57983.1 ater|EAU33228.1 atub|OJI83099.1 aver|OJJ04418.1 awen|OJJ30285.1 azon|OJJ43974.1

Group4539: abra|OJJ78062.1 acar|OOF98792.1 acla|EAW06621.1 afis|EAW16144.1 afla|EED48686.1 afum|EAL88773.1 agla|OJJ86192.1 akaw|GAA85931.1 aluc|OJZ91133.1 anid|CBF79590.1 anig|EHA19832.1 aory|EIT73255.1 asyd|OJJ57981.1 ater|EAU33229.1 atub|OJI83143.1 aver|OJJ04419.1 awen|OJJ30283.1 azon|OJJ43973.1

Group4540: abra|OJJ78073.1 acar|OOF98790.1 acla|EAW15356.1 afis|EAW20356.1 afla|EED46131.1 afum|EAL85473.1 agla|OJJ81763.1 akaw|GAA85921.1 aluc|OJZ91146.1 anid|CBF74973.1 anig|EHA19844.1 aory|EIT76365.1 asyd|OJJ60088.1 ater|EAU34035.1 atub|OJI83160.1 aver|OJJ04175.1 awen|OJJ38942.1 azon|OJJ45059.1

Group4541: abra|OJJ78074.1 acar|OOF98789.1 afis|EAW16145.1 afla|EED48687.1 agla|OJJ86198.1 akaw|GAA85920.1 aluc|OJZ91147.1 anid|CBF79583.1 anig|EHA19845.1 aory|EIT73058.1 asyd|OJJ57977.1 ater|EAU33230.1 atub|OJI83161.1 aver|OJJ04423.1 awen|OJJ30281.1 azon|OJJ43972.1 acla|EAW06620.1 afum|EAL88774.1

Group4542: abra|OJJ78078.1 acar|OOF98786.1 acla|EAW06611.1 afis|EAW16204.1 afla|EED48697.1 afum|EAL88832.1 agla|OJJ86215.1 akaw|GAA85916.1 aluc|OJZ91150.1 anid|CBF79563.1 anig|EHA19848.1 aory|EIT73216.1 asyd|OJJ57913.1 ater|EAU33246.1 atub|OJI83164.1 aver|OJJ04480.1 awen|OJJ30268.1 azon|OJJ43964.1

Group4543: abra|OJJ78083.1 acar|OOF98783.1 acla|EAW06613.1 afis|EAW16177.1 afla|EED54377.1 afum|EAL88810.1 agla|OJJ86213.1 akaw|GAA85909.1 aluc|OJZ91157.1 anid|CBF79566.1 anig|EHA19853.1 aory|EIT75854.1 asyd|OJJ57917.1 ater|EAU33243.1 atub|OJI83172.1 aver|OJJ04474.1 awen|OJJ30272.1 azon|OJJ43966.1

Group4545: abra|OJJ78110.1 acar|OOF98756.1 acla|EAW06606.1 afis|EAW16217.1 afla|EED48711.1 afum|EAL88844.1 agla|OJJ86227.1 akaw|GAA85896.1 aluc|OJZ91175.1 anid|CBF79538.1 anig|EHA19867.1 aory|EIT73084.1 asyd|OJJ57899.1 ater|EAU33261.1 atub|OJI83188.1 aver|OJJ04496.1 awen|OJJ30255.1 azon|OJJ43955.1

Group4546: abra|OJJ78112.1 acar|OOF98755.1 acla|EAW06605.1 afis|EAW16218.1 afla|EED48712.1 afum|EAL88846.1 agla|OJJ86225.1 akaw|GAA85895.1 aluc|OJZ91177.1 anid|CBF79536.1 anig|EHA19868.1 aory|EIT73123.1 asyd|OJJ57898.1 ater|EAU33262.1 atub|OJI83189.1 aver|OJJ04497.1 awen|OJJ30254.1 azon|OJJ43957.1

Group4548: abra|OJJ78189.1 acar|OOF98670.1 acla|EAW06798.1 afis|EAW16347.1 afla|EED51752.1 afum|EAL88972.1 agla|OJJ88241.1 akaw|GAA85840.1 aluc|OJZ91260.1 anid|CBF89212.1 anig|EHA19948.1 aory|EIT80660.1 asyd|OJJ62523.1 ater|EAU36051.1 atub|OJI83249.1 aver|OJI96642.1 awen|OJJ37750.1 azon|OJJ47558.1

Group4550: abra|OJJ78206.1 acar|OOF98648.1 acla|EAW11242.1 afis|EAW18261.1 afla|EED48720.1 afum|EAL87625.1 agla|OJJ88165.1 akaw|GAA85820.1 aluc|OJZ91279.1 anid|CBF79529.1 anig|EHA19964.1 aory|EIT73088.1 asyd|OJJ57894.1 ater|EAU33269.1 atub|OJI83269.1 aver|OJJ04501.1 awen|OJJ30635.1 azon|OJJ47983.1

Group4551: abra|OJJ78213.1 acar|OOF98642.1 acla|EAW11237.1 afis|EAW18268.1 afla|EED48733.1 afum|EAL87633.1 agla|OJJ88159.1 akaw|GAA85814.1 aluc|OJZ91285.1 anid|CBF79516.1 anig|EHA19970.1 aory|EIT73048.1 asyd|OJJ57888.1 ater|EAU33275.1 atub|OJI83275.1 aver|OJJ04506.1 awen|OJJ30642.1 azon|OJJ47973.1

Group4552: abra|OJJ78216.1 acar|OOF98640.1 acla|EAW11234.1 afis|EAW18271.1 afla|EED48737.1 afum|EAL87636.1 agla|OJJ88156.1 akaw|GAA85811.1 aluc|OJZ91288.1 anid|CBF79510.1 anig|EHA19973.1 aory|EIT73125.1 asyd|OJJ57885.1 ater|EAU33277.1 atub|OJI83278.1 aver|OJJ04509.1 awen|OJJ30644.1 azon|OJJ47971.1

Group4553: abra|OJJ78217.1 acar|OOF98639.1 acla|EAW11232.1 afis|EAW18272.1 afla|EED48738.1 afum|EAL87637.1 agla|OJJ88155.1 akaw|GAA85807.1 aluc|OJZ91291.1 anid|CBF79508.1 anig|EHA19974.1 aory|EIT73235.1 asyd|OJJ57884.1 ater|EAU33278.1 atub|OJI83282.1 aver|OJJ04510.1 awen|OJJ30645.1 azon|OJJ47970.1

Group4554: abra|OJJ78218.1 acar|OOF98638.1 acla|EAW11231.1 afis|EAW18273.1 afla|EED48739.1 afum|EAL87638.1 agla|OJJ88154.1 akaw|GAA85806.1 aluc|OJZ91292.1 anid|CBF79507.1 anig|EHA19975.1 aory|EIT73224.1 asyd|OJJ57883.1 ater|EAU33279.1 atub|OJI83283.1 aver|OJJ04511.1 awen|OJJ30646.1 azon|OJJ47969.1

Group4555: abra|OJJ78220.1 acar|OOF98636.1 acla|EAW11229.1 afis|EAW18274.1 afla|EED48741.1 afum|EAL87640.1 agla|OJJ88152.1 akaw|GAA85805.1 aluc|OJZ91294.1 anid|CBF79503.1 anig|EHA19977.1 aory|EIT73281.1 asyd|OJJ57881.1 ater|EAU33280.1 atub|OJI83285.1 aver|OJJ04513.1 awen|OJJ30647.1 azon|OJJ47968.1

Group4556: abra|OJJ78221.1 acar|OOF98635.1 acla|EAW11228.1 afis|EAW18275.1 afla|EED48742.1 afum|EAL87641.1 agla|OJJ88151.1 akaw|GAA85804.1 aluc|OJZ91296.1 anid|CBF79499.1 anig|EHA19978.1 aory|EIT73223.1 asyd|OJJ57880.1 ater|EAU33281.1 atub|OJI83286.1 aver|OJJ04514.1 awen|OJJ30648.1 azon|OJJ47967.1

Group4557: abra|OJJ78226.1 acar|OOF98632.1 acla|EAW09284.1 afis|EAW21863.1 afla|EED46448.1 afum|EAL92877.2 agla|OJJ84270.1 akaw|GAA85799.1 aluc|OJZ91301.1 anid|CBF76755.1 anig|EHA19983.1 aory|EIT72377.1 asyd|OJJ61262.1 ater|EAU30861.1 atub|OJI83292.1 aver|OJJ00588.1 awen|OJJ34153.1 azon|OJJ43475.1

Group4558: abra|OJJ78229.1 acar|OOF98630.1 acla|EAW09280.1 afis|EAW21867.1 afla|EED46442.1 afum|EAL92881.1 agla|OJJ84268.1 akaw|GAA85795.1 aluc|OJZ91304.1 anid|CBF76763.1 anig|EHA19986.1 aory|EIT72329.1 asyd|OJJ61265.1 ater|EAU30864.1 atub|OJI83295.1 aver|OJJ00592.1 awen|OJJ34156.1 azon|OJJ43473.1

Group4559: abra|OJJ78230.1 acar|OOF98629.1 acla|EAW09279.1 afis|EAW21868.1 afla|EED46441.1 afum|EAL92882.1 agla|OJJ84267.1 akaw|GAA85794.1 aluc|OJZ91305.1 anid|CBF76765.1 anig|EHA19987.1 aory|EIT72366.1 asyd|OJJ61266.1 ater|EAU30866.1 atub|OJI83296.1 aver|OJJ00593.1 awen|OJJ34157.1 azon|OJJ43472.1

Group4560: abra|OJJ78231.1 acar|OOF98628.1 acla|EAW09278.1 afis|EAW21869.1 afla|EED46440.1 afum|EAL92883.1 agla|OJJ84264.1 akaw|GAA85793.1 aluc|OJZ91306.1 anid|CBF76767.1 anig|EHA19988.1 aory|EIT72428.1 asyd|OJJ61267.1 ater|EAU30867.1 atub|OJI83297.1 aver|OJJ00594.1 awen|OJJ34158.1 azon|OJJ43471.1

Group4561: abra|OJJ78233.1 acar|OOF98627.1 acla|EAW09277.1 afis|EAW21870.1 afla|EED46438.1 afum|EAL92884.1 agla|OJJ84262.1 akaw|GAA85792.1 aluc|OJZ91307.1 anid|CBF76769.1 anig|EHA19991.1 aory|EIT72393.1 asyd|OJJ61268.1 ater|EAU30868.1 atub|OJI83300.1 aver|OJJ00597.1 awen|OJJ34159.1 azon|OJJ43470.1

Group4562: abra|OJJ78234.1 acar|OOF98626.1 acla|EAW09276.1 afis|EAW21871.1 afla|EED46437.1 afum|EAL92885.1 agla|OJJ84261.1 akaw|GAA85791.1 aluc|OJZ91308.1 anid|CBF76771.1 anig|EHA19992.1

aory|EIT72319.1 asyd|OJJ61269.1 ater|EAU30869.1 atub|OJI83301.1 aver|OJJ00598.1 awen|OJJ34160.1 azon|OJJ43469.1

Group4563: abra|OJJ78237.1 acar|OOF98623.1 acla|EAW09273.1 afis|EAW21875.1 afla|EED46433.1 afum|EAL92889.2 agla|OJJ84257.1 akaw|GAA85788.1 aluc|OJZ91311.1 anid|CBF76776.1 anig|EHA19995.1 aory|EIT72389.1 asyd|OJJ61273.1 ater|EAU30874.1 atub|OJI83304.1 aver|OJJ00601.1 awen|OJJ34164.1 azon|OJJ43466.1

Group4564: abra|OJJ78238.1 acar|OOF98622.1 acla|EAW09272.1 afis|EAW21876.1 afla|EED46432.1 afum|EAL92890.1 agla|OJJ84256.1 akaw|GAA85787.1 aluc|OJZ91312.1 anid|CBF76778.1 anig|EHA19996.1 aory|EIT72368.1 asyd|OJJ61274.1 ater|EAU30875.1 atub|OJI83305.1 aver|OJJ00602.1 awen|OJJ34165.1 azon|OJJ43465.1

Group4565: abra|OJJ78239.1 acar|OOF98621.1 acla|EAW09271.1 afis|EAW21877.1 afla|EED46431.1 afum|EAL92891.2 agla|OJJ84255.1 akaw|GAA85786.1 aluc|OJZ91313.1 anid|CBF76780.1 anig|EHA19997.1 aory|EIT72420.1 asyd|OJJ61275.1 ater|EAU30876.1 atub|OJI83306.1 aver|OJJ00603.1 awen|OJJ34166.1 azon|OJJ43464.1

Group4566: abra|OJJ78241.1 acar|OOF98619.1 acla|EAW09269.1 afis|EAW21879.1 afla|EED46429.1 afum|EAL92893.1 agla|OJJ84253.1 akaw|GAA85783.1 aluc|OJZ91315.1 anid|CBF76784.1 anig|EHA19998.1 aory|EIT72355.1 asyd|OJJ61277.1 ater|EAU30877.1 atub|OJI83309.1 aver|OJJ00605.1 awen|OJJ34168.1 azon|OJJ43462.1

Group4567: abra|OJJ78242.1 acar|OOF98618.1 acla|EAW09268.1 afis|EAW21880.1 afla|EED46428.1 afum|EAL92894.2 agla|OJJ84252.1 akaw|GAA85782.1 aluc|OJZ91316.1 anid|CBF76786.1 anig|EHA19999.1 aory|EIT72330.1 asyd|OJJ61278.1 ater|EAU30878.1 atub|OJI83310.1 aver|OJJ00606.1 awen|OJJ34169.1 azon|OJJ43461.1

Group4568: abra|OJJ78244.1 acar|OOF98616.1 acla|EAW09265.1 afis|EAW21883.1 afla|EED46424.1 afum|EAL92897.1 agla|OJJ84251.1 akaw|GAA85780.1 aluc|OJZ91318.1 anid|CBF76788.1 anig|EHA20001.1 aory|EIT72383.1 asyd|OJJ61280.1 ater|EAU30881.1 atub|OJI83312.1 aver|OJJ00607.1 awen|OJJ34172.1 azon|OJJ43460.1

Group4569: abra|OJJ78247.1 acar|OOF98615.1 acla|EAW09263.1 afis|EAW21884.1 afla|EED46423.1 afum|EAL92898.1 agla|OJJ84249.1 akaw|GAA85779.1 aluc|OJZ91321.1 anid|CBF76790.1 anig|EHA20003.1 aory|EIT72443.1 asyd|OJJ61281.1 ater|EAU30882.1 atub|OJI83314.1 aver|OJJ00608.1 awen|OJJ34173.1 azon|OJJ43459.1

Group4570: abra|OJJ78249.1 acar|OOF98614.1 acla|EAW09262.1 afis|EAW21885.1 afla|EED46422.1 afum|EAL92899.1 agla|OJJ84248.1 akaw|GAA85778.1 aluc|OJZ91322.1 anid|CBF76792.1 anig|EHA20004.1 aory|EIT72353.1 asyd|OJJ61282.1 ater|EAU30883.1 atub|OJI83315.1 aver|OJJ00609.1 awen|OJJ34174.1 azon|OJJ43458.1

Group4571: abra|OJJ78250.1 acar|OOF98612.1 acla|EAW09261.1 afis|EAW21886.1 afla|EED46421.1 afum|EAL92900.1 agla|OJJ84247.1 akaw|GAA85777.1 aluc|OJZ91323.1 anid|CBF76794.1 anig|EHA20005.1 aory|EIT72442.1 asyd|OJJ61283.1 ater|EAU30884.1 atub|OJI83316.1 aver|OJJ00610.1 awen|OJJ34175.1 azon|OJJ43457.1

Group4572: abra|OJJ78252.1 acar|OOF98611.1 acla|EAW09259.1 afis|EAW21888.1 afla|EED46419.1 afum|EAL92902.1 agla|OJJ84245.1 akaw|GAA85775.1 aluc|OJZ91325.1 anid|CBF76798.1 anig|EHA20007.1 aory|EIT72406.1 asyd|OJJ61285.1 ater|EAU30885.1 atub|OJI83318.1 aver|OJJ00611.1 awen|OJJ34177.1 azon|OJJ43455.1

Group4573: abra|OJJ78253.1 acar|OOF98610.1 acla|EAW09258.1 afis|EAW21889.1 afla|EED46418.1 afum|EAL92903.1 agla|OJJ84244.1 akaw|GAA85774.1 aluc|OJZ91326.1 anid|CBF76800.1 anig|EHA20008.1 aory|EIT72432.1 asyd|OJJ61286.1 ater|EAU30886.1 atub|OJI83319.1 aver|OJJ00612.1 awen|OJJ34178.1 azon|OJJ43454.1

Group4574: abra|OJJ78255.1 acar|OOF98607.1 acla|EAW09254.1 afis|EAW21893.1 afla|EED46415.1 afum|EAL92907.1 agla|OJJ84239.1 akaw|GAA85771.1 aluc|OJZ91329.1 anid|CBF76806.1 anig|EHA20011.1 aory|EIT72313.1 asyd|OJJ61289.1 ater|EAU30889.1 atub|OJI83322.1 aver|OJJ00615.1 awen|OJJ34182.1 azon|OJJ43450.1

Group4575: abra|OJJ78256.1 acar|OOF98606.1 acla|EAW09253.1 afis|EAW21894.1 afla|EED46414.1 afum|EAL92908.1 agla|OJJ84238.1 akaw|GAA85770.1 aluc|OJZ91330.1 anid|CBF76808.1 anig|EHA20012.1 aory|EIT72378.1 asyd|OJJ61290.1 ater|EAU30890.1 atub|OJI83323.1 aver|OJJ00616.1 awen|OJJ34183.1 azon|OJJ43449.1

Group4576: abra|OJJ78260.1 acar|OOF98600.1 acla|EAW09250.1 afis|EAW21897.1 afla|EED46411.1 afum|EAL92910.1 agla|OJJ84227.1 akaw|GAA85765.1 aluc|OJZ91335.1 anid|CBF76814.1 anig|EHA20016.1 aory|EIT72440.1 asyd|OJJ61293.1 ater|EAU30892.1 atub|OJI83327.1 aver|OJJ00619.1 awen|OJJ34186.1 azon|OJJ43447.1

Group4577: abra|OJJ78261.1 acar|OOF98599.1 acla|EAW09249.1 afis|EAW21898.1 afla|EED46408.1 afum|EAL92911.2 agla|OJJ84228.1 akaw|GAA85764.1 aluc|OJZ91336.1 anid|CBF76816.1 anig|EHA20017.1 aory|EIT72346.1 asyd|OJJ61294.1 ater|EAU30893.1 atub|OJI83328.1 aver|OJJ00620.1 awen|OJJ34187.1 azon|OJJ43446.1

Group4578: abra|OJJ78263.1 acar|OOF98598.1 acla|EAW09248.1 afis|EAW21899.1 afla|EED46407.1 afum|EAL92912.1 agla|OJJ84229.1 akaw|GAA85762.1 aluc|OJZ91338.1 anid|CBF76818.1 anig|EHA20018.1 aory|EIT72328.1 asyd|OJJ61295.1 ater|EAU30894.1 atub|OJI83330.1 aver|OJJ00621.1 awen|OJJ34188.1 azon|OJJ43445.1

Group4579: abra|OJJ78264.1 acar|OOF98597.1 acla|EAW09247.1 afis|EAW21900.1 afla|EED46406.1 afum|EAL92913.1 agla|OJJ84230.1 akaw|GAA85761.1 aluc|OJZ91339.1 anid|CBF76820.1 anig|EHA20019.1 aory|EIT72352.1 asyd|OJJ61296.1 ater|EAU30895.1 atub|OJI83331.1 aver|OJJ00622.1 awen|OJJ34189.1 azon|OJJ43444.1

Group4580: abra|OJJ78266.1 acar|OOF98596.1 acla|EAW09246.1 afis|EAW21901.1 afla|EED46405.1 afum|EAL92914.1 agla|OJJ84231.1 akaw|GAA85760.1 aluc|OJZ91340.1 anid|CBF76822.1 anig|EHA20020.1 aory|EIT72444.1 asyd|OJJ61297.1 ater|EAU30896.1 atub|OJI83332.1 aver|OJJ00623.1 awen|OJJ34190.1 azon|OJJ43443.1

Group4581: abra|OJJ78267.1 acar|OOF98595.1 acla|EAW09245.1 afis|EAW21902.1 afla|EED46403.1 afum|EAL92915.1 agla|OJJ84232.1 akaw|GAA85759.1 aluc|OJZ91341.1 anid|CBF76824.1 anig|EHA20021.1 aory|EIT72339.1 asyd|OJJ61298.1 ater|EAU30897.1 atub|OJI83333.1 aver|OJJ00624.1 awen|OJJ34191.1 azon|OJJ43442.1

Group4582: abra|OJJ78268.1 acar|OOF98594.1 acla|EAW09244.1 afis|EAW21903.1 afla|EED46402.1 afum|EAL92916.1 agla|OJJ84233.1 akaw|GAA85758.1 aluc|OJZ91342.1 anid|CBF76826.1 anig|EHA20022.1 aory|EIT72439.1 asyd|OJJ61299.1 ater|EAU30898.1 atub|OJI83334.1 aver|OJJ00625.1 awen|OJJ34192.1 azon|OJJ43441.1

Group4583: abra|OJJ78275.1 acar|OOF98592.1 acla|EAW09239.1 afis|EAW21909.1 afla|EED46641.1 afum|EAL92921.1 agla|OJJ84222.1 akaw|GAA85751.1 aluc|OJZ91350.1 anid|CBF76837.1 anig|EHA20028.1 aory|EIT77855.1 asyd|OJJ61305.1 ater|EAU30909.1 atub|OJI83342.1 aver|OJJ00630.1 awen|OJJ34199.1 azon|OJJ43435.1

Group4584: abra|OJJ78277.1 acar|OOF98590.1 acla|EAW09237.1 afis|EAW21911.1 afla|EED46639.1 afum|EAL92923.1 agla|OJJ84220.1 akaw|GAA85749.1 aluc|OJZ91352.1 anid|CBF76842.1 anig|EHA20030.1 aory|EIT77915.1 asyd|OJJ61307.1 ater|EAU30911.1 atub|OJI83344.1 aver|OJJ00632.1 awen|OJJ34201.1 azon|OJJ43433.1

Group4585: abra|OJJ78278.1 acar|OOF98588.1 acla|EAW09236.1 afis|EAW21912.1 afla|EED46638.1 afum|EAL92924.1 agla|OJJ84219.1 akaw|GAA85748.1 aluc|OJZ91353.1 anid|CBF76844.1 anig|EHA20031.1 aory|EIT77829.1 asyd|OJJ61308.1 ater|EAU30912.1 atub|OJI83345.1 aver|OJJ00633.1 awen|OJJ34202.1 azon|OJJ43432.1

Group4586: abra|OJJ78280.1 acar|OOF98589.1 acla|EAW09235.1 afis|EAW21913.1 afla|EED46637.1 afum|EAL92925.1 agla|OJJ84210.1 akaw|GAA85747.1 aluc|OJZ91354.1 anid|CBF76846.1 anig|EHA20032.1 aory|EIT77951.1 asyd|OJJ61309.1 ater|EAU30914.1 atub|OJI83347.1 aver|OJJ00634.1 awen|OJJ34203.1 azon|OJJ43431.1

Group4587: abra|OJJ78282.1 acar|OOF98586.1 acla|EAW09233.1 afis|EAW21915.1 afla|EED46634.1 afum|EAL92927.1 agla|OJJ84208.1 akaw|GAA85745.1 aluc|OJZ91356.1 anid|CBF76850.1 anig|EHA20034.1 aory|EIT77867.1 asyd|OJJ61311.1 ater|EAU30916.1 atub|OJI83349.1 aver|OJJ00636.1 awen|OJJ34205.1 azon|OJJ43429.1

Group4588: abra|OJJ78284.1 acar|OOF98584.1 acla|EAW09231.1 afis|EAW21917.1 afla|EED46632.1 afum|EAL92929.1 agla|OJJ84206.1 akaw|GAA85743.1 aluc|OJZ91358.1 anid|CBF76854.1 anig|EHA20036.1 aory|EIT77843.1 asyd|OJJ61313.1 ater|EAU30919.1 atub|OJI83351.1 aver|OJJ00638.1 awen|OJJ34207.1 azon|OJJ43427.1

Group4589: abra|OJJ78285.1 acar|OOF98583.1 acla|EAW09230.1 afis|EAW21918.1 afum|EAL92930.1 agla|OJJ84205.1 akaw|GAA85742.1 aluc|OJZ91359.1 anid|CBF76856.1 anig|EHA20037.1 asyd|OJJ61314.1 ater|EAU30920.1 atub|OJI83352.1 aver|OJJ00639.1 awen|OJJ34208.1 azon|OJJ43426.1 afla|EED46631.1 aory|EIT77816.1

Group4590: abra|OJJ78287.1 acar|OOF98581.1 acla|EAW09228.1 afis|EAW21920.1 afla|EED46630.1 afum|EAL92932.1 agla|OJJ84203.1 akaw|GAA85740.1 aluc|OJZ91361.1 anid|CBF76860.1 anig|EHA20039.1 aory|EIT77823.1 asyd|OJJ61316.1 ater|EAU30925.1 atub|OJI83354.1 aver|OJJ00641.1 awen|OJJ34210.1 azon|OJJ43424.1

Group4591: abra|OJJ78288.1 acar|OOF98579.1 acla|EAW09227.1 afis|EAW21921.1 afla|EED46629.1 afum|EAL92934.1 agla|OJJ84202.1 akaw|GAA85739.1 aluc|OJZ91362.1 anid|CBF76862.1 anig|EHA20040.1

aory|EIT77918.1 asyd|OJJ61317.1 ater|EAU30926.1 atub|OJI83355.1 aver|OJJ00642.1 awen|OJJ34211.1 azon|OJJ43423.1

Group4592: abra|OJJ78289.1 acar|OOF98580.1 acla|EAW09226.1 afis|EAW21922.1 afla|EED46628.1 afum|EAL92935.1 agla|OJJ84201.1 akaw|GAA85738.1 aluc|OJZ91363.1 anid|CBF76864.1 anig|EHA20041.1 aory|EIT77831.1 asyd|OJJ61318.1 ater|EAU30927.1 atub|OJI83356.1 aver|OJJ00643.1 awen|OJJ34212.1 azon|OJJ43422.1

Group4593: abra|OJJ78290.1 acar|OOF98578.1 acla|EAW09225.1 afis|EAW21923.1 afla|EED46627.1 afum|EAL92936.1 agla|OJJ84200.1 akaw|GAA85737.1 aluc|OJZ91364.1 anid|CBF76866.1 anig|EHA20042.1 aory|EIT77896.1 asyd|OJJ61319.1 ater|EAU30928.1 atub|OJI83357.1 aver|OJJ00644.1 awen|OJJ34213.1 azon|OJJ43421.1

Group4594: abra|OJJ78292.1 acar|OOF98576.1 afis|EAW21924.1 afla|EED46615.1 afum|EAL92937.1 akaw|GAA85735.1 aluc|OJZ91366.1 anig|EHA20044.1 aory|EIT77890.1 ater|EAU30929.1 atub|OJI83359.1 aver|OJJ00647.1 azon|OJJ43420.1 acla|EAW09224.1 agla|OJJ84199.1 anid|CBF76872.1 asyd|OJJ61322.1 awen|OJJ34214.1

Group4595: abra|OJJ78293.1 acar|OOF98575.1 acla|EAW09223.1 afis|EAW21925.1 afla|EED46616.1 afum|EAL92938.1 agla|OJJ84198.1 akaw|GAA85734.1 aluc|OJZ91367.1 anid|CBF76882.1 anig|EHA20045.1 aory|EIT77869.1 asyd|OJJ61327.1 ater|EAU30930.1 atub|OJI83360.1 aver|OJJ00652.1 awen|OJJ34215.1 azon|OJJ43419.1

Group4596: abra|OJJ78298.1 acar|OOF98571.1 acla|EAW09218.1 afis|EAW21930.1 afla|EED46623.1 afum|EAL92942.1 agla|OJJ84191.1 akaw|GAA85729.1 aluc|OJZ91372.1 anid|CBF76884.1 anig|EHA20050.1 aory|EIT77852.1 asyd|OJJ61328.1 ater|EAU30936.1 atub|OJI83365.1 aver|OJJ00654.1 awen|OJJ34220.1 azon|OJJ43414.1

Group4597: abra|OJJ78299.1 acar|OOF98567.1 acla|EAW09216.1 afis|EAW21931.1 afla|EED46626.1 afum|EAL92943.2 agla|OJJ84190.1 akaw|GAA85727.1 aluc|OJZ91376.1 anid|CBF76885.1 anig|EHA20051.1 aory|EIT77815.1 asyd|OJJ61330.1 ater|EAU30937.1 atub|OJI83366.1 aver|OJJ00655.1 awen|OJJ34222.1 azon|OJJ43413.1

Group4598: abra|OJJ78301.1 acar|OOF98565.1 acla|EAW09214.1 afis|EAW21933.1 afla|EED46353.1 afum|EAL92946.1 agla|OJJ84187.1 akaw|GAA85725.1 aluc|OJZ91378.1 anid|CBF76887.1 anig|EHA20053.1 aory|EIT81186.1 asyd|OJJ61333.1 ater|EAU30938.1 atub|OJI83371.1 aver|OJJ00657.1 awen|OJJ34224.1 azon|OJJ43411.1

Group4599: abra|OJJ78302.1 acar|OOF98564.1 acla|EAW09213.1 afis|EAW21934.1 afla|EED46354.1 afum|EAL92947.1 agla|OJJ84186.1 akaw|GAA85724.1 aluc|OJZ91379.1 anid|CBF76889.1 anig|EHA20054.1 aory|EIT81211.1 asyd|OJJ61334.1 ater|EAU30939.1 atub|OJI83372.1 aver|OJJ00658.1 awen|OJJ34225.1 azon|OJJ43410.1

Group4600: abra|OJJ78308.1 acar|OOF98558.1 acla|EAW09212.1 afis|EAW21935.1 afla|EED46652.1 afum|EAL92948.2 agla|OJJ84185.1 akaw|GAA85721.1 aluc|OJZ91383.1 anid|CBF76899.1 anig|EHA20060.1 aory|EIT77952.1 asyd|OJJ61338.1 ater|EAU30941.1 atub|OJI83376.1 aver|OJJ00662.1 awen|OJJ34226.1 azon|OJJ43409.1

Group4601: abra|OJJ78319.1 acar|OOF98556.1 acla|EAW09206.1 afis|EAW21941.1 afla|EED46658.1 afum|EAL92955.1 agla|OJJ84182.1 akaw|GAA85711.1 aluc|OJZ91392.1 anid|CBF84078.1 anig|EHA20070.1 aory|EIT77820.1 asyd|OJJ61342.1 ater|EAU30946.1 atub|OJI83385.1 aver|OJJ00666.1 awen|OJJ34232.1 azon|OJJ43403.1

Group4602: abra|OJJ78323.1 acar|OOF98551.1 acla|EAW09202.1 afis|EAW21945.1 afla|EED46662.1 afum|EAL92959.1 agla|OJJ84179.1 akaw|GAA85705.1 aluc|OJZ91398.1 anid|CBF84050.1 anig|EHA20075.1 aory|EIT77962.1 asyd|OJJ61343.1 ater|EAU30993.1 atub|OJI83390.1 aver|OJJ00667.1 awen|OJJ34236.1 azon|OJJ43400.1

Group4603: abra|OJJ78325.1 acar|OOF98550.1 acla|EAW09201.1 afis|EAW21946.1 afla|EED46663.1 afum|EAL92960.1 agla|OJJ84174.1 akaw|GAA85704.1 aluc|OJZ91399.1 anid|CBF84059.1 anig|EHA20076.1 aory|EIT77808.1 asyd|OJJ61345.1 ater|EAU30994.1 atub|OJI83391.1 aver|OJJ00670.1 awen|OJJ34237.1 azon|OJJ43399.1

Group4604: abra|OJJ78327.1 acar|OOF98548.1 acla|EAW09199.1 afis|EAW21948.1 afla|EED46666.1 afum|EAL92962.1 agla|OJJ84172.1 akaw|GAA85702.1 aluc|OJZ91401.1 anid|CBF84062.1 anig|EHA20078.1 aory|EIT77953.1 asyd|OJJ61347.1 ater|EAU30996.1 atub|OJI83393.1 aver|OJJ00674.1 awen|OJJ34240.1 azon|OJJ43397.1

Group4605: abra|OJJ78328.1 acar|OOF98547.1 acla|EAW09198.1 afis|EAW21949.1 afum|EAL92963.1 agla|OJJ84171.1 akaw|GAA85701.1 aluc|OJZ91402.1 anid|CBF84064.1 anig|EHA20079.1 aory|EIT77849.1 asyd|OJJ61348.1 ater|EAU30997.1 atub|OJI83395.1 aver|OJJ00675.1 awen|OJJ34241.1 azon|OJJ43396.1 afla|EED46667.1

Group4606: abra|OJJ78329.1 acar|OOF98546.1 acla|EAW09197.1 afis|EAW21950.1 afla|EED46668.1 afum|EAL92964.1 agla|OJJ84170.1 akaw|GAA85700.1 aluc|OJZ91403.1 anid|CBF84066.1 anig|EHA20080.1 aory|EIT77926.1 asyd|OJJ61349.1 ater|EAU30998.1 atub|OJI83396.1 aver|OJJ00676.1 awen|OJJ34242.1 azon|OJJ43395.1

Group4607: abra|OJJ78334.1 acar|OOF98544.1 acla|EAW09194.1 afis|EAW21952.1 afla|EED46669.1 afum|EAL92966.1 agla|OJJ84159.1 akaw|GAA85696.1 aluc|OJZ91407.1 anid|CBF84069.1 anig|EHA20083.1 aory|EIT77916.1 asyd|OJJ61351.1 ater|EAU31000.1 atub|OJI83400.1 aver|OJJ00678.1 awen|OJJ34244.1 azon|OJJ43394.1

Group4608: abra|OJJ78337.1 acar|OOF98543.1 acla|EAW09193.1 afis|EAW21953.1 afla|EED46671.1 afum|EAL92967.1 agla|OJJ84160.1 akaw|GAA85695.1 aluc|OJZ91409.1 anid|CBF84070.1 anig|EHA20084.1 aory|EIT77807.1 asyd|OJJ61352.1 ater|EAU31001.1 atub|OJI83403.1 aver|OJJ00679.1 awen|OJJ34245.1 azon|OJJ43393.1

Group4609: abra|OJJ78339.1 acar|OOF98541.1 acla|EAW09192.1 afis|EAW21954.1 afla|EED46672.1 afum|EAL92968.1 agla|OJJ84161.1 akaw|GAA85694.1 aluc|OJZ91411.1 anid|CBF84074.1 anig|EHA20085.1 aory|EIT77806.1 asyd|OJJ61353.1 ater|EAU31060.1 atub|OJI83404.1 aver|OJJ00680.1 awen|OJJ34246.1 azon|OJJ43392.1

Group4610: abra|OJJ78345.1 acar|OOF90269.1 acla|EAW09189.1 afis|EAW21957.1 afla|EED46676.1 afum|EAL92971.1 agla|OJJ84164.1 akaw|GAA85688.1 aluc|OJZ91418.1 anid|CBF84037.1 anig|EHA20090.1 aory|EIT77863.1 asyd|OJJ61362.1 ater|EAU31063.1 atub|OJI83412.1 aver|OJJ00689.1 awen|OJJ34249.1 azon|OJJ43389.1

Group4611: abra|OJJ78346.1 acar|OOF90268.1 acla|EAW09188.1 afis|EAW21958.1 afla|EED46678.1 afum|EAL92972.1 agla|OJJ84158.1 akaw|GAA85687.1 aluc|OJZ91420.1 anid|CBF84041.1 anig|EHA20091.1 aory|EIT77880.1 asyd|OJJ61360.1 ater|EAU31078.1 atub|OJI83413.1 aver|OJJ00687.1 awen|OJJ34251.1 azon|OJJ43387.1

Group4612: abra|OJJ78347.1 acar|OOF90267.1 acla|EAW09187.1 afis|EAW21959.1 afla|EED46679.1 afum|EAL92973.1 agla|OJJ84157.1 akaw|GAA85686.1 aluc|OJZ91421.1 anid|CBF84043.1 anig|EHA20092.1 aory|EIT77956.1 asyd|OJJ61359.1 ater|EAU31079.1 atub|OJI83414.1 aver|OJJ00686.1 awen|OJJ34252.1 azon|OJJ43386.1

Group4613: abra|OJJ78351.1 acar|OOF98536.1 acla|EAW09184.1 afis|EAW21962.1 afla|EED46683.1 afum|EAL92976.1 agla|OJJ84141.1 akaw|GAA85683.1 aluc|OJZ91425.1 anid|CBF82742.1 anig|EHA20095.1 aory|EIT77876.1 asyd|OJJ61622.1 ater|EAU31082.1 atub|OJI83418.1 aver|OJJ00946.1 awen|OJJ34256.1 azon|OJJ43383.1

Group4614: abra|OJJ78353.1 acar|OOF98535.1 acla|EAW09182.1 afis|EAW21964.1 afla|EED46685.1 afum|EAL92978.2 agla|OJJ84139.1 akaw|GAA85681.1 aluc|OJZ91428.1 anid|CBF82738.1 anig|EHA20097.1 aory|EIT77960.1 asyd|OJJ61624.1 ater|EAU31084.1 atub|OJI83421.1 aver|OJJ00949.1 awen|OJJ34258.1 azon|OJJ43381.1

Group4615: abra|OJJ78354.1 acar|OOF98534.1 acla|EAW09181.1 afis|EAW21965.1 afla|EED46686.1 afum|EAL92979.1 agla|OJJ84138.1 akaw|GAA85680.1 aluc|OJZ91429.1 anid|CBF82736.1 anig|EHA20098.1 aory|EIT77854.1 asyd|OJJ61625.1 ater|EAU31085.1 atub|OJI83422.1 aver|OJJ00950.1 awen|OJJ34259.1 azon|OJJ43378.1

Group4616: abra|OJJ78356.1 acar|OOF98533.1 acla|EAW09178.1 afis|EAW21968.1 afla|EED46689.1 afum|EAL92982.1 agla|OJJ84131.1 akaw|GAA85677.1 aluc|OJZ91431.1 anid|CBF82731.1 anig|EHA20100.1 aory|EIT77885.1 asyd|OJJ61628.1 ater|EAU31088.1 atub|OJI83424.1 aver|OJJ00952.1 awen|OJJ34261.1 azon|OJJ43376.1

Group4617: abra|OJJ78363.1 acar|OOF98524.1 acla|EAW09172.1 afis|EAW21974.1 afla|EED46697.1 afum|EAL92989.1 agla|OJJ84128.1 akaw|GAA85670.1 aluc|OJZ91438.1 anid|CBF82716.1 anig|EHA20103.1 aory|EIT77925.1 asyd|OJJ61637.1 ater|EAU31092.1 atub|OJI83432.1 aver|OJJ00960.1 awen|OJJ34267.1 azon|OJJ43369.1

Group4618: abra|OJJ78365.1 acar|OOF98521.1 acla|EAW09170.1 afis|EAW21976.1 afla|EED46699.1 afum|EAL92991.1 agla|OJJ84125.1 akaw|GAA85668.1 aluc|OJZ91442.1 anid|CBF82720.1 anig|EHA20106.1 aory|EIT77848.1 asyd|OJJ61635.1 ater|EAU31094.1 atub|OJI83436.1 aver|OJJ00958.1 awen|OJJ34269.1 azon|OJJ43368.1

Group4619: abra|OJJ78366.1 acar|OOF98520.1 acla|EAW09169.1 afis|EAW21977.1 afla|EED46700.1 afum|EAL92992.1 agla|OJJ84124.1 akaw|GAA85667.1 aluc|OJZ91443.1 anid|CBF82722.1 anig|EHA20107.1 aory|EIT77809.1 asyd|OJJ61634.1 ater|EAU31095.1 atub|OJI83437.1 aver|OJJ00957.1 awen|OJJ34270.1 azon|OJJ43366.1

Group4620: abra|OJJ78368.1 acar|OOF98518.1 acla|EAW09167.1 afis|EAW21979.1 afla|EED46702.1 afum|EAL92994.1 agla|OJJ84120.1 akaw|GAA85665.1 aluc|OJZ91445.1 anid|CBF82711.1 anig|EHA20109.1

aory|EIT77847.1 asyd|OJJ61640.1 ater|EAU31097.1 atub|OJI83439.1 aver|OJJ00963.1 awen|OJJ34272.1 azon|OJJ43364.1

Group4621: abra|OJJ78369.1 acar|OOF90130.1 acla|EAW09166.1 afis|EAW21980.1 afla|EED46704.1 afum|EAL92995.1 agla|OJJ84119.1 akaw|GAA85664.1 aluc|OJZ91446.1 anid|CBF82709.1 anig|EHA20110.1 aory|EIT77845.1 asyd|OJJ61641.1 ater|EAU31100.1 atub|OJI83440.1 aver|OJJ00964.1 awen|OJJ34274.1 azon|OJJ43363.1

Group4622: abra|OJJ78371.1 acar|OOF90128.1 acla|EAW09164.1 afis|EAW21982.1 afla|EED46706.1 afum|EAL92997.1 agla|OJJ84118.1 akaw|GAA85662.1 aluc|OJZ91448.1 anid|CBF82705.1 anig|EHA20112.1 aory|EIT77874.1 asyd|OJJ61643.1 ater|EAU31101.1 atub|OJI83441.1 aver|OJJ00965.1 awen|OJJ34276.1 azon|OJJ43361.1

Group4623: abra|OJJ78372.1 acar|OOF90125.1 acla|EAW09163.1 afis|EAW21983.1 afla|EED46707.1 afum|EAL92998.1 agla|OJJ84115.1 akaw|GAA85661.1 aluc|OJZ91449.1 anid|CBF82703.1 anig|EHA20113.1 aory|EIT77859.1 asyd|OJJ61644.1 ater|EAU31102.1 atub|OJI83442.1 aver|OJJ00966.1 awen|OJJ34280.1 azon|OJJ43360.1

Group4624: abra|OJJ78374.1 acar|OOF98515.1 acla|EAW09161.1 afis|EAW21985.1 afla|EED46709.1 afum|EAL93000.1 agla|OJJ84113.1 akaw|GAA85659.1 aluc|OJZ91451.1 anid|CBF82698.1 anig|EHA20115.1 aory|EIT77948.1 asyd|OJJ61646.1 ater|EAU31104.1 atub|OJI83444.1 aver|OJJ00968.1 awen|OJJ34282.1 azon|OJJ43358.1

Group4625: abra|OJJ78375.1 acar|OOF98514.1 acla|EAW09160.1 afis|EAW21986.1 afla|EED46710.1 afum|EAL93002.1 agla|OJJ84112.1 akaw|GAA85658.1 aluc|OJZ91452.1 anid|CBF82696.1 anig|EHA20116.1 aory|EIT77888.1 asyd|OJJ61647.1 ater|EAU31106.1 atub|OJI83445.1 aver|OJJ00969.1 awen|OJJ34283.1 azon|OJJ43357.1

Group4626: abra|OJJ78378.1 acar|OOF98510.1 acla|EAW09155.1 afis|EAW21991.1 afla|EED46715.1 afum|EAL93006.2 agla|OJJ84107.1 akaw|GAA85654.1 aluc|OJZ91457.1 anid|CBF82687.1 anig|EHA20120.1 aory|EIT77887.1 asyd|OJJ61652.1 ater|EAU31110.1 atub|OJI83450.1 aver|OJJ00973.1 awen|OJJ34287.1 azon|OJJ43351.1

Group4627: abra|OJJ78380.1 acar|OOF98508.1 acla|EAW09153.1 afis|EAW21993.1 afla|EED46717.1 afum|EAL93008.1 agla|OJJ84104.1 akaw|GAA85652.1 aluc|OJZ91459.1 anid|CBF82683.1 anig|EHA20122.1 aory|EIT77842.1 asyd|OJJ61655.1 ater|EAU31112.1 atub|OJI83452.1 aver|OJJ00976.1 awen|OJJ34289.1 azon|OJJ43349.1

Group4628: abra|OJJ78381.1 acar|OOF98507.1 acla|EAW09152.1 afis|EAW21994.1 afla|EED46718.1 afum|EAL93009.1 agla|OJJ84103.1 akaw|GAA85651.1 aluc|OJZ91460.1 anid|CBF82681.1 anig|EHA20123.1 aory|EIT77902.1 asyd|OJJ61656.1 ater|EAU31113.1 atub|OJI83453.1 aver|OJJ00977.1 awen|OJJ34290.1 azon|OJJ43348.1

Group4629: abra|OJJ78382.1 acar|OOF98506.1 acla|EAW09151.1 afis|EAW21995.1 afla|EED46719.1 afum|EAL93010.1 agla|OJJ84102.1 akaw|GAA85650.1 aluc|OJZ91461.1 anid|CBF82679.1 anig|EHA20124.1 aory|EIT77906.1 asyd|OJJ61657.1 ater|EAU31114.1 atub|OJI83454.1 aver|OJJ00978.1 awen|OJJ34291.1 azon|OJJ43347.1

Group4630: abra|OJJ78383.1 acar|OOF98505.1 acla|EAW09150.1 afis|EAW21996.1 afla|EED46720.1 afum|EAL93011.1 agla|OJJ84101.1 akaw|GAA85649.1 aluc|OJZ91462.1 anid|CBF82677.1 anig|EHA20125.1 aory|EIT77812.1 asyd|OJJ61658.1 ater|EAU31115.1 atub|OJI83455.1 aver|OJJ00979.1 awen|OJJ34292.1 azon|OJJ43346.1

Group4631: abra|OJJ78384.1 acar|OOF98504.1 acla|EAW09149.1 afis|EAW21997.1 afla|EED46721.1 afum|EAL93012.1 agla|OJJ84100.1 akaw|GAA85648.1 aluc|OJZ91463.1 anid|CBF82675.1 anig|EHA20126.1 aory|EIT77875.1 asyd|OJJ61659.1 ater|EAU31116.1 atub|OJI83456.1 aver|OJJ00980.1 awen|OJJ34293.1 azon|OJJ43345.1

Group4632: abra|OJJ78385.1 acar|OOF98503.1 acla|EAW09148.1 afis|EAW21998.1 afla|EED46722.1 afum|EAL93013.1 agla|OJJ84099.1 akaw|GAA85647.1 aluc|OJZ91464.1 anid|CBF82674.1 anig|EHA20127.1 aory|EIT77891.1 asyd|OJJ61660.1 ater|EAU31117.1 atub|OJI83457.1 aver|OJJ00981.1 awen|OJJ34294.1 azon|OJJ43344.1

Group4633: abra|OJJ78386.1 acar|OOF98502.1 acla|EAW09147.1 afis|EAW21999.1 afla|EED46723.1 afum|EAL93014.1 agla|OJJ84098.1 akaw|GAA85646.1 aluc|OJZ91465.1 anid|CBF82672.1 anig|EHA20128.1 aory|EIT77907.1 asyd|OJJ61661.1 ater|EAU31118.1 atub|OJI83458.1 aver|OJJ00982.1 awen|OJJ34295.1 azon|OJJ43343.1

Group4634: abra|OJJ78388.1 acar|OOF98500.1 acla|EAW09145.1 afis|EAW22002.1 afla|EED46727.1 afum|EAL93017.1 agla|OJJ84096.1 akaw|GAA85644.1 aluc|OJZ91467.1 anid|CBF82666.1 anig|EHA20129.1 aory|EIT77844.1 asyd|OJJ61664.1 ater|EAU31121.1 atub|OJI83460.1 aver|OJJ00985.1 awen|OJJ34297.1 azon|OJJ43341.1

**Additional data file 3:** *A. fumigatus* and *A. fischeri* orthology analysis (Inparanoid output).

####################################
Group summary

one_to_one: 8536
one_to_many: 10
many_to_one: 14
many_to_many: 2

Total groups of orthologs: 8562
------------------------------------
8581 in-paralogs from *A. fumigatus*
8579 in-paralogs from *A. fischeri*
####################################

---

Group of orthologs #1. Best score 14205 bits
Score difference with first non-orthologous sequence - *A. fumigatus*:11599 *A. fischeri*:12414
afum|EAL90366.1 100.00% afis|EAW22836.1 100.00%
Bootstrap support for afum|EAL90366.1 as seed ortholog is 100%.
Bootstrap support for afis|EAW22836.1 as seed ortholog is 100%.

---

Group of orthologs #2. Best score 11940 bits
Score difference with first non-orthologous sequence - *A. fumigatus*:11940 *A. fischeri*:11940
afum|EAL93103.1 100.00% afis|EAW18419.1 100.00%
Bootstrap support for afum|EAL93103.1 as seed ortholog is 100%.
Bootstrap support for afis|EAW18419.1 as seed ortholog is 100%.

---

Group of orthologs #3. Best score 11375 bits
Score difference with first non-orthologous sequence - *A. fumigatus*:11375 *A. fischeri*:11375
afum|EAL93509.1 100.00% afis|EAW18777.1 100.00%
Bootstrap support for afum|EAL93509.1 as seed ortholog is 100%.
Bootstrap support for afis|EAW18777.1 as seed ortholog is 100%.

---

Group of orthologs #4. Best score 10648 bits
Score difference with first non-orthologous sequence - *A. fumigatus*:10648 *A. fischeri*:10078
afum|EAL91050.1 100.00% afis|EAW22140.1 100.00%
Bootstrap support for afum|EAL91050.1 as seed ortholog is 100%.
Bootstrap support for afis|EAW22140.1 as seed ortholog is 100%.

---

Group of orthologs #5. Best score 10326 bits
Score difference with first non-orthologous sequence - *A. fumigatus*:10326 *A. fischeri*:10326
afum|EAL91432.1 100.00% afis|EAW17619.1 100.00%
Bootstrap support for afum|EAL91432.1 as seed ortholog is 100%.
Bootstrap support for afis|EAW17619.1 as seed ortholog is 100%.

---

Group of orthologs #6. Best score 9512 bits
Score difference with first non-orthologous sequence - *A. fumigatus*:9512 *A. fischeri*:9512
afum|EAL89711.1 100.00% afis|EAW25043.1 100.00%
Bootstrap support for afum|EAL89711.1 as seed ortholog is 100%.
Bootstrap support for afis|EAW25043.1 as seed ortholog is 100%.

---

Group of orthologs #7. Best score 9397 bits
Score difference with first non-orthologous sequence - *A. fumigatus*:9397 *A. fischeri*:9397
afum|EAL86047.1 100.00% afis|EAW24375.1 100.00%

Bootstrap support for afum|EAL86047.1 as seed ortholog is 100%.
Bootstrap support for afis|EAW24375.1 as seed ortholog is 100%.

---

Group of orthologs #8. Best score 7433 bits
Score difference with first non-orthologous sequence - *A. fumigatus*:7433 *A. fischeri*:7433
afum|EAL89636.1 100.00% afis|EAW24976.1 100.00%
Bootstrap support for afum|EAL89636.1 as seed ortholog is 100%.
Bootstrap support for afis|EAW24976.1 as seed ortholog is 100%.

---

Group of orthologs #9. Best score 7236 bits
Score difference with first non-orthologous sequence - *A. fumigatus*:7236 *A. fischeri*:7236
afum|EAL93647.1 100.00% afis|EAW18916.1 100.00%
Bootstrap support for afum|EAL93647.1 as seed ortholog is 100%.
Bootstrap support for afis|EAW18916.1 as seed ortholog is 100%.

---

Group of orthologs #10. Best score 6785 bits
Score difference with first non-orthologous sequence - *A. fumigatus*:6785 *A. fischeri*:6785
afum|EAL93940.1 100.00% afis|EAW19210.1 100.00%
Bootstrap support for afum|EAL93940.1 as seed ortholog is 100%.
Bootstrap support for afis|EAW19210.1 as seed ortholog is 100%.

---

Group of orthologs #11. Best score 6674 bits
Score difference with first non-orthologous sequence - *A. fumigatus*:6674 *A. fischeri*:6674
afum|EAL92789.1 100.00% afis|EAW21759.1 100.00%
Bootstrap support for afum|EAL92789.1 as seed ortholog is 100%.
Bootstrap support for afis|EAW21759.1 as seed ortholog is 100%.

---

Group of orthologs #12. Best score 6583 bits
Score difference with first non-orthologous sequence - *A. fumigatus*:6583 *A. fischeri*:6583
afum|EAL87400.1 100.00% afis|EAW23970.1 100.00%
Bootstrap support for afum|EAL87400.1 as seed ortholog is 100%.
Bootstrap support for afis|EAW23970.1 as seed ortholog is 100%.

---

Group of orthologs #13. Best score 6522 bits
Score difference with first non-orthologous sequence - *A. fumigatus*:6522 *A. fischeri*:6522
afum|EAL89387.1 100.00% afis|EAW24741.1 100.00%
Bootstrap support for afum|EAL89387.1 as seed ortholog is 100%.
Bootstrap support for afis|EAW24741.1 as seed ortholog is 100%.

---

Group of orthologs #14. Best score 6417 bits
Score difference with first non-orthologous sequence - *A. fumigatus*:6417 *A. fischeri*:6417
afum|EAL91349.1 100.00% afis|EAW17535.1 100.00%
Bootstrap support for afum|EAL91349.1 as seed ortholog is 100%.
Bootstrap support for afis|EAW17535.1 as seed ortholog is 100%.

---

Group of orthologs #15. Best score 6185 bits
Score difference with first non-orthologous sequence - *A. fumigatus*:6185 *A. fischeri*:6185
afum|EAL88949.1 100.00% afis|EAW16325.1 100.00%
Bootstrap support for afum|EAL88949.1 as seed ortholog is 100%.
Bootstrap support for afis|EAW16325.1 as seed ortholog is 100%.

---

Group of orthologs #16. Best score 6172 bits
Score difference with first non-orthologous sequence - *A. fumigatus*:6172 *A. fischeri*:6172
afum|EAL93093.2 100.00% afis|EAW18413.1 100.00%
Bootstrap support for afum|EAL93093.2 as seed ortholog is 100%.
Bootstrap support for afis|EAW18413.1 as seed ortholog is 100%.

Group of orthologs #17. Best score 5983 bits
Score difference with first non-orthologous sequence - *A. fumigatus*:5983 *A. fischeri*:5983
afum|EAL88675.1 100.00% afis|EAW16043.1 100.00%
Bootstrap support for afum|EAL88675.1 as seed ortholog is 100%.
Bootstrap support for afis|EAW16043.1 as seed ortholog is 100%.

Group of orthologs #18. Best score 5929 bits
Score difference with first non-orthologous sequence - *A. fumigatus*:5002 *A. fischeri*:4763
afum|EAL87227.2 100.00% afis|EAW23793.1 100.00%
Bootstrap support for afum|EAL87227.2 as seed ortholog is 100%.
Bootstrap support for afis|EAW23793.1 as seed ortholog is 100%.

Group of orthologs #19. Best score 5911 bits
Score difference with first non-orthologous sequence - *A. fumigatus*:5911 *A. fischeri*:5911
afum|EAL86374.1 100.00% afis|EAW16823.1 100.00%
Bootstrap support for afum|EAL86374.1 as seed ortholog is 100%.
Bootstrap support for afis|EAW16823.1 as seed ortholog is 100%.

Group of orthologs #20. Best score 5871 bits
Score difference with first non-orthologous sequence - *A. fumigatus*:5871 *A. fischeri*:5871
afum|EAL93422.1 100.00% afis|EAW18700.1 100.00%
Bootstrap support for afum|EAL93422.1 as seed ortholog is 100%.
Bootstrap support for afis|EAW18700.1 as seed ortholog is 100%.

Group of orthologs #21. Best score 5870 bits
Score difference with first non-orthologous sequence - *A. fumigatus*:5870 *A. fischeri*:5870
afum|EBA27443.1 100.00% afis|EAW18066.1 100.00%
Bootstrap support for afum|EBA27443.1 as seed ortholog is 100%.
Bootstrap support for afis|EAW18066.1 as seed ortholog is 100%.

Group of orthologs #22. Best score 5866 bits
Score difference with first non-orthologous sequence - *A. fumigatus*:5866 *A. fischeri*:5866
afum|EAL90874.1 100.00% afis|EAW22316.1 100.00%
Bootstrap support for afum|EAL90874.1 as seed ortholog is 100%.
Bootstrap support for afis|EAW22316.1 as seed ortholog is 100%.

Group of orthologs #23. Best score 5864 bits
Score difference with first non-orthologous sequence - *A. fumigatus*:5864 *A. fischeri*:5864
afum|EAL91679.1 100.00% afis|EAW17871.1 100.00%
Bootstrap support for afum|EAL91679.1 as seed ortholog is 100%.
Bootstrap support for afis|EAW17871.1 as seed ortholog is 100%.

Group of orthologs #24. Best score 5859 bits
Score difference with first non-orthologous sequence - *A. fumigatus*:5859 *A. fischeri*:5859
afum|EAL93140.1 100.00% afis|EAW18455.1 100.00%
Bootstrap support for afum|EAL93140.1 as seed ortholog is 100%.
Bootstrap support for afis|EAW18455.1 as seed ortholog is 100%.

Group of orthologs #25. Best score 5799 bits
Score difference with first non-orthologous sequence - *A. fumigatus*:5799 *A. fischeri*:5799
afum|EAL88599.1 100.00% afis|EAW15966.1 100.00%
Bootstrap support for afum|EAL88599.1 as seed ortholog is 100%.
Bootstrap support for afis|EAW15966.1 as seed ortholog is 100%.

Group of orthologs #26. Best score 5759 bits

Score difference with first non-orthologous sequence - *A. fumigatus*:4173   *A. fischeri*:3893
afum|EAL92117.1 100.00% afis|EAW21094.1 100.00%
Bootstrap support for afum|EAL92117.1 as seed ortholog is 100%.
Bootstrap support for afis|EAW21094.1 as seed ortholog is 100%.

---

Group of orthologs #27. Best score 5743 bits
Score difference with first non-orthologous sequence - *A. fumigatus*:5743   *A. fischeri*:5743
afum|EAL93322.1 100.00% afis|EAW18612.1 100.00%
Bootstrap support for afum|EAL93322.1 as seed ortholog is 100%.
Bootstrap support for afis|EAW18612.1 as seed ortholog is 100%.

---

Group of orthologs #28. Best score 5634 bits
Score difference with first non-orthologous sequence - *A. fumigatus*:5634   *A. fischeri*:5634
afum|EAL91729.1 100.00% afis|EAW17918.1 100.00%
Bootstrap support for afum|EAL91729.1 as seed ortholog is 100%.
Bootstrap support for afis|EAW17918.1 as seed ortholog is 100%.

---

Group of orthologs #29. Best score 5572 bits
Score difference with first non-orthologous sequence - *A. fumigatus*:4603   *A. fischeri*:4524
afum|EAL86424.2 100.00% afis|EAW16886.1 100.00%
Bootstrap support for afum|EAL86424.2 as seed ortholog is 100%.
Bootstrap support for afis|EAW16886.1 as seed ortholog is 100%.

---

Group of orthologs #30. Best score 5531 bits
Score difference with first non-orthologous sequence - *A. fumigatus*:5531   *A. fischeri*:4010
afum|EAL92328.1 100.00% afis|EAW21312.1 100.00%
Bootstrap support for afum|EAL92328.1 as seed ortholog is 100%.
Bootstrap support for afis|EAW21312.1 as seed ortholog is 100%.

---

Group of orthologs #31. Best score 5498 bits
Score difference with first non-orthologous sequence - *A. fumigatus*:5498   *A. fischeri*:5498
afum|EAL93163.1 100.00% afis|EAW18476.1 100.00%
Bootstrap support for afum|EAL93163.1 as seed ortholog is 100%.
Bootstrap support for afis|EAW18476.1 as seed ortholog is 100%.

---

Group of orthologs #32. Best score 5455 bits
Score difference with first non-orthologous sequence - *A. fumigatus*:5455   *A. fischeri*:5455
afum|EAL92557.1 100.00% afis|EAW21537.1 100.00%
Bootstrap support for afum|EAL92557.1 as seed ortholog is 100%.
Bootstrap support for afis|EAW21537.1 as seed ortholog is 100%.

---

Group of orthologs #33. Best score 5388 bits
Score difference with first non-orthologous sequence - *A. fumigatus*:5388   *A. fischeri*:5388
afum|EAL92625.1 100.00% afis|EAW21606.1 100.00%
Bootstrap support for afum|EAL92625.1 as seed ortholog is 100%.
Bootstrap support for afis|EAW21606.1 as seed ortholog is 100%.

---

Group of orthologs #34. Best score 5339 bits
Score difference with first non-orthologous sequence - *A. fumigatus*:4350   *A. fischeri*:4671
afum|EAL89230.2 100.00% afis|EAW20415.1 100.00%
Bootstrap support for afum|EAL89230.2 as seed ortholog is 100%.
Bootstrap support for afis|EAW20415.1 as seed ortholog is 100%.

---

Group of orthologs #35. Best score 5327 bits
Score difference with first non-orthologous sequence - *A. fumigatus*:5327   *A. fischeri*:5327
afum|EAL85242.1 100.00% afis|EAW20124.1 100.00%

Bootstrap support for afum|EAL85242.1 as seed ortholog is 100%.
Bootstrap support for afis|EAW20124.1 as seed ortholog is 100%.

---

Group of orthologs #36. Best score 5312 bits
Score difference with first non-orthologous sequence - *A. fumigatus*:5312 *A. fischeri*:5312
afum|EAL88977.1 100.00% afis|EAW16352.1 100.00%
Bootstrap support for afum|EAL88977.1 as seed ortholog is 100%.
Bootstrap support for afis|EAW16352.1 as seed ortholog is 100%.

---

Group of orthologs #37. Best score 5179 bits
Score difference with first non-orthologous sequence - *A. fumigatus*:5179 *A. fischeri*:5179
afum|EAL88447.1 100.00% afis|EAW23034.1 100.00%
Bootstrap support for afum|EAL88447.1 as seed ortholog is 100%.
Bootstrap support for afis|EAW23034.1 as seed ortholog is 100%.

---

Group of orthologs #38. Best score 5163 bits
Score difference with first non-orthologous sequence - *A. fumigatus*:5163 *A. fischeri*:4549
afum|EAL92059.1 100.00% afis|EAW21021.1 100.00%
Bootstrap support for afum|EAL92059.1 as seed ortholog is 100%.
Bootstrap support for afis|EAW21021.1 as seed ortholog is 100%.

---

Group of orthologs #39. Best score 5140 bits
Score difference with first non-orthologous sequence - *A. fumigatus*:5140 *A. fischeri*:5140
afum|EAL88693.1 100.00% afis|EAW16060.1 100.00%
Bootstrap support for afum|EAL88693.1 as seed ortholog is 100%.
Bootstrap support for afis|EAW16060.1 as seed ortholog is 100%.

---

Group of orthologs #40. Best score 5109 bits
Score difference with first non-orthologous sequence - *A. fumigatus*:5109 *A. fischeri*:5109
afum|EAL88703.2 100.00% afis|EAW16071.1 100.00%
Bootstrap support for afum|EAL88703.2 as seed ortholog is 100%.
Bootstrap support for afis|EAW16071.1 as seed ortholog is 100%.

---

Group of orthologs #41. Best score 5097 bits
Score difference with first non-orthologous sequence - *A. fumigatus*:5097 *A. fischeri*:3680
afum|EAL94057.1 100.00% afis|EAW19338.1 100.00%
Bootstrap support for afum|EAL94057.1 as seed ortholog is 100%.
Bootstrap support for afis|EAW19338.1 as seed ortholog is 100%.

---

Group of orthologs #42. Best score 5084 bits
Score difference with first non-orthologous sequence - *A. fumigatus*:5084 *A. fischeri*:5084
afum|EAL93587.1 100.00% afis|EAW18856.1 100.00%
Bootstrap support for afum|EAL93587.1 as seed ortholog is 100%.
Bootstrap support for afis|EAW18856.1 as seed ortholog is 100%.

---

Group of orthologs #43. Best score 5040 bits
Score difference with first non-orthologous sequence - *A. fumigatus*:5040 *A. fischeri*:5040
afum|EAL86701.1 100.00% afis|EAW17283.1 100.00%
Bootstrap support for afum|EAL86701.1 as seed ortholog is 100%.
Bootstrap support for afis|EAW17283.1 as seed ortholog is 100%.

---

Group of orthologs #44. Best score 5025 bits
Score difference with first non-orthologous sequence - *A. fumigatus*:5025 *A. fischeri*:5025
afum|EAL90438.1 100.00% afis|EAW22761.1 100.00%
Bootstrap support for afum|EAL90438.1 as seed ortholog is 100%.
Bootstrap support for afis|EAW22761.1 as seed ortholog is 100%.

---

Group of orthologs #45. Best score 4986 bits
Score difference with first non-orthologous sequence - *A. fumigatus*:4986 *A. fischeri*:4568
afum|EAL92291.2 100.00% afis|EAW21276.1 100.00%
Bootstrap support for afum|EAL92291.2 as seed ortholog is 100%.
Bootstrap support for afis|EAW21276.1 as seed ortholog is 100%.

---

Group of orthologs #46. Best score 4921 bits
Score difference with first non-orthologous sequence - *A. fumigatus*:4921 *A. fischeri*:4546
afum|EAL88817.1 100.00% afis|EAW16186.1 100.00%
Bootstrap support for afum|EAL88817.1 as seed ortholog is 100%.
Bootstrap support for afis|EAW16186.1 as seed ortholog is 100%.

---

Group of orthologs #47. Best score 4900 bits
Score difference with first non-orthologous sequence - *A. fumigatus*:4900 *A. fischeri*:4900
afum|EAL85833.1 100.00% afis|EAW15490.1 100.00%
Bootstrap support for afum|EAL85833.1 as seed ortholog is 100%.
Bootstrap support for afis|EAW15490.1 as seed ortholog is 100%.

---

Group of orthologs #48. Best score 4899 bits
Score difference with first non-orthologous sequence - *A. fumigatus*:4899 *A. fischeri*:4899
afum|EAL88144.1 100.00% afis|EAW23334.1 100.00%
Bootstrap support for afum|EAL88144.1 as seed ortholog is 100%.
Bootstrap support for afis|EAW23334.1 as seed ortholog is 100%.

---

Group of orthologs #49. Best score 4897 bits
Score difference with first non-orthologous sequence - *A. fumigatus*:4897 *A. fischeri*:4897
afum|EAL93796.1 100.00% afis|EAW19067.1 100.00%
Bootstrap support for afum|EAL93796.1 as seed ortholog is 100%.
Bootstrap support for afis|EAW19067.1 as seed ortholog is 100%.

---

Group of orthologs #50. Best score 4859 bits
Score difference with first non-orthologous sequence - *A. fumigatus*:4859 *A. fischeri*:4256
afum|EAL86616.2 100.00% afis|EAW17190.1 100.00%
Bootstrap support for afum|EAL86616.2 as seed ortholog is 100%.
Bootstrap support for afis|EAW17190.1 as seed ortholog is 100%.

---

Group of orthologs #51. Best score 4833 bits
Score difference with first non-orthologous sequence - *A. fumigatus*:4833 *A. fischeri*:4833
afum|EAL86131.1 100.00% afis|EAW24461.1 100.00%
Bootstrap support for afum|EAL86131.1 as seed ortholog is 100%.
Bootstrap support for afis|EAW24461.1 as seed ortholog is 100%.

---

Group of orthologs #52. Best score 4831 bits
Score difference with first non-orthologous sequence - *A. fumigatus*:4831 *A. fischeri*:4831
afum|EAL92973.1 100.00% afis|EAW21959.1 100.00%
Bootstrap support for afum|EAL92973.1 as seed ortholog is 100%.
Bootstrap support for afis|EAW21959.1 as seed ortholog is 100%.

---

Group of orthologs #53. Best score 4781 bits
Score difference with first non-orthologous sequence - *A. fumigatus*:4781 *A. fischeri*:4781
afum|EAL93208.1 100.00% afis|EAW18513.1 100.00%
Bootstrap support for afum|EAL93208.1 as seed ortholog is 100%.
Bootstrap support for afis|EAW18513.1 as seed ortholog is 100%.

---

Group of orthologs #54. Best score 4771 bits

Score difference with first non-orthologous sequence - *A. fumigatus*:4771   *A. fischeri*:4771
afum|EAL84455.1 100.00% afis|EAW20605.1 100.00%
Bootstrap support for afum|EAL84455.1 as seed ortholog is 100%.
Bootstrap support for afis|EAW20605.1 as seed ortholog is 100%.

---

Group of orthologs #55. Best score 4762 bits
Score difference with first non-orthologous sequence - *A. fumigatus*:4762   *A. fischeri*:4762
afum|EAL85631.1 100.00% afis|EAW15717.1 100.00%
Bootstrap support for afum|EAL85631.1 as seed ortholog is 100%.
Bootstrap support for afis|EAW15717.1 as seed ortholog is 100%.

---

Group of orthologs #56. Best score 4759 bits
Score difference with first non-orthologous sequence - *A. fumigatus*:4759   *A. fischeri*:4759
afum|EAL90472.1 100.00% afis|EAW22724.1 100.00%
Bootstrap support for afum|EAL90472.1 as seed ortholog is 100%.
Bootstrap support for afis|EAW22724.1 as seed ortholog is 100%.

---

Group of orthologs #57. Best score 4753 bits
Score difference with first non-orthologous sequence - *A. fumigatus*:4753   *A. fischeri*:4753
afum|EAL89662.1 100.00% afis|EAW25032.1 100.00%
Bootstrap support for afum|EAL89662.1 as seed ortholog is 100%.
Bootstrap support for afis|EAW25032.1 as seed ortholog is 100%.

---

Group of orthologs #58. Best score 4751 bits
Score difference with first non-orthologous sequence - *A. fumigatus*:4751   *A. fischeri*:4751
afum|EAL89159.1 100.00% afis|EAW16561.1 100.00%
Bootstrap support for afum|EAL89159.1 as seed ortholog is 100%.
Bootstrap support for afis|EAW16561.1 as seed ortholog is 100%.

---

Group of orthologs #59. Best score 4742 bits
Score difference with first non-orthologous sequence - *A. fumigatus*:4742   *A. fischeri*:4742
afum|EAL92302.1 100.00% afis|EAW21285.1 100.00%
Bootstrap support for afum|EAL92302.1 as seed ortholog is 100%.
Bootstrap support for afis|EAW21285.1 as seed ortholog is 100%.

---

Group of orthologs #60. Best score 4729 bits
Score difference with first non-orthologous sequence - *A. fumigatus*:4729   *A. fischeri*:4729
afum|EAL91630.1 100.00% afis|EAW17820.1 100.00%
Bootstrap support for afum|EAL91630.1 as seed ortholog is 100%.
Bootstrap support for afis|EAW17820.1 as seed ortholog is 100%.

---

Group of orthologs #61. Best score 4716 bits
Score difference with first non-orthologous sequence - *A. fumigatus*:4247   *A. fischeri*:3701
afum|EAL86624.1 100.00% afis|EAW17196.1 100.00%
Bootstrap support for afum|EAL86624.1 as seed ortholog is 100%.
Bootstrap support for afis|EAW17196.1 as seed ortholog is 100%.

---

Group of orthologs #62. Best score 4707 bits
Score difference with first non-orthologous sequence - *A. fumigatus*:4707   *A. fischeri*:4707
afum|EAL89080.1 100.00% afis|EAW16486.1 100.00%
Bootstrap support for afum|EAL89080.1 as seed ortholog is 100%.
Bootstrap support for afis|EAW16486.1 as seed ortholog is 100%.

---

Group of orthologs #63. Best score 4704 bits
Score difference with first non-orthologous sequence - *A. fumigatus*:4704   *A. fischeri*:4704
afum|EAL86902.1 100.00% afis|EAW19620.1 100.00%

Bootstrap support for afum|EAL86902.1 as seed ortholog is 100%.
Bootstrap support for afis|EAW19620.1 as seed ortholog is 100%.

---

Group of orthologs #64. Best score 4703 bits
Score difference with first non-orthologous sequence - *A. fumigatus*:4211 *A. fischeri*:3368
afum|EAL85149.1 100.00% afis|EAW19749.1 100.00%
Bootstrap support for afum|EAL85149.1 as seed ortholog is 100%.
Bootstrap support for afis|EAW19749.1 as seed ortholog is 100%.

---

Group of orthologs #65. Best score 4676 bits
Score difference with first non-orthologous sequence - *A. fumigatus*:4676 *A. fischeri*:4676
afum|EAL84624.2 100.00% afis|EAW20424.1 100.00%
Bootstrap support for afum|EAL84624.2 as seed ortholog is 100%.
Bootstrap support for afis|EAW20424.1 as seed ortholog is 100%.

---

Group of orthologs #66. Best score 4655 bits
Score difference with first non-orthologous sequence - *A. fumigatus*:4655 *A. fischeri*:4655
afum|EAL89683.1 100.00% afis|EAW25013.1 100.00%
Bootstrap support for afum|EAL89683.1 as seed ortholog is 100%.
Bootstrap support for afis|EAW25013.1 as seed ortholog is 100%.

---

Group of orthologs #67. Best score 4641 bits
Score difference with first non-orthologous sequence - *A. fumigatus*:4641 *A. fischeri*:4641
afum|EAL89151.2 100.00% afis|EAW16553.1 100.00%
Bootstrap support for afum|EAL89151.2 as seed ortholog is 100%.
Bootstrap support for afis|EAW16553.1 as seed ortholog is 100%.

---

Group of orthologs #68. Best score 4635 bits
Score difference with first non-orthologous sequence - *A. fumigatus*:4635 *A. fischeri*:2814
afum|EAL87273.1 100.00% afis|EAW23842.1 100.00%
Bootstrap support for afum|EAL87273.1 as seed ortholog is 100%.
Bootstrap support for afis|EAW23842.1 as seed ortholog is 100%.

---

Group of orthologs #69. Best score 4622 bits
Score difference with first non-orthologous sequence - *A. fumigatus*:4622 *A. fischeri*:4622
afum|EAL88344.1 100.00% afis|EAW23137.1 100.00%
Bootstrap support for afum|EAL88344.1 as seed ortholog is 100%.
Bootstrap support for afis|EAW23137.1 as seed ortholog is 100%.

---

Group of orthologs #70. Best score 4506 bits
Score difference with first non-orthologous sequence - *A. fumigatus*:3735 *A. fischeri*:2592
afum|EAL85129.1 100.00% afis|EAW19771.1 100.00%
Bootstrap support for afum|EAL85129.1 as seed ortholog is 100%.
Bootstrap support for afis|EAW19771.1 as seed ortholog is 100%.

---

Group of orthologs #71. Best score 4479 bits
Score difference with first non-orthologous sequence - *A. fumigatus*:4479 *A. fischeri*:4479
afum|EAL90832.1 100.00% afis|EAW22359.1 100.00%
Bootstrap support for afum|EAL90832.1 as seed ortholog is 100%.
Bootstrap support for afis|EAW22359.1 as seed ortholog is 100%.

---

Group of orthologs #72. Best score 4468 bits
Score difference with first non-orthologous sequence - *A. fumigatus*:4468 *A. fischeri*:4468
afum|EAL92330.1 100.00% afis|EAW21313.1 100.00%
Bootstrap support for afum|EAL92330.1 as seed ortholog is 100%.
Bootstrap support for afis|EAW21313.1 as seed ortholog is 100%.

Group of orthologs #73. Best score 4462 bits
Score difference with first non-orthologous sequence - *A. fumigatus*:4462 *A. fischeri*:4462
afum|EAL88295.1 100.00% afis|EAW23188.1 100.00%
Bootstrap support for afum|EAL88295.1 as seed ortholog is 100%.
Bootstrap support for afis|EAW23188.1 as seed ortholog is 100%.

---

Group of orthologs #74. Best score 4451 bits
Score difference with first non-orthologous sequence - *A. fumigatus*:4451 *A. fischeri*:4451
afum|EAL86700.1 100.00% afis|EAW17282.1 100.00%
Bootstrap support for afum|EAL86700.1 as seed ortholog is 100%.
Bootstrap support for afis|EAW17282.1 as seed ortholog is 100%.

---

Group of orthologs #75. Best score 4438 bits
Score difference with first non-orthologous sequence - *A. fumigatus*:4438 *A. fischeri*:4438
afum|EAL87610.1 100.00% afis|EAW18245.1 100.00%
Bootstrap support for afum|EAL87610.1 as seed ortholog is 100%.
Bootstrap support for afis|EAW18245.1 as seed ortholog is 100%.

---

Group of orthologs #76. Best score 4419 bits
Score difference with first non-orthologous sequence - *A. fumigatus*:4419 *A. fischeri*:4151
afum|EAL87861.1 100.00% afis|EAW23605.1 100.00%
Bootstrap support for afum|EAL87861.1 as seed ortholog is 100%.
Bootstrap support for afis|EAW23605.1 as seed ortholog is 100%.

---

Group of orthologs #77. Best score 4407 bits
Score difference with first non-orthologous sequence - *A. fumigatus*:4407 *A. fischeri*:4407
afum|EAL87012.1 100.00% afis|EAW19483.1 100.00%
Bootstrap support for afum|EAL87012.1 as seed ortholog is 100%.
Bootstrap support for afis|EAW19483.1 as seed ortholog is 100%.

---

Group of orthologs #78. Best score 4400 bits
Score difference with first non-orthologous sequence - *A. fumigatus*:4400 *A. fischeri*:4400
afum|EAL93639.1 100.00% afis|EAW18907.1 100.00%
Bootstrap support for afum|EAL93639.1 as seed ortholog is 100%.
Bootstrap support for afis|EAW18907.1 as seed ortholog is 100%.

---

Group of orthologs #79. Best score 4394 bits
Score difference with first non-orthologous sequence - *A. fumigatus*:4394 *A. fischeri*:4394
afum|EAL87631.1 100.00% afis|EAW18266.1 100.00%
Bootstrap support for afum|EAL87631.1 as seed ortholog is 100%.
Bootstrap support for afis|EAW18266.1 as seed ortholog is 100%.

---

Group of orthologs #80. Best score 4392 bits
Score difference with first non-orthologous sequence - *A. fumigatus*:4392 *A. fischeri*:4392
afum|EBA27403.1 100.00% afis|EAW22870.1 100.00%
Bootstrap support for afum|EBA27403.1 as seed ortholog is 100%.
Bootstrap support for afis|EAW22870.1 as seed ortholog is 100%.

---

Group of orthologs #81. Best score 4385 bits
Score difference with first non-orthologous sequence - *A. fumigatus*:4385 *A. fischeri*:4385
afum|EAL91853.1 100.00% afis|EAW18042.1 100.00%
Bootstrap support for afum|EAL91853.1 as seed ortholog is 100%.
Bootstrap support for afis|EAW18042.1 as seed ortholog is 100%.

---

Group of orthologs #82. Best score 4315 bits

Score difference with first non-orthologous sequence - *A. fumigatus*:4315   *A. fischeri*:4315
afum|EAL93000.1 100.00% afis|EAW21985.1 100.00%
Bootstrap support for afum|EAL93000.1 as seed ortholog is 100%.
Bootstrap support for afis|EAW21985.1 as seed ortholog is 100%.

---

Group of orthologs #83. Best score 4305 bits
Score difference with first non-orthologous sequence - *A. fumigatus*:4305   *A. fischeri*:4305
afum|EBA27508.1 100.00% afis|EAW19212.1 100.00%
Bootstrap support for afum|EBA27508.1 as seed ortholog is 100%.
Bootstrap support for afis|EAW19212.1 as seed ortholog is 100%.

---

Group of orthologs #84. Best score 4275 bits
Score difference with first non-orthologous sequence - *A. fumigatus*:4275   *A. fischeri*:4275
afum|EAL90799.1 100.00% afis|EAW22392.1 100.00%
Bootstrap support for afum|EAL90799.1 as seed ortholog is 100%.
Bootstrap support for afis|EAW22392.1 as seed ortholog is 100%.

---

Group of orthologs #85. Best score 4256 bits
Score difference with first non-orthologous sequence - *A. fumigatus*:4256   *A. fischeri*:4256
afum|EAL93638.1 100.00% afis|EAW18906.1 100.00%
Bootstrap support for afum|EAL93638.1 as seed ortholog is 100%.
Bootstrap support for afis|EAW18906.1 as seed ortholog is 100%.

---

Group of orthologs #86. Best score 4236 bits
Score difference with first non-orthologous sequence - *A. fumigatus*:4236   *A. fischeri*:4236
afum|EAL85643.1 100.00% afis|EAW15704.1 100.00%
Bootstrap support for afum|EAL85643.1 as seed ortholog is 100%.
Bootstrap support for afis|EAW15704.1 as seed ortholog is 100%.

---

Group of orthologs #87. Best score 4202 bits
Score difference with first non-orthologous sequence - *A. fumigatus*:4202   *A. fischeri*:4202
afum|EAL88739.2 100.00% afis|EAW16107.1 100.00%
Bootstrap support for afum|EAL88739.2 as seed ortholog is 100%.
Bootstrap support for afis|EAW16107.1 as seed ortholog is 100%.

---

Group of orthologs #88. Best score 4184 bits
Score difference with first non-orthologous sequence - *A. fumigatus*:4184   *A. fischeri*:4184
afum|EAL85962.1 100.00% afis|EAW24294.1 100.00%
Bootstrap support for afum|EAL85962.1 as seed ortholog is 100%.
Bootstrap support for afis|EAW24294.1 as seed ortholog is 100%.

---

Group of orthologs #89. Best score 4183 bits
Score difference with first non-orthologous sequence - *A. fumigatus*:4183   *A. fischeri*:2404
afum|EAL84875.1 100.00% afis|EAW20700.1 100.00%
Bootstrap support for afum|EAL84875.1 as seed ortholog is 100%.
Bootstrap support for afis|EAW20700.1 as seed ortholog is 100%.

---

Group of orthologs #90. Best score 4175 bits
Score difference with first non-orthologous sequence - *A. fumigatus*:4175   *A. fischeri*:4175
afum|EAL93901.1 100.00% afis|EAW19171.1 100.00%
Bootstrap support for afum|EAL93901.1 as seed ortholog is 100%.
Bootstrap support for afis|EAW19171.1 as seed ortholog is 100%.

---

Group of orthologs #91. Best score 4165 bits
Score difference with first non-orthologous sequence - *A. fumigatus*:4165   *A. fischeri*:4165
afum|EAL90642.1 100.00% afis|EAW22553.1 100.00%

Bootstrap support for afum|EAL90642.1 as seed ortholog is 100%.
Bootstrap support for afis|EAW22553.1 as seed ortholog is 100%.

---

Group of orthologs #92. Best score 4157 bits
Score difference with first non-orthologous sequence - *A. fumigatus*:4157 *A. fischeri*:4157
afum|EAL88235.1 100.00% afis|EAW23246.1 100.00%
Bootstrap support for afum|EAL88235.1 as seed ortholog is 100%.
Bootstrap support for afis|EAW23246.1 as seed ortholog is 100%.

---

Group of orthologs #93. Best score 4131 bits
Score difference with first non-orthologous sequence - *A. fumigatus*:4131 *A. fischeri*:4131
afum|EAL85546.1 100.00% afis|EAW15804.1 100.00%
Bootstrap support for afum|EAL85546.1 as seed ortholog is 100%.
Bootstrap support for afis|EAW15804.1 as seed ortholog is 100%.

---

Group of orthologs #94. Best score 4113 bits
Score difference with first non-orthologous sequence - *A. fumigatus*:4113 *A. fischeri*:4113
afum|EAL88889.1 100.00% afis|EAW16268.1 100.00%
Bootstrap support for afum|EAL88889.1 as seed ortholog is 100%.
Bootstrap support for afis|EAW16268.1 as seed ortholog is 100%.

---

Group of orthologs #95. Best score 4097 bits
Score difference with first non-orthologous sequence - *A. fumigatus*:4097 *A. fischeri*:4097
afum|EAL87638.1 100.00% afis|EAW18273.1 100.00%
Bootstrap support for afum|EAL87638.1 as seed ortholog is 100%.
Bootstrap support for afis|EAW18273.1 as seed ortholog is 100%.

---

Group of orthologs #96. Best score 4094 bits
Score difference with first non-orthologous sequence - *A. fumigatus*:4094 *A. fischeri*:1469
afum|EAL89339.1 100.00% afis|EAW24697.1 100.00%
Bootstrap support for afum|EAL89339.1 as seed ortholog is 100%.
Bootstrap support for afis|EAW24697.1 as seed ortholog is 100%.

---

Group of orthologs #97. Best score 4086 bits
Score difference with first non-orthologous sequence - *A. fumigatus*:4086 *A. fischeri*:4086
afum|EAL88508.1 100.00% afis|EAW15874.1 100.00%
Bootstrap support for afum|EAL88508.1 as seed ortholog is 100%.
Bootstrap support for afis|EAW15874.1 as seed ortholog is 100%.

---

Group of orthologs #98. Best score 4084 bits
Score difference with first non-orthologous sequence - *A. fumigatus*:4084 *A. fischeri*:4084
afum|EAL91969.1 100.00% afis|EAW24115.1 100.00%
Bootstrap support for afum|EAL91969.1 as seed ortholog is 100%.
Bootstrap support for afis|EAW24115.1 as seed ortholog is 100%.

---

Group of orthologs #99. Best score 4083 bits
Score difference with first non-orthologous sequence - *A. fumigatus*:4083 *A. fischeri*:4083
afum|EAL89175.1 100.00% afis|EAW16579.1 100.00%
Bootstrap support for afum|EAL89175.1 as seed ortholog is 100%.
Bootstrap support for afis|EAW16579.1 as seed ortholog is 100%.

---

Group of orthologs #100. Best score 4063 bits
Score difference with first non-orthologous sequence - *A. fumigatus*:4063 *A. fischeri*:4063
afum|EAL94019.1 100.00% afis|EAW19300.1 100.00%
Bootstrap support for afum|EAL94019.1 as seed ortholog is 100%.
Bootstrap support for afis|EAW19300.1 as seed ortholog is 100%.

---

Group of orthologs #101. Best score 4049 bits
Score difference with first non-orthologous sequence - *A. fumigatus*:4049   *A. fischeri*:4049
afum|EAL85557.1 100.00% afis|EAW15793.1 100.00%
Bootstrap support for afum|EAL85557.1 as seed ortholog is 100%.
Bootstrap support for afis|EAW15793.1 as seed ortholog is 100%.

---

Group of orthologs #102. Best score 4031 bits
Score difference with first non-orthologous sequence - *A. fumigatus*:4031   *A. fischeri*:2093
afum|EAL84397.1 100.00% afis|EAW25724.1 100.00%
Bootstrap support for afum|EAL84397.1 as seed ortholog is 100%.
Bootstrap support for afis|EAW25724.1 as seed ortholog is 100%.

---

Group of orthologs #103. Best score 4025 bits
Score difference with first non-orthologous sequence - *A. fumigatus*:4025   *A. fischeri*:4025
afum|EBA27477.1 100.00% afis|EAW21862.1 100.00%
Bootstrap support for afum|EBA27477.1 as seed ortholog is 100%.
Bootstrap support for afis|EAW21862.1 as seed ortholog is 100%.

---

Group of orthologs #104. Best score 4017 bits
Score difference with first non-orthologous sequence - *A. fumigatus*:4017   *A. fischeri*:4017
afum|EAL90842.1 100.00% afis|EAW22350.1 100.00%
Bootstrap support for afum|EAL90842.1 as seed ortholog is 100%.
Bootstrap support for afis|EAW22350.1 as seed ortholog is 100%.

---

Group of orthologs #105. Best score 4015 bits
Score difference with first non-orthologous sequence - *A. fumigatus*:4015   *A. fischeri*:4015
afum|EAL90722.1 100.00% afis|EAW22469.1 100.00%
Bootstrap support for afum|EAL90722.1 as seed ortholog is 100%.
Bootstrap support for afis|EAW22469.1 as seed ortholog is 100%.

---

Group of orthologs #106. Best score 4002 bits
Score difference with first non-orthologous sequence - *A. fumigatus*:4002   *A. fischeri*:4002
afum|EAL91935.1 100.00% afis|EAW24080.1 100.00%
Bootstrap support for afum|EAL91935.1 as seed ortholog is 100%.
Bootstrap support for afis|EAW24080.1 as seed ortholog is 100%.

---

Group of orthologs #107. Best score 3995 bits
Score difference with first non-orthologous sequence - *A. fumigatus*:3995   *A. fischeri*:3995
afum|EAL85196.1 100.00% afis|EAW20078.1 100.00%
Bootstrap support for afum|EAL85196.1 as seed ortholog is 100%.
Bootstrap support for afis|EAW20078.1 as seed ortholog is 100%.

---

Group of orthologs #108. Best score 3983 bits
Score difference with first non-orthologous sequence - *A. fumigatus*:3983   *A. fischeri*:3983
afum|EAL91297.1 100.00% afis|EAW17468.1 100.00%
Bootstrap support for afum|EAL91297.1 as seed ortholog is 100%.
Bootstrap support for afis|EAW17468.1 as seed ortholog is 100%.

---

Group of orthologs #109. Best score 3948 bits
Score difference with first non-orthologous sequence - *A. fumigatus*:3948   *A. fischeri*:3948
afum|EAL90017.1 100.00% afis|EAW25355.1 100.00%
Bootstrap support for afum|EAL90017.1 as seed ortholog is 100%.
Bootstrap support for afis|EAW25355.1 as seed ortholog is 100%.

---

Group of orthologs #110. Best score 3945 bits

Score difference with first non-orthologous sequence - *A. fumigatus*:3945 *A. fischeri*:3945
afum|EAL84261.1 100.00% afis|EAW25543.1 100.00%
Bootstrap support for afum|EAL84261.1 as seed ortholog is 100%.
Bootstrap support for afis|EAW25543.1 as seed ortholog is 100%.

---

Group of orthologs #111. Best score 3937 bits
Score difference with first non-orthologous sequence - *A. fumigatus*:3937 *A. fischeri*:3937
afum|EAL84657.1 100.00% afis|EAW20976.1 100.00%
Bootstrap support for afum|EAL84657.1 as seed ortholog is 100%.
Bootstrap support for afis|EAW20976.1 as seed ortholog is 100%.

---

Group of orthologs #112. Best score 3937 bits
Score difference with first non-orthologous sequence - *A. fumigatus*:3937 *A. fischeri*:3937
afum|EAL87094.1 100.00% afis|EAW19400.1 100.00%
Bootstrap support for afum|EAL87094.1 as seed ortholog is 100%.
Bootstrap support for afis|EAW19400.1 as seed ortholog is 100%.

---

Group of orthologs #113. Best score 3936 bits
Score difference with first non-orthologous sequence - *A. fumigatus*:3936 *A. fischeri*:3936
afum|EAL88050.1 100.00% afis|EAW23419.1 100.00%
Bootstrap support for afum|EAL88050.1 as seed ortholog is 100%.
Bootstrap support for afis|EAW23419.1 as seed ortholog is 100%.

---

Group of orthologs #114. Best score 3909 bits
Score difference with first non-orthologous sequence - *A. fumigatus*:3651 *A. fischeri*:3611
afum|EAL84933.1 100.00% afis|EAW19983.1 100.00%
Bootstrap support for afum|EAL84933.1 as seed ortholog is 100%.
Bootstrap support for afis|EAW19983.1 as seed ortholog is 100%.

---

Group of orthologs #115. Best score 3892 bits
Score difference with first non-orthologous sequence - *A. fumigatus*:3892 *A. fischeri*:3892
afum|EAL89207.1 100.00% afis|EAW16611.1 100.00%
Bootstrap support for afum|EAL89207.1 as seed ortholog is 100%.
Bootstrap support for afis|EAW16611.1 as seed ortholog is 100%.

---

Group of orthologs #116. Best score 3861 bits
Score difference with first non-orthologous sequence - *A. fumigatus*:3861 *A. fischeri*:3861
afum|EAL93767.1 100.00% afis|EAW19038.1 100.00%
Bootstrap support for afum|EAL93767.1 as seed ortholog is 100%.
Bootstrap support for afis|EAW19038.1 as seed ortholog is 100%.

---

Group of orthologs #117. Best score 3858 bits
Score difference with first non-orthologous sequence - *A. fumigatus*:3858 *A. fischeri*:3858
afum|EAL92473.1 100.00% afis|EAW21454.1 100.00%
Bootstrap support for afum|EAL92473.1 as seed ortholog is 100%.
Bootstrap support for afis|EAW21454.1 as seed ortholog is 100%.

---

Group of orthologs #118. Best score 3841 bits
Score difference with first non-orthologous sequence - *A. fumigatus*:3841 *A. fischeri*:3841
afum|EAL87459.1 100.00% afis|EAW24028.1 100.00%
Bootstrap support for afum|EAL87459.1 as seed ortholog is 100%.
Bootstrap support for afis|EAW24028.1 as seed ortholog is 100%.

---

Group of orthologs #119. Best score 3840 bits
Score difference with first non-orthologous sequence - *A. fumigatus*:3840 *A. fischeri*:3840
afum|EAL93708.1 100.00% afis|EAW18974.1 100.00%

Bootstrap support for afum|EAL93708.1 as seed ortholog is 100%.
Bootstrap support for afis|EAW18974.1 as seed ortholog is 100%.

---

Group of orthologs #120. Best score 3832 bits
Score difference with first non-orthologous sequence - *A. fumigatus*:3832 *A. fischeri*:3832
afum|EAL91827.1 100.00% afis|EAW18015.1 100.00%
Bootstrap support for afum|EAL91827.1 as seed ortholog is 100%.
Bootstrap support for afis|EAW18015.1 as seed ortholog is 100%.

---

Group of orthologs #121. Best score 3814 bits
Score difference with first non-orthologous sequence - *A. fumigatus*:3389 *A. fischeri*:3305
afum|EAL91682.1 100.00% afis|EAW17874.1 100.00%
Bootstrap support for afum|EAL91682.1 as seed ortholog is 100%.
Bootstrap support for afis|EAW17874.1 as seed ortholog is 100%.

---

Group of orthologs #122. Best score 3813 bits
Score difference with first non-orthologous sequence - *A. fumigatus*:3813 *A. fischeri*:3813
afum|EAL89461.1 100.00% afis|EAW24817.1 100.00%
Bootstrap support for afum|EAL89461.1 as seed ortholog is 100%.
Bootstrap support for afis|EAW24817.1 as seed ortholog is 100%.

---

Group of orthologs #123. Best score 3804 bits
Score difference with first non-orthologous sequence - *A. fumigatus*:3804 *A. fischeri*:3804
afum|EAL89903.1 100.00% afis|EAW25241.1 100.00%
Bootstrap support for afum|EAL89903.1 as seed ortholog is 100%.
Bootstrap support for afis|EAW25241.1 as seed ortholog is 100%.

---

Group of orthologs #124. Best score 3796 bits
Score difference with first non-orthologous sequence - *A. fumigatus*:3796 *A. fischeri*:3796
afum|EAL92890.1 100.00% afis|EAW21876.1 100.00%
Bootstrap support for afum|EAL92890.1 as seed ortholog is 100%.
Bootstrap support for afis|EAW21876.1 as seed ortholog is 100%.

---

Group of orthologs #125. Best score 3796 bits
Score difference with first non-orthologous sequence - *A. fumigatus*:3796 *A. fischeri*:3796
afum|EAL92993.1 100.00% afis|EAW21978.1 100.00%
Bootstrap support for afum|EAL92993.1 as seed ortholog is 100%.
Bootstrap support for afis|EAW21978.1 as seed ortholog is 100%.

---

Group of orthologs #126. Best score 3787 bits
Score difference with first non-orthologous sequence - *A. fumigatus*:3787 *A. fischeri*:3787
afum|EAL91920.1 100.00% afis|EAW24065.1 100.00%
Bootstrap support for afum|EAL91920.1 as seed ortholog is 100%.
Bootstrap support for afis|EAW24065.1 as seed ortholog is 100%.

---

Group of orthologs #127. Best score 3758 bits
Score difference with first non-orthologous sequence - *A. fumigatus*:3758 *A. fischeri*:3758
afum|EAL88965.1 100.00% afis|EAW16340.1 100.00%
Bootstrap support for afum|EAL88965.1 as seed ortholog is 100%.
Bootstrap support for afis|EAW16340.1 as seed ortholog is 100%.

---

Group of orthologs #128. Best score 3751 bits
Score difference with first non-orthologous sequence - *A. fumigatus*:3751 *A. fischeri*:3751
afum|EAL90706.1 100.00% afis|EAW22487.1 100.00%
Bootstrap support for afum|EAL90706.1 as seed ortholog is 100%.
Bootstrap support for afis|EAW22487.1 as seed ortholog is 100%.

Group of orthologs #129. Best score 3749 bits
Score difference with first non-orthologous sequence - *A. fumigatus*:3749 *A. fischeri*:3749
afum|EAL91423.1 100.00% afis|EAW17610.1 100.00%
Bootstrap support for afum|EAL91423.1 as seed ortholog is 100%.
Bootstrap support for afis|EAW17610.1 as seed ortholog is 100%.

Group of orthologs #130. Best score 3737 bits
Score difference with first non-orthologous sequence - *A. fumigatus*:3737 *A. fischeri*:3737
afum|EAL93245.1 100.00% afis|EAW18545.1 100.00%
Bootstrap support for afum|EAL93245.1 as seed ortholog is 100%.
Bootstrap support for afis|EAW18545.1 as seed ortholog is 100%.

Group of orthologs #131. Best score 3724 bits
Score difference with first non-orthologous sequence - *A. fumigatus*:3724 *A. fischeri*:3724
afum|EAL91221.2 100.00% afis|EAW17387.1 100.00%
Bootstrap support for afum|EAL91221.2 as seed ortholog is 100%.
Bootstrap support for afis|EAW17387.1 as seed ortholog is 100%.

Group of orthologs #132. Best score 3712 bits
Score difference with first non-orthologous sequence - *A. fumigatus*:3712 *A. fischeri*:3712
afum|EAL87569.1 100.00% afis|EAW18204.1 100.00%
Bootstrap support for afum|EAL87569.1 as seed ortholog is 100%.
Bootstrap support for afis|EAW18204.1 as seed ortholog is 100%.

Group of orthologs #133. Best score 3704 bits
Score difference with first non-orthologous sequence - *A. fumigatus*:3704 *A. fischeri*:3704
afum|EAL90827.1 100.00% afis|EAW22364.1 100.00%
Bootstrap support for afum|EAL90827.1 as seed ortholog is 100%.
Bootstrap support for afis|EAW22364.1 as seed ortholog is 100%.

Group of orthologs #134. Best score 3703 bits
Score difference with first non-orthologous sequence - *A. fumigatus*:3703 *A. fischeri*:3703
afum|EAL88239.1 100.00% afis|EAW23243.1 100.00%
Bootstrap support for afum|EAL88239.1 as seed ortholog is 100%.
Bootstrap support for afis|EAW23243.1 as seed ortholog is 100%.

Group of orthologs #135. Best score 3694 bits
Score difference with first non-orthologous sequence - *A. fumigatus*:3694 *A. fischeri*:3694
afum|EAL93161.1 100.00% afis|EAW18474.1 100.00%
Bootstrap support for afum|EAL93161.1 as seed ortholog is 100%.
Bootstrap support for afis|EAW18474.1 as seed ortholog is 100%.

Group of orthologs #136. Best score 3684 bits
Score difference with first non-orthologous sequence - *A. fumigatus*:3684 *A. fischeri*:3684
afum|EAL85320.1 100.00% afis|EAW20200.1 100.00%
Bootstrap support for afum|EAL85320.1 as seed ortholog is 100%.
Bootstrap support for afis|EAW20200.1 as seed ortholog is 100%.

Group of orthologs #137. Best score 3674 bits
Score difference with first non-orthologous sequence - *A. fumigatus*:3674 *A. fischeri*:3674
afum|EBA27495.1 100.00% afis|EAW18891.1 100.00%
Bootstrap support for afum|EBA27495.1 as seed ortholog is 100%.
Bootstrap support for afis|EAW18891.1 as seed ortholog is 100%.

Group of orthologs #138. Best score 3673 bits

Score difference with first non-orthologous sequence - *A. fumigatus*:3673 *A. fischeri*:3673
afum|EAL93713.2 100.00% afis|EAW18977.1 100.00%
Bootstrap support for afum|EAL93713.2 as seed ortholog is 100%.
Bootstrap support for afis|EAW18977.1 as seed ortholog is 100%.

---

Group of orthologs #139. Best score 3672 bits
Score difference with first non-orthologous sequence - *A. fumigatus*:3672 *A. fischeri*:3672
afum|EAL92943.2 100.00% afis|EAW21931.1 100.00%
Bootstrap support for afum|EAL92943.2 as seed ortholog is 100%.
Bootstrap support for afis|EAW21931.1 as seed ortholog is 100%.

---

Group of orthologs #140. Best score 3670 bits
Score difference with first non-orthologous sequence - *A. fumigatus*:3670 *A. fischeri*:2800
afum|EAL84837.1 100.00% afis|EAW20787.1 100.00%
Bootstrap support for afum|EAL84837.1 as seed ortholog is 100%.
Bootstrap support for afis|EAW20787.1 as seed ortholog is 100%.

---

Group of orthologs #141. Best score 3664 bits
Score difference with first non-orthologous sequence - *A. fumigatus*:3664 *A. fischeri*:3664
afum|EAL90724.1 100.00% afis|EAW22467.1 100.00%
Bootstrap support for afum|EAL90724.1 as seed ortholog is 100%.
Bootstrap support for afis|EAW22467.1 as seed ortholog is 100%.

---

Group of orthologs #142. Best score 3662 bits
Score difference with first non-orthologous sequence - *A. fumigatus*:3662 *A. fischeri*:3662
afum|EAL85639.1 100.00% afis|EAW15708.1 100.00%
Bootstrap support for afum|EAL85639.1 as seed ortholog is 100%.
Bootstrap support for afis|EAW15708.1 as seed ortholog is 100%.

---

Group of orthologs #143. Best score 3658 bits
Score difference with first non-orthologous sequence - *A. fumigatus*:3658 *A. fischeri*:3658
afum|EAL92340.1 100.00% afis|EAW21323.1 100.00%
Bootstrap support for afum|EAL92340.1 as seed ortholog is 100%.
Bootstrap support for afis|EAW21323.1 as seed ortholog is 100%.

---

Group of orthologs #144. Best score 3656 bits
Score difference with first non-orthologous sequence - *A. fumigatus*:3656 *A. fischeri*:3656
afum|EAL88733.1 100.00% afis|EAW16101.1 100.00%
Bootstrap support for afum|EAL88733.1 as seed ortholog is 100%.
Bootstrap support for afis|EAW16101.1 as seed ortholog is 100%.

---

Group of orthologs #145. Best score 3655 bits
Score difference with first non-orthologous sequence - *A. fumigatus*:3655 *A. fischeri*:2550
afum|EAL85604.1 100.00% afis|EAW15744.1 100.00%
Bootstrap support for afum|EAL85604.1 as seed ortholog is 100%.
Bootstrap support for afis|EAW15744.1 as seed ortholog is 100%.

---

Group of orthologs #146. Best score 3651 bits
Score difference with first non-orthologous sequence - *A. fumigatus*:3651 *A. fischeri*:3651
afum|EAL87595.1 100.00% afis|EAW18228.1 100.00%
Bootstrap support for afum|EAL87595.1 as seed ortholog is 100%.
Bootstrap support for afis|EAW18228.1 as seed ortholog is 100%.

---

Group of orthologs #147. Best score 3651 bits
Score difference with first non-orthologous sequence - *A. fumigatus*:3651 *A. fischeri*:3651
afum|EAL87926.1 100.00% afis|EAW23540.1 100.00%

Bootstrap support for afum|EAL87926.1 as seed ortholog is 100%.
Bootstrap support for afis|EAW23540.1 as seed ortholog is 100%.

---

Group of orthologs #148. Best score 3636 bits
Score difference with first non-orthologous sequence - *A. fumigatus*:3636 *A. fischeri*:3636
afum|EAL85832.1 100.00% afis|EAW15491.1 100.00%
Bootstrap support for afum|EAL85832.1 as seed ortholog is 100%.
Bootstrap support for afis|EAW15491.1 as seed ortholog is 100%.

---

Group of orthologs #149. Best score 3633 bits
Score difference with first non-orthologous sequence - *A. fumigatus*:3633 *A. fischeri*:3633
afum|EAL91653.1 100.00% afis|EAW17844.1 100.00%
Bootstrap support for afum|EAL91653.1 as seed ortholog is 100%.
Bootstrap support for afis|EAW17844.1 as seed ortholog is 100%.

---

Group of orthologs #150. Best score 3624 bits
Score difference with first non-orthologous sequence - *A. fumigatus*:3624 *A. fischeri*:3624
afum|EAL89586.1 100.00% afis|EAW24930.1 100.00%
Bootstrap support for afum|EAL89586.1 as seed ortholog is 100%.
Bootstrap support for afis|EAW24930.1 as seed ortholog is 100%.

---

Group of orthologs #151. Best score 3622 bits
Score difference with first non-orthologous sequence - *A. fumigatus*:3622 *A. fischeri*:3622
afum|EAL88318.1 100.00% afis|EAW23166.1 100.00%
Bootstrap support for afum|EAL88318.1 as seed ortholog is 100%.
Bootstrap support for afis|EAW23166.1 as seed ortholog is 100%.

---

Group of orthologs #152. Best score 3618 bits
Score difference with first non-orthologous sequence - *A. fumigatus*:3618 *A. fischeri*:3618
afum|EAL90357.1 100.00% afis|EAW22845.1 100.00%
Bootstrap support for afum|EAL90357.1 as seed ortholog is 100%.
Bootstrap support for afis|EAW22845.1 as seed ortholog is 100%.

---

Group of orthologs #153. Best score 3611 bits
Score difference with first non-orthologous sequence - *A. fumigatus*:3611 *A. fischeri*:1776
afum|EAL93809.1 100.00% afis|EAW19080.1 100.00%
Bootstrap support for afum|EAL93809.1 as seed ortholog is 100%.
Bootstrap support for afis|EAW19080.1 as seed ortholog is 100%.

---

Group of orthologs #154. Best score 3608 bits
Score difference with first non-orthologous sequence - *A. fumigatus*:3608 *A. fischeri*:3608
afum|EAL91348.1 100.00% afis|EAW17534.1 100.00%
Bootstrap support for afum|EAL91348.1 as seed ortholog is 100%.
Bootstrap support for afis|EAW17534.1 as seed ortholog is 100%.

---

Group of orthologs #155. Best score 3606 bits
Score difference with first non-orthologous sequence - *A. fumigatus*:3606 *A. fischeri*:3606
afum|EAL91424.1 100.00% afis|EAW17611.1 100.00%
Bootstrap support for afum|EAL91424.1 as seed ortholog is 100%.
Bootstrap support for afis|EAW17611.1 as seed ortholog is 100%.

---

Group of orthologs #156. Best score 3600 bits
Score difference with first non-orthologous sequence - *A. fumigatus*:3600 *A. fischeri*:3600
afum|EAL84710.1 100.00% afis|EAW20924.1 100.00%
Bootstrap support for afum|EAL84710.1 as seed ortholog is 100%.
Bootstrap support for afis|EAW20924.1 as seed ortholog is 100%.

Group of orthologs #157. Best score 3600 bits
Score difference with first non-orthologous sequence - *A. fumigatus*:3203 *A. fischeri*:3092
afum|EAL91801.1 100.00% afis|EAW17991.1 100.00%
Bootstrap support for afum|EAL91801.1 as seed ortholog is 100%.
Bootstrap support for afis|EAW17991.1 as seed ortholog is 100%.

Group of orthologs #158. Best score 3599 bits
Score difference with first non-orthologous sequence - *A. fumigatus*:3599 *A. fischeri*:3599
afum|EAL84891.1 100.00% afis|EAW20026.1 100.00%
Bootstrap support for afum|EAL84891.1 as seed ortholog is 100%.
Bootstrap support for afis|EAW20026.1 as seed ortholog is 100%.

Group of orthologs #159. Best score 3582 bits
Score difference with first non-orthologous sequence - *A. fumigatus*:3582 *A. fischeri*:3582
afum|EAL87349.1 100.00% afis|EAW23924.1 100.00%
Bootstrap support for afum|EAL87349.1 as seed ortholog is 100%.
Bootstrap support for afis|EAW23924.1 as seed ortholog is 100%.

Group of orthologs #160. Best score 3581 bits
Score difference with first non-orthologous sequence - *A. fumigatus*:3581 *A. fischeri*:1671
afum|EAL90765.1 100.00% afis|EAW22426.1 100.00%
Bootstrap support for afum|EAL90765.1 as seed ortholog is 100%.
Bootstrap support for afis|EAW22426.1 as seed ortholog is 100%.

Group of orthologs #161. Best score 3579 bits
Score difference with first non-orthologous sequence - *A. fumigatus*:3579 *A. fischeri*:3579
afum|EAL93856.1 100.00% afis|EAW19124.1 100.00%
Bootstrap support for afum|EAL93856.1 as seed ortholog is 100%.
Bootstrap support for afis|EAW19124.1 as seed ortholog is 100%.

Group of orthologs #162. Best score 3562 bits
Score difference with first non-orthologous sequence - *A. fumigatus*:3562 *A. fischeri*:3562
afum|EAL89572.2 100.00% afis|EAW24917.1 100.00%
Bootstrap support for afum|EAL89572.2 as seed ortholog is 100%.
Bootstrap support for afis|EAW24917.1 as seed ortholog is 100%.

Group of orthologs #163. Best score 3541 bits
Score difference with first non-orthologous sequence - *A. fumigatus*:3541 *A. fischeri*:3541
afum|EAL85935.1 100.00% afis|EAW24267.1 100.00%
Bootstrap support for afum|EAL85935.1 as seed ortholog is 100%.
Bootstrap support for afis|EAW24267.1 as seed ortholog is 100%.

Group of orthologs #164. Best score 3539 bits
Score difference with first non-orthologous sequence - *A. fumigatus*:3539 *A. fischeri*:3539
afum|EAL87589.2 100.00% afis|EAW18223.1 100.00%
Bootstrap support for afum|EAL87589.2 as seed ortholog is 100%.
Bootstrap support for afis|EAW18223.1 as seed ortholog is 100%.

Group of orthologs #165. Best score 3535 bits
Score difference with first non-orthologous sequence - *A. fumigatus*:3535 *A. fischeri*:3535
afum|EAL89767.1 100.00% afis|EAW25096.1 100.00%
Bootstrap support for afum|EAL89767.1 as seed ortholog is 100%.
Bootstrap support for afis|EAW25096.1 as seed ortholog is 100%.

Group of orthologs #166. Best score 3528 bits

Score difference with first non-orthologous sequence - *A. fumigatus*:3528 *A. fischeri*:3528
afum|EAL88418.1 100.00% afis|EAW23061.1 100.00%
Bootstrap support for afum|EAL88418.1 as seed ortholog is 100%.
Bootstrap support for afis|EAW23061.1 as seed ortholog is 100%.

---

Group of orthologs #167. Best score 3512 bits
Score difference with first non-orthologous sequence - *A. fumigatus*:3512 *A. fischeri*:3512
afum|EAL87348.1 100.00% afis|EAW23923.1 100.00%
Bootstrap support for afum|EAL87348.1 as seed ortholog is 100%.
Bootstrap support for afis|EAW23923.1 as seed ortholog is 100%.

---

Group of orthologs #168. Best score 3510 bits
Score difference with first non-orthologous sequence - *A. fumigatus*:3510 *A. fischeri*:3510
afum|EAL93368.1 100.00% afis|EAW18652.1 100.00%
Bootstrap support for afum|EAL93368.1 as seed ortholog is 100%.
Bootstrap support for afis|EAW18652.1 as seed ortholog is 100%.

---

Group of orthologs #169. Best score 3508 bits
Score difference with first non-orthologous sequence - *A. fumigatus*:3508 *A. fischeri*:3508
afum|EAL87508.1 100.00% afis|EAW18150.1 100.00%
Bootstrap support for afum|EAL87508.1 as seed ortholog is 100%.
Bootstrap support for afis|EAW18150.1 as seed ortholog is 100%.

---

Group of orthologs #170. Best score 3508 bits
Score difference with first non-orthologous sequence - *A. fumigatus*:3508 *A. fischeri*:2433
afum|EAL91655.1 100.00% afis|EAW17846.1 100.00%
Bootstrap support for afum|EAL91655.1 as seed ortholog is 100%.
Bootstrap support for afis|EAW17846.1 as seed ortholog is 100%.

---

Group of orthologs #171. Best score 3505 bits
Score difference with first non-orthologous sequence - *A. fumigatus*:3505 *A. fischeri*:3505
afum|EAL89449.1 100.00% afis|EAW24807.1 100.00%
Bootstrap support for afum|EAL89449.1 as seed ortholog is 100%.
Bootstrap support for afis|EAW24807.1 as seed ortholog is 100%.

---

Group of orthologs #172. Best score 3497 bits
Score difference with first non-orthologous sequence - *A. fumigatus*:3497 *A. fischeri*:3497
afum|EAL84448.1 100.00% afis|EAW20613.1 100.00%
Bootstrap support for afum|EAL84448.1 as seed ortholog is 100%.
Bootstrap support for afis|EAW20613.1 as seed ortholog is 100%.

---

Group of orthologs #173. Best score 3497 bits
Score difference with first non-orthologous sequence - *A. fumigatus*:3497 *A. fischeri*:3497
afum|EAL92151.1 100.00% afis|EAW21133.1 100.00%
Bootstrap support for afum|EAL92151.1 as seed ortholog is 100%.
Bootstrap support for afis|EAW21133.1 as seed ortholog is 100%.

---

Group of orthologs #174. Best score 3495 bits
Score difference with first non-orthologous sequence - *A. fumigatus*:3495 *A. fischeri*:3495
afum|EAL90031.1 100.00% afis|EAW25371.1 100.00%
Bootstrap support for afum|EAL90031.1 as seed ortholog is 100%.
Bootstrap support for afis|EAW25371.1 as seed ortholog is 100%.

---

Group of orthologs #175. Best score 3487 bits
Score difference with first non-orthologous sequence - *A. fumigatus*:3487 *A. fischeri*:3487
afum|EAL87083.1 100.00% afis|EAW19411.1 100.00%

Bootstrap support for afum|EAL87083.1 as seed ortholog is 100%.
Bootstrap support for afis|EAW19411.1 as seed ortholog is 100%.

---

Group of orthologs #176. Best score 3485 bits
Score difference with first non-orthologous sequence - *A. fumigatus*:3485 *A. fischeri*:3485
afum|EAL88146.2 100.00% afis|EAW23332.1 100.00%
Bootstrap support for afum|EAL88146.2 as seed ortholog is 100%.
Bootstrap support for afis|EAW23332.1 as seed ortholog is 100%.

---

Group of orthologs #177. Best score 3483 bits
Score difference with first non-orthologous sequence - *A. fumigatus*:3483 *A. fischeri*:1933
afum|EAL92840.1 100.00% afis|EAW21824.1 100.00%
Bootstrap support for afum|EAL92840.1 as seed ortholog is 100%.
Bootstrap support for afis|EAW21824.1 as seed ortholog is 100%.

---

Group of orthologs #178. Best score 3482 bits
Score difference with first non-orthologous sequence - *A. fumigatus*:3482 *A. fischeri*:2322
afum|EAL88655.1 100.00% afis|EAW16026.1 100.00%
Bootstrap support for afum|EAL88655.1 as seed ortholog is 100%.
Bootstrap support for afis|EAW16026.1 as seed ortholog is 100%.

---

Group of orthologs #179. Best score 3480 bits
Score difference with first non-orthologous sequence - *A. fumigatus*:3480 *A. fischeri*:3480
afum|EAL91433.2 100.00% afis|EAW17620.1 100.00%
Bootstrap support for afum|EAL91433.2 as seed ortholog is 100%.
Bootstrap support for afis|EAW17620.1 as seed ortholog is 100%.

---

Group of orthologs #180. Best score 3479 bits
Score difference with first non-orthologous sequence - *A. fumigatus*:3479 *A. fischeri*:3479
afum|EAL87333.1 100.00% afis|EAW23905.1 100.00%
Bootstrap support for afum|EAL87333.1 as seed ortholog is 100%.
Bootstrap support for afis|EAW23905.1 as seed ortholog is 100%.

---

Group of orthologs #181. Best score 3465 bits
Score difference with first non-orthologous sequence - *A. fumigatus*:3465 *A. fischeri*:3465
afum|EBA27410.1 100.00% afis|EAW22631.1 100.00%
Bootstrap support for afum|EBA27410.1 as seed ortholog is 100%.
Bootstrap support for afis|EAW22631.1 as seed ortholog is 100%.

---

Group of orthologs #182. Best score 3463 bits
Score difference with first non-orthologous sequence - *A. fumigatus*:3463 *A. fischeri*:3463
afum|EAL93984.1 100.00% afis|EAW19257.1 100.00%
Bootstrap support for afum|EAL93984.1 as seed ortholog is 100%.
Bootstrap support for afis|EAW19257.1 as seed ortholog is 100%.

---

Group of orthologs #183. Best score 3449 bits
Score difference with first non-orthologous sequence - *A. fumigatus*:3449 *A. fischeri*:3449
afum|EAL92994.1 100.00% afis|EAW21979.1 100.00%
Bootstrap support for afum|EAL92994.1 as seed ortholog is 100%.
Bootstrap support for afis|EAW21979.1 as seed ortholog is 100%.

---

Group of orthologs #184. Best score 3447 bits
Score difference with first non-orthologous sequence - *A. fumigatus*:3447 *A. fischeri*:3447
afum|EAL92431.1 100.00% afis|EAW21416.1 100.00%
Bootstrap support for afum|EAL92431.1 as seed ortholog is 100%.
Bootstrap support for afis|EAW21416.1 as seed ortholog is 100%.

Group of orthologs #185. Best score 3443 bits
Score difference with first non-orthologous sequence - *A. fumigatus*:3443 *A. fischeri*:1633
afum|EAL86223.1 100.00% afis|EAW24562.1 100.00%
Bootstrap support for afum|EAL86223.1 as seed ortholog is 100%.
Bootstrap support for afis|EAW24562.1 as seed ortholog is 100%.

---

Group of orthologs #186. Best score 3431 bits
Score difference with first non-orthologous sequence - *A. fumigatus*:3258 *A. fischeri*:3431
afum|EAL88509.2 100.00% afis|EAW15875.1 100.00%
Bootstrap support for afum|EAL88509.2 as seed ortholog is 100%.
Bootstrap support for afis|EAW15875.1 as seed ortholog is 100%.

---

Group of orthologs #187. Best score 3430 bits
Score difference with first non-orthologous sequence - *A. fumigatus*:3430 *A. fischeri*:3430
afum|EAL85798.1 100.00% afis|EAW15530.1 100.00%
Bootstrap support for afum|EAL85798.1 as seed ortholog is 100%.
Bootstrap support for afis|EAW15530.1 as seed ortholog is 100%.

---

Group of orthologs #188. Best score 3425 bits
Score difference with first non-orthologous sequence - *A. fumigatus*:3080 *A. fischeri*:3017
afum|EAL91018.1 100.00% afis|EAW22171.1 100.00%
Bootstrap support for afum|EAL91018.1 as seed ortholog is 100%.
Bootstrap support for afis|EAW22171.1 as seed ortholog is 100%.

---

Group of orthologs #189. Best score 3424 bits
Score difference with first non-orthologous sequence - *A. fumigatus*:3424 *A. fischeri*:3424
afum|EAL92038.1 100.00% afis|EAW24176.1 100.00%
Bootstrap support for afum|EAL92038.1 as seed ortholog is 100%.
Bootstrap support for afis|EAW24176.1 as seed ortholog is 100%.

---

Group of orthologs #190. Best score 3420 bits
Score difference with first non-orthologous sequence - *A. fumigatus*:3420 *A. fischeri*:3420
afum|EAL85993.1 100.00% afis|EAW24323.1 100.00%
Bootstrap support for afum|EAL85993.1 as seed ortholog is 100%.
Bootstrap support for afis|EAW24323.1 as seed ortholog is 100%.

---

Group of orthologs #191. Best score 3420 bits
Score difference with first non-orthologous sequence - *A. fumigatus*:3420 *A. fischeri*:3420
afum|EAL93092.1 100.00% afis|EAW18412.1 100.00%
Bootstrap support for afum|EAL93092.1 as seed ortholog is 100%.
Bootstrap support for afis|EAW18412.1 as seed ortholog is 100%.

---

Group of orthologs #192. Best score 3419 bits
Score difference with first non-orthologous sequence - *A. fumigatus*:3419 *A. fischeri*:3419
afum|EAL92977.1 100.00% afis|EAW21963.1 100.00%
Bootstrap support for afum|EAL92977.1 as seed ortholog is 100%.
Bootstrap support for afis|EAW21963.1 as seed ortholog is 100%.

---

Group of orthologs #193. Best score 3416 bits
Score difference with first non-orthologous sequence - *A. fumigatus*:3416 *A. fischeri*:1873
afum|EAL86078.1 100.00% afis|EAW24409.1 100.00%
Bootstrap support for afum|EAL86078.1 as seed ortholog is 100%.
Bootstrap support for afis|EAW24409.1 as seed ortholog is 100%.

---

Group of orthologs #194. Best score 3411 bits

Score difference with first non-orthologous sequence - *A. fumigatus*:3411   *A. fischeri*:3176
afum|EAL89667.1 100.00% afis|EAW25028.1 100.00%
Bootstrap support for afum|EAL89667.1 as seed ortholog is 100%.
Bootstrap support for afis|EAW25028.1 as seed ortholog is 100%.

---

Group of orthologs #195. Best score 3403 bits
Score difference with first non-orthologous sequence - *A. fumigatus*:3403   *A. fischeri*:3403
afum|EAL92776.1 100.00% afis|EAW21743.1 100.00%
Bootstrap support for afum|EAL92776.1 as seed ortholog is 100%.
Bootstrap support for afis|EAW21743.1 as seed ortholog is 100%.

---

Group of orthologs #196. Best score 3378 bits
Score difference with first non-orthologous sequence - *A. fumigatus*:3378   *A. fischeri*:3378
afum|EAL87585.1 100.00% afis|EAW18219.1 100.00%
Bootstrap support for afum|EAL87585.1 as seed ortholog is 100%.
Bootstrap support for afis|EAW18219.1 as seed ortholog is 100%.

---

Group of orthologs #197. Best score 3377 bits
Score difference with first non-orthologous sequence - *A. fumigatus*:3377   *A. fischeri*:1251
afum|EAL86423.2 100.00% afis|EAW16885.1 100.00%
Bootstrap support for afum|EAL86423.2 as seed ortholog is 100%.
Bootstrap support for afis|EAW16885.1 as seed ortholog is 100%.

---

Group of orthologs #198. Best score 3376 bits
Score difference with first non-orthologous sequence - *A. fumigatus*:3023   *A. fischeri*:2999
afum|EAL87248.1 100.00% afis|EAW23813.1 100.00%
Bootstrap support for afum|EAL87248.1 as seed ortholog is 100%.
Bootstrap support for afis|EAW23813.1 as seed ortholog is 100%.

---

Group of orthologs #199. Best score 3362 bits
Score difference with first non-orthologous sequence - *A. fumigatus*:3362   *A. fischeri*:3362
afum|EAL91962.1 100.00% afis|EAW24108.1 100.00%
Bootstrap support for afum|EAL91962.1 as seed ortholog is 100%.
Bootstrap support for afis|EAW24108.1 as seed ortholog is 100%.

---

Group of orthologs #200. Best score 3359 bits
Score difference with first non-orthologous sequence - *A. fumigatus*:3359   *A. fischeri*:3359
afum|EAL87314.2 100.00% afis|EAW23887.1 100.00%
Bootstrap support for afum|EAL87314.2 as seed ortholog is 100%.
Bootstrap support for afis|EAW23887.1 as seed ortholog is 100%.

---

Group of orthologs #201. Best score 3356 bits
Score difference with first non-orthologous sequence - *A. fumigatus*:3356   *A. fischeri*:1461
afum|EAL91073.1 100.00% afis|EAW22112.1 100.00%
Bootstrap support for afum|EAL91073.1 as seed ortholog is 100%.
Bootstrap support for afis|EAW22112.1 as seed ortholog is 100%.

---

Group of orthologs #202. Best score 3351 bits
Score difference with first non-orthologous sequence - *A. fumigatus*:3351   *A. fischeri*:3351
afum|EAL84582.1 100.00% afis|EAW20471.1 100.00%
Bootstrap support for afum|EAL84582.1 as seed ortholog is 100%.
Bootstrap support for afis|EAW20471.1 as seed ortholog is 100%.

---

Group of orthologs #203. Best score 3351 bits
Score difference with first non-orthologous sequence - *A. fumigatus*:3351   *A. fischeri*:3154
afum|EAL85691.1 100.00% afis|EAW22082.1 100.00%

Bootstrap support for afum|EAL85691.1 as seed ortholog is 100%.
Bootstrap support for afis|EAW22082.1 as seed ortholog is 100%.

---

Group of orthologs #204. Best score 3346 bits
Score difference with first non-orthologous sequence - *A. fumigatus*:3346 *A. fischeri*:3346
afum|EAL87688.1 100.00% afis|EAW18321.1 100.00%
Bootstrap support for afum|EAL87688.1 as seed ortholog is 100%.
Bootstrap support for afis|EAW18321.1 as seed ortholog is 100%.

---

Group of orthologs #205. Best score 3345 bits
Score difference with first non-orthologous sequence - *A. fumigatus*:3014 *A. fischeri*:3000
afum|EAL91783.1 100.00% afis|EAW17974.1 100.00%
Bootstrap support for afum|EAL91783.1 as seed ortholog is 100%.
Bootstrap support for afis|EAW17974.1 as seed ortholog is 100%.

---

Group of orthologs #206. Best score 3337 bits
Score difference with first non-orthologous sequence - *A. fumigatus*:2977 *A. fischeri*:2954
afum|EAL91555.1 100.00% afis|EAW17741.1 100.00%
Bootstrap support for afum|EAL91555.1 as seed ortholog is 100%.
Bootstrap support for afis|EAW17741.1 as seed ortholog is 100%.

---

Group of orthologs #207. Best score 3335 bits
Score difference with first non-orthologous sequence - *A. fumigatus*:3335 *A. fischeri*:2018
afum|EAL90415.1 100.00% afis|EAW22784.1 100.00%
Bootstrap support for afum|EAL90415.1 as seed ortholog is 100%.
Bootstrap support for afis|EAW22784.1 as seed ortholog is 100%.

---

Group of orthologs #208. Best score 3332 bits
Score difference with first non-orthologous sequence - *A. fumigatus*:3332 *A. fischeri*:3332
afum|EAL90652.1 100.00% afis|EAW22542.1 100.00%
Bootstrap support for afum|EAL90652.1 as seed ortholog is 100%.
Bootstrap support for afis|EAW22542.1 as seed ortholog is 100%.

---

Group of orthologs #209. Best score 3331 bits
Score difference with first non-orthologous sequence - *A. fumigatus*:3331 *A. fischeri*:3331
afum|EBA27486.1 100.00% afis|EAW18485.1 100.00%
Bootstrap support for afum|EBA27486.1 as seed ortholog is 100%.
Bootstrap support for afis|EAW18485.1 as seed ortholog is 100%.

---

Group of orthologs #210. Best score 3325 bits
Score difference with first non-orthologous sequence - *A. fumigatus*:3325 *A. fischeri*:3325
afum|EAL90586.1 100.00% afis|EAW22612.1 100.00%
Bootstrap support for afum|EAL90586.1 as seed ortholog is 100%.
Bootstrap support for afis|EAW22612.1 as seed ortholog is 100%.

---

Group of orthologs #211. Best score 3323 bits
Score difference with first non-orthologous sequence - *A. fumigatus*:2480 *A. fischeri*:3323
afum|EAL85910.2 100.00% afis|EAW24242.1 100.00%
Bootstrap support for afum|EAL85910.2 as seed ortholog is 100%.
Bootstrap support for afis|EAW24242.1 as seed ortholog is 100%.

---

Group of orthologs #212. Best score 3323 bits
Score difference with first non-orthologous sequence - *A. fumigatus*:3323 *A. fischeri*:3323
afum|EAL86032.1 100.00% afis|EAW24362.1 100.00%
Bootstrap support for afum|EAL86032.1 as seed ortholog is 100%.
Bootstrap support for afis|EAW24362.1 as seed ortholog is 100%.

---

Group of orthologs #213. Best score 3321 bits
Score difference with first non-orthologous sequence - *A. fumigatus*:3321 *A. fischeri*:3321
afum|EAL92697.1 100.00% afis|EAW21662.1 100.00%
Bootstrap support for afum|EAL92697.1 as seed ortholog is 100%.
Bootstrap support for afis|EAW21662.1 as seed ortholog is 100%.

---

Group of orthologs #214. Best score 3319 bits
Score difference with first non-orthologous sequence - *A. fumigatus*:3319 *A. fischeri*:3319
afum|EAL89913.1 100.00% afis|EAW25251.1 100.00%
Bootstrap support for afum|EAL89913.1 as seed ortholog is 100%.
Bootstrap support for afis|EAW25251.1 as seed ortholog is 100%.

---

Group of orthologs #215. Best score 3318 bits
Score difference with first non-orthologous sequence - *A. fumigatus*:3318 *A. fischeri*:3318
afum|EAL88494.1 100.00% afis|EAW15858.1 100.00%
Bootstrap support for afum|EAL88494.1 as seed ortholog is 100%.
Bootstrap support for afis|EAW15858.1 as seed ortholog is 100%.

---

Group of orthologs #216. Best score 3317 bits
Score difference with first non-orthologous sequence - *A. fumigatus*:3317 *A. fischeri*:3317
afum|EAL85556.1 100.00% afis|EAW15794.1 100.00%
Bootstrap support for afum|EAL85556.1 as seed ortholog is 100%.
Bootstrap support for afis|EAW15794.1 as seed ortholog is 100%.

---

Group of orthologs #217. Best score 3312 bits
Score difference with first non-orthologous sequence - *A. fumigatus*:3312 *A. fischeri*:3312
afum|EAL90469.1 100.00% afis|EAW22726.1 100.00%
Bootstrap support for afum|EAL90469.1 as seed ortholog is 100%.
Bootstrap support for afis|EAW22726.1 as seed ortholog is 100%.

---

Group of orthologs #218. Best score 3296 bits
Score difference with first non-orthologous sequence - *A. fumigatus*:3296 *A. fischeri*:3296
afum|EAL87319.1 100.00% afis|EAW23891.1 100.00%
Bootstrap support for afum|EAL87319.1 as seed ortholog is 100%.
Bootstrap support for afis|EAW23891.1 as seed ortholog is 100%.

---

Group of orthologs #219. Best score 3294 bits
Score difference with first non-orthologous sequence - *A. fumigatus*:3294 *A. fischeri*:2339
afum|EAL84904.1 100.00% afis|EAW20014.1 100.00%
Bootstrap support for afum|EAL84904.1 as seed ortholog is 100%.
Bootstrap support for afis|EAW20014.1 as seed ortholog is 100%.

---

Group of orthologs #220. Best score 3284 bits
Score difference with first non-orthologous sequence - *A. fumigatus*:3284 *A. fischeri*:3284
afum|EAL90421.1 100.00% afis|EAW22777.1 100.00%
Bootstrap support for afum|EAL90421.1 as seed ortholog is 100%.
Bootstrap support for afis|EAW22777.1 as seed ortholog is 100%.

---

Group of orthologs #221. Best score 3283 bits
Score difference with first non-orthologous sequence - *A. fumigatus*:2353 *A. fischeri*:2271
afum|EAL90367.2 100.00% afis|EAW22835.1 100.00%
Bootstrap support for afum|EAL90367.2 as seed ortholog is 100%.
Bootstrap support for afis|EAW22835.1 as seed ortholog is 100%.

---

Group of orthologs #222. Best score 3278 bits

Score difference with first non-orthologous sequence - *A. fumigatus*:3278 *A. fischeri*:3278
afum|EAL93678.1 100.00% afis|EAW18945.1 100.00%
Bootstrap support for afum|EAL93678.1 as seed ortholog is 100%.
Bootstrap support for afis|EAW18945.1 as seed ortholog is 100%.

---

Group of orthologs #223. Best score 3268 bits
Score difference with first non-orthologous sequence - *A. fumigatus*:3268 *A. fischeri*:3268
afum|EAL89495.1 100.00% afis|EAW24849.1 100.00%
Bootstrap support for afum|EAL89495.1 as seed ortholog is 100%.
Bootstrap support for afis|EAW24849.1 as seed ortholog is 100%.

---

Group of orthologs #224. Best score 3259 bits
Score difference with first non-orthologous sequence - *A. fumigatus*:3259 *A. fischeri*:3259
afum|EAL90587.1 100.00% afis|EAW22611.1 100.00%
Bootstrap support for afum|EAL90587.1 as seed ortholog is 100%.
Bootstrap support for afis|EAW22611.1 as seed ortholog is 100%.

---

Group of orthologs #225. Best score 3251 bits
Score difference with first non-orthologous sequence - *A. fumigatus*:3251 *A. fischeri*:3251
afum|EAL88253.1 100.00% afis|EAW23230.1 100.00%
Bootstrap support for afum|EAL88253.1 as seed ortholog is 100%.
Bootstrap support for afis|EAW23230.1 as seed ortholog is 100%.

---

Group of orthologs #226. Best score 3245 bits
Score difference with first non-orthologous sequence - *A. fumigatus*:3245 *A. fischeri*:3245
afum|EAL84659.1 100.00% afis|EAW20974.1 100.00%
Bootstrap support for afum|EAL84659.1 as seed ortholog is 100%.
Bootstrap support for afis|EAW20974.1 as seed ortholog is 100%.

---

Group of orthologs #227. Best score 3244 bits
Score difference with first non-orthologous sequence - *A. fumigatus*:3244 *A. fischeri*:3244
afum|EAL85640.1 100.00% afis|EAW15707.1 100.00%
Bootstrap support for afum|EAL85640.1 as seed ortholog is 100%.
Bootstrap support for afis|EAW15707.1 as seed ortholog is 100%.

---

Group of orthologs #228. Best score 3243 bits
Score difference with first non-orthologous sequence - *A. fumigatus*:3243 *A. fischeri*:3243
afum|EAL87572.1 100.00% afis|EAW18207.1 100.00%
Bootstrap support for afum|EAL87572.1 as seed ortholog is 100%.
Bootstrap support for afis|EAW18207.1 as seed ortholog is 100%.

---

Group of orthologs #229. Best score 3243 bits
Score difference with first non-orthologous sequence - *A. fumigatus*:3243 *A. fischeri*:3243
afum|EAL93181.1 100.00% afis|EAW18491.1 100.00%
Bootstrap support for afum|EAL93181.1 as seed ortholog is 100%.
Bootstrap support for afis|EAW18491.1 as seed ortholog is 100%.

---

Group of orthologs #230. Best score 3243 bits
Score difference with first non-orthologous sequence - *A. fumigatus*:3243 *A. fischeri*:3243
afum|EAL87405.1 100.00% afis|EAW23975.1 100.00%
Bootstrap support for afum|EAL87405.1 as seed ortholog is 100%.
Bootstrap support for afis|EAW23975.1 as seed ortholog is 100%.

---

Group of orthologs #231. Best score 3242 bits
Score difference with first non-orthologous sequence - *A. fumigatus*:3242 *A. fischeri*:3242
afum|EAL87643.1 100.00% afis|EAW18277.1 100.00%

Bootstrap support for afum|EAL87643.1 as seed ortholog is 100%.
Bootstrap support for afis|EAW18277.1 as seed ortholog is 100%.

---

Group of orthologs #232. Best score 3240 bits
Score difference with first non-orthologous sequence - *A. fumigatus*:2872 *A. fischeri*:2782
afum|EAL89872.1 100.00% afis|EAW25211.1 100.00%
Bootstrap support for afum|EAL89872.1 as seed ortholog is 100%.
Bootstrap support for afis|EAW25211.1 as seed ortholog is 100%.

---

Group of orthologs #233. Best score 3238 bits
Score difference with first non-orthologous sequence - *A. fumigatus*:3238 *A. fischeri*:1622
afum|EAL89669.1 100.00% afis|EAW25026.1 100.00%
Bootstrap support for afum|EAL89669.1 as seed ortholog is 100%.
Bootstrap support for afis|EAW25026.1 as seed ortholog is 100%.

---

Group of orthologs #234. Best score 3233 bits
Score difference with first non-orthologous sequence - *A. fumigatus*:3233 *A. fischeri*:3233
afum|EAL86870.1 100.00% afis|EAW19652.1 100.00%
Bootstrap support for afum|EAL86870.1 as seed ortholog is 100%.
Bootstrap support for afis|EAW19652.1 as seed ortholog is 100%.

---

Group of orthologs #235. Best score 3226 bits
Score difference with first non-orthologous sequence - *A. fumigatus*:3226 *A. fischeri*:3226
afum|EAL93858.1 100.00% afis|EAW19126.1 100.00%
Bootstrap support for afum|EAL93858.1 as seed ortholog is 100%.
Bootstrap support for afis|EAW19126.1 as seed ortholog is 100%.

---

Group of orthologs #236. Best score 3226 bits
Score difference with first non-orthologous sequence - *A. fumigatus*:3226 *A. fischeri*:3226
afum|EAL90602.1 100.00% afis|EAW22596.1 100.00%
Bootstrap support for afum|EAL90602.1 as seed ortholog is 100%.
Bootstrap support for afis|EAW22596.1 as seed ortholog is 100%.

---

Group of orthologs #237. Best score 3226 bits
Score difference with first non-orthologous sequence - *A. fumigatus*:3226 *A. fischeri*:3226
afum|EAL92889.2 100.00% afis|EAW21875.1 100.00%
Bootstrap support for afum|EAL92889.2 as seed ortholog is 100%.
Bootstrap support for afis|EAW21875.1 as seed ortholog is 100%.

---

Group of orthologs #238. Best score 3220 bits
Score difference with first non-orthologous sequence - *A. fumigatus*:3220 *A. fischeri*:1817
afum|EAL84314.2 100.00% afis|EAW25626.1 100.00%
Bootstrap support for afum|EAL84314.2 as seed ortholog is 100%.
Bootstrap support for afis|EAW25626.1 as seed ortholog is 100%.

---

Group of orthologs #239. Best score 3218 bits
Score difference with first non-orthologous sequence - *A. fumigatus*:3218 *A. fischeri*:3218
afum|EAL93143.1 100.00% afis|EAW18458.1 100.00%
Bootstrap support for afum|EAL93143.1 as seed ortholog is 100%.
Bootstrap support for afis|EAW18458.1 as seed ortholog is 100%.

---

Group of orthologs #240. Best score 3200 bits
Score difference with first non-orthologous sequence - *A. fumigatus*:3200 *A. fischeri*:3200
afum|EAL85173.2 100.00% afis|EAW20053.1 100.00%
Bootstrap support for afum|EAL85173.2 as seed ortholog is 100%.
Bootstrap support for afis|EAW20053.1 as seed ortholog is 100%.

Group of orthologs #241. Best score 3199 bits
Score difference with first non-orthologous sequence - *A. fumigatus*:3199 *A. fischeri*:3199
afum|EAL85016.1 100.00% afis|EAW19882.1 100.00%
Bootstrap support for afum|EAL85016.1 as seed ortholog is 100%.
Bootstrap support for afis|EAW19882.1 as seed ortholog is 100%.

Group of orthologs #242. Best score 3186 bits
Score difference with first non-orthologous sequence - *A. fumigatus*:3186 *A. fischeri*:3186
afum|EAL89147.1 100.00% afis|EAW16549.1 100.00%
Bootstrap support for afum|EAL89147.1 as seed ortholog is 100%.
Bootstrap support for afis|EAW16549.1 as seed ortholog is 100%.

Group of orthologs #243. Best score 3167 bits
Score difference with first non-orthologous sequence - *A. fumigatus*:3167 *A. fischeri*:3167
afum|EAL93640.1 100.00% afis|EAW18908.1 100.00%
Bootstrap support for afum|EAL93640.1 as seed ortholog is 100%.
Bootstrap support for afis|EAW18908.1 as seed ortholog is 100%.

Group of orthologs #244. Best score 3151 bits
Score difference with first non-orthologous sequence - *A. fumigatus*:3151 *A. fischeri*:3151
afum|EAL84438.1 100.00% afis|EAW20629.1 100.00%
Bootstrap support for afum|EAL84438.1 as seed ortholog is 100%.
Bootstrap support for afis|EAW20629.1 as seed ortholog is 100%.

Group of orthologs #245. Best score 3149 bits
Score difference with first non-orthologous sequence - *A. fumigatus*:3149 *A. fischeri*:3149
afum|EAL89164.1 100.00% afis|EAW16567.1 100.00%
Bootstrap support for afum|EAL89164.1 as seed ortholog is 100%.
Bootstrap support for afis|EAW16567.1 as seed ortholog is 100%.

Group of orthologs #246. Best score 3147 bits
Score difference with first non-orthologous sequence - *A. fumigatus*:3147 *A. fischeri*:3147
afum|EAL93911.1 100.00% afis|EAW19181.1 100.00%
Bootstrap support for afum|EAL93911.1 as seed ortholog is 100%.
Bootstrap support for afis|EAW19181.1 as seed ortholog is 100%.

Group of orthologs #247. Best score 3131 bits
Score difference with first non-orthologous sequence - *A. fumigatus*:3131 *A. fischeri*:3131
afum|EAL87451.1 100.00% afis|EAW24021.1 100.00%
Bootstrap support for afum|EAL87451.1 as seed ortholog is 100%.
Bootstrap support for afis|EAW24021.1 as seed ortholog is 100%.

Group of orthologs #248. Best score 3124 bits
Score difference with first non-orthologous sequence - *A. fumigatus*:3124 *A. fischeri*:3124
afum|EAL91292.1 100.00% afis|EAW17463.1 100.00%
Bootstrap support for afum|EAL91292.1 as seed ortholog is 100%.
Bootstrap support for afis|EAW17463.1 as seed ortholog is 100%.

Group of orthologs #249. Best score 3117 bits
Score difference with first non-orthologous sequence - *A. fumigatus*:3117 *A. fischeri*:3117
afum|EAL93341.1 100.00% afis|EAW18629.1 100.00%
Bootstrap support for afum|EAL93341.1 as seed ortholog is 100%.
Bootstrap support for afis|EAW18629.1 as seed ortholog is 100%.

Group of orthologs #250. Best score 3111 bits

Score difference with first non-orthologous sequence - *A. fumigatus*:3111 *A. fischeri*:3111
afum|EAL87538.1 100.00% afis|EAW18177.1 100.00%
Bootstrap support for afum|EAL87538.1 as seed ortholog is 100%.
Bootstrap support for afis|EAW18177.1 as seed ortholog is 100%.

---

Group of orthologs #251. Best score 3110 bits
Score difference with first non-orthologous sequence - *A. fumigatus*:3110 *A. fischeri*:3110
afum|EAL91658.1 100.00% afis|EAW17849.1 100.00%
Bootstrap support for afum|EAL91658.1 as seed ortholog is 100%.
Bootstrap support for afis|EAW17849.1 as seed ortholog is 100%.

---

Group of orthologs #252. Best score 3101 bits
Score difference with first non-orthologous sequence - *A. fumigatus*:3101 *A. fischeri*:3101
afum|EAL90368.1 100.00% afis|EAW22834.1 100.00%
Bootstrap support for afum|EAL90368.1 as seed ortholog is 100%.
Bootstrap support for afis|EAW22834.1 as seed ortholog is 100%.

---

Group of orthologs #253. Best score 3101 bits
Score difference with first non-orthologous sequence - *A. fumigatus*:2811 *A. fischeri*:2741
afum|EAL92361.1 100.00% afis|EAW21344.1 100.00%
Bootstrap support for afum|EAL92361.1 as seed ortholog is 100%.
Bootstrap support for afis|EAW21344.1 as seed ortholog is 100%.

---

Group of orthologs #254. Best score 3095 bits
Score difference with first non-orthologous sequence - *A. fumigatus*:3095 *A. fischeri*:3095
afum|EAL92207.1 100.00% afis|EAW21185.1 100.00%
Bootstrap support for afum|EAL92207.1 as seed ortholog is 100%.
Bootstrap support for afis|EAW21185.1 as seed ortholog is 100%.

---

Group of orthologs #255. Best score 3094 bits
Score difference with first non-orthologous sequence - *A. fumigatus*:3094 *A. fischeri*:3094
afum|EAL85569.1 100.00% afis|EAW15781.1 100.00%
Bootstrap support for afum|EAL85569.1 as seed ortholog is 100%.
Bootstrap support for afis|EAW15781.1 as seed ortholog is 100%.

---

Group of orthologs #256. Best score 3094 bits
Score difference with first non-orthologous sequence - *A. fumigatus*:3094 *A. fischeri*:3094
afum|EAL87564.1 100.00% afis|EAW18199.1 100.00%
Bootstrap support for afum|EAL87564.1 as seed ortholog is 100%.
Bootstrap support for afis|EAW18199.1 as seed ortholog is 100%.

---

Group of orthologs #257. Best score 3087 bits
Score difference with first non-orthologous sequence - *A. fumigatus*:2208 *A. fischeri*:2861
afum|EAL89365.2 100.00% afis|EAW19928.1 100.00%
Bootstrap support for afum|EAL89365.2 as seed ortholog is 100%.
Bootstrap support for afis|EAW19928.1 as seed ortholog is 100%.

---

Group of orthologs #258. Best score 3085 bits
Score difference with first non-orthologous sequence - *A. fumigatus*:2907 *A. fischeri*:1211
afum|EAL91987.1 100.00% afis|EAW24131.1 100.00%
Bootstrap support for afum|EAL91987.1 as seed ortholog is 100%.
Bootstrap support for afis|EAW24131.1 as seed ortholog is 100%.

---

Group of orthologs #259. Best score 3080 bits
Score difference with first non-orthologous sequence - *A. fumigatus*:3080 *A. fischeri*:3080
afum|EAL87250.1 100.00% afis|EAW23815.1 100.00%

Bootstrap support for afum|EAL87250.1 as seed ortholog is 100%.
Bootstrap support for afis|EAW23815.1 as seed ortholog is 100%.

---

Group of orthologs #260. Best score 3073 bits
Score difference with first non-orthologous sequence - *A. fumigatus*:3073 *A. fischeri*:3073
afum|EAL87279.1 100.00% afis|EAW23851.1 100.00%
Bootstrap support for afum|EAL87279.1 as seed ortholog is 100%.
Bootstrap support for afis|EAW23851.1 as seed ortholog is 100%.

---

Group of orthologs #261. Best score 3073 bits
Score difference with first non-orthologous sequence - *A. fumigatus*:3073 *A. fischeri*:3073
afum|EAL93295.2 100.00% afis|EAW18588.1 100.00%
Bootstrap support for afum|EAL93295.2 as seed ortholog is 100%.
Bootstrap support for afis|EAW18588.1 as seed ortholog is 100%.

---

Group of orthologs #262. Best score 3070 bits
Score difference with first non-orthologous sequence - *A. fumigatus*:3070 *A. fischeri*:3070
afum|EAL84441.1 100.00% afis|EAW20627.1 100.00%
Bootstrap support for afum|EAL84441.1 as seed ortholog is 100%.
Bootstrap support for afis|EAW20627.1 as seed ortholog is 100%.

---

Group of orthologs #263. Best score 3069 bits
Score difference with first non-orthologous sequence - *A. fumigatus*:3069 *A. fischeri*:2790
afum|EAL89269.1 100.00% afis|EAW16665.1 100.00%
Bootstrap support for afum|EAL89269.1 as seed ortholog is 100%.
Bootstrap support for afis|EAW16665.1 as seed ortholog is 100%.

---

Group of orthologs #264. Best score 3068 bits
Score difference with first non-orthologous sequence - *A. fumigatus*:3068 *A. fischeri*:3068
afum|EAL90316.1 100.00% afis|EAW22883.1 100.00%
Bootstrap support for afum|EAL90316.1 as seed ortholog is 100%.
Bootstrap support for afis|EAW22883.1 as seed ortholog is 100%.

---

Group of orthologs #265. Best score 3064 bits
Score difference with first non-orthologous sequence - *A. fumigatus*:3064 *A. fischeri*:2087
afum|EAL86806.1 100.00% afis|EAW19720.1 100.00%
Bootstrap support for afum|EAL86806.1 as seed ortholog is 100%.
Bootstrap support for afis|EAW19720.1 as seed ortholog is 100%.

---

Group of orthologs #266. Best score 3064 bits
Score difference with first non-orthologous sequence - *A. fumigatus*:3064 *A. fischeri*:3064
afum|EAL93162.1 100.00% afis|EAW18475.1 100.00%
Bootstrap support for afum|EAL93162.1 as seed ortholog is 100%.
Bootstrap support for afis|EAW18475.1 as seed ortholog is 100%.

---

Group of orthologs #267. Best score 3059 bits
Score difference with first non-orthologous sequence - *A. fumigatus*:3059 *A. fischeri*:3059
afum|EAL93591.1 100.00% afis|EAW18859.1 100.00%
Bootstrap support for afum|EAL93591.1 as seed ortholog is 100%.
Bootstrap support for afis|EAW18859.1 as seed ortholog is 100%.

---

Group of orthologs #268. Best score 3053 bits
Score difference with first non-orthologous sequence - *A. fumigatus*:3053 *A. fischeri*:3053
afum|EAL85904.1 100.00% afis|EAW24235.1 100.00%
Bootstrap support for afum|EAL85904.1 as seed ortholog is 100%.
Bootstrap support for afis|EAW24235.1 as seed ortholog is 100%.

---

Group of orthologs #269. Best score 3050 bits
Score difference with first non-orthologous sequence - *A. fumigatus*:3050 *A. fischeri*:3050
afum|EAL84702.1 100.00% afis|EAW20932.1 100.00%
Bootstrap support for afum|EAL84702.1 as seed ortholog is 100%.
Bootstrap support for afis|EAW20932.1 as seed ortholog is 100%.

---

Group of orthologs #270. Best score 3046 bits
Score difference with first non-orthologous sequence - *A. fumigatus*:3046 *A. fischeri*:3046
afum|EAL90701.1 100.00% afis|EAW22492.1 100.00%
Bootstrap support for afum|EAL90701.1 as seed ortholog is 100%.
Bootstrap support for afis|EAW22492.1 as seed ortholog is 100%.

---

Group of orthologs #271. Best score 3038 bits
Score difference with first non-orthologous sequence - *A. fumigatus*:3038 *A. fischeri*:3038
afum|EAL86129.1 100.00% afis|EAW24459.1 100.00%
Bootstrap support for afum|EAL86129.1 as seed ortholog is 100%.
Bootstrap support for afis|EAW24459.1 as seed ortholog is 100%.

---

Group of orthologs #272. Best score 3032 bits
Score difference with first non-orthologous sequence - *A. fumigatus*:3032 *A. fischeri*:3032
afum|EAL89074.1 100.00% afis|EAW16480.1 100.00%
Bootstrap support for afum|EAL89074.1 as seed ortholog is 100%.
Bootstrap support for afis|EAW16480.1 as seed ortholog is 100%.

---

Group of orthologs #273. Best score 3024 bits
Score difference with first non-orthologous sequence - *A. fumigatus*:3024 *A. fischeri*:2726
afum|EAL92613.1 100.00% afis|EAW21595.1 100.00%
Bootstrap support for afum|EAL92613.1 as seed ortholog is 100%.
Bootstrap support for afis|EAW21595.1 as seed ortholog is 100%.

---

Group of orthologs #274. Best score 3021 bits
Score difference with first non-orthologous sequence - *A. fumigatus*:3021 *A. fischeri*:3021
afum|EAL86709.1 100.00% afis|EAW17289.1 100.00%
Bootstrap support for afum|EAL86709.1 as seed ortholog is 100%.
Bootstrap support for afis|EAW17289.1 as seed ortholog is 100%.

---

Group of orthologs #275. Best score 3021 bits
Score difference with first non-orthologous sequence - *A. fumigatus*:3021 *A. fischeri*:3021
afum|EAL85329.1 100.00% afis|EAW20212.1 100.00%
Bootstrap support for afum|EAL85329.1 as seed ortholog is 100%.
Bootstrap support for afis|EAW20212.1 as seed ortholog is 100%.

---

Group of orthologs #276. Best score 3020 bits
Score difference with first non-orthologous sequence - *A. fumigatus*:3020 *A. fischeri*:3020
afum|EAL85965.1 100.00% afis|EAW24297.1 100.00%
Bootstrap support for afum|EAL85965.1 as seed ortholog is 100%.
Bootstrap support for afis|EAW24297.1 as seed ortholog is 100%.

---

Group of orthologs #277. Best score 3018 bits
Score difference with first non-orthologous sequence - *A. fumigatus*:3018 *A. fischeri*:3018
afum|EAL89526.1 100.00% afis|EAW24876.1 100.00%
Bootstrap support for afum|EAL89526.1 as seed ortholog is 100%.
Bootstrap support for afis|EAW24876.1 as seed ortholog is 100%.

---

Group of orthologs #278. Best score 3013 bits

Score difference with first non-orthologous sequence - *A. fumigatus*:2825   *A. fischeri*:2179
afum|EAL86625.1    100.00%    afis|EAW17197.1    100.00%
Bootstrap support for afum|EAL86625.1 as seed ortholog is 100%.
Bootstrap support for afis|EAW17197.1 as seed ortholog is 100%.

---

Group of orthologs #279. Best score 3008 bits
Score difference with first non-orthologous sequence - *A. fumigatus*:3008   *A. fischeri*:3008
afum|EAL88681.1    100.00%    afis|EAW16048.1    100.00%
Bootstrap support for afum|EAL88681.1 as seed ortholog is 100%.
Bootstrap support for afis|EAW16048.1 as seed ortholog is 100%.

---

Group of orthologs #280. Best score 3006 bits
Score difference with first non-orthologous sequence - *A. fumigatus*:3006   *A. fischeri*:3006
afum|EAL93922.1    100.00%    afis|EAW19191.1    100.00%
Bootstrap support for afum|EAL93922.1 as seed ortholog is 100%.
Bootstrap support for afis|EAW19191.1 as seed ortholog is 100%.

---

Group of orthologs #281. Best score 2996 bits
Score difference with first non-orthologous sequence - *A. fumigatus*:2996   *A. fischeri*:2996
afum|EAL88105.1    100.00%    afis|EAW23368.1    100.00%
Bootstrap support for afum|EAL88105.1 as seed ortholog is 100%.
Bootstrap support for afis|EAW23368.1 as seed ortholog is 100%.

---

Group of orthologs #282. Best score 2993 bits
Score difference with first non-orthologous sequence - *A. fumigatus*:2993   *A. fischeri*:2993
afum|EAL93475.1    100.00%    afis|EAW18749.1    100.00%
Bootstrap support for afum|EAL93475.1 as seed ortholog is 100%.
Bootstrap support for afis|EAW18749.1 as seed ortholog is 100%.

---

Group of orthologs #283. Best score 2991 bits
Score difference with first non-orthologous sequence - *A. fumigatus*:2991   *A. fischeri*:2702
afum|EAL91592.1    100.00%    afis|EAW17784.1    100.00%
Bootstrap support for afum|EAL91592.1 as seed ortholog is 100%.
Bootstrap support for afis|EAW17784.1 as seed ortholog is 100%.

---

Group of orthologs #284. Best score 2989 bits
Score difference with first non-orthologous sequence - *A. fumigatus*:2989   *A. fischeri*:2989
afum|EAL88849.1    100.00%    afis|EAW16221.1    100.00%
Bootstrap support for afum|EAL88849.1 as seed ortholog is 100%.
Bootstrap support for afis|EAW16221.1 as seed ortholog is 100%.

---

Group of orthologs #285. Best score 2983 bits
Score difference with first non-orthologous sequence - *A. fumigatus*:2983   *A. fischeri*:2983
afum|EAL92501.1    100.00%    afis|EAW21481.1    100.00%
Bootstrap support for afum|EAL92501.1 as seed ortholog is 100%.
Bootstrap support for afis|EAW21481.1 as seed ortholog is 100%.

---

Group of orthologs #286. Best score 2982 bits
Score difference with first non-orthologous sequence - *A. fumigatus*:2982   *A. fischeri*:2982
afum|EAL90113.1    100.00%    afis|EAW25454.1    100.00%
Bootstrap support for afum|EAL90113.1 as seed ortholog is 100%.
Bootstrap support for afis|EAW25454.1 as seed ortholog is 100%.

---

Group of orthologs #287. Best score 2978 bits
Score difference with first non-orthologous sequence - *A. fumigatus*:2978   *A. fischeri*:2978
afum|EAL89680.1    100.00%    afis|EAW25016.1    100.00%

Bootstrap support for afum|EAL89680.1 as seed ortholog is 100%.
Bootstrap support for afis|EAW25016.1 as seed ortholog is 100%.

---

Group of orthologs #288. Best score 2975 bits
Score difference with first non-orthologous sequence - *A. fumigatus*:2975 *A. fischeri*:2975
afum|EAL91540.1 100.00% afis|EAW17725.1 100.00%
Bootstrap support for afum|EAL91540.1 as seed ortholog is 100%.
Bootstrap support for afis|EAW17725.1 as seed ortholog is 100%.

---

Group of orthologs #289. Best score 2972 bits
Score difference with first non-orthologous sequence - *A. fumigatus*:2972 *A. fischeri*:2972
afum|EAL88506.1 100.00% afis|EAW15872.1 100.00%
Bootstrap support for afum|EAL88506.1 as seed ortholog is 100%.
Bootstrap support for afis|EAW15872.1 as seed ortholog is 100%.

---

Group of orthologs #290. Best score 2966 bits
Score difference with first non-orthologous sequence - *A. fumigatus*:2966 *A. fischeri*:2966
afum|EAL85644.1 100.00% afis|EAW15703.1 100.00%
Bootstrap support for afum|EAL85644.1 as seed ortholog is 100%.
Bootstrap support for afis|EAW15703.1 as seed ortholog is 100%.

---

Group of orthologs #291. Best score 2966 bits
Score difference with first non-orthologous sequence - *A. fumigatus*:2966 *A. fischeri*:2966
afum|EAL91717.2 100.00% afis|EAW17908.1 100.00%
Bootstrap support for afum|EAL91717.2 as seed ortholog is 100%.
Bootstrap support for afis|EAW17908.1 as seed ortholog is 100%.

---

Group of orthologs #292. Best score 2965 bits
Score difference with first non-orthologous sequence - *A. fumigatus*:2965 *A. fischeri*:2965
afum|EAL93070.1 100.00% afis|EAW18394.1 100.00%
Bootstrap support for afum|EAL93070.1 as seed ortholog is 100%.
Bootstrap support for afis|EAW18394.1 as seed ortholog is 100%.

---

Group of orthologs #293. Best score 2963 bits
Score difference with first non-orthologous sequence - *A. fumigatus*:2963 *A. fischeri*:2963
afum|EAL92461.2 100.00% afis|EAW21443.1 100.00%
Bootstrap support for afum|EAL92461.2 as seed ortholog is 100%.
Bootstrap support for afis|EAW21443.1 as seed ortholog is 100%.

---

Group of orthologs #294. Best score 2962 bits
Score difference with first non-orthologous sequence - *A. fumigatus*:2962 *A. fischeri*:2962
afum|EAL93673.1 100.00% afis|EAW18940.1 100.00%
Bootstrap support for afum|EAL93673.1 as seed ortholog is 100%.
Bootstrap support for afis|EAW18940.1 as seed ortholog is 100%.

---

Group of orthologs #295. Best score 2957 bits
Score difference with first non-orthologous sequence - *A. fumigatus*:2957 *A. fischeri*:2957
afum|EAL92750.1 100.00% afis|EAW21716.1 100.00%
Bootstrap support for afum|EAL92750.1 as seed ortholog is 100%.
Bootstrap support for afis|EAW21716.1 as seed ortholog is 100%.

---

Group of orthologs #296. Best score 2956 bits
Score difference with first non-orthologous sequence - *A. fumigatus*:2956 *A. fischeri*:2956
afum|EAL89204.1 100.00% afis|EAW16608.1 100.00%
Bootstrap support for afum|EAL89204.1 as seed ortholog is 100%.
Bootstrap support for afis|EAW16608.1 as seed ortholog is 100%.

---
Group of orthologs #297. Best score 2952 bits
Score difference with first non-orthologous sequence - *A. fumigatus*:2952 *A. fischeri*:2952
afum|EAL91525.1 100.00% afis|EAW17710.1 100.00%
Bootstrap support for afum|EAL91525.1 as seed ortholog is 100%.
Bootstrap support for afis|EAW17710.1 as seed ortholog is 100%.

---
Group of orthologs #298. Best score 2951 bits
Score difference with first non-orthologous sequence - *A. fumigatus*:2951 *A. fischeri*:2951
afum|EAL93327.1 100.00% afis|EAW18617.1 100.00%
Bootstrap support for afum|EAL93327.1 as seed ortholog is 100%.
Bootstrap support for afis|EAW18617.1 as seed ortholog is 100%.

---
Group of orthologs #299. Best score 2950 bits
Score difference with first non-orthologous sequence - *A. fumigatus*:2950 *A. fischeri*:2950
afum|EAL91886.1 100.00% afis|EAW18074.1 100.00%
Bootstrap support for afum|EAL91886.1 as seed ortholog is 100%.
Bootstrap support for afis|EAW18074.1 as seed ortholog is 100%.

---
Group of orthologs #300. Best score 2944 bits
Score difference with first non-orthologous sequence - *A. fumigatus*:2944 *A. fischeri*:2944
afum|EAL85809.1 100.00% afis|EAW15520.1 100.00%
Bootstrap support for afum|EAL85809.1 as seed ortholog is 100%.
Bootstrap support for afis|EAW15520.1 as seed ortholog is 100%.

---
Group of orthologs #301. Best score 2939 bits
Score difference with first non-orthologous sequence - *A. fumigatus*:2750 *A. fischeri*:1304
afum|EAL89381.1 100.00% afis|EAW24734.1 100.00%
Bootstrap support for afum|EAL89381.1 as seed ortholog is 100%.
Bootstrap support for afis|EAW24734.1 as seed ortholog is 100%.

---
Group of orthologs #302. Best score 2938 bits
Score difference with first non-orthologous sequence - *A. fumigatus*:2938 *A. fischeri*:2938
afum|EAL91744.1 100.00% afis|EAW17933.1 100.00%
Bootstrap support for afum|EAL91744.1 as seed ortholog is 100%.
Bootstrap support for afis|EAW17933.1 as seed ortholog is 100%.

---
Group of orthologs #303. Best score 2936 bits
Score difference with first non-orthologous sequence - *A. fumigatus*:2936 *A. fischeri*:2936
afum|EAL93800.1 100.00% afis|EAW19072.1 100.00%
Bootstrap support for afum|EAL93800.1 as seed ortholog is 100%.
Bootstrap support for afis|EAW19072.1 as seed ortholog is 100%.

---
Group of orthologs #304. Best score 2935 bits
Score difference with first non-orthologous sequence - *A. fumigatus*:2935 *A. fischeri*:2935
afum|EAL92254.1 100.00% afis|EAW21241.1 100.00%
Bootstrap support for afum|EAL92254.1 as seed ortholog is 100%.
Bootstrap support for afis|EAW21241.1 as seed ortholog is 100%.

---
Group of orthologs #305. Best score 2929 bits
Score difference with first non-orthologous sequence - *A. fumigatus*:2929 *A. fischeri*:2929
afum|EAL87966.1 100.00% afis|EAW23497.1 100.00%
Bootstrap support for afum|EAL87966.1 as seed ortholog is 100%.
Bootstrap support for afis|EAW23497.1 as seed ortholog is 100%.

---
Group of orthologs #306. Best score 2925 bits

Score difference with first non-orthologous sequence - *A. fumigatus*:2781 *A. fischeri*:2118
afum|EAL86649.1 100.00% afis|EAW17227.1 100.00%
Bootstrap support for afum|EAL86649.1 as seed ortholog is 100%.
Bootstrap support for afis|EAW17227.1 as seed ortholog is 100%.

---

Group of orthologs #307. Best score 2919 bits
Score difference with first non-orthologous sequence - *A. fumigatus*:2919 *A. fischeri*:2919
afum|EAL88403.1 100.00% afis|EAW23076.1 100.00%
Bootstrap support for afum|EAL88403.1 as seed ortholog is 100%.
Bootstrap support for afis|EAW23076.1 as seed ortholog is 100%.

---

Group of orthologs #308. Best score 2919 bits
Score difference with first non-orthologous sequence - *A. fumigatus*:2779 *A. fischeri*:2477
afum|EAL89906.1 100.00% afis|EAW25244.1 100.00%
Bootstrap support for afum|EAL89906.1 as seed ortholog is 100%.
Bootstrap support for afis|EAW25244.1 as seed ortholog is 100%.

---

Group of orthologs #309. Best score 2916 bits
Score difference with first non-orthologous sequence - *A. fumigatus*:2916 *A. fischeri*:2916
afum|EAL92694.1 100.00% afis|EAW21659.1 100.00%
Bootstrap support for afum|EAL92694.1 as seed ortholog is 100%.
Bootstrap support for afis|EAW21659.1 as seed ortholog is 100%.

---

Group of orthologs #310. Best score 2914 bits
Score difference with first non-orthologous sequence - *A. fumigatus*:2776 *A. fischeri*:2117
afum|EAL90601.1 100.00% afis|EAW22597.1 100.00%
Bootstrap support for afum|EAL90601.1 as seed ortholog is 100%.
Bootstrap support for afis|EAW22597.1 as seed ortholog is 100%.

---

Group of orthologs #311. Best score 2914 bits
Score difference with first non-orthologous sequence - *A. fumigatus*:2914 *A. fischeri*:2914
afum|EAL92422.1 100.00% afis|EAW21408.1 100.00%
Bootstrap support for afum|EAL92422.1 as seed ortholog is 100%.
Bootstrap support for afis|EAW21408.1 as seed ortholog is 100%.

---

Group of orthologs #312. Best score 2911 bits
Score difference with first non-orthologous sequence - *A. fumigatus*:2911 *A. fischeri*:2911
afum|EAL88486.1 100.00% afis|EAW15850.1 100.00%
Bootstrap support for afum|EAL88486.1 as seed ortholog is 100%.
Bootstrap support for afis|EAW15850.1 as seed ortholog is 100%.

---

Group of orthologs #313. Best score 2907 bits
Score difference with first non-orthologous sequence - *A. fumigatus*:2907 *A. fischeri*:2907
afum|EBA27439.1 100.00% afis|EAW17917.1 100.00%
Bootstrap support for afum|EBA27439.1 as seed ortholog is 100%.
Bootstrap support for afis|EAW17917.1 as seed ortholog is 100%.

---

Group of orthologs #314. Best score 2900 bits
Score difference with first non-orthologous sequence - *A. fumigatus*:2900 *A. fischeri*:2900
afum|EAL88260.1 100.00% afis|EAW23224.1 100.00%
Bootstrap support for afum|EAL88260.1 as seed ortholog is 100%.
Bootstrap support for afis|EAW23224.1 as seed ortholog is 100%.

---

Group of orthologs #315. Best score 2899 bits
Score difference with first non-orthologous sequence - *A. fumigatus*:2461 *A. fischeri*:2517
afum|EAL89388.1 100.00% afis|EAW24742.1 100.00%

Bootstrap support for afum|EAL89388.1 as seed ortholog is 100%.
Bootstrap support for afis|EAW24742.1 as seed ortholog is 100%.

---

Group of orthologs #316. Best score 2898 bits
Score difference with first non-orthologous sequence - *A. fumigatus*:2898 *A. fischeri*:2898
afum|EAL86090.1 100.00% afis|EAW24422.1 100.00%
Bootstrap support for afum|EAL86090.1 as seed ortholog is 100%.
Bootstrap support for afis|EAW24422.1 as seed ortholog is 100%.

---

Group of orthologs #317. Best score 2897 bits
Score difference with first non-orthologous sequence - *A. fumigatus*:2897 *A. fischeri*:2897
afum|EAL86946.1 100.00% afis|EAW19571.1 100.00%
Bootstrap support for afum|EAL86946.1 as seed ortholog is 100%.
Bootstrap support for afis|EAW19571.1 as seed ortholog is 100%.

---

Group of orthologs #318. Best score 2897 bits
Score difference with first non-orthologous sequence - *A. fumigatus*:2604 *A. fischeri*:2897
afum|EAL91355.1 100.00% afis|EAW17541.1 100.00%
Bootstrap support for afum|EAL91355.1 as seed ortholog is 100%.
Bootstrap support for afis|EAW17541.1 as seed ortholog is 100%.

---

Group of orthologs #319. Best score 2897 bits
Score difference with first non-orthologous sequence - *A. fumigatus*:2897 *A. fischeri*:2897
afum|EAL87958.1 100.00% afis|EAW23503.1 100.00%
Bootstrap support for afum|EAL87958.1 as seed ortholog is 100%.
Bootstrap support for afis|EAW23503.1 as seed ortholog is 100%.

---

Group of orthologs #320. Best score 2894 bits
Score difference with first non-orthologous sequence - *A. fumigatus*:2894 *A. fischeri*:2894
afum|EAL85968.1 100.00% afis|EAW24300.1 100.00%
Bootstrap support for afum|EAL85968.1 as seed ortholog is 100%.
Bootstrap support for afis|EAW24300.1 as seed ortholog is 100%.

---

Group of orthologs #321. Best score 2891 bits
Score difference with first non-orthologous sequence - *A. fumigatus*:2891 *A. fischeri*:2891
afum|EAL87288.1 100.00% afis|EAW23860.1 100.00%
Bootstrap support for afum|EAL87288.1 as seed ortholog is 100%.
Bootstrap support for afis|EAW23860.1 as seed ortholog is 100%.

---

Group of orthologs #322. Best score 2891 bits
Score difference with first non-orthologous sequence - *A. fumigatus*:2891 *A. fischeri*:2891
afum|EAL88130.1 100.00% afis|EAW23346.1 100.00%
Bootstrap support for afum|EAL88130.1 as seed ortholog is 100%.
Bootstrap support for afis|EAW23346.1 as seed ortholog is 100%.

---

Group of orthologs #323. Best score 2887 bits
Score difference with first non-orthologous sequence - *A. fumigatus*:2887 *A. fischeri*:2887
afum|EAL90074.1 100.00% afis|EAW25415.1 100.00%
Bootstrap support for afum|EAL90074.1 as seed ortholog is 100%.
Bootstrap support for afis|EAW25415.1 as seed ortholog is 100%.

---

Group of orthologs #324. Best score 2885 bits
Score difference with first non-orthologous sequence - *A. fumigatus*:2885 *A. fischeri*:2885
afum|EAL85326.1 100.00% afis|EAW20209.1 100.00%
Bootstrap support for afum|EAL85326.1 as seed ortholog is 100%.
Bootstrap support for afis|EAW20209.1 as seed ortholog is 100%.

Group of orthologs #325. Best score 2885 bits
Score difference with first non-orthologous sequence - *A. fumigatus*:2885 *A. fischeri*:2885
afum|EAL93203.1 100.00% afis|EAW18508.1 100.00%
Bootstrap support for afum|EAL93203.1 as seed ortholog is 100%.
Bootstrap support for afis|EAW18508.1 as seed ortholog is 100%.

Group of orthologs #326. Best score 2882 bits
Score difference with first non-orthologous sequence - *A. fumigatus*:2882 *A. fischeri*:2882
afum|EAL88664.1 100.00% afis|EAW16033.1 100.00%
Bootstrap support for afum|EAL88664.1 as seed ortholog is 100%.
Bootstrap support for afis|EAW16033.1 as seed ortholog is 100%.

Group of orthologs #327. Best score 2880 bits
Score difference with first non-orthologous sequence - *A. fumigatus*:2880 *A. fischeri*:2880
afum|EAL88775.1 100.00% afis|EAW16146.1 100.00%
Bootstrap support for afum|EAL88775.1 as seed ortholog is 100%.
Bootstrap support for afis|EAW16146.1 as seed ortholog is 100%.

Group of orthologs #328. Best score 2877 bits
Score difference with first non-orthologous sequence - *A. fumigatus*:549 *A. fischeri*:2877
afum|EAL92855.1 100.00% afis|EAW21843.1 100.00%
Bootstrap support for afum|EAL92855.1 as seed ortholog is 100%.
Bootstrap support for afis|EAW21843.1 as seed ortholog is 100%.

Group of orthologs #329. Best score 2877 bits
Score difference with first non-orthologous sequence - *A. fumigatus*:2877 *A. fischeri*:2877
afum|EAL89957.1 100.00% afis|EAW25294.1 100.00%
Bootstrap support for afum|EAL89957.1 as seed ortholog is 100%.
Bootstrap support for afis|EAW25294.1 as seed ortholog is 100%.

Group of orthologs #330. Best score 2876 bits
Score difference with first non-orthologous sequence - *A. fumigatus*:2876 *A. fischeri*:1609
afum|EAL85730.1 100.00% afis|EAW15603.1 100.00%
Bootstrap support for afum|EAL85730.1 as seed ortholog is 100%.
Bootstrap support for afis|EAW15603.1 as seed ortholog is 100%.

Group of orthologs #331. Best score 2873 bits
Score difference with first non-orthologous sequence - *A. fumigatus*:2716 *A. fischeri*:1669
afum|EAL85692.1 100.00% afis|EAW22083.1 100.00%
Bootstrap support for afum|EAL85692.1 as seed ortholog is 100%.
Bootstrap support for afis|EAW22083.1 as seed ortholog is 100%.

Group of orthologs #332. Best score 2866 bits
Score difference with first non-orthologous sequence - *A. fumigatus*:2866 *A. fischeri*:1751
afum|EAL89841.1 100.00% afis|EAW25168.1 100.00%
Bootstrap support for afum|EAL89841.1 as seed ortholog is 100%.
Bootstrap support for afis|EAW25168.1 as seed ortholog is 100%.

Group of orthologs #333. Best score 2864 bits
Score difference with first non-orthologous sequence - *A. fumigatus*:2864 *A. fischeri*:2864
afum|EAL84533.1 100.00% afis|EAW20524.1 100.00%
Bootstrap support for afum|EAL84533.1 as seed ortholog is 100%.
Bootstrap support for afis|EAW20524.1 as seed ortholog is 100%.

Group of orthologs #334. Best score 2863 bits

Score difference with first non-orthologous sequence - *A. fumigatus*:2863   *A. fischeri*:2863
afum|EAL94018.1 100.00% afis|EAW19299.1 100.00%
Bootstrap support for afum|EAL94018.1 as seed ortholog is 100%.
Bootstrap support for afis|EAW19299.1 as seed ortholog is 100%.

---

Group of orthologs #335. Best score 2863 bits
Score difference with first non-orthologous sequence - *A. fumigatus*:2863   *A. fischeri*:2863
afum|EAL93013.1 100.00% afis|EAW21998.1 100.00%
Bootstrap support for afum|EAL93013.1 as seed ortholog is 100%.
Bootstrap support for afis|EAW21998.1 as seed ortholog is 100%.

---

Group of orthologs #336. Best score 2855 bits
Score difference with first non-orthologous sequence - *A. fumigatus*:2855   *A. fischeri*:2855
afum|EAL90016.1 100.00% afis|EAW25354.1 100.00%
Bootstrap support for afum|EAL90016.1 as seed ortholog is 100%.
Bootstrap support for afis|EAW25354.1 as seed ortholog is 100%.

---

Group of orthologs #337. Best score 2850 bits
Score difference with first non-orthologous sequence - *A. fumigatus*:2850   *A. fischeri*:2850
afum|EAL89179.1 100.00% afis|EAW16584.1 100.00%
Bootstrap support for afum|EAL89179.1 as seed ortholog is 100%.
Bootstrap support for afis|EAW16584.1 as seed ortholog is 100%.

---

Group of orthologs #338. Best score 2848 bits
Score difference with first non-orthologous sequence - *A. fumigatus*:2848   *A. fischeri*:2848
afum|EAL93652.1 100.00% afis|EAW18921.1 100.00%
Bootstrap support for afum|EAL93652.1 as seed ortholog is 100%.
Bootstrap support for afis|EAW18921.1 as seed ortholog is 100%.

---

Group of orthologs #339. Best score 2847 bits
Score difference with first non-orthologous sequence - *A. fumigatus*:2847   *A. fischeri*:2847
afum|EAL91325.1 100.00% afis|EAW17502.1 100.00%
Bootstrap support for afum|EAL91325.1 as seed ortholog is 100%.
Bootstrap support for afis|EAW17502.1 as seed ortholog is 100%.

---

Group of orthologs #340. Best score 2845 bits
Score difference with first non-orthologous sequence - *A. fumigatus*:2659   *A. fischeri*:1183
afum|EAL86560.1 100.00% afis|EAW17072.1 100.00%
Bootstrap support for afum|EAL86560.1 as seed ortholog is 100%.
Bootstrap support for afis|EAW17072.1 as seed ortholog is 100%.

---

Group of orthologs #341. Best score 2845 bits
Score difference with first non-orthologous sequence - *A. fumigatus*:2845   *A. fischeri*:2845
afum|EAL87001.1 100.00% afis|EAW19493.1 100.00%
Bootstrap support for afum|EAL87001.1 as seed ortholog is 100%.
Bootstrap support for afis|EAW19493.1 as seed ortholog is 100%.

---

Group of orthologs #342. Best score 2844 bits
Score difference with first non-orthologous sequence - *A. fumigatus*:2844   *A. fischeri*:2844
afum|EAL90758.1 100.00% afis|EAW22433.1 100.00%
Bootstrap support for afum|EAL90758.1 as seed ortholog is 100%.
Bootstrap support for afis|EAW22433.1 as seed ortholog is 100%.

---

Group of orthologs #343. Best score 2843 bits
Score difference with first non-orthologous sequence - *A. fumigatus*:2843   *A. fischeri*:2843
afum|EAL87633.1 100.00% afis|EAW18268.1 100.00%

Bootstrap support for afum|EAL87633.1 as seed ortholog is 100%.
Bootstrap support for afis|EAW18268.1 as seed ortholog is 100%.

---

Group of orthologs #344. Best score 2842 bits
Score difference with first non-orthologous sequence - *A. fumigatus*:2842 *A. fischeri*:2842
afum|EAL85607.1 100.00% afis|EAW15741.1 100.00%
Bootstrap support for afum|EAL85607.1 as seed ortholog is 100%.
Bootstrap support for afis|EAW15741.1 as seed ortholog is 100%.

---

Group of orthologs #345. Best score 2842 bits
Score difference with first non-orthologous sequence - *A. fumigatus*:2842 *A. fischeri*:2842
afum|EAL93310.1 100.00% afis|EAW18602.1 100.00%
Bootstrap support for afum|EAL93310.1 as seed ortholog is 100%.
Bootstrap support for afis|EAW18602.1 as seed ortholog is 100%.

---

Group of orthologs #346. Best score 2841 bits
Score difference with first non-orthologous sequence - *A. fumigatus*:2841 *A. fischeri*:2841
afum|EAL89067.1 100.00% afis|EAW16473.1 100.00%
Bootstrap support for afum|EAL89067.1 as seed ortholog is 100%.
Bootstrap support for afis|EAW16473.1 as seed ortholog is 100%.

---

Group of orthologs #347. Best score 2838 bits
Score difference with first non-orthologous sequence - *A. fumigatus*:2838 *A. fischeri*:2838
afum|EAL85713.2 100.00% afis|EAW15621.1 100.00%
Bootstrap support for afum|EAL85713.2 as seed ortholog is 100%.
Bootstrap support for afis|EAW15621.1 as seed ortholog is 100%.

---

Group of orthologs #348. Best score 2838 bits
Score difference with first non-orthologous sequence - *A. fumigatus*:2838 *A. fischeri*:2838
afum|EAL84688.1 100.00% afis|EAW20946.1 100.00%
Bootstrap support for afum|EAL84688.1 as seed ortholog is 100%.
Bootstrap support for afis|EAW20946.1 as seed ortholog is 100%.

---

Group of orthologs #349. Best score 2831 bits
Score difference with first non-orthologous sequence - *A. fumigatus*:2831 *A. fischeri*:2831
afum|EAL87013.2 100.00% afis|EAW19482.1 100.00%
Bootstrap support for afum|EAL87013.2 as seed ortholog is 100%.
Bootstrap support for afis|EAW19482.1 as seed ortholog is 100%.

---

Group of orthologs #350. Best score 2830 bits
Score difference with first non-orthologous sequence - *A. fumigatus*:2119 *A. fischeri*:2830
afum|EAL89167.1 100.00% afis|EAW16570.1 100.00%
Bootstrap support for afum|EAL89167.1 as seed ortholog is 100%.
Bootstrap support for afis|EAW16570.1 as seed ortholog is 100%.

---

Group of orthologs #351. Best score 2827 bits
Score difference with first non-orthologous sequence - *A. fumigatus*:2827 *A. fischeri*:2827
afum|EAL90233.1 100.00% afis|EAW22964.1 100.00%
Bootstrap support for afum|EAL90233.1 as seed ortholog is 100%.
Bootstrap support for afis|EAW22964.1 as seed ortholog is 100%.

---

Group of orthologs #352. Best score 2820 bits
Score difference with first non-orthologous sequence - *A. fumigatus*:2068 *A. fischeri*:2079
afum|EAL86268.1 100.00% afis|EAW22323.1 100.00%
afis|EAW21065.1 60.37%
Bootstrap support for afum|EAL86268.1 as seed ortholog is 100%.

Bootstrap support for afis|EAW22323.1 as seed ortholog is 100%.

---

Group of orthologs #353. Best score 2820 bits
Score difference with first non-orthologous sequence - *A. fumigatus*:2820 *A. fischeri*:2820
afum|EAL87652.1 100.00% afis|EAW18286.1 100.00%
Bootstrap support for afum|EAL87652.1 as seed ortholog is 100%.
Bootstrap support for afis|EAW18286.1 as seed ortholog is 100%.

---

Group of orthologs #354. Best score 2819 bits
Score difference with first non-orthologous sequence - *A. fumigatus*:2819 *A. fischeri*:2819
afum|EAL91453.1 100.00% afis|EAW17638.1 100.00%
Bootstrap support for afum|EAL91453.1 as seed ortholog is 100%.
Bootstrap support for afis|EAW17638.1 as seed ortholog is 100%.

---

Group of orthologs #355. Best score 2819 bits
Score difference with first non-orthologous sequence - *A. fumigatus*:2819 *A. fischeri*:2819
afum|EAL92481.1 100.00% afis|EAW21462.1 100.00%
Bootstrap support for afum|EAL92481.1 as seed ortholog is 100%.
Bootstrap support for afis|EAW21462.1 as seed ortholog is 100%.

---

Group of orthologs #356. Best score 2819 bits
Score difference with first non-orthologous sequence - *A. fumigatus*:2819 *A. fischeri*:2819
afum|EAL92007.1 100.00% afis|EAW24152.1 100.00%
Bootstrap support for afum|EAL92007.1 as seed ortholog is 100%.
Bootstrap support for afis|EAW24152.1 as seed ortholog is 100%.

---

Group of orthologs #357. Best score 2814 bits
Score difference with first non-orthologous sequence - *A. fumigatus*:2814 *A. fischeri*:2814
afum|EAL93703.1 100.00% afis|EAW18970.1 100.00%
Bootstrap support for afum|EAL93703.1 as seed ortholog is 100%.
Bootstrap support for afis|EAW18970.1 as seed ortholog is 100%.

---

Group of orthologs #358. Best score 2813 bits
Score difference with first non-orthologous sequence - *A. fumigatus*:2813 *A. fischeri*:2813
afum|EAL93905.1 100.00% afis|EAW19175.1 100.00%
Bootstrap support for afum|EAL93905.1 as seed ortholog is 100%.
Bootstrap support for afis|EAW19175.1 as seed ortholog is 100%.

---

Group of orthologs #359. Best score 2812 bits
Score difference with first non-orthologous sequence - *A. fumigatus*:2604 *A. fischeri*:2812
afum|EAL93631.1 100.00% afis|EAW18899.1 100.00%
Bootstrap support for afum|EAL93631.1 as seed ortholog is 100%.
Bootstrap support for afis|EAW18899.1 as seed ortholog is 100%.

---

Group of orthologs #360. Best score 2811 bits
Score difference with first non-orthologous sequence - *A. fumigatus*:2811 *A. fischeri*:2811
afum|EAL87031.2 100.00% afis|EAW19465.1 100.00%
Bootstrap support for afum|EAL87031.2 as seed ortholog is 100%.
Bootstrap support for afis|EAW19465.1 as seed ortholog is 100%.

---

Group of orthologs #361. Best score 2810 bits
Score difference with first non-orthologous sequence - *A. fumigatus*:2810 *A. fischeri*:2154
afum|EAL93746.1 100.00% afis|EAW19012.1 100.00%
Bootstrap support for afum|EAL93746.1 as seed ortholog is 100%.
Bootstrap support for afis|EAW19012.1 as seed ortholog is 100%.

---

Group of orthologs #362. Best score 2809 bits
Score difference with first non-orthologous sequence - *A. fumigatus*:2429 *A. fischeri*:2809
afum|EAL84386.2 100.00% afis|EAW25714.1 100.00%
Bootstrap support for afum|EAL84386.2 as seed ortholog is 100%.
Bootstrap support for afis|EAW25714.1 as seed ortholog is 100%.

---

Group of orthologs #363. Best score 2805 bits
Score difference with first non-orthologous sequence - *A. fumigatus*:2805 *A. fischeri*:2805
afum|EAL85262.1 100.00% afis|EAW20147.1 100.00%
Bootstrap support for afum|EAL85262.1 as seed ortholog is 100%.
Bootstrap support for afis|EAW20147.1 as seed ortholog is 100%.

---

Group of orthologs #364. Best score 2803 bits
Score difference with first non-orthologous sequence - *A. fumigatus*:2803 *A. fischeri*:2803
afum|EAL90622.2 100.00% afis|EAW22576.1 100.00%
Bootstrap support for afum|EAL90622.2 as seed ortholog is 100%.
Bootstrap support for afis|EAW22576.1 as seed ortholog is 100%.

---

Group of orthologs #365. Best score 2802 bits
Score difference with first non-orthologous sequence - *A. fumigatus*:2802 *A. fischeri*:2802
afum|EAL84646.1 100.00% afis|EAW20984.1 100.00%
Bootstrap support for afum|EAL84646.1 as seed ortholog is 100%.
Bootstrap support for afis|EAW20984.1 as seed ortholog is 100%.

---

Group of orthologs #366. Best score 2800 bits
Score difference with first non-orthologous sequence - *A. fumigatus*:2800 *A. fischeri*:2800
afum|EAL85221.1 100.00% afis|EAW20104.1 100.00%
Bootstrap support for afum|EAL85221.1 as seed ortholog is 100%.
Bootstrap support for afis|EAW20104.1 as seed ortholog is 100%.

---

Group of orthologs #367. Best score 2797 bits
Score difference with first non-orthologous sequence - *A. fumigatus*:2797 *A. fischeri*:2797
afum|EAL87559.1 100.00% afis|EAW18193.1 100.00%
Bootstrap support for afum|EAL87559.1 as seed ortholog is 100%.
Bootstrap support for afis|EAW18193.1 as seed ortholog is 100%.

---

Group of orthologs #368. Best score 2793 bits
Score difference with first non-orthologous sequence - *A. fumigatus*:2793 *A. fischeri*:2793
afum|EAL90333.1 100.00% afis|EAW22866.1 100.00%
Bootstrap support for afum|EAL90333.1 as seed ortholog is 100%.
Bootstrap support for afis|EAW22866.1 as seed ortholog is 100%.

---

Group of orthologs #369. Best score 2782 bits
Score difference with first non-orthologous sequence - *A. fumigatus*:2782 *A. fischeri*:2782
afum|EAL84839.2 100.00% afis|EAW20783.1 100.00%
Bootstrap support for afum|EAL84839.2 as seed ortholog is 100%.
Bootstrap support for afis|EAW20783.1 as seed ortholog is 100%.

---

Group of orthologs #370. Best score 2778 bits
Score difference with first non-orthologous sequence - *A. fumigatus*:2778 *A. fischeri*:2778
afum|EAL93302.1 100.00% afis|EAW18594.1 100.00%
Bootstrap support for afum|EAL93302.1 as seed ortholog is 100%.
Bootstrap support for afis|EAW18594.1 as seed ortholog is 100%.

---

Group of orthologs #371. Best score 2777 bits
Score difference with first non-orthologous sequence - *A. fumigatus*:2777 *A. fischeri*:2777

afum|EAL92579.1 100.00% afis|EAW21560.1 100.00%
Bootstrap support for afum|EAL92579.1 as seed ortholog is 100%.
Bootstrap support for afis|EAW21560.1 as seed ortholog is 100%.

---

Group of orthologs #372. Best score 2774 bits
Score difference with first non-orthologous sequence - *A. fumigatus*:2774 *A. fischeri*:2774
afum|EAL90010.2 100.00% afis|EAW25347.1 100.00%
Bootstrap support for afum|EAL90010.2 as seed ortholog is 100%.
Bootstrap support for afis|EAW25347.1 as seed ortholog is 100%.

---

Group of orthologs #373. Best score 2773 bits
Score difference with first non-orthologous sequence - *A. fumigatus*:2773 *A. fischeri*:2773
afum|EAL86937.1 100.00% afis|EAW19581.1 100.00%
Bootstrap support for afum|EAL86937.1 as seed ortholog is 100%.
Bootstrap support for afis|EAW19581.1 as seed ortholog is 100%.

---

Group of orthologs #374. Best score 2772 bits
Score difference with first non-orthologous sequence - *A. fumigatus*:2772 *A. fischeri*:2772
afum|EAL91794.1 100.00% afis|EAW17985.1 100.00%
Bootstrap support for afum|EAL91794.1 as seed ortholog is 100%.
Bootstrap support for afis|EAW17985.1 as seed ortholog is 100%.

---

Group of orthologs #375. Best score 2771 bits
Score difference with first non-orthologous sequence - *A. fumigatus*:2771 *A. fischeri*:2771
afum|EAL85199.1 100.00% afis|EAW20080.1 100.00%
Bootstrap support for afum|EAL85199.1 as seed ortholog is 100%.
Bootstrap support for afis|EAW20080.1 as seed ortholog is 100%.

---

Group of orthologs #376. Best score 2771 bits
Score difference with first non-orthologous sequence - *A. fumigatus*:2771 *A. fischeri*:2771
afum|EAL92546.1 100.00% afis|EAW21525.1 100.00%
Bootstrap support for afum|EAL92546.1 as seed ortholog is 100%.
Bootstrap support for afis|EAW21525.1 as seed ortholog is 100%.

---

Group of orthologs #377. Best score 2766 bits
Score difference with first non-orthologous sequence - *A. fumigatus*:2766 *A. fischeri*:2766
afum|EAL92644.1 100.00% afis|EAW21610.1 100.00%
Bootstrap support for afum|EAL92644.1 as seed ortholog is 100%.
Bootstrap support for afis|EAW21610.1 as seed ortholog is 100%.

---

Group of orthologs #378. Best score 2762 bits
Score difference with first non-orthologous sequence - *A. fumigatus*:2481 *A. fischeri*:2762
afum|EAL92309.1 100.00% afis|EAW21292.1 100.00%
Bootstrap support for afum|EAL92309.1 as seed ortholog is 100%.
Bootstrap support for afis|EAW21292.1 as seed ortholog is 100%.

---

Group of orthologs #379. Best score 2761 bits
Score difference with first non-orthologous sequence - *A. fumigatus*:2761 *A. fischeri*:2761
afum|EAL90131.1 100.00% afis|EAW25467.1 100.00%
Bootstrap support for afum|EAL90131.1 as seed ortholog is 100%.
Bootstrap support for afis|EAW25467.1 as seed ortholog is 100%.

---

Group of orthologs #380. Best score 2759 bits
Score difference with first non-orthologous sequence - *A. fumigatus*:2759 *A. fischeri*:2759
afum|EAL86296.2 100.00% afis|EAW16722.1 100.00%
Bootstrap support for afum|EAL86296.2 as seed ortholog is 100%.

Bootstrap support for afis|EAW16722.1 as seed ortholog is 100%.

---

Group of orthologs #381. Best score 2755 bits
Score difference with first non-orthologous sequence - *A. fumigatus*:2755 *A. fischeri*:2755
afum|EAL92825.2 100.00% afis|EAW21800.1 100.00%
Bootstrap support for afum|EAL92825.2 as seed ortholog is 100%.
Bootstrap support for afis|EAW21800.1 as seed ortholog is 100%.

---

Group of orthologs #382. Best score 2753 bits
Score difference with first non-orthologous sequence - *A. fumigatus*:2753 *A. fischeri*:2753
afum|EAL93377.1 100.00% afis|EAW18659.1 100.00%
Bootstrap support for afum|EAL93377.1 as seed ortholog is 100%.
Bootstrap support for afis|EAW18659.1 as seed ortholog is 100%.

---

Group of orthologs #383. Best score 2752 bits
Score difference with first non-orthologous sequence - *A. fumigatus*:2752 *A. fischeri*:2752
afum|EBA27334.1 100.00% afis|EAW23363.1 100.00%
Bootstrap support for afum|EBA27334.1 as seed ortholog is 100%.
Bootstrap support for afis|EAW23363.1 as seed ortholog is 100%.

---

Group of orthologs #384. Best score 2749 bits
Score difference with first non-orthologous sequence - *A. fumigatus*:2749 *A. fischeri*:2749
afum|EAL93314.1 100.00% afis|EAW18605.1 100.00%
Bootstrap support for afum|EAL93314.1 as seed ortholog is 100%.
Bootstrap support for afis|EAW18605.1 as seed ortholog is 100%.

---

Group of orthologs #385. Best score 2747 bits
Score difference with first non-orthologous sequence - *A. fumigatus*:2747 *A. fischeri*:2747
afum|EAL87884.1 100.00% afis|EAW23581.1 100.00%
Bootstrap support for afum|EAL87884.1 as seed ortholog is 100%.
Bootstrap support for afis|EAW23581.1 as seed ortholog is 100%.

---

Group of orthologs #386. Best score 2740 bits
Score difference with first non-orthologous sequence - *A. fumigatus*:2740 *A. fischeri*:2740
afum|EAL84754.1 100.00% afis|EAW20878.1 100.00%
Bootstrap support for afum|EAL84754.1 as seed ortholog is 100%.
Bootstrap support for afis|EAW20878.1 as seed ortholog is 100%.

---

Group of orthologs #387. Best score 2740 bits
Score difference with first non-orthologous sequence - *A. fumigatus*:2740 *A. fischeri*:2740
afum|EAL93773.1 100.00% afis|EAW19044.1 100.00%
Bootstrap support for afum|EAL93773.1 as seed ortholog is 100%.
Bootstrap support for afis|EAW19044.1 as seed ortholog is 100%.

---

Group of orthologs #388. Best score 2740 bits
Score difference with first non-orthologous sequence - *A. fumigatus*:2740 *A. fischeri*:2740
afum|EAL90881.1 100.00% afis|EAW22309.1 100.00%
Bootstrap support for afum|EAL90881.1 as seed ortholog is 100%.
Bootstrap support for afis|EAW22309.1 as seed ortholog is 100%.

---

Group of orthologs #389. Best score 2739 bits
Score difference with first non-orthologous sequence - *A. fumigatus*:2739 *A. fischeri*:2739
afum|EAL86103.1 100.00% afis|EAW24435.1 100.00%
Bootstrap support for afum|EAL86103.1 as seed ortholog is 100%.
Bootstrap support for afis|EAW24435.1 as seed ortholog is 100%.

---

Group of orthologs #390. Best score 2735 bits
Score difference with first non-orthologous sequence - *A. fumigatus*:2735 *A. fischeri*:2735
afum|EAL88224.1 100.00% afis|EAW23257.1 100.00%
Bootstrap support for afum|EAL88224.1 as seed ortholog is 100%.
Bootstrap support for afis|EAW23257.1 as seed ortholog is 100%.

---

Group of orthologs #391. Best score 2734 bits
Score difference with first non-orthologous sequence - *A. fumigatus*:2734 *A. fischeri*:2734
afum|EAL89103.1 100.00% afis|EAW16508.1 100.00%
Bootstrap support for afum|EAL89103.1 as seed ortholog is 100%.
Bootstrap support for afis|EAW16508.1 as seed ortholog is 100%.

---

Group of orthologs #392. Best score 2734 bits
Score difference with first non-orthologous sequence - *A. fumigatus*:2734 *A. fischeri*:1410
afum|EAL92512.1 100.00% afis|EAW21493.1 100.00%
Bootstrap support for afum|EAL92512.1 as seed ortholog is 100%.
Bootstrap support for afis|EAW21493.1 as seed ortholog is 100%.

---

Group of orthologs #393. Best score 2734 bits
Score difference with first non-orthologous sequence - *A. fumigatus*:2734 *A. fischeri*:2734
afum|EAL89706.1 100.00% afis|EAW25038.1 100.00%
Bootstrap support for afum|EAL89706.1 as seed ortholog is 100%.
Bootstrap support for afis|EAW25038.1 as seed ortholog is 100%.

---

Group of orthologs #394. Best score 2729 bits
Score difference with first non-orthologous sequence - *A. fumigatus*:2729 *A. fischeri*:2729
afum|EAL86917.1 100.00% afis|EAW19601.1 100.00%
Bootstrap support for afum|EAL86917.1 as seed ortholog is 100%.
Bootstrap support for afis|EAW19601.1 as seed ortholog is 100%.

---

Group of orthologs #395. Best score 2727 bits
Score difference with first non-orthologous sequence - *A. fumigatus*:568 *A. fischeri*:378
afum|EAL93101.1 100.00% afis|EAW18417.1 100.00%
Bootstrap support for afum|EAL93101.1 as seed ortholog is 100%.
Bootstrap support for afis|EAW18417.1 as seed ortholog is 100%.

---

Group of orthologs #396. Best score 2721 bits
Score difference with first non-orthologous sequence - *A. fumigatus*:2721 *A. fischeri*:2721
afum|EAL84586.2 100.00% afis|EAW20466.1 100.00%
Bootstrap support for afum|EAL84586.2 as seed ortholog is 100%.
Bootstrap support for afis|EAW20466.1 as seed ortholog is 100%.

---

Group of orthologs #397. Best score 2715 bits
Score difference with first non-orthologous sequence - *A. fumigatus*:2715 *A. fischeri*:2715
afum|EAL91416.1 100.00% afis|EAW17603.1 100.00%
Bootstrap support for afum|EAL91416.1 as seed ortholog is 100%.
Bootstrap support for afis|EAW17603.1 as seed ortholog is 100%.

---

Group of orthologs #398. Best score 2714 bits
Score difference with first non-orthologous sequence - *A. fumigatus*:2714 *A. fischeri*:2714
afum|EAL85980.1 100.00% afis|EAW24312.1 100.00%
Bootstrap support for afum|EAL85980.1 as seed ortholog is 100%.
Bootstrap support for afis|EAW24312.1 as seed ortholog is 100%.

---

Group of orthologs #399. Best score 2713 bits
Score difference with first non-orthologous sequence - *A. fumigatus*:2713 *A. fischeri*:2713

afum|EAL85768.1 100.00% afis|EAW15565.1 100.00%
Bootstrap support for afum|EAL85768.1 as seed ortholog is 100%.
Bootstrap support for afis|EAW15565.1 as seed ortholog is 100%.

---

Group of orthologs #400. Best score 2712 bits
Score difference with first non-orthologous sequence - *A. fumigatus*:2712 *A. fischeri*:2712
afum|EAL91798.1 100.00% afis|EAW17989.1 100.00%
Bootstrap support for afum|EAL91798.1 as seed ortholog is 100%.
Bootstrap support for afis|EAW17989.1 as seed ortholog is 100%.

---

Group of orthologs #401. Best score 2710 bits
Score difference with first non-orthologous sequence - *A. fumigatus*:2710 *A. fischeri*:2710
afum|EAL85769.1 100.00% afis|EAW15563.1 100.00%
Bootstrap support for afum|EAL85769.1 as seed ortholog is 100%.
Bootstrap support for afis|EAW15563.1 as seed ortholog is 100%.

---

Group of orthologs #402. Best score 2705 bits
Score difference with first non-orthologous sequence - *A. fumigatus*:2705 *A. fischeri*:2705
afum|EAL88175.1 100.00% afis|EAW23303.1 100.00%
Bootstrap support for afum|EAL88175.1 as seed ortholog is 100%.
Bootstrap support for afis|EAW23303.1 as seed ortholog is 100%.

---

Group of orthologs #403. Best score 2703 bits
Score difference with first non-orthologous sequence - *A. fumigatus*:2703 *A. fischeri*:2703
afum|EAL90140.1 100.00% afis|EAW25477.1 100.00%
Bootstrap support for afum|EAL90140.1 as seed ortholog is 100%.
Bootstrap support for afis|EAW25477.1 as seed ortholog is 100%.

---

Group of orthologs #404. Best score 2695 bits
Score difference with first non-orthologous sequence - *A. fumigatus*:2695 *A. fischeri*:2695
afum|EAL89473.1 100.00% afis|EAW24828.1 100.00%
Bootstrap support for afum|EAL89473.1 as seed ortholog is 100%.
Bootstrap support for afis|EAW24828.1 as seed ortholog is 100%.

---

Group of orthologs #405. Best score 2693 bits
Score difference with first non-orthologous sequence - *A. fumigatus*:2693 *A. fischeri*:2693
afum|EAL93951.2 100.00% afis|EAW19218.1 100.00%
Bootstrap support for afum|EAL93951.2 as seed ortholog is 100%.
Bootstrap support for afis|EAW19218.1 as seed ortholog is 100%.

---

Group of orthologs #406. Best score 2690 bits
Score difference with first non-orthologous sequence - *A. fumigatus*:2690 *A. fischeri*:2690
afum|EAL91852.1 100.00% afis|EAW18041.1 100.00%
Bootstrap support for afum|EAL91852.1 as seed ortholog is 100%.
Bootstrap support for afis|EAW18041.1 as seed ortholog is 100%.

---

Group of orthologs #407. Best score 2689 bits
Score difference with first non-orthologous sequence - *A. fumigatus*:2689 *A. fischeri*:1755
afum|EAL84400.2 100.00% afis|EAW25729.1 100.00%
Bootstrap support for afum|EAL84400.2 as seed ortholog is 100%.
Bootstrap support for afis|EAW25729.1 as seed ortholog is 100%.

---

Group of orthologs #408. Best score 2689 bits
Score difference with first non-orthologous sequence - *A. fumigatus*:2689 *A. fischeri*:2689
afum|EAL92261.1 100.00% afis|EAW21248.1 100.00%
Bootstrap support for afum|EAL92261.1 as seed ortholog is 100%.

Bootstrap support for afis|EAW21248.1 as seed ortholog is 100%.

---

Group of orthologs #409. Best score 2689 bits
Score difference with first non-orthologous sequence - *A. fumigatus*:2689 *A. fischeri*:2689
afum|EAL89940.1 100.00% afis|EAW25277.1 100.00%
Bootstrap support for afum|EAL89940.1 as seed ortholog is 100%.
Bootstrap support for afis|EAW25277.1 as seed ortholog is 100%.

---

Group of orthologs #410. Best score 2688 bits
Score difference with first non-orthologous sequence - *A. fumigatus*:2688 *A. fischeri*:1747
afum|EAL92660.1 100.00% afis|EAW21624.1 100.00%
Bootstrap support for afum|EAL92660.1 as seed ortholog is 100%.
Bootstrap support for afis|EAW21624.1 as seed ortholog is 100%.

---

Group of orthologs #411. Best score 2687 bits
Score difference with first non-orthologous sequence - *A. fumigatus*:2687 *A. fischeri*:2687
afum|EAL89174.1 100.00% afis|EAW16578.1 100.00%
Bootstrap support for afum|EAL89174.1 as seed ortholog is 100%.
Bootstrap support for afis|EAW16578.1 as seed ortholog is 100%.

---

Group of orthologs #412. Best score 2687 bits
Score difference with first non-orthologous sequence - *A. fumigatus*:2687 *A. fischeri*:2687
afum|EAL90812.1 100.00% afis|EAW22379.1 100.00%
Bootstrap support for afum|EAL90812.1 as seed ortholog is 100%.
Bootstrap support for afis|EAW22379.1 as seed ortholog is 100%.

---

Group of orthologs #413. Best score 2686 bits
Score difference with first non-orthologous sequence - *A. fumigatus*:2686 *A. fischeri*:2686
afum|EAL84576.1 100.00% afis|EAW20478.1 100.00%
Bootstrap support for afum|EAL84576.1 as seed ortholog is 100%.
Bootstrap support for afis|EAW20478.1 as seed ortholog is 100%.

---

Group of orthologs #414. Best score 2685 bits
Score difference with first non-orthologous sequence - *A. fumigatus*:2685 *A. fischeri*:2685
afum|EAL87566.1 100.00% afis|EAW18201.1 100.00%
Bootstrap support for afum|EAL87566.1 as seed ortholog is 100%.
Bootstrap support for afis|EAW18201.1 as seed ortholog is 100%.

---

Group of orthologs #415. Best score 2685 bits
Score difference with first non-orthologous sequence - *A. fumigatus*:2685 *A. fischeri*:2685
afum|EAL92517.1 100.00% afis|EAW21498.1 100.00%
Bootstrap support for afum|EAL92517.1 as seed ortholog is 100%.
Bootstrap support for afis|EAW21498.1 as seed ortholog is 100%.

---

Group of orthologs #416. Best score 2684 bits
Score difference with first non-orthologous sequence - *A. fumigatus*:2684 *A. fischeri*:2684
afum|EAL92736.1 100.00% afis|EAW21702.1 100.00%
Bootstrap support for afum|EAL92736.1 as seed ortholog is 100%.
Bootstrap support for afis|EAW21702.1 as seed ortholog is 100%.

---

Group of orthologs #417. Best score 2683 bits
Score difference with first non-orthologous sequence - *A. fumigatus*:2329 *A. fischeri*:2683
afum|EAL89531.1 100.00% afis|EAW24881.1 100.00%
Bootstrap support for afum|EAL89531.1 as seed ortholog is 100%.
Bootstrap support for afis|EAW24881.1 as seed ortholog is 100%.

---

Group of orthologs #418. Best score 2677 bits
Score difference with first non-orthologous sequence - *A. fumigatus*:2677 *A. fischeri*:2677
afum|EAL88946.1 100.00% afis|EAW16323.1 100.00%
Bootstrap support for afum|EAL88946.1 as seed ortholog is 100%.
Bootstrap support for afis|EAW16323.1 as seed ortholog is 100%.

---

Group of orthologs #419. Best score 2674 bits
Score difference with first non-orthologous sequence - *A. fumigatus*:2674 *A. fischeri*:2674
afum|EAL93068.1 100.00% afis|EAW18392.1 100.00%
Bootstrap support for afum|EAL93068.1 as seed ortholog is 100%.
Bootstrap support for afis|EAW18392.1 as seed ortholog is 100%.

---

Group of orthologs #420. Best score 2673 bits
Score difference with first non-orthologous sequence - *A. fumigatus*:2673 *A. fischeri*:2673
afum|EAL87091.1 100.00% afis|EAW19404.1 100.00%
Bootstrap support for afum|EAL87091.1 as seed ortholog is 100%.
Bootstrap support for afis|EAW19404.1 as seed ortholog is 100%.

---

Group of orthologs #421. Best score 2671 bits
Score difference with first non-orthologous sequence - *A. fumigatus*:2671 *A. fischeri*:2671
afum|EAL90657.1 100.00% afis|EAW22537.1 100.00%
Bootstrap support for afum|EAL90657.1 as seed ortholog is 100%.
Bootstrap support for afis|EAW22537.1 as seed ortholog is 100%.

---

Group of orthologs #422. Best score 2669 bits
Score difference with first non-orthologous sequence - *A. fumigatus*:2669 *A. fischeri*:2669
afum|EAL84734.2 100.00% afis|EAW20898.1 100.00%
Bootstrap support for afum|EAL84734.2 as seed ortholog is 100%.
Bootstrap support for afis|EAW20898.1 as seed ortholog is 100%.

---

Group of orthologs #423. Best score 2665 bits
Score difference with first non-orthologous sequence - *A. fumigatus*:2665 *A. fischeri*:2665
afum|EAL87102.1 100.00% afis|EAW19392.1 100.00%
Bootstrap support for afum|EAL87102.1 as seed ortholog is 100%.
Bootstrap support for afis|EAW19392.1 as seed ortholog is 100%.

---

Group of orthologs #424. Best score 2664 bits
Score difference with first non-orthologous sequence - *A. fumigatus*:2664 *A. fischeri*:2664
afum|EAL90616.1 100.00% afis|EAW22583.1 100.00%
Bootstrap support for afum|EAL90616.1 as seed ortholog is 100%.
Bootstrap support for afis|EAW22583.1 as seed ortholog is 100%.

---

Group of orthologs #425. Best score 2655 bits
Score difference with first non-orthologous sequence - *A. fumigatus*:2655 *A. fischeri*:2655
afum|EAL84700.1 100.00% afis|EAW20934.1 100.00%
Bootstrap support for afum|EAL84700.1 as seed ortholog is 100%.
Bootstrap support for afis|EAW20934.1 as seed ortholog is 100%.

---

Group of orthologs #426. Best score 2654 bits
Score difference with first non-orthologous sequence - *A. fumigatus*:2654 *A. fischeri*:2654
afum|EAL89659.2 100.00% afis|EAW25035.1 100.00%
Bootstrap support for afum|EAL89659.2 as seed ortholog is 100%.
Bootstrap support for afis|EAW25035.1 as seed ortholog is 100%.

---

Group of orthologs #427. Best score 2652 bits
Score difference with first non-orthologous sequence - *A. fumigatus*:2459 *A. fischeri*:1810

afum|EAL84844.1 100.00% afis|EAW20776.1 100.00%
Bootstrap support for afum|EAL84844.1 as seed ortholog is 100%.
Bootstrap support for afis|EAW20776.1 as seed ortholog is 100%.

---

Group of orthologs #428. Best score 2648 bits
Score difference with first non-orthologous sequence - *A. fumigatus*:2307 *A. fischeri*:2648
afum|EAL88772.1 100.00% afis|EAW16143.1 100.00%
Bootstrap support for afum|EAL88772.1 as seed ortholog is 100%.
Bootstrap support for afis|EAW16143.1 as seed ortholog is 100%.

---

Group of orthologs #429. Best score 2645 bits
Score difference with first non-orthologous sequence - *A. fumigatus*:2645 *A. fischeri*:2645
afum|EAL88583.1 100.00% afis|EAW15950.1 100.00%
Bootstrap support for afum|EAL88583.1 as seed ortholog is 100%.
Bootstrap support for afis|EAW15950.1 as seed ortholog is 100%.

---

Group of orthologs #430. Best score 2644 bits
Score difference with first non-orthologous sequence - *A. fumigatus*:2644 *A. fischeri*:2644
afum|EAL86928.1 100.00% afis|EAW19590.1 100.00%
Bootstrap support for afum|EAL86928.1 as seed ortholog is 100%.
Bootstrap support for afis|EAW19590.1 as seed ortholog is 100%.

---

Group of orthologs #431. Best score 2643 bits
Score difference with first non-orthologous sequence - *A. fumigatus*:2643 *A. fischeri*:2643
afum|EAL89447.1 100.00% afis|EAW24805.1 100.00%
Bootstrap support for afum|EAL89447.1 as seed ortholog is 100%.
Bootstrap support for afis|EAW24805.1 as seed ortholog is 100%.

---

Group of orthologs #432. Best score 2640 bits
Score difference with first non-orthologous sequence - *A. fumigatus*:2640 *A. fischeri*:2640
afum|EAL88603.1 100.00% afis|EAW15971.1 100.00%
Bootstrap support for afum|EAL88603.1 as seed ortholog is 100%.
Bootstrap support for afis|EAW15971.1 as seed ortholog is 100%.

---

Group of orthologs #433. Best score 2640 bits
Score difference with first non-orthologous sequence - *A. fumigatus*:2640 *A. fischeri*:2640
afum|EAL88708.1 100.00% afis|EAW16076.1 100.00%
Bootstrap support for afum|EAL88708.1 as seed ortholog is 100%.
Bootstrap support for afis|EAW16076.1 as seed ortholog is 100%.

---

Group of orthologs #434. Best score 2640 bits
Score difference with first non-orthologous sequence - *A. fumigatus*:2640 *A. fischeri*:2640
afum|EAL90638.1 100.00% afis|EAW22557.1 100.00%
Bootstrap support for afum|EAL90638.1 as seed ortholog is 100%.
Bootstrap support for afis|EAW22557.1 as seed ortholog is 100%.

---

Group of orthologs #435. Best score 2640 bits
Score difference with first non-orthologous sequence - *A. fumigatus*:2640 *A. fischeri*:2640
afum|EAL92756.1 100.00% afis|EAW21723.1 100.00%
Bootstrap support for afum|EAL92756.1 as seed ortholog is 100%.
Bootstrap support for afis|EAW21723.1 as seed ortholog is 100%.

---

Group of orthologs #436. Best score 2639 bits
Score difference with first non-orthologous sequence - *A. fumigatus*:2288 *A. fischeri*:2639
afum|EAL84352.1 100.00% afis|EAW25682.1 100.00%
Bootstrap support for afum|EAL84352.1 as seed ortholog is 100%.

Bootstrap support for afis|EAW25682.1 as seed ortholog is 100%.

---

Group of orthologs #437. Best score 2634 bits
Score difference with first non-orthologous sequence - *A. fumigatus*:2634   *A. fischeri*:2634
afum|EAL87464.1 100.00% afis|EAW24033.1 100.00%
Bootstrap support for afum|EAL87464.1 as seed ortholog is 100%.
Bootstrap support for afis|EAW24033.1 as seed ortholog is 100%.

---

Group of orthologs #438. Best score 2631 bits
Score difference with first non-orthologous sequence - *A. fumigatus*:2631   *A. fischeri*:2631
afum|EAL88586.1 100.00% afis|EAW15953.1 100.00%
Bootstrap support for afum|EAL88586.1 as seed ortholog is 100%.
Bootstrap support for afis|EAW15953.1 as seed ortholog is 100%.

---

Group of orthologs #439. Best score 2631 bits
Score difference with first non-orthologous sequence - *A. fumigatus*:2631   *A. fischeri*:2631
afum|EAL93045.1 100.00% afis|EAW18370.1 100.00%
Bootstrap support for afum|EAL93045.1 as seed ortholog is 100%.
Bootstrap support for afis|EAW18370.1 as seed ortholog is 100%.

---

Group of orthologs #440. Best score 2630 bits
Score difference with first non-orthologous sequence - *A. fumigatus*:2630   *A. fischeri*:2630
afum|EAL85681.1 100.00% afis|EAW15667.1 100.00%
Bootstrap support for afum|EAL85681.1 as seed ortholog is 100%.
Bootstrap support for afis|EAW15667.1 as seed ortholog is 100%.

---

Group of orthologs #441. Best score 2629 bits
Score difference with first non-orthologous sequence - *A. fumigatus*:2629   *A. fischeri*:2629
afum|EAL85989.1 100.00% afis|EAW24319.1 100.00%
Bootstrap support for afum|EAL85989.1 as seed ortholog is 100%.
Bootstrap support for afis|EAW24319.1 as seed ortholog is 100%.

---

Group of orthologs #442. Best score 2626 bits
Score difference with first non-orthologous sequence - *A. fumigatus*:2626   *A. fischeri*:2626
afum|EAL84615.1 100.00% afis|EAW20437.1 100.00%
Bootstrap support for afum|EAL84615.1 as seed ortholog is 100%.
Bootstrap support for afis|EAW20437.1 as seed ortholog is 100%.

---

Group of orthologs #443. Best score 2626 bits
Score difference with first non-orthologous sequence - *A. fumigatus*:2626   *A. fischeri*:2626
afum|EAL86087.1 100.00% afis|EAW24418.1 100.00%
Bootstrap support for afum|EAL86087.1 as seed ortholog is 100%.
Bootstrap support for afis|EAW24418.1 as seed ortholog is 100%.

---

Group of orthologs #444. Best score 2626 bits
Score difference with first non-orthologous sequence - *A. fumigatus*:2626   *A. fischeri*:2626
afum|EAL93010.1 100.00% afis|EAW21995.1 100.00%
Bootstrap support for afum|EAL93010.1 as seed ortholog is 100%.
Bootstrap support for afis|EAW21995.1 as seed ortholog is 100%.

---

Group of orthologs #445. Best score 2624 bits
Score difference with first non-orthologous sequence - *A. fumigatus*:2624   *A. fischeri*:2624
afum|EAL91524.2 100.00% afis|EAW17709.1 100.00%
Bootstrap support for afum|EAL91524.2 as seed ortholog is 100%.
Bootstrap support for afis|EAW17709.1 as seed ortholog is 100%.

---

Group of orthologs #446. Best score 2622 bits
Score difference with first non-orthologous sequence - *A. fumigatus*:2622 *A. fischeri*:2622
afum|EAL84733.1 100.00% afis|EAW20899.1 100.00%
Bootstrap support for afum|EAL84733.1 as seed ortholog is 100%.
Bootstrap support for afis|EAW20899.1 as seed ortholog is 100%.

---

Group of orthologs #447. Best score 2622 bits
Score difference with first non-orthologous sequence - *A. fumigatus*:2622 *A. fischeri*:2622
afum|EAL93959.1 100.00% afis|EAW19230.1 100.00%
Bootstrap support for afum|EAL93959.1 as seed ortholog is 100%.
Bootstrap support for afis|EAW19230.1 as seed ortholog is 100%.

---

Group of orthologs #448. Best score 2618 bits
Score difference with first non-orthologous sequence - *A. fumigatus*:2618 *A. fischeri*:2618
afum|EAL90202.1 100.00% afis|EAW25539.1 100.00%
Bootstrap support for afum|EAL90202.1 as seed ortholog is 100%.
Bootstrap support for afis|EAW25539.1 as seed ortholog is 100%.

---

Group of orthologs #449. Best score 2617 bits
Score difference with first non-orthologous sequence - *A. fumigatus*:2617 *A. fischeri*:2617
afum|EAL87691.1 100.00% afis|EAW18325.1 100.00%
Bootstrap support for afum|EAL87691.1 as seed ortholog is 100%.
Bootstrap support for afis|EAW18325.1 as seed ortholog is 100%.

---

Group of orthologs #450. Best score 2615 bits
Score difference with first non-orthologous sequence - *A. fumigatus*:2615 *A. fischeri*:2615
afum|EAL93066.1 100.00% afis|EAW18391.1 100.00%
Bootstrap support for afum|EAL93066.1 as seed ortholog is 100%.
Bootstrap support for afis|EAW18391.1 as seed ortholog is 100%.

---

Group of orthologs #451. Best score 2614 bits
Score difference with first non-orthologous sequence - *A. fumigatus*:2614 *A. fischeri*:2614
afum|EAL85200.1 100.00% afis|EAW20081.1 100.00%
Bootstrap support for afum|EAL85200.1 as seed ortholog is 100%.
Bootstrap support for afis|EAW20081.1 as seed ortholog is 100%.

---

Group of orthologs #452. Best score 2613 bits
Score difference with first non-orthologous sequence - *A. fumigatus*:2613 *A. fischeri*:2613
afum|EAL89986.1 100.00% afis|EAW25325.1 100.00%
Bootstrap support for afum|EAL89986.1 as seed ortholog is 100%.
Bootstrap support for afis|EAW25325.1 as seed ortholog is 100%.

---

Group of orthologs #453. Best score 2612 bits
Score difference with first non-orthologous sequence - *A. fumigatus*:2612 *A. fischeri*:2612
afum|EAL85722.1 100.00% afis|EAW15613.1 100.00%
Bootstrap support for afum|EAL85722.1 as seed ortholog is 100%.
Bootstrap support for afis|EAW15613.1 as seed ortholog is 100%.

---

Group of orthologs #454. Best score 2612 bits
Score difference with first non-orthologous sequence - *A. fumigatus*:2612 *A. fischeri*:2028
afum|EAL84566.1 100.00% afis|EAW20489.1 100.00%
Bootstrap support for afum|EAL84566.1 as seed ortholog is 100%.
Bootstrap support for afis|EAW20489.1 as seed ortholog is 100%.

---

Group of orthologs #455. Best score 2611 bits
Score difference with first non-orthologous sequence - *A. fumigatus*:2611 *A. fischeri*:2611

afum|EAL88390.1 100.00% afis|EAW23089.1 100.00%
Bootstrap support for afum|EAL88390.1 as seed ortholog is 100%.
Bootstrap support for afis|EAW23089.1 as seed ortholog is 100%.

---

Group of orthologs #456. Best score 2611 bits
Score difference with first non-orthologous sequence - *A. fumigatus*:2611 *A. fischeri*:1685
afum|EAL92371.2 100.00% afis|EAW21356.1 100.00%
Bootstrap support for afum|EAL92371.2 as seed ortholog is 100%.
Bootstrap support for afis|EAW21356.1 as seed ortholog is 100%.

---

Group of orthologs #457. Best score 2609 bits
Score difference with first non-orthologous sequence - *A. fumigatus*:2609 *A. fischeri*:2609
afum|EAL92040.1 100.00% afis|EAW24178.1 100.00%
Bootstrap support for afum|EAL92040.1 as seed ortholog is 100%.
Bootstrap support for afis|EAW24178.1 as seed ortholog is 100%.

---

Group of orthologs #458. Best score 2608 bits
Score difference with first non-orthologous sequence - *A. fumigatus*:2608 *A. fischeri*:2608
afum|EAL93468.1 100.00% afis|EAW18742.1 100.00%
Bootstrap support for afum|EAL93468.1 as seed ortholog is 100%.
Bootstrap support for afis|EAW18742.1 as seed ortholog is 100%.

---

Group of orthologs #459. Best score 2607 bits
Score difference with first non-orthologous sequence - *A. fumigatus*:2607 *A. fischeri*:2607
afum|EAL85331.1 100.00% afis|EAW20214.1 100.00%
Bootstrap support for afum|EAL85331.1 as seed ortholog is 100%.
Bootstrap support for afis|EAW20214.1 as seed ortholog is 100%.

---

Group of orthologs #460. Best score 2605 bits
Score difference with first non-orthologous sequence - *A. fumigatus*:2605 *A. fischeri*:2605
afum|EAL85302.2 100.00% afis|EAW20184.1 100.00%
Bootstrap support for afum|EAL85302.2 as seed ortholog is 100%.
Bootstrap support for afis|EAW20184.1 as seed ortholog is 100%.

---

Group of orthologs #461. Best score 2604 bits
Score difference with first non-orthologous sequence - *A. fumigatus*:2604 *A. fischeri*:2604
afum|EAL93406.1 100.00% afis|EAW18685.1 100.00%
Bootstrap support for afum|EAL93406.1 as seed ortholog is 100%.
Bootstrap support for afis|EAW18685.1 as seed ortholog is 100%.

---

Group of orthologs #462. Best score 2602 bits
Score difference with first non-orthologous sequence - *A. fumigatus*:2602 *A. fischeri*:2602
afum|EAL88850.1 100.00% afis|EAW16222.1 100.00%
Bootstrap support for afum|EAL88850.1 as seed ortholog is 100%.
Bootstrap support for afis|EAW16222.1 as seed ortholog is 100%.

---

Group of orthologs #463. Best score 2601 bits
Score difference with first non-orthologous sequence - *A. fumigatus*:2601 *A. fischeri*:2601
afum|EAL89627.1 100.00% afis|EAW24968.1 100.00%
Bootstrap support for afum|EAL89627.1 as seed ortholog is 100%.
Bootstrap support for afis|EAW24968.1 as seed ortholog is 100%.

---

Group of orthologs #464. Best score 2598 bits
Score difference with first non-orthologous sequence - *A. fumigatus*:2598 *A. fischeri*:2598
afum|EAL91940.1 100.00% afis|EAW24085.1 100.00%
Bootstrap support for afum|EAL91940.1 as seed ortholog is 100%.

Bootstrap support for afis|EAW24085.1 as seed ortholog is 100%.

---

Group of orthologs #465. Best score 2594 bits
Score difference with first non-orthologous sequence - *A. fumigatus*:2594 *A. fischeri*:2594
afum|EAL90793.1 100.00% afis|EAW22398.1 100.00%
Bootstrap support for afum|EAL90793.1 as seed ortholog is 100%.
Bootstrap support for afis|EAW22398.1 as seed ortholog is 100%.

---

Group of orthologs #466. Best score 2593 bits
Score difference with first non-orthologous sequence - *A. fumigatus*:2593 *A. fischeri*:2593
afum|EAL90435.1 100.00% afis|EAW22764.1 100.00%
Bootstrap support for afum|EAL90435.1 as seed ortholog is 100%.
Bootstrap support for afis|EAW22764.1 as seed ortholog is 100%.

---

Group of orthologs #467. Best score 2592 bits
Score difference with first non-orthologous sequence - *A. fumigatus*:2592 *A. fischeri*:2592
afum|EAL88711.1 100.00% afis|EAW16079.1 100.00%
Bootstrap support for afum|EAL88711.1 as seed ortholog is 100%.
Bootstrap support for afis|EAW16079.1 as seed ortholog is 100%.

---

Group of orthologs #468. Best score 2589 bits
Score difference with first non-orthologous sequence - *A. fumigatus*:2589 *A. fischeri*:2589
afum|EAL90072.1 100.00% afis|EAW25413.1 100.00%
Bootstrap support for afum|EAL90072.1 as seed ortholog is 100%.
Bootstrap support for afis|EAW25413.1 as seed ortholog is 100%.

---

Group of orthologs #469. Best score 2588 bits
Score difference with first non-orthologous sequence - *A. fumigatus*:2588 *A. fischeri*:2588
afum|EAL90208.1 100.00% afis|EAW22993.1 100.00%
Bootstrap support for afum|EAL90208.1 as seed ortholog is 100%.
Bootstrap support for afis|EAW22993.1 as seed ortholog is 100%.

---

Group of orthologs #470. Best score 2587 bits
Score difference with first non-orthologous sequence - *A. fumigatus*:2587 *A. fischeri*:2587
afum|EAL88209.1 100.00% afis|EAW23270.1 100.00%
Bootstrap support for afum|EAL88209.1 as seed ortholog is 100%.
Bootstrap support for afis|EAW23270.1 as seed ortholog is 100%.

---

Group of orthologs #471. Best score 2587 bits
Score difference with first non-orthologous sequence - *A. fumigatus*:2587 *A. fischeri*:2587
afum|EAL90705.1 100.00% afis|EAW22488.1 100.00%
Bootstrap support for afum|EAL90705.1 as seed ortholog is 100%.
Bootstrap support for afis|EAW22488.1 as seed ortholog is 100%.

---

Group of orthologs #472. Best score 2580 bits
Score difference with first non-orthologous sequence - *A. fumigatus*:2580 *A. fischeri*:2580
afum|EAL85248.1 100.00% afis|EAW20132.1 100.00%
Bootstrap support for afum|EAL85248.1 as seed ortholog is 100%.
Bootstrap support for afis|EAW20132.1 as seed ortholog is 100%.

---

Group of orthologs #473. Best score 2580 bits
Score difference with first non-orthologous sequence - *A. fumigatus*:2580 *A. fischeri*:2580
afum|EAL93196.2 100.00% afis|EAW18503.1 100.00%
Bootstrap support for afum|EAL93196.2 as seed ortholog is 100%.
Bootstrap support for afis|EAW18503.1 as seed ortholog is 100%.

---

Group of orthologs #474. Best score 2578 bits
Score difference with first non-orthologous sequence - *A. fumigatus*:2578 *A. fischeri*:2578
afum|EAL88604.1 100.00% afis|EAW15972.1 100.00%
Bootstrap support for afum|EAL88604.1 as seed ortholog is 100%.
Bootstrap support for afis|EAW15972.1 as seed ortholog is 100%.

---

Group of orthologs #475. Best score 2578 bits
Score difference with first non-orthologous sequence - *A. fumigatus*:2578 *A. fischeri*:2578
afum|EAL88088.1 100.00% afis|EAW23383.1 100.00%
Bootstrap support for afum|EAL88088.1 as seed ortholog is 100%.
Bootstrap support for afis|EAW23383.1 as seed ortholog is 100%.

---

Group of orthologs #476. Best score 2568 bits
Score difference with first non-orthologous sequence - *A. fumigatus*:2568 *A. fischeri*:2568
afum|EAL87930.1 100.00% afis|EAW23536.1 100.00%
Bootstrap support for afum|EAL87930.1 as seed ortholog is 100%.
Bootstrap support for afis|EAW23536.1 as seed ortholog is 100%.

---

Group of orthologs #477. Best score 2566 bits
Score difference with first non-orthologous sequence - *A. fumigatus*:2566 *A. fischeri*:2566
afum|EAL87024.1 100.00% afis|EAW19472.1 100.00%
Bootstrap support for afum|EAL87024.1 as seed ortholog is 100%.
Bootstrap support for afis|EAW19472.1 as seed ortholog is 100%.

---

Group of orthologs #478. Best score 2566 bits
Score difference with first non-orthologous sequence - *A. fumigatus*:2566 *A. fischeri*:2566
afum|EAL91351.1 100.00% afis|EAW17537.1 100.00%
Bootstrap support for afum|EAL91351.1 as seed ortholog is 100%.
Bootstrap support for afis|EAW17537.1 as seed ortholog is 100%.

---

Group of orthologs #479. Best score 2565 bits
Score difference with first non-orthologous sequence - *A. fumigatus*:2565 *A. fischeri*:2565
afum|EAL93014.1 100.00% afis|EAW21999.1 100.00%
Bootstrap support for afum|EAL93014.1 as seed ortholog is 100%.
Bootstrap support for afis|EAW21999.1 as seed ortholog is 100%.

---

Group of orthologs #480. Best score 2563 bits
Score difference with first non-orthologous sequence - *A. fumigatus*:2563 *A. fischeri*:2563
afum|EAL90703.1 100.00% afis|EAW22490.1 100.00%
Bootstrap support for afum|EAL90703.1 as seed ortholog is 100%.
Bootstrap support for afis|EAW22490.1 as seed ortholog is 100%.

---

Group of orthologs #481. Best score 2562 bits
Score difference with first non-orthologous sequence - *A. fumigatus*:2562 *A. fischeri*:2562
afum|EAL89143.1 100.00% afis|EAW16545.1 100.00%
Bootstrap support for afum|EAL89143.1 as seed ortholog is 100%.
Bootstrap support for afis|EAW16545.1 as seed ortholog is 100%.

---

Group of orthologs #482. Best score 2561 bits
Score difference with first non-orthologous sequence - *A. fumigatus*:2561 *A. fischeri*:2561
afum|EAL86513.1 100.00% afis|EAW16989.1 100.00%
Bootstrap support for afum|EAL86513.1 as seed ortholog is 100%.
Bootstrap support for afis|EAW16989.1 as seed ortholog is 100%.

---

Group of orthologs #483. Best score 2561 bits
Score difference with first non-orthologous sequence - *A. fumigatus*:2561 *A. fischeri*:2561

afum|EAL88011.1 100.00% afis|EAW23455.1 100.00%
Bootstrap support for afum|EAL88011.1 as seed ortholog is 100%.
Bootstrap support for afis|EAW23455.1 as seed ortholog is 100%.

---

Group of orthologs #484. Best score 2560 bits
Score difference with first non-orthologous sequence - *A. fumigatus*:2560 *A. fischeri*:2325
afum|EAL87015.1 100.00% afis|EAW19480.1 100.00%
Bootstrap support for afum|EAL87015.1 as seed ortholog is 100%.
Bootstrap support for afis|EAW19480.1 as seed ortholog is 100%.

---

Group of orthologs #485. Best score 2559 bits
Score difference with first non-orthologous sequence - *A. fumigatus*:2559 *A. fischeri*:2559
afum|EAL90054.1 100.00% afis|EAW25395.1 100.00%
Bootstrap support for afum|EAL90054.1 as seed ortholog is 100%.
Bootstrap support for afis|EAW25395.1 as seed ortholog is 100%.

---

Group of orthologs #486. Best score 2558 bits
Score difference with first non-orthologous sequence - *A. fumigatus*:2558 *A. fischeri*:2431
afum|EAL84665.1 100.00% afis|EAW20969.1 100.00%
Bootstrap support for afum|EAL84665.1 as seed ortholog is 100%.
Bootstrap support for afis|EAW20969.1 as seed ortholog is 100%.

---

Group of orthologs #487. Best score 2558 bits
Score difference with first non-orthologous sequence - *A. fumigatus*:2558 *A. fischeri*:2558
afum|EAL89493.1 100.00% afis|EAW24847.1 100.00%
Bootstrap support for afum|EAL89493.1 as seed ortholog is 100%.
Bootstrap support for afis|EAW24847.1 as seed ortholog is 100%.

---

Group of orthologs #488. Best score 2557 bits
Score difference with first non-orthologous sequence - *A. fumigatus*:2557 *A. fischeri*:2557
afum|EAL90782.1 100.00% afis|EAW22410.1 100.00%
Bootstrap support for afum|EAL90782.1 as seed ortholog is 100%.
Bootstrap support for afis|EAW22410.1 as seed ortholog is 100%.

---

Group of orthologs #489. Best score 2553 bits
Score difference with first non-orthologous sequence - *A. fumigatus*:2553 *A. fischeri*:2553
afum|EAL90403.1 100.00% afis|EAW22796.1 100.00%
Bootstrap support for afum|EAL90403.1 as seed ortholog is 100%.
Bootstrap support for afis|EAW22796.1 as seed ortholog is 100%.

---

Group of orthologs #490. Best score 2553 bits
Score difference with first non-orthologous sequence - *A. fumigatus*:2553 *A. fischeri*:1567
afum|EAL89712.1 100.00% afis|EAW25044.1 100.00%
Bootstrap support for afum|EAL89712.1 as seed ortholog is 100%.
Bootstrap support for afis|EAW25044.1 as seed ortholog is 100%.

---

Group of orthologs #491. Best score 2552 bits
Score difference with first non-orthologous sequence - *A. fumigatus*:2552 *A. fischeri*:2552
afum|EAL94013.1 100.00% afis|EAW19294.1 100.00%
Bootstrap support for afum|EAL94013.1 as seed ortholog is 100%.
Bootstrap support for afis|EAW19294.1 as seed ortholog is 100%.

---

Group of orthologs #492. Best score 2551 bits
Score difference with first non-orthologous sequence - *A. fumigatus*:2551 *A. fischeri*:2551
afum|EAL90975.1 100.00% afis|EAW22219.1 100.00%
Bootstrap support for afum|EAL90975.1 as seed ortholog is 100%.

Bootstrap support for afis|EAW22219.1 as seed ortholog is 100%.

---

Group of orthologs #493. Best score 2548 bits
Score difference with first non-orthologous sequence - *A. fumigatus*:2548 *A. fischeri*:2548
afum|EAL93213.1 100.00% afis|EAW18518.1 100.00%
Bootstrap support for afum|EAL93213.1 as seed ortholog is 100%.
Bootstrap support for afis|EAW18518.1 as seed ortholog is 100%.

---

Group of orthologs #494. Best score 2547 bits
Score difference with first non-orthologous sequence - *A. fumigatus*:2547 *A. fischeri*:2547
afum|EAL84525.1 100.00% afis|EAW20532.1 100.00%
Bootstrap support for afum|EAL84525.1 as seed ortholog is 100%.
Bootstrap support for afis|EAW20532.1 as seed ortholog is 100%.

---

Group of orthologs #495. Best score 2544 bits
Score difference with first non-orthologous sequence - *A. fumigatus*:2544 *A. fischeri*:2544
afum|EAL91264.1 100.00% afis|EAW17437.1 100.00%
Bootstrap support for afum|EAL91264.1 as seed ortholog is 100%.
Bootstrap support for afis|EAW17437.1 as seed ortholog is 100%.

---

Group of orthologs #496. Best score 2540 bits
Score difference with first non-orthologous sequence - *A. fumigatus*:2540 *A. fischeri*:2540
afum|EAL93296.2 100.00% afis|EAW18589.1 100.00%
Bootstrap support for afum|EAL93296.2 as seed ortholog is 100%.
Bootstrap support for afis|EAW18589.1 as seed ortholog is 100%.

---

Group of orthologs #497. Best score 2539 bits
Score difference with first non-orthologous sequence - *A. fumigatus*:2539 *A. fischeri*:2539
afum|EAL85615.2 100.00% afis|EAW15732.1 100.00%
Bootstrap support for afum|EAL85615.2 as seed ortholog is 100%.
Bootstrap support for afis|EAW15732.1 as seed ortholog is 100%.

---

Group of orthologs #498. Best score 2534 bits
Score difference with first non-orthologous sequence - *A. fumigatus*:2390 *A. fischeri*:2396
afum|EAL92384.1 100.00% afis|EAW21367.1 100.00%
Bootstrap support for afum|EAL92384.1 as seed ortholog is 100%.
Bootstrap support for afis|EAW21367.1 as seed ortholog is 100%.

---

Group of orthologs #499. Best score 2532 bits
Score difference with first non-orthologous sequence - *A. fumigatus*:2532 *A. fischeri*:2532
afum|EAL93546.1 100.00% afis|EAW18810.1 100.00%
Bootstrap support for afum|EAL93546.1 as seed ortholog is 100%.
Bootstrap support for afis|EAW18810.1 as seed ortholog is 100%.

---

Group of orthologs #500. Best score 2520 bits
Score difference with first non-orthologous sequence - *A. fumigatus*:2520 *A. fischeri*:2520
afum|EAL90877.1 100.00% afis|EAW22313.1 100.00%
Bootstrap support for afum|EAL90877.1 as seed ortholog is 100%.
Bootstrap support for afis|EAW22313.1 as seed ortholog is 100%.

---

Group of orthologs #501. Best score 2518 bits
Score difference with first non-orthologous sequence - *A. fumigatus*:2518 *A. fischeri*:2518
afum|EAL89938.1 100.00% afis|EAW25275.1 100.00%
Bootstrap support for afum|EAL89938.1 as seed ortholog is 100%.
Bootstrap support for afis|EAW25275.1 as seed ortholog is 100%.

---

Group of orthologs #502. Best score 2517 bits
Score difference with first non-orthologous sequence - *A. fumigatus*:2517 *A. fischeri*:2517
afum|EAL88242.1 100.00% afis|EAW23240.1 100.00%
Bootstrap support for afum|EAL88242.1 as seed ortholog is 100%.
Bootstrap support for afis|EAW23240.1 as seed ortholog is 100%.

---

Group of orthologs #503. Best score 2515 bits
Score difference with first non-orthologous sequence - *A. fumigatus*:2515 *A. fischeri*:2232
afum|EAL85670.1 100.00% afis|EAW15677.1 100.00%
Bootstrap support for afum|EAL85670.1 as seed ortholog is 100%.
Bootstrap support for afis|EAW15677.1 as seed ortholog is 100%.

---

Group of orthologs #504. Best score 2514 bits
Score difference with first non-orthologous sequence - *A. fumigatus*:2514 *A. fischeri*:2514
afum|EAL90308.1 100.00% afis|EAW22890.1 100.00%
Bootstrap support for afum|EAL90308.1 as seed ortholog is 100%.
Bootstrap support for afis|EAW22890.1 as seed ortholog is 100%.

---

Group of orthologs #505. Best score 2514 bits
Score difference with first non-orthologous sequence - *A. fumigatus*:2514 *A. fischeri*:2514
afum|EAL92700.1 100.00% afis|EAW21665.1 100.00%
Bootstrap support for afum|EAL92700.1 as seed ortholog is 100%.
Bootstrap support for afis|EAW21665.1 as seed ortholog is 100%.

---

Group of orthologs #506. Best score 2513 bits
Score difference with first non-orthologous sequence - *A. fumigatus*:2513 *A. fischeri*:1141
afum|EAL89306.1 100.00% afis|EAW17294.1 100.00%
afis|EAW19388.1 9.12%
Bootstrap support for afum|EAL89306.1 as seed ortholog is 100%.
Bootstrap support for afis|EAW17294.1 as seed ortholog is 100%.

---

Group of orthologs #507. Best score 2512 bits
Score difference with first non-orthologous sequence - *A. fumigatus*:2512 *A. fischeri*:2512
afum|EAL86026.2 100.00% afis|EAW24356.1 100.00%
Bootstrap support for afum|EAL86026.2 as seed ortholog is 100%.
Bootstrap support for afis|EAW24356.1 as seed ortholog is 100%.

---

Group of orthologs #508. Best score 2512 bits
Score difference with first non-orthologous sequence - *A. fumigatus*:2512 *A. fischeri*:2512
afum|EAL88017.1 100.00% afis|EAW23448.1 100.00%
Bootstrap support for afum|EAL88017.1 as seed ortholog is 100%.
Bootstrap support for afis|EAW23448.1 as seed ortholog is 100%.

---

Group of orthologs #509. Best score 2511 bits
Score difference with first non-orthologous sequence - *A. fumigatus*:2511 *A. fischeri*:2511
afum|EAL90413.1 100.00% afis|EAW22786.1 100.00%
Bootstrap support for afum|EAL90413.1 as seed ortholog is 100%.
Bootstrap support for afis|EAW22786.1 as seed ortholog is 100%.

---

Group of orthologs #510. Best score 2511 bits
Score difference with first non-orthologous sequence - *A. fumigatus*:2511 *A. fischeri*:2511
afum|EAL89400.1 100.00% afis|EAW24758.1 100.00%
Bootstrap support for afum|EAL89400.1 as seed ortholog is 100%.
Bootstrap support for afis|EAW24758.1 as seed ortholog is 100%.

---

Group of orthologs #511. Best score 2510 bits

Score difference with first non-orthologous sequence - *A. fumigatus*:2510   *A. fischeri*:2510
afum|EAL89996.1 100.00% afis|EAW25333.1 100.00%
Bootstrap support for afum|EAL89996.1 as seed ortholog is 100%.
Bootstrap support for afis|EAW25333.1 as seed ortholog is 100%.

---

Group of orthologs #512. Best score 2509 bits
Score difference with first non-orthologous sequence - *A. fumigatus*:2509   *A. fischeri*:2509
afum|EAL86876.1 100.00% afis|EAW19646.1 100.00%
Bootstrap support for afum|EAL86876.1 as seed ortholog is 100%.
Bootstrap support for afis|EAW19646.1 as seed ortholog is 100%.

---

Group of orthologs #513. Best score 2507 bits
Score difference with first non-orthologous sequence - *A. fumigatus*:2507   *A. fischeri*:2507
afum|EAL90600.1 100.00% afis|EAW22598.1 100.00%
Bootstrap support for afum|EAL90600.1 as seed ortholog is 100%.
Bootstrap support for afis|EAW22598.1 as seed ortholog is 100%.

---

Group of orthologs #514. Best score 2504 bits
Score difference with first non-orthologous sequence - *A. fumigatus*:2504   *A. fischeri*:2504
afum|EAL88717.1 100.00% afis|EAW16085.1 100.00%
Bootstrap support for afum|EAL88717.1 as seed ortholog is 100%.
Bootstrap support for afis|EAW16085.1 as seed ortholog is 100%.

---

Group of orthologs #515. Best score 2504 bits
Score difference with first non-orthologous sequence - *A. fumigatus*:2504   *A. fischeri*:2504
afum|EAL87432.1 100.00% afis|EAW24002.1 100.00%
Bootstrap support for afum|EAL87432.1 as seed ortholog is 100%.
Bootstrap support for afis|EAW24002.1 as seed ortholog is 100%.

---

Group of orthologs #516. Best score 2502 bits
Score difference with first non-orthologous sequence - *A. fumigatus*:2502   *A. fischeri*:2502
afum|EAL90311.1 100.00% afis|EAW22887.1 100.00%
Bootstrap support for afum|EAL90311.1 as seed ortholog is 100%.
Bootstrap support for afis|EAW22887.1 as seed ortholog is 100%.

---

Group of orthologs #517. Best score 2500 bits
Score difference with first non-orthologous sequence - *A. fumigatus*:2500   *A. fischeri*:2500
afum|EAL87709.1 100.00% afis|EAW18345.1 100.00%
Bootstrap support for afum|EAL87709.1 as seed ortholog is 100%.
Bootstrap support for afis|EAW18345.1 as seed ortholog is 100%.

---

Group of orthologs #518. Best score 2500 bits
Score difference with first non-orthologous sequence - *A. fumigatus*:2246   *A. fischeri*:2500
afum|EAL91285.1 100.00% afis|EAW17457.1 100.00%
Bootstrap support for afum|EAL91285.1 as seed ortholog is 100%.
Bootstrap support for afis|EAW17457.1 as seed ortholog is 100%.

---

Group of orthologs #519. Best score 2496 bits
Score difference with first non-orthologous sequence - *A. fumigatus*:2496   *A. fischeri*:2496
afum|EAL86905.1 100.00% afis|EAW19617.1 100.00%
Bootstrap support for afum|EAL86905.1 as seed ortholog is 100%.
Bootstrap support for afis|EAW19617.1 as seed ortholog is 100%.

---

Group of orthologs #520. Best score 2496 bits
Score difference with first non-orthologous sequence - *A. fumigatus*:2496   *A. fischeri*:2496
afum|EAL91812.1 100.00% afis|EAW18001.1 100.00%

Bootstrap support for afum|EAL91812.1 as seed ortholog is 100%.
Bootstrap support for afis|EAW18001.1 as seed ortholog is 100%.

---

Group of orthologs #521. Best score 2492 bits
Score difference with first non-orthologous sequence - *A. fumigatus*:2492 *A. fischeri*:2492
afum|EAL87601.2 100.00% afis|EAW18233.1 100.00%
Bootstrap support for afum|EAL87601.2 as seed ortholog is 100%.
Bootstrap support for afis|EAW18233.1 as seed ortholog is 100%.

---

Group of orthologs #522. Best score 2490 bits
Score difference with first non-orthologous sequence - *A. fumigatus*:2490 *A. fischeri*:2490
afum|EAL88985.1 100.00% afis|EAW16360.1 100.00%
Bootstrap support for afum|EAL88985.1 as seed ortholog is 100%.
Bootstrap support for afis|EAW16360.1 as seed ortholog is 100%.

---

Group of orthologs #523. Best score 2490 bits
Score difference with first non-orthologous sequence - *A. fumigatus*:2490 *A. fischeri*:2490
afum|EAL94015.1 100.00% afis|EAW19296.1 100.00%
Bootstrap support for afum|EAL94015.1 as seed ortholog is 100%.
Bootstrap support for afis|EAW19296.1 as seed ortholog is 100%.

---

Group of orthologs #524. Best score 2488 bits
Score difference with first non-orthologous sequence - *A. fumigatus*:2488 *A. fischeri*:2488
afum|EAL91164.1 100.00% afis|EAW17327.1 100.00%
Bootstrap support for afum|EAL91164.1 as seed ortholog is 100%.
Bootstrap support for afis|EAW17327.1 as seed ortholog is 100%.

---

Group of orthologs #525. Best score 2484 bits
Score difference with first non-orthologous sequence - *A. fumigatus*:2484 *A. fischeri*:2484
afum|EAL84620.1 100.00% afis|EAW20430.1 100.00%
Bootstrap support for afum|EAL84620.1 as seed ortholog is 100%.
Bootstrap support for afis|EAW20430.1 as seed ortholog is 100%.

---

Group of orthologs #526. Best score 2483 bits
Score difference with first non-orthologous sequence - *A. fumigatus*:2483 *A. fischeri*:2483
afum|EAL90236.1 100.00% afis|EAW22961.1 100.00%
Bootstrap support for afum|EAL90236.1 as seed ortholog is 100%.
Bootstrap support for afis|EAW22961.1 as seed ortholog is 100%.

---

Group of orthologs #527. Best score 2482 bits
Score difference with first non-orthologous sequence - *A. fumigatus*:2482 *A. fischeri*:2482
afum|EAL88109.1 100.00% afis|EAW23364.1 100.00%
Bootstrap support for afum|EAL88109.1 as seed ortholog is 100%.
Bootstrap support for afis|EAW23364.1 as seed ortholog is 100%.

---

Group of orthologs #528. Best score 2481 bits
Score difference with first non-orthologous sequence - *A. fumigatus*:2481 *A. fischeri*:2481
afum|EAL91382.1 100.00% afis|EAW17569.1 100.00%
Bootstrap support for afum|EAL91382.1 as seed ortholog is 100%.
Bootstrap support for afis|EAW17569.1 as seed ortholog is 100%.

---

Group of orthologs #529. Best score 2476 bits
Score difference with first non-orthologous sequence - *A. fumigatus*:2476 *A. fischeri*:2476
afum|EAL88071.1 100.00% afis|EAW23399.1 100.00%
Bootstrap support for afum|EAL88071.1 as seed ortholog is 100%.
Bootstrap support for afis|EAW23399.1 as seed ortholog is 100%.

Group of orthologs #530. Best score 2476 bits
Score difference with first non-orthologous sequence - *A. fumigatus*:2476 *A. fischeri*:2476
afum|EAL90026.1 100.00% afis|EAW25364.1 100.00%
Bootstrap support for afum|EAL90026.1 as seed ortholog is 100%.
Bootstrap support for afis|EAW25364.1 as seed ortholog is 100%.

---

Group of orthologs #531. Best score 2475 bits
Score difference with first non-orthologous sequence - *A. fumigatus*:2475 *A. fischeri*:2475
afum|EAL91426.1 100.00% afis|EAW17613.1 100.00%
Bootstrap support for afum|EAL91426.1 as seed ortholog is 100%.
Bootstrap support for afis|EAW17613.1 as seed ortholog is 100%.

---

Group of orthologs #532. Best score 2475 bits
Score difference with first non-orthologous sequence - *A. fumigatus*:2475 *A. fischeri*:2475
afum|EAL93516.2 100.00% afis|EAW18784.1 100.00%
Bootstrap support for afum|EAL93516.2 as seed ortholog is 100%.
Bootstrap support for afis|EAW18784.1 as seed ortholog is 100%.

---

Group of orthologs #533. Best score 2473 bits
Score difference with first non-orthologous sequence - *A. fumigatus*:2473 *A. fischeri*:2473
afum|EAL87963.1 100.00% afis|EAW23499.1 100.00%
Bootstrap support for afum|EAL87963.1 as seed ortholog is 100%.
Bootstrap support for afis|EAW23499.1 as seed ortholog is 100%.

---

Group of orthologs #534. Best score 2471 bits
Score difference with first non-orthologous sequence - *A. fumigatus*:2471 *A. fischeri*:2471
afum|EAL93160.2 100.00% afis|EAW18473.1 100.00%
Bootstrap support for afum|EAL93160.2 as seed ortholog is 100%.
Bootstrap support for afis|EAW18473.1 as seed ortholog is 100%.

---

Group of orthologs #535. Best score 2468 bits
Score difference with first non-orthologous sequence - *A. fumigatus*:2468 *A. fischeri*:2468
afum|EAL88231.2 100.00% afis|EAW23250.1 100.00%
Bootstrap support for afum|EAL88231.2 as seed ortholog is 100%.
Bootstrap support for afis|EAW23250.1 as seed ortholog is 100%.

---

Group of orthologs #536. Best score 2467 bits
Score difference with first non-orthologous sequence - *A. fumigatus*:2467 *A. fischeri*:2467
afum|EAL92953.1 100.00% afis|EAW21940.1 100.00%
Bootstrap support for afum|EAL92953.1 as seed ortholog is 100%.
Bootstrap support for afis|EAW21940.1 as seed ortholog is 100%.

---

Group of orthologs #537. Best score 2464 bits
Score difference with first non-orthologous sequence - *A. fumigatus*:2464 *A. fischeri*:2464
afum|EAL89203.1 100.00% afis|EAW16606.1 100.00%
Bootstrap support for afum|EAL89203.1 as seed ortholog is 100%.
Bootstrap support for afis|EAW16606.1 as seed ortholog is 100%.

---

Group of orthologs #538. Best score 2464 bits
Score difference with first non-orthologous sequence - *A. fumigatus*:2464 *A. fischeri*:2464
afum|EAL93963.1 100.00% afis|EAW19235.1 100.00%
Bootstrap support for afum|EAL93963.1 as seed ortholog is 100%.
Bootstrap support for afis|EAW19235.1 as seed ortholog is 100%.

---

Group of orthologs #539. Best score 2464 bits

Score difference with first non-orthologous sequence - *A. fumigatus*:2464   *A. fischeri*:2464
afum|EAL92611.1 100.00% afis|EAW21593.1 100.00%
Bootstrap support for afum|EAL92611.1 as seed ortholog is 100%.
Bootstrap support for afis|EAW21593.1 as seed ortholog is 100%.

---

Group of orthologs #540. Best score 2464 bits
Score difference with first non-orthologous sequence - *A. fumigatus*:2464   *A. fischeri*:2464
afum|EAL89568.1 100.00% afis|EAW24914.1 100.00%
Bootstrap support for afum|EAL89568.1 as seed ortholog is 100%.
Bootstrap support for afis|EAW24914.1 as seed ortholog is 100%.

---

Group of orthologs #541. Best score 2463 bits
Score difference with first non-orthologous sequence - *A. fumigatus*:2463   *A. fischeri*:2463
afum|EAL91407.1 100.00% afis|EAW17594.1 100.00%
Bootstrap support for afum|EAL91407.1 as seed ortholog is 100%.
Bootstrap support for afis|EAW17594.1 as seed ortholog is 100%.

---

Group of orthologs #542. Best score 2463 bits
Score difference with first non-orthologous sequence - *A. fumigatus*:2463   *A. fischeri*:2463
afum|EAL93873.1 100.00% afis|EAW19142.1 100.00%
Bootstrap support for afum|EAL93873.1 as seed ortholog is 100%.
Bootstrap support for afis|EAW19142.1 as seed ortholog is 100%.

---

Group of orthologs #543. Best score 2461 bits
Score difference with first non-orthologous sequence - *A. fumigatus*:2461   *A. fischeri*:2191
afum|EAL87677.1 100.00% afis|EAW18311.1 100.00%
Bootstrap support for afum|EAL87677.1 as seed ortholog is 100%.
Bootstrap support for afis|EAW18311.1 as seed ortholog is 100%.

---

Group of orthologs #544. Best score 2461 bits
Score difference with first non-orthologous sequence - *A. fumigatus*:2461   *A. fischeri*:2169
afum|EAL87230.1 100.00% afis|EAW23796.1 100.00%
Bootstrap support for afum|EAL87230.1 as seed ortholog is 100%.
Bootstrap support for afis|EAW23796.1 as seed ortholog is 100%.

---

Group of orthologs #545. Best score 2460 bits
Score difference with first non-orthologous sequence - *A. fumigatus*:2460   *A. fischeri*:2460
afum|EAL90702.1 100.00% afis|EAW22491.1 100.00%
Bootstrap support for afum|EAL90702.1 as seed ortholog is 100%.
Bootstrap support for afis|EAW22491.1 as seed ortholog is 100%.

---

Group of orthologs #546. Best score 2460 bits
Score difference with first non-orthologous sequence - *A. fumigatus*:2276   *A. fischeri*:2460
afum|EBA27302.1 100.00% afis|EAW23728.1 100.00%
Bootstrap support for afum|EBA27302.1 as seed ortholog is 100%.
Bootstrap support for afis|EAW23728.1 as seed ortholog is 100%.

---

Group of orthologs #547. Best score 2458 bits
Score difference with first non-orthologous sequence - *A. fumigatus*:2458   *A. fischeri*:2458
afum|EAL85649.1 100.00% afis|EAW15699.1 100.00%
Bootstrap support for afum|EAL85649.1 as seed ortholog is 100%.
Bootstrap support for afis|EAW15699.1 as seed ortholog is 100%.

---

Group of orthologs #548. Best score 2458 bits
Score difference with first non-orthologous sequence - *A. fumigatus*:2458   *A. fischeri*:2458
afum|EAL93535.1 100.00% afis|EAW18799.1 100.00%

Bootstrap support for afum|EAL93535.1 as seed ortholog is 100%.
Bootstrap support for afis|EAW18799.1 as seed ortholog is 100%.

---

Group of orthologs #549. Best score 2455 bits
Score difference with first non-orthologous sequence - *A. fumigatus*:2455 *A. fischeri*:2455
afum|EAL93644.1 100.00% afis|EAW18913.1 100.00%
Bootstrap support for afum|EAL93644.1 as seed ortholog is 100%.
Bootstrap support for afis|EAW18913.1 as seed ortholog is 100%.

---

Group of orthologs #550. Best score 2453 bits
Score difference with first non-orthologous sequence - *A. fumigatus*:2453 *A. fischeri*:2453
afum|EAL91563.1 100.00% afis|EAW17749.1 100.00%
Bootstrap support for afum|EAL91563.1 as seed ortholog is 100%.
Bootstrap support for afis|EAW17749.1 as seed ortholog is 100%.

---

Group of orthologs #551. Best score 2450 bits
Score difference with first non-orthologous sequence - *A. fumigatus*:2450 *A. fischeri*:2450
afum|EAL88968.1 100.00% afis|EAW16343.1 100.00%
Bootstrap support for afum|EAL88968.1 as seed ortholog is 100%.
Bootstrap support for afis|EAW16343.1 as seed ortholog is 100%.

---

Group of orthologs #552. Best score 2449 bits
Score difference with first non-orthologous sequence - *A. fumigatus*:2449 *A. fischeri*:2449
afum|EAL90999.1 100.00% afis|EAW22192.1 100.00%
Bootstrap support for afum|EAL90999.1 as seed ortholog is 100%.
Bootstrap support for afis|EAW22192.1 as seed ortholog is 100%.

---

Group of orthologs #553. Best score 2447 bits
Score difference with first non-orthologous sequence - *A. fumigatus*:2447 *A. fischeri*:2447
afum|EAL88600.1 100.00% afis|EAW15967.1 100.00%
Bootstrap support for afum|EAL88600.1 as seed ortholog is 100%.
Bootstrap support for afis|EAW15967.1 as seed ortholog is 100%.

---

Group of orthologs #554. Best score 2446 bits
Score difference with first non-orthologous sequence - *A. fumigatus*:2446 *A. fischeri*:2446
afum|EAL85260.1 100.00% afis|EAW20145.1 100.00%
Bootstrap support for afum|EAL85260.1 as seed ortholog is 100%.
Bootstrap support for afis|EAW20145.1 as seed ortholog is 100%.

---

Group of orthologs #555. Best score 2445 bits
Score difference with first non-orthologous sequence - *A. fumigatus*:2445 *A. fischeri*:2445
afum|EAL84739.1 100.00% afis|EAW20894.1 100.00%
Bootstrap support for afum|EAL84739.1 as seed ortholog is 100%.
Bootstrap support for afis|EAW20894.1 as seed ortholog is 100%.

---

Group of orthologs #556. Best score 2445 bits
Score difference with first non-orthologous sequence - *A. fumigatus*:2445 *A. fischeri*:2445
afum|EAL91013.1 100.00% afis|EAW22176.1 100.00%
Bootstrap support for afum|EAL91013.1 as seed ortholog is 100%.
Bootstrap support for afis|EAW22176.1 as seed ortholog is 100%.

---

Group of orthologs #557. Best score 2445 bits
Score difference with first non-orthologous sequence - *A. fumigatus*:2445 *A. fischeri*:2445
afum|EAL92479.1 100.00% afis|EAW21460.1 100.00%
Bootstrap support for afum|EAL92479.1 as seed ortholog is 100%.
Bootstrap support for afis|EAW21460.1 as seed ortholog is 100%.

Group of orthologs #558. Best score 2441 bits
Score difference with first non-orthologous sequence - *A. fumigatus*:2441 *A. fischeri*:2441
afum|EAL91419.1 100.00% afis|EAW17606.1 100.00%
Bootstrap support for afum|EAL91419.1 as seed ortholog is 100%.
Bootstrap support for afis|EAW17606.1 as seed ortholog is 100%.

---

Group of orthologs #559. Best score 2440 bits
Score difference with first non-orthologous sequence - *A. fumigatus*:2440 *A. fischeri*:2440
afum|EAL93787.1 100.00% afis|EAW19058.1 100.00%
Bootstrap support for afum|EAL93787.1 as seed ortholog is 100%.
Bootstrap support for afis|EAW19058.1 as seed ortholog is 100%.

---

Group of orthologs #560. Best score 2435 bits
Score difference with first non-orthologous sequence - *A. fumigatus*:2435 *A. fischeri*:2435
afum|EAL89515.1 100.00% afis|EAW24865.1 100.00%
Bootstrap support for afum|EAL89515.1 as seed ortholog is 100%.
Bootstrap support for afis|EAW24865.1 as seed ortholog is 100%.

---

Group of orthologs #561. Best score 2434 bits
Score difference with first non-orthologous sequence - *A. fumigatus*:2434 *A. fischeri*:2434
afum|EAL88743.1 100.00% afis|EAW16112.1 100.00%
Bootstrap support for afum|EAL88743.1 as seed ortholog is 100%.
Bootstrap support for afis|EAW16112.1 as seed ortholog is 100%.

---

Group of orthologs #562. Best score 2434 bits
Score difference with first non-orthologous sequence - *A. fumigatus*:2434 *A. fischeri*:2434
afum|EAL88480.1 100.00% afis|EAW23000.1 100.00%
Bootstrap support for afum|EAL88480.1 as seed ortholog is 100%.
Bootstrap support for afis|EAW23000.1 as seed ortholog is 100%.

---

Group of orthologs #563. Best score 2433 bits
Score difference with first non-orthologous sequence - *A. fumigatus*:2433 *A. fischeri*:2433
afum|EAL93201.1 100.00% afis|EAW18507.1 100.00%
Bootstrap support for afum|EAL93201.1 as seed ortholog is 100%.
Bootstrap support for afis|EAW18507.1 as seed ortholog is 100%.

---

Group of orthologs #564. Best score 2431 bits
Score difference with first non-orthologous sequence - *A. fumigatus*:2431 *A. fischeri*:2431
afum|EAL90611.1 100.00% afis|EAW22588.1 100.00%
Bootstrap support for afum|EAL90611.1 as seed ortholog is 100%.
Bootstrap support for afis|EAW22588.1 as seed ortholog is 100%.

---

Group of orthologs #565. Best score 2429 bits
Score difference with first non-orthologous sequence - *A. fumigatus*:2429 *A. fischeri*:2429
afum|EAL84752.1 100.00% afis|EAW20880.1 100.00%
Bootstrap support for afum|EAL84752.1 as seed ortholog is 100%.
Bootstrap support for afis|EAW20880.1 as seed ortholog is 100%.

---

Group of orthologs #566. Best score 2428 bits
Score difference with first non-orthologous sequence - *A. fumigatus*:2428 *A. fischeri*:2428
afum|EAL93440.1 100.00% afis|EAW18715.1 100.00%
Bootstrap support for afum|EAL93440.1 as seed ortholog is 100%.
Bootstrap support for afis|EAW18715.1 as seed ortholog is 100%.

---

Group of orthologs #567. Best score 2418 bits

Score difference with first non-orthologous sequence - *A. fumigatus*:2418   *A. fischeri*:2418
afum|EAL90704.1 100.00% afis|EAW22489.1 100.00%
Bootstrap support for afum|EAL90704.1 as seed ortholog is 100%.
Bootstrap support for afis|EAW22489.1 as seed ortholog is 100%.

---

Group of orthologs #568. Best score 2417 bits
Score difference with first non-orthologous sequence - *A. fumigatus*:2417   *A. fischeri*:2417
afum|EAL88554.1 100.00% afis|EAW15918.1 100.00%
Bootstrap support for afum|EAL88554.1 as seed ortholog is 100%.
Bootstrap support for afis|EAW15918.1 as seed ortholog is 100%.

---

Group of orthologs #569. Best score 2416 bits
Score difference with first non-orthologous sequence - *A. fumigatus*:2416   *A. fischeri*:1300
afum|EAL84790.1 100.00% afis|EAW20842.1 100.00%
Bootstrap support for afum|EAL84790.1 as seed ortholog is 100%.
Bootstrap support for afis|EAW20842.1 as seed ortholog is 100%.

---

Group of orthologs #570. Best score 2414 bits
Score difference with first non-orthologous sequence - *A. fumigatus*:2414   *A. fischeri*:2414
afum|EAL90293.1 100.00% afis|EAW22904.1 100.00%
Bootstrap support for afum|EAL90293.1 as seed ortholog is 100%.
Bootstrap support for afis|EAW22904.1 as seed ortholog is 100%.

---

Group of orthologs #571. Best score 2413 bits
Score difference with first non-orthologous sequence - *A. fumigatus*:2413   *A. fischeri*:2279
afum|EAL88812.1 100.00% afis|EAW16180.1 100.00%
Bootstrap support for afum|EAL88812.1 as seed ortholog is 100%.
Bootstrap support for afis|EAW16180.1 as seed ortholog is 100%.

---

Group of orthologs #572. Best score 2409 bits
Score difference with first non-orthologous sequence - *A. fumigatus*:2228   *A. fischeri*:1821
afum|EAL85682.1 100.00% afis|EAW15666.1 100.00%
Bootstrap support for afum|EAL85682.1 as seed ortholog is 100%.
Bootstrap support for afis|EAW15666.1 as seed ortholog is 100%.

---

Group of orthologs #573. Best score 2409 bits
Score difference with first non-orthologous sequence - *A. fumigatus*:2409   *A. fischeri*:2409
afum|EAL90749.1 100.00% afis|EAW22442.1 100.00%
Bootstrap support for afum|EAL90749.1 as seed ortholog is 100%.
Bootstrap support for afis|EAW22442.1 as seed ortholog is 100%.

---

Group of orthologs #574. Best score 2409 bits
Score difference with first non-orthologous sequence - *A. fumigatus*:2409   *A. fischeri*:2409
afum|EAL92181.1 100.00% afis|EAW21162.1 100.00%
Bootstrap support for afum|EAL92181.1 as seed ortholog is 100%.
Bootstrap support for afis|EAW21162.1 as seed ortholog is 100%.

---

Group of orthologs #575. Best score 2407 bits
Score difference with first non-orthologous sequence - *A. fumigatus*:2407   *A. fischeri*:2407
afum|EAL85544.1 100.00% afis|EAW15807.1 100.00%
Bootstrap support for afum|EAL85544.1 as seed ortholog is 100%.
Bootstrap support for afis|EAW15807.1 as seed ortholog is 100%.

---

Group of orthologs #576. Best score 2404 bits
Score difference with first non-orthologous sequence - *A. fumigatus*:2404   *A. fischeri*:2226
afum|EAL85951.1 100.00% afis|EAW24282.1 100.00%

Bootstrap support for afum|EAL85951.1 as seed ortholog is 100%.
Bootstrap support for afis|EAW24282.1 as seed ortholog is 100%.

---

Group of orthologs #577. Best score 2400 bits
Score difference with first non-orthologous sequence - *A. fumigatus*:2400 *A. fischeri*:2400
afum|EAL86893.1 100.00% afis|EAW19629.1 100.00%
Bootstrap support for afum|EAL86893.1 as seed ortholog is 100%.
Bootstrap support for afis|EAW19629.1 as seed ortholog is 100%.

---

Group of orthologs #578. Best score 2399 bits
Score difference with first non-orthologous sequence - *A. fumigatus*:2399 *A. fischeri*:2399
afum|EAL87469.1 100.00% afis|EAW18112.1 100.00%
Bootstrap support for afum|EAL87469.1 as seed ortholog is 100%.
Bootstrap support for afis|EAW18112.1 as seed ortholog is 100%.

---

Group of orthologs #579. Best score 2399 bits
Score difference with first non-orthologous sequence - *A. fumigatus*:2399 *A. fischeri*:2399
afum|EAL88015.1 100.00% afis|EAW23451.1 100.00%
Bootstrap support for afum|EAL88015.1 as seed ortholog is 100%.
Bootstrap support for afis|EAW23451.1 as seed ortholog is 100%.

---

Group of orthologs #580. Best score 2398 bits
Score difference with first non-orthologous sequence - *A. fumigatus*:2398 *A. fischeri*:2398
afum|EAL88305.1 100.00% afis|EAW23178.1 100.00%
Bootstrap support for afum|EAL88305.1 as seed ortholog is 100%.
Bootstrap support for afis|EAW23178.1 as seed ortholog is 100%.

---

Group of orthologs #581. Best score 2394 bits
Score difference with first non-orthologous sequence - *A. fumigatus*:2394 *A. fischeri*:2394
afum|EAL88888.1 100.00% afis|EAW16267.1 100.00%
Bootstrap support for afum|EAL88888.1 as seed ortholog is 100%.
Bootstrap support for afis|EAW16267.1 as seed ortholog is 100%.

---

Group of orthologs #582. Best score 2394 bits
Score difference with first non-orthologous sequence - *A. fumigatus*:2394 *A. fischeri*:2394
afum|EAL85301.1 100.00% afis|EAW20183.1 100.00%
Bootstrap support for afum|EAL85301.1 as seed ortholog is 100%.
Bootstrap support for afis|EAW20183.1 as seed ortholog is 100%.

---

Group of orthologs #583. Best score 2394 bits
Score difference with first non-orthologous sequence - *A. fumigatus*:2394 *A. fischeri*:2394
afum|EAL93884.1 100.00% afis|EAW19153.1 100.00%
Bootstrap support for afum|EAL93884.1 as seed ortholog is 100%.
Bootstrap support for afis|EAW19153.1 as seed ortholog is 100%.

---

Group of orthologs #584. Best score 2392 bits
Score difference with first non-orthologous sequence - *A. fumigatus*:2392 *A. fischeri*:2392
afum|EAL90893.1 100.00% afis|EAW22288.1 100.00%
Bootstrap support for afum|EAL90893.1 as seed ortholog is 100%.
Bootstrap support for afis|EAW22288.1 as seed ortholog is 100%.

---

Group of orthologs #585. Best score 2392 bits
Score difference with first non-orthologous sequence - *A. fumigatus*:2392 *A. fischeri*:2392
afum|EAL90292.1 100.00% afis|EAW22905.1 100.00%
Bootstrap support for afum|EAL90292.1 as seed ortholog is 100%.
Bootstrap support for afis|EAW22905.1 as seed ortholog is 100%.

---

Group of orthologs #586. Best score 2391 bits
Score difference with first non-orthologous sequence - *A. fumigatus*:2391 *A. fischeri*:2391
afum|EAL91792.1 100.00% afis|EAW17983.1 100.00%
Bootstrap support for afum|EAL91792.1 as seed ortholog is 100%.
Bootstrap support for afis|EAW17983.1 as seed ortholog is 100%.

---

Group of orthologs #587. Best score 2390 bits
Score difference with first non-orthologous sequence - *A. fumigatus*:2390 *A. fischeri*:2390
afum|EAL87696.1 100.00% afis|EAW18331.1 100.00%
Bootstrap support for afum|EAL87696.1 as seed ortholog is 100%.
Bootstrap support for afis|EAW18331.1 as seed ortholog is 100%.

---

Group of orthologs #588. Best score 2386 bits
Score difference with first non-orthologous sequence - *A. fumigatus*:2386 *A. fischeri*:2386
afum|EAL87412.1 100.00% afis|EAW23982.1 100.00%
Bootstrap support for afum|EAL87412.1 as seed ortholog is 100%.
Bootstrap support for afis|EAW23982.1 as seed ortholog is 100%.

---

Group of orthologs #589. Best score 2385 bits
Score difference with first non-orthologous sequence - *A. fumigatus*:2385 *A. fischeri*:2385
afum|EAL92443.1 100.00% afis|EAW21428.1 100.00%
Bootstrap support for afum|EAL92443.1 as seed ortholog is 100%.
Bootstrap support for afis|EAW21428.1 as seed ortholog is 100%.

---

Group of orthologs #590. Best score 2385 bits
Score difference with first non-orthologous sequence - *A. fumigatus*:2385 *A. fischeri*:2385
afum|EBA27407.1 100.00% afis|EAW22807.1 100.00%
Bootstrap support for afum|EBA27407.1 as seed ortholog is 100%.
Bootstrap support for afis|EAW22807.1 as seed ortholog is 100%.

---

Group of orthologs #591. Best score 2384 bits
Score difference with first non-orthologous sequence - *A. fumigatus*:2384 *A. fischeri*:2290
afum|EAL88707.1 100.00% afis|EAW16075.1 100.00%
Bootstrap support for afum|EAL88707.1 as seed ortholog is 100%.
Bootstrap support for afis|EAW16075.1 as seed ortholog is 100%.

---

Group of orthologs #592. Best score 2384 bits
Score difference with first non-orthologous sequence - *A. fumigatus*:2384 *A. fischeri*:2384
afum|EAL89699.1 100.00% afis|EAW24999.1 100.00%
Bootstrap support for afum|EAL89699.1 as seed ortholog is 100%.
Bootstrap support for afis|EAW24999.1 as seed ortholog is 100%.

---

Group of orthologs #593. Best score 2382 bits
Score difference with first non-orthologous sequence - *A. fumigatus*:2382 *A. fischeri*:2382
afum|EAL92609.1 100.00% afis|EAW21591.1 100.00%
Bootstrap support for afum|EAL92609.1 as seed ortholog is 100%.
Bootstrap support for afis|EAW21591.1 as seed ortholog is 100%.

---

Group of orthologs #594. Best score 2381 bits
Score difference with first non-orthologous sequence - *A. fumigatus*:2381 *A. fischeri*:2381
afum|EAL87922.1 100.00% afis|EAW23544.1 100.00%
Bootstrap support for afum|EAL87922.1 as seed ortholog is 100%.
Bootstrap support for afis|EAW23544.1 as seed ortholog is 100%.

---

Group of orthologs #595. Best score 2373 bits

Score difference with first non-orthologous sequence - *A. fumigatus*:2373   *A. fischeri*:2373
afum|EAL87004.1 100.00% afis|EAW19490.1 100.00%
Bootstrap support for afum|EAL87004.1 as seed ortholog is 100%.
Bootstrap support for afis|EAW19490.1 as seed ortholog is 100%.

---

Group of orthologs #596. Best score 2373 bits
Score difference with first non-orthologous sequence - *A. fumigatus*:2373   *A. fischeri*:1985
afum|EAL87141.1 100.00% afis|EAW23700.1 100.00%
Bootstrap support for afum|EAL87141.1 as seed ortholog is 100%.
Bootstrap support for afis|EAW23700.1 as seed ortholog is 100%.

---

Group of orthologs #597. Best score 2372 bits
Score difference with first non-orthologous sequence - *A. fumigatus*:2372   *A. fischeri*:2372
afum|EAL88520.1 100.00% afis|EAW15886.1 100.00%
Bootstrap support for afum|EAL88520.1 as seed ortholog is 100%.
Bootstrap support for afis|EAW15886.1 as seed ortholog is 100%.

---

Group of orthologs #598. Best score 2372 bits
Score difference with first non-orthologous sequence - *A. fumigatus*:2372   *A. fischeri*:1046
afum|EAL84546.1 100.00% afis|EAW20510.1 100.00%
Bootstrap support for afum|EAL84546.1 as seed ortholog is 100%.
Bootstrap support for afis|EAW20510.1 as seed ortholog is 100%.

---

Group of orthologs #599. Best score 2372 bits
Score difference with first non-orthologous sequence - *A. fumigatus*:2372   *A. fischeri*:2372
afum|EAL88234.2 100.00% afis|EAW23247.1 100.00%
Bootstrap support for afum|EAL88234.2 as seed ortholog is 100%.
Bootstrap support for afis|EAW23247.1 as seed ortholog is 100%.

---

Group of orthologs #600. Best score 2372 bits
Score difference with first non-orthologous sequence - *A. fumigatus*:2372   *A. fischeri*:2372
afum|EAL92477.1 100.00% afis|EAW21458.1 100.00%
Bootstrap support for afum|EAL92477.1 as seed ortholog is 100%.
Bootstrap support for afis|EAW21458.1 as seed ortholog is 100%.

---

Group of orthologs #601. Best score 2371 bits
Score difference with first non-orthologous sequence - *A. fumigatus*:2371   *A. fischeri*:2371
afum|EAL92565.1 100.00% afis|EAW21545.1 100.00%
Bootstrap support for afum|EAL92565.1 as seed ortholog is 100%.
Bootstrap support for afis|EAW21545.1 as seed ortholog is 100%.

---

Group of orthologs #602. Best score 2369 bits
Score difference with first non-orthologous sequence - *A. fumigatus*:2369   *A. fischeri*:2369
afum|EAL91807.2 100.00% afis|EAW17997.1 100.00%
Bootstrap support for afum|EAL91807.2 as seed ortholog is 100%.
Bootstrap support for afis|EAW17997.1 as seed ortholog is 100%.

---

Group of orthologs #603. Best score 2368 bits
Score difference with first non-orthologous sequence - *A. fumigatus*:2368   *A. fischeri*:2368
afum|EAL91669.2 100.00% afis|EAW17861.1 100.00%
Bootstrap support for afum|EAL91669.2 as seed ortholog is 100%.
Bootstrap support for afis|EAW17861.1 as seed ortholog is 100%.

---

Group of orthologs #604. Best score 2368 bits
Score difference with first non-orthologous sequence - *A. fumigatus*:2368   *A. fischeri*:2368
afum|EAL93530.1 100.00% afis|EAW18794.1 100.00%

Bootstrap support for afum|EAL93530.1 as seed ortholog is 100%.
Bootstrap support for afis|EAW18794.1 as seed ortholog is 100%.

---

Group of orthologs #605. Best score 2368 bits
Score difference with first non-orthologous sequence - *A. fumigatus*:2368 *A. fischeri*:831
afum|EAL89773.1 100.00% afis|EAW25102.1 100.00%
Bootstrap support for afum|EAL89773.1 as seed ortholog is 100%.
Bootstrap support for afis|EAW25102.1 as seed ortholog is 100%.

---

Group of orthologs #606. Best score 2364 bits
Score difference with first non-orthologous sequence - *A. fumigatus*:2364 *A. fischeri*:2364
afum|EAL87533.1 100.00% afis|EAW18171.1 100.00%
Bootstrap support for afum|EAL87533.1 as seed ortholog is 100%.
Bootstrap support for afis|EAW18171.1 as seed ortholog is 100%.

---

Group of orthologs #607. Best score 2363 bits
Score difference with first non-orthologous sequence - *A. fumigatus*:2363 *A. fischeri*:2363
afum|EAL89628.1 100.00% afis|EAW24969.1 100.00%
Bootstrap support for afum|EAL89628.1 as seed ortholog is 100%.
Bootstrap support for afis|EAW24969.1 as seed ortholog is 100%.

---

Group of orthologs #608. Best score 2362 bits
Score difference with first non-orthologous sequence - *A. fumigatus*:2362 *A. fischeri*:2362
afum|EAL91215.1 100.00% afis|EAW17380.1 100.00%
Bootstrap support for afum|EAL91215.1 as seed ortholog is 100%.
Bootstrap support for afis|EAW17380.1 as seed ortholog is 100%.

---

Group of orthologs #609. Best score 2361 bits
Score difference with first non-orthologous sequence - *A. fumigatus*:2361 *A. fischeri*:2361
afum|EAL86322.1 100.00% afis|EAW16747.1 100.00%
Bootstrap support for afum|EAL86322.1 as seed ortholog is 100%.
Bootstrap support for afis|EAW16747.1 as seed ortholog is 100%.

---

Group of orthologs #610. Best score 2360 bits
Score difference with first non-orthologous sequence - *A. fumigatus*:2360 *A. fischeri*:2360
afum|EAL85168.2 100.00% afis|EAW20047.1 100.00%
Bootstrap support for afum|EAL85168.2 as seed ortholog is 100%.
Bootstrap support for afis|EAW20047.1 as seed ortholog is 100%.

---

Group of orthologs #611. Best score 2360 bits
Score difference with first non-orthologous sequence - *A. fumigatus*:2360 *A. fischeri*:2360
afum|EAL93636.1 100.00% afis|EAW18904.1 100.00%
Bootstrap support for afum|EAL93636.1 as seed ortholog is 100%.
Bootstrap support for afis|EAW18904.1 as seed ortholog is 100%.

---

Group of orthologs #612. Best score 2358 bits
Score difference with first non-orthologous sequence - *A. fumigatus*:2358 *A. fischeri*:2293
afum|EAL86551.2 100.00% afis|EAW17054.1 100.00%
Bootstrap support for afum|EAL86551.2 as seed ortholog is 100%.
Bootstrap support for afis|EAW17054.1 as seed ortholog is 100%.

---

Group of orthologs #613. Best score 2356 bits
Score difference with first non-orthologous sequence - *A. fumigatus*:2356 *A. fischeri*:2356
afum|EAL84393.2 100.00% afis|EAW25718.1 100.00%
Bootstrap support for afum|EAL84393.2 as seed ortholog is 100%.
Bootstrap support for afis|EAW25718.1 as seed ortholog is 100%.

---

Group of orthologs #614. Best score 2354 bits
Score difference with first non-orthologous sequence - *A. fumigatus*:2354 *A. fischeri*:2354
afum|EAL85243.1 100.00% afis|EAW20125.1 100.00%
Bootstrap support for afum|EAL85243.1 as seed ortholog is 100%.
Bootstrap support for afis|EAW20125.1 as seed ortholog is 100%.

---

Group of orthologs #615. Best score 2352 bits
Score difference with first non-orthologous sequence - *A. fumigatus*:2352 *A. fischeri*:2352
afum|EAL86053.1 100.00% afis|EAW24381.1 100.00%
Bootstrap support for afum|EAL86053.1 as seed ortholog is 100%.
Bootstrap support for afis|EAW24381.1 as seed ortholog is 100%.

---

Group of orthologs #616. Best score 2349 bits
Score difference with first non-orthologous sequence - *A. fumigatus*:2349 *A. fischeri*:2061
afum|EAL89843.1 100.00% afis|EAW25170.1 100.00%
Bootstrap support for afum|EAL89843.1 as seed ortholog is 100%.
Bootstrap support for afis|EAW25170.1 as seed ortholog is 100%.

---

Group of orthologs #617. Best score 2348 bits
Score difference with first non-orthologous sequence - *A. fumigatus*:2348 *A. fischeri*:2075
afum|EAL85956.1 100.00% afis|EAW24287.1 100.00%
Bootstrap support for afum|EAL85956.1 as seed ortholog is 100%.
Bootstrap support for afis|EAW24287.1 as seed ortholog is 100%.

---

Group of orthologs #618. Best score 2347 bits
Score difference with first non-orthologous sequence - *A. fumigatus*:2347 *A. fischeri*:2347
afum|EAL84365.1 100.00% afis|EAW21006.1 100.00%
Bootstrap support for afum|EAL84365.1 as seed ortholog is 100%.
Bootstrap support for afis|EAW21006.1 as seed ortholog is 100%.

---

Group of orthologs #619. Best score 2343 bits
Score difference with first non-orthologous sequence - *A. fumigatus*:2343 *A. fischeri*:2343
afum|EAL87485.1 100.00% afis|EAW18127.1 100.00%
Bootstrap support for afum|EAL87485.1 as seed ortholog is 100%.
Bootstrap support for afis|EAW18127.1 as seed ortholog is 100%.

---

Group of orthologs #620. Best score 2343 bits
Score difference with first non-orthologous sequence - *A. fumigatus*:2343 *A. fischeri*:2343
afum|EAL93470.1 100.00% afis|EAW18744.1 100.00%
Bootstrap support for afum|EAL93470.1 as seed ortholog is 100%.
Bootstrap support for afis|EAW18744.1 as seed ortholog is 100%.

---

Group of orthologs #621. Best score 2343 bits
Score difference with first non-orthologous sequence - *A. fumigatus*:2343 *A. fischeri*:2343
afum|EAL92189.2 100.00% afis|EAW21170.1 100.00%
Bootstrap support for afum|EAL92189.2 as seed ortholog is 100%.
Bootstrap support for afis|EAW21170.1 as seed ortholog is 100%.

---

Group of orthologs #622. Best score 2342 bits
Score difference with first non-orthologous sequence - *A. fumigatus*:2342 *A. fischeri*:2182
afum|EAL92809.1 100.00% afis|EAW21780.1 100.00%
Bootstrap support for afum|EAL92809.1 as seed ortholog is 100%.
Bootstrap support for afis|EAW21780.1 as seed ortholog is 100%.

---

Group of orthologs #623. Best score 2336 bits

Score difference with first non-orthologous sequence - *A. fumigatus*:2336   *A. fischeri*:2336
afum|EAL85844.1 100.00% afis|EAW15477.1 100.00%
Bootstrap support for afum|EAL85844.1 as seed ortholog is 100%.
Bootstrap support for afis|EAW15477.1 as seed ortholog is 100%.

---

Group of orthologs #624. Best score 2334 bits
Score difference with first non-orthologous sequence - *A. fumigatus*:988   *A. fischeri*:1457
afum|EAL85113.2 100.00% afis|EAW16804.1 100.00%
Bootstrap support for afum|EAL85113.2 as seed ortholog is 100%.
Bootstrap support for afis|EAW16804.1 as seed ortholog is 100%.

---

Group of orthologs #625. Best score 2332 bits
Score difference with first non-orthologous sequence - *A. fumigatus*:2332   *A. fischeri*:2332
afum|EAL87382.1 100.00% afis|EAW23954.1 100.00%
Bootstrap support for afum|EAL87382.1 as seed ortholog is 100%.
Bootstrap support for afis|EAW23954.1 as seed ortholog is 100%.

---

Group of orthologs #626. Best score 2324 bits
Score difference with first non-orthologous sequence - *A. fumigatus*:2324   *A. fischeri*:2324
afum|EAL87224.1 100.00% afis|EAW23790.1 100.00%
Bootstrap support for afum|EAL87224.1 as seed ortholog is 100%.
Bootstrap support for afis|EAW23790.1 as seed ortholog is 100%.

---

Group of orthologs #627. Best score 2321 bits
Score difference with first non-orthologous sequence - *A. fumigatus*:2321   *A. fischeri*:2321
afum|EBA27405.1 100.00% afis|EAW22820.1 100.00%
Bootstrap support for afum|EBA27405.1 as seed ortholog is 100%.
Bootstrap support for afis|EAW22820.1 as seed ortholog is 100%.

---

Group of orthologs #628. Best score 2320 bits
Score difference with first non-orthologous sequence - *A. fumigatus*:2320   *A. fischeri*:2320
afum|EAL92278.1 100.00% afis|EAW21263.1 100.00%
Bootstrap support for afum|EAL92278.1 as seed ortholog is 100%.
Bootstrap support for afis|EAW21263.1 as seed ortholog is 100%.

---

Group of orthologs #629. Best score 2319 bits
Score difference with first non-orthologous sequence - *A. fumigatus*:2319   *A. fischeri*:2319
afum|EAL87623.1 100.00% afis|EAW18260.1 100.00%
Bootstrap support for afum|EAL87623.1 as seed ortholog is 100%.
Bootstrap support for afis|EAW18260.1 as seed ortholog is 100%.

---

Group of orthologs #630. Best score 2319 bits
Score difference with first non-orthologous sequence - *A. fumigatus*:2319   *A. fischeri*:2319
afum|EAL88415.1 100.00% afis|EAW23064.1 100.00%
Bootstrap support for afum|EAL88415.1 as seed ortholog is 100%.
Bootstrap support for afis|EAW23064.1 as seed ortholog is 100%.

---

Group of orthologs #631. Best score 2319 bits
Score difference with first non-orthologous sequence - *A. fumigatus*:2319   *A. fischeri*:2319
afum|EAL91994.1 100.00% afis|EAW24139.1 100.00%
Bootstrap support for afum|EAL91994.1 as seed ortholog is 100%.
Bootstrap support for afis|EAW24139.1 as seed ortholog is 100%.

---

Group of orthologs #632. Best score 2318 bits
Score difference with first non-orthologous sequence - *A. fumigatus*:2318   *A. fischeri*:2318
afum|EAL92726.1 100.00% afis|EAW21692.1 100.00%

Bootstrap support for afum|EAL92726.1 as seed ortholog is 100%.
Bootstrap support for afis|EAW21692.1 as seed ortholog is 100%.

---

Group of orthologs #633. Best score 2317 bits
Score difference with first non-orthologous sequence - *A. fumigatus*:2317 *A. fischeri*:2317
afum|EAL91408.1 100.00% afis|EAW17595.1 100.00%
Bootstrap support for afum|EAL91408.1 as seed ortholog is 100%.
Bootstrap support for afis|EAW17595.1 as seed ortholog is 100%.

---

Group of orthologs #634. Best score 2317 bits
Score difference with first non-orthologous sequence - *A. fumigatus*:2317 *A. fischeri*:2317
afum|EBA27217.1 100.00% afis|EAW19848.1 100.00%
Bootstrap support for afum|EBA27217.1 as seed ortholog is 100%.
Bootstrap support for afis|EAW19848.1 as seed ortholog is 100%.

---

Group of orthologs #635. Best score 2316 bits
Score difference with first non-orthologous sequence - *A. fumigatus*:2316 *A. fischeri*:2316
afum|EAL91617.1 100.00% afis|EAW17809.1 100.00%
Bootstrap support for afum|EAL91617.1 as seed ortholog is 100%.
Bootstrap support for afis|EAW17809.1 as seed ortholog is 100%.

---

Group of orthologs #636. Best score 2314 bits
Score difference with first non-orthologous sequence - *A. fumigatus*:2314 *A. fischeri*:2314
afum|EAL87385.1 100.00% afis|EAW23957.1 100.00%
Bootstrap support for afum|EAL87385.1 as seed ortholog is 100%.
Bootstrap support for afis|EAW23957.1 as seed ortholog is 100%.

---

Group of orthologs #637. Best score 2306 bits
Score difference with first non-orthologous sequence - *A. fumigatus*:2306 *A. fischeri*:2306
afum|EAL92998.1 100.00% afis|EAW21983.1 100.00%
Bootstrap support for afum|EAL92998.1 as seed ortholog is 100%.
Bootstrap support for afis|EAW21983.1 as seed ortholog is 100%.

---

Group of orthologs #638. Best score 2305 bits
Score difference with first non-orthologous sequence - *A. fumigatus*:2305 *A. fischeri*:2305
afum|EBA27293.1 100.00% afis|EAW19625.1 100.00%
Bootstrap support for afum|EBA27293.1 as seed ortholog is 100%.
Bootstrap support for afis|EAW19625.1 as seed ortholog is 100%.

---

Group of orthologs #639. Best score 2304 bits
Score difference with first non-orthologous sequence - *A. fumigatus*:2304 *A. fischeri*:2304
afum|EAL93817.1 100.00% afis|EAW19087.1 100.00%
Bootstrap support for afum|EAL93817.1 as seed ortholog is 100%.
Bootstrap support for afis|EAW19087.1 as seed ortholog is 100%.

---

Group of orthologs #640. Best score 2303 bits
Score difference with first non-orthologous sequence - *A. fumigatus*:2303 *A. fischeri*:2303
afum|EAL87487.1 100.00% afis|EAW18130.1 100.00%
Bootstrap support for afum|EAL87487.1 as seed ortholog is 100%.
Bootstrap support for afis|EAW18130.1 as seed ortholog is 100%.

---

Group of orthologs #641. Best score 2303 bits
Score difference with first non-orthologous sequence - *A. fumigatus*:2303 *A. fischeri*:2303
afum|EAL92203.1 100.00% afis|EAW21182.1 100.00%
Bootstrap support for afum|EAL92203.1 as seed ortholog is 100%.
Bootstrap support for afis|EAW21182.1 as seed ortholog is 100%.

---
Group of orthologs #642. Best score 2302 bits
Score difference with first non-orthologous sequence - *A. fumigatus*:2302 *A. fischeri*:2302
afum|EAL85157.1 100.00% afis|EAW20036.1 100.00%
Bootstrap support for afum|EAL85157.1 as seed ortholog is 100%.
Bootstrap support for afis|EAW20036.1 as seed ortholog is 100%.

---
Group of orthologs #643. Best score 2302 bits
Score difference with first non-orthologous sequence - *A. fumigatus*:2302 *A. fischeri*:2302
afum|EAL91519.2 100.00% afis|EAW17704.1 100.00%
Bootstrap support for afum|EAL91519.2 as seed ortholog is 100%.
Bootstrap support for afis|EAW17704.1 as seed ortholog is 100%.

---
Group of orthologs #644. Best score 2300 bits
Score difference with first non-orthologous sequence - *A. fumigatus*:2300 *A. fischeri*:2300
afum|EBA27469.1 100.00% afis|EAW21652.1 100.00%
Bootstrap support for afum|EBA27469.1 as seed ortholog is 100%.
Bootstrap support for afis|EAW21652.1 as seed ortholog is 100%.

---
Group of orthologs #645. Best score 2299 bits
Score difference with first non-orthologous sequence - *A. fumigatus*:2299 *A. fischeri*:2299
afum|EAL92923.1 100.00% afis|EAW21911.1 100.00%
Bootstrap support for afum|EAL92923.1 as seed ortholog is 100%.
Bootstrap support for afis|EAW21911.1 as seed ortholog is 100%.

---
Group of orthologs #646. Best score 2298 bits
Score difference with first non-orthologous sequence - *A. fumigatus*:2298 *A. fischeri*:2298
afum|EAL88285.1 100.00% afis|EAW23198.1 100.00%
Bootstrap support for afum|EAL88285.1 as seed ortholog is 100%.
Bootstrap support for afis|EAW23198.1 as seed ortholog is 100%.

---
Group of orthologs #647. Best score 2298 bits
Score difference with first non-orthologous sequence - *A. fumigatus*:2298 *A. fischeri*:2298
afum|EAL92415.1 100.00% afis|EAW21401.1 100.00%
Bootstrap support for afum|EAL92415.1 as seed ortholog is 100%.
Bootstrap support for afis|EAW21401.1 as seed ortholog is 100%.

---
Group of orthologs #648. Best score 2298 bits
Score difference with first non-orthologous sequence - *A. fumigatus*:2298 *A. fischeri*:2298
afum|EAL91953.2 100.00% afis|EAW24099.1 100.00%
Bootstrap support for afum|EAL91953.2 as seed ortholog is 100%.
Bootstrap support for afis|EAW24099.1 as seed ortholog is 100%.

---
Group of orthologs #649. Best score 2296 bits
Score difference with first non-orthologous sequence - *A. fumigatus*:2296 *A. fischeri*:2296
afum|EAL91950.1 100.00% afis|EAW24096.1 100.00%
Bootstrap support for afum|EAL91950.1 as seed ortholog is 100%.
Bootstrap support for afis|EAW24096.1 as seed ortholog is 100%.

---
Group of orthologs #650. Best score 2295 bits
Score difference with first non-orthologous sequence - *A. fumigatus*:2295 *A. fischeri*:2295
afum|EAL84592.1 100.00% afis|EAW20461.1 100.00%
Bootstrap support for afum|EAL84592.1 as seed ortholog is 100%.
Bootstrap support for afis|EAW20461.1 as seed ortholog is 100%.

---
Group of orthologs #651. Best score 2294 bits

Score difference with first non-orthologous sequence - *A. fumigatus*:2294 *A. fischeri*:2294
afum|EAL92449.1 100.00% afis|EAW21434.1 100.00%
Bootstrap support for afum|EAL92449.1 as seed ortholog is 100%.
Bootstrap support for afis|EAW21434.1 as seed ortholog is 100%.

---

Group of orthologs #652. Best score 2294 bits
Score difference with first non-orthologous sequence - *A. fumigatus*:2294 *A. fischeri*:2294
afum|EAL92607.1 100.00% afis|EAW21589.1 100.00%
Bootstrap support for afum|EAL92607.1 as seed ortholog is 100%.
Bootstrap support for afis|EAW21589.1 as seed ortholog is 100%.

---

Group of orthologs #653. Best score 2291 bits
Score difference with first non-orthologous sequence - *A. fumigatus*:2291 *A. fischeri*:2291
afum|EAL93851.1 100.00% afis|EAW19119.1 100.00%
Bootstrap support for afum|EAL93851.1 as seed ortholog is 100%.
Bootstrap support for afis|EAW19119.1 as seed ortholog is 100%.

---

Group of orthologs #654. Best score 2291 bits
Score difference with first non-orthologous sequence - *A. fumigatus*:2291 *A. fischeri*:2291
afum|EAL89909.1 100.00% afis|EAW25247.1 100.00%
Bootstrap support for afum|EAL89909.1 as seed ortholog is 100%.
Bootstrap support for afis|EAW25247.1 as seed ortholog is 100%.

---

Group of orthologs #655. Best score 2288 bits
Score difference with first non-orthologous sequence - *A. fumigatus*:2288 *A. fischeri*:2288
afum|EAL85787.1 100.00% afis|EAW15544.1 100.00%
Bootstrap support for afum|EAL85787.1 as seed ortholog is 100%.
Bootstrap support for afis|EAW15544.1 as seed ortholog is 100%.

---

Group of orthologs #656. Best score 2288 bits
Score difference with first non-orthologous sequence - *A. fumigatus*:2288 *A. fischeri*:2288
afum|EAL90867.1 100.00% afis|EAW22325.1 100.00%
Bootstrap support for afum|EAL90867.1 as seed ortholog is 100%.
Bootstrap support for afis|EAW22325.1 as seed ortholog is 100%.

---

Group of orthologs #657. Best score 2286 bits
Score difference with first non-orthologous sequence - *A. fumigatus*:2286 *A. fischeri*:2286
afum|EAL92556.2 100.00% afis|EAW21536.1 100.00%
Bootstrap support for afum|EAL92556.2 as seed ortholog is 100%.
Bootstrap support for afis|EAW21536.1 as seed ortholog is 100%.

---

Group of orthologs #658. Best score 2285 bits
Score difference with first non-orthologous sequence - *A. fumigatus*:2285 *A. fischeri*:2285
afum|EAL90987.2 100.00% afis|EAW22206.1 100.00%
Bootstrap support for afum|EAL90987.2 as seed ortholog is 100%.
Bootstrap support for afis|EAW22206.1 as seed ortholog is 100%.

---

Group of orthologs #659. Best score 2281 bits
Score difference with first non-orthologous sequence - *A. fumigatus*:2281 *A. fischeri*:2281
afum|EAL87882.1 100.00% afis|EAW23582.1 100.00%
Bootstrap support for afum|EAL87882.1 as seed ortholog is 100%.
Bootstrap support for afis|EAW23582.1 as seed ortholog is 100%.

---

Group of orthologs #660. Best score 2278 bits
Score difference with first non-orthologous sequence - *A. fumigatus*:2278 *A. fischeri*:2278
afum|EAL90221.1 100.00% afis|EAW22976.1 100.00%

Bootstrap support for afum|EAL90221.1 as seed ortholog is 100%.
Bootstrap support for afis|EAW22976.1 as seed ortholog is 100%.

---

Group of orthologs #661. Best score 2276 bits
Score difference with first non-orthologous sequence - *A. fumigatus*:2276 *A. fischeri*:2276
afum|EAL85866.1 100.00% afis|EAW24194.1 100.00%
Bootstrap support for afum|EAL85866.1 as seed ortholog is 100%.
Bootstrap support for afis|EAW24194.1 as seed ortholog is 100%.

---

Group of orthologs #662. Best score 2274 bits
Score difference with first non-orthologous sequence - *A. fumigatus*:2274 *A. fischeri*:2274
afum|EAL85549.2 100.00% afis|EAW15801.1 100.00%
Bootstrap support for afum|EAL85549.2 as seed ortholog is 100%.
Bootstrap support for afis|EAW15801.1 as seed ortholog is 100%.

---

Group of orthologs #663. Best score 2272 bits
Score difference with first non-orthologous sequence - *A. fumigatus*:2272 *A. fischeri*:2272
afum|EAL87969.1 100.00% afis|EAW23494.1 100.00%
Bootstrap support for afum|EAL87969.1 as seed ortholog is 100%.
Bootstrap support for afis|EAW23494.1 as seed ortholog is 100%.

---

Group of orthologs #664. Best score 2272 bits
Score difference with first non-orthologous sequence - *A. fumigatus*:2272 *A. fischeri*:2272
afum|EAL93765.2 100.00% afis|EAW19036.1 100.00%
Bootstrap support for afum|EAL93765.2 as seed ortholog is 100%.
Bootstrap support for afis|EAW19036.1 as seed ortholog is 100%.

---

Group of orthologs #665. Best score 2271 bits
Score difference with first non-orthologous sequence - *A. fumigatus*:2271 *A. fischeri*:2271
afum|EBA27315.1 100.00% afis|EAW18169.1 100.00%
Bootstrap support for afum|EBA27315.1 as seed ortholog is 100%.
Bootstrap support for afis|EAW18169.1 as seed ortholog is 100%.

---

Group of orthologs #666. Best score 2269 bits
Score difference with first non-orthologous sequence - *A. fumigatus*:2026 *A. fischeri*:2269
afum|EAL84317.1 100.00% afis|EAW25629.1 100.00%
Bootstrap support for afum|EAL84317.1 as seed ortholog is 100%.
Bootstrap support for afis|EAW25629.1 as seed ortholog is 100%.

---

Group of orthologs #667. Best score 2266 bits
Score difference with first non-orthologous sequence - *A. fumigatus*:2266 *A. fischeri*:2266
afum|EAL85638.1 100.00% afis|EAW15709.1 100.00%
Bootstrap support for afum|EAL85638.1 as seed ortholog is 100%.
Bootstrap support for afis|EAW15709.1 as seed ortholog is 100%.

---

Group of orthologs #668. Best score 2265 bits
Score difference with first non-orthologous sequence - *A. fumigatus*:2265 *A. fischeri*:2265
afum|EAL91819.1 100.00% afis|EAW18007.1 100.00%
Bootstrap support for afum|EAL91819.1 as seed ortholog is 100%.
Bootstrap support for afis|EAW18007.1 as seed ortholog is 100%.

---

Group of orthologs #669. Best score 2265 bits
Score difference with first non-orthologous sequence - *A. fumigatus*:2265 *A. fischeri*:2265
afum|EAL92978.2 100.00% afis|EAW21964.1 100.00%
Bootstrap support for afum|EAL92978.2 as seed ortholog is 100%.
Bootstrap support for afis|EAW21964.1 as seed ortholog is 100%.

---

Group of orthologs #670. Best score 2263 bits
Score difference with first non-orthologous sequence - *A. fumigatus*:2263 *A. fischeri*:2263
afum|EAL93977.1 100.00% afis|EAW19248.1 100.00%
Bootstrap support for afum|EAL93977.1 as seed ortholog is 100%.
Bootstrap support for afis|EAW19248.1 as seed ortholog is 100%.

---

Group of orthologs #671. Best score 2262 bits
Score difference with first non-orthologous sequence - *A. fumigatus*:2262 *A. fischeri*:2262
afum|EAL85611.1 100.00% afis|EAW15737.1 100.00%
Bootstrap support for afum|EAL85611.1 as seed ortholog is 100%.
Bootstrap support for afis|EAW15737.1 as seed ortholog is 100%.

---

Group of orthologs #672. Best score 2262 bits
Score difference with first non-orthologous sequence - *A. fumigatus*:2262 *A. fischeri*:2262
afum|EAL87980.1 100.00% afis|EAW23483.1 100.00%
Bootstrap support for afum|EAL87980.1 as seed ortholog is 100%.
Bootstrap support for afis|EAW23483.1 as seed ortholog is 100%.

---

Group of orthologs #673. Best score 2260 bits
Score difference with first non-orthologous sequence - *A. fumigatus*:2260 *A. fischeri*:2029
afum|EAL86228.1 100.00% afis|EAW24565.1 100.00%
Bootstrap support for afum|EAL86228.1 as seed ortholog is 100%.
Bootstrap support for afis|EAW24565.1 as seed ortholog is 100%.

---

Group of orthologs #674. Best score 2259 bits
Score difference with first non-orthologous sequence - *A. fumigatus*:2259 *A. fischeri*:2259
afum|EAL90918.1 100.00% afis|EAW22262.1 100.00%
Bootstrap support for afum|EAL90918.1 as seed ortholog is 100%.
Bootstrap support for afis|EAW22262.1 as seed ortholog is 100%.

---

Group of orthologs #675. Best score 2259 bits
Score difference with first non-orthologous sequence - *A. fumigatus*:2259 *A. fischeri*:2259
afum|EAL89963.2 100.00% afis|EAW25300.1 100.00%
Bootstrap support for afum|EAL89963.2 as seed ortholog is 100%.
Bootstrap support for afis|EAW25300.1 as seed ortholog is 100%.

---

Group of orthologs #676. Best score 2256 bits
Score difference with first non-orthologous sequence - *A. fumigatus*:2256 *A. fischeri*:2256
afum|EAL91210.1 100.00% afis|EAW17374.1 100.00%
Bootstrap support for afum|EAL91210.1 as seed ortholog is 100%.
Bootstrap support for afis|EAW17374.1 as seed ortholog is 100%.

---

Group of orthologs #677. Best score 2256 bits
Score difference with first non-orthologous sequence - *A. fumigatus*:2256 *A. fischeri*:2256
afum|EAL90450.1 100.00% afis|EAW22749.1 100.00%
Bootstrap support for afum|EAL90450.1 as seed ortholog is 100%.
Bootstrap support for afis|EAW22749.1 as seed ortholog is 100%.

---

Group of orthologs #678. Best score 2255 bits
Score difference with first non-orthologous sequence - *A. fumigatus*:2255 *A. fischeri*:2255
afum|EAL88976.1 100.00% afis|EAW16351.1 100.00%
Bootstrap support for afum|EAL88976.1 as seed ortholog is 100%.
Bootstrap support for afis|EAW16351.1 as seed ortholog is 100%.

---

Group of orthologs #679. Best score 2254 bits

Score difference with first non-orthologous sequence - *A. fumigatus*:2254   *A. fischeri*:2254
afum|EAL87374.1    100.00%    afis|EAW23948.1    100.00%
Bootstrap support for afum|EAL87374.1 as seed ortholog is 100%.
Bootstrap support for afis|EAW23948.1 as seed ortholog is 100%.

---

Group of orthologs #680. Best score 2253 bits
Score difference with first non-orthologous sequence - *A. fumigatus*:2253   *A. fischeri*:1901
afum|EAL88529.1    100.00%    afis|EAW15894.1    100.00%
Bootstrap support for afum|EAL88529.1 as seed ortholog is 100%.
Bootstrap support for afis|EAW15894.1 as seed ortholog is 100%.

---

Group of orthologs #681. Best score 2253 bits
Score difference with first non-orthologous sequence - *A. fumigatus*:2253   *A. fischeri*:2253
afum|EAL90033.1    100.00%    afis|EAW25373.1    100.00%
Bootstrap support for afum|EAL90033.1 as seed ortholog is 100%.
Bootstrap support for afis|EAW25373.1 as seed ortholog is 100%.

---

Group of orthologs #682. Best score 2252 bits
Score difference with first non-orthologous sequence - *A. fumigatus*:2252   *A. fischeri*:2252
afum|EAL92216.1    100.00%    afis|EAW21205.1    100.00%
Bootstrap support for afum|EAL92216.1 as seed ortholog is 100%.
Bootstrap support for afis|EAW21205.1 as seed ortholog is 100%.

---

Group of orthologs #683. Best score 2251 bits
Score difference with first non-orthologous sequence - *A. fumigatus*:2251   *A. fischeri*:2251
afum|EBA27384.1    100.00%    afis|EAW24972.1    100.00%
Bootstrap support for afum|EBA27384.1 as seed ortholog is 100%.
Bootstrap support for afis|EAW24972.1 as seed ortholog is 100%.

---

Group of orthologs #684. Best score 2250 bits
Score difference with first non-orthologous sequence - *A. fumigatus*:2250   *A. fischeri*:2250
afum|EAL92520.1    100.00%    afis|EAW21501.1    100.00%
Bootstrap support for afum|EAL92520.1 as seed ortholog is 100%.
Bootstrap support for afis|EAW21501.1 as seed ortholog is 100%.

---

Group of orthologs #685. Best score 2250 bits
Score difference with first non-orthologous sequence - *A. fumigatus*:2250   *A. fischeri*:2250
afum|EAL89549.1    100.00%    afis|EAW24895.1    100.00%
Bootstrap support for afum|EAL89549.1 as seed ortholog is 100%.
Bootstrap support for afis|EAW24895.1 as seed ortholog is 100%.

---

Group of orthologs #686. Best score 2245 bits
Score difference with first non-orthologous sequence - *A. fumigatus*:2245   *A. fischeri*:2245
afum|EAL89075.1    100.00%    afis|EAW16481.1    100.00%
Bootstrap support for afum|EAL89075.1 as seed ortholog is 100%.
Bootstrap support for afis|EAW16481.1 as seed ortholog is 100%.

---

Group of orthologs #687. Best score 2244 bits
Score difference with first non-orthologous sequence - *A. fumigatus*:2244   *A. fischeri*:2244
afum|EAL86875.1    100.00%    afis|EAW19647.1    100.00%
Bootstrap support for afum|EAL86875.1 as seed ortholog is 100%.
Bootstrap support for afis|EAW19647.1 as seed ortholog is 100%.

---

Group of orthologs #688. Best score 2244 bits
Score difference with first non-orthologous sequence - *A. fumigatus*:2244   *A. fischeri*:2244
afum|EAL88308.1    100.00%    afis|EAW23175.1    100.00%

Bootstrap support for afum|EAL88308.1 as seed ortholog is 100%.
Bootstrap support for afis|EAW23175.1 as seed ortholog is 100%.

---

Group of orthologs #689. Best score 2242 bits
Score difference with first non-orthologous sequence - *A. fumigatus*:2242 *A. fischeri*:2242
afum|EAL84578.1 100.00% afis|EAW20475.1 100.00%
Bootstrap support for afum|EAL84578.1 as seed ortholog is 100%.
Bootstrap support for afis|EAW20475.1 as seed ortholog is 100%.

---

Group of orthologs #690. Best score 2242 bits
Score difference with first non-orthologous sequence - *A. fumigatus*:2242 *A. fischeri*:2242
afum|EAL93007.1 100.00% afis|EAW21992.1 100.00%
Bootstrap support for afum|EAL93007.1 as seed ortholog is 100%.
Bootstrap support for afis|EAW21992.1 as seed ortholog is 100%.

---

Group of orthologs #691. Best score 2239 bits
Score difference with first non-orthologous sequence - *A. fumigatus*:2239 *A. fischeri*:2239
afum|EAL90424.1 100.00% afis|EAW22774.1 100.00%
Bootstrap support for afum|EAL90424.1 as seed ortholog is 100%.
Bootstrap support for afis|EAW22774.1 as seed ortholog is 100%.

---

Group of orthologs #692. Best score 2238 bits
Score difference with first non-orthologous sequence - *A. fumigatus*:2238 *A. fischeri*:2238
afum|EAL89123.2 100.00% afis|EAW16527.1 100.00%
Bootstrap support for afum|EAL89123.2 as seed ortholog is 100%.
Bootstrap support for afis|EAW16527.1 as seed ortholog is 100%.

---

Group of orthologs #693. Best score 2237 bits
Score difference with first non-orthologous sequence - *A. fumigatus*:2237 *A. fischeri*:2237
afum|EAL90836.1 100.00% afis|EAW22355.1 100.00%
Bootstrap support for afum|EAL90836.1 as seed ortholog is 100%.
Bootstrap support for afis|EAW22355.1 as seed ortholog is 100%.

---

Group of orthologs #694. Best score 2237 bits
Score difference with first non-orthologous sequence - *A. fumigatus*:2237 *A. fischeri*:886
afum|EAL92146.1 100.00% afis|EAW21123.1 100.00%
Bootstrap support for afum|EAL92146.1 as seed ortholog is 100%.
Bootstrap support for afis|EAW21123.1 as seed ortholog is 100%.

---

Group of orthologs #695. Best score 2237 bits
Score difference with first non-orthologous sequence - *A. fumigatus*:2237 *A. fischeri*:2237
afum|EAL92712.1 100.00% afis|EAW21678.1 100.00%
Bootstrap support for afum|EAL92712.1 as seed ortholog is 100%.
Bootstrap support for afis|EAW21678.1 as seed ortholog is 100%.

---

Group of orthologs #696. Best score 2237 bits
Score difference with first non-orthologous sequence - *A. fumigatus*:2237 *A. fischeri*:2237
afum|EAL90062.1 100.00% afis|EAW25403.1 100.00%
Bootstrap support for afum|EAL90062.1 as seed ortholog is 100%.
Bootstrap support for afis|EAW25403.1 as seed ortholog is 100%.

---

Group of orthologs #697. Best score 2236 bits
Score difference with first non-orthologous sequence - *A. fumigatus*:2236 *A. fischeri*:2236
afum|EAL88669.1 100.00% afis|EAW16038.1 100.00%
Bootstrap support for afum|EAL88669.1 as seed ortholog is 100%.
Bootstrap support for afis|EAW16038.1 as seed ortholog is 100%.

---

Group of orthologs #698. Best score 2234 bits
Score difference with first non-orthologous sequence - *A. fumigatus*:2234 *A. fischeri*:2234
afum|EAL88611.1 100.00% afis|EAW15979.1 100.00%
Bootstrap support for afum|EAL88611.1 as seed ortholog is 100%.
Bootstrap support for afis|EAW15979.1 as seed ortholog is 100%.

---

Group of orthologs #699. Best score 2234 bits
Score difference with first non-orthologous sequence - *A. fumigatus*:2234 *A. fischeri*:2234
afum|EAL93669.2 100.00% afis|EAW18936.1 100.00%
Bootstrap support for afum|EAL93669.2 as seed ortholog is 100%.
Bootstrap support for afis|EAW18936.1 as seed ortholog is 100%.

---

Group of orthologs #700. Best score 2234 bits
Score difference with first non-orthologous sequence - *A. fumigatus*:2234 *A. fischeri*:1484
afum|EAL90777.1 100.00% afis|EAW22415.1 100.00%
Bootstrap support for afum|EAL90777.1 as seed ortholog is 100%.
Bootstrap support for afis|EAW22415.1 as seed ortholog is 100%.

---

Group of orthologs #701. Best score 2233 bits
Score difference with first non-orthologous sequence - *A. fumigatus*:2233 *A. fischeri*:2233
afum|EAL84534.1 100.00% afis|EAW20523.1 100.00%
Bootstrap support for afum|EAL84534.1 as seed ortholog is 100%.
Bootstrap support for afis|EAW20523.1 as seed ortholog is 100%.

---

Group of orthologs #702. Best score 2233 bits
Score difference with first non-orthologous sequence - *A. fumigatus*:2233 *A. fischeri*:2233
afum|EAL85175.2 100.00% afis|EAW20055.1 100.00%
Bootstrap support for afum|EAL85175.2 as seed ortholog is 100%.
Bootstrap support for afis|EAW20055.1 as seed ortholog is 100%.

---

Group of orthologs #703. Best score 2232 bits
Score difference with first non-orthologous sequence - *A. fumigatus*:2232 *A. fischeri*:2232
afum|EAL90184.1 100.00% afis|EAW25519.1 100.00%
Bootstrap support for afum|EAL90184.1 as seed ortholog is 100%.
Bootstrap support for afis|EAW25519.1 as seed ortholog is 100%.

---

Group of orthologs #704. Best score 2231 bits
Score difference with first non-orthologous sequence - *A. fumigatus*:2231 *A. fischeri*:1583
afum|EAL87284.1 100.00% afis|EAW23856.1 100.00%
Bootstrap support for afum|EAL87284.1 as seed ortholog is 100%.
Bootstrap support for afis|EAW23856.1 as seed ortholog is 100%.

---

Group of orthologs #705. Best score 2229 bits
Score difference with first non-orthologous sequence - *A. fumigatus*:2229 *A. fischeri*:2229
afum|EAL89194.1 100.00% afis|EAW16598.1 100.00%
Bootstrap support for afum|EAL89194.1 as seed ortholog is 100%.
Bootstrap support for afis|EAW16598.1 as seed ortholog is 100%.

---

Group of orthologs #706. Best score 2229 bits
Score difference with first non-orthologous sequence - *A. fumigatus*:2229 *A. fischeri*:2229
afum|EAL87570.1 100.00% afis|EAW18205.1 100.00%
Bootstrap support for afum|EAL87570.1 as seed ortholog is 100%.
Bootstrap support for afis|EAW18205.1 as seed ortholog is 100%.

---

Group of orthologs #707. Best score 2229 bits

Score difference with first non-orthologous sequence - *A. fumigatus*:2229 *A. fischeri*:2229
afum|EAL93894.1 100.00% afis|EAW19164.1 100.00%
Bootstrap support for afum|EAL93894.1 as seed ortholog is 100%.
Bootstrap support for afis|EAW19164.1 as seed ortholog is 100%.

---

Group of orthologs #708. Best score 2225 bits
Score difference with first non-orthologous sequence - *A. fumigatus*:2225 *A. fischeri*:2225
afum|EAL85435.2 100.00% afis|EAW20307.1 100.00%
Bootstrap support for afum|EAL85435.2 as seed ortholog is 100%.
Bootstrap support for afis|EAW20307.1 as seed ortholog is 100%.

---

Group of orthologs #709. Best score 2225 bits
Score difference with first non-orthologous sequence - *A. fumigatus*:2225 *A. fischeri*:2225
afum|EAL88363.1 100.00% afis|EAW23117.1 100.00%
Bootstrap support for afum|EAL88363.1 as seed ortholog is 100%.
Bootstrap support for afis|EAW23117.1 as seed ortholog is 100%.

---

Group of orthologs #710. Best score 2221 bits
Score difference with first non-orthologous sequence - *A. fumigatus*:2221 *A. fischeri*:2221
afum|EAL87076.1 100.00% afis|EAW19418.1 100.00%
Bootstrap support for afum|EAL87076.1 as seed ortholog is 100%.
Bootstrap support for afis|EAW19418.1 as seed ortholog is 100%.

---

Group of orthologs #711. Best score 2221 bits
Score difference with first non-orthologous sequence - *A. fumigatus*:2221 *A. fischeri*:2221
afum|EAL88138.1 100.00% afis|EAW23340.1 100.00%
Bootstrap support for afum|EAL88138.1 as seed ortholog is 100%.
Bootstrap support for afis|EAW23340.1 as seed ortholog is 100%.

---

Group of orthologs #712. Best score 2221 bits
Score difference with first non-orthologous sequence - *A. fumigatus*:2221 *A. fischeri*:2221
afum|EAL93795.1 100.00% afis|EAW19066.1 100.00%
Bootstrap support for afum|EAL93795.1 as seed ortholog is 100%.
Bootstrap support for afis|EAW19066.1 as seed ortholog is 100%.

---

Group of orthologs #713. Best score 2221 bits
Score difference with first non-orthologous sequence - *A. fumigatus*:2221 *A. fischeri*:2221
afum|EAL93980.1 100.00% afis|EAW19251.1 100.00%
Bootstrap support for afum|EAL93980.1 as seed ortholog is 100%.
Bootstrap support for afis|EAW19251.1 as seed ortholog is 100%.

---

Group of orthologs #714. Best score 2221 bits
Score difference with first non-orthologous sequence - *A. fumigatus*:2221 *A. fischeri*:2221
afum|EAL92596.1 100.00% afis|EAW21578.1 100.00%
Bootstrap support for afum|EAL92596.1 as seed ortholog is 100%.
Bootstrap support for afis|EAW21578.1 as seed ortholog is 100%.

---

Group of orthologs #715. Best score 2221 bits
Score difference with first non-orthologous sequence - *A. fumigatus*:2221 *A. fischeri*:2221
afum|EAL89882.1 100.00% afis|EAW25221.1 100.00%
Bootstrap support for afum|EAL89882.1 as seed ortholog is 100%.
Bootstrap support for afis|EAW25221.1 as seed ortholog is 100%.

---

Group of orthologs #716. Best score 2220 bits
Score difference with first non-orthologous sequence - *A. fumigatus*:1926 *A. fischeri*:2220
afum|EAL87088.1 100.00% afis|EAW19406.1 100.00%

Bootstrap support for afum|EAL87088.1 as seed ortholog is 100%.
Bootstrap support for afis|EAW19406.1 as seed ortholog is 100%.

---

Group of orthologs #717. Best score 2218 bits
Score difference with first non-orthologous sequence - *A. fumigatus*:2218 *A. fischeri*:2218
afum|EAL87082.1 100.00% afis|EAW19412.1 100.00%
Bootstrap support for afum|EAL87082.1 as seed ortholog is 100%.
Bootstrap support for afis|EAW19412.1 as seed ortholog is 100%.

---

Group of orthologs #718. Best score 2218 bits
Score difference with first non-orthologous sequence - *A. fumigatus*:2218 *A. fischeri*:2218
afum|EAL93585.1 100.00% afis|EAW18854.1 100.00%
Bootstrap support for afum|EAL93585.1 as seed ortholog is 100%.
Bootstrap support for afis|EAW18854.1 as seed ortholog is 100%.

---

Group of orthologs #719. Best score 2212 bits
Score difference with first non-orthologous sequence - *A. fumigatus*:2212 *A. fischeri*:2212
afum|EAL88133.1 100.00% afis|EAW23344.1 100.00%
Bootstrap support for afum|EAL88133.1 as seed ortholog is 100%.
Bootstrap support for afis|EAW23344.1 as seed ortholog is 100%.

---

Group of orthologs #720. Best score 2212 bits
Score difference with first non-orthologous sequence - *A. fumigatus*:2212 *A. fischeri*:2212
afum|EAL92434.1 100.00% afis|EAW21419.1 100.00%
Bootstrap support for afum|EAL92434.1 as seed ortholog is 100%.
Bootstrap support for afis|EAW21419.1 as seed ortholog is 100%.

---

Group of orthologs #721. Best score 2211 bits
Score difference with first non-orthologous sequence - *A. fumigatus*:2211 *A. fischeri*:2211
afum|EAL91842.1 100.00% afis|EAW18031.1 100.00%
Bootstrap support for afum|EAL91842.1 as seed ortholog is 100%.
Bootstrap support for afis|EAW18031.1 as seed ortholog is 100%.

---

Group of orthologs #722. Best score 2211 bits
Score difference with first non-orthologous sequence - *A. fumigatus*:2211 *A. fischeri*:2211
afum|EAL87465.1 100.00% afis|EAW24034.1 100.00%
Bootstrap support for afum|EAL87465.1 as seed ortholog is 100%.
Bootstrap support for afis|EAW24034.1 as seed ortholog is 100%.

---

Group of orthologs #723. Best score 2210 bits
Score difference with first non-orthologous sequence - *A. fumigatus*:2210 *A. fischeri*:2210
afum|EAL91992.1 100.00% afis|EAW24137.1 100.00%
Bootstrap support for afum|EAL91992.1 as seed ortholog is 100%.
Bootstrap support for afis|EAW24137.1 as seed ortholog is 100%.

---

Group of orthologs #724. Best score 2209 bits
Score difference with first non-orthologous sequence - *A. fumigatus*:2209 *A. fischeri*:2209
afum|EAL88966.1 100.00% afis|EAW16341.1 100.00%
Bootstrap support for afum|EAL88966.1 as seed ortholog is 100%.
Bootstrap support for afis|EAW16341.1 as seed ortholog is 100%.

---

Group of orthologs #725. Best score 2209 bits
Score difference with first non-orthologous sequence - *A. fumigatus*:2209 *A. fischeri*:2209
afum|EAL90491.1 100.00% afis|EAW22701.1 100.00%
Bootstrap support for afum|EAL90491.1 as seed ortholog is 100%.
Bootstrap support for afis|EAW22701.1 as seed ortholog is 100%.

Group of orthologs #726. Best score 2208 bits
Score difference with first non-orthologous sequence - *A. fumigatus*:2208 *A. fischeri*:2208
afum|EAL92409.1 100.00% afis|EAW21395.1 100.00%
Bootstrap support for afum|EAL92409.1 as seed ortholog is 100%.
Bootstrap support for afis|EAW21395.1 as seed ortholog is 100%.

Group of orthologs #727. Best score 2207 bits
Score difference with first non-orthologous sequence - *A. fumigatus*:2207 *A. fischeri*:2207
afum|EAL91674.1 100.00% afis|EAW17866.1 100.00%
Bootstrap support for afum|EAL91674.1 as seed ortholog is 100%.
Bootstrap support for afis|EAW17866.1 as seed ortholog is 100%.

Group of orthologs #728. Best score 2206 bits
Score difference with first non-orthologous sequence - *A. fumigatus*:2206 *A. fischeri*:2206
afum|EAL84447.1 100.00% afis|EAW20614.1 100.00%
Bootstrap support for afum|EAL84447.1 as seed ortholog is 100%.
Bootstrap support for afis|EAW20614.1 as seed ortholog is 100%.

Group of orthologs #729. Best score 2205 bits
Score difference with first non-orthologous sequence - *A. fumigatus*:2205 *A. fischeri*:2205
afum|EAL88735.1 100.00% afis|EAW16103.1 100.00%
Bootstrap support for afum|EAL88735.1 as seed ortholog is 100%.
Bootstrap support for afis|EAW16103.1 as seed ortholog is 100%.

Group of orthologs #730. Best score 2203 bits
Score difference with first non-orthologous sequence - *A. fumigatus*:2203 *A. fischeri*:2203
afum|EAL85292.1 100.00% afis|EAW20174.1 100.00%
Bootstrap support for afum|EAL85292.1 as seed ortholog is 100%.
Bootstrap support for afis|EAW20174.1 as seed ortholog is 100%.

Group of orthologs #731. Best score 2203 bits
Score difference with first non-orthologous sequence - *A. fumigatus*:2203 *A. fischeri*:2203
afum|EAL92604.1 100.00% afis|EAW21586.1 100.00%
Bootstrap support for afum|EAL92604.1 as seed ortholog is 100%.
Bootstrap support for afis|EAW21586.1 as seed ortholog is 100%.

Group of orthologs #732. Best score 2200 bits
Score difference with first non-orthologous sequence - *A. fumigatus*:2200 *A. fischeri*:2200
afum|EAL87030.1 100.00% afis|EAW19466.1 100.00%
Bootstrap support for afum|EAL87030.1 as seed ortholog is 100%.
Bootstrap support for afis|EAW19466.1 as seed ortholog is 100%.

Group of orthologs #733. Best score 2200 bits
Score difference with first non-orthologous sequence - *A. fumigatus*:2200 *A. fischeri*:2200
afum|EAL92516.1 100.00% afis|EAW21497.1 100.00%
Bootstrap support for afum|EAL92516.1 as seed ortholog is 100%.
Bootstrap support for afis|EAW21497.1 as seed ortholog is 100%.

Group of orthologs #734. Best score 2197 bits
Score difference with first non-orthologous sequence - *A. fumigatus*:2197 *A. fischeri*:2197
afum|EAL88094.1 100.00% afis|EAW23377.1 100.00%
Bootstrap support for afum|EAL88094.1 as seed ortholog is 100%.
Bootstrap support for afis|EAW23377.1 as seed ortholog is 100%.

Group of orthologs #735. Best score 2197 bits

Score difference with first non-orthologous sequence - *A. fumigatus*:2197   *A. fischeri*:2197
afum|EAL93416.1 100.00% afis|EAW18694.1 100.00%
Bootstrap support for afum|EAL93416.1 as seed ortholog is 100%.
Bootstrap support for afis|EAW18694.1 as seed ortholog is 100%.

---

Group of orthologs #736. Best score 2193 bits
Score difference with first non-orthologous sequence - *A. fumigatus*:2193   *A. fischeri*:2193
afum|EAL93478.1 100.00% afis|EAW18751.1 100.00%
Bootstrap support for afum|EAL93478.1 as seed ortholog is 100%.
Bootstrap support for afis|EAW18751.1 as seed ortholog is 100%.

---

Group of orthologs #737. Best score 2191 bits
Score difference with first non-orthologous sequence - *A. fumigatus*:2191   *A. fischeri*:2191
afum|EAL88710.1 100.00% afis|EAW16078.1 100.00%
Bootstrap support for afum|EAL88710.1 as seed ortholog is 100%.
Bootstrap support for afis|EAW16078.1 as seed ortholog is 100%.

---

Group of orthologs #738. Best score 2191 bits
Score difference with first non-orthologous sequence - *A. fumigatus*:2191   *A. fischeri*:2191
afum|EAL89647.1 100.00% afis|EAW24987.1 100.00%
Bootstrap support for afum|EAL89647.1 as seed ortholog is 100%.
Bootstrap support for afis|EAW24987.1 as seed ortholog is 100%.

---

Group of orthologs #739. Best score 2190 bits
Score difference with first non-orthologous sequence - *A. fumigatus*:2190   *A. fischeri*:2190
afum|EAL90843.1 100.00% afis|EAW22349.1 100.00%
Bootstrap support for afum|EAL90843.1 as seed ortholog is 100%.
Bootstrap support for afis|EAW22349.1 as seed ortholog is 100%.

---

Group of orthologs #740. Best score 2184 bits
Score difference with first non-orthologous sequence - *A. fumigatus*:2184   *A. fischeri*:2184
afum|EAL88237.1 100.00% afis|EAW23245.1 100.00%
Bootstrap support for afum|EAL88237.1 as seed ortholog is 100%.
Bootstrap support for afis|EAW23245.1 as seed ortholog is 100%.

---

Group of orthologs #741. Best score 2184 bits
Score difference with first non-orthologous sequence - *A. fumigatus*:1863   *A. fischeri*:2184
afum|EAL91936.1 100.00% afis|EAW24081.1 100.00%
Bootstrap support for afum|EAL91936.1 as seed ortholog is 100%.
Bootstrap support for afis|EAW24081.1 as seed ortholog is 100%.

---

Group of orthologs #742. Best score 2181 bits
Score difference with first non-orthologous sequence - *A. fumigatus*:2181   *A. fischeri*:2181
afum|EAL87099.1 100.00% afis|EAW19395.1 100.00%
Bootstrap support for afum|EAL87099.1 as seed ortholog is 100%.
Bootstrap support for afis|EAW19395.1 as seed ortholog is 100%.

---

Group of orthologs #743. Best score 2181 bits
Score difference with first non-orthologous sequence - *A. fumigatus*:2181   *A. fischeri*:2181
afum|EAL93460.1 100.00% afis|EAW18735.1 100.00%
Bootstrap support for afum|EAL93460.1 as seed ortholog is 100%.
Bootstrap support for afis|EAW18735.1 as seed ortholog is 100%.

---

Group of orthologs #744. Best score 2181 bits
Score difference with first non-orthologous sequence - *A. fumigatus*:2181   *A. fischeri*:1227
afum|EAL92480.1 100.00% afis|EAW21461.1 100.00%

Bootstrap support for afum|EAL92480.1 as seed ortholog is 100%.
Bootstrap support for afis|EAW21461.1 as seed ortholog is 100%.

---

Group of orthologs #745. Best score 2180 bits
Score difference with first non-orthologous sequence - *A. fumigatus*:2180 *A. fischeri*:2180
afum|EAL91821.1 100.00% afis|EAW18009.1 100.00%
Bootstrap support for afum|EAL91821.1 as seed ortholog is 100%.
Bootstrap support for afis|EAW18009.1 as seed ortholog is 100%.

---

Group of orthologs #746. Best score 2180 bits
Score difference with first non-orthologous sequence - *A. fumigatus*:2180 *A. fischeri*:2180
afum|EAL90858.1 100.00% afis|EAW22334.1 100.00%
Bootstrap support for afum|EAL90858.1 as seed ortholog is 100%.
Bootstrap support for afis|EAW22334.1 as seed ortholog is 100%.

---

Group of orthologs #747. Best score 2179 bits
Score difference with first non-orthologous sequence - *A. fumigatus*:2179 *A. fischeri*:2179
afum|EAL87362.1 100.00% afis|EAW23937.1 100.00%
Bootstrap support for afum|EAL87362.1 as seed ortholog is 100%.
Bootstrap support for afis|EAW23937.1 as seed ortholog is 100%.

---

Group of orthologs #748. Best score 2179 bits
Score difference with first non-orthologous sequence - *A. fumigatus*:2179 *A. fischeri*:2179
afum|EAL92790.2 100.00% afis|EAW21760.1 100.00%
Bootstrap support for afum|EAL92790.2 as seed ortholog is 100%.
Bootstrap support for afis|EAW21760.1 as seed ortholog is 100%.

---

Group of orthologs #749. Best score 2178 bits
Score difference with first non-orthologous sequence - *A. fumigatus*:2178 *A. fischeri*:2178
afum|EAL93859.1 100.00% afis|EAW19127.1 100.00%
Bootstrap support for afum|EAL93859.1 as seed ortholog is 100%.
Bootstrap support for afis|EAW19127.1 as seed ortholog is 100%.

---

Group of orthologs #750. Best score 2177 bits
Score difference with first non-orthologous sequence - *A. fumigatus*:2177 *A. fischeri*:2177
afum|EBA27429.1 100.00% afis|EAW17449.1 100.00%
Bootstrap support for afum|EBA27429.1 as seed ortholog is 100%.
Bootstrap support for afis|EAW17449.1 as seed ortholog is 100%.

---

Group of orthologs #751. Best score 2175 bits
Score difference with first non-orthologous sequence - *A. fumigatus*:2175 *A. fischeri*:2175
afum|EAL92955.1 100.00% afis|EAW21941.1 100.00%
Bootstrap support for afum|EAL92955.1 as seed ortholog is 100%.
Bootstrap support for afis|EAW21941.1 as seed ortholog is 100%.

---

Group of orthologs #752. Best score 2173 bits
Score difference with first non-orthologous sequence - *A. fumigatus*:2173 *A. fischeri*:2173
afum|EAL88891.1 100.00% afis|EAW16270.1 100.00%
Bootstrap support for afum|EAL88891.1 as seed ortholog is 100%.
Bootstrap support for afis|EAW16270.1 as seed ortholog is 100%.

---

Group of orthologs #753. Best score 2171 bits
Score difference with first non-orthologous sequence - *A. fumigatus*:2171 *A. fischeri*:2171
afum|EAL92619.1 100.00% afis|EAW21600.1 100.00%
Bootstrap support for afum|EAL92619.1 as seed ortholog is 100%.
Bootstrap support for afis|EAW21600.1 as seed ortholog is 100%.

---

Group of orthologs #754. Best score 2170 bits
Score difference with first non-orthologous sequence - *A. fumigatus*:2170 *A. fischeri*:2170
afum|EAL91954.1 100.00% afis|EAW24100.1 100.00%
Bootstrap support for afum|EAL91954.1 as seed ortholog is 100%.
Bootstrap support for afis|EAW24100.1 as seed ortholog is 100%.

---

Group of orthologs #755. Best score 2169 bits
Score difference with first non-orthologous sequence - *A. fumigatus*:2169 *A. fischeri*:2169
afum|EAL89138.1 100.00% afis|EAW16541.1 100.00%
Bootstrap support for afum|EAL89138.1 as seed ortholog is 100%.
Bootstrap support for afis|EAW16541.1 as seed ortholog is 100%.

---

Group of orthologs #756. Best score 2168 bits
Score difference with first non-orthologous sequence - *A. fumigatus*:2168 *A. fischeri*:2168
afum|EAL91732.1 100.00% afis|EAW17921.1 100.00%
Bootstrap support for afum|EAL91732.1 as seed ortholog is 100%.
Bootstrap support for afis|EAW17921.1 as seed ortholog is 100%.

---

Group of orthologs #757. Best score 2167 bits
Score difference with first non-orthologous sequence - *A. fumigatus*:2167 *A. fischeri*:2167
afum|EAL91882.1 100.00% afis|EAW18070.1 100.00%
Bootstrap support for afum|EAL91882.1 as seed ortholog is 100%.
Bootstrap support for afis|EAW18070.1 as seed ortholog is 100%.

---

Group of orthologs #758. Best score 2166 bits
Score difference with first non-orthologous sequence - *A. fumigatus*:2166 *A. fischeri*:2003
afum|EAL87959.1 100.00% afis|EAW23502.1 100.00%
Bootstrap support for afum|EAL87959.1 as seed ortholog is 100%.
Bootstrap support for afis|EAW23502.1 as seed ortholog is 100%.

---

Group of orthologs #759. Best score 2166 bits
Score difference with first non-orthologous sequence - *A. fumigatus*:2166 *A. fischeri*:2166
afum|EAL93253.1 100.00% afis|EAW18553.1 100.00%
Bootstrap support for afum|EAL93253.1 as seed ortholog is 100%.
Bootstrap support for afis|EAW18553.1 as seed ortholog is 100%.

---

Group of orthologs #760. Best score 2165 bits
Score difference with first non-orthologous sequence - *A. fumigatus*:2165 *A. fischeri*:2165
afum|EAL90750.1 100.00% afis|EAW22441.1 100.00%
Bootstrap support for afum|EAL90750.1 as seed ortholog is 100%.
Bootstrap support for afis|EAW22441.1 as seed ortholog is 100%.

---

Group of orthologs #761. Best score 2165 bits
Score difference with first non-orthologous sequence - *A. fumigatus*:2165 *A. fischeri*:2165
afum|EAL90041.1 100.00% afis|EAW25383.1 100.00%
Bootstrap support for afum|EAL90041.1 as seed ortholog is 100%.
Bootstrap support for afis|EAW25383.1 as seed ortholog is 100%.

---

Group of orthologs #762. Best score 2163 bits
Score difference with first non-orthologous sequence - *A. fumigatus*:2163 *A. fischeri*:2163
afum|EAL87698.1 100.00% afis|EAW18333.1 100.00%
Bootstrap support for afum|EAL87698.1 as seed ortholog is 100%.
Bootstrap support for afis|EAW18333.1 as seed ortholog is 100%.

---

Group of orthologs #763. Best score 2162 bits

Score difference with first non-orthologous sequence - *A. fumigatus*:2162   *A. fischeri*:2162
afum|EAL85747.1 100.00% afis|EAW15585.1 100.00%
Bootstrap support for afum|EAL85747.1 as seed ortholog is 100%.
Bootstrap support for afis|EAW15585.1 as seed ortholog is 100%.

---

Group of orthologs #764. Best score 2159 bits
Score difference with first non-orthologous sequence - *A. fumigatus*:2159   *A. fischeri*:2159
afum|EAL85191.1 100.00% afis|EAW20073.1 100.00%
Bootstrap support for afum|EAL85191.1 as seed ortholog is 100%.
Bootstrap support for afis|EAW20073.1 as seed ortholog is 100%.

---

Group of orthologs #765. Best score 2157 bits
Score difference with first non-orthologous sequence - *A. fumigatus*:2157   *A. fischeri*:2157
afum|EAL90525.1 100.00% afis|EAW22667.1 100.00%
Bootstrap support for afum|EAL90525.1 as seed ortholog is 100%.
Bootstrap support for afis|EAW22667.1 as seed ortholog is 100%.

---

Group of orthologs #766. Best score 2156 bits
Score difference with first non-orthologous sequence - *A. fumigatus*:2156   *A. fischeri*:2156
afum|EAL86923.1 100.00% afis|EAW19595.1 100.00%
Bootstrap support for afum|EAL86923.1 as seed ortholog is 100%.
Bootstrap support for afis|EAW19595.1 as seed ortholog is 100%.

---

Group of orthologs #767. Best score 2156 bits
Score difference with first non-orthologous sequence - *A. fumigatus*:2156   *A. fischeri*:2156
afum|EAL93019.1 100.00% afis|EAW22004.1 100.00%
Bootstrap support for afum|EAL93019.1 as seed ortholog is 100%.
Bootstrap support for afis|EAW22004.1 as seed ortholog is 100%.

---

Group of orthologs #768. Best score 2153 bits
Score difference with first non-orthologous sequence - *A. fumigatus*:2153   *A. fischeri*:2153
afum|EAL84539.1 100.00% afis|EAW20518.1 100.00%
Bootstrap support for afum|EAL84539.1 as seed ortholog is 100%.
Bootstrap support for afis|EAW20518.1 as seed ortholog is 100%.

---

Group of orthologs #769. Best score 2153 bits
Score difference with first non-orthologous sequence - *A. fumigatus*:2153   *A. fischeri*:2153
afum|EAL90917.1 100.00% afis|EAW22263.1 100.00%
Bootstrap support for afum|EAL90917.1 as seed ortholog is 100%.
Bootstrap support for afis|EAW22263.1 as seed ortholog is 100%.

---

Group of orthologs #770. Best score 2153 bits
Score difference with first non-orthologous sequence - *A. fumigatus*:2153   *A. fischeri*:674
afum|EAL89049.1 100.00% afis|EAW25605.1 100.00%
Bootstrap support for afum|EAL89049.1 as seed ortholog is 100%.
Bootstrap support for afis|EAW25605.1 as seed ortholog is 99%.

---

Group of orthologs #771. Best score 2149 bits
Score difference with first non-orthologous sequence - *A. fumigatus*:2149   *A. fischeri*:2149
afum|EAL91437.1 100.00% afis|EAW17623.1 100.00%
Bootstrap support for afum|EAL91437.1 as seed ortholog is 100%.
Bootstrap support for afis|EAW17623.1 as seed ortholog is 100%.

---

Group of orthologs #772. Best score 2148 bits
Score difference with first non-orthologous sequence - *A. fumigatus*:2148   *A. fischeri*:2148
afum|EAL92067.2 100.00% afis|EAW21029.1 100.00%

Bootstrap support for afum|EAL92067.2 as seed ortholog is 100%.
Bootstrap support for afis|EAW21029.1 as seed ortholog is 100%.

---

Group of orthologs #773. Best score 2146 bits
Score difference with first non-orthologous sequence - *A. fumigatus*:2146 *A. fischeri*:2146
afum|EAL85609.1 100.00% afis|EAW15739.1 100.00%
Bootstrap support for afum|EAL85609.1 as seed ortholog is 100%.
Bootstrap support for afis|EAW15739.1 as seed ortholog is 100%.

---

Group of orthologs #774. Best score 2144 bits
Score difference with first non-orthologous sequence - *A. fumigatus*:2144 *A. fischeri*:2144
afum|EAL93186.1 100.00% afis|EAW18496.1 100.00%
Bootstrap support for afum|EAL93186.1 as seed ortholog is 100%.
Bootstrap support for afis|EAW18496.1 as seed ortholog is 100%.

---

Group of orthologs #775. Best score 2143 bits
Score difference with first non-orthologous sequence - *A. fumigatus*:2143 *A. fischeri*:2143
afum|EAL85996.1 100.00% afis|EAW24326.1 100.00%
Bootstrap support for afum|EAL85996.1 as seed ortholog is 100%.
Bootstrap support for afis|EAW24326.1 as seed ortholog is 100%.

---

Group of orthologs #776. Best score 2143 bits
Score difference with first non-orthologous sequence - *A. fumigatus*:2143 *A. fischeri*:2143
afum|EAL88404.1 100.00% afis|EAW23075.1 100.00%
Bootstrap support for afum|EAL88404.1 as seed ortholog is 100%.
Bootstrap support for afis|EAW23075.1 as seed ortholog is 100%.

---

Group of orthologs #777. Best score 2143 bits
Score difference with first non-orthologous sequence - *A. fumigatus*:2143 *A. fischeri*:2143
afum|EBA27333.1 100.00% afis|EAW23422.1 100.00%
Bootstrap support for afum|EBA27333.1 as seed ortholog is 100%.
Bootstrap support for afis|EAW23422.1 as seed ortholog is 100%.

---

Group of orthologs #778. Best score 2142 bits
Score difference with first non-orthologous sequence - *A. fumigatus*:2142 *A. fischeri*:2142
afum|EAL87353.1 100.00% afis|EAW23928.1 100.00%
Bootstrap support for afum|EAL87353.1 as seed ortholog is 100%.
Bootstrap support for afis|EAW23928.1 as seed ortholog is 100%.

---

Group of orthologs #779. Best score 2138 bits
Score difference with first non-orthologous sequence - *A. fumigatus*:2138 *A. fischeri*:2138
afum|EAL88566.1 100.00% afis|EAW15930.1 100.00%
Bootstrap support for afum|EAL88566.1 as seed ortholog is 100%.
Bootstrap support for afis|EAW15930.1 as seed ortholog is 100%.

---

Group of orthologs #780. Best score 2138 bits
Score difference with first non-orthologous sequence - *A. fumigatus*:2138 *A. fischeri*:2138
afum|EAL90624.2 100.00% afis|EAW22574.1 100.00%
Bootstrap support for afum|EAL90624.2 as seed ortholog is 100%.
Bootstrap support for afis|EAW22574.1 as seed ortholog is 100%.

---

Group of orthologs #781. Best score 2137 bits
Score difference with first non-orthologous sequence - *A. fumigatus*:2137 *A. fischeri*:2137
afum|EAL89932.1 100.00% afis|EAW25269.1 100.00%
Bootstrap support for afum|EAL89932.1 as seed ortholog is 100%.
Bootstrap support for afis|EAW25269.1 as seed ortholog is 100%.

---

Group of orthologs #782. Best score 2136 bits
Score difference with first non-orthologous sequence - *A. fumigatus*:1948 *A. fischeri*:1500
afum|EAL88289.1 100.00% afis|EAW23194.1 100.00%
Bootstrap support for afum|EAL88289.1 as seed ortholog is 100%.
Bootstrap support for afis|EAW23194.1 as seed ortholog is 100%.

---

Group of orthologs #783. Best score 2135 bits
Score difference with first non-orthologous sequence - *A. fumigatus*:2135 *A. fischeri*:2135
afum|EAL90781.1 100.00% afis|EAW22411.1 100.00%
Bootstrap support for afum|EAL90781.1 as seed ortholog is 100%.
Bootstrap support for afis|EAW22411.1 as seed ortholog is 100%.

---

Group of orthologs #784. Best score 2132 bits
Score difference with first non-orthologous sequence - *A. fumigatus*:2132 *A. fischeri*:2132
afum|EAL89129.1 100.00% afis|EAW16532.1 100.00%
Bootstrap support for afum|EAL89129.1 as seed ortholog is 100%.
Bootstrap support for afis|EAW16532.1 as seed ortholog is 100%.

---

Group of orthologs #785. Best score 2129 bits
Score difference with first non-orthologous sequence - *A. fumigatus*:2129 *A. fischeri*:2129
afum|EAL90390.1 100.00% afis|EAW22811.1 100.00%
Bootstrap support for afum|EAL90390.1 as seed ortholog is 100%.
Bootstrap support for afis|EAW22811.1 as seed ortholog is 100%.

---

Group of orthologs #786. Best score 2128 bits
Score difference with first non-orthologous sequence - *A. fumigatus*:2128 *A. fischeri*:2128
afum|EAL88527.1 100.00% afis|EAW15892.1 100.00%
Bootstrap support for afum|EAL88527.1 as seed ortholog is 100%.
Bootstrap support for afis|EAW15892.1 as seed ortholog is 100%.

---

Group of orthologs #787. Best score 2128 bits
Score difference with first non-orthologous sequence - *A. fumigatus*:2128 *A. fischeri*:2128
afum|EAL86968.1 100.00% afis|EAW19539.1 100.00%
Bootstrap support for afum|EAL86968.1 as seed ortholog is 100%.
Bootstrap support for afis|EAW19539.1 as seed ortholog is 100%.

---

Group of orthologs #788. Best score 2127 bits
Score difference with first non-orthologous sequence - *A. fumigatus*:2127 *A. fischeri*:2127
afum|EAL91445.1 100.00% afis|EAW17630.1 100.00%
Bootstrap support for afum|EAL91445.1 as seed ortholog is 100%.
Bootstrap support for afis|EAW17630.1 as seed ortholog is 100%.

---

Group of orthologs #789. Best score 2125 bits
Score difference with first non-orthologous sequence - *A. fumigatus*:2125 *A. fischeri*:2125
afum|EAL88293.1 100.00% afis|EAW23190.1 100.00%
Bootstrap support for afum|EAL88293.1 as seed ortholog is 100%.
Bootstrap support for afis|EAW23190.1 as seed ortholog is 100%.

---

Group of orthologs #790. Best score 2124 bits
Score difference with first non-orthologous sequence - *A. fumigatus*:2124 *A. fischeri*:884
afum|EAL86742.1 100.00% afis|EAW20667.1 100.00%
Bootstrap support for afum|EAL86742.1 as seed ortholog is 100%.
Bootstrap support for afis|EAW20667.1 as seed ortholog is 100%.

---

Group of orthologs #791. Best score 2123 bits

Score difference with first non-orthologous sequence - *A. fumigatus*:2123   *A. fischeri*:2123
afum|EAL90615.1 100.00% afis|EAW22584.1 100.00%
Bootstrap support for afum|EAL90615.1 as seed ortholog is 100%.
Bootstrap support for afis|EAW22584.1 as seed ortholog is 100%.

---

Group of orthologs #792. Best score 2122 bits
Score difference with first non-orthologous sequence - *A. fumigatus*:2122   *A. fischeri*:2122
afum|EAL85842.1 100.00% afis|EAW15479.1 100.00%
Bootstrap support for afum|EAL85842.1 as seed ortholog is 100%.
Bootstrap support for afis|EAW15479.1 as seed ortholog is 100%.

---

Group of orthologs #793. Best score 2121 bits
Score difference with first non-orthologous sequence - *A. fumigatus*:2121   *A. fischeri*:2121
afum|EAL88062.1 100.00% afis|EAW23408.1 100.00%
Bootstrap support for afum|EAL88062.1 as seed ortholog is 100%.
Bootstrap support for afis|EAW23408.1 as seed ortholog is 100%.

---

Group of orthologs #794. Best score 2120 bits
Score difference with first non-orthologous sequence - *A. fumigatus*:2120   *A. fischeri*:1578
afum|EAL90953.1 100.00% afis|EAW22239.1 100.00%
Bootstrap support for afum|EAL90953.1 as seed ortholog is 100%.
Bootstrap support for afis|EAW22239.1 as seed ortholog is 100%.

---

Group of orthologs #795. Best score 2119 bits
Score difference with first non-orthologous sequence - *A. fumigatus*:2119   *A. fischeri*:2119
afum|EAL86530.1 100.00% afis|EAW17010.1 100.00%
Bootstrap support for afum|EAL86530.1 as seed ortholog is 100%.
Bootstrap support for afis|EAW17010.1 as seed ortholog is 100%.

---

Group of orthologs #796. Best score 2119 bits
Score difference with first non-orthologous sequence - *A. fumigatus*:2119   *A. fischeri*:2119
afum|EAL87496.1 100.00% afis|EAW18139.1 100.00%
Bootstrap support for afum|EAL87496.1 as seed ortholog is 100%.
Bootstrap support for afis|EAW18139.1 as seed ortholog is 100%.

---

Group of orthologs #797. Best score 2119 bits
Score difference with first non-orthologous sequence - *A. fumigatus*:2119   *A. fischeri*:2119
afum|EAL89648.1 100.00% afis|EAW24988.1 100.00%
Bootstrap support for afum|EAL89648.1 as seed ortholog is 100%.
Bootstrap support for afis|EAW24988.1 as seed ortholog is 100%.

---

Group of orthologs #798. Best score 2118 bits
Score difference with first non-orthologous sequence - *A. fumigatus*:2118   *A. fischeri*:2118
afum|EAL85966.1 100.00% afis|EAW24298.1 100.00%
Bootstrap support for afum|EAL85966.1 as seed ortholog is 100%.
Bootstrap support for afis|EAW24298.1 as seed ortholog is 100%.

---

Group of orthologs #799. Best score 2118 bits
Score difference with first non-orthologous sequence - *A. fumigatus*:2118   *A. fischeri*:2118
afum|EAL88178.1 100.00% afis|EAW23301.1 100.00%
Bootstrap support for afum|EAL88178.1 as seed ortholog is 100%.
Bootstrap support for afis|EAW23301.1 as seed ortholog is 100%.

---

Group of orthologs #800. Best score 2116 bits
Score difference with first non-orthologous sequence - *A. fumigatus*:2116   *A. fischeri*:2116
afum|EAL93437.1 100.00% afis|EAW18712.1 100.00%

Bootstrap support for afum|EAL93437.1 as seed ortholog is 100%.
Bootstrap support for afis|EAW18712.1 as seed ortholog is 100%.

---

Group of orthologs #801. Best score 2115 bits
Score difference with first non-orthologous sequence - *A. fumigatus*:2115 *A. fischeri*:2115
afum|EAL91440.1 100.00% afis|EAW17626.1 100.00%
Bootstrap support for afum|EAL91440.1 as seed ortholog is 100%.
Bootstrap support for afis|EAW17626.1 as seed ortholog is 100%.

---

Group of orthologs #802. Best score 2113 bits
Score difference with first non-orthologous sequence - *A. fumigatus*:2021 *A. fischeri*:1859
afum|EAL84730.1 100.00% afis|EAW20901.1 100.00%
Bootstrap support for afum|EAL84730.1 as seed ortholog is 100%.
Bootstrap support for afis|EAW20901.1 as seed ortholog is 100%.

---

Group of orthologs #803. Best score 2113 bits
Score difference with first non-orthologous sequence - *A. fumigatus*:2113 *A. fischeri*:2113
afum|EBA27321.1 100.00% afis|EAW18239.1 100.00%
Bootstrap support for afum|EBA27321.1 as seed ortholog is 100%.
Bootstrap support for afis|EAW18239.1 as seed ortholog is 100%.

---

Group of orthologs #804. Best score 2112 bits
Score difference with first non-orthologous sequence - *A. fumigatus*:2112 *A. fischeri*:1359
afum|EAL89297.1 100.00% afis|EAW16700.1 100.00%
Bootstrap support for afum|EAL89297.1 as seed ortholog is 100%.
Bootstrap support for afis|EAW16700.1 as seed ortholog is 100%.

---

Group of orthologs #805. Best score 2112 bits
Score difference with first non-orthologous sequence - *A. fumigatus*:2112 *A. fischeri*:2112
afum|EAL93105.1 100.00% afis|EAW18421.1 100.00%
Bootstrap support for afum|EAL93105.1 as seed ortholog is 100%.
Bootstrap support for afis|EAW18421.1 as seed ortholog is 100%.

---

Group of orthologs #806. Best score 2112 bits
Score difference with first non-orthologous sequence - *A. fumigatus*:2112 *A. fischeri*:2112
afum|EAL90614.1 100.00% afis|EAW22585.1 100.00%
Bootstrap support for afum|EAL90614.1 as seed ortholog is 100%.
Bootstrap support for afis|EAW22585.1 as seed ortholog is 100%.

---

Group of orthologs #807. Best score 2112 bits
Score difference with first non-orthologous sequence - *A. fumigatus*:2112 *A. fischeri*:2112
afum|EAL89629.1 100.00% afis|EAW24970.1 100.00%
Bootstrap support for afum|EAL89629.1 as seed ortholog is 100%.
Bootstrap support for afis|EAW24970.1 as seed ortholog is 100%.

---

Group of orthologs #808. Best score 2111 bits
Score difference with first non-orthologous sequence - *A. fumigatus*:2111 *A. fischeri*:2111
afum|EAL92865.1 100.00% afis|EAW21852.1 100.00%
Bootstrap support for afum|EAL92865.1 as seed ortholog is 100%.
Bootstrap support for afis|EAW21852.1 as seed ortholog is 100%.

---

Group of orthologs #809. Best score 2110 bits
Score difference with first non-orthologous sequence - *A. fumigatus*:2110 *A. fischeri*:2110
afum|EAL89884.1 100.00% afis|EAW25223.1 100.00%
Bootstrap support for afum|EAL89884.1 as seed ortholog is 100%.
Bootstrap support for afis|EAW25223.1 as seed ortholog is 100%.

Group of orthologs #810. Best score 2109 bits
Score difference with first non-orthologous sequence - *A. fumigatus*:2109 *A. fischeri*:1901
afum|EAL91411.1 100.00% afis|EAW17598.1 100.00%
Bootstrap support for afum|EAL91411.1 as seed ortholog is 100%.
Bootstrap support for afis|EAW17598.1 as seed ortholog is 100%.

---

Group of orthologs #811. Best score 2108 bits
Score difference with first non-orthologous sequence - *A. fumigatus*:2108 *A. fischeri*:2108
afum|EAL94058.1 100.00% afis|EAW19339.1 100.00%
Bootstrap support for afum|EAL94058.1 as seed ortholog is 100%.
Bootstrap support for afis|EAW19339.1 as seed ortholog is 100%.

---

Group of orthologs #812. Best score 2108 bits
Score difference with first non-orthologous sequence - *A. fumigatus*:2108 *A. fischeri*:2108
afum|EBA27387.1 100.00% afis|EAW25008.1 100.00%
Bootstrap support for afum|EBA27387.1 as seed ortholog is 100%.
Bootstrap support for afis|EAW25008.1 as seed ortholog is 100%.

---

Group of orthologs #813. Best score 2107 bits
Score difference with first non-orthologous sequence - *A. fumigatus*:2107 *A. fischeri*:2107
afum|EAL92500.1 100.00% afis|EAW21480.1 100.00%
Bootstrap support for afum|EAL92500.1 as seed ortholog is 100%.
Bootstrap support for afis|EAW21480.1 as seed ortholog is 100%.

---

Group of orthologs #814. Best score 2106 bits
Score difference with first non-orthologous sequence - *A. fumigatus*:2106 *A. fischeri*:2106
afum|EAL85812.1 100.00% afis|EAW15517.1 100.00%
Bootstrap support for afum|EAL85812.1 as seed ortholog is 100%.
Bootstrap support for afis|EAW15517.1 as seed ortholog is 100%.

---

Group of orthologs #815. Best score 2106 bits
Score difference with first non-orthologous sequence - *A. fumigatus*:2106 *A. fischeri*:2106
afum|EAL91638.1 100.00% afis|EAW17828.1 100.00%
Bootstrap support for afum|EAL91638.1 as seed ortholog is 100%.
Bootstrap support for afis|EAW17828.1 as seed ortholog is 100%.

---

Group of orthologs #816. Best score 2105 bits
Score difference with first non-orthologous sequence - *A. fumigatus*:2105 *A. fischeri*:2105
afum|EAL90048.2 100.00% afis|EAW25389.1 100.00%
Bootstrap support for afum|EAL90048.2 as seed ortholog is 100%.
Bootstrap support for afis|EAW25389.1 as seed ortholog is 100%.

---

Group of orthologs #817. Best score 2104 bits
Score difference with first non-orthologous sequence - *A. fumigatus*:2104 *A. fischeri*:2104
afum|EAL87414.1 100.00% afis|EAW23985.1 100.00%
Bootstrap support for afum|EAL87414.1 as seed ortholog is 100%.
Bootstrap support for afis|EAW23985.1 as seed ortholog is 100%.

---

Group of orthologs #818. Best score 2104 bits
Score difference with first non-orthologous sequence - *A. fumigatus*:2104 *A. fischeri*:2104
afum|EAL88104.2 100.00% afis|EAW23369.1 100.00%
Bootstrap support for afum|EAL88104.2 as seed ortholog is 100%.
Bootstrap support for afis|EAW23369.1 as seed ortholog is 100%.

---

Group of orthologs #819. Best score 2103 bits

Score difference with first non-orthologous sequence - *A. fumigatus*:2103   *A. fischeri*:2103
afum|EAL93467.1    100.00%    afis|EAW18741.1    100.00%
Bootstrap support for afum|EAL93467.1 as seed ortholog is 100%.
Bootstrap support for afis|EAW18741.1 as seed ortholog is 100%.

---

Group of orthologs #820. Best score 2103 bits
Score difference with first non-orthologous sequence - *A. fumigatus*:2103   *A. fischeri*:2103
afum|EBA27297.1    100.00%    afis|EAW19500.1    100.00%
Bootstrap support for afum|EBA27297.1 as seed ortholog is 100%.
Bootstrap support for afis|EAW19500.1 as seed ortholog is 100%.

---

Group of orthologs #821. Best score 2102 bits
Score difference with first non-orthologous sequence - *A. fumigatus*:2102   *A. fischeri*:2102
afum|EAL87081.1    100.00%    afis|EAW19413.1    100.00%
Bootstrap support for afum|EAL87081.1 as seed ortholog is 100%.
Bootstrap support for afis|EAW19413.1 as seed ortholog is 100%.

---

Group of orthologs #822. Best score 2102 bits
Score difference with first non-orthologous sequence - *A. fumigatus*:2102   *A. fischeri*:2102
afum|EAL91493.1    100.00%    afis|EAW17679.1    100.00%
Bootstrap support for afum|EAL91493.1 as seed ortholog is 100%.
Bootstrap support for afis|EAW17679.1 as seed ortholog is 100%.

---

Group of orthologs #823. Best score 2102 bits
Score difference with first non-orthologous sequence - *A. fumigatus*:2102   *A. fischeri*:2102
afum|EAL87365.1    100.00%    afis|EAW23940.1    100.00%
Bootstrap support for afum|EAL87365.1 as seed ortholog is 100%.
Bootstrap support for afis|EAW23940.1 as seed ortholog is 100%.

---

Group of orthologs #824. Best score 2099 bits
Score difference with first non-orthologous sequence - *A. fumigatus*:2099   *A. fischeri*:2099
afum|EAL86703.1    100.00%    afis|EAW17285.1    100.00%
Bootstrap support for afum|EAL86703.1 as seed ortholog is 100%.
Bootstrap support for afis|EAW17285.1 as seed ortholog is 100%.

---

Group of orthologs #825. Best score 2097 bits
Score difference with first non-orthologous sequence - *A. fumigatus*:2097   *A. fischeri*:2097
afum|EAL85872.1    100.00%    afis|EAW24202.1    100.00%
Bootstrap support for afum|EAL85872.1 as seed ortholog is 100%.
Bootstrap support for afis|EAW24202.1 as seed ortholog is 100%.

---

Group of orthologs #826. Best score 2097 bits
Score difference with first non-orthologous sequence - *A. fumigatus*:2097   *A. fischeri*:2097
afum|EAL90876.1    100.00%    afis|EAW22314.1    100.00%
Bootstrap support for afum|EAL90876.1 as seed ortholog is 100%.
Bootstrap support for afis|EAW22314.1 as seed ortholog is 100%.

---

Group of orthologs #827. Best score 2096 bits
Score difference with first non-orthologous sequence - *A. fumigatus*:2096   *A. fischeri*:1768
afum|EAL85182.2    100.00%    afis|EAW20062.1    100.00%
Bootstrap support for afum|EAL85182.2 as seed ortholog is 100%.
Bootstrap support for afis|EAW20062.1 as seed ortholog is 100%.

---

Group of orthologs #828. Best score 2096 bits
Score difference with first non-orthologous sequence - *A. fumigatus*:2096   *A. fischeri*:2096
afum|EAL91676.1    100.00%    afis|EAW17868.1    100.00%

Bootstrap support for afum|EAL91676.1 as seed ortholog is 100%.
Bootstrap support for afis|EAW17868.1 as seed ortholog is 100%.

---

Group of orthologs #829. Best score 2096 bits
Score difference with first non-orthologous sequence - *A. fumigatus*:2096 *A. fischeri*:2096
afum|EAL93659.1 100.00% afis|EAW18928.1 100.00%
Bootstrap support for afum|EAL93659.1 as seed ortholog is 100%.
Bootstrap support for afis|EAW18928.1 as seed ortholog is 100%.

---

Group of orthologs #830. Best score 2096 bits
Score difference with first non-orthologous sequence - *A. fumigatus*:2096 *A. fischeri*:2096
afum|EAL92435.1 100.00% afis|EAW21420.1 100.00%
Bootstrap support for afum|EAL92435.1 as seed ortholog is 100%.
Bootstrap support for afis|EAW21420.1 as seed ortholog is 100%.

---

Group of orthologs #831. Best score 2095 bits
Score difference with first non-orthologous sequence - *A. fumigatus*:1859 *A. fischeri*:1512
afum|EBA27423.1 100.00% afis|EAW17324.1 100.00%
Bootstrap support for afum|EBA27423.1 as seed ortholog is 100%.
Bootstrap support for afis|EAW17324.1 as seed ortholog is 100%.

---

Group of orthologs #832. Best score 2095 bits
Score difference with first non-orthologous sequence - *A. fumigatus*:2095 *A. fischeri*:2095
afum|EAL94016.1 100.00% afis|EAW19297.1 100.00%
Bootstrap support for afum|EAL94016.1 as seed ortholog is 100%.
Bootstrap support for afis|EAW19297.1 as seed ortholog is 100%.

---

Group of orthologs #833. Best score 2095 bits
Score difference with first non-orthologous sequence - *A. fumigatus*:2095 *A. fischeri*:2095
afum|EAL90125.1 100.00% afis|EAW25462.1 100.00%
Bootstrap support for afum|EAL90125.1 as seed ortholog is 100%.
Bootstrap support for afis|EAW25462.1 as seed ortholog is 100%.

---

Group of orthologs #834. Best score 2093 bits
Score difference with first non-orthologous sequence - *A. fumigatus*:2093 *A. fischeri*:2093
afum|EAL84762.1 100.00% afis|EAW20872.1 100.00%
Bootstrap support for afum|EAL84762.1 as seed ortholog is 100%.
Bootstrap support for afis|EAW20872.1 as seed ortholog is 100%.

---

Group of orthologs #835. Best score 2093 bits
Score difference with first non-orthologous sequence - *A. fumigatus*:2093 *A. fischeri*:2093
afum|EAL91690.1 100.00% afis|EAW17882.1 100.00%
Bootstrap support for afum|EAL91690.1 as seed ortholog is 100%.
Bootstrap support for afis|EAW17882.1 as seed ortholog is 100%.

---

Group of orthologs #836. Best score 2093 bits
Score difference with first non-orthologous sequence - *A. fumigatus*:2093 *A. fischeri*:2093
afum|EAL93152.1 100.00% afis|EAW18466.1 100.00%
Bootstrap support for afum|EAL93152.1 as seed ortholog is 100%.
Bootstrap support for afis|EAW18466.1 as seed ortholog is 100%.

---

Group of orthologs #837. Best score 2092 bits
Score difference with first non-orthologous sequence - *A. fumigatus*:2092 *A. fischeri*:2092
afum|EAL90884.1 100.00% afis|EAW22297.1 100.00%
Bootstrap support for afum|EAL90884.1 as seed ortholog is 100%.
Bootstrap support for afis|EAW22297.1 as seed ortholog is 100%.

---

Group of orthologs #838. Best score 2091 bits
Score difference with first non-orthologous sequence - *A. fumigatus*:2091 *A. fischeri*:2091
afum|EAL90180.1 100.00% afis|EAW25515.1 100.00%
Bootstrap support for afum|EAL90180.1 as seed ortholog is 100%.
Bootstrap support for afis|EAW25515.1 as seed ortholog is 100%.

---

Group of orthologs #839. Best score 2089 bits
Score difference with first non-orthologous sequence - *A. fumigatus*:2089 *A. fischeri*:2089
afum|EAL88909.1 100.00% afis|EAW16287.1 100.00%
Bootstrap support for afum|EAL88909.1 as seed ortholog is 100%.
Bootstrap support for afis|EAW16287.1 as seed ortholog is 100%.

---

Group of orthologs #840. Best score 2089 bits
Score difference with first non-orthologous sequence - *A. fumigatus*:2089 *A. fischeri*:2089
afum|EBA27345.1 100.00% afis|EAW23092.1 100.00%
Bootstrap support for afum|EBA27345.1 as seed ortholog is 100%.
Bootstrap support for afis|EAW23092.1 as seed ortholog is 100%.

---

Group of orthologs #841. Best score 2088 bits
Score difference with first non-orthologous sequence - *A. fumigatus*:2088 *A. fischeri*:2088
afum|EAL87086.1 100.00% afis|EAW19408.1 100.00%
Bootstrap support for afum|EAL87086.1 as seed ortholog is 100%.
Bootstrap support for afis|EAW19408.1 as seed ortholog is 100%.

---

Group of orthologs #842. Best score 2088 bits
Score difference with first non-orthologous sequence - *A. fumigatus*:2088 *A. fischeri*:2088
afum|EAL90119.1 100.00% afis|EAW25459.1 100.00%
Bootstrap support for afum|EAL90119.1 as seed ortholog is 100%.
Bootstrap support for afis|EAW25459.1 as seed ortholog is 100%.

---

Group of orthologs #843. Best score 2086 bits
Score difference with first non-orthologous sequence - *A. fumigatus*:2086 *A. fischeri*:2086
afum|EAL85575.1 100.00% afis|EAW15774.1 100.00%
Bootstrap support for afum|EAL85575.1 as seed ortholog is 100%.
Bootstrap support for afis|EAW15774.1 as seed ortholog is 100%.

---

Group of orthologs #844. Best score 2086 bits
Score difference with first non-orthologous sequence - *A. fumigatus*:2086 *A. fischeri*:2086
afum|EAL94000.1 100.00% afis|EAW19274.1 100.00%
Bootstrap support for afum|EAL94000.1 as seed ortholog is 100%.
Bootstrap support for afis|EAW19274.1 as seed ortholog is 100%.

---

Group of orthologs #845. Best score 2085 bits
Score difference with first non-orthologous sequence - *A. fumigatus*:2085 *A. fischeri*:2085
afum|EAL90900.1 100.00% afis|EAW22281.1 100.00%
Bootstrap support for afum|EAL90900.1 as seed ortholog is 100%.
Bootstrap support for afis|EAW22281.1 as seed ortholog is 100%.

---

Group of orthologs #846. Best score 2084 bits
Score difference with first non-orthologous sequence - *A. fumigatus*:2084 *A. fischeri*:2084
afum|EAL87613.1 100.00% afis|EAW18248.1 100.00%
Bootstrap support for afum|EAL87613.1 as seed ortholog is 100%.
Bootstrap support for afis|EAW18248.1 as seed ortholog is 100%.

---

Group of orthologs #847. Best score 2084 bits

Score difference with first non-orthologous sequence - *A. fumigatus*:2084   *A. fischeri*:2084
afum|EAL88351.1 100.00% afis|EAW23129.1 100.00%
Bootstrap support for afum|EAL88351.1 as seed ortholog is 100%.
Bootstrap support for afis|EAW23129.1 as seed ortholog is 100%.

---

Group of orthologs #848. Best score 2082 bits
Score difference with first non-orthologous sequence - *A. fumigatus*:2082   *A. fischeri*:2082
afum|EAL91815.2 100.00% afis|EAW18004.1 100.00%
Bootstrap support for afum|EAL91815.2 as seed ortholog is 100%.
Bootstrap support for afis|EAW18004.1 as seed ortholog is 100%.

---

Group of orthologs #849. Best score 2082 bits
Score difference with first non-orthologous sequence - *A. fumigatus*:2082   *A. fischeri*:2082
afum|EAL87433.1 100.00% afis|EAW24003.1 100.00%
Bootstrap support for afum|EAL87433.1 as seed ortholog is 100%.
Bootstrap support for afis|EAW24003.1 as seed ortholog is 100%.

---

Group of orthologs #850. Best score 2081 bits
Score difference with first non-orthologous sequence - *A. fumigatus*:2081   *A. fischeri*:2081
afum|EAL89223.1 100.00% afis|EAW16613.1 100.00%
Bootstrap support for afum|EAL89223.1 as seed ortholog is 100%.
Bootstrap support for afis|EAW16613.1 as seed ortholog is 100%.

---

Group of orthologs #851. Best score 2081 bits
Score difference with first non-orthologous sequence - *A. fumigatus*:2081   *A. fischeri*:2081
afum|EAL84303.1 100.00% afis|EAW25597.1 100.00%
Bootstrap support for afum|EAL84303.1 as seed ortholog is 100%.
Bootstrap support for afis|EAW25597.1 as seed ortholog is 100%.

---

Group of orthologs #852. Best score 2081 bits
Score difference with first non-orthologous sequence - *A. fumigatus*:2081   *A. fischeri*:2081
afum|EAL92545.1 100.00% afis|EAW21524.1 100.00%
Bootstrap support for afum|EAL92545.1 as seed ortholog is 100%.
Bootstrap support for afis|EAW21524.1 as seed ortholog is 100%.

---

Group of orthologs #853. Best score 2080 bits
Score difference with first non-orthologous sequence - *A. fumigatus*:2080   *A. fischeri*:2080
afum|EAL88750.1 100.00% afis|EAW16119.1 100.00%
Bootstrap support for afum|EAL88750.1 as seed ortholog is 100%.
Bootstrap support for afis|EAW16119.1 as seed ortholog is 100%.

---

Group of orthologs #854. Best score 2080 bits
Score difference with first non-orthologous sequence - *A. fumigatus*:2080   *A. fischeri*:2080
afum|EAL84545.2 100.00% afis|EAW20511.1 100.00%
Bootstrap support for afum|EAL84545.2 as seed ortholog is 100%.
Bootstrap support for afis|EAW20511.1 as seed ortholog is 100%.

---

Group of orthologs #855. Best score 2077 bits
Score difference with first non-orthologous sequence - *A. fumigatus*:2077   *A. fischeri*:1531
afum|EAL89076.1 100.00% afis|EAW16482.1 100.00%
Bootstrap support for afum|EAL89076.1 as seed ortholog is 100%.
Bootstrap support for afis|EAW16482.1 as seed ortholog is 100%.

---

Group of orthologs #856. Best score 2076 bits
Score difference with first non-orthologous sequence - *A. fumigatus*:2076   *A. fischeri*:1594
afum|EAL92566.1 100.00% afis|EAW21546.1 100.00%

Bootstrap support for afum|EAL92566.1 as seed ortholog is 100%.
Bootstrap support for afis|EAW21546.1 as seed ortholog is 100%.

---

Group of orthologs #857. Best score 2076 bits
Score difference with first non-orthologous sequence - *A. fumigatus*:2076 *A. fischeri*:2076
afum|EAL92960.1 100.00% afis|EAW21946.1 100.00%
Bootstrap support for afum|EAL92960.1 as seed ortholog is 100%.
Bootstrap support for afis|EAW21946.1 as seed ortholog is 100%.

---

Group of orthologs #858. Best score 2074 bits
Score difference with first non-orthologous sequence - *A. fumigatus*:2074 *A. fischeri*:2074
afum|EAL90548.1 100.00% afis|EAW22644.1 100.00%
Bootstrap support for afum|EAL90548.1 as seed ortholog is 100%.
Bootstrap support for afis|EAW22644.1 as seed ortholog is 100%.

---

Group of orthologs #859. Best score 2073 bits
Score difference with first non-orthologous sequence - *A. fumigatus*:2073 *A. fischeri*:2073
afum|EAL87106.1 100.00% afis|EAW19389.1 100.00%
Bootstrap support for afum|EAL87106.1 as seed ortholog is 100%.
Bootstrap support for afis|EAW19389.1 as seed ortholog is 100%.

---

Group of orthologs #860. Best score 2073 bits
Score difference with first non-orthologous sequence - *A. fumigatus*:2073 *A. fischeri*:2073
afum|EAL90677.1 100.00% afis|EAW22516.1 100.00%
Bootstrap support for afum|EAL90677.1 as seed ortholog is 100%.
Bootstrap support for afis|EAW22516.1 as seed ortholog is 100%.

---

Group of orthologs #861. Best score 2072 bits
Score difference with first non-orthologous sequence - *A. fumigatus*:2072 *A. fischeri*:2072
afum|EAL89661.1 100.00% afis|EAW25033.1 100.00%
Bootstrap support for afum|EAL89661.1 as seed ortholog is 100%.
Bootstrap support for afis|EAW25033.1 as seed ortholog is 100%.

---

Group of orthologs #862. Best score 2071 bits
Score difference with first non-orthologous sequence - *A. fumigatus*:2071 *A. fischeri*:2071
afum|EAL87021.1 100.00% afis|EAW19475.1 100.00%
Bootstrap support for afum|EAL87021.1 as seed ortholog is 100%.
Bootstrap support for afis|EAW19475.1 as seed ortholog is 100%.

---

Group of orthologs #863. Best score 2071 bits
Score difference with first non-orthologous sequence - *A. fumigatus*:2071 *A. fischeri*:2071
afum|EAL91381.1 100.00% afis|EAW17567.1 100.00%
Bootstrap support for afum|EAL91381.1 as seed ortholog is 100%.
Bootstrap support for afis|EAW17567.1 as seed ortholog is 100%.

---

Group of orthologs #864. Best score 2070 bits
Score difference with first non-orthologous sequence - *A. fumigatus*:2070 *A. fischeri*:2070
afum|EAL92774.1 100.00% afis|EAW21741.1 100.00%
Bootstrap support for afum|EAL92774.1 as seed ortholog is 100%.
Bootstrap support for afis|EAW21741.1 as seed ortholog is 100%.

---

Group of orthologs #865. Best score 2069 bits
Score difference with first non-orthologous sequence - *A. fumigatus*:2069 *A. fischeri*:2069
afum|EAL90862.1 100.00% afis|EAW22330.1 100.00%
Bootstrap support for afum|EAL90862.1 as seed ortholog is 100%.
Bootstrap support for afis|EAW22330.1 as seed ortholog is 100%.

---

Group of orthologs #866. Best score 2068 bits
Score difference with first non-orthologous sequence - *A. fumigatus*:1962 *A. fischeri*:1761
afum|EAL85291.1 100.00% afis|EAW20173.1 100.00%
Bootstrap support for afum|EAL85291.1 as seed ortholog is 100%.
Bootstrap support for afis|EAW20173.1 as seed ortholog is 100%.

---

Group of orthologs #867. Best score 2067 bits
Score difference with first non-orthologous sequence - *A. fumigatus*:2067 *A. fischeri*:2067
afum|EAL91451.2 100.00% afis|EAW17636.1 100.00%
Bootstrap support for afum|EAL91451.2 as seed ortholog is 100%.
Bootstrap support for afis|EAW17636.1 as seed ortholog is 100%.

---

Group of orthologs #868. Best score 2066 bits
Score difference with first non-orthologous sequence - *A. fumigatus*:2066 *A. fischeri*:1948
afum|EAL90655.1 100.00% afis|EAW22539.1 100.00%
Bootstrap support for afum|EAL90655.1 as seed ortholog is 100%.
Bootstrap support for afis|EAW22539.1 as seed ortholog is 100%.

---

Group of orthologs #869. Best score 2066 bits
Score difference with first non-orthologous sequence - *A. fumigatus*:2066 *A. fischeri*:1980
afum|EAL90490.1 100.00% afis|EAW22702.1 100.00%
Bootstrap support for afum|EAL90490.1 as seed ortholog is 100%.
Bootstrap support for afis|EAW22702.1 as seed ortholog is 100%.

---

Group of orthologs #870. Best score 2065 bits
Score difference with first non-orthologous sequence - *A. fumigatus*:2065 *A. fischeri*:2065
afum|EAL91517.1 100.00% afis|EAW17702.1 100.00%
Bootstrap support for afum|EAL91517.1 as seed ortholog is 100%.
Bootstrap support for afis|EAW17702.1 as seed ortholog is 100%.

---

Group of orthologs #871. Best score 2063 bits
Score difference with first non-orthologous sequence - *A. fumigatus*:2063 *A. fischeri*:2063
afum|EAL88425.1 100.00% afis|EAW23055.1 100.00%
Bootstrap support for afum|EAL88425.1 as seed ortholog is 100%.
Bootstrap support for afis|EAW23055.1 as seed ortholog is 100%.

---

Group of orthologs #872. Best score 2062 bits
Score difference with first non-orthologous sequence - *A. fumigatus*:2062 *A. fischeri*:2062
afum|EAL87406.1 100.00% afis|EAW23976.1 100.00%
Bootstrap support for afum|EAL87406.1 as seed ortholog is 100%.
Bootstrap support for afis|EAW23976.1 as seed ortholog is 100%.

---

Group of orthologs #873. Best score 2062 bits
Score difference with first non-orthologous sequence - *A. fumigatus*:2062 *A. fischeri*:2062
afum|EAL87917.2 100.00% afis|EAW23549.1 100.00%
Bootstrap support for afum|EAL87917.2 as seed ortholog is 100%.
Bootstrap support for afis|EAW23549.1 as seed ortholog is 100%.

---

Group of orthologs #874. Best score 2062 bits
Score difference with first non-orthologous sequence - *A. fumigatus*:2062 *A. fischeri*:2062
afum|EAL93356.1 100.00% afis|EAW18642.1 100.00%
Bootstrap support for afum|EAL93356.1 as seed ortholog is 100%.
Bootstrap support for afis|EAW18642.1 as seed ortholog is 100%.

---

Group of orthologs #875. Best score 2061 bits

Score difference with first non-orthologous sequence - *A. fumigatus*:2061   *A. fischeri*:2061
afum|EAL91347.1 100.00% afis|EAW17533.1 100.00%
Bootstrap support for afum|EAL91347.1 as seed ortholog is 100%.
Bootstrap support for afis|EAW17533.1 as seed ortholog is 100%.

---

Group of orthologs #876. Best score 2061 bits
Score difference with first non-orthologous sequence - *A. fumigatus*:2061   *A. fischeri*:2061
afum|EAL89881.1 100.00% afis|EAW25220.1 100.00%
Bootstrap support for afum|EAL89881.1 as seed ortholog is 100%.
Bootstrap support for afis|EAW25220.1 as seed ortholog is 100%.

---

Group of orthologs #877. Best score 2058 bits
Score difference with first non-orthologous sequence - *A. fumigatus*:2058   *A. fischeri*:2058
afum|EAL87276.2 100.00% afis|EAW23848.1 100.00%
Bootstrap support for afum|EAL87276.2 as seed ortholog is 100%.
Bootstrap support for afis|EAW23848.1 as seed ortholog is 100%.

---

Group of orthologs #878. Best score 2058 bits
Score difference with first non-orthologous sequence - *A. fumigatus*:2058   *A. fischeri*:2058
afum|EAL94091.1 100.00% afis|EAW19373.1 100.00%
Bootstrap support for afum|EAL94091.1 as seed ortholog is 100%.
Bootstrap support for afis|EAW19373.1 as seed ortholog is 100%.

---

Group of orthologs #879. Best score 2058 bits
Score difference with first non-orthologous sequence - *A. fumigatus*:2058   *A. fischeri*:2058
afum|EAL92888.1 100.00% afis|EAW21874.1 100.00%
Bootstrap support for afum|EAL92888.1 as seed ortholog is 100%.
Bootstrap support for afis|EAW21874.1 as seed ortholog is 100%.

---

Group of orthologs #880. Best score 2056 bits
Score difference with first non-orthologous sequence - *A. fumigatus*:1823   *A. fischeri*:1485
afum|EAL88721.1 100.00% afis|EAW16089.1 100.00%
Bootstrap support for afum|EAL88721.1 as seed ortholog is 100%.
Bootstrap support for afis|EAW16089.1 as seed ortholog is 100%.

---

Group of orthologs #881. Best score 2056 bits
Score difference with first non-orthologous sequence - *A. fumigatus*:2056   *A. fischeri*:661
afum|EAL86225.1 100.00% afis|EAW24564.1 100.00%
Bootstrap support for afum|EAL86225.1 as seed ortholog is 100%.
Bootstrap support for afis|EAW24564.1 as seed ortholog is 100%.

---

Group of orthologs #882. Best score 2055 bits
Score difference with first non-orthologous sequence - *A. fumigatus*:2055   *A. fischeri*:2055
afum|EAL85089.1 100.00% afis|EAW19804.1 100.00%
Bootstrap support for afum|EAL85089.1 as seed ortholog is 100%.
Bootstrap support for afis|EAW19804.1 as seed ortholog is 100%.

---

Group of orthologs #883. Best score 2055 bits
Score difference with first non-orthologous sequence - *A. fumigatus*:2055   *A. fischeri*:2055
afum|EAL87441.1 100.00% afis|EAW24011.1 100.00%
Bootstrap support for afum|EAL87441.1 as seed ortholog is 100%.
Bootstrap support for afis|EAW24011.1 as seed ortholog is 100%.

---

Group of orthologs #884. Best score 2054 bits
Score difference with first non-orthologous sequence - *A. fumigatus*:2054   *A. fischeri*:2054
afum|EAL86909.1 100.00% afis|EAW19613.1 100.00%

Bootstrap support for afum|EAL86909.1 as seed ortholog is 100%.
Bootstrap support for afis|EAW19613.1 as seed ortholog is 100%.

---

Group of orthologs #885. Best score 2053 bits
Score difference with first non-orthologous sequence - *A. fumigatus*:2053 *A. fischeri*:2053
afum|EAL84379.1 100.00% afis|EAW25706.1 100.00%
Bootstrap support for afum|EAL84379.1 as seed ortholog is 100%.
Bootstrap support for afis|EAW25706.1 as seed ortholog is 100%.

---

Group of orthologs #886. Best score 2052 bits
Score difference with first non-orthologous sequence - *A. fumigatus*:2052 *A. fischeri*:2052
afum|EAL86913.1 100.00% afis|EAW19609.1 100.00%
Bootstrap support for afum|EAL86913.1 as seed ortholog is 100%.
Bootstrap support for afis|EAW19609.1 as seed ortholog is 100%.

---

Group of orthologs #887. Best score 2050 bits
Score difference with first non-orthologous sequence - *A. fumigatus*:2050 *A. fischeri*:2050
afum|EAL93104.1 100.00% afis|EAW18420.1 100.00%
Bootstrap support for afum|EAL93104.1 as seed ortholog is 100%.
Bootstrap support for afis|EAW18420.1 as seed ortholog is 100%.

---

Group of orthologs #888. Best score 2049 bits
Score difference with first non-orthologous sequence - *A. fumigatus*:2049 *A. fischeri*:2049
afum|EAL87579.1 100.00% afis|EAW18213.1 100.00%
Bootstrap support for afum|EAL87579.1 as seed ortholog is 100%.
Bootstrap support for afis|EAW18213.1 as seed ortholog is 100%.

---

Group of orthologs #889. Best score 2049 bits
Score difference with first non-orthologous sequence - *A. fumigatus*:2049 *A. fischeri*:2049
afum|EAL89562.1 100.00% afis|EAW24909.1 100.00%
Bootstrap support for afum|EAL89562.1 as seed ortholog is 100%.
Bootstrap support for afis|EAW24909.1 as seed ortholog is 100%.

---

Group of orthologs #890. Best score 2049 bits
Score difference with first non-orthologous sequence - *A. fumigatus*:2049 *A. fischeri*:2049
afum|EAL89979.1 100.00% afis|EAW25318.1 100.00%
Bootstrap support for afum|EAL89979.1 as seed ortholog is 100%.
Bootstrap support for afis|EAW25318.1 as seed ortholog is 100%.

---

Group of orthologs #891. Best score 2049 bits
Score difference with first non-orthologous sequence - *A. fumigatus*:2049 *A. fischeri*:2049
afum|EAL90171.1 100.00% afis|EAW25505.1 100.00%
Bootstrap support for afum|EAL90171.1 as seed ortholog is 100%.
Bootstrap support for afis|EAW25505.1 as seed ortholog is 100%.

---

Group of orthologs #892. Best score 2048 bits
Score difference with first non-orthologous sequence - *A. fumigatus*:2048 *A. fischeri*:2048
afum|EAL84535.1 100.00% afis|EAW20522.1 100.00%
Bootstrap support for afum|EAL84535.1 as seed ortholog is 100%.
Bootstrap support for afis|EAW20522.1 as seed ortholog is 100%.

---

Group of orthologs #893. Best score 2048 bits
Score difference with first non-orthologous sequence - *A. fumigatus*:2048 *A. fischeri*:2048
afum|EAL91543.1 100.00% afis|EAW17728.1 100.00%
Bootstrap support for afum|EAL91543.1 as seed ortholog is 100%.
Bootstrap support for afis|EAW17728.1 as seed ortholog is 100%.

---

Group of orthologs #894. Best score 2047 bits
Score difference with first non-orthologous sequence - *A. fumigatus*:2047 *A. fischeri*:2047
afum|EAL87871.1 100.00% afis|EAW23596.1 100.00%
Bootstrap support for afum|EAL87871.1 as seed ortholog is 100%.
Bootstrap support for afis|EAW23596.1 as seed ortholog is 100%.

---

Group of orthologs #895. Best score 2046 bits
Score difference with first non-orthologous sequence - *A. fumigatus*:2046 *A. fischeri*:2046
afum|EAL88367.1 100.00% afis|EAW23112.1 100.00%
Bootstrap support for afum|EAL88367.1 as seed ortholog is 100%.
Bootstrap support for afis|EAW23112.1 as seed ortholog is 100%.

---

Group of orthologs #896. Best score 2045 bits
Score difference with first non-orthologous sequence - *A. fumigatus*:2045 *A. fischeri*:2045
afum|EAL89258.1 100.00% afis|EAW16654.1 100.00%
Bootstrap support for afum|EAL89258.1 as seed ortholog is 100%.
Bootstrap support for afis|EAW16654.1 as seed ortholog is 100%.

---

Group of orthologs #897. Best score 2043 bits
Score difference with first non-orthologous sequence - *A. fumigatus*:2043 *A. fischeri*:2043
afum|EAL93106.1 100.00% afis|EAW18422.1 100.00%
Bootstrap support for afum|EAL93106.1 as seed ortholog is 100%.
Bootstrap support for afis|EAW18422.1 as seed ortholog is 100%.

---

Group of orthologs #898. Best score 2042 bits
Score difference with first non-orthologous sequence - *A. fumigatus*:2042 *A. fischeri*:2042
afum|EAL90256.1 100.00% afis|EAW22940.1 100.00%
Bootstrap support for afum|EAL90256.1 as seed ortholog is 100%.
Bootstrap support for afis|EAW22940.1 as seed ortholog is 100%.

---

Group of orthologs #899. Best score 2042 bits
Score difference with first non-orthologous sequence - *A. fumigatus*:2042 *A. fischeri*:2042
afum|EAL92380.1 100.00% afis|EAW21364.1 100.00%
Bootstrap support for afum|EAL92380.1 as seed ortholog is 100%.
Bootstrap support for afis|EAW21364.1 as seed ortholog is 100%.

---

Group of orthologs #900. Best score 2041 bits
Score difference with first non-orthologous sequence - *A. fumigatus*:2041 *A. fischeri*:2041
afum|EAL91010.1 100.00% afis|EAW22179.1 100.00%
Bootstrap support for afum|EAL91010.1 as seed ortholog is 100%.
Bootstrap support for afis|EAW22179.1 as seed ortholog is 100%.

---

Group of orthologs #901. Best score 2040 bits
Score difference with first non-orthologous sequence - *A. fumigatus*:2040 *A. fischeri*:2040
afum|EAL86911.1 100.00% afis|EAW19611.1 100.00%
Bootstrap support for afum|EAL86911.1 as seed ortholog is 100%.
Bootstrap support for afis|EAW19611.1 as seed ortholog is 100%.

---

Group of orthologs #902. Best score 2040 bits
Score difference with first non-orthologous sequence - *A. fumigatus*:2040 *A. fischeri*:2040
afum|EAL87326.1 100.00% afis|EAW23899.1 100.00%
Bootstrap support for afum|EAL87326.1 as seed ortholog is 100%.
Bootstrap support for afis|EAW23899.1 as seed ortholog is 100%.

---

Group of orthologs #903. Best score 2040 bits

Score difference with first non-orthologous sequence - *A. fumigatus*:2040   *A. fischeri*:2040
afum|EAL89728.1 100.00% afis|EAW25060.1 100.00%
Bootstrap support for afum|EAL89728.1 as seed ortholog is 100%.
Bootstrap support for afis|EAW25060.1 as seed ortholog is 100%.

---

Group of orthologs #904. Best score 2039 bits
Score difference with first non-orthologous sequence - *A. fumigatus*:2039   *A. fischeri*:2039
afum|EAL85453.1 100.00% afis|EAW20327.1 100.00%
Bootstrap support for afum|EAL85453.1 as seed ortholog is 100%.
Bootstrap support for afis|EAW20327.1 as seed ortholog is 100%.

---

Group of orthologs #905. Best score 2039 bits
Score difference with first non-orthologous sequence - *A. fumigatus*:2039   *A. fischeri*:2039
afum|EAL91559.1 100.00% afis|EAW17747.1 100.00%
Bootstrap support for afum|EAL91559.1 as seed ortholog is 100%.
Bootstrap support for afis|EAW17747.1 as seed ortholog is 100%.

---

Group of orthologs #906. Best score 2039 bits
Score difference with first non-orthologous sequence - *A. fumigatus*:2039   *A. fischeri*:2039
afum|EAL93537.1 100.00% afis|EAW18801.1 100.00%
Bootstrap support for afum|EAL93537.1 as seed ortholog is 100%.
Bootstrap support for afis|EAW18801.1 as seed ortholog is 100%.

---

Group of orthologs #907. Best score 2039 bits
Score difference with first non-orthologous sequence - *A. fumigatus*:2039   *A. fischeri*:2039
afum|EAL89537.1 100.00% afis|EAW24886.1 100.00%
Bootstrap support for afum|EAL89537.1 as seed ortholog is 100%.
Bootstrap support for afis|EAW24886.1 as seed ortholog is 100%.

---

Group of orthologs #908. Best score 2037 bits
Score difference with first non-orthologous sequence - *A. fumigatus*:2037   *A. fischeri*:2037
afum|EAL93862.1 100.00% afis|EAW19130.1 100.00%
Bootstrap support for afum|EAL93862.1 as seed ortholog is 100%.
Bootstrap support for afis|EAW19130.1 as seed ortholog is 100%.

---

Group of orthologs #909. Best score 2037 bits
Score difference with first non-orthologous sequence - *A. fumigatus*:2037   *A. fischeri*:2037
afum|EAL92338.1 100.00% afis|EAW21321.1 100.00%
Bootstrap support for afum|EAL92338.1 as seed ortholog is 100%.
Bootstrap support for afis|EAW21321.1 as seed ortholog is 100%.

---

Group of orthologs #910. Best score 2037 bits
Score difference with first non-orthologous sequence - *A. fumigatus*:2037   *A. fischeri*:2037
afum|EAL89379.1 100.00% afis|EAW24732.1 100.00%
Bootstrap support for afum|EAL89379.1 as seed ortholog is 100%.
Bootstrap support for afis|EAW24732.1 as seed ortholog is 100%.

---

Group of orthologs #911. Best score 2036 bits
Score difference with first non-orthologous sequence - *A. fumigatus*:2036   *A. fischeri*:2036
afum|EAL93515.1 100.00% afis|EAW18783.1 100.00%
Bootstrap support for afum|EAL93515.1 as seed ortholog is 100%.
Bootstrap support for afis|EAW18783.1 as seed ortholog is 100%.

---

Group of orthologs #912. Best score 2033 bits
Score difference with first non-orthologous sequence - *A. fumigatus*:2033   *A. fischeri*:2033
afum|EAL87037.1 100.00% afis|EAW19459.1 100.00%

Bootstrap support for afum|EAL87037.1 as seed ortholog is 100%.
Bootstrap support for afis|EAW19459.1 as seed ortholog is 100%.

---

Group of orthologs #913. Best score 2033 bits
Score difference with first non-orthologous sequence - *A. fumigatus*:1840 *A. fischeri*:2033
afum|EAL90633.1 100.00% afis|EAW22562.1 100.00%
Bootstrap support for afum|EAL90633.1 as seed ortholog is 100%.
Bootstrap support for afis|EAW22562.1 as seed ortholog is 100%.

---

Group of orthologs #914. Best score 2032 bits
Score difference with first non-orthologous sequence - *A. fumigatus*:2032 *A. fischeri*:2032
afum|EAL88774.1 100.00% afis|EAW16145.1 100.00%
Bootstrap support for afum|EAL88774.1 as seed ortholog is 100%.
Bootstrap support for afis|EAW16145.1 as seed ortholog is 100%.

---

Group of orthologs #915. Best score 2032 bits
Score difference with first non-orthologous sequence - *A. fumigatus*:2032 *A. fischeri*:2032
afum|EAL89693.1 100.00% afis|EAW25005.1 100.00%
Bootstrap support for afum|EAL89693.1 as seed ortholog is 100%.
Bootstrap support for afis|EAW25005.1 as seed ortholog is 100%.

---

Group of orthologs #916. Best score 2031 bits
Score difference with first non-orthologous sequence - *A. fumigatus*:2031 *A. fischeri*:2031
afum|EAL88550.2 100.00% afis|EAW15915.1 100.00%
Bootstrap support for afum|EAL88550.2 as seed ortholog is 100%.
Bootstrap support for afis|EAW15915.1 as seed ortholog is 100%.

---

Group of orthologs #917. Best score 2029 bits
Score difference with first non-orthologous sequence - *A. fumigatus*:2029 *A. fischeri*:2029
afum|EAL87658.1 100.00% afis|EAW18292.1 100.00%
Bootstrap support for afum|EAL87658.1 as seed ortholog is 100%.
Bootstrap support for afis|EAW18292.1 as seed ortholog is 100%.

---

Group of orthologs #918. Best score 2029 bits
Score difference with first non-orthologous sequence - *A. fumigatus*:2029 *A. fischeri*:2029
afum|EAL89731.2 100.00% afis|EAW25062.1 100.00%
Bootstrap support for afum|EAL89731.2 as seed ortholog is 100%.
Bootstrap support for afis|EAW25062.1 as seed ortholog is 100%.

---

Group of orthologs #919. Best score 2028 bits
Score difference with first non-orthologous sequence - *A. fumigatus*:2028 *A. fischeri*:2028
afum|EAL88181.2 100.00% afis|EAW23298.1 100.00%
Bootstrap support for afum|EAL88181.2 as seed ortholog is 100%.
Bootstrap support for afis|EAW23298.1 as seed ortholog is 100%.

---

Group of orthologs #920. Best score 2027 bits
Score difference with first non-orthologous sequence - *A. fumigatus*:2027 *A. fischeri*:2027
afum|EAL86919.1 100.00% afis|EAW19599.1 100.00%
Bootstrap support for afum|EAL86919.1 as seed ortholog is 100%.
Bootstrap support for afis|EAW19599.1 as seed ortholog is 100%.

---

Group of orthologs #921. Best score 2027 bits
Score difference with first non-orthologous sequence - *A. fumigatus*:2027 *A. fischeri*:1836
afum|EAL92854.1 100.00% afis|EAW21842.1 100.00%
Bootstrap support for afum|EAL92854.1 as seed ortholog is 100%.
Bootstrap support for afis|EAW21842.1 as seed ortholog is 100%.

Group of orthologs #922. Best score 2025 bits
Score difference with first non-orthologous sequence - *A. fumigatus*:2025 *A. fischeri*:641
afum|EAL88803.1 100.00% afis|EAW16169.1 100.00%
Bootstrap support for afum|EAL88803.1 as seed ortholog is 100%.
Bootstrap support for afis|EAW16169.1 as seed ortholog is 100%.

---

Group of orthologs #923. Best score 2025 bits
Score difference with first non-orthologous sequence - *A. fumigatus*:2025 *A. fischeri*:2025
afum|EAL91422.1 100.00% afis|EAW17609.1 100.00%
Bootstrap support for afum|EAL91422.1 as seed ortholog is 100%.
Bootstrap support for afis|EAW17609.1 as seed ortholog is 100%.

---

Group of orthologs #924. Best score 2025 bits
Score difference with first non-orthologous sequence - *A. fumigatus*:2025 *A. fischeri*:2025
afum|EAL90788.1 100.00% afis|EAW22403.1 100.00%
Bootstrap support for afum|EAL90788.1 as seed ortholog is 100%.
Bootstrap support for afis|EAW22403.1 as seed ortholog is 100%.

---

Group of orthologs #925. Best score 2024 bits
Score difference with first non-orthologous sequence - *A. fumigatus*:2024 *A. fischeri*:2024
afum|EAL89496.1 100.00% afis|EAW24850.1 100.00%
Bootstrap support for afum|EAL89496.1 as seed ortholog is 100%.
Bootstrap support for afis|EAW24850.1 as seed ortholog is 100%.

---

Group of orthologs #926. Best score 2023 bits
Score difference with first non-orthologous sequence - *A. fumigatus*:2023 *A. fischeri*:2023
afum|EAL93426.1 100.00% afis|EAW18703.1 100.00%
Bootstrap support for afum|EAL93426.1 as seed ortholog is 100%.
Bootstrap support for afis|EAW18703.1 as seed ortholog is 100%.

---

Group of orthologs #927. Best score 2022 bits
Score difference with first non-orthologous sequence - *A. fumigatus*:2022 *A. fischeri*:2022
afum|EAL88441.1 100.00% afis|EAW23040.1 100.00%
Bootstrap support for afum|EAL88441.1 as seed ortholog is 100%.
Bootstrap support for afis|EAW23040.1 as seed ortholog is 100%.

---

Group of orthologs #928. Best score 2021 bits
Score difference with first non-orthologous sequence - *A. fumigatus*:2021 *A. fischeri*:2021
afum|EAL85327.1 100.00% afis|EAW20210.1 100.00%
Bootstrap support for afum|EAL85327.1 as seed ortholog is 100%.
Bootstrap support for afis|EAW20210.1 as seed ortholog is 100%.

---

Group of orthologs #929. Best score 2021 bits
Score difference with first non-orthologous sequence - *A. fumigatus*:2021 *A. fischeri*:2021
afum|EAL92192.1 100.00% afis|EAW21172.1 100.00%
Bootstrap support for afum|EAL92192.1 as seed ortholog is 100%.
Bootstrap support for afis|EAW21172.1 as seed ortholog is 100%.

---

Group of orthologs #930. Best score 2020 bits
Score difference with first non-orthologous sequence - *A. fumigatus*:2020 *A. fischeri*:2020
afum|EAL92185.1 100.00% afis|EAW21166.1 100.00%
Bootstrap support for afum|EAL92185.1 as seed ortholog is 100%.
Bootstrap support for afis|EAW21166.1 as seed ortholog is 100%.

---

Group of orthologs #931. Best score 2019 bits

Score difference with first non-orthologous sequence - *A. fumigatus*:2019 *A. fischeri*:2019
afum|EAL92336.1 100.00% afis|EAW21319.1 100.00%
Bootstrap support for afum|EAL92336.1 as seed ortholog is 100%.
Bootstrap support for afis|EAW21319.1 as seed ortholog is 100%.

---

Group of orthologs #932. Best score 2018 bits
Score difference with first non-orthologous sequence - *A. fumigatus*:1916 *A. fischeri*:2018
afum|EAL90683.1 100.00% afis|EAW22510.1 100.00%
Bootstrap support for afum|EAL90683.1 as seed ortholog is 100%.
Bootstrap support for afis|EAW22510.1 as seed ortholog is 100%.

---

Group of orthologs #933. Best score 2016 bits
Score difference with first non-orthologous sequence - *A. fumigatus*:2016 *A. fischeri*:2016
afum|EAL88213.1 100.00% afis|EAW23266.1 100.00%
Bootstrap support for afum|EAL88213.1 as seed ortholog is 100%.
Bootstrap support for afis|EAW23266.1 as seed ortholog is 100%.

---

Group of orthologs #934. Best score 2016 bits
Score difference with first non-orthologous sequence - *A. fumigatus*:2016 *A. fischeri*:2016
afum|EAL89534.1 100.00% afis|EAW24884.1 100.00%
Bootstrap support for afum|EAL89534.1 as seed ortholog is 100%.
Bootstrap support for afis|EAW24884.1 as seed ortholog is 100%.

---

Group of orthologs #935. Best score 2014 bits
Score difference with first non-orthologous sequence - *A. fumigatus*:2014 *A. fischeri*:2014
afum|EAL85619.1 100.00% afis|EAW15728.1 100.00%
Bootstrap support for afum|EAL85619.1 as seed ortholog is 100%.
Bootstrap support for afis|EAW15728.1 as seed ortholog is 100%.

---

Group of orthologs #936. Best score 2014 bits
Score difference with first non-orthologous sequence - *A. fumigatus*:2014 *A. fischeri*:2014
afum|EAL92004.1 100.00% afis|EAW24149.1 100.00%
Bootstrap support for afum|EAL92004.1 as seed ortholog is 100%.
Bootstrap support for afis|EAW24149.1 as seed ortholog is 100%.

---

Group of orthologs #937. Best score 2013 bits
Score difference with first non-orthologous sequence - *A. fumigatus*:2013 *A. fischeri*:2013
afum|EAL88221.1 100.00% afis|EAW23260.1 100.00%
Bootstrap support for afum|EAL88221.1 as seed ortholog is 100%.
Bootstrap support for afis|EAW23260.1 as seed ortholog is 100%.

---

Group of orthologs #938. Best score 2011 bits
Score difference with first non-orthologous sequence - *A. fumigatus*:2011 *A. fischeri*:2011
afum|EAL85194.1 100.00% afis|EAW20076.1 100.00%
Bootstrap support for afum|EAL85194.1 as seed ortholog is 100%.
Bootstrap support for afis|EAW20076.1 as seed ortholog is 100%.

---

Group of orthologs #939. Best score 2011 bits
Score difference with first non-orthologous sequence - *A. fumigatus*:2011 *A. fischeri*:1854
afum|EAL88797.1 100.00% afis|EAW20720.1 100.00%
Bootstrap support for afum|EAL88797.1 as seed ortholog is 100%.
Bootstrap support for afis|EAW20720.1 as seed ortholog is 100%.

---

Group of orthologs #940. Best score 2011 bits
Score difference with first non-orthologous sequence - *A. fumigatus*:2011 *A. fischeri*:2011
afum|EAL88122.2 100.00% afis|EAW23354.1 100.00%

Bootstrap support for afum|EAL88122.2 as seed ortholog is 100%.
Bootstrap support for afis|EAW23354.1 as seed ortholog is 100%.

---

Group of orthologs #941. Best score 2010 bits
Score difference with first non-orthologous sequence - *A. fumigatus*:2010 *A. fischeri*:2010
afum|EAL91181.1 100.00% afis|EAW17337.1 100.00%
Bootstrap support for afum|EAL91181.1 as seed ortholog is 100%.
Bootstrap support for afis|EAW17337.1 as seed ortholog is 100%.

---

Group of orthologs #942. Best score 2009 bits
Score difference with first non-orthologous sequence - *A. fumigatus*:2009 *A. fischeri*:1932
afum|EBA27220.1 100.00% afis|EAW19769.1 100.00%
Bootstrap support for afum|EBA27220.1 as seed ortholog is 100%.
Bootstrap support for afis|EAW19769.1 as seed ortholog is 100%.

---

Group of orthologs #943. Best score 2008 bits
Score difference with first non-orthologous sequence - *A. fumigatus*:2008 *A. fischeri*:2008
afum|EAL89553.1 100.00% afis|EAW24899.1 100.00%
Bootstrap support for afum|EAL89553.1 as seed ortholog is 100%.
Bootstrap support for afis|EAW24899.1 as seed ortholog is 100%.

---

Group of orthologs #944. Best score 2006 bits
Score difference with first non-orthologous sequence - *A. fumigatus*:2006 *A. fischeri*:2006
afum|EAL88650.1 100.00% afis|EAW16021.1 100.00%
Bootstrap support for afum|EAL88650.1 as seed ortholog is 100%.
Bootstrap support for afis|EAW16021.1 as seed ortholog is 100%.

---

Group of orthologs #945. Best score 2006 bits
Score difference with first non-orthologous sequence - *A. fumigatus*:2006 *A. fischeri*:2006
afum|EAL85166.1 100.00% afis|EAW20045.1 100.00%
Bootstrap support for afum|EAL85166.1 as seed ortholog is 100%.
Bootstrap support for afis|EAW20045.1 as seed ortholog is 100%.

---

Group of orthologs #946. Best score 2006 bits
Score difference with first non-orthologous sequence - *A. fumigatus*:2006 *A. fischeri*:2006
afum|EAL87925.1 100.00% afis|EAW23541.1 100.00%
Bootstrap support for afum|EAL87925.1 as seed ortholog is 100%.
Bootstrap support for afis|EAW23541.1 as seed ortholog is 100%.

---

Group of orthologs #947. Best score 2005 bits
Score difference with first non-orthologous sequence - *A. fumigatus*:2005 *A. fischeri*:2005
afum|EAL90739.1 100.00% afis|EAW22452.1 100.00%
Bootstrap support for afum|EAL90739.1 as seed ortholog is 100%.
Bootstrap support for afis|EAW22452.1 as seed ortholog is 100%.

---

Group of orthologs #948. Best score 2004 bits
Score difference with first non-orthologous sequence - *A. fumigatus*:2004 *A. fischeri*:2004
afum|EAL93390.1 100.00% afis|EAW18671.1 100.00%
Bootstrap support for afum|EAL93390.1 as seed ortholog is 100%.
Bootstrap support for afis|EAW18671.1 as seed ortholog is 100%.

---

Group of orthologs #949. Best score 2004 bits
Score difference with first non-orthologous sequence - *A. fumigatus*:2004 *A. fischeri*:2004
afum|EAL93643.1 100.00% afis|EAW18912.1 100.00%
Bootstrap support for afum|EAL93643.1 as seed ortholog is 100%.
Bootstrap support for afis|EAW18912.1 as seed ortholog is 100%.

---

Group of orthologs #950. Best score 2003 bits
Score difference with first non-orthologous sequence - *A. fumigatus*:2003 *A. fischeri*:2003
afum|EAL86951.1 100.00% afis|EAW19560.1 100.00%
Bootstrap support for afum|EAL86951.1 as seed ortholog is 100%.
Bootstrap support for afis|EAW19560.1 as seed ortholog is 100%.

---

Group of orthologs #951. Best score 2003 bits
Score difference with first non-orthologous sequence - *A. fumigatus*:2003 *A. fischeri*:2003
afum|EAL91845.1 100.00% afis|EAW18034.1 100.00%
Bootstrap support for afum|EAL91845.1 as seed ortholog is 100%.
Bootstrap support for afis|EAW18034.1 as seed ortholog is 100%.

---

Group of orthologs #952. Best score 2002 bits
Score difference with first non-orthologous sequence - *A. fumigatus*:2002 *A. fischeri*:2002
afum|EAL91684.1 100.00% afis|EAW17876.1 100.00%
Bootstrap support for afum|EAL91684.1 as seed ortholog is 100%.
Bootstrap support for afis|EAW17876.1 as seed ortholog is 100%.

---

Group of orthologs #953. Best score 2002 bits
Score difference with first non-orthologous sequence - *A. fumigatus*:2002 *A. fischeri*:2002
afum|EAL92490.1 100.00% afis|EAW21471.1 100.00%
Bootstrap support for afum|EAL92490.1 as seed ortholog is 100%.
Bootstrap support for afis|EAW21471.1 as seed ortholog is 100%.

---

Group of orthologs #954. Best score 2000 bits
Score difference with first non-orthologous sequence - *A. fumigatus*:2000 *A. fischeri*:2000
afum|EBA27413.1 100.00% afis|EAW22391.1 100.00%
Bootstrap support for afum|EBA27413.1 as seed ortholog is 100%.
Bootstrap support for afis|EAW22391.1 as seed ortholog is 100%.

---

Group of orthologs #955. Best score 1999 bits
Score difference with first non-orthologous sequence - *A. fumigatus*:1999 *A. fischeri*:1999
afum|EAL89069.1 100.00% afis|EAW16475.1 100.00%
Bootstrap support for afum|EAL89069.1 as seed ortholog is 100%.
Bootstrap support for afis|EAW16475.1 as seed ortholog is 100%.

---

Group of orthologs #956. Best score 1999 bits
Score difference with first non-orthologous sequence - *A. fumigatus*:1999 *A. fischeri*:1999
afum|EAL90305.2 100.00% afis|EAW22893.1 100.00%
Bootstrap support for afum|EAL90305.2 as seed ortholog is 100%.
Bootstrap support for afis|EAW22893.1 as seed ortholog is 100%.

---

Group of orthologs #957. Best score 1998 bits
Score difference with first non-orthologous sequence - *A. fumigatus*:1221 *A. fischeri*:1472
afum|EAL91877.1 100.00% afis|EAW18065.1 100.00%
Bootstrap support for afum|EAL91877.1 as seed ortholog is 100%.
Bootstrap support for afis|EAW18065.1 as seed ortholog is 100%.

---

Group of orthologs #958. Best score 1997 bits
Score difference with first non-orthologous sequence - *A. fumigatus*:1997 *A. fischeri*:1997
afum|EAL86002.1 100.00% afis|EAW24332.1 100.00%
Bootstrap support for afum|EAL86002.1 as seed ortholog is 100%.
Bootstrap support for afis|EAW24332.1 as seed ortholog is 100%.

---

Group of orthologs #959. Best score 1997 bits

Score difference with first non-orthologous sequence - *A. fumigatus*:1997 *A. fischeri*:1997
afum|EAL87204.1 100.00% afis|EAW23770.1 100.00%
Bootstrap support for afum|EAL87204.1 as seed ortholog is 100%.
Bootstrap support for afis|EAW23770.1 as seed ortholog is 100%.

---

Group of orthologs #960. Best score 1997 bits
Score difference with first non-orthologous sequence - *A. fumigatus*:1997 *A. fischeri*:1997
afum|EAL87332.1 100.00% afis|EAW23904.1 100.00%
Bootstrap support for afum|EAL87332.1 as seed ortholog is 100%.
Bootstrap support for afis|EAW23904.1 as seed ortholog is 100%.

---

Group of orthologs #961. Best score 1997 bits
Score difference with first non-orthologous sequence - *A. fumigatus*:1997 *A. fischeri*:1997
afum|EAL93999.2 100.00% afis|EAW19273.1 100.00%
Bootstrap support for afum|EAL93999.2 as seed ortholog is 100%.
Bootstrap support for afis|EAW19273.1 as seed ortholog is 100%.

---

Group of orthologs #962. Best score 1997 bits
Score difference with first non-orthologous sequence - *A. fumigatus*:1997 *A. fischeri*:1997
afum|EAL92316.1 100.00% afis|EAW21299.1 100.00%
Bootstrap support for afum|EAL92316.1 as seed ortholog is 100%.
Bootstrap support for afis|EAW21299.1 as seed ortholog is 100%.

---

Group of orthologs #963. Best score 1995 bits
Score difference with first non-orthologous sequence - *A. fumigatus*:1995 *A. fischeri*:1995
afum|EAL90298.1 100.00% afis|EAW22899.1 100.00%
Bootstrap support for afum|EAL90298.1 as seed ortholog is 100%.
Bootstrap support for afis|EAW22899.1 as seed ortholog is 100%.

---

Group of orthologs #964. Best score 1994 bits
Score difference with first non-orthologous sequence - *A. fumigatus*:1994 *A. fischeri*:1994
afum|EAL90146.1 100.00% afis|EAW25481.1 100.00%
Bootstrap support for afum|EAL90146.1 as seed ortholog is 100%.
Bootstrap support for afis|EAW25481.1 as seed ortholog is 100%.

---

Group of orthologs #965. Best score 1993 bits
Score difference with first non-orthologous sequence - *A. fumigatus*:1993 *A. fischeri*:1993
afum|EAL92766.1 100.00% afis|EAW21733.1 100.00%
Bootstrap support for afum|EAL92766.1 as seed ortholog is 100%.
Bootstrap support for afis|EAW21733.1 as seed ortholog is 100%.

---

Group of orthologs #966. Best score 1993 bits
Score difference with first non-orthologous sequence - *A. fumigatus*:1993 *A. fischeri*:1993
afum|EAL89601.1 100.00% afis|EAW24944.1 100.00%
Bootstrap support for afum|EAL89601.1 as seed ortholog is 100%.
Bootstrap support for afis|EAW24944.1 as seed ortholog is 100%.

---

Group of orthologs #967. Best score 1992 bits
Score difference with first non-orthologous sequence - *A. fumigatus*:1992 *A. fischeri*:1992
afum|EAL92775.1 100.00% afis|EAW21742.1 100.00%
Bootstrap support for afum|EAL92775.1 as seed ortholog is 100%.
Bootstrap support for afis|EAW21742.1 as seed ortholog is 100%.

---

Group of orthologs #968. Best score 1991 bits
Score difference with first non-orthologous sequence - *A. fumigatus*:1991 *A. fischeri*:1991
afum|EAL88521.2 100.00% afis|EAW15887.1 100.00%

Bootstrap support for afum|EAL88521.2 as seed ortholog is 100%.
Bootstrap support for afis|EAW15887.1 as seed ortholog is 100%.

---

Group of orthologs #969. Best score 1991 bits
Score difference with first non-orthologous sequence - *A. fumigatus*:1991 *A. fischeri*:1991
afum|EAL88958.1 100.00% afis|EAW16333.1 100.00%
Bootstrap support for afum|EAL88958.1 as seed ortholog is 100%.
Bootstrap support for afis|EAW16333.1 as seed ortholog is 100%.

---

Group of orthologs #970. Best score 1989 bits
Score difference with first non-orthologous sequence - *A. fumigatus*:1989 *A. fischeri*:1581
afum|EAL90406.1 100.00% afis|EAW22793.1 100.00%
Bootstrap support for afum|EAL90406.1 as seed ortholog is 100%.
Bootstrap support for afis|EAW22793.1 as seed ortholog is 100%.

---

Group of orthologs #971. Best score 1988 bits
Score difference with first non-orthologous sequence - *A. fumigatus*:1988 *A. fischeri*:1988
afum|EAL87562.2 100.00% afis|EAW18191.1 100.00%
Bootstrap support for afum|EAL87562.2 as seed ortholog is 100%.
Bootstrap support for afis|EAW18191.1 as seed ortholog is 100%.

---

Group of orthologs #972. Best score 1988 bits
Score difference with first non-orthologous sequence - *A. fumigatus*:1988 *A. fischeri*:1988
afum|EAL84637.1 100.00% afis|EAW20992.1 100.00%
Bootstrap support for afum|EAL84637.1 as seed ortholog is 100%.
Bootstrap support for afis|EAW20992.1 as seed ortholog is 100%.

---

Group of orthologs #973. Best score 1988 bits
Score difference with first non-orthologous sequence - *A. fumigatus*:1988 *A. fischeri*:1988
afum|EAL90565.1 100.00% afis|EAW22630.1 100.00%
Bootstrap support for afum|EAL90565.1 as seed ortholog is 100%.
Bootstrap support for afis|EAW22630.1 as seed ortholog is 100%.

---

Group of orthologs #974. Best score 1988 bits
Score difference with first non-orthologous sequence - *A. fumigatus*:1988 *A. fischeri*:1988
afum|EAL90396.1 100.00% afis|EAW22803.1 100.00%
Bootstrap support for afum|EAL90396.1 as seed ortholog is 100%.
Bootstrap support for afis|EAW22803.1 as seed ortholog is 100%.

---

Group of orthologs #975. Best score 1987 bits
Score difference with first non-orthologous sequence - *A. fumigatus*:1618 *A. fischeri*:1987
afum|EAL90494.1 100.00% afis|EAW22698.1 100.00%
Bootstrap support for afum|EAL90494.1 as seed ortholog is 100%.
Bootstrap support for afis|EAW22698.1 as seed ortholog is 100%.

---

Group of orthologs #976. Best score 1986 bits
Score difference with first non-orthologous sequence - *A. fumigatus*:1986 *A. fischeri*:1986
afum|EAL86427.2 100.00% afis|EAW16889.1 100.00%
Bootstrap support for afum|EAL86427.2 as seed ortholog is 100%.
Bootstrap support for afis|EAW16889.1 as seed ortholog is 100%.

---

Group of orthologs #977. Best score 1984 bits
Score difference with first non-orthologous sequence - *A. fumigatus*:1984 *A. fischeri*:1984
afum|EAL92424.1 100.00% afis|EAW21410.1 100.00%
Bootstrap support for afum|EAL92424.1 as seed ortholog is 100%.
Bootstrap support for afis|EAW21410.1 as seed ortholog is 100%.

Group of orthologs #978. Best score 1983 bits
Score difference with first non-orthologous sequence - *A. fumigatus*:1983 *A. fischeri*:1983
afum|EAL93648.1 100.00% afis|EAW18917.1 100.00%
Bootstrap support for afum|EAL93648.1 as seed ortholog is 100%.
Bootstrap support for afis|EAW18917.1 as seed ortholog is 100%.

Group of orthologs #979. Best score 1982 bits
Score difference with first non-orthologous sequence - *A. fumigatus*:1982 *A. fischeri*:1982
afum|EAL90279.1 100.00% afis|EAW22918.1 100.00%
Bootstrap support for afum|EAL90279.1 as seed ortholog is 100%.
Bootstrap support for afis|EAW22918.1 as seed ortholog is 100%.

Group of orthologs #980. Best score 1982 bits
Score difference with first non-orthologous sequence - *A. fumigatus*:1929 *A. fischeri*:1982
afum|EAL89575.1 100.00% afis|EAW24920.1 100.00%
Bootstrap support for afum|EAL89575.1 as seed ortholog is 100%.
Bootstrap support for afis|EAW24920.1 as seed ortholog is 100%.

Group of orthologs #981. Best score 1981 bits
Score difference with first non-orthologous sequence - *A. fumigatus*:1981 *A. fischeri*:1981
afum|EAL88189.1 100.00% afis|EAW23290.1 100.00%
Bootstrap support for afum|EAL88189.1 as seed ortholog is 100%.
Bootstrap support for afis|EAW23290.1 as seed ortholog is 100%.

Group of orthologs #982. Best score 1979 bits
Score difference with first non-orthologous sequence - *A. fumigatus*:1979 *A. fischeri*:1979
afum|EAL85875.1 100.00% afis|EAW24205.1 100.00%
Bootstrap support for afum|EAL85875.1 as seed ortholog is 100%.
Bootstrap support for afis|EAW24205.1 as seed ortholog is 100%.

Group of orthologs #983. Best score 1979 bits
Score difference with first non-orthologous sequence - *A. fumigatus*:1979 *A. fischeri*:1979
afum|EAL89733.1 100.00% afis|EAW25064.1 100.00%
Bootstrap support for afum|EAL89733.1 as seed ortholog is 100%.
Bootstrap support for afis|EAW25064.1 as seed ortholog is 100%.

Group of orthologs #984. Best score 1978 bits
Score difference with first non-orthologous sequence - *A. fumigatus*:1978 *A. fischeri*:1978
afum|EAL91464.1 100.00% afis|EAW17648.1 100.00%
Bootstrap support for afum|EAL91464.1 as seed ortholog is 100%.
Bootstrap support for afis|EAW17648.1 as seed ortholog is 100%.

Group of orthologs #985. Best score 1977 bits
Score difference with first non-orthologous sequence - *A. fumigatus*:1133 *A. fischeri*:1469
afum|EAL89285.1 100.00% afis|EAW16681.1 100.00%
Bootstrap support for afum|EAL89285.1 as seed ortholog is 100%.
Bootstrap support for afis|EAW16681.1 as seed ortholog is 100%.

Group of orthologs #986. Best score 1977 bits
Score difference with first non-orthologous sequence - *A. fumigatus*:1977 *A. fischeri*:1977
afum|EAL93881.1 100.00% afis|EAW19150.1 100.00%
Bootstrap support for afum|EAL93881.1 as seed ortholog is 100%.
Bootstrap support for afis|EAW19150.1 as seed ortholog is 100%.

Group of orthologs #987. Best score 1977 bits

Score difference with first non-orthologous sequence - *A. fumigatus*:1977 *A. fischeri*:1977
afum|EAL92861.2 100.00% afis|EAW21848.1 100.00%
Bootstrap support for afum|EAL92861.2 as seed ortholog is 100%.
Bootstrap support for afis|EAW21848.1 as seed ortholog is 100%.

---

Group of orthologs #988. Best score 1976 bits
Score difference with first non-orthologous sequence - *A. fumigatus*:1976 *A. fischeri*:1976
afum|EAL91372.1 100.00% afis|EAW17558.1 100.00%
Bootstrap support for afum|EAL91372.1 as seed ortholog is 100%.
Bootstrap support for afis|EAW17558.1 as seed ortholog is 100%.

---

Group of orthologs #989. Best score 1976 bits
Score difference with first non-orthologous sequence - *A. fumigatus*:1976 *A. fischeri*:1976
afum|EAL92301.1 100.00% afis|EAW21284.1 100.00%
Bootstrap support for afum|EAL92301.1 as seed ortholog is 100%.
Bootstrap support for afis|EAW21284.1 as seed ortholog is 100%.

---

Group of orthologs #990. Best score 1975 bits
Score difference with first non-orthologous sequence - *A. fumigatus*:1975 *A. fischeri*:1975
afum|EAL86321.1 100.00% afis|EAW16746.1 100.00%
Bootstrap support for afum|EAL86321.1 as seed ortholog is 100%.
Bootstrap support for afis|EAW16746.1 as seed ortholog is 100%.

---

Group of orthologs #991. Best score 1975 bits
Score difference with first non-orthologous sequence - *A. fumigatus*:1975 *A. fischeri*:1975
afum|EAL85410.1 100.00% afis|EAW20276.1 100.00%
Bootstrap support for afum|EAL85410.1 as seed ortholog is 100%.
Bootstrap support for afis|EAW20276.1 as seed ortholog is 100%.

---

Group of orthologs #992. Best score 1975 bits
Score difference with first non-orthologous sequence - *A. fumigatus*:1975 *A. fischeri*:1975
afum|EAL85973.1 100.00% afis|EAW24305.1 100.00%
Bootstrap support for afum|EAL85973.1 as seed ortholog is 100%.
Bootstrap support for afis|EAW24305.1 as seed ortholog is 100%.

---

Group of orthologs #993. Best score 1975 bits
Score difference with first non-orthologous sequence - *A. fumigatus*:1975 *A. fischeri*:1975
afum|EAL90128.2 100.00% afis|EAW25465.1 100.00%
Bootstrap support for afum|EAL90128.2 as seed ortholog is 100%.
Bootstrap support for afis|EAW25465.1 as seed ortholog is 100%.

---

Group of orthologs #994. Best score 1974 bits
Score difference with first non-orthologous sequence - *A. fumigatus*:1974 *A. fischeri*:1974
afum|EAL85645.1 100.00% afis|EAW15702.1 100.00%
Bootstrap support for afum|EAL85645.1 as seed ortholog is 100%.
Bootstrap support for afis|EAW15702.1 as seed ortholog is 100%.

---

Group of orthologs #995. Best score 1974 bits
Score difference with first non-orthologous sequence - *A. fumigatus*:1201 *A. fischeri*:1495
afum|EAL89033.1 100.00% afis|EAW16409.1 100.00%
Bootstrap support for afum|EAL89033.1 as seed ortholog is 100%.
Bootstrap support for afis|EAW16409.1 as seed ortholog is 100%.

---

Group of orthologs #996. Best score 1974 bits
Score difference with first non-orthologous sequence - *A. fumigatus*:1863 *A. fischeri*:1888
afum|EAL93921.1 100.00% afis|EAW19189.1 100.00%

Bootstrap support for afum|EAL93921.1 as seed ortholog is 100%.
Bootstrap support for afis|EAW19189.1 as seed ortholog is 100%.

---

Group of orthologs #997. Best score 1974 bits
Score difference with first non-orthologous sequence - *A. fumigatus*:1974 *A. fischeri*:1974
afum|EBA27208.1 100.00% afis|EAW20802.1 100.00%
Bootstrap support for afum|EBA27208.1 as seed ortholog is 100%.
Bootstrap support for afis|EAW20802.1 as seed ortholog is 100%.

---

Group of orthologs #998. Best score 1974 bits
Score difference with first non-orthologous sequence - *A. fumigatus*:1974 *A. fischeri*:1974
afum|EAL92984.1 100.00% afis|EAW21970.1 100.00%
Bootstrap support for afum|EAL92984.1 as seed ortholog is 100%.
Bootstrap support for afis|EAW21970.1 as seed ortholog is 100%.

---

Group of orthologs #999. Best score 1973 bits
Score difference with first non-orthologous sequence - *A. fumigatus*:1973 *A. fischeri*:1973
afum|EAL84668.1 100.00% afis|EAW20965.1 100.00%
Bootstrap support for afum|EAL84668.1 as seed ortholog is 100%.
Bootstrap support for afis|EAW20965.1 as seed ortholog is 100%.

---

Group of orthologs #1000. Best score 1972 bits
Score difference with first non-orthologous sequence - *A. fumigatus*:1972 *A. fischeri*:1214
afum|EAL85004.1 100.00% afis|EAW19900.1 100.00%
Bootstrap support for afum|EAL85004.1 as seed ortholog is 100%.
Bootstrap support for afis|EAW19900.1 as seed ortholog is 100%.

---

Group of orthologs #1001. Best score 1972 bits
Score difference with first non-orthologous sequence - *A. fumigatus*:1972 *A. fischeri*:1972
afum|EBA27491.1 100.00% afis|EAW18663.1 100.00%
Bootstrap support for afum|EBA27491.1 as seed ortholog is 100%.
Bootstrap support for afis|EAW18663.1 as seed ortholog is 100%.

---

Group of orthologs #1002. Best score 1970 bits
Score difference with first non-orthologous sequence - *A. fumigatus*:1970 *A. fischeri*:1970
afum|EAL90139.1 100.00% afis|EAW25476.1 100.00%
Bootstrap support for afum|EAL90139.1 as seed ortholog is 100%.
Bootstrap support for afis|EAW25476.1 as seed ortholog is 100%.

---

Group of orthologs #1003. Best score 1970 bits
Score difference with first non-orthologous sequence - *A. fumigatus*:1970 *A. fischeri*:1970
afum|EBA27300.1 100.00% afis|EAW23683.1 100.00%
Bootstrap support for afum|EBA27300.1 as seed ortholog is 100%.
Bootstrap support for afis|EAW23683.1 as seed ortholog is 100%.

---

Group of orthologs #1004. Best score 1968 bits
Score difference with first non-orthologous sequence - *A. fumigatus*:1968 *A. fischeri*:1968
afum|EAL92919.2 100.00% afis|EAW21907.1 100.00%
Bootstrap support for afum|EAL92919.2 as seed ortholog is 100%.
Bootstrap support for afis|EAW21907.1 as seed ortholog is 100%.

---

Group of orthologs #1005. Best score 1967 bits
Score difference with first non-orthologous sequence - *A. fumigatus*:1967 *A. fischeri*:1967
afum|EAL91759.1 100.00% afis|EAW17949.1 100.00%
Bootstrap support for afum|EAL91759.1 as seed ortholog is 100%.
Bootstrap support for afis|EAW17949.1 as seed ortholog is 100%.

---

Group of orthologs #1006. Best score 1967 bits
Score difference with first non-orthologous sequence - *A. fumigatus*:1967 *A. fischeri*:1967
afum|EAL87445.1 100.00% afis|EAW24015.1 100.00%
Bootstrap support for afum|EAL87445.1 as seed ortholog is 100%.
Bootstrap support for afis|EAW24015.1 as seed ortholog is 100%.

---

Group of orthologs #1007. Best score 1966 bits
Score difference with first non-orthologous sequence - *A. fumigatus*:1966 *A. fischeri*:1966
afum|EAL91092.1 100.00% afis|EAW22078.1 100.00%
Bootstrap support for afum|EAL91092.1 as seed ortholog is 100%.
Bootstrap support for afis|EAW22078.1 as seed ortholog is 100%.

---

Group of orthologs #1008. Best score 1966 bits
Score difference with first non-orthologous sequence - *A. fumigatus*:1966 *A. fischeri*:1966
afum|EAL89650.1 100.00% afis|EAW24990.1 100.00%
Bootstrap support for afum|EAL89650.1 as seed ortholog is 100%.
Bootstrap support for afis|EAW24990.1 as seed ortholog is 100%.

---

Group of orthologs #1009. Best score 1965 bits
Score difference with first non-orthologous sequence - *A. fumigatus*:1402 *A. fischeri*:1491
afum|EAL86307.1 100.00% afis|EAW16731.1 100.00%
Bootstrap support for afum|EAL86307.1 as seed ortholog is 100%.
Bootstrap support for afis|EAW16731.1 as seed ortholog is 100%.

---

Group of orthologs #1010. Best score 1963 bits
Score difference with first non-orthologous sequence - *A. fumigatus*:1963 *A. fischeri*:1963
afum|EAL89734.1 100.00% afis|EAW25065.1 100.00%
Bootstrap support for afum|EAL89734.1 as seed ortholog is 100%.
Bootstrap support for afis|EAW25065.1 as seed ortholog is 100%.

---

Group of orthologs #1011. Best score 1962 bits
Score difference with first non-orthologous sequence - *A. fumigatus*:1962 *A. fischeri*:1962
afum|EAL93491.1 100.00% afis|EAW18760.1 100.00%
Bootstrap support for afum|EAL93491.1 as seed ortholog is 100%.
Bootstrap support for afis|EAW18760.1 as seed ortholog is 100%.

---

Group of orthologs #1012. Best score 1961 bits
Score difference with first non-orthologous sequence - *A. fumigatus*:1961 *A. fischeri*:1961
afum|EBA27357.1 100.00% afis|EAW16148.1 100.00%
Bootstrap support for afum|EBA27357.1 as seed ortholog is 100%.
Bootstrap support for afis|EAW16148.1 as seed ortholog is 100%.

---

Group of orthologs #1013. Best score 1958 bits
Score difference with first non-orthologous sequence - *A. fumigatus*:1958 *A. fischeri*:1958
afum|EAL87344.1 100.00% afis|EAW23918.1 100.00%
Bootstrap support for afum|EAL87344.1 as seed ortholog is 100%.
Bootstrap support for afis|EAW23918.1 as seed ortholog is 100%.

---

Group of orthologs #1014. Best score 1958 bits
Score difference with first non-orthologous sequence - *A. fumigatus*:1958 *A. fischeri*:1958
afum|EAL94007.1 100.00% afis|EAW19281.1 100.00%
Bootstrap support for afum|EAL94007.1 as seed ortholog is 100%.
Bootstrap support for afis|EAW19281.1 as seed ortholog is 100%.

---

Group of orthologs #1015. Best score 1957 bits

Score difference with first non-orthologous sequence - *A. fumigatus*:1957 *A. fischeri*:1957
afum|EAL93683.1 100.00% afis|EAW18949.1 100.00%
Bootstrap support for afum|EAL93683.1 as seed ortholog is 100%.
Bootstrap support for afis|EAW18949.1 as seed ortholog is 100%.

---

Group of orthologs #1016. Best score 1956 bits
Score difference with first non-orthologous sequence - *A. fumigatus*:1956 *A. fischeri*:1956
afum|EAL84483.1 100.00% afis|EAW20581.1 100.00%
Bootstrap support for afum|EAL84483.1 as seed ortholog is 100%.
Bootstrap support for afis|EAW20581.1 as seed ortholog is 100%.

---

Group of orthologs #1017. Best score 1954 bits
Score difference with first non-orthologous sequence - *A. fumigatus*:1954 *A. fischeri*:1954
afum|EAL93197.1 100.00% afis|EAW18504.1 100.00%
Bootstrap support for afum|EAL93197.1 as seed ortholog is 100%.
Bootstrap support for afis|EAW18504.1 as seed ortholog is 100%.

---

Group of orthologs #1018. Best score 1953 bits
Score difference with first non-orthologous sequence - *A. fumigatus*:1953 *A. fischeri*:1953
afum|EAL86031.1 100.00% afis|EAW24361.1 100.00%
Bootstrap support for afum|EAL86031.1 as seed ortholog is 100%.
Bootstrap support for afis|EAW24361.1 as seed ortholog is 100%.

---

Group of orthologs #1019. Best score 1951 bits
Score difference with first non-orthologous sequence - *A. fumigatus*:1951 *A. fischeri*:1951
afum|EAL92467.1 100.00% afis|EAW21448.1 100.00%
Bootstrap support for afum|EAL92467.1 as seed ortholog is 100%.
Bootstrap support for afis|EAW21448.1 as seed ortholog is 100%.

---

Group of orthologs #1020. Best score 1950 bits
Score difference with first non-orthologous sequence - *A. fumigatus*:1950 *A. fischeri*:1950
afum|EAL91831.1 100.00% afis|EAW18019.1 100.00%
Bootstrap support for afum|EAL91831.1 as seed ortholog is 100%.
Bootstrap support for afis|EAW18019.1 as seed ortholog is 100%.

---

Group of orthologs #1021. Best score 1949 bits
Score difference with first non-orthologous sequence - *A. fumigatus*:1949 *A. fischeri*:783
afum|EAL88616.1 100.00% afis|EAW15984.1 100.00%
Bootstrap support for afum|EAL88616.1 as seed ortholog is 100%.
Bootstrap support for afis|EAW15984.1 as seed ortholog is 100%.

---

Group of orthologs #1022. Best score 1949 bits
Score difference with first non-orthologous sequence - *A. fumigatus*:1949 *A. fischeri*:1949
afum|EAL88158.1 100.00% afis|EAW23318.1 100.00%
Bootstrap support for afum|EAL88158.1 as seed ortholog is 100%.
Bootstrap support for afis|EAW23318.1 as seed ortholog is 100%.

---

Group of orthologs #1023. Best score 1948 bits
Score difference with first non-orthologous sequence - *A. fumigatus*:1948 *A. fischeri*:1948
afum|EAL91531.1 100.00% afis|EAW17716.1 100.00%
Bootstrap support for afum|EAL91531.1 as seed ortholog is 100%.
Bootstrap support for afis|EAW17716.1 as seed ortholog is 100%.

---

Group of orthologs #1024. Best score 1947 bits
Score difference with first non-orthologous sequence - *A. fumigatus*:1947 *A. fischeri*:1947
afum|EAL92440.1 100.00% afis|EAW21425.1 100.00%

Bootstrap support for afum|EAL92440.1 as seed ortholog is 100%.
Bootstrap support for afis|EAW21425.1 as seed ortholog is 100%.

---

Group of orthologs #1025. Best score 1947 bits
Score difference with first non-orthologous sequence - *A. fumigatus*:1947 *A. fischeri*:1947
afum|EAL92917.1 100.00% afis|EAW21904.1 100.00%
Bootstrap support for afum|EAL92917.1 as seed ortholog is 100%.
Bootstrap support for afis|EAW21904.1 as seed ortholog is 100%.

---

Group of orthologs #1026. Best score 1946 bits
Score difference with first non-orthologous sequence - *A. fumigatus*:1946 *A. fischeri*:1946
afum|EAL86894.1 100.00% afis|EAW19627.1 100.00%
Bootstrap support for afum|EAL86894.1 as seed ortholog is 100%.
Bootstrap support for afis|EAW19627.1 as seed ortholog is 100%.

---

Group of orthologs #1027. Best score 1946 bits
Score difference with first non-orthologous sequence - *A. fumigatus*:1946 *A. fischeri*:1946
afum|EAL92536.1 100.00% afis|EAW21516.1 100.00%
Bootstrap support for afum|EAL92536.1 as seed ortholog is 100%.
Bootstrap support for afis|EAW21516.1 as seed ortholog is 100%.

---

Group of orthologs #1028. Best score 1945 bits
Score difference with first non-orthologous sequence - *A. fumigatus*:1945 *A. fischeri*:1945
afum|EAL87680.1 100.00% afis|EAW18313.1 100.00%
Bootstrap support for afum|EAL87680.1 as seed ortholog is 100%.
Bootstrap support for afis|EAW18313.1 as seed ortholog is 100%.

---

Group of orthologs #1029. Best score 1945 bits
Score difference with first non-orthologous sequence - *A. fumigatus*:1945 *A. fischeri*:1945
afum|EAL91404.1 100.00% afis|EAW17591.1 100.00%
Bootstrap support for afum|EAL91404.1 as seed ortholog is 100%.
Bootstrap support for afis|EAW17591.1 as seed ortholog is 100%.

---

Group of orthologs #1030. Best score 1945 bits
Score difference with first non-orthologous sequence - *A. fumigatus*:1945 *A. fischeri*:1945
afum|EAL84368.2 100.00% afis|EAW25694.1 100.00%
Bootstrap support for afum|EAL84368.2 as seed ortholog is 100%.
Bootstrap support for afis|EAW25694.1 as seed ortholog is 100%.

---

Group of orthologs #1031. Best score 1945 bits
Score difference with first non-orthologous sequence - *A. fumigatus*:1945 *A. fischeri*:1945
afum|EAL87135.1 100.00% afis|EAW23690.1 100.00%
Bootstrap support for afum|EAL87135.1 as seed ortholog is 100%.
Bootstrap support for afis|EAW23690.1 as seed ortholog is 100%.

---

Group of orthologs #1032. Best score 1945 bits
Score difference with first non-orthologous sequence - *A. fumigatus*:1945 *A. fischeri*:1945
afum|EAL88194.1 100.00% afis|EAW23285.1 100.00%
Bootstrap support for afum|EAL88194.1 as seed ortholog is 100%.
Bootstrap support for afis|EAW23285.1 as seed ortholog is 100%.

---

Group of orthologs #1033. Best score 1945 bits
Score difference with first non-orthologous sequence - *A. fumigatus*:1945 *A. fischeri*:1945
afum|EAL91955.1 100.00% afis|EAW24101.1 100.00%
Bootstrap support for afum|EAL91955.1 as seed ortholog is 100%.
Bootstrap support for afis|EAW24101.1 as seed ortholog is 100%.

Group of orthologs #1034. Best score 1944 bits
Score difference with first non-orthologous sequence - *A. fumigatus*:1944 *A. fischeri*:1944
afum|EAL93983.1 100.00% afis|EAW19254.1 100.00%
Bootstrap support for afum|EAL93983.1 as seed ortholog is 100%.
Bootstrap support for afis|EAW19254.1 as seed ortholog is 100%.

Group of orthologs #1035. Best score 1942 bits
Score difference with first non-orthologous sequence - *A. fumigatus*:1942 *A. fischeri*:1942
afum|EAL91640.1 100.00% afis|EAW17830.1 100.00%
Bootstrap support for afum|EAL91640.1 as seed ortholog is 100%.
Bootstrap support for afis|EAW17830.1 as seed ortholog is 100%.

Group of orthologs #1036. Best score 1939 bits
Score difference with first non-orthologous sequence - *A. fumigatus*:1939 *A. fischeri*:1939
afum|EAL91662.1 100.00% afis|EAW17854.1 100.00%
Bootstrap support for afum|EAL91662.1 as seed ortholog is 100%.
Bootstrap support for afis|EAW17854.1 as seed ortholog is 100%.

Group of orthologs #1037. Best score 1939 bits
Score difference with first non-orthologous sequence - *A. fumigatus*:1939 *A. fischeri*:1939
afum|EAL87295.1 100.00% afis|EAW23867.1 100.00%
Bootstrap support for afum|EAL87295.1 as seed ortholog is 100%.
Bootstrap support for afis|EAW23867.1 as seed ortholog is 100%.

Group of orthologs #1038. Best score 1939 bits
Score difference with first non-orthologous sequence - *A. fumigatus*:1939 *A. fischeri*:1939
afum|EAL92822.2 100.00% afis|EAW21797.1 100.00%
Bootstrap support for afum|EAL92822.2 as seed ortholog is 100%.
Bootstrap support for afis|EAW21797.1 as seed ortholog is 100%.

Group of orthologs #1039. Best score 1938 bits
Score difference with first non-orthologous sequence - *A. fumigatus*:1938 *A. fischeri*:1422
afum|EAL88648.1 100.00% afis|EAW16019.1 100.00%
Bootstrap support for afum|EAL88648.1 as seed ortholog is 100%.
Bootstrap support for afis|EAW16019.1 as seed ortholog is 100%.

Group of orthologs #1040. Best score 1938 bits
Score difference with first non-orthologous sequence - *A. fumigatus*:1828 *A. fischeri*:1320
afum|EAL88730.1 100.00% afis|EAW16098.1 100.00%
Bootstrap support for afum|EAL88730.1 as seed ortholog is 100%.
Bootstrap support for afis|EAW16098.1 as seed ortholog is 100%.

Group of orthologs #1041. Best score 1938 bits
Score difference with first non-orthologous sequence - *A. fumigatus*:1938 *A. fischeri*:1938
afum|EAL87836.1 100.00% afis|EAW23624.1 100.00%
Bootstrap support for afum|EAL87836.1 as seed ortholog is 100%.
Bootstrap support for afis|EAW23624.1 as seed ortholog is 100%.

Group of orthologs #1042. Best score 1938 bits
Score difference with first non-orthologous sequence - *A. fumigatus*:1938 *A. fischeri*:1938
afum|EAL93532.1 100.00% afis|EAW18796.1 100.00%
Bootstrap support for afum|EAL93532.1 as seed ortholog is 100%.
Bootstrap support for afis|EAW18796.1 as seed ortholog is 100%.

Group of orthologs #1043. Best score 1937 bits

Score difference with first non-orthologous sequence - *A. fumigatus*:1937   *A. fischeri*:1937
afum|EAL90538.1 100.00% afis|EAW22654.1 100.00%
Bootstrap support for afum|EAL90538.1 as seed ortholog is 100%.
Bootstrap support for afis|EAW22654.1 as seed ortholog is 100%.

---

Group of orthologs #1044. Best score 1936 bits
Score difference with first non-orthologous sequence - *A. fumigatus*:1936   *A. fischeri*:1936
afum|EAL88444.1 100.00% afis|EAW23037.1 100.00%
Bootstrap support for afum|EAL88444.1 as seed ortholog is 100%.
Bootstrap support for afis|EAW23037.1 as seed ortholog is 100%.

---

Group of orthologs #1045. Best score 1936 bits
Score difference with first non-orthologous sequence - *A. fumigatus*:1936   *A. fischeri*:1936
afum|EBA27467.1 100.00% afis|EAW21569.1 100.00%
Bootstrap support for afum|EBA27467.1 as seed ortholog is 100%.
Bootstrap support for afis|EAW21569.1 as seed ortholog is 100%.

---

Group of orthologs #1046. Best score 1934 bits
Score difference with first non-orthologous sequence - *A. fumigatus*:1934   *A. fischeri*:1934
afum|EAL86943.1 100.00% afis|EAW19574.1 100.00%
Bootstrap support for afum|EAL86943.1 as seed ortholog is 100%.
Bootstrap support for afis|EAW19574.1 as seed ortholog is 100%.

---

Group of orthologs #1047. Best score 1934 bits
Score difference with first non-orthologous sequence - *A. fumigatus*:1934   *A. fischeri*:1934
afum|EAL93719.1 100.00% afis|EAW18983.1 100.00%
Bootstrap support for afum|EAL93719.1 as seed ortholog is 100%.
Bootstrap support for afis|EAW18983.1 as seed ortholog is 100%.

---

Group of orthologs #1048. Best score 1934 bits
Score difference with first non-orthologous sequence - *A. fumigatus*:1934   *A. fischeri*:1934
afum|EAL89566.2 100.00% afis|EAW24912.1 100.00%
Bootstrap support for afum|EAL89566.2 as seed ortholog is 100%.
Bootstrap support for afis|EAW24912.1 as seed ortholog is 100%.

---

Group of orthologs #1049. Best score 1933 bits
Score difference with first non-orthologous sequence - *A. fumigatus*:1780   *A. fischeri*:1933
afum|EAL89130.2 100.00% afis|EAW16533.1 100.00%
Bootstrap support for afum|EAL89130.2 as seed ortholog is 100%.
Bootstrap support for afis|EAW16533.1 as seed ortholog is 100%.

---

Group of orthologs #1050. Best score 1933 bits
Score difference with first non-orthologous sequence - *A. fumigatus*:1933   *A. fischeri*:1933
afum|EBA27309.1 100.00% afis|EAW23889.1 100.00%
Bootstrap support for afum|EBA27309.1 as seed ortholog is 100%.
Bootstrap support for afis|EAW23889.1 as seed ortholog is 100%.

---

Group of orthologs #1051. Best score 1932 bits
Score difference with first non-orthologous sequence - *A. fumigatus*:1932   *A. fischeri*:1932
afum|EAL85621.1 100.00% afis|EAW15726.1 100.00%
Bootstrap support for afum|EAL85621.1 as seed ortholog is 100%.
Bootstrap support for afis|EAW15726.1 as seed ortholog is 100%.

---

Group of orthologs #1052. Best score 1931 bits
Score difference with first non-orthologous sequence - *A. fumigatus*:1931   *A. fischeri*:1931
afum|EAL84890.1 100.00% afis|EAW20027.1 100.00%

Bootstrap support for afum|EAL84890.1 as seed ortholog is 100%.
Bootstrap support for afis|EAW20027.1 as seed ortholog is 100%.

---

Group of orthologs #1053. Best score 1930 bits
Score difference with first non-orthologous sequence - *A. fumigatus*:1930 *A. fischeri*:1930
afum|EAL84899.2 100.00% afis|EAW20018.1 100.00%
Bootstrap support for afum|EAL84899.2 as seed ortholog is 100%.
Bootstrap support for afis|EAW20018.1 as seed ortholog is 100%.

---

Group of orthologs #1054. Best score 1929 bits
Score difference with first non-orthologous sequence - *A. fumigatus*:1929 *A. fischeri*:1297
afum|EAL89165.1 100.00% afis|EAW16568.1 100.00%
Bootstrap support for afum|EAL89165.1 as seed ortholog is 100%.
Bootstrap support for afis|EAW16568.1 as seed ortholog is 100%.

---

Group of orthologs #1055. Best score 1929 bits
Score difference with first non-orthologous sequence - *A. fumigatus*:1929 *A. fischeri*:1521
afum|EAL87000.1 100.00% afis|EAW19494.1 100.00%
Bootstrap support for afum|EAL87000.1 as seed ortholog is 100%.
Bootstrap support for afis|EAW19494.1 as seed ortholog is 100%.

---

Group of orthologs #1056. Best score 1929 bits
Score difference with first non-orthologous sequence - *A. fumigatus*:1929 *A. fischeri*:1929
afum|EAL91429.1 100.00% afis|EAW17616.1 100.00%
Bootstrap support for afum|EAL91429.1 as seed ortholog is 100%.
Bootstrap support for afis|EAW17616.1 as seed ortholog is 100%.

---

Group of orthologs #1057. Best score 1929 bits
Score difference with first non-orthologous sequence - *A. fumigatus*:1929 *A. fischeri*:1929
afum|EAL88392.1 100.00% afis|EAW23087.1 100.00%
Bootstrap support for afum|EAL88392.1 as seed ortholog is 100%.
Bootstrap support for afis|EAW23087.1 as seed ortholog is 100%.

---

Group of orthologs #1058. Best score 1929 bits
Score difference with first non-orthologous sequence - *A. fumigatus*:1929 *A. fischeri*:1929
afum|EAL93571.1 100.00% afis|EAW18837.1 100.00%
Bootstrap support for afum|EAL93571.1 as seed ortholog is 100%.
Bootstrap support for afis|EAW18837.1 as seed ortholog is 100%.

---

Group of orthologs #1059. Best score 1929 bits
Score difference with first non-orthologous sequence - *A. fumigatus*:1929 *A. fischeri*:1929
afum|EAL92988.1 100.00% afis|EAW21973.1 100.00%
Bootstrap support for afum|EAL92988.1 as seed ortholog is 100%.
Bootstrap support for afis|EAW21973.1 as seed ortholog is 100%.

---

Group of orthologs #1060. Best score 1928 bits
Score difference with first non-orthologous sequence - *A. fumigatus*:1928 *A. fischeri*:1928
afum|EAL91475.1 100.00% afis|EAW17660.1 100.00%
Bootstrap support for afum|EAL91475.1 as seed ortholog is 100%.
Bootstrap support for afis|EAW17660.1 as seed ortholog is 100%.

---

Group of orthologs #1061. Best score 1928 bits
Score difference with first non-orthologous sequence - *A. fumigatus*:1928 *A. fischeri*:681
afum|EAL88166.1 100.00% afis|EAW23311.1 100.00%
Bootstrap support for afum|EAL88166.1 as seed ortholog is 100%.
Bootstrap support for afis|EAW23311.1 as seed ortholog is 100%.

---

Group of orthologs #1062. Best score 1927 bits
Score difference with first non-orthologous sequence - *A. fumigatus*:1927 *A. fischeri*:1927
afum|EAL92703.1 100.00% afis|EAW21668.1 100.00%
Bootstrap support for afum|EAL92703.1 as seed ortholog is 100%.
Bootstrap support for afis|EAW21668.1 as seed ortholog is 100%.

---

Group of orthologs #1063. Best score 1926 bits
Score difference with first non-orthologous sequence - *A. fumigatus*:1926 *A. fischeri*:1926
afum|EAL92529.1 100.00% afis|EAW21509.1 100.00%
Bootstrap support for afum|EAL92529.1 as seed ortholog is 100%.
Bootstrap support for afis|EAW21509.1 as seed ortholog is 100%.

---

Group of orthologs #1064. Best score 1925 bits
Score difference with first non-orthologous sequence - *A. fumigatus*:1925 *A. fischeri*:1925
afum|EAL88057.1 100.00% afis|EAW23412.1 100.00%
Bootstrap support for afum|EAL88057.1 as seed ortholog is 100%.
Bootstrap support for afis|EAW23412.1 as seed ortholog is 100%.

---

Group of orthologs #1065. Best score 1925 bits
Score difference with first non-orthologous sequence - *A. fumigatus*:1925 *A. fischeri*:1925
afum|EAL89885.1 100.00% afis|EAW25224.1 100.00%
Bootstrap support for afum|EAL89885.1 as seed ortholog is 100%.
Bootstrap support for afis|EAW25224.1 as seed ortholog is 100%.

---

Group of orthologs #1066. Best score 1924 bits
Score difference with first non-orthologous sequence - *A. fumigatus*:1740 *A. fischeri*:1327
afum|EAL84958.1 100.00% afis|EAW19956.1 100.00%
Bootstrap support for afum|EAL84958.1 as seed ortholog is 100%.
Bootstrap support for afis|EAW19956.1 as seed ortholog is 100%.

---

Group of orthologs #1067. Best score 1924 bits
Score difference with first non-orthologous sequence - *A. fumigatus*:1924 *A. fischeri*:1924
afum|EAL92138.1 100.00% afis|EAW21115.1 100.00%
Bootstrap support for afum|EAL92138.1 as seed ortholog is 100%.
Bootstrap support for afis|EAW21115.1 as seed ortholog is 100%.

---

Group of orthologs #1068. Best score 1922 bits
Score difference with first non-orthologous sequence - *A. fumigatus*:1922 *A. fischeri*:1922
afum|EAL88715.1 100.00% afis|EAW16083.1 100.00%
Bootstrap support for afum|EAL88715.1 as seed ortholog is 100%.
Bootstrap support for afis|EAW16083.1 as seed ortholog is 100%.

---

Group of orthologs #1069. Best score 1920 bits
Score difference with first non-orthologous sequence - *A. fumigatus*:1920 *A. fischeri*:1920
afum|EAL85567.1 100.00% afis|EAW15783.1 100.00%
Bootstrap support for afum|EAL85567.1 as seed ortholog is 100%.
Bootstrap support for afis|EAW15783.1 as seed ortholog is 100%.

---

Group of orthologs #1070. Best score 1920 bits
Score difference with first non-orthologous sequence - *A. fumigatus*:1920 *A. fischeri*:1920
afum|EAL87198.1 100.00% afis|EAW23765.1 100.00%
Bootstrap support for afum|EAL87198.1 as seed ortholog is 100%.
Bootstrap support for afis|EAW23765.1 as seed ortholog is 100%.

---

Group of orthologs #1071. Best score 1920 bits

Score difference with first non-orthologous sequence - *A. fumigatus*:1920 *A. fischeri*:1920
afum|EAL93819.1 100.00% afis|EAW19089.1 100.00%
Bootstrap support for afum|EAL93819.1 as seed ortholog is 100%.
Bootstrap support for afis|EAW19089.1 as seed ortholog is 100%.

---

Group of orthologs #1072. Best score 1920 bits
Score difference with first non-orthologous sequence - *A. fumigatus*:1920 *A. fischeri*:1920
afum|EAL91982.1 100.00% afis|EAW24126.1 100.00%
Bootstrap support for afum|EAL91982.1 as seed ortholog is 100%.
Bootstrap support for afis|EAW24126.1 as seed ortholog is 100%.

---

Group of orthologs #1073. Best score 1919 bits
Score difference with first non-orthologous sequence - *A. fumigatus*:1919 *A. fischeri*:1919
afum|EAL93200.1 100.00% afis|EAW18506.1 100.00%
Bootstrap support for afum|EAL93200.1 as seed ortholog is 100%.
Bootstrap support for afis|EAW18506.1 as seed ortholog is 100%.

---

Group of orthologs #1074. Best score 1918 bits
Score difference with first non-orthologous sequence - *A. fumigatus*:1918 *A. fischeri*:1918
afum|EAL85187.1 100.00% afis|EAW20069.1 100.00%
Bootstrap support for afum|EAL85187.1 as seed ortholog is 100%.
Bootstrap support for afis|EAW20069.1 as seed ortholog is 100%.

---

Group of orthologs #1075. Best score 1917 bits
Score difference with first non-orthologous sequence - *A. fumigatus*:1499 *A. fischeri*:1723
afum|EAL88092.1 100.00% afis|EAW23379.1 100.00%
Bootstrap support for afum|EAL88092.1 as seed ortholog is 100%.
Bootstrap support for afis|EAW23379.1 as seed ortholog is 100%.

---

Group of orthologs #1076. Best score 1917 bits
Score difference with first non-orthologous sequence - *A. fumigatus*:1917 *A. fischeri*:1917
afum|EAL90568.1 100.00% afis|EAW22628.1 100.00%
Bootstrap support for afum|EAL90568.1 as seed ortholog is 100%.
Bootstrap support for afis|EAW22628.1 as seed ortholog is 100%.

---

Group of orthologs #1077. Best score 1917 bits
Score difference with first non-orthologous sequence - *A. fumigatus*:1917 *A. fischeri*:1917
afum|EAL92920.1 100.00% afis|EAW21908.1 100.00%
Bootstrap support for afum|EAL92920.1 as seed ortholog is 100%.
Bootstrap support for afis|EAW21908.1 as seed ortholog is 100%.

---

Group of orthologs #1078. Best score 1916 bits
Score difference with first non-orthologous sequence - *A. fumigatus*:1916 *A. fischeri*:1916
afum|EAL92975.1 100.00% afis|EAW21961.1 100.00%
Bootstrap support for afum|EAL92975.1 as seed ortholog is 100%.
Bootstrap support for afis|EAW21961.1 as seed ortholog is 100%.

---

Group of orthologs #1079. Best score 1914 bits
Score difference with first non-orthologous sequence - *A. fumigatus*:1914 *A. fischeri*:1914
afum|EAL85219.1 100.00% afis|EAW20102.1 100.00%
Bootstrap support for afum|EAL85219.1 as seed ortholog is 100%.
Bootstrap support for afis|EAW20102.1 as seed ortholog is 100%.

---

Group of orthologs #1080. Best score 1914 bits
Score difference with first non-orthologous sequence - *A. fumigatus*:1914 *A. fischeri*:1914
afum|EAL85911.1 100.00% afis|EAW24243.1 100.00%

Bootstrap support for afum|EAL85911.1 as seed ortholog is 100%.
Bootstrap support for afis|EAW24243.1 as seed ortholog is 100%.

---

Group of orthologs #1081. Best score 1913 bits
Score difference with first non-orthologous sequence - *A. fumigatus*:1913 *A. fischeri*:1913
afum|EAL87267.1 100.00% afis|EAW23836.1 100.00%
Bootstrap support for afum|EAL87267.1 as seed ortholog is 100%.
Bootstrap support for afis|EAW23836.1 as seed ortholog is 100%.

---

Group of orthologs #1082. Best score 1913 bits
Score difference with first non-orthologous sequence - *A. fumigatus*:1913 *A. fischeri*:1913
afum|EAL88203.1 100.00% afis|EAW23276.1 100.00%
Bootstrap support for afum|EAL88203.1 as seed ortholog is 100%.
Bootstrap support for afis|EAW23276.1 as seed ortholog is 100%.

---

Group of orthologs #1083. Best score 1913 bits
Score difference with first non-orthologous sequence - *A. fumigatus*:1913 *A. fischeri*:1913
afum|EAL93821.1 100.00% afis|EAW19091.1 100.00%
Bootstrap support for afum|EAL93821.1 as seed ortholog is 100%.
Bootstrap support for afis|EAW19091.1 as seed ortholog is 100%.

---

Group of orthologs #1084. Best score 1911 bits
Score difference with first non-orthologous sequence - *A. fumigatus*:1911 *A. fischeri*:1911
afum|EAL86114.1 100.00% afis|EAW24445.1 100.00%
Bootstrap support for afum|EAL86114.1 as seed ortholog is 100%.
Bootstrap support for afis|EAW24445.1 as seed ortholog is 100%.

---

Group of orthologs #1085. Best score 1911 bits
Score difference with first non-orthologous sequence - *A. fumigatus*:1911 *A. fischeri*:1911
afum|EAL91024.1 100.00% afis|EAW22167.1 100.00%
Bootstrap support for afum|EAL91024.1 as seed ortholog is 100%.
Bootstrap support for afis|EAW22167.1 as seed ortholog is 100%.

---

Group of orthologs #1086. Best score 1911 bits
Score difference with first non-orthologous sequence - *A. fumigatus*:1911 *A. fischeri*:1911
afum|EAL89977.2 100.00% afis|EAW25316.1 100.00%
Bootstrap support for afum|EAL89977.2 as seed ortholog is 100%.
Bootstrap support for afis|EAW25316.1 as seed ortholog is 100%.

---

Group of orthologs #1087. Best score 1909 bits
Score difference with first non-orthologous sequence - *A. fumigatus*:1909 *A. fischeri*:1909
afum|EAL85558.1 100.00% afis|EAW15792.1 100.00%
Bootstrap support for afum|EAL85558.1 as seed ortholog is 100%.
Bootstrap support for afis|EAW15792.1 as seed ortholog is 100%.

---

Group of orthologs #1088. Best score 1908 bits
Score difference with first non-orthologous sequence - *A. fumigatus*:1908 *A. fischeri*:1908
afum|EAL89937.1 100.00% afis|EAW25274.1 100.00%
Bootstrap support for afum|EAL89937.1 as seed ortholog is 100%.
Bootstrap support for afis|EAW25274.1 as seed ortholog is 100%.

---

Group of orthologs #1089. Best score 1907 bits
Score difference with first non-orthologous sequence - *A. fumigatus*:1907 *A. fischeri*:1907
afum|EAL84323.1 100.00% afis|EAW25635.1 100.00%
Bootstrap support for afum|EAL84323.1 as seed ortholog is 100%.
Bootstrap support for afis|EAW25635.1 as seed ortholog is 100%.

Group of orthologs #1090. Best score 1907 bits
Score difference with first non-orthologous sequence - *A. fumigatus*:1907 *A. fischeri*:1907
afum|EAL88052.2 100.00% afis|EAW23417.1 100.00%
Bootstrap support for afum|EAL88052.2 as seed ortholog is 100%.
Bootstrap support for afis|EAW23417.1 as seed ortholog is 100%.

Group of orthologs #1091. Best score 1906 bits
Score difference with first non-orthologous sequence - *A. fumigatus*:1906 *A. fischeri*:1906
afum|EAL93581.1 100.00% afis|EAW18850.1 100.00%
Bootstrap support for afum|EAL93581.1 as seed ortholog is 100%.
Bootstrap support for afis|EAW18850.1 as seed ortholog is 100%.

Group of orthologs #1092. Best score 1905 bits
Score difference with first non-orthologous sequence - *A. fumigatus*:1905 *A. fischeri*:1905
afum|EAL88642.1 100.00% afis|EAW16012.1 100.00%
Bootstrap support for afum|EAL88642.1 as seed ortholog is 100%.
Bootstrap support for afis|EAW16012.1 as seed ortholog is 100%.

Group of orthologs #1093. Best score 1904 bits
Score difference with first non-orthologous sequence - *A. fumigatus*:1904 *A. fischeri*:1904
afum|EAL86710.1 100.00% afis|EAW17290.1 100.00%
Bootstrap support for afum|EAL86710.1 as seed ortholog is 100%.
Bootstrap support for afis|EAW17290.1 as seed ortholog is 100%.

Group of orthologs #1094. Best score 1904 bits
Score difference with first non-orthologous sequence - *A. fumigatus*:1904 *A. fischeri*:1534
afum|EAL89214.1 100.00% afis|EAW16623.1 100.00%
Bootstrap support for afum|EAL89214.1 as seed ortholog is 100%.
Bootstrap support for afis|EAW16623.1 as seed ortholog is 100%.

Group of orthologs #1095. Best score 1904 bits
Score difference with first non-orthologous sequence - *A. fumigatus*:1904 *A. fischeri*:1904
afum|EAL86061.1 100.00% afis|EAW24389.1 100.00%
Bootstrap support for afum|EAL86061.1 as seed ortholog is 100%.
Bootstrap support for afis|EAW24389.1 as seed ortholog is 100%.

Group of orthologs #1096. Best score 1903 bits
Score difference with first non-orthologous sequence - *A. fumigatus*:1903 *A. fischeri*:1903
afum|EAL91308.1 100.00% afis|EAW17479.1 100.00%
Bootstrap support for afum|EAL91308.1 as seed ortholog is 100%.
Bootstrap support for afis|EAW17479.1 as seed ortholog is 100%.

Group of orthologs #1097. Best score 1903 bits
Score difference with first non-orthologous sequence - *A. fumigatus*:1903 *A. fischeri*:1903
afum|EAL90249.1 100.00% afis|EAW22948.1 100.00%
Bootstrap support for afum|EAL90249.1 as seed ortholog is 100%.
Bootstrap support for afis|EAW22948.1 as seed ortholog is 100%.

Group of orthologs #1098. Best score 1903 bits
Score difference with first non-orthologous sequence - *A. fumigatus*:1903 *A. fischeri*:1903
afum|EAL92296.1 100.00% afis|EAW21280.1 100.00%
Bootstrap support for afum|EAL92296.1 as seed ortholog is 100%.
Bootstrap support for afis|EAW21280.1 as seed ortholog is 100%.

Group of orthologs #1099. Best score 1902 bits

Score difference with first non-orthologous sequence - *A. fumigatus*:1902 *A. fischeri*:1902
afum|EAL85530.1 100.00% afis|EAW15821.1 100.00%
Bootstrap support for afum|EAL85530.1 as seed ortholog is 100%.
Bootstrap support for afis|EAW15821.1 as seed ortholog is 100%.

---

Group of orthologs #1100. Best score 1902 bits
Score difference with first non-orthologous sequence - *A. fumigatus*:1902 *A. fischeri*:1902
afum|EAL89184.1 100.00% afis|EAW16589.1 100.00%
Bootstrap support for afum|EAL89184.1 as seed ortholog is 100%.
Bootstrap support for afis|EAW16589.1 as seed ortholog is 100%.

---

Group of orthologs #1101. Best score 1902 bits
Score difference with first non-orthologous sequence - *A. fumigatus*:1902 *A. fischeri*:1902
afum|EAL94036.1 100.00% afis|EAW19319.1 100.00%
Bootstrap support for afum|EAL94036.1 as seed ortholog is 100%.
Bootstrap support for afis|EAW19319.1 as seed ortholog is 100%.

---

Group of orthologs #1102. Best score 1902 bits
Score difference with first non-orthologous sequence - *A. fumigatus*:1902 *A. fischeri*:1902
afum|EAL89888.1 100.00% afis|EAW25227.1 100.00%
Bootstrap support for afum|EAL89888.1 as seed ortholog is 100%.
Bootstrap support for afis|EAW25227.1 as seed ortholog is 100%.

---

Group of orthologs #1103. Best score 1899 bits
Score difference with first non-orthologous sequence - *A. fumigatus*:1899 *A. fischeri*:1899
afum|EAL85924.1 100.00% afis|EAW24256.1 100.00%
Bootstrap support for afum|EAL85924.1 as seed ortholog is 100%.
Bootstrap support for afis|EAW24256.1 as seed ortholog is 100%.

---

Group of orthologs #1104. Best score 1899 bits
Score difference with first non-orthologous sequence - *A. fumigatus*:1899 *A. fischeri*:1899
afum|EAL89704.1 100.00% afis|EAW24997.1 100.00%
Bootstrap support for afum|EAL89704.1 as seed ortholog is 100%.
Bootstrap support for afis|EAW24997.1 as seed ortholog is 100%.

---

Group of orthologs #1105. Best score 1898 bits
Score difference with first non-orthologous sequence - *A. fumigatus*:1898 *A. fischeri*:1898
afum|EAL89133.1 100.00% afis|EAW16536.1 100.00%
Bootstrap support for afum|EAL89133.1 as seed ortholog is 100%.
Bootstrap support for afis|EAW16536.1 as seed ortholog is 100%.

---

Group of orthologs #1106. Best score 1898 bits
Score difference with first non-orthologous sequence - *A. fumigatus*:1898 *A. fischeri*:1898
afum|EAL91452.1 100.00% afis|EAW17637.1 100.00%
Bootstrap support for afum|EAL91452.1 as seed ortholog is 100%.
Bootstrap support for afis|EAW17637.1 as seed ortholog is 100%.

---

Group of orthologs #1107. Best score 1898 bits
Score difference with first non-orthologous sequence - *A. fumigatus*:1898 *A. fischeri*:1898
afum|EAL87913.1 100.00% afis|EAW23553.1 100.00%
Bootstrap support for afum|EAL87913.1 as seed ortholog is 100%.
Bootstrap support for afis|EAW23553.1 as seed ortholog is 100%.

---

Group of orthologs #1108. Best score 1896 bits
Score difference with first non-orthologous sequence - *A. fumigatus*:1896 *A. fischeri*:1896
afum|EAL88886.1 100.00% afis|EAW16265.1 100.00%

Bootstrap support for afum|EAL88886.1 as seed ortholog is 100%.
Bootstrap support for afis|EAW16265.1 as seed ortholog is 100%.

---

Group of orthologs #1109. Best score 1896 bits
Score difference with first non-orthologous sequence - *A. fumigatus*:1896 *A. fischeri*:1896
afum|EAL87041.1 100.00% afis|EAW19455.1 100.00%
Bootstrap support for afum|EAL87041.1 as seed ortholog is 100%.
Bootstrap support for afis|EAW19455.1 as seed ortholog is 100%.

---

Group of orthologs #1110. Best score 1896 bits
Score difference with first non-orthologous sequence - *A. fumigatus*:1896 *A. fischeri*:1896
afum|EAL86079.1 100.00% afis|EAW24410.1 100.00%
Bootstrap support for afum|EAL86079.1 as seed ortholog is 100%.
Bootstrap support for afis|EAW24410.1 as seed ortholog is 100%.

---

Group of orthologs #1111. Best score 1895 bits
Score difference with first non-orthologous sequence - *A. fumigatus*:1895 *A. fischeri*:1895
afum|EAL87645.1 100.00% afis|EAW18279.1 100.00%
Bootstrap support for afum|EAL87645.1 as seed ortholog is 100%.
Bootstrap support for afis|EAW18279.1 as seed ortholog is 100%.

---

Group of orthologs #1112. Best score 1895 bits
Score difference with first non-orthologous sequence - *A. fumigatus*:1895 *A. fischeri*:1895
afum|EAL86126.1 100.00% afis|EAW24456.1 100.00%
Bootstrap support for afum|EAL86126.1 as seed ortholog is 100%.
Bootstrap support for afis|EAW24456.1 as seed ortholog is 100%.

---

Group of orthologs #1113. Best score 1895 bits
Score difference with first non-orthologous sequence - *A. fumigatus*:1895 *A. fischeri*:1895
afum|EAL93219.1 100.00% afis|EAW18524.1 100.00%
Bootstrap support for afum|EAL93219.1 as seed ortholog is 100%.
Bootstrap support for afis|EAW18524.1 as seed ortholog is 100%.

---

Group of orthologs #1114. Best score 1894 bits
Score difference with first non-orthologous sequence - *A. fumigatus*:1814 *A. fischeri*:1480
afum|EAL84713.1 100.00% afis|EAW20921.1 100.00%
Bootstrap support for afum|EAL84713.1 as seed ortholog is 100%.
Bootstrap support for afis|EAW20921.1 as seed ortholog is 100%.

---

Group of orthologs #1115. Best score 1893 bits
Score difference with first non-orthologous sequence - *A. fumigatus*:1893 *A. fischeri*:1893
afum|EAL92405.1 100.00% afis|EAW21391.1 100.00%
Bootstrap support for afum|EAL92405.1 as seed ortholog is 100%.
Bootstrap support for afis|EAW21391.1 as seed ortholog is 100%.

---

Group of orthologs #1116. Best score 1889 bits
Score difference with first non-orthologous sequence - *A. fumigatus*:1140 *A. fischeri*:1533
afum|EAL87814.1 100.00% afis|EAW16772.1 100.00%
Bootstrap support for afum|EAL87814.1 as seed ortholog is 100%.
Bootstrap support for afis|EAW16772.1 as seed ortholog is 100%.

---

Group of orthologs #1117. Best score 1888 bits
Score difference with first non-orthologous sequence - *A. fumigatus*:1888 *A. fischeri*:1888
afum|EAL91522.2 100.00% afis|EAW17708.1 100.00%
Bootstrap support for afum|EAL91522.2 as seed ortholog is 100%.
Bootstrap support for afis|EAW17708.1 as seed ortholog is 100%.

---

Group of orthologs #1118. Best score 1887 bits
Score difference with first non-orthologous sequence - *A. fumigatus*:1887 *A. fischeri*:1887
afum|EAL85777.1 100.00% afis|EAW15555.1 100.00%
Bootstrap support for afum|EAL85777.1 as seed ortholog is 100%.
Bootstrap support for afis|EAW15555.1 as seed ortholog is 100%.

---

Group of orthologs #1119. Best score 1886 bits
Score difference with first non-orthologous sequence - *A. fumigatus*:1886 *A. fischeri*:1886
afum|EAL92468.1 100.00% afis|EAW21449.1 100.00%
Bootstrap support for afum|EAL92468.1 as seed ortholog is 100%.
Bootstrap support for afis|EAW21449.1 as seed ortholog is 100%.

---

Group of orthologs #1120. Best score 1885 bits
Score difference with first non-orthologous sequence - *A. fumigatus*:1885 *A. fischeri*:1885
afum|EAL84885.1 100.00% afis|EAW20031.1 100.00%
Bootstrap support for afum|EAL84885.1 as seed ortholog is 100%.
Bootstrap support for afis|EAW20031.1 as seed ortholog is 100%.

---

Group of orthologs #1121. Best score 1885 bits
Score difference with first non-orthologous sequence - *A. fumigatus*:1885 *A. fischeri*:1885
afum|EAL92696.1 100.00% afis|EAW21661.1 100.00%
Bootstrap support for afum|EAL92696.1 as seed ortholog is 100%.
Bootstrap support for afis|EAW21661.1 as seed ortholog is 100%.

---

Group of orthologs #1122. Best score 1885 bits
Score difference with first non-orthologous sequence - *A. fumigatus*:1885 *A. fischeri*:1885
afum|EAL89607.1 100.00% afis|EAW24950.1 100.00%
Bootstrap support for afum|EAL89607.1 as seed ortholog is 100%.
Bootstrap support for afis|EAW24950.1 as seed ortholog is 100%.

---

Group of orthologs #1123. Best score 1884 bits
Score difference with first non-orthologous sequence - *A. fumigatus*:1884 *A. fischeri*:1704
afum|EAL94039.1 100.00% afis|EAW19321.1 100.00%
Bootstrap support for afum|EAL94039.1 as seed ortholog is 100%.
Bootstrap support for afis|EAW19321.1 as seed ortholog is 100%.

---

Group of orthologs #1124. Best score 1883 bits
Score difference with first non-orthologous sequence - *A. fumigatus*:1883 *A. fischeri*:1883
afum|EAL93309.1 100.00% afis|EAW18601.1 100.00%
Bootstrap support for afum|EAL93309.1 as seed ortholog is 100%.
Bootstrap support for afis|EAW18601.1 as seed ortholog is 100%.

---

Group of orthologs #1125. Best score 1883 bits
Score difference with first non-orthologous sequence - *A. fumigatus*:1883 *A. fischeri*:1883
afum|EAL90830.1 100.00% afis|EAW22361.1 100.00%
Bootstrap support for afum|EAL90830.1 as seed ortholog is 100%.
Bootstrap support for afis|EAW22361.1 as seed ortholog is 100%.

---

Group of orthologs #1126. Best score 1881 bits
Score difference with first non-orthologous sequence - *A. fumigatus*:1881 *A. fischeri*:1881
afum|EAL90020.1 100.00% afis|EAW25358.1 100.00%
Bootstrap support for afum|EAL90020.1 as seed ortholog is 100%.
Bootstrap support for afis|EAW25358.1 as seed ortholog is 100%.

---

Group of orthologs #1127. Best score 1879 bits

Score difference with first non-orthologous sequence - *A. fumigatus*:1879   *A. fischeri*:1879
afum|EAL88939.1 100.00% afis|EAW16316.1 100.00%
Bootstrap support for afum|EAL88939.1 as seed ortholog is 100%.
Bootstrap support for afis|EAW16316.1 as seed ortholog is 100%.

---

Group of orthologs #1128. Best score 1879 bits
Score difference with first non-orthologous sequence - *A. fumigatus*:1879   *A. fischeri*:1879
afum|EAL93508.1 100.00% afis|EAW18776.1 100.00%
Bootstrap support for afum|EAL93508.1 as seed ortholog is 100%.
Bootstrap support for afis|EAW18776.1 as seed ortholog is 100%.

---

Group of orthologs #1129. Best score 1878 bits
Score difference with first non-orthologous sequence - *A. fumigatus*:1878   *A. fischeri*:1878
afum|EAL88195.1 100.00% afis|EAW23284.1 100.00%
Bootstrap support for afum|EAL88195.1 as seed ortholog is 100%.
Bootstrap support for afis|EAW23284.1 as seed ortholog is 100%.

---

Group of orthologs #1130. Best score 1878 bits
Score difference with first non-orthologous sequence - *A. fumigatus*:1878   *A. fischeri*:1878
afum|EAL87952.1 100.00% afis|EAW23509.1 100.00%
Bootstrap support for afum|EAL87952.1 as seed ortholog is 100%.
Bootstrap support for afis|EAW23509.1 as seed ortholog is 100%.

---

Group of orthologs #1131. Best score 1878 bits
Score difference with first non-orthologous sequence - *A. fumigatus*:1878   *A. fischeri*:1878
afum|EAL92813.1 100.00% afis|EAW21786.1 100.00%
Bootstrap support for afum|EAL92813.1 as seed ortholog is 100%.
Bootstrap support for afis|EAW21786.1 as seed ortholog is 100%.

---

Group of orthologs #1132. Best score 1877 bits
Score difference with first non-orthologous sequence - *A. fumigatus*:1877   *A. fischeri*:1877
afum|EAL86595.1 100.00% afis|EAW17237.1 100.00%
Bootstrap support for afum|EAL86595.1 as seed ortholog is 100%.
Bootstrap support for afis|EAW17237.1 as seed ortholog is 100%.

---

Group of orthologs #1133. Best score 1874 bits
Score difference with first non-orthologous sequence - *A. fumigatus*:1793   *A. fischeri*:1732
afum|EAL88097.1 100.00% afis|EAW23374.1 100.00%
Bootstrap support for afum|EAL88097.1 as seed ortholog is 100%.
Bootstrap support for afis|EAW23374.1 as seed ortholog is 100%.

---

Group of orthologs #1134. Best score 1873 bits
Score difference with first non-orthologous sequence - *A. fumigatus*:1873   *A. fischeri*:1873
afum|EAL91776.1 100.00% afis|EAW17967.1 100.00%
Bootstrap support for afum|EAL91776.1 as seed ortholog is 100%.
Bootstrap support for afis|EAW17967.1 as seed ortholog is 100%.

---

Group of orthologs #1135. Best score 1873 bits
Score difference with first non-orthologous sequence - *A. fumigatus*:1873   *A. fischeri*:1873
afum|EAL93924.1 100.00% afis|EAW19193.1 100.00%
Bootstrap support for afum|EAL93924.1 as seed ortholog is 100%.
Bootstrap support for afis|EAW19193.1 as seed ortholog is 100%.

---

Group of orthologs #1136. Best score 1872 bits
Score difference with first non-orthologous sequence - *A. fumigatus*:1872   *A. fischeri*:1872
afum|EAL86984.1 100.00% afis|EAW19512.1 100.00%

Bootstrap support for afum|EAL86984.1 as seed ortholog is 100%.
Bootstrap support for afis|EAW19512.1 as seed ortholog is 100%.

---

Group of orthologs #1137. Best score 1872 bits
Score difference with first non-orthologous sequence - *A. fumigatus*:1872 *A. fischeri*:1872
afum|EBA27298.1 100.00% afis|EAW19499.1 100.00%
Bootstrap support for afum|EBA27298.1 as seed ortholog is 100%.
Bootstrap support for afis|EAW19499.1 as seed ortholog is 100%.

---

Group of orthologs #1138. Best score 1872 bits
Score difference with first non-orthologous sequence - *A. fumigatus*:1872 *A. fischeri*:1872
afum|EAL92874.1 100.00% afis|EAW21861.1 100.00%
Bootstrap support for afum|EAL92874.1 as seed ortholog is 100%.
Bootstrap support for afis|EAW21861.1 as seed ortholog is 100%.

---

Group of orthologs #1139. Best score 1870 bits
Score difference with first non-orthologous sequence - *A. fumigatus*:1870 *A. fischeri*:1870
afum|EAL88327.2 100.00% afis|EAW23154.1 100.00%
Bootstrap support for afum|EAL88327.2 as seed ortholog is 100%.
Bootstrap support for afis|EAW23154.1 as seed ortholog is 100%.

---

Group of orthologs #1140. Best score 1870 bits
Score difference with first non-orthologous sequence - *A. fumigatus*:1870 *A. fischeri*:1870
afum|EBA27318.1 100.00% afis|EAW18194.1 100.00%
Bootstrap support for afum|EBA27318.1 as seed ortholog is 100%.
Bootstrap support for afis|EAW18194.1 as seed ortholog is 100%.

---

Group of orthologs #1141. Best score 1869 bits
Score difference with first non-orthologous sequence - *A. fumigatus*:1869 *A. fischeri*:1869
afum|EAL91403.1 100.00% afis|EAW17590.1 100.00%
Bootstrap support for afum|EAL91403.1 as seed ortholog is 100%.
Bootstrap support for afis|EAW17590.1 as seed ortholog is 100%.

---

Group of orthologs #1142. Best score 1869 bits
Score difference with first non-orthologous sequence - *A. fumigatus*:1869 *A. fischeri*:1727
afum|EAL94059.1 100.00% afis|EAW19340.1 100.00%
Bootstrap support for afum|EAL94059.1 as seed ortholog is 100%.
Bootstrap support for afis|EAW19340.1 as seed ortholog is 100%.

---

Group of orthologs #1143. Best score 1869 bits
Score difference with first non-orthologous sequence - *A. fumigatus*:1869 *A. fischeri*:1251
afum|EBA27254.1 100.00% afis|EAW24364.1 100.00%
Bootstrap support for afum|EBA27254.1 as seed ortholog is 100%.
Bootstrap support for afis|EAW24364.1 as seed ortholog is 100%.

---

Group of orthologs #1144. Best score 1865 bits
Score difference with first non-orthologous sequence - *A. fumigatus*:1865 *A. fischeri*:1865
afum|EAL84898.1 100.00% afis|EAW20019.1 100.00%
Bootstrap support for afum|EAL84898.1 as seed ortholog is 100%.
Bootstrap support for afis|EAW20019.1 as seed ortholog is 100%.

---

Group of orthologs #1145. Best score 1865 bits
Score difference with first non-orthologous sequence - *A. fumigatus*:1865 *A. fischeri*:1865
afum|EAL93632.1 100.00% afis|EAW18900.1 100.00%
Bootstrap support for afum|EAL93632.1 as seed ortholog is 100%.
Bootstrap support for afis|EAW18900.1 as seed ortholog is 100%.

---

Group of orthologs #1146. Best score 1865 bits
Score difference with first non-orthologous sequence - *A. fumigatus*:1865 *A. fischeri*:1865
afum|EAL90052.1 100.00% afis|EAW25393.1 100.00%
Bootstrap support for afum|EAL90052.1 as seed ortholog is 100%.
Bootstrap support for afis|EAW25393.1 as seed ortholog is 100%.

---

Group of orthologs #1147. Best score 1864 bits
Score difference with first non-orthologous sequence - *A. fumigatus*:1864 *A. fischeri*:1864
afum|EAL93421.1 100.00% afis|EAW18699.1 100.00%
Bootstrap support for afum|EAL93421.1 as seed ortholog is 100%.
Bootstrap support for afis|EAW18699.1 as seed ortholog is 100%.

---

Group of orthologs #1148. Best score 1864 bits
Score difference with first non-orthologous sequence - *A. fumigatus*:1864 *A. fischeri*:1864
afum|EAL90445.1 100.00% afis|EAW22754.1 100.00%
Bootstrap support for afum|EAL90445.1 as seed ortholog is 100%.
Bootstrap support for afis|EAW22754.1 as seed ortholog is 100%.

---

Group of orthologs #1149. Best score 1863 bits
Score difference with first non-orthologous sequence - *A. fumigatus*:1863 *A. fischeri*:1760
afum|EAL87632.1 100.00% afis|EAW18267.1 100.00%
Bootstrap support for afum|EAL87632.1 as seed ortholog is 100%.
Bootstrap support for afis|EAW18267.1 as seed ortholog is 100%.

---

Group of orthologs #1150. Best score 1863 bits
Score difference with first non-orthologous sequence - *A. fumigatus*:1863 *A. fischeri*:1863
afum|EAL91649.1 100.00% afis|EAW17839.1 100.00%
Bootstrap support for afum|EAL91649.1 as seed ortholog is 100%.
Bootstrap support for afis|EAW17839.1 as seed ortholog is 100%.

---

Group of orthologs #1151. Best score 1862 bits
Score difference with first non-orthologous sequence - *A. fumigatus*:1862 *A. fischeri*:1862
afum|EAL88457.1 100.00% afis|EAW23025.1 100.00%
Bootstrap support for afum|EAL88457.1 as seed ortholog is 100%.
Bootstrap support for afis|EAW23025.1 as seed ortholog is 100%.

---

Group of orthologs #1152. Best score 1862 bits
Score difference with first non-orthologous sequence - *A. fumigatus*:1862 *A. fischeri*:1862
afum|EAL89754.1 100.00% afis|EAW25085.1 100.00%
Bootstrap support for afum|EAL89754.1 as seed ortholog is 100%.
Bootstrap support for afis|EAW25085.1 as seed ortholog is 100%.

---

Group of orthologs #1153. Best score 1861 bits
Score difference with first non-orthologous sequence - *A. fumigatus*:1861 *A. fischeri*:1861
afum|EAL86001.1 100.00% afis|EAW24331.1 100.00%
Bootstrap support for afum|EAL86001.1 as seed ortholog is 100%.
Bootstrap support for afis|EAW24331.1 as seed ortholog is 100%.

---

Group of orthologs #1154. Best score 1861 bits
Score difference with first non-orthologous sequence - *A. fumigatus*:1861 *A. fischeri*:1861
afum|EAL88207.1 100.00% afis|EAW23272.1 100.00%
Bootstrap support for afum|EAL88207.1 as seed ortholog is 100%.
Bootstrap support for afis|EAW23272.1 as seed ortholog is 100%.

---

Group of orthologs #1155. Best score 1860 bits

Score difference with first non-orthologous sequence - *A. fumigatus*:1860   *A. fischeri*:1860
afum|EAL88086.2   100.00%   afis|EAW23385.1   100.00%
Bootstrap support for afum|EAL88086.2 as seed ortholog is 100%.
Bootstrap support for afis|EAW23385.1 as seed ortholog is 100%.

---

Group of orthologs #1156. Best score 1860 bits
Score difference with first non-orthologous sequence - *A. fumigatus*:1860   *A. fischeri*:1860
afum|EAL87912.2   100.00%   afis|EAW23554.1   100.00%
Bootstrap support for afum|EAL87912.2 as seed ortholog is 100%.
Bootstrap support for afis|EAW23554.1 as seed ortholog is 100%.

---

Group of orthologs #1157. Best score 1859 bits
Score difference with first non-orthologous sequence - *A. fumigatus*:1720   *A. fischeri*:1859
afum|EAL86489.1   100.00%   afis|EAW16962.1   100.00%
Bootstrap support for afum|EAL86489.1 as seed ortholog is 100%.
Bootstrap support for afis|EAW16962.1 as seed ortholog is 100%.

---

Group of orthologs #1158. Best score 1859 bits
Score difference with first non-orthologous sequence - *A. fumigatus*:1859   *A. fischeri*:1859
afum|EAL90723.1   100.00%   afis|EAW22468.1   100.00%
Bootstrap support for afum|EAL90723.1 as seed ortholog is 100%.
Bootstrap support for afis|EAW22468.1 as seed ortholog is 100%.

---

Group of orthologs #1159. Best score 1858 bits
Score difference with first non-orthologous sequence - *A. fumigatus*:1858   *A. fischeri*:1858
afum|EAL85346.1   100.00%   afis|EAW20229.1   100.00%
Bootstrap support for afum|EAL85346.1 as seed ortholog is 100%.
Bootstrap support for afis|EAW20229.1 as seed ortholog is 100%.

---

Group of orthologs #1160. Best score 1857 bits
Score difference with first non-orthologous sequence - *A. fumigatus*:1857   *A. fischeri*:1857
afum|EAL91710.1   100.00%   afis|EAW17902.1   100.00%
Bootstrap support for afum|EAL91710.1 as seed ortholog is 100%.
Bootstrap support for afis|EAW17902.1 as seed ortholog is 100%.

---

Group of orthologs #1161. Best score 1857 bits
Score difference with first non-orthologous sequence - *A. fumigatus*:1857   *A. fischeri*:1857
afum|EAL85930.1   100.00%   afis|EAW24262.1   100.00%
Bootstrap support for afum|EAL85930.1 as seed ortholog is 100%.
Bootstrap support for afis|EAW24262.1 as seed ortholog is 100%.

---

Group of orthologs #1162. Best score 1855 bits
Score difference with first non-orthologous sequence - *A. fumigatus*:1855   *A. fischeri*:1855
afum|EAL87939.1   100.00%   afis|EAW23526.1   100.00%
Bootstrap support for afum|EAL87939.1 as seed ortholog is 100%.
Bootstrap support for afis|EAW23526.1 as seed ortholog is 100%.

---

Group of orthologs #1163. Best score 1855 bits
Score difference with first non-orthologous sequence - *A. fumigatus*:1855   *A. fischeri*:1855
afum|EAL93551.1   100.00%   afis|EAW18815.1   100.00%
Bootstrap support for afum|EAL93551.1 as seed ortholog is 100%.
Bootstrap support for afis|EAW18815.1 as seed ortholog is 100%.

---

Group of orthologs #1164. Best score 1854 bits
Score difference with first non-orthologous sequence - *A. fumigatus*:1719   *A. fischeri*:1701
afum|EAL87216.1   100.00%   afis|EAW23782.1   100.00%

Bootstrap support for afum|EAL87216.1 as seed ortholog is 100%.
Bootstrap support for afis|EAW23782.1 as seed ortholog is 100%.

---

Group of orthologs #1165. Best score 1854 bits
Score difference with first non-orthologous sequence - *A. fumigatus*:1854 *A. fischeri*:1854
afum|EAL93041.1 100.00% afis|EAW18367.1 100.00%
Bootstrap support for afum|EAL93041.1 as seed ortholog is 100%.
Bootstrap support for afis|EAW18367.1 as seed ortholog is 100%.

---

Group of orthologs #1166. Best score 1854 bits
Score difference with first non-orthologous sequence - *A. fumigatus*:1854 *A. fischeri*:1854
afum|EAL92008.1 100.00% afis|EAW24153.1 100.00%
Bootstrap support for afum|EAL92008.1 as seed ortholog is 100%.
Bootstrap support for afis|EAW24153.1 as seed ortholog is 100%.

---

Group of orthologs #1167. Best score 1851 bits
Score difference with first non-orthologous sequence - *A. fumigatus*:1851 *A. fischeri*:1851
afum|EAL85030.1 100.00% afis|EAW19866.1 100.00%
Bootstrap support for afum|EAL85030.1 as seed ortholog is 100%.
Bootstrap support for afis|EAW19866.1 as seed ortholog is 100%.

---

Group of orthologs #1168. Best score 1851 bits
Score difference with first non-orthologous sequence - *A. fumigatus*:1851 *A. fischeri*:1851
afum|EAL91947.2 100.00% afis|EAW24092.1 100.00%
Bootstrap support for afum|EAL91947.2 as seed ortholog is 100%.
Bootstrap support for afis|EAW24092.1 as seed ortholog is 100%.

---

Group of orthologs #1169. Best score 1850 bits
Score difference with first non-orthologous sequence - *A. fumigatus*:1850 *A. fischeri*:1850
afum|EAL91289.1 100.00% afis|EAW17461.1 100.00%
Bootstrap support for afum|EAL91289.1 as seed ortholog is 100%.
Bootstrap support for afis|EAW17461.1 as seed ortholog is 100%.

---

Group of orthologs #1170. Best score 1850 bits
Score difference with first non-orthologous sequence - *A. fumigatus*:1850 *A. fischeri*:1850
afum|EAL92456.1 100.00% afis|EAW21440.1 100.00%
Bootstrap support for afum|EAL92456.1 as seed ortholog is 100%.
Bootstrap support for afis|EAW21440.1 as seed ortholog is 100%.

---

Group of orthologs #1171. Best score 1849 bits
Score difference with first non-orthologous sequence - *A. fumigatus*:1849 *A. fischeri*:1849
afum|EAL85744.1 100.00% afis|EAW15588.1 100.00%
Bootstrap support for afum|EAL85744.1 as seed ortholog is 100%.
Bootstrap support for afis|EAW15588.1 as seed ortholog is 100%.

---

Group of orthologs #1172. Best score 1849 bits
Score difference with first non-orthologous sequence - *A. fumigatus*:1849 *A. fischeri*:1849
afum|EAL89855.1 100.00% afis|EAW25181.1 100.00%
Bootstrap support for afum|EAL89855.1 as seed ortholog is 100%.
Bootstrap support for afis|EAW25181.1 as seed ortholog is 100%.

---

Group of orthologs #1173. Best score 1849 bits
Score difference with first non-orthologous sequence - *A. fumigatus*:1849 *A. fischeri*:1849
afum|EAL90144.1 100.00% afis|EAW25479.1 100.00%
Bootstrap support for afum|EAL90144.1 as seed ortholog is 100%.
Bootstrap support for afis|EAW25479.1 as seed ortholog is 100%.

Group of orthologs #1174. Best score 1848 bits
Score difference with first non-orthologous sequence - *A. fumigatus*:1848 *A. fischeri*:1848
afum|EAL85001.1 100.00% afis|EAW19906.1 100.00%
Bootstrap support for afum|EAL85001.1 as seed ortholog is 100%.
Bootstrap support for afis|EAW19906.1 as seed ortholog is 100%.

---

Group of orthologs #1175. Best score 1848 bits
Score difference with first non-orthologous sequence - *A. fumigatus*:1848 *A. fischeri*:1848
afum|EAL84612.1 100.00% afis|EAW20440.1 100.00%
Bootstrap support for afum|EAL84612.1 as seed ortholog is 100%.
Bootstrap support for afis|EAW20440.1 as seed ortholog is 100%.

---

Group of orthologs #1176. Best score 1848 bits
Score difference with first non-orthologous sequence - *A. fumigatus*:1848 *A. fischeri*:1848
afum|EAL88019.1 100.00% afis|EAW23445.1 100.00%
Bootstrap support for afum|EAL88019.1 as seed ortholog is 100%.
Bootstrap support for afis|EAW23445.1 as seed ortholog is 100%.

---

Group of orthologs #1177. Best score 1847 bits
Score difference with first non-orthologous sequence - *A. fumigatus*:1847 *A. fischeri*:1847
afum|EAL86572.1 100.00% afis|EAW17098.1 100.00%
Bootstrap support for afum|EAL86572.1 as seed ortholog is 100%.
Bootstrap support for afis|EAW17098.1 as seed ortholog is 100%.

---

Group of orthologs #1178. Best score 1847 bits
Score difference with first non-orthologous sequence - *A. fumigatus*:1847 *A. fischeri*:1847
afum|EAL89922.1 100.00% afis|EAW25260.1 100.00%
Bootstrap support for afum|EAL89922.1 as seed ortholog is 100%.
Bootstrap support for afis|EAW25260.1 as seed ortholog is 100%.

---

Group of orthologs #1179. Best score 1846 bits
Score difference with first non-orthologous sequence - *A. fumigatus*:1846 *A. fischeri*:1846
afum|EAL88986.1 100.00% afis|EAW16361.1 100.00%
Bootstrap support for afum|EAL88986.1 as seed ortholog is 100%.
Bootstrap support for afis|EAW16361.1 as seed ortholog is 100%.

---

Group of orthologs #1180. Best score 1846 bits
Score difference with first non-orthologous sequence - *A. fumigatus*:1846 *A. fischeri*:1846
afum|EAL88474.1 100.00% afis|EAW23006.1 100.00%
Bootstrap support for afum|EAL88474.1 as seed ortholog is 100%.
Bootstrap support for afis|EAW23006.1 as seed ortholog is 100%.

---

Group of orthologs #1181. Best score 1845 bits
Score difference with first non-orthologous sequence - *A. fumigatus*:1845 *A. fischeri*:1845
afum|EAL84555.1 100.00% afis|EAW20500.1 100.00%
Bootstrap support for afum|EAL84555.1 as seed ortholog is 100%.
Bootstrap support for afis|EAW20500.1 as seed ortholog is 100%.

---

Group of orthologs #1182. Best score 1845 bits
Score difference with first non-orthologous sequence - *A. fumigatus*:1845 *A. fischeri*:1845
afum|EAL87509.1 100.00% afis|EAW18151.1 100.00%
Bootstrap support for afum|EAL87509.1 as seed ortholog is 100%.
Bootstrap support for afis|EAW18151.1 as seed ortholog is 100%.

---

Group of orthologs #1183. Best score 1845 bits

Score difference with first non-orthologous sequence - *A. fumigatus*:1845 *A. fischeri*:1845
afum|EAL90854.1 100.00% afis|EAW22338.1 100.00%
Bootstrap support for afum|EAL90854.1 as seed ortholog is 100%.
Bootstrap support for afis|EAW22338.1 as seed ortholog is 100%.

---

Group of orthologs #1184. Best score 1844 bits
Score difference with first non-orthologous sequence - *A. fumigatus*:1844 *A. fischeri*:1844
afum|EAL91450.1 100.00% afis|EAW17635.1 100.00%
Bootstrap support for afum|EAL91450.1 as seed ortholog is 100%.
Bootstrap support for afis|EAW17635.1 as seed ortholog is 100%.

---

Group of orthologs #1185. Best score 1844 bits
Score difference with first non-orthologous sequence - *A. fumigatus*:1844 *A. fischeri*:1844
afum|EAL90188.2 100.00% afis|EAW25523.1 100.00%
Bootstrap support for afum|EAL90188.2 as seed ortholog is 100%.
Bootstrap support for afis|EAW25523.1 as seed ortholog is 100%.

---

Group of orthologs #1186. Best score 1843 bits
Score difference with first non-orthologous sequence - *A. fumigatus*:1843 *A. fischeri*:1843
afum|EAL85724.1 100.00% afis|EAW15610.1 100.00%
Bootstrap support for afum|EAL85724.1 as seed ortholog is 100%.
Bootstrap support for afis|EAW15610.1 as seed ortholog is 100%.

---

Group of orthologs #1187. Best score 1842 bits
Score difference with first non-orthologous sequence - *A. fumigatus*:1842 *A. fischeri*:1842
afum|EAL88633.1 100.00% afis|EAW16000.1 100.00%
Bootstrap support for afum|EAL88633.1 as seed ortholog is 100%.
Bootstrap support for afis|EAW16000.1 as seed ortholog is 100%.

---

Group of orthologs #1188. Best score 1841 bits
Score difference with first non-orthologous sequence - *A. fumigatus*:1664 *A. fischeri*:1601
afum|EAL91888.1 100.00% afis|EAW18076.1 100.00%
Bootstrap support for afum|EAL91888.1 as seed ortholog is 100%.
Bootstrap support for afis|EAW18076.1 as seed ortholog is 100%.

---

Group of orthologs #1189. Best score 1841 bits
Score difference with first non-orthologous sequence - *A. fumigatus*:1841 *A. fischeri*:1841
afum|EAL92591.2 100.00% afis|EAW21573.1 100.00%
Bootstrap support for afum|EAL92591.2 as seed ortholog is 100%.
Bootstrap support for afis|EAW21573.1 as seed ortholog is 100%.

---

Group of orthologs #1190. Best score 1840 bits
Score difference with first non-orthologous sequence - *A. fumigatus*:1840 *A. fischeri*:1425
afum|EAL84484.1 100.00% afis|EAW20580.1 100.00%
Bootstrap support for afum|EAL84484.1 as seed ortholog is 100%.
Bootstrap support for afis|EAW20580.1 as seed ortholog is 100%.

---

Group of orthologs #1191. Best score 1840 bits
Score difference with first non-orthologous sequence - *A. fumigatus*:1840 *A. fischeri*:1840
afum|EAL89635.1 100.00% afis|EAW24975.1 100.00%
Bootstrap support for afum|EAL89635.1 as seed ortholog is 100%.
Bootstrap support for afis|EAW24975.1 as seed ortholog is 100%.

---

Group of orthologs #1192. Best score 1839 bits
Score difference with first non-orthologous sequence - *A. fumigatus*:1839 *A. fischeri*:1839
afum|EAL89874.1 100.00% afis|EAW25213.1 100.00%

Bootstrap support for afum|EAL89874.1 as seed ortholog is 100%.
Bootstrap support for afis|EAW25213.1 as seed ortholog is 100%.

---

Group of orthologs #1193. Best score 1838 bits
Score difference with first non-orthologous sequence - *A. fumigatus*:1704 *A. fischeri*:1838
afum|EAL85898.1 100.00% afis|EAW24229.1 100.00%
Bootstrap support for afum|EAL85898.1 as seed ortholog is 100%.
Bootstrap support for afis|EAW24229.1 as seed ortholog is 100%.

---

Group of orthologs #1194. Best score 1838 bits
Score difference with first non-orthologous sequence - *A. fumigatus*:1838 *A. fischeri*:1241
afum|EAL86075.1 100.00% afis|EAW24404.1 100.00%
Bootstrap support for afum|EAL86075.1 as seed ortholog is 100%.
Bootstrap support for afis|EAW24404.1 as seed ortholog is 100%.

---

Group of orthologs #1195. Best score 1838 bits
Score difference with first non-orthologous sequence - *A. fumigatus*:1838 *A. fischeri*:1838
afum|EAL90025.1 100.00% afis|EAW25363.1 100.00%
Bootstrap support for afum|EAL90025.1 as seed ortholog is 100%.
Bootstrap support for afis|EAW25363.1 as seed ortholog is 100%.

---

Group of orthologs #1196. Best score 1837 bits
Score difference with first non-orthologous sequence - *A. fumigatus*:1837 *A. fischeri*:1837
afum|EAL85673.1 100.00% afis|EAW15674.1 100.00%
Bootstrap support for afum|EAL85673.1 as seed ortholog is 100%.
Bootstrap support for afis|EAW15674.1 as seed ortholog is 100%.

---

Group of orthologs #1197. Best score 1835 bits
Score difference with first non-orthologous sequence - *A. fumigatus*:1835 *A. fischeri*:1835
afum|EAL90269.1 100.00% afis|EAW22928.1 100.00%
Bootstrap support for afum|EAL90269.1 as seed ortholog is 100%.
Bootstrap support for afis|EAW22928.1 as seed ortholog is 100%.

---

Group of orthologs #1198. Best score 1835 bits
Score difference with first non-orthologous sequence - *A. fumigatus*:1835 *A. fischeri*:1835
afum|EAL92553.1 100.00% afis|EAW21532.1 100.00%
Bootstrap support for afum|EAL92553.1 as seed ortholog is 100%.
Bootstrap support for afis|EAW21532.1 as seed ortholog is 100%.

---

Group of orthologs #1199. Best score 1834 bits
Score difference with first non-orthologous sequence - *A. fumigatus*:1834 *A. fischeri*:1202
afum|EAL89039.1 100.00% afis|EAW16415.1 100.00%
Bootstrap support for afum|EAL89039.1 as seed ortholog is 100%.
Bootstrap support for afis|EAW16415.1 as seed ortholog is 100%.

---

Group of orthologs #1200. Best score 1833 bits
Score difference with first non-orthologous sequence - *A. fumigatus*:1833 *A. fischeri*:1833
afum|EBA27274.1 100.00% afis|EAW16942.1 100.00%
Bootstrap support for afum|EBA27274.1 as seed ortholog is 100%.
Bootstrap support for afis|EAW16942.1 as seed ortholog is 100%.

---

Group of orthologs #1201. Best score 1831 bits
Score difference with first non-orthologous sequence - *A. fumigatus*:1831 *A. fischeri*:1831
afum|EAL88954.1 100.00% afis|EAW16330.1 100.00%
Bootstrap support for afum|EAL88954.1 as seed ortholog is 100%.
Bootstrap support for afis|EAW16330.1 as seed ortholog is 100%.

---

Group of orthologs #1202. Best score 1831 bits
Score difference with first non-orthologous sequence - *A. fumigatus*:1831 *A. fischeri*:1831
afum|EAL90306.1 100.00% afis|EAW22892.1 100.00%
Bootstrap support for afum|EAL90306.1 as seed ortholog is 100%.
Bootstrap support for afis|EAW22892.1 as seed ortholog is 100%.

---

Group of orthologs #1203. Best score 1830 bits
Score difference with first non-orthologous sequence - *A. fumigatus*:1830 *A. fischeri*:1830
afum|EAL88996.1 100.00% afis|EAW16371.1 100.00%
Bootstrap support for afum|EAL88996.1 as seed ortholog is 100%.
Bootstrap support for afis|EAW16371.1 as seed ortholog is 100%.

---

Group of orthologs #1204. Best score 1830 bits
Score difference with first non-orthologous sequence - *A. fumigatus*:1830 *A. fischeri*:1830
afum|EAL87897.2 100.00% afis|EAW23570.1 100.00%
Bootstrap support for afum|EAL87897.2 as seed ortholog is 100%.
Bootstrap support for afis|EAW23570.1 as seed ortholog is 100%.

---

Group of orthologs #1205. Best score 1829 bits
Score difference with first non-orthologous sequence - *A. fumigatus*:1829 *A. fischeri*:1829
afum|EAL89430.1 100.00% afis|EAW24790.1 100.00%
Bootstrap support for afum|EAL89430.1 as seed ortholog is 100%.
Bootstrap support for afis|EAW24790.1 as seed ortholog is 100%.

---

Group of orthologs #1206. Best score 1828 bits
Score difference with first non-orthologous sequence - *A. fumigatus*:1828 *A. fischeri*:1828
afum|EAL89639.1 100.00% afis|EAW24979.1 100.00%
Bootstrap support for afum|EAL89639.1 as seed ortholog is 100%.
Bootstrap support for afis|EAW24979.1 as seed ortholog is 100%.

---

Group of orthologs #1207. Best score 1827 bits
Score difference with first non-orthologous sequence - *A. fumigatus*:1827 *A. fischeri*:1827
afum|EAL88856.1 100.00% afis|EAW16233.1 100.00%
Bootstrap support for afum|EAL88856.1 as seed ortholog is 100%.
Bootstrap support for afis|EAW16233.1 as seed ortholog is 100%.

---

Group of orthologs #1208. Best score 1827 bits
Score difference with first non-orthologous sequence - *A. fumigatus*:1827 *A. fischeri*:1827
afum|EAL90915.1 100.00% afis|EAW22265.1 100.00%
Bootstrap support for afum|EAL90915.1 as seed ortholog is 100%.
Bootstrap support for afis|EAW22265.1 as seed ortholog is 100%.

---

Group of orthologs #1209. Best score 1826 bits
Score difference with first non-orthologous sequence - *A. fumigatus*:1826 *A. fischeri*:1826
afum|EAL90961.1 100.00% afis|EAW22232.1 100.00%
Bootstrap support for afum|EAL90961.1 as seed ortholog is 100%.
Bootstrap support for afis|EAW22232.1 as seed ortholog is 100%.

---

Group of orthologs #1210. Best score 1825 bits
Score difference with first non-orthologous sequence - *A. fumigatus*:1825 *A. fischeri*:1825
afum|EAL91231.1 100.00% afis|EAW17393.1 100.00%
Bootstrap support for afum|EAL91231.1 as seed ortholog is 100%.
Bootstrap support for afis|EAW17393.1 as seed ortholog is 100%.

---

Group of orthologs #1211. Best score 1825 bits

Score difference with first non-orthologous sequence - *A. fumigatus*:1825 *A. fischeri*:1825
afum|EAL88004.1 100.00% afis|EAW23462.1 100.00%
Bootstrap support for afum|EAL88004.1 as seed ortholog is 100%.
Bootstrap support for afis|EAW23462.1 as seed ortholog is 100%.

---

Group of orthologs #1212. Best score 1824 bits
Score difference with first non-orthologous sequence - *A. fumigatus*:1824 *A. fischeri*:1438
afum|EAL92821.1 100.00% afis|EAW21796.1 100.00%
Bootstrap support for afum|EAL92821.1 as seed ortholog is 100%.
Bootstrap support for afis|EAW21796.1 as seed ortholog is 100%.

---

Group of orthologs #1213. Best score 1823 bits
Score difference with first non-orthologous sequence - *A. fumigatus*:1823 *A. fischeri*:1823
afum|EAL86120.1 100.00% afis|EAW24451.1 100.00%
Bootstrap support for afum|EAL86120.1 as seed ortholog is 100%.
Bootstrap support for afis|EAW24451.1 as seed ortholog is 100%.

---

Group of orthologs #1214. Best score 1823 bits
Score difference with first non-orthologous sequence - *A. fumigatus*:1823 *A. fischeri*:1823
afum|EAL88401.1 100.00% afis|EAW23078.1 100.00%
Bootstrap support for afum|EAL88401.1 as seed ortholog is 100%.
Bootstrap support for afis|EAW23078.1 as seed ortholog is 100%.

---

Group of orthologs #1215. Best score 1823 bits
Score difference with first non-orthologous sequence - *A. fumigatus*:1823 *A. fischeri*:1823
afum|EAL90444.2 100.00% afis|EAW22755.1 100.00%
Bootstrap support for afum|EAL90444.2 as seed ortholog is 100%.
Bootstrap support for afis|EAW22755.1 as seed ortholog is 100%.

---

Group of orthologs #1216. Best score 1821 bits
Score difference with first non-orthologous sequence - *A. fumigatus*:1821 *A. fischeri*:1821
afum|EAL85650.1 100.00% afis|EAW15698.1 100.00%
Bootstrap support for afum|EAL85650.1 as seed ortholog is 100%.
Bootstrap support for afis|EAW15698.1 as seed ortholog is 100%.

---

Group of orthologs #1217. Best score 1821 bits
Score difference with first non-orthologous sequence - *A. fumigatus*:1821 *A. fischeri*:1821
afum|EAL90373.2 100.00% afis|EAW22829.1 100.00%
Bootstrap support for afum|EAL90373.2 as seed ortholog is 100%.
Bootstrap support for afis|EAW22829.1 as seed ortholog is 100%.

---

Group of orthologs #1218. Best score 1819 bits
Score difference with first non-orthologous sequence - *A. fumigatus*:1819 *A. fischeri*:1819
afum|EAL89096.1 100.00% afis|EAW16501.1 100.00%
Bootstrap support for afum|EAL89096.1 as seed ortholog is 100%.
Bootstrap support for afis|EAW16501.1 as seed ortholog is 100%.

---

Group of orthologs #1219. Best score 1819 bits
Score difference with first non-orthologous sequence - *A. fumigatus*:1819 *A. fischeri*:1819
afum|EAL88358.1 100.00% afis|EAW23122.1 100.00%
Bootstrap support for afum|EAL88358.1 as seed ortholog is 100%.
Bootstrap support for afis|EAW23122.1 as seed ortholog is 100%.

---

Group of orthologs #1220. Best score 1819 bits
Score difference with first non-orthologous sequence - *A. fumigatus*:1819 *A. fischeri*:1819
afum|EAL93584.1 100.00% afis|EAW18853.1 100.00%

Bootstrap support for afum|EAL93584.1 as seed ortholog is 100%.
Bootstrap support for afis|EAW18853.1 as seed ortholog is 100%.

---

Group of orthologs #1221. Best score 1819 bits
Score difference with first non-orthologous sequence - *A. fumigatus*:1819 *A. fischeri*:1819
afum|EAL92258.1 100.00% afis|EAW21245.1 100.00%
Bootstrap support for afum|EAL92258.1 as seed ortholog is 100%.
Bootstrap support for afis|EAW21245.1 as seed ortholog is 100%.

---

Group of orthologs #1222. Best score 1819 bits
Score difference with first non-orthologous sequence - *A. fumigatus*:1819 *A. fischeri*:1819
afum|EAL92465.2 100.00% afis|EAW21446.1 100.00%
Bootstrap support for afum|EAL92465.2 as seed ortholog is 100%.
Bootstrap support for afis|EAW21446.1 as seed ortholog is 100%.

---

Group of orthologs #1223. Best score 1818 bits
Score difference with first non-orthologous sequence - *A. fumigatus*:1818 *A. fischeri*:1818
afum|EAL88129.1 100.00% afis|EAW23347.1 100.00%
Bootstrap support for afum|EAL88129.1 as seed ortholog is 100%.
Bootstrap support for afis|EAW23347.1 as seed ortholog is 100%.

---

Group of orthologs #1224. Best score 1816 bits
Score difference with first non-orthologous sequence - *A. fumigatus*:1816 *A. fischeri*:1816
afum|EAL91951.2 100.00% afis|EAW24097.1 100.00%
Bootstrap support for afum|EAL91951.2 as seed ortholog is 100%.
Bootstrap support for afis|EAW24097.1 as seed ortholog is 100%.

---

Group of orthologs #1225. Best score 1813 bits
Score difference with first non-orthologous sequence - *A. fumigatus*:1813 *A. fischeri*:1813
afum|EAL87366.1 100.00% afis|EAW23941.1 100.00%
Bootstrap support for afum|EAL87366.1 as seed ortholog is 100%.
Bootstrap support for afis|EAW23941.1 as seed ortholog is 100%.

---

Group of orthologs #1226. Best score 1813 bits
Score difference with first non-orthologous sequence - *A. fumigatus*:1813 *A. fischeri*:1813
afum|EAL89944.1 100.00% afis|EAW25281.1 100.00%
Bootstrap support for afum|EAL89944.1 as seed ortholog is 100%.
Bootstrap support for afis|EAW25281.1 as seed ortholog is 100%.

---

Group of orthologs #1227. Best score 1812 bits
Score difference with first non-orthologous sequence - *A. fumigatus*:1812 *A. fischeri*:1812
afum|EAL87915.1 100.00% afis|EAW23551.1 100.00%
Bootstrap support for afum|EAL87915.1 as seed ortholog is 100%.
Bootstrap support for afis|EAW23551.1 as seed ortholog is 100%.

---

Group of orthologs #1228. Best score 1812 bits
Score difference with first non-orthologous sequence - *A. fumigatus*:1291 *A. fischeri*:1812
afum|EBA27314.1 100.00% afis|EAW18160.1 100.00%
Bootstrap support for afum|EBA27314.1 as seed ortholog is 100%.
Bootstrap support for afis|EAW18160.1 as seed ortholog is 100%.

---

Group of orthologs #1229. Best score 1812 bits
Score difference with first non-orthologous sequence - *A. fumigatus*:1812 *A. fischeri*:1812
afum|EAL90237.1 100.00% afis|EAW22960.1 100.00%
Bootstrap support for afum|EAL90237.1 as seed ortholog is 100%.
Bootstrap support for afis|EAW22960.1 as seed ortholog is 100%.

---

Group of orthologs #1230. Best score 1812 bits
Score difference with first non-orthologous sequence - *A. fumigatus*:1812 *A. fischeri*:1812
afum|EAL89739.1 100.00% afis|EAW25070.1 100.00%
Bootstrap support for afum|EAL89739.1 as seed ortholog is 100%.
Bootstrap support for afis|EAW25070.1 as seed ortholog is 100%.

---

Group of orthologs #1231. Best score 1810 bits
Score difference with first non-orthologous sequence - *A. fumigatus*:1810 *A. fischeri*:1810
afum|EAL88077.1 100.00% afis|EAW23393.1 100.00%
Bootstrap support for afum|EAL88077.1 as seed ortholog is 100%.
Bootstrap support for afis|EAW23393.1 as seed ortholog is 100%.

---

Group of orthologs #1232. Best score 1810 bits
Score difference with first non-orthologous sequence - *A. fumigatus*:1810 *A. fischeri*:1810
afum|EAL93872.1 100.00% afis|EAW19141.1 100.00%
Bootstrap support for afum|EAL93872.1 as seed ortholog is 100%.
Bootstrap support for afis|EAW19141.1 as seed ortholog is 100%.

---

Group of orthologs #1233. Best score 1808 bits
Score difference with first non-orthologous sequence - *A. fumigatus*:1808 *A. fischeri*:1808
afum|EAL92464.2 100.00% afis|EAW21445.1 100.00%
Bootstrap support for afum|EAL92464.2 as seed ortholog is 100%.
Bootstrap support for afis|EAW21445.1 as seed ortholog is 100%.

---

Group of orthologs #1234. Best score 1808 bits
Score difference with first non-orthologous sequence - *A. fumigatus*:1808 *A. fischeri*:1808
afum|EAL89823.1 100.00% afis|EAW25148.1 100.00%
Bootstrap support for afum|EAL89823.1 as seed ortholog is 100%.
Bootstrap support for afis|EAW25148.1 as seed ortholog is 100%.

---

Group of orthologs #1235. Best score 1807 bits
Score difference with first non-orthologous sequence - *A. fumigatus*:972 *A. fischeri*:1567
afum|EAL86491.1 100.00% afis|EAW16964.1 100.00%
Bootstrap support for afum|EAL86491.1 as seed ortholog is 100%.
Bootstrap support for afis|EAW16964.1 as seed ortholog is 100%.

---

Group of orthologs #1236. Best score 1806 bits
Score difference with first non-orthologous sequence - *A. fumigatus*:1806 *A. fischeri*:1806
afum|EAL91480.2 100.00% afis|EAW17666.1 100.00%
Bootstrap support for afum|EAL91480.2 as seed ortholog is 100%.
Bootstrap support for afis|EAW17666.1 as seed ortholog is 100%.

---

Group of orthologs #1237. Best score 1804 bits
Score difference with first non-orthologous sequence - *A. fumigatus*:1804 *A. fischeri*:1804
afum|EAL92559.2 100.00% afis|EAW21539.1 100.00%
Bootstrap support for afum|EAL92559.2 as seed ortholog is 100%.
Bootstrap support for afis|EAW21539.1 as seed ortholog is 100%.

---

Group of orthologs #1238. Best score 1804 bits
Score difference with first non-orthologous sequence - *A. fumigatus*:1646 *A. fischeri*:1804
afum|EAL89318.1 100.00% afis|EAW24652.1 100.00%
Bootstrap support for afum|EAL89318.1 as seed ortholog is 100%.
Bootstrap support for afis|EAW24652.1 as seed ortholog is 100%.

---

Group of orthologs #1239. Best score 1804 bits

Score difference with first non-orthologous sequence - *A. fumigatus*:1804 *A. fischeri*:1804
afum|EAL89423.1 100.00% afis|EAW24784.1 100.00%
Bootstrap support for afum|EAL89423.1 as seed ortholog is 100%.
Bootstrap support for afis|EAW24784.1 as seed ortholog is 100%.

---

Group of orthologs #1240. Best score 1803 bits
Score difference with first non-orthologous sequence - *A. fumigatus*:1803 *A. fischeri*:1803
afum|EAL84450.1 100.00% afis|EAW20611.1 100.00%
Bootstrap support for afum|EAL84450.1 as seed ortholog is 100%.
Bootstrap support for afis|EAW20611.1 as seed ortholog is 100%.

---

Group of orthologs #1241. Best score 1803 bits
Score difference with first non-orthologous sequence - *A. fumigatus*:1803 *A. fischeri*:1803
afum|EAL85252.1 100.00% afis|EAW20137.1 100.00%
Bootstrap support for afum|EAL85252.1 as seed ortholog is 100%.
Bootstrap support for afis|EAW20137.1 as seed ortholog is 100%.

---

Group of orthologs #1242. Best score 1803 bits
Score difference with first non-orthologous sequence - *A. fumigatus*:1604 *A. fischeri*:1109
afum|EAL91070.1 100.00% afis|EAW22115.1 100.00%
Bootstrap support for afum|EAL91070.1 as seed ortholog is 100%.
Bootstrap support for afis|EAW22115.1 as seed ortholog is 100%.

---

Group of orthologs #1243. Best score 1802 bits
Score difference with first non-orthologous sequence - *A. fumigatus*:1802 *A. fischeri*:1802
afum|EAL88546.2 100.00% afis|EAW15911.1 100.00%
Bootstrap support for afum|EAL88546.2 as seed ortholog is 100%.
Bootstrap support for afis|EAW15911.1 as seed ortholog is 100%.

---

Group of orthologs #1244. Best score 1802 bits
Score difference with first non-orthologous sequence - *A. fumigatus*:1802 *A. fischeri*:1802
afum|EAL89828.1 100.00% afis|EAW25153.1 100.00%
Bootstrap support for afum|EAL89828.1 as seed ortholog is 100%.
Bootstrap support for afis|EAW25153.1 as seed ortholog is 100%.

---

Group of orthologs #1245. Best score 1801 bits
Score difference with first non-orthologous sequence - *A. fumigatus*:1626 *A. fischeri*:1604
afum|EAL84954.1 100.00% afis|EAW19961.1 100.00%
Bootstrap support for afum|EAL84954.1 as seed ortholog is 100%.
Bootstrap support for afis|EAW19961.1 as seed ortholog is 100%.

---

Group of orthologs #1246. Best score 1801 bits
Score difference with first non-orthologous sequence - *A. fumigatus*:1801 *A. fischeri*:1801
afum|EAL91889.1 100.00% afis|EAW18077.1 100.00%
Bootstrap support for afum|EAL91889.1 as seed ortholog is 100%.
Bootstrap support for afis|EAW18077.1 as seed ortholog is 100%.

---

Group of orthologs #1247. Best score 1801 bits
Score difference with first non-orthologous sequence - *A. fumigatus*:1801 *A. fischeri*:1801
afum|EAL93723.1 100.00% afis|EAW18987.1 100.00%
Bootstrap support for afum|EAL93723.1 as seed ortholog is 100%.
Bootstrap support for afis|EAW18987.1 as seed ortholog is 100%.

---

Group of orthologs #1248. Best score 1800 bits
Score difference with first non-orthologous sequence - *A. fumigatus*:1800 *A. fischeri*:1800
afum|EAL85212.1 100.00% afis|EAW20094.1 100.00%

Bootstrap support for afum|EAL85212.1 as seed ortholog is 100%.
Bootstrap support for afis|EAW20094.1 as seed ortholog is 100%.

---

Group of orthologs #1249. Best score 1800 bits
Score difference with first non-orthologous sequence - *A. fumigatus*:1800 *A. fischeri*:1800
afum|EAL90375.2 100.00% afis|EAW22825.1 100.00%
Bootstrap support for afum|EAL90375.2 as seed ortholog is 100%.
Bootstrap support for afis|EAW22825.1 as seed ortholog is 100%.

---

Group of orthologs #1250. Best score 1799 bits
Score difference with first non-orthologous sequence - *A. fumigatus*:1381 *A. fischeri*:1698
afum|EAL88706.2 100.00% afis|EAW16074.1 100.00%
Bootstrap support for afum|EAL88706.2 as seed ortholog is 100%.
Bootstrap support for afis|EAW16074.1 as seed ortholog is 100%.

---

Group of orthologs #1251. Best score 1797 bits
Score difference with first non-orthologous sequence - *A. fumigatus*:1797 *A. fischeri*:1797
afum|EAL88673.1 100.00% afis|EAW16041.1 100.00%
Bootstrap support for afum|EAL88673.1 as seed ortholog is 100%.
Bootstrap support for afis|EAW16041.1 as seed ortholog is 100%.

---

Group of orthologs #1252. Best score 1797 bits
Score difference with first non-orthologous sequence - *A. fumigatus*:1797 *A. fischeri*:1797
afum|EAL90792.1 100.00% afis|EAW22399.1 100.00%
Bootstrap support for afum|EAL90792.1 as seed ortholog is 100%.
Bootstrap support for afis|EAW22399.1 as seed ortholog is 100%.

---

Group of orthologs #1253. Best score 1796 bits
Score difference with first non-orthologous sequence - *A. fumigatus*:1796 *A. fischeri*:1796
afum|EAL91396.1 100.00% afis|EAW17583.1 100.00%
Bootstrap support for afum|EAL91396.1 as seed ortholog is 100%.
Bootstrap support for afis|EAW17583.1 as seed ortholog is 100%.

---

Group of orthologs #1254. Best score 1796 bits
Score difference with first non-orthologous sequence - *A. fumigatus*:1796 *A. fischeri*:1796
afum|EAL92836.1 100.00% afis|EAW21820.1 100.00%
Bootstrap support for afum|EAL92836.1 as seed ortholog is 100%.
Bootstrap support for afis|EAW21820.1 as seed ortholog is 100%.

---

Group of orthologs #1255. Best score 1794 bits
Score difference with first non-orthologous sequence - *A. fumigatus*:1794 *A. fischeri*:1794
afum|EAL93790.1 100.00% afis|EAW19061.1 100.00%
Bootstrap support for afum|EAL93790.1 as seed ortholog is 100%.
Bootstrap support for afis|EAW19061.1 as seed ortholog is 100%.

---

Group of orthologs #1256. Best score 1794 bits
Score difference with first non-orthologous sequence - *A. fumigatus*:1794 *A. fischeri*:1794
afum|EAL92812.1 100.00% afis|EAW21785.1 100.00%
Bootstrap support for afum|EAL92812.1 as seed ortholog is 100%.
Bootstrap support for afis|EAW21785.1 as seed ortholog is 100%.

---

Group of orthologs #1257. Best score 1792 bits
Score difference with first non-orthologous sequence - *A. fumigatus*:1792 *A. fischeri*:1792
afum|EAL91478.1 100.00% afis|EAW17664.1 100.00%
Bootstrap support for afum|EAL91478.1 as seed ortholog is 100%.
Bootstrap support for afis|EAW17664.1 as seed ortholog is 100%.

Group of orthologs #1258. Best score 1791 bits
Score difference with first non-orthologous sequence - *A. fumigatus*:1791 *A. fischeri*:1791
afum|EAL91505.1 100.00% afis|EAW17691.1 100.00%
Bootstrap support for afum|EAL91505.1 as seed ortholog is 100%.
Bootstrap support for afis|EAW17691.1 as seed ortholog is 100%.

Group of orthologs #1259. Best score 1790 bits
Score difference with first non-orthologous sequence - *A. fumigatus*:1790 *A. fischeri*:1790
afum|EAL87014.1 100.00% afis|EAW19481.1 100.00%
Bootstrap support for afum|EAL87014.1 as seed ortholog is 100%.
Bootstrap support for afis|EAW19481.1 as seed ortholog is 100%.

Group of orthologs #1260. Best score 1790 bits
Score difference with first non-orthologous sequence - *A. fumigatus*:1790 *A. fischeri*:1790
afum|EAL91041.1 100.00% afis|EAW22149.1 100.00%
Bootstrap support for afum|EAL91041.1 as seed ortholog is 100%.
Bootstrap support for afis|EAW22149.1 as seed ortholog is 100%.

Group of orthologs #1261. Best score 1790 bits
Score difference with first non-orthologous sequence - *A. fumigatus*:1790 *A. fischeri*:1790
afum|EAL89480.1 100.00% afis|EAW24835.1 100.00%
Bootstrap support for afum|EAL89480.1 as seed ortholog is 100%.
Bootstrap support for afis|EAW24835.1 as seed ortholog is 100%.

Group of orthologs #1262. Best score 1789 bits
Score difference with first non-orthologous sequence - *A. fumigatus*:1789 *A. fischeri*:1789
afum|EAL90291.1 100.00% afis|EAW22906.1 100.00%
Bootstrap support for afum|EAL90291.1 as seed ortholog is 100%.
Bootstrap support for afis|EAW22906.1 as seed ortholog is 100%.

Group of orthologs #1263. Best score 1788 bits
Score difference with first non-orthologous sequence - *A. fumigatus*:1788 *A. fischeri*:1788
afum|EAL84801.1 100.00% afis|EAW20831.1 100.00%
Bootstrap support for afum|EAL84801.1 as seed ortholog is 100%.
Bootstrap support for afis|EAW20831.1 as seed ortholog is 100%.

Group of orthologs #1264. Best score 1788 bits
Score difference with first non-orthologous sequence - *A. fumigatus*:1788 *A. fischeri*:1788
afum|EAL92273.1 100.00% afis|EAW21258.1 100.00%
Bootstrap support for afum|EAL92273.1 as seed ortholog is 100%.
Bootstrap support for afis|EAW21258.1 as seed ortholog is 100%.

Group of orthologs #1265. Best score 1788 bits
Score difference with first non-orthologous sequence - *A. fumigatus*:1667 *A. fischeri*:1610
afum|EAL92289.2 100.00% afis|EAW21274.1 100.00%
Bootstrap support for afum|EAL92289.2 as seed ortholog is 100%.
Bootstrap support for afis|EAW21274.1 as seed ortholog is 100%.

Group of orthologs #1266. Best score 1788 bits
Score difference with first non-orthologous sequence - *A. fumigatus*:1788 *A. fischeri*:1788
afum|EAL89491.1 100.00% afis|EAW24845.1 100.00%
Bootstrap support for afum|EAL89491.1 as seed ortholog is 100%.
Bootstrap support for afis|EAW24845.1 as seed ortholog is 100%.

Group of orthologs #1267. Best score 1787 bits

Score difference with first non-orthologous sequence - *A. fumigatus*:1787   *A. fischeri*:1787
afum|EAL88304.1 100.00% afis|EAW23179.1 100.00%
Bootstrap support for afum|EAL88304.1 as seed ortholog is 100%.
Bootstrap support for afis|EAW23179.1 as seed ortholog is 100%.

---

Group of orthologs #1268. Best score 1785 bits
Score difference with first non-orthologous sequence - *A. fumigatus*:1785   *A. fischeri*:1785
afum|EAL85652.2 100.00% afis|EAW15696.1 100.00%
Bootstrap support for afum|EAL85652.2 as seed ortholog is 100%.
Bootstrap support for afis|EAW15696.1 as seed ortholog is 100%.

---

Group of orthologs #1269. Best score 1785 bits
Score difference with first non-orthologous sequence - *A. fumigatus*:1785   *A. fischeri*:1785
afum|EAL93966.1 100.00% afis|EAW19238.1 100.00%
Bootstrap support for afum|EAL93966.1 as seed ortholog is 100%.
Bootstrap support for afis|EAW19238.1 as seed ortholog is 100%.

---

Group of orthologs #1270. Best score 1785 bits
Score difference with first non-orthologous sequence - *A. fumigatus*:1785   *A. fischeri*:1785
afum|EAL90707.1 100.00% afis|EAW22486.1 100.00%
Bootstrap support for afum|EAL90707.1 as seed ortholog is 100%.
Bootstrap support for afis|EAW22486.1 as seed ortholog is 100%.

---

Group of orthologs #1271. Best score 1785 bits
Score difference with first non-orthologous sequence - *A. fumigatus*:1785   *A. fischeri*:1547
afum|EAL92667.1 100.00% afis|EAW21634.1 100.00%
Bootstrap support for afum|EAL92667.1 as seed ortholog is 100%.
Bootstrap support for afis|EAW21634.1 as seed ortholog is 100%.

---

Group of orthologs #1272. Best score 1785 bits
Score difference with first non-orthologous sequence - *A. fumigatus*:1785   *A. fischeri*:1785
afum|EAL90090.1 100.00% afis|EAW25432.1 100.00%
Bootstrap support for afum|EAL90090.1 as seed ortholog is 100%.
Bootstrap support for afis|EAW25432.1 as seed ortholog is 100%.

---

Group of orthologs #1273. Best score 1784 bits
Score difference with first non-orthologous sequence - *A. fumigatus*:1784   *A. fischeri*:1208
afum|EAL87630.1 100.00% afis|EAW18265.1 100.00%
Bootstrap support for afum|EAL87630.1 as seed ortholog is 100%.
Bootstrap support for afis|EAW18265.1 as seed ortholog is 100%.

---

Group of orthologs #1274. Best score 1784 bits
Score difference with first non-orthologous sequence - *A. fumigatus*:1784   *A. fischeri*:1784
afum|EAL91939.1 100.00% afis|EAW24084.1 100.00%
Bootstrap support for afum|EAL91939.1 as seed ortholog is 100%.
Bootstrap support for afis|EAW24084.1 as seed ortholog is 100%.

---

Group of orthologs #1275. Best score 1783 bits
Score difference with first non-orthologous sequence - *A. fumigatus*:1783   *A. fischeri*:1783
afum|EAL90388.2 100.00% afis|EAW22813.1 100.00%
Bootstrap support for afum|EAL90388.2 as seed ortholog is 100%.
Bootstrap support for afis|EAW22813.1 as seed ortholog is 100%.

---

Group of orthologs #1276. Best score 1782 bits
Score difference with first non-orthologous sequence - *A. fumigatus*:1782   *A. fischeri*:1060
afum|EAL85241.2 100.00% afis|EAW20123.1 100.00%

Bootstrap support for afum|EAL85241.2 as seed ortholog is 100%.
Bootstrap support for afis|EAW20123.1 as seed ortholog is 100%.

---

Group of orthologs #1277. Best score 1782 bits
Score difference with first non-orthologous sequence - *A. fumigatus*:1782 *A. fischeri*:1782
afum|EAL92324.1 100.00% afis|EAW21308.1 100.00%
Bootstrap support for afum|EAL92324.1 as seed ortholog is 100%.
Bootstrap support for afis|EAW21308.1 as seed ortholog is 100%.

---

Group of orthologs #1278. Best score 1781 bits
Score difference with first non-orthologous sequence - *A. fumigatus*:1781 *A. fischeri*:1781
afum|EAL85822.1 100.00% afis|EAW15502.1 100.00%
Bootstrap support for afum|EAL85822.1 as seed ortholog is 100%.
Bootstrap support for afis|EAW15502.1 as seed ortholog is 100%.

---

Group of orthologs #1279. Best score 1780 bits
Score difference with first non-orthologous sequence - *A. fumigatus*:1780 *A. fischeri*:1780
afum|EAL87988.1 100.00% afis|EAW23477.1 100.00%
Bootstrap support for afum|EAL87988.1 as seed ortholog is 100%.
Bootstrap support for afis|EAW23477.1 as seed ortholog is 100%.

---

Group of orthologs #1280. Best score 1780 bits
Score difference with first non-orthologous sequence - *A. fumigatus*:1780 *A. fischeri*:1780
afum|EAL92332.1 100.00% afis|EAW21315.1 100.00%
Bootstrap support for afum|EAL92332.1 as seed ortholog is 100%.
Bootstrap support for afis|EAW21315.1 as seed ortholog is 100%.

---

Group of orthologs #1281. Best score 1779 bits
Score difference with first non-orthologous sequence - *A. fumigatus*:1779 *A. fischeri*:1779
afum|EAL91256.1 100.00% afis|EAW17428.1 100.00%
Bootstrap support for afum|EAL91256.1 as seed ortholog is 100%.
Bootstrap support for afis|EAW17428.1 as seed ortholog is 100%.

---

Group of orthologs #1282. Best score 1778 bits
Score difference with first non-orthologous sequence - *A. fumigatus*:1778 *A. fischeri*:1778
afum|EAL93649.2 100.00% afis|EAW18918.1 100.00%
Bootstrap support for afum|EAL93649.2 as seed ortholog is 100%.
Bootstrap support for afis|EAW18918.1 as seed ortholog is 100%.

---

Group of orthologs #1283. Best score 1778 bits
Score difference with first non-orthologous sequence - *A. fumigatus*:1778 *A. fischeri*:1778
afum|EAL90851.1 100.00% afis|EAW22341.1 100.00%
Bootstrap support for afum|EAL90851.1 as seed ortholog is 100%.
Bootstrap support for afis|EAW22341.1 as seed ortholog is 100%.

---

Group of orthologs #1284. Best score 1777 bits
Score difference with first non-orthologous sequence - *A. fumigatus*:1777 *A. fischeri*:1777
afum|EAL93354.2 100.00% afis|EAW18640.1 100.00%
Bootstrap support for afum|EAL93354.2 as seed ortholog is 100%.
Bootstrap support for afis|EAW18640.1 as seed ortholog is 100%.

---

Group of orthologs #1285. Best score 1777 bits
Score difference with first non-orthologous sequence - *A. fumigatus*:1777 *A. fischeri*:1777
afum|EAL90179.1 100.00% afis|EAW25514.1 100.00%
Bootstrap support for afum|EAL90179.1 as seed ortholog is 100%.
Bootstrap support for afis|EAW25514.1 as seed ortholog is 100%.

Group of orthologs #1286. Best score 1776 bits
Score difference with first non-orthologous sequence - *A. fumigatus*:1776 *A. fischeri*:1776
afum|EAL91469.1 100.00% afis|EAW17654.1 100.00%
Bootstrap support for afum|EAL91469.1 as seed ortholog is 100%.
Bootstrap support for afis|EAW17654.1 as seed ortholog is 100%.

Group of orthologs #1287. Best score 1776 bits
Score difference with first non-orthologous sequence - *A. fumigatus*:1776 *A. fischeri*:1776
afum|EAL92593.1 100.00% afis|EAW21575.1 100.00%
Bootstrap support for afum|EAL92593.1 as seed ortholog is 100%.
Bootstrap support for afis|EAW21575.1 as seed ortholog is 100%.

Group of orthologs #1288. Best score 1775 bits
Score difference with first non-orthologous sequence - *A. fumigatus*:1775 *A. fischeri*:1775
afum|EAL88921.1 100.00% afis|EAW16299.1 100.00%
Bootstrap support for afum|EAL88921.1 as seed ortholog is 100%.
Bootstrap support for afis|EAW16299.1 as seed ortholog is 100%.

Group of orthologs #1289. Best score 1775 bits
Score difference with first non-orthologous sequence - *A. fumigatus*:1775 *A. fischeri*:1775
afum|EAL89880.1 100.00% afis|EAW25219.1 100.00%
Bootstrap support for afum|EAL89880.1 as seed ortholog is 100%.
Bootstrap support for afis|EAW25219.1 as seed ortholog is 100%.

Group of orthologs #1290. Best score 1774 bits
Score difference with first non-orthologous sequence - *A. fumigatus*:1774 *A. fischeri*:1774
afum|EAL90997.1 100.00% afis|EAW22194.1 100.00%
Bootstrap support for afum|EAL90997.1 as seed ortholog is 100%.
Bootstrap support for afis|EAW22194.1 as seed ortholog is 100%.

Group of orthologs #1291. Best score 1773 bits
Score difference with first non-orthologous sequence - *A. fumigatus*:1773 *A. fischeri*:1075
afum|EAL84245.1 100.00% afis|EAW15402.1 100.00%
Bootstrap support for afum|EAL84245.1 as seed ortholog is 100%.
Bootstrap support for afis|EAW15402.1 as seed ortholog is 100%.

Group of orthologs #1292. Best score 1773 bits
Score difference with first non-orthologous sequence - *A. fumigatus*:1773 *A. fischeri*:1773
afum|EAL91507.1 100.00% afis|EAW17693.1 100.00%
Bootstrap support for afum|EAL91507.1 as seed ortholog is 100%.
Bootstrap support for afis|EAW17693.1 as seed ortholog is 100%.

Group of orthologs #1293. Best score 1769 bits
Score difference with first non-orthologous sequence - *A. fumigatus*:1769 *A. fischeri*:1769
afum|EAL88887.1 100.00% afis|EAW16266.1 100.00%
Bootstrap support for afum|EAL88887.1 as seed ortholog is 100%.
Bootstrap support for afis|EAW16266.1 as seed ortholog is 100%.

Group of orthologs #1294. Best score 1768 bits
Score difference with first non-orthologous sequence - *A. fumigatus*:1768 *A. fischeri*:1768
afum|EAL86185.1 100.00% afis|EAW24519.1 100.00%
Bootstrap support for afum|EAL86185.1 as seed ortholog is 100%.
Bootstrap support for afis|EAW24519.1 as seed ortholog is 100%.

Group of orthologs #1295. Best score 1768 bits

Score difference with first non-orthologous sequence - *A. fumigatus*:1768 *A. fischeri*:1768
afum|EAL93608.1 100.00% afis|EAW18876.1 100.00%
Bootstrap support for afum|EAL93608.1 as seed ortholog is 100%.
Bootstrap support for afis|EAW18876.1 as seed ortholog is 100%.

---

Group of orthologs #1296. Best score 1768 bits
Score difference with first non-orthologous sequence - *A. fumigatus*:1653 *A. fischeri*:1768
afum|EAL89589.1 100.00% afis|EAW24933.1 100.00%
Bootstrap support for afum|EAL89589.1 as seed ortholog is 100%.
Bootstrap support for afis|EAW24933.1 as seed ortholog is 100%.

---

Group of orthologs #1297. Best score 1768 bits
Score difference with first non-orthologous sequence - *A. fumigatus*:1768 *A. fischeri*:1768
afum|EAL89655.1 100.00% afis|EAW24994.1 100.00%
Bootstrap support for afum|EAL89655.1 as seed ortholog is 100%.
Bootstrap support for afis|EAW24994.1 as seed ortholog is 100%.

---

Group of orthologs #1298. Best score 1767 bits
Score difference with first non-orthologous sequence - *A. fumigatus*:1767 *A. fischeri*:1767
afum|EAL90011.1 100.00% afis|EAW25348.1 100.00%
Bootstrap support for afum|EAL90011.1 as seed ortholog is 100%.
Bootstrap support for afis|EAW25348.1 as seed ortholog is 100%.

---

Group of orthologs #1299. Best score 1766 bits
Score difference with first non-orthologous sequence - *A. fumigatus*:1766 *A. fischeri*:1766
afum|EAL88741.1 100.00% afis|EAW16110.1 100.00%
Bootstrap support for afum|EAL88741.1 as seed ortholog is 100%.
Bootstrap support for afis|EAW16110.1 as seed ortholog is 100%.

---

Group of orthologs #1300. Best score 1766 bits
Score difference with first non-orthologous sequence - *A. fumigatus*:1766 *A. fischeri*:1766
afum|EAL88975.1 100.00% afis|EAW16350.1 100.00%
Bootstrap support for afum|EAL88975.1 as seed ortholog is 100%.
Bootstrap support for afis|EAW16350.1 as seed ortholog is 100%.

---

Group of orthologs #1301. Best score 1766 bits
Score difference with first non-orthologous sequence - *A. fumigatus*:1766 *A. fischeri*:1766
afum|EAL84540.1 100.00% afis|EAW20517.1 100.00%
Bootstrap support for afum|EAL84540.1 as seed ortholog is 100%.
Bootstrap support for afis|EAW20517.1 as seed ortholog is 100%.

---

Group of orthologs #1302. Best score 1766 bits
Score difference with first non-orthologous sequence - *A. fumigatus*:1766 *A. fischeri*:1766
afum|EAL88435.1 100.00% afis|EAW23046.1 100.00%
Bootstrap support for afum|EAL88435.1 as seed ortholog is 100%.
Bootstrap support for afis|EAW23046.1 as seed ortholog is 100%.

---

Group of orthologs #1303. Best score 1766 bits
Score difference with first non-orthologous sequence - *A. fumigatus*:1766 *A. fischeri*:1766
afum|EAL92387.1 100.00% afis|EAW21370.1 100.00%
Bootstrap support for afum|EAL92387.1 as seed ortholog is 100%.
Bootstrap support for afis|EAW21370.1 as seed ortholog is 100%.

---

Group of orthologs #1304. Best score 1765 bits
Score difference with first non-orthologous sequence - *A. fumigatus*:1765 *A. fischeri*:1765
afum|EAL84445.1 100.00% afis|EAW20616.1 100.00%

Bootstrap support for afum|EAL84445.1 as seed ortholog is 100%.
Bootstrap support for afis|EAW20616.1 as seed ortholog is 100%.

---

Group of orthologs #1305. Best score 1765 bits
Score difference with first non-orthologous sequence - *A. fumigatus*:1549 *A. fischeri*:1120
afum|EAL86858.1 100.00% afis|EAW19665.1 100.00%
Bootstrap support for afum|EAL86858.1 as seed ortholog is 100%.
Bootstrap support for afis|EAW19665.1 as seed ortholog is 100%.

---

Group of orthologs #1306. Best score 1765 bits
Score difference with first non-orthologous sequence - *A. fumigatus*:1765 *A. fischeri*:1443
afum|EAL93199.1 100.00% afis|EAW18505.1 100.00%
Bootstrap support for afum|EAL93199.1 as seed ortholog is 100%.
Bootstrap support for afis|EAW18505.1 as seed ortholog is 100%.

---

Group of orthologs #1307. Best score 1762 bits
Score difference with first non-orthologous sequence - *A. fumigatus*:1762 *A. fischeri*:1762
afum|EAL85019.1 100.00% afis|EAW19879.1 100.00%
Bootstrap support for afum|EAL85019.1 as seed ortholog is 100%.
Bootstrap support for afis|EAW19879.1 as seed ortholog is 100%.

---

Group of orthologs #1308. Best score 1762 bits
Score difference with first non-orthologous sequence - *A. fumigatus*:1762 *A. fischeri*:1762
afum|EAL92543.1 100.00% afis|EAW21522.1 100.00%
Bootstrap support for afum|EAL92543.1 as seed ortholog is 100%.
Bootstrap support for afis|EAW21522.1 as seed ortholog is 100%.

---

Group of orthologs #1309. Best score 1762 bits
Score difference with first non-orthologous sequence - *A. fumigatus*:1762 *A. fischeri*:1762
afum|EAL92681.1 100.00% afis|EAW21647.1 100.00%
Bootstrap support for afum|EAL92681.1 as seed ortholog is 100%.
Bootstrap support for afis|EAW21647.1 as seed ortholog is 100%.

---

Group of orthologs #1310. Best score 1760 bits
Score difference with first non-orthologous sequence - *A. fumigatus*:1760 *A. fischeri*:1760
afum|EAL93886.1 100.00% afis|EAW19155.1 100.00%
Bootstrap support for afum|EAL93886.1 as seed ortholog is 100%.
Bootstrap support for afis|EAW19155.1 as seed ortholog is 100%.

---

Group of orthologs #1311. Best score 1760 bits
Score difference with first non-orthologous sequence - *A. fumigatus*:1760 *A. fischeri*:1760
afum|EAL90641.1 100.00% afis|EAW22554.1 100.00%
Bootstrap support for afum|EAL90641.1 as seed ortholog is 100%.
Bootstrap support for afis|EAW22554.1 as seed ortholog is 100%.

---

Group of orthologs #1312. Best score 1760 bits
Score difference with first non-orthologous sequence - *A. fumigatus*:1760 *A. fischeri*:1760
afum|EAL92824.2 100.00% afis|EAW21799.1 100.00%
Bootstrap support for afum|EAL92824.2 as seed ortholog is 100%.
Bootstrap support for afis|EAW21799.1 as seed ortholog is 100%.

---

Group of orthologs #1313. Best score 1759 bits
Score difference with first non-orthologous sequence - *A. fumigatus*:1759 *A. fischeri*:1759
afum|EAL92310.1 100.00% afis|EAW21293.1 100.00%
Bootstrap support for afum|EAL92310.1 as seed ortholog is 100%.
Bootstrap support for afis|EAW21293.1 as seed ortholog is 100%.

Group of orthologs #1314. Best score 1758 bits
Score difference with first non-orthologous sequence - *A. fumigatus*:1758 *A. fischeri*:1758
afum|EAL89170.2 100.00% afis|EAW16573.1 100.00%
Bootstrap support for afum|EAL89170.2 as seed ortholog is 100%.
Bootstrap support for afis|EAW16573.1 as seed ortholog is 100%.

Group of orthologs #1315. Best score 1757 bits
Score difference with first non-orthologous sequence - *A. fumigatus*:1757 *A. fischeri*:1044
afum|EAL91554.1 100.00% afis|EAW17740.1 100.00%
Bootstrap support for afum|EAL91554.1 as seed ortholog is 100%.
Bootstrap support for afis|EAW17740.1 as seed ortholog is 100%.

Group of orthologs #1316. Best score 1756 bits
Score difference with first non-orthologous sequence - *A. fumigatus*:1756 *A. fischeri*:1756
afum|EAL88315.1 100.00% afis|EAW23169.1 100.00%
Bootstrap support for afum|EAL88315.1 as seed ortholog is 100%.
Bootstrap support for afis|EAW23169.1 as seed ortholog is 100%.

Group of orthologs #1317. Best score 1754 bits
Score difference with first non-orthologous sequence - *A. fumigatus*:1754 *A. fischeri*:1754
afum|EAL85990.1 100.00% afis|EAW24320.1 100.00%
Bootstrap support for afum|EAL85990.1 as seed ortholog is 100%.
Bootstrap support for afis|EAW24320.1 as seed ortholog is 100%.

Group of orthologs #1318. Best score 1754 bits
Score difference with first non-orthologous sequence - *A. fumigatus*:1754 *A. fischeri*:1754
afum|EAL87218.2 100.00% afis|EAW23784.1 100.00%
Bootstrap support for afum|EAL87218.2 as seed ortholog is 100%.
Bootstrap support for afis|EAW23784.1 as seed ortholog is 100%.

Group of orthologs #1319. Best score 1753 bits
Score difference with first non-orthologous sequence - *A. fumigatus*:1753 *A. fischeri*:1753
afum|EAL86445.1 100.00% afis|EAW16909.1 100.00%
Bootstrap support for afum|EAL86445.1 as seed ortholog is 100%.
Bootstrap support for afis|EAW16909.1 as seed ortholog is 100%.

Group of orthologs #1320. Best score 1753 bits
Score difference with first non-orthologous sequence - *A. fumigatus*:1753 *A. fischeri*:1753
afum|EAL86038.2 100.00% afis|EAW24366.1 100.00%
Bootstrap support for afum|EAL86038.2 as seed ortholog is 100%.
Bootstrap support for afis|EAW24366.1 as seed ortholog is 100%.

Group of orthologs #1321. Best score 1752 bits
Score difference with first non-orthologous sequence - *A. fumigatus*:1752 *A. fischeri*:1752
afum|EAL86644.1 100.00% afis|EAW17219.1 100.00%
Bootstrap support for afum|EAL86644.1 as seed ortholog is 100%.
Bootstrap support for afis|EAW17219.1 as seed ortholog is 100%.

Group of orthologs #1322. Best score 1752 bits
Score difference with first non-orthologous sequence - *A. fumigatus*:1752 *A. fischeri*:1752
afum|EAL87606.1 100.00% afis|EAW18242.1 100.00%
Bootstrap support for afum|EAL87606.1 as seed ortholog is 100%.
Bootstrap support for afis|EAW18242.1 as seed ortholog is 100%.

Group of orthologs #1323. Best score 1751 bits

Score difference with first non-orthologous sequence - *A. fumigatus*:1751 *A. fischeri*:1751
afum|EAL88873.1 100.00% afis|EAW16245.1 100.00%
Bootstrap support for afum|EAL88873.1 as seed ortholog is 100%.
Bootstrap support for afis|EAW16245.1 as seed ortholog is 100%.

---

Group of orthologs #1324. Best score 1750 bits
Score difference with first non-orthologous sequence - *A. fumigatus*:1750 *A. fischeri*:1750
afum|EAL93543.1 100.00% afis|EAW18807.1 100.00%
Bootstrap support for afum|EAL93543.1 as seed ortholog is 100%.
Bootstrap support for afis|EAW18807.1 as seed ortholog is 100%.

---

Group of orthologs #1325. Best score 1750 bits
Score difference with first non-orthologous sequence - *A. fumigatus*:1750 *A. fischeri*:1750
afum|EAL90756.1 100.00% afis|EAW22435.1 100.00%
Bootstrap support for afum|EAL90756.1 as seed ortholog is 100%.
Bootstrap support for afis|EAW22435.1 as seed ortholog is 100%.

---

Group of orthologs #1326. Best score 1750 bits
Score difference with first non-orthologous sequence - *A. fumigatus*:1750 *A. fischeri*:1750
afum|EAL92472.1 100.00% afis|EAW21453.1 100.00%
Bootstrap support for afum|EAL92472.1 as seed ortholog is 100%.
Bootstrap support for afis|EAW21453.1 as seed ortholog is 100%.

---

Group of orthologs #1327. Best score 1749 bits
Score difference with first non-orthologous sequence - *A. fumigatus*:1749 *A. fischeri*:1749
afum|EAL91279.1 100.00% afis|EAW17451.1 100.00%
Bootstrap support for afum|EAL91279.1 as seed ortholog is 100%.
Bootstrap support for afis|EAW17451.1 as seed ortholog is 100%.

---

Group of orthologs #1328. Best score 1749 bits
Score difference with first non-orthologous sequence - *A. fumigatus*:1749 *A. fischeri*:1749
afum|EAL87443.1 100.00% afis|EAW24013.1 100.00%
Bootstrap support for afum|EAL87443.1 as seed ortholog is 100%.
Bootstrap support for afis|EAW24013.1 as seed ortholog is 100%.

---

Group of orthologs #1329. Best score 1749 bits
Score difference with first non-orthologous sequence - *A. fumigatus*:1749 *A. fischeri*:1749
afum|EAL93805.2 100.00% afis|EAW19077.1 100.00%
Bootstrap support for afum|EAL93805.2 as seed ortholog is 100%.
Bootstrap support for afis|EAW19077.1 as seed ortholog is 100%.

---

Group of orthologs #1330. Best score 1749 bits
Score difference with first non-orthologous sequence - *A. fumigatus*:1749 *A. fischeri*:1749
afum|EAL90808.1 100.00% afis|EAW22383.1 100.00%
Bootstrap support for afum|EAL90808.1 as seed ortholog is 100%.
Bootstrap support for afis|EAW22383.1 as seed ortholog is 100%.

---

Group of orthologs #1331. Best score 1748 bits
Score difference with first non-orthologous sequence - *A. fumigatus*:1748 *A. fischeri*:1748
afum|EAL88208.1 100.00% afis|EAW23271.1 100.00%
Bootstrap support for afum|EAL88208.1 as seed ortholog is 100%.
Bootstrap support for afis|EAW23271.1 as seed ortholog is 100%.

---

Group of orthologs #1332. Best score 1747 bits
Score difference with first non-orthologous sequence - *A. fumigatus*:1747 *A. fischeri*:1747
afum|EAL85183.1 100.00% afis|EAW20063.1 100.00%

Bootstrap support for afum|EAL85183.1 as seed ortholog is 100%.
Bootstrap support for afis|EAW20063.1 as seed ortholog is 100%.

---

Group of orthologs #1333. Best score 1747 bits
Score difference with first non-orthologous sequence - *A. fumigatus*:1747 *A. fischeri*:1747
afum|EAL93004.1 100.00% afis|EAW21988.1 100.00%
Bootstrap support for afum|EAL93004.1 as seed ortholog is 100%.
Bootstrap support for afis|EAW21988.1 as seed ortholog is 100%.

---

Group of orthologs #1334. Best score 1747 bits
Score difference with first non-orthologous sequence - *A. fumigatus*:1747 *A. fischeri*:1747
afum|EAL89800.1 100.00% afis|EAW25129.1 100.00%
Bootstrap support for afum|EAL89800.1 as seed ortholog is 100%.
Bootstrap support for afis|EAW25129.1 as seed ortholog is 100%.

---

Group of orthologs #1335. Best score 1746 bits
Score difference with first non-orthologous sequence - *A. fumigatus*:1356 *A. fischeri*:1286
afum|EAL86540.1 100.00% afis|EAW17018.1 100.00%
Bootstrap support for afum|EAL86540.1 as seed ortholog is 100%.
Bootstrap support for afis|EAW17018.1 as seed ortholog is 100%.

---

Group of orthologs #1336. Best score 1746 bits
Score difference with first non-orthologous sequence - *A. fumigatus*:1746 *A. fischeri*:1746
afum|EAL90899.1 100.00% afis|EAW22282.1 100.00%
Bootstrap support for afum|EAL90899.1 as seed ortholog is 100%.
Bootstrap support for afis|EAW22282.1 as seed ortholog is 100%.

---

Group of orthologs #1337. Best score 1744 bits
Score difference with first non-orthologous sequence - *A. fumigatus*:1744 *A. fischeri*:1744
afum|EAL87671.1 100.00% afis|EAW18305.1 100.00%
Bootstrap support for afum|EAL87671.1 as seed ortholog is 100%.
Bootstrap support for afis|EAW18305.1 as seed ortholog is 100%.

---

Group of orthologs #1338. Best score 1742 bits
Score difference with first non-orthologous sequence - *A. fumigatus*:1742 *A. fischeri*:1742
afum|EAL87583.1 100.00% afis|EAW18217.1 100.00%
Bootstrap support for afum|EAL87583.1 as seed ortholog is 100%.
Bootstrap support for afis|EAW18217.1 as seed ortholog is 100%.

---

Group of orthologs #1339. Best score 1742 bits
Score difference with first non-orthologous sequence - *A. fumigatus*:1742 *A. fischeri*:1742
afum|EAL90694.1 100.00% afis|EAW22499.1 100.00%
Bootstrap support for afum|EAL90694.1 as seed ortholog is 100%.
Bootstrap support for afis|EAW22499.1 as seed ortholog is 100%.

---

Group of orthologs #1340. Best score 1741 bits
Score difference with first non-orthologous sequence - *A. fumigatus*:1741 *A. fischeri*:1741
afum|EAL86920.1 100.00% afis|EAW19598.1 100.00%
Bootstrap support for afum|EAL86920.1 as seed ortholog is 100%.
Bootstrap support for afis|EAW19598.1 as seed ortholog is 100%.

---

Group of orthologs #1341. Best score 1741 bits
Score difference with first non-orthologous sequence - *A. fumigatus*:1741 *A. fischeri*:1741
afum|EAL89544.1 100.00% afis|EAW24891.1 100.00%
Bootstrap support for afum|EAL89544.1 as seed ortholog is 100%.
Bootstrap support for afis|EAW24891.1 as seed ortholog is 100%.

Group of orthologs #1342. Best score 1740 bits
Score difference with first non-orthologous sequence - *A. fumigatus*:1740 *A. fischeri*:1740
afum|EAL86879.1 100.00% afis|EAW19643.1 100.00%
Bootstrap support for afum|EAL86879.1 as seed ortholog is 100%.
Bootstrap support for afis|EAW19643.1 as seed ortholog is 100%.

Group of orthologs #1343. Best score 1740 bits
Score difference with first non-orthologous sequence - *A. fumigatus*:1740 *A. fischeri*:1740
afum|EAL93961.1 100.00% afis|EAW19232.1 100.00%
Bootstrap support for afum|EAL93961.1 as seed ortholog is 100%.
Bootstrap support for afis|EAW19232.1 as seed ortholog is 100%.

Group of orthologs #1344. Best score 1739 bits
Score difference with first non-orthologous sequence - *A. fumigatus*:1739 *A. fischeri*:1739
afum|EAL87567.1 100.00% afis|EAW18202.1 100.00%
Bootstrap support for afum|EAL87567.1 as seed ortholog is 100%.
Bootstrap support for afis|EAW18202.1 as seed ortholog is 100%.

Group of orthologs #1345. Best score 1739 bits
Score difference with first non-orthologous sequence - *A. fumigatus*:1739 *A. fischeri*:1739
afum|EAL86128.1 100.00% afis|EAW24458.1 100.00%
Bootstrap support for afum|EAL86128.1 as seed ortholog is 100%.
Bootstrap support for afis|EAW24458.1 as seed ortholog is 100%.

Group of orthologs #1346. Best score 1739 bits
Score difference with first non-orthologous sequence - *A. fumigatus*:1739 *A. fischeri*:1739
afum|EAL92963.1 100.00% afis|EAW21949.1 100.00%
Bootstrap support for afum|EAL92963.1 as seed ortholog is 100%.
Bootstrap support for afis|EAW21949.1 as seed ortholog is 100%.

Group of orthologs #1347. Best score 1738 bits
Score difference with first non-orthologous sequence - *A. fumigatus*:1738 *A. fischeri*:1738
afum|EAL91526.1 100.00% afis|EAW17711.1 100.00%
Bootstrap support for afum|EAL91526.1 as seed ortholog is 100%.
Bootstrap support for afis|EAW17711.1 as seed ortholog is 100%.

Group of orthologs #1348. Best score 1738 bits
Score difference with first non-orthologous sequence - *A. fumigatus*:1738 *A. fischeri*:1738
afum|EAL93394.1 100.00% afis|EAW18674.1 100.00%
Bootstrap support for afum|EAL93394.1 as seed ortholog is 100%.
Bootstrap support for afis|EAW18674.1 as seed ortholog is 100%.

Group of orthologs #1349. Best score 1738 bits
Score difference with first non-orthologous sequence - *A. fumigatus*:1738 *A. fischeri*:1738
afum|EAL92948.2 100.00% afis|EAW21935.1 100.00%
Bootstrap support for afum|EAL92948.2 as seed ortholog is 100%.
Bootstrap support for afis|EAW21935.1 as seed ortholog is 100%.

Group of orthologs #1350. Best score 1737 bits
Score difference with first non-orthologous sequence - *A. fumigatus*:1737 *A. fischeri*:1737
afum|EAL86512.1 100.00% afis|EAW16988.1 100.00%
Bootstrap support for afum|EAL86512.1 as seed ortholog is 100%.
Bootstrap support for afis|EAW16988.1 as seed ortholog is 100%.

Group of orthologs #1351. Best score 1737 bits

Score difference with first non-orthologous sequence - *A. fumigatus*:1737   *A. fischeri*:1737
afum|EAL88410.1   100.00%   afis|EAW23069.1   100.00%
Bootstrap support for afum|EAL88410.1 as seed ortholog is 100%.
Bootstrap support for afis|EAW23069.1 as seed ortholog is 100%.

---

Group of orthologs #1352. Best score 1737 bits
Score difference with first non-orthologous sequence - *A. fumigatus*:1737   *A. fischeri*:1737
afum|EAL90064.1   100.00%   afis|EAW25405.1   100.00%
Bootstrap support for afum|EAL90064.1 as seed ortholog is 100%.
Bootstrap support for afis|EAW25405.1 as seed ortholog is 100%.

---

Group of orthologs #1353. Best score 1735 bits
Score difference with first non-orthologous sequence - *A. fumigatus*:1735   *A. fischeri*:1735
afum|EAL88596.1   100.00%   afis|EAW15963.1   100.00%
Bootstrap support for afum|EAL88596.1 as seed ortholog is 100%.
Bootstrap support for afis|EAW15963.1 as seed ortholog is 100%.

---

Group of orthologs #1354. Best score 1735 bits
Score difference with first non-orthologous sequence - *A. fumigatus*:1735   *A. fischeri*:1735
afum|EAL85325.1   100.00%   afis|EAW20208.1   100.00%
Bootstrap support for afum|EAL85325.1 as seed ortholog is 100%.
Bootstrap support for afis|EAW20208.1 as seed ortholog is 100%.

---

Group of orthologs #1355. Best score 1735 bits
Score difference with first non-orthologous sequence - *A. fumigatus*:1735   *A. fischeri*:1735
afum|EAL84641.1   100.00%   afis|EAW20988.1   100.00%
Bootstrap support for afum|EAL84641.1 as seed ortholog is 100%.
Bootstrap support for afis|EAW20988.1 as seed ortholog is 100%.

---

Group of orthologs #1356. Best score 1735 bits
Score difference with first non-orthologous sequence - *A. fumigatus*:1735   *A. fischeri*:1735
afum|EAL86056.1   100.00%   afis|EAW24384.1   100.00%
Bootstrap support for afum|EAL86056.1 as seed ortholog is 100%.
Bootstrap support for afis|EAW24384.1 as seed ortholog is 100%.

---

Group of orthologs #1357. Best score 1734 bits
Score difference with first non-orthologous sequence - *A. fumigatus*:1734   *A. fischeri*:1734
afum|EAL90250.1   100.00%   afis|EAW22947.1   100.00%
Bootstrap support for afum|EAL90250.1 as seed ortholog is 100%.
Bootstrap support for afis|EAW22947.1 as seed ortholog is 100%.

---

Group of orthologs #1358. Best score 1733 bits
Score difference with first non-orthologous sequence - *A. fumigatus*:1502   *A. fischeri*:1583
afum|EAL92856.1   100.00%   afis|EAW21844.1   100.00%
Bootstrap support for afum|EAL92856.1 as seed ortholog is 100%.
Bootstrap support for afis|EAW21844.1 as seed ortholog is 100%.

---

Group of orthologs #1359. Best score 1732 bits
Score difference with first non-orthologous sequence - *A. fumigatus*:1732   *A. fischeri*:1732
afum|EAL87710.1   100.00%   afis|EAW18346.1   100.00%
Bootstrap support for afum|EAL87710.1 as seed ortholog is 100%.
Bootstrap support for afis|EAW18346.1 as seed ortholog is 100%.

---

Group of orthologs #1360. Best score 1731 bits
Score difference with first non-orthologous sequence - *A. fumigatus*:1731   *A. fischeri*:1459
afum|EAL84705.1   100.00%   afis|EAW20929.1   100.00%

Bootstrap support for afum|EAL84705.1 as seed ortholog is 100%.
Bootstrap support for afis|EAW20929.1 as seed ortholog is 100%.

---

Group of orthologs #1361. Best score 1731 bits
Score difference with first non-orthologous sequence - *A. fumigatus*:1624 *A. fischeri*:1622
afum|EAL93120.1 100.00% afis|EAW18438.1 100.00%
Bootstrap support for afum|EAL93120.1 as seed ortholog is 100%.
Bootstrap support for afis|EAW18438.1 as seed ortholog is 100%.

---

Group of orthologs #1362. Best score 1731 bits
Score difference with first non-orthologous sequence - *A. fumigatus*:1731 *A. fischeri*:1731
afum|EAL89503.2 100.00% afis|EAW24855.1 100.00%
Bootstrap support for afum|EAL89503.2 as seed ortholog is 100%.
Bootstrap support for afis|EAW24855.1 as seed ortholog is 100%.

---

Group of orthologs #1363. Best score 1730 bits
Score difference with first non-orthologous sequence - *A. fumigatus*:1730 *A. fischeri*:1730
afum|EAL88082.1 100.00% afis|EAW23389.1 100.00%
Bootstrap support for afum|EAL88082.1 as seed ortholog is 100%.
Bootstrap support for afis|EAW23389.1 as seed ortholog is 100%.

---

Group of orthologs #1364. Best score 1729 bits
Score difference with first non-orthologous sequence - *A. fumigatus*:1729 *A. fischeri*:1729
afum|EAL94014.1 100.00% afis|EAW19295.1 100.00%
Bootstrap support for afum|EAL94014.1 as seed ortholog is 100%.
Bootstrap support for afis|EAW19295.1 as seed ortholog is 100%.

---

Group of orthologs #1365. Best score 1728 bits
Score difference with first non-orthologous sequence - *A. fumigatus*:1728 *A. fischeri*:1728
afum|EAL87575.2 100.00% afis|EAW18209.1 100.00%
Bootstrap support for afum|EAL87575.2 as seed ortholog is 100%.
Bootstrap support for afis|EAW18209.1 as seed ortholog is 100%.

---

Group of orthologs #1366. Best score 1728 bits
Score difference with first non-orthologous sequence - *A. fumigatus*:1728 *A. fischeri*:1728
afum|EAL85950.1 100.00% afis|EAW24281.1 100.00%
Bootstrap support for afum|EAL85950.1 as seed ortholog is 100%.
Bootstrap support for afis|EAW24281.1 as seed ortholog is 100%.

---

Group of orthologs #1367. Best score 1728 bits
Score difference with first non-orthologous sequence - *A. fumigatus*:1582 *A. fischeri*:1728
afum|EAL86122.1 100.00% afis|EAW24453.1 100.00%
Bootstrap support for afum|EAL86122.1 as seed ortholog is 100%.
Bootstrap support for afis|EAW24453.1 as seed ortholog is 100%.

---

Group of orthologs #1368. Best score 1728 bits
Score difference with first non-orthologous sequence - *A. fumigatus*:1728 *A. fischeri*:1728
afum|EAL88424.1 100.00% afis|EAW23056.1 100.00%
Bootstrap support for afum|EAL88424.1 as seed ortholog is 100%.
Bootstrap support for afis|EAW23056.1 as seed ortholog is 100%.

---

Group of orthologs #1369. Best score 1728 bits
Score difference with first non-orthologous sequence - *A. fumigatus*:1728 *A. fischeri*:1728
afum|EAL93005.2 100.00% afis|EAW21989.1 100.00%
Bootstrap support for afum|EAL93005.2 as seed ortholog is 100%.
Bootstrap support for afis|EAW21989.1 as seed ortholog is 100%.

Group of orthologs #1370. Best score 1727 bits
Score difference with first non-orthologous sequence - *A. fumigatus*:1727 *A. fischeri*:1727
afum|EAL91115.2 100.00% afis|EAW15512.1 100.00%
afum|EAL90946.1 87.50%
Bootstrap support for afum|EAL91115.2 as seed ortholog is 100%.
Bootstrap support for afis|EAW15512.1 as seed ortholog is 100%.

Group of orthologs #1371. Best score 1727 bits
Score difference with first non-orthologous sequence - *A. fumigatus*:1727 *A. fischeri*:1518
afum|EAL88534.1 100.00% afis|EAW15899.1 100.00%
Bootstrap support for afum|EAL88534.1 as seed ortholog is 100%.
Bootstrap support for afis|EAW15899.1 as seed ortholog is 100%.

Group of orthologs #1372. Best score 1727 bits
Score difference with first non-orthologous sequence - *A. fumigatus*:1618 *A. fischeri*:1727
afum|EAL88865.1 100.00% afis|EAW16243.1 100.00%
Bootstrap support for afum|EAL88865.1 as seed ortholog is 100%.
Bootstrap support for afis|EAW16243.1 as seed ortholog is 100%.

Group of orthologs #1373. Best score 1727 bits
Score difference with first non-orthologous sequence - *A. fumigatus*:1727 *A. fischeri*:1727
afum|EAL87383.1 100.00% afis|EAW23955.1 100.00%
Bootstrap support for afum|EAL87383.1 as seed ortholog is 100%.
Bootstrap support for afis|EAW23955.1 as seed ortholog is 100%.

Group of orthologs #1374. Best score 1727 bits
Score difference with first non-orthologous sequence - *A. fumigatus*:1727 *A. fischeri*:1727
afum|EAL93237.1 100.00% afis|EAW18539.1 100.00%
Bootstrap support for afum|EAL93237.1 as seed ortholog is 100%.
Bootstrap support for afis|EAW18539.1 as seed ortholog is 100%.

Group of orthologs #1375. Best score 1726 bits
Score difference with first non-orthologous sequence - *A. fumigatus*:1726 *A. fischeri*:1726
afum|EAL87535.1 100.00% afis|EAW18174.1 100.00%
Bootstrap support for afum|EAL87535.1 as seed ortholog is 100%.
Bootstrap support for afis|EAW18174.1 as seed ortholog is 100%.

Group of orthologs #1376. Best score 1726 bits
Score difference with first non-orthologous sequence - *A. fumigatus*:1726 *A. fischeri*:1726
afum|EAL88135.1 100.00% afis|EAW23343.1 100.00%
Bootstrap support for afum|EAL88135.1 as seed ortholog is 100%.
Bootstrap support for afis|EAW23343.1 as seed ortholog is 100%.

Group of orthologs #1377. Best score 1725 bits
Score difference with first non-orthologous sequence - *A. fumigatus*:1725 *A. fischeri*:1725
afum|EAL85886.1 100.00% afis|EAW24217.1 100.00%
Bootstrap support for afum|EAL85886.1 as seed ortholog is 100%.
Bootstrap support for afis|EAW24217.1 as seed ortholog is 100%.

Group of orthologs #1378. Best score 1723 bits
Score difference with first non-orthologous sequence - *A. fumigatus*:1723 *A. fischeri*:1723
afum|EAL88704.1 100.00% afis|EAW16072.1 100.00%
Bootstrap support for afum|EAL88704.1 as seed ortholog is 100%.
Bootstrap support for afis|EAW16072.1 as seed ortholog is 100%.

Group of orthologs #1379. Best score 1723 bits
Score difference with first non-orthologous sequence - *A. fumigatus*:1723 *A. fischeri*:1723
afum|EAL93275.1 100.00% afis|EAW18572.1 100.00%
Bootstrap support for afum|EAL93275.1 as seed ortholog is 100%.
Bootstrap support for afis|EAW18572.1 as seed ortholog is 100%.

---

Group of orthologs #1380. Best score 1722 bits
Score difference with first non-orthologous sequence - *A. fumigatus*:1722 *A. fischeri*:1722
afum|EAL85834.1 100.00% afis|EAW15489.1 100.00%
Bootstrap support for afum|EAL85834.1 as seed ortholog is 100%.
Bootstrap support for afis|EAW15489.1 as seed ortholog is 100%.

---

Group of orthologs #1381. Best score 1722 bits
Score difference with first non-orthologous sequence - *A. fumigatus*:1722 *A. fischeri*:1722
afum|EAL85568.1 100.00% afis|EAW15782.1 100.00%
Bootstrap support for afum|EAL85568.1 as seed ortholog is 100%.
Bootstrap support for afis|EAW15782.1 as seed ortholog is 100%.

---

Group of orthologs #1382. Best score 1722 bits
Score difference with first non-orthologous sequence - *A. fumigatus*:1722 *A. fischeri*:1722
afum|EAL85887.1 100.00% afis|EAW24218.1 100.00%
Bootstrap support for afum|EAL85887.1 as seed ortholog is 100%.
Bootstrap support for afis|EAW24218.1 as seed ortholog is 100%.

---

Group of orthologs #1383. Best score 1721 bits
Score difference with first non-orthologous sequence - *A. fumigatus*:1721 *A. fischeri*:1721
afum|EAL85055.2 100.00% afis|EAW19838.1 100.00%
Bootstrap support for afum|EAL85055.2 as seed ortholog is 100%.
Bootstrap support for afis|EAW19838.1 as seed ortholog is 100%.

---

Group of orthologs #1384. Best score 1721 bits
Score difference with first non-orthologous sequence - *A. fumigatus*:1721 *A. fischeri*:1721
afum|EAL85474.1 100.00% afis|EAW20357.1 100.00%
Bootstrap support for afum|EAL85474.1 as seed ortholog is 100%.
Bootstrap support for afis|EAW20357.1 as seed ortholog is 100%.

---

Group of orthologs #1385. Best score 1720 bits
Score difference with first non-orthologous sequence - *A. fumigatus*:1720 *A. fischeri*:1720
afum|EAL84884.1 100.00% afis|EAW20032.1 100.00%
Bootstrap support for afum|EAL84884.1 as seed ortholog is 100%.
Bootstrap support for afis|EAW20032.1 as seed ortholog is 100%.

---

Group of orthologs #1386. Best score 1719 bits
Score difference with first non-orthologous sequence - *A. fumigatus*:1591 *A. fischeri*:1633
afum|EAL91057.2 100.00% afis|EAW22133.1 100.00%
Bootstrap support for afum|EAL91057.2 as seed ortholog is 100%.
Bootstrap support for afis|EAW22133.1 as seed ortholog is 100%.

---

Group of orthologs #1387. Best score 1718 bits
Score difference with first non-orthologous sequence - *A. fumigatus*:1718 *A. fischeri*:1718
afum|EAL88477.1 100.00% afis|EAW23003.1 100.00%
Bootstrap support for afum|EAL88477.1 as seed ortholog is 100%.
Bootstrap support for afis|EAW23003.1 as seed ortholog is 100%.

---

Group of orthologs #1388. Best score 1717 bits
Score difference with first non-orthologous sequence - *A. fumigatus*:1717 *A. fischeri*:1717

afum|EAL85058.1 100.00% afis|EAW19834.1 100.00%
Bootstrap support for afum|EAL85058.1 as seed ortholog is 100%.
Bootstrap support for afis|EAW19834.1 as seed ortholog is 100%.

---

Group of orthologs #1389. Best score 1717 bits
Score difference with first non-orthologous sequence - *A. fumigatus*:1717 *A. fischeri*:1717
afum|EAL84806.1 100.00% afis|EAW20823.1 100.00%
Bootstrap support for afum|EAL84806.1 as seed ortholog is 100%.
Bootstrap support for afis|EAW20823.1 as seed ortholog is 100%.

---

Group of orthologs #1390. Best score 1716 bits
Score difference with first non-orthologous sequence - *A. fumigatus*:1716 *A. fischeri*:1716
afum|EAL87571.1 100.00% afis|EAW18206.1 100.00%
Bootstrap support for afum|EAL87571.1 as seed ortholog is 100%.
Bootstrap support for afis|EAW18206.1 as seed ortholog is 100%.

---

Group of orthologs #1391. Best score 1716 bits
Score difference with first non-orthologous sequence - *A. fumigatus*:1716 *A. fischeri*:1716
afum|EAL85497.1 100.00% afis|EAW20398.1 100.00%
Bootstrap support for afum|EAL85497.1 as seed ortholog is 100%.
Bootstrap support for afis|EAW20398.1 as seed ortholog is 100%.

---

Group of orthologs #1392. Best score 1716 bits
Score difference with first non-orthologous sequence - *A. fumigatus*:1716 *A. fischeri*:1716
afum|EAL91552.1 100.00% afis|EAW17737.1 100.00%
Bootstrap support for afum|EAL91552.1 as seed ortholog is 100%.
Bootstrap support for afis|EAW17737.1 as seed ortholog is 100%.

---

Group of orthologs #1393. Best score 1716 bits
Score difference with first non-orthologous sequence - *A. fumigatus*:1716 *A. fischeri*:1716
afum|EAL92157.2 100.00% afis|EAW21138.1 100.00%
Bootstrap support for afum|EAL92157.2 as seed ortholog is 100%.
Bootstrap support for afis|EAW21138.1 as seed ortholog is 100%.

---

Group of orthologs #1394. Best score 1716 bits
Score difference with first non-orthologous sequence - *A. fumigatus*:1716 *A. fischeri*:1716
afum|EAL92237.1 100.00% afis|EAW21224.1 100.00%
Bootstrap support for afum|EAL92237.1 as seed ortholog is 100%.
Bootstrap support for afis|EAW21224.1 as seed ortholog is 100%.

---

Group of orthologs #1395. Best score 1715 bits
Score difference with first non-orthologous sequence - *A. fumigatus*:1715 *A. fischeri*:1715
afum|EAL89609.1 100.00% afis|EAW24952.1 100.00%
Bootstrap support for afum|EAL89609.1 as seed ortholog is 100%.
Bootstrap support for afis|EAW24952.1 as seed ortholog is 100%.

---

Group of orthologs #1396. Best score 1714 bits
Score difference with first non-orthologous sequence - *A. fumigatus*:1714 *A. fischeri*:1714
afum|EAL86052.1 100.00% afis|EAW24380.1 100.00%
Bootstrap support for afum|EAL86052.1 as seed ortholog is 100%.
Bootstrap support for afis|EAW24380.1 as seed ortholog is 100%.

---

Group of orthologs #1397. Best score 1714 bits
Score difference with first non-orthologous sequence - *A. fumigatus*:1714 *A. fischeri*:1637
afum|EAL87440.1 100.00% afis|EAW24010.1 100.00%
Bootstrap support for afum|EAL87440.1 as seed ortholog is 100%.

Bootstrap support for afis|EAW24010.1 as seed ortholog is 100%.

---

Group of orthologs #1398. Best score 1714 bits
Score difference with first non-orthologous sequence - *A. fumigatus*:1714 *A. fischeri*:1714
afum|EAL92505.1 100.00% afis|EAW21485.1 100.00%
Bootstrap support for afum|EAL92505.1 as seed ortholog is 100%.
Bootstrap support for afis|EAW21485.1 as seed ortholog is 100%.

---

Group of orthologs #1399. Best score 1713 bits
Score difference with first non-orthologous sequence - *A. fumigatus*:1713 *A. fischeri*:1713
afum|EAL84723.1 100.00% afis|EAW20911.1 100.00%
Bootstrap support for afum|EAL84723.1 as seed ortholog is 100%.
Bootstrap support for afis|EAW20911.1 as seed ortholog is 100%.

---

Group of orthologs #1400. Best score 1713 bits
Score difference with first non-orthologous sequence - *A. fumigatus*:1713 *A. fischeri*:1713
afum|EAL93380.1 100.00% afis|EAW18661.1 100.00%
Bootstrap support for afum|EAL93380.1 as seed ortholog is 100%.
Bootstrap support for afis|EAW18661.1 as seed ortholog is 100%.

---

Group of orthologs #1401. Best score 1712 bits
Score difference with first non-orthologous sequence - *A. fumigatus*:1712 *A. fischeri*:1712
afum|EAL86273.1 100.00% afis|EAW20691.1 100.00%
Bootstrap support for afum|EAL86273.1 as seed ortholog is 100%.
Bootstrap support for afis|EAW20691.1 as seed ortholog is 100%.

---

Group of orthologs #1402. Best score 1712 bits
Score difference with first non-orthologous sequence - *A. fumigatus*:1712 *A. fischeri*:1712
afum|EAL87339.1 100.00% afis|EAW23910.1 100.00%
Bootstrap support for afum|EAL87339.1 as seed ortholog is 100%.
Bootstrap support for afis|EAW23910.1 as seed ortholog is 100%.

---

Group of orthologs #1403. Best score 1711 bits
Score difference with first non-orthologous sequence - *A. fumigatus*:1711 *A. fischeri*:1711
afum|EAL88981.1 100.00% afis|EAW16356.1 100.00%
Bootstrap support for afum|EAL88981.1 as seed ortholog is 100%.
Bootstrap support for afis|EAW16356.1 as seed ortholog is 100%.

---

Group of orthologs #1404. Best score 1711 bits
Score difference with first non-orthologous sequence - *A. fumigatus*:1711 *A. fischeri*:1711
afum|EAL93900.1 100.00% afis|EAW19170.1 100.00%
Bootstrap support for afum|EAL93900.1 as seed ortholog is 100%.
Bootstrap support for afis|EAW19170.1 as seed ortholog is 100%.

---

Group of orthologs #1405. Best score 1710 bits
Score difference with first non-orthologous sequence - *A. fumigatus*:1710 *A. fischeri*:1250
afum|EAL87190.2 100.00% afis|EAW23754.1 100.00%
Bootstrap support for afum|EAL87190.2 as seed ortholog is 100%.
Bootstrap support for afis|EAW23754.1 as seed ortholog is 100%.

---

Group of orthologs #1406. Best score 1710 bits
Score difference with first non-orthologous sequence - *A. fumigatus*:1710 *A. fischeri*:1710
afum|EAL93085.1 100.00% afis|EAW18405.1 100.00%
Bootstrap support for afum|EAL93085.1 as seed ortholog is 100%.
Bootstrap support for afis|EAW18405.1 as seed ortholog is 100%.

---

Group of orthologs #1407. Best score 1710 bits
Score difference with first non-orthologous sequence - *A. fumigatus*:1710 *A. fischeri*:1710
afum|EAL93125.1 100.00% afis|EAW18442.1 100.00%
Bootstrap support for afum|EAL93125.1 as seed ortholog is 100%.
Bootstrap support for afis|EAW18442.1 as seed ortholog is 100%.

---

Group of orthologs #1408. Best score 1710 bits
Score difference with first non-orthologous sequence - *A. fumigatus*:1710 *A. fischeri*:1710
afum|EAL88280.1 100.00% afis|EAW23203.1 100.00%
Bootstrap support for afum|EAL88280.1 as seed ortholog is 100%.
Bootstrap support for afis|EAW23203.1 as seed ortholog is 100%.

---

Group of orthologs #1409. Best score 1710 bits
Score difference with first non-orthologous sequence - *A. fumigatus*:1710 *A. fischeri*:1710
afum|EAL90161.1 100.00% afis|EAW25495.1 100.00%
Bootstrap support for afum|EAL90161.1 as seed ortholog is 100%.
Bootstrap support for afis|EAW25495.1 as seed ortholog is 100%.

---

Group of orthologs #1410. Best score 1709 bits
Score difference with first non-orthologous sequence - *A. fumigatus*:1709 *A. fischeri*:1709
afum|EAL89043.1 100.00% afis|EAW16442.1 100.00%
Bootstrap support for afum|EAL89043.1 as seed ortholog is 100%.
Bootstrap support for afis|EAW16442.1 as seed ortholog is 100%.

---

Group of orthologs #1411. Best score 1709 bits
Score difference with first non-orthologous sequence - *A. fumigatus*:1559 *A. fischeri*:1579
afum|EAL84877.1 100.00% afis|EAW20702.1 100.00%
Bootstrap support for afum|EAL84877.1 as seed ortholog is 100%.
Bootstrap support for afis|EAW20702.1 as seed ortholog is 100%.

---

Group of orthologs #1412. Best score 1709 bits
Score difference with first non-orthologous sequence - *A. fumigatus*:1589 *A. fischeri*:1592
afum|EAL84763.1 100.00% afis|EAW20871.1 100.00%
Bootstrap support for afum|EAL84763.1 as seed ortholog is 100%.
Bootstrap support for afis|EAW20871.1 as seed ortholog is 100%.

---

Group of orthologs #1413. Best score 1709 bits
Score difference with first non-orthologous sequence - *A. fumigatus*:1709 *A. fischeri*:1709
afum|EAL88396.1 100.00% afis|EAW23083.1 100.00%
Bootstrap support for afum|EAL88396.1 as seed ortholog is 100%.
Bootstrap support for afis|EAW23083.1 as seed ortholog is 100%.

---

Group of orthologs #1414. Best score 1709 bits
Score difference with first non-orthologous sequence - *A. fumigatus*:1709 *A. fischeri*:1709
afum|EAL88432.1 100.00% afis|EAW23048.1 100.00%
Bootstrap support for afum|EAL88432.1 as seed ortholog is 100%.
Bootstrap support for afis|EAW23048.1 as seed ortholog is 100%.

---

Group of orthologs #1415. Best score 1708 bits
Score difference with first non-orthologous sequence - *A. fumigatus*:1708 *A. fischeri*:1708
afum|EAL89670.1 100.00% afis|EAW25025.1 100.00%
Bootstrap support for afum|EAL89670.1 as seed ortholog is 100%.
Bootstrap support for afis|EAW25025.1 as seed ortholog is 100%.

---

Group of orthologs #1416. Best score 1706 bits
Score difference with first non-orthologous sequence - *A. fumigatus*:1706 *A. fischeri*:1706

afum|EAL84682.1 100.00% afis|EAW20952.1 100.00%
Bootstrap support for afum|EAL84682.1 as seed ortholog is 100%.
Bootstrap support for afis|EAW20952.1 as seed ortholog is 100%.

---

Group of orthologs #1417. Best score 1706 bits
Score difference with first non-orthologous sequence - *A. fumigatus*:1706 *A. fischeri*:1706
afum|EAL88204.1 100.00% afis|EAW23275.1 100.00%
Bootstrap support for afum|EAL88204.1 as seed ortholog is 100%.
Bootstrap support for afis|EAW23275.1 as seed ortholog is 100%.

---

Group of orthologs #1418. Best score 1706 bits
Score difference with first non-orthologous sequence - *A. fumigatus*:1706 *A. fischeri*:1706
afum|EAL90219.1 100.00% afis|EAW22978.1 100.00%
Bootstrap support for afum|EAL90219.1 as seed ortholog is 100%.
Bootstrap support for afis|EAW22978.1 as seed ortholog is 100%.

---

Group of orthologs #1419. Best score 1705 bits
Score difference with first non-orthologous sequence - *A. fumigatus*:1705 *A. fischeri*:521
afum|EAL93619.1 100.00% afis|EAW18886.1 100.00%
Bootstrap support for afum|EAL93619.1 as seed ortholog is 100%.
Bootstrap support for afis|EAW18886.1 as seed ortholog is 100%.

---

Group of orthologs #1420. Best score 1703 bits
Score difference with first non-orthologous sequence - *A. fumigatus*:1703 *A. fischeri*:1703
afum|EAL85552.1 100.00% afis|EAW15798.1 100.00%
Bootstrap support for afum|EAL85552.1 as seed ortholog is 100%.
Bootstrap support for afis|EAW15798.1 as seed ortholog is 100%.

---

Group of orthologs #1421. Best score 1703 bits
Score difference with first non-orthologous sequence - *A. fumigatus*:1618 *A. fischeri*:1703
afum|EAL90516.1 100.00% afis|EAW22676.1 100.00%
Bootstrap support for afum|EAL90516.1 as seed ortholog is 100%.
Bootstrap support for afis|EAW22676.1 as seed ortholog is 100%.

---

Group of orthologs #1422. Best score 1702 bits
Score difference with first non-orthologous sequence - *A. fumigatus*:1702 *A. fischeri*:1702
afum|EAL85215.1 100.00% afis|EAW20097.1 100.00%
Bootstrap support for afum|EAL85215.1 as seed ortholog is 100%.
Bootstrap support for afis|EAW20097.1 as seed ortholog is 100%.

---

Group of orthologs #1423. Best score 1702 bits
Score difference with first non-orthologous sequence - *A. fumigatus*:1702 *A. fischeri*:1702
afum|EAL91301.1 100.00% afis|EAW17472.1 100.00%
Bootstrap support for afum|EAL91301.1 as seed ortholog is 100%.
Bootstrap support for afis|EAW17472.1 as seed ortholog is 100%.

---

Group of orthologs #1424. Best score 1702 bits
Score difference with first non-orthologous sequence - *A. fumigatus*:1702 *A. fischeri*:1702
afum|EBA27341.1 100.00% afis|EAW23208.1 100.00%
Bootstrap support for afum|EBA27341.1 as seed ortholog is 100%.
Bootstrap support for afis|EAW23208.1 as seed ortholog is 100%.

---

Group of orthologs #1425. Best score 1701 bits
Score difference with first non-orthologous sequence - *A. fumigatus*:1701 *A. fischeri*:1701
afum|EAL85878.1 100.00% afis|EAW24208.1 100.00%
Bootstrap support for afum|EAL85878.1 as seed ortholog is 100%.

Bootstrap support for afis|EAW24208.1 as seed ortholog is 100%.

---

Group of orthologs #1426. Best score 1700 bits
Score difference with first non-orthologous sequence - *A. fumigatus*:1700 *A. fischeri*:1700
afum|EAL84341.1 100.00% afis|EAW25669.1 100.00%
Bootstrap support for afum|EAL84341.1 as seed ortholog is 100%.
Bootstrap support for afis|EAW25669.1 as seed ortholog is 100%.

---

Group of orthologs #1427. Best score 1699 bits
Score difference with first non-orthologous sequence - *A. fumigatus*:1699 *A. fischeri*:1699
afum|EAL92263.1 100.00% afis|EAW21250.1 100.00%
Bootstrap support for afum|EAL92263.1 as seed ortholog is 100%.
Bootstrap support for afis|EAW21250.1 as seed ortholog is 100%.

---

Group of orthologs #1428. Best score 1699 bits
Score difference with first non-orthologous sequence - *A. fumigatus*:1699 *A. fischeri*:1699
afum|EAL92033.1 100.00% afis|EAW24171.1 100.00%
Bootstrap support for afum|EAL92033.1 as seed ortholog is 100%.
Bootstrap support for afis|EAW24171.1 as seed ortholog is 100%.

---

Group of orthologs #1429. Best score 1697 bits
Score difference with first non-orthologous sequence - *A. fumigatus*:1697 *A. fischeri*:1697
afum|EAL91509.1 100.00% afis|EAW17695.1 100.00%
Bootstrap support for afum|EAL91509.1 as seed ortholog is 100%.
Bootstrap support for afis|EAW17695.1 as seed ortholog is 100%.

---

Group of orthologs #1430. Best score 1696 bits
Score difference with first non-orthologous sequence - *A. fumigatus*:1696 *A. fischeri*:1696
afum|EAL87971.1 100.00% afis|EAW23492.1 100.00%
Bootstrap support for afum|EAL87971.1 as seed ortholog is 100%.
Bootstrap support for afis|EAW23492.1 as seed ortholog is 100%.

---

Group of orthologs #1431. Best score 1694 bits
Score difference with first non-orthologous sequence - *A. fumigatus*:1694 *A. fischeri*:1694
afum|EAL84272.1 100.00% afis|EAW25549.1 100.00%
Bootstrap support for afum|EAL84272.1 as seed ortholog is 100%.
Bootstrap support for afis|EAW25549.1 as seed ortholog is 100%.

---

Group of orthologs #1432. Best score 1694 bits
Score difference with first non-orthologous sequence - *A. fumigatus*:1694 *A. fischeri*:1694
afum|EAL85958.1 100.00% afis|EAW24290.1 100.00%
Bootstrap support for afum|EAL85958.1 as seed ortholog is 100%.
Bootstrap support for afis|EAW24290.1 as seed ortholog is 100%.

---

Group of orthologs #1433. Best score 1694 bits
Score difference with first non-orthologous sequence - *A. fumigatus*:1694 *A. fischeri*:1647
afum|EAL92771.1 100.00% afis|EAW21737.1 100.00%
Bootstrap support for afum|EAL92771.1 as seed ortholog is 100%.
Bootstrap support for afis|EAW21737.1 as seed ortholog is 100%.

---

Group of orthologs #1434. Best score 1693 bits
Score difference with first non-orthologous sequence - *A. fumigatus*:1693 *A. fischeri*:1693
afum|EAL85210.1 100.00% afis|EAW20091.1 100.00%
Bootstrap support for afum|EAL85210.1 as seed ortholog is 100%.
Bootstrap support for afis|EAW20091.1 as seed ortholog is 100%.

---

Group of orthologs #1435. Best score 1692 bits
Score difference with first non-orthologous sequence - *A. fumigatus*:1692 *A. fischeri*:1692
afum|EAL85602.1 100.00% afis|EAW15746.1 100.00%
Bootstrap support for afum|EAL85602.1 as seed ortholog is 100%.
Bootstrap support for afis|EAW15746.1 as seed ortholog is 100%.

---

Group of orthologs #1436. Best score 1692 bits
Score difference with first non-orthologous sequence - *A. fumigatus*:1692 *A. fischeri*:1692
afum|EAL90210.1 100.00% afis|EAW22990.1 100.00%
Bootstrap support for afum|EAL90210.1 as seed ortholog is 100%.
Bootstrap support for afis|EAW22990.1 as seed ortholog is 100%.

---

Group of orthologs #1437. Best score 1691 bits
Score difference with first non-orthologous sequence - *A. fumigatus*:1691 *A. fischeri*:1691
afum|EAL87701.1 100.00% afis|EAW18337.1 100.00%
Bootstrap support for afum|EAL87701.1 as seed ortholog is 100%.
Bootstrap support for afis|EAW18337.1 as seed ortholog is 100%.

---

Group of orthologs #1438. Best score 1691 bits
Score difference with first non-orthologous sequence - *A. fumigatus*:1691 *A. fischeri*:1691
afum|EAL87940.1 100.00% afis|EAW23525.1 100.00%
Bootstrap support for afum|EAL87940.1 as seed ortholog is 100%.
Bootstrap support for afis|EAW23525.1 as seed ortholog is 100%.

---

Group of orthologs #1439. Best score 1691 bits
Score difference with first non-orthologous sequence - *A. fumigatus*:1691 *A. fischeri*:1691
afum|EAL90592.1 100.00% afis|EAW22606.1 100.00%
Bootstrap support for afum|EAL90592.1 as seed ortholog is 100%.
Bootstrap support for afis|EAW22606.1 as seed ortholog is 100%.

---

Group of orthologs #1440. Best score 1691 bits
Score difference with first non-orthologous sequence - *A. fumigatus*:1691 *A. fischeri*:1691
afum|EAL90593.1 100.00% afis|EAW22605.1 100.00%
Bootstrap support for afum|EAL90593.1 as seed ortholog is 100%.
Bootstrap support for afis|EAW22605.1 as seed ortholog is 100%.

---

Group of orthologs #1441. Best score 1690 bits
Score difference with first non-orthologous sequence - *A. fumigatus*:1690 *A. fischeri*:1690
afum|EAL91503.1 100.00% afis|EAW17689.1 100.00%
Bootstrap support for afum|EAL91503.1 as seed ortholog is 100%.
Bootstrap support for afis|EAW17689.1 as seed ortholog is 100%.

---

Group of orthologs #1442. Best score 1690 bits
Score difference with first non-orthologous sequence - *A. fumigatus*:1690 *A. fischeri*:1690
afum|EAL91004.1 100.00% afis|EAW22185.1 100.00%
Bootstrap support for afum|EAL91004.1 as seed ortholog is 100%.
Bootstrap support for afis|EAW22185.1 as seed ortholog is 100%.

---

Group of orthologs #1443. Best score 1690 bits
Score difference with first non-orthologous sequence - *A. fumigatus*:1690 *A. fischeri*:1690
afum|EAL90957.1 100.00% afis|EAW22236.1 100.00%
Bootstrap support for afum|EAL90957.1 as seed ortholog is 100%.
Bootstrap support for afis|EAW22236.1 as seed ortholog is 100%.

---

Group of orthologs #1444. Best score 1688 bits
Score difference with first non-orthologous sequence - *A. fumigatus*:1688 *A. fischeri*:1688

afum|EAL85772.1 100.00% afis|EAW15560.1 100.00%
Bootstrap support for afum|EAL85772.1 as seed ortholog is 100%.
Bootstrap support for afis|EAW15560.1 as seed ortholog is 100%.

---

Group of orthologs #1445. Best score 1688 bits
Score difference with first non-orthologous sequence - *A. fumigatus*:1114 *A. fischeri*:1688
afum|EAL91860.1 100.00% afis|EAW18048.1 100.00%
Bootstrap support for afum|EAL91860.1 as seed ortholog is 100%.
Bootstrap support for afis|EAW18048.1 as seed ortholog is 100%.

---

Group of orthologs #1446. Best score 1688 bits
Score difference with first non-orthologous sequence - *A. fumigatus*:1688 *A. fischeri*:1688
afum|EAL93721.1 100.00% afis|EAW18985.1 100.00%
Bootstrap support for afum|EAL93721.1 as seed ortholog is 100%.
Bootstrap support for afis|EAW18985.1 as seed ortholog is 100%.

---

Group of orthologs #1447. Best score 1688 bits
Score difference with first non-orthologous sequence - *A. fumigatus*:1595 *A. fischeri*:1281
afum|EAL90913.1 100.00% afis|EAW22267.1 100.00%
Bootstrap support for afum|EAL90913.1 as seed ortholog is 100%.
Bootstrap support for afis|EAW22267.1 as seed ortholog is 100%.

---

Group of orthologs #1448. Best score 1688 bits
Score difference with first non-orthologous sequence - *A. fumigatus*:1688 *A. fischeri*:1688
afum|EAL89519.1 100.00% afis|EAW24869.1 100.00%
Bootstrap support for afum|EAL89519.1 as seed ortholog is 100%.
Bootstrap support for afis|EAW24869.1 as seed ortholog is 100%.

---

Group of orthologs #1449. Best score 1687 bits
Score difference with first non-orthologous sequence - *A. fumigatus*:1687 *A. fischeri*:1687
afum|EAL85737.2 100.00% afis|EAW15594.1 100.00%
Bootstrap support for afum|EAL85737.2 as seed ortholog is 100%.
Bootstrap support for afis|EAW15594.1 as seed ortholog is 100%.

---

Group of orthologs #1450. Best score 1687 bits
Score difference with first non-orthologous sequence - *A. fumigatus*:1687 *A. fischeri*:1048
afum|EAL86575.1 100.00% afis|EAW17111.1 100.00%
Bootstrap support for afum|EAL86575.1 as seed ortholog is 100%.
Bootstrap support for afis|EAW17111.1 as seed ortholog is 100%.

---

Group of orthologs #1451. Best score 1687 bits
Score difference with first non-orthologous sequence - *A. fumigatus*:1687 *A. fischeri*:1687
afum|EAL84482.1 100.00% afis|EAW20582.1 100.00%
Bootstrap support for afum|EAL84482.1 as seed ortholog is 100%.
Bootstrap support for afis|EAW20582.1 as seed ortholog is 100%.

---

Group of orthologs #1452. Best score 1686 bits
Score difference with first non-orthologous sequence - *A. fumigatus*:1686 *A. fischeri*:1686
afum|EAL87417.2 100.00% afis|EAW23988.1 100.00%
Bootstrap support for afum|EAL87417.2 as seed ortholog is 100%.
Bootstrap support for afis|EAW23988.1 as seed ortholog is 100%.

---

Group of orthologs #1453. Best score 1684 bits
Score difference with first non-orthologous sequence - *A. fumigatus*:1684 *A. fischeri*:1684
afum|EAL86060.1 100.00% afis|EAW24388.1 100.00%
Bootstrap support for afum|EAL86060.1 as seed ortholog is 100%.

Bootstrap support for afis|EAW24388.1 as seed ortholog is 100%.

---

Group of orthologs #1454. Best score 1684 bits
Score difference with first non-orthologous sequence - *A. fumigatus*:1684 *A. fischeri*:1684
afum|EAL93413.2 100.00% afis|EAW18692.1 100.00%
Bootstrap support for afum|EAL93413.2 as seed ortholog is 100%.
Bootstrap support for afis|EAW18692.1 as seed ortholog is 100%.

---

Group of orthologs #1455. Best score 1681 bits
Score difference with first non-orthologous sequence - *A. fumigatus*:1681 *A. fischeri*:1681
afum|EAL90690.1 100.00% afis|EAW22503.1 100.00%
Bootstrap support for afum|EAL90690.1 as seed ortholog is 100%.
Bootstrap support for afis|EAW22503.1 as seed ortholog is 100%.

---

Group of orthologs #1456. Best score 1681 bits
Score difference with first non-orthologous sequence - *A. fumigatus*:1681 *A. fischeri*:1681
afum|EAL90290.1 100.00% afis|EAW22907.1 100.00%
Bootstrap support for afum|EAL90290.1 as seed ortholog is 100%.
Bootstrap support for afis|EAW22907.1 as seed ortholog is 100%.

---

Group of orthologs #1457. Best score 1681 bits
Score difference with first non-orthologous sequence - *A. fumigatus*:1681 *A. fischeri*:1681
afum|EAL90364.2 100.00% afis|EAW22838.1 100.00%
Bootstrap support for afum|EAL90364.2 as seed ortholog is 100%.
Bootstrap support for afis|EAW22838.1 as seed ortholog is 100%.

---

Group of orthologs #1458. Best score 1681 bits
Score difference with first non-orthologous sequence - *A. fumigatus*:1681 *A. fischeri*:1681
afum|EAL89676.1 100.00% afis|EAW25019.1 100.00%
Bootstrap support for afum|EAL89676.1 as seed ortholog is 100%.
Bootstrap support for afis|EAW25019.1 as seed ortholog is 100%.

---

Group of orthologs #1459. Best score 1680 bits
Score difference with first non-orthologous sequence - *A. fumigatus*:1680 *A. fischeri*:1680
afum|EAL87653.1 100.00% afis|EAW18287.1 100.00%
Bootstrap support for afum|EAL87653.1 as seed ortholog is 100%.
Bootstrap support for afis|EAW18287.1 as seed ortholog is 100%.

---

Group of orthologs #1460. Best score 1679 bits
Score difference with first non-orthologous sequence - *A. fumigatus*:1679 *A. fischeri*:1679
afum|EAL87373.1 100.00% afis|EAW23947.1 100.00%
Bootstrap support for afum|EAL87373.1 as seed ortholog is 100%.
Bootstrap support for afis|EAW23947.1 as seed ortholog is 100%.

---

Group of orthologs #1461. Best score 1679 bits
Score difference with first non-orthologous sequence - *A. fumigatus*:1679 *A. fischeri*:1679
afum|EAL90360.1 100.00% afis|EAW22843.1 100.00%
Bootstrap support for afum|EAL90360.1 as seed ortholog is 100%.
Bootstrap support for afis|EAW22843.1 as seed ortholog is 100%.

---

Group of orthologs #1462. Best score 1678 bits
Score difference with first non-orthologous sequence - *A. fumigatus*:1678 *A. fischeri*:1678
afum|EAL93228.1 100.00% afis|EAW18531.1 100.00%
Bootstrap support for afum|EAL93228.1 as seed ortholog is 100%.
Bootstrap support for afis|EAW18531.1 as seed ortholog is 100%.

---

Group of orthologs #1463. Best score 1678 bits
Score difference with first non-orthologous sequence - *A. fumigatus*:1678 *A. fischeri*:1678
afum|EAL93384.1 100.00% afis|EAW18666.1 100.00%
Bootstrap support for afum|EAL93384.1 as seed ortholog is 100%.
Bootstrap support for afis|EAW18666.1 as seed ortholog is 100%.

---

Group of orthologs #1464. Best score 1678 bits
Score difference with first non-orthologous sequence - *A. fumigatus*:1587 *A. fischeri*:675
afum|EAL92550.1 100.00% afis|EAW21529.1 100.00%
Bootstrap support for afum|EAL92550.1 as seed ortholog is 100%.
Bootstrap support for afis|EAW21529.1 as seed ortholog is 100%.

---

Group of orthologs #1465. Best score 1678 bits
Score difference with first non-orthologous sequence - *A. fumigatus*:1678 *A. fischeri*:1678
afum|EBA27225.1 100.00% afis|EAW20135.1 100.00%
Bootstrap support for afum|EBA27225.1 as seed ortholog is 100%.
Bootstrap support for afis|EAW20135.1 as seed ortholog is 100%.

---

Group of orthologs #1466. Best score 1678 bits
Score difference with first non-orthologous sequence - *A. fumigatus*:1678 *A. fischeri*:1678
afum|EAL89752.1 100.00% afis|EAW25083.1 100.00%
Bootstrap support for afum|EAL89752.1 as seed ortholog is 100%.
Bootstrap support for afis|EAW25083.1 as seed ortholog is 100%.

---

Group of orthologs #1467. Best score 1677 bits
Score difference with first non-orthologous sequence - *A. fumigatus*:1677 *A. fischeri*:1677
afum|EAL93158.1 100.00% afis|EAW18471.1 100.00%
Bootstrap support for afum|EAL93158.1 as seed ortholog is 100%.
Bootstrap support for afis|EAW18471.1 as seed ortholog is 100%.

---

Group of orthologs #1468. Best score 1677 bits
Score difference with first non-orthologous sequence - *A. fumigatus*:1677 *A. fischeri*:1677
afum|EAL90623.1 100.00% afis|EAW22575.1 100.00%
Bootstrap support for afum|EAL90623.1 as seed ortholog is 100%.
Bootstrap support for afis|EAW22575.1 as seed ortholog is 100%.

---

Group of orthologs #1469. Best score 1676 bits
Score difference with first non-orthologous sequence - *A. fumigatus*:1676 *A. fischeri*:1676
afum|EAL90348.1 100.00% afis|EAW22852.1 100.00%
Bootstrap support for afum|EAL90348.1 as seed ortholog is 100%.
Bootstrap support for afis|EAW22852.1 as seed ortholog is 100%.

---

Group of orthologs #1470. Best score 1675 bits
Score difference with first non-orthologous sequence - *A. fumigatus*:1675 *A. fischeri*:1675
afum|EAL88496.1 100.00% afis|EAW15861.1 100.00%
Bootstrap support for afum|EAL88496.1 as seed ortholog is 100%.
Bootstrap support for afis|EAW15861.1 as seed ortholog is 100%.

---

Group of orthologs #1471. Best score 1675 bits
Score difference with first non-orthologous sequence - *A. fumigatus*:1675 *A. fischeri*:1675
afum|EAL93771.1 100.00% afis|EAW19042.1 100.00%
Bootstrap support for afum|EAL93771.1 as seed ortholog is 100%.
Bootstrap support for afis|EAW19042.1 as seed ortholog is 100%.

---

Group of orthologs #1472. Best score 1675 bits
Score difference with first non-orthologous sequence - *A. fumigatus*:1675 *A. fischeri*:1675

afum|EAL90282.1 100.00% afis|EAW22915.1 100.00%
Bootstrap support for afum|EAL90282.1 as seed ortholog is 100%.
Bootstrap support for afis|EAW22915.1 as seed ortholog is 100%.

---

Group of orthologs #1473. Best score 1675 bits
Score difference with first non-orthologous sequence - *A. fumigatus*:1675 *A. fischeri*:1675
afum|EAL89459.1 100.00% afis|EAW24815.1 100.00%
Bootstrap support for afum|EAL89459.1 as seed ortholog is 100%.
Bootstrap support for afis|EAW24815.1 as seed ortholog is 100%.

---

Group of orthologs #1474. Best score 1674 bits
Score difference with first non-orthologous sequence - *A. fumigatus*:1674 *A. fischeri*:1674
afum|EAL87057.1 100.00% afis|EAW19439.1 100.00%
Bootstrap support for afum|EAL87057.1 as seed ortholog is 100%.
Bootstrap support for afis|EAW19439.1 as seed ortholog is 100%.

---

Group of orthologs #1475. Best score 1674 bits
Score difference with first non-orthologous sequence - *A. fumigatus*:1674 *A. fischeri*:1674
afum|EAL93064.1 100.00% afis|EAW18389.1 100.00%
Bootstrap support for afum|EAL93064.1 as seed ortholog is 100%.
Bootstrap support for afis|EAW18389.1 as seed ortholog is 100%.

---

Group of orthologs #1476. Best score 1673 bits
Score difference with first non-orthologous sequence - *A. fumigatus*:1673 *A. fischeri*:1673
afum|EAL87359.1 100.00% afis|EAW23934.1 100.00%
Bootstrap support for afum|EAL87359.1 as seed ortholog is 100%.
Bootstrap support for afis|EAW23934.1 as seed ortholog is 100%.

---

Group of orthologs #1477. Best score 1672 bits
Score difference with first non-orthologous sequence - *A. fumigatus*:1672 *A. fischeri*:1672
afum|EAL93443.2 100.00% afis|EAW18719.1 100.00%
Bootstrap support for afum|EAL93443.2 as seed ortholog is 100%.
Bootstrap support for afis|EAW18719.1 as seed ortholog is 100%.

---

Group of orthologs #1478. Best score 1671 bits
Score difference with first non-orthologous sequence - *A. fumigatus*:1671 *A. fischeri*:1671
afum|EAL88842.1 100.00% afis|EAW16215.1 100.00%
Bootstrap support for afum|EAL88842.1 as seed ortholog is 100%.
Bootstrap support for afis|EAW16215.1 as seed ortholog is 100%.

---

Group of orthologs #1479. Best score 1671 bits
Score difference with first non-orthologous sequence - *A. fumigatus*:1671 *A. fischeri*:1671
afum|EAL84273.1 100.00% afis|EAW25550.1 100.00%
Bootstrap support for afum|EAL84273.1 as seed ortholog is 100%.
Bootstrap support for afis|EAW25550.1 as seed ortholog is 100%.

---

Group of orthologs #1480. Best score 1671 bits
Score difference with first non-orthologous sequence - *A. fumigatus*:1671 *A. fischeri*:1671
afum|EAL91002.1 100.00% afis|EAW22188.1 100.00%
Bootstrap support for afum|EAL91002.1 as seed ortholog is 100%.
Bootstrap support for afis|EAW22188.1 as seed ortholog is 100%.

---

Group of orthologs #1481. Best score 1670 bits
Score difference with first non-orthologous sequence - *A. fumigatus*:1670 *A. fischeri*:1670
afum|EAL89440.1 100.00% afis|EAW24798.1 100.00%
Bootstrap support for afum|EAL89440.1 as seed ortholog is 100%.

Bootstrap support for afis|EAW24798.1 as seed ortholog is 100%.

---

Group of orthologs #1482. Best score 1669 bits
Score difference with first non-orthologous sequence - *A. fumigatus*:1669 *A. fischeri*:1064
afum|EAL86763.1 100.00% afis|EAW16767.1 100.00%
Bootstrap support for afum|EAL86763.1 as seed ortholog is 100%.
Bootstrap support for afis|EAW16767.1 as seed ortholog is 100%.

---

Group of orthologs #1483. Best score 1669 bits
Score difference with first non-orthologous sequence - *A. fumigatus*:1669 *A. fischeri*:1669
afum|EAL88487.1 100.00% afis|EAW15851.1 100.00%
Bootstrap support for afum|EAL88487.1 as seed ortholog is 100%.
Bootstrap support for afis|EAW15851.1 as seed ortholog is 100%.

---

Group of orthologs #1484. Best score 1669 bits
Score difference with first non-orthologous sequence - *A. fumigatus*:1669 *A. fischeri*:1669
afum|EAL90400.1 100.00% afis|EAW22799.1 100.00%
Bootstrap support for afum|EAL90400.1 as seed ortholog is 100%.
Bootstrap support for afis|EAW22799.1 as seed ortholog is 100%.

---

Group of orthologs #1485. Best score 1669 bits
Score difference with first non-orthologous sequence - *A. fumigatus*:1669 *A. fischeri*:1669
afum|EAL92390.1 100.00% afis|EAW21373.1 100.00%
Bootstrap support for afum|EAL92390.1 as seed ortholog is 100%.
Bootstrap support for afis|EAW21373.1 as seed ortholog is 100%.

---

Group of orthologs #1486. Best score 1669 bits
Score difference with first non-orthologous sequence - *A. fumigatus*:1669 *A. fischeri*:1669
afum|EAL89707.1 100.00% afis|EAW25039.1 100.00%
Bootstrap support for afum|EAL89707.1 as seed ortholog is 100%.
Bootstrap support for afis|EAW25039.1 as seed ortholog is 100%.

---

Group of orthologs #1487. Best score 1668 bits
Score difference with first non-orthologous sequence - *A. fumigatus*:671 *A. fischeri*:1668
afum|EAL93075.1 100.00% afis|EAW18398.1 100.00%
Bootstrap support for afum|EAL93075.1 as seed ortholog is 100%.
Bootstrap support for afis|EAW18398.1 as seed ortholog is 100%.

---

Group of orthologs #1488. Best score 1668 bits
Score difference with first non-orthologous sequence - *A. fumigatus*:1159 *A. fischeri*:1668
afum|EAL92373.1 100.00% afis|EAW21358.1 100.00%
Bootstrap support for afum|EAL92373.1 as seed ortholog is 100%.
Bootstrap support for afis|EAW21358.1 as seed ortholog is 100%.

---

Group of orthologs #1489. Best score 1667 bits
Score difference with first non-orthologous sequence - *A. fumigatus*:1667 *A. fischeri*:1667
afum|EAL87536.1 100.00% afis|EAW18175.1 100.00%
Bootstrap support for afum|EAL87536.1 as seed ortholog is 100%.
Bootstrap support for afis|EAW18175.1 as seed ortholog is 100%.

---

Group of orthologs #1490. Best score 1667 bits
Score difference with first non-orthologous sequence - *A. fumigatus*:1667 *A. fischeri*:1667
afum|EAL92156.1 100.00% afis|EAW21137.1 100.00%
Bootstrap support for afum|EAL92156.1 as seed ortholog is 100%.
Bootstrap support for afis|EAW21137.1 as seed ortholog is 100%.

---

Group of orthologs #1491. Best score 1666 bits
Score difference with first non-orthologous sequence - *A. fumigatus*:1666 *A. fischeri*:1666
afum|EAL89093.1 100.00% afis|EAW16498.1 100.00%
Bootstrap support for afum|EAL89093.1 as seed ortholog is 100%.
Bootstrap support for afis|EAW16498.1 as seed ortholog is 100%.

---

Group of orthologs #1492. Best score 1666 bits
Score difference with first non-orthologous sequence - *A. fumigatus*:1291 *A. fischeri*:1666
afum|EAL87052.1 100.00% afis|EAW19443.1 100.00%
Bootstrap support for afum|EAL87052.1 as seed ortholog is 100%.
Bootstrap support for afis|EAW19443.1 as seed ortholog is 100%.

---

Group of orthologs #1493. Best score 1666 bits
Score difference with first non-orthologous sequence - *A. fumigatus*:1666 *A. fischeri*:1666
afum|EAL86949.1 100.00% afis|EAW19563.1 100.00%
Bootstrap support for afum|EAL86949.1 as seed ortholog is 100%.
Bootstrap support for afis|EAW19563.1 as seed ortholog is 100%.

---

Group of orthologs #1494. Best score 1666 bits
Score difference with first non-orthologous sequence - *A. fumigatus*:1666 *A. fischeri*:1666
afum|EAL90971.1 100.00% afis|EAW22223.1 100.00%
Bootstrap support for afum|EAL90971.1 as seed ortholog is 100%.
Bootstrap support for afis|EAW22223.1 as seed ortholog is 100%.

---

Group of orthologs #1495. Best score 1666 bits
Score difference with first non-orthologous sequence - *A. fumigatus*:1666 *A. fischeri*:1666
afum|EAL92864.1 100.00% afis|EAW21851.1 100.00%
Bootstrap support for afum|EAL92864.1 as seed ortholog is 100%.
Bootstrap support for afis|EAW21851.1 as seed ortholog is 100%.

---

Group of orthologs #1496. Best score 1665 bits
Score difference with first non-orthologous sequence - *A. fumigatus*:1665 *A. fischeri*:1665
afum|EAL86304.2 100.00% afis|EAW16729.1 100.00%
Bootstrap support for afum|EAL86304.2 as seed ortholog is 100%.
Bootstrap support for afis|EAW16729.1 as seed ortholog is 100%.

---

Group of orthologs #1497. Best score 1665 bits
Score difference with first non-orthologous sequence - *A. fumigatus*:1665 *A. fischeri*:1665
afum|EAL88634.1 100.00% afis|EAW16002.1 100.00%
Bootstrap support for afum|EAL88634.1 as seed ortholog is 100%.
Bootstrap support for afis|EAW16002.1 as seed ortholog is 100%.

---

Group of orthologs #1498. Best score 1665 bits
Score difference with first non-orthologous sequence - *A. fumigatus*:1665 *A. fischeri*:1305
afum|EAL88828.1 100.00% afis|EAW16201.1 100.00%
Bootstrap support for afum|EAL88828.1 as seed ortholog is 100%.
Bootstrap support for afis|EAW16201.1 as seed ortholog is 100%.

---

Group of orthologs #1499. Best score 1665 bits
Score difference with first non-orthologous sequence - *A. fumigatus*:1665 *A. fischeri*:1665
afum|EAL87947.2 100.00% afis|EAW23516.1 100.00%
Bootstrap support for afum|EAL87947.2 as seed ortholog is 100%.
Bootstrap support for afis|EAW23516.1 as seed ortholog is 100%.

---

Group of orthologs #1500. Best score 1665 bits
Score difference with first non-orthologous sequence - *A. fumigatus*:1665 *A. fischeri*:1665

afum|EAL93706.1 100.00% afis|EAW18973.1 100.00%
Bootstrap support for afum|EAL93706.1 as seed ortholog is 100%.
Bootstrap support for afis|EAW18973.1 as seed ortholog is 100%.

---

Group of orthologs #1501. Best score 1665 bits
Score difference with first non-orthologous sequence - *A. fumigatus*:1665 *A. fischeri*:1665
afum|EAL92539.1 100.00% afis|EAW21518.1 100.00%
Bootstrap support for afum|EAL92539.1 as seed ortholog is 100%.
Bootstrap support for afis|EAW21518.1 as seed ortholog is 100%.

---

Group of orthologs #1502. Best score 1665 bits
Score difference with first non-orthologous sequence - *A. fumigatus*:1665 *A. fischeri*:1665
afum|EAL89854.1 100.00% afis|EAW25180.1 100.00%
Bootstrap support for afum|EAL89854.1 as seed ortholog is 100%.
Bootstrap support for afis|EAW25180.1 as seed ortholog is 100%.

---

Group of orthologs #1503. Best score 1664 bits
Score difference with first non-orthologous sequence - *A. fumigatus*:1275 *A. fischeri*:1025
afum|EAL85490.1 100.00% afis|EAW20376.1 100.00%
Bootstrap support for afum|EAL85490.1 as seed ortholog is 100%.
Bootstrap support for afis|EAW20376.1 as seed ortholog is 100%.

---

Group of orthologs #1504. Best score 1664 bits
Score difference with first non-orthologous sequence - *A. fumigatus*:1664 *A. fischeri*:379
afum|EAL90682.1 100.00% afis|EAW22511.1 100.00%
Bootstrap support for afum|EAL90682.1 as seed ortholog is 100%.
Bootstrap support for afis|EAW22511.1 as seed ortholog is 100%.

---

Group of orthologs #1505. Best score 1663 bits
Score difference with first non-orthologous sequence - *A. fumigatus*:1663 *A. fischeri*:1663
afum|EAL90172.1 100.00% afis|EAW25506.1 100.00%
Bootstrap support for afum|EAL90172.1 as seed ortholog is 100%.
Bootstrap support for afis|EAW25506.1 as seed ortholog is 100%.

---

Group of orthologs #1506. Best score 1662 bits
Score difference with first non-orthologous sequence - *A. fumigatus*:1662 *A. fischeri*:1662
afum|EAL93913.1 100.00% afis|EAW19183.1 100.00%
Bootstrap support for afum|EAL93913.1 as seed ortholog is 100%.
Bootstrap support for afis|EAW19183.1 as seed ortholog is 100%.

---

Group of orthologs #1507. Best score 1662 bits
Score difference with first non-orthologous sequence - *A. fumigatus*:1662 *A. fischeri*:1662
afum|EAL90386.1 100.00% afis|EAW22815.1 100.00%
Bootstrap support for afum|EAL90386.1 as seed ortholog is 100%.
Bootstrap support for afis|EAW22815.1 as seed ortholog is 100%.

---

Group of orthologs #1508. Best score 1661 bits
Score difference with first non-orthologous sequence - *A. fumigatus*:1661 *A. fischeri*:1661
afum|EAL85629.2 100.00% afis|EAW15719.1 100.00%
Bootstrap support for afum|EAL85629.2 as seed ortholog is 100%.
Bootstrap support for afis|EAW15719.1 as seed ortholog is 100%.

---

Group of orthologs #1509. Best score 1661 bits
Score difference with first non-orthologous sequence - *A. fumigatus*:1661 *A. fischeri*:1661
afum|EAL89144.1 100.00% afis|EAW16546.1 100.00%
Bootstrap support for afum|EAL89144.1 as seed ortholog is 100%.

Bootstrap support for afis|EAW16546.1 as seed ortholog is 100%.

---

Group of orthologs #1510. Best score 1661 bits
Score difference with first non-orthologous sequence - *A. fumigatus*:1661 *A. fischeri*:1661
afum|EAL93286.1 100.00% afis|EAW18582.1 100.00%
Bootstrap support for afum|EAL93286.1 as seed ortholog is 100%.
Bootstrap support for afis|EAW18582.1 as seed ortholog is 100%.

---

Group of orthologs #1511. Best score 1661 bits
Score difference with first non-orthologous sequence - *A. fumigatus*:1661 *A. fischeri*:1661
afum|EAL93610.1 100.00% afis|EAW18878.1 100.00%
Bootstrap support for afum|EAL93610.1 as seed ortholog is 100%.
Bootstrap support for afis|EAW18878.1 as seed ortholog is 100%.

---

Group of orthologs #1512. Best score 1661 bits
Score difference with first non-orthologous sequence - *A. fumigatus*:1661 *A. fischeri*:1661
afum|EAL92386.1 100.00% afis|EAW21369.1 100.00%
Bootstrap support for afum|EAL92386.1 as seed ortholog is 100%.
Bootstrap support for afis|EAW21369.1 as seed ortholog is 100%.

---

Group of orthologs #1513. Best score 1660 bits
Score difference with first non-orthologous sequence - *A. fumigatus*:1660 *A. fischeri*:1660
afum|EAL85888.1 100.00% afis|EAW24219.1 100.00%
Bootstrap support for afum|EAL85888.1 as seed ortholog is 100%.
Bootstrap support for afis|EAW24219.1 as seed ortholog is 100%.

---

Group of orthologs #1514. Best score 1660 bits
Score difference with first non-orthologous sequence - *A. fumigatus*:1660 *A. fischeri*:1660
afum|EAL87163.1 100.00% afis|EAW23723.1 100.00%
Bootstrap support for afum|EAL87163.1 as seed ortholog is 100%.
Bootstrap support for afis|EAW23723.1 as seed ortholog is 100%.

---

Group of orthologs #1515. Best score 1660 bits
Score difference with first non-orthologous sequence - *A. fumigatus*:1660 *A. fischeri*:1660
afum|EAL90349.1 100.00% afis|EAW22851.1 100.00%
Bootstrap support for afum|EAL90349.1 as seed ortholog is 100%.
Bootstrap support for afis|EAW22851.1 as seed ortholog is 100%.

---

Group of orthologs #1516. Best score 1660 bits
Score difference with first non-orthologous sequence - *A. fumigatus*:1660 *A. fischeri*:1660
afum|EAL90242.1 100.00% afis|EAW22955.1 100.00%
Bootstrap support for afum|EAL90242.1 as seed ortholog is 100%.
Bootstrap support for afis|EAW22955.1 as seed ortholog is 100%.

---

Group of orthologs #1517. Best score 1659 bits
Score difference with first non-orthologous sequence - *A. fumigatus*:1659 *A. fischeri*:1503
afum|EAL88947.1 100.00% afis|EAW16324.1 100.00%
Bootstrap support for afum|EAL88947.1 as seed ortholog is 100%.
Bootstrap support for afis|EAW16324.1 as seed ortholog is 100%.

---

Group of orthologs #1518. Best score 1659 bits
Score difference with first non-orthologous sequence - *A. fumigatus*:1659 *A. fischeri*:1659
afum|EAL89127.1 100.00% afis|EAW16530.1 100.00%
Bootstrap support for afum|EAL89127.1 as seed ortholog is 100%.
Bootstrap support for afis|EAW16530.1 as seed ortholog is 100%.

---

Group of orthologs #1519. Best score 1659 bits
Score difference with first non-orthologous sequence - *A. fumigatus*:1659 *A. fischeri*:1659
afum|EAL84683.2 100.00% afis|EAW20951.1 100.00%
Bootstrap support for afum|EAL84683.2 as seed ortholog is 100%.
Bootstrap support for afis|EAW20951.1 as seed ortholog is 100%.

---

Group of orthologs #1520. Best score 1659 bits
Score difference with first non-orthologous sequence - *A. fumigatus*:1659 *A. fischeri*:1659
afum|EAL86068.1 100.00% afis|EAW24396.1 100.00%
Bootstrap support for afum|EAL86068.1 as seed ortholog is 100%.
Bootstrap support for afis|EAW24396.1 as seed ortholog is 100%.

---

Group of orthologs #1521. Best score 1659 bits
Score difference with first non-orthologous sequence - *A. fumigatus*:1659 *A. fischeri*:1659
afum|EAL90628.1 100.00% afis|EAW22570.1 100.00%
Bootstrap support for afum|EAL90628.1 as seed ortholog is 100%.
Bootstrap support for afis|EAW22570.1 as seed ortholog is 100%.

---

Group of orthologs #1522. Best score 1658 bits
Score difference with first non-orthologous sequence - *A. fumigatus*:1658 *A. fischeri*:1658
afum|EAL85510.1 100.00% afis|EAW15841.1 100.00%
Bootstrap support for afum|EAL85510.1 as seed ortholog is 100%.
Bootstrap support for afis|EAW15841.1 as seed ortholog is 100%.

---

Group of orthologs #1523. Best score 1657 bits
Score difference with first non-orthologous sequence - *A. fumigatus*:1657 *A. fischeri*:1657
afum|EAL87674.1 100.00% afis|EAW18308.1 100.00%
Bootstrap support for afum|EAL87674.1 as seed ortholog is 100%.
Bootstrap support for afis|EAW18308.1 as seed ortholog is 100%.

---

Group of orthologs #1524. Best score 1657 bits
Score difference with first non-orthologous sequence - *A. fumigatus*:1657 *A. fischeri*:1561
afum|EAL86792.1 100.00% afis|EAW19737.1 100.00%
Bootstrap support for afum|EAL86792.1 as seed ortholog is 100%.
Bootstrap support for afis|EAW19737.1 as seed ortholog is 100%.

---

Group of orthologs #1525. Best score 1656 bits
Score difference with first non-orthologous sequence - *A. fumigatus*:1656 *A. fischeri*:1656
afum|EAL93374.1 100.00% afis|EAW18657.1 100.00%
Bootstrap support for afum|EAL93374.1 as seed ortholog is 100%.
Bootstrap support for afis|EAW18657.1 as seed ortholog is 100%.

---

Group of orthologs #1526. Best score 1655 bits
Score difference with first non-orthologous sequence - *A. fumigatus*:1655 *A. fischeri*:1655
afum|EAL88925.1 100.00% afis|EAW16303.1 100.00%
Bootstrap support for afum|EAL88925.1 as seed ortholog is 100%.
Bootstrap support for afis|EAW16303.1 as seed ortholog is 100%.

---

Group of orthologs #1527. Best score 1654 bits
Score difference with first non-orthologous sequence - *A. fumigatus*:1654 *A. fischeri*:1654
afum|EAL93620.1 100.00% afis|EAW18887.1 100.00%
Bootstrap support for afum|EAL93620.1 as seed ortholog is 100%.
Bootstrap support for afis|EAW18887.1 as seed ortholog is 100%.

---

Group of orthologs #1528. Best score 1653 bits
Score difference with first non-orthologous sequence - *A. fumigatus*:1653 *A. fischeri*:1653

afum|EAL91708.1 100.00% afis|EAW17900.1 100.00%
Bootstrap support for afum|EAL91708.1 as seed ortholog is 100%.
Bootstrap support for afis|EAW17900.1 as seed ortholog is 100%.

---

Group of orthologs #1529. Best score 1652 bits
Score difference with first non-orthologous sequence - *A. fumigatus*:1652 *A. fischeri*:1652
afum|EAL86067.1 100.00% afis|EAW24395.1 100.00%
Bootstrap support for afum|EAL86067.1 as seed ortholog is 100%.
Bootstrap support for afis|EAW24395.1 as seed ortholog is 100%.

---

Group of orthologs #1530. Best score 1652 bits
Score difference with first non-orthologous sequence - *A. fumigatus*:1652 *A. fischeri*:1471
afum|EAL93091.1 100.00% afis|EAW18411.1 100.00%
Bootstrap support for afum|EAL93091.1 as seed ortholog is 100%.
Bootstrap support for afis|EAW18411.1 as seed ortholog is 100%.

---

Group of orthologs #1531. Best score 1652 bits
Score difference with first non-orthologous sequence - *A. fumigatus*:1395 *A. fischeri*:1574
afum|EAL92894.2 100.00% afis|EAW21880.1 100.00%
Bootstrap support for afum|EAL92894.2 as seed ortholog is 100%.
Bootstrap support for afis|EAW21880.1 as seed ortholog is 100%.

---

Group of orthologs #1532. Best score 1651 bits
Score difference with first non-orthologous sequence - *A. fumigatus*:1651 *A. fischeri*:1442
afum|EAL86480.1 100.00% afis|EAW16952.1 100.00%
Bootstrap support for afum|EAL86480.1 as seed ortholog is 100%.
Bootstrap support for afis|EAW16952.1 as seed ortholog is 100%.

---

Group of orthologs #1533. Best score 1651 bits
Score difference with first non-orthologous sequence - *A. fumigatus*:1651 *A. fischeri*:1028
afum|EAL86791.1 100.00% afis|EAW19738.1 100.00%
Bootstrap support for afum|EAL86791.1 as seed ortholog is 100%.
Bootstrap support for afis|EAW19738.1 as seed ortholog is 100%.

---

Group of orthologs #1534. Best score 1651 bits
Score difference with first non-orthologous sequence - *A. fumigatus*:1651 *A. fischeri*:1651
afum|EAL93043.1 100.00% afis|EAW18368.1 100.00%
Bootstrap support for afum|EAL93043.1 as seed ortholog is 100%.
Bootstrap support for afis|EAW18368.1 as seed ortholog is 100%.

---

Group of orthologs #1535. Best score 1651 bits
Score difference with first non-orthologous sequence - *A. fumigatus*:1651 *A. fischeri*:1651
afum|EAL93852.1 100.00% afis|EAW19120.1 100.00%
Bootstrap support for afum|EAL93852.1 as seed ortholog is 100%.
Bootstrap support for afis|EAW19120.1 as seed ortholog is 100%.

---

Group of orthologs #1536. Best score 1650 bits
Score difference with first non-orthologous sequence - *A. fumigatus*:1650 *A. fischeri*:1527
afum|EAL84712.1 100.00% afis|EAW20922.1 100.00%
Bootstrap support for afum|EAL84712.1 as seed ortholog is 100%.
Bootstrap support for afis|EAW20922.1 as seed ortholog is 100%.

---

Group of orthologs #1537. Best score 1650 bits
Score difference with first non-orthologous sequence - *A. fumigatus*:1650 *A. fischeri*:1650
afum|EAL91721.1 100.00% afis|EAW17912.1 100.00%
Bootstrap support for afum|EAL91721.1 as seed ortholog is 100%.

Bootstrap support for afis|EAW17912.1 as seed ortholog is 100%.

---

Group of orthologs #1538. Best score 1650 bits
Score difference with first non-orthologous sequence - *A. fumigatus*:1650 *A. fischeri*:1650
afum|EAL92983.1 100.00% afis|EAW21969.1 100.00%
Bootstrap support for afum|EAL92983.1 as seed ortholog is 100%.
Bootstrap support for afis|EAW21969.1 as seed ortholog is 100%.

---

Group of orthologs #1539. Best score 1649 bits
Score difference with first non-orthologous sequence - *A. fumigatus*:1649 *A. fischeri*:1649
afum|EAL90825.1 100.00% afis|EAW22367.1 100.00%
Bootstrap support for afum|EAL90825.1 as seed ortholog is 100%.
Bootstrap support for afis|EAW22367.1 as seed ortholog is 100%.

---

Group of orthologs #1540. Best score 1649 bits
Score difference with first non-orthologous sequence - *A. fumigatus*:1649 *A. fischeri*:1649
afum|EAL90021.1 100.00% afis|EAW25359.1 100.00%
Bootstrap support for afum|EAL90021.1 as seed ortholog is 100%.
Bootstrap support for afis|EAW25359.1 as seed ortholog is 100%.

---

Group of orthologs #1541. Best score 1648 bits
Score difference with first non-orthologous sequence - *A. fumigatus*:1648 *A. fischeri*:1648
afum|EAL87634.1 100.00% afis|EAW18269.1 100.00%
Bootstrap support for afum|EAL87634.1 as seed ortholog is 100%.
Bootstrap support for afis|EAW18269.1 as seed ortholog is 100%.

---

Group of orthologs #1542. Best score 1648 bits
Score difference with first non-orthologous sequence - *A. fumigatus*:1648 *A. fischeri*:1648
afum|EAL90729.1 100.00% afis|EAW22462.1 100.00%
Bootstrap support for afum|EAL90729.1 as seed ortholog is 100%.
Bootstrap support for afis|EAW22462.1 as seed ortholog is 100%.

---

Group of orthologs #1543. Best score 1647 bits
Score difference with first non-orthologous sequence - *A. fumigatus*:1647 *A. fischeri*:1647
afum|EAL88884.1 100.00% afis|EAW16263.1 100.00%
Bootstrap support for afum|EAL88884.1 as seed ortholog is 100%.
Bootstrap support for afis|EAW16263.1 as seed ortholog is 100%.

---

Group of orthologs #1544. Best score 1647 bits
Score difference with first non-orthologous sequence - *A. fumigatus*:1647 *A. fischeri*:1647
afum|EAL87590.1 100.00% afis|EAW18224.1 100.00%
Bootstrap support for afum|EAL87590.1 as seed ortholog is 100%.
Bootstrap support for afis|EAW18224.1 as seed ortholog is 100%.

---

Group of orthologs #1545. Best score 1647 bits
Score difference with first non-orthologous sequence - *A. fumigatus*:1550 *A. fischeri*:1647
afum|EAL91190.1 100.00% afis|EAW17346.1 100.00%
Bootstrap support for afum|EAL91190.1 as seed ortholog is 100%.
Bootstrap support for afis|EAW17346.1 as seed ortholog is 100%.

---

Group of orthologs #1546. Best score 1647 bits
Score difference with first non-orthologous sequence - *A. fumigatus*:1647 *A. fischeri*:1647
afum|EAL93680.1 100.00% afis|EAW18946.1 100.00%
Bootstrap support for afum|EAL93680.1 as seed ortholog is 100%.
Bootstrap support for afis|EAW18946.1 as seed ortholog is 100%.

---

Group of orthologs #1547. Best score 1646 bits
Score difference with first non-orthologous sequence - *A. fumigatus*:1646 *A. fischeri*:1646
afum|EAL85029.1 100.00% afis|EAW19868.1 100.00%
Bootstrap support for afum|EAL85029.1 as seed ortholog is 100%.
Bootstrap support for afis|EAW19868.1 as seed ortholog is 100%.

---

Group of orthologs #1548. Best score 1646 bits
Score difference with first non-orthologous sequence - *A. fumigatus*:1646 *A. fischeri*:1646
afum|EAL85245.1 100.00% afis|EAW20127.1 100.00%
Bootstrap support for afum|EAL85245.1 as seed ortholog is 100%.
Bootstrap support for afis|EAW20127.1 as seed ortholog is 100%.

---

Group of orthologs #1549. Best score 1646 bits
Score difference with first non-orthologous sequence - *A. fumigatus*:1646 *A. fischeri*:1646
afum|EAL92104.2 100.00% afis|EAW21083.1 100.00%
Bootstrap support for afum|EAL92104.2 as seed ortholog is 100%.
Bootstrap support for afis|EAW21083.1 as seed ortholog is 100%.

---

Group of orthologs #1550. Best score 1645 bits
Score difference with first non-orthologous sequence - *A. fumigatus*:1645 *A. fischeri*:1645
afum|EAL86518.2 100.00% afis|EAW16994.1 100.00%
Bootstrap support for afum|EAL86518.2 as seed ortholog is 100%.
Bootstrap support for afis|EAW16994.1 as seed ortholog is 100%.

---

Group of orthologs #1551. Best score 1645 bits
Score difference with first non-orthologous sequence - *A. fumigatus*:1645 *A. fischeri*:1645
afum|EAL84766.2 100.00% afis|EAW20868.1 100.00%
Bootstrap support for afum|EAL84766.2 as seed ortholog is 100%.
Bootstrap support for afis|EAW20868.1 as seed ortholog is 100%.

---

Group of orthologs #1552. Best score 1645 bits
Score difference with first non-orthologous sequence - *A. fumigatus*:1645 *A. fischeri*:1645
afum|EAL87038.1 100.00% afis|EAW19458.1 100.00%
Bootstrap support for afum|EAL87038.1 as seed ortholog is 100%.
Bootstrap support for afis|EAW19458.1 as seed ortholog is 100%.

---

Group of orthologs #1553. Best score 1645 bits
Score difference with first non-orthologous sequence - *A. fumigatus*:1645 *A. fischeri*:1645
afum|EAL90693.1 100.00% afis|EAW22500.1 100.00%
Bootstrap support for afum|EAL90693.1 as seed ortholog is 100%.
Bootstrap support for afis|EAW22500.1 as seed ortholog is 100%.

---

Group of orthologs #1554. Best score 1643 bits
Score difference with first non-orthologous sequence - *A. fumigatus*:1643 *A. fischeri*:1643
afum|EAL89087.1 100.00% afis|EAW16493.1 100.00%
Bootstrap support for afum|EAL89087.1 as seed ortholog is 100%.
Bootstrap support for afis|EAW16493.1 as seed ortholog is 100%.

---

Group of orthologs #1555. Best score 1643 bits
Score difference with first non-orthologous sequence - *A. fumigatus*:1643 *A. fischeri*:1643
afum|EAL93329.1 100.00% afis|EAW18619.1 100.00%
Bootstrap support for afum|EAL93329.1 as seed ortholog is 100%.
Bootstrap support for afis|EAW18619.1 as seed ortholog is 100%.

---

Group of orthologs #1556. Best score 1643 bits
Score difference with first non-orthologous sequence - *A. fumigatus*:1296 *A. fischeri*:1643

afum|EAL92129.1 100.00% afis|EAW21108.1 100.00%
Bootstrap support for afum|EAL92129.1 as seed ortholog is 100%.
Bootstrap support for afis|EAW21108.1 as seed ortholog is 100%.

---

Group of orthologs #1557. Best score 1641 bits
Score difference with first non-orthologous sequence - *A. fumigatus*:1641 *A. fischeri*:1641
afum|EAL84607.1 100.00% afis|EAW20446.1 100.00%
Bootstrap support for afum|EAL84607.1 as seed ortholog is 100%.
Bootstrap support for afis|EAW20446.1 as seed ortholog is 100%.

---

Group of orthologs #1558. Best score 1641 bits
Score difference with first non-orthologous sequence - *A. fumigatus*:1641 *A. fischeri*:1641
afum|EAL87436.1 100.00% afis|EAW24006.1 100.00%
Bootstrap support for afum|EAL87436.1 as seed ortholog is 100%.
Bootstrap support for afis|EAW24006.1 as seed ortholog is 100%.

---

Group of orthologs #1559. Best score 1641 bits
Score difference with first non-orthologous sequence - *A. fumigatus*:1641 *A. fischeri*:1641
afum|EAL92971.1 100.00% afis|EAW21957.1 100.00%
Bootstrap support for afum|EAL92971.1 as seed ortholog is 100%.
Bootstrap support for afis|EAW21957.1 as seed ortholog is 100%.

---

Group of orthologs #1560. Best score 1641 bits
Score difference with first non-orthologous sequence - *A. fumigatus*:1641 *A. fischeri*:1641
afum|EAL90141.1 100.00% afis|EAW25478.1 100.00%
Bootstrap support for afum|EAL90141.1 as seed ortholog is 100%.
Bootstrap support for afis|EAW25478.1 as seed ortholog is 100%.

---

Group of orthologs #1561. Best score 1640 bits
Score difference with first non-orthologous sequence - *A. fumigatus*:1640 *A. fischeri*:1640
afum|EAL85053.1 100.00% afis|EAW19842.1 100.00%
Bootstrap support for afum|EAL85053.1 as seed ortholog is 100%.
Bootstrap support for afis|EAW19842.1 as seed ortholog is 100%.

---

Group of orthologs #1562. Best score 1640 bits
Score difference with first non-orthologous sequence - *A. fumigatus*:1640 *A. fischeri*:1640
afum|EAL90806.1 100.00% afis|EAW22384.1 100.00%
Bootstrap support for afum|EAL90806.1 as seed ortholog is 100%.
Bootstrap support for afis|EAW22384.1 as seed ortholog is 100%.

---

Group of orthologs #1563. Best score 1640 bits
Score difference with first non-orthologous sequence - *A. fumigatus*:1640 *A. fischeri*:1640
afum|EAL92563.1 100.00% afis|EAW21543.1 100.00%
Bootstrap support for afum|EAL92563.1 as seed ortholog is 100%.
Bootstrap support for afis|EAW21543.1 as seed ortholog is 100%.

---

Group of orthologs #1564. Best score 1640 bits
Score difference with first non-orthologous sequence - *A. fumigatus*:1640 *A. fischeri*:1640
afum|EAL89894.1 100.00% afis|EAW25233.1 100.00%
Bootstrap support for afum|EAL89894.1 as seed ortholog is 100%.
Bootstrap support for afis|EAW25233.1 as seed ortholog is 100%.

---

Group of orthologs #1565. Best score 1639 bits
Score difference with first non-orthologous sequence - *A. fumigatus*:1639 *A. fischeri*:1639
afum|EAL89253.1 100.00% afis|EAW16649.1 100.00%
Bootstrap support for afum|EAL89253.1 as seed ortholog is 100%.

Bootstrap support for afis|EAW16649.1 as seed ortholog is 100%.

---

Group of orthologs #1566. Best score 1639 bits
Score difference with first non-orthologous sequence - *A. fumigatus*:1639 *A. fischeri*:1639
afum|EAL86008.1 100.00% afis|EAW24337.1 100.00%
Bootstrap support for afum|EAL86008.1 as seed ortholog is 100%.
Bootstrap support for afis|EAW24337.1 as seed ortholog is 100%.

---

Group of orthologs #1567. Best score 1639 bits
Score difference with first non-orthologous sequence - *A. fumigatus*:854 *A. fischeri*:919
afum|EAL89817.2 100.00% afis|EAW25144.1 100.00%
Bootstrap support for afum|EAL89817.2 as seed ortholog is 100%.
Bootstrap support for afis|EAW25144.1 as seed ortholog is 100%.

---

Group of orthologs #1568. Best score 1638 bits
Score difference with first non-orthologous sequence - *A. fumigatus*:1638 *A. fischeri*:1638
afum|EAL91442.1 100.00% afis|EAW17628.1 100.00%
Bootstrap support for afum|EAL91442.1 as seed ortholog is 100%.
Bootstrap support for afis|EAW17628.1 as seed ortholog is 100%.

---

Group of orthologs #1569. Best score 1638 bits
Score difference with first non-orthologous sequence - *A. fumigatus*:1638 *A. fischeri*:1638
afum|EAL88302.1 100.00% afis|EAW23181.1 100.00%
Bootstrap support for afum|EAL88302.1 as seed ortholog is 100%.
Bootstrap support for afis|EAW23181.1 as seed ortholog is 100%.

---

Group of orthologs #1570. Best score 1638 bits
Score difference with first non-orthologous sequence - *A. fumigatus*:1638 *A. fischeri*:1638
afum|EAL87924.1 100.00% afis|EAW23542.1 100.00%
Bootstrap support for afum|EAL87924.1 as seed ortholog is 100%.
Bootstrap support for afis|EAW23542.1 as seed ortholog is 100%.

---

Group of orthologs #1571. Best score 1637 bits
Score difference with first non-orthologous sequence - *A. fumigatus*:1637 *A. fischeri*:1637
afum|EAL88684.1 100.00% afis|EAW16052.1 100.00%
Bootstrap support for afum|EAL88684.1 as seed ortholog is 100%.
Bootstrap support for afis|EAW16052.1 as seed ortholog is 100%.

---

Group of orthologs #1572. Best score 1637 bits
Score difference with first non-orthologous sequence - *A. fumigatus*:1637 *A. fischeri*:1259
afum|EAL88026.1 100.00% afis|EAW23436.1 100.00%
Bootstrap support for afum|EAL88026.1 as seed ortholog is 100%.
Bootstrap support for afis|EAW23436.1 as seed ortholog is 100%.

---

Group of orthologs #1573. Best score 1637 bits
Score difference with first non-orthologous sequence - *A. fumigatus*:1637 *A. fischeri*:1637
afum|EAL91973.1 100.00% afis|EAW24118.1 100.00%
Bootstrap support for afum|EAL91973.1 as seed ortholog is 100%.
Bootstrap support for afis|EAW24118.1 as seed ortholog is 100%.

---

Group of orthologs #1574. Best score 1636 bits
Score difference with first non-orthologous sequence - *A. fumigatus*:1636 *A. fischeri*:1636
afum|EAL86699.1 100.00% afis|EAW17281.1 100.00%
Bootstrap support for afum|EAL86699.1 as seed ortholog is 100%.
Bootstrap support for afis|EAW17281.1 as seed ortholog is 100%.

---

Group of orthologs #1575. Best score 1636 bits
Score difference with first non-orthologous sequence - *A. fumigatus*:1636 *A. fischeri*:1636
afum|EAL85092.1 100.00% afis|EAW19798.1 100.00%
Bootstrap support for afum|EAL85092.1 as seed ortholog is 100%.
Bootstrap support for afis|EAW19798.1 as seed ortholog is 100%.

---

Group of orthologs #1576. Best score 1636 bits
Score difference with first non-orthologous sequence - *A. fumigatus*:1279 *A. fischeri*:1636
afum|EAL90676.1 100.00% afis|EAW22517.1 100.00%
Bootstrap support for afum|EAL90676.1 as seed ortholog is 100%.
Bootstrap support for afis|EAW22517.1 as seed ortholog is 100%.

---

Group of orthologs #1577. Best score 1635 bits
Score difference with first non-orthologous sequence - *A. fumigatus*:1635 *A. fischeri*:1635
afum|EAL85837.1 100.00% afis|EAW15486.1 100.00%
Bootstrap support for afum|EAL85837.1 as seed ortholog is 100%.
Bootstrap support for afis|EAW15486.1 as seed ortholog is 100%.

---

Group of orthologs #1578. Best score 1635 bits
Score difference with first non-orthologous sequence - *A. fumigatus*:1635 *A. fischeri*:1560
afum|EAL93364.1 100.00% afis|EAW18648.1 100.00%
Bootstrap support for afum|EAL93364.1 as seed ortholog is 100%.
Bootstrap support for afis|EAW18648.1 as seed ortholog is 100%.

---

Group of orthologs #1579. Best score 1634 bits
Score difference with first non-orthologous sequence - *A. fumigatus*:1634 *A. fischeri*:1634
afum|EAL87724.1 100.00% afis|EAW18360.1 100.00%
Bootstrap support for afum|EAL87724.1 as seed ortholog is 100%.
Bootstrap support for afis|EAW18360.1 as seed ortholog is 100%.

---

Group of orthologs #1580. Best score 1634 bits
Score difference with first non-orthologous sequence - *A. fumigatus*:1634 *A. fischeri*:1634
afum|EAL90517.1 100.00% afis|EAW22675.1 100.00%
Bootstrap support for afum|EAL90517.1 as seed ortholog is 100%.
Bootstrap support for afis|EAW22675.1 as seed ortholog is 100%.

---

Group of orthologs #1581. Best score 1634 bits
Score difference with first non-orthologous sequence - *A. fumigatus*:1634 *A. fischeri*:1634
afum|EAL91975.1 100.00% afis|EAW24120.1 100.00%
Bootstrap support for afum|EAL91975.1 as seed ortholog is 100%.
Bootstrap support for afis|EAW24120.1 as seed ortholog is 100%.

---

Group of orthologs #1582. Best score 1633 bits
Score difference with first non-orthologous sequence - *A. fumigatus*:1633 *A. fischeri*:1633
afum|EAL88475.1 100.00% afis|EAW23005.1 100.00%
Bootstrap support for afum|EAL88475.1 as seed ortholog is 100%.
Bootstrap support for afis|EAW23005.1 as seed ortholog is 100%.

---

Group of orthologs #1583. Best score 1633 bits
Score difference with first non-orthologous sequence - *A. fumigatus*:1633 *A. fischeri*:1633
afum|EAL93481.1 100.00% afis|EAW18754.1 100.00%
Bootstrap support for afum|EAL93481.1 as seed ortholog is 100%.
Bootstrap support for afis|EAW18754.1 as seed ortholog is 100%.

---

Group of orthologs #1584. Best score 1633 bits
Score difference with first non-orthologous sequence - *A. fumigatus*:1633 *A. fischeri*:1633

afum|EAL90834.1 100.00% afis|EAW22357.1 100.00%
Bootstrap support for afum|EAL90834.1 as seed ortholog is 100%.
Bootstrap support for afis|EAW22357.1 as seed ortholog is 100%.

---

Group of orthologs #1585. Best score 1632 bits
Score difference with first non-orthologous sequence - *A. fumigatus*:1632 *A. fischeri*:1545
afum|EAL86938.1 100.00% afis|EAW19580.1 100.00%
Bootstrap support for afum|EAL86938.1 as seed ortholog is 100%.
Bootstrap support for afis|EAW19580.1 as seed ortholog is 100%.

---

Group of orthologs #1586. Best score 1631 bits
Score difference with first non-orthologous sequence - *A. fumigatus*:1631 *A. fischeri*:1631
afum|EAL85596.1 100.00% afis|EAW15753.1 100.00%
Bootstrap support for afum|EAL85596.1 as seed ortholog is 100%.
Bootstrap support for afis|EAW15753.1 as seed ortholog is 100%.

---

Group of orthologs #1587. Best score 1631 bits
Score difference with first non-orthologous sequence - *A. fumigatus*:1631 *A. fischeri*:1631
afum|EAL84888.1 100.00% afis|EAW20029.1 100.00%
Bootstrap support for afum|EAL84888.1 as seed ortholog is 100%.
Bootstrap support for afis|EAW20029.1 as seed ortholog is 100%.

---

Group of orthologs #1588. Best score 1631 bits
Score difference with first non-orthologous sequence - *A. fumigatus*:1631 *A. fischeri*:1631
afum|EAL91278.2 100.00% afis|EAW17450.1 100.00%
Bootstrap support for afum|EAL91278.2 as seed ortholog is 100%.
Bootstrap support for afis|EAW17450.1 as seed ortholog is 100%.

---

Group of orthologs #1589. Best score 1631 bits
Score difference with first non-orthologous sequence - *A. fumigatus*:1631 *A. fischeri*:1631
afum|EAL91921.1 100.00% afis|EAW24066.1 100.00%
Bootstrap support for afum|EAL91921.1 as seed ortholog is 100%.
Bootstrap support for afis|EAW24066.1 as seed ortholog is 100%.

---

Group of orthologs #1590. Best score 1629 bits
Score difference with first non-orthologous sequence - *A. fumigatus*:1629 *A. fischeri*:1629
afum|EAL89215.1 100.00% afis|EAW16622.1 100.00%
Bootstrap support for afum|EAL89215.1 as seed ortholog is 100%.
Bootstrap support for afis|EAW16622.1 as seed ortholog is 100%.

---

Group of orthologs #1591. Best score 1629 bits
Score difference with first non-orthologous sequence - *A. fumigatus*:1432 *A. fischeri*:1513
afum|EAL84833.1 100.00% afis|EAW20796.1 100.00%
Bootstrap support for afum|EAL84833.1 as seed ortholog is 100%.
Bootstrap support for afis|EAW20796.1 as seed ortholog is 100%.

---

Group of orthologs #1592. Best score 1629 bits
Score difference with first non-orthologous sequence - *A. fumigatus*:1629 *A. fischeri*:1629
afum|EAL91280.1 100.00% afis|EAW17452.1 100.00%
Bootstrap support for afum|EAL91280.1 as seed ortholog is 100%.
Bootstrap support for afis|EAW17452.1 as seed ortholog is 100%.

---

Group of orthologs #1593. Best score 1628 bits
Score difference with first non-orthologous sequence - *A. fumigatus*:1262 *A. fischeri*:1101
afum|EAL87255.1 100.00% afis|EAW23820.1 100.00%
Bootstrap support for afum|EAL87255.1 as seed ortholog is 100%.

Bootstrap support for afis|EAW23820.1 as seed ortholog is 100%.

---

Group of orthologs #1594. Best score 1628 bits
Score difference with first non-orthologous sequence - *A. fumigatus*:1628 *A. fischeri*:1628
afum|EAL90757.1 100.00% afis|EAW22434.1 100.00%
Bootstrap support for afum|EAL90757.1 as seed ortholog is 100%.
Bootstrap support for afis|EAW22434.1 as seed ortholog is 100%.

---

Group of orthologs #1595. Best score 1628 bits
Score difference with first non-orthologous sequence - *A. fumigatus*:1628 *A. fischeri*:1314
afum|EAL90530.1 100.00% afis|EAW22662.1 100.00%
Bootstrap support for afum|EAL90530.1 as seed ortholog is 100%.
Bootstrap support for afis|EAW22662.1 as seed ortholog is 100%.

---

Group of orthologs #1596. Best score 1627 bits
Score difference with first non-orthologous sequence - *A. fumigatus*:1271 *A. fischeri*:811
afum|EAL86253.1 100.00% afis|EAW24627.1 100.00%
Bootstrap support for afum|EAL86253.1 as seed ortholog is 100%.
Bootstrap support for afis|EAW24627.1 as seed ortholog is 100%.

---

Group of orthologs #1597. Best score 1627 bits
Score difference with first non-orthologous sequence - *A. fumigatus*:1627 *A. fischeri*:1627
afum|EBA27180.1 100.00% afis|EAW20583.1 100.00%
Bootstrap support for afum|EBA27180.1 as seed ortholog is 100%.
Bootstrap support for afis|EAW20583.1 as seed ortholog is 100%.

---

Group of orthologs #1598. Best score 1626 bits
Score difference with first non-orthologous sequence - *A. fumigatus*:1626 *A. fischeri*:1626
afum|EAL88875.1 100.00% afis|EAW16247.1 100.00%
Bootstrap support for afum|EAL88875.1 as seed ortholog is 100%.
Bootstrap support for afis|EAW16247.1 as seed ortholog is 100%.

---

Group of orthologs #1599. Best score 1624 bits
Score difference with first non-orthologous sequence - *A. fumigatus*:1123 *A. fischeri*:1624
afum|EAL88829.2 100.00% afis|EAW21809.1 100.00%
Bootstrap support for afum|EAL88829.2 as seed ortholog is 100%.
Bootstrap support for afis|EAW21809.1 as seed ortholog is 100%.

---

Group of orthologs #1600. Best score 1624 bits
Score difference with first non-orthologous sequence - *A. fumigatus*:1417 *A. fischeri*:1580
afum|EAL89364.1 100.00% afis|EAW24716.1 100.00%
Bootstrap support for afum|EAL89364.1 as seed ortholog is 100%.
Bootstrap support for afis|EAW24716.1 as seed ortholog is 100%.

---

Group of orthologs #1601. Best score 1624 bits
Score difference with first non-orthologous sequence - *A. fumigatus*:1624 *A. fischeri*:1624
afum|EAL92788.2 100.00% afis|EAW21758.1 100.00%
Bootstrap support for afum|EAL92788.2 as seed ortholog is 100%.
Bootstrap support for afis|EAW21758.1 as seed ortholog is 100%.

---

Group of orthologs #1602. Best score 1624 bits
Score difference with first non-orthologous sequence - *A. fumigatus*:1490 *A. fischeri*:1501
afum|EAL91949.1 100.00% afis|EAW24095.1 100.00%
Bootstrap support for afum|EAL91949.1 as seed ortholog is 100%.
Bootstrap support for afis|EAW24095.1 as seed ortholog is 100%.

---

Group of orthologs #1603. Best score 1623 bits
Score difference with first non-orthologous sequence - *A. fumigatus*:1623 *A. fischeri*:1623
afum|EAL85868.1 100.00% afis|EAW24196.1 100.00%
Bootstrap support for afum|EAL85868.1 as seed ortholog is 100%.
Bootstrap support for afis|EAW24196.1 as seed ortholog is 100%.

---

Group of orthologs #1604. Best score 1622 bits
Score difference with first non-orthologous sequence - *A. fumigatus*:1622 *A. fischeri*:1622
afum|EAL88188.1 100.00% afis|EAW23291.1 100.00%
Bootstrap support for afum|EAL88188.1 as seed ortholog is 100%.
Bootstrap support for afis|EAW23291.1 as seed ortholog is 100%.

---

Group of orthologs #1605. Best score 1622 bits
Score difference with first non-orthologous sequence - *A. fumigatus*:1622 *A. fischeri*:1622
afum|EAL94001.1 100.00% afis|EAW19275.1 100.00%
Bootstrap support for afum|EAL94001.1 as seed ortholog is 100%.
Bootstrap support for afis|EAW19275.1 as seed ortholog is 100%.

---

Group of orthologs #1606. Best score 1621 bits
Score difference with first non-orthologous sequence - *A. fumigatus*:1621 *A. fischeri*:1506
afum|EAL85753.2 100.00% afis|EAW15581.1 100.00%
Bootstrap support for afum|EAL85753.2 as seed ortholog is 100%.
Bootstrap support for afis|EAW15581.1 as seed ortholog is 100%.

---

Group of orthologs #1607. Best score 1621 bits
Score difference with first non-orthologous sequence - *A. fumigatus*:1621 *A. fischeri*:1621
afum|EAL89082.1 100.00% afis|EAW16488.1 100.00%
Bootstrap support for afum|EAL89082.1 as seed ortholog is 100%.
Bootstrap support for afis|EAW16488.1 as seed ortholog is 100%.

---

Group of orthologs #1608. Best score 1621 bits
Score difference with first non-orthologous sequence - *A. fumigatus*:1541 *A. fischeri*:910
afum|EAL86238.1 100.00% afis|EAW24608.1 100.00%
Bootstrap support for afum|EAL86238.1 as seed ortholog is 100%.
Bootstrap support for afis|EAW24608.1 as seed ortholog is 100%.

---

Group of orthologs #1609. Best score 1621 bits
Score difference with first non-orthologous sequence - *A. fumigatus*:1621 *A. fischeri*:1621
afum|EAL90434.1 100.00% afis|EAW22765.1 100.00%
Bootstrap support for afum|EAL90434.1 as seed ortholog is 100%.
Bootstrap support for afis|EAW22765.1 as seed ortholog is 100%.

---

Group of orthologs #1610. Best score 1619 bits
Score difference with first non-orthologous sequence - *A. fumigatus*:1619 *A. fischeri*:1619
afum|EAL92303.1 100.00% afis|EAW21286.1 100.00%
Bootstrap support for afum|EAL92303.1 as seed ortholog is 100%.
Bootstrap support for afis|EAW21286.1 as seed ortholog is 100%.

---

Group of orthologs #1611. Best score 1619 bits
Score difference with first non-orthologous sequence - *A. fumigatus*:1619 *A. fischeri*:1619
afum|EAL89615.2 100.00% afis|EAW24957.1 100.00%
Bootstrap support for afum|EAL89615.2 as seed ortholog is 100%.
Bootstrap support for afis|EAW24957.1 as seed ortholog is 100%.

---

Group of orthologs #1612. Best score 1618 bits
Score difference with first non-orthologous sequence - *A. fumigatus*:1243 *A. fischeri*:1618

afum|EAL86351.1 100.00% afis|EAW16780.1 100.00%
Bootstrap support for afum|EAL86351.1 as seed ortholog is 100%.
Bootstrap support for afis|EAW16780.1 as seed ortholog is 100%.

---

Group of orthologs #1613. Best score 1618 bits
Score difference with first non-orthologous sequence - *A. fumigatus*:1618 *A. fischeri*:1618
afum|EAL85871.1 100.00% afis|EAW24199.1 100.00%
Bootstrap support for afum|EAL85871.1 as seed ortholog is 100%.
Bootstrap support for afis|EAW24199.1 as seed ortholog is 100%.

---

Group of orthologs #1614. Best score 1618 bits
Score difference with first non-orthologous sequence - *A. fumigatus*:1618 *A. fischeri*:1618
afum|EAL88324.1 100.00% afis|EAW23159.1 100.00%
Bootstrap support for afum|EAL88324.1 as seed ortholog is 100%.
Bootstrap support for afis|EAW23159.1 as seed ortholog is 100%.

---

Group of orthologs #1615. Best score 1618 bits
Score difference with first non-orthologous sequence - *A. fumigatus*:1618 *A. fischeri*:1618
afum|EAL92305.1 100.00% afis|EAW21288.1 100.00%
Bootstrap support for afum|EAL92305.1 as seed ortholog is 100%.
Bootstrap support for afis|EAW21288.1 as seed ortholog is 100%.

---

Group of orthologs #1616. Best score 1618 bits
Score difference with first non-orthologous sequence - *A. fumigatus*:1618 *A. fischeri*:1618
afum|EAL89748.1 100.00% afis|EAW25079.1 100.00%
Bootstrap support for afum|EAL89748.1 as seed ortholog is 100%.
Bootstrap support for afis|EAW25079.1 as seed ortholog is 100%.

---

Group of orthologs #1617. Best score 1618 bits
Score difference with first non-orthologous sequence - *A. fumigatus*:1618 *A. fischeri*:1510
afum|EAL89860.2 100.00% afis|EAW25186.1 100.00%
Bootstrap support for afum|EAL89860.2 as seed ortholog is 100%.
Bootstrap support for afis|EAW25186.1 as seed ortholog is 100%.

---

Group of orthologs #1618. Best score 1617 bits
Score difference with first non-orthologous sequence - *A. fumigatus*:1617 *A. fischeri*:1617
afum|EAL87428.1 100.00% afis|EAW23998.1 100.00%
Bootstrap support for afum|EAL87428.1 as seed ortholog is 100%.
Bootstrap support for afis|EAW23998.1 as seed ortholog is 100%.

---

Group of orthologs #1619. Best score 1616 bits
Score difference with first non-orthologous sequence - *A. fumigatus*:1616 *A. fischeri*:1616
afum|EAL86576.1 100.00% afis|EAW17112.1 100.00%
Bootstrap support for afum|EAL86576.1 as seed ortholog is 100%.
Bootstrap support for afis|EAW17112.1 as seed ortholog is 100%.

---

Group of orthologs #1620. Best score 1616 bits
Score difference with first non-orthologous sequence - *A. fumigatus*:1616 *A. fischeri*:1616
afum|EBA27513.1 100.00% afis|EAW19327.1 100.00%
Bootstrap support for afum|EBA27513.1 as seed ortholog is 100%.
Bootstrap support for afis|EAW19327.1 as seed ortholog is 100%.

---

Group of orthologs #1621. Best score 1615 bits
Score difference with first non-orthologous sequence - *A. fumigatus*:1615 *A. fischeri*:1615
afum|EAL87289.1 100.00% afis|EAW23861.1 100.00%
Bootstrap support for afum|EAL87289.1 as seed ortholog is 100%.

Bootstrap support for afis|EAW23861.1 as seed ortholog is 100%.

---

Group of orthologs #1622. Best score 1615 bits
Score difference with first non-orthologous sequence - *A. fumigatus*:1615 *A. fischeri*:1615
afum|EAL87311.1 100.00% afis|EAW23883.1 100.00%
Bootstrap support for afum|EAL87311.1 as seed ortholog is 100%.
Bootstrap support for afis|EAW23883.1 as seed ortholog is 100%.

---

Group of orthologs #1623. Best score 1615 bits
Score difference with first non-orthologous sequence - *A. fumigatus*:1615 *A. fischeri*:1615
afum|EAL92396.1 100.00% afis|EAW21381.1 100.00%
Bootstrap support for afum|EAL92396.1 as seed ortholog is 100%.
Bootstrap support for afis|EAW21381.1 as seed ortholog is 100%.

---

Group of orthologs #1624. Best score 1614 bits
Score difference with first non-orthologous sequence - *A. fumigatus*:1614 *A. fischeri*:1614
afum|EAL85234.1 100.00% afis|EAW20117.1 100.00%
Bootstrap support for afum|EAL85234.1 as seed ortholog is 100%.
Bootstrap support for afis|EAW20117.1 as seed ortholog is 100%.

---

Group of orthologs #1625. Best score 1614 bits
Score difference with first non-orthologous sequence - *A. fumigatus*:1614 *A. fischeri*:1614
afum|EAL93412.1 100.00% afis|EAW18691.1 100.00%
Bootstrap support for afum|EAL93412.1 as seed ortholog is 100%.
Bootstrap support for afis|EAW18691.1 as seed ortholog is 100%.

---

Group of orthologs #1626. Best score 1614 bits
Score difference with first non-orthologous sequence - *A. fumigatus*:1247 *A. fischeri*:1614
afum|EAL89682.1 100.00% afis|EAW25014.1 100.00%
Bootstrap support for afum|EAL89682.1 as seed ortholog is 100%.
Bootstrap support for afis|EAW25014.1 as seed ortholog is 100%.

---

Group of orthologs #1627. Best score 1613 bits
Score difference with first non-orthologous sequence - *A. fumigatus*:1613 *A. fischeri*:1613
afum|EAL92595.1 100.00% afis|EAW21576.1 100.00%
Bootstrap support for afum|EAL92595.1 as seed ortholog is 100%.
Bootstrap support for afis|EAW21576.1 as seed ortholog is 100%.

---

Group of orthologs #1628. Best score 1613 bits
Score difference with first non-orthologous sequence - *A. fumigatus*:1613 *A. fischeri*:1613
afum|EAL92820.1 100.00% afis|EAW21795.1 100.00%
Bootstrap support for afum|EAL92820.1 as seed ortholog is 100%.
Bootstrap support for afis|EAW21795.1 as seed ortholog is 100%.

---

Group of orthologs #1629. Best score 1613 bits
Score difference with first non-orthologous sequence - *A. fumigatus*:1613 *A. fischeri*:1613
afum|EAL92940.1 100.00% afis|EAW21927.1 100.00%
Bootstrap support for afum|EAL92940.1 as seed ortholog is 100%.
Bootstrap support for afis|EAW21927.1 as seed ortholog is 100%.

---

Group of orthologs #1630. Best score 1612 bits
Score difference with first non-orthologous sequence - *A. fumigatus*:1553 *A. fischeri*:1612
afum|EAL85757.1 100.00% afis|EAW15577.1 100.00%
Bootstrap support for afum|EAL85757.1 as seed ortholog is 100%.
Bootstrap support for afis|EAW15577.1 as seed ortholog is 100%.

---

Group of orthologs #1631. Best score 1612 bits
Score difference with first non-orthologous sequence - *A. fumigatus*:1612 *A. fischeri*:1612
afum|EAL85205.1 100.00% afis|EAW20086.1 100.00%
Bootstrap support for afum|EAL85205.1 as seed ortholog is 100%.
Bootstrap support for afis|EAW20086.1 as seed ortholog is 100%.

---

Group of orthologs #1632. Best score 1612 bits
Score difference with first non-orthologous sequence - *A. fumigatus*:1612 *A. fischeri*:1612
afum|EAL88170.1 100.00% afis|EAW23307.1 100.00%
Bootstrap support for afum|EAL88170.1 as seed ortholog is 100%.
Bootstrap support for afis|EAW23307.1 as seed ortholog is 100%.

---

Group of orthologs #1633. Best score 1612 bits
Score difference with first non-orthologous sequence - *A. fumigatus*:1612 *A. fischeri*:1494
afum|EAL90950.1 100.00% afis|EAW22243.1 100.00%
Bootstrap support for afum|EAL90950.1 as seed ortholog is 100%.
Bootstrap support for afis|EAW22243.1 as seed ortholog is 100%.

---

Group of orthologs #1634. Best score 1612 bits
Score difference with first non-orthologous sequence - *A. fumigatus*:1612 *A. fischeri*:1612
afum|EAL90399.1 100.00% afis|EAW22800.1 100.00%
Bootstrap support for afum|EAL90399.1 as seed ortholog is 100%.
Bootstrap support for afis|EAW22800.1 as seed ortholog is 100%.

---

Group of orthologs #1635. Best score 1610 bits
Score difference with first non-orthologous sequence - *A. fumigatus*:1610 *A. fischeri*:1610
afum|EAL88552.1 100.00% afis|EAW15917.1 100.00%
Bootstrap support for afum|EAL88552.1 as seed ortholog is 100%.
Bootstrap support for afis|EAW15917.1 as seed ortholog is 100%.

---

Group of orthologs #1636. Best score 1610 bits
Score difference with first non-orthologous sequence - *A. fumigatus*:1610 *A. fischeri*:1610
afum|EAL90520.1 100.00% afis|EAW22672.1 100.00%
Bootstrap support for afum|EAL90520.1 as seed ortholog is 100%.
Bootstrap support for afis|EAW22672.1 as seed ortholog is 100%.

---

Group of orthologs #1637. Best score 1609 bits
Score difference with first non-orthologous sequence - *A. fumigatus*:1609 *A. fischeri*:1609
afum|EAL87716.1 100.00% afis|EAW18353.1 100.00%
Bootstrap support for afum|EAL87716.1 as seed ortholog is 100%.
Bootstrap support for afis|EAW18353.1 as seed ortholog is 100%.

---

Group of orthologs #1638. Best score 1609 bits
Score difference with first non-orthologous sequence - *A. fumigatus*:1609 *A. fischeri*:1609
afum|EAL92363.1 100.00% afis|EAW21346.1 100.00%
Bootstrap support for afum|EAL92363.1 as seed ortholog is 100%.
Bootstrap support for afis|EAW21346.1 as seed ortholog is 100%.

---

Group of orthologs #1639. Best score 1608 bits
Score difference with first non-orthologous sequence - *A. fumigatus*:1483 *A. fischeri*:1493
afum|EAL85499.2 100.00% afis|EAW20400.1 100.00%
Bootstrap support for afum|EAL85499.2 as seed ortholog is 100%.
Bootstrap support for afis|EAW20400.1 as seed ortholog is 100%.

---

Group of orthologs #1640. Best score 1608 bits
Score difference with first non-orthologous sequence - *A. fumigatus*:1608 *A. fischeri*:1608

afum|EAL92493.1 100.00% afis|EAW21474.1 100.00%
Bootstrap support for afum|EAL92493.1 as seed ortholog is 100%.
Bootstrap support for afis|EAW21474.1 as seed ortholog is 100%.

---

Group of orthologs #1641. Best score 1607 bits
Score difference with first non-orthologous sequence - *A. fumigatus*:1607 *A. fischeri*:1607
afum|EAL89686.1 100.00% afis|EAW25010.1 100.00%
Bootstrap support for afum|EAL89686.1 as seed ortholog is 100%.
Bootstrap support for afis|EAW25010.1 as seed ortholog is 100%.

---

Group of orthologs #1642. Best score 1606 bits
Score difference with first non-orthologous sequence - *A. fumigatus*:1606 *A. fischeri*:1606
afum|EAL88982.1 100.00% afis|EAW16357.1 100.00%
Bootstrap support for afum|EAL88982.1 as seed ortholog is 100%.
Bootstrap support for afis|EAW16357.1 as seed ortholog is 100%.

---

Group of orthologs #1643. Best score 1606 bits
Score difference with first non-orthologous sequence - *A. fumigatus*:1606 *A. fischeri*:1606
afum|EAL93696.1 100.00% afis|EAW18964.1 100.00%
Bootstrap support for afum|EAL93696.1 as seed ortholog is 100%.
Bootstrap support for afis|EAW18964.1 as seed ortholog is 100%.

---

Group of orthologs #1644. Best score 1606 bits
Score difference with first non-orthologous sequence - *A. fumigatus*:1606 *A. fischeri*:1606
afum|EAL91053.1 100.00% afis|EAW22137.1 100.00%
Bootstrap support for afum|EAL91053.1 as seed ortholog is 100%.
Bootstrap support for afis|EAW22137.1 as seed ortholog is 100%.

---

Group of orthologs #1645. Best score 1605 bits
Score difference with first non-orthologous sequence - *A. fumigatus*:1605 *A. fischeri*:1605
afum|EAL92470.1 100.00% afis|EAW21451.1 100.00%
Bootstrap support for afum|EAL92470.1 as seed ortholog is 100%.
Bootstrap support for afis|EAW21451.1 as seed ortholog is 100%.

---

Group of orthologs #1646. Best score 1604 bits
Score difference with first non-orthologous sequence - *A. fumigatus*:1509 *A. fischeri*:1171
afum|EAL85460.1 100.00% afis|EAW20343.1 100.00%
Bootstrap support for afum|EAL85460.1 as seed ortholog is 100%.
Bootstrap support for afis|EAW20343.1 as seed ortholog is 100%.

---

Group of orthologs #1647. Best score 1604 bits
Score difference with first non-orthologous sequence - *A. fumigatus*:1604 *A. fischeri*:1604
afum|EAL86963.1 100.00% afis|EAW19546.1 100.00%
Bootstrap support for afum|EAL86963.1 as seed ortholog is 100%.
Bootstrap support for afis|EAW19546.1 as seed ortholog is 100%.

---

Group of orthologs #1648. Best score 1604 bits
Score difference with first non-orthologous sequence - *A. fumigatus*:1604 *A. fischeri*:1604
afum|EAL93752.1 100.00% afis|EAW19021.1 100.00%
Bootstrap support for afum|EAL93752.1 as seed ortholog is 100%.
Bootstrap support for afis|EAW19021.1 as seed ortholog is 100%.

---

Group of orthologs #1649. Best score 1604 bits
Score difference with first non-orthologous sequence - *A. fumigatus*:1604 *A. fischeri*:1604
afum|EAL89969.1 100.00% afis|EAW25306.1 100.00%
Bootstrap support for afum|EAL89969.1 as seed ortholog is 100%.

Bootstrap support for afis|EAW25306.1 as seed ortholog is 100%.

---

Group of orthologs #1650. Best score 1603 bits
Score difference with first non-orthologous sequence - *A. fumigatus*:1547 *A. fischeri*:1039
afum|EAL89057.2 100.00% afis|EAW16461.1 100.00%
Bootstrap support for afum|EAL89057.2 as seed ortholog is 100%.
Bootstrap support for afis|EAW16461.1 as seed ortholog is 100%.

---

Group of orthologs #1651. Best score 1603 bits
Score difference with first non-orthologous sequence - *A. fumigatus*:1603 *A. fischeri*:1603
afum|EAL90220.2 100.00% afis|EAW22977.1 100.00%
Bootstrap support for afum|EAL90220.2 as seed ortholog is 100%.
Bootstrap support for afis|EAW22977.1 as seed ortholog is 100%.

---

Group of orthologs #1652. Best score 1603 bits
Score difference with first non-orthologous sequence - *A. fumigatus*:1603 *A. fischeri*:1603
afum|EAL92153.1 100.00% afis|EAW21134.1 100.00%
Bootstrap support for afum|EAL92153.1 as seed ortholog is 100%.
Bootstrap support for afis|EAW21134.1 as seed ortholog is 100%.

---

Group of orthologs #1653. Best score 1601 bits
Score difference with first non-orthologous sequence - *A. fumigatus*:1601 *A. fischeri*:1601
afum|EAL84883.1 100.00% afis|EAW20033.1 100.00%
Bootstrap support for afum|EAL84883.1 as seed ortholog is 100%.
Bootstrap support for afis|EAW20033.1 as seed ortholog is 100%.

---

Group of orthologs #1654. Best score 1601 bits
Score difference with first non-orthologous sequence - *A. fumigatus*:1601 *A. fischeri*:1601
afum|EAL87637.1 100.00% afis|EAW18272.1 100.00%
Bootstrap support for afum|EAL87637.1 as seed ortholog is 100%.
Bootstrap support for afis|EAW18272.1 as seed ortholog is 100%.

---

Group of orthologs #1655. Best score 1601 bits
Score difference with first non-orthologous sequence - *A. fumigatus*:769 *A. fischeri*:1601
afum|EAL86157.1 100.00% afis|EAW24488.1 100.00%
Bootstrap support for afum|EAL86157.1 as seed ortholog is 100%.
Bootstrap support for afis|EAW24488.1 as seed ortholog is 100%.

---

Group of orthologs #1656. Best score 1600 bits
Score difference with first non-orthologous sequence - *A. fumigatus*:1600 *A. fischeri*:1600
afum|EAL85786.1 100.00% afis|EAW15546.1 100.00%
Bootstrap support for afum|EAL85786.1 as seed ortholog is 100%.
Bootstrap support for afis|EAW15546.1 as seed ortholog is 100%.

---

Group of orthologs #1657. Best score 1600 bits
Score difference with first non-orthologous sequence - *A. fumigatus*:1529 *A. fischeri*:1600
afum|EAL91008.1 100.00% afis|EAW22181.1 100.00%
Bootstrap support for afum|EAL91008.1 as seed ortholog is 100%.
Bootstrap support for afis|EAW22181.1 as seed ortholog is 100%.

---

Group of orthologs #1658. Best score 1597 bits
Score difference with first non-orthologous sequence - *A. fumigatus*:1597 *A. fischeri*:1479
afum|EAL89271.2 100.00% afis|EAW16667.1 100.00%
Bootstrap support for afum|EAL89271.2 as seed ortholog is 100%.
Bootstrap support for afis|EAW16667.1 as seed ortholog is 100%.

---

Group of orthologs #1659. Best score 1597 bits
Score difference with first non-orthologous sequence - *A. fumigatus*:1597   *A. fischeri*:1597
afum|EAL89603.1 100.00% afis|EAW24946.1 100.00%
Bootstrap support for afum|EAL89603.1 as seed ortholog is 100%.
Bootstrap support for afis|EAW24946.1 as seed ortholog is 100%.

---

Group of orthologs #1660. Best score 1596 bits
Score difference with first non-orthologous sequence - *A. fumigatus*:1596   *A. fischeri*:1596
afum|EAL91761.1 100.00% afis|EAW17951.1 100.00%
Bootstrap support for afum|EAL91761.1 as seed ortholog is 100%.
Bootstrap support for afis|EAW17951.1 as seed ortholog is 100%.

---

Group of orthologs #1661. Best score 1596 bits
Score difference with first non-orthologous sequence - *A. fumigatus*:1596   *A. fischeri*:1596
afum|EAL88470.1 100.00% afis|EAW23011.1 100.00%
Bootstrap support for afum|EAL88470.1 as seed ortholog is 100%.
Bootstrap support for afis|EAW23011.1 as seed ortholog is 100%.

---

Group of orthologs #1662. Best score 1596 bits
Score difference with first non-orthologous sequence - *A. fumigatus*:1596   *A. fischeri*:1596
afum|EAL93617.1 100.00% afis|EAW18884.1 100.00%
Bootstrap support for afum|EAL93617.1 as seed ortholog is 100%.
Bootstrap support for afis|EAW18884.1 as seed ortholog is 100%.

---

Group of orthologs #1663. Best score 1596 bits
Score difference with first non-orthologous sequence - *A. fumigatus*:1596   *A. fischeri*:1341
afum|EAL93928.1 100.00% afis|EAW19196.1 100.00%
Bootstrap support for afum|EAL93928.1 as seed ortholog is 100%.
Bootstrap support for afis|EAW19196.1 as seed ortholog is 100%.

---

Group of orthologs #1664. Best score 1596 bits
Score difference with first non-orthologous sequence - *A. fumigatus*:1596   *A. fischeri*:1596
afum|EAL90509.1 100.00% afis|EAW22683.1 100.00%
Bootstrap support for afum|EAL90509.1 as seed ortholog is 100%.
Bootstrap support for afis|EAW22683.1 as seed ortholog is 100%.

---

Group of orthologs #1665. Best score 1593 bits
Score difference with first non-orthologous sequence - *A. fumigatus*:1593   *A. fischeri*:1593
afum|EAL93897.1 100.00% afis|EAW19167.1 100.00%
Bootstrap support for afum|EAL93897.1 as seed ortholog is 100%.
Bootstrap support for afis|EAW19167.1 as seed ortholog is 100%.

---

Group of orthologs #1666. Best score 1593 bits
Score difference with first non-orthologous sequence - *A. fumigatus*:1593   *A. fischeri*:1593
afum|EAL90720.1 100.00% afis|EAW22471.1 100.00%
Bootstrap support for afum|EAL90720.1 as seed ortholog is 100%.
Bootstrap support for afis|EAW22471.1 as seed ortholog is 100%.

---

Group of orthologs #1667. Best score 1592 bits
Score difference with first non-orthologous sequence - *A. fumigatus*:919   *A. fischeri*:1592
afum|EAL84865.1 100.00% afis|EAW20752.1 100.00%
Bootstrap support for afum|EAL84865.1 as seed ortholog is 100%.
Bootstrap support for afis|EAW20752.1 as seed ortholog is 100%.

---

Group of orthologs #1668. Best score 1591 bits
Score difference with first non-orthologous sequence - *A. fumigatus*:1591   *A. fischeri*:1591

afum|EAL91944.1 100.00% afis|EAW24089.1 100.00%
Bootstrap support for afum|EAL91944.1 as seed ortholog is 100%.
Bootstrap support for afis|EAW24089.1 as seed ortholog is 100%.

---

Group of orthologs #1669. Best score 1591 bits
Score difference with first non-orthologous sequence - *A. fumigatus*:1591 *A. fischeri*:1591
afum|EBA27343.1 100.00% afis|EAW23165.1 100.00%
Bootstrap support for afum|EBA27343.1 as seed ortholog is 100%.
Bootstrap support for afis|EAW23165.1 as seed ortholog is 100%.

---

Group of orthologs #1670. Best score 1590 bits
Score difference with first non-orthologous sequence - *A. fumigatus*:1590 *A. fischeri*:1590
afum|EAL88119.2 100.00% afis|EAW23357.1 100.00%
Bootstrap support for afum|EAL88119.2 as seed ortholog is 100%.
Bootstrap support for afis|EAW23357.1 as seed ortholog is 100%.

---

Group of orthologs #1671. Best score 1590 bits
Score difference with first non-orthologous sequence - *A. fumigatus*:1590 *A. fischeri*:1590
afum|EAL87995.1 100.00% afis|EAW23471.1 100.00%
Bootstrap support for afum|EAL87995.1 as seed ortholog is 100%.
Bootstrap support for afis|EAW23471.1 as seed ortholog is 100%.

---

Group of orthologs #1672. Best score 1590 bits
Score difference with first non-orthologous sequence - *A. fumigatus*:1590 *A. fischeri*:1590
afum|EAL93715.1 100.00% afis|EAW18979.1 100.00%
Bootstrap support for afum|EAL93715.1 as seed ortholog is 100%.
Bootstrap support for afis|EAW18979.1 as seed ortholog is 100%.

---

Group of orthologs #1673. Best score 1590 bits
Score difference with first non-orthologous sequence - *A. fumigatus*:1590 *A. fischeri*:1590
afum|EAL90768.2 100.00% afis|EAW22423.1 100.00%
Bootstrap support for afum|EAL90768.2 as seed ortholog is 100%.
Bootstrap support for afis|EAW22423.1 as seed ortholog is 100%.

---

Group of orthologs #1674. Best score 1589 bits
Score difference with first non-orthologous sequence - *A. fumigatus*:1589 *A. fischeri*:1589
afum|EAL87693.1 100.00% afis|EAW18327.1 100.00%
Bootstrap support for afum|EAL87693.1 as seed ortholog is 100%.
Bootstrap support for afis|EAW18327.1 as seed ortholog is 100%.

---

Group of orthologs #1675. Best score 1589 bits
Score difference with first non-orthologous sequence - *A. fumigatus*:1589 *A. fischeri*:1589
afum|EAL93950.1 100.00% afis|EAW19217.1 100.00%
Bootstrap support for afum|EAL93950.1 as seed ortholog is 100%.
Bootstrap support for afis|EAW19217.1 as seed ortholog is 100%.

---

Group of orthologs #1676. Best score 1589 bits
Score difference with first non-orthologous sequence - *A. fumigatus*:1589 *A. fischeri*:1589
afum|EAL90923.1 100.00% afis|EAW22255.1 100.00%
Bootstrap support for afum|EAL90923.1 as seed ortholog is 100%.
Bootstrap support for afis|EAW22255.1 as seed ortholog is 100%.

---

Group of orthologs #1677. Best score 1589 bits
Score difference with first non-orthologous sequence - *A. fumigatus*:1589 *A. fischeri*:1589
afum|EAL92011.1 100.00% afis|EAW24156.1 100.00%
Bootstrap support for afum|EAL92011.1 as seed ortholog is 100%.

Bootstrap support for afis|EAW24156.1 as seed ortholog is 100%.

---

Group of orthologs #1678. Best score 1588 bits
Score difference with first non-orthologous sequence - *A. fumigatus*:1588 *A. fischeri*:1588
afum|EAL89267.1 100.00% afis|EAW16663.1 100.00%
Bootstrap support for afum|EAL89267.1 as seed ortholog is 100%.
Bootstrap support for afis|EAW16663.1 as seed ortholog is 100%.

---

Group of orthologs #1679. Best score 1588 bits
Score difference with first non-orthologous sequence - *A. fumigatus*:1588 *A. fischeri*:1588
afum|EAL92877.2 100.00% afis|EAW21863.1 100.00%
Bootstrap support for afum|EAL92877.2 as seed ortholog is 100%.
Bootstrap support for afis|EAW21863.1 as seed ortholog is 100%.

---

Group of orthologs #1680. Best score 1587 bits
Score difference with first non-orthologous sequence - *A. fumigatus*:1538 *A. fischeri*:1125
afum|EAL86559.2 100.00% afis|EAW17071.1 100.00%
Bootstrap support for afum|EAL86559.2 as seed ortholog is 100%.
Bootstrap support for afis|EAW17071.1 as seed ortholog is 100%.

---

Group of orthologs #1681. Best score 1587 bits
Score difference with first non-orthologous sequence - *A. fumigatus*:1587 *A. fischeri*:1587
afum|EAL86244.1 100.00% afis|EAW24615.1 100.00%
Bootstrap support for afum|EAL86244.1 as seed ortholog is 100%.
Bootstrap support for afis|EAW24615.1 as seed ortholog is 100%.

---

Group of orthologs #1682. Best score 1587 bits
Score difference with first non-orthologous sequence - *A. fumigatus*:1587 *A. fischeri*:1587
afum|EAL88198.2 100.00% afis|EAW23281.1 100.00%
Bootstrap support for afum|EAL88198.2 as seed ortholog is 100%.
Bootstrap support for afis|EAW23281.1 as seed ortholog is 100%.

---

Group of orthologs #1683. Best score 1587 bits
Score difference with first non-orthologous sequence - *A. fumigatus*:1587 *A. fischeri*:1587
afum|EAL90230.1 100.00% afis|EAW22967.1 100.00%
Bootstrap support for afum|EAL90230.1 as seed ortholog is 100%.
Bootstrap support for afis|EAW22967.1 as seed ortholog is 100%.

---

Group of orthologs #1684. Best score 1587 bits
Score difference with first non-orthologous sequence - *A. fumigatus*:1587 *A. fischeri*:1587
afum|EAL92172.1 100.00% afis|EAW21154.1 100.00%
Bootstrap support for afum|EAL92172.1 as seed ortholog is 100%.
Bootstrap support for afis|EAW21154.1 as seed ortholog is 100%.

---

Group of orthologs #1685. Best score 1586 bits
Score difference with first non-orthologous sequence - *A. fumigatus*:1586 *A. fischeri*:1586
afum|EAL92194.1 100.00% afis|EAW21174.1 100.00%
Bootstrap support for afum|EAL92194.1 as seed ortholog is 100%.
Bootstrap support for afis|EAW21174.1 as seed ortholog is 100%.

---

Group of orthologs #1686. Best score 1585 bits
Score difference with first non-orthologous sequence - *A. fumigatus*:1585 *A. fischeri*:1585
afum|EAL85539.1 100.00% afis|EAW15812.1 100.00%
Bootstrap support for afum|EAL85539.1 as seed ortholog is 100%.
Bootstrap support for afis|EAW15812.1 as seed ortholog is 100%.

---

Group of orthologs #1687. Best score 1584 bits
Score difference with first non-orthologous sequence - *A. fumigatus*:1492 *A. fischeri*:1584
afum|EAL87628.1 100.00% afis|EAW18264.1 100.00%
Bootstrap support for afum|EAL87628.1 as seed ortholog is 100%.
Bootstrap support for afis|EAW18264.1 as seed ortholog is 100%.

---

Group of orthologs #1688. Best score 1584 bits
Score difference with first non-orthologous sequence - *A. fumigatus*:1584 *A. fischeri*:1584
afum|EAL90921.1 100.00% afis|EAW22259.1 100.00%
Bootstrap support for afum|EAL90921.1 as seed ortholog is 100%.
Bootstrap support for afis|EAW22259.1 as seed ortholog is 100%.

---

Group of orthologs #1689. Best score 1583 bits
Score difference with first non-orthologous sequence - *A. fumigatus*:1583 *A. fischeri*:1583
afum|EAL91388.1 100.00% afis|EAW17575.1 100.00%
Bootstrap support for afum|EAL91388.1 as seed ortholog is 100%.
Bootstrap support for afis|EAW17575.1 as seed ortholog is 100%.

---

Group of orthologs #1690. Best score 1583 bits
Score difference with first non-orthologous sequence - *A. fumigatus*:1583 *A. fischeri*:1583
afum|EAL93251.1 100.00% afis|EAW18551.1 100.00%
Bootstrap support for afum|EAL93251.1 as seed ortholog is 100%.
Bootstrap support for afis|EAW18551.1 as seed ortholog is 100%.

---

Group of orthologs #1691. Best score 1582 bits
Score difference with first non-orthologous sequence - *A. fumigatus*:1582 *A. fischeri*:1582
afum|EAL92794.1 100.00% afis|EAW21764.1 100.00%
Bootstrap support for afum|EAL92794.1 as seed ortholog is 100%.
Bootstrap support for afis|EAW21764.1 as seed ortholog is 100%.

---

Group of orthologs #1692. Best score 1582 bits
Score difference with first non-orthologous sequence - *A. fumigatus*:1582 *A. fischeri*:1582
afum|EAL92811.1 100.00% afis|EAW21784.1 100.00%
Bootstrap support for afum|EAL92811.1 as seed ortholog is 100%.
Bootstrap support for afis|EAW21784.1 as seed ortholog is 100%.

---

Group of orthologs #1693. Best score 1581 bits
Score difference with first non-orthologous sequence - *A. fumigatus*:1581 *A. fischeri*:1581
afum|EAL93500.2 100.00% afis|EAW18769.1 100.00%
Bootstrap support for afum|EAL93500.2 as seed ortholog is 100%.
Bootstrap support for afis|EAW18769.1 as seed ortholog is 100%.

---

Group of orthologs #1694. Best score 1581 bits
Score difference with first non-orthologous sequence - *A. fumigatus*:1581 *A. fischeri*:1581
afum|EAL90868.1 100.00% afis|EAW22324.1 100.00%
Bootstrap support for afum|EAL90868.1 as seed ortholog is 100%.
Bootstrap support for afis|EAW22324.1 as seed ortholog is 100%.

---

Group of orthologs #1695. Best score 1581 bits
Score difference with first non-orthologous sequence - *A. fumigatus*:1581 *A. fischeri*:1581
afum|EAL89788.1 100.00% afis|EAW25117.1 100.00%
Bootstrap support for afum|EAL89788.1 as seed ortholog is 100%.
Bootstrap support for afis|EAW25117.1 as seed ortholog is 100%.

---

Group of orthologs #1696. Best score 1581 bits
Score difference with first non-orthologous sequence - *A. fumigatus*:1581 *A. fischeri*:1581

afum|EAL90167.1 100.00% afis|EAW25501.1 100.00%
Bootstrap support for afum|EAL90167.1 as seed ortholog is 100%.
Bootstrap support for afis|EAW25501.1 as seed ortholog is 100%.

---

Group of orthologs #1697. Best score 1579 bits
Score difference with first non-orthologous sequence - *A. fumigatus*:1579 *A. fischeri*:1579
afum|EAL91865.1 100.00% afis|EAW18054.1 100.00%
Bootstrap support for afum|EAL91865.1 as seed ortholog is 100%.
Bootstrap support for afis|EAW18054.1 as seed ortholog is 100%.

---

Group of orthologs #1698. Best score 1579 bits
Score difference with first non-orthologous sequence - *A. fumigatus*:1579 *A. fischeri*:1579
afum|EAL88459.1 100.00% afis|EAW23023.1 100.00%
Bootstrap support for afum|EAL88459.1 as seed ortholog is 100%.
Bootstrap support for afis|EAW23023.1 as seed ortholog is 100%.

---

Group of orthologs #1699. Best score 1579 bits
Score difference with first non-orthologous sequence - *A. fumigatus*:1579 *A. fischeri*:1579
afum|EAL93495.1 100.00% afis|EAW18763.1 100.00%
Bootstrap support for afum|EAL93495.1 as seed ortholog is 100%.
Bootstrap support for afis|EAW18763.1 as seed ortholog is 100%.

---

Group of orthologs #1700. Best score 1579 bits
Score difference with first non-orthologous sequence - *A. fumigatus*:1579 *A. fischeri*:1579
afum|EAL89420.1 100.00% afis|EAW24781.1 100.00%
Bootstrap support for afum|EAL89420.1 as seed ortholog is 100%.
Bootstrap support for afis|EAW24781.1 as seed ortholog is 100%.

---

Group of orthologs #1701. Best score 1578 bits
Score difference with first non-orthologous sequence - *A. fumigatus*:1578 *A. fischeri*:1578
afum|EAL89154.1 100.00% afis|EAW16556.1 100.00%
Bootstrap support for afum|EAL89154.1 as seed ortholog is 100%.
Bootstrap support for afis|EAW16556.1 as seed ortholog is 100%.

---

Group of orthologs #1702. Best score 1578 bits
Score difference with first non-orthologous sequence - *A. fumigatus*:1366 *A. fischeri*:1316
afum|EBA27239.1 100.00% afis|EAW15597.1 100.00%
Bootstrap support for afum|EBA27239.1 as seed ortholog is 100%.
Bootstrap support for afis|EAW15597.1 as seed ortholog is 100%.

---

Group of orthologs #1703. Best score 1577 bits
Score difference with first non-orthologous sequence - *A. fumigatus*:1577 *A. fischeri*:1577
afum|EAL91527.1 100.00% afis|EAW17712.1 100.00%
Bootstrap support for afum|EAL91527.1 as seed ortholog is 100%.
Bootstrap support for afis|EAW17712.1 as seed ortholog is 100%.

---

Group of orthologs #1704. Best score 1577 bits
Score difference with first non-orthologous sequence - *A. fumigatus*:1577 *A. fischeri*:1577
afum|EAL87207.1 100.00% afis|EAW23773.1 100.00%
Bootstrap support for afum|EAL87207.1 as seed ortholog is 100%.
Bootstrap support for afis|EAW23773.1 as seed ortholog is 100%.

---

Group of orthologs #1705. Best score 1576 bits
Score difference with first non-orthologous sequence - *A. fumigatus*:1576 *A. fischeri*:1576
afum|EAL87556.1 100.00% afis|EAW18195.1 100.00%
Bootstrap support for afum|EAL87556.1 as seed ortholog is 100%.

Bootstrap support for afis|EAW18195.1 as seed ortholog is 100%.

---

Group of orthologs #1706. Best score 1576 bits
Score difference with first non-orthologous sequence - *A. fumigatus*:1576 *A. fischeri*:1576
afum|EAL93137.1 100.00% afis|EAW18452.1 100.00%
Bootstrap support for afum|EAL93137.1 as seed ortholog is 100%.
Bootstrap support for afis|EAW18452.1 as seed ortholog is 100%.

---

Group of orthologs #1707. Best score 1576 bits
Score difference with first non-orthologous sequence - *A. fumigatus*:1576 *A. fischeri*:1202
afum|EAL93814.1 100.00% afis|EAW19084.1 100.00%
Bootstrap support for afum|EAL93814.1 as seed ortholog is 100%.
Bootstrap support for afis|EAW19084.1 as seed ortholog is 100%.

---

Group of orthologs #1708. Best score 1576 bits
Score difference with first non-orthologous sequence - *A. fumigatus*:1576 *A. fischeri*:1576
afum|EAL92183.1 100.00% afis|EAW21164.1 100.00%
Bootstrap support for afum|EAL92183.1 as seed ortholog is 100%.
Bootstrap support for afis|EAW21164.1 as seed ortholog is 100%.

---

Group of orthologs #1709. Best score 1574 bits
Score difference with first non-orthologous sequence - *A. fumigatus*:1574 *A. fischeri*:1574
afum|EAL91449.1 100.00% afis|EAW17634.1 100.00%
Bootstrap support for afum|EAL91449.1 as seed ortholog is 100%.
Bootstrap support for afis|EAW17634.1 as seed ortholog is 100%.

---

Group of orthologs #1710. Best score 1574 bits
Score difference with first non-orthologous sequence - *A. fumigatus*:1574 *A. fischeri*:1574
afum|EAL88167.1 100.00% afis|EAW23310.1 100.00%
Bootstrap support for afum|EAL88167.1 as seed ortholog is 100%.
Bootstrap support for afis|EAW23310.1 as seed ortholog is 100%.

---

Group of orthologs #1711. Best score 1574 bits
Score difference with first non-orthologous sequence - *A. fumigatus*:1574 *A. fischeri*:1574
afum|EAL89746.1 100.00% afis|EAW25077.1 100.00%
Bootstrap support for afum|EAL89746.1 as seed ortholog is 100%.
Bootstrap support for afis|EAW25077.1 as seed ortholog is 100%.

---

Group of orthologs #1712. Best score 1573 bits
Score difference with first non-orthologous sequence - *A. fumigatus*:1573 *A. fischeri*:1573
afum|EAL86918.1 100.00% afis|EAW19600.1 100.00%
Bootstrap support for afum|EAL86918.1 as seed ortholog is 100%.
Bootstrap support for afis|EAW19600.1 as seed ortholog is 100%.

---

Group of orthologs #1713. Best score 1573 bits
Score difference with first non-orthologous sequence - *A. fumigatus*:1573 *A. fischeri*:1573
afum|EAL90760.1 100.00% afis|EAW22431.1 100.00%
Bootstrap support for afum|EAL90760.1 as seed ortholog is 100%.
Bootstrap support for afis|EAW22431.1 as seed ortholog is 100%.

---

Group of orthologs #1714. Best score 1572 bits
Score difference with first non-orthologous sequence - *A. fumigatus*:1572 *A. fischeri*:1572
afum|EAL88622.1 100.00% afis|EAW15990.1 100.00%
Bootstrap support for afum|EAL88622.1 as seed ortholog is 100%.
Bootstrap support for afis|EAW15990.1 as seed ortholog is 100%.

---

Group of orthologs #1715. Best score 1571 bits
Score difference with first non-orthologous sequence - *A. fumigatus*:1571 *A. fischeri*:1571
afum|EAL91793.1 100.00% afis|EAW17984.1 100.00%
Bootstrap support for afum|EAL91793.1 as seed ortholog is 100%.
Bootstrap support for afis|EAW17984.1 as seed ortholog is 100%.

---

Group of orthologs #1716. Best score 1571 bits
Score difference with first non-orthologous sequence - *A. fumigatus*:1571 *A. fischeri*:1571
afum|EAL93087.1 100.00% afis|EAW18407.1 100.00%
Bootstrap support for afum|EAL93087.1 as seed ortholog is 100%.
Bootstrap support for afis|EAW18407.1 as seed ortholog is 100%.

---

Group of orthologs #1717. Best score 1571 bits
Score difference with first non-orthologous sequence - *A. fumigatus*:1571 *A. fischeri*:1571
afum|EAL90359.1 100.00% afis|EAW22844.1 100.00%
Bootstrap support for afum|EAL90359.1 as seed ortholog is 100%.
Bootstrap support for afis|EAW22844.1 as seed ortholog is 100%.

---

Group of orthologs #1718. Best score 1569 bits
Score difference with first non-orthologous sequence - *A. fumigatus*:1569 *A. fischeri*:1569
afum|EAL91311.1 100.00% afis|EAW17489.1 100.00%
Bootstrap support for afum|EAL91311.1 as seed ortholog is 100%.
Bootstrap support for afis|EAW17489.1 as seed ortholog is 100%.

---

Group of orthologs #1719. Best score 1569 bits
Score difference with first non-orthologous sequence - *A. fumigatus*:1569 *A. fischeri*:1569
afum|EAL92196.1 100.00% afis|EAW21176.1 100.00%
Bootstrap support for afum|EAL92196.1 as seed ortholog is 100%.
Bootstrap support for afis|EAW21176.1 as seed ortholog is 100%.

---

Group of orthologs #1720. Best score 1569 bits
Score difference with first non-orthologous sequence - *A. fumigatus*:1569 *A. fischeri*:812
afum|EAL92370.2 100.00% afis|EAW21355.1 100.00%
Bootstrap support for afum|EAL92370.2 as seed ortholog is 100%.
Bootstrap support for afis|EAW21355.1 as seed ortholog is 100%.

---

Group of orthologs #1721. Best score 1569 bits
Score difference with first non-orthologous sequence - *A. fumigatus*:1569 *A. fischeri*:1569
afum|EAL90198.1 100.00% afis|EAW25534.1 100.00%
Bootstrap support for afum|EAL90198.1 as seed ortholog is 100%.
Bootstrap support for afis|EAW25534.1 as seed ortholog is 100%.

---

Group of orthologs #1722. Best score 1568 bits
Score difference with first non-orthologous sequence - *A. fumigatus*:1568 *A. fischeri*:1568
afum|EAL88389.1 100.00% afis|EAW23090.1 100.00%
Bootstrap support for afum|EAL88389.1 as seed ortholog is 100%.
Bootstrap support for afis|EAW23090.1 as seed ortholog is 100%.

---

Group of orthologs #1723. Best score 1568 bits
Score difference with first non-orthologous sequence - *A. fumigatus*:1568 *A. fischeri*:1568
afum|EAL89749.1 100.00% afis|EAW25080.1 100.00%
Bootstrap support for afum|EAL89749.1 as seed ortholog is 100%.
Bootstrap support for afis|EAW25080.1 as seed ortholog is 100%.

---

Group of orthologs #1724. Best score 1567 bits
Score difference with first non-orthologous sequence - *A. fumigatus*:1567 *A. fischeri*:1567

afum|EAL88661.1 100.00% afis|EAW16030.1 100.00%
Bootstrap support for afum|EAL88661.1 as seed ortholog is 100%.
Bootstrap support for afis|EAW16030.1 as seed ortholog is 100%.

---

Group of orthologs #1725. Best score 1567 bits
Score difference with first non-orthologous sequence - *A. fumigatus*:1567 *A. fischeri*:1567
afum|EAL84887.1 100.00% afis|EAW20030.1 100.00%
Bootstrap support for afum|EAL84887.1 as seed ortholog is 100%.
Bootstrap support for afis|EAW20030.1 as seed ortholog is 100%.

---

Group of orthologs #1726. Best score 1566 bits
Score difference with first non-orthologous sequence - *A. fumigatus*:1566 *A. fischeri*:1566
afum|EAL86048.1 100.00% afis|EAW24376.1 100.00%
Bootstrap support for afum|EAL86048.1 as seed ortholog is 100%.
Bootstrap support for afis|EAW24376.1 as seed ortholog is 100%.

---

Group of orthologs #1727. Best score 1566 bits
Score difference with first non-orthologous sequence - *A. fumigatus*:1566 *A. fischeri*:1566
afum|EAL93547.1 100.00% afis|EAW18811.1 100.00%
Bootstrap support for afum|EAL93547.1 as seed ortholog is 100%.
Bootstrap support for afis|EAW18811.1 as seed ortholog is 100%.

---

Group of orthologs #1728. Best score 1566 bits
Score difference with first non-orthologous sequence - *A. fumigatus*:1566 *A. fischeri*:1566
afum|EAL90266.1 100.00% afis|EAW22931.1 100.00%
Bootstrap support for afum|EAL90266.1 as seed ortholog is 100%.
Bootstrap support for afis|EAW22931.1 as seed ortholog is 100%.

---

Group of orthologs #1729. Best score 1565 bits
Score difference with first non-orthologous sequence - *A. fumigatus*:1565 *A. fischeri*:1565
afum|EAL85719.1 100.00% afis|EAW15616.1 100.00%
Bootstrap support for afum|EAL85719.1 as seed ortholog is 100%.
Bootstrap support for afis|EAW15616.1 as seed ortholog is 100%.

---

Group of orthologs #1730. Best score 1565 bits
Score difference with first non-orthologous sequence - *A. fumigatus*:1565 *A. fischeri*:1414
afum|EAL86674.1 100.00% afis|EAW17256.1 100.00%
Bootstrap support for afum|EAL86674.1 as seed ortholog is 100%.
Bootstrap support for afis|EAW17256.1 as seed ortholog is 100%.

---

Group of orthologs #1731. Best score 1565 bits
Score difference with first non-orthologous sequence - *A. fumigatus*:1565 *A. fischeri*:1565
afum|EAL91864.1 100.00% afis|EAW18053.1 100.00%
Bootstrap support for afum|EAL91864.1 as seed ortholog is 100%.
Bootstrap support for afis|EAW18053.1 as seed ortholog is 100%.

---

Group of orthologs #1732. Best score 1565 bits
Score difference with first non-orthologous sequence - *A. fumigatus*:997 *A. fischeri*:1565
afum|EAL89591.1 100.00% afis|EAW24934.1 100.00%
Bootstrap support for afum|EAL89591.1 as seed ortholog is 100%.
Bootstrap support for afis|EAW24934.1 as seed ortholog is 100%.

---

Group of orthologs #1733. Best score 1565 bits
Score difference with first non-orthologous sequence - *A. fumigatus*:1565 *A. fischeri*:1565
afum|EAL92002.1 100.00% afis|EAW24147.1 100.00%
Bootstrap support for afum|EAL92002.1 as seed ortholog is 100%.

Bootstrap support for afis|EAW24147.1 as seed ortholog is 100%.

---

Group of orthologs #1734. Best score 1564 bits
Score difference with first non-orthologous sequence - *A. fumigatus*:1564 *A. fischeri*:1494
afum|EAL85755.1 100.00% afis|EAW15579.1 100.00%
Bootstrap support for afum|EAL85755.1 as seed ortholog is 100%.
Bootstrap support for afis|EAW15579.1 as seed ortholog is 100%.

---

Group of orthologs #1735. Best score 1564 bits
Score difference with first non-orthologous sequence - *A. fumigatus*:1564 *A. fischeri*:1564
afum|EAL87027.1 100.00% afis|EAW19469.1 100.00%
Bootstrap support for afum|EAL87027.1 as seed ortholog is 100%.
Bootstrap support for afis|EAW19469.1 as seed ortholog is 100%.

---

Group of orthologs #1736. Best score 1564 bits
Score difference with first non-orthologous sequence - *A. fumigatus*:1564 *A. fischeri*:1564
afum|EAL89983.1 100.00% afis|EAW25322.1 100.00%
Bootstrap support for afum|EAL89983.1 as seed ortholog is 100%.
Bootstrap support for afis|EAW25322.1 as seed ortholog is 100%.

---

Group of orthologs #1737. Best score 1563 bits
Score difference with first non-orthologous sequence - *A. fumigatus*:1563 *A. fischeri*:1563
afum|EAL88222.1 100.00% afis|EAW23259.1 100.00%
Bootstrap support for afum|EAL88222.1 as seed ortholog is 100%.
Bootstrap support for afis|EAW23259.1 as seed ortholog is 100%.

---

Group of orthologs #1738. Best score 1563 bits
Score difference with first non-orthologous sequence - *A. fumigatus*:1563 *A. fischeri*:1563
afum|EAL90216.1 100.00% afis|EAW22981.1 100.00%
Bootstrap support for afum|EAL90216.1 as seed ortholog is 100%.
Bootstrap support for afis|EAW22981.1 as seed ortholog is 100%.

---

Group of orthologs #1739. Best score 1562 bits
Score difference with first non-orthologous sequence - *A. fumigatus*:1562 *A. fischeri*:1562
afum|EAL92614.1 100.00% afis|EAW21596.1 100.00%
Bootstrap support for afum|EAL92614.1 as seed ortholog is 100%.
Bootstrap support for afis|EAW21596.1 as seed ortholog is 100%.

---

Group of orthologs #1740. Best score 1561 bits
Score difference with first non-orthologous sequence - *A. fumigatus*:1561 *A. fischeri*:1561
afum|EAL88665.1 100.00% afis|EAW16034.1 100.00%
Bootstrap support for afum|EAL88665.1 as seed ortholog is 100%.
Bootstrap support for afis|EAW16034.1 as seed ortholog is 100%.

---

Group of orthologs #1741. Best score 1561 bits
Score difference with first non-orthologous sequence - *A. fumigatus*:695 *A. fischeri*:1561
afum|EAL91515.2 100.00% afis|EAW17700.1 100.00%
Bootstrap support for afum|EAL91515.2 as seed ortholog is 100%.
Bootstrap support for afis|EAW17700.1 as seed ortholog is 100%.

---

Group of orthologs #1742. Best score 1559 bits
Score difference with first non-orthologous sequence - *A. fumigatus*:1559 *A. fischeri*:1279
afum|EAL86889.1 100.00% afis|EAW19633.1 100.00%
Bootstrap support for afum|EAL86889.1 as seed ortholog is 100%.
Bootstrap support for afis|EAW19633.1 as seed ortholog is 100%.

---

Group of orthologs #1743. Best score 1558 bits
Score difference with first non-orthologous sequence - *A. fumigatus*:1558 *A. fischeri*:1558
afum|EAL90261.1 100.00% afis|EAW22935.1 100.00%
Bootstrap support for afum|EAL90261.1 as seed ortholog is 100%.
Bootstrap support for afis|EAW22935.1 as seed ortholog is 100%.

---

Group of orthologs #1744. Best score 1558 bits
Score difference with first non-orthologous sequence - *A. fumigatus*:1558 *A. fischeri*:1558
afum|EAL92209.1 100.00% afis|EAW21187.1 100.00%
Bootstrap support for afum|EAL92209.1 as seed ortholog is 100%.
Bootstrap support for afis|EAW21187.1 as seed ortholog is 100%.

---

Group of orthologs #1745. Best score 1557 bits
Score difference with first non-orthologous sequence - *A. fumigatus*:1557 *A. fischeri*:1557
afum|EAL87079.1 100.00% afis|EAW19415.1 100.00%
Bootstrap support for afum|EAL87079.1 as seed ortholog is 100%.
Bootstrap support for afis|EAW19415.1 as seed ortholog is 100%.

---

Group of orthologs #1746. Best score 1557 bits
Score difference with first non-orthologous sequence - *A. fumigatus*:1557 *A. fischeri*:1557
afum|EAL90223.1 100.00% afis|EAW22974.1 100.00%
Bootstrap support for afum|EAL90223.1 as seed ortholog is 100%.
Bootstrap support for afis|EAW22974.1 as seed ortholog is 100%.

---

Group of orthologs #1747. Best score 1555 bits
Score difference with first non-orthologous sequence - *A. fumigatus*:1555 *A. fischeri*:1555
afum|EAL90372.2 100.00% afis|EAW22830.1 100.00%
Bootstrap support for afum|EAL90372.2 as seed ortholog is 100%.
Bootstrap support for afis|EAW22830.1 as seed ortholog is 100%.

---

Group of orthologs #1748. Best score 1554 bits
Score difference with first non-orthologous sequence - *A. fumigatus*:1554 *A. fischeri*:1554
afum|EAL88582.1 100.00% afis|EAW15949.1 100.00%
Bootstrap support for afum|EAL88582.1 as seed ortholog is 100%.
Bootstrap support for afis|EAW15949.1 as seed ortholog is 100%.

---

Group of orthologs #1749. Best score 1554 bits
Score difference with first non-orthologous sequence - *A. fumigatus*:1554 *A. fischeri*:1554
afum|EAL90831.1 100.00% afis|EAW22360.1 100.00%
Bootstrap support for afum|EAL90831.1 as seed ortholog is 100%.
Bootstrap support for afis|EAW22360.1 as seed ortholog is 100%.

---

Group of orthologs #1750. Best score 1554 bits
Score difference with first non-orthologous sequence - *A. fumigatus*:1554 *A. fischeri*:1554
afum|EAL90068.1 100.00% afis|EAW25409.1 100.00%
Bootstrap support for afum|EAL90068.1 as seed ortholog is 100%.
Bootstrap support for afis|EAW25409.1 as seed ortholog is 100%.

---

Group of orthologs #1751. Best score 1553 bits
Score difference with first non-orthologous sequence - *A. fumigatus*:1553 *A. fischeri*:1553
afum|EAL85193.2 100.00% afis|EAW20075.1 100.00%
Bootstrap support for afum|EAL85193.2 as seed ortholog is 100%.
Bootstrap support for afis|EAW20075.1 as seed ortholog is 100%.

---

Group of orthologs #1752. Best score 1553 bits
Score difference with first non-orthologous sequence - *A. fumigatus*:1553 *A. fischeri*:1553

afum|EAL89124.1 100.00% afis|EAW16528.1 100.00%
Bootstrap support for afum|EAL89124.1 as seed ortholog is 100%.
Bootstrap support for afis|EAW16528.1 as seed ortholog is 100%.

---

Group of orthologs #1753. Best score 1552 bits
Score difference with first non-orthologous sequence - *A. fumigatus*:1552 *A. fischeri*:1480
afum|EAL86598.1 100.00% afis|EAW17161.1 100.00%
Bootstrap support for afum|EAL86598.1 as seed ortholog is 100%.
Bootstrap support for afis|EAW17161.1 as seed ortholog is 100%.

---

Group of orthologs #1754. Best score 1552 bits
Score difference with first non-orthologous sequence - *A. fumigatus*:1552 *A. fischeri*:1552
afum|EAL85066.1 100.00% afis|EAW19826.1 100.00%
Bootstrap support for afum|EAL85066.1 as seed ortholog is 100%.
Bootstrap support for afis|EAW19826.1 as seed ortholog is 100%.

---

Group of orthologs #1755. Best score 1552 bits
Score difference with first non-orthologous sequence - *A. fumigatus*:1552 *A. fischeri*:1552
afum|EAL88892.1 100.00% afis|EAW16271.1 100.00%
Bootstrap support for afum|EAL88892.1 as seed ortholog is 100%.
Bootstrap support for afis|EAW16271.1 as seed ortholog is 100%.

---

Group of orthologs #1756. Best score 1552 bits
Score difference with first non-orthologous sequence - *A. fumigatus*:1552 *A. fischeri*:1552
afum|EAL87233.1 100.00% afis|EAW23798.1 100.00%
Bootstrap support for afum|EAL87233.1 as seed ortholog is 100%.
Bootstrap support for afis|EAW23798.1 as seed ortholog is 100%.

---

Group of orthologs #1757. Best score 1552 bits
Score difference with first non-orthologous sequence - *A. fumigatus*:1552 *A. fischeri*:1552
afum|EAL88066.1 100.00% afis|EAW23404.1 100.00%
Bootstrap support for afum|EAL88066.1 as seed ortholog is 100%.
Bootstrap support for afis|EAW23404.1 as seed ortholog is 100%.

---

Group of orthologs #1758. Best score 1552 bits
Score difference with first non-orthologous sequence - *A. fumigatus*:1552 *A. fischeri*:1552
afum|EAL93357.1 100.00% afis|EAW18643.1 100.00%
Bootstrap support for afum|EAL93357.1 as seed ortholog is 100%.
Bootstrap support for afis|EAW18643.1 as seed ortholog is 100%.

---

Group of orthologs #1759. Best score 1552 bits
Score difference with first non-orthologous sequence - *A. fumigatus*:1552 *A. fischeri*:1552
afum|EAL90580.1 100.00% afis|EAW22617.1 100.00%
Bootstrap support for afum|EAL90580.1 as seed ortholog is 100%.
Bootstrap support for afis|EAW22617.1 as seed ortholog is 100%.

---

Group of orthologs #1760. Best score 1552 bits
Score difference with first non-orthologous sequence - *A. fumigatus*:1552 *A. fischeri*:1552
afum|EAL90609.1 100.00% afis|EAW22590.1 100.00%
Bootstrap support for afum|EAL90609.1 as seed ortholog is 100%.
Bootstrap support for afis|EAW22590.1 as seed ortholog is 100%.

---

Group of orthologs #1761. Best score 1550 bits
Score difference with first non-orthologous sequence - *A. fumigatus*:1550 *A. fischeri*:1550
afum|EAL85456.1 100.00% afis|EAW20329.1 100.00%
Bootstrap support for afum|EAL85456.1 as seed ortholog is 100%.

Bootstrap support for afis|EAW20329.1 as seed ortholog is 100%.

---

Group of orthologs #1762. Best score 1549 bits
Score difference with first non-orthologous sequence - *A. fumigatus*:1549 *A. fischeri*:1549
afum|EAL91768.1 100.00% afis|EAW17957.1 100.00%
Bootstrap support for afum|EAL91768.1 as seed ortholog is 100%.
Bootstrap support for afis|EAW17957.1 as seed ortholog is 100%.

---

Group of orthologs #1763. Best score 1549 bits
Score difference with first non-orthologous sequence - *A. fumigatus*:1549 *A. fischeri*:1549
afum|EAL91835.1 100.00% afis|EAW18023.1 100.00%
Bootstrap support for afum|EAL91835.1 as seed ortholog is 100%.
Bootstrap support for afis|EAW18023.1 as seed ortholog is 100%.

---

Group of orthologs #1764. Best score 1549 bits
Score difference with first non-orthologous sequence - *A. fumigatus*:1549 *A. fischeri*:1549
afum|EAL93579.1 100.00% afis|EAW18847.1 100.00%
Bootstrap support for afum|EAL93579.1 as seed ortholog is 100%.
Bootstrap support for afis|EAW18847.1 as seed ortholog is 100%.

---

Group of orthologs #1765. Best score 1547 bits
Score difference with first non-orthologous sequence - *A. fumigatus*:1547 *A. fischeri*:1547
afum|EAL89261.1 100.00% afis|EAW16657.1 100.00%
Bootstrap support for afum|EAL89261.1 as seed ortholog is 100%.
Bootstrap support for afis|EAW16657.1 as seed ortholog is 100%.

---

Group of orthologs #1766. Best score 1547 bits
Score difference with first non-orthologous sequence - *A. fumigatus*:1547 *A. fischeri*:1547
afum|EAL91699.1 100.00% afis|EAW17890.1 100.00%
Bootstrap support for afum|EAL91699.1 as seed ortholog is 100%.
Bootstrap support for afis|EAW17890.1 as seed ortholog is 100%.

---

Group of orthologs #1767. Best score 1547 bits
Score difference with first non-orthologous sequence - *A. fumigatus*:1272 *A. fischeri*:1140
afum|EAL90698.1 100.00% afis|EAW22495.1 100.00%
Bootstrap support for afum|EAL90698.1 as seed ortholog is 100%.
Bootstrap support for afis|EAW22495.1 as seed ortholog is 100%.

---

Group of orthologs #1768. Best score 1546 bits
Score difference with first non-orthologous sequence - *A. fumigatus*:1546 *A. fischeri*:1546
afum|EAL88768.1 100.00% afis|EAW16139.1 100.00%
Bootstrap support for afum|EAL88768.1 as seed ortholog is 100%.
Bootstrap support for afis|EAW16139.1 as seed ortholog is 100%.

---

Group of orthologs #1769. Best score 1546 bits
Score difference with first non-orthologous sequence - *A. fumigatus*:1546 *A. fischeri*:1546
afum|EAL86088.1 100.00% afis|EAW24419.1 100.00%
Bootstrap support for afum|EAL86088.1 as seed ortholog is 100%.
Bootstrap support for afis|EAW24419.1 as seed ortholog is 100%.

---

Group of orthologs #1770. Best score 1546 bits
Score difference with first non-orthologous sequence - *A. fumigatus*:1546 *A. fischeri*:1546
afum|EBA27500.1 100.00% afis|EAW18925.1 100.00%
Bootstrap support for afum|EBA27500.1 as seed ortholog is 100%.
Bootstrap support for afis|EAW18925.1 as seed ortholog is 100%.

---

Group of orthologs #1771. Best score 1545 bits
Score difference with first non-orthologous sequence - *A. fumigatus*:1545 *A. fischeri*:1545
afum|EAL88640.1 100.00% afis|EAW16010.1 100.00%
Bootstrap support for afum|EAL88640.1 as seed ortholog is 100%.
Bootstrap support for afis|EAW16010.1 as seed ortholog is 100%.

---

Group of orthologs #1772. Best score 1545 bits
Score difference with first non-orthologous sequence - *A. fumigatus*:1545 *A. fischeri*:1545
afum|EAL88013.1 100.00% afis|EAW23453.1 100.00%
Bootstrap support for afum|EAL88013.1 as seed ortholog is 100%.
Bootstrap support for afis|EAW23453.1 as seed ortholog is 100%.

---

Group of orthologs #1773. Best score 1545 bits
Score difference with first non-orthologous sequence - *A. fumigatus*:1545 *A. fischeri*:1545
afum|EAL93385.1 100.00% afis|EAW18667.1 100.00%
Bootstrap support for afum|EAL93385.1 as seed ortholog is 100%.
Bootstrap support for afis|EAW18667.1 as seed ortholog is 100%.

---

Group of orthologs #1774. Best score 1545 bits
Score difference with first non-orthologous sequence - *A. fumigatus*:1545 *A. fischeri*:1545
afum|EAL92673.1 100.00% afis|EAW21639.1 100.00%
Bootstrap support for afum|EAL92673.1 as seed ortholog is 100%.
Bootstrap support for afis|EAW21639.1 as seed ortholog is 100%.

---

Group of orthologs #1775. Best score 1544 bits
Score difference with first non-orthologous sequence - *A. fumigatus*:1337 *A. fischeri*:1384
afum|EAL91095.1 100.00% afis|EAW22075.1 100.00%
Bootstrap support for afum|EAL91095.1 as seed ortholog is 100%.
Bootstrap support for afis|EAW22075.1 as seed ortholog is 100%.

---

Group of orthologs #1776. Best score 1543 bits
Score difference with first non-orthologous sequence - *A. fumigatus*:1543 *A. fischeri*:1543
afum|EAL92274.1 100.00% afis|EAW21259.1 100.00%
Bootstrap support for afum|EAL92274.1 as seed ortholog is 100%.
Bootstrap support for afis|EAW21259.1 as seed ortholog is 100%.

---

Group of orthologs #1777. Best score 1542 bits
Score difference with first non-orthologous sequence - *A. fumigatus*:1542 *A. fischeri*:1542
afum|EAL85920.2 100.00% afis|EAW24251.1 100.00%
Bootstrap support for afum|EAL85920.2 as seed ortholog is 100%.
Bootstrap support for afis|EAW24251.1 as seed ortholog is 100%.

---

Group of orthologs #1778. Best score 1541 bits
Score difference with first non-orthologous sequence - *A. fumigatus*:1391 *A. fischeri*:1541
afum|EAL84369.1 100.00% afis|EAW25695.1 100.00%
Bootstrap support for afum|EAL84369.1 as seed ortholog is 100%.
Bootstrap support for afis|EAW25695.1 as seed ortholog is 100%.

---

Group of orthologs #1779. Best score 1540 bits
Score difference with first non-orthologous sequence - *A. fumigatus*:1540 *A. fischeri*:1540
afum|EAL90521.1 100.00% afis|EAW22671.1 100.00%
Bootstrap support for afum|EAL90521.1 as seed ortholog is 100%.
Bootstrap support for afis|EAW22671.1 as seed ortholog is 100%.

---

Group of orthologs #1780. Best score 1540 bits
Score difference with first non-orthologous sequence - *A. fumigatus*:1540 *A. fischeri*:1540

afum|EAL89622.1 100.00% afis|EAW24964.1 100.00%
Bootstrap support for afum|EAL89622.1 as seed ortholog is 100%.
Bootstrap support for afis|EAW24964.1 as seed ortholog is 100%.

---

Group of orthologs #1781. Best score 1539 bits
Score difference with first non-orthologous sequence - *A. fumigatus*:1539 *A. fischeri*:1539
afum|EAL85783.1 100.00% afis|EAW15549.1 100.00%
Bootstrap support for afum|EAL85783.1 as seed ortholog is 100%.
Bootstrap support for afis|EAW15549.1 as seed ortholog is 100%.

---

Group of orthologs #1782. Best score 1539 bits
Score difference with first non-orthologous sequence - *A. fumigatus*:1539 *A. fischeri*:1539
afum|EAL89104.2 100.00% afis|EAW16509.1 100.00%
Bootstrap support for afum|EAL89104.2 as seed ortholog is 100%.
Bootstrap support for afis|EAW16509.1 as seed ortholog is 100%.

---

Group of orthologs #1783. Best score 1539 bits
Score difference with first non-orthologous sequence - *A. fumigatus*:1539 *A. fischeri*:1539
afum|EAL91824.1 100.00% afis|EAW18012.1 100.00%
Bootstrap support for afum|EAL91824.1 as seed ortholog is 100%.
Bootstrap support for afis|EAW18012.1 as seed ortholog is 100%.

---

Group of orthologs #1784. Best score 1539 bits
Score difference with first non-orthologous sequence - *A. fumigatus*:1539 *A. fischeri*:1539
afum|EAL93040.1 100.00% afis|EAW18366.1 100.00%
Bootstrap support for afum|EAL93040.1 as seed ortholog is 100%.
Bootstrap support for afis|EAW18366.1 as seed ortholog is 100%.

---

Group of orthologs #1785. Best score 1538 bits
Score difference with first non-orthologous sequence - *A. fumigatus*:1313 *A. fischeri*:1208
afum|EAL93849.1 100.00% afis|EAW19117.1 100.00%
Bootstrap support for afum|EAL93849.1 as seed ortholog is 100%.
Bootstrap support for afis|EAW19117.1 as seed ortholog is 100%.

---

Group of orthologs #1786. Best score 1537 bits
Score difference with first non-orthologous sequence - *A. fumigatus*:1537 *A. fischeri*:1537
afum|EAL85601.1 100.00% afis|EAW15748.1 100.00%
Bootstrap support for afum|EAL85601.1 as seed ortholog is 100%.
Bootstrap support for afis|EAW15748.1 as seed ortholog is 100%.

---

Group of orthologs #1787. Best score 1537 bits
Score difference with first non-orthologous sequence - *A. fumigatus*:1537 *A. fischeri*:1537
afum|EAL88379.1 100.00% afis|EAW23099.1 100.00%
Bootstrap support for afum|EAL88379.1 as seed ortholog is 100%.
Bootstrap support for afis|EAW23099.1 as seed ortholog is 100%.

---

Group of orthologs #1788. Best score 1537 bits
Score difference with first non-orthologous sequence - *A. fumigatus*:1537 *A. fischeri*:1537
afum|EAL91085.1 100.00% afis|EAW22106.1 100.00%
Bootstrap support for afum|EAL91085.1 as seed ortholog is 100%.
Bootstrap support for afis|EAW22106.1 as seed ortholog is 100%.

---

Group of orthologs #1789. Best score 1536 bits
Score difference with first non-orthologous sequence - *A. fumigatus*:1536 *A. fischeri*:1536
afum|EAL85065.1 100.00% afis|EAW19827.1 100.00%
Bootstrap support for afum|EAL85065.1 as seed ortholog is 100%.

Bootstrap support for afis|EAW19827.1 as seed ortholog is 100%.

---

Group of orthologs #1790. Best score 1536 bits
Score difference with first non-orthologous sequence - *A. fumigatus*:1536 *A. fischeri*:1536
afum|EAL93456.1 100.00% afis|EAW18731.1 100.00%
Bootstrap support for afum|EAL93456.1 as seed ortholog is 100%.
Bootstrap support for afis|EAW18731.1 as seed ortholog is 100%.

---

Group of orthologs #1791. Best score 1536 bits
Score difference with first non-orthologous sequence - *A. fumigatus*:1536 *A. fischeri*:1536
afum|EAL92413.1 100.00% afis|EAW21399.1 100.00%
Bootstrap support for afum|EAL92413.1 as seed ortholog is 100%.
Bootstrap support for afis|EAW21399.1 as seed ortholog is 100%.

---

Group of orthologs #1792. Best score 1535 bits
Score difference with first non-orthologous sequence - *A. fumigatus*:1535 *A. fischeri*:1535
afum|EAL86686.1 100.00% afis|EAW17269.1 100.00%
Bootstrap support for afum|EAL86686.1 as seed ortholog is 100%.
Bootstrap support for afis|EAW17269.1 as seed ortholog is 100%.

---

Group of orthologs #1793. Best score 1535 bits
Score difference with first non-orthologous sequence - *A. fumigatus*:1535 *A. fischeri*:1535
afum|EAL88876.1 100.00% afis|EAW16248.1 100.00%
Bootstrap support for afum|EAL88876.1 as seed ortholog is 100%.
Bootstrap support for afis|EAW16248.1 as seed ortholog is 100%.

---

Group of orthologs #1794. Best score 1535 bits
Score difference with first non-orthologous sequence - *A. fumigatus*:1535 *A. fischeri*:1535
afum|EAL91948.1 100.00% afis|EAW24093.1 100.00%
Bootstrap support for afum|EAL91948.1 as seed ortholog is 100%.
Bootstrap support for afis|EAW24093.1 as seed ortholog is 100%.

---

Group of orthologs #1795. Best score 1534 bits
Score difference with first non-orthologous sequence - *A. fumigatus*:1534 *A. fischeri*:1534
afum|EAL91867.1 100.00% afis|EAW18056.1 100.00%
Bootstrap support for afum|EAL91867.1 as seed ortholog is 100%.
Bootstrap support for afis|EAW18056.1 as seed ortholog is 100%.

---

Group of orthologs #1796. Best score 1534 bits
Score difference with first non-orthologous sequence - *A. fumigatus*:1534 *A. fischeri*:1534
afum|EAL93290.1 100.00% afis|EAW18586.1 100.00%
Bootstrap support for afum|EAL93290.1 as seed ortholog is 100%.
Bootstrap support for afis|EAW18586.1 as seed ortholog is 100%.

---

Group of orthologs #1797. Best score 1534 bits
Score difference with first non-orthologous sequence - *A. fumigatus*:1534 *A. fischeri*:1534
afum|EAL93698.2 100.00% afis|EAW18966.1 100.00%
Bootstrap support for afum|EAL93698.2 as seed ortholog is 100%.
Bootstrap support for afis|EAW18966.1 as seed ortholog is 100%.

---

Group of orthologs #1798. Best score 1534 bits
Score difference with first non-orthologous sequence - *A. fumigatus*:1534 *A. fischeri*:1534
afum|EAL92509.1 100.00% afis|EAW21490.1 100.00%
Bootstrap support for afum|EAL92509.1 as seed ortholog is 100%.
Bootstrap support for afis|EAW21490.1 as seed ortholog is 100%.

---

Group of orthologs #1799. Best score 1533 bits
Score difference with first non-orthologous sequence - *A. fumigatus*:1533 *A. fischeri*:1533
afum|EAL85628.1 100.00% afis|EAW15720.1 100.00%
Bootstrap support for afum|EAL85628.1 as seed ortholog is 100%.
Bootstrap support for afis|EAW15720.1 as seed ortholog is 100%.

---

Group of orthologs #1800. Best score 1533 bits
Score difference with first non-orthologous sequence - *A. fumigatus*:1533 *A. fischeri*:1533
afum|EAL86016.1 100.00% afis|EAW24346.1 100.00%
Bootstrap support for afum|EAL86016.1 as seed ortholog is 100%.
Bootstrap support for afis|EAW24346.1 as seed ortholog is 100%.

---

Group of orthologs #1801. Best score 1533 bits
Score difference with first non-orthologous sequence - *A. fumigatus*:1533 *A. fischeri*:1533
afum|EAL88275.1 100.00% afis|EAW23209.1 100.00%
Bootstrap support for afum|EAL88275.1 as seed ortholog is 100%.
Bootstrap support for afis|EAW23209.1 as seed ortholog is 100%.

---

Group of orthologs #1802. Best score 1531 bits
Score difference with first non-orthologous sequence - *A. fumigatus*:1531 *A. fischeri*:1531
afum|EAL93670.1 100.00% afis|EAW18937.1 100.00%
Bootstrap support for afum|EAL93670.1 as seed ortholog is 100%.
Bootstrap support for afis|EAW18937.1 as seed ortholog is 100%.

---

Group of orthologs #1803. Best score 1531 bits
Score difference with first non-orthologous sequence - *A. fumigatus*:1531 *A. fischeri*:1531
afum|EAL92045.1 100.00% afis|EAW24185.1 100.00%
Bootstrap support for afum|EAL92045.1 as seed ortholog is 100%.
Bootstrap support for afis|EAW24185.1 as seed ortholog is 100%.

---

Group of orthologs #1804. Best score 1530 bits
Score difference with first non-orthologous sequence - *A. fumigatus*:1530 *A. fischeri*:1530
afum|EAL90627.1 100.00% afis|EAW22571.1 100.00%
Bootstrap support for afum|EAL90627.1 as seed ortholog is 100%.
Bootstrap support for afis|EAW22571.1 as seed ortholog is 100%.

---

Group of orthologs #1805. Best score 1530 bits
Score difference with first non-orthologous sequence - *A. fumigatus*:1530 *A. fischeri*:1530
afum|EAL92514.1 100.00% afis|EAW21495.1 100.00%
Bootstrap support for afum|EAL92514.1 as seed ortholog is 100%.
Bootstrap support for afis|EAW21495.1 as seed ortholog is 100%.

---

Group of orthologs #1806. Best score 1529 bits
Score difference with first non-orthologous sequence - *A. fumigatus*:1529 *A. fischeri*:1529
afum|EAL84580.2 100.00% afis|EAW20472.1 100.00%
Bootstrap support for afum|EAL84580.2 as seed ortholog is 100%.
Bootstrap support for afis|EAW20472.1 as seed ortholog is 100%.

---

Group of orthologs #1807. Best score 1526 bits
Score difference with first non-orthologous sequence - *A. fumigatus*:1526 *A. fischeri*:1526
afum|EAL92307.1 100.00% afis|EAW21290.1 100.00%
Bootstrap support for afum|EAL92307.1 as seed ortholog is 100%.
Bootstrap support for afis|EAW21290.1 as seed ortholog is 100%.

---

Group of orthologs #1808. Best score 1526 bits
Score difference with first non-orthologous sequence - *A. fumigatus*:1526 *A. fischeri*:1526

afum|EAL91996.1 100.00% afis|EAW24141.1 100.00%
Bootstrap support for afum|EAL91996.1 as seed ortholog is 100%.
Bootstrap support for afis|EAW24141.1 as seed ortholog is 100%.

---

Group of orthologs #1809. Best score 1525 bits
Score difference with first non-orthologous sequence - *A. fumigatus*:1525 *A. fischeri*:1437
afum|EAL86271.1 100.00% afis|EAW20695.1 100.00%
Bootstrap support for afum|EAL86271.1 as seed ortholog is 100%.
Bootstrap support for afis|EAW20695.1 as seed ortholog is 100%.

---

Group of orthologs #1810. Best score 1525 bits
Score difference with first non-orthologous sequence - *A. fumigatus*:1525 *A. fischeri*:1525
afum|EBA27442.1 100.00% afis|EAW18058.1 100.00%
Bootstrap support for afum|EBA27442.1 as seed ortholog is 100%.
Bootstrap support for afis|EAW18058.1 as seed ortholog is 100%.

---

Group of orthologs #1811. Best score 1525 bits
Score difference with first non-orthologous sequence - *A. fumigatus*:1525 *A. fischeri*:1525
afum|EBA27184.1 100.00% afis|EAW20554.1 100.00%
Bootstrap support for afum|EBA27184.1 as seed ortholog is 100%.
Bootstrap support for afis|EAW20554.1 as seed ortholog is 100%.

---

Group of orthologs #1812. Best score 1525 bits
Score difference with first non-orthologous sequence - *A. fumigatus*:1525 *A. fischeri*:1525
afum|EAL89721.1 100.00% afis|EAW25053.1 100.00%
Bootstrap support for afum|EAL89721.1 as seed ortholog is 100%.
Bootstrap support for afis|EAW25053.1 as seed ortholog is 100%.

---

Group of orthologs #1813. Best score 1525 bits
Score difference with first non-orthologous sequence - *A. fumigatus*:1525 *A. fischeri*:1525
afum|EAL91968.1 100.00% afis|EAW24114.1 100.00%
Bootstrap support for afum|EAL91968.1 as seed ortholog is 100%.
Bootstrap support for afis|EAW24114.1 as seed ortholog is 100%.

---

Group of orthologs #1814. Best score 1524 bits
Score difference with first non-orthologous sequence - *A. fumigatus*:1524 *A. fischeri*:1524
afum|EAL93455.2 100.00% afis|EAW18730.1 100.00%
Bootstrap support for afum|EAL93455.2 as seed ortholog is 100%.
Bootstrap support for afis|EAW18730.1 as seed ortholog is 100%.

---

Group of orthologs #1815. Best score 1524 bits
Score difference with first non-orthologous sequence - *A. fumigatus*:1524 *A. fischeri*:1524
afum|EAL93711.2 100.00% afis|EAW18976.1 100.00%
Bootstrap support for afum|EAL93711.2 as seed ortholog is 100%.
Bootstrap support for afis|EAW18976.1 as seed ortholog is 100%.

---

Group of orthologs #1816. Best score 1524 bits
Score difference with first non-orthologous sequence - *A. fumigatus*:1524 *A. fischeri*:1524
afum|EAL92725.1 100.00% afis|EAW21691.1 100.00%
Bootstrap support for afum|EAL92725.1 as seed ortholog is 100%.
Bootstrap support for afis|EAW21691.1 as seed ortholog is 100%.

---

Group of orthologs #1817. Best score 1524 bits
Score difference with first non-orthologous sequence - *A. fumigatus*:1524 *A. fischeri*:1524
afum|EAL92782.1 100.00% afis|EAW21752.1 100.00%
Bootstrap support for afum|EAL92782.1 as seed ortholog is 100%.

Bootstrap support for afis|EAW21752.1 as seed ortholog is 100%.

---

Group of orthologs #1818. Best score 1524 bits
Score difference with first non-orthologous sequence - *A. fumigatus*:1524 *A. fischeri*:1524
afum|EAL92972.1 100.00% afis|EAW21958.1 100.00%
Bootstrap support for afum|EAL92972.1 as seed ortholog is 100%.
Bootstrap support for afis|EAW21958.1 as seed ortholog is 100%.

---

Group of orthologs #1819. Best score 1523 bits
Score difference with first non-orthologous sequence - *A. fumigatus*:1523 *A. fischeri*:1523
afum|EAL87713.1 100.00% afis|EAW18350.1 100.00%
Bootstrap support for afum|EAL87713.1 as seed ortholog is 100%.
Bootstrap support for afis|EAW18350.1 as seed ortholog is 100%.

---

Group of orthologs #1820. Best score 1523 bits
Score difference with first non-orthologous sequence - *A. fumigatus*:1523 *A. fischeri*:1523
afum|EAL92242.1 100.00% afis|EAW21230.1 100.00%
Bootstrap support for afum|EAL92242.1 as seed ortholog is 100%.
Bootstrap support for afis|EAW21230.1 as seed ortholog is 100%.

---

Group of orthologs #1821. Best score 1522 bits
Score difference with first non-orthologous sequence - *A. fumigatus*:1522 *A. fischeri*:1522
afum|EAL85881.1 100.00% afis|EAW24212.1 100.00%
Bootstrap support for afum|EAL85881.1 as seed ortholog is 100%.
Bootstrap support for afis|EAW24212.1 as seed ortholog is 100%.

---

Group of orthologs #1822. Best score 1522 bits
Score difference with first non-orthologous sequence - *A. fumigatus*:1522 *A. fischeri*:1522
afum|EAL93084.1 100.00% afis|EAW18404.1 100.00%
Bootstrap support for afum|EAL93084.1 as seed ortholog is 100%.
Bootstrap support for afis|EAW18404.1 as seed ortholog is 100%.

---

Group of orthologs #1823. Best score 1522 bits
Score difference with first non-orthologous sequence - *A. fumigatus*:1522 *A. fischeri*:1522
afum|EAL93812.1 100.00% afis|EAW19083.1 100.00%
Bootstrap support for afum|EAL93812.1 as seed ortholog is 100%.
Bootstrap support for afis|EAW19083.1 as seed ortholog is 100%.

---

Group of orthologs #1824. Best score 1521 bits
Score difference with first non-orthologous sequence - *A. fumigatus*:1521 *A. fischeri*:1521
afum|EAL88296.1 100.00% afis|EAW23187.1 100.00%
Bootstrap support for afum|EAL88296.1 as seed ortholog is 100%.
Bootstrap support for afis|EAW23187.1 as seed ortholog is 100%.

---

Group of orthologs #1825. Best score 1521 bits
Score difference with first non-orthologous sequence - *A. fumigatus*:1521 *A. fischeri*:1521
afum|EAL92925.1 100.00% afis|EAW21913.1 100.00%
Bootstrap support for afum|EAL92925.1 as seed ortholog is 100%.
Bootstrap support for afis|EAW21913.1 as seed ortholog is 100%.

---

Group of orthologs #1826. Best score 1520 bits
Score difference with first non-orthologous sequence - *A. fumigatus*:1520 *A. fischeri*:1520
afum|EAL90476.1 100.00% afis|EAW22718.1 100.00%
Bootstrap support for afum|EAL90476.1 as seed ortholog is 100%.
Bootstrap support for afis|EAW22718.1 as seed ortholog is 100%.

---

Group of orthologs #1827. Best score 1519 bits
Score difference with first non-orthologous sequence - *A. fumigatus*:1519 *A. fischeri*:1519
afum|EAL85310.1 100.00% afis|EAW20191.1 100.00%
Bootstrap support for afum|EAL85310.1 as seed ortholog is 100%.
Bootstrap support for afis|EAW20191.1 as seed ortholog is 100%.

---

Group of orthologs #1828. Best score 1518 bits
Score difference with first non-orthologous sequence - *A. fumigatus*:1518 *A. fischeri*:1518
afum|EAL85654.1 100.00% afis|EAW15694.1 100.00%
Bootstrap support for afum|EAL85654.1 as seed ortholog is 100%.
Bootstrap support for afis|EAW15694.1 as seed ortholog is 100%.

---

Group of orthologs #1829. Best score 1518 bits
Score difference with first non-orthologous sequence - *A. fumigatus*:1518 *A. fischeri*:1518
afum|EAL89265.1 100.00% afis|EAW16661.1 100.00%
Bootstrap support for afum|EAL89265.1 as seed ortholog is 100%.
Bootstrap support for afis|EAW16661.1 as seed ortholog is 100%.

---

Group of orthologs #1830. Best score 1518 bits
Score difference with first non-orthologous sequence - *A. fumigatus*:1518 *A. fischeri*:1518
afum|EAL87257.1 100.00% afis|EAW23824.1 100.00%
Bootstrap support for afum|EAL87257.1 as seed ortholog is 100%.
Bootstrap support for afis|EAW23824.1 as seed ortholog is 100%.

---

Group of orthologs #1831. Best score 1518 bits
Score difference with first non-orthologous sequence - *A. fumigatus*:1518 *A. fischeri*:1368
afum|EAL93929.2 100.00% afis|EAW19197.1 100.00%
Bootstrap support for afum|EAL93929.2 as seed ortholog is 100%.
Bootstrap support for afis|EAW19197.1 as seed ortholog is 100%.

---

Group of orthologs #1832. Best score 1518 bits
Score difference with first non-orthologous sequence - *A. fumigatus*:1518 *A. fischeri*:1518
afum|EAL93986.1 100.00% afis|EAW19259.1 100.00%
Bootstrap support for afum|EAL93986.1 as seed ortholog is 100%.
Bootstrap support for afis|EAW19259.1 as seed ortholog is 100%.

---

Group of orthologs #1833. Best score 1518 bits
Score difference with first non-orthologous sequence - *A. fumigatus*:1518 *A. fischeri*:1167
afum|EAL90124.1 100.00% afis|EAW25461.1 100.00%
Bootstrap support for afum|EAL90124.1 as seed ortholog is 100%.
Bootstrap support for afis|EAW25461.1 as seed ortholog is 100%.

---

Group of orthologs #1834. Best score 1516 bits
Score difference with first non-orthologous sequence - *A. fumigatus*:1516 *A. fischeri*:1516
afum|EAL91322.1 100.00% afis|EAW17499.1 100.00%
Bootstrap support for afum|EAL91322.1 as seed ortholog is 100%.
Bootstrap support for afis|EAW17499.1 as seed ortholog is 100%.

---

Group of orthologs #1835. Best score 1516 bits
Score difference with first non-orthologous sequence - *A. fumigatus*:1516 *A. fischeri*:1516
afum|EAL93425.1 100.00% afis|EAW18702.1 100.00%
Bootstrap support for afum|EAL93425.1 as seed ortholog is 100%.
Bootstrap support for afis|EAW18702.1 as seed ortholog is 100%.

---

Group of orthologs #1836. Best score 1516 bits
Score difference with first non-orthologous sequence - *A. fumigatus*:1516 *A. fischeri*:1516

afum|EAL90078.1 100.00% afis|EAW25419.1 100.00%
Bootstrap support for afum|EAL90078.1 as seed ortholog is 100%.
Bootstrap support for afis|EAW25419.1 as seed ortholog is 100%.

---

Group of orthologs #1837. Best score 1514 bits
Score difference with first non-orthologous sequence - *A. fumigatus*:1514 *A. fischeri*:1514
afum|EAL88452.1 100.00% afis|EAW23030.1 100.00%
Bootstrap support for afum|EAL88452.1 as seed ortholog is 100%.
Bootstrap support for afis|EAW23030.1 as seed ortholog is 100%.

---

Group of orthologs #1838. Best score 1513 bits
Score difference with first non-orthologous sequence - *A. fumigatus*:1513 *A. fischeri*:1513
afum|EAL91460.1 100.00% afis|EAW17645.1 100.00%
Bootstrap support for afum|EAL91460.1 as seed ortholog is 100%.
Bootstrap support for afis|EAW17645.1 as seed ortholog is 100%.

---

Group of orthologs #1839. Best score 1513 bits
Score difference with first non-orthologous sequence - *A. fumigatus*:1513 *A. fischeri*:1513
afum|EAL89623.1 100.00% afis|EAW24965.1 100.00%
Bootstrap support for afum|EAL89623.1 as seed ortholog is 100%.
Bootstrap support for afis|EAW24965.1 as seed ortholog is 100%.

---

Group of orthologs #1840. Best score 1512 bits
Score difference with first non-orthologous sequence - *A. fumigatus*:1512 *A. fischeri*:1512
afum|EAL88960.1 100.00% afis|EAW16335.1 100.00%
Bootstrap support for afum|EAL88960.1 as seed ortholog is 100%.
Bootstrap support for afis|EAW16335.1 as seed ortholog is 100%.

---

Group of orthologs #1841. Best score 1512 bits
Score difference with first non-orthologous sequence - *A. fumigatus*:1512 *A. fischeri*:1512
afum|EAL87568.1 100.00% afis|EAW18203.1 100.00%
Bootstrap support for afum|EAL87568.1 as seed ortholog is 100%.
Bootstrap support for afis|EAW18203.1 as seed ortholog is 100%.

---

Group of orthologs #1842. Best score 1512 bits
Score difference with first non-orthologous sequence - *A. fumigatus*:1512 *A. fischeri*:1512
afum|EAL90443.1 100.00% afis|EAW22756.1 100.00%
Bootstrap support for afum|EAL90443.1 as seed ortholog is 100%.
Bootstrap support for afis|EAW22756.1 as seed ortholog is 100%.

---

Group of orthologs #1843. Best score 1512 bits
Score difference with first non-orthologous sequence - *A. fumigatus*:1512 *A. fischeri*:1512
afum|EAL92349.1 100.00% afis|EAW21331.1 100.00%
Bootstrap support for afum|EAL92349.1 as seed ortholog is 100%.
Bootstrap support for afis|EAW21331.1 as seed ortholog is 100%.

---

Group of orthologs #1844. Best score 1512 bits
Score difference with first non-orthologous sequence - *A. fumigatus*:1512 *A. fischeri*:1512
afum|EAL92599.2 100.00% afis|EAW21581.1 100.00%
Bootstrap support for afum|EAL92599.2 as seed ortholog is 100%.
Bootstrap support for afis|EAW21581.1 as seed ortholog is 100%.

---

Group of orthologs #1845. Best score 1511 bits
Score difference with first non-orthologous sequence - *A. fumigatus*:1511 *A. fischeri*:1511
afum|EAL85526.1 100.00% afis|EAW15825.1 100.00%
Bootstrap support for afum|EAL85526.1 as seed ortholog is 100%.

Bootstrap support for afis|EAW15825.1 as seed ortholog is 100%.

---

Group of orthologs #1846. Best score 1511 bits
Score difference with first non-orthologous sequence - *A. fumigatus*:1511 *A. fischeri*:1511
afum|EAL91288.1 100.00% afis|EAW17460.1 100.00%
Bootstrap support for afum|EAL91288.1 as seed ortholog is 100%.
Bootstrap support for afis|EAW17460.1 as seed ortholog is 100%.

---

Group of orthologs #1847. Best score 1511 bits
Score difference with first non-orthologous sequence - *A. fumigatus*:1511 *A. fischeri*:1511
afum|EAL89778.1 100.00% afis|EAW25107.1 100.00%
Bootstrap support for afum|EAL89778.1 as seed ortholog is 100%.
Bootstrap support for afis|EAW25107.1 as seed ortholog is 100%.

---

Group of orthologs #1848. Best score 1510 bits
Score difference with first non-orthologous sequence - *A. fumigatus*:1510 *A. fischeri*:1510
afum|EAL92358.1 100.00% afis|EAW21341.1 100.00%
Bootstrap support for afum|EAL92358.1 as seed ortholog is 100%.
Bootstrap support for afis|EAW21341.1 as seed ortholog is 100%.

---

Group of orthologs #1849. Best score 1509 bits
Score difference with first non-orthologous sequence - *A. fumigatus*:1509 *A. fischeri*:1509
afum|EAL85665.1 100.00% afis|EAW15683.1 100.00%
Bootstrap support for afum|EAL85665.1 as seed ortholog is 100%.
Bootstrap support for afis|EAW15683.1 as seed ortholog is 100%.

---

Group of orthologs #1850. Best score 1509 bits
Score difference with first non-orthologous sequence - *A. fumigatus*:1509 *A. fischeri*:1509
afum|EAL85211.1 100.00% afis|EAW20092.1 100.00%
Bootstrap support for afum|EAL85211.1 as seed ortholog is 100%.
Bootstrap support for afis|EAW20092.1 as seed ortholog is 100%.

---

Group of orthologs #1851. Best score 1509 bits
Score difference with first non-orthologous sequence - *A. fumigatus*:1509 *A. fischeri*:1509
afum|EAL89268.1 100.00% afis|EAW16664.1 100.00%
Bootstrap support for afum|EAL89268.1 as seed ortholog is 100%.
Bootstrap support for afis|EAW16664.1 as seed ortholog is 100%.

---

Group of orthologs #1852. Best score 1508 bits
Score difference with first non-orthologous sequence - *A. fumigatus*:1508 *A. fischeri*:1508
afum|EAL85617.1 100.00% afis|EAW15730.1 100.00%
Bootstrap support for afum|EAL85617.1 as seed ortholog is 100%.
Bootstrap support for afis|EAW15730.1 as seed ortholog is 100%.

---

Group of orthologs #1853. Best score 1508 bits
Score difference with first non-orthologous sequence - *A. fumigatus*:1508 *A. fischeri*:1508
afum|EAL87446.1 100.00% afis|EAW24016.1 100.00%
Bootstrap support for afum|EAL87446.1 as seed ortholog is 100%.
Bootstrap support for afis|EAW24016.1 as seed ortholog is 100%.

---

Group of orthologs #1854. Best score 1507 bits
Score difference with first non-orthologous sequence - *A. fumigatus*:1507 *A. fischeri*:1507
afum|EAL88128.1 100.00% afis|EAW23348.1 100.00%
Bootstrap support for afum|EAL88128.1 as seed ortholog is 100%.
Bootstrap support for afis|EAW23348.1 as seed ortholog is 100%.

---

Group of orthologs #1855. Best score 1507 bits
Score difference with first non-orthologous sequence - *A. fumigatus*:1507   *A. fischeri*:1507
afum|EAL93431.1    100.00%    afis|EAW18708.1    100.00%
Bootstrap support for afum|EAL93431.1 as seed ortholog is 100%.
Bootstrap support for afis|EAW18708.1 as seed ortholog is 100%.

---

Group of orthologs #1856. Best score 1507 bits
Score difference with first non-orthologous sequence - *A. fumigatus*:1507   *A. fischeri*:1507
afum|EBA27488.1    100.00%    afis|EAW18606.1    100.00%
Bootstrap support for afum|EBA27488.1 as seed ortholog is 100%.
Bootstrap support for afis|EAW18606.1 as seed ortholog is 100%.

---

Group of orthologs #1857. Best score 1507 bits
Score difference with first non-orthologous sequence - *A. fumigatus*:1507   *A. fischeri*:1507
afum|EAL92675.1    100.00%    afis|EAW21641.1    100.00%
Bootstrap support for afum|EAL92675.1 as seed ortholog is 100%.
Bootstrap support for afis|EAW21641.1 as seed ortholog is 100%.

---

Group of orthologs #1858. Best score 1506 bits
Score difference with first non-orthologous sequence - *A. fumigatus*:1347   *A. fischeri*:1415
afum|EAL89054.1    100.00%    afis|EAW16458.1    100.00%
Bootstrap support for afum|EAL89054.1 as seed ortholog is 100%.
Bootstrap support for afis|EAW16458.1 as seed ortholog is 100%.

---

Group of orthologs #1859. Best score 1506 bits
Score difference with first non-orthologous sequence - *A. fumigatus*:1506   *A. fischeri*:1506
afum|EAL87887.1    100.00%    afis|EAW23578.1    100.00%
Bootstrap support for afum|EAL87887.1 as seed ortholog is 100%.
Bootstrap support for afis|EAW23578.1 as seed ortholog is 100%.

---

Group of orthologs #1860. Best score 1506 bits
Score difference with first non-orthologous sequence - *A. fumigatus*:1506   *A. fischeri*:1506
afum|EAL93754.1    100.00%    afis|EAW19023.1    100.00%
Bootstrap support for afum|EAL93754.1 as seed ortholog is 100%.
Bootstrap support for afis|EAW19023.1 as seed ortholog is 100%.

---

Group of orthologs #1861. Best score 1506 bits
Score difference with first non-orthologous sequence - *A. fumigatus*:1506   *A. fischeri*:1506
afum|EAL92959.1    100.00%    afis|EAW21945.1    100.00%
Bootstrap support for afum|EAL92959.1 as seed ortholog is 100%.
Bootstrap support for afis|EAW21945.1 as seed ortholog is 100%.

---

Group of orthologs #1862. Best score 1506 bits
Score difference with first non-orthologous sequence - *A. fumigatus*:1506   *A. fischeri*:1506
afum|EAL89914.1    100.00%    afis|EAW25252.1    100.00%
Bootstrap support for afum|EAL89914.1 as seed ortholog is 100%.
Bootstrap support for afis|EAW25252.1 as seed ortholog is 100%.

---

Group of orthologs #1863. Best score 1505 bits
Score difference with first non-orthologous sequence - *A. fumigatus*:1505   *A. fischeri*:1505
afum|EAL90589.1    100.00%    afis|EAW22609.1    100.00%
Bootstrap support for afum|EAL90589.1 as seed ortholog is 100%.
Bootstrap support for afis|EAW22609.1 as seed ortholog is 100%.

---

Group of orthologs #1864. Best score 1505 bits
Score difference with first non-orthologous sequence - *A. fumigatus*:1505   *A. fischeri*:1505

afum|EAL92341.1 100.00% afis|EAW21324.1 100.00%
Bootstrap support for afum|EAL92341.1 as seed ortholog is 100%.
Bootstrap support for afis|EAW21324.1 as seed ortholog is 100%.

---

Group of orthologs #1865. Best score 1504 bits
Score difference with first non-orthologous sequence - *A. fumigatus*:1345 *A. fischeri*:1327
afum|EAL85662.1 100.00% afis|EAW15686.1 100.00%
Bootstrap support for afum|EAL85662.1 as seed ortholog is 100%.
Bootstrap support for afis|EAW15686.1 as seed ortholog is 100%.

---

Group of orthologs #1866. Best score 1503 bits
Score difference with first non-orthologous sequence - *A. fumigatus*:1503 *A. fischeri*:1503
afum|EAL91715.1 100.00% afis|EAW17906.1 100.00%
Bootstrap support for afum|EAL91715.1 as seed ortholog is 100%.
Bootstrap support for afis|EAW17906.1 as seed ortholog is 100%.

---

Group of orthologs #1867. Best score 1503 bits
Score difference with first non-orthologous sequence - *A. fumigatus*:1503 *A. fischeri*:1503
afum|EAL93215.1 100.00% afis|EAW18520.1 100.00%
Bootstrap support for afum|EAL93215.1 as seed ortholog is 100%.
Bootstrap support for afis|EAW18520.1 as seed ortholog is 100%.

---

Group of orthologs #1868. Best score 1502 bits
Score difference with first non-orthologous sequence - *A. fumigatus*:1502 *A. fischeri*:1502
afum|EAL86569.1 100.00% afis|EAW17092.1 100.00%
Bootstrap support for afum|EAL86569.1 as seed ortholog is 100%.
Bootstrap support for afis|EAW17092.1 as seed ortholog is 100%.

---

Group of orthologs #1869. Best score 1502 bits
Score difference with first non-orthologous sequence - *A. fumigatus*:1502 *A. fischeri*:1502
afum|EAL85317.1 100.00% afis|EAW20197.1 100.00%
Bootstrap support for afum|EAL85317.1 as seed ortholog is 100%.
Bootstrap support for afis|EAW20197.1 as seed ortholog is 100%.

---

Group of orthologs #1870. Best score 1502 bits
Score difference with first non-orthologous sequence - *A. fumigatus*:1502 *A. fischeri*:1502
afum|EAL87597.1 100.00% afis|EAW18230.1 100.00%
Bootstrap support for afum|EAL87597.1 as seed ortholog is 100%.
Bootstrap support for afis|EAW18230.1 as seed ortholog is 100%.

---

Group of orthologs #1871. Best score 1502 bits
Score difference with first non-orthologous sequence - *A. fumigatus*:1502 *A. fischeri*:1502
afum|EAL85468.1 100.00% afis|EAW20350.1 100.00%
Bootstrap support for afum|EAL85468.1 as seed ortholog is 100%.
Bootstrap support for afis|EAW20350.1 as seed ortholog is 100%.

---

Group of orthologs #1872. Best score 1502 bits
Score difference with first non-orthologous sequence - *A. fumigatus*:1502 *A. fischeri*:1502
afum|EAL89671.1 100.00% afis|EAW25024.1 100.00%
Bootstrap support for afum|EAL89671.1 as seed ortholog is 100%.
Bootstrap support for afis|EAW25024.1 as seed ortholog is 100%.

---

Group of orthologs #1873. Best score 1502 bits
Score difference with first non-orthologous sequence - *A. fumigatus*:1502 *A. fischeri*:1502
afum|EBA27454.1 100.00% afis|EAW21222.1 100.00%
Bootstrap support for afum|EBA27454.1 as seed ortholog is 100%.

Bootstrap support for afis|EAW21222.1 as seed ortholog is 100%.

---

Group of orthologs #1874. Best score 1501 bits
Score difference with first non-orthologous sequence - *A. fumigatus*:1501 *A. fischeri*:1424
afum|EAL85676.2 100.00% afis|EAW15671.1 100.00%
Bootstrap support for afum|EAL85676.2 as seed ortholog is 100%.
Bootstrap support for afis|EAW15671.1 as seed ortholog is 100%.

---

Group of orthologs #1875. Best score 1501 bits
Score difference with first non-orthologous sequence - *A. fumigatus*:1501 *A. fischeri*:1501
afum|EAL91765.1 100.00% afis|EAW17954.1 100.00%
Bootstrap support for afum|EAL91765.1 as seed ortholog is 100%.
Bootstrap support for afis|EAW17954.1 as seed ortholog is 100%.

---

Group of orthologs #1876. Best score 1501 bits
Score difference with first non-orthologous sequence - *A. fumigatus*:1501 *A. fischeri*:1501
afum|EAL92256.1 100.00% afis|EAW21243.1 100.00%
Bootstrap support for afum|EAL92256.1 as seed ortholog is 100%.
Bootstrap support for afis|EAW21243.1 as seed ortholog is 100%.

---

Group of orthologs #1877. Best score 1501 bits
Score difference with first non-orthologous sequence - *A. fumigatus*:1501 *A. fischeri*:1501
afum|EAL89468.1 100.00% afis|EAW24824.1 100.00%
Bootstrap support for afum|EAL89468.1 as seed ortholog is 100%.
Bootstrap support for afis|EAW24824.1 as seed ortholog is 100%.

---

Group of orthologs #1878. Best score 1500 bits
Score difference with first non-orthologous sequence - *A. fumigatus*:1500 *A. fischeri*:1500
afum|EAL86702.1 100.00% afis|EAW17284.1 100.00%
Bootstrap support for afum|EAL86702.1 as seed ortholog is 100%.
Bootstrap support for afis|EAW17284.1 as seed ortholog is 100%.

---

Group of orthologs #1879. Best score 1500 bits
Score difference with first non-orthologous sequence - *A. fumigatus*:1500 *A. fischeri*:1500
afum|EAL88760.1 100.00% afis|EAW16130.1 100.00%
Bootstrap support for afum|EAL88760.1 as seed ortholog is 100%.
Bootstrap support for afis|EAW16130.1 as seed ortholog is 100%.

---

Group of orthologs #1880. Best score 1500 bits
Score difference with first non-orthologous sequence - *A. fumigatus*:1435 *A. fischeri*:1500
afum|EAL88852.1 100.00% afis|EAW16229.1 100.00%
Bootstrap support for afum|EAL88852.1 as seed ortholog is 100%.
Bootstrap support for afis|EAW16229.1 as seed ortholog is 100%.

---

Group of orthologs #1881. Best score 1500 bits
Score difference with first non-orthologous sequence - *A. fumigatus*:1500 *A. fischeri*:1500
afum|EAL87252.1 100.00% afis|EAW23817.1 100.00%
Bootstrap support for afum|EAL87252.1 as seed ortholog is 100%.
Bootstrap support for afis|EAW23817.1 as seed ortholog is 100%.

---

Group of orthologs #1882. Best score 1499 bits
Score difference with first non-orthologous sequence - *A. fumigatus*:1073 *A. fischeri*:1499
afum|EAL91863.1 100.00% afis|EAW18052.1 100.00%
Bootstrap support for afum|EAL91863.1 as seed ortholog is 100%.
Bootstrap support for afis|EAW18052.1 as seed ortholog is 100%.

---

Group of orthologs #1883. Best score 1499 bits
Score difference with first non-orthologous sequence - *A. fumigatus*:1499 *A. fischeri*:1499
afum|EAL87304.1 100.00% afis|EAW23876.1 100.00%
Bootstrap support for afum|EAL87304.1 as seed ortholog is 100%.
Bootstrap support for afis|EAW23876.1 as seed ortholog is 100%.

---

Group of orthologs #1884. Best score 1498 bits
Score difference with first non-orthologous sequence - *A. fumigatus*:1498 *A. fischeri*:1498
afum|EAL86692.1 100.00% afis|EAW17274.1 100.00%
Bootstrap support for afum|EAL86692.1 as seed ortholog is 100%.
Bootstrap support for afis|EAW17274.1 as seed ortholog is 100%.

---

Group of orthologs #1885. Best score 1496 bits
Score difference with first non-orthologous sequence - *A. fumigatus*:1496 *A. fischeri*:1496
afum|EAL87689.2 100.00% afis|EAW18322.1 100.00%
Bootstrap support for afum|EAL87689.2 as seed ortholog is 100%.
Bootstrap support for afis|EAW18322.1 as seed ortholog is 100%.

---

Group of orthologs #1886. Best score 1496 bits
Score difference with first non-orthologous sequence - *A. fumigatus*:1496 *A. fischeri*:1496
afum|EBA27367.1 100.00% afis|EAW16497.1 100.00%
Bootstrap support for afum|EBA27367.1 as seed ortholog is 100%.
Bootstrap support for afis|EAW16497.1 as seed ortholog is 100%.

---

Group of orthologs #1887. Best score 1496 bits
Score difference with first non-orthologous sequence - *A. fumigatus*:1496 *A. fischeri*:1496
afum|EAL93912.1 100.00% afis|EAW19182.1 100.00%
Bootstrap support for afum|EAL93912.1 as seed ortholog is 100%.
Bootstrap support for afis|EAW19182.1 as seed ortholog is 100%.

---

Group of orthologs #1888. Best score 1496 bits
Score difference with first non-orthologous sequence - *A. fumigatus*:1496 *A. fischeri*:1315
afum|EAL94048.2 100.00% afis|EAW19331.1 100.00%
Bootstrap support for afum|EAL94048.2 as seed ortholog is 100%.
Bootstrap support for afis|EAW19331.1 as seed ortholog is 100%.

---

Group of orthologs #1889. Best score 1496 bits
Score difference with first non-orthologous sequence - *A. fumigatus*:1496 *A. fischeri*:1496
afum|EAL90803.1 100.00% afis|EAW22387.1 100.00%
Bootstrap support for afum|EAL90803.1 as seed ortholog is 100%.
Bootstrap support for afis|EAW22387.1 as seed ortholog is 100%.

---

Group of orthologs #1890. Best score 1495 bits
Score difference with first non-orthologous sequence - *A. fumigatus*:1495 *A. fischeri*:1495
afum|EAL84250.1 100.00% afis|EAW15398.1 100.00%
Bootstrap support for afum|EAL84250.1 as seed ortholog is 100%.
Bootstrap support for afis|EAW15398.1 as seed ortholog is 100%.

---

Group of orthologs #1891. Best score 1494 bits
Score difference with first non-orthologous sequence - *A. fumigatus*:1494 *A. fischeri*:1494
afum|EAL92957.1 100.00% afis|EAW21943.1 100.00%
Bootstrap support for afum|EAL92957.1 as seed ortholog is 100%.
Bootstrap support for afis|EAW21943.1 as seed ortholog is 100%.

---

Group of orthologs #1892. Best score 1493 bits
Score difference with first non-orthologous sequence - *A. fumigatus*:1493 *A. fischeri*:1493

afum|EAL85223.1 100.00% afis|EAW20106.1 100.00%
Bootstrap support for afum|EAL85223.1 as seed ortholog is 100%.
Bootstrap support for afis|EAW20106.1 as seed ortholog is 100%.

---

Group of orthologs #1893. Best score 1493 bits
Score difference with first non-orthologous sequence - *A. fumigatus*:1493 *A. fischeri*:1493
afum|EAL89213.1 100.00% afis|EAW16624.1 100.00%
Bootstrap support for afum|EAL89213.1 as seed ortholog is 100%.
Bootstrap support for afis|EAW16624.1 as seed ortholog is 100%.

---

Group of orthologs #1894. Best score 1493 bits
Score difference with first non-orthologous sequence - *A. fumigatus*:1493 *A. fischeri*:1493
afum|EAL90804.1 100.00% afis|EAW22386.1 100.00%
Bootstrap support for afum|EAL90804.1 as seed ortholog is 100%.
Bootstrap support for afis|EAW22386.1 as seed ortholog is 100%.

---

Group of orthologs #1895. Best score 1492 bits
Score difference with first non-orthologous sequence - *A. fumigatus*:1492 *A. fischeri*:1492
afum|EAL87139.1 100.00% afis|EAW23695.1 100.00%
Bootstrap support for afum|EAL87139.1 as seed ortholog is 100%.
Bootstrap support for afis|EAW23695.1 as seed ortholog is 100%.

---

Group of orthologs #1896. Best score 1491 bits
Score difference with first non-orthologous sequence - *A. fumigatus*:1491 *A. fischeri*:1491
afum|EAL84628.1 100.00% afis|EAW20422.1 100.00%
Bootstrap support for afum|EAL84628.1 as seed ortholog is 100%.
Bootstrap support for afis|EAW20422.1 as seed ortholog is 100%.

---

Group of orthologs #1897. Best score 1490 bits
Score difference with first non-orthologous sequence - *A. fumigatus*:1490 *A. fischeri*:1490
afum|EAL85273.1 100.00% afis|EAW20157.1 100.00%
Bootstrap support for afum|EAL85273.1 as seed ortholog is 100%.
Bootstrap support for afis|EAW20157.1 as seed ortholog is 100%.

---

Group of orthologs #1898. Best score 1490 bits
Score difference with first non-orthologous sequence - *A. fumigatus*:1490 *A. fischeri*:1490
afum|EAL88269.1 100.00% afis|EAW23215.1 100.00%
Bootstrap support for afum|EAL88269.1 as seed ortholog is 100%.
Bootstrap support for afis|EAW23215.1 as seed ortholog is 100%.

---

Group of orthologs #1899. Best score 1490 bits
Score difference with first non-orthologous sequence - *A. fumigatus*:1490 *A. fischeri*:1490
afum|EAL90035.1 100.00% afis|EAW25376.1 100.00%
Bootstrap support for afum|EAL90035.1 as seed ortholog is 100%.
Bootstrap support for afis|EAW25376.1 as seed ortholog is 100%.

---

Group of orthologs #1900. Best score 1489 bits
Score difference with first non-orthologous sequence - *A. fumigatus*:1489 *A. fischeri*:1489
afum|EAL88722.1 100.00% afis|EAW16090.1 100.00%
Bootstrap support for afum|EAL88722.1 as seed ortholog is 100%.
Bootstrap support for afis|EAW16090.1 as seed ortholog is 100%.

---

Group of orthologs #1901. Best score 1487 bits
Score difference with first non-orthologous sequence - *A. fumigatus*:1487 *A. fischeri*:1487
afum|EAL88843.1 100.00% afis|EAW16216.1 100.00%
Bootstrap support for afum|EAL88843.1 as seed ortholog is 100%.

Bootstrap support for afis|EAW16216.1 as seed ortholog is 100%.

---

Group of orthologs #1902. Best score 1487 bits
Score difference with first non-orthologous sequence - *A. fumigatus*:1487 *A. fischeri*:1487
afum|EAL90479.1 100.00% afis|EAW22715.1 100.00%
Bootstrap support for afum|EAL90479.1 as seed ortholog is 100%.
Bootstrap support for afis|EAW22715.1 as seed ortholog is 100%.

---

Group of orthologs #1903. Best score 1487 bits
Score difference with first non-orthologous sequence - *A. fumigatus*:1487 *A. fischeri*:1487
afum|EAL90037.1 100.00% afis|EAW25378.1 100.00%
Bootstrap support for afum|EAL90037.1 as seed ortholog is 100%.
Bootstrap support for afis|EAW25378.1 as seed ortholog is 100%.

---

Group of orthologs #1904. Best score 1487 bits
Score difference with first non-orthologous sequence - *A. fumigatus*:1487 *A. fischeri*:1487
afum|EAL90182.1 100.00% afis|EAW25517.1 100.00%
Bootstrap support for afum|EAL90182.1 as seed ortholog is 100%.
Bootstrap support for afis|EAW25517.1 as seed ortholog is 100%.

---

Group of orthologs #1905. Best score 1486 bits
Score difference with first non-orthologous sequence - *A. fumigatus*:1486 *A. fischeri*:1486
afum|EAL85540.1 100.00% afis|EAW15811.1 100.00%
Bootstrap support for afum|EAL85540.1 as seed ortholog is 100%.
Bootstrap support for afis|EAW15811.1 as seed ortholog is 100%.

---

Group of orthologs #1906. Best score 1486 bits
Score difference with first non-orthologous sequence - *A. fumigatus*:1486 *A. fischeri*:1486
afum|EAL93973.1 100.00% afis|EAW19244.1 100.00%
Bootstrap support for afum|EAL93973.1 as seed ortholog is 100%.
Bootstrap support for afis|EAW19244.1 as seed ortholog is 100%.

---

Group of orthologs #1907. Best score 1486 bits
Score difference with first non-orthologous sequence - *A. fumigatus*:1486 *A. fischeri*:1486
afum|EAL92381.1 100.00% afis|EAW21365.1 100.00%
Bootstrap support for afum|EAL92381.1 as seed ortholog is 100%.
Bootstrap support for afis|EAW21365.1 as seed ortholog is 100%.

---

Group of orthologs #1908. Best score 1485 bits
Score difference with first non-orthologous sequence - *A. fumigatus*:1485 *A. fischeri*:1485
afum|EAL91885.1 100.00% afis|EAW18073.1 100.00%
Bootstrap support for afum|EAL91885.1 as seed ortholog is 100%.
Bootstrap support for afis|EAW18073.1 as seed ortholog is 100%.

---

Group of orthologs #1909. Best score 1484 bits
Score difference with first non-orthologous sequence - *A. fumigatus*:1484 *A. fischeri*:1484
afum|EAL84549.1 100.00% afis|EAW20507.1 100.00%
Bootstrap support for afum|EAL84549.1 as seed ortholog is 100%.
Bootstrap support for afis|EAW20507.1 as seed ortholog is 100%.

---

Group of orthologs #1910. Best score 1484 bits
Score difference with first non-orthologous sequence - *A. fumigatus*:1484 *A. fischeri*:1484
afum|EAL89770.1 100.00% afis|EAW25099.1 100.00%
Bootstrap support for afum|EAL89770.1 as seed ortholog is 100%.
Bootstrap support for afis|EAW25099.1 as seed ortholog is 100%.

---

Group of orthologs #1911. Best score 1483 bits
Score difference with first non-orthologous sequence - *A. fumigatus*:1483 *A. fischeri*:1483
afum|EAL91359.1 100.00% afis|EAW17545.1 100.00%
Bootstrap support for afum|EAL91359.1 as seed ortholog is 100%.
Bootstrap support for afis|EAW17545.1 as seed ortholog is 100%.

---

Group of orthologs #1912. Best score 1483 bits
Score difference with first non-orthologous sequence - *A. fumigatus*:1333 *A. fischeri*:1435
afum|EAL92317.1 100.00% afis|EAW21300.1 100.00%
Bootstrap support for afum|EAL92317.1 as seed ortholog is 100%.
Bootstrap support for afis|EAW21300.1 as seed ortholog is 100%.

---

Group of orthologs #1913. Best score 1483 bits
Score difference with first non-orthologous sequence - *A. fumigatus*:1483 *A. fischeri*:1483
afum|EAL89768.1 100.00% afis|EAW25097.1 100.00%
Bootstrap support for afum|EAL89768.1 as seed ortholog is 100%.
Bootstrap support for afis|EAW25097.1 as seed ortholog is 100%.

---

Group of orthologs #1914. Best score 1482 bits
Score difference with first non-orthologous sequence - *A. fumigatus*:1482 *A. fischeri*:1482
afum|EAL91802.1 100.00% afis|EAW17992.1 100.00%
Bootstrap support for afum|EAL91802.1 as seed ortholog is 100%.
Bootstrap support for afis|EAW17992.1 as seed ortholog is 100%.

---

Group of orthologs #1915. Best score 1482 bits
Score difference with first non-orthologous sequence - *A. fumigatus*:1482 *A. fischeri*:1482
afum|EAL93731.1 100.00% afis|EAW18996.1 100.00%
Bootstrap support for afum|EAL93731.1 as seed ortholog is 100%.
Bootstrap support for afis|EAW18996.1 as seed ortholog is 100%.

---

Group of orthologs #1916. Best score 1482 bits
Score difference with first non-orthologous sequence - *A. fumigatus*:1482 *A. fischeri*:1482
afum|EAL90819.1 100.00% afis|EAW22373.1 100.00%
Bootstrap support for afum|EAL90819.1 as seed ortholog is 100%.
Bootstrap support for afis|EAW22373.1 as seed ortholog is 100%.

---

Group of orthologs #1917. Best score 1481 bits
Score difference with first non-orthologous sequence - *A. fumigatus*:1481 *A. fischeri*:1481
afum|EAL93165.1 100.00% afis|EAW18477.1 100.00%
Bootstrap support for afum|EAL93165.1 as seed ortholog is 100%.
Bootstrap support for afis|EAW18477.1 as seed ortholog is 100%.

---

Group of orthologs #1918. Best score 1481 bits
Score difference with first non-orthologous sequence - *A. fumigatus*:1481 *A. fischeri*:1481
afum|EAL89618.1 100.00% afis|EAW24960.1 100.00%
Bootstrap support for afum|EAL89618.1 as seed ortholog is 100%.
Bootstrap support for afis|EAW24960.1 as seed ortholog is 100%.

---

Group of orthologs #1919. Best score 1481 bits
Score difference with first non-orthologous sequence - *A. fumigatus*:1481 *A. fischeri*:1481
afum|EBA27400.1 100.00% afis|EAW22992.1 100.00%
Bootstrap support for afum|EBA27400.1 as seed ortholog is 100%.
Bootstrap support for afis|EAW22992.1 as seed ortholog is 100%.

---

Group of orthologs #1920. Best score 1480 bits
Score difference with first non-orthologous sequence - *A. fumigatus*:1480 *A. fischeri*:1149

afum|EAL86652.1 100.00% afis|EAW17232.1 100.00%
Bootstrap support for afum|EAL86652.1 as seed ortholog is 100%.
Bootstrap support for afis|EAW17232.1 as seed ortholog is 100%.

---

Group of orthologs #1921. Best score 1480 bits
Score difference with first non-orthologous sequence - *A. fumigatus*:1480 *A. fischeri*:1480
afum|EAL84324.1 100.00% afis|EAW25636.1 100.00%
Bootstrap support for afum|EAL84324.1 as seed ortholog is 100%.
Bootstrap support for afis|EAW25636.1 as seed ortholog is 100%.

---

Group of orthologs #1922. Best score 1480 bits
Score difference with first non-orthologous sequence - *A. fumigatus*:1480 *A. fischeri*:1480
afum|EAL93180.1 100.00% afis|EAW18490.1 100.00%
Bootstrap support for afum|EAL93180.1 as seed ortholog is 100%.
Bootstrap support for afis|EAW18490.1 as seed ortholog is 100%.

---

Group of orthologs #1923. Best score 1480 bits
Score difference with first non-orthologous sequence - *A. fumigatus*:1424 *A. fischeri*:1480
afum|EAL92128.1 100.00% afis|EAW21107.1 100.00%
Bootstrap support for afum|EAL92128.1 as seed ortholog is 100%.
Bootstrap support for afis|EAW21107.1 as seed ortholog is 100%.

---

Group of orthologs #1924. Best score 1479 bits
Score difference with first non-orthologous sequence - *A. fumigatus*:1479 *A. fischeri*:1479
afum|EAL93898.1 100.00% afis|EAW19168.1 100.00%
Bootstrap support for afum|EAL93898.1 as seed ortholog is 100%.
Bootstrap support for afis|EAW19168.1 as seed ortholog is 100%.

---

Group of orthologs #1925. Best score 1478 bits
Score difference with first non-orthologous sequence - *A. fumigatus*:1478 *A. fischeri*:1478
afum|EAL87684.1 100.00% afis|EAW18317.1 100.00%
Bootstrap support for afum|EAL87684.1 as seed ortholog is 100%.
Bootstrap support for afis|EAW18317.1 as seed ortholog is 100%.

---

Group of orthologs #1926. Best score 1478 bits
Score difference with first non-orthologous sequence - *A. fumigatus*:1478 *A. fischeri*:1478
afum|EAL91894.1 100.00% afis|EAW18082.1 100.00%
Bootstrap support for afum|EAL91894.1 as seed ortholog is 100%.
Bootstrap support for afis|EAW18082.1 as seed ortholog is 100%.

---

Group of orthologs #1927. Best score 1478 bits
Score difference with first non-orthologous sequence - *A. fumigatus*:1478 *A. fischeri*:1478
afum|EAL87263.1 100.00% afis|EAW23831.1 100.00%
Bootstrap support for afum|EAL87263.1 as seed ortholog is 100%.
Bootstrap support for afis|EAW23831.1 as seed ortholog is 100%.

---

Group of orthologs #1928. Best score 1478 bits
Score difference with first non-orthologous sequence - *A. fumigatus*:1478 *A. fischeri*:1478
afum|EAL93697.2 100.00% afis|EAW18965.1 100.00%
Bootstrap support for afum|EAL93697.2 as seed ortholog is 100%.
Bootstrap support for afis|EAW18965.1 as seed ortholog is 100%.

---

Group of orthologs #1929. Best score 1478 bits
Score difference with first non-orthologous sequence - *A. fumigatus*:1478 *A. fischeri*:1478
afum|EAL90382.1 100.00% afis|EAW22819.1 100.00%
Bootstrap support for afum|EAL90382.1 as seed ortholog is 100%.

Bootstrap support for afis|EAW22819.1 as seed ortholog is 100%.

---

Group of orthologs #1930. Best score 1477 bits
Score difference with first non-orthologous sequence - *A. fumigatus*:1477 *A. fischeri*:1477
afum|EAL88625.1 100.00% afis|EAW15993.1 100.00%
Bootstrap support for afum|EAL88625.1 as seed ortholog is 100%.
Bootstrap support for afis|EAW15993.1 as seed ortholog is 100%.

---

Group of orthologs #1931. Best score 1477 bits
Score difference with first non-orthologous sequence - *A. fumigatus*:1263 *A. fischeri*:1124
afum|EAL84913.1 100.00% afis|EAW20005.1 100.00%
Bootstrap support for afum|EAL84913.1 as seed ortholog is 100%.
Bootstrap support for afis|EAW20005.1 as seed ortholog is 100%.

---

Group of orthologs #1932. Best score 1477 bits
Score difference with first non-orthologous sequence - *A. fumigatus*:1477 *A. fischeri*:1477
afum|EAL86837.2 100.00% afis|EAW19689.1 100.00%
Bootstrap support for afum|EAL86837.2 as seed ortholog is 100%.
Bootstrap support for afis|EAW19689.1 as seed ortholog is 100%.

---

Group of orthologs #1933. Best score 1477 bits
Score difference with first non-orthologous sequence - *A. fumigatus*:1477 *A. fischeri*:1477
afum|EAL93050.1 100.00% afis|EAW18375.1 100.00%
Bootstrap support for afum|EAL93050.1 as seed ortholog is 100%.
Bootstrap support for afis|EAW18375.1 as seed ortholog is 100%.

---

Group of orthologs #1934. Best score 1477 bits
Score difference with first non-orthologous sequence - *A. fumigatus*:1477 *A. fischeri*:1477
afum|EAL93489.1 100.00% afis|EAW18759.1 100.00%
Bootstrap support for afum|EAL93489.1 as seed ortholog is 100%.
Bootstrap support for afis|EAW18759.1 as seed ortholog is 100%.

---

Group of orthologs #1935. Best score 1477 bits
Score difference with first non-orthologous sequence - *A. fumigatus*:1477 *A. fischeri*:1477
afum|EAL90253.1 100.00% afis|EAW22944.1 100.00%
Bootstrap support for afum|EAL90253.1 as seed ortholog is 100%.
Bootstrap support for afis|EAW22944.1 as seed ortholog is 100%.

---

Group of orthologs #1936. Best score 1477 bits
Score difference with first non-orthologous sequence - *A. fumigatus*:1477 *A. fischeri*:1477
afum|EAL92881.1 100.00% afis|EAW21867.1 100.00%
Bootstrap support for afum|EAL92881.1 as seed ortholog is 100%.
Bootstrap support for afis|EAW21867.1 as seed ortholog is 100%.

---

Group of orthologs #1937. Best score 1477 bits
Score difference with first non-orthologous sequence - *A. fumigatus*:1477 *A. fischeri*:1477
afum|EAL92921.1 100.00% afis|EAW21909.1 100.00%
Bootstrap support for afum|EAL92921.1 as seed ortholog is 100%.
Bootstrap support for afis|EAW21909.1 as seed ortholog is 100%.

---

Group of orthologs #1938. Best score 1476 bits
Score difference with first non-orthologous sequence - *A. fumigatus*:1476 *A. fischeri*:1476
afum|EAL86220.1 100.00% afis|EAW24556.1 100.00%
Bootstrap support for afum|EAL86220.1 as seed ortholog is 100%.
Bootstrap support for afis|EAW24556.1 as seed ortholog is 100%.

---

Group of orthologs #1939. Best score 1476 bits
Score difference with first non-orthologous sequence - *A. fumigatus*:1476 *A. fischeri*:1476
afum|EAL88210.2 100.00% afis|EAW23269.1 100.00%
Bootstrap support for afum|EAL88210.2 as seed ortholog is 100%.
Bootstrap support for afis|EAW23269.1 as seed ortholog is 100%.

---

Group of orthologs #1940. Best score 1476 bits
Score difference with first non-orthologous sequence - *A. fumigatus*:1476 *A. fischeri*:1476
afum|EAL90972.1 100.00% afis|EAW22222.1 100.00%
Bootstrap support for afum|EAL90972.1 as seed ortholog is 100%.
Bootstrap support for afis|EAW22222.1 as seed ortholog is 100%.

---

Group of orthologs #1941. Best score 1476 bits
Score difference with first non-orthologous sequence - *A. fumigatus*:1476 *A. fischeri*:1476
afum|EAL89465.1 100.00% afis|EAW24821.1 100.00%
Bootstrap support for afum|EAL89465.1 as seed ortholog is 100%.
Bootstrap support for afis|EAW24821.1 as seed ortholog is 100%.

---

Group of orthologs #1942. Best score 1476 bits
Score difference with first non-orthologous sequence - *A. fumigatus*:1322 *A. fischeri*:1083
afum|EAL90166.1 100.00% afis|EAW25500.1 100.00%
Bootstrap support for afum|EAL90166.1 as seed ortholog is 100%.
Bootstrap support for afis|EAW25500.1 as seed ortholog is 100%.

---

Group of orthologs #1943. Best score 1475 bits
Score difference with first non-orthologous sequence - *A. fumigatus*:1475 *A. fischeri*:1475
afum|EAL85268.1 100.00% afis|EAW20153.1 100.00%
Bootstrap support for afum|EAL85268.1 as seed ortholog is 100%.
Bootstrap support for afis|EAW20153.1 as seed ortholog is 100%.

---

Group of orthologs #1944. Best score 1475 bits
Score difference with first non-orthologous sequence - *A. fumigatus*:1399 *A. fischeri*:1384
afum|EAL85459.1 100.00% afis|EAW20338.1 100.00%
Bootstrap support for afum|EAL85459.1 as seed ortholog is 100%.
Bootstrap support for afis|EAW20338.1 as seed ortholog is 100%.

---

Group of orthologs #1945. Best score 1475 bits
Score difference with first non-orthologous sequence - *A. fumigatus*:1475 *A. fischeri*:1411
afum|EAL91321.1 100.00% afis|EAW17498.1 100.00%
Bootstrap support for afum|EAL91321.1 as seed ortholog is 100%.
Bootstrap support for afis|EAW17498.1 as seed ortholog is 100%.

---

Group of orthologs #1946. Best score 1475 bits
Score difference with first non-orthologous sequence - *A. fumigatus*:1475 *A. fischeri*:1475
afum|EAL86049.1 100.00% afis|EAW24377.1 100.00%
Bootstrap support for afum|EAL86049.1 as seed ortholog is 100%.
Bootstrap support for afis|EAW24377.1 as seed ortholog is 100%.

---

Group of orthologs #1947. Best score 1475 bits
Score difference with first non-orthologous sequence - *A. fumigatus*:1475 *A. fischeri*:1475
afum|EAL88467.2 100.00% afis|EAW23014.1 100.00%
Bootstrap support for afum|EAL88467.2 as seed ortholog is 100%.
Bootstrap support for afis|EAW23014.1 as seed ortholog is 100%.

---

Group of orthologs #1948. Best score 1474 bits
Score difference with first non-orthologous sequence - *A. fumigatus*:1474 *A. fischeri*:1474

afum|EAL91604.2 100.00% afis|EAW17796.1 100.00%
Bootstrap support for afum|EAL91604.2 as seed ortholog is 100%.
Bootstrap support for afis|EAW17796.1 as seed ortholog is 100%.

---

Group of orthologs #1949. Best score 1474 bits
Score difference with first non-orthologous sequence - *A. fumigatus*:1474 *A. fischeri*:1474
afum|EAL85970.1 100.00% afis|EAW24302.1 100.00%
Bootstrap support for afum|EAL85970.1 as seed ortholog is 100%.
Bootstrap support for afis|EAW24302.1 as seed ortholog is 100%.

---

Group of orthologs #1950. Best score 1474 bits
Score difference with first non-orthologous sequence - *A. fumigatus*:1474 *A. fischeri*:1351
afum|EAL86137.2 100.00% afis|EAW24466.1 100.00%
Bootstrap support for afum|EAL86137.2 as seed ortholog is 100%.
Bootstrap support for afis|EAW24466.1 as seed ortholog is 100%.

---

Group of orthologs #1951. Best score 1474 bits
Score difference with first non-orthologous sequence - *A. fumigatus*:1474 *A. fischeri*:1474
afum|EAL87442.1 100.00% afis|EAW24012.1 100.00%
Bootstrap support for afum|EAL87442.1 as seed ortholog is 100%.
Bootstrap support for afis|EAW24012.1 as seed ortholog is 100%.

---

Group of orthologs #1952. Best score 1474 bits
Score difference with first non-orthologous sequence - *A. fumigatus*:1474 *A. fischeri*:1474
afum|EAL92249.1 100.00% afis|EAW21237.1 100.00%
Bootstrap support for afum|EAL92249.1 as seed ortholog is 100%.
Bootstrap support for afis|EAW21237.1 as seed ortholog is 100%.

---

Group of orthologs #1953. Best score 1474 bits
Score difference with first non-orthologous sequence - *A. fumigatus*:1474 *A. fischeri*:1474
afum|EAL89625.2 100.00% afis|EAW24967.1 100.00%
Bootstrap support for afum|EAL89625.2 as seed ortholog is 100%.
Bootstrap support for afis|EAW24967.1 as seed ortholog is 100%.

---

Group of orthologs #1954. Best score 1474 bits
Score difference with first non-orthologous sequence - *A. fumigatus*:1474 *A. fischeri*:1474
afum|EAL92911.2 100.00% afis|EAW21898.1 100.00%
Bootstrap support for afum|EAL92911.2 as seed ortholog is 100%.
Bootstrap support for afis|EAW21898.1 as seed ortholog is 100%.

---

Group of orthologs #1955. Best score 1473 bits
Score difference with first non-orthologous sequence - *A. fumigatus*:1473 *A. fischeri*:1473
afum|EAL88617.1 100.00% afis|EAW15985.1 100.00%
Bootstrap support for afum|EAL88617.1 as seed ortholog is 100%.
Bootstrap support for afis|EAW15985.1 as seed ortholog is 100%.

---

Group of orthologs #1956. Best score 1473 bits
Score difference with first non-orthologous sequence - *A. fumigatus*:1473 *A. fischeri*:1473
afum|EAL87069.1 100.00% afis|EAW19425.1 100.00%
Bootstrap support for afum|EAL87069.1 as seed ortholog is 100%.
Bootstrap support for afis|EAW19425.1 as seed ortholog is 100%.

---

Group of orthologs #1957. Best score 1473 bits
Score difference with first non-orthologous sequence - *A. fumigatus*:1261 *A. fischeri*:1331
afum|EAL86983.1 100.00% afis|EAW19513.1 100.00%
Bootstrap support for afum|EAL86983.1 as seed ortholog is 100%.

Bootstrap support for afis|EAW19513.1 as seed ortholog is 100%.

---

Group of orthologs #1958. Best score 1473 bits
Score difference with first non-orthologous sequence - *A. fumigatus*:1473 *A. fischeri*:1473
afum|EAL92352.1 100.00% afis|EAW21334.1 100.00%
Bootstrap support for afum|EAL92352.1 as seed ortholog is 100%.
Bootstrap support for afis|EAW21334.1 as seed ortholog is 100%.

---

Group of orthologs #1959. Best score 1473 bits
Score difference with first non-orthologous sequence - *A. fumigatus*:1473 *A. fischeri*:1473
afum|EAL92748.1 100.00% afis|EAW21713.1 100.00%
Bootstrap support for afum|EAL92748.1 as seed ortholog is 100%.
Bootstrap support for afis|EAW21713.1 as seed ortholog is 100%.

---

Group of orthologs #1960. Best score 1473 bits
Score difference with first non-orthologous sequence - *A. fumigatus*:1473 *A. fischeri*:1473
afum|EAL89935.1 100.00% afis|EAW25272.1 100.00%
Bootstrap support for afum|EAL89935.1 as seed ortholog is 100%.
Bootstrap support for afis|EAW25272.1 as seed ortholog is 100%.

---

Group of orthologs #1961. Best score 1472 bits
Score difference with first non-orthologous sequence - *A. fumigatus*:1472 *A. fischeri*:1472
afum|EAL84621.1 100.00% afis|EAW20429.1 100.00%
Bootstrap support for afum|EAL84621.1 as seed ortholog is 100%.
Bootstrap support for afis|EAW20429.1 as seed ortholog is 100%.

---

Group of orthologs #1962. Best score 1472 bits
Score difference with first non-orthologous sequence - *A. fumigatus*:1404 *A. fischeri*:1365
afum|EAL90766.1 100.00% afis|EAW22425.1 100.00%
Bootstrap support for afum|EAL90766.1 as seed ortholog is 100%.
Bootstrap support for afis|EAW22425.1 as seed ortholog is 100%.

---

Group of orthologs #1963. Best score 1471 bits
Score difference with first non-orthologous sequence - *A. fumigatus*:1471 *A. fischeri*:1471
afum|EAL93554.1 100.00% afis|EAW18818.1 100.00%
Bootstrap support for afum|EAL93554.1 as seed ortholog is 100%.
Bootstrap support for afis|EAW18818.1 as seed ortholog is 100%.

---

Group of orthologs #1964. Best score 1471 bits
Score difference with first non-orthologous sequence - *A. fumigatus*:1292 *A. fischeri*:1252
afum|EAL92081.1 100.00% afis|EAW21045.1 100.00%
Bootstrap support for afum|EAL92081.1 as seed ortholog is 100%.
Bootstrap support for afis|EAW21045.1 as seed ortholog is 100%.

---

Group of orthologs #1965. Best score 1470 bits
Score difference with first non-orthologous sequence - *A. fumigatus*:1470 *A. fischeri*:1470
afum|EAL91293.1 100.00% afis|EAW17464.1 100.00%
Bootstrap support for afum|EAL91293.1 as seed ortholog is 100%.
Bootstrap support for afis|EAW17464.1 as seed ortholog is 100%.

---

Group of orthologs #1966. Best score 1470 bits
Score difference with first non-orthologous sequence - *A. fumigatus*:1470 *A. fischeri*:1470
afum|EAL91754.1 100.00% afis|EAW17943.1 100.00%
Bootstrap support for afum|EAL91754.1 as seed ortholog is 100%.
Bootstrap support for afis|EAW17943.1 as seed ortholog is 100%.

---

Group of orthologs #1967. Best score 1470 bits
Score difference with first non-orthologous sequence - *A. fumigatus*:1470 *A. fischeri*:1470
afum|EAL86110.1 100.00% afis|EAW24442.1 100.00%
Bootstrap support for afum|EAL86110.1 as seed ortholog is 100%.
Bootstrap support for afis|EAW24442.1 as seed ortholog is 100%.

---

Group of orthologs #1968. Best score 1470 bits
Score difference with first non-orthologous sequence - *A. fumigatus*:1470 *A. fischeri*:1470
afum|EAL93145.1 100.00% afis|EAW18459.1 100.00%
Bootstrap support for afum|EAL93145.1 as seed ortholog is 100%.
Bootstrap support for afis|EAW18459.1 as seed ortholog is 100%.

---

Group of orthologs #1969. Best score 1470 bits
Score difference with first non-orthologous sequence - *A. fumigatus*:1470 *A. fischeri*:1470
afum|EAL91974.1 100.00% afis|EAW24119.1 100.00%
Bootstrap support for afum|EAL91974.1 as seed ortholog is 100%.
Bootstrap support for afis|EAW24119.1 as seed ortholog is 100%.

---

Group of orthologs #1970. Best score 1469 bits
Score difference with first non-orthologous sequence - *A. fumigatus*:1469 *A. fischeri*:1469
afum|EAL84318.1 100.00% afis|EAW25630.1 100.00%
Bootstrap support for afum|EAL84318.1 as seed ortholog is 100%.
Bootstrap support for afis|EAW25630.1 as seed ortholog is 100%.

---

Group of orthologs #1971. Best score 1469 bits
Score difference with first non-orthologous sequence - *A. fumigatus*:1469 *A. fischeri*:1469
afum|EAL93867.1 100.00% afis|EAW19135.1 100.00%
Bootstrap support for afum|EAL93867.1 as seed ortholog is 100%.
Bootstrap support for afis|EAW19135.1 as seed ortholog is 100%.

---

Group of orthologs #1972. Best score 1469 bits
Score difference with first non-orthologous sequence - *A. fumigatus*:1145 *A. fischeri*:1096
afum|EAL90712.1 100.00% afis|EAW22482.1 100.00%
Bootstrap support for afum|EAL90712.1 as seed ortholog is 100%.
Bootstrap support for afis|EAW22482.1 as seed ortholog is 100%.

---

Group of orthologs #1973. Best score 1469 bits
Score difference with first non-orthologous sequence - *A. fumigatus*:1373 *A. fischeri*:1074
afum|EAL92404.2 100.00% afis|EAW21390.1 100.00%
Bootstrap support for afum|EAL92404.2 as seed ortholog is 100%.
Bootstrap support for afis|EAW21390.1 as seed ortholog is 100%.

---

Group of orthologs #1974. Best score 1468 bits
Score difference with first non-orthologous sequence - *A. fumigatus*:1468 *A. fischeri*:1468
afum|EAL89064.1 100.00% afis|EAW16470.1 100.00%
Bootstrap support for afum|EAL89064.1 as seed ortholog is 100%.
Bootstrap support for afis|EAW16470.1 as seed ortholog is 100%.

---

Group of orthologs #1975. Best score 1467 bits
Score difference with first non-orthologous sequence - *A. fumigatus*:1467 *A. fischeri*:1467
afum|EAL84765.1 100.00% afis|EAW20869.1 100.00%
Bootstrap support for afum|EAL84765.1 as seed ortholog is 100%.
Bootstrap support for afis|EAW20869.1 as seed ortholog is 100%.

---

Group of orthologs #1976. Best score 1467 bits
Score difference with first non-orthologous sequence - *A. fumigatus*:1467 *A. fischeri*:1467

afum|EAL87424.1 100.00% afis|EAW23994.1 100.00%
Bootstrap support for afum|EAL87424.1 as seed ortholog is 100%.
Bootstrap support for afis|EAW23994.1 as seed ortholog is 100%.

---

Group of orthologs #1977. Best score 1467 bits
Score difference with first non-orthologous sequence - *A. fumigatus*:1467 *A. fischeri*:1467
afum|EAL93925.2 100.00% afis|EAW19194.1 100.00%
Bootstrap support for afum|EAL93925.2 as seed ortholog is 100%.
Bootstrap support for afis|EAW19194.1 as seed ortholog is 100%.

---

Group of orthologs #1978. Best score 1466 bits
Score difference with first non-orthologous sequence - *A. fumigatus*:1370 *A. fischeri*:1185
afum|EAL84905.1 100.00% afis|EAW20013.1 100.00%
Bootstrap support for afum|EAL84905.1 as seed ortholog is 100%.
Bootstrap support for afis|EAW20013.1 as seed ortholog is 100%.

---

Group of orthologs #1979. Best score 1466 bits
Score difference with first non-orthologous sequence - *A. fumigatus*:1466 *A. fischeri*:1466
afum|EAL89158.1 100.00% afis|EAW16560.1 100.00%
Bootstrap support for afum|EAL89158.1 as seed ortholog is 100%.
Bootstrap support for afis|EAW16560.1 as seed ortholog is 100%.

---

Group of orthologs #1980. Best score 1466 bits
Score difference with first non-orthologous sequence - *A. fumigatus*:1466 *A. fischeri*:1466
afum|EAL89181.2 100.00% afis|EAW16586.1 100.00%
Bootstrap support for afum|EAL89181.2 as seed ortholog is 100%.
Bootstrap support for afis|EAW16586.1 as seed ortholog is 100%.

---

Group of orthologs #1981. Best score 1466 bits
Score difference with first non-orthologous sequence - *A. fumigatus*:1466 *A. fischeri*:1466
afum|EAL91294.1 100.00% afis|EAW17465.1 100.00%
Bootstrap support for afum|EAL91294.1 as seed ortholog is 100%.
Bootstrap support for afis|EAW17465.1 as seed ortholog is 100%.

---

Group of orthologs #1982. Best score 1466 bits
Score difference with first non-orthologous sequence - *A. fumigatus*:1466 *A. fischeri*:1466
afum|EAL91390.1 100.00% afis|EAW17577.1 100.00%
Bootstrap support for afum|EAL91390.1 as seed ortholog is 100%.
Bootstrap support for afis|EAW17577.1 as seed ortholog is 100%.

---

Group of orthologs #1983. Best score 1466 bits
Score difference with first non-orthologous sequence - *A. fumigatus*:1466 *A. fischeri*:1466
afum|EAL86171.1 100.00% afis|EAW24501.1 100.00%
Bootstrap support for afum|EAL86171.1 as seed ortholog is 100%.
Bootstrap support for afis|EAW24501.1 as seed ortholog is 100%.

---

Group of orthologs #1984. Best score 1464 bits
Score difference with first non-orthologous sequence - *A. fumigatus*:1464 *A. fischeri*:1464
afum|EAL85771.1 100.00% afis|EAW15561.1 100.00%
Bootstrap support for afum|EAL85771.1 as seed ortholog is 100%.
Bootstrap support for afis|EAW15561.1 as seed ortholog is 100%.

---

Group of orthologs #1985. Best score 1464 bits
Score difference with first non-orthologous sequence - *A. fumigatus*:1464 *A. fischeri*:1464
afum|EAL85633.1 100.00% afis|EAW15715.1 100.00%
Bootstrap support for afum|EAL85633.1 as seed ortholog is 100%.

Bootstrap support for afis|EAW15715.1 as seed ortholog is 100%.

---

Group of orthologs #1986. Best score 1464 bits
Score difference with first non-orthologous sequence - *A. fumigatus*:1464 *A. fischeri*:1464
afum|EAL88890.1 100.00% afis|EAW16269.1 100.00%
Bootstrap support for afum|EAL88890.1 as seed ortholog is 100%.
Bootstrap support for afis|EAW16269.1 as seed ortholog is 100%.

---

Group of orthologs #1987. Best score 1464 bits
Score difference with first non-orthologous sequence - *A. fumigatus*:1243 *A. fischeri*:1105
afum|EAL90554.2 100.00% afis|EAW22638.1 100.00%
Bootstrap support for afum|EAL90554.2 as seed ortholog is 100%.
Bootstrap support for afis|EAW22638.1 as seed ortholog is 100%.

---

Group of orthologs #1988. Best score 1463 bits
Score difference with first non-orthologous sequence - *A. fumigatus*:1463 *A. fischeri*:1463
afum|EAL88915.1 100.00% afis|EAW16293.1 100.00%
Bootstrap support for afum|EAL88915.1 as seed ortholog is 100%.
Bootstrap support for afis|EAW16293.1 as seed ortholog is 100%.

---

Group of orthologs #1989. Best score 1463 bits
Score difference with first non-orthologous sequence - *A. fumigatus*:1422 *A. fischeri*:1400
afum|EAL93657.1 100.00% afis|EAW18926.1 100.00%
Bootstrap support for afum|EAL93657.1 as seed ortholog is 100%.
Bootstrap support for afis|EAW18926.1 as seed ortholog is 100%.

---

Group of orthologs #1990. Best score 1463 bits
Score difference with first non-orthologous sequence - *A. fumigatus*:1463 *A. fischeri*:1463
afum|EAL89444.1 100.00% afis|EAW24803.1 100.00%
Bootstrap support for afum|EAL89444.1 as seed ortholog is 100%.
Bootstrap support for afis|EAW24803.1 as seed ortholog is 100%.

---

Group of orthologs #1991. Best score 1462 bits
Score difference with first non-orthologous sequence - *A. fumigatus*:1462 *A. fischeri*:1462
afum|EAL86695.1 100.00% afis|EAW17277.1 100.00%
Bootstrap support for afum|EAL86695.1 as seed ortholog is 100%.
Bootstrap support for afis|EAW17277.1 as seed ortholog is 100%.

---

Group of orthologs #1992. Best score 1462 bits
Score difference with first non-orthologous sequence - *A. fumigatus*:1462 *A. fischeri*:1462
afum|EAL92571.1 100.00% afis|EAW21551.1 100.00%
Bootstrap support for afum|EAL92571.1 as seed ortholog is 100%.
Bootstrap support for afis|EAW21551.1 as seed ortholog is 100%.

---

Group of orthologs #1993. Best score 1462 bits
Score difference with first non-orthologous sequence - *A. fumigatus*:1462 *A. fischeri*:1462
afum|EAL92010.1 100.00% afis|EAW24155.1 100.00%
Bootstrap support for afum|EAL92010.1 as seed ortholog is 100%.
Bootstrap support for afis|EAW24155.1 as seed ortholog is 100%.

---

Group of orthologs #1994. Best score 1461 bits
Score difference with first non-orthologous sequence - *A. fumigatus*:1461 *A. fischeri*:1461
afum|EAL85099.1 100.00% afis|EAW19792.1 100.00%
Bootstrap support for afum|EAL85099.1 as seed ortholog is 100%.
Bootstrap support for afis|EAW19792.1 as seed ortholog is 100%.

---

Group of orthologs #1995. Best score 1461 bits
Score difference with first non-orthologous sequence - *A. fumigatus*:1461 *A. fischeri*:1461
afum|EAL84966.1 100.00% afis|EAW19949.1 100.00%
Bootstrap support for afum|EAL84966.1 as seed ortholog is 100%.
Bootstrap support for afis|EAW19949.1 as seed ortholog is 100%.

---

Group of orthologs #1996. Best score 1461 bits
Score difference with first non-orthologous sequence - *A. fumigatus*:1461 *A. fischeri*:1461
afum|EAL87578.1 100.00% afis|EAW18212.1 100.00%
Bootstrap support for afum|EAL87578.1 as seed ortholog is 100%.
Bootstrap support for afis|EAW18212.1 as seed ortholog is 100%.

---

Group of orthologs #1997. Best score 1461 bits
Score difference with first non-orthologous sequence - *A. fumigatus*:1461 *A. fischeri*:1461
afum|EAL90890.1 100.00% afis|EAW22291.1 100.00%
Bootstrap support for afum|EAL90890.1 as seed ortholog is 100%.
Bootstrap support for afis|EAW22291.1 as seed ortholog is 100%.

---

Group of orthologs #1998. Best score 1460 bits
Score difference with first non-orthologous sequence - *A. fumigatus*:1460 *A. fischeri*:1460
afum|EAL90974.1 100.00% afis|EAW22220.1 100.00%
Bootstrap support for afum|EAL90974.1 as seed ortholog is 100%.
Bootstrap support for afis|EAW22220.1 as seed ortholog is 100%.

---

Group of orthologs #1999. Best score 1460 bits
Score difference with first non-orthologous sequence - *A. fumigatus*:1460 *A. fischeri*:1372
afum|EAL90493.1 100.00% afis|EAW22699.1 100.00%
Bootstrap support for afum|EAL90493.1 as seed ortholog is 100%.
Bootstrap support for afis|EAW22699.1 as seed ortholog is 100%.

---

Group of orthologs #2000. Best score 1459 bits
Score difference with first non-orthologous sequence - *A. fumigatus*:1459 *A. fischeri*:1459
afum|EAL85269.1 100.00% afis|EAW20154.1 100.00%
Bootstrap support for afum|EAL85269.1 as seed ortholog is 100%.
Bootstrap support for afis|EAW20154.1 as seed ortholog is 100%.

---

Group of orthologs #2001. Best score 1459 bits
Score difference with first non-orthologous sequence - *A. fumigatus*:1459 *A. fischeri*:1459
afum|EAL87704.1 100.00% afis|EAW18341.1 100.00%
Bootstrap support for afum|EAL87704.1 as seed ortholog is 100%.
Bootstrap support for afis|EAW18341.1 as seed ortholog is 100%.

---

Group of orthologs #2002. Best score 1459 bits
Score difference with first non-orthologous sequence - *A. fumigatus*:1459 *A. fischeri*:1459
afum|EAL91872.1 100.00% afis|EAW18061.1 100.00%
Bootstrap support for afum|EAL91872.1 as seed ortholog is 100%.
Bootstrap support for afis|EAW18061.1 as seed ortholog is 100%.

---

Group of orthologs #2003. Best score 1459 bits
Score difference with first non-orthologous sequence - *A. fumigatus*:1459 *A. fischeri*:1459
afum|EAL91064.1 100.00% afis|EAW22123.1 100.00%
Bootstrap support for afum|EAL91064.1 as seed ortholog is 100%.
Bootstrap support for afis|EAW22123.1 as seed ortholog is 100%.

---

Group of orthologs #2004. Best score 1459 bits
Score difference with first non-orthologous sequence - *A. fumigatus*:1459 *A. fischeri*:1459

afum|EAL92497.1 100.00% afis|EAW21477.1 100.00%
Bootstrap support for afum|EAL92497.1 as seed ortholog is 100%.
Bootstrap support for afis|EAW21477.1 as seed ortholog is 100%.

---

Group of orthologs #2005. Best score 1458 bits
Score difference with first non-orthologous sequence - *A. fumigatus*:1458 *A. fischeri*:1458
afum|EAL93722.1 100.00% afis|EAW18986.1 100.00%
Bootstrap support for afum|EAL93722.1 as seed ortholog is 100%.
Bootstrap support for afis|EAW18986.1 as seed ortholog is 100%.

---

Group of orthologs #2006. Best score 1457 bits
Score difference with first non-orthologous sequence - *A. fumigatus*:1457 *A. fischeri*:1457
afum|EAL87264.2 100.00% afis|EAW23832.1 100.00%
Bootstrap support for afum|EAL87264.2 as seed ortholog is 100%.
Bootstrap support for afis|EAW23832.1 as seed ortholog is 100%.

---

Group of orthologs #2007. Best score 1457 bits
Score difference with first non-orthologous sequence - *A. fumigatus*:1457 *A. fischeri*:1457
afum|EAL90730.1 100.00% afis|EAW22461.1 100.00%
Bootstrap support for afum|EAL90730.1 as seed ortholog is 100%.
Bootstrap support for afis|EAW22461.1 as seed ortholog is 100%.

---

Group of orthologs #2008. Best score 1456 bits
Score difference with first non-orthologous sequence - *A. fumigatus*:1456 *A. fischeri*:1456
afum|EAL84832.1 100.00% afis|EAW20797.1 100.00%
Bootstrap support for afum|EAL84832.1 as seed ortholog is 100%.
Bootstrap support for afis|EAW20797.1 as seed ortholog is 100%.

---

Group of orthologs #2009. Best score 1456 bits
Score difference with first non-orthologous sequence - *A. fumigatus*:1456 *A. fischeri*:1456
afum|EAL88272.1 100.00% afis|EAW23212.1 100.00%
Bootstrap support for afum|EAL88272.1 as seed ortholog is 100%.
Bootstrap support for afis|EAW23212.1 as seed ortholog is 100%.

---

Group of orthologs #2010. Best score 1455 bits
Score difference with first non-orthologous sequence - *A. fumigatus*:1455 *A. fischeri*:1455
afum|EAL88951.1 100.00% afis|EAW16327.1 100.00%
Bootstrap support for afum|EAL88951.1 as seed ortholog is 100%.
Bootstrap support for afis|EAW16327.1 as seed ortholog is 100%.

---

Group of orthologs #2011. Best score 1455 bits
Score difference with first non-orthologous sequence - *A. fumigatus*:1455 *A. fischeri*:1455
afum|EAL87505.1 100.00% afis|EAW18147.1 100.00%
Bootstrap support for afum|EAL87505.1 as seed ortholog is 100%.
Bootstrap support for afis|EAW18147.1 as seed ortholog is 100%.

---

Group of orthologs #2012. Best score 1455 bits
Score difference with first non-orthologous sequence - *A. fumigatus*:1286 *A. fischeri*:938
afum|EAL92769.1 100.00% afis|EAW21736.1 100.00%
Bootstrap support for afum|EAL92769.1 as seed ortholog is 100%.
Bootstrap support for afis|EAW21736.1 as seed ortholog is 100%.

---

Group of orthologs #2013. Best score 1454 bits
Score difference with first non-orthologous sequence - *A. fumigatus*:1454 *A. fischeri*:1454
afum|EAL91666.1 100.00% afis|EAW17858.1 100.00%
Bootstrap support for afum|EAL91666.1 as seed ortholog is 100%.

Bootstrap support for afis|EAW17858.1 as seed ortholog is 100%.

---

Group of orthologs #2014. Best score 1454 bits
Score difference with first non-orthologous sequence - *A. fumigatus*:1454 *A. fischeri*:1454
afum|EAL93693.1 100.00% afis|EAW18960.1 100.00%
Bootstrap support for afum|EAL93693.1 as seed ortholog is 100%.
Bootstrap support for afis|EAW18960.1 as seed ortholog is 100%.

---

Group of orthologs #2015. Best score 1453 bits
Score difference with first non-orthologous sequence - *A. fumigatus*:1222 *A. fischeri*:1453
afum|EAL85524.1 100.00% afis|EAW15827.1 100.00%
Bootstrap support for afum|EAL85524.1 as seed ortholog is 100%.
Bootstrap support for afis|EAW15827.1 as seed ortholog is 100%.

---

Group of orthologs #2016. Best score 1453 bits
Score difference with first non-orthologous sequence - *A. fumigatus*:1453 *A. fischeri*:1453
afum|EAL85934.1 100.00% afis|EAW24266.1 100.00%
Bootstrap support for afum|EAL85934.1 as seed ortholog is 100%.
Bootstrap support for afis|EAW24266.1 as seed ortholog is 100%.

---

Group of orthologs #2017. Best score 1452 bits
Score difference with first non-orthologous sequence - *A. fumigatus*:1452 *A. fischeri*:1452
afum|EAL84235.1 100.00% afis|EAW15411.1 100.00%
Bootstrap support for afum|EAL84235.1 as seed ortholog is 100%.
Bootstrap support for afis|EAW15411.1 as seed ortholog is 100%.

---

Group of orthologs #2018. Best score 1452 bits
Score difference with first non-orthologous sequence - *A. fumigatus*:1452 *A. fischeri*:1452
afum|EAL87576.2 100.00% afis|EAW18210.1 100.00%
Bootstrap support for afum|EAL87576.2 as seed ortholog is 100%.
Bootstrap support for afis|EAW18210.1 as seed ortholog is 100%.

---

Group of orthologs #2019. Best score 1452 bits
Score difference with first non-orthologous sequence - *A. fumigatus*:1452 *A. fischeri*:1221
afum|EAL91180.1 100.00% afis|EAW17336.1 100.00%
Bootstrap support for afum|EAL91180.1 as seed ortholog is 100%.
Bootstrap support for afis|EAW17336.1 as seed ortholog is 100%.

---

Group of orthologs #2020. Best score 1452 bits
Score difference with first non-orthologous sequence - *A. fumigatus*:1452 *A. fischeri*:1452
afum|EAL88472.1 100.00% afis|EAW23008.1 100.00%
Bootstrap support for afum|EAL88472.1 as seed ortholog is 100%.
Bootstrap support for afis|EAW23008.1 as seed ortholog is 100%.

---

Group of orthologs #2021. Best score 1452 bits
Score difference with first non-orthologous sequence - *A. fumigatus*:1452 *A. fischeri*:1452
afum|EAL93445.1 100.00% afis|EAW18721.1 100.00%
Bootstrap support for afum|EAL93445.1 as seed ortholog is 100%.
Bootstrap support for afis|EAW18721.1 as seed ortholog is 100%.

---

Group of orthologs #2022. Best score 1452 bits
Score difference with first non-orthologous sequence - *A. fumigatus*:1452 *A. fischeri*:1452
afum|EAL92833.1 100.00% afis|EAW21817.1 100.00%
Bootstrap support for afum|EAL92833.1 as seed ortholog is 100%.
Bootstrap support for afis|EAW21817.1 as seed ortholog is 100%.

---

Group of orthologs #2023. Best score 1452 bits
Score difference with first non-orthologous sequence - *A. fumigatus*:1452 *A. fischeri*:1452
afum|EAL89879.1 100.00% afis|EAW25218.1 100.00%
Bootstrap support for afum|EAL89879.1 as seed ortholog is 100%.
Bootstrap support for afis|EAW25218.1 as seed ortholog is 100%.

---

Group of orthologs #2024. Best score 1451 bits
Score difference with first non-orthologous sequence - *A. fumigatus*:1451 *A. fischeri*:1451
afum|EAL88844.1 100.00% afis|EAW16217.1 100.00%
Bootstrap support for afum|EAL88844.1 as seed ortholog is 100%.
Bootstrap support for afis|EAW16217.1 as seed ortholog is 100%.

---

Group of orthologs #2025. Best score 1451 bits
Score difference with first non-orthologous sequence - *A. fumigatus*:1451 *A. fischeri*:1451
afum|EAL84498.1 100.00% afis|EAW20563.1 100.00%
Bootstrap support for afum|EAL84498.1 as seed ortholog is 100%.
Bootstrap support for afis|EAW20563.1 as seed ortholog is 100%.

---

Group of orthologs #2026. Best score 1451 bits
Score difference with first non-orthologous sequence - *A. fumigatus*:1451 *A. fischeri*:1451
afum|EAL84689.2 100.00% afis|EAW20945.1 100.00%
Bootstrap support for afum|EAL84689.2 as seed ortholog is 100%.
Bootstrap support for afis|EAW20945.1 as seed ortholog is 100%.

---

Group of orthologs #2027. Best score 1450 bits
Score difference with first non-orthologous sequence - *A. fumigatus*:996 *A. fischeri*:980
afum|EAL89031.1 100.00% afis|EAW16392.1 100.00%
Bootstrap support for afum|EAL89031.1 as seed ortholog is 100%.
Bootstrap support for afis|EAW16392.1 as seed ortholog is 100%.

---

Group of orthologs #2028. Best score 1450 bits
Score difference with first non-orthologous sequence - *A. fumigatus*:1450 *A. fischeri*:1450
afum|EAL86805.2 100.00% afis|EAW19721.1 100.00%
Bootstrap support for afum|EAL86805.2 as seed ortholog is 100%.
Bootstrap support for afis|EAW19721.1 as seed ortholog is 100%.

---

Group of orthologs #2029. Best score 1450 bits
Score difference with first non-orthologous sequence - *A. fumigatus*:1450 *A. fischeri*:1450
afum|EAL93747.2 100.00% afis|EAW19013.1 100.00%
Bootstrap support for afum|EAL93747.2 as seed ortholog is 100%.
Bootstrap support for afis|EAW19013.1 as seed ortholog is 100%.

---

Group of orthologs #2030. Best score 1450 bits
Score difference with first non-orthologous sequence - *A. fumigatus*:1450 *A. fischeri*:1450
afum|EAL92178.1 100.00% afis|EAW21160.1 100.00%
Bootstrap support for afum|EAL92178.1 as seed ortholog is 100%.
Bootstrap support for afis|EAW21160.1 as seed ortholog is 100%.

---

Group of orthologs #2031. Best score 1450 bits
Score difference with first non-orthologous sequence - *A. fumigatus*:1450 *A. fischeri*:1450
afum|EAL89772.1 100.00% afis|EAW25101.1 100.00%
Bootstrap support for afum|EAL89772.1 as seed ortholog is 100%.
Bootstrap support for afis|EAW25101.1 as seed ortholog is 100%.

---

Group of orthologs #2032. Best score 1450 bits
Score difference with first non-orthologous sequence - *A. fumigatus*:1450 *A. fischeri*:1450

afum|EAL89886.1 100.00% afis|EAW25225.1 100.00%
Bootstrap support for afum|EAL89886.1 as seed ortholog is 100%.
Bootstrap support for afis|EAW25225.1 as seed ortholog is 100%.

---

Group of orthologs #2033. Best score 1449 bits
Score difference with first non-orthologous sequence - *A. fumigatus*:1449 *A. fischeri*:1449
afum|EAL86514.1 100.00% afis|EAW16990.1 100.00%
Bootstrap support for afum|EAL86514.1 as seed ortholog is 100%.
Bootstrap support for afis|EAW16990.1 as seed ortholog is 100%.

---

Group of orthologs #2034. Best score 1449 bits
Score difference with first non-orthologous sequence - *A. fumigatus*:1449 *A. fischeri*:1377
afum|EAL89182.1 100.00% afis|EAW16587.1 100.00%
Bootstrap support for afum|EAL89182.1 as seed ortholog is 100%.
Bootstrap support for afis|EAW16587.1 as seed ortholog is 100%.

---

Group of orthologs #2035. Best score 1449 bits
Score difference with first non-orthologous sequence - *A. fumigatus*:1449 *A. fischeri*:1449
afum|EAL88273.1 100.00% afis|EAW23211.1 100.00%
Bootstrap support for afum|EAL88273.1 as seed ortholog is 100%.
Bootstrap support for afis|EAW23211.1 as seed ortholog is 100%.

---

Group of orthologs #2036. Best score 1449 bits
Score difference with first non-orthologous sequence - *A. fumigatus*:1449 *A. fischeri*:1449
afum|EAL90691.1 100.00% afis|EAW22502.1 100.00%
Bootstrap support for afum|EAL90691.1 as seed ortholog is 100%.
Bootstrap support for afis|EAW22502.1 as seed ortholog is 100%.

---

Group of orthologs #2037. Best score 1449 bits
Score difference with first non-orthologous sequence - *A. fumigatus*:1449 *A. fischeri*:1449
afum|EAL90410.2 100.00% afis|EAW22789.1 100.00%
Bootstrap support for afum|EAL90410.2 as seed ortholog is 100%.
Bootstrap support for afis|EAW22789.1 as seed ortholog is 100%.

---

Group of orthologs #2038. Best score 1449 bits
Score difference with first non-orthologous sequence - *A. fumigatus*:1449 *A. fischeri*:1449
afum|EAL89865.1 100.00% afis|EAW25202.1 100.00%
Bootstrap support for afum|EAL89865.1 as seed ortholog is 100%.
Bootstrap support for afis|EAW25202.1 as seed ortholog is 100%.

---

Group of orthologs #2039. Best score 1448 bits
Score difference with first non-orthologous sequence - *A. fumigatus*:1448 *A. fischeri*:1448
afum|EAL89293.1 100.00% afis|EAW16696.1 100.00%
Bootstrap support for afum|EAL89293.1 as seed ortholog is 100%.
Bootstrap support for afis|EAW16696.1 as seed ortholog is 100%.

---

Group of orthologs #2040. Best score 1447 bits
Score difference with first non-orthologous sequence - *A. fumigatus*:1447 *A. fischeri*:1447
afum|EAL90717.1 100.00% afis|EAW22474.1 100.00%
Bootstrap support for afum|EAL90717.1 as seed ortholog is 100%.
Bootstrap support for afis|EAW22474.1 as seed ortholog is 100%.

---

Group of orthologs #2041. Best score 1447 bits
Score difference with first non-orthologous sequence - *A. fumigatus*:1285 *A. fischeri*:1015
afum|EAL90464.2 100.00% afis|EAW22731.1 100.00%
Bootstrap support for afum|EAL90464.2 as seed ortholog is 100%.

Bootstrap support for afis|EAW22731.1 as seed ortholog is 100%.

---

Group of orthologs #2042. Best score 1446 bits
Score difference with first non-orthologous sequence - *A. fumigatus*:1446 *A. fischeri*:1446
afum|EAL91284.1 100.00% afis|EAW17456.1 100.00%
Bootstrap support for afum|EAL91284.1 as seed ortholog is 100%.
Bootstrap support for afis|EAW17456.1 as seed ortholog is 100%.

---

Group of orthologs #2043. Best score 1445 bits
Score difference with first non-orthologous sequence - *A. fumigatus*:1445 *A. fischeri*:1445
afum|EAL84560.2 100.00% afis|EAW20494.1 100.00%
Bootstrap support for afum|EAL84560.2 as seed ortholog is 100%.
Bootstrap support for afis|EAW20494.1 as seed ortholog is 100%.

---

Group of orthologs #2044. Best score 1445 bits
Score difference with first non-orthologous sequence - *A. fumigatus*:1445 *A. fischeri*:1445
afum|EAL91357.1 100.00% afis|EAW17543.1 100.00%
Bootstrap support for afum|EAL91357.1 as seed ortholog is 100%.
Bootstrap support for afis|EAW17543.1 as seed ortholog is 100%.

---

Group of orthologs #2045. Best score 1445 bits
Score difference with first non-orthologous sequence - *A. fumigatus*:1445 *A. fischeri*:1445
afum|EAL91724.1 100.00% afis|EAW17914.1 100.00%
Bootstrap support for afum|EAL91724.1 as seed ortholog is 100%.
Bootstrap support for afis|EAW17914.1 as seed ortholog is 100%.

---

Group of orthologs #2046. Best score 1445 bits
Score difference with first non-orthologous sequence - *A. fumigatus*:1150 *A. fischeri*:1045
afum|EAL87112.1 100.00% afis|EAW23667.1 100.00%
Bootstrap support for afum|EAL87112.1 as seed ortholog is 100%.
Bootstrap support for afis|EAW23667.1 as seed ortholog is 100%.

---

Group of orthologs #2047. Best score 1445 bits
Score difference with first non-orthologous sequence - *A. fumigatus*:1127 *A. fischeri*:691
afum|EAL87168.1 100.00% afis|EAW23730.1 100.00%
Bootstrap support for afum|EAL87168.1 as seed ortholog is 100%.
Bootstrap support for afis|EAW23730.1 as seed ortholog is 100%.

---

Group of orthologs #2048. Best score 1445 bits
Score difference with first non-orthologous sequence - *A. fumigatus*:1445 *A. fischeri*:1445
afum|EAL87408.1 100.00% afis|EAW23978.1 100.00%
Bootstrap support for afum|EAL87408.1 as seed ortholog is 100%.
Bootstrap support for afis|EAW23978.1 as seed ortholog is 100%.

---

Group of orthologs #2049. Best score 1445 bits
Score difference with first non-orthologous sequence - *A. fumigatus*:1200 *A. fischeri*:1445
afum|EAL88241.1 100.00% afis|EAW23241.1 100.00%
Bootstrap support for afum|EAL88241.1 as seed ortholog is 100%.
Bootstrap support for afis|EAW23241.1 as seed ortholog is 100%.

---

Group of orthologs #2050. Best score 1444 bits
Score difference with first non-orthologous sequence - *A. fumigatus*:1444 *A. fischeri*:1444
afum|EAL89116.1 100.00% afis|EAW16521.1 100.00%
Bootstrap support for afum|EAL89116.1 as seed ortholog is 100%.
Bootstrap support for afis|EAW16521.1 as seed ortholog is 100%.

---

Group of orthologs #2051. Best score 1444 bits
Score difference with first non-orthologous sequence - *A. fumigatus*:951 *A. fischeri*:650
afum|EAL87170.1 100.00% afis|EAW23731.1 100.00%
Bootstrap support for afum|EAL87170.1 as seed ortholog is 100%.
Bootstrap support for afis|EAW23731.1 as seed ortholog is 100%.

---

Group of orthologs #2052. Best score 1444 bits
Score difference with first non-orthologous sequence - *A. fumigatus*:1444 *A. fischeri*:1444
afum|EAL93123.1 100.00% afis|EAW18441.1 100.00%
Bootstrap support for afum|EAL93123.1 as seed ortholog is 100%.
Bootstrap support for afis|EAW18441.1 as seed ortholog is 100%.

---

Group of orthologs #2053. Best score 1444 bits
Score difference with first non-orthologous sequence - *A. fumigatus*:1444 *A. fischeri*:1444
afum|EAL93147.1 100.00% afis|EAW18461.1 100.00%
Bootstrap support for afum|EAL93147.1 as seed ortholog is 100%.
Bootstrap support for afis|EAW18461.1 as seed ortholog is 100%.

---

Group of orthologs #2054. Best score 1444 bits
Score difference with first non-orthologous sequence - *A. fumigatus*:1444 *A. fischeri*:1444
afum|EAL87878.1 100.00% afis|EAW23586.1 100.00%
Bootstrap support for afum|EAL87878.1 as seed ortholog is 100%.
Bootstrap support for afis|EAW23586.1 as seed ortholog is 100%.

---

Group of orthologs #2055. Best score 1444 bits
Score difference with first non-orthologous sequence - *A. fumigatus*:1444 *A. fischeri*:1444
afum|EBA27487.1 100.00% afis|EAW18527.1 100.00%
Bootstrap support for afum|EBA27487.1 as seed ortholog is 100%.
Bootstrap support for afis|EAW18527.1 as seed ortholog is 100%.

---

Group of orthologs #2056. Best score 1443 bits
Score difference with first non-orthologous sequence - *A. fumigatus*:1344 *A. fischeri*:1217
afum|EAL84492.1 100.00% afis|EAW20573.1 100.00%
Bootstrap support for afum|EAL84492.1 as seed ortholog is 100%.
Bootstrap support for afis|EAW20573.1 as seed ortholog is 100%.

---

Group of orthologs #2057. Best score 1443 bits
Score difference with first non-orthologous sequence - *A. fumigatus*:1388 *A. fischeri*:1374
afum|EAL86999.1 100.00% afis|EAW19495.1 100.00%
Bootstrap support for afum|EAL86999.1 as seed ortholog is 100%.
Bootstrap support for afis|EAW19495.1 as seed ortholog is 100%.

---

Group of orthologs #2058. Best score 1443 bits
Score difference with first non-orthologous sequence - *A. fumigatus*:1443 *A. fischeri*:1443
afum|EAL88252.1 100.00% afis|EAW23231.1 100.00%
Bootstrap support for afum|EAL88252.1 as seed ortholog is 100%.
Bootstrap support for afis|EAW23231.1 as seed ortholog is 100%.

---

Group of orthologs #2059. Best score 1443 bits
Score difference with first non-orthologous sequence - *A. fumigatus*:1443 *A. fischeri*:1443
afum|EAL89668.1 100.00% afis|EAW25027.1 100.00%
Bootstrap support for afum|EAL89668.1 as seed ortholog is 100%.
Bootstrap support for afis|EAW25027.1 as seed ortholog is 100%.

---

Group of orthologs #2060. Best score 1443 bits
Score difference with first non-orthologous sequence - *A. fumigatus*:1443 *A. fischeri*:1443

afum|EAL89766.2 100.00% afis|EAW25095.1 100.00%
Bootstrap support for afum|EAL89766.2 as seed ortholog is 100%.
Bootstrap support for afis|EAW25095.1 as seed ortholog is 100%.

---

Group of orthologs #2061. Best score 1442 bits
Score difference with first non-orthologous sequence - *A. fumigatus*:1442 *A. fischeri*:1442
afum|EAL85541.1 100.00% afis|EAW15810.1 100.00%
Bootstrap support for afum|EAL85541.1 as seed ortholog is 100%.
Bootstrap support for afis|EAW15810.1 as seed ortholog is 100%.

---

Group of orthologs #2062. Best score 1442 bits
Score difference with first non-orthologous sequence - *A. fumigatus*:1442 *A. fischeri*:1442
afum|EAL88510.1 100.00% afis|EAW15876.1 100.00%
Bootstrap support for afum|EAL88510.1 as seed ortholog is 100%.
Bootstrap support for afis|EAW15876.1 as seed ortholog is 100%.

---

Group of orthologs #2063. Best score 1442 bits
Score difference with first non-orthologous sequence - *A. fumigatus*:1442 *A. fischeri*:1442
afum|EAL87560.1 100.00% afis|EAW18192.1 100.00%
Bootstrap support for afum|EAL87560.1 as seed ortholog is 100%.
Bootstrap support for afis|EAW18192.1 as seed ortholog is 100%.

---

Group of orthologs #2064. Best score 1442 bits
Score difference with first non-orthologous sequence - *A. fumigatus*:1442 *A. fischeri*:1442
afum|EAL90350.1 100.00% afis|EAW22850.1 100.00%
Bootstrap support for afum|EAL90350.1 as seed ortholog is 100%.
Bootstrap support for afis|EAW22850.1 as seed ortholog is 100%.

---

Group of orthologs #2065. Best score 1442 bits
Score difference with first non-orthologous sequence - *A. fumigatus*:1442 *A. fischeri*:1169
afum|EAL92365.1 100.00% afis|EAW21348.1 100.00%
Bootstrap support for afum|EAL92365.1 as seed ortholog is 100%.
Bootstrap support for afis|EAW21348.1 as seed ortholog is 100%.

---

Group of orthologs #2066. Best score 1441 bits
Score difference with first non-orthologous sequence - *A. fumigatus*:1441 *A. fischeri*:1441
afum|EAL87736.1 100.00% afis|EAW17067.1 100.00%
Bootstrap support for afum|EAL87736.1 as seed ortholog is 100%.
Bootstrap support for afis|EAW17067.1 as seed ortholog is 100%.

---

Group of orthologs #2067. Best score 1441 bits
Score difference with first non-orthologous sequence - *A. fumigatus*:1441 *A. fischeri*:1441
afum|EAL93907.1 100.00% afis|EAW19177.1 100.00%
Bootstrap support for afum|EAL93907.1 as seed ortholog is 100%.
Bootstrap support for afis|EAW19177.1 as seed ortholog is 100%.

---

Group of orthologs #2068. Best score 1441 bits
Score difference with first non-orthologous sequence - *A. fumigatus*:1441 *A. fischeri*:1381
afum|EAL89866.2 100.00% afis|EAW25205.1 100.00%
Bootstrap support for afum|EAL89866.2 as seed ortholog is 100%.
Bootstrap support for afis|EAW25205.1 as seed ortholog is 100%.

---

Group of orthologs #2069. Best score 1440 bits
Score difference with first non-orthologous sequence - *A. fumigatus*:1440 *A. fischeri*:1440
afum|EAL90523.1 100.00% afis|EAW22669.1 100.00%
Bootstrap support for afum|EAL90523.1 as seed ortholog is 100%.

Bootstrap support for afis|EAW22669.1 as seed ortholog is 100%.

---

Group of orthologs #2070. Best score 1440 bits
Score difference with first non-orthologous sequence - *A. fumigatus*:1440   *A. fischeri*:1440
afum|EAL89955.1 100.00% afis|EAW25292.1 100.00%
Bootstrap support for afum|EAL89955.1 as seed ortholog is 100%.
Bootstrap support for afis|EAW25292.1 as seed ortholog is 100%.

---

Group of orthologs #2071. Best score 1440 bits
Score difference with first non-orthologous sequence - *A. fumigatus*:1440   *A. fischeri*:1440
afum|EAL89980.2 100.00% afis|EAW25319.1 100.00%
Bootstrap support for afum|EAL89980.2 as seed ortholog is 100%.
Bootstrap support for afis|EAW25319.1 as seed ortholog is 100%.

---

Group of orthologs #2072. Best score 1439 bits
Score difference with first non-orthologous sequence - *A. fumigatus*:1357   *A. fischeri*:1113
afum|EAL88578.2 100.00% afis|EAW15944.1 100.00%
Bootstrap support for afum|EAL88578.2 as seed ortholog is 100%.
Bootstrap support for afis|EAW15944.1 as seed ortholog is 100%.

---

Group of orthologs #2073. Best score 1439 bits
Score difference with first non-orthologous sequence - *A. fumigatus*:1362   *A. fischeri*:1439
afum|EAL85467.1 100.00% afis|EAW20347.1 100.00%
Bootstrap support for afum|EAL85467.1 as seed ortholog is 100%.
Bootstrap support for afis|EAW20347.1 as seed ortholog is 100%.

---

Group of orthologs #2074. Best score 1439 bits
Score difference with first non-orthologous sequence - *A. fumigatus*:1439   *A. fischeri*:1439
afum|EAL91045.1 100.00% afis|EAW22145.1 100.00%
Bootstrap support for afum|EAL91045.1 as seed ortholog is 100%.
Bootstrap support for afis|EAW22145.1 as seed ortholog is 100%.

---

Group of orthologs #2075. Best score 1438 bits
Score difference with first non-orthologous sequence - *A. fumigatus*:1033   *A. fischeri*:862
afum|EAL85734.1 100.00% afis|EAW15598.1 100.00%
Bootstrap support for afum|EAL85734.1 as seed ortholog is 100%.
Bootstrap support for afis|EAW15598.1 as seed ortholog is 100%.

---

Group of orthologs #2076. Best score 1438 bits
Score difference with first non-orthologous sequence - *A. fumigatus*:1290   *A. fischeri*:1230
afum|EAL86567.1 100.00% afis|EAW17089.1 100.00%
Bootstrap support for afum|EAL86567.1 as seed ortholog is 100%.
Bootstrap support for afis|EAW17089.1 as seed ortholog is 100%.

---

Group of orthologs #2077. Best score 1438 bits
Score difference with first non-orthologous sequence - *A. fumigatus*:1438   *A. fischeri*:1438
afum|EAL88148.1 100.00% afis|EAW23331.1 100.00%
Bootstrap support for afum|EAL88148.1 as seed ortholog is 100%.
Bootstrap support for afis|EAW23331.1 as seed ortholog is 100%.

---

Group of orthologs #2078. Best score 1438 bits
Score difference with first non-orthologous sequence - *A. fumigatus*:1237   *A. fischeri*:1237
afum|EAL89507.1 100.00% afis|EAW24859.1 100.00%
Bootstrap support for afum|EAL89507.1 as seed ortholog is 100%.
Bootstrap support for afis|EAW24859.1 as seed ortholog is 100%.

---

Group of orthologs #2079. Best score 1437 bits
Score difference with first non-orthologous sequence - *A. fumigatus*:1294 *A. fischeri*:1259
afum|EAL91502.1 100.00% afis|EAW17688.1 100.00%
Bootstrap support for afum|EAL91502.1 as seed ortholog is 100%.
Bootstrap support for afis|EAW17688.1 as seed ortholog is 100%.

---

Group of orthologs #2080. Best score 1436 bits
Score difference with first non-orthologous sequence - *A. fumigatus*:1436 *A. fischeri*:1306
afum|EAL91182.1 100.00% afis|EAW17338.1 100.00%
Bootstrap support for afum|EAL91182.1 as seed ortholog is 100%.
Bootstrap support for afis|EAW17338.1 as seed ortholog is 100%.

---

Group of orthologs #2081. Best score 1436 bits
Score difference with first non-orthologous sequence - *A. fumigatus*:1436 *A. fischeri*:1436
afum|EAL88397.1 100.00% afis|EAW23082.1 100.00%
Bootstrap support for afum|EAL88397.1 as seed ortholog is 100%.
Bootstrap support for afis|EAW23082.1 as seed ortholog is 100%.

---

Group of orthologs #2082. Best score 1436 bits
Score difference with first non-orthologous sequence - *A. fumigatus*:1436 *A. fischeri*:1436
afum|EAL93258.1 100.00% afis|EAW18558.1 100.00%
Bootstrap support for afum|EAL93258.1 as seed ortholog is 100%.
Bootstrap support for afis|EAW18558.1 as seed ortholog is 100%.

---

Group of orthologs #2083. Best score 1436 bits
Score difference with first non-orthologous sequence - *A. fumigatus*:1436 *A. fischeri*:1436
afum|EAL93403.1 100.00% afis|EAW18682.1 100.00%
Bootstrap support for afum|EAL93403.1 as seed ortholog is 100%.
Bootstrap support for afis|EAW18682.1 as seed ortholog is 100%.

---

Group of orthologs #2084. Best score 1436 bits
Score difference with first non-orthologous sequence - *A. fumigatus*:1436 *A. fischeri*:1436
afum|EBA27485.1 100.00% afis|EAW18484.1 100.00%
Bootstrap support for afum|EBA27485.1 as seed ortholog is 100%.
Bootstrap support for afis|EAW18484.1 as seed ortholog is 100%.

---

Group of orthologs #2085. Best score 1436 bits
Score difference with first non-orthologous sequence - *A. fumigatus*:1436 *A. fischeri*:1436
afum|EAL93965.2 100.00% afis|EAW19237.1 100.00%
Bootstrap support for afum|EAL93965.2 as seed ortholog is 100%.
Bootstrap support for afis|EAW19237.1 as seed ortholog is 100%.

---

Group of orthologs #2086. Best score 1436 bits
Score difference with first non-orthologous sequence - *A. fumigatus*:1436 *A. fischeri*:1436
afum|EAL91032.1 100.00% afis|EAW22158.1 100.00%
Bootstrap support for afum|EAL91032.1 as seed ortholog is 100%.
Bootstrap support for afis|EAW22158.1 as seed ortholog is 100%.

---

Group of orthologs #2087. Best score 1436 bits
Score difference with first non-orthologous sequence - *A. fumigatus*:1436 *A. fischeri*:1436
afum|EAL89460.2 100.00% afis|EAW24816.1 100.00%
Bootstrap support for afum|EAL89460.2 as seed ortholog is 100%.
Bootstrap support for afis|EAW24816.1 as seed ortholog is 100%.

---

Group of orthologs #2088. Best score 1436 bits
Score difference with first non-orthologous sequence - *A. fumigatus*:1436 *A. fischeri*:1436

afum|EAL92950.2 100.00% afis|EAW21937.1 100.00%
Bootstrap support for afum|EAL92950.2 as seed ortholog is 100%.
Bootstrap support for afis|EAW21937.1 as seed ortholog is 100%.

---

Group of orthologs #2089. Best score 1435 bits
Score difference with first non-orthologous sequence - *A. fumigatus*:1169 *A. fischeri*:912
afum|EAL92724.1 100.00% afis|EAW21690.1 100.00%
Bootstrap support for afum|EAL92724.1 as seed ortholog is 100%.
Bootstrap support for afis|EAW21690.1 as seed ortholog is 100%.

---

Group of orthologs #2090. Best score 1434 bits
Score difference with first non-orthologous sequence - *A. fumigatus*:1327 *A. fischeri*:1296
afum|EAL85288.1 100.00% afis|EAW20170.1 100.00%
Bootstrap support for afum|EAL85288.1 as seed ortholog is 100%.
Bootstrap support for afis|EAW20170.1 as seed ortholog is 100%.

---

Group of orthologs #2091. Best score 1434 bits
Score difference with first non-orthologous sequence - *A. fumigatus*:1434 *A. fischeri*:1434
afum|EAL87287.1 100.00% afis|EAW23859.1 100.00%
Bootstrap support for afum|EAL87287.1 as seed ortholog is 100%.
Bootstrap support for afis|EAW23859.1 as seed ortholog is 100%.

---

Group of orthologs #2092. Best score 1434 bits
Score difference with first non-orthologous sequence - *A. fumigatus*:1131 *A. fischeri*:1065
afum|EAL91937.1 100.00% afis|EAW24082.1 100.00%
Bootstrap support for afum|EAL91937.1 as seed ortholog is 100%.
Bootstrap support for afis|EAW24082.1 as seed ortholog is 100%.

---

Group of orthologs #2093. Best score 1433 bits
Score difference with first non-orthologous sequence - *A. fumigatus*:1433 *A. fischeri*:1433
afum|EAL85803.1 100.00% afis|EAW15525.1 100.00%
Bootstrap support for afum|EAL85803.1 as seed ortholog is 100%.
Bootstrap support for afis|EAW15525.1 as seed ortholog is 100%.

---

Group of orthologs #2094. Best score 1433 bits
Score difference with first non-orthologous sequence - *A. fumigatus*:1268 *A. fischeri*:1239
afum|EAL85140.1 100.00% afis|EAW19757.1 100.00%
Bootstrap support for afum|EAL85140.1 as seed ortholog is 100%.
Bootstrap support for afis|EAW19757.1 as seed ortholog is 100%.

---

Group of orthologs #2095. Best score 1433 bits
Score difference with first non-orthologous sequence - *A. fumigatus*:1433 *A. fischeri*:1433
afum|EAL87676.1 100.00% afis|EAW18310.1 100.00%
Bootstrap support for afum|EAL87676.1 as seed ortholog is 100%.
Bootstrap support for afis|EAW18310.1 as seed ortholog is 100%.

---

Group of orthologs #2096. Best score 1433 bits
Score difference with first non-orthologous sequence - *A. fumigatus*:1433 *A. fischeri*:1433
afum|EAL88352.1 100.00% afis|EAW23128.1 100.00%
Bootstrap support for afum|EAL88352.1 as seed ortholog is 100%.
Bootstrap support for afis|EAW23128.1 as seed ortholog is 100%.

---

Group of orthologs #2097. Best score 1433 bits
Score difference with first non-orthologous sequence - *A. fumigatus*:1433 *A. fischeri*:1433
afum|EAL89435.1 100.00% afis|EAW24793.1 100.00%
Bootstrap support for afum|EAL89435.1 as seed ortholog is 100%.

Bootstrap support for afis|EAW24793.1 as seed ortholog is 100%.

---

Group of orthologs #2098. Best score 1432 bits
Score difference with first non-orthologous sequence - *A. fumigatus*:1432 *A. fischeri*:1432
afum|EAL85162.1 100.00% afis|EAW20042.1 100.00%
Bootstrap support for afum|EAL85162.1 as seed ortholog is 100%.
Bootstrap support for afis|EAW20042.1 as seed ortholog is 100%.

---

Group of orthologs #2099. Best score 1432 bits
Score difference with first non-orthologous sequence - *A. fumigatus*:1133 *A. fischeri*:1432
afum|EAL84861.1 100.00% afis|EAW20759.1 100.00%
Bootstrap support for afum|EAL84861.1 as seed ortholog is 100%.
Bootstrap support for afis|EAW20759.1 as seed ortholog is 100%.

---

Group of orthologs #2100. Best score 1432 bits
Score difference with first non-orthologous sequence - *A. fumigatus*:1263 *A. fischeri*:999
afum|EAL93797.1 100.00% afis|EAW19068.1 100.00%
Bootstrap support for afum|EAL93797.1 as seed ortholog is 100%.
Bootstrap support for afis|EAW19068.1 as seed ortholog is 100%.

---

Group of orthologs #2101. Best score 1432 bits
Score difference with first non-orthologous sequence - *A. fumigatus*:1432 *A. fischeri*:1432
afum|EAL89985.1 100.00% afis|EAW25324.1 100.00%
Bootstrap support for afum|EAL89985.1 as seed ortholog is 100%.
Bootstrap support for afis|EAW25324.1 as seed ortholog is 100%.

---

Group of orthologs #2102. Best score 1431 bits
Score difference with first non-orthologous sequence - *A. fumigatus*:1431 *A. fischeri*:1243
afum|EAL85927.2 100.00% afis|EAW24259.1 100.00%
Bootstrap support for afum|EAL85927.2 as seed ortholog is 100%.
Bootstrap support for afis|EAW24259.1 as seed ortholog is 100%.

---

Group of orthologs #2103. Best score 1431 bits
Score difference with first non-orthologous sequence - *A. fumigatus*:1431 *A. fischeri*:1087
afum|EAL89346.1 100.00% afis|EAW24704.1 100.00%
Bootstrap support for afum|EAL89346.1 as seed ortholog is 100%.
Bootstrap support for afis|EAW24704.1 as seed ortholog is 100%.

---

Group of orthologs #2104. Best score 1430 bits
Score difference with first non-orthologous sequence - *A. fumigatus*:1430 *A. fischeri*:1430
afum|EAL88752.1 100.00% afis|EAW16121.1 100.00%
Bootstrap support for afum|EAL88752.1 as seed ortholog is 100%.
Bootstrap support for afis|EAW16121.1 as seed ortholog is 100%.

---

Group of orthologs #2105. Best score 1429 bits
Score difference with first non-orthologous sequence - *A. fumigatus*:1429 *A. fischeri*:1244
afum|EAL86418.1 100.00% afis|EAW16882.1 100.00%
Bootstrap support for afum|EAL86418.1 as seed ortholog is 100%.
Bootstrap support for afis|EAW16882.1 as seed ortholog is 100%.

---

Group of orthologs #2106. Best score 1429 bits
Score difference with first non-orthologous sequence - *A. fumigatus*:1429 *A. fischeri*:1429
afum|EAL84767.1 100.00% afis|EAW20867.1 100.00%
Bootstrap support for afum|EAL84767.1 as seed ortholog is 100%.
Bootstrap support for afis|EAW20867.1 as seed ortholog is 100%.

---

Group of orthologs #2107. Best score 1428 bits
Score difference with first non-orthologous sequence - *A. fumigatus*:1428 *A. fischeri*:1428
afum|EAL89278.1 100.00% afis|EAW16673.1 100.00%
Bootstrap support for afum|EAL89278.1 as seed ortholog is 100%.
Bootstrap support for afis|EAW16673.1 as seed ortholog is 100%.

---

Group of orthologs #2108. Best score 1428 bits
Score difference with first non-orthologous sequence - *A. fumigatus*:1428 *A. fischeri*:1428
afum|EAL88278.1 100.00% afis|EAW23205.1 100.00%
Bootstrap support for afum|EAL88278.1 as seed ortholog is 100%.
Bootstrap support for afis|EAW23205.1 as seed ortholog is 100%.

---

Group of orthologs #2109. Best score 1428 bits
Score difference with first non-orthologous sequence - *A. fumigatus*:1108 *A. fischeri*:1044
afum|EAL93452.1 100.00% afis|EAW18727.1 100.00%
Bootstrap support for afum|EAL93452.1 as seed ortholog is 100%.
Bootstrap support for afis|EAW18727.1 as seed ortholog is 100%.

---

Group of orthologs #2110. Best score 1428 bits
Score difference with first non-orthologous sequence - *A. fumigatus*:1428 *A. fischeri*:1428
afum|EAL90286.1 100.00% afis|EAW22911.1 100.00%
Bootstrap support for afum|EAL90286.1 as seed ortholog is 100%.
Bootstrap support for afis|EAW22911.1 as seed ortholog is 100%.

---

Group of orthologs #2111. Best score 1428 bits
Score difference with first non-orthologous sequence - *A. fumigatus*:1428 *A. fischeri*:1428
afum|EBA27416.1 100.00% afis|EAW22365.1 100.00%
Bootstrap support for afum|EBA27416.1 as seed ortholog is 100%.
Bootstrap support for afis|EAW22365.1 as seed ortholog is 100%.

---

Group of orthologs #2112. Best score 1427 bits
Score difference with first non-orthologous sequence - *A. fumigatus*:1221 *A. fischeri*:1274
afum|EAL85727.1 100.00% afis|EAW15606.1 100.00%
Bootstrap support for afum|EAL85727.1 as seed ortholog is 100%.
Bootstrap support for afis|EAW15606.1 as seed ortholog is 100%.

---

Group of orthologs #2113. Best score 1427 bits
Score difference with first non-orthologous sequence - *A. fumigatus*:1427 *A. fischeri*:1427
afum|EAL84616.1 100.00% afis|EAW20436.1 100.00%
Bootstrap support for afum|EAL84616.1 as seed ortholog is 100%.
Bootstrap support for afis|EAW20436.1 as seed ortholog is 100%.

---

Group of orthologs #2114. Best score 1427 bits
Score difference with first non-orthologous sequence - *A. fumigatus*:1427 *A. fischeri*:1427
afum|EAL92123.1 100.00% afis|EAW21102.1 100.00%
Bootstrap support for afum|EAL92123.1 as seed ortholog is 100%.
Bootstrap support for afis|EAW21102.1 as seed ortholog is 100%.

---

Group of orthologs #2115. Best score 1427 bits
Score difference with first non-orthologous sequence - *A. fumigatus*:1427 *A. fischeri*:1427
afum|EAL90635.1 100.00% afis|EAW22560.1 100.00%
Bootstrap support for afum|EAL90635.1 as seed ortholog is 100%.
Bootstrap support for afis|EAW22560.1 as seed ortholog is 100%.

---

Group of orthologs #2116. Best score 1426 bits
Score difference with first non-orthologous sequence - *A. fumigatus*:1302 *A. fischeri*:1426

afum|EAL92057.1 100.00% afis|EAW21019.1 100.00%
Bootstrap support for afum|EAL92057.1 as seed ortholog is 100%.
Bootstrap support for afis|EAW21019.1 as seed ortholog is 100%.

---

Group of orthologs #2117. Best score 1425 bits
Score difference with first non-orthologous sequence - *A. fumigatus*:1425 *A. fischeri*:1425
afum|EAL85748.2 100.00% afis|EAW15584.1 100.00%
Bootstrap support for afum|EAL85748.2 as seed ortholog is 100%.
Bootstrap support for afis|EAW15584.1 as seed ortholog is 100%.

---

Group of orthologs #2118. Best score 1425 bits
Score difference with first non-orthologous sequence - *A. fumigatus*:1224 *A. fischeri*:1425
afum|EAL84496.1 100.00% afis|EAW20565.1 100.00%
Bootstrap support for afum|EAL84496.1 as seed ortholog is 100%.
Bootstrap support for afis|EAW20565.1 as seed ortholog is 100%.

---

Group of orthologs #2119. Best score 1425 bits
Score difference with first non-orthologous sequence - *A. fumigatus*:1425 *A. fischeri*:1425
afum|EAL86817.2 100.00% afis|EAW19710.1 100.00%
Bootstrap support for afum|EAL86817.2 as seed ortholog is 100%.
Bootstrap support for afis|EAW19710.1 as seed ortholog is 100%.

---

Group of orthologs #2120. Best score 1425 bits
Score difference with first non-orthologous sequence - *A. fumigatus*:1425 *A. fischeri*:1425
afum|EAL86006.1 100.00% afis|EAW24336.1 100.00%
Bootstrap support for afum|EAL86006.1 as seed ortholog is 100%.
Bootstrap support for afis|EAW24336.1 as seed ortholog is 100%.

---

Group of orthologs #2121. Best score 1425 bits
Score difference with first non-orthologous sequence - *A. fumigatus*:1287 *A. fischeri*:1279
afum|EAL92499.1 100.00% afis|EAW21479.1 100.00%
Bootstrap support for afum|EAL92499.1 as seed ortholog is 100%.
Bootstrap support for afis|EAW21479.1 as seed ortholog is 100%.

---

Group of orthologs #2122. Best score 1424 bits
Score difference with first non-orthologous sequence - *A. fumigatus*:1424 *A. fischeri*:1424
afum|EAL84882.1 100.00% afis|EAW20034.1 100.00%
Bootstrap support for afum|EAL84882.1 as seed ortholog is 100%.
Bootstrap support for afis|EAW20034.1 as seed ortholog is 100%.

---

Group of orthologs #2123. Best score 1423 bits
Score difference with first non-orthologous sequence - *A. fumigatus*:1423 *A. fischeri*:1423
afum|EAL86431.1 100.00% afis|EAW16893.1 100.00%
Bootstrap support for afum|EAL86431.1 as seed ortholog is 100%.
Bootstrap support for afis|EAW16893.1 as seed ortholog is 100%.

---

Group of orthologs #2124. Best score 1423 bits
Score difference with first non-orthologous sequence - *A. fumigatus*:1423 *A. fischeri*:1423
afum|EAL88940.1 100.00% afis|EAW16317.1 100.00%
Bootstrap support for afum|EAL88940.1 as seed ortholog is 100%.
Bootstrap support for afis|EAW16317.1 as seed ortholog is 100%.

---

Group of orthologs #2125. Best score 1423 bits
Score difference with first non-orthologous sequence - *A. fumigatus*:1423 *A. fischeri*:1423
afum|EAL91077.1 100.00% afis|EAW22107.1 100.00%
Bootstrap support for afum|EAL91077.1 as seed ortholog is 100%.

Bootstrap support for afis|EAW22107.1 as seed ortholog is 100%.

---

Group of orthologs #2126. Best score 1423 bits
Score difference with first non-orthologous sequence - *A. fumigatus*:1423 *A. fischeri*:1423
afum|EAL92268.1 100.00% afis|EAW21253.1 100.00%
Bootstrap support for afum|EAL92268.1 as seed ortholog is 100%.
Bootstrap support for afis|EAW21253.1 as seed ortholog is 100%.

---

Group of orthologs #2127. Best score 1423 bits
Score difference with first non-orthologous sequence - *A. fumigatus*:1423 *A. fischeri*:1423
afum|EAL92643.1 100.00% afis|EAW21609.1 100.00%
Bootstrap support for afum|EAL92643.1 as seed ortholog is 100%.
Bootstrap support for afis|EAW21609.1 as seed ortholog is 100%.

---

Group of orthologs #2128. Best score 1423 bits
Score difference with first non-orthologous sequence - *A. fumigatus*:1423 *A. fischeri*:1423
afum|EAL89875.1 100.00% afis|EAW25214.1 100.00%
Bootstrap support for afum|EAL89875.1 as seed ortholog is 100%.
Bootstrap support for afis|EAW25214.1 as seed ortholog is 100%.

---

Group of orthologs #2129. Best score 1422 bits
Score difference with first non-orthologous sequence - *A. fumigatus*:1192 *A. fischeri*:1236
afum|EAL84462.1 100.00% afis|EAW20599.1 100.00%
Bootstrap support for afum|EAL84462.1 as seed ortholog is 100%.
Bootstrap support for afis|EAW20599.1 as seed ortholog is 100%.

---

Group of orthologs #2130. Best score 1421 bits
Score difference with first non-orthologous sequence - *A. fumigatus*:1421 *A. fischeri*:1421
afum|EAL92437.1 100.00% afis|EAW21422.1 100.00%
Bootstrap support for afum|EAL92437.1 as seed ortholog is 100%.
Bootstrap support for afis|EAW21422.1 as seed ortholog is 100%.

---

Group of orthologs #2131. Best score 1421 bits
Score difference with first non-orthologous sequence - *A. fumigatus*:1421 *A. fischeri*:1421
afum|EBA27196.1 100.00% afis|EAW20455.1 100.00%
Bootstrap support for afum|EBA27196.1 as seed ortholog is 100%.
Bootstrap support for afis|EAW20455.1 as seed ortholog is 100%.

---

Group of orthologs #2132. Best score 1421 bits
Score difference with first non-orthologous sequence - *A. fumigatus*:1421 *A. fischeri*:1421
afum|EAL93030.1 100.00% afis|EAW22015.1 100.00%
Bootstrap support for afum|EAL93030.1 as seed ortholog is 100%.
Bootstrap support for afis|EAW22015.1 as seed ortholog is 100%.

---

Group of orthologs #2133. Best score 1420 bits
Score difference with first non-orthologous sequence - *A. fumigatus*:1420 *A. fischeri*:1316
afum|EAL84726.1 100.00% afis|EAW20907.1 100.00%
Bootstrap support for afum|EAL84726.1 as seed ortholog is 100%.
Bootstrap support for afis|EAW20907.1 as seed ortholog is 100%.

---

Group of orthologs #2134. Best score 1420 bits
Score difference with first non-orthologous sequence - *A. fumigatus*:1420 *A. fischeri*:1420
afum|EAL91177.1 100.00% afis|EAW17334.1 100.00%
Bootstrap support for afum|EAL91177.1 as seed ortholog is 100%.
Bootstrap support for afis|EAW17334.1 as seed ortholog is 100%.

---

Group of orthologs #2135. Best score 1420 bits
Score difference with first non-orthologous sequence - *A. fumigatus*:1420 *A. fischeri*:1420
afum|EAL90919.1 100.00% afis|EAW22261.1 100.00%
Bootstrap support for afum|EAL90919.1 as seed ortholog is 100%.
Bootstrap support for afis|EAW22261.1 as seed ortholog is 100%.

---

Group of orthologs #2136. Best score 1420 bits
Score difference with first non-orthologous sequence - *A. fumigatus*:1420 *A. fischeri*:1278
afum|EAL94050.2 100.00% afis|EAW19333.1 100.00%
Bootstrap support for afum|EAL94050.2 as seed ortholog is 100%.
Bootstrap support for afis|EAW19333.1 as seed ortholog is 100%.

---

Group of orthologs #2137. Best score 1420 bits
Score difference with first non-orthologous sequence - *A. fumigatus*:1420 *A. fischeri*:1239
afum|EAL94051.2 100.00% afis|EAW19334.1 100.00%
Bootstrap support for afum|EAL94051.2 as seed ortholog is 100%.
Bootstrap support for afis|EAW19334.1 as seed ortholog is 100%.

---

Group of orthologs #2138. Best score 1419 bits
Score difference with first non-orthologous sequence - *A. fumigatus*:1196 *A. fischeri*:1419
afum|EAL86434.1 100.00% afis|EAW16899.1 100.00%
Bootstrap support for afum|EAL86434.1 as seed ortholog is 100%.
Bootstrap support for afis|EAW16899.1 as seed ortholog is 100%.

---

Group of orthologs #2139. Best score 1418 bits
Score difference with first non-orthologous sequence - *A. fumigatus*:1418 *A. fischeri*:1106
afum|EAL84263.1 100.00% afis|EAW25545.1 100.00%
Bootstrap support for afum|EAL84263.1 as seed ortholog is 100%.
Bootstrap support for afis|EAW25545.1 as seed ortholog is 100%.

---

Group of orthologs #2140. Best score 1418 bits
Score difference with first non-orthologous sequence - *A. fumigatus*:1418 *A. fischeri*:1418
afum|EAL92586.1 100.00% afis|EAW21567.1 100.00%
Bootstrap support for afum|EAL92586.1 as seed ortholog is 100%.
Bootstrap support for afis|EAW21567.1 as seed ortholog is 100%.

---

Group of orthologs #2141. Best score 1417 bits
Score difference with first non-orthologous sequence - *A. fumigatus*:1417 *A. fischeri*:1417
afum|EAL89113.1 100.00% afis|EAW16518.1 100.00%
Bootstrap support for afum|EAL89113.1 as seed ortholog is 100%.
Bootstrap support for afis|EAW16518.1 as seed ortholog is 100%.

---

Group of orthologs #2142. Best score 1417 bits
Score difference with first non-orthologous sequence - *A. fumigatus*:1417 *A. fischeri*:1417
afum|EAL86950.1 100.00% afis|EAW19562.1 100.00%
Bootstrap support for afum|EAL86950.1 as seed ortholog is 100%.
Bootstrap support for afis|EAW19562.1 as seed ortholog is 100%.

---

Group of orthologs #2143. Best score 1417 bits
Score difference with first non-orthologous sequence - *A. fumigatus*:1417 *A. fischeri*:1417
afum|EAL85442.1 100.00% afis|EAW22025.1 100.00%
Bootstrap support for afum|EAL85442.1 as seed ortholog is 100%.
Bootstrap support for afis|EAW22025.1 as seed ortholog is 100%.

---

Group of orthologs #2144. Best score 1417 bits
Score difference with first non-orthologous sequence - *A. fumigatus*:1417 *A. fischeri*:1417

afum|EAL91839.1 100.00% afis|EAW18028.1 100.00%
Bootstrap support for afum|EAL91839.1 as seed ortholog is 100%.
Bootstrap support for afis|EAW18028.1 as seed ortholog is 100%.

---

Group of orthologs #2145. Best score 1417 bits
Score difference with first non-orthologous sequence - *A. fumigatus*:1417 *A. fischeri*:1417
afum|EAL90721.1 100.00% afis|EAW22470.1 100.00%
Bootstrap support for afum|EAL90721.1 as seed ortholog is 100%.
Bootstrap support for afis|EAW22470.1 as seed ortholog is 100%.

---

Group of orthologs #2146. Best score 1417 bits
Score difference with first non-orthologous sequence - *A. fumigatus*:1417 *A. fischeri*:1417
afum|EAL89501.1 100.00% afis|EAW24854.1 100.00%
Bootstrap support for afum|EAL89501.1 as seed ortholog is 100%.
Bootstrap support for afis|EAW24854.1 as seed ortholog is 100%.

---

Group of orthologs #2147. Best score 1416 bits
Score difference with first non-orthologous sequence - *A. fumigatus*:1416 *A. fischeri*:1324
afum|EAL84741.1 100.00% afis|EAW20892.1 100.00%
Bootstrap support for afum|EAL84741.1 as seed ortholog is 100%.
Bootstrap support for afis|EAW20892.1 as seed ortholog is 100%.

---

Group of orthologs #2148. Best score 1416 bits
Score difference with first non-orthologous sequence - *A. fumigatus*:1416 *A. fischeri*:1416
afum|EAL87381.1 100.00% afis|EAW23953.1 100.00%
Bootstrap support for afum|EAL87381.1 as seed ortholog is 100%.
Bootstrap support for afis|EAW23953.1 as seed ortholog is 100%.

---

Group of orthologs #2149. Best score 1416 bits
Score difference with first non-orthologous sequence - *A. fumigatus*:1416 *A. fischeri*:1416
afum|EAL92402.1 100.00% afis|EAW21387.1 100.00%
Bootstrap support for afum|EAL92402.1 as seed ortholog is 100%.
Bootstrap support for afis|EAW21387.1 as seed ortholog is 100%.

---

Group of orthologs #2150. Best score 1415 bits
Score difference with first non-orthologous sequence - *A. fumigatus*:1415 *A. fischeri*:1415
afum|EAL89157.1 100.00% afis|EAW16559.1 100.00%
Bootstrap support for afum|EAL89157.1 as seed ortholog is 100%.
Bootstrap support for afis|EAW16559.1 as seed ortholog is 100%.

---

Group of orthologs #2151. Best score 1415 bits
Score difference with first non-orthologous sequence - *A. fumigatus*:1415 *A. fischeri*:1415
afum|EAL90376.1 100.00% afis|EAW22824.1 100.00%
Bootstrap support for afum|EAL90376.1 as seed ortholog is 100%.
Bootstrap support for afis|EAW22824.1 as seed ortholog is 100%.

---

Group of orthologs #2152. Best score 1414 bits
Score difference with first non-orthologous sequence - *A. fumigatus*:1084 *A. fischeri*:1128
afum|EAL86141.1 100.00% afis|EAW24469.1 100.00%
Bootstrap support for afum|EAL86141.1 as seed ortholog is 100%.
Bootstrap support for afis|EAW24469.1 as seed ortholog is 100%.

---

Group of orthologs #2153. Best score 1414 bits
Score difference with first non-orthologous sequence - *A. fumigatus*:1141 *A. fischeri*:1414
afum|EBA27340.1 100.00% afis|EAW23213.1 100.00%
Bootstrap support for afum|EBA27340.1 as seed ortholog is 100%.

Bootstrap support for afis|EAW23213.1 as seed ortholog is 100%.

---

Group of orthologs #2154. Best score 1413 bits
Score difference with first non-orthologous sequence - *A. fumigatus*:1176 *A. fischeri*:797
afum|EAL86343.2 100.00% afis|EAW16774.1 100.00%
Bootstrap support for afum|EAL86343.2 as seed ortholog is 100%.
Bootstrap support for afis|EAW16774.1 as seed ortholog is 100%.

---

Group of orthologs #2155. Best score 1413 bits
Score difference with first non-orthologous sequence - *A. fumigatus*:1413 *A. fischeri*:1335
afum|EAL87664.2 100.00% afis|EAW18298.1 100.00%
Bootstrap support for afum|EAL87664.2 as seed ortholog is 100%.
Bootstrap support for afis|EAW18298.1 as seed ortholog is 100%.

---

Group of orthologs #2156. Best score 1413 bits
Score difference with first non-orthologous sequence - *A. fumigatus*:1413 *A. fischeri*:1413
afum|EAL91461.1 100.00% afis|EAW17646.1 100.00%
Bootstrap support for afum|EAL91461.1 as seed ortholog is 100%.
Bootstrap support for afis|EAW17646.1 as seed ortholog is 100%.

---

Group of orthologs #2157. Best score 1413 bits
Score difference with first non-orthologous sequence - *A. fumigatus*:1057 *A. fischeri*:1077
afum|EAL90181.1 100.00% afis|EAW25516.1 100.00%
Bootstrap support for afum|EAL90181.1 as seed ortholog is 100%.
Bootstrap support for afis|EAW25516.1 as seed ortholog is 100%.

---

Group of orthologs #2158. Best score 1412 bits
Score difference with first non-orthologous sequence - *A. fumigatus*:1412 *A. fischeri*:1412
afum|EAL88585.1 100.00% afis|EAW15952.1 100.00%
Bootstrap support for afum|EAL88585.1 as seed ortholog is 100%.
Bootstrap support for afis|EAW15952.1 as seed ortholog is 100%.

---

Group of orthologs #2159. Best score 1412 bits
Score difference with first non-orthologous sequence - *A. fumigatus*:1412 *A. fischeri*:1412
afum|EAL88901.1 100.00% afis|EAW16280.1 100.00%
Bootstrap support for afum|EAL88901.1 as seed ortholog is 100%.
Bootstrap support for afis|EAW16280.1 as seed ortholog is 100%.

---

Group of orthologs #2160. Best score 1412 bits
Score difference with first non-orthologous sequence - *A. fumigatus*:1412 *A. fischeri*:1412
afum|EAL87499.1 100.00% afis|EAW18142.1 100.00%
Bootstrap support for afum|EAL87499.1 as seed ortholog is 100%.
Bootstrap support for afis|EAW18142.1 as seed ortholog is 100%.

---

Group of orthologs #2161. Best score 1412 bits
Score difference with first non-orthologous sequence - *A. fumigatus*:1412 *A. fischeri*:1412
afum|EAL87040.1 100.00% afis|EAW19456.1 100.00%
Bootstrap support for afum|EAL87040.1 as seed ortholog is 100%.
Bootstrap support for afis|EAW19456.1 as seed ortholog is 100%.

---

Group of orthologs #2162. Best score 1412 bits
Score difference with first non-orthologous sequence - *A. fumigatus*:1412 *A. fischeri*:1412
afum|EAL91980.1 100.00% afis|EAW24124.1 100.00%
Bootstrap support for afum|EAL91980.1 as seed ortholog is 100%.
Bootstrap support for afis|EAW24124.1 as seed ortholog is 100%.

---

Group of orthologs #2163. Best score 1412 bits
Score difference with first non-orthologous sequence - *A. fumigatus*:1412 *A. fischeri*:1412
afum|EAL92003.1 100.00% afis|EAW24148.1 100.00%
Bootstrap support for afum|EAL92003.1 as seed ortholog is 100%.
Bootstrap support for afis|EAW24148.1 as seed ortholog is 100%.

---

Group of orthologs #2164. Best score 1411 bits
Score difference with first non-orthologous sequence - *A. fumigatus*:1411 *A. fischeri*:1411
afum|EAL85554.1 100.00% afis|EAW15796.1 100.00%
Bootstrap support for afum|EAL85554.1 as seed ortholog is 100%.
Bootstrap support for afis|EAW15796.1 as seed ortholog is 100%.

---

Group of orthologs #2165. Best score 1411 bits
Score difference with first non-orthologous sequence - *A. fumigatus*:1411 *A. fischeri*:1411
afum|EAL84543.2 100.00% afis|EAW20514.1 100.00%
Bootstrap support for afum|EAL84543.2 as seed ortholog is 100%.
Bootstrap support for afis|EAW20514.1 as seed ortholog is 100%.

---

Group of orthologs #2166. Best score 1411 bits
Score difference with first non-orthologous sequence - *A. fumigatus*:1411 *A. fischeri*:1411
afum|EAL91890.1 100.00% afis|EAW18078.1 100.00%
Bootstrap support for afum|EAL91890.1 as seed ortholog is 100%.
Bootstrap support for afis|EAW18078.1 as seed ortholog is 100%.

---

Group of orthologs #2167. Best score 1411 bits
Score difference with first non-orthologous sequence - *A. fumigatus*:1411 *A. fischeri*:1411
afum|EAL92119.1 100.00% afis|EAW21096.1 100.00%
Bootstrap support for afum|EAL92119.1 as seed ortholog is 100%.
Bootstrap support for afis|EAW21096.1 as seed ortholog is 100%.

---

Group of orthologs #2168. Best score 1411 bits
Score difference with first non-orthologous sequence - *A. fumigatus*:1411 *A. fischeri*:1411
afum|EAL92803.1 100.00% afis|EAW21774.1 100.00%
Bootstrap support for afum|EAL92803.1 as seed ortholog is 100%.
Bootstrap support for afis|EAW21774.1 as seed ortholog is 100%.

---

Group of orthologs #2169. Best score 1411 bits
Score difference with first non-orthologous sequence - *A. fumigatus*:1411 *A. fischeri*:1411
afum|EAL92916.1 100.00% afis|EAW21903.1 100.00%
Bootstrap support for afum|EAL92916.1 as seed ortholog is 100%.
Bootstrap support for afis|EAW21903.1 as seed ortholog is 100%.

---

Group of orthologs #2170. Best score 1411 bits
Score difference with first non-orthologous sequence - *A. fumigatus*:1411 *A. fischeri*:804
afum|EAL89781.1 100.00% afis|EAW25110.1 100.00%
Bootstrap support for afum|EAL89781.1 as seed ortholog is 100%.
Bootstrap support for afis|EAW25110.1 as seed ortholog is 100%.

---

Group of orthologs #2171. Best score 1411 bits
Score difference with first non-orthologous sequence - *A. fumigatus*:1411 *A. fischeri*:1411
afum|EBA27457.1 100.00% afis|EAW21269.1 100.00%
Bootstrap support for afum|EBA27457.1 as seed ortholog is 100%.
Bootstrap support for afis|EAW21269.1 as seed ortholog is 100%.

---

Group of orthologs #2172. Best score 1411 bits
Score difference with first non-orthologous sequence - *A. fumigatus*:1411 *A. fischeri*:1411

afum|EAL90060.1 100.00% afis|EAW25401.1 100.00%
Bootstrap support for afum|EAL90060.1 as seed ortholog is 100%.
Bootstrap support for afis|EAW25401.1 as seed ortholog is 100%.

---

Group of orthologs #2173. Best score 1410 bits
Score difference with first non-orthologous sequence - *A. fumigatus*:1410 *A. fischeri*:1410
afum|EAL85094.1 100.00% afis|EAW19796.1 100.00%
Bootstrap support for afum|EAL85094.1 as seed ortholog is 100%.
Bootstrap support for afis|EAW19796.1 as seed ortholog is 100%.

---

Group of orthologs #2174. Best score 1410 bits
Score difference with first non-orthologous sequence - *A. fumigatus*:1410 *A. fischeri*:1410
afum|EAL86000.1 100.00% afis|EAW24330.1 100.00%
Bootstrap support for afum|EAL86000.1 as seed ortholog is 100%.
Bootstrap support for afis|EAW24330.1 as seed ortholog is 100%.

---

Group of orthologs #2175. Best score 1410 bits
Score difference with first non-orthologous sequence - *A. fumigatus*:1410 *A. fischeri*:1410
afum|EAL89791.1 100.00% afis|EAW25120.1 100.00%
Bootstrap support for afum|EAL89791.1 as seed ortholog is 100%.
Bootstrap support for afis|EAW25120.1 as seed ortholog is 100%.

---

Group of orthologs #2176. Best score 1410 bits
Score difference with first non-orthologous sequence - *A. fumigatus*:1410 *A. fischeri*:1410
afum|EAL90079.1 100.00% afis|EAW25420.1 100.00%
Bootstrap support for afum|EAL90079.1 as seed ortholog is 100%.
Bootstrap support for afis|EAW25420.1 as seed ortholog is 100%.

---

Group of orthologs #2177. Best score 1409 bits
Score difference with first non-orthologous sequence - *A. fumigatus*:1409 *A. fischeri*:1409
afum|EAL85885.1 100.00% afis|EAW24216.1 100.00%
Bootstrap support for afum|EAL85885.1 as seed ortholog is 100%.
Bootstrap support for afis|EAW24216.1 as seed ortholog is 100%.

---

Group of orthologs #2178. Best score 1409 bits
Score difference with first non-orthologous sequence - *A. fumigatus*:1409 *A. fischeri*:1409
afum|EAL93761.1 100.00% afis|EAW19032.1 100.00%
Bootstrap support for afum|EAL93761.1 as seed ortholog is 100%.
Bootstrap support for afis|EAW19032.1 as seed ortholog is 100%.

---

Group of orthologs #2179. Best score 1408 bits
Score difference with first non-orthologous sequence - *A. fumigatus*:1408 *A. fischeri*:1408
afum|EAL85664.1 100.00% afis|EAW15684.1 100.00%
Bootstrap support for afum|EAL85664.1 as seed ortholog is 100%.
Bootstrap support for afis|EAW15684.1 as seed ortholog is 100%.

---

Group of orthologs #2180. Best score 1408 bits
Score difference with first non-orthologous sequence - *A. fumigatus*:1408 *A. fischeri*:1408
afum|EAL93834.1 100.00% afis|EAW19102.1 100.00%
Bootstrap support for afum|EAL93834.1 as seed ortholog is 100%.
Bootstrap support for afis|EAW19102.1 as seed ortholog is 100%.

---

Group of orthologs #2181. Best score 1408 bits
Score difference with first non-orthologous sequence - *A. fumigatus*:1408 *A. fischeri*:1311
afum|EAL91049.1 100.00% afis|EAW22141.1 100.00%
Bootstrap support for afum|EAL91049.1 as seed ortholog is 100%.

Bootstrap support for afis|EAW22141.1 as seed ortholog is 100%.

---

Group of orthologs #2182. Best score 1407 bits
Score difference with first non-orthologous sequence - *A. fumigatus*:355 *A. fischeri*:1407
afum|EAL85174.2 100.00% afis|EAW20054.1 100.00%
Bootstrap support for afum|EAL85174.2 as seed ortholog is 100%.
Bootstrap support for afis|EAW20054.1 as seed ortholog is 100%.

---

Group of orthologs #2183. Best score 1407 bits
Score difference with first non-orthologous sequence - *A. fumigatus*:1407 *A. fischeri*:1407
afum|EAL86058.1 100.00% afis|EAW24386.1 100.00%
Bootstrap support for afum|EAL86058.1 as seed ortholog is 100%.
Bootstrap support for afis|EAW24386.1 as seed ortholog is 100%.

---

Group of orthologs #2184. Best score 1405 bits
Score difference with first non-orthologous sequence - *A. fumigatus*:1405 *A. fischeri*:1405
afum|EAL88662.1 100.00% afis|EAW16031.1 100.00%
Bootstrap support for afum|EAL88662.1 as seed ortholog is 100%.
Bootstrap support for afis|EAW16031.1 as seed ortholog is 100%.

---

Group of orthologs #2185. Best score 1405 bits
Score difference with first non-orthologous sequence - *A. fumigatus*:1405 *A. fischeri*:1405
afum|EAL88695.1 100.00% afis|EAW16063.1 100.00%
Bootstrap support for afum|EAL88695.1 as seed ortholog is 100%.
Bootstrap support for afis|EAW16063.1 as seed ortholog is 100%.

---

Group of orthologs #2186. Best score 1405 bits
Score difference with first non-orthologous sequence - *A. fumigatus*:1405 *A. fischeri*:1405
afum|EAL88314.1 100.00% afis|EAW23170.1 100.00%
Bootstrap support for afum|EAL88314.1 as seed ortholog is 100%.
Bootstrap support for afis|EAW23170.1 as seed ortholog is 100%.

---

Group of orthologs #2187. Best score 1405 bits
Score difference with first non-orthologous sequence - *A. fumigatus*:1405 *A. fischeri*:1405
afum|EAL88202.1 100.00% afis|EAW23277.1 100.00%
Bootstrap support for afum|EAL88202.1 as seed ortholog is 100%.
Bootstrap support for afis|EAW23277.1 as seed ortholog is 100%.

---

Group of orthologs #2188. Best score 1405 bits
Score difference with first non-orthologous sequence - *A. fumigatus*:1405 *A. fischeri*:1405
afum|EAL93806.2 100.00% afis|EAW19078.1 100.00%
Bootstrap support for afum|EAL93806.2 as seed ortholog is 100%.
Bootstrap support for afis|EAW19078.1 as seed ortholog is 100%.

---

Group of orthologs #2189. Best score 1404 bits
Score difference with first non-orthologous sequence - *A. fumigatus*:1404 *A. fischeri*:1404
afum|EAL88987.1 100.00% afis|EAW16362.1 100.00%
Bootstrap support for afum|EAL88987.1 as seed ortholog is 100%.
Bootstrap support for afis|EAW16362.1 as seed ortholog is 100%.

---

Group of orthologs #2190. Best score 1404 bits
Score difference with first non-orthologous sequence - *A. fumigatus*:1404 *A. fischeri*:1404
afum|EAL92837.1 100.00% afis|EAW21821.1 100.00%
Bootstrap support for afum|EAL92837.1 as seed ortholog is 100%.
Bootstrap support for afis|EAW21821.1 as seed ortholog is 100%.

---

Group of orthologs #2191. Best score 1404 bits
Score difference with first non-orthologous sequence - *A. fumigatus*:1404 *A. fischeri*:1404
afum|EAL90018.1 100.00% afis|EAW25356.1 100.00%
Bootstrap support for afum|EAL90018.1 as seed ortholog is 100%.
Bootstrap support for afis|EAW25356.1 as seed ortholog is 100%.

---

Group of orthologs #2192. Best score 1403 bits
Score difference with first non-orthologous sequence - *A. fumigatus*:1403 *A. fischeri*:1058
afum|EAL88969.1 100.00% afis|EAW16344.1 100.00%
Bootstrap support for afum|EAL88969.1 as seed ortholog is 100%.
Bootstrap support for afis|EAW16344.1 as seed ortholog is 100%.

---

Group of orthologs #2193. Best score 1403 bits
Score difference with first non-orthologous sequence - *A. fumigatus*:1403 *A. fischeri*:1074
afum|EAL85409.1 100.00% afis|EAW20275.1 100.00%
Bootstrap support for afum|EAL85409.1 as seed ortholog is 100%.
Bootstrap support for afis|EAW20275.1 as seed ortholog is 100%.

---

Group of orthologs #2194. Best score 1402 bits
Score difference with first non-orthologous sequence - *A. fumigatus*:1402 *A. fischeri*:1175
afum|EAL85855.1 100.00% afis|EAW15467.1 100.00%
Bootstrap support for afum|EAL85855.1 as seed ortholog is 100%.
Bootstrap support for afis|EAW15467.1 as seed ortholog is 100%.

---

Group of orthologs #2195. Best score 1402 bits
Score difference with first non-orthologous sequence - *A. fumigatus*:1402 *A. fischeri*:1119
afum|EAL85448.1 100.00% afis|EAW20323.1 100.00%
Bootstrap support for afum|EAL85448.1 as seed ortholog is 100%.
Bootstrap support for afis|EAW20323.1 as seed ortholog is 100%.

---

Group of orthologs #2196. Best score 1402 bits
Score difference with first non-orthologous sequence - *A. fumigatus*:1307 *A. fischeri*:1402
afum|EAL87656.1 100.00% afis|EAW18291.1 100.00%
Bootstrap support for afum|EAL87656.1 as seed ortholog is 100%.
Bootstrap support for afis|EAW18291.1 as seed ortholog is 100%.

---

Group of orthologs #2197. Best score 1402 bits
Score difference with first non-orthologous sequence - *A. fumigatus*:1402 *A. fischeri*:1402
afum|EAL84328.2 100.00% afis|EAW25641.1 100.00%
Bootstrap support for afum|EAL84328.2 as seed ortholog is 100%.
Bootstrap support for afis|EAW25641.1 as seed ortholog is 100%.

---

Group of orthologs #2198. Best score 1402 bits
Score difference with first non-orthologous sequence - *A. fumigatus*:1402 *A. fischeri*:1402
afum|EAL87269.1 100.00% afis|EAW23838.1 100.00%
Bootstrap support for afum|EAL87269.1 as seed ortholog is 100%.
Bootstrap support for afis|EAW23838.1 as seed ortholog is 100%.

---

Group of orthologs #2199. Best score 1402 bits
Score difference with first non-orthologous sequence - *A. fumigatus*:1402 *A. fischeri*:398
afum|EAL93716.2 100.00% afis|EAW18980.1 100.00%
Bootstrap support for afum|EAL93716.2 as seed ortholog is 100%.
Bootstrap support for afis|EAW18980.1 as seed ortholog is 100%.

---

Group of orthologs #2200. Best score 1401 bits
Score difference with first non-orthologous sequence - *A. fumigatus*:1264 *A. fischeri*:1146

afum|EAL85736.2 100.00% afis|EAW15595.1 100.00%
Bootstrap support for afum|EAL85736.2 as seed ortholog is 100%.
Bootstrap support for afis|EAW15595.1 as seed ortholog is 100%.

---

Group of orthologs #2201. Best score 1401 bits
Score difference with first non-orthologous sequence - *A. fumigatus*:1401 *A. fischeri*:1317
afum|EAL86626.2 100.00% afis|EAW17198.1 100.00%
Bootstrap support for afum|EAL86626.2 as seed ortholog is 100%.
Bootstrap support for afis|EAW17198.1 as seed ortholog is 100%.

---

Group of orthologs #2202. Best score 1401 bits
Score difference with first non-orthologous sequence - *A. fumigatus*:1401 *A. fischeri*:1401
afum|EAL87511.1 100.00% afis|EAW18152.1 100.00%
Bootstrap support for afum|EAL87511.1 as seed ortholog is 100%.
Bootstrap support for afis|EAW18152.1 as seed ortholog is 100%.

---

Group of orthologs #2203. Best score 1401 bits
Score difference with first non-orthologous sequence - *A. fumigatus*:584 *A. fischeri*:1401
afum|EAL84638.1 100.00% afis|EAW20991.1 100.00%
Bootstrap support for afum|EAL84638.1 as seed ortholog is 100%.
Bootstrap support for afis|EAW20991.1 as seed ortholog is 100%.

---

Group of orthologs #2204. Best score 1401 bits
Score difference with first non-orthologous sequence - *A. fumigatus*:1401 *A. fischeri*:1401
afum|EAL93604.1 100.00% afis|EAW18872.1 100.00%
Bootstrap support for afum|EAL93604.1 as seed ortholog is 100%.
Bootstrap support for afis|EAW18872.1 as seed ortholog is 100%.

---

Group of orthologs #2205. Best score 1401 bits
Score difference with first non-orthologous sequence - *A. fumigatus*:1401 *A. fischeri*:1401
afum|EAL92103.1 100.00% afis|EAW21082.1 100.00%
Bootstrap support for afum|EAL92103.1 as seed ortholog is 100%.
Bootstrap support for afis|EAW21082.1 as seed ortholog is 100%.

---

Group of orthologs #2206. Best score 1401 bits
Score difference with first non-orthologous sequence - *A. fumigatus*:1401 *A. fischeri*:1401
afum|EAL89912.1 100.00% afis|EAW25250.1 100.00%
Bootstrap support for afum|EAL89912.1 as seed ortholog is 100%.
Bootstrap support for afis|EAW25250.1 as seed ortholog is 100%.

---

Group of orthologs #2207. Best score 1400 bits
Score difference with first non-orthologous sequence - *A. fumigatus*:1203 *A. fischeri*:1093
afum|EAL88478.1 100.00% afis|EAW23002.1 100.00%
Bootstrap support for afum|EAL88478.1 as seed ortholog is 100%.
Bootstrap support for afis|EAW23002.1 as seed ortholog is 100%.

---

Group of orthologs #2208. Best score 1400 bits
Score difference with first non-orthologous sequence - *A. fumigatus*:1400 *A. fischeri*:1400
afum|EAL89970.1 100.00% afis|EAW25307.1 100.00%
Bootstrap support for afum|EAL89970.1 as seed ortholog is 100%.
Bootstrap support for afis|EAW25307.1 as seed ortholog is 100%.

---

Group of orthologs #2209. Best score 1399 bits
Score difference with first non-orthologous sequence - *A. fumigatus*:1292 *A. fischeri*:1318
afum|EAL85799.1 100.00% afis|EAW15529.1 100.00%
Bootstrap support for afum|EAL85799.1 as seed ortholog is 100%.

Bootstrap support for afis|EAW15529.1 as seed ortholog is 100%.

---

Group of orthologs #2210. Best score 1399 bits
Score difference with first non-orthologous sequence - *A. fumigatus*:29 *A. fischeri*:1399
afum|EAL89358.1 100.00% afis|EAW18029.1 100.00%
Bootstrap support for afum|EAL89358.1 as seed ortholog is 79%.
Bootstrap support for afis|EAW18029.1 as seed ortholog is 100%.

---

Group of orthologs #2211. Best score 1399 bits
Score difference with first non-orthologous sequence - *A. fumigatus*:1399 *A. fischeri*:1399
afum|EAL84283.1 100.00% afis|EAW25559.1 100.00%
Bootstrap support for afum|EAL84283.1 as seed ortholog is 100%.
Bootstrap support for afis|EAW25559.1 as seed ortholog is 100%.

---

Group of orthologs #2212. Best score 1398 bits
Score difference with first non-orthologous sequence - *A. fumigatus*:1087 *A. fischeri*:1398
afum|EAL87042.1 100.00% afis|EAW19454.1 100.00%
Bootstrap support for afum|EAL87042.1 as seed ortholog is 100%.
Bootstrap support for afis|EAW19454.1 as seed ortholog is 100%.

---

Group of orthologs #2213. Best score 1398 bits
Score difference with first non-orthologous sequence - *A. fumigatus*:1398 *A. fischeri*:1398
afum|EAL90581.1 100.00% afis|EAW22616.1 100.00%
Bootstrap support for afum|EAL90581.1 as seed ortholog is 100%.
Bootstrap support for afis|EAW22616.1 as seed ortholog is 100%.

---

Group of orthologs #2214. Best score 1398 bits
Score difference with first non-orthologous sequence - *A. fumigatus*:1398 *A. fischeri*:1225
afum|EAL91915.1 100.00% afis|EAW24060.1 100.00%
Bootstrap support for afum|EAL91915.1 as seed ortholog is 100%.
Bootstrap support for afis|EAW24060.1 as seed ortholog is 100%.

---

Group of orthologs #2215. Best score 1397 bits
Score difference with first non-orthologous sequence - *A. fumigatus*:1397 *A. fischeri*:1397
afum|EAL87310.1 100.00% afis|EAW23882.1 100.00%
Bootstrap support for afum|EAL87310.1 as seed ortholog is 100%.
Bootstrap support for afis|EAW23882.1 as seed ortholog is 100%.

---

Group of orthologs #2216. Best score 1396 bits
Score difference with first non-orthologous sequence - *A. fumigatus*:1396 *A. fischeri*:1396
afum|EAL88718.1 100.00% afis|EAW16086.1 100.00%
Bootstrap support for afum|EAL88718.1 as seed ortholog is 100%.
Bootstrap support for afis|EAW16086.1 as seed ortholog is 100%.

---

Group of orthologs #2217. Best score 1396 bits
Score difference with first non-orthologous sequence - *A. fumigatus*:1396 *A. fischeri*:1396
afum|EAL85281.1 100.00% afis|EAW20163.1 100.00%
Bootstrap support for afum|EAL85281.1 as seed ortholog is 100%.
Bootstrap support for afis|EAW20163.1 as seed ortholog is 100%.

---

Group of orthologs #2218. Best score 1395 bits
Score difference with first non-orthologous sequence - *A. fumigatus*:1395 *A. fischeri*:1395
afum|EAL88698.1 100.00% afis|EAW16066.1 100.00%
Bootstrap support for afum|EAL88698.1 as seed ortholog is 100%.
Bootstrap support for afis|EAW16066.1 as seed ortholog is 100%.

---

Group of orthologs #2219. Best score 1395 bits
Score difference with first non-orthologous sequence - *A. fumigatus*:1010 *A. fischeri*:870
afum|EAL84745.1 100.00% afis|EAW20888.1 100.00%
Bootstrap support for afum|EAL84745.1 as seed ortholog is 100%.
Bootstrap support for afis|EAW20888.1 as seed ortholog is 100%.

---

Group of orthologs #2220. Best score 1395 bits
Score difference with first non-orthologous sequence - *A. fumigatus*:1395 *A. fischeri*:1395
afum|EAL84634.1 100.00% afis|EAW20994.1 100.00%
Bootstrap support for afum|EAL84634.1 as seed ortholog is 100%.
Bootstrap support for afis|EAW20994.1 as seed ortholog is 100%.

---

Group of orthologs #2221. Best score 1395 bits
Score difference with first non-orthologous sequence - *A. fumigatus*:1395 *A. fischeri*:1395
afum|EAL90839.1 100.00% afis|EAW22353.1 100.00%
Bootstrap support for afum|EAL90839.1 as seed ortholog is 100%.
Bootstrap support for afis|EAW22353.1 as seed ortholog is 100%.

---

Group of orthologs #2222. Best score 1395 bits
Score difference with first non-orthologous sequence - *A. fumigatus*:1395 *A. fischeri*:1395
afum|EAL92193.1 100.00% afis|EAW21173.1 100.00%
Bootstrap support for afum|EAL92193.1 as seed ortholog is 100%.
Bootstrap support for afis|EAW21173.1 as seed ortholog is 100%.

---

Group of orthologs #2223. Best score 1394 bits
Score difference with first non-orthologous sequence - *A. fumigatus*:1394 *A. fischeri*:1394
afum|EAL86174.1 100.00% afis|EAW24506.1 100.00%
Bootstrap support for afum|EAL86174.1 as seed ortholog is 100%.
Bootstrap support for afis|EAW24506.1 as seed ortholog is 100%.

---

Group of orthologs #2224. Best score 1394 bits
Score difference with first non-orthologous sequence - *A. fumigatus*:1323 *A. fischeri*:1394
afum|EAL90969.1 100.00% afis|EAW22225.1 100.00%
Bootstrap support for afum|EAL90969.1 as seed ortholog is 100%.
Bootstrap support for afis|EAW22225.1 as seed ortholog is 100%.

---

Group of orthologs #2225. Best score 1394 bits
Score difference with first non-orthologous sequence - *A. fumigatus*:1394 *A. fischeri*:1394
afum|EAL90240.1 100.00% afis|EAW22957.1 100.00%
Bootstrap support for afum|EAL90240.1 as seed ortholog is 100%.
Bootstrap support for afis|EAW22957.1 as seed ortholog is 100%.

---

Group of orthologs #2226. Best score 1393 bits
Score difference with first non-orthologous sequence - *A. fumigatus*:1214 *A. fischeri*:1202
afum|EAL84421.1 100.00% afis|EAW20651.1 100.00%
Bootstrap support for afum|EAL84421.1 as seed ortholog is 100%.
Bootstrap support for afis|EAW20651.1 as seed ortholog is 100%.

---

Group of orthologs #2227. Best score 1393 bits
Score difference with first non-orthologous sequence - *A. fumigatus*:1393 *A. fischeri*:1393
afum|EAL87708.1 100.00% afis|EAW18344.1 100.00%
Bootstrap support for afum|EAL87708.1 as seed ortholog is 100%.
Bootstrap support for afis|EAW18344.1 as seed ortholog is 100%.

---

Group of orthologs #2228. Best score 1393 bits
Score difference with first non-orthologous sequence - *A. fumigatus*:1393 *A. fischeri*:1393

afum|EAL88169.1 100.00% afis|EAW23308.1 100.00%
Bootstrap support for afum|EAL88169.1 as seed ortholog is 100%.
Bootstrap support for afis|EAW23308.1 as seed ortholog is 100%.

---

Group of orthologs #2229. Best score 1392 bits
Score difference with first non-orthologous sequence - *A. fumigatus*:1392 *A. fischeri*:1392
afum|EAL88309.1 100.00% afis|EAW23174.1 100.00%
Bootstrap support for afum|EAL88309.1 as seed ortholog is 100%.
Bootstrap support for afis|EAW23174.1 as seed ortholog is 100%.

---

Group of orthologs #2230. Best score 1392 bits
Score difference with first non-orthologous sequence - *A. fumigatus*:1392 *A. fischeri*:1392
afum|EAL92077.2 100.00% afis|EAW21038.1 100.00%
Bootstrap support for afum|EAL92077.2 as seed ortholog is 100%.
Bootstrap support for afis|EAW21038.1 as seed ortholog is 100%.

---

Group of orthologs #2231. Best score 1392 bits
Score difference with first non-orthologous sequence - *A. fumigatus*:1392 *A. fischeri*:1392
afum|EAL92841.1 100.00% afis|EAW21826.1 100.00%
Bootstrap support for afum|EAL92841.1 as seed ortholog is 100%.
Bootstrap support for afis|EAW21826.1 as seed ortholog is 100%.

---

Group of orthologs #2232. Best score 1392 bits
Score difference with first non-orthologous sequence - *A. fumigatus*:1392 *A. fischeri*:1392
afum|EAL92915.1 100.00% afis|EAW21902.1 100.00%
Bootstrap support for afum|EAL92915.1 as seed ortholog is 100%.
Bootstrap support for afis|EAW21902.1 as seed ortholog is 100%.

---

Group of orthologs #2233. Best score 1391 bits
Score difference with first non-orthologous sequence - *A. fumigatus*:1310 *A. fischeri*:1391
afum|EAL86834.1 100.00% afis|EAW19692.1 100.00%
Bootstrap support for afum|EAL86834.1 as seed ortholog is 100%.
Bootstrap support for afis|EAW19692.1 as seed ortholog is 100%.

---

Group of orthologs #2234. Best score 1391 bits
Score difference with first non-orthologous sequence - *A. fumigatus*:1391 *A. fischeri*:1063
afum|EAL91600.1 100.00% afis|EAW17791.1 100.00%
Bootstrap support for afum|EAL91600.1 as seed ortholog is 100%.
Bootstrap support for afis|EAW17791.1 as seed ortholog is 100%.

---

Group of orthologs #2235. Best score 1391 bits
Score difference with first non-orthologous sequence - *A. fumigatus*:1391 *A. fischeri*:1391
afum|EAL89870.1 100.00% afis|EAW25209.1 100.00%
Bootstrap support for afum|EAL89870.1 as seed ortholog is 100%.
Bootstrap support for afis|EAW25209.1 as seed ortholog is 100%.

---

Group of orthologs #2236. Best score 1390 bits
Score difference with first non-orthologous sequence - *A. fumigatus*:1390 *A. fischeri*:1390
afum|EAL88773.1 100.00% afis|EAW16144.1 100.00%
Bootstrap support for afum|EAL88773.1 as seed ortholog is 100%.
Bootstrap support for afis|EAW16144.1 as seed ortholog is 100%.

---

Group of orthologs #2237. Best score 1390 bits
Score difference with first non-orthologous sequence - *A. fumigatus*:1390 *A. fischeri*:1390
afum|EAL89509.1 100.00% afis|EAW24860.1 100.00%
Bootstrap support for afum|EAL89509.1 as seed ortholog is 100%.

Bootstrap support for afis|EAW24860.1 as seed ortholog is 100%.

---

Group of orthologs #2238. Best score 1389 bits
Score difference with first non-orthologous sequence - *A. fumigatus*:1389 *A. fischeri*:1389
afum|EAL85545.1 100.00% afis|EAW15805.1 100.00%
Bootstrap support for afum|EAL85545.1 as seed ortholog is 100%.
Bootstrap support for afis|EAW15805.1 as seed ortholog is 100%.

---

Group of orthologs #2239. Best score 1389 bits
Score difference with first non-orthologous sequence - *A. fumigatus*:1389 *A. fischeri*:1389
afum|EAL88643.1 100.00% afis|EAW16014.1 100.00%
Bootstrap support for afum|EAL88643.1 as seed ortholog is 100%.
Bootstrap support for afis|EAW16014.1 as seed ortholog is 100%.

---

Group of orthologs #2240. Best score 1389 bits
Score difference with first non-orthologous sequence - *A. fumigatus*:1389 *A. fischeri*:1389
afum|EAL86881.1 100.00% afis|EAW19641.1 100.00%
Bootstrap support for afum|EAL86881.1 as seed ortholog is 100%.
Bootstrap support for afis|EAW19641.1 as seed ortholog is 100%.

---

Group of orthologs #2241. Best score 1388 bits
Score difference with first non-orthologous sequence - *A. fumigatus*:1388 *A. fischeri*:1388
afum|EAL84725.1 100.00% afis|EAW20908.1 100.00%
Bootstrap support for afum|EAL84725.1 as seed ortholog is 100%.
Bootstrap support for afis|EAW20908.1 as seed ortholog is 100%.

---

Group of orthologs #2242. Best score 1388 bits
Score difference with first non-orthologous sequence - *A. fumigatus*:1388 *A. fischeri*:1388
afum|EAL90909.1 100.00% afis|EAW22271.1 100.00%
Bootstrap support for afum|EAL90909.1 as seed ortholog is 100%.
Bootstrap support for afis|EAW22271.1 as seed ortholog is 100%.

---

Group of orthologs #2243. Best score 1388 bits
Score difference with first non-orthologous sequence - *A. fumigatus*:1388 *A. fischeri*:1388
afum|EAL92574.1 100.00% afis|EAW21554.1 100.00%
Bootstrap support for afum|EAL92574.1 as seed ortholog is 100%.
Bootstrap support for afis|EAW21554.1 as seed ortholog is 100%.

---

Group of orthologs #2244. Best score 1388 bits
Score difference with first non-orthologous sequence - *A. fumigatus*:1388 *A. fischeri*:1388
afum|EBA27465.1 100.00% afis|EAW21555.1 100.00%
Bootstrap support for afum|EBA27465.1 as seed ortholog is 100%.
Bootstrap support for afis|EAW21555.1 as seed ortholog is 100%.

---

Group of orthologs #2245. Best score 1387 bits
Score difference with first non-orthologous sequence - *A. fumigatus*:1387 *A. fischeri*:1387
afum|EAL85226.1 100.00% afis|EAW20109.1 100.00%
Bootstrap support for afum|EAL85226.1 as seed ortholog is 100%.
Bootstrap support for afis|EAW20109.1 as seed ortholog is 100%.

---

Group of orthologs #2246. Best score 1387 bits
Score difference with first non-orthologous sequence - *A. fumigatus*:1387 *A. fischeri*:1136
afum|EAL88442.1 100.00% afis|EAW23039.1 100.00%
Bootstrap support for afum|EAL88442.1 as seed ortholog is 100%.
Bootstrap support for afis|EAW23039.1 as seed ortholog is 100%.

---

Group of orthologs #2247. Best score 1387 bits
Score difference with first non-orthologous sequence - *A. fumigatus*:1387 *A. fischeri*:1387
afum|EAL90995.1 100.00% afis|EAW22196.1 100.00%
Bootstrap support for afum|EAL90995.1 as seed ortholog is 100%.
Bootstrap support for afis|EAW22196.1 as seed ortholog is 100%.

---

Group of orthologs #2248. Best score 1387 bits
Score difference with first non-orthologous sequence - *A. fumigatus*:1387 *A. fischeri*:1387
afum|EAL90864.1 100.00% afis|EAW22328.1 100.00%
Bootstrap support for afum|EAL90864.1 as seed ortholog is 100%.
Bootstrap support for afis|EAW22328.1 as seed ortholog is 100%.

---

Group of orthologs #2249. Best score 1386 bits
Score difference with first non-orthologous sequence - *A. fumigatus*:1323 *A. fischeri*:1250
afum|EAL91363.1 100.00% afis|EAW17549.1 100.00%
Bootstrap support for afum|EAL91363.1 as seed ortholog is 100%.
Bootstrap support for afis|EAW17549.1 as seed ortholog is 100%.

---

Group of orthologs #2250. Best score 1386 bits
Score difference with first non-orthologous sequence - *A. fumigatus*:1386 *A. fischeri*:1386
afum|EAL91618.1 100.00% afis|EAW17810.1 100.00%
Bootstrap support for afum|EAL91618.1 as seed ortholog is 100%.
Bootstrap support for afis|EAW17810.1 as seed ortholog is 100%.

---

Group of orthologs #2251. Best score 1386 bits
Score difference with first non-orthologous sequence - *A. fumigatus*:1386 *A. fischeri*:1386
afum|EAL93875.1 100.00% afis|EAW19144.1 100.00%
Bootstrap support for afum|EAL93875.1 as seed ortholog is 100%.
Bootstrap support for afis|EAW19144.1 as seed ortholog is 100%.

---

Group of orthologs #2252. Best score 1386 bits
Score difference with first non-orthologous sequence - *A. fumigatus*:1386 *A. fischeri*:1386
afum|EAL90515.1 100.00% afis|EAW22677.1 100.00%
Bootstrap support for afum|EAL90515.1 as seed ortholog is 100%.
Bootstrap support for afis|EAW22677.1 as seed ortholog is 100%.

---

Group of orthologs #2253. Best score 1385 bits
Score difference with first non-orthologous sequence - *A. fumigatus*:1385 *A. fischeri*:1385
afum|EAL85037.1 100.00% afis|EAW19858.1 100.00%
Bootstrap support for afum|EAL85037.1 as seed ortholog is 100%.
Bootstrap support for afis|EAW19858.1 as seed ortholog is 100%.

---

Group of orthologs #2254. Best score 1385 bits
Score difference with first non-orthologous sequence - *A. fumigatus*:1385 *A. fischeri*:1385
afum|EAL84425.1 100.00% afis|EAW20648.1 100.00%
Bootstrap support for afum|EAL84425.1 as seed ortholog is 100%.
Bootstrap support for afis|EAW20648.1 as seed ortholog is 100%.

---

Group of orthologs #2255. Best score 1385 bits
Score difference with first non-orthologous sequence - *A. fumigatus*:1094 *A. fischeri*:1173
afum|EAL84357.1 100.00% afis|EAW25688.1 100.00%
Bootstrap support for afum|EAL84357.1 as seed ortholog is 100%.
Bootstrap support for afis|EAW25688.1 as seed ortholog is 100%.

---

Group of orthologs #2256. Best score 1385 bits
Score difference with first non-orthologous sequence - *A. fumigatus*:1385 *A. fischeri*:1385

afum|EAL86045.1 100.00% afis|EAW24373.1 100.00%
Bootstrap support for afum|EAL86045.1 as seed ortholog is 100%.
Bootstrap support for afis|EAW24373.1 as seed ortholog is 100%.

---

Group of orthologs #2257. Best score 1385 bits
Score difference with first non-orthologous sequence - *A. fumigatus*:1385 *A. fischeri*:1385
afum|EAL88190.1 100.00% afis|EAW23289.1 100.00%
Bootstrap support for afum|EAL88190.1 as seed ortholog is 100%.
Bootstrap support for afis|EAW23289.1 as seed ortholog is 100%.

---

Group of orthologs #2258. Best score 1385 bits
Score difference with first non-orthologous sequence - *A. fumigatus*:1385 *A. fischeri*:1385
afum|EAL90584.1 100.00% afis|EAW22614.1 100.00%
Bootstrap support for afum|EAL90584.1 as seed ortholog is 100%.
Bootstrap support for afis|EAW22614.1 as seed ortholog is 100%.

---

Group of orthologs #2259. Best score 1385 bits
Score difference with first non-orthologous sequence - *A. fumigatus*:1385 *A. fischeri*:1385
afum|EAL89310.1 100.00% afis|EAW24660.1 100.00%
Bootstrap support for afum|EAL89310.1 as seed ortholog is 100%.
Bootstrap support for afis|EAW24660.1 as seed ortholog is 100%.

---

Group of orthologs #2260. Best score 1384 bits
Score difference with first non-orthologous sequence - *A. fumigatus*:1384 *A. fischeri*:1384
afum|EAL85587.1 100.00% afis|EAW15761.1 100.00%
Bootstrap support for afum|EAL85587.1 as seed ortholog is 100%.
Bootstrap support for afis|EAW15761.1 as seed ortholog is 100%.

---

Group of orthologs #2261. Best score 1384 bits
Score difference with first non-orthologous sequence - *A. fumigatus*:1384 *A. fischeri*:502
afum|EAL84916.1 100.00% afis|EAW20003.1 100.00%
Bootstrap support for afum|EAL84916.1 as seed ortholog is 100%.
Bootstrap support for afis|EAW20003.1 as seed ortholog is 100%.

---

Group of orthologs #2262. Best score 1384 bits
Score difference with first non-orthologous sequence - *A. fumigatus*:1384 *A. fischeri*:1384
afum|EAL92308.1 100.00% afis|EAW21291.1 100.00%
Bootstrap support for afum|EAL92308.1 as seed ortholog is 100%.
Bootstrap support for afis|EAW21291.1 as seed ortholog is 100%.

---

Group of orthologs #2263. Best score 1383 bits
Score difference with first non-orthologous sequence - *A. fumigatus*:1220 *A. fischeri*:1192
afum|EAL84776.1 100.00% afis|EAW20859.1 100.00%
Bootstrap support for afum|EAL84776.1 as seed ortholog is 100%.
Bootstrap support for afis|EAW20859.1 as seed ortholog is 100%.

---

Group of orthologs #2264. Best score 1383 bits
Score difference with first non-orthologous sequence - *A. fumigatus*:1383 *A. fischeri*:1383
afum|EAL91691.1 100.00% afis|EAW17883.1 100.00%
Bootstrap support for afum|EAL91691.1 as seed ortholog is 100%.
Bootstrap support for afis|EAW17883.1 as seed ortholog is 100%.

---

Group of orthologs #2265. Best score 1383 bits
Score difference with first non-orthologous sequence - *A. fumigatus*:1383 *A. fischeri*:1383
afum|EAL87363.1 100.00% afis|EAW23938.1 100.00%
Bootstrap support for afum|EAL87363.1 as seed ortholog is 100%.

Bootstrap support for afis|EAW23938.1 as seed ortholog is 100%.

---

Group of orthologs #2266. Best score 1383 bits
Score difference with first non-orthologous sequence - *A. fumigatus*:1383 *A. fischeri*:862
afum|EAL88049.1 100.00% afis|EAW23420.1 100.00%
Bootstrap support for afum|EAL88049.1 as seed ortholog is 100%.
Bootstrap support for afis|EAW23420.1 as seed ortholog is 100%.

---

Group of orthologs #2267. Best score 1382 bits
Score difference with first non-orthologous sequence - *A. fumigatus*:1382 *A. fischeri*:1382
afum|EAL86935.2 100.00% afis|EAW19583.1 100.00%
Bootstrap support for afum|EAL86935.2 as seed ortholog is 100%.
Bootstrap support for afis|EAW19583.1 as seed ortholog is 100%.

---

Group of orthologs #2268. Best score 1382 bits
Score difference with first non-orthologous sequence - *A. fumigatus*:1382 *A. fischeri*:1382
afum|EAL88310.1 100.00% afis|EAW23173.1 100.00%
Bootstrap support for afum|EAL88310.1 as seed ortholog is 100%.
Bootstrap support for afis|EAW23173.1 as seed ortholog is 100%.

---

Group of orthologs #2269. Best score 1382 bits
Score difference with first non-orthologous sequence - *A. fumigatus*:1382 *A. fischeri*:1382
afum|EAL89394.1 100.00% afis|EAW24747.1 100.00%
Bootstrap support for afum|EAL89394.1 as seed ortholog is 100%.
Bootstrap support for afis|EAW24747.1 as seed ortholog is 100%.

---

Group of orthologs #2270. Best score 1381 bits
Score difference with first non-orthologous sequence - *A. fumigatus*:1381 *A. fischeri*:1155
afum|EAL86822.1 100.00% afis|EAW19704.1 100.00%
Bootstrap support for afum|EAL86822.1 as seed ortholog is 100%.
Bootstrap support for afis|EAW19704.1 as seed ortholog is 100%.

---

Group of orthologs #2271. Best score 1381 bits
Score difference with first non-orthologous sequence - *A. fumigatus*:1142 *A. fischeri*:1078
afum|EAL92830.1 100.00% afis|EAW21804.1 100.00%
Bootstrap support for afum|EAL92830.1 as seed ortholog is 100%.
Bootstrap support for afis|EAW21804.1 as seed ortholog is 100%.

---

Group of orthologs #2272. Best score 1380 bits
Score difference with first non-orthologous sequence - *A. fumigatus*:1380 *A. fischeri*:942
afum|EAL84550.1 100.00% afis|EAW20506.1 100.00%
Bootstrap support for afum|EAL84550.1 as seed ortholog is 100%.
Bootstrap support for afis|EAW20506.1 as seed ortholog is 100%.

---

Group of orthologs #2273. Best score 1380 bits
Score difference with first non-orthologous sequence - *A. fumigatus*:1249 *A. fischeri*:646
afum|EAL89216.2 100.00% afis|EAW16621.1 100.00%
Bootstrap support for afum|EAL89216.2 as seed ortholog is 100%.
Bootstrap support for afis|EAW16621.1 as seed ortholog is 100%.

---

Group of orthologs #2274. Best score 1379 bits
Score difference with first non-orthologous sequence - *A. fumigatus*:1379 *A. fischeri*:1379
afum|EAL89097.1 100.00% afis|EAW16502.1 100.00%
Bootstrap support for afum|EAL89097.1 as seed ortholog is 100%.
Bootstrap support for afis|EAW16502.1 as seed ortholog is 100%.

---

Group of orthologs #2275. Best score 1379 bits
Score difference with first non-orthologous sequence - *A. fumigatus*:1379 *A. fischeri*:1379
afum|EAL91281.2 100.00% afis|EAW17453.1 100.00%
Bootstrap support for afum|EAL91281.2 as seed ortholog is 100%.
Bootstrap support for afis|EAW17453.1 as seed ortholog is 100%.

---

Group of orthologs #2276. Best score 1379 bits
Score difference with first non-orthologous sequence - *A. fumigatus*:1379 *A. fischeri*:1379
afum|EAL93175.1 100.00% afis|EAW18486.1 100.00%
Bootstrap support for afum|EAL93175.1 as seed ortholog is 100%.
Bootstrap support for afis|EAW18486.1 as seed ortholog is 100%.

---

Group of orthologs #2277. Best score 1379 bits
Score difference with first non-orthologous sequence - *A. fumigatus*:789 *A. fischeri*:1221
afum|EAL90667.1 100.00% afis|EAW22527.1 100.00%
Bootstrap support for afum|EAL90667.1 as seed ortholog is 100%.
Bootstrap support for afis|EAW22527.1 as seed ortholog is 100%.

---

Group of orthologs #2278. Best score 1378 bits
Score difference with first non-orthologous sequence - *A. fumigatus*:1378 *A. fischeri*:1378
afum|EAL91274.1 100.00% afis|EAW17447.1 100.00%
Bootstrap support for afum|EAL91274.1 as seed ortholog is 100%.
Bootstrap support for afis|EAW17447.1 as seed ortholog is 100%.

---

Group of orthologs #2279. Best score 1378 bits
Score difference with first non-orthologous sequence - *A. fumigatus*:1256 *A. fischeri*:1228
afum|EAL88212.1 100.00% afis|EAW23267.1 100.00%
Bootstrap support for afum|EAL88212.1 as seed ortholog is 100%.
Bootstrap support for afis|EAW23267.1 as seed ortholog is 100%.

---

Group of orthologs #2280. Best score 1378 bits
Score difference with first non-orthologous sequence - *A. fumigatus*:1378 *A. fischeri*:1378
afum|EAL93692.1 100.00% afis|EAW18959.1 100.00%
Bootstrap support for afum|EAL93692.1 as seed ortholog is 100%.
Bootstrap support for afis|EAW18959.1 as seed ortholog is 100%.

---

Group of orthologs #2281. Best score 1378 bits
Score difference with first non-orthologous sequence - *A. fumigatus*:1378 *A. fischeri*:1378
afum|EAL93915.1 100.00% afis|EAW19184.1 100.00%
Bootstrap support for afum|EAL93915.1 as seed ortholog is 100%.
Bootstrap support for afis|EAW19184.1 as seed ortholog is 100%.

---

Group of orthologs #2282. Best score 1377 bits
Score difference with first non-orthologous sequence - *A. fumigatus*:1267 *A. fischeri*:1248
afum|EAL88726.1 100.00% afis|EAW16094.1 100.00%
Bootstrap support for afum|EAL88726.1 as seed ortholog is 100%.
Bootstrap support for afis|EAW16094.1 as seed ortholog is 100%.

---

Group of orthologs #2283. Best score 1377 bits
Score difference with first non-orthologous sequence - *A. fumigatus*:1377 *A. fischeri*:1377
afum|EAL91843.1 100.00% afis|EAW18032.1 100.00%
Bootstrap support for afum|EAL91843.1 as seed ortholog is 100%.
Bootstrap support for afis|EAW18032.1 as seed ortholog is 100%.

---

Group of orthologs #2284. Best score 1377 bits
Score difference with first non-orthologous sequence - *A. fumigatus*:1377 *A. fischeri*:1377

afum|EAL87309.1 100.00% afis|EAW23881.1 100.00%
Bootstrap support for afum|EAL87309.1 as seed ortholog is 100%.
Bootstrap support for afis|EAW23881.1 as seed ortholog is 100%.

---

Group of orthologs #2285. Best score 1377 bits
Score difference with first non-orthologous sequence - *A. fumigatus*:1377 *A. fischeri*:1377
afum|EAL92471.1 100.00% afis|EAW21452.1 100.00%
Bootstrap support for afum|EAL92471.1 as seed ortholog is 100%.
Bootstrap support for afis|EAW21452.1 as seed ortholog is 100%.

---

Group of orthologs #2286. Best score 1376 bits
Score difference with first non-orthologous sequence - *A. fumigatus*:1376 *A. fischeri*:1376
afum|EAL87679.1 100.00% afis|EAW18312.1 100.00%
Bootstrap support for afum|EAL87679.1 as seed ortholog is 100%.
Bootstrap support for afis|EAW18312.1 as seed ortholog is 100%.

---

Group of orthologs #2287. Best score 1376 bits
Score difference with first non-orthologous sequence - *A. fumigatus*:1376 *A. fischeri*:1376
afum|EAL91803.1 100.00% afis|EAW17993.1 100.00%
Bootstrap support for afum|EAL91803.1 as seed ortholog is 100%.
Bootstrap support for afis|EAW17993.1 as seed ortholog is 100%.

---

Group of orthologs #2288. Best score 1376 bits
Score difference with first non-orthologous sequence - *A. fumigatus*:1376 *A. fischeri*:1376
afum|EAL86119.1 100.00% afis|EAW24450.1 100.00%
Bootstrap support for afum|EAL86119.1 as seed ortholog is 100%.
Bootstrap support for afis|EAW24450.1 as seed ortholog is 100%.

---

Group of orthologs #2289. Best score 1376 bits
Score difference with first non-orthologous sequence - *A. fumigatus*:1376 *A. fischeri*:1376
afum|EAL93142.1 100.00% afis|EAW18457.1 100.00%
Bootstrap support for afum|EAL93142.1 as seed ortholog is 100%.
Bootstrap support for afis|EAW18457.1 as seed ortholog is 100%.

---

Group of orthologs #2290. Best score 1376 bits
Score difference with first non-orthologous sequence - *A. fumigatus*:1376 *A. fischeri*:1376
afum|EAL93553.1 100.00% afis|EAW18817.1 100.00%
Bootstrap support for afum|EAL93553.1 as seed ortholog is 100%.
Bootstrap support for afis|EAW18817.1 as seed ortholog is 100%.

---

Group of orthologs #2291. Best score 1375 bits
Score difference with first non-orthologous sequence - *A. fumigatus*:1375 *A. fischeri*:1375
afum|EAL93492.2 100.00% afis|EAW18761.1 100.00%
Bootstrap support for afum|EAL93492.2 as seed ortholog is 100%.
Bootstrap support for afis|EAW18761.1 as seed ortholog is 100%.

---

Group of orthologs #2292. Best score 1375 bits
Score difference with first non-orthologous sequence - *A. fumigatus*:1375 *A. fischeri*:1235
afum|EAL90895.1 100.00% afis|EAW22286.1 100.00%
Bootstrap support for afum|EAL90895.1 as seed ortholog is 100%.
Bootstrap support for afis|EAW22286.1 as seed ortholog is 100%.

---

Group of orthologs #2293. Best score 1375 bits
Score difference with first non-orthologous sequence - *A. fumigatus*:1375 *A. fischeri*:1375
afum|EAL92661.1 100.00% afis|EAW21625.1 100.00%
Bootstrap support for afum|EAL92661.1 as seed ortholog is 100%.

Bootstrap support for afis|EAW21625.1 as seed ortholog is 100%.

---

Group of orthologs #2294. Best score 1375 bits
Score difference with first non-orthologous sequence - *A. fumigatus*:1375 *A. fischeri*:1375
afum|EBA27195.1 100.00% afis|EAW20458.1 100.00%
Bootstrap support for afum|EBA27195.1 as seed ortholog is 100%.
Bootstrap support for afis|EAW20458.1 as seed ortholog is 100%.

---

Group of orthologs #2295. Best score 1375 bits
Score difference with first non-orthologous sequence - *A. fumigatus*:1375 *A. fischeri*:1375
afum|EAL89597.1 100.00% afis|EAW24940.1 100.00%
Bootstrap support for afum|EAL89597.1 as seed ortholog is 100%.
Bootstrap support for afis|EAW24940.1 as seed ortholog is 100%.

---

Group of orthologs #2296. Best score 1374 bits
Score difference with first non-orthologous sequence - *A. fumigatus*:1374 *A. fischeri*:1374
afum|EAL88573.1 100.00% afis|EAW15937.1 100.00%
Bootstrap support for afum|EAL88573.1 as seed ortholog is 100%.
Bootstrap support for afis|EAW15937.1 as seed ortholog is 100%.

---

Group of orthologs #2297. Best score 1374 bits
Score difference with first non-orthologous sequence - *A. fumigatus*:1374 *A. fischeri*:1374
afum|EAL91314.1 100.00% afis|EAW17492.1 100.00%
Bootstrap support for afum|EAL91314.1 as seed ortholog is 100%.
Bootstrap support for afis|EAW17492.1 as seed ortholog is 100%.

---

Group of orthologs #2298. Best score 1373 bits
Score difference with first non-orthologous sequence - *A. fumigatus*:1166 *A. fischeri*:1309
afum|EAL84191.1 100.00% afis|EAW15454.1 100.00%
Bootstrap support for afum|EAL84191.1 as seed ortholog is 100%.
Bootstrap support for afis|EAW15454.1 as seed ortholog is 100%.

---

Group of orthologs #2299. Best score 1373 bits
Score difference with first non-orthologous sequence - *A. fumigatus*:1373 *A. fischeri*:1373
afum|EAL88654.1 100.00% afis|EAW16025.1 100.00%
Bootstrap support for afum|EAL88654.1 as seed ortholog is 100%.
Bootstrap support for afis|EAW16025.1 as seed ortholog is 100%.

---

Group of orthologs #2300. Best score 1373 bits
Score difference with first non-orthologous sequence - *A. fumigatus*:1069 *A. fischeri*:1074
afum|EAL85049.1 100.00% afis|EAW19845.1 100.00%
Bootstrap support for afum|EAL85049.1 as seed ortholog is 100%.
Bootstrap support for afis|EAW19845.1 as seed ortholog is 100%.

---

Group of orthologs #2301. Best score 1373 bits
Score difference with first non-orthologous sequence - *A. fumigatus*:1373 *A. fischeri*:1373
afum|EAL91705.1 100.00% afis|EAW17897.1 100.00%
Bootstrap support for afum|EAL91705.1 as seed ortholog is 100%.
Bootstrap support for afis|EAW17897.1 as seed ortholog is 100%.

---

Group of orthologs #2302. Best score 1373 bits
Score difference with first non-orthologous sequence - *A. fumigatus*:1373 *A. fischeri*:1373
afum|EAL91881.2 100.00% afis|EAW18069.1 100.00%
Bootstrap support for afum|EAL91881.2 as seed ortholog is 100%.
Bootstrap support for afis|EAW18069.1 as seed ortholog is 100%.

---

Group of orthologs #2303. Best score 1373 bits
Score difference with first non-orthologous sequence - *A. fumigatus*:1287 *A. fischeri*:1315
afum|EAL94076.1 100.00% afis|EAW19356.1 100.00%
Bootstrap support for afum|EAL94076.1 as seed ortholog is 100%.
Bootstrap support for afis|EAW19356.1 as seed ortholog is 100%.

---

Group of orthologs #2304. Best score 1372 bits
Score difference with first non-orthologous sequence - *A. fumigatus*:1244 *A. fischeri*:1225
afum|EAL86470.1 100.00% afis|EAW16943.1 100.00%
Bootstrap support for afum|EAL86470.1 as seed ortholog is 100%.
Bootstrap support for afis|EAW16943.1 as seed ortholog is 100%.

---

Group of orthologs #2305. Best score 1372 bits
Score difference with first non-orthologous sequence - *A. fumigatus*:1372 *A. fischeri*:1372
afum|EAL88106.1 100.00% afis|EAW23367.1 100.00%
Bootstrap support for afum|EAL88106.1 as seed ortholog is 100%.
Bootstrap support for afis|EAW23367.1 as seed ortholog is 100%.

---

Group of orthologs #2306. Best score 1372 bits
Score difference with first non-orthologous sequence - *A. fumigatus*:1124 *A. fischeri*:1310
afum|EAL92474.1 100.00% afis|EAW21455.1 100.00%
Bootstrap support for afum|EAL92474.1 as seed ortholog is 100%.
Bootstrap support for afis|EAW21455.1 as seed ortholog is 100%.

---

Group of orthologs #2307. Best score 1372 bits
Score difference with first non-orthologous sequence - *A. fumigatus*:1372 *A. fischeri*:1372
afum|EAL90051.1 100.00% afis|EAW25392.1 100.00%
Bootstrap support for afum|EAL90051.1 as seed ortholog is 100%.
Bootstrap support for afis|EAW25392.1 as seed ortholog is 100%.

---

Group of orthologs #2308. Best score 1371 bits
Score difference with first non-orthologous sequence - *A. fumigatus*:1297 *A. fischeri*:1318
afum|EAL89013.2 100.00% afis|EAW16385.1 100.00%
Bootstrap support for afum|EAL89013.2 as seed ortholog is 100%.
Bootstrap support for afis|EAW16385.1 as seed ortholog is 100%.

---

Group of orthologs #2309. Best score 1371 bits
Score difference with first non-orthologous sequence - *A. fumigatus*:1371 *A. fischeri*:1371
afum|EAL92244.1 100.00% afis|EAW21232.1 100.00%
Bootstrap support for afum|EAL92244.1 as seed ortholog is 100%.
Bootstrap support for afis|EAW21232.1 as seed ortholog is 100%.

---

Group of orthologs #2310. Best score 1370 bits
Score difference with first non-orthologous sequence - *A. fumigatus*:1370 *A. fischeri*:1132
afum|EAL85850.1 100.00% afis|EAW15471.1 100.00%
Bootstrap support for afum|EAL85850.1 as seed ortholog is 100%.
Bootstrap support for afis|EAW15471.1 as seed ortholog is 100%.

---

Group of orthologs #2311. Best score 1370 bits
Score difference with first non-orthologous sequence - *A. fumigatus*:1370 *A. fischeri*:1370
afum|EAL86391.1 100.00% afis|EAW16845.1 100.00%
Bootstrap support for afum|EAL86391.1 as seed ortholog is 100%.
Bootstrap support for afis|EAW16845.1 as seed ortholog is 100%.

---

Group of orthologs #2312. Best score 1370 bits
Score difference with first non-orthologous sequence - *A. fumigatus*:1370 *A. fischeri*:1323

afum|EAL89173.1 100.00% afis|EAW16577.1 100.00%
Bootstrap support for afum|EAL89173.1 as seed ortholog is 100%.
Bootstrap support for afis|EAW16577.1 as seed ortholog is 100%.

---

Group of orthologs #2313. Best score 1370 bits
Score difference with first non-orthologous sequence - *A. fumigatus*:1370 *A. fischeri*:598
afum|EAL93446.1 100.00% afis|EAW18722.1 100.00%
Bootstrap support for afum|EAL93446.1 as seed ortholog is 100%.
Bootstrap support for afis|EAW18722.1 as seed ortholog is 100%.

---

Group of orthologs #2314. Best score 1370 bits
Score difference with first non-orthologous sequence - *A. fumigatus*:1370 *A. fischeri*:1370
afum|EAL92150.1 100.00% afis|EAW21132.1 100.00%
Bootstrap support for afum|EAL92150.1 as seed ortholog is 100%.
Bootstrap support for afis|EAW21132.1 as seed ortholog is 100%.

---

Group of orthologs #2315. Best score 1369 bits
Score difference with first non-orthologous sequence - *A. fumigatus*:1369 *A. fischeri*:1369
afum|EAL88748.1 100.00% afis|EAW16117.1 100.00%
Bootstrap support for afum|EAL88748.1 as seed ortholog is 100%.
Bootstrap support for afis|EAW16117.1 as seed ortholog is 100%.

---

Group of orthologs #2316. Best score 1369 bits
Score difference with first non-orthologous sequence - *A. fumigatus*:1369 *A. fischeri*:1369
afum|EAL89050.1 100.00% afis|EAW16455.1 100.00%
Bootstrap support for afum|EAL89050.1 as seed ortholog is 100%.
Bootstrap support for afis|EAW16455.1 as seed ortholog is 100%.

---

Group of orthologs #2317. Best score 1369 bits
Score difference with first non-orthologous sequence - *A. fumigatus*:1369 *A. fischeri*:1369
afum|EAL91254.1 100.00% afis|EAW17426.1 100.00%
Bootstrap support for afum|EAL91254.1 as seed ortholog is 100%.
Bootstrap support for afis|EAW17426.1 as seed ortholog is 100%.

---

Group of orthologs #2318. Best score 1369 bits
Score difference with first non-orthologous sequence - *A. fumigatus*:1257 *A. fischeri*:1369
afum|EAL93351.1 100.00% afis|EAW18638.1 100.00%
Bootstrap support for afum|EAL93351.1 as seed ortholog is 100%.
Bootstrap support for afis|EAW18638.1 as seed ortholog is 100%.

---

Group of orthologs #2319. Best score 1369 bits
Score difference with first non-orthologous sequence - *A. fumigatus*:1369 *A. fischeri*:1369
afum|EAL93395.2 100.00% afis|EAW18675.1 100.00%
Bootstrap support for afum|EAL93395.2 as seed ortholog is 100%.
Bootstrap support for afis|EAW18675.1 as seed ortholog is 100%.

---

Group of orthologs #2320. Best score 1368 bits
Score difference with first non-orthologous sequence - *A. fumigatus*:1368 *A. fischeri*:1368
afum|EAL85752.1 100.00% afis|EAW15582.1 100.00%
Bootstrap support for afum|EAL85752.1 as seed ortholog is 100%.
Bootstrap support for afis|EAW15582.1 as seed ortholog is 100%.

---

Group of orthologs #2321. Best score 1368 bits
Score difference with first non-orthologous sequence - *A. fumigatus*:1368 *A. fischeri*:1368
afum|EAL89148.1 100.00% afis|EAW16550.1 100.00%
Bootstrap support for afum|EAL89148.1 as seed ortholog is 100%.

Bootstrap support for afis|EAW16550.1 as seed ortholog is 100%.

---

Group of orthologs #2322. Best score 1367 bits
Score difference with first non-orthologous sequence - *A. fumigatus*:1185 *A. fischeri*:1367
afum|EAL86554.1 100.00% afis|EAW17058.1 100.00%
Bootstrap support for afum|EAL86554.1 as seed ortholog is 100%.
Bootstrap support for afis|EAW17058.1 as seed ortholog is 100%.

---

Group of orthologs #2323. Best score 1367 bits
Score difference with first non-orthologous sequence - *A. fumigatus*:1367 *A. fischeri*:1138
afum|EAL84969.1 100.00% afis|EAW19946.1 100.00%
Bootstrap support for afum|EAL84969.1 as seed ortholog is 100%.
Bootstrap support for afis|EAW19946.1 as seed ortholog is 100%.

---

Group of orthologs #2324. Best score 1367 bits
Score difference with first non-orthologous sequence - *A. fumigatus*:1367 *A. fischeri*:1367
afum|EAL84285.2 100.00% afis|EAW25575.1 100.00%
Bootstrap support for afum|EAL84285.2 as seed ortholog is 100%.
Bootstrap support for afis|EAW25575.1 as seed ortholog is 100%.

---

Group of orthologs #2325. Best score 1367 bits
Score difference with first non-orthologous sequence - *A. fumigatus*:1367 *A. fischeri*:1367
afum|EBA27283.1 100.00% afis|EAW17242.1 100.00%
Bootstrap support for afum|EBA27283.1 as seed ortholog is 100%.
Bootstrap support for afis|EAW17242.1 as seed ortholog is 100%.

---

Group of orthologs #2326. Best score 1367 bits
Score difference with first non-orthologous sequence - *A. fumigatus*:1367 *A. fischeri*:1367
afum|EAL88226.1 100.00% afis|EAW23255.1 100.00%
Bootstrap support for afum|EAL88226.1 as seed ortholog is 100%.
Bootstrap support for afis|EAW23255.1 as seed ortholog is 100%.

---

Group of orthologs #2327. Best score 1367 bits
Score difference with first non-orthologous sequence - *A. fumigatus*:1367 *A. fischeri*:1367
afum|EAL93294.2 100.00% afis|EAW18587.1 100.00%
Bootstrap support for afum|EAL93294.2 as seed ortholog is 100%.
Bootstrap support for afis|EAW18587.1 as seed ortholog is 100%.

---

Group of orthologs #2328. Best score 1367 bits
Score difference with first non-orthologous sequence - *A. fumigatus*:1367 *A. fischeri*:1367
afum|EAL90719.1 100.00% afis|EAW22472.1 100.00%
Bootstrap support for afum|EAL90719.1 as seed ortholog is 100%.
Bootstrap support for afis|EAW22472.1 as seed ortholog is 100%.

---

Group of orthologs #2329. Best score 1366 bits
Score difference with first non-orthologous sequence - *A. fumigatus*:1366 *A. fischeri*:1366
afum|EAL94010.1 100.00% afis|EAW19284.1 100.00%
Bootstrap support for afum|EAL94010.1 as seed ortholog is 100%.
Bootstrap support for afis|EAW19284.1 as seed ortholog is 100%.

---

Group of orthologs #2330. Best score 1366 bits
Score difference with first non-orthologous sequence - *A. fumigatus*:1366 *A. fischeri*:1366
afum|EAL89741.1 100.00% afis|EAW25072.1 100.00%
Bootstrap support for afum|EAL89741.1 as seed ortholog is 100%.
Bootstrap support for afis|EAW25072.1 as seed ortholog is 100%.

---

Group of orthologs #2331. Best score 1365 bits
Score difference with first non-orthologous sequence - *A. fumigatus*:1365 *A. fischeri*:1365
afum|EAL84703.2 100.00% afis|EAW20931.1 100.00%
Bootstrap support for afum|EAL84703.2 as seed ortholog is 100%.
Bootstrap support for afis|EAW20931.1 as seed ortholog is 100%.

---

Group of orthologs #2332. Best score 1365 bits
Score difference with first non-orthologous sequence - *A. fumigatus*:1308 *A. fischeri*:1269
afum|EAL92219.1 100.00% afis|EAW21208.1 100.00%
Bootstrap support for afum|EAL92219.1 as seed ortholog is 100%.
Bootstrap support for afis|EAW21208.1 as seed ortholog is 100%.

---

Group of orthologs #2333. Best score 1364 bits
Score difference with first non-orthologous sequence - *A. fumigatus*:1364 *A. fischeri*:1364
afum|EAL85785.1 100.00% afis|EAW15547.1 100.00%
Bootstrap support for afum|EAL85785.1 as seed ortholog is 100%.
Bootstrap support for afis|EAW15547.1 as seed ortholog is 100%.

---

Group of orthologs #2334. Best score 1364 bits
Score difference with first non-orthologous sequence - *A. fumigatus*:1364 *A. fischeri*:1364
afum|EAL84651.2 100.00% afis|EAW20980.1 100.00%
Bootstrap support for afum|EAL84651.2 as seed ortholog is 100%.
Bootstrap support for afis|EAW20980.1 as seed ortholog is 100%.

---

Group of orthologs #2335. Best score 1364 bits
Score difference with first non-orthologous sequence - *A. fumigatus*:1364 *A. fischeri*:1364
afum|EAL85976.1 100.00% afis|EAW24308.1 100.00%
Bootstrap support for afum|EAL85976.1 as seed ortholog is 100%.
Bootstrap support for afis|EAW24308.1 as seed ortholog is 100%.

---

Group of orthologs #2336. Best score 1364 bits
Score difference with first non-orthologous sequence - *A. fumigatus*:1364 *A. fischeri*:1364
afum|EAL85985.1 100.00% afis|EAW24317.1 100.00%
Bootstrap support for afum|EAL85985.1 as seed ortholog is 100%.
Bootstrap support for afis|EAW24317.1 as seed ortholog is 100%.

---

Group of orthologs #2337. Best score 1364 bits
Score difference with first non-orthologous sequence - *A. fumigatus*:1364 *A. fischeri*:1364
afum|EAL93218.1 100.00% afis|EAW18523.1 100.00%
Bootstrap support for afum|EAL93218.1 as seed ortholog is 100%.
Bootstrap support for afis|EAW18523.1 as seed ortholog is 100%.

---

Group of orthologs #2338. Best score 1364 bits
Score difference with first non-orthologous sequence - *A. fumigatus*:1364 *A. fischeri*:1364
afum|EAL93775.1 100.00% afis|EAW19046.1 100.00%
Bootstrap support for afum|EAL93775.1 as seed ortholog is 100%.
Bootstrap support for afis|EAW19046.1 as seed ortholog is 100%.

---

Group of orthologs #2339. Best score 1364 bits
Score difference with first non-orthologous sequence - *A. fumigatus*:1364 *A. fischeri*:1364
afum|EAL91009.1 100.00% afis|EAW22180.1 100.00%
Bootstrap support for afum|EAL91009.1 as seed ortholog is 100%.
Bootstrap support for afis|EAW22180.1 as seed ortholog is 100%.

---

Group of orthologs #2340. Best score 1364 bits
Score difference with first non-orthologous sequence - *A. fumigatus*:1364 *A. fischeri*:1364

afum|EAL94062.1 100.00% afis|EAW19343.1 100.00%
Bootstrap support for afum|EAL94062.1 as seed ortholog is 100%.
Bootstrap support for afis|EAW19343.1 as seed ortholog is 100%.

---

Group of orthologs #2341. Best score 1363 bits
Score difference with first non-orthologous sequence - *A. fumigatus*:1130 *A. fischeri*:1124
afum|EAL84798.1 100.00% afis|EAW20834.1 100.00%
Bootstrap support for afum|EAL84798.1 as seed ortholog is 100%.
Bootstrap support for afis|EAW20834.1 as seed ortholog is 100%.

---

Group of orthologs #2342. Best score 1363 bits
Score difference with first non-orthologous sequence - *A. fumigatus*:1016 *A. fischeri*:1071
afum|EAL86169.1 100.00% afis|EAW24499.1 100.00%
Bootstrap support for afum|EAL86169.1 as seed ortholog is 100%.
Bootstrap support for afis|EAW24499.1 as seed ortholog is 100%.

---

Group of orthologs #2343. Best score 1363 bits
Score difference with first non-orthologous sequence - *A. fumigatus*:1363 *A. fischeri*:1291
afum|EAL93613.1 100.00% afis|EAW18881.1 100.00%
Bootstrap support for afum|EAL93613.1 as seed ortholog is 100%.
Bootstrap support for afis|EAW18881.1 as seed ortholog is 100%.

---

Group of orthologs #2344. Best score 1362 bits
Score difference with first non-orthologous sequence - *A. fumigatus*:1362 *A. fischeri*:1362
afum|EAL87872.1 100.00% afis|EAW23595.1 100.00%
Bootstrap support for afum|EAL87872.1 as seed ortholog is 100%.
Bootstrap support for afis|EAW23595.1 as seed ortholog is 100%.

---

Group of orthologs #2345. Best score 1361 bits
Score difference with first non-orthologous sequence - *A. fumigatus*:1154 *A. fischeri*:1163
afum|EAL86440.1 100.00% afis|EAW16904.1 100.00%
Bootstrap support for afum|EAL86440.1 as seed ortholog is 100%.
Bootstrap support for afis|EAW16904.1 as seed ortholog is 100%.

---

Group of orthologs #2346. Best score 1361 bits
Score difference with first non-orthologous sequence - *A. fumigatus*:1361 *A. fischeri*:1229
afum|EAL91855.2 100.00% afis|EAW18044.1 100.00%
Bootstrap support for afum|EAL91855.2 as seed ortholog is 100%.
Bootstrap support for afis|EAW18044.1 as seed ortholog is 100%.

---

Group of orthologs #2347. Best score 1361 bits
Score difference with first non-orthologous sequence - *A. fumigatus*:1361 *A. fischeri*:1361
afum|EAL89638.1 100.00% afis|EAW24978.1 100.00%
Bootstrap support for afum|EAL89638.1 as seed ortholog is 100%.
Bootstrap support for afis|EAW24978.1 as seed ortholog is 100%.

---

Group of orthologs #2348. Best score 1360 bits
Score difference with first non-orthologous sequence - *A. fumigatus*:1360 *A. fischeri*:1360
afum|EAL85085.1 100.00% afis|EAW19807.1 100.00%
Bootstrap support for afum|EAL85085.1 as seed ortholog is 100%.
Bootstrap support for afis|EAW19807.1 as seed ortholog is 100%.

---

Group of orthologs #2349. Best score 1360 bits
Score difference with first non-orthologous sequence - *A. fumigatus*:1360 *A. fischeri*:1360
afum|EAL86070.1 100.00% afis|EAW24399.1 100.00%
Bootstrap support for afum|EAL86070.1 as seed ortholog is 100%.

Bootstrap support for afis|EAW24399.1 as seed ortholog is 100%.

---

Group of orthologs #2350. Best score 1360 bits
Score difference with first non-orthologous sequence - *A. fumigatus*:1360 *A. fischeri*:1360
afum|EAL87134.1 100.00% afis|EAW23688.1 100.00%
Bootstrap support for afum|EAL87134.1 as seed ortholog is 100%.
Bootstrap support for afis|EAW23688.1 as seed ortholog is 100%.

---

Group of orthologs #2351. Best score 1360 bits
Score difference with first non-orthologous sequence - *A. fumigatus*:1360 *A. fischeri*:1360
afum|EAL90880.1 100.00% afis|EAW22310.1 100.00%
Bootstrap support for afum|EAL90880.1 as seed ortholog is 100%.
Bootstrap support for afis|EAW22310.1 as seed ortholog is 100%.

---

Group of orthologs #2352. Best score 1359 bits
Score difference with first non-orthologous sequence - *A. fumigatus*:1165 *A. fischeri*:1359
afum|EAL89192.1 100.00% afis|EAW16596.1 100.00%
Bootstrap support for afum|EAL89192.1 as seed ortholog is 100%.
Bootstrap support for afis|EAW16596.1 as seed ortholog is 100%.

---

Group of orthologs #2353. Best score 1359 bits
Score difference with first non-orthologous sequence - *A. fumigatus*:1359 *A. fischeri*:1359
afum|EAL90898.1 100.00% afis|EAW22283.1 100.00%
Bootstrap support for afum|EAL90898.1 as seed ortholog is 100%.
Bootstrap support for afis|EAW22283.1 as seed ortholog is 100%.

---

Group of orthologs #2354. Best score 1359 bits
Score difference with first non-orthologous sequence - *A. fumigatus*:1359 *A. fischeri*:1359
afum|EAL90859.1 100.00% afis|EAW22333.1 100.00%
Bootstrap support for afum|EAL90859.1 as seed ortholog is 100%.
Bootstrap support for afis|EAW22333.1 as seed ortholog is 100%.

---

Group of orthologs #2355. Best score 1358 bits
Score difference with first non-orthologous sequence - *A. fumigatus*:1358 *A. fischeri*:1358
afum|EAL91262.1 100.00% afis|EAW17435.1 100.00%
Bootstrap support for afum|EAL91262.1 as seed ortholog is 100%.
Bootstrap support for afis|EAW17435.1 as seed ortholog is 100%.

---

Group of orthologs #2356. Best score 1358 bits
Score difference with first non-orthologous sequence - *A. fumigatus*:1358 *A. fischeri*:1358
afum|EAL90653.1 100.00% afis|EAW22541.1 100.00%
Bootstrap support for afum|EAL90653.1 as seed ortholog is 100%.
Bootstrap support for afis|EAW22541.1 as seed ortholog is 100%.

---

Group of orthologs #2357. Best score 1357 bits
Score difference with first non-orthologous sequence - *A. fumigatus*:1039 *A. fischeri*:1357
afum|EAL87118.1 100.00% afis|EAW23675.1 100.00%
Bootstrap support for afum|EAL87118.1 as seed ortholog is 100%.
Bootstrap support for afis|EAW23675.1 as seed ortholog is 100%.

---

Group of orthologs #2358. Best score 1357 bits
Score difference with first non-orthologous sequence - *A. fumigatus*:1357 *A. fischeri*:1357
afum|EAL93225.1 100.00% afis|EAW18528.1 100.00%
Bootstrap support for afum|EAL93225.1 as seed ortholog is 100%.
Bootstrap support for afis|EAW18528.1 as seed ortholog is 100%.

---

Group of orthologs #2359. Best score 1357 bits
Score difference with first non-orthologous sequence - *A. fumigatus*:1357 *A. fischeri*:1357
afum|EAL93947.1 100.00% afis|EAW19214.1 100.00%
Bootstrap support for afum|EAL93947.1 as seed ortholog is 100%.
Bootstrap support for afis|EAW19214.1 as seed ortholog is 100%.

---

Group of orthologs #2360. Best score 1357 bits
Score difference with first non-orthologous sequence - *A. fumigatus*:1357 *A. fischeri*:1357
afum|EAL90829.1 100.00% afis|EAW22362.1 100.00%
Bootstrap support for afum|EAL90829.1 as seed ortholog is 100%.
Bootstrap support for afis|EAW22362.1 as seed ortholog is 100%.

---

Group of orthologs #2361. Best score 1357 bits
Score difference with first non-orthologous sequence - *A. fumigatus*:1357 *A. fischeri*:1357
afum|EAL92910.1 100.00% afis|EAW21897.1 100.00%
Bootstrap support for afum|EAL92910.1 as seed ortholog is 100%.
Bootstrap support for afis|EAW21897.1 as seed ortholog is 100%.

---

Group of orthologs #2362. Best score 1357 bits
Score difference with first non-orthologous sequence - *A. fumigatus*:1357 *A. fischeri*:1357
afum|EAL90091.1 100.00% afis|EAW25433.1 100.00%
Bootstrap support for afum|EAL90091.1 as seed ortholog is 100%.
Bootstrap support for afis|EAW25433.1 as seed ortholog is 100%.

---

Group of orthologs #2363. Best score 1356 bits
Score difference with first non-orthologous sequence - *A. fumigatus*:1356 *A. fischeri*:467
afum|EAL84562.1 100.00% afis|EAW20492.1 100.00%
Bootstrap support for afum|EAL84562.1 as seed ortholog is 100%.
Bootstrap support for afis|EAW20492.1 as seed ortholog is 100%.

---

Group of orthologs #2364. Best score 1356 bits
Score difference with first non-orthologous sequence - *A. fumigatus*:1356 *A. fischeri*:1356
afum|EAL91751.1 100.00% afis|EAW17940.1 100.00%
Bootstrap support for afum|EAL91751.1 as seed ortholog is 100%.
Bootstrap support for afis|EAW17940.1 as seed ortholog is 100%.

---

Group of orthologs #2365. Best score 1356 bits
Score difference with first non-orthologous sequence - *A. fumigatus*:1356 *A. fischeri*:1356
afum|EAL93207.1 100.00% afis|EAW18512.1 100.00%
Bootstrap support for afum|EAL93207.1 as seed ortholog is 100%.
Bootstrap support for afis|EAW18512.1 as seed ortholog is 100%.

---

Group of orthologs #2366. Best score 1356 bits
Score difference with first non-orthologous sequence - *A. fumigatus*:1356 *A. fischeri*:1356
afum|EBA27432.1 100.00% afis|EAW17571.1 100.00%
Bootstrap support for afum|EBA27432.1 as seed ortholog is 100%.
Bootstrap support for afis|EAW17571.1 as seed ortholog is 100%.

---

Group of orthologs #2367. Best score 1356 bits
Score difference with first non-orthologous sequence - *A. fumigatus*:1356 *A. fischeri*:1356
afum|EAL93461.1 100.00% afis|EAW18736.1 100.00%
Bootstrap support for afum|EAL93461.1 as seed ortholog is 100%.
Bootstrap support for afis|EAW18736.1 as seed ortholog is 100%.

---

Group of orthologs #2368. Best score 1355 bits
Score difference with first non-orthologous sequence - *A. fumigatus*:1060 *A. fischeri*:1355

afum|EAL86439.1 100.00% afis|EAW16903.1 100.00%
Bootstrap support for afum|EAL86439.1 as seed ortholog is 100%.
Bootstrap support for afis|EAW16903.1 as seed ortholog is 100%.

---

Group of orthologs #2369. Best score 1355 bits
Score difference with first non-orthologous sequence - *A. fumigatus*:1355 *A. fischeri*:1355
afum|EAL84708.1 100.00% afis|EAW20926.1 100.00%
Bootstrap support for afum|EAL84708.1 as seed ortholog is 100%.
Bootstrap support for afis|EAW20926.1 as seed ortholog is 100%.

---

Group of orthologs #2370. Best score 1355 bits
Score difference with first non-orthologous sequence - *A. fumigatus*:1047 *A. fischeri*:1051
afum|EAL86815.1 100.00% afis|EAW19712.1 100.00%
Bootstrap support for afum|EAL86815.1 as seed ortholog is 100%.
Bootstrap support for afis|EAW19712.1 as seed ortholog is 100%.

---

Group of orthologs #2371. Best score 1355 bits
Score difference with first non-orthologous sequence - *A. fumigatus*:1355 *A. fischeri*:1355
afum|EAL88247.1 100.00% afis|EAW23235.1 100.00%
Bootstrap support for afum|EAL88247.1 as seed ortholog is 100%.
Bootstrap support for afis|EAW23235.1 as seed ortholog is 100%.

---

Group of orthologs #2372. Best score 1354 bits
Score difference with first non-orthologous sequence - *A. fumigatus*:1354 *A. fischeri*:1354
afum|EAL91698.1 100.00% afis|EAW17889.1 100.00%
Bootstrap support for afum|EAL91698.1 as seed ortholog is 100%.
Bootstrap support for afis|EAW17889.1 as seed ortholog is 100%.

---

Group of orthologs #2373. Best score 1354 bits
Score difference with first non-orthologous sequence - *A. fumigatus*:1354 *A. fischeri*:1354
afum|EAL90570.1 100.00% afis|EAW22626.1 100.00%
Bootstrap support for afum|EAL90570.1 as seed ortholog is 100%.
Bootstrap support for afis|EAW22626.1 as seed ortholog is 100%.

---

Group of orthologs #2374. Best score 1354 bits
Score difference with first non-orthologous sequence - *A. fumigatus*:1354 *A. fischeri*:1243
afum|EAL92659.1 100.00% afis|EAW21623.1 100.00%
Bootstrap support for afum|EAL92659.1 as seed ortholog is 100%.
Bootstrap support for afis|EAW21623.1 as seed ortholog is 100%.

---

Group of orthologs #2375. Best score 1354 bits
Score difference with first non-orthologous sequence - *A. fumigatus*:770 *A. fischeri*:1354
afum|EAL92886.1 100.00% afis|EAW21872.1 100.00%
Bootstrap support for afum|EAL92886.1 as seed ortholog is 100%.
Bootstrap support for afis|EAW21872.1 as seed ortholog is 100%.

---

Group of orthologs #2376. Best score 1353 bits
Score difference with first non-orthologous sequence - *A. fumigatus*:1353 *A. fischeri*:1175
afum|EAL84983.1 100.00% afis|EAW19926.1 100.00%
Bootstrap support for afum|EAL84983.1 as seed ortholog is 100%.
Bootstrap support for afis|EAW19926.1 as seed ortholog is 100%.

---

Group of orthologs #2377. Best score 1353 bits
Score difference with first non-orthologous sequence - *A. fumigatus*:1353 *A. fischeri*:1353
afum|EAL92966.1 100.00% afis|EAW21952.1 100.00%
Bootstrap support for afum|EAL92966.1 as seed ortholog is 100%.

Bootstrap support for afis|EAW21952.1 as seed ortholog is 100%.

---

Group of orthologs #2378. Best score 1353 bits
Score difference with first non-orthologous sequence - *A. fumigatus*:1353 *A. fischeri*:1353
afum|EAL90050.1 100.00% afis|EAW25391.1 100.00%
Bootstrap support for afum|EAL90050.1 as seed ortholog is 100%.
Bootstrap support for afis|EAW25391.1 as seed ortholog is 100%.

---

Group of orthologs #2379. Best score 1352 bits
Score difference with first non-orthologous sequence - *A. fumigatus*:1352 *A. fischeri*:1352
afum|EAL91714.1 100.00% afis|EAW17905.1 100.00%
Bootstrap support for afum|EAL91714.1 as seed ortholog is 100%.
Bootstrap support for afis|EAW17905.1 as seed ortholog is 100%.

---

Group of orthologs #2380. Best score 1352 bits
Score difference with first non-orthologous sequence - *A. fumigatus*:1352 *A. fischeri*:1352
afum|EAL85863.1 100.00% afis|EAW24190.1 100.00%
Bootstrap support for afum|EAL85863.1 as seed ortholog is 100%.
Bootstrap support for afis|EAW24190.1 as seed ortholog is 100%.

---

Group of orthologs #2381. Best score 1352 bits
Score difference with first non-orthologous sequence - *A. fumigatus*:1352 *A. fischeri*:1261
afum|EAL86134.1 100.00% afis|EAW24463.1 100.00%
Bootstrap support for afum|EAL86134.1 as seed ortholog is 100%.
Bootstrap support for afis|EAW24463.1 as seed ortholog is 100%.

---

Group of orthologs #2382. Best score 1352 bits
Score difference with first non-orthologous sequence - *A. fumigatus*:1352 *A. fischeri*:1352
afum|EAL91072.1 100.00% afis|EAW22113.1 100.00%
Bootstrap support for afum|EAL91072.1 as seed ortholog is 100%.
Bootstrap support for afis|EAW22113.1 as seed ortholog is 100%.

---

Group of orthologs #2383. Best score 1352 bits
Score difference with first non-orthologous sequence - *A. fumigatus*:1242 *A. fischeri*:943
afum|EAL89328.1 100.00% afis|EAW24687.1 100.00%
Bootstrap support for afum|EAL89328.1 as seed ortholog is 100%.
Bootstrap support for afis|EAW24687.1 as seed ortholog is 100%.

---

Group of orthologs #2384. Best score 1351 bits
Score difference with first non-orthologous sequence - *A. fumigatus*:1351 *A. fischeri*:1218
afum|EAL86585.1 100.00% afis|EAW17124.1 100.00%
Bootstrap support for afum|EAL86585.1 as seed ortholog is 100%.
Bootstrap support for afis|EAW17124.1 as seed ortholog is 100%.

---

Group of orthologs #2385. Best score 1351 bits
Score difference with first non-orthologous sequence - *A. fumigatus*:1351 *A. fischeri*:1351
afum|EAL85457.1 100.00% afis|EAW20331.1 100.00%
Bootstrap support for afum|EAL85457.1 as seed ortholog is 100%.
Bootstrap support for afis|EAW20331.1 as seed ortholog is 100%.

---

Group of orthologs #2386. Best score 1351 bits
Score difference with first non-orthologous sequence - *A. fumigatus*:1351 *A. fischeri*:1351
afum|EAL87025.1 100.00% afis|EAW19471.1 100.00%
Bootstrap support for afum|EAL87025.1 as seed ortholog is 100%.
Bootstrap support for afis|EAW19471.1 as seed ortholog is 100%.

---

Group of orthologs #2387. Best score 1351 bits
Score difference with first non-orthologous sequence - *A. fumigatus*:1351 *A. fischeri*:1351
afum|EAL91346.1 100.00% afis|EAW17532.1 100.00%
Bootstrap support for afum|EAL91346.1 as seed ortholog is 100%.
Bootstrap support for afis|EAW17532.1 as seed ortholog is 100%.

---

Group of orthologs #2388. Best score 1351 bits
Score difference with first non-orthologous sequence - *A. fumigatus*:1028 *A. fischeri*:1042
afum|EAL84375.1 100.00% afis|EAW25702.1 100.00%
Bootstrap support for afum|EAL84375.1 as seed ortholog is 100%.
Bootstrap support for afis|EAW25702.1 as seed ortholog is 100%.

---

Group of orthologs #2389. Best score 1351 bits
Score difference with first non-orthologous sequence - *A. fumigatus*:1351 *A. fischeri*:1269
afum|EAL93402.1 100.00% afis|EAW18681.1 100.00%
Bootstrap support for afum|EAL93402.1 as seed ortholog is 100%.
Bootstrap support for afis|EAW18681.1 as seed ortholog is 100%.

---

Group of orthologs #2390. Best score 1351 bits
Score difference with first non-orthologous sequence - *A. fumigatus*:1351 *A. fischeri*:1351
afum|EAL90244.1 100.00% afis|EAW22953.1 100.00%
Bootstrap support for afum|EAL90244.1 as seed ortholog is 100%.
Bootstrap support for afis|EAW22953.1 as seed ortholog is 100%.

---

Group of orthologs #2391. Best score 1350 bits
Score difference with first non-orthologous sequence - *A. fumigatus*:1350 *A. fischeri*:1350
afum|EAL89954.1 100.00% afis|EAW25291.1 100.00%
Bootstrap support for afum|EAL89954.1 as seed ortholog is 100%.
Bootstrap support for afis|EAW25291.1 as seed ortholog is 100%.

---

Group of orthologs #2392. Best score 1349 bits
Score difference with first non-orthologous sequence - *A. fumigatus*:1349 *A. fischeri*:1349
afum|EAL85666.1 100.00% afis|EAW15682.1 100.00%
Bootstrap support for afum|EAL85666.1 as seed ortholog is 100%.
Bootstrap support for afis|EAW15682.1 as seed ortholog is 100%.

---

Group of orthologs #2393. Best score 1349 bits
Score difference with first non-orthologous sequence - *A. fumigatus*:1349 *A. fischeri*:1349
afum|EAL91258.1 100.00% afis|EAW17430.1 100.00%
Bootstrap support for afum|EAL91258.1 as seed ortholog is 100%.
Bootstrap support for afis|EAW17430.1 as seed ortholog is 100%.

---

Group of orthologs #2394. Best score 1349 bits
Score difference with first non-orthologous sequence - *A. fumigatus*:1349 *A. fischeri*:1349
afum|EAL88337.2 100.00% afis|EAW23144.1 100.00%
Bootstrap support for afum|EAL88337.2 as seed ortholog is 100%.
Bootstrap support for afis|EAW23144.1 as seed ortholog is 100%.

---

Group of orthologs #2395. Best score 1348 bits
Score difference with first non-orthologous sequence - *A. fumigatus*:1348 *A. fischeri*:1348
afum|EAL85582.1 100.00% afis|EAW15766.1 100.00%
Bootstrap support for afum|EAL85582.1 as seed ortholog is 100%.
Bootstrap support for afis|EAW15766.1 as seed ortholog is 100%.

---

Group of orthologs #2396. Best score 1348 bits
Score difference with first non-orthologous sequence - *A. fumigatus*:1348 *A. fischeri*:1348

afum|EAL84993.2 100.00% afis|EAW19913.1 100.00%
Bootstrap support for afum|EAL84993.2 as seed ortholog is 100%.
Bootstrap support for afis|EAW19913.1 as seed ortholog is 100%.

---

Group of orthologs #2397. Best score 1348 bits
Score difference with first non-orthologous sequence - *A. fumigatus*:1292 *A. fischeri*:1348
afum|EAL86975.1 100.00% afis|EAW19531.1 100.00%
Bootstrap support for afum|EAL86975.1 as seed ortholog is 100%.
Bootstrap support for afis|EAW19531.1 as seed ortholog is 100%.

---

Group of orthologs #2398. Best score 1348 bits
Score difference with first non-orthologous sequence - *A. fumigatus*:1348 *A. fischeri*:1348
afum|EAL91850.1 100.00% afis|EAW18039.1 100.00%
Bootstrap support for afum|EAL91850.1 as seed ortholog is 100%.
Bootstrap support for afis|EAW18039.1 as seed ortholog is 100%.

---

Group of orthologs #2399. Best score 1348 bits
Score difference with first non-orthologous sequence - *A. fumigatus*:1348 *A. fischeri*:1348
afum|EAL88365.1 100.00% afis|EAW23115.1 100.00%
Bootstrap support for afum|EAL88365.1 as seed ortholog is 100%.
Bootstrap support for afis|EAW23115.1 as seed ortholog is 100%.

---

Group of orthologs #2400. Best score 1348 bits
Score difference with first non-orthologous sequence - *A. fumigatus*:1258 *A. fischeri*:1213
afum|EAL93311.1 100.00% afis|EAW18603.1 100.00%
Bootstrap support for afum|EAL93311.1 as seed ortholog is 100%.
Bootstrap support for afis|EAW18603.1 as seed ortholog is 100%.

---

Group of orthologs #2401. Best score 1348 bits
Score difference with first non-orthologous sequence - *A. fumigatus*:918 *A. fischeri*:959
afum|EBA27325.1 100.00% afis|EAW23630.1 100.00%
Bootstrap support for afum|EBA27325.1 as seed ortholog is 100%.
Bootstrap support for afis|EAW23630.1 as seed ortholog is 100%.

---

Group of orthologs #2402. Best score 1347 bits
Score difference with first non-orthologous sequence - *A. fumigatus*:1347 *A. fischeri*:1347
afum|EBA27303.1 100.00% afis|EAW23760.1 100.00%
Bootstrap support for afum|EBA27303.1 as seed ortholog is 100%.
Bootstrap support for afis|EAW23760.1 as seed ortholog is 100%.

---

Group of orthologs #2403. Best score 1346 bits
Score difference with first non-orthologous sequence - *A. fumigatus*:1280 *A. fischeri*:1346
afum|EAL86680.1 100.00% afis|EAW17263.1 100.00%
Bootstrap support for afum|EAL86680.1 as seed ortholog is 100%.
Bootstrap support for afis|EAW17263.1 as seed ortholog is 100%.

---

Group of orthologs #2404. Best score 1346 bits
Score difference with first non-orthologous sequence - *A. fumigatus*:1143 *A. fischeri*:1346
afum|EAL86801.1 100.00% afis|EAW19725.1 100.00%
Bootstrap support for afum|EAL86801.1 as seed ortholog is 100%.
Bootstrap support for afis|EAW19725.1 as seed ortholog is 100%.

---

Group of orthologs #2405. Best score 1346 bits
Score difference with first non-orthologous sequence - *A. fumigatus*:1346 *A. fischeri*:1346
afum|EAL88423.2 100.00% afis|EAW23057.1 100.00%
Bootstrap support for afum|EAL88423.2 as seed ortholog is 100%.

Bootstrap support for afis|EAW23057.1 as seed ortholog is 100%.

---

Group of orthologs #2406. Best score 1346 bits
Score difference with first non-orthologous sequence - *A. fumigatus*:604 *A. fischeri*:1346
afum|EAL93846.1 100.00% afis|EAW19114.1 100.00%
Bootstrap support for afum|EAL93846.1 as seed ortholog is 100%.
Bootstrap support for afis|EAW19114.1 as seed ortholog is 100%.

---

Group of orthologs #2407. Best score 1345 bits
Score difference with first non-orthologous sequence - *A. fumigatus*:1345 *A. fischeri*:1345
afum|EAL86121.1 100.00% afis|EAW24452.1 100.00%
Bootstrap support for afum|EAL86121.1 as seed ortholog is 100%.
Bootstrap support for afis|EAW24452.1 as seed ortholog is 100%.

---

Group of orthologs #2408. Best score 1345 bits
Score difference with first non-orthologous sequence - *A. fumigatus*:1194 *A. fischeri*:1139
afum|EAL91061.1 100.00% afis|EAW22127.1 100.00%
Bootstrap support for afum|EAL91061.1 as seed ortholog is 100%.
Bootstrap support for afis|EAW22127.1 as seed ortholog is 100%.

---

Group of orthologs #2409. Best score 1345 bits
Score difference with first non-orthologous sequence - *A. fumigatus*:1228 *A. fischeri*:1345
afum|EAL89324.1 100.00% afis|EAW24671.1 100.00%
Bootstrap support for afum|EAL89324.1 as seed ortholog is 100%.
Bootstrap support for afis|EAW24671.1 as seed ortholog is 100%.

---

Group of orthologs #2410. Best score 1344 bits
Score difference with first non-orthologous sequence - *A. fumigatus*:948 *A. fischeri*:898
afum|EAL85677.1 100.00% afis|EAW15670.1 100.00%
Bootstrap support for afum|EAL85677.1 as seed ortholog is 100%.
Bootstrap support for afis|EAW15670.1 as seed ortholog is 100%.

---

Group of orthologs #2411. Best score 1344 bits
Score difference with first non-orthologous sequence - *A. fumigatus*:1344 *A. fischeri*:1344
afum|EAL86012.1 100.00% afis|EAW24341.1 100.00%
Bootstrap support for afum|EAL86012.1 as seed ortholog is 100%.
Bootstrap support for afis|EAW24341.1 as seed ortholog is 100%.

---

Group of orthologs #2412. Best score 1344 bits
Score difference with first non-orthologous sequence - *A. fumigatus*:1200 *A. fischeri*:860
afum|EBA27373.1 100.00% afis|EAW16675.1 100.00%
Bootstrap support for afum|EBA27373.1 as seed ortholog is 100%.
Bootstrap support for afis|EAW16675.1 as seed ortholog is 100%.

---

Group of orthologs #2413. Best score 1344 bits
Score difference with first non-orthologous sequence - *A. fumigatus*:1300 *A. fischeri*:1344
afum|EAL87129.1 100.00% afis|EAW23684.1 100.00%
Bootstrap support for afum|EAL87129.1 as seed ortholog is 100%.
Bootstrap support for afis|EAW23684.1 as seed ortholog is 100%.

---

Group of orthologs #2414. Best score 1344 bits
Score difference with first non-orthologous sequence - *A. fumigatus*:1344 *A. fischeri*:1344
afum|EAL90524.1 100.00% afis|EAW22668.1 100.00%
Bootstrap support for afum|EAL90524.1 as seed ortholog is 100%.
Bootstrap support for afis|EAW22668.1 as seed ortholog is 100%.

---

Group of orthologs #2415. Best score 1344 bits
Score difference with first non-orthologous sequence - *A. fumigatus*:1344 *A. fischeri*:1344
afum|EAL89893.1 100.00% afis|EAW25232.1 100.00%
Bootstrap support for afum|EAL89893.1 as seed ortholog is 100%.
Bootstrap support for afis|EAW25232.1 as seed ortholog is 100%.

---

Group of orthologs #2416. Best score 1343 bits
Score difference with first non-orthologous sequence - *A. fumigatus*:1343 *A. fischeri*:1343
afum|EAL84523.1 100.00% afis|EAW20535.1 100.00%
Bootstrap support for afum|EAL84523.1 as seed ortholog is 100%.
Bootstrap support for afis|EAW20535.1 as seed ortholog is 100%.

---

Group of orthologs #2417. Best score 1343 bits
Score difference with first non-orthologous sequence - *A. fumigatus*:1343 *A. fischeri*:1283
afum|EAL84510.1 100.00% afis|EAW20550.1 100.00%
Bootstrap support for afum|EAL84510.1 as seed ortholog is 100%.
Bootstrap support for afis|EAW20550.1 as seed ortholog is 100%.

---

Group of orthologs #2418. Best score 1343 bits
Score difference with first non-orthologous sequence - *A. fumigatus*:1343 *A. fischeri*:1343
afum|EAL89522.1 100.00% afis|EAW24872.1 100.00%
Bootstrap support for afum|EAL89522.1 as seed ortholog is 100%.
Bootstrap support for afis|EAW24872.1 as seed ortholog is 100%.

---

Group of orthologs #2419. Best score 1343 bits
Score difference with first non-orthologous sequence - *A. fumigatus*:1343 *A. fischeri*:1343
afum|EAL89533.1 100.00% afis|EAW24883.1 100.00%
Bootstrap support for afum|EAL89533.1 as seed ortholog is 100%.
Bootstrap support for afis|EAW24883.1 as seed ortholog is 100%.

---

Group of orthologs #2420. Best score 1342 bits
Score difference with first non-orthologous sequence - *A. fumigatus*:1342 *A. fischeri*:1342
afum|EAL91441.1 100.00% afis|EAW17627.1 100.00%
Bootstrap support for afum|EAL91441.1 as seed ortholog is 100%.
Bootstrap support for afis|EAW17627.1 as seed ortholog is 100%.

---

Group of orthologs #2421. Best score 1342 bits
Score difference with first non-orthologous sequence - *A. fumigatus*:1342 *A. fischeri*:1342
afum|EAL86115.1 100.00% afis|EAW24446.1 100.00%
Bootstrap support for afum|EAL86115.1 as seed ortholog is 100%.
Bootstrap support for afis|EAW24446.1 as seed ortholog is 100%.

---

Group of orthologs #2422. Best score 1342 bits
Score difference with first non-orthologous sequence - *A. fumigatus*:1342 *A. fischeri*:1342
afum|EAL92908.1 100.00% afis|EAW21894.1 100.00%
Bootstrap support for afum|EAL92908.1 as seed ortholog is 100%.
Bootstrap support for afis|EAW21894.1 as seed ortholog is 100%.

---

Group of orthologs #2423. Best score 1341 bits
Score difference with first non-orthologous sequence - *A. fumigatus*:1341 *A. fischeri*:1341
afum|EAL88636.1 100.00% afis|EAW16004.1 100.00%
Bootstrap support for afum|EAL88636.1 as seed ortholog is 100%.
Bootstrap support for afis|EAW16004.1 as seed ortholog is 100%.

---

Group of orthologs #2424. Best score 1341 bits
Score difference with first non-orthologous sequence - *A. fumigatus*:65 *A. fischeri*:1341

afum|EAL85181.2 100.00% afis|EAW20061.1 100.00%
Bootstrap support for afum|EAL85181.2 as seed ortholog is 98%.
Bootstrap support for afis|EAW20061.1 as seed ortholog is 100%.

---

Group of orthologs #2425. Best score 1341 bits
Score difference with first non-orthologous sequence - *A. fumigatus*:1341 *A. fischeri*:1005
afum|EAL85348.1 100.00% afis|EAW20231.1 100.00%
Bootstrap support for afum|EAL85348.1 as seed ortholog is 100%.
Bootstrap support for afis|EAW20231.1 as seed ortholog is 100%.

---

Group of orthologs #2426. Best score 1341 bits
Score difference with first non-orthologous sequence - *A. fumigatus*:1341 *A. fischeri*:1341
afum|EAL86092.1 100.00% afis|EAW24424.1 100.00%
Bootstrap support for afum|EAL86092.1 as seed ortholog is 100%.
Bootstrap support for afis|EAW24424.1 as seed ortholog is 100%.

---

Group of orthologs #2427. Best score 1341 bits
Score difference with first non-orthologous sequence - *A. fumigatus*:960 *A. fischeri*:1341
afum|EAL93176.1 100.00% afis|EAW18487.1 100.00%
Bootstrap support for afum|EAL93176.1 as seed ortholog is 100%.
Bootstrap support for afis|EAW18487.1 as seed ortholog is 100%.

---

Group of orthologs #2428. Best score 1341 bits
Score difference with first non-orthologous sequence - *A. fumigatus*:1341 *A. fischeri*:1341
afum|EAL93906.1 100.00% afis|EAW19176.1 100.00%
Bootstrap support for afum|EAL93906.1 as seed ortholog is 100%.
Bootstrap support for afis|EAW19176.1 as seed ortholog is 100%.

---

Group of orthologs #2429. Best score 1340 bits
Score difference with first non-orthologous sequence - *A. fumigatus*:1340 *A. fischeri*:1272
afum|EAL86646.1 100.00% afis|EAW17224.1 100.00%
Bootstrap support for afum|EAL86646.1 as seed ortholog is 100%.
Bootstrap support for afis|EAW17224.1 as seed ortholog is 100%.

---

Group of orthologs #2430. Best score 1340 bits
Score difference with first non-orthologous sequence - *A. fumigatus*:1340 *A. fischeri*:1340
afum|EAL86940.2 100.00% afis|EAW19578.1 100.00%
Bootstrap support for afum|EAL86940.2 as seed ortholog is 100%.
Bootstrap support for afis|EAW19578.1 as seed ortholog is 100%.

---

Group of orthologs #2431. Best score 1340 bits
Score difference with first non-orthologous sequence - *A. fumigatus*:1080 *A. fischeri*:950
afum|EAL85690.1 100.00% afis|EAW22081.1 100.00%
Bootstrap support for afum|EAL85690.1 as seed ortholog is 100%.
Bootstrap support for afis|EAW22081.1 as seed ortholog is 100%.

---

Group of orthologs #2432. Best score 1340 bits
Score difference with first non-orthologous sequence - *A. fumigatus*:1340 *A. fischeri*:1057
afum|EAL91627.1 100.00% afis|EAW17817.1 100.00%
Bootstrap support for afum|EAL91627.1 as seed ortholog is 100%.
Bootstrap support for afis|EAW17817.1 as seed ortholog is 100%.

---

Group of orthologs #2433. Best score 1340 bits
Score difference with first non-orthologous sequence - *A. fumigatus*:1340 *A. fischeri*:1340
afum|EAL94030.1 100.00% afis|EAW19314.1 100.00%
Bootstrap support for afum|EAL94030.1 as seed ortholog is 100%.

Bootstrap support for afis|EAW19314.1 as seed ortholog is 100%.

---

Group of orthologs #2434. Best score 1340 bits
Score difference with first non-orthologous sequence - *A. fumigatus*:1340 *A. fischeri*:1340
afum|EAL92513.1 100.00% afis|EAW21494.1 100.00%
Bootstrap support for afum|EAL92513.1 as seed ortholog is 100%.
Bootstrap support for afis|EAW21494.1 as seed ortholog is 100%.

---

Group of orthologs #2435. Best score 1339 bits
Score difference with first non-orthologous sequence - *A. fumigatus*:1339 *A. fischeri*:1339
afum|EAL85801.1 100.00% afis|EAW15527.1 100.00%
Bootstrap support for afum|EAL85801.1 as seed ortholog is 100%.
Bootstrap support for afis|EAW15527.1 as seed ortholog is 100%.

---

Group of orthologs #2436. Best score 1339 bits
Score difference with first non-orthologous sequence - *A. fumigatus*:1244 *A. fischeri*:1238
afum|EAL86641.1 100.00% afis|EAW17215.1 100.00%
Bootstrap support for afum|EAL86641.1 as seed ortholog is 100%.
Bootstrap support for afis|EAW17215.1 as seed ortholog is 100%.

---

Group of orthologs #2437. Best score 1339 bits
Score difference with first non-orthologous sequence - *A. fumigatus*:1339 *A. fischeri*:1339
afum|EAL88565.1 100.00% afis|EAW15929.1 100.00%
Bootstrap support for afum|EAL88565.1 as seed ortholog is 100%.
Bootstrap support for afis|EAW15929.1 as seed ortholog is 100%.

---

Group of orthologs #2438. Best score 1339 bits
Score difference with first non-orthologous sequence - *A. fumigatus*:1339 *A. fischeri*:1339
afum|EAL84768.1 100.00% afis|EAW20866.1 100.00%
Bootstrap support for afum|EAL84768.1 as seed ortholog is 100%.
Bootstrap support for afis|EAW20866.1 as seed ortholog is 100%.

---

Group of orthologs #2439. Best score 1339 bits
Score difference with first non-orthologous sequence - *A. fumigatus*:1339 *A. fischeri*:1154
afum|EAL91405.1 100.00% afis|EAW17592.1 100.00%
Bootstrap support for afum|EAL91405.1 as seed ortholog is 100%.
Bootstrap support for afis|EAW17592.1 as seed ortholog is 100%.

---

Group of orthologs #2440. Best score 1339 bits
Score difference with first non-orthologous sequence - *A. fumigatus*:1063 *A. fischeri*:695
afum|EAL88355.1 100.00% afis|EAW23125.1 100.00%
Bootstrap support for afum|EAL88355.1 as seed ortholog is 100%.
Bootstrap support for afis|EAW23125.1 as seed ortholog is 100%.

---

Group of orthologs #2441. Best score 1339 bits
Score difference with first non-orthologous sequence - *A. fumigatus*:1339 *A. fischeri*:1339
afum|EAL89545.1 100.00% afis|EAW24892.1 100.00%
Bootstrap support for afum|EAL89545.1 as seed ortholog is 100%.
Bootstrap support for afis|EAW24892.1 as seed ortholog is 100%.

---

Group of orthologs #2442. Best score 1338 bits
Score difference with first non-orthologous sequence - *A. fumigatus*:1338 *A. fischeri*:1338
afum|EAL84454.1 100.00% afis|EAW20606.1 100.00%
Bootstrap support for afum|EAL84454.1 as seed ortholog is 100%.
Bootstrap support for afis|EAW20606.1 as seed ortholog is 100%.

---

Group of orthologs #2443. Best score 1338 bits
Score difference with first non-orthologous sequence - *A. fumigatus*:1338 *A. fischeri*:1338
afum|EAL87087.1 100.00% afis|EAW19407.1 100.00%
Bootstrap support for afum|EAL87087.1 as seed ortholog is 100%.
Bootstrap support for afis|EAW19407.1 as seed ortholog is 100%.

---

Group of orthologs #2444. Best score 1337 bits
Score difference with first non-orthologous sequence - *A. fumigatus*:1337 *A. fischeri*:1337
afum|EAL92400.1 100.00% afis|EAW21385.1 100.00%
Bootstrap support for afum|EAL92400.1 as seed ortholog is 100%.
Bootstrap support for afis|EAW21385.1 as seed ortholog is 100%.

---

Group of orthologs #2445. Best score 1337 bits
Score difference with first non-orthologous sequence - *A. fumigatus*:1337 *A. fischeri*:1337
afum|EAL92469.1 100.00% afis|EAW21450.1 100.00%
Bootstrap support for afum|EAL92469.1 as seed ortholog is 100%.
Bootstrap support for afis|EAW21450.1 as seed ortholog is 100%.

---

Group of orthologs #2446. Best score 1337 bits
Score difference with first non-orthologous sequence - *A. fumigatus*:1337 *A. fischeri*:1337
afum|EAL89974.1 100.00% afis|EAW25313.1 100.00%
Bootstrap support for afum|EAL89974.1 as seed ortholog is 100%.
Bootstrap support for afis|EAW25313.1 as seed ortholog is 100%.

---

Group of orthologs #2447. Best score 1336 bits
Score difference with first non-orthologous sequence - *A. fumigatus*:1264 *A. fischeri*:1240
afum|EAL86024.2 100.00% afis|EAW24353.1 100.00%
Bootstrap support for afum|EAL86024.2 as seed ortholog is 100%.
Bootstrap support for afis|EAW24353.1 as seed ortholog is 100%.

---

Group of orthologs #2448. Best score 1336 bits
Score difference with first non-orthologous sequence - *A. fumigatus*:1336 *A. fischeri*:1336
afum|EAL93563.1 100.00% afis|EAW18829.1 100.00%
Bootstrap support for afum|EAL93563.1 as seed ortholog is 100%.
Bootstrap support for afis|EAW18829.1 as seed ortholog is 100%.

---

Group of orthologs #2449. Best score 1335 bits
Score difference with first non-orthologous sequence - *A. fumigatus*:1335 *A. fischeri*:1335
afum|EAL84631.1 100.00% afis|EAW20997.1 100.00%
Bootstrap support for afum|EAL84631.1 as seed ortholog is 100%.
Bootstrap support for afis|EAW20997.1 as seed ortholog is 100%.

---

Group of orthologs #2450. Best score 1335 bits
Score difference with first non-orthologous sequence - *A. fumigatus*:1335 *A. fischeri*:1335
afum|EAL91665.1 100.00% afis|EAW17857.1 100.00%
Bootstrap support for afum|EAL91665.1 as seed ortholog is 100%.
Bootstrap support for afis|EAW17857.1 as seed ortholog is 100%.

---

Group of orthologs #2451. Best score 1335 bits
Score difference with first non-orthologous sequence - *A. fumigatus*:1335 *A. fischeri*:1335
afum|EAL91838.1 100.00% afis|EAW18027.1 100.00%
Bootstrap support for afum|EAL91838.1 as seed ortholog is 100%.
Bootstrap support for afis|EAW18027.1 as seed ortholog is 100%.

---

Group of orthologs #2452. Best score 1335 bits
Score difference with first non-orthologous sequence - *A. fumigatus*:1188 *A. fischeri*:823

afum|EAL88065.2 100.00% afis|EAW23405.1 100.00%
Bootstrap support for afum|EAL88065.2 as seed ortholog is 100%.
Bootstrap support for afis|EAW23405.1 as seed ortholog is 100%.

---

Group of orthologs #2453. Best score 1335 bits
Score difference with first non-orthologous sequence - *A. fumigatus*:1335 *A. fischeri*:1335
afum|EAL93704.1 100.00% afis|EAW18971.1 100.00%
Bootstrap support for afum|EAL93704.1 as seed ortholog is 100%.
Bootstrap support for afis|EAW18971.1 as seed ortholog is 100%.

---

Group of orthologs #2454. Best score 1335 bits
Score difference with first non-orthologous sequence - *A. fumigatus*:1335 *A. fischeri*:1335
afum|EAL90878.1 100.00% afis|EAW22312.1 100.00%
Bootstrap support for afum|EAL90878.1 as seed ortholog is 100%.
Bootstrap support for afis|EAW22312.1 as seed ortholog is 100%.

---

Group of orthologs #2455. Best score 1333 bits
Score difference with first non-orthologous sequence - *A. fumigatus*:1145 *A. fischeri*:894
afum|EAL87611.1 100.00% afis|EAW18246.1 100.00%
Bootstrap support for afum|EAL87611.1 as seed ortholog is 100%.
Bootstrap support for afis|EAW18246.1 as seed ortholog is 100%.

---

Group of orthologs #2456. Best score 1333 bits
Score difference with first non-orthologous sequence - *A. fumigatus*:1333 *A. fischeri*:1333
afum|EAL88400.1 100.00% afis|EAW23079.1 100.00%
Bootstrap support for afum|EAL88400.1 as seed ortholog is 100%.
Bootstrap support for afis|EAW23079.1 as seed ortholog is 100%.

---

Group of orthologs #2457. Best score 1333 bits
Score difference with first non-orthologous sequence - *A. fumigatus*:1333 *A. fischeri*:1333
afum|EAL91958.1 100.00% afis|EAW24104.1 100.00%
Bootstrap support for afum|EAL91958.1 as seed ortholog is 100%.
Bootstrap support for afis|EAW24104.1 as seed ortholog is 100%.

---

Group of orthologs #2458. Best score 1332 bits
Score difference with first non-orthologous sequence - *A. fumigatus*:1332 *A. fischeri*:1332
afum|EAL85653.1 100.00% afis|EAW15695.1 100.00%
Bootstrap support for afum|EAL85653.1 as seed ortholog is 100%.
Bootstrap support for afis|EAW15695.1 as seed ortholog is 100%.

---

Group of orthologs #2459. Best score 1332 bits
Score difference with first non-orthologous sequence - *A. fumigatus*:386 *A. fischeri*:1332
afum|EAL86308.1 100.00% afis|EAW16734.1 100.00%
Bootstrap support for afum|EAL86308.1 as seed ortholog is 100%.
Bootstrap support for afis|EAW16734.1 as seed ortholog is 100%.

---

Group of orthologs #2460. Best score 1332 bits
Score difference with first non-orthologous sequence - *A. fumigatus*:1332 *A. fischeri*:504
afum|EAL86188.1 100.00% afis|EAW24523.1 100.00%
Bootstrap support for afum|EAL86188.1 as seed ortholog is 100%.
Bootstrap support for afis|EAW24523.1 as seed ortholog is 100%.

---

Group of orthologs #2461. Best score 1332 bits
Score difference with first non-orthologous sequence - *A. fumigatus*:1332 *A. fischeri*:1332
afum|EAL88382.2 100.00% afis|EAW23096.1 100.00%
Bootstrap support for afum|EAL88382.2 as seed ortholog is 100%.

Bootstrap support for afis|EAW23096.1 as seed ortholog is 100%.

---

Group of orthologs #2462. Best score 1332 bits
Score difference with first non-orthologous sequence - *A. fumigatus*:1332 *A. fischeri*:1332
afum|EAL88320.1 100.00% afis|EAW23163.1 100.00%
Bootstrap support for afum|EAL88320.1 as seed ortholog is 100%.
Bootstrap support for afis|EAW23163.1 as seed ortholog is 100%.

---

Group of orthologs #2463. Best score 1332 bits
Score difference with first non-orthologous sequence - *A. fumigatus*:1332 *A. fischeri*:1332
afum|EAL90904.1 100.00% afis|EAW22277.1 100.00%
Bootstrap support for afum|EAL90904.1 as seed ortholog is 100%.
Bootstrap support for afis|EAW22277.1 as seed ortholog is 100%.

---

Group of orthologs #2464. Best score 1332 bits
Score difference with first non-orthologous sequence - *A. fumigatus*:1332 *A. fischeri*:1175
afum|EAL92226.1 100.00% afis|EAW21215.1 100.00%
Bootstrap support for afum|EAL92226.1 as seed ortholog is 100%.
Bootstrap support for afis|EAW21215.1 as seed ortholog is 100%.

---

Group of orthologs #2465. Best score 1331 bits
Score difference with first non-orthologous sequence - *A. fumigatus*:1331 *A. fischeri*:1331
afum|EAL88676.1 100.00% afis|EAW16044.1 100.00%
Bootstrap support for afum|EAL88676.1 as seed ortholog is 100%.
Bootstrap support for afis|EAW16044.1 as seed ortholog is 100%.

---

Group of orthologs #2466. Best score 1330 bits
Score difference with first non-orthologous sequence - *A. fumigatus*:1330 *A. fischeri*:1330
afum|EAL84516.1 100.00% afis|EAW20542.1 100.00%
Bootstrap support for afum|EAL84516.1 as seed ortholog is 100%.
Bootstrap support for afis|EAW20542.1 as seed ortholog is 100%.

---

Group of orthologs #2467. Best score 1330 bits
Score difference with first non-orthologous sequence - *A. fumigatus*:1330 *A. fischeri*:1330
afum|EAL84724.2 100.00% afis|EAW20910.1 100.00%
Bootstrap support for afum|EAL84724.2 as seed ortholog is 100%.
Bootstrap support for afis|EAW20910.1 as seed ortholog is 100%.

---

Group of orthologs #2468. Best score 1330 bits
Score difference with first non-orthologous sequence - *A. fumigatus*:1330 *A. fischeri*:1330
afum|EAL93274.1 100.00% afis|EAW18571.1 100.00%
Bootstrap support for afum|EAL93274.1 as seed ortholog is 100%.
Bootstrap support for afis|EAW18571.1 as seed ortholog is 100%.

---

Group of orthologs #2469. Best score 1330 bits
Score difference with first non-orthologous sequence - *A. fumigatus*:1330 *A. fischeri*:1154
afum|EAL90643.1 100.00% afis|EAW22552.1 100.00%
Bootstrap support for afum|EAL90643.1 as seed ortholog is 100%.
Bootstrap support for afis|EAW22552.1 as seed ortholog is 100%.

---

Group of orthologs #2470. Best score 1329 bits
Score difference with first non-orthologous sequence - *A. fumigatus*:1329 *A. fischeri*:1329
afum|EAL85553.2 100.00% afis|EAW15797.1 100.00%
Bootstrap support for afum|EAL85553.2 as seed ortholog is 100%.
Bootstrap support for afis|EAW15797.1 as seed ortholog is 100%.

---

Group of orthologs #2471. Best score 1329 bits
Score difference with first non-orthologous sequence - *A. fumigatus*:1329 *A. fischeri*:1017
afum|EAL86453.1 100.00% afis|EAW16927.1 100.00%
Bootstrap support for afum|EAL86453.1 as seed ortholog is 100%.
Bootstrap support for afis|EAW16927.1 as seed ortholog is 100%.

---

Group of orthologs #2472. Best score 1329 bits
Score difference with first non-orthologous sequence - *A. fumigatus*:1164 *A. fischeri*:1040
afum|EAL84915.1 100.00% afis|EAW20004.1 100.00%
Bootstrap support for afum|EAL84915.1 as seed ortholog is 100%.
Bootstrap support for afis|EAW20004.1 as seed ortholog is 100%.

---

Group of orthologs #2473. Best score 1329 bits
Score difference with first non-orthologous sequence - *A. fumigatus*:1122 *A. fischeri*:1126
afum|EAL85178.1 100.00% afis|EAW20058.1 100.00%
Bootstrap support for afum|EAL85178.1 as seed ortholog is 100%.
Bootstrap support for afis|EAW20058.1 as seed ortholog is 100%.

---

Group of orthologs #2474. Best score 1328 bits
Score difference with first non-orthologous sequence - *A. fumigatus*:1328 *A. fischeri*:1328
afum|EAL91626.1 100.00% afis|EAW17816.1 100.00%
Bootstrap support for afum|EAL91626.1 as seed ortholog is 100%.
Bootstrap support for afis|EAW17816.1 as seed ortholog is 100%.

---

Group of orthologs #2475. Best score 1328 bits
Score difference with first non-orthologous sequence - *A. fumigatus*:1328 *A. fischeri*:1328
afum|EAL86077.1 100.00% afis|EAW24408.1 100.00%
Bootstrap support for afum|EAL86077.1 as seed ortholog is 100%.
Bootstrap support for afis|EAW24408.1 as seed ortholog is 100%.

---

Group of orthologs #2476. Best score 1328 bits
Score difference with first non-orthologous sequence - *A. fumigatus*:1328 *A. fischeri*:1328
afum|EAL87434.1 100.00% afis|EAW24004.1 100.00%
Bootstrap support for afum|EAL87434.1 as seed ortholog is 100%.
Bootstrap support for afis|EAW24004.1 as seed ortholog is 100%.

---

Group of orthologs #2477. Best score 1328 bits
Score difference with first non-orthologous sequence - *A. fumigatus*:1328 *A. fischeri*:940
afum|EAL93386.1 100.00% afis|EAW18668.1 100.00%
Bootstrap support for afum|EAL93386.1 as seed ortholog is 100%.
Bootstrap support for afis|EAW18668.1 as seed ortholog is 100%.

---

Group of orthologs #2478. Best score 1328 bits
Score difference with first non-orthologous sequence - *A. fumigatus*:1328 *A. fischeri*:1328
afum|EAL92367.1 100.00% afis|EAW21352.1 100.00%
Bootstrap support for afum|EAL92367.1 as seed ortholog is 100%.
Bootstrap support for afis|EAW21352.1 as seed ortholog is 100%.

---

Group of orthologs #2479. Best score 1327 bits
Score difference with first non-orthologous sequence - *A. fumigatus*:1327 *A. fischeri*:1327
afum|EAL85600.1 100.00% afis|EAW15749.1 100.00%
Bootstrap support for afum|EAL85600.1 as seed ortholog is 100%.
Bootstrap support for afis|EAW15749.1 as seed ortholog is 100%.

---

Group of orthologs #2480. Best score 1327 bits
Score difference with first non-orthologous sequence - *A. fumigatus*:1327 *A. fischeri*:1327

afum|EAL91520.2 100.00% afis|EAW17705.1 100.00%
Bootstrap support for afum|EAL91520.2 as seed ortholog is 100%.
Bootstrap support for afis|EAW17705.1 as seed ortholog is 100%.

---

Group of orthologs #2481. Best score 1327 bits
Score difference with first non-orthologous sequence - *A. fumigatus*:995 *A. fischeri*:1327
afum|EAL85879.1 100.00% afis|EAW24209.1 100.00%
Bootstrap support for afum|EAL85879.1 as seed ortholog is 100%.
Bootstrap support for afis|EAW24209.1 as seed ortholog is 100%.

---

Group of orthologs #2482. Best score 1327 bits
Score difference with first non-orthologous sequence - *A. fumigatus*:1164 *A. fischeri*:1327
afum|EAL86025.2 100.00% afis|EAW24354.1 100.00%
Bootstrap support for afum|EAL86025.2 as seed ortholog is 100%.
Bootstrap support for afis|EAW24354.1 as seed ortholog is 100%.

---

Group of orthologs #2483. Best score 1327 bits
Score difference with first non-orthologous sequence - *A. fumigatus*:1327 *A. fischeri*:1327
afum|EAL89450.2 100.00% afis|EAW24808.1 100.00%
Bootstrap support for afum|EAL89450.2 as seed ortholog is 100%.
Bootstrap support for afis|EAW24808.1 as seed ortholog is 100%.

---

Group of orthologs #2484. Best score 1327 bits
Score difference with first non-orthologous sequence - *A. fumigatus*:1327 *A. fischeri*:1327
afum|EAL89895.1 100.00% afis|EAW25234.1 100.00%
Bootstrap support for afum|EAL89895.1 as seed ortholog is 100%.
Bootstrap support for afis|EAW25234.1 as seed ortholog is 100%.

---

Group of orthologs #2485. Best score 1326 bits
Score difference with first non-orthologous sequence - *A. fumigatus*:1061 *A. fischeri*:1227
afum|EAL84908.1 100.00% afis|EAW20010.1 100.00%
Bootstrap support for afum|EAL84908.1 as seed ortholog is 100%.
Bootstrap support for afis|EAW20010.1 as seed ortholog is 100%.

---

Group of orthologs #2486. Best score 1326 bits
Score difference with first non-orthologous sequence - *A. fumigatus*:1326 *A. fischeri*:1326
afum|EAL91597.1 100.00% afis|EAW17788.1 100.00%
Bootstrap support for afum|EAL91597.1 as seed ortholog is 100%.
Bootstrap support for afis|EAW17788.1 as seed ortholog is 100%.

---

Group of orthologs #2487. Best score 1326 bits
Score difference with first non-orthologous sequence - *A. fumigatus*:1326 *A. fischeri*:989
afum|EAL87957.1 100.00% afis|EAW23504.1 100.00%
Bootstrap support for afum|EAL87957.1 as seed ortholog is 100%.
Bootstrap support for afis|EAW23504.1 as seed ortholog is 100%.

---

Group of orthologs #2488. Best score 1326 bits
Score difference with first non-orthologous sequence - *A. fumigatus*:1326 *A. fischeri*:1326
afum|EAL87929.1 100.00% afis|EAW23537.1 100.00%
Bootstrap support for afum|EAL87929.1 as seed ortholog is 100%.
Bootstrap support for afis|EAW23537.1 as seed ortholog is 100%.

---

Group of orthologs #2489. Best score 1326 bits
Score difference with first non-orthologous sequence - *A. fumigatus*:1326 *A. fischeri*:1326
afum|EAL92849.1 100.00% afis|EAW21834.1 100.00%
Bootstrap support for afum|EAL92849.1 as seed ortholog is 100%.

Bootstrap support for afis|EAW21834.1 as seed ortholog is 100%.

---

Group of orthologs #2490. Best score 1325 bits
Score difference with first non-orthologous sequence - *A. fumigatus*:1325 *A. fischeri*:1325
afum|EAL87329.1 100.00% afis|EAW23902.1 100.00%
Bootstrap support for afum|EAL87329.1 as seed ortholog is 100%.
Bootstrap support for afis|EAW23902.1 as seed ortholog is 100%.

---

Group of orthologs #2491. Best score 1325 bits
Score difference with first non-orthologous sequence - *A. fumigatus*:1182 *A. fischeri*:1106
afum|EAL92115.1 100.00% afis|EAW21092.1 100.00%
Bootstrap support for afum|EAL92115.1 as seed ortholog is 100%.
Bootstrap support for afis|EAW21092.1 as seed ortholog is 100%.

---

Group of orthologs #2492. Best score 1325 bits
Score difference with first non-orthologous sequence - *A. fumigatus*:1325 *A. fischeri*:1325
afum|EAL89691.2 100.00% afis|EAW25006.1 100.00%
Bootstrap support for afum|EAL89691.2 as seed ortholog is 100%.
Bootstrap support for afis|EAW25006.1 as seed ortholog is 100%.

---

Group of orthologs #2493. Best score 1324 bits
Score difference with first non-orthologous sequence - *A. fumigatus*:1107 *A. fischeri*:1187
afum|EAL88366.1 100.00% afis|EAW23113.1 100.00%
Bootstrap support for afum|EAL88366.1 as seed ortholog is 100%.
Bootstrap support for afis|EAW23113.1 as seed ortholog is 100%.

---

Group of orthologs #2494. Best score 1324 bits
Score difference with first non-orthologous sequence - *A. fumigatus*:1324 *A. fischeri*:1324
afum|EAL93789.1 100.00% afis|EAW19060.1 100.00%
Bootstrap support for afum|EAL93789.1 as seed ortholog is 100%.
Bootstrap support for afis|EAW19060.1 as seed ortholog is 100%.

---

Group of orthologs #2495. Best score 1324 bits
Score difference with first non-orthologous sequence - *A. fumigatus*:1324 *A. fischeri*:1171
afum|EAL92212.1 100.00% afis|EAW21202.1 100.00%
Bootstrap support for afum|EAL92212.1 as seed ortholog is 100%.
Bootstrap support for afis|EAW21202.1 as seed ortholog is 100%.

---

Group of orthologs #2496. Best score 1324 bits
Score difference with first non-orthologous sequence - *A. fumigatus*:1324 *A. fischeri*:1324
afum|EAL89939.1 100.00% afis|EAW25276.1 100.00%
Bootstrap support for afum|EAL89939.1 as seed ortholog is 100%.
Bootstrap support for afis|EAW25276.1 as seed ortholog is 100%.

---

Group of orthologs #2497. Best score 1324 bits
Score difference with first non-orthologous sequence - *A. fumigatus*:1223 *A. fischeri*:1197
afum|EBA27381.1 100.00% afis|EAW24765.1 100.00%
Bootstrap support for afum|EBA27381.1 as seed ortholog is 100%.
Bootstrap support for afis|EAW24765.1 as seed ortholog is 100%.

---

Group of orthologs #2498. Best score 1323 bits
Score difference with first non-orthologous sequence - *A. fumigatus*:1323 *A. fischeri*:1323
afum|EAL88560.2 100.00% afis|EAW15923.1 100.00%
Bootstrap support for afum|EAL88560.2 as seed ortholog is 100%.
Bootstrap support for afis|EAW15923.1 as seed ortholog is 100%.

---

Group of orthologs #2499. Best score 1323 bits
Score difference with first non-orthologous sequence - *A. fumigatus*:1323 *A. fischeri*:1323
afum|EAL89504.1 100.00% afis|EAW24856.1 100.00%
Bootstrap support for afum|EAL89504.1 as seed ortholog is 100%.
Bootstrap support for afis|EAW24856.1 as seed ortholog is 100%.

---

Group of orthologs #2500. Best score 1322 bits
Score difference with first non-orthologous sequence - *A. fumigatus*:1322 *A. fischeri*:1322
afum|EAL85289.1 100.00% afis|EAW20171.1 100.00%
Bootstrap support for afum|EAL85289.1 as seed ortholog is 100%.
Bootstrap support for afis|EAW20171.1 as seed ortholog is 100%.

---

Group of orthologs #2501. Best score 1322 bits
Score difference with first non-orthologous sequence - *A. fumigatus*:1322 *A. fischeri*:1322
afum|EAL89470.1 100.00% afis|EAW24826.1 100.00%
Bootstrap support for afum|EAL89470.1 as seed ortholog is 100%.
Bootstrap support for afis|EAW24826.1 as seed ortholog is 100%.

---

Group of orthologs #2502. Best score 1321 bits
Score difference with first non-orthologous sequence - *A. fumigatus*:1321 *A. fischeri*:1321
afum|EAL84584.1 100.00% afis|EAW20468.1 100.00%
Bootstrap support for afum|EAL84584.1 as seed ortholog is 100%.
Bootstrap support for afis|EAW20468.1 as seed ortholog is 100%.

---

Group of orthologs #2503. Best score 1321 bits
Score difference with first non-orthologous sequence - *A. fumigatus*:1110 *A. fischeri*:1148
afum|EAL93718.1 100.00% afis|EAW18982.1 100.00%
Bootstrap support for afum|EAL93718.1 as seed ortholog is 100%.
Bootstrap support for afis|EAW18982.1 as seed ortholog is 100%.

---

Group of orthologs #2504. Best score 1321 bits
Score difference with first non-orthologous sequence - *A. fumigatus*:1321 *A. fischeri*:1321
afum|EAL93899.1 100.00% afis|EAW19169.1 100.00%
Bootstrap support for afum|EAL93899.1 as seed ortholog is 100%.
Bootstrap support for afis|EAW19169.1 as seed ortholog is 100%.

---

Group of orthologs #2505. Best score 1321 bits
Score difference with first non-orthologous sequence - *A. fumigatus*:1321 *A. fischeri*:1321
afum|EBA27212.1 100.00% afis|EAW19979.1 100.00%
Bootstrap support for afum|EBA27212.1 as seed ortholog is 100%.
Bootstrap support for afis|EAW19979.1 as seed ortholog is 100%.

---

Group of orthologs #2506. Best score 1321 bits
Score difference with first non-orthologous sequence - *A. fumigatus*:1321 *A. fischeri*:1321
afum|EAL89792.1 100.00% afis|EAW25121.1 100.00%
Bootstrap support for afum|EAL89792.1 as seed ortholog is 100%.
Bootstrap support for afis|EAW25121.1 as seed ortholog is 100%.

---

Group of orthologs #2507. Best score 1320 bits
Score difference with first non-orthologous sequence - *A. fumigatus*:1059 *A. fischeri*:1320
afum|EAL88956.1 100.00% afis|EAW16331.1 100.00%
Bootstrap support for afum|EAL88956.1 as seed ortholog is 100%.
Bootstrap support for afis|EAW16331.1 as seed ortholog is 100%.

---

Group of orthologs #2508. Best score 1320 bits
Score difference with first non-orthologous sequence - *A. fumigatus*:1234 *A. fischeri*:1320

afum|EAL91599.1 100.00% afis|EAW17790.1 100.00%
Bootstrap support for afum|EAL91599.1 as seed ortholog is 100%.
Bootstrap support for afis|EAW17790.1 as seed ortholog is 100%.

---

Group of orthologs #2509. Best score 1320 bits
Score difference with first non-orthologous sequence - *A. fumigatus*:1320 *A. fischeri*:1320
afum|EAL92618.1 100.00% afis|EAW21599.1 100.00%
Bootstrap support for afum|EAL92618.1 as seed ortholog is 100%.
Bootstrap support for afis|EAW21599.1 as seed ortholog is 100%.

---

Group of orthologs #2510. Best score 1320 bits
Score difference with first non-orthologous sequence - *A. fumigatus*:1320 *A. fischeri*:1320
afum|EBA27203.1 100.00% afis|EAW20900.1 100.00%
Bootstrap support for afum|EBA27203.1 as seed ortholog is 100%.
Bootstrap support for afis|EAW20900.1 as seed ortholog is 100%.

---

Group of orthologs #2511. Best score 1319 bits
Score difference with first non-orthologous sequence - *A. fumigatus*:1319 *A. fischeri*:1319
afum|EAL91425.1 100.00% afis|EAW17612.1 100.00%
Bootstrap support for afum|EAL91425.1 as seed ortholog is 100%.
Bootstrap support for afis|EAW17612.1 as seed ortholog is 100%.

---

Group of orthologs #2512. Best score 1319 bits
Score difference with first non-orthologous sequence - *A. fumigatus*:1319 *A. fischeri*:1319
afum|EAL91462.1 100.00% afis|EAW17647.1 100.00%
Bootstrap support for afum|EAL91462.1 as seed ortholog is 100%.
Bootstrap support for afis|EAW17647.1 as seed ortholog is 100%.

---

Group of orthologs #2513. Best score 1319 bits
Score difference with first non-orthologous sequence - *A. fumigatus*:760 *A. fischeri*:1120
afum|EAL92781.2 100.00% afis|EAW21750.1 100.00%
Bootstrap support for afum|EAL92781.2 as seed ortholog is 100%.
Bootstrap support for afis|EAW21750.1 as seed ortholog is 100%.

---

Group of orthologs #2514. Best score 1318 bits
Score difference with first non-orthologous sequence - *A. fumigatus*:1318 *A. fischeri*:1318
afum|EAL93770.1 100.00% afis|EAW19041.1 100.00%
Bootstrap support for afum|EAL93770.1 as seed ortholog is 100%.
Bootstrap support for afis|EAW19041.1 as seed ortholog is 100%.

---

Group of orthologs #2515. Best score 1318 bits
Score difference with first non-orthologous sequence - *A. fumigatus*:1133 *A. fischeri*:819
afum|EAL89427.1 100.00% afis|EAW24788.1 100.00%
Bootstrap support for afum|EAL89427.1 as seed ortholog is 100%.
Bootstrap support for afis|EAW24788.1 as seed ortholog is 100%.

---

Group of orthologs #2516. Best score 1317 bits
Score difference with first non-orthologous sequence - *A. fumigatus*:1317 *A. fischeri*:1317
afum|EAL88991.1 100.00% afis|EAW16366.1 100.00%
Bootstrap support for afum|EAL88991.1 as seed ortholog is 100%.
Bootstrap support for afis|EAW16366.1 as seed ortholog is 100%.

---

Group of orthologs #2517. Best score 1317 bits
Score difference with first non-orthologous sequence - *A. fumigatus*:1128 *A. fischeri*:1317
afum|EAL84394.2 100.00% afis|EAW25719.1 100.00%
Bootstrap support for afum|EAL84394.2 as seed ortholog is 100%.

Bootstrap support for afis|EAW25719.1 as seed ortholog is 100%.

---

Group of orthologs #2518. Best score 1317 bits
Score difference with first non-orthologous sequence - *A. fumigatus*:1048 *A. fischeri*:1317
afum|EAL87356.1 100.00% afis|EAW23931.1 100.00%
Bootstrap support for afum|EAL87356.1 as seed ortholog is 100%.
Bootstrap support for afis|EAW23931.1 as seed ortholog is 100%.

---

Group of orthologs #2519. Best score 1317 bits
Score difference with first non-orthologous sequence - *A. fumigatus*:1317 *A. fischeri*:1250
afum|EAL91998.1 100.00% afis|EAW24143.1 100.00%
Bootstrap support for afum|EAL91998.1 as seed ortholog is 100%.
Bootstrap support for afis|EAW24143.1 as seed ortholog is 100%.

---

Group of orthologs #2520. Best score 1316 bits
Score difference with first non-orthologous sequence - *A. fumigatus*:1316 *A. fischeri*:1048
afum|EAL89217.1 100.00% afis|EAW16620.1 100.00%
Bootstrap support for afum|EAL89217.1 as seed ortholog is 100%.
Bootstrap support for afis|EAW16620.1 as seed ortholog is 100%.

---

Group of orthologs #2521. Best score 1316 bits
Score difference with first non-orthologous sequence - *A. fumigatus*:1220 *A. fischeri*:1316
afum|EAL87695.1 100.00% afis|EAW18330.1 100.00%
Bootstrap support for afum|EAL87695.1 as seed ortholog is 100%.
Bootstrap support for afis|EAW18330.1 as seed ortholog is 100%.

---

Group of orthologs #2522. Best score 1316 bits
Score difference with first non-orthologous sequence - *A. fumigatus*:1316 *A. fischeri*:1222
afum|EAL89372.1 100.00% afis|EAW24723.1 100.00%
Bootstrap support for afum|EAL89372.1 as seed ortholog is 100%.
Bootstrap support for afis|EAW24723.1 as seed ortholog is 100%.

---

Group of orthologs #2523. Best score 1315 bits
Score difference with first non-orthologous sequence - *A. fumigatus*:1315 *A. fischeri*:1315
afum|EAL88593.1 100.00% afis|EAW15960.1 100.00%
Bootstrap support for afum|EAL88593.1 as seed ortholog is 100%.
Bootstrap support for afis|EAW15960.1 as seed ortholog is 100%.

---

Group of orthologs #2524. Best score 1315 bits
Score difference with first non-orthologous sequence - *A. fumigatus*:960 *A. fischeri*:1031
afum|EAL91624.1 100.00% afis|EAW17815.1 100.00%
Bootstrap support for afum|EAL91624.1 as seed ortholog is 100%.
Bootstrap support for afis|EAW17815.1 as seed ortholog is 100%.

---

Group of orthologs #2525. Best score 1315 bits
Score difference with first non-orthologous sequence - *A. fumigatus*:1315 *A. fischeri*:1315
afum|EAL92752.1 100.00% afis|EAW21719.1 100.00%
Bootstrap support for afum|EAL92752.1 as seed ortholog is 100%.
Bootstrap support for afis|EAW21719.1 as seed ortholog is 100%.

---

Group of orthologs #2526. Best score 1315 bits
Score difference with first non-orthologous sequence - *A. fumigatus*:1040 *A. fischeri*:873
afum|EAL89485.1 100.00% afis|EAW24839.1 100.00%
Bootstrap support for afum|EAL89485.1 as seed ortholog is 100%.
Bootstrap support for afis|EAW24839.1 as seed ortholog is 100%.

---

Group of orthologs #2527. Best score 1315 bits
Score difference with first non-orthologous sequence - *A. fumigatus*:1315 *A. fischeri*:1315
afum|EAL89936.1 100.00% afis|EAW25273.1 100.00%
Bootstrap support for afum|EAL89936.1 as seed ortholog is 100%.
Bootstrap support for afis|EAW25273.1 as seed ortholog is 100%.

---

Group of orthologs #2528. Best score 1314 bits
Score difference with first non-orthologous sequence - *A. fumigatus*:1314 *A. fischeri*:1314
afum|EAL84561.1 100.00% afis|EAW20493.1 100.00%
Bootstrap support for afum|EAL84561.1 as seed ortholog is 100%.
Bootstrap support for afis|EAW20493.1 as seed ortholog is 100%.

---

Group of orthologs #2529. Best score 1314 bits
Score difference with first non-orthologous sequence - *A. fumigatus*:1241 *A. fischeri*:1078
afum|EAL84320.1 100.00% afis|EAW25632.1 100.00%
Bootstrap support for afum|EAL84320.1 as seed ortholog is 100%.
Bootstrap support for afis|EAW25632.1 as seed ortholog is 100%.

---

Group of orthologs #2530. Best score 1314 bits
Score difference with first non-orthologous sequence - *A. fumigatus*:863 *A. fischeri*:1164
afum|EAL87274.1 100.00% afis|EAW23843.1 100.00%
Bootstrap support for afum|EAL87274.1 as seed ortholog is 100%.
Bootstrap support for afis|EAW23843.1 as seed ortholog is 100%.

---

Group of orthologs #2531. Best score 1314 bits
Score difference with first non-orthologous sequence - *A. fumigatus*:1314 *A. fischeri*:1314
afum|EAL88012.1 100.00% afis|EAW23454.1 100.00%
Bootstrap support for afum|EAL88012.1 as seed ortholog is 100%.
Bootstrap support for afis|EAW23454.1 as seed ortholog is 100%.

---

Group of orthologs #2532. Best score 1314 bits
Score difference with first non-orthologous sequence - *A. fumigatus*:928 *A. fischeri*:974
afum|EAL93985.1 100.00% afis|EAW19258.1 100.00%
Bootstrap support for afum|EAL93985.1 as seed ortholog is 100%.
Bootstrap support for afis|EAW19258.1 as seed ortholog is 100%.

---

Group of orthologs #2533. Best score 1313 bits
Score difference with first non-orthologous sequence - *A. fumigatus*:1261 *A. fischeri*:1313
afum|EAL84687.1 100.00% afis|EAW20947.1 100.00%
Bootstrap support for afum|EAL84687.1 as seed ortholog is 100%.
Bootstrap support for afis|EAW20947.1 as seed ortholog is 100%.

---

Group of orthologs #2534. Best score 1313 bits
Score difference with first non-orthologous sequence - *A. fumigatus*:1313 *A. fischeri*:1313
afum|EAL91302.1 100.00% afis|EAW17473.1 100.00%
Bootstrap support for afum|EAL91302.1 as seed ortholog is 100%.
Bootstrap support for afis|EAW17473.1 as seed ortholog is 100%.

---

Group of orthologs #2535. Best score 1313 bits
Score difference with first non-orthologous sequence - *A. fumigatus*:1313 *A. fischeri*:1313
afum|EAL92318.1 100.00% afis|EAW21301.1 100.00%
Bootstrap support for afum|EAL92318.1 as seed ortholog is 100%.
Bootstrap support for afis|EAW21301.1 as seed ortholog is 100%.

---

Group of orthologs #2536. Best score 1313 bits
Score difference with first non-orthologous sequence - *A. fumigatus*:1313 *A. fischeri*:1313

afum|EAL89442.1 100.00% afis|EAW24801.1 100.00%
Bootstrap support for afum|EAL89442.1 as seed ortholog is 100%.
Bootstrap support for afis|EAW24801.1 as seed ortholog is 100%.

---

Group of orthologs #2537. Best score 1313 bits
Score difference with first non-orthologous sequence - *A. fumigatus*:1313 *A. fischeri*:1313
afum|EAL89814.1 100.00% afis|EAW25142.1 100.00%
Bootstrap support for afum|EAL89814.1 as seed ortholog is 100%.
Bootstrap support for afis|EAW25142.1 as seed ortholog is 100%.

---

Group of orthologs #2538. Best score 1313 bits
Score difference with first non-orthologous sequence - *A. fumigatus*:964 *A. fischeri*:864
afum|EAL89929.1 100.00% afis|EAW25266.1 100.00%
Bootstrap support for afum|EAL89929.1 as seed ortholog is 100%.
Bootstrap support for afis|EAW25266.1 as seed ortholog is 100%.

---

Group of orthologs #2539. Best score 1312 bits
Score difference with first non-orthologous sequence - *A. fumigatus*:1312 *A. fischeri*:1312
afum|EAL87714.1 100.00% afis|EAW18351.1 100.00%
Bootstrap support for afum|EAL87714.1 as seed ortholog is 100%.
Bootstrap support for afis|EAW18351.1 as seed ortholog is 100%.

---

Group of orthologs #2540. Best score 1312 bits
Score difference with first non-orthologous sequence - *A. fumigatus*:1312 *A. fischeri*:1312
afum|EAL91485.1 100.00% afis|EAW17671.1 100.00%
Bootstrap support for afum|EAL91485.1 as seed ortholog is 100%.
Bootstrap support for afis|EAW17671.1 as seed ortholog is 100%.

---

Group of orthologs #2541. Best score 1312 bits
Score difference with first non-orthologous sequence - *A. fumigatus*:1127 *A. fischeri*:1031
afum|EAL84335.1 100.00% afis|EAW25663.1 100.00%
Bootstrap support for afum|EAL84335.1 as seed ortholog is 100%.
Bootstrap support for afis|EAW25663.1 as seed ortholog is 100%.

---

Group of orthologs #2542. Best score 1312 bits
Score difference with first non-orthologous sequence - *A. fumigatus*:1312 *A. fischeri*:1312
afum|EAL91000.1 100.00% afis|EAW22191.1 100.00%
Bootstrap support for afum|EAL91000.1 as seed ortholog is 100%.
Bootstrap support for afis|EAW22191.1 as seed ortholog is 100%.

---

Group of orthologs #2543. Best score 1312 bits
Score difference with first non-orthologous sequence - *A. fumigatus*:1312 *A. fischeri*:1312
afum|EAL92685.2 100.00% afis|EAW21651.1 100.00%
Bootstrap support for afum|EAL92685.2 as seed ortholog is 100%.
Bootstrap support for afis|EAW21651.1 as seed ortholog is 100%.

---

Group of orthologs #2544. Best score 1311 bits
Score difference with first non-orthologous sequence - *A. fumigatus*:1311 *A. fischeri*:1311
afum|EAL84619.1 100.00% afis|EAW20432.1 100.00%
Bootstrap support for afum|EAL84619.1 as seed ortholog is 100%.
Bootstrap support for afis|EAW20432.1 as seed ortholog is 100%.

---

Group of orthologs #2545. Best score 1311 bits
Score difference with first non-orthologous sequence - *A. fumigatus*:1311 *A. fischeri*:507
afum|EAL84797.1 100.00% afis|EAW20837.1 100.00%
Bootstrap support for afum|EAL84797.1 as seed ortholog is 100%.

Bootstrap support for afis|EAW20837.1 as seed ortholog is 100%.

---

Group of orthologs #2546. Best score 1311 bits
Score difference with first non-orthologous sequence - *A. fumigatus*:1311 *A. fischeri*:1311
afum|EAL91252.1 100.00% afis|EAW17424.1 100.00%
Bootstrap support for afum|EAL91252.1 as seed ortholog is 100%.
Bootstrap support for afis|EAW17424.1 as seed ortholog is 100%.

---

Group of orthologs #2547. Best score 1311 bits
Score difference with first non-orthologous sequence - *A. fumigatus*:1311 *A. fischeri*:1311
afum|EAL86234.1 100.00% afis|EAW24571.1 100.00%
Bootstrap support for afum|EAL86234.1 as seed ortholog is 100%.
Bootstrap support for afis|EAW24571.1 as seed ortholog is 100%.

---

Group of orthologs #2548. Best score 1311 bits
Score difference with first non-orthologous sequence - *A. fumigatus*:1311 *A. fischeri*:1311
afum|EAL92063.1 100.00% afis|EAW21024.1 100.00%
Bootstrap support for afum|EAL92063.1 as seed ortholog is 100%.
Bootstrap support for afis|EAW21024.1 as seed ortholog is 100%.

---

Group of orthologs #2549. Best score 1311 bits
Score difference with first non-orthologous sequence - *A. fumigatus*:1311 *A. fischeri*:1311
afum|EAL90784.1 100.00% afis|EAW22408.1 100.00%
Bootstrap support for afum|EAL90784.1 as seed ortholog is 100%.
Bootstrap support for afis|EAW22408.1 as seed ortholog is 100%.

---

Group of orthologs #2550. Best score 1310 bits
Score difference with first non-orthologous sequence - *A. fumigatus*:1310 *A. fischeri*:1310
afum|EAL87489.1 100.00% afis|EAW18132.1 100.00%
Bootstrap support for afum|EAL87489.1 as seed ortholog is 100%.
Bootstrap support for afis|EAW18132.1 as seed ortholog is 100%.

---

Group of orthologs #2551. Best score 1310 bits
Score difference with first non-orthologous sequence - *A. fumigatus*:1310 *A. fischeri*:1310
afum|EAL87497.1 100.00% afis|EAW18140.1 100.00%
Bootstrap support for afum|EAL87497.1 as seed ortholog is 100%.
Bootstrap support for afis|EAW18140.1 as seed ortholog is 100%.

---

Group of orthologs #2552. Best score 1310 bits
Score difference with first non-orthologous sequence - *A. fumigatus*:1310 *A. fischeri*:1310
afum|EAL88426.1 100.00% afis|EAW23054.1 100.00%
Bootstrap support for afum|EAL88426.1 as seed ortholog is 100%.
Bootstrap support for afis|EAW23054.1 as seed ortholog is 100%.

---

Group of orthologs #2553. Best score 1310 bits
Score difference with first non-orthologous sequence - *A. fumigatus*:1310 *A. fischeri*:1310
afum|EBA27452.1 100.00% afis|EAW21171.1 100.00%
Bootstrap support for afum|EBA27452.1 as seed ortholog is 100%.
Bootstrap support for afis|EAW21171.1 as seed ortholog is 100%.

---

Group of orthologs #2554. Best score 1309 bits
Score difference with first non-orthologous sequence - *A. fumigatus*:1309 *A. fischeri*:1309
afum|EAL93108.1 100.00% afis|EAW18427.1 100.00%
Bootstrap support for afum|EAL93108.1 as seed ortholog is 100%.
Bootstrap support for afis|EAW18427.1 as seed ortholog is 100%.

---

Group of orthologs #2555. Best score 1309 bits
Score difference with first non-orthologous sequence - *A. fumigatus*:1309 *A. fischeri*:1309
afum|EAL87975.1 100.00% afis|EAW23488.1 100.00%
Bootstrap support for afum|EAL87975.1 as seed ortholog is 100%.
Bootstrap support for afis|EAW23488.1 as seed ortholog is 100%.

---

Group of orthologs #2556. Best score 1309 bits
Score difference with first non-orthologous sequence - *A. fumigatus*:1309 *A. fischeri*:1309
afum|EAL91028.1 100.00% afis|EAW22163.1 100.00%
Bootstrap support for afum|EAL91028.1 as seed ortholog is 100%.
Bootstrap support for afis|EAW22163.1 as seed ortholog is 100%.

---

Group of orthologs #2557. Best score 1308 bits
Score difference with first non-orthologous sequence - *A. fumigatus*:1308 *A. fischeri*:1308
afum|EAL88686.1 100.00% afis|EAW16054.1 100.00%
Bootstrap support for afum|EAL88686.1 as seed ortholog is 100%.
Bootstrap support for afis|EAW16054.1 as seed ortholog is 100%.

---

Group of orthologs #2558. Best score 1308 bits
Score difference with first non-orthologous sequence - *A. fumigatus*:1308 *A. fischeri*:1308
afum|EAL89002.1 100.00% afis|EAW16377.1 100.00%
Bootstrap support for afum|EAL89002.1 as seed ortholog is 100%.
Bootstrap support for afis|EAW16377.1 as seed ortholog is 100%.

---

Group of orthologs #2559. Best score 1308 bits
Score difference with first non-orthologous sequence - *A. fumigatus*:1308 *A. fischeri*:1308
afum|EAL93370.1 100.00% afis|EAW18654.1 100.00%
Bootstrap support for afum|EAL93370.1 as seed ortholog is 100%.
Bootstrap support for afis|EAW18654.1 as seed ortholog is 100%.

---

Group of orthologs #2560. Best score 1308 bits
Score difference with first non-orthologous sequence - *A. fumigatus*:1308 *A. fischeri*:1308
afum|EAL92502.1 100.00% afis|EAW21482.1 100.00%
Bootstrap support for afum|EAL92502.1 as seed ortholog is 100%.
Bootstrap support for afis|EAW21482.1 as seed ortholog is 100%.

---

Group of orthologs #2561. Best score 1308 bits
Score difference with first non-orthologous sequence - *A. fumigatus*:958 *A. fischeri*:1164
afum|EAL89684.1 100.00% afis|EAW25012.1 100.00%
Bootstrap support for afum|EAL89684.1 as seed ortholog is 100%.
Bootstrap support for afis|EAW25012.1 as seed ortholog is 100%.

---

Group of orthologs #2562. Best score 1308 bits
Score difference with first non-orthologous sequence - *A. fumigatus*:1308 *A. fischeri*:1308
afum|EAL93006.2 100.00% afis|EAW21991.1 100.00%
Bootstrap support for afum|EAL93006.2 as seed ortholog is 100%.
Bootstrap support for afis|EAW21991.1 as seed ortholog is 100%.

---

Group of orthologs #2563. Best score 1307 bits
Score difference with first non-orthologous sequence - *A. fumigatus*:1307 *A. fischeri*:1307
afum|EAL84216.1 100.00% afis|EAW15427.1 100.00%
Bootstrap support for afum|EAL84216.1 as seed ortholog is 100%.
Bootstrap support for afis|EAW15427.1 as seed ortholog is 100%.

---

Group of orthologs #2564. Best score 1307 bits
Score difference with first non-orthologous sequence - *A. fumigatus*:1307 *A. fischeri*:1307

afum|EAL85839.1 100.00% afis|EAW15483.1 100.00%
Bootstrap support for afum|EAL85839.1 as seed ortholog is 100%.
Bootstrap support for afis|EAW15483.1 as seed ortholog is 100%.

---

Group of orthologs #2565. Best score 1307 bits
Score difference with first non-orthologous sequence - *A. fumigatus*:1307 *A. fischeri*:1307
afum|EAL84779.1 100.00% afis|EAW20853.1 100.00%
Bootstrap support for afum|EAL84779.1 as seed ortholog is 100%.
Bootstrap support for afis|EAW20853.1 as seed ortholog is 100%.

---

Group of orthologs #2566. Best score 1307 bits
Score difference with first non-orthologous sequence - *A. fumigatus*:1307 *A. fischeri*:1307
afum|EAL84727.1 100.00% afis|EAW20906.1 100.00%
Bootstrap support for afum|EAL84727.1 as seed ortholog is 100%.
Bootstrap support for afis|EAW20906.1 as seed ortholog is 100%.

---

Group of orthologs #2567. Best score 1307 bits
Score difference with first non-orthologous sequence - *A. fumigatus*:1307 *A. fischeri*:1244
afum|EAL86866.1 100.00% afis|EAW19657.1 100.00%
Bootstrap support for afum|EAL86866.1 as seed ortholog is 100%.
Bootstrap support for afis|EAW19657.1 as seed ortholog is 100%.

---

Group of orthologs #2568. Best score 1307 bits
Score difference with first non-orthologous sequence - *A. fumigatus*:1307 *A. fischeri*:1307
afum|EAL85901.2 100.00% afis|EAW24232.1 100.00%
Bootstrap support for afum|EAL85901.2 as seed ortholog is 100%.
Bootstrap support for afis|EAW24232.1 as seed ortholog is 100%.

---

Group of orthologs #2569. Best score 1307 bits
Score difference with first non-orthologous sequence - *A. fumigatus*:949 *A. fischeri*:995
afum|EAL90365.2 100.00% afis|EAW22837.1 100.00%
Bootstrap support for afum|EAL90365.2 as seed ortholog is 100%.
Bootstrap support for afis|EAW22837.1 as seed ortholog is 100%.

---

Group of orthologs #2570. Best score 1307 bits
Score difference with first non-orthologous sequence - *A. fumigatus*:1141 *A. fischeri*:877
afum|EAL92198.1 100.00% afis|EAW21178.1 100.00%
Bootstrap support for afum|EAL92198.1 as seed ortholog is 100%.
Bootstrap support for afis|EAW21178.1 as seed ortholog is 100%.

---

Group of orthologs #2571. Best score 1306 bits
Score difference with first non-orthologous sequence - *A. fumigatus*:1306 *A. fischeri*:1306
afum|EAL85685.1 100.00% afis|EAW15663.1 100.00%
Bootstrap support for afum|EAL85685.1 as seed ortholog is 100%.
Bootstrap support for afis|EAW15663.1 as seed ortholog is 100%.

---

Group of orthologs #2572. Best score 1306 bits
Score difference with first non-orthologous sequence - *A. fumigatus*:1306 *A. fischeri*:1306
afum|EAL85296.1 100.00% afis|EAW20178.1 100.00%
Bootstrap support for afum|EAL85296.1 as seed ortholog is 100%.
Bootstrap support for afis|EAW20178.1 as seed ortholog is 100%.

---

Group of orthologs #2573. Best score 1306 bits
Score difference with first non-orthologous sequence - *A. fumigatus*:1306 *A. fischeri*:1306
afum|EAL91887.1 100.00% afis|EAW18075.1 100.00%
Bootstrap support for afum|EAL91887.1 as seed ortholog is 100%.

Bootstrap support for afis|EAW18075.1 as seed ortholog is 100%.

---

Group of orthologs #2574. Best score 1306 bits
Score difference with first non-orthologous sequence - *A. fumigatus*:1101 *A. fischeri*:1152
afum|EAL93146.1 100.00% afis|EAW18460.1 100.00%
Bootstrap support for afum|EAL93146.1 as seed ortholog is 100%.
Bootstrap support for afis|EAW18460.1 as seed ortholog is 100%.

---

Group of orthologs #2575. Best score 1305 bits
Score difference with first non-orthologous sequence - *A. fumigatus*:1305 *A. fischeri*:917
afum|EAL86833.1 100.00% afis|EAW19693.1 100.00%
Bootstrap support for afum|EAL86833.1 as seed ortholog is 100%.
Bootstrap support for afis|EAW19693.1 as seed ortholog is 100%.

---

Group of orthologs #2576. Best score 1305 bits
Score difference with first non-orthologous sequence - *A. fumigatus*:1305 *A. fischeri*:1305
afum|EAL89677.1 100.00% afis|EAW25018.1 100.00%
Bootstrap support for afum|EAL89677.1 as seed ortholog is 100%.
Bootstrap support for afis|EAW25018.1 as seed ortholog is 100%.

---

Group of orthologs #2577. Best score 1304 bits
Score difference with first non-orthologous sequence - *A. fumigatus*:1304 *A. fischeri*:1304
afum|EAL91282.1 100.00% afis|EAW17454.1 100.00%
Bootstrap support for afum|EAL91282.1 as seed ortholog is 100%.
Bootstrap support for afis|EAW17454.1 as seed ortholog is 100%.

---

Group of orthologs #2578. Best score 1304 bits
Score difference with first non-orthologous sequence - *A. fumigatus*:1304 *A. fischeri*:1304
afum|EAL93220.1 100.00% afis|EAW18525.1 100.00%
Bootstrap support for afum|EAL93220.1 as seed ortholog is 100%.
Bootstrap support for afis|EAW18525.1 as seed ortholog is 100%.

---

Group of orthologs #2579. Best score 1304 bits
Score difference with first non-orthologous sequence - *A. fumigatus*:1304 *A. fischeri*:733
afum|EAL94046.1 100.00% afis|EAW19329.1 100.00%
Bootstrap support for afum|EAL94046.1 as seed ortholog is 100%.
Bootstrap support for afis|EAW19329.1 as seed ortholog is 100%.

---

Group of orthologs #2580. Best score 1304 bits
Score difference with first non-orthologous sequence - *A. fumigatus*:1220 *A. fischeri*:1144
afum|EAL89484.1 100.00% afis|EAW24838.1 100.00%
Bootstrap support for afum|EAL89484.1 as seed ortholog is 100%.
Bootstrap support for afis|EAW24838.1 as seed ortholog is 100%.

---

Group of orthologs #2581. Best score 1303 bits
Score difference with first non-orthologous sequence - *A. fumigatus*:1303 *A. fischeri*:1303
afum|EAL93964.1 100.00% afis|EAW19236.1 100.00%
Bootstrap support for afum|EAL93964.1 as seed ortholog is 100%.
Bootstrap support for afis|EAW19236.1 as seed ortholog is 100%.

---

Group of orthologs #2582. Best score 1302 bits
Score difference with first non-orthologous sequence - *A. fumigatus*:1302 *A. fischeri*:579
afum|EAL88067.1 100.00% afis|EAW23403.1 100.00%
Bootstrap support for afum|EAL88067.1 as seed ortholog is 100%.
Bootstrap support for afis|EAW23403.1 as seed ortholog is 100%.

---

Group of orthologs #2583. Best score 1301 bits
Score difference with first non-orthologous sequence - *A. fumigatus*:915 *A. fischeri*:1301
afum|EAL84785.1 100.00% afis|EAW20848.1 100.00%
Bootstrap support for afum|EAL84785.1 as seed ortholog is 100%.
Bootstrap support for afis|EAW20848.1 as seed ortholog is 100%.

---

Group of orthologs #2584. Best score 1301 bits
Score difference with first non-orthologous sequence - *A. fumigatus*:1301 *A. fischeri*:1223
afum|EAL86861.1 100.00% afis|EAW19661.1 100.00%
Bootstrap support for afum|EAL86861.1 as seed ortholog is 100%.
Bootstrap support for afis|EAW19661.1 as seed ortholog is 100%.

---

Group of orthologs #2585. Best score 1301 bits
Score difference with first non-orthologous sequence - *A. fumigatus*:1301 *A. fischeri*:1301
afum|EAL92866.1 100.00% afis|EAW21853.1 100.00%
Bootstrap support for afum|EAL92866.1 as seed ortholog is 100%.
Bootstrap support for afis|EAW21853.1 as seed ortholog is 100%.

---

Group of orthologs #2586. Best score 1300 bits
Score difference with first non-orthologous sequence - *A. fumigatus*:543 *A. fischeri*:819
afum|EAL86373.1 100.00% afis|EAW16822.1 100.00%
Bootstrap support for afum|EAL86373.1 as seed ortholog is 100%.
Bootstrap support for afis|EAW16822.1 as seed ortholog is 100%.

---

Group of orthologs #2587. Best score 1300 bits
Score difference with first non-orthologous sequence - *A. fumigatus*:1300 *A. fischeri*:1300
afum|EAL91787.1 100.00% afis|EAW17978.1 100.00%
Bootstrap support for afum|EAL91787.1 as seed ortholog is 100%.
Bootstrap support for afis|EAW17978.1 as seed ortholog is 100%.

---

Group of orthologs #2588. Best score 1300 bits
Score difference with first non-orthologous sequence - *A. fumigatus*:1300 *A. fischeri*:1300
afum|EAL90663.1 100.00% afis|EAW22531.1 100.00%
Bootstrap support for afum|EAL90663.1 as seed ortholog is 100%.
Bootstrap support for afis|EAW22531.1 as seed ortholog is 100%.

---

Group of orthologs #2589. Best score 1300 bits
Score difference with first non-orthologous sequence - *A. fumigatus*:1300 *A. fischeri*:1300
afum|EAL90458.2 100.00% afis|EAW22737.1 100.00%
Bootstrap support for afum|EAL90458.2 as seed ortholog is 100%.
Bootstrap support for afis|EAW22737.1 as seed ortholog is 100%.

---

Group of orthologs #2590. Best score 1299 bits
Score difference with first non-orthologous sequence - *A. fumigatus*:1183 *A. fischeri*:1299
afum|EAL85228.2 100.00% afis|EAW20111.1 100.00%
Bootstrap support for afum|EAL85228.2 as seed ortholog is 100%.
Bootstrap support for afis|EAW20111.1 as seed ortholog is 100%.

---

Group of orthologs #2591. Best score 1299 bits
Score difference with first non-orthologous sequence - *A. fumigatus*:1299 *A. fischeri*:1299
afum|EAL87542.1 100.00% afis|EAW18181.1 100.00%
Bootstrap support for afum|EAL87542.1 as seed ortholog is 100%.
Bootstrap support for afis|EAW18181.1 as seed ortholog is 100%.

---

Group of orthologs #2592. Best score 1299 bits
Score difference with first non-orthologous sequence - *A. fumigatus*:1299 *A. fischeri*:1139

afum|EAL86991.2 100.00% afis|EAW19502.1 100.00%
Bootstrap support for afum|EAL86991.2 as seed ortholog is 100%.
Bootstrap support for afis|EAW19502.1 as seed ortholog is 100%.

---

Group of orthologs #2593. Best score 1299 bits
Score difference with first non-orthologous sequence - *A. fumigatus*:1299 *A. fischeri*:1040
afum|EAL85693.1 100.00% afis|EAW22084.1 100.00%
Bootstrap support for afum|EAL85693.1 as seed ortholog is 100%.
Bootstrap support for afis|EAW22084.1 as seed ortholog is 100%.

---

Group of orthologs #2594. Best score 1299 bits
Score difference with first non-orthologous sequence - *A. fumigatus*:1299 *A. fischeri*:1112
afum|EBA27236.1 100.00% afis|EAW15735.1 100.00%
Bootstrap support for afum|EBA27236.1 as seed ortholog is 100%.
Bootstrap support for afis|EAW15735.1 as seed ortholog is 100%.

---

Group of orthologs #2595. Best score 1299 bits
Score difference with first non-orthologous sequence - *A. fumigatus*:1299 *A. fischeri*:1299
afum|EAL88383.1 100.00% afis|EAW23095.1 100.00%
Bootstrap support for afum|EAL88383.1 as seed ortholog is 100%.
Bootstrap support for afis|EAW23095.1 as seed ortholog is 100%.

---

Group of orthologs #2596. Best score 1298 bits
Score difference with first non-orthologous sequence - *A. fumigatus*:1152 *A. fischeri*:1058
afum|EAL85853.1 100.00% afis|EAW15469.1 100.00%
Bootstrap support for afum|EAL85853.1 as seed ortholog is 100%.
Bootstrap support for afis|EAW15469.1 as seed ortholog is 100%.

---

Group of orthologs #2597. Best score 1298 bits
Score difference with first non-orthologous sequence - *A. fumigatus*:1298 *A. fischeri*:1298
afum|EAL91901.1 100.00% afis|EAW18092.1 100.00%
Bootstrap support for afum|EAL91901.1 as seed ortholog is 100%.
Bootstrap support for afis|EAW18092.1 as seed ortholog is 100%.

---

Group of orthologs #2598. Best score 1298 bits
Score difference with first non-orthologous sequence - *A. fumigatus*:1298 *A. fischeri*:1298
afum|EAL93948.1 100.00% afis|EAW19215.1 100.00%
Bootstrap support for afum|EAL93948.1 as seed ortholog is 100%.
Bootstrap support for afis|EAW19215.1 as seed ortholog is 100%.

---

Group of orthologs #2599. Best score 1298 bits
Score difference with first non-orthologous sequence - *A. fumigatus*:1069 *A. fischeri*:989
afum|EAL92061.1 100.00% afis|EAW21022.1 100.00%
Bootstrap support for afum|EAL92061.1 as seed ortholog is 100%.
Bootstrap support for afis|EAW21022.1 as seed ortholog is 100%.

---

Group of orthologs #2600. Best score 1298 bits
Score difference with first non-orthologous sequence - *A. fumigatus*:1298 *A. fischeri*:1298
afum|EAL92976.1 100.00% afis|EAW21962.1 100.00%
Bootstrap support for afum|EAL92976.1 as seed ortholog is 100%.
Bootstrap support for afis|EAW21962.1 as seed ortholog is 100%.

---

Group of orthologs #2601. Best score 1297 bits
Score difference with first non-orthologous sequence - *A. fumigatus*:1297 *A. fischeri*:978
afum|EAL85034.1 100.00% afis|EAW19861.1 100.00%
Bootstrap support for afum|EAL85034.1 as seed ortholog is 100%.

Bootstrap support for afis|EAW19861.1 as seed ortholog is 100%.

---

Group of orthologs #2602. Best score 1297 bits
Score difference with first non-orthologous sequence - *A. fumigatus*:1297 *A. fischeri*:1190
afum|EAL84729.2 100.00% afis|EAW20902.1 100.00%
Bootstrap support for afum|EAL84729.2 as seed ortholog is 100%.
Bootstrap support for afis|EAW20902.1 as seed ortholog is 100%.

---

Group of orthologs #2603. Best score 1297 bits
Score difference with first non-orthologous sequence - *A. fumigatus*:1297 *A. fischeri*:1297
afum|EAL93625.1 100.00% afis|EAW18893.1 100.00%
Bootstrap support for afum|EAL93625.1 as seed ortholog is 100%.
Bootstrap support for afis|EAW18893.1 as seed ortholog is 100%.

---

Group of orthologs #2604. Best score 1297 bits
Score difference with first non-orthologous sequence - *A. fumigatus*:1297 *A. fischeri*:1297
afum|EAL90879.1 100.00% afis|EAW22311.1 100.00%
Bootstrap support for afum|EAL90879.1 as seed ortholog is 100%.
Bootstrap support for afis|EAW22311.1 as seed ortholog is 100%.

---

Group of orthologs #2605. Best score 1296 bits
Score difference with first non-orthologous sequence - *A. fumigatus*:1296 *A. fischeri*:1296
afum|EAL89100.1 100.00% afis|EAW16505.1 100.00%
Bootstrap support for afum|EAL89100.1 as seed ortholog is 100%.
Bootstrap support for afis|EAW16505.1 as seed ortholog is 100%.

---

Group of orthologs #2606. Best score 1296 bits
Score difference with first non-orthologous sequence - *A. fumigatus*:1296 *A. fischeri*:1296
afum|EAL93934.1 100.00% afis|EAW19204.1 100.00%
Bootstrap support for afum|EAL93934.1 as seed ortholog is 100%.
Bootstrap support for afis|EAW19204.1 as seed ortholog is 100%.

---

Group of orthologs #2607. Best score 1296 bits
Score difference with first non-orthologous sequence - *A. fumigatus*:1296 *A. fischeri*:1296
afum|EAL90922.1 100.00% afis|EAW22256.1 100.00%
Bootstrap support for afum|EAL90922.1 as seed ortholog is 100%.
Bootstrap support for afis|EAW22256.1 as seed ortholog is 100%.

---

Group of orthologs #2608. Best score 1295 bits
Score difference with first non-orthologous sequence - *A. fumigatus*:1295 *A. fischeri*:1295
afum|EAL91471.1 100.00% afis|EAW17656.1 100.00%
Bootstrap support for afum|EAL91471.1 as seed ortholog is 100%.
Bootstrap support for afis|EAW17656.1 as seed ortholog is 100%.

---

Group of orthologs #2609. Best score 1295 bits
Score difference with first non-orthologous sequence - *A. fumigatus*:1024 *A. fischeri*:1015
afum|EAL91091.2 100.00% afis|EAW22098.1 100.00%
Bootstrap support for afum|EAL91091.2 as seed ortholog is 100%.
Bootstrap support for afis|EAW22098.1 as seed ortholog is 100%.

---

Group of orthologs #2610. Best score 1295 bits
Score difference with first non-orthologous sequence - *A. fumigatus*:1295 *A. fischeri*:1095
afum|EAL90778.1 100.00% afis|EAW22414.1 100.00%
Bootstrap support for afum|EAL90778.1 as seed ortholog is 100%.
Bootstrap support for afis|EAW22414.1 as seed ortholog is 100%.

---

Group of orthologs #2611. Best score 1295 bits
Score difference with first non-orthologous sequence - *A. fumigatus*:1295 *A. fischeri*:1295
afum|EAL90243.1 100.00% afis|EAW22954.1 100.00%
Bootstrap support for afum|EAL90243.1 as seed ortholog is 100%.
Bootstrap support for afis|EAW22954.1 as seed ortholog is 100%.

---

Group of orthologs #2612. Best score 1294 bits
Score difference with first non-orthologous sequence - *A. fumigatus*:1294 *A. fischeri*:1294
afum|EAL85184.1 100.00% afis|EAW20064.1 100.00%
Bootstrap support for afum|EAL85184.1 as seed ortholog is 100%.
Bootstrap support for afis|EAW20064.1 as seed ortholog is 100%.

---

Group of orthologs #2613. Best score 1294 bits
Score difference with first non-orthologous sequence - *A. fumigatus*:1294 *A. fischeri*:1294
afum|EAL85321.1 100.00% afis|EAW20201.1 100.00%
Bootstrap support for afum|EAL85321.1 as seed ortholog is 100%.
Bootstrap support for afis|EAW20201.1 as seed ortholog is 100%.

---

Group of orthologs #2614. Best score 1294 bits
Score difference with first non-orthologous sequence - *A. fumigatus*:1294 *A. fischeri*:1294
afum|EAL87343.1 100.00% afis|EAW23916.1 100.00%
Bootstrap support for afum|EAL87343.1 as seed ortholog is 100%.
Bootstrap support for afis|EAW23916.1 as seed ortholog is 100%.

---

Group of orthologs #2615. Best score 1293 bits
Score difference with first non-orthologous sequence - *A. fumigatus*:1293 *A. fischeri*:1293
afum|EAL85538.1 100.00% afis|EAW15813.1 100.00%
Bootstrap support for afum|EAL85538.1 as seed ortholog is 100%.
Bootstrap support for afis|EAW15813.1 as seed ortholog is 100%.

---

Group of orthologs #2616. Best score 1293 bits
Score difference with first non-orthologous sequence - *A. fumigatus*:1293 *A. fischeri*:1293
afum|EAL85198.1 100.00% afis|EAW20079.1 100.00%
Bootstrap support for afum|EAL85198.1 as seed ortholog is 100%.
Bootstrap support for afis|EAW20079.1 as seed ortholog is 100%.

---

Group of orthologs #2617. Best score 1293 bits
Score difference with first non-orthologous sequence - *A. fumigatus*:1293 *A. fischeri*:797
afum|EAL85334.1 100.00% afis|EAW20219.1 100.00%
Bootstrap support for afum|EAL85334.1 as seed ortholog is 100%.
Bootstrap support for afis|EAW20219.1 as seed ortholog is 100%.

---

Group of orthologs #2618. Best score 1293 bits
Score difference with first non-orthologous sequence - *A. fumigatus*:1293 *A. fischeri*:1293
afum|EAL85484.1 100.00% afis|EAW20370.1 100.00%
Bootstrap support for afum|EAL85484.1 as seed ortholog is 100%.
Bootstrap support for afis|EAW20370.1 as seed ortholog is 100%.

---

Group of orthologs #2619. Best score 1293 bits
Score difference with first non-orthologous sequence - *A. fumigatus*:1293 *A. fischeri*:1293
afum|EAL87066.1 100.00% afis|EAW19428.1 100.00%
Bootstrap support for afum|EAL87066.1 as seed ortholog is 100%.
Bootstrap support for afis|EAW19428.1 as seed ortholog is 100%.

---

Group of orthologs #2620. Best score 1293 bits
Score difference with first non-orthologous sequence - *A. fumigatus*:1293 *A. fischeri*:1293

afum|EAL92286.1 100.00% afis|EAW21271.1 100.00%
Bootstrap support for afum|EAL92286.1 as seed ortholog is 100%.
Bootstrap support for afis|EAW21271.1 as seed ortholog is 100%.

---

Group of orthologs #2621. Best score 1292 bits
Score difference with first non-orthologous sequence - *A. fumigatus*:644 *A. fischeri*:1292
afum|EAL85087.1 100.00% afis|EAW19806.1 100.00%
Bootstrap support for afum|EAL85087.1 as seed ortholog is 100%.
Bootstrap support for afis|EAW19806.1 as seed ortholog is 100%.

---

Group of orthologs #2622. Best score 1292 bits
Score difference with first non-orthologous sequence - *A. fumigatus*:1292 *A. fischeri*:1292
afum|EAL86964.1 100.00% afis|EAW19545.1 100.00%
Bootstrap support for afum|EAL86964.1 as seed ortholog is 100%.
Bootstrap support for afis|EAW19545.1 as seed ortholog is 100%.

---

Group of orthologs #2623. Best score 1292 bits
Score difference with first non-orthologous sequence - *A. fumigatus*:1292 *A. fischeri*:1292
afum|EAL87403.1 100.00% afis|EAW23973.1 100.00%
Bootstrap support for afum|EAL87403.1 as seed ortholog is 100%.
Bootstrap support for afis|EAW23973.1 as seed ortholog is 100%.

---

Group of orthologs #2624. Best score 1292 bits
Score difference with first non-orthologous sequence - *A. fumigatus*:1292 *A. fischeri*:1292
afum|EAL87460.1 100.00% afis|EAW24029.1 100.00%
Bootstrap support for afum|EAL87460.1 as seed ortholog is 100%.
Bootstrap support for afis|EAW24029.1 as seed ortholog is 100%.

---

Group of orthologs #2625. Best score 1292 bits
Score difference with first non-orthologous sequence - *A. fumigatus*:1292 *A. fischeri*:1292
afum|EAL90908.1 100.00% afis|EAW22273.1 100.00%
Bootstrap support for afum|EAL90908.1 as seed ortholog is 100%.
Bootstrap support for afis|EAW22273.1 as seed ortholog is 100%.

---

Group of orthologs #2626. Best score 1291 bits
Score difference with first non-orthologous sequence - *A. fumigatus*:1142 *A. fischeri*:1054
afum|EAL84967.1 100.00% afis|EAW19948.1 100.00%
Bootstrap support for afum|EAL84967.1 as seed ortholog is 100%.
Bootstrap support for afis|EAW19948.1 as seed ortholog is 100%.

---

Group of orthologs #2627. Best score 1291 bits
Score difference with first non-orthologous sequence - *A. fumigatus*:1291 *A. fischeri*:1291
afum|EAL84695.1 100.00% afis|EAW20939.1 100.00%
Bootstrap support for afum|EAL84695.1 as seed ortholog is 100%.
Bootstrap support for afis|EAW20939.1 as seed ortholog is 100%.

---

Group of orthologs #2628. Best score 1291 bits
Score difference with first non-orthologous sequence - *A. fumigatus*:1180 *A. fischeri*:1175
afum|EAL91166.1 100.00% afis|EAW17329.1 100.00%
Bootstrap support for afum|EAL91166.1 as seed ortholog is 100%.
Bootstrap support for afis|EAW17329.1 as seed ortholog is 100%.

---

Group of orthologs #2629. Best score 1291 bits
Score difference with first non-orthologous sequence - *A. fumigatus*:1166 *A. fischeri*:860
afum|EAL90486.1 100.00% afis|EAW22706.1 100.00%
Bootstrap support for afum|EAL90486.1 as seed ortholog is 100%.

Bootstrap support for afis|EAW22706.1 as seed ortholog is 100%.

---

Group of orthologs #2630. Best score 1291 bits
Score difference with first non-orthologous sequence - *A. fumigatus*:1291 *A. fischeri*:1190
afum|EBA27198.1 100.00% afis|EAW20444.1 100.00%
Bootstrap support for afum|EBA27198.1 as seed ortholog is 100%.
Bootstrap support for afis|EAW20444.1 as seed ortholog is 100%.

---

Group of orthologs #2631. Best score 1290 bits
Score difference with first non-orthologous sequence - *A. fumigatus*:1290 *A. fischeri*:1290
afum|EAL88910.1 100.00% afis|EAW16288.1 100.00%
Bootstrap support for afum|EAL88910.1 as seed ortholog is 100%.
Bootstrap support for afis|EAW16288.1 as seed ortholog is 100%.

---

Group of orthologs #2632. Best score 1290 bits
Score difference with first non-orthologous sequence - *A. fumigatus*:1290 *A. fischeri*:1290
afum|EAL84622.1 100.00% afis|EAW20428.1 100.00%
Bootstrap support for afum|EAL84622.1 as seed ortholog is 100%.
Bootstrap support for afis|EAW20428.1 as seed ortholog is 100%.

---

Group of orthologs #2633. Best score 1290 bits
Score difference with first non-orthologous sequence - *A. fumigatus*:1290 *A. fischeri*:1290
afum|EAL87275.1 100.00% afis|EAW23844.1 100.00%
Bootstrap support for afum|EAL87275.1 as seed ortholog is 100%.
Bootstrap support for afis|EAW23844.1 as seed ortholog is 100%.

---

Group of orthologs #2634. Best score 1290 bits
Score difference with first non-orthologous sequence - *A. fumigatus*:1290 *A. fischeri*:1290
afum|EAL87448.1 100.00% afis|EAW24018.1 100.00%
Bootstrap support for afum|EAL87448.1 as seed ortholog is 100%.
Bootstrap support for afis|EAW24018.1 as seed ortholog is 100%.

---

Group of orthologs #2635. Best score 1289 bits
Score difference with first non-orthologous sequence - *A. fumigatus*:1289 *A. fischeri*:1289
afum|EAL92280.1 100.00% afis|EAW21265.1 100.00%
Bootstrap support for afum|EAL92280.1 as seed ortholog is 100%.
Bootstrap support for afis|EAW21265.1 as seed ortholog is 100%.

---

Group of orthologs #2636. Best score 1288 bits
Score difference with first non-orthologous sequence - *A. fumigatus*:1288 *A. fischeri*:1288
afum|EAL85827.1 100.00% afis|EAW15496.1 100.00%
Bootstrap support for afum|EAL85827.1 as seed ortholog is 100%.
Bootstrap support for afis|EAW15496.1 as seed ortholog is 100%.

---

Group of orthologs #2637. Best score 1288 bits
Score difference with first non-orthologous sequence - *A. fumigatus*:1288 *A. fischeri*:1288
afum|EAL85792.1 100.00% afis|EAW15538.1 100.00%
Bootstrap support for afum|EAL85792.1 as seed ortholog is 100%.
Bootstrap support for afis|EAW15538.1 as seed ortholog is 100%.

---

Group of orthologs #2638. Best score 1288 bits
Score difference with first non-orthologous sequence - *A. fumigatus*:1288 *A. fischeri*:1137
afum|EAL89295.1 100.00% afis|EAW16698.1 100.00%
Bootstrap support for afum|EAL89295.1 as seed ortholog is 100%.
Bootstrap support for afis|EAW16698.1 as seed ortholog is 100%.

---

Group of orthologs #2639. Best score 1288 bits
Score difference with first non-orthologous sequence - *A. fumigatus*:1288 *A. fischeri*:1288
afum|EAL93098.1 100.00% afis|EAW18416.1 100.00%
Bootstrap support for afum|EAL93098.1 as seed ortholog is 100%.
Bootstrap support for afis|EAW18416.1 as seed ortholog is 100%.

---

Group of orthologs #2640. Best score 1288 bits
Score difference with first non-orthologous sequence - *A. fumigatus*:773 *A. fischeri*:910
afum|EAL93641.1 100.00% afis|EAW18909.1 100.00%
Bootstrap support for afum|EAL93641.1 as seed ortholog is 100%.
Bootstrap support for afis|EAW18909.1 as seed ortholog is 100%.

---

Group of orthologs #2641. Best score 1288 bits
Score difference with first non-orthologous sequence - *A. fumigatus*:1288 *A. fischeri*:1288
afum|EAL93658.1 100.00% afis|EAW18927.1 100.00%
Bootstrap support for afum|EAL93658.1 as seed ortholog is 100%.
Bootstrap support for afis|EAW18927.1 as seed ortholog is 100%.

---

Group of orthologs #2642. Best score 1287 bits
Score difference with first non-orthologous sequence - *A. fumigatus*:1287 *A. fischeri*:1287
afum|EAL91443.1 100.00% afis|EAW17629.1 100.00%
Bootstrap support for afum|EAL91443.1 as seed ortholog is 100%.
Bootstrap support for afis|EAW17629.1 as seed ortholog is 100%.

---

Group of orthologs #2643. Best score 1287 bits
Score difference with first non-orthologous sequence - *A. fumigatus*:1077 *A. fischeri*:1033
afum|EAL91570.1 100.00% afis|EAW17760.1 100.00%
Bootstrap support for afum|EAL91570.1 as seed ortholog is 100%.
Bootstrap support for afis|EAW17760.1 as seed ortholog is 100%.

---

Group of orthologs #2644. Best score 1287 bits
Score difference with first non-orthologous sequence - *A. fumigatus*:1287 *A. fischeri*:1287
afum|EAL91650.1 100.00% afis|EAW17841.1 100.00%
Bootstrap support for afum|EAL91650.1 as seed ortholog is 100%.
Bootstrap support for afis|EAW17841.1 as seed ortholog is 100%.

---

Group of orthologs #2645. Best score 1287 bits
Score difference with first non-orthologous sequence - *A. fumigatus*:1287 *A. fischeri*:1287
afum|EAL92577.1 100.00% afis|EAW21558.1 100.00%
Bootstrap support for afum|EAL92577.1 as seed ortholog is 100%.
Bootstrap support for afis|EAW21558.1 as seed ortholog is 100%.

---

Group of orthologs #2646. Best score 1286 bits
Score difference with first non-orthologous sequence - *A. fumigatus*:1286 *A. fischeri*:1286
afum|EAL85869.1 100.00% afis|EAW24197.1 100.00%
Bootstrap support for afum|EAL85869.1 as seed ortholog is 100%.
Bootstrap support for afis|EAW24197.1 as seed ortholog is 100%.

---

Group of orthologs #2647. Best score 1286 bits
Score difference with first non-orthologous sequence - *A. fumigatus*:1286 *A. fischeri*:1286
afum|EAL92259.1 100.00% afis|EAW21246.1 100.00%
Bootstrap support for afum|EAL92259.1 as seed ortholog is 100%.
Bootstrap support for afis|EAW21246.1 as seed ortholog is 100%.

---

Group of orthologs #2648. Best score 1285 bits
Score difference with first non-orthologous sequence - *A. fumigatus*:1121 *A. fischeri*:1285

afum|EAL88689.1 100.00% afis|EAW16057.1 100.00%
Bootstrap support for afum|EAL88689.1 as seed ortholog is 100%.
Bootstrap support for afis|EAW16057.1 as seed ortholog is 100%.

---

Group of orthologs #2649. Best score 1285 bits
Score difference with first non-orthologous sequence - *A. fumigatus*:1134 *A. fischeri*:1196
afum|EAL91096.1 100.00% afis|EAW22074.1 100.00%
Bootstrap support for afum|EAL91096.1 as seed ortholog is 100%.
Bootstrap support for afis|EAW22074.1 as seed ortholog is 100%.

---

Group of orthologs #2650. Best score 1285 bits
Score difference with first non-orthologous sequence - *A. fumigatus*:1285 *A. fischeri*:1285
afum|EAL89883.1 100.00% afis|EAW25222.1 100.00%
Bootstrap support for afum|EAL89883.1 as seed ortholog is 100%.
Bootstrap support for afis|EAW25222.1 as seed ortholog is 100%.

---

Group of orthologs #2651. Best score 1284 bits
Score difference with first non-orthologous sequence - *A. fumigatus*:1284 *A. fischeri*:1284
afum|EAL87415.2 100.00% afis|EAW23986.1 100.00%
Bootstrap support for afum|EAL87415.2 as seed ortholog is 100%.
Bootstrap support for afis|EAW23986.1 as seed ortholog is 100%.

---

Group of orthologs #2652. Best score 1284 bits
Score difference with first non-orthologous sequence - *A. fumigatus*:1284 *A. fischeri*:1284
afum|EBA27307.1 100.00% afis|EAW23885.1 100.00%
Bootstrap support for afum|EBA27307.1 as seed ortholog is 100%.
Bootstrap support for afis|EAW23885.1 as seed ortholog is 100%.

---

Group of orthologs #2653. Best score 1283 bits
Score difference with first non-orthologous sequence - *A. fumigatus*:1283 *A. fischeri*:1283
afum|EAL84955.1 100.00% afis|EAW19960.1 100.00%
Bootstrap support for afum|EAL84955.1 as seed ortholog is 100%.
Bootstrap support for afis|EAW19960.1 as seed ortholog is 100%.

---

Group of orthologs #2654. Best score 1283 bits
Score difference with first non-orthologous sequence - *A. fumigatus*:1283 *A. fischeri*:1283
afum|EAL84528.1 100.00% afis|EAW20529.1 100.00%
Bootstrap support for afum|EAL84528.1 as seed ortholog is 100%.
Bootstrap support for afis|EAW20529.1 as seed ortholog is 100%.

---

Group of orthologs #2655. Best score 1282 bits
Score difference with first non-orthologous sequence - *A. fumigatus*:1282 *A. fischeri*:653
afum|EAL84526.1 100.00% afis|EAW20531.1 100.00%
Bootstrap support for afum|EAL84526.1 as seed ortholog is 100%.
Bootstrap support for afis|EAW20531.1 as seed ortholog is 100%.

---

Group of orthologs #2656. Best score 1282 bits
Score difference with first non-orthologous sequence - *A. fumigatus*:1282 *A. fischeri*:1282
afum|EAL89077.1 100.00% afis|EAW16483.1 100.00%
Bootstrap support for afum|EAL89077.1 as seed ortholog is 100%.
Bootstrap support for afis|EAW16483.1 as seed ortholog is 100%.

---

Group of orthologs #2657. Best score 1282 bits
Score difference with first non-orthologous sequence - *A. fumigatus*:1282 *A. fischeri*:1282
afum|EAL87090.1 100.00% afis|EAW19405.1 100.00%
Bootstrap support for afum|EAL87090.1 as seed ortholog is 100%.

Bootstrap support for afis|EAW19405.1 as seed ortholog is 100%.

---

Group of orthologs #2658. Best score 1282 bits
Score difference with first non-orthologous sequence - *A. fumigatus*:1282 *A. fischeri*:1282
afum|EAL88107.1 100.00% afis|EAW23366.1 100.00%
Bootstrap support for afum|EAL88107.1 as seed ortholog is 100%.
Bootstrap support for afis|EAW23366.1 as seed ortholog is 100%.

---

Group of orthologs #2659. Best score 1282 bits
Score difference with first non-orthologous sequence - *A. fumigatus*:1282 *A. fischeri*:1282
afum|EAL93469.1 100.00% afis|EAW18743.1 100.00%
Bootstrap support for afum|EAL93469.1 as seed ortholog is 100%.
Bootstrap support for afis|EAW18743.1 as seed ortholog is 100%.

---

Group of orthologs #2660. Best score 1282 bits
Score difference with first non-orthologous sequence - *A. fumigatus*:1282 *A. fischeri*:1282
afum|EAL93736.1 100.00% afis|EAW19002.1 100.00%
Bootstrap support for afum|EAL93736.1 as seed ortholog is 100%.
Bootstrap support for afis|EAW19002.1 as seed ortholog is 100%.

---

Group of orthologs #2661. Best score 1281 bits
Score difference with first non-orthologous sequence - *A. fumigatus*:1281 *A. fischeri*:1281
afum|EAL85307.1 100.00% afis|EAW20188.1 100.00%
Bootstrap support for afum|EAL85307.1 as seed ortholog is 100%.
Bootstrap support for afis|EAW20188.1 as seed ortholog is 100%.

---

Group of orthologs #2662. Best score 1281 bits
Score difference with first non-orthologous sequence - *A. fumigatus*:1281 *A. fischeri*:1281
afum|EAL84647.2 100.00% afis|EAW20983.1 100.00%
Bootstrap support for afum|EAL84647.2 as seed ortholog is 100%.
Bootstrap support for afis|EAW20983.1 as seed ortholog is 100%.

---

Group of orthologs #2663. Best score 1281 bits
Score difference with first non-orthologous sequence - *A. fumigatus*:1281 *A. fischeri*:1209
afum|EAL91752.1 100.00% afis|EAW17941.1 100.00%
Bootstrap support for afum|EAL91752.1 as seed ortholog is 100%.
Bootstrap support for afis|EAW17941.1 as seed ortholog is 100%.

---

Group of orthologs #2664. Best score 1281 bits
Score difference with first non-orthologous sequence - *A. fumigatus*:1281 *A. fischeri*:1281
afum|EAL85889.1 100.00% afis|EAW24220.1 100.00%
Bootstrap support for afum|EAL85889.1 as seed ortholog is 100%.
Bootstrap support for afis|EAW24220.1 as seed ortholog is 100%.

---

Group of orthologs #2665. Best score 1281 bits
Score difference with first non-orthologous sequence - *A. fumigatus*:1281 *A. fischeri*:1281
afum|EAL93078.1 100.00% afis|EAW18400.1 100.00%
Bootstrap support for afum|EAL93078.1 as seed ortholog is 100%.
Bootstrap support for afis|EAW18400.1 as seed ortholog is 100%.

---

Group of orthologs #2666. Best score 1281 bits
Score difference with first non-orthologous sequence - *A. fumigatus*:1281 *A. fischeri*:1281
afum|EAL89990.1 100.00% afis|EAW25329.1 100.00%
Bootstrap support for afum|EAL89990.1 as seed ortholog is 100%.
Bootstrap support for afis|EAW25329.1 as seed ortholog is 100%.

---

Group of orthologs #2667. Best score 1280 bits
Score difference with first non-orthologous sequence - *A. fumigatus*:1280 *A. fischeri*:1280
afum|EAL85324.1 100.00% afis|EAW20206.1 100.00%
Bootstrap support for afum|EAL85324.1 as seed ortholog is 100%.
Bootstrap support for afis|EAW20206.1 as seed ortholog is 100%.

---

Group of orthologs #2668. Best score 1280 bits
Score difference with first non-orthologous sequence - *A. fumigatus*:1280 *A. fischeri*:667
afum|EAL84403.1 100.00% afis|EAW25732.1 100.00%
Bootstrap support for afum|EAL84403.1 as seed ortholog is 100%.
Bootstrap support for afis|EAW25732.1 as seed ortholog is 100%.

---

Group of orthologs #2669. Best score 1280 bits
Score difference with first non-orthologous sequence - *A. fumigatus*:1280 *A. fischeri*:1280
afum|EAL92860.1 100.00% afis|EAW21847.1 100.00%
Bootstrap support for afum|EAL92860.1 as seed ortholog is 100%.
Bootstrap support for afis|EAW21847.1 as seed ortholog is 100%.

---

Group of orthologs #2670. Best score 1279 bits
Score difference with first non-orthologous sequence - *A. fumigatus*:1149 *A. fischeri*:922
afum|EAL86325.1 100.00% afis|EAW16753.1 100.00%
Bootstrap support for afum|EAL86325.1 as seed ortholog is 100%.
Bootstrap support for afis|EAW16753.1 as seed ortholog is 100%.

---

Group of orthologs #2671. Best score 1279 bits
Score difference with first non-orthologous sequence - *A. fumigatus*:1139 *A. fischeri*:1184
afum|EAL88522.2 100.00% afis|EAW15888.1 100.00%
Bootstrap support for afum|EAL88522.2 as seed ortholog is 100%.
Bootstrap support for afis|EAW15888.1 as seed ortholog is 100%.

---

Group of orthologs #2672. Best score 1279 bits
Score difference with first non-orthologous sequence - *A. fumigatus*:1279 *A. fischeri*:1279
afum|EAL88663.1 100.00% afis|EAW16032.1 100.00%
Bootstrap support for afum|EAL88663.1 as seed ortholog is 100%.
Bootstrap support for afis|EAW16032.1 as seed ortholog is 100%.

---

Group of orthologs #2673. Best score 1279 bits
Score difference with first non-orthologous sequence - *A. fumigatus*:1279 *A. fischeri*:355
afum|EAL89006.1 100.00% afis|EAW16381.1 100.00%
Bootstrap support for afum|EAL89006.1 as seed ortholog is 100%.
Bootstrap support for afis|EAW16381.1 as seed ortholog is 100%.

---

Group of orthologs #2674. Best score 1279 bits
Score difference with first non-orthologous sequence - *A. fumigatus*:1279 *A. fischeri*:1279
afum|EAL87723.1 100.00% afis|EAW18359.1 100.00%
Bootstrap support for afum|EAL87723.1 as seed ortholog is 100%.
Bootstrap support for afis|EAW18359.1 as seed ortholog is 100%.

---

Group of orthologs #2675. Best score 1279 bits
Score difference with first non-orthologous sequence - *A. fumigatus*:1138 *A. fischeri*:1034
afum|EAL91165.1 100.00% afis|EAW17328.1 100.00%
Bootstrap support for afum|EAL91165.1 as seed ortholog is 100%.
Bootstrap support for afis|EAW17328.1 as seed ortholog is 100%.

---

Group of orthologs #2676. Best score 1279 bits
Score difference with first non-orthologous sequence - *A. fumigatus*:1279 *A. fischeri*:1279

afum|EAL91397.1 100.00% afis|EAW17584.1 100.00%
Bootstrap support for afum|EAL91397.1 as seed ortholog is 100%.
Bootstrap support for afis|EAW17584.1 as seed ortholog is 100%.

---

Group of orthologs #2677. Best score 1279 bits
Score difference with first non-orthologous sequence - *A. fumigatus*:1279 *A. fischeri*:1279
afum|EAL93971.2 100.00% afis|EAW19242.1 100.00%
Bootstrap support for afum|EAL93971.2 as seed ortholog is 100%.
Bootstrap support for afis|EAW19242.1 as seed ortholog is 100%.

---

Group of orthologs #2678. Best score 1279 bits
Score difference with first non-orthologous sequence - *A. fumigatus*:1279 *A. fischeri*:1279
afum|EAL93974.1 100.00% afis|EAW19245.1 100.00%
Bootstrap support for afum|EAL93974.1 as seed ortholog is 100%.
Bootstrap support for afis|EAW19245.1 as seed ortholog is 100%.

---

Group of orthologs #2679. Best score 1279 bits
Score difference with first non-orthologous sequence - *A. fumigatus*:1279 *A. fischeri*:1279
afum|EAL92765.1 100.00% afis|EAW21732.1 100.00%
Bootstrap support for afum|EAL92765.1 as seed ortholog is 100%.
Bootstrap support for afis|EAW21732.1 as seed ortholog is 100%.

---

Group of orthologs #2680. Best score 1279 bits
Score difference with first non-orthologous sequence - *A. fumigatus*:820 *A. fischeri*:1279
afum|EAL90163.1 100.00% afis|EAW25497.1 100.00%
Bootstrap support for afum|EAL90163.1 as seed ortholog is 100%.
Bootstrap support for afis|EAW25497.1 as seed ortholog is 100%.

---

Group of orthologs #2681. Best score 1279 bits
Score difference with first non-orthologous sequence - *A. fumigatus*:1279 *A. fischeri*:1279
afum|EAL91989.2 100.00% afis|EAW24134.1 100.00%
Bootstrap support for afum|EAL91989.2 as seed ortholog is 100%.
Bootstrap support for afis|EAW24134.1 as seed ortholog is 100%.

---

Group of orthologs #2682. Best score 1278 bits
Score difference with first non-orthologous sequence - *A. fumigatus*:1278 *A. fischeri*:1278
afum|EAL86394.1 100.00% afis|EAW16848.1 100.00%
Bootstrap support for afum|EAL86394.1 as seed ortholog is 100%.
Bootstrap support for afis|EAW16848.1 as seed ortholog is 100%.

---

Group of orthologs #2683. Best score 1278 bits
Score difference with first non-orthologous sequence - *A. fumigatus*:1278 *A. fischeri*:1278
afum|EAL88653.1 100.00% afis|EAW16024.1 100.00%
Bootstrap support for afum|EAL88653.1 as seed ortholog is 100%.
Bootstrap support for afis|EAW16024.1 as seed ortholog is 100%.

---

Group of orthologs #2684. Best score 1278 bits
Score difference with first non-orthologous sequence - *A. fumigatus*:940 *A. fischeri*:847
afum|EAL85074.1 100.00% afis|EAW19817.1 100.00%
Bootstrap support for afum|EAL85074.1 as seed ortholog is 100%.
Bootstrap support for afis|EAW19817.1 as seed ortholog is 100%.

---

Group of orthologs #2685. Best score 1278 bits
Score difference with first non-orthologous sequence - *A. fumigatus*:1278 *A. fischeri*:1278
afum|EAL87051.1 100.00% afis|EAW19444.1 100.00%
Bootstrap support for afum|EAL87051.1 as seed ortholog is 100%.

Bootstrap support for afis|EAW19444.1 as seed ortholog is 100%.

---

Group of orthologs #2686. Best score 1277 bits
Score difference with first non-orthologous sequence - *A. fumigatus*:1111 *A. fischeri*:1073
afum|EAL85147.2 100.00% afis|EAW19750.1 100.00%
Bootstrap support for afum|EAL85147.2 as seed ortholog is 100%.
Bootstrap support for afis|EAW19750.1 as seed ortholog is 100%.

---

Group of orthologs #2687. Best score 1277 bits
Score difference with first non-orthologous sequence - *A. fumigatus*:1277 *A. fischeri*:1277
afum|EAL91739.1 100.00% afis|EAW17928.1 100.00%
Bootstrap support for afum|EAL91739.1 as seed ortholog is 100%.
Bootstrap support for afis|EAW17928.1 as seed ortholog is 100%.

---

Group of orthologs #2688. Best score 1277 bits
Score difference with first non-orthologous sequence - *A. fumigatus*:1277 *A. fischeri*:1277
afum|EAL89467.1 100.00% afis|EAW24823.1 100.00%
Bootstrap support for afum|EAL89467.1 as seed ortholog is 100%.
Bootstrap support for afis|EAW24823.1 as seed ortholog is 100%.

---

Group of orthologs #2689. Best score 1276 bits
Score difference with first non-orthologous sequence - *A. fumigatus*:1276 *A. fischeri*:1276
afum|EAL84557.1 100.00% afis|EAW20497.1 100.00%
Bootstrap support for afum|EAL84557.1 as seed ortholog is 100%.
Bootstrap support for afis|EAW20497.1 as seed ortholog is 100%.

---

Group of orthologs #2690. Best score 1276 bits
Score difference with first non-orthologous sequence - *A. fumigatus*:1276 *A. fischeri*:1276
afum|EAL91312.1 100.00% afis|EAW17490.1 100.00%
Bootstrap support for afum|EAL91312.1 as seed ortholog is 100%.
Bootstrap support for afis|EAW17490.1 as seed ortholog is 100%.

---

Group of orthologs #2691. Best score 1276 bits
Score difference with first non-orthologous sequence - *A. fumigatus*:1178 *A. fischeri*:1276
afum|EAL91686.1 100.00% afis|EAW17878.1 100.00%
Bootstrap support for afum|EAL91686.1 as seed ortholog is 100%.
Bootstrap support for afis|EAW17878.1 as seed ortholog is 100%.

---

Group of orthologs #2692. Best score 1276 bits
Score difference with first non-orthologous sequence - *A. fumigatus*:1276 *A. fischeri*:1153
afum|EAL89229.1 100.00% afis|EAW20416.1 100.00%
Bootstrap support for afum|EAL89229.1 as seed ortholog is 100%.
Bootstrap support for afis|EAW20416.1 as seed ortholog is 100%.

---

Group of orthologs #2693. Best score 1276 bits
Score difference with first non-orthologous sequence - *A. fumigatus*:1276 *A. fischeri*:1276
afum|EAL93522.2 100.00% afis|EAW18788.1 100.00%
Bootstrap support for afum|EAL93522.2 as seed ortholog is 100%.
Bootstrap support for afis|EAW18788.1 as seed ortholog is 100%.

---

Group of orthologs #2694. Best score 1276 bits
Score difference with first non-orthologous sequence - *A. fumigatus*:1276 *A. fischeri*:1276
afum|EAL92419.1 100.00% afis|EAW21405.1 100.00%
Bootstrap support for afum|EAL92419.1 as seed ortholog is 100%.
Bootstrap support for afis|EAW21405.1 as seed ortholog is 100%.

---

Group of orthologs #2695. Best score 1276 bits
Score difference with first non-orthologous sequence - *A. fumigatus*:1276 *A. fischeri*:1276
afum|EAL90061.1 100.00% afis|EAW25402.1 100.00%
Bootstrap support for afum|EAL90061.1 as seed ortholog is 100%.
Bootstrap support for afis|EAW25402.1 as seed ortholog is 100%.

---

Group of orthologs #2696. Best score 1275 bits
Score difference with first non-orthologous sequence - *A. fumigatus*:1275 *A. fischeri*:1275
afum|EAL84558.1 100.00% afis|EAW20496.1 100.00%
Bootstrap support for afum|EAL84558.1 as seed ortholog is 100%.
Bootstrap support for afis|EAW20496.1 as seed ortholog is 100%.

---

Group of orthologs #2697. Best score 1275 bits
Score difference with first non-orthologous sequence - *A. fumigatus*:1275 *A. fischeri*:856
afum|EAL85486.1 100.00% afis|EAW20372.1 100.00%
Bootstrap support for afum|EAL85486.1 as seed ortholog is 100%.
Bootstrap support for afis|EAW20372.1 as seed ortholog is 100%.

---

Group of orthologs #2698. Best score 1275 bits
Score difference with first non-orthologous sequence - *A. fumigatus*:1275 *A. fischeri*:1275
afum|EAL88408.1 100.00% afis|EAW23071.1 100.00%
Bootstrap support for afum|EAL88408.1 as seed ortholog is 100%.
Bootstrap support for afis|EAW23071.1 as seed ortholog is 100%.

---

Group of orthologs #2699. Best score 1275 bits
Score difference with first non-orthologous sequence - *A. fumigatus*:1275 *A. fischeri*:1275
afum|EAL90783.1 100.00% afis|EAW22409.1 100.00%
Bootstrap support for afum|EAL90783.1 as seed ortholog is 100%.
Bootstrap support for afis|EAW22409.1 as seed ortholog is 100%.

---

Group of orthologs #2700. Best score 1274 bits
Score difference with first non-orthologous sequence - *A. fumigatus*:1274 *A. fischeri*:1274
afum|EAL84329.1 100.00% afis|EAW25642.1 100.00%
Bootstrap support for afum|EAL84329.1 as seed ortholog is 100%.
Bootstrap support for afis|EAW25642.1 as seed ortholog is 100%.

---

Group of orthologs #2701. Best score 1274 bits
Score difference with first non-orthologous sequence - *A. fumigatus*:1155 *A. fischeri*:1077
afum|EAL92325.1 100.00% afis|EAW21309.1 100.00%
Bootstrap support for afum|EAL92325.1 as seed ortholog is 100%.
Bootstrap support for afis|EAW21309.1 as seed ortholog is 100%.

---

Group of orthologs #2702. Best score 1273 bits
Score difference with first non-orthologous sequence - *A. fumigatus*:1273 *A. fischeri*:1273
afum|EAL87659.1 100.00% afis|EAW18293.1 100.00%
Bootstrap support for afum|EAL87659.1 as seed ortholog is 100%.
Bootstrap support for afis|EAW18293.1 as seed ortholog is 100%.

---

Group of orthologs #2703. Best score 1273 bits
Score difference with first non-orthologous sequence - *A. fumigatus*:1044 *A. fischeri*:427
afum|EAL91537.1 100.00% afis|EAW17722.1 100.00%
Bootstrap support for afum|EAL91537.1 as seed ortholog is 100%.
Bootstrap support for afis|EAW17722.1 as seed ortholog is 100%.

---

Group of orthologs #2704. Best score 1273 bits
Score difference with first non-orthologous sequence - *A. fumigatus*:1273 *A. fischeri*:1273

afum|EAL87199.1 100.00% afis|EAW23766.1 100.00%
Bootstrap support for afum|EAL87199.1 as seed ortholog is 100%.
Bootstrap support for afis|EAW23766.1 as seed ortholog is 100%.

---

Group of orthologs #2705. Best score 1273 bits
Score difference with first non-orthologous sequence - *A. fumigatus*:1273 *A. fischeri*:1273
afum|EAL87206.1 100.00% afis|EAW23772.1 100.00%
Bootstrap support for afum|EAL87206.1 as seed ortholog is 100%.
Bootstrap support for afis|EAW23772.1 as seed ortholog is 100%.

---

Group of orthologs #2706. Best score 1273 bits
Score difference with first non-orthologous sequence - *A. fumigatus*:1273 *A. fischeri*:1273
afum|EAL93582.1 100.00% afis|EAW18851.1 100.00%
Bootstrap support for afum|EAL93582.1 as seed ortholog is 100%.
Bootstrap support for afis|EAW18851.1 as seed ortholog is 100%.

---

Group of orthologs #2707. Best score 1272 bits
Score difference with first non-orthologous sequence - *A. fumigatus*:1067 *A. fischeri*:981
afum|EAL85706.1 100.00% afis|EAW15628.1 100.00%
Bootstrap support for afum|EAL85706.1 as seed ortholog is 100%.
Bootstrap support for afis|EAW15628.1 as seed ortholog is 100%.

---

Group of orthologs #2708. Best score 1272 bits
Score difference with first non-orthologous sequence - *A. fumigatus*:1272 *A. fischeri*:1272
afum|EAL91647.1 100.00% afis|EAW17837.1 100.00%
Bootstrap support for afum|EAL91647.1 as seed ortholog is 100%.
Bootstrap support for afis|EAW17837.1 as seed ortholog is 100%.

---

Group of orthologs #2709. Best score 1272 bits
Score difference with first non-orthologous sequence - *A. fumigatus*:1272 *A. fischeri*:1272
afum|EAL92622.1 100.00% afis|EAW21603.1 100.00%
Bootstrap support for afum|EAL92622.1 as seed ortholog is 100%.
Bootstrap support for afis|EAW21603.1 as seed ortholog is 100%.

---

Group of orthologs #2710. Best score 1272 bits
Score difference with first non-orthologous sequence - *A. fumigatus*:1272 *A. fischeri*:1272
afum|EAL90009.1 100.00% afis|EAW25346.1 100.00%
Bootstrap support for afum|EAL90009.1 as seed ortholog is 100%.
Bootstrap support for afis|EAW25346.1 as seed ortholog is 100%.

---

Group of orthologs #2711. Best score 1272 bits
Score difference with first non-orthologous sequence - *A. fumigatus*:1272 *A. fischeri*:1272
afum|EAL91942.1 100.00% afis|EAW24087.1 100.00%
Bootstrap support for afum|EAL91942.1 as seed ortholog is 100%.
Bootstrap support for afis|EAW24087.1 as seed ortholog is 100%.

---

Group of orthologs #2712. Best score 1271 bits
Score difference with first non-orthologous sequence - *A. fumigatus*:1112 *A. fischeri*:946
afum|EAL85818.1 100.00% afis|EAW15506.1 100.00%
Bootstrap support for afum|EAL85818.1 as seed ortholog is 100%.
Bootstrap support for afis|EAW15506.1 as seed ortholog is 100%.

---

Group of orthologs #2713. Best score 1271 bits
Score difference with first non-orthologous sequence - *A. fumigatus*:1271 *A. fischeri*:1271
afum|EAL89094.1 100.00% afis|EAW16499.1 100.00%
Bootstrap support for afum|EAL89094.1 as seed ortholog is 100%.

Bootstrap support for afis|EAW16499.1 as seed ortholog is 100%.

---

Group of orthologs #2714. Best score 1271 bits
Score difference with first non-orthologous sequence - *A. fumigatus*:1271 *A. fischeri*:1271
afum|EAL91457.1 100.00% afis|EAW17642.1 100.00%
Bootstrap support for afum|EAL91457.1 as seed ortholog is 100%.
Bootstrap support for afis|EAW17642.1 as seed ortholog is 100%.

---

Group of orthologs #2715. Best score 1271 bits
Score difference with first non-orthologous sequence - *A. fumigatus*:1025 *A. fischeri*:932
afum|EAL84305.1 100.00% afis|EAW25600.1 100.00%
Bootstrap support for afum|EAL84305.1 as seed ortholog is 100%.
Bootstrap support for afis|EAW25600.1 as seed ortholog is 100%.

---

Group of orthologs #2716. Best score 1271 bits
Score difference with first non-orthologous sequence - *A. fumigatus*:737 *A. fischeri*:742
afum|EAL93089.1 100.00% afis|EAW18409.1 100.00%
Bootstrap support for afum|EAL93089.1 as seed ortholog is 100%.
Bootstrap support for afis|EAW18409.1 as seed ortholog is 100%.

---

Group of orthologs #2717. Best score 1271 bits
Score difference with first non-orthologous sequence - *A. fumigatus*:1271 *A. fischeri*:1271
afum|EAL87962.2 100.00% afis|EAW23500.1 100.00%
Bootstrap support for afum|EAL87962.2 as seed ortholog is 100%.
Bootstrap support for afis|EAW23500.1 as seed ortholog is 100%.

---

Group of orthologs #2718. Best score 1271 bits
Score difference with first non-orthologous sequence - *A. fumigatus*:1271 *A. fischeri*:1271
afum|EAL92399.1 100.00% afis|EAW21384.1 100.00%
Bootstrap support for afum|EAL92399.1 as seed ortholog is 100%.
Bootstrap support for afis|EAW21384.1 as seed ortholog is 100%.

---

Group of orthologs #2719. Best score 1271 bits
Score difference with first non-orthologous sequence - *A. fumigatus*:1271 *A. fischeri*:1271
afum|EAL89511.1 100.00% afis|EAW24862.1 100.00%
Bootstrap support for afum|EAL89511.1 as seed ortholog is 100%.
Bootstrap support for afis|EAW24862.1 as seed ortholog is 100%.

---

Group of orthologs #2720. Best score 1270 bits
Score difference with first non-orthologous sequence - *A. fumigatus*:1270 *A. fischeri*:1224
afum|EAL87096.1 100.00% afis|EAW19398.1 100.00%
Bootstrap support for afum|EAL87096.1 as seed ortholog is 100%.
Bootstrap support for afis|EAW19398.1 as seed ortholog is 100%.

---

Group of orthologs #2721. Best score 1270 bits
Score difference with first non-orthologous sequence - *A. fumigatus*:1116 *A. fischeri*:1270
afum|EAL86081.1 100.00% afis|EAW24412.1 100.00%
Bootstrap support for afum|EAL86081.1 as seed ortholog is 100%.
Bootstrap support for afis|EAW24412.1 as seed ortholog is 100%.

---

Group of orthologs #2722. Best score 1270 bits
Score difference with first non-orthologous sequence - *A. fumigatus*:1270 *A. fischeri*:1270
afum|EAL91938.1 100.00% afis|EAW24083.1 100.00%
Bootstrap support for afum|EAL91938.1 as seed ortholog is 100%.
Bootstrap support for afis|EAW24083.1 as seed ortholog is 100%.

---

Group of orthologs #2723. Best score 1269 bits
Score difference with first non-orthologous sequence - *A. fumigatus*:1269 *A. fischeri*:1269
afum|EAL85715.1 100.00% afis|EAW15620.1 100.00%
Bootstrap support for afum|EAL85715.1 as seed ortholog is 100%.
Bootstrap support for afis|EAW15620.1 as seed ortholog is 100%.

---

Group of orthologs #2724. Best score 1269 bits
Score difference with first non-orthologous sequence - *A. fumigatus*:1269 *A. fischeri*:1269
afum|EAL91361.1 100.00% afis|EAW17547.1 100.00%
Bootstrap support for afum|EAL91361.1 as seed ortholog is 100%.
Bootstrap support for afis|EAW17547.1 as seed ortholog is 100%.

---

Group of orthologs #2725. Best score 1269 bits
Score difference with first non-orthologous sequence - *A. fumigatus*:1269 *A. fischeri*:1269
afum|EAL88064.1 100.00% afis|EAW23406.1 100.00%
Bootstrap support for afum|EAL88064.1 as seed ortholog is 100%.
Bootstrap support for afis|EAW23406.1 as seed ortholog is 100%.

---

Group of orthologs #2726. Best score 1269 bits
Score difference with first non-orthologous sequence - *A. fumigatus*:1269 *A. fischeri*:1269
afum|EAL87456.1 100.00% afis|EAW24025.1 100.00%
Bootstrap support for afum|EAL87456.1 as seed ortholog is 100%.
Bootstrap support for afis|EAW24025.1 as seed ortholog is 100%.

---

Group of orthologs #2727. Best score 1269 bits
Score difference with first non-orthologous sequence - *A. fumigatus*:1111 *A. fischeri*:1062
afum|EAL89334.1 100.00% afis|EAW24692.1 100.00%
Bootstrap support for afum|EAL89334.1 as seed ortholog is 100%.
Bootstrap support for afis|EAW24692.1 as seed ortholog is 100%.

---

Group of orthologs #2728. Best score 1269 bits
Score difference with first non-orthologous sequence - *A. fumigatus*:1269 *A. fischeri*:1269
afum|EAL92753.1 100.00% afis|EAW21720.1 100.00%
Bootstrap support for afum|EAL92753.1 as seed ortholog is 100%.
Bootstrap support for afis|EAW21720.1 as seed ortholog is 100%.

---

Group of orthologs #2729. Best score 1269 bits
Score difference with first non-orthologous sequence - *A. fumigatus*:1269 *A. fischeri*:1269
afum|EAL89605.1 100.00% afis|EAW24948.1 100.00%
Bootstrap support for afum|EAL89605.1 as seed ortholog is 100%.
Bootstrap support for afis|EAW24948.1 as seed ortholog is 100%.

---

Group of orthologs #2730. Best score 1269 bits
Score difference with first non-orthologous sequence - *A. fumigatus*:1269 *A. fischeri*:1269
afum|EAL89673.1 100.00% afis|EAW25022.1 100.00%
Bootstrap support for afum|EAL89673.1 as seed ortholog is 100%.
Bootstrap support for afis|EAW25022.1 as seed ortholog is 100%.

---

Group of orthologs #2731. Best score 1268 bits
Score difference with first non-orthologous sequence - *A. fumigatus*:1268 *A. fischeri*:1268
afum|EAL88639.1 100.00% afis|EAW16009.1 100.00%
Bootstrap support for afum|EAL88639.1 as seed ortholog is 100%.
Bootstrap support for afis|EAW16009.1 as seed ortholog is 100%.

---

Group of orthologs #2732. Best score 1268 bits
Score difference with first non-orthologous sequence - *A. fumigatus*:1268 *A. fischeri*:1268

afum|EAL91376.1 100.00% afis|EAW17562.1 100.00%
Bootstrap support for afum|EAL91376.1 as seed ortholog is 100%.
Bootstrap support for afis|EAW17562.1 as seed ortholog is 100%.

---

Group of orthologs #2733. Best score 1268 bits
Score difference with first non-orthologous sequence - *A. fumigatus*:1268 *A. fischeri*:1268
afum|EAL87223.2 100.00% afis|EAW23789.1 100.00%
Bootstrap support for afum|EAL87223.2 as seed ortholog is 100%.
Bootstrap support for afis|EAW23789.1 as seed ortholog is 100%.

---

Group of orthologs #2734. Best score 1267 bits
Score difference with first non-orthologous sequence - *A. fumigatus*:1198 *A. fischeri*:838
afum|EAL88551.2 100.00% afis|EAW15916.1 100.00%
Bootstrap support for afum|EAL88551.2 as seed ortholog is 100%.
Bootstrap support for afis|EAW15916.1 as seed ortholog is 100%.

---

Group of orthologs #2735. Best score 1267 bits
Score difference with first non-orthologous sequence - *A. fumigatus*:1267 *A. fischeri*:1267
afum|EAL84277.1 100.00% afis|EAW25553.1 100.00%
Bootstrap support for afum|EAL84277.1 as seed ortholog is 100%.
Bootstrap support for afis|EAW25553.1 as seed ortholog is 100%.

---

Group of orthologs #2736. Best score 1267 bits
Score difference with first non-orthologous sequence - *A. fumigatus*:1267 *A. fischeri*:1267
afum|EAL86254.1 100.00% afis|EAW24632.1 100.00%
Bootstrap support for afum|EAL86254.1 as seed ortholog is 100%.
Bootstrap support for afis|EAW24632.1 as seed ortholog is 100%.

---

Group of orthologs #2737. Best score 1267 bits
Score difference with first non-orthologous sequence - *A. fumigatus*:1267 *A. fischeri*:1267
afum|EAL88332.1 100.00% afis|EAW23149.1 100.00%
Bootstrap support for afum|EAL88332.1 as seed ortholog is 100%.
Bootstrap support for afis|EAW23149.1 as seed ortholog is 100%.

---

Group of orthologs #2738. Best score 1266 bits
Score difference with first non-orthologous sequence - *A. fumigatus*:1266 *A. fischeri*:1266
afum|EAL84542.1 100.00% afis|EAW20515.1 100.00%
Bootstrap support for afum|EAL84542.1 as seed ortholog is 100%.
Bootstrap support for afis|EAW20515.1 as seed ortholog is 100%.

---

Group of orthologs #2739. Best score 1266 bits
Score difference with first non-orthologous sequence - *A. fumigatus*:1135 *A. fischeri*:1266
afum|EAL84486.2 100.00% afis|EAW20578.1 100.00%
Bootstrap support for afum|EAL84486.2 as seed ortholog is 100%.
Bootstrap support for afis|EAW20578.1 as seed ortholog is 100%.

---

Group of orthologs #2740. Best score 1266 bits
Score difference with first non-orthologous sequence - *A. fumigatus*:1266 *A. fischeri*:1266
afum|EAL87534.1 100.00% afis|EAW18172.1 100.00%
Bootstrap support for afum|EAL87534.1 as seed ortholog is 100%.
Bootstrap support for afis|EAW18172.1 as seed ortholog is 100%.

---

Group of orthologs #2741. Best score 1266 bits
Score difference with first non-orthologous sequence - *A. fumigatus*:1266 *A. fischeri*:922
afum|EAL89292.1 100.00% afis|EAW16694.1 100.00%
Bootstrap support for afum|EAL89292.1 as seed ortholog is 100%.

Bootstrap support for afis|EAW16694.1 as seed ortholog is 100%.

---

Group of orthologs #2742. Best score 1266 bits
Score difference with first non-orthologous sequence - *A. fumigatus*:1113 *A. fischeri*:808
afum|EAL92895.1 100.00% afis|EAW21881.1 100.00%
Bootstrap support for afum|EAL92895.1 as seed ortholog is 100%.
Bootstrap support for afis|EAW21881.1 as seed ortholog is 100%.

---

Group of orthologs #2743. Best score 1265 bits
Score difference with first non-orthologous sequence - *A. fumigatus*:1265 *A. fischeri*:1265
afum|EAL85709.1 100.00% afis|EAW15625.1 100.00%
Bootstrap support for afum|EAL85709.1 as seed ortholog is 100%.
Bootstrap support for afis|EAW15625.1 as seed ortholog is 100%.

---

Group of orthologs #2744. Best score 1265 bits
Score difference with first non-orthologous sequence - *A. fumigatus*:940 *A. fischeri*:914
afum|EAL86556.1 100.00% afis|EAW17060.1 100.00%
Bootstrap support for afum|EAL86556.1 as seed ortholog is 100%.
Bootstrap support for afis|EAW17060.1 as seed ortholog is 100%.

---

Group of orthologs #2745. Best score 1265 bits
Score difference with first non-orthologous sequence - *A. fumigatus*:1087 *A. fischeri*:983
afum|EAL89286.1 100.00% afis|EAW16682.1 100.00%
Bootstrap support for afum|EAL89286.1 as seed ortholog is 100%.
Bootstrap support for afis|EAW16682.1 as seed ortholog is 100%.

---

Group of orthologs #2746. Best score 1265 bits
Score difference with first non-orthologous sequence - *A. fumigatus*:1265 *A. fischeri*:1265
afum|EAL90608.1 100.00% afis|EAW22591.1 100.00%
Bootstrap support for afum|EAL90608.1 as seed ortholog is 100%.
Bootstrap support for afis|EAW22591.1 as seed ortholog is 100%.

---

Group of orthologs #2747. Best score 1264 bits
Score difference with first non-orthologous sequence - *A. fumigatus*:1264 *A. fischeri*:1264
afum|EAL88644.1 100.00% afis|EAW16015.1 100.00%
Bootstrap support for afum|EAL88644.1 as seed ortholog is 100%.
Bootstrap support for afis|EAW16015.1 as seed ortholog is 100%.

---

Group of orthologs #2748. Best score 1264 bits
Score difference with first non-orthologous sequence - *A. fumigatus*:1264 *A. fischeri*:1264
afum|EAL92017.1 100.00% afis|EAW24161.1 100.00%
Bootstrap support for afum|EAL92017.1 as seed ortholog is 100%.
Bootstrap support for afis|EAW24161.1 as seed ortholog is 100%.

---

Group of orthologs #2749. Best score 1263 bits
Score difference with first non-orthologous sequence - *A. fumigatus*:1263 *A. fischeri*:1263
afum|EAL89073.1 100.00% afis|EAW16479.1 100.00%
Bootstrap support for afum|EAL89073.1 as seed ortholog is 100%.
Bootstrap support for afis|EAW16479.1 as seed ortholog is 100%.

---

Group of orthologs #2750. Best score 1263 bits
Score difference with first non-orthologous sequence - *A. fumigatus*:1096 *A. fischeri*:1047
afum|EAL84286.1 100.00% afis|EAW25576.1 100.00%
Bootstrap support for afum|EAL84286.1 as seed ortholog is 100%.
Bootstrap support for afis|EAW25576.1 as seed ortholog is 100%.

---

Group of orthologs #2751. Best score 1263 bits
Score difference with first non-orthologous sequence - *A. fumigatus*:1263 *A. fischeri*:1263
afum|EAL86083.1 100.00% afis|EAW24414.1 100.00%
Bootstrap support for afum|EAL86083.1 as seed ortholog is 100%.
Bootstrap support for afis|EAW24414.1 as seed ortholog is 100%.

---

Group of orthologs #2752. Best score 1263 bits
Score difference with first non-orthologous sequence - *A. fumigatus*:1263 *A. fischeri*:1263
afum|EAL90098.1 100.00% afis|EAW25440.1 100.00%
Bootstrap support for afum|EAL90098.1 as seed ortholog is 100%.
Bootstrap support for afis|EAW25440.1 as seed ortholog is 100%.

---

Group of orthologs #2753. Best score 1262 bits
Score difference with first non-orthologous sequence - *A. fumigatus*:1262 *A. fischeri*:1262
afum|EAL86934.1 100.00% afis|EAW19584.1 100.00%
Bootstrap support for afum|EAL86934.1 as seed ortholog is 100%.
Bootstrap support for afis|EAW19584.1 as seed ortholog is 100%.

---

Group of orthologs #2754. Best score 1262 bits
Score difference with first non-orthologous sequence - *A. fumigatus*:1262 *A. fischeri*:1262
afum|EAL90982.1 100.00% afis|EAW22212.1 100.00%
Bootstrap support for afum|EAL90982.1 as seed ortholog is 100%.
Bootstrap support for afis|EAW22212.1 as seed ortholog is 100%.

---

Group of orthologs #2755. Best score 1261 bits
Score difference with first non-orthologous sequence - *A. fumigatus*:1261 *A. fischeri*:1261
afum|EAL86673.2 100.00% afis|EAW17255.1 100.00%
Bootstrap support for afum|EAL86673.2 as seed ortholog is 100%.
Bootstrap support for afis|EAW17255.1 as seed ortholog is 100%.

---

Group of orthologs #2756. Best score 1261 bits
Score difference with first non-orthologous sequence - *A. fumigatus*:1261 *A. fischeri*:1261
afum|EAL93751.1 100.00% afis|EAW19020.1 100.00%
Bootstrap support for afum|EAL93751.1 as seed ortholog is 100%.
Bootstrap support for afis|EAW19020.1 as seed ortholog is 100%.

---

Group of orthologs #2757. Best score 1261 bits
Score difference with first non-orthologous sequence - *A. fumigatus*:1261 *A. fischeri*:1147
afum|EAL94088.1 100.00% afis|EAW19369.1 100.00%
Bootstrap support for afum|EAL94088.1 as seed ortholog is 100%.
Bootstrap support for afis|EAW19369.1 as seed ortholog is 100%.

---

Group of orthologs #2758. Best score 1260 bits
Score difference with first non-orthologous sequence - *A. fumigatus*:825 *A. fischeri*:956
afum|EAL88697.1 100.00% afis|EAW16065.1 100.00%
Bootstrap support for afum|EAL88697.1 as seed ortholog is 100%.
Bootstrap support for afis|EAW16065.1 as seed ortholog is 100%.

---

Group of orthologs #2759. Best score 1260 bits
Score difference with first non-orthologous sequence - *A. fumigatus*:1260 *A. fischeri*:1260
afum|EAL87855.2 100.00% afis|EAW23609.1 100.00%
Bootstrap support for afum|EAL87855.2 as seed ortholog is 100%.
Bootstrap support for afis|EAW23609.1 as seed ortholog is 100%.

---

Group of orthologs #2760. Best score 1260 bits
Score difference with first non-orthologous sequence - *A. fumigatus*:1260 *A. fischeri*:1260

afum|EAL90949.1 100.00% afis|EAW22244.1 100.00%
Bootstrap support for afum|EAL90949.1 as seed ortholog is 100%.
Bootstrap support for afis|EAW22244.1 as seed ortholog is 100%.

---

Group of orthologs #2761. Best score 1260 bits
Score difference with first non-orthologous sequence - *A. fumigatus*:1260 *A. fischeri*:1260
afum|EAL90740.1 100.00% afis|EAW22451.1 100.00%
Bootstrap support for afum|EAL90740.1 as seed ortholog is 100%.
Bootstrap support for afis|EAW22451.1 as seed ortholog is 100%.

---

Group of orthologs #2762. Best score 1259 bits
Score difference with first non-orthologous sequence - *A. fumigatus*:1259 *A. fischeri*:1259
afum|EAL88833.1 100.00% afis|EAW16205.1 100.00%
Bootstrap support for afum|EAL88833.1 as seed ortholog is 100%.
Bootstrap support for afis|EAW16205.1 as seed ortholog is 100%.

---

Group of orthologs #2763. Best score 1259 bits
Score difference with first non-orthologous sequence - *A. fumigatus*:1259 *A. fischeri*:1259
afum|EAL93342.1 100.00% afis|EAW18630.1 100.00%
Bootstrap support for afum|EAL93342.1 as seed ortholog is 100%.
Bootstrap support for afis|EAW18630.1 as seed ortholog is 100%.

---

Group of orthologs #2764. Best score 1259 bits
Score difference with first non-orthologous sequence - *A. fumigatus*:1259 *A. fischeri*:1259
afum|EAL89390.1 100.00% afis|EAW24744.1 100.00%
Bootstrap support for afum|EAL89390.1 as seed ortholog is 100%.
Bootstrap support for afis|EAW24744.1 as seed ortholog is 100%.

---

Group of orthologs #2765. Best score 1259 bits
Score difference with first non-orthologous sequence - *A. fumigatus*:571 *A. fischeri*:451
afum|EAL89971.1 100.00% afis|EAW25308.1 100.00%
Bootstrap support for afum|EAL89971.1 as seed ortholog is 100%.
Bootstrap support for afis|EAW25308.1 as seed ortholog is 100%.

---

Group of orthologs #2766. Best score 1259 bits
Score difference with first non-orthologous sequence - *A. fumigatus*:1259 *A. fischeri*:1163
afum|EAL90015.1 100.00% afis|EAW25353.1 100.00%
Bootstrap support for afum|EAL90015.1 as seed ortholog is 100%.
Bootstrap support for afis|EAW25353.1 as seed ortholog is 100%.

---

Group of orthologs #2767. Best score 1258 bits
Score difference with first non-orthologous sequence - *A. fumigatus*:899 *A. fischeri*:681
afum|EAL93236.1 100.00% afis|EAW18538.1 100.00%
Bootstrap support for afum|EAL93236.1 as seed ortholog is 100%.
Bootstrap support for afis|EAW18538.1 as seed ortholog is 100%.

---

Group of orthologs #2768. Best score 1258 bits
Score difference with first non-orthologous sequence - *A. fumigatus*:1258 *A. fischeri*:1258
afum|EAL91039.1 100.00% afis|EAW22151.1 100.00%
Bootstrap support for afum|EAL91039.1 as seed ortholog is 100%.
Bootstrap support for afis|EAW22151.1 as seed ortholog is 100%.

---

Group of orthologs #2769. Best score 1258 bits
Score difference with first non-orthologous sequence - *A. fumigatus*:1258 *A. fischeri*:1258
afum|EAL89809.2 100.00% afis|EAW25138.1 100.00%
Bootstrap support for afum|EAL89809.2 as seed ortholog is 100%.

Bootstrap support for afis|EAW25138.1 as seed ortholog is 100%.

---

Group of orthologs #2770. Best score 1257 bits
Score difference with first non-orthologous sequence - *A. fumigatus*:1078 *A. fischeri*:1189
afum|EAL89226.2 100.00% afis|EAW16630.1 100.00%
Bootstrap support for afum|EAL89226.2 as seed ortholog is 100%.
Bootstrap support for afis|EAW16630.1 as seed ortholog is 100%.

---

Group of orthologs #2771. Best score 1257 bits
Score difference with first non-orthologous sequence - *A. fumigatus*:1257 *A. fischeri*:1257
afum|EAL87039.1 100.00% afis|EAW19457.1 100.00%
Bootstrap support for afum|EAL87039.1 as seed ortholog is 100%.
Bootstrap support for afis|EAW19457.1 as seed ortholog is 100%.

---

Group of orthologs #2772. Best score 1257 bits
Score difference with first non-orthologous sequence - *A. fumigatus*:1128 *A. fischeri*:1166
afum|EAL86794.1 100.00% afis|EAW19735.1 100.00%
Bootstrap support for afum|EAL86794.1 as seed ortholog is 100%.
Bootstrap support for afis|EAW19735.1 as seed ortholog is 100%.

---

Group of orthologs #2773. Best score 1257 bits
Score difference with first non-orthologous sequence - *A. fumigatus*:1257 *A. fischeri*:1257
afum|EAL91773.1 100.00% afis|EAW17963.1 100.00%
Bootstrap support for afum|EAL91773.1 as seed ortholog is 100%.
Bootstrap support for afis|EAW17963.1 as seed ortholog is 100%.

---

Group of orthologs #2774. Best score 1257 bits
Score difference with first non-orthologous sequence - *A. fumigatus*:840 *A. fischeri*:463
afum|EAL86150.1 100.00% afis|EAW24479.1 100.00%
Bootstrap support for afum|EAL86150.1 as seed ortholog is 100%.
Bootstrap support for afis|EAW24479.1 as seed ortholog is 100%.

---

Group of orthologs #2775. Best score 1257 bits
Score difference with first non-orthologous sequence - *A. fumigatus*:1257 *A. fischeri*:1257
afum|EAL93597.1 100.00% afis|EAW18865.1 100.00%
Bootstrap support for afum|EAL93597.1 as seed ortholog is 100%.
Bootstrap support for afis|EAW18865.1 as seed ortholog is 100%.

---

Group of orthologs #2776. Best score 1256 bits
Score difference with first non-orthologous sequence - *A. fumigatus*:1256 *A. fischeri*:1256
afum|EAL87002.1 100.00% afis|EAW19492.1 100.00%
Bootstrap support for afum|EAL87002.1 as seed ortholog is 100%.
Bootstrap support for afis|EAW19492.1 as seed ortholog is 100%.

---

Group of orthologs #2777. Best score 1256 bits
Score difference with first non-orthologous sequence - *A. fumigatus*:1042 *A. fischeri*:774
afum|EAL91170.2 100.00% afis|EAW17333.1 100.00%
Bootstrap support for afum|EAL91170.2 as seed ortholog is 100%.
Bootstrap support for afis|EAW17333.1 as seed ortholog is 100%.

---

Group of orthologs #2778. Best score 1256 bits
Score difference with first non-orthologous sequence - *A. fumigatus*:1256 *A. fischeri*:1256
afum|EAL93303.1 100.00% afis|EAW18595.1 100.00%
Bootstrap support for afum|EAL93303.1 as seed ortholog is 100%.
Bootstrap support for afis|EAW18595.1 as seed ortholog is 100%.

---

Group of orthologs #2779. Best score 1256 bits
Score difference with first non-orthologous sequence - *A. fumigatus*:1022 *A. fischeri*:1256
afum|EAL90708.1 100.00% afis|EAW22485.1 100.00%
Bootstrap support for afum|EAL90708.1 as seed ortholog is 100%.
Bootstrap support for afis|EAW22485.1 as seed ortholog is 100%.

---

Group of orthologs #2780. Best score 1255 bits
Score difference with first non-orthologous sequence - *A. fumigatus*:1255 *A. fischeri*:1255
afum|EAL85819.1 100.00% afis|EAW15505.1 100.00%
Bootstrap support for afum|EAL85819.1 as seed ortholog is 100%.
Bootstrap support for afis|EAW15505.1 as seed ortholog is 100%.

---

Group of orthologs #2781. Best score 1255 bits
Score difference with first non-orthologous sequence - *A. fumigatus*:1255 *A. fischeri*:1255
afum|EAL88598.1 100.00% afis|EAW15965.1 100.00%
Bootstrap support for afum|EAL88598.1 as seed ortholog is 100%.
Bootstrap support for afis|EAW15965.1 as seed ortholog is 100%.

---

Group of orthologs #2782. Best score 1255 bits
Score difference with first non-orthologous sequence - *A. fumigatus*:1255 *A. fischeri*:1255
afum|EAL89135.1 100.00% afis|EAW16538.1 100.00%
Bootstrap support for afum|EAL89135.1 as seed ortholog is 100%.
Bootstrap support for afis|EAW16538.1 as seed ortholog is 100%.

---

Group of orthologs #2783. Best score 1255 bits
Score difference with first non-orthologous sequence - *A. fumigatus*:1255 *A. fischeri*:1255
afum|EAL87429.1 100.00% afis|EAW23999.1 100.00%
Bootstrap support for afum|EAL87429.1 as seed ortholog is 100%.
Bootstrap support for afis|EAW23999.1 as seed ortholog is 100%.

---

Group of orthologs #2784. Best score 1255 bits
Score difference with first non-orthologous sequence - *A. fumigatus*:1255 *A. fischeri*:1255
afum|EAL93396.1 100.00% afis|EAW18676.1 100.00%
Bootstrap support for afum|EAL93396.1 as seed ortholog is 100%.
Bootstrap support for afis|EAW18676.1 as seed ortholog is 100%.

---

Group of orthologs #2785. Best score 1255 bits
Score difference with first non-orthologous sequence - *A. fumigatus*:1167 *A. fischeri*:458
afum|EAL93753.1 100.00% afis|EAW19022.1 100.00%
Bootstrap support for afum|EAL93753.1 as seed ortholog is 100%.
Bootstrap support for afis|EAW19022.1 as seed ortholog is 100%.

---

Group of orthologs #2786. Best score 1254 bits
Score difference with first non-orthologous sequence - *A. fumigatus*:1254 *A. fischeri*:1185
afum|EAL86315.1 100.00% afis|EAW16740.1 100.00%
Bootstrap support for afum|EAL86315.1 as seed ortholog is 100%.
Bootstrap support for afis|EAW16740.1 as seed ortholog is 100%.

---

Group of orthologs #2787. Best score 1254 bits
Score difference with first non-orthologous sequence - *A. fumigatus*:1254 *A. fischeri*:1254
afum|EAL88724.1 100.00% afis|EAW16092.1 100.00%
Bootstrap support for afum|EAL88724.1 as seed ortholog is 100%.
Bootstrap support for afis|EAW16092.1 as seed ortholog is 100%.

---

Group of orthologs #2788. Best score 1254 bits
Score difference with first non-orthologous sequence - *A. fumigatus*:1078 *A. fischeri*:1254

afum|EAL84500.1 100.00% afis|EAW20561.1 100.00%
Bootstrap support for afum|EAL84500.1 as seed ortholog is 100%.
Bootstrap support for afis|EAW20561.1 as seed ortholog is 100%.

---

Group of orthologs #2789. Best score 1254 bits
Score difference with first non-orthologous sequence - *A. fumigatus*:1254 *A. fischeri*:1254
afum|EAL93705.1 100.00% afis|EAW18972.1 100.00%
Bootstrap support for afum|EAL93705.1 as seed ortholog is 100%.
Bootstrap support for afis|EAW18972.1 as seed ortholog is 100%.

---

Group of orthologs #2790. Best score 1254 bits
Score difference with first non-orthologous sequence - *A. fumigatus*:1117 *A. fischeri*:1075
afum|EAL93810.1 100.00% afis|EAW19081.1 100.00%
Bootstrap support for afum|EAL93810.1 as seed ortholog is 100%.
Bootstrap support for afis|EAW19081.1 as seed ortholog is 100%.

---

Group of orthologs #2791. Best score 1253 bits
Score difference with first non-orthologous sequence - *A. fumigatus*:139 *A. fischeri*:1050
afum|EAL91800.1 100.00% afis|EAW17990.1 100.00%
afis|EAW20711.1 40.96%
Bootstrap support for afum|EAL91800.1 as seed ortholog is 99%.
Bootstrap support for afis|EAW17990.1 as seed ortholog is 100%.

---

Group of orthologs #2792. Best score 1253 bits
Score difference with first non-orthologous sequence - *A. fumigatus*:1253 *A. fischeri*:1253
afum|EAL85688.2 100.00% afis|EAW15658.1 100.00%
Bootstrap support for afum|EAL85688.2 as seed ortholog is 100%.
Bootstrap support for afis|EAW15658.1 as seed ortholog is 100%.

---

Group of orthologs #2793. Best score 1253 bits
Score difference with first non-orthologous sequence - *A. fumigatus*:1253 *A. fischeri*:1253
afum|EAL87827.2 100.00% afis|EAW23633.1 100.00%
Bootstrap support for afum|EAL87827.2 as seed ortholog is 100%.
Bootstrap support for afis|EAW23633.1 as seed ortholog is 100%.

---

Group of orthologs #2794. Best score 1253 bits
Score difference with first non-orthologous sequence - *A. fumigatus*:1253 *A. fischeri*:1253
afum|EAL92652.1 100.00% afis|EAW21617.1 100.00%
Bootstrap support for afum|EAL92652.1 as seed ortholog is 100%.
Bootstrap support for afis|EAW21617.1 as seed ortholog is 100%.

---

Group of orthologs #2795. Best score 1252 bits
Score difference with first non-orthologous sequence - *A. fumigatus*:1058 *A. fischeri*:1252
afum|EAL88754.1 100.00% afis|EAW16124.1 100.00%
Bootstrap support for afum|EAL88754.1 as seed ortholog is 100%.
Bootstrap support for afis|EAW16124.1 as seed ortholog is 100%.

---

Group of orthologs #2796. Best score 1252 bits
Score difference with first non-orthologous sequence - *A. fumigatus*:1252 *A. fischeri*:1252
afum|EAL88244.1 100.00% afis|EAW23238.1 100.00%
Bootstrap support for afum|EAL88244.1 as seed ortholog is 100%.
Bootstrap support for afis|EAW23238.1 as seed ortholog is 100%.

---

Group of orthologs #2797. Best score 1251 bits
Score difference with first non-orthologous sequence - *A. fumigatus*:1251 *A. fischeri*:1109
afum|EAL85044.1 100.00% afis|EAW19849.1 100.00%

Bootstrap support for afum|EAL85044.1 as seed ortholog is 100%.
Bootstrap support for afis|EAW19849.1 as seed ortholog is 100%.

---

Group of orthologs #2798. Best score 1251 bits
Score difference with first non-orthologous sequence - *A. fumigatus*:1251 *A. fischeri*:1251
afum|EAL87008.1 100.00% afis|EAW19486.1 100.00%
Bootstrap support for afum|EAL87008.1 as seed ortholog is 100%.
Bootstrap support for afis|EAW19486.1 as seed ortholog is 100%.

---

Group of orthologs #2799. Best score 1251 bits
Score difference with first non-orthologous sequence - *A. fumigatus*:944 *A. fischeri*:1251
afum|EAL86230.1 100.00% afis|EAW24567.1 100.00%
Bootstrap support for afum|EAL86230.1 as seed ortholog is 100%.
Bootstrap support for afis|EAW24567.1 as seed ortholog is 100%.

---

Group of orthologs #2800. Best score 1251 bits
Score difference with first non-orthologous sequence - *A. fumigatus*:1141 *A. fischeri*:1251
afum|EAL93135.1 100.00% afis|EAW18450.1 100.00%
Bootstrap support for afum|EAL93135.1 as seed ortholog is 100%.
Bootstrap support for afis|EAW18450.1 as seed ortholog is 100%.

---

Group of orthologs #2801. Best score 1251 bits
Score difference with first non-orthologous sequence - *A. fumigatus*:1094 *A. fischeri*:1251
afum|EAL90241.1 100.00% afis|EAW22956.1 100.00%
Bootstrap support for afum|EAL90241.1 as seed ortholog is 100%.
Bootstrap support for afis|EAW22956.1 as seed ortholog is 100%.

---

Group of orthologs #2802. Best score 1251 bits
Score difference with first non-orthologous sequence - *A. fumigatus*:1251 *A. fischeri*:1251
afum|EAL92705.1 100.00% afis|EAW21670.1 100.00%
Bootstrap support for afum|EAL92705.1 as seed ortholog is 100%.
Bootstrap support for afis|EAW21670.1 as seed ortholog is 100%.

---

Group of orthologs #2803. Best score 1250 bits
Score difference with first non-orthologous sequence - *A. fumigatus*:1020 *A. fischeri*:414
afum|EAL84459.1 100.00% afis|EAW20602.1 100.00%
Bootstrap support for afum|EAL84459.1 as seed ortholog is 100%.
Bootstrap support for afis|EAW20602.1 as seed ortholog is 100%.

---

Group of orthologs #2804. Best score 1250 bits
Score difference with first non-orthologous sequence - *A. fumigatus*:1250 *A. fischeri*:1250
afum|EAL87342.1 100.00% afis|EAW23913.1 100.00%
Bootstrap support for afum|EAL87342.1 as seed ortholog is 100%.
Bootstrap support for afis|EAW23913.1 as seed ortholog is 100%.

---

Group of orthologs #2805. Best score 1250 bits
Score difference with first non-orthologous sequence - *A. fumigatus*:1250 *A. fischeri*:1250
afum|EAL88021.1 100.00% afis|EAW23442.1 100.00%
Bootstrap support for afum|EAL88021.1 as seed ortholog is 100%.
Bootstrap support for afis|EAW23442.1 as seed ortholog is 100%.

---

Group of orthologs #2806. Best score 1250 bits
Score difference with first non-orthologous sequence - *A. fumigatus*:1250 *A. fischeri*:1250
afum|EAL93350.1 100.00% afis|EAW18637.1 100.00%
Bootstrap support for afum|EAL93350.1 as seed ortholog is 100%.
Bootstrap support for afis|EAW18637.1 as seed ortholog is 100%.

---

Group of orthologs #2807. Best score 1250 bits
Score difference with first non-orthologous sequence - *A. fumigatus*:798 *A. fischeri*:1056
afum|EAL91046.1 100.00% afis|EAW22144.1 100.00%
Bootstrap support for afum|EAL91046.1 as seed ortholog is 100%.
Bootstrap support for afis|EAW22144.1 as seed ortholog is 100%.

---

Group of orthologs #2808. Best score 1249 bits
Score difference with first non-orthologous sequence - *A. fumigatus*:1106 *A. fischeri*:1119
afum|EAL90466.1 100.00% afis|EAW22729.1 100.00%
Bootstrap support for afum|EAL90466.1 as seed ortholog is 100%.
Bootstrap support for afis|EAW22729.1 as seed ortholog is 100%.

---

Group of orthologs #2809. Best score 1249 bits
Score difference with first non-orthologous sequence - *A. fumigatus*:1249 *A. fischeri*:1249
afum|EAL90471.1 100.00% afis|EAW22725.1 100.00%
Bootstrap support for afum|EAL90471.1 as seed ortholog is 100%.
Bootstrap support for afis|EAW22725.1 as seed ortholog is 100%.

---

Group of orthologs #2810. Best score 1248 bits
Score difference with first non-orthologous sequence - *A. fumigatus*:1107 *A. fischeri*:731
afum|EAL85732.1 100.00% afis|EAW15601.1 100.00%
Bootstrap support for afum|EAL85732.1 as seed ortholog is 100%.
Bootstrap support for afis|EAW15601.1 as seed ortholog is 100%.

---

Group of orthologs #2811. Best score 1248 bits
Score difference with first non-orthologous sequence - *A. fumigatus*:1248 *A. fischeri*:1248
afum|EAL88342.1 100.00% afis|EAW23139.1 100.00%
Bootstrap support for afum|EAL88342.1 as seed ortholog is 100%.
Bootstrap support for afis|EAW23139.1 as seed ortholog is 100%.

---

Group of orthologs #2812. Best score 1248 bits
Score difference with first non-orthologous sequence - *A. fumigatus*:1248 *A. fischeri*:1248
afum|EBA27409.1 100.00% afis|EAW22692.1 100.00%
Bootstrap support for afum|EBA27409.1 as seed ortholog is 100%.
Bootstrap support for afis|EAW22692.1 as seed ortholog is 100%.

---

Group of orthologs #2813. Best score 1247 bits
Score difference with first non-orthologous sequence - *A. fumigatus*:1247 *A. fischeri*:1247
afum|EAL91857.1 100.00% afis|EAW18045.1 100.00%
Bootstrap support for afum|EAL91857.1 as seed ortholog is 100%.
Bootstrap support for afis|EAW18045.1 as seed ortholog is 100%.

---

Group of orthologs #2814. Best score 1247 bits
Score difference with first non-orthologous sequence - *A. fumigatus*:982 *A. fischeri*:1247
afum|EAL84297.2 100.00% afis|EAW25589.1 100.00%
Bootstrap support for afum|EAL84297.2 as seed ortholog is 100%.
Bootstrap support for afis|EAW25589.1 as seed ortholog is 100%.

---

Group of orthologs #2815. Best score 1247 bits
Score difference with first non-orthologous sequence - *A. fumigatus*:1247 *A. fischeri*:1247
afum|EAL93059.1 100.00% afis|EAW18384.1 100.00%
Bootstrap support for afum|EAL93059.1 as seed ortholog is 100%.
Bootstrap support for afis|EAW18384.1 as seed ortholog is 100%.

---

Group of orthologs #2816. Best score 1247 bits

Score difference with first non-orthologous sequence - *A. fumigatus*:1115 *A. fischeri*:1040
afum|EAL93119.1 100.00% afis|EAW18437.1 100.00%
Bootstrap support for afum|EAL93119.1 as seed ortholog is 100%.
Bootstrap support for afis|EAW18437.1 as seed ortholog is 100%.

---

Group of orthologs #2817. Best score 1247 bits
Score difference with first non-orthologous sequence - *A. fumigatus*:1247 *A. fischeri*:1247
afum|EAL93190.1 100.00% afis|EAW18499.1 100.00%
Bootstrap support for afum|EAL93190.1 as seed ortholog is 100%.
Bootstrap support for afis|EAW18499.1 as seed ortholog is 100%.

---

Group of orthologs #2818. Best score 1247 bits
Score difference with first non-orthologous sequence - *A. fumigatus*:967 *A. fischeri*:1112
afum|EAL87838.2 100.00% afis|EAW23622.1 100.00%
Bootstrap support for afum|EAL87838.2 as seed ortholog is 100%.
Bootstrap support for afis|EAW23622.1 as seed ortholog is 100%.

---

Group of orthologs #2819. Best score 1247 bits
Score difference with first non-orthologous sequence - *A. fumigatus*:860 *A. fischeri*:1164
afum|EAL92111.1 100.00% afis|EAW21088.1 100.00%
Bootstrap support for afum|EAL92111.1 as seed ortholog is 100%.
Bootstrap support for afis|EAW21088.1 as seed ortholog is 100%.

---

Group of orthologs #2820. Best score 1247 bits
Score difference with first non-orthologous sequence - *A. fumigatus*:747 *A. fischeri*:716
afum|EAL90594.1 100.00% afis|EAW22604.1 100.00%
Bootstrap support for afum|EAL90594.1 as seed ortholog is 100%.
Bootstrap support for afis|EAW22604.1 as seed ortholog is 100%.

---

Group of orthologs #2821. Best score 1246 bits
Score difference with first non-orthologous sequence - *A. fumigatus*:1246 *A. fischeri*:1246
afum|EAL85515.1 100.00% afis|EAW15836.1 100.00%
Bootstrap support for afum|EAL85515.1 as seed ortholog is 100%.
Bootstrap support for afis|EAW15836.1 as seed ortholog is 100%.

---

Group of orthologs #2822. Best score 1246 bits
Score difference with first non-orthologous sequence - *A. fumigatus*:1246 *A. fischeri*:1246
afum|EAL86509.1 100.00% afis|EAW16982.1 100.00%
Bootstrap support for afum|EAL86509.1 as seed ortholog is 100%.
Bootstrap support for afis|EAW16982.1 as seed ortholog is 100%.

---

Group of orthologs #2823. Best score 1246 bits
Score difference with first non-orthologous sequence - *A. fumigatus*:1246 *A. fischeri*:1246
afum|EAL88638.2 100.00% afis|EAW16006.1 100.00%
Bootstrap support for afum|EAL88638.2 as seed ortholog is 100%.
Bootstrap support for afis|EAW16006.1 as seed ortholog is 100%.

---

Group of orthologs #2824. Best score 1246 bits
Score difference with first non-orthologous sequence - *A. fumigatus*:1246 *A. fischeri*:1246
afum|EAL88938.1 100.00% afis|EAW16315.1 100.00%
Bootstrap support for afum|EAL88938.1 as seed ortholog is 100%.
Bootstrap support for afis|EAW16315.1 as seed ortholog is 100%.

---

Group of orthologs #2825. Best score 1246 bits
Score difference with first non-orthologous sequence - *A. fumigatus*:1246 *A. fischeri*:1246
afum|EAL86156.1 100.00% afis|EAW24487.1 100.00%

Bootstrap support for afum|EAL86156.1 as seed ortholog is 100%.
Bootstrap support for afis|EAW24487.1 as seed ortholog is 100%.

---

Group of orthologs #2826. Best score 1246 bits
Score difference with first non-orthologous sequence - *A. fumigatus*:1246 *A. fischeri*:1246
afum|EAL93505.1 100.00% afis|EAW18773.1 100.00%
Bootstrap support for afum|EAL93505.1 as seed ortholog is 100%.
Bootstrap support for afis|EAW18773.1 as seed ortholog is 100%.

---

Group of orthologs #2827. Best score 1246 bits
Score difference with first non-orthologous sequence - *A. fumigatus*:1246 *A. fischeri*:1246
afum|EAL92112.1 100.00% afis|EAW21089.1 100.00%
Bootstrap support for afum|EAL92112.1 as seed ortholog is 100%.
Bootstrap support for afis|EAW21089.1 as seed ortholog is 100%.

---

Group of orthologs #2828. Best score 1246 bits
Score difference with first non-orthologous sequence - *A. fumigatus*:1246 *A. fischeri*:1246
afum|EAL89755.1 100.00% afis|EAW25086.1 100.00%
Bootstrap support for afum|EAL89755.1 as seed ortholog is 100%.
Bootstrap support for afis|EAW25086.1 as seed ortholog is 100%.

---

Group of orthologs #2829. Best score 1245 bits
Score difference with first non-orthologous sequence - *A. fumigatus*:1245 *A. fischeri*:1245
afum|EAL89081.1 100.00% afis|EAW16487.1 100.00%
Bootstrap support for afum|EAL89081.1 as seed ortholog is 100%.
Bootstrap support for afis|EAW16487.1 as seed ortholog is 100%.

---

Group of orthologs #2830. Best score 1245 bits
Score difference with first non-orthologous sequence - *A. fumigatus*:1079 *A. fischeri*:1005
afum|EAL86021.1 100.00% afis|EAW24351.1 100.00%
Bootstrap support for afum|EAL86021.1 as seed ortholog is 100%.
Bootstrap support for afis|EAW24351.1 as seed ortholog is 100%.

---

Group of orthologs #2831. Best score 1245 bits
Score difference with first non-orthologous sequence - *A. fumigatus*:771 *A. fischeri*:1094
afum|EAL86152.1 100.00% afis|EAW24483.1 100.00%
Bootstrap support for afum|EAL86152.1 as seed ortholog is 100%.
Bootstrap support for afis|EAW24483.1 as seed ortholog is 100%.

---

Group of orthologs #2832. Best score 1245 bits
Score difference with first non-orthologous sequence - *A. fumigatus*:1085 *A. fischeri*:1135
afum|EBA27224.1 100.00% afis|EAW20120.1 100.00%
Bootstrap support for afum|EBA27224.1 as seed ortholog is 100%.
Bootstrap support for afis|EAW20120.1 as seed ortholog is 100%.

---

Group of orthologs #2833. Best score 1244 bits
Score difference with first non-orthologous sequence - *A. fumigatus*:1244 *A. fischeri*:1244
afum|EAL87513.1 100.00% afis|EAW18154.1 100.00%
Bootstrap support for afum|EAL87513.1 as seed ortholog is 100%.
Bootstrap support for afis|EAW18154.1 as seed ortholog is 100%.

---

Group of orthologs #2834. Best score 1244 bits
Score difference with first non-orthologous sequence - *A. fumigatus*:1244 *A. fischeri*:1015
afum|EAL85365.2 100.00% afis|EAW20247.1 100.00%
Bootstrap support for afum|EAL85365.2 as seed ortholog is 100%.
Bootstrap support for afis|EAW20247.1 as seed ortholog is 100%.

---

Group of orthologs #2835. Best score 1244 bits
Score difference with first non-orthologous sequence - *A. fumigatus*:1131 *A. fischeri*:1135
afum|EAL86133.1 100.00% afis|EAW24462.1 100.00%
Bootstrap support for afum|EAL86133.1 as seed ortholog is 100%.
Bootstrap support for afis|EAW24462.1 as seed ortholog is 100%.

---

Group of orthologs #2836. Best score 1244 bits
Score difference with first non-orthologous sequence - *A. fumigatus*:1244 *A. fischeri*:1244
afum|EAL88292.1 100.00% afis|EAW23191.1 100.00%
Bootstrap support for afum|EAL88292.1 as seed ortholog is 100%.
Bootstrap support for afis|EAW23191.1 as seed ortholog is 100%.

---

Group of orthologs #2837. Best score 1244 bits
Score difference with first non-orthologous sequence - *A. fumigatus*:1244 *A. fischeri*:1244
afum|EAL90296.1 100.00% afis|EAW22901.1 100.00%
Bootstrap support for afum|EAL90296.1 as seed ortholog is 100%.
Bootstrap support for afis|EAW22901.1 as seed ortholog is 100%.

---

Group of orthologs #2838. Best score 1244 bits
Score difference with first non-orthologous sequence - *A. fumigatus*:1244 *A. fischeri*:1244
afum|EAL92334.1 100.00% afis|EAW21317.1 100.00%
Bootstrap support for afum|EAL92334.1 as seed ortholog is 100%.
Bootstrap support for afis|EAW21317.1 as seed ortholog is 100%.

---

Group of orthologs #2839. Best score 1244 bits
Score difference with first non-orthologous sequence - *A. fumigatus*:1244 *A. fischeri*:1244
afum|EAL92398.1 100.00% afis|EAW21383.1 100.00%
Bootstrap support for afum|EAL92398.1 as seed ortholog is 100%.
Bootstrap support for afis|EAW21383.1 as seed ortholog is 100%.

---

Group of orthologs #2840. Best score 1244 bits
Score difference with first non-orthologous sequence - *A. fumigatus*:1244 *A. fischeri*:1244
afum|EAL92792.1 100.00% afis|EAW21762.1 100.00%
Bootstrap support for afum|EAL92792.1 as seed ortholog is 100%.
Bootstrap support for afis|EAW21762.1 as seed ortholog is 100%.

---

Group of orthologs #2841. Best score 1243 bits
Score difference with first non-orthologous sequence - *A. fumigatus*:1243 *A. fischeri*:1243
afum|EAL84819.1 100.00% afis|EAW20809.1 100.00%
Bootstrap support for afum|EAL84819.1 as seed ortholog is 100%.
Bootstrap support for afis|EAW20809.1 as seed ortholog is 100%.

---

Group of orthologs #2842. Best score 1243 bits
Score difference with first non-orthologous sequence - *A. fumigatus*:948 *A. fischeri*:1089
afum|EAL91654.1 100.00% afis|EAW17845.1 100.00%
Bootstrap support for afum|EAL91654.1 as seed ortholog is 100%.
Bootstrap support for afis|EAW17845.1 as seed ortholog is 100%.

---

Group of orthologs #2843. Best score 1243 bits
Score difference with first non-orthologous sequence - *A. fumigatus*:1243 *A. fischeri*:1243
afum|EAL88101.1 100.00% afis|EAW23371.1 100.00%
Bootstrap support for afum|EAL88101.1 as seed ortholog is 100%.
Bootstrap support for afis|EAW23371.1 as seed ortholog is 100%.

---

Group of orthologs #2844. Best score 1243 bits

Score difference with first non-orthologous sequence - *A. fumigatus*:1243   *A. fischeri*:1243
afum|EAL93995.1 100.00% afis|EAW19268.1 100.00%
Bootstrap support for afum|EAL93995.1 as seed ortholog is 100%.
Bootstrap support for afis|EAW19268.1 as seed ortholog is 100%.

---

Group of orthologs #2845. Best score 1242 bits
Score difference with first non-orthologous sequence - *A. fumigatus*:1242   *A. fischeri*:1242
afum|EAL84527.1 100.00% afis|EAW20530.1 100.00%
Bootstrap support for afum|EAL84527.1 as seed ortholog is 100%.
Bootstrap support for afis|EAW20530.1 as seed ortholog is 100%.

---

Group of orthologs #2846. Best score 1242 bits
Score difference with first non-orthologous sequence - *A. fumigatus*:1040   *A. fischeri*:774
afum|EAL88999.1 100.00% afis|EAW16375.1 100.00%
Bootstrap support for afum|EAL88999.1 as seed ortholog is 100%.
Bootstrap support for afis|EAW16375.1 as seed ortholog is 100%.

---

Group of orthologs #2847. Best score 1242 bits
Score difference with first non-orthologous sequence - *A. fumigatus*:1242   *A. fischeri*:1242
afum|EAL91859.1 100.00% afis|EAW18047.1 100.00%
Bootstrap support for afum|EAL91859.1 as seed ortholog is 100%.
Bootstrap support for afis|EAW18047.1 as seed ortholog is 100%.

---

Group of orthologs #2848. Best score 1241 bits
Score difference with first non-orthologous sequence - *A. fumigatus*:1241   *A. fischeri*:1165
afum|EAL89264.1 100.00% afis|EAW16660.1 100.00%
Bootstrap support for afum|EAL89264.1 as seed ortholog is 100%.
Bootstrap support for afis|EAW16660.1 as seed ortholog is 100%.

---

Group of orthologs #2849. Best score 1241 bits
Score difference with first non-orthologous sequence - *A. fumigatus*:1241   *A. fischeri*:1241
afum|EAL87397.1 100.00% afis|EAW23967.1 100.00%
Bootstrap support for afum|EAL87397.1 as seed ortholog is 100%.
Bootstrap support for afis|EAW23967.1 as seed ortholog is 100%.

---

Group of orthologs #2850. Best score 1241 bits
Score difference with first non-orthologous sequence - *A. fumigatus*:1241   *A. fischeri*:1045
afum|EAL93820.1 100.00% afis|EAW19090.1 100.00%
Bootstrap support for afum|EAL93820.1 as seed ortholog is 100%.
Bootstrap support for afis|EAW19090.1 as seed ortholog is 100%.

---

Group of orthologs #2851. Best score 1241 bits
Score difference with first non-orthologous sequence - *A. fumigatus*:1241   *A. fischeri*:1241
afum|EAL90820.2 100.00% afis|EAW22372.1 100.00%
Bootstrap support for afum|EAL90820.2 as seed ortholog is 100%.
Bootstrap support for afis|EAW22372.1 as seed ortholog is 100%.

---

Group of orthologs #2852. Best score 1241 bits
Score difference with first non-orthologous sequence - *A. fumigatus*:1241   *A. fischeri*:1241
afum|EAL90361.1 100.00% afis|EAW22842.1 100.00%
Bootstrap support for afum|EAL90361.1 as seed ortholog is 100%.
Bootstrap support for afis|EAW22842.1 as seed ortholog is 100%.

---

Group of orthologs #2853. Best score 1241 bits
Score difference with first non-orthologous sequence - *A. fumigatus*:1130   *A. fischeri*:1144
afum|EAL90209.1 100.00% afis|EAW22991.1 100.00%

Bootstrap support for afum|EAL90209.1 as seed ortholog is 100%.
Bootstrap support for afis|EAW22991.1 as seed ortholog is 100%.

---

Group of orthologs #2854. Best score 1241 bits
Score difference with first non-orthologous sequence - *A. fumigatus*:1241 *A. fischeri*:1241
afum|EAL90029.2 100.00% afis|EAW25369.1 100.00%
Bootstrap support for afum|EAL90029.2 as seed ortholog is 100%.
Bootstrap support for afis|EAW25369.1 as seed ortholog is 100%.

---

Group of orthologs #2855. Best score 1241 bits
Score difference with first non-orthologous sequence - *A. fumigatus*:1241 *A. fischeri*:1241
afum|EAL90174.1 100.00% afis|EAW25508.1 100.00%
Bootstrap support for afum|EAL90174.1 as seed ortholog is 100%.
Bootstrap support for afis|EAW25508.1 as seed ortholog is 100%.

---

Group of orthologs #2856. Best score 1240 bits
Score difference with first non-orthologous sequence - *A. fumigatus*:1033 *A. fischeri*:1160
afum|EAL87911.1 100.00% afis|EAW23555.1 100.00%
Bootstrap support for afum|EAL87911.1 as seed ortholog is 100%.
Bootstrap support for afis|EAW23555.1 as seed ortholog is 100%.

---

Group of orthologs #2857. Best score 1240 bits
Score difference with first non-orthologous sequence - *A. fumigatus*:962 *A. fischeri*:813
afum|EAL93250.1 100.00% afis|EAW18550.1 100.00%
Bootstrap support for afum|EAL93250.1 as seed ortholog is 100%.
Bootstrap support for afis|EAW18550.1 as seed ortholog is 100%.

---

Group of orthologs #2858. Best score 1240 bits
Score difference with first non-orthologous sequence - *A. fumigatus*:1240 *A. fischeri*:1184
afum|EAL90099.1 100.00% afis|EAW25441.1 100.00%
Bootstrap support for afum|EAL90099.1 as seed ortholog is 100%.
Bootstrap support for afis|EAW25441.1 as seed ortholog is 100%.

---

Group of orthologs #2859. Best score 1239 bits
Score difference with first non-orthologous sequence - *A. fumigatus*:1239 *A. fischeri*:1239
afum|EAL84613.1 100.00% afis|EAW20439.1 100.00%
Bootstrap support for afum|EAL84613.1 as seed ortholog is 100%.
Bootstrap support for afis|EAW20439.1 as seed ortholog is 100%.

---

Group of orthologs #2860. Best score 1239 bits
Score difference with first non-orthologous sequence - *A. fumigatus*:971 *A. fischeri*:1104
afum|EAL84435.1 100.00% afis|EAW20633.1 100.00%
Bootstrap support for afum|EAL84435.1 as seed ortholog is 100%.
Bootstrap support for afis|EAW20633.1 as seed ortholog is 100%.

---

Group of orthologs #2861. Best score 1239 bits
Score difference with first non-orthologous sequence - *A. fumigatus*:1239 *A. fischeri*:1239
afum|EAL93774.1 100.00% afis|EAW19045.1 100.00%
Bootstrap support for afum|EAL93774.1 as seed ortholog is 100%.
Bootstrap support for afis|EAW19045.1 as seed ortholog is 100%.

---

Group of orthologs #2862. Best score 1239 bits
Score difference with first non-orthologous sequence - *A. fumigatus*:1066 *A. fischeri*:1239
afum|EAL90561.1 100.00% afis|EAW22632.1 100.00%
Bootstrap support for afum|EAL90561.1 as seed ortholog is 100%.
Bootstrap support for afis|EAW22632.1 as seed ortholog is 100%.

---

Group of orthologs #2863. Best score 1239 bits
Score difference with first non-orthologous sequence - *A. fumigatus*:1189 *A. fischeri*:1083
afum|EAL90178.1 100.00% afis|EAW25513.1 100.00%
Bootstrap support for afum|EAL90178.1 as seed ortholog is 100%.
Bootstrap support for afis|EAW25513.1 as seed ortholog is 100%.

---

Group of orthologs #2864. Best score 1238 bits
Score difference with first non-orthologous sequence - *A. fumigatus*:1238 *A. fischeri*:1238
afum|EAL84749.1 100.00% afis|EAW20883.1 100.00%
Bootstrap support for afum|EAL84749.1 as seed ortholog is 100%.
Bootstrap support for afis|EAW20883.1 as seed ortholog is 100%.

---

Group of orthologs #2865. Best score 1238 bits
Score difference with first non-orthologous sequence - *A. fumigatus*:1238 *A. fischeri*:1238
afum|EAL92314.1 100.00% afis|EAW21297.1 100.00%
Bootstrap support for afum|EAL92314.1 as seed ortholog is 100%.
Bootstrap support for afis|EAW21297.1 as seed ortholog is 100%.

---

Group of orthologs #2866. Best score 1237 bits
Score difference with first non-orthologous sequence - *A. fumigatus*:1237 *A. fischeri*:1237
afum|EAL88658.1 100.00% afis|EAW16028.1 100.00%
Bootstrap support for afum|EAL88658.1 as seed ortholog is 100%.
Bootstrap support for afis|EAW16028.1 as seed ortholog is 100%.

---

Group of orthologs #2867. Best score 1237 bits
Score difference with first non-orthologous sequence - *A. fumigatus*:1237 *A. fischeri*:1237
afum|EAL88990.1 100.00% afis|EAW16365.1 100.00%
Bootstrap support for afum|EAL88990.1 as seed ortholog is 100%.
Bootstrap support for afis|EAW16365.1 as seed ortholog is 100%.

---

Group of orthologs #2868. Best score 1237 bits
Score difference with first non-orthologous sequence - *A. fumigatus*:1237 *A. fischeri*:1237
afum|EAL92164.1 100.00% afis|EAW21145.1 100.00%
Bootstrap support for afum|EAL92164.1 as seed ortholog is 100%.
Bootstrap support for afis|EAW21145.1 as seed ortholog is 100%.

---

Group of orthologs #2869. Best score 1236 bits
Score difference with first non-orthologous sequence - *A. fumigatus*:627 *A. fischeri*:709
afum|EAL88822.1 100.00% afis|EAW16191.1 100.00%
Bootstrap support for afum|EAL88822.1 as seed ortholog is 100%.
Bootstrap support for afis|EAW16191.1 as seed ortholog is 100%.

---

Group of orthologs #2870. Best score 1236 bits
Score difference with first non-orthologous sequence - *A. fumigatus*:1028 *A. fischeri*:978
afum|EAL85050.1 100.00% afis|EAW19844.1 100.00%
Bootstrap support for afum|EAL85050.1 as seed ortholog is 100%.
Bootstrap support for afis|EAW19844.1 as seed ortholog is 100%.

---

Group of orthologs #2871. Best score 1236 bits
Score difference with first non-orthologous sequence - *A. fumigatus*:1236 *A. fischeri*:1236
afum|EAL84610.2 100.00% afis|EAW20442.1 100.00%
Bootstrap support for afum|EAL84610.2 as seed ortholog is 100%.
Bootstrap support for afis|EAW20442.1 as seed ortholog is 100%.

---

Group of orthologs #2872. Best score 1236 bits

Score difference with first non-orthologous sequence - *A. fumigatus*:1236 *A. fischeri*:1236
afum|EAL91934.1 100.00% afis|EAW24079.1 100.00%
Bootstrap support for afum|EAL91934.1 as seed ortholog is 100%.
Bootstrap support for afis|EAW24079.1 as seed ortholog is 100%.

---

Group of orthologs #2873. Best score 1235 bits
Score difference with first non-orthologous sequence - *A. fumigatus*:1053 *A. fischeri*:1235
afum|EAL87222.1 100.00% afis|EAW23788.1 100.00%
Bootstrap support for afum|EAL87222.1 as seed ortholog is 100%.
Bootstrap support for afis|EAW23788.1 as seed ortholog is 100%.

---

Group of orthologs #2874. Best score 1235 bits
Score difference with first non-orthologous sequence - *A. fumigatus*:1235 *A. fischeri*:1235
afum|EAL87453.1 100.00% afis|EAW24022.1 100.00%
Bootstrap support for afum|EAL87453.1 as seed ortholog is 100%.
Bootstrap support for afis|EAW24022.1 as seed ortholog is 100%.

---

Group of orthologs #2875. Best score 1235 bits
Score difference with first non-orthologous sequence - *A. fumigatus*:1235 *A. fischeri*:1235
afum|EAL88042.1 100.00% afis|EAW24585.1 100.00%
Bootstrap support for afum|EAL88042.1 as seed ortholog is 100%.
Bootstrap support for afis|EAW24585.1 as seed ortholog is 100%.

---

Group of orthologs #2876. Best score 1235 bits
Score difference with first non-orthologous sequence - *A. fumigatus*:1235 *A. fischeri*:1235
afum|EAL90764.1 100.00% afis|EAW22427.1 100.00%
Bootstrap support for afum|EAL90764.1 as seed ortholog is 100%.
Bootstrap support for afis|EAW22427.1 as seed ortholog is 100%.

---

Group of orthologs #2877. Best score 1234 bits
Score difference with first non-orthologous sequence - *A. fumigatus*:1080 *A. fischeri*:1234
afum|EAL85083.1 100.00% afis|EAW19809.1 100.00%
Bootstrap support for afum|EAL85083.1 as seed ortholog is 100%.
Bootstrap support for afis|EAW19809.1 as seed ortholog is 100%.

---

Group of orthologs #2878. Best score 1234 bits
Score difference with first non-orthologous sequence - *A. fumigatus*:1234 *A. fischeri*:1234
afum|EAL84559.1 100.00% afis|EAW20495.1 100.00%
Bootstrap support for afum|EAL84559.1 as seed ortholog is 100%.
Bootstrap support for afis|EAW20495.1 as seed ortholog is 100%.

---

Group of orthologs #2879. Best score 1234 bits
Score difference with first non-orthologous sequence - *A. fumigatus*:1234 *A. fischeri*:1234
afum|EAL89066.1 100.00% afis|EAW16472.1 100.00%
Bootstrap support for afum|EAL89066.1 as seed ortholog is 100%.
Bootstrap support for afis|EAW16472.1 as seed ortholog is 100%.

---

Group of orthologs #2880. Best score 1234 bits
Score difference with first non-orthologous sequence - *A. fumigatus*:1234 *A. fischeri*:1234
afum|EAL87059.1 100.00% afis|EAW19436.1 100.00%
Bootstrap support for afum|EAL87059.1 as seed ortholog is 100%.
Bootstrap support for afis|EAW19436.1 as seed ortholog is 100%.

---

Group of orthologs #2881. Best score 1234 bits
Score difference with first non-orthologous sequence - *A. fumigatus*:1234 *A. fischeri*:1234
afum|EAL85919.1 100.00% afis|EAW24250.1 100.00%

Bootstrap support for afum|EAL85919.1 as seed ortholog is 100%.
Bootstrap support for afis|EAW24250.1 as seed ortholog is 100%.

---

Group of orthologs #2882. Best score 1234 bits
Score difference with first non-orthologous sequence - *A. fumigatus*:1234 *A. fischeri*:1234
afum|EAL92084.1 100.00% afis|EAW21048.1 100.00%
Bootstrap support for afum|EAL92084.1 as seed ortholog is 100%.
Bootstrap support for afis|EAW21048.1 as seed ortholog is 100%.

---

Group of orthologs #2883. Best score 1234 bits
Score difference with first non-orthologous sequence - *A. fumigatus*:1234 *A. fischeri*:1234
afum|EAL90319.1 100.00% afis|EAW22880.1 100.00%
Bootstrap support for afum|EAL90319.1 as seed ortholog is 100%.
Bootstrap support for afis|EAW22880.1 as seed ortholog is 100%.

---

Group of orthologs #2884. Best score 1234 bits
Score difference with first non-orthologous sequence - *A. fumigatus*:1112 *A. fischeri*:1099
afum|EAL89443.1 100.00% afis|EAW24802.1 100.00%
Bootstrap support for afum|EAL89443.1 as seed ortholog is 100%.
Bootstrap support for afis|EAW24802.1 as seed ortholog is 100%.

---

Group of orthologs #2885. Best score 1234 bits
Score difference with first non-orthologous sequence - *A. fumigatus*:1234 *A. fischeri*:1234
afum|EAL90005.2 100.00% afis|EAW25342.1 100.00%
Bootstrap support for afum|EAL90005.2 as seed ortholog is 100%.
Bootstrap support for afis|EAW25342.1 as seed ortholog is 100%.

---

Group of orthologs #2886. Best score 1233 bits
Score difference with first non-orthologous sequence - *A. fumigatus*:1030 *A. fischeri*:957
afum|EAL86487.2 100.00% afis|EAW16961.1 100.00%
Bootstrap support for afum|EAL86487.2 as seed ortholog is 100%.
Bootstrap support for afis|EAW16961.1 as seed ortholog is 100%.

---

Group of orthologs #2887. Best score 1233 bits
Score difference with first non-orthologous sequence - *A. fumigatus*:1233 *A. fischeri*:1140
afum|EAL86840.1 100.00% afis|EAW19682.1 100.00%
Bootstrap support for afum|EAL86840.1 as seed ortholog is 100%.
Bootstrap support for afis|EAW19682.1 as seed ortholog is 100%.

---

Group of orthologs #2888. Best score 1233 bits
Score difference with first non-orthologous sequence - *A. fumigatus*:1233 *A. fischeri*:1233
afum|EAL92851.1 100.00% afis|EAW21837.1 100.00%
Bootstrap support for afum|EAL92851.1 as seed ortholog is 100%.
Bootstrap support for afis|EAW21837.1 as seed ortholog is 100%.

---

Group of orthologs #2889. Best score 1232 bits
Score difference with first non-orthologous sequence - *A. fumigatus*:1232 *A. fischeri*:1232
afum|EAL88589.1 100.00% afis|EAW15956.1 100.00%
Bootstrap support for afum|EAL88589.1 as seed ortholog is 100%.
Bootstrap support for afis|EAW15956.1 as seed ortholog is 100%.

---

Group of orthologs #2890. Best score 1232 bits
Score difference with first non-orthologous sequence - *A. fumigatus*:1232 *A. fischeri*:1232
afum|EAL88692.1 100.00% afis|EAW16059.1 100.00%
Bootstrap support for afum|EAL88692.1 as seed ortholog is 100%.
Bootstrap support for afis|EAW16059.1 as seed ortholog is 100%.

Group of orthologs #2891. Best score 1232 bits
Score difference with first non-orthologous sequence - *A. fumigatus*:1232 *A. fischeri*:1232
afum|EAL89098.1 100.00% afis|EAW16503.1 100.00%
Bootstrap support for afum|EAL89098.1 as seed ortholog is 100%.
Bootstrap support for afis|EAW16503.1 as seed ortholog is 100%.

Group of orthologs #2892. Best score 1232 bits
Score difference with first non-orthologous sequence - *A. fumigatus*:1163 *A. fischeri*:1135
afum|EAL91074.1 100.00% afis|EAW22110.1 100.00%
Bootstrap support for afum|EAL91074.1 as seed ortholog is 100%.
Bootstrap support for afis|EAW22110.1 as seed ortholog is 100%.

Group of orthologs #2893. Best score 1232 bits
Score difference with first non-orthologous sequence - *A. fumigatus*:1232 *A. fischeri*:1232
afum|EAL90545.1 100.00% afis|EAW22647.1 100.00%
Bootstrap support for afum|EAL90545.1 as seed ortholog is 100%.
Bootstrap support for afis|EAW22647.1 as seed ortholog is 100%.

Group of orthologs #2894. Best score 1232 bits
Score difference with first non-orthologous sequence - *A. fumigatus*:1232 *A. fischeri*:1232
afum|EAL92492.1 100.00% afis|EAW21473.1 100.00%
Bootstrap support for afum|EAL92492.1 as seed ortholog is 100%.
Bootstrap support for afis|EAW21473.1 as seed ortholog is 100%.

Group of orthologs #2895. Best score 1232 bits
Score difference with first non-orthologous sequence - *A. fumigatus*:1232 *A. fischeri*:1232
afum|EAL89685.1 100.00% afis|EAW25011.1 100.00%
Bootstrap support for afum|EAL89685.1 as seed ortholog is 100%.
Bootstrap support for afis|EAW25011.1 as seed ortholog is 100%.

Group of orthologs #2896. Best score 1231 bits
Score difference with first non-orthologous sequence - *A. fumigatus*:1231 *A. fischeri*:1231
afum|EAL85076.1 100.00% afis|EAW19815.1 100.00%
Bootstrap support for afum|EAL85076.1 as seed ortholog is 100%.
Bootstrap support for afis|EAW19815.1 as seed ortholog is 100%.

Group of orthologs #2897. Best score 1231 bits
Score difference with first non-orthologous sequence - *A. fumigatus*:869 *A. fischeri*:1231
afum|EAL86153.1 100.00% afis|EAW24484.1 100.00%
Bootstrap support for afum|EAL86153.1 as seed ortholog is 100%.
Bootstrap support for afis|EAW24484.1 as seed ortholog is 100%.

Group of orthologs #2898. Best score 1230 bits
Score difference with first non-orthologous sequence - *A. fumigatus*:1230 *A. fischeri*:1230
afum|EAL84907.1 100.00% afis|EAW20011.1 100.00%
Bootstrap support for afum|EAL84907.1 as seed ortholog is 100%.
Bootstrap support for afis|EAW20011.1 as seed ortholog is 100%.

Group of orthologs #2899. Best score 1230 bits
Score difference with first non-orthologous sequence - *A. fumigatus*:986 *A. fischeri*:947
afum|EAL88993.1 100.00% afis|EAW16368.1 100.00%
Bootstrap support for afum|EAL88993.1 as seed ortholog is 100%.
Bootstrap support for afis|EAW16368.1 as seed ortholog is 100%.

Group of orthologs #2900. Best score 1230 bits

Score difference with first non-orthologous sequence - *A. fumigatus*:1067   *A. fischeri*:1230
afum|EAL89277.1 100.00% afis|EAW16672.1 100.00%
Bootstrap support for afum|EAL89277.1 as seed ortholog is 100%.
Bootstrap support for afis|EAW16672.1 as seed ortholog is 100%.

---

Group of orthologs #2901. Best score 1230 bits
Score difference with first non-orthologous sequence - *A. fumigatus*:1230   *A. fischeri*:1230
afum|EAL87662.1 100.00% afis|EAW18296.1 100.00%
Bootstrap support for afum|EAL87662.1 as seed ortholog is 100%.
Bootstrap support for afis|EAW18296.1 as seed ortholog is 100%.

---

Group of orthologs #2902. Best score 1230 bits
Score difference with first non-orthologous sequence - *A. fumigatus*:1230   *A. fischeri*:1230
afum|EAL86927.1 100.00% afis|EAW19591.1 100.00%
Bootstrap support for afum|EAL86927.1 as seed ortholog is 100%.
Bootstrap support for afis|EAW19591.1 as seed ortholog is 100%.

---

Group of orthologs #2903. Best score 1230 bits
Score difference with first non-orthologous sequence - *A. fumigatus*:1230   *A. fischeri*:1230
afum|EAL91233.1 100.00% afis|EAW17395.1 100.00%
Bootstrap support for afum|EAL91233.1 as seed ortholog is 100%.
Bootstrap support for afis|EAW17395.1 as seed ortholog is 100%.

---

Group of orthologs #2904. Best score 1230 bits
Score difference with first non-orthologous sequence - *A. fumigatus*:1230   *A. fischeri*:1230
afum|EAL91749.1 100.00% afis|EAW17938.1 100.00%
Bootstrap support for afum|EAL91749.1 as seed ortholog is 100%.
Bootstrap support for afis|EAW17938.1 as seed ortholog is 100%.

---

Group of orthologs #2905. Best score 1230 bits
Score difference with first non-orthologous sequence - *A. fumigatus*:1230   *A. fischeri*:1230
afum|EAL93408.2 100.00% afis|EAW18687.1 100.00%
Bootstrap support for afum|EAL93408.2 as seed ortholog is 100%.
Bootstrap support for afis|EAW18687.1 as seed ortholog is 100%.

---

Group of orthologs #2906. Best score 1230 bits
Score difference with first non-orthologous sequence - *A. fumigatus*:1230   *A. fischeri*:1230
afum|EAL92678.1 100.00% afis|EAW21644.1 100.00%
Bootstrap support for afum|EAL92678.1 as seed ortholog is 100%.
Bootstrap support for afis|EAW21644.1 as seed ortholog is 100%.

---

Group of orthologs #2907. Best score 1229 bits
Score difference with first non-orthologous sequence - *A. fumigatus*:1050   *A. fischeri*:1229
afum|EAL84220.1 100.00% afis|EAW15424.1 100.00%
Bootstrap support for afum|EAL84220.1 as seed ortholog is 100%.
Bootstrap support for afis|EAW15424.1 as seed ortholog is 100%.

---

Group of orthologs #2908. Best score 1229 bits
Score difference with first non-orthologous sequence - *A. fumigatus*:1229   *A. fischeri*:1229
afum|EAL84897.1 100.00% afis|EAW20020.1 100.00%
Bootstrap support for afum|EAL84897.1 as seed ortholog is 100%.
Bootstrap support for afis|EAW20020.1 as seed ortholog is 100%.

---

Group of orthologs #2909. Best score 1229 bits
Score difference with first non-orthologous sequence - *A. fumigatus*:929   *A. fischeri*:1229
afum|EAL86210.1 100.00% afis|EAW24546.1 100.00%

Bootstrap support for afum|EAL86210.1 as seed ortholog is 100%.
Bootstrap support for afis|EAW24546.1 as seed ortholog is 100%.

---

Group of orthologs #2910. Best score 1229 bits
Score difference with first non-orthologous sequence - *A. fumigatus*:1110 *A. fischeri*:1140
afum|EAL87948.1 100.00% afis|EAW23515.1 100.00%
Bootstrap support for afum|EAL87948.1 as seed ortholog is 100%.
Bootstrap support for afis|EAW23515.1 as seed ortholog is 100%.

---

Group of orthologs #2911. Best score 1229 bits
Score difference with first non-orthologous sequence - *A. fumigatus*:1229 *A. fischeri*:1229
afum|EAL92269.1 100.00% afis|EAW21254.1 100.00%
Bootstrap support for afum|EAL92269.1 as seed ortholog is 100%.
Bootstrap support for afis|EAW21254.1 as seed ortholog is 100%.

---

Group of orthologs #2912. Best score 1228 bits
Score difference with first non-orthologous sequence - *A. fumigatus*:1228 *A. fischeri*:1228
afum|EAL88053.2 100.00% afis|EAW23416.1 100.00%
Bootstrap support for afum|EAL88053.2 as seed ortholog is 100%.
Bootstrap support for afis|EAW23416.1 as seed ortholog is 100%.

---

Group of orthologs #2913. Best score 1228 bits
Score difference with first non-orthologous sequence - *A. fumigatus*:1228 *A. fischeri*:1228
afum|EAL92389.1 100.00% afis|EAW21372.1 100.00%
Bootstrap support for afum|EAL92389.1 as seed ortholog is 100%.
Bootstrap support for afis|EAW21372.1 as seed ortholog is 100%.

---

Group of orthologs #2914. Best score 1227 bits
Score difference with first non-orthologous sequence - *A. fumigatus*:1227 *A. fischeri*:1227
afum|EAL89187.1 100.00% afis|EAW16591.1 100.00%
Bootstrap support for afum|EAL89187.1 as seed ortholog is 100%.
Bootstrap support for afis|EAW16591.1 as seed ortholog is 100%.

---

Group of orthologs #2915. Best score 1227 bits
Score difference with first non-orthologous sequence - *A. fumigatus*:993 *A. fischeri*:858
afum|EAL86961.1 100.00% afis|EAW19548.1 100.00%
Bootstrap support for afum|EAL86961.1 as seed ortholog is 100%.
Bootstrap support for afis|EAW19548.1 as seed ortholog is 100%.

---

Group of orthologs #2916. Best score 1227 bits
Score difference with first non-orthologous sequence - *A. fumigatus*:796 *A. fischeri*:807
afum|EAL87905.1 100.00% afis|EAW23563.1 100.00%
Bootstrap support for afum|EAL87905.1 as seed ortholog is 100%.
Bootstrap support for afis|EAW23563.1 as seed ortholog is 100%.

---

Group of orthologs #2917. Best score 1227 bits
Score difference with first non-orthologous sequence - *A. fumigatus*:1227 *A. fischeri*:1227
afum|EAL92802.1 100.00% afis|EAW21773.1 100.00%
Bootstrap support for afum|EAL92802.1 as seed ortholog is 100%.
Bootstrap support for afis|EAW21773.1 as seed ortholog is 100%.

---

Group of orthologs #2918. Best score 1226 bits
Score difference with first non-orthologous sequence - *A. fumigatus*:1226 *A. fischeri*:1226
afum|EAL91646.1 100.00% afis|EAW17836.1 100.00%
Bootstrap support for afum|EAL91646.1 as seed ortholog is 100%.
Bootstrap support for afis|EAW17836.1 as seed ortholog is 100%.

---

Group of orthologs #2919. Best score 1226 bits
Score difference with first non-orthologous sequence - *A. fumigatus*:1226 *A. fischeri*:1226
afum|EAL91738.1 100.00% afis|EAW17927.1 100.00%
Bootstrap support for afum|EAL91738.1 as seed ortholog is 100%.
Bootstrap support for afis|EAW17927.1 as seed ortholog is 100%.

---

Group of orthologs #2920. Best score 1226 bits
Score difference with first non-orthologous sequence - *A. fumigatus*:1226 *A. fischeri*:966
afum|EAL84284.1 100.00% afis|EAW25560.1 100.00%
Bootstrap support for afum|EAL84284.1 as seed ortholog is 100%.
Bootstrap support for afis|EAW25560.1 as seed ortholog is 100%.

---

Group of orthologs #2921. Best score 1226 bits
Score difference with first non-orthologous sequence - *A. fumigatus*:1226 *A. fischeri*:1226
afum|EAL92608.1 100.00% afis|EAW21590.1 100.00%
Bootstrap support for afum|EAL92608.1 as seed ortholog is 100%.
Bootstrap support for afis|EAW21590.1 as seed ortholog is 100%.

---

Group of orthologs #2922. Best score 1225 bits
Score difference with first non-orthologous sequence - *A. fumigatus*:957 *A. fischeri*:1078
afum|EAL86490.1 100.00% afis|EAW16963.1 100.00%
Bootstrap support for afum|EAL86490.1 as seed ortholog is 100%.
Bootstrap support for afis|EAW16963.1 as seed ortholog is 100%.

---

Group of orthologs #2923. Best score 1225 bits
Score difference with first non-orthologous sequence - *A. fumigatus*:34 *A. fischeri*:1225
afum|EAL85180.1 100.00% afis|EAW20060.1 100.00%
Bootstrap support for afum|EAL85180.1 as seed ortholog is 91%.
Bootstrap support for afis|EAW20060.1 as seed ortholog is 100%.

---

Group of orthologs #2924. Best score 1225 bits
Score difference with first non-orthologous sequence - *A. fumigatus*:1225 *A. fischeri*:1225
afum|EAL84676.1 100.00% afis|EAW20957.1 100.00%
Bootstrap support for afum|EAL84676.1 as seed ortholog is 100%.
Bootstrap support for afis|EAW20957.1 as seed ortholog is 100%.

---

Group of orthologs #2925. Best score 1225 bits
Score difference with first non-orthologous sequence - *A. fumigatus*:1225 *A. fischeri*:1225
afum|EAL93359.1 100.00% afis|EAW18644.1 100.00%
Bootstrap support for afum|EAL93359.1 as seed ortholog is 100%.
Bootstrap support for afis|EAW18644.1 as seed ortholog is 100%.

---

Group of orthologs #2926. Best score 1225 bits
Score difference with first non-orthologous sequence - *A. fumigatus*:1074 *A. fischeri*:1225
afum|EAL90861.1 100.00% afis|EAW22331.1 100.00%
Bootstrap support for afum|EAL90861.1 as seed ortholog is 100%.
Bootstrap support for afis|EAW22331.1 as seed ortholog is 100%.

---

Group of orthologs #2927. Best score 1225 bits
Score difference with first non-orthologous sequence - *A. fumigatus*:1225 *A. fischeri*:1225
afum|EAL90275.1 100.00% afis|EAW22922.1 100.00%
Bootstrap support for afum|EAL90275.1 as seed ortholog is 100%.
Bootstrap support for afis|EAW22922.1 as seed ortholog is 100%.

---

Group of orthologs #2928. Best score 1225 bits

Score difference with first non-orthologous sequence - *A. fumigatus*:1225 *A. fischeri*:1067
afum|EAL89317.1 100.00% afis|EAW24653.1 100.00%
Bootstrap support for afum|EAL89317.1 as seed ortholog is 100%.
Bootstrap support for afis|EAW24653.1 as seed ortholog is 100%.

---

Group of orthologs #2929. Best score 1225 bits
Score difference with first non-orthologous sequence - *A. fumigatus*:1225 *A. fischeri*:1225
afum|EAL89674.1 100.00% afis|EAW25021.1 100.00%
Bootstrap support for afum|EAL89674.1 as seed ortholog is 100%.
Bootstrap support for afis|EAW25021.1 as seed ortholog is 100%.

---

Group of orthologs #2930. Best score 1223 bits
Score difference with first non-orthologous sequence - *A. fumigatus*:1223 *A. fischeri*:1223
afum|EAL85209.1 100.00% afis|EAW20090.1 100.00%
Bootstrap support for afum|EAL85209.1 as seed ortholog is 100%.
Bootstrap support for afis|EAW20090.1 as seed ortholog is 100%.

---

Group of orthologs #2931. Best score 1223 bits
Score difference with first non-orthologous sequence - *A. fumigatus*:1223 *A. fischeri*:1223
afum|EAL87626.1 100.00% afis|EAW18262.1 100.00%
Bootstrap support for afum|EAL87626.1 as seed ortholog is 100%.
Bootstrap support for afis|EAW18262.1 as seed ortholog is 100%.

---

Group of orthologs #2932. Best score 1223 bits
Score difference with first non-orthologous sequence - *A. fumigatus*:1223 *A. fischeri*:1223
afum|EAL91326.1 100.00% afis|EAW17504.1 100.00%
Bootstrap support for afum|EAL91326.1 as seed ortholog is 100%.
Bootstrap support for afis|EAW17504.1 as seed ortholog is 100%.

---

Group of orthologs #2933. Best score 1223 bits
Score difference with first non-orthologous sequence - *A. fumigatus*:1223 *A. fischeri*:1223
afum|EAL91551.1 100.00% afis|EAW17736.1 100.00%
Bootstrap support for afum|EAL91551.1 as seed ortholog is 100%.
Bootstrap support for afis|EAW17736.1 as seed ortholog is 100%.

---

Group of orthologs #2934. Best score 1223 bits
Score difference with first non-orthologous sequence - *A. fumigatus*:1223 *A. fischeri*:1223
afum|EAL91639.1 100.00% afis|EAW17829.1 100.00%
Bootstrap support for afum|EAL91639.1 as seed ortholog is 100%.
Bootstrap support for afis|EAW17829.1 as seed ortholog is 100%.

---

Group of orthologs #2935. Best score 1223 bits
Score difference with first non-orthologous sequence - *A. fumigatus*:1096 *A. fischeri*:1095
afum|EAL93187.1 100.00% afis|EAW18497.1 100.00%
Bootstrap support for afum|EAL93187.1 as seed ortholog is 100%.
Bootstrap support for afis|EAW18497.1 as seed ortholog is 100%.

---

Group of orthologs #2936. Best score 1223 bits
Score difference with first non-orthologous sequence - *A. fumigatus*:1223 *A. fischeri*:1223
afum|EAL92186.1 100.00% afis|EAW21167.1 100.00%
Bootstrap support for afum|EAL92186.1 as seed ortholog is 100%.
Bootstrap support for afis|EAW21167.1 as seed ortholog is 100%.

---

Group of orthologs #2937. Best score 1223 bits
Score difference with first non-orthologous sequence - *A. fumigatus*:1223 *A. fischeri*:1223
afum|EAL92199.1 100.00% afis|EAW21179.1 100.00%

Bootstrap support for afum|EAL92199.1 as seed ortholog is 100%.
Bootstrap support for afis|EAW21179.1 as seed ortholog is 100%.

---

Group of orthologs #2938. Best score 1223 bits
Score difference with first non-orthologous sequence - *A. fumigatus*:1223 *A. fischeri*:1223
afum|EAL92729.1 100.00% afis|EAW21695.1 100.00%
Bootstrap support for afum|EAL92729.1 as seed ortholog is 100%.
Bootstrap support for afis|EAW21695.1 as seed ortholog is 100%.

---

Group of orthologs #2939. Best score 1223 bits
Score difference with first non-orthologous sequence - *A. fumigatus*:1223 *A. fischeri*:1223
afum|EAL93023.1 100.00% afis|EAW22008.1 100.00%
Bootstrap support for afum|EAL93023.1 as seed ortholog is 100%.
Bootstrap support for afis|EAW22008.1 as seed ortholog is 100%.

---

Group of orthologs #2940. Best score 1222 bits
Score difference with first non-orthologous sequence - *A. fumigatus*:996 *A. fischeri*:949
afum|EAL86562.1 100.00% afis|EAW17076.1 100.00%
Bootstrap support for afum|EAL86562.1 as seed ortholog is 100%.
Bootstrap support for afis|EAW17076.1 as seed ortholog is 100%.

---

Group of orthologs #2941. Best score 1222 bits
Score difference with first non-orthologous sequence - *A. fumigatus*:832 *A. fischeri*:1137
afum|EAL87523.1 100.00% afis|EAW18163.1 100.00%
Bootstrap support for afum|EAL87523.1 as seed ortholog is 100%.
Bootstrap support for afis|EAW18163.1 as seed ortholog is 100%.

---

Group of orthologs #2942. Best score 1222 bits
Score difference with first non-orthologous sequence - *A. fumigatus*:855 *A. fischeri*:1222
afum|EAL85421.1 100.00% afis|EAW20290.1 100.00%
Bootstrap support for afum|EAL85421.1 as seed ortholog is 100%.
Bootstrap support for afis|EAW20290.1 as seed ortholog is 100%.

---

Group of orthologs #2943. Best score 1222 bits
Score difference with first non-orthologous sequence - *A. fumigatus*:1222 *A. fischeri*:1222
afum|EAL87706.1 100.00% afis|EAW18343.1 100.00%
Bootstrap support for afum|EAL87706.1 as seed ortholog is 100%.
Bootstrap support for afis|EAW18343.1 as seed ortholog is 100%.

---

Group of orthologs #2944. Best score 1222 bits
Score difference with first non-orthologous sequence - *A. fumigatus*:1222 *A. fischeri*:1222
afum|EAL90914.1 100.00% afis|EAW22266.1 100.00%
Bootstrap support for afum|EAL90914.1 as seed ortholog is 100%.
Bootstrap support for afis|EAW22266.1 as seed ortholog is 100%.

---

Group of orthologs #2945. Best score 1222 bits
Score difference with first non-orthologous sequence - *A. fumigatus*:1222 *A. fischeri*:1222
afum|EAL92551.1 100.00% afis|EAW21530.1 100.00%
Bootstrap support for afum|EAL92551.1 as seed ortholog is 100%.
Bootstrap support for afis|EAW21530.1 as seed ortholog is 100%.

---

Group of orthologs #2946. Best score 1222 bits
Score difference with first non-orthologous sequence - *A. fumigatus*:1222 *A. fischeri*:1222
afum|EAL89633.1 100.00% afis|EAW24973.1 100.00%
Bootstrap support for afum|EAL89633.1 as seed ortholog is 100%.
Bootstrap support for afis|EAW24973.1 as seed ortholog is 100%.

Group of orthologs #2947. Best score 1221 bits
Score difference with first non-orthologous sequence - *A. fumigatus*:1221 *A. fischeri*:1138
afum|EAL88571.1 100.00% afis|EAW15935.1 100.00%
Bootstrap support for afum|EAL88571.1 as seed ortholog is 100%.
Bootstrap support for afis|EAW15935.1 as seed ortholog is 100%.

Group of orthologs #2948. Best score 1221 bits
Score difference with first non-orthologous sequence - *A. fumigatus*:1221 *A. fischeri*:897
afum|EAL88818.2 100.00% afis|EAW16187.1 100.00%
Bootstrap support for afum|EAL88818.2 as seed ortholog is 100%.
Bootstrap support for afis|EAW16187.1 as seed ortholog is 100%.

Group of orthologs #2949. Best score 1221 bits
Score difference with first non-orthologous sequence - *A. fumigatus*:1221 *A. fischeri*:1221
afum|EAL89012.1 100.00% afis|EAW16384.1 100.00%
Bootstrap support for afum|EAL89012.1 as seed ortholog is 100%.
Bootstrap support for afis|EAW16384.1 as seed ortholog is 100%.

Group of orthologs #2950. Best score 1221 bits
Score difference with first non-orthologous sequence - *A. fumigatus*:1030 *A. fischeri*:999
afum|EAL87308.1 100.00% afis|EAW23880.1 100.00%
Bootstrap support for afum|EAL87308.1 as seed ortholog is 100%.
Bootstrap support for afis|EAW23880.1 as seed ortholog is 100%.

Group of orthologs #2951. Best score 1221 bits
Score difference with first non-orthologous sequence - *A. fumigatus*:1118 *A. fischeri*:1105
afum|EAL92132.1 100.00% afis|EAW21110.1 100.00%
Bootstrap support for afum|EAL92132.1 as seed ortholog is 100%.
Bootstrap support for afis|EAW21110.1 as seed ortholog is 100%.

Group of orthologs #2952. Best score 1221 bits
Score difference with first non-orthologous sequence - *A. fumigatus*:1016 *A. fischeri*:1176
afum|EAL94082.1 100.00% afis|EAW19362.1 100.00%
Bootstrap support for afum|EAL94082.1 as seed ortholog is 100%.
Bootstrap support for afis|EAW19362.1 as seed ortholog is 100%.

Group of orthologs #2953. Best score 1220 bits
Score difference with first non-orthologous sequence - *A. fumigatus*:1220 *A. fischeri*:1048
afum|EAL86609.1 100.00% afis|EAW17179.1 100.00%
Bootstrap support for afum|EAL86609.1 as seed ortholog is 100%.
Bootstrap support for afis|EAW17179.1 as seed ortholog is 100%.

Group of orthologs #2954. Best score 1220 bits
Score difference with first non-orthologous sequence - *A. fumigatus*:1167 *A. fischeri*:1028
afum|EAL93088.1 100.00% afis|EAW18408.1 100.00%
Bootstrap support for afum|EAL93088.1 as seed ortholog is 100%.
Bootstrap support for afis|EAW18408.1 as seed ortholog is 100%.

Group of orthologs #2955. Best score 1220 bits
Score difference with first non-orthologous sequence - *A. fumigatus*:1220 *A. fischeri*:1220
afum|EAL93241.1 100.00% afis|EAW18541.1 100.00%
Bootstrap support for afum|EAL93241.1 as seed ortholog is 100%.
Bootstrap support for afis|EAW18541.1 as seed ortholog is 100%.

Group of orthologs #2956. Best score 1220 bits

Score difference with first non-orthologous sequence - *A. fumigatus*:1220   *A. fischeri*:1220
afum|EAL93453.1 100.00% afis|EAW18728.1 100.00%
Bootstrap support for afum|EAL93453.1 as seed ortholog is 100%.
Bootstrap support for afis|EAW18728.1 as seed ortholog is 100%.

---

Group of orthologs #2957. Best score 1220 bits
Score difference with first non-orthologous sequence - *A. fumigatus*:1220   *A. fischeri*:1220
afum|EAL90822.2 100.00% afis|EAW22370.1 100.00%
Bootstrap support for afum|EAL90822.2 as seed ortholog is 100%.
Bootstrap support for afis|EAW22370.1 as seed ortholog is 100%.

---

Group of orthologs #2958. Best score 1220 bits
Score difference with first non-orthologous sequence - *A. fumigatus*:1220   *A. fischeri*:1220
afum|EAL92319.1 100.00% afis|EAW21302.1 100.00%
Bootstrap support for afum|EAL92319.1 as seed ortholog is 100%.
Bootstrap support for afis|EAW21302.1 as seed ortholog is 100%.

---

Group of orthologs #2959. Best score 1220 bits
Score difference with first non-orthologous sequence - *A. fumigatus*:1220   *A. fischeri*:1220
afum|EAL89500.1 100.00% afis|EAW24853.1 100.00%
Bootstrap support for afum|EAL89500.1 as seed ortholog is 100%.
Bootstrap support for afis|EAW24853.1 as seed ortholog is 100%.

---

Group of orthologs #2960. Best score 1219 bits
Score difference with first non-orthologous sequence - *A. fumigatus*:1028   *A. fischeri*:875
afum|EAL86627.2 100.00% afis|EAW17200.1 100.00%
Bootstrap support for afum|EAL86627.2 as seed ortholog is 100%.
Bootstrap support for afis|EAW17200.1 as seed ortholog is 100%.

---

Group of orthologs #2961. Best score 1219 bits
Score difference with first non-orthologous sequence - *A. fumigatus*:1219   *A. fischeri*:1219
afum|EAL85090.1 100.00% afis|EAW19800.1 100.00%
Bootstrap support for afum|EAL85090.1 as seed ortholog is 100%.
Bootstrap support for afis|EAW19800.1 as seed ortholog is 100%.

---

Group of orthologs #2962. Best score 1219 bits
Score difference with first non-orthologous sequence - *A. fumigatus*:1219   *A. fischeri*:1219
afum|EAL84579.1 100.00% afis|EAW20473.1 100.00%
Bootstrap support for afum|EAL84579.1 as seed ortholog is 100%.
Bootstrap support for afis|EAW20473.1 as seed ortholog is 100%.

---

Group of orthologs #2963. Best score 1219 bits
Score difference with first non-orthologous sequence - *A. fumigatus*:1219   *A. fischeri*:1219
afum|EAL89111.2 100.00% afis|EAW16516.1 100.00%
Bootstrap support for afum|EAL89111.2 as seed ortholog is 100%.
Bootstrap support for afis|EAW16516.1 as seed ortholog is 100%.

---

Group of orthologs #2964. Best score 1219 bits
Score difference with first non-orthologous sequence - *A. fumigatus*:1219   *A. fischeri*:1219
afum|EAL86082.1 100.00% afis|EAW24413.1 100.00%
Bootstrap support for afum|EAL86082.1 as seed ortholog is 100%.
Bootstrap support for afis|EAW24413.1 as seed ortholog is 100%.

---

Group of orthologs #2965. Best score 1219 bits
Score difference with first non-orthologous sequence - *A. fumigatus*:974   *A. fischeri*:1219
afum|EAL87235.2 100.00% afis|EAW23801.1 100.00%

Bootstrap support for afum|EAL87235.2 as seed ortholog is 100%.
Bootstrap support for afis|EAW23801.1 as seed ortholog is 100%.

---

Group of orthologs #2966. Best score 1219 bits
Score difference with first non-orthologous sequence - *A. fumigatus*:1219 *A. fischeri*:1219
afum|EAL90968.1 100.00% afis|EAW22226.1 100.00%
Bootstrap support for afum|EAL90968.1 as seed ortholog is 100%.
Bootstrap support for afis|EAW22226.1 as seed ortholog is 100%.

---

Group of orthologs #2967. Best score 1219 bits
Score difference with first non-orthologous sequence - *A. fumigatus*:1219 *A. fischeri*:1219
afum|EAL90235.1 100.00% afis|EAW22962.1 100.00%
Bootstrap support for afum|EAL90235.1 as seed ortholog is 100%.
Bootstrap support for afis|EAW22962.1 as seed ortholog is 100%.

---

Group of orthologs #2968. Best score 1218 bits
Score difference with first non-orthologous sequence - *A. fumigatus*:1218 *A. fischeri*:1218
afum|EAL86582.1 100.00% afis|EAW17121.1 100.00%
Bootstrap support for afum|EAL86582.1 as seed ortholog is 100%.
Bootstrap support for afis|EAW17121.1 as seed ortholog is 100%.

---

Group of orthologs #2969. Best score 1218 bits
Score difference with first non-orthologous sequence - *A. fumigatus*:1218 *A. fischeri*:1218
afum|EAL88607.1 100.00% afis|EAW15975.1 100.00%
Bootstrap support for afum|EAL88607.1 as seed ortholog is 100%.
Bootstrap support for afis|EAW15975.1 as seed ortholog is 100%.

---

Group of orthologs #2970. Best score 1218 bits
Score difference with first non-orthologous sequence - *A. fumigatus*:1218 *A. fischeri*:1218
afum|EAL89298.1 100.00% afis|EAW16707.1 100.00%
Bootstrap support for afum|EAL89298.1 as seed ortholog is 100%.
Bootstrap support for afis|EAW16707.1 as seed ortholog is 100%.

---

Group of orthologs #2971. Best score 1218 bits
Score difference with first non-orthologous sequence - *A. fumigatus*:1218 *A. fischeri*:1218
afum|EAL93259.1 100.00% afis|EAW18559.1 100.00%
Bootstrap support for afum|EAL93259.1 as seed ortholog is 100%.
Bootstrap support for afis|EAW18559.1 as seed ortholog is 100%.

---

Group of orthologs #2972. Best score 1218 bits
Score difference with first non-orthologous sequence - *A. fumigatus*:1218 *A. fischeri*:1218
afum|EAL90896.1 100.00% afis|EAW22285.1 100.00%
Bootstrap support for afum|EAL90896.1 as seed ortholog is 100%.
Bootstrap support for afis|EAW22285.1 as seed ortholog is 100%.

---

Group of orthologs #2973. Best score 1218 bits
Score difference with first non-orthologous sequence - *A. fumigatus*:1218 *A. fischeri*:1218
afum|EAL90617.1 100.00% afis|EAW22582.1 100.00%
Bootstrap support for afum|EAL90617.1 as seed ortholog is 100%.
Bootstrap support for afis|EAW22582.1 as seed ortholog is 100%.

---

Group of orthologs #2974. Best score 1218 bits
Score difference with first non-orthologous sequence - *A. fumigatus*:1218 *A. fischeri*:1218
afum|EAL89548.1 100.00% afis|EAW24894.1 100.00%
Bootstrap support for afum|EAL89548.1 as seed ortholog is 100%.
Bootstrap support for afis|EAW24894.1 as seed ortholog is 100%.

---

Group of orthologs #2975. Best score 1217 bits
Score difference with first non-orthologous sequence - *A. fumigatus*:937 *A. fischeri*:808
afum|EAL86573.1 100.00% afis|EAW17109.1 100.00%
Bootstrap support for afum|EAL86573.1 as seed ortholog is 100%.
Bootstrap support for afis|EAW17109.1 as seed ortholog is 100%.

---

Group of orthologs #2976. Best score 1217 bits
Score difference with first non-orthologous sequence - *A. fumigatus*:1217 *A. fischeri*:1217
afum|EAL90970.1 100.00% afis|EAW22224.1 100.00%
Bootstrap support for afum|EAL90970.1 as seed ortholog is 100%.
Bootstrap support for afis|EAW22224.1 as seed ortholog is 100%.

---

Group of orthologs #2977. Best score 1217 bits
Score difference with first non-orthologous sequence - *A. fumigatus*:1126 *A. fischeri*:1016
afum|EAL90511.1 100.00% afis|EAW22681.1 100.00%
Bootstrap support for afum|EAL90511.1 as seed ortholog is 100%.
Bootstrap support for afis|EAW22681.1 as seed ortholog is 100%.

---

Group of orthologs #2978. Best score 1217 bits
Score difference with first non-orthologous sequence - *A. fumigatus*:1217 *A. fischeri*:873
afum|EAL90462.1 100.00% afis|EAW22733.1 100.00%
Bootstrap support for afum|EAL90462.1 as seed ortholog is 100%.
Bootstrap support for afis|EAW22733.1 as seed ortholog is 100%.

---

Group of orthologs #2979. Best score 1216 bits
Score difference with first non-orthologous sequence - *A. fumigatus*:1216 *A. fischeri*:1216
afum|EAL91830.1 100.00% afis|EAW18018.1 100.00%
Bootstrap support for afum|EAL91830.1 as seed ortholog is 100%.
Bootstrap support for afis|EAW18018.1 as seed ortholog is 100%.

---

Group of orthologs #2980. Best score 1216 bits
Score difference with first non-orthologous sequence - *A. fumigatus*:880 *A. fischeri*:1069
afum|EAL93131.1 100.00% afis|EAW18446.1 100.00%
Bootstrap support for afum|EAL93131.1 as seed ortholog is 100%.
Bootstrap support for afis|EAW18446.1 as seed ortholog is 100%.

---

Group of orthologs #2981. Best score 1216 bits
Score difference with first non-orthologous sequence - *A. fumigatus*:1216 *A. fischeri*:1216
afum|EAL93301.1 100.00% afis|EAW18593.1 100.00%
Bootstrap support for afum|EAL93301.1 as seed ortholog is 100%.
Bootstrap support for afis|EAW18593.1 as seed ortholog is 100%.

---

Group of orthologs #2982. Best score 1216 bits
Score difference with first non-orthologous sequence - *A. fumigatus*:1216 *A. fischeri*:1216
afum|EAL92759.1 100.00% afis|EAW21726.1 100.00%
Bootstrap support for afum|EAL92759.1 as seed ortholog is 100%.
Bootstrap support for afis|EAW21726.1 as seed ortholog is 100%.

---

Group of orthologs #2983. Best score 1216 bits
Score difference with first non-orthologous sequence - *A. fumigatus*:1216 *A. fischeri*:150
afum|EAL89552.2 100.00% afis|EAW24898.1 100.00%
Bootstrap support for afum|EAL89552.2 as seed ortholog is 100%.
Bootstrap support for afis|EAW24898.1 as seed ortholog is 100%.

---

Group of orthologs #2984. Best score 1215 bits

Score difference with first non-orthologous sequence - *A. fumigatus*:807 *A. fischeri*:1215
afum|EAL91605.2 100.00% afis|EAW17797.1 100.00%
Bootstrap support for afum|EAL91605.2 as seed ortholog is 100%.
Bootstrap support for afis|EAW17797.1 as seed ortholog is 100%.

---

Group of orthologs #2985. Best score 1215 bits
Score difference with first non-orthologous sequence - *A. fumigatus*:1215 *A. fischeri*:1215
afum|EAL87297.1 100.00% afis|EAW23869.1 100.00%
Bootstrap support for afum|EAL87297.1 as seed ortholog is 100%.
Bootstrap support for afis|EAW23869.1 as seed ortholog is 100%.

---

Group of orthologs #2986. Best score 1215 bits
Score difference with first non-orthologous sequence - *A. fumigatus*:1215 *A. fischeri*:1215
afum|EAL90454.1 100.00% afis|EAW22741.1 100.00%
Bootstrap support for afum|EAL90454.1 as seed ortholog is 100%.
Bootstrap support for afis|EAW22741.1 as seed ortholog is 100%.

---

Group of orthologs #2987. Best score 1214 bits
Score difference with first non-orthologous sequence - *A. fumigatus*:1214 *A. fischeri*:1214
afum|EAL88262.1 100.00% afis|EAW23222.1 100.00%
Bootstrap support for afum|EAL88262.1 as seed ortholog is 100%.
Bootstrap support for afis|EAW23222.1 as seed ortholog is 100%.

---

Group of orthologs #2988. Best score 1214 bits
Score difference with first non-orthologous sequence - *A. fumigatus*:1214 *A. fischeri*:1214
afum|EAL93432.1 100.00% afis|EAW18709.1 100.00%
Bootstrap support for afum|EAL93432.1 as seed ortholog is 100%.
Bootstrap support for afis|EAW18709.1 as seed ortholog is 100%.

---

Group of orthologs #2989. Best score 1214 bits
Score difference with first non-orthologous sequence - *A. fumigatus*:1214 *A. fischeri*:1214
afum|EAL93439.1 100.00% afis|EAW18714.1 100.00%
Bootstrap support for afum|EAL93439.1 as seed ortholog is 100%.
Bootstrap support for afis|EAW18714.1 as seed ortholog is 100%.

---

Group of orthologs #2990. Best score 1214 bits
Score difference with first non-orthologous sequence - *A. fumigatus*:1214 *A. fischeri*:1066
afum|EAL89314.2 100.00% afis|EAW24656.1 100.00%
Bootstrap support for afum|EAL89314.2 as seed ortholog is 100%.
Bootstrap support for afis|EAW24656.1 as seed ortholog is 100%.

---

Group of orthologs #2991. Best score 1214 bits
Score difference with first non-orthologous sequence - *A. fumigatus*:1214 *A. fischeri*:1214
afum|EAL92691.1 100.00% afis|EAW21656.1 100.00%
Bootstrap support for afum|EAL92691.1 as seed ortholog is 100%.
Bootstrap support for afis|EAW21656.1 as seed ortholog is 100%.

---

Group of orthologs #2992. Best score 1213 bits
Score difference with first non-orthologous sequence - *A. fumigatus*:1213 *A. fischeri*:1213
afum|EAL87475.1 100.00% afis|EAW18117.1 100.00%
Bootstrap support for afum|EAL87475.1 as seed ortholog is 100%.
Bootstrap support for afis|EAW18117.1 as seed ortholog is 100%.

---

Group of orthologs #2993. Best score 1213 bits
Score difference with first non-orthologous sequence - *A. fumigatus*:924 *A. fischeri*:1213
afum|EAL91553.1 100.00% afis|EAW17738.1 100.00%

Bootstrap support for afum|EAL91553.1 as seed ortholog is 100%.
Bootstrap support for afis|EAW17738.1 as seed ortholog is 100%.

---

Group of orthologs #2994. Best score 1213 bits
Score difference with first non-orthologous sequence - *A. fumigatus*:1213 *A. fischeri*:1213
afum|EAL85907.1 100.00% afis|EAW24239.1 100.00%
Bootstrap support for afum|EAL85907.1 as seed ortholog is 100%.
Bootstrap support for afis|EAW24239.1 as seed ortholog is 100%.

---

Group of orthologs #2995. Best score 1213 bits
Score difference with first non-orthologous sequence - *A. fumigatus*:1213 *A. fischeri*:1067
afum|EAL86149.1 100.00% afis|EAW24476.1 100.00%
Bootstrap support for afum|EAL86149.1 as seed ortholog is 100%.
Bootstrap support for afis|EAW24476.1 as seed ortholog is 100%.

---

Group of orthologs #2996. Best score 1213 bits
Score difference with first non-orthologous sequence - *A. fumigatus*:1213 *A. fischeri*:1213
afum|EAL93281.1 100.00% afis|EAW18578.1 100.00%
Bootstrap support for afum|EAL93281.1 as seed ortholog is 100%.
Bootstrap support for afis|EAW18578.1 as seed ortholog is 100%.

---

Group of orthologs #2997. Best score 1213 bits
Score difference with first non-orthologous sequence - *A. fumigatus*:1213 *A. fischeri*:1213
afum|EAL93618.1 100.00% afis|EAW18885.1 100.00%
Bootstrap support for afum|EAL93618.1 as seed ortholog is 100%.
Bootstrap support for afis|EAW18885.1 as seed ortholog is 100%.

---

Group of orthologs #2998. Best score 1213 bits
Score difference with first non-orthologous sequence - *A. fumigatus*:1213 *A. fischeri*:1213
afum|EAL93020.1 100.00% afis|EAW22005.1 100.00%
Bootstrap support for afum|EAL93020.1 as seed ortholog is 100%.
Bootstrap support for afis|EAW22005.1 as seed ortholog is 100%.

---

Group of orthologs #2999. Best score 1212 bits
Score difference with first non-orthologous sequence - *A. fumigatus*:1212 *A. fischeri*:1212
afum|EAL87103.1 100.00% afis|EAW19391.1 100.00%
Bootstrap support for afum|EAL87103.1 as seed ortholog is 100%.
Bootstrap support for afis|EAW19391.1 as seed ortholog is 100%.

---

Group of orthologs #3000. Best score 1212 bits
Score difference with first non-orthologous sequence - *A. fumigatus*:1005 *A. fischeri*:1120
afum|EAL87301.1 100.00% afis|EAW23873.1 100.00%
Bootstrap support for afum|EAL87301.1 as seed ortholog is 100%.
Bootstrap support for afis|EAW23873.1 as seed ortholog is 100%.

---

Group of orthologs #3001. Best score 1212 bits
Score difference with first non-orthologous sequence - *A. fumigatus*:1212 *A. fischeri*:1212
afum|EAL87322.1 100.00% afis|EAW23894.1 100.00%
Bootstrap support for afum|EAL87322.1 as seed ortholog is 100%.
Bootstrap support for afis|EAW23894.1 as seed ortholog is 100%.

---

Group of orthologs #3002. Best score 1212 bits
Score difference with first non-orthologous sequence - *A. fumigatus*:1212 *A. fischeri*:1212
afum|EAL92843.1 100.00% afis|EAW21828.1 100.00%
Bootstrap support for afum|EAL92843.1 as seed ortholog is 100%.
Bootstrap support for afis|EAW21828.1 as seed ortholog is 100%.

---

Group of orthologs #3003. Best score 1211 bits
Score difference with first non-orthologous sequence - *A. fumigatus*:926 *A. fischeri*:762
afum|EAL84529.1 100.00% afis|EAW20528.1 100.00%
Bootstrap support for afum|EAL84529.1 as seed ortholog is 100%.
Bootstrap support for afis|EAW20528.1 as seed ortholog is 100%.

---

Group of orthologs #3004. Best score 1211 bits
Score difference with first non-orthologous sequence - *A. fumigatus*:997 *A. fischeri*:976
afum|EAL84541.1 100.00% afis|EAW20516.1 100.00%
Bootstrap support for afum|EAL84541.1 as seed ortholog is 100%.
Bootstrap support for afis|EAW20516.1 as seed ortholog is 100%.

---

Group of orthologs #3005. Best score 1211 bits
Score difference with first non-orthologous sequence - *A. fumigatus*:1211 *A. fischeri*:1211
afum|EAL85253.1 100.00% afis|EAW20138.1 100.00%
Bootstrap support for afum|EAL85253.1 as seed ortholog is 100%.
Bootstrap support for afis|EAW20138.1 as seed ortholog is 100%.

---

Group of orthologs #3006. Best score 1211 bits
Score difference with first non-orthologous sequence - *A. fumigatus*:1211 *A. fischeri*:1211
afum|EAL87307.1 100.00% afis|EAW23879.1 100.00%
Bootstrap support for afum|EAL87307.1 as seed ortholog is 100%.
Bootstrap support for afis|EAW23879.1 as seed ortholog is 100%.

---

Group of orthologs #3007. Best score 1211 bits
Score difference with first non-orthologous sequence - *A. fumigatus*:1211 *A. fischeri*:1211
afum|EAL87886.1 100.00% afis|EAW23579.1 100.00%
Bootstrap support for afum|EAL87886.1 as seed ortholog is 100%.
Bootstrap support for afis|EAW23579.1 as seed ortholog is 100%.

---

Group of orthologs #3008. Best score 1211 bits
Score difference with first non-orthologous sequence - *A. fumigatus*:1211 *A. fischeri*:1211
afum|EAL87439.1 100.00% afis|EAW24009.1 100.00%
Bootstrap support for afum|EAL87439.1 as seed ortholog is 100%.
Bootstrap support for afis|EAW24009.1 as seed ortholog is 100%.

---

Group of orthologs #3009. Best score 1211 bits
Score difference with first non-orthologous sequence - *A. fumigatus*:1048 *A. fischeri*:957
afum|EAL90998.1 100.00% afis|EAW22193.1 100.00%
Bootstrap support for afum|EAL90998.1 as seed ortholog is 100%.
Bootstrap support for afis|EAW22193.1 as seed ortholog is 100%.

---

Group of orthologs #3010. Best score 1211 bits
Score difference with first non-orthologous sequence - *A. fumigatus*:1211 *A. fischeri*:1211
afum|EAL90347.1 100.00% afis|EAW22853.1 100.00%
Bootstrap support for afum|EAL90347.1 as seed ortholog is 100%.
Bootstrap support for afis|EAW22853.1 as seed ortholog is 100%.

---

Group of orthologs #3011. Best score 1211 bits
Score difference with first non-orthologous sequence - *A. fumigatus*:1055 *A. fischeri*:1050
afum|EAL92407.1 100.00% afis|EAW21393.1 100.00%
Bootstrap support for afum|EAL92407.1 as seed ortholog is 100%.
Bootstrap support for afis|EAW21393.1 as seed ortholog is 100%.

---

Group of orthologs #3012. Best score 1211 bits

Score difference with first non-orthologous sequence - *A. fumigatus*:1211 *A. fischeri*:1211
afum|EAL92898.1 100.00% afis|EAW21884.1 100.00%
Bootstrap support for afum|EAL92898.1 as seed ortholog is 100%.
Bootstrap support for afis|EAW21884.1 as seed ortholog is 100%.

---

Group of orthologs #3013. Best score 1210 bits
Score difference with first non-orthologous sequence - *A. fumigatus*:1210 *A. fischeri*:1210
afum|EAL88882.1 100.00% afis|EAW16261.1 100.00%
Bootstrap support for afum|EAL88882.1 as seed ortholog is 100%.
Bootstrap support for afis|EAW16261.1 as seed ortholog is 100%.

---

Group of orthologs #3014. Best score 1210 bits
Score difference with first non-orthologous sequence - *A. fumigatus*:1003 *A. fischeri*:775
afum|EAL87892.2 100.00% afis|EAW23573.1 100.00%
Bootstrap support for afum|EAL87892.2 as seed ortholog is 100%.
Bootstrap support for afis|EAW23573.1 as seed ortholog is 100%.

---

Group of orthologs #3015. Best score 1210 bits
Score difference with first non-orthologous sequence - *A. fumigatus*:1210 *A. fischeri*:1210
afum|EAL94009.1 100.00% afis|EAW19283.1 100.00%
Bootstrap support for afum|EAL94009.1 as seed ortholog is 100%.
Bootstrap support for afis|EAW19283.1 as seed ortholog is 100%.

---

Group of orthologs #3016. Best score 1209 bits
Score difference with first non-orthologous sequence - *A. fumigatus*:1209 *A. fischeri*:1209
afum|EAL86672.2 100.00% afis|EAW17253.1 100.00%
Bootstrap support for afum|EAL86672.2 as seed ortholog is 100%.
Bootstrap support for afis|EAW17253.1 as seed ortholog is 100%.

---

Group of orthologs #3017. Best score 1209 bits
Score difference with first non-orthologous sequence - *A. fumigatus*:1209 *A. fischeri*:1115
afum|EAL84770.1 100.00% afis|EAW20865.1 100.00%
Bootstrap support for afum|EAL84770.1 as seed ortholog is 100%.
Bootstrap support for afis|EAW20865.1 as seed ortholog is 100%.

---

Group of orthologs #3018. Best score 1209 bits
Score difference with first non-orthologous sequence - *A. fumigatus*:1209 *A. fischeri*:1209
afum|EAL91637.1 100.00% afis|EAW17827.1 100.00%
Bootstrap support for afum|EAL91637.1 as seed ortholog is 100%.
Bootstrap support for afis|EAW17827.1 as seed ortholog is 100%.

---

Group of orthologs #3019. Best score 1209 bits
Score difference with first non-orthologous sequence - *A. fumigatus*:1209 *A. fischeri*:580
afum|EAL87192.1 100.00% afis|EAW23756.1 100.00%
Bootstrap support for afum|EAL87192.1 as seed ortholog is 100%.
Bootstrap support for afis|EAW23756.1 as seed ortholog is 100%.

---

Group of orthologs #3020. Best score 1209 bits
Score difference with first non-orthologous sequence - *A. fumigatus*:1209 *A. fischeri*:1209
afum|EAL88371.1 100.00% afis|EAW23108.1 100.00%
Bootstrap support for afum|EAL88371.1 as seed ortholog is 100%.
Bootstrap support for afis|EAW23108.1 as seed ortholog is 100%.

---

Group of orthologs #3021. Best score 1209 bits
Score difference with first non-orthologous sequence - *A. fumigatus*:1209 *A. fischeri*:1209
afum|EAL88316.1 100.00% afis|EAW23168.1 100.00%

Bootstrap support for afum|EAL88316.1 as seed ortholog is 100%.
Bootstrap support for afis|EAW23168.1 as seed ortholog is 100%.

---

Group of orthologs #3022. Best score 1209 bits
Score difference with first non-orthologous sequence - *A. fumigatus*:1056 *A. fischeri*:927
afum|EAL92170.1 100.00% afis|EAW21151.1 100.00%
Bootstrap support for afum|EAL92170.1 as seed ortholog is 100%.
Bootstrap support for afis|EAW21151.1 as seed ortholog is 100%.

---

Group of orthologs #3023. Best score 1208 bits
Score difference with first non-orthologous sequence - *A. fumigatus*:1208 *A. fischeri*:1208
afum|EAL85222.1 100.00% afis|EAW20105.1 100.00%
Bootstrap support for afum|EAL85222.1 as seed ortholog is 100%.
Bootstrap support for afis|EAW20105.1 as seed ortholog is 100%.

---

Group of orthologs #3024. Best score 1208 bits
Score difference with first non-orthologous sequence - *A. fumigatus*:1208 *A. fischeri*:1208
afum|EAL91217.1 100.00% afis|EAW17382.1 100.00%
Bootstrap support for afum|EAL91217.1 as seed ortholog is 100%.
Bootstrap support for afis|EAW17382.1 as seed ortholog is 100%.

---

Group of orthologs #3025. Best score 1208 bits
Score difference with first non-orthologous sequence - *A. fumigatus*:1119 *A. fischeri*:934
afum|EAL91320.1 100.00% afis|EAW17497.1 100.00%
Bootstrap support for afum|EAL91320.1 as seed ortholog is 100%.
Bootstrap support for afis|EAW17497.1 as seed ortholog is 100%.

---

Group of orthologs #3026. Best score 1208 bits
Score difference with first non-orthologous sequence - *A. fumigatus*:1208 *A. fischeri*:1208
afum|EAL86062.1 100.00% afis|EAW24390.1 100.00%
Bootstrap support for afum|EAL86062.1 as seed ortholog is 100%.
Bootstrap support for afis|EAW24390.1 as seed ortholog is 100%.

---

Group of orthologs #3027. Best score 1208 bits
Score difference with first non-orthologous sequence - *A. fumigatus*:981 *A. fischeri*:1055
afum|EAL90801.1 100.00% afis|EAW22389.1 100.00%
Bootstrap support for afum|EAL90801.1 as seed ortholog is 100%.
Bootstrap support for afis|EAW22389.1 as seed ortholog is 100%.

---

Group of orthologs #3028. Best score 1207 bits
Score difference with first non-orthologous sequence - *A. fumigatus*:1103 *A. fischeri*:1075
afum|EBA27218.1 100.00% afis|EAW19822.1 100.00%
Bootstrap support for afum|EBA27218.1 as seed ortholog is 100%.
Bootstrap support for afis|EAW19822.1 as seed ortholog is 100%.

---

Group of orthologs #3029. Best score 1207 bits
Score difference with first non-orthologous sequence - *A. fumigatus*:1207 *A. fischeri*:1207
afum|EAL89558.1 100.00% afis|EAW24905.1 100.00%
Bootstrap support for afum|EAL89558.1 as seed ortholog is 100%.
Bootstrap support for afis|EAW24905.1 as seed ortholog is 100%.

---

Group of orthologs #3030. Best score 1206 bits
Score difference with first non-orthologous sequence - *A. fumigatus*:1003 *A. fischeri*:1110
afum|EAL87492.1 100.00% afis|EAW18135.1 100.00%
Bootstrap support for afum|EAL87492.1 as seed ortholog is 100%.
Bootstrap support for afis|EAW18135.1 as seed ortholog is 100%.

Group of orthologs #3031. Best score 1206 bits
Score difference with first non-orthologous sequence - *A. fumigatus*:1206 *A. fischeri*:1206
afum|EAL93383.1 100.00% afis|EAW18665.1 100.00%
Bootstrap support for afum|EAL93383.1 as seed ortholog is 100%.
Bootstrap support for afis|EAW18665.1 as seed ortholog is 100%.

Group of orthologs #3032. Best score 1206 bits
Score difference with first non-orthologous sequence - *A. fumigatus*:1206 *A. fischeri*:1206
afum|EAL90802.1 100.00% afis|EAW22388.1 100.00%
Bootstrap support for afum|EAL90802.1 as seed ortholog is 100%.
Bootstrap support for afis|EAW22388.1 as seed ortholog is 100%.

Group of orthologs #3033. Best score 1206 bits
Score difference with first non-orthologous sequence - *A. fumigatus*:1206 *A. fischeri*:1206
afum|EAL92567.1 100.00% afis|EAW21547.1 100.00%
Bootstrap support for afum|EAL92567.1 as seed ortholog is 100%.
Bootstrap support for afis|EAW21547.1 as seed ortholog is 100%.

Group of orthologs #3034. Best score 1206 bits
Score difference with first non-orthologous sequence - *A. fumigatus*:1206 *A. fischeri*:1206
afum|EAL90104.1 100.00% afis|EAW25446.1 100.00%
Bootstrap support for afum|EAL90104.1 as seed ortholog is 100%.
Bootstrap support for afis|EAW25446.1 as seed ortholog is 100%.

Group of orthologs #3035. Best score 1205 bits
Score difference with first non-orthologous sequence - *A. fumigatus*:1051 *A. fischeri*:842
afum|EAL86168.1 100.00% afis|EAW24498.1 100.00%
Bootstrap support for afum|EAL86168.1 as seed ortholog is 100%.
Bootstrap support for afis|EAW24498.1 as seed ortholog is 100%.

Group of orthologs #3036. Best score 1205 bits
Score difference with first non-orthologous sequence - *A. fumigatus*:1205 *A. fischeri*:1205
afum|EAL90742.1 100.00% afis|EAW22449.1 100.00%
Bootstrap support for afum|EAL90742.1 as seed ortholog is 100%.
Bootstrap support for afis|EAW22449.1 as seed ortholog is 100%.

Group of orthologs #3037. Best score 1205 bits
Score difference with first non-orthologous sequence - *A. fumigatus*:1066 *A. fischeri*:979
afum|EAL89840.1 100.00% afis|EAW25167.1 100.00%
Bootstrap support for afum|EAL89840.1 as seed ortholog is 100%.
Bootstrap support for afis|EAW25167.1 as seed ortholog is 100%.

Group of orthologs #3038. Best score 1205 bits
Score difference with first non-orthologous sequence - *A. fumigatus*:934 *A. fischeri*:921
afum|EAL89982.1 100.00% afis|EAW25321.1 100.00%
Bootstrap support for afum|EAL89982.1 as seed ortholog is 100%.
Bootstrap support for afis|EAW25321.1 as seed ortholog is 100%.

Group of orthologs #3039. Best score 1204 bits
Score difference with first non-orthologous sequence - *A. fumigatus*:975 *A. fischeri*:1060
afum|EAL86466.1 100.00% afis|EAW16939.1 100.00%
Bootstrap support for afum|EAL86466.1 as seed ortholog is 100%.
Bootstrap support for afis|EAW16939.1 as seed ortholog is 100%.

Group of orthologs #3040. Best score 1204 bits

Score difference with first non-orthologous sequence - *A. fumigatus*:988   *A. fischeri*:1028
afum|EAL85190.1 100.00% afis|EAW20072.1 100.00%
Bootstrap support for afum|EAL85190.1 as seed ortholog is 100%.
Bootstrap support for afis|EAW20072.1 as seed ortholog is 100%.

---

Group of orthologs #3041. Best score 1204 bits
Score difference with first non-orthologous sequence - *A. fumigatus*:1204   *A. fischeri*:1204
afum|EAL85319.1 100.00% afis|EAW20199.1 100.00%
Bootstrap support for afum|EAL85319.1 as seed ortholog is 100%.
Bootstrap support for afis|EAW20199.1 as seed ortholog is 100%.

---

Group of orthologs #3042. Best score 1204 bits
Score difference with first non-orthologous sequence - *A. fumigatus*:1012   *A. fischeri*:909
afum|EAL90967.1 100.00% afis|EAW22227.1 100.00%
Bootstrap support for afum|EAL90967.1 as seed ortholog is 100%.
Bootstrap support for afis|EAW22227.1 as seed ortholog is 100%.

---

Group of orthologs #3043. Best score 1204 bits
Score difference with first non-orthologous sequence - *A. fumigatus*:1204   *A. fischeri*:1204
afum|EAL89350.1 100.00% afis|EAW24708.1 100.00%
Bootstrap support for afum|EAL89350.1 as seed ortholog is 100%.
Bootstrap support for afis|EAW24708.1 as seed ortholog is 100%.

---

Group of orthologs #3044. Best score 1203 bits
Score difference with first non-orthologous sequence - *A. fumigatus*:861   *A. fischeri*:768
afum|EAL88988.1 100.00% afis|EAW16363.1 100.00%
Bootstrap support for afum|EAL88988.1 as seed ortholog is 100%.
Bootstrap support for afis|EAW16363.1 as seed ortholog is 100%.

---

Group of orthologs #3045. Best score 1203 bits
Score difference with first non-orthologous sequence - *A. fumigatus*:1203   *A. fischeri*:1203
afum|EAL93497.1 100.00% afis|EAW18765.1 100.00%
Bootstrap support for afum|EAL93497.1 as seed ortholog is 100%.
Bootstrap support for afis|EAW18765.1 as seed ortholog is 100%.

---

Group of orthologs #3046. Best score 1203 bits
Score difference with first non-orthologous sequence - *A. fumigatus*:1203   *A. fischeri*:949
afum|EAL92287.1 100.00% afis|EAW21272.1 100.00%
Bootstrap support for afum|EAL92287.1 as seed ortholog is 100%.
Bootstrap support for afis|EAW21272.1 as seed ortholog is 100%.

---

Group of orthologs #3047. Best score 1203 bits
Score difference with first non-orthologous sequence - *A. fumigatus*:1203   *A. fischeri*:1203
afum|EAL91965.1 100.00% afis|EAW24111.1 100.00%
Bootstrap support for afum|EAL91965.1 as seed ortholog is 100%.
Bootstrap support for afis|EAW24111.1 as seed ortholog is 100%.

---

Group of orthologs #3048. Best score 1202 bits
Score difference with first non-orthologous sequence - *A. fumigatus*:1202   *A. fischeri*:1202
afum|EAL84234.1 100.00% afis|EAW15412.1 100.00%
Bootstrap support for afum|EAL84234.1 as seed ortholog is 100%.
Bootstrap support for afis|EAW15412.1 as seed ortholog is 100%.

---

Group of orthologs #3049. Best score 1202 bits
Score difference with first non-orthologous sequence - *A. fumigatus*:1202   *A. fischeri*:1202
afum|EAL88931.1 100.00% afis|EAW16309.1 100.00%

Bootstrap support for afum|EAL88931.1 as seed ortholog is 100%.
Bootstrap support for afis|EAW16309.1 as seed ortholog is 100%.

---

Group of orthologs #3050. Best score 1202 bits
Score difference with first non-orthologous sequence - *A. fumigatus*:1202 *A. fischeri*:1202
afum|EAL86916.1 100.00% afis|EAW19602.1 100.00%
Bootstrap support for afum|EAL86916.1 as seed ortholog is 100%.
Bootstrap support for afis|EAW19602.1 as seed ortholog is 100%.

---

Group of orthologs #3051. Best score 1202 bits
Score difference with first non-orthologous sequence - *A. fumigatus*:1202 *A. fischeri*:1202
afum|EAL92323.1 100.00% afis|EAW21307.1 100.00%
Bootstrap support for afum|EAL92323.1 as seed ortholog is 100%.
Bootstrap support for afis|EAW21307.1 as seed ortholog is 100%.

---

Group of orthologs #3052. Best score 1202 bits
Score difference with first non-orthologous sequence - *A. fumigatus*:1202 *A. fischeri*:1057
afum|EAL89316.1 100.00% afis|EAW24654.1 100.00%
Bootstrap support for afum|EAL89316.1 as seed ortholog is 100%.
Bootstrap support for afis|EAW24654.1 as seed ortholog is 100%.

---

Group of orthologs #3053. Best score 1201 bits
Score difference with first non-orthologous sequence - *A. fumigatus*:1201 *A. fischeri*:1201
afum|EAL85591.1 100.00% afis|EAW15757.1 100.00%
Bootstrap support for afum|EAL85591.1 as seed ortholog is 100%.
Bootstrap support for afis|EAW15757.1 as seed ortholog is 100%.

---

Group of orthologs #3054. Best score 1201 bits
Score difference with first non-orthologous sequence - *A. fumigatus*:1201 *A. fischeri*:1201
afum|EAL93269.2 100.00% afis|EAW18567.1 100.00%
Bootstrap support for afum|EAL93269.2 as seed ortholog is 100%.
Bootstrap support for afis|EAW18567.1 as seed ortholog is 100%.

---

Group of orthologs #3055. Best score 1201 bits
Score difference with first non-orthologous sequence - *A. fumigatus*:1201 *A. fischeri*:1201
afum|EAL92578.1 100.00% afis|EAW21559.1 100.00%
Bootstrap support for afum|EAL92578.1 as seed ortholog is 100%.
Bootstrap support for afis|EAW21559.1 as seed ortholog is 100%.

---

Group of orthologs #3056. Best score 1200 bits
Score difference with first non-orthologous sequence - *A. fumigatus*:1039 *A. fischeri*:1200
afum|EAL84460.1 100.00% afis|EAW20601.1 100.00%
Bootstrap support for afum|EAL84460.1 as seed ortholog is 100%.
Bootstrap support for afis|EAW20601.1 as seed ortholog is 100%.

---

Group of orthologs #3057. Best score 1200 bits
Score difference with first non-orthologous sequence - *A. fumigatus*:1200 *A. fischeri*:1200
afum|EAL88384.2 100.00% afis|EAW23094.1 100.00%
Bootstrap support for afum|EAL88384.2 as seed ortholog is 100%.
Bootstrap support for afis|EAW23094.1 as seed ortholog is 100%.

---

Group of orthologs #3058. Best score 1200 bits
Score difference with first non-orthologous sequence - *A. fumigatus*:1200 *A. fischeri*:1200
afum|EAL88123.1 100.00% afis|EAW23353.1 100.00%
Bootstrap support for afum|EAL88123.1 as seed ortholog is 100%.
Bootstrap support for afis|EAW23353.1 as seed ortholog is 100%.

---

Group of orthologs #3059. Best score 1200 bits
Score difference with first non-orthologous sequence - *A. fumigatus*:1200 *A. fischeri*:1200
afum|EAL93555.1 100.00% afis|EAW18819.1 100.00%
Bootstrap support for afum|EAL93555.1 as seed ortholog is 100%.
Bootstrap support for afis|EAW18819.1 as seed ortholog is 100%.

---

Group of orthologs #3060. Best score 1200 bits
Score difference with first non-orthologous sequence - *A. fumigatus*:1200 *A. fischeri*:1200
afum|EAL90416.1 100.00% afis|EAW22783.1 100.00%
Bootstrap support for afum|EAL90416.1 as seed ortholog is 100%.
Bootstrap support for afis|EAW22783.1 as seed ortholog is 100%.

---

Group of orthologs #3061. Best score 1200 bits
Score difference with first non-orthologous sequence - *A. fumigatus*:1200 *A. fischeri*:1200
afum|EAL89850.1 100.00% afis|EAW25177.1 100.00%
Bootstrap support for afum|EAL89850.1 as seed ortholog is 100%.
Bootstrap support for afis|EAW25177.1 as seed ortholog is 100%.

---

Group of orthologs #3062. Best score 1200 bits
Score difference with first non-orthologous sequence - *A. fumigatus*:1200 *A. fischeri*:522
afum|EAL91927.1 100.00% afis|EAW24071.1 100.00%
Bootstrap support for afum|EAL91927.1 as seed ortholog is 100%.
Bootstrap support for afis|EAW24071.1 as seed ortholog is 100%.

---

Group of orthologs #3063. Best score 1199 bits
Score difference with first non-orthologous sequence - *A. fumigatus*:863 *A. fischeri*:909
afum|EAL88685.1 100.00% afis|EAW16053.1 100.00%
Bootstrap support for afum|EAL88685.1 as seed ortholog is 100%.
Bootstrap support for afis|EAW16053.1 as seed ortholog is 100%.

---

Group of orthologs #3064. Best score 1199 bits
Score difference with first non-orthologous sequence - *A. fumigatus*:881 *A. fischeri*:1080
afum|EAL85491.1 100.00% afis|EAW20377.1 100.00%
Bootstrap support for afum|EAL85491.1 as seed ortholog is 100%.
Bootstrap support for afis|EAW20377.1 as seed ortholog is 100%.

---

Group of orthologs #3065. Best score 1199 bits
Score difference with first non-orthologous sequence - *A. fumigatus*:1086 *A. fischeri*:1053
afum|EAL93046.1 100.00% afis|EAW18371.1 100.00%
Bootstrap support for afum|EAL93046.1 as seed ortholog is 100%.
Bootstrap support for afis|EAW18371.1 as seed ortholog is 100%.

---

Group of orthologs #3066. Best score 1199 bits
Score difference with first non-orthologous sequence - *A. fumigatus*:867 *A. fischeri*:649
afum|EAL87881.1 100.00% afis|EAW23583.1 100.00%
Bootstrap support for afum|EAL87881.1 as seed ortholog is 100%.
Bootstrap support for afis|EAW23583.1 as seed ortholog is 100%.

---

Group of orthologs #3067. Best score 1199 bits
Score difference with first non-orthologous sequence - *A. fumigatus*:1199 *A. fischeri*:1199
afum|EAL90744.1 100.00% afis|EAW22447.1 100.00%
Bootstrap support for afum|EAL90744.1 as seed ortholog is 100%.
Bootstrap support for afis|EAW22447.1 as seed ortholog is 100%.

---

Group of orthologs #3068. Best score 1198 bits

Score difference with first non-orthologous sequence - *A. fumigatus*:891   *A. fischeri*:1089
afum|EAL85857.1 100.00% afis|EAW15465.1 100.00%
Bootstrap support for afum|EAL85857.1 as seed ortholog is 100%.
Bootstrap support for afis|EAW15465.1 as seed ortholog is 100%.

---

Group of orthologs #3069. Best score 1198 bits
Score difference with first non-orthologous sequence - *A. fumigatus*:1198   *A. fischeri*:1198
afum|EAL91603.1 100.00% afis|EAW17795.1 100.00%
Bootstrap support for afum|EAL91603.1 as seed ortholog is 100%.
Bootstrap support for afis|EAW17795.1 as seed ortholog is 100%.

---

Group of orthologs #3070. Best score 1198 bits
Score difference with first non-orthologous sequence - *A. fumigatus*:1120   *A. fischeri*:1198
afum|EAL91629.1 100.00% afis|EAW17819.1 100.00%
Bootstrap support for afum|EAL91629.1 as seed ortholog is 100%.
Bootstrap support for afis|EAW17819.1 as seed ortholog is 100%.

---

Group of orthologs #3071. Best score 1198 bits
Score difference with first non-orthologous sequence - *A. fumigatus*:1198   *A. fischeri*:1198
afum|EAL85971.1 100.00% afis|EAW24303.1 100.00%
Bootstrap support for afum|EAL85971.1 as seed ortholog is 100%.
Bootstrap support for afis|EAW24303.1 as seed ortholog is 100%.

---

Group of orthologs #3072. Best score 1197 bits
Score difference with first non-orthologous sequence - *A. fumigatus*:880   *A. fischeri*:1067
afum|EAL84775.2 100.00% afis|EAW20860.1 100.00%
Bootstrap support for afum|EAL84775.2 as seed ortholog is 100%.
Bootstrap support for afis|EAW20860.1 as seed ortholog is 100%.

---

Group of orthologs #3073. Best score 1197 bits
Score difference with first non-orthologous sequence - *A. fumigatus*:1197   *A. fischeri*:1197
afum|EAL87600.1 100.00% afis|EAW18232.1 100.00%
Bootstrap support for afum|EAL87600.1 as seed ortholog is 100%.
Bootstrap support for afis|EAW18232.1 as seed ortholog is 100%.

---

Group of orthologs #3074. Best score 1197 bits
Score difference with first non-orthologous sequence - *A. fumigatus*:891   *A. fischeri*:543
afum|EAL87675.1 100.00% afis|EAW18309.1 100.00%
Bootstrap support for afum|EAL87675.1 as seed ortholog is 100%.
Bootstrap support for afis|EAW18309.1 as seed ortholog is 100%.

---

Group of orthologs #3075. Best score 1197 bits
Score difference with first non-orthologous sequence - *A. fumigatus*:1197   *A. fischeri*:1197
afum|EAL86106.1 100.00% afis|EAW24438.1 100.00%
Bootstrap support for afum|EAL86106.1 as seed ortholog is 100%.
Bootstrap support for afis|EAW24438.1 as seed ortholog is 100%.

---

Group of orthologs #3076. Best score 1197 bits
Score difference with first non-orthologous sequence - *A. fumigatus*:1197   *A. fischeri*:1197
afum|EBA27498.1 100.00% afis|EAW18915.1 100.00%
Bootstrap support for afum|EBA27498.1 as seed ortholog is 100%.
Bootstrap support for afis|EAW18915.1 as seed ortholog is 100%.

---

Group of orthologs #3077. Best score 1197 bits
Score difference with first non-orthologous sequence - *A. fumigatus*:1197   *A. fischeri*:1197
afum|EAL90313.1 100.00% afis|EAW22885.1 100.00%

Bootstrap support for afum|EAL90313.1 as seed ortholog is 100%.
Bootstrap support for afis|EAW22885.1 as seed ortholog is 100%.

---

Group of orthologs #3078. Best score 1197 bits
Score difference with first non-orthologous sequence - *A. fumigatus*:1197 *A. fischeri*:1197
afum|EAL92283.1 100.00% afis|EAW21268.1 100.00%
Bootstrap support for afum|EAL92283.1 as seed ortholog is 100%.
Bootstrap support for afis|EAW21268.1 as seed ortholog is 100%.

---

Group of orthologs #3079. Best score 1197 bits
Score difference with first non-orthologous sequence - *A. fumigatus*:1197 *A. fischeri*:1197
afum|EAL92819.1 100.00% afis|EAW21794.1 100.00%
Bootstrap support for afum|EAL92819.1 as seed ortholog is 100%.
Bootstrap support for afis|EAW21794.1 as seed ortholog is 100%.

---

Group of orthologs #3080. Best score 1197 bits
Score difference with first non-orthologous sequence - *A. fumigatus*:744 *A. fischeri*:446
afum|EAL92853.1 100.00% afis|EAW21841.1 100.00%
Bootstrap support for afum|EAL92853.1 as seed ortholog is 100%.
Bootstrap support for afis|EAW21841.1 as seed ortholog is 100%.

---

Group of orthologs #3081. Best score 1197 bits
Score difference with first non-orthologous sequence - *A. fumigatus*:983 *A. fischeri*:384
afum|EAL91966.1 100.00% afis|EAW24112.1 100.00%
Bootstrap support for afum|EAL91966.1 as seed ortholog is 100%.
Bootstrap support for afis|EAW24112.1 as seed ortholog is 100%.

---

Group of orthologs #3082. Best score 1196 bits
Score difference with first non-orthologous sequence - *A. fumigatus*:1149 *A. fischeri*:1003
afum|EAL89045.1 100.00% afis|EAW16444.1 100.00%
Bootstrap support for afum|EAL89045.1 as seed ortholog is 100%.
Bootstrap support for afis|EAW16444.1 as seed ortholog is 100%.

---

Group of orthologs #3083. Best score 1196 bits
Score difference with first non-orthologous sequence - *A. fumigatus*:1196 *A. fischeri*:1196
afum|EAL84693.1 100.00% afis|EAW20941.1 100.00%
Bootstrap support for afum|EAL84693.1 as seed ortholog is 100%.
Bootstrap support for afis|EAW20941.1 as seed ortholog is 100%.

---

Group of orthologs #3084. Best score 1196 bits
Score difference with first non-orthologous sequence - *A. fumigatus*:982 *A. fischeri*:933
afum|EAL87449.1 100.00% afis|EAW24019.1 100.00%
Bootstrap support for afum|EAL87449.1 as seed ortholog is 100%.
Bootstrap support for afis|EAW24019.1 as seed ortholog is 100%.

---

Group of orthologs #3085. Best score 1196 bits
Score difference with first non-orthologous sequence - *A. fumigatus*:1011 *A. fischeri*:1196
afum|EAL92432.2 100.00% afis|EAW21417.1 100.00%
Bootstrap support for afum|EAL92432.2 as seed ortholog is 100%.
Bootstrap support for afis|EAW21417.1 as seed ortholog is 100%.

---

Group of orthologs #3086. Best score 1196 bits
Score difference with first non-orthologous sequence - *A. fumigatus*:1196 *A. fischeri*:1196
afum|EAL92621.1 100.00% afis|EAW21602.1 100.00%
Bootstrap support for afum|EAL92621.1 as seed ortholog is 100%.
Bootstrap support for afis|EAW21602.1 as seed ortholog is 100%.

Group of orthologs #3087. Best score 1195 bits
Score difference with first non-orthologous sequence - *A. fumigatus*:1034 *A. fischeri*:980
afum|EAL86420.1 100.00% afis|EAW16884.1 100.00%
Bootstrap support for afum|EAL86420.1 as seed ortholog is 100%.
Bootstrap support for afis|EAW16884.1 as seed ortholog is 100%.

Group of orthologs #3088. Best score 1195 bits
Score difference with first non-orthologous sequence - *A. fumigatus*:1195 *A. fischeri*:1195
afum|EAL88155.1 100.00% afis|EAW23324.1 100.00%
Bootstrap support for afum|EAL88155.1 as seed ortholog is 100%.
Bootstrap support for afis|EAW23324.1 as seed ortholog is 100%.

Group of orthologs #3089. Best score 1194 bits
Score difference with first non-orthologous sequence - *A. fumigatus*:1194 *A. fischeri*:1194
afum|EAL85826.1 100.00% afis|EAW15497.1 100.00%
Bootstrap support for afum|EAL85826.1 as seed ortholog is 100%.
Bootstrap support for afis|EAW15497.1 as seed ortholog is 100%.

Group of orthologs #3090. Best score 1194 bits
Score difference with first non-orthologous sequence - *A. fumigatus*:889 *A. fischeri*:264
afum|EAL86664.2 100.00% afis|EAW17245.1 100.00%
Bootstrap support for afum|EAL86664.2 as seed ortholog is 100%.
Bootstrap support for afis|EAW17245.1 as seed ortholog is 100%.

Group of orthologs #3091. Best score 1194 bits
Score difference with first non-orthologous sequence - *A. fumigatus*:1194 *A. fischeri*:1194
afum|EAL88942.1 100.00% afis|EAW16318.1 100.00%
Bootstrap support for afum|EAL88942.1 as seed ortholog is 100%.
Bootstrap support for afis|EAW16318.1 as seed ortholog is 100%.

Group of orthologs #3092. Best score 1194 bits
Score difference with first non-orthologous sequence - *A. fumigatus*:1194 *A. fischeri*:874
afum|EAL84413.1 100.00% afis|EAW20685.1 100.00%
Bootstrap support for afum|EAL84413.1 as seed ortholog is 100%.
Bootstrap support for afis|EAW20685.1 as seed ortholog is 100%.

Group of orthologs #3093. Best score 1194 bits
Score difference with first non-orthologous sequence - *A. fumigatus*:1194 *A. fischeri*:1194
afum|EAL89079.1 100.00% afis|EAW16485.1 100.00%
Bootstrap support for afum|EAL89079.1 as seed ortholog is 100%.
Bootstrap support for afis|EAW16485.1 as seed ortholog is 100%.

Group of orthologs #3094. Best score 1194 bits
Score difference with first non-orthologous sequence - *A. fumigatus*:1194 *A. fischeri*:1194
afum|EAL87473.1 100.00% afis|EAW18116.1 100.00%
Bootstrap support for afum|EAL87473.1 as seed ortholog is 100%.
Bootstrap support for afis|EAW18116.1 as seed ortholog is 100%.

Group of orthologs #3095. Best score 1194 bits
Score difference with first non-orthologous sequence - *A. fumigatus*:1194 *A. fischeri*:1194
afum|EAL85366.1 100.00% afis|EAW20248.1 100.00%
Bootstrap support for afum|EAL85366.1 as seed ortholog is 100%.
Bootstrap support for afis|EAW20248.1 as seed ortholog is 100%.

Group of orthologs #3096. Best score 1194 bits

Score difference with first non-orthologous sequence - *A. fumigatus*:1194 *A. fischeri*:1194
afum|EAL87080.2 100.00% afis|EAW19414.1 100.00%
Bootstrap support for afum|EAL87080.2 as seed ortholog is 100%.
Bootstrap support for afis|EAW19414.1 as seed ortholog is 100%.

---

Group of orthologs #3097. Best score 1194 bits
Score difference with first non-orthologous sequence - *A. fumigatus*:1194 *A. fischeri*:1194
afum|EAL84381.1 100.00% afis|EAW25708.1 100.00%
Bootstrap support for afum|EAL84381.1 as seed ortholog is 100%.
Bootstrap support for afis|EAW25708.1 as seed ortholog is 100%.

---

Group of orthologs #3098. Best score 1194 bits
Score difference with first non-orthologous sequence - *A. fumigatus*:1194 *A. fischeri*:1194
afum|EAL86180.1 100.00% afis|EAW24514.1 100.00%
Bootstrap support for afum|EAL86180.1 as seed ortholog is 100%.
Bootstrap support for afis|EAW24514.1 as seed ortholog is 100%.

---

Group of orthologs #3099. Best score 1194 bits
Score difference with first non-orthologous sequence - *A. fumigatus*:1194 *A. fischeri*:1103
afum|EAL94077.1 100.00% afis|EAW19357.1 100.00%
Bootstrap support for afum|EAL94077.1 as seed ortholog is 100%.
Bootstrap support for afis|EAW19357.1 as seed ortholog is 100%.

---

Group of orthologs #3100. Best score 1194 bits
Score difference with first non-orthologous sequence - *A. fumigatus*:1116 *A. fischeri*:1050
afum|EAL89312.1 100.00% afis|EAW24658.1 100.00%
Bootstrap support for afum|EAL89312.1 as seed ortholog is 100%.
Bootstrap support for afis|EAW24658.1 as seed ortholog is 100%.

---

Group of orthologs #3101. Best score 1194 bits
Score difference with first non-orthologous sequence - *A. fumigatus*:1125 *A. fischeri*:1045
afum|EAL89411.1 100.00% afis|EAW24770.1 100.00%
Bootstrap support for afum|EAL89411.1 as seed ortholog is 100%.
Bootstrap support for afis|EAW24770.1 as seed ortholog is 100%.

---

Group of orthologs #3102. Best score 1194 bits
Score difference with first non-orthologous sequence - *A. fumigatus*:1194 *A. fischeri*:1194
afum|EAL92863.2 100.00% afis|EAW21850.1 100.00%
Bootstrap support for afum|EAL92863.2 as seed ortholog is 100%.
Bootstrap support for afis|EAW21850.1 as seed ortholog is 100%.

---

Group of orthologs #3103. Best score 1194 bits
Score difference with first non-orthologous sequence - *A. fumigatus*:1194 *A. fischeri*:1194
afum|EAL89981.1 100.00% afis|EAW25320.1 100.00%
Bootstrap support for afum|EAL89981.1 as seed ortholog is 100%.
Bootstrap support for afis|EAW25320.1 as seed ortholog is 100%.

---

Group of orthologs #3104. Best score 1193 bits
Score difference with first non-orthologous sequence - *A. fumigatus*:1049 *A. fischeri*:969
afum|EAL88824.1 100.00% afis|EAW16193.1 100.00%
Bootstrap support for afum|EAL88824.1 as seed ortholog is 100%.
Bootstrap support for afis|EAW16193.1 as seed ortholog is 100%.

---

Group of orthologs #3105. Best score 1193 bits
Score difference with first non-orthologous sequence - *A. fumigatus*:1193 *A. fischeri*:1193
afum|EAL87670.1 100.00% afis|EAW18304.1 100.00%

Bootstrap support for afum|EAL87670.1 as seed ortholog is 100%.
Bootstrap support for afis|EAW18304.1 as seed ortholog is 100%.

---

Group of orthologs #3106. Best score 1193 bits
Score difference with first non-orthologous sequence - *A. fumigatus*:1193 *A. fischeri*:1193
afum|EAL93784.1 100.00% afis|EAW19055.1 100.00%
Bootstrap support for afum|EAL93784.1 as seed ortholog is 100%.
Bootstrap support for afis|EAW19055.1 as seed ortholog is 100%.

---

Group of orthologs #3107. Best score 1193 bits
Score difference with first non-orthologous sequence - *A. fumigatus*:1193 *A. fischeri*:1193
afum|EAL92448.2 100.00% afis|EAW21433.1 100.00%
Bootstrap support for afum|EAL92448.2 as seed ortholog is 100%.
Bootstrap support for afis|EAW21433.1 as seed ortholog is 100%.

---

Group of orthologs #3108. Best score 1193 bits
Score difference with first non-orthologous sequence - *A. fumigatus*:1193 *A. fischeri*:1193
afum|EAL89498.1 100.00% afis|EAW24851.1 100.00%
Bootstrap support for afum|EAL89498.1 as seed ortholog is 100%.
Bootstrap support for afis|EAW24851.1 as seed ortholog is 100%.

---

Group of orthologs #3109. Best score 1192 bits
Score difference with first non-orthologous sequence - *A. fumigatus*:914 *A. fischeri*:850
afum|EAL86471.1 100.00% afis|EAW16944.1 100.00%
Bootstrap support for afum|EAL86471.1 as seed ortholog is 100%.
Bootstrap support for afis|EAW16944.1 as seed ortholog is 100%.

---

Group of orthologs #3110. Best score 1192 bits
Score difference with first non-orthologous sequence - *A. fumigatus*:1192 *A. fischeri*:1192
afum|EAL88581.1 100.00% afis|EAW15948.1 100.00%
Bootstrap support for afum|EAL88581.1 as seed ortholog is 100%.
Bootstrap support for afis|EAW15948.1 as seed ortholog is 100%.

---

Group of orthologs #3111. Best score 1192 bits
Score difference with first non-orthologous sequence - *A. fumigatus*:1192 *A. fischeri*:1192
afum|EAL88957.1 100.00% afis|EAW16332.1 100.00%
Bootstrap support for afum|EAL88957.1 as seed ortholog is 100%.
Bootstrap support for afis|EAW16332.1 as seed ortholog is 100%.

---

Group of orthologs #3112. Best score 1192 bits
Score difference with first non-orthologous sequence - *A. fumigatus*:26 *A. fischeri*:917
afum|EAL85177.1 100.00% afis|EAW20057.1 100.00%
Bootstrap support for afum|EAL85177.1 as seed ortholog is 87%.
Bootstrap support for afis|EAW20057.1 as seed ortholog is 100%.

---

Group of orthologs #3113. Best score 1192 bits
Score difference with first non-orthologous sequence - *A. fumigatus*:1192 *A. fischeri*:1044
afum|EAL91192.2 100.00% afis|EAW17349.1 100.00%
Bootstrap support for afum|EAL91192.2 as seed ortholog is 100%.
Bootstrap support for afis|EAW17349.1 as seed ortholog is 100%.

---

Group of orthologs #3114. Best score 1192 bits
Score difference with first non-orthologous sequence - *A. fumigatus*:1192 *A. fischeri*:1192
afum|EAL91307.1 100.00% afis|EAW17478.1 100.00%
Bootstrap support for afum|EAL91307.1 as seed ortholog is 100%.
Bootstrap support for afis|EAW17478.1 as seed ortholog is 100%.

Group of orthologs #3115. Best score 1192 bits
Score difference with first non-orthologous sequence - *A. fumigatus*:1192 *A. fischeri*:768
afum|EAL91608.1 100.00% afis|EAW17800.1 100.00%
Bootstrap support for afum|EAL91608.1 as seed ortholog is 100%.
Bootstrap support for afis|EAW17800.1 as seed ortholog is 100%.

Group of orthologs #3116. Best score 1192 bits
Score difference with first non-orthologous sequence - *A. fumigatus*:1050 *A. fischeri*:934
afum|EAL84319.1 100.00% afis|EAW25631.1 100.00%
Bootstrap support for afum|EAL84319.1 as seed ortholog is 100%.
Bootstrap support for afis|EAW25631.1 as seed ortholog is 100%.

Group of orthologs #3117. Best score 1192 bits
Score difference with first non-orthologous sequence - *A. fumigatus*:1089 *A. fischeri*:1007
afum|EAL91054.1 100.00% afis|EAW22136.1 100.00%
Bootstrap support for afum|EAL91054.1 as seed ortholog is 100%.
Bootstrap support for afis|EAW22136.1 as seed ortholog is 100%.

Group of orthologs #3118. Best score 1191 bits
Score difference with first non-orthologous sequence - *A. fumigatus*:1191 *A. fischeri*:1191
afum|EAL88630.1 100.00% afis|EAW15998.1 100.00%
Bootstrap support for afum|EAL88630.1 as seed ortholog is 100%.
Bootstrap support for afis|EAW15998.1 as seed ortholog is 100%.

Group of orthologs #3119. Best score 1191 bits
Score difference with first non-orthologous sequence - *A. fumigatus*:1191 *A. fischeri*:1191
afum|EAL85033.1 100.00% afis|EAW19862.1 100.00%
Bootstrap support for afum|EAL85033.1 as seed ortholog is 100%.
Bootstrap support for afis|EAW19862.1 as seed ortholog is 100%.

Group of orthologs #3120. Best score 1191 bits
Score difference with first non-orthologous sequence - *A. fumigatus*:1011 *A. fischeri*:986
afum|EAL84361.1 100.00% afis|EAW25691.1 100.00%
Bootstrap support for afum|EAL84361.1 as seed ortholog is 100%.
Bootstrap support for afis|EAW25691.1 as seed ortholog is 100%.

Group of orthologs #3121. Best score 1191 bits
Score difference with first non-orthologous sequence - *A. fumigatus*:1191 *A. fischeri*:1191
afum|EAL88282.1 100.00% afis|EAW23201.1 100.00%
Bootstrap support for afum|EAL88282.1 as seed ortholog is 100%.
Bootstrap support for afis|EAW23201.1 as seed ortholog is 100%.

Group of orthologs #3122. Best score 1191 bits
Score difference with first non-orthologous sequence - *A. fumigatus*:1191 *A. fischeri*:1191
afum|EAL88080.1 100.00% afis|EAW23391.1 100.00%
Bootstrap support for afum|EAL88080.1 as seed ortholog is 100%.
Bootstrap support for afis|EAW23391.1 as seed ortholog is 100%.

Group of orthologs #3123. Best score 1191 bits
Score difference with first non-orthologous sequence - *A. fumigatus*:803 *A. fischeri*:1078
afum|EAL89567.1 100.00% afis|EAW24913.1 100.00%
Bootstrap support for afum|EAL89567.1 as seed ortholog is 100%.
Bootstrap support for afis|EAW24913.1 as seed ortholog is 100%.

Group of orthologs #3124. Best score 1191 bits

Score difference with first non-orthologous sequence - *A. fumigatus*:1191 *A. fischeri*:1191
afum|EBA27336.1 100.00% afis|EAW23359.1 100.00%
Bootstrap support for afum|EBA27336.1 as seed ortholog is 100%.
Bootstrap support for afis|EAW23359.1 as seed ortholog is 100%.

---

Group of orthologs #3125. Best score 1190 bits
Score difference with first non-orthologous sequence - *A. fumigatus*:1190 *A. fischeri*:1190
afum|EAL85108.1 100.00% afis|EAW19784.1 100.00%
Bootstrap support for afum|EAL85108.1 as seed ortholog is 100%.
Bootstrap support for afis|EAW19784.1 as seed ortholog is 100%.

---

Group of orthologs #3126. Best score 1190 bits
Score difference with first non-orthologous sequence - *A. fumigatus*:1190 *A. fischeri*:1190
afum|EAL91967.1 100.00% afis|EAW24113.1 100.00%
Bootstrap support for afum|EAL91967.1 as seed ortholog is 100%.
Bootstrap support for afis|EAW24113.1 as seed ortholog is 100%.

---

Group of orthologs #3127. Best score 1189 bits
Score difference with first non-orthologous sequence - *A. fumigatus*:1189 *A. fischeri*:1189
afum|EAL88628.2 100.00% afis|EAW15996.1 100.00%
Bootstrap support for afum|EAL88628.2 as seed ortholog is 100%.
Bootstrap support for afis|EAW15996.1 as seed ortholog is 100%.

---

Group of orthologs #3128. Best score 1189 bits
Score difference with first non-orthologous sequence - *A. fumigatus*:1189 *A. fischeri*:1022
afum|EAL88980.1 100.00% afis|EAW16355.1 100.00%
Bootstrap support for afum|EAL88980.1 as seed ortholog is 100%.
Bootstrap support for afis|EAW16355.1 as seed ortholog is 100%.

---

Group of orthologs #3129. Best score 1189 bits
Score difference with first non-orthologous sequence - *A. fumigatus*:992 *A. fischeri*:889
afum|EAL93838.1 100.00% afis|EAW19107.1 100.00%
Bootstrap support for afum|EAL93838.1 as seed ortholog is 100%.
Bootstrap support for afis|EAW19107.1 as seed ortholog is 100%.

---

Group of orthologs #3130. Best score 1189 bits
Score difference with first non-orthologous sequence - *A. fumigatus*:1099 *A. fischeri*:1012
afum|EAL90951.1 100.00% afis|EAW22242.1 100.00%
Bootstrap support for afum|EAL90951.1 as seed ortholog is 100%.
Bootstrap support for afis|EAW22242.1 as seed ortholog is 100%.

---

Group of orthologs #3131. Best score 1188 bits
Score difference with first non-orthologous sequence - *A. fumigatus*:1188 *A. fischeri*:1188
afum|EAL88769.2 100.00% afis|EAW16140.1 100.00%
Bootstrap support for afum|EAL88769.2 as seed ortholog is 100%.
Bootstrap support for afis|EAW16140.1 as seed ortholog is 100%.

---

Group of orthologs #3132. Best score 1188 bits
Score difference with first non-orthologous sequence - *A. fumigatus*:1188 *A. fischeri*:1188
afum|EAL85914.1 100.00% afis|EAW24246.1 100.00%
Bootstrap support for afum|EAL85914.1 as seed ortholog is 100%.
Bootstrap support for afis|EAW24246.1 as seed ortholog is 100%.

---

Group of orthologs #3133. Best score 1188 bits
Score difference with first non-orthologous sequence - *A. fumigatus*:1188 *A. fischeri*:1188
afum|EAL85939.1 100.00% afis|EAW24271.1 100.00%

Bootstrap support for afum|EAL85939.1 as seed ortholog is 100%.
Bootstrap support for afis|EAW24271.1 as seed ortholog is 100%.

---

Group of orthologs #3134. Best score 1188 bits
Score difference with first non-orthologous sequence - *A. fumigatus*:923 *A. fischeri*:968
afum|EAL93870.1 100.00% afis|EAW19138.1 100.00%
Bootstrap support for afum|EAL93870.1 as seed ortholog is 100%.
Bootstrap support for afis|EAW19138.1 as seed ortholog is 100%.

---

Group of orthologs #3135. Best score 1188 bits
Score difference with first non-orthologous sequence - *A. fumigatus*:937 *A. fischeri*:1188
afum|EAL92777.1 100.00% afis|EAW21744.1 100.00%
Bootstrap support for afum|EAL92777.1 as seed ortholog is 100%.
Bootstrap support for afis|EAW21744.1 as seed ortholog is 100%.

---

Group of orthologs #3136. Best score 1188 bits
Score difference with first non-orthologous sequence - *A. fumigatus*:1046 *A. fischeri*:1188
afum|EAL90086.1 100.00% afis|EAW25427.1 100.00%
Bootstrap support for afum|EAL90086.1 as seed ortholog is 100%.
Bootstrap support for afis|EAW25427.1 as seed ortholog is 100%.

---

Group of orthologs #3137. Best score 1187 bits
Score difference with first non-orthologous sequence - *A. fumigatus*:1187 *A. fischeri*:1187
afum|EAL88696.1 100.00% afis|EAW16064.1 100.00%
Bootstrap support for afum|EAL88696.1 as seed ortholog is 100%.
Bootstrap support for afis|EAW16064.1 as seed ortholog is 100%.

---

Group of orthologs #3138. Best score 1187 bits
Score difference with first non-orthologous sequence - *A. fumigatus*:1187 *A. fischeri*:1187
afum|EAL84547.1 100.00% afis|EAW20509.1 100.00%
Bootstrap support for afum|EAL84547.1 as seed ortholog is 100%.
Bootstrap support for afis|EAW20509.1 as seed ortholog is 100%.

---

Group of orthologs #3139. Best score 1187 bits
Score difference with first non-orthologous sequence - *A. fumigatus*:587 *A. fischeri*:1187
afum|EAL85419.1 100.00% afis|EAW20287.1 100.00%
Bootstrap support for afum|EAL85419.1 as seed ortholog is 100%.
Bootstrap support for afis|EAW20287.1 as seed ortholog is 100%.

---

Group of orthologs #3140. Best score 1187 bits
Score difference with first non-orthologous sequence - *A. fumigatus*:1187 *A. fischeri*:1187
afum|EAL91350.1 100.00% afis|EAW17536.1 100.00%
Bootstrap support for afum|EAL91350.1 as seed ortholog is 100%.
Bootstrap support for afis|EAW17536.1 as seed ortholog is 100%.

---

Group of orthologs #3141. Best score 1187 bits
Score difference with first non-orthologous sequence - *A. fumigatus*:1187 *A. fischeri*:1187
afum|EAL93577.1 100.00% afis|EAW18846.1 100.00%
Bootstrap support for afum|EAL93577.1 as seed ortholog is 100%.
Bootstrap support for afis|EAW18846.1 as seed ortholog is 100%.

---

Group of orthologs #3142. Best score 1187 bits
Score difference with first non-orthologous sequence - *A. fumigatus*:1187 *A. fischeri*:1187
afum|EAL90871.1 100.00% afis|EAW22319.1 100.00%
Bootstrap support for afum|EAL90871.1 as seed ortholog is 100%.
Bootstrap support for afis|EAW22319.1 as seed ortholog is 100%.

---

Group of orthologs #3143. Best score 1186 bits
Score difference with first non-orthologous sequence - *A. fumigatus*:1186 *A. fischeri*:1186
afum|EAL88688.1 100.00% afis|EAW16056.1 100.00%
Bootstrap support for afum|EAL88688.1 as seed ortholog is 100%.
Bootstrap support for afis|EAW16056.1 as seed ortholog is 100%.

---

Group of orthologs #3144. Best score 1186 bits
Score difference with first non-orthologous sequence - *A. fumigatus*:1186 *A. fischeri*:1186
afum|EAL90838.1 100.00% afis|EAW22354.1 100.00%
Bootstrap support for afum|EAL90838.1 as seed ortholog is 100%.
Bootstrap support for afis|EAW22354.1 as seed ortholog is 100%.

---

Group of orthologs #3145. Best score 1186 bits
Score difference with first non-orthologous sequence - *A. fumigatus*:1186 *A. fischeri*:1186
afum|EAL92549.1 100.00% afis|EAW21528.1 100.00%
Bootstrap support for afum|EAL92549.1 as seed ortholog is 100%.
Bootstrap support for afis|EAW21528.1 as seed ortholog is 100%.

---

Group of orthologs #3146. Best score 1186 bits
Score difference with first non-orthologous sequence - *A. fumigatus*:1186 *A. fischeri*:1186
afum|EAL90045.1 100.00% afis|EAW25386.1 100.00%
Bootstrap support for afum|EAL90045.1 as seed ortholog is 100%.
Bootstrap support for afis|EAW25386.1 as seed ortholog is 100%.

---

Group of orthologs #3147. Best score 1185 bits
Score difference with first non-orthologous sequence - *A. fumigatus*:843 *A. fischeri*:1020
afum|EAL86476.1 100.00% afis|EAW16949.1 100.00%
Bootstrap support for afum|EAL86476.1 as seed ortholog is 100%.
Bootstrap support for afis|EAW16949.1 as seed ortholog is 100%.

---

Group of orthologs #3148. Best score 1185 bits
Score difference with first non-orthologous sequence - *A. fumigatus*:1185 *A. fischeri*:1185
afum|EAL87029.1 100.00% afis|EAW19467.1 100.00%
Bootstrap support for afum|EAL87029.1 as seed ortholog is 100%.
Bootstrap support for afis|EAW19467.1 as seed ortholog is 100%.

---

Group of orthologs #3149. Best score 1185 bits
Score difference with first non-orthologous sequence - *A. fumigatus*:1185 *A. fischeri*:1185
afum|EAL91402.1 100.00% afis|EAW17589.1 100.00%
Bootstrap support for afum|EAL91402.1 as seed ortholog is 100%.
Bootstrap support for afis|EAW17589.1 as seed ortholog is 100%.

---

Group of orthologs #3150. Best score 1185 bits
Score difference with first non-orthologous sequence - *A. fumigatus*:1185 *A. fischeri*:1185
afum|EAL93065.1 100.00% afis|EAW18390.1 100.00%
Bootstrap support for afum|EAL93065.1 as seed ortholog is 100%.
Bootstrap support for afis|EAW18390.1 as seed ortholog is 100%.

---

Group of orthologs #3151. Best score 1185 bits
Score difference with first non-orthologous sequence - *A. fumigatus*:984 *A. fischeri*:1034
afum|EAL90404.1 100.00% afis|EAW22795.1 100.00%
Bootstrap support for afum|EAL90404.1 as seed ortholog is 100%.
Bootstrap support for afis|EAW22795.1 as seed ortholog is 100%.

---

Group of orthologs #3152. Best score 1185 bits

Score difference with first non-orthologous sequence - *A. fumigatus*:1185 *A. fischeri*:1185
afum|EAL92163.1 100.00% afis|EAW21144.1 100.00%
Bootstrap support for afum|EAL92163.1 as seed ortholog is 100%.
Bootstrap support for afis|EAW21144.1 as seed ortholog is 100%.

---

Group of orthologs #3153. Best score 1184 bits
Score difference with first non-orthologous sequence - *A. fumigatus*:1184 *A. fischeri*:1184
afum|EAL88893.1 100.00% afis|EAW16272.1 100.00%
Bootstrap support for afum|EAL88893.1 as seed ortholog is 100%.
Bootstrap support for afis|EAW16272.1 as seed ortholog is 100%.

---

Group of orthologs #3154. Best score 1184 bits
Score difference with first non-orthologous sequence - *A. fumigatus*:1184 *A. fischeri*:1184
afum|EAL84972.1 100.00% afis|EAW19940.1 100.00%
Bootstrap support for afum|EAL84972.1 as seed ortholog is 100%.
Bootstrap support for afis|EAW19940.1 as seed ortholog is 100%.

---

Group of orthologs #3155. Best score 1184 bits
Score difference with first non-orthologous sequence - *A. fumigatus*:1184 *A. fischeri*:1184
afum|EAL87682.1 100.00% afis|EAW18315.1 100.00%
Bootstrap support for afum|EAL87682.1 as seed ortholog is 100%.
Bootstrap support for afis|EAW18315.1 as seed ortholog is 100%.

---

Group of orthologs #3156. Best score 1184 bits
Score difference with first non-orthologous sequence - *A. fumigatus*:732 *A. fischeri*:814
afum|EAL91169.1 100.00% afis|EAW17332.1 100.00%
Bootstrap support for afum|EAL91169.1 as seed ortholog is 100%.
Bootstrap support for afis|EAW17332.1 as seed ortholog is 100%.

---

Group of orthologs #3157. Best score 1184 bits
Score difference with first non-orthologous sequence - *A. fumigatus*:1184 *A. fischeri*:1184
afum|EAL91680.1 100.00% afis|EAW17872.1 100.00%
Bootstrap support for afum|EAL91680.1 as seed ortholog is 100%.
Bootstrap support for afis|EAW17872.1 as seed ortholog is 100%.

---

Group of orthologs #3158. Best score 1184 bits
Score difference with first non-orthologous sequence - *A. fumigatus*:1041 *A. fischeri*:939
afum|EAL87183.1 100.00% afis|EAW23749.1 100.00%
Bootstrap support for afum|EAL87183.1 as seed ortholog is 100%.
Bootstrap support for afis|EAW23749.1 as seed ortholog is 100%.

---

Group of orthologs #3159. Best score 1183 bits
Score difference with first non-orthologous sequence - *A. fumigatus*:1183 *A. fischeri*:1183
afum|EAL88620.1 100.00% afis|EAW15988.1 100.00%
Bootstrap support for afum|EAL88620.1 as seed ortholog is 100%.
Bootstrap support for afis|EAW15988.1 as seed ortholog is 100%.

---

Group of orthologs #3160. Best score 1183 bits
Score difference with first non-orthologous sequence - *A. fumigatus*:1183 *A. fischeri*:1183
afum|EAL91659.1 100.00% afis|EAW17850.1 100.00%
Bootstrap support for afum|EAL91659.1 as seed ortholog is 100%.
Bootstrap support for afis|EAW17850.1 as seed ortholog is 100%.

---

Group of orthologs #3161. Best score 1182 bits
Score difference with first non-orthologous sequence - *A. fumigatus*:1182 *A. fischeri*:1182
afum|EAL84218.1 100.00% afis|EAW15426.1 100.00%

Bootstrap support for afum|EAL84218.1 as seed ortholog is 100%.
Bootstrap support for afis|EAW15426.1 as seed ortholog is 100%.

---

Group of orthologs #3162. Best score 1182 bits
Score difference with first non-orthologous sequence - *A. fumigatus*:1182 *A. fischeri*:1035
afum|EAL84911.1 100.00% afis|EAW20007.1 100.00%
Bootstrap support for afum|EAL84911.1 as seed ortholog is 100%.
Bootstrap support for afis|EAW20007.1 as seed ortholog is 100%.

---

Group of orthologs #3163. Best score 1182 bits
Score difference with first non-orthologous sequence - *A. fumigatus*:1182 *A. fischeri*:1182
afum|EAL87607.1 100.00% afis|EAW18243.1 100.00%
Bootstrap support for afum|EAL87607.1 as seed ortholog is 100%.
Bootstrap support for afis|EAW18243.1 as seed ortholog is 100%.

---

Group of orthologs #3164. Best score 1182 bits
Score difference with first non-orthologous sequence - *A. fumigatus*:1182 *A. fischeri*:1182
afum|EAL91875.1 100.00% afis|EAW18063.1 100.00%
Bootstrap support for afum|EAL91875.1 as seed ortholog is 100%.
Bootstrap support for afis|EAW18063.1 as seed ortholog is 100%.

---

Group of orthologs #3165. Best score 1182 bits
Score difference with first non-orthologous sequence - *A. fumigatus*:1182 *A. fischeri*:1182
afum|EAL93192.1 100.00% afis|EAW18501.1 100.00%
Bootstrap support for afum|EAL93192.1 as seed ortholog is 100%.
Bootstrap support for afis|EAW18501.1 as seed ortholog is 100%.

---

Group of orthologs #3166. Best score 1181 bits
Score difference with first non-orthologous sequence - *A. fumigatus*:1181 *A. fischeri*:1181
afum|EAL84221.1 100.00% afis|EAW15423.1 100.00%
Bootstrap support for afum|EAL84221.1 as seed ortholog is 100%.
Bootstrap support for afis|EAW15423.1 as seed ortholog is 100%.

---

Group of orthologs #3167. Best score 1181 bits
Score difference with first non-orthologous sequence - *A. fumigatus*:1181 *A. fischeri*:1181
afum|EAL86691.1 100.00% afis|EAW17273.1 100.00%
Bootstrap support for afum|EAL86691.1 as seed ortholog is 100%.
Bootstrap support for afis|EAW17273.1 as seed ortholog is 100%.

---

Group of orthologs #3168. Best score 1181 bits
Score difference with first non-orthologous sequence - *A. fumigatus*:1181 *A. fischeri*:1181
afum|EAL84556.1 100.00% afis|EAW20498.1 100.00%
Bootstrap support for afum|EAL84556.1 as seed ortholog is 100%.
Bootstrap support for afis|EAW20498.1 as seed ortholog is 100%.

---

Group of orthologs #3169. Best score 1181 bits
Score difference with first non-orthologous sequence - *A. fumigatus*:299 *A. fischeri*:1181
afum|EAL85171.1 100.00% afis|EAW20050.1 100.00%
Bootstrap support for afum|EAL85171.1 as seed ortholog is 100%.
Bootstrap support for afis|EAW20050.1 as seed ortholog is 100%.

---

Group of orthologs #3170. Best score 1181 bits
Score difference with first non-orthologous sequence - *A. fumigatus*:1181 *A. fischeri*:1181
afum|EAL85284.1 100.00% afis|EAW20166.1 100.00%
Bootstrap support for afum|EAL85284.1 as seed ortholog is 100%.
Bootstrap support for afis|EAW20166.1 as seed ortholog is 100%.

Group of orthologs #3171. Best score 1181 bits
Score difference with first non-orthologous sequence - *A. fumigatus*:1181 *A. fischeri*:1181
afum|EAL84706.1 100.00% afis|EAW20928.1 100.00%
Bootstrap support for afum|EAL84706.1 as seed ortholog is 100%.
Bootstrap support for afis|EAW20928.1 as seed ortholog is 100%.

Group of orthologs #3172. Best score 1181 bits
Score difference with first non-orthologous sequence - *A. fumigatus*:1119 *A. fischeri*:1132
afum|EAL85436.1 100.00% afis|EAW20313.1 100.00%
Bootstrap support for afum|EAL85436.1 as seed ortholog is 100%.
Bootstrap support for afis|EAW20313.1 as seed ortholog is 100%.

Group of orthologs #3173. Best score 1181 bits
Score difference with first non-orthologous sequence - *A. fumigatus*:1064 *A. fischeri*:1030
afum|EAL84292.1 100.00% afis|EAW25582.1 100.00%
Bootstrap support for afum|EAL84292.1 as seed ortholog is 100%.
Bootstrap support for afis|EAW25582.1 as seed ortholog is 100%.

Group of orthologs #3174. Best score 1181 bits
Score difference with first non-orthologous sequence - *A. fumigatus*:1035 *A. fischeri*:546
afum|EAL93549.1 100.00% afis|EAW18813.1 100.00%
Bootstrap support for afum|EAL93549.1 as seed ortholog is 100%.
Bootstrap support for afis|EAW18813.1 as seed ortholog is 100%.

Group of orthologs #3175. Best score 1180 bits
Score difference with first non-orthologous sequence - *A. fumigatus*:806 *A. fischeri*:781
afum|EAL84937.1 100.00% afis|EAW19973.1 100.00%
Bootstrap support for afum|EAL84937.1 as seed ortholog is 100%.
Bootstrap support for afis|EAW19973.1 as seed ortholog is 100%.

Group of orthologs #3176. Best score 1180 bits
Score difference with first non-orthologous sequence - *A. fumigatus*:1180 *A. fischeri*:1180
afum|EAL93938.2 100.00% afis|EAW19208.1 100.00%
Bootstrap support for afum|EAL93938.2 as seed ortholog is 100%.
Bootstrap support for afis|EAW19208.1 as seed ortholog is 100%.

Group of orthologs #3177. Best score 1180 bits
Score difference with first non-orthologous sequence - *A. fumigatus*:1180 *A. fischeri*:1124
afum|EAL89646.1 100.00% afis|EAW24986.1 100.00%
Bootstrap support for afum|EAL89646.1 as seed ortholog is 100%.
Bootstrap support for afis|EAW24986.1 as seed ortholog is 100%.

Group of orthologs #3178. Best score 1180 bits
Score difference with first non-orthologous sequence - *A. fumigatus*:1180 *A. fischeri*:1180
afum|EAL89953.1 100.00% afis|EAW25290.1 100.00%
Bootstrap support for afum|EAL89953.1 as seed ortholog is 100%.
Bootstrap support for afis|EAW25290.1 as seed ortholog is 100%.

Group of orthologs #3179. Best score 1179 bits
Score difference with first non-orthologous sequence - *A. fumigatus*:1179 *A. fischeri*:1179
afum|EAL86436.1 100.00% afis|EAW16901.1 100.00%
Bootstrap support for afum|EAL86436.1 as seed ortholog is 100%.
Bootstrap support for afis|EAW16901.1 as seed ortholog is 100%.

Group of orthologs #3180. Best score 1179 bits

Score difference with first non-orthologous sequence - *A. fumigatus*:1179 *A. fischeri*:1179
afum|EAL85172.1 100.00% afis|EAW20052.1 100.00%
Bootstrap support for afum|EAL85172.1 as seed ortholog is 100%.
Bootstrap support for afis|EAW20052.1 as seed ortholog is 100%.

---

Group of orthologs #3181. Best score 1179 bits
Score difference with first non-orthologous sequence - *A. fumigatus*:1179 *A. fischeri*:1179
afum|EAL87545.1 100.00% afis|EAW18183.1 100.00%
Bootstrap support for afum|EAL87545.1 as seed ortholog is 100%.
Bootstrap support for afis|EAW18183.1 as seed ortholog is 100%.

---

Group of orthologs #3182. Best score 1179 bits
Score difference with first non-orthologous sequence - *A. fumigatus*:1179 *A. fischeri*:1179
afum|EAL84670.1 100.00% afis|EAW20963.1 100.00%
Bootstrap support for afum|EAL84670.1 as seed ortholog is 100%.
Bootstrap support for afis|EAW20963.1 as seed ortholog is 100%.

---

Group of orthologs #3183. Best score 1179 bits
Score difference with first non-orthologous sequence - *A. fumigatus*:1179 *A. fischeri*:957
afum|EAL91069.1 100.00% afis|EAW22117.1 100.00%
Bootstrap support for afum|EAL91069.1 as seed ortholog is 100%.
Bootstrap support for afis|EAW22117.1 as seed ortholog is 100%.

---

Group of orthologs #3184. Best score 1179 bits
Score difference with first non-orthologous sequence - *A. fumigatus*:1179 *A. fischeri*:1179
afum|EAL92706.1 100.00% afis|EAW21671.1 100.00%
Bootstrap support for afum|EAL92706.1 as seed ortholog is 100%.
Bootstrap support for afis|EAW21671.1 as seed ortholog is 100%.

---

Group of orthologs #3185. Best score 1179 bits
Score difference with first non-orthologous sequence - *A. fumigatus*:1179 *A. fischeri*:1179
afum|EAL89664.1 100.00% afis|EAW25031.1 100.00%
Bootstrap support for afum|EAL89664.1 as seed ortholog is 100%.
Bootstrap support for afis|EAW25031.1 as seed ortholog is 100%.

---

Group of orthologs #3186. Best score 1179 bits
Score difference with first non-orthologous sequence - *A. fumigatus*:1179 *A. fischeri*:1179
afum|EAL92999.1 100.00% afis|EAW21984.1 100.00%
Bootstrap support for afum|EAL92999.1 as seed ortholog is 100%.
Bootstrap support for afis|EAW21984.1 as seed ortholog is 100%.

---

Group of orthologs #3187. Best score 1178 bits
Score difference with first non-orthologous sequence - *A. fumigatus*:1178 *A. fischeri*:1178
afum|EAL88728.1 100.00% afis|EAW16096.1 100.00%
Bootstrap support for afum|EAL88728.1 as seed ortholog is 100%.
Bootstrap support for afis|EAW16096.1 as seed ortholog is 100%.

---

Group of orthologs #3188. Best score 1178 bits
Score difference with first non-orthologous sequence - *A. fumigatus*:1019 *A. fischeri*:970
afum|EAL88761.1 100.00% afis|EAW16131.1 100.00%
Bootstrap support for afum|EAL88761.1 as seed ortholog is 100%.
Bootstrap support for afis|EAW16131.1 as seed ortholog is 100%.

---

Group of orthologs #3189. Best score 1178 bits
Score difference with first non-orthologous sequence - *A. fumigatus*:1178 *A. fischeri*:1178
afum|EAL89176.1 100.00% afis|EAW16580.1 100.00%

Bootstrap support for afum|EAL89176.1 as seed ortholog is 100%.
Bootstrap support for afis|EAW16580.1 as seed ortholog is 100%.

---

Group of orthologs #3190. Best score 1178 bits
Score difference with first non-orthologous sequence - *A. fumigatus*:1178 *A. fischeri*:1178
afum|EAL87996.1 100.00% afis|EAW23470.1 100.00%
Bootstrap support for afum|EAL87996.1 as seed ortholog is 100%.
Bootstrap support for afis|EAW23470.1 as seed ortholog is 100%.

---

Group of orthologs #3191. Best score 1178 bits
Score difference with first non-orthologous sequence - *A. fumigatus*:829 *A. fischeri*:412
afum|EAL87859.1 100.00% afis|EAW23606.1 100.00%
Bootstrap support for afum|EAL87859.1 as seed ortholog is 100%.
Bootstrap support for afis|EAW23606.1 as seed ortholog is 100%.

---

Group of orthologs #3192. Best score 1178 bits
Score difference with first non-orthologous sequence - *A. fumigatus*:1178 *A. fischeri*:1178
afum|EAL90456.1 100.00% afis|EAW22739.1 100.00%
Bootstrap support for afum|EAL90456.1 as seed ortholog is 100%.
Bootstrap support for afis|EAW22739.1 as seed ortholog is 100%.

---

Group of orthologs #3193. Best score 1178 bits
Score difference with first non-orthologous sequence - *A. fumigatus*:1178 *A. fischeri*:1178
afum|EAL92617.1 100.00% afis|EAW21598.1 100.00%
Bootstrap support for afum|EAL92617.1 as seed ortholog is 100%.
Bootstrap support for afis|EAW21598.1 as seed ortholog is 100%.

---

Group of orthologs #3194. Best score 1177 bits
Score difference with first non-orthologous sequence - *A. fumigatus*:1177 *A. fischeri*:1177
afum|EAL85669.1 100.00% afis|EAW15679.1 100.00%
Bootstrap support for afum|EAL85669.1 as seed ortholog is 100%.
Bootstrap support for afis|EAW15679.1 as seed ortholog is 100%.

---

Group of orthologs #3195. Best score 1177 bits
Score difference with first non-orthologous sequence - *A. fumigatus*:1177 *A. fischeri*:1177
afum|EAL84661.1 100.00% afis|EAW20972.1 100.00%
Bootstrap support for afum|EAL84661.1 as seed ortholog is 100%.
Bootstrap support for afis|EAW20972.1 as seed ortholog is 100%.

---

Group of orthologs #3196. Best score 1177 bits
Score difference with first non-orthologous sequence - *A. fumigatus*:1177 *A. fischeri*:1177
afum|EAL86941.2 100.00% afis|EAW19577.1 100.00%
Bootstrap support for afum|EAL86941.2 as seed ortholog is 100%.
Bootstrap support for afis|EAW19577.1 as seed ortholog is 100%.

---

Group of orthologs #3197. Best score 1177 bits
Score difference with first non-orthologous sequence - *A. fumigatus*:1177 *A. fischeri*:1177
afum|EAL93369.1 100.00% afis|EAW18653.1 100.00%
Bootstrap support for afum|EAL93369.1 as seed ortholog is 100%.
Bootstrap support for afis|EAW18653.1 as seed ortholog is 100%.

---

Group of orthologs #3198. Best score 1177 bits
Score difference with first non-orthologous sequence - *A. fumigatus*:1177 *A. fischeri*:1177
afum|EAL89580.1 100.00% afis|EAW24925.1 100.00%
Bootstrap support for afum|EAL89580.1 as seed ortholog is 100%.
Bootstrap support for afis|EAW24925.1 as seed ortholog is 100%.

---

Group of orthologs #3199. Best score 1176 bits
Score difference with first non-orthologous sequence - *A. fumigatus*:986 *A. fischeri*:1103
afum|EAL86367.1 100.00% afis|EAW16816.1 100.00%
Bootstrap support for afum|EAL86367.1 as seed ortholog is 100%.
Bootstrap support for afis|EAW16816.1 as seed ortholog is 100%.

---

Group of orthologs #3200. Best score 1176 bits
Score difference with first non-orthologous sequence - *A. fumigatus*:1176 *A. fischeri*:1176
afum|EAL88719.1 100.00% afis|EAW16087.1 100.00%
Bootstrap support for afum|EAL88719.1 as seed ortholog is 100%.
Bootstrap support for afis|EAW16087.1 as seed ortholog is 100%.

---

Group of orthologs #3201. Best score 1176 bits
Score difference with first non-orthologous sequence - *A. fumigatus*:1176 *A. fischeri*:1176
afum|EAL91898.1 100.00% afis|EAW18086.1 100.00%
Bootstrap support for afum|EAL91898.1 as seed ortholog is 100%.
Bootstrap support for afis|EAW18086.1 as seed ortholog is 100%.

---

Group of orthologs #3202. Best score 1176 bits
Score difference with first non-orthologous sequence - *A. fumigatus*:1176 *A. fischeri*:1176
afum|EAL86010.1 100.00% afis|EAW24339.1 100.00%
Bootstrap support for afum|EAL86010.1 as seed ortholog is 100%.
Bootstrap support for afis|EAW24339.1 as seed ortholog is 100%.

---

Group of orthologs #3203. Best score 1176 bits
Score difference with first non-orthologous sequence - *A. fumigatus*:1176 *A. fischeri*:1176
afum|EAL91025.1 100.00% afis|EAW22166.1 100.00%
Bootstrap support for afum|EAL91025.1 as seed ortholog is 100%.
Bootstrap support for afis|EAW22166.1 as seed ortholog is 100%.

---

Group of orthologs #3204. Best score 1176 bits
Score difference with first non-orthologous sequence - *A. fumigatus*:1176 *A. fischeri*:1176
afum|EAL90258.1 100.00% afis|EAW22938.1 100.00%
Bootstrap support for afum|EAL90258.1 as seed ortholog is 100%.
Bootstrap support for afis|EAW22938.1 as seed ortholog is 100%.

---

Group of orthologs #3205. Best score 1176 bits
Score difference with first non-orthologous sequence - *A. fumigatus*:1176 *A. fischeri*:1176
afum|EAL92773.1 100.00% afis|EAW21739.1 100.00%
Bootstrap support for afum|EAL92773.1 as seed ortholog is 100%.
Bootstrap support for afis|EAW21739.1 as seed ortholog is 100%.

---

Group of orthologs #3206. Best score 1176 bits
Score difference with first non-orthologous sequence - *A. fumigatus*:1176 *A. fischeri*:1176
afum|EAL90030.1 100.00% afis|EAW25370.1 100.00%
Bootstrap support for afum|EAL90030.1 as seed ortholog is 100%.
Bootstrap support for afis|EAW25370.1 as seed ortholog is 100%.

---

Group of orthologs #3207. Best score 1175 bits
Score difference with first non-orthologous sequence - *A. fumigatus*:1175 *A. fischeri*:1175
afum|EAL85186.1 100.00% afis|EAW20068.1 100.00%
Bootstrap support for afum|EAL85186.1 as seed ortholog is 100%.
Bootstrap support for afis|EAW20068.1 as seed ortholog is 100%.

---

Group of orthologs #3208. Best score 1175 bits

Score difference with first non-orthologous sequence - *A. fumigatus*:1104 *A. fischeri*:1071
afum|EAL91562.1 100.00% afis|EAW17748.1 100.00%
Bootstrap support for afum|EAL91562.1 as seed ortholog is 100%.
Bootstrap support for afis|EAW17748.1 as seed ortholog is 100%.

---

Group of orthologs #3209. Best score 1175 bits
Score difference with first non-orthologous sequence - *A. fumigatus*:1175 *A. fischeri*:925
afum|EAL92114.1 100.00% afis|EAW21091.1 100.00%
Bootstrap support for afum|EAL92114.1 as seed ortholog is 100%.
Bootstrap support for afis|EAW21091.1 as seed ortholog is 100%.

---

Group of orthologs #3210. Best score 1175 bits
Score difference with first non-orthologous sequence - *A. fumigatus*:1175 *A. fischeri*:1175
afum|EAL92670.1 100.00% afis|EAW21636.1 100.00%
Bootstrap support for afum|EAL92670.1 as seed ortholog is 100%.
Bootstrap support for afis|EAW21636.1 as seed ortholog is 100%.

---

Group of orthologs #3211. Best score 1175 bits
Score difference with first non-orthologous sequence - *A. fumigatus*:1175 *A. fischeri*:1175
afum|EAL91943.1 100.00% afis|EAW24088.1 100.00%
Bootstrap support for afum|EAL91943.1 as seed ortholog is 100%.
Bootstrap support for afis|EAW24088.1 as seed ortholog is 100%.

---

Group of orthologs #3212. Best score 1174 bits
Score difference with first non-orthologous sequence - *A. fumigatus*:833 *A. fischeri*:832
afum|EAL88666.2 100.00% afis|EAW16035.1 100.00%
Bootstrap support for afum|EAL88666.2 as seed ortholog is 100%.
Bootstrap support for afis|EAW16035.1 as seed ortholog is 100%.

---

Group of orthologs #3213. Best score 1174 bits
Score difference with first non-orthologous sequence - *A. fumigatus*:1174 *A. fischeri*:1174
afum|EAL89266.1 100.00% afis|EAW16662.1 100.00%
Bootstrap support for afum|EAL89266.1 as seed ortholog is 100%.
Bootstrap support for afis|EAW16662.1 as seed ortholog is 100%.

---

Group of orthologs #3214. Best score 1174 bits
Score difference with first non-orthologous sequence - *A. fumigatus*:1174 *A. fischeri*:1174
afum|EAL93738.1 100.00% afis|EAW19004.1 100.00%
Bootstrap support for afum|EAL93738.1 as seed ortholog is 100%.
Bootstrap support for afis|EAW19004.1 as seed ortholog is 100%.

---

Group of orthologs #3215. Best score 1174 bits
Score difference with first non-orthologous sequence - *A. fumigatus*:1007 *A. fischeri*:546
afum|EAL89790.1 100.00% afis|EAW25119.1 100.00%
Bootstrap support for afum|EAL89790.1 as seed ortholog is 100%.
Bootstrap support for afis|EAW25119.1 as seed ortholog is 100%.

---

Group of orthologs #3216. Best score 1173 bits
Score difference with first non-orthologous sequence - *A. fumigatus*:1173 *A. fischeri*:1173
afum|EAL85686.1 100.00% afis|EAW15662.1 100.00%
Bootstrap support for afum|EAL85686.1 as seed ortholog is 100%.
Bootstrap support for afis|EAW15662.1 as seed ortholog is 100%.

---

Group of orthologs #3217. Best score 1172 bits
Score difference with first non-orthologous sequence - *A. fumigatus*:1172 *A. fischeri*:1172
afum|EAL89296.1 100.00% afis|EAW16699.1 100.00%

Bootstrap support for afum|EAL89296.1 as seed ortholog is 100%.
Bootstrap support for afis|EAW16699.1 as seed ortholog is 100%.

---

Group of orthologs #3218. Best score 1172 bits
Score difference with first non-orthologous sequence - *A. fumigatus*:948 *A. fischeri*:770
afum|EAL86860.2 100.00% afis|EAW19662.1 100.00%
Bootstrap support for afum|EAL86860.2 as seed ortholog is 100%.
Bootstrap support for afis|EAW19662.1 as seed ortholog is 100%.

---

Group of orthologs #3219. Best score 1172 bits
Score difference with first non-orthologous sequence - *A. fumigatus*:1172 *A. fischeri*:1172
afum|EAL93514.1 100.00% afis|EAW18782.1 100.00%
Bootstrap support for afum|EAL93514.1 as seed ortholog is 100%.
Bootstrap support for afis|EAW18782.1 as seed ortholog is 100%.

---

Group of orthologs #3220. Best score 1172 bits
Score difference with first non-orthologous sequence - *A. fumigatus*:1172 *A. fischeri*:1172
afum|EAL89487.1 100.00% afis|EAW24841.1 100.00%
Bootstrap support for afum|EAL89487.1 as seed ortholog is 100%.
Bootstrap support for afis|EAW24841.1 as seed ortholog is 100%.

---

Group of orthologs #3221. Best score 1172 bits
Score difference with first non-orthologous sequence - *A. fumigatus*:1172 *A. fischeri*:1172
afum|EAL93015.1 100.00% afis|EAW22000.1 100.00%
Bootstrap support for afum|EAL93015.1 as seed ortholog is 100%.
Bootstrap support for afis|EAW22000.1 as seed ortholog is 100%.

---

Group of orthologs #3222. Best score 1171 bits
Score difference with first non-orthologous sequence - *A. fumigatus*:1171 *A. fischeri*:1171
afum|EAL88916.1 100.00% afis|EAW16294.1 100.00%
Bootstrap support for afum|EAL88916.1 as seed ortholog is 100%.
Bootstrap support for afis|EAW16294.1 as seed ortholog is 100%.

---

Group of orthologs #3223. Best score 1171 bits
Score difference with first non-orthologous sequence - *A. fumigatus*:918 *A. fischeri*:226
afum|EAL84311.1 100.00% afis|EAW25623.1 100.00%
Bootstrap support for afum|EAL84311.1 as seed ortholog is 100%.
Bootstrap support for afis|EAW25623.1 as seed ortholog is 100%.

---

Group of orthologs #3224. Best score 1171 bits
Score difference with first non-orthologous sequence - *A. fumigatus*:1171 *A. fischeri*:1171
afum|EAL91985.1 100.00% afis|EAW24129.1 100.00%
Bootstrap support for afum|EAL91985.1 as seed ortholog is 100%.
Bootstrap support for afis|EAW24129.1 as seed ortholog is 100%.

---

Group of orthologs #3225. Best score 1170 bits
Score difference with first non-orthologous sequence - *A. fumigatus*:1170 *A. fischeri*:1170
afum|EAL86482.2 100.00% afis|EAW16955.1 100.00%
Bootstrap support for afum|EAL86482.2 as seed ortholog is 100%.
Bootstrap support for afis|EAW16955.1 as seed ortholog is 100%.

---

Group of orthologs #3226. Best score 1170 bits
Score difference with first non-orthologous sequence - *A. fumigatus*:1170 *A. fischeri*:1031
afum|EAL85230.1 100.00% afis|EAW20113.1 100.00%
Bootstrap support for afum|EAL85230.1 as seed ortholog is 100%.
Bootstrap support for afis|EAW20113.1 as seed ortholog is 100%.

Group of orthologs #3227. Best score 1170 bits
Score difference with first non-orthologous sequence - *A. fumigatus*:1170 *A. fischeri*:497
afum|EAL84738.2 100.00% afis|EAW20895.1 100.00%
Bootstrap support for afum|EAL84738.2 as seed ortholog is 100%.
Bootstrap support for afis|EAW20895.1 as seed ortholog is 100%.

---

Group of orthologs #3228. Best score 1170 bits
Score difference with first non-orthologous sequence - *A. fumigatus*:1170 *A. fischeri*:1170
afum|EAL91694.1 100.00% afis|EAW17886.1 100.00%
Bootstrap support for afum|EAL91694.1 as seed ortholog is 100%.
Bootstrap support for afis|EAW17886.1 as seed ortholog is 100%.

---

Group of orthologs #3229. Best score 1170 bits
Score difference with first non-orthologous sequence - *A. fumigatus*:972 *A. fischeri*:1170
afum|EAL93811.1 100.00% afis|EAW19082.1 100.00%
Bootstrap support for afum|EAL93811.1 as seed ortholog is 100%.
Bootstrap support for afis|EAW19082.1 as seed ortholog is 100%.

---

Group of orthologs #3230. Best score 1170 bits
Score difference with first non-orthologous sequence - *A. fumigatus*:1170 *A. fischeri*:1067
afum|EAL90991.1 100.00% afis|EAW22201.1 100.00%
Bootstrap support for afum|EAL90991.1 as seed ortholog is 100%.
Bootstrap support for afis|EAW22201.1 as seed ortholog is 100%.

---

Group of orthologs #3231. Best score 1170 bits
Score difference with first non-orthologous sequence - *A. fumigatus*:1170 *A. fischeri*:1170
afum|EAL92658.1 100.00% afis|EAW21622.1 100.00%
Bootstrap support for afum|EAL92658.1 as seed ortholog is 100%.
Bootstrap support for afis|EAW21622.1 as seed ortholog is 100%.

---

Group of orthologs #3232. Best score 1169 bits
Score difference with first non-orthologous sequence - *A. fumigatus*:1169 *A. fischeri*:989
afum|EAL85858.1 100.00% afis|EAW15463.1 100.00%
Bootstrap support for afum|EAL85858.1 as seed ortholog is 100%.
Bootstrap support for afis|EAW15463.1 as seed ortholog is 100%.

---

Group of orthologs #3233. Best score 1169 bits
Score difference with first non-orthologous sequence - *A. fumigatus*:1169 *A. fischeri*:1169
afum|EAL85710.1 100.00% afis|EAW15624.1 100.00%
Bootstrap support for afum|EAL85710.1 as seed ortholog is 100%.
Bootstrap support for afis|EAW15624.1 as seed ortholog is 100%.

---

Group of orthologs #3234. Best score 1169 bits
Score difference with first non-orthologous sequence - *A. fumigatus*:1169 *A. fischeri*:1169
afum|EAL85595.1 100.00% afis|EAW15754.1 100.00%
Bootstrap support for afum|EAL85595.1 as seed ortholog is 100%.
Bootstrap support for afis|EAW15754.1 as seed ortholog is 100%.

---

Group of orthologs #3235. Best score 1169 bits
Score difference with first non-orthologous sequence - *A. fumigatus*:1003 *A. fischeri*:960
afum|EAL85144.1 100.00% afis|EAW19753.1 100.00%
Bootstrap support for afum|EAL85144.1 as seed ortholog is 100%.
Bootstrap support for afis|EAW19753.1 as seed ortholog is 100%.

---

Group of orthologs #3236. Best score 1169 bits

Score difference with first non-orthologous sequence - *A. fumigatus*:1169 *A. fischeri*:1169
afum|EAL84599.1 100.00% afis|EAW20454.1 100.00%
Bootstrap support for afum|EAL84599.1 as seed ortholog is 100%.
Bootstrap support for afis|EAW20454.1 as seed ortholog is 100%.

---

Group of orthologs #3237. Best score 1169 bits
Score difference with first non-orthologous sequence - *A. fumigatus*:1169 *A. fischeri*:1169
afum|EAL89155.1 100.00% afis|EAW16557.1 100.00%
Bootstrap support for afum|EAL89155.1 as seed ortholog is 100%.
Bootstrap support for afis|EAW16557.1 as seed ortholog is 100%.

---

Group of orthologs #3238. Best score 1169 bits
Score difference with first non-orthologous sequence - *A. fumigatus*:1169 *A. fischeri*:1169
afum|EAL85472.1 100.00% afis|EAW20355.1 100.00%
Bootstrap support for afum|EAL85472.1 as seed ortholog is 100%.
Bootstrap support for afis|EAW20355.1 as seed ortholog is 100%.

---

Group of orthologs #3239. Best score 1169 bits
Score difference with first non-orthologous sequence - *A. fumigatus*:1169 *A. fischeri*:1169
afum|EAL91362.1 100.00% afis|EAW17548.1 100.00%
Bootstrap support for afum|EAL91362.1 as seed ortholog is 100%.
Bootstrap support for afis|EAW17548.1 as seed ortholog is 100%.

---

Group of orthologs #3240. Best score 1169 bits
Score difference with first non-orthologous sequence - *A. fumigatus*:1169 *A. fischeri*:1169
afum|EAL90903.1 100.00% afis|EAW22278.1 100.00%
Bootstrap support for afum|EAL90903.1 as seed ortholog is 100%.
Bootstrap support for afis|EAW22278.1 as seed ortholog is 100%.

---

Group of orthologs #3241. Best score 1168 bits
Score difference with first non-orthologous sequence - *A. fumigatus*:1168 *A. fischeri*:996
afum|EAL85695.1 100.00% afis|EAW22086.1 100.00%
Bootstrap support for afum|EAL85695.1 as seed ortholog is 100%.
Bootstrap support for afis|EAW22086.1 as seed ortholog is 100%.

---

Group of orthologs #3242. Best score 1168 bits
Score difference with first non-orthologous sequence - *A. fumigatus*:1168 *A. fischeri*:1168
afum|EAL86246.1 100.00% afis|EAW24618.1 100.00%
Bootstrap support for afum|EAL86246.1 as seed ortholog is 100%.
Bootstrap support for afis|EAW24618.1 as seed ortholog is 100%.

---

Group of orthologs #3243. Best score 1168 bits
Score difference with first non-orthologous sequence - *A. fumigatus*:1168 *A. fischeri*:1168
afum|EBA27317.1 100.00% afis|EAW18184.1 100.00%
Bootstrap support for afum|EBA27317.1 as seed ortholog is 100%.
Bootstrap support for afis|EAW18184.1 as seed ortholog is 100%.

---

Group of orthologs #3244. Best score 1168 bits
Score difference with first non-orthologous sequence - *A. fumigatus*:888 *A. fischeri*:865
afum|EAL94043.1 100.00% afis|EAW19325.1 100.00%
Bootstrap support for afum|EAL94043.1 as seed ortholog is 100%.
Bootstrap support for afis|EAW19325.1 as seed ortholog is 100%.

---

Group of orthologs #3245. Best score 1167 bits
Score difference with first non-orthologous sequence - *A. fumigatus*:1167 *A. fischeri*:1167
afum|EAL85816.1 100.00% afis|EAW15515.1 100.00%

Bootstrap support for afum|EAL85816.1 as seed ortholog is 100%.
Bootstrap support for afis|EAW15515.1 as seed ortholog is 100%.

---

Group of orthologs #3246. Best score 1167 bits
Score difference with first non-orthologous sequence - *A. fumigatus*:1167 *A. fischeri*:1167
afum|EAL87644.1 100.00% afis|EAW18278.1 100.00%
Bootstrap support for afum|EAL87644.1 as seed ortholog is 100%.
Bootstrap support for afis|EAW18278.1 as seed ortholog is 100%.

---

Group of orthologs #3247. Best score 1166 bits
Score difference with first non-orthologous sequence - *A. fumigatus*:974 *A. fischeri*:1000
afum|EAL85093.1 100.00% afis|EAW19797.1 100.00%
Bootstrap support for afum|EAL85093.1 as seed ortholog is 100%.
Bootstrap support for afis|EAW19797.1 as seed ortholog is 100%.

---

Group of orthologs #3248. Best score 1166 bits
Score difference with first non-orthologous sequence - *A. fumigatus*:1166 *A. fischeri*:630
afum|EAL86910.1 100.00% afis|EAW19612.1 100.00%
Bootstrap support for afum|EAL86910.1 as seed ortholog is 100%.
Bootstrap support for afis|EAW19612.1 as seed ortholog is 100%.

---

Group of orthologs #3249. Best score 1166 bits
Score difference with first non-orthologous sequence - *A. fumigatus*:1166 *A. fischeri*:1166
afum|EAL91409.1 100.00% afis|EAW17596.1 100.00%
Bootstrap support for afum|EAL91409.1 as seed ortholog is 100%.
Bootstrap support for afis|EAW17596.1 as seed ortholog is 100%.

---

Group of orthologs #3250. Best score 1166 bits
Score difference with first non-orthologous sequence - *A. fumigatus*:970 *A. fischeri*:1166
afum|EAL86011.1 100.00% afis|EAW24340.1 100.00%
Bootstrap support for afum|EAL86011.1 as seed ortholog is 100%.
Bootstrap support for afis|EAW24340.1 as seed ortholog is 100%.

---

Group of orthologs #3251. Best score 1166 bits
Score difference with first non-orthologous sequence - *A. fumigatus*:1166 *A. fischeri*:1166
afum|EAL93782.1 100.00% afis|EAW19053.1 100.00%
Bootstrap support for afum|EAL93782.1 as seed ortholog is 100%.
Bootstrap support for afis|EAW19053.1 as seed ortholog is 100%.

---

Group of orthologs #3252. Best score 1166 bits
Score difference with first non-orthologous sequence - *A. fumigatus*:1166 *A. fischeri*:976
afum|EAL90962.1 100.00% afis|EAW22231.1 100.00%
Bootstrap support for afum|EAL90962.1 as seed ortholog is 100%.
Bootstrap support for afis|EAW22231.1 as seed ortholog is 100%.

---

Group of orthologs #3253. Best score 1166 bits
Score difference with first non-orthologous sequence - *A. fumigatus*:1166 *A. fischeri*:1166
afum|EAL92747.1 100.00% afis|EAW21712.1 100.00%
Bootstrap support for afum|EAL92747.1 as seed ortholog is 100%.
Bootstrap support for afis|EAW21712.1 as seed ortholog is 100%.

---

Group of orthologs #3254. Best score 1165 bits
Score difference with first non-orthologous sequence - *A. fumigatus*:1165 *A. fischeri*:1165
afum|EAL89038.1 100.00% afis|EAW16414.1 100.00%
Bootstrap support for afum|EAL89038.1 as seed ortholog is 100%.
Bootstrap support for afis|EAW16414.1 as seed ortholog is 100%.

Group of orthologs #3255. Best score 1165 bits
Score difference with first non-orthologous sequence - *A. fumigatus*:1165 *A. fischeri*:1165
afum|EAL93333.1 100.00% afis|EAW18623.1 100.00%
Bootstrap support for afum|EAL93333.1 as seed ortholog is 100%.
Bootstrap support for afis|EAW18623.1 as seed ortholog is 100%.

Group of orthologs #3256. Best score 1165 bits
Score difference with first non-orthologous sequence - *A. fumigatus*:1165 *A. fischeri*:1165
afum|EAL89593.1 100.00% afis|EAW24936.1 100.00%
Bootstrap support for afum|EAL89593.1 as seed ortholog is 100%.
Bootstrap support for afis|EAW24936.1 as seed ortholog is 100%.

Group of orthologs #3257. Best score 1164 bits
Score difference with first non-orthologous sequence - *A. fumigatus*:1164 *A. fischeri*:1164
afum|EAL88377.1 100.00% afis|EAW23101.1 100.00%
Bootstrap support for afum|EAL88377.1 as seed ortholog is 100%.
Bootstrap support for afis|EAW23101.1 as seed ortholog is 100%.

Group of orthologs #3258. Best score 1164 bits
Score difference with first non-orthologous sequence - *A. fumigatus*:1164 *A. fischeri*:1102
afum|EAL89327.1 100.00% afis|EAW24686.1 100.00%
Bootstrap support for afum|EAL89327.1 as seed ortholog is 100%.
Bootstrap support for afis|EAW24686.1 as seed ortholog is 100%.

Group of orthologs #3259. Best score 1163 bits
Score difference with first non-orthologous sequence - *A. fumigatus*:1163 *A. fischeri*:1163
afum|EAL84711.1 100.00% afis|EAW20923.1 100.00%
Bootstrap support for afum|EAL84711.1 as seed ortholog is 100%.
Bootstrap support for afis|EAW20923.1 as seed ortholog is 100%.

Group of orthologs #3260. Best score 1163 bits
Score difference with first non-orthologous sequence - *A. fumigatus*:937 *A. fischeri*:817
afum|EAL86034.1 100.00% afis|EAW24363.1 100.00%
Bootstrap support for afum|EAL86034.1 as seed ortholog is 100%.
Bootstrap support for afis|EAW24363.1 as seed ortholog is 100%.

Group of orthologs #3261. Best score 1163 bits
Score difference with first non-orthologous sequence - *A. fumigatus*:1163 *A. fischeri*:1078
afum|EAL86042.1 100.00% afis|EAW24370.1 100.00%
Bootstrap support for afum|EAL86042.1 as seed ortholog is 100%.
Bootstrap support for afis|EAW24370.1 as seed ortholog is 100%.

Group of orthologs #3262. Best score 1163 bits
Score difference with first non-orthologous sequence - *A. fumigatus*:846 *A. fischeri*:739
afum|EAL87171.1 100.00% afis|EAW23733.1 100.00%
Bootstrap support for afum|EAL87171.1 as seed ortholog is 100%.
Bootstrap support for afis|EAW23733.1 as seed ortholog is 100%.

Group of orthologs #3263. Best score 1163 bits
Score difference with first non-orthologous sequence - *A. fumigatus*:1163 *A. fischeri*:1163
afum|EAL88087.1 100.00% afis|EAW23384.1 100.00%
Bootstrap support for afum|EAL88087.1 as seed ortholog is 100%.
Bootstrap support for afis|EAW23384.1 as seed ortholog is 100%.

Group of orthologs #3264. Best score 1163 bits

Score difference with first non-orthologous sequence - *A. fumigatus*:1163   *A. fischeri*:1163
afum|EAL87837.1 100.00% afis|EAW23623.1 100.00%
Bootstrap support for afum|EAL87837.1 as seed ortholog is 100%.
Bootstrap support for afis|EAW23623.1 as seed ortholog is 100%.

---

Group of orthologs #3265. Best score 1163 bits
Score difference with first non-orthologous sequence - *A. fumigatus*:1163   *A. fischeri*:1163
afum|EAL93527.1 100.00% afis|EAW18792.1 100.00%
Bootstrap support for afum|EAL93527.1 as seed ortholog is 100%.
Bootstrap support for afis|EAW18792.1 as seed ortholog is 100%.

---

Group of orthologs #3266. Best score 1162 bits
Score difference with first non-orthologous sequence - *A. fumigatus*:352   *A. fischeri*:1162
afum|EAL89177.1 100.00% afis|EAW16582.1 100.00%
Bootstrap support for afum|EAL89177.1 as seed ortholog is 100%.
Bootstrap support for afis|EAW16582.1 as seed ortholog is 100%.

---

Group of orthologs #3267. Best score 1162 bits
Score difference with first non-orthologous sequence - *A. fumigatus*:838   *A. fischeri*:736
afum|EAL84793.1 100.00% afis|EAW20840.1 100.00%
Bootstrap support for afum|EAL84793.1 as seed ortholog is 100%.
Bootstrap support for afis|EAW20840.1 as seed ortholog is 100%.

---

Group of orthologs #3268. Best score 1162 bits
Score difference with first non-orthologous sequence - *A. fumigatus*:850   *A. fischeri*:750
afum|EAL89300.1 100.00% afis|EAW16712.1 100.00%
Bootstrap support for afum|EAL89300.1 as seed ortholog is 100%.
Bootstrap support for afis|EAW16712.1 as seed ortholog is 100%.

---

Group of orthologs #3269. Best score 1162 bits
Score difference with first non-orthologous sequence - *A. fumigatus*:1021   *A. fischeri*:797
afum|EAL85995.1 100.00% afis|EAW24325.1 100.00%
Bootstrap support for afum|EAL85995.1 as seed ortholog is 100%.
Bootstrap support for afis|EAW24325.1 as seed ortholog is 100%.

---

Group of orthologs #3270. Best score 1162 bits
Score difference with first non-orthologous sequence - *A. fumigatus*:1162   *A. fischeri*:1162
afum|EAL88359.1 100.00% afis|EAW23121.1 100.00%
Bootstrap support for afum|EAL88359.1 as seed ortholog is 100%.
Bootstrap support for afis|EAW23121.1 as seed ortholog is 100%.

---

Group of orthologs #3271. Best score 1162 bits
Score difference with first non-orthologous sequence - *A. fumigatus*:1162   *A. fischeri*:1162
afum|EAL93972.1 100.00% afis|EAW19243.1 100.00%
Bootstrap support for afum|EAL93972.1 as seed ortholog is 100%.
Bootstrap support for afis|EAW19243.1 as seed ortholog is 100%.

---

Group of orthologs #3272. Best score 1162 bits
Score difference with first non-orthologous sequence - *A. fumigatus*:1162   *A. fischeri*:1162
afum|EAL92533.1 100.00% afis|EAW21513.1 100.00%
Bootstrap support for afum|EAL92533.1 as seed ortholog is 100%.
Bootstrap support for afis|EAW21513.1 as seed ortholog is 100%.

---

Group of orthologs #3273. Best score 1161 bits
Score difference with first non-orthologous sequence - *A. fumigatus*:988   *A. fischeri*:904
afum|EAL86450.1 100.00% afis|EAW16925.1 100.00%

Bootstrap support for afum|EAL86450.1 as seed ortholog is 100%.
Bootstrap support for afis|EAW16925.1 as seed ortholog is 100%.

---

Group of orthologs #3274. Best score 1161 bits
Score difference with first non-orthologous sequence - *A. fumigatus*:1161 *A. fischeri*:717
afum|EAL88027.1 100.00% afis|EAW16155.1 100.00%
Bootstrap support for afum|EAL88027.1 as seed ortholog is 100%.
Bootstrap support for afis|EAW16155.1 as seed ortholog is 100%.

---

Group of orthologs #3275. Best score 1161 bits
Score difference with first non-orthologous sequence - *A. fumigatus*:1047 *A. fischeri*:1161
afum|EAL86538.1 100.00% afis|EAW20678.1 100.00%
Bootstrap support for afum|EAL86538.1 as seed ortholog is 100%.
Bootstrap support for afis|EAW20678.1 as seed ortholog is 100%.

---

Group of orthologs #3276. Best score 1161 bits
Score difference with first non-orthologous sequence - *A. fumigatus*:1161 *A. fischeri*:1161
afum|EAL91415.1 100.00% afis|EAW17602.1 100.00%
Bootstrap support for afum|EAL91415.1 as seed ortholog is 100%.
Bootstrap support for afis|EAW17602.1 as seed ortholog is 100%.

---

Group of orthologs #3277. Best score 1161 bits
Score difference with first non-orthologous sequence - *A. fumigatus*:1161 *A. fischeri*:1161
afum|EAL87120.1 100.00% afis|EAW23677.1 100.00%
Bootstrap support for afum|EAL87120.1 as seed ortholog is 100%.
Bootstrap support for afis|EAW23677.1 as seed ortholog is 100%.

---

Group of orthologs #3278. Best score 1161 bits
Score difference with first non-orthologous sequence - *A. fumigatus*:1161 *A. fischeri*:1161
afum|EAL89403.2 100.00% afis|EAW24761.1 100.00%
Bootstrap support for afum|EAL89403.2 as seed ortholog is 100%.
Bootstrap support for afis|EAW24761.1 as seed ortholog is 100%.

---

Group of orthologs #3279. Best score 1160 bits
Score difference with first non-orthologous sequence - *A. fumigatus*:774 *A. fischeri*:921
afum|EAL85316.1 100.00% afis|EAW20196.1 100.00%
Bootstrap support for afum|EAL85316.1 as seed ortholog is 100%.
Bootstrap support for afis|EAW20196.1 as seed ortholog is 100%.

---

Group of orthologs #3280. Best score 1160 bits
Score difference with first non-orthologous sequence - *A. fumigatus*:1160 *A. fischeri*:1160
afum|EAL91299.1 100.00% afis|EAW17470.1 100.00%
Bootstrap support for afum|EAL91299.1 as seed ortholog is 100%.
Bootstrap support for afis|EAW17470.1 as seed ortholog is 100%.

---

Group of orthologs #3281. Best score 1160 bits
Score difference with first non-orthologous sequence - *A. fumigatus*:1160 *A. fischeri*:1160
afum|EAL85915.1 100.00% afis|EAW24247.1 100.00%
Bootstrap support for afum|EAL85915.1 as seed ortholog is 100%.
Bootstrap support for afis|EAW24247.1 as seed ortholog is 100%.

---

Group of orthologs #3282. Best score 1160 bits
Score difference with first non-orthologous sequence - *A. fumigatus*:1160 *A. fischeri*:1160
afum|EAL87219.2 100.00% afis|EAW23785.1 100.00%
Bootstrap support for afum|EAL87219.2 as seed ortholog is 100%.
Bootstrap support for afis|EAW23785.1 as seed ortholog is 100%.

Group of orthologs #3283. Best score 1160 bits
Score difference with first non-orthologous sequence - *A. fumigatus*:1160 *A. fischeri*:1160
afum|EAL87977.1 100.00% afis|EAW23486.1 100.00%
Bootstrap support for afum|EAL87977.1 as seed ortholog is 100%.
Bootstrap support for afis|EAW23486.1 as seed ortholog is 100%.

Group of orthologs #3284. Best score 1160 bits
Score difference with first non-orthologous sequence - *A. fumigatus*:1160 *A. fischeri*:1160
afum|EAL90814.1 100.00% afis|EAW22378.1 100.00%
Bootstrap support for afum|EAL90814.1 as seed ortholog is 100%.
Bootstrap support for afis|EAW22378.1 as seed ortholog is 100%.

Group of orthologs #3285. Best score 1160 bits
Score difference with first non-orthologous sequence - *A. fumigatus*:1160 *A. fischeri*:1160
afum|EBA27189.1 100.00% afis|EAW20538.1 100.00%
Bootstrap support for afum|EBA27189.1 as seed ortholog is 100%.
Bootstrap support for afis|EAW20538.1 as seed ortholog is 100%.

Group of orthologs #3286. Best score 1160 bits
Score difference with first non-orthologous sequence - *A. fumigatus*:1160 *A. fischeri*:1160
afum|EAL89620.1 100.00% afis|EAW24962.1 100.00%
Bootstrap support for afum|EAL89620.1 as seed ortholog is 100%.
Bootstrap support for afis|EAW24962.1 as seed ortholog is 100%.

Group of orthologs #3287. Best score 1160 bits
Score difference with first non-orthologous sequence - *A. fumigatus*:1160 *A. fischeri*:1160
afum|EAL92023.1 100.00% afis|EAW24166.1 100.00%
Bootstrap support for afum|EAL92023.1 as seed ortholog is 100%.
Bootstrap support for afis|EAW24166.1 as seed ortholog is 100%.

Group of orthologs #3288. Best score 1160 bits
Score difference with first non-orthologous sequence - *A. fumigatus*:1160 *A. fischeri*:1160
afum|EBA27171.1 100.00% afis|EAW25557.1 100.00%
Bootstrap support for afum|EBA27171.1 as seed ortholog is 100%.
Bootstrap support for afis|EAW25557.1 as seed ortholog is 100%.

Group of orthologs #3289. Best score 1159 bits
Score difference with first non-orthologous sequence - *A. fumigatus*:1159 *A. fischeri*:1159
afum|EAL85511.2 100.00% afis|EAW15840.1 100.00%
Bootstrap support for afum|EAL85511.2 as seed ortholog is 100%.
Bootstrap support for afis|EAW15840.1 as seed ortholog is 100%.

Group of orthologs #3290. Best score 1159 bits
Score difference with first non-orthologous sequence - *A. fumigatus*:663 *A. fischeri*:729
afum|EAL86395.1 100.00% afis|EAW16849.1 100.00%
Bootstrap support for afum|EAL86395.1 as seed ortholog is 100%.
Bootstrap support for afis|EAW16849.1 as seed ortholog is 100%.

Group of orthologs #3291. Best score 1159 bits
Score difference with first non-orthologous sequence - *A. fumigatus*:1159 *A. fischeri*:1159
afum|EAL88629.1 100.00% afis|EAW15997.1 100.00%
Bootstrap support for afum|EAL88629.1 as seed ortholog is 100%.
Bootstrap support for afis|EAW15997.1 as seed ortholog is 100%.

Group of orthologs #3292. Best score 1159 bits

Score difference with first non-orthologous sequence - *A. fumigatus*:1159 *A. fischeri*:1159
afum|EAL93488.2 100.00% afis|EAW18758.1 100.00%
Bootstrap support for afum|EAL93488.2 as seed ortholog is 100%.
Bootstrap support for afis|EAW18758.1 as seed ortholog is 100%.

---

Group of orthologs #3293. Best score 1159 bits
Score difference with first non-orthologous sequence - *A. fumigatus*:1159 *A. fischeri*:1159
afum|EAL90912.1 100.00% afis|EAW22268.1 100.00%
Bootstrap support for afum|EAL90912.1 as seed ortholog is 100%.
Bootstrap support for afis|EAW22268.1 as seed ortholog is 100%.

---

Group of orthologs #3294. Best score 1158 bits
Score difference with first non-orthologous sequence - *A. fumigatus*:1158 *A. fischeri*:1158
afum|EAL85580.1 100.00% afis|EAW15768.1 100.00%
Bootstrap support for afum|EAL85580.1 as seed ortholog is 100%.
Bootstrap support for afis|EAW15768.1 as seed ortholog is 100%.

---

Group of orthologs #3295. Best score 1158 bits
Score difference with first non-orthologous sequence - *A. fumigatus*:1158 *A. fischeri*:454
afum|EAL89034.1 100.00% afis|EAW16410.1 100.00%
Bootstrap support for afum|EAL89034.1 as seed ortholog is 100%.
Bootstrap support for afis|EAW16410.1 as seed ortholog is 100%.

---

Group of orthologs #3296. Best score 1158 bits
Score difference with first non-orthologous sequence - *A. fumigatus*:1158 *A. fischeri*:1158
afum|EAL91673.1 100.00% afis|EAW17865.1 100.00%
Bootstrap support for afum|EAL91673.1 as seed ortholog is 100%.
Bootstrap support for afis|EAW17865.1 as seed ortholog is 100%.

---

Group of orthologs #3297. Best score 1158 bits
Score difference with first non-orthologous sequence - *A. fumigatus*:1158 *A. fischeri*:1158
afum|EAL93724.1 100.00% afis|EAW18988.1 100.00%
Bootstrap support for afum|EAL93724.1 as seed ortholog is 100%.
Bootstrap support for afis|EAW18988.1 as seed ortholog is 100%.

---

Group of orthologs #3298. Best score 1157 bits
Score difference with first non-orthologous sequence - *A. fumigatus*:1157 *A. fischeri*:1157
afum|EAL92082.1 100.00% afis|EAW21046.1 100.00%
Bootstrap support for afum|EAL92082.1 as seed ortholog is 100%.
Bootstrap support for afis|EAW21046.1 as seed ortholog is 100%.

---

Group of orthologs #3299. Best score 1157 bits
Score difference with first non-orthologous sequence - *A. fumigatus*:1012 *A. fischeri*:1050
afum|EAL89410.1 100.00% afis|EAW24769.1 100.00%
Bootstrap support for afum|EAL89410.1 as seed ortholog is 100%.
Bootstrap support for afis|EAW24769.1 as seed ortholog is 100%.

---

Group of orthologs #3300. Best score 1156 bits
Score difference with first non-orthologous sequence - *A. fumigatus*:949 *A. fischeri*:1156
afum|EAL85259.1 100.00% afis|EAW20144.1 100.00%
Bootstrap support for afum|EAL85259.1 as seed ortholog is 100%.
Bootstrap support for afis|EAW20144.1 as seed ortholog is 100%.

---

Group of orthologs #3301. Best score 1156 bits
Score difference with first non-orthologous sequence - *A. fumigatus*:1156 *A. fischeri*:1156
afum|EAL85967.1 100.00% afis|EAW24299.1 100.00%

Bootstrap support for afum|EAL85967.1 as seed ortholog is 100%.
Bootstrap support for afis|EAW24299.1 as seed ortholog is 100%.

---

Group of orthologs #3302. Best score 1156 bits
Score difference with first non-orthologous sequence - *A. fumigatus*:1156 *A. fischeri*:1156
afum|EAL92425.1 100.00% afis|EAW21411.1 100.00%
Bootstrap support for afum|EAL92425.1 as seed ortholog is 100%.
Bootstrap support for afis|EAW21411.1 as seed ortholog is 100%.

---

Group of orthologs #3303. Best score 1156 bits
Score difference with first non-orthologous sequence - *A. fumigatus*:1156 *A. fischeri*:1156
afum|EAL89776.1 100.00% afis|EAW25105.1 100.00%
Bootstrap support for afum|EAL89776.1 as seed ortholog is 100%.
Bootstrap support for afis|EAW25105.1 as seed ortholog is 100%.

---

Group of orthologs #3304. Best score 1155 bits
Score difference with first non-orthologous sequence - *A. fumigatus*:1155 *A. fischeri*:835
afum|EAL84720.1 100.00% afis|EAW20914.1 100.00%
Bootstrap support for afum|EAL84720.1 as seed ortholog is 100%.
Bootstrap support for afis|EAW20914.1 as seed ortholog is 100%.

---

Group of orthologs #3305. Best score 1155 bits
Score difference with first non-orthologous sequence - *A. fumigatus*:1155 *A. fischeri*:1155
afum|EAL91722.1 100.00% afis|EAW17913.1 100.00%
Bootstrap support for afum|EAL91722.1 as seed ortholog is 100%.
Bootstrap support for afis|EAW17913.1 as seed ortholog is 100%.

---

Group of orthologs #3306. Best score 1155 bits
Score difference with first non-orthologous sequence - *A. fumigatus*:1155 *A. fischeri*:1155
afum|EAL92891.2 100.00% afis|EAW21877.1 100.00%
Bootstrap support for afum|EAL92891.2 as seed ortholog is 100%.
Bootstrap support for afis|EAW21877.1 as seed ortholog is 100%.

---

Group of orthologs #3307. Best score 1155 bits
Score difference with first non-orthologous sequence - *A. fumigatus*:1155 *A. fischeri*:1155
afum|EAL92935.1 100.00% afis|EAW21922.1 100.00%
Bootstrap support for afum|EAL92935.1 as seed ortholog is 100%.
Bootstrap support for afis|EAW21922.1 as seed ortholog is 100%.

---

Group of orthologs #3308. Best score 1155 bits
Score difference with first non-orthologous sequence - *A. fumigatus*:1155 *A. fischeri*:1155
afum|EAL89787.1 100.00% afis|EAW25116.1 100.00%
Bootstrap support for afum|EAL89787.1 as seed ortholog is 100%.
Bootstrap support for afis|EAW25116.1 as seed ortholog is 100%.

---

Group of orthologs #3309. Best score 1154 bits
Score difference with first non-orthologous sequence - *A. fumigatus*:939 *A. fischeri*:792
afum|EAL85766.1 100.00% afis|EAW15567.1 100.00%
Bootstrap support for afum|EAL85766.1 as seed ortholog is 100%.
Bootstrap support for afis|EAW15567.1 as seed ortholog is 100%.

---

Group of orthologs #3310. Best score 1154 bits
Score difference with first non-orthologous sequence - *A. fumigatus*:1154 *A. fischeri*:1154
afum|EAL84633.1 100.00% afis|EAW20995.1 100.00%
Bootstrap support for afum|EAL84633.1 as seed ortholog is 100%.
Bootstrap support for afis|EAW20995.1 as seed ortholog is 100%.

---

Group of orthologs #3311. Best score 1154 bits
Score difference with first non-orthologous sequence - *A. fumigatus*:1154 *A. fischeri*:1154
afum|EAL91375.1 100.00% afis|EAW17561.1 100.00%
Bootstrap support for afum|EAL91375.1 as seed ortholog is 100%.
Bootstrap support for afis|EAW17561.1 as seed ortholog is 100%.

---

Group of orthologs #3312. Best score 1154 bits
Score difference with first non-orthologous sequence - *A. fumigatus*:1154 *A. fischeri*:1154
afum|EAL92924.1 100.00% afis|EAW21912.1 100.00%
Bootstrap support for afum|EAL92924.1 as seed ortholog is 100%.
Bootstrap support for afis|EAW21912.1 as seed ortholog is 100%.

---

Group of orthologs #3313. Best score 1154 bits
Score difference with first non-orthologous sequence - *A. fumigatus*:1154 *A. fischeri*:1154
afum|EAL92974.1 100.00% afis|EAW21960.1 100.00%
Bootstrap support for afum|EAL92974.1 as seed ortholog is 100%.
Bootstrap support for afis|EAW21960.1 as seed ortholog is 100%.

---

Group of orthologs #3314. Best score 1154 bits
Score difference with first non-orthologous sequence - *A. fumigatus*:1154 *A. fischeri*:1154
afum|EAL89771.1 100.00% afis|EAW25100.1 100.00%
Bootstrap support for afum|EAL89771.1 as seed ortholog is 100%.
Bootstrap support for afis|EAW25100.1 as seed ortholog is 100%.

---

Group of orthologs #3315. Best score 1153 bits
Score difference with first non-orthologous sequence - *A. fumigatus*:1153 *A. fischeri*:1059
afum|EAL86419.1 100.00% afis|EAW16883.1 100.00%
Bootstrap support for afum|EAL86419.1 as seed ortholog is 100%.
Bootstrap support for afis|EAW16883.1 as seed ortholog is 100%.

---

Group of orthologs #3316. Best score 1153 bits
Score difference with first non-orthologous sequence - *A. fumigatus*:772 *A. fischeri*:536
afum|EAL86435.1 100.00% afis|EAW16900.1 100.00%
Bootstrap support for afum|EAL86435.1 as seed ortholog is 100%.
Bootstrap support for afis|EAW16900.1 as seed ortholog is 100%.

---

Group of orthologs #3317. Best score 1153 bits
Score difference with first non-orthologous sequence - *A. fumigatus*:1153 *A. fischeri*:1153
afum|EAL86835.1 100.00% afis|EAW19691.1 100.00%
Bootstrap support for afum|EAL86835.1 as seed ortholog is 100%.
Bootstrap support for afis|EAW19691.1 as seed ortholog is 100%.

---

Group of orthologs #3318. Best score 1153 bits
Score difference with first non-orthologous sequence - *A. fumigatus*:1002 *A. fischeri*:929
afum|EAL91287.1 100.00% afis|EAW17459.1 100.00%
Bootstrap support for afum|EAL91287.1 as seed ortholog is 100%.
Bootstrap support for afis|EAW17459.1 as seed ortholog is 100%.

---

Group of orthologs #3319. Best score 1153 bits
Score difference with first non-orthologous sequence - *A. fumigatus*:1153 *A. fischeri*:1153
afum|EAL90212.1 100.00% afis|EAW22987.1 100.00%
Bootstrap support for afum|EAL90212.1 as seed ortholog is 100%.
Bootstrap support for afis|EAW22987.1 as seed ortholog is 100%.

---

Group of orthologs #3320. Best score 1153 bits

Score difference with first non-orthologous sequence - *A. fumigatus*:1153   *A. fischeri*:1153
afum|EBA27214.1 100.00% afis|EAW19924.1 100.00%
Bootstrap support for afum|EBA27214.1 as seed ortholog is 100%.
Bootstrap support for afis|EAW19924.1 as seed ortholog is 100%.

---

Group of orthologs #3321. Best score 1153 bits
Score difference with first non-orthologous sequence - *A. fumigatus*:1153   *A. fischeri*:1153
afum|EAL92043.1 100.00% afis|EAW24183.1 100.00%
Bootstrap support for afum|EAL92043.1 as seed ortholog is 100%.
Bootstrap support for afis|EAW24183.1 as seed ortholog is 100%.

---

Group of orthologs #3322. Best score 1152 bits
Score difference with first non-orthologous sequence - *A. fumigatus*:981   *A. fischeri*:617
afum|EAL92382.2 100.00% afis|EAW21366.1 100.00%
Bootstrap support for afum|EAL92382.2 as seed ortholog is 100%.
Bootstrap support for afis|EAW21366.1 as seed ortholog is 100%.

---

Group of orthologs #3323. Best score 1152 bits
Score difference with first non-orthologous sequence - *A. fumigatus*:1152   *A. fischeri*:1152
afum|EAL89651.1 100.00% afis|EAW24991.1 100.00%
Bootstrap support for afum|EAL89651.1 as seed ortholog is 100%.
Bootstrap support for afis|EAW24991.1 as seed ortholog is 100%.

---

Group of orthologs #3324. Best score 1152 bits
Score difference with first non-orthologous sequence - *A. fumigatus*:1019   *A. fischeri*:1152
afum|EAL91972.2 100.00% afis|EAW24117.1 100.00%
Bootstrap support for afum|EAL91972.2 as seed ortholog is 100%.
Bootstrap support for afis|EAW24117.1 as seed ortholog is 100%.

---

Group of orthologs #3325. Best score 1151 bits
Score difference with first non-orthologous sequence - *A. fumigatus*:1027   *A. fischeri*:998
afum|EAL86675.1 100.00% afis|EAW17258.1 100.00%
Bootstrap support for afum|EAL86675.1 as seed ortholog is 100%.
Bootstrap support for afis|EAW17258.1 as seed ortholog is 100%.

---

Group of orthologs #3326. Best score 1151 bits
Score difference with first non-orthologous sequence - *A. fumigatus*:1151   *A. fischeri*:1151
afum|EBA27207.1 100.00% afis|EAW20815.1 100.00%
Bootstrap support for afum|EBA27207.1 as seed ortholog is 100%.
Bootstrap support for afis|EAW20815.1 as seed ortholog is 100%.

---

Group of orthologs #3327. Best score 1150 bits
Score difference with first non-orthologous sequence - *A. fumigatus*:1150   *A. fischeri*:1150
afum|EAL85287.1 100.00% afis|EAW20169.1 100.00%
Bootstrap support for afum|EAL85287.1 as seed ortholog is 100%.
Bootstrap support for afis|EAW20169.1 as seed ortholog is 100%.

---

Group of orthologs #3328. Best score 1150 bits
Score difference with first non-orthologous sequence - *A. fumigatus*:1150   *A. fischeri*:1150
afum|EAL87591.1 100.00% afis|EAW18225.1 100.00%
Bootstrap support for afum|EAL87591.1 as seed ortholog is 100%.
Bootstrap support for afis|EAW18225.1 as seed ortholog is 100%.

---

Group of orthologs #3329. Best score 1150 bits
Score difference with first non-orthologous sequence - *A. fumigatus*:1150   *A. fischeri*:1150
afum|EAL93786.1 100.00% afis|EAW19057.1 100.00%

Bootstrap support for afum|EAL93786.1 as seed ortholog is 100%.
Bootstrap support for afis|EAW19057.1 as seed ortholog is 100%.

---

Group of orthologs #3330. Best score 1150 bits
Score difference with first non-orthologous sequence - *A. fumigatus*:1150 *A. fischeri*:1150
afum|EAL90752.1 100.00% afis|EAW22439.1 100.00%
Bootstrap support for afum|EAL90752.1 as seed ortholog is 100%.
Bootstrap support for afis|EAW22439.1 as seed ortholog is 100%.

---

Group of orthologs #3331. Best score 1150 bits
Score difference with first non-orthologous sequence - *A. fumigatus*:1150 *A. fischeri*:1150
afum|EAL90585.1 100.00% afis|EAW22613.1 100.00%
Bootstrap support for afum|EAL90585.1 as seed ortholog is 100%.
Bootstrap support for afis|EAW22613.1 as seed ortholog is 100%.

---

Group of orthologs #3332. Best score 1149 bits
Score difference with first non-orthologous sequence - *A. fumigatus*:864 *A. fischeri*:892
afum|EAL85145.1 100.00% afis|EAW19752.1 100.00%
Bootstrap support for afum|EAL85145.1 as seed ortholog is 100%.
Bootstrap support for afis|EAW19752.1 as seed ortholog is 100%.

---

Group of orthologs #3333. Best score 1149 bits
Score difference with first non-orthologous sequence - *A. fumigatus*:1149 *A. fischeri*:1149
afum|EAL87495.1 100.00% afis|EAW18138.1 100.00%
Bootstrap support for afum|EAL87495.1 as seed ortholog is 100%.
Bootstrap support for afis|EAW18138.1 as seed ortholog is 100%.

---

Group of orthologs #3334. Best score 1149 bits
Score difference with first non-orthologous sequence - *A. fumigatus*:1149 *A. fischeri*:258
afum|EAL84338.1 100.00% afis|EAW25668.1 100.00%
Bootstrap support for afum|EAL84338.1 as seed ortholog is 100%.
Bootstrap support for afis|EAW25668.1 as seed ortholog is 100%.

---

Group of orthologs #3335. Best score 1149 bits
Score difference with first non-orthologous sequence - *A. fumigatus*:1149 *A. fischeri*:1149
afum|EAL85905.1 100.00% afis|EAW24236.1 100.00%
Bootstrap support for afum|EAL85905.1 as seed ortholog is 100%.
Bootstrap support for afis|EAW24236.1 as seed ortholog is 100%.

---

Group of orthologs #3336. Best score 1148 bits
Score difference with first non-orthologous sequence - *A. fumigatus*:1148 *A. fischeri*:1148
afum|EAL86348.2 100.00% afis|EAW16778.1 100.00%
Bootstrap support for afum|EAL86348.2 as seed ortholog is 100%.
Bootstrap support for afis|EAW16778.1 as seed ortholog is 100%.

---

Group of orthologs #3337. Best score 1148 bits
Score difference with first non-orthologous sequence - *A. fumigatus*:1148 *A. fischeri*:1148
afum|EAL91546.1 100.00% afis|EAW17731.1 100.00%
Bootstrap support for afum|EAL91546.1 as seed ortholog is 100%.
Bootstrap support for afis|EAW17731.1 as seed ortholog is 100%.

---

Group of orthologs #3338. Best score 1148 bits
Score difference with first non-orthologous sequence - *A. fumigatus*:1148 *A. fischeri*:1030
afum|EAL92188.2 100.00% afis|EAW21169.1 100.00%
Bootstrap support for afum|EAL92188.2 as seed ortholog is 100%.
Bootstrap support for afis|EAW21169.1 as seed ortholog is 100%.

---

Group of orthologs #3339. Best score 1148 bits
Score difference with first non-orthologous sequence - *A. fumigatus*:1148 *A. fischeri*:1148
afum|EAL92034.1 100.00% afis|EAW24172.1 100.00%
Bootstrap support for afum|EAL92034.1 as seed ortholog is 100%.
Bootstrap support for afis|EAW24172.1 as seed ortholog is 100%.

---

Group of orthologs #3340. Best score 1147 bits
Score difference with first non-orthologous sequence - *A. fumigatus*:1147 *A. fischeri*:1147
afum|EAL88897.1 100.00% afis|EAW16276.1 100.00%
Bootstrap support for afum|EAL88897.1 as seed ortholog is 100%.
Bootstrap support for afis|EAW16276.1 as seed ortholog is 100%.

---

Group of orthologs #3341. Best score 1147 bits
Score difference with first non-orthologous sequence - *A. fumigatus*:629 *A. fischeri*:1147
afum|EAL85367.1 100.00% afis|EAW20249.1 100.00%
Bootstrap support for afum|EAL85367.1 as seed ortholog is 100%.
Bootstrap support for afis|EAW20249.1 as seed ortholog is 100%.

---

Group of orthologs #3342. Best score 1147 bits
Score difference with first non-orthologous sequence - *A. fumigatus*:1147 *A. fischeri*:350
afum|EAL86880.1 100.00% afis|EAW19642.1 100.00%
Bootstrap support for afum|EAL86880.1 as seed ortholog is 100%.
Bootstrap support for afis|EAW19642.1 as seed ortholog is 100%.

---

Group of orthologs #3343. Best score 1147 bits
Score difference with first non-orthologous sequence - *A. fumigatus*:1147 *A. fischeri*:1147
afum|EAL85891.2 100.00% afis|EAW24222.1 100.00%
Bootstrap support for afum|EAL85891.2 as seed ortholog is 100%.
Bootstrap support for afis|EAW24222.1 as seed ortholog is 100%.

---

Group of orthologs #3344. Best score 1147 bits
Score difference with first non-orthologous sequence - *A. fumigatus*:1147 *A. fischeri*:1147
afum|EAL93666.2 100.00% afis|EAW18933.1 100.00%
Bootstrap support for afum|EAL93666.2 as seed ortholog is 100%.
Bootstrap support for afis|EAW18933.1 as seed ortholog is 100%.

---

Group of orthologs #3345. Best score 1147 bits
Score difference with first non-orthologous sequence - *A. fumigatus*:846 *A. fischeri*:1028
afum|EAL93829.1 100.00% afis|EAW19097.1 100.00%
Bootstrap support for afum|EAL93829.1 as seed ortholog is 100%.
Bootstrap support for afis|EAW19097.1 as seed ortholog is 100%.

---

Group of orthologs #3346. Best score 1146 bits
Score difference with first non-orthologous sequence - *A. fumigatus*:1146 *A. fischeri*:1146
afum|EAL87617.1 100.00% afis|EAW18254.1 100.00%
Bootstrap support for afum|EAL87617.1 as seed ortholog is 100%.
Bootstrap support for afis|EAW18254.1 as seed ortholog is 100%.

---

Group of orthologs #3347. Best score 1146 bits
Score difference with first non-orthologous sequence - *A. fumigatus*:1146 *A. fischeri*:1146
afum|EAL93267.2 100.00% afis|EAW18565.1 100.00%
Bootstrap support for afum|EAL93267.2 as seed ortholog is 100%.
Bootstrap support for afis|EAW18565.1 as seed ortholog is 100%.

---

Group of orthologs #3348. Best score 1146 bits

Score difference with first non-orthologous sequence - *A. fumigatus*:1146   *A. fischeri*:1146
afum|EAL91075.1 100.00% afis|EAW22109.1 100.00%
Bootstrap support for afum|EAL91075.1 as seed ortholog is 100%.
Bootstrap support for afis|EAW22109.1 as seed ortholog is 100%.

---

Group of orthologs #3349. Best score 1146 bits
Score difference with first non-orthologous sequence - *A. fumigatus*:1146   *A. fischeri*:958
afum|EAL90787.1 100.00% afis|EAW22404.1 100.00%
Bootstrap support for afum|EAL90787.1 as seed ortholog is 100%.
Bootstrap support for afis|EAW22404.1 as seed ortholog is 100%.

---

Group of orthologs #3350. Best score 1145 bits
Score difference with first non-orthologous sequence - *A. fumigatus*:1145   *A. fischeri*:1145
afum|EAL87722.1 100.00% afis|EAW18358.1 100.00%
Bootstrap support for afum|EAL87722.1 as seed ortholog is 100%.
Bootstrap support for afis|EAW18358.1 as seed ortholog is 100%.

---

Group of orthologs #3351. Best score 1145 bits
Score difference with first non-orthologous sequence - *A. fumigatus*:1145   *A. fischeri*:1145
afum|EAL85882.1 100.00% afis|EAW24213.1 100.00%
Bootstrap support for afum|EAL85882.1 as seed ortholog is 100%.
Bootstrap support for afis|EAW24213.1 as seed ortholog is 100%.

---

Group of orthologs #3352. Best score 1145 bits
Score difference with first non-orthologous sequence - *A. fumigatus*:894   *A. fischeri*:846
afum|EAL92997.1 100.00% afis|EAW21982.1 100.00%
Bootstrap support for afum|EAL92997.1 as seed ortholog is 100%.
Bootstrap support for afis|EAW21982.1 as seed ortholog is 100%.

---

Group of orthologs #3353. Best score 1144 bits
Score difference with first non-orthologous sequence - *A. fumigatus*:1083   *A. fischeri*:769
afum|EAL86952.1 100.00% afis|EAW19559.1 100.00%
Bootstrap support for afum|EAL86952.1 as seed ortholog is 100%.
Bootstrap support for afis|EAW19559.1 as seed ortholog is 100%.

---

Group of orthologs #3354. Best score 1144 bits
Score difference with first non-orthologous sequence - *A. fumigatus*:1144   *A. fischeri*:1144
afum|EAL88417.1 100.00% afis|EAW23062.1 100.00%
Bootstrap support for afum|EAL88417.1 as seed ortholog is 100%.
Bootstrap support for afis|EAW23062.1 as seed ortholog is 100%.

---

Group of orthologs #3355. Best score 1144 bits
Score difference with first non-orthologous sequence - *A. fumigatus*:1144   *A. fischeri*:1144
afum|EBA27330.1 100.00% afis|EAW23456.1 100.00%
Bootstrap support for afum|EBA27330.1 as seed ortholog is 100%.
Bootstrap support for afis|EAW23456.1 as seed ortholog is 100%.

---

Group of orthologs #3356. Best score 1143 bits
Score difference with first non-orthologous sequence - *A. fumigatus*:881   *A. fischeri*:903
afum|EAL86606.1 100.00% afis|EAW17176.1 100.00%
Bootstrap support for afum|EAL86606.1 as seed ortholog is 100%.
Bootstrap support for afis|EAW17176.1 as seed ortholog is 100%.

---

Group of orthologs #3357. Best score 1143 bits
Score difference with first non-orthologous sequence - *A. fumigatus*:1143   *A. fischeri*:1143
afum|EAL86622.1 100.00% afis|EAW17194.1 100.00%

Bootstrap support for afum|EAL86622.1 as seed ortholog is 100%.
Bootstrap support for afis|EAW17194.1 as seed ortholog is 100%.

---

Group of orthologs #3358. Best score 1143 bits
Score difference with first non-orthologous sequence - *A. fumigatus*:1143 *A. fischeri*:1143
afum|EAL86685.1 100.00% afis|EAW17268.1 100.00%
Bootstrap support for afum|EAL86685.1 as seed ortholog is 100%.
Bootstrap support for afis|EAW17268.1 as seed ortholog is 100%.

---

Group of orthologs #3359. Best score 1143 bits
Score difference with first non-orthologous sequence - *A. fumigatus*:1143 *A. fischeri*:1062
afum|EAL87835.1 100.00% afis|EAW23625.1 100.00%
Bootstrap support for afum|EAL87835.1 as seed ortholog is 100%.
Bootstrap support for afis|EAW23625.1 as seed ortholog is 100%.

---

Group of orthologs #3360. Best score 1143 bits
Score difference with first non-orthologous sequence - *A. fumigatus*:1143 *A. fischeri*:893
afum|EAL93266.1 100.00% afis|EAW18564.1 100.00%
Bootstrap support for afum|EAL93266.1 as seed ortholog is 100%.
Bootstrap support for afis|EAW18564.1 as seed ortholog is 100%.

---

Group of orthologs #3361. Best score 1143 bits
Score difference with first non-orthologous sequence - *A. fumigatus*:957 *A. fischeri*:1079
afum|EAL89371.1 100.00% afis|EAW24722.1 100.00%
Bootstrap support for afum|EAL89371.1 as seed ortholog is 100%.
Bootstrap support for afis|EAW24722.1 as seed ortholog is 100%.

---

Group of orthologs #3362. Best score 1142 bits
Score difference with first non-orthologous sequence - *A. fumigatus*:1142 *A. fischeri*:1142
afum|EAL84772.1 100.00% afis|EAW20863.1 100.00%
Bootstrap support for afum|EAL84772.1 as seed ortholog is 100%.
Bootstrap support for afis|EAW20863.1 as seed ortholog is 100%.

---

Group of orthologs #3363. Best score 1142 bits
Score difference with first non-orthologous sequence - *A. fumigatus*:1142 *A. fischeri*:1142
afum|EAL88360.1 100.00% afis|EAW23120.1 100.00%
Bootstrap support for afum|EAL88360.1 as seed ortholog is 100%.
Bootstrap support for afis|EAW23120.1 as seed ortholog is 100%.

---

Group of orthologs #3364. Best score 1142 bits
Score difference with first non-orthologous sequence - *A. fumigatus*:1027 *A. fischeri*:999
afum|EAL90654.2 100.00% afis|EAW22540.1 100.00%
Bootstrap support for afum|EAL90654.2 as seed ortholog is 100%.
Bootstrap support for afis|EAW22540.1 as seed ortholog is 100%.

---

Group of orthologs #3365. Best score 1142 bits
Score difference with first non-orthologous sequence - *A. fumigatus*:1142 *A. fischeri*:1142
afum|EAL90405.1 100.00% afis|EAW22794.1 100.00%
Bootstrap support for afum|EAL90405.1 as seed ortholog is 100%.
Bootstrap support for afis|EAW22794.1 as seed ortholog is 100%.

---

Group of orthologs #3366. Best score 1142 bits
Score difference with first non-orthologous sequence - *A. fumigatus*:762 *A. fischeri*:819
afum|EBA27371.1 100.00% afis|EAW20414.1 100.00%
Bootstrap support for afum|EBA27371.1 as seed ortholog is 100%.
Bootstrap support for afis|EAW20414.1 as seed ortholog is 100%.

Group of orthologs #3367. Best score 1141 bits
Score difference with first non-orthologous sequence - *A. fumigatus*:1141 *A. fischeri*:1141
afum|EAL89188.1 100.00% afis|EAW16592.1 100.00%
Bootstrap support for afum|EAL89188.1 as seed ortholog is 100%.
Bootstrap support for afis|EAW16592.1 as seed ortholog is 100%.

Group of orthologs #3368. Best score 1141 bits
Score difference with first non-orthologous sequence - *A. fumigatus*:1141 *A. fischeri*:1141
afum|EAL87587.1 100.00% afis|EAW18221.1 100.00%
Bootstrap support for afum|EAL87587.1 as seed ortholog is 100%.
Bootstrap support for afis|EAW18221.1 as seed ortholog is 100%.

Group of orthologs #3369. Best score 1141 bits
Score difference with first non-orthologous sequence - *A. fumigatus*:1141 *A. fischeri*:1141
afum|EAL89291.1 100.00% afis|EAW16693.1 100.00%
Bootstrap support for afum|EAL89291.1 as seed ortholog is 100%.
Bootstrap support for afis|EAW16693.1 as seed ortholog is 100%.

Group of orthologs #3370. Best score 1141 bits
Score difference with first non-orthologous sequence - *A. fumigatus*:1141 *A. fischeri*:1141
afum|EAL91196.1 100.00% afis|EAW17359.1 100.00%
Bootstrap support for afum|EAL91196.1 as seed ortholog is 100%.
Bootstrap support for afis|EAW17359.1 as seed ortholog is 100%.

Group of orthologs #3371. Best score 1141 bits
Score difference with first non-orthologous sequence - *A. fumigatus*:1141 *A. fischeri*:1141
afum|EAL86027.1 100.00% afis|EAW24357.1 100.00%
Bootstrap support for afum|EAL86027.1 as seed ortholog is 100%.
Bootstrap support for afis|EAW24357.1 as seed ortholog is 100%.

Group of orthologs #3372. Best score 1141 bits
Score difference with first non-orthologous sequence - *A. fumigatus*:1141 *A. fischeri*:667
afum|EAL88283.1 100.00% afis|EAW23200.1 100.00%
Bootstrap support for afum|EAL88283.1 as seed ortholog is 100%.
Bootstrap support for afis|EAW23200.1 as seed ortholog is 100%.

Group of orthologs #3373. Best score 1141 bits
Score difference with first non-orthologous sequence - *A. fumigatus*:1141 *A. fischeri*:1141
afum|EAL90425.1 100.00% afis|EAW22773.1 100.00%
Bootstrap support for afum|EAL90425.1 as seed ortholog is 100%.
Bootstrap support for afis|EAW22773.1 as seed ortholog is 100%.

Group of orthologs #3374. Best score 1141 bits
Score difference with first non-orthologous sequence - *A. fumigatus*:950 *A. fischeri*:952
afum|EAL89614.2 100.00% afis|EAW24956.1 100.00%
Bootstrap support for afum|EAL89614.2 as seed ortholog is 100%.
Bootstrap support for afis|EAW24956.1 as seed ortholog is 100%.

Group of orthologs #3375. Best score 1141 bits
Score difference with first non-orthologous sequence - *A. fumigatus*:1141 *A. fischeri*:1141
afum|EAL92907.1 100.00% afis|EAW21893.1 100.00%
Bootstrap support for afum|EAL92907.1 as seed ortholog is 100%.
Bootstrap support for afis|EAW21893.1 as seed ortholog is 100%.

Group of orthologs #3376. Best score 1141 bits

Score difference with first non-orthologous sequence - *A. fumigatus*:1141   *A. fischeri*:1141
afum|EAL89847.1 100.00% afis|EAW25174.1 100.00%
Bootstrap support for afum|EAL89847.1 as seed ortholog is 100%.
Bootstrap support for afis|EAW25174.1 as seed ortholog is 100%.

---

Group of orthologs #3377. Best score 1140 bits
Score difference with first non-orthologous sequence - *A. fumigatus*:1140   *A. fischeri*:1140
afum|EAL86586.1 100.00% afis|EAW17126.1 100.00%
Bootstrap support for afum|EAL86586.1 as seed ortholog is 100%.
Bootstrap support for afis|EAW17126.1 as seed ortholog is 100%.

---

Group of orthologs #3378. Best score 1140 bits
Score difference with first non-orthologous sequence - *A. fumigatus*:1140   *A. fischeri*:1140
afum|EAL88971.1 100.00% afis|EAW16346.1 100.00%
Bootstrap support for afum|EAL88971.1 as seed ortholog is 100%.
Bootstrap support for afis|EAW16346.1 as seed ortholog is 100%.

---

Group of orthologs #3379. Best score 1140 bits
Score difference with first non-orthologous sequence - *A. fumigatus*:1140   *A. fischeri*:1140
afum|EAL84392.2 100.00% afis|EAW25717.1 100.00%
Bootstrap support for afum|EAL84392.2 as seed ortholog is 100%.
Bootstrap support for afis|EAW25717.1 as seed ortholog is 100%.

---

Group of orthologs #3380. Best score 1140 bits
Score difference with first non-orthologous sequence - *A. fumigatus*:1140   *A. fischeri*:1140
afum|EAL93831.1 100.00% afis|EAW19099.1 100.00%
Bootstrap support for afum|EAL93831.1 as seed ortholog is 100%.
Bootstrap support for afis|EAW19099.1 as seed ortholog is 100%.

---

Group of orthologs #3381. Best score 1139 bits
Score difference with first non-orthologous sequence - *A. fumigatus*:1139   *A. fischeri*:1139
afum|EAL88592.1 100.00% afis|EAW15959.1 100.00%
Bootstrap support for afum|EAL88592.1 as seed ortholog is 100%.
Bootstrap support for afis|EAW15959.1 as seed ortholog is 100%.

---

Group of orthologs #3382. Best score 1139 bits
Score difference with first non-orthologous sequence - *A. fumigatus*:877   *A. fischeri*:746
afum|EAL85217.1 100.00% afis|EAW20100.1 100.00%
Bootstrap support for afum|EAL85217.1 as seed ortholog is 100%.
Bootstrap support for afis|EAW20100.1 as seed ortholog is 100%.

---

Group of orthologs #3383. Best score 1139 bits
Score difference with first non-orthologous sequence - *A. fumigatus*:1139   *A. fischeri*:1139
afum|EAL87516.1 100.00% afis|EAW18157.1 100.00%
Bootstrap support for afum|EAL87516.1 as seed ortholog is 100%.
Bootstrap support for afis|EAW18157.1 as seed ortholog is 100%.

---

Group of orthologs #3384. Best score 1139 bits
Score difference with first non-orthologous sequence - *A. fumigatus*:1139   *A. fischeri*:1139
afum|EAL89848.1 100.00% afis|EAW25175.1 100.00%
Bootstrap support for afum|EAL89848.1 as seed ortholog is 100%.
Bootstrap support for afis|EAW25175.1 as seed ortholog is 100%.

---

Group of orthologs #3385. Best score 1139 bits
Score difference with first non-orthologous sequence - *A. fumigatus*:1139   *A. fischeri*:1139
afum|EAL90112.1 100.00% afis|EAW25453.1 100.00%

Bootstrap support for afum|EAL90112.1 as seed ortholog is 100%.
Bootstrap support for afis|EAW25453.1 as seed ortholog is 100%.

---

Group of orthologs #3386. Best score 1138 bits
Score difference with first non-orthologous sequence - *A. fumigatus*:890 *A. fischeri*:942
afum|EAL84203.1 100.00% afis|EAW15444.1 100.00%
Bootstrap support for afum|EAL84203.1 as seed ortholog is 100%.
Bootstrap support for afis|EAW15444.1 as seed ortholog is 100%.

---

Group of orthologs #3387. Best score 1138 bits
Score difference with first non-orthologous sequence - *A. fumigatus*:811 *A. fischeri*:1138
afum|EAL91153.1 100.00% afis|EAW17317.1 100.00%
Bootstrap support for afum|EAL91153.1 as seed ortholog is 100%.
Bootstrap support for afis|EAW17317.1 as seed ortholog is 100%.

---

Group of orthologs #3388. Best score 1138 bits
Score difference with first non-orthologous sequence - *A. fumigatus*:1138 *A. fischeri*:1138
afum|EAL91538.1 100.00% afis|EAW17723.1 100.00%
Bootstrap support for afum|EAL91538.1 as seed ortholog is 100%.
Bootstrap support for afis|EAW17723.1 as seed ortholog is 100%.

---

Group of orthologs #3389. Best score 1138 bits
Score difference with first non-orthologous sequence - *A. fumigatus*:1138 *A. fischeri*:1138
afum|EAL88068.1 100.00% afis|EAW23402.1 100.00%
Bootstrap support for afum|EAL88068.1 as seed ortholog is 100%.
Bootstrap support for afis|EAW23402.1 as seed ortholog is 100%.

---

Group of orthologs #3390. Best score 1138 bits
Score difference with first non-orthologous sequence - *A. fumigatus*:1138 *A. fischeri*:964
afum|EAL93541.1 100.00% afis|EAW18805.1 100.00%
Bootstrap support for afum|EAL93541.1 as seed ortholog is 100%.
Bootstrap support for afis|EAW18805.1 as seed ortholog is 100%.

---

Group of orthologs #3391. Best score 1138 bits
Score difference with first non-orthologous sequence - *A. fumigatus*:923 *A. fischeri*:913
afum|EAL94070.1 100.00% afis|EAW19351.1 100.00%
Bootstrap support for afum|EAL94070.1 as seed ortholog is 100%.
Bootstrap support for afis|EAW19351.1 as seed ortholog is 100%.

---

Group of orthologs #3392. Best score 1138 bits
Score difference with first non-orthologous sequence - *A. fumigatus*:968 *A. fischeri*:1138
afum|EAL89899.1 100.00% afis|EAW25238.1 100.00%
Bootstrap support for afum|EAL89899.1 as seed ortholog is 100%.
Bootstrap support for afis|EAW25238.1 as seed ortholog is 100%.

---

Group of orthologs #3393. Best score 1138 bits
Score difference with first non-orthologous sequence - *A. fumigatus*:1048 *A. fischeri*:1034
afum|EAL90103.1 100.00% afis|EAW25445.1 100.00%
Bootstrap support for afum|EAL90103.1 as seed ortholog is 100%.
Bootstrap support for afis|EAW25445.1 as seed ortholog is 100%.

---

Group of orthologs #3394. Best score 1137 bits
Score difference with first non-orthologous sequence - *A. fumigatus*:959 *A. fischeri*:817
afum|EAL84195.2 100.00% afis|EAW15451.1 100.00%
Bootstrap support for afum|EAL84195.2 as seed ortholog is 100%.
Bootstrap support for afis|EAW15451.1 as seed ortholog is 100%.

---

Group of orthologs #3395. Best score 1137 bits
Score difference with first non-orthologous sequence - *A. fumigatus*:1137 *A. fischeri*:1137
afum|EAL84771.1 100.00% afis|EAW20864.1 100.00%
Bootstrap support for afum|EAL84771.1 as seed ortholog is 100%.
Bootstrap support for afis|EAW20864.1 as seed ortholog is 100%.

---

Group of orthologs #3396. Best score 1137 bits
Score difference with first non-orthologous sequence - *A. fumigatus*:1137 *A. fischeri*:1137
afum|EAL86887.1 100.00% afis|EAW19635.1 100.00%
Bootstrap support for afum|EAL86887.1 as seed ortholog is 100%.
Bootstrap support for afis|EAW19635.1 as seed ortholog is 100%.

---

Group of orthologs #3397. Best score 1137 bits
Score difference with first non-orthologous sequence - *A. fumigatus*:1137 *A. fischeri*:1137
afum|EAL91871.1 100.00% afis|EAW18060.1 100.00%
Bootstrap support for afum|EAL91871.1 as seed ortholog is 100%.
Bootstrap support for afis|EAW18060.1 as seed ortholog is 100%.

---

Group of orthologs #3398. Best score 1137 bits
Score difference with first non-orthologous sequence - *A. fumigatus*:1137 *A. fischeri*:1137
afum|EAL93134.1 100.00% afis|EAW18449.1 100.00%
Bootstrap support for afum|EAL93134.1 as seed ortholog is 100%.
Bootstrap support for afis|EAW18449.1 as seed ortholog is 100%.

---

Group of orthologs #3399. Best score 1137 bits
Score difference with first non-orthologous sequence - *A. fumigatus*:842 *A. fischeri*:843
afum|EBA27372.1 100.00% afis|EAW20413.1 100.00%
Bootstrap support for afum|EBA27372.1 as seed ortholog is 100%.
Bootstrap support for afis|EAW20413.1 as seed ortholog is 100%.

---

Group of orthologs #3400. Best score 1137 bits
Score difference with first non-orthologous sequence - *A. fumigatus*:1137 *A. fischeri*:1137
afum|EAL89738.1 100.00% afis|EAW25069.1 100.00%
Bootstrap support for afum|EAL89738.1 as seed ortholog is 100%.
Bootstrap support for afis|EAW25069.1 as seed ortholog is 100%.

---

Group of orthologs #3401. Best score 1136 bits
Score difference with first non-orthologous sequence - *A. fumigatus*:1136 *A. fischeri*:1136
afum|EAL86448.1 100.00% afis|EAW16921.1 100.00%
Bootstrap support for afum|EAL86448.1 as seed ortholog is 100%.
Bootstrap support for afis|EAW16921.1 as seed ortholog is 100%.

---

Group of orthologs #3402. Best score 1136 bits
Score difference with first non-orthologous sequence - *A. fumigatus*:1136 *A. fischeri*:1136
afum|EAL84522.1 100.00% afis|EAW20536.1 100.00%
Bootstrap support for afum|EAL84522.1 as seed ortholog is 100%.
Bootstrap support for afis|EAW20536.1 as seed ortholog is 100%.

---

Group of orthologs #3403. Best score 1136 bits
Score difference with first non-orthologous sequence - *A. fumigatus*:993 *A. fischeri*:831
afum|EAL87549.1 100.00% afis|EAW18186.1 100.00%
Bootstrap support for afum|EAL87549.1 as seed ortholog is 100%.
Bootstrap support for afis|EAW18186.1 as seed ortholog is 100%.

---

Group of orthologs #3404. Best score 1136 bits

Score difference with first non-orthologous sequence - *A. fumigatus*:1136 *A. fischeri*:1136
afum|EAL91785.1 100.00% afis|EAW17976.1 100.00%
Bootstrap support for afum|EAL91785.1 as seed ortholog is 100%.
Bootstrap support for afis|EAW17976.1 as seed ortholog is 100%.

---

Group of orthologs #3405. Best score 1136 bits
Score difference with first non-orthologous sequence - *A. fumigatus*:574 *A. fischeri*:512
afum|EAL86044.1 100.00% afis|EAW24372.1 100.00%
Bootstrap support for afum|EAL86044.1 as seed ortholog is 100%.
Bootstrap support for afis|EAW24372.1 as seed ortholog is 100%.

---

Group of orthologs #3406. Best score 1136 bits
Score difference with first non-orthologous sequence - *A. fumigatus*:914 *A. fischeri*:860
afum|EAL86195.1 100.00% afis|EAW24530.1 100.00%
Bootstrap support for afum|EAL86195.1 as seed ortholog is 100%.
Bootstrap support for afis|EAW24530.1 as seed ortholog is 100%.

---

Group of orthologs #3407. Best score 1136 bits
Score difference with first non-orthologous sequence - *A. fumigatus*:1002 *A. fischeri*:1136
afum|EAL88430.1 100.00% afis|EAW23050.1 100.00%
Bootstrap support for afum|EAL88430.1 as seed ortholog is 100%.
Bootstrap support for afis|EAW23050.1 as seed ortholog is 100%.

---

Group of orthologs #3408. Best score 1136 bits
Score difference with first non-orthologous sequence - *A. fumigatus*:1136 *A. fischeri*:1136
afum|EAL93465.1 100.00% afis|EAW18739.1 100.00%
Bootstrap support for afum|EAL93465.1 as seed ortholog is 100%.
Bootstrap support for afis|EAW18739.1 as seed ortholog is 100%.

---

Group of orthologs #3409. Best score 1136 bits
Score difference with first non-orthologous sequence - *A. fumigatus*:1057 *A. fischeri*:1136
afum|EAL90024.1 100.00% afis|EAW25362.1 100.00%
Bootstrap support for afum|EAL90024.1 as seed ortholog is 100%.
Bootstrap support for afis|EAW25362.1 as seed ortholog is 100%.

---

Group of orthologs #3410. Best score 1136 bits
Score difference with first non-orthologous sequence - *A. fumigatus*:1136 *A. fischeri*:1136
afum|EAL90089.1 100.00% afis|EAW25431.1 100.00%
Bootstrap support for afum|EAL90089.1 as seed ortholog is 100%.
Bootstrap support for afis|EAW25431.1 as seed ortholog is 100%.

---

Group of orthologs #3411. Best score 1135 bits
Score difference with first non-orthologous sequence - *A. fumigatus*:823 *A. fischeri*:955
afum|EAL91688.1 100.00% afis|EAW17880.1 100.00%
Bootstrap support for afum|EAL91688.1 as seed ortholog is 100%.
Bootstrap support for afis|EAW17880.1 as seed ortholog is 100%.

---

Group of orthologs #3412. Best score 1135 bits
Score difference with first non-orthologous sequence - *A. fumigatus*:1135 *A. fischeri*:1135
afum|EAL91784.1 100.00% afis|EAW17975.1 100.00%
Bootstrap support for afum|EAL91784.1 as seed ortholog is 100%.
Bootstrap support for afis|EAW17975.1 as seed ortholog is 100%.

---

Group of orthologs #3413. Best score 1135 bits
Score difference with first non-orthologous sequence - *A. fumigatus*:1067 *A. fischeri*:949
afum|EBA27281.1 100.00% afis|EAW17185.1 100.00%

Bootstrap support for afum|EBA27281.1 as seed ortholog is 100%.
Bootstrap support for afis|EAW17185.1 as seed ortholog is 100%.

---

Group of orthologs #3414. Best score 1135 bits
Score difference with first non-orthologous sequence - *A. fumigatus*:1135 *A. fischeri*:1135
afum|EAL89337.1 100.00% afis|EAW24695.1 100.00%
Bootstrap support for afum|EAL89337.1 as seed ortholog is 100%.
Bootstrap support for afis|EAW24695.1 as seed ortholog is 100%.

---

Group of orthologs #3415. Best score 1134 bits
Score difference with first non-orthologous sequence - *A. fumigatus*:1134 *A. fischeri*:1134
afum|EAL85847.1 100.00% afis|EAW15474.1 100.00%
Bootstrap support for afum|EAL85847.1 as seed ortholog is 100%.
Bootstrap support for afis|EAW15474.1 as seed ortholog is 100%.

---

Group of orthologs #3416. Best score 1134 bits
Score difference with first non-orthologous sequence - *A. fumigatus*:920 *A. fischeri*:892
afum|EAL85091.1 100.00% afis|EAW19799.1 100.00%
Bootstrap support for afum|EAL85091.1 as seed ortholog is 100%.
Bootstrap support for afis|EAW19799.1 as seed ortholog is 100%.

---

Group of orthologs #3417. Best score 1134 bits
Score difference with first non-orthologous sequence - *A. fumigatus*:1134 *A. fischeri*:1134
afum|EAL91820.1 100.00% afis|EAW18008.1 100.00%
Bootstrap support for afum|EAL91820.1 as seed ortholog is 100%.
Bootstrap support for afis|EAW18008.1 as seed ortholog is 100%.

---

Group of orthologs #3418. Best score 1134 bits
Score difference with first non-orthologous sequence - *A. fumigatus*:1134 *A. fischeri*:1134
afum|EAL87209.1 100.00% afis|EAW23775.1 100.00%
Bootstrap support for afum|EAL87209.1 as seed ortholog is 100%.
Bootstrap support for afis|EAW23775.1 as seed ortholog is 100%.

---

Group of orthologs #3419. Best score 1134 bits
Score difference with first non-orthologous sequence - *A. fumigatus*:1134 *A. fischeri*:1134
afum|EAL93414.1 100.00% afis|EAW18693.1 100.00%
Bootstrap support for afum|EAL93414.1 as seed ortholog is 100%.
Bootstrap support for afis|EAW18693.1 as seed ortholog is 100%.

---

Group of orthologs #3420. Best score 1134 bits
Score difference with first non-orthologous sequence - *A. fumigatus*:1134 *A. fischeri*:1134
afum|EAL92764.1 100.00% afis|EAW21731.1 100.00%
Bootstrap support for afum|EAL92764.1 as seed ortholog is 100%.
Bootstrap support for afis|EAW21731.1 as seed ortholog is 100%.

---

Group of orthologs #3421. Best score 1134 bits
Score difference with first non-orthologous sequence - *A. fumigatus*:1134 *A. fischeri*:1134
afum|EAL89757.1 100.00% afis|EAW25088.1 100.00%
Bootstrap support for afum|EAL89757.1 as seed ortholog is 100%.
Bootstrap support for afis|EAW25088.1 as seed ortholog is 100%.

---

Group of orthologs #3422. Best score 1134 bits
Score difference with first non-orthologous sequence - *A. fumigatus*:1134 *A. fischeri*:1134
afum|EAL89812.1 100.00% afis|EAW25140.1 100.00%
Bootstrap support for afum|EAL89812.1 as seed ortholog is 100%.
Bootstrap support for afis|EAW25140.1 as seed ortholog is 100%.

Group of orthologs #3423. Best score 1134 bits
Score difference with first non-orthologous sequence - *A. fumigatus*:1134 *A. fischeri*:1134
afum|EAL91925.1 100.00% afis|EAW24069.1 100.00%
Bootstrap support for afum|EAL91925.1 as seed ortholog is 100%.
Bootstrap support for afis|EAW24069.1 as seed ortholog is 100%.

Group of orthologs #3424. Best score 1133 bits
Score difference with first non-orthologous sequence - *A. fumigatus*:1133 *A. fischeri*:1133
afum|EAL90823.1 100.00% afis|EAW22369.1 100.00%
Bootstrap support for afum|EAL90823.1 as seed ortholog is 100%.
Bootstrap support for afis|EAW22369.1 as seed ortholog is 100%.

Group of orthologs #3425. Best score 1133 bits
Score difference with first non-orthologous sequence - *A. fumigatus*:1133 *A. fischeri*:1133
afum|EAL92649.1 100.00% afis|EAW21615.1 100.00%
Bootstrap support for afum|EAL92649.1 as seed ortholog is 100%.
Bootstrap support for afis|EAW21615.1 as seed ortholog is 100%.

Group of orthologs #3426. Best score 1133 bits
Score difference with first non-orthologous sequence - *A. fumigatus*:1133 *A. fischeri*:1133
afum|EAL89927.1 100.00% afis|EAW25264.1 100.00%
Bootstrap support for afum|EAL89927.1 as seed ortholog is 100%.
Bootstrap support for afis|EAW25264.1 as seed ortholog is 100%.

Group of orthologs #3427. Best score 1133 bits
Score difference with first non-orthologous sequence - *A. fumigatus*:1133 *A. fischeri*:1133
afum|EAL91929.1 100.00% afis|EAW24073.1 100.00%
Bootstrap support for afum|EAL91929.1 as seed ortholog is 100%.
Bootstrap support for afis|EAW24073.1 as seed ortholog is 100%.

Group of orthologs #3428. Best score 1132 bits
Score difference with first non-orthologous sequence - *A. fumigatus*:1132 *A. fischeri*:1132
afum|EAL84467.1 100.00% afis|EAW20594.1 100.00%
Bootstrap support for afum|EAL84467.1 as seed ortholog is 100%.
Bootstrap support for afis|EAW20594.1 as seed ortholog is 100%.

Group of orthologs #3429. Best score 1132 bits
Score difference with first non-orthologous sequence - *A. fumigatus*:1132 *A. fischeri*:1132
afum|EAL89239.1 100.00% afis|EAW16637.1 100.00%
Bootstrap support for afum|EAL89239.1 as seed ortholog is 100%.
Bootstrap support for afis|EAW16637.1 as seed ortholog is 100%.

Group of orthologs #3430. Best score 1132 bits
Score difference with first non-orthologous sequence - *A. fumigatus*:1132 *A. fischeri*:1132
afum|EAL91353.1 100.00% afis|EAW17539.1 100.00%
Bootstrap support for afum|EAL91353.1 as seed ortholog is 100%.
Bootstrap support for afis|EAW17539.1 as seed ortholog is 100%.

Group of orthologs #3431. Best score 1132 bits
Score difference with first non-orthologous sequence - *A. fumigatus*:1132 *A. fischeri*:1132
afum|EAL93061.1 100.00% afis|EAW18386.1 100.00%
Bootstrap support for afum|EAL93061.1 as seed ortholog is 100%.
Bootstrap support for afis|EAW18386.1 as seed ortholog is 100%.

Group of orthologs #3432. Best score 1132 bits

Score difference with first non-orthologous sequence - *A. fumigatus*:1132   *A. fischeri*:1132
afum|EAL90762.1 100.00% afis|EAW22429.1 100.00%
Bootstrap support for afum|EAL90762.1 as seed ortholog is 100%.
Bootstrap support for afis|EAW22429.1 as seed ortholog is 100%.

---

Group of orthologs #3433. Best score 1131 bits
Score difference with first non-orthologous sequence - *A. fumigatus*:1131   *A. fischeri*:1131
afum|EAL89180.1 100.00% afis|EAW16585.1 100.00%
Bootstrap support for afum|EAL89180.1 as seed ortholog is 100%.
Bootstrap support for afis|EAW16585.1 as seed ortholog is 100%.

---

Group of orthologs #3434. Best score 1131 bits
Score difference with first non-orthologous sequence - *A. fumigatus*:815   *A. fischeri*:1131
afum|EAL84848.1 100.00% afis|EAW20771.1 100.00%
Bootstrap support for afum|EAL84848.1 as seed ortholog is 100%.
Bootstrap support for afis|EAW20771.1 as seed ortholog is 100%.

---

Group of orthologs #3435. Best score 1131 bits
Score difference with first non-orthologous sequence - *A. fumigatus*:1131   *A. fischeri*:1131
afum|EAL87370.1 100.00% afis|EAW23945.1 100.00%
Bootstrap support for afum|EAL87370.1 as seed ortholog is 100%.
Bootstrap support for afis|EAW23945.1 as seed ortholog is 100%.

---

Group of orthologs #3436. Best score 1131 bits
Score difference with first non-orthologous sequence - *A. fumigatus*:1131   *A. fischeri*:1131
afum|EAL87404.1 100.00% afis|EAW23974.1 100.00%
Bootstrap support for afum|EAL87404.1 as seed ortholog is 100%.
Bootstrap support for afis|EAW23974.1 as seed ortholog is 100%.

---

Group of orthologs #3437. Best score 1131 bits
Score difference with first non-orthologous sequence - *A. fumigatus*:1131   *A. fischeri*:841
afum|EAL87447.1 100.00% afis|EAW24017.1 100.00%
Bootstrap support for afum|EAL87447.1 as seed ortholog is 100%.
Bootstrap support for afis|EAW24017.1 as seed ortholog is 100%.

---

Group of orthologs #3438. Best score 1131 bits
Score difference with first non-orthologous sequence - *A. fumigatus*:1131   *A. fischeri*:1131
afum|EAL93331.1 100.00% afis|EAW18621.1 100.00%
Bootstrap support for afum|EAL93331.1 as seed ortholog is 100%.
Bootstrap support for afis|EAW18621.1 as seed ortholog is 100%.

---

Group of orthologs #3439. Best score 1131 bits
Score difference with first non-orthologous sequence - *A. fumigatus*:1131   *A. fischeri*:1131
afum|EAL90924.1 100.00% afis|EAW22254.1 100.00%
Bootstrap support for afum|EAL90924.1 as seed ortholog is 100%.
Bootstrap support for afis|EAW22254.1 as seed ortholog is 100%.

---

Group of orthologs #3440. Best score 1131 bits
Score difference with first non-orthologous sequence - *A. fumigatus*:1131   *A. fischeri*:1131
afum|EAL90818.1 100.00% afis|EAW22374.1 100.00%
Bootstrap support for afum|EAL90818.1 as seed ortholog is 100%.
Bootstrap support for afis|EAW22374.1 as seed ortholog is 100%.

---

Group of orthologs #3441. Best score 1131 bits
Score difference with first non-orthologous sequence - *A. fumigatus*:954   *A. fischeri*:879
afum|EAL92663.1 100.00% afis|EAW21628.1 100.00%

Bootstrap support for afum|EAL92663.1 as seed ortholog is 100%.
Bootstrap support for afis|EAW21628.1 as seed ortholog is 100%.

---

Group of orthologs #3442. Best score 1130 bits
Score difference with first non-orthologous sequence - *A. fumigatus*:1130 *A. fischeri*:1130
afum|EAL86382.2 100.00% afis|EAW16835.1 100.00%
Bootstrap support for afum|EAL86382.2 as seed ortholog is 100%.
Bootstrap support for afis|EAW16835.1 as seed ortholog is 100%.

---

Group of orthologs #3443. Best score 1130 bits
Score difference with first non-orthologous sequence - *A. fumigatus*:1130 *A. fischeri*:731
afum|EAL85042.1 100.00% afis|EAW19851.1 100.00%
Bootstrap support for afum|EAL85042.1 as seed ortholog is 100%.
Bootstrap support for afis|EAW19851.1 as seed ortholog is 100%.

---

Group of orthologs #3444. Best score 1130 bits
Score difference with first non-orthologous sequence - *A. fumigatus*:1130 *A. fischeri*:1130
afum|EAL87540.1 100.00% afis|EAW18179.1 100.00%
Bootstrap support for afum|EAL87540.1 as seed ortholog is 100%.
Bootstrap support for afis|EAW18179.1 as seed ortholog is 100%.

---

Group of orthologs #3445. Best score 1130 bits
Score difference with first non-orthologous sequence - *A. fumigatus*:1015 *A. fischeri*:1038
afum|EAL89270.1 100.00% afis|EAW16666.1 100.00%
Bootstrap support for afum|EAL89270.1 as seed ortholog is 100%.
Bootstrap support for afis|EAW16666.1 as seed ortholog is 100%.

---

Group of orthologs #3446. Best score 1130 bits
Score difference with first non-orthologous sequence - *A. fumigatus*:956 *A. fischeri*:1130
afum|EAL85488.1 100.00% afis|EAW20374.1 100.00%
Bootstrap support for afum|EAL85488.1 as seed ortholog is 100%.
Bootstrap support for afis|EAW20374.1 as seed ortholog is 100%.

---

Group of orthologs #3447. Best score 1130 bits
Score difference with first non-orthologous sequence - *A. fumigatus*:1130 *A. fischeri*:1130
afum|EAL88323.1 100.00% afis|EAW23160.1 100.00%
Bootstrap support for afum|EAL88323.1 as seed ortholog is 100%.
Bootstrap support for afis|EAW23160.1 as seed ortholog is 100%.

---

Group of orthologs #3448. Best score 1130 bits
Score difference with first non-orthologous sequence - *A. fumigatus*:1130 *A. fischeri*:1130
afum|EAL88218.1 100.00% afis|EAW23262.1 100.00%
Bootstrap support for afum|EAL88218.1 as seed ortholog is 100%.
Bootstrap support for afis|EAW23262.1 as seed ortholog is 100%.

---

Group of orthologs #3449. Best score 1130 bits
Score difference with first non-orthologous sequence - *A. fumigatus*:1130 *A. fischeri*:1051
afum|EAL90553.2 100.00% afis|EAW22639.1 100.00%
Bootstrap support for afum|EAL90553.2 as seed ortholog is 100%.
Bootstrap support for afis|EAW22639.1 as seed ortholog is 100%.

---

Group of orthologs #3450. Best score 1130 bits
Score difference with first non-orthologous sequence - *A. fumigatus*:986 *A. fischeri*:815
afum|EAL90572.1 100.00% afis|EAW22624.1 100.00%
Bootstrap support for afum|EAL90572.1 as seed ortholog is 100%.
Bootstrap support for afis|EAW22624.1 as seed ortholog is 100%.

---

Group of orthologs #3451. Best score 1130 bits
Score difference with first non-orthologous sequence - *A. fumigatus*:872 *A. fischeri*:835
afum|EAL92311.1 100.00% afis|EAW21294.1 100.00%
Bootstrap support for afum|EAL92311.1 as seed ortholog is 100%.
Bootstrap support for afis|EAW21294.1 as seed ortholog is 100%.

---

Group of orthologs #3452. Best score 1130 bits
Score difference with first non-orthologous sequence - *A. fumigatus*:1130 *A. fischeri*:1130
afum|EAL89422.1 100.00% afis|EAW24783.1 100.00%
Bootstrap support for afum|EAL89422.1 as seed ortholog is 100%.
Bootstrap support for afis|EAW24783.1 as seed ortholog is 100%.

---

Group of orthologs #3453. Best score 1130 bits
Score difference with first non-orthologous sequence - *A. fumigatus*:1130 *A. fischeri*:1130
afum|EAL92913.1 100.00% afis|EAW21900.1 100.00%
Bootstrap support for afum|EAL92913.1 as seed ortholog is 100%.
Bootstrap support for afis|EAW21900.1 as seed ortholog is 100%.

---

Group of orthologs #3454. Best score 1130 bits
Score difference with first non-orthologous sequence - *A. fumigatus*:1130 *A. fischeri*:1130
afum|EAL92979.1 100.00% afis|EAW21965.1 100.00%
Bootstrap support for afum|EAL92979.1 as seed ortholog is 100%.
Bootstrap support for afis|EAW21965.1 as seed ortholog is 100%.

---

Group of orthologs #3455. Best score 1129 bits
Score difference with first non-orthologous sequence - *A. fumigatus*:1129 *A. fischeri*:1129
afum|EAL85593.1 100.00% afis|EAW15756.1 100.00%
Bootstrap support for afum|EAL85593.1 as seed ortholog is 100%.
Bootstrap support for afis|EAW15756.1 as seed ortholog is 100%.

---

Group of orthologs #3456. Best score 1129 bits
Score difference with first non-orthologous sequence - *A. fumigatus*:981 *A. fischeri*:1129
afum|EAL86900.1 100.00% afis|EAW19621.1 100.00%
Bootstrap support for afum|EAL86900.1 as seed ortholog is 100%.
Bootstrap support for afis|EAW19621.1 as seed ortholog is 100%.

---

Group of orthologs #3457. Best score 1129 bits
Score difference with first non-orthologous sequence - *A. fumigatus*:1129 *A. fischeri*:1129
afum|EAL86043.1 100.00% afis|EAW24371.1 100.00%
Bootstrap support for afum|EAL86043.1 as seed ortholog is 100%.
Bootstrap support for afis|EAW24371.1 as seed ortholog is 100%.

---

Group of orthologs #3458. Best score 1129 bits
Score difference with first non-orthologous sequence - *A. fumigatus*:1129 *A. fischeri*:1129
afum|EAL91924.1 100.00% afis|EAW24068.1 100.00%
Bootstrap support for afum|EAL91924.1 as seed ortholog is 100%.
Bootstrap support for afis|EAW24068.1 as seed ortholog is 100%.

---

Group of orthologs #3459. Best score 1128 bits
Score difference with first non-orthologous sequence - *A. fumigatus*:1128 *A. fischeri*:1128
afum|EAL85293.1 100.00% afis|EAW20175.1 100.00%
Bootstrap support for afum|EAL85293.1 as seed ortholog is 100%.
Bootstrap support for afis|EAW20175.1 as seed ortholog is 100%.

---

Group of orthologs #3460. Best score 1128 bits

Score difference with first non-orthologous sequence - *A. fumigatus*:1128   *A. fischeri*:1128
afum|EAL87291.1 100.00% afis|EAW23863.1 100.00%
Bootstrap support for afum|EAL87291.1 as seed ortholog is 100%.
Bootstrap support for afis|EAW23863.1 as seed ortholog is 100%.

---

Group of orthologs #3461. Best score 1128 bits
Score difference with first non-orthologous sequence - *A. fumigatus*:1128   *A. fischeri*:1128
afum|EAL88025.1 100.00% afis|EAW23437.1 100.00%
Bootstrap support for afum|EAL88025.1 as seed ortholog is 100%.
Bootstrap support for afis|EAW23437.1 as seed ortholog is 100%.

---

Group of orthologs #3462. Best score 1128 bits
Score difference with first non-orthologous sequence - *A. fumigatus*:1128   *A. fischeri*:1128
afum|EAL88103.2 100.00% afis|EAW23370.1 100.00%
Bootstrap support for afum|EAL88103.2 as seed ortholog is 100%.
Bootstrap support for afis|EAW23370.1 as seed ortholog is 100%.

---

Group of orthologs #3463. Best score 1128 bits
Score difference with first non-orthologous sequence - *A. fumigatus*:929   *A. fischeri*:933
afum|EAL92290.2 100.00% afis|EAW21275.1 100.00%
Bootstrap support for afum|EAL92290.2 as seed ortholog is 100%.
Bootstrap support for afis|EAW21275.1 as seed ortholog is 100%.

---

Group of orthologs #3464. Best score 1128 bits
Score difference with first non-orthologous sequence - *A. fumigatus*:1128   *A. fischeri*:1128
afum|EAL89804.1 100.00% afis|EAW25133.1 100.00%
Bootstrap support for afum|EAL89804.1 as seed ortholog is 100%.
Bootstrap support for afis|EAW25133.1 as seed ortholog is 100%.

---

Group of orthologs #3465. Best score 1128 bits
Score difference with first non-orthologous sequence - *A. fumigatus*:1128   *A. fischeri*:1128
afum|EAL89933.1 100.00% afis|EAW25270.1 100.00%
Bootstrap support for afum|EAL89933.1 as seed ortholog is 100%.
Bootstrap support for afis|EAW25270.1 as seed ortholog is 100%.

---

Group of orthologs #3466. Best score 1127 bits
Score difference with first non-orthologous sequence - *A. fumigatus*:1127   *A. fischeri*:1022
afum|EAL88928.1 100.00% afis|EAW16306.1 100.00%
Bootstrap support for afum|EAL88928.1 as seed ortholog is 100%.
Bootstrap support for afis|EAW16306.1 as seed ortholog is 100%.

---

Group of orthologs #3467. Best score 1127 bits
Score difference with first non-orthologous sequence - *A. fumigatus*:1127   *A. fischeri*:1127
afum|EAL85256.1 100.00% afis|EAW20141.1 100.00%
Bootstrap support for afum|EAL85256.1 as seed ortholog is 100%.
Bootstrap support for afis|EAW20141.1 as seed ortholog is 100%.

---

Group of orthologs #3468. Best score 1127 bits
Score difference with first non-orthologous sequence - *A. fumigatus*:1127   *A. fischeri*:1005
afum|EAL91183.1 100.00% afis|EAW17339.1 100.00%
Bootstrap support for afum|EAL91183.1 as seed ortholog is 100%.
Bootstrap support for afis|EAW17339.1 as seed ortholog is 100%.

---

Group of orthologs #3469. Best score 1127 bits
Score difference with first non-orthologous sequence - *A. fumigatus*:1127   *A. fischeri*:1127
afum|EAL91736.1 100.00% afis|EAW17925.1 100.00%

Bootstrap support for afum|EAL91736.1 as seed ortholog is 100%.
Bootstrap support for afis|EAW17925.1 as seed ortholog is 100%.

---

Group of orthologs #3470. Best score 1127 bits
Score difference with first non-orthologous sequence - *A. fumigatus*:1127 *A. fischeri*:1127
afum|EAL91891.1 100.00% afis|EAW18079.1 100.00%
Bootstrap support for afum|EAL91891.1 as seed ortholog is 100%.
Bootstrap support for afis|EAW18079.1 as seed ortholog is 100%.

---

Group of orthologs #3471. Best score 1127 bits
Score difference with first non-orthologous sequence - *A. fumigatus*:732 *A. fischeri*:879
afum|EAL88074.1 100.00% afis|EAW23396.1 100.00%
Bootstrap support for afum|EAL88074.1 as seed ortholog is 100%.
Bootstrap support for afis|EAW23396.1 as seed ortholog is 100%.

---

Group of orthologs #3472. Best score 1127 bits
Score difference with first non-orthologous sequence - *A. fumigatus*:1127 *A. fischeri*:1127
afum|EAL93634.1 100.00% afis|EAW18902.1 100.00%
Bootstrap support for afum|EAL93634.1 as seed ortholog is 100%.
Bootstrap support for afis|EAW18902.1 as seed ortholog is 100%.

---

Group of orthologs #3473. Best score 1126 bits
Score difference with first non-orthologous sequence - *A. fumigatus*:1126 *A. fischeri*:810
afum|EAL86784.2 100.00% afis|EAW19742.1 100.00%
Bootstrap support for afum|EAL86784.2 as seed ortholog is 100%.
Bootstrap support for afis|EAW19742.1 as seed ortholog is 100%.

---

Group of orthologs #3474. Best score 1126 bits
Score difference with first non-orthologous sequence - *A. fumigatus*:1126 *A. fischeri*:1126
afum|EAL91260.1 100.00% afis|EAW17432.1 100.00%
Bootstrap support for afum|EAL91260.1 as seed ortholog is 100%.
Bootstrap support for afis|EAW17432.1 as seed ortholog is 100%.

---

Group of orthologs #3475. Best score 1126 bits
Score difference with first non-orthologous sequence - *A. fumigatus*:1126 *A. fischeri*:1126
afum|EAL91829.1 100.00% afis|EAW18017.1 100.00%
Bootstrap support for afum|EAL91829.1 as seed ortholog is 100%.
Bootstrap support for afis|EAW18017.1 as seed ortholog is 100%.

---

Group of orthologs #3476. Best score 1125 bits
Score difference with first non-orthologous sequence - *A. fumigatus*:1125 *A. fischeri*:1125
afum|EAL88528.1 100.00% afis|EAW15893.1 100.00%
Bootstrap support for afum|EAL88528.1 as seed ortholog is 100%.
Bootstrap support for afis|EAW15893.1 as seed ortholog is 100%.

---

Group of orthologs #3477. Best score 1125 bits
Score difference with first non-orthologous sequence - *A. fumigatus*:1125 *A. fischeri*:1125
afum|EAL88727.1 100.00% afis|EAW16095.1 100.00%
Bootstrap support for afum|EAL88727.1 as seed ortholog is 100%.
Bootstrap support for afis|EAW16095.1 as seed ortholog is 100%.

---

Group of orthologs #3478. Best score 1125 bits
Score difference with first non-orthologous sequence - *A. fumigatus*:1125 *A. fischeri*:1125
afum|EAL87494.1 100.00% afis|EAW18137.1 100.00%
Bootstrap support for afum|EAL87494.1 as seed ortholog is 100%.
Bootstrap support for afis|EAW18137.1 as seed ortholog is 100%.

Group of orthologs #3479. Best score 1125 bits
Score difference with first non-orthologous sequence - *A. fumigatus*:990 *A. fischeri*:842
afum|EAL86136.1 100.00% afis|EAW24465.1 100.00%
Bootstrap support for afum|EAL86136.1 as seed ortholog is 100%.
Bootstrap support for afis|EAW24465.1 as seed ortholog is 100%.

---

Group of orthologs #3480. Best score 1125 bits
Score difference with first non-orthologous sequence - *A. fumigatus*:1125 *A. fischeri*:1125
afum|EAL86144.1 100.00% afis|EAW24471.1 100.00%
Bootstrap support for afum|EAL86144.1 as seed ortholog is 100%.
Bootstrap support for afis|EAW24471.1 as seed ortholog is 100%.

---

Group of orthologs #3481. Best score 1125 bits
Score difference with first non-orthologous sequence - *A. fumigatus*:1125 *A. fischeri*:1125
afum|EAL92078.1 100.00% afis|EAW21039.1 100.00%
Bootstrap support for afum|EAL92078.1 as seed ortholog is 100%.
Bootstrap support for afis|EAW21039.1 as seed ortholog is 100%.

---

Group of orthologs #3482. Best score 1125 bits
Score difference with first non-orthologous sequence - *A. fumigatus*:1125 *A. fischeri*:954
afum|EAL89351.1 100.00% afis|EAW24710.1 100.00%
Bootstrap support for afum|EAL89351.1 as seed ortholog is 100%.
Bootstrap support for afis|EAW24710.1 as seed ortholog is 100%.

---

Group of orthologs #3483. Best score 1125 bits
Score difference with first non-orthologous sequence - *A. fumigatus*:1125 *A. fischeri*:1125
afum|EAL92680.1 100.00% afis|EAW21646.1 100.00%
Bootstrap support for afum|EAL92680.1 as seed ortholog is 100%.
Bootstrap support for afis|EAW21646.1 as seed ortholog is 100%.

---

Group of orthologs #3484. Best score 1124 bits
Score difference with first non-orthologous sequence - *A. fumigatus*:1124 *A. fischeri*:1124
afum|EAL84294.1 100.00% afis|EAW25587.1 100.00%
Bootstrap support for afum|EAL84294.1 as seed ortholog is 100%.
Bootstrap support for afis|EAW25587.1 as seed ortholog is 100%.

---

Group of orthologs #3485. Best score 1124 bits
Score difference with first non-orthologous sequence - *A. fumigatus*:1124 *A. fischeri*:1124
afum|EAL91914.1 100.00% afis|EAW24059.1 100.00%
Bootstrap support for afum|EAL91914.1 as seed ortholog is 100%.
Bootstrap support for afis|EAW24059.1 as seed ortholog is 100%.

---

Group of orthologs #3486. Best score 1123 bits
Score difference with first non-orthologous sequence - *A. fumigatus*:790 *A. fischeri*:680
afum|EAL86982.1 100.00% afis|EAW19514.1 100.00%
Bootstrap support for afum|EAL86982.1 as seed ortholog is 100%.
Bootstrap support for afis|EAW19514.1 as seed ortholog is 100%.

---

Group of orthologs #3487. Best score 1123 bits
Score difference with first non-orthologous sequence - *A. fumigatus*:1123 *A. fischeri*:1123
afum|EAL85999.1 100.00% afis|EAW24329.1 100.00%
Bootstrap support for afum|EAL85999.1 as seed ortholog is 100%.
Bootstrap support for afis|EAW24329.1 as seed ortholog is 100%.

---

Group of orthologs #3488. Best score 1123 bits

Score difference with first non-orthologous sequence - *A. fumigatus*:1123 *A. fischeri*:1123
afum|EAL90395.1 100.00% afis|EAW22804.1 100.00%
Bootstrap support for afum|EAL90395.1 as seed ortholog is 100%.
Bootstrap support for afis|EAW22804.1 as seed ortholog is 100%.

---

Group of orthologs #3489. Best score 1122 bits
Score difference with first non-orthologous sequence - *A. fumigatus*:1122 *A. fischeri*:1122
afum|EAL88536.1 100.00% afis|EAW15901.1 100.00%
Bootstrap support for afum|EAL88536.1 as seed ortholog is 100%.
Bootstrap support for afis|EAW15901.1 as seed ortholog is 100%.

---

Group of orthologs #3490. Best score 1122 bits
Score difference with first non-orthologous sequence - *A. fumigatus*:1122 *A. fischeri*:1122
afum|EAL88623.1 100.00% afis|EAW15991.1 100.00%
Bootstrap support for afum|EAL88623.1 as seed ortholog is 100%.
Bootstrap support for afis|EAW15991.1 as seed ortholog is 100%.

---

Group of orthologs #3491. Best score 1122 bits
Score difference with first non-orthologous sequence - *A. fumigatus*:1122 *A. fischeri*:1122
afum|EAL88930.1 100.00% afis|EAW16308.1 100.00%
Bootstrap support for afum|EAL88930.1 as seed ortholog is 100%.
Bootstrap support for afis|EAW16308.1 as seed ortholog is 100%.

---

Group of orthologs #3492. Best score 1122 bits
Score difference with first non-orthologous sequence - *A. fumigatus*:1029 *A. fischeri*:1018
afum|EAL84326.1 100.00% afis|EAW25639.1 100.00%
Bootstrap support for afum|EAL84326.1 as seed ortholog is 100%.
Bootstrap support for afis|EAW25639.1 as seed ortholog is 100%.

---

Group of orthologs #3493. Best score 1121 bits
Score difference with first non-orthologous sequence - *A. fumigatus*:1121 *A. fischeri*:1121
afum|EAL85598.1 100.00% afis|EAW15751.1 100.00%
Bootstrap support for afum|EAL85598.1 as seed ortholog is 100%.
Bootstrap support for afis|EAW15751.1 as seed ortholog is 100%.

---

Group of orthologs #3494. Best score 1121 bits
Score difference with first non-orthologous sequence - *A. fumigatus*:1121 *A. fischeri*:1121
afum|EAL92337.1 100.00% afis|EAW21320.1 100.00%
Bootstrap support for afum|EAL92337.1 as seed ortholog is 100%.
Bootstrap support for afis|EAW21320.1 as seed ortholog is 100%.

---

Group of orthologs #3495. Best score 1121 bits
Score difference with first non-orthologous sequence - *A. fumigatus*:1121 *A. fischeri*:1121
afum|EAL89448.1 100.00% afis|EAW24806.1 100.00%
Bootstrap support for afum|EAL89448.1 as seed ortholog is 100%.
Bootstrap support for afis|EAW24806.1 as seed ortholog is 100%.

---

Group of orthologs #3496. Best score 1121 bits
Score difference with first non-orthologous sequence - *A. fumigatus*:1121 *A. fischeri*:1121
afum|EAL89831.1 100.00% afis|EAW25156.1 100.00%
Bootstrap support for afum|EAL89831.1 as seed ortholog is 100%.
Bootstrap support for afis|EAW25156.1 as seed ortholog is 100%.

---

Group of orthologs #3497. Best score 1120 bits
Score difference with first non-orthologous sequence - *A. fumigatus*:1120 *A. fischeri*:1120
afum|EAL87702.1 100.00% afis|EAW18339.1 100.00%

Bootstrap support for afum|EAL87702.1 as seed ortholog is 100%.
Bootstrap support for afis|EAW18339.1 as seed ortholog is 100%.

---

Group of orthologs #3498. Best score 1120 bits
Score difference with first non-orthologous sequence - *A. fumigatus*:1120 *A. fischeri*:1120
afum|EAL86904.1 100.00% afis|EAW19618.1 100.00%
Bootstrap support for afum|EAL86904.1 as seed ortholog is 100%.
Bootstrap support for afis|EAW19618.1 as seed ortholog is 100%.

---

Group of orthologs #3499. Best score 1120 bits
Score difference with first non-orthologous sequence - *A. fumigatus*:972 *A. fischeri*:1120
afum|EAL90398.1 100.00% afis|EAW22801.1 100.00%
Bootstrap support for afum|EAL90398.1 as seed ortholog is 100%.
Bootstrap support for afis|EAW22801.1 as seed ortholog is 100%.

---

Group of orthologs #3500. Best score 1119 bits
Score difference with first non-orthologous sequence - *A. fumigatus*:1119 *A. fischeri*:1119
afum|EAL85576.1 100.00% afis|EAW15772.1 100.00%
Bootstrap support for afum|EAL85576.1 as seed ortholog is 100%.
Bootstrap support for afis|EAW15772.1 as seed ortholog is 100%.

---

Group of orthologs #3501. Best score 1119 bits
Score difference with first non-orthologous sequence - *A. fumigatus*:729 *A. fischeri*:779
afum|EAL86486.1 100.00% afis|EAW16959.1 100.00%
Bootstrap support for afum|EAL86486.1 as seed ortholog is 100%.
Bootstrap support for afis|EAW16959.1 as seed ortholog is 100%.

---

Group of orthologs #3502. Best score 1119 bits
Score difference with first non-orthologous sequence - *A. fumigatus*:1119 *A. fischeri*:1119
afum|EAL88409.1 100.00% afis|EAW23070.1 100.00%
Bootstrap support for afum|EAL88409.1 as seed ortholog is 100%.
Bootstrap support for afis|EAW23070.1 as seed ortholog is 100%.

---

Group of orthologs #3503. Best score 1119 bits
Score difference with first non-orthologous sequence - *A. fumigatus*:1119 *A. fischeri*:1119
afum|EAL88443.1 100.00% afis|EAW23038.1 100.00%
Bootstrap support for afum|EAL88443.1 as seed ortholog is 100%.
Bootstrap support for afis|EAW23038.1 as seed ortholog is 100%.

---

Group of orthologs #3504. Best score 1119 bits
Score difference with first non-orthologous sequence - *A. fumigatus*:1119 *A. fischeri*:1119
afum|EAL92359.1 100.00% afis|EAW21342.1 100.00%
Bootstrap support for afum|EAL92359.1 as seed ortholog is 100%.
Bootstrap support for afis|EAW21342.1 as seed ortholog is 100%.

---

Group of orthologs #3505. Best score 1118 bits
Score difference with first non-orthologous sequence - *A. fumigatus*:1118 *A. fischeri*:1118
afum|EAL86962.1 100.00% afis|EAW19547.1 100.00%
Bootstrap support for afum|EAL86962.1 as seed ortholog is 100%.
Bootstrap support for afis|EAW19547.1 as seed ortholog is 100%.

---

Group of orthologs #3506. Best score 1118 bits
Score difference with first non-orthologous sequence - *A. fumigatus*:1118 *A. fischeri*:1118
afum|EAL91703.1 100.00% afis|EAW17895.1 100.00%
Bootstrap support for afum|EAL91703.1 as seed ortholog is 100%.
Bootstrap support for afis|EAW17895.1 as seed ortholog is 100%.

Group of orthologs #3507. Best score 1118 bits
Score difference with first non-orthologous sequence - *A. fumigatus*:1118 *A. fischeri*:1118
afum|EAL88356.1 100.00% afis|EAW23124.1 100.00%
Bootstrap support for afum|EAL88356.1 as seed ortholog is 100%.
Bootstrap support for afis|EAW23124.1 as seed ortholog is 100%.

Group of orthologs #3508. Best score 1117 bits
Score difference with first non-orthologous sequence - *A. fumigatus*:1117 *A. fischeri*:1117
afum|EAL88024.1 100.00% afis|EAW23438.1 100.00%
Bootstrap support for afum|EAL88024.1 as seed ortholog is 100%.
Bootstrap support for afis|EAW23438.1 as seed ortholog is 100%.

Group of orthologs #3509. Best score 1117 bits
Score difference with first non-orthologous sequence - *A. fumigatus*:1117 *A. fischeri*:1117
afum|EAL90151.1 100.00% afis|EAW25486.1 100.00%
Bootstrap support for afum|EAL90151.1 as seed ortholog is 100%.
Bootstrap support for afis|EAW25486.1 as seed ortholog is 100%.

Group of orthologs #3510. Best score 1116 bits
Score difference with first non-orthologous sequence - *A. fumigatus*:1116 *A. fischeri*:1116
afum|EAL86793.1 100.00% afis|EAW19736.1 100.00%
Bootstrap support for afum|EAL86793.1 as seed ortholog is 100%.
Bootstrap support for afis|EAW19736.1 as seed ortholog is 100%.

Group of orthologs #3511. Best score 1116 bits
Score difference with first non-orthologous sequence - *A. fumigatus*:1116 *A. fischeri*:1116
afum|EAL91406.1 100.00% afis|EAW17593.1 100.00%
Bootstrap support for afum|EAL91406.1 as seed ortholog is 100%.
Bootstrap support for afis|EAW17593.1 as seed ortholog is 100%.

Group of orthologs #3512. Best score 1116 bits
Score difference with first non-orthologous sequence - *A. fumigatus*:1116 *A. fischeri*:1116
afum|EAL88297.1 100.00% afis|EAW23186.1 100.00%
Bootstrap support for afum|EAL88297.1 as seed ortholog is 100%.
Bootstrap support for afis|EAW23186.1 as seed ortholog is 100%.

Group of orthologs #3513. Best score 1116 bits
Score difference with first non-orthologous sequence - *A. fumigatus*:1116 *A. fischeri*:1116
afum|EAL93324.1 100.00% afis|EAW18614.1 100.00%
Bootstrap support for afum|EAL93324.1 as seed ortholog is 100%.
Bootstrap support for afis|EAW18614.1 as seed ortholog is 100%.

Group of orthologs #3514. Best score 1116 bits
Score difference with first non-orthologous sequence - *A. fumigatus*:940 *A. fischeri*:1116
afum|EAL93417.1 100.00% afis|EAW18695.1 100.00%
Bootstrap support for afum|EAL93417.1 as seed ortholog is 100%.
Bootstrap support for afis|EAW18695.1 as seed ortholog is 100%.

Group of orthologs #3515. Best score 1116 bits
Score difference with first non-orthologous sequence - *A. fumigatus*:1116 *A. fischeri*:1116
afum|EAL92672.2 100.00% afis|EAW21638.1 100.00%
Bootstrap support for afum|EAL92672.2 as seed ortholog is 100%.
Bootstrap support for afis|EAW21638.1 as seed ortholog is 100%.

Group of orthologs #3516. Best score 1116 bits

Score difference with first non-orthologous sequence - *A. fumigatus*:1116   *A. fischeri*:1116
afum|EAL92969.1   100.00%   afis|EAW21955.1   100.00%
Bootstrap support for afum|EAL92969.1 as seed ortholog is 100%.
Bootstrap support for afis|EAW21955.1 as seed ortholog is 100%.

---

Group of orthologs #3517. Best score 1116 bits
Score difference with first non-orthologous sequence - *A. fumigatus*:1116   *A. fischeri*:1116
afum|EAL89742.1   100.00%   afis|EAW25073.1   100.00%
Bootstrap support for afum|EAL89742.1 as seed ortholog is 100%.
Bootstrap support for afis|EAW25073.1 as seed ortholog is 100%.

---

Group of orthologs #3518. Best score 1115 bits
Score difference with first non-orthologous sequence - *A. fumigatus*:1115   *A. fischeri*:1115
afum|EAL85680.1   100.00%   afis|EAW15668.1   100.00%
Bootstrap support for afum|EAL85680.1 as seed ortholog is 100%.
Bootstrap support for afis|EAW15668.1 as seed ortholog is 100%.

---

Group of orthologs #3519. Best score 1115 bits
Score difference with first non-orthologous sequence - *A. fumigatus*:1115   *A. fischeri*:1115
afum|EAL90863.1   100.00%   afis|EAW22329.1   100.00%
Bootstrap support for afum|EAL90863.1 as seed ortholog is 100%.
Bootstrap support for afis|EAW22329.1 as seed ortholog is 100%.

---

Group of orthologs #3520. Best score 1115 bits
Score difference with first non-orthologous sequence - *A. fumigatus*:1115   *A. fischeri*:1115
afum|EAL90496.1   100.00%   afis|EAW22696.1   100.00%
Bootstrap support for afum|EAL90496.1 as seed ortholog is 100%.
Bootstrap support for afis|EAW22696.1 as seed ortholog is 100%.

---

Group of orthologs #3521. Best score 1114 bits
Score difference with first non-orthologous sequence - *A. fumigatus*:1039   *A. fischeri*:1114
afum|EAL85846.1   100.00%   afis|EAW15475.1   100.00%
Bootstrap support for afum|EAL85846.1 as seed ortholog is 100%.
Bootstrap support for afis|EAW15475.1 as seed ortholog is 100%.

---

Group of orthologs #3522. Best score 1114 bits
Score difference with first non-orthologous sequence - *A. fumigatus*:1114   *A. fischeri*:1114
afum|EAL87967.1   100.00%   afis|EAW23496.1   100.00%
Bootstrap support for afum|EAL87967.1 as seed ortholog is 100%.
Bootstrap support for afis|EAW23496.1 as seed ortholog is 100%.

---

Group of orthologs #3523. Best score 1114 bits
Score difference with first non-orthologous sequence - *A. fumigatus*:1114   *A. fischeri*:1114
afum|EAL90247.1   100.00%   afis|EAW22950.1   100.00%
Bootstrap support for afum|EAL90247.1 as seed ortholog is 100%.
Bootstrap support for afis|EAW22950.1 as seed ortholog is 100%.

---

Group of orthologs #3524. Best score 1114 bits
Score difference with first non-orthologous sequence - *A. fumigatus*:1015   *A. fischeri*:1114
afum|EAL92368.1   100.00%   afis|EAW21353.1   100.00%
Bootstrap support for afum|EAL92368.1 as seed ortholog is 100%.
Bootstrap support for afis|EAW21353.1 as seed ortholog is 100%.

---

Group of orthologs #3525. Best score 1113 bits
Score difference with first non-orthologous sequence - *A. fumigatus*:852   *A. fischeri*:688
afum|EAL84572.1   100.00%   afis|EAW20482.1   100.00%

Bootstrap support for afum|EAL84572.1 as seed ortholog is 100%.
Bootstrap support for afis|EAW20482.1 as seed ortholog is 100%.

---

Group of orthologs #3526. Best score 1113 bits
Score difference with first non-orthologous sequence - *A. fumigatus*:1113 *A. fischeri*:1113
afum|EAL89107.2 100.00% afis|EAW16512.1 100.00%
Bootstrap support for afum|EAL89107.2 as seed ortholog is 100%.
Bootstrap support for afis|EAW16512.1 as seed ortholog is 100%.

---

Group of orthologs #3527. Best score 1113 bits
Score difference with first non-orthologous sequence - *A. fumigatus*:1113 *A. fischeri*:1113
afum|EAL87983.1 100.00% afis|EAW23481.1 100.00%
Bootstrap support for afum|EAL87983.1 as seed ortholog is 100%.
Bootstrap support for afis|EAW23481.1 as seed ortholog is 100%.

---

Group of orthologs #3528. Best score 1113 bits
Score difference with first non-orthologous sequence - *A. fumigatus*:1113 *A. fischeri*:1113
afum|EAL92118.1 100.00% afis|EAW21095.1 100.00%
Bootstrap support for afum|EAL92118.1 as seed ortholog is 100%.
Bootstrap support for afis|EAW21095.1 as seed ortholog is 100%.

---

Group of orthologs #3529. Best score 1113 bits
Score difference with first non-orthologous sequence - *A. fumigatus*:933 *A. fischeri*:728
afum|EAL92366.1 100.00% afis|EAW21351.1 100.00%
Bootstrap support for afum|EAL92366.1 as seed ortholog is 100%.
Bootstrap support for afis|EAW21351.1 as seed ortholog is 100%.

---

Group of orthologs #3530. Best score 1112 bits
Score difference with first non-orthologous sequence - *A. fumigatus*:869 *A. fischeri*:727
afum|EAL86306.2 100.00% afis|EAW16730.1 100.00%
Bootstrap support for afum|EAL86306.2 as seed ortholog is 100%.
Bootstrap support for afis|EAW16730.1 as seed ortholog is 100%.

---

Group of orthologs #3531. Best score 1112 bits
Score difference with first non-orthologous sequence - *A. fumigatus*:847 *A. fischeri*:808
afum|EAL84778.1 100.00% afis|EAW20856.1 100.00%
Bootstrap support for afum|EAL84778.1 as seed ortholog is 100%.
Bootstrap support for afis|EAW20856.1 as seed ortholog is 100%.

---

Group of orthologs #3532. Best score 1112 bits
Score difference with first non-orthologous sequence - *A. fumigatus*:1112 *A. fischeri*:1112
afum|EAL91476.2 100.00% afis|EAW17662.1 100.00%
Bootstrap support for afum|EAL91476.2 as seed ortholog is 100%.
Bootstrap support for afis|EAW17662.1 as seed ortholog is 100%.

---

Group of orthologs #3533. Best score 1112 bits
Score difference with first non-orthologous sequence - *A. fumigatus*:1112 *A. fischeri*:1004
afum|EAL84287.2 100.00% afis|EAW25577.1 100.00%
Bootstrap support for afum|EAL84287.2 as seed ortholog is 100%.
Bootstrap support for afis|EAW25577.1 as seed ortholog is 100%.

---

Group of orthologs #3534. Best score 1112 bits
Score difference with first non-orthologous sequence - *A. fumigatus*:782 *A. fischeri*:1112
afum|EAL86184.1 100.00% afis|EAW24518.1 100.00%
Bootstrap support for afum|EAL86184.1 as seed ortholog is 100%.
Bootstrap support for afis|EAW24518.1 as seed ortholog is 100%.

---

Group of orthologs #3535. Best score 1112 bits
Score difference with first non-orthologous sequence - *A. fumigatus*:1112 *A. fischeri*:1112
afum|EAL93048.1 100.00% afis|EAW18373.1 100.00%
Bootstrap support for afum|EAL93048.1 as seed ortholog is 100%.
Bootstrap support for afis|EAW18373.1 as seed ortholog is 100%.

---

Group of orthologs #3536. Best score 1112 bits
Score difference with first non-orthologous sequence - *A. fumigatus*:901 *A. fischeri*:902
afum|EAL93863.1 100.00% afis|EAW19131.1 100.00%
Bootstrap support for afum|EAL93863.1 as seed ortholog is 100%.
Bootstrap support for afis|EAW19131.1 as seed ortholog is 100%.

---

Group of orthologs #3537. Best score 1112 bits
Score difference with first non-orthologous sequence - *A. fumigatus*:1112 *A. fischeri*:1112
afum|EAL91086.1 100.00% afis|EAW22105.1 100.00%
Bootstrap support for afum|EAL91086.1 as seed ortholog is 100%.
Bootstrap support for afis|EAW22105.1 as seed ortholog is 100%.

---

Group of orthologs #3538. Best score 1111 bits
Score difference with first non-orthologous sequence - *A. fumigatus*:1111 *A. fischeri*:1111
afum|EAL86679.1 100.00% afis|EAW17262.1 100.00%
Bootstrap support for afum|EAL86679.1 as seed ortholog is 100%.
Bootstrap support for afis|EAW17262.1 as seed ortholog is 100%.

---

Group of orthologs #3539. Best score 1111 bits
Score difference with first non-orthologous sequence - *A. fumigatus*:902 *A. fischeri*:1111
afum|EAL84950.2 100.00% afis|EAW19968.1 100.00%
Bootstrap support for afum|EAL84950.2 as seed ortholog is 100%.
Bootstrap support for afis|EAW19968.1 as seed ortholog is 100%.

---

Group of orthologs #3540. Best score 1110 bits
Score difference with first non-orthologous sequence - *A. fumigatus*:1110 *A. fischeri*:1110
afum|EAL88631.1 100.00% afis|EAW15999.1 100.00%
Bootstrap support for afum|EAL88631.1 as seed ortholog is 100%.
Bootstrap support for afis|EAW15999.1 as seed ortholog is 100%.

---

Group of orthologs #3541. Best score 1110 bits
Score difference with first non-orthologous sequence - *A. fumigatus*:1110 *A. fischeri*:939
afum|EAL88979.1 100.00% afis|EAW16354.1 100.00%
Bootstrap support for afum|EAL88979.1 as seed ortholog is 100%.
Bootstrap support for afis|EAW16354.1 as seed ortholog is 100%.

---

Group of orthologs #3542. Best score 1110 bits
Score difference with first non-orthologous sequence - *A. fumigatus*:1110 *A. fischeri*:1110
afum|EAL89145.1 100.00% afis|EAW16547.1 100.00%
Bootstrap support for afum|EAL89145.1 as seed ortholog is 100%.
Bootstrap support for afis|EAW16547.1 as seed ortholog is 100%.

---

Group of orthologs #3543. Best score 1110 bits
Score difference with first non-orthologous sequence - *A. fumigatus*:1110 *A. fischeri*:1110
afum|EAL85959.1 100.00% afis|EAW24291.1 100.00%
Bootstrap support for afum|EAL85959.1 as seed ortholog is 100%.
Bootstrap support for afis|EAW24291.1 as seed ortholog is 100%.

---

Group of orthologs #3544. Best score 1110 bits

Score difference with first non-orthologous sequence - *A. fumigatus*:1110 *A. fischeri*:1110
afum|EAL87953.1 100.00% afis|EAW23508.1 100.00%
Bootstrap support for afum|EAL87953.1 as seed ortholog is 100%.
Bootstrap support for afis|EAW23508.1 as seed ortholog is 100%.

---

Group of orthologs #3545. Best score 1110 bits
Score difference with first non-orthologous sequence - *A. fumigatus*:689 *A. fischeri*:785
afum|EAL87885.1 100.00% afis|EAW23580.1 100.00%
Bootstrap support for afum|EAL87885.1 as seed ortholog is 100%.
Bootstrap support for afis|EAW23580.1 as seed ortholog is 100%.

---

Group of orthologs #3546. Best score 1110 bits
Score difference with first non-orthologous sequence - *A. fumigatus*:1110 *A. fischeri*:758
afum|EAL93735.1 100.00% afis|EAW19000.1 100.00%
Bootstrap support for afum|EAL93735.1 as seed ortholog is 100%.
Bootstrap support for afis|EAW19000.1 as seed ortholog is 100%.

---

Group of orthologs #3547. Best score 1110 bits
Score difference with first non-orthologous sequence - *A. fumigatus*:1110 *A. fischeri*:1110
afum|EBA27499.1 100.00% afis|EAW18924.1 100.00%
Bootstrap support for afum|EBA27499.1 as seed ortholog is 100%.
Bootstrap support for afis|EAW18924.1 as seed ortholog is 100%.

---

Group of orthologs #3548. Best score 1110 bits
Score difference with first non-orthologous sequence - *A. fumigatus*:1110 *A. fischeri*:1110
afum|EAL92360.1 100.00% afis|EAW21343.1 100.00%
Bootstrap support for afum|EAL92360.1 as seed ortholog is 100%.
Bootstrap support for afis|EAW21343.1 as seed ortholog is 100%.

---

Group of orthologs #3549. Best score 1110 bits
Score difference with first non-orthologous sequence - *A. fumigatus*:1110 *A. fischeri*:1110
afum|EAL92515.1 100.00% afis|EAW21496.1 100.00%
Bootstrap support for afum|EAL92515.1 as seed ortholog is 100%.
Bootstrap support for afis|EAW21496.1 as seed ortholog is 100%.

---

Group of orthologs #3550. Best score 1109 bits
Score difference with first non-orthologous sequence - *A. fumigatus*:334 *A. fischeri*:516
afum|EAL86318.1 100.00% afis|EAW16743.1 100.00%
Bootstrap support for afum|EAL86318.1 as seed ortholog is 100%.
Bootstrap support for afis|EAW16743.1 as seed ortholog is 100%.

---

Group of orthologs #3551. Best score 1109 bits
Score difference with first non-orthologous sequence - *A. fumigatus*:1109 *A. fischeri*:346
afum|EAL88562.1 100.00% afis|EAW15926.1 100.00%
Bootstrap support for afum|EAL88562.1 as seed ortholog is 100%.
Bootstrap support for afis|EAW15926.1 as seed ortholog is 100%.

---

Group of orthologs #3552. Best score 1109 bits
Score difference with first non-orthologous sequence - *A. fumigatus*:764 *A. fischeri*:1109
afum|EAL85373.1 100.00% afis|EAW20265.1 100.00%
Bootstrap support for afum|EAL85373.1 as seed ortholog is 100%.
Bootstrap support for afis|EAW20265.1 as seed ortholog is 100%.

---

Group of orthologs #3553. Best score 1109 bits
Score difference with first non-orthologous sequence - *A. fumigatus*:1109 *A. fischeri*:1109
afum|EAL88350.2 100.00% afis|EAW23131.1 100.00%

Bootstrap support for afum|EAL88350.2 as seed ortholog is 100%.
Bootstrap support for afis|EAW23131.1 as seed ortholog is 100%.

---

Group of orthologs #3554. Best score 1109 bits
Score difference with first non-orthologous sequence - *A. fumigatus*:1109 *A. fischeri*:1109
afum|EAL90028.2 100.00% afis|EAW25368.1 100.00%
Bootstrap support for afum|EAL90028.2 as seed ortholog is 100%.
Bootstrap support for afis|EAW25368.1 as seed ortholog is 100%.

---

Group of orthologs #3555. Best score 1108 bits
Score difference with first non-orthologous sequence - *A. fumigatus*:1108 *A. fischeri*:1108
afum|EAL86892.1 100.00% afis|EAW19630.1 100.00%
Bootstrap support for afum|EAL86892.1 as seed ortholog is 100%.
Bootstrap support for afis|EAW19630.1 as seed ortholog is 100%.

---

Group of orthologs #3556. Best score 1108 bits
Score difference with first non-orthologous sequence - *A. fumigatus*:907 *A. fischeri*:703
afum|EAL90510.1 100.00% afis|EAW22682.1 100.00%
Bootstrap support for afum|EAL90510.1 as seed ortholog is 100%.
Bootstrap support for afis|EAW22682.1 as seed ortholog is 100%.

---

Group of orthologs #3557. Best score 1108 bits
Score difference with first non-orthologous sequence - *A. fumigatus*:1108 *A. fischeri*:1108
afum|EAL92217.1 100.00% afis|EAW21206.1 100.00%
Bootstrap support for afum|EAL92217.1 as seed ortholog is 100%.
Bootstrap support for afis|EAW21206.1 as seed ortholog is 100%.

---

Group of orthologs #3558. Best score 1108 bits
Score difference with first non-orthologous sequence - *A. fumigatus*:1108 *A. fischeri*:1108
afum|EAL92616.1 100.00% afis|EAW21597.1 100.00%
Bootstrap support for afum|EAL92616.1 as seed ortholog is 100%.
Bootstrap support for afis|EAW21597.1 as seed ortholog is 100%.

---

Group of orthologs #3559. Best score 1107 bits
Score difference with first non-orthologous sequence - *A. fumigatus*:1107 *A. fischeri*:1107
afum|EAL84593.1 100.00% afis|EAW20460.1 100.00%
Bootstrap support for afum|EAL84593.1 as seed ortholog is 100%.
Bootstrap support for afis|EAW20460.1 as seed ortholog is 100%.

---

Group of orthologs #3560. Best score 1107 bits
Score difference with first non-orthologous sequence - *A. fumigatus*:824 *A. fischeri*:890
afum|EAL85480.2 100.00% afis|EAW20364.1 100.00%
Bootstrap support for afum|EAL85480.2 as seed ortholog is 100%.
Bootstrap support for afis|EAW20364.1 as seed ortholog is 100%.

---

Group of orthologs #3561. Best score 1107 bits
Score difference with first non-orthologous sequence - *A. fumigatus*:1107 *A. fischeri*:1032
afum|EAL91696.1 100.00% afis|EAW17887.1 100.00%
Bootstrap support for afum|EAL91696.1 as seed ortholog is 100%.
Bootstrap support for afis|EAW17887.1 as seed ortholog is 100%.

---

Group of orthologs #3562. Best score 1107 bits
Score difference with first non-orthologous sequence - *A. fumigatus*:1107 *A. fischeri*:600
afum|EAL84290.1 100.00% afis|EAW25579.1 100.00%
Bootstrap support for afum|EAL84290.1 as seed ortholog is 100%.
Bootstrap support for afis|EAW25579.1 as seed ortholog is 100%.

Group of orthologs #3563. Best score 1106 bits
Score difference with first non-orthologous sequence - *A. fumigatus*:1106 *A. fischeri*:1106
afum|EAL84466.1 100.00% afis|EAW20595.1 100.00%
Bootstrap support for afum|EAL84466.1 as seed ortholog is 100%.
Bootstrap support for afis|EAW20595.1 as seed ortholog is 100%.

Group of orthologs #3564. Best score 1106 bits
Score difference with first non-orthologous sequence - *A. fumigatus*:853 *A. fischeri*:835
afum|EAL84431.1 100.00% afis|EAW20637.1 100.00%
Bootstrap support for afum|EAL84431.1 as seed ortholog is 100%.
Bootstrap support for afis|EAW20637.1 as seed ortholog is 100%.

Group of orthologs #3565. Best score 1106 bits
Score difference with first non-orthologous sequence - *A. fumigatus*:1106 *A. fischeri*:1106
afum|EAL86981.2 100.00% afis|EAW19515.1 100.00%
Bootstrap support for afum|EAL86981.2 as seed ortholog is 100%.
Bootstrap support for afis|EAW19515.1 as seed ortholog is 100%.

Group of orthologs #3566. Best score 1106 bits
Score difference with first non-orthologous sequence - *A. fumigatus*:1106 *A. fischeri*:1106
afum|EAL91158.1 100.00% afis|EAW17322.1 100.00%
Bootstrap support for afum|EAL91158.1 as seed ortholog is 100%.
Bootstrap support for afis|EAW17322.1 as seed ortholog is 100%.

Group of orthologs #3567. Best score 1106 bits
Score difference with first non-orthologous sequence - *A. fumigatus*:1040 *A. fischeri*:662
afum|EAL87965.1 100.00% afis|EAW23498.1 100.00%
Bootstrap support for afum|EAL87965.1 as seed ortholog is 100%.
Bootstrap support for afis|EAW23498.1 as seed ortholog is 100%.

Group of orthologs #3568. Best score 1106 bits
Score difference with first non-orthologous sequence - *A. fumigatus*:1106 *A. fischeri*:1106
afum|EAL92100.1 100.00% afis|EAW21077.1 100.00%
Bootstrap support for afum|EAL92100.1 as seed ortholog is 100%.
Bootstrap support for afis|EAW21077.1 as seed ortholog is 100%.

Group of orthologs #3569. Best score 1106 bits
Score difference with first non-orthologous sequence - *A. fumigatus*:1106 *A. fischeri*:1106
afum|EAL92755.1 100.00% afis|EAW21722.1 100.00%
Bootstrap support for afum|EAL92755.1 as seed ortholog is 100%.
Bootstrap support for afis|EAW21722.1 as seed ortholog is 100%.

Group of orthologs #3570. Best score 1106 bits
Score difference with first non-orthologous sequence - *A. fumigatus*:1106 *A. fischeri*:1106
afum|EAL89910.1 100.00% afis|EAW25248.1 100.00%
Bootstrap support for afum|EAL89910.1 as seed ortholog is 100%.
Bootstrap support for afis|EAW25248.1 as seed ortholog is 100%.

Group of orthologs #3571. Best score 1106 bits
Score difference with first non-orthologous sequence - *A. fumigatus*:1106 *A. fischeri*:1106
afum|EAL92042.1 100.00% afis|EAW24182.1 100.00%
Bootstrap support for afum|EAL92042.1 as seed ortholog is 100%.
Bootstrap support for afis|EAW24182.1 as seed ortholog is 100%.

Group of orthologs #3572. Best score 1105 bits

Score difference with first non-orthologous sequence - *A. fumigatus*:961   *A. fischeri*:886
afum|EAL85079.1   100.00%   afis|EAW19812.1   100.00%
Bootstrap support for afum|EAL85079.1 as seed ortholog is 100%.
Bootstrap support for afis|EAW19812.1 as seed ortholog is 100%.

---

Group of orthologs #3573. Best score 1105 bits
Score difference with first non-orthologous sequence - *A. fumigatus*:1105   *A. fischeri*:1105
afum|EAL89084.1   100.00%   afis|EAW16490.1   100.00%
Bootstrap support for afum|EAL89084.1 as seed ortholog is 100%.
Bootstrap support for afis|EAW16490.1 as seed ortholog is 100%.

---

Group of orthologs #3574. Best score 1105 bits
Score difference with first non-orthologous sequence - *A. fumigatus*:1105   *A. fischeri*:1105
afum|EAL91548.1   100.00%   afis|EAW17733.1   100.00%
Bootstrap support for afum|EAL91548.1 as seed ortholog is 100%.
Bootstrap support for afis|EAW17733.1 as seed ortholog is 100%.

---

Group of orthologs #3575. Best score 1105 bits
Score difference with first non-orthologous sequence - *A. fumigatus*:844   *A. fischeri*:910
afum|EAL84356.1   100.00%   afis|EAW25687.1   100.00%
Bootstrap support for afum|EAL84356.1 as seed ortholog is 100%.
Bootstrap support for afis|EAW25687.1 as seed ortholog is 100%.

---

Group of orthologs #3576. Best score 1105 bits
Score difference with first non-orthologous sequence - *A. fumigatus*:1105   *A. fischeri*:1105
afum|EAL90148.1   100.00%   afis|EAW25483.1   100.00%
Bootstrap support for afum|EAL90148.1 as seed ortholog is 100%.
Bootstrap support for afis|EAW25483.1 as seed ortholog is 100%.

---

Group of orthologs #3577. Best score 1104 bits
Score difference with first non-orthologous sequence - *A. fumigatus*:1104   *A. fischeri*:906
afum|EAL86561.1   100.00%   afis|EAW17073.1   100.00%
Bootstrap support for afum|EAL86561.1 as seed ortholog is 100%.
Bootstrap support for afis|EAW17073.1 as seed ortholog is 100%.

---

Group of orthologs #3578. Best score 1104 bits
Score difference with first non-orthologous sequence - *A. fumigatus*:1104   *A. fischeri*:1104
afum|EAL86250.1   100.00%   afis|EAW24622.1   100.00%
Bootstrap support for afum|EAL86250.1 as seed ortholog is 100%.
Bootstrap support for afis|EAW24622.1 as seed ortholog is 100%.

---

Group of orthologs #3579. Best score 1104 bits
Score difference with first non-orthologous sequence - *A. fumigatus*:1104   *A. fischeri*:1104
afum|EAL87844.1   100.00%   afis|EAW23618.1   100.00%
Bootstrap support for afum|EAL87844.1 as seed ortholog is 100%.
Bootstrap support for afis|EAW23618.1 as seed ortholog is 100%.

---

Group of orthologs #3580. Best score 1103 bits
Score difference with first non-orthologous sequence - *A. fumigatus*:926   *A. fischeri*:968
afum|EAL88587.1   100.00%   afis|EAW15954.1   100.00%
Bootstrap support for afum|EAL88587.1 as seed ortholog is 100%.
Bootstrap support for afis|EAW15954.1 as seed ortholog is 100%.

---

Group of orthologs #3581. Best score 1103 bits
Score difference with first non-orthologous sequence - *A. fumigatus*:1103   *A. fischeri*:1103
afum|EAL84979.1   100.00%   afis|EAW19929.1   100.00%

Bootstrap support for afum|EAL84979.1 as seed ortholog is 100%.
Bootstrap support for afis|EAW19929.1 as seed ortholog is 100%.

---

Group of orthologs #3582. Best score 1103 bits
Score difference with first non-orthologous sequence - *A. fumigatus*:1103 *A. fischeri*:1103
afum|EAL84604.1 100.00% afis|EAW20448.1 100.00%
Bootstrap support for afum|EAL84604.1 as seed ortholog is 100%.
Bootstrap support for afis|EAW20448.1 as seed ortholog is 100%.

---

Group of orthologs #3583. Best score 1103 bits
Score difference with first non-orthologous sequence - *A. fumigatus*:1103 *A. fischeri*:1103
afum|EAL85312.2 100.00% afis|EAW20192.1 100.00%
Bootstrap support for afum|EAL85312.2 as seed ortholog is 100%.
Bootstrap support for afis|EAW20192.1 as seed ortholog is 100%.

---

Group of orthologs #3584. Best score 1103 bits
Score difference with first non-orthologous sequence - *A. fumigatus*:1103 *A. fischeri*:1103
afum|EAL87187.2 100.00% afis|EAW23750.1 100.00%
Bootstrap support for afum|EAL87187.2 as seed ortholog is 100%.
Bootstrap support for afis|EAW23750.1 as seed ortholog is 100%.

---

Group of orthologs #3585. Best score 1103 bits
Score difference with first non-orthologous sequence - *A. fumigatus*:1103 *A. fischeri*:1103
afum|EAL93032.2 100.00% afis|EAW18361.1 100.00%
Bootstrap support for afum|EAL93032.2 as seed ortholog is 100%.
Bootstrap support for afis|EAW18361.1 as seed ortholog is 100%.

---

Group of orthologs #3586. Best score 1103 bits
Score difference with first non-orthologous sequence - *A. fumigatus*:1103 *A. fischeri*:1103
afum|EAL92554.1 100.00% afis|EAW21533.1 100.00%
Bootstrap support for afum|EAL92554.1 as seed ortholog is 100%.
Bootstrap support for afis|EAW21533.1 as seed ortholog is 100%.

---

Group of orthologs #3587. Best score 1103 bits
Score difference with first non-orthologous sequence - *A. fumigatus*:1103 *A. fischeri*:1103
afum|EAL90126.1 100.00% afis|EAW25463.1 100.00%
Bootstrap support for afum|EAL90126.1 as seed ortholog is 100%.
Bootstrap support for afis|EAW25463.1 as seed ortholog is 100%.

---

Group of orthologs #3588. Best score 1102 bits
Score difference with first non-orthologous sequence - *A. fumigatus*:1102 *A. fischeri*:1102
afum|EAL85588.1 100.00% afis|EAW15760.1 100.00%
Bootstrap support for afum|EAL85588.1 as seed ortholog is 100%.
Bootstrap support for afis|EAW15760.1 as seed ortholog is 100%.

---

Group of orthologs #3589. Best score 1102 bits
Score difference with first non-orthologous sequence - *A. fumigatus*:1102 *A. fischeri*:1102
afum|EAL89070.1 100.00% afis|EAW16476.1 100.00%
Bootstrap support for afum|EAL89070.1 as seed ortholog is 100%.
Bootstrap support for afis|EAW16476.1 as seed ortholog is 100%.

---

Group of orthologs #3590. Best score 1102 bits
Score difference with first non-orthologous sequence - *A. fumigatus*:1102 *A. fischeri*:1102
afum|EAL86996.1 100.00% afis|EAW19497.1 100.00%
Bootstrap support for afum|EAL86996.1 as seed ortholog is 100%.
Bootstrap support for afis|EAW19497.1 as seed ortholog is 100%.

---

Group of orthologs #3591. Best score 1102 bits
Score difference with first non-orthologous sequence - *A. fumigatus*:1102 *A. fischeri*:1102
afum|EAL93794.1 100.00% afis|EAW19065.1 100.00%
Bootstrap support for afum|EAL93794.1 as seed ortholog is 100%.
Bootstrap support for afis|EAW19065.1 as seed ortholog is 100%.

---

Group of orthologs #3592. Best score 1102 bits
Score difference with first non-orthologous sequence - *A. fumigatus*:1102 *A. fischeri*:1102
afum|EAL92131.1 100.00% afis|EAW21109.1 100.00%
Bootstrap support for afum|EAL92131.1 as seed ortholog is 100%.
Bootstrap support for afis|EAW21109.1 as seed ortholog is 100%.

---

Group of orthologs #3593. Best score 1102 bits
Score difference with first non-orthologous sequence - *A. fumigatus*:1102 *A. fischeri*:1102
afum|EAL90776.1 100.00% afis|EAW22416.1 100.00%
Bootstrap support for afum|EAL90776.1 as seed ortholog is 100%.
Bootstrap support for afis|EAW22416.1 as seed ortholog is 100%.

---

Group of orthologs #3594. Best score 1102 bits
Score difference with first non-orthologous sequence - *A. fumigatus*:1102 *A. fischeri*:1102
afum|EAL91932.1 100.00% afis|EAW24077.1 100.00%
Bootstrap support for afum|EAL91932.1 as seed ortholog is 100%.
Bootstrap support for afis|EAW24077.1 as seed ortholog is 100%.

---

Group of orthologs #3595. Best score 1101 bits
Score difference with first non-orthologous sequence - *A. fumigatus*:1101 *A. fischeri*:1101
afum|EAL88984.1 100.00% afis|EAW16359.1 100.00%
Bootstrap support for afum|EAL88984.1 as seed ortholog is 100%.
Bootstrap support for afis|EAW16359.1 as seed ortholog is 100%.

---

Group of orthologs #3596. Best score 1101 bits
Score difference with first non-orthologous sequence - *A. fumigatus*:1101 *A. fischeri*:1101
afum|EAL84590.1 100.00% afis|EAW20463.1 100.00%
Bootstrap support for afum|EAL84590.1 as seed ortholog is 100%.
Bootstrap support for afis|EAW20463.1 as seed ortholog is 100%.

---

Group of orthologs #3597. Best score 1101 bits
Score difference with first non-orthologous sequence - *A. fumigatus*:1101 *A. fischeri*:1101
afum|EAL93263.1 100.00% afis|EAW18562.1 100.00%
Bootstrap support for afum|EAL93263.1 as seed ortholog is 100%.
Bootstrap support for afis|EAW18562.1 as seed ortholog is 100%.

---

Group of orthologs #3598. Best score 1101 bits
Score difference with first non-orthologous sequence - *A. fumigatus*:827 *A. fischeri*:717
afum|EAL89987.1 100.00% afis|EAW25326.1 100.00%
Bootstrap support for afum|EAL89987.1 as seed ortholog is 100%.
Bootstrap support for afis|EAW25326.1 as seed ortholog is 100%.

---

Group of orthologs #3599. Best score 1100 bits
Score difference with first non-orthologous sequence - *A. fumigatus*:1100 *A. fischeri*:1100
afum|EAL84583.1 100.00% afis|EAW20469.1 100.00%
Bootstrap support for afum|EAL84583.1 as seed ortholog is 100%.
Bootstrap support for afis|EAW20469.1 as seed ortholog is 100%.

---

Group of orthologs #3600. Best score 1100 bits

Score difference with first non-orthologous sequence - *A. fumigatus*:717 *A. fischeri*:1100
afum|EAL85912.1 100.00% afis|EAW24244.1 100.00%
Bootstrap support for afum|EAL85912.1 as seed ortholog is 100%.
Bootstrap support for afis|EAW24244.1 as seed ortholog is 100%.

---

Group of orthologs #3601. Best score 1100 bits
Score difference with first non-orthologous sequence - *A. fumigatus*:1100 *A. fischeri*:1100
afum|EAL85925.1 100.00% afis|EAW24257.1 100.00%
Bootstrap support for afum|EAL85925.1 as seed ortholog is 100%.
Bootstrap support for afis|EAW24257.1 as seed ortholog is 100%.

---

Group of orthologs #3602. Best score 1100 bits
Score difference with first non-orthologous sequence - *A. fumigatus*:1100 *A. fischeri*:1100
afum|EAL88328.1 100.00% afis|EAW23153.1 100.00%
Bootstrap support for afum|EAL88328.1 as seed ortholog is 100%.
Bootstrap support for afis|EAW23153.1 as seed ortholog is 100%.

---

Group of orthologs #3603. Best score 1100 bits
Score difference with first non-orthologous sequence - *A. fumigatus*:1100 *A. fischeri*:1100
afum|EAL88162.1 100.00% afis|EAW23315.1 100.00%
Bootstrap support for afum|EAL88162.1 as seed ortholog is 100%.
Bootstrap support for afis|EAW23315.1 as seed ortholog is 100%.

---

Group of orthologs #3604. Best score 1100 bits
Score difference with first non-orthologous sequence - *A. fumigatus*:1100 *A. fischeri*:763
afum|EAL93260.1 100.00% afis|EAW18560.1 100.00%
Bootstrap support for afum|EAL93260.1 as seed ortholog is 100%.
Bootstrap support for afis|EAW18560.1 as seed ortholog is 100%.

---

Group of orthologs #3605. Best score 1100 bits
Score difference with first non-orthologous sequence - *A. fumigatus*:1100 *A. fischeri*:1100
afum|EAL93338.1 100.00% afis|EAW18626.1 100.00%
Bootstrap support for afum|EAL93338.1 as seed ortholog is 100%.
Bootstrap support for afis|EAW18626.1 as seed ortholog is 100%.

---

Group of orthologs #3606. Best score 1100 bits
Score difference with first non-orthologous sequence - *A. fumigatus*:1100 *A. fischeri*:1100
afum|EAL90539.1 100.00% afis|EAW22653.1 100.00%
Bootstrap support for afum|EAL90539.1 as seed ortholog is 100%.
Bootstrap support for afis|EAW22653.1 as seed ortholog is 100%.

---

Group of orthologs #3607. Best score 1100 bits
Score difference with first non-orthologous sequence - *A. fumigatus*:1000 *A. fischeri*:838
afum|EAL90320.1 100.00% afis|EAW22879.1 100.00%
Bootstrap support for afum|EAL90320.1 as seed ortholog is 100%.
Bootstrap support for afis|EAW22879.1 as seed ortholog is 100%.

---

Group of orthologs #3608. Best score 1100 bits
Score difference with first non-orthologous sequence - *A. fumigatus*:1100 *A. fischeri*:1100
afum|EAL92850.1 100.00% afis|EAW21835.1 100.00%
Bootstrap support for afum|EAL92850.1 as seed ortholog is 100%.
Bootstrap support for afis|EAW21835.1 as seed ortholog is 100%.

---

Group of orthologs #3609. Best score 1100 bits
Score difference with first non-orthologous sequence - *A. fumigatus*:1100 *A. fischeri*:1100
afum|EBA27468.1 100.00% afis|EAW21570.1 100.00%

Bootstrap support for afum|EBA27468.1 as seed ortholog is 100%.
Bootstrap support for afis|EAW21570.1 as seed ortholog is 100%.

---

Group of orthologs #3610. Best score 1099 bits
Score difference with first non-orthologous sequence - *A. fumigatus*:700 *A. fischeri*:1099
afum|EAL91076.2 100.00% afis|EAW22108.1 100.00%
Bootstrap support for afum|EAL91076.2 as seed ortholog is 100%.
Bootstrap support for afis|EAW22108.1 as seed ortholog is 100%.

---

Group of orthologs #3611. Best score 1098 bits
Score difference with first non-orthologous sequence - *A. fumigatus*:1098 *A. fischeri*:1098
afum|EAL87699.1 100.00% afis|EAW18334.1 100.00%
Bootstrap support for afum|EAL87699.1 as seed ortholog is 100%.
Bootstrap support for afis|EAW18334.1 as seed ortholog is 100%.

---

Group of orthologs #3612. Best score 1098 bits
Score difference with first non-orthologous sequence - *A. fumigatus*:1098 *A. fischeri*:1098
afum|EAL91675.1 100.00% afis|EAW17867.1 100.00%
Bootstrap support for afum|EAL91675.1 as seed ortholog is 100%.
Bootstrap support for afis|EAW17867.1 as seed ortholog is 100%.

---

Group of orthologs #3613. Best score 1098 bits
Score difference with first non-orthologous sequence - *A. fumigatus*:1098 *A. fischeri*:1098
afum|EAL87315.1 100.00% afis|EAW23888.1 100.00%
Bootstrap support for afum|EAL87315.1 as seed ortholog is 100%.
Bootstrap support for afis|EAW23888.1 as seed ortholog is 100%.

---

Group of orthologs #3614. Best score 1097 bits
Score difference with first non-orthologous sequence - *A. fumigatus*:1097 *A. fischeri*:842
afum|EAL88872.1 100.00% afis|EAW16244.1 100.00%
Bootstrap support for afum|EAL88872.1 as seed ortholog is 100%.
Bootstrap support for afis|EAW16244.1 as seed ortholog is 100%.

---

Group of orthologs #3615. Best score 1097 bits
Score difference with first non-orthologous sequence - *A. fumigatus*:940 *A. fischeri*:950
afum|EAL85349.1 100.00% afis|EAW20232.1 100.00%
Bootstrap support for afum|EAL85349.1 as seed ortholog is 100%.
Bootstrap support for afis|EAW20232.1 as seed ortholog is 100%.

---

Group of orthologs #3616. Best score 1097 bits
Score difference with first non-orthologous sequence - *A. fumigatus*:907 *A. fischeri*:952
afum|EAL85404.1 100.00% afis|EAW20270.1 100.00%
Bootstrap support for afum|EAL85404.1 as seed ortholog is 100%.
Bootstrap support for afis|EAW20270.1 as seed ortholog is 100%.

---

Group of orthologs #3617. Best score 1097 bits
Score difference with first non-orthologous sequence - *A. fumigatus*:1097 *A. fischeri*:1097
afum|EAL91265.1 100.00% afis|EAW17439.1 100.00%
Bootstrap support for afum|EAL91265.1 as seed ortholog is 100%.
Bootstrap support for afis|EAW17439.1 as seed ortholog is 100%.

---

Group of orthologs #3618. Best score 1097 bits
Score difference with first non-orthologous sequence - *A. fumigatus*:1097 *A. fischeri*:1097
afum|EAL91702.1 100.00% afis|EAW17893.1 100.00%
Bootstrap support for afum|EAL91702.1 as seed ortholog is 100%.
Bootstrap support for afis|EAW17893.1 as seed ortholog is 100%.

---

Group of orthologs #3619. Best score 1097 bits
Score difference with first non-orthologous sequence - *A. fumigatus*:975 *A. fischeri*:887
afum|EAL93109.1 100.00% afis|EAW18429.1 100.00%
Bootstrap support for afum|EAL93109.1 as seed ortholog is 100%.
Bootstrap support for afis|EAW18429.1 as seed ortholog is 100%.

---

Group of orthologs #3620. Best score 1097 bits
Score difference with first non-orthologous sequence - *A. fumigatus*:1097 *A. fischeri*:1097
afum|EAL93410.1 100.00% afis|EAW18689.1 100.00%
Bootstrap support for afum|EAL93410.1 as seed ortholog is 100%.
Bootstrap support for afis|EAW18689.1 as seed ortholog is 100%.

---

Group of orthologs #3621. Best score 1097 bits
Score difference with first non-orthologous sequence - *A. fumigatus*:1097 *A. fischeri*:1097
afum|EAL92072.2 100.00% afis|EAW21034.1 100.00%
Bootstrap support for afum|EAL92072.2 as seed ortholog is 100%.
Bootstrap support for afis|EAW21034.1 as seed ortholog is 100%.

---

Group of orthologs #3622. Best score 1097 bits
Score difference with first non-orthologous sequence - *A. fumigatus*:1097 *A. fischeri*:1097
afum|EAL90660.1 100.00% afis|EAW22534.1 100.00%
Bootstrap support for afum|EAL90660.1 as seed ortholog is 100%.
Bootstrap support for afis|EAW22534.1 as seed ortholog is 100%.

---

Group of orthologs #3623. Best score 1097 bits
Score difference with first non-orthologous sequence - *A. fumigatus*:1097 *A. fischeri*:1097
afum|EAL92009.1 100.00% afis|EAW24154.1 100.00%
Bootstrap support for afum|EAL92009.1 as seed ortholog is 100%.
Bootstrap support for afis|EAW24154.1 as seed ortholog is 100%.

---

Group of orthologs #3624. Best score 1096 bits
Score difference with first non-orthologous sequence - *A. fumigatus*:1096 *A. fischeri*:1096
afum|EAL86797.1 100.00% afis|EAW19731.1 100.00%
Bootstrap support for afum|EAL86797.1 as seed ortholog is 100%.
Bootstrap support for afis|EAW19731.1 as seed ortholog is 100%.

---

Group of orthologs #3625. Best score 1096 bits
Score difference with first non-orthologous sequence - *A. fumigatus*:1096 *A. fischeri*:1096
afum|EAL85897.1 100.00% afis|EAW24228.1 100.00%
Bootstrap support for afum|EAL85897.1 as seed ortholog is 100%.
Bootstrap support for afis|EAW24228.1 as seed ortholog is 100%.

---

Group of orthologs #3626. Best score 1096 bits
Score difference with first non-orthologous sequence - *A. fumigatus*:1096 *A. fischeri*:1096
afum|EAL88156.2 100.00% afis|EAW23320.1 100.00%
Bootstrap support for afum|EAL88156.2 as seed ortholog is 100%.
Bootstrap support for afis|EAW23320.1 as seed ortholog is 100%.

---

Group of orthologs #3627. Best score 1096 bits
Score difference with first non-orthologous sequence - *A. fumigatus*:1096 *A. fischeri*:1096
afum|EAL93931.1 100.00% afis|EAW19200.1 100.00%
Bootstrap support for afum|EAL93931.1 as seed ortholog is 100%.
Bootstrap support for afis|EAW19200.1 as seed ortholog is 100%.

---

Group of orthologs #3628. Best score 1096 bits

Score difference with first non-orthologous sequence - *A. fumigatus*:1096 *A. fischeri*:1096
afum|EAL91917.1 100.00% afis|EAW24062.1 100.00%
Bootstrap support for afum|EAL91917.1 as seed ortholog is 100%.
Bootstrap support for afis|EAW24062.1 as seed ortholog is 100%.

---

Group of orthologs #3629. Best score 1096 bits
Score difference with first non-orthologous sequence - *A. fumigatus*:1096 *A. fischeri*:1096
afum|EBA27339.1 100.00% afis|EAW23214.1 100.00%
Bootstrap support for afum|EBA27339.1 as seed ortholog is 100%.
Bootstrap support for afis|EAW23214.1 as seed ortholog is 100%.

---

Group of orthologs #3630. Best score 1095 bits
Score difference with first non-orthologous sequence - *A. fumigatus*:1095 *A. fischeri*:1095
afum|EAL91556.1 100.00% afis|EAW17744.1 100.00%
Bootstrap support for afum|EAL91556.1 as seed ortholog is 100%.
Bootstrap support for afis|EAW17744.1 as seed ortholog is 100%.

---

Group of orthologs #3631. Best score 1095 bits
Score difference with first non-orthologous sequence - *A. fumigatus*:1095 *A. fischeri*:1095
afum|EAL93684.1 100.00% afis|EAW18950.1 100.00%
Bootstrap support for afum|EAL93684.1 as seed ortholog is 100%.
Bootstrap support for afis|EAW18950.1 as seed ortholog is 100%.

---

Group of orthologs #3632. Best score 1095 bits
Score difference with first non-orthologous sequence - *A. fumigatus*:1095 *A. fischeri*:1095
afum|EAL90066.1 100.00% afis|EAW25407.1 100.00%
Bootstrap support for afum|EAL90066.1 as seed ortholog is 100%.
Bootstrap support for afis|EAW25407.1 as seed ortholog is 100%.

---

Group of orthologs #3633. Best score 1094 bits
Score difference with first non-orthologous sequence - *A. fumigatus*:1094 *A. fischeri*:1094
afum|EAL85532.1 100.00% afis|EAW15819.1 100.00%
Bootstrap support for afum|EAL85532.1 as seed ortholog is 100%.
Bootstrap support for afis|EAW15819.1 as seed ortholog is 100%.

---

Group of orthologs #3634. Best score 1094 bits
Score difference with first non-orthologous sequence - *A. fumigatus*:1094 *A. fischeri*:421
afum|EAL86473.1 100.00% afis|EAW16946.1 100.00%
Bootstrap support for afum|EAL86473.1 as seed ortholog is 100%.
Bootstrap support for afis|EAW16946.1 as seed ortholog is 100%.

---

Group of orthologs #3635. Best score 1094 bits
Score difference with first non-orthologous sequence - *A. fumigatus*:1094 *A. fischeri*:1094
afum|EAL88542.1 100.00% afis|EAW15907.1 100.00%
Bootstrap support for afum|EAL88542.1 as seed ortholog is 100%.
Bootstrap support for afis|EAW15907.1 as seed ortholog is 100%.

---

Group of orthologs #3636. Best score 1094 bits
Score difference with first non-orthologous sequence - *A. fumigatus*:1094 *A. fischeri*:1094
afum|EAL85432.1 100.00% afis|EAW20302.1 100.00%
Bootstrap support for afum|EAL85432.1 as seed ortholog is 100%.
Bootstrap support for afis|EAW20302.1 as seed ortholog is 100%.

---

Group of orthologs #3637. Best score 1094 bits
Score difference with first non-orthologous sequence - *A. fumigatus*:1094 *A. fischeri*:1094
afum|EAL91822.1 100.00% afis|EAW18010.1 100.00%

Bootstrap support for afum|EAL91822.1 as seed ortholog is 100%.
Bootstrap support for afis|EAW18010.1 as seed ortholog is 100%.

---

Group of orthologs #3638. Best score 1094 bits
Score difference with first non-orthologous sequence - *A. fumigatus*:1094 *A. fischeri*:1094
afum|EAL90726.1 100.00% afis|EAW22465.1 100.00%
Bootstrap support for afum|EAL90726.1 as seed ortholog is 100%.
Bootstrap support for afis|EAW22465.1 as seed ortholog is 100%.

---

Group of orthologs #3639. Best score 1094 bits
Score difference with first non-orthologous sequence - *A. fumigatus*:1094 *A. fischeri*:1094
afum|EAL90418.1 100.00% afis|EAW22781.1 100.00%
Bootstrap support for afum|EAL90418.1 as seed ortholog is 100%.
Bootstrap support for afis|EAW22781.1 as seed ortholog is 100%.

---

Group of orthologs #3640. Best score 1094 bits
Score difference with first non-orthologous sequence - *A. fumigatus*:989 *A. fischeri*:929
afum|EAL92288.1 100.00% afis|EAW21273.1 100.00%
Bootstrap support for afum|EAL92288.1 as seed ortholog is 100%.
Bootstrap support for afis|EAW21273.1 as seed ortholog is 100%.

---

Group of orthologs #3641. Best score 1093 bits
Score difference with first non-orthologous sequence - *A. fumigatus*:1093 *A. fischeri*:1093
afum|EAL88904.1 100.00% afis|EAW16283.1 100.00%
Bootstrap support for afum|EAL88904.1 as seed ortholog is 100%.
Bootstrap support for afis|EAW16283.1 as seed ortholog is 100%.

---

Group of orthologs #3642. Best score 1093 bits
Score difference with first non-orthologous sequence - *A. fumigatus*:901 *A. fischeri*:876
afum|EAL84518.2 100.00% afis|EAW20540.1 100.00%
Bootstrap support for afum|EAL84518.2 as seed ortholog is 100%.
Bootstrap support for afis|EAW20540.1 as seed ortholog is 100%.

---

Group of orthologs #3643. Best score 1093 bits
Score difference with first non-orthologous sequence - *A. fumigatus*:1093 *A. fischeri*:1093
afum|EAL89072.1 100.00% afis|EAW16478.1 100.00%
Bootstrap support for afum|EAL89072.1 as seed ortholog is 100%.
Bootstrap support for afis|EAW16478.1 as seed ortholog is 100%.

---

Group of orthologs #3644. Best score 1093 bits
Score difference with first non-orthologous sequence - *A. fumigatus*:1093 *A. fischeri*:1093
afum|EAL89109.1 100.00% afis|EAW16514.1 100.00%
Bootstrap support for afum|EAL89109.1 as seed ortholog is 100%.
Bootstrap support for afis|EAW16514.1 as seed ortholog is 100%.

---

Group of orthologs #3645. Best score 1093 bits
Score difference with first non-orthologous sequence - *A. fumigatus*:1093 *A. fischeri*:1093
afum|EAL84856.1 100.00% afis|EAW20763.1 100.00%
Bootstrap support for afum|EAL84856.1 as seed ortholog is 100%.
Bootstrap support for afis|EAW20763.1 as seed ortholog is 100%.

---

Group of orthologs #3646. Best score 1093 bits
Score difference with first non-orthologous sequence - *A. fumigatus*:1093 *A. fischeri*:1093
afum|EAL85481.2 100.00% afis|EAW20365.1 100.00%
Bootstrap support for afum|EAL85481.2 as seed ortholog is 100%.
Bootstrap support for afis|EAW20365.1 as seed ortholog is 100%.

---

Group of orthologs #3647. Best score 1093 bits
Score difference with first non-orthologous sequence - *A. fumigatus*:1093 *A. fischeri*:1093
afum|EAL85964.1 100.00% afis|EAW24296.1 100.00%
Bootstrap support for afum|EAL85964.1 as seed ortholog is 100%.
Bootstrap support for afis|EAW24296.1 as seed ortholog is 100%.

---

Group of orthologs #3648. Best score 1093 bits
Score difference with first non-orthologous sequence - *A. fumigatus*:1093 *A. fischeri*:1093
afum|EAL88333.1 100.00% afis|EAW23148.1 100.00%
Bootstrap support for afum|EAL88333.1 as seed ortholog is 100%.
Bootstrap support for afis|EAW23148.1 as seed ortholog is 100%.

---

Group of orthologs #3649. Best score 1093 bits
Score difference with first non-orthologous sequence - *A. fumigatus*:1018 *A. fischeri*:1093
afum|EAL88340.1 100.00% afis|EAW23141.1 100.00%
Bootstrap support for afum|EAL88340.1 as seed ortholog is 100%.
Bootstrap support for afis|EAW23141.1 as seed ortholog is 100%.

---

Group of orthologs #3650. Best score 1093 bits
Score difference with first non-orthologous sequence - *A. fumigatus*:1093 *A. fischeri*:1093
afum|EAL88154.1 100.00% afis|EAW23325.1 100.00%
Bootstrap support for afum|EAL88154.1 as seed ortholog is 100%.
Bootstrap support for afis|EAW23325.1 as seed ortholog is 100%.

---

Group of orthologs #3651. Best score 1093 bits
Score difference with first non-orthologous sequence - *A. fumigatus*:1093 *A. fischeri*:1093
afum|EAL93279.1 100.00% afis|EAW18576.1 100.00%
Bootstrap support for afum|EAL93279.1 as seed ortholog is 100%.
Bootstrap support for afis|EAW18576.1 as seed ortholog is 100%.

---

Group of orthologs #3652. Best score 1092 bits
Score difference with first non-orthologous sequence - *A. fumigatus*:1092 *A. fischeri*:1092
afum|EAL84903.1 100.00% afis|EAW20015.1 100.00%
Bootstrap support for afum|EAL84903.1 as seed ortholog is 100%.
Bootstrap support for afis|EAW20015.1 as seed ortholog is 100%.

---

Group of orthologs #3653. Best score 1092 bits
Score difference with first non-orthologous sequence - *A. fumigatus*:1092 *A. fischeri*:1092
afum|EAL92264.1 100.00% afis|EAW21251.1 100.00%
Bootstrap support for afum|EAL92264.1 as seed ortholog is 100%.
Bootstrap support for afis|EAW21251.1 as seed ortholog is 100%.

---

Group of orthologs #3654. Best score 1091 bits
Score difference with first non-orthologous sequence - *A. fumigatus*:1091 *A. fischeri*:1091
afum|EAL87544.1 100.00% afis|EAW18182.1 100.00%
Bootstrap support for afum|EAL87544.1 as seed ortholog is 100%.
Bootstrap support for afis|EAW18182.1 as seed ortholog is 100%.

---

Group of orthologs #3655. Best score 1091 bits
Score difference with first non-orthologous sequence - *A. fumigatus*:1091 *A. fischeri*:1091
afum|EAL87072.1 100.00% afis|EAW19422.1 100.00%
Bootstrap support for afum|EAL87072.1 as seed ortholog is 100%.
Bootstrap support for afis|EAW19422.1 as seed ortholog is 100%.

---

Group of orthologs #3656. Best score 1091 bits

Score difference with first non-orthologous sequence - *A. fumigatus*:1091   *A. fischeri*:1091
afum|EAL87335.1 100.00% afis|EAW23907.1 100.00%
Bootstrap support for afum|EAL87335.1 as seed ortholog is 100%.
Bootstrap support for afis|EAW23907.1 as seed ortholog is 100%.

---

Group of orthologs #3657. Best score 1091 bits
Score difference with first non-orthologous sequence - *A. fumigatus*:1091   *A. fischeri*:1025
afum|EAL90084.1 100.00% afis|EAW25425.1 100.00%
Bootstrap support for afum|EAL90084.1 as seed ortholog is 100%.
Bootstrap support for afis|EAW25425.1 as seed ortholog is 100%.

---

Group of orthologs #3658. Best score 1090 bits
Score difference with first non-orthologous sequence - *A. fumigatus*:1090   *A. fischeri*:1090
afum|EAL85286.1 100.00% afis|EAW20168.1 100.00%
Bootstrap support for afum|EAL85286.1 as seed ortholog is 100%.
Bootstrap support for afis|EAW20168.1 as seed ortholog is 100%.

---

Group of orthologs #3659. Best score 1090 bits
Score difference with first non-orthologous sequence - *A. fumigatus*:1090   *A. fischeri*:1090
afum|EAL84866.1 100.00% afis|EAW20748.1 100.00%
Bootstrap support for afum|EAL84866.1 as seed ortholog is 100%.
Bootstrap support for afis|EAW20748.1 as seed ortholog is 100%.

---

Group of orthologs #3660. Best score 1090 bits
Score difference with first non-orthologous sequence - *A. fumigatus*:1090   *A. fischeri*:1090
afum|EAL91270.1 100.00% afis|EAW17444.1 100.00%
Bootstrap support for afum|EAL91270.1 as seed ortholog is 100%.
Bootstrap support for afis|EAW17444.1 as seed ortholog is 100%.

---

Group of orthologs #3661. Best score 1090 bits
Score difference with first non-orthologous sequence - *A. fumigatus*:1003   *A. fischeri*:897
afum|EAL89233.1 100.00% afis|EAW20411.1 100.00%
Bootstrap support for afum|EAL89233.1 as seed ortholog is 100%.
Bootstrap support for afis|EAW20411.1 as seed ortholog is 100%.

---

Group of orthologs #3662. Best score 1090 bits
Score difference with first non-orthologous sequence - *A. fumigatus*:923   *A. fischeri*:596
afum|EAL94029.1 100.00% afis|EAW19313.1 100.00%
Bootstrap support for afum|EAL94029.1 as seed ortholog is 100%.
Bootstrap support for afis|EAW19313.1 as seed ortholog is 100%.

---

Group of orthologs #3663. Best score 1090 bits
Score difference with first non-orthologous sequence - *A. fumigatus*:1090   *A. fischeri*:1090
afum|EAL90222.1 100.00% afis|EAW22975.1 100.00%
Bootstrap support for afum|EAL90222.1 as seed ortholog is 100%.
Bootstrap support for afis|EAW22975.1 as seed ortholog is 100%.

---

Group of orthologs #3664. Best score 1090 bits
Score difference with first non-orthologous sequence - *A. fumigatus*:1090   *A. fischeri*:1090
afum|EAL89313.1 100.00% afis|EAW24657.1 100.00%
Bootstrap support for afum|EAL89313.1 as seed ortholog is 100%.
Bootstrap support for afis|EAW24657.1 as seed ortholog is 100%.

---

Group of orthologs #3665. Best score 1090 bits
Score difference with first non-orthologous sequence - *A. fumigatus*:1090   *A. fischeri*:1090
afum|EAL92600.1 100.00% afis|EAW21582.1 100.00%

Bootstrap support for afum|EAL92600.1 as seed ortholog is 100%.
Bootstrap support for afis|EAW21582.1 as seed ortholog is 100%.

---

Group of orthologs #3666. Best score 1090 bits
Score difference with first non-orthologous sequence - *A. fumigatus*:1090 *A. fischeri*:398
afum|EAL89873.1 100.00% afis|EAW25212.1 100.00%
Bootstrap support for afum|EAL89873.1 as seed ortholog is 100%.
Bootstrap support for afis|EAW25212.1 as seed ortholog is 100%.

---

Group of orthologs #3667. Best score 1089 bits
Score difference with first non-orthologous sequence - *A. fumigatus*:1089 *A. fischeri*:1089
afum|EAL85020.1 100.00% afis|EAW19877.1 100.00%
Bootstrap support for afum|EAL85020.1 as seed ortholog is 100%.
Bootstrap support for afis|EAW19877.1 as seed ortholog is 100%.

---

Group of orthologs #3668. Best score 1089 bits
Score difference with first non-orthologous sequence - *A. fumigatus*:1089 *A. fischeri*:1089
afum|EAL91557.1 100.00% afis|EAW17745.1 100.00%
Bootstrap support for afum|EAL91557.1 as seed ortholog is 100%.
Bootstrap support for afis|EAW17745.1 as seed ortholog is 100%.

---

Group of orthologs #3669. Best score 1089 bits
Score difference with first non-orthologous sequence - *A. fumigatus*:1089 *A. fischeri*:1089
afum|EAL93047.1 100.00% afis|EAW18372.1 100.00%
Bootstrap support for afum|EAL93047.1 as seed ortholog is 100%.
Bootstrap support for afis|EAW18372.1 as seed ortholog is 100%.

---

Group of orthologs #3670. Best score 1089 bits
Score difference with first non-orthologous sequence - *A. fumigatus*:1089 *A. fischeri*:1089
afum|EAL93170.1 100.00% afis|EAW18481.1 100.00%
Bootstrap support for afum|EAL93170.1 as seed ortholog is 100%.
Bootstrap support for afis|EAW18481.1 as seed ortholog is 100%.

---

Group of orthologs #3671. Best score 1089 bits
Score difference with first non-orthologous sequence - *A. fumigatus*:1089 *A. fischeri*:1089
afum|EAL90192.1 100.00% afis|EAW25527.1 100.00%
Bootstrap support for afum|EAL90192.1 as seed ortholog is 100%.
Bootstrap support for afis|EAW25527.1 as seed ortholog is 100%.

---

Group of orthologs #3672. Best score 1088 bits
Score difference with first non-orthologous sequence - *A. fumigatus*:1088 *A. fischeri*:1088
afum|EAL88734.1 100.00% afis|EAW16102.1 100.00%
Bootstrap support for afum|EAL88734.1 as seed ortholog is 100%.
Bootstrap support for afis|EAW16102.1 as seed ortholog is 100%.

---

Group of orthologs #3673. Best score 1088 bits
Score difference with first non-orthologous sequence - *A. fumigatus*:1088 *A. fischeri*:1088
afum|EAL87143.1 100.00% afis|EAW23702.1 100.00%
Bootstrap support for afum|EAL87143.1 as seed ortholog is 100%.
Bootstrap support for afis|EAW23702.1 as seed ortholog is 100%.

---

Group of orthologs #3674. Best score 1088 bits
Score difference with first non-orthologous sequence - *A. fumigatus*:1088 *A. fischeri*:1088
afum|EAL87954.1 100.00% afis|EAW23507.1 100.00%
Bootstrap support for afum|EAL87954.1 as seed ortholog is 100%.
Bootstrap support for afis|EAW23507.1 as seed ortholog is 100%.

Group of orthologs #3675. Best score 1088 bits
Score difference with first non-orthologous sequence - *A. fumigatus*:1088 *A. fischeri*:1088
afum|EAL93542.1 100.00% afis|EAW18806.1 100.00%
Bootstrap support for afum|EAL93542.1 as seed ortholog is 100%.
Bootstrap support for afis|EAW18806.1 as seed ortholog is 100%.

Group of orthologs #3676. Best score 1088 bits
Score difference with first non-orthologous sequence - *A. fumigatus*:1088 *A. fischeri*:1088
afum|EAL89336.1 100.00% afis|EAW24694.1 100.00%
Bootstrap support for afum|EAL89336.1 as seed ortholog is 100%.
Bootstrap support for afis|EAW24694.1 as seed ortholog is 100%.

Group of orthologs #3677. Best score 1088 bits
Score difference with first non-orthologous sequence - *A. fumigatus*:1088 *A. fischeri*:1088
afum|EAL89401.1 100.00% afis|EAW24759.1 100.00%
Bootstrap support for afum|EAL89401.1 as seed ortholog is 100%.
Bootstrap support for afis|EAW24759.1 as seed ortholog is 100%.

Group of orthologs #3678. Best score 1087 bits
Score difference with first non-orthologous sequence - *A. fumigatus*:1087 *A. fischeri*:1087
afum|EAL87554.1 100.00% afis|EAW18197.1 100.00%
Bootstrap support for afum|EAL87554.1 as seed ortholog is 100%.
Bootstrap support for afis|EAW18197.1 as seed ortholog is 100%.

Group of orthologs #3679. Best score 1087 bits
Score difference with first non-orthologous sequence - *A. fumigatus*:1087 *A. fischeri*:1087
afum|EAL91189.2 100.00% afis|EAW17345.1 100.00%
Bootstrap support for afum|EAL91189.2 as seed ortholog is 100%.
Bootstrap support for afis|EAW17345.1 as seed ortholog is 100%.

Group of orthologs #3680. Best score 1087 bits
Score difference with first non-orthologous sequence - *A. fumigatus*:1000 *A. fischeri*:1087
afum|EAL85992.1 100.00% afis|EAW24322.1 100.00%
Bootstrap support for afum|EAL85992.1 as seed ortholog is 100%.
Bootstrap support for afis|EAW24322.1 as seed ortholog is 100%.

Group of orthologs #3681. Best score 1087 bits
Score difference with first non-orthologous sequence - *A. fumigatus*:1087 *A. fischeri*:1087
afum|EAL87868.1 100.00% afis|EAW23598.1 100.00%
Bootstrap support for afum|EAL87868.1 as seed ortholog is 100%.
Bootstrap support for afis|EAW23598.1 as seed ortholog is 100%.

Group of orthologs #3682. Best score 1087 bits
Score difference with first non-orthologous sequence - *A. fumigatus*:1087 *A. fischeri*:1087
afum|EAL89331.1 100.00% afis|EAW24690.1 100.00%
Bootstrap support for afum|EAL89331.1 as seed ortholog is 100%.
Bootstrap support for afis|EAW24690.1 as seed ortholog is 100%.

Group of orthologs #3683. Best score 1087 bits
Score difference with first non-orthologous sequence - *A. fumigatus*:1087 *A. fischeri*:1087
afum|EAL89672.1 100.00% afis|EAW25023.1 100.00%
Bootstrap support for afum|EAL89672.1 as seed ortholog is 100%.
Bootstrap support for afis|EAW25023.1 as seed ortholog is 100%.

Group of orthologs #3684. Best score 1086 bits

Score difference with first non-orthologous sequence - *A. fumigatus*:1086   *A. fischeri*:1086
afum|EAL85590.1   100.00%   afis|EAW15758.1   100.00%
Bootstrap support for afum|EAL85590.1 as seed ortholog is 100%.
Bootstrap support for afis|EAW15758.1 as seed ortholog is 100%.

---

Group of orthologs #3685. Best score 1086 bits
Score difference with first non-orthologous sequence - *A. fumigatus*:1086   *A. fischeri*:1086
afum|EAL84677.1   100.00%   afis|EAW20956.1   100.00%
Bootstrap support for afum|EAL84677.1 as seed ortholog is 100%.
Bootstrap support for afis|EAW20956.1 as seed ortholog is 100%.

---

Group of orthologs #3686. Best score 1085 bits
Score difference with first non-orthologous sequence - *A. fumigatus*:1085   *A. fischeri*:1085
afum|EAL87056.1   100.00%   afis|EAW19440.1   100.00%
Bootstrap support for afum|EAL87056.1 as seed ortholog is 100%.
Bootstrap support for afis|EAW19440.1 as seed ortholog is 100%.

---

Group of orthologs #3687. Best score 1085 bits
Score difference with first non-orthologous sequence - *A. fumigatus*:554   *A. fischeri*:632
afum|EAL92453.2   100.00%   afis|EAW21438.1   100.00%
Bootstrap support for afum|EAL92453.2 as seed ortholog is 100%.
Bootstrap support for afis|EAW21438.1 as seed ortholog is 100%.

---

Group of orthologs #3688. Best score 1084 bits
Score difference with first non-orthologous sequence - *A. fumigatus*:1084   *A. fischeri*:638
afum|EAL85859.1   100.00%   afis|EAW15460.1   100.00%
Bootstrap support for afum|EAL85859.1 as seed ortholog is 100%.
Bootstrap support for afis|EAW15460.1 as seed ortholog is 100%.

---

Group of orthologs #3689. Best score 1084 bits
Score difference with first non-orthologous sequence - *A. fumigatus*:1084   *A. fischeri*:1084
afum|EAL86693.1   100.00%   afis|EAW17275.1   100.00%
Bootstrap support for afum|EAL86693.1 as seed ortholog is 100%.
Bootstrap support for afis|EAW17275.1 as seed ortholog is 100%.

---

Group of orthologs #3690. Best score 1084 bits
Score difference with first non-orthologous sequence - *A. fumigatus*:1011   *A. fischeri*:995
afum|EAL84874.1   100.00%   afis|EAW20699.1   100.00%
Bootstrap support for afum|EAL84874.1 as seed ortholog is 100%.
Bootstrap support for afis|EAW20699.1 as seed ortholog is 100%.

---

Group of orthologs #3691. Best score 1084 bits
Score difference with first non-orthologous sequence - *A. fumigatus*:1084   *A. fischeri*:1084
afum|EAL91295.1   100.00%   afis|EAW17466.1   100.00%
Bootstrap support for afum|EAL91295.1 as seed ortholog is 100%.
Bootstrap support for afis|EAW17466.1 as seed ortholog is 100%.

---

Group of orthologs #3692. Best score 1084 bits
Score difference with first non-orthologous sequence - *A. fumigatus*:1084   *A. fischeri*:1084
afum|EAL91786.1   100.00%   afis|EAW17977.1   100.00%
Bootstrap support for afum|EAL91786.1 as seed ortholog is 100%.
Bootstrap support for afis|EAW17977.1 as seed ortholog is 100%.

---

Group of orthologs #3693. Best score 1084 bits
Score difference with first non-orthologous sequence - *A. fumigatus*:1084   *A. fischeri*:1084
afum|EAL90558.2   100.00%   afis|EAW22635.1   100.00%

Bootstrap support for afum|EAL90558.2 as seed ortholog is 100%.
Bootstrap support for afis|EAW22635.1 as seed ortholog is 100%.

---

Group of orthologs #3694. Best score 1084 bits
Score difference with first non-orthologous sequence - *A. fumigatus*:1084 *A. fischeri*:1084
afum|EAL89657.1 100.00% afis|EAW25037.1 100.00%
Bootstrap support for afum|EAL89657.1 as seed ortholog is 100%.
Bootstrap support for afis|EAW25037.1 as seed ortholog is 100%.

---

Group of orthologs #3695. Best score 1083 bits
Score difference with first non-orthologous sequence - *A. fumigatus*:1083 *A. fischeri*:1083
afum|EAL86116.1 100.00% afis|EAW24447.1 100.00%
Bootstrap support for afum|EAL86116.1 as seed ortholog is 100%.
Bootstrap support for afis|EAW24447.1 as seed ortholog is 100%.

---

Group of orthologs #3696. Best score 1083 bits
Score difference with first non-orthologous sequence - *A. fumigatus*:549 *A. fischeri*:583
afum|EAL87116.1 100.00% afis|EAW23673.1 100.00%
Bootstrap support for afum|EAL87116.1 as seed ortholog is 100%.
Bootstrap support for afis|EAW23673.1 as seed ortholog is 100%.

---

Group of orthologs #3697. Best score 1083 bits
Score difference with first non-orthologous sequence - *A. fumigatus*:1083 *A. fischeri*:1083
afum|EAL87220.2 100.00% afis|EAW23786.1 100.00%
Bootstrap support for afum|EAL87220.2 as seed ortholog is 100%.
Bootstrap support for afis|EAW23786.1 as seed ortholog is 100%.

---

Group of orthologs #3698. Best score 1083 bits
Score difference with first non-orthologous sequence - *A. fumigatus*:831 *A. fischeri*:1083
afum|EAL92847.1 100.00% afis|EAW21832.1 100.00%
Bootstrap support for afum|EAL92847.1 as seed ortholog is 100%.
Bootstrap support for afis|EAW21832.1 as seed ortholog is 100%.

---

Group of orthologs #3699. Best score 1082 bits
Score difference with first non-orthologous sequence - *A. fumigatus*:1082 *A. fischeri*:715
afum|EAL85504.1 100.00% afis|EAW15847.1 100.00%
Bootstrap support for afum|EAL85504.1 as seed ortholog is 100%.
Bootstrap support for afis|EAW15847.1 as seed ortholog is 100%.

---

Group of orthologs #3700. Best score 1082 bits
Score difference with first non-orthologous sequence - *A. fumigatus*:1082 *A. fischeri*:1082
afum|EAL88223.1 100.00% afis|EAW23258.1 100.00%
Bootstrap support for afum|EAL88223.1 as seed ortholog is 100%.
Bootstrap support for afis|EAW23258.1 as seed ortholog is 100%.

---

Group of orthologs #3701. Best score 1082 bits
Score difference with first non-orthologous sequence - *A. fumigatus*:1082 *A. fischeri*:908
afum|EAL93742.1 100.00% afis|EAW19008.1 100.00%
Bootstrap support for afum|EAL93742.1 as seed ortholog is 100%.
Bootstrap support for afis|EAW19008.1 as seed ortholog is 100%.

---

Group of orthologs #3702. Best score 1082 bits
Score difference with first non-orthologous sequence - *A. fumigatus*:939 *A. fischeri*:1082
afum|EAL92086.1 100.00% afis|EAW21050.1 100.00%
Bootstrap support for afum|EAL92086.1 as seed ortholog is 100%.
Bootstrap support for afis|EAW21050.1 as seed ortholog is 100%.

---

Group of orthologs #3703. Best score 1082 bits
Score difference with first non-orthologous sequence - *A. fumigatus*:1082 *A. fischeri*:1082
afum|EAL90905.1 100.00% afis|EAW22276.1 100.00%
Bootstrap support for afum|EAL90905.1 as seed ortholog is 100%.
Bootstrap support for afis|EAW22276.1 as seed ortholog is 100%.

---

Group of orthologs #3704. Best score 1082 bits
Score difference with first non-orthologous sequence - *A. fumigatus*:1082 *A. fischeri*:1082
afum|EAL89716.1 100.00% afis|EAW25048.1 100.00%
Bootstrap support for afum|EAL89716.1 as seed ortholog is 100%.
Bootstrap support for afis|EAW25048.1 as seed ortholog is 100%.

---

Group of orthologs #3705. Best score 1082 bits
Score difference with first non-orthologous sequence - *A. fumigatus*:1082 *A. fischeri*:997
afum|EAL93009.1 100.00% afis|EAW21994.1 100.00%
Bootstrap support for afum|EAL93009.1 as seed ortholog is 100%.
Bootstrap support for afis|EAW21994.1 as seed ortholog is 100%.

---

Group of orthologs #3706. Best score 1081 bits
Score difference with first non-orthologous sequence - *A. fumigatus*:1081 *A. fischeri*:396
afum|EAL85407.1 100.00% afis|EAW20273.1 100.00%
Bootstrap support for afum|EAL85407.1 as seed ortholog is 100%.
Bootstrap support for afis|EAW20273.1 as seed ortholog is 100%.

---

Group of orthologs #3707. Best score 1081 bits
Score difference with first non-orthologous sequence - *A. fumigatus*:1081 *A. fischeri*:1081
afum|EAL91467.1 100.00% afis|EAW17652.1 100.00%
Bootstrap support for afum|EAL91467.1 as seed ortholog is 100%.
Bootstrap support for afis|EAW17652.1 as seed ortholog is 100%.

---

Group of orthologs #3708. Best score 1081 bits
Score difference with first non-orthologous sequence - *A. fumigatus*:1081 *A. fischeri*:1081
afum|EAL87148.1 100.00% afis|EAW23704.1 100.00%
Bootstrap support for afum|EAL87148.1 as seed ortholog is 100%.
Bootstrap support for afis|EAW23704.1 as seed ortholog is 100%.

---

Group of orthologs #3709. Best score 1081 bits
Score difference with first non-orthologous sequence - *A. fumigatus*:1081 *A. fischeri*:1081
afum|EAL92436.1 100.00% afis|EAW21421.1 100.00%
Bootstrap support for afum|EAL92436.1 as seed ortholog is 100%.
Bootstrap support for afis|EAW21421.1 as seed ortholog is 100%.

---

Group of orthologs #3710. Best score 1081 bits
Score difference with first non-orthologous sequence - *A. fumigatus*:1081 *A. fischeri*:1081
afum|EAL92648.1 100.00% afis|EAW21614.1 100.00%
Bootstrap support for afum|EAL92648.1 as seed ortholog is 100%.
Bootstrap support for afis|EAW21614.1 as seed ortholog is 100%.

---

Group of orthologs #3711. Best score 1081 bits
Score difference with first non-orthologous sequence - *A. fumigatus*:945 *A. fischeri*:880
afum|EAL89404.1 100.00% afis|EAW24762.1 100.00%
Bootstrap support for afum|EAL89404.1 as seed ortholog is 100%.
Bootstrap support for afis|EAW24762.1 as seed ortholog is 100%.

---

Group of orthologs #3712. Best score 1080 bits

Score difference with first non-orthologous sequence - *A. fumigatus*:1080   *A. fischeri*:1080
afum|EAL88671.1 100.00% afis|EAW16039.1 100.00%
Bootstrap support for afum|EAL88671.1 as seed ortholog is 100%.
Bootstrap support for afis|EAW16039.1 as seed ortholog is 100%.

---

Group of orthologs #3713. Best score 1080 bits
Score difference with first non-orthologous sequence - *A. fumigatus*:1080   *A. fischeri*:1080
afum|EAL87529.2 100.00% afis|EAW18168.1 100.00%
Bootstrap support for afum|EAL87529.2 as seed ortholog is 100%.
Bootstrap support for afis|EAW18168.1 as seed ortholog is 100%.

---

Group of orthologs #3714. Best score 1080 bits
Score difference with first non-orthologous sequence - *A. fumigatus*:1080   *A. fischeri*:1080
afum|EAL93233.1 100.00% afis|EAW18535.1 100.00%
Bootstrap support for afum|EAL93233.1 as seed ortholog is 100%.
Bootstrap support for afis|EAW18535.1 as seed ortholog is 100%.

---

Group of orthologs #3715. Best score 1080 bits
Score difference with first non-orthologous sequence - *A. fumigatus*:1080   *A. fischeri*:1080
afum|EAL92585.1 100.00% afis|EAW21566.1 100.00%
Bootstrap support for afum|EAL92585.1 as seed ortholog is 100%.
Bootstrap support for afis|EAW21566.1 as seed ortholog is 100%.

---

Group of orthologs #3716. Best score 1080 bits
Score difference with first non-orthologous sequence - *A. fumigatus*:1080   *A. fischeri*:881
afum|EAL89380.1 100.00% afis|EAW24733.1 100.00%
Bootstrap support for afum|EAL89380.1 as seed ortholog is 100%.
Bootstrap support for afis|EAW24733.1 as seed ortholog is 100%.

---

Group of orthologs #3717. Best score 1079 bits
Score difference with first non-orthologous sequence - *A. fumigatus*:1079   *A. fischeri*:1079
afum|EAL90384.1 100.00% afis|EAW22817.1 100.00%
Bootstrap support for afum|EAL90384.1 as seed ortholog is 100%.
Bootstrap support for afis|EAW22817.1 as seed ortholog is 100%.

---

Group of orthologs #3718. Best score 1079 bits
Score difference with first non-orthologous sequence - *A. fumigatus*:1079   *A. fischeri*:1079
afum|EAL89561.1 100.00% afis|EAW24908.1 100.00%
Bootstrap support for afum|EAL89561.1 as seed ortholog is 100%.
Bootstrap support for afis|EAW24908.1 as seed ortholog is 100%.

---

Group of orthologs #3719. Best score 1078 bits
Score difference with first non-orthologous sequence - *A. fumigatus*:1078   *A. fischeri*:1078
afum|EAL84514.1 100.00% afis|EAW20544.1 100.00%
Bootstrap support for afum|EAL84514.1 as seed ortholog is 100%.
Bootstrap support for afis|EAW20544.1 as seed ortholog is 100%.

---

Group of orthologs #3720. Best score 1078 bits
Score difference with first non-orthologous sequence - *A. fumigatus*:1078   *A. fischeri*:1078
afum|EAL87635.1 100.00% afis|EAW18270.1 100.00%
Bootstrap support for afum|EAL87635.1 as seed ortholog is 100%.
Bootstrap support for afis|EAW18270.1 as seed ortholog is 100%.

---

Group of orthologs #3721. Best score 1078 bits
Score difference with first non-orthologous sequence - *A. fumigatus*:1078   *A. fischeri*:963
afum|EAL86789.1 100.00% afis|EAW19740.1 100.00%

Bootstrap support for afum|EAL86789.1 as seed ortholog is 100%.
Bootstrap support for afis|EAW19740.1 as seed ortholog is 100%.

---

Group of orthologs #3722. Best score 1078 bits
Score difference with first non-orthologous sequence - *A. fumigatus*:877 *A. fischeri*:967
afum|EAL89235.1 100.00% afis|EAW20409.1 100.00%
Bootstrap support for afum|EAL89235.1 as seed ortholog is 100%.
Bootstrap support for afis|EAW20409.1 as seed ortholog is 100%.

---

Group of orthologs #3723. Best score 1078 bits
Score difference with first non-orthologous sequence - *A. fumigatus*:1078 *A. fischeri*:1078
afum|EAL85981.1 100.00% afis|EAW24313.1 100.00%
Bootstrap support for afum|EAL85981.1 as seed ortholog is 100%.
Bootstrap support for afis|EAW24313.1 as seed ortholog is 100%.

---

Group of orthologs #3724. Best score 1078 bits
Score difference with first non-orthologous sequence - *A. fumigatus*:1078 *A. fischeri*:1078
afum|EAL93289.1 100.00% afis|EAW18585.1 100.00%
Bootstrap support for afum|EAL93289.1 as seed ortholog is 100%.
Bootstrap support for afis|EAW18585.1 as seed ortholog is 100%.

---

Group of orthologs #3725. Best score 1078 bits
Score difference with first non-orthologous sequence - *A. fumigatus*:1078 *A. fischeri*:1078
afum|EAL89505.2 100.00% afis|EAW24857.1 100.00%
Bootstrap support for afum|EAL89505.2 as seed ortholog is 100%.
Bootstrap support for afis|EAW24857.1 as seed ortholog is 100%.

---

Group of orthologs #3726. Best score 1077 bits
Score difference with first non-orthologous sequence - *A. fumigatus*:988 *A. fischeri*:1077
afum|EAL91205.2 100.00% afis|EAW17368.1 100.00%
Bootstrap support for afum|EAL91205.2 as seed ortholog is 100%.
Bootstrap support for afis|EAW17368.1 as seed ortholog is 100%.

---

Group of orthologs #3727. Best score 1077 bits
Score difference with first non-orthologous sequence - *A. fumigatus*:1077 *A. fischeri*:1077
afum|EAL91365.1 100.00% afis|EAW17551.1 100.00%
Bootstrap support for afum|EAL91365.1 as seed ortholog is 100%.
Bootstrap support for afis|EAW17551.1 as seed ortholog is 100%.

---

Group of orthologs #3728. Best score 1077 bits
Score difference with first non-orthologous sequence - *A. fumigatus*:1077 *A. fischeri*:1077
afum|EAL91598.1 100.00% afis|EAW17789.1 100.00%
Bootstrap support for afum|EAL91598.1 as seed ortholog is 100%.
Bootstrap support for afis|EAW17789.1 as seed ortholog is 100%.

---

Group of orthologs #3729. Best score 1077 bits
Score difference with first non-orthologous sequence - *A. fumigatus*:1077 *A. fischeri*:1077
afum|EAL91087.1 100.00% afis|EAW22104.1 100.00%
Bootstrap support for afum|EAL91087.1 as seed ortholog is 100%.
Bootstrap support for afis|EAW22104.1 as seed ortholog is 100%.

---

Group of orthologs #3730. Best score 1077 bits
Score difference with first non-orthologous sequence - *A. fumigatus*:1077 *A. fischeri*:1077
afum|EAL90661.1 100.00% afis|EAW22533.1 100.00%
Bootstrap support for afum|EAL90661.1 as seed ortholog is 100%.
Bootstrap support for afis|EAW22533.1 as seed ortholog is 100%.

---
Group of orthologs #3731. Best score 1077 bits
Score difference with first non-orthologous sequence - *A. fumigatus*:1077 *A. fischeri*:1077
afum|EBA27509.1 100.00% afis|EAW19239.1 100.00%
Bootstrap support for afum|EBA27509.1 as seed ortholog is 100%.
Bootstrap support for afis|EAW19239.1 as seed ortholog is 100%.

---
Group of orthologs #3732. Best score 1076 bits
Score difference with first non-orthologous sequence - *A. fumigatus*:834 *A. fischeri*:693
afum|EAL86303.2 100.00% afis|EAW16728.1 100.00%
Bootstrap support for afum|EAL86303.2 as seed ortholog is 100%.
Bootstrap support for afis|EAW16728.1 as seed ortholog is 100%.

---
Group of orthologs #3733. Best score 1076 bits
Score difference with first non-orthologous sequence - *A. fumigatus*:1076 *A. fischeri*:1076
afum|EAL90988.1 100.00% afis|EAW22205.1 100.00%
Bootstrap support for afum|EAL90988.1 as seed ortholog is 100%.
Bootstrap support for afis|EAW22205.1 as seed ortholog is 100%.

---
Group of orthologs #3734. Best score 1076 bits
Score difference with first non-orthologous sequence - *A. fumigatus*:1076 *A. fischeri*:1076
afum|EAL89577.2 100.00% afis|EAW24922.1 100.00%
Bootstrap support for afum|EAL89577.2 as seed ortholog is 100%.
Bootstrap support for afis|EAW24922.1 as seed ortholog is 100%.

---
Group of orthologs #3735. Best score 1075 bits
Score difference with first non-orthologous sequence - *A. fumigatus*:1075 *A. fischeri*:1075
afum|EAL84233.1 100.00% afis|EAW15413.1 100.00%
Bootstrap support for afum|EAL84233.1 as seed ortholog is 100%.
Bootstrap support for afis|EAW15413.1 as seed ortholog is 100%.

---
Group of orthologs #3736. Best score 1075 bits
Score difference with first non-orthologous sequence - *A. fumigatus*:1075 *A. fischeri*:735
afum|EAL86353.1 100.00% afis|EAW16782.1 100.00%
Bootstrap support for afum|EAL86353.1 as seed ortholog is 100%.
Bootstrap support for afis|EAW16782.1 as seed ortholog is 100%.

---
Group of orthologs #3737. Best score 1075 bits
Score difference with first non-orthologous sequence - *A. fumigatus*:842 *A. fischeri*:1075
afum|EAL84470.1 100.00% afis|EAW20591.1 100.00%
Bootstrap support for afum|EAL84470.1 as seed ortholog is 100%.
Bootstrap support for afis|EAW20591.1 as seed ortholog is 100%.

---
Group of orthologs #3738. Best score 1075 bits
Score difference with first non-orthologous sequence - *A. fumigatus*:1075 *A. fischeri*:1075
afum|EAL91456.1 100.00% afis|EAW17641.1 100.00%
Bootstrap support for afum|EAL91456.1 as seed ortholog is 100%.
Bootstrap support for afis|EAW17641.1 as seed ortholog is 100%.

---
Group of orthologs #3739. Best score 1075 bits
Score difference with first non-orthologous sequence - *A. fumigatus*:1075 *A. fischeri*:1075
afum|EAL93682.1 100.00% afis|EAW18948.1 100.00%
Bootstrap support for afum|EAL93682.1 as seed ortholog is 100%.
Bootstrap support for afis|EAW18948.1 as seed ortholog is 100%.

---
Group of orthologs #3740. Best score 1075 bits

Score difference with first non-orthologous sequence - *A. fumigatus*:1075   *A. fischeri*:1075
afum|EAL90513.1   100.00%   afis|EAW22679.1   100.00%
Bootstrap support for afum|EAL90513.1 as seed ortholog is 100%.
Bootstrap support for afis|EAW22679.1 as seed ortholog is 100%.

---

Group of orthologs #3741. Best score 1074 bits
Score difference with first non-orthologous sequence - *A. fumigatus*:1074   *A. fischeri*:1074
afum|EAL86447.1   100.00%   afis|EAW16911.1   100.00%
Bootstrap support for afum|EAL86447.1 as seed ortholog is 100%.
Bootstrap support for afis|EAW16911.1 as seed ortholog is 100%.

---

Group of orthologs #3742. Best score 1074 bits
Score difference with first non-orthologous sequence - *A. fumigatus*:785   *A. fischeri*:791
afum|EAL85142.1   100.00%   afis|EAW19755.1   100.00%
Bootstrap support for afum|EAL85142.1 as seed ortholog is 100%.
Bootstrap support for afis|EAW19755.1 as seed ortholog is 100%.

---

Group of orthologs #3743. Best score 1074 bits
Score difference with first non-orthologous sequence - *A. fumigatus*:1074   *A. fischeri*:1074
afum|EAL87593.2   100.00%   afis|EAW18226.1   100.00%
Bootstrap support for afum|EAL87593.2 as seed ortholog is 100%.
Bootstrap support for afis|EAW18226.1 as seed ortholog is 100%.

---

Group of orthologs #3744. Best score 1074 bits
Score difference with first non-orthologous sequence - *A. fumigatus*:1074   *A. fischeri*:1074
afum|EAL87215.1   100.00%   afis|EAW23781.1   100.00%
Bootstrap support for afum|EAL87215.1 as seed ortholog is 100%.
Bootstrap support for afis|EAW23781.1 as seed ortholog is 100%.

---

Group of orthologs #3745. Best score 1074 bits
Score difference with first non-orthologous sequence - *A. fumigatus*:1074   *A. fischeri*:1074
afum|EAL87854.1   100.00%   afis|EAW23610.1   100.00%
Bootstrap support for afum|EAL87854.1 as seed ortholog is 100%.
Bootstrap support for afis|EAW23610.1 as seed ortholog is 100%.

---

Group of orthologs #3746. Best score 1074 bits
Score difference with first non-orthologous sequence - *A. fumigatus*:1074   *A. fischeri*:1074
afum|EAL93807.1   100.00%   afis|EAW19079.1   100.00%
Bootstrap support for afum|EAL93807.1 as seed ortholog is 100%.
Bootstrap support for afis|EAW19079.1 as seed ortholog is 100%.

---

Group of orthologs #3747. Best score 1073 bits
Score difference with first non-orthologous sequence - *A. fumigatus*:1073   *A. fischeri*:1073
afum|EAL85283.1   100.00%   afis|EAW20165.1   100.00%
Bootstrap support for afum|EAL85283.1 as seed ortholog is 100%.
Bootstrap support for afis|EAW20165.1 as seed ortholog is 100%.

---

Group of orthologs #3748. Best score 1073 bits
Score difference with first non-orthologous sequence - *A. fumigatus*:1073   *A. fischeri*:1073
afum|EAL91162.1   100.00%   afis|EAW17325.1   100.00%
Bootstrap support for afum|EAL91162.1 as seed ortholog is 100%.
Bootstrap support for afis|EAW17325.1 as seed ortholog is 100%.

---

Group of orthologs #3749. Best score 1073 bits
Score difference with first non-orthologous sequence - *A. fumigatus*:1073   *A. fischeri*:1073
afum|EAL93372.1   100.00%   afis|EAW18656.1   100.00%

Bootstrap support for afum|EAL93372.1 as seed ortholog is 100%.
Bootstrap support for afis|EAW18656.1 as seed ortholog is 100%.

---

Group of orthologs #3750. Best score 1072 bits
Score difference with first non-orthologous sequence - *A. fumigatus*:975 *A. fischeri*:870
afum|EAL89241.1 100.00% afis|EAW16639.1 100.00%
Bootstrap support for afum|EAL89241.1 as seed ortholog is 100%.
Bootstrap support for afis|EAW16639.1 as seed ortholog is 100%.

---

Group of orthologs #3751. Best score 1072 bits
Score difference with first non-orthologous sequence - *A. fumigatus*:1072 *A. fischeri*:1072
afum|EAL84717.1 100.00% afis|EAW20917.1 100.00%
Bootstrap support for afum|EAL84717.1 as seed ortholog is 100%.
Bootstrap support for afis|EAW20917.1 as seed ortholog is 100%.

---

Group of orthologs #3752. Best score 1072 bits
Score difference with first non-orthologous sequence - *A. fumigatus*:1072 *A. fischeri*:1072
afum|EAL91412.2 100.00% afis|EAW17599.1 100.00%
Bootstrap support for afum|EAL91412.2 as seed ortholog is 100%.
Bootstrap support for afis|EAW17599.1 as seed ortholog is 100%.

---

Group of orthologs #3753. Best score 1072 bits
Score difference with first non-orthologous sequence - *A. fumigatus*:1072 *A. fischeri*:1072
afum|EAL87272.1 100.00% afis|EAW23841.1 100.00%
Bootstrap support for afum|EAL87272.1 as seed ortholog is 100%.
Bootstrap support for afis|EAW23841.1 as seed ortholog is 100%.

---

Group of orthologs #3754. Best score 1072 bits
Score difference with first non-orthologous sequence - *A. fumigatus*:1072 *A. fischeri*:1072
afum|EAL88243.1 100.00% afis|EAW23239.1 100.00%
Bootstrap support for afum|EAL88243.1 as seed ortholog is 100%.
Bootstrap support for afis|EAW23239.1 as seed ortholog is 100%.

---

Group of orthologs #3755. Best score 1072 bits
Score difference with first non-orthologous sequence - *A. fumigatus*:1072 *A. fischeri*:1072
afum|EAL93427.2 100.00% afis|EAW18704.1 100.00%
Bootstrap support for afum|EAL93427.2 as seed ortholog is 100%.
Bootstrap support for afis|EAW18704.1 as seed ortholog is 100%.

---

Group of orthologs #3756. Best score 1072 bits
Score difference with first non-orthologous sequence - *A. fumigatus*:1072 *A. fischeri*:1072
afum|EAL93538.1 100.00% afis|EAW18802.1 100.00%
Bootstrap support for afum|EAL93538.1 as seed ortholog is 100%.
Bootstrap support for afis|EAW18802.1 as seed ortholog is 100%.

---

Group of orthologs #3757. Best score 1072 bits
Score difference with first non-orthologous sequence - *A. fumigatus*:517 *A. fischeri*:521
afum|EAL94045.1 100.00% afis|EAW19328.1 100.00%
Bootstrap support for afum|EAL94045.1 as seed ortholog is 100%.
Bootstrap support for afis|EAW19328.1 as seed ortholog is 100%.

---

Group of orthologs #3758. Best score 1072 bits
Score difference with first non-orthologous sequence - *A. fumigatus*:1072 *A. fischeri*:1072
afum|EAL90323.1 100.00% afis|EAW22876.1 100.00%
Bootstrap support for afum|EAL90323.1 as seed ortholog is 100%.
Bootstrap support for afis|EAW22876.1 as seed ortholog is 100%.

Group of orthologs #3759. Best score 1072 bits
Score difference with first non-orthologous sequence - *A. fumigatus*:1072 *A. fischeri*:1072
afum|EAL92350.1 100.00% afis|EAW21332.1 100.00%
Bootstrap support for afum|EAL92350.1 as seed ortholog is 100%.
Bootstrap support for afis|EAW21332.1 as seed ortholog is 100%.

---

Group of orthologs #3760. Best score 1072 bits
Score difference with first non-orthologous sequence - *A. fumigatus*:1072 *A. fischeri*:1072
afum|EAL92016.1 100.00% afis|EAW24160.1 100.00%
Bootstrap support for afum|EAL92016.1 as seed ortholog is 100%.
Bootstrap support for afis|EAW24160.1 as seed ortholog is 100%.

---

Group of orthologs #3761. Best score 1071 bits
Score difference with first non-orthologous sequence - *A. fumigatus*:1071 *A. fischeri*:1071
afum|EAL86803.1 100.00% afis|EAW19723.1 100.00%
Bootstrap support for afum|EAL86803.1 as seed ortholog is 100%.
Bootstrap support for afis|EAW19723.1 as seed ortholog is 100%.

---

Group of orthologs #3762. Best score 1071 bits
Score difference with first non-orthologous sequence - *A. fumigatus*:1071 *A. fischeri*:1071
afum|EAL91336.1 100.00% afis|EAW17518.1 100.00%
Bootstrap support for afum|EAL91336.1 as seed ortholog is 100%.
Bootstrap support for afis|EAW17518.1 as seed ortholog is 100%.

---

Group of orthologs #3763. Best score 1071 bits
Score difference with first non-orthologous sequence - *A. fumigatus*:1071 *A. fischeri*:904
afum|EAL87300.1 100.00% afis|EAW23872.1 100.00%
Bootstrap support for afum|EAL87300.1 as seed ortholog is 100%.
Bootstrap support for afis|EAW23872.1 as seed ortholog is 100%.

---

Group of orthologs #3764. Best score 1071 bits
Score difference with first non-orthologous sequence - *A. fumigatus*:1071 *A. fischeri*:1071
afum|EAL93484.1 100.00% afis|EAW18756.1 100.00%
Bootstrap support for afum|EAL93484.1 as seed ortholog is 100%.
Bootstrap support for afis|EAW18756.1 as seed ortholog is 100%.

---

Group of orthologs #3765. Best score 1071 bits
Score difference with first non-orthologous sequence - *A. fumigatus*:1071 *A. fischeri*:1071
afum|EAL90684.1 100.00% afis|EAW22509.1 100.00%
Bootstrap support for afum|EAL90684.1 as seed ortholog is 100%.
Bootstrap support for afis|EAW22509.1 as seed ortholog is 100%.

---

Group of orthologs #3766. Best score 1071 bits
Score difference with first non-orthologous sequence - *A. fumigatus*:943 *A. fischeri*:892
afum|EAL90137.1 100.00% afis|EAW25474.1 100.00%
Bootstrap support for afum|EAL90137.1 as seed ortholog is 100%.
Bootstrap support for afis|EAW25474.1 as seed ortholog is 100%.

---

Group of orthologs #3767. Best score 1070 bits
Score difference with first non-orthologous sequence - *A. fumigatus*:1070 *A. fischeri*:1070
afum|EAL85784.1 100.00% afis|EAW15548.1 100.00%
Bootstrap support for afum|EAL85784.1 as seed ortholog is 100%.
Bootstrap support for afis|EAW15548.1 as seed ortholog is 100%.

---

Group of orthologs #3768. Best score 1070 bits

Score difference with first non-orthologous sequence - *A. fumigatus*:777   *A. fischeri*:551
afum|EAL85729.1 100.00% afis|EAW15604.1 100.00%
Bootstrap support for afum|EAL85729.1 as seed ortholog is 100%.
Bootstrap support for afis|EAW15604.1 as seed ortholog is 100%.

---

Group of orthologs #3769. Best score 1070 bits
Score difference with first non-orthologous sequence - *A. fumigatus*:1070   *A. fischeri*:1070
afum|EAL87462.1 100.00% afis|EAW24031.1 100.00%
Bootstrap support for afum|EAL87462.1 as seed ortholog is 100%.
Bootstrap support for afis|EAW24031.1 as seed ortholog is 100%.

---

Group of orthologs #3770. Best score 1070 bits
Score difference with first non-orthologous sequence - *A. fumigatus*:1070   *A. fischeri*:1070
afum|EAL90846.1 100.00% afis|EAW22346.1 100.00%
Bootstrap support for afum|EAL90846.1 as seed ortholog is 100%.
Bootstrap support for afis|EAW22346.1 as seed ortholog is 100%.

---

Group of orthologs #3771. Best score 1069 bits
Score difference with first non-orthologous sequence - *A. fumigatus*:1069   *A. fischeri*:1069
afum|EAL84468.1 100.00% afis|EAW20593.1 100.00%
Bootstrap support for afum|EAL84468.1 as seed ortholog is 100%.
Bootstrap support for afis|EAW20593.1 as seed ortholog is 100%.

---

Group of orthologs #3772. Best score 1069 bits
Score difference with first non-orthologous sequence - *A. fumigatus*:1069   *A. fischeri*:1069
afum|EAL93169.1 100.00% afis|EAW18480.1 100.00%
Bootstrap support for afum|EAL93169.1 as seed ortholog is 100%.
Bootstrap support for afis|EAW18480.1 as seed ortholog is 100%.

---

Group of orthologs #3773. Best score 1068 bits
Score difference with first non-orthologous sequence - *A. fumigatus*:1068   *A. fischeri*:1068
afum|EAL89238.1 100.00% afis|EAW16636.1 100.00%
Bootstrap support for afum|EAL89238.1 as seed ortholog is 100%.
Bootstrap support for afis|EAW16636.1 as seed ortholog is 100%.

---

Group of orthologs #3774. Best score 1068 bits
Score difference with first non-orthologous sequence - *A. fumigatus*:1068   *A. fischeri*:1068
afum|EAL89254.1 100.00% afis|EAW16650.1 100.00%
Bootstrap support for afum|EAL89254.1 as seed ortholog is 100%.
Bootstrap support for afis|EAW16650.1 as seed ortholog is 100%.

---

Group of orthologs #3775. Best score 1068 bits
Score difference with first non-orthologous sequence - *A. fumigatus*:1068   *A. fischeri*:523
afum|EAL93248.1 100.00% afis|EAW18548.1 100.00%
Bootstrap support for afum|EAL93248.1 as seed ortholog is 100%.
Bootstrap support for afis|EAW18548.1 as seed ortholog is 100%.

---

Group of orthologs #3776. Best score 1068 bits
Score difference with first non-orthologous sequence - *A. fumigatus*:1068   *A. fischeri*:1068
afum|EAL91931.2 100.00% afis|EAW24076.1 100.00%
Bootstrap support for afum|EAL91931.2 as seed ortholog is 100%.
Bootstrap support for afis|EAW24076.1 as seed ortholog is 100%.

---

Group of orthologs #3777. Best score 1067 bits
Score difference with first non-orthologous sequence - *A. fumigatus*:987   *A. fischeri*:933
afum|EAL85913.2 100.00% afis|EAW24245.1 100.00%

Bootstrap support for afum|EAL85913.2 as seed ortholog is 100%.
Bootstrap support for afis|EAW24245.1 as seed ortholog is 100%.

---

Group of orthologs #3778. Best score 1066 bits
Score difference with first non-orthologous sequence - *A. fumigatus*:1066 *A. fischeri*:1066
afum|EAL86863.1 100.00% afis|EAW19660.1 100.00%
Bootstrap support for afum|EAL86863.1 as seed ortholog is 100%.
Bootstrap support for afis|EAW19660.1 as seed ortholog is 100%.

---

Group of orthologs #3779. Best score 1066 bits
Score difference with first non-orthologous sequence - *A. fumigatus*:1066 *A. fischeri*:1066
afum|EAL90341.1 100.00% afis|EAW22859.1 100.00%
Bootstrap support for afum|EAL90341.1 as seed ortholog is 100%.
Bootstrap support for afis|EAW22859.1 as seed ortholog is 100%.

---

Group of orthologs #3780. Best score 1065 bits
Score difference with first non-orthologous sequence - *A. fumigatus*:1065 *A. fischeri*:1065
afum|EAL91854.1 100.00% afis|EAW18043.1 100.00%
Bootstrap support for afum|EAL91854.1 as seed ortholog is 100%.
Bootstrap support for afis|EAW18043.1 as seed ortholog is 100%.

---

Group of orthologs #3781. Best score 1065 bits
Score difference with first non-orthologous sequence - *A. fumigatus*:1065 *A. fischeri*:731
afum|EAL88127.1 100.00% afis|EAW23349.1 100.00%
Bootstrap support for afum|EAL88127.1 as seed ortholog is 100%.
Bootstrap support for afis|EAW23349.1 as seed ortholog is 100%.

---

Group of orthologs #3782. Best score 1064 bits
Score difference with first non-orthologous sequence - *A. fumigatus*:930 *A. fischeri*:977
afum|EAL87618.1 100.00% afis|EAW18255.1 100.00%
Bootstrap support for afum|EAL87618.1 as seed ortholog is 100%.
Bootstrap support for afis|EAW18255.1 as seed ortholog is 100%.

---

Group of orthologs #3783. Best score 1064 bits
Score difference with first non-orthologous sequence - *A. fumigatus*:1064 *A. fischeri*:1064
afum|EAL93880.1 100.00% afis|EAW19149.1 100.00%
Bootstrap support for afum|EAL93880.1 as seed ortholog is 100%.
Bootstrap support for afis|EAW19149.1 as seed ortholog is 100%.

---

Group of orthologs #3784. Best score 1064 bits
Score difference with first non-orthologous sequence - *A. fumigatus*:843 *A. fischeri*:775
afum|EAL92293.2 100.00% afis|EAW21278.1 100.00%
Bootstrap support for afum|EAL92293.2 as seed ortholog is 100%.
Bootstrap support for afis|EAW21278.1 as seed ortholog is 100%.

---

Group of orthologs #3785. Best score 1064 bits
Score difference with first non-orthologous sequence - *A. fumigatus*:1064 *A. fischeri*:1064
afum|EAL92445.1 100.00% afis|EAW21430.1 100.00%
Bootstrap support for afum|EAL92445.1 as seed ortholog is 100%.
Bootstrap support for afis|EAW21430.1 as seed ortholog is 100%.

---

Group of orthologs #3786. Best score 1063 bits
Score difference with first non-orthologous sequence - *A. fumigatus*:1063 *A. fischeri*:959
afum|EAL86814.1 100.00% afis|EAW19713.1 100.00%
Bootstrap support for afum|EAL86814.1 as seed ortholog is 100%.
Bootstrap support for afis|EAW19713.1 as seed ortholog is 100%.

Group of orthologs #3787. Best score 1063 bits
Score difference with first non-orthologous sequence - *A. fumigatus*:1063 *A. fischeri*:1063
afum|EAL92793.1 100.00% afis|EAW21763.1 100.00%
Bootstrap support for afum|EAL92793.1 as seed ortholog is 100%.
Bootstrap support for afis|EAW21763.1 as seed ortholog is 100%.

Group of orthologs #3788. Best score 1062 bits
Score difference with first non-orthologous sequence - *A. fumigatus*:1062 *A. fischeri*:1062
afum|EAL85625.2 100.00% afis|EAW15722.1 100.00%
Bootstrap support for afum|EAL85625.2 as seed ortholog is 100%.
Bootstrap support for afis|EAW15722.1 as seed ortholog is 100%.

Group of orthologs #3789. Best score 1062 bits
Score difference with first non-orthologous sequence - *A. fumigatus*:1062 *A. fischeri*:1062
afum|EAL84963.1 100.00% afis|EAW19952.1 100.00%
Bootstrap support for afum|EAL84963.1 as seed ortholog is 100%.
Bootstrap support for afis|EAW19952.1 as seed ortholog is 100%.

Group of orthologs #3790. Best score 1062 bits
Score difference with first non-orthologous sequence - *A. fumigatus*:1062 *A. fischeri*:1062
afum|EAL89206.1 100.00% afis|EAW16610.1 100.00%
Bootstrap support for afum|EAL89206.1 as seed ortholog is 100%.
Bootstrap support for afis|EAW16610.1 as seed ortholog is 100%.

Group of orthologs #3791. Best score 1062 bits
Score difference with first non-orthologous sequence - *A. fumigatus*:751 *A. fischeri*:1062
afum|EAL85350.1 100.00% afis|EAW20233.1 100.00%
Bootstrap support for afum|EAL85350.1 as seed ortholog is 100%.
Bootstrap support for afis|EAW20233.1 as seed ortholog is 100%.

Group of orthologs #3792. Best score 1062 bits
Score difference with first non-orthologous sequence - *A. fumigatus*:1062 *A. fischeri*:1062
afum|EAL87249.1 100.00% afis|EAW23814.1 100.00%
Bootstrap support for afum|EAL87249.1 as seed ortholog is 100%.
Bootstrap support for afis|EAW23814.1 as seed ortholog is 100%.

Group of orthologs #3793. Best score 1062 bits
Score difference with first non-orthologous sequence - *A. fumigatus*:1062 *A. fischeri*:1062
afum|EAL93626.1 100.00% afis|EAW18894.1 100.00%
Bootstrap support for afum|EAL93626.1 as seed ortholog is 100%.
Bootstrap support for afis|EAW18894.1 as seed ortholog is 100%.

Group of orthologs #3794. Best score 1062 bits
Score difference with first non-orthologous sequence - *A. fumigatus*:1062 *A. fischeri*:1062
afum|EAL90459.1 100.00% afis|EAW22736.1 100.00%
Bootstrap support for afum|EAL90459.1 as seed ortholog is 100%.
Bootstrap support for afis|EAW22736.1 as seed ortholog is 100%.

Group of orthologs #3795. Best score 1062 bits
Score difference with first non-orthologous sequence - *A. fumigatus*:1062 *A. fischeri*:1062
afum|EAL90226.1 100.00% afis|EAW22971.1 100.00%
Bootstrap support for afum|EAL90226.1 as seed ortholog is 100%.
Bootstrap support for afis|EAW22971.1 as seed ortholog is 100%.

Group of orthologs #3796. Best score 1061 bits

Score difference with first non-orthologous sequence - *A. fumigatus*:1061   *A. fischeri*:938
afum|EAL93845.1 100.00% afis|EAW19113.1 100.00%
Bootstrap support for afum|EAL93845.1 as seed ortholog is 100%.
Bootstrap support for afis|EAW19113.1 as seed ortholog is 100%.

---

Group of orthologs #3797. Best score 1060 bits
Score difference with first non-orthologous sequence - *A. fumigatus*:1060   *A. fischeri*:1060
afum|EAL90006.1 100.00% afis|EAW25343.1 100.00%
Bootstrap support for afum|EAL90006.1 as seed ortholog is 100%.
Bootstrap support for afis|EAW25343.1 as seed ortholog is 100%.

---

Group of orthologs #3798. Best score 1059 bits
Score difference with first non-orthologous sequence - *A. fumigatus*:1059   *A. fischeri*:1059
afum|EAL88626.1 100.00% afis|EAW15994.1 100.00%
Bootstrap support for afum|EAL88626.1 as seed ortholog is 100%.
Bootstrap support for afis|EAW15994.1 as seed ortholog is 100%.

---

Group of orthologs #3799. Best score 1059 bits
Score difference with first non-orthologous sequence - *A. fumigatus*:1059   *A. fischeri*:881
afum|EAL88855.1 100.00% afis|EAW16232.1 100.00%
Bootstrap support for afum|EAL88855.1 as seed ortholog is 100%.
Bootstrap support for afis|EAW16232.1 as seed ortholog is 100%.

---

Group of orthologs #3800. Best score 1059 bits
Score difference with first non-orthologous sequence - *A. fumigatus*:1059   *A. fischeri*:1059
afum|EAL87537.1 100.00% afis|EAW18176.1 100.00%
Bootstrap support for afum|EAL87537.1 as seed ortholog is 100%.
Bootstrap support for afis|EAW18176.1 as seed ortholog is 100%.

---

Group of orthologs #3801. Best score 1059 bits
Score difference with first non-orthologous sequence - *A. fumigatus*:1059   *A. fischeri*:1059
afum|EAL91504.1 100.00% afis|EAW17690.1 100.00%
Bootstrap support for afum|EAL91504.1 as seed ortholog is 100%.
Bootstrap support for afis|EAW17690.1 as seed ortholog is 100%.

---

Group of orthologs #3802. Best score 1059 bits
Score difference with first non-orthologous sequence - *A. fumigatus*:1059   *A. fischeri*:1059
afum|EAL93501.2 100.00% afis|EAW18770.1 100.00%
Bootstrap support for afum|EAL93501.2 as seed ortholog is 100%.
Bootstrap support for afis|EAW18770.1 as seed ortholog is 100%.

---

Group of orthologs #3803. Best score 1059 bits
Score difference with first non-orthologous sequence - *A. fumigatus*:983   *A. fischeri*:797
afum|EBA27209.1 100.00% afis|EAW19985.1 100.00%
Bootstrap support for afum|EBA27209.1 as seed ortholog is 100%.
Bootstrap support for afis|EAW19985.1 as seed ortholog is 100%.

---

Group of orthologs #3804. Best score 1059 bits
Score difference with first non-orthologous sequence - *A. fumigatus*:1059   *A. fischeri*:1059
afum|EAL92576.1 100.00% afis|EAW21557.1 100.00%
Bootstrap support for afum|EAL92576.1 as seed ortholog is 100%.
Bootstrap support for afis|EAW21557.1 as seed ortholog is 100%.

---

Group of orthologs #3805. Best score 1059 bits
Score difference with first non-orthologous sequence - *A. fumigatus*:1059   *A. fischeri*:987
afum|EBA27192.1 100.00% afis|EAW20512.1 100.00%

Bootstrap support for afum|EBA27192.1 as seed ortholog is 100%.
Bootstrap support for afis|EAW20512.1 as seed ortholog is 100%.

---

Group of orthologs #3806. Best score 1059 bits
Score difference with first non-orthologous sequence - *A. fumigatus*:1059 *A. fischeri*:1059
afum|EAL89642.1 100.00% afis|EAW24982.1 100.00%
Bootstrap support for afum|EAL89642.1 as seed ortholog is 100%.
Bootstrap support for afis|EAW24982.1 as seed ortholog is 100%.

---

Group of orthologs #3807. Best score 1059 bits
Score difference with first non-orthologous sequence - *A. fumigatus*:1059 *A. fischeri*:968
afum|EAL89926.2 100.00% afis|EAW25263.1 100.00%
Bootstrap support for afum|EAL89926.2 as seed ortholog is 100%.
Bootstrap support for afis|EAW25263.1 as seed ortholog is 100%.

---

Group of orthologs #3808. Best score 1058 bits
Score difference with first non-orthologous sequence - *A. fumigatus*:1058 *A. fischeri*:1058
afum|EAL86682.1 100.00% afis|EAW17265.1 100.00%
Bootstrap support for afum|EAL86682.1 as seed ortholog is 100%.
Bootstrap support for afis|EAW17265.1 as seed ortholog is 100%.

---

Group of orthologs #3809. Best score 1058 bits
Score difference with first non-orthologous sequence - *A. fumigatus*:1058 *A. fischeri*:1058
afum|EAL84671.1 100.00% afis|EAW20962.1 100.00%
Bootstrap support for afum|EAL84671.1 as seed ortholog is 100%.
Bootstrap support for afis|EAW20962.1 as seed ortholog is 100%.

---

Group of orthologs #3810. Best score 1058 bits
Score difference with first non-orthologous sequence - *A. fumigatus*:1058 *A. fischeri*:1058
afum|EAL91671.1 100.00% afis|EAW17863.1 100.00%
Bootstrap support for afum|EAL91671.1 as seed ortholog is 100%.
Bootstrap support for afis|EAW17863.1 as seed ortholog is 100%.

---

Group of orthologs #3811. Best score 1058 bits
Score difference with first non-orthologous sequence - *A. fumigatus*:1058 *A. fischeri*:1058
afum|EAL87409.1 100.00% afis|EAW23979.1 100.00%
Bootstrap support for afum|EAL87409.1 as seed ortholog is 100%.
Bootstrap support for afis|EAW23979.1 as seed ortholog is 100%.

---

Group of orthologs #3812. Best score 1058 bits
Score difference with first non-orthologous sequence - *A. fumigatus*:1058 *A. fischeri*:1058
afum|EAL87846.1 100.00% afis|EAW23616.1 100.00%
Bootstrap support for afum|EAL87846.1 as seed ortholog is 100%.
Bootstrap support for afis|EAW23616.1 as seed ortholog is 100%.

---

Group of orthologs #3813. Best score 1058 bits
Score difference with first non-orthologous sequence - *A. fumigatus*:1058 *A. fischeri*:1058
afum|EAL93891.1 100.00% afis|EAW19161.1 100.00%
Bootstrap support for afum|EAL93891.1 as seed ortholog is 100%.
Bootstrap support for afis|EAW19161.1 as seed ortholog is 100%.

---

Group of orthologs #3814. Best score 1058 bits
Score difference with first non-orthologous sequence - *A. fumigatus*:1058 *A. fischeri*:1058
afum|EAL91033.1 100.00% afis|EAW22157.1 100.00%
Bootstrap support for afum|EAL91033.1 as seed ortholog is 100%.
Bootstrap support for afis|EAW22157.1 as seed ortholog is 100%.

Group of orthologs #3815. Best score 1058 bits
Score difference with first non-orthologous sequence - *A. fumigatus*:1058 *A. fischeri*:1058
afum|EAL90981.2 100.00% afis|EAW22213.1 100.00%
Bootstrap support for afum|EAL90981.2 as seed ortholog is 100%.
Bootstrap support for afis|EAW22213.1 as seed ortholog is 100%.

Group of orthologs #3816. Best score 1058 bits
Score difference with first non-orthologous sequence - *A. fumigatus*:1058 *A. fischeri*:1058
afum|EAL92737.1 100.00% afis|EAW21703.1 100.00%
Bootstrap support for afum|EAL92737.1 as seed ortholog is 100%.
Bootstrap support for afis|EAW21703.1 as seed ortholog is 100%.

Group of orthologs #3817. Best score 1058 bits
Score difference with first non-orthologous sequence - *A. fumigatus*:1058 *A. fischeri*:1011
afum|EAL91945.1 100.00% afis|EAW24090.1 100.00%
Bootstrap support for afum|EAL91945.1 as seed ortholog is 100%.
Bootstrap support for afis|EAW24090.1 as seed ortholog is 100%.

Group of orthologs #3818. Best score 1057 bits
Score difference with first non-orthologous sequence - *A. fumigatus*:1057 *A. fischeri*:1057
afum|EAL85661.1 100.00% afis|EAW15687.1 100.00%
Bootstrap support for afum|EAL85661.1 as seed ortholog is 100%.
Bootstrap support for afis|EAW15687.1 as seed ortholog is 100%.

Group of orthologs #3819. Best score 1057 bits
Score difference with first non-orthologous sequence - *A. fumigatus*:1057 *A. fischeri*:1057
afum|EAL85663.1 100.00% afis|EAW15685.1 100.00%
Bootstrap support for afum|EAL85663.1 as seed ortholog is 100%.
Bootstrap support for afis|EAW15685.1 as seed ortholog is 100%.

Group of orthologs #3820. Best score 1057 bits
Score difference with first non-orthologous sequence - *A. fumigatus*:502 *A. fischeri*:900
afum|EAL87749.2 100.00% afis|EAW20715.1 100.00%
Bootstrap support for afum|EAL87749.2 as seed ortholog is 100%.
Bootstrap support for afis|EAW20715.1 as seed ortholog is 100%.

Group of orthologs #3821. Best score 1057 bits
Score difference with first non-orthologous sequence - *A. fumigatus*:1057 *A. fischeri*:1057
afum|EAL93340.1 100.00% afis|EAW18628.1 100.00%
Bootstrap support for afum|EAL93340.1 as seed ortholog is 100%.
Bootstrap support for afis|EAW18628.1 as seed ortholog is 100%.

Group of orthologs #3822. Best score 1057 bits
Score difference with first non-orthologous sequence - *A. fumigatus*:1057 *A. fischeri*:1057
afum|EAL89602.1 100.00% afis|EAW24945.1 100.00%
Bootstrap support for afum|EAL89602.1 as seed ortholog is 100%.
Bootstrap support for afis|EAW24945.1 as seed ortholog is 100%.

Group of orthologs #3823. Best score 1057 bits
Score difference with first non-orthologous sequence - *A. fumigatus*:1057 *A. fischeri*:1057
afum|EAL90134.1 100.00% afis|EAW25470.1 100.00%
Bootstrap support for afum|EAL90134.1 as seed ortholog is 100%.
Bootstrap support for afis|EAW25470.1 as seed ortholog is 100%.

Group of orthologs #3824. Best score 1056 bits

Score difference with first non-orthologous sequence - *A. fumigatus*:1056 *A. fischeri*:1056
afum|EAL88896.1 100.00% afis|EAW16275.1 100.00%
Bootstrap support for afum|EAL88896.1 as seed ortholog is 100%.
Bootstrap support for afis|EAW16275.1 as seed ortholog is 100%.

---

Group of orthologs #3825. Best score 1056 bits
Score difference with first non-orthologous sequence - *A. fumigatus*:1056 *A. fischeri*:1056
afum|EAL87672.1 100.00% afis|EAW18306.1 100.00%
Bootstrap support for afum|EAL87672.1 as seed ortholog is 100%.
Bootstrap support for afis|EAW18306.1 as seed ortholog is 100%.

---

Group of orthologs #3826. Best score 1056 bits
Score difference with first non-orthologous sequence - *A. fumigatus*:1056 *A. fischeri*:1056
afum|EAL87266.1 100.00% afis|EAW23835.1 100.00%
Bootstrap support for afum|EAL87266.1 as seed ortholog is 100%.
Bootstrap support for afis|EAW23835.1 as seed ortholog is 100%.

---

Group of orthologs #3827. Best score 1056 bits
Score difference with first non-orthologous sequence - *A. fumigatus*:882 *A. fischeri*:793
afum|EAL93348.2 100.00% afis|EAW18635.1 100.00%
Bootstrap support for afum|EAL93348.2 as seed ortholog is 100%.
Bootstrap support for afis|EAW18635.1 as seed ortholog is 100%.

---

Group of orthologs #3828. Best score 1056 bits
Score difference with first non-orthologous sequence - *A. fumigatus*:1056 *A. fischeri*:897
afum|EAL92127.1 100.00% afis|EAW21106.1 100.00%
Bootstrap support for afum|EAL92127.1 as seed ortholog is 100%.
Bootstrap support for afis|EAW21106.1 as seed ortholog is 100%.

---

Group of orthologs #3829. Best score 1056 bits
Score difference with first non-orthologous sequence - *A. fumigatus*:1056 *A. fischeri*:1056
afum|EAL90556.2 100.00% afis|EAW22636.1 100.00%
Bootstrap support for afum|EAL90556.2 as seed ortholog is 100%.
Bootstrap support for afis|EAW22636.1 as seed ortholog is 100%.

---

Group of orthologs #3830. Best score 1055 bits
Score difference with first non-orthologous sequence - *A. fumigatus*:1055 *A. fischeri*:1055
afum|EAL85848.1 100.00% afis|EAW15473.1 100.00%
Bootstrap support for afum|EAL85848.1 as seed ortholog is 100%.
Bootstrap support for afis|EAW15473.1 as seed ortholog is 100%.

---

Group of orthologs #3831. Best score 1055 bits
Score difference with first non-orthologous sequence - *A. fumigatus*:1055 *A. fischeri*:1055
afum|EAL85796.2 100.00% afis|EAW15532.1 100.00%
Bootstrap support for afum|EAL85796.2 as seed ortholog is 100%.
Bootstrap support for afis|EAW15532.1 as seed ortholog is 100%.

---

Group of orthologs #3832. Best score 1055 bits
Score difference with first non-orthologous sequence - *A. fumigatus*:1055 *A. fischeri*:1055
afum|EAL85523.1 100.00% afis|EAW15828.1 100.00%
Bootstrap support for afum|EAL85523.1 as seed ortholog is 100%.
Bootstrap support for afis|EAW15828.1 as seed ortholog is 100%.

---

Group of orthologs #3833. Best score 1055 bits
Score difference with first non-orthologous sequence - *A. fumigatus*:1055 *A. fischeri*:1055
afum|EAL88543.1 100.00% afis|EAW15908.1 100.00%

Bootstrap support for afum|EAL88543.1 as seed ortholog is 100%.
Bootstrap support for afis|EAW15908.1 as seed ortholog is 100%.

---

Group of orthologs #3834. Best score 1055 bits
Score difference with first non-orthologous sequence - *A. fumigatus*:1055 *A. fischeri*:354
afum|EBA27496.1 100.00% afis|EAW18892.1 100.00%
Bootstrap support for afum|EBA27496.1 as seed ortholog is 100%.
Bootstrap support for afis|EAW18892.1 as seed ortholog is 100%.

---

Group of orthologs #3835. Best score 1055 bits
Score difference with first non-orthologous sequence - *A. fumigatus*:1055 *A. fischeri*:1055
afum|EAL90532.1 100.00% afis|EAW22660.1 100.00%
Bootstrap support for afum|EAL90532.1 as seed ortholog is 100%.
Bootstrap support for afis|EAW22660.1 as seed ortholog is 100%.

---

Group of orthologs #3836. Best score 1055 bits
Score difference with first non-orthologous sequence - *A. fumigatus*:1055 *A. fischeri*:1055
afum|EAL90460.1 100.00% afis|EAW22735.1 100.00%
Bootstrap support for afum|EAL90460.1 as seed ortholog is 100%.
Bootstrap support for afis|EAW22735.1 as seed ortholog is 100%.

---

Group of orthologs #3837. Best score 1054 bits
Score difference with first non-orthologous sequence - *A. fumigatus*:1054 *A. fischeri*:1054
afum|EAL91369.1 100.00% afis|EAW17555.1 100.00%
Bootstrap support for afum|EAL91369.1 as seed ortholog is 100%.
Bootstrap support for afis|EAW17555.1 as seed ortholog is 100%.

---

Group of orthologs #3838. Best score 1054 bits
Score difference with first non-orthologous sequence - *A. fumigatus*:1054 *A. fischeri*:1054
afum|EAL86158.1 100.00% afis|EAW24489.1 100.00%
Bootstrap support for afum|EAL86158.1 as seed ortholog is 100%.
Bootstrap support for afis|EAW24489.1 as seed ortholog is 100%.

---

Group of orthologs #3839. Best score 1054 bits
Score difference with first non-orthologous sequence - *A. fumigatus*:1054 *A. fischeri*:1054
afum|EAL93018.1 100.00% afis|EAW22003.1 100.00%
Bootstrap support for afum|EAL93018.1 as seed ortholog is 100%.
Bootstrap support for afis|EAW22003.1 as seed ortholog is 100%.

---

Group of orthologs #3840. Best score 1053 bits
Score difference with first non-orthologous sequence - *A. fumigatus*:1053 *A. fischeri*:1053
afum|EAL85282.1 100.00% afis|EAW20164.1 100.00%
Bootstrap support for afum|EAL85282.1 as seed ortholog is 100%.
Bootstrap support for afis|EAW20164.1 as seed ortholog is 100%.

---

Group of orthologs #3841. Best score 1053 bits
Score difference with first non-orthologous sequence - *A. fumigatus*:1053 *A. fischeri*:1053
afum|EAL87068.1 100.00% afis|EAW19426.1 100.00%
Bootstrap support for afum|EAL87068.1 as seed ortholog is 100%.
Bootstrap support for afis|EAW19426.1 as seed ortholog is 100%.

---

Group of orthologs #3842. Best score 1053 bits
Score difference with first non-orthologous sequence - *A. fumigatus*:1053 *A. fischeri*:1053
afum|EAL85884.1 100.00% afis|EAW24215.1 100.00%
Bootstrap support for afum|EAL85884.1 as seed ortholog is 100%.
Bootstrap support for afis|EAW24215.1 as seed ortholog is 100%.

---

Group of orthologs #3843. Best score 1053 bits
Score difference with first non-orthologous sequence - *A. fumigatus*:1053 *A. fischeri*:1053
afum|EAL88230.1 100.00% afis|EAW23251.1 100.00%
Bootstrap support for afum|EAL88230.1 as seed ortholog is 100%.
Bootstrap support for afis|EAW23251.1 as seed ortholog is 100%.

---

Group of orthologs #3844. Best score 1053 bits
Score difference with first non-orthologous sequence - *A. fumigatus*:1053 *A. fischeri*:1053
afum|EAL93635.1 100.00% afis|EAW18903.1 100.00%
Bootstrap support for afum|EAL93635.1 as seed ortholog is 100%.
Bootstrap support for afis|EAW18903.1 as seed ortholog is 100%.

---

Group of orthologs #3845. Best score 1053 bits
Score difference with first non-orthologous sequence - *A. fumigatus*:1053 *A. fischeri*:1053
afum|EAL92690.1 100.00% afis|EAW21655.1 100.00%
Bootstrap support for afum|EAL92690.1 as seed ortholog is 100%.
Bootstrap support for afis|EAW21655.1 as seed ortholog is 100%.

---

Group of orthologs #3846. Best score 1053 bits
Score difference with first non-orthologous sequence - *A. fumigatus*:1053 *A. fischeri*:1053
afum|EAL90008.2 100.00% afis|EAW25345.1 100.00%
Bootstrap support for afum|EAL90008.2 as seed ortholog is 100%.
Bootstrap support for afis|EAW25345.1 as seed ortholog is 100%.

---

Group of orthologs #3847. Best score 1052 bits
Score difference with first non-orthologous sequence - *A. fumigatus*:1052 *A. fischeri*:1052
afum|EAL86640.1 100.00% afis|EAW17213.1 100.00%
Bootstrap support for afum|EAL86640.1 as seed ortholog is 100%.
Bootstrap support for afis|EAW17213.1 as seed ortholog is 100%.

---

Group of orthologs #3848. Best score 1052 bits
Score difference with first non-orthologous sequence - *A. fumigatus*:1052 *A. fischeri*:1052
afum|EAL91811.1 100.00% afis|EAW18000.1 100.00%
Bootstrap support for afum|EAL91811.1 as seed ortholog is 100%.
Bootstrap support for afis|EAW18000.1 as seed ortholog is 100%.

---

Group of orthologs #3849. Best score 1051 bits
Score difference with first non-orthologous sequence - *A. fumigatus*:1051 *A. fischeri*:1051
afum|EAL85482.1 100.00% afis|EAW20366.1 100.00%
Bootstrap support for afum|EAL85482.1 as seed ortholog is 100%.
Bootstrap support for afis|EAW20366.1 as seed ortholog is 100%.

---

Group of orthologs #3850. Best score 1051 bits
Score difference with first non-orthologous sequence - *A. fumigatus*:1051 *A. fischeri*:1051
afum|EAL89527.1 100.00% afis|EAW24877.1 100.00%
Bootstrap support for afum|EAL89527.1 as seed ortholog is 100%.
Bootstrap support for afis|EAW24877.1 as seed ortholog is 100%.

---

Group of orthologs #3851. Best score 1050 bits
Score difference with first non-orthologous sequence - *A. fumigatus*:1050 *A. fischeri*:1050
afum|EAL88151.1 100.00% afis|EAW23328.1 100.00%
Bootstrap support for afum|EAL88151.1 as seed ortholog is 100%.
Bootstrap support for afis|EAW23328.1 as seed ortholog is 100%.

---

Group of orthologs #3852. Best score 1050 bits

Score difference with first non-orthologous sequence - *A. fumigatus*:1050 *A. fischeri*:1050
afum|EAL93655.1 100.00% afis|EAW18923.1 100.00%
Bootstrap support for afum|EAL93655.1 as seed ortholog is 100%.
Bootstrap support for afis|EAW18923.1 as seed ortholog is 100%.

---

Group of orthologs #3853. Best score 1050 bits
Score difference with first non-orthologous sequence - *A. fumigatus*:1050 *A. fischeri*:926
afum|EAL91011.1 100.00% afis|EAW22178.1 100.00%
Bootstrap support for afum|EAL91011.1 as seed ortholog is 100%.
Bootstrap support for afis|EAW22178.1 as seed ortholog is 100%.

---

Group of orthologs #3854. Best score 1049 bits
Score difference with first non-orthologous sequence - *A. fumigatus*:1049 *A. fischeri*:967
afum|EAL85054.2 100.00% afis|EAW19839.1 100.00%
Bootstrap support for afum|EAL85054.2 as seed ortholog is 100%.
Bootstrap support for afis|EAW19839.1 as seed ortholog is 100%.

---

Group of orthologs #3855. Best score 1049 bits
Score difference with first non-orthologous sequence - *A. fumigatus*:1049 *A. fischeri*:1049
afum|EAL84289.1 100.00% afis|EAW25578.1 100.00%
Bootstrap support for afum|EAL84289.1 as seed ortholog is 100%.
Bootstrap support for afis|EAW25578.1 as seed ortholog is 100%.

---

Group of orthologs #3856. Best score 1049 bits
Score difference with first non-orthologous sequence - *A. fumigatus*:1049 *A. fischeri*:1049
afum|EAL90810.1 100.00% afis|EAW22381.1 100.00%
Bootstrap support for afum|EAL90810.1 as seed ortholog is 100%.
Bootstrap support for afis|EAW22381.1 as seed ortholog is 100%.

---

Group of orthologs #3857. Best score 1048 bits
Score difference with first non-orthologous sequence - *A. fumigatus*:1048 *A. fischeri*:1048
afum|EAL85776.1 100.00% afis|EAW15556.1 100.00%
Bootstrap support for afum|EAL85776.1 as seed ortholog is 100%.
Bootstrap support for afis|EAW15556.1 as seed ortholog is 100%.

---

Group of orthologs #3858. Best score 1048 bits
Score difference with first non-orthologous sequence - *A. fumigatus*:1048 *A. fischeri*:1048
afum|EAL85347.2 100.00% afis|EAW20230.1 100.00%
Bootstrap support for afum|EAL85347.2 as seed ortholog is 100%.
Bootstrap support for afis|EAW20230.1 as seed ortholog is 100%.

---

Group of orthologs #3859. Best score 1048 bits
Score difference with first non-orthologous sequence - *A. fumigatus*:906 *A. fischeri*:422
afum|EAL86183.1 100.00% afis|EAW24517.1 100.00%
Bootstrap support for afum|EAL86183.1 as seed ortholog is 100%.
Bootstrap support for afis|EAW24517.1 as seed ortholog is 100%.

---

Group of orthologs #3860. Best score 1048 bits
Score difference with first non-orthologous sequence - *A. fumigatus*:1048 *A. fischeri*:1048
afum|EAL88411.1 100.00% afis|EAW23068.1 100.00%
Bootstrap support for afum|EAL88411.1 as seed ortholog is 100%.
Bootstrap support for afis|EAW23068.1 as seed ortholog is 100%.

---

Group of orthologs #3861. Best score 1048 bits
Score difference with first non-orthologous sequence - *A. fumigatus*:1048 *A. fischeri*:1048
afum|EAL87946.1 100.00% afis|EAW23518.1 100.00%

Bootstrap support for afum|EAL87946.1 as seed ortholog is 100%.
Bootstrap support for afis|EAW23518.1 as seed ortholog is 100%.

---

Group of orthologs #3862. Best score 1048 bits
Score difference with first non-orthologous sequence - *A. fumigatus*:1048 *A. fischeri*:1048
afum|EAL90994.1 100.00% afis|EAW22197.1 100.00%
Bootstrap support for afum|EAL90994.1 as seed ortholog is 100%.
Bootstrap support for afis|EAW22197.1 as seed ortholog is 100%.

---

Group of orthologs #3863. Best score 1048 bits
Score difference with first non-orthologous sequence - *A. fumigatus*:1048 *A. fischeri*:1048
afum|EAL92645.1 100.00% afis|EAW21611.1 100.00%
Bootstrap support for afum|EAL92645.1 as seed ortholog is 100%.
Bootstrap support for afis|EAW21611.1 as seed ortholog is 100%.

---

Group of orthologs #3864. Best score 1047 bits
Score difference with first non-orthologous sequence - *A. fumigatus*:852 *A. fischeri*:1047
afum|EAL89137.1 100.00% afis|EAW16540.1 100.00%
Bootstrap support for afum|EAL89137.1 as seed ortholog is 100%.
Bootstrap support for afis|EAW16540.1 as seed ortholog is 100%.

---

Group of orthologs #3865. Best score 1047 bits
Score difference with first non-orthologous sequence - *A. fumigatus*:1047 *A. fischeri*:632
afum|EAL89190.1 100.00% afis|EAW16594.1 100.00%
Bootstrap support for afum|EAL89190.1 as seed ortholog is 100%.
Bootstrap support for afis|EAW16594.1 as seed ortholog is 100%.

---

Group of orthologs #3866. Best score 1047 bits
Score difference with first non-orthologous sequence - *A. fumigatus*:1047 *A. fischeri*:861
afum|EAL86810.1 100.00% afis|EAW19717.1 100.00%
Bootstrap support for afum|EAL86810.1 as seed ortholog is 100%.
Bootstrap support for afis|EAW19717.1 as seed ortholog is 100%.

---

Group of orthologs #3867. Best score 1047 bits
Score difference with first non-orthologous sequence - *A. fumigatus*:1047 *A. fischeri*:1047
afum|EAL89793.1 100.00% afis|EAW25122.1 100.00%
Bootstrap support for afum|EAL89793.1 as seed ortholog is 100%.
Bootstrap support for afis|EAW25122.1 as seed ortholog is 100%.

---

Group of orthologs #3868. Best score 1046 bits
Score difference with first non-orthologous sequence - *A. fumigatus*:1046 *A. fischeri*:1046
afum|EAL86528.2 100.00% afis|EAW17009.1 100.00%
Bootstrap support for afum|EAL86528.2 as seed ortholog is 100%.
Bootstrap support for afis|EAW17009.1 as seed ortholog is 100%.

---

Group of orthologs #3869. Best score 1046 bits
Score difference with first non-orthologous sequence - *A. fumigatus*:1046 *A. fischeri*:1046
afum|EAL86965.1 100.00% afis|EAW19544.1 100.00%
Bootstrap support for afum|EAL86965.1 as seed ortholog is 100%.
Bootstrap support for afis|EAW19544.1 as seed ortholog is 100%.

---

Group of orthologs #3870. Best score 1046 bits
Score difference with first non-orthologous sequence - *A. fumigatus*:1046 *A. fischeri*:1046
afum|EAL93115.1 100.00% afis|EAW18434.1 100.00%
Bootstrap support for afum|EAL93115.1 as seed ortholog is 100%.
Bootstrap support for afis|EAW18434.1 as seed ortholog is 100%.

---

Group of orthologs #3871. Best score 1046 bits
Score difference with first non-orthologous sequence - *A. fumigatus*:1046 *A. fischeri*:1046
afum|EAL88083.1 100.00% afis|EAW23388.1 100.00%
Bootstrap support for afum|EAL88083.1 as seed ortholog is 100%.
Bootstrap support for afis|EAW23388.1 as seed ortholog is 100%.

---

Group of orthologs #3872. Best score 1046 bits
Score difference with first non-orthologous sequence - *A. fumigatus*:1046 *A. fischeri*:1046
afum|EAL93328.1 100.00% afis|EAW18618.1 100.00%
Bootstrap support for afum|EAL93328.1 as seed ortholog is 100%.
Bootstrap support for afis|EAW18618.1 as seed ortholog is 100%.

---

Group of orthologs #3873. Best score 1046 bits
Score difference with first non-orthologous sequence - *A. fumigatus*:1046 *A. fischeri*:1046
afum|EAL90547.1 100.00% afis|EAW22645.1 100.00%
Bootstrap support for afum|EAL90547.1 as seed ortholog is 100%.
Bootstrap support for afis|EAW22645.1 as seed ortholog is 100%.

---

Group of orthologs #3874. Best score 1046 bits
Score difference with first non-orthologous sequence - *A. fumigatus*:1046 *A. fischeri*:1046
afum|EAL90288.1 100.00% afis|EAW22909.1 100.00%
Bootstrap support for afum|EAL90288.1 as seed ortholog is 100%.
Bootstrap support for afis|EAW22909.1 as seed ortholog is 100%.

---

Group of orthologs #3875. Best score 1046 bits
Score difference with first non-orthologous sequence - *A. fumigatus*:1046 *A. fischeri*:1046
afum|EAL92956.1 100.00% afis|EAW21942.1 100.00%
Bootstrap support for afum|EAL92956.1 as seed ortholog is 100%.
Bootstrap support for afis|EAW21942.1 as seed ortholog is 100%.

---

Group of orthologs #3876. Best score 1045 bits
Score difference with first non-orthologous sequence - *A. fumigatus*:1045 *A. fischeri*:1045
afum|EAL88709.1 100.00% afis|EAW16077.1 100.00%
Bootstrap support for afum|EAL88709.1 as seed ortholog is 100%.
Bootstrap support for afis|EAW16077.1 as seed ortholog is 100%.

---

Group of orthologs #3877. Best score 1045 bits
Score difference with first non-orthologous sequence - *A. fumigatus*:1045 *A. fischeri*:1045
afum|EAL91447.1 100.00% afis|EAW17632.1 100.00%
Bootstrap support for afum|EAL91447.1 as seed ortholog is 100%.
Bootstrap support for afis|EAW17632.1 as seed ortholog is 100%.

---

Group of orthologs #3878. Best score 1045 bits
Score difference with first non-orthologous sequence - *A. fumigatus*:1045 *A. fischeri*:1045
afum|EAL91823.1 100.00% afis|EAW18011.1 100.00%
Bootstrap support for afum|EAL91823.1 as seed ortholog is 100%.
Bootstrap support for afis|EAW18011.1 as seed ortholog is 100%.

---

Group of orthologs #3879. Best score 1045 bits
Score difference with first non-orthologous sequence - *A. fumigatus*:1045 *A. fischeri*:1045
afum|EAL93141.1 100.00% afis|EAW18456.1 100.00%
Bootstrap support for afum|EAL93141.1 as seed ortholog is 100%.
Bootstrap support for afis|EAW18456.1 as seed ortholog is 100%.

---

Group of orthologs #3880. Best score 1045 bits

Score difference with first non-orthologous sequence - *A. fumigatus*:1045 *A. fischeri*:1045
afum|EAL88069.1 100.00% afis|EAW23401.1 100.00%
Bootstrap support for afum|EAL88069.1 as seed ortholog is 100%.
Bootstrap support for afis|EAW23401.1 as seed ortholog is 100%.

---

Group of orthologs #3881. Best score 1045 bits
Score difference with first non-orthologous sequence - *A. fumigatus*:939 *A. fischeri*:1045
afum|EAL91097.1 100.00% afis|EAW22073.1 100.00%
Bootstrap support for afum|EAL91097.1 as seed ortholog is 100%.
Bootstrap support for afis|EAW22073.1 as seed ortholog is 100%.

---

Group of orthologs #3882. Best score 1045 bits
Score difference with first non-orthologous sequence - *A. fumigatus*:939 *A. fischeri*:813
afum|EAL89332.1 100.00% afis|EAW24691.1 100.00%
Bootstrap support for afum|EAL89332.1 as seed ortholog is 100%.
Bootstrap support for afis|EAW24691.1 as seed ortholog is 100%.

---

Group of orthologs #3883. Best score 1045 bits
Score difference with first non-orthologous sequence - *A. fumigatus*:1045 *A. fischeri*:937
afum|EAL92709.1 100.00% afis|EAW21675.1 100.00%
Bootstrap support for afum|EAL92709.1 as seed ortholog is 100%.
Bootstrap support for afis|EAW21675.1 as seed ortholog is 100%.

---

Group of orthologs #3884. Best score 1044 bits
Score difference with first non-orthologous sequence - *A. fumigatus*:1044 *A. fischeri*:1044
afum|EAL85227.1 100.00% afis|EAW20110.1 100.00%
Bootstrap support for afum|EAL85227.1 as seed ortholog is 100%.
Bootstrap support for afis|EAW20110.1 as seed ortholog is 100%.

---

Group of orthologs #3885. Best score 1044 bits
Score difference with first non-orthologous sequence - *A. fumigatus*:890 *A. fischeri*:813
afum|EAL91844.1 100.00% afis|EAW18033.1 100.00%
Bootstrap support for afum|EAL91844.1 as seed ortholog is 100%.
Bootstrap support for afis|EAW18033.1 as seed ortholog is 100%.

---

Group of orthologs #3886. Best score 1044 bits
Score difference with first non-orthologous sequence - *A. fumigatus*:713 *A. fischeri*:628
afum|EAL91847.1 100.00% afis|EAW18036.1 100.00%
Bootstrap support for afum|EAL91847.1 as seed ortholog is 100%.
Bootstrap support for afis|EAW18036.1 as seed ortholog is 100%.

---

Group of orthologs #3887. Best score 1044 bits
Score difference with first non-orthologous sequence - *A. fumigatus*:1044 *A. fischeri*:1044
afum|EAL85938.1 100.00% afis|EAW24270.1 100.00%
Bootstrap support for afum|EAL85938.1 as seed ortholog is 100%.
Bootstrap support for afis|EAW24270.1 as seed ortholog is 100%.

---

Group of orthologs #3888. Best score 1044 bits
Score difference with first non-orthologous sequence - *A. fumigatus*:1044 *A. fischeri*:1044
afum|EAL88406.1 100.00% afis|EAW23073.1 100.00%
Bootstrap support for afum|EAL88406.1 as seed ortholog is 100%.
Bootstrap support for afis|EAW23073.1 as seed ortholog is 100%.

---

Group of orthologs #3889. Best score 1044 bits
Score difference with first non-orthologous sequence - *A. fumigatus*:1044 *A. fischeri*:1044
afum|EAL88006.1 100.00% afis|EAW23460.1 100.00%

Bootstrap support for afum|EAL88006.1 as seed ortholog is 100%.
Bootstrap support for afis|EAW23460.1 as seed ortholog is 100%.

---

Group of orthologs #3890. Best score 1044 bits
Score difference with first non-orthologous sequence - *A. fumigatus*:1044 *A. fischeri*:1044
afum|EAL90436.2 100.00% afis|EAW22763.1 100.00%
Bootstrap support for afum|EAL90436.2 as seed ortholog is 100%.
Bootstrap support for afis|EAW22763.1 as seed ortholog is 100%.

---

Group of orthologs #3891. Best score 1044 bits
Score difference with first non-orthologous sequence - *A. fumigatus*:1044 *A. fischeri*:1044
afum|EAL92897.1 100.00% afis|EAW21883.1 100.00%
Bootstrap support for afum|EAL92897.1 as seed ortholog is 100%.
Bootstrap support for afis|EAW21883.1 as seed ortholog is 100%.

---

Group of orthologs #3892. Best score 1043 bits
Score difference with first non-orthologous sequence - *A. fumigatus*:908 *A. fischeri*:867
afum|EAL85216.1 100.00% afis|EAW20098.1 100.00%
Bootstrap support for afum|EAL85216.1 as seed ortholog is 100%.
Bootstrap support for afis|EAW20098.1 as seed ortholog is 100%.

---

Group of orthologs #3893. Best score 1043 bits
Score difference with first non-orthologous sequence - *A. fumigatus*:1043 *A. fischeri*:662
afum|EAL91204.1 100.00% afis|EAW17367.1 100.00%
Bootstrap support for afum|EAL91204.1 as seed ortholog is 100%.
Bootstrap support for afis|EAW17367.1 as seed ortholog is 100%.

---

Group of orthologs #3894. Best score 1043 bits
Score difference with first non-orthologous sequence - *A. fumigatus*:1043 *A. fischeri*:1043
afum|EAL91733.1 100.00% afis|EAW17922.1 100.00%
Bootstrap support for afum|EAL91733.1 as seed ortholog is 100%.
Bootstrap support for afis|EAW17922.1 as seed ortholog is 100%.

---

Group of orthologs #3895. Best score 1043 bits
Score difference with first non-orthologous sequence - *A. fumigatus*:1043 *A. fischeri*:1043
afum|EAL91817.1 100.00% afis|EAW18005.1 100.00%
Bootstrap support for afum|EAL91817.1 as seed ortholog is 100%.
Bootstrap support for afis|EAW18005.1 as seed ortholog is 100%.

---

Group of orthologs #3896. Best score 1043 bits
Score difference with first non-orthologous sequence - *A. fumigatus*:722 *A. fischeri*:615
afum|EAL93436.2 100.00% afis|EAW17486.1 100.00%
Bootstrap support for afum|EAL93436.2 as seed ortholog is 100%.
Bootstrap support for afis|EAW17486.1 as seed ortholog is 100%.

---

Group of orthologs #3897. Best score 1043 bits
Score difference with first non-orthologous sequence - *A. fumigatus*:1043 *A. fischeri*:1043
afum|EAL87369.1 100.00% afis|EAW23944.1 100.00%
Bootstrap support for afum|EAL87369.1 as seed ortholog is 100%.
Bootstrap support for afis|EAW23944.1 as seed ortholog is 100%.

---

Group of orthologs #3898. Best score 1043 bits
Score difference with first non-orthologous sequence - *A. fumigatus*:1043 *A. fischeri*:1043
afum|EAL88225.1 100.00% afis|EAW23256.1 100.00%
Bootstrap support for afum|EAL88225.1 as seed ortholog is 100%.
Bootstrap support for afis|EAW23256.1 as seed ortholog is 100%.

Group of orthologs #3899. Best score 1043 bits
Score difference with first non-orthologous sequence - *A. fumigatus*:1043 *A. fischeri*:1043
afum|EAL90678.1 100.00% afis|EAW22515.1 100.00%
Bootstrap support for afum|EAL90678.1 as seed ortholog is 100%.
Bootstrap support for afis|EAW22515.1 as seed ortholog is 100%.

---

Group of orthologs #3900. Best score 1043 bits
Score difference with first non-orthologous sequence - *A. fumigatus*:1043 *A. fischeri*:1043
afum|EAL90666.2 100.00% afis|EAW22528.1 100.00%
Bootstrap support for afum|EAL90666.2 as seed ortholog is 100%.
Bootstrap support for afis|EAW22528.1 as seed ortholog is 100%.

---

Group of orthologs #3901. Best score 1043 bits
Score difference with first non-orthologous sequence - *A. fumigatus*:1043 *A. fischeri*:1043
afum|EAL90651.1 100.00% afis|EAW22544.1 100.00%
Bootstrap support for afum|EAL90651.1 as seed ortholog is 100%.
Bootstrap support for afis|EAW22544.1 as seed ortholog is 100%.

---

Group of orthologs #3902. Best score 1043 bits
Score difference with first non-orthologous sequence - *A. fumigatus*:1043 *A. fischeri*:1043
afum|EAL92597.1 100.00% afis|EAW21579.1 100.00%
Bootstrap support for afum|EAL92597.1 as seed ortholog is 100%.
Bootstrap support for afis|EAW21579.1 as seed ortholog is 100%.

---

Group of orthologs #3903. Best score 1043 bits
Score difference with first non-orthologous sequence - *A. fumigatus*:1043 *A. fischeri*:866
afum|EAL89421.1 100.00% afis|EAW24782.1 100.00%
Bootstrap support for afum|EAL89421.1 as seed ortholog is 100%.
Bootstrap support for afis|EAW24782.1 as seed ortholog is 100%.

---

Group of orthologs #3904. Best score 1043 bits
Score difference with first non-orthologous sequence - *A. fumigatus*:1043 *A. fischeri*:279
afum|EAL89439.1 100.00% afis|EAW24797.1 100.00%
Bootstrap support for afum|EAL89439.1 as seed ortholog is 100%.
Bootstrap support for afis|EAW24797.1 as seed ortholog is 100%.

---

Group of orthologs #3905. Best score 1042 bits
Score difference with first non-orthologous sequence - *A. fumigatus*:1042 *A. fischeri*:750
afum|EAL84906.1 100.00% afis|EAW20012.1 100.00%
Bootstrap support for afum|EAL84906.1 as seed ortholog is 100%.
Bootstrap support for afis|EAW20012.1 as seed ortholog is 100%.

---

Group of orthologs #3906. Best score 1042 bits
Score difference with first non-orthologous sequence - *A. fumigatus*:1042 *A. fischeri*:1042
afum|EAL93033.1 100.00% afis|EAW18362.1 100.00%
Bootstrap support for afum|EAL93033.1 as seed ortholog is 100%.
Bootstrap support for afis|EAW18362.1 as seed ortholog is 100%.

---

Group of orthologs #3907. Best score 1042 bits
Score difference with first non-orthologous sequence - *A. fumigatus*:843 *A. fischeri*:821
afum|EAL93572.1 100.00% afis|EAW18838.1 100.00%
Bootstrap support for afum|EAL93572.1 as seed ortholog is 100%.
Bootstrap support for afis|EAW18838.1 as seed ortholog is 100%.

---

Group of orthologs #3908. Best score 1042 bits

Score difference with first non-orthologous sequence - *A. fumigatus*:1042 *A. fischeri*:1042
afum|EAL90551.1 100.00% afis|EAW22641.1 100.00%
Bootstrap support for afum|EAL90551.1 as seed ortholog is 100%.
Bootstrap support for afis|EAW22641.1 as seed ortholog is 100%.

---

Group of orthologs #3909. Best score 1042 bits
Score difference with first non-orthologous sequence - *A. fumigatus*:1042 *A. fischeri*:1042
afum|EAL92796.1 100.00% afis|EAW21766.1 100.00%
Bootstrap support for afum|EAL92796.1 as seed ortholog is 100%.
Bootstrap support for afis|EAW21766.1 as seed ortholog is 100%.

---

Group of orthologs #3910. Best score 1041 bits
Score difference with first non-orthologous sequence - *A. fumigatus*:1041 *A. fischeri*:1041
afum|EAL87478.1 100.00% afis|EAW18120.1 100.00%
Bootstrap support for afum|EAL87478.1 as seed ortholog is 100%.
Bootstrap support for afis|EAW18120.1 as seed ortholog is 100%.

---

Group of orthologs #3911. Best score 1041 bits
Score difference with first non-orthologous sequence - *A. fumigatus*:1041 *A. fischeri*:1041
afum|EAL91007.1 100.00% afis|EAW22182.1 100.00%
Bootstrap support for afum|EAL91007.1 as seed ortholog is 100%.
Bootstrap support for afis|EAW22182.1 as seed ortholog is 100%.

---

Group of orthologs #3912. Best score 1040 bits
Score difference with first non-orthologous sequence - *A. fumigatus*:1040 *A. fischeri*:1040
afum|EAL93276.1 100.00% afis|EAW18573.1 100.00%
Bootstrap support for afum|EAL93276.1 as seed ortholog is 100%.
Bootstrap support for afis|EAW18573.1 as seed ortholog is 100%.

---

Group of orthologs #3913. Best score 1040 bits
Score difference with first non-orthologous sequence - *A. fumigatus*:1040 *A. fischeri*:1040
afum|EAL93472.1 100.00% afis|EAW18746.1 100.00%
Bootstrap support for afum|EAL93472.1 as seed ortholog is 100%.
Bootstrap support for afis|EAW18746.1 as seed ortholog is 100%.

---

Group of orthologs #3914. Best score 1040 bits
Score difference with first non-orthologous sequence - *A. fumigatus*:1040 *A. fischeri*:766
afum|EAL92569.1 100.00% afis|EAW21549.1 100.00%
Bootstrap support for afum|EAL92569.1 as seed ortholog is 100%.
Bootstrap support for afis|EAW21549.1 as seed ortholog is 100%.

---

Group of orthologs #3915. Best score 1040 bits
Score difference with first non-orthologous sequence - *A. fumigatus*:1040 *A. fischeri*:1040
afum|EAL92848.1 100.00% afis|EAW21833.1 100.00%
Bootstrap support for afum|EAL92848.1 as seed ortholog is 100%.
Bootstrap support for afis|EAW21833.1 as seed ortholog is 100%.

---

Group of orthologs #3916. Best score 1040 bits
Score difference with first non-orthologous sequence - *A. fumigatus*:1040 *A. fischeri*:1040
afum|EAL92945.2 100.00% afis|EAW21932.1 100.00%
Bootstrap support for afum|EAL92945.2 as seed ortholog is 100%.
Bootstrap support for afis|EAW21932.1 as seed ortholog is 100%.

---

Group of orthologs #3917. Best score 1039 bits
Score difference with first non-orthologous sequence - *A. fumigatus*:1039 *A. fischeri*:1039
afum|EAL87517.1 100.00% afis|EAW18158.1 100.00%

Bootstrap support for afum|EAL87517.1 as seed ortholog is 100%.
Bootstrap support for afis|EAW18158.1 as seed ortholog is 100%.

---

Group of orthologs #3918. Best score 1039 bits
Score difference with first non-orthologous sequence - *A. fumigatus*:1039 *A. fischeri*:1039
afum|EAL93246.1 100.00% afis|EAW18546.1 100.00%
Bootstrap support for afum|EAL93246.1 as seed ortholog is 100%.
Bootstrap support for afis|EAW18546.1 as seed ortholog is 100%.

---

Group of orthologs #3919. Best score 1039 bits
Score difference with first non-orthologous sequence - *A. fumigatus*:1039 *A. fischeri*:1039
afum|EAL90780.1 100.00% afis|EAW22412.1 100.00%
Bootstrap support for afum|EAL90780.1 as seed ortholog is 100%.
Bootstrap support for afis|EAW22412.1 as seed ortholog is 100%.

---

Group of orthologs #3920. Best score 1039 bits
Score difference with first non-orthologous sequence - *A. fumigatus*:1039 *A. fischeri*:1039
afum|EAL90274.1 100.00% afis|EAW22923.1 100.00%
Bootstrap support for afum|EAL90274.1 as seed ortholog is 100%.
Bootstrap support for afis|EAW22923.1 as seed ortholog is 100%.

---

Group of orthologs #3921. Best score 1038 bits
Score difference with first non-orthologous sequence - *A. fumigatus*:1038 *A. fischeri*:1038
afum|EAL84508.1 100.00% afis|EAW20553.1 100.00%
Bootstrap support for afum|EAL84508.1 as seed ortholog is 100%.
Bootstrap support for afis|EAW20553.1 as seed ortholog is 100%.

---

Group of orthologs #3922. Best score 1038 bits
Score difference with first non-orthologous sequence - *A. fumigatus*:1038 *A. fischeri*:1038
afum|EAL85225.1 100.00% afis|EAW20108.1 100.00%
Bootstrap support for afum|EAL85225.1 as seed ortholog is 100%.
Bootstrap support for afis|EAW20108.1 as seed ortholog is 100%.

---

Group of orthologs #3923. Best score 1038 bits
Score difference with first non-orthologous sequence - *A. fumigatus*:1038 *A. fischeri*:1038
afum|EAL88334.1 100.00% afis|EAW23147.1 100.00%
Bootstrap support for afum|EAL88334.1 as seed ortholog is 100%.
Bootstrap support for afis|EAW23147.1 as seed ortholog is 100%.

---

Group of orthologs #3924. Best score 1038 bits
Score difference with first non-orthologous sequence - *A. fumigatus*:1038 *A. fischeri*:1038
afum|EBA27226.1 100.00% afis|EAW20136.1 100.00%
Bootstrap support for afum|EBA27226.1 as seed ortholog is 100%.
Bootstrap support for afis|EAW20136.1 as seed ortholog is 100%.

---

Group of orthologs #3925. Best score 1037 bits
Score difference with first non-orthologous sequence - *A. fumigatus*:1037 *A. fischeri*:1037
afum|EAL84600.1 100.00% afis|EAW20453.1 100.00%
Bootstrap support for afum|EAL84600.1 as seed ortholog is 100%.
Bootstrap support for afis|EAW20453.1 as seed ortholog is 100%.

---

Group of orthologs #3926. Best score 1037 bits
Score difference with first non-orthologous sequence - *A. fumigatus*:852 *A. fischeri*:876
afum|EAL87292.1 100.00% afis|EAW23864.1 100.00%
Bootstrap support for afum|EAL87292.1 as seed ortholog is 100%.
Bootstrap support for afis|EAW23864.1 as seed ortholog is 100%.

---

Group of orthologs #3927. Best score 1037 bits
Score difference with first non-orthologous sequence - *A. fumigatus*:1037 *A. fischeri*:1037
afum|EAL93382.1 100.00% afis|EAW18664.1 100.00%
Bootstrap support for afum|EAL93382.1 as seed ortholog is 100%.
Bootstrap support for afis|EAW18664.1 as seed ortholog is 100%.

---

Group of orthologs #3928. Best score 1037 bits
Score difference with first non-orthologous sequence - *A. fumigatus*:731 *A. fischeri*:732
afum|EAL93544.1 100.00% afis|EAW18808.1 100.00%
Bootstrap support for afum|EAL93544.1 as seed ortholog is 100%.
Bootstrap support for afis|EAW18808.1 as seed ortholog is 100%.

---

Group of orthologs #3929. Best score 1037 bits
Score difference with first non-orthologous sequence - *A. fumigatus*:1037 *A. fischeri*:1037
afum|EAL90483.1 100.00% afis|EAW22709.1 100.00%
Bootstrap support for afum|EAL90483.1 as seed ortholog is 100%.
Bootstrap support for afis|EAW22709.1 as seed ortholog is 100%.

---

Group of orthologs #3930. Best score 1037 bits
Score difference with first non-orthologous sequence - *A. fumigatus*:1037 *A. fischeri*:1037
afum|EAL90417.1 100.00% afis|EAW22782.1 100.00%
Bootstrap support for afum|EAL90417.1 as seed ortholog is 100%.
Bootstrap support for afis|EAW22782.1 as seed ortholog is 100%.

---

Group of orthologs #3931. Best score 1037 bits
Score difference with first non-orthologous sequence - *A. fumigatus*:1037 *A. fischeri*:1037
afum|EAL89997.1 100.00% afis|EAW25334.1 100.00%
Bootstrap support for afum|EAL89997.1 as seed ortholog is 100%.
Bootstrap support for afis|EAW25334.1 as seed ortholog is 100%.

---

Group of orthologs #3932. Best score 1036 bits
Score difference with first non-orthologous sequence - *A. fumigatus*:369 *A. fischeri*:421
afum|EAL85027.1 100.00% afis|EAW19870.1 100.00%
Bootstrap support for afum|EAL85027.1 as seed ortholog is 100%.
Bootstrap support for afis|EAW19870.1 as seed ortholog is 100%.

---

Group of orthologs #3933. Best score 1036 bits
Score difference with first non-orthologous sequence - *A. fumigatus*:1036 *A. fischeri*:1036
afum|EAL91508.1 100.00% afis|EAW17694.1 100.00%
Bootstrap support for afum|EAL91508.1 as seed ortholog is 100%.
Bootstrap support for afis|EAW17694.1 as seed ortholog is 100%.

---

Group of orthologs #3934. Best score 1036 bits
Score difference with first non-orthologous sequence - *A. fumigatus*:970 *A. fischeri*:1036
afum|EAL87176.1 100.00% afis|EAW23739.1 100.00%
Bootstrap support for afum|EAL87176.1 as seed ortholog is 100%.
Bootstrap support for afis|EAW23739.1 as seed ortholog is 100%.

---

Group of orthologs #3935. Best score 1036 bits
Score difference with first non-orthologous sequence - *A. fumigatus*:1036 *A. fischeri*:1036
afum|EAL88336.1 100.00% afis|EAW23145.1 100.00%
Bootstrap support for afum|EAL88336.1 as seed ortholog is 100%.
Bootstrap support for afis|EAW23145.1 as seed ortholog is 100%.

---

Group of orthologs #3936. Best score 1036 bits

Score difference with first non-orthologous sequence - *A. fumigatus*:1036   *A. fischeri*:1036
afum|EAL90157.1 100.00% afis|EAW25492.1 100.00%
Bootstrap support for afum|EAL90157.1 as seed ortholog is 100%.
Bootstrap support for afis|EAW25492.1 as seed ortholog is 100%.

---

Group of orthologs #3937. Best score 1035 bits
Score difference with first non-orthologous sequence - *A. fumigatus*:1035   *A. fischeri*:1035
afum|EAL93962.1 100.00% afis|EAW19233.1 100.00%
Bootstrap support for afum|EAL93962.1 as seed ortholog is 100%.
Bootstrap support for afis|EAW19233.1 as seed ortholog is 100%.

---

Group of orthologs #3938. Best score 1035 bits
Score difference with first non-orthologous sequence - *A. fumigatus*:1035   *A. fischeri*:1035
afum|EAL90402.1 100.00% afis|EAW22797.1 100.00%
Bootstrap support for afum|EAL90402.1 as seed ortholog is 100%.
Bootstrap support for afis|EAW22797.1 as seed ortholog is 100%.

---

Group of orthologs #3939. Best score 1035 bits
Score difference with first non-orthologous sequence - *A. fumigatus*:1035   *A. fischeri*:1035
afum|EAL89649.1 100.00% afis|EAW24989.1 100.00%
Bootstrap support for afum|EAL89649.1 as seed ortholog is 100%.
Bootstrap support for afis|EAW24989.1 as seed ortholog is 100%.

---

Group of orthologs #3940. Best score 1035 bits
Score difference with first non-orthologous sequence - *A. fumigatus*:1035   *A. fischeri*:772
afum|EAL89747.1 100.00% afis|EAW25078.1 100.00%
Bootstrap support for afum|EAL89747.1 as seed ortholog is 100%.
Bootstrap support for afis|EAW25078.1 as seed ortholog is 100%.

---

Group of orthologs #3941. Best score 1034 bits
Score difference with first non-orthologous sequence - *A. fumigatus*:1034   *A. fischeri*:905
afum|EAL86671.2 100.00% afis|EAW17252.1 100.00%
Bootstrap support for afum|EAL86671.2 as seed ortholog is 100%.
Bootstrap support for afis|EAW17252.1 as seed ortholog is 100%.

---

Group of orthologs #3942. Best score 1034 bits
Score difference with first non-orthologous sequence - *A. fumigatus*:1034   *A. fischeri*:1034
afum|EAL88532.1 100.00% afis|EAW15897.1 100.00%
Bootstrap support for afum|EAL88532.1 as seed ortholog is 100%.
Bootstrap support for afis|EAW15897.1 as seed ortholog is 100%.

---

Group of orthologs #3943. Best score 1034 bits
Score difference with first non-orthologous sequence - *A. fumigatus*:875   *A. fischeri*:851
afum|EAL84524.1 100.00% afis|EAW20534.1 100.00%
Bootstrap support for afum|EAL84524.1 as seed ortholog is 100%.
Bootstrap support for afis|EAW20534.1 as seed ortholog is 100%.

---

Group of orthologs #3944. Best score 1034 bits
Score difference with first non-orthologous sequence - *A. fumigatus*:1034   *A. fischeri*:1034
afum|EAL91805.1 100.00% afis|EAW17995.1 100.00%
Bootstrap support for afum|EAL91805.1 as seed ortholog is 100%.
Bootstrap support for afis|EAW17995.1 as seed ortholog is 100%.

---

Group of orthologs #3945. Best score 1034 bits
Score difference with first non-orthologous sequence - *A. fumigatus*:731   *A. fischeri*:692
afum|EAL93768.1 100.00% afis|EAW19039.1 100.00%

Bootstrap support for afum|EAL93768.1 as seed ortholog is 100%.
Bootstrap support for afis|EAW19039.1 as seed ortholog is 100%.

---

Group of orthologs #3946. Best score 1034 bits
Score difference with first non-orthologous sequence - *A. fumigatus*:1034 *A. fischeri*:1034
afum|EAL92351.1 100.00% afis|EAW21333.1 100.00%
Bootstrap support for afum|EAL92351.1 as seed ortholog is 100%.
Bootstrap support for afis|EAW21333.1 as seed ortholog is 100%.

---

Group of orthologs #3947. Best score 1034 bits
Score difference with first non-orthologous sequence - *A. fumigatus*:1034 *A. fischeri*:1034
afum|EAL92647.1 100.00% afis|EAW21613.1 100.00%
Bootstrap support for afum|EAL92647.1 as seed ortholog is 100%.
Bootstrap support for afis|EAW21613.1 as seed ortholog is 100%.

---

Group of orthologs #3948. Best score 1034 bits
Score difference with first non-orthologous sequence - *A. fumigatus*:1034 *A. fischeri*:1034
afum|EAL90095.1 100.00% afis|EAW25437.1 100.00%
Bootstrap support for afum|EAL90095.1 as seed ortholog is 100%.
Bootstrap support for afis|EAW25437.1 as seed ortholog is 100%.

---

Group of orthologs #3949. Best score 1033 bits
Score difference with first non-orthologous sequence - *A. fumigatus*:1033 *A. fischeri*:1033
afum|EAL84917.1 100.00% afis|EAW20002.1 100.00%
Bootstrap support for afum|EAL84917.1 as seed ortholog is 100%.
Bootstrap support for afis|EAW20002.1 as seed ortholog is 100%.

---

Group of orthologs #3950. Best score 1033 bits
Score difference with first non-orthologous sequence - *A. fumigatus*:1033 *A. fischeri*:725
afum|EAL85478.2 100.00% afis|EAW20362.1 100.00%
Bootstrap support for afum|EAL85478.2 as seed ortholog is 100%.
Bootstrap support for afis|EAW20362.1 as seed ortholog is 100%.

---

Group of orthologs #3951. Best score 1033 bits
Score difference with first non-orthologous sequence - *A. fumigatus*:918 *A. fischeri*:1033
afum|EAL86159.1 100.00% afis|EAW24490.1 100.00%
Bootstrap support for afum|EAL86159.1 as seed ortholog is 100%.
Bootstrap support for afis|EAW24490.1 as seed ortholog is 100%.

---

Group of orthologs #3952. Best score 1033 bits
Score difference with first non-orthologous sequence - *A. fumigatus*:1033 *A. fischeri*:1033
afum|EAL90816.1 100.00% afis|EAW22376.1 100.00%
Bootstrap support for afum|EAL90816.1 as seed ortholog is 100%.
Bootstrap support for afis|EAW22376.1 as seed ortholog is 100%.

---

Group of orthologs #3953. Best score 1033 bits
Score difference with first non-orthologous sequence - *A. fumigatus*:1033 *A. fischeri*:1033
afum|EAL89941.1 100.00% afis|EAW25278.1 100.00%
Bootstrap support for afum|EAL89941.1 as seed ortholog is 100%.
Bootstrap support for afis|EAW25278.1 as seed ortholog is 100%.

---

Group of orthologs #3954. Best score 1032 bits
Score difference with first non-orthologous sequence - *A. fumigatus*:1032 *A. fischeri*:1032
afum|EAL86687.1 100.00% afis|EAW17270.1 100.00%
Bootstrap support for afum|EAL86687.1 as seed ortholog is 100%.
Bootstrap support for afis|EAW17270.1 as seed ortholog is 100%.

---

Group of orthologs #3955. Best score 1032 bits
Score difference with first non-orthologous sequence - *A. fumigatus*:1032 *A. fischeri*:959
afum|EAL84951.2 100.00% afis|EAW19969.1 100.00%
Bootstrap support for afum|EAL84951.2 as seed ortholog is 100%.
Bootstrap support for afis|EAW19969.1 as seed ortholog is 100%.

---

Group of orthologs #3956. Best score 1032 bits
Score difference with first non-orthologous sequence - *A. fumigatus*:1032 *A. fischeri*:1032
afum|EAL93221.1 100.00% afis|EAW18526.1 100.00%
Bootstrap support for afum|EAL93221.1 as seed ortholog is 100%.
Bootstrap support for afis|EAW18526.1 as seed ortholog is 100%.

---

Group of orthologs #3957. Best score 1032 bits
Score difference with first non-orthologous sequence - *A. fumigatus*:1032 *A. fischeri*:1032
afum|EAL90420.1 100.00% afis|EAW22778.1 100.00%
Bootstrap support for afum|EAL90420.1 as seed ortholog is 100%.
Bootstrap support for afis|EAW22778.1 as seed ortholog is 100%.

---

Group of orthologs #3958. Best score 1031 bits
Score difference with first non-orthologous sequence - *A. fumigatus*:1031 *A. fischeri*:1031
afum|EAL84998.1 100.00% afis|EAW19908.1 100.00%
Bootstrap support for afum|EAL84998.1 as seed ortholog is 100%.
Bootstrap support for afis|EAW19908.1 as seed ortholog is 100%.

---

Group of orthologs #3959. Best score 1031 bits
Score difference with first non-orthologous sequence - *A. fumigatus*:1031 *A. fischeri*:747
afum|EAL84721.1 100.00% afis|EAW20913.1 100.00%
Bootstrap support for afum|EAL84721.1 as seed ortholog is 100%.
Bootstrap support for afis|EAW20913.1 as seed ortholog is 100%.

---

Group of orthologs #3960. Best score 1031 bits
Score difference with first non-orthologous sequence - *A. fumigatus*:1031 *A. fischeri*:1031
afum|EAL86091.1 100.00% afis|EAW24423.1 100.00%
Bootstrap support for afum|EAL86091.1 as seed ortholog is 100%.
Bootstrap support for afis|EAW24423.1 as seed ortholog is 100%.

---

Group of orthologs #3961. Best score 1031 bits
Score difference with first non-orthologous sequence - *A. fumigatus*:1031 *A. fischeri*:1031
afum|EAL93927.2 100.00% afis|EAW19195.1 100.00%
Bootstrap support for afum|EAL93927.2 as seed ortholog is 100%.
Bootstrap support for afis|EAW19195.1 as seed ortholog is 100%.

---

Group of orthologs #3962. Best score 1031 bits
Score difference with first non-orthologous sequence - *A. fumigatus*:1031 *A. fischeri*:1031
afum|EAL90911.1 100.00% afis|EAW22269.1 100.00%
Bootstrap support for afum|EAL90911.1 as seed ortholog is 100%.
Bootstrap support for afis|EAW22269.1 as seed ortholog is 100%.

---

Group of orthologs #3963. Best score 1030 bits
Score difference with first non-orthologous sequence - *A. fumigatus*:1030 *A. fischeri*:1030
afum|EAL86821.1 100.00% afis|EAW19705.1 100.00%
Bootstrap support for afum|EAL86821.1 as seed ortholog is 100%.
Bootstrap support for afis|EAW19705.1 as seed ortholog is 100%.

---

Group of orthologs #3964. Best score 1030 bits

Score difference with first non-orthologous sequence - *A. fumigatus*:1030 *A. fischeri*:1030
afum|EAL91575.1 100.00% afis|EAW17765.1 100.00%
Bootstrap support for afum|EAL91575.1 as seed ortholog is 100%.
Bootstrap support for afis|EAW17765.1 as seed ortholog is 100%.

---

Group of orthologs #3965. Best score 1030 bits
Score difference with first non-orthologous sequence - *A. fumigatus*:1030 *A. fischeri*:1030
afum|EAL93321.1 100.00% afis|EAW18611.1 100.00%
Bootstrap support for afum|EAL93321.1 as seed ortholog is 100%.
Bootstrap support for afis|EAW18611.1 as seed ortholog is 100%.

---

Group of orthologs #3966. Best score 1029 bits
Score difference with first non-orthologous sequence - *A. fumigatus*:1029 *A. fischeri*:1029
afum|EAL86743.1 100.00% afis|EAW16809.1 100.00%
Bootstrap support for afum|EAL86743.1 as seed ortholog is 100%.
Bootstrap support for afis|EAW16809.1 as seed ortholog is 100%.

---

Group of orthologs #3967. Best score 1029 bits
Score difference with first non-orthologous sequence - *A. fumigatus*:1029 *A. fischeri*:1029
afum|EAL87669.1 100.00% afis|EAW18303.1 100.00%
Bootstrap support for afum|EAL87669.1 as seed ortholog is 100%.
Bootstrap support for afis|EAW18303.1 as seed ortholog is 100%.

---

Group of orthologs #3968. Best score 1029 bits
Score difference with first non-orthologous sequence - *A. fumigatus*:1029 *A. fischeri*:1029
afum|EAL91216.1 100.00% afis|EAW17381.1 100.00%
Bootstrap support for afum|EAL91216.1 as seed ortholog is 100%.
Bootstrap support for afis|EAW17381.1 as seed ortholog is 100%.

---

Group of orthologs #3969. Best score 1029 bits
Score difference with first non-orthologous sequence - *A. fumigatus*:1029 *A. fischeri*:1029
afum|EAL91893.1 100.00% afis|EAW18081.1 100.00%
Bootstrap support for afum|EAL91893.1 as seed ortholog is 100%.
Bootstrap support for afis|EAW18081.1 as seed ortholog is 100%.

---

Group of orthologs #3970. Best score 1029 bits
Score difference with first non-orthologous sequence - *A. fumigatus*:1029 *A. fischeri*:968
afum|EAL92785.1 100.00% afis|EAW21755.1 100.00%
Bootstrap support for afum|EAL92785.1 as seed ortholog is 100%.
Bootstrap support for afis|EAW21755.1 as seed ortholog is 100%.

---

Group of orthologs #3971. Best score 1029 bits
Score difference with first non-orthologous sequence - *A. fumigatus*:1029 *A. fischeri*:1029
afum|EAL92026.1 100.00% afis|EAW24168.1 100.00%
Bootstrap support for afum|EAL92026.1 as seed ortholog is 100%.
Bootstrap support for afis|EAW24168.1 as seed ortholog is 100%.

---

Group of orthologs #3972. Best score 1028 bits
Score difference with first non-orthologous sequence - *A. fumigatus*:1028 *A. fischeri*:1028
afum|EAL86474.1 100.00% afis|EAW16947.1 100.00%
Bootstrap support for afum|EAL86474.1 as seed ortholog is 100%.
Bootstrap support for afis|EAW16947.1 as seed ortholog is 100%.

---

Group of orthologs #3973. Best score 1028 bits
Score difference with first non-orthologous sequence - *A. fumigatus*:1028 *A. fischeri*:1028
afum|EAL88731.1 100.00% afis|EAW16099.1 100.00%

Bootstrap support for afum|EAL88731.1 as seed ortholog is 100%.
Bootstrap support for afis|EAW16099.1 as seed ortholog is 100%.

---

Group of orthologs #3974. Best score 1028 bits
Score difference with first non-orthologous sequence - *A. fumigatus*:816 *A. fischeri*:933
afum|EAL85500.2 100.00% afis|EAW20401.1 100.00%
Bootstrap support for afum|EAL85500.2 as seed ortholog is 100%.
Bootstrap support for afis|EAW20401.1 as seed ortholog is 100%.

---

Group of orthologs #3975. Best score 1028 bits
Score difference with first non-orthologous sequence - *A. fumigatus*:1028 *A. fischeri*:1028
afum|EAL93126.1 100.00% afis|EAW18443.1 100.00%
Bootstrap support for afum|EAL93126.1 as seed ortholog is 100%.
Bootstrap support for afis|EAW18443.1 as seed ortholog is 100%.

---

Group of orthologs #3976. Best score 1028 bits
Score difference with first non-orthologous sequence - *A. fumigatus*:1028 *A. fischeri*:1028
afum|EAL87876.1 100.00% afis|EAW23590.1 100.00%
Bootstrap support for afum|EAL87876.1 as seed ortholog is 100%.
Bootstrap support for afis|EAW23590.1 as seed ortholog is 100%.

---

Group of orthologs #3977. Best score 1028 bits
Score difference with first non-orthologous sequence - *A. fumigatus*:1028 *A. fischeri*:1028
afum|EAL93566.1 100.00% afis|EAW18832.1 100.00%
Bootstrap support for afum|EAL93566.1 as seed ortholog is 100%.
Bootstrap support for afis|EAW18832.1 as seed ortholog is 100%.

---

Group of orthologs #3978. Best score 1028 bits
Score difference with first non-orthologous sequence - *A. fumigatus*:1028 *A. fischeri*:1028
afum|EAL92246.1 100.00% afis|EAW21234.1 100.00%
Bootstrap support for afum|EAL92246.1 as seed ortholog is 100%.
Bootstrap support for afis|EAW21234.1 as seed ortholog is 100%.

---

Group of orthologs #3979. Best score 1027 bits
Score difference with first non-orthologous sequence - *A. fumigatus*:901 *A. fischeri*:1027
afum|EAL86830.1 100.00% afis|EAW19697.1 100.00%
Bootstrap support for afum|EAL86830.1 as seed ortholog is 100%.
Bootstrap support for afis|EAW19697.1 as seed ortholog is 100%.

---

Group of orthologs #3980. Best score 1027 bits
Score difference with first non-orthologous sequence - *A. fumigatus*:1027 *A. fischeri*:940
afum|EAL91529.1 100.00% afis|EAW17714.1 100.00%
Bootstrap support for afum|EAL91529.1 as seed ortholog is 100%.
Bootstrap support for afis|EAW17714.1 as seed ortholog is 100%.

---

Group of orthologs #3981. Best score 1027 bits
Score difference with first non-orthologous sequence - *A. fumigatus*:1027 *A. fischeri*:1027
afum|EAL89595.1 100.00% afis|EAW24938.1 100.00%
Bootstrap support for afum|EAL89595.1 as seed ortholog is 100%.
Bootstrap support for afis|EAW24938.1 as seed ortholog is 100%.

---

Group of orthologs #3982. Best score 1026 bits
Score difference with first non-orthologous sequence - *A. fumigatus*:819 *A. fischeri*:535
afum|EAL85105.2 100.00% afis|EAW19789.1 100.00%
Bootstrap support for afum|EAL85105.2 as seed ortholog is 100%.
Bootstrap support for afis|EAW19789.1 as seed ortholog is 100%.

Group of orthologs #3983. Best score 1026 bits
Score difference with first non-orthologous sequence - *A. fumigatus*:1026 *A. fischeri*:1026
afum|EAL87299.1 100.00% afis|EAW23871.1 100.00%
Bootstrap support for afum|EAL87299.1 as seed ortholog is 100%.
Bootstrap support for afis|EAW23871.1 as seed ortholog is 100%.

Group of orthologs #3984. Best score 1026 bits
Score difference with first non-orthologous sequence - *A. fumigatus*:1026 *A. fischeri*:1026
afum|EAL88439.1 100.00% afis|EAW23042.1 100.00%
Bootstrap support for afum|EAL88439.1 as seed ortholog is 100%.
Bootstrap support for afis|EAW23042.1 as seed ortholog is 100%.

Group of orthologs #3985. Best score 1026 bits
Score difference with first non-orthologous sequence - *A. fumigatus*:1026 *A. fischeri*:1026
afum|EAL91015.1 100.00% afis|EAW22174.1 100.00%
Bootstrap support for afum|EAL91015.1 as seed ortholog is 100%.
Bootstrap support for afis|EAW22174.1 as seed ortholog is 100%.

Group of orthologs #3986. Best score 1025 bits
Score difference with first non-orthologous sequence - *A. fumigatus*:1025 *A. fischeri*:1025
afum|EAL84418.2 100.00% afis|EAW20679.1 100.00%
Bootstrap support for afum|EAL84418.2 as seed ortholog is 100%.
Bootstrap support for afis|EAW20679.1 as seed ortholog is 100%.

Group of orthologs #3987. Best score 1025 bits
Score difference with first non-orthologous sequence - *A. fumigatus*:1025 *A. fischeri*:1025
afum|EAL87522.1 100.00% afis|EAW18162.1 100.00%
Bootstrap support for afum|EAL87522.1 as seed ortholog is 100%.
Bootstrap support for afis|EAW18162.1 as seed ortholog is 100%.

Group of orthologs #3988. Best score 1025 bits
Score difference with first non-orthologous sequence - *A. fumigatus*:1025 *A. fischeri*:1025
afum|EAL93668.2 100.00% afis|EAW18935.1 100.00%
Bootstrap support for afum|EAL93668.2 as seed ortholog is 100%.
Bootstrap support for afis|EAW18935.1 as seed ortholog is 100%.

Group of orthologs #3989. Best score 1024 bits
Score difference with first non-orthologous sequence - *A. fumigatus*:1024 *A. fischeri*:1024
afum|EAL88597.1 100.00% afis|EAW15964.1 100.00%
Bootstrap support for afum|EAL88597.1 as seed ortholog is 100%.
Bootstrap support for afis|EAW15964.1 as seed ortholog is 100%.

Group of orthologs #3990. Best score 1024 bits
Score difference with first non-orthologous sequence - *A. fumigatus*:1024 *A. fischeri*:1024
afum|EAL85266.1 100.00% afis|EAW20151.1 100.00%
Bootstrap support for afum|EAL85266.1 as seed ortholog is 100%.
Bootstrap support for afis|EAW20151.1 as seed ortholog is 100%.

Group of orthologs #3991. Best score 1024 bits
Score difference with first non-orthologous sequence - *A. fumigatus*:917 *A. fischeri*:1024
afum|EAL87526.1 100.00% afis|EAW18166.1 100.00%
Bootstrap support for afum|EAL87526.1 as seed ortholog is 100%.
Bootstrap support for afis|EAW18166.1 as seed ortholog is 100%.

Group of orthologs #3992. Best score 1024 bits

Score difference with first non-orthologous sequence - *A. fumigatus*:793   *A. fischeri*:771
afum|EAL84821.1   100.00%   afis|EAW20807.1   100.00%
Bootstrap support for afum|EAL84821.1 as seed ortholog is 100%.
Bootstrap support for afis|EAW20807.1 as seed ortholog is 100%.

---

Group of orthologs #3993. Best score 1024 bits
Score difference with first non-orthologous sequence - *A. fumigatus*:1024   *A. fischeri*:1024
afum|EAL86992.2   100.00%   afis|EAW19501.1   100.00%
Bootstrap support for afum|EAL86992.2 as seed ortholog is 100%.
Bootstrap support for afis|EAW19501.1 as seed ortholog is 100%.

---

Group of orthologs #3994. Best score 1024 bits
Score difference with first non-orthologous sequence - *A. fumigatus*:1024   *A. fischeri*:646
afum|EAL86838.1   100.00%   afis|EAW19686.1   100.00%
Bootstrap support for afum|EAL86838.1 as seed ortholog is 100%.
Bootstrap support for afis|EAW19686.1 as seed ortholog is 100%.

---

Group of orthologs #3995. Best score 1024 bits
Score difference with first non-orthologous sequence - *A. fumigatus*:1024   *A. fischeri*:1024
afum|EAL85909.2   100.00%   afis|EAW24241.1   100.00%
Bootstrap support for afum|EAL85909.2 as seed ortholog is 100%.
Bootstrap support for afis|EAW24241.1 as seed ortholog is 100%.

---

Group of orthologs #3996. Best score 1024 bits
Score difference with first non-orthologous sequence - *A. fumigatus*:1024   *A. fischeri*:1024
afum|EAL93179.1   100.00%   afis|EAW18489.1   100.00%
Bootstrap support for afum|EAL93179.1 as seed ortholog is 100%.
Bootstrap support for afis|EAW18489.1 as seed ortholog is 100%.

---

Group of orthologs #3997. Best score 1024 bits
Score difference with first non-orthologous sequence - *A. fumigatus*:1024   *A. fischeri*:927
afum|EAL92292.1   100.00%   afis|EAW21277.1   100.00%
Bootstrap support for afum|EAL92292.1 as seed ortholog is 100%.
Bootstrap support for afis|EAW21277.1 as seed ortholog is 100%.

---

Group of orthologs #3998. Best score 1024 bits
Score difference with first non-orthologous sequence - *A. fumigatus*:1024   *A. fischeri*:1024
afum|EAL92964.1   100.00%   afis|EAW21950.1   100.00%
Bootstrap support for afum|EAL92964.1 as seed ortholog is 100%.
Bootstrap support for afis|EAW21950.1 as seed ortholog is 100%.

---

Group of orthologs #3999. Best score 1024 bits
Score difference with first non-orthologous sequence - *A. fumigatus*:1024   *A. fischeri*:1024
afum|EAL89962.2   100.00%   afis|EAW25299.1   100.00%
Bootstrap support for afum|EAL89962.2 as seed ortholog is 100%.
Bootstrap support for afis|EAW25299.1 as seed ortholog is 100%.

---

Group of orthologs #4000. Best score 1023 bits
Score difference with first non-orthologous sequence - *A. fumigatus*:1023   *A. fischeri*:421
afum|EAL86472.2   100.00%   afis|EAW16945.1   100.00%
Bootstrap support for afum|EAL86472.2 as seed ortholog is 100%.
Bootstrap support for afis|EAW16945.1 as seed ortholog is 100%.

---

Group of orthologs #4001. Best score 1023 bits
Score difference with first non-orthologous sequence - *A. fumigatus*:764   *A. fischeri*:587
afum|EAL86614.1   100.00%   afis|EAW17188.1   100.00%

Bootstrap support for afum|EAL86614.1 as seed ortholog is 100%.
Bootstrap support for afis|EAW17188.1 as seed ortholog is 100%.

---

Group of orthologs #4002. Best score 1023 bits
Score difference with first non-orthologous sequence - *A. fumigatus*:1023 *A. fischeri*:1023
afum|EAL87681.1 100.00% afis|EAW18314.1 100.00%
Bootstrap support for afum|EAL87681.1 as seed ortholog is 100%.
Bootstrap support for afis|EAW18314.1 as seed ortholog is 100%.

---

Group of orthologs #4003. Best score 1023 bits
Score difference with first non-orthologous sequence - *A. fumigatus*:1023 *A. fischeri*:1023
afum|EAL87050.1 100.00% afis|EAW19445.1 100.00%
Bootstrap support for afum|EAL87050.1 as seed ortholog is 100%.
Bootstrap support for afis|EAW19445.1 as seed ortholog is 100%.

---

Group of orthologs #4004. Best score 1023 bits
Score difference with first non-orthologous sequence - *A. fumigatus*:815 *A. fischeri*:1023
afum|EAL91186.1 100.00% afis|EAW17341.1 100.00%
Bootstrap support for afum|EAL91186.1 as seed ortholog is 100%.
Bootstrap support for afis|EAW17341.1 as seed ortholog is 100%.

---

Group of orthologs #4005. Best score 1023 bits
Score difference with first non-orthologous sequence - *A. fumigatus*:1023 *A. fischeri*:1023
afum|EAL87918.2 100.00% afis|EAW23548.1 100.00%
Bootstrap support for afum|EAL87918.2 as seed ortholog is 100%.
Bootstrap support for afis|EAW23548.1 as seed ortholog is 100%.

---

Group of orthologs #4006. Best score 1023 bits
Score difference with first non-orthologous sequence - *A. fumigatus*:230 *A. fischeri*:207
afum|EBA27475.1 100.00% afis|EAW21838.1 100.00%
Bootstrap support for afum|EBA27475.1 as seed ortholog is 100%.
Bootstrap support for afis|EAW21838.1 as seed ortholog is 100%.

---

Group of orthologs #4007. Best score 1022 bits
Score difference with first non-orthologous sequence - *A. fumigatus*:801 *A. fischeri*:783
afum|EAL84952.1 100.00% afis|EAW19970.1 100.00%
Bootstrap support for afum|EAL84952.1 as seed ortholog is 100%.
Bootstrap support for afis|EAW19970.1 as seed ortholog is 100%.

---

Group of orthologs #4008. Best score 1022 bits
Score difference with first non-orthologous sequence - *A. fumigatus*:1022 *A. fischeri*:1022
afum|EAL87711.1 100.00% afis|EAW18347.1 100.00%
Bootstrap support for afum|EAL87711.1 as seed ortholog is 100%.
Bootstrap support for afis|EAW18347.1 as seed ortholog is 100%.

---

Group of orthologs #4009. Best score 1022 bits
Score difference with first non-orthologous sequence - *A. fumigatus*:740 *A. fischeri*:879
afum|EAL92049.1 100.00% afis|EAW19748.1 100.00%
Bootstrap support for afum|EAL92049.1 as seed ortholog is 100%.
Bootstrap support for afis|EAW19748.1 as seed ortholog is 100%.

---

Group of orthologs #4010. Best score 1022 bits
Score difference with first non-orthologous sequence - *A. fumigatus*:1022 *A. fischeri*:1022
afum|EAL93671.1 100.00% afis|EAW18938.1 100.00%
Bootstrap support for afum|EAL93671.1 as seed ortholog is 100%.
Bootstrap support for afis|EAW18938.1 as seed ortholog is 100%.

Group of orthologs #4011. Best score 1022 bits
Score difference with first non-orthologous sequence - *A. fumigatus*:1022 *A. fischeri*:845
afum|EBA27258.1 100.00% afis|EAW24508.1 100.00%
Bootstrap support for afum|EBA27258.1 as seed ortholog is 100%.
Bootstrap support for afis|EAW24508.1 as seed ortholog is 100%.

Group of orthologs #4012. Best score 1021 bits
Score difference with first non-orthologous sequence - *A. fumigatus*:1021 *A. fischeri*:1021
afum|EAL85179.1 100.00% afis|EAW20059.1 100.00%
afum|EAL87759.1 20.00%
Bootstrap support for afum|EAL85179.1 as seed ortholog is 100%.
Bootstrap support for afis|EAW20059.1 as seed ortholog is 100%.

Group of orthologs #4013. Best score 1021 bits
Score difference with first non-orthologous sequence - *A. fumigatus*:1021 *A. fischeri*:1021
afum|EAL86388.1 100.00% afis|EAW16841.1 100.00%
Bootstrap support for afum|EAL86388.1 as seed ortholog is 100%.
Bootstrap support for afis|EAW16841.1 as seed ortholog is 100%.

Group of orthologs #4014. Best score 1021 bits
Score difference with first non-orthologous sequence - *A. fumigatus*:1021 *A. fischeri*:1021
afum|EAL85235.1 100.00% afis|EAW20118.1 100.00%
Bootstrap support for afum|EAL85235.1 as seed ortholog is 100%.
Bootstrap support for afis|EAW20118.1 as seed ortholog is 100%.

Group of orthologs #4015. Best score 1021 bits
Score difference with first non-orthologous sequence - *A. fumigatus*:1021 *A. fischeri*:1021
afum|EAL87646.1 100.00% afis|EAW18280.1 100.00%
Bootstrap support for afum|EAL87646.1 as seed ortholog is 100%.
Bootstrap support for afis|EAW18280.1 as seed ortholog is 100%.

Group of orthologs #4016. Best score 1021 bits
Score difference with first non-orthologous sequence - *A. fumigatus*:1021 *A. fischeri*:1021
afum|EAL87651.1 100.00% afis|EAW18285.1 100.00%
Bootstrap support for afum|EAL87651.1 as seed ortholog is 100%.
Bootstrap support for afis|EAW18285.1 as seed ortholog is 100%.

Group of orthologs #4017. Best score 1021 bits
Score difference with first non-orthologous sequence - *A. fumigatus*:1021 *A. fischeri*:1021
afum|EAL87700.1 100.00% afis|EAW18336.1 100.00%
Bootstrap support for afum|EAL87700.1 as seed ortholog is 100%.
Bootstrap support for afis|EAW18336.1 as seed ortholog is 100%.

Group of orthologs #4018. Best score 1021 bits
Score difference with first non-orthologous sequence - *A. fumigatus*:1021 *A. fischeri*:1021
afum|EAL91309.1 100.00% afis|EAW17481.1 100.00%
Bootstrap support for afum|EAL91309.1 as seed ortholog is 100%.
Bootstrap support for afis|EAW17481.1 as seed ortholog is 100%.

Group of orthologs #4019. Best score 1021 bits
Score difference with first non-orthologous sequence - *A. fumigatus*:1021 *A. fischeri*:1021
afum|EAL91370.1 100.00% afis|EAW17556.1 100.00%
Bootstrap support for afum|EAL91370.1 as seed ortholog is 100%.
Bootstrap support for afis|EAW17556.1 as seed ortholog is 100%.

Group of orthologs #4020. Best score 1021 bits
Score difference with first non-orthologous sequence - *A. fumigatus*:1021 *A. fischeri*:1021
afum|EAL93212.1 100.00% afis|EAW18517.1 100.00%
Bootstrap support for afum|EAL93212.1 as seed ortholog is 100%.
Bootstrap support for afis|EAW18517.1 as seed ortholog is 100%.

---

Group of orthologs #4021. Best score 1021 bits
Score difference with first non-orthologous sequence - *A. fumigatus*:1021 *A. fischeri*:1021
afum|EBA27492.1 100.00% afis|EAW18716.1 100.00%
Bootstrap support for afum|EBA27492.1 as seed ortholog is 100%.
Bootstrap support for afis|EAW18716.1 as seed ortholog is 100%.

---

Group of orthologs #4022. Best score 1021 bits
Score difference with first non-orthologous sequence - *A. fumigatus*:1021 *A. fischeri*:1021
afum|EAL90966.1 100.00% afis|EAW22228.1 100.00%
Bootstrap support for afum|EAL90966.1 as seed ortholog is 100%.
Bootstrap support for afis|EAW22228.1 as seed ortholog is 100%.

---

Group of orthologs #4023. Best score 1021 bits
Score difference with first non-orthologous sequence - *A. fumigatus*:1021 *A. fischeri*:1021
afum|EAL90094.1 100.00% afis|EAW25436.1 100.00%
Bootstrap support for afum|EAL90094.1 as seed ortholog is 100%.
Bootstrap support for afis|EAW25436.1 as seed ortholog is 100%.

---

Group of orthologs #4024. Best score 1021 bits
Score difference with first non-orthologous sequence - *A. fumigatus*:1021 *A. fischeri*:1021
afum|EAL90136.2 100.00% afis|EAW25473.1 100.00%
Bootstrap support for afum|EAL90136.2 as seed ortholog is 100%.
Bootstrap support for afis|EAW25473.1 as seed ortholog is 100%.

---

Group of orthologs #4025. Best score 1020 bits
Score difference with first non-orthologous sequence - *A. fumigatus*:1020 *A. fischeri*:1020
afum|EAL91328.2 100.00% afis|EAW17506.1 100.00%
Bootstrap support for afum|EAL91328.2 as seed ortholog is 100%.
Bootstrap support for afis|EAW17506.1 as seed ortholog is 100%.

---

Group of orthologs #4026. Best score 1020 bits
Score difference with first non-orthologous sequence - *A. fumigatus*:1020 *A. fischeri*:1020
afum|EAL91635.1 100.00% afis|EAW17825.1 100.00%
Bootstrap support for afum|EAL91635.1 as seed ortholog is 100%.
Bootstrap support for afis|EAW17825.1 as seed ortholog is 100%.

---

Group of orthologs #4027. Best score 1020 bits
Score difference with first non-orthologous sequence - *A. fumigatus*:1020 *A. fischeri*:1020
afum|EAL87132.1 100.00% afis|EAW23687.1 100.00%
Bootstrap support for afum|EAL87132.1 as seed ortholog is 100%.
Bootstrap support for afis|EAW23687.1 as seed ortholog is 100%.

---

Group of orthologs #4028. Best score 1020 bits
Score difference with first non-orthologous sequence - *A. fumigatus*:1020 *A. fischeri*:1020
afum|EAL90461.1 100.00% afis|EAW22734.1 100.00%
Bootstrap support for afum|EAL90461.1 as seed ortholog is 100%.
Bootstrap support for afis|EAW22734.1 as seed ortholog is 100%.

---

Group of orthologs #4029. Best score 1020 bits
Score difference with first non-orthologous sequence - *A. fumigatus*:741 *A. fischeri*:737

afum|EAL91923.1 100.00% afis|EAW24067.1 100.00%
Bootstrap support for afum|EAL91923.1 as seed ortholog is 100%.
Bootstrap support for afis|EAW24067.1 as seed ortholog is 100%.

---

Group of orthologs #4030. Best score 1019 bits
Score difference with first non-orthologous sequence - *A. fumigatus*:1019 *A. fischeri*:1019
afum|EAL91238.1 100.00% afis|EAW17400.1 100.00%
Bootstrap support for afum|EAL91238.1 as seed ortholog is 100%.
Bootstrap support for afis|EAW17400.1 as seed ortholog is 100%.

---

Group of orthologs #4031. Best score 1019 bits
Score difference with first non-orthologous sequence - *A. fumigatus*:1019 *A. fischeri*:1019
afum|EAL86186.1 100.00% afis|EAW24521.1 100.00%
Bootstrap support for afum|EAL86186.1 as seed ortholog is 100%.
Bootstrap support for afis|EAW24521.1 as seed ortholog is 100%.

---

Group of orthologs #4032. Best score 1019 bits
Score difference with first non-orthologous sequence - *A. fumigatus*:1019 *A. fischeri*:878
afum|EAL93836.1 100.00% afis|EAW19104.1 100.00%
Bootstrap support for afum|EAL93836.1 as seed ortholog is 100%.
Bootstrap support for afis|EAW19104.1 as seed ortholog is 100%.

---

Group of orthologs #4033. Best score 1019 bits
Score difference with first non-orthologous sequence - *A. fumigatus*:1019 *A. fischeri*:1019
afum|EAL91094.1 100.00% afis|EAW22076.1 100.00%
Bootstrap support for afum|EAL91094.1 as seed ortholog is 100%.
Bootstrap support for afis|EAW22076.1 as seed ortholog is 100%.

---

Group of orthologs #4034. Best score 1019 bits
Score difference with first non-orthologous sequence - *A. fumigatus*:1019 *A. fischeri*:1019
afum|EAL89311.1 100.00% afis|EAW24659.1 100.00%
Bootstrap support for afum|EAL89311.1 as seed ortholog is 100%.
Bootstrap support for afis|EAW24659.1 as seed ortholog is 100%.

---

Group of orthologs #4035. Best score 1018 bits
Score difference with first non-orthologous sequence - *A. fumigatus*:766 *A. fischeri*:1018
afum|EAL84213.1 100.00% afis|EAW15430.1 100.00%
Bootstrap support for afum|EAL84213.1 as seed ortholog is 100%.
Bootstrap support for afis|EAW15430.1 as seed ortholog is 100%.

---

Group of orthologs #4036. Best score 1018 bits
Score difference with first non-orthologous sequence - *A. fumigatus*:1018 *A. fischeri*:1018
afum|EAL91446.1 100.00% afis|EAW17631.1 100.00%
Bootstrap support for afum|EAL91446.1 as seed ortholog is 100%.
Bootstrap support for afis|EAW17631.1 as seed ortholog is 100%.

---

Group of orthologs #4037. Best score 1018 bits
Score difference with first non-orthologous sequence - *A. fumigatus*:1018 *A. fischeri*:1018
afum|EAL91532.1 100.00% afis|EAW17717.1 100.00%
Bootstrap support for afum|EAL91532.1 as seed ortholog is 100%.
Bootstrap support for afis|EAW17717.1 as seed ortholog is 100%.

---

Group of orthologs #4038. Best score 1018 bits
Score difference with first non-orthologous sequence - *A. fumigatus*:1018 *A. fischeri*:1018
afum|EAL91711.1 100.00% afis|EAW17903.1 100.00%
Bootstrap support for afum|EAL91711.1 as seed ortholog is 100%.

Bootstrap support for afis|EAW17903.1 as seed ortholog is 100%.

---

Group of orthologs #4039. Best score 1018 bits
Score difference with first non-orthologous sequence - *A. fumigatus*:1018 *A. fischeri*:1018
afum|EAL84362.1 100.00% afis|EAW25692.1 100.00%
Bootstrap support for afum|EAL84362.1 as seed ortholog is 100%.
Bootstrap support for afis|EAW25692.1 as seed ortholog is 100%.

---

Group of orthologs #4040. Best score 1018 bits
Score difference with first non-orthologous sequence - *A. fumigatus*:1018 *A. fischeri*:1018
afum|EAL90875.1 100.00% afis|EAW22315.1 100.00%
Bootstrap support for afum|EAL90875.1 as seed ortholog is 100%.
Bootstrap support for afis|EAW22315.1 as seed ortholog is 100%.

---

Group of orthologs #4041. Best score 1018 bits
Score difference with first non-orthologous sequence - *A. fumigatus*:1018 *A. fischeri*:1018
afum|EAL90671.1 100.00% afis|EAW22523.1 100.00%
Bootstrap support for afum|EAL90671.1 as seed ortholog is 100%.
Bootstrap support for afis|EAW22523.1 as seed ortholog is 100%.

---

Group of orthologs #4042. Best score 1016 bits
Score difference with first non-orthologous sequence - *A. fumigatus*:1016 *A. fischeri*:1016
afum|EAL85789.1 100.00% afis|EAW15541.1 100.00%
Bootstrap support for afum|EAL85789.1 as seed ortholog is 100%.
Bootstrap support for afis|EAW15541.1 as seed ortholog is 100%.

---

Group of orthologs #4043. Best score 1016 bits
Score difference with first non-orthologous sequence - *A. fumigatus*:1016 *A. fischeri*:1016
afum|EAL89059.1 100.00% afis|EAW16463.1 100.00%
Bootstrap support for afum|EAL89059.1 as seed ortholog is 100%.
Bootstrap support for afis|EAW16463.1 as seed ortholog is 100%.

---

Group of orthologs #4044. Best score 1016 bits
Score difference with first non-orthologous sequence - *A. fumigatus*:1016 *A. fischeri*:1016
afum|EAL86882.1 100.00% afis|EAW19640.1 100.00%
Bootstrap support for afum|EAL86882.1 as seed ortholog is 100%.
Bootstrap support for afis|EAW19640.1 as seed ortholog is 100%.

---

Group of orthologs #4045. Best score 1015 bits
Score difference with first non-orthologous sequence - *A. fumigatus*:1015 *A. fischeri*:1015
afum|EAL85723.1 100.00% afis|EAW15612.1 100.00%
Bootstrap support for afum|EAL85723.1 as seed ortholog is 100%.
Bootstrap support for afis|EAW15612.1 as seed ortholog is 100%.

---

Group of orthologs #4046. Best score 1015 bits
Score difference with first non-orthologous sequence - *A. fumigatus*:1015 *A. fischeri*:877
afum|EAL88819.2 100.00% afis|EAW16188.1 100.00%
Bootstrap support for afum|EAL88819.2 as seed ortholog is 100%.
Bootstrap support for afis|EAW16188.1 as seed ortholog is 100%.

---

Group of orthologs #4047. Best score 1015 bits
Score difference with first non-orthologous sequence - *A. fumigatus*:816 *A. fischeri*:1015
afum|EAL91340.1 100.00% afis|EAW17525.1 100.00%
Bootstrap support for afum|EAL91340.1 as seed ortholog is 100%.
Bootstrap support for afis|EAW17525.1 as seed ortholog is 100%.

---

Group of orthologs #4048. Best score 1015 bits
Score difference with first non-orthologous sequence - *A. fumigatus*:1015 *A. fischeri*:1015
afum|EAL86030.1 100.00% afis|EAW24360.1 100.00%
Bootstrap support for afum|EAL86030.1 as seed ortholog is 100%.
Bootstrap support for afis|EAW24360.1 as seed ortholog is 100%.

---

Group of orthologs #4049. Best score 1015 bits
Score difference with first non-orthologous sequence - *A. fumigatus*:944 *A. fischeri*:861
afum|EAL87115.1 100.00% afis|EAW23670.1 100.00%
Bootstrap support for afum|EAL87115.1 as seed ortholog is 100%.
Bootstrap support for afis|EAW23670.1 as seed ortholog is 100%.

---

Group of orthologs #4050. Best score 1015 bits
Score difference with first non-orthologous sequence - *A. fumigatus*:1015 *A. fischeri*:1015
afum|EAL88405.1 100.00% afis|EAW23074.1 100.00%
Bootstrap support for afum|EAL88405.1 as seed ortholog is 100%.
Bootstrap support for afis|EAW23074.1 as seed ortholog is 100%.

---

Group of orthologs #4051. Best score 1015 bits
Score difference with first non-orthologous sequence - *A. fumigatus*:1015 *A. fischeri*:1015
afum|EAL88381.2 100.00% afis|EAW23097.1 100.00%
Bootstrap support for afum|EAL88381.2 as seed ortholog is 100%.
Bootstrap support for afis|EAW23097.1 as seed ortholog is 100%.

---

Group of orthologs #4052. Best score 1015 bits
Score difference with first non-orthologous sequence - *A. fumigatus*:1015 *A. fischeri*:1015
afum|EAL90674.1 100.00% afis|EAW22519.1 100.00%
Bootstrap support for afum|EAL90674.1 as seed ortholog is 100%.
Bootstrap support for afis|EAW22519.1 as seed ortholog is 100%.

---

Group of orthologs #4053. Best score 1015 bits
Score difference with first non-orthologous sequence - *A. fumigatus*:1015 *A. fischeri*:815
afum|EAL94084.2 100.00% afis|EAW19365.1 100.00%
Bootstrap support for afum|EAL94084.2 as seed ortholog is 100%.
Bootstrap support for afis|EAW19365.1 as seed ortholog is 100%.

---

Group of orthologs #4054. Best score 1014 bits
Score difference with first non-orthologous sequence - *A. fumigatus*:1014 *A. fischeri*:1014
afum|EAL86552.2 100.00% afis|EAW17055.1 100.00%
Bootstrap support for afum|EAL86552.2 as seed ortholog is 100%.
Bootstrap support for afis|EAW17055.1 as seed ortholog is 100%.

---

Group of orthologs #4055. Best score 1014 bits
Score difference with first non-orthologous sequence - *A. fumigatus*:1014 *A. fischeri*:1014
afum|EAL85125.1 100.00% afis|EAW19774.1 100.00%
Bootstrap support for afum|EAL85125.1 as seed ortholog is 100%.
Bootstrap support for afis|EAW19774.1 as seed ortholog is 100%.

---

Group of orthologs #4056. Best score 1014 bits
Score difference with first non-orthologous sequence - *A. fumigatus*:1014 *A. fischeri*:1014
afum|EAL87982.1 100.00% afis|EAW23482.1 100.00%
Bootstrap support for afum|EAL87982.1 as seed ortholog is 100%.
Bootstrap support for afis|EAW23482.1 as seed ortholog is 100%.

---

Group of orthologs #4057. Best score 1014 bits
Score difference with first non-orthologous sequence - *A. fumigatus*:1014 *A. fischeri*:1014

afum|EAL88007.2 100.00% afis|EAW23459.1 100.00%
Bootstrap support for afum|EAL88007.2 as seed ortholog is 100%.
Bootstrap support for afis|EAW23459.1 as seed ortholog is 100%.

---

Group of orthologs #4058. Best score 1014 bits
Score difference with first non-orthologous sequence - *A. fumigatus*:1014 *A. fischeri*:1014
afum|EAL88085.1 100.00% afis|EAW23386.1 100.00%
Bootstrap support for afum|EAL88085.1 as seed ortholog is 100%.
Bootstrap support for afis|EAW23386.1 as seed ortholog is 100%.

---

Group of orthologs #4059. Best score 1014 bits
Score difference with first non-orthologous sequence - *A. fumigatus*:1014 *A. fischeri*:1014
afum|EAL93651.2 100.00% afis|EAW18920.1 100.00%
Bootstrap support for afum|EAL93651.2 as seed ortholog is 100%.
Bootstrap support for afis|EAW18920.1 as seed ortholog is 100%.

---

Group of orthologs #4060. Best score 1013 bits
Score difference with first non-orthologous sequence - *A. fumigatus*:1013 *A. fischeri*:1013
afum|EAL85906.1 100.00% afis|EAW24238.1 100.00%
Bootstrap support for afum|EAL85906.1 as seed ortholog is 100%.
Bootstrap support for afis|EAW24238.1 as seed ortholog is 100%.

---

Group of orthologs #4061. Best score 1013 bits
Score difference with first non-orthologous sequence - *A. fumigatus*:1013 *A. fischeri*:1013
afum|EAL87357.1 100.00% afis|EAW23932.1 100.00%
Bootstrap support for afum|EAL87357.1 as seed ortholog is 100%.
Bootstrap support for afis|EAW23932.1 as seed ortholog is 100%.

---

Group of orthologs #4062. Best score 1013 bits
Score difference with first non-orthologous sequence - *A. fumigatus*:1013 *A. fischeri*:1013
afum|EAL88186.1 100.00% afis|EAW23293.1 100.00%
Bootstrap support for afum|EAL88186.1 as seed ortholog is 100%.
Bootstrap support for afis|EAW23293.1 as seed ortholog is 100%.

---

Group of orthologs #4063. Best score 1013 bits
Score difference with first non-orthologous sequence - *A. fumigatus*:1013 *A. fischeri*:1013
afum|EAL88215.1 100.00% afis|EAW23265.1 100.00%
Bootstrap support for afum|EAL88215.1 as seed ortholog is 100%.
Bootstrap support for afis|EAW23265.1 as seed ortholog is 100%.

---

Group of orthologs #4064. Best score 1013 bits
Score difference with first non-orthologous sequence - *A. fumigatus*:1013 *A. fischeri*:1013
afum|EAL92531.1 100.00% afis|EAW21511.1 100.00%
Bootstrap support for afum|EAL92531.1 as seed ortholog is 100%.
Bootstrap support for afis|EAW21511.1 as seed ortholog is 100%.

---

Group of orthologs #4065. Best score 1013 bits
Score difference with first non-orthologous sequence - *A. fumigatus*:1013 *A. fischeri*:1013
afum|EAL92006.1 100.00% afis|EAW24151.1 100.00%
Bootstrap support for afum|EAL92006.1 as seed ortholog is 100%.
Bootstrap support for afis|EAW24151.1 as seed ortholog is 100%.

---

Group of orthologs #4066. Best score 1012 bits
Score difference with first non-orthologous sequence - *A. fumigatus*:1012 *A. fischeri*:1012
afum|EAL85543.1 100.00% afis|EAW15808.1 100.00%
Bootstrap support for afum|EAL85543.1 as seed ortholog is 100%.

Bootstrap support for afis|EAW15808.1 as seed ortholog is 100%.

---

Group of orthologs #4067. Best score 1012 bits
Score difference with first non-orthologous sequence - *A. fumigatus*:1012 *A. fischeri*:1012
afum|EAL89198.1 100.00% afis|EAW16602.1 100.00%
Bootstrap support for afum|EAL89198.1 as seed ortholog is 100%.
Bootstrap support for afis|EAW16602.1 as seed ortholog is 100%.

---

Group of orthologs #4068. Best score 1012 bits
Score difference with first non-orthologous sequence - *A. fumigatus*:1012 *A. fischeri*:942
afum|EAL91706.1 100.00% afis|EAW17898.1 100.00%
Bootstrap support for afum|EAL91706.1 as seed ortholog is 100%.
Bootstrap support for afis|EAW17898.1 as seed ortholog is 100%.

---

Group of orthologs #4069. Best score 1012 bits
Score difference with first non-orthologous sequence - *A. fumigatus*:1012 *A. fischeri*:664
afum|EAL86066.1 100.00% afis|EAW24394.1 100.00%
Bootstrap support for afum|EAL86066.1 as seed ortholog is 100%.
Bootstrap support for afis|EAW24394.1 as seed ortholog is 100%.

---

Group of orthologs #4070. Best score 1012 bits
Score difference with first non-orthologous sequence - *A. fumigatus*:1012 *A. fischeri*:1012
afum|EAL91016.1 100.00% afis|EAW22173.1 100.00%
Bootstrap support for afum|EAL91016.1 as seed ortholog is 100%.
Bootstrap support for afis|EAW22173.1 as seed ortholog is 100%.

---

Group of orthologs #4071. Best score 1012 bits
Score difference with first non-orthologous sequence - *A. fumigatus*:1012 *A. fischeri*:1012
afum|EAL90518.1 100.00% afis|EAW22674.1 100.00%
Bootstrap support for afum|EAL90518.1 as seed ortholog is 100%.
Bootstrap support for afis|EAW22674.1 as seed ortholog is 100%.

---

Group of orthologs #4072. Best score 1011 bits
Score difference with first non-orthologous sequence - *A. fumigatus*:1011 *A. fischeri*:1011
afum|EAL84974.1 100.00% afis|EAW19936.1 100.00%
Bootstrap support for afum|EAL84974.1 as seed ortholog is 100%.
Bootstrap support for afis|EAW19936.1 as seed ortholog is 100%.

---

Group of orthologs #4073. Best score 1011 bits
Score difference with first non-orthologous sequence - *A. fumigatus*:1011 *A. fischeri*:1011
afum|EAL87486.1 100.00% afis|EAW18129.1 100.00%
Bootstrap support for afum|EAL87486.1 as seed ortholog is 100%.
Bootstrap support for afis|EAW18129.1 as seed ortholog is 100%.

---

Group of orthologs #4074. Best score 1011 bits
Score difference with first non-orthologous sequence - *A. fumigatus*:1011 *A. fischeri*:1011
afum|EAL92408.1 100.00% afis|EAW21394.1 100.00%
Bootstrap support for afum|EAL92408.1 as seed ortholog is 100%.
Bootstrap support for afis|EAW21394.1 as seed ortholog is 100%.

---

Group of orthologs #4075. Best score 1010 bits
Score difference with first non-orthologous sequence - *A. fumigatus*:1010 *A. fischeri*:1010
afum|EAL88317.2 100.00% afis|EAW23167.1 100.00%
Bootstrap support for afum|EAL88317.2 as seed ortholog is 100%.
Bootstrap support for afis|EAW23167.1 as seed ortholog is 100%.

---

Group of orthologs #4076. Best score 1010 bits
Score difference with first non-orthologous sequence - *A. fumigatus*:1010 *A. fischeri*:1010
afum|EAL88136.2 100.00% afis|EAW23342.1 100.00%
Bootstrap support for afum|EAL88136.2 as seed ortholog is 100%.
Bootstrap support for afis|EAW23342.1 as seed ortholog is 100%.

---

Group of orthologs #4077. Best score 1010 bits
Score difference with first non-orthologous sequence - *A. fumigatus*:1010 *A. fischeri*:1010
afum|EAL92403.2 100.00% afis|EAW21388.1 100.00%
Bootstrap support for afum|EAL92403.2 as seed ortholog is 100%.
Bootstrap support for afis|EAW21388.1 as seed ortholog is 100%.

---

Group of orthologs #4078. Best score 1009 bits
Score difference with first non-orthologous sequence - *A. fumigatus*:1009 *A. fischeri*:1009
afum|EAL87866.1 100.00% afis|EAW23601.1 100.00%
Bootstrap support for afum|EAL87866.1 as seed ortholog is 100%.
Bootstrap support for afis|EAW23601.1 as seed ortholog is 100%.

---

Group of orthologs #4079. Best score 1009 bits
Score difference with first non-orthologous sequence - *A. fumigatus*:1009 *A. fischeri*:1009
afum|EAL93969.1 100.00% afis|EAW19240.1 100.00%
Bootstrap support for afum|EAL93969.1 as seed ortholog is 100%.
Bootstrap support for afis|EAW19240.1 as seed ortholog is 100%.

---

Group of orthologs #4080. Best score 1009 bits
Score difference with first non-orthologous sequence - *A. fumigatus*:1009 *A. fischeri*:1009
afum|EAL90599.1 100.00% afis|EAW22599.1 100.00%
Bootstrap support for afum|EAL90599.1 as seed ortholog is 100%.
Bootstrap support for afis|EAW22599.1 as seed ortholog is 100%.

---

Group of orthologs #4081. Best score 1009 bits
Score difference with first non-orthologous sequence - *A. fumigatus*:1009 *A. fischeri*:1009
afum|EAL90452.1 100.00% afis|EAW22744.1 100.00%
Bootstrap support for afum|EAL90452.1 as seed ortholog is 100%.
Bootstrap support for afis|EAW22744.1 as seed ortholog is 100%.

---

Group of orthologs #4082. Best score 1008 bits
Score difference with first non-orthologous sequence - *A. fumigatus*:1008 *A. fischeri*:1008
afum|EAL85067.1 100.00% afis|EAW19825.1 100.00%
Bootstrap support for afum|EAL85067.1 as seed ortholog is 100%.
Bootstrap support for afis|EAW19825.1 as seed ortholog is 100%.

---

Group of orthologs #4083. Best score 1008 bits
Score difference with first non-orthologous sequence - *A. fumigatus*:1008 *A. fischeri*:1008
afum|EAL89256.1 100.00% afis|EAW16652.1 100.00%
Bootstrap support for afum|EAL89256.1 as seed ortholog is 100%.
Bootstrap support for afis|EAW16652.1 as seed ortholog is 100%.

---

Group of orthologs #4084. Best score 1008 bits
Score difference with first non-orthologous sequence - *A. fumigatus*:1008 *A. fischeri*:925
afum|EAL91147.1 100.00% afis|EAW17312.1 100.00%
Bootstrap support for afum|EAL91147.1 as seed ortholog is 100%.
Bootstrap support for afis|EAW17312.1 as seed ortholog is 100%.

---

Group of orthologs #4085. Best score 1008 bits
Score difference with first non-orthologous sequence - *A. fumigatus*:1008 *A. fischeri*:1008

afum|EAL93919.1 100.00% afis|EAW19188.1 100.00%
Bootstrap support for afum|EAL93919.1 as seed ortholog is 100%.
Bootstrap support for afis|EAW19188.1 as seed ortholog is 100%.

---

Group of orthologs #4086. Best score 1008 bits
Score difference with first non-orthologous sequence - *A. fumigatus*:1008 *A. fischeri*:516
afum|EAL89345.1 100.00% afis|EAW24703.1 100.00%
Bootstrap support for afum|EAL89345.1 as seed ortholog is 100%.
Bootstrap support for afis|EAW24703.1 as seed ortholog is 100%.

---

Group of orthologs #4087. Best score 1007 bits
Score difference with first non-orthologous sequence - *A. fumigatus*:1007 *A. fischeri*:641
afum|EAL85790.1 100.00% afis|EAW15540.1 100.00%
Bootstrap support for afum|EAL85790.1 as seed ortholog is 100%.
Bootstrap support for afis|EAW15540.1 as seed ortholog is 100%.

---

Group of orthologs #4088. Best score 1007 bits
Score difference with first non-orthologous sequence - *A. fumigatus*:713 *A. fischeri*:788
afum|EAL86352.1 100.00% afis|EAW16781.1 100.00%
Bootstrap support for afum|EAL86352.1 as seed ortholog is 100%.
Bootstrap support for afis|EAW16781.1 as seed ortholog is 100%.

---

Group of orthologs #4089. Best score 1007 bits
Score difference with first non-orthologous sequence - *A. fumigatus*:1007 *A. fischeri*:1007
afum|EAL88742.1 100.00% afis|EAW16111.1 100.00%
Bootstrap support for afum|EAL88742.1 as seed ortholog is 100%.
Bootstrap support for afis|EAW16111.1 as seed ortholog is 100%.

---

Group of orthologs #4090. Best score 1007 bits
Score difference with first non-orthologous sequence - *A. fumigatus*:1007 *A. fischeri*:1007
afum|EAL84707.1 100.00% afis|EAW20927.1 100.00%
Bootstrap support for afum|EAL84707.1 as seed ortholog is 100%.
Bootstrap support for afis|EAW20927.1 as seed ortholog is 100%.

---

Group of orthologs #4091. Best score 1007 bits
Score difference with first non-orthologous sequence - *A. fumigatus*:1007 *A. fischeri*:1007
afum|EAL87032.1 100.00% afis|EAW19464.1 100.00%
Bootstrap support for afum|EAL87032.1 as seed ortholog is 100%.
Bootstrap support for afis|EAW19464.1 as seed ortholog is 100%.

---

Group of orthologs #4092. Best score 1007 bits
Score difference with first non-orthologous sequence - *A. fumigatus*:1007 *A. fischeri*:1007
afum|EAL87271.1 100.00% afis|EAW23840.1 100.00%
Bootstrap support for afum|EAL87271.1 as seed ortholog is 100%.
Bootstrap support for afis|EAW23840.1 as seed ortholog is 100%.

---

Group of orthologs #4093. Best score 1007 bits
Score difference with first non-orthologous sequence - *A. fumigatus*:597 *A. fischeri*:550
afum|EAL93930.1 100.00% afis|EAW19198.1 100.00%
Bootstrap support for afum|EAL93930.1 as seed ortholog is 100%.
Bootstrap support for afis|EAW19198.1 as seed ortholog is 100%.

---

Group of orthologs #4094. Best score 1006 bits
Score difference with first non-orthologous sequence - *A. fumigatus*:1006 *A. fischeri*:1006
afum|EAL86355.1 100.00% afis|EAW16785.1 100.00%
Bootstrap support for afum|EAL86355.1 as seed ortholog is 100%.

Bootstrap support for afis|EAW16785.1 as seed ortholog is 100%.

---

Group of orthologs #4095. Best score 1006 bits
Score difference with first non-orthologous sequence - *A. fumigatus*:1006 *A. fischeri*:1006
afum|EAL85159.1 100.00% afis|EAW20038.1 100.00%
Bootstrap support for afum|EAL85159.1 as seed ortholog is 100%.
Bootstrap support for afis|EAW20038.1 as seed ortholog is 100%.

---

Group of orthologs #4096. Best score 1006 bits
Score difference with first non-orthologous sequence - *A. fumigatus*:1006 *A. fischeri*:1006
afum|EAL87666.1 100.00% afis|EAW18300.1 100.00%
Bootstrap support for afum|EAL87666.1 as seed ortholog is 100%.
Bootstrap support for afis|EAW18300.1 as seed ortholog is 100%.

---

Group of orthologs #4097. Best score 1006 bits
Score difference with first non-orthologous sequence - *A. fumigatus*:1006 *A. fischeri*:1006
afum|EAL86820.1 100.00% afis|EAW19706.1 100.00%
Bootstrap support for afum|EAL86820.1 as seed ortholog is 100%.
Bootstrap support for afis|EAW19706.1 as seed ortholog is 100%.

---

Group of orthologs #4098. Best score 1006 bits
Score difference with first non-orthologous sequence - *A. fumigatus*:1006 *A. fischeri*:1006
afum|EBA27244.1 100.00% afis|EAW15522.1 100.00%
Bootstrap support for afum|EBA27244.1 as seed ortholog is 100%.
Bootstrap support for afis|EAW15522.1 as seed ortholog is 100%.

---

Group of orthologs #4099. Best score 1006 bits
Score difference with first non-orthologous sequence - *A. fumigatus*:1006 *A. fischeri*:1006
afum|EAL91777.1 100.00% afis|EAW17968.1 100.00%
Bootstrap support for afum|EAL91777.1 as seed ortholog is 100%.
Bootstrap support for afis|EAW17968.1 as seed ortholog is 100%.

---

Group of orthologs #4100. Best score 1006 bits
Score difference with first non-orthologous sequence - *A. fumigatus*:1006 *A. fischeri*:895
afum|EAL86040.1 100.00% afis|EAW24368.1 100.00%
Bootstrap support for afum|EAL86040.1 as seed ortholog is 100%.
Bootstrap support for afis|EAW24368.1 as seed ortholog is 100%.

---

Group of orthologs #4101. Best score 1006 bits
Score difference with first non-orthologous sequence - *A. fumigatus*:1006 *A. fischeri*:1006
afum|EBA27506.1 100.00% afis|EAW19158.1 100.00%
Bootstrap support for afum|EBA27506.1 as seed ortholog is 100%.
Bootstrap support for afis|EAW19158.1 as seed ortholog is 100%.

---

Group of orthologs #4102. Best score 1006 bits
Score difference with first non-orthologous sequence - *A. fumigatus*:1006 *A. fischeri*:1006
afum|EAL89890.2 100.00% afis|EAW25229.1 100.00%
Bootstrap support for afum|EAL89890.2 as seed ortholog is 100%.
Bootstrap support for afis|EAW25229.1 as seed ortholog is 100%.

---

Group of orthologs #4103. Best score 1005 bits
Score difference with first non-orthologous sequence - *A. fumigatus*:773 *A. fischeri*:566
afum|EAL94033.1 100.00% afis|EAW19316.1 100.00%
Bootstrap support for afum|EAL94033.1 as seed ortholog is 100%.
Bootstrap support for afis|EAW19316.1 as seed ortholog is 100%.

---

Group of orthologs #4104. Best score 1005 bits
Score difference with first non-orthologous sequence - *A. fumigatus*:1005 *A. fischeri*:1005
afum|EAL90307.1 100.00% afis|EAW22891.1 100.00%
Bootstrap support for afum|EAL90307.1 as seed ortholog is 100%.
Bootstrap support for afis|EAW22891.1 as seed ortholog is 100%.

---

Group of orthologs #4105. Best score 1005 bits
Score difference with first non-orthologous sequence - *A. fumigatus*:665 *A. fischeri*:1005
afum|EAL90083.1 100.00% afis|EAW25424.1 100.00%
Bootstrap support for afum|EAL90083.1 as seed ortholog is 100%.
Bootstrap support for afis|EAW25424.1 as seed ortholog is 100%.

---

Group of orthologs #4106. Best score 1004 bits
Score difference with first non-orthologous sequence - *A. fumigatus*:1004 *A. fischeri*:1004
afum|EAL85712.1 100.00% afis|EAW15622.1 100.00%
Bootstrap support for afum|EAL85712.1 as seed ortholog is 100%.
Bootstrap support for afis|EAW15622.1 as seed ortholog is 100%.

---

Group of orthologs #4107. Best score 1004 bits
Score difference with first non-orthologous sequence - *A. fumigatus*:1004 *A. fischeri*:1004
afum|EAL87697.1 100.00% afis|EAW18332.1 100.00%
Bootstrap support for afum|EAL87697.1 as seed ortholog is 100%.
Bootstrap support for afis|EAW18332.1 as seed ortholog is 100%.

---

Group of orthologs #4108. Best score 1004 bits
Score difference with first non-orthologous sequence - *A. fumigatus*:1004 *A. fischeri*:1004
afum|EAL87210.1 100.00% afis|EAW23776.1 100.00%
Bootstrap support for afum|EAL87210.1 as seed ortholog is 100%.
Bootstrap support for afis|EAW23776.1 as seed ortholog is 100%.

---

Group of orthologs #4109. Best score 1004 bits
Score difference with first non-orthologous sequence - *A. fumigatus*:1004 *A. fischeri*:1004
afum|EAL93525.1 100.00% afis|EAW18790.1 100.00%
Bootstrap support for afum|EAL93525.1 as seed ortholog is 100%.
Bootstrap support for afis|EAW18790.1 as seed ortholog is 100%.

---

Group of orthologs #4110. Best score 1004 bits
Score difference with first non-orthologous sequence - *A. fumigatus*:1004 *A. fischeri*:1004
afum|EAL89805.1 100.00% afis|EAW25134.1 100.00%
Bootstrap support for afum|EAL89805.1 as seed ortholog is 100%.
Bootstrap support for afis|EAW25134.1 as seed ortholog is 100%.

---

Group of orthologs #4111. Best score 1003 bits
Score difference with first non-orthologous sequence - *A. fumigatus*:909 *A. fischeri*:1003
afum|EAL85408.1 100.00% afis|EAW20274.1 100.00%
Bootstrap support for afum|EAL85408.1 as seed ortholog is 100%.
Bootstrap support for afis|EAW20274.1 as seed ortholog is 100%.

---

Group of orthologs #4112. Best score 1003 bits
Score difference with first non-orthologous sequence - *A. fumigatus*:792 *A. fischeri*:1003
afum|EAL91657.1 100.00% afis|EAW17848.1 100.00%
Bootstrap support for afum|EAL91657.1 as seed ortholog is 100%.
Bootstrap support for afis|EAW17848.1 as seed ortholog is 100%.

---

Group of orthologs #4113. Best score 1003 bits
Score difference with first non-orthologous sequence - *A. fumigatus*:1003 *A. fischeri*:1003

afum|EAL87927.1 100.00% afis|EAW23539.1 100.00%
Bootstrap support for afum|EAL87927.1 as seed ortholog is 100%.
Bootstrap support for afis|EAW23539.1 as seed ortholog is 100%.

---

Group of orthologs #4114. Best score 1003 bits
Score difference with first non-orthologous sequence - *A. fumigatus*:1003 *A. fischeri*:1003
afum|EAL91961.1 100.00% afis|EAW24107.1 100.00%
Bootstrap support for afum|EAL91961.1 as seed ortholog is 100%.
Bootstrap support for afis|EAW24107.1 as seed ortholog is 100%.

---

Group of orthologs #4115. Best score 1002 bits
Score difference with first non-orthologous sequence - *A. fumigatus*:1002 *A. fischeri*:1002
afum|EAL86689.1 100.00% afis|EAW17272.1 100.00%
Bootstrap support for afum|EAL86689.1 as seed ortholog is 100%.
Bootstrap support for afis|EAW17272.1 as seed ortholog is 100%.

---

Group of orthologs #4116. Best score 1002 bits
Score difference with first non-orthologous sequence - *A. fumigatus*:1002 *A. fischeri*:1002
afum|EAL84548.1 100.00% afis|EAW20508.1 100.00%
Bootstrap support for afum|EAL84548.1 as seed ortholog is 100%.
Bootstrap support for afis|EAW20508.1 as seed ortholog is 100%.

---

Group of orthologs #4117. Best score 1002 bits
Score difference with first non-orthologous sequence - *A. fumigatus*:1002 *A. fischeri*:1002
afum|EAL84753.1 100.00% afis|EAW20879.1 100.00%
Bootstrap support for afum|EAL84753.1 as seed ortholog is 100%.
Bootstrap support for afis|EAW20879.1 as seed ortholog is 100%.

---

Group of orthologs #4118. Best score 1002 bits
Score difference with first non-orthologous sequence - *A. fumigatus*:1002 *A. fischeri*:1002
afum|EAL93994.1 100.00% afis|EAW19267.1 100.00%
Bootstrap support for afum|EAL93994.1 as seed ortholog is 100%.
Bootstrap support for afis|EAW19267.1 as seed ortholog is 100%.

---

Group of orthologs #4119. Best score 1002 bits
Score difference with first non-orthologous sequence - *A. fumigatus*:1002 *A. fischeri*:243
afum|EAL89335.1 100.00% afis|EAW24693.1 100.00%
Bootstrap support for afum|EAL89335.1 as seed ortholog is 100%.
Bootstrap support for afis|EAW24693.1 as seed ortholog is 100%.

---

Group of orthologs #4120. Best score 1001 bits
Score difference with first non-orthologous sequence - *A. fumigatus*:1001 *A. fischeri*:1001
afum|EAL88899.1 100.00% afis|EAW16278.1 100.00%
Bootstrap support for afum|EAL88899.1 as seed ortholog is 100%.
Bootstrap support for afis|EAW16278.1 as seed ortholog is 100%.

---

Group of orthologs #4121. Best score 1001 bits
Score difference with first non-orthologous sequence - *A. fumigatus*:1001 *A. fischeri*:1001
afum|EAL91753.1 100.00% afis|EAW17942.1 100.00%
Bootstrap support for afum|EAL91753.1 as seed ortholog is 100%.
Bootstrap support for afis|EAW17942.1 as seed ortholog is 100%.

---

Group of orthologs #4122. Best score 1001 bits
Score difference with first non-orthologous sequence - *A. fumigatus*:1001 *A. fischeri*:1001
afum|EAL87380.1 100.00% afis|EAW23952.1 100.00%
Bootstrap support for afum|EAL87380.1 as seed ortholog is 100%.

Bootstrap support for afis|EAW23952.1 as seed ortholog is 100%.

---

Group of orthologs #4123. Best score 1001 bits
Score difference with first non-orthologous sequence - *A. fumigatus*:1001 *A. fischeri*:1001
afum|EAL87903.1 100.00% afis|EAW23565.1 100.00%
Bootstrap support for afum|EAL87903.1 as seed ortholog is 100%.
Bootstrap support for afis|EAW23565.1 as seed ortholog is 100%.

---

Group of orthologs #4124. Best score 1001 bits
Score difference with first non-orthologous sequence - *A. fumigatus*:1001 *A. fischeri*:828
afum|EAL93743.1 100.00% afis|EAW19009.1 100.00%
Bootstrap support for afum|EAL93743.1 as seed ortholog is 100%.
Bootstrap support for afis|EAW19009.1 as seed ortholog is 100%.

---

Group of orthologs #4125. Best score 1001 bits
Score difference with first non-orthologous sequence - *A. fumigatus*:1001 *A. fischeri*:1001
afum|EAL92391.1 100.00% afis|EAW21374.1 100.00%
Bootstrap support for afum|EAL92391.1 as seed ortholog is 100%.
Bootstrap support for afis|EAW21374.1 as seed ortholog is 100%.

---

Group of orthologs #4126. Best score 1000 bits
Score difference with first non-orthologous sequence - *A. fumigatus*:1000 *A. fischeri*:1000
afum|EAL85634.1 100.00% afis|EAW15714.1 100.00%
Bootstrap support for afum|EAL85634.1 as seed ortholog is 100%.
Bootstrap support for afis|EAW15714.1 as seed ortholog is 100%.

---

Group of orthologs #4127. Best score 1000 bits
Score difference with first non-orthologous sequence - *A. fumigatus*:1000 *A. fischeri*:1000
afum|EAL87525.2 100.00% afis|EAW18165.1 100.00%
Bootstrap support for afum|EAL87525.2 as seed ortholog is 100%.
Bootstrap support for afis|EAW18165.1 as seed ortholog is 100%.

---

Group of orthologs #4128. Best score 1000 bits
Score difference with first non-orthologous sequence - *A. fumigatus*:139 *A. fischeri*:604
afum|EAL90432.1 100.00% afis|EAW16867.1 100.00%
Bootstrap support for afum|EAL90432.1 as seed ortholog is 99%.
Bootstrap support for afis|EAW16867.1 as seed ortholog is 100%.

---

Group of orthologs #4129. Best score 1000 bits
Score difference with first non-orthologous sequence - *A. fumigatus*:1000 *A. fischeri*:1000
afum|EAL93204.1 100.00% afis|EAW18509.1 100.00%
Bootstrap support for afum|EAL93204.1 as seed ortholog is 100%.
Bootstrap support for afis|EAW18509.1 as seed ortholog is 100%.

---

Group of orthologs #4130. Best score 1000 bits
Score difference with first non-orthologous sequence - *A. fumigatus*:1000 *A. fischeri*:1000
afum|EAL88438.1 100.00% afis|EAW23043.1 100.00%
Bootstrap support for afum|EAL88438.1 as seed ortholog is 100%.
Bootstrap support for afis|EAW23043.1 as seed ortholog is 100%.

---

Group of orthologs #4131. Best score 1000 bits
Score difference with first non-orthologous sequence - *A. fumigatus*:1000 *A. fischeri*:1000
afum|EAL88182.1 100.00% afis|EAW23296.1 100.00%
Bootstrap support for afum|EAL88182.1 as seed ortholog is 100%.
Bootstrap support for afis|EAW23296.1 as seed ortholog is 100%.

---

Group of orthologs #4132. Best score 1000 bits
Score difference with first non-orthologous sequence - *A. fumigatus*:1000 *A. fischeri*:1000
afum|EAL93799.1 100.00% afis|EAW19071.1 100.00%
Bootstrap support for afum|EAL93799.1 as seed ortholog is 100%.
Bootstrap support for afis|EAW19071.1 as seed ortholog is 100%.

---

Group of orthologs #4133. Best score 1000 bits
Score difference with first non-orthologous sequence - *A. fumigatus*:1000 *A. fischeri*:1000
afum|EAL92528.1 100.00% afis|EAW21508.1 100.00%
Bootstrap support for afum|EAL92528.1 as seed ortholog is 100%.
Bootstrap support for afis|EAW21508.1 as seed ortholog is 100%.

---

Group of orthologs #4134. Best score 1000 bits
Score difference with first non-orthologous sequence - *A. fumigatus*:1000 *A. fischeri*:1000
afum|EAL89492.1 100.00% afis|EAW24846.1 100.00%
Bootstrap support for afum|EAL89492.1 as seed ortholog is 100%.
Bootstrap support for afis|EAW24846.1 as seed ortholog is 100%.

---

Group of orthologs #4135. Best score 1000 bits
Score difference with first non-orthologous sequence - *A. fumigatus*:1000 *A. fischeri*:332
afum|EAL90047.1 100.00% afis|EAW25388.1 100.00%
Bootstrap support for afum|EAL90047.1 as seed ortholog is 100%.
Bootstrap support for afis|EAW25388.1 as seed ortholog is 100%.

---

Group of orthologs #4136. Best score 999 bits
Score difference with first non-orthologous sequence - *A. fumigatus*:999 *A. fischeri*:442
afum|EAL85740.1 100.00% afis|EAW15592.1 100.00%
Bootstrap support for afum|EAL85740.1 as seed ortholog is 100%.
Bootstrap support for afis|EAW15592.1 as seed ortholog is 100%.

---

Group of orthologs #4137. Best score 999 bits
Score difference with first non-orthologous sequence - *A. fumigatus*:999 *A. fischeri*:999
afum|EAL88588.1 100.00% afis|EAW15955.1 100.00%
Bootstrap support for afum|EAL88588.1 as seed ortholog is 100%.
Bootstrap support for afis|EAW15955.1 as seed ortholog is 100%.

---

Group of orthologs #4138. Best score 999 bits
Score difference with first non-orthologous sequence - *A. fumigatus*:999 *A. fischeri*:826
afum|EAL84879.1 100.00% afis|EAW20729.1 100.00%
Bootstrap support for afum|EAL84879.1 as seed ortholog is 100%.
Bootstrap support for afis|EAW20729.1 as seed ortholog is 100%.

---

Group of orthologs #4139. Best score 999 bits
Score difference with first non-orthologous sequence - *A. fumigatus*:999 *A. fischeri*:999
afum|EAL87667.2 100.00% afis|EAW18301.1 100.00%
Bootstrap support for afum|EAL87667.2 as seed ortholog is 100%.
Bootstrap support for afis|EAW18301.1 as seed ortholog is 100%.

---

Group of orthologs #4140. Best score 999 bits
Score difference with first non-orthologous sequence - *A. fumigatus*:999 *A. fischeri*:999
afum|EAL86947.1 100.00% afis|EAW19569.1 100.00%
Bootstrap support for afum|EAL86947.1 as seed ortholog is 100%.
Bootstrap support for afis|EAW19569.1 as seed ortholog is 100%.

---

Group of orthologs #4141. Best score 999 bits
Score difference with first non-orthologous sequence - *A. fumigatus*:999 *A. fischeri*:999

afum|EAL87111.1 100.00% afis|EAW23666.1 100.00%
Bootstrap support for afum|EAL87111.1 as seed ortholog is 100%.
Bootstrap support for afis|EAW23666.1 as seed ortholog is 100%.

---

Group of orthologs #4142. Best score 999 bits
Score difference with first non-orthologous sequence - *A. fumigatus*:999 *A. fischeri*:999
afum|EAL87181.1 100.00% afis|EAW23746.1 100.00%
Bootstrap support for afum|EAL87181.1 as seed ortholog is 100%.
Bootstrap support for afis|EAW23746.1 as seed ortholog is 100%.

---

Group of orthologs #4143. Best score 999 bits
Score difference with first non-orthologous sequence - *A. fumigatus*:999 *A. fischeri*:999
afum|EAL92147.1 100.00% afis|EAW21124.1 100.00%
Bootstrap support for afum|EAL92147.1 as seed ortholog is 100%.
Bootstrap support for afis|EAW21124.1 as seed ortholog is 100%.

---

Group of orthologs #4144. Best score 999 bits
Score difference with first non-orthologous sequence - *A. fumigatus*:999 *A. fischeri*:999
afum|EAL92169.1 100.00% afis|EAW21150.1 100.00%
Bootstrap support for afum|EAL92169.1 as seed ortholog is 100%.
Bootstrap support for afis|EAW21150.1 as seed ortholog is 100%.

---

Group of orthologs #4145. Best score 999 bits
Score difference with first non-orthologous sequence - *A. fumigatus*:911 *A. fischeri*:999
afum|EAL92260.1 100.00% afis|EAW21247.1 100.00%
Bootstrap support for afum|EAL92260.1 as seed ortholog is 100%.
Bootstrap support for afis|EAW21247.1 as seed ortholog is 100%.

---

Group of orthologs #4146. Best score 999 bits
Score difference with first non-orthologous sequence - *A. fumigatus*:999 *A. fischeri*:999
afum|EAL91906.2 100.00% afis|EAW24046.1 100.00%
Bootstrap support for afum|EAL91906.2 as seed ortholog is 100%.
Bootstrap support for afis|EAW24046.1 as seed ortholog is 100%.

---

Group of orthologs #4147. Best score 999 bits
Score difference with first non-orthologous sequence - *A. fumigatus*:999 *A. fischeri*:999
afum|EAL91919.1 100.00% afis|EAW24064.1 100.00%
Bootstrap support for afum|EAL91919.1 as seed ortholog is 100%.
Bootstrap support for afis|EAW24064.1 as seed ortholog is 100%.

---

Group of orthologs #4148. Best score 998 bits
Score difference with first non-orthologous sequence - *A. fumigatus*:998 *A. fischeri*:998
afum|EAL85583.1 100.00% afis|EAW15765.1 100.00%
Bootstrap support for afum|EAL85583.1 as seed ortholog is 100%.
Bootstrap support for afis|EAW15765.1 as seed ortholog is 100%.

---

Group of orthologs #4149. Best score 998 bits
Score difference with first non-orthologous sequence - *A. fumigatus*:998 *A. fischeri*:998
afum|EAL91458.1 100.00% afis|EAW17643.1 100.00%
Bootstrap support for afum|EAL91458.1 as seed ortholog is 100%.
Bootstrap support for afis|EAW17643.1 as seed ortholog is 100%.

---

Group of orthologs #4150. Best score 997 bits
Score difference with first non-orthologous sequence - *A. fumigatus*:997 *A. fischeri*:997
afum|EAL88535.1 100.00% afis|EAW15900.1 100.00%
Bootstrap support for afum|EAL88535.1 as seed ortholog is 100%.

Bootstrap support for afis|EAW15900.1 as seed ortholog is 100%.

---

Group of orthologs #4151. Best score 997 bits
Score difference with first non-orthologous sequence - *A. fumigatus*:997 *A. fischeri*:997
afum|EAL87668.2 100.00% afis|EAW18302.1 100.00%
Bootstrap support for afum|EAL87668.2 as seed ortholog is 100%.
Bootstrap support for afis|EAW18302.1 as seed ortholog is 100%.

---

Group of orthologs #4152. Best score 997 bits
Score difference with first non-orthologous sequence - *A. fumigatus*:997 *A. fischeri*:803
afum|EAL86990.1 100.00% afis|EAW19503.1 100.00%
Bootstrap support for afum|EAL86990.1 as seed ortholog is 100%.
Bootstrap support for afis|EAW19503.1 as seed ortholog is 100%.

---

Group of orthologs #4153. Best score 997 bits
Score difference with first non-orthologous sequence - *A. fumigatus*:997 *A. fischeri*:997
afum|EBA27427.1 100.00% afis|EAW17386.1 100.00%
Bootstrap support for afum|EBA27427.1 as seed ortholog is 100%.
Bootstrap support for afis|EAW17386.1 as seed ortholog is 100%.

---

Group of orthologs #4154. Best score 997 bits
Score difference with first non-orthologous sequence - *A. fumigatus*:997 *A. fischeri*:997
afum|EAL93755.1 100.00% afis|EAW19024.1 100.00%
Bootstrap support for afum|EAL93755.1 as seed ortholog is 100%.
Bootstrap support for afis|EAW19024.1 as seed ortholog is 100%.

---

Group of orthologs #4155. Best score 997 bits
Score difference with first non-orthologous sequence - *A. fumigatus*:997 *A. fischeri*:997
afum|EAL92683.1 100.00% afis|EAW21649.1 100.00%
Bootstrap support for afum|EAL92683.1 as seed ortholog is 100%.
Bootstrap support for afis|EAW21649.1 as seed ortholog is 100%.

---

Group of orthologs #4156. Best score 996 bits
Score difference with first non-orthologous sequence - *A. fumigatus*:996 *A. fischeri*:996
afum|EAL88737.1 100.00% afis|EAW16105.1 100.00%
Bootstrap support for afum|EAL88737.1 as seed ortholog is 100%.
Bootstrap support for afis|EAW16105.1 as seed ortholog is 100%.

---

Group of orthologs #4157. Best score 996 bits
Score difference with first non-orthologous sequence - *A. fumigatus*:996 *A. fischeri*:996
afum|EAL84750.1 100.00% afis|EAW20882.1 100.00%
Bootstrap support for afum|EAL84750.1 as seed ortholog is 100%.
Bootstrap support for afis|EAW20882.1 as seed ortholog is 100%.

---

Group of orthologs #4158. Best score 996 bits
Score difference with first non-orthologous sequence - *A. fumigatus*:996 *A. fischeri*:996
afum|EAL91770.1 100.00% afis|EAW17959.1 100.00%
Bootstrap support for afum|EAL91770.1 as seed ortholog is 100%.
Bootstrap support for afis|EAW17959.1 as seed ortholog is 100%.

---

Group of orthologs #4159. Best score 996 bits
Score difference with first non-orthologous sequence - *A. fumigatus*:996 *A. fischeri*:855
afum|EAL88149.2 100.00% afis|EAW23330.1 100.00%
Bootstrap support for afum|EAL88149.2 as seed ortholog is 100%.
Bootstrap support for afis|EAW23330.1 as seed ortholog is 100%.

---

Group of orthologs #4160. Best score 996 bits
Score difference with first non-orthologous sequence - *A. fumigatus*:996 *A. fischeri*:996
afum|EAL92173.1 100.00% afis|EAW21155.1 100.00%
Bootstrap support for afum|EAL92173.1 as seed ortholog is 100%.
Bootstrap support for afis|EAW21155.1 as seed ortholog is 100%.

---

Group of orthologs #4161. Best score 996 bits
Score difference with first non-orthologous sequence - *A. fumigatus*:893 *A. fischeri*:920
afum|EAL89408.2 100.00% afis|EAW24766.1 100.00%
Bootstrap support for afum|EAL89408.2 as seed ortholog is 100%.
Bootstrap support for afis|EAW24766.1 as seed ortholog is 100%.

---

Group of orthologs #4162. Best score 996 bits
Score difference with first non-orthologous sequence - *A. fumigatus*:996 *A. fischeri*:996
afum|EAL89576.1 100.00% afis|EAW24921.1 100.00%
Bootstrap support for afum|EAL89576.1 as seed ortholog is 100%.
Bootstrap support for afis|EAW24921.1 as seed ortholog is 100%.

---

Group of orthologs #4163. Best score 996 bits
Score difference with first non-orthologous sequence - *A. fumigatus*:996 *A. fischeri*:996
afum|EAL89978.2 100.00% afis|EAW25317.1 100.00%
Bootstrap support for afum|EAL89978.2 as seed ortholog is 100%.
Bootstrap support for afis|EAW25317.1 as seed ortholog is 100%.

---

Group of orthologs #4164. Best score 996 bits
Score difference with first non-orthologous sequence - *A. fumigatus*:996 *A. fischeri*:996
afum|EAL90115.2 100.00% afis|EAW25456.1 100.00%
Bootstrap support for afum|EAL90115.2 as seed ortholog is 100%.
Bootstrap support for afis|EAW25456.1 as seed ortholog is 100%.

---

Group of orthologs #4165. Best score 995 bits
Score difference with first non-orthologous sequence - *A. fumigatus*:995 *A. fischeri*:876
afum|EAL85107.2 100.00% afis|EAW19787.1 100.00%
Bootstrap support for afum|EAL85107.2 as seed ortholog is 100%.
Bootstrap support for afis|EAW19787.1 as seed ortholog is 100%.

---

Group of orthologs #4166. Best score 995 bits
Score difference with first non-orthologous sequence - *A. fumigatus*:995 *A. fischeri*:821
afum|EAL91737.1 100.00% afis|EAW17926.1 100.00%
Bootstrap support for afum|EAL91737.1 as seed ortholog is 100%.
Bootstrap support for afis|EAW17926.1 as seed ortholog is 100%.

---

Group of orthologs #4167. Best score 995 bits
Score difference with first non-orthologous sequence - *A. fumigatus*:767 *A. fischeri*:535
afum|EAL88228.1 100.00% afis|EAW23253.1 100.00%
Bootstrap support for afum|EAL88228.1 as seed ortholog is 100%.
Bootstrap support for afis|EAW23253.1 as seed ortholog is 100%.

---

Group of orthologs #4168. Best score 995 bits
Score difference with first non-orthologous sequence - *A. fumigatus*:995 *A. fischeri*:927
afum|EAL93892.2 100.00% afis|EAW19162.1 100.00%
Bootstrap support for afum|EAL93892.2 as seed ortholog is 100%.
Bootstrap support for afis|EAW19162.1 as seed ortholog is 100%.

---

Group of orthologs #4169. Best score 995 bits
Score difference with first non-orthologous sequence - *A. fumigatus*:995 *A. fischeri*:995

afum|EAL92142.2 100.00% afis|EAW21118.1 100.00%
Bootstrap support for afum|EAL92142.2 as seed ortholog is 100%.
Bootstrap support for afis|EAW21118.1 as seed ortholog is 100%.

---

Group of orthologs #4170. Best score 995 bits
Score difference with first non-orthologous sequence - *A. fumigatus*:995 *A. fischeri*:995
afum|EAL89695.1 100.00% afis|EAW25003.1 100.00%
Bootstrap support for afum|EAL89695.1 as seed ortholog is 100%.
Bootstrap support for afis|EAW25003.1 as seed ortholog is 100%.

---

Group of orthologs #4171. Best score 994 bits
Score difference with first non-orthologous sequence - *A. fumigatus*:994 *A. fischeri*:994
afum|EAL93195.1 100.00% afis|EAW18502.1 100.00%
Bootstrap support for afum|EAL93195.1 as seed ortholog is 100%.
Bootstrap support for afis|EAW18502.1 as seed ortholog is 100%.

---

Group of orthologs #4172. Best score 994 bits
Score difference with first non-orthologous sequence - *A. fumigatus*:994 *A. fischeri*:994
afum|EAL92001.1 100.00% afis|EAW24146.1 100.00%
Bootstrap support for afum|EAL92001.1 as seed ortholog is 100%.
Bootstrap support for afis|EAW24146.1 as seed ortholog is 100%.

---

Group of orthologs #4173. Best score 993 bits
Score difference with first non-orthologous sequence - *A. fumigatus*:993 *A. fischeri*:993
afum|EAL86311.1 100.00% afis|EAW16736.1 100.00%
Bootstrap support for afum|EAL86311.1 as seed ortholog is 100%.
Bootstrap support for afis|EAW16736.1 as seed ortholog is 100%.

---

Group of orthologs #4174. Best score 993 bits
Score difference with first non-orthologous sequence - *A. fumigatus*:993 *A. fischeri*:993
afum|EAL89065.1 100.00% afis|EAW16471.1 100.00%
Bootstrap support for afum|EAL89065.1 as seed ortholog is 100%.
Bootstrap support for afis|EAW16471.1 as seed ortholog is 100%.

---

Group of orthologs #4175. Best score 993 bits
Score difference with first non-orthologous sequence - *A. fumigatus*:993 *A. fischeri*:993
afum|EAL91791.1 100.00% afis|EAW17982.1 100.00%
Bootstrap support for afum|EAL91791.1 as seed ortholog is 100%.
Bootstrap support for afis|EAW17982.1 as seed ortholog is 100%.

---

Group of orthologs #4176. Best score 993 bits
Score difference with first non-orthologous sequence - *A. fumigatus*:993 *A. fischeri*:993
afum|EAL88322.1 100.00% afis|EAW23161.1 100.00%
Bootstrap support for afum|EAL88322.1 as seed ortholog is 100%.
Bootstrap support for afis|EAW23161.1 as seed ortholog is 100%.

---

Group of orthologs #4177. Best score 993 bits
Score difference with first non-orthologous sequence - *A. fumigatus*:993 *A. fischeri*:993
afum|EAL92484.1 100.00% afis|EAW21464.1 100.00%
Bootstrap support for afum|EAL92484.1 as seed ortholog is 100%.
Bootstrap support for afis|EAW21464.1 as seed ortholog is 100%.

---

Group of orthologs #4178. Best score 992 bits
Score difference with first non-orthologous sequence - *A. fumigatus*:992 *A. fischeri*:992
afum|EAL84515.1 100.00% afis|EAW20543.1 100.00%
Bootstrap support for afum|EAL84515.1 as seed ortholog is 100%.

Bootstrap support for afis|EAW20543.1 as seed ortholog is 100%.

---

Group of orthologs #4179. Best score 992 bits
Score difference with first non-orthologous sequence - *A. fumigatus*:992 *A. fischeri*:992
afum|EAL89183.1 100.00% afis|EAW16588.1 100.00%
Bootstrap support for afum|EAL89183.1 as seed ortholog is 100%.
Bootstrap support for afis|EAW16588.1 as seed ortholog is 100%.

---

Group of orthologs #4180. Best score 992 bits
Score difference with first non-orthologous sequence - *A. fumigatus*:910 *A. fischeri*:992
afum|EAL89287.1 100.00% afis|EAW16683.1 100.00%
Bootstrap support for afum|EAL89287.1 as seed ortholog is 100%.
Bootstrap support for afis|EAW16683.1 as seed ortholog is 100%.

---

Group of orthologs #4181. Best score 992 bits
Score difference with first non-orthologous sequence - *A. fumigatus*:992 *A. fischeri*:992
afum|EAL86970.2 100.00% afis|EAW19536.1 100.00%
Bootstrap support for afum|EAL86970.2 as seed ortholog is 100%.
Bootstrap support for afis|EAW19536.1 as seed ortholog is 100%.

---

Group of orthologs #4182. Best score 992 bits
Score difference with first non-orthologous sequence - *A. fumigatus*:992 *A. fischeri*:992
afum|EAL91472.1 100.00% afis|EAW17657.1 100.00%
Bootstrap support for afum|EAL91472.1 as seed ortholog is 100%.
Bootstrap support for afis|EAW17657.1 as seed ortholog is 100%.

---

Group of orthologs #4183. Best score 992 bits
Score difference with first non-orthologous sequence - *A. fumigatus*:992 *A. fischeri*:992
afum|EAL87261.1 100.00% afis|EAW23829.1 100.00%
Bootstrap support for afum|EAL87261.1 as seed ortholog is 100%.
Bootstrap support for afis|EAW23829.1 as seed ortholog is 100%.

---

Group of orthologs #4184. Best score 992 bits
Score difference with first non-orthologous sequence - *A. fumigatus*:992 *A. fischeri*:992
afum|EAL88265.1 100.00% afis|EAW23219.1 100.00%
Bootstrap support for afum|EAL88265.1 as seed ortholog is 100%.
Bootstrap support for afis|EAW23219.1 as seed ortholog is 100%.

---

Group of orthologs #4185. Best score 992 bits
Score difference with first non-orthologous sequence - *A. fumigatus*:992 *A. fischeri*:992
afum|EAL92711.1 100.00% afis|EAW21677.1 100.00%
Bootstrap support for afum|EAL92711.1 as seed ortholog is 100%.
Bootstrap support for afis|EAW21677.1 as seed ortholog is 100%.

---

Group of orthologs #4186. Best score 991 bits
Score difference with first non-orthologous sequence - *A. fumigatus*:991 *A. fischeri*:844
afum|EAL85041.1 100.00% afis|EAW19854.1 100.00%
afis|EAW24644.1 6.03%
Bootstrap support for afum|EAL85041.1 as seed ortholog is 100%.
Bootstrap support for afis|EAW19854.1 as seed ortholog is 100%.

---

Group of orthologs #4187. Best score 991 bits
Score difference with first non-orthologous sequence - *A. fumigatus*:991 *A. fischeri*:991
afum|EAL85765.1 100.00% afis|EAW15568.1 100.00%
Bootstrap support for afum|EAL85765.1 as seed ortholog is 100%.
Bootstrap support for afis|EAW15568.1 as seed ortholog is 100%.

---

Group of orthologs #4188. Best score 991 bits
Score difference with first non-orthologous sequence - *A. fumigatus*:991 *A. fischeri*:782
afum|EAL86251.1 100.00% afis|EAW24623.1 100.00%
Bootstrap support for afum|EAL86251.1 as seed ortholog is 100%.
Bootstrap support for afis|EAW24623.1 as seed ortholog is 100%.

---

Group of orthologs #4189. Best score 991 bits
Score difference with first non-orthologous sequence - *A. fumigatus*:991 *A. fischeri*:991
afum|EAL93798.1 100.00% afis|EAW19070.1 100.00%
Bootstrap support for afum|EAL93798.1 as seed ortholog is 100%.
Bootstrap support for afis|EAW19070.1 as seed ortholog is 100%.

---

Group of orthologs #4190. Best score 991 bits
Score difference with first non-orthologous sequence - *A. fumigatus*:991 *A. fischeri*:991
afum|EAL91048.2 100.00% afis|EAW22142.1 100.00%
Bootstrap support for afum|EAL91048.2 as seed ortholog is 100%.
Bootstrap support for afis|EAW22142.1 as seed ortholog is 100%.

---

Group of orthologs #4191. Best score 990 bits
Score difference with first non-orthologous sequence - *A. fumigatus*:990 *A. fischeri*:497
afum|EAL85513.1 100.00% afis|EAW15838.1 100.00%
Bootstrap support for afum|EAL85513.1 as seed ortholog is 100%.
Bootstrap support for afis|EAW15838.1 as seed ortholog is 100%.

---

Group of orthologs #4192. Best score 990 bits
Score difference with first non-orthologous sequence - *A. fumigatus*:905 *A. fischeri*:584
afum|EAL86483.2 100.00% afis|EAW16956.1 100.00%
Bootstrap support for afum|EAL86483.2 as seed ortholog is 100%.
Bootstrap support for afis|EAW16956.1 as seed ortholog is 100%.

---

Group of orthologs #4193. Best score 990 bits
Score difference with first non-orthologous sequence - *A. fumigatus*:990 *A. fischeri*:871
afum|EAL88838.1 100.00% afis|EAW16211.1 100.00%
Bootstrap support for afum|EAL88838.1 as seed ortholog is 100%.
Bootstrap support for afis|EAW16211.1 as seed ortholog is 100%.

---

Group of orthologs #4194. Best score 990 bits
Score difference with first non-orthologous sequence - *A. fumigatus*:832 *A. fischeri*:736
afum|EAL89035.1 100.00% afis|EAW16411.1 100.00%
Bootstrap support for afum|EAL89035.1 as seed ortholog is 100%.
Bootstrap support for afis|EAW16411.1 as seed ortholog is 100%.

---

Group of orthologs #4195. Best score 990 bits
Score difference with first non-orthologous sequence - *A. fumigatus*:990 *A. fischeri*:990
afum|EAL85170.2 100.00% afis|EAW20049.1 100.00%
Bootstrap support for afum|EAL85170.2 as seed ortholog is 100%.
Bootstrap support for afis|EAW20049.1 as seed ortholog is 100%.

---

Group of orthologs #4196. Best score 990 bits
Score difference with first non-orthologous sequence - *A. fumigatus*:990 *A. fischeri*:990
afum|EAL90498.1 100.00% afis|EAW22695.1 100.00%
Bootstrap support for afum|EAL90498.1 as seed ortholog is 100%.
Bootstrap support for afis|EAW22695.1 as seed ortholog is 100%.

---

Group of orthologs #4197. Best score 990 bits

Score difference with first non-orthologous sequence - *A. fumigatus*:990 *A. fischeri*:876
afum|EBA27456.1 100.00% afis|EAW21226.1 100.00%
Bootstrap support for afum|EBA27456.1 as seed ortholog is 100%.
Bootstrap support for afis|EAW21226.1 as seed ortholog is 100%.

---

Group of orthologs #4198. Best score 990 bits
Score difference with first non-orthologous sequence - *A. fumigatus*:990 *A. fischeri*:990
afum|EAL90170.1 100.00% afis|EAW25504.1 100.00%
Bootstrap support for afum|EAL90170.1 as seed ortholog is 100%.
Bootstrap support for afis|EAW25504.1 as seed ortholog is 100%.

---

Group of orthologs #4199. Best score 989 bits
Score difference with first non-orthologous sequence - *A. fumigatus*:989 *A. fischeri*:989
afum|EAL88649.1 100.00% afis|EAW16020.1 100.00%
Bootstrap support for afum|EAL88649.1 as seed ortholog is 100%.
Bootstrap support for afis|EAW16020.1 as seed ortholog is 100%.

---

Group of orthologs #4200. Best score 989 bits
Score difference with first non-orthologous sequence - *A. fumigatus*:989 *A. fischeri*:989
afum|EAL84322.1 100.00% afis|EAW25634.1 100.00%
Bootstrap support for afum|EAL84322.1 as seed ortholog is 100%.
Bootstrap support for afis|EAW25634.1 as seed ortholog is 100%.

---

Group of orthologs #4201. Best score 989 bits
Score difference with first non-orthologous sequence - *A. fumigatus*:701 *A. fischeri*:640
afum|EAL86205.2 100.00% afis|EAW24543.1 100.00%
Bootstrap support for afum|EAL86205.2 as seed ortholog is 100%.
Bootstrap support for afis|EAW24543.1 as seed ortholog is 100%.

---

Group of orthologs #4202. Best score 989 bits
Score difference with first non-orthologous sequence - *A. fumigatus*:791 *A. fischeri*:815
afum|EAL87843.1 100.00% afis|EAW23619.1 100.00%
Bootstrap support for afum|EAL87843.1 as seed ortholog is 100%.
Bootstrap support for afis|EAW23619.1 as seed ortholog is 100%.

---

Group of orthologs #4203. Best score 989 bits
Score difference with first non-orthologous sequence - *A. fumigatus*:989 *A. fischeri*:989
afum|EAL94052.1 100.00% afis|EAW19335.1 100.00%
Bootstrap support for afum|EAL94052.1 as seed ortholog is 100%.
Bootstrap support for afis|EAW19335.1 as seed ortholog is 100%.

---

Group of orthologs #4204. Best score 989 bits
Score difference with first non-orthologous sequence - *A. fumigatus*:989 *A. fischeri*:989
afum|EAL92458.2 100.00% afis|EAW21441.1 100.00%
Bootstrap support for afum|EAL92458.2 as seed ortholog is 100%.
Bootstrap support for afis|EAW21441.1 as seed ortholog is 100%.

---

Group of orthologs #4205. Best score 989 bits
Score difference with first non-orthologous sequence - *A. fumigatus*:989 *A. fischeri*:989
afum|EBA27388.1 100.00% afis|EAW25193.1 100.00%
Bootstrap support for afum|EBA27388.1 as seed ortholog is 100%.
Bootstrap support for afis|EAW25193.1 as seed ortholog is 100%.

---

Group of orthologs #4206. Best score 989 bits
Score difference with first non-orthologous sequence - *A. fumigatus*:989 *A. fischeri*:989
afum|EBA27395.1 100.00% afis|EAW25512.1 100.00%

Bootstrap support for afum|EBA27395.1 as seed ortholog is 100%.
Bootstrap support for afis|EAW25512.1 as seed ortholog is 100%.

---

Group of orthologs #4207. Best score 988 bits
Score difference with first non-orthologous sequence - *A. fumigatus*:988 *A. fischeri*:988
afum|EAL85805.1 100.00% afis|EAW15523.1 100.00%
Bootstrap support for afum|EAL85805.1 as seed ortholog is 100%.
Bootstrap support for afis|EAW15523.1 as seed ortholog is 100%.

---

Group of orthologs #4208. Best score 988 bits
Score difference with first non-orthologous sequence - *A. fumigatus*:730 *A. fischeri*:554
afum|EAL88978.1 100.00% afis|EAW16353.1 100.00%
Bootstrap support for afum|EAL88978.1 as seed ortholog is 100%.
Bootstrap support for afis|EAW16353.1 as seed ortholog is 100%.

---

Group of orthologs #4209. Best score 988 bits
Score difference with first non-orthologous sequence - *A. fumigatus*:988 *A. fischeri*:988
afum|EAL84342.1 100.00% afis|EAW25670.1 100.00%
Bootstrap support for afum|EAL84342.1 as seed ortholog is 100%.
Bootstrap support for afis|EAW25670.1 as seed ortholog is 100%.

---

Group of orthologs #4210. Best score 988 bits
Score difference with first non-orthologous sequence - *A. fumigatus*:988 *A. fischeri*:988
afum|EAL86004.1 100.00% afis|EAW24334.1 100.00%
Bootstrap support for afum|EAL86004.1 as seed ortholog is 100%.
Bootstrap support for afis|EAW24334.1 as seed ortholog is 100%.

---

Group of orthologs #4211. Best score 988 bits
Score difference with first non-orthologous sequence - *A. fumigatus*:988 *A. fischeri*:988
afum|EAL88482.1 100.00% afis|EAW22998.1 100.00%
Bootstrap support for afum|EAL88482.1 as seed ortholog is 100%.
Bootstrap support for afis|EAW22998.1 as seed ortholog is 100%.

---

Group of orthologs #4212. Best score 988 bits
Score difference with first non-orthologous sequence - *A. fumigatus*:988 *A. fischeri*:988
afum|EAL93792.1 100.00% afis|EAW19063.1 100.00%
Bootstrap support for afum|EAL93792.1 as seed ortholog is 100%.
Bootstrap support for afis|EAW19063.1 as seed ortholog is 100%.

---

Group of orthologs #4213. Best score 988 bits
Score difference with first non-orthologous sequence - *A. fumigatus*:988 *A. fischeri*:988
afum|EAL92442.1 100.00% afis|EAW21427.1 100.00%
Bootstrap support for afum|EAL92442.1 as seed ortholog is 100%.
Bootstrap support for afis|EAW21427.1 as seed ortholog is 100%.

---

Group of orthologs #4214. Best score 987 bits
Score difference with first non-orthologous sequence - *A. fumigatus*:987 *A. fischeri*:987
afum|EAL85548.1 100.00% afis|EAW15802.1 100.00%
Bootstrap support for afum|EAL85548.1 as seed ortholog is 100%.
Bootstrap support for afis|EAW15802.1 as seed ortholog is 100%.

---

Group of orthologs #4215. Best score 987 bits
Score difference with first non-orthologous sequence - *A. fumigatus*:987 *A. fischeri*:987
afum|EAL89276.1 100.00% afis|EAW16671.1 100.00%
Bootstrap support for afum|EAL89276.1 as seed ortholog is 100%.
Bootstrap support for afis|EAW16671.1 as seed ortholog is 100%.

Group of orthologs #4216. Best score 987 bits
Score difference with first non-orthologous sequence - *A. fumigatus*:987 *A. fischeri*:937
afum|EAL91168.1 100.00% afis|EAW17331.1 100.00%
Bootstrap support for afum|EAL91168.1 as seed ortholog is 100%.
Bootstrap support for afis|EAW17331.1 as seed ortholog is 100%.

Group of orthologs #4217. Best score 987 bits
Score difference with first non-orthologous sequence - *A. fumigatus*:987 *A. fischeri*:987
afum|EAL88341.1 100.00% afis|EAW23140.1 100.00%
Bootstrap support for afum|EAL88341.1 as seed ortholog is 100%.
Bootstrap support for afis|EAW23140.1 as seed ortholog is 100%.

Group of orthologs #4218. Best score 987 bits
Score difference with first non-orthologous sequence - *A. fumigatus*:987 *A. fischeri*:987
afum|EAL88020.1 100.00% afis|EAW23444.1 100.00%
Bootstrap support for afum|EAL88020.1 as seed ortholog is 100%.
Bootstrap support for afis|EAW23444.1 as seed ortholog is 100%.

Group of orthologs #4219. Best score 987 bits
Score difference with first non-orthologous sequence - *A. fumigatus*:987 *A. fischeri*:987
afum|EAL93430.1 100.00% afis|EAW18707.1 100.00%
Bootstrap support for afum|EAL93430.1 as seed ortholog is 100%.
Bootstrap support for afis|EAW18707.1 as seed ortholog is 100%.

Group of orthologs #4220. Best score 987 bits
Score difference with first non-orthologous sequence - *A. fumigatus*:987 *A. fischeri*:987
afum|EAL92488.1 100.00% afis|EAW21469.1 100.00%
Bootstrap support for afum|EAL92488.1 as seed ortholog is 100%.
Bootstrap support for afis|EAW21469.1 as seed ortholog is 100%.

Group of orthologs #4221. Best score 987 bits
Score difference with first non-orthologous sequence - *A. fumigatus*:987 *A. fischeri*:987
afum|EAL92581.1 100.00% afis|EAW21562.1 100.00%
Bootstrap support for afum|EAL92581.1 as seed ortholog is 100%.
Bootstrap support for afis|EAW21562.1 as seed ortholog is 100%.

Group of orthologs #4222. Best score 987 bits
Score difference with first non-orthologous sequence - *A. fumigatus*:790 *A. fischeri*:987
afum|EAL92751.1 100.00% afis|EAW21717.1 100.00%
Bootstrap support for afum|EAL92751.1 as seed ortholog is 100%.
Bootstrap support for afis|EAW21717.1 as seed ortholog is 100%.

Group of orthologs #4223. Best score 987 bits
Score difference with first non-orthologous sequence - *A. fumigatus*:741 *A. fischeri*:987
afum|EAL92929.1 100.00% afis|EAW21917.1 100.00%
Bootstrap support for afum|EAL92929.1 as seed ortholog is 100%.
Bootstrap support for afis|EAW21917.1 as seed ortholog is 100%.

Group of orthologs #4224. Best score 987 bits
Score difference with first non-orthologous sequence - *A. fumigatus*:987 *A. fischeri*:987
afum|EBA27466.1 100.00% afis|EAW21556.1 100.00%
Bootstrap support for afum|EBA27466.1 as seed ortholog is 100%.
Bootstrap support for afis|EAW21556.1 as seed ortholog is 100%.

Group of orthologs #4225. Best score 986 bits

Score difference with first non-orthologous sequence - *A. fumigatus*:886   *A. fischeri*:986
afum|EAL86399.1 100.00% afis|EAW16856.1 100.00%
Bootstrap support for afum|EAL86399.1 as seed ortholog is 100%.
Bootstrap support for afis|EAW16856.1 as seed ortholog is 100%.

---

Group of orthologs #4226. Best score 986 bits
Score difference with first non-orthologous sequence - *A. fumigatus*:986   *A. fischeri*:986
afum|EAL87661.2 100.00% afis|EAW18295.1 100.00%
Bootstrap support for afum|EAL87661.2 as seed ortholog is 100%.
Bootstrap support for afis|EAW18295.1 as seed ortholog is 100%.

---

Group of orthologs #4227. Best score 986 bits
Score difference with first non-orthologous sequence - *A. fumigatus*:986   *A. fischeri*:986
afum|EAL91833.1 100.00% afis|EAW18021.1 100.00%
Bootstrap support for afum|EAL91833.1 as seed ortholog is 100%.
Bootstrap support for afis|EAW18021.1 as seed ortholog is 100%.

---

Group of orthologs #4228. Best score 986 bits
Score difference with first non-orthologous sequence - *A. fumigatus*:459   *A. fischeri*:986
afum|EAL84372.1 100.00% afis|EAW25697.1 100.00%
Bootstrap support for afum|EAL84372.1 as seed ortholog is 100%.
Bootstrap support for afis|EAW25697.1 as seed ortholog is 100%.

---

Group of orthologs #4229. Best score 985 bits
Score difference with first non-orthologous sequence - *A. fumigatus*:985   *A. fischeri*:985
afum|EAL85823.1 100.00% afis|EAW15500.1 100.00%
Bootstrap support for afum|EAL85823.1 as seed ortholog is 100%.
Bootstrap support for afis|EAW15500.1 as seed ortholog is 100%.

---

Group of orthologs #4230. Best score 985 bits
Score difference with first non-orthologous sequence - *A. fumigatus*:985   *A. fischeri*:985
afum|EAL88153.1 100.00% afis|EAW23326.1 100.00%
Bootstrap support for afum|EAL88153.1 as seed ortholog is 100%.
Bootstrap support for afis|EAW23326.1 as seed ortholog is 100%.

---

Group of orthologs #4231. Best score 985 bits
Score difference with first non-orthologous sequence - *A. fumigatus*:985   *A. fischeri*:985
afum|EAL92582.1 100.00% afis|EAW21563.1 100.00%
Bootstrap support for afum|EAL92582.1 as seed ortholog is 100%.
Bootstrap support for afis|EAW21563.1 as seed ortholog is 100%.

---

Group of orthologs #4232. Best score 984 bits
Score difference with first non-orthologous sequence - *A. fumigatus*:742   *A. fischeri*:800
afum|EAL88922.1 100.00% afis|EAW16300.1 100.00%
Bootstrap support for afum|EAL88922.1 as seed ortholog is 100%.
Bootstrap support for afis|EAW16300.1 as seed ortholog is 100%.

---

Group of orthologs #4233. Best score 984 bits
Score difference with first non-orthologous sequence - *A. fumigatus*:984   *A. fischeri*:849
afum|EAL85203.1 100.00% afis|EAW20084.1 100.00%
Bootstrap support for afum|EAL85203.1 as seed ortholog is 100%.
Bootstrap support for afis|EAW20084.1 as seed ortholog is 100%.

---

Group of orthologs #4234. Best score 984 bits
Score difference with first non-orthologous sequence - *A. fumigatus*:984   *A. fischeri*:984
afum|EAL92688.1 100.00% afis|EAW21653.1 100.00%

Bootstrap support for afum|EAL92688.1 as seed ortholog is 100%.
Bootstrap support for afis|EAW21653.1 as seed ortholog is 100%.

---

Group of orthologs #4235. Best score 984 bits
Score difference with first non-orthologous sequence - *A. fumigatus*:984 *A. fischeri*:984
afum|EAL92879.1 100.00% afis|EAW21865.1 100.00%
Bootstrap support for afum|EAL92879.1 as seed ortholog is 100%.
Bootstrap support for afis|EAW21865.1 as seed ortholog is 100%.

---

Group of orthologs #4236. Best score 983 bits
Score difference with first non-orthologous sequence - *A. fumigatus*:983 *A. fischeri*:983
afum|EAL85760.1 100.00% afis|EAW15574.1 100.00%
Bootstrap support for afum|EAL85760.1 as seed ortholog is 100%.
Bootstrap support for afis|EAW15574.1 as seed ortholog is 100%.

---

Group of orthologs #4237. Best score 983 bits
Score difference with first non-orthologous sequence - *A. fumigatus*:983 *A. fischeri*:983
afum|EAL86522.2 100.00% afis|EAW16997.1 100.00%
Bootstrap support for afum|EAL86522.2 as seed ortholog is 100%.
Bootstrap support for afis|EAW16997.1 as seed ortholog is 100%.

---

Group of orthologs #4238. Best score 983 bits
Score difference with first non-orthologous sequence - *A. fumigatus*:983 *A. fischeri*:983
afum|EAL85969.1 100.00% afis|EAW24301.1 100.00%
Bootstrap support for afum|EAL85969.1 as seed ortholog is 100%.
Bootstrap support for afis|EAW24301.1 as seed ortholog is 100%.

---

Group of orthologs #4239. Best score 983 bits
Score difference with first non-orthologous sequence - *A. fumigatus*:983 *A. fischeri*:983
afum|EAL86170.1 100.00% afis|EAW24500.1 100.00%
Bootstrap support for afum|EAL86170.1 as seed ortholog is 100%.
Bootstrap support for afis|EAW24500.1 as seed ortholog is 100%.

---

Group of orthologs #4240. Best score 983 bits
Score difference with first non-orthologous sequence - *A. fumigatus*:983 *A. fischeri*:983
afum|EAL87834.1 100.00% afis|EAW23626.1 100.00%
Bootstrap support for afum|EAL87834.1 as seed ortholog is 100%.
Bootstrap support for afis|EAW23626.1 as seed ortholog is 100%.

---

Group of orthologs #4241. Best score 983 bits
Score difference with first non-orthologous sequence - *A. fumigatus*:549 *A. fischeri*:430
afum|EAL89775.1 100.00% afis|EAW25104.1 100.00%
Bootstrap support for afum|EAL89775.1 as seed ortholog is 100%.
Bootstrap support for afis|EAW25104.1 as seed ortholog is 100%.

---

Group of orthologs #4242. Best score 982 bits
Score difference with first non-orthologous sequence - *A. fumigatus*:982 *A. fischeri*:982
afum|EAL94086.1 100.00% afis|EAW19367.1 100.00%
Bootstrap support for afum|EAL94086.1 as seed ortholog is 100%.
Bootstrap support for afis|EAW19367.1 as seed ortholog is 100%.

---

Group of orthologs #4243. Best score 982 bits
Score difference with first non-orthologous sequence - *A. fumigatus*:982 *A. fischeri*:982
afum|EAL92450.1 100.00% afis|EAW21435.1 100.00%
Bootstrap support for afum|EAL92450.1 as seed ortholog is 100%.
Bootstrap support for afis|EAW21435.1 as seed ortholog is 100%.

Group of orthologs #4244. Best score 982 bits
Score difference with first non-orthologous sequence - *A. fumigatus*:982 *A. fischeri*:982
afum|EAL89968.2 100.00% afis|EAW25305.1 100.00%
Bootstrap support for afum|EAL89968.2 as seed ortholog is 100%.
Bootstrap support for afis|EAW25305.1 as seed ortholog is 100%.

---

Group of orthologs #4245. Best score 981 bits
Score difference with first non-orthologous sequence - *A. fumigatus*:981 *A. fischeri*:981
afum|EAL88540.1 100.00% afis|EAW15905.1 100.00%
Bootstrap support for afum|EAL88540.1 as seed ortholog is 100%.
Bootstrap support for afis|EAW15905.1 as seed ortholog is 100%.

---

Group of orthologs #4246. Best score 981 bits
Score difference with first non-orthologous sequence - *A. fumigatus*:981 *A. fischeri*:908
afum|EAL89146.1 100.00% afis|EAW16548.1 100.00%
Bootstrap support for afum|EAL89146.1 as seed ortholog is 100%.
Bootstrap support for afis|EAW16548.1 as seed ortholog is 100%.

---

Group of orthologs #4247. Best score 981 bits
Score difference with first non-orthologous sequence - *A. fumigatus*:981 *A. fischeri*:981
afum|EAL91334.1 100.00% afis|EAW17516.1 100.00%
Bootstrap support for afum|EAL91334.1 as seed ortholog is 100%.
Bootstrap support for afis|EAW17516.1 as seed ortholog is 100%.

---

Group of orthologs #4248. Best score 981 bits
Score difference with first non-orthologous sequence - *A. fumigatus*:981 *A. fischeri*:981
afum|EAL88427.1 100.00% afis|EAW23053.1 100.00%
Bootstrap support for afum|EAL88427.1 as seed ortholog is 100%.
Bootstrap support for afis|EAW23053.1 as seed ortholog is 100%.

---

Group of orthologs #4249. Best score 981 bits
Score difference with first non-orthologous sequence - *A. fumigatus*:981 *A. fischeri*:913
afum|EAL92778.1 100.00% afis|EAW21746.1 100.00%
Bootstrap support for afum|EAL92778.1 as seed ortholog is 100%.
Bootstrap support for afis|EAW21746.1 as seed ortholog is 100%.

---

Group of orthologs #4250. Best score 981 bits
Score difference with first non-orthologous sequence - *A. fumigatus*:981 *A. fischeri*:981
afum|EBA27471.1 100.00% afis|EAW21747.1 100.00%
Bootstrap support for afum|EBA27471.1 as seed ortholog is 100%.
Bootstrap support for afis|EAW21747.1 as seed ortholog is 100%.

---

Group of orthologs #4251. Best score 980 bits
Score difference with first non-orthologous sequence - *A. fumigatus*:980 *A. fischeri*:980
afum|EAL86365.1 100.00% afis|EAW16796.1 100.00%
Bootstrap support for afum|EAL86365.1 as seed ortholog is 100%.
Bootstrap support for afis|EAW16796.1 as seed ortholog is 100%.

---

Group of orthologs #4252. Best score 980 bits
Score difference with first non-orthologous sequence - *A. fumigatus*:678 *A. fischeri*:980
afum|EAL91237.1 100.00% afis|EAW17399.1 100.00%
Bootstrap support for afum|EAL91237.1 as seed ortholog is 100%.
Bootstrap support for afis|EAW17399.1 as seed ortholog is 100%.

---

Group of orthologs #4253. Best score 980 bits

Score difference with first non-orthologous sequence - *A. fumigatus*:980 *A. fischeri*:884
afum|EAL88072.1 100.00% afis|EAW23398.1 100.00%
Bootstrap support for afum|EAL88072.1 as seed ortholog is 100%.
Bootstrap support for afis|EAW23398.1 as seed ortholog is 100%.

---

Group of orthologs #4254. Best score 980 bits
Score difference with first non-orthologous sequence - *A. fumigatus*:980 *A. fischeri*:980
afum|EAL93763.1 100.00% afis|EAW19034.1 100.00%
Bootstrap support for afum|EAL93763.1 as seed ortholog is 100%.
Bootstrap support for afis|EAW19034.1 as seed ortholog is 100%.

---

Group of orthologs #4255. Best score 980 bits
Score difference with first non-orthologous sequence - *A. fumigatus*:980 *A. fischeri*:980
afum|EAL90507.1 100.00% afis|EAW22686.1 100.00%
Bootstrap support for afum|EAL90507.1 as seed ortholog is 100%.
Bootstrap support for afis|EAW22686.1 as seed ortholog is 100%.

---

Group of orthologs #4256. Best score 980 bits
Score difference with first non-orthologous sequence - *A. fumigatus*:980 *A. fischeri*:980
afum|EAL90228.1 100.00% afis|EAW22969.1 100.00%
Bootstrap support for afum|EAL90228.1 as seed ortholog is 100%.
Bootstrap support for afis|EAW22969.1 as seed ortholog is 100%.

---

Group of orthologs #4257. Best score 980 bits
Score difference with first non-orthologous sequence - *A. fumigatus*:980 *A. fischeri*:980
afum|EAL89908.1 100.00% afis|EAW25246.1 100.00%
Bootstrap support for afum|EAL89908.1 as seed ortholog is 100%.
Bootstrap support for afis|EAW25246.1 as seed ortholog is 100%.

---

Group of orthologs #4258. Best score 979 bits
Score difference with first non-orthologous sequence - *A. fumigatus*:917 *A. fischeri*:979
afum|EAL84511.1 100.00% afis|EAW20548.1 100.00%
Bootstrap support for afum|EAL84511.1 as seed ortholog is 100%.
Bootstrap support for afis|EAW20548.1 as seed ortholog is 100%.

---

Group of orthologs #4259. Best score 979 bits
Score difference with first non-orthologous sequence - *A. fumigatus*:979 *A. fischeri*:979
afum|EAL93663.1 100.00% afis|EAW18931.1 100.00%
Bootstrap support for afum|EAL93663.1 as seed ortholog is 100%.
Bootstrap support for afis|EAW18931.1 as seed ortholog is 100%.

---

Group of orthologs #4260. Best score 979 bits
Score difference with first non-orthologous sequence - *A. fumigatus*:979 *A. fischeri*:979
afum|EAL90741.1 100.00% afis|EAW22450.1 100.00%
Bootstrap support for afum|EAL90741.1 as seed ortholog is 100%.
Bootstrap support for afis|EAW22450.1 as seed ortholog is 100%.

---

Group of orthologs #4261. Best score 979 bits
Score difference with first non-orthologous sequence - *A. fumigatus*:979 *A. fischeri*:979
afum|EBA27338.1 100.00% afis|EAW23317.1 100.00%
Bootstrap support for afum|EBA27338.1 as seed ortholog is 100%.
Bootstrap support for afis|EAW23317.1 as seed ortholog is 100%.

---

Group of orthologs #4262. Best score 978 bits
Score difference with first non-orthologous sequence - *A. fumigatus*:978 *A. fischeri*:978
afum|EAL86867.1 100.00% afis|EAW19655.1 100.00%

Bootstrap support for afum|EAL86867.1 as seed ortholog is 100%.
Bootstrap support for afis|EAW19655.1 as seed ortholog is 100%.

---

Group of orthologs #4263. Best score 978 bits
Score difference with first non-orthologous sequence - *A. fumigatus*:978 *A. fischeri*:978
afum|EAL91017.1 100.00% afis|EAW22172.1 100.00%
Bootstrap support for afum|EAL91017.1 as seed ortholog is 100%.
Bootstrap support for afis|EAW22172.1 as seed ortholog is 100%.

---

Group of orthologs #4264. Best score 978 bits
Score difference with first non-orthologous sequence - *A. fumigatus*:978 *A. fischeri*:978
afum|EAL90034.1 100.00% afis|EAW25375.1 100.00%
Bootstrap support for afum|EAL90034.1 as seed ortholog is 100%.
Bootstrap support for afis|EAW25375.1 as seed ortholog is 100%.

---

Group of orthologs #4265. Best score 977 bits
Score difference with first non-orthologous sequence - *A. fumigatus*:977 *A. fischeri*:931
afum|EAL88815.1 100.00% afis|EAW16184.1 100.00%
Bootstrap support for afum|EAL88815.1 as seed ortholog is 100%.
Bootstrap support for afis|EAW16184.1 as seed ortholog is 100%.

---

Group of orthologs #4266. Best score 977 bits
Score difference with first non-orthologous sequence - *A. fumigatus*:977 *A. fischeri*:803
afum|EAL86831.1 100.00% afis|EAW19696.1 100.00%
Bootstrap support for afum|EAL86831.1 as seed ortholog is 100%.
Bootstrap support for afis|EAW19696.1 as seed ortholog is 100%.

---

Group of orthologs #4267. Best score 977 bits
Score difference with first non-orthologous sequence - *A. fumigatus*:977 *A. fischeri*:917
afum|EAL86193.1 100.00% afis|EAW24528.1 100.00%
Bootstrap support for afum|EAL86193.1 as seed ortholog is 100%.
Bootstrap support for afis|EAW24528.1 as seed ortholog is 100%.

---

Group of orthologs #4268. Best score 977 bits
Score difference with first non-orthologous sequence - *A. fumigatus*:977 *A. fischeri*:977
afum|EAL93376.1 100.00% afis|EAW18660.1 100.00%
Bootstrap support for afum|EAL93376.1 as seed ortholog is 100%.
Bootstrap support for afis|EAW18660.1 as seed ortholog is 100%.

---

Group of orthologs #4269. Best score 977 bits
Score difference with first non-orthologous sequence - *A. fumigatus*:977 *A. fischeri*:977
afum|EAL90634.1 100.00% afis|EAW22561.1 100.00%
Bootstrap support for afum|EAL90634.1 as seed ortholog is 100%.
Bootstrap support for afis|EAW22561.1 as seed ortholog is 100%.

---

Group of orthologs #4270. Best score 977 bits
Score difference with first non-orthologous sequence - *A. fumigatus*:977 *A. fischeri*:977
afum|EAL90110.1 100.00% afis|EAW25451.1 100.00%
Bootstrap support for afum|EAL90110.1 as seed ortholog is 100%.
Bootstrap support for afis|EAW25451.1 as seed ortholog is 100%.

---

Group of orthologs #4271. Best score 976 bits
Score difference with first non-orthologous sequence - *A. fumigatus*:976 *A. fischeri*:976
afum|EAL86926.1 100.00% afis|EAW19592.1 100.00%
Bootstrap support for afum|EAL86926.1 as seed ortholog is 100%.
Bootstrap support for afis|EAW19592.1 as seed ortholog is 100%.

Group of orthologs #4272. Best score 976 bits
Score difference with first non-orthologous sequence - *A. fumigatus*:976 *A. fischeri*:976
afum|EAL91200.1 100.00% afis|EAW17363.1 100.00%
Bootstrap support for afum|EAL91200.1 as seed ortholog is 100%.
Bootstrap support for afis|EAW17363.1 as seed ortholog is 100%.

Group of orthologs #4273. Best score 976 bits
Score difference with first non-orthologous sequence - *A. fumigatus*:976 *A. fischeri*:976
afum|EAL91779.1 100.00% afis|EAW17970.1 100.00%
Bootstrap support for afum|EAL91779.1 as seed ortholog is 100%.
Bootstrap support for afis|EAW17970.1 as seed ortholog is 100%.

Group of orthologs #4274. Best score 976 bits
Score difference with first non-orthologous sequence - *A. fumigatus*:976 *A. fischeri*:553
afum|EAL86017.1 100.00% afis|EAW24347.1 100.00%
Bootstrap support for afum|EAL86017.1 as seed ortholog is 100%.
Bootstrap support for afis|EAW24347.1 as seed ortholog is 100%.

Group of orthologs #4275. Best score 976 bits
Score difference with first non-orthologous sequence - *A. fumigatus*:976 *A. fischeri*:976
afum|EAL89946.1 100.00% afis|EAW25283.1 100.00%
Bootstrap support for afum|EAL89946.1 as seed ortholog is 100%.
Bootstrap support for afis|EAW25283.1 as seed ortholog is 100%.

Group of orthologs #4276. Best score 976 bits
Score difference with first non-orthologous sequence - *A. fumigatus*:976 *A. fischeri*:976
afum|EAL90082.1 100.00% afis|EAW25423.1 100.00%
Bootstrap support for afum|EAL90082.1 as seed ortholog is 100%.
Bootstrap support for afis|EAW25423.1 as seed ortholog is 100%.

Group of orthologs #4277. Best score 976 bits
Score difference with first non-orthologous sequence - *A. fumigatus*:976 *A. fischeri*:879
afum|EAL90114.1 100.00% afis|EAW25455.1 100.00%
Bootstrap support for afum|EAL90114.1 as seed ortholog is 100%.
Bootstrap support for afis|EAW25455.1 as seed ortholog is 100%.

Group of orthologs #4278. Best score 975 bits
Score difference with first non-orthologous sequence - *A. fumigatus*:975 *A. fischeri*:975
afum|EAL85668.1 100.00% afis|EAW15680.1 100.00%
Bootstrap support for afum|EAL85668.1 as seed ortholog is 100%.
Bootstrap support for afis|EAW15680.1 as seed ortholog is 100%.

Group of orthologs #4279. Best score 975 bits
Score difference with first non-orthologous sequence - *A. fumigatus*:975 *A. fischeri*:975
afum|EAL93694.1 100.00% afis|EAW18962.1 100.00%
Bootstrap support for afum|EAL93694.1 as seed ortholog is 100%.
Bootstrap support for afis|EAW18962.1 as seed ortholog is 100%.

Group of orthologs #4280. Best score 974 bits
Score difference with first non-orthologous sequence - *A. fumigatus*:746 *A. fischeri*:815
afum|EAL86443.2 100.00% afis|EAW16906.1 100.00%
Bootstrap support for afum|EAL86443.2 as seed ortholog is 100%.
Bootstrap support for afis|EAW16906.1 as seed ortholog is 100%.

Group of orthologs #4281. Best score 974 bits

Score difference with first non-orthologous sequence - *A. fumigatus*:974 *A. fischeri*:974
afum|EAL86678.1 100.00% afis|EAW17261.1 100.00%
Bootstrap support for afum|EAL86678.1 as seed ortholog is 100%.
Bootstrap support for afis|EAW17261.1 as seed ortholog is 100%.

---

Group of orthologs #4282. Best score 974 bits
Score difference with first non-orthologous sequence - *A. fumigatus*:974 *A. fischeri*:974
afum|EAL88563.1 100.00% afis|EAW15927.1 100.00%
Bootstrap support for afum|EAL88563.1 as seed ortholog is 100%.
Bootstrap support for afis|EAW15927.1 as seed ortholog is 100%.

---

Group of orthologs #4283. Best score 974 bits
Score difference with first non-orthologous sequence - *A. fumigatus*:974 *A. fischeri*:610
afum|EAL84740.1 100.00% afis|EAW20893.1 100.00%
Bootstrap support for afum|EAL84740.1 as seed ortholog is 100%.
Bootstrap support for afis|EAW20893.1 as seed ortholog is 100%.

---

Group of orthologs #4284. Best score 974 bits
Score difference with first non-orthologous sequence - *A. fumigatus*:974 *A. fischeri*:974
afum|EAL91883.1 100.00% afis|EAW18071.1 100.00%
Bootstrap support for afum|EAL91883.1 as seed ortholog is 100%.
Bootstrap support for afis|EAW18071.1 as seed ortholog is 100%.

---

Group of orthologs #4285. Best score 974 bits
Score difference with first non-orthologous sequence - *A. fumigatus*:974 *A. fischeri*:974
afum|EAL89726.1 100.00% afis|EAW25058.1 100.00%
Bootstrap support for afum|EAL89726.1 as seed ortholog is 100%.
Bootstrap support for afis|EAW25058.1 as seed ortholog is 100%.

---

Group of orthologs #4286. Best score 973 bits
Score difference with first non-orthologous sequence - *A. fumigatus*:973 *A. fischeri*:973
afum|EAL85446.1 100.00% afis|EAW20318.1 100.00%
Bootstrap support for afum|EAL85446.1 as seed ortholog is 100%.
Bootstrap support for afis|EAW20318.1 as seed ortholog is 100%.

---

Group of orthologs #4287. Best score 973 bits
Score difference with first non-orthologous sequence - *A. fumigatus*:973 *A. fischeri*:973
afum|EAL91607.1 100.00% afis|EAW17799.1 100.00%
Bootstrap support for afum|EAL91607.1 as seed ortholog is 100%.
Bootstrap support for afis|EAW17799.1 as seed ortholog is 100%.

---

Group of orthologs #4288. Best score 973 bits
Score difference with first non-orthologous sequence - *A. fumigatus*:973 *A. fischeri*:973
afum|EAL90597.1 100.00% afis|EAW22601.1 100.00%
Bootstrap support for afum|EAL90597.1 as seed ortholog is 100%.
Bootstrap support for afis|EAW22601.1 as seed ortholog is 100%.

---

Group of orthologs #4289. Best score 973 bits
Score difference with first non-orthologous sequence - *A. fumigatus*:973 *A. fischeri*:973
afum|EAL89478.1 100.00% afis|EAW24833.1 100.00%
Bootstrap support for afum|EAL89478.1 as seed ortholog is 100%.
Bootstrap support for afis|EAW24833.1 as seed ortholog is 100%.

---

Group of orthologs #4290. Best score 972 bits
Score difference with first non-orthologous sequence - *A. fumigatus*:972 *A. fischeri*:972
afum|EAL85314.1 100.00% afis|EAW20194.1 100.00%

Bootstrap support for afum|EAL85314.1 as seed ortholog is 100%.
Bootstrap support for afis|EAW20194.1 as seed ortholog is 100%.

---

Group of orthologs #4291. Best score 972 bits
Score difference with first non-orthologous sequence - *A. fumigatus*:972 *A. fischeri*:639
afum|EAL84304.1 100.00% afis|EAW25598.1 100.00%
Bootstrap support for afum|EAL84304.1 as seed ortholog is 100%.
Bootstrap support for afis|EAW25598.1 as seed ortholog is 100%.

---

Group of orthologs #4292. Best score 972 bits
Score difference with first non-orthologous sequence - *A. fumigatus*:972 *A. fischeri*:972
afum|EBA27273.1 100.00% afis|EAW16941.1 100.00%
Bootstrap support for afum|EBA27273.1 as seed ortholog is 100%.
Bootstrap support for afis|EAW16941.1 as seed ortholog is 100%.

---

Group of orthologs #4293. Best score 972 bits
Score difference with first non-orthologous sequence - *A. fumigatus*:972 *A. fischeri*:972
afum|EAL90189.1 100.00% afis|EAW25524.1 100.00%
Bootstrap support for afum|EAL90189.1 as seed ortholog is 100%.
Bootstrap support for afis|EAW25524.1 as seed ortholog is 100%.

---

Group of orthologs #4294. Best score 971 bits
Score difference with first non-orthologous sequence - *A. fumigatus*:870 *A. fischeri*:884
afum|EAL86429.1 100.00% afis|EAW16891.1 100.00%
Bootstrap support for afum|EAL86429.1 as seed ortholog is 100%.
Bootstrap support for afis|EAW16891.1 as seed ortholog is 100%.

---

Group of orthologs #4295. Best score 971 bits
Score difference with first non-orthologous sequence - *A. fumigatus*:971 *A. fischeri*:927
afum|EAL86694.1 100.00% afis|EAW17276.1 100.00%
Bootstrap support for afum|EAL86694.1 as seed ortholog is 100%.
Bootstrap support for afis|EAW17276.1 as seed ortholog is 100%.

---

Group of orthologs #4296. Best score 971 bits
Score difference with first non-orthologous sequence - *A. fumigatus*:729 *A. fischeri*:971
afum|EAL88561.1 100.00% afis|EAW15925.1 100.00%
Bootstrap support for afum|EAL88561.1 as seed ortholog is 100%.
Bootstrap support for afis|EAW15925.1 as seed ortholog is 100%.

---

Group of orthologs #4297. Best score 971 bits
Score difference with first non-orthologous sequence - *A. fumigatus*:971 *A. fischeri*:971
afum|EAL88814.1 100.00% afis|EAW16183.1 100.00%
Bootstrap support for afum|EAL88814.1 as seed ortholog is 100%.
Bootstrap support for afis|EAW16183.1 as seed ortholog is 100%.

---

Group of orthologs #4298. Best score 971 bits
Score difference with first non-orthologous sequence - *A. fumigatus*:971 *A. fischeri*:971
afum|EAL91379.1 100.00% afis|EAW17565.1 100.00%
Bootstrap support for afum|EAL91379.1 as seed ortholog is 100%.
Bootstrap support for afis|EAW17565.1 as seed ortholog is 100%.

---

Group of orthologs #4299. Best score 971 bits
Score difference with first non-orthologous sequence - *A. fumigatus*:971 *A. fischeri*:922
afum|EAL84355.1 100.00% afis|EAW25686.1 100.00%
Bootstrap support for afum|EAL84355.1 as seed ortholog is 100%.
Bootstrap support for afis|EAW25686.1 as seed ortholog is 100%.

Group of orthologs #4300. Best score 971 bits
Score difference with first non-orthologous sequence - *A. fumigatus*:971 *A. fischeri*:971
afum|EAL88303.1 100.00% afis|EAW23180.1 100.00%
Bootstrap support for afum|EAL88303.1 as seed ortholog is 100%.
Bootstrap support for afis|EAW23180.1 as seed ortholog is 100%.

---

Group of orthologs #4301. Best score 971 bits
Score difference with first non-orthologous sequence - *A. fumigatus*:971 *A. fischeri*:971
afum|EAL93337.1 100.00% afis|EAW18625.1 100.00%
Bootstrap support for afum|EAL93337.1 as seed ortholog is 100%.
Bootstrap support for afis|EAW18625.1 as seed ortholog is 100%.

---

Group of orthologs #4302. Best score 971 bits
Score difference with first non-orthologous sequence - *A. fumigatus*:971 *A. fischeri*:971
afum|EAL93442.2 100.00% afis|EAW18718.1 100.00%
Bootstrap support for afum|EAL93442.2 as seed ortholog is 100%.
Bootstrap support for afis|EAW18718.1 as seed ortholog is 100%.

---

Group of orthologs #4303. Best score 971 bits
Score difference with first non-orthologous sequence - *A. fumigatus*:971 *A. fischeri*:971
afum|EAL93887.1 100.00% afis|EAW19156.1 100.00%
Bootstrap support for afum|EAL93887.1 as seed ortholog is 100%.
Bootstrap support for afis|EAW19156.1 as seed ortholog is 100%.

---

Group of orthologs #4304. Best score 971 bits
Score difference with first non-orthologous sequence - *A. fumigatus*:971 *A. fischeri*:971
afum|EAL93978.1 100.00% afis|EAW19249.1 100.00%
Bootstrap support for afum|EAL93978.1 as seed ortholog is 100%.
Bootstrap support for afis|EAW19249.1 as seed ortholog is 100%.

---

Group of orthologs #4305. Best score 970 bits
Score difference with first non-orthologous sequence - *A. fumigatus*:970 *A. fischeri*:970
afum|EAL85689.1 100.00% afis|EAW15657.1 100.00%
Bootstrap support for afum|EAL85689.1 as seed ortholog is 100%.
Bootstrap support for afis|EAW15657.1 as seed ortholog is 100%.

---

Group of orthologs #4306. Best score 970 bits
Score difference with first non-orthologous sequence - *A. fumigatus*:970 *A. fischeri*:970
afum|EAL85647.2 100.00% afis|EAW15701.1 100.00%
Bootstrap support for afum|EAL85647.2 as seed ortholog is 100%.
Bootstrap support for afis|EAW15701.1 as seed ortholog is 100%.

---

Group of orthologs #4307. Best score 970 bits
Score difference with first non-orthologous sequence - *A. fumigatus*:970 *A. fischeri*:970
afum|EAL91398.2 100.00% afis|EAW17585.1 100.00%
Bootstrap support for afum|EAL91398.2 as seed ortholog is 100%.
Bootstrap support for afis|EAW17585.1 as seed ortholog is 100%.

---

Group of orthologs #4308. Best score 970 bits
Score difference with first non-orthologous sequence - *A. fumigatus*:970 *A. fischeri*:970
afum|EAL89523.1 100.00% afis|EAW24873.1 100.00%
Bootstrap support for afum|EAL89523.1 as seed ortholog is 100%.
Bootstrap support for afis|EAW24873.1 as seed ortholog is 100%.

---

Group of orthologs #4309. Best score 970 bits

Score difference with first non-orthologous sequence - *A. fumigatus*:970 *A. fischeri*:970
afum|EBA27386.1 100.00% afis|EAW24985.1 100.00%
Bootstrap support for afum|EBA27386.1 as seed ortholog is 100%.
Bootstrap support for afis|EAW24985.1 as seed ortholog is 100%.

---

Group of orthologs #4310. Best score 969 bits
Score difference with first non-orthologous sequence - *A. fumigatus*:969 *A. fischeri*:969
afum|EAL87514.1 100.00% afis|EAW18155.1 100.00%
Bootstrap support for afum|EAL87514.1 as seed ortholog is 100%.
Bootstrap support for afis|EAW18155.1 as seed ortholog is 100%.

---

Group of orthologs #4311. Best score 969 bits
Score difference with first non-orthologous sequence - *A. fumigatus*:969 *A. fischeri*:969
afum|EBA27504.1 100.00% afis|EAW19096.1 100.00%
Bootstrap support for afum|EBA27504.1 as seed ortholog is 100%.
Bootstrap support for afis|EAW19096.1 as seed ortholog is 100%.

---

Group of orthologs #4312. Best score 969 bits
Score difference with first non-orthologous sequence - *A. fumigatus*:969 *A. fischeri*:969
afum|EAL89826.2 100.00% afis|EAW25151.1 100.00%
Bootstrap support for afum|EAL89826.2 as seed ortholog is 100%.
Bootstrap support for afis|EAW25151.1 as seed ortholog is 100%.

---

Group of orthologs #4313. Best score 968 bits
Score difference with first non-orthologous sequence - *A. fumigatus*:968 *A. fischeri*:920
afum|EAL86405.2 100.00% afis|EAW16866.1 100.00%
Bootstrap support for afum|EAL86405.2 as seed ortholog is 100%.
Bootstrap support for afis|EAW16866.1 as seed ortholog is 100%.

---

Group of orthologs #4314. Best score 968 bits
Score difference with first non-orthologous sequence - *A. fumigatus*:686 *A. fischeri*:752
afum|EAL87527.2 100.00% afis|EAW18167.1 100.00%
Bootstrap support for afum|EAL87527.2 as seed ortholog is 100%.
Bootstrap support for afis|EAW18167.1 as seed ortholog is 100%.

---

Group of orthologs #4315. Best score 968 bits
Score difference with first non-orthologous sequence - *A. fumigatus*:968 *A. fischeri*:968
afum|EAL86240.1 100.00% afis|EAW24610.1 100.00%
Bootstrap support for afum|EAL86240.1 as seed ortholog is 100%.
Bootstrap support for afis|EAW24610.1 as seed ortholog is 100%.

---

Group of orthologs #4316. Best score 968 bits
Score difference with first non-orthologous sequence - *A. fumigatus*:968 *A. fischeri*:968
afum|EAL90528.1 100.00% afis|EAW22664.1 100.00%
Bootstrap support for afum|EAL90528.1 as seed ortholog is 100%.
Bootstrap support for afis|EAW22664.1 as seed ortholog is 100%.

---

Group of orthologs #4317. Best score 968 bits
Score difference with first non-orthologous sequence - *A. fumigatus*:968 *A. fischeri*:968
afum|EAL90340.1 100.00% afis|EAW22860.1 100.00%
Bootstrap support for afum|EAL90340.1 as seed ortholog is 100%.
Bootstrap support for afis|EAW22860.1 as seed ortholog is 100%.

---

Group of orthologs #4318. Best score 968 bits
Score difference with first non-orthologous sequence - *A. fumigatus*:968 *A. fischeri*:968
afum|EAL90363.1 100.00% afis|EAW22840.1 100.00%

Bootstrap support for afum|EAL90363.1 as seed ortholog is 100%.
Bootstrap support for afis|EAW22840.1 as seed ortholog is 100%.

---

Group of orthologs #4319. Best score 968 bits
Score difference with first non-orthologous sequence - *A. fumigatus*:968 *A. fischeri*:968
afum|EAL89414.2 100.00% afis|EAW24772.1 100.00%
Bootstrap support for afum|EAL89414.2 as seed ortholog is 100%.
Bootstrap support for afis|EAW24772.1 as seed ortholog is 100%.

---

Group of orthologs #4320. Best score 968 bits
Score difference with first non-orthologous sequence - *A. fumigatus*:968 *A. fischeri*:968
afum|EAL92805.1 100.00% afis|EAW21776.1 100.00%
Bootstrap support for afum|EAL92805.1 as seed ortholog is 100%.
Bootstrap support for afis|EAW21776.1 as seed ortholog is 100%.

---

Group of orthologs #4321. Best score 968 bits
Score difference with first non-orthologous sequence - *A. fumigatus*:968 *A. fischeri*:968
afum|EAL92846.1 100.00% afis|EAW21831.1 100.00%
Bootstrap support for afum|EAL92846.1 as seed ortholog is 100%.
Bootstrap support for afis|EAW21831.1 as seed ortholog is 100%.

---

Group of orthologs #4322. Best score 968 bits
Score difference with first non-orthologous sequence - *A. fumigatus*:968 *A. fischeri*:968
afum|EAL89750.1 100.00% afis|EAW25081.1 100.00%
Bootstrap support for afum|EAL89750.1 as seed ortholog is 100%.
Bootstrap support for afis|EAW25081.1 as seed ortholog is 100%.

---

Group of orthologs #4323. Best score 968 bits
Score difference with first non-orthologous sequence - *A. fumigatus*:968 *A. fischeri*:968
afum|EAL89785.1 100.00% afis|EAW25114.1 100.00%
Bootstrap support for afum|EAL89785.1 as seed ortholog is 100%.
Bootstrap support for afis|EAW25114.1 as seed ortholog is 100%.

---

Group of orthologs #4324. Best score 968 bits
Score difference with first non-orthologous sequence - *A. fumigatus*:868 *A. fischeri*:968
afum|EBA27264.1 100.00% afis|EAW21064.1 100.00%
Bootstrap support for afum|EBA27264.1 as seed ortholog is 100%.
Bootstrap support for afis|EAW21064.1 as seed ortholog is 100%.

---

Group of orthologs #4325. Best score 967 bits
Score difference with first non-orthologous sequence - *A. fumigatus*:967 *A. fischeri*:967
afum|EAL85751.1 100.00% afis|EAW15583.1 100.00%
Bootstrap support for afum|EAL85751.1 as seed ortholog is 100%.
Bootstrap support for afis|EAW15583.1 as seed ortholog is 100%.

---

Group of orthologs #4326. Best score 967 bits
Score difference with first non-orthologous sequence - *A. fumigatus*:967 *A. fischeri*:967
afum|EAL87393.1 100.00% afis|EAW23963.1 100.00%
Bootstrap support for afum|EAL87393.1 as seed ortholog is 100%.
Bootstrap support for afis|EAW23963.1 as seed ortholog is 100%.

---

Group of orthologs #4327. Best score 967 bits
Score difference with first non-orthologous sequence - *A. fumigatus*:967 *A. fischeri*:967
afum|EAL90468.2 100.00% afis|EAW22727.1 100.00%
Bootstrap support for afum|EAL90468.2 as seed ortholog is 100%.
Bootstrap support for afis|EAW22727.1 as seed ortholog is 100%.

Group of orthologs #4328. Best score 967 bits
Score difference with first non-orthologous sequence - *A. fumigatus*:967 *A. fischeri*:967
afum|EAL90260.1 100.00% afis|EAW22936.1 100.00%
Bootstrap support for afum|EAL90260.1 as seed ortholog is 100%.
Bootstrap support for afis|EAW22936.1 as seed ortholog is 100%.

Group of orthologs #4329. Best score 967 bits
Score difference with first non-orthologous sequence - *A. fumigatus*:967 *A. fischeri*:967
afum|EAL92377.1 100.00% afis|EAW21361.1 100.00%
Bootstrap support for afum|EAL92377.1 as seed ortholog is 100%.
Bootstrap support for afis|EAW21361.1 as seed ortholog is 100%.

Group of orthologs #4330. Best score 967 bits
Score difference with first non-orthologous sequence - *A. fumigatus*:967 *A. fischeri*:967
afum|EAL92912.1 100.00% afis|EAW21899.1 100.00%
Bootstrap support for afum|EAL92912.1 as seed ortholog is 100%.
Bootstrap support for afis|EAW21899.1 as seed ortholog is 100%.

Group of orthologs #4331. Best score 966 bits
Score difference with first non-orthologous sequence - *A. fumigatus*:966 *A. fischeri*:966
afum|EAL88641.1 100.00% afis|EAW16011.1 100.00%
Bootstrap support for afum|EAL88641.1 as seed ortholog is 100%.
Bootstrap support for afis|EAW16011.1 as seed ortholog is 100%.

Group of orthologs #4332. Best score 966 bits
Score difference with first non-orthologous sequence - *A. fumigatus*:966 *A. fischeri*:966
afum|EAL87577.1 100.00% afis|EAW18211.1 100.00%
Bootstrap support for afum|EAL87577.1 as seed ortholog is 100%.
Bootstrap support for afis|EAW18211.1 as seed ortholog is 100%.

Group of orthologs #4333. Best score 966 bits
Score difference with first non-orthologous sequence - *A. fumigatus*:966 *A. fischeri*:708
afum|EAL87098.1 100.00% afis|EAW19396.1 100.00%
Bootstrap support for afum|EAL87098.1 as seed ortholog is 100%.
Bootstrap support for afis|EAW19396.1 as seed ortholog is 100%.

Group of orthologs #4334. Best score 966 bits
Score difference with first non-orthologous sequence - *A. fumigatus*:966 *A. fischeri*:966
afum|EAL91466.1 100.00% afis|EAW17651.1 100.00%
Bootstrap support for afum|EAL91466.1 as seed ortholog is 100%.
Bootstrap support for afis|EAW17651.1 as seed ortholog is 100%.

Group of orthologs #4335. Best score 966 bits
Score difference with first non-orthologous sequence - *A. fumigatus*:966 *A. fischeri*:826
afum|EAL89236.1 100.00% afis|EAW20408.1 100.00%
Bootstrap support for afum|EAL89236.1 as seed ortholog is 100%.
Bootstrap support for afis|EAW20408.1 as seed ortholog is 100%.

Group of orthologs #4336. Best score 965 bits
Score difference with first non-orthologous sequence - *A. fumigatus*:965 *A. fischeri*:965
afum|EAL84440.1 100.00% afis|EAW20626.1 100.00%
Bootstrap support for afum|EAL84440.1 as seed ortholog is 100%.
Bootstrap support for afis|EAW20626.1 as seed ortholog is 100%.

Group of orthologs #4337. Best score 965 bits

Score difference with first non-orthologous sequence - *A. fumigatus*:965   *A. fischeri*:965
afum|EAL85232.1 100.00% afis|EAW20115.1 100.00%
Bootstrap support for afum|EAL85232.1 as seed ortholog is 100%.
Bootstrap support for afis|EAW20115.1 as seed ortholog is 100%.

---

Group of orthologs #4338. Best score 965 bits
Score difference with first non-orthologous sequence - *A. fumigatus*:965   *A. fischeri*:965
afum|EAL91836.1 100.00% afis|EAW18024.1 100.00%
Bootstrap support for afum|EAL91836.1 as seed ortholog is 100%.
Bootstrap support for afis|EAW18024.1 as seed ortholog is 100%.

---

Group of orthologs #4339. Best score 965 bits
Score difference with first non-orthologous sequence - *A. fumigatus*:965   *A. fischeri*:965
afum|EAL85942.1 100.00% afis|EAW24274.1 100.00%
Bootstrap support for afum|EAL85942.1 as seed ortholog is 100%.
Bootstrap support for afis|EAW24274.1 as seed ortholog is 100%.

---

Group of orthologs #4340. Best score 965 bits
Score difference with first non-orthologous sequence - *A. fumigatus*:965   *A. fischeri*:965
afum|EAL88436.1 100.00% afis|EAW23045.1 100.00%
Bootstrap support for afum|EAL88436.1 as seed ortholog is 100%.
Bootstrap support for afis|EAW23045.1 as seed ortholog is 100%.

---

Group of orthologs #4341. Best score 965 bits
Score difference with first non-orthologous sequence - *A. fumigatus*:965   *A. fischeri*:965
afum|EAL88120.2 100.00% afis|EAW23356.1 100.00%
Bootstrap support for afum|EAL88120.2 as seed ortholog is 100%.
Bootstrap support for afis|EAW23356.1 as seed ortholog is 100%.

---

Group of orthologs #4342. Best score 965 bits
Score difference with first non-orthologous sequence - *A. fumigatus*:965   *A. fischeri*:965
afum|EAL88255.1 100.00% afis|EAW23228.1 100.00%
Bootstrap support for afum|EAL88255.1 as seed ortholog is 100%.
Bootstrap support for afis|EAW23228.1 as seed ortholog is 100%.

---

Group of orthologs #4343. Best score 965 bits
Score difference with first non-orthologous sequence - *A. fumigatus*:965   *A. fischeri*:965
afum|EAL90948.1 100.00% afis|EAW22245.1 100.00%
Bootstrap support for afum|EAL90948.1 as seed ortholog is 100%.
Bootstrap support for afis|EAW22245.1 as seed ortholog is 100%.

---

Group of orthologs #4344. Best score 965 bits
Score difference with first non-orthologous sequence - *A. fumigatus*:965   *A. fischeri*:965
afum|EAL92165.1 100.00% afis|EAW21146.1 100.00%
Bootstrap support for afum|EAL92165.1 as seed ortholog is 100%.
Bootstrap support for afis|EAW21146.1 as seed ortholog is 100%.

---

Group of orthologs #4345. Best score 964 bits
Score difference with first non-orthologous sequence - *A. fumigatus*:964   *A. fischeri*:650
afum|EAL86377.1 100.00% afis|EAW16829.1 100.00%
Bootstrap support for afum|EAL86377.1 as seed ortholog is 100%.
Bootstrap support for afis|EAW16829.1 as seed ortholog is 100%.

---

Group of orthologs #4346. Best score 964 bits
Score difference with first non-orthologous sequence - *A. fumigatus*:964   *A. fischeri*:964
afum|EAL84932.1 100.00% afis|EAW19984.1 100.00%

Bootstrap support for afum|EAL84932.1 as seed ortholog is 100%.
Bootstrap support for afis|EAW19984.1 as seed ortholog is 100%.

---

Group of orthologs #4347. Best score 964 bits
Score difference with first non-orthologous sequence - *A. fumigatus*:964 *A. fischeri*:964
afum|EAL89011.1 100.00% afis|EAW16383.1 100.00%
Bootstrap support for afum|EAL89011.1 as seed ortholog is 100%.
Bootstrap support for afis|EAW16383.1 as seed ortholog is 100%.

---

Group of orthologs #4348. Best score 964 bits
Score difference with first non-orthologous sequence - *A. fumigatus*:835 *A. fischeri*:800
afum|EAL86243.1 100.00% afis|EAW24613.1 100.00%
Bootstrap support for afum|EAL86243.1 as seed ortholog is 100%.
Bootstrap support for afis|EAW24613.1 as seed ortholog is 100%.

---

Group of orthologs #4349. Best score 964 bits
Score difference with first non-orthologous sequence - *A. fumigatus*:964 *A. fischeri*:964
afum|EAL92177.1 100.00% afis|EAW21159.1 100.00%
Bootstrap support for afum|EAL92177.1 as seed ortholog is 100%.
Bootstrap support for afis|EAW21159.1 as seed ortholog is 100%.

---

Group of orthologs #4350. Best score 963 bits
Score difference with first non-orthologous sequence - *A. fumigatus*:963 *A. fischeri*:963
afum|EAL85632.1 100.00% afis|EAW15716.1 100.00%
Bootstrap support for afum|EAL85632.1 as seed ortholog is 100%.
Bootstrap support for afis|EAW15716.1 as seed ortholog is 100%.

---

Group of orthologs #4351. Best score 963 bits
Score difference with first non-orthologous sequence - *A. fumigatus*:963 *A. fischeri*:963
afum|EAL87062.1 100.00% afis|EAW19433.1 100.00%
Bootstrap support for afum|EAL87062.1 as seed ortholog is 100%.
Bootstrap support for afis|EAW19433.1 as seed ortholog is 100%.

---

Group of orthologs #4352. Best score 963 bits
Score difference with first non-orthologous sequence - *A. fumigatus*:963 *A. fischeri*:963
afum|EAL92410.1 100.00% afis|EAW21396.1 100.00%
Bootstrap support for afum|EAL92410.1 as seed ortholog is 100%.
Bootstrap support for afis|EAW21396.1 as seed ortholog is 100%.

---

Group of orthologs #4353. Best score 962 bits
Score difference with first non-orthologous sequence - *A. fumigatus*:859 *A. fischeri*:878
afum|EAL91300.1 100.00% afis|EAW17471.1 100.00%
Bootstrap support for afum|EAL91300.1 as seed ortholog is 100%.
Bootstrap support for afis|EAW17471.1 as seed ortholog is 100%.

---

Group of orthologs #4354. Best score 962 bits
Score difference with first non-orthologous sequence - *A. fumigatus*:962 *A. fischeri*:962
afum|EAL92091.1 100.00% afis|EAW21055.1 100.00%
Bootstrap support for afum|EAL92091.1 as seed ortholog is 100%.
Bootstrap support for afis|EAW21055.1 as seed ortholog is 100%.

---

Group of orthologs #4355. Best score 962 bits
Score difference with first non-orthologous sequence - *A. fumigatus*:962 *A. fischeri*:962
afum|EAL91059.1 100.00% afis|EAW22131.1 100.00%
Bootstrap support for afum|EAL91059.1 as seed ortholog is 100%.
Bootstrap support for afis|EAW22131.1 as seed ortholog is 100%.

Group of orthologs #4356. Best score 962 bits
Score difference with first non-orthologous sequence - *A. fumigatus*:962 *A. fischeri*:962
afum|EAL90857.2 100.00% afis|EAW22335.1 100.00%
Bootstrap support for afum|EAL90857.2 as seed ortholog is 100%.
Bootstrap support for afis|EAW22335.1 as seed ortholog is 100%.

---

Group of orthologs #4357. Best score 962 bits
Score difference with first non-orthologous sequence - *A. fumigatus*:962 *A. fischeri*:962
afum|EAL90662.1 100.00% afis|EAW22532.1 100.00%
Bootstrap support for afum|EAL90662.1 as seed ortholog is 100%.
Bootstrap support for afis|EAW22532.1 as seed ortholog is 100%.

---

Group of orthologs #4358. Best score 961 bits
Score difference with first non-orthologous sequence - *A. fumigatus*:961 *A. fischeri*:961
afum|EAL85255.1 100.00% afis|EAW20140.1 100.00%
Bootstrap support for afum|EAL85255.1 as seed ortholog is 100%.
Bootstrap support for afis|EAW20140.1 as seed ortholog is 100%.

---

Group of orthologs #4359. Best score 961 bits
Score difference with first non-orthologous sequence - *A. fumigatus*:961 *A. fischeri*:961
afum|EAL85696.1 100.00% afis|EAW22087.1 100.00%
Bootstrap support for afum|EAL85696.1 as seed ortholog is 100%.
Bootstrap support for afis|EAW22087.1 as seed ortholog is 100%.

---

Group of orthologs #4360. Best score 961 bits
Score difference with first non-orthologous sequence - *A. fumigatus*:961 *A. fischeri*:961
afum|EAL91716.2 100.00% afis|EAW17907.1 100.00%
Bootstrap support for afum|EAL91716.2 as seed ortholog is 100%.
Bootstrap support for afis|EAW17907.1 as seed ortholog is 100%.

---

Group of orthologs #4361. Best score 961 bits
Score difference with first non-orthologous sequence - *A. fumigatus*:961 *A. fischeri*:961
afum|EAL87355.1 100.00% afis|EAW23930.1 100.00%
Bootstrap support for afum|EAL87355.1 as seed ortholog is 100%.
Bootstrap support for afis|EAW23930.1 as seed ortholog is 100%.

---

Group of orthologs #4362. Best score 961 bits
Score difference with first non-orthologous sequence - *A. fumigatus*:961 *A. fischeri*:961
afum|EAL93172.1 100.00% afis|EAW18483.1 100.00%
Bootstrap support for afum|EAL93172.1 as seed ortholog is 100%.
Bootstrap support for afis|EAW18483.1 as seed ortholog is 100%.

---

Group of orthologs #4363. Best score 961 bits
Score difference with first non-orthologous sequence - *A. fumigatus*:961 *A. fischeri*:961
afum|EAL88479.2 100.00% afis|EAW23001.1 100.00%
Bootstrap support for afum|EAL88479.2 as seed ortholog is 100%.
Bootstrap support for afis|EAW23001.1 as seed ortholog is 100%.

---

Group of orthologs #4364. Best score 961 bits
Score difference with first non-orthologous sequence - *A. fumigatus*:961 *A. fischeri*:961
afum|EAL88378.2 100.00% afis|EAW23100.1 100.00%
Bootstrap support for afum|EAL88378.2 as seed ortholog is 100%.
Bootstrap support for afis|EAW23100.1 as seed ortholog is 100%.

---

Group of orthologs #4365. Best score 961 bits

Score difference with first non-orthologous sequence - *A. fumigatus*:961 *A. fischeri*:961
afum|EAL93257.1 100.00% afis|EAW18557.1 100.00%
Bootstrap support for afum|EAL93257.1 as seed ortholog is 100%.
Bootstrap support for afis|EAW18557.1 as seed ortholog is 100%.

---

Group of orthologs #4366. Best score 961 bits
Score difference with first non-orthologous sequence - *A. fumigatus*:961 *A. fischeri*:961
afum|EAL93552.1 100.00% afis|EAW18816.1 100.00%
Bootstrap support for afum|EAL93552.1 as seed ortholog is 100%.
Bootstrap support for afis|EAW18816.1 as seed ortholog is 100%.

---

Group of orthologs #4367. Best score 961 bits
Score difference with first non-orthologous sequence - *A. fumigatus*:961 *A. fischeri*:961
afum|EAL94041.1 100.00% afis|EAW19323.1 100.00%
Bootstrap support for afum|EAL94041.1 as seed ortholog is 100%.
Bootstrap support for afis|EAW19323.1 as seed ortholog is 100%.

---

Group of orthologs #4368. Best score 961 bits
Score difference with first non-orthologous sequence - *A. fumigatus*:961 *A. fischeri*:961
afum|EAL90805.1 100.00% afis|EAW22385.1 100.00%
Bootstrap support for afum|EAL90805.1 as seed ortholog is 100%.
Bootstrap support for afis|EAW22385.1 as seed ortholog is 100%.

---

Group of orthologs #4369. Best score 961 bits
Score difference with first non-orthologous sequence - *A. fumigatus*:730 *A. fischeri*:652
afum|EAL90771.1 100.00% afis|EAW22421.1 100.00%
Bootstrap support for afum|EAL90771.1 as seed ortholog is 100%.
Bootstrap support for afis|EAW22421.1 as seed ortholog is 100%.

---

Group of orthologs #4370. Best score 961 bits
Score difference with first non-orthologous sequence - *A. fumigatus*:961 *A. fischeri*:961
afum|EAL90576.1 100.00% afis|EAW22620.1 100.00%
Bootstrap support for afum|EAL90576.1 as seed ortholog is 100%.
Bootstrap support for afis|EAW22620.1 as seed ortholog is 100%.

---

Group of orthologs #4371. Best score 961 bits
Score difference with first non-orthologous sequence - *A. fumigatus*:842 *A. fischeri*:961
afum|EAL90285.1 100.00% afis|EAW22912.1 100.00%
Bootstrap support for afum|EAL90285.1 as seed ortholog is 100%.
Bootstrap support for afis|EAW22912.1 as seed ortholog is 100%.

---

Group of orthologs #4372. Best score 960 bits
Score difference with first non-orthologous sequence - *A. fumigatus*:960 *A. fischeri*:960
afum|EAL85641.1 100.00% afis|EAW15706.1 100.00%
Bootstrap support for afum|EAL85641.1 as seed ortholog is 100%.
Bootstrap support for afis|EAW15706.1 as seed ortholog is 100%.

---

Group of orthologs #4373. Best score 960 bits
Score difference with first non-orthologous sequence - *A. fumigatus*:960 *A. fischeri*:960
afum|EAL84513.1 100.00% afis|EAW20546.1 100.00%
Bootstrap support for afum|EAL84513.1 as seed ortholog is 100%.
Bootstrap support for afis|EAW20546.1 as seed ortholog is 100%.

---

Group of orthologs #4374. Best score 960 bits
Score difference with first non-orthologous sequence - *A. fumigatus*:960 *A. fischeri*:960
afum|EAL93981.1 100.00% afis|EAW19252.1 100.00%

Bootstrap support for afum|EAL93981.1 as seed ortholog is 100%.
Bootstrap support for afis|EAW19252.1 as seed ortholog is 100%.

---

Group of orthologs #4375. Best score 960 bits
Score difference with first non-orthologous sequence - *A. fumigatus*:960 *A. fischeri*:960
afum|EAL90257.1 100.00% afis|EAW22939.1 100.00%
Bootstrap support for afum|EAL90257.1 as seed ortholog is 100%.
Bootstrap support for afis|EAW22939.1 as seed ortholog is 100%.

---

Group of orthologs #4376. Best score 960 bits
Score difference with first non-orthologous sequence - *A. fumigatus*:960 *A. fischeri*:960
afum|EAL89656.1 100.00% afis|EAW24995.1 100.00%
Bootstrap support for afum|EAL89656.1 as seed ortholog is 100%.
Bootstrap support for afis|EAW24995.1 as seed ortholog is 100%.

---

Group of orthologs #4377. Best score 959 bits
Score difference with first non-orthologous sequence - *A. fumigatus*:828 *A. fischeri*:866
afum|EAL85369.1 100.00% afis|EAW20251.1 100.00%
Bootstrap support for afum|EAL85369.1 as seed ortholog is 100%.
Bootstrap support for afis|EAW20251.1 as seed ortholog is 100%.

---

Group of orthologs #4378. Best score 959 bits
Score difference with first non-orthologous sequence - *A. fumigatus*:959 *A. fischeri*:959
afum|EAL84385.1 100.00% afis|EAW25713.1 100.00%
Bootstrap support for afum|EAL84385.1 as seed ortholog is 100%.
Bootstrap support for afis|EAW25713.1 as seed ortholog is 100%.

---

Group of orthologs #4379. Best score 959 bits
Score difference with first non-orthologous sequence - *A. fumigatus*:959 *A. fischeri*:421
afum|EAL84398.1 100.00% afis|EAW25725.1 100.00%
Bootstrap support for afum|EAL84398.1 as seed ortholog is 100%.
Bootstrap support for afis|EAW25725.1 as seed ortholog is 100%.

---

Group of orthologs #4380. Best score 959 bits
Score difference with first non-orthologous sequence - *A. fumigatus*:959 *A. fischeri*:513
afum|EAL86194.1 100.00% afis|EAW24529.1 100.00%
Bootstrap support for afum|EAL86194.1 as seed ortholog is 100%.
Bootstrap support for afis|EAW24529.1 as seed ortholog is 100%.

---

Group of orthologs #4381. Best score 959 bits
Score difference with first non-orthologous sequence - *A. fumigatus*:959 *A. fischeri*:959
afum|EAL88412.1 100.00% afis|EAW23067.1 100.00%
Bootstrap support for afum|EAL88412.1 as seed ortholog is 100%.
Bootstrap support for afis|EAW23067.1 as seed ortholog is 100%.

---

Group of orthologs #4382. Best score 959 bits
Score difference with first non-orthologous sequence - *A. fumigatus*:959 *A. fischeri*:832
afum|EAL93335.1 100.00% afis|EAW18624.1 100.00%
Bootstrap support for afum|EAL93335.1 as seed ortholog is 100%.
Bootstrap support for afis|EAW18624.1 as seed ortholog is 100%.

---

Group of orthologs #4383. Best score 959 bits
Score difference with first non-orthologous sequence - *A. fumigatus*:959 *A. fischeri*:959
afum|EAL91058.1 100.00% afis|EAW22132.1 100.00%
Bootstrap support for afum|EAL91058.1 as seed ortholog is 100%.
Bootstrap support for afis|EAW22132.1 as seed ortholog is 100%.

---

Group of orthologs #4384. Best score 959 bits
Score difference with first non-orthologous sequence - *A. fumigatus*:959 *A. fischeri*:959
afum|EAL92460.1 100.00% afis|EAW21442.1 100.00%
Bootstrap support for afum|EAL92460.1 as seed ortholog is 100%.
Bootstrap support for afis|EAW21442.1 as seed ortholog is 100%.

---

Group of orthologs #4385. Best score 959 bits
Score difference with first non-orthologous sequence - *A. fumigatus*:959 *A. fischeri*:959
afum|EAL89329.1 100.00% afis|EAW24688.1 100.00%
Bootstrap support for afum|EAL89329.1 as seed ortholog is 100%.
Bootstrap support for afis|EAW24688.1 as seed ortholog is 100%.

---

Group of orthologs #4386. Best score 959 bits
Score difference with first non-orthologous sequence - *A. fumigatus*:868 *A. fischeri*:959
afum|EAL89780.2 100.00% afis|EAW25109.1 100.00%
Bootstrap support for afum|EAL89780.2 as seed ortholog is 100%.
Bootstrap support for afis|EAW25109.1 as seed ortholog is 100%.

---

Group of orthologs #4387. Best score 958 bits
Score difference with first non-orthologous sequence - *A. fumigatus*:958 *A. fischeri*:958
afum|EAL88811.1 100.00% afis|EAW16178.1 100.00%
Bootstrap support for afum|EAL88811.1 as seed ortholog is 100%.
Bootstrap support for afis|EAW16178.1 as seed ortholog is 100%.

---

Group of orthologs #4388. Best score 958 bits
Score difference with first non-orthologous sequence - *A. fumigatus*:958 *A. fischeri*:373
afum|EAL84505.1 100.00% afis|EAW20556.1 100.00%
Bootstrap support for afum|EAL84505.1 as seed ortholog is 100%.
Bootstrap support for afis|EAW20556.1 as seed ortholog is 100%.

---

Group of orthologs #4389. Best score 958 bits
Score difference with first non-orthologous sequence - *A. fumigatus*:702 *A. fischeri*:699
afum|EAL84852.1 100.00% afis|EAW20767.1 100.00%
Bootstrap support for afum|EAL84852.1 as seed ortholog is 100%.
Bootstrap support for afis|EAW20767.1 as seed ortholog is 100%.

---

Group of orthologs #4390. Best score 958 bits
Score difference with first non-orthologous sequence - *A. fumigatus*:958 *A. fischeri*:958
afum|EAL86828.1 100.00% afis|EAW19699.1 100.00%
Bootstrap support for afum|EAL86828.1 as seed ortholog is 100%.
Bootstrap support for afis|EAW19699.1 as seed ortholog is 100%.

---

Group of orthologs #4391. Best score 958 bits
Score difference with first non-orthologous sequence - *A. fumigatus*:958 *A. fischeri*:958
afum|EAL86076.1 100.00% afis|EAW24405.1 100.00%
Bootstrap support for afum|EAL86076.1 as seed ortholog is 100%.
Bootstrap support for afis|EAW24405.1 as seed ortholog is 100%.

---

Group of orthologs #4392. Best score 958 bits
Score difference with first non-orthologous sequence - *A. fumigatus*:958 *A. fischeri*:958
afum|EAL93486.1 100.00% afis|EAW18757.1 100.00%
Bootstrap support for afum|EAL93486.1 as seed ortholog is 100%.
Bootstrap support for afis|EAW18757.1 as seed ortholog is 100%.

---

Group of orthologs #4393. Best score 958 bits

Score difference with first non-orthologous sequence - *A. fumigatus*:958 *A. fischeri*:958
afum|EAL90844.1 100.00% afis|EAW22348.1 100.00%
Bootstrap support for afum|EAL90844.1 as seed ortholog is 100%.
Bootstrap support for afis|EAW22348.1 as seed ortholog is 100%.

---

Group of orthologs #4394. Best score 958 bits
Score difference with first non-orthologous sequence - *A. fumigatus*:958 *A. fischeri*:958
afum|EAL89610.1 100.00% afis|EAW24953.1 100.00%
Bootstrap support for afum|EAL89610.1 as seed ortholog is 100%.
Bootstrap support for afis|EAW24953.1 as seed ortholog is 100%.

---

Group of orthologs #4395. Best score 957 bits
Score difference with first non-orthologous sequence - *A. fumigatus*:957 *A. fischeri*:957
afum|EAL85487.1 100.00% afis|EAW20373.1 100.00%
Bootstrap support for afum|EAL85487.1 as seed ortholog is 100%.
Bootstrap support for afis|EAW20373.1 as seed ortholog is 100%.

---

Group of orthologs #4396. Best score 957 bits
Score difference with first non-orthologous sequence - *A. fumigatus*:957 *A. fischeri*:957
afum|EAL91417.1 100.00% afis|EAW17604.1 100.00%
Bootstrap support for afum|EAL91417.1 as seed ortholog is 100%.
Bootstrap support for afis|EAW17604.1 as seed ortholog is 100%.

---

Group of orthologs #4397. Best score 957 bits
Score difference with first non-orthologous sequence - *A. fumigatus*:957 *A. fischeri*:957
afum|EAL87999.1 100.00% afis|EAW23467.1 100.00%
Bootstrap support for afum|EAL87999.1 as seed ortholog is 100%.
Bootstrap support for afis|EAW23467.1 as seed ortholog is 100%.

---

Group of orthologs #4398. Best score 956 bits
Score difference with first non-orthologous sequence - *A. fumigatus*:956 *A. fischeri*:956
afum|EAL89299.1 100.00% afis|EAW16711.1 100.00%
Bootstrap support for afum|EAL89299.1 as seed ortholog is 100%.
Bootstrap support for afis|EAW16711.1 as seed ortholog is 100%.

---

Group of orthologs #4399. Best score 956 bits
Score difference with first non-orthologous sequence - *A. fumigatus*:956 *A. fischeri*:956
afum|EAL91876.1 100.00% afis|EAW18064.1 100.00%
Bootstrap support for afum|EAL91876.1 as seed ortholog is 100%.
Bootstrap support for afis|EAW18064.1 as seed ortholog is 100%.

---

Group of orthologs #4400. Best score 956 bits
Score difference with first non-orthologous sequence - *A. fumigatus*:956 *A. fischeri*:956
afum|EAL87119.1 100.00% afis|EAW23676.1 100.00%
Bootstrap support for afum|EAL87119.1 as seed ortholog is 100%.
Bootstrap support for afis|EAW23676.1 as seed ortholog is 100%.

---

Group of orthologs #4401. Best score 956 bits
Score difference with first non-orthologous sequence - *A. fumigatus*:956 *A. fischeri*:956
afum|EAL92218.1 100.00% afis|EAW21207.1 100.00%
Bootstrap support for afum|EAL92218.1 as seed ortholog is 100%.
Bootstrap support for afis|EAW21207.1 as seed ortholog is 100%.

---

Group of orthologs #4402. Best score 956 bits
Score difference with first non-orthologous sequence - *A. fumigatus*:956 *A. fischeri*:897
afum|EAL92626.1 100.00% afis|EAW21607.1 100.00%

Bootstrap support for afum|EAL92626.1 as seed ortholog is 100%.
Bootstrap support for afis|EAW21607.1 as seed ortholog is 100%.

---

Group of orthologs #4403. Best score 955 bits
Score difference with first non-orthologous sequence - *A. fumigatus*:955 *A. fischeri*:798
afum|EAL85527.1 100.00% afis|EAW15824.1 100.00%
Bootstrap support for afum|EAL85527.1 as seed ortholog is 100%.
Bootstrap support for afis|EAW15824.1 as seed ortholog is 100%.

---

Group of orthologs #4404. Best score 955 bits
Score difference with first non-orthologous sequence - *A. fumigatus*:701 *A. fischeri*:770
afum|EAL86384.1 100.00% afis|EAW16836.1 100.00%
Bootstrap support for afum|EAL86384.1 as seed ortholog is 100%.
Bootstrap support for afis|EAW16836.1 as seed ortholog is 100%.

---

Group of orthologs #4405. Best score 955 bits
Score difference with first non-orthologous sequence - *A. fumigatus*:886 *A. fischeri*:833
afum|EAL86437.1 100.00% afis|EAW16902.1 100.00%
Bootstrap support for afum|EAL86437.1 as seed ortholog is 100%.
Bootstrap support for afis|EAW16902.1 as seed ortholog is 100%.

---

Group of orthologs #4406. Best score 955 bits
Score difference with first non-orthologous sequence - *A. fumigatus*:955 *A. fischeri*:955
afum|EAL88992.2 100.00% afis|EAW16367.1 100.00%
Bootstrap support for afum|EAL88992.2 as seed ortholog is 100%.
Bootstrap support for afis|EAW16367.1 as seed ortholog is 100%.

---

Group of orthologs #4407. Best score 955 bits
Score difference with first non-orthologous sequence - *A. fumigatus*:538 *A. fischeri*:955
afum|EAL84490.1 100.00% afis|EAW20575.1 100.00%
Bootstrap support for afum|EAL84490.1 as seed ortholog is 100%.
Bootstrap support for afis|EAW20575.1 as seed ortholog is 100%.

---

Group of orthologs #4408. Best score 955 bits
Score difference with first non-orthologous sequence - *A. fumigatus*:692 *A. fischeri*:694
afum|EAL91584.2 100.00% afis|EAW17776.1 100.00%
Bootstrap support for afum|EAL91584.2 as seed ortholog is 100%.
Bootstrap support for afis|EAW17776.1 as seed ortholog is 100%.

---

Group of orthologs #4409. Best score 955 bits
Score difference with first non-orthologous sequence - *A. fumigatus*:882 *A. fischeri*:955
afum|EAL93053.1 100.00% afis|EAW18378.1 100.00%
Bootstrap support for afum|EAL93053.1 as seed ortholog is 100%.
Bootstrap support for afis|EAW18378.1 as seed ortholog is 100%.

---

Group of orthologs #4410. Best score 955 bits
Score difference with first non-orthologous sequence - *A. fumigatus*:955 *A. fischeri*:955
afum|EAL88171.1 100.00% afis|EAW23306.1 100.00%
Bootstrap support for afum|EAL88171.1 as seed ortholog is 100%.
Bootstrap support for afis|EAW23306.1 as seed ortholog is 100%.

---

Group of orthologs #4411. Best score 955 bits
Score difference with first non-orthologous sequence - *A. fumigatus*:955 *A. fischeri*:955
afum|EAL93479.1 100.00% afis|EAW18752.1 100.00%
Bootstrap support for afum|EAL93479.1 as seed ortholog is 100%.
Bootstrap support for afis|EAW18752.1 as seed ortholog is 100%.

---

Group of orthologs #4412. Best score 955 bits
Score difference with first non-orthologous sequence - *A. fumigatus*:955 *A. fischeri*:955
afum|EAL89520.1 100.00% afis|EAW24870.1 100.00%
Bootstrap support for afum|EAL89520.1 as seed ortholog is 100%.
Bootstrap support for afis|EAW24870.1 as seed ortholog is 100%.

---

Group of orthologs #4413. Best score 955 bits
Score difference with first non-orthologous sequence - *A. fumigatus*:955 *A. fischeri*:955
afum|EAL89934.1 100.00% afis|EAW25271.1 100.00%
Bootstrap support for afum|EAL89934.1 as seed ortholog is 100%.
Bootstrap support for afis|EAW25271.1 as seed ortholog is 100%.

---

Group of orthologs #4414. Best score 955 bits
Score difference with first non-orthologous sequence - *A. fumigatus*:955 *A. fischeri*:955
afum|EAL92021.1 100.00% afis|EAW24164.1 100.00%
Bootstrap support for afum|EAL92021.1 as seed ortholog is 100%.
Bootstrap support for afis|EAW24164.1 as seed ortholog is 100%.

---

Group of orthologs #4415. Best score 954 bits
Score difference with first non-orthologous sequence - *A. fumigatus*:954 *A. fischeri*:954
afum|EAL88758.1 100.00% afis|EAW16128.1 100.00%
Bootstrap support for afum|EAL88758.1 as seed ortholog is 100%.
Bootstrap support for afis|EAW16128.1 as seed ortholog is 100%.

---

Group of orthologs #4416. Best score 954 bits
Score difference with first non-orthologous sequence - *A. fumigatus*:954 *A. fischeri*:954
afum|EAL92139.1 100.00% afis|EAW21116.1 100.00%
Bootstrap support for afum|EAL92139.1 as seed ortholog is 100%.
Bootstrap support for afis|EAW21116.1 as seed ortholog is 100%.

---

Group of orthologs #4417. Best score 954 bits
Score difference with first non-orthologous sequence - *A. fumigatus*:668 *A. fischeri*:954
afum|EAL89779.1 100.00% afis|EAW25108.1 100.00%
Bootstrap support for afum|EAL89779.1 as seed ortholog is 100%.
Bootstrap support for afis|EAW25108.1 as seed ortholog is 100%.

---

Group of orthologs #4418. Best score 953 bits
Score difference with first non-orthologous sequence - *A. fumigatus*:953 *A. fischeri*:953
afum|EAL84686.1 100.00% afis|EAW20948.1 100.00%
Bootstrap support for afum|EAL84686.1 as seed ortholog is 100%.
Bootstrap support for afis|EAW20948.1 as seed ortholog is 100%.

---

Group of orthologs #4419. Best score 953 bits
Score difference with first non-orthologous sequence - *A. fumigatus*:953 *A. fischeri*:953
afum|EAL88416.1 100.00% afis|EAW23063.1 100.00%
Bootstrap support for afum|EAL88416.1 as seed ortholog is 100%.
Bootstrap support for afis|EAW23063.1 as seed ortholog is 100%.

---

Group of orthologs #4420. Best score 953 bits
Score difference with first non-orthologous sequence - *A. fumigatus*:953 *A. fischeri*:953
afum|EAL92069.1 100.00% afis|EAW21030.1 100.00%
Bootstrap support for afum|EAL92069.1 as seed ortholog is 100%.
Bootstrap support for afis|EAW21030.1 as seed ortholog is 100%.

---

Group of orthologs #4421. Best score 953 bits

Score difference with first non-orthologous sequence - *A. fumigatus*:953 *A. fischeri*:853
afum|EAL92441.1 100.00% afis|EAW21426.1 100.00%
Bootstrap support for afum|EAL92441.1 as seed ortholog is 100%.
Bootstrap support for afis|EAW21426.1 as seed ortholog is 100%.

---

Group of orthologs #4422. Best score 952 bits
Score difference with first non-orthologous sequence - *A. fumigatus*:952 *A. fischeri*:952
afum|EAL84876.1 100.00% afis|EAW20701.1 100.00%
Bootstrap support for afum|EAL84876.1 as seed ortholog is 100%.
Bootstrap support for afis|EAW20701.1 as seed ortholog is 100%.

---

Group of orthologs #4423. Best score 952 bits
Score difference with first non-orthologous sequence - *A. fumigatus*:952 *A. fischeri*:952
afum|EAL93458.1 100.00% afis|EAW18733.1 100.00%
Bootstrap support for afum|EAL93458.1 as seed ortholog is 100%.
Bootstrap support for afis|EAW18733.1 as seed ortholog is 100%.

---

Group of orthologs #4424. Best score 952 bits
Score difference with first non-orthologous sequence - *A. fumigatus*:952 *A. fischeri*:952
afum|EAL93466.1 100.00% afis|EAW18740.1 100.00%
Bootstrap support for afum|EAL93466.1 as seed ortholog is 100%.
Bootstrap support for afis|EAW18740.1 as seed ortholog is 100%.

---

Group of orthologs #4425. Best score 952 bits
Score difference with first non-orthologous sequence - *A. fumigatus*:952 *A. fischeri*:952
afum|EAL90254.1 100.00% afis|EAW22943.1 100.00%
Bootstrap support for afum|EAL90254.1 as seed ortholog is 100%.
Bootstrap support for afis|EAW22943.1 as seed ortholog is 100%.

---

Group of orthologs #4426. Best score 952 bits
Score difference with first non-orthologous sequence - *A. fumigatus*:952 *A. fischeri*:952
afum|EAL92552.1 100.00% afis|EAW21531.1 100.00%
Bootstrap support for afum|EAL92552.1 as seed ortholog is 100%.
Bootstrap support for afis|EAW21531.1 as seed ortholog is 100%.

---

Group of orthologs #4427. Best score 951 bits
Score difference with first non-orthologous sequence - *A. fumigatus*:951 *A. fischeri*:951
afum|EAL85733.1 100.00% afis|EAW15599.1 100.00%
Bootstrap support for afum|EAL85733.1 as seed ortholog is 100%.
Bootstrap support for afis|EAW15599.1 as seed ortholog is 100%.

---

Group of orthologs #4428. Best score 951 bits
Score difference with first non-orthologous sequence - *A. fumigatus*:577 *A. fischeri*:525
afum|EAL86605.1 100.00% afis|EAW17175.1 100.00%
Bootstrap support for afum|EAL86605.1 as seed ortholog is 100%.
Bootstrap support for afis|EAW17175.1 as seed ortholog is 100%.

---

Group of orthologs #4429. Best score 951 bits
Score difference with first non-orthologous sequence - *A. fumigatus*:951 *A. fischeri*:951
afum|EAL84759.1 100.00% afis|EAW20875.1 100.00%
Bootstrap support for afum|EAL84759.1 as seed ortholog is 100%.
Bootstrap support for afis|EAW20875.1 as seed ortholog is 100%.

---

Group of orthologs #4430. Best score 951 bits
Score difference with first non-orthologous sequence - *A. fumigatus*:951 *A. fischeri*:951
afum|EAL87191.1 100.00% afis|EAW23755.1 100.00%

Bootstrap support for afum|EAL87191.1 as seed ortholog is 100%.
Bootstrap support for afis|EAW23755.1 as seed ortholog is 100%.

---

Group of orthologs #4431. Best score 951 bits
Score difference with first non-orthologous sequence - *A. fumigatus*:951 *A. fischeri*:785
afum|EAL90797.1 100.00% afis|EAW22394.1 100.00%
Bootstrap support for afum|EAL90797.1 as seed ortholog is 100%.
Bootstrap support for afis|EAW22394.1 as seed ortholog is 100%.

---

Group of orthologs #4432. Best score 951 bits
Score difference with first non-orthologous sequence - *A. fumigatus*:951 *A. fischeri*:951
afum|EAL92738.1 100.00% afis|EAW21704.1 100.00%
Bootstrap support for afum|EAL92738.1 as seed ortholog is 100%.
Bootstrap support for afis|EAW21704.1 as seed ortholog is 100%.

---

Group of orthologs #4433. Best score 951 bits
Score difference with first non-orthologous sequence - *A. fumigatus*:951 *A. fischeri*:951
afum|EAL90075.1 100.00% afis|EAW25416.1 100.00%
Bootstrap support for afum|EAL90075.1 as seed ortholog is 100%.
Bootstrap support for afis|EAW25416.1 as seed ortholog is 100%.

---

Group of orthologs #4434. Best score 951 bits
Score difference with first non-orthologous sequence - *A. fumigatus*:951 *A. fischeri*:951
afum|EAL92039.1 100.00% afis|EAW24177.1 100.00%
Bootstrap support for afum|EAL92039.1 as seed ortholog is 100%.
Bootstrap support for afis|EAW24177.1 as seed ortholog is 100%.

---

Group of orthologs #4435. Best score 950 bits
Score difference with first non-orthologous sequence - *A. fumigatus*:950 *A. fischeri*:950
afum|EAL92808.1 100.00% afis|EAW21779.1 100.00%
Bootstrap support for afum|EAL92808.1 as seed ortholog is 100%.
Bootstrap support for afis|EAW21779.1 as seed ortholog is 100%.

---

Group of orthologs #4436. Best score 949 bits
Score difference with first non-orthologous sequence - *A. fumigatus*:949 *A. fischeri*:877
afum|EAL91718.1 100.00% afis|EAW17909.1 100.00%
Bootstrap support for afum|EAL91718.1 as seed ortholog is 100%.
Bootstrap support for afis|EAW17909.1 as seed ortholog is 100%.

---

Group of orthologs #4437. Best score 949 bits
Score difference with first non-orthologous sequence - *A. fumigatus*:949 *A. fischeri*:949
afum|EAL91742.1 100.00% afis|EAW17931.1 100.00%
Bootstrap support for afum|EAL91742.1 as seed ortholog is 100%.
Bootstrap support for afis|EAW17931.1 as seed ortholog is 100%.

---

Group of orthologs #4438. Best score 949 bits
Score difference with first non-orthologous sequence - *A. fumigatus*:949 *A. fischeri*:949
afum|EAL93121.2 100.00% afis|EAW18439.1 100.00%
Bootstrap support for afum|EAL93121.2 as seed ortholog is 100%.
Bootstrap support for afis|EAW18439.1 as seed ortholog is 100%.

---

Group of orthologs #4439. Best score 949 bits
Score difference with first non-orthologous sequence - *A. fumigatus*:949 *A. fischeri*:949
afum|EAL89477.1 100.00% afis|EAW24832.1 100.00%
Bootstrap support for afum|EAL89477.1 as seed ortholog is 100%.
Bootstrap support for afis|EAW24832.1 as seed ortholog is 100%.

---

Group of orthologs #4440. Best score 949 bits
Score difference with first non-orthologous sequence - *A. fumigatus*:949 *A. fischeri*:949
afum|EAL89579.1 100.00% afis|EAW24924.1 100.00%
Bootstrap support for afum|EAL89579.1 as seed ortholog is 100%.
Bootstrap support for afis|EAW24924.1 as seed ortholog is 100%.

---

Group of orthologs #4441. Best score 949 bits
Score difference with first non-orthologous sequence - *A. fumigatus*:949 *A. fischeri*:949
afum|EAL89868.1 100.00% afis|EAW25207.1 100.00%
Bootstrap support for afum|EAL89868.1 as seed ortholog is 100%.
Bootstrap support for afis|EAW25207.1 as seed ortholog is 100%.

---

Group of orthologs #4442. Best score 948 bits
Score difference with first non-orthologous sequence - *A. fumigatus*:948 *A. fischeri*:948
afum|EAL84228.1 100.00% afis|EAW15417.1 100.00%
Bootstrap support for afum|EAL84228.1 as seed ortholog is 100%.
Bootstrap support for afis|EAW15417.1 as seed ortholog is 100%.

---

Group of orthologs #4443. Best score 948 bits
Score difference with first non-orthologous sequence - *A. fumigatus*:948 *A. fischeri*:948
afum|EAL91497.1 100.00% afis|EAW17683.1 100.00%
Bootstrap support for afum|EAL91497.1 as seed ortholog is 100%.
Bootstrap support for afis|EAW17683.1 as seed ortholog is 100%.

---

Group of orthologs #4444. Best score 948 bits
Score difference with first non-orthologous sequence - *A. fumigatus*:948 *A. fischeri*:948
afum|EAL91619.1 100.00% afis|EAW17811.1 100.00%
Bootstrap support for afum|EAL91619.1 as seed ortholog is 100%.
Bootstrap support for afis|EAW17811.1 as seed ortholog is 100%.

---

Group of orthologs #4445. Best score 948 bits
Score difference with first non-orthologous sequence - *A. fumigatus*:948 *A. fischeri*:948
afum|EBA27436.1 100.00% afis|EAW17672.1 100.00%
Bootstrap support for afum|EBA27436.1 as seed ortholog is 100%.
Bootstrap support for afis|EAW17672.1 as seed ortholog is 100%.

---

Group of orthologs #4446. Best score 948 bits
Score difference with first non-orthologous sequence - *A. fumigatus*:948 *A. fischeri*:948
afum|EAL90356.1 100.00% afis|EAW22846.1 100.00%
Bootstrap support for afum|EAL90356.1 as seed ortholog is 100%.
Bootstrap support for afis|EAW22846.1 as seed ortholog is 100%.

---

Group of orthologs #4447. Best score 947 bits
Score difference with first non-orthologous sequence - *A. fumigatus*:947 *A. fischeri*:947
afum|EAL89228.1 100.00% afis|EAW16632.1 100.00%
Bootstrap support for afum|EAL89228.1 as seed ortholog is 100%.
Bootstrap support for afis|EAW16632.1 as seed ortholog is 100%.

---

Group of orthologs #4448. Best score 947 bits
Score difference with first non-orthologous sequence - *A. fumigatus*:687 *A. fischeri*:947
afum|EBA27170.1 100.00% afis|EAW15420.1 100.00%
Bootstrap support for afum|EBA27170.1 as seed ortholog is 100%.
Bootstrap support for afis|EAW15420.1 as seed ortholog is 100%.

---

Group of orthologs #4449. Best score 947 bits

Score difference with first non-orthologous sequence - *A. fumigatus*:947   *A. fischeri*:947
afum|EAL88112.1 100.00% afis|EAW23362.1 100.00%
Bootstrap support for afum|EAL88112.1 as seed ortholog is 100%.
Bootstrap support for afis|EAW23362.1 as seed ortholog is 100%.

---

Group of orthologs #4450. Best score 947 bits
Score difference with first non-orthologous sequence - *A. fumigatus*:947   *A. fischeri*:947
afum|EAL87832.1 100.00% afis|EAW23628.1 100.00%
Bootstrap support for afum|EAL87832.1 as seed ortholog is 100%.
Bootstrap support for afis|EAW23628.1 as seed ortholog is 100%.

---

Group of orthologs #4451. Best score 947 bits
Score difference with first non-orthologous sequence - *A. fumigatus*:947   *A. fischeri*:947
afum|EAL90065.1 100.00% afis|EAW25406.1 100.00%
Bootstrap support for afum|EAL90065.1 as seed ortholog is 100%.
Bootstrap support for afis|EAW25406.1 as seed ortholog is 100%.

---

Group of orthologs #4452. Best score 946 bits
Score difference with first non-orthologous sequence - *A. fumigatus*:946   *A. fischeri*:946
afum|EAL88615.1 100.00% afis|EAW15983.1 100.00%
Bootstrap support for afum|EAL88615.1 as seed ortholog is 100%.
Bootstrap support for afis|EAW15983.1 as seed ortholog is 100%.

---

Group of orthologs #4453. Best score 946 bits
Score difference with first non-orthologous sequence - *A. fumigatus*:946   *A. fischeri*:946
afum|EAL89199.1 100.00% afis|EAW16603.1 100.00%
Bootstrap support for afum|EAL89199.1 as seed ortholog is 100%.
Bootstrap support for afis|EAW16603.1 as seed ortholog is 100%.

---

Group of orthologs #4454. Best score 946 bits
Score difference with first non-orthologous sequence - *A. fumigatus*:946   *A. fischeri*:946
afum|EAL85945.1 100.00% afis|EAW24276.1 100.00%
Bootstrap support for afum|EAL85945.1 as seed ortholog is 100%.
Bootstrap support for afis|EAW24276.1 as seed ortholog is 100%.

---

Group of orthologs #4455. Best score 946 bits
Score difference with first non-orthologous sequence - *A. fumigatus*:946   *A. fischeri*:946
afum|EAL88263.1 100.00% afis|EAW23221.1 100.00%
Bootstrap support for afum|EAL88263.1 as seed ortholog is 100%.
Bootstrap support for afis|EAW23221.1 as seed ortholog is 100%.

---

Group of orthologs #4456. Best score 946 bits
Score difference with first non-orthologous sequence - *A. fumigatus*:946   *A. fischeri*:946
afum|EAL90798.1 100.00% afis|EAW22393.1 100.00%
Bootstrap support for afum|EAL90798.1 as seed ortholog is 100%.
Bootstrap support for afis|EAW22393.1 as seed ortholog is 100%.

---

Group of orthologs #4457. Best score 946 bits
Score difference with first non-orthologous sequence - *A. fumigatus*:946   *A. fischeri*:946
afum|EAL92707.1 100.00% afis|EAW21672.1 100.00%
Bootstrap support for afum|EAL92707.1 as seed ortholog is 100%.
Bootstrap support for afis|EAW21672.1 as seed ortholog is 100%.

---

Group of orthologs #4458. Best score 946 bits
Score difference with first non-orthologous sequence - *A. fumigatus*:946   *A. fischeri*:946
afum|EBA27348.1 100.00% afis|EAW23010.1 100.00%

Bootstrap support for afum|EBA27348.1 as seed ortholog is 100%.
Bootstrap support for afis|EAW23010.1 as seed ortholog is 100%.

---

Group of orthologs #4459. Best score 945 bits
Score difference with first non-orthologous sequence - *A. fumigatus*:945 *A. fischeri*:945
afum|EAL93558.1 100.00% afis|EAW18821.1 100.00%
Bootstrap support for afum|EAL93558.1 as seed ortholog is 100%.
Bootstrap support for afis|EAW18821.1 as seed ortholog is 100%.

---

Group of orthologs #4460. Best score 945 bits
Score difference with first non-orthologous sequence - *A. fumigatus*:890 *A. fischeri*:945
afum|EAL90394.1 100.00% afis|EAW22805.1 100.00%
Bootstrap support for afum|EAL90394.1 as seed ortholog is 100%.
Bootstrap support for afis|EAW22805.1 as seed ortholog is 100%.

---

Group of orthologs #4461. Best score 945 bits
Score difference with first non-orthologous sequence - *A. fumigatus*:945 *A. fischeri*:945
afum|EAL89518.1 100.00% afis|EAW24868.1 100.00%
Bootstrap support for afum|EAL89518.1 as seed ortholog is 100%.
Bootstrap support for afis|EAW24868.1 as seed ortholog is 100%.

---

Group of orthologs #4462. Best score 945 bits
Score difference with first non-orthologous sequence - *A. fumigatus*:945 *A. fischeri*:945
afum|EAL90183.1 100.00% afis|EAW25518.1 100.00%
Bootstrap support for afum|EAL90183.1 as seed ortholog is 100%.
Bootstrap support for afis|EAW25518.1 as seed ortholog is 100%.

---

Group of orthologs #4463. Best score 944 bits
Score difference with first non-orthologous sequence - *A. fumigatus*:944 *A. fischeri*:944
afum|EAL84735.1 100.00% afis|EAW20897.1 100.00%
Bootstrap support for afum|EAL84735.1 as seed ortholog is 100%.
Bootstrap support for afis|EAW20897.1 as seed ortholog is 100%.

---

Group of orthologs #4464. Best score 944 bits
Score difference with first non-orthologous sequence - *A. fumigatus*:944 *A. fischeri*:944
afum|EAL87046.1 100.00% afis|EAW19450.1 100.00%
Bootstrap support for afum|EAL87046.1 as seed ortholog is 100%.
Bootstrap support for afis|EAW19450.1 as seed ortholog is 100%.

---

Group of orthologs #4465. Best score 944 bits
Score difference with first non-orthologous sequence - *A. fumigatus*:944 *A. fischeri*:944
afum|EAL87294.1 100.00% afis|EAW23866.1 100.00%
Bootstrap support for afum|EAL87294.1 as seed ortholog is 100%.
Bootstrap support for afis|EAW23866.1 as seed ortholog is 100%.

---

Group of orthologs #4466. Best score 944 bits
Score difference with first non-orthologous sequence - *A. fumigatus*:944 *A. fischeri*:944
afum|EAL92801.1 100.00% afis|EAW21772.1 100.00%
Bootstrap support for afum|EAL92801.1 as seed ortholog is 100%.
Bootstrap support for afis|EAW21772.1 as seed ortholog is 100%.

---

Group of orthologs #4467. Best score 944 bits
Score difference with first non-orthologous sequence - *A. fumigatus*:944 *A. fischeri*:944
afum|EAL92902.1 100.00% afis|EAW21888.1 100.00%
Bootstrap support for afum|EAL92902.1 as seed ortholog is 100%.
Bootstrap support for afis|EAW21888.1 as seed ortholog is 100%.

---

Group of orthologs #4468. Best score 943 bits
Score difference with first non-orthologous sequence - *A. fumigatus*:943 *A. fischeri*:943
afum|EAL86454.1 100.00% afis|EAW16929.1 100.00%
Bootstrap support for afum|EAL86454.1 as seed ortholog is 100%.
Bootstrap support for afis|EAW16929.1 as seed ortholog is 100%.

---

Group of orthologs #4469. Best score 943 bits
Score difference with first non-orthologous sequence - *A. fumigatus*:943 *A. fischeri*:943
afum|EAL84805.1 100.00% afis|EAW20824.1 100.00%
Bootstrap support for afum|EAL84805.1 as seed ortholog is 100%.
Bootstrap support for afis|EAW20824.1 as seed ortholog is 100%.

---

Group of orthologs #4470. Best score 943 bits
Score difference with first non-orthologous sequence - *A. fumigatus*:943 *A. fischeri*:943
afum|EAL87036.1 100.00% afis|EAW19460.1 100.00%
Bootstrap support for afum|EAL87036.1 as seed ortholog is 100%.
Bootstrap support for afis|EAW19460.1 as seed ortholog is 100%.

---

Group of orthologs #4471. Best score 943 bits
Score difference with first non-orthologous sequence - *A. fumigatus*:943 *A. fischeri*:943
afum|EAL91542.1 100.00% afis|EAW17727.1 100.00%
Bootstrap support for afum|EAL91542.1 as seed ortholog is 100%.
Bootstrap support for afis|EAW17727.1 as seed ortholog is 100%.

---

Group of orthologs #4472. Best score 943 bits
Score difference with first non-orthologous sequence - *A. fumigatus*:943 *A. fischeri*:943
afum|EAL87121.1 100.00% afis|EAW23678.1 100.00%
Bootstrap support for afum|EAL87121.1 as seed ortholog is 100%.
Bootstrap support for afis|EAW23678.1 as seed ortholog is 100%.

---

Group of orthologs #4473. Best score 943 bits
Score difference with first non-orthologous sequence - *A. fumigatus*:943 *A. fischeri*:943
afum|EAL89409.1 100.00% afis|EAW24768.1 100.00%
Bootstrap support for afum|EAL89409.1 as seed ortholog is 100%.
Bootstrap support for afis|EAW24768.1 as seed ortholog is 100%.

---

Group of orthologs #4474. Best score 942 bits
Score difference with first non-orthologous sequence - *A. fumigatus*:942 *A. fischeri*:942
afum|EAL85672.1 100.00% afis|EAW15675.1 100.00%
Bootstrap support for afum|EAL85672.1 as seed ortholog is 100%.
Bootstrap support for afis|EAW15675.1 as seed ortholog is 100%.

---

Group of orthologs #4475. Best score 942 bits
Score difference with first non-orthologous sequence - *A. fumigatus*:781 *A. fischeri*:792
afum|EAL86613.2 100.00% afis|EAW17187.1 100.00%
Bootstrap support for afum|EAL86613.2 as seed ortholog is 100%.
Bootstrap support for afis|EAW17187.1 as seed ortholog is 100%.

---

Group of orthologs #4476. Best score 942 bits
Score difference with first non-orthologous sequence - *A. fumigatus*:886 *A. fischeri*:545
afum|EAL84862.1 100.00% afis|EAW20757.1 100.00%
Bootstrap support for afum|EAL84862.1 as seed ortholog is 100%.
Bootstrap support for afis|EAW20757.1 as seed ortholog is 100%.

---

Group of orthologs #4477. Best score 942 bits

Score difference with first non-orthologous sequence - *A. fumigatus*:942 *A. fischeri*:942
afum|EAL91383.1 100.00% afis|EAW17570.1 100.00%
Bootstrap support for afum|EAL91383.1 as seed ortholog is 100%.
Bootstrap support for afis|EAW17570.1 as seed ortholog is 100%.

---

Group of orthologs #4478. Best score 942 bits
Score difference with first non-orthologous sequence - *A. fumigatus*:942 *A. fischeri*:942
afum|EAL91414.1 100.00% afis|EAW17600.1 100.00%
Bootstrap support for afum|EAL91414.1 as seed ortholog is 100%.
Bootstrap support for afis|EAW17600.1 as seed ortholog is 100%.

---

Group of orthologs #4479. Best score 942 bits
Score difference with first non-orthologous sequence - *A. fumigatus*:942 *A. fischeri*:942
afum|EAL85937.1 100.00% afis|EAW24269.1 100.00%
Bootstrap support for afum|EAL85937.1 as seed ortholog is 100%.
Bootstrap support for afis|EAW24269.1 as seed ortholog is 100%.

---

Group of orthologs #4480. Best score 942 bits
Score difference with first non-orthologous sequence - *A. fumigatus*:942 *A. fischeri*:942
afum|EAL93114.1 100.00% afis|EAW18433.1 100.00%
Bootstrap support for afum|EAL93114.1 as seed ortholog is 100%.
Bootstrap support for afis|EAW18433.1 as seed ortholog is 100%.

---

Group of orthologs #4481. Best score 942 bits
Score difference with first non-orthologous sequence - *A. fumigatus*:942 *A. fischeri*:942
afum|EAL93850.1 100.00% afis|EAW19118.1 100.00%
Bootstrap support for afum|EAL93850.1 as seed ortholog is 100%.
Bootstrap support for afis|EAW19118.1 as seed ortholog is 100%.

---

Group of orthologs #4482. Best score 942 bits
Score difference with first non-orthologous sequence - *A. fumigatus*:942 *A. fischeri*:942
afum|EAL90419.1 100.00% afis|EAW22780.1 100.00%
Bootstrap support for afum|EAL90419.1 as seed ortholog is 100%.
Bootstrap support for afis|EAW22780.1 as seed ortholog is 100%.

---

Group of orthologs #4483. Best score 942 bits
Score difference with first non-orthologous sequence - *A. fumigatus*:942 *A. fischeri*:942
afum|EAL89582.1 100.00% afis|EAW24926.1 100.00%
Bootstrap support for afum|EAL89582.1 as seed ortholog is 100%.
Bootstrap support for afis|EAW24926.1 as seed ortholog is 100%.

---

Group of orthologs #4484. Best score 941 bits
Score difference with first non-orthologous sequence - *A. fumigatus*:488 *A. fischeri*:941
afum|EAL85767.2 100.00% afis|EAW15566.1 100.00%
Bootstrap support for afum|EAL85767.2 as seed ortholog is 100%.
Bootstrap support for afis|EAW15566.1 as seed ortholog is 100%.

---

Group of orthologs #4485. Best score 941 bits
Score difference with first non-orthologous sequence - *A. fumigatus*:941 *A. fischeri*:941
afum|EAL85675.1 100.00% afis|EAW15673.1 100.00%
Bootstrap support for afum|EAL85675.1 as seed ortholog is 100%.
Bootstrap support for afis|EAW15673.1 as seed ortholog is 100%.

---

Group of orthologs #4486. Best score 941 bits
Score difference with first non-orthologous sequence - *A. fumigatus*:941 *A. fischeri*:941
afum|EAL86796.1 100.00% afis|EAW19732.1 100.00%

Bootstrap support for afum|EAL86796.1 as seed ortholog is 100%.
Bootstrap support for afis|EAW19732.1 as seed ortholog is 100%.

---

Group of orthologs #4487. Best score 941 bits
Score difference with first non-orthologous sequence - *A. fumigatus*:941 *A. fischeri*:941
afum|EAL91218.1 100.00% afis|EAW17383.1 100.00%
Bootstrap support for afum|EAL91218.1 as seed ortholog is 100%.
Bootstrap support for afis|EAW17383.1 as seed ortholog is 100%.

---

Group of orthologs #4488. Best score 941 bits
Score difference with first non-orthologous sequence - *A. fumigatus*:941 *A. fischeri*:941
afum|EAL93510.1 100.00% afis|EAW18778.1 100.00%
Bootstrap support for afum|EAL93510.1 as seed ortholog is 100%.
Bootstrap support for afis|EAW18778.1 as seed ortholog is 100%.

---

Group of orthologs #4489. Best score 941 bits
Score difference with first non-orthologous sequence - *A. fumigatus*:941 *A. fischeri*:941
afum|EAL90665.1 100.00% afis|EAW22529.1 100.00%
Bootstrap support for afum|EAL90665.1 as seed ortholog is 100%.
Bootstrap support for afis|EAW22529.1 as seed ortholog is 100%.

---

Group of orthologs #4490. Best score 941 bits
Score difference with first non-orthologous sequence - *A. fumigatus*:941 *A. fischeri*:941
afum|EAL92294.1 100.00% afis|EAW21279.1 100.00%
Bootstrap support for afum|EAL92294.1 as seed ortholog is 100%.
Bootstrap support for afis|EAW21279.1 as seed ortholog is 100%.

---

Group of orthologs #4491. Best score 941 bits
Score difference with first non-orthologous sequence - *A. fumigatus*:941 *A. fischeri*:941
afum|EAL92760.1 100.00% afis|EAW21727.1 100.00%
Bootstrap support for afum|EAL92760.1 as seed ortholog is 100%.
Bootstrap support for afis|EAW21727.1 as seed ortholog is 100%.

---

Group of orthologs #4492. Best score 940 bits
Score difference with first non-orthologous sequence - *A. fumigatus*:823 *A. fischeri*:813
afum|EAL88861.1 100.00% afis|EAW16240.1 100.00%
Bootstrap support for afum|EAL88861.1 as seed ortholog is 100%.
Bootstrap support for afis|EAW16240.1 as seed ortholog is 100%.

---

Group of orthologs #4493. Best score 940 bits
Score difference with first non-orthologous sequence - *A. fumigatus*:940 *A. fischeri*:698
afum|EAL89802.1 100.00% afis|EAW25131.1 100.00%
Bootstrap support for afum|EAL89802.1 as seed ortholog is 100%.
Bootstrap support for afis|EAW25131.1 as seed ortholog is 100%.

---

Group of orthologs #4494. Best score 939 bits
Score difference with first non-orthologous sequence - *A. fumigatus*:939 *A. fischeri*:708
afum|EAL86463.1 100.00% afis|EAW16938.1 100.00%
Bootstrap support for afum|EAL86463.1 as seed ortholog is 100%.
Bootstrap support for afis|EAW16938.1 as seed ortholog is 100%.

---

Group of orthologs #4495. Best score 939 bits
Score difference with first non-orthologous sequence - *A. fumigatus*:939 *A. fischeri*:883
afum|EAL86615.1 100.00% afis|EAW17189.1 100.00%
Bootstrap support for afum|EAL86615.1 as seed ortholog is 100%.
Bootstrap support for afis|EAW17189.1 as seed ortholog is 100%.

---

Group of orthologs #4496. Best score 939 bits
Score difference with first non-orthologous sequence - *A. fumigatus*:939 *A. fischeri*:939
afum|EAL87061.1 100.00% afis|EAW19434.1 100.00%
Bootstrap support for afum|EAL87061.1 as seed ortholog is 100%.
Bootstrap support for afis|EAW19434.1 as seed ortholog is 100%.

---

Group of orthologs #4497. Best score 939 bits
Score difference with first non-orthologous sequence - *A. fumigatus*:939 *A. fischeri*:939
afum|EAL88460.1 100.00% afis|EAW23022.1 100.00%
Bootstrap support for afum|EAL88460.1 as seed ortholog is 100%.
Bootstrap support for afis|EAW23022.1 as seed ortholog is 100%.

---

Group of orthologs #4498. Best score 939 bits
Score difference with first non-orthologous sequence - *A. fumigatus*:939 *A. fischeri*:939
afum|EBA27497.1 100.00% afis|EAW18914.1 100.00%
Bootstrap support for afum|EBA27497.1 as seed ortholog is 100%.
Bootstrap support for afis|EAW18914.1 as seed ortholog is 100%.

---

Group of orthologs #4499. Best score 939 bits
Score difference with first non-orthologous sequence - *A. fumigatus*:939 *A. fischeri*:939
afum|EAL92968.1 100.00% afis|EAW21954.1 100.00%
Bootstrap support for afum|EAL92968.1 as seed ortholog is 100%.
Bootstrap support for afis|EAW21954.1 as seed ortholog is 100%.

---

Group of orthologs #4500. Best score 938 bits
Score difference with first non-orthologous sequence - *A. fumigatus*:938 *A. fischeri*:938
afum|EAL85529.1 100.00% afis|EAW15822.1 100.00%
Bootstrap support for afum|EAL85529.1 as seed ortholog is 100%.
Bootstrap support for afis|EAW15822.1 as seed ortholog is 100%.

---

Group of orthologs #4501. Best score 938 bits
Score difference with first non-orthologous sequence - *A. fumigatus*:938 *A. fischeri*:649
afum|EAL86390.1 100.00% afis|EAW16844.1 100.00%
Bootstrap support for afum|EAL86390.1 as seed ortholog is 100%.
Bootstrap support for afis|EAW16844.1 as seed ortholog is 100%.

---

Group of orthologs #4502. Best score 938 bits
Score difference with first non-orthologous sequence - *A. fumigatus*:938 *A. fischeri*:938
afum|EAL84764.1 100.00% afis|EAW20870.1 100.00%
Bootstrap support for afum|EAL84764.1 as seed ortholog is 100%.
Bootstrap support for afis|EAW20870.1 as seed ortholog is 100%.

---

Group of orthologs #4503. Best score 938 bits
Score difference with first non-orthologous sequence - *A. fumigatus*:938 *A. fischeri*:938
afum|EAL92282.1 100.00% afis|EAW21267.1 100.00%
Bootstrap support for afum|EAL92282.1 as seed ortholog is 100%.
Bootstrap support for afis|EAW21267.1 as seed ortholog is 100%.

---

Group of orthologs #4504. Best score 937 bits
Score difference with first non-orthologous sequence - *A. fumigatus*:937 *A. fischeri*:937
afum|EAL91316.1 100.00% afis|EAW17494.1 100.00%
Bootstrap support for afum|EAL91316.1 as seed ortholog is 100%.
Bootstrap support for afis|EAW17494.1 as seed ortholog is 100%.

---

Group of orthologs #4505. Best score 937 bits

Score difference with first non-orthologous sequence - *A. fumigatus*:937 *A. fischeri*:937
afum|EAL93130.1 100.00% afis|EAW18445.1 100.00%
Bootstrap support for afum|EAL93130.1 as seed ortholog is 100%.
Bootstrap support for afis|EAW18445.1 as seed ortholog is 100%.

---

Group of orthologs #4506. Best score 937 bits
Score difference with first non-orthologous sequence - *A. fumigatus*:496 *A. fischeri*:706
afum|EAL90955.1 100.00% afis|EAW22238.1 100.00%
Bootstrap support for afum|EAL90955.1 as seed ortholog is 100%.
Bootstrap support for afis|EAW22238.1 as seed ortholog is 100%.

---

Group of orthologs #4507. Best score 937 bits
Score difference with first non-orthologous sequence - *A. fumigatus*:937 *A. fischeri*:937
afum|EAL89374.1 100.00% afis|EAW24726.1 100.00%
Bootstrap support for afum|EAL89374.1 as seed ortholog is 100%.
Bootstrap support for afis|EAW24726.1 as seed ortholog is 100%.

---

Group of orthologs #4508. Best score 936 bits
Score difference with first non-orthologous sequence - *A. fumigatus*:787 *A. fischeri*:787
afum|EAL87627.1 100.00% afis|EAW18263.1 100.00%
Bootstrap support for afum|EAL87627.1 as seed ortholog is 100%.
Bootstrap support for afis|EAW18263.1 as seed ortholog is 100%.

---

Group of orthologs #4509. Best score 936 bits
Score difference with first non-orthologous sequence - *A. fumigatus*:936 *A. fischeri*:936
afum|EAL88307.2 100.00% afis|EAW23176.1 100.00%
Bootstrap support for afum|EAL88307.2 as seed ortholog is 100%.
Bootstrap support for afis|EAW23176.1 as seed ortholog is 100%.

---

Group of orthologs #4510. Best score 936 bits
Score difference with first non-orthologous sequence - *A. fumigatus*:936 *A. fischeri*:936
afum|EAL88099.1 100.00% afis|EAW23373.1 100.00%
Bootstrap support for afum|EAL88099.1 as seed ortholog is 100%.
Bootstrap support for afis|EAW23373.1 as seed ortholog is 100%.

---

Group of orthologs #4511. Best score 936 bits
Score difference with first non-orthologous sequence - *A. fumigatus*:936 *A. fischeri*:936
afum|EAL93450.1 100.00% afis|EAW18725.1 100.00%
Bootstrap support for afum|EAL93450.1 as seed ortholog is 100%.
Bootstrap support for afis|EAW18725.1 as seed ortholog is 100%.

---

Group of orthologs #4512. Best score 936 bits
Score difference with first non-orthologous sequence - *A. fumigatus*:936 *A. fischeri*:936
afum|EAL89454.1 100.00% afis|EAW24811.1 100.00%
Bootstrap support for afum|EAL89454.1 as seed ortholog is 100%.
Bootstrap support for afis|EAW24811.1 as seed ortholog is 100%.

---

Group of orthologs #4513. Best score 936 bits
Score difference with first non-orthologous sequence - *A. fumigatus*:936 *A. fischeri*:936
afum|EAL89554.1 100.00% afis|EAW24900.1 100.00%
Bootstrap support for afum|EAL89554.1 as seed ortholog is 100%.
Bootstrap support for afis|EAW24900.1 as seed ortholog is 100%.

---

Group of orthologs #4514. Best score 935 bits
Score difference with first non-orthologous sequence - *A. fumigatus*:935 *A. fischeri*:935
afum|EAL85005.1 100.00% afis|EAW19896.1 100.00%

Bootstrap support for afum|EAL85005.1 as seed ortholog is 100%.
Bootstrap support for afis|EAW19896.1 as seed ortholog is 100%.

---

Group of orthologs #4515. Best score 935 bits
Score difference with first non-orthologous sequence - *A. fumigatus*:935   *A. fischeri*:755
afum|EAL84953.1 100.00% afis|EAW19971.1 100.00%
Bootstrap support for afum|EAL84953.1 as seed ortholog is 100%.
Bootstrap support for afis|EAW19971.1 as seed ortholog is 100%.

---

Group of orthologs #4516. Best score 935 bits
Score difference with first non-orthologous sequence - *A. fumigatus*:935   *A. fischeri*:935
afum|EAL86065.1 100.00% afis|EAW24393.1 100.00%
Bootstrap support for afum|EAL86065.1 as seed ortholog is 100%.
Bootstrap support for afis|EAW24393.1 as seed ortholog is 100%.

---

Group of orthologs #4517. Best score 935 bits
Score difference with first non-orthologous sequence - *A. fumigatus*:935   *A. fischeri*:935
afum|EAL87131.1 100.00% afis|EAW23686.1 100.00%
Bootstrap support for afum|EAL87131.1 as seed ortholog is 100%.
Bootstrap support for afis|EAW23686.1 as seed ortholog is 100%.

---

Group of orthologs #4518. Best score 935 bits
Score difference with first non-orthologous sequence - *A. fumigatus*:935   *A. fischeri*:935
afum|EAL87955.1 100.00% afis|EAW23506.1 100.00%
Bootstrap support for afum|EAL87955.1 as seed ortholog is 100%.
Bootstrap support for afis|EAW23506.1 as seed ortholog is 100%.

---

Group of orthologs #4519. Best score 935 bits
Score difference with first non-orthologous sequence - *A. fumigatus*:935   *A. fischeri*:935
afum|EAL93695.1 100.00% afis|EAW18963.1 100.00%
Bootstrap support for afum|EAL93695.1 as seed ortholog is 100%.
Bootstrap support for afis|EAW18963.1 as seed ortholog is 100%.

---

Group of orthologs #4520. Best score 934 bits
Score difference with first non-orthologous sequence - *A. fumigatus*:934   *A. fischeri*:934
afum|EAL88613.1 100.00% afis|EAW15981.1 100.00%
Bootstrap support for afum|EAL88613.1 as seed ortholog is 100%.
Bootstrap support for afis|EAW15981.1 as seed ortholog is 100%.

---

Group of orthologs #4521. Best score 934 bits
Score difference with first non-orthologous sequence - *A. fumigatus*:934   *A. fischeri*:265
afum|EAL89005.1 100.00% afis|EAW16380.1 100.00%
Bootstrap support for afum|EAL89005.1 as seed ortholog is 100%.
Bootstrap support for afis|EAW16380.1 as seed ortholog is 100%.

---

Group of orthologs #4522. Best score 934 bits
Score difference with first non-orthologous sequence - *A. fumigatus*:934   *A. fischeri*:934
afum|EAL89252.1 100.00% afis|EAW16648.1 100.00%
Bootstrap support for afum|EAL89252.1 as seed ortholog is 100%.
Bootstrap support for afis|EAW16648.1 as seed ortholog is 100%.

---

Group of orthologs #4523. Best score 934 bits
Score difference with first non-orthologous sequence - *A. fumigatus*:934   *A. fischeri*:934
afum|EAL91272.1 100.00% afis|EAW17446.1 100.00%
Bootstrap support for afum|EAL91272.1 as seed ortholog is 100%.
Bootstrap support for afis|EAW17446.1 as seed ortholog is 100%.

Group of orthologs #4524. Best score 934 bits
Score difference with first non-orthologous sequence - *A. fumigatus*:934 *A. fischeri*:661
afum|EAL86632.1 100.00% afis|EAW23637.1 100.00%
Bootstrap support for afum|EAL86632.1 as seed ortholog is 100%.
Bootstrap support for afis|EAW23637.1 as seed ortholog is 100%.

Group of orthologs #4525. Best score 934 bits
Score difference with first non-orthologous sequence - *A. fumigatus*:934 *A. fischeri*:934
afum|EAL85997.1 100.00% afis|EAW24327.1 100.00%
Bootstrap support for afum|EAL85997.1 as seed ortholog is 100%.
Bootstrap support for afis|EAW24327.1 as seed ortholog is 100%.

Group of orthologs #4526. Best score 934 bits
Score difference with first non-orthologous sequence - *A. fumigatus*:934 *A. fischeri*:934
afum|EAL90591.1 100.00% afis|EAW22607.1 100.00%
Bootstrap support for afum|EAL90591.1 as seed ortholog is 100%.
Bootstrap support for afis|EAW22607.1 as seed ortholog is 100%.

Group of orthologs #4527. Best score 934 bits
Score difference with first non-orthologous sequence - *A. fumigatus*:455 *A. fischeri*:934
afum|EAL94074.2 100.00% afis|EAW19354.1 100.00%
Bootstrap support for afum|EAL94074.2 as seed ortholog is 100%.
Bootstrap support for afis|EAW19354.1 as seed ortholog is 100%.

Group of orthologs #4528. Best score 933 bits
Score difference with first non-orthologous sequence - *A. fumigatus*:933 *A. fischeri*:933
afum|EAL88621.1 100.00% afis|EAW15989.1 100.00%
Bootstrap support for afum|EAL88621.1 as seed ortholog is 100%.
Bootstrap support for afis|EAW15989.1 as seed ortholog is 100%.

Group of orthologs #4529. Best score 933 bits
Score difference with first non-orthologous sequence - *A. fumigatus*:933 *A. fischeri*:933
afum|EAL85231.1 100.00% afis|EAW20114.1 100.00%
Bootstrap support for afum|EAL85231.1 as seed ortholog is 100%.
Bootstrap support for afis|EAW20114.1 as seed ortholog is 100%.

Group of orthologs #4530. Best score 933 bits
Score difference with first non-orthologous sequence - *A. fumigatus*:933 *A. fischeri*:933
afum|EAL86955.2 100.00% afis|EAW19556.1 100.00%
Bootstrap support for afum|EAL86955.2 as seed ortholog is 100%.
Bootstrap support for afis|EAW19556.1 as seed ortholog is 100%.

Group of orthologs #4531. Best score 933 bits
Score difference with first non-orthologous sequence - *A. fumigatus*:933 *A. fischeri*:933
afum|EAL91268.1 100.00% afis|EAW17442.1 100.00%
Bootstrap support for afum|EAL91268.1 as seed ortholog is 100%.
Bootstrap support for afis|EAW17442.1 as seed ortholog is 100%.

Group of orthologs #4532. Best score 933 bits
Score difference with first non-orthologous sequence - *A. fumigatus*:933 *A. fischeri*:634
afum|EAL85972.1 100.00% afis|EAW24304.1 100.00%
Bootstrap support for afum|EAL85972.1 as seed ortholog is 100%.
Bootstrap support for afis|EAW24304.1 as seed ortholog is 100%.

Group of orthologs #4533. Best score 933 bits

Score difference with first non-orthologous sequence - *A. fumigatus*:933 *A. fischeri*:933
afum|EAL93953.1 100.00% afis|EAW19220.1 100.00%
Bootstrap support for afum|EAL93953.1 as seed ortholog is 100%.
Bootstrap support for afis|EAW19220.1 as seed ortholog is 100%.

---

Group of orthologs #4534. Best score 933 bits
Score difference with first non-orthologous sequence - *A. fumigatus*:933 *A. fischeri*:933
afum|EAL90542.1 100.00% afis|EAW22650.1 100.00%
Bootstrap support for afum|EAL90542.1 as seed ortholog is 100%.
Bootstrap support for afis|EAW22650.1 as seed ortholog is 100%.

---

Group of orthologs #4535. Best score 933 bits
Score difference with first non-orthologous sequence - *A. fumigatus*:933 *A. fischeri*:933
afum|EAL90309.1 100.00% afis|EAW22889.1 100.00%
Bootstrap support for afum|EAL90309.1 as seed ortholog is 100%.
Bootstrap support for afis|EAW22889.1 as seed ortholog is 100%.

---

Group of orthologs #4536. Best score 933 bits
Score difference with first non-orthologous sequence - *A. fumigatus*:464 *A. fischeri*:807
afum|EAL92780.2 100.00% afis|EAW21749.1 100.00%
Bootstrap support for afum|EAL92780.2 as seed ortholog is 100%.
Bootstrap support for afis|EAW21749.1 as seed ortholog is 100%.

---

Group of orthologs #4537. Best score 932 bits
Score difference with first non-orthologous sequence - *A. fumigatus*:398 *A. fischeri*:390
afum|EAL85095.2 100.00% afis|EAW19795.1 100.00%
Bootstrap support for afum|EAL85095.2 as seed ortholog is 100%.
Bootstrap support for afis|EAW19795.1 as seed ortholog is 100%.

---

Group of orthologs #4538. Best score 932 bits
Score difference with first non-orthologous sequence - *A. fumigatus*:932 *A. fischeri*:932
afum|EAL85169.1 100.00% afis|EAW20048.1 100.00%
Bootstrap support for afum|EAL85169.1 as seed ortholog is 100%.
Bootstrap support for afis|EAW20048.1 as seed ortholog is 100%.

---

Group of orthologs #4539. Best score 932 bits
Score difference with first non-orthologous sequence - *A. fumigatus*:932 *A. fischeri*:932
afum|EAL89275.1 100.00% afis|EAW16670.1 100.00%
Bootstrap support for afum|EAL89275.1 as seed ortholog is 100%.
Bootstrap support for afis|EAW16670.1 as seed ortholog is 100%.

---

Group of orthologs #4540. Best score 932 bits
Score difference with first non-orthologous sequence - *A. fumigatus*:579 *A. fischeri*:932
afum|EAL86117.1 100.00% afis|EAW24448.1 100.00%
Bootstrap support for afum|EAL86117.1 as seed ortholog is 100%.
Bootstrap support for afis|EAW24448.1 as seed ortholog is 100%.

---

Group of orthologs #4541. Best score 932 bits
Score difference with first non-orthologous sequence - *A. fumigatus*:932 *A. fischeri*:932
afum|EAL88343.1 100.00% afis|EAW23138.1 100.00%
Bootstrap support for afum|EAL88343.1 as seed ortholog is 100%.
Bootstrap support for afis|EAW23138.1 as seed ortholog is 100%.

---

Group of orthologs #4542. Best score 932 bits
Score difference with first non-orthologous sequence - *A. fumigatus*:932 *A. fischeri*:932
afum|EAL90686.1 100.00% afis|EAW22507.1 100.00%

Bootstrap support for afum|EAL90686.1 as seed ortholog is 100%.
Bootstrap support for afis|EAW22507.1 as seed ortholog is 100%.

---

Group of orthologs #4543. Best score 931 bits
Score difference with first non-orthologous sequence - *A. fumigatus*:931 *A. fischeri*:931
afum|EAL88533.1 100.00% afis|EAW15898.1 100.00%
Bootstrap support for afum|EAL88533.1 as seed ortholog is 100%.
Bootstrap support for afis|EAW15898.1 as seed ortholog is 100%.

---

Group of orthologs #4544. Best score 931 bits
Score difference with first non-orthologous sequence - *A. fumigatus*:931 *A. fischeri*:931
afum|EAL87614.1 100.00% afis|EAW18249.1 100.00%
Bootstrap support for afum|EAL87614.1 as seed ortholog is 100%.
Bootstrap support for afis|EAW18249.1 as seed ortholog is 100%.

---

Group of orthologs #4545. Best score 931 bits
Score difference with first non-orthologous sequence - *A. fumigatus*:931 *A. fischeri*:931
afum|EAL85463.1 100.00% afis|EAW20341.1 100.00%
Bootstrap support for afum|EAL85463.1 as seed ortholog is 100%.
Bootstrap support for afis|EAW20341.1 as seed ortholog is 100%.

---

Group of orthologs #4546. Best score 931 bits
Score difference with first non-orthologous sequence - *A. fumigatus*:931 *A. fischeri*:931
afum|EAL93730.1 100.00% afis|EAW18995.1 100.00%
Bootstrap support for afum|EAL93730.1 as seed ortholog is 100%.
Bootstrap support for afis|EAW18995.1 as seed ortholog is 100%.

---

Group of orthologs #4547. Best score 931 bits
Score difference with first non-orthologous sequence - *A. fumigatus*:854 *A. fischeri*:814
afum|EAL89368.2 100.00% afis|EAW24719.1 100.00%
Bootstrap support for afum|EAL89368.2 as seed ortholog is 100%.
Bootstrap support for afis|EAW24719.1 as seed ortholog is 100%.

---

Group of orthologs #4548. Best score 931 bits
Score difference with first non-orthologous sequence - *A. fumigatus*:931 *A. fischeri*:931
afum|EAL92858.1 100.00% afis|EAW21846.1 100.00%
Bootstrap support for afum|EAL92858.1 as seed ortholog is 100%.
Bootstrap support for afis|EAW21846.1 as seed ortholog is 100%.

---

Group of orthologs #4549. Best score 930 bits
Score difference with first non-orthologous sequence - *A. fumigatus*:930 *A. fischeri*:930
afum|EAL91399.1 100.00% afis|EAW17586.1 100.00%
Bootstrap support for afum|EAL91399.1 as seed ortholog is 100%.
Bootstrap support for afis|EAW17586.1 as seed ortholog is 100%.

---

Group of orthologs #4550. Best score 930 bits
Score difference with first non-orthologous sequence - *A. fumigatus*:930 *A. fischeri*:930
afum|EAL87906.1 100.00% afis|EAW23562.1 100.00%
Bootstrap support for afum|EAL87906.1 as seed ortholog is 100%.
Bootstrap support for afis|EAW23562.1 as seed ortholog is 100%.

---

Group of orthologs #4551. Best score 930 bits
Score difference with first non-orthologous sequence - *A. fumigatus*:930 *A. fischeri*:930
afum|EAL93367.1 100.00% afis|EAW18651.1 100.00%
Bootstrap support for afum|EAL93367.1 as seed ortholog is 100%.
Bootstrap support for afis|EAW18651.1 as seed ortholog is 100%.

Group of orthologs #4552. Best score 930 bits
Score difference with first non-orthologous sequence - *A. fumigatus*:930 *A. fischeri*:930
afum|EAL94005.1 100.00% afis|EAW19279.1 100.00%
Bootstrap support for afum|EAL94005.1 as seed ortholog is 100%.
Bootstrap support for afis|EAW19279.1 as seed ortholog is 100%.

Group of orthologs #4553. Best score 930 bits
Score difference with first non-orthologous sequence - *A. fumigatus*:930 *A. fischeri*:930
afum|EAL90504.1 100.00% afis|EAW22689.1 100.00%
Bootstrap support for afum|EAL90504.1 as seed ortholog is 100%.
Bootstrap support for afis|EAW22689.1 as seed ortholog is 100%.

Group of orthologs #4554. Best score 930 bits
Score difference with first non-orthologous sequence - *A. fumigatus*:930 *A. fischeri*:930
afum|EAL90408.1 100.00% afis|EAW22791.1 100.00%
Bootstrap support for afum|EAL90408.1 as seed ortholog is 100%.
Bootstrap support for afis|EAW22791.1 as seed ortholog is 100%.

Group of orthologs #4555. Best score 930 bits
Score difference with first non-orthologous sequence - *A. fumigatus*:930 *A. fischeri*:930
afum|EAL92313.1 100.00% afis|EAW21296.1 100.00%
Bootstrap support for afum|EAL92313.1 as seed ortholog is 100%.
Bootstrap support for afis|EAW21296.1 as seed ortholog is 100%.

Group of orthologs #4556. Best score 929 bits
Score difference with first non-orthologous sequence - *A. fumigatus*:929 *A. fischeri*:929
afum|EAL93231.1 100.00% afis|EAW18533.1 100.00%
Bootstrap support for afum|EAL93231.1 as seed ortholog is 100%.
Bootstrap support for afis|EAW18533.1 as seed ortholog is 100%.

Group of orthologs #4557. Best score 929 bits
Score difference with first non-orthologous sequence - *A. fumigatus*:929 *A. fischeri*:929
afum|EAL90619.2 100.00% afis|EAW22580.1 100.00%
Bootstrap support for afum|EAL90619.2 as seed ortholog is 100%.
Bootstrap support for afis|EAW22580.1 as seed ortholog is 100%.

Group of orthologs #4558. Best score 928 bits
Score difference with first non-orthologous sequence - *A. fumigatus*:928 *A. fischeri*:928
afum|EAL84512.1 100.00% afis|EAW20547.1 100.00%
Bootstrap support for afum|EAL84512.1 as seed ortholog is 100%.
Bootstrap support for afis|EAW20547.1 as seed ortholog is 100%.

Group of orthologs #4559. Best score 928 bits
Score difference with first non-orthologous sequence - *A. fumigatus*:928 *A. fischeri*:928
afum|EAL85313.1 100.00% afis|EAW20193.1 100.00%
Bootstrap support for afum|EAL85313.1 as seed ortholog is 100%.
Bootstrap support for afis|EAW20193.1 as seed ortholog is 100%.

Group of orthologs #4560. Best score 928 bits
Score difference with first non-orthologous sequence - *A. fumigatus*:928 *A. fischeri*:928
afum|EAL93418.1 100.00% afis|EAW18696.1 100.00%
Bootstrap support for afum|EAL93418.1 as seed ortholog is 100%.
Bootstrap support for afis|EAW18696.1 as seed ortholog is 100%.

Group of orthologs #4561. Best score 928 bits

Score difference with first non-orthologous sequence - *A. fumigatus*:928   *A. fischeri*:928
afum|EAL92903.1   100.00%   afis|EAW21889.1   100.00%
Bootstrap support for afum|EAL92903.1 as seed ortholog is 100%.
Bootstrap support for afis|EAW21889.1 as seed ortholog is 100%.

---

Group of orthologs #4562. Best score 927 bits
Score difference with first non-orthologous sequence - *A. fumigatus*:712   *A. fischeri*:534
afum|EAL88929.1   100.00%   afis|EAW16307.1   100.00%
Bootstrap support for afum|EAL88929.1 as seed ortholog is 100%.
Bootstrap support for afis|EAW16307.1 as seed ortholog is 100%.

---

Group of orthologs #4563. Best score 927 bits
Score difference with first non-orthologous sequence - *A. fumigatus*:927   *A. fischeri*:927
afum|EAL87616.1   100.00%   afis|EAW18253.1   100.00%
Bootstrap support for afum|EAL87616.1 as seed ortholog is 100%.
Bootstrap support for afis|EAW18253.1 as seed ortholog is 100%.

---

Group of orthologs #4564. Best score 927 bits
Score difference with first non-orthologous sequence - *A. fumigatus*:927   *A. fischeri*:927
afum|EAL93607.1   100.00%   afis|EAW18875.1   100.00%
Bootstrap support for afum|EAL93607.1 as seed ortholog is 100%.
Bootstrap support for afis|EAW18875.1 as seed ortholog is 100%.

---

Group of orthologs #4565. Best score 927 bits
Score difference with first non-orthologous sequence - *A. fumigatus*:927   *A. fischeri*:927
afum|EAL93762.2   100.00%   afis|EAW19033.1   100.00%
Bootstrap support for afum|EAL93762.2 as seed ortholog is 100%.
Bootstrap support for afis|EAW19033.1 as seed ortholog is 100%.

---

Group of orthologs #4566. Best score 927 bits
Score difference with first non-orthologous sequence - *A. fumigatus*:927   *A. fischeri*:927
afum|EAL93970.2   100.00%   afis|EAW19241.1   100.00%
Bootstrap support for afum|EAL93970.2 as seed ortholog is 100%.
Bootstrap support for afis|EAW19241.1 as seed ortholog is 100%.

---

Group of orthologs #4567. Best score 927 bits
Score difference with first non-orthologous sequence - *A. fumigatus*:927   *A. fischeri*:927
afum|EAL89613.1   100.00%   afis|EAW24955.1   100.00%
Bootstrap support for afum|EAL89613.1 as seed ortholog is 100%.
Bootstrap support for afis|EAW24955.1 as seed ortholog is 100%.

---

Group of orthologs #4568. Best score 926 bits
Score difference with first non-orthologous sequence - *A. fumigatus*:926   *A. fischeri*:926
afum|EAL88502.1   100.00%   afis|EAW15868.1   100.00%
Bootstrap support for afum|EAL88502.1 as seed ortholog is 100%.
Bootstrap support for afis|EAW15868.1 as seed ortholog is 100%.

---

Group of orthologs #4569. Best score 926 bits
Score difference with first non-orthologous sequence - *A. fumigatus*:859   *A. fischeri*:734
afum|EAL89001.1   100.00%   afis|EAW16376.1   100.00%
Bootstrap support for afum|EAL89001.1 as seed ortholog is 100%.
Bootstrap support for afis|EAW16376.1 as seed ortholog is 100%.

---

Group of orthologs #4570. Best score 926 bits
Score difference with first non-orthologous sequence - *A. fumigatus*:926   *A. fischeri*:717
afum|EAL84871.1   100.00%   afis|EAW20697.1   100.00%

Bootstrap support for afum|EAL84871.1 as seed ortholog is 100%.
Bootstrap support for afis|EAW20697.1 as seed ortholog is 100%.

---

Group of orthologs #4571. Best score 926 bits
Score difference with first non-orthologous sequence - *A. fumigatus*:926 *A. fischeri*:926
afum|EAL86925.1 100.00% afis|EAW19593.1 100.00%
Bootstrap support for afum|EAL86925.1 as seed ortholog is 100%.
Bootstrap support for afis|EAW19593.1 as seed ortholog is 100%.

---

Group of orthologs #4572. Best score 926 bits
Score difference with first non-orthologous sequence - *A. fumigatus*:926 *A. fischeri*:926
afum|EAL91235.1 100.00% afis|EAW17397.1 100.00%
Bootstrap support for afum|EAL91235.1 as seed ortholog is 100%.
Bootstrap support for afis|EAW17397.1 as seed ortholog is 100%.

---

Group of orthologs #4573. Best score 926 bits
Score difference with first non-orthologous sequence - *A. fumigatus*:688 *A. fischeri*:803
afum|EAL84363.1 100.00% afis|EAW25693.1 100.00%
Bootstrap support for afum|EAL84363.1 as seed ortholog is 100%.
Bootstrap support for afis|EAW25693.1 as seed ortholog is 100%.

---

Group of orthologs #4574. Best score 926 bits
Score difference with first non-orthologous sequence - *A. fumigatus*:807 *A. fischeri*:926
afum|EAL87225.1 100.00% afis|EAW23791.1 100.00%
Bootstrap support for afum|EAL87225.1 as seed ortholog is 100%.
Bootstrap support for afis|EAW23791.1 as seed ortholog is 100%.

---

Group of orthologs #4575. Best score 925 bits
Score difference with first non-orthologous sequence - *A. fumigatus*:925 *A. fischeri*:925
afum|EAL86986.1 100.00% afis|EAW19508.1 100.00%
Bootstrap support for afum|EAL86986.1 as seed ortholog is 100%.
Bootstrap support for afis|EAW19508.1 as seed ortholog is 100%.

---

Group of orthologs #4576. Best score 925 bits
Score difference with first non-orthologous sequence - *A. fumigatus*:925 *A. fischeri*:925
afum|EAL91354.1 100.00% afis|EAW17540.1 100.00%
Bootstrap support for afum|EAL91354.1 as seed ortholog is 100%.
Bootstrap support for afis|EAW17540.1 as seed ortholog is 100%.

---

Group of orthologs #4577. Best score 925 bits
Score difference with first non-orthologous sequence - *A. fumigatus*:925 *A. fischeri*:925
afum|EAL87180.1 100.00% afis|EAW23744.1 100.00%
Bootstrap support for afum|EAL87180.1 as seed ortholog is 100%.
Bootstrap support for afis|EAW23744.1 as seed ortholog is 100%.

---

Group of orthologs #4578. Best score 925 bits
Score difference with first non-orthologous sequence - *A. fumigatus*:925 *A. fischeri*:491
afum|EAL87193.2 100.00% afis|EAW23759.1 100.00%
Bootstrap support for afum|EAL87193.2 as seed ortholog is 100%.
Bootstrap support for afis|EAW23759.1 as seed ortholog is 100%.

---

Group of orthologs #4579. Best score 924 bits
Score difference with first non-orthologous sequence - *A. fumigatus*:924 *A. fischeri*:924
afum|EAL85542.1 100.00% afis|EAW15809.1 100.00%
Bootstrap support for afum|EAL85542.1 as seed ortholog is 100%.
Bootstrap support for afis|EAW15809.1 as seed ortholog is 100%.

Group of orthologs #4580. Best score 924 bits
Score difference with first non-orthologous sequence - *A. fumigatus*:924 *A. fischeri*:924
afum|EAL88515.1 100.00% afis|EAW15881.1 100.00%
Bootstrap support for afum|EAL88515.1 as seed ortholog is 100%.
Bootstrap support for afis|EAW15881.1 as seed ortholog is 100%.

Group of orthologs #4581. Best score 924 bits
Score difference with first non-orthologous sequence - *A. fumigatus*:814 *A. fischeri*:823
afum|EAL85462.1 100.00% afis|EAW20342.1 100.00%
Bootstrap support for afum|EAL85462.1 as seed ortholog is 100%.
Bootstrap support for afis|EAW20342.1 as seed ortholog is 100%.

Group of orthologs #4582. Best score 924 bits
Score difference with first non-orthologous sequence - *A. fumigatus*:924 *A. fischeri*:924
afum|EAL91454.1 100.00% afis|EAW17639.1 100.00%
Bootstrap support for afum|EAL91454.1 as seed ortholog is 100%.
Bootstrap support for afis|EAW17639.1 as seed ortholog is 100%.

Group of orthologs #4583. Best score 924 bits
Score difference with first non-orthologous sequence - *A. fumigatus*:924 *A. fischeri*:924
afum|EAL90032.1 100.00% afis|EAW25372.1 100.00%
Bootstrap support for afum|EAL90032.1 as seed ortholog is 100%.
Bootstrap support for afis|EAW25372.1 as seed ortholog is 100%.

Group of orthologs #4584. Best score 923 bits
Score difference with first non-orthologous sequence - *A. fumigatus*:851 *A. fischeri*:809
afum|EAL85070.1 100.00% afis|EAW19823.1 100.00%
Bootstrap support for afum|EAL85070.1 as seed ortholog is 100%.
Bootstrap support for afis|EAW19823.1 as seed ortholog is 100%.

Group of orthologs #4585. Best score 923 bits
Score difference with first non-orthologous sequence - *A. fumigatus*:923 *A. fischeri*:923
afum|EAL89119.1 100.00% afis|EAW16524.1 100.00%
Bootstrap support for afum|EAL89119.1 as seed ortholog is 100%.
Bootstrap support for afis|EAW16524.1 as seed ortholog is 100%.

Group of orthologs #4586. Best score 923 bits
Score difference with first non-orthologous sequence - *A. fumigatus*:923 *A. fischeri*:923
afum|EAL91163.1 100.00% afis|EAW17326.1 100.00%
Bootstrap support for afum|EAL91163.1 as seed ortholog is 100%.
Bootstrap support for afis|EAW17326.1 as seed ortholog is 100%.

Group of orthologs #4587. Best score 923 bits
Score difference with first non-orthologous sequence - *A. fumigatus*:923 *A. fischeri*:923
afum|EAL91586.1 100.00% afis|EAW17777.1 100.00%
Bootstrap support for afum|EAL91586.1 as seed ortholog is 100%.
Bootstrap support for afis|EAW17777.1 as seed ortholog is 100%.

Group of orthologs #4588. Best score 923 bits
Score difference with first non-orthologous sequence - *A. fumigatus*:923 *A. fischeri*:923
afum|EAL93757.1 100.00% afis|EAW19026.1 100.00%
Bootstrap support for afum|EAL93757.1 as seed ortholog is 100%.
Bootstrap support for afis|EAW19026.1 as seed ortholog is 100%.

Group of orthologs #4589. Best score 923 bits

Score difference with first non-orthologous sequence - *A. fumigatus*:923   *A. fischeri*:923
afum|EAL92873.1 100.00% afis|EAW21860.1 100.00%
Bootstrap support for afum|EAL92873.1 as seed ortholog is 100%.
Bootstrap support for afis|EAW21860.1 as seed ortholog is 100%.

---

Group of orthologs #4590. Best score 922 bits
Score difference with first non-orthologous sequence - *A. fumigatus*:922   *A. fischeri*:922
afum|EAL85843.1 100.00% afis|EAW15478.1 100.00%
Bootstrap support for afum|EAL85843.1 as seed ortholog is 100%.
Bootstrap support for afis|EAW15478.1 as seed ortholog is 100%.

---

Group of orthologs #4591. Best score 922 bits
Score difference with first non-orthologous sequence - *A. fumigatus*:922   *A. fischeri*:922
afum|EAL86459.1 100.00% afis|EAW16934.1 100.00%
Bootstrap support for afum|EAL86459.1 as seed ortholog is 100%.
Bootstrap support for afis|EAW16934.1 as seed ortholog is 100%.

---

Group of orthologs #4592. Best score 922 bits
Score difference with first non-orthologous sequence - *A. fumigatus*:922   *A. fischeri*:922
afum|EAL84614.1 100.00% afis|EAW20438.1 100.00%
Bootstrap support for afum|EAL84614.1 as seed ortholog is 100%.
Bootstrap support for afis|EAW20438.1 as seed ortholog is 100%.

---

Group of orthologs #4593. Best score 922 bits
Score difference with first non-orthologous sequence - *A. fumigatus*:922   *A. fischeri*:922
afum|EBA27365.1 100.00% afis|EAW16400.1 100.00%
Bootstrap support for afum|EBA27365.1 as seed ortholog is 100%.
Bootstrap support for afis|EAW16400.1 as seed ortholog is 100%.

---

Group of orthologs #4594. Best score 922 bits
Score difference with first non-orthologous sequence - *A. fumigatus*:922   *A. fischeri*:922
afum|EAL88469.1 100.00% afis|EAW23012.1 100.00%
Bootstrap support for afum|EAL88469.1 as seed ortholog is 100%.
Bootstrap support for afis|EAW23012.1 as seed ortholog is 100%.

---

Group of orthologs #4595. Best score 922 bits
Score difference with first non-orthologous sequence - *A. fumigatus*:922   *A. fischeri*:922
afum|EAL93590.2 100.00% afis|EAW18858.1 100.00%
Bootstrap support for afum|EAL93590.2 as seed ortholog is 100%.
Bootstrap support for afis|EAW18858.1 as seed ortholog is 100%.

---

Group of orthologs #4596. Best score 922 bits
Score difference with first non-orthologous sequence - *A. fumigatus*:922   *A. fischeri*:922
afum|EAL90508.1 100.00% afis|EAW22685.1 100.00%
Bootstrap support for afum|EAL90508.1 as seed ortholog is 100%.
Bootstrap support for afis|EAW22685.1 as seed ortholog is 100%.

---

Group of orthologs #4597. Best score 922 bits
Score difference with first non-orthologous sequence - *A. fumigatus*:922   *A. fischeri*:922
afum|EAL90607.1 100.00% afis|EAW22592.1 100.00%
Bootstrap support for afum|EAL90607.1 as seed ortholog is 100%.
Bootstrap support for afis|EAW22592.1 as seed ortholog is 100%.

---

Group of orthologs #4598. Best score 922 bits
Score difference with first non-orthologous sequence - *A. fumigatus*:922   *A. fischeri*:922
afum|EAL90391.1 100.00% afis|EAW22810.1 100.00%

Bootstrap support for afum|EAL90391.1 as seed ortholog is 100%.
Bootstrap support for afis|EAW22810.1 as seed ortholog is 100%.

---

Group of orthologs #4599. Best score 922 bits
Score difference with first non-orthologous sequence - *A. fumigatus*:922 *A. fischeri*:535
afum|EAL92682.2 100.00% afis|EAW21648.1 100.00%
Bootstrap support for afum|EAL92682.2 as seed ortholog is 100%.
Bootstrap support for afis|EAW21648.1 as seed ortholog is 100%.

---

Group of orthologs #4600. Best score 921 bits
Score difference with first non-orthologous sequence - *A. fumigatus*:921 *A. fischeri*:921
afum|EAL84229.2 100.00% afis|EAW15416.1 100.00%
Bootstrap support for afum|EAL84229.2 as seed ortholog is 100%.
Bootstrap support for afis|EAW15416.1 as seed ortholog is 100%.

---

Group of orthologs #4601. Best score 921 bits
Score difference with first non-orthologous sequence - *A. fumigatus*:921 *A. fischeri*:921
afum|EAL86972.1 100.00% afis|EAW19534.1 100.00%
Bootstrap support for afum|EAL86972.1 as seed ortholog is 100%.
Bootstrap support for afis|EAW19534.1 as seed ortholog is 100%.

---

Group of orthologs #4602. Best score 921 bits
Score difference with first non-orthologous sequence - *A. fumigatus*:921 *A. fischeri*:921
afum|EAL91368.1 100.00% afis|EAW17554.1 100.00%
Bootstrap support for afum|EAL91368.1 as seed ortholog is 100%.
Bootstrap support for afis|EAW17554.1 as seed ortholog is 100%.

---

Group of orthologs #4603. Best score 921 bits
Score difference with first non-orthologous sequence - *A. fumigatus*:921 *A. fischeri*:921
afum|EAL86063.1 100.00% afis|EAW24391.1 100.00%
Bootstrap support for afum|EAL86063.1 as seed ortholog is 100%.
Bootstrap support for afis|EAW24391.1 as seed ortholog is 100%.

---

Group of orthologs #4604. Best score 921 bits
Score difference with first non-orthologous sequence - *A. fumigatus*:921 *A. fischeri*:921
afum|EAL87450.1 100.00% afis|EAW24020.1 100.00%
Bootstrap support for afum|EAL87450.1 as seed ortholog is 100%.
Bootstrap support for afis|EAW24020.1 as seed ortholog is 100%.

---

Group of orthologs #4605. Best score 921 bits
Score difference with first non-orthologous sequence - *A. fumigatus*:921 *A. fischeri*:921
afum|EAL93667.2 100.00% afis|EAW18934.1 100.00%
Bootstrap support for afum|EAL93667.2 as seed ortholog is 100%.
Bootstrap support for afis|EAW18934.1 as seed ortholog is 100%.

---

Group of orthologs #4606. Best score 921 bits
Score difference with first non-orthologous sequence - *A. fumigatus*:921 *A. fischeri*:921
afum|EAL90331.1 100.00% afis|EAW22868.1 100.00%
Bootstrap support for afum|EAL90331.1 as seed ortholog is 100%.
Bootstrap support for afis|EAW22868.1 as seed ortholog is 100%.

---

Group of orthologs #4607. Best score 921 bits
Score difference with first non-orthologous sequence - *A. fumigatus*:921 *A. fischeri*:611
afum|EAL89340.1 100.00% afis|EAW24698.1 100.00%
Bootstrap support for afum|EAL89340.1 as seed ortholog is 100%.
Bootstrap support for afis|EAW24698.1 as seed ortholog is 100%.

Group of orthologs #4608. Best score 921 bits
Score difference with first non-orthologous sequence - *A. fumigatus*:921 *A. fischeri*:921
afum|EAL92939.1 100.00% afis|EAW21926.1 100.00%
Bootstrap support for afum|EAL92939.1 as seed ortholog is 100%.
Bootstrap support for afis|EAW21926.1 as seed ortholog is 100%.

Group of orthologs #4609. Best score 920 bits
Score difference with first non-orthologous sequence - *A. fumigatus*:920 *A. fischeri*:920
afum|EAL85932.1 100.00% afis|EAW24264.1 100.00%
Bootstrap support for afum|EAL85932.1 as seed ortholog is 100%.
Bootstrap support for afis|EAW24264.1 as seed ortholog is 100%.

Group of orthologs #4610. Best score 920 bits
Score difference with first non-orthologous sequence - *A. fumigatus*:920 *A. fischeri*:920
afum|EAL93122.1 100.00% afis|EAW18440.1 100.00%
Bootstrap support for afum|EAL93122.1 as seed ortholog is 100%.
Bootstrap support for afis|EAW18440.1 as seed ortholog is 100%.

Group of orthologs #4611. Best score 920 bits
Score difference with first non-orthologous sequence - *A. fumigatus*:920 *A. fischeri*:879
afum|EAL88009.1 100.00% afis|EAW23458.1 100.00%
Bootstrap support for afum|EAL88009.1 as seed ortholog is 100%.
Bootstrap support for afis|EAW23458.1 as seed ortholog is 100%.

Group of orthologs #4612. Best score 920 bits
Score difference with first non-orthologous sequence - *A. fumigatus*:920 *A. fischeri*:920
afum|EAL87857.1 100.00% afis|EAW23608.1 100.00%
Bootstrap support for afum|EAL87857.1 as seed ortholog is 100%.
Bootstrap support for afis|EAW23608.1 as seed ortholog is 100%.

Group of orthologs #4613. Best score 920 bits
Score difference with first non-orthologous sequence - *A. fumigatus*:380 *A. fischeri*:483
afum|EAL93420.1 100.00% afis|EAW18698.1 100.00%
Bootstrap support for afum|EAL93420.1 as seed ortholog is 100%.
Bootstrap support for afis|EAW18698.1 as seed ortholog is 100%.

Group of orthologs #4614. Best score 920 bits
Score difference with first non-orthologous sequence - *A. fumigatus*:920 *A. fischeri*:920
afum|EAL93766.1 100.00% afis|EAW19037.1 100.00%
Bootstrap support for afum|EAL93766.1 as seed ortholog is 100%.
Bootstrap support for afis|EAW19037.1 as seed ortholog is 100%.

Group of orthologs #4615. Best score 919 bits
Score difference with first non-orthologous sequence - *A. fumigatus*:919 *A. fischeri*:919
afum|EAL88702.1 100.00% afis|EAW16070.1 100.00%
Bootstrap support for afum|EAL88702.1 as seed ortholog is 100%.
Bootstrap support for afis|EAW16070.1 as seed ortholog is 100%.

Group of orthologs #4616. Best score 919 bits
Score difference with first non-orthologous sequence - *A. fumigatus*:919 *A. fischeri*:919
afum|EAL88456.1 100.00% afis|EAW23026.1 100.00%
Bootstrap support for afum|EAL88456.1 as seed ortholog is 100%.
Bootstrap support for afis|EAW23026.1 as seed ortholog is 100%.

Group of orthologs #4617. Best score 919 bits

Score difference with first non-orthologous sequence - *A. fumigatus*:919 *A. fischeri*:919
afum|EAL87842.1 100.00% afis|EAW23620.1 100.00%
Bootstrap support for afum|EAL87842.1 as seed ortholog is 100%.
Bootstrap support for afis|EAW23620.1 as seed ortholog is 100%.

---

Group of orthologs #4618. Best score 919 bits
Score difference with first non-orthologous sequence - *A. fumigatus*:919 *A. fischeri*:919
afum|EAL90725.1 100.00% afis|EAW22466.1 100.00%
Bootstrap support for afum|EAL90725.1 as seed ortholog is 100%.
Bootstrap support for afis|EAW22466.1 as seed ortholog is 100%.

---

Group of orthologs #4619. Best score 919 bits
Score difference with first non-orthologous sequence - *A. fumigatus*:919 *A. fischeri*:919
afum|EAL90276.1 100.00% afis|EAW22921.1 100.00%
Bootstrap support for afum|EAL90276.1 as seed ortholog is 100%.
Bootstrap support for afis|EAW22921.1 as seed ortholog is 100%.

---

Group of orthologs #4620. Best score 919 bits
Score difference with first non-orthologous sequence - *A. fumigatus*:831 *A. fischeri*:875
afum|EAL89574.1 100.00% afis|EAW24919.1 100.00%
Bootstrap support for afum|EAL89574.1 as seed ortholog is 100%.
Bootstrap support for afis|EAW24919.1 as seed ortholog is 100%.

---

Group of orthologs #4621. Best score 919 bits
Score difference with first non-orthologous sequence - *A. fumigatus*:919 *A. fischeri*:919
afum|EAL90100.1 100.00% afis|EAW25442.1 100.00%
Bootstrap support for afum|EAL90100.1 as seed ortholog is 100%.
Bootstrap support for afis|EAW25442.1 as seed ortholog is 100%.

---

Group of orthologs #4622. Best score 918 bits
Score difference with first non-orthologous sequence - *A. fumigatus*:828 *A. fischeri*:918
afum|EAL88505.1 100.00% afis|EAW15871.1 100.00%
Bootstrap support for afum|EAL88505.1 as seed ortholog is 100%.
Bootstrap support for afis|EAW15871.1 as seed ortholog is 100%.

---

Group of orthologs #4623. Best score 918 bits
Score difference with first non-orthologous sequence - *A. fumigatus*:663 *A. fischeri*:733
afum|EAL85141.1 100.00% afis|EAW19756.1 100.00%
Bootstrap support for afum|EAL85141.1 as seed ortholog is 100%.
Bootstrap support for afis|EAW19756.1 as seed ortholog is 100%.

---

Group of orthologs #4624. Best score 918 bits
Score difference with first non-orthologous sequence - *A. fumigatus*:918 *A. fischeri*:857
afum|EAL91286.1 100.00% afis|EAW17458.1 100.00%
Bootstrap support for afum|EAL91286.1 as seed ortholog is 100%.
Bootstrap support for afis|EAW17458.1 as seed ortholog is 100%.

---

Group of orthologs #4625. Best score 918 bits
Score difference with first non-orthologous sequence - *A. fumigatus*:918 *A. fischeri*:918
afum|EAL92987.1 100.00% afis|EAW21972.1 100.00%
Bootstrap support for afum|EAL92987.1 as seed ortholog is 100%.
Bootstrap support for afis|EAW21972.1 as seed ortholog is 100%.

---

Group of orthologs #4626. Best score 918 bits
Score difference with first non-orthologous sequence - *A. fumigatus*:918 *A. fischeri*:918
afum|EAL89960.1 100.00% afis|EAW25297.1 100.00%

Bootstrap support for afum|EAL89960.1 as seed ortholog is 100%.
Bootstrap support for afis|EAW25297.1 as seed ortholog is 100%.

---

Group of orthologs #4627. Best score 917 bits
Score difference with first non-orthologous sequence - *A. fumigatus*:917 *A. fischeri*:917
afum|EAL84644.1 100.00% afis|EAW20986.1 100.00%
Bootstrap support for afum|EAL84644.1 as seed ortholog is 100%.
Bootstrap support for afis|EAW20986.1 as seed ortholog is 100%.

---

Group of orthologs #4628. Best score 917 bits
Score difference with first non-orthologous sequence - *A. fumigatus*:917 *A. fischeri*:917
afum|EAL91214.1 100.00% afis|EAW17379.1 100.00%
Bootstrap support for afum|EAL91214.1 as seed ortholog is 100%.
Bootstrap support for afis|EAW17379.1 as seed ortholog is 100%.

---

Group of orthologs #4629. Best score 917 bits
Score difference with first non-orthologous sequence - *A. fumigatus*:917 *A. fischeri*:917
afum|EAL84325.1 100.00% afis|EAW25638.1 100.00%
Bootstrap support for afum|EAL84325.1 as seed ortholog is 100%.
Bootstrap support for afis|EAW25638.1 as seed ortholog is 100%.

---

Group of orthologs #4630. Best score 917 bits
Score difference with first non-orthologous sequence - *A. fumigatus*:917 *A. fischeri*:917
afum|EAL90916.1 100.00% afis|EAW22264.1 100.00%
Bootstrap support for afum|EAL90916.1 as seed ortholog is 100%.
Bootstrap support for afis|EAW22264.1 as seed ortholog is 100%.

---

Group of orthologs #4631. Best score 916 bits
Score difference with first non-orthologous sequence - *A. fumigatus*:916 *A. fischeri*:495
afum|EAL85880.1 100.00% afis|EAW24211.1 100.00%
Bootstrap support for afum|EAL85880.1 as seed ortholog is 100%.
Bootstrap support for afis|EAW24211.1 as seed ortholog is 100%.

---

Group of orthologs #4632. Best score 916 bits
Score difference with first non-orthologous sequence - *A. fumigatus*:916 *A. fischeri*:916
afum|EAL88266.1 100.00% afis|EAW23218.1 100.00%
Bootstrap support for afum|EAL88266.1 as seed ortholog is 100%.
Bootstrap support for afis|EAW23218.1 as seed ortholog is 100%.

---

Group of orthologs #4633. Best score 916 bits
Score difference with first non-orthologous sequence - *A. fumigatus*:804 *A. fischeri*:916
afum|EAL93026.1 100.00% afis|EAW22011.1 100.00%
Bootstrap support for afum|EAL93026.1 as seed ortholog is 100%.
Bootstrap support for afis|EAW22011.1 as seed ortholog is 100%.

---

Group of orthologs #4634. Best score 915 bits
Score difference with first non-orthologous sequence - *A. fumigatus*:915 *A. fischeri*:915
afum|EAL86400.1 100.00% afis|EAW16857.1 100.00%
Bootstrap support for afum|EAL86400.1 as seed ortholog is 100%.
Bootstrap support for afis|EAW16857.1 as seed ortholog is 100%.

---

Group of orthologs #4635. Best score 915 bits
Score difference with first non-orthologous sequence - *A. fumigatus*:722 *A. fischeri*:637
afum|EAL86211.1 100.00% afis|EAW24547.1 100.00%
Bootstrap support for afum|EAL86211.1 as seed ortholog is 100%.
Bootstrap support for afis|EAW24547.1 as seed ortholog is 100%.

---
Group of orthologs #4636. Best score 915 bits
Score difference with first non-orthologous sequence - *A. fumigatus*:915 *A. fischeri*:915
afum|EAL92098.1 100.00% afis|EAW21075.1 100.00%
Bootstrap support for afum|EAL92098.1 as seed ortholog is 100%.
Bootstrap support for afis|EAW21075.1 as seed ortholog is 100%.

---
Group of orthologs #4637. Best score 914 bits
Score difference with first non-orthologous sequence - *A. fumigatus*:914 *A. fischeri*:826
afum|EAL86807.2 100.00% afis|EAW19719.1 100.00%
Bootstrap support for afum|EAL86807.2 as seed ortholog is 100%.
Bootstrap support for afis|EAW19719.1 as seed ortholog is 100%.

---
Group of orthologs #4638. Best score 913 bits
Score difference with first non-orthologous sequence - *A. fumigatus*:913 *A. fischeri*:913
afum|EAL84434.1 100.00% afis|EAW20634.1 100.00%
Bootstrap support for afum|EAL84434.1 as seed ortholog is 100%.
Bootstrap support for afis|EAW20634.1 as seed ortholog is 100%.

---
Group of orthologs #4639. Best score 913 bits
Score difference with first non-orthologous sequence - *A. fumigatus*:913 *A. fischeri*:913
afum|EAL91541.1 100.00% afis|EAW17726.1 100.00%
Bootstrap support for afum|EAL91541.1 as seed ortholog is 100%.
Bootstrap support for afis|EAW17726.1 as seed ortholog is 100%.

---
Group of orthologs #4640. Best score 913 bits
Score difference with first non-orthologous sequence - *A. fumigatus*:913 *A. fischeri*:913
afum|EAL91870.1 100.00% afis|EAW18059.1 100.00%
Bootstrap support for afum|EAL91870.1 as seed ortholog is 100%.
Bootstrap support for afis|EAW18059.1 as seed ortholog is 100%.

---
Group of orthologs #4641. Best score 912 bits
Score difference with first non-orthologous sequence - *A. fumigatus*:912 *A. fischeri*:912
afum|EAL86064.1 100.00% afis|EAW24392.1 100.00%
Bootstrap support for afum|EAL86064.1 as seed ortholog is 100%.
Bootstrap support for afis|EAW24392.1 as seed ortholog is 100%.

---
Group of orthologs #4642. Best score 912 bits
Score difference with first non-orthologous sequence - *A. fumigatus*:912 *A. fischeri*:912
afum|EAL88338.1 100.00% afis|EAW23143.1 100.00%
Bootstrap support for afum|EAL88338.1 as seed ortholog is 100%.
Bootstrap support for afis|EAW23143.1 as seed ortholog is 100%.

---
Group of orthologs #4643. Best score 912 bits
Score difference with first non-orthologous sequence - *A. fumigatus*:912 *A. fischeri*:912
afum|EAL93361.1 100.00% afis|EAW18646.1 100.00%
Bootstrap support for afum|EAL93361.1 as seed ortholog is 100%.
Bootstrap support for afis|EAW18646.1 as seed ortholog is 100%.

---
Group of orthologs #4644. Best score 912 bits
Score difference with first non-orthologous sequence - *A. fumigatus*:912 *A. fischeri*:912
afum|EAL92744.1 100.00% afis|EAW21709.1 100.00%
Bootstrap support for afum|EAL92744.1 as seed ortholog is 100%.
Bootstrap support for afis|EAW21709.1 as seed ortholog is 100%.

---
Group of orthologs #4645. Best score 912 bits

Score difference with first non-orthologous sequence - *A. fumigatus*:912 *A. fischeri*:912
afum|EBA27344.1 100.00% afis|EAW23164.1 100.00%
Bootstrap support for afum|EBA27344.1 as seed ortholog is 100%.
Bootstrap support for afis|EAW23164.1 as seed ortholog is 100%.

---

Group of orthologs #4646. Best score 911 bits
Score difference with first non-orthologous sequence - *A. fumigatus*:911 *A. fischeri*:911
afum|EAL89117.1 100.00% afis|EAW16522.1 100.00%
Bootstrap support for afum|EAL89117.1 as seed ortholog is 100%.
Bootstrap support for afis|EAW16522.1 as seed ortholog is 100%.

---

Group of orthologs #4647. Best score 911 bits
Score difference with first non-orthologous sequence - *A. fumigatus*:797 *A. fischeri*:860
afum|EAL91392.1 100.00% afis|EAW17579.1 100.00%
Bootstrap support for afum|EAL91392.1 as seed ortholog is 100%.
Bootstrap support for afis|EAW17579.1 as seed ortholog is 100%.

---

Group of orthologs #4648. Best score 911 bits
Score difference with first non-orthologous sequence - *A. fumigatus*:911 *A. fischeri*:481
afum|EAL85952.1 100.00% afis|EAW24283.1 100.00%
Bootstrap support for afum|EAL85952.1 as seed ortholog is 100%.
Bootstrap support for afis|EAW24283.1 as seed ortholog is 100%.

---

Group of orthologs #4649. Best score 911 bits
Score difference with first non-orthologous sequence - *A. fumigatus*:911 *A. fischeri*:911
afum|EAL87243.1 100.00% afis|EAW23809.1 100.00%
Bootstrap support for afum|EAL87243.1 as seed ortholog is 100%.
Bootstrap support for afis|EAW23809.1 as seed ortholog is 100%.

---

Group of orthologs #4650. Best score 911 bits
Score difference with first non-orthologous sequence - *A. fumigatus*:911 *A. fischeri*:911
afum|EAL93097.1 100.00% afis|EAW18415.1 100.00%
Bootstrap support for afum|EAL93097.1 as seed ortholog is 100%.
Bootstrap support for afis|EAW18415.1 as seed ortholog is 100%.

---

Group of orthologs #4651. Best score 911 bits
Score difference with first non-orthologous sequence - *A. fumigatus*:911 *A. fischeri*:911
afum|EAL88391.1 100.00% afis|EAW23088.1 100.00%
Bootstrap support for afum|EAL88391.1 as seed ortholog is 100%.
Bootstrap support for afis|EAW23088.1 as seed ortholog is 100%.

---

Group of orthologs #4652. Best score 911 bits
Score difference with first non-orthologous sequence - *A. fumigatus*:911 *A. fischeri*:911
afum|EBA27505.1 100.00% afis|EAW19157.1 100.00%
Bootstrap support for afum|EBA27505.1 as seed ortholog is 100%.
Bootstrap support for afis|EAW19157.1 as seed ortholog is 100%.

---

Group of orthologs #4653. Best score 911 bits
Score difference with first non-orthologous sequence - *A. fumigatus*:911 *A. fischeri*:911
afum|EAL93031.1 100.00% afis|EAW22016.1 100.00%
Bootstrap support for afum|EAL93031.1 as seed ortholog is 100%.
Bootstrap support for afis|EAW22016.1 as seed ortholog is 100%.

---

Group of orthologs #4654. Best score 910 bits
Score difference with first non-orthologous sequence - *A. fumigatus*:910 *A. fischeri*:910
afum|EAL85038.1 100.00% afis|EAW19857.1 100.00%

Bootstrap support for afum|EAL85038.1 as seed ortholog is 100%.
Bootstrap support for afis|EAW19857.1 as seed ortholog is 100%.

---

Group of orthologs #4655. Best score 910 bits
Score difference with first non-orthologous sequence - *A. fumigatus*:910 *A. fischeri*:910
afum|EAL84461.1 100.00% afis|EAW20600.1 100.00%
Bootstrap support for afum|EAL84461.1 as seed ortholog is 100%.
Bootstrap support for afis|EAW20600.1 as seed ortholog is 100%.

---

Group of orthologs #4656. Best score 910 bits
Score difference with first non-orthologous sequence - *A. fumigatus*:910 *A. fischeri*:803
afum|EAL87350.1 100.00% afis|EAW23925.1 100.00%
Bootstrap support for afum|EAL87350.1 as seed ortholog is 100%.
Bootstrap support for afis|EAW23925.1 as seed ortholog is 100%.

---

Group of orthologs #4657. Best score 910 bits
Score difference with first non-orthologous sequence - *A. fumigatus*:910 *A. fischeri*:910
afum|EAL88448.1 100.00% afis|EAW23033.1 100.00%
Bootstrap support for afum|EAL88448.1 as seed ortholog is 100%.
Bootstrap support for afis|EAW23033.1 as seed ortholog is 100%.

---

Group of orthologs #4658. Best score 910 bits
Score difference with first non-orthologous sequence - *A. fumigatus*:910 *A. fischeri*:861
afum|EAL89385.1 100.00% afis|EAW24738.1 100.00%
Bootstrap support for afum|EAL89385.1 as seed ortholog is 100%.
Bootstrap support for afis|EAW24738.1 as seed ortholog is 100%.

---

Group of orthologs #4659. Best score 910 bits
Score difference with first non-orthologous sequence - *A. fumigatus*:910 *A. fischeri*:910
afum|EAL89452.1 100.00% afis|EAW24809.1 100.00%
Bootstrap support for afum|EAL89452.1 as seed ortholog is 100%.
Bootstrap support for afis|EAW24809.1 as seed ortholog is 100%.

---

Group of orthologs #4660. Best score 910 bits
Score difference with first non-orthologous sequence - *A. fumigatus*:910 *A. fischeri*:910
afum|EAL89821.1 100.00% afis|EAW25146.1 100.00%
Bootstrap support for afum|EAL89821.1 as seed ortholog is 100%.
Bootstrap support for afis|EAW25146.1 as seed ortholog is 100%.

---

Group of orthologs #4661. Best score 910 bits
Score difference with first non-orthologous sequence - *A. fumigatus*:910 *A. fischeri*:910
afum|EAL90055.1 100.00% afis|EAW25396.1 100.00%
Bootstrap support for afum|EAL90055.1 as seed ortholog is 100%.
Bootstrap support for afis|EAW25396.1 as seed ortholog is 100%.

---

Group of orthologs #4662. Best score 909 bits
Score difference with first non-orthologous sequence - *A. fumigatus*:909 *A. fischeri*:909
afum|EAL84655.1 100.00% afis|EAW20978.1 100.00%
Bootstrap support for afum|EAL84655.1 as seed ortholog is 100%.
Bootstrap support for afis|EAW20978.1 as seed ortholog is 100%.

---

Group of orthologs #4663. Best score 909 bits
Score difference with first non-orthologous sequence - *A. fumigatus*:909 *A. fischeri*:718
afum|EAL90612.1 100.00% afis|EAW22587.1 100.00%
Bootstrap support for afum|EAL90612.1 as seed ortholog is 100%.
Bootstrap support for afis|EAW22587.1 as seed ortholog is 100%.

---

Group of orthologs #4664. Best score 908 bits
Score difference with first non-orthologous sequence - *A. fumigatus*:908 *A. fischeri*:908
afum|EAL85821.1 100.00% afis|EAW15503.1 100.00%
Bootstrap support for afum|EAL85821.1 as seed ortholog is 100%.
Bootstrap support for afis|EAW15503.1 as seed ortholog is 100%.

---

Group of orthologs #4665. Best score 908 bits
Score difference with first non-orthologous sequence - *A. fumigatus*:835 *A. fischeri*:799
afum|EAL84696.1 100.00% afis|EAW20938.1 100.00%
Bootstrap support for afum|EAL84696.1 as seed ortholog is 100%.
Bootstrap support for afis|EAW20938.1 as seed ortholog is 100%.

---

Group of orthologs #4666. Best score 908 bits
Score difference with first non-orthologous sequence - *A. fumigatus*:908 *A. fischeri*:908
afum|EAL89052.1 100.00% afis|EAW17167.1 100.00%
Bootstrap support for afum|EAL89052.1 as seed ortholog is 100%.
Bootstrap support for afis|EAW17167.1 as seed ortholog is 100%.

---

Group of orthologs #4667. Best score 908 bits
Score difference with first non-orthologous sequence - *A. fumigatus*:908 *A. fischeri*:908
afum|EAL91455.1 100.00% afis|EAW17640.1 100.00%
Bootstrap support for afum|EAL91455.1 as seed ortholog is 100%.
Bootstrap support for afis|EAW17640.1 as seed ortholog is 100%.

---

Group of orthologs #4668. Best score 908 bits
Score difference with first non-orthologous sequence - *A. fumigatus*:908 *A. fischeri*:908
afum|EAL91775.1 100.00% afis|EAW17966.1 100.00%
Bootstrap support for afum|EAL91775.1 as seed ortholog is 100%.
Bootstrap support for afis|EAW17966.1 as seed ortholog is 100%.

---

Group of orthologs #4669. Best score 908 bits
Score difference with first non-orthologous sequence - *A. fumigatus*:908 *A. fischeri*:908
afum|EAL87920.1 100.00% afis|EAW23546.1 100.00%
Bootstrap support for afum|EAL87920.1 as seed ortholog is 100%.
Bootstrap support for afis|EAW23546.1 as seed ortholog is 100%.

---

Group of orthologs #4670. Best score 908 bits
Score difference with first non-orthologous sequence - *A. fumigatus*:908 *A. fischeri*:908
afum|EAL93419.1 100.00% afis|EAW18697.1 100.00%
Bootstrap support for afum|EAL93419.1 as seed ortholog is 100%.
Bootstrap support for afis|EAW18697.1 as seed ortholog is 100%.

---

Group of orthologs #4671. Best score 907 bits
Score difference with first non-orthologous sequence - *A. fumigatus*:907 *A. fischeri*:907
afum|EAL85555.1 100.00% afis|EAW15795.1 100.00%
Bootstrap support for afum|EAL85555.1 as seed ortholog is 100%.
Bootstrap support for afis|EAW15795.1 as seed ortholog is 100%.

---

Group of orthologs #4672. Best score 907 bits
Score difference with first non-orthologous sequence - *A. fumigatus*:907 *A. fischeri*:907
afum|EAL86645.1 100.00% afis|EAW17223.1 100.00%
Bootstrap support for afum|EAL86645.1 as seed ortholog is 100%.
Bootstrap support for afis|EAW17223.1 as seed ortholog is 100%.

---

Group of orthologs #4673. Best score 907 bits

Score difference with first non-orthologous sequence - *A. fumigatus*:907 *A. fischeri*:907
afum|EAL84444.1 100.00% afis|EAW20617.1 100.00%
Bootstrap support for afum|EAL84444.1 as seed ortholog is 100%.
Bootstrap support for afis|EAW20617.1 as seed ortholog is 100%.

---

Group of orthologs #4674. Best score 907 bits
Score difference with first non-orthologous sequence - *A. fumigatus*:708 *A. fischeri*:590
afum|EAL84423.1 100.00% afis|EAW20650.1 100.00%
Bootstrap support for afum|EAL84423.1 as seed ortholog is 100%.
Bootstrap support for afis|EAW20650.1 as seed ortholog is 100%.

---

Group of orthologs #4675. Best score 907 bits
Score difference with first non-orthologous sequence - *A. fumigatus*:907 *A. fischeri*:907
afum|EAL89240.1 100.00% afis|EAW16638.1 100.00%
Bootstrap support for afum|EAL89240.1 as seed ortholog is 100%.
Bootstrap support for afis|EAW16638.1 as seed ortholog is 100%.

---

Group of orthologs #4676. Best score 907 bits
Score difference with first non-orthologous sequence - *A. fumigatus*:907 *A. fischeri*:907
afum|EAL91566.1 100.00% afis|EAW17753.1 100.00%
Bootstrap support for afum|EAL91566.1 as seed ortholog is 100%.
Bootstrap support for afis|EAW17753.1 as seed ortholog is 100%.

---

Group of orthologs #4677. Best score 907 bits
Score difference with first non-orthologous sequence - *A. fumigatus*:907 *A. fischeri*:907
afum|EAL90870.1 100.00% afis|EAW22321.1 100.00%
Bootstrap support for afum|EAL90870.1 as seed ortholog is 100%.
Bootstrap support for afis|EAW22321.1 as seed ortholog is 100%.

---

Group of orthologs #4678. Best score 907 bits
Score difference with first non-orthologous sequence - *A. fumigatus*:907 *A. fischeri*:907
afum|EAL92257.1 100.00% afis|EAW21244.1 100.00%
Bootstrap support for afum|EAL92257.1 as seed ortholog is 100%.
Bootstrap support for afis|EAW21244.1 as seed ortholog is 100%.

---

Group of orthologs #4679. Best score 907 bits
Score difference with first non-orthologous sequence - *A. fumigatus*:907 *A. fischeri*:756
afum|EBA27190.1 100.00% afis|EAW20537.1 100.00%
Bootstrap support for afum|EBA27190.1 as seed ortholog is 100%.
Bootstrap support for afis|EAW20537.1 as seed ortholog is 100%.

---

Group of orthologs #4680. Best score 907 bits
Score difference with first non-orthologous sequence - *A. fumigatus*:907 *A. fischeri*:907
afum|EAL92980.1 100.00% afis|EAW21966.1 100.00%
Bootstrap support for afum|EAL92980.1 as seed ortholog is 100%.
Bootstrap support for afis|EAW21966.1 as seed ortholog is 100%.

---

Group of orthologs #4681. Best score 907 bits
Score difference with first non-orthologous sequence - *A. fumigatus*:907 *A. fischeri*:833
afum|EAL89897.1 100.00% afis|EAW25236.1 100.00%
Bootstrap support for afum|EAL89897.1 as seed ortholog is 100%.
Bootstrap support for afis|EAW25236.1 as seed ortholog is 100%.

---

Group of orthologs #4682. Best score 906 bits
Score difference with first non-orthologous sequence - *A. fumigatus*:906 *A. fischeri*:906
afum|EAL86696.1 100.00% afis|EAW17278.1 100.00%

Bootstrap support for afum|EAL86696.1 as seed ortholog is 100%.
Bootstrap support for afis|EAW17278.1 as seed ortholog is 100%.

---

Group of orthologs #4683. Best score 906 bits
Score difference with first non-orthologous sequence - *A. fumigatus*:906 *A. fischeri*:906
afum|EAL85961.1 100.00% afis|EAW24293.1 100.00%
Bootstrap support for afum|EAL85961.1 as seed ortholog is 100%.
Bootstrap support for afis|EAW24293.1 as seed ortholog is 100%.

---

Group of orthologs #4684. Best score 905 bits
Score difference with first non-orthologous sequence - *A. fumigatus*:905 *A. fischeri*:710
afum|EAL86368.2 100.00% afis|EAW16817.1 100.00%
Bootstrap support for afum|EAL86368.2 as seed ortholog is 100%.
Bootstrap support for afis|EAW16817.1 as seed ortholog is 100%.

---

Group of orthologs #4685. Best score 905 bits
Score difference with first non-orthologous sequence - *A. fumigatus*:905 *A. fischeri*:905
afum|EAL86604.2 100.00% afis|EAW17174.1 100.00%
Bootstrap support for afum|EAL86604.2 as seed ortholog is 100%.
Bootstrap support for afis|EAW17174.1 as seed ortholog is 100%.

---

Group of orthologs #4686. Best score 905 bits
Score difference with first non-orthologous sequence - *A. fumigatus*:905 *A. fischeri*:905
afum|EAL91623.1 100.00% afis|EAW17814.1 100.00%
Bootstrap support for afum|EAL91623.1 as seed ortholog is 100%.
Bootstrap support for afis|EAW17814.1 as seed ortholog is 100%.

---

Group of orthologs #4687. Best score 905 bits
Score difference with first non-orthologous sequence - *A. fumigatus*:905 *A. fischeri*:905
afum|EAL86130.1 100.00% afis|EAW24460.1 100.00%
Bootstrap support for afum|EAL86130.1 as seed ortholog is 100%.
Bootstrap support for afis|EAW24460.1 as seed ortholog is 100%.

---

Group of orthologs #4688. Best score 905 bits
Score difference with first non-orthologous sequence - *A. fumigatus*:905 *A. fischeri*:905
afum|EAL92141.1 100.00% afis|EAW21117.1 100.00%
Bootstrap support for afum|EAL92141.1 as seed ortholog is 100%.
Bootstrap support for afis|EAW21117.1 as seed ortholog is 100%.

---

Group of orthologs #4689. Best score 905 bits
Score difference with first non-orthologous sequence - *A. fumigatus*:905 *A. fischeri*:905
afum|EAL90204.1 100.00% afis|EAW22995.1 100.00%
Bootstrap support for afum|EAL90204.1 as seed ortholog is 100%.
Bootstrap support for afis|EAW22995.1 as seed ortholog is 100%.

---

Group of orthologs #4690. Best score 905 bits
Score difference with first non-orthologous sequence - *A. fumigatus*:905 *A. fischeri*:760
afum|EAL94080.1 100.00% afis|EAW19360.1 100.00%
Bootstrap support for afum|EAL94080.1 as seed ortholog is 100%.
Bootstrap support for afis|EAW19360.1 as seed ortholog is 100%.

---

Group of orthologs #4691. Best score 905 bits
Score difference with first non-orthologous sequence - *A. fumigatus*:705 *A. fischeri*:905
afum|EAL92746.1 100.00% afis|EAW21711.1 100.00%
Bootstrap support for afum|EAL92746.1 as seed ortholog is 100%.
Bootstrap support for afis|EAW21711.1 as seed ortholog is 100%.

---
Group of orthologs #4692. Best score 904 bits
Score difference with first non-orthologous sequence - *A. fumigatus*:904 *A. fischeri*:904
afum|EAL85618.1 100.00% afis|EAW15729.1 100.00%
Bootstrap support for afum|EAL85618.1 as seed ortholog is 100%.
Bootstrap support for afis|EAW15729.1 as seed ortholog is 100%.

---
Group of orthologs #4693. Best score 904 bits
Score difference with first non-orthologous sequence - *A. fumigatus*:904 *A. fischeri*:904
afum|EAL84964.1 100.00% afis|EAW19951.1 100.00%
Bootstrap support for afum|EAL84964.1 as seed ortholog is 100%.
Bootstrap support for afis|EAW19951.1 as seed ortholog is 100%.

---
Group of orthologs #4694. Best score 904 bits
Score difference with first non-orthologous sequence - *A. fumigatus*:738 *A. fischeri*:675
afum|EAL85416.1 100.00% afis|EAW20282.1 100.00%
Bootstrap support for afum|EAL85416.1 as seed ortholog is 100%.
Bootstrap support for afis|EAW20282.1 as seed ortholog is 100%.

---
Group of orthologs #4695. Best score 904 bits
Score difference with first non-orthologous sequence - *A. fumigatus*:904 *A. fischeri*:904
afum|EAL86932.1 100.00% afis|EAW19586.1 100.00%
Bootstrap support for afum|EAL86932.1 as seed ortholog is 100%.
Bootstrap support for afis|EAW19586.1 as seed ortholog is 100%.

---
Group of orthologs #4696. Best score 904 bits
Score difference with first non-orthologous sequence - *A. fumigatus*:904 *A. fischeri*:904
afum|EAL91431.1 100.00% afis|EAW17618.1 100.00%
Bootstrap support for afum|EAL91431.1 as seed ortholog is 100%.
Bootstrap support for afis|EAW17618.1 as seed ortholog is 100%.

---
Group of orthologs #4697. Best score 904 bits
Score difference with first non-orthologous sequence - *A. fumigatus*:904 *A. fischeri*:904
afum|EAL85963.1 100.00% afis|EAW24295.1 100.00%
Bootstrap support for afum|EAL85963.1 as seed ortholog is 100%.
Bootstrap support for afis|EAW24295.1 as seed ortholog is 100%.

---
Group of orthologs #4698. Best score 904 bits
Score difference with first non-orthologous sequence - *A. fumigatus*:904 *A. fischeri*:724
afum|EAL87202.1 100.00% afis|EAW23768.1 100.00%
Bootstrap support for afum|EAL87202.1 as seed ortholog is 100%.
Bootstrap support for afis|EAW23768.1 as seed ortholog is 100%.

---
Group of orthologs #4699. Best score 904 bits
Score difference with first non-orthologous sequence - *A. fumigatus*:904 *A. fischeri*:904
afum|EAL88000.1 100.00% afis|EAW23466.1 100.00%
Bootstrap support for afum|EAL88000.1 as seed ortholog is 100%.
Bootstrap support for afis|EAW23466.1 as seed ortholog is 100%.

---
Group of orthologs #4700. Best score 903 bits
Score difference with first non-orthologous sequence - *A. fumigatus*:903 *A. fischeri*:680
afum|EAL84239.1 100.00% afis|EAW15408.1 100.00%
Bootstrap support for afum|EAL84239.1 as seed ortholog is 100%.
Bootstrap support for afis|EAW15408.1 as seed ortholog is 100%.

---
Group of orthologs #4701. Best score 903 bits

Score difference with first non-orthologous sequence - *A. fumigatus*:903 *A. fischeri*:903
afum|EAL88483.1 100.00% afis|EAW22997.1 100.00%
Bootstrap support for afum|EAL88483.1 as seed ortholog is 100%.
Bootstrap support for afis|EAW22997.1 as seed ortholog is 100%.

---

Group of orthologs #4702. Best score 902 bits
Score difference with first non-orthologous sequence - *A. fumigatus*:902 *A. fischeri*:796
afum|EAL84226.1 100.00% afis|EAW15418.1 100.00%
Bootstrap support for afum|EAL84226.1 as seed ortholog is 100%.
Bootstrap support for afis|EAW15418.1 as seed ortholog is 100%.

---

Group of orthologs #4703. Best score 902 bits
Score difference with first non-orthologous sequence - *A. fumigatus*:902 *A. fischeri*:902
afum|EAL88753.1 100.00% afis|EAW16123.1 100.00%
Bootstrap support for afum|EAL88753.1 as seed ortholog is 100%.
Bootstrap support for afis|EAW16123.1 as seed ortholog is 100%.

---

Group of orthologs #4704. Best score 902 bits
Score difference with first non-orthologous sequence - *A. fumigatus*:902 *A. fischeri*:902
afum|EAL91167.2 100.00% afis|EAW17330.1 100.00%
Bootstrap support for afum|EAL91167.2 as seed ortholog is 100%.
Bootstrap support for afis|EAW17330.1 as seed ortholog is 100%.

---

Group of orthologs #4705. Best score 902 bits
Score difference with first non-orthologous sequence - *A. fumigatus*:902 *A. fischeri*:902
afum|EAL91851.1 100.00% afis|EAW18040.1 100.00%
Bootstrap support for afum|EAL91851.1 as seed ortholog is 100%.
Bootstrap support for afis|EAW18040.1 as seed ortholog is 100%.

---

Group of orthologs #4706. Best score 902 bits
Score difference with first non-orthologous sequence - *A. fumigatus*:902 *A. fischeri*:902
afum|EAL87880.1 100.00% afis|EAW23584.1 100.00%
Bootstrap support for afum|EAL87880.1 as seed ortholog is 100%.
Bootstrap support for afis|EAW23584.1 as seed ortholog is 100%.

---

Group of orthologs #4707. Best score 902 bits
Score difference with first non-orthologous sequence - *A. fumigatus*:902 *A. fischeri*:834
afum|EAL92053.1 100.00% afis|EAW21015.1 100.00%
Bootstrap support for afum|EAL92053.1 as seed ortholog is 100%.
Bootstrap support for afis|EAW21015.1 as seed ortholog is 100%.

---

Group of orthologs #4708. Best score 902 bits
Score difference with first non-orthologous sequence - *A. fumigatus*:902 *A. fischeri*:902
afum|EAL90910.1 100.00% afis|EAW22270.1 100.00%
Bootstrap support for afum|EAL90910.1 as seed ortholog is 100%.
Bootstrap support for afis|EAW22270.1 as seed ortholog is 100%.

---

Group of orthologs #4709. Best score 902 bits
Score difference with first non-orthologous sequence - *A. fumigatus*:902 *A. fischeri*:902
afum|EAL90106.1 100.00% afis|EAW25448.1 100.00%
Bootstrap support for afum|EAL90106.1 as seed ortholog is 100%.
Bootstrap support for afis|EAW25448.1 as seed ortholog is 100%.

---

Group of orthologs #4710. Best score 901 bits
Score difference with first non-orthologous sequence - *A. fumigatus*:901 *A. fischeri*:901
afum|EAL88061.2 100.00% afis|EAW23409.1 100.00%

Bootstrap support for afum|EAL88061.2 as seed ortholog is 100%.
Bootstrap support for afis|EAW23409.1 as seed ortholog is 100%.

---

Group of orthologs #4711. Best score 901 bits
Score difference with first non-orthologous sequence - *A. fumigatus*:901 *A. fischeri*:575
afum|EAL90648.1 100.00% afis|EAW22547.1 100.00%
Bootstrap support for afum|EAL90648.1 as seed ortholog is 100%.
Bootstrap support for afis|EAW22547.1 as seed ortholog is 100%.

---

Group of orthologs #4712. Best score 901 bits
Score difference with first non-orthologous sequence - *A. fumigatus*:901 *A. fischeri*:901
afum|EAL92412.1 100.00% afis|EAW21398.1 100.00%
Bootstrap support for afum|EAL92412.1 as seed ortholog is 100%.
Bootstrap support for afis|EAW21398.1 as seed ortholog is 100%.

---

Group of orthologs #4713. Best score 900 bits
Score difference with first non-orthologous sequence - *A. fumigatus*:900 *A. fischeri*:900
afum|EAL89200.1 100.00% afis|EAW16604.1 100.00%
Bootstrap support for afum|EAL89200.1 as seed ortholog is 100%.
Bootstrap support for afis|EAW16604.1 as seed ortholog is 100%.

---

Group of orthologs #4714. Best score 900 bits
Score difference with first non-orthologous sequence - *A. fumigatus*:900 *A. fischeri*:598
afum|EAL84309.2 100.00% afis|EAW25621.1 100.00%
Bootstrap support for afum|EAL84309.2 as seed ortholog is 100%.
Bootstrap support for afis|EAW25621.1 as seed ortholog is 100%.

---

Group of orthologs #4715. Best score 900 bits
Score difference with first non-orthologous sequence - *A. fumigatus*:900 *A. fischeri*:900
afum|EAL85894.1 100.00% afis|EAW24226.1 100.00%
Bootstrap support for afum|EAL85894.1 as seed ortholog is 100%.
Bootstrap support for afis|EAW24226.1 as seed ortholog is 100%.

---

Group of orthologs #4716. Best score 900 bits
Score difference with first non-orthologous sequence - *A. fumigatus*:900 *A. fischeri*:900
afum|EAL90053.1 100.00% afis|EAW25394.1 100.00%
Bootstrap support for afum|EAL90053.1 as seed ortholog is 100%.
Bootstrap support for afis|EAW25394.1 as seed ortholog is 100%.

---

Group of orthologs #4717. Best score 899 bits
Score difference with first non-orthologous sequence - *A. fumigatus*:899 *A. fischeri*:899
afum|EAL86362.1 100.00% afis|EAW16793.1 100.00%
Bootstrap support for afum|EAL86362.1 as seed ortholog is 100%.
Bootstrap support for afis|EAW16793.1 as seed ortholog is 100%.

---

Group of orthologs #4718. Best score 899 bits
Score difference with first non-orthologous sequence - *A. fumigatus*:820 *A. fischeri*:657
afum|EAL86461.1 100.00% afis|EAW16936.1 100.00%
Bootstrap support for afum|EAL86461.1 as seed ortholog is 100%.
Bootstrap support for afis|EAW16936.1 as seed ortholog is 100%.

---

Group of orthologs #4719. Best score 899 bits
Score difference with first non-orthologous sequence - *A. fumigatus*:899 *A. fischeri*:899
afum|EAL88504.1 100.00% afis|EAW15870.1 100.00%
Bootstrap support for afum|EAL88504.1 as seed ortholog is 100%.
Bootstrap support for afis|EAW15870.1 as seed ortholog is 100%.

Group of orthologs #4720. Best score 899 bits
Score difference with first non-orthologous sequence - *A. fumigatus*:899 *A. fischeri*:745
afum|EAL88816.2 100.00% afis|EAW16185.1 100.00%
Bootstrap support for afum|EAL88816.2 as seed ortholog is 100%.
Bootstrap support for afis|EAW16185.1 as seed ortholog is 100%.

Group of orthologs #4721. Best score 899 bits
Score difference with first non-orthologous sequence - *A. fumigatus*:899 *A. fischeri*:899
afum|EAL84570.1 100.00% afis|EAW20485.1 100.00%
Bootstrap support for afum|EAL84570.1 as seed ortholog is 100%.
Bootstrap support for afis|EAW20485.1 as seed ortholog is 100%.

Group of orthologs #4722. Best score 899 bits
Score difference with first non-orthologous sequence - *A. fumigatus*:899 *A. fischeri*:899
afum|EAL89053.1 100.00% afis|EAW16456.1 100.00%
Bootstrap support for afum|EAL89053.1 as seed ortholog is 100%.
Bootstrap support for afis|EAW16456.1 as seed ortholog is 100%.

Group of orthologs #4723. Best score 899 bits
Score difference with first non-orthologous sequence - *A. fumigatus*:899 *A. fischeri*:899
afum|EAL84820.1 100.00% afis|EAW20808.1 100.00%
Bootstrap support for afum|EAL84820.1 as seed ortholog is 100%.
Bootstrap support for afis|EAW20808.1 as seed ortholog is 100%.

Group of orthologs #4724. Best score 899 bits
Score difference with first non-orthologous sequence - *A. fumigatus*:899 *A. fischeri*:899
afum|EAL90506.1 100.00% afis|EAW22687.1 100.00%
Bootstrap support for afum|EAL90506.1 as seed ortholog is 100%.
Bootstrap support for afis|EAW22687.1 as seed ortholog is 100%.

Group of orthologs #4725. Best score 899 bits
Score difference with first non-orthologous sequence - *A. fumigatus*:899 *A. fischeri*:899
afum|EAL90231.1 100.00% afis|EAW22966.1 100.00%
Bootstrap support for afum|EAL90231.1 as seed ortholog is 100%.
Bootstrap support for afis|EAW22966.1 as seed ortholog is 100%.

Group of orthologs #4726. Best score 899 bits
Score difference with first non-orthologous sequence - *A. fumigatus*:899 *A. fischeri*:899
afum|EAL92327.1 100.00% afis|EAW21311.1 100.00%
Bootstrap support for afum|EAL92327.1 as seed ortholog is 100%.
Bootstrap support for afis|EAW21311.1 as seed ortholog is 100%.

Group of orthologs #4727. Best score 898 bits
Score difference with first non-orthologous sequence - *A. fumigatus*:898 *A. fischeri*:690
afum|EAL89009.2 100.00% afis|EAW20386.1 100.00%
Bootstrap support for afum|EAL89009.2 as seed ortholog is 100%.
Bootstrap support for afis|EAW20386.1 as seed ortholog is 100%.

Group of orthologs #4728. Best score 898 bits
Score difference with first non-orthologous sequence - *A. fumigatus*:898 *A. fischeri*:898
afum|EAL88380.2 100.00% afis|EAW23098.1 100.00%
Bootstrap support for afum|EAL88380.2 as seed ortholog is 100%.
Bootstrap support for afis|EAW23098.1 as seed ortholog is 100%.

Group of orthologs #4729. Best score 898 bits

Score difference with first non-orthologous sequence - *A. fumigatus*:898   *A. fischeri*:898
afum|EAL87909.1   100.00%   afis|EAW23557.1   100.00%
Bootstrap support for afum|EAL87909.1 as seed ortholog is 100%.
Bootstrap support for afis|EAW23557.1 as seed ortholog is 100%.

---

Group of orthologs #4730. Best score 898 bits
Score difference with first non-orthologous sequence - *A. fumigatus*:898   *A. fischeri*:898
afum|EAL92335.1   100.00%   afis|EAW21318.1   100.00%
Bootstrap support for afum|EAL92335.1 as seed ortholog is 100%.
Bootstrap support for afis|EAW21318.1 as seed ortholog is 100%.

---

Group of orthologs #4731. Best score 897 bits
Score difference with first non-orthologous sequence - *A. fumigatus*:897   *A. fischeri*:897
afum|EAL93166.1   100.00%   afis|EAW18478.1   100.00%
Bootstrap support for afum|EAL93166.1 as seed ortholog is 100%.
Bootstrap support for afis|EAW18478.1 as seed ortholog is 100%.

---

Group of orthologs #4732. Best score 897 bits
Score difference with first non-orthologous sequence - *A. fumigatus*:897   *A. fischeri*:897
afum|EAL88217.1   100.00%   afis|EAW23263.1   100.00%
Bootstrap support for afum|EAL88217.1 as seed ortholog is 100%.
Bootstrap support for afis|EAW23263.1 as seed ortholog is 100%.

---

Group of orthologs #4733. Best score 897 bits
Score difference with first non-orthologous sequence - *A. fumigatus*:814   *A. fischeri*:721
afum|EAL90626.1   100.00%   afis|EAW22572.1   100.00%
Bootstrap support for afum|EAL90626.1 as seed ortholog is 100%.
Bootstrap support for afis|EAW22572.1 as seed ortholog is 100%.

---

Group of orthologs #4734. Best score 896 bits
Score difference with first non-orthologous sequence - *A. fumigatus*:896   *A. fischeri*:896
afum|EAL86653.1   100.00%   afis|EAW17236.1   100.00%
Bootstrap support for afum|EAL86653.1 as seed ortholog is 100%.
Bootstrap support for afis|EAW17236.1 as seed ortholog is 100%.

---

Group of orthologs #4735. Best score 896 bits
Score difference with first non-orthologous sequence - *A. fumigatus*:896   *A. fischeri*:896
afum|EAL89030.2   100.00%   afis|EAW16397.1   100.00%
Bootstrap support for afum|EAL89030.2 as seed ortholog is 100%.
Bootstrap support for afis|EAW16397.1 as seed ortholog is 100%.

---

Group of orthologs #4736. Best score 896 bits
Score difference with first non-orthologous sequence - *A. fumigatus*:896   *A. fischeri*:849
afum|EAL91530.1   100.00%   afis|EAW17715.1   100.00%
Bootstrap support for afum|EAL91530.1 as seed ortholog is 100%.
Bootstrap support for afis|EAW17715.1 as seed ortholog is 100%.

---

Group of orthologs #4737. Best score 896 bits
Score difference with first non-orthologous sequence - *A. fumigatus*:896   *A. fischeri*:896
afum|EAL92487.1   100.00%   afis|EAW21467.1   100.00%
Bootstrap support for afum|EAL92487.1 as seed ortholog is 100%.
Bootstrap support for afis|EAW21467.1 as seed ortholog is 100%.

---

Group of orthologs #4738. Best score 896 bits
Score difference with first non-orthologous sequence - *A. fumigatus*:896   *A. fischeri*:896
afum|EAL91983.1   100.00%   afis|EAW24127.1   100.00%

Bootstrap support for afum|EAL91983.1 as seed ortholog is 100%.
Bootstrap support for afis|EAW24127.1 as seed ortholog is 100%.

---

Group of orthologs #4739. Best score 896 bits
Score difference with first non-orthologous sequence - *A. fumigatus*:896 *A. fischeri*:468
afum|EAL92020.1 100.00% afis|EAW24163.1 100.00%
Bootstrap support for afum|EAL92020.1 as seed ortholog is 100%.
Bootstrap support for afis|EAW24163.1 as seed ortholog is 100%.

---

Group of orthologs #4740. Best score 895 bits
Score difference with first non-orthologous sequence - *A. fumigatus*:895 *A. fischeri*:895
afum|EAL84573.1 100.00% afis|EAW20481.1 100.00%
Bootstrap support for afum|EAL84573.1 as seed ortholog is 100%.
Bootstrap support for afis|EAW20481.1 as seed ortholog is 100%.

---

Group of orthologs #4741. Best score 895 bits
Score difference with first non-orthologous sequence - *A. fumigatus*:895 *A. fischeri*:447
afum|EAL84493.1 100.00% afis|EAW20569.1 100.00%
Bootstrap support for afum|EAL84493.1 as seed ortholog is 100%.
Bootstrap support for afis|EAW20569.1 as seed ortholog is 100%.

---

Group of orthologs #4742. Best score 895 bits
Score difference with first non-orthologous sequence - *A. fumigatus*:895 *A. fischeri*:895
afum|EAL85264.1 100.00% afis|EAW20149.1 100.00%
Bootstrap support for afum|EAL85264.1 as seed ortholog is 100%.
Bootstrap support for afis|EAW20149.1 as seed ortholog is 100%.

---

Group of orthologs #4743. Best score 895 bits
Score difference with first non-orthologous sequence - *A. fumigatus*:895 *A. fischeri*:835
afum|EAL87017.1 100.00% afis|EAW19479.1 100.00%
Bootstrap support for afum|EAL87017.1 as seed ortholog is 100%.
Bootstrap support for afis|EAW19479.1 as seed ortholog is 100%.

---

Group of orthologs #4744. Best score 895 bits
Score difference with first non-orthologous sequence - *A. fumigatus*:895 *A. fischeri*:895
afum|EAL87045.1 100.00% afis|EAW19451.1 100.00%
Bootstrap support for afum|EAL87045.1 as seed ortholog is 100%.
Bootstrap support for afis|EAW19451.1 as seed ortholog is 100%.

---

Group of orthologs #4745. Best score 895 bits
Score difference with first non-orthologous sequence - *A. fumigatus*:895 *A. fischeri*:895
afum|EAL84274.1 100.00% afis|EAW25551.1 100.00%
Bootstrap support for afum|EAL84274.1 as seed ortholog is 100%.
Bootstrap support for afis|EAW25551.1 as seed ortholog is 100%.

---

Group of orthologs #4746. Best score 895 bits
Score difference with first non-orthologous sequence - *A. fumigatus*:895 *A. fischeri*:895
afum|EAL93060.1 100.00% afis|EAW18385.1 100.00%
Bootstrap support for afum|EAL93060.1 as seed ortholog is 100%.
Bootstrap support for afis|EAW18385.1 as seed ortholog is 100%.

---

Group of orthologs #4747. Best score 895 bits
Score difference with first non-orthologous sequence - *A. fumigatus*:895 *A. fischeri*:895
afum|EAL93278.1 100.00% afis|EAW18575.1 100.00%
Bootstrap support for afum|EAL93278.1 as seed ortholog is 100%.
Bootstrap support for afis|EAW18575.1 as seed ortholog is 100%.

---

Group of orthologs #4748. Best score 895 bits
Score difference with first non-orthologous sequence - *A. fumigatus*:895 *A. fischeri*:895
afum|EAL90227.2 100.00% afis|EAW22970.1 100.00%
Bootstrap support for afum|EAL90227.2 as seed ortholog is 100%.
Bootstrap support for afis|EAW22970.1 as seed ortholog is 100%.

---

Group of orthologs #4749. Best score 895 bits
Score difference with first non-orthologous sequence - *A. fumigatus*:895 *A. fischeri*:895
afum|EAL90168.1 100.00% afis|EAW25502.1 100.00%
Bootstrap support for afum|EAL90168.1 as seed ortholog is 100%.
Bootstrap support for afis|EAW25502.1 as seed ortholog is 100%.

---

Group of orthologs #4750. Best score 894 bits
Score difference with first non-orthologous sequence - *A. fumigatus*:894 *A. fischeri*:894
afum|EAL86467.1 100.00% afis|EAW16940.1 100.00%
Bootstrap support for afum|EAL86467.1 as seed ortholog is 100%.
Bootstrap support for afis|EAW16940.1 as seed ortholog is 100%.

---

Group of orthologs #4751. Best score 894 bits
Score difference with first non-orthologous sequence - *A. fumigatus*:894 *A. fischeri*:894
afum|EAL87160.2 100.00% afis|EAW23715.1 100.00%
Bootstrap support for afum|EAL87160.2 as seed ortholog is 100%.
Bootstrap support for afis|EAW23715.1 as seed ortholog is 100%.

---

Group of orthologs #4752. Best score 894 bits
Score difference with first non-orthologous sequence - *A. fumigatus*:894 *A. fischeri*:894
afum|EAL92702.1 100.00% afis|EAW21667.1 100.00%
Bootstrap support for afum|EAL92702.1 as seed ortholog is 100%.
Bootstrap support for afis|EAW21667.1 as seed ortholog is 100%.

---

Group of orthologs #4753. Best score 894 bits
Score difference with first non-orthologous sequence - *A. fumigatus*:894 *A. fischeri*:894
afum|EAL89481.2 100.00% afis|EAW24836.1 100.00%
Bootstrap support for afum|EAL89481.2 as seed ortholog is 100%.
Bootstrap support for afis|EAW24836.1 as seed ortholog is 100%.

---

Group of orthologs #4754. Best score 893 bits
Score difference with first non-orthologous sequence - *A. fumigatus*:893 *A. fischeri*:893
afum|EAL86401.1 100.00% afis|EAW16858.1 100.00%
Bootstrap support for afum|EAL86401.1 as seed ortholog is 100%.
Bootstrap support for afis|EAW16858.1 as seed ortholog is 100%.

---

Group of orthologs #4755. Best score 893 bits
Score difference with first non-orthologous sequence - *A. fumigatus*:454 *A. fischeri*:893
afum|EAL87500.1 100.00% afis|EAW18143.1 100.00%
Bootstrap support for afum|EAL87500.1 as seed ortholog is 100%.
Bootstrap support for afis|EAW18143.1 as seed ortholog is 100%.

---

Group of orthologs #4756. Best score 893 bits
Score difference with first non-orthologous sequence - *A. fumigatus*:893 *A. fischeri*:893
afum|EAL91760.1 100.00% afis|EAW17950.1 100.00%
Bootstrap support for afum|EAL91760.1 as seed ortholog is 100%.
Bootstrap support for afis|EAW17950.1 as seed ortholog is 100%.

---

Group of orthologs #4757. Best score 893 bits

Score difference with first non-orthologous sequence - *A. fumigatus*:893 *A. fischeri*:893
afum|EAL93744.1 100.00% afis|EAW19010.1 100.00%
Bootstrap support for afum|EAL93744.1 as seed ortholog is 100%.
Bootstrap support for afis|EAW19010.1 as seed ortholog is 100%.

---

Group of orthologs #4758. Best score 893 bits
Score difference with first non-orthologous sequence - *A. fumigatus*:893 *A. fischeri*:893
afum|EAL93976.1 100.00% afis|EAW19247.1 100.00%
Bootstrap support for afum|EAL93976.1 as seed ortholog is 100%.
Bootstrap support for afis|EAW19247.1 as seed ortholog is 100%.

---

Group of orthologs #4759. Best score 893 bits
Score difference with first non-orthologous sequence - *A. fumigatus*:893 *A. fischeri*:893
afum|EAL90852.1 100.00% afis|EAW22340.1 100.00%
Bootstrap support for afum|EAL90852.1 as seed ortholog is 100%.
Bootstrap support for afis|EAW22340.1 as seed ortholog is 100%.

---

Group of orthologs #4760. Best score 893 bits
Score difference with first non-orthologous sequence - *A. fumigatus*:893 *A. fischeri*:893
afum|EAL90976.1 100.00% afis|EAW22218.1 100.00%
Bootstrap support for afum|EAL90976.1 as seed ortholog is 100%.
Bootstrap support for afis|EAW22218.1 as seed ortholog is 100%.

---

Group of orthologs #4761. Best score 893 bits
Score difference with first non-orthologous sequence - *A. fumigatus*:893 *A. fischeri*:893
afum|EAL89863.2 100.00% afis|EAW25200.1 100.00%
Bootstrap support for afum|EAL89863.2 as seed ortholog is 100%.
Bootstrap support for afis|EAW25200.1 as seed ortholog is 100%.

---

Group of orthologs #4762. Best score 892 bits
Score difference with first non-orthologous sequence - *A. fumigatus*:892 *A. fischeri*:892
afum|EAL89003.1 100.00% afis|EAW16378.1 100.00%
Bootstrap support for afum|EAL89003.1 as seed ortholog is 100%.
Bootstrap support for afis|EAW16378.1 as seed ortholog is 100%.

---

Group of orthologs #4763. Best score 892 bits
Score difference with first non-orthologous sequence - *A. fumigatus*:892 *A. fischeri*:892
afum|EAL89027.1 100.00% afis|EAW16399.1 100.00%
Bootstrap support for afum|EAL89027.1 as seed ortholog is 100%.
Bootstrap support for afis|EAW16399.1 as seed ortholog is 100%.

---

Group of orthologs #4764. Best score 892 bits
Score difference with first non-orthologous sequence - *A. fumigatus*:892 *A. fischeri*:892
afum|EAL85218.1 100.00% afis|EAW20101.1 100.00%
Bootstrap support for afum|EAL85218.1 as seed ortholog is 100%.
Bootstrap support for afis|EAW20101.1 as seed ortholog is 100%.

---

Group of orthologs #4765. Best score 892 bits
Score difference with first non-orthologous sequence - *A. fumigatus*:892 *A. fischeri*:892
afum|EAL86829.1 100.00% afis|EAW19698.1 100.00%
Bootstrap support for afum|EAL86829.1 as seed ortholog is 100%.
Bootstrap support for afis|EAW19698.1 as seed ortholog is 100%.

---

Group of orthologs #4766. Best score 892 bits
Score difference with first non-orthologous sequence - *A. fumigatus*:722 *A. fischeri*:773
afum|EAL91613.1 100.00% afis|EAW17805.1 100.00%

Bootstrap support for afum|EAL91613.1 as seed ortholog is 100%.
Bootstrap support for afis|EAW17805.1 as seed ortholog is 100%.

---

Group of orthologs #4767. Best score 892 bits
Score difference with first non-orthologous sequence - *A. fumigatus*:892 *A. fischeri*:892
afum|EAL91902.1 100.00% afis|EAW18094.1 100.00%
Bootstrap support for afum|EAL91902.1 as seed ortholog is 100%.
Bootstrap support for afis|EAW18094.1 as seed ortholog is 100%.

---

Group of orthologs #4768. Best score 892 bits
Score difference with first non-orthologous sequence - *A. fumigatus*:892 *A. fischeri*:892
afum|EAL86086.1 100.00% afis|EAW24417.1 100.00%
Bootstrap support for afum|EAL86086.1 as seed ortholog is 100%.
Bootstrap support for afis|EAW24417.1 as seed ortholog is 100%.

---

Group of orthologs #4769. Best score 892 bits
Score difference with first non-orthologous sequence - *A. fumigatus*:892 *A. fischeri*:892
afum|EAL93079.1 100.00% afis|EAW18401.1 100.00%
Bootstrap support for afum|EAL93079.1 as seed ortholog is 100%.
Bootstrap support for afis|EAW18401.1 as seed ortholog is 100%.

---

Group of orthologs #4770. Best score 892 bits
Score difference with first non-orthologous sequence - *A. fumigatus*:892 *A. fischeri*:892
afum|EAL88298.1 100.00% afis|EAW23185.1 100.00%
Bootstrap support for afum|EAL88298.1 as seed ortholog is 100%.
Bootstrap support for afis|EAW23185.1 as seed ortholog is 100%.

---

Group of orthologs #4771. Best score 892 bits
Score difference with first non-orthologous sequence - *A. fumigatus*:892 *A. fischeri*:892
afum|EAL87894.1 100.00% afis|EAW23572.1 100.00%
Bootstrap support for afum|EAL87894.1 as seed ortholog is 100%.
Bootstrap support for afis|EAW23572.1 as seed ortholog is 100%.

---

Group of orthologs #4772. Best score 892 bits
Score difference with first non-orthologous sequence - *A. fumigatus*:892 *A. fischeri*:294
afum|EAL93960.1 100.00% afis|EAW19231.1 100.00%
Bootstrap support for afum|EAL93960.1 as seed ortholog is 100%.
Bootstrap support for afis|EAW19231.1 as seed ortholog is 100%.

---

Group of orthologs #4773. Best score 891 bits
Score difference with first non-orthologous sequence - *A. fumigatus*:891 *A. fischeri*:891
afum|EAL87179.1 100.00% afis|EAW23742.1 100.00%
Bootstrap support for afum|EAL87179.1 as seed ortholog is 100%.
Bootstrap support for afis|EAW23742.1 as seed ortholog is 100%.

---

Group of orthologs #4774. Best score 891 bits
Score difference with first non-orthologous sequence - *A. fumigatus*:891 *A. fischeri*:891
afum|EAL87358.1 100.00% afis|EAW23933.1 100.00%
Bootstrap support for afum|EAL87358.1 as seed ortholog is 100%.
Bootstrap support for afis|EAW23933.1 as seed ortholog is 100%.

---

Group of orthologs #4775. Best score 891 bits
Score difference with first non-orthologous sequence - *A. fumigatus*:891 *A. fischeri*:891
afum|EAL92035.1 100.00% afis|EAW24173.1 100.00%
Bootstrap support for afum|EAL92035.1 as seed ortholog is 100%.
Bootstrap support for afis|EAW24173.1 as seed ortholog is 100%.

Group of orthologs #4776. Best score 890 bits
Score difference with first non-orthologous sequence - *A. fumigatus*:890 *A. fischeri*:890
afum|EAL87092.1 100.00% afis|EAW19403.1 100.00%
Bootstrap support for afum|EAL87092.1 as seed ortholog is 100%.
Bootstrap support for afis|EAW19403.1 as seed ortholog is 100%.

Group of orthologs #4777. Best score 890 bits
Score difference with first non-orthologous sequence - *A. fumigatus*:890 *A. fischeri*:890
afum|EAL88248.1 100.00% afis|EAW23234.1 100.00%
Bootstrap support for afum|EAL88248.1 as seed ortholog is 100%.
Bootstrap support for afis|EAW23234.1 as seed ortholog is 100%.

Group of orthologs #4778. Best score 890 bits
Score difference with first non-orthologous sequence - *A. fumigatus*:890 *A. fischeri*:890
afum|EAL87976.1 100.00% afis|EAW23487.1 100.00%
Bootstrap support for afum|EAL87976.1 as seed ortholog is 100%.
Bootstrap support for afis|EAW23487.1 as seed ortholog is 100%.

Group of orthologs #4779. Best score 890 bits
Score difference with first non-orthologous sequence - *A. fumigatus*:890 *A. fischeri*:890
afum|EAL90070.1 100.00% afis|EAW25411.1 100.00%
Bootstrap support for afum|EAL90070.1 as seed ortholog is 100%.
Bootstrap support for afis|EAW25411.1 as seed ortholog is 100%.

Group of orthologs #4780. Best score 889 bits
Score difference with first non-orthologous sequence - *A. fumigatus*:889 *A. fischeri*:889
afum|EAL87491.1 100.00% afis|EAW18134.1 100.00%
Bootstrap support for afum|EAL87491.1 as seed ortholog is 100%.
Bootstrap support for afis|EAW18134.1 as seed ortholog is 100%.

Group of orthologs #4781. Best score 889 bits
Score difference with first non-orthologous sequence - *A. fumigatus*:889 *A. fischeri*:842
afum|EAL87524.1 100.00% afis|EAW18164.1 100.00%
Bootstrap support for afum|EAL87524.1 as seed ortholog is 100%.
Bootstrap support for afis|EAW18164.1 as seed ortholog is 100%.

Group of orthologs #4782. Best score 889 bits
Score difference with first non-orthologous sequence - *A. fumigatus*:178 *A. fischeri*:889
afum|EAL85364.1 100.00% afis|EAW20246.1 100.00%
Bootstrap support for afum|EAL85364.1 as seed ortholog is 100%.
Bootstrap support for afis|EAW20246.1 as seed ortholog is 100%.

Group of orthologs #4783. Best score 889 bits
Score difference with first non-orthologous sequence - *A. fumigatus*:889 *A. fischeri*:889
afum|EAL85415.1 100.00% afis|EAW20281.1 100.00%
Bootstrap support for afum|EAL85415.1 as seed ortholog is 100%.
Bootstrap support for afis|EAW20281.1 as seed ortholog is 100%.

Group of orthologs #4784. Best score 889 bits
Score difference with first non-orthologous sequence - *A. fumigatus*:889 *A. fischeri*:889
afum|EAL91329.1 100.00% afis|EAW17507.1 100.00%
Bootstrap support for afum|EAL91329.1 as seed ortholog is 100%.
Bootstrap support for afis|EAW17507.1 as seed ortholog is 100%.

Group of orthologs #4785. Best score 889 bits

Score difference with first non-orthologous sequence - *A. fumigatus*:889 *A. fischeri*:889
afum|EAL85896.1 100.00% afis|EAW24227.1 100.00%
Bootstrap support for afum|EAL85896.1 as seed ortholog is 100%.
Bootstrap support for afis|EAW24227.1 as seed ortholog is 100%.

---

Group of orthologs #4786. Best score 889 bits
Score difference with first non-orthologous sequence - *A. fumigatus*:889 *A. fischeri*:889
afum|EAL88407.1 100.00% afis|EAW23072.1 100.00%
Bootstrap support for afum|EAL88407.1 as seed ortholog is 100%.
Bootstrap support for afis|EAW23072.1 as seed ortholog is 100%.

---

Group of orthologs #4787. Best score 889 bits
Score difference with first non-orthologous sequence - *A. fumigatus*:889 *A. fischeri*:889
afum|EAL93601.1 100.00% afis|EAW18869.1 100.00%
Bootstrap support for afum|EAL93601.1 as seed ortholog is 100%.
Bootstrap support for afis|EAW18869.1 as seed ortholog is 100%.

---

Group of orthologs #4788. Best score 889 bits
Score difference with first non-orthologous sequence - *A. fumigatus*:889 *A. fischeri*:889
afum|EAL90477.1 100.00% afis|EAW22717.1 100.00%
Bootstrap support for afum|EAL90477.1 as seed ortholog is 100%.
Bootstrap support for afis|EAW22717.1 as seed ortholog is 100%.

---

Group of orthologs #4789. Best score 889 bits
Score difference with first non-orthologous sequence - *A. fumigatus*:889 *A. fischeri*:889
afum|EAL90156.1 100.00% afis|EAW25491.1 100.00%
Bootstrap support for afum|EAL90156.1 as seed ortholog is 100%.
Bootstrap support for afis|EAW25491.1 as seed ortholog is 100%.

---

Group of orthologs #4790. Best score 888 bits
Score difference with first non-orthologous sequence - *A. fumigatus*:888 *A. fischeri*:888
afum|EAL89262.1 100.00% afis|EAW16658.1 100.00%
Bootstrap support for afum|EAL89262.1 as seed ortholog is 100%.
Bootstrap support for afis|EAW16658.1 as seed ortholog is 100%.

---

Group of orthologs #4791. Best score 888 bits
Score difference with first non-orthologous sequence - *A. fumigatus*:888 *A. fischeri*:888
afum|EAL93672.1 100.00% afis|EAW18939.1 100.00%
Bootstrap support for afum|EAL93672.1 as seed ortholog is 100%.
Bootstrap support for afis|EAW18939.1 as seed ortholog is 100%.

---

Group of orthologs #4792. Best score 888 bits
Score difference with first non-orthologous sequence - *A. fumigatus*:888 *A. fischeri*:888
afum|EAL91029.1 100.00% afis|EAW22162.1 100.00%
Bootstrap support for afum|EAL91029.1 as seed ortholog is 100%.
Bootstrap support for afis|EAW22162.1 as seed ortholog is 100%.

---

Group of orthologs #4793. Best score 888 bits
Score difference with first non-orthologous sequence - *A. fumigatus*:714 *A. fischeri*:888
afum|EAL92610.1 100.00% afis|EAW21592.1 100.00%
Bootstrap support for afum|EAL92610.1 as seed ortholog is 100%.
Bootstrap support for afis|EAW21592.1 as seed ortholog is 100%.

---

Group of orthologs #4794. Best score 888 bits
Score difference with first non-orthologous sequence - *A. fumigatus*:888 *A. fischeri*:888
afum|EAL89528.1 100.00% afis|EAW24878.1 100.00%

Bootstrap support for afum|EAL89528.1 as seed ortholog is 100%.
Bootstrap support for afis|EAW24878.1 as seed ortholog is 100%.

---

Group of orthologs #4795. Best score 888 bits
Score difference with first non-orthologous sequence - *A. fumigatus*:888 *A. fischeri*:888
afum|EAL89658.1 100.00% afis|EAW25036.1 100.00%
Bootstrap support for afum|EAL89658.1 as seed ortholog is 100%.
Bootstrap support for afis|EAW25036.1 as seed ortholog is 100%.

---

Group of orthologs #4796. Best score 887 bits
Score difference with first non-orthologous sequence - *A. fumigatus*:887 *A. fischeri*:887
afum|EAL88570.1 100.00% afis|EAW15934.1 100.00%
Bootstrap support for afum|EAL88570.1 as seed ortholog is 100%.
Bootstrap support for afis|EAW15934.1 as seed ortholog is 100%.

---

Group of orthologs #4797. Best score 887 bits
Score difference with first non-orthologous sequence - *A. fumigatus*:810 *A. fischeri*:776
afum|EAL84925.1 100.00% afis|EAW19990.1 100.00%
Bootstrap support for afum|EAL84925.1 as seed ortholog is 100%.
Bootstrap support for afis|EAW19990.1 as seed ortholog is 100%.

---

Group of orthologs #4798. Best score 887 bits
Score difference with first non-orthologous sequence - *A. fumigatus*:887 *A. fischeri*:887
afum|EAL85399.1 100.00% afis|EAW20625.1 100.00%
Bootstrap support for afum|EAL85399.1 as seed ortholog is 100%.
Bootstrap support for afis|EAW20625.1 as seed ortholog is 100%.

---

Group of orthologs #4799. Best score 887 bits
Score difference with first non-orthologous sequence - *A. fumigatus*:887 *A. fischeri*:887
afum|EAL91377.1 100.00% afis|EAW17563.1 100.00%
Bootstrap support for afum|EAL91377.1 as seed ortholog is 100%.
Bootstrap support for afis|EAW17563.1 as seed ortholog is 100%.

---

Group of orthologs #4800. Best score 887 bits
Score difference with first non-orthologous sequence - *A. fumigatus*:887 *A. fischeri*:887
afum|EAL86172.1 100.00% afis|EAW24504.1 100.00%
Bootstrap support for afum|EAL86172.1 as seed ortholog is 100%.
Bootstrap support for afis|EAW24504.1 as seed ortholog is 100%.

---

Group of orthologs #4801. Best score 887 bits
Score difference with first non-orthologous sequence - *A. fumigatus*:887 *A. fischeri*:887
afum|EAL93273.1 100.00% afis|EAW18570.1 100.00%
Bootstrap support for afum|EAL93273.1 as seed ortholog is 100%.
Bootstrap support for afis|EAW18570.1 as seed ortholog is 100%.

---

Group of orthologs #4802. Best score 887 bits
Score difference with first non-orthologous sequence - *A. fumigatus*:887 *A. fischeri*:887
afum|EAL93860.1 100.00% afis|EAW19128.1 100.00%
Bootstrap support for afum|EAL93860.1 as seed ortholog is 100%.
Bootstrap support for afis|EAW19128.1 as seed ortholog is 100%.

---

Group of orthologs #4803. Best score 887 bits
Score difference with first non-orthologous sequence - *A. fumigatus*:887 *A. fischeri*:887
afum|EAL92657.2 100.00% afis|EAW21621.1 100.00%
Bootstrap support for afum|EAL92657.2 as seed ortholog is 100%.
Bootstrap support for afis|EAW21621.1 as seed ortholog is 100%.

Group of orthologs #4804. Best score 887 bits
Score difference with first non-orthologous sequence - *A. fumigatus*:887 *A. fischeri*:887
afum|EBA27228.1 100.00% afis|EAW21814.1 100.00%
Bootstrap support for afum|EBA27228.1 as seed ortholog is 100%.
Bootstrap support for afis|EAW21814.1 as seed ortholog is 100%.

Group of orthologs #4805. Best score 886 bits
Score difference with first non-orthologous sequence - *A. fumigatus*:886 *A. fischeri*:736
afum|EAL86381.1 100.00% afis|EAW16834.1 100.00%
Bootstrap support for afum|EAL86381.1 as seed ortholog is 100%.
Bootstrap support for afis|EAW16834.1 as seed ortholog is 100%.

Group of orthologs #4806. Best score 886 bits
Score difference with first non-orthologous sequence - *A. fumigatus*:886 *A. fischeri*:817
afum|EAL87095.1 100.00% afis|EAW19399.1 100.00%
Bootstrap support for afum|EAL87095.1 as seed ortholog is 100%.
Bootstrap support for afis|EAW19399.1 as seed ortholog is 100%.

Group of orthologs #4807. Best score 886 bits
Score difference with first non-orthologous sequence - *A. fumigatus*:886 *A. fischeri*:886
afum|EAL87923.2 100.00% afis|EAW23543.1 100.00%
Bootstrap support for afum|EAL87923.2 as seed ortholog is 100%.
Bootstrap support for afis|EAW23543.1 as seed ortholog is 100%.

Group of orthologs #4808. Best score 886 bits
Score difference with first non-orthologous sequence - *A. fumigatus*:886 *A. fischeri*:886
afum|EAL93499.1 100.00% afis|EAW18768.1 100.00%
Bootstrap support for afum|EAL93499.1 as seed ortholog is 100%.
Bootstrap support for afis|EAW18768.1 as seed ortholog is 100%.

Group of orthologs #4809. Best score 886 bits
Score difference with first non-orthologous sequence - *A. fumigatus*:886 *A. fischeri*:886
afum|EAL90527.1 100.00% afis|EAW22665.1 100.00%
Bootstrap support for afum|EAL90527.1 as seed ortholog is 100%.
Bootstrap support for afis|EAW22665.1 as seed ortholog is 100%.

Group of orthologs #4810. Best score 886 bits
Score difference with first non-orthologous sequence - *A. fumigatus*:886 *A. fischeri*:886
afum|EAL89876.1 100.00% afis|EAW25215.1 100.00%
Bootstrap support for afum|EAL89876.1 as seed ortholog is 100%.
Bootstrap support for afis|EAW25215.1 as seed ortholog is 100%.

Group of orthologs #4811. Best score 885 bits
Score difference with first non-orthologous sequence - *A. fumigatus*:885 *A. fischeri*:885
afum|EAL90668.1 100.00% afis|EAW22526.1 100.00%
Bootstrap support for afum|EAL90668.1 as seed ortholog is 100%.
Bootstrap support for afis|EAW22526.1 as seed ortholog is 100%.

Group of orthologs #4812. Best score 884 bits
Score difference with first non-orthologous sequence - *A. fumigatus*:884 *A. fischeri*:884
afum|EAL86816.2 100.00% afis|EAW19711.1 100.00%
Bootstrap support for afum|EAL86816.2 as seed ortholog is 100%.
Bootstrap support for afis|EAW19711.1 as seed ortholog is 100%.

Group of orthologs #4813. Best score 884 bits

Score difference with first non-orthologous sequence - *A. fumigatus*:884 *A. fischeri*:884
afum|EAL91862.1 100.00% afis|EAW18051.1 100.00%
Bootstrap support for afum|EAL91862.1 as seed ortholog is 100%.
Bootstrap support for afis|EAW18051.1 as seed ortholog is 100%.

---

Group of orthologs #4814. Best score 884 bits
Score difference with first non-orthologous sequence - *A. fumigatus*:884 *A. fischeri*:884
afum|EAL84384.1 100.00% afis|EAW25711.1 100.00%
Bootstrap support for afum|EAL84384.1 as seed ortholog is 100%.
Bootstrap support for afis|EAW25711.1 as seed ortholog is 100%.

---

Group of orthologs #4815. Best score 884 bits
Score difference with first non-orthologous sequence - *A. fumigatus*:884 *A. fischeri*:884
afum|EAL87167.1 100.00% afis|EAW23729.1 100.00%
Bootstrap support for afum|EAL87167.1 as seed ortholog is 100%.
Bootstrap support for afis|EAW23729.1 as seed ortholog is 100%.

---

Group of orthologs #4816. Best score 884 bits
Score difference with first non-orthologous sequence - *A. fumigatus*:884 *A. fischeri*:842
afum|EAL88375.1 100.00% afis|EAW23104.1 100.00%
Bootstrap support for afum|EAL88375.1 as seed ortholog is 100%.
Bootstrap support for afis|EAW23104.1 as seed ortholog is 100%.

---

Group of orthologs #4817. Best score 884 bits
Score difference with first non-orthologous sequence - *A. fumigatus*:884 *A. fischeri*:884
afum|EAL93284.1 100.00% afis|EAW18580.1 100.00%
Bootstrap support for afum|EAL93284.1 as seed ortholog is 100%.
Bootstrap support for afis|EAW18580.1 as seed ortholog is 100%.

---

Group of orthologs #4818. Best score 884 bits
Score difference with first non-orthologous sequence - *A. fumigatus*:884 *A. fischeri*:884
afum|EAL93586.1 100.00% afis|EAW18855.1 100.00%
Bootstrap support for afum|EAL93586.1 as seed ortholog is 100%.
Bootstrap support for afis|EAW18855.1 as seed ortholog is 100%.

---

Group of orthologs #4819. Best score 883 bits
Score difference with first non-orthologous sequence - *A. fumigatus*:259 *A. fischeri*:547
afum|EAL84968.2 100.00% afis|EAW19947.1 100.00%
Bootstrap support for afum|EAL84968.2 as seed ortholog is 100%.
Bootstrap support for afis|EAW19947.1 as seed ortholog is 100%.

---

Group of orthologs #4820. Best score 883 bits
Score difference with first non-orthologous sequence - *A. fumigatus*:883 *A. fischeri*:883
afum|EAL88918.1 100.00% afis|EAW16296.1 100.00%
Bootstrap support for afum|EAL88918.1 as seed ortholog is 100%.
Bootstrap support for afis|EAW16296.1 as seed ortholog is 100%.

---

Group of orthologs #4821. Best score 883 bits
Score difference with first non-orthologous sequence - *A. fumigatus*:883 *A. fischeri*:883
afum|EAL86003.1 100.00% afis|EAW24333.1 100.00%
Bootstrap support for afum|EAL86003.1 as seed ortholog is 100%.
Bootstrap support for afis|EAW24333.1 as seed ortholog is 100%.

---

Group of orthologs #4822. Best score 883 bits
Score difference with first non-orthologous sequence - *A. fumigatus*:883 *A. fischeri*:691
afum|EAL86145.1 100.00% afis|EAW24472.1 100.00%

Bootstrap support for afum|EAL86145.1 as seed ortholog is 100%.
Bootstrap support for afis|EAW24472.1 as seed ortholog is 100%.

---

Group of orthologs #4823. Best score 883 bits
Score difference with first non-orthologous sequence - *A. fumigatus*:883 *A. fischeri*:883
afum|EAL88288.2 100.00% afis|EAW23195.1 100.00%
Bootstrap support for afum|EAL88288.2 as seed ortholog is 100%.
Bootstrap support for afis|EAW23195.1 as seed ortholog is 100%.

---

Group of orthologs #4824. Best score 883 bits
Score difference with first non-orthologous sequence - *A. fumigatus*:883 *A. fischeri*:883
afum|EAL90080.2 100.00% afis|EAW25421.1 100.00%
Bootstrap support for afum|EAL90080.2 as seed ortholog is 100%.
Bootstrap support for afis|EAW25421.1 as seed ortholog is 100%.

---

Group of orthologs #4825. Best score 882 bits
Score difference with first non-orthologous sequence - *A. fumigatus*:818 *A. fischeri*:648
afum|EAL84864.1 100.00% afis|EAW20753.1 100.00%
Bootstrap support for afum|EAL84864.1 as seed ortholog is 100%.
Bootstrap support for afis|EAW20753.1 as seed ortholog is 100%.

---

Group of orthologs #4826. Best score 882 bits
Score difference with first non-orthologous sequence - *A. fumigatus*:882 *A. fischeri*:882
afum|EAL89259.1 100.00% afis|EAW16655.1 100.00%
Bootstrap support for afum|EAL89259.1 as seed ortholog is 100%.
Bootstrap support for afis|EAW16655.1 as seed ortholog is 100%.

---

Group of orthologs #4827. Best score 882 bits
Score difference with first non-orthologous sequence - *A. fumigatus*:882 *A. fischeri*:882
afum|EAL93661.1 100.00% afis|EAW18930.1 100.00%
Bootstrap support for afum|EAL93661.1 as seed ortholog is 100%.
Bootstrap support for afis|EAW18930.1 as seed ortholog is 100%.

---

Group of orthologs #4828. Best score 881 bits
Score difference with first non-orthologous sequence - *A. fumigatus*:881 *A. fischeri*:789
afum|EAL85679.1 100.00% afis|EAW15669.1 100.00%
Bootstrap support for afum|EAL85679.1 as seed ortholog is 100%.
Bootstrap support for afis|EAW15669.1 as seed ortholog is 100%.

---

Group of orthologs #4829. Best score 881 bits
Score difference with first non-orthologous sequence - *A. fumigatus*:881 *A. fischeri*:795
afum|EAL85572.1 100.00% afis|EAW15777.1 100.00%
Bootstrap support for afum|EAL85572.1 as seed ortholog is 100%.
Bootstrap support for afis|EAW15777.1 as seed ortholog is 100%.

---

Group of orthologs #4830. Best score 881 bits
Score difference with first non-orthologous sequence - *A. fumigatus*:881 *A. fischeri*:881
afum|EAL88541.1 100.00% afis|EAW15906.1 100.00%
Bootstrap support for afum|EAL88541.1 as seed ortholog is 100%.
Bootstrap support for afis|EAW15906.1 as seed ortholog is 100%.

---

Group of orthologs #4831. Best score 881 bits
Score difference with first non-orthologous sequence - *A. fumigatus*:881 *A. fischeri*:636
afum|EAL85420.1 100.00% afis|EAW20289.1 100.00%
Bootstrap support for afum|EAL85420.1 as seed ortholog is 100%.
Bootstrap support for afis|EAW20289.1 as seed ortholog is 100%.

Group of orthologs #4832. Best score 881 bits
Score difference with first non-orthologous sequence - *A. fumigatus*:881 *A. fischeri*:881
afum|EAL85940.2 100.00% afis|EAW24272.1 100.00%
Bootstrap support for afum|EAL85940.2 as seed ortholog is 100%.
Bootstrap support for afis|EAW24272.1 as seed ortholog is 100%.

---

Group of orthologs #4833. Best score 881 bits
Score difference with first non-orthologous sequence - *A. fumigatus*:881 *A. fischeri*:881
afum|EAL87302.1 100.00% afis|EAW23874.1 100.00%
Bootstrap support for afum|EAL87302.1 as seed ortholog is 100%.
Bootstrap support for afis|EAW23874.1 as seed ortholog is 100%.

---

Group of orthologs #4834. Best score 881 bits
Score difference with first non-orthologous sequence - *A. fumigatus*:881 *A. fischeri*:881
afum|EAL89538.2 100.00% afis|EAW24887.1 100.00%
Bootstrap support for afum|EAL89538.2 as seed ortholog is 100%.
Bootstrap support for afis|EAW24887.1 as seed ortholog is 100%.

---

Group of orthologs #4835. Best score 881 bits
Score difference with first non-orthologous sequence - *A. fumigatus*:881 *A. fischeri*:881
afum|EAL89988.1 100.00% afis|EAW25327.1 100.00%
Bootstrap support for afum|EAL89988.1 as seed ortholog is 100%.
Bootstrap support for afis|EAW25327.1 as seed ortholog is 100%.

---

Group of orthologs #4836. Best score 880 bits
Score difference with first non-orthologous sequence - *A. fumigatus*:880 *A. fischeri*:880
afum|EAL85791.1 100.00% afis|EAW15539.1 100.00%
Bootstrap support for afum|EAL85791.1 as seed ortholog is 100%.
Bootstrap support for afis|EAW15539.1 as seed ortholog is 100%.

---

Group of orthologs #4837. Best score 880 bits
Score difference with first non-orthologous sequence - *A. fumigatus*:880 *A. fischeri*:880
afum|EAL88713.1 100.00% afis|EAW16081.1 100.00%
Bootstrap support for afum|EAL88713.1 as seed ortholog is 100%.
Bootstrap support for afis|EAW16081.1 as seed ortholog is 100%.

---

Group of orthologs #4838. Best score 880 bits
Score difference with first non-orthologous sequence - *A. fumigatus*:880 *A. fischeri*:880
afum|EAL86969.2 100.00% afis|EAW19537.1 100.00%
Bootstrap support for afum|EAL86969.2 as seed ortholog is 100%.
Bootstrap support for afis|EAW19537.1 as seed ortholog is 100%.

---

Group of orthologs #4839. Best score 880 bits
Score difference with first non-orthologous sequence - *A. fumigatus*:880 *A. fischeri*:880
afum|EAL90259.1 100.00% afis|EAW22937.1 100.00%
Bootstrap support for afum|EAL90259.1 as seed ortholog is 100%.
Bootstrap support for afis|EAW22937.1 as seed ortholog is 100%.

---

Group of orthologs #4840. Best score 880 bits
Score difference with first non-orthologous sequence - *A. fumigatus*:880 *A. fischeri*:880
afum|EAL89896.2 100.00% afis|EAW25235.1 100.00%
Bootstrap support for afum|EAL89896.2 as seed ortholog is 100%.
Bootstrap support for afis|EAW25235.1 as seed ortholog is 100%.

---

Group of orthologs #4841. Best score 879 bits

Score difference with first non-orthologous sequence - *A. fumigatus*:718   *A. fischeri*:879
afum|EAL88568.1 100.00% afis|EAW15932.1 100.00%
Bootstrap support for afum|EAL88568.1 as seed ortholog is 100%.
Bootstrap support for afis|EAW15932.1 as seed ortholog is 100%.

---

Group of orthologs #4842. Best score 879 bits
Score difference with first non-orthologous sequence - *A. fumigatus*:879   *A. fischeri*:879
afum|EAL84694.1 100.00% afis|EAW20940.1 100.00%
Bootstrap support for afum|EAL84694.1 as seed ortholog is 100%.
Bootstrap support for afis|EAW20940.1 as seed ortholog is 100%.

---

Group of orthologs #4843. Best score 879 bits
Score difference with first non-orthologous sequence - *A. fumigatus*:879   *A. fischeri*:879
afum|EAL87361.1 100.00% afis|EAW23936.1 100.00%
Bootstrap support for afum|EAL87361.1 as seed ortholog is 100%.
Bootstrap support for afis|EAW23936.1 as seed ortholog is 100%.

---

Group of orthologs #4844. Best score 878 bits
Score difference with first non-orthologous sequence - *A. fumigatus*:878   *A. fischeri*:878
afum|EAL91315.1 100.00% afis|EAW17493.1 100.00%
Bootstrap support for afum|EAL91315.1 as seed ortholog is 100%.
Bootstrap support for afis|EAW17493.1 as seed ortholog is 100%.

---

Group of orthologs #4845. Best score 878 bits
Score difference with first non-orthologous sequence - *A. fumigatus*:878   *A. fischeri*:878
afum|EAL90571.1 100.00% afis|EAW22625.1 100.00%
Bootstrap support for afum|EAL90571.1 as seed ortholog is 100%.
Bootstrap support for afis|EAW22625.1 as seed ortholog is 100%.

---

Group of orthologs #4846. Best score 878 bits
Score difference with first non-orthologous sequence - *A. fumigatus*:804   *A. fischeri*:676
afum|EAL92187.1 100.00% afis|EAW21168.1 100.00%
Bootstrap support for afum|EAL92187.1 as seed ortholog is 100%.
Bootstrap support for afis|EAW21168.1 as seed ortholog is 100%.

---

Group of orthologs #4847. Best score 878 bits
Score difference with first non-orthologous sequence - *A. fumigatus*:878   *A. fischeri*:878
afum|EAL92558.2 100.00% afis|EAW21538.1 100.00%
Bootstrap support for afum|EAL92558.2 as seed ortholog is 100%.
Bootstrap support for afis|EAW21538.1 as seed ortholog is 100%.

---

Group of orthologs #4848. Best score 878 bits
Score difference with first non-orthologous sequence - *A. fumigatus*:878   *A. fischeri*:878
afum|EAL92982.1 100.00% afis|EAW21968.1 100.00%
Bootstrap support for afum|EAL92982.1 as seed ortholog is 100%.
Bootstrap support for afis|EAW21968.1 as seed ortholog is 100%.

---

Group of orthologs #4849. Best score 878 bits
Score difference with first non-orthologous sequence - *A. fumigatus*:878   *A. fischeri*:878
afum|EAL89891.1 100.00% afis|EAW25230.1 100.00%
Bootstrap support for afum|EAL89891.1 as seed ortholog is 100%.
Bootstrap support for afis|EAW25230.1 as seed ortholog is 100%.

---

Group of orthologs #4850. Best score 878 bits
Score difference with first non-orthologous sequence - *A. fumigatus*:878   *A. fischeri*:878
afum|EAL90069.1 100.00% afis|EAW25410.1 100.00%

Bootstrap support for afum|EAL90069.1 as seed ortholog is 100%.
Bootstrap support for afis|EAW25410.1 as seed ortholog is 100%.

---

Group of orthologs #4851. Best score 877 bits
Score difference with first non-orthologous sequence - *A. fumigatus*:877 *A. fischeri*:877
afum|EAL85533.1 100.00% afis|EAW15818.1 100.00%
Bootstrap support for afum|EAL85533.1 as seed ortholog is 100%.
Bootstrap support for afis|EAW15818.1 as seed ortholog is 100%.

---

Group of orthologs #4852. Best score 877 bits
Score difference with first non-orthologous sequence - *A. fumigatus*:877 *A. fischeri*:877
afum|EAL85521.1 100.00% afis|EAW15831.1 100.00%
Bootstrap support for afum|EAL85521.1 as seed ortholog is 100%.
Bootstrap support for afis|EAW15831.1 as seed ortholog is 100%.

---

Group of orthologs #4853. Best score 877 bits
Score difference with first non-orthologous sequence - *A. fumigatus*:877 *A. fischeri*:738
afum|EAL89197.1 100.00% afis|EAW16601.1 100.00%
Bootstrap support for afum|EAL89197.1 as seed ortholog is 100%.
Bootstrap support for afis|EAW16601.1 as seed ortholog is 100%.

---

Group of orthologs #4854. Best score 877 bits
Score difference with first non-orthologous sequence - *A. fumigatus*:877 *A. fischeri*:877
afum|EAL85870.1 100.00% afis|EAW24198.1 100.00%
Bootstrap support for afum|EAL85870.1 as seed ortholog is 100%.
Bootstrap support for afis|EAW24198.1 as seed ortholog is 100%.

---

Group of orthologs #4855. Best score 877 bits
Score difference with first non-orthologous sequence - *A. fumigatus*:877 *A. fischeri*:877
afum|EAL93801.1 100.00% afis|EAW19073.1 100.00%
Bootstrap support for afum|EAL93801.1 as seed ortholog is 100%.
Bootstrap support for afis|EAW19073.1 as seed ortholog is 100%.

---

Group of orthologs #4856. Best score 877 bits
Score difference with first non-orthologous sequence - *A. fumigatus*:877 *A. fischeri*:877
afum|EAL92406.1 100.00% afis|EAW21392.1 100.00%
Bootstrap support for afum|EAL92406.1 as seed ortholog is 100%.
Bootstrap support for afis|EAW21392.1 as seed ortholog is 100%.

---

Group of orthologs #4857. Best score 877 bits
Score difference with first non-orthologous sequence - *A. fumigatus*:877 *A. fischeri*:877
afum|EAL89994.1 100.00% afis|EAW25332.1 100.00%
Bootstrap support for afum|EAL89994.1 as seed ortholog is 100%.
Bootstrap support for afis|EAW25332.1 as seed ortholog is 100%.

---

Group of orthologs #4858. Best score 876 bits
Score difference with first non-orthologous sequence - *A. fumigatus*:876 *A. fischeri*:876
afum|EAL85829.1 100.00% afis|EAW15494.1 100.00%
Bootstrap support for afum|EAL85829.1 as seed ortholog is 100%.
Bootstrap support for afis|EAW15494.1 as seed ortholog is 100%.

---

Group of orthologs #4859. Best score 876 bits
Score difference with first non-orthologous sequence - *A. fumigatus*:876 *A. fischeri*:496
afum|EAL86376.1 100.00% afis|EAW16827.1 100.00%
Bootstrap support for afum|EAL86376.1 as seed ortholog is 100%.
Bootstrap support for afis|EAW16827.1 as seed ortholog is 100%.

Group of orthologs #4860. Best score 876 bits
Score difference with first non-orthologous sequence - *A. fumigatus*:876 *A. fischeri*:819
afum|EAL86013.1 100.00% afis|EAW24342.1 100.00%
Bootstrap support for afum|EAL86013.1 as seed ortholog is 100%.
Bootstrap support for afis|EAW24342.1 as seed ortholog is 100%.

Group of orthologs #4861. Best score 876 bits
Score difference with first non-orthologous sequence - *A. fumigatus*:876 *A. fischeri*:876
afum|EAL89494.1 100.00% afis|EAW24848.1 100.00%
Bootstrap support for afum|EAL89494.1 as seed ortholog is 100%.
Bootstrap support for afis|EAW24848.1 as seed ortholog is 100%.

Group of orthologs #4862. Best score 875 bits
Score difference with first non-orthologous sequence - *A. fumigatus*:875 *A. fischeri*:875
afum|EAL86346.1 100.00% afis|EAW16777.1 100.00%
Bootstrap support for afum|EAL86346.1 as seed ortholog is 100%.
Bootstrap support for afis|EAW16777.1 as seed ortholog is 100%.

Group of orthologs #4863. Best score 875 bits
Score difference with first non-orthologous sequence - *A. fumigatus*:875 *A. fischeri*:875
afum|EAL87063.1 100.00% afis|EAW19431.1 100.00%
Bootstrap support for afum|EAL87063.1 as seed ortholog is 100%.
Bootstrap support for afis|EAW19431.1 as seed ortholog is 100%.

Group of orthologs #4864. Best score 875 bits
Score difference with first non-orthologous sequence - *A. fumigatus*:875 *A. fischeri*:875
afum|EAL87932.1 100.00% afis|EAW23534.1 100.00%
Bootstrap support for afum|EAL87932.1 as seed ortholog is 100%.
Bootstrap support for afis|EAW23534.1 as seed ortholog is 100%.

Group of orthologs #4865. Best score 875 bits
Score difference with first non-orthologous sequence - *A. fumigatus*:875 *A. fischeri*:746
afum|EAL92099.1 100.00% afis|EAW21076.1 100.00%
Bootstrap support for afum|EAL92099.1 as seed ortholog is 100%.
Bootstrap support for afis|EAW21076.1 as seed ortholog is 100%.

Group of orthologs #4866. Best score 875 bits
Score difference with first non-orthologous sequence - *A. fumigatus*:875 *A. fischeri*:875
afum|EAL91991.1 100.00% afis|EAW24136.1 100.00%
Bootstrap support for afum|EAL91991.1 as seed ortholog is 100%.
Bootstrap support for afis|EAW24136.1 as seed ortholog is 100%.

Group of orthologs #4867. Best score 874 bits
Score difference with first non-orthologous sequence - *A. fumigatus*:874 *A. fischeri*:874
afum|EAL85642.1 100.00% afis|EAW15705.1 100.00%
Bootstrap support for afum|EAL85642.1 as seed ortholog is 100%.
Bootstrap support for afis|EAW15705.1 as seed ortholog is 100%.

Group of orthologs #4868. Best score 874 bits
Score difference with first non-orthologous sequence - *A. fumigatus*:723 *A. fischeri*:822
afum|EAL84666.1 100.00% afis|EAW20967.1 100.00%
Bootstrap support for afum|EAL84666.1 as seed ortholog is 100%.
Bootstrap support for afis|EAW20967.1 as seed ortholog is 100%.

Group of orthologs #4869. Best score 874 bits

Score difference with first non-orthologous sequence - *A. fumigatus*:874 *A. fischeri*:874
afum|EAL87705.1 100.00% afis|EAW18342.1 100.00%
Bootstrap support for afum|EAL87705.1 as seed ortholog is 100%.
Bootstrap support for afis|EAW18342.1 as seed ortholog is 100%.

---

Group of orthologs #4870. Best score 874 bits
Score difference with first non-orthologous sequence - *A. fumigatus*:874 *A. fischeri*:874
afum|EAL91810.1 100.00% afis|EAW17999.1 100.00%
Bootstrap support for afum|EAL91810.1 as seed ortholog is 100%.
Bootstrap support for afis|EAW17999.1 as seed ortholog is 100%.

---

Group of orthologs #4871. Best score 874 bits
Score difference with first non-orthologous sequence - *A. fumigatus*:874 *A. fischeri*:874
afum|EAL87354.1 100.00% afis|EAW23929.1 100.00%
Bootstrap support for afum|EAL87354.1 as seed ortholog is 100%.
Bootstrap support for afis|EAW23929.1 as seed ortholog is 100%.

---

Group of orthologs #4872. Best score 874 bits
Score difference with first non-orthologous sequence - *A. fumigatus*:874 *A. fischeri*:874
afum|EAL93739.1 100.00% afis|EAW19005.1 100.00%
Bootstrap support for afum|EAL93739.1 as seed ortholog is 100%.
Bootstrap support for afis|EAW19005.1 as seed ortholog is 100%.

---

Group of orthologs #4873. Best score 874 bits
Score difference with first non-orthologous sequence - *A. fumigatus*:874 *A. fischeri*:874
afum|EAL92312.1 100.00% afis|EAW21295.1 100.00%
Bootstrap support for afum|EAL92312.1 as seed ortholog is 100%.
Bootstrap support for afis|EAW21295.1 as seed ortholog is 100%.

---

Group of orthologs #4874. Best score 874 bits
Score difference with first non-orthologous sequence - *A. fumigatus*:874 *A. fischeri*:542
afum|EAL89402.1 100.00% afis|EAW24760.1 100.00%
Bootstrap support for afum|EAL89402.1 as seed ortholog is 100%.
Bootstrap support for afis|EAW24760.1 as seed ortholog is 100%.

---

Group of orthologs #4875. Best score 874 bits
Score difference with first non-orthologous sequence - *A. fumigatus*:107 *A. fischeri*:874
afum|EAL89592.1 100.00% afis|EAW24935.1 100.00%
Bootstrap support for afum|EAL89592.1 as seed ortholog is 99%.
Bootstrap support for afis|EAW24935.1 as seed ortholog is 100%.

---

Group of orthologs #4876. Best score 874 bits
Score difference with first non-orthologous sequence - *A. fumigatus*:874 *A. fischeri*:874
afum|EAL92885.1 100.00% afis|EAW21871.1 100.00%
Bootstrap support for afum|EAL92885.1 as seed ortholog is 100%.
Bootstrap support for afis|EAW21871.1 as seed ortholog is 100%.

---

Group of orthologs #4877. Best score 873 bits
Score difference with first non-orthologous sequence - *A. fumigatus*:873 *A. fischeri*:873
afum|EAL84987.2 100.00% afis|EAW19922.1 100.00%
Bootstrap support for afum|EAL84987.2 as seed ortholog is 100%.
Bootstrap support for afis|EAW19922.1 as seed ortholog is 100%.

---

Group of orthologs #4878. Best score 873 bits
Score difference with first non-orthologous sequence - *A. fumigatus*:255 *A. fischeri*:103
afum|EAL88796.1 100.00% afis|EAW20719.1 100.00%

Bootstrap support for afum|EAL88796.1 as seed ortholog is 100%.
Bootstrap support for afis|EAW20719.1 as seed ortholog is 99%.

---

Group of orthologs #4879. Best score 873 bits
Score difference with first non-orthologous sequence - *A. fumigatus*:873 *A. fischeri*:873
afum|EAL89599.1 100.00% afis|EAW24942.1 100.00%
Bootstrap support for afum|EAL89599.1 as seed ortholog is 100%.
Bootstrap support for afis|EAW24942.1 as seed ortholog is 100%.

---

Group of orthologs #4880. Best score 872 bits
Score difference with first non-orthologous sequence - *A. fumigatus*:872 *A. fischeri*:872
afum|EAL86804.2 100.00% afis|EAW19722.1 100.00%
Bootstrap support for afum|EAL86804.2 as seed ortholog is 100%.
Bootstrap support for afis|EAW19722.1 as seed ortholog is 100%.

---

Group of orthologs #4881. Best score 872 bits
Score difference with first non-orthologous sequence - *A. fumigatus*:872 *A. fischeri*:872
afum|EAL93365.1 100.00% afis|EAW18649.1 100.00%
Bootstrap support for afum|EAL93365.1 as seed ortholog is 100%.
Bootstrap support for afis|EAW18649.1 as seed ortholog is 100%.

---

Group of orthologs #4882. Best score 871 bits
Score difference with first non-orthologous sequence - *A. fumigatus*:871 *A. fischeri*:871
afum|EAL85525.1 100.00% afis|EAW15826.1 100.00%
Bootstrap support for afum|EAL85525.1 as seed ortholog is 100%.
Bootstrap support for afis|EAW15826.1 as seed ortholog is 100%.

---

Group of orthologs #4883. Best score 871 bits
Score difference with first non-orthologous sequence - *A. fumigatus*:871 *A. fischeri*:871
afum|EAL84722.1 100.00% afis|EAW20912.1 100.00%
Bootstrap support for afum|EAL84722.1 as seed ortholog is 100%.
Bootstrap support for afis|EAW20912.1 as seed ortholog is 100%.

---

Group of orthologs #4884. Best score 871 bits
Score difference with first non-orthologous sequence - *A. fumigatus*:871 *A. fischeri*:56
afum|EAL86274.1 100.00% afis|EAW20692.1 100.00%
Bootstrap support for afum|EAL86274.1 as seed ortholog is 100%.
Bootstrap support for afis|EAW20692.1 as seed ortholog is 95%.

---

Group of orthologs #4885. Best score 871 bits
Score difference with first non-orthologous sequence - *A. fumigatus*:871 *A. fischeri*:871
afum|EAL92245.1 100.00% afis|EAW21233.1 100.00%
Bootstrap support for afum|EAL92245.1 as seed ortholog is 100%.
Bootstrap support for afis|EAW21233.1 as seed ortholog is 100%.

---

Group of orthologs #4886. Best score 871 bits
Score difference with first non-orthologous sequence - *A. fumigatus*:871 *A. fischeri*:871
afum|EAL89869.1 100.00% afis|EAW25208.1 100.00%
Bootstrap support for afum|EAL89869.1 as seed ortholog is 100%.
Bootstrap support for afis|EAW25208.1 as seed ortholog is 100%.

---

Group of orthologs #4887. Best score 870 bits
Score difference with first non-orthologous sequence - *A. fumigatus*:870 *A. fischeri*:718
afum|EAL84210.1 100.00% afis|EAW15434.1 100.00%
Bootstrap support for afum|EAL84210.1 as seed ortholog is 100%.
Bootstrap support for afis|EAW15434.1 as seed ortholog is 100%.

Group of orthologs #4888. Best score 870 bits
Score difference with first non-orthologous sequence - *A. fumigatus*:870 *A. fischeri*:870
afum|EAL89088.1 100.00% afis|EAW16494.1 100.00%
Bootstrap support for afum|EAL89088.1 as seed ortholog is 100%.
Bootstrap support for afis|EAW16494.1 as seed ortholog is 100%.

Group of orthologs #4889. Best score 870 bits
Score difference with first non-orthologous sequence - *A. fumigatus*:870 *A. fischeri*:707
afum|EAL89122.1 100.00% afis|EAW16526.1 100.00%
Bootstrap support for afum|EAL89122.1 as seed ortholog is 100%.
Bootstrap support for afis|EAW16526.1 as seed ortholog is 100%.

Group of orthologs #4890. Best score 870 bits
Score difference with first non-orthologous sequence - *A. fumigatus*:870 *A. fischeri*:582
afum|EAL85339.1 100.00% afis|EAW20224.1 100.00%
Bootstrap support for afum|EAL85339.1 as seed ortholog is 100%.
Bootstrap support for afis|EAW20224.1 as seed ortholog is 100%.

Group of orthologs #4891. Best score 870 bits
Score difference with first non-orthologous sequence - *A. fumigatus*:672 *A. fischeri*:562
afum|EAL84814.1 100.00% afis|EAW20814.1 100.00%
Bootstrap support for afum|EAL84814.1 as seed ortholog is 100%.
Bootstrap support for afis|EAW20814.1 as seed ortholog is 100%.

Group of orthologs #4892. Best score 870 bits
Score difference with first non-orthologous sequence - *A. fumigatus*:870 *A. fischeri*:870
afum|EBA27349.1 100.00% afis|EAW15849.1 100.00%
Bootstrap support for afum|EBA27349.1 as seed ortholog is 100%.
Bootstrap support for afis|EAW15849.1 as seed ortholog is 100%.

Group of orthologs #4893. Best score 870 bits
Score difference with first non-orthologous sequence - *A. fumigatus*:870 *A. fischeri*:713
afum|EAL90906.1 100.00% afis|EAW22275.1 100.00%
Bootstrap support for afum|EAL90906.1 as seed ortholog is 100%.
Bootstrap support for afis|EAW22275.1 as seed ortholog is 100%.

Group of orthologs #4894. Best score 869 bits
Score difference with first non-orthologous sequence - *A. fumigatus*:805 *A. fischeri*:628
afum|EAL87033.1 100.00% afis|EAW19463.1 100.00%
Bootstrap support for afum|EAL87033.1 as seed ortholog is 100%.
Bootstrap support for afis|EAW19463.1 as seed ortholog is 100%.

Group of orthologs #4895. Best score 869 bits
Score difference with first non-orthologous sequence - *A. fumigatus*:869 *A. fischeri*:869
afum|EAL90336.1 100.00% afis|EAW22864.1 100.00%
Bootstrap support for afum|EAL90336.1 as seed ortholog is 100%.
Bootstrap support for afis|EAW22864.1 as seed ortholog is 100%.

Group of orthologs #4896. Best score 869 bits
Score difference with first non-orthologous sequence - *A. fumigatus*:869 *A. fischeri*:869
afum|EAL94092.1 100.00% afis|EAW19378.1 100.00%
Bootstrap support for afum|EAL94092.1 as seed ortholog is 100%.
Bootstrap support for afis|EAW19378.1 as seed ortholog is 100%.

Group of orthologs #4897. Best score 868 bits

Score difference with first non-orthologous sequence - *A. fumigatus*:385   *A. fischeri*:868
afum|EAL85817.1   100.00%   afis|EAW15514.1   100.00%
Bootstrap support for afum|EAL85817.1 as seed ortholog is 100%.
Bootstrap support for afis|EAW15514.1 as seed ortholog is 100%.

---

Group of orthologs #4898. Best score 868 bits
Score difference with first non-orthologous sequence - *A. fumigatus*:868   *A. fischeri*:868
afum|EAL86515.1   100.00%   afis|EAW16991.1   100.00%
Bootstrap support for afum|EAL86515.1 as seed ortholog is 100%.
Bootstrap support for afis|EAW16991.1 as seed ortholog is 100%.

---

Group of orthologs #4899. Best score 868 bits
Score difference with first non-orthologous sequence - *A. fumigatus*:868   *A. fischeri*:780
afum|EAL84479.1   100.00%   afis|EAW20586.1   100.00%
Bootstrap support for afum|EAL84479.1 as seed ortholog is 100%.
Bootstrap support for afis|EAW20586.1 as seed ortholog is 100%.

---

Group of orthologs #4900. Best score 868 bits
Score difference with first non-orthologous sequence - *A. fumigatus*:868   *A. fischeri*:868
afum|EAL89101.1   100.00%   afis|EAW16506.1   100.00%
Bootstrap support for afum|EAL89101.1 as seed ortholog is 100%.
Bootstrap support for afis|EAW16506.1 as seed ortholog is 100%.

---

Group of orthologs #4901. Best score 868 bits
Score difference with first non-orthologous sequence - *A. fumigatus*:868   *A. fischeri*:868
afum|EAL87077.1   100.00%   afis|EAW19417.1   100.00%
Bootstrap support for afum|EAL87077.1 as seed ortholog is 100%.
Bootstrap support for afis|EAW19417.1 as seed ortholog is 100%.

---

Group of orthologs #4902. Best score 868 bits
Score difference with first non-orthologous sequence - *A. fumigatus*:868   *A. fischeri*:868
afum|EAL86085.2   100.00%   afis|EAW24416.1   100.00%
Bootstrap support for afum|EAL86085.2 as seed ortholog is 100%.
Bootstrap support for afis|EAW24416.1 as seed ortholog is 100%.

---

Group of orthologs #4903. Best score 868 bits
Score difference with first non-orthologous sequence - *A. fumigatus*:868   *A. fischeri*:220
afum|EAL88126.1   100.00%   afis|EAW23350.1   100.00%
Bootstrap support for afum|EAL88126.1 as seed ortholog is 100%.
Bootstrap support for afis|EAW23350.1 as seed ortholog is 100%.

---

Group of orthologs #4904. Best score 868 bits
Score difference with first non-orthologous sequence - *A. fumigatus*:735   *A. fischeri*:422
afum|EAL93564.1   100.00%   afis|EAW18830.1   100.00%
Bootstrap support for afum|EAL93564.1 as seed ortholog is 100%.
Bootstrap support for afis|EAW18830.1 as seed ortholog is 100%.

---

Group of orthologs #4905. Best score 868 bits
Score difference with first non-orthologous sequence - *A. fumigatus*:713   *A. fischeri*:868
afum|EAL90848.1   100.00%   afis|EAW22343.1   100.00%
Bootstrap support for afum|EAL90848.1 as seed ortholog is 100%.
Bootstrap support for afis|EAW22343.1 as seed ortholog is 100%.

---

Group of orthologs #4906. Best score 868 bits
Score difference with first non-orthologous sequence - *A. fumigatus*:868   *A. fischeri*:868
afum|EAL90377.1   100.00%   afis|EAW22823.1   100.00%

Bootstrap support for afum|EAL90377.1 as seed ortholog is 100%.
Bootstrap support for afis|EAW22823.1 as seed ortholog is 100%.

---

Group of orthologs #4907. Best score 868 bits
Score difference with first non-orthologous sequence - *A. fumigatus*:868 *A. fischeri*:868
afum|EAL92343.1 100.00% afis|EAW21325.1 100.00%
Bootstrap support for afum|EAL92343.1 as seed ortholog is 100%.
Bootstrap support for afis|EAW21325.1 as seed ortholog is 100%.

---

Group of orthologs #4908. Best score 868 bits
Score difference with first non-orthologous sequence - *A. fumigatus*:868 *A. fischeri*:868
afum|EAL92491.1 100.00% afis|EAW21472.1 100.00%
Bootstrap support for afum|EAL92491.1 as seed ortholog is 100%.
Bootstrap support for afis|EAW21472.1 as seed ortholog is 100%.

---

Group of orthologs #4909. Best score 868 bits
Score difference with first non-orthologous sequence - *A. fumigatus*:868 *A. fischeri*:868
afum|EAL89641.1 100.00% afis|EAW24981.1 100.00%
Bootstrap support for afum|EAL89641.1 as seed ortholog is 100%.
Bootstrap support for afis|EAW24981.1 as seed ortholog is 100%.

---

Group of orthologs #4910. Best score 868 bits
Score difference with first non-orthologous sequence - *A. fumigatus*:868 *A. fischeri*:868
afum|EAL90203.1 100.00% afis|EAW25540.1 100.00%
Bootstrap support for afum|EAL90203.1 as seed ortholog is 100%.
Bootstrap support for afis|EAW25540.1 as seed ortholog is 100%.

---

Group of orthologs #4911. Best score 867 bits
Score difference with first non-orthologous sequence - *A. fumigatus*:770 *A. fischeri*:783
afum|EAL86309.1 100.00% afis|EAW16735.1 100.00%
Bootstrap support for afum|EAL86309.1 as seed ortholog is 100%.
Bootstrap support for afis|EAW16735.1 as seed ortholog is 100%.

---

Group of orthologs #4912. Best score 867 bits
Score difference with first non-orthologous sequence - *A. fumigatus*:600 *A. fischeri*:681
afum|EAL84922.1 100.00% afis|EAW19995.1 100.00%
Bootstrap support for afum|EAL84922.1 as seed ortholog is 100%.
Bootstrap support for afis|EAW19995.1 as seed ortholog is 100%.

---

Group of orthologs #4913. Best score 867 bits
Score difference with first non-orthologous sequence - *A. fumigatus*:867 *A. fischeri*:867
afum|EAL94047.1 100.00% afis|EAW19330.1 100.00%
Bootstrap support for afum|EAL94047.1 as seed ortholog is 100%.
Bootstrap support for afis|EAW19330.1 as seed ortholog is 100%.

---

Group of orthologs #4914. Best score 866 bits
Score difference with first non-orthologous sequence - *A. fumigatus*:866 *A. fischeri*:866
afum|EAL91371.1 100.00% afis|EAW17557.1 100.00%
Bootstrap support for afum|EAL91371.1 as seed ortholog is 100%.
Bootstrap support for afis|EAW17557.1 as seed ortholog is 100%.

---

Group of orthologs #4915. Best score 866 bits
Score difference with first non-orthologous sequence - *A. fumigatus*:866 *A. fischeri*:808
afum|EAL91656.1 100.00% afis|EAW17847.1 100.00%
Bootstrap support for afum|EAL91656.1 as seed ortholog is 100%.
Bootstrap support for afis|EAW17847.1 as seed ortholog is 100%.

Group of orthologs #4916. Best score 866 bits
Score difference with first non-orthologous sequence - *A. fumigatus*:709 *A. fischeri*:517
afum|EAL87172.1 100.00% afis|EAW23734.1 100.00%
Bootstrap support for afum|EAL87172.1 as seed ortholog is 100%.
Bootstrap support for afis|EAW23734.1 as seed ortholog is 100%.

---

Group of orthologs #4917. Best score 866 bits
Score difference with first non-orthologous sequence - *A. fumigatus*:866 *A. fischeri*:866
afum|EAL87251.1 100.00% afis|EAW23816.1 100.00%
Bootstrap support for afum|EAL87251.1 as seed ortholog is 100%.
Bootstrap support for afis|EAW23816.1 as seed ortholog is 100%.

---

Group of orthologs #4918. Best score 866 bits
Score difference with first non-orthologous sequence - *A. fumigatus*:866 *A. fischeri*:866
afum|EAL90625.1 100.00% afis|EAW22573.1 100.00%
Bootstrap support for afum|EAL90625.1 as seed ortholog is 100%.
Bootstrap support for afis|EAW22573.1 as seed ortholog is 100%.

---

Group of orthologs #4919. Best score 866 bits
Score difference with first non-orthologous sequence - *A. fumigatus*:866 *A. fischeri*:866
afum|EAL89729.1 100.00% afis|EAW25061.1 100.00%
Bootstrap support for afum|EAL89729.1 as seed ortholog is 100%.
Bootstrap support for afis|EAW25061.1 as seed ortholog is 100%.

---

Group of orthologs #4920. Best score 866 bits
Score difference with first non-orthologous sequence - *A. fumigatus*:866 *A. fischeri*:866
afum|EAL89769.1 100.00% afis|EAW25098.1 100.00%
Bootstrap support for afum|EAL89769.1 as seed ortholog is 100%.
Bootstrap support for afis|EAW25098.1 as seed ortholog is 100%.

---

Group of orthologs #4921. Best score 866 bits
Score difference with first non-orthologous sequence - *A. fumigatus*:866 *A. fischeri*:866
afum|EBA27385.1 100.00% afis|EAW24984.1 100.00%
Bootstrap support for afum|EBA27385.1 as seed ortholog is 100%.
Bootstrap support for afis|EAW24984.1 as seed ortholog is 100%.

---

Group of orthologs #4922. Best score 865 bits
Score difference with first non-orthologous sequence - *A. fumigatus*:765 *A. fischeri*:742
afum|EAL86555.1 100.00% afis|EAW17059.1 100.00%
Bootstrap support for afum|EAL86555.1 as seed ortholog is 100%.
Bootstrap support for afis|EAW17059.1 as seed ortholog is 100%.

---

Group of orthologs #4923. Best score 865 bits
Score difference with first non-orthologous sequence - *A. fumigatus*:865 *A. fischeri*:865
afum|EAL88964.1 100.00% afis|EAW16339.1 100.00%
Bootstrap support for afum|EAL88964.1 as seed ortholog is 100%.
Bootstrap support for afis|EAW16339.1 as seed ortholog is 100%.

---

Group of orthologs #4924. Best score 865 bits
Score difference with first non-orthologous sequence - *A. fumigatus*:865 *A. fischeri*:865
afum|EAL91539.1 100.00% afis|EAW17724.1 100.00%
Bootstrap support for afum|EAL91539.1 as seed ortholog is 100%.
Bootstrap support for afis|EAW17724.1 as seed ortholog is 100%.

---

Group of orthologs #4925. Best score 865 bits

Score difference with first non-orthologous sequence - *A. fumigatus*:865 *A. fischeri*:865
afum|EAL86051.1 100.00% afis|EAW24379.1 100.00%
Bootstrap support for afum|EAL86051.1 as seed ortholog is 100%.
Bootstrap support for afis|EAW24379.1 as seed ortholog is 100%.

---

Group of orthologs #4926. Best score 865 bits
Score difference with first non-orthologous sequence - *A. fumigatus*:865 *A. fischeri*:865
afum|EAL88376.1 100.00% afis|EAW23103.1 100.00%
Bootstrap support for afum|EAL88376.1 as seed ortholog is 100%.
Bootstrap support for afis|EAW23103.1 as seed ortholog is 100%.

---

Group of orthologs #4927. Best score 865 bits
Score difference with first non-orthologous sequence - *A. fumigatus*:865 *A. fischeri*:865
afum|EAL87961.1 100.00% afis|EAW23501.1 100.00%
Bootstrap support for afum|EAL87961.1 as seed ortholog is 100%.
Bootstrap support for afis|EAW23501.1 as seed ortholog is 100%.

---

Group of orthologs #4928. Best score 865 bits
Score difference with first non-orthologous sequence - *A. fumigatus*:813 *A. fischeri*:721
afum|EAL93622.1 100.00% afis|EAW18889.1 100.00%
Bootstrap support for afum|EAL93622.1 as seed ortholog is 100%.
Bootstrap support for afis|EAW18889.1 as seed ortholog is 100%.

---

Group of orthologs #4929. Best score 865 bits
Score difference with first non-orthologous sequence - *A. fumigatus*:865 *A. fischeri*:865
afum|EAL90560.1 100.00% afis|EAW22633.1 100.00%
Bootstrap support for afum|EAL90560.1 as seed ortholog is 100%.
Bootstrap support for afis|EAW22633.1 as seed ortholog is 100%.

---

Group of orthologs #4930. Best score 864 bits
Score difference with first non-orthologous sequence - *A. fumigatus*:864 *A. fischeri*:864
afum|EAL88544.1 100.00% afis|EAW15909.1 100.00%
Bootstrap support for afum|EAL88544.1 as seed ortholog is 100%.
Bootstrap support for afis|EAW15909.1 as seed ortholog is 100%.

---

Group of orthologs #4931. Best score 864 bits
Score difference with first non-orthologous sequence - *A. fumigatus*:625 *A. fischeri*:195
afum|EAL85417.1 100.00% afis|EAW20283.1 100.00%
Bootstrap support for afum|EAL85417.1 as seed ortholog is 100%.
Bootstrap support for afis|EAW20283.1 as seed ortholog is 100%.

---

Group of orthologs #4932. Best score 864 bits
Score difference with first non-orthologous sequence - *A. fumigatus*:864 *A. fischeri*:671
afum|EAL85450.1 100.00% afis|EAW20325.1 100.00%
Bootstrap support for afum|EAL85450.1 as seed ortholog is 100%.
Bootstrap support for afis|EAW20325.1 as seed ortholog is 100%.

---

Group of orthologs #4933. Best score 864 bits
Score difference with first non-orthologous sequence - *A. fumigatus*:864 *A. fischeri*:864
afum|EAL86979.1 100.00% afis|EAW19518.1 100.00%
Bootstrap support for afum|EAL86979.1 as seed ortholog is 100%.
Bootstrap support for afis|EAW19518.1 as seed ortholog is 100%.

---

Group of orthologs #4934. Best score 864 bits
Score difference with first non-orthologous sequence - *A. fumigatus*:864 *A. fischeri*:864
afum|EAL91387.1 100.00% afis|EAW17574.1 100.00%

Bootstrap support for afum|EAL91387.1 as seed ortholog is 100%.
Bootstrap support for afis|EAW17574.1 as seed ortholog is 100%.

---

Group of orthologs #4935. Best score 864 bits
Score difference with first non-orthologous sequence - *A. fumigatus*:864 *A. fischeri*:864
afum|EAL93307.1 100.00% afis|EAW18599.1 100.00%
Bootstrap support for afum|EAL93307.1 as seed ortholog is 100%.
Bootstrap support for afis|EAW18599.1 as seed ortholog is 100%.

---

Group of orthologs #4936. Best score 864 bits
Score difference with first non-orthologous sequence - *A. fumigatus*:479 *A. fischeri*:784
afum|EBA27290.1 100.00% afis|EAW19708.1 100.00%
Bootstrap support for afum|EBA27290.1 as seed ortholog is 100%.
Bootstrap support for afis|EAW19708.1 as seed ortholog is 100%.

---

Group of orthologs #4937. Best score 864 bits
Score difference with first non-orthologous sequence - *A. fumigatus*:864 *A. fischeri*:864
afum|EAL92962.1 100.00% afis|EAW21948.1 100.00%
Bootstrap support for afum|EAL92962.1 as seed ortholog is 100%.
Bootstrap support for afis|EAW21948.1 as seed ortholog is 100%.

---

Group of orthologs #4938. Best score 863 bits
Score difference with first non-orthologous sequence - *A. fumigatus*:863 *A. fischeri*:863
afum|EAL84846.1 100.00% afis|EAW20773.1 100.00%
Bootstrap support for afum|EAL84846.1 as seed ortholog is 100%.
Bootstrap support for afis|EAW20773.1 as seed ortholog is 100%.

---

Group of orthologs #4939. Best score 863 bits
Score difference with first non-orthologous sequence - *A. fumigatus*:863 *A. fischeri*:863
afum|EAL91323.1 100.00% afis|EAW17500.1 100.00%
Bootstrap support for afum|EAL91323.1 as seed ortholog is 100%.
Bootstrap support for afis|EAW17500.1 as seed ortholog is 100%.

---

Group of orthologs #4940. Best score 863 bits
Score difference with first non-orthologous sequence - *A. fumigatus*:863 *A. fischeri*:863
afum|EAL91896.1 100.00% afis|EAW18084.1 100.00%
Bootstrap support for afum|EAL91896.1 as seed ortholog is 100%.
Bootstrap support for afis|EAW18084.1 as seed ortholog is 100%.

---

Group of orthologs #4941. Best score 863 bits
Score difference with first non-orthologous sequence - *A. fumigatus*:863 *A. fischeri*:863
afum|EAL87298.1 100.00% afis|EAW23870.1 100.00%
Bootstrap support for afum|EAL87298.1 as seed ortholog is 100%.
Bootstrap support for afis|EAW23870.1 as seed ortholog is 100%.

---

Group of orthologs #4942. Best score 863 bits
Score difference with first non-orthologous sequence - *A. fumigatus*:863 *A. fischeri*:863
afum|EAL87364.1 100.00% afis|EAW23939.1 100.00%
Bootstrap support for afum|EAL87364.1 as seed ortholog is 100%.
Bootstrap support for afis|EAW23939.1 as seed ortholog is 100%.

---

Group of orthologs #4943. Best score 863 bits
Score difference with first non-orthologous sequence - *A. fumigatus*:863 *A. fischeri*:863
afum|EAL93903.1 100.00% afis|EAW19173.1 100.00%
Bootstrap support for afum|EAL93903.1 as seed ortholog is 100%.
Bootstrap support for afis|EAW19173.1 as seed ortholog is 100%.

Group of orthologs #4944. Best score 863 bits
Score difference with first non-orthologous sequence - *A. fumigatus*:863 *A. fischeri*:863
afum|EAL90646.1 100.00% afis|EAW22549.1 100.00%
Bootstrap support for afum|EAL90646.1 as seed ortholog is 100%.
Bootstrap support for afis|EAW22549.1 as seed ortholog is 100%.

Group of orthologs #4945. Best score 863 bits
Score difference with first non-orthologous sequence - *A. fumigatus*:863 *A. fischeri*:863
afum|EBA27250.1 100.00% afis|EAW24224.1 100.00%
Bootstrap support for afum|EBA27250.1 as seed ortholog is 100%.
Bootstrap support for afis|EAW24224.1 as seed ortholog is 100%.

Group of orthologs #4946. Best score 862 bits
Score difference with first non-orthologous sequence - *A. fumigatus*:862 *A. fischeri*:862
afum|EAL86498.1 100.00% afis|EAW16970.1 100.00%
Bootstrap support for afum|EAL86498.1 as seed ortholog is 100%.
Bootstrap support for afis|EAW16970.1 as seed ortholog is 100%.

Group of orthologs #4947. Best score 862 bits
Score difference with first non-orthologous sequence - *A. fumigatus*:862 *A. fischeri*:862
afum|EAL85933.1 100.00% afis|EAW24265.1 100.00%
Bootstrap support for afum|EAL85933.1 as seed ortholog is 100%.
Bootstrap support for afis|EAW24265.1 as seed ortholog is 100%.

Group of orthologs #4948. Best score 862 bits
Score difference with first non-orthologous sequence - *A. fumigatus*:862 *A. fischeri*:746
afum|EAL89901.1 100.00% afis|EAW25240.1 100.00%
Bootstrap support for afum|EAL89901.1 as seed ortholog is 100%.
Bootstrap support for afis|EAW25240.1 as seed ortholog is 100%.

Group of orthologs #4949. Best score 862 bits
Score difference with first non-orthologous sequence - *A. fumigatus*:862 *A. fischeri*:862
afum|EAL91990.1 100.00% afis|EAW24135.1 100.00%
Bootstrap support for afum|EAL91990.1 as seed ortholog is 100%.
Bootstrap support for afis|EAW24135.1 as seed ortholog is 100%.

Group of orthologs #4950. Best score 861 bits
Score difference with first non-orthologous sequence - *A. fumigatus*:861 *A. fischeri*:861
afum|EAL85655.1 100.00% afis|EAW15693.1 100.00%
Bootstrap support for afum|EAL85655.1 as seed ortholog is 100%.
Bootstrap support for afis|EAW15693.1 as seed ortholog is 100%.

Group of orthologs #4951. Best score 861 bits
Score difference with first non-orthologous sequence - *A. fumigatus*:669 *A. fischeri*:861
afum|EAL88725.1 100.00% afis|EAW16093.1 100.00%
Bootstrap support for afum|EAL88725.1 as seed ortholog is 100%.
Bootstrap support for afis|EAW16093.1 as seed ortholog is 100%.

Group of orthologs #4952. Best score 861 bits
Score difference with first non-orthologous sequence - *A. fumigatus*:861 *A. fischeri*:502
afum|EAL88841.1 100.00% afis|EAW16214.1 100.00%
Bootstrap support for afum|EAL88841.1 as seed ortholog is 100%.
Bootstrap support for afis|EAW16214.1 as seed ortholog is 100%.

Group of orthologs #4953. Best score 861 bits

Score difference with first non-orthologous sequence - *A. fumigatus*:861 *A. fischeri*:861
afum|EAL84786.1 100.00% afis|EAW20847.1 100.00%
Bootstrap support for afum|EAL84786.1 as seed ortholog is 100%.
Bootstrap support for afis|EAW20847.1 as seed ortholog is 100%.

---

Group of orthologs #4954. Best score 861 bits
Score difference with first non-orthologous sequence - *A. fumigatus*:861 *A. fischeri*:861
afum|EAL86123.1 100.00% afis|EAW24454.1 100.00%
Bootstrap support for afum|EAL86123.1 as seed ortholog is 100%.
Bootstrap support for afis|EAW24454.1 as seed ortholog is 100%.

---

Group of orthologs #4955. Best score 861 bits
Score difference with first non-orthologous sequence - *A. fumigatus*:861 *A. fischeri*:548
afum|EAL91052.1 100.00% afis|EAW22138.1 100.00%
Bootstrap support for afum|EAL91052.1 as seed ortholog is 100%.
Bootstrap support for afis|EAW22138.1 as seed ortholog is 100%.

---

Group of orthologs #4956. Best score 861 bits
Score difference with first non-orthologous sequence - *A. fumigatus*:861 *A. fischeri*:861
afum|EAL90985.1 100.00% afis|EAW22209.1 100.00%
Bootstrap support for afum|EAL90985.1 as seed ortholog is 100%.
Bootstrap support for afis|EAW22209.1 as seed ortholog is 100%.

---

Group of orthologs #4957. Best score 861 bits
Score difference with first non-orthologous sequence - *A. fumigatus*:861 *A. fischeri*:861
afum|EAL92378.1 100.00% afis|EAW21362.1 100.00%
Bootstrap support for afum|EAL92378.1 as seed ortholog is 100%.
Bootstrap support for afis|EAW21362.1 as seed ortholog is 100%.

---

Group of orthologs #4958. Best score 861 bits
Score difference with first non-orthologous sequence - *A. fumigatus*:861 *A. fischeri*:861
afum|EAL92414.1 100.00% afis|EAW21400.1 100.00%
Bootstrap support for afum|EAL92414.1 as seed ortholog is 100%.
Bootstrap support for afis|EAW21400.1 as seed ortholog is 100%.

---

Group of orthologs #4959. Best score 860 bits
Score difference with first non-orthologous sequence - *A. fumigatus*:860 *A. fischeri*:300
afum|EAL85052.1 100.00% afis|EAW19843.1 100.00%
Bootstrap support for afum|EAL85052.1 as seed ortholog is 100%.
Bootstrap support for afis|EAW19843.1 as seed ortholog is 100%.

---

Group of orthologs #4960. Best score 860 bits
Score difference with first non-orthologous sequence - *A. fumigatus*:860 *A. fischeri*:860
afum|EAL85189.1 100.00% afis|EAW20071.1 100.00%
Bootstrap support for afum|EAL85189.1 as seed ortholog is 100%.
Bootstrap support for afis|EAW20071.1 as seed ortholog is 100%.

---

Group of orthologs #4961. Best score 860 bits
Score difference with first non-orthologous sequence - *A. fumigatus*:860 *A. fischeri*:860
afum|EAL84755.1 100.00% afis|EAW20877.1 100.00%
Bootstrap support for afum|EAL84755.1 as seed ortholog is 100%.
Bootstrap support for afis|EAW20877.1 as seed ortholog is 100%.

---

Group of orthologs #4962. Best score 860 bits
Score difference with first non-orthologous sequence - *A. fumigatus*:860 *A. fischeri*:860
afum|EAL91781.1 100.00% afis|EAW17972.1 100.00%

Bootstrap support for afum|EAL91781.1 as seed ortholog is 100%.
Bootstrap support for afis|EAW17972.1 as seed ortholog is 100%.

---

Group of orthologs #4963. Best score 860 bits
Score difference with first non-orthologous sequence - *A. fumigatus*:807 *A. fischeri*:860
afum|EAL88339.1 100.00% afis|EAW23142.1 100.00%
Bootstrap support for afum|EAL88339.1 as seed ortholog is 100%.
Bootstrap support for afis|EAW23142.1 as seed ortholog is 100%.

---

Group of orthologs #4964. Best score 860 bits
Score difference with first non-orthologous sequence - *A. fumigatus*:860 *A. fischeri*:860
afum|EAL92418.1 100.00% afis|EAW21404.1 100.00%
Bootstrap support for afum|EAL92418.1 as seed ortholog is 100%.
Bootstrap support for afis|EAW21404.1 as seed ortholog is 100%.

---

Group of orthologs #4965. Best score 860 bits
Score difference with first non-orthologous sequence - *A. fumigatus*:860 *A. fischeri*:860
afum|EAL89375.2 100.00% afis|EAW24727.1 100.00%
Bootstrap support for afum|EAL89375.2 as seed ortholog is 100%.
Bootstrap support for afis|EAW24727.1 as seed ortholog is 100%.

---

Group of orthologs #4966. Best score 860 bits
Score difference with first non-orthologous sequence - *A. fumigatus*:860 *A. fischeri*:860
afum|EAL90153.1 100.00% afis|EAW25488.1 100.00%
Bootstrap support for afum|EAL90153.1 as seed ortholog is 100%.
Bootstrap support for afis|EAW25488.1 as seed ortholog is 100%.

---

Group of orthologs #4967. Best score 860 bits
Score difference with first non-orthologous sequence - *A. fumigatus*:860 *A. fischeri*:860
afum|EAL92012.2 100.00% afis|EAW24157.1 100.00%
Bootstrap support for afum|EAL92012.2 as seed ortholog is 100%.
Bootstrap support for afis|EAW24157.1 as seed ortholog is 100%.

---

Group of orthologs #4968. Best score 859 bits
Score difference with first non-orthologous sequence - *A. fumigatus*:859 *A. fischeri*:859
afum|EAL87685.1 100.00% afis|EAW18318.1 100.00%
Bootstrap support for afum|EAL87685.1 as seed ortholog is 100%.
Bootstrap support for afis|EAW18318.1 as seed ortholog is 100%.

---

Group of orthologs #4969. Best score 859 bits
Score difference with first non-orthologous sequence - *A. fumigatus*:859 *A. fischeri*:859
afum|EAL91741.1 100.00% afis|EAW17930.1 100.00%
Bootstrap support for afum|EAL91741.1 as seed ortholog is 100%.
Bootstrap support for afis|EAW17930.1 as seed ortholog is 100%.

---

Group of orthologs #4970. Best score 859 bits
Score difference with first non-orthologous sequence - *A. fumigatus*:859 *A. fischeri*:859
afum|EAL85926.2 100.00% afis|EAW24258.1 100.00%
Bootstrap support for afum|EAL85926.2 as seed ortholog is 100%.
Bootstrap support for afis|EAW24258.1 as seed ortholog is 100%.

---

Group of orthologs #4971. Best score 859 bits
Score difference with first non-orthologous sequence - *A. fumigatus*:859 *A. fischeri*:859
afum|EBA27368.1 100.00% afis|EAW16575.1 100.00%
Bootstrap support for afum|EBA27368.1 as seed ortholog is 100%.
Bootstrap support for afis|EAW16575.1 as seed ortholog is 100%.

Group of orthologs #4972. Best score 859 bits
Score difference with first non-orthologous sequence - *A. fumigatus*:631 *A. fischeri*:859
afum|EAL87874.1 100.00% afis|EAW23592.1 100.00%
Bootstrap support for afum|EAL87874.1 as seed ortholog is 100%.
Bootstrap support for afis|EAW23592.1 as seed ortholog is 100%.

---

Group of orthologs #4973. Best score 858 bits
Score difference with first non-orthologous sequence - *A. fumigatus*:858 *A. fischeri*:763
afum|EAL91621.1 100.00% afis|EAW17812.1 100.00%
Bootstrap support for afum|EAL91621.1 as seed ortholog is 100%.
Bootstrap support for afis|EAW17812.1 as seed ortholog is 100%.

---

Group of orthologs #4974. Best score 858 bits
Score difference with first non-orthologous sequence - *A. fumigatus*:858 *A. fischeri*:858
afum|EAL87990.1 100.00% afis|EAW23475.1 100.00%
Bootstrap support for afum|EAL87990.1 as seed ortholog is 100%.
Bootstrap support for afis|EAW23475.1 as seed ortholog is 100%.

---

Group of orthologs #4975. Best score 858 bits
Score difference with first non-orthologous sequence - *A. fumigatus*:858 *A. fischeri*:858
afum|EAL89457.1 100.00% afis|EAW24813.1 100.00%
Bootstrap support for afum|EAL89457.1 as seed ortholog is 100%.
Bootstrap support for afis|EAW24813.1 as seed ortholog is 100%.

---

Group of orthologs #4976. Best score 858 bits
Score difference with first non-orthologous sequence - *A. fumigatus*:858 *A. fischeri*:858
afum|EAL92741.1 100.00% afis|EAW21706.1 100.00%
Bootstrap support for afum|EAL92741.1 as seed ortholog is 100%.
Bootstrap support for afis|EAW21706.1 as seed ortholog is 100%.

---

Group of orthologs #4977. Best score 857 bits
Score difference with first non-orthologous sequence - *A. fumigatus*:857 *A. fischeri*:543
afum|EAL85731.1 100.00% afis|EAW15602.1 100.00%
Bootstrap support for afum|EAL85731.1 as seed ortholog is 100%.
Bootstrap support for afis|EAW15602.1 as seed ortholog is 100%.

---

Group of orthologs #4978. Best score 857 bits
Score difference with first non-orthologous sequence - *A. fumigatus*:857 *A. fischeri*:857
afum|EAL88657.1 100.00% afis|EAW16027.1 100.00%
Bootstrap support for afum|EAL88657.1 as seed ortholog is 100%.
Bootstrap support for afis|EAW16027.1 as seed ortholog is 100%.

---

Group of orthologs #4979. Best score 857 bits
Score difference with first non-orthologous sequence - *A. fumigatus*:729 *A. fischeri*:676
afum|EAL88827.1 100.00% afis|EAW16200.1 100.00%
Bootstrap support for afum|EAL88827.1 as seed ortholog is 100%.
Bootstrap support for afis|EAW16200.1 as seed ortholog is 100%.

---

Group of orthologs #4980. Best score 857 bits
Score difference with first non-orthologous sequence - *A. fumigatus*:857 *A. fischeri*:857
afum|EAL84709.1 100.00% afis|EAW20925.1 100.00%
Bootstrap support for afum|EAL84709.1 as seed ortholog is 100%.
Bootstrap support for afis|EAW20925.1 as seed ortholog is 100%.

---

Group of orthologs #4981. Best score 857 bits

Score difference with first non-orthologous sequence - *A. fumigatus*:857 *A. fischeri*:857
afum|EAL92417.1 100.00% afis|EAW21403.1 100.00%
Bootstrap support for afum|EAL92417.1 as seed ortholog is 100%.
Bootstrap support for afis|EAW21403.1 as seed ortholog is 100%.

---

Group of orthologs #4982. Best score 856 bits
Score difference with first non-orthologous sequence - *A. fumigatus*:856 *A. fischeri*:856
afum|EAL85062.1 100.00% afis|EAW19830.1 100.00%
Bootstrap support for afum|EAL85062.1 as seed ortholog is 100%.
Bootstrap support for afis|EAW19830.1 as seed ortholog is 100%.

---

Group of orthologs #4983. Best score 856 bits
Score difference with first non-orthologous sequence - *A. fumigatus*:856 *A. fischeri*:507
afum|EAL84971.1 100.00% afis|EAW19942.1 100.00%
Bootstrap support for afum|EAL84971.1 as seed ortholog is 100%.
Bootstrap support for afis|EAW19942.1 as seed ortholog is 100%.

---

Group of orthologs #4984. Best score 856 bits
Score difference with first non-orthologous sequence - *A. fumigatus*:856 *A. fischeri*:856
afum|EAL89016.2 100.00% afis|EAW16390.1 100.00%
Bootstrap support for afum|EAL89016.2 as seed ortholog is 100%.
Bootstrap support for afis|EAW16390.1 as seed ortholog is 100%.

---

Group of orthologs #4985. Best score 856 bits
Score difference with first non-orthologous sequence - *A. fumigatus*:704 *A. fischeri*:697
afum|EAL84818.1 100.00% afis|EAW20810.1 100.00%
Bootstrap support for afum|EAL84818.1 as seed ortholog is 100%.
Bootstrap support for afis|EAW20810.1 as seed ortholog is 100%.

---

Group of orthologs #4986. Best score 856 bits
Score difference with first non-orthologous sequence - *A. fumigatus*:856 *A. fischeri*:856
afum|EAL87375.1 100.00% afis|EAW23949.1 100.00%
Bootstrap support for afum|EAL87375.1 as seed ortholog is 100%.
Bootstrap support for afis|EAW23949.1 as seed ortholog is 100%.

---

Group of orthologs #4987. Best score 856 bits
Score difference with first non-orthologous sequence - *A. fumigatus*:856 *A. fischeri*:856
afum|EAL88429.1 100.00% afis|EAW23051.1 100.00%
Bootstrap support for afum|EAL88429.1 as seed ortholog is 100%.
Bootstrap support for afis|EAW23051.1 as seed ortholog is 100%.

---

Group of orthologs #4988. Best score 856 bits
Score difference with first non-orthologous sequence - *A. fumigatus*:856 *A. fischeri*:856
afum|EAL92519.1 100.00% afis|EAW21500.1 100.00%
Bootstrap support for afum|EAL92519.1 as seed ortholog is 100%.
Bootstrap support for afis|EAW21500.1 as seed ortholog is 100%.

---

Group of orthologs #4989. Best score 855 bits
Score difference with first non-orthologous sequence - *A. fumigatus*:855 *A. fischeri*:855
afum|EAL85528.1 100.00% afis|EAW15823.1 100.00%
Bootstrap support for afum|EAL85528.1 as seed ortholog is 100%.
Bootstrap support for afis|EAW15823.1 as seed ortholog is 100%.

---

Group of orthologs #4990. Best score 855 bits
Score difference with first non-orthologous sequence - *A. fumigatus*:855 *A. fischeri*:855
afum|EAL86412.1 100.00% afis|EAW16875.1 100.00%

Bootstrap support for afum|EAL86412.1 as seed ortholog is 100%.
Bootstrap support for afis|EAW16875.1 as seed ortholog is 100%.

---

Group of orthologs #4991. Best score 855 bits
Score difference with first non-orthologous sequence - *A. fumigatus*:855 *A. fischeri*:855
afum|EAL87551.1 100.00% afis|EAW18188.1 100.00%
Bootstrap support for afum|EAL87551.1 as seed ortholog is 100%.
Bootstrap support for afis|EAW18188.1 as seed ortholog is 100%.

---

Group of orthologs #4992. Best score 855 bits
Score difference with first non-orthologous sequence - *A. fumigatus*:855 *A. fischeri*:855
afum|EAL86818.1 100.00% afis|EAW19709.1 100.00%
Bootstrap support for afum|EAL86818.1 as seed ortholog is 100%.
Bootstrap support for afis|EAW19709.1 as seed ortholog is 100%.

---

Group of orthologs #4993. Best score 855 bits
Score difference with first non-orthologous sequence - *A. fumigatus*:855 *A. fischeri*:855
afum|EAL87323.1 100.00% afis|EAW23895.1 100.00%
Bootstrap support for afum|EAL87323.1 as seed ortholog is 100%.
Bootstrap support for afis|EAW23895.1 as seed ortholog is 100%.

---

Group of orthologs #4994. Best score 855 bits
Score difference with first non-orthologous sequence - *A. fumigatus*:855 *A. fischeri*:855
afum|EAL93433.1 100.00% afis|EAW18710.1 100.00%
Bootstrap support for afum|EAL93433.1 as seed ortholog is 100%.
Bootstrap support for afis|EAW18710.1 as seed ortholog is 100%.

---

Group of orthologs #4995. Best score 855 bits
Score difference with first non-orthologous sequence - *A. fumigatus*:855 *A. fischeri*:316
afum|EAL93788.1 100.00% afis|EAW19059.1 100.00%
Bootstrap support for afum|EAL93788.1 as seed ortholog is 100%.
Bootstrap support for afis|EAW19059.1 as seed ortholog is 100%.

---

Group of orthologs #4996. Best score 855 bits
Score difference with first non-orthologous sequence - *A. fumigatus*:855 *A. fischeri*:582
afum|EAL92125.1 100.00% afis|EAW21104.1 100.00%
Bootstrap support for afum|EAL92125.1 as seed ortholog is 100%.
Bootstrap support for afis|EAW21104.1 as seed ortholog is 100%.

---

Group of orthologs #4997. Best score 855 bits
Score difference with first non-orthologous sequence - *A. fumigatus*:403 *A. fischeri*:855
afum|EAL90484.1 100.00% afis|EAW22708.1 100.00%
Bootstrap support for afum|EAL90484.1 as seed ortholog is 100%.
Bootstrap support for afis|EAW22708.1 as seed ortholog is 100%.

---

Group of orthologs #4998. Best score 855 bits
Score difference with first non-orthologous sequence - *A. fumigatus*:855 *A. fischeri*:855
afum|EAL91984.1 100.00% afis|EAW24128.1 100.00%
Bootstrap support for afum|EAL91984.1 as seed ortholog is 100%.
Bootstrap support for afis|EAW24128.1 as seed ortholog is 100%.

---

Group of orthologs #4999. Best score 854 bits
Score difference with first non-orthologous sequence - *A. fumigatus*:854 *A. fischeri*:854
afum|EAL88537.1 100.00% afis|EAW15902.1 100.00%
Bootstrap support for afum|EAL88537.1 as seed ortholog is 100%.
Bootstrap support for afis|EAW15902.1 as seed ortholog is 100%.

---

Group of orthologs #5000. Best score 854 bits
Score difference with first non-orthologous sequence - *A. fumigatus*:854 *A. fischeri*:854
afum|EAL84416.2 100.00% afis|EAW20680.1 100.00%
Bootstrap support for afum|EAL84416.2 as seed ortholog is 100%.
Bootstrap support for afis|EAW20680.1 as seed ortholog is 100%.

---

Group of orthologs #5001. Best score 854 bits
Score difference with first non-orthologous sequence - *A. fumigatus*:764 *A. fischeri*:854
afum|EAL90535.1 100.00% afis|EAW22657.1 100.00%
Bootstrap support for afum|EAL90535.1 as seed ortholog is 100%.
Bootstrap support for afis|EAW22657.1 as seed ortholog is 100%.

---

Group of orthologs #5002. Best score 853 bits
Score difference with first non-orthologous sequence - *A. fumigatus*:853 *A. fischeri*:853
afum|EAL85589.1 100.00% afis|EAW15759.1 100.00%
Bootstrap support for afum|EAL85589.1 as seed ortholog is 100%.
Bootstrap support for afis|EAW15759.1 as seed ortholog is 100%.

---

Group of orthologs #5003. Best score 853 bits
Score difference with first non-orthologous sequence - *A. fumigatus*:853 *A. fischeri*:537
afum|EAL85344.1 100.00% afis|EAW20227.1 100.00%
Bootstrap support for afum|EAL85344.1 as seed ortholog is 100%.
Bootstrap support for afis|EAW20227.1 as seed ortholog is 100%.

---

Group of orthologs #5004. Best score 853 bits
Score difference with first non-orthologous sequence - *A. fumigatus*:853 *A. fischeri*:853
afum|EAL91651.1 100.00% afis|EAW17842.1 100.00%
Bootstrap support for afum|EAL91651.1 as seed ortholog is 100%.
Bootstrap support for afis|EAW17842.1 as seed ortholog is 100%.

---

Group of orthologs #5005. Best score 853 bits
Score difference with first non-orthologous sequence - *A. fumigatus*:853 *A. fischeri*:853
afum|EAL87226.1 100.00% afis|EAW23792.1 100.00%
Bootstrap support for afum|EAL87226.1 as seed ortholog is 100%.
Bootstrap support for afis|EAW23792.1 as seed ortholog is 100%.

---

Group of orthologs #5006. Best score 852 bits
Score difference with first non-orthologous sequence - *A. fumigatus*:852 *A. fischeri*:852
afum|EAL88187.1 100.00% afis|EAW23292.1 100.00%
Bootstrap support for afum|EAL88187.1 as seed ortholog is 100%.
Bootstrap support for afis|EAW23292.1 as seed ortholog is 100%.

---

Group of orthologs #5007. Best score 852 bits
Score difference with first non-orthologous sequence - *A. fumigatus*:852 *A. fischeri*:852
afum|EAL93676.1 100.00% afis|EAW18943.1 100.00%
Bootstrap support for afum|EAL93676.1 as seed ortholog is 100%.
Bootstrap support for afis|EAW18943.1 as seed ortholog is 100%.

---

Group of orthologs #5008. Best score 852 bits
Score difference with first non-orthologous sequence - *A. fumigatus*:852 *A. fischeri*:852
afum|EAL92155.1 100.00% afis|EAW21136.1 100.00%
Bootstrap support for afum|EAL92155.1 as seed ortholog is 100%.
Bootstrap support for afis|EAW21136.1 as seed ortholog is 100%.

---

Group of orthologs #5009. Best score 852 bits

Score difference with first non-orthologous sequence - *A. fumigatus*:852   *A. fischeri*:687
afum|EAL89341.2 100.00% afis|EAW24699.1 100.00%
Bootstrap support for afum|EAL89341.2 as seed ortholog is 100%.
Bootstrap support for afis|EAW24699.1 as seed ortholog is 100%.

---

Group of orthologs #5010. Best score 852 bits
Score difference with first non-orthologous sequence - *A. fumigatus*:852   *A. fischeri*:852
afum|EAL93002.1 100.00% afis|EAW21986.1 100.00%
Bootstrap support for afum|EAL93002.1 as seed ortholog is 100%.
Bootstrap support for afis|EAW21986.1 as seed ortholog is 100%.

---

Group of orthologs #5011. Best score 851 bits
Score difference with first non-orthologous sequence - *A. fumigatus*:851   *A. fischeri*:736
afum|EAL87831.1 100.00% afis|EAW23629.1 100.00%
Bootstrap support for afum|EAL87831.1 as seed ortholog is 100%.
Bootstrap support for afis|EAW23629.1 as seed ortholog is 100%.

---

Group of orthologs #5012. Best score 851 bits
Score difference with first non-orthologous sequence - *A. fumigatus*:851   *A. fischeri*:851
afum|EAL89950.1 100.00% afis|EAW25287.1 100.00%
Bootstrap support for afum|EAL89950.1 as seed ortholog is 100%.
Bootstrap support for afis|EAW25287.1 as seed ortholog is 100%.

---

Group of orthologs #5013. Best score 850 bits
Score difference with first non-orthologous sequence - *A. fumigatus*:850   *A. fischeri*:850
afum|EAL86508.1 100.00% afis|EAW16981.1 100.00%
Bootstrap support for afum|EAL86508.1 as seed ortholog is 100%.
Bootstrap support for afis|EAW16981.1 as seed ortholog is 100%.

---

Group of orthologs #5014. Best score 850 bits
Score difference with first non-orthologous sequence - *A. fumigatus*:850   *A. fischeri*:850
afum|EAL84910.1 100.00% afis|EAW20008.1 100.00%
Bootstrap support for afum|EAL84910.1 as seed ortholog is 100%.
Bootstrap support for afis|EAW20008.1 as seed ortholog is 100%.

---

Group of orthologs #5015. Best score 850 bits
Score difference with first non-orthologous sequence - *A. fumigatus*:748   *A. fischeri*:850
afum|EAL90569.1 100.00% afis|EAW22627.1 100.00%
Bootstrap support for afum|EAL90569.1 as seed ortholog is 100%.
Bootstrap support for afis|EAW22627.1 as seed ortholog is 100%.

---

Group of orthologs #5016. Best score 850 bits
Score difference with first non-orthologous sequence - *A. fumigatus*:850   *A. fischeri*:850
afum|EAL92423.1 100.00% afis|EAW21409.1 100.00%
Bootstrap support for afum|EAL92423.1 as seed ortholog is 100%.
Bootstrap support for afis|EAW21409.1 as seed ortholog is 100%.

---

Group of orthologs #5017. Best score 850 bits
Score difference with first non-orthologous sequence - *A. fumigatus*:850   *A. fischeri*:850
afum|EAL89525.1 100.00% afis|EAW24875.1 100.00%
Bootstrap support for afum|EAL89525.1 as seed ortholog is 100%.
Bootstrap support for afis|EAW24875.1 as seed ortholog is 100%.

---

Group of orthologs #5018. Best score 850 bits
Score difference with first non-orthologous sequence - *A. fumigatus*:850   *A. fischeri*:850
afum|EAL89529.1 100.00% afis|EAW24879.1 100.00%

Bootstrap support for afum|EAL89529.1 as seed ortholog is 100%.
Bootstrap support for afis|EAW24879.1 as seed ortholog is 100%.

---

Group of orthologs #5019. Best score 850 bits
Score difference with first non-orthologous sequence - *A. fumigatus*:850 *A. fischeri*:850
afum|EAL89587.1 100.00% afis|EAW24931.1 100.00%
Bootstrap support for afum|EAL89587.1 as seed ortholog is 100%.
Bootstrap support for afis|EAW24931.1 as seed ortholog is 100%.

---

Group of orthologs #5020. Best score 850 bits
Score difference with first non-orthologous sequence - *A. fumigatus*:850 *A. fischeri*:850
afum|EAL90059.1 100.00% afis|EAW25400.1 100.00%
Bootstrap support for afum|EAL90059.1 as seed ortholog is 100%.
Bootstrap support for afis|EAW25400.1 as seed ortholog is 100%.

---

Group of orthologs #5021. Best score 849 bits
Score difference with first non-orthologous sequence - *A. fumigatus*:768 *A. fischeri*:793
afum|EAL84225.1 100.00% afis|EAW15419.1 100.00%
Bootstrap support for afum|EAL84225.1 as seed ortholog is 100%.
Bootstrap support for afis|EAW15419.1 as seed ortholog is 100%.

---

Group of orthologs #5022. Best score 849 bits
Score difference with first non-orthologous sequence - *A. fumigatus*:782 *A. fischeri*:721
afum|EAL86859.2 100.00% afis|EAW19663.1 100.00%
Bootstrap support for afum|EAL86859.2 as seed ortholog is 100%.
Bootstrap support for afis|EAW19663.1 as seed ortholog is 100%.

---

Group of orthologs #5023. Best score 849 bits
Score difference with first non-orthologous sequence - *A. fumigatus*:849 *A. fischeri*:849
afum|EAL92573.1 100.00% afis|EAW21553.1 100.00%
Bootstrap support for afum|EAL92573.1 as seed ortholog is 100%.
Bootstrap support for afis|EAW21553.1 as seed ortholog is 100%.

---

Group of orthologs #5024. Best score 849 bits
Score difference with first non-orthologous sequence - *A. fumigatus*:849 *A. fischeri*:849
afum|EAL91918.1 100.00% afis|EAW24063.1 100.00%
Bootstrap support for afum|EAL91918.1 as seed ortholog is 100%.
Bootstrap support for afis|EAW24063.1 as seed ortholog is 100%.

---

Group of orthologs #5025. Best score 848 bits
Score difference with first non-orthologous sequence - *A. fumigatus*:848 *A. fischeri*:848
afum|EAL91118.1 100.00% afis|EAW20299.1 100.00%
afum|EAL90938.1 36.36%
Bootstrap support for afum|EAL91118.1 as seed ortholog is 100%.
Bootstrap support for afis|EAW20299.1 as seed ortholog is 100%.

---

Group of orthologs #5026. Best score 848 bits
Score difference with first non-orthologous sequence - *A. fumigatus*:848 *A. fischeri*:669
afum|EAL86599.1 100.00% afis|EAW17162.1 100.00%
Bootstrap support for afum|EAL86599.1 as seed ortholog is 100%.
Bootstrap support for afis|EAW17162.1 as seed ortholog is 100%.

---

Group of orthologs #5027. Best score 848 bits
Score difference with first non-orthologous sequence - *A. fumigatus*:848 *A. fischeri*:848
afum|EAL89255.1 100.00% afis|EAW16651.1 100.00%
Bootstrap support for afum|EAL89255.1 as seed ortholog is 100%.

Bootstrap support for afis|EAW16651.1 as seed ortholog is 100%.

---

Group of orthologs #5028. Best score 848 bits
Score difference with first non-orthologous sequence - *A. fumigatus*:848 *A. fischeri*:801
afum|EAL84823.1 100.00% afis|EAW20805.1 100.00%
Bootstrap support for afum|EAL84823.1 as seed ortholog is 100%.
Bootstrap support for afis|EAW20805.1 as seed ortholog is 100%.

---

Group of orthologs #5029. Best score 848 bits
Score difference with first non-orthologous sequence - *A. fumigatus*:848 *A. fischeri*:848
afum|EAL87599.1 100.00% afis|EAW18231.1 100.00%
Bootstrap support for afum|EAL87599.1 as seed ortholog is 100%.
Bootstrap support for afis|EAW18231.1 as seed ortholog is 100%.

---

Group of orthologs #5030. Best score 848 bits
Score difference with first non-orthologous sequence - *A. fumigatus*:848 *A. fischeri*:848
afum|EAL85987.1 100.00% afis|EAW24318.1 100.00%
Bootstrap support for afum|EAL85987.1 as seed ortholog is 100%.
Bootstrap support for afis|EAW24318.1 as seed ortholog is 100%.

---

Group of orthologs #5031. Best score 848 bits
Score difference with first non-orthologous sequence - *A. fumigatus*:848 *A. fischeri*:848
afum|EAL93211.1 100.00% afis|EAW18516.1 100.00%
Bootstrap support for afum|EAL93211.1 as seed ortholog is 100%.
Bootstrap support for afis|EAW18516.1 as seed ortholog is 100%.

---

Group of orthologs #5032. Best score 848 bits
Score difference with first non-orthologous sequence - *A. fumigatus*:848 *A. fischeri*:848
afum|EAL88330.2 100.00% afis|EAW23151.1 100.00%
Bootstrap support for afum|EAL88330.2 as seed ortholog is 100%.
Bootstrap support for afis|EAW23151.1 as seed ortholog is 100%.

---

Group of orthologs #5033. Best score 848 bits
Score difference with first non-orthologous sequence - *A. fumigatus*:848 *A. fischeri*:848
afum|EAL89472.1 100.00% afis|EAW24827.1 100.00%
Bootstrap support for afum|EAL89472.1 as seed ortholog is 100%.
Bootstrap support for afis|EAW24827.1 as seed ortholog is 100%.

---

Group of orthologs #5034. Best score 848 bits
Score difference with first non-orthologous sequence - *A. fumigatus*:848 *A. fischeri*:848
afum|EAL90101.1 100.00% afis|EAW25443.1 100.00%
Bootstrap support for afum|EAL90101.1 as seed ortholog is 100%.
Bootstrap support for afis|EAW25443.1 as seed ortholog is 100%.

---

Group of orthologs #5035. Best score 847 bits
Score difference with first non-orthologous sequence - *A. fumigatus*:847 *A. fischeri*:847
afum|EAL86790.1 100.00% afis|EAW19739.1 100.00%
Bootstrap support for afum|EAL86790.1 as seed ortholog is 100%.
Bootstrap support for afis|EAW19739.1 as seed ortholog is 100%.

---

Group of orthologs #5036. Best score 847 bits
Score difference with first non-orthologous sequence - *A. fumigatus*:847 *A. fischeri*:682
afum|EAL91224.1 100.00% afis|EAW17388.1 100.00%
Bootstrap support for afum|EAL91224.1 as seed ortholog is 100%.
Bootstrap support for afis|EAW17388.1 as seed ortholog is 100%.

---

Group of orthologs #5037. Best score 847 bits
Score difference with first non-orthologous sequence - *A. fumigatus*:847 *A. fischeri*:847
afum|EAL90302.1 100.00% afis|EAW22896.1 100.00%
Bootstrap support for afum|EAL90302.1 as seed ortholog is 100%.
Bootstrap support for afis|EAW22896.1 as seed ortholog is 100%.

---

Group of orthologs #5038. Best score 847 bits
Score difference with first non-orthologous sequence - *A. fumigatus*:847 *A. fischeri*:847
afum|EAL94069.1 100.00% afis|EAW19349.1 100.00%
Bootstrap support for afum|EAL94069.1 as seed ortholog is 100%.
Bootstrap support for afis|EAW19349.1 as seed ortholog is 100%.

---

Group of orthologs #5039. Best score 847 bits
Score difference with first non-orthologous sequence - *A. fumigatus*:847 *A. fischeri*:847
afum|EAL89998.1 100.00% afis|EAW25335.1 100.00%
Bootstrap support for afum|EAL89998.1 as seed ortholog is 100%.
Bootstrap support for afis|EAW25335.1 as seed ortholog is 100%.

---

Group of orthologs #5040. Best score 847 bits
Score difference with first non-orthologous sequence - *A. fumigatus*:847 *A. fischeri*:847
afum|EAL90007.1 100.00% afis|EAW25344.1 100.00%
Bootstrap support for afum|EAL90007.1 as seed ortholog is 100%.
Bootstrap support for afis|EAW25344.1 as seed ortholog is 100%.

---

Group of orthologs #5041. Best score 846 bits
Score difference with first non-orthologous sequence - *A. fumigatus*:846 *A. fischeri*:846
afum|EAL88837.1 100.00% afis|EAW16210.1 100.00%
Bootstrap support for afum|EAL88837.1 as seed ortholog is 100%.
Bootstrap support for afis|EAW16210.1 as seed ortholog is 100%.

---

Group of orthologs #5042. Best score 846 bits
Score difference with first non-orthologous sequence - *A. fumigatus*:846 *A. fischeri*:661
afum|EAL88874.1 100.00% afis|EAW16246.1 100.00%
Bootstrap support for afum|EAL88874.1 as seed ortholog is 100%.
Bootstrap support for afis|EAW16246.1 as seed ortholog is 100%.

---

Group of orthologs #5043. Best score 846 bits
Score difference with first non-orthologous sequence - *A. fumigatus*:846 *A. fischeri*:846
afum|EAL89078.1 100.00% afis|EAW16484.1 100.00%
Bootstrap support for afum|EAL89078.1 as seed ortholog is 100%.
Bootstrap support for afis|EAW16484.1 as seed ortholog is 100%.

---

Group of orthologs #5044. Best score 846 bits
Score difference with first non-orthologous sequence - *A. fumigatus*:846 *A. fischeri*:846
afum|EAL91692.1 100.00% afis|EAW17884.1 100.00%
Bootstrap support for afum|EAL91692.1 as seed ortholog is 100%.
Bootstrap support for afis|EAW17884.1 as seed ortholog is 100%.

---

Group of orthologs #5045. Best score 846 bits
Score difference with first non-orthologous sequence - *A. fumigatus*:846 *A. fischeri*:846
afum|EAL93893.1 100.00% afis|EAW19163.1 100.00%
Bootstrap support for afum|EAL93893.1 as seed ortholog is 100%.
Bootstrap support for afis|EAW19163.1 as seed ortholog is 100%.

---

Group of orthologs #5046. Best score 846 bits
Score difference with first non-orthologous sequence - *A. fumigatus*:846 *A. fischeri*:332

afum|EAL93904.1 100.00% afis|EAW19174.1 100.00%
Bootstrap support for afum|EAL93904.1 as seed ortholog is 100%.
Bootstrap support for afis|EAW19174.1 as seed ortholog is 100%.

---

Group of orthologs #5047. Best score 846 bits
Score difference with first non-orthologous sequence - *A. fumigatus*:846 *A. fischeri*:846
afum|EAL90849.1 100.00% afis|EAW22342.1 100.00%
Bootstrap support for afum|EAL90849.1 as seed ortholog is 100%.
Bootstrap support for afis|EAW22342.1 as seed ortholog is 100%.

---

Group of orthologs #5048. Best score 845 bits
Score difference with first non-orthologous sequence - *A. fumigatus*:845 *A. fischeri*:845
afum|EAL88564.1 100.00% afis|EAW15928.1 100.00%
Bootstrap support for afum|EAL88564.1 as seed ortholog is 100%.
Bootstrap support for afis|EAW15928.1 as seed ortholog is 100%.

---

Group of orthologs #5049. Best score 845 bits
Score difference with first non-orthologous sequence - *A. fumigatus*:845 *A. fischeri*:845
afum|EAL88919.1 100.00% afis|EAW16297.1 100.00%
Bootstrap support for afum|EAL88919.1 as seed ortholog is 100%.
Bootstrap support for afis|EAW16297.1 as seed ortholog is 100%.

---

Group of orthologs #5050. Best score 845 bits
Score difference with first non-orthologous sequence - *A. fumigatus*:589 *A. fischeri*:768
afum|EAL85338.1 100.00% afis|EAW20223.1 100.00%
Bootstrap support for afum|EAL85338.1 as seed ortholog is 100%.
Bootstrap support for afis|EAW20223.1 as seed ortholog is 100%.

---

Group of orthologs #5051. Best score 845 bits
Score difference with first non-orthologous sequence - *A. fumigatus*:789 *A. fischeri*:723
afum|EAL91244.2 100.00% afis|EAW17410.1 100.00%
Bootstrap support for afum|EAL91244.2 as seed ortholog is 100%.
Bootstrap support for afis|EAW17410.1 as seed ortholog is 100%.

---

Group of orthologs #5052. Best score 845 bits
Score difference with first non-orthologous sequence - *A. fumigatus*:845 *A. fischeri*:694
afum|EAL91689.2 100.00% afis|EAW17881.1 100.00%
Bootstrap support for afum|EAL91689.2 as seed ortholog is 100%.
Bootstrap support for afis|EAW17881.1 as seed ortholog is 100%.

---

Group of orthologs #5053. Best score 845 bits
Score difference with first non-orthologous sequence - *A. fumigatus*:845 *A. fischeri*:845
afum|EAL88264.1 100.00% afis|EAW23220.1 100.00%
Bootstrap support for afum|EAL88264.1 as seed ortholog is 100%.
Bootstrap support for afis|EAW23220.1 as seed ortholog is 100%.

---

Group of orthologs #5054. Best score 845 bits
Score difference with first non-orthologous sequence - *A. fumigatus*:845 *A. fischeri*:466
afum|EAL90588.1 100.00% afis|EAW22610.1 100.00%
Bootstrap support for afum|EAL90588.1 as seed ortholog is 100%.
Bootstrap support for afis|EAW22610.1 as seed ortholog is 100%.

---

Group of orthologs #5055. Best score 845 bits
Score difference with first non-orthologous sequence - *A. fumigatus*:845 *A. fischeri*:845
afum|EAL92362.1 100.00% afis|EAW21345.1 100.00%
Bootstrap support for afum|EAL92362.1 as seed ortholog is 100%.

Bootstrap support for afis|EAW21345.1 as seed ortholog is 100%.

---

Group of orthologs #5056. Best score 845 bits
Score difference with first non-orthologous sequence - *A. fumigatus*:845 *A. fischeri*:845
afum|EAL89660.1 100.00% afis|EAW25034.1 100.00%
Bootstrap support for afum|EAL89660.1 as seed ortholog is 100%.
Bootstrap support for afis|EAW25034.1 as seed ortholog is 100%.

---

Group of orthologs #5057. Best score 844 bits
Score difference with first non-orthologous sequence - *A. fumigatus*:844 *A. fischeri*:844
afum|EAL86326.1 100.00% afis|EAW16754.1 100.00%
Bootstrap support for afum|EAL86326.1 as seed ortholog is 100%.
Bootstrap support for afis|EAW16754.1 as seed ortholog is 100%.

---

Group of orthologs #5058. Best score 844 bits
Score difference with first non-orthologous sequence - *A. fumigatus*:844 *A. fischeri*:844
afum|EAL86014.1 100.00% afis|EAW24343.1 100.00%
Bootstrap support for afum|EAL86014.1 as seed ortholog is 100%.
Bootstrap support for afis|EAW24343.1 as seed ortholog is 100%.

---

Group of orthologs #5059. Best score 844 bits
Score difference with first non-orthologous sequence - *A. fumigatus*:844 *A. fischeri*:844
afum|EAL88374.1 100.00% afis|EAW23105.1 100.00%
Bootstrap support for afum|EAL88374.1 as seed ortholog is 100%.
Bootstrap support for afis|EAW23105.1 as seed ortholog is 100%.

---

Group of orthologs #5060. Best score 844 bits
Score difference with first non-orthologous sequence - *A. fumigatus*:844 *A. fischeri*:844
afum|EAL93861.1 100.00% afis|EAW19129.1 100.00%
Bootstrap support for afum|EAL93861.1 as seed ortholog is 100%.
Bootstrap support for afis|EAW19129.1 as seed ortholog is 100%.

---

Group of orthologs #5061. Best score 844 bits
Score difference with first non-orthologous sequence - *A. fumigatus*:844 *A. fischeri*:779
afum|EAL90485.1 100.00% afis|EAW22707.1 100.00%
Bootstrap support for afum|EAL90485.1 as seed ortholog is 100%.
Bootstrap support for afis|EAW22707.1 as seed ortholog is 100%.

---

Group of orthologs #5062. Best score 843 bits
Score difference with first non-orthologous sequence - *A. fumigatus*:843 *A. fischeri*:843
afum|EAL91386.1 100.00% afis|EAW17572.1 100.00%
Bootstrap support for afum|EAL91386.1 as seed ortholog is 100%.
Bootstrap support for afis|EAW17572.1 as seed ortholog is 100%.

---

Group of orthologs #5063. Best score 843 bits
Score difference with first non-orthologous sequence - *A. fumigatus*:800 *A. fischeri*:678
afum|EAL93102.1 100.00% afis|EAW18418.1 100.00%
Bootstrap support for afum|EAL93102.1 as seed ortholog is 100%.
Bootstrap support for afis|EAW18418.1 as seed ortholog is 100%.

---

Group of orthologs #5064. Best score 843 bits
Score difference with first non-orthologous sequence - *A. fumigatus*:843 *A. fischeri*:746
afum|EAL93280.1 100.00% afis|EAW18577.1 100.00%
Bootstrap support for afum|EAL93280.1 as seed ortholog is 100%.
Bootstrap support for afis|EAW18577.1 as seed ortholog is 100%.

---

Group of orthologs #5065. Best score 843 bits
Score difference with first non-orthologous sequence - *A. fumigatus*:843 *A. fischeri*:843
afum|EAL90947.1 100.00% afis|EAW22246.1 100.00%
Bootstrap support for afum|EAL90947.1 as seed ortholog is 100%.
Bootstrap support for afis|EAW22246.1 as seed ortholog is 100%.

---

Group of orthologs #5066. Best score 842 bits
Score difference with first non-orthologous sequence - *A. fumigatus*:842 *A. fischeri*:842
afum|EAL85566.1 100.00% afis|EAW15784.1 100.00%
Bootstrap support for afum|EAL85566.1 as seed ortholog is 100%.
Bootstrap support for afis|EAW15784.1 as seed ortholog is 100%.

---

Group of orthologs #5067. Best score 842 bits
Score difference with first non-orthologous sequence - *A. fumigatus*:842 *A. fischeri*:842
afum|EAL88924.1 100.00% afis|EAW16302.1 100.00%
Bootstrap support for afum|EAL88924.1 as seed ortholog is 100%.
Bootstrap support for afis|EAW16302.1 as seed ortholog is 100%.

---

Group of orthologs #5068. Best score 842 bits
Score difference with first non-orthologous sequence - *A. fumigatus*:842 *A. fischeri*:842
afum|EAL84574.1 100.00% afis|EAW20480.1 100.00%
Bootstrap support for afum|EAL84574.1 as seed ortholog is 100%.
Bootstrap support for afis|EAW20480.1 as seed ortholog is 100%.

---

Group of orthologs #5069. Best score 842 bits
Score difference with first non-orthologous sequence - *A. fumigatus*:842 *A. fischeri*:842
afum|EAL91339.2 100.00% afis|EAW17524.1 100.00%
Bootstrap support for afum|EAL91339.2 as seed ortholog is 100%.
Bootstrap support for afis|EAW17524.1 as seed ortholog is 100%.

---

Group of orthologs #5070. Best score 842 bits
Score difference with first non-orthologous sequence - *A. fumigatus*:842 *A. fischeri*:842
afum|EAL87399.1 100.00% afis|EAW23969.1 100.00%
Bootstrap support for afum|EAL87399.1 as seed ortholog is 100%.
Bootstrap support for afis|EAW23969.1 as seed ortholog is 100%.

---

Group of orthologs #5071. Best score 842 bits
Score difference with first non-orthologous sequence - *A. fumigatus*:842 *A. fischeri*:842
afum|EAL88468.1 100.00% afis|EAW23013.1 100.00%
Bootstrap support for afum|EAL88468.1 as seed ortholog is 100%.
Bootstrap support for afis|EAW23013.1 as seed ortholog is 100%.

---

Group of orthologs #5072. Best score 842 bits
Score difference with first non-orthologous sequence - *A. fumigatus*:713 *A. fischeri*:716
afum|EAL88173.1 100.00% afis|EAW23304.1 100.00%
Bootstrap support for afum|EAL88173.1 as seed ortholog is 100%.
Bootstrap support for afis|EAW23304.1 as seed ortholog is 100%.

---

Group of orthologs #5073. Best score 842 bits
Score difference with first non-orthologous sequence - *A. fumigatus*:842 *A. fischeri*:842
afum|EAL88003.1 100.00% afis|EAW23463.1 100.00%
Bootstrap support for afum|EAL88003.1 as seed ortholog is 100%.
Bootstrap support for afis|EAW23463.1 as seed ortholog is 100%.

---

Group of orthologs #5074. Best score 842 bits
Score difference with first non-orthologous sequence - *A. fumigatus*:842 *A. fischeri*:842

afum|EAL89565.1 100.00% afis|EAW24911.1 100.00%
Bootstrap support for afum|EAL89565.1 as seed ortholog is 100%.
Bootstrap support for afis|EAW24911.1 as seed ortholog is 100%.

---

Group of orthologs #5075. Best score 841 bits
Score difference with first non-orthologous sequence - *A. fumigatus*:841 *A. fischeri*:841
afum|EAL87280.1 100.00% afis|EAW23852.1 100.00%
Bootstrap support for afum|EAL87280.1 as seed ortholog is 100%.
Bootstrap support for afis|EAW23852.1 as seed ortholog is 100%.

---

Group of orthologs #5076. Best score 841 bits
Score difference with first non-orthologous sequence - *A. fumigatus*:841 *A. fischeri*:841
afum|EAL93262.1 100.00% afis|EAW18561.1 100.00%
Bootstrap support for afum|EAL93262.1 as seed ortholog is 100%.
Bootstrap support for afis|EAW18561.1 as seed ortholog is 100%.

---

Group of orthologs #5077. Best score 841 bits
Score difference with first non-orthologous sequence - *A. fumigatus*:841 *A. fischeri*:841
afum|EAL92124.1 100.00% afis|EAW21103.1 100.00%
Bootstrap support for afum|EAL92124.1 as seed ortholog is 100%.
Bootstrap support for afis|EAW21103.1 as seed ortholog is 100%.

---

Group of orthologs #5078. Best score 841 bits
Score difference with first non-orthologous sequence - *A. fumigatus*:763 *A. fischeri*:744
afum|EAL90455.1 100.00% afis|EAW22740.1 100.00%
Bootstrap support for afum|EAL90455.1 as seed ortholog is 100%.
Bootstrap support for afis|EAW22740.1 as seed ortholog is 100%.

---

Group of orthologs #5079. Best score 841 bits
Score difference with first non-orthologous sequence - *A. fumigatus*:841 *A. fischeri*:841
afum|EBA27458.1 100.00% afis|EAW21270.1 100.00%
Bootstrap support for afum|EBA27458.1 as seed ortholog is 100%.
Bootstrap support for afis|EAW21270.1 as seed ortholog is 100%.

---

Group of orthologs #5080. Best score 840 bits
Score difference with first non-orthologous sequence - *A. fumigatus*:785 *A. fischeri*:751
afum|EAL91044.1 100.00% afis|EAW22146.1 100.00%
Bootstrap support for afum|EAL91044.1 as seed ortholog is 100%.
Bootstrap support for afis|EAW22146.1 as seed ortholog is 100%.

---

Group of orthologs #5081. Best score 839 bits
Score difference with first non-orthologous sequence - *A. fumigatus*:839 *A. fischeri*:839
afum|EAL87058.1 100.00% afis|EAW19438.1 100.00%
Bootstrap support for afum|EAL87058.1 as seed ortholog is 100%.
Bootstrap support for afis|EAW19438.1 as seed ortholog is 100%.

---

Group of orthologs #5082. Best score 839 bits
Score difference with first non-orthologous sequence - *A. fumigatus*:706 *A. fischeri*:716
afum|EAL85694.1 100.00% afis|EAW22085.1 100.00%
Bootstrap support for afum|EAL85694.1 as seed ortholog is 100%.
Bootstrap support for afis|EAW22085.1 as seed ortholog is 100%.

---

Group of orthologs #5083. Best score 839 bits
Score difference with first non-orthologous sequence - *A. fumigatus*:839 *A. fischeri*:839
afum|EAL85899.2 100.00% afis|EAW24230.1 100.00%
Bootstrap support for afum|EAL85899.2 as seed ortholog is 100%.

Bootstrap support for afis|EAW24230.1 as seed ortholog is 100%.

---

Group of orthologs #5084. Best score 839 bits
Score difference with first non-orthologous sequence - *A. fumigatus*:839 *A. fischeri*:839
afum|EAL91001.1 100.00% afis|EAW22190.1 100.00%
Bootstrap support for afum|EAL91001.1 as seed ortholog is 100%.
Bootstrap support for afis|EAW22190.1 as seed ortholog is 100%.

---

Group of orthologs #5085. Best score 839 bits
Score difference with first non-orthologous sequence - *A. fumigatus*:839 *A. fischeri*:839
afum|EAL90369.1 100.00% afis|EAW22833.1 100.00%
Bootstrap support for afum|EAL90369.1 as seed ortholog is 100%.
Bootstrap support for afis|EAW22833.1 as seed ortholog is 100%.

---

Group of orthologs #5086. Best score 839 bits
Score difference with first non-orthologous sequence - *A. fumigatus*:839 *A. fischeri*:839
afum|EAL92733.1 100.00% afis|EAW21699.1 100.00%
Bootstrap support for afum|EAL92733.1 as seed ortholog is 100%.
Bootstrap support for afis|EAW21699.1 as seed ortholog is 100%.

---

Group of orthologs #5087. Best score 839 bits
Score difference with first non-orthologous sequence - *A. fumigatus*:839 *A. fischeri*:839
afum|EAL89678.1 100.00% afis|EAW25017.1 100.00%
Bootstrap support for afum|EAL89678.1 as seed ortholog is 100%.
Bootstrap support for afis|EAW25017.1 as seed ortholog is 100%.

---

Group of orthologs #5088. Best score 838 bits
Score difference with first non-orthologous sequence - *A. fumigatus*:838 *A. fischeri*:838
afum|EAL88652.1 100.00% afis|EAW16023.1 100.00%
Bootstrap support for afum|EAL88652.1 as seed ortholog is 100%.
Bootstrap support for afis|EAW16023.1 as seed ortholog is 100%.

---

Group of orthologs #5089. Best score 838 bits
Score difference with first non-orthologous sequence - *A. fumigatus*:775 *A. fischeri*:838
afum|EAL87019.1 100.00% afis|EAW19477.1 100.00%
Bootstrap support for afum|EAL87019.1 as seed ortholog is 100%.
Bootstrap support for afis|EAW19477.1 as seed ortholog is 100%.

---

Group of orthologs #5090. Best score 838 bits
Score difference with first non-orthologous sequence - *A. fumigatus*:838 *A. fischeri*:538
afum|EAL86888.1 100.00% afis|EAW19634.1 100.00%
Bootstrap support for afum|EAL86888.1 as seed ortholog is 100%.
Bootstrap support for afis|EAW19634.1 as seed ortholog is 100%.

---

Group of orthologs #5091. Best score 838 bits
Score difference with first non-orthologous sequence - *A. fumigatus*:838 *A. fischeri*:838
afum|EAL92090.1 100.00% afis|EAW21054.1 100.00%
Bootstrap support for afum|EAL92090.1 as seed ortholog is 100%.
Bootstrap support for afis|EAW21054.1 as seed ortholog is 100%.

---

Group of orthologs #5092. Best score 838 bits
Score difference with first non-orthologous sequence - *A. fumigatus*:838 *A. fischeri*:838
afum|EAL90214.1 100.00% afis|EAW22985.1 100.00%
Bootstrap support for afum|EAL90214.1 as seed ortholog is 100%.
Bootstrap support for afis|EAW22985.1 as seed ortholog is 100%.

---

Group of orthologs #5093. Best score 837 bits
Score difference with first non-orthologous sequence - *A. fumigatus*:837 *A. fischeri*:837
afum|EAL84889.1 100.00% afis|EAW20028.1 100.00%
Bootstrap support for afum|EAL84889.1 as seed ortholog is 100%.
Bootstrap support for afis|EAW20028.1 as seed ortholog is 100%.

---

Group of orthologs #5094. Best score 837 bits
Score difference with first non-orthologous sequence - *A. fumigatus*:837 *A. fischeri*:692
afum|EAL89248.1 100.00% afis|EAW16644.1 100.00%
Bootstrap support for afum|EAL89248.1 as seed ortholog is 100%.
Bootstrap support for afis|EAW16644.1 as seed ortholog is 100%.

---

Group of orthologs #5095. Best score 837 bits
Score difference with first non-orthologous sequence - *A. fumigatus*:584 *A. fischeri*:723
afum|EBA27444.1 100.00% afis|EAW18067.1 100.00%
Bootstrap support for afum|EBA27444.1 as seed ortholog is 100%.
Bootstrap support for afis|EAW18067.1 as seed ortholog is 100%.

---

Group of orthologs #5096. Best score 837 bits
Score difference with first non-orthologous sequence - *A. fumigatus*:837 *A. fischeri*:837
afum|EAL90326.1 100.00% afis|EAW22872.1 100.00%
Bootstrap support for afum|EAL90326.1 as seed ortholog is 100%.
Bootstrap support for afis|EAW22872.1 as seed ortholog is 100%.

---

Group of orthologs #5097. Best score 836 bits
Score difference with first non-orthologous sequence - *A. fumigatus*:672 *A. fischeri*:836
afum|EAL85536.1 100.00% afis|EAW15815.1 100.00%
Bootstrap support for afum|EAL85536.1 as seed ortholog is 100%.
Bootstrap support for afis|EAW15815.1 as seed ortholog is 100%.

---

Group of orthologs #5098. Best score 836 bits
Score difference with first non-orthologous sequence - *A. fumigatus*:836 *A. fischeri*:836
afum|EAL87047.1 100.00% afis|EAW19449.1 100.00%
Bootstrap support for afum|EAL87047.1 as seed ortholog is 100%.
Bootstrap support for afis|EAW19449.1 as seed ortholog is 100%.

---

Group of orthologs #5099. Best score 836 bits
Score difference with first non-orthologous sequence - *A. fumigatus*:523 *A. fischeri*:721
afum|EAL86869.1 100.00% afis|EAW19653.1 100.00%
Bootstrap support for afum|EAL86869.1 as seed ortholog is 100%.
Bootstrap support for afis|EAW19653.1 as seed ortholog is 100%.

---

Group of orthologs #5100. Best score 836 bits
Score difference with first non-orthologous sequence - *A. fumigatus*:836 *A. fischeri*:836
afum|EAL91439.1 100.00% afis|EAW17625.1 100.00%
Bootstrap support for afum|EAL91439.1 as seed ortholog is 100%.
Bootstrap support for afis|EAW17625.1 as seed ortholog is 100%.

---

Group of orthologs #5101. Best score 836 bits
Score difference with first non-orthologous sequence - *A. fumigatus*:836 *A. fischeri*:836
afum|EAL91536.1 100.00% afis|EAW17721.1 100.00%
Bootstrap support for afum|EAL91536.1 as seed ortholog is 100%.
Bootstrap support for afis|EAW17721.1 as seed ortholog is 100%.

---

Group of orthologs #5102. Best score 836 bits
Score difference with first non-orthologous sequence - *A. fumigatus*:836 *A. fischeri*:836

afum|EAL91884.1 100.00% afis|EAW18072.1 100.00%
Bootstrap support for afum|EAL91884.1 as seed ortholog is 100%.
Bootstrap support for afis|EAW18072.1 as seed ortholog is 100%.

---

Group of orthologs #5103. Best score 836 bits
Score difference with first non-orthologous sequence - *A. fumigatus*:567 *A. fischeri*:558
afum|EAL87265.1 100.00% afis|EAW23834.1 100.00%
Bootstrap support for afum|EAL87265.1 as seed ortholog is 100%.
Bootstrap support for afis|EAW23834.1 as seed ortholog is 100%.

---

Group of orthologs #5104. Best score 836 bits
Score difference with first non-orthologous sequence - *A. fumigatus*:836 *A. fischeri*:836
afum|EAL92065.1 100.00% afis|EAW21027.1 100.00%
Bootstrap support for afum|EAL92065.1 as seed ortholog is 100%.
Bootstrap support for afis|EAW21027.1 as seed ortholog is 100%.

---

Group of orthologs #5105. Best score 836 bits
Score difference with first non-orthologous sequence - *A. fumigatus*:836 *A. fischeri*:742
afum|EAL90669.1 100.00% afis|EAW22525.1 100.00%
Bootstrap support for afum|EAL90669.1 as seed ortholog is 100%.
Bootstrap support for afis|EAW22525.1 as seed ortholog is 100%.

---

Group of orthologs #5106. Best score 836 bits
Score difference with first non-orthologous sequence - *A. fumigatus*:836 *A. fischeri*:836
afum|EAL90605.1 100.00% afis|EAW22594.1 100.00%
Bootstrap support for afum|EAL90605.1 as seed ortholog is 100%.
Bootstrap support for afis|EAW22594.1 as seed ortholog is 100%.

---

Group of orthologs #5107. Best score 836 bits
Score difference with first non-orthologous sequence - *A. fumigatus*:836 *A. fischeri*:836
afum|EAL92620.1 100.00% afis|EAW21601.1 100.00%
Bootstrap support for afum|EAL92620.1 as seed ortholog is 100%.
Bootstrap support for afis|EAW21601.1 as seed ortholog is 100%.

---

Group of orthologs #5108. Best score 836 bits
Score difference with first non-orthologous sequence - *A. fumigatus*:836 *A. fischeri*:836
afum|EAL92946.1 100.00% afis|EAW21933.1 100.00%
Bootstrap support for afum|EAL92946.1 as seed ortholog is 100%.
Bootstrap support for afis|EAW21933.1 as seed ortholog is 100%.

---

Group of orthologs #5109. Best score 836 bits
Score difference with first non-orthologous sequence - *A. fumigatus*:836 *A. fischeri*:836
afum|EAL89762.2 100.00% afis|EAW25092.1 100.00%
Bootstrap support for afum|EAL89762.2 as seed ortholog is 100%.
Bootstrap support for afis|EAW25092.1 as seed ortholog is 100%.

---

Group of orthologs #5110. Best score 835 bits
Score difference with first non-orthologous sequence - *A. fumigatus*:667 *A. fischeri*:612
afum|EAL84996.2 100.00% afis|EAW19910.1 100.00%
Bootstrap support for afum|EAL84996.2 as seed ortholog is 100%.
Bootstrap support for afis|EAW19910.1 as seed ortholog is 100%.

---

Group of orthologs #5111. Best score 835 bits
Score difference with first non-orthologous sequence - *A. fumigatus*:835 *A. fischeri*:835
afum|EAL88912.1 100.00% afis|EAW16290.1 100.00%
Bootstrap support for afum|EAL88912.1 as seed ortholog is 100%.

Bootstrap support for afis|EAW16290.1 as seed ortholog is 100%.

---

Group of orthologs #5112. Best score 835 bits
Score difference with first non-orthologous sequence - *A. fumigatus*:835 *A. fischeri*:835
afum|EAL88997.1 100.00% afis|EAW16373.1 100.00%
Bootstrap support for afum|EAL88997.1 as seed ortholog is 100%.
Bootstrap support for afis|EAW16373.1 as seed ortholog is 100%.

---

Group of orthologs #5113. Best score 835 bits
Score difference with first non-orthologous sequence - *A. fumigatus*:835 *A. fischeri*:730
afum|EAL85315.1 100.00% afis|EAW20195.1 100.00%
Bootstrap support for afum|EAL85315.1 as seed ortholog is 100%.
Bootstrap support for afis|EAW20195.1 as seed ortholog is 100%.

---

Group of orthologs #5114. Best score 835 bits
Score difference with first non-orthologous sequence - *A. fumigatus*:835 *A. fischeri*:835
afum|EAL88393.1 100.00% afis|EAW23086.1 100.00%
Bootstrap support for afum|EAL88393.1 as seed ortholog is 100%.
Bootstrap support for afis|EAW23086.1 as seed ortholog is 100%.

---

Group of orthologs #5115. Best score 835 bits
Score difference with first non-orthologous sequence - *A. fumigatus*:835 *A. fischeri*:835
afum|EAL88267.1 100.00% afis|EAW23217.1 100.00%
Bootstrap support for afum|EAL88267.1 as seed ortholog is 100%.
Bootstrap support for afis|EAW23217.1 as seed ortholog is 100%.

---

Group of orthologs #5116. Best score 835 bits
Score difference with first non-orthologous sequence - *A. fumigatus*:835 *A. fischeri*:835
afum|EAL87970.1 100.00% afis|EAW23493.1 100.00%
Bootstrap support for afum|EAL87970.1 as seed ortholog is 100%.
Bootstrap support for afis|EAW23493.1 as seed ortholog is 100%.

---

Group of orthologs #5117. Best score 834 bits
Score difference with first non-orthologous sequence - *A. fumigatus*:834 *A. fischeri*:834
afum|EAL85534.1 100.00% afis|EAW15817.1 100.00%
Bootstrap support for afum|EAL85534.1 as seed ortholog is 100%.
Bootstrap support for afis|EAW15817.1 as seed ortholog is 100%.

---

Group of orthologs #5118. Best score 834 bits
Score difference with first non-orthologous sequence - *A. fumigatus*:834 *A. fischeri*:834
afum|EAL85202.1 100.00% afis|EAW20083.1 100.00%
Bootstrap support for afum|EAL85202.1 as seed ortholog is 100%.
Bootstrap support for afis|EAW20083.1 as seed ortholog is 100%.

---

Group of orthologs #5119. Best score 834 bits
Score difference with first non-orthologous sequence - *A. fumigatus*:834 *A. fischeri*:581
afum|EAL87195.1 100.00% afis|EAW23762.1 100.00%
Bootstrap support for afum|EAL87195.1 as seed ortholog is 100%.
Bootstrap support for afis|EAW23762.1 as seed ortholog is 100%.

---

Group of orthologs #5120. Best score 834 bits
Score difference with first non-orthologous sequence - *A. fumigatus*:834 *A. fischeri*:834
afum|EAL93062.1 100.00% afis|EAW18387.1 100.00%
Bootstrap support for afum|EAL93062.1 as seed ortholog is 100%.
Bootstrap support for afis|EAW18387.1 as seed ortholog is 100%.

---

Group of orthologs #5121. Best score 833 bits
Score difference with first non-orthologous sequence - *A. fumigatus*:710 *A. fischeri*:618
afum|EAL85908.1 100.00% afis|EAW24240.1 100.00%
Bootstrap support for afum|EAL85908.1 as seed ortholog is 100%.
Bootstrap support for afis|EAW24240.1 as seed ortholog is 100%.

---

Group of orthologs #5122. Best score 833 bits
Score difference with first non-orthologous sequence - *A. fumigatus*:833 *A. fischeri*:833
afum|EAL93055.1 100.00% afis|EAW18380.1 100.00%
Bootstrap support for afum|EAL93055.1 as seed ortholog is 100%.
Bootstrap support for afis|EAW18380.1 as seed ortholog is 100%.

---

Group of orthologs #5123. Best score 833 bits
Score difference with first non-orthologous sequence - *A. fumigatus*:833 *A. fischeri*:833
afum|EAL87933.1 100.00% afis|EAW23533.1 100.00%
Bootstrap support for afum|EAL87933.1 as seed ortholog is 100%.
Bootstrap support for afis|EAW23533.1 as seed ortholog is 100%.

---

Group of orthologs #5124. Best score 833 bits
Score difference with first non-orthologous sequence - *A. fumigatus*:833 *A. fischeri*:833
afum|EAL90885.1 100.00% afis|EAW22296.1 100.00%
Bootstrap support for afum|EAL90885.1 as seed ortholog is 100%.
Bootstrap support for afis|EAW22296.1 as seed ortholog is 100%.

---

Group of orthologs #5125. Best score 833 bits
Score difference with first non-orthologous sequence - *A. fumigatus*:833 *A. fischeri*:833
afum|EAL92671.1 100.00% afis|EAW21637.1 100.00%
Bootstrap support for afum|EAL92671.1 as seed ortholog is 100%.
Bootstrap support for afis|EAW21637.1 as seed ortholog is 100%.

---

Group of orthologs #5126. Best score 832 bits
Score difference with first non-orthologous sequence - *A. fumigatus*:832 *A. fischeri*:832
afum|EAL84591.1 100.00% afis|EAW20462.1 100.00%
Bootstrap support for afum|EAL84591.1 as seed ortholog is 100%.
Bootstrap support for afis|EAW20462.1 as seed ortholog is 100%.

---

Group of orthologs #5127. Best score 832 bits
Score difference with first non-orthologous sequence - *A. fumigatus*:832 *A. fischeri*:832
afum|EAL85318.1 100.00% afis|EAW20198.1 100.00%
Bootstrap support for afum|EAL85318.1 as seed ortholog is 100%.
Bootstrap support for afis|EAW20198.1 as seed ortholog is 100%.

---

Group of orthologs #5128. Best score 832 bits
Score difference with first non-orthologous sequence - *A. fumigatus*:645 *A. fischeri*:832
afum|EAL84849.1 100.00% afis|EAW20770.1 100.00%
Bootstrap support for afum|EAL84849.1 as seed ortholog is 100%.
Bootstrap support for afis|EAW20770.1 as seed ortholog is 100%.

---

Group of orthologs #5129. Best score 832 bits
Score difference with first non-orthologous sequence - *A. fumigatus*:832 *A. fischeri*:832
afum|EAL87044.1 100.00% afis|EAW19452.1 100.00%
Bootstrap support for afum|EAL87044.1 as seed ortholog is 100%.
Bootstrap support for afis|EAW19452.1 as seed ortholog is 100%.

---

Group of orthologs #5130. Best score 832 bits
Score difference with first non-orthologous sequence - *A. fumigatus*:832 *A. fischeri*:832

afum|EAL91283.1 100.00% afis|EAW17455.1 100.00%
Bootstrap support for afum|EAL91283.1 as seed ortholog is 100%.
Bootstrap support for afis|EAW17455.1 as seed ortholog is 100%.

---

Group of orthologs #5131. Best score 832 bits
Score difference with first non-orthologous sequence - *A. fumigatus*:832 *A. fischeri*:832
afum|EAL86028.1 100.00% afis|EAW24358.1 100.00%
Bootstrap support for afum|EAL86028.1 as seed ortholog is 100%.
Bootstrap support for afis|EAW24358.1 as seed ortholog is 100%.

---

Group of orthologs #5132. Best score 832 bits
Score difference with first non-orthologous sequence - *A. fumigatus*:832 *A. fischeri*:832
afum|EAL87877.1 100.00% afis|EAW23587.1 100.00%
Bootstrap support for afum|EAL87877.1 as seed ortholog is 100%.
Bootstrap support for afis|EAW23587.1 as seed ortholog is 100%.

---

Group of orthologs #5133. Best score 832 bits
Score difference with first non-orthologous sequence - *A. fumigatus*:832 *A. fischeri*:471
afum|EAL93319.1 100.00% afis|EAW18609.1 100.00%
Bootstrap support for afum|EAL93319.1 as seed ortholog is 100%.
Bootstrap support for afis|EAW18609.1 as seed ortholog is 100%.

---

Group of orthologs #5134. Best score 831 bits
Score difference with first non-orthologous sequence - *A. fumigatus*:831 *A. fischeri*:831
afum|EAL84571.1 100.00% afis|EAW20484.1 100.00%
Bootstrap support for afum|EAL84571.1 as seed ortholog is 100%.
Bootstrap support for afis|EAW20484.1 as seed ortholog is 100%.

---

Group of orthologs #5135. Best score 831 bits
Score difference with first non-orthologous sequence - *A. fumigatus*:831 *A. fischeri*:831
afum|EAL85929.1 100.00% afis|EAW24261.1 100.00%
Bootstrap support for afum|EAL85929.1 as seed ortholog is 100%.
Bootstrap support for afis|EAW24261.1 as seed ortholog is 100%.

---

Group of orthologs #5136. Best score 831 bits
Score difference with first non-orthologous sequence - *A. fumigatus*:831 *A. fischeri*:831
afum|EAL88402.1 100.00% afis|EAW23077.1 100.00%
Bootstrap support for afum|EAL88402.1 as seed ortholog is 100%.
Bootstrap support for afis|EAW23077.1 as seed ortholog is 100%.

---

Group of orthologs #5137. Best score 831 bits
Score difference with first non-orthologous sequence - *A. fumigatus*:831 *A. fischeri*:831
afum|EAL93874.1 100.00% afis|EAW19143.1 100.00%
Bootstrap support for afum|EAL93874.1 as seed ortholog is 100%.
Bootstrap support for afis|EAW19143.1 as seed ortholog is 100%.

---

Group of orthologs #5138. Best score 831 bits
Score difference with first non-orthologous sequence - *A. fumigatus*:831 *A. fischeri*:646
afum|EAL94040.1 100.00% afis|EAW19322.1 100.00%
Bootstrap support for afum|EAL94040.1 as seed ortholog is 100%.
Bootstrap support for afis|EAW19322.1 as seed ortholog is 100%.

---

Group of orthologs #5139. Best score 831 bits
Score difference with first non-orthologous sequence - *A. fumigatus*:542 *A. fischeri*:697
afum|EBA27245.1 100.00% afis|EAW25746.1 100.00%
Bootstrap support for afum|EBA27245.1 as seed ortholog is 100%.

Bootstrap support for afis|EAW25746.1 as seed ortholog is 100%.

---

Group of orthologs #5140. Best score 830 bits
Score difference with first non-orthologous sequence - *A. fumigatus*:830 *A. fischeri*:830
afum|EAL88605.1 100.00% afis|EAW15973.1 100.00%
Bootstrap support for afum|EAL88605.1 as seed ortholog is 100%.
Bootstrap support for afis|EAW15973.1 as seed ortholog is 100%.

---

Group of orthologs #5141. Best score 830 bits
Score difference with first non-orthologous sequence - *A. fumigatus*:830 *A. fischeri*:830
afum|EAL88757.1 100.00% afis|EAW16127.1 100.00%
Bootstrap support for afum|EAL88757.1 as seed ortholog is 100%.
Bootstrap support for afis|EAW16127.1 as seed ortholog is 100%.

---

Group of orthologs #5142. Best score 830 bits
Score difference with first non-orthologous sequence - *A. fumigatus*:830 *A. fischeri*:830
afum|EAL84568.1 100.00% afis|EAW20487.1 100.00%
Bootstrap support for afum|EAL84568.1 as seed ortholog is 100%.
Bootstrap support for afis|EAW20487.1 as seed ortholog is 100%.

---

Group of orthologs #5143. Best score 830 bits
Score difference with first non-orthologous sequence - *A. fumigatus*:761 *A. fischeri*:745
afum|EAL84799.1 100.00% afis|EAW20833.1 100.00%
Bootstrap support for afum|EAL84799.1 as seed ortholog is 100%.
Bootstrap support for afis|EAW20833.1 as seed ortholog is 100%.

---

Group of orthologs #5144. Best score 830 bits
Score difference with first non-orthologous sequence - *A. fumigatus*:830 *A. fischeri*:830
afum|EAL84681.2 100.00% afis|EAW20953.1 100.00%
Bootstrap support for afum|EAL84681.2 as seed ortholog is 100%.
Bootstrap support for afis|EAW20953.1 as seed ortholog is 100%.

---

Group of orthologs #5145. Best score 830 bits
Score difference with first non-orthologous sequence - *A. fumigatus*:319 *A. fischeri*:711
afum|EAL86967.1 100.00% afis|EAW19541.1 100.00%
Bootstrap support for afum|EAL86967.1 as seed ortholog is 100%.
Bootstrap support for afis|EAW19541.1 as seed ortholog is 100%.

---

Group of orthologs #5146. Best score 830 bits
Score difference with first non-orthologous sequence - *A. fumigatus*:830 *A. fischeri*:830
afum|EAL91434.1 100.00% afis|EAW17621.1 100.00%
Bootstrap support for afum|EAL91434.1 as seed ortholog is 100%.
Bootstrap support for afis|EAW17621.1 as seed ortholog is 100%.

---

Group of orthologs #5147. Best score 830 bits
Score difference with first non-orthologous sequence - *A. fumigatus*:830 *A. fischeri*:830
afum|EAL91474.1 100.00% afis|EAW17659.1 100.00%
Bootstrap support for afum|EAL91474.1 as seed ortholog is 100%.
Bootstrap support for afis|EAW17659.1 as seed ortholog is 100%.

---

Group of orthologs #5148. Best score 830 bits
Score difference with first non-orthologous sequence - *A. fumigatus*:830 *A. fischeri*:830
afum|EAL89419.1 100.00% afis|EAW24780.1 100.00%
Bootstrap support for afum|EAL89419.1 as seed ortholog is 100%.
Bootstrap support for afis|EAW24780.1 as seed ortholog is 100%.

---

Group of orthologs #5149. Best score 830 bits
Score difference with first non-orthologous sequence - *A. fumigatus*:830 *A. fischeri*:830
afum|EBA27197.1 100.00% afis|EAW20445.1 100.00%
Bootstrap support for afum|EBA27197.1 as seed ortholog is 100%.
Bootstrap support for afis|EAW20445.1 as seed ortholog is 100%.

---

Group of orthologs #5150. Best score 830 bits
Score difference with first non-orthologous sequence - *A. fumigatus*:830 *A. fischeri*:830
afum|EAL90176.2 100.00% afis|EAW25510.1 100.00%
Bootstrap support for afum|EAL90176.2 as seed ortholog is 100%.
Bootstrap support for afis|EAW25510.1 as seed ortholog is 100%.

---

Group of orthologs #5151. Best score 829 bits
Score difference with first non-orthologous sequence - *A. fumigatus*:829 *A. fischeri*:710
afum|EAL87174.1 100.00% afis|EAW23736.1 100.00%
Bootstrap support for afum|EAL87174.1 as seed ortholog is 100%.
Bootstrap support for afis|EAW23736.1 as seed ortholog is 100%.

---

Group of orthologs #5152. Best score 828 bits
Score difference with first non-orthologous sequence - *A. fumigatus*:828 *A. fischeri*:697
afum|EAL85249.1 100.00% afis|EAW20133.1 100.00%
Bootstrap support for afum|EAL85249.1 as seed ortholog is 100%.
Bootstrap support for afis|EAW20133.1 as seed ortholog is 100%.

---

Group of orthologs #5153. Best score 828 bits
Score difference with first non-orthologous sequence - *A. fumigatus*:828 *A. fischeri*:828
afum|EAL91481.1 100.00% afis|EAW17667.1 100.00%
Bootstrap support for afum|EAL91481.1 as seed ortholog is 100%.
Bootstrap support for afis|EAW17667.1 as seed ortholog is 100%.

---

Group of orthologs #5154. Best score 828 bits
Score difference with first non-orthologous sequence - *A. fumigatus*:828 *A. fischeri*:828
afum|EAL86073.1 100.00% afis|EAW24402.1 100.00%
Bootstrap support for afum|EAL86073.1 as seed ortholog is 100%.
Bootstrap support for afis|EAW24402.1 as seed ortholog is 100%.

---

Group of orthologs #5155. Best score 828 bits
Score difference with first non-orthologous sequence - *A. fumigatus*:828 *A. fischeri*:828
afum|EAL87867.1 100.00% afis|EAW23600.1 100.00%
Bootstrap support for afum|EAL87867.1 as seed ortholog is 100%.
Bootstrap support for afis|EAW23600.1 as seed ortholog is 100%.

---

Group of orthologs #5156. Best score 828 bits
Score difference with first non-orthologous sequence - *A. fumigatus*:828 *A. fischeri*:828
afum|EAL93511.1 100.00% afis|EAW18779.1 100.00%
Bootstrap support for afum|EAL93511.1 as seed ortholog is 100%.
Bootstrap support for afis|EAW18779.1 as seed ortholog is 100%.

---

Group of orthologs #5157. Best score 828 bits
Score difference with first non-orthologous sequence - *A. fumigatus*:828 *A. fischeri*:751
afum|EAL90463.1 100.00% afis|EAW22732.1 100.00%
Bootstrap support for afum|EAL90463.1 as seed ortholog is 100%.
Bootstrap support for afis|EAW22732.1 as seed ortholog is 100%.

---

Group of orthologs #5158. Best score 828 bits
Score difference with first non-orthologous sequence - *A. fumigatus*:828 *A. fischeri*:828

afum|EAL89532.1 100.00% afis|EAW24882.1 100.00%
Bootstrap support for afum|EAL89532.1 as seed ortholog is 100%.
Bootstrap support for afis|EAW24882.1 as seed ortholog is 100%.

---

Group of orthologs #5159. Best score 827 bits
Score difference with first non-orthologous sequence - *A. fumigatus*:827 *A. fischeri*:827
afum|EAL84787.1 100.00% afis|EAW20846.1 100.00%
Bootstrap support for afum|EAL84787.1 as seed ortholog is 100%.
Bootstrap support for afis|EAW20846.1 as seed ortholog is 100%.

---

Group of orthologs #5160. Best score 827 bits
Score difference with first non-orthologous sequence - *A. fumigatus*:753 *A. fischeri*:543
afum|EAL84792.1 100.00% afis|EAW20841.1 100.00%
Bootstrap support for afum|EAL84792.1 as seed ortholog is 100%.
Bootstrap support for afis|EAW20841.1 as seed ortholog is 100%.

---

Group of orthologs #5161. Best score 827 bits
Score difference with first non-orthologous sequence - *A. fumigatus*:827 *A. fischeri*:827
afum|EAL84757.1 100.00% afis|EAW20876.1 100.00%
Bootstrap support for afum|EAL84757.1 as seed ortholog is 100%.
Bootstrap support for afis|EAW20876.1 as seed ortholog is 100%.

---

Group of orthologs #5162. Best score 827 bits
Score difference with first non-orthologous sequence - *A. fumigatus*:827 *A. fischeri*:827
afum|EAL91477.2 100.00% afis|EAW17663.1 100.00%
Bootstrap support for afum|EAL91477.2 as seed ortholog is 100%.
Bootstrap support for afis|EAW17663.1 as seed ortholog is 100%.

---

Group of orthologs #5163. Best score 827 bits
Score difference with first non-orthologous sequence - *A. fumigatus*:827 *A. fischeri*:827
afum|EAL92240.1 100.00% afis|EAW21228.1 100.00%
Bootstrap support for afum|EAL92240.1 as seed ortholog is 100%.
Bootstrap support for afis|EAW21228.1 as seed ortholog is 100%.

---

Group of orthologs #5164. Best score 827 bits
Score difference with first non-orthologous sequence - *A. fumigatus*:827 *A. fischeri*:827
afum|EAL92754.1 100.00% afis|EAW21721.1 100.00%
Bootstrap support for afum|EAL92754.1 as seed ortholog is 100%.
Bootstrap support for afis|EAW21721.1 as seed ortholog is 100%.

---

Group of orthologs #5165. Best score 826 bits
Score difference with first non-orthologous sequence - *A. fumigatus*:826 *A. fischeri*:826
afum|EAL85606.1 100.00% afis|EAW15742.1 100.00%
Bootstrap support for afum|EAL85606.1 as seed ortholog is 100%.
Bootstrap support for afis|EAW15742.1 as seed ortholog is 100%.

---

Group of orthologs #5166. Best score 826 bits
Score difference with first non-orthologous sequence - *A. fumigatus*:826 *A. fischeri*:826
afum|EAL86392.1 100.00% afis|EAW16846.1 100.00%
Bootstrap support for afum|EAL86392.1 as seed ortholog is 100%.
Bootstrap support for afis|EAW16846.1 as seed ortholog is 100%.

---

Group of orthologs #5167. Best score 826 bits
Score difference with first non-orthologous sequence - *A. fumigatus*:826 *A. fischeri*:826
afum|EAL88913.1 100.00% afis|EAW16291.1 100.00%
Bootstrap support for afum|EAL88913.1 as seed ortholog is 100%.

Bootstrap support for afis|EAW16291.1 as seed ortholog is 100%.

---

Group of orthologs #5168. Best score 826 bits
Score difference with first non-orthologous sequence - *A. fumigatus*:826 *A. fischeri*:826
afum|EAL87480.1 100.00% afis|EAW18122.1 100.00%
Bootstrap support for afum|EAL87480.1 as seed ortholog is 100%.
Bootstrap support for afis|EAW18122.1 as seed ortholog is 100%.

---

Group of orthologs #5169. Best score 826 bits
Score difference with first non-orthologous sequence - *A. fumigatus*:826 *A. fischeri*:826
afum|EAL84685.1 100.00% afis|EAW20949.1 100.00%
Bootstrap support for afum|EAL84685.1 as seed ortholog is 100%.
Bootstrap support for afis|EAW20949.1 as seed ortholog is 100%.

---

Group of orthologs #5170. Best score 826 bits
Score difference with first non-orthologous sequence - *A. fumigatus*:826 *A. fischeri*:826
afum|EAL87005.1 100.00% afis|EAW19489.1 100.00%
Bootstrap support for afum|EAL87005.1 as seed ortholog is 100%.
Bootstrap support for afis|EAW19489.1 as seed ortholog is 100%.

---

Group of orthologs #5171. Best score 826 bits
Score difference with first non-orthologous sequence - *A. fumigatus*:826 *A. fischeri*:826
afum|EAL86059.1 100.00% afis|EAW24387.1 100.00%
Bootstrap support for afum|EAL86059.1 as seed ortholog is 100%.
Bootstrap support for afis|EAW24387.1 as seed ortholog is 100%.

---

Group of orthologs #5172. Best score 826 bits
Score difference with first non-orthologous sequence - *A. fumigatus*:826 *A. fischeri*:826
afum|EAL87978.2 100.00% afis|EAW23485.1 100.00%
Bootstrap support for afum|EAL87978.2 as seed ortholog is 100%.
Bootstrap support for afis|EAW23485.1 as seed ortholog is 100%.

---

Group of orthologs #5173. Best score 826 bits
Score difference with first non-orthologous sequence - *A. fumigatus*:826 *A. fischeri*:826
afum|EAL90487.1 100.00% afis|EAW22705.1 100.00%
Bootstrap support for afum|EAL90487.1 as seed ortholog is 100%.
Bootstrap support for afis|EAW22705.1 as seed ortholog is 100%.

---

Group of orthologs #5174. Best score 826 bits
Score difference with first non-orthologous sequence - *A. fumigatus*:826 *A. fischeri*:826
afum|EAL92562.1 100.00% afis|EAW21542.1 100.00%
Bootstrap support for afum|EAL92562.1 as seed ortholog is 100%.
Bootstrap support for afis|EAW21542.1 as seed ortholog is 100%.

---

Group of orthologs #5175. Best score 826 bits
Score difference with first non-orthologous sequence - *A. fumigatus*:764 *A. fischeri*:826
afum|EAL92713.1 100.00% afis|EAW21679.1 100.00%
Bootstrap support for afum|EAL92713.1 as seed ortholog is 100%.
Bootstrap support for afis|EAW21679.1 as seed ortholog is 100%.

---

Group of orthologs #5176. Best score 826 bits
Score difference with first non-orthologous sequence - *A. fumigatus*:826 *A. fischeri*:826
afum|EBA27448.1 100.00% afis|EAW21120.1 100.00%
Bootstrap support for afum|EBA27448.1 as seed ortholog is 100%.
Bootstrap support for afis|EAW21120.1 as seed ortholog is 100%.

---

Group of orthologs #5177. Best score 826 bits
Score difference with first non-orthologous sequence - *A. fumigatus*:826 *A. fischeri*:826
afum|EBA27453.1 100.00% afis|EAW21221.1 100.00%
Bootstrap support for afum|EBA27453.1 as seed ortholog is 100%.
Bootstrap support for afis|EAW21221.1 as seed ortholog is 100%.

---

Group of orthologs #5178. Best score 825 bits
Score difference with first non-orthologous sequence - *A. fumigatus*:825 *A. fischeri*:825
afum|EAL86621.1 100.00% afis|EAW17193.1 100.00%
Bootstrap support for afum|EAL86621.1 as seed ortholog is 100%.
Bootstrap support for afis|EAW17193.1 as seed ortholog is 100%.

---

Group of orthologs #5179. Best score 825 bits
Score difference with first non-orthologous sequence - *A. fumigatus*:825 *A. fischeri*:825
afum|EAL85304.2 100.00% afis|EAW20185.1 100.00%
Bootstrap support for afum|EAL85304.2 as seed ortholog is 100%.
Bootstrap support for afis|EAW20185.1 as seed ortholog is 100%.

---

Group of orthologs #5180. Best score 825 bits
Score difference with first non-orthologous sequence - *A. fumigatus*:825 *A. fischeri*:825
afum|EAL87504.1 100.00% afis|EAW18146.1 100.00%
Bootstrap support for afum|EAL87504.1 as seed ortholog is 100%.
Bootstrap support for afis|EAW18146.1 as seed ortholog is 100%.

---

Group of orthologs #5181. Best score 825 bits
Score difference with first non-orthologous sequence - *A. fumigatus*:825 *A. fischeri*:825
afum|EAL87162.1 100.00% afis|EAW23722.1 100.00%
Bootstrap support for afum|EAL87162.1 as seed ortholog is 100%.
Bootstrap support for afis|EAW23722.1 as seed ortholog is 100%.

---

Group of orthologs #5182. Best score 825 bits
Score difference with first non-orthologous sequence - *A. fumigatus*:825 *A. fischeri*:825
afum|EAL88137.2 100.00% afis|EAW23341.1 100.00%
Bootstrap support for afum|EAL88137.2 as seed ortholog is 100%.
Bootstrap support for afis|EAW23341.1 as seed ortholog is 100%.

---

Group of orthologs #5183. Best score 825 bits
Score difference with first non-orthologous sequence - *A. fumigatus*:825 *A. fischeri*:825
afum|EAL93975.1 100.00% afis|EAW19246.1 100.00%
Bootstrap support for afum|EAL93975.1 as seed ortholog is 100%.
Bootstrap support for afis|EAW19246.1 as seed ortholog is 100%.

---

Group of orthologs #5184. Best score 825 bits
Score difference with first non-orthologous sequence - *A. fumigatus*:825 *A. fischeri*:825
afum|EAL90889.1 100.00% afis|EAW22292.1 100.00%
Bootstrap support for afum|EAL90889.1 as seed ortholog is 100%.
Bootstrap support for afis|EAW22292.1 as seed ortholog is 100%.

---

Group of orthologs #5185. Best score 825 bits
Score difference with first non-orthologous sequence - *A. fumigatus*:637 *A. fischeri*:462
afum|EAL91047.1 100.00% afis|EAW22143.1 100.00%
Bootstrap support for afum|EAL91047.1 as seed ortholog is 100%.
Bootstrap support for afis|EAW22143.1 as seed ortholog is 100%.

---

Group of orthologs #5186. Best score 824 bits
Score difference with first non-orthologous sequence - *A. fumigatus*:824 *A. fischeri*:824

afum|EAL84788.1 100.00% afis|EAW20845.1 100.00%
Bootstrap support for afum|EAL84788.1 as seed ortholog is 100%.
Bootstrap support for afis|EAW20845.1 as seed ortholog is 100%.

---

Group of orthologs #5187. Best score 824 bits
Score difference with first non-orthologous sequence - *A. fumigatus*:824 *A. fischeri*:824
afum|EAL86939.1 100.00% afis|EAW19579.1 100.00%
Bootstrap support for afum|EAL86939.1 as seed ortholog is 100%.
Bootstrap support for afis|EAW19579.1 as seed ortholog is 100%.

---

Group of orthologs #5188. Best score 824 bits
Score difference with first non-orthologous sequence - *A. fumigatus*:824 *A. fischeri*:824
afum|EAL86931.1 100.00% afis|EAW19587.1 100.00%
Bootstrap support for afum|EAL86931.1 as seed ortholog is 100%.
Bootstrap support for afis|EAW19587.1 as seed ortholog is 100%.

---

Group of orthologs #5189. Best score 824 bits
Score difference with first non-orthologous sequence - *A. fumigatus*:698 *A. fischeri*:669
afum|EAL91846.1 100.00% afis|EAW18035.1 100.00%
Bootstrap support for afum|EAL91846.1 as seed ortholog is 100%.
Bootstrap support for afis|EAW18035.1 as seed ortholog is 100%.

---

Group of orthologs #5190. Best score 824 bits
Score difference with first non-orthologous sequence - *A. fumigatus*:824 *A. fischeri*:824
afum|EAL93996.1 100.00% afis|EAW19269.1 100.00%
Bootstrap support for afum|EAL93996.1 as seed ortholog is 100%.
Bootstrap support for afis|EAW19269.1 as seed ortholog is 100%.

---

Group of orthologs #5191. Best score 824 bits
Score difference with first non-orthologous sequence - *A. fumigatus*:824 *A. fischeri*:824
afum|EAL90422.1 100.00% afis|EAW22776.1 100.00%
Bootstrap support for afum|EAL90422.1 as seed ortholog is 100%.
Bootstrap support for afis|EAW22776.1 as seed ortholog is 100%.

---

Group of orthologs #5192. Best score 824 bits
Score difference with first non-orthologous sequence - *A. fumigatus*:824 *A. fischeri*:824
afum|EBA27415.1 100.00% afis|EAW22366.1 100.00%
Bootstrap support for afum|EBA27415.1 as seed ortholog is 100%.
Bootstrap support for afis|EAW22366.1 as seed ortholog is 100%.

---

Group of orthologs #5193. Best score 823 bits
Score difference with first non-orthologous sequence - *A. fumigatus*:823 *A. fischeri*:823
afum|EAL85060.1 100.00% afis|EAW19832.1 100.00%
Bootstrap support for afum|EAL85060.1 as seed ortholog is 100%.
Bootstrap support for afis|EAW19832.1 as seed ortholog is 100%.

---

Group of orthologs #5194. Best score 823 bits
Score difference with first non-orthologous sequence - *A. fumigatus*:823 *A. fischeri*:747
afum|EAL85010.1 100.00% afis|EAW19888.1 100.00%
Bootstrap support for afum|EAL85010.1 as seed ortholog is 100%.
Bootstrap support for afis|EAW19888.1 as seed ortholog is 100%.

---

Group of orthologs #5195. Best score 823 bits
Score difference with first non-orthologous sequence - *A. fumigatus*:823 *A. fischeri*:823
afum|EAL84464.1 100.00% afis|EAW20597.1 100.00%
Bootstrap support for afum|EAL84464.1 as seed ortholog is 100%.

Bootstrap support for afis|EAW20597.1 as seed ortholog is 100%.

---

Group of orthologs #5196. Best score 823 bits
Score difference with first non-orthologous sequence - *A. fumigatus*:823 *A. fischeri*:823
afum|EAL87482.1 100.00% afis|EAW18124.1 100.00%
Bootstrap support for afum|EAL87482.1 as seed ortholog is 100%.
Bootstrap support for afis|EAW18124.1 as seed ortholog is 100%.

---

Group of orthologs #5197. Best score 823 bits
Score difference with first non-orthologous sequence - *A. fumigatus*:823 *A. fischeri*:823
afum|EAL86908.1 100.00% afis|EAW19614.1 100.00%
Bootstrap support for afum|EAL86908.1 as seed ortholog is 100%.
Bootstrap support for afis|EAW19614.1 as seed ortholog is 100%.

---

Group of orthologs #5198. Best score 823 bits
Score difference with first non-orthologous sequence - *A. fumigatus*:823 *A. fischeri*:823
afum|EAL91225.1 100.00% afis|EAW17389.1 100.00%
Bootstrap support for afum|EAL91225.1 as seed ortholog is 100%.
Bootstrap support for afis|EAW17389.1 as seed ortholog is 100%.

---

Group of orthologs #5199. Best score 823 bits
Score difference with first non-orthologous sequence - *A. fumigatus*:823 *A. fischeri*:823
afum|EAL86057.1 100.00% afis|EAW24385.1 100.00%
Bootstrap support for afum|EAL86057.1 as seed ortholog is 100%.
Bootstrap support for afis|EAW24385.1 as seed ortholog is 100%.

---

Group of orthologs #5200. Best score 823 bits
Score difference with first non-orthologous sequence - *A. fumigatus*:823 *A. fischeri*:823
afum|EAL88331.1 100.00% afis|EAW23150.1 100.00%
Bootstrap support for afum|EAL88331.1 as seed ortholog is 100%.
Bootstrap support for afis|EAW23150.1 as seed ortholog is 100%.

---

Group of orthologs #5201. Best score 823 bits
Score difference with first non-orthologous sequence - *A. fumigatus*:635 *A. fischeri*:823
afum|EAL89389.1 100.00% afis|EAW24743.1 100.00%
Bootstrap support for afum|EAL89389.1 as seed ortholog is 100%.
Bootstrap support for afis|EAW24743.1 as seed ortholog is 100%.

---

Group of orthologs #5202. Best score 822 bits
Score difference with first non-orthologous sequence - *A. fumigatus*:822 *A. fischeri*:569
afum|EAL84970.1 100.00% afis|EAW19943.1 100.00%
Bootstrap support for afum|EAL84970.1 as seed ortholog is 100%.
Bootstrap support for afis|EAW19943.1 as seed ortholog is 100%.

---

Group of orthologs #5203. Best score 822 bits
Score difference with first non-orthologous sequence - *A. fumigatus*:663 *A. fischeri*:681
afum|EAL91494.1 100.00% afis|EAW17680.1 100.00%
Bootstrap support for afum|EAL91494.1 as seed ortholog is 100%.
Bootstrap support for afis|EAW17680.1 as seed ortholog is 100%.

---

Group of orthologs #5204. Best score 822 bits
Score difference with first non-orthologous sequence - *A. fumigatus*:822 *A. fischeri*:822
afum|EAL88414.1 100.00% afis|EAW23065.1 100.00%
Bootstrap support for afum|EAL88414.1 as seed ortholog is 100%.
Bootstrap support for afis|EAW23065.1 as seed ortholog is 100%.

---

Group of orthologs #5205. Best score 822 bits
Score difference with first non-orthologous sequence - *A. fumigatus*:822 *A. fischeri*:822
afum|EAL88279.1 100.00% afis|EAW23204.1 100.00%
Bootstrap support for afum|EAL88279.1 as seed ortholog is 100%.
Bootstrap support for afis|EAW23204.1 as seed ortholog is 100%.

---

Group of orthologs #5206. Best score 822 bits
Score difference with first non-orthologous sequence - *A. fumigatus*:822 *A. fischeri*:822
afum|EAL90262.1 100.00% afis|EAW22934.1 100.00%
Bootstrap support for afum|EAL90262.1 as seed ortholog is 100%.
Bootstrap support for afis|EAW22934.1 as seed ortholog is 100%.

---

Group of orthologs #5207. Best score 822 bits
Score difference with first non-orthologous sequence - *A. fumigatus*:822 *A. fischeri*:720
afum|EAL92393.1 100.00% afis|EAW21376.1 100.00%
Bootstrap support for afum|EAL92393.1 as seed ortholog is 100%.
Bootstrap support for afis|EAW21376.1 as seed ortholog is 100%.

---

Group of orthologs #5208. Best score 822 bits
Score difference with first non-orthologous sequence - *A. fumigatus*:822 *A. fischeri*:822
afum|EAL89573.1 100.00% afis|EAW24918.1 100.00%
Bootstrap support for afum|EAL89573.1 as seed ortholog is 100%.
Bootstrap support for afis|EAW24918.1 as seed ortholog is 100%.

---

Group of orthologs #5209. Best score 821 bits
Score difference with first non-orthologous sequence - *A. fumigatus*:821 *A. fischeri*:638
afum|EAL88594.1 100.00% afis|EAW15961.1 100.00%
Bootstrap support for afum|EAL88594.1 as seed ortholog is 100%.
Bootstrap support for afis|EAW15961.1 as seed ortholog is 100%.

---

Group of orthologs #5210. Best score 821 bits
Score difference with first non-orthologous sequence - *A. fumigatus*:821 *A. fischeri*:821
afum|EAL84912.1 100.00% afis|EAW20006.1 100.00%
Bootstrap support for afum|EAL84912.1 as seed ortholog is 100%.
Bootstrap support for afis|EAW20006.1 as seed ortholog is 100%.

---

Group of orthologs #5211. Best score 821 bits
Score difference with first non-orthologous sequence - *A. fumigatus*:821 *A. fischeri*:821
afum|EAL87054.1 100.00% afis|EAW19442.1 100.00%
Bootstrap support for afum|EAL87054.1 as seed ortholog is 100%.
Bootstrap support for afis|EAW19442.1 as seed ortholog is 100%.

---

Group of orthologs #5212. Best score 821 bits
Score difference with first non-orthologous sequence - *A. fumigatus*:821 *A. fischeri*:821
afum|EAL87201.1 100.00% afis|EAW23767.1 100.00%
Bootstrap support for afum|EAL87201.1 as seed ortholog is 100%.
Bootstrap support for afis|EAW23767.1 as seed ortholog is 100%.

---

Group of orthologs #5213. Best score 821 bits
Score difference with first non-orthologous sequence - *A. fumigatus*:821 *A. fischeri*:821
afum|EAL87253.1 100.00% afis|EAW23818.1 100.00%
Bootstrap support for afum|EAL87253.1 as seed ortholog is 100%.
Bootstrap support for afis|EAW23818.1 as seed ortholog is 100%.

---

Group of orthologs #5214. Best score 821 bits
Score difference with first non-orthologous sequence - *A. fumigatus*:821 *A. fischeri*:821

afum|EAL87386.1 100.00% afis|EAW23958.1 100.00%
Bootstrap support for afum|EAL87386.1 as seed ortholog is 100%.
Bootstrap support for afis|EAW23958.1 as seed ortholog is 100%.

---

Group of orthologs #5215. Best score 821 bits
Score difference with first non-orthologous sequence - *A. fumigatus*:821 *A. fischeri*:821
afum|EAL88462.1 100.00% afis|EAW23020.1 100.00%
Bootstrap support for afum|EAL88462.1 as seed ortholog is 100%.
Bootstrap support for afis|EAW23020.1 as seed ortholog is 100%.

---

Group of orthologs #5216. Best score 821 bits
Score difference with first non-orthologous sequence - *A. fumigatus*:821 *A. fischeri*:821
afum|EAL89751.1 100.00% afis|EAW25082.1 100.00%
Bootstrap support for afum|EAL89751.1 as seed ortholog is 100%.
Bootstrap support for afis|EAW25082.1 as seed ortholog is 100%.

---

Group of orthologs #5217. Best score 821 bits
Score difference with first non-orthologous sequence - *A. fumigatus*:821 *A. fischeri*:821
afum|EAL90108.1 100.00% afis|EAW25449.1 100.00%
Bootstrap support for afum|EAL90108.1 as seed ortholog is 100%.
Bootstrap support for afis|EAW25449.1 as seed ortholog is 100%.

---

Group of orthologs #5218. Best score 820 bits
Score difference with first non-orthologous sequence - *A. fumigatus*:820 *A. fischeri*:694
afum|EAL88491.1 100.00% afis|EAW15855.1 100.00%
Bootstrap support for afum|EAL88491.1 as seed ortholog is 100%.
Bootstrap support for afis|EAW15855.1 as seed ortholog is 100%.

---

Group of orthologs #5219. Best score 820 bits
Score difference with first non-orthologous sequence - *A. fumigatus*:820 *A. fischeri*:820
afum|EAL85285.1 100.00% afis|EAW20167.1 100.00%
Bootstrap support for afum|EAL85285.1 as seed ortholog is 100%.
Bootstrap support for afis|EAW20167.1 as seed ortholog is 100%.

---

Group of orthologs #5220. Best score 820 bits
Score difference with first non-orthologous sequence - *A. fumigatus*:820 *A. fischeri*:820
afum|EAL93171.1 100.00% afis|EAW18482.1 100.00%
Bootstrap support for afum|EAL93171.1 as seed ortholog is 100%.
Bootstrap support for afis|EAW18482.1 as seed ortholog is 100%.

---

Group of orthologs #5221. Best score 820 bits
Score difference with first non-orthologous sequence - *A. fumigatus*:820 *A. fischeri*:731
afum|EAL91023.1 100.00% afis|EAW22168.1 100.00%
Bootstrap support for afum|EAL91023.1 as seed ortholog is 100%.
Bootstrap support for afis|EAW22168.1 as seed ortholog is 100%.

---

Group of orthologs #5222. Best score 820 bits
Score difference with first non-orthologous sequence - *A. fumigatus*:820 *A. fischeri*:820
afum|EAL90990.1 100.00% afis|EAW22202.1 100.00%
Bootstrap support for afum|EAL90990.1 as seed ortholog is 100%.
Bootstrap support for afis|EAW22202.1 as seed ortholog is 100%.

---

Group of orthologs #5223. Best score 820 bits
Score difference with first non-orthologous sequence - *A. fumigatus*:820 *A. fischeri*:820
afum|EAL89370.1 100.00% afis|EAW24721.1 100.00%
Bootstrap support for afum|EAL89370.1 as seed ortholog is 100%.

Bootstrap support for afis|EAW24721.1 as seed ortholog is 100%.

---

Group of orthologs #5224. Best score 820 bits
Score difference with first non-orthologous sequence - *A. fumigatus*:820 *A. fischeri*:820
afum|EAL90038.1 100.00% afis|EAW25379.1 100.00%
Bootstrap support for afum|EAL90038.1 as seed ortholog is 100%.
Bootstrap support for afis|EAW25379.1 as seed ortholog is 100%.

---

Group of orthologs #5225. Best score 818 bits
Score difference with first non-orthologous sequence - *A. fumigatus*:818 *A. fischeri*:818
afum|EAL85860.1 100.00% afis|EAW15459.1 100.00%
Bootstrap support for afum|EAL85860.1 as seed ortholog is 100%.
Bootstrap support for afis|EAW15459.1 as seed ortholog is 100%.

---

Group of orthologs #5226. Best score 818 bits
Score difference with first non-orthologous sequence - *A. fumigatus*:818 *A. fischeri*:818
afum|EAL93699.1 100.00% afis|EAW18967.1 100.00%
Bootstrap support for afum|EAL93699.1 as seed ortholog is 100%.
Bootstrap support for afis|EAW18967.1 as seed ortholog is 100%.

---

Group of orthologs #5227. Best score 818 bits
Score difference with first non-orthologous sequence - *A. fumigatus*:818 *A. fischeri*:818
afum|EAL90596.1 100.00% afis|EAW22602.1 100.00%
Bootstrap support for afum|EAL90596.1 as seed ortholog is 100%.
Bootstrap support for afis|EAW22602.1 as seed ortholog is 100%.

---

Group of orthologs #5228. Best score 818 bits
Score difference with first non-orthologous sequence - *A. fumigatus*:818 *A. fischeri*:818
afum|EAL92814.1 100.00% afis|EAW21787.1 100.00%
Bootstrap support for afum|EAL92814.1 as seed ortholog is 100%.
Bootstrap support for afis|EAW21787.1 as seed ortholog is 100%.

---

Group of orthologs #5229. Best score 817 bits
Score difference with first non-orthologous sequence - *A. fumigatus*:732 *A. fischeri*:517
afum|EAL85026.1 100.00% afis|EAW19871.1 100.00%
Bootstrap support for afum|EAL85026.1 as seed ortholog is 100%.
Bootstrap support for afis|EAW19871.1 as seed ortholog is 100%.

---

Group of orthologs #5230. Best score 817 bits
Score difference with first non-orthologous sequence - *A. fumigatus*:817 *A. fischeri*:700
afum|EAL84977.2 100.00% afis|EAW19932.1 100.00%
Bootstrap support for afum|EAL84977.2 as seed ortholog is 100%.
Bootstrap support for afis|EAW19932.1 as seed ortholog is 100%.

---

Group of orthologs #5231. Best score 817 bits
Score difference with first non-orthologous sequence - *A. fumigatus*:817 *A. fischeri*:817
afum|EAL84645.1 100.00% afis|EAW20985.1 100.00%
Bootstrap support for afum|EAL84645.1 as seed ortholog is 100%.
Bootstrap support for afis|EAW20985.1 as seed ortholog is 100%.

---

Group of orthologs #5232. Best score 817 bits
Score difference with first non-orthologous sequence - *A. fumigatus*:713 *A. fischeri*:722
afum|EAL84654.1 100.00% afis|EAW20979.1 100.00%
Bootstrap support for afum|EAL84654.1 as seed ortholog is 100%.
Bootstrap support for afis|EAW20979.1 as seed ortholog is 100%.

---

Group of orthologs #5233. Best score 817 bits
Score difference with first non-orthologous sequence - *A. fumigatus*:817 *A. fischeri*:817
afum|EAL87186.2 100.00% afis|EAW20267.1 100.00%
Bootstrap support for afum|EAL87186.2 as seed ortholog is 100%.
Bootstrap support for afis|EAW20267.1 as seed ortholog is 100%.

---

Group of orthologs #5234. Best score 817 bits
Score difference with first non-orthologous sequence - *A. fumigatus*:817 *A. fischeri*:817
afum|EAL93155.1 100.00% afis|EAW18469.1 100.00%
Bootstrap support for afum|EAL93155.1 as seed ortholog is 100%.
Bootstrap support for afis|EAW18469.1 as seed ortholog is 100%.

---

Group of orthologs #5235. Best score 817 bits
Score difference with first non-orthologous sequence - *A. fumigatus*:817 *A. fischeri*:817
afum|EAL87372.1 100.00% afis|EAW23946.1 100.00%
Bootstrap support for afum|EAL87372.1 as seed ortholog is 100%.
Bootstrap support for afis|EAW23946.1 as seed ortholog is 100%.

---

Group of orthologs #5236. Best score 817 bits
Score difference with first non-orthologous sequence - *A. fumigatus*:817 *A. fischeri*:817
afum|EAL88191.1 100.00% afis|EAW23288.1 100.00%
Bootstrap support for afum|EAL88191.1 as seed ortholog is 100%.
Bootstrap support for afis|EAW23288.1 as seed ortholog is 100%.

---

Group of orthologs #5237. Best score 817 bits
Score difference with first non-orthologous sequence - *A. fumigatus*:817 *A. fischeri*:817
afum|EAL89555.1 100.00% afis|EAW24901.1 100.00%
Bootstrap support for afum|EAL89555.1 as seed ortholog is 100%.
Bootstrap support for afis|EAW24901.1 as seed ortholog is 100%.

---

Group of orthologs #5238. Best score 817 bits
Score difference with first non-orthologous sequence - *A. fumigatus*:735 *A. fischeri*:547
afum|EAL91971.1 100.00% afis|EAW24116.1 100.00%
Bootstrap support for afum|EAL91971.1 as seed ortholog is 100%.
Bootstrap support for afis|EAW24116.1 as seed ortholog is 100%.

---

Group of orthologs #5239. Best score 816 bits
Score difference with first non-orthologous sequence - *A. fumigatus*:816 *A. fischeri*:816
afum|EAL88646.1 100.00% afis|EAW16017.1 100.00%
Bootstrap support for afum|EAL88646.1 as seed ortholog is 100%.
Bootstrap support for afis|EAW16017.1 as seed ortholog is 100%.

---

Group of orthologs #5240. Best score 816 bits
Score difference with first non-orthologous sequence - *A. fumigatus*:816 *A. fischeri*:816
afum|EAL88699.1 100.00% afis|EAW16067.1 100.00%
Bootstrap support for afum|EAL88699.1 as seed ortholog is 100%.
Bootstrap support for afis|EAW16067.1 as seed ortholog is 100%.

---

Group of orthologs #5241. Best score 816 bits
Score difference with first non-orthologous sequence - *A. fumigatus*:816 *A. fischeri*:816
afum|EAL85121.1 100.00% afis|EAW19776.1 100.00%
Bootstrap support for afum|EAL85121.1 as seed ortholog is 100%.
Bootstrap support for afis|EAW19776.1 as seed ortholog is 100%.

---

Group of orthologs #5242. Best score 816 bits
Score difference with first non-orthologous sequence - *A. fumigatus*:816 *A. fischeri*:816

afum|EAL85498.1 100.00% afis|EAW20399.1 100.00%
Bootstrap support for afum|EAL85498.1 as seed ortholog is 100%.
Bootstrap support for afis|EAW20399.1 as seed ortholog is 100%.

---

Group of orthologs #5243. Best score 816 bits
Score difference with first non-orthologous sequence - *A. fumigatus*:738 *A. fischeri*:816
afum|EAL91063.1 100.00% afis|EAW22125.1 100.00%
Bootstrap support for afum|EAL91063.1 as seed ortholog is 100%.
Bootstrap support for afis|EAW22125.1 as seed ortholog is 100%.

---

Group of orthologs #5244. Best score 816 bits
Score difference with first non-orthologous sequence - *A. fumigatus*:816 *A. fischeri*:816
afum|EAL90019.1 100.00% afis|EAW25357.1 100.00%
Bootstrap support for afum|EAL90019.1 as seed ortholog is 100%.
Bootstrap support for afis|EAW25357.1 as seed ortholog is 100%.

---

Group of orthologs #5245. Best score 815 bits
Score difference with first non-orthologous sequence - *A. fumigatus*:815 *A. fischeri*:815
afum|EAL85239.1 100.00% afis|EAW20121.1 100.00%
Bootstrap support for afum|EAL85239.1 as seed ortholog is 100%.
Bootstrap support for afis|EAW20121.1 as seed ortholog is 100%.

---

Group of orthologs #5246. Best score 815 bits
Score difference with first non-orthologous sequence - *A. fumigatus*:764 *A. fischeri*:697
afum|EAL91367.1 100.00% afis|EAW17553.1 100.00%
Bootstrap support for afum|EAL91367.1 as seed ortholog is 100%.
Bootstrap support for afis|EAW17553.1 as seed ortholog is 100%.

---

Group of orthologs #5247. Best score 815 bits
Score difference with first non-orthologous sequence - *A. fumigatus*:815 *A. fischeri*:815
afum|EAL85991.1 100.00% afis|EAW24321.1 100.00%
Bootstrap support for afum|EAL85991.1 as seed ortholog is 100%.
Bootstrap support for afis|EAW24321.1 as seed ortholog is 100%.

---

Group of orthologs #5248. Best score 815 bits
Score difference with first non-orthologous sequence - *A. fumigatus*:697 *A. fischeri*:307
afum|EAL87376.1 100.00% afis|EAW23950.1 100.00%
Bootstrap support for afum|EAL87376.1 as seed ortholog is 100%.
Bootstrap support for afis|EAW23950.1 as seed ortholog is 100%.

---

Group of orthologs #5249. Best score 815 bits
Score difference with first non-orthologous sequence - *A. fumigatus*:815 *A. fischeri*:815
afum|EAL90387.1 100.00% afis|EAW22814.1 100.00%
Bootstrap support for afum|EAL90387.1 as seed ortholog is 100%.
Bootstrap support for afis|EAW22814.1 as seed ortholog is 100%.

---

Group of orthologs #5250. Best score 814 bits
Score difference with first non-orthologous sequence - *A. fumigatus*:814 *A. fischeri*:814
afum|EAL88767.1 100.00% afis|EAW16138.1 100.00%
Bootstrap support for afum|EAL88767.1 as seed ortholog is 100%.
Bootstrap support for afis|EAW16138.1 as seed ortholog is 100%.

---

Group of orthologs #5251. Best score 814 bits
Score difference with first non-orthologous sequence - *A. fumigatus*:814 *A. fischeri*:814
afum|EAL85345.2 100.00% afis|EAW20228.1 100.00%
Bootstrap support for afum|EAL85345.2 as seed ortholog is 100%.

Bootstrap support for afis|EAW20228.1 as seed ortholog is 100%.

---

Group of orthologs #5252. Best score 814 bits
Score difference with first non-orthologous sequence - *A. fumigatus*:814 *A. fischeri*:814
afum|EAL91797.1 100.00% afis|EAW17988.1 100.00%
Bootstrap support for afum|EAL91797.1 as seed ortholog is 100%.
Bootstrap support for afis|EAW17988.1 as seed ortholog is 100%.

---

Group of orthologs #5253. Best score 814 bits
Score difference with first non-orthologous sequence - *A. fumigatus*:814 *A. fischeri*:814
afum|EAL92590.1 100.00% afis|EAW21572.1 100.00%
Bootstrap support for afum|EAL92590.1 as seed ortholog is 100%.
Bootstrap support for afis|EAW21572.1 as seed ortholog is 100%.

---

Group of orthologs #5254. Best score 813 bits
Score difference with first non-orthologous sequence - *A. fumigatus*:813 *A. fischeri*:700
afum|EAL89251.1 100.00% afis|EAW16647.1 100.00%
Bootstrap support for afum|EAL89251.1 as seed ortholog is 100%.
Bootstrap support for afis|EAW16647.1 as seed ortholog is 100%.

---

Group of orthologs #5255. Best score 813 bits
Score difference with first non-orthologous sequence - *A. fumigatus*:813 *A. fischeri*:813
afum|EAL88058.1 100.00% afis|EAW23411.1 100.00%
Bootstrap support for afum|EAL88058.1 as seed ortholog is 100%.
Bootstrap support for afis|EAW23411.1 as seed ortholog is 100%.

---

Group of orthologs #5256. Best score 813 bits
Score difference with first non-orthologous sequence - *A. fumigatus*:813 *A. fischeri*:813
afum|EAL93764.2 100.00% afis|EAW19035.1 100.00%
Bootstrap support for afum|EAL93764.2 as seed ortholog is 100%.
Bootstrap support for afis|EAW19035.1 as seed ortholog is 100%.

---

Group of orthologs #5257. Best score 813 bits
Score difference with first non-orthologous sequence - *A. fumigatus*:813 *A. fischeri*:768
afum|EAL92829.2 100.00% afis|EAW21803.1 100.00%
Bootstrap support for afum|EAL92829.2 as seed ortholog is 100%.
Bootstrap support for afis|EAW21803.1 as seed ortholog is 100%.

---

Group of orthologs #5258. Best score 813 bits
Score difference with first non-orthologous sequence - *A. fumigatus*:813 *A. fischeri*:700
afum|EAL89993.1 100.00% afis|EAW25331.1 100.00%
Bootstrap support for afum|EAL89993.1 as seed ortholog is 100%.
Bootstrap support for afis|EAW25331.1 as seed ortholog is 100%.

---

Group of orthologs #5259. Best score 812 bits
Score difference with first non-orthologous sequence - *A. fumigatus*:812 *A. fischeri*:294
afum|EAL86669.1 100.00% afis|EAW17250.1 100.00%
Bootstrap support for afum|EAL86669.1 as seed ortholog is 100%.
Bootstrap support for afis|EAW17250.1 as seed ortholog is 100%.

---

Group of orthologs #5260. Best score 812 bits
Score difference with first non-orthologous sequence - *A. fumigatus*:812 *A. fischeri*:812
afum|EAL85097.1 100.00% afis|EAW19793.1 100.00%
Bootstrap support for afum|EAL85097.1 as seed ortholog is 100%.
Bootstrap support for afis|EAW19793.1 as seed ortholog is 100%.

---

Group of orthologs #5261. Best score 812 bits
Score difference with first non-orthologous sequence - *A. fumigatus*:812 *A. fischeri*:812
afum|EAL84530.1 100.00% afis|EAW20527.1 100.00%
Bootstrap support for afum|EAL84530.1 as seed ortholog is 100%.
Bootstrap support for afis|EAW20527.1 as seed ortholog is 100%.

---

Group of orthologs #5262. Best score 812 bits
Score difference with first non-orthologous sequence - *A. fumigatus*:812 *A. fischeri*:649
afum|EAL91622.1 100.00% afis|EAW17813.1 100.00%
Bootstrap support for afum|EAL91622.1 as seed ortholog is 100%.
Bootstrap support for afis|EAW17813.1 as seed ortholog is 100%.

---

Group of orthologs #5263. Best score 812 bits
Score difference with first non-orthologous sequence - *A. fumigatus*:812 *A. fischeri*:486
afum|EAL86221.1 100.00% afis|EAW24558.1 100.00%
Bootstrap support for afum|EAL86221.1 as seed ortholog is 100%.
Bootstrap support for afis|EAW24558.1 as seed ortholog is 100%.

---

Group of orthologs #5264. Best score 811 bits
Score difference with first non-orthologous sequence - *A. fumigatus*:811 *A. fischeri*:811
afum|EAL87070.1 100.00% afis|EAW19424.1 100.00%
Bootstrap support for afum|EAL87070.1 as seed ortholog is 100%.
Bootstrap support for afis|EAW19424.1 as seed ortholog is 100%.

---

Group of orthologs #5265. Best score 811 bits
Score difference with first non-orthologous sequence - *A. fumigatus*:811 *A. fischeri*:811
afum|EAL86924.1 100.00% afis|EAW19594.1 100.00%
Bootstrap support for afum|EAL86924.1 as seed ortholog is 100%.
Bootstrap support for afis|EAW19594.1 as seed ortholog is 100%.

---

Group of orthologs #5266. Best score 811 bits
Score difference with first non-orthologous sequence - *A. fumigatus*:811 *A. fischeri*:321
afum|EAL91148.1 100.00% afis|EAW17313.1 100.00%
Bootstrap support for afum|EAL91148.1 as seed ortholog is 100%.
Bootstrap support for afis|EAW17313.1 as seed ortholog is 100%.

---

Group of orthologs #5267. Best score 811 bits
Score difference with first non-orthologous sequence - *A. fumigatus*:811 *A. fischeri*:811
afum|EAL91211.1 100.00% afis|EAW17375.1 100.00%
Bootstrap support for afum|EAL91211.1 as seed ortholog is 100%.
Bootstrap support for afis|EAW17375.1 as seed ortholog is 100%.

---

Group of orthologs #5268. Best score 811 bits
Score difference with first non-orthologous sequence - *A. fumigatus*:811 *A. fischeri*:811
afum|EAL91564.1 100.00% afis|EAW17751.1 100.00%
Bootstrap support for afum|EAL91564.1 as seed ortholog is 100%.
Bootstrap support for afis|EAW17751.1 as seed ortholog is 100%.

---

Group of orthologs #5269. Best score 811 bits
Score difference with first non-orthologous sequence - *A. fumigatus*:811 *A. fischeri*:724
afum|EAL87890.1 100.00% afis|EAW23575.1 100.00%
Bootstrap support for afum|EAL87890.1 as seed ortholog is 100%.
Bootstrap support for afis|EAW23575.1 as seed ortholog is 100%.

---

Group of orthologs #5270. Best score 811 bits
Score difference with first non-orthologous sequence - *A. fumigatus*:811 *A. fischeri*:744

afum|EAL93448.1 100.00% afis|EAW18723.1 100.00%
Bootstrap support for afum|EAL93448.1 as seed ortholog is 100%.
Bootstrap support for afis|EAW18723.1 as seed ortholog is 100%.

---

Group of orthologs #5271. Best score 811 bits
Score difference with first non-orthologous sequence - *A. fumigatus*:692 *A. fischeri*:385
afum|EAL93890.2 100.00% afis|EAW19160.1 100.00%
Bootstrap support for afum|EAL93890.2 as seed ortholog is 100%.
Bootstrap support for afis|EAW19160.1 as seed ortholog is 100%.

---

Group of orthologs #5272. Best score 811 bits
Score difference with first non-orthologous sequence - *A. fumigatus*:811 *A. fischeri*:811
afum|EAL92506.1 100.00% afis|EAW21486.1 100.00%
Bootstrap support for afum|EAL92506.1 as seed ortholog is 100%.
Bootstrap support for afis|EAW21486.1 as seed ortholog is 100%.

---

Group of orthologs #5273. Best score 811 bits
Score difference with first non-orthologous sequence - *A. fumigatus*:811 *A. fischeri*:811
afum|EAL89594.1 100.00% afis|EAW24937.1 100.00%
Bootstrap support for afum|EAL89594.1 as seed ortholog is 100%.
Bootstrap support for afis|EAW24937.1 as seed ortholog is 100%.

---

Group of orthologs #5274. Best score 811 bits
Score difference with first non-orthologous sequence - *A. fumigatus*:811 *A. fischeri*:811
afum|EAL89640.1 100.00% afis|EAW24980.1 100.00%
Bootstrap support for afum|EAL89640.1 as seed ortholog is 100%.
Bootstrap support for afis|EAW24980.1 as seed ortholog is 100%.

---

Group of orthologs #5275. Best score 811 bits
Score difference with first non-orthologous sequence - *A. fumigatus*:811 *A. fischeri*:811
afum|EAL91956.1 100.00% afis|EAW24102.1 100.00%
Bootstrap support for afum|EAL91956.1 as seed ortholog is 100%.
Bootstrap support for afis|EAW24102.1 as seed ortholog is 100%.

---

Group of orthologs #5276. Best score 810 bits
Score difference with first non-orthologous sequence - *A. fumigatus*:810 *A. fischeri*:810
afum|EAL85820.1 100.00% afis|EAW15504.1 100.00%
Bootstrap support for afum|EAL85820.1 as seed ortholog is 100%.
Bootstrap support for afis|EAW15504.1 as seed ortholog is 100%.

---

Group of orthologs #5277. Best score 810 bits
Score difference with first non-orthologous sequence - *A. fumigatus*:810 *A. fischeri*:810
afum|EAL89191.1 100.00% afis|EAW16595.1 100.00%
Bootstrap support for afum|EAL89191.1 as seed ortholog is 100%.
Bootstrap support for afis|EAW16595.1 as seed ortholog is 100%.

---

Group of orthologs #5278. Best score 810 bits
Score difference with first non-orthologous sequence - *A. fumigatus*:810 *A. fischeri*:810
afum|EAL86884.1 100.00% afis|EAW19638.1 100.00%
Bootstrap support for afum|EAL86884.1 as seed ortholog is 100%.
Bootstrap support for afis|EAW19638.1 as seed ortholog is 100%.

---

Group of orthologs #5279. Best score 810 bits
Score difference with first non-orthologous sequence - *A. fumigatus*:810 *A. fischeri*:810
afum|EAL88206.1 100.00% afis|EAW23273.1 100.00%
Bootstrap support for afum|EAL88206.1 as seed ortholog is 100%.

Bootstrap support for afis|EAW23273.1 as seed ortholog is 100%.

---

Group of orthologs #5280. Best score 810 bits
Score difference with first non-orthologous sequence - *A. fumigatus*:810 *A. fischeri*:810
afum|EAL90526.1 100.00% afis|EAW22666.1 100.00%
Bootstrap support for afum|EAL90526.1 as seed ortholog is 100%.
Bootstrap support for afis|EAW22666.1 as seed ortholog is 100%.

---

Group of orthologs #5281. Best score 810 bits
Score difference with first non-orthologous sequence - *A. fumigatus*:810 *A. fischeri*:810
afum|EAL92345.1 100.00% afis|EAW21327.1 100.00%
Bootstrap support for afum|EAL92345.1 as seed ortholog is 100%.
Bootstrap support for afis|EAW21327.1 as seed ortholog is 100%.

---

Group of orthologs #5282. Best score 810 bits
Score difference with first non-orthologous sequence - *A. fumigatus*:810 *A. fischeri*:810
afum|EAL92446.2 100.00% afis|EAW21431.1 100.00%
Bootstrap support for afum|EAL92446.2 as seed ortholog is 100%.
Bootstrap support for afis|EAW21431.1 as seed ortholog is 100%.

---

Group of orthologs #5283. Best score 809 bits
Score difference with first non-orthologous sequence - *A. fumigatus*:809 *A. fischeri*:809
afum|EAL85770.1 100.00% afis|EAW15562.1 100.00%
Bootstrap support for afum|EAL85770.1 as seed ortholog is 100%.
Bootstrap support for afis|EAW15562.1 as seed ortholog is 100%.

---

Group of orthologs #5284. Best score 809 bits
Score difference with first non-orthologous sequence - *A. fumigatus*:809 *A. fischeri*:809
afum|EAL85867.1 100.00% afis|EAW24195.1 100.00%
Bootstrap support for afum|EAL85867.1 as seed ortholog is 100%.
Bootstrap support for afis|EAW24195.1 as seed ortholog is 100%.

---

Group of orthologs #5285. Best score 809 bits
Score difference with first non-orthologous sequence - *A. fumigatus*:747 *A. fischeri*:605
afum|EAL87334.1 100.00% afis|EAW23906.1 100.00%
Bootstrap support for afum|EAL87334.1 as seed ortholog is 100%.
Bootstrap support for afis|EAW23906.1 as seed ortholog is 100%.

---

Group of orthologs #5286. Best score 809 bits
Score difference with first non-orthologous sequence - *A. fumigatus*:809 *A. fischeri*:649
afum|EAL90442.2 100.00% afis|EAW22757.1 100.00%
Bootstrap support for afum|EAL90442.2 as seed ortholog is 100%.
Bootstrap support for afis|EAW22757.1 as seed ortholog is 100%.

---

Group of orthologs #5287. Best score 809 bits
Score difference with first non-orthologous sequence - *A. fumigatus*:809 *A. fischeri*:809
afum|EAL92416.1 100.00% afis|EAW21402.1 100.00%
Bootstrap support for afum|EAL92416.1 as seed ortholog is 100%.
Bootstrap support for afis|EAW21402.1 as seed ortholog is 100%.

---

Group of orthologs #5288. Best score 808 bits
Score difference with first non-orthologous sequence - *A. fumigatus*:808 *A. fischeri*:808
afum|EAL85659.1 100.00% afis|EAW15689.1 100.00%
Bootstrap support for afum|EAL85659.1 as seed ortholog is 100%.
Bootstrap support for afis|EAW15689.1 as seed ortholog is 100%.

---

Group of orthologs #5289. Best score 808 bits
Score difference with first non-orthologous sequence - *A. fumigatus*:808 *A. fischeri*:721
afum|EAL88860.1 100.00% afis|EAW16239.1 100.00%
Bootstrap support for afum|EAL88860.1 as seed ortholog is 100%.
Bootstrap support for afis|EAW16239.1 as seed ortholog is 100%.

---

Group of orthologs #5290. Best score 808 bits
Score difference with first non-orthologous sequence - *A. fumigatus*:808 *A. fischeri*:808
afum|EAL84679.1 100.00% afis|EAW20955.1 100.00%
Bootstrap support for afum|EAL84679.1 as seed ortholog is 100%.
Bootstrap support for afis|EAW20955.1 as seed ortholog is 100%.

---

Group of orthologs #5291. Best score 808 bits
Score difference with first non-orthologous sequence - *A. fumigatus*:808 *A. fischeri*:724
afum|EBA27424.1 100.00% afis|EAW17342.1 100.00%
Bootstrap support for afum|EBA27424.1 as seed ortholog is 100%.
Bootstrap support for afis|EAW17342.1 as seed ortholog is 100%.

---

Group of orthologs #5292. Best score 808 bits
Score difference with first non-orthologous sequence - *A. fumigatus*:808 *A. fischeri*:808
afum|EAL88440.1 100.00% afis|EAW23041.1 100.00%
Bootstrap support for afum|EAL88440.1 as seed ortholog is 100%.
Bootstrap support for afis|EAW23041.1 as seed ortholog is 100%.

---

Group of orthologs #5293. Best score 808 bits
Score difference with first non-orthologous sequence - *A. fumigatus*:808 *A. fischeri*:808
afum|EAL93923.2 100.00% afis|EAW19192.1 100.00%
Bootstrap support for afum|EAL93923.2 as seed ortholog is 100%.
Bootstrap support for afis|EAW19192.1 as seed ortholog is 100%.

---

Group of orthologs #5294. Best score 808 bits
Score difference with first non-orthologous sequence - *A. fumigatus*:808 *A. fischeri*:808
afum|EAL89942.1 100.00% afis|EAW25279.1 100.00%
Bootstrap support for afum|EAL89942.1 as seed ortholog is 100%.
Bootstrap support for afis|EAW25279.1 as seed ortholog is 100%.

---

Group of orthologs #5295. Best score 807 bits
Score difference with first non-orthologous sequence - *A. fumigatus*:807 *A. fischeri*:807
afum|EAL85581.1 100.00% afis|EAW15767.1 100.00%
Bootstrap support for afum|EAL85581.1 as seed ortholog is 100%.
Bootstrap support for afis|EAW15767.1 as seed ortholog is 100%.

---

Group of orthologs #5296. Best score 807 bits
Score difference with first non-orthologous sequence - *A. fumigatus*:807 *A. fischeri*:807
afum|EAL88526.2 100.00% afis|EAW15891.1 100.00%
Bootstrap support for afum|EAL88526.2 as seed ortholog is 100%.
Bootstrap support for afis|EAW15891.1 as seed ortholog is 100%.

---

Group of orthologs #5297. Best score 807 bits
Score difference with first non-orthologous sequence - *A. fumigatus*:807 *A. fischeri*:807
afum|EAL88729.1 100.00% afis|EAW16097.1 100.00%
Bootstrap support for afum|EAL88729.1 as seed ortholog is 100%.
Bootstrap support for afis|EAW16097.1 as seed ortholog is 100%.

---

Group of orthologs #5298. Best score 807 bits
Score difference with first non-orthologous sequence - *A. fumigatus*:807 *A. fischeri*:807

afum|EAL87075.1 100.00% afis|EAW19419.1 100.00%
Bootstrap support for afum|EAL87075.1 as seed ortholog is 100%.
Bootstrap support for afis|EAW19419.1 as seed ortholog is 100%.

---

Group of orthologs #5299. Best score 807 bits
Score difference with first non-orthologous sequence - *A. fumigatus*:807 *A. fischeri*:807
afum|EAL86854.2 100.00% afis|EAW19669.1 100.00%
Bootstrap support for afum|EAL86854.2 as seed ortholog is 100%.
Bootstrap support for afis|EAW19669.1 as seed ortholog is 100%.

---

Group of orthologs #5300. Best score 807 bits
Score difference with first non-orthologous sequence - *A. fumigatus*:807 *A. fischeri*:648
afum|EAL92201.1 100.00% afis|EAW21180.1 100.00%
Bootstrap support for afum|EAL92201.1 as seed ortholog is 100%.
Bootstrap support for afis|EAW21180.1 as seed ortholog is 100%.

---

Group of orthologs #5301. Best score 807 bits
Score difference with first non-orthologous sequence - *A. fumigatus*:757 *A. fischeri*:539
afum|EAL89964.1 100.00% afis|EAW25301.1 100.00%
Bootstrap support for afum|EAL89964.1 as seed ortholog is 100%.
Bootstrap support for afis|EAW25301.1 as seed ortholog is 100%.

---

Group of orthologs #5302. Best score 807 bits
Score difference with first non-orthologous sequence - *A. fumigatus*:807 *A. fischeri*:807
afum|EBA27342.1 100.00% afis|EAW23207.1 100.00%
Bootstrap support for afum|EBA27342.1 as seed ortholog is 100%.
Bootstrap support for afis|EAW23207.1 as seed ortholog is 100%.

---

Group of orthologs #5303. Best score 806 bits
Score difference with first non-orthologous sequence - *A. fumigatus*:689 *A. fischeri*:668
afum|EAL86317.2 100.00% afis|EAW16742.1 100.00%
Bootstrap support for afum|EAL86317.2 as seed ortholog is 100%.
Bootstrap support for afis|EAW16742.1 as seed ortholog is 100%.

---

Group of orthologs #5304. Best score 806 bits
Score difference with first non-orthologous sequence - *A. fumigatus*:806 *A. fischeri*:806
afum|EAL86361.1 100.00% afis|EAW16792.1 100.00%
Bootstrap support for afum|EAL86361.1 as seed ortholog is 100%.
Bootstrap support for afis|EAW16792.1 as seed ortholog is 100%.

---

Group of orthologs #5305. Best score 806 bits
Score difference with first non-orthologous sequence - *A. fumigatus*:806 *A. fischeri*:806
afum|EAL86020.2 100.00% afis|EAW24350.1 100.00%
Bootstrap support for afum|EAL86020.2 as seed ortholog is 100%.
Bootstrap support for afis|EAW24350.1 as seed ortholog is 100%.

---

Group of orthologs #5306. Best score 806 bits
Score difference with first non-orthologous sequence - *A. fumigatus*:806 *A. fischeri*:737
afum|EAL93371.1 100.00% afis|EAW18655.1 100.00%
Bootstrap support for afum|EAL93371.1 as seed ortholog is 100%.
Bootstrap support for afis|EAW18655.1 as seed ortholog is 100%.

---

Group of orthologs #5307. Best score 806 bits
Score difference with first non-orthologous sequence - *A. fumigatus*:806 *A. fischeri*:806
afum|EAL93717.1 100.00% afis|EAW18981.1 100.00%
Bootstrap support for afum|EAL93717.1 as seed ortholog is 100%.

Bootstrap support for afis|EAW18981.1 as seed ortholog is 100%.

---

Group of orthologs #5308. Best score 806 bits
Score difference with first non-orthologous sequence - *A. fumigatus*:806 *A. fischeri*:626
afum|EAL88040.1 100.00% afis|EAW24586.1 100.00%
Bootstrap support for afum|EAL88040.1 as seed ortholog is 100%.
Bootstrap support for afis|EAW24586.1 as seed ortholog is 100%.

---

Group of orthologs #5309. Best score 806 bits
Score difference with first non-orthologous sequence - *A. fumigatus*:806 *A. fischeri*:806
afum|EAL93008.1 100.00% afis|EAW21993.1 100.00%
Bootstrap support for afum|EAL93008.1 as seed ortholog is 100%.
Bootstrap support for afis|EAW21993.1 as seed ortholog is 100%.

---

Group of orthologs #5310. Best score 805 bits
Score difference with first non-orthologous sequence - *A. fumigatus*:805 *A. fischeri*:805
afum|EAL91374.1 100.00% afis|EAW17560.1 100.00%
Bootstrap support for afum|EAL91374.1 as seed ortholog is 100%.
Bootstrap support for afis|EAW17560.1 as seed ortholog is 100%.

---

Group of orthologs #5311. Best score 805 bits
Score difference with first non-orthologous sequence - *A. fumigatus*:805 *A. fischeri*:805
afum|EAL91636.1 100.00% afis|EAW17826.1 100.00%
Bootstrap support for afum|EAL91636.1 as seed ortholog is 100%.
Bootstrap support for afis|EAW17826.1 as seed ortholog is 100%.

---

Group of orthologs #5312. Best score 805 bits
Score difference with first non-orthologous sequence - *A. fumigatus*:805 *A. fischeri*:805
afum|EAL91725.2 100.00% afis|EAW17915.1 100.00%
Bootstrap support for afum|EAL91725.2 as seed ortholog is 100%.
Bootstrap support for afis|EAW17915.1 as seed ortholog is 100%.

---

Group of orthologs #5313. Best score 805 bits
Score difference with first non-orthologous sequence - *A. fumigatus*:805 *A. fischeri*:805
afum|EAL86050.1 100.00% afis|EAW24378.1 100.00%
Bootstrap support for afum|EAL86050.1 as seed ortholog is 100%.
Bootstrap support for afis|EAW24378.1 as seed ortholog is 100%.

---

Group of orthologs #5314. Best score 805 bits
Score difference with first non-orthologous sequence - *A. fumigatus*:805 *A. fischeri*:805
afum|EAL87437.1 100.00% afis|EAW24007.1 100.00%
Bootstrap support for afum|EAL87437.1 as seed ortholog is 100%.
Bootstrap support for afis|EAW24007.1 as seed ortholog is 100%.

---

Group of orthologs #5315. Best score 805 bits
Score difference with first non-orthologous sequence - *A. fumigatus*:805 *A. fischeri*:805
afum|EAL94023.1 100.00% afis|EAW19305.1 100.00%
Bootstrap support for afum|EAL94023.1 as seed ortholog is 100%.
Bootstrap support for afis|EAW19305.1 as seed ortholog is 100%.

---

Group of orthologs #5316. Best score 804 bits
Score difference with first non-orthologous sequence - *A. fumigatus*:804 *A. fischeri*:804
afum|EAL87762.1 100.00% afis|EAW20056.1 100.00%
afum|EAL85176.2 97.44%
Bootstrap support for afum|EAL87762.1 as seed ortholog is 100%.
Bootstrap support for afis|EAW20056.1 as seed ortholog is 100%.

Group of orthologs #5317. Best score 804 bits
Score difference with first non-orthologous sequence - *A. fumigatus*:804 *A. fischeri*:519
afum|EAL85836.1 100.00% afis|EAW15487.1 100.00%
Bootstrap support for afum|EAL85836.1 as seed ortholog is 100%.
Bootstrap support for afis|EAW15487.1 as seed ortholog is 100%.

---

Group of orthologs #5318. Best score 804 bits
Score difference with first non-orthologous sequence - *A. fumigatus*:804 *A. fischeri*:720
afum|EAL86363.1 100.00% afis|EAW16794.1 100.00%
Bootstrap support for afum|EAL86363.1 as seed ortholog is 100%.
Bootstrap support for afis|EAW16794.1 as seed ortholog is 100%.

---

Group of orthologs #5319. Best score 804 bits
Score difference with first non-orthologous sequence - *A. fumigatus*:804 *A. fischeri*:804
afum|EAL84643.1 100.00% afis|EAW20987.1 100.00%
Bootstrap support for afum|EAL84643.1 as seed ortholog is 100%.
Bootstrap support for afis|EAW20987.1 as seed ortholog is 100%.

---

Group of orthologs #5320. Best score 804 bits
Score difference with first non-orthologous sequence - *A. fumigatus*:804 *A. fischeri*:804
afum|EAL91826.1 100.00% afis|EAW18014.1 100.00%
Bootstrap support for afum|EAL91826.1 as seed ortholog is 100%.
Bootstrap support for afis|EAW18014.1 as seed ortholog is 100%.

---

Group of orthologs #5321. Best score 804 bits
Score difference with first non-orthologous sequence - *A. fumigatus*:804 *A. fischeri*:804
afum|EAL93506.1 100.00% afis|EAW18774.1 100.00%
Bootstrap support for afum|EAL93506.1 as seed ortholog is 100%.
Bootstrap support for afis|EAW18774.1 as seed ortholog is 100%.

---

Group of orthologs #5322. Best score 804 bits
Score difference with first non-orthologous sequence - *A. fumigatus*:804 *A. fischeri*:804
afum|EAL90327.1 100.00% afis|EAW22871.1 100.00%
Bootstrap support for afum|EAL90327.1 as seed ortholog is 100%.
Bootstrap support for afis|EAW22871.1 as seed ortholog is 100%.

---

Group of orthologs #5323. Best score 803 bits
Score difference with first non-orthologous sequence - *A. fumigatus*:673 *A. fischeri*:512
afum|EAL88877.1 100.00% afis|EAW16249.1 100.00%
Bootstrap support for afum|EAL88877.1 as seed ortholog is 100%.
Bootstrap support for afis|EAW16249.1 as seed ortholog is 100%.

---

Group of orthologs #5324. Best score 803 bits
Score difference with first non-orthologous sequence - *A. fumigatus*:803 *A. fischeri*:803
afum|EAL89161.1 100.00% afis|EAW16564.1 100.00%
Bootstrap support for afum|EAL89161.1 as seed ortholog is 100%.
Bootstrap support for afis|EAW16564.1 as seed ortholog is 100%.

---

Group of orthologs #5325. Best score 803 bits
Score difference with first non-orthologous sequence - *A. fumigatus*:803 *A. fischeri*:803
afum|EAL86886.1 100.00% afis|EAW19636.1 100.00%
Bootstrap support for afum|EAL86886.1 as seed ortholog is 100%.
Bootstrap support for afis|EAW19636.1 as seed ortholog is 100%.

---

Group of orthologs #5326. Best score 803 bits

Score difference with first non-orthologous sequence - *A. fumigatus*:803 *A. fischeri*:803
afum|EAL88368.1 100.00% afis|EAW23111.1 100.00%
Bootstrap support for afum|EAL88368.1 as seed ortholog is 100%.
Bootstrap support for afis|EAW23111.1 as seed ortholog is 100%.

---

Group of orthologs #5327. Best score 803 bits
Score difference with first non-orthologous sequence - *A. fumigatus*:803 *A. fischeri*:803
afum|EAL88100.1 100.00% afis|EAW23372.1 100.00%
Bootstrap support for afum|EAL88100.1 as seed ortholog is 100%.
Bootstrap support for afis|EAW23372.1 as seed ortholog is 100%.

---

Group of orthologs #5328. Best score 803 bits
Score difference with first non-orthologous sequence - *A. fumigatus*:803 *A. fischeri*:803
afum|EAL92795.1 100.00% afis|EAW21765.1 100.00%
Bootstrap support for afum|EAL92795.1 as seed ortholog is 100%.
Bootstrap support for afis|EAW21765.1 as seed ortholog is 100%.

---

Group of orthologs #5329. Best score 802 bits
Score difference with first non-orthologous sequence - *A. fumigatus*:802 *A. fischeri*:713
afum|EAL86503.1 100.00% afis|EAW16978.1 100.00%
Bootstrap support for afum|EAL86503.1 as seed ortholog is 100%.
Bootstrap support for afis|EAW16978.1 as seed ortholog is 100%.

---

Group of orthologs #5330. Best score 802 bits
Score difference with first non-orthologous sequence - *A. fumigatus*:802 *A. fischeri*:802
afum|EAL88963.1 100.00% afis|EAW16337.1 100.00%
Bootstrap support for afum|EAL88963.1 as seed ortholog is 100%.
Bootstrap support for afis|EAW16337.1 as seed ortholog is 100%.

---

Group of orthologs #5331. Best score 802 bits
Score difference with first non-orthologous sequence - *A. fumigatus*:802 *A. fischeri*:802
afum|EAL91506.1 100.00% afis|EAW17692.1 100.00%
Bootstrap support for afum|EAL91506.1 as seed ortholog is 100%.
Bootstrap support for afis|EAW17692.1 as seed ortholog is 100%.

---

Group of orthologs #5332. Best score 802 bits
Score difference with first non-orthologous sequence - *A. fumigatus*:747 *A. fischeri*:802
afum|EAL91610.1 100.00% afis|EAW17802.1 100.00%
Bootstrap support for afum|EAL91610.1 as seed ortholog is 100%.
Bootstrap support for afis|EAW17802.1 as seed ortholog is 100%.

---

Group of orthologs #5333. Best score 802 bits
Score difference with first non-orthologous sequence - *A. fumigatus*:802 *A. fischeri*:802
afum|EAL91818.1 100.00% afis|EAW18006.1 100.00%
Bootstrap support for afum|EAL91818.1 as seed ortholog is 100%.
Bootstrap support for afis|EAW18006.1 as seed ortholog is 100%.

---

Group of orthologs #5334. Best score 802 bits
Score difference with first non-orthologous sequence - *A. fumigatus*:802 *A. fischeri*:802
afum|EAL85861.1 100.00% afis|EAW24188.1 100.00%
Bootstrap support for afum|EAL85861.1 as seed ortholog is 100%.
Bootstrap support for afis|EAW24188.1 as seed ortholog is 100%.

---

Group of orthologs #5335. Best score 802 bits
Score difference with first non-orthologous sequence - *A. fumigatus*:655 *A. fischeri*:629
afum|EAL87239.1 100.00% afis|EAW23806.1 100.00%

Bootstrap support for afum|EAL87239.1 as seed ortholog is 100%.
Bootstrap support for afis|EAW23806.1 as seed ortholog is 100%.

---

Group of orthologs #5336. Best score 802 bits
Score difference with first non-orthologous sequence - *A. fumigatus*:573 *A. fischeri*:432
afum|EAL93769.1 100.00% afis|EAW19040.1 100.00%
Bootstrap support for afum|EAL93769.1 as seed ortholog is 100%.
Bootstrap support for afis|EAW19040.1 as seed ortholog is 100%.

---

Group of orthologs #5337. Best score 802 bits
Score difference with first non-orthologous sequence - *A. fumigatus*:802 *A. fischeri*:802
afum|EAL91952.1 100.00% afis|EAW24098.1 100.00%
Bootstrap support for afum|EAL91952.1 as seed ortholog is 100%.
Bootstrap support for afis|EAW24098.1 as seed ortholog is 100%.

---

Group of orthologs #5338. Best score 801 bits
Score difference with first non-orthologous sequence - *A. fumigatus*:801 *A. fischeri*:369
afum|EAL84207.1 100.00% afis|EAW15438.1 100.00%
Bootstrap support for afum|EAL84207.1 as seed ortholog is 100%.
Bootstrap support for afis|EAW15438.1 as seed ortholog is 100%.

---

Group of orthologs #5339. Best score 801 bits
Score difference with first non-orthologous sequence - *A. fumigatus*:756 *A. fischeri*:801
afum|EAL85267.1 100.00% afis|EAW20152.1 100.00%
Bootstrap support for afum|EAL85267.1 as seed ortholog is 100%.
Bootstrap support for afis|EAW20152.1 as seed ortholog is 100%.

---

Group of orthologs #5340. Best score 801 bits
Score difference with first non-orthologous sequence - *A. fumigatus*:801 *A. fischeri*:801
afum|EAL86985.1 100.00% afis|EAW19510.1 100.00%
Bootstrap support for afum|EAL86985.1 as seed ortholog is 100%.
Bootstrap support for afis|EAW19510.1 as seed ortholog is 100%.

---

Group of orthologs #5341. Best score 801 bits
Score difference with first non-orthologous sequence - *A. fumigatus*:391 *A. fischeri*:289
afum|EAL93843.2 100.00% afis|EAW19112.1 100.00%
Bootstrap support for afum|EAL93843.2 as seed ortholog is 100%.
Bootstrap support for afis|EAW19112.1 as seed ortholog is 100%.

---

Group of orthologs #5342. Best score 801 bits
Score difference with first non-orthologous sequence - *A. fumigatus*:801 *A. fischeri*:801
afum|EAL90759.1 100.00% afis|EAW22432.1 100.00%
Bootstrap support for afum|EAL90759.1 as seed ortholog is 100%.
Bootstrap support for afis|EAW22432.1 as seed ortholog is 100%.

---

Group of orthologs #5343. Best score 801 bits
Score difference with first non-orthologous sequence - *A. fumigatus*:801 *A. fischeri*:801
afum|EAL90549.2 100.00% afis|EAW22643.1 100.00%
Bootstrap support for afum|EAL90549.2 as seed ortholog is 100%.
Bootstrap support for afis|EAW22643.1 as seed ortholog is 100%.

---

Group of orthologs #5344. Best score 801 bits
Score difference with first non-orthologous sequence - *A. fumigatus*:801 *A. fischeri*:801
afum|EAL94063.1 100.00% afis|EAW19344.1 100.00%
Bootstrap support for afum|EAL94063.1 as seed ortholog is 100%.
Bootstrap support for afis|EAW19344.1 as seed ortholog is 100%.

Group of orthologs #5345. Best score 800 bits
Score difference with first non-orthologous sequence - *A. fumigatus*:800 *A. fischeri*:800
afum|EAL85247.1 100.00% afis|EAW20131.1 100.00%
Bootstrap support for afum|EAL85247.1 as seed ortholog is 100%.
Bootstrap support for afis|EAW20131.1 as seed ortholog is 100%.

Group of orthologs #5346. Best score 800 bits
Score difference with first non-orthologous sequence - *A. fumigatus*:675 *A. fischeri*:758
afum|EAL86197.1 100.00% afis|EAW24531.1 100.00%
Bootstrap support for afum|EAL86197.1 as seed ortholog is 100%.
Bootstrap support for afis|EAW24531.1 as seed ortholog is 100%.

Group of orthologs #5347. Best score 799 bits
Score difference with first non-orthologous sequence - *A. fumigatus*:799 *A. fischeri*:799
afum|EAL86449.1 100.00% afis|EAW16923.1 100.00%
Bootstrap support for afum|EAL86449.1 as seed ortholog is 100%.
Bootstrap support for afis|EAW16923.1 as seed ortholog is 100%.

Group of orthologs #5348. Best score 799 bits
Score difference with first non-orthologous sequence - *A. fumigatus*:799 *A. fischeri*:635
afum|EAL91482.1 100.00% afis|EAW17668.1 100.00%
Bootstrap support for afum|EAL91482.1 as seed ortholog is 100%.
Bootstrap support for afis|EAW17668.1 as seed ortholog is 100%.

Group of orthologs #5349. Best score 799 bits
Score difference with first non-orthologous sequence - *A. fumigatus*:799 *A. fischeri*:799
afum|EAL93117.1 100.00% afis|EAW18436.1 100.00%
Bootstrap support for afum|EAL93117.1 as seed ortholog is 100%.
Bootstrap support for afis|EAW18436.1 as seed ortholog is 100%.

Group of orthologs #5350. Best score 799 bits
Score difference with first non-orthologous sequence - *A. fumigatus*:746 *A. fischeri*:701
afum|EAL87921.2 100.00% afis|EAW23545.1 100.00%
Bootstrap support for afum|EAL87921.2 as seed ortholog is 100%.
Bootstrap support for afis|EAW23545.1 as seed ortholog is 100%.

Group of orthologs #5351. Best score 799 bits
Score difference with first non-orthologous sequence - *A. fumigatus*:799 *A. fischeri*:799
afum|EAL92503.1 100.00% afis|EAW21483.1 100.00%
Bootstrap support for afum|EAL92503.1 as seed ortholog is 100%.
Bootstrap support for afis|EAW21483.1 as seed ortholog is 100%.

Group of orthologs #5352. Best score 798 bits
Score difference with first non-orthologous sequence - *A. fumigatus*:647 *A. fischeri*:644
afum|EAL85702.1 100.00% afis|EAW15637.1 100.00%
Bootstrap support for afum|EAL85702.1 as seed ortholog is 100%.
Bootstrap support for afis|EAW15637.1 as seed ortholog is 100%.

Group of orthologs #5353. Best score 798 bits
Score difference with first non-orthologous sequence - *A. fumigatus*:798 *A. fischeri*:798
afum|EAL84810.1 100.00% afis|EAW20819.1 100.00%
Bootstrap support for afum|EAL84810.1 as seed ortholog is 100%.
Bootstrap support for afis|EAW20819.1 as seed ortholog is 100%.

Group of orthologs #5354. Best score 798 bits

Score difference with first non-orthologous sequence - *A. fumigatus*:798 *A. fischeri*:798
afum|EAL91491.1 100.00% afis|EAW17678.1 100.00%
Bootstrap support for afum|EAL91491.1 as seed ortholog is 100%.
Bootstrap support for afis|EAW17678.1 as seed ortholog is 100%.

---

Group of orthologs #5355. Best score 798 bits
Score difference with first non-orthologous sequence - *A. fumigatus*:696 *A. fischeri*:443
afum|EAL85893.1 100.00% afis|EAW24225.1 100.00%
Bootstrap support for afum|EAL85893.1 as seed ortholog is 100%.
Bootstrap support for afis|EAW24225.1 as seed ortholog is 100%.

---

Group of orthologs #5356. Best score 798 bits
Score difference with first non-orthologous sequence - *A. fumigatus*:798 *A. fischeri*:798
afum|EAL87419.1 100.00% afis|EAW23989.1 100.00%
Bootstrap support for afum|EAL87419.1 as seed ortholog is 100%.
Bootstrap support for afis|EAW23989.1 as seed ortholog is 100%.

---

Group of orthologs #5357. Best score 798 bits
Score difference with first non-orthologous sequence - *A. fumigatus*:751 *A. fischeri*:654
afum|EAL87879.1 100.00% afis|EAW23585.1 100.00%
Bootstrap support for afum|EAL87879.1 as seed ortholog is 100%.
Bootstrap support for afis|EAW23585.1 as seed ortholog is 100%.

---

Group of orthologs #5358. Best score 798 bits
Score difference with first non-orthologous sequence - *A. fumigatus*:798 *A. fischeri*:798
afum|EAL93570.1 100.00% afis|EAW18834.1 100.00%
Bootstrap support for afum|EAL93570.1 as seed ortholog is 100%.
Bootstrap support for afis|EAW18834.1 as seed ortholog is 100%.

---

Group of orthologs #5359. Best score 798 bits
Score difference with first non-orthologous sequence - *A. fumigatus*:798 *A. fischeri*:798
afum|EAL90763.1 100.00% afis|EAW22428.1 100.00%
Bootstrap support for afum|EAL90763.1 as seed ortholog is 100%.
Bootstrap support for afis|EAW22428.1 as seed ortholog is 100%.

---

Group of orthologs #5360. Best score 798 bits
Score difference with first non-orthologous sequence - *A. fumigatus*:798 *A. fischeri*:706
afum|EAL90465.1 100.00% afis|EAW22730.1 100.00%
Bootstrap support for afum|EAL90465.1 as seed ortholog is 100%.
Bootstrap support for afis|EAW22730.1 as seed ortholog is 100%.

---

Group of orthologs #5361. Best score 798 bits
Score difference with first non-orthologous sequence - *A. fumigatus*:798 *A. fischeri*:719
afum|EAL90441.1 100.00% afis|EAW22758.1 100.00%
Bootstrap support for afum|EAL90441.1 as seed ortholog is 100%.
Bootstrap support for afis|EAW22758.1 as seed ortholog is 100%.

---

Group of orthologs #5362. Best score 798 bits
Score difference with first non-orthologous sequence - *A. fumigatus*:798 *A. fischeri*:746
afum|EBA27507.1 100.00% afis|EAW19159.1 100.00%
Bootstrap support for afum|EBA27507.1 as seed ortholog is 100%.
Bootstrap support for afis|EAW19159.1 as seed ortholog is 100%.

---

Group of orthologs #5363. Best score 798 bits
Score difference with first non-orthologous sequence - *A. fumigatus*:798 *A. fischeri*:798
afum|EAL92642.1 100.00% afis|EAW21608.1 100.00%

Bootstrap support for afum|EAL92642.1 as seed ortholog is 100%.
Bootstrap support for afis|EAW21608.1 as seed ortholog is 100%.

---

Group of orthologs #5364. Best score 798 bits
Score difference with first non-orthologous sequence - *A. fumigatus*:680 *A. fischeri*:544
afum|EAL89619.1 100.00% afis|EAW24961.1 100.00%
Bootstrap support for afum|EAL89619.1 as seed ortholog is 100%.
Bootstrap support for afis|EAW24961.1 as seed ortholog is 100%.

---

Group of orthologs #5365. Best score 797 bits
Score difference with first non-orthologous sequence - *A. fumigatus*:797 *A. fischeri*:797
afum|EAL87588.1 100.00% afis|EAW18222.1 100.00%
Bootstrap support for afum|EAL87588.1 as seed ortholog is 100%.
Bootstrap support for afis|EAW18222.1 as seed ortholog is 100%.

---

Group of orthologs #5366. Best score 797 bits
Score difference with first non-orthologous sequence - *A. fumigatus*:797 *A. fischeri*:797
afum|EAL93603.1 100.00% afis|EAW18871.1 100.00%
Bootstrap support for afum|EAL93603.1 as seed ortholog is 100%.
Bootstrap support for afis|EAW18871.1 as seed ortholog is 100%.

---

Group of orthologs #5367. Best score 797 bits
Score difference with first non-orthologous sequence - *A. fumigatus*:797 *A. fischeri*:645
afum|EAL93957.1 100.00% afis|EAW19226.1 100.00%
Bootstrap support for afum|EAL93957.1 as seed ortholog is 100%.
Bootstrap support for afis|EAW19226.1 as seed ortholog is 100%.

---

Group of orthologs #5368. Best score 797 bits
Score difference with first non-orthologous sequence - *A. fumigatus*:797 *A. fischeri*:797
afum|EAL90978.1 100.00% afis|EAW22216.1 100.00%
Bootstrap support for afum|EAL90978.1 as seed ortholog is 100%.
Bootstrap support for afis|EAW22216.1 as seed ortholog is 100%.

---

Group of orthologs #5369. Best score 796 bits
Score difference with first non-orthologous sequence - *A. fumigatus*:796 *A. fischeri*:796
afum|EAL88687.1 100.00% afis|EAW16055.1 100.00%
Bootstrap support for afum|EAL88687.1 as seed ortholog is 100%.
Bootstrap support for afis|EAW16055.1 as seed ortholog is 100%.

---

Group of orthologs #5370. Best score 796 bits
Score difference with first non-orthologous sequence - *A. fumigatus*:796 *A. fischeri*:796
afum|EAL86107.1 100.00% afis|EAW24439.1 100.00%
Bootstrap support for afum|EAL86107.1 as seed ortholog is 100%.
Bootstrap support for afis|EAW24439.1 as seed ortholog is 100%.

---

Group of orthologs #5371. Best score 796 bits
Score difference with first non-orthologous sequence - *A. fumigatus*:796 *A. fischeri*:796
afum|EAL87130.1 100.00% afis|EAW23685.1 100.00%
Bootstrap support for afum|EAL87130.1 as seed ortholog is 100%.
Bootstrap support for afis|EAW23685.1 as seed ortholog is 100%.

---

Group of orthologs #5372. Best score 796 bits
Score difference with first non-orthologous sequence - *A. fumigatus*:796 *A. fischeri*:649
afum|EAL93988.1 100.00% afis|EAW19261.1 100.00%
Bootstrap support for afum|EAL93988.1 as seed ortholog is 100%.
Bootstrap support for afis|EAW19261.1 as seed ortholog is 100%.

Group of orthologs #5373. Best score 796 bits
Score difference with first non-orthologous sequence - *A. fumigatus*:796 *A. fischeri*:796
afum|EAL92606.1 100.00% afis|EAW21588.1 100.00%
Bootstrap support for afum|EAL92606.1 as seed ortholog is 100%.
Bootstrap support for afis|EAW21588.1 as seed ortholog is 100%.

Group of orthologs #5374. Best score 796 bits
Score difference with first non-orthologous sequence - *A. fumigatus*:796 *A. fischeri*:796
afum|EAL89608.1 100.00% afis|EAW24951.1 100.00%
Bootstrap support for afum|EAL89608.1 as seed ortholog is 100%.
Bootstrap support for afis|EAW24951.1 as seed ortholog is 100%.

Group of orthologs #5375. Best score 795 bits
Score difference with first non-orthologous sequence - *A. fumigatus*:795 *A. fischeri*:701
afum|EAL85743.1 100.00% afis|EAW15589.1 100.00%
Bootstrap support for afum|EAL85743.1 as seed ortholog is 100%.
Bootstrap support for afis|EAW15589.1 as seed ortholog is 100%.

Group of orthologs #5376. Best score 795 bits
Score difference with first non-orthologous sequence - *A. fumigatus*:795 *A. fischeri*:795
afum|EAL85614.1 100.00% afis|EAW15733.1 100.00%
Bootstrap support for afum|EAL85614.1 as seed ortholog is 100%.
Bootstrap support for afis|EAW15733.1 as seed ortholog is 100%.

Group of orthologs #5377. Best score 795 bits
Score difference with first non-orthologous sequence - *A. fumigatus*:795 *A. fischeri*:795
afum|EAL86698.1 100.00% afis|EAW17280.1 100.00%
Bootstrap support for afum|EAL86698.1 as seed ortholog is 100%.
Bootstrap support for afis|EAW17280.1 as seed ortholog is 100%.

Group of orthologs #5378. Best score 795 bits
Score difference with first non-orthologous sequence - *A. fumigatus*:795 *A. fischeri*:795
afum|EAL85207.1 100.00% afis|EAW20088.1 100.00%
Bootstrap support for afum|EAL85207.1 as seed ortholog is 100%.
Bootstrap support for afis|EAW20088.1 as seed ortholog is 100%.

Group of orthologs #5379. Best score 795 bits
Score difference with first non-orthologous sequence - *A. fumigatus*:722 *A. fischeri*:500
afum|EAL91146.1 100.00% afis|EAW17311.1 100.00%
Bootstrap support for afum|EAL91146.1 as seed ortholog is 100%.
Bootstrap support for afis|EAW17311.1 as seed ortholog is 100%.

Group of orthologs #5380. Best score 795 bits
Score difference with first non-orthologous sequence - *A. fumigatus*:795 *A. fischeri*:795
afum|EAL90959.1 100.00% afis|EAW22234.1 100.00%
Bootstrap support for afum|EAL90959.1 as seed ortholog is 100%.
Bootstrap support for afis|EAW22234.1 as seed ortholog is 100%.

Group of orthologs #5381. Best score 795 bits
Score difference with first non-orthologous sequence - *A. fumigatus*:795 *A. fischeri*:795
afum|EAL90533.1 100.00% afis|EAW22659.1 100.00%
Bootstrap support for afum|EAL90533.1 as seed ortholog is 100%.
Bootstrap support for afis|EAW22659.1 as seed ortholog is 100%.

Group of orthologs #5382. Best score 795 bits

Score difference with first non-orthologous sequence - *A. fumigatus*:795 *A. fischeri*:795
afum|EAL92798.1 100.00% afis|EAW21769.1 100.00%
Bootstrap support for afum|EAL92798.1 as seed ortholog is 100%.
Bootstrap support for afis|EAW21769.1 as seed ortholog is 100%.

---

Group of orthologs #5383. Best score 794 bits
Score difference with first non-orthologous sequence - *A. fumigatus*:794 *A. fischeri*:794
afum|EAL88677.1 100.00% afis|EAW16045.1 100.00%
Bootstrap support for afum|EAL88677.1 as seed ortholog is 100%.
Bootstrap support for afis|EAW16045.1 as seed ortholog is 100%.

---

Group of orthologs #5384. Best score 794 bits
Score difference with first non-orthologous sequence - *A. fumigatus*:794 *A. fischeri*:611
afum|EAL87085.1 100.00% afis|EAW19409.1 100.00%
Bootstrap support for afum|EAL87085.1 as seed ortholog is 100%.
Bootstrap support for afis|EAW19409.1 as seed ortholog is 100%.

---

Group of orthologs #5385. Best score 794 bits
Score difference with first non-orthologous sequence - *A. fumigatus*:794 *A. fischeri*:794
afum|EAL86898.1 100.00% afis|EAW19624.1 100.00%
Bootstrap support for afum|EAL86898.1 as seed ortholog is 100%.
Bootstrap support for afis|EAW19624.1 as seed ortholog is 100%.

---

Group of orthologs #5386. Best score 794 bits
Score difference with first non-orthologous sequence - *A. fumigatus*:794 *A. fischeri*:794
afum|EAL91337.1 100.00% afis|EAW17519.1 100.00%
Bootstrap support for afum|EAL91337.1 as seed ortholog is 100%.
Bootstrap support for afis|EAW17519.1 as seed ortholog is 100%.

---

Group of orthologs #5387. Best score 794 bits
Score difference with first non-orthologous sequence - *A. fumigatus*:750 *A. fischeri*:693
afum|EAL91580.1 100.00% afis|EAW17771.1 100.00%
Bootstrap support for afum|EAL91580.1 as seed ortholog is 100%.
Bootstrap support for afis|EAW17771.1 as seed ortholog is 100%.

---

Group of orthologs #5388. Best score 794 bits
Score difference with first non-orthologous sequence - *A. fumigatus*:794 *A. fischeri*:794
afum|EAL86154.1 100.00% afis|EAW24485.1 100.00%
Bootstrap support for afum|EAL86154.1 as seed ortholog is 100%.
Bootstrap support for afis|EAW24485.1 as seed ortholog is 100%.

---

Group of orthologs #5389. Best score 794 bits
Score difference with first non-orthologous sequence - *A. fumigatus*:794 *A. fischeri*:794
afum|EAL89722.1 100.00% afis|EAW25054.1 100.00%
Bootstrap support for afum|EAL89722.1 as seed ortholog is 100%.
Bootstrap support for afis|EAW25054.1 as seed ortholog is 100%.

---

Group of orthologs #5390. Best score 793 bits
Score difference with first non-orthologous sequence - *A. fumigatus*:793 *A. fischeri*:793
afum|EAL85716.1 100.00% afis|EAW15619.1 100.00%
Bootstrap support for afum|EAL85716.1 as seed ortholog is 100%.
Bootstrap support for afis|EAW15619.1 as seed ortholog is 100%.

---

Group of orthologs #5391. Best score 793 bits
Score difference with first non-orthologous sequence - *A. fumigatus*:793 *A. fischeri*:793
afum|EBA27355.1 100.00% afis|EAW16037.1 100.00%

Bootstrap support for afum|EBA27355.1 as seed ortholog is 100%.
Bootstrap support for afis|EAW16037.1 as seed ortholog is 100%.

---

Group of orthologs #5392. Best score 793 bits
Score difference with first non-orthologous sequence - *A. fumigatus*:793 *A. fischeri*:713
afum|EAL90772.1 100.00% afis|EAW22420.1 100.00%
Bootstrap support for afum|EAL90772.1 as seed ortholog is 100%.
Bootstrap support for afis|EAW22420.1 as seed ortholog is 100%.

---

Group of orthologs #5393. Best score 793 bits
Score difference with first non-orthologous sequence - *A. fumigatus*:793 *A. fischeri*:793
afum|EBA27347.1 100.00% afis|EAW23018.1 100.00%
Bootstrap support for afum|EBA27347.1 as seed ortholog is 100%.
Bootstrap support for afis|EAW23018.1 as seed ortholog is 100%.

---

Group of orthologs #5394. Best score 792 bits
Score difference with first non-orthologous sequence - *A. fumigatus*:792 *A. fischeri*:792
afum|EAL88610.1 100.00% afis|EAW15978.1 100.00%
Bootstrap support for afum|EAL88610.1 as seed ortholog is 100%.
Bootstrap support for afis|EAW15978.1 as seed ortholog is 100%.

---

Group of orthologs #5395. Best score 792 bits
Score difference with first non-orthologous sequence - *A. fumigatus*:728 *A. fischeri*:387
afum|EAL84822.1 100.00% afis|EAW20806.1 100.00%
Bootstrap support for afum|EAL84822.1 as seed ortholog is 100%.
Bootstrap support for afis|EAW20806.1 as seed ortholog is 100%.

---

Group of orthologs #5396. Best score 792 bits
Score difference with first non-orthologous sequence - *A. fumigatus*:792 *A. fischeri*:792
afum|EAL93512.1 100.00% afis|EAW18780.1 100.00%
Bootstrap support for afum|EAL93512.1 as seed ortholog is 100%.
Bootstrap support for afis|EAW18780.1 as seed ortholog is 100%.

---

Group of orthologs #5397. Best score 792 bits
Score difference with first non-orthologous sequence - *A. fumigatus*:792 *A. fischeri*:792
afum|EAL93855.1 100.00% afis|EAW19123.1 100.00%
Bootstrap support for afum|EAL93855.1 as seed ortholog is 100%.
Bootstrap support for afis|EAW19123.1 as seed ortholog is 100%.

---

Group of orthologs #5398. Best score 791 bits
Score difference with first non-orthologous sequence - *A. fumigatus*:791 *A. fischeri*:791
afum|EAL85537.1 100.00% afis|EAW15814.1 100.00%
Bootstrap support for afum|EAL85537.1 as seed ortholog is 100%.
Bootstrap support for afis|EAW15814.1 as seed ortholog is 100%.

---

Group of orthologs #5399. Best score 791 bits
Score difference with first non-orthologous sequence - *A. fumigatus*:791 *A. fischeri*:791
afum|EAL86517.1 100.00% afis|EAW16993.1 100.00%
Bootstrap support for afum|EAL86517.1 as seed ortholog is 100%.
Bootstrap support for afis|EAW16993.1 as seed ortholog is 100%.

---

Group of orthologs #5400. Best score 791 bits
Score difference with first non-orthologous sequence - *A. fumigatus*:791 *A. fischeri*:791
afum|EAL86727.1 100.00% afis|EAW17155.1 100.00%
Bootstrap support for afum|EAL86727.1 as seed ortholog is 100%.
Bootstrap support for afis|EAW17155.1 as seed ortholog is 100%.

Group of orthologs #5401. Best score 791 bits
Score difference with first non-orthologous sequence - *A. fumigatus*:791 *A. fischeri*:791
afum|EAL87715.1 100.00% afis|EAW18352.1 100.00%
Bootstrap support for afum|EAL87715.1 as seed ortholog is 100%.
Bootstrap support for afis|EAW18352.1 as seed ortholog is 100%.

Group of orthologs #5402. Best score 791 bits
Score difference with first non-orthologous sequence - *A. fumigatus*:715 *A. fischeri*:728
afum|EAL86175.1 100.00% afis|EAW24507.1 100.00%
Bootstrap support for afum|EAL86175.1 as seed ortholog is 100%.
Bootstrap support for afis|EAW24507.1 as seed ortholog is 100%.

Group of orthologs #5403. Best score 791 bits
Score difference with first non-orthologous sequence - *A. fumigatus*:791 *A. fischeri*:791
afum|EBA27425.1 100.00% afis|EAW17343.1 100.00%
Bootstrap support for afum|EBA27425.1 as seed ortholog is 100%.
Bootstrap support for afis|EAW17343.1 as seed ortholog is 100%.

Group of orthologs #5404. Best score 791 bits
Score difference with first non-orthologous sequence - *A. fumigatus*:791 *A. fischeri*:791
afum|EAL87919.1 100.00% afis|EAW23547.1 100.00%
Bootstrap support for afum|EAL87919.1 as seed ortholog is 100%.
Bootstrap support for afis|EAW23547.1 as seed ortholog is 100%.

Group of orthologs #5405. Best score 791 bits
Score difference with first non-orthologous sequence - *A. fumigatus*:791 *A. fischeri*:791
afum|EAL87895.2 100.00% afis|EAW23571.1 100.00%
Bootstrap support for afum|EAL87895.2 as seed ortholog is 100%.
Bootstrap support for afis|EAW23571.1 as seed ortholog is 100%.

Group of orthologs #5406. Best score 791 bits
Score difference with first non-orthologous sequence - *A. fumigatus*:791 *A. fischeri*:791
afum|EAL91034.1 100.00% afis|EAW22156.1 100.00%
Bootstrap support for afum|EAL91034.1 as seed ortholog is 100%.
Bootstrap support for afis|EAW22156.1 as seed ortholog is 100%.

Group of orthologs #5407. Best score 791 bits
Score difference with first non-orthologous sequence - *A. fumigatus*:791 *A. fischeri*:791
afum|EAL92241.2 100.00% afis|EAW21229.1 100.00%
Bootstrap support for afum|EAL92241.2 as seed ortholog is 100%.
Bootstrap support for afis|EAW21229.1 as seed ortholog is 100%.

Group of orthologs #5408. Best score 791 bits
Score difference with first non-orthologous sequence - *A. fumigatus*:791 *A. fischeri*:791
afum|EAL92510.1 100.00% afis|EAW21491.1 100.00%
Bootstrap support for afum|EAL92510.1 as seed ortholog is 100%.
Bootstrap support for afis|EAW21491.1 as seed ortholog is 100%.

Group of orthologs #5409. Best score 791 bits
Score difference with first non-orthologous sequence - *A. fumigatus*:791 *A. fischeri*:791
afum|EAL89878.1 100.00% afis|EAW25217.1 100.00%
Bootstrap support for afum|EAL89878.1 as seed ortholog is 100%.
Bootstrap support for afis|EAW25217.1 as seed ortholog is 100%.

Group of orthologs #5410. Best score 790 bits

Score difference with first non-orthologous sequence - *A. fumigatus*:790 *A. fischeri*:790
afum|EAL85192.1 100.00% afis|EAW20074.1 100.00%
Bootstrap support for afum|EAL85192.1 as seed ortholog is 100%.
Bootstrap support for afis|EAW20074.1 as seed ortholog is 100%.

---

Group of orthologs #5411. Best score 790 bits
Score difference with first non-orthologous sequence - *A. fumigatus*:790 *A. fischeri*:790
afum|EAL89195.1 100.00% afis|EAW16599.1 100.00%
Bootstrap support for afum|EAL89195.1 as seed ortholog is 100%.
Bootstrap support for afis|EAW16599.1 as seed ortholog is 100%.

---

Group of orthologs #5412. Best score 790 bits
Score difference with first non-orthologous sequence - *A. fumigatus*:723 *A. fischeri*:690
afum|EAL91389.1 100.00% afis|EAW17576.1 100.00%
Bootstrap support for afum|EAL91389.1 as seed ortholog is 100%.
Bootstrap support for afis|EAW17576.1 as seed ortholog is 100%.

---

Group of orthologs #5413. Best score 790 bits
Score difference with first non-orthologous sequence - *A. fumigatus*:790 *A. fischeri*:790
afum|EAL90312.1 100.00% afis|EAW22886.1 100.00%
Bootstrap support for afum|EAL90312.1 as seed ortholog is 100%.
Bootstrap support for afis|EAW22886.1 as seed ortholog is 100%.

---

Group of orthologs #5414. Best score 790 bits
Score difference with first non-orthologous sequence - *A. fumigatus*:790 *A. fischeri*:790
afum|EAL91981.1 100.00% afis|EAW24125.1 100.00%
Bootstrap support for afum|EAL91981.1 as seed ortholog is 100%.
Bootstrap support for afis|EAW24125.1 as seed ortholog is 100%.

---

Group of orthologs #5415. Best score 789 bits
Score difference with first non-orthologous sequence - *A. fumigatus*:789 *A. fischeri*:689
afum|EAL87541.1 100.00% afis|EAW18180.1 100.00%
Bootstrap support for afum|EAL87541.1 as seed ortholog is 100%.
Bootstrap support for afis|EAW18180.1 as seed ortholog is 100%.

---

Group of orthologs #5416. Best score 789 bits
Score difference with first non-orthologous sequence - *A. fumigatus*:789 *A. fischeri*:789
afum|EAL87619.1 100.00% afis|EAW18256.1 100.00%
Bootstrap support for afum|EAL87619.1 as seed ortholog is 100%.
Bootstrap support for afis|EAW18256.1 as seed ortholog is 100%.

---

Group of orthologs #5417. Best score 789 bits
Score difference with first non-orthologous sequence - *A. fumigatus*:789 *A. fischeri*:595
afum|EAL85444.1 100.00% afis|EAW20316.1 100.00%
Bootstrap support for afum|EAL85444.1 as seed ortholog is 100%.
Bootstrap support for afis|EAW20316.1 as seed ortholog is 100%.

---

Group of orthologs #5418. Best score 789 bits
Score difference with first non-orthologous sequence - *A. fumigatus*:719 *A. fischeri*:676
afum|EAL91468.1 100.00% afis|EAW17653.1 100.00%
Bootstrap support for afum|EAL91468.1 as seed ortholog is 100%.
Bootstrap support for afis|EAW17653.1 as seed ortholog is 100%.

---

Group of orthologs #5419. Best score 789 bits
Score difference with first non-orthologous sequence - *A. fumigatus*:789 *A. fischeri*:789
afum|EAL88287.1 100.00% afis|EAW23196.1 100.00%

Bootstrap support for afum|EAL88287.1 as seed ortholog is 100%.
Bootstrap support for afis|EAW23196.1 as seed ortholog is 100%.

---

Group of orthologs #5420. Best score 789 bits
Score difference with first non-orthologous sequence - *A. fumigatus*:789 *A. fischeri*:789
afum|EAL93243.1 100.00% afis|EAW18543.1 100.00%
Bootstrap support for afum|EAL93243.1 as seed ortholog is 100%.
Bootstrap support for afis|EAW18543.1 as seed ortholog is 100%.

---

Group of orthologs #5421. Best score 789 bits
Score difference with first non-orthologous sequence - *A. fumigatus*:789 *A. fischeri*:789
afum|EAL93710.1 100.00% afis|EAW18975.1 100.00%
Bootstrap support for afum|EAL93710.1 as seed ortholog is 100%.
Bootstrap support for afis|EAW18975.1 as seed ortholog is 100%.

---

Group of orthologs #5422. Best score 789 bits
Score difference with first non-orthologous sequence - *A. fumigatus*:789 *A. fischeri*:789
afum|EAL92716.2 100.00% afis|EAW21682.1 100.00%
Bootstrap support for afum|EAL92716.2 as seed ortholog is 100%.
Bootstrap support for afis|EAW21682.1 as seed ortholog is 100%.

---

Group of orthologs #5423. Best score 789 bits
Score difference with first non-orthologous sequence - *A. fumigatus*:789 *A. fischeri*:789
afum|EAL89961.1 100.00% afis|EAW25298.1 100.00%
Bootstrap support for afum|EAL89961.1 as seed ortholog is 100%.
Bootstrap support for afis|EAW25298.1 as seed ortholog is 100%.

---

Group of orthologs #5424. Best score 788 bits
Score difference with first non-orthologous sequence - *A. fumigatus*:714 *A. fischeri*:676
afum|EAL88635.2 100.00% afis|EAW16003.1 100.00%
Bootstrap support for afum|EAL88635.2 as seed ortholog is 100%.
Bootstrap support for afis|EAW16003.1 as seed ortholog is 100%.

---

Group of orthologs #5425. Best score 788 bits
Score difference with first non-orthologous sequence - *A. fumigatus*:788 *A. fischeri*:788
afum|EAL88647.1 100.00% afis|EAW16018.1 100.00%
Bootstrap support for afum|EAL88647.1 as seed ortholog is 100%.
Bootstrap support for afis|EAW16018.1 as seed ortholog is 100%.

---

Group of orthologs #5426. Best score 788 bits
Score difference with first non-orthologous sequence - *A. fumigatus*:788 *A. fischeri*:788
afum|EAL85276.1 100.00% afis|EAW20159.1 100.00%
Bootstrap support for afum|EAL85276.1 as seed ortholog is 100%.
Bootstrap support for afis|EAW20159.1 as seed ortholog is 100%.

---

Group of orthologs #5427. Best score 788 bits
Score difference with first non-orthologous sequence - *A. fumigatus*:788 *A. fischeri*:788
afum|EAL84858.1 100.00% afis|EAW20761.1 100.00%
Bootstrap support for afum|EAL84858.1 as seed ortholog is 100%.
Bootstrap support for afis|EAW20761.1 as seed ortholog is 100%.

---

Group of orthologs #5428. Best score 788 bits
Score difference with first non-orthologous sequence - *A. fumigatus*:788 *A. fischeri*:788
afum|EAL84293.1 100.00% afis|EAW25583.1 100.00%
Bootstrap support for afum|EAL84293.1 as seed ortholog is 100%.
Bootstrap support for afis|EAW25583.1 as seed ortholog is 100%.

---
Group of orthologs #5429. Best score 788 bits
Score difference with first non-orthologous sequence - *A. fumigatus*:788 *A. fischeri*:788
afum|EAL93902.1 100.00% afis|EAW19172.1 100.00%
Bootstrap support for afum|EAL93902.1 as seed ortholog is 100%.
Bootstrap support for afis|EAW19172.1 as seed ortholog is 100%.

---
Group of orthologs #5430. Best score 787 bits
Score difference with first non-orthologous sequence - *A. fumigatus*:787 *A. fischeri*:787
afum|EAL85840.1 100.00% afis|EAW15482.1 100.00%
Bootstrap support for afum|EAL85840.1 as seed ortholog is 100%.
Bootstrap support for afis|EAW15482.1 as seed ortholog is 100%.

---
Group of orthologs #5431. Best score 787 bits
Score difference with first non-orthologous sequence - *A. fumigatus*:787 *A. fischeri*:657
afum|EAL86510.1 100.00% afis|EAW16983.1 100.00%
Bootstrap support for afum|EAL86510.1 as seed ortholog is 100%.
Bootstrap support for afis|EAW16983.1 as seed ortholog is 100%.

---
Group of orthologs #5432. Best score 787 bits
Score difference with first non-orthologous sequence - *A. fumigatus*:787 *A. fischeri*:787
afum|EAL88056.1 100.00% afis|EAW23413.1 100.00%
Bootstrap support for afum|EAL88056.1 as seed ortholog is 100%.
Bootstrap support for afis|EAW23413.1 as seed ortholog is 100%.

---
Group of orthologs #5433. Best score 787 bits
Score difference with first non-orthologous sequence - *A. fumigatus*:787 *A. fischeri*:787
afum|EAL92438.1 100.00% afis|EAW21423.1 100.00%
Bootstrap support for afum|EAL92438.1 as seed ortholog is 100%.
Bootstrap support for afis|EAW21423.1 as seed ortholog is 100%.

---
Group of orthologs #5434. Best score 787 bits
Score difference with first non-orthologous sequence - *A. fumigatus*:787 *A. fischeri*:787
afum|EAL89434.1 100.00% afis|EAW24792.1 100.00%
Bootstrap support for afum|EAL89434.1 as seed ortholog is 100%.
Bootstrap support for afis|EAW24792.1 as seed ortholog is 100%.

---
Group of orthologs #5435. Best score 786 bits
Score difference with first non-orthologous sequence - *A. fumigatus*:786 *A. fischeri*:657
afum|EAL84236.1 100.00% afis|EAW15410.1 100.00%
Bootstrap support for afum|EAL84236.1 as seed ortholog is 100%.
Bootstrap support for afis|EAW15410.1 as seed ortholog is 100%.

---
Group of orthologs #5436. Best score 786 bits
Score difference with first non-orthologous sequence - *A. fumigatus*:738 *A. fischeri*:693
afum|EAL86403.2 100.00% afis|EAW16864.1 100.00%
Bootstrap support for afum|EAL86403.2 as seed ortholog is 100%.
Bootstrap support for afis|EAW16864.1 as seed ortholog is 100%.

---
Group of orthologs #5437. Best score 786 bits
Score difference with first non-orthologous sequence - *A. fumigatus*:363 *A. fischeri*:503
afum|EAL86497.1 100.00% afis|EAW16969.1 100.00%
Bootstrap support for afum|EAL86497.1 as seed ortholog is 100%.
Bootstrap support for afis|EAW16969.1 as seed ortholog is 100%.

---
Group of orthologs #5438. Best score 786 bits

Score difference with first non-orthologous sequence - *A. fumigatus*:786 *A. fischeri*:786
afum|EAL88517.1 100.00% afis|EAW15883.1 100.00%
Bootstrap support for afum|EAL88517.1 as seed ortholog is 100%.
Bootstrap support for afis|EAW15883.1 as seed ortholog is 100%.

---

Group of orthologs #5439. Best score 786 bits
Score difference with first non-orthologous sequence - *A. fumigatus*:786 *A. fischeri*:786
afum|EAL86826.1 100.00% afis|EAW19701.1 100.00%
Bootstrap support for afum|EAL86826.1 as seed ortholog is 100%.
Bootstrap support for afis|EAW19701.1 as seed ortholog is 100%.

---

Group of orthologs #5440. Best score 786 bits
Score difference with first non-orthologous sequence - *A. fumigatus*:786 *A. fischeri*:671
afum|EAL92116.1 100.00% afis|EAW21093.1 100.00%
Bootstrap support for afum|EAL92116.1 as seed ortholog is 100%.
Bootstrap support for afis|EAW21093.1 as seed ortholog is 100%.

---

Group of orthologs #5441. Best score 786 bits
Score difference with first non-orthologous sequence - *A. fumigatus*:786 *A. fischeri*:786
afum|EAL92369.1 100.00% afis|EAW21354.1 100.00%
Bootstrap support for afum|EAL92369.1 as seed ortholog is 100%.
Bootstrap support for afis|EAW21354.1 as seed ortholog is 100%.

---

Group of orthologs #5442. Best score 786 bits
Score difference with first non-orthologous sequence - *A. fumigatus*:646 *A. fischeri*:658
afum|EAL89665.1 100.00% afis|EAW25030.1 100.00%
Bootstrap support for afum|EAL89665.1 as seed ortholog is 100%.
Bootstrap support for afis|EAW25030.1 as seed ortholog is 100%.

---

Group of orthologs #5443. Best score 785 bits
Score difference with first non-orthologous sequence - *A. fumigatus*:732 *A. fischeri*:585
afum|EAL87553.1 100.00% afis|EAW18189.1 100.00%
Bootstrap support for afum|EAL87553.1 as seed ortholog is 100%.
Bootstrap support for afis|EAW18189.1 as seed ortholog is 100%.

---

Group of orthologs #5444. Best score 785 bits
Score difference with first non-orthologous sequence - *A. fumigatus*:785 *A. fischeri*:785
afum|EAL87164.1 100.00% afis|EAW23725.1 100.00%
Bootstrap support for afum|EAL87164.1 as seed ortholog is 100%.
Bootstrap support for afis|EAW23725.1 as seed ortholog is 100%.

---

Group of orthologs #5445. Best score 785 bits
Score difference with first non-orthologous sequence - *A. fumigatus*:704 *A. fischeri*:471
afum|EAL92392.1 100.00% afis|EAW21375.1 100.00%
Bootstrap support for afum|EAL92392.1 as seed ortholog is 100%.
Bootstrap support for afis|EAW21375.1 as seed ortholog is 100%.

---

Group of orthologs #5446. Best score 785 bits
Score difference with first non-orthologous sequence - *A. fumigatus*:785 *A. fischeri*:785
afum|EBA27223.1 100.00% afis|EAW20039.1 100.00%
Bootstrap support for afum|EBA27223.1 as seed ortholog is 100%.
Bootstrap support for afis|EAW20039.1 as seed ortholog is 100%.

---

Group of orthologs #5447. Best score 785 bits
Score difference with first non-orthologous sequence - *A. fumigatus*:785 *A. fischeri*:785
afum|EAL92692.1 100.00% afis|EAW21657.1 100.00%

Bootstrap support for afum|EAL92692.1 as seed ortholog is 100%.
Bootstrap support for afis|EAW21657.1 as seed ortholog is 100%.

---

Group of orthologs #5448. Best score 785 bits
Score difference with first non-orthologous sequence - *A. fumigatus*:785 *A. fischeri*:785
afum|EAL89488.1 100.00% afis|EAW24842.1 100.00%
Bootstrap support for afum|EAL89488.1 as seed ortholog is 100%.
Bootstrap support for afis|EAW24842.1 as seed ortholog is 100%.

---

Group of orthologs #5449. Best score 784 bits
Score difference with first non-orthologous sequence - *A. fumigatus*:784 *A. fischeri*:784
afum|EAL85146.1 100.00% afis|EAW19751.1 100.00%
Bootstrap support for afum|EAL85146.1 as seed ortholog is 100%.
Bootstrap support for afis|EAW19751.1 as seed ortholog is 100%.

---

Group of orthologs #5450. Best score 784 bits
Score difference with first non-orthologous sequence - *A. fumigatus*:784 *A. fischeri*:512
afum|EAL91435.1 100.00% afis|EAW17622.1 100.00%
Bootstrap support for afum|EAL91435.1 as seed ortholog is 100%.
Bootstrap support for afis|EAW17622.1 as seed ortholog is 100%.

---

Group of orthologs #5451. Best score 784 bits
Score difference with first non-orthologous sequence - *A. fumigatus*:784 *A. fischeri*:784
afum|EAL91062.1 100.00% afis|EAW22126.1 100.00%
Bootstrap support for afum|EAL91062.1 as seed ortholog is 100%.
Bootstrap support for afis|EAW22126.1 as seed ortholog is 100%.

---

Group of orthologs #5452. Best score 784 bits
Score difference with first non-orthologous sequence - *A. fumigatus*:784 *A. fischeri*:784
afum|EAL92180.1 100.00% afis|EAW21161.1 100.00%
Bootstrap support for afum|EAL92180.1 as seed ortholog is 100%.
Bootstrap support for afis|EAW21161.1 as seed ortholog is 100%.

---

Group of orthologs #5453. Best score 783 bits
Score difference with first non-orthologous sequence - *A. fumigatus*:783 *A. fischeri*:783
afum|EAL86354.1 100.00% afis|EAW16783.1 100.00%
Bootstrap support for afum|EAL86354.1 as seed ortholog is 100%.
Bootstrap support for afis|EAW16783.1 as seed ortholog is 100%.

---

Group of orthologs #5454. Best score 783 bits
Score difference with first non-orthologous sequence - *A. fumigatus*:783 *A. fischeri*:570
afum|EAL85012.1 100.00% afis|EAW19885.1 100.00%
Bootstrap support for afum|EAL85012.1 as seed ortholog is 100%.
Bootstrap support for afis|EAW19885.1 as seed ortholog is 100%.

---

Group of orthologs #5455. Best score 783 bits
Score difference with first non-orthologous sequence - *A. fumigatus*:783 *A. fischeri*:783
afum|EAL84893.1 100.00% afis|EAW20024.1 100.00%
Bootstrap support for afum|EAL84893.1 as seed ortholog is 100%.
Bootstrap support for afis|EAW20024.1 as seed ortholog is 100%.

---

Group of orthologs #5456. Best score 783 bits
Score difference with first non-orthologous sequence - *A. fumigatus*:783 *A. fischeri*:783
afum|EAL84605.1 100.00% afis|EAW20447.1 100.00%
Bootstrap support for afum|EAL84605.1 as seed ortholog is 100%.
Bootstrap support for afis|EAW20447.1 as seed ortholog is 100%.

---

Group of orthologs #5457. Best score 783 bits
Score difference with first non-orthologous sequence - *A. fumigatus*:783 *A. fischeri*:783

| | | | |
|---|---|---|---|
| afum\|EAL91707.1 | 100.00% | afis\|EAW17899.1 | 100.00% |

Bootstrap support for afum|EAL91707.1 as seed ortholog is 100%.
Bootstrap support for afis|EAW17899.1 as seed ortholog is 100%.

---

Group of orthologs #5458. Best score 783 bits
Score difference with first non-orthologous sequence - *A. fumigatus*:783 *A. fischeri*:783

| | | | |
|---|---|---|---|
| afum\|EAL85921.2 | 100.00% | afis\|EAW24252.1 | 100.00% |

Bootstrap support for afum|EAL85921.2 as seed ortholog is 100%.
Bootstrap support for afis|EAW24252.1 as seed ortholog is 100%.

---

Group of orthologs #5459. Best score 783 bits
Score difference with first non-orthologous sequence - *A. fumigatus*:783 *A. fischeri*:519

| | | | |
|---|---|---|---|
| afum\|EAL88145.1 | 100.00% | afis\|EAW23333.1 | 100.00% |

Bootstrap support for afum|EAL88145.1 as seed ortholog is 100%.
Bootstrap support for afis|EAW23333.1 as seed ortholog is 100%.

---

Group of orthologs #5460. Best score 783 bits
Score difference with first non-orthologous sequence - *A. fumigatus*:783 *A. fischeri*:783

| | | | |
|---|---|---|---|
| afum\|EAL93740.1 | 100.00% | afis\|EAW19006.1 | 100.00% |

Bootstrap support for afum|EAL93740.1 as seed ortholog is 100%.
Bootstrap support for afis|EAW19006.1 as seed ortholog is 100%.

---

Group of orthologs #5461. Best score 783 bits
Score difference with first non-orthologous sequence - *A. fumigatus*:783 *A. fischeri*:783

| | | | |
|---|---|---|---|
| afum\|EAL91005.1 | 100.00% | afis\|EAW22184.1 | 100.00% |

Bootstrap support for afum|EAL91005.1 as seed ortholog is 100%.
Bootstrap support for afis|EAW22184.1 as seed ortholog is 100%.

---

Group of orthologs #5462. Best score 783 bits
Score difference with first non-orthologous sequence - *A. fumigatus*:783 *A. fischeri*:783

| | | | |
|---|---|---|---|
| afum\|EAL92394.1 | 100.00% | afis\|EAW21379.1 | 100.00% |

Bootstrap support for afum|EAL92394.1 as seed ortholog is 100%.
Bootstrap support for afis|EAW21379.1 as seed ortholog is 100%.

---

Group of orthologs #5463. Best score 783 bits
Score difference with first non-orthologous sequence - *A. fumigatus*:783 *A. fischeri*:783

| | | | |
|---|---|---|---|
| afum\|EAL90165.1 | 100.00% | afis\|EAW25499.1 | 100.00% |

Bootstrap support for afum|EAL90165.1 as seed ortholog is 100%.
Bootstrap support for afis|EAW25499.1 as seed ortholog is 100%.

---

Group of orthologs #5464. Best score 782 bits
Score difference with first non-orthologous sequence - *A. fumigatus*:782 *A. fischeri*:782

| | | | |
|---|---|---|---|
| afum\|EAL85599.1 | 100.00% | afis\|EAW15750.1 | 100.00% |

Bootstrap support for afum|EAL85599.1 as seed ortholog is 100%.
Bootstrap support for afis|EAW15750.1 as seed ortholog is 100%.

---

Group of orthologs #5465. Best score 782 bits
Score difference with first non-orthologous sequence - *A. fumigatus*:527 *A. fischeri*:507

| | | | |
|---|---|---|---|
| afum\|EAL88863.1 | 100.00% | afis\|EAW16241.1 | 100.00% |

Bootstrap support for afum|EAL88863.1 as seed ortholog is 100%.
Bootstrap support for afis|EAW16241.1 as seed ortholog is 100%.

---

Group of orthologs #5466. Best score 782 bits

Score difference with first non-orthologous sequence - *A. fumigatus*:653 *A. fischeri*:653
afum|EAL91199.1 100.00% afis|EAW17362.1 100.00%
Bootstrap support for afum|EAL91199.1 as seed ortholog is 100%.
Bootstrap support for afis|EAW17362.1 as seed ortholog is 100%.

---

Group of orthologs #5467. Best score 782 bits
Score difference with first non-orthologous sequence - *A. fumigatus*:782 *A. fischeri*:782
afum|EAL88164.1 100.00% afis|EAW23313.1 100.00%
Bootstrap support for afum|EAL88164.1 as seed ortholog is 100%.
Bootstrap support for afis|EAW23313.1 as seed ortholog is 100%.

---

Group of orthologs #5468. Best score 782 bits
Score difference with first non-orthologous sequence - *A. fumigatus*:782 *A. fischeri*:782
afum|EAL93660.1 100.00% afis|EAW18929.1 100.00%
Bootstrap support for afum|EAL93660.1 as seed ortholog is 100%.
Bootstrap support for afis|EAW18929.1 as seed ortholog is 100%.

---

Group of orthologs #5469. Best score 782 bits
Score difference with first non-orthologous sequence - *A. fumigatus*:782 *A. fischeri*:782
afum|EAL93869.1 100.00% afis|EAW19137.1 100.00%
Bootstrap support for afum|EAL93869.1 as seed ortholog is 100%.
Bootstrap support for afis|EAW19137.1 as seed ortholog is 100%.

---

Group of orthologs #5470. Best score 782 bits
Score difference with first non-orthologous sequence - *A. fumigatus*:782 *A. fischeri*:782
afum|EAL90426.1 100.00% afis|EAW22772.1 100.00%
Bootstrap support for afum|EAL90426.1 as seed ortholog is 100%.
Bootstrap support for afis|EAW22772.1 as seed ortholog is 100%.

---

Group of orthologs #5471. Best score 782 bits
Score difference with first non-orthologous sequence - *A. fumigatus*:631 *A. fischeri*:782
afum|EAL89476.1 100.00% afis|EAW24831.1 100.00%
Bootstrap support for afum|EAL89476.1 as seed ortholog is 100%.
Bootstrap support for afis|EAW24831.1 as seed ortholog is 100%.

---

Group of orthologs #5472. Best score 781 bits
Score difference with first non-orthologous sequence - *A. fumigatus*:781 *A. fischeri*:781
afum|EAL88284.1 100.00% afis|EAW23199.1 100.00%
Bootstrap support for afum|EAL88284.1 as seed ortholog is 100%.
Bootstrap support for afis|EAW23199.1 as seed ortholog is 100%.

---

Group of orthologs #5473. Best score 781 bits
Score difference with first non-orthologous sequence - *A. fumigatus*:741 *A. fischeri*:576
afum|EAL90925.1 100.00% afis|EAW22249.1 100.00%
Bootstrap support for afum|EAL90925.1 as seed ortholog is 100%.
Bootstrap support for afis|EAW22249.1 as seed ortholog is 100%.

---

Group of orthologs #5474. Best score 781 bits
Score difference with first non-orthologous sequence - *A. fumigatus*:781 *A. fischeri*:781
afum|EAL90317.1 100.00% afis|EAW22882.1 100.00%
Bootstrap support for afum|EAL90317.1 as seed ortholog is 100%.
Bootstrap support for afis|EAW22882.1 as seed ortholog is 100%.

---

Group of orthologs #5475. Best score 780 bits
Score difference with first non-orthologous sequence - *A. fumigatus*:780 *A. fischeri*:780
afum|EAL88559.1 100.00% afis|EAW15922.1 100.00%

Bootstrap support for afum|EAL88559.1 as seed ortholog is 100%.
Bootstrap support for afis|EAW15922.1 as seed ortholog is 100%.

---

Group of orthologs #5476. Best score 780 bits
Score difference with first non-orthologous sequence - *A. fumigatus*:780 *A. fischeri*:780
afum|EAL84961.1 100.00% afis|EAW19954.1 100.00%
Bootstrap support for afum|EAL84961.1 as seed ortholog is 100%.
Bootstrap support for afis|EAW19954.1 as seed ortholog is 100%.

---

Group of orthologs #5477. Best score 780 bits
Score difference with first non-orthologous sequence - *A. fumigatus*:780 *A. fischeri*:293
afum|EAL84928.1 100.00% afis|EAW19987.1 100.00%
Bootstrap support for afum|EAL84928.1 as seed ortholog is 100%.
Bootstrap support for afis|EAW19987.1 as seed ortholog is 100%.

---

Group of orthologs #5478. Best score 780 bits
Score difference with first non-orthologous sequence - *A. fumigatus*:677 *A. fischeri*:655
afum|EAL87140.1 100.00% afis|EAW23699.1 100.00%
Bootstrap support for afum|EAL87140.1 as seed ortholog is 100%.
Bootstrap support for afis|EAW23699.1 as seed ortholog is 100%.

---

Group of orthologs #5479. Best score 780 bits
Score difference with first non-orthologous sequence - *A. fumigatus*:780 *A. fischeri*:780
afum|EAL92229.1 100.00% afis|EAW21217.1 100.00%
Bootstrap support for afum|EAL92229.1 as seed ortholog is 100%.
Bootstrap support for afis|EAW21217.1 as seed ortholog is 100%.

---

Group of orthologs #5480. Best score 780 bits
Score difference with first non-orthologous sequence - *A. fumigatus*:701 *A. fischeri*:700
afum|EAL92300.2 100.00% afis|EAW21283.1 100.00%
Bootstrap support for afum|EAL92300.2 as seed ortholog is 100%.
Bootstrap support for afis|EAW21283.1 as seed ortholog is 100%.

---

Group of orthologs #5481. Best score 779 bits
Score difference with first non-orthologous sequence - *A. fumigatus*:779 *A. fischeri*:779
afum|EAL86524.1 100.00% afis|EAW16999.1 100.00%
Bootstrap support for afum|EAL86524.1 as seed ortholog is 100%.
Bootstrap support for afis|EAW16999.1 as seed ortholog is 100%.

---

Group of orthologs #5482. Best score 779 bits
Score difference with first non-orthologous sequence - *A. fumigatus*:779 *A. fischeri*:779
afum|EAL88905.1 100.00% afis|EAW16284.1 100.00%
Bootstrap support for afum|EAL88905.1 as seed ortholog is 100%.
Bootstrap support for afis|EAW16284.1 as seed ortholog is 100%.

---

Group of orthologs #5483. Best score 779 bits
Score difference with first non-orthologous sequence - *A. fumigatus*:779 *A. fischeri*:779
afum|EAL87654.2 100.00% afis|EAW18288.1 100.00%
Bootstrap support for afum|EAL87654.2 as seed ortholog is 100%.
Bootstrap support for afis|EAW18288.1 as seed ortholog is 100%.

---

Group of orthologs #5484. Best score 779 bits
Score difference with first non-orthologous sequence - *A. fumigatus*:779 *A. fischeri*:779
afum|EAL86945.1 100.00% afis|EAW19572.1 100.00%
Bootstrap support for afum|EAL86945.1 as seed ortholog is 100%.
Bootstrap support for afis|EAW19572.1 as seed ortholog is 100%.

Group of orthologs #5485. Best score 779 bits
Score difference with first non-orthologous sequence - *A. fumigatus*:779 *A. fischeri*:701
afum|EAL87875.1 100.00% afis|EAW23591.1 100.00%
Bootstrap support for afum|EAL87875.1 as seed ortholog is 100%.
Bootstrap support for afis|EAW23591.1 as seed ortholog is 100%.

Group of orthologs #5486. Best score 779 bits
Score difference with first non-orthologous sequence - *A. fumigatus*:779 *A. fischeri*:570
afum|EAL92524.1 100.00% afis|EAW21505.1 100.00%
Bootstrap support for afum|EAL92524.1 as seed ortholog is 100%.
Bootstrap support for afis|EAW21505.1 as seed ortholog is 100%.

Group of orthologs #5487. Best score 779 bits
Score difference with first non-orthologous sequence - *A. fumigatus*:668 *A. fischeri*:648
afum|EAL89621.1 100.00% afis|EAW24963.1 100.00%
Bootstrap support for afum|EAL89621.1 as seed ortholog is 100%.
Bootstrap support for afis|EAW24963.1 as seed ortholog is 100%.

Group of orthologs #5488. Best score 779 bits
Score difference with first non-orthologous sequence - *A. fumigatus*:779 *A. fischeri*:779
afum|EAL92941.1 100.00% afis|EAW21928.1 100.00%
Bootstrap support for afum|EAL92941.1 as seed ortholog is 100%.
Bootstrap support for afis|EAW21928.1 as seed ortholog is 100%.

Group of orthologs #5489. Best score 779 bits
Score difference with first non-orthologous sequence - *A. fumigatus*:779 *A. fischeri*:779
afum|EAL92991.1 100.00% afis|EAW21976.1 100.00%
Bootstrap support for afum|EAL92991.1 as seed ortholog is 100%.
Bootstrap support for afis|EAW21976.1 as seed ortholog is 100%.

Group of orthologs #5490. Best score 778 bits
Score difference with first non-orthologous sequence - *A. fumigatus*:778 *A. fischeri*:778
afum|EAL85032.1 100.00% afis|EAW19863.1 100.00%
Bootstrap support for afum|EAL85032.1 as seed ortholog is 100%.
Bootstrap support for afis|EAW19863.1 as seed ortholog is 100%.

Group of orthologs #5491. Best score 778 bits
Score difference with first non-orthologous sequence - *A. fumigatus*:778 *A. fischeri*:778
afum|EAL88952.1 100.00% afis|EAW16328.1 100.00%
Bootstrap support for afum|EAL88952.1 as seed ortholog is 100%.
Bootstrap support for afis|EAW16328.1 as seed ortholog is 100%.

Group of orthologs #5492. Best score 778 bits
Score difference with first non-orthologous sequence - *A. fumigatus*:778 *A. fischeri*:778
afum|EAL91687.1 100.00% afis|EAW17879.1 100.00%
Bootstrap support for afum|EAL91687.1 as seed ortholog is 100%.
Bootstrap support for afis|EAW17879.1 as seed ortholog is 100%.

Group of orthologs #5493. Best score 778 bits
Score difference with first non-orthologous sequence - *A. fumigatus*:778 *A. fischeri*:532
afum|EAL84350.1 100.00% afis|EAW25680.1 100.00%
Bootstrap support for afum|EAL84350.1 as seed ortholog is 100%.
Bootstrap support for afis|EAW25680.1 as seed ortholog is 100%.

Group of orthologs #5494. Best score 778 bits

Score difference with first non-orthologous sequence - *A. fumigatus*:778 *A. fischeri*:778
afum|EAL90071.1 100.00% afis|EAW25412.1 100.00%
Bootstrap support for afum|EAL90071.1 as seed ortholog is 100%.
Bootstrap support for afis|EAW25412.1 as seed ortholog is 100%.

---

Group of orthologs #5495. Best score 777 bits
Score difference with first non-orthologous sequence - *A. fumigatus*:350 *A. fischeri*:420
afum|EAL84639.1 100.00% afis|EAW20990.1 100.00%
Bootstrap support for afum|EAL84639.1 as seed ortholog is 100%.
Bootstrap support for afis|EAW20990.1 as seed ortholog is 100%.

---

Group of orthologs #5496. Best score 777 bits
Score difference with first non-orthologous sequence - *A. fumigatus*:777 *A. fischeri*:777
afum|EAL87649.1 100.00% afis|EAW18283.1 100.00%
Bootstrap support for afum|EAL87649.1 as seed ortholog is 100%.
Bootstrap support for afis|EAW18283.1 as seed ortholog is 100%.

---

Group of orthologs #5497. Best score 777 bits
Score difference with first non-orthologous sequence - *A. fumigatus*:777 *A. fischeri*:777
afum|EAL88199.1 100.00% afis|EAW23280.1 100.00%
Bootstrap support for afum|EAL88199.1 as seed ortholog is 100%.
Bootstrap support for afis|EAW23280.1 as seed ortholog is 100%.

---

Group of orthologs #5498. Best score 777 bits
Score difference with first non-orthologous sequence - *A. fumigatus*:777 *A. fischeri*:777
afum|EAL93318.1 100.00% afis|EAW18608.1 100.00%
Bootstrap support for afum|EAL93318.1 as seed ortholog is 100%.
Bootstrap support for afis|EAW18608.1 as seed ortholog is 100%.

---

Group of orthologs #5499. Best score 777 bits
Score difference with first non-orthologous sequence - *A. fumigatus*:777 *A. fischeri*:777
afum|EAL90550.2 100.00% afis|EAW22642.1 100.00%
Bootstrap support for afum|EAL90550.2 as seed ortholog is 100%.
Bootstrap support for afis|EAW22642.1 as seed ortholog is 100%.

---

Group of orthologs #5500. Best score 777 bits
Score difference with first non-orthologous sequence - *A. fumigatus*:777 *A. fischeri*:777
afum|EBA27455.1 100.00% afis|EAW21225.1 100.00%
Bootstrap support for afum|EBA27455.1 as seed ortholog is 100%.
Bootstrap support for afis|EAW21225.1 as seed ortholog is 100%.

---

Group of orthologs #5501. Best score 776 bits
Score difference with first non-orthologous sequence - *A. fumigatus*:776 *A. fischeri*:776
afum|EAL89246.1 100.00% afis|EAW16643.1 100.00%
Bootstrap support for afum|EAL89246.1 as seed ortholog is 100%.
Bootstrap support for afis|EAW16643.1 as seed ortholog is 100%.

---

Group of orthologs #5502. Best score 776 bits
Score difference with first non-orthologous sequence - *A. fumigatus*:776 *A. fischeri*:776
afum|EAL91499.1 100.00% afis|EAW17685.1 100.00%
Bootstrap support for afum|EAL91499.1 as seed ortholog is 100%.
Bootstrap support for afis|EAW17685.1 as seed ortholog is 100%.

---

Group of orthologs #5503. Best score 776 bits
Score difference with first non-orthologous sequence - *A. fumigatus*:776 *A. fischeri*:776
afum|EAL86200.1 100.00% afis|EAW24535.1 100.00%

Bootstrap support for afum|EAL86200.1 as seed ortholog is 100%.
Bootstrap support for afis|EAW24535.1 as seed ortholog is 100%.

---

Group of orthologs #5504. Best score 776 bits
Score difference with first non-orthologous sequence - *A. fumigatus*:776 *A. fischeri*:776
afum|EAL88002.1 100.00% afis|EAW23464.1 100.00%
Bootstrap support for afum|EAL88002.1 as seed ortholog is 100%.
Bootstrap support for afis|EAW23464.1 as seed ortholog is 100%.

---

Group of orthologs #5505. Best score 776 bits
Score difference with first non-orthologous sequence - *A. fumigatus*:776 *A. fischeri*:776
afum|EAL93653.1 100.00% afis|EAW18922.1 100.00%
Bootstrap support for afum|EAL93653.1 as seed ortholog is 100%.
Bootstrap support for afis|EAW18922.1 as seed ortholog is 100%.

---

Group of orthologs #5506. Best score 776 bits
Score difference with first non-orthologous sequence - *A. fumigatus*:776 *A. fischeri*:776
afum|EAL93979.1 100.00% afis|EAW19250.1 100.00%
Bootstrap support for afum|EAL93979.1 as seed ortholog is 100%.
Bootstrap support for afis|EAW19250.1 as seed ortholog is 100%.

---

Group of orthologs #5507. Best score 776 bits
Score difference with first non-orthologous sequence - *A. fumigatus*:776 *A. fischeri*:776
afum|EAL92427.1 100.00% afis|EAW21413.1 100.00%
Bootstrap support for afum|EAL92427.1 as seed ortholog is 100%.
Bootstrap support for afis|EAW21413.1 as seed ortholog is 100%.

---

Group of orthologs #5508. Best score 776 bits
Score difference with first non-orthologous sequence - *A. fumigatus*:776 *A. fischeri*:776
afum|EAL89705.1 100.00% afis|EAW24996.1 100.00%
Bootstrap support for afum|EAL89705.1 as seed ortholog is 100%.
Bootstrap support for afis|EAW24996.1 as seed ortholog is 100%.

---

Group of orthologs #5509. Best score 775 bits
Score difference with first non-orthologous sequence - *A. fumigatus*:653 *A. fischeri*:552
afum|EAL84395.2 100.00% afis|EAW25721.1 100.00%
Bootstrap support for afum|EAL84395.2 as seed ortholog is 100%.
Bootstrap support for afis|EAW25721.1 as seed ortholog is 100%.

---

Group of orthologs #5510. Best score 775 bits
Score difference with first non-orthologous sequence - *A. fumigatus*:775 *A. fischeri*:638
afum|EAL90692.1 100.00% afis|EAW22501.1 100.00%
Bootstrap support for afum|EAL90692.1 as seed ortholog is 100%.
Bootstrap support for afis|EAW22501.1 as seed ortholog is 100%.

---

Group of orthologs #5511. Best score 775 bits
Score difference with first non-orthologous sequence - *A. fumigatus*:775 *A. fischeri*:775
afum|EAL92728.1 100.00% afis|EAW21694.1 100.00%
Bootstrap support for afum|EAL92728.1 as seed ortholog is 100%.
Bootstrap support for afis|EAW21694.1 as seed ortholog is 100%.

---

Group of orthologs #5512. Best score 774 bits
Score difference with first non-orthologous sequence - *A. fumigatus*:774 *A. fischeri*:774
afum|EAL84189.1 100.00% afis|EAW15456.1 100.00%
Bootstrap support for afum|EAL84189.1 as seed ortholog is 100%.
Bootstrap support for afis|EAW15456.1 as seed ortholog is 100%.

---

Group of orthologs #5513. Best score 774 bits
Score difference with first non-orthologous sequence - *A. fumigatus*:774 *A. fischeri*:774
afum|EAL85559.1 100.00% afis|EAW15791.1 100.00%
Bootstrap support for afum|EAL85559.1 as seed ortholog is 100%.
Bootstrap support for afis|EAW15791.1 as seed ortholog is 100%.

---

Group of orthologs #5514. Best score 774 bits
Score difference with first non-orthologous sequence - *A. fumigatus*:774 *A. fischeri*:774
afum|EAL88878.1 100.00% afis|EAW16257.1 100.00%
Bootstrap support for afum|EAL88878.1 as seed ortholog is 100%.
Bootstrap support for afis|EAW16257.1 as seed ortholog is 100%.

---

Group of orthologs #5515. Best score 774 bits
Score difference with first non-orthologous sequence - *A. fumigatus*:774 *A. fischeri*:774
afum|EAL84377.1 100.00% afis|EAW25704.1 100.00%
Bootstrap support for afum|EAL84377.1 as seed ortholog is 100%.
Bootstrap support for afis|EAW25704.1 as seed ortholog is 100%.

---

Group of orthologs #5516. Best score 773 bits
Score difference with first non-orthologous sequence - *A. fumigatus*:773 *A. fischeri*:773
afum|EAL86587.1 100.00% afis|EAW17127.1 100.00%
Bootstrap support for afum|EAL86587.1 as seed ortholog is 100%.
Bootstrap support for afis|EAW17127.1 as seed ortholog is 100%.

---

Group of orthologs #5517. Best score 773 bits
Score difference with first non-orthologous sequence - *A. fumigatus*:773 *A. fischeri*:488
afum|EAL88809.2 100.00% afis|EAW16176.1 100.00%
Bootstrap support for afum|EAL88809.2 as seed ortholog is 100%.
Bootstrap support for afis|EAW16176.1 as seed ortholog is 100%.

---

Group of orthologs #5518. Best score 773 bits
Score difference with first non-orthologous sequence - *A. fumigatus*:620 *A. fischeri*:773
afum|EAL84938.1 100.00% afis|EAW19972.1 100.00%
Bootstrap support for afum|EAL84938.1 as seed ortholog is 100%.
Bootstrap support for afis|EAW19972.1 as seed ortholog is 100%.

---

Group of orthologs #5519. Best score 773 bits
Score difference with first non-orthologous sequence - *A. fumigatus*:773 *A. fischeri*:773
afum|EAL91330.1 100.00% afis|EAW17511.1 100.00%
Bootstrap support for afum|EAL91330.1 as seed ortholog is 100%.
Bootstrap support for afis|EAW17511.1 as seed ortholog is 100%.

---

Group of orthologs #5520. Best score 773 bits
Score difference with first non-orthologous sequence - *A. fumigatus*:620 *A. fischeri*:773
afum|EAL87152.1 100.00% afis|EAW23708.1 100.00%
Bootstrap support for afum|EAL87152.1 as seed ortholog is 100%.
Bootstrap support for afis|EAW23708.1 as seed ortholog is 100%.

---

Group of orthologs #5521. Best score 773 bits
Score difference with first non-orthologous sequence - *A. fumigatus*:773 *A. fischeri*:773
afum|EAL88326.1 100.00% afis|EAW23157.1 100.00%
Bootstrap support for afum|EAL88326.1 as seed ortholog is 100%.
Bootstrap support for afis|EAW23157.1 as seed ortholog is 100%.

---

Group of orthologs #5522. Best score 773 bits

Score difference with first non-orthologous sequence - *A. fumigatus*:773   *A. fischeri*:773
afum|EAL93815.1   100.00%   afis|EAW19085.1   100.00%
Bootstrap support for afum|EAL93815.1 as seed ortholog is 100%.
Bootstrap support for afis|EAW19085.1 as seed ortholog is 100%.

---

Group of orthologs #5523. Best score 773 bits
Score difference with first non-orthologous sequence - *A. fumigatus*:773   *A. fischeri*:773
afum|EAL93822.1   100.00%   afis|EAW19092.1   100.00%
Bootstrap support for afum|EAL93822.1 as seed ortholog is 100%.
Bootstrap support for afis|EAW19092.1 as seed ortholog is 100%.

---

Group of orthologs #5524. Best score 773 bits
Score difference with first non-orthologous sequence - *A. fumigatus*:773   *A. fischeri*:773
afum|EAL90488.1   100.00%   afis|EAW22704.1   100.00%
Bootstrap support for afum|EAL90488.1 as seed ortholog is 100%.
Bootstrap support for afis|EAW22704.1 as seed ortholog is 100%.

---

Group of orthologs #5525. Best score 773 bits
Score difference with first non-orthologous sequence - *A. fumigatus*:773   *A. fischeri*:773
afum|EAL90482.1   100.00%   afis|EAW22712.1   100.00%
Bootstrap support for afum|EAL90482.1 as seed ortholog is 100%.
Bootstrap support for afis|EAW22712.1 as seed ortholog is 100%.

---

Group of orthologs #5526. Best score 773 bits
Score difference with first non-orthologous sequence - *A. fumigatus*:773   *A. fischeri*:773
afum|EAL92379.1   100.00%   afis|EAW21363.1   100.00%
Bootstrap support for afum|EAL92379.1 as seed ortholog is 100%.
Bootstrap support for afis|EAW21363.1 as seed ortholog is 100%.

---

Group of orthologs #5527. Best score 773 bits
Score difference with first non-orthologous sequence - *A. fumigatus*:773   *A. fischeri*:773
afum|EAL89330.1   100.00%   afis|EAW24689.1   100.00%
Bootstrap support for afum|EAL89330.1 as seed ortholog is 100%.
Bootstrap support for afis|EAW24689.1 as seed ortholog is 100%.

---

Group of orthologs #5528. Best score 773 bits
Score difference with first non-orthologous sequence - *A. fumigatus*:773   *A. fischeri*:692
afum|EAL89560.1   100.00%   afis|EAW24907.1   100.00%
Bootstrap support for afum|EAL89560.1 as seed ortholog is 100%.
Bootstrap support for afis|EAW24907.1 as seed ortholog is 100%.

---

Group of orthologs #5529. Best score 773 bits
Score difference with first non-orthologous sequence - *A. fumigatus*:773   *A. fischeri*:773
afum|EAL89703.1   100.00%   afis|EAW24998.1   100.00%
Bootstrap support for afum|EAL89703.1 as seed ortholog is 100%.
Bootstrap support for afis|EAW24998.1 as seed ortholog is 100%.

---

Group of orthologs #5530. Best score 773 bits
Score difference with first non-orthologous sequence - *A. fumigatus*:773   *A. fischeri*:773
afum|EAL89904.1   100.00%   afis|EAW25242.1   100.00%
Bootstrap support for afum|EAL89904.1 as seed ortholog is 100%.
Bootstrap support for afis|EAW25242.1 as seed ortholog is 100%.

---

Group of orthologs #5531. Best score 773 bits
Score difference with first non-orthologous sequence - *A. fumigatus*:773   *A. fischeri*:773
afum|EAL90067.1   100.00%   afis|EAW25408.1   100.00%

Bootstrap support for afum|EAL90067.1 as seed ortholog is 100%.
Bootstrap support for afis|EAW25408.1 as seed ortholog is 100%.

---

Group of orthologs #5532. Best score 772 bits
Score difference with first non-orthologous sequence - *A. fumigatus*:675 *A. fischeri*:543
afum|EAL85762.1 100.00% afis|EAW15571.1 100.00%
Bootstrap support for afum|EAL85762.1 as seed ortholog is 100%.
Bootstrap support for afis|EAW15571.1 as seed ortholog is 100%.

---

Group of orthologs #5533. Best score 772 bits
Score difference with first non-orthologous sequence - *A. fumigatus*:722 *A. fischeri*:636
afum|EAL84909.1 100.00% afis|EAW20009.1 100.00%
Bootstrap support for afum|EAL84909.1 as seed ortholog is 100%.
Bootstrap support for afis|EAW20009.1 as seed ortholog is 100%.

---

Group of orthologs #5534. Best score 772 bits
Score difference with first non-orthologous sequence - *A. fumigatus*:772 *A. fischeri*:772
afum|EAL91319.1 100.00% afis|EAW17496.1 100.00%
Bootstrap support for afum|EAL91319.1 as seed ortholog is 100%.
Bootstrap support for afis|EAW17496.1 as seed ortholog is 100%.

---

Group of orthologs #5535. Best score 772 bits
Score difference with first non-orthologous sequence - *A. fumigatus*:772 *A. fischeri*:652
afum|EAL93323.1 100.00% afis|EAW18613.1 100.00%
Bootstrap support for afum|EAL93323.1 as seed ortholog is 100%.
Bootstrap support for afis|EAW18613.1 as seed ortholog is 100%.

---

Group of orthologs #5536. Best score 772 bits
Score difference with first non-orthologous sequence - *A. fumigatus*:772 *A. fischeri*:772
afum|EAL92321.1 100.00% afis|EAW21305.1 100.00%
Bootstrap support for afum|EAL92321.1 as seed ortholog is 100%.
Bootstrap support for afis|EAW21305.1 as seed ortholog is 100%.

---

Group of orthologs #5537. Best score 771 bits
Score difference with first non-orthologous sequence - *A. fumigatus*:771 *A. fischeri*:771
afum|EAL85518.1 100.00% afis|EAW15834.1 100.00%
Bootstrap support for afum|EAL85518.1 as seed ortholog is 100%.
Bootstrap support for afis|EAW15834.1 as seed ortholog is 100%.

---

Group of orthologs #5538. Best score 771 bits
Score difference with first non-orthologous sequence - *A. fumigatus*:771 *A. fischeri*:771
afum|EAL88900.1 100.00% afis|EAW16279.1 100.00%
Bootstrap support for afum|EAL88900.1 as seed ortholog is 100%.
Bootstrap support for afis|EAW16279.1 as seed ortholog is 100%.

---

Group of orthologs #5539. Best score 771 bits
Score difference with first non-orthologous sequence - *A. fumigatus*:771 *A. fischeri*:771
afum|EAL84777.1 100.00% afis|EAW20858.1 100.00%
Bootstrap support for afum|EAL84777.1 as seed ortholog is 100%.
Bootstrap support for afis|EAW20858.1 as seed ortholog is 100%.

---

Group of orthologs #5540. Best score 771 bits
Score difference with first non-orthologous sequence - *A. fumigatus*:771 *A. fischeri*:771
afum|EAL84690.2 100.00% afis|EAW20944.1 100.00%
Bootstrap support for afum|EAL84690.2 as seed ortholog is 100%.
Bootstrap support for afis|EAW20944.1 as seed ortholog is 100%.

---
Group of orthologs #5541. Best score 771 bits
Score difference with first non-orthologous sequence - *A. fumigatus*:771 *A. fischeri*:584
afum|EAL86140.1 100.00% afis|EAW24468.1 100.00%
Bootstrap support for afum|EAL86140.1 as seed ortholog is 100%.
Bootstrap support for afis|EAW24468.1 as seed ortholog is 100%.

---
Group of orthologs #5542. Best score 771 bits
Score difference with first non-orthologous sequence - *A. fumigatus*:771 *A. fischeri*:771
afum|EAL87124.1 100.00% afis|EAW23681.1 100.00%
Bootstrap support for afum|EAL87124.1 as seed ortholog is 100%.
Bootstrap support for afis|EAW23681.1 as seed ortholog is 100%.

---
Group of orthologs #5543. Best score 771 bits
Score difference with first non-orthologous sequence - *A. fumigatus*:771 *A. fischeri*:718
afum|EAL89325.1 100.00% afis|EAW24685.1 100.00%
Bootstrap support for afum|EAL89325.1 as seed ortholog is 100%.
Bootstrap support for afis|EAW24685.1 as seed ortholog is 100%.

---
Group of orthologs #5544. Best score 771 bits
Score difference with first non-orthologous sequence - *A. fumigatus*:771 *A. fischeri*:771
afum|EAL90160.1 100.00% afis|EAW25494.1 100.00%
Bootstrap support for afum|EAL90160.1 as seed ortholog is 100%.
Bootstrap support for afis|EAW25494.1 as seed ortholog is 100%.

---
Group of orthologs #5545. Best score 770 bits
Score difference with first non-orthologous sequence - *A. fumigatus*:702 *A. fischeri*:658
afum|EAL84214.1 100.00% afis|EAW15429.1 100.00%
Bootstrap support for afum|EAL84214.1 as seed ortholog is 100%.
Bootstrap support for afis|EAW15429.1 as seed ortholog is 100%.

---
Group of orthologs #5546. Best score 770 bits
Score difference with first non-orthologous sequence - *A. fumigatus*:770 *A. fischeri*:527
afum|EAL88825.1 100.00% afis|EAW16194.1 100.00%
Bootstrap support for afum|EAL88825.1 as seed ortholog is 100%.
Bootstrap support for afis|EAW16194.1 as seed ortholog is 100%.

---
Group of orthologs #5547. Best score 770 bits
Score difference with first non-orthologous sequence - *A. fumigatus*:770 *A. fischeri*:770
afum|EAL84564.1 100.00% afis|EAW20490.1 100.00%
Bootstrap support for afum|EAL84564.1 as seed ortholog is 100%.
Bootstrap support for afis|EAW20490.1 as seed ortholog is 100%.

---
Group of orthologs #5548. Best score 770 bits
Score difference with first non-orthologous sequence - *A. fumigatus*:770 *A. fischeri*:770
afum|EAL87498.1 100.00% afis|EAW18141.1 100.00%
Bootstrap support for afum|EAL87498.1 as seed ortholog is 100%.
Bootstrap support for afis|EAW18141.1 as seed ortholog is 100%.

---
Group of orthologs #5549. Best score 770 bits
Score difference with first non-orthologous sequence - *A. fumigatus*:722 *A. fischeri*:694
afum|EAL91188.1 100.00% afis|EAW17344.1 100.00%
Bootstrap support for afum|EAL91188.1 as seed ortholog is 100%.
Bootstrap support for afis|EAW17344.1 as seed ortholog is 100%.

---
Group of orthologs #5550. Best score 770 bits

Score difference with first non-orthologous sequence - *A. fumigatus*:770 *A. fischeri*:770
afum|EAL86104.1 100.00% afis|EAW24436.1 100.00%
Bootstrap support for afum|EAL86104.1 as seed ortholog is 100%.
Bootstrap support for afis|EAW24436.1 as seed ortholog is 100%.

---

Group of orthologs #5551. Best score 770 bits
Score difference with first non-orthologous sequence - *A. fumigatus*:770 *A. fischeri*:569
afum|EAL87221.1 100.00% afis|EAW23787.1 100.00%
Bootstrap support for afum|EAL87221.1 as seed ortholog is 100%.
Bootstrap support for afis|EAW23787.1 as seed ortholog is 100%.

---

Group of orthologs #5552. Best score 770 bits
Score difference with first non-orthologous sequence - *A. fumigatus*:770 *A. fischeri*:770
afum|EAL88347.1 100.00% afis|EAW23134.1 100.00%
Bootstrap support for afum|EAL88347.1 as seed ortholog is 100%.
Bootstrap support for afis|EAW23134.1 as seed ortholog is 100%.

---

Group of orthologs #5553. Best score 770 bits
Score difference with first non-orthologous sequence - *A. fumigatus*:770 *A. fischeri*:770
afum|EAL88143.1 100.00% afis|EAW23335.1 100.00%
Bootstrap support for afum|EAL88143.1 as seed ortholog is 100%.
Bootstrap support for afis|EAW23335.1 as seed ortholog is 100%.

---

Group of orthologs #5554. Best score 770 bits
Score difference with first non-orthologous sequence - *A. fumigatus*:770 *A. fischeri*:549
afum|EAL87430.1 100.00% afis|EAW24000.1 100.00%
Bootstrap support for afum|EAL87430.1 as seed ortholog is 100%.
Bootstrap support for afis|EAW24000.1 as seed ortholog is 100%.

---

Group of orthologs #5555. Best score 770 bits
Score difference with first non-orthologous sequence - *A. fumigatus*:770 *A. fischeri*:770
afum|EAL92211.1 100.00% afis|EAW21191.1 100.00%
Bootstrap support for afum|EAL92211.1 as seed ortholog is 100%.
Bootstrap support for afis|EAW21191.1 as seed ortholog is 100%.

---

Group of orthologs #5556. Best score 770 bits
Score difference with first non-orthologous sequence - *A. fumigatus*:770 *A. fischeri*:648
afum|EAL89398.1 100.00% afis|EAW24755.1 100.00%
Bootstrap support for afum|EAL89398.1 as seed ortholog is 100%.
Bootstrap support for afis|EAW24755.1 as seed ortholog is 100%.

---

Group of orthologs #5557. Best score 770 bits
Score difference with first non-orthologous sequence - *A. fumigatus*:770 *A. fischeri*:770
afum|EAL92704.1 100.00% afis|EAW21669.1 100.00%
Bootstrap support for afum|EAL92704.1 as seed ortholog is 100%.
Bootstrap support for afis|EAW21669.1 as seed ortholog is 100%.

---

Group of orthologs #5558. Best score 769 bits
Score difference with first non-orthologous sequence - *A. fumigatus*:677 *A. fischeri*:621
afum|EAL85501.1 100.00% afis|EAW20402.1 100.00%
Bootstrap support for afum|EAL85501.1 as seed ortholog is 100%.
Bootstrap support for afis|EAW20402.1 as seed ortholog is 100%.

---

Group of orthologs #5559. Best score 769 bits
Score difference with first non-orthologous sequence - *A. fumigatus*:769 *A. fischeri*:769
afum|EAL93178.1 100.00% afis|EAW18488.1 100.00%

Bootstrap support for afum|EAL93178.1 as seed ortholog is 100%.
Bootstrap support for afis|EAW18488.1 as seed ortholog is 100%.

---

Group of orthologs #5560. Best score 769 bits
Score difference with first non-orthologous sequence - *A. fumigatus*:769 *A. fischeri*:769
afum|EAL92385.2 100.00% afis|EAW21368.1 100.00%
Bootstrap support for afum|EAL92385.2 as seed ortholog is 100%.
Bootstrap support for afis|EAW21368.1 as seed ortholog is 100%.

---

Group of orthologs #5561. Best score 769 bits
Score difference with first non-orthologous sequence - *A. fumigatus*:769 *A. fischeri*:769
afum|EAL89418.1 100.00% afis|EAW24778.1 100.00%
Bootstrap support for afum|EAL89418.1 as seed ortholog is 100%.
Bootstrap support for afis|EAW24778.1 as seed ortholog is 100%.

---

Group of orthologs #5562. Best score 769 bits
Score difference with first non-orthologous sequence - *A. fumigatus*:769 *A. fischeri*:769
afum|EAL92932.1 100.00% afis|EAW21920.1 100.00%
Bootstrap support for afum|EAL92932.1 as seed ortholog is 100%.
Bootstrap support for afis|EAW21920.1 as seed ortholog is 100%.

---

Group of orthologs #5563. Best score 768 bits
Score difference with first non-orthologous sequence - *A. fumigatus*:768 *A. fischeri*:558
afum|EAL87582.1 100.00% afis|EAW18216.1 100.00%
Bootstrap support for afum|EAL87582.1 as seed ortholog is 100%.
Bootstrap support for afis|EAW18216.1 as seed ortholog is 100%.

---

Group of orthologs #5564. Best score 768 bits
Score difference with first non-orthologous sequence - *A. fumigatus*:768 *A. fischeri*:768
afum|EAL87640.1 100.00% afis|EAW18274.1 100.00%
Bootstrap support for afum|EAL87640.1 as seed ortholog is 100%.
Bootstrap support for afis|EAW18274.1 as seed ortholog is 100%.

---

Group of orthologs #5565. Best score 768 bits
Score difference with first non-orthologous sequence - *A. fumigatus*:768 *A. fischeri*:768
afum|EAL87979.1 100.00% afis|EAW23484.1 100.00%
Bootstrap support for afum|EAL87979.1 as seed ortholog is 100%.
Bootstrap support for afis|EAW23484.1 as seed ortholog is 100%.

---

Group of orthologs #5566. Best score 768 bits
Score difference with first non-orthologous sequence - *A. fumigatus*:768 *A. fischeri*:768
afum|EAL93288.1 100.00% afis|EAW18583.1 100.00%
Bootstrap support for afum|EAL93288.1 as seed ortholog is 100%.
Bootstrap support for afis|EAW18583.1 as seed ortholog is 100%.

---

Group of orthologs #5567. Best score 768 bits
Score difference with first non-orthologous sequence - *A. fumigatus*:768 *A. fischeri*:768
afum|EAL92088.2 100.00% afis|EAW21052.1 100.00%
Bootstrap support for afum|EAL92088.2 as seed ortholog is 100%.
Bootstrap support for afis|EAW21052.1 as seed ortholog is 100%.

---

Group of orthologs #5568. Best score 768 bits
Score difference with first non-orthologous sequence - *A. fumigatus*:768 *A. fischeri*:768
afum|EAL90735.1 100.00% afis|EAW22456.1 100.00%
Bootstrap support for afum|EAL90735.1 as seed ortholog is 100%.
Bootstrap support for afis|EAW22456.1 as seed ortholog is 100%.

Group of orthologs #5569. Best score 767 bits
Score difference with first non-orthologous sequence - *A. fumigatus*:767 *A. fischeri*:767
afum|EAL84965.1 100.00% afis|EAW19950.1 100.00%
Bootstrap support for afum|EAL84965.1 as seed ortholog is 100%.
Bootstrap support for afis|EAW19950.1 as seed ortholog is 100%.

Group of orthologs #5570. Best score 767 bits
Score difference with first non-orthologous sequence - *A. fumigatus*:767 *A. fischeri*:767
afum|EAL84611.1 100.00% afis|EAW20441.1 100.00%
Bootstrap support for afum|EAL84611.1 as seed ortholog is 100%.
Bootstrap support for afis|EAW20441.1 as seed ortholog is 100%.

Group of orthologs #5571. Best score 767 bits
Score difference with first non-orthologous sequence - *A. fumigatus*:679 *A. fischeri*:336
afum|EAL87839.1 100.00% afis|EAW17484.1 100.00%
Bootstrap support for afum|EAL87839.1 as seed ortholog is 100%.
Bootstrap support for afis|EAW17484.1 as seed ortholog is 100%.

Group of orthologs #5572. Best score 767 bits
Score difference with first non-orthologous sequence - *A. fumigatus*:767 *A. fischeri*:452
afum|EAL84878.1 100.00% afis|EAW20703.1 100.00%
Bootstrap support for afum|EAL84878.1 as seed ortholog is 100%.
Bootstrap support for afis|EAW20703.1 as seed ortholog is 100%.

Group of orthologs #5573. Best score 767 bits
Score difference with first non-orthologous sequence - *A. fumigatus*:767 *A. fischeri*:767
afum|EAL84800.1 100.00% afis|EAW20832.1 100.00%
Bootstrap support for afum|EAL84800.1 as seed ortholog is 100%.
Bootstrap support for afis|EAW20832.1 as seed ortholog is 100%.

Group of orthologs #5574. Best score 767 bits
Score difference with first non-orthologous sequence - *A. fumigatus*:697 *A. fischeri*:767
afum|EAL85998.1 100.00% afis|EAW24328.1 100.00%
Bootstrap support for afum|EAL85998.1 as seed ortholog is 100%.
Bootstrap support for afis|EAW24328.1 as seed ortholog is 100%.

Group of orthologs #5575. Best score 767 bits
Score difference with first non-orthologous sequence - *A. fumigatus*:767 *A. fischeri*:767
afum|EAL87973.2 100.00% afis|EAW23490.1 100.00%
Bootstrap support for afum|EAL87973.2 as seed ortholog is 100%.
Bootstrap support for afis|EAW23490.1 as seed ortholog is 100%.

Group of orthologs #5576. Best score 767 bits
Score difference with first non-orthologous sequence - *A. fumigatus*:767 *A. fischeri*:767
afum|EAL93589.2 100.00% afis|EAW18857.1 100.00%
Bootstrap support for afum|EAL93589.2 as seed ortholog is 100%.
Bootstrap support for afis|EAW18857.1 as seed ortholog is 100%.

Group of orthologs #5577. Best score 767 bits
Score difference with first non-orthologous sequence - *A. fumigatus*:767 *A. fischeri*:767
afum|EAL93599.1 100.00% afis|EAW18867.1 100.00%
Bootstrap support for afum|EAL93599.1 as seed ortholog is 100%.
Bootstrap support for afis|EAW18867.1 as seed ortholog is 100%.

Group of orthologs #5578. Best score 767 bits

Score difference with first non-orthologous sequence - *A. fumigatus*:767 *A. fischeri*:767
afum|EAL90865.1 100.00% afis|EAW22327.1 100.00%
Bootstrap support for afum|EAL90865.1 as seed ortholog is 100%.
Bootstrap support for afis|EAW22327.1 as seed ortholog is 100%.

---

Group of orthologs #5579. Best score 767 bits
Score difference with first non-orthologous sequence - *A. fumigatus*:767 *A. fischeri*:767
afum|EAL90566.1 100.00% afis|EAW22629.1 100.00%
Bootstrap support for afum|EAL90566.1 as seed ortholog is 100%.
Bootstrap support for afis|EAW22629.1 as seed ortholog is 100%.

---

Group of orthologs #5580. Best score 767 bits
Score difference with first non-orthologous sequence - *A. fumigatus*:767 *A. fischeri*:767
afum|EAL90439.1 100.00% afis|EAW22760.1 100.00%
Bootstrap support for afum|EAL90439.1 as seed ortholog is 100%.
Bootstrap support for afis|EAW22760.1 as seed ortholog is 100%.

---

Group of orthologs #5581. Best score 767 bits
Score difference with first non-orthologous sequence - *A. fumigatus*:767 *A. fischeri*:767
afum|EAL90281.1 100.00% afis|EAW22916.1 100.00%
Bootstrap support for afum|EAL90281.1 as seed ortholog is 100%.
Bootstrap support for afis|EAW22916.1 as seed ortholog is 100%.

---

Group of orthologs #5582. Best score 766 bits
Score difference with first non-orthologous sequence - *A. fumigatus*:766 *A. fischeri*:766
afum|EAL88832.1 100.00% afis|EAW16204.1 100.00%
Bootstrap support for afum|EAL88832.1 as seed ortholog is 100%.
Bootstrap support for afis|EAW16204.1 as seed ortholog is 100%.

---

Group of orthologs #5583. Best score 766 bits
Score difference with first non-orthologous sequence - *A. fumigatus*:671 *A. fischeri*:628
afum|EAL91709.1 100.00% afis|EAW17901.1 100.00%
Bootstrap support for afum|EAL91709.1 as seed ortholog is 100%.
Bootstrap support for afis|EAW17901.1 as seed ortholog is 100%.

---

Group of orthologs #5584. Best score 766 bits
Score difference with first non-orthologous sequence - *A. fumigatus*:766 *A. fischeri*:766
afum|EAL88179.1 100.00% afis|EAW23300.1 100.00%
Bootstrap support for afum|EAL88179.1 as seed ortholog is 100%.
Bootstrap support for afis|EAW23300.1 as seed ortholog is 100%.

---

Group of orthologs #5585. Best score 766 bits
Score difference with first non-orthologous sequence - *A. fumigatus*:634 *A. fischeri*:499
afum|EAL93317.1 100.00% afis|EAW18607.1 100.00%
Bootstrap support for afum|EAL93317.1 as seed ortholog is 100%.
Bootstrap support for afis|EAW18607.1 as seed ortholog is 100%.

---

Group of orthologs #5586. Best score 766 bits
Score difference with first non-orthologous sequence - *A. fumigatus*:766 *A. fischeri*:766
afum|EAL90920.1 100.00% afis|EAW22260.1 100.00%
Bootstrap support for afum|EAL90920.1 as seed ortholog is 100%.
Bootstrap support for afis|EAW22260.1 as seed ortholog is 100%.

---

Group of orthologs #5587. Best score 766 bits
Score difference with first non-orthologous sequence - *A. fumigatus*:766 *A. fischeri*:766
afum|EAL90529.1 100.00% afis|EAW22663.1 100.00%

Bootstrap support for afum|EAL90529.1 as seed ortholog is 100%.
Bootstrap support for afis|EAW22663.1 as seed ortholog is 100%.

---

Group of orthologs #5588. Best score 766 bits
Score difference with first non-orthologous sequence - *A. fumigatus*:766 *A. fischeri*:766
afum|EAL89598.1 100.00% afis|EAW24941.1 100.00%
Bootstrap support for afum|EAL89598.1 as seed ortholog is 100%.
Bootstrap support for afis|EAW24941.1 as seed ortholog is 100%.

---

Group of orthologs #5589. Best score 765 bits
Score difference with first non-orthologous sequence - *A. fumigatus*:765 *A. fischeri*:765
afum|EAL84457.1 100.00% afis|EAW20604.1 100.00%
Bootstrap support for afum|EAL84457.1 as seed ortholog is 100%.
Bootstrap support for afis|EAW20604.1 as seed ortholog is 100%.

---

Group of orthologs #5590. Best score 765 bits
Score difference with first non-orthologous sequence - *A. fumigatus*:765 *A. fischeri*:765
afum|EAL91813.1 100.00% afis|EAW18002.1 100.00%
Bootstrap support for afum|EAL91813.1 as seed ortholog is 100%.
Bootstrap support for afis|EAW18002.1 as seed ortholog is 100%.

---

Group of orthologs #5591. Best score 765 bits
Score difference with first non-orthologous sequence - *A. fumigatus*:482 *A. fischeri*:659
afum|EAL87177.1 100.00% afis|EAW23740.1 100.00%
Bootstrap support for afum|EAL87177.1 as seed ortholog is 100%.
Bootstrap support for afis|EAW23740.1 as seed ortholog is 100%.

---

Group of orthologs #5592. Best score 765 bits
Score difference with first non-orthologous sequence - *A. fumigatus*:765 *A. fischeri*:765
afum|EAL88395.1 100.00% afis|EAW23084.1 100.00%
Bootstrap support for afum|EAL88395.1 as seed ortholog is 100%.
Bootstrap support for afis|EAW23084.1 as seed ortholog is 100%.

---

Group of orthologs #5593. Best score 764 bits
Score difference with first non-orthologous sequence - *A. fumigatus*:480 *A. fischeri*:764
afum|EAL84241.1 100.00% afis|EAW15405.1 100.00%
Bootstrap support for afum|EAL84241.1 as seed ortholog is 100%.
Bootstrap support for afis|EAW15405.1 as seed ortholog is 100%.

---

Group of orthologs #5594. Best score 764 bits
Score difference with first non-orthologous sequence - *A. fumigatus*:764 *A. fischeri*:712
afum|EAL84202.1 100.00% afis|EAW15445.1 100.00%
Bootstrap support for afum|EAL84202.1 as seed ortholog is 100%.
Bootstrap support for afis|EAW15445.1 as seed ortholog is 100%.

---

Group of orthologs #5595. Best score 764 bits
Score difference with first non-orthologous sequence - *A. fumigatus*:764 *A. fischeri*:764
afum|EAL84575.1 100.00% afis|EAW20479.1 100.00%
Bootstrap support for afum|EAL84575.1 as seed ortholog is 100%.
Bootstrap support for afis|EAW20479.1 as seed ortholog is 100%.

---

Group of orthologs #5596. Best score 764 bits
Score difference with first non-orthologous sequence - *A. fumigatus*:764 *A. fischeri*:764
afum|EAL89227.1 100.00% afis|EAW16631.1 100.00%
Bootstrap support for afum|EAL89227.1 as seed ortholog is 100%.
Bootstrap support for afis|EAW16631.1 as seed ortholog is 100%.

Group of orthologs #5597. Best score 764 bits
Score difference with first non-orthologous sequence - *A. fumigatus*:764 *A. fischeri*:764
afum|EAL86783.1 100.00% afis|EAW19743.1 100.00%
Bootstrap support for afum|EAL86783.1 as seed ortholog is 100%.
Bootstrap support for afis|EAW19743.1 as seed ortholog is 100%.

Group of orthologs #5598. Best score 764 bits
Score difference with first non-orthologous sequence - *A. fumigatus*:764 *A. fischeri*:764
afum|EAL91745.1 100.00% afis|EAW17934.1 100.00%
Bootstrap support for afum|EAL91745.1 as seed ortholog is 100%.
Bootstrap support for afis|EAW17934.1 as seed ortholog is 100%.

Group of orthologs #5599. Best score 764 bits
Score difference with first non-orthologous sequence - *A. fumigatus*:764 *A. fischeri*:764
afum|EAL86233.1 100.00% afis|EAW24570.1 100.00%
Bootstrap support for afum|EAL86233.1 as seed ortholog is 100%.
Bootstrap support for afis|EAW24570.1 as seed ortholog is 100%.

Group of orthologs #5600. Best score 764 bits
Score difference with first non-orthologous sequence - *A. fumigatus*:764 *A. fischeri*:764
afum|EAL93477.1 100.00% afis|EAW18750.1 100.00%
Bootstrap support for afum|EAL93477.1 as seed ortholog is 100%.
Bootstrap support for afis|EAW18750.1 as seed ortholog is 100%.

Group of orthologs #5601. Best score 764 bits
Score difference with first non-orthologous sequence - *A. fumigatus*:764 *A. fischeri*:764
afum|EAL93895.1 100.00% afis|EAW19165.1 100.00%
Bootstrap support for afum|EAL93895.1 as seed ortholog is 100%.
Bootstrap support for afis|EAW19165.1 as seed ortholog is 100%.

Group of orthologs #5602. Best score 764 bits
Score difference with first non-orthologous sequence - *A. fumigatus*:764 *A. fischeri*:764
afum|EAL90856.1 100.00% afis|EAW22336.1 100.00%
Bootstrap support for afum|EAL90856.1 as seed ortholog is 100%.
Bootstrap support for afis|EAW22336.1 as seed ortholog is 100%.

Group of orthologs #5603. Best score 764 bits
Score difference with first non-orthologous sequence - *A. fumigatus*:764 *A. fischeri*:764
afum|EAL92523.1 100.00% afis|EAW21504.1 100.00%
Bootstrap support for afum|EAL92523.1 as seed ortholog is 100%.
Bootstrap support for afis|EAW21504.1 as seed ortholog is 100%.

Group of orthologs #5604. Best score 764 bits
Score difference with first non-orthologous sequence - *A. fumigatus*:764 *A. fischeri*:764
afum|EAL92723.2 100.00% afis|EAW21689.1 100.00%
Bootstrap support for afum|EAL92723.2 as seed ortholog is 100%.
Bootstrap support for afis|EAW21689.1 as seed ortholog is 100%.

Group of orthologs #5605. Best score 764 bits
Score difference with first non-orthologous sequence - *A. fumigatus*:764 *A. fischeri*:764
afum|EAL93016.1 100.00% afis|EAW22001.1 100.00%
Bootstrap support for afum|EAL93016.1 as seed ortholog is 100%.
Bootstrap support for afis|EAW22001.1 as seed ortholog is 100%.

Group of orthologs #5606. Best score 763 bits

Score difference with first non-orthologous sequence - *A. fumigatus*:500 *A. fischeri*:301
afum|EAL87548.1 100.00% afis|EAW18185.1 100.00%
Bootstrap support for afum|EAL87548.1 as seed ortholog is 100%.
Bootstrap support for afis|EAW18185.1 as seed ortholog is 100%.

---

Group of orthologs #5607. Best score 763 bits
Score difference with first non-orthologous sequence - *A. fumigatus*:763 *A. fischeri*:208
afum|EAL91641.1 100.00% afis|EAW17831.1 100.00%
Bootstrap support for afum|EAL91641.1 as seed ortholog is 100%.
Bootstrap support for afis|EAW17831.1 as seed ortholog is 100%.

---

Group of orthologs #5608. Best score 763 bits
Score difference with first non-orthologous sequence - *A. fumigatus*:763 *A. fischeri*:763
afum|EAL91663.1 100.00% afis|EAW17855.1 100.00%
Bootstrap support for afum|EAL91663.1 as seed ortholog is 100%.
Bootstrap support for afis|EAW17855.1 as seed ortholog is 100%.

---

Group of orthologs #5609. Best score 763 bits
Score difference with first non-orthologous sequence - *A. fumigatus*:763 *A. fischeri*:763
afum|EAL88033.1 100.00% afis|EAW23423.1 100.00%
Bootstrap support for afum|EAL88033.1 as seed ortholog is 100%.
Bootstrap support for afis|EAW23423.1 as seed ortholog is 100%.

---

Group of orthologs #5610. Best score 763 bits
Score difference with first non-orthologous sequence - *A. fumigatus*:763 *A. fischeri*:763
afum|EAL90645.1 100.00% afis|EAW22550.1 100.00%
Bootstrap support for afum|EAL90645.1 as seed ortholog is 100%.
Bootstrap support for afis|EAW22550.1 as seed ortholog is 100%.

---

Group of orthologs #5611. Best score 762 bits
Score difference with first non-orthologous sequence - *A. fumigatus*:580 *A. fischeri*:544
afum|EAL85025.1 100.00% afis|EAW19874.1 100.00%
Bootstrap support for afum|EAL85025.1 as seed ortholog is 100%.
Bootstrap support for afis|EAW19874.1 as seed ortholog is 100%.

---

Group of orthologs #5612. Best score 762 bits
Score difference with first non-orthologous sequence - *A. fumigatus*:762 *A. fischeri*:762
afum|EAL86147.1 100.00% afis|EAW24475.1 100.00%
Bootstrap support for afum|EAL86147.1 as seed ortholog is 100%.
Bootstrap support for afis|EAW24475.1 as seed ortholog is 100%.

---

Group of orthologs #5613. Best score 762 bits
Score difference with first non-orthologous sequence - *A. fumigatus*:762 *A. fischeri*:599
afum|EAL86218.1 100.00% afis|EAW24553.1 100.00%
Bootstrap support for afum|EAL86218.1 as seed ortholog is 100%.
Bootstrap support for afis|EAW24553.1 as seed ortholog is 100%.

---

Group of orthologs #5614. Best score 762 bits
Score difference with first non-orthologous sequence - *A. fumigatus*:762 *A. fischeri*:762
afum|EAL92126.1 100.00% afis|EAW21105.1 100.00%
Bootstrap support for afum|EAL92126.1 as seed ortholog is 100%.
Bootstrap support for afis|EAW21105.1 as seed ortholog is 100%.

---

Group of orthologs #5615. Best score 762 bits
Score difference with first non-orthologous sequence - *A. fumigatus*:762 *A. fischeri*:762
afum|EAL90845.1 100.00% afis|EAW22347.1 100.00%

Bootstrap support for afum|EAL90845.1 as seed ortholog is 100%.
Bootstrap support for afis|EAW22347.1 as seed ortholog is 100%.

---

Group of orthologs #5616. Best score 762 bits
Score difference with first non-orthologous sequence - *A. fumigatus*:762 *A. fischeri*:762
afum|EAL89710.1 100.00% afis|EAW25042.1 100.00%
Bootstrap support for afum|EAL89710.1 as seed ortholog is 100%.
Bootstrap support for afis|EAW25042.1 as seed ortholog is 100%.

---

Group of orthologs #5617. Best score 761 bits
Score difference with first non-orthologous sequence - *A. fumigatus*:761 *A. fischeri*:761
afum|EAL88736.1 100.00% afis|EAW16104.1 100.00%
Bootstrap support for afum|EAL88736.1 as seed ortholog is 100%.
Bootstrap support for afis|EAW16104.1 as seed ortholog is 100%.

---

Group of orthologs #5618. Best score 761 bits
Score difference with first non-orthologous sequence - *A. fumigatus*:761 *A. fischeri*:644
afum|EAL88805.1 100.00% afis|EAW16173.1 100.00%
Bootstrap support for afum|EAL88805.1 as seed ortholog is 100%.
Bootstrap support for afis|EAW16173.1 as seed ortholog is 100%.

---

Group of orthologs #5619. Best score 761 bits
Score difference with first non-orthologous sequence - *A. fumigatus*:395 *A. fischeri*:519
afum|EAL85138.1 100.00% afis|EAW19764.1 100.00%
Bootstrap support for afum|EAL85138.1 as seed ortholog is 100%.
Bootstrap support for afis|EAW19764.1 as seed ortholog is 100%.

---

Group of orthologs #5620. Best score 760 bits
Score difference with first non-orthologous sequence - *A. fumigatus*:462 *A. fischeri*:248
afum|EAL88847.1 100.00% afis|EAW16219.1 100.00%
Bootstrap support for afum|EAL88847.1 as seed ortholog is 100%.
Bootstrap support for afis|EAW16219.1 as seed ortholog is 100%.

---

Group of orthologs #5621. Best score 760 bits
Score difference with first non-orthologous sequence - *A. fumigatus*:679 *A. fischeri*:760
afum|EAL84815.1 100.00% afis|EAW20813.1 100.00%
Bootstrap support for afum|EAL84815.1 as seed ortholog is 100%.
Bootstrap support for afis|EAW20813.1 as seed ortholog is 100%.

---

Group of orthologs #5622. Best score 760 bits
Score difference with first non-orthologous sequence - *A. fumigatus*:760 *A. fischeri*:760
afum|EAL84382.1 100.00% afis|EAW25709.1 100.00%
Bootstrap support for afum|EAL84382.1 as seed ortholog is 100%.
Bootstrap support for afis|EAW25709.1 as seed ortholog is 100%.

---

Group of orthologs #5623. Best score 760 bits
Score difference with first non-orthologous sequence - *A. fumigatus*:760 *A. fischeri*:760
afum|EAL85954.1 100.00% afis|EAW24285.1 100.00%
Bootstrap support for afum|EAL85954.1 as seed ortholog is 100%.
Bootstrap support for afis|EAW24285.1 as seed ortholog is 100%.

---

Group of orthologs #5624. Best score 760 bits
Score difference with first non-orthologous sequence - *A. fumigatus*:760 *A. fischeri*:760
afum|EAL86191.1 100.00% afis|EAW24527.1 100.00%
Bootstrap support for afum|EAL86191.1 as seed ortholog is 100%.
Bootstrap support for afis|EAW24527.1 as seed ortholog is 100%.

Group of orthologs #5625. Best score 760 bits
Score difference with first non-orthologous sequence - *A. fumigatus*:760 *A. fischeri*:552
afum|EAL93111.2 100.00% afis|EAW18431.1 100.00%
Bootstrap support for afum|EAL93111.2 as seed ortholog is 100%.
Bootstrap support for afis|EAW18431.1 as seed ortholog is 100%.

Group of orthologs #5626. Best score 760 bits
Score difference with first non-orthologous sequence - *A. fumigatus*:760 *A. fischeri*:672
afum|EAL92083.1 100.00% afis|EAW21047.1 100.00%
Bootstrap support for afum|EAL92083.1 as seed ortholog is 100%.
Bootstrap support for afis|EAW21047.1 as seed ortholog is 100%.

Group of orthologs #5627. Best score 760 bits
Score difference with first non-orthologous sequence - *A. fumigatus*:760 *A. fischeri*:760
afum|EAL92542.1 100.00% afis|EAW21521.1 100.00%
Bootstrap support for afum|EAL92542.1 as seed ortholog is 100%.
Bootstrap support for afis|EAW21521.1 as seed ortholog is 100%.

Group of orthologs #5628. Best score 759 bits
Score difference with first non-orthologous sequence - *A. fumigatus*:759 *A. fischeri*:759
afum|EAL85507.1 100.00% afis|EAW15844.1 100.00%
Bootstrap support for afum|EAL85507.1 as seed ortholog is 100%.
Bootstrap support for afis|EAW15844.1 as seed ortholog is 100%.

Group of orthologs #5629. Best score 759 bits
Score difference with first non-orthologous sequence - *A. fumigatus*:759 *A. fischeri*:759
afum|EAL88723.1 100.00% afis|EAW16091.1 100.00%
Bootstrap support for afum|EAL88723.1 as seed ortholog is 100%.
Bootstrap support for afis|EAW16091.1 as seed ortholog is 100%.

Group of orthologs #5630. Best score 759 bits
Score difference with first non-orthologous sequence - *A. fumigatus*:759 *A. fischeri*:759
afum|EAL85082.1 100.00% afis|EAW19810.1 100.00%
Bootstrap support for afum|EAL85082.1 as seed ortholog is 100%.
Bootstrap support for afis|EAW19810.1 as seed ortholog is 100%.

Group of orthologs #5631. Best score 759 bits
Score difference with first non-orthologous sequence - *A. fumigatus*:759 *A. fischeri*:467
afum|EAL84975.2 100.00% afis|EAW19935.1 100.00%
Bootstrap support for afum|EAL84975.2 as seed ortholog is 100%.
Bootstrap support for afis|EAW19935.1 as seed ortholog is 100%.

Group of orthologs #5632. Best score 759 bits
Score difference with first non-orthologous sequence - *A. fumigatus*:759 *A. fischeri*:759
afum|EAL85405.2 100.00% afis|EAW20271.1 100.00%
Bootstrap support for afum|EAL85405.2 as seed ortholog is 100%.
Bootstrap support for afis|EAW20271.1 as seed ortholog is 100%.

Group of orthologs #5633. Best score 759 bits
Score difference with first non-orthologous sequence - *A. fumigatus*:759 *A. fischeri*:759
afum|EAL91678.1 100.00% afis|EAW17870.1 100.00%
Bootstrap support for afum|EAL91678.1 as seed ortholog is 100%.
Bootstrap support for afis|EAW17870.1 as seed ortholog is 100%.

Group of orthologs #5634. Best score 759 bits

Score difference with first non-orthologous sequence - *A. fumigatus*:759 *A. fischeri*:759
afum|EAL89898.1 100.00% afis|EAW25237.1 100.00%
Bootstrap support for afum|EAL89898.1 as seed ortholog is 100%.
Bootstrap support for afis|EAW25237.1 as seed ortholog is 100%.

---

Group of orthologs #5635. Best score 758 bits
Score difference with first non-orthologous sequence - *A. fumigatus*:758 *A. fischeri*:758
afum|EAL87282.1 100.00% afis|EAW23854.1 100.00%
Bootstrap support for afum|EAL87282.1 as seed ortholog is 100%.
Bootstrap support for afis|EAW23854.1 as seed ortholog is 100%.

---

Group of orthologs #5636. Best score 758 bits
Score difference with first non-orthologous sequence - *A. fumigatus*:670 *A. fischeri*:670
afum|EAL90199.1 100.00% afis|EAW25535.1 100.00%
Bootstrap support for afum|EAL90199.1 as seed ortholog is 100%.
Bootstrap support for afis|EAW25535.1 as seed ortholog is 100%.

---

Group of orthologs #5637. Best score 757 bits
Score difference with first non-orthologous sequence - *A. fumigatus*:757 *A. fischeri*:503
afum|EAL86331.1 100.00% afis|EAW16759.1 100.00%
Bootstrap support for afum|EAL86331.1 as seed ortholog is 100%.
Bootstrap support for afis|EAW16759.1 as seed ortholog is 100%.

---

Group of orthologs #5638. Best score 757 bits
Score difference with first non-orthologous sequence - *A. fumigatus*:757 *A. fischeri*:757
afum|EAL86334.1 100.00% afis|EAW16763.1 100.00%
Bootstrap support for afum|EAL86334.1 as seed ortholog is 100%.
Bootstrap support for afis|EAW16763.1 as seed ortholog is 100%.

---

Group of orthologs #5639. Best score 757 bits
Score difference with first non-orthologous sequence - *A. fumigatus*:757 *A. fischeri*:757
afum|EAL86663.2 100.00% afis|EAW17243.1 100.00%
Bootstrap support for afum|EAL86663.2 as seed ortholog is 100%.
Bootstrap support for afis|EAW17243.1 as seed ortholog is 100%.

---

Group of orthologs #5640. Best score 757 bits
Score difference with first non-orthologous sequence - *A. fumigatus*:757 *A. fischeri*:757
afum|EAL84995.1 100.00% afis|EAW19912.1 100.00%
Bootstrap support for afum|EAL84995.1 as seed ortholog is 100%.
Bootstrap support for afis|EAW19912.1 as seed ortholog is 100%.

---

Group of orthologs #5641. Best score 757 bits
Score difference with first non-orthologous sequence - *A. fumigatus*:757 *A. fischeri*:757
afum|EAL87281.1 100.00% afis|EAW23853.1 100.00%
Bootstrap support for afum|EAL87281.1 as seed ortholog is 100%.
Bootstrap support for afis|EAW23853.1 as seed ortholog is 100%.

---

Group of orthologs #5642. Best score 757 bits
Score difference with first non-orthologous sequence - *A. fumigatus*:757 *A. fischeri*:757
afum|EAL92266.1 100.00% afis|EAW21252.1 100.00%
Bootstrap support for afum|EAL92266.1 as seed ortholog is 100%.
Bootstrap support for afis|EAW21252.1 as seed ortholog is 100%.

---

Group of orthologs #5643. Best score 757 bits
Score difference with first non-orthologous sequence - *A. fumigatus*:757 *A. fischeri*:757
afum|EAL92893.1 100.00% afis|EAW21879.1 100.00%

Bootstrap support for afum|EAL92893.1 as seed ortholog is 100%.
Bootstrap support for afis|EAW21879.1 as seed ortholog is 100%.

---

Group of orthologs #5644. Best score 757 bits
Score difference with first non-orthologous sequence - *A. fumigatus*:757 *A. fischeri*:757
afum|EAL92914.1 100.00% afis|EAW21901.1 100.00%
Bootstrap support for afum|EAL92914.1 as seed ortholog is 100%.
Bootstrap support for afis|EAW21901.1 as seed ortholog is 100%.

---

Group of orthologs #5645. Best score 757 bits
Score difference with first non-orthologous sequence - *A. fumigatus*:757 *A. fischeri*:757
afum|EAL92022.1 100.00% afis|EAW24165.1 100.00%
Bootstrap support for afum|EAL92022.1 as seed ortholog is 100%.
Bootstrap support for afis|EAW24165.1 as seed ortholog is 100%.

---

Group of orthologs #5646. Best score 756 bits
Score difference with first non-orthologous sequence - *A. fumigatus*:756 *A. fischeri*:256
afum|EAL86357.1 100.00% afis|EAW16787.1 100.00%
Bootstrap support for afum|EAL86357.1 as seed ortholog is 100%.
Bootstrap support for afis|EAW16787.1 as seed ortholog is 100%.

---

Group of orthologs #5647. Best score 756 bits
Score difference with first non-orthologous sequence - *A. fumigatus*:520 *A. fischeri*:704
afum|EAL87501.1 100.00% afis|EAW18144.1 100.00%
Bootstrap support for afum|EAL87501.1 as seed ortholog is 100%.
Bootstrap support for afis|EAW18144.1 as seed ortholog is 100%.

---

Group of orthologs #5648. Best score 756 bits
Score difference with first non-orthologous sequence - *A. fumigatus*:756 *A. fischeri*:756
afum|EAL89490.1 100.00% afis|EAW24844.1 100.00%
Bootstrap support for afum|EAL89490.1 as seed ortholog is 100%.
Bootstrap support for afis|EAW24844.1 as seed ortholog is 100%.

---

Group of orthologs #5649. Best score 755 bits
Score difference with first non-orthologous sequence - *A. fumigatus*:755 *A. fischeri*:755
afum|EAL84554.1 100.00% afis|EAW20501.1 100.00%
Bootstrap support for afum|EAL84554.1 as seed ortholog is 100%.
Bootstrap support for afis|EAW20501.1 as seed ortholog is 100%.

---

Group of orthologs #5650. Best score 755 bits
Score difference with first non-orthologous sequence - *A. fumigatus*:639 *A. fischeri*:619
afum|EAL84348.1 100.00% afis|EAW25678.1 100.00%
Bootstrap support for afum|EAL84348.1 as seed ortholog is 100%.
Bootstrap support for afis|EAW25678.1 as seed ortholog is 100%.

---

Group of orthologs #5651. Best score 755 bits
Score difference with first non-orthologous sequence - *A. fumigatus*:755 *A. fischeri*:755
afum|EAL93226.1 100.00% afis|EAW18529.1 100.00%
Bootstrap support for afum|EAL93226.1 as seed ortholog is 100%.
Bootstrap support for afis|EAW18529.1 as seed ortholog is 100%.

---

Group of orthologs #5652. Best score 755 bits
Score difference with first non-orthologous sequence - *A. fumigatus*:755 *A. fischeri*:755
afum|EAL90774.1 100.00% afis|EAW22418.1 100.00%
Bootstrap support for afum|EAL90774.1 as seed ortholog is 100%.
Bootstrap support for afis|EAW22418.1 as seed ortholog is 100%.

Group of orthologs #5653. Best score 755 bits
Score difference with first non-orthologous sequence - *A. fumigatus*:755 *A. fischeri*:755
afum|EAL89714.1 100.00% afis|EAW25046.1 100.00%
Bootstrap support for afum|EAL89714.1 as seed ortholog is 100%.
Bootstrap support for afis|EAW25046.1 as seed ortholog is 100%.

---

Group of orthologs #5654. Best score 755 bits
Score difference with first non-orthologous sequence - *A. fumigatus*:669 *A. fischeri*:569
afum|EAL89918.1 100.00% afis|EAW25257.1 100.00%
Bootstrap support for afum|EAL89918.1 as seed ortholog is 100%.
Bootstrap support for afis|EAW25257.1 as seed ortholog is 100%.

---

Group of orthologs #5655. Best score 754 bits
Score difference with first non-orthologous sequence - *A. fumigatus*:427 *A. fischeri*:561
afum|EAL86432.1 100.00% afis|EAW16895.1 100.00%
Bootstrap support for afum|EAL86432.1 as seed ortholog is 100%.
Bootstrap support for afis|EAW16895.1 as seed ortholog is 100%.

---

Group of orthologs #5656. Best score 754 bits
Score difference with first non-orthologous sequence - *A. fumigatus*:754 *A. fischeri*:754
afum|EAL88771.1 100.00% afis|EAW16142.1 100.00%
Bootstrap support for afum|EAL88771.1 as seed ortholog is 100%.
Bootstrap support for afis|EAW16142.1 as seed ortholog is 100%.

---

Group of orthologs #5657. Best score 754 bits
Score difference with first non-orthologous sequence - *A. fumigatus*:754 *A. fischeri*:754
afum|EAL87989.2 100.00% afis|EAW23476.1 100.00%
Bootstrap support for afum|EAL87989.2 as seed ortholog is 100%.
Bootstrap support for afis|EAW23476.1 as seed ortholog is 100%.

---

Group of orthologs #5658. Best score 754 bits
Score difference with first non-orthologous sequence - *A. fumigatus*:754 *A. fischeri*:754
afum|EAL87455.1 100.00% afis|EAW24024.1 100.00%
Bootstrap support for afum|EAL87455.1 as seed ortholog is 100%.
Bootstrap support for afis|EAW24024.1 as seed ortholog is 100%.

---

Group of orthologs #5659. Best score 754 bits
Score difference with first non-orthologous sequence - *A. fumigatus*:754 *A. fischeri*:754
afum|EAL92135.1 100.00% afis|EAW21112.1 100.00%
Bootstrap support for afum|EAL92135.1 as seed ortholog is 100%.
Bootstrap support for afis|EAW21112.1 as seed ortholog is 100%.

---

Group of orthologs #5660. Best score 754 bits
Score difference with first non-orthologous sequence - *A. fumigatus*:754 *A. fischeri*:754
afum|EAL90345.1 100.00% afis|EAW22855.1 100.00%
Bootstrap support for afum|EAL90345.1 as seed ortholog is 100%.
Bootstrap support for afis|EAW22855.1 as seed ortholog is 100%.

---

Group of orthologs #5661. Best score 754 bits
Score difference with first non-orthologous sequence - *A. fumigatus*:754 *A. fischeri*:754
afum|EAL92275.1 100.00% afis|EAW21260.1 100.00%
Bootstrap support for afum|EAL92275.1 as seed ortholog is 100%.
Bootstrap support for afis|EAW21260.1 as seed ortholog is 100%.

---

Group of orthologs #5662. Best score 754 bits

Score difference with first non-orthologous sequence - *A. fumigatus*:754 *A. fischeri*:754
afum|EAL92306.1 100.00% afis|EAW21289.1 100.00%
Bootstrap support for afum|EAL92306.1 as seed ortholog is 100%.
Bootstrap support for afis|EAW21289.1 as seed ortholog is 100%.

---

Group of orthologs #5663. Best score 754 bits
Score difference with first non-orthologous sequence - *A. fumigatus*:754 *A. fischeri*:754
afum|EAL89437.1 100.00% afis|EAW24795.1 100.00%
Bootstrap support for afum|EAL89437.1 as seed ortholog is 100%.
Bootstrap support for afis|EAW24795.1 as seed ortholog is 100%.

---

Group of orthologs #5664. Best score 753 bits
Score difference with first non-orthologous sequence - *A. fumigatus*:753 *A. fischeri*:753
afum|EAL84429.1 100.00% afis|EAW20641.1 100.00%
Bootstrap support for afum|EAL84429.1 as seed ortholog is 100%.
Bootstrap support for afis|EAW20641.1 as seed ortholog is 100%.

---

Group of orthologs #5665. Best score 753 bits
Score difference with first non-orthologous sequence - *A. fumigatus*:753 *A. fischeri*:753
afum|EAL89142.2 100.00% afis|EAW16544.1 100.00%
Bootstrap support for afum|EAL89142.2 as seed ortholog is 100%.
Bootstrap support for afis|EAW16544.1 as seed ortholog is 100%.

---

Group of orthologs #5666. Best score 753 bits
Score difference with first non-orthologous sequence - *A. fumigatus*:753 *A. fischeri*:753
afum|EAL87483.1 100.00% afis|EAW18125.1 100.00%
Bootstrap support for afum|EAL87483.1 as seed ortholog is 100%.
Bootstrap support for afis|EAW18125.1 as seed ortholog is 100%.

---

Group of orthologs #5667. Best score 753 bits
Score difference with first non-orthologous sequence - *A. fumigatus*:753 *A. fischeri*:753
afum|EAL87006.1 100.00% afis|EAW19488.1 100.00%
Bootstrap support for afum|EAL87006.1 as seed ortholog is 100%.
Bootstrap support for afis|EAW19488.1 as seed ortholog is 100%.

---

Group of orthologs #5668. Best score 753 bits
Score difference with first non-orthologous sequence - *A. fumigatus*:753 *A. fischeri*:753
afum|EAL91814.2 100.00% afis|EAW18003.1 100.00%
Bootstrap support for afum|EAL91814.2 as seed ortholog is 100%.
Bootstrap support for afis|EAW18003.1 as seed ortholog is 100%.

---

Group of orthologs #5669. Best score 753 bits
Score difference with first non-orthologous sequence - *A. fumigatus*:753 *A. fischeri*:753
afum|EAL87900.1 100.00% afis|EAW23567.1 100.00%
Bootstrap support for afum|EAL87900.1 as seed ortholog is 100%.
Bootstrap support for afis|EAW23567.1 as seed ortholog is 100%.

---

Group of orthologs #5670. Best score 753 bits
Score difference with first non-orthologous sequence - *A. fumigatus*:753 *A. fischeri*:753
afum|EAL94006.1 100.00% afis|EAW19280.1 100.00%
Bootstrap support for afum|EAL94006.1 as seed ortholog is 100%.
Bootstrap support for afis|EAW19280.1 as seed ortholog is 100%.

---

Group of orthologs #5671. Best score 753 bits
Score difference with first non-orthologous sequence - *A. fumigatus*:753 *A. fischeri*:753
afum|EAL90604.1 100.00% afis|EAW22595.1 100.00%

Bootstrap support for afum|EAL90604.1 as seed ortholog is 100%.
Bootstrap support for afis|EAW22595.1 as seed ortholog is 100%.

---

Group of orthologs #5672. Best score 753 bits
Score difference with first non-orthologous sequence - *A. fumigatus*:753 *A. fischeri*:753
afum|EAL92184.1 100.00% afis|EAW21165.1 100.00%
Bootstrap support for afum|EAL92184.1 as seed ortholog is 100%.
Bootstrap support for afis|EAW21165.1 as seed ortholog is 100%.

---

Group of orthologs #5673. Best score 753 bits
Score difference with first non-orthologous sequence - *A. fumigatus*:753 *A. fischeri*:753
afum|EAL92845.1 100.00% afis|EAW21830.1 100.00%
Bootstrap support for afum|EAL92845.1 as seed ortholog is 100%.
Bootstrap support for afis|EAW21830.1 as seed ortholog is 100%.

---

Group of orthologs #5674. Best score 753 bits
Score difference with first non-orthologous sequence - *A. fumigatus*:753 *A. fischeri*:753
afum|EAL91911.1 100.00% afis|EAW25195.1 100.00%
Bootstrap support for afum|EAL91911.1 as seed ortholog is 100%.
Bootstrap support for afis|EAW25195.1 as seed ortholog is 100%.

---

Group of orthologs #5675. Best score 752 bits
Score difference with first non-orthologous sequence - *A. fumigatus*:642 *A. fischeri*:617
afum|EAL86344.1 100.00% afis|EAW16775.1 100.00%
Bootstrap support for afum|EAL86344.1 as seed ortholog is 100%.
Bootstrap support for afis|EAW16775.1 as seed ortholog is 100%.

---

Group of orthologs #5676. Best score 752 bits
Score difference with first non-orthologous sequence - *A. fumigatus*:752 *A. fischeri*:752
afum|EAL87477.1 100.00% afis|EAW18119.1 100.00%
Bootstrap support for afum|EAL87477.1 as seed ortholog is 100%.
Bootstrap support for afis|EAW18119.1 as seed ortholog is 100%.

---

Group of orthologs #5677. Best score 752 bits
Score difference with first non-orthologous sequence - *A. fumigatus*:674 *A. fischeri*:655
afum|EAL89218.1 100.00% afis|EAW16619.1 100.00%
Bootstrap support for afum|EAL89218.1 as seed ortholog is 100%.
Bootstrap support for afis|EAW16619.1 as seed ortholog is 100%.

---

Group of orthologs #5678. Best score 752 bits
Score difference with first non-orthologous sequence - *A. fumigatus*:752 *A. fischeri*:752
afum|EAL87242.1 100.00% afis|EAW23808.1 100.00%
Bootstrap support for afum|EAL87242.1 as seed ortholog is 100%.
Bootstrap support for afis|EAW23808.1 as seed ortholog is 100%.

---

Group of orthologs #5679. Best score 752 bits
Score difference with first non-orthologous sequence - *A. fumigatus*:752 *A. fischeri*:752
afum|EAL93326.1 100.00% afis|EAW18616.1 100.00%
Bootstrap support for afum|EAL93326.1 as seed ortholog is 100%.
Bootstrap support for afis|EAW18616.1 as seed ortholog is 100%.

---

Group of orthologs #5680. Best score 752 bits
Score difference with first non-orthologous sequence - *A. fumigatus*:752 *A. fischeri*:631
afum|EAL93745.2 100.00% afis|EAW19011.1 100.00%
Bootstrap support for afum|EAL93745.2 as seed ortholog is 100%.
Bootstrap support for afis|EAW19011.1 as seed ortholog is 100%.

---
Group of orthologs #5681. Best score 752 bits
Score difference with first non-orthologous sequence - *A. fumigatus*:752 *A. fischeri*:752
afum|EAL92428.1 100.00% afis|EAW21414.1 100.00%
Bootstrap support for afum|EAL92428.1 as seed ortholog is 100%.
Bootstrap support for afis|EAW21414.1 as seed ortholog is 100%.

---
Group of orthologs #5682. Best score 751 bits
Score difference with first non-orthologous sequence - *A. fumigatus*:751 *A. fischeri*:751
afum|EAL87506.1 100.00% afis|EAW18148.1 100.00%
Bootstrap support for afum|EAL87506.1 as seed ortholog is 100%.
Bootstrap support for afis|EAW18148.1 as seed ortholog is 100%.

---
Group of orthologs #5683. Best score 751 bits
Score difference with first non-orthologous sequence - *A. fumigatus*:751 *A. fischeri*:751
afum|EAL84715.1 100.00% afis|EAW20919.1 100.00%
Bootstrap support for afum|EAL84715.1 as seed ortholog is 100%.
Bootstrap support for afis|EAW20919.1 as seed ortholog is 100%.

---
Group of orthologs #5684. Best score 751 bits
Score difference with first non-orthologous sequence - *A. fumigatus*:751 *A. fischeri*:751
afum|EAL91514.1 100.00% afis|EAW17699.1 100.00%
Bootstrap support for afum|EAL91514.1 as seed ortholog is 100%.
Bootstrap support for afis|EAW17699.1 as seed ortholog is 100%.

---
Group of orthologs #5685. Best score 751 bits
Score difference with first non-orthologous sequence - *A. fumigatus*:656 *A. fischeri*:544
afum|EAL88193.1 100.00% afis|EAW23286.1 100.00%
Bootstrap support for afum|EAL88193.1 as seed ortholog is 100%.
Bootstrap support for afis|EAW23286.1 as seed ortholog is 100%.

---
Group of orthologs #5686. Best score 751 bits
Score difference with first non-orthologous sequence - *A. fumigatus*:751 *A. fischeri*:751
afum|EAL93992.1 100.00% afis|EAW19265.1 100.00%
Bootstrap support for afum|EAL93992.1 as seed ortholog is 100%.
Bootstrap support for afis|EAW19265.1 as seed ortholog is 100%.

---
Group of orthologs #5687. Best score 751 bits
Score difference with first non-orthologous sequence - *A. fumigatus*:751 *A. fischeri*:751
afum|EAL90754.1 100.00% afis|EAW22437.1 100.00%
Bootstrap support for afum|EAL90754.1 as seed ortholog is 100%.
Bootstrap support for afis|EAW22437.1 as seed ortholog is 100%.

---
Group of orthologs #5688. Best score 751 bits
Score difference with first non-orthologous sequence - *A. fumigatus*:751 *A. fischeri*:751
afum|EAL90287.1 100.00% afis|EAW22910.1 100.00%
Bootstrap support for afum|EAL90287.1 as seed ortholog is 100%.
Bootstrap support for afis|EAW22910.1 as seed ortholog is 100%.

---
Group of orthologs #5689. Best score 751 bits
Score difference with first non-orthologous sequence - *A. fumigatus*:751 *A. fischeri*:686
afum|EAL89535.2 100.00% afis|EAW24885.1 100.00%
Bootstrap support for afum|EAL89535.2 as seed ortholog is 100%.
Bootstrap support for afis|EAW24885.1 as seed ortholog is 100%.

---
Group of orthologs #5690. Best score 751 bits

Score difference with first non-orthologous sequence - *A. fumigatus*:751   *A. fischeri*:751
afum|EAL92981.1 100.00% afis|EAW21967.1 100.00%
Bootstrap support for afum|EAL92981.1 as seed ortholog is 100%.
Bootstrap support for afis|EAW21967.1 as seed ortholog is 100%.

---

Group of orthologs #5691. Best score 751 bits
Score difference with first non-orthologous sequence - *A. fumigatus*:751   *A. fischeri*:751
afum|EAL89799.1 100.00% afis|EAW25128.1 100.00%
Bootstrap support for afum|EAL89799.1 as seed ortholog is 100%.
Bootstrap support for afis|EAW25128.1 as seed ortholog is 100%.

---

Group of orthologs #5692. Best score 750 bits
Score difference with first non-orthologous sequence - *A. fumigatus*:750   *A. fischeri*:750
afum|EAL89120.1 100.00% afis|EAW16525.1 100.00%
Bootstrap support for afum|EAL89120.1 as seed ortholog is 100%.
Bootstrap support for afis|EAW16525.1 as seed ortholog is 100%.

---

Group of orthologs #5693. Best score 750 bits
Score difference with first non-orthologous sequence - *A. fumigatus*:750   *A. fischeri*:750
afum|EAL85953.1 100.00% afis|EAW24284.1 100.00%
Bootstrap support for afum|EAL85953.1 as seed ortholog is 100%.
Bootstrap support for afis|EAW24284.1 as seed ortholog is 100%.

---

Group of orthologs #5694. Best score 750 bits
Score difference with first non-orthologous sequence - *A. fumigatus*:750   *A. fischeri*:750
afum|EAL88299.1 100.00% afis|EAW23184.1 100.00%
Bootstrap support for afum|EAL88299.1 as seed ortholog is 100%.
Bootstrap support for afis|EAW23184.1 as seed ortholog is 100%.

---

Group of orthologs #5695. Best score 750 bits
Score difference with first non-orthologous sequence - *A. fumigatus*:750   *A. fischeri*:542
afum|EAL92205.1 100.00% afis|EAW21184.1 100.00%
Bootstrap support for afum|EAL92205.1 as seed ortholog is 100%.
Bootstrap support for afis|EAW21184.1 as seed ortholog is 100%.

---

Group of orthologs #5696. Best score 750 bits
Score difference with first non-orthologous sequence - *A. fumigatus*:750   *A. fischeri*:750
afum|EAL92210.1 100.00% afis|EAW21188.1 100.00%
Bootstrap support for afum|EAL92210.1 as seed ortholog is 100%.
Bootstrap support for afis|EAW21188.1 as seed ortholog is 100%.

---

Group of orthologs #5697. Best score 749 bits
Score difference with first non-orthologous sequence - *A. fumigatus*:749   *A. fischeri*:749
afum|EAL93633.1 100.00% afis|EAW18901.1 100.00%
Bootstrap support for afum|EAL93633.1 as seed ortholog is 100%.
Bootstrap support for afis|EAW18901.1 as seed ortholog is 100%.

---

Group of orthologs #5698. Best score 749 bits
Score difference with first non-orthologous sequence - *A. fumigatus*:749   *A. fischeri*:749
afum|EAL90522.1 100.00% afis|EAW22670.1 100.00%
Bootstrap support for afum|EAL90522.1 as seed ortholog is 100%.
Bootstrap support for afis|EAW22670.1 as seed ortholog is 100%.

---

Group of orthologs #5699. Best score 749 bits
Score difference with first non-orthologous sequence - *A. fumigatus*:749   *A. fischeri*:539
afum|EAL90355.1 100.00% afis|EAW22847.1 100.00%

Bootstrap support for afum|EAL90355.1 as seed ortholog is 100%.
Bootstrap support for afis|EAW22847.1 as seed ortholog is 100%.

---

Group of orthologs #5700. Best score 749 bits
Score difference with first non-orthologous sequence - *A. fumigatus*:536 *A. fischeri*:293
afum|EAL92662.1 100.00% afis|EAW21627.1 100.00%
Bootstrap support for afum|EAL92662.1 as seed ortholog is 100%.
Bootstrap support for afis|EAW21627.1 as seed ortholog is 100%.

---

Group of orthologs #5701. Best score 749 bits
Score difference with first non-orthologous sequence - *A. fumigatus*:749 *A. fischeri*:749
afum|EAL92857.2 100.00% afis|EAW21845.1 100.00%
Bootstrap support for afum|EAL92857.2 as seed ortholog is 100%.
Bootstrap support for afis|EAW21845.1 as seed ortholog is 100%.

---

Group of orthologs #5702. Best score 749 bits
Score difference with first non-orthologous sequence - *A. fumigatus*:749 *A. fischeri*:749
afum|EAL92992.1 100.00% afis|EAW21977.1 100.00%
Bootstrap support for afum|EAL92992.1 as seed ortholog is 100%.
Bootstrap support for afis|EAW21977.1 as seed ortholog is 100%.

---

Group of orthologs #5703. Best score 748 bits
Score difference with first non-orthologous sequence - *A. fumigatus*:748 *A. fischeri*:748
afum|EAL87625.1 100.00% afis|EAW18261.1 100.00%
Bootstrap support for afum|EAL87625.1 as seed ortholog is 100%.
Bootstrap support for afis|EAW18261.1 as seed ortholog is 100%.

---

Group of orthologs #5704. Best score 748 bits
Score difference with first non-orthologous sequence - *A. fumigatus*:748 *A. fischeri*:684
afum|EAL86055.1 100.00% afis|EAW24383.1 100.00%
Bootstrap support for afum|EAL86055.1 as seed ortholog is 100%.
Bootstrap support for afis|EAW24383.1 as seed ortholog is 100%.

---

Group of orthologs #5705. Best score 748 bits
Score difference with first non-orthologous sequence - *A. fumigatus*:748 *A. fischeri*:748
afum|EAL93523.1 100.00% afis|EAW18789.1 100.00%
Bootstrap support for afum|EAL93523.1 as seed ortholog is 100%.
Bootstrap support for afis|EAW18789.1 as seed ortholog is 100%.

---

Group of orthologs #5706. Best score 748 bits
Score difference with first non-orthologous sequence - *A. fumigatus*:748 *A. fischeri*:748
afum|EAL93616.1 100.00% afis|EAW18883.1 100.00%
Bootstrap support for afum|EAL93616.1 as seed ortholog is 100%.
Bootstrap support for afis|EAW18883.1 as seed ortholog is 100%.

---

Group of orthologs #5707. Best score 748 bits
Score difference with first non-orthologous sequence - *A. fumigatus*:748 *A. fischeri*:748
afum|EAL93857.1 100.00% afis|EAW19125.1 100.00%
Bootstrap support for afum|EAL93857.1 as seed ortholog is 100%.
Bootstrap support for afis|EAW19125.1 as seed ortholog is 100%.

---

Group of orthologs #5708. Best score 748 bits
Score difference with first non-orthologous sequence - *A. fumigatus*:748 *A. fischeri*:748
afum|EAL90753.1 100.00% afis|EAW22438.1 100.00%
Bootstrap support for afum|EAL90753.1 as seed ortholog is 100%.
Bootstrap support for afis|EAW22438.1 as seed ortholog is 100%.

---
Group of orthologs #5709. Best score 748 bits
Score difference with first non-orthologous sequence - *A. fumigatus*:597 *A. fischeri*:456
afum|EAL90303.1 100.00% afis|EAW22895.1 100.00%
Bootstrap support for afum|EAL90303.1 as seed ortholog is 100%.
Bootstrap support for afis|EAW22895.1 as seed ortholog is 100%.

---
Group of orthologs #5710. Best score 748 bits
Score difference with first non-orthologous sequence - *A. fumigatus*:748 *A. fischeri*:423
afum|EAL94083.1 100.00% afis|EAW19364.1 100.00%
Bootstrap support for afum|EAL94083.1 as seed ortholog is 100%.
Bootstrap support for afis|EAW19364.1 as seed ortholog is 100%.

---
Group of orthologs #5711. Best score 747 bits
Score difference with first non-orthologous sequence - *A. fumigatus*:747 *A. fischeri*:747
afum|EAL86643.2 100.00% afis|EAW17218.1 100.00%
Bootstrap support for afum|EAL86643.2 as seed ortholog is 100%.
Bootstrap support for afis|EAW17218.1 as seed ortholog is 100%.

---
Group of orthologs #5712. Best score 747 bits
Score difference with first non-orthologous sequence - *A. fumigatus*:747 *A. fischeri*:651
afum|EAL84924.1 100.00% afis|EAW19993.1 100.00%
Bootstrap support for afum|EAL84924.1 as seed ortholog is 100%.
Bootstrap support for afis|EAW19993.1 as seed ortholog is 100%.

---
Group of orthologs #5713. Best score 747 bits
Score difference with first non-orthologous sequence - *A. fumigatus*:747 *A. fischeri*:747
afum|EAL91040.1 100.00% afis|EAW22150.1 100.00%
Bootstrap support for afum|EAL91040.1 as seed ortholog is 100%.
Bootstrap support for afis|EAW22150.1 as seed ortholog is 100%.

---
Group of orthologs #5714. Best score 747 bits
Score difference with first non-orthologous sequence - *A. fumigatus*:747 *A. fischeri*:747
afum|EAL90986.2 100.00% afis|EAW22207.1 100.00%
Bootstrap support for afum|EAL90986.2 as seed ortholog is 100%.
Bootstrap support for afis|EAW22207.1 as seed ortholog is 100%.

---
Group of orthologs #5715. Best score 747 bits
Score difference with first non-orthologous sequence - *A. fumigatus*:747 *A. fischeri*:747
afum|EAL90248.1 100.00% afis|EAW22949.1 100.00%
Bootstrap support for afum|EAL90248.1 as seed ortholog is 100%.
Bootstrap support for afis|EAW22949.1 as seed ortholog is 100%.

---
Group of orthologs #5716. Best score 746 bits
Score difference with first non-orthologous sequence - *A. fumigatus*:746 *A. fischeri*:746
afum|EAL84990.1 100.00% afis|EAW19918.1 100.00%
Bootstrap support for afum|EAL84990.1 as seed ortholog is 100%.
Bootstrap support for afis|EAW19918.1 as seed ortholog is 100%.

---
Group of orthologs #5717. Best score 746 bits
Score difference with first non-orthologous sequence - *A. fumigatus*:746 *A. fischeri*:746
afum|EAL88168.1 100.00% afis|EAW23309.1 100.00%
Bootstrap support for afum|EAL88168.1 as seed ortholog is 100%.
Bootstrap support for afis|EAW23309.1 as seed ortholog is 100%.

---
Group of orthologs #5718. Best score 745 bits

Score difference with first non-orthologous sequence - *A. fumigatus*:745 *A. fischeri*:745
afum|EAL86602.1 100.00% afis|EAW17170.1 100.00%
Bootstrap support for afum|EAL86602.1 as seed ortholog is 100%.
Bootstrap support for afis|EAW17170.1 as seed ortholog is 100%.

---

Group of orthologs #5719. Best score 745 bits
Score difference with first non-orthologous sequence - *A. fumigatus*:745 *A. fischeri*:745
afum|EAL88580.1 100.00% afis|EAW15947.1 100.00%
Bootstrap support for afum|EAL88580.1 as seed ortholog is 100%.
Bootstrap support for afis|EAW15947.1 as seed ortholog is 100%.

---

Group of orthologs #5720. Best score 745 bits
Score difference with first non-orthologous sequence - *A. fumigatus*:664 *A. fischeri*:117
afum|EAL88651.1 100.00% afis|EAW16022.1 100.00%
Bootstrap support for afum|EAL88651.1 as seed ortholog is 100%.
Bootstrap support for afis|EAW16022.1 as seed ortholog is 99%.

---

Group of orthologs #5721. Best score 745 bits
Score difference with first non-orthologous sequence - *A. fumigatus*:745 *A. fischeri*:745
afum|EAL85100.2 100.00% afis|EAW19791.1 100.00%
Bootstrap support for afum|EAL85100.2 as seed ortholog is 100%.
Bootstrap support for afis|EAW19791.1 as seed ortholog is 100%.

---

Group of orthologs #5722. Best score 745 bits
Score difference with first non-orthologous sequence - *A. fumigatus*:745 *A. fischeri*:745
afum|EAL85160.1 100.00% afis|EAW20040.1 100.00%
Bootstrap support for afum|EAL85160.1 as seed ortholog is 100%.
Bootstrap support for afis|EAW20040.1 as seed ortholog is 100%.

---

Group of orthologs #5723. Best score 745 bits
Score difference with first non-orthologous sequence - *A. fumigatus*:745 *A. fischeri*:516
afum|EAL86242.1 100.00% afis|EAW24612.1 100.00%
Bootstrap support for afum|EAL86242.1 as seed ortholog is 100%.
Bootstrap support for afis|EAW24612.1 as seed ortholog is 100%.

---

Group of orthologs #5724. Best score 745 bits
Score difference with first non-orthologous sequence - *A. fumigatus*:745 *A. fischeri*:745
afum|EAL87410.1 100.00% afis|EAW23980.1 100.00%
Bootstrap support for afum|EAL87410.1 as seed ortholog is 100%.
Bootstrap support for afis|EAW23980.1 as seed ortholog is 100%.

---

Group of orthologs #5725. Best score 745 bits
Score difference with first non-orthologous sequence - *A. fumigatus*:745 *A. fischeri*:745
afum|EAL90478.1 100.00% afis|EAW22716.1 100.00%
Bootstrap support for afum|EAL90478.1 as seed ortholog is 100%.
Bootstrap support for afis|EAW22716.1 as seed ortholog is 100%.

---

Group of orthologs #5726. Best score 745 bits
Score difference with first non-orthologous sequence - *A. fumigatus*:745 *A. fischeri*:745
afum|EAL89811.1 100.00% afis|EAW25139.1 100.00%
Bootstrap support for afum|EAL89811.1 as seed ortholog is 100%.
Bootstrap support for afis|EAW25139.1 as seed ortholog is 100%.

---

Group of orthologs #5727. Best score 744 bits
Score difference with first non-orthologous sequence - *A. fumigatus*:744 *A. fischeri*:744
afum|EAL88879.1 100.00% afis|EAW16258.1 100.00%

Bootstrap support for afum|EAL88879.1 as seed ortholog is 100%.
Bootstrap support for afis|EAW16258.1 as seed ortholog is 100%.

---

Group of orthologs #5728. Best score 744 bits
Score difference with first non-orthologous sequence - *A. fumigatus*:744 *A. fischeri*:744
afum|EAL84552.1 100.00% afis|EAW20504.1 100.00%
Bootstrap support for afum|EAL84552.1 as seed ortholog is 100%.
Bootstrap support for afis|EAW20504.1 as seed ortholog is 100%.

---

Group of orthologs #5729. Best score 744 bits
Score difference with first non-orthologous sequence - *A. fumigatus*:744 *A. fischeri*:744
afum|EAL85946.1 100.00% afis|EAW24277.1 100.00%
Bootstrap support for afum|EAL85946.1 as seed ortholog is 100%.
Bootstrap support for afis|EAW24277.1 as seed ortholog is 100%.

---

Group of orthologs #5730. Best score 744 bits
Score difference with first non-orthologous sequence - *A. fumigatus*:744 *A. fischeri*:744
afum|EAL93882.1 100.00% afis|EAW19151.1 100.00%
Bootstrap support for afum|EAL93882.1 as seed ortholog is 100%.
Bootstrap support for afis|EAW19151.1 as seed ortholog is 100%.

---

Group of orthologs #5731. Best score 744 bits
Score difference with first non-orthologous sequence - *A. fumigatus*:744 *A. fischeri*:356
afum|EAL89349.1 100.00% afis|EAW24707.1 100.00%
Bootstrap support for afum|EAL89349.1 as seed ortholog is 100%.
Bootstrap support for afis|EAW24707.1 as seed ortholog is 100%.

---

Group of orthologs #5732. Best score 743 bits
Score difference with first non-orthologous sequence - *A. fumigatus*:743 *A. fischeri*:743
afum|EAL85584.1 100.00% afis|EAW15764.1 100.00%
Bootstrap support for afum|EAL85584.1 as seed ortholog is 100%.
Bootstrap support for afis|EAW15764.1 as seed ortholog is 100%.

---

Group of orthologs #5733. Best score 743 bits
Score difference with first non-orthologous sequence - *A. fumigatus*:743 *A. fischeri*:743
afum|EAL88624.2 100.00% afis|EAW15992.1 100.00%
Bootstrap support for afum|EAL88624.2 as seed ortholog is 100%.
Bootstrap support for afis|EAW15992.1 as seed ortholog is 100%.

---

Group of orthologs #5734. Best score 743 bits
Score difference with first non-orthologous sequence - *A. fumigatus*:743 *A. fischeri*:743
afum|EAL84927.1 100.00% afis|EAW19988.1 100.00%
Bootstrap support for afum|EAL84927.1 as seed ortholog is 100%.
Bootstrap support for afis|EAW19988.1 as seed ortholog is 100%.

---

Group of orthologs #5735. Best score 743 bits
Score difference with first non-orthologous sequence - *A. fumigatus*:743 *A. fischeri*:743
afum|EAL91338.1 100.00% afis|EAW17523.1 100.00%
Bootstrap support for afum|EAL91338.1 as seed ortholog is 100%.
Bootstrap support for afis|EAW17523.1 as seed ortholog is 100%.

---

Group of orthologs #5736. Best score 743 bits
Score difference with first non-orthologous sequence - *A. fumigatus*:743 *A. fischeri*:743
afum|EAL91512.1 100.00% afis|EAW17698.1 100.00%
Bootstrap support for afum|EAL91512.1 as seed ortholog is 100%.
Bootstrap support for afis|EAW17698.1 as seed ortholog is 100%.

---
Group of orthologs #5737. Best score 742 bits
Score difference with first non-orthologous sequence - *A. fumigatus*:742 *A. fischeri*:742
afum|EAL84838.2 100.00% afis|EAW20786.1 100.00%
Bootstrap support for afum|EAL84838.2 as seed ortholog is 100%.
Bootstrap support for afis|EAW20786.1 as seed ortholog is 100%.

---
Group of orthologs #5738. Best score 742 bits
Score difference with first non-orthologous sequence - *A. fumigatus*:742 *A. fischeri*:742
afum|EAL87719.1 100.00% afis|EAW18356.1 100.00%
Bootstrap support for afum|EAL87719.1 as seed ortholog is 100%.
Bootstrap support for afis|EAW18356.1 as seed ortholog is 100%.

---
Group of orthologs #5739. Best score 742 bits
Score difference with first non-orthologous sequence - *A. fumigatus*:742 *A. fischeri*:742
afum|EAL91567.1 100.00% afis|EAW17754.1 100.00%
Bootstrap support for afum|EAL91567.1 as seed ortholog is 100%.
Bootstrap support for afis|EAW17754.1 as seed ortholog is 100%.

---
Group of orthologs #5740. Best score 742 bits
Score difference with first non-orthologous sequence - *A. fumigatus*:612 *A. fischeri*:421
afum|EAL89010.1 100.00% afis|EAW20387.1 100.00%
Bootstrap support for afum|EAL89010.1 as seed ortholog is 100%.
Bootstrap support for afis|EAW20387.1 as seed ortholog is 100%.

---
Group of orthologs #5741. Best score 742 bits
Score difference with first non-orthologous sequence - *A. fumigatus*:742 *A. fischeri*:742
afum|EAL86237.1 100.00% afis|EAW24607.1 100.00%
Bootstrap support for afum|EAL86237.1 as seed ortholog is 100%.
Bootstrap support for afis|EAW24607.1 as seed ortholog is 100%.

---
Group of orthologs #5742. Best score 742 bits
Score difference with first non-orthologous sequence - *A. fumigatus*:742 *A. fischeri*:742
afum|EAL93205.1 100.00% afis|EAW18510.1 100.00%
Bootstrap support for afum|EAL93205.1 as seed ortholog is 100%.
Bootstrap support for afis|EAW18510.1 as seed ortholog is 100%.

---
Group of orthologs #5743. Best score 742 bits
Score difference with first non-orthologous sequence - *A. fumigatus*:742 *A. fischeri*:742
afum|EAL90996.1 100.00% afis|EAW22195.1 100.00%
Bootstrap support for afum|EAL90996.1 as seed ortholog is 100%.
Bootstrap support for afis|EAW22195.1 as seed ortholog is 100%.

---
Group of orthologs #5744. Best score 742 bits
Score difference with first non-orthologous sequence - *A. fumigatus*:742 *A. fischeri*:742
afum|EAL90664.1 100.00% afis|EAW22530.1 100.00%
Bootstrap support for afum|EAL90664.1 as seed ortholog is 100%.
Bootstrap support for afis|EAW22530.1 as seed ortholog is 100%.

---
Group of orthologs #5745. Best score 742 bits
Score difference with first non-orthologous sequence - *A. fumigatus*:742 *A. fischeri*:686
afum|EAL90583.1 100.00% afis|EAW22615.1 100.00%
Bootstrap support for afum|EAL90583.1 as seed ortholog is 100%.
Bootstrap support for afis|EAW22615.1 as seed ortholog is 100%.

---
Group of orthologs #5746. Best score 742 bits

Score difference with first non-orthologous sequence - *A. fumigatus*:742 *A. fischeri*:456
afum|EAL92221.1 100.00% afis|EAW21210.1 100.00%
Bootstrap support for afum|EAL92221.1 as seed ortholog is 100%.
Bootstrap support for afis|EAW21210.1 as seed ortholog is 100%.

---

Group of orthologs #5747. Best score 742 bits
Score difference with first non-orthologous sequence - *A. fumigatus*:742 *A. fischeri*:742
afum|EAL92900.1 100.00% afis|EAW21886.1 100.00%
Bootstrap support for afum|EAL92900.1 as seed ortholog is 100%.
Bootstrap support for afis|EAW21886.1 as seed ortholog is 100%.

---

Group of orthologs #5748. Best score 742 bits
Score difference with first non-orthologous sequence - *A. fumigatus*:742 *A. fischeri*:742
afum|EAL89846.1 100.00% afis|EAW25172.1 100.00%
Bootstrap support for afum|EAL89846.1 as seed ortholog is 100%.
Bootstrap support for afis|EAW25172.1 as seed ortholog is 100%.

---

Group of orthologs #5749. Best score 741 bits
Score difference with first non-orthologous sequence - *A. fumigatus*:741 *A. fischeri*:741
afum|EAL85551.1 100.00% afis|EAW15799.1 100.00%
Bootstrap support for afum|EAL85551.1 as seed ortholog is 100%.
Bootstrap support for afis|EAW15799.1 as seed ortholog is 100%.

---

Group of orthologs #5750. Best score 741 bits
Score difference with first non-orthologous sequence - *A. fumigatus*:741 *A. fischeri*:741
afum|EAL90897.1 100.00% afis|EAW22284.1 100.00%
Bootstrap support for afum|EAL90897.1 as seed ortholog is 100%.
Bootstrap support for afis|EAW22284.1 as seed ortholog is 100%.

---

Group of orthologs #5751. Best score 741 bits
Score difference with first non-orthologous sequence - *A. fumigatus*:672 *A. fischeri*:648
afum|EAL92225.1 100.00% afis|EAW21214.1 100.00%
Bootstrap support for afum|EAL92225.1 as seed ortholog is 100%.
Bootstrap support for afis|EAW21214.1 as seed ortholog is 100%.

---

Group of orthologs #5752. Best score 740 bits
Score difference with first non-orthologous sequence - *A. fumigatus*:640 *A. fischeri*:740
afum|EAL91766.1 100.00% afis|EAW17955.1 100.00%
Bootstrap support for afum|EAL91766.1 as seed ortholog is 100%.
Bootstrap support for afis|EAW17955.1 as seed ortholog is 100%.

---

Group of orthologs #5753. Best score 740 bits
Score difference with first non-orthologous sequence - *A. fumigatus*:583 *A. fischeri*:589
afum|EAL92120.1 100.00% afis|EAW21097.1 100.00%
Bootstrap support for afum|EAL92120.1 as seed ortholog is 100%.
Bootstrap support for afis|EAW21097.1 as seed ortholog is 100%.

---

Group of orthologs #5754. Best score 740 bits
Score difference with first non-orthologous sequence - *A. fumigatus*:740 *A. fischeri*:740
afum|EAL91051.1 100.00% afis|EAW22139.1 100.00%
Bootstrap support for afum|EAL91051.1 as seed ortholog is 100%.
Bootstrap support for afis|EAW22139.1 as seed ortholog is 100%.

---

Group of orthologs #5755. Best score 740 bits
Score difference with first non-orthologous sequence - *A. fumigatus*:740 *A. fischeri*:740
afum|EAL92742.1 100.00% afis|EAW21707.1 100.00%

Bootstrap support for afum|EAL92742.1 as seed ortholog is 100%.
Bootstrap support for afis|EAW21707.1 as seed ortholog is 100%.

---

Group of orthologs #5756. Best score 739 bits
Score difference with first non-orthologous sequence - *A. fumigatus*:739 *A. fischeri*:739
afum|EAL85721.1 100.00% afis|EAW15614.1 100.00%
Bootstrap support for afum|EAL85721.1 as seed ortholog is 100%.
Bootstrap support for afis|EAW15614.1 as seed ortholog is 100%.

---

Group of orthologs #5757. Best score 739 bits
Score difference with first non-orthologous sequence - *A. fumigatus*:739 *A. fischeri*:40
afum|EAL84427.1 100.00% afis|EAW20646.1 100.00%
Bootstrap support for afum|EAL84427.1 as seed ortholog is 100%.
Bootstrap support for afis|EAW20646.1 as seed ortholog is 94%.

---

Group of orthologs #5758. Best score 739 bits
Score difference with first non-orthologous sequence - *A. fumigatus*:739 *A. fischeri*:739
afum|EAL87484.1 100.00% afis|EAW18126.1 100.00%
Bootstrap support for afum|EAL87484.1 as seed ortholog is 100%.
Bootstrap support for afis|EAW18126.1 as seed ortholog is 100%.

---

Group of orthologs #5759. Best score 739 bits
Score difference with first non-orthologous sequence - *A. fumigatus*:739 *A. fischeri*:739
afum|EAL86005.1 100.00% afis|EAW24335.1 100.00%
Bootstrap support for afum|EAL86005.1 as seed ortholog is 100%.
Bootstrap support for afis|EAW24335.1 as seed ortholog is 100%.

---

Group of orthologs #5760. Best score 739 bits
Score difference with first non-orthologous sequence - *A. fumigatus*:739 *A. fischeri*:621
afum|EAL87293.1 100.00% afis|EAW23865.1 100.00%
Bootstrap support for afum|EAL87293.1 as seed ortholog is 100%.
Bootstrap support for afis|EAW23865.1 as seed ortholog is 100%.

---

Group of orthologs #5761. Best score 739 bits
Score difference with first non-orthologous sequence - *A. fumigatus*:739 *A. fischeri*:739
afum|EAL90718.1 100.00% afis|EAW22473.1 100.00%
Bootstrap support for afum|EAL90718.1 as seed ortholog is 100%.
Bootstrap support for afis|EAW22473.1 as seed ortholog is 100%.

---

Group of orthologs #5762. Best score 739 bits
Score difference with first non-orthologous sequence - *A. fumigatus*:739 *A. fischeri*:739
afum|EAL90246.1 100.00% afis|EAW22951.1 100.00%
Bootstrap support for afum|EAL90246.1 as seed ortholog is 100%.
Bootstrap support for afis|EAW22951.1 as seed ortholog is 100%.

---

Group of orthologs #5763. Best score 738 bits
Score difference with first non-orthologous sequence - *A. fumigatus*:738 *A. fischeri*:738
afum|EAL88576.1 100.00% afis|EAW15942.1 100.00%
Bootstrap support for afum|EAL88576.1 as seed ortholog is 100%.
Bootstrap support for afis|EAW15942.1 as seed ortholog is 100%.

---

Group of orthologs #5764. Best score 738 bits
Score difference with first non-orthologous sequence - *A. fumigatus*:738 *A. fischeri*:738
afum|EAL84673.1 100.00% afis|EAW20960.1 100.00%
Bootstrap support for afum|EAL84673.1 as seed ortholog is 100%.
Bootstrap support for afis|EAW20960.1 as seed ortholog is 100%.

---
Group of orthologs #5765. Best score 738 bits
Score difference with first non-orthologous sequence - *A. fumigatus*:663 *A. fischeri*:609
afum|EAL86987.1 100.00% afis|EAW19507.1 100.00%
Bootstrap support for afum|EAL86987.1 as seed ortholog is 100%.
Bootstrap support for afis|EAW19507.1 as seed ortholog is 100%.

---
Group of orthologs #5766. Best score 738 bits
Score difference with first non-orthologous sequence - *A. fumigatus*:738 *A. fischeri*:738
afum|EAL93306.1 100.00% afis|EAW18598.1 100.00%
Bootstrap support for afum|EAL93306.1 as seed ortholog is 100%.
Bootstrap support for afis|EAW18598.1 as seed ortholog is 100%.

---
Group of orthologs #5767. Best score 738 bits
Score difference with first non-orthologous sequence - *A. fumigatus*:738 *A. fischeri*:738
afum|EAL89416.1 100.00% afis|EAW24775.1 100.00%
Bootstrap support for afum|EAL89416.1 as seed ortholog is 100%.
Bootstrap support for afis|EAW24775.1 as seed ortholog is 100%.

---
Group of orthologs #5768. Best score 738 bits
Score difference with first non-orthologous sequence - *A. fumigatus*:738 *A. fischeri*:738
afum|EAL89524.1 100.00% afis|EAW24874.1 100.00%
Bootstrap support for afum|EAL89524.1 as seed ortholog is 100%.
Bootstrap support for afis|EAW24874.1 as seed ortholog is 100%.

---
Group of orthologs #5769. Best score 738 bits
Score difference with first non-orthologous sequence - *A. fumigatus*:738 *A. fischeri*:738
afum|EAL89713.1 100.00% afis|EAW25045.1 100.00%
Bootstrap support for afum|EAL89713.1 as seed ortholog is 100%.
Bootstrap support for afis|EAW25045.1 as seed ortholog is 100%.

---
Group of orthologs #5770. Best score 737 bits
Score difference with first non-orthologous sequence - *A. fumigatus*:656 *A. fischeri*:737
afum|EAL84200.1 100.00% afis|EAW15446.1 100.00%
Bootstrap support for afum|EAL84200.1 as seed ortholog is 100%.
Bootstrap support for afis|EAW15446.1 as seed ortholog is 100%.

---
Group of orthologs #5771. Best score 737 bits
Score difference with first non-orthologous sequence - *A. fumigatus*:737 *A. fischeri*:737
afum|EAL86360.1 100.00% afis|EAW16791.1 100.00%
Bootstrap support for afum|EAL86360.1 as seed ortholog is 100%.
Bootstrap support for afis|EAW16791.1 as seed ortholog is 100%.

---
Group of orthologs #5772. Best score 737 bits
Score difference with first non-orthologous sequence - *A. fumigatus*:737 *A. fischeri*:737
afum|EAL88700.2 100.00% afis|EAW16068.1 100.00%
Bootstrap support for afum|EAL88700.2 as seed ortholog is 100%.
Bootstrap support for afis|EAW16068.1 as seed ortholog is 100%.

---
Group of orthologs #5773. Best score 737 bits
Score difference with first non-orthologous sequence - *A. fumigatus*:635 *A. fischeri*:393
afum|EAL85271.1 100.00% afis|EAW20156.1 100.00%
Bootstrap support for afum|EAL85271.1 as seed ortholog is 100%.
Bootstrap support for afis|EAW20156.1 as seed ortholog is 100%.

---
Group of orthologs #5774. Best score 737 bits

Score difference with first non-orthologous sequence - *A. fumigatus*:737   *A. fischeri*:737
afum|EAL88481.1 100.00% afis|EAW22999.1 100.00%
Bootstrap support for afum|EAL88481.1 as seed ortholog is 100%.
Bootstrap support for afis|EAW22999.1 as seed ortholog is 100%.

---

Group of orthologs #5775. Best score 737 bits
Score difference with first non-orthologous sequence - *A. fumigatus*:737   *A. fischeri*:737
afum|EAL88240.1 100.00% afis|EAW23242.1 100.00%
Bootstrap support for afum|EAL88240.1 as seed ortholog is 100%.
Bootstrap support for afis|EAW23242.1 as seed ortholog is 100%.

---

Group of orthologs #5776. Best score 737 bits
Score difference with first non-orthologous sequence - *A. fumigatus*:737   *A. fischeri*:737
afum|EAL92227.2 100.00% afis|EAW21216.1 100.00%
Bootstrap support for afum|EAL92227.2 as seed ortholog is 100%.
Bootstrap support for afis|EAW21216.1 as seed ortholog is 100%.

---

Group of orthologs #5777. Best score 736 bits
Score difference with first non-orthologous sequence - *A. fumigatus*:736   *A. fischeri*:736
afum|EAL85063.1 100.00% afis|EAW19829.1 100.00%
Bootstrap support for afum|EAL85063.1 as seed ortholog is 100%.
Bootstrap support for afis|EAW19829.1 as seed ortholog is 100%.

---

Group of orthologs #5778. Best score 736 bits
Score difference with first non-orthologous sequence - *A. fumigatus*:736   *A. fischeri*:736
afum|EAL85261.1 100.00% afis|EAW20146.1 100.00%
Bootstrap support for afum|EAL85261.1 as seed ortholog is 100%.
Bootstrap support for afis|EAW20146.1 as seed ortholog is 100%.

---

Group of orthologs #5779. Best score 736 bits
Score difference with first non-orthologous sequence - *A. fumigatus*:736   *A. fischeri*:674
afum|EAL84828.1 100.00% afis|EAW20801.1 100.00%
Bootstrap support for afum|EAL84828.1 as seed ortholog is 100%.
Bootstrap support for afis|EAW20801.1 as seed ortholog is 100%.

---

Group of orthologs #5780. Best score 736 bits
Score difference with first non-orthologous sequence - *A. fumigatus*:736   *A. fischeri*:656
afum|EAL86102.1 100.00% afis|EAW24434.1 100.00%
Bootstrap support for afum|EAL86102.1 as seed ortholog is 100%.
Bootstrap support for afis|EAW24434.1 as seed ortholog is 100%.

---

Group of orthologs #5781. Best score 736 bits
Score difference with first non-orthologous sequence - *A. fumigatus*:736   *A. fischeri*:736
afum|EAL87241.1 100.00% afis|EAW23807.1 100.00%
Bootstrap support for afum|EAL87241.1 as seed ortholog is 100%.
Bootstrap support for afis|EAW23807.1 as seed ortholog is 100%.

---

Group of orthologs #5782. Best score 736 bits
Score difference with first non-orthologous sequence - *A. fumigatus*:736   *A. fischeri*:736
afum|EAL93700.2 100.00% afis|EAW18968.1 100.00%
Bootstrap support for afum|EAL93700.2 as seed ortholog is 100%.
Bootstrap support for afis|EAW18968.1 as seed ortholog is 100%.

---

Group of orthologs #5783. Best score 736 bits
Score difference with first non-orthologous sequence - *A. fumigatus*:736   *A. fischeri*:736
afum|EAL92761.1 100.00% afis|EAW21728.1 100.00%

Bootstrap support for afum|EAL92761.1 as seed ortholog is 100%.
Bootstrap support for afis|EAW21728.1 as seed ortholog is 100%.

---

Group of orthologs #5784. Best score 735 bits
Score difference with first non-orthologous sequence - *A. fumigatus*:735 *A. fischeri*:735
afum|EAL84829.1 100.00% afis|EAW20800.1 100.00%
Bootstrap support for afum|EAL84829.1 as seed ortholog is 100%.
Bootstrap support for afis|EAW20800.1 as seed ortholog is 100%.

---

Group of orthologs #5785. Best score 735 bits
Score difference with first non-orthologous sequence - *A. fumigatus*:735 *A. fischeri*:735
afum|EAL91789.1 100.00% afis|EAW17980.1 100.00%
Bootstrap support for afum|EAL91789.1 as seed ortholog is 100%.
Bootstrap support for afis|EAW17980.1 as seed ortholog is 100%.

---

Group of orthologs #5786. Best score 735 bits
Score difference with first non-orthologous sequence - *A. fumigatus*:735 *A. fischeri*:735
afum|EAL91795.1 100.00% afis|EAW17986.1 100.00%
Bootstrap support for afum|EAL91795.1 as seed ortholog is 100%.
Bootstrap support for afis|EAW17986.1 as seed ortholog is 100%.

---

Group of orthologs #5787. Best score 735 bits
Score difference with first non-orthologous sequence - *A. fumigatus*:735 *A. fischeri*:735
afum|EAL93320.1 100.00% afis|EAW18610.1 100.00%
Bootstrap support for afum|EAL93320.1 as seed ortholog is 100%.
Bootstrap support for afis|EAW18610.1 as seed ortholog is 100%.

---

Group of orthologs #5788. Best score 735 bits
Score difference with first non-orthologous sequence - *A. fumigatus*:735 *A. fischeri*:735
afum|EAL90630.1 100.00% afis|EAW22568.1 100.00%
Bootstrap support for afum|EAL90630.1 as seed ortholog is 100%.
Bootstrap support for afis|EAW22568.1 as seed ortholog is 100%.

---

Group of orthologs #5789. Best score 735 bits
Score difference with first non-orthologous sequence - *A. fumigatus*:735 *A. fischeri*:735
afum|EAL92236.1 100.00% afis|EAW21223.1 100.00%
Bootstrap support for afum|EAL92236.1 as seed ortholog is 100%.
Bootstrap support for afis|EAW21223.1 as seed ortholog is 100%.

---

Group of orthologs #5790. Best score 735 bits
Score difference with first non-orthologous sequence - *A. fumigatus*:735 *A. fischeri*:735
afum|EAL92252.1 100.00% afis|EAW21239.1 100.00%
Bootstrap support for afum|EAL92252.1 as seed ortholog is 100%.
Bootstrap support for afis|EAW21239.1 as seed ortholog is 100%.

---

Group of orthologs #5791. Best score 735 bits
Score difference with first non-orthologous sequence - *A. fumigatus*:735 *A. fischeri*:476
afum|EAL92272.1 100.00% afis|EAW21257.1 100.00%
Bootstrap support for afum|EAL92272.1 as seed ortholog is 100%.
Bootstrap support for afis|EAW21257.1 as seed ortholog is 100%.

---

Group of orthologs #5792. Best score 734 bits
Score difference with first non-orthologous sequence - *A. fumigatus*:734 *A. fischeri*:734
afum|EAL84212.1 100.00% afis|EAW15431.1 100.00%
Bootstrap support for afum|EAL84212.1 as seed ortholog is 100%.
Bootstrap support for afis|EAW15431.1 as seed ortholog is 100%.

---

Group of orthologs #5793. Best score 734 bits
Score difference with first non-orthologous sequence - *A. fumigatus*:734 *A. fischeri*:734
afum|EAL88579.1 100.00% afis|EAW15945.1 100.00%
Bootstrap support for afum|EAL88579.1 as seed ortholog is 100%.
Bootstrap support for afis|EAW15945.1 as seed ortholog is 100%.

---

Group of orthologs #5794. Best score 734 bits
Score difference with first non-orthologous sequence - *A. fumigatus*:734 *A. fischeri*:734
afum|EAL88883.1 100.00% afis|EAW16262.1 100.00%
Bootstrap support for afum|EAL88883.1 as seed ortholog is 100%.
Bootstrap support for afis|EAW16262.1 as seed ortholog is 100%.

---

Group of orthologs #5795. Best score 734 bits
Score difference with first non-orthologous sequence - *A. fumigatus*:734 *A. fischeri*:555
afum|EAL84585.1 100.00% afis|EAW20467.1 100.00%
Bootstrap support for afum|EAL84585.1 as seed ortholog is 100%.
Bootstrap support for afis|EAW20467.1 as seed ortholog is 100%.

---

Group of orthologs #5796. Best score 734 bits
Score difference with first non-orthologous sequence - *A. fumigatus*:679 *A. fischeri*:634
afum|EAL93058.1 100.00% afis|EAW18383.1 100.00%
Bootstrap support for afum|EAL93058.1 as seed ortholog is 100%.
Bootstrap support for afis|EAW18383.1 as seed ortholog is 100%.

---

Group of orthologs #5797. Best score 734 bits
Score difference with first non-orthologous sequence - *A. fumigatus*:734 *A. fischeri*:734
afum|EAL93991.1 100.00% afis|EAW19264.1 100.00%
Bootstrap support for afum|EAL93991.1 as seed ortholog is 100%.
Bootstrap support for afis|EAW19264.1 as seed ortholog is 100%.

---

Group of orthologs #5798. Best score 734 bits
Score difference with first non-orthologous sequence - *A. fumigatus*:734 *A. fischeri*:734
afum|EAL90324.1 100.00% afis|EAW22874.1 100.00%
Bootstrap support for afum|EAL90324.1 as seed ortholog is 100%.
Bootstrap support for afis|EAW22874.1 as seed ortholog is 100%.

---

Group of orthologs #5799. Best score 734 bits
Score difference with first non-orthologous sequence - *A. fumigatus*:734 *A. fischeri*:734
afum|EAL89719.1 100.00% afis|EAW25051.1 100.00%
Bootstrap support for afum|EAL89719.1 as seed ortholog is 100%.
Bootstrap support for afis|EAW25051.1 as seed ortholog is 100%.

---

Group of orthologs #5800. Best score 733 bits
Score difference with first non-orthologous sequence - *A. fumigatus*:733 *A. fischeri*:733
afum|EAL85250.1 100.00% afis|EAW20134.1 100.00%
Bootstrap support for afum|EAL85250.1 as seed ortholog is 100%.
Bootstrap support for afis|EAW20134.1 as seed ortholog is 100%.

---

Group of orthologs #5801. Best score 733 bits
Score difference with first non-orthologous sequence - *A. fumigatus*:733 *A. fischeri*:733
afum|EAL87026.1 100.00% afis|EAW19470.1 100.00%
Bootstrap support for afum|EAL87026.1 as seed ortholog is 100%.
Bootstrap support for afis|EAW19470.1 as seed ortholog is 100%.

---

Group of orthologs #5802. Best score 733 bits

Score difference with first non-orthologous sequence - *A. fumigatus*:733 *A. fischeri*:733
afum|EAL90716.1 100.00% afis|EAW22475.1 100.00%
Bootstrap support for afum|EAL90716.1 as seed ortholog is 100%.
Bootstrap support for afis|EAW22475.1 as seed ortholog is 100%.

---

Group of orthologs #5803. Best score 733 bits
Score difference with first non-orthologous sequence - *A. fumigatus*:733 *A. fischeri*:733
afum|EAL90344.1 100.00% afis|EAW22856.1 100.00%
Bootstrap support for afum|EAL90344.1 as seed ortholog is 100%.
Bootstrap support for afis|EAW22856.1 as seed ortholog is 100%.

---

Group of orthologs #5804. Best score 732 bits
Score difference with first non-orthologous sequence - *A. fumigatus*:732 *A. fischeri*:732
afum|EAL85061.1 100.00% afis|EAW19831.1 100.00%
Bootstrap support for afum|EAL85061.1 as seed ortholog is 100%.
Bootstrap support for afis|EAW19831.1 as seed ortholog is 100%.

---

Group of orthologs #5805. Best score 732 bits
Score difference with first non-orthologous sequence - *A. fumigatus*:732 *A. fischeri*:732
afum|EAL88923.1 100.00% afis|EAW16301.1 100.00%
Bootstrap support for afum|EAL88923.1 as seed ortholog is 100%.
Bootstrap support for afis|EAW16301.1 as seed ortholog is 100%.

---

Group of orthologs #5806. Best score 732 bits
Score difference with first non-orthologous sequence - *A. fumigatus*:732 *A. fischeri*:549
afum|EAL84501.1 100.00% afis|EAW20560.1 100.00%
Bootstrap support for afum|EAL84501.1 as seed ortholog is 100%.
Bootstrap support for afis|EAW20560.1 as seed ortholog is 100%.

---

Group of orthologs #5807. Best score 732 bits
Score difference with first non-orthologous sequence - *A. fumigatus*:732 *A. fischeri*:732
afum|EAL88238.1 100.00% afis|EAW23244.1 100.00%
Bootstrap support for afum|EAL88238.1 as seed ortholog is 100%.
Bootstrap support for afis|EAW23244.1 as seed ortholog is 100%.

---

Group of orthologs #5808. Best score 732 bits
Score difference with first non-orthologous sequence - *A. fumigatus*:732 *A. fischeri*:469
afum|EAL92110.2 100.00% afis|EAW21087.1 100.00%
Bootstrap support for afum|EAL92110.2 as seed ortholog is 100%.
Bootstrap support for afis|EAW21087.1 as seed ortholog is 100%.

---

Group of orthologs #5809. Best score 731 bits
Score difference with first non-orthologous sequence - *A. fumigatus*:731 *A. fischeri*:731
afum|EAL85585.1 100.00% afis|EAW15763.1 100.00%
Bootstrap support for afum|EAL85585.1 as seed ortholog is 100%.
Bootstrap support for afis|EAW15763.1 as seed ortholog is 100%.

---

Group of orthologs #5810. Best score 731 bits
Score difference with first non-orthologous sequence - *A. fumigatus*:731 *A. fischeri*:731
afum|EAL87615.1 100.00% afis|EAW18250.1 100.00%
Bootstrap support for afum|EAL87615.1 as seed ortholog is 100%.
Bootstrap support for afis|EAW18250.1 as seed ortholog is 100%.

---

Group of orthologs #5811. Best score 731 bits
Score difference with first non-orthologous sequence - *A. fumigatus*:731 *A. fischeri*:731
afum|EBA27201.1 100.00% afis|EAW20968.1 100.00%

Bootstrap support for afum|EBA27201.1 as seed ortholog is 100%.
Bootstrap support for afis|EAW20968.1 as seed ortholog is 100%.

---

Group of orthologs #5812. Best score 730 bits
Score difference with first non-orthologous sequence - *A. fumigatus*:730 *A. fischeri*:730
afum|EAL85220.1 100.00% afis|EAW20103.1 100.00%
Bootstrap support for afum|EAL85220.1 as seed ortholog is 100%.
Bootstrap support for afis|EAW20103.1 as seed ortholog is 100%.

---

Group of orthologs #5813. Best score 730 bits
Score difference with first non-orthologous sequence - *A. fumigatus*:730 *A. fischeri*:730
afum|EAL91496.1 100.00% afis|EAW17682.1 100.00%
Bootstrap support for afum|EAL91496.1 as seed ortholog is 100%.
Bootstrap support for afis|EAW17682.1 as seed ortholog is 100%.

---

Group of orthologs #5814. Best score 730 bits
Score difference with first non-orthologous sequence - *A. fumigatus*:730 *A. fischeri*:730
afum|EAL93049.1 100.00% afis|EAW18374.1 100.00%
Bootstrap support for afum|EAL93049.1 as seed ortholog is 100%.
Bootstrap support for afis|EAW18374.1 as seed ortholog is 100%.

---

Group of orthologs #5815. Best score 730 bits
Score difference with first non-orthologous sequence - *A. fumigatus*:730 *A. fischeri*:730
afum|EAL88453.2 100.00% afis|EAW23028.1 100.00%
Bootstrap support for afum|EAL88453.2 as seed ortholog is 100%.
Bootstrap support for afis|EAW23028.1 as seed ortholog is 100%.

---

Group of orthologs #5816. Best score 730 bits
Score difference with first non-orthologous sequence - *A. fumigatus*:730 *A. fischeri*:730
afum|EAL93750.1 100.00% afis|EAW19018.1 100.00%
Bootstrap support for afum|EAL93750.1 as seed ortholog is 100%.
Bootstrap support for afis|EAW19018.1 as seed ortholog is 100%.

---

Group of orthologs #5817. Best score 730 bits
Score difference with first non-orthologous sequence - *A. fumigatus*:730 *A. fischeri*:730
afum|EAL90636.1 100.00% afis|EAW22559.1 100.00%
Bootstrap support for afum|EAL90636.1 as seed ortholog is 100%.
Bootstrap support for afis|EAW22559.1 as seed ortholog is 100%.

---

Group of orthologs #5818. Best score 730 bits
Score difference with first non-orthologous sequence - *A. fumigatus*:599 *A. fischeri*:477
afum|EAL90322.1 100.00% afis|EAW22877.1 100.00%
Bootstrap support for afum|EAL90322.1 as seed ortholog is 100%.
Bootstrap support for afis|EAW22877.1 as seed ortholog is 100%.

---

Group of orthologs #5819. Best score 730 bits
Score difference with first non-orthologous sequence - *A. fumigatus*:730 *A. fischeri*:730
afum|EAL92787.1 100.00% afis|EAW21757.1 100.00%
Bootstrap support for afum|EAL92787.1 as seed ortholog is 100%.
Bootstrap support for afis|EAW21757.1 as seed ortholog is 100%.

---

Group of orthologs #5820. Best score 730 bits
Score difference with first non-orthologous sequence - *A. fumigatus*:730 *A. fischeri*:730
afum|EAL89530.1 100.00% afis|EAW24880.1 100.00%
Bootstrap support for afum|EAL89530.1 as seed ortholog is 100%.
Bootstrap support for afis|EAW24880.1 as seed ortholog is 100%.

Group of orthologs #5821. Best score 730 bits
Score difference with first non-orthologous sequence - *A. fumigatus*:730 *A. fischeri*:730
afum|EAL89637.1 100.00% afis|EAW24977.1 100.00%
Bootstrap support for afum|EAL89637.1 as seed ortholog is 100%.
Bootstrap support for afis|EAW24977.1 as seed ortholog is 100%.

---

Group of orthologs #5822. Best score 730 bits
Score difference with first non-orthologous sequence - *A. fumigatus*:730 *A. fischeri*:730
afum|EAL92937.1 100.00% afis|EAW21924.1 100.00%
Bootstrap support for afum|EAL92937.1 as seed ortholog is 100%.
Bootstrap support for afis|EAW21924.1 as seed ortholog is 100%.

---

Group of orthologs #5823. Best score 730 bits
Score difference with first non-orthologous sequence - *A. fumigatus*:730 *A. fischeri*:730
afum|EBA27308.1 100.00% afis|EAW23886.1 100.00%
Bootstrap support for afum|EBA27308.1 as seed ortholog is 100%.
Bootstrap support for afis|EAW23886.1 as seed ortholog is 100%.

---

Group of orthologs #5824. Best score 729 bits
Score difference with first non-orthologous sequence - *A. fumigatus*:729 *A. fischeri*:729
afum|EAL84588.1 100.00% afis|EAW20465.1 100.00%
Bootstrap support for afum|EAL84588.1 as seed ortholog is 100%.
Bootstrap support for afis|EAW20465.1 as seed ortholog is 100%.

---

Group of orthologs #5825. Best score 729 bits
Score difference with first non-orthologous sequence - *A. fumigatus*:729 *A. fischeri*:729
afum|EAL91428.1 100.00% afis|EAW17615.1 100.00%
Bootstrap support for afum|EAL91428.1 as seed ortholog is 100%.
Bootstrap support for afis|EAW17615.1 as seed ortholog is 100%.

---

Group of orthologs #5826. Best score 729 bits
Score difference with first non-orthologous sequence - *A. fumigatus*:729 *A. fischeri*:729
afum|EAL85876.1 100.00% afis|EAW24206.1 100.00%
Bootstrap support for afum|EAL85876.1 as seed ortholog is 100%.
Bootstrap support for afis|EAW24206.1 as seed ortholog is 100%.

---

Group of orthologs #5827. Best score 729 bits
Score difference with first non-orthologous sequence - *A. fumigatus*:729 *A. fischeri*:729
afum|EAL93252.1 100.00% afis|EAW18552.1 100.00%
Bootstrap support for afum|EAL93252.1 as seed ortholog is 100%.
Bootstrap support for afis|EAW18552.1 as seed ortholog is 100%.

---

Group of orthologs #5828. Best score 729 bits
Score difference with first non-orthologous sequence - *A. fumigatus*:729 *A. fischeri*:729
afum|EAL87463.1 100.00% afis|EAW24032.1 100.00%
Bootstrap support for afum|EAL87463.1 as seed ortholog is 100%.
Bootstrap support for afis|EAW24032.1 as seed ortholog is 100%.

---

Group of orthologs #5829. Best score 729 bits
Score difference with first non-orthologous sequence - *A. fumigatus*:729 *A. fischeri*:729
afum|EAL93793.2 100.00% afis|EAW19064.1 100.00%
Bootstrap support for afum|EAL93793.2 as seed ortholog is 100%.
Bootstrap support for afis|EAW19064.1 as seed ortholog is 100%.

---

Group of orthologs #5830. Best score 729 bits

Score difference with first non-orthologous sequence - *A. fumigatus*:729 *A. fischeri*:729
afum|EAL92695.1 100.00% afis|EAW21660.1 100.00%
Bootstrap support for afum|EAL92695.1 as seed ortholog is 100%.
Bootstrap support for afis|EAW21660.1 as seed ortholog is 100%.

---

Group of orthologs #5831. Best score 729 bits
Score difference with first non-orthologous sequence - *A. fumigatus*:729 *A. fischeri*:729
afum|EAL89559.1 100.00% afis|EAW24906.1 100.00%
Bootstrap support for afum|EAL89559.1 as seed ortholog is 100%.
Bootstrap support for afis|EAW24906.1 as seed ortholog is 100%.

---

Group of orthologs #5832. Best score 729 bits
Score difference with first non-orthologous sequence - *A. fumigatus*:729 *A. fischeri*:729
afum|EAL92835.1 100.00% afis|EAW21819.1 100.00%
Bootstrap support for afum|EAL92835.1 as seed ortholog is 100%.
Bootstrap support for afis|EAW21819.1 as seed ortholog is 100%.

---

Group of orthologs #5833. Best score 728 bits
Score difference with first non-orthologous sequence - *A. fumigatus*:728 *A. fischeri*:728
afum|EAL86312.1 100.00% afis|EAW16737.1 100.00%
Bootstrap support for afum|EAL86312.1 as seed ortholog is 100%.
Bootstrap support for afis|EAW16737.1 as seed ortholog is 100%.

---

Group of orthologs #5834. Best score 728 bits
Score difference with first non-orthologous sequence - *A. fumigatus*:728 *A. fischeri*:728
afum|EAL86568.1 100.00% afis|EAW17090.1 100.00%
Bootstrap support for afum|EAL86568.1 as seed ortholog is 100%.
Bootstrap support for afis|EAW17090.1 as seed ortholog is 100%.

---

Group of orthologs #5835. Best score 728 bits
Score difference with first non-orthologous sequence - *A. fumigatus*:728 *A. fischeri*:728
afum|EAL93206.1 100.00% afis|EAW18511.1 100.00%
Bootstrap support for afum|EAL93206.1 as seed ortholog is 100%.
Bootstrap support for afis|EAW18511.1 as seed ortholog is 100%.

---

Group of orthologs #5836. Best score 728 bits
Score difference with first non-orthologous sequence - *A. fumigatus*:728 *A. fischeri*:728
afum|EAL90728.1 100.00% afis|EAW22463.1 100.00%
Bootstrap support for afum|EAL90728.1 as seed ortholog is 100%.
Bootstrap support for afis|EAW22463.1 as seed ortholog is 100%.

---

Group of orthologs #5837. Best score 728 bits
Score difference with first non-orthologous sequence - *A. fumigatus*:728 *A. fischeri*:728
afum|EAL92869.1 100.00% afis|EAW21856.1 100.00%
Bootstrap support for afum|EAL92869.1 as seed ortholog is 100%.
Bootstrap support for afis|EAW21856.1 as seed ortholog is 100%.

---

Group of orthologs #5838. Best score 727 bits
Score difference with first non-orthologous sequence - *A. fumigatus*:496 *A. fischeri*:637
afum|EAL88810.1 100.00% afis|EAW16177.1 100.00%
Bootstrap support for afum|EAL88810.1 as seed ortholog is 100%.
Bootstrap support for afis|EAW16177.1 as seed ortholog is 100%.

---

Group of orthologs #5839. Best score 727 bits
Score difference with first non-orthologous sequence - *A. fumigatus*:727 *A. fischeri*:727
afum|EAL86948.1 100.00% afis|EAW19564.1 100.00%

Bootstrap support for afum|EAL86948.1 as seed ortholog is 100%.
Bootstrap support for afis|EAW19564.1 as seed ortholog is 100%.

---

Group of orthologs #5840. Best score 727 bits
Score difference with first non-orthologous sequence - *A. fumigatus*:727 *A. fischeri*:727
afum|EAL87175.1 100.00% afis|EAW23738.1 100.00%
Bootstrap support for afum|EAL87175.1 as seed ortholog is 100%.
Bootstrap support for afis|EAW23738.1 as seed ortholog is 100%.

---

Group of orthologs #5841. Best score 727 bits
Score difference with first non-orthologous sequence - *A. fumigatus*:727 *A. fischeri*:627
afum|EAL88031.1 100.00% afis|EAW23430.1 100.00%
Bootstrap support for afum|EAL88031.1 as seed ortholog is 100%.
Bootstrap support for afis|EAW23430.1 as seed ortholog is 100%.

---

Group of orthologs #5842. Best score 727 bits
Score difference with first non-orthologous sequence - *A. fumigatus*:727 *A. fischeri*:727
afum|EAL93347.1 100.00% afis|EAW18634.1 100.00%
Bootstrap support for afum|EAL93347.1 as seed ortholog is 100%.
Bootstrap support for afis|EAW18634.1 as seed ortholog is 100%.

---

Group of orthologs #5843. Best score 727 bits
Score difference with first non-orthologous sequence - *A. fumigatus*:727 *A. fischeri*:727
afum|EAL93375.1 100.00% afis|EAW18658.1 100.00%
Bootstrap support for afum|EAL93375.1 as seed ortholog is 100%.
Bootstrap support for afis|EAW18658.1 as seed ortholog is 100%.

---

Group of orthologs #5844. Best score 727 bits
Score difference with first non-orthologous sequence - *A. fumigatus*:727 *A. fischeri*:237
afum|EAL89338.1 100.00% afis|EAW24696.1 100.00%
Bootstrap support for afum|EAL89338.1 as seed ortholog is 100%.
Bootstrap support for afis|EAW24696.1 as seed ortholog is 100%.

---

Group of orthologs #5845. Best score 727 bits
Score difference with first non-orthologous sequence - *A. fumigatus*:727 *A. fischeri*:605
afum|EAL89697.1 100.00% afis|EAW25001.1 100.00%
Bootstrap support for afum|EAL89697.1 as seed ortholog is 100%.
Bootstrap support for afis|EAW25001.1 as seed ortholog is 100%.

---

Group of orthologs #5846. Best score 726 bits
Score difference with first non-orthologous sequence - *A. fumigatus*:726 *A. fischeri*:280
afum|EAL88602.1 100.00% afis|EAW15969.1 100.00%
Bootstrap support for afum|EAL88602.1 as seed ortholog is 100%.
Bootstrap support for afis|EAW15969.1 as seed ortholog is 100%.

---

Group of orthologs #5847. Best score 726 bits
Score difference with first non-orthologous sequence - *A. fumigatus*:649 *A. fischeri*:655
afum|EAL91672.1 100.00% afis|EAW17864.1 100.00%
Bootstrap support for afum|EAL91672.1 as seed ortholog is 100%.
Bootstrap support for afis|EAW17864.1 as seed ortholog is 100%.

---

Group of orthologs #5848. Best score 726 bits
Score difference with first non-orthologous sequence - *A. fumigatus*:726 *A. fischeri*:726
afum|EAL88219.1 100.00% afis|EAW23261.1 100.00%
Bootstrap support for afum|EAL88219.1 as seed ortholog is 100%.
Bootstrap support for afis|EAW23261.1 as seed ortholog is 100%.

Group of orthologs #5849. Best score 726 bits
Score difference with first non-orthologous sequence - *A. fumigatus*:625 *A. fischeri*:525
afum|EAL93871.2 100.00% afis|EAW19139.1 100.00%
Bootstrap support for afum|EAL93871.2 as seed ortholog is 100%.
Bootstrap support for afis|EAW19139.1 as seed ortholog is 100%.

Group of orthologs #5850. Best score 726 bits
Score difference with first non-orthologous sequence - *A. fumigatus*:726 *A. fischeri*:726
afum|EAL90505.1 100.00% afis|EAW22688.1 100.00%
Bootstrap support for afum|EAL90505.1 as seed ortholog is 100%.
Bootstrap support for afis|EAW22688.1 as seed ortholog is 100%.

Group of orthologs #5851. Best score 726 bits
Score difference with first non-orthologous sequence - *A. fumigatus*:726 *A. fischeri*:726
afum|EAL90433.1 100.00% afis|EAW22766.1 100.00%
Bootstrap support for afum|EAL90433.1 as seed ortholog is 100%.
Bootstrap support for afis|EAW22766.1 as seed ortholog is 100%.

Group of orthologs #5852. Best score 726 bits
Score difference with first non-orthologous sequence - *A. fumigatus*:726 *A. fischeri*:661
afum|EAL94081.1 100.00% afis|EAW19361.1 100.00%
Bootstrap support for afum|EAL94081.1 as seed ortholog is 100%.
Bootstrap support for afis|EAW19361.1 as seed ortholog is 100%.

Group of orthologs #5853. Best score 725 bits
Score difference with first non-orthologous sequence - *A. fumigatus*:725 *A. fischeri*:725
afum|EAL88574.1 100.00% afis|EAW15939.1 100.00%
Bootstrap support for afum|EAL88574.1 as seed ortholog is 100%.
Bootstrap support for afis|EAW15939.1 as seed ortholog is 100%.

Group of orthologs #5854. Best score 725 bits
Score difference with first non-orthologous sequence - *A. fumigatus*:725 *A. fischeri*:725
afum|EAL84282.1 100.00% afis|EAW25558.1 100.00%
Bootstrap support for afum|EAL84282.1 as seed ortholog is 100%.
Bootstrap support for afis|EAW25558.1 as seed ortholog is 100%.

Group of orthologs #5855. Best score 725 bits
Score difference with first non-orthologous sequence - *A. fumigatus*:725 *A. fischeri*:725
afum|EAL84376.1 100.00% afis|EAW25703.1 100.00%
Bootstrap support for afum|EAL84376.1 as seed ortholog is 100%.
Bootstrap support for afis|EAW25703.1 as seed ortholog is 100%.

Group of orthologs #5856. Best score 725 bits
Score difference with first non-orthologous sequence - *A. fumigatus*:725 *A. fischeri*:725
afum|EAL86124.1 100.00% afis|EAW24455.1 100.00%
Bootstrap support for afum|EAL86124.1 as seed ortholog is 100%.
Bootstrap support for afis|EAW24455.1 as seed ortholog is 100%.

Group of orthologs #5857. Best score 725 bits
Score difference with first non-orthologous sequence - *A. fumigatus*:725 *A. fischeri*:725
afum|EAL88165.1 100.00% afis|EAW23312.1 100.00%
Bootstrap support for afum|EAL88165.1 as seed ortholog is 100%.
Bootstrap support for afis|EAW23312.1 as seed ortholog is 100%.

Group of orthologs #5858. Best score 725 bits

Score difference with first non-orthologous sequence - *A. fumigatus*:725 *A. fischeri*:725
afum|EAL87850.1 100.00% afis|EAW23613.1 100.00%
Bootstrap support for afum|EAL87850.1 as seed ortholog is 100%.
Bootstrap support for afis|EAW23613.1 as seed ortholog is 100%.

---

Group of orthologs #5859. Best score 725 bits
Score difference with first non-orthologous sequence - *A. fumigatus*:725 *A. fischeri*:725
afum|EBA27493.1 100.00% afis|EAW18717.1 100.00%
Bootstrap support for afum|EBA27493.1 as seed ortholog is 100%.
Bootstrap support for afis|EAW18717.1 as seed ortholog is 100%.

---

Group of orthologs #5860. Best score 725 bits
Score difference with first non-orthologous sequence - *A. fumigatus*:725 *A. fischeri*:725
afum|EAL92601.2 100.00% afis|EAW21583.1 100.00%
Bootstrap support for afum|EAL92601.2 as seed ortholog is 100%.
Bootstrap support for afis|EAW21583.1 as seed ortholog is 100%.

---

Group of orthologs #5861. Best score 724 bits
Score difference with first non-orthologous sequence - *A. fumigatus*:724 *A. fischeri*:724
afum|EAL88682.1 100.00% afis|EAW16049.1 100.00%
Bootstrap support for afum|EAL88682.1 as seed ortholog is 100%.
Bootstrap support for afis|EAW16049.1 as seed ortholog is 100%.

---

Group of orthologs #5862. Best score 723 bits
Score difference with first non-orthologous sequence - *A. fumigatus*:723 *A. fischeri*:723
afum|EAL85941.2 100.00% afis|EAW24273.1 100.00%
Bootstrap support for afum|EAL85941.2 as seed ortholog is 100%.
Bootstrap support for afis|EAW24273.1 as seed ortholog is 100%.

---

Group of orthologs #5863. Best score 723 bits
Score difference with first non-orthologous sequence - *A. fumigatus*:723 *A. fischeri*:503
afum|EAL87149.1 100.00% afis|EAW23705.1 100.00%
Bootstrap support for afum|EAL87149.1 as seed ortholog is 100%.
Bootstrap support for afis|EAW23705.1 as seed ortholog is 100%.

---

Group of orthologs #5864. Best score 723 bits
Score difference with first non-orthologous sequence - *A. fumigatus*:723 *A. fischeri*:723
afum|EAL90473.1 100.00% afis|EAW22723.1 100.00%
Bootstrap support for afum|EAL90473.1 as seed ortholog is 100%.
Bootstrap support for afis|EAW22723.1 as seed ortholog is 100%.

---

Group of orthologs #5865. Best score 723 bits
Score difference with first non-orthologous sequence - *A. fumigatus*:723 *A. fischeri*:723
afum|EBA27191.1 100.00% afis|EAW20513.1 100.00%
Bootstrap support for afum|EBA27191.1 as seed ortholog is 100%.
Bootstrap support for afis|EAW20513.1 as seed ortholog is 100%.

---

Group of orthologs #5866. Best score 723 bits
Score difference with first non-orthologous sequence - *A. fumigatus*:723 *A. fischeri*:723
afum|EAL89784.1 100.00% afis|EAW25113.1 100.00%
Bootstrap support for afum|EAL89784.1 as seed ortholog is 100%.
Bootstrap support for afis|EAW25113.1 as seed ortholog is 100%.

---

Group of orthologs #5867. Best score 723 bits
Score difference with first non-orthologous sequence - *A. fumigatus*:723 *A. fischeri*:723
afum|EAL92005.1 100.00% afis|EAW24150.1 100.00%

Bootstrap support for afum|EAL92005.1 as seed ortholog is 100%.
Bootstrap support for afis|EAW24150.1 as seed ortholog is 100%.

---

Group of orthologs #5868. Best score 722 bits
Score difference with first non-orthologous sequence - *A. fumigatus*:722 *A. fischeri*:722
afum|EAL85358.1 100.00% afis|EAW20240.1 100.00%
Bootstrap support for afum|EAL85358.1 as seed ortholog is 100%.
Bootstrap support for afis|EAW20240.1 as seed ortholog is 100%.

---

Group of orthologs #5869. Best score 722 bits
Score difference with first non-orthologous sequence - *A. fumigatus*:603 *A. fischeri*:549
afum|EAL92722.1 100.00% afis|EAW21688.1 100.00%
Bootstrap support for afum|EAL92722.1 as seed ortholog is 100%.
Bootstrap support for afis|EAW21688.1 as seed ortholog is 100%.

---

Group of orthologs #5870. Best score 722 bits
Score difference with first non-orthologous sequence - *A. fumigatus*:659 *A. fischeri*:516
afum|EAL92041.1 100.00% afis|EAW24180.1 100.00%
Bootstrap support for afum|EAL92041.1 as seed ortholog is 100%.
Bootstrap support for afis|EAW24180.1 as seed ortholog is 100%.

---

Group of orthologs #5871. Best score 721 bits
Score difference with first non-orthologous sequence - *A. fumigatus*:721 *A. fischeri*:721
afum|EAL84945.2 100.00% afis|EAW19964.1 100.00%
Bootstrap support for afum|EAL84945.2 as seed ortholog is 100%.
Bootstrap support for afis|EAW19964.1 as seed ortholog is 100%.

---

Group of orthologs #5872. Best score 721 bits
Score difference with first non-orthologous sequence - *A. fumigatus*:553 *A. fischeri*:721
afum|EAL87476.1 100.00% afis|EAW18118.1 100.00%
Bootstrap support for afum|EAL87476.1 as seed ortholog is 100%.
Bootstrap support for afis|EAW18118.1 as seed ortholog is 100%.

---

Group of orthologs #5873. Best score 721 bits
Score difference with first non-orthologous sequence - *A. fumigatus*:721 *A. fischeri*:721
afum|EAL85370.1 100.00% afis|EAW20252.1 100.00%
Bootstrap support for afum|EAL85370.1 as seed ortholog is 100%.
Bootstrap support for afis|EAW20252.1 as seed ortholog is 100%.

---

Group of orthologs #5874. Best score 721 bits
Score difference with first non-orthologous sequence - *A. fumigatus*:721 *A. fischeri*:721
afum|EAL92495.1 100.00% afis|EAW21475.1 100.00%
Bootstrap support for afum|EAL92495.1 as seed ortholog is 100%.
Bootstrap support for afis|EAW21475.1 as seed ortholog is 100%.

---

Group of orthologs #5875. Best score 721 bits
Score difference with first non-orthologous sequence - *A. fumigatus*:721 *A. fischeri*:721
afum|EAL94037.1 100.00% afis|EAW20260.1 100.00%
Bootstrap support for afum|EAL94037.1 as seed ortholog is 100%.
Bootstrap support for afis|EAW20260.1 as seed ortholog is 100%.

---

Group of orthologs #5876. Best score 721 bits
Score difference with first non-orthologous sequence - *A. fumigatus*:721 *A. fischeri*:644
afum|EAL92990.1 100.00% afis|EAW21975.1 100.00%
Bootstrap support for afum|EAL92990.1 as seed ortholog is 100%.
Bootstrap support for afis|EAW21975.1 as seed ortholog is 100%.

Group of orthologs #5877. Best score 721 bits
Score difference with first non-orthologous sequence - *A. fumigatus*:721 *A. fischeri*:721
afum|EAL90173.2 100.00% afis|EAW25507.1 100.00%
Bootstrap support for afum|EAL90173.2 as seed ortholog is 100%.
Bootstrap support for afis|EAW25507.1 as seed ortholog is 100%.

Group of orthologs #5878. Best score 720 bits
Score difference with first non-orthologous sequence - *A. fumigatus*:720 *A. fischeri*:720
afum|EAL88511.1 100.00% afis|EAW15877.1 100.00%
Bootstrap support for afum|EAL88511.1 as seed ortholog is 100%.
Bootstrap support for afis|EAW15877.1 as seed ortholog is 100%.

Group of orthologs #5879. Best score 720 bits
Score difference with first non-orthologous sequence - *A. fumigatus*:720 *A. fischeri*:720
afum|EAL89105.1 100.00% afis|EAW16510.1 100.00%
Bootstrap support for afum|EAL89105.1 as seed ortholog is 100%.
Bootstrap support for afis|EAW16510.1 as seed ortholog is 100%.

Group of orthologs #5880. Best score 720 bits
Score difference with first non-orthologous sequence - *A. fumigatus*:639 *A. fischeri*:653
afum|EAL89212.2 100.00% afis|EAW16625.1 100.00%
Bootstrap support for afum|EAL89212.2 as seed ortholog is 100%.
Bootstrap support for afis|EAW16625.1 as seed ortholog is 100%.

Group of orthologs #5881. Best score 720 bits
Score difference with first non-orthologous sequence - *A. fumigatus*:720 *A. fischeri*:720
afum|EAL87197.1 100.00% afis|EAW23764.1 100.00%
Bootstrap support for afum|EAL87197.1 as seed ortholog is 100%.
Bootstrap support for afis|EAW23764.1 as seed ortholog is 100%.

Group of orthologs #5882. Best score 720 bits
Score difference with first non-orthologous sequence - *A. fumigatus*:720 *A. fischeri*:720
afum|EAL88124.2 100.00% afis|EAW23352.1 100.00%
Bootstrap support for afum|EAL88124.2 as seed ortholog is 100%.
Bootstrap support for afis|EAW23352.1 as seed ortholog is 100%.

Group of orthologs #5883. Best score 720 bits
Score difference with first non-orthologous sequence - *A. fumigatus*:720 *A. fischeri*:548
afum|EAL87908.1 100.00% afis|EAW23558.1 100.00%
Bootstrap support for afum|EAL87908.1 as seed ortholog is 100%.
Bootstrap support for afis|EAW23558.1 as seed ortholog is 100%.

Group of orthologs #5884. Best score 720 bits
Score difference with first non-orthologous sequence - *A. fumigatus*:720 *A. fischeri*:720
afum|EAL93629.1 100.00% afis|EAW18897.1 100.00%
Bootstrap support for afum|EAL93629.1 as seed ortholog is 100%.
Bootstrap support for afis|EAW18897.1 as seed ortholog is 100%.

Group of orthologs #5885. Best score 720 bits
Score difference with first non-orthologous sequence - *A. fumigatus*:678 *A. fischeri*:648
afum|EAL90992.1 100.00% afis|EAW22200.1 100.00%
Bootstrap support for afum|EAL90992.1 as seed ortholog is 100%.
Bootstrap support for afis|EAW22200.1 as seed ortholog is 100%.

Group of orthologs #5886. Best score 720 bits

Score difference with first non-orthologous sequence - *A. fumigatus*:641 *A. fischeri*:600
afum|EAL94087.1 100.00% afis|EAW19368.1 100.00%
Bootstrap support for afum|EAL94087.1 as seed ortholog is 100%.
Bootstrap support for afis|EAW19368.1 as seed ortholog is 100%.

---

Group of orthologs #5887. Best score 720 bits
Score difference with first non-orthologous sequence - *A. fumigatus*:720 *A. fischeri*:720
afum|EAL92947.1 100.00% afis|EAW21934.1 100.00%
Bootstrap support for afum|EAL92947.1 as seed ortholog is 100%.
Bootstrap support for afis|EAW21934.1 as seed ortholog is 100%.

---

Group of orthologs #5888. Best score 720 bits
Score difference with first non-orthologous sequence - *A. fumigatus*:720 *A. fischeri*:459
afum|EAL90087.1 100.00% afis|EAW25428.1 100.00%
Bootstrap support for afum|EAL90087.1 as seed ortholog is 100%.
Bootstrap support for afis|EAW25428.1 as seed ortholog is 100%.

---

Group of orthologs #5889. Best score 719 bits
Score difference with first non-orthologous sequence - *A. fumigatus*:719 *A. fischeri*:719
afum|EAL84494.1 100.00% afis|EAW20568.1 100.00%
Bootstrap support for afum|EAL84494.1 as seed ortholog is 100%.
Bootstrap support for afis|EAW20568.1 as seed ortholog is 100%.

---

Group of orthologs #5890. Best score 719 bits
Score difference with first non-orthologous sequence - *A. fumigatus*:719 *A. fischeri*:719
afum|EAL87502.1 100.00% afis|EAW18145.1 100.00%
Bootstrap support for afum|EAL87502.1 as seed ortholog is 100%.
Bootstrap support for afis|EAW18145.1 as seed ortholog is 100%.

---

Group of orthologs #5891. Best score 719 bits
Score difference with first non-orthologous sequence - *A. fumigatus*:719 *A. fischeri*:719
afum|EAL85449.1 100.00% afis|EAW20324.1 100.00%
Bootstrap support for afum|EAL85449.1 as seed ortholog is 100%.
Bootstrap support for afis|EAW20324.1 as seed ortholog is 100%.

---

Group of orthologs #5892. Best score 719 bits
Score difference with first non-orthologous sequence - *A. fumigatus*:719 *A. fischeri*:719
afum|EAL87402.1 100.00% afis|EAW23972.1 100.00%
Bootstrap support for afum|EAL87402.1 as seed ortholog is 100%.
Bootstrap support for afis|EAW23972.1 as seed ortholog is 100%.

---

Group of orthologs #5893. Best score 719 bits
Score difference with first non-orthologous sequence - *A. fumigatus*:625 *A. fischeri*:613
afum|EAL90731.1 100.00% afis|EAW22460.1 100.00%
Bootstrap support for afum|EAL90731.1 as seed ortholog is 100%.
Bootstrap support for afis|EAW22460.1 as seed ortholog is 100%.

---

Group of orthologs #5894. Best score 718 bits
Score difference with first non-orthologous sequence - *A. fumigatus*:718 *A. fischeri*:718
afum|EAL86415.1 100.00% afis|EAW16878.1 100.00%
Bootstrap support for afum|EAL86415.1 as seed ortholog is 100%.
Bootstrap support for afis|EAW16878.1 as seed ortholog is 100%.

---

Group of orthologs #5895. Best score 718 bits
Score difference with first non-orthologous sequence - *A. fumigatus*:718 *A. fischeri*:718
afum|EAL89272.1 100.00% afis|EAW16668.1 100.00%

Bootstrap support for afum|EAL89272.1 as seed ortholog is 100%.
Bootstrap support for afis|EAW16668.1 as seed ortholog is 100%.

---

Group of orthologs #5896. Best score 718 bits
Score difference with first non-orthologous sequence - *A. fumigatus*:718 *A. fischeri*:718
afum|EAL87105.1 100.00% afis|EAW19390.1 100.00%
Bootstrap support for afum|EAL87105.1 as seed ortholog is 100%.
Bootstrap support for afis|EAW19390.1 as seed ortholog is 100%.

---

Group of orthologs #5897. Best score 718 bits
Score difference with first non-orthologous sequence - *A. fumigatus*:718 *A. fischeri*:718
afum|EAL91571.1 100.00% afis|EAW17761.1 100.00%
Bootstrap support for afum|EAL91571.1 as seed ortholog is 100%.
Bootstrap support for afis|EAW17761.1 as seed ortholog is 100%.

---

Group of orthologs #5898. Best score 718 bits
Score difference with first non-orthologous sequence - *A. fumigatus*:718 *A. fischeri*:718
afum|EAL87244.1 100.00% afis|EAW23810.1 100.00%
Bootstrap support for afum|EAL87244.1 as seed ortholog is 100%.
Bootstrap support for afis|EAW23810.1 as seed ortholog is 100%.

---

Group of orthologs #5899. Best score 718 bits
Score difference with first non-orthologous sequence - *A. fumigatus*:718 *A. fischeri*:718
afum|EBA27440.1 100.00% afis|EAW17965.1 100.00%
Bootstrap support for afum|EBA27440.1 as seed ortholog is 100%.
Bootstrap support for afis|EAW17965.1 as seed ortholog is 100%.

---

Group of orthologs #5900. Best score 718 bits
Score difference with first non-orthologous sequence - *A. fumigatus*:718 *A. fischeri*:718
afum|EAL93533.1 100.00% afis|EAW18797.1 100.00%
Bootstrap support for afum|EAL93533.1 as seed ortholog is 100%.
Bootstrap support for afis|EAW18797.1 as seed ortholog is 100%.

---

Group of orthologs #5901. Best score 718 bits
Score difference with first non-orthologous sequence - *A. fumigatus*:718 *A. fischeri*:718
afum|EAL92806.1 100.00% afis|EAW21777.1 100.00%
Bootstrap support for afum|EAL92806.1 as seed ortholog is 100%.
Bootstrap support for afis|EAW21777.1 as seed ortholog is 100%.

---

Group of orthologs #5902. Best score 718 bits
Score difference with first non-orthologous sequence - *A. fumigatus*:718 *A. fischeri*:718
afum|EAL89836.1 100.00% afis|EAW25161.1 100.00%
Bootstrap support for afum|EAL89836.1 as seed ortholog is 100%.
Bootstrap support for afis|EAW25161.1 as seed ortholog is 100%.

---

Group of orthologs #5903. Best score 717 bits
Score difference with first non-orthologous sequence - *A. fumigatus*:640 *A. fischeri*:613
afum|EAL88902.1 100.00% afis|EAW16281.1 100.00%
Bootstrap support for afum|EAL88902.1 as seed ortholog is 100%.
Bootstrap support for afis|EAW16281.1 as seed ortholog is 100%.

---

Group of orthologs #5904. Best score 717 bits
Score difference with first non-orthologous sequence - *A. fumigatus*:717 *A. fischeri*:717
afum|EAL88933.1 100.00% afis|EAW16311.1 100.00%
Bootstrap support for afum|EAL88933.1 as seed ortholog is 100%.
Bootstrap support for afis|EAW16311.1 as seed ortholog is 100%.

Group of orthologs #5905. Best score 717 bits
Score difference with first non-orthologous sequence - *A. fumigatus*:717 *A. fischeri*:717
afum|EAL84563.1 100.00% afis|EAW20491.1 100.00%
Bootstrap support for afum|EAL84563.1 as seed ortholog is 100%.
Bootstrap support for afis|EAW20491.1 as seed ortholog is 100%.

---

Group of orthologs #5906. Best score 717 bits
Score difference with first non-orthologous sequence - *A. fumigatus*:717 *A. fischeri*:717
afum|EAL87470.1 100.00% afis|EAW18113.1 100.00%
Bootstrap support for afum|EAL87470.1 as seed ortholog is 100%.
Bootstrap support for afis|EAW18113.1 as seed ortholog is 100%.

---

Group of orthologs #5907. Best score 716 bits
Score difference with first non-orthologous sequence - *A. fumigatus*:716 *A. fischeri*:716
afum|EAL88595.1 100.00% afis|EAW15962.1 100.00%
Bootstrap support for afum|EAL88595.1 as seed ortholog is 100%.
Bootstrap support for afis|EAW15962.1 as seed ortholog is 100%.

---

Group of orthologs #5908. Best score 716 bits
Score difference with first non-orthologous sequence - *A. fumigatus*:716 *A. fischeri*:716
afum|EAL87647.1 100.00% afis|EAW18281.1 100.00%
Bootstrap support for afum|EAL87647.1 as seed ortholog is 100%.
Bootstrap support for afis|EAW18281.1 as seed ortholog is 100%.

---

Group of orthologs #5909. Best score 716 bits
Score difference with first non-orthologous sequence - *A. fumigatus*:716 *A. fischeri*:716
afum|EAL86808.1 100.00% afis|EAW19718.1 100.00%
Bootstrap support for afum|EAL86808.1 as seed ortholog is 100%.
Bootstrap support for afis|EAW19718.1 as seed ortholog is 100%.

---

Group of orthologs #5910. Best score 716 bits
Score difference with first non-orthologous sequence - *A. fumigatus*:716 *A. fischeri*:716
afum|EAL91483.1 100.00% afis|EAW17669.1 100.00%
Bootstrap support for afum|EAL91483.1 as seed ortholog is 100%.
Bootstrap support for afis|EAW17669.1 as seed ortholog is 100%.

---

Group of orthologs #5911. Best score 716 bits
Score difference with first non-orthologous sequence - *A. fumigatus*:716 *A. fischeri*:716
afum|EAL87268.1 100.00% afis|EAW23837.1 100.00%
Bootstrap support for afum|EAL87268.1 as seed ortholog is 100%.
Bootstrap support for afis|EAW23837.1 as seed ortholog is 100%.

---

Group of orthologs #5912. Best score 716 bits
Score difference with first non-orthologous sequence - *A. fumigatus*:716 *A. fischeri*:716
afum|EAL88364.1 100.00% afis|EAW23116.1 100.00%
Bootstrap support for afum|EAL88364.1 as seed ortholog is 100%.
Bootstrap support for afis|EAW23116.1 as seed ortholog is 100%.

---

Group of orthologs #5913. Best score 716 bits
Score difference with first non-orthologous sequence - *A. fumigatus*:716 *A. fischeri*:716
afum|EAL93249.1 100.00% afis|EAW18549.1 100.00%
Bootstrap support for afum|EAL93249.1 as seed ortholog is 100%.
Bootstrap support for afis|EAW18549.1 as seed ortholog is 100%.

---

Group of orthologs #5914. Best score 715 bits

Score difference with first non-orthologous sequence - *A. fumigatus*:715 *A. fischeri*:715
afum|EAL86458.1 100.00% afis|EAW16933.1 100.00%
Bootstrap support for afum|EAL86458.1 as seed ortholog is 100%.
Bootstrap support for afis|EAW16933.1 as seed ortholog is 100%.

---

Group of orthologs #5915. Best score 715 bits
Score difference with first non-orthologous sequence - *A. fumigatus*:715 *A. fischeri*:715
afum|EAL85994.1 100.00% afis|EAW24324.1 100.00%
Bootstrap support for afum|EAL85994.1 as seed ortholog is 100%.
Bootstrap support for afis|EAW24324.1 as seed ortholog is 100%.

---

Group of orthologs #5916. Best score 715 bits
Score difference with first non-orthologous sequence - *A. fumigatus*:715 *A. fischeri*:715
afum|EAL90791.1 100.00% afis|EAW22400.1 100.00%
Bootstrap support for afum|EAL90791.1 as seed ortholog is 100%.
Bootstrap support for afis|EAW22400.1 as seed ortholog is 100%.

---

Group of orthologs #5917. Best score 715 bits
Score difference with first non-orthologous sequence - *A. fumigatus*:715 *A. fischeri*:715
afum|EAL90681.1 100.00% afis|EAW22512.1 100.00%
Bootstrap support for afum|EAL90681.1 as seed ortholog is 100%.
Bootstrap support for afis|EAW22512.1 as seed ortholog is 100%.

---

Group of orthologs #5918. Best score 715 bits
Score difference with first non-orthologous sequence - *A. fumigatus*:715 *A. fischeri*:715
afum|EAL92786.1 100.00% afis|EAW21756.1 100.00%
Bootstrap support for afum|EAL92786.1 as seed ortholog is 100%.
Bootstrap support for afis|EAW21756.1 as seed ortholog is 100%.

---

Group of orthologs #5919. Best score 714 bits
Score difference with first non-orthologous sequence - *A. fumigatus*:571 *A. fischeri*:714
afum|EAL88950.1 100.00% afis|EAW16326.1 100.00%
Bootstrap support for afum|EAL88950.1 as seed ortholog is 100%.
Bootstrap support for afis|EAW16326.1 as seed ortholog is 100%.

---

Group of orthologs #5920. Best score 714 bits
Score difference with first non-orthologous sequence - *A. fumigatus*:714 *A. fischeri*:714
afum|EAL84857.1 100.00% afis|EAW20762.1 100.00%
Bootstrap support for afum|EAL84857.1 as seed ortholog is 100%.
Bootstrap support for afis|EAW20762.1 as seed ortholog is 100%.

---

Group of orthologs #5921. Best score 714 bits
Score difference with first non-orthologous sequence - *A. fumigatus*:714 *A. fischeri*:714
afum|EAL85883.1 100.00% afis|EAW24214.1 100.00%
Bootstrap support for afum|EAL85883.1 as seed ortholog is 100%.
Bootstrap support for afis|EAW24214.1 as seed ortholog is 100%.

---

Group of orthologs #5922. Best score 714 bits
Score difference with first non-orthologous sequence - *A. fumigatus*:714 *A. fischeri*:365
afum|EAL84396.1 100.00% afis|EAW25723.1 100.00%
Bootstrap support for afum|EAL84396.1 as seed ortholog is 100%.
Bootstrap support for afis|EAW25723.1 as seed ortholog is 100%.

---

Group of orthologs #5923. Best score 714 bits
Score difference with first non-orthologous sequence - *A. fumigatus*:714 *A. fischeri*:714
afum|EAL93737.1 100.00% afis|EAW19003.1 100.00%

Bootstrap support for afum|EAL93737.1 as seed ortholog is 100%.
Bootstrap support for afis|EAW19003.1 as seed ortholog is 100%.

---

Group of orthologs #5924. Best score 714 bits
Score difference with first non-orthologous sequence - *A. fumigatus*:714 *A. fischeri*:714
afum|EAL91090.1 100.00% afis|EAW22099.1 100.00%
Bootstrap support for afum|EAL91090.1 as seed ortholog is 100%.
Bootstrap support for afis|EAW22099.1 as seed ortholog is 100%.

---

Group of orthologs #5925. Best score 714 bits
Score difference with first non-orthologous sequence - *A. fumigatus*:714 *A. fischeri*:714
afum|EAL90540.1 100.00% afis|EAW22652.1 100.00%
Bootstrap support for afum|EAL90540.1 as seed ortholog is 100%.
Bootstrap support for afis|EAW22652.1 as seed ortholog is 100%.

---

Group of orthologs #5926. Best score 714 bits
Score difference with first non-orthologous sequence - *A. fumigatus*:714 *A. fischeri*:714
afum|EAL92202.1 100.00% afis|EAW21181.1 100.00%
Bootstrap support for afum|EAL92202.1 as seed ortholog is 100%.
Bootstrap support for afis|EAW21181.1 as seed ortholog is 100%.

---

Group of orthologs #5927. Best score 713 bits
Score difference with first non-orthologous sequence - *A. fumigatus*:713 *A. fischeri*:713
afum|EAL91242.1 100.00% afis|EAW17407.1 100.00%
Bootstrap support for afum|EAL91242.1 as seed ortholog is 100%.
Bootstrap support for afis|EAW17407.1 as seed ortholog is 100%.

---

Group of orthologs #5928. Best score 713 bits
Score difference with first non-orthologous sequence - *A. fumigatus*:713 *A. fischeri*:713
afum|EAL91360.1 100.00% afis|EAW17546.1 100.00%
Bootstrap support for afum|EAL91360.1 as seed ortholog is 100%.
Bootstrap support for afis|EAW17546.1 as seed ortholog is 100%.

---

Group of orthologs #5929. Best score 713 bits
Score difference with first non-orthologous sequence - *A. fumigatus*:713 *A. fischeri*:713
afum|EAL87351.1 100.00% afis|EAW23926.1 100.00%
Bootstrap support for afum|EAL87351.1 as seed ortholog is 100%.
Bootstrap support for afis|EAW23926.1 as seed ortholog is 100%.

---

Group of orthologs #5930. Best score 713 bits
Score difference with first non-orthologous sequence - *A. fumigatus*:713 *A. fischeri*:713
afum|EAL88078.1 100.00% afis|EAW23392.1 100.00%
Bootstrap support for afum|EAL88078.1 as seed ortholog is 100%.
Bootstrap support for afis|EAW23392.1 as seed ortholog is 100%.

---

Group of orthologs #5931. Best score 713 bits
Score difference with first non-orthologous sequence - *A. fumigatus*:713 *A. fischeri*:713
afum|EAL91037.1 100.00% afis|EAW22153.1 100.00%
Bootstrap support for afum|EAL91037.1 as seed ortholog is 100%.
Bootstrap support for afis|EAW22153.1 as seed ortholog is 100%.

---

Group of orthologs #5932. Best score 713 bits
Score difference with first non-orthologous sequence - *A. fumigatus*:713 *A. fischeri*:713
afum|EAL90595.1 100.00% afis|EAW22603.1 100.00%
Bootstrap support for afum|EAL90595.1 as seed ortholog is 100%.
Bootstrap support for afis|EAW22603.1 as seed ortholog is 100%.

Group of orthologs #5933. Best score 713 bits
Score difference with first non-orthologous sequence - *A. fumigatus*:713 *A. fischeri*:713
afum|EAL92433.1 100.00% afis|EAW21418.1 100.00%
Bootstrap support for afum|EAL92433.1 as seed ortholog is 100%.
Bootstrap support for afis|EAW21418.1 as seed ortholog is 100%.

---

Group of orthologs #5934. Best score 713 bits
Score difference with first non-orthologous sequence - *A. fumigatus*:713 *A. fischeri*:713
afum|EAL92677.1 100.00% afis|EAW21643.1 100.00%
Bootstrap support for afum|EAL92677.1 as seed ortholog is 100%.
Bootstrap support for afis|EAW21643.1 as seed ortholog is 100%.

---

Group of orthologs #5935. Best score 713 bits
Score difference with first non-orthologous sequence - *A. fumigatus*:713 *A. fischeri*:713
afum|EAL92745.1 100.00% afis|EAW21710.1 100.00%
Bootstrap support for afum|EAL92745.1 as seed ortholog is 100%.
Bootstrap support for afis|EAW21710.1 as seed ortholog is 100%.

---

Group of orthologs #5936. Best score 713 bits
Score difference with first non-orthologous sequence - *A. fumigatus*:713 *A. fischeri*:713
afum|EAL92906.1 100.00% afis|EAW21892.1 100.00%
Bootstrap support for afum|EAL92906.1 as seed ortholog is 100%.
Bootstrap support for afis|EAW21892.1 as seed ortholog is 100%.

---

Group of orthologs #5937. Best score 713 bits
Score difference with first non-orthologous sequence - *A. fumigatus*:713 *A. fischeri*:713
afum|EAL92934.1 100.00% afis|EAW21921.1 100.00%
Bootstrap support for afum|EAL92934.1 as seed ortholog is 100%.
Bootstrap support for afis|EAW21921.1 as seed ortholog is 100%.

---

Group of orthologs #5938. Best score 713 bits
Score difference with first non-orthologous sequence - *A. fumigatus*:713 *A. fischeri*:713
afum|EAL90092.1 100.00% afis|EAW25434.1 100.00%
Bootstrap support for afum|EAL90092.1 as seed ortholog is 100%.
Bootstrap support for afis|EAW25434.1 as seed ortholog is 100%.

---

Group of orthologs #5939. Best score 712 bits
Score difference with first non-orthologous sequence - *A. fumigatus*:712 *A. fischeri*:534
afum|EAL86411.1 100.00% afis|EAW16873.1 100.00%
Bootstrap support for afum|EAL86411.1 as seed ortholog is 100%.
Bootstrap support for afis|EAW16873.1 as seed ortholog is 100%.

---

Group of orthologs #5940. Best score 712 bits
Score difference with first non-orthologous sequence - *A. fumigatus*:712 *A. fischeri*:712
afum|EAL84873.1 100.00% afis|EAW20698.1 100.00%
Bootstrap support for afum|EAL84873.1 as seed ortholog is 100%.
Bootstrap support for afis|EAW20698.1 as seed ortholog is 100%.

---

Group of orthologs #5941. Best score 712 bits
Score difference with first non-orthologous sequence - *A. fumigatus*:712 *A. fischeri*:712
afum|EAL91259.1 100.00% afis|EAW17431.1 100.00%
Bootstrap support for afum|EAL91259.1 as seed ortholog is 100%.
Bootstrap support for afis|EAW17431.1 as seed ortholog is 100%.

---

Group of orthologs #5942. Best score 712 bits

Score difference with first non-orthologous sequence - *A. fumigatus*:712   *A. fischeri*:712
afum|EAL87150.1 100.00% afis|EAW23706.1 100.00%
Bootstrap support for afum|EAL87150.1 as seed ortholog is 100%.
Bootstrap support for afis|EAW23706.1 as seed ortholog is 100%.

---

Group of orthologs #5943. Best score 712 bits
Score difference with first non-orthologous sequence - *A. fumigatus*:712   *A. fischeri*:712
afum|EAL88055.1 100.00% afis|EAW23414.1 100.00%
Bootstrap support for afum|EAL88055.1 as seed ortholog is 100%.
Bootstrap support for afis|EAW23414.1 as seed ortholog is 100%.

---

Group of orthologs #5944. Best score 712 bits
Score difference with first non-orthologous sequence - *A. fumigatus*:712   *A. fischeri*:712
afum|EAL92253.1 100.00% afis|EAW21240.1 100.00%
Bootstrap support for afum|EAL92253.1 as seed ortholog is 100%.
Bootstrap support for afis|EAW21240.1 as seed ortholog is 100%.

---

Group of orthologs #5945. Best score 711 bits
Score difference with first non-orthologous sequence - *A. fumigatus*:633   *A. fischeri*:577
afum|EAL91661.1 100.00% afis|EAW17853.1 100.00%
Bootstrap support for afum|EAL91661.1 as seed ortholog is 100%.
Bootstrap support for afis|EAW17853.1 as seed ortholog is 100%.

---

Group of orthologs #5946. Best score 711 bits
Score difference with first non-orthologous sequence - *A. fumigatus*:711   *A. fischeri*:711
afum|EAL93502.1 100.00% afis|EAW18771.1 100.00%
Bootstrap support for afum|EAL93502.1 as seed ortholog is 100%.
Bootstrap support for afis|EAW18771.1 as seed ortholog is 100%.

---

Group of orthologs #5947. Best score 711 bits
Score difference with first non-orthologous sequence - *A. fumigatus*:711   *A. fischeri*:711
afum|EAL90695.1 100.00% afis|EAW22498.1 100.00%
Bootstrap support for afum|EAL90695.1 as seed ortholog is 100%.
Bootstrap support for afis|EAW22498.1 as seed ortholog is 100%.

---

Group of orthologs #5948. Best score 711 bits
Score difference with first non-orthologous sequence - *A. fumigatus*:711   *A. fischeri*:711
afum|EAL90268.1 100.00% afis|EAW22929.1 100.00%
Bootstrap support for afum|EAL90268.1 as seed ortholog is 100%.
Bootstrap support for afis|EAW22929.1 as seed ortholog is 100%.

---

Group of orthologs #5949. Best score 711 bits
Score difference with first non-orthologous sequence - *A. fumigatus*:608   *A. fischeri*:570
afum|EAL92504.1 100.00% afis|EAW21484.1 100.00%
Bootstrap support for afum|EAL92504.1 as seed ortholog is 100%.
Bootstrap support for afis|EAW21484.1 as seed ortholog is 100%.

---

Group of orthologs #5950. Best score 711 bits
Score difference with first non-orthologous sequence - *A. fumigatus*:711   *A. fischeri*:711
afum|EAL93021.2 100.00% afis|EAW22006.1 100.00%
Bootstrap support for afum|EAL93021.2 as seed ortholog is 100%.
Bootstrap support for afis|EAW22006.1 as seed ortholog is 100%.

---

Group of orthologs #5951. Best score 710 bits
Score difference with first non-orthologous sequence - *A. fumigatus*:710   *A. fischeri*:710
afum|EAL84215.1 100.00% afis|EAW15428.1 100.00%

Bootstrap support for afum|EAL84215.1 as seed ortholog is 100%.
Bootstrap support for afis|EAW15428.1 as seed ortholog is 100%.

---

Group of orthologs #5952. Best score 710 bits
Score difference with first non-orthologous sequence - *A. fumigatus*:710 *A. fischeri*:710
afum|EAL86564.2 100.00% afis|EAW17087.1 100.00%
Bootstrap support for afum|EAL86564.2 as seed ortholog is 100%.
Bootstrap support for afis|EAW17087.1 as seed ortholog is 100%.

---

Group of orthologs #5953. Best score 710 bits
Score difference with first non-orthologous sequence - *A. fumigatus*:710 *A. fischeri*:710
afum|EAL84488.2 100.00% afis|EAW20577.1 100.00%
Bootstrap support for afum|EAL84488.2 as seed ortholog is 100%.
Bootstrap support for afis|EAW20577.1 as seed ortholog is 100%.

---

Group of orthologs #5954. Best score 710 bits
Score difference with first non-orthologous sequence - *A. fumigatus*:308 *A. fischeri*:130
afum|EAL85274.1 100.00% afis|EAW20158.1 100.00%
Bootstrap support for afum|EAL85274.1 as seed ortholog is 100%.
Bootstrap support for afis|EAW20158.1 as seed ortholog is 99%.

---

Group of orthologs #5955. Best score 710 bits
Score difference with first non-orthologous sequence - *A. fumigatus*:710 *A. fischeri*:710
afum|EAL87580.1 100.00% afis|EAW18214.1 100.00%
Bootstrap support for afum|EAL87580.1 as seed ortholog is 100%.
Bootstrap support for afis|EAW18214.1 as seed ortholog is 100%.

---

Group of orthologs #5956. Best score 710 bits
Score difference with first non-orthologous sequence - *A. fumigatus*:612 *A. fischeri*:421
afum|EAL84698.1 100.00% afis|EAW20936.1 100.00%
Bootstrap support for afum|EAL84698.1 as seed ortholog is 100%.
Bootstrap support for afis|EAW20936.1 as seed ortholog is 100%.

---

Group of orthologs #5957. Best score 710 bits
Score difference with first non-orthologous sequence - *A. fumigatus*:710 *A. fischeri*:710
afum|EAL90639.1 100.00% afis|EAW22556.1 100.00%
Bootstrap support for afum|EAL90639.1 as seed ortholog is 100%.
Bootstrap support for afis|EAW22556.1 as seed ortholog is 100%.

---

Group of orthologs #5958. Best score 710 bits
Score difference with first non-orthologous sequence - *A. fumigatus*:710 *A. fischeri*:710
afum|EAL90264.1 100.00% afis|EAW22932.1 100.00%
Bootstrap support for afum|EAL90264.1 as seed ortholog is 100%.
Bootstrap support for afis|EAW22932.1 as seed ortholog is 100%.

---

Group of orthologs #5959. Best score 709 bits
Score difference with first non-orthologous sequence - *A. fumigatus*:709 *A. fischeri*:709
afum|EAL84495.1 100.00% afis|EAW20566.1 100.00%
Bootstrap support for afum|EAL84495.1 as seed ortholog is 100%.
Bootstrap support for afis|EAW20566.1 as seed ortholog is 100%.

---

Group of orthologs #5960. Best score 709 bits
Score difference with first non-orthologous sequence - *A. fumigatus*:709 *A. fischeri*:709
afum|EAL89475.1 100.00% afis|EAW24830.1 100.00%
Bootstrap support for afum|EAL89475.1 as seed ortholog is 100%.
Bootstrap support for afis|EAW24830.1 as seed ortholog is 100%.

Group of orthologs #5961. Best score 709 bits
Score difference with first non-orthologous sequence - *A. fumigatus*:709 *A. fischeri*:709
afum|EAL89808.2 100.00% afis|EAW25137.1 100.00%
Bootstrap support for afum|EAL89808.2 as seed ortholog is 100%.
Bootstrap support for afis|EAW25137.1 as seed ortholog is 100%.

Group of orthologs #5962. Best score 708 bits
Score difference with first non-orthologous sequence - *A. fumigatus*:708 *A. fischeri*:708
afum|EAL85522.2 100.00% afis|EAW15829.1 100.00%
Bootstrap support for afum|EAL85522.2 as seed ortholog is 100%.
Bootstrap support for afis|EAW15829.1 as seed ortholog is 100%.

Group of orthologs #5963. Best score 708 bits
Score difference with first non-orthologous sequence - *A. fumigatus*:539 *A. fischeri*:548
afum|EAL87189.1 100.00% afis|EAW23752.1 100.00%
Bootstrap support for afum|EAL87189.1 as seed ortholog is 100%.
Bootstrap support for afis|EAW23752.1 as seed ortholog is 100%.

Group of orthologs #5964. Best score 708 bits
Score difference with first non-orthologous sequence - *A. fumigatus*:708 *A. fischeri*:241
afum|EAL92085.1 100.00% afis|EAW21049.1 100.00%
Bootstrap support for afum|EAL92085.1 as seed ortholog is 100%.
Bootstrap support for afis|EAW21049.1 as seed ortholog is 100%.

Group of orthologs #5965. Best score 708 bits
Score difference with first non-orthologous sequence - *A. fumigatus*:708 *A. fischeri*:708
afum|EAL90332.1 100.00% afis|EAW22867.1 100.00%
Bootstrap support for afum|EAL90332.1 as seed ortholog is 100%.
Bootstrap support for afis|EAW22867.1 as seed ortholog is 100%.

Group of orthologs #5966. Best score 707 bits
Score difference with first non-orthologous sequence - *A. fumigatus*:707 *A. fischeri*:544
afum|EAL85852.1 100.00% afis|EAW15470.1 100.00%
Bootstrap support for afum|EAL85852.1 as seed ortholog is 100%.
Bootstrap support for afis|EAW15470.1 as seed ortholog is 100%.

Group of orthologs #5967. Best score 707 bits
Score difference with first non-orthologous sequence - *A. fumigatus*:707 *A. fischeri*:707
afum|EAL89242.1 100.00% afis|EAW16640.1 100.00%
Bootstrap support for afum|EAL89242.1 as seed ortholog is 100%.
Bootstrap support for afis|EAW16640.1 as seed ortholog is 100%.

Group of orthologs #5968. Best score 707 bits
Score difference with first non-orthologous sequence - *A. fumigatus*:707 *A. fischeri*:565
afum|EAL86836.1 100.00% afis|EAW19690.1 100.00%
Bootstrap support for afum|EAL86836.1 as seed ortholog is 100%.
Bootstrap support for afis|EAW19690.1 as seed ortholog is 100%.

Group of orthologs #5969. Best score 707 bits
Score difference with first non-orthologous sequence - *A. fumigatus*:707 *A. fischeri*:707
afum|EAL89234.2 100.00% afis|EAW20410.1 100.00%
Bootstrap support for afum|EAL89234.2 as seed ortholog is 100%.
Bootstrap support for afis|EAW20410.1 as seed ortholog is 100%.

Group of orthologs #5970. Best score 707 bits

Score difference with first non-orthologous sequence - *A. fumigatus*:707 *A. fischeri*:707
afum|EAL91825.1 100.00% afis|EAW18013.1 100.00%
Bootstrap support for afum|EAL91825.1 as seed ortholog is 100%.
Bootstrap support for afis|EAW18013.1 as seed ortholog is 100%.

---

Group of orthologs #5971. Best score 707 bits
Score difference with first non-orthologous sequence - *A. fumigatus*:707 *A. fischeri*:707
afum|EAL86080.1 100.00% afis|EAW24411.1 100.00%
Bootstrap support for afum|EAL86080.1 as seed ortholog is 100%.
Bootstrap support for afis|EAW24411.1 as seed ortholog is 100%.

---

Group of orthologs #5972. Best score 707 bits
Score difference with first non-orthologous sequence - *A. fumigatus*:707 *A. fischeri*:707
afum|EAL90135.1 100.00% afis|EAW25471.1 100.00%
Bootstrap support for afum|EAL90135.1 as seed ortholog is 100%.
Bootstrap support for afis|EAW25471.1 as seed ortholog is 100%.

---

Group of orthologs #5973. Best score 706 bits
Score difference with first non-orthologous sequence - *A. fumigatus*:706 *A. fischeri*:706
afum|EAL91194.1 100.00% afis|EAW17351.1 100.00%
Bootstrap support for afum|EAL91194.1 as seed ortholog is 100%.
Bootstrap support for afis|EAW17351.1 as seed ortholog is 100%.

---

Group of orthologs #5974. Best score 706 bits
Score difference with first non-orthologous sequence - *A. fumigatus*:706 *A. fischeri*:706
afum|EAL91782.1 100.00% afis|EAW17973.1 100.00%
Bootstrap support for afum|EAL91782.1 as seed ortholog is 100%.
Bootstrap support for afis|EAW17973.1 as seed ortholog is 100%.

---

Group of orthologs #5975. Best score 706 bits
Score difference with first non-orthologous sequence - *A. fumigatus*:706 *A. fischeri*:706
afum|EAL88113.1 100.00% afis|EAW23361.1 100.00%
Bootstrap support for afum|EAL88113.1 as seed ortholog is 100%.
Bootstrap support for afis|EAW23361.1 as seed ortholog is 100%.

---

Group of orthologs #5976. Best score 706 bits
Score difference with first non-orthologous sequence - *A. fumigatus*:706 *A. fischeri*:706
afum|EAL92842.1 100.00% afis|EAW21827.1 100.00%
Bootstrap support for afum|EAL92842.1 as seed ortholog is 100%.
Bootstrap support for afis|EAW21827.1 as seed ortholog is 100%.

---

Group of orthologs #5977. Best score 705 bits
Score difference with first non-orthologous sequence - *A. fumigatus*:705 *A. fischeri*:705
afum|EAL85720.1 100.00% afis|EAW15615.1 100.00%
Bootstrap support for afum|EAL85720.1 as seed ortholog is 100%.
Bootstrap support for afis|EAW15615.1 as seed ortholog is 100%.

---

Group of orthologs #5978. Best score 705 bits
Score difference with first non-orthologous sequence - *A. fumigatus*:705 *A. fischeri*:705
afum|EAL86324.1 100.00% afis|EAW16752.1 100.00%
Bootstrap support for afum|EAL86324.1 as seed ortholog is 100%.
Bootstrap support for afis|EAW16752.1 as seed ortholog is 100%.

---

Group of orthologs #5979. Best score 705 bits
Score difference with first non-orthologous sequence - *A. fumigatus*:437 *A. fischeri*:627
afum|EAL86387.1 100.00% afis|EAW16840.1 100.00%

Bootstrap support for afum|EAL86387.1 as seed ortholog is 100%.
Bootstrap support for afis|EAW16840.1 as seed ortholog is 100%.

---

Group of orthologs #5980. Best score 705 bits
Score difference with first non-orthologous sequence - *A. fumigatus*:705 *A. fischeri*:705
afum|EAL88738.1 100.00% afis|EAW16106.1 100.00%
Bootstrap support for afum|EAL88738.1 as seed ortholog is 100%.
Bootstrap support for afis|EAW16106.1 as seed ortholog is 100%.

---

Group of orthologs #5981. Best score 705 bits
Score difference with first non-orthologous sequence - *A. fumigatus*:705 *A. fischeri*:705
afum|EAL85309.1 100.00% afis|EAW20190.1 100.00%
Bootstrap support for afum|EAL85309.1 as seed ortholog is 100%.
Bootstrap support for afis|EAW20190.1 as seed ortholog is 100%.

---

Group of orthologs #5982. Best score 705 bits
Score difference with first non-orthologous sequence - *A. fumigatus*:312 *A. fischeri*:705
afum|EAL91587.1 100.00% afis|EAW17778.1 100.00%
Bootstrap support for afum|EAL91587.1 as seed ortholog is 100%.
Bootstrap support for afis|EAW17778.1 as seed ortholog is 100%.

---

Group of orthologs #5983. Best score 705 bits
Score difference with first non-orthologous sequence - *A. fumigatus*:705 *A. fischeri*:705
afum|EAL86247.1 100.00% afis|EAW24619.1 100.00%
Bootstrap support for afum|EAL86247.1 as seed ortholog is 100%.
Bootstrap support for afis|EAW24619.1 as seed ortholog is 100%.

---

Group of orthologs #5984. Best score 705 bits
Score difference with first non-orthologous sequence - *A. fumigatus*:705 *A. fischeri*:705
afum|EAL89436.1 100.00% afis|EAW24794.1 100.00%
Bootstrap support for afum|EAL89436.1 as seed ortholog is 100%.
Bootstrap support for afis|EAW24794.1 as seed ortholog is 100%.

---

Group of orthologs #5985. Best score 704 bits
Score difference with first non-orthologous sequence - *A. fumigatus*:704 *A. fischeri*:704
afum|EAL86592.1 100.00% afis|EAW17132.1 100.00%
Bootstrap support for afum|EAL86592.1 as seed ortholog is 100%.
Bootstrap support for afis|EAW17132.1 as seed ortholog is 100%.

---

Group of orthologs #5986. Best score 704 bits
Score difference with first non-orthologous sequence - *A. fumigatus*:704 *A. fischeri*:704
afum|EAL88720.1 100.00% afis|EAW16088.1 100.00%
Bootstrap support for afum|EAL88720.1 as seed ortholog is 100%.
Bootstrap support for afis|EAW16088.1 as seed ortholog is 100%.

---

Group of orthologs #5987. Best score 704 bits
Score difference with first non-orthologous sequence - *A. fumigatus*:704 *A. fischeri*:704
afum|EAL85208.1 100.00% afis|EAW20089.1 100.00%
Bootstrap support for afum|EAL85208.1 as seed ortholog is 100%.
Bootstrap support for afis|EAW20089.1 as seed ortholog is 100%.

---

Group of orthologs #5988. Best score 704 bits
Score difference with first non-orthologous sequence - *A. fumigatus*:704 *A. fischeri*:526
afum|EAL87621.1 100.00% afis|EAW18258.1 100.00%
Bootstrap support for afum|EAL87621.1 as seed ortholog is 100%.
Bootstrap support for afis|EAW18258.1 as seed ortholog is 100%.

Group of orthologs #5989. Best score 704 bits
Score difference with first non-orthologous sequence - *A. fumigatus*:704 *A. fischeri*:704
afum|EAL91697.1 100.00% afis|EAW17888.1 100.00%
Bootstrap support for afum|EAL91697.1 as seed ortholog is 100%.
Bootstrap support for afis|EAW17888.1 as seed ortholog is 100%.

---

Group of orthologs #5990. Best score 704 bits
Score difference with first non-orthologous sequence - *A. fumigatus*:704 *A. fischeri*:704
afum|EAL87869.1 100.00% afis|EAW23597.1 100.00%
Bootstrap support for afum|EAL87869.1 as seed ortholog is 100%.
Bootstrap support for afis|EAW23597.1 as seed ortholog is 100%.

---

Group of orthologs #5991. Best score 704 bits
Score difference with first non-orthologous sequence - *A. fumigatus*:704 *A. fischeri*:704
afum|EAL93282.1 100.00% afis|EAW18579.1 100.00%
Bootstrap support for afum|EAL93282.1 as seed ortholog is 100%.
Bootstrap support for afis|EAW18579.1 as seed ortholog is 100%.

---

Group of orthologs #5992. Best score 704 bits
Score difference with first non-orthologous sequence - *A. fumigatus*:704 *A. fischeri*:704
afum|EAL92087.1 100.00% afis|EAW21051.1 100.00%
Bootstrap support for afum|EAL92087.1 as seed ortholog is 100%.
Bootstrap support for afis|EAW21051.1 as seed ortholog is 100%.

---

Group of orthologs #5993. Best score 704 bits
Score difference with first non-orthologous sequence - *A. fumigatus*:704 *A. fischeri*:704
afum|EAL90649.1 100.00% afis|EAW22546.1 100.00%
Bootstrap support for afum|EAL90649.1 as seed ortholog is 100%.
Bootstrap support for afis|EAW22546.1 as seed ortholog is 100%.

---

Group of orthologs #5994. Best score 704 bits
Score difference with first non-orthologous sequence - *A. fumigatus*:704 *A. fischeri*:704
afum|EAL89753.1 100.00% afis|EAW25084.1 100.00%
Bootstrap support for afum|EAL89753.1 as seed ortholog is 100%.
Bootstrap support for afis|EAW25084.1 as seed ortholog is 100%.

---

Group of orthologs #5995. Best score 704 bits
Score difference with first non-orthologous sequence - *A. fumigatus*:704 *A. fischeri*:704
afum|EAL90076.1 100.00% afis|EAW25417.1 100.00%
Bootstrap support for afum|EAL90076.1 as seed ortholog is 100%.
Bootstrap support for afis|EAW25417.1 as seed ortholog is 100%.

---

Group of orthologs #5996. Best score 704 bits
Score difference with first non-orthologous sequence - *A. fumigatus*:704 *A. fischeri*:704
afum|EAL90201.1 100.00% afis|EAW25538.1 100.00%
Bootstrap support for afum|EAL90201.1 as seed ortholog is 100%.
Bootstrap support for afis|EAW25538.1 as seed ortholog is 100%.

---

Group of orthologs #5997. Best score 703 bits
Score difference with first non-orthologous sequence - *A. fumigatus*:703 *A. fischeri*:703
afum|EAL87388.1 100.00% afis|EAW23959.1 100.00%
Bootstrap support for afum|EAL87388.1 as seed ortholog is 100%.
Bootstrap support for afis|EAW23959.1 as seed ortholog is 100%.

---

Group of orthologs #5998. Best score 703 bits

Score difference with first non-orthologous sequence - *A. fumigatus*:703   *A. fischeri*:703
afum|EAL93595.1 100.00% afis|EAW18863.1 100.00%
Bootstrap support for afum|EAL93595.1 as seed ortholog is 100%.
Bootstrap support for afis|EAW18863.1 as seed ortholog is 100%.

---

Group of orthologs #5999. Best score 703 bits
Score difference with first non-orthologous sequence - *A. fumigatus*:703   *A. fischeri*:703
afum|EAL90902.1 100.00% afis|EAW22279.1 100.00%
Bootstrap support for afum|EAL90902.1 as seed ortholog is 100%.
Bootstrap support for afis|EAW22279.1 as seed ortholog is 100%.

---

Group of orthologs #6000. Best score 703 bits
Score difference with first non-orthologous sequence - *A. fumigatus*:703   *A. fischeri*:703
afum|EAL92160.1 100.00% afis|EAW21141.1 100.00%
Bootstrap support for afum|EAL92160.1 as seed ortholog is 100%.
Bootstrap support for afis|EAW21141.1 as seed ortholog is 100%.

---

Group of orthologs #6001. Best score 703 bits
Score difference with first non-orthologous sequence - *A. fumigatus*:703   *A. fischeri*:661
afum|EAL90043.1 100.00% afis|EAW25384.1 100.00%
Bootstrap support for afum|EAL90043.1 as seed ortholog is 100%.
Bootstrap support for afis|EAW25384.1 as seed ortholog is 100%.

---

Group of orthologs #6002. Best score 702 bits
Score difference with first non-orthologous sequence - *A. fumigatus*:702   *A. fischeri*:702
afum|EAL86462.1 100.00% afis|EAW16937.1 100.00%
Bootstrap support for afum|EAL86462.1 as seed ortholog is 100%.
Bootstrap support for afis|EAW16937.1 as seed ortholog is 100%.

---

Group of orthologs #6003. Best score 702 bits
Score difference with first non-orthologous sequence - *A. fumigatus*:702   *A. fischeri*:702
afum|EAL87384.1 100.00% afis|EAW23956.1 100.00%
Bootstrap support for afum|EAL87384.1 as seed ortholog is 100%.
Bootstrap support for afis|EAW23956.1 as seed ortholog is 100%.

---

Group of orthologs #6004. Best score 702 bits
Score difference with first non-orthologous sequence - *A. fumigatus*:524   *A. fischeri*:527
afum|EAL87991.2 100.00% afis|EAW23474.1 100.00%
Bootstrap support for afum|EAL87991.2 as seed ortholog is 100%.
Bootstrap support for afis|EAW23474.1 as seed ortholog is 100%.

---

Group of orthologs #6005. Best score 702 bits
Score difference with first non-orthologous sequence - *A. fumigatus*:702   *A. fischeri*:401
afum|EAL93841.2 100.00% afis|EAW19110.1 100.00%
Bootstrap support for afum|EAL93841.2 as seed ortholog is 100%.
Bootstrap support for afis|EAW19110.1 as seed ortholog is 100%.

---

Group of orthologs #6006. Best score 702 bits
Score difference with first non-orthologous sequence - *A. fumigatus*:702   *A. fischeri*:702
afum|EAL92096.1 100.00% afis|EAW21073.1 100.00%
Bootstrap support for afum|EAL92096.1 as seed ortholog is 100%.
Bootstrap support for afis|EAW21073.1 as seed ortholog is 100%.

---

Group of orthologs #6007. Best score 702 bits
Score difference with first non-orthologous sequence - *A. fumigatus*:702   *A. fischeri*:702
afum|EAL90883.1 100.00% afis|EAW22298.1 100.00%

Bootstrap support for afum|EAL90883.1 as seed ortholog is 100%.
Bootstrap support for afis|EAW22298.1 as seed ortholog is 100%.

---

Group of orthologs #6008. Best score 702 bits
Score difference with first non-orthologous sequence - *A. fumigatus*:702 *A. fischeri*:702
afum|EBA27394.1 100.00% afis|EAW25380.1 100.00%
Bootstrap support for afum|EBA27394.1 as seed ortholog is 100%.
Bootstrap support for afis|EAW25380.1 as seed ortholog is 100%.

---

Group of orthologs #6009. Best score 701 bits
Score difference with first non-orthologous sequence - *A. fumigatus*:701 *A. fischeri*:532
afum|EAL85841.1 100.00% afis|EAW15481.1 100.00%
Bootstrap support for afum|EAL85841.1 as seed ortholog is 100%.
Bootstrap support for afis|EAW15481.1 as seed ortholog is 100%.

---

Group of orthologs #6010. Best score 701 bits
Score difference with first non-orthologous sequence - *A. fumigatus*:601 *A. fischeri*:595
afum|EAL88967.1 100.00% afis|EAW16342.1 100.00%
Bootstrap support for afum|EAL88967.1 as seed ortholog is 100%.
Bootstrap support for afis|EAW16342.1 as seed ortholog is 100%.

---

Group of orthologs #6011. Best score 701 bits
Score difference with first non-orthologous sequence - *A. fumigatus*:701 *A. fischeri*:701
afum|EAL89152.1 100.00% afis|EAW16554.1 100.00%
Bootstrap support for afum|EAL89152.1 as seed ortholog is 100%.
Bootstrap support for afis|EAW16554.1 as seed ortholog is 100%.

---

Group of orthologs #6012. Best score 701 bits
Score difference with first non-orthologous sequence - *A. fumigatus*:701 *A. fischeri*:701
afum|EAL86957.1 100.00% afis|EAW19552.1 100.00%
Bootstrap support for afum|EAL86957.1 as seed ortholog is 100%.
Bootstrap support for afis|EAW19552.1 as seed ortholog is 100%.

---

Group of orthologs #6013. Best score 701 bits
Score difference with first non-orthologous sequence - *A. fumigatus*:701 *A. fischeri*:581
afum|EAL86958.1 100.00% afis|EAW19551.1 100.00%
Bootstrap support for afum|EAL86958.1 as seed ortholog is 100%.
Bootstrap support for afis|EAW19551.1 as seed ortholog is 100%.

---

Group of orthologs #6014. Best score 701 bits
Score difference with first non-orthologous sequence - *A. fumigatus*:701 *A. fischeri*:701
afum|EAL92584.1 100.00% afis|EAW21565.1 100.00%
Bootstrap support for afum|EAL92584.1 as seed ortholog is 100%.
Bootstrap support for afis|EAW21565.1 as seed ortholog is 100%.

---

Group of orthologs #6015. Best score 700 bits
Score difference with first non-orthologous sequence - *A. fumigatus*:700 *A. fischeri*:700
afum|EAL85824.1 100.00% afis|EAW15499.1 100.00%
Bootstrap support for afum|EAL85824.1 as seed ortholog is 100%.
Bootstrap support for afis|EAW15499.1 as seed ortholog is 100%.

---

Group of orthologs #6016. Best score 700 bits
Score difference with first non-orthologous sequence - *A. fumigatus*:625 *A. fischeri*:50
afum|EAL86477.2 100.00% afis|EAW16950.1 100.00%
Bootstrap support for afum|EAL86477.2 as seed ortholog is 100%.
Bootstrap support for afis|EAW16950.1 as seed ortholog is 96%.

Group of orthologs #6017. Best score 700 bits
Score difference with first non-orthologous sequence - *A. fumigatus*:700 *A. fischeri*:700
afum|EAL86501.1 100.00% afis|EAW17526.1 100.00%
Bootstrap support for afum|EAL86501.1 as seed ortholog is 100%.
Bootstrap support for afis|EAW17526.1 as seed ortholog is 100%.

Group of orthologs #6018. Best score 700 bits
Score difference with first non-orthologous sequence - *A. fumigatus*:700 *A. fischeri*:700
afum|EAL91298.1 100.00% afis|EAW17469.1 100.00%
Bootstrap support for afum|EAL91298.1 as seed ortholog is 100%.
Bootstrap support for afis|EAW17469.1 as seed ortholog is 100%.

Group of orthologs #6019. Best score 700 bits
Score difference with first non-orthologous sequence - *A. fumigatus*:700 *A. fischeri*:700
afum|EAL90817.1 100.00% afis|EAW22375.1 100.00%
Bootstrap support for afum|EAL90817.1 as seed ortholog is 100%.
Bootstrap support for afis|EAW22375.1 as seed ortholog is 100%.

Group of orthologs #6020. Best score 700 bits
Score difference with first non-orthologous sequence - *A. fumigatus*:700 *A. fischeri*:543
afum|EAL92270.1 100.00% afis|EAW21255.1 100.00%
Bootstrap support for afum|EAL92270.1 as seed ortholog is 100%.
Bootstrap support for afis|EAW21255.1 as seed ortholog is 100%.

Group of orthologs #6021. Best score 699 bits
Score difference with first non-orthologous sequence - *A. fumigatus*:699 *A. fischeri*:699
afum|EAL86416.1 100.00% afis|EAW16879.1 100.00%
Bootstrap support for afum|EAL86416.1 as seed ortholog is 100%.
Bootstrap support for afis|EAW16879.1 as seed ortholog is 100%.

Group of orthologs #6022. Best score 699 bits
Score difference with first non-orthologous sequence - *A. fumigatus*:699 *A. fischeri*:699
afum|EAL84569.1 100.00% afis|EAW20486.1 100.00%
Bootstrap support for afum|EAL84569.1 as seed ortholog is 100%.
Bootstrap support for afis|EAW20486.1 as seed ortholog is 100%.

Group of orthologs #6023. Best score 699 bits
Score difference with first non-orthologous sequence - *A. fumigatus*:699 *A. fischeri*:699
afum|EAL85164.2 100.00% afis|EAW20044.1 100.00%
Bootstrap support for afum|EAL85164.2 as seed ortholog is 100%.
Bootstrap support for afis|EAW20044.1 as seed ortholog is 100%.

Group of orthologs #6024. Best score 699 bits
Score difference with first non-orthologous sequence - *A. fumigatus*:699 *A. fischeri*:699
afum|EAL91616.1 100.00% afis|EAW17808.1 100.00%
Bootstrap support for afum|EAL91616.1 as seed ortholog is 100%.
Bootstrap support for afis|EAW17808.1 as seed ortholog is 100%.

Group of orthologs #6025. Best score 699 bits
Score difference with first non-orthologous sequence - *A. fumigatus*:699 *A. fischeri*:699
afum|EAL86101.1 100.00% afis|EAW24433.1 100.00%
Bootstrap support for afum|EAL86101.1 as seed ortholog is 100%.
Bootstrap support for afis|EAW24433.1 as seed ortholog is 100%.

Group of orthologs #6026. Best score 699 bits

Score difference with first non-orthologous sequence - *A. fumigatus*:699   *A. fischeri*:699
afum|EAL93057.1 100.00% afis|EAW18382.1 100.00%
Bootstrap support for afum|EAL93057.1 as seed ortholog is 100%.
Bootstrap support for afis|EAW18382.1 as seed ortholog is 100%.

---

Group of orthologs #6027. Best score 698 bits
Score difference with first non-orthologous sequence - *A. fumigatus*:698   *A. fischeri*:698
afum|EAL89007.1 100.00% afis|EAW16382.1 100.00%
Bootstrap support for afum|EAL89007.1 as seed ortholog is 100%.
Bootstrap support for afis|EAW16382.1 as seed ortholog is 100%.

---

Group of orthologs #6028. Best score 698 bits
Score difference with first non-orthologous sequence - *A. fumigatus*:598   *A. fischeri*:698
afum|EAL89162.1 100.00% afis|EAW16565.1 100.00%
Bootstrap support for afum|EAL89162.1 as seed ortholog is 100%.
Bootstrap support for afis|EAW16565.1 as seed ortholog is 100%.

---

Group of orthologs #6029. Best score 698 bits
Score difference with first non-orthologous sequence - *A. fumigatus*:698   *A. fischeri*:698
afum|EAL87493.1 100.00% afis|EAW18136.1 100.00%
Bootstrap support for afum|EAL87493.1 as seed ortholog is 100%.
Bootstrap support for afis|EAW18136.1 as seed ortholog is 100%.

---

Group of orthologs #6030. Best score 698 bits
Score difference with first non-orthologous sequence - *A. fumigatus*:698   *A. fischeri*:698
afum|EAL87020.1 100.00% afis|EAW19476.1 100.00%
Bootstrap support for afum|EAL87020.1 as seed ortholog is 100%.
Bootstrap support for afis|EAW19476.1 as seed ortholog is 100%.

---

Group of orthologs #6031. Best score 698 bits
Score difference with first non-orthologous sequence - *A. fumigatus*:698   *A. fischeri*:698
afum|EAL86855.1 100.00% afis|EAW19668.1 100.00%
Bootstrap support for afum|EAL86855.1 as seed ortholog is 100%.
Bootstrap support for afis|EAW19668.1 as seed ortholog is 100%.

---

Group of orthologs #6032. Best score 698 bits
Score difference with first non-orthologous sequence - *A. fumigatus*:698   *A. fischeri*:698
afum|EBA27354.1 100.00% afis|EAW16036.1 100.00%
Bootstrap support for afum|EBA27354.1 as seed ortholog is 100%.
Bootstrap support for afis|EAW16036.1 as seed ortholog is 100%.

---

Group of orthologs #6033. Best score 698 bits
Score difference with first non-orthologous sequence - *A. fumigatus*:698   *A. fischeri*:698
afum|EAL88250.1 100.00% afis|EAW23232.1 100.00%
Bootstrap support for afum|EAL88250.1 as seed ortholog is 100%.
Bootstrap support for afis|EAW23232.1 as seed ortholog is 100%.

---

Group of orthologs #6034. Best score 698 bits
Score difference with first non-orthologous sequence - *A. fumigatus*:698   *A. fischeri*:698
afum|EAL87945.2 100.00% afis|EAW23519.1 100.00%
Bootstrap support for afum|EAL87945.2 as seed ortholog is 100%.
Bootstrap support for afis|EAW23519.1 as seed ortholog is 100%.

---

Group of orthologs #6035. Best score 698 bits
Score difference with first non-orthologous sequence - *A. fumigatus*:698   *A. fischeri*:614
afum|EAL93387.1 100.00% afis|EAW18669.1 100.00%

Bootstrap support for afum|EAL93387.1 as seed ortholog is 100%.
Bootstrap support for afis|EAW18669.1 as seed ortholog is 100%.

---

Group of orthologs #6036. Best score 698 bits
Score difference with first non-orthologous sequence - *A. fumigatus*:614 *A. fischeri*:698
afum|EAL93519.1 100.00% afis|EAW18786.1 100.00%
Bootstrap support for afum|EAL93519.1 as seed ortholog is 100%.
Bootstrap support for afis|EAW18786.1 as seed ortholog is 100%.

---

Group of orthologs #6037. Best score 698 bits
Score difference with first non-orthologous sequence - *A. fumigatus*:698 *A. fischeri*:698
afum|EAL90650.1 100.00% afis|EAW22545.1 100.00%
Bootstrap support for afum|EAL90650.1 as seed ortholog is 100%.
Bootstrap support for afis|EAW22545.1 as seed ortholog is 100%.

---

Group of orthologs #6038. Best score 698 bits
Score difference with first non-orthologous sequence - *A. fumigatus*:698 *A. fischeri*:698
afum|EAL92534.1 100.00% afis|EAW21514.1 100.00%
Bootstrap support for afum|EAL92534.1 as seed ortholog is 100%.
Bootstrap support for afis|EAW21514.1 as seed ortholog is 100%.

---

Group of orthologs #6039. Best score 697 bits
Score difference with first non-orthologous sequence - *A. fumigatus*:697 *A. fischeri*:697
afum|EAL84458.1 100.00% afis|EAW20603.1 100.00%
Bootstrap support for afum|EAL84458.1 as seed ortholog is 100%.
Bootstrap support for afis|EAW20603.1 as seed ortholog is 100%.

---

Group of orthologs #6040. Best score 697 bits
Score difference with first non-orthologous sequence - *A. fumigatus*:697 *A. fischeri*:637
afum|EAL87532.1 100.00% afis|EAW18170.1 100.00%
Bootstrap support for afum|EAL87532.1 as seed ortholog is 100%.
Bootstrap support for afis|EAW18170.1 as seed ortholog is 100%.

---

Group of orthologs #6041. Best score 697 bits
Score difference with first non-orthologous sequence - *A. fumigatus*:621 *A. fischeri*:597
afum|EAL86181.1 100.00% afis|EAW24515.1 100.00%
Bootstrap support for afum|EAL86181.1 as seed ortholog is 100%.
Bootstrap support for afis|EAW24515.1 as seed ortholog is 100%.

---

Group of orthologs #6042. Best score 696 bits
Score difference with first non-orthologous sequence - *A. fumigatus*:696 *A. fischeri*:696
afum|EAL84567.1 100.00% afis|EAW20488.1 100.00%
Bootstrap support for afum|EAL84567.1 as seed ortholog is 100%.
Bootstrap support for afis|EAW20488.1 as seed ortholog is 100%.

---

Group of orthologs #6043. Best score 696 bits
Score difference with first non-orthologous sequence - *A. fumigatus*:696 *A. fischeri*:696
afum|EAL87010.1 100.00% afis|EAW19485.1 100.00%
Bootstrap support for afum|EAL87010.1 as seed ortholog is 100%.
Bootstrap support for afis|EAW19485.1 as seed ortholog is 100%.

---

Group of orthologs #6044. Best score 696 bits
Score difference with first non-orthologous sequence - *A. fumigatus*:696 *A. fischeri*:696
afum|EAL91767.1 100.00% afis|EAW17956.1 100.00%
Bootstrap support for afum|EAL91767.1 as seed ortholog is 100%.
Bootstrap support for afis|EAW17956.1 as seed ortholog is 100%.

---

Group of orthologs #6045. Best score 696 bits
Score difference with first non-orthologous sequence - *A. fumigatus*:696 *A. fischeri*:696

| | | | |
|---|---|---|---|
| afum\|EAL85948.1 | 100.00% | afis\|EAW24279.1 | 100.00% |

Bootstrap support for afum|EAL85948.1 as seed ortholog is 100%.
Bootstrap support for afis|EAW24279.1 as seed ortholog is 100%.

---

Group of orthologs #6046. Best score 696 bits
Score difference with first non-orthologous sequence - *A. fumigatus*:696 *A. fischeri*:696

| | | | |
|---|---|---|---|
| afum\|EAL90796.1 | 100.00% | afis\|EAW22395.1 | 100.00% |

Bootstrap support for afum|EAL90796.1 as seed ortholog is 100%.
Bootstrap support for afis|EAW22395.1 as seed ortholog is 100%.

---

Group of orthologs #6047. Best score 695 bits
Score difference with first non-orthologous sequence - *A. fumigatus*:695 *A. fischeri*:695

| | | | |
|---|---|---|---|
| afum\|EAL84199.2 | 100.00% | afis\|EAW15447.1 | 100.00% |

Bootstrap support for afum|EAL84199.2 as seed ortholog is 100%.
Bootstrap support for afis|EAW15447.1 as seed ortholog is 100%.

---

Group of orthologs #6048. Best score 695 bits
Score difference with first non-orthologous sequence - *A. fumigatus*:695 *A. fischeri*:695

| | | | |
|---|---|---|---|
| afum\|EAL84436.1 | 100.00% | afis\|EAW20631.1 | 100.00% |

Bootstrap support for afum|EAL84436.1 as seed ortholog is 100%.
Bootstrap support for afis|EAW20631.1 as seed ortholog is 100%.

---

Group of orthologs #6049. Best score 695 bits
Score difference with first non-orthologous sequence - *A. fumigatus*:566 *A. fischeri*:519

| | | | |
|---|---|---|---|
| afum\|EAL91612.1 | 100.00% | afis\|EAW17804.1 | 100.00% |

Bootstrap support for afum|EAL91612.1 as seed ortholog is 100%.
Bootstrap support for afis|EAW17804.1 as seed ortholog is 100%.

---

Group of orthologs #6050. Best score 695 bits
Score difference with first non-orthologous sequence - *A. fumigatus*:695 *A. fischeri*:695

| | | | |
|---|---|---|---|
| afum\|EBA27277.1 | 100.00% | afis\|EAW16987.1 | 100.00% |

Bootstrap support for afum|EBA27277.1 as seed ortholog is 100%.
Bootstrap support for afis|EAW16987.1 as seed ortholog is 100%.

---

Group of orthologs #6051. Best score 695 bits
Score difference with first non-orthologous sequence - *A. fumigatus*:695 *A. fischeri*:695

| | | | |
|---|---|---|---|
| afum\|EAL93182.1 | 100.00% | afis\|EAW18492.1 | 100.00% |

Bootstrap support for afum|EAL93182.1 as seed ortholog is 100%.
Bootstrap support for afis|EAW18492.1 as seed ortholog is 100%.

---

Group of orthologs #6052. Best score 694 bits
Score difference with first non-orthologous sequence - *A. fumigatus*:694 *A. fischeri*:449

| | | | |
|---|---|---|---|
| afum\|EAL89134.1 | 100.00% | afis\|EAW16537.1 | 100.00% |

Bootstrap support for afum|EAL89134.1 as seed ortholog is 100%.
Bootstrap support for afis|EAW16537.1 as seed ortholog is 100%.

---

Group of orthologs #6053. Best score 694 bits
Score difference with first non-orthologous sequence - *A. fumigatus*:694 *A. fischeri*:694

| | | | |
|---|---|---|---|
| afum\|EAL87596.1 | 100.00% | afis\|EAW18229.1 | 100.00% |

Bootstrap support for afum|EAL87596.1 as seed ortholog is 100%.
Bootstrap support for afis|EAW18229.1 as seed ortholog is 100%.

---

Group of orthologs #6054. Best score 694 bits

Score difference with first non-orthologous sequence - *A. fumigatus*:694   *A. fischeri*:694
afum|EAL91582.1 100.00% afis|EAW17773.1 100.00%
Bootstrap support for afum|EAL91582.1 as seed ortholog is 100%.
Bootstrap support for afis|EAW17773.1 as seed ortholog is 100%.

---

Group of orthologs #6055. Best score 694 bits
Score difference with first non-orthologous sequence - *A. fumigatus*:694   *A. fischeri*:694
afum|EAL91642.1 100.00% afis|EAW17832.1 100.00%
Bootstrap support for afum|EAL91642.1 as seed ortholog is 100%.
Bootstrap support for afis|EAW17832.1 as seed ortholog is 100%.

---

Group of orthologs #6056. Best score 694 bits
Score difference with first non-orthologous sequence - *A. fumigatus*:694   *A. fischeri*:585
afum|EAL84358.1 100.00% afis|EAW25689.1 100.00%
Bootstrap support for afum|EAL84358.1 as seed ortholog is 100%.
Bootstrap support for afis|EAW25689.1 as seed ortholog is 100%.

---

Group of orthologs #6057. Best score 694 bits
Score difference with first non-orthologous sequence - *A. fumigatus*:694   *A. fischeri*:694
afum|EAL88063.2 100.00% afis|EAW23407.1 100.00%
Bootstrap support for afum|EAL88063.2 as seed ortholog is 100%.
Bootstrap support for afis|EAW23407.1 as seed ortholog is 100%.

---

Group of orthologs #6058. Best score 694 bits
Score difference with first non-orthologous sequence - *A. fumigatus*:694   *A. fischeri*:694
afum|EAL92107.2 100.00% afis|EAW21085.1 100.00%
Bootstrap support for afum|EAL92107.2 as seed ortholog is 100%.
Bootstrap support for afis|EAW21085.1 as seed ortholog is 100%.

---

Group of orthologs #6059. Best score 694 bits
Score difference with first non-orthologous sequence - *A. fumigatus*:694   *A. fischeri*:694
afum|EAL89789.1 100.00% afis|EAW25118.1 100.00%
Bootstrap support for afum|EAL89789.1 as seed ortholog is 100%.
Bootstrap support for afis|EAW25118.1 as seed ortholog is 100%.

---

Group of orthologs #6060. Best score 693 bits
Score difference with first non-orthologous sequence - *A. fumigatus*:613   *A. fischeri*:618
afum|EAL87018.1 100.00% afis|EAW19478.1 100.00%
Bootstrap support for afum|EAL87018.1 as seed ortholog is 100%.
Bootstrap support for afis|EAW19478.1 as seed ortholog is 100%.

---

Group of orthologs #6061. Best score 693 bits
Score difference with first non-orthologous sequence - *A. fumigatus*:693   *A. fischeri*:592
afum|EAL86959.1 100.00% afis|EAW19550.1 100.00%
Bootstrap support for afum|EAL86959.1 as seed ortholog is 100%.
Bootstrap support for afis|EAW19550.1 as seed ortholog is 100%.

---

Group of orthologs #6062. Best score 693 bits
Score difference with first non-orthologous sequence - *A. fumigatus*:693   *A. fischeri*:693
afum|EAL87208.1 100.00% afis|EAW23774.1 100.00%
Bootstrap support for afum|EAL87208.1 as seed ortholog is 100%.
Bootstrap support for afis|EAW23774.1 as seed ortholog is 100%.

---

Group of orthologs #6063. Best score 693 bits
Score difference with first non-orthologous sequence - *A. fumigatus*:693   *A. fischeri*:693
afum|EAL90392.1 100.00% afis|EAW22809.1 100.00%

Bootstrap support for afum|EAL90392.1 as seed ortholog is 100%.
Bootstrap support for afis|EAW22809.1 as seed ortholog is 100%.

---

Group of orthologs #6064. Best score 692 bits
Score difference with first non-orthologous sequence - *A. fumigatus*:692 *A. fischeri*:692
afum|EAL88751.1 100.00% afis|EAW16120.1 100.00%
Bootstrap support for afum|EAL88751.1 as seed ortholog is 100%.
Bootstrap support for afis|EAW16120.1 as seed ortholog is 100%.

---

Group of orthologs #6065. Best score 692 bits
Score difference with first non-orthologous sequence - *A. fumigatus*:692 *A. fischeri*:494
afum|EAL85128.1 100.00% afis|EAW19772.1 100.00%
Bootstrap support for afum|EAL85128.1 as seed ortholog is 100%.
Bootstrap support for afis|EAW19772.1 as seed ortholog is 100%.

---

Group of orthologs #6066. Best score 692 bits
Score difference with first non-orthologous sequence - *A. fumigatus*:692 *A. fischeri*:692
afum|EAL87060.1 100.00% afis|EAW19435.1 100.00%
Bootstrap support for afum|EAL87060.1 as seed ortholog is 100%.
Bootstrap support for afis|EAW19435.1 as seed ortholog is 100%.

---

Group of orthologs #6067. Best score 692 bits
Score difference with first non-orthologous sequence - *A. fumigatus*:692 *A. fischeri*:692
afum|EAL91757.1 100.00% afis|EAW17947.1 100.00%
Bootstrap support for afum|EAL91757.1 as seed ortholog is 100%.
Bootstrap support for afis|EAW17947.1 as seed ortholog is 100%.

---

Group of orthologs #6068. Best score 692 bits
Score difference with first non-orthologous sequence - *A. fumigatus*:692 *A. fischeri*:692
afum|EAL87888.1 100.00% afis|EAW23577.1 100.00%
Bootstrap support for afum|EAL87888.1 as seed ortholog is 100%.
Bootstrap support for afis|EAW23577.1 as seed ortholog is 100%.

---

Group of orthologs #6069. Best score 692 bits
Score difference with first non-orthologous sequence - *A. fumigatus*:692 *A. fischeri*:692
afum|EAL93936.1 100.00% afis|EAW19206.1 100.00%
Bootstrap support for afum|EAL93936.1 as seed ortholog is 100%.
Bootstrap support for afis|EAW19206.1 as seed ortholog is 100%.

---

Group of orthologs #6070. Best score 692 bits
Score difference with first non-orthologous sequence - *A. fumigatus*:692 *A. fischeri*:692
afum|EAL92986.1 100.00% afis|EAW21971.1 100.00%
Bootstrap support for afum|EAL92986.1 as seed ortholog is 100%.
Bootstrap support for afis|EAW21971.1 as seed ortholog is 100%.

---

Group of orthologs #6071. Best score 691 bits
Score difference with first non-orthologous sequence - *A. fumigatus*:691 *A. fischeri*:691
afum|EAL85838.1 100.00% afis|EAW15484.1 100.00%
Bootstrap support for afum|EAL85838.1 as seed ortholog is 100%.
Bootstrap support for afis|EAW15484.1 as seed ortholog is 100%.

---

Group of orthologs #6072. Best score 691 bits
Score difference with first non-orthologous sequence - *A. fumigatus*:691 *A. fischeri*:691
afum|EAL88936.1 100.00% afis|EAW16314.1 100.00%
Bootstrap support for afum|EAL88936.1 as seed ortholog is 100%.
Bootstrap support for afis|EAW16314.1 as seed ortholog is 100%.

Group of orthologs #6073. Best score 691 bits
Score difference with first non-orthologous sequence - *A. fumigatus*:691 *A. fischeri*:691
afum|EAL91489.1 100.00% afis|EAW17676.1 100.00%
Bootstrap support for afum|EAL91489.1 as seed ortholog is 100%.
Bootstrap support for afis|EAW17676.1 as seed ortholog is 100%.

---

Group of orthologs #6074. Best score 691 bits
Score difference with first non-orthologous sequence - *A. fumigatus*:691 *A. fischeri*:446
afum|EAL91895.1 100.00% afis|EAW18083.1 100.00%
Bootstrap support for afum|EAL91895.1 as seed ortholog is 100%.
Bootstrap support for afis|EAW18083.1 as seed ortholog is 100%.

---

Group of orthologs #6075. Best score 691 bits
Score difference with first non-orthologous sequence - *A. fumigatus*:599 *A. fischeri*:607
afum|EAL87457.1 100.00% afis|EAW24026.1 100.00%
Bootstrap support for afum|EAL87457.1 as seed ortholog is 100%.
Bootstrap support for afis|EAW24026.1 as seed ortholog is 100%.

---

Group of orthologs #6076. Best score 691 bits
Score difference with first non-orthologous sequence - *A. fumigatus*:691 *A. fischeri*:691
afum|EAL93910.1 100.00% afis|EAW19180.1 100.00%
Bootstrap support for afum|EAL93910.1 as seed ortholog is 100%.
Bootstrap support for afis|EAW19180.1 as seed ortholog is 100%.

---

Group of orthologs #6077. Best score 691 bits
Score difference with first non-orthologous sequence - *A. fumigatus*:691 *A. fischeri*:691
afum|EAL92489.1 100.00% afis|EAW21470.1 100.00%
Bootstrap support for afum|EAL92489.1 as seed ortholog is 100%.
Bootstrap support for afis|EAW21470.1 as seed ortholog is 100%.

---

Group of orthologs #6078. Best score 691 bits
Score difference with first non-orthologous sequence - *A. fumigatus*:691 *A. fischeri*:691
afum|EAL92669.1 100.00% afis|EAW21635.1 100.00%
Bootstrap support for afum|EAL92669.1 as seed ortholog is 100%.
Bootstrap support for afis|EAW21635.1 as seed ortholog is 100%.

---

Group of orthologs #6079. Best score 691 bits
Score difference with first non-orthologous sequence - *A. fumigatus*:691 *A. fischeri*:691
afum|EAL90096.1 100.00% afis|EAW25438.1 100.00%
Bootstrap support for afum|EAL90096.1 as seed ortholog is 100%.
Bootstrap support for afis|EAW25438.1 as seed ortholog is 100%.

---

Group of orthologs #6080. Best score 691 bits
Score difference with first non-orthologous sequence - *A. fumigatus*:691 *A. fischeri*:691
afum|EAL90186.2 100.00% afis|EAW25521.1 100.00%
Bootstrap support for afum|EAL90186.2 as seed ortholog is 100%.
Bootstrap support for afis|EAW25521.1 as seed ortholog is 100%.

---

Group of orthologs #6081. Best score 690 bits
Score difference with first non-orthologous sequence - *A. fumigatus*:690 *A. fischeri*:690
afum|EAL85297.1 100.00% afis|EAW20179.1 100.00%
Bootstrap support for afum|EAL85297.1 as seed ortholog is 100%.
Bootstrap support for afis|EAW20179.1 as seed ortholog is 100%.

---

Group of orthologs #6082. Best score 690 bits

Score difference with first non-orthologous sequence - *A. fumigatus*:638 *A. fischeri*:646
afum|EAL85423.1 100.00% afis|EAW20291.1 100.00%
Bootstrap support for afum|EAL85423.1 as seed ortholog is 100%.
Bootstrap support for afis|EAW20291.1 as seed ortholog is 100%.

---

Group of orthologs #6083. Best score 690 bits
Score difference with first non-orthologous sequence - *A. fumigatus*:690 *A. fischeri*:690
afum|EAL91574.1 100.00% afis|EAW17764.1 100.00%
Bootstrap support for afum|EAL91574.1 as seed ortholog is 100%.
Bootstrap support for afis|EAW17764.1 as seed ortholog is 100%.

---

Group of orthologs #6084. Best score 690 bits
Score difference with first non-orthologous sequence - *A. fumigatus*:690 *A. fischeri*:690
afum|EAL92970.2 100.00% afis|EAW21956.1 100.00%
Bootstrap support for afum|EAL92970.2 as seed ortholog is 100%.
Bootstrap support for afis|EAW21956.1 as seed ortholog is 100%.

---

Group of orthologs #6085. Best score 690 bits
Score difference with first non-orthologous sequence - *A. fumigatus*:690 *A. fischeri*:690
afum|EAL89743.1 100.00% afis|EAW25074.1 100.00%
Bootstrap support for afum|EAL89743.1 as seed ortholog is 100%.
Bootstrap support for afis|EAW25074.1 as seed ortholog is 100%.

---

Group of orthologs #6086. Best score 690 bits
Score difference with first non-orthologous sequence - *A. fumigatus*:690 *A. fischeri*:690
afum|EAL89892.1 100.00% afis|EAW25231.1 100.00%
Bootstrap support for afum|EAL89892.1 as seed ortholog is 100%.
Bootstrap support for afis|EAW25231.1 as seed ortholog is 100%.

---

Group of orthologs #6087. Best score 689 bits
Score difference with first non-orthologous sequence - *A. fumigatus*:689 *A. fischeri*:689
afum|EAL84312.1 100.00% afis|EAW25624.1 100.00%
Bootstrap support for afum|EAL84312.1 as seed ortholog is 100%.
Bootstrap support for afis|EAW25624.1 as seed ortholog is 100%.

---

Group of orthologs #6088. Best score 689 bits
Score difference with first non-orthologous sequence - *A. fumigatus*:435 *A. fischeri*:689
afum|EAL86100.1 100.00% afis|EAW24432.1 100.00%
Bootstrap support for afum|EAL86100.1 as seed ortholog is 100%.
Bootstrap support for afis|EAW24432.1 as seed ortholog is 100%.

---

Group of orthologs #6089. Best score 689 bits
Score difference with first non-orthologous sequence - *A. fumigatus*:689 *A. fischeri*:689
afum|EAL93156.1 100.00% afis|EAW18470.1 100.00%
Bootstrap support for afum|EAL93156.1 as seed ortholog is 100%.
Bootstrap support for afis|EAW18470.1 as seed ortholog is 100%.

---

Group of orthologs #6090. Best score 689 bits
Score difference with first non-orthologous sequence - *A. fumigatus*:689 *A. fischeri*:689
afum|EAL92844.1 100.00% afis|EAW21829.1 100.00%
Bootstrap support for afum|EAL92844.1 as seed ortholog is 100%.
Bootstrap support for afis|EAW21829.1 as seed ortholog is 100%.

---

Group of orthologs #6091. Best score 688 bits
Score difference with first non-orthologous sequence - *A. fumigatus*:688 *A. fischeri*:688
afum|EAL86428.2 100.00% afis|EAW16890.1 100.00%

Bootstrap support for afum|EAL86428.2 as seed ortholog is 100%.
Bootstrap support for afis|EAW16890.1 as seed ortholog is 100%.

---

Group of orthologs #6092. Best score 688 bits
Score difference with first non-orthologous sequence - *A. fumigatus*:546 *A. fischeri*:541
afum|EAL86446.1 100.00% afis|EAW16910.1 100.00%
Bootstrap support for afum|EAL86446.1 as seed ortholog is 100%.
Bootstrap support for afis|EAW16910.1 as seed ortholog is 100%.

---

Group of orthologs #6093. Best score 688 bits
Score difference with first non-orthologous sequence - *A. fumigatus*:688 *A. fischeri*:688
afum|EAL88614.1 100.00% afis|EAW15982.1 100.00%
Bootstrap support for afum|EAL88614.1 as seed ortholog is 100%.
Bootstrap support for afis|EAW15982.1 as seed ortholog is 100%.

---

Group of orthologs #6094. Best score 688 bits
Score difference with first non-orthologous sequence - *A. fumigatus*:688 *A. fischeri*:688
afum|EAL84863.1 100.00% afis|EAW20755.1 100.00%
Bootstrap support for afum|EAL84863.1 as seed ortholog is 100%.
Bootstrap support for afis|EAW20755.1 as seed ortholog is 100%.

---

Group of orthologs #6095. Best score 688 bits
Score difference with first non-orthologous sequence - *A. fumigatus*:688 *A. fischeri*:688
afum|EAL84383.1 100.00% afis|EAW25710.1 100.00%
Bootstrap support for afum|EAL84383.1 as seed ortholog is 100%.
Bootstrap support for afis|EAW25710.1 as seed ortholog is 100%.

---

Group of orthologs #6096. Best score 688 bits
Score difference with first non-orthologous sequence - *A. fumigatus*:550 *A. fischeri*:589
afum|EAL86202.1 100.00% afis|EAW24538.1 100.00%
Bootstrap support for afum|EAL86202.1 as seed ortholog is 100%.
Bootstrap support for afis|EAW24538.1 as seed ortholog is 100%.

---

Group of orthologs #6097. Best score 688 bits
Score difference with first non-orthologous sequence - *A. fumigatus*:688 *A. fischeri*:688
afum|EAL88420.1 100.00% afis|EAW23059.1 100.00%
Bootstrap support for afum|EAL88420.1 as seed ortholog is 100%.
Bootstrap support for afis|EAW23059.1 as seed ortholog is 100%.

---

Group of orthologs #6098. Best score 688 bits
Score difference with first non-orthologous sequence - *A. fumigatus*:688 *A. fischeri*:688
afum|EAL87873.1 100.00% afis|EAW23594.1 100.00%
Bootstrap support for afum|EAL87873.1 as seed ortholog is 100%.
Bootstrap support for afis|EAW23594.1 as seed ortholog is 100%.

---

Group of orthologs #6099. Best score 688 bits
Score difference with first non-orthologous sequence - *A. fumigatus*:688 *A. fischeri*:453
afum|EAL93883.1 100.00% afis|EAW19152.1 100.00%
Bootstrap support for afum|EAL93883.1 as seed ortholog is 100%.
Bootstrap support for afis|EAW19152.1 as seed ortholog is 100%.

---

Group of orthologs #6100. Best score 688 bits
Score difference with first non-orthologous sequence - *A. fumigatus*:688 *A. fischeri*:688
afum|EAL90325.2 100.00% afis|EAW22873.1 100.00%
Bootstrap support for afum|EAL90325.2 as seed ortholog is 100%.
Bootstrap support for afis|EAW22873.1 as seed ortholog is 100%.

Group of orthologs #6101. Best score 688 bits
Score difference with first non-orthologous sequence - *A. fumigatus*:688 *A. fischeri*:688
afum|EAL90295.1 100.00% afis|EAW22902.1 100.00%
Bootstrap support for afum|EAL90295.1 as seed ortholog is 100%.
Bootstrap support for afis|EAW22902.1 as seed ortholog is 100%.

Group of orthologs #6102. Best score 688 bits
Score difference with first non-orthologous sequence - *A. fumigatus*:688 *A. fischeri*:688
afum|EAL89930.1 100.00% afis|EAW25267.1 100.00%
Bootstrap support for afum|EAL89930.1 as seed ortholog is 100%.
Bootstrap support for afis|EAW25267.1 as seed ortholog is 100%.

Group of orthologs #6103. Best score 688 bits
Score difference with first non-orthologous sequence - *A. fumigatus*:688 *A. fischeri*:688
afum|EAL89952.1 100.00% afis|EAW25289.1 100.00%
Bootstrap support for afum|EAL89952.1 as seed ortholog is 100%.
Bootstrap support for afis|EAW25289.1 as seed ortholog is 100%.

Group of orthologs #6104. Best score 687 bits
Score difference with first non-orthologous sequence - *A. fumigatus*:687 *A. fischeri*:687
afum|EAL84674.1 100.00% afis|EAW20959.1 100.00%
Bootstrap support for afum|EAL84674.1 as seed ortholog is 100%.
Bootstrap support for afis|EAW20959.1 as seed ortholog is 100%.

Group of orthologs #6105. Best score 687 bits
Score difference with first non-orthologous sequence - *A. fumigatus*:687 *A. fischeri*:687
afum|EAL91572.1 100.00% afis|EAW17762.1 100.00%
Bootstrap support for afum|EAL91572.1 as seed ortholog is 100%.
Bootstrap support for afis|EAW17762.1 as seed ortholog is 100%.

Group of orthologs #6106. Best score 687 bits
Score difference with first non-orthologous sequence - *A. fumigatus*:687 *A. fischeri*:514
afum|EAL88473.1 100.00% afis|EAW23007.1 100.00%
Bootstrap support for afum|EAL88473.1 as seed ortholog is 100%.
Bootstrap support for afis|EAW23007.1 as seed ortholog is 100%.

Group of orthologs #6107. Best score 687 bits
Score difference with first non-orthologous sequence - *A. fumigatus*:687 *A. fischeri*:687
afum|EAL93580.1 100.00% afis|EAW18849.1 100.00%
Bootstrap support for afum|EAL93580.1 as seed ortholog is 100%.
Bootstrap support for afis|EAW18849.1 as seed ortholog is 100%.

Group of orthologs #6108. Best score 686 bits
Score difference with first non-orthologous sequence - *A. fumigatus*:92 *A. fischeri*:686
afum|EAL85475.1 100.00% afis|EAW20358.1 100.00%
Bootstrap support for afum|EAL85475.1 as seed ortholog is 99%.
Bootstrap support for afis|EAW20358.1 as seed ortholog is 100%.

Group of orthologs #6109. Best score 686 bits
Score difference with first non-orthologous sequence - *A. fumigatus*:686 *A. fischeri*:686
afum|EAL85917.2 100.00% afis|EAW24249.1 100.00%
Bootstrap support for afum|EAL85917.2 as seed ortholog is 100%.
Bootstrap support for afis|EAW24249.1 as seed ortholog is 100%.

Group of orthologs #6110. Best score 686 bits

Score difference with first non-orthologous sequence - *A. fumigatus*:686   *A. fischeri*:686
afum|EAL93138.1 100.00% afis|EAW18453.1 100.00%
Bootstrap support for afum|EAL93138.1 as seed ortholog is 100%.
Bootstrap support for afis|EAW18453.1 as seed ortholog is 100%.

---

Group of orthologs #6111. Best score 686 bits
Score difference with first non-orthologous sequence - *A. fumigatus*:686   *A. fischeri*:686
afum|EAL93011.1 100.00% afis|EAW21996.1 100.00%
Bootstrap support for afum|EAL93011.1 as seed ortholog is 100%.
Bootstrap support for afis|EAW21996.1 as seed ortholog is 100%.

---

Group of orthologs #6112. Best score 685 bits
Score difference with first non-orthologous sequence - *A. fumigatus*:685   *A. fischeri*:685
afum|EAL85550.2 100.00% afis|EAW15800.1 100.00%
Bootstrap support for afum|EAL85550.2 as seed ortholog is 100%.
Bootstrap support for afis|EAW15800.1 as seed ortholog is 100%.

---

Group of orthologs #6113. Best score 685 bits
Score difference with first non-orthologous sequence - *A. fumigatus*:685   *A. fischeri*:685
afum|EAL84855.1 100.00% afis|EAW20764.1 100.00%
Bootstrap support for afum|EAL84855.1 as seed ortholog is 100%.
Bootstrap support for afis|EAW20764.1 as seed ortholog is 100%.

---

Group of orthologs #6114. Best score 685 bits
Score difference with first non-orthologous sequence - *A. fumigatus*:685   *A. fischeri*:685
afum|EAL86891.1 100.00% afis|EAW19631.1 100.00%
Bootstrap support for afum|EAL86891.1 as seed ortholog is 100%.
Bootstrap support for afis|EAW19631.1 as seed ortholog is 100%.

---

Group of orthologs #6115. Best score 685 bits
Score difference with first non-orthologous sequence - *A. fumigatus*:685   *A. fischeri*:685
afum|EAL91364.1 100.00% afis|EAW17550.1 100.00%
Bootstrap support for afum|EAL91364.1 as seed ortholog is 100%.
Bootstrap support for afis|EAW17550.1 as seed ortholog is 100%.

---

Group of orthologs #6116. Best score 685 bits
Score difference with first non-orthologous sequence - *A. fumigatus*:615   *A. fischeri*:633
afum|EAL87182.1 100.00% afis|EAW23747.1 100.00%
Bootstrap support for afum|EAL87182.1 as seed ortholog is 100%.
Bootstrap support for afis|EAW23747.1 as seed ortholog is 100%.

---

Group of orthologs #6117. Best score 685 bits
Score difference with first non-orthologous sequence - *A. fumigatus*:685   *A. fischeri*:685
afum|EAL88258.1 100.00% afis|EAW23225.1 100.00%
Bootstrap support for afum|EAL88258.1 as seed ortholog is 100%.
Bootstrap support for afis|EAW23225.1 as seed ortholog is 100%.

---

Group of orthologs #6118. Best score 685 bits
Score difference with first non-orthologous sequence - *A. fumigatus*:685   *A. fischeri*:685
afum|EAL93982.1 100.00% afis|EAW19253.1 100.00%
Bootstrap support for afum|EAL93982.1 as seed ortholog is 100%.
Bootstrap support for afis|EAW19253.1 as seed ortholog is 100%.

---

Group of orthologs #6119. Best score 685 bits
Score difference with first non-orthologous sequence - *A. fumigatus*:685   *A. fischeri*:685
afum|EAL92717.1 100.00% afis|EAW21683.1 100.00%

Bootstrap support for afum|EAL92717.1 as seed ortholog is 100%.
Bootstrap support for afis|EAW21683.1 as seed ortholog is 100%.

---

Group of orthologs #6120. Best score 685 bits
Score difference with first non-orthologous sequence - *A. fumigatus*:685 *A. fischeri*:685
afum|EAL92831.1 100.00% afis|EAW21815.1 100.00%
Bootstrap support for afum|EAL92831.1 as seed ortholog is 100%.
Bootstrap support for afis|EAW21815.1 as seed ortholog is 100%.

---

Group of orthologs #6121. Best score 684 bits
Score difference with first non-orthologous sequence - *A. fumigatus*:684 *A. fischeri*:684
afum|EAL84845.1 100.00% afis|EAW20775.1 100.00%
Bootstrap support for afum|EAL84845.1 as seed ortholog is 100%.
Bootstrap support for afis|EAW20775.1 as seed ortholog is 100%.

---

Group of orthologs #6122. Best score 684 bits
Score difference with first non-orthologous sequence - *A. fumigatus*:684 *A. fischeri*:684
afum|EAL91748.1 100.00% afis|EAW17937.1 100.00%
Bootstrap support for afum|EAL91748.1 as seed ortholog is 100%.
Bootstrap support for afis|EAW17937.1 as seed ortholog is 100%.

---

Group of orthologs #6123. Best score 684 bits
Score difference with first non-orthologous sequence - *A. fumigatus*:684 *A. fischeri*:684
afum|EAL93399.1 100.00% afis|EAW18679.1 100.00%
Bootstrap support for afum|EAL93399.1 as seed ortholog is 100%.
Bootstrap support for afis|EAW18679.1 as seed ortholog is 100%.

---

Group of orthologs #6124. Best score 683 bits
Score difference with first non-orthologous sequence - *A. fumigatus*:683 *A. fischeri*:683
afum|EAL84900.1 100.00% afis|EAW20017.1 100.00%
Bootstrap support for afum|EAL84900.1 as seed ortholog is 100%.
Bootstrap support for afis|EAW20017.1 as seed ortholog is 100%.

---

Group of orthologs #6125. Best score 682 bits
Score difference with first non-orthologous sequence - *A. fumigatus*:682 *A. fischeri*:682
afum|EAL85804.1 100.00% afis|EAW15524.1 100.00%
Bootstrap support for afum|EAL85804.1 as seed ortholog is 100%.
Bootstrap support for afis|EAW15524.1 as seed ortholog is 100%.

---

Group of orthologs #6126. Best score 682 bits
Score difference with first non-orthologous sequence - *A. fumigatus*:682 *A. fischeri*:682
afum|EAL85773.1 100.00% afis|EAW15559.1 100.00%
Bootstrap support for afum|EAL85773.1 as seed ortholog is 100%.
Bootstrap support for afis|EAW15559.1 as seed ortholog is 100%.

---

Group of orthologs #6127. Best score 682 bits
Score difference with first non-orthologous sequence - *A. fumigatus*:682 *A. fischeri*:682
afum|EAL84982.2 100.00% afis|EAW19927.1 100.00%
Bootstrap support for afum|EAL84982.2 as seed ortholog is 100%.
Bootstrap support for afis|EAW19927.1 as seed ortholog is 100%.

---

Group of orthologs #6128. Best score 682 bits
Score difference with first non-orthologous sequence - *A. fumigatus*:682 *A. fischeri*:682
afum|EAL87022.1 100.00% afis|EAW19474.1 100.00%
Bootstrap support for afum|EAL87022.1 as seed ortholog is 100%.
Bootstrap support for afis|EAW19474.1 as seed ortholog is 100%.

Group of orthologs #6129. Best score 682 bits
Score difference with first non-orthologous sequence - *A. fumigatus*:682 *A. fischeri*:682
afum|EAL87173.1 100.00% afis|EAW23735.1 100.00%
Bootstrap support for afum|EAL87173.1 as seed ortholog is 100%.
Bootstrap support for afis|EAW23735.1 as seed ortholog is 100%.

Group of orthologs #6130. Best score 682 bits
Score difference with first non-orthologous sequence - *A. fumigatus*:682 *A. fischeri*:682
afum|EAL93343.2 100.00% afis|EAW18631.1 100.00%
Bootstrap support for afum|EAL93343.2 as seed ortholog is 100%.
Bootstrap support for afis|EAW18631.1 as seed ortholog is 100%.

Group of orthologs #6131. Best score 682 bits
Score difference with first non-orthologous sequence - *A. fumigatus*:682 *A. fischeri*:682
afum|EAL89624.2 100.00% afis|EAW24966.1 100.00%
Bootstrap support for afum|EAL89624.2 as seed ortholog is 100%.
Bootstrap support for afis|EAW24966.1 as seed ortholog is 100%.

Group of orthologs #6132. Best score 682 bits
Score difference with first non-orthologous sequence - *A. fumigatus*:682 *A. fischeri*:682
afum|EAL89830.1 100.00% afis|EAW25155.1 100.00%
Bootstrap support for afum|EAL89830.1 as seed ortholog is 100%.
Bootstrap support for afis|EAW25155.1 as seed ortholog is 100%.

Group of orthologs #6133. Best score 681 bits
Score difference with first non-orthologous sequence - *A. fumigatus*:681 *A. fischeri*:681
afum|EAL84237.1 100.00% afis|EAW15409.1 100.00%
Bootstrap support for afum|EAL84237.1 as seed ortholog is 100%.
Bootstrap support for afis|EAW15409.1 as seed ortholog is 100%.

Group of orthologs #6134. Best score 681 bits
Score difference with first non-orthologous sequence - *A. fumigatus*:619 *A. fischeri*:681
afum|EAL84198.2 100.00% afis|EAW15448.1 100.00%
Bootstrap support for afum|EAL84198.2 as seed ortholog is 100%.
Bootstrap support for afis|EAW15448.1 as seed ortholog is 100%.

Group of orthologs #6135. Best score 681 bits
Score difference with first non-orthologous sequence - *A. fumigatus*:681 *A. fischeri*:356
afum|EAL86864.1 100.00% afis|EAW19659.1 100.00%
Bootstrap support for afum|EAL86864.1 as seed ortholog is 100%.
Bootstrap support for afis|EAW19659.1 as seed ortholog is 100%.

Group of orthologs #6136. Best score 681 bits
Score difference with first non-orthologous sequence - *A. fumigatus*:428 *A. fischeri*:398
afum|EAL87949.1 100.00% afis|EAW23514.1 100.00%
Bootstrap support for afum|EAL87949.1 as seed ortholog is 100%.
Bootstrap support for afis|EAW23514.1 as seed ortholog is 100%.

Group of orthologs #6137. Best score 681 bits
Score difference with first non-orthologous sequence - *A. fumigatus*:681 *A. fischeri*:681
afum|EAL93990.1 100.00% afis|EAW19263.1 100.00%
Bootstrap support for afum|EAL93990.1 as seed ortholog is 100%.
Bootstrap support for afis|EAW19263.1 as seed ortholog is 100%.

Group of orthologs #6138. Best score 680 bits

Score difference with first non-orthologous sequence - *A. fumigatus*:680 *A. fischeri*:363
afum|EAL85143.1 100.00% afis|EAW19754.1 100.00%
Bootstrap support for afum|EAL85143.1 as seed ortholog is 100%.
Bootstrap support for afis|EAW19754.1 as seed ortholog is 100%.

---

Group of orthologs #6139. Best score 680 bits
Score difference with first non-orthologous sequence - *A. fumigatus*:680 *A. fischeri*:680
afum|EAL89290.1 100.00% afis|EAW16690.1 100.00%
Bootstrap support for afum|EAL89290.1 as seed ortholog is 100%.
Bootstrap support for afis|EAW16690.1 as seed ortholog is 100%.

---

Group of orthologs #6140. Best score 680 bits
Score difference with first non-orthologous sequence - *A. fumigatus*:680 *A. fischeri*:680
afum|EAL89782.2 100.00% afis|EAW25111.1 100.00%
Bootstrap support for afum|EAL89782.2 as seed ortholog is 100%.
Bootstrap support for afis|EAW25111.1 as seed ortholog is 100%.

---

Group of orthologs #6141. Best score 679 bits
Score difference with first non-orthologous sequence - *A. fumigatus*:679 *A. fischeri*:679
afum|EAL86668.1 100.00% afis|EAW17249.1 100.00%
Bootstrap support for afum|EAL86668.1 as seed ortholog is 100%.
Bootstrap support for afis|EAW17249.1 as seed ortholog is 100%.

---

Group of orthologs #6142. Best score 679 bits
Score difference with first non-orthologous sequence - *A. fumigatus*:679 *A. fischeri*:679
afum|EAL88601.1 100.00% afis|EAW15968.1 100.00%
Bootstrap support for afum|EAL88601.1 as seed ortholog is 100%.
Bootstrap support for afis|EAW15968.1 as seed ortholog is 100%.

---

Group of orthologs #6143. Best score 679 bits
Score difference with first non-orthologous sequence - *A. fumigatus*:588 *A. fischeri*:679
afum|EAL88823.1 100.00% afis|EAW16192.1 100.00%
Bootstrap support for afum|EAL88823.1 as seed ortholog is 100%.
Bootstrap support for afis|EAW16192.1 as seed ortholog is 100%.

---

Group of orthologs #6144. Best score 679 bits
Score difference with first non-orthologous sequence - *A. fumigatus*:609 *A. fischeri*:591
afum|EAL89185.1 100.00% afis|EAW16590.1 100.00%
Bootstrap support for afum|EAL89185.1 as seed ortholog is 100%.
Bootstrap support for afis|EAW16590.1 as seed ortholog is 100%.

---

Group of orthologs #6145. Best score 679 bits
Score difference with first non-orthologous sequence - *A. fumigatus*:378 *A. fischeri*:679
afum|EAL91583.1 100.00% afis|EAW17775.1 100.00%
Bootstrap support for afum|EAL91583.1 as seed ortholog is 100%.
Bootstrap support for afis|EAW17775.1 as seed ortholog is 100%.

---

Group of orthologs #6146. Best score 679 bits
Score difference with first non-orthologous sequence - *A. fumigatus*:679 *A. fischeri*:679
afum|EAL87318.1 100.00% afis|EAW23890.1 100.00%
Bootstrap support for afum|EAL87318.1 as seed ortholog is 100%.
Bootstrap support for afis|EAW23890.1 as seed ortholog is 100%.

---

Group of orthologs #6147. Best score 679 bits
Score difference with first non-orthologous sequence - *A. fumigatus*:679 *A. fischeri*:679
afum|EAL87914.2 100.00% afis|EAW23552.1 100.00%

Bootstrap support for afum|EAL87914.2 as seed ortholog is 100%.
Bootstrap support for afis|EAW23552.1 as seed ortholog is 100%.

---

Group of orthologs #6148. Best score 679 bits
Score difference with first non-orthologous sequence - *A. fumigatus*:679 *A. fischeri*:629
afum|EAL90977.1 100.00% afis|EAW22217.1 100.00%
Bootstrap support for afum|EAL90977.1 as seed ortholog is 100%.
Bootstrap support for afis|EAW22217.1 as seed ortholog is 100%.

---

Group of orthologs #6149. Best score 679 bits
Score difference with first non-orthologous sequence - *A. fumigatus*:679 *A. fischeri*:679
afum|EAL90789.1 100.00% afis|EAW22402.1 100.00%
Bootstrap support for afum|EAL90789.1 as seed ortholog is 100%.
Bootstrap support for afis|EAW22402.1 as seed ortholog is 100%.

---

Group of orthologs #6150. Best score 679 bits
Score difference with first non-orthologous sequence - *A. fumigatus*:679 *A. fischeri*:679
afum|EAL92478.1 100.00% afis|EAW21459.1 100.00%
Bootstrap support for afum|EAL92478.1 as seed ortholog is 100%.
Bootstrap support for afis|EAW21459.1 as seed ortholog is 100%.

---

Group of orthologs #6151. Best score 679 bits
Score difference with first non-orthologous sequence - *A. fumigatus*:679 *A. fischeri*:679
afum|EAL89373.1 100.00% afis|EAW24724.1 100.00%
Bootstrap support for afum|EAL89373.1 as seed ortholog is 100%.
Bootstrap support for afis|EAW24724.1 as seed ortholog is 100%.

---

Group of orthologs #6152. Best score 678 bits
Score difference with first non-orthologous sequence - *A. fumigatus*:678 *A. fischeri*:678
afum|EAL88609.1 100.00% afis|EAW15977.1 100.00%
Bootstrap support for afum|EAL88609.1 as seed ortholog is 100%.
Bootstrap support for afis|EAW15977.1 as seed ortholog is 100%.

---

Group of orthologs #6153. Best score 678 bits
Score difference with first non-orthologous sequence - *A. fumigatus*:678 *A. fischeri*:678
afum|EAL88959.1 100.00% afis|EAW16334.1 100.00%
Bootstrap support for afum|EAL88959.1 as seed ortholog is 100%.
Bootstrap support for afis|EAW16334.1 as seed ortholog is 100%.

---

Group of orthologs #6154. Best score 678 bits
Score difference with first non-orthologous sequence - *A. fumigatus*:678 *A. fischeri*:678
afum|EAL89095.1 100.00% afis|EAW16500.1 100.00%
Bootstrap support for afum|EAL89095.1 as seed ortholog is 100%.
Bootstrap support for afis|EAW16500.1 as seed ortholog is 100%.

---

Group of orthologs #6155. Best score 678 bits
Score difference with first non-orthologous sequence - *A. fumigatus*:678 *A. fischeri*:678
afum|EAL84850.1 100.00% afis|EAW20768.1 100.00%
Bootstrap support for afum|EAL84850.1 as seed ortholog is 100%.
Bootstrap support for afis|EAW20768.1 as seed ortholog is 100%.

---

Group of orthologs #6156. Best score 678 bits
Score difference with first non-orthologous sequence - *A. fumigatus*:678 *A. fischeri*:678
afum|EAL86929.1 100.00% afis|EAW19589.1 100.00%
Bootstrap support for afum|EAL86929.1 as seed ortholog is 100%.
Bootstrap support for afis|EAW19589.1 as seed ortholog is 100%.

Group of orthologs #6157. Best score 678 bits
Score difference with first non-orthologous sequence - *A. fumigatus*:678 *A. fischeri*:678
afum|EAL89606.1 100.00% afis|EAW24949.1 100.00%
Bootstrap support for afum|EAL89606.1 as seed ortholog is 100%.
Bootstrap support for afis|EAW24949.1 as seed ortholog is 100%.

Group of orthologs #6158. Best score 677 bits
Score difference with first non-orthologous sequence - *A. fumigatus*:677 *A. fischeri*:677
afum|EAL87985.1 100.00% afis|EAW20747.1 100.00%
Bootstrap support for afum|EAL87985.1 as seed ortholog is 100%.
Bootstrap support for afis|EAW20747.1 as seed ortholog is 100%.

Group of orthologs #6159. Best score 677 bits
Score difference with first non-orthologous sequence - *A. fumigatus*:677 *A. fischeri*:677
afum|EAL87205.1 100.00% afis|EAW23771.1 100.00%
Bootstrap support for afum|EAL87205.1 as seed ortholog is 100%.
Bootstrap support for afis|EAW23771.1 as seed ortholog is 100%.

Group of orthologs #6160. Best score 677 bits
Score difference with first non-orthologous sequence - *A. fumigatus*:677 *A. fischeri*:677
afum|EAL92927.1 100.00% afis|EAW21915.1 100.00%
Bootstrap support for afum|EAL92927.1 as seed ortholog is 100%.
Bootstrap support for afis|EAW21915.1 as seed ortholog is 100%.

Group of orthologs #6161. Best score 676 bits
Score difference with first non-orthologous sequence - *A. fumigatus*:676 *A. fischeri*:676
afum|EAL88683.1 100.00% afis|EAW16050.1 100.00%
Bootstrap support for afum|EAL88683.1 as seed ortholog is 100%.
Bootstrap support for afis|EAW16050.1 as seed ortholog is 100%.

Group of orthologs #6162. Best score 676 bits
Score difference with first non-orthologous sequence - *A. fumigatus*:676 *A. fischeri*:566
afum|EAL84989.2 100.00% afis|EAW19919.1 100.00%
Bootstrap support for afum|EAL84989.2 as seed ortholog is 100%.
Bootstrap support for afis|EAW19919.1 as seed ortholog is 100%.

Group of orthologs #6163. Best score 676 bits
Score difference with first non-orthologous sequence - *A. fumigatus*:587 *A. fischeri*:592
afum|EAL88932.1 100.00% afis|EAW16310.1 100.00%
Bootstrap support for afum|EAL88932.1 as seed ortholog is 100%.
Bootstrap support for afis|EAW16310.1 as seed ortholog is 100%.

Group of orthologs #6164. Best score 676 bits
Score difference with first non-orthologous sequence - *A. fumigatus*:676 *A. fischeri*:514
afum|EAL87539.1 100.00% afis|EAW18178.1 100.00%
Bootstrap support for afum|EAL87539.1 as seed ortholog is 100%.
Bootstrap support for afis|EAW18178.1 as seed ortholog is 100%.

Group of orthologs #6165. Best score 676 bits
Score difference with first non-orthologous sequence - *A. fumigatus*:676 *A. fischeri*:676
afum|EAL84859.1 100.00% afis|EAW20760.1 100.00%
Bootstrap support for afum|EAL84859.1 as seed ortholog is 100%.
Bootstrap support for afis|EAW20760.1 as seed ortholog is 100%.

Group of orthologs #6166. Best score 676 bits

Score difference with first non-orthologous sequence - *A. fumigatus*:676 *A. fischeri*:676
afum|EAL91579.1 100.00% afis|EAW17770.1 100.00%
Bootstrap support for afum|EAL91579.1 as seed ortholog is 100%.
Bootstrap support for afis|EAW17770.1 as seed ortholog is 100%.

---

Group of orthologs #6167. Best score 676 bits
Score difference with first non-orthologous sequence - *A. fumigatus*:676 *A. fischeri*:676
afum|EAL90873.1 100.00% afis|EAW22317.1 100.00%
Bootstrap support for afum|EAL90873.1 as seed ortholog is 100%.
Bootstrap support for afis|EAW22317.1 as seed ortholog is 100%.

---

Group of orthologs #6168. Best score 676 bits
Score difference with first non-orthologous sequence - *A. fumigatus*:676 *A. fischeri*:676
afum|EAL89516.1 100.00% afis|EAW24866.1 100.00%
Bootstrap support for afum|EAL89516.1 as seed ortholog is 100%.
Bootstrap support for afis|EAW24866.1 as seed ortholog is 100%.

---

Group of orthologs #6169. Best score 675 bits
Score difference with first non-orthologous sequence - *A. fumigatus*:675 *A. fischeri*:675
afum|EAL85106.2 100.00% afis|EAW19788.1 100.00%
Bootstrap support for afum|EAL85106.2 as seed ortholog is 100%.
Bootstrap support for afis|EAW19788.1 as seed ortholog is 100%.

---

Group of orthologs #6170. Best score 675 bits
Score difference with first non-orthologous sequence - *A. fumigatus*:675 *A. fischeri*:675
afum|EAL85028.1 100.00% afis|EAW19869.1 100.00%
Bootstrap support for afum|EAL85028.1 as seed ortholog is 100%.
Bootstrap support for afis|EAW19869.1 as seed ortholog is 100%.

---

Group of orthologs #6171. Best score 675 bits
Score difference with first non-orthologous sequence - *A. fumigatus*:574 *A. fischeri*:675
afum|EAL84918.1 100.00% afis|EAW20001.1 100.00%
Bootstrap support for afum|EAL84918.1 as seed ortholog is 100%.
Bootstrap support for afis|EAW20001.1 as seed ortholog is 100%.

---

Group of orthologs #6172. Best score 675 bits
Score difference with first non-orthologous sequence - *A. fumigatus*:675 *A. fischeri*:675
afum|EAL91804.1 100.00% afis|EAW17994.1 100.00%
Bootstrap support for afum|EAL91804.1 as seed ortholog is 100%.
Bootstrap support for afis|EAW17994.1 as seed ortholog is 100%.

---

Group of orthologs #6173. Best score 675 bits
Score difference with first non-orthologous sequence - *A. fumigatus*:220 *A. fischeri*:352
afum|EAL84310.1 100.00% afis|EAW25622.1 100.00%
Bootstrap support for afum|EAL84310.1 as seed ortholog is 100%.
Bootstrap support for afis|EAW25622.1 as seed ortholog is 100%.

---

Group of orthologs #6174. Best score 675 bits
Score difference with first non-orthologous sequence - *A. fumigatus*:675 *A. fischeri*:609
afum|EAL90860.1 100.00% afis|EAW22332.1 100.00%
Bootstrap support for afum|EAL90860.1 as seed ortholog is 100%.
Bootstrap support for afis|EAW22332.1 as seed ortholog is 100%.

---

Group of orthologs #6175. Best score 675 bits
Score difference with first non-orthologous sequence - *A. fumigatus*:675 *A. fischeri*:675
afum|EAL92572.2 100.00% afis|EAW21552.1 100.00%

Bootstrap support for afum|EAL92572.2 as seed ortholog is 100%.
Bootstrap support for afis|EAW21552.1 as seed ortholog is 100%.

---

Group of orthologs #6176. Best score 674 bits
Score difference with first non-orthologous sequence - *A. fumigatus*:674 *A. fischeri*:674
afum|EAL84531.1 100.00% afis|EAW20526.1 100.00%
Bootstrap support for afum|EAL84531.1 as seed ortholog is 100%.
Bootstrap support for afis|EAW20526.1 as seed ortholog is 100%.

---

Group of orthologs #6177. Best score 674 bits
Score difference with first non-orthologous sequence - *A. fumigatus*:674 *A. fischeri*:674
afum|EAL89071.1 100.00% afis|EAW16477.1 100.00%
Bootstrap support for afum|EAL89071.1 as seed ortholog is 100%.
Bootstrap support for afis|EAW16477.1 as seed ortholog is 100%.

---

Group of orthologs #6178. Best score 674 bits
Score difference with first non-orthologous sequence - *A. fumigatus*:674 *A. fischeri*:674
afum|EAL93825.1 100.00% afis|EAW19094.1 100.00%
Bootstrap support for afum|EAL93825.1 as seed ortholog is 100%.
Bootstrap support for afis|EAW19094.1 as seed ortholog is 100%.

---

Group of orthologs #6179. Best score 674 bits
Score difference with first non-orthologous sequence - *A. fumigatus*:674 *A. fischeri*:674
afum|EBA27213.1 100.00% afis|EAW19976.1 100.00%
Bootstrap support for afum|EBA27213.1 as seed ortholog is 100%.
Bootstrap support for afis|EAW19976.1 as seed ortholog is 100%.

---

Group of orthologs #6180. Best score 673 bits
Score difference with first non-orthologous sequence - *A. fumigatus*:585 *A. fischeri*:578
afum|EAL85687.1 100.00% afis|EAW15661.1 100.00%
Bootstrap support for afum|EAL85687.1 as seed ortholog is 100%.
Bootstrap support for afis|EAW15661.1 as seed ortholog is 100%.

---

Group of orthologs #6181. Best score 673 bits
Score difference with first non-orthologous sequence - *A. fumigatus*:673 *A. fischeri*:673
afum|EAL88606.1 100.00% afis|EAW15974.1 100.00%
Bootstrap support for afum|EAL88606.1 as seed ortholog is 100%.
Bootstrap support for afis|EAW15974.1 as seed ortholog is 100%.

---

Group of orthologs #6182. Best score 673 bits
Score difference with first non-orthologous sequence - *A. fumigatus*:673 *A. fischeri*:673
afum|EAL85024.1 100.00% afis|EAW19875.1 100.00%
Bootstrap support for afum|EAL85024.1 as seed ortholog is 100%.
Bootstrap support for afis|EAW19875.1 as seed ortholog is 100%.

---

Group of orthologs #6183. Best score 673 bits
Score difference with first non-orthologous sequence - *A. fumigatus*:673 *A. fischeri*:673
afum|EAL89086.1 100.00% afis|EAW16492.1 100.00%
Bootstrap support for afum|EAL89086.1 as seed ortholog is 100%.
Bootstrap support for afis|EAW16492.1 as seed ortholog is 100%.

---

Group of orthologs #6184. Best score 673 bits
Score difference with first non-orthologous sequence - *A. fumigatus*:673 *A. fischeri*:673
afum|EAL91465.1 100.00% afis|EAW17649.1 100.00%
Bootstrap support for afum|EAL91465.1 as seed ortholog is 100%.
Bootstrap support for afis|EAW17649.1 as seed ortholog is 100%.

---

Group of orthologs #6185. Best score 673 bits
Score difference with first non-orthologous sequence - *A. fumigatus*:673 *A. fischeri*:495
afum|EAL91769.1 100.00% afis|EAW17958.1 100.00%
Bootstrap support for afum|EAL91769.1 as seed ortholog is 100%.
Bootstrap support for afis|EAW17958.1 as seed ortholog is 100%.

---

Group of orthologs #6186. Best score 673 bits
Score difference with first non-orthologous sequence - *A. fumigatus*:625 *A. fischeri*:376
afum|EAL86069.1 100.00% afis|EAW24398.1 100.00%
Bootstrap support for afum|EAL86069.1 as seed ortholog is 100%.
Bootstrap support for afis|EAW24398.1 as seed ortholog is 100%.

---

Group of orthologs #6187. Best score 673 bits
Score difference with first non-orthologous sequence - *A. fumigatus*:673 *A. fischeri*:673
afum|EAL86135.1 100.00% afis|EAW24464.1 100.00%
Bootstrap support for afum|EAL86135.1 as seed ortholog is 100%.
Bootstrap support for afis|EAW24464.1 as seed ortholog is 100%.

---

Group of orthologs #6188. Best score 673 bits
Score difference with first non-orthologous sequence - *A. fumigatus*:673 *A. fischeri*:673
afum|EAL87321.1 100.00% afis|EAW23893.1 100.00%
Bootstrap support for afum|EAL87321.1 as seed ortholog is 100%.
Bootstrap support for afis|EAW23893.1 as seed ortholog is 100%.

---

Group of orthologs #6189. Best score 673 bits
Score difference with first non-orthologous sequence - *A. fumigatus*:673 *A. fischeri*:673
afum|EAL93946.1 100.00% afis|EAW19213.1 100.00%
Bootstrap support for afum|EAL93946.1 as seed ortholog is 100%.
Bootstrap support for afis|EAW19213.1 as seed ortholog is 100%.

---

Group of orthologs #6190. Best score 673 bits
Score difference with first non-orthologous sequence - *A. fumigatus*:673 *A. fischeri*:673
afum|EAL90621.1 100.00% afis|EAW22577.1 100.00%
Bootstrap support for afum|EAL90621.1 as seed ortholog is 100%.
Bootstrap support for afis|EAW22577.1 as seed ortholog is 100%.

---

Group of orthologs #6191. Best score 673 bits
Score difference with first non-orthologous sequence - *A. fumigatus*:673 *A. fischeri*:673
afum|EAL89681.1 100.00% afis|EAW25015.1 100.00%
Bootstrap support for afum|EAL89681.1 as seed ortholog is 100%.
Bootstrap support for afis|EAW25015.1 as seed ortholog is 100%.

---

Group of orthologs #6192. Best score 673 bits
Score difference with first non-orthologous sequence - *A. fumigatus*:673 *A. fischeri*:673
afum|EAL93012.1 100.00% afis|EAW21997.1 100.00%
Bootstrap support for afum|EAL93012.1 as seed ortholog is 100%.
Bootstrap support for afis|EAW21997.1 as seed ortholog is 100%.

---

Group of orthologs #6193. Best score 673 bits
Score difference with first non-orthologous sequence - *A. fumigatus*:581 *A. fischeri*:454
afum|EAL89916.1 100.00% afis|EAW25253.1 100.00%
Bootstrap support for afum|EAL89916.1 as seed ortholog is 100%.
Bootstrap support for afis|EAW25253.1 as seed ortholog is 100%.

---

Group of orthologs #6194. Best score 672 bits

Score difference with first non-orthologous sequence - *A. fumigatus*:672 *A. fischeri*:672
afum|EAL86339.2 100.00% afis|EAW16771.1 100.00%
Bootstrap support for afum|EAL86339.2 as seed ortholog is 100%.
Bootstrap support for afis|EAW16771.1 as seed ortholog is 100%.

---

Group of orthologs #6195. Best score 672 bits
Score difference with first non-orthologous sequence - *A. fumigatus*:672 *A. fischeri*:672
afum|EAL88926.1 100.00% afis|EAW16304.1 100.00%
Bootstrap support for afum|EAL88926.1 as seed ortholog is 100%.
Bootstrap support for afis|EAW16304.1 as seed ortholog is 100%.

---

Group of orthologs #6196. Best score 672 bits
Score difference with first non-orthologous sequence - *A. fumigatus*:672 *A. fischeri*:672
afum|EAL85188.1 100.00% afis|EAW20070.1 100.00%
Bootstrap support for afum|EAL85188.1 as seed ortholog is 100%.
Bootstrap support for afis|EAW20070.1 as seed ortholog is 100%.

---

Group of orthologs #6197. Best score 672 bits
Score difference with first non-orthologous sequence - *A. fumigatus*:672 *A. fischeri*:672
afum|EAL87690.2 100.00% afis|EAW18323.1 100.00%
Bootstrap support for afum|EAL87690.2 as seed ortholog is 100%.
Bootstrap support for afis|EAW18323.1 as seed ortholog is 100%.

---

Group of orthologs #6198. Best score 672 bits
Score difference with first non-orthologous sequence - *A. fumigatus*:672 *A. fischeri*:672
afum|EAL87078.1 100.00% afis|EAW19416.1 100.00%
Bootstrap support for afum|EAL87078.1 as seed ortholog is 100%.
Bootstrap support for afis|EAW19416.1 as seed ortholog is 100%.

---

Group of orthologs #6199. Best score 672 bits
Score difference with first non-orthologous sequence - *A. fumigatus*:672 *A. fischeri*:589
afum|EAL86846.1 100.00% afis|EAW19676.1 100.00%
Bootstrap support for afum|EAL86846.1 as seed ortholog is 100%.
Bootstrap support for afis|EAW19676.1 as seed ortholog is 100%.

---

Group of orthologs #6200. Best score 672 bits
Score difference with first non-orthologous sequence - *A. fumigatus*:672 *A. fischeri*:525
afum|EAL91206.2 100.00% afis|EAW17369.1 100.00%
Bootstrap support for afum|EAL91206.2 as seed ortholog is 100%.
Bootstrap support for afis|EAW17369.1 as seed ortholog is 100%.

---

Group of orthologs #6201. Best score 672 bits
Score difference with first non-orthologous sequence - *A. fumigatus*:672 *A. fischeri*:672
afum|EAL86099.1 100.00% afis|EAW24431.1 100.00%
Bootstrap support for afum|EAL86099.1 as seed ortholog is 100%.
Bootstrap support for afis|EAW24431.1 as seed ortholog is 100%.

---

Group of orthologs #6202. Best score 672 bits
Score difference with first non-orthologous sequence - *A. fumigatus*:672 *A. fischeri*:672
afum|EAL90656.2 100.00% afis|EAW22538.1 100.00%
Bootstrap support for afum|EAL90656.2 as seed ortholog is 100%.
Bootstrap support for afis|EAW22538.1 as seed ortholog is 100%.

---

Group of orthologs #6203. Best score 672 bits
Score difference with first non-orthologous sequence - *A. fumigatus*:672 *A. fischeri*:672
afum|EBA27299.1 100.00% afis|EAW19448.1 100.00%

Bootstrap support for afum|EBA27299.1 as seed ortholog is 100%.
Bootstrap support for afis|EAW19448.1 as seed ortholog is 100%.

---

Group of orthologs #6204. Best score 672 bits
Score difference with first non-orthologous sequence - *A. fumigatus*:580 *A. fischeri*:605
afum|EAL92527.1 100.00% afis|EAW21507.1 100.00%
Bootstrap support for afum|EAL92527.1 as seed ortholog is 100%.
Bootstrap support for afis|EAW21507.1 as seed ortholog is 100%.

---

Group of orthologs #6205. Best score 671 bits
Score difference with first non-orthologous sequence - *A. fumigatus*:671 *A. fischeri*:671
afum|EAL91700.1 100.00% afis|EAW17891.1 100.00%
Bootstrap support for afum|EAL91700.1 as seed ortholog is 100%.
Bootstrap support for afis|EAW17891.1 as seed ortholog is 100%.

---

Group of orthologs #6206. Best score 671 bits
Score difference with first non-orthologous sequence - *A. fumigatus*:671 *A. fischeri*:671
afum|EAL88073.1 100.00% afis|EAW23397.1 100.00%
Bootstrap support for afum|EAL88073.1 as seed ortholog is 100%.
Bootstrap support for afis|EAW23397.1 as seed ortholog is 100%.

---

Group of orthologs #6207. Best score 671 bits
Score difference with first non-orthologous sequence - *A. fumigatus*:671 *A. fischeri*:671
afum|EAL93677.1 100.00% afis|EAW18944.1 100.00%
Bootstrap support for afum|EAL93677.1 as seed ortholog is 100%.
Bootstrap support for afis|EAW18944.1 as seed ortholog is 100%.

---

Group of orthologs #6208. Best score 671 bits
Score difference with first non-orthologous sequence - *A. fumigatus*:671 *A. fischeri*:671
afum|EAL93989.1 100.00% afis|EAW19262.1 100.00%
Bootstrap support for afum|EAL93989.1 as seed ortholog is 100%.
Bootstrap support for afis|EAW19262.1 as seed ortholog is 100%.

---

Group of orthologs #6209. Best score 671 bits
Score difference with first non-orthologous sequence - *A. fumigatus*:488 *A. fischeri*:303
afum|EAL90338.1 100.00% afis|EAW22862.1 100.00%
Bootstrap support for afum|EAL90338.1 as seed ortholog is 100%.
Bootstrap support for afis|EAW22862.1 as seed ortholog is 100%.

---

Group of orthologs #6210. Best score 671 bits
Score difference with first non-orthologous sequence - *A. fumigatus*:671 *A. fischeri*:671
afum|EAL92561.1 100.00% afis|EAW21541.1 100.00%
Bootstrap support for afum|EAL92561.1 as seed ortholog is 100%.
Bootstrap support for afis|EAW21541.1 as seed ortholog is 100%.

---

Group of orthologs #6211. Best score 671 bits
Score difference with first non-orthologous sequence - *A. fumigatus*:589 *A. fischeri*:671
afum|EAL89907.2 100.00% afis|EAW25245.1 100.00%
Bootstrap support for afum|EAL89907.2 as seed ortholog is 100%.
Bootstrap support for afis|EAW25245.1 as seed ortholog is 100%.

---

Group of orthologs #6212. Best score 671 bits
Score difference with first non-orthologous sequence - *A. fumigatus*:671 *A. fischeri*:671
afum|EAL90147.1 100.00% afis|EAW25482.1 100.00%
Bootstrap support for afum|EAL90147.1 as seed ortholog is 100%.
Bootstrap support for afis|EAW25482.1 as seed ortholog is 100%.

---

Group of orthologs #6213. Best score 670 bits
Score difference with first non-orthologous sequence - *A. fumigatus*:670 *A. fischeri*:670

| | | | |
|---|---|---|---|
| afum\|EAL85081.1 | 100.00% | afis\|EAW19811.1 | 100.00% |

Bootstrap support for afum|EAL85081.1 as seed ortholog is 100%.
Bootstrap support for afis|EAW19811.1 as seed ortholog is 100%.

---

Group of orthologs #6214. Best score 670 bits
Score difference with first non-orthologous sequence - *A. fumigatus*:670 *A. fischeri*:670

| | | | |
|---|---|---|---|
| afum\|EAL86933.1 | 100.00% | afis\|EAW19585.1 | 100.00% |

Bootstrap support for afum|EAL86933.1 as seed ortholog is 100%.
Bootstrap support for afis|EAW19585.1 as seed ortholog is 100%.

---

Group of orthologs #6215. Best score 670 bits
Score difference with first non-orthologous sequence - *A. fumigatus*:670 *A. fischeri*:670

| | | | |
|---|---|---|---|
| afum\|EAL93600.1 | 100.00% | afis\|EAW18868.1 | 100.00% |

Bootstrap support for afum|EAL93600.1 as seed ortholog is 100%.
Bootstrap support for afis|EAW18868.1 as seed ortholog is 100%.

---

Group of orthologs #6216. Best score 670 bits
Score difference with first non-orthologous sequence - *A. fumigatus*:670 *A. fischeri*:670

| | | | |
|---|---|---|---|
| afum\|EAL91030.1 | 100.00% | afis\|EAW22161.1 | 100.00% |

Bootstrap support for afum|EAL91030.1 as seed ortholog is 100%.
Bootstrap support for afis|EAW22161.1 as seed ortholog is 100%.

---

Group of orthologs #6217. Best score 670 bits
Score difference with first non-orthologous sequence - *A. fumigatus*:670 *A. fischeri*:670

| | | | |
|---|---|---|---|
| afum\|EAL90734.1 | 100.00% | afis\|EAW22457.1 | 100.00% |

Bootstrap support for afum|EAL90734.1 as seed ortholog is 100%.
Bootstrap support for afis|EAW22457.1 as seed ortholog is 100%.

---

Group of orthologs #6218. Best score 670 bits
Score difference with first non-orthologous sequence - *A. fumigatus*:670 *A. fischeri*:670

| | | | |
|---|---|---|---|
| afum\|EAL92699.1 | 100.00% | afis\|EAW21664.1 | 100.00% |

Bootstrap support for afum|EAL92699.1 as seed ortholog is 100%.
Bootstrap support for afis|EAW21664.1 as seed ortholog is 100%.

---

Group of orthologs #6219. Best score 669 bits
Score difference with first non-orthologous sequence - *A. fumigatus*:570 *A. fischeri*:415

| | | | |
|---|---|---|---|
| afum\|EAL86574.1 | 100.00% | afis\|EAW17110.1 | 100.00% |

Bootstrap support for afum|EAL86574.1 as seed ortholog is 100%.
Bootstrap support for afis|EAW17110.1 as seed ortholog is 100%.

---

Group of orthologs #6220. Best score 669 bits
Score difference with first non-orthologous sequence - *A. fumigatus*:669 *A. fischeri*:147

| | | | |
|---|---|---|---|
| afum\|EAL89915.1 | 100.00% | afis\|EAW16226.1 | 100.00% |

Bootstrap support for afum|EAL89915.1 as seed ortholog is 100%.
Bootstrap support for afis|EAW16226.1 as seed ortholog is 100%.

---

Group of orthologs #6221. Best score 669 bits
Score difference with first non-orthologous sequence - *A. fumigatus*:669 *A. fischeri*:669

| | | | |
|---|---|---|---|
| afum\|EAL87101.1 | 100.00% | afis\|EAW19393.1 | 100.00% |

Bootstrap support for afum|EAL87101.1 as seed ortholog is 100%.
Bootstrap support for afis|EAW19393.1 as seed ortholog is 100%.

---

Group of orthologs #6222. Best score 669 bits

Score difference with first non-orthologous sequence - *A. fumigatus*:669 *A. fischeri*:669
afum|EAL93404.1 100.00% afis|EAW18683.1 100.00%
Bootstrap support for afum|EAL93404.1 as seed ortholog is 100%.
Bootstrap support for afis|EAW18683.1 as seed ortholog is 100%.

---

Group of orthologs #6223. Best score 669 bits
Score difference with first non-orthologous sequence - *A. fumigatus*:669 *A. fischeri*:669
afum|EAL93444.1 100.00% afis|EAW18720.1 100.00%
Bootstrap support for afum|EAL93444.1 as seed ortholog is 100%.
Bootstrap support for afis|EAW18720.1 as seed ortholog is 100%.

---

Group of orthologs #6224. Best score 669 bits
Score difference with first non-orthologous sequence - *A. fumigatus*:669 *A. fischeri*:669
afum|EAL90534.1 100.00% afis|EAW22658.1 100.00%
Bootstrap support for afum|EAL90534.1 as seed ortholog is 100%.
Bootstrap support for afis|EAW22658.1 as seed ortholog is 100%.

---

Group of orthologs #6225. Best score 669 bits
Score difference with first non-orthologous sequence - *A. fumigatus*:669 *A. fischeri*:669
afum|EAL92168.1 100.00% afis|EAW21149.1 100.00%
Bootstrap support for afum|EAL92168.1 as seed ortholog is 100%.
Bootstrap support for afis|EAW21149.1 as seed ortholog is 100%.

---

Group of orthologs #6226. Best score 669 bits
Score difference with first non-orthologous sequence - *A. fumigatus*:669 *A. fischeri*:669
afum|EAL89735.2 100.00% afis|EAW25066.1 100.00%
Bootstrap support for afum|EAL89735.2 as seed ortholog is 100%.
Bootstrap support for afis|EAW25066.1 as seed ortholog is 100%.

---

Group of orthologs #6227. Best score 668 bits
Score difference with first non-orthologous sequence - *A. fumigatus*:668 *A. fischeri*:668
afum|EAL88618.1 100.00% afis|EAW15986.1 100.00%
Bootstrap support for afum|EAL88618.1 as seed ortholog is 100%.
Bootstrap support for afis|EAW15986.1 as seed ortholog is 100%.

---

Group of orthologs #6228. Best score 668 bits
Score difference with first non-orthologous sequence - *A. fumigatus*:668 *A. fischeri*:668
afum|EAL91378.1 100.00% afis|EAW17564.1 100.00%
Bootstrap support for afum|EAL91378.1 as seed ortholog is 100%.
Bootstrap support for afis|EAW17564.1 as seed ortholog is 100%.

---

Group of orthologs #6229. Best score 668 bits
Score difference with first non-orthologous sequence - *A. fumigatus*:668 *A. fischeri*:668
afum|EAL86094.1 100.00% afis|EAW24426.1 100.00%
Bootstrap support for afum|EAL86094.1 as seed ortholog is 100%.
Bootstrap support for afis|EAW24426.1 as seed ortholog is 100%.

---

Group of orthologs #6230. Best score 668 bits
Score difference with first non-orthologous sequence - *A. fumigatus*:668 *A. fischeri*:668
afum|EAL93885.1 100.00% afis|EAW19154.1 100.00%
Bootstrap support for afum|EAL93885.1 as seed ortholog is 100%.
Bootstrap support for afis|EAW19154.1 as seed ortholog is 100%.

---

Group of orthologs #6231. Best score 668 bits
Score difference with first non-orthologous sequence - *A. fumigatus*:668 *A. fischeri*:668
afum|EAL90097.1 100.00% afis|EAW25439.1 100.00%

Bootstrap support for afum|EAL90097.1 as seed ortholog is 100%.
Bootstrap support for afis|EAW25439.1 as seed ortholog is 100%.

---

Group of orthologs #6232. Best score 667 bits
Score difference with first non-orthologous sequence - *A. fumigatus*:667 *A. fischeri*:667
afum|EAL91393.1 100.00% afis|EAW17580.1 100.00%
Bootstrap support for afum|EAL91393.1 as seed ortholog is 100%.
Bootstrap support for afis|EAW17580.1 as seed ortholog is 100%.

---

Group of orthologs #6233. Best score 667 bits
Score difference with first non-orthologous sequence - *A. fumigatus*:667 *A. fischeri*:667
afum|EAL91644.1 100.00% afis|EAW17834.1 100.00%
Bootstrap support for afum|EAL91644.1 as seed ortholog is 100%.
Bootstrap support for afis|EAW17834.1 as seed ortholog is 100%.

---

Group of orthologs #6234. Best score 667 bits
Score difference with first non-orthologous sequence - *A. fumigatus*:667 *A. fischeri*:667
afum|EAL87123.1 100.00% afis|EAW23680.1 100.00%
Bootstrap support for afum|EAL87123.1 as seed ortholog is 100%.
Bootstrap support for afis|EAW23680.1 as seed ortholog is 100%.

---

Group of orthologs #6235. Best score 667 bits
Score difference with first non-orthologous sequence - *A. fumigatus*:667 *A. fischeri*:600
afum|EAL93749.1 100.00% afis|EAW19016.1 100.00%
Bootstrap support for afum|EAL93749.1 as seed ortholog is 100%.
Bootstrap support for afis|EAW19016.1 as seed ortholog is 100%.

---

Group of orthologs #6236. Best score 667 bits
Score difference with first non-orthologous sequence - *A. fumigatus*:667 *A. fischeri*:521
afum|EAL90575.1 100.00% afis|EAW22621.1 100.00%
Bootstrap support for afum|EAL90575.1 as seed ortholog is 100%.
Bootstrap support for afis|EAW22621.1 as seed ortholog is 100%.

---

Group of orthologs #6237. Best score 666 bits
Score difference with first non-orthologous sequence - *A. fumigatus*:666 *A. fischeri*:666
afum|EAL86684.1 100.00% afis|EAW17267.1 100.00%
Bootstrap support for afum|EAL86684.1 as seed ortholog is 100%.
Bootstrap support for afis|EAW17267.1 as seed ortholog is 100%.

---

Group of orthologs #6238. Best score 666 bits
Score difference with first non-orthologous sequence - *A. fumigatus*:666 *A. fischeri*:666
afum|EAL88846.1 100.00% afis|EAW16218.1 100.00%
Bootstrap support for afum|EAL88846.1 as seed ortholog is 100%.
Bootstrap support for afis|EAW16218.1 as seed ortholog is 100%.

---

Group of orthologs #6239. Best score 666 bits
Score difference with first non-orthologous sequence - *A. fumigatus*:666 *A. fischeri*:666
afum|EAL91734.1 100.00% afis|EAW17923.1 100.00%
Bootstrap support for afum|EAL91734.1 as seed ortholog is 100%.
Bootstrap support for afis|EAW17923.1 as seed ortholog is 100%.

---

Group of orthologs #6240. Best score 666 bits
Score difference with first non-orthologous sequence - *A. fumigatus*:666 *A. fischeri*:666
afum|EAL86113.1 100.00% afis|EAW24444.1 100.00%
Bootstrap support for afum|EAL86113.1 as seed ortholog is 100%.
Bootstrap support for afis|EAW24444.1 as seed ortholog is 100%.

---

Group of orthologs #6241. Best score 666 bits
Score difference with first non-orthologous sequence - *A. fumigatus*:666 *A. fischeri*:666
afum|EAL93454.1 100.00% afis|EAW18729.1 100.00%
Bootstrap support for afum|EAL93454.1 as seed ortholog is 100%.
Bootstrap support for afis|EAW18729.1 as seed ortholog is 100%.

---

Group of orthologs #6242. Best score 666 bits
Score difference with first non-orthologous sequence - *A. fumigatus*:305 *A. fischeri*:666
afum|EAL90746.1 100.00% afis|EAW22445.1 100.00%
Bootstrap support for afum|EAL90746.1 as seed ortholog is 100%.
Bootstrap support for afis|EAW22445.1 as seed ortholog is 100%.

---

Group of orthologs #6243. Best score 666 bits
Score difference with first non-orthologous sequence - *A. fumigatus*:666 *A. fischeri*:666
afum|EAL90775.1 100.00% afis|EAW22417.1 100.00%
Bootstrap support for afum|EAL90775.1 as seed ortholog is 100%.
Bootstrap support for afis|EAW22417.1 as seed ortholog is 100%.

---

Group of orthologs #6244. Best score 665 bits
Score difference with first non-orthologous sequence - *A. fumigatus*:665 *A. fischeri*:665
afum|EAL86648.1 100.00% afis|EAW17226.1 100.00%
Bootstrap support for afum|EAL86648.1 as seed ortholog is 100%.
Bootstrap support for afis|EAW17226.1 as seed ortholog is 100%.

---

Group of orthologs #6245. Best score 665 bits
Score difference with first non-orthologous sequence - *A. fumigatus*:665 *A. fischeri*:665
afum|EAL91755.1 100.00% afis|EAW17944.1 100.00%
Bootstrap support for afum|EAL91755.1 as seed ortholog is 100%.
Bootstrap support for afis|EAW17944.1 as seed ortholog is 100%.

---

Group of orthologs #6246. Best score 665 bits
Score difference with first non-orthologous sequence - *A. fumigatus*:665 *A. fischeri*:665
afum|EAL93268.1 100.00% afis|EAW18566.1 100.00%
Bootstrap support for afum|EAL93268.1 as seed ortholog is 100%.
Bootstrap support for afis|EAW18566.1 as seed ortholog is 100%.

---

Group of orthologs #6247. Best score 665 bits
Score difference with first non-orthologous sequence - *A. fumigatus*:665 *A. fischeri*:665
afum|EAL93802.1 100.00% afis|EAW19074.1 100.00%
Bootstrap support for afum|EAL93802.1 as seed ortholog is 100%.
Bootstrap support for afis|EAW19074.1 as seed ortholog is 100%.

---

Group of orthologs #6248. Best score 664 bits
Score difference with first non-orthologous sequence - *A. fumigatus*:664 *A. fischeri*:664
afum|EAL91144.1 100.00% afis|EAW17309.1 100.00%
Bootstrap support for afum|EAL91144.1 as seed ortholog is 100%.
Bootstrap support for afis|EAW17309.1 as seed ortholog is 100%.

---

Group of orthologs #6249. Best score 664 bits
Score difference with first non-orthologous sequence - *A. fumigatus*:664 *A. fischeri*:664
afum|EAL85936.1 100.00% afis|EAW24268.1 100.00%
Bootstrap support for afum|EAL85936.1 as seed ortholog is 100%.
Bootstrap support for afis|EAW24268.1 as seed ortholog is 100%.

---

Group of orthologs #6250. Best score 664 bits

Score difference with first non-orthologous sequence - *A. fumigatus*:664   *A. fischeri*:664
afum|EAL93209.1 100.00% afis|EAW18514.1 100.00%
Bootstrap support for afum|EAL93209.1 as seed ortholog is 100%.
Bootstrap support for afis|EAW18514.1 as seed ortholog is 100%.

---

Group of orthologs #6251. Best score 664 bits
Score difference with first non-orthologous sequence - *A. fumigatus*:664   *A. fischeri*:664
afum|EAL92058.1 100.00% afis|EAW21020.1 100.00%
Bootstrap support for afum|EAL92058.1 as seed ortholog is 100%.
Bootstrap support for afis|EAW21020.1 as seed ortholog is 100%.

---

Group of orthologs #6252. Best score 664 bits
Score difference with first non-orthologous sequence - *A. fumigatus*:664   *A. fischeri*:664
afum|EBA27199.1 100.00% afis|EAW20443.1 100.00%
Bootstrap support for afum|EBA27199.1 as seed ortholog is 100%.
Bootstrap support for afis|EAW20443.1 as seed ortholog is 100%.

---

Group of orthologs #6253. Best score 664 bits
Score difference with first non-orthologous sequence - *A. fumigatus*:664   *A. fischeri*:664
afum|EAL89616.1 100.00% afis|EAW24958.1 100.00%
Bootstrap support for afum|EAL89616.1 as seed ortholog is 100%.
Bootstrap support for afis|EAW24958.1 as seed ortholog is 100%.

---

Group of orthologs #6254. Best score 664 bits
Score difference with first non-orthologous sequence - *A. fumigatus*:664   *A. fischeri*:664
afum|EAL89921.1 100.00% afis|EAW25259.1 100.00%
Bootstrap support for afum|EAL89921.1 as seed ortholog is 100%.
Bootstrap support for afis|EAW25259.1 as seed ortholog is 100%.

---

Group of orthologs #6255. Best score 664 bits
Score difference with first non-orthologous sequence - *A. fumigatus*:664   *A. fischeri*:664
afum|EAL90057.1 100.00% afis|EAW25398.1 100.00%
Bootstrap support for afum|EAL90057.1 as seed ortholog is 100%.
Bootstrap support for afis|EAW25398.1 as seed ortholog is 100%.

---

Group of orthologs #6256. Best score 663 bits
Score difference with first non-orthologous sequence - *A. fumigatus*:663   *A. fischeri*:663
afum|EAL88935.1 100.00% afis|EAW16313.1 100.00%
Bootstrap support for afum|EAL88935.1 as seed ortholog is 100%.
Bootstrap support for afis|EAW16313.1 as seed ortholog is 100%.

---

Group of orthologs #6257. Best score 663 bits
Score difference with first non-orthologous sequence - *A. fumigatus*:576   *A. fischeri*:557
afum|EAL84517.2 100.00% afis|EAW20541.1 100.00%
Bootstrap support for afum|EAL84517.2 as seed ortholog is 100%.
Bootstrap support for afis|EAW20541.1 as seed ortholog is 100%.

---

Group of orthologs #6258. Best score 663 bits
Score difference with first non-orthologous sequence - *A. fumigatus*:663   *A. fischeri*:663
afum|EAL87584.1 100.00% afis|EAW18218.1 100.00%
Bootstrap support for afum|EAL87584.1 as seed ortholog is 100%.
Bootstrap support for afis|EAW18218.1 as seed ortholog is 100%.

---

Group of orthologs #6259. Best score 663 bits
Score difference with first non-orthologous sequence - *A. fumigatus*:663   *A. fischeri*:498
afum|EAL84354.1 100.00% afis|EAW25685.1 100.00%

Bootstrap support for afum|EAL84354.1 as seed ortholog is 100%.
Bootstrap support for afis|EAW25685.1 as seed ortholog is 100%.

---

Group of orthologs #6260. Best score 663 bits
Score difference with first non-orthologous sequence - *A. fumigatus*:663 *A. fischeri*:663
afum|EAL91056.1 100.00% afis|EAW22134.1 100.00%
Bootstrap support for afum|EAL91056.1 as seed ortholog is 100%.
Bootstrap support for afis|EAW22134.1 as seed ortholog is 100%.

---

Group of orthologs #6261. Best score 663 bits
Score difference with first non-orthologous sequence - *A. fumigatus*:663 *A. fischeri*:663
afum|EAL90637.1 100.00% afis|EAW22558.1 100.00%
Bootstrap support for afum|EAL90637.1 as seed ortholog is 100%.
Bootstrap support for afis|EAW22558.1 as seed ortholog is 100%.

---

Group of orthologs #6262. Best score 663 bits
Score difference with first non-orthologous sequence - *A. fumigatus*:663 *A. fischeri*:558
afum|EAL91928.1 100.00% afis|EAW24072.1 100.00%
Bootstrap support for afum|EAL91928.1 as seed ortholog is 100%.
Bootstrap support for afis|EAW24072.1 as seed ortholog is 100%.

---

Group of orthologs #6263. Best score 663 bits
Score difference with first non-orthologous sequence - *A. fumigatus*:663 *A. fischeri*:663
afum|EBA27380.1 100.00% afis|EAW24740.1 100.00%
Bootstrap support for afum|EBA27380.1 as seed ortholog is 100%.
Bootstrap support for afis|EAW24740.1 as seed ortholog is 100%.

---

Group of orthologs #6264. Best score 662 bits
Score difference with first non-orthologous sequence - *A. fumigatus*:662 *A. fischeri*:662
afum|EAL93247.1 100.00% afis|EAW18547.1 100.00%
Bootstrap support for afum|EAL93247.1 as seed ortholog is 100%.
Bootstrap support for afis|EAW18547.1 as seed ortholog is 100%.

---

Group of orthologs #6265. Best score 661 bits
Score difference with first non-orthologous sequence - *A. fumigatus*:552 *A. fischeri*:430
afum|EAL87550.1 100.00% afis|EAW18187.1 100.00%
Bootstrap support for afum|EAL87550.1 as seed ortholog is 100%.
Bootstrap support for afis|EAW18187.1 as seed ortholog is 100%.

---

Group of orthologs #6266. Best score 661 bits
Score difference with first non-orthologous sequence - *A. fumigatus*:452 *A. fischeri*:661
afum|EAL86548.1 100.00% afis|EAW19903.1 100.00%
Bootstrap support for afum|EAL86548.1 as seed ortholog is 100%.
Bootstrap support for afis|EAW19903.1 as seed ortholog is 100%.

---

Group of orthologs #6267. Best score 661 bits
Score difference with first non-orthologous sequence - *A. fumigatus*:661 *A. fischeri*:661
afum|EAL85903.1 100.00% afis|EAW24234.1 100.00%
Bootstrap support for afum|EAL85903.1 as seed ortholog is 100%.
Bootstrap support for afis|EAW24234.1 as seed ortholog is 100%.

---

Group of orthologs #6268. Best score 661 bits
Score difference with first non-orthologous sequence - *A. fumigatus*:661 *A. fischeri*:661
afum|EAL86046.1 100.00% afis|EAW24374.1 100.00%
Bootstrap support for afum|EAL86046.1 as seed ortholog is 100%.
Bootstrap support for afis|EAW24374.1 as seed ortholog is 100%.

Group of orthologs #6269. Best score 661 bits
Score difference with first non-orthologous sequence - *A. fumigatus*:661 *A. fischeri*:661
afum|EAL87438.1 100.00% afis|EAW24008.1 100.00%
Bootstrap support for afum|EAL87438.1 as seed ortholog is 100%.
Bootstrap support for afis|EAW24008.1 as seed ortholog is 100%.

---

Group of orthologs #6270. Best score 660 bits
Score difference with first non-orthologous sequence - *A. fumigatus*:660 *A. fischeri*:570
afum|EAL85496.1 100.00% afis|EAW20385.1 100.00%
Bootstrap support for afum|EAL85496.1 as seed ortholog is 100%.
Bootstrap support for afis|EAW20385.1 as seed ortholog is 100%.

---

Group of orthologs #6271. Best score 660 bits
Score difference with first non-orthologous sequence - *A. fumigatus*:660 *A. fischeri*:660
afum|EAL86178.1 100.00% afis|EAW24509.1 100.00%
Bootstrap support for afum|EAL86178.1 as seed ortholog is 100%.
Bootstrap support for afis|EAW24509.1 as seed ortholog is 100%.

---

Group of orthologs #6272. Best score 660 bits
Score difference with first non-orthologous sequence - *A. fumigatus*:660 *A. fischeri*:660
afum|EAL90342.1 100.00% afis|EAW22858.1 100.00%
Bootstrap support for afum|EAL90342.1 as seed ortholog is 100%.
Bootstrap support for afis|EAW22858.1 as seed ortholog is 100%.

---

Group of orthologs #6273. Best score 660 bits
Score difference with first non-orthologous sequence - *A. fumigatus*:660 *A. fischeri*:660
afum|EAL92476.1 100.00% afis|EAW21457.1 100.00%
Bootstrap support for afum|EAL92476.1 as seed ortholog is 100%.
Bootstrap support for afis|EAW21457.1 as seed ortholog is 100%.

---

Group of orthologs #6274. Best score 659 bits
Score difference with first non-orthologous sequence - *A. fumigatus*:599 *A. fischeri*:598
afum|EAL86330.1 100.00% afis|EAW16758.1 100.00%
Bootstrap support for afum|EAL86330.1 as seed ortholog is 100%.
Bootstrap support for afis|EAW16758.1 as seed ortholog is 100%.

---

Group of orthologs #6275. Best score 659 bits
Score difference with first non-orthologous sequence - *A. fumigatus*:501 *A. fischeri*:409
afum|EAL86683.1 100.00% afis|EAW17266.1 100.00%
Bootstrap support for afum|EAL86683.1 as seed ortholog is 100%.
Bootstrap support for afis|EAW17266.1 as seed ortholog is 100%.

---

Group of orthologs #6276. Best score 659 bits
Score difference with first non-orthologous sequence - *A. fumigatus*:659 *A. fischeri*:659
afum|EAL88755.1 100.00% afis|EAW16125.1 100.00%
Bootstrap support for afum|EAL88755.1 as seed ortholog is 100%.
Bootstrap support for afis|EAW16125.1 as seed ortholog is 100%.

---

Group of orthologs #6277. Best score 659 bits
Score difference with first non-orthologous sequence - *A. fumigatus*:659 *A. fischeri*:659
afum|EAL84761.1 100.00% afis|EAW20873.1 100.00%
Bootstrap support for afum|EAL84761.1 as seed ortholog is 100%.
Bootstrap support for afis|EAW20873.1 as seed ortholog is 100%.

---

Group of orthologs #6278. Best score 659 bits

Score difference with first non-orthologous sequence - *A. fumigatus*:659 *A. fischeri*:475
afum|EAL90577.1 100.00% afis|EAW22619.1 100.00%
Bootstrap support for afum|EAL90577.1 as seed ortholog is 100%.
Bootstrap support for afis|EAW22619.1 as seed ortholog is 100%.

---

Group of orthologs #6279. Best score 658 bits
Score difference with first non-orthologous sequence - *A. fumigatus*:658 *A. fischeri*:658
afum|EAL86262.1 100.00% afis|EAW15656.1 100.00%
Bootstrap support for afum|EAL86262.1 as seed ortholog is 100%.
Bootstrap support for afis|EAW15656.1 as seed ortholog is 100%.

---

Group of orthologs #6280. Best score 658 bits
Score difference with first non-orthologous sequence - *A. fumigatus*:658 *A. fischeri*:658
afum|EAL89131.1 100.00% afis|EAW16534.1 100.00%
Bootstrap support for afum|EAL89131.1 as seed ortholog is 100%.
Bootstrap support for afis|EAW16534.1 as seed ortholog is 100%.

---

Group of orthologs #6281. Best score 658 bits
Score difference with first non-orthologous sequence - *A. fumigatus*:658 *A. fischeri*:658
afum|EAL84743.1 100.00% afis|EAW20890.1 100.00%
Bootstrap support for afum|EAL84743.1 as seed ortholog is 100%.
Bootstrap support for afis|EAW20890.1 as seed ortholog is 100%.

---

Group of orthologs #6282. Best score 658 bits
Score difference with first non-orthologous sequence - *A. fumigatus*:658 *A. fischeri*:658
afum|EAL85947.1 100.00% afis|EAW24278.1 100.00%
Bootstrap support for afum|EAL85947.1 as seed ortholog is 100%.
Bootstrap support for afis|EAW24278.1 as seed ortholog is 100%.

---

Group of orthologs #6283. Best score 658 bits
Score difference with first non-orthologous sequence - *A. fumigatus*:658 *A. fischeri*:658
afum|EAL89463.1 100.00% afis|EAW24819.1 100.00%
Bootstrap support for afum|EAL89463.1 as seed ortholog is 100%.
Bootstrap support for afis|EAW24819.1 as seed ortholog is 100%.

---

Group of orthologs #6284. Best score 658 bits
Score difference with first non-orthologous sequence - *A. fumigatus*:658 *A. fischeri*:658
afum|EAL92818.1 100.00% afis|EAW21793.1 100.00%
Bootstrap support for afum|EAL92818.1 as seed ortholog is 100%.
Bootstrap support for afis|EAW21793.1 as seed ortholog is 100%.

---

Group of orthologs #6285. Best score 657 bits
Score difference with first non-orthologous sequence - *A. fumigatus*:657 *A. fischeri*:657
afum|EAL85531.1 100.00% afis|EAW15820.1 100.00%
Bootstrap support for afum|EAL85531.1 as seed ortholog is 100%.
Bootstrap support for afis|EAW15820.1 as seed ortholog is 100%.

---

Group of orthologs #6286. Best score 657 bits
Score difference with first non-orthologous sequence - *A. fumigatus*:657 *A. fischeri*:657
afum|EAL87260.1 100.00% afis|EAW23828.1 100.00%
Bootstrap support for afum|EAL87260.1 as seed ortholog is 100%.
Bootstrap support for afis|EAW23828.1 as seed ortholog is 100%.

---

Group of orthologs #6287. Best score 657 bits
Score difference with first non-orthologous sequence - *A. fumigatus*:657 *A. fischeri*:657
afum|EAL90679.1 100.00% afis|EAW22514.1 100.00%

Bootstrap support for afum|EAL90679.1 as seed ortholog is 100%.
Bootstrap support for afis|EAW22514.1 as seed ortholog is 100%.

---

Group of orthologs #6288. Best score 657 bits
Score difference with first non-orthologous sequence - *A. fumigatus*:657 *A. fischeri*:657
afum|EAL90339.1 100.00% afis|EAW22861.1 100.00%
Bootstrap support for afum|EAL90339.1 as seed ortholog is 100%.
Bootstrap support for afis|EAW22861.1 as seed ortholog is 100%.

---

Group of orthologs #6289. Best score 657 bits
Score difference with first non-orthologous sequence - *A. fumigatus*:657 *A. fischeri*:657
afum|EAL92767.1 100.00% afis|EAW21734.1 100.00%
Bootstrap support for afum|EAL92767.1 as seed ortholog is 100%.
Bootstrap support for afis|EAW21734.1 as seed ortholog is 100%.

---

Group of orthologs #6290. Best score 657 bits
Score difference with first non-orthologous sequence - *A. fumigatus*:657 *A. fischeri*:657
afum|EAL89797.1 100.00% afis|EAW25126.1 100.00%
Bootstrap support for afum|EAL89797.1 as seed ortholog is 100%.
Bootstrap support for afis|EAW25126.1 as seed ortholog is 100%.

---

Group of orthologs #6291. Best score 656 bits
Score difference with first non-orthologous sequence - *A. fumigatus*:656 *A. fischeri*:656
afum|EAL84424.1 100.00% afis|EAW20649.1 100.00%
Bootstrap support for afum|EAL84424.1 as seed ortholog is 100%.
Bootstrap support for afis|EAW20649.1 as seed ortholog is 100%.

---

Group of orthologs #6292. Best score 656 bits
Score difference with first non-orthologous sequence - *A. fumigatus*:449 *A. fischeri*:308
afum|EAL89024.1 100.00% afis|EAW16402.1 100.00%
Bootstrap support for afum|EAL89024.1 as seed ortholog is 100%.
Bootstrap support for afis|EAW16402.1 as seed ortholog is 100%.

---

Group of orthologs #6293. Best score 656 bits
Score difference with first non-orthologous sequence - *A. fumigatus*:656 *A. fischeri*:571
afum|EAL85477.1 100.00% afis|EAW20360.1 100.00%
Bootstrap support for afum|EAL85477.1 as seed ortholog is 100%.
Bootstrap support for afis|EAW20360.1 as seed ortholog is 100%.

---

Group of orthologs #6294. Best score 656 bits
Score difference with first non-orthologous sequence - *A. fumigatus*:656 *A. fischeri*:656
afum|EAL91731.1 100.00% afis|EAW17920.1 100.00%
Bootstrap support for afum|EAL91731.1 as seed ortholog is 100%.
Bootstrap support for afis|EAW17920.1 as seed ortholog is 100%.

---

Group of orthologs #6295. Best score 656 bits
Score difference with first non-orthologous sequence - *A. fumigatus*:656 *A. fischeri*:516
afum|EBA27280.1 100.00% afis|EAW17184.1 100.00%
Bootstrap support for afum|EBA27280.1 as seed ortholog is 100%.
Bootstrap support for afis|EAW17184.1 as seed ortholog is 100%.

---

Group of orthologs #6296. Best score 656 bits
Score difference with first non-orthologous sequence - *A. fumigatus*:656 *A. fischeri*:656
afum|EAL92159.1 100.00% afis|EAW21140.1 100.00%
Bootstrap support for afum|EAL92159.1 as seed ortholog is 100%.
Bootstrap support for afis|EAW21140.1 as seed ortholog is 100%.

Group of orthologs #6297. Best score 655 bits
Score difference with first non-orthologous sequence - *A. fumigatus*:595 *A. fischeri*:571
afum|EAL87889.1 100.00% afis|EAW23576.1 100.00%
Bootstrap support for afum|EAL87889.1 as seed ortholog is 100%.
Bootstrap support for afis|EAW23576.1 as seed ortholog is 100%.

---

Group of orthologs #6298. Best score 655 bits
Score difference with first non-orthologous sequence - *A. fumigatus*:586 *A. fischeri*:655
afum|EAL93956.1 100.00% afis|EAW19225.1 100.00%
Bootstrap support for afum|EAL93956.1 as seed ortholog is 100%.
Bootstrap support for afis|EAW19225.1 as seed ortholog is 100%.

---

Group of orthologs #6299. Best score 655 bits
Score difference with first non-orthologous sequence - *A. fumigatus*:655 *A. fischeri*:655
afum|EBA27179.1 100.00% afis|EAW20584.1 100.00%
Bootstrap support for afum|EBA27179.1 as seed ortholog is 100%.
Bootstrap support for afis|EAW20584.1 as seed ortholog is 100%.

---

Group of orthologs #6300. Best score 654 bits
Score difference with first non-orthologous sequence - *A. fumigatus*:654 *A. fischeri*:654
afum|EAL85708.1 100.00% afis|EAW15626.1 100.00%
Bootstrap support for afum|EAL85708.1 as seed ortholog is 100%.
Bootstrap support for afis|EAW15626.1 as seed ortholog is 100%.

---

Group of orthologs #6301. Best score 654 bits
Score difference with first non-orthologous sequence - *A. fumigatus*:654 *A. fischeri*:654
afum|EAL85612.2 100.00% afis|EAW15736.1 100.00%
Bootstrap support for afum|EAL85612.2 as seed ortholog is 100%.
Bootstrap support for afis|EAW15736.1 as seed ortholog is 100%.

---

Group of orthologs #6302. Best score 654 bits
Score difference with first non-orthologous sequence - *A. fumigatus*:654 *A. fischeri*:654
afum|EAL88489.1 100.00% afis|EAW15853.1 100.00%
Bootstrap support for afum|EAL88489.1 as seed ortholog is 100%.
Bootstrap support for afis|EAW15853.1 as seed ortholog is 100%.

---

Group of orthologs #6303. Best score 654 bits
Score difference with first non-orthologous sequence - *A. fumigatus*:654 *A. fischeri*:654
afum|EAL88740.1 100.00% afis|EAW16108.1 100.00%
Bootstrap support for afum|EAL88740.1 as seed ortholog is 100%.
Bootstrap support for afis|EAW16108.1 as seed ortholog is 100%.

---

Group of orthologs #6304. Best score 654 bits
Score difference with first non-orthologous sequence - *A. fumigatus*:654 *A. fischeri*:654
afum|EAL85949.1 100.00% afis|EAW24280.1 100.00%
Bootstrap support for afum|EAL85949.1 as seed ortholog is 100%.
Bootstrap support for afis|EAW24280.1 as seed ortholog is 100%.

---

Group of orthologs #6305. Best score 654 bits
Score difference with first non-orthologous sequence - *A. fumigatus*:654 *A. fischeri*:654
afum|EAL88399.1 100.00% afis|EAW23080.1 100.00%
Bootstrap support for afum|EAL88399.1 as seed ortholog is 100%.
Bootstrap support for afis|EAW23080.1 as seed ortholog is 100%.

---

Group of orthologs #6306. Best score 654 bits

Score difference with first non-orthologous sequence - *A. fumigatus*:654 *A. fischeri*:654
afum|EAL88161.2 100.00% afis|EAW23316.1 100.00%
Bootstrap support for afum|EAL88161.2 as seed ortholog is 100%.
Bootstrap support for afis|EAW23316.1 as seed ortholog is 100%.

---

Group of orthologs #6307. Best score 654 bits
Score difference with first non-orthologous sequence - *A. fumigatus*:654 *A. fischeri*:654
afum|EAL87984.1 100.00% afis|EAW23480.1 100.00%
Bootstrap support for afum|EAL87984.1 as seed ortholog is 100%.
Bootstrap support for afis|EAW23480.1 as seed ortholog is 100%.

---

Group of orthologs #6308. Best score 654 bits
Score difference with first non-orthologous sequence - *A. fumigatus*:654 *A. fischeri*:654
afum|EAL93397.1 100.00% afis|EAW18677.1 100.00%
Bootstrap support for afum|EAL93397.1 as seed ortholog is 100%.
Bootstrap support for afis|EAW18677.1 as seed ortholog is 100%.

---

Group of orthologs #6309. Best score 654 bits
Score difference with first non-orthologous sequence - *A. fumigatus*:654 *A. fischeri*:600
afum|EAL92136.1 100.00% afis|EAW21113.1 100.00%
Bootstrap support for afum|EAL92136.1 as seed ortholog is 100%.
Bootstrap support for afis|EAW21113.1 as seed ortholog is 100%.

---

Group of orthologs #6310. Best score 653 bits
Score difference with first non-orthologous sequence - *A. fumigatus*:592 *A. fischeri*:574
afum|EAL85701.2 100.00% afis|EAW15638.1 100.00%
Bootstrap support for afum|EAL85701.2 as seed ortholog is 100%.
Bootstrap support for afis|EAW15638.1 as seed ortholog is 100%.

---

Group of orthologs #6311. Best score 653 bits
Score difference with first non-orthologous sequence - *A. fumigatus*:653 *A. fischeri*:653
afum|EAL85333.1 100.00% afis|EAW20218.1 100.00%
Bootstrap support for afum|EAL85333.1 as seed ortholog is 100%.
Bootstrap support for afis|EAW20218.1 as seed ortholog is 100%.

---

Group of orthologs #6312. Best score 653 bits
Score difference with first non-orthologous sequence - *A. fumigatus*:653 *A. fischeri*:653
afum|EAL84299.2 100.00% afis|EAW25592.1 100.00%
Bootstrap support for afum|EAL84299.2 as seed ortholog is 100%.
Bootstrap support for afis|EAW25592.1 as seed ortholog is 100%.

---

Group of orthologs #6313. Best score 653 bits
Score difference with first non-orthologous sequence - *A. fumigatus*:653 *A. fischeri*:653
afum|EAL93132.1 100.00% afis|EAW18447.1 100.00%
Bootstrap support for afum|EAL93132.1 as seed ortholog is 100%.
Bootstrap support for afis|EAW18447.1 as seed ortholog is 100%.

---

Group of orthologs #6314. Best score 653 bits
Score difference with first non-orthologous sequence - *A. fumigatus*:653 *A. fischeri*:653
afum|EAL87993.1 100.00% afis|EAW23473.1 100.00%
Bootstrap support for afum|EAL87993.1 as seed ortholog is 100%.
Bootstrap support for afis|EAW23473.1 as seed ortholog is 100%.

---

Group of orthologs #6315. Best score 653 bits
Score difference with first non-orthologous sequence - *A. fumigatus*:653 *A. fischeri*:653
afum|EAL93559.1 100.00% afis|EAW18822.1 100.00%

Bootstrap support for afum|EAL93559.1 as seed ortholog is 100%.
Bootstrap support for afis|EAW18822.1 as seed ortholog is 100%.

---

Group of orthologs #6316. Best score 652 bits
Score difference with first non-orthologous sequence - *A. fumigatus*:652 *A. fischeri*:652
afum|EAL88172.1 100.00% afis|EAW23305.1 100.00%
Bootstrap support for afum|EAL88172.1 as seed ortholog is 100%.
Bootstrap support for afis|EAW23305.1 as seed ortholog is 100%.

---

Group of orthologs #6317. Best score 652 bits
Score difference with first non-orthologous sequence - *A. fumigatus*:652 *A. fischeri*:113
afum|EAL93583.1 100.00% afis|EAW18852.1 100.00%
Bootstrap support for afum|EAL93583.1 as seed ortholog is 100%.
Bootstrap support for afis|EAW18852.1 as seed ortholog is 100%.

---

Group of orthologs #6318. Best score 652 bits
Score difference with first non-orthologous sequence - *A. fumigatus*:652 *A. fischeri*:652
afum|EBA27502.1 100.00% afis|EAW19015.1 100.00%
Bootstrap support for afum|EBA27502.1 as seed ortholog is 100%.
Bootstrap support for afis|EAW19015.1 as seed ortholog is 100%.

---

Group of orthologs #6319. Best score 652 bits
Score difference with first non-orthologous sequence - *A. fumigatus*:652 *A. fischeri*:652
afum|EAL92684.1 100.00% afis|EAW21650.1 100.00%
Bootstrap support for afum|EAL92684.1 as seed ortholog is 100%.
Bootstrap support for afis|EAW21650.1 as seed ortholog is 100%.

---

Group of orthologs #6320. Best score 651 bits
Score difference with first non-orthologous sequence - *A. fumigatus*:651 *A. fischeri*:651
afum|EAL88732.1 100.00% afis|EAW16100.1 100.00%
Bootstrap support for afum|EAL88732.1 as seed ortholog is 100%.
Bootstrap support for afis|EAW16100.1 as seed ortholog is 100%.

---

Group of orthologs #6321. Best score 651 bits
Score difference with first non-orthologous sequence - *A. fumigatus*:651 *A. fischeri*:651
afum|EAL87093.1 100.00% afis|EAW19402.1 100.00%
Bootstrap support for afum|EAL87093.1 as seed ortholog is 100%.
Bootstrap support for afis|EAW19402.1 as seed ortholog is 100%.

---

Group of orthologs #6322. Best score 651 bits
Score difference with first non-orthologous sequence - *A. fumigatus*:651 *A. fischeri*:651
afum|EAL93548.1 100.00% afis|EAW18812.1 100.00%
Bootstrap support for afum|EAL93548.1 as seed ortholog is 100%.
Bootstrap support for afis|EAW18812.1 as seed ortholog is 100%.

---

Group of orthologs #6323. Best score 651 bits
Score difference with first non-orthologous sequence - *A. fumigatus*:651 *A. fischeri*:651
afum|EAL93691.1 100.00% afis|EAW18958.1 100.00%
Bootstrap support for afum|EAL93691.1 as seed ortholog is 100%.
Bootstrap support for afis|EAW18958.1 as seed ortholog is 100%.

---

Group of orthologs #6324. Best score 651 bits
Score difference with first non-orthologous sequence - *A. fumigatus*:651 *A. fischeri*:462
afum|EAL90815.1 100.00% afis|EAW22377.1 100.00%
Bootstrap support for afum|EAL90815.1 as seed ortholog is 100%.
Bootstrap support for afis|EAW22377.1 as seed ortholog is 100%.

Group of orthologs #6325. Best score 651 bits
Score difference with first non-orthologous sequence - *A. fumigatus*:651 *A. fischeri*:651
afum|EAL89426.1 100.00% afis|EAW24787.1 100.00%
Bootstrap support for afum|EAL89426.1 as seed ortholog is 100%.
Bootstrap support for afis|EAW24787.1 as seed ortholog is 100%.

---

Group of orthologs #6326. Best score 651 bits
Score difference with first non-orthologous sequence - *A. fumigatus*:651 *A. fischeri*:651
afum|EAL92816.1 100.00% afis|EAW21791.1 100.00%
Bootstrap support for afum|EAL92816.1 as seed ortholog is 100%.
Bootstrap support for afis|EAW21791.1 as seed ortholog is 100%.

---

Group of orthologs #6327. Best score 651 bits
Score difference with first non-orthologous sequence - *A. fumigatus*:651 *A. fischeri*:651
afum|EAL89837.1 100.00% afis|EAW25163.1 100.00%
Bootstrap support for afum|EAL89837.1 as seed ortholog is 100%.
Bootstrap support for afis|EAW25163.1 as seed ortholog is 100%.

---

Group of orthologs #6328. Best score 650 bits
Score difference with first non-orthologous sequence - *A. fumigatus*:650 *A. fischeri*:650
afum|EAL89115.1 100.00% afis|EAW16520.1 100.00%
Bootstrap support for afum|EAL89115.1 as seed ortholog is 100%.
Bootstrap support for afis|EAW16520.1 as seed ortholog is 100%.

---

Group of orthologs #6329. Best score 650 bits
Score difference with first non-orthologous sequence - *A. fumigatus*:650 *A. fischeri*:650
afum|EAL85299.1 100.00% afis|EAW20181.1 100.00%
Bootstrap support for afum|EAL85299.1 as seed ortholog is 100%.
Bootstrap support for afis|EAW20181.1 as seed ortholog is 100%.

---

Group of orthologs #6330. Best score 650 bits
Score difference with first non-orthologous sequence - *A. fumigatus*:650 *A. fischeri*:650
afum|EAL86956.1 100.00% afis|EAW19553.1 100.00%
Bootstrap support for afum|EAL86956.1 as seed ortholog is 100%.
Bootstrap support for afis|EAW19553.1 as seed ortholog is 100%.

---

Group of orthologs #6331. Best score 650 bits
Score difference with first non-orthologous sequence - *A. fumigatus*:650 *A. fischeri*:650
afum|EAL91142.1 100.00% afis|EAW17306.1 100.00%
Bootstrap support for afum|EAL91142.1 as seed ortholog is 100%.
Bootstrap support for afis|EAW17306.1 as seed ortholog is 100%.

---

Group of orthologs #6332. Best score 650 bits
Score difference with first non-orthologous sequence - *A. fumigatus*:650 *A. fischeri*:650
afum|EAL84295.2 100.00% afis|EAW25588.1 100.00%
Bootstrap support for afum|EAL84295.2 as seed ortholog is 100%.
Bootstrap support for afis|EAW25588.1 as seed ortholog is 100%.

---

Group of orthologs #6333. Best score 650 bits
Score difference with first non-orthologous sequence - *A. fumigatus*:571 *A. fischeri*:540
afum|EAL93840.1 100.00% afis|EAW19109.1 100.00%
Bootstrap support for afum|EAL93840.1 as seed ortholog is 100%.
Bootstrap support for afis|EAW19109.1 as seed ortholog is 100%.

---

Group of orthologs #6334. Best score 650 bits

Score difference with first non-orthologous sequence - *A. fumigatus*:650 *A. fischeri*:650
afum|EAL93842.2 100.00% afis|EAW19111.1 100.00%
Bootstrap support for afum|EAL93842.2 as seed ortholog is 100%.
Bootstrap support for afis|EAW19111.1 as seed ortholog is 100%.

---

Group of orthologs #6335. Best score 650 bits
Score difference with first non-orthologous sequence - *A. fumigatus*:650 *A. fischeri*:650
afum|EBA27511.1 100.00% afis|EAW19271.1 100.00%
Bootstrap support for afum|EBA27511.1 as seed ortholog is 100%.
Bootstrap support for afis|EAW19271.1 as seed ortholog is 100%.

---

Group of orthologs #6336. Best score 650 bits
Score difference with first non-orthologous sequence - *A. fumigatus*:650 *A. fischeri*:650
afum|EAL91916.1 100.00% afis|EAW24061.1 100.00%
Bootstrap support for afum|EAL91916.1 as seed ortholog is 100%.
Bootstrap support for afis|EAW24061.1 as seed ortholog is 100%.

---

Group of orthologs #6337. Best score 649 bits
Score difference with first non-orthologous sequence - *A. fumigatus*:407 *A. fischeri*:293
afum|EAL86386.1 100.00% afis|EAW16839.1 100.00%
Bootstrap support for afum|EAL86386.1 as seed ortholog is 100%.
Bootstrap support for afis|EAW16839.1 as seed ortholog is 100%.

---

Group of orthologs #6338. Best score 649 bits
Score difference with first non-orthologous sequence - *A. fumigatus*:552 *A. fischeri*:514
afum|EAL84402.2 100.00% afis|EAW25731.1 100.00%
Bootstrap support for afum|EAL84402.2 as seed ortholog is 100%.
Bootstrap support for afis|EAW25731.1 as seed ortholog is 100%.

---

Group of orthologs #6339. Best score 649 bits
Score difference with first non-orthologous sequence - *A. fumigatus*:649 *A. fischeri*:649
afum|EAL93037.1 100.00% afis|EAW18364.1 100.00%
Bootstrap support for afum|EAL93037.1 as seed ortholog is 100%.
Bootstrap support for afis|EAW18364.1 as seed ortholog is 100%.

---

Group of orthologs #6340. Best score 649 bits
Score difference with first non-orthologous sequence - *A. fumigatus*:649 *A. fischeri*:649
afum|EAL92568.1 100.00% afis|EAW21548.1 100.00%
Bootstrap support for afum|EAL92568.1 as seed ortholog is 100%.
Bootstrap support for afis|EAW21548.1 as seed ortholog is 100%.

---

Group of orthologs #6341. Best score 649 bits
Score difference with first non-orthologous sequence - *A. fumigatus*:649 *A. fischeri*:649
afum|EAL89433.1 100.00% afis|EAW24791.1 100.00%
Bootstrap support for afum|EAL89433.1 as seed ortholog is 100%.
Bootstrap support for afis|EAW24791.1 as seed ortholog is 100%.

---

Group of orthologs #6342. Best score 649 bits
Score difference with first non-orthologous sequence - *A. fumigatus*:649 *A. fischeri*:649
afum|EAL90133.1 100.00% afis|EAW25469.1 100.00%
Bootstrap support for afum|EAL90133.1 as seed ortholog is 100%.
Bootstrap support for afis|EAW25469.1 as seed ortholog is 100%.

---

Group of orthologs #6343. Best score 648 bits
Score difference with first non-orthologous sequence - *A. fumigatus*:648 *A. fischeri*:648
afum|EAL85671.1 100.00% afis|EAW15676.1 100.00%

Bootstrap support for afum|EAL85671.1 as seed ortholog is 100%.
Bootstrap support for afis|EAW15676.1 as seed ortholog is 100%.

---

Group of orthologs #6344. Best score 648 bits
Score difference with first non-orthologous sequence - *A. fumigatus*:579 *A. fischeri*:582
afum|EAL86313.2 100.00% afis|EAW16738.1 100.00%
Bootstrap support for afum|EAL86313.2 as seed ortholog is 100%.
Bootstrap support for afis|EAW16738.1 as seed ortholog is 100%.

---

Group of orthologs #6345. Best score 648 bits
Score difference with first non-orthologous sequence - *A. fumigatus*:648 *A. fischeri*:648
afum|EAL87100.1 100.00% afis|EAW19394.1 100.00%
Bootstrap support for afum|EAL87100.1 as seed ortholog is 100%.
Bootstrap support for afis|EAW19394.1 as seed ortholog is 100%.

---

Group of orthologs #6346. Best score 648 bits
Score difference with first non-orthologous sequence - *A. fumigatus*:405 *A. fischeri*:567
afum|EAL86841.1 100.00% afis|EAW19681.1 100.00%
Bootstrap support for afum|EAL86841.1 as seed ortholog is 100%.
Bootstrap support for afis|EAW19681.1 as seed ortholog is 100%.

---

Group of orthologs #6347. Best score 648 bits
Score difference with first non-orthologous sequence - *A. fumigatus*:574 *A. fischeri*:648
afum|EAL90177.1 100.00% afis|EAW25511.1 100.00%
Bootstrap support for afum|EAL90177.1 as seed ortholog is 100%.
Bootstrap support for afis|EAW25511.1 as seed ortholog is 100%.

---

Group of orthologs #6348. Best score 647 bits
Score difference with first non-orthologous sequence - *A. fumigatus*:325 *A. fischeri*:556
afum|EAL85667.1 100.00% afis|EAW15681.1 100.00%
Bootstrap support for afum|EAL85667.1 as seed ortholog is 100%.
Bootstrap support for afis|EAW15681.1 as seed ortholog is 100%.

---

Group of orthologs #6349. Best score 647 bits
Score difference with first non-orthologous sequence - *A. fumigatus*:647 *A. fischeri*:135
afum|EAL84480.1 100.00% afis|EAW20585.1 100.00%
Bootstrap support for afum|EAL84480.1 as seed ortholog is 100%.
Bootstrap support for afis|EAW20585.1 as seed ortholog is 100%.

---

Group of orthologs #6350. Best score 647 bits
Score difference with first non-orthologous sequence - *A. fumigatus*:647 *A. fischeri*:647
afum|EAL84701.1 100.00% afis|EAW20933.1 100.00%
Bootstrap support for afum|EAL84701.1 as seed ortholog is 100%.
Bootstrap support for afis|EAW20933.1 as seed ortholog is 100%.

---

Group of orthologs #6351. Best score 647 bits
Score difference with first non-orthologous sequence - *A. fumigatus*:647 *A. fischeri*:647
afum|EAL85473.1 100.00% afis|EAW20356.1 100.00%
Bootstrap support for afum|EAL85473.1 as seed ortholog is 100%.
Bootstrap support for afis|EAW20356.1 as seed ortholog is 100%.

---

Group of orthologs #6352. Best score 647 bits
Score difference with first non-orthologous sequence - *A. fumigatus*:422 *A. fischeri*:355
afum|EAL86173.1 100.00% afis|EAW24505.1 100.00%
Bootstrap support for afum|EAL86173.1 as seed ortholog is 100%.
Bootstrap support for afis|EAW24505.1 as seed ortholog is 100%.

---

Group of orthologs #6353. Best score 647 bits
Score difference with first non-orthologous sequence - *A. fumigatus*:647 *A. fischeri*:647
afum|EAL93791.1 100.00% afis|EAW19062.1 100.00%
Bootstrap support for afum|EAL93791.1 as seed ortholog is 100%.
Bootstrap support for afis|EAW19062.1 as seed ortholog is 100%.

---

Group of orthologs #6354. Best score 647 bits
Score difference with first non-orthologous sequence - *A. fumigatus*:647 *A. fischeri*:500
afum|EAL90872.2 100.00% afis|EAW22318.1 100.00%
Bootstrap support for afum|EAL90872.2 as seed ortholog is 100%.
Bootstrap support for afis|EAW22318.1 as seed ortholog is 100%.

---

Group of orthologs #6355. Best score 647 bits
Score difference with first non-orthologous sequence - *A. fumigatus*:647 *A. fischeri*:647
afum|EAL90795.1 100.00% afis|EAW22396.1 100.00%
Bootstrap support for afum|EAL90795.1 as seed ortholog is 100%.
Bootstrap support for afis|EAW22396.1 as seed ortholog is 100%.

---

Group of orthologs #6356. Best score 647 bits
Score difference with first non-orthologous sequence - *A. fumigatus*:202 *A. fischeri*:647
afum|EAL91907.2 100.00% afis|EAW24053.1 100.00%
Bootstrap support for afum|EAL91907.2 as seed ortholog is 100%.
Bootstrap support for afis|EAW24053.1 as seed ortholog is 100%.

---

Group of orthologs #6357. Best score 646 bits
Score difference with first non-orthologous sequence - *A. fumigatus*:646 *A. fischeri*:646
afum|EAL84988.1 100.00% afis|EAW19921.1 100.00%
Bootstrap support for afum|EAL84988.1 as seed ortholog is 100%.
Bootstrap support for afis|EAW19921.1 as seed ortholog is 100%.

---

Group of orthologs #6358. Best score 646 bits
Score difference with first non-orthologous sequence - *A. fumigatus*:646 *A. fischeri*:646
afum|EAL89205.1 100.00% afis|EAW16609.1 100.00%
Bootstrap support for afum|EAL89205.1 as seed ortholog is 100%.
Bootstrap support for afis|EAW16609.1 as seed ortholog is 100%.

---

Group of orthologs #6359. Best score 646 bits
Score difference with first non-orthologous sequence - *A. fumigatus*:646 *A. fischeri*:646
afum|EAL87331.1 100.00% afis|EAW23903.1 100.00%
Bootstrap support for afum|EAL87331.1 as seed ortholog is 100%.
Bootstrap support for afis|EAW23903.1 as seed ortholog is 100%.

---

Group of orthologs #6360. Best score 646 bits
Score difference with first non-orthologous sequence - *A. fumigatus*:646 *A. fischeri*:646
afum|EAL88466.1 100.00% afis|EAW23015.1 100.00%
Bootstrap support for afum|EAL88466.1 as seed ortholog is 100%.
Bootstrap support for afis|EAW23015.1 as seed ortholog is 100%.

---

Group of orthologs #6361. Best score 646 bits
Score difference with first non-orthologous sequence - *A. fumigatus*:646 *A. fischeri*:646
afum|EAL90952.1 100.00% afis|EAW22241.1 100.00%
Bootstrap support for afum|EAL90952.1 as seed ortholog is 100%.
Bootstrap support for afis|EAW22241.1 as seed ortholog is 100%.

---

Group of orthologs #6362. Best score 646 bits

Score difference with first non-orthologous sequence - *A. fumigatus*:646 *A. fischeri*:646
afum|EAL89446.1 100.00% afis|EAW24804.1 100.00%
Bootstrap support for afum|EAL89446.1 as seed ortholog is 100%.
Bootstrap support for afis|EAW24804.1 as seed ortholog is 100%.

---

Group of orthologs #6363. Best score 645 bits
Score difference with first non-orthologous sequence - *A. fumigatus*:645 *A. fischeri*:645
afum|EAL88854.1 100.00% afis|EAW16231.1 100.00%
Bootstrap support for afum|EAL88854.1 as seed ortholog is 100%.
Bootstrap support for afis|EAW16231.1 as seed ortholog is 100%.

---

Group of orthologs #6364. Best score 645 bits
Score difference with first non-orthologous sequence - *A. fumigatus*:645 *A. fischeri*:645
afum|EAL89026.1 100.00% afis|EAW16401.1 100.00%
Bootstrap support for afum|EAL89026.1 as seed ortholog is 100%.
Bootstrap support for afis|EAW16401.1 as seed ortholog is 100%.

---

Group of orthologs #6365. Best score 645 bits
Score difference with first non-orthologous sequence - *A. fumigatus*:645 *A. fischeri*:645
afum|EAL86885.1 100.00% afis|EAW19637.1 100.00%
Bootstrap support for afum|EAL86885.1 as seed ortholog is 100%.
Bootstrap support for afis|EAW19637.1 as seed ortholog is 100%.

---

Group of orthologs #6366. Best score 645 bits
Score difference with first non-orthologous sequence - *A. fumigatus*:645 *A. fischeri*:645
afum|EAL88385.1 100.00% afis|EAW23093.1 100.00%
Bootstrap support for afum|EAL88385.1 as seed ortholog is 100%.
Bootstrap support for afis|EAW23093.1 as seed ortholog is 100%.

---

Group of orthologs #6367. Best score 645 bits
Score difference with first non-orthologous sequence - *A. fumigatus*:645 *A. fischeri*:645
afum|EAL88091.1 100.00% afis|EAW23380.1 100.00%
Bootstrap support for afum|EAL88091.1 as seed ortholog is 100%.
Bootstrap support for afis|EAW23380.1 as seed ortholog is 100%.

---

Group of orthologs #6368. Best score 644 bits
Score difference with first non-orthologous sequence - *A. fumigatus*:644 *A. fischeri*:644
afum|EAL88962.1 100.00% afis|EAW16336.1 100.00%
Bootstrap support for afum|EAL88962.1 as seed ortholog is 100%.
Bootstrap support for afis|EAW16336.1 as seed ortholog is 100%.

---

Group of orthologs #6369. Best score 644 bits
Score difference with first non-orthologous sequence - *A. fumigatus*:589 *A. fischeri*:569
afum|EAL86998.1 100.00% afis|EAW19496.1 100.00%
Bootstrap support for afum|EAL86998.1 as seed ortholog is 100%.
Bootstrap support for afis|EAW19496.1 as seed ortholog is 100%.

---

Group of orthologs #6370. Best score 644 bits
Score difference with first non-orthologous sequence - *A. fumigatus*:644 *A. fischeri*:644
afum|EAL87262.1 100.00% afis|EAW23830.1 100.00%
Bootstrap support for afum|EAL87262.1 as seed ortholog is 100%.
Bootstrap support for afis|EAW23830.1 as seed ortholog is 100%.

---

Group of orthologs #6371. Best score 644 bits
Score difference with first non-orthologous sequence - *A. fumigatus*:644 *A. fischeri*:644
afum|EAL88421.1 100.00% afis|EAW23058.1 100.00%

Bootstrap support for afum|EAL88421.1 as seed ortholog is 100%.
Bootstrap support for afis|EAW23058.1 as seed ortholog is 100%.

---

Group of orthologs #6372. Best score 644 bits
Score difference with first non-orthologous sequence - *A. fumigatus*:644 *A. fischeri*:644
afum|EAL89321.1 100.00% afis|EAW22771.1 100.00%
Bootstrap support for afum|EAL89321.1 as seed ortholog is 100%.
Bootstrap support for afis|EAW22771.1 as seed ortholog is 100%.

---

Group of orthologs #6373. Best score 644 bits
Score difference with first non-orthologous sequence - *A. fumigatus*:538 *A. fischeri*:541
afum|EAL90785.1 100.00% afis|EAW22407.1 100.00%
Bootstrap support for afum|EAL90785.1 as seed ortholog is 100%.
Bootstrap support for afis|EAW22407.1 as seed ortholog is 100%.

---

Group of orthologs #6374. Best score 644 bits
Score difference with first non-orthologous sequence - *A. fumigatus*:644 *A. fischeri*:566
afum|EAL90480.1 100.00% afis|EAW22714.1 100.00%
Bootstrap support for afum|EAL90480.1 as seed ortholog is 100%.
Bootstrap support for afis|EAW22714.1 as seed ortholog is 100%.

---

Group of orthologs #6375. Best score 644 bits
Score difference with first non-orthologous sequence - *A. fumigatus*:644 *A. fischeri*:644
afum|EAL92666.2 100.00% afis|EAW21633.1 100.00%
Bootstrap support for afum|EAL92666.2 as seed ortholog is 100%.
Bootstrap support for afis|EAW21633.1 as seed ortholog is 100%.

---

Group of orthologs #6376. Best score 644 bits
Score difference with first non-orthologous sequence - *A. fumigatus*:644 *A. fischeri*:644
afum|EAL89486.1 100.00% afis|EAW24840.1 100.00%
Bootstrap support for afum|EAL89486.1 as seed ortholog is 100%.
Bootstrap support for afis|EAW24840.1 as seed ortholog is 100%.

---

Group of orthologs #6377. Best score 644 bits
Score difference with first non-orthologous sequence - *A. fumigatus*:644 *A. fischeri*:644
afum|EAL92951.2 100.00% afis|EAW22060.1 100.00%
Bootstrap support for afum|EAL92951.2 as seed ortholog is 100%.
Bootstrap support for afis|EAW22060.1 as seed ortholog is 100%.

---

Group of orthologs #6378. Best score 643 bits
Score difference with first non-orthologous sequence - *A. fumigatus*:643 *A. fischeri*:585
afum|EAL86623.1 100.00% afis|EAW17195.1 100.00%
Bootstrap support for afum|EAL86623.1 as seed ortholog is 100%.
Bootstrap support for afis|EAW17195.1 as seed ortholog is 100%.

---

Group of orthologs #6379. Best score 643 bits
Score difference with first non-orthologous sequence - *A. fumigatus*:583 *A. fischeri*:557
afum|EAL91849.1 100.00% afis|EAW18038.1 100.00%
Bootstrap support for afum|EAL91849.1 as seed ortholog is 100%.
Bootstrap support for afis|EAW18038.1 as seed ortholog is 100%.

---

Group of orthologs #6380. Best score 643 bits
Score difference with first non-orthologous sequence - *A. fumigatus*:643 *A. fischeri*:643
afum|EAL93256.1 100.00% afis|EAW18556.1 100.00%
Bootstrap support for afum|EAL93256.1 as seed ortholog is 100%.
Bootstrap support for afis|EAW18556.1 as seed ortholog is 100%.

---
Group of orthologs #6381. Best score 643 bits
Score difference with first non-orthologous sequence - *A. fumigatus*:643 *A. fischeri*:643
afum|EAL90632.1 100.00% afis|EAW22563.1 100.00%
Bootstrap support for afum|EAL90632.1 as seed ortholog is 100%.
Bootstrap support for afis|EAW22563.1 as seed ortholog is 100%.

---
Group of orthologs #6382. Best score 642 bits
Score difference with first non-orthologous sequence - *A. fumigatus*:642 *A. fischeri*:642
afum|EAL86676.1 100.00% afis|EAW17259.1 100.00%
Bootstrap support for afum|EAL86676.1 as seed ortholog is 100%.
Bootstrap support for afis|EAW17259.1 as seed ortholog is 100%.

---
Group of orthologs #6383. Best score 642 bits
Score difference with first non-orthologous sequence - *A. fumigatus*:642 *A. fischeri*:642
afum|EAL91670.2 100.00% afis|EAW17862.1 100.00%
Bootstrap support for afum|EAL91670.2 as seed ortholog is 100%.
Bootstrap support for afis|EAW17862.1 as seed ortholog is 100%.

---
Group of orthologs #6384. Best score 642 bits
Score difference with first non-orthologous sequence - *A. fumigatus*:554 *A. fischeri*:382
afum|EBA27174.1 100.00% afis|EAW25637.1 100.00%
Bootstrap support for afum|EBA27174.1 as seed ortholog is 100%.
Bootstrap support for afis|EAW25637.1 as seed ortholog is 100%.

---
Group of orthologs #6385. Best score 641 bits
Score difference with first non-orthologous sequence - *A. fumigatus*:641 *A. fischeri*:641
afum|EAL88538.1 100.00% afis|EAW15903.1 100.00%
Bootstrap support for afum|EAL88538.1 as seed ortholog is 100%.
Bootstrap support for afis|EAW15903.1 as seed ortholog is 100%.

---
Group of orthologs #6386. Best score 641 bits
Score difference with first non-orthologous sequence - *A. fumigatus*:641 *A. fischeri*:350
afum|EAL85493.1 100.00% afis|EAW20380.1 100.00%
Bootstrap support for afum|EAL85493.1 as seed ortholog is 100%.
Bootstrap support for afis|EAW20380.1 as seed ortholog is 100%.

---
Group of orthologs #6387. Best score 641 bits
Score difference with first non-orthologous sequence - *A. fumigatus*:641 *A. fischeri*:641
afum|EAL86142.1 100.00% afis|EAW24470.1 100.00%
Bootstrap support for afum|EAL86142.1 as seed ortholog is 100%.
Bootstrap support for afis|EAW24470.1 as seed ortholog is 100%.

---
Group of orthologs #6388. Best score 641 bits
Score difference with first non-orthologous sequence - *A. fumigatus*:641 *A. fischeri*:641
afum|EBA27276.1 100.00% afis|EAW16985.1 100.00%
Bootstrap support for afum|EBA27276.1 as seed ortholog is 100%.
Bootstrap support for afis|EAW16985.1 as seed ortholog is 100%.

---
Group of orthologs #6389. Best score 641 bits
Score difference with first non-orthologous sequence - *A. fumigatus*:641 *A. fischeri*:641
afum|EAL92109.1 100.00% afis|EAW21086.1 100.00%
Bootstrap support for afum|EAL92109.1 as seed ortholog is 100%.
Bootstrap support for afis|EAW21086.1 as seed ortholog is 100%.

---
Group of orthologs #6390. Best score 640 bits

Score difference with first non-orthologous sequence - *A. fumigatus*:640 *A. fischeri*:640
afum|EAL87468.1 100.00% afis|EAW18111.1 100.00%
Bootstrap support for afum|EAL87468.1 as seed ortholog is 100%.
Bootstrap support for afis|EAW18111.1 as seed ortholog is 100%.

---

Group of orthologs #6391. Best score 640 bits
Score difference with first non-orthologous sequence - *A. fumigatus*:598 *A. fischeri*:543
afum|EAL91366.1 100.00% afis|EAW17552.1 100.00%
Bootstrap support for afum|EAL91366.1 as seed ortholog is 100%.
Bootstrap support for afis|EAW17552.1 as seed ortholog is 100%.

---

Group of orthologs #6392. Best score 640 bits
Score difference with first non-orthologous sequence - *A. fumigatus*:640 *A. fischeri*:640
afum|EAL86098.1 100.00% afis|EAW24430.1 100.00%
Bootstrap support for afum|EAL86098.1 as seed ortholog is 100%.
Bootstrap support for afis|EAW24430.1 as seed ortholog is 100%.

---

Group of orthologs #6393. Best score 640 bits
Score difference with first non-orthologous sequence - *A. fumigatus*:319 *A. fischeri*:140
afum|EAL87858.1 100.00% afis|EAW23607.1 100.00%
Bootstrap support for afum|EAL87858.1 as seed ortholog is 100%.
Bootstrap support for afis|EAW23607.1 as seed ortholog is 100%.

---

Group of orthologs #6394. Best score 640 bits
Score difference with first non-orthologous sequence - *A. fumigatus*:569 *A. fischeri*:553
afum|EAL90727.1 100.00% afis|EAW22464.1 100.00%
Bootstrap support for afum|EAL90727.1 as seed ortholog is 100%.
Bootstrap support for afis|EAW22464.1 as seed ortholog is 100%.

---

Group of orthologs #6395. Best score 639 bits
Score difference with first non-orthologous sequence - *A. fumigatus*:639 *A. fischeri*:639
afum|EAL85279.1 100.00% afis|EAW20161.1 100.00%
Bootstrap support for afum|EAL85279.1 as seed ortholog is 100%.
Bootstrap support for afis|EAW20161.1 as seed ortholog is 100%.

---

Group of orthologs #6396. Best score 639 bits
Score difference with first non-orthologous sequence - *A. fumigatus*:564 *A. fischeri*:554
afum|EAL89250.1 100.00% afis|EAW16646.1 100.00%
Bootstrap support for afum|EAL89250.1 as seed ortholog is 100%.
Bootstrap support for afis|EAW16646.1 as seed ortholog is 100%.

---

Group of orthologs #6397. Best score 639 bits
Score difference with first non-orthologous sequence - *A. fumigatus*:639 *A. fischeri*:639
afum|EAL93080.1 100.00% afis|EAW18402.1 100.00%
Bootstrap support for afum|EAL93080.1 as seed ortholog is 100%.
Bootstrap support for afis|EAW18402.1 as seed ortholog is 100%.

---

Group of orthologs #6398. Best score 639 bits
Score difference with first non-orthologous sequence - *A. fumigatus*:639 *A. fischeri*:639
afum|EAL93539.1 100.00% afis|EAW18803.1 100.00%
Bootstrap support for afum|EAL93539.1 as seed ortholog is 100%.
Bootstrap support for afis|EAW18803.1 as seed ortholog is 100%.

---

Group of orthologs #6399. Best score 639 bits
Score difference with first non-orthologous sequence - *A. fumigatus*:639 *A. fischeri*:639
afum|EAL90631.1 100.00% afis|EAW22567.1 100.00%

Bootstrap support for afum|EAL90631.1 as seed ortholog is 100%.
Bootstrap support for afis|EAW22567.1 as seed ortholog is 100%.

---

Group of orthologs #6400. Best score 639 bits
Score difference with first non-orthologous sequence - *A. fumigatus*:639 *A. fischeri*:639
afum|EAL91913.1 100.00% afis|EAW24057.1 100.00%
Bootstrap support for afum|EAL91913.1 as seed ortholog is 100%.
Bootstrap support for afis|EAW24057.1 as seed ortholog is 100%.

---

Group of orthologs #6401. Best score 638 bits
Score difference with first non-orthologous sequence - *A. fumigatus*:638 *A. fischeri*:638
afum|EAL84432.1 100.00% afis|EAW20636.1 100.00%
Bootstrap support for afum|EAL84432.1 as seed ortholog is 100%.
Bootstrap support for afis|EAW20636.1 as seed ortholog is 100%.

---

Group of orthologs #6402. Best score 638 bits
Score difference with first non-orthologous sequence - *A. fumigatus*:638 *A. fischeri*:436
afum|EAL87113.1 100.00% afis|EAW23668.1 100.00%
Bootstrap support for afum|EAL87113.1 as seed ortholog is 100%.
Bootstrap support for afis|EAW23668.1 as seed ortholog is 100%.

---

Group of orthologs #6403. Best score 638 bits
Score difference with first non-orthologous sequence - *A. fumigatus*:638 *A. fischeri*:638
afum|EAL88361.1 100.00% afis|EAW23119.1 100.00%
Bootstrap support for afum|EAL88361.1 as seed ortholog is 100%.
Bootstrap support for afis|EAW23119.1 as seed ortholog is 100%.

---

Group of orthologs #6404. Best score 638 bits
Score difference with first non-orthologous sequence - *A. fumigatus*:638 *A. fischeri*:566
afum|EAL93360.1 100.00% afis|EAW18645.1 100.00%
Bootstrap support for afum|EAL93360.1 as seed ortholog is 100%.
Bootstrap support for afis|EAW18645.1 as seed ortholog is 100%.

---

Group of orthologs #6405. Best score 638 bits
Score difference with first non-orthologous sequence - *A. fumigatus*:638 *A. fischeri*:638
afum|EAL92884.1 100.00% afis|EAW21870.1 100.00%
Bootstrap support for afum|EAL92884.1 as seed ortholog is 100%.
Bootstrap support for afis|EAW21870.1 as seed ortholog is 100%.

---

Group of orthologs #6406. Best score 637 bits
Score difference with first non-orthologous sequence - *A. fumigatus*:637 *A. fischeri*:637
afum|EAL85630.2 100.00% afis|EAW15718.1 100.00%
Bootstrap support for afum|EAL85630.2 as seed ortholog is 100%.
Bootstrap support for afis|EAW15718.1 as seed ortholog is 100%.

---

Group of orthologs #6407. Best score 637 bits
Score difference with first non-orthologous sequence - *A. fumigatus*:637 *A. fischeri*:637
afum|EAL88927.1 100.00% afis|EAW16305.1 100.00%
Bootstrap support for afum|EAL88927.1 as seed ortholog is 100%.
Bootstrap support for afis|EAW16305.1 as seed ortholog is 100%.

---

Group of orthologs #6408. Best score 637 bits
Score difference with first non-orthologous sequence - *A. fumigatus*:637 *A. fischeri*:637
afum|EAL88362.1 100.00% afis|EAW23118.1 100.00%
Bootstrap support for afum|EAL88362.1 as seed ortholog is 100%.
Bootstrap support for afis|EAW23118.1 as seed ortholog is 100%.

Group of orthologs #6409. Best score 637 bits
Score difference with first non-orthologous sequence - *A. fumigatus*:637 *A. fischeri*:637
afum|EAL88329.1 100.00% afis|EAW23152.1 100.00%
Bootstrap support for afum|EAL88329.1 as seed ortholog is 100%.
Bootstrap support for afis|EAW23152.1 as seed ortholog is 100%.

Group of orthologs #6410. Best score 637 bits
Score difference with first non-orthologous sequence - *A. fumigatus*:637 *A. fischeri*:637
afum|EAL91012.1 100.00% afis|EAW22177.1 100.00%
Bootstrap support for afum|EAL91012.1 as seed ortholog is 100%.
Bootstrap support for afis|EAW22177.1 as seed ortholog is 100%.

Group of orthologs #6411. Best score 637 bits
Score difference with first non-orthologous sequence - *A. fumigatus*:637 *A. fischeri*:637
afum|EBA27470.1 100.00% afis|EAW21715.1 100.00%
Bootstrap support for afum|EBA27470.1 as seed ortholog is 100%.
Bootstrap support for afis|EAW21715.1 as seed ortholog is 100%.

Group of orthologs #6412. Best score 636 bits
Score difference with first non-orthologous sequence - *A. fumigatus*:636 *A. fischeri*:636
afum|EAL88672.1 100.00% afis|EAW16040.1 100.00%
Bootstrap support for afum|EAL88672.1 as seed ortholog is 100%.
Bootstrap support for afis|EAW16040.1 as seed ortholog is 100%.

Group of orthologs #6413. Best score 636 bits
Score difference with first non-orthologous sequence - *A. fumigatus*:636 *A. fischeri*:636
afum|EAL88690.2 100.00% afis|EAW16058.1 100.00%
Bootstrap support for afum|EAL88690.2 as seed ortholog is 100%.
Bootstrap support for afis|EAW16058.1 as seed ortholog is 100%.

Group of orthologs #6414. Best score 636 bits
Score difference with first non-orthologous sequence - *A. fumigatus*:636 *A. fischeri*:636
afum|EAL86189.1 100.00% afis|EAW24524.1 100.00%
Bootstrap support for afum|EAL86189.1 as seed ortholog is 100%.
Bootstrap support for afis|EAW24524.1 as seed ortholog is 100%.

Group of orthologs #6415. Best score 636 bits
Score difference with first non-orthologous sequence - *A. fumigatus*:636 *A. fischeri*:636
afum|EAL88274.1 100.00% afis|EAW23210.1 100.00%
Bootstrap support for afum|EAL88274.1 as seed ortholog is 100%.
Bootstrap support for afis|EAW23210.1 as seed ortholog is 100%.

Group of orthologs #6416. Best score 636 bits
Score difference with first non-orthologous sequence - *A. fumigatus*:636 *A. fischeri*:636
afum|EAL91089.1 100.00% afis|EAW22100.1 100.00%
Bootstrap support for afum|EAL91089.1 as seed ortholog is 100%.
Bootstrap support for afis|EAW22100.1 as seed ortholog is 100%.

Group of orthologs #6417. Best score 636 bits
Score difference with first non-orthologous sequence - *A. fumigatus*:636 *A. fischeri*:636
afum|EAL90502.1 100.00% afis|EAW22690.1 100.00%
Bootstrap support for afum|EAL90502.1 as seed ortholog is 100%.
Bootstrap support for afis|EAW22690.1 as seed ortholog is 100%.

Group of orthologs #6418. Best score 636 bits

Score difference with first non-orthologous sequence - *A. fumigatus*:636   *A. fischeri*:636
afum|EAL92589.1 100.00% afis|EAW21571.1 100.00%
Bootstrap support for afum|EAL92589.1 as seed ortholog is 100%.
Bootstrap support for afis|EAW21571.1 as seed ortholog is 100%.

---

Group of orthologs #6419. Best score 636 bits
Score difference with first non-orthologous sequence - *A. fumigatus*:636   *A. fischeri*:636
afum|EAL92870.1 100.00% afis|EAW21857.1 100.00%
Bootstrap support for afum|EAL92870.1 as seed ortholog is 100%.
Bootstrap support for afis|EAW21857.1 as seed ortholog is 100%.

---

Group of orthologs #6420. Best score 636 bits
Score difference with first non-orthologous sequence - *A. fumigatus*:636   *A. fischeri*:636
afum|EAL89675.1 100.00% afis|EAW25020.1 100.00%
Bootstrap support for afum|EAL89675.1 as seed ortholog is 100%.
Bootstrap support for afis|EAW25020.1 as seed ortholog is 100%.

---

Group of orthologs #6421. Best score 636 bits
Score difference with first non-orthologous sequence - *A. fumigatus*:636   *A. fischeri*:636
afum|EAL90193.2 100.00% afis|EAW25528.1 100.00%
Bootstrap support for afum|EAL90193.2 as seed ortholog is 100%.
Bootstrap support for afis|EAW25528.1 as seed ortholog is 100%.

---

Group of orthologs #6422. Best score 635 bits
Score difference with first non-orthologous sequence - *A. fumigatus*:635   *A. fischeri*:635
afum|EAL86327.1 100.00% afis|EAW16755.1 100.00%
Bootstrap support for afum|EAL86327.1 as seed ortholog is 100%.
Bootstrap support for afis|EAW16755.1 as seed ortholog is 100%.

---

Group of orthologs #6423. Best score 635 bits
Score difference with first non-orthologous sequence - *A. fumigatus*:635   *A. fischeri*:635
afum|EAL88911.1 100.00% afis|EAW16289.1 100.00%
Bootstrap support for afum|EAL88911.1 as seed ortholog is 100%.
Bootstrap support for afis|EAW16289.1 as seed ortholog is 100%.

---

Group of orthologs #6424. Best score 635 bits
Score difference with first non-orthologous sequence - *A. fumigatus*:635   *A. fischeri*:635
afum|EAL93451.1 100.00% afis|EAW18726.1 100.00%
Bootstrap support for afum|EAL93451.1 as seed ortholog is 100%.
Bootstrap support for afis|EAW18726.1 as seed ortholog is 100%.

---

Group of orthologs #6425. Best score 635 bits
Score difference with first non-orthologous sequence - *A. fumigatus*:635   *A. fischeri*:635
afum|EAL93606.1 100.00% afis|EAW18874.1 100.00%
Bootstrap support for afum|EAL93606.1 as seed ortholog is 100%.
Bootstrap support for afis|EAW18874.1 as seed ortholog is 100%.

---

Group of orthologs #6426. Best score 635 bits
Score difference with first non-orthologous sequence - *A. fumigatus*:635   *A. fischeri*:635
afum|EAL93779.1 100.00% afis|EAW19050.1 100.00%
Bootstrap support for afum|EAL93779.1 as seed ortholog is 100%.
Bootstrap support for afis|EAW19050.1 as seed ortholog is 100%.

---

Group of orthologs #6427. Best score 635 bits
Score difference with first non-orthologous sequence - *A. fumigatus*:635   *A. fischeri*:635
afum|EAL91065.1 100.00% afis|EAW22122.1 100.00%

Bootstrap support for afum|EAL91065.1 as seed ortholog is 100%.
Bootstrap support for afis|EAW22122.1 as seed ortholog is 100%.

---

Group of orthologs #6428. Best score 635 bits
Score difference with first non-orthologous sequence - *A. fumigatus*:635 *A. fischeri*:635
afum|EAL89839.1 100.00% afis|EAW25166.1 100.00%
Bootstrap support for afum|EAL89839.1 as seed ortholog is 100%.
Bootstrap support for afis|EAW25166.1 as seed ortholog is 100%.

---

Group of orthologs #6429. Best score 634 bits
Score difference with first non-orthologous sequence - *A. fumigatus*:439 *A. fischeri*:345
afum|EAL88806.1 100.00% afis|EAW16174.1 100.00%
Bootstrap support for afum|EAL88806.1 as seed ortholog is 100%.
Bootstrap support for afis|EAW16174.1 as seed ortholog is 100%.

---

Group of orthologs #6430. Best score 634 bits
Score difference with first non-orthologous sequence - *A. fumigatus*:634 *A. fischeri*:634
afum|EAL84426.1 100.00% afis|EAW20647.1 100.00%
Bootstrap support for afum|EAL84426.1 as seed ortholog is 100%.
Bootstrap support for afis|EAW20647.1 as seed ortholog is 100%.

---

Group of orthologs #6431. Best score 634 bits
Score difference with first non-orthologous sequence - *A. fumigatus*:634 *A. fischeri*:257
afum|EAL91897.1 100.00% afis|EAW18085.1 100.00%
Bootstrap support for afum|EAL91897.1 as seed ortholog is 100%.
Bootstrap support for afis|EAW18085.1 as seed ortholog is 100%.

---

Group of orthologs #6432. Best score 634 bits
Score difference with first non-orthologous sequence - *A. fumigatus*:634 *A. fischeri*:634
afum|EAL86179.1 100.00% afis|EAW24510.1 100.00%
Bootstrap support for afum|EAL86179.1 as seed ortholog is 100%.
Bootstrap support for afis|EAW24510.1 as seed ortholog is 100%.

---

Group of orthologs #6433. Best score 634 bits
Score difference with first non-orthologous sequence - *A. fumigatus*:634 *A. fischeri*:634
afum|EAL87956.1 100.00% afis|EAW23505.1 100.00%
Bootstrap support for afum|EAL87956.1 as seed ortholog is 100%.
Bootstrap support for afis|EAW23505.1 as seed ortholog is 100%.

---

Group of orthologs #6434. Best score 633 bits
Score difference with first non-orthologous sequence - *A. fumigatus*:633 *A. fischeri*:633
afum|EAL85011.1 100.00% afis|EAW19886.1 100.00%
Bootstrap support for afum|EAL85011.1 as seed ortholog is 100%.
Bootstrap support for afis|EAW19886.1 as seed ortholog is 100%.

---

Group of orthologs #6435. Best score 633 bits
Score difference with first non-orthologous sequence - *A. fumigatus*:633 *A. fischeri*:633
afum|EAL88934.1 100.00% afis|EAW16312.1 100.00%
Bootstrap support for afum|EAL88934.1 as seed ortholog is 100%.
Bootstrap support for afis|EAW16312.1 as seed ortholog is 100%.

---

Group of orthologs #6436. Best score 633 bits
Score difference with first non-orthologous sequence - *A. fumigatus*:633 *A. fischeri*:633
afum|EAL84672.1 100.00% afis|EAW20961.1 100.00%
Bootstrap support for afum|EAL84672.1 as seed ortholog is 100%.
Bootstrap support for afis|EAW20961.1 as seed ortholog is 100%.

Group of orthologs #6437. Best score 633 bits
Score difference with first non-orthologous sequence - *A. fumigatus*:633 *A. fischeri*:633
afum|EAL88018.1 100.00% afis|EAW23447.1 100.00%
Bootstrap support for afum|EAL88018.1 as seed ortholog is 100%.
Bootstrap support for afis|EAW23447.1 as seed ortholog is 100%.

Group of orthologs #6438. Best score 633 bits
Score difference with first non-orthologous sequence - *A. fumigatus*:633 *A. fischeri*:432
afum|EAL93391.1 100.00% afis|EAW18672.1 100.00%
Bootstrap support for afum|EAL93391.1 as seed ortholog is 100%.
Bootstrap support for afis|EAW18672.1 as seed ortholog is 100%.

Group of orthologs #6439. Best score 633 bits
Score difference with first non-orthologous sequence - *A. fumigatus*:633 *A. fischeri*:633
afum|EAL90688.1 100.00% afis|EAW22505.1 100.00%
Bootstrap support for afum|EAL90688.1 as seed ortholog is 100%.
Bootstrap support for afis|EAW22505.1 as seed ortholog is 100%.

Group of orthologs #6440. Best score 633 bits
Score difference with first non-orthologous sequence - *A. fumigatus*:633 *A. fischeri*:633
afum|EBA27329.1 100.00% afis|EAW23457.1 100.00%
Bootstrap support for afum|EBA27329.1 as seed ortholog is 100%.
Bootstrap support for afis|EAW23457.1 as seed ortholog is 100%.

Group of orthologs #6441. Best score 632 bits
Score difference with first non-orthologous sequence - *A. fumigatus*:632 *A. fischeri*:632
afum|EAL86320.1 100.00% afis|EAW16745.1 100.00%
Bootstrap support for afum|EAL86320.1 as seed ortholog is 100%.
Bootstrap support for afis|EAW16745.1 as seed ortholog is 100%.

Group of orthologs #6442. Best score 632 bits
Score difference with first non-orthologous sequence - *A. fumigatus*:632 *A. fischeri*:632
afum|EAL88518.1 100.00% afis|EAW15884.1 100.00%
Bootstrap support for afum|EAL88518.1 as seed ortholog is 100%.
Bootstrap support for afis|EAW15884.1 as seed ortholog is 100%.

Group of orthologs #6443. Best score 632 bits
Score difference with first non-orthologous sequence - *A. fumigatus*:632 *A. fischeri*:632
afum|EAL84577.1 100.00% afis|EAW20476.1 100.00%
Bootstrap support for afum|EAL84577.1 as seed ortholog is 100%.
Bootstrap support for afis|EAW20476.1 as seed ortholog is 100%.

Group of orthologs #6444. Best score 632 bits
Score difference with first non-orthologous sequence - *A. fumigatus*:632 *A. fischeri*:632
afum|EAL92388.1 100.00% afis|EAW21371.1 100.00%
Bootstrap support for afum|EAL92388.1 as seed ortholog is 100%.
Bootstrap support for afis|EAW21371.1 as seed ortholog is 100%.

Group of orthologs #6445. Best score 632 bits
Score difference with first non-orthologous sequence - *A. fumigatus*:632 *A. fischeri*:632
afum|EAL90077.1 100.00% afis|EAW25418.1 100.00%
Bootstrap support for afum|EAL90077.1 as seed ortholog is 100%.
Bootstrap support for afis|EAW25418.1 as seed ortholog is 100%.

Group of orthologs #6446. Best score 631 bits

Score difference with first non-orthologous sequence - *A. fumigatus*:518   *A. fischeri*:571
afum|EAL85739.2 100.00% afis|EAW15593.1 100.00%
Bootstrap support for afum|EAL85739.2 as seed ortholog is 100%.
Bootstrap support for afis|EAW15593.1 as seed ortholog is 100%.

---

Group of orthologs #6447. Best score 631 bits
Score difference with first non-orthologous sequence - *A. fumigatus*:631   *A. fischeri*:383
afum|EAL85635.1 100.00% afis|EAW15713.1 100.00%
Bootstrap support for afum|EAL85635.1 as seed ortholog is 100%.
Bootstrap support for afis|EAW15713.1 as seed ortholog is 100%.

---

Group of orthologs #6448. Best score 631 bits
Score difference with first non-orthologous sequence - *A. fumigatus*:61   *A. fischeri*:208
afum|EAL89046.1 100.00% afis|EAW16449.1 100.00%
Bootstrap support for afum|EAL89046.1 as seed ortholog is 76%.
Bootstrap support for afis|EAW16449.1 as seed ortholog is 99%.

---

Group of orthologs #6449. Best score 631 bits
Score difference with first non-orthologous sequence - *A. fumigatus*:631   *A. fischeri*:631
afum|EAL91303.1 100.00% afis|EAW17474.1 100.00%
Bootstrap support for afum|EAL91303.1 as seed ortholog is 100%.
Bootstrap support for afis|EAW17474.1 as seed ortholog is 100%.

---

Group of orthologs #6450. Best score 631 bits
Score difference with first non-orthologous sequence - *A. fumigatus*:631   *A. fischeri*:631
afum|EAL93998.1 100.00% afis|EAW19272.1 100.00%
Bootstrap support for afum|EAL93998.1 as seed ortholog is 100%.
Bootstrap support for afis|EAW19272.1 as seed ortholog is 100%.

---

Group of orthologs #6451. Best score 631 bits
Score difference with first non-orthologous sequence - *A. fumigatus*:631   *A. fischeri*:631
afum|EAL89584.2 100.00% afis|EAW24928.1 100.00%
Bootstrap support for afum|EAL89584.2 as seed ortholog is 100%.
Bootstrap support for afis|EAW24928.1 as seed ortholog is 100%.

---

Group of orthologs #6452. Best score 630 bits
Score difference with first non-orthologous sequence - *A. fumigatus*:630   *A. fischeri*:630
afum|EAL89304.1 100.00% afis|EAW25774.1 100.00%
afis|EAW23657.1 99.38%
afis|EAW24635.1 99.11%
afis|EAW15379.1 23.79%
afis|EAW23656.1 20.92%
Bootstrap support for afum|EAL89304.1 as seed ortholog is 100%.
Bootstrap support for afis|EAW25774.1 as seed ortholog is 100%.

---

Group of orthologs #6453. Best score 630 bits
Score difference with first non-orthologous sequence - *A. fumigatus*:630   *A. fischeri*:630
afum|EAL85931.1 100.00% afis|EAW24263.1 100.00%
Bootstrap support for afum|EAL85931.1 as seed ortholog is 100%.
Bootstrap support for afis|EAW24263.1 as seed ortholog is 100%.

---

Group of orthologs #6454. Best score 630 bits
Score difference with first non-orthologous sequence - *A. fumigatus*:590   *A. fischeri*:461
afum|EAL92498.1 100.00% afis|EAW21478.1 100.00%
Bootstrap support for afum|EAL92498.1 as seed ortholog is 100%.
Bootstrap support for afis|EAW21478.1 as seed ortholog is 100%.

---

Group of orthologs #6455. Best score 630 bits
Score difference with first non-orthologous sequence - *A. fumigatus*:630 *A. fischeri*:630
afum|EAL92936.1 100.00% afis|EAW21923.1 100.00%
Bootstrap support for afum|EAL92936.1 as seed ortholog is 100%.
Bootstrap support for afis|EAW21923.1 as seed ortholog is 100%.

---

Group of orthologs #6456. Best score 629 bits
Score difference with first non-orthologous sequence - *A. fumigatus*:629 *A. fischeri*:629
afum|EAL85078.1 100.00% afis|EAW19813.1 100.00%
Bootstrap support for afum|EAL85078.1 as seed ortholog is 100%.
Bootstrap support for afis|EAW19813.1 as seed ortholog is 100%.

---

Group of orthologs #6457. Best score 629 bits
Score difference with first non-orthologous sequence - *A. fumigatus*:629 *A. fischeri*:629
afum|EAL85195.1 100.00% afis|EAW20077.1 100.00%
Bootstrap support for afum|EAL85195.1 as seed ortholog is 100%.
Bootstrap support for afis|EAW20077.1 as seed ortholog is 100%.

---

Group of orthologs #6458. Best score 629 bits
Score difference with first non-orthologous sequence - *A. fumigatus*:629 *A. fischeri*:373
afum|EAL84774.1 100.00% afis|EAW20861.1 100.00%
Bootstrap support for afum|EAL84774.1 as seed ortholog is 100%.
Bootstrap support for afis|EAW20861.1 as seed ortholog is 100%.

---

Group of orthologs #6459. Best score 629 bits
Score difference with first non-orthologous sequence - *A. fumigatus*:629 *A. fischeri*:629
afum|EBA27479.1 100.00% afis|EAW21896.1 100.00%
Bootstrap support for afum|EBA27479.1 as seed ortholog is 100%.
Bootstrap support for afis|EAW21896.1 as seed ortholog is 100%.

---

Group of orthologs #6460. Best score 628 bits
Score difference with first non-orthologous sequence - *A. fumigatus*:519 *A. fischeri*:493
afum|EAL86375.1 100.00% afis|EAW16826.1 100.00%
Bootstrap support for afum|EAL86375.1 as seed ortholog is 100%.
Bootstrap support for afis|EAW16826.1 as seed ortholog is 100%.

---

Group of orthologs #6461. Best score 628 bits
Score difference with first non-orthologous sequence - *A. fumigatus*:628 *A. fischeri*:628
afum|EAL88498.1 100.00% afis|EAW15863.1 100.00%
Bootstrap support for afum|EAL88498.1 as seed ortholog is 100%.
Bootstrap support for afis|EAW15863.1 as seed ortholog is 100%.

---

Group of orthologs #6462. Best score 628 bits
Score difference with first non-orthologous sequence - *A. fumigatus*:628 *A. fischeri*:628
afum|EAL91568.1 100.00% afis|EAW17755.1 100.00%
Bootstrap support for afum|EAL91568.1 as seed ortholog is 100%.
Bootstrap support for afis|EAW17755.1 as seed ortholog is 100%.

---

Group of orthologs #6463. Best score 628 bits
Score difference with first non-orthologous sequence - *A. fumigatus*:628 *A. fischeri*:628
afum|EAL93188.1 100.00% afis|EAW18498.1 100.00%
Bootstrap support for afum|EAL93188.1 as seed ortholog is 100%.
Bootstrap support for afis|EAW18498.1 as seed ortholog is 100%.

---

Group of orthologs #6464. Best score 628 bits

Score difference with first non-orthologous sequence - *A. fumigatus*:628 *A. fischeri*:291
afum|EAL90866.1 100.00% afis|EAW22326.1 100.00%
Bootstrap support for afum|EAL90866.1 as seed ortholog is 100%.
Bootstrap support for afis|EAW22326.1 as seed ortholog is 100%.

---

Group of orthologs #6465. Best score 627 bits
Score difference with first non-orthologous sequence - *A. fumigatus*:627 *A. fischeri*:627
afum|EAL85800.1 100.00% afis|EAW15528.1 100.00%
Bootstrap support for afum|EAL85800.1 as seed ortholog is 100%.
Bootstrap support for afis|EAW15528.1 as seed ortholog is 100%.

---

Group of orthologs #6466. Best score 627 bits
Score difference with first non-orthologous sequence - *A. fumigatus*:627 *A. fischeri*:627
afum|EAL88531.1 100.00% afis|EAW15896.1 100.00%
Bootstrap support for afum|EAL88531.1 as seed ortholog is 100%.
Bootstrap support for afis|EAW15896.1 as seed ortholog is 100%.

---

Group of orthologs #6467. Best score 627 bits
Score difference with first non-orthologous sequence - *A. fumigatus*:627 *A. fischeri*:627
afum|EAL92071.1 100.00% afis|EAW21033.1 100.00%
Bootstrap support for afum|EAL92071.1 as seed ortholog is 100%.
Bootstrap support for afis|EAW21033.1 as seed ortholog is 100%.

---

Group of orthologs #6468. Best score 627 bits
Score difference with first non-orthologous sequence - *A. fumigatus*:627 *A. fischeri*:627
afum|EAL90886.1 100.00% afis|EAW22295.1 100.00%
Bootstrap support for afum|EAL90886.1 as seed ortholog is 100%.
Bootstrap support for afis|EAW22295.1 as seed ortholog is 100%.

---

Group of orthologs #6469. Best score 627 bits
Score difference with first non-orthologous sequence - *A. fumigatus*:627 *A. fischeri*:627
afum|EAL90882.1 100.00% afis|EAW22308.1 100.00%
Bootstrap support for afum|EAL90882.1 as seed ortholog is 100%.
Bootstrap support for afis|EAW22308.1 as seed ortholog is 100%.

---

Group of orthologs #6470. Best score 627 bits
Score difference with first non-orthologous sequence - *A. fumigatus*:627 *A. fischeri*:627
afum|EAL89289.1 100.00% afis|EAW25571.1 100.00%
Bootstrap support for afum|EAL89289.1 as seed ortholog is 100%.
Bootstrap support for afis|EAW25571.1 as seed ortholog is 100%.

---

Group of orthologs #6471. Best score 626 bits
Score difference with first non-orthologous sequence - *A. fumigatus*:626 *A. fischeri*:626
afum|EAL88821.2 100.00% afis|EAW16190.1 100.00%
Bootstrap support for afum|EAL88821.2 as seed ortholog is 100%.
Bootstrap support for afis|EAW16190.1 as seed ortholog is 100%.

---

Group of orthologs #6472. Best score 626 bits
Score difference with first non-orthologous sequence - *A. fumigatus*:626 *A. fischeri*:455
afum|EAL91213.1 100.00% afis|EAW17378.1 100.00%
Bootstrap support for afum|EAL91213.1 as seed ortholog is 100%.
Bootstrap support for afis|EAW17378.1 as seed ortholog is 100%.

---

Group of orthologs #6473. Best score 626 bits
Score difference with first non-orthologous sequence - *A. fumigatus*:626 *A. fischeri*:626
afum|EAL91304.1 100.00% afis|EAW17475.1 100.00%

Bootstrap support for afum|EAL91304.1 as seed ortholog is 100%.
Bootstrap support for afis|EAW17475.1 as seed ortholog is 100%.

---

Group of orthologs #6474. Best score 626 bits
Score difference with first non-orthologous sequence - *A. fumigatus*:626 *A. fischeri*:626
afum|EAL93526.1 100.00% afis|EAW18791.1 100.00%
Bootstrap support for afum|EAL93526.1 as seed ortholog is 100%.
Bootstrap support for afis|EAW18791.1 as seed ortholog is 100%.

---

Group of orthologs #6475. Best score 626 bits
Score difference with first non-orthologous sequence - *A. fumigatus*:626 *A. fischeri*:626
afum|EAL93776.1 100.00% afis|EAW19047.1 100.00%
Bootstrap support for afum|EAL93776.1 as seed ortholog is 100%.
Bootstrap support for afis|EAW19047.1 as seed ortholog is 100%.

---

Group of orthologs #6476. Best score 625 bits
Score difference with first non-orthologous sequence - *A. fumigatus*:625 *A. fischeri*:625
afum|EAL86516.1 100.00% afis|EAW16992.1 100.00%
Bootstrap support for afum|EAL86516.1 as seed ortholog is 100%.
Bootstrap support for afis|EAW16992.1 as seed ortholog is 100%.

---

Group of orthologs #6477. Best score 625 bits
Score difference with first non-orthologous sequence - *A. fumigatus*:625 *A. fischeri*:625
afum|EAL84602.1 100.00% afis|EAW20451.1 100.00%
Bootstrap support for afum|EAL84602.1 as seed ortholog is 100%.
Bootstrap support for afis|EAW20451.1 as seed ortholog is 100%.

---

Group of orthologs #6478. Best score 625 bits
Score difference with first non-orthologous sequence - *A. fumigatus*:625 *A. fischeri*:625
afum|EAL91236.1 100.00% afis|EAW17398.1 100.00%
Bootstrap support for afum|EAL91236.1 as seed ortholog is 100%.
Bootstrap support for afis|EAW17398.1 as seed ortholog is 100%.

---

Group of orthologs #6479. Best score 625 bits
Score difference with first non-orthologous sequence - *A. fumigatus*:625 *A. fischeri*:625
afum|EAL91806.2 100.00% afis|EAW17996.1 100.00%
Bootstrap support for afum|EAL91806.2 as seed ortholog is 100%.
Bootstrap support for afis|EAW17996.1 as seed ortholog is 100%.

---

Group of orthologs #6480. Best score 625 bits
Score difference with first non-orthologous sequence - *A. fumigatus*:625 *A. fischeri*:625
afum|EAL93531.1 100.00% afis|EAW18795.1 100.00%
Bootstrap support for afum|EAL93531.1 as seed ortholog is 100%.
Bootstrap support for afis|EAW18795.1 as seed ortholog is 100%.

---

Group of orthologs #6481. Best score 625 bits
Score difference with first non-orthologous sequence - *A. fumigatus*:403 *A. fischeri*:249
afum|EAL94012.2 100.00% afis|EAW19978.1 100.00%
Bootstrap support for afum|EAL94012.2 as seed ortholog is 100%.
Bootstrap support for afis|EAW19978.1 as seed ortholog is 100%.

---

Group of orthologs #6482. Best score 625 bits
Score difference with first non-orthologous sequence - *A. fumigatus*:625 *A. fischeri*:625
afum|EAL92721.2 100.00% afis|EAW21687.1 100.00%
Bootstrap support for afum|EAL92721.2 as seed ortholog is 100%.
Bootstrap support for afis|EAW21687.1 as seed ortholog is 100%.

Group of orthologs #6483. Best score 625 bits
Score difference with first non-orthologous sequence - *A. fumigatus*:625 *A. fischeri*:625
afum|EAL92731.1 100.00% afis|EAW21697.1 100.00%
Bootstrap support for afum|EAL92731.1 as seed ortholog is 100%.
Bootstrap support for afis|EAW21697.1 as seed ortholog is 100%.

Group of orthologs #6484. Best score 624 bits
Score difference with first non-orthologous sequence - *A. fumigatus*:624 *A. fischeri*:467
afum|EAL86329.1 100.00% afis|EAW16757.1 100.00%
Bootstrap support for afum|EAL86329.1 as seed ortholog is 100%.
Bootstrap support for afis|EAW16757.1 as seed ortholog is 100%.

Group of orthologs #6485. Best score 624 bits
Score difference with first non-orthologous sequence - *A. fumigatus*:624 *A. fischeri*:624
afum|EAL86578.1 100.00% afis|EAW17116.1 100.00%
Bootstrap support for afum|EAL86578.1 as seed ortholog is 100%.
Bootstrap support for afis|EAW17116.1 as seed ortholog is 100%.

Group of orthologs #6486. Best score 624 bits
Score difference with first non-orthologous sequence - *A. fumigatus*:624 *A. fischeri*:549
afum|EAL85119.2 100.00% afis|EAW19779.1 100.00%
Bootstrap support for afum|EAL85119.2 as seed ortholog is 100%.
Bootstrap support for afis|EAW19779.1 as seed ortholog is 100%.

Group of orthologs #6487. Best score 624 bits
Score difference with first non-orthologous sequence - *A. fumigatus*:624 *A. fischeri*:624
afum|EAL84537.1 100.00% afis|EAW20520.1 100.00%
Bootstrap support for afum|EAL84537.1 as seed ortholog is 100%.
Bootstrap support for afis|EAW20520.1 as seed ortholog is 100%.

Group of orthologs #6488. Best score 624 bits
Score difference with first non-orthologous sequence - *A. fumigatus*:624 *A. fischeri*:536
afum|EAL89055.1 100.00% afis|EAW16459.1 100.00%
Bootstrap support for afum|EAL89055.1 as seed ortholog is 100%.
Bootstrap support for afis|EAW16459.1 as seed ortholog is 100%.

Group of orthologs #6489. Best score 624 bits
Score difference with first non-orthologous sequence - *A. fumigatus*:499 *A. fischeri*:510
afum|EAL85236.2 100.00% afis|EAW20119.1 100.00%
Bootstrap support for afum|EAL85236.2 as seed ortholog is 100%.
Bootstrap support for afis|EAW20119.1 as seed ortholog is 100%.

Group of orthologs #6490. Best score 624 bits
Score difference with first non-orthologous sequence - *A. fumigatus*:624 *A. fischeri*:624
afum|EAL91291.1 100.00% afis|EAW17462.1 100.00%
Bootstrap support for afum|EAL91291.1 as seed ortholog is 100%.
Bootstrap support for afis|EAW17462.1 as seed ortholog is 100%.

Group of orthologs #6491. Best score 624 bits
Score difference with first non-orthologous sequence - *A. fumigatus*:624 *A. fischeri*:624
afum|EAL87426.1 100.00% afis|EAW23996.1 100.00%
Bootstrap support for afum|EAL87426.1 as seed ortholog is 100%.
Bootstrap support for afis|EAW23996.1 as seed ortholog is 100%.

Group of orthologs #6492. Best score 624 bits

Score difference with first non-orthologous sequence - *A. fumigatus*:624 *A. fischeri*:624
afum|EAL93312.1 100.00% afis|EAW18604.1 100.00%
Bootstrap support for afum|EAL93312.1 as seed ortholog is 100%.
Bootstrap support for afis|EAW18604.1 as seed ortholog is 100%.

---

Group of orthologs #6493. Best score 624 bits
Score difference with first non-orthologous sequence - *A. fumigatus*:624 *A. fischeri*:624
afum|EBA27441.1 100.00% afis|EAW18057.1 100.00%
Bootstrap support for afum|EBA27441.1 as seed ortholog is 100%.
Bootstrap support for afis|EAW18057.1 as seed ortholog is 100%.

---

Group of orthologs #6494. Best score 624 bits
Score difference with first non-orthologous sequence - *A. fumigatus*:624 *A. fischeri*:624
afum|EAL94021.1 100.00% afis|EAW19303.1 100.00%
Bootstrap support for afum|EAL94021.1 as seed ortholog is 100%.
Bootstrap support for afis|EAW19303.1 as seed ortholog is 100%.

---

Group of orthologs #6495. Best score 624 bits
Score difference with first non-orthologous sequence - *A. fumigatus*:624 *A. fischeri*:624
afum|EAL90887.1 100.00% afis|EAW22294.1 100.00%
Bootstrap support for afum|EAL90887.1 as seed ortholog is 100%.
Bootstrap support for afis|EAW22294.1 as seed ortholog is 100%.

---

Group of orthologs #6496. Best score 624 bits
Score difference with first non-orthologous sequence - *A. fumigatus*:503 *A. fischeri*:391
afum|EAL90767.1 100.00% afis|EAW22424.1 100.00%
Bootstrap support for afum|EAL90767.1 as seed ortholog is 100%.
Bootstrap support for afis|EAW22424.1 as seed ortholog is 100%.

---

Group of orthologs #6497. Best score 623 bits
Score difference with first non-orthologous sequence - *A. fumigatus*:623 *A. fischeri*:623
afum|EAL85845.1 100.00% afis|EAW15476.1 100.00%
Bootstrap support for afum|EAL85845.1 as seed ortholog is 100%.
Bootstrap support for afis|EAW15476.1 as seed ortholog is 100%.

---

Group of orthologs #6498. Best score 623 bits
Score difference with first non-orthologous sequence - *A. fumigatus*:623 *A. fischeri*:623
afum|EAL84935.1 100.00% afis|EAW19980.1 100.00%
Bootstrap support for afum|EAL84935.1 as seed ortholog is 100%.
Bootstrap support for afis|EAW19980.1 as seed ortholog is 100%.

---

Group of orthologs #6499. Best score 623 bits
Score difference with first non-orthologous sequence - *A. fumigatus*:623 *A. fischeri*:623
afum|EAL85330.1 100.00% afis|EAW20213.1 100.00%
Bootstrap support for afum|EAL85330.1 as seed ortholog is 100%.
Bootstrap support for afis|EAW20213.1 as seed ortholog is 100%.

---

Group of orthologs #6500. Best score 623 bits
Score difference with first non-orthologous sequence - *A. fumigatus*:529 *A. fischeri*:527
afum|EAL89438.1 100.00% afis|EAW24796.1 100.00%
Bootstrap support for afum|EAL89438.1 as seed ortholog is 100%.
Bootstrap support for afis|EAW24796.1 as seed ortholog is 100%.

---

Group of orthologs #6501. Best score 623 bits
Score difference with first non-orthologous sequence - *A. fumigatus*:623 *A. fischeri*:623
afum|EAL89949.1 100.00% afis|EAW25286.1 100.00%

Bootstrap support for afum|EAL89949.1 as seed ortholog is 100%.
Bootstrap support for afis|EAW25286.1 as seed ortholog is 100%.

---

Group of orthologs #6502. Best score 622 bits
Score difference with first non-orthologous sequence - *A. fumigatus*:622 *A. fischeri*:622
afum|EAL85573.1 100.00% afis|EAW15776.1 100.00%
Bootstrap support for afum|EAL85573.1 as seed ortholog is 100%.
Bootstrap support for afis|EAW15776.1 as seed ortholog is 100%.

---

Group of orthologs #6503. Best score 622 bits
Score difference with first non-orthologous sequence - *A. fumigatus*:622 *A. fischeri*:622
afum|EAL84595.1 100.00% afis|EAW20457.1 100.00%
Bootstrap support for afum|EAL84595.1 as seed ortholog is 100%.
Bootstrap support for afis|EAW20457.1 as seed ortholog is 100%.

---

Group of orthologs #6504. Best score 622 bits
Score difference with first non-orthologous sequence - *A. fumigatus*:525 *A. fischeri*:507
afum|EAL87035.1 100.00% afis|EAW19461.1 100.00%
Bootstrap support for afum|EAL87035.1 as seed ortholog is 100%.
Bootstrap support for afis|EAW19461.1 as seed ortholog is 100%.

---

Group of orthologs #6505. Best score 622 bits
Score difference with first non-orthologous sequence - *A. fumigatus*:622 *A. fischeri*:622
afum|EAL87347.1 100.00% afis|EAW23922.1 100.00%
Bootstrap support for afum|EAL87347.1 as seed ortholog is 100%.
Bootstrap support for afis|EAW23922.1 as seed ortholog is 100%.

---

Group of orthologs #6506. Best score 622 bits
Score difference with first non-orthologous sequence - *A. fumigatus*:622 *A. fischeri*:622
afum|EBA27414.1 100.00% afis|EAW22390.1 100.00%
Bootstrap support for afum|EBA27414.1 as seed ortholog is 100%.
Bootstrap support for afis|EAW22390.1 as seed ortholog is 100%.

---

Group of orthologs #6507. Best score 621 bits
Score difference with first non-orthologous sequence - *A. fumigatus*:621 *A. fischeri*:621
afum|EAL88499.1 100.00% afis|EAW15865.1 100.00%
Bootstrap support for afum|EAL88499.1 as seed ortholog is 100%.
Bootstrap support for afis|EAW15865.1 as seed ortholog is 100%.

---

Group of orthologs #6508. Best score 621 bits
Score difference with first non-orthologous sequence - *A. fumigatus*:621 *A. fischeri*:621
afum|EAL91511.1 100.00% afis|EAW17697.1 100.00%
Bootstrap support for afum|EAL91511.1 as seed ortholog is 100%.
Bootstrap support for afis|EAW17697.1 as seed ortholog is 100%.

---

Group of orthologs #6509. Best score 621 bits
Score difference with first non-orthologous sequence - *A. fumigatus*:555 *A. fischeri*:510
afum|EAL87229.2 100.00% afis|EAW23795.1 100.00%
Bootstrap support for afum|EAL87229.2 as seed ortholog is 100%.
Bootstrap support for afis|EAW23795.1 as seed ortholog is 100%.

---

Group of orthologs #6510. Best score 621 bits
Score difference with first non-orthologous sequence - *A. fumigatus*:621 *A. fischeri*:621
afum|EAL92570.1 100.00% afis|EAW21550.1 100.00%
Bootstrap support for afum|EAL92570.1 as seed ortholog is 100%.
Bootstrap support for afis|EAW21550.1 as seed ortholog is 100%.

Group of orthologs #6511. Best score 621 bits
Score difference with first non-orthologous sequence - *A. fumigatus*:621 *A. fischeri*:621
afum|EAL89462.1 100.00% afis|EAW24818.1 100.00%
Bootstrap support for afum|EAL89462.1 as seed ortholog is 100%.
Bootstrap support for afis|EAW24818.1 as seed ortholog is 100%.

---

Group of orthologs #6512. Best score 620 bits
Score difference with first non-orthologous sequence - *A. fumigatus*:620 *A. fischeri*:620
afum|EAL85622.1 100.00% afis|EAW15725.1 100.00%
Bootstrap support for afum|EAL85622.1 as seed ortholog is 100%.
Bootstrap support for afis|EAW15725.1 as seed ortholog is 100%.

---

Group of orthologs #6513. Best score 620 bits
Score difference with first non-orthologous sequence - *A. fumigatus*:620 *A. fischeri*:620
afum|EAL85561.1 100.00% afis|EAW15789.1 100.00%
Bootstrap support for afum|EAL85561.1 as seed ortholog is 100%.
Bootstrap support for afis|EAW15789.1 as seed ortholog is 100%.

---

Group of orthologs #6514. Best score 620 bits
Score difference with first non-orthologous sequence - *A. fumigatus*:620 *A. fischeri*:620
afum|EAL86704.1 100.00% afis|EAW17286.1 100.00%
Bootstrap support for afum|EAL86704.1 as seed ortholog is 100%.
Bootstrap support for afis|EAW17286.1 as seed ortholog is 100%.

---

Group of orthologs #6515. Best score 620 bits
Score difference with first non-orthologous sequence - *A. fumigatus*:620 *A. fischeri*:517
afum|EAL84978.1 100.00% afis|EAW19931.1 100.00%
Bootstrap support for afum|EAL84978.1 as seed ortholog is 100%.
Bootstrap support for afis|EAW19931.1 as seed ortholog is 100%.

---

Group of orthologs #6516. Best score 620 bits
Score difference with first non-orthologous sequence - *A. fumigatus*:620 *A. fischeri*:620
afum|EAL91201.1 100.00% afis|EAW17364.1 100.00%
Bootstrap support for afum|EAL91201.1 as seed ortholog is 100%.
Bootstrap support for afis|EAW17364.1 as seed ortholog is 100%.

---

Group of orthologs #6517. Best score 620 bits
Score difference with first non-orthologous sequence - *A. fumigatus*:620 *A. fischeri*:620
afum|EAL91780.1 100.00% afis|EAW17971.1 100.00%
Bootstrap support for afum|EAL91780.1 as seed ortholog is 100%.
Bootstrap support for afis|EAW17971.1 as seed ortholog is 100%.

---

Group of orthologs #6518. Best score 620 bits
Score difference with first non-orthologous sequence - *A. fumigatus*:620 *A. fischeri*:620
afum|EAL92148.1 100.00% afis|EAW21130.1 100.00%
Bootstrap support for afum|EAL92148.1 as seed ortholog is 100%.
Bootstrap support for afis|EAW21130.1 as seed ortholog is 100%.

---

Group of orthologs #6519. Best score 620 bits
Score difference with first non-orthologous sequence - *A. fumigatus*:620 *A. fischeri*:620
afum|EAL92623.1 100.00% afis|EAW21604.1 100.00%
Bootstrap support for afum|EAL92623.1 as seed ortholog is 100%.
Bootstrap support for afis|EAW21604.1 as seed ortholog is 100%.

---

Group of orthologs #6520. Best score 620 bits

Score difference with first non-orthologous sequence - *A. fumigatus*:620 *A. fischeri*:620
afum|EAL89514.1 100.00% afis|EAW24864.1 100.00%
Bootstrap support for afum|EAL89514.1 as seed ortholog is 100%.
Bootstrap support for afis|EAW24864.1 as seed ortholog is 100%.

---

Group of orthologs #6521. Best score 620 bits
Score difference with first non-orthologous sequence - *A. fumigatus*:620 *A. fischeri*:620
afum|EAL92967.1 100.00% afis|EAW21953.1 100.00%
Bootstrap support for afum|EAL92967.1 as seed ortholog is 100%.
Bootstrap support for afis|EAW21953.1 as seed ortholog is 100%.

---

Group of orthologs #6522. Best score 620 bits
Score difference with first non-orthologous sequence - *A. fumigatus*:620 *A. fischeri*:620
afum|EAL89928.1 100.00% afis|EAW25265.1 100.00%
Bootstrap support for afum|EAL89928.1 as seed ortholog is 100%.
Bootstrap support for afis|EAW25265.1 as seed ortholog is 100%.

---

Group of orthologs #6523. Best score 619 bits
Score difference with first non-orthologous sequence - *A. fumigatus*:619 *A. fischeri*:444
afum|EAL86581.1 100.00% afis|EAW17119.1 100.00%
Bootstrap support for afum|EAL86581.1 as seed ortholog is 100%.
Bootstrap support for afis|EAW17119.1 as seed ortholog is 100%.

---

Group of orthologs #6524. Best score 619 bits
Score difference with first non-orthologous sequence - *A. fumigatus*:440 *A. fischeri*:329
afum|EAL86612.1 100.00% afis|EAW17186.1 100.00%
Bootstrap support for afum|EAL86612.1 as seed ortholog is 100%.
Bootstrap support for afis|EAW17186.1 as seed ortholog is 100%.

---

Group of orthologs #6525. Best score 619 bits
Score difference with first non-orthologous sequence - *A. fumigatus*:524 *A. fischeri*:518
afum|EAL89221.1 100.00% afis|EAW16615.1 100.00%
Bootstrap support for afum|EAL89221.1 as seed ortholog is 100%.
Bootstrap support for afis|EAW16615.1 as seed ortholog is 100%.

---

Group of orthologs #6526. Best score 619 bits
Score difference with first non-orthologous sequence - *A. fumigatus*:619 *A. fischeri*:619
afum|EAL84748.1 100.00% afis|EAW20885.1 100.00%
Bootstrap support for afum|EAL84748.1 as seed ortholog is 100%.
Bootstrap support for afis|EAW20885.1 as seed ortholog is 100%.

---

Group of orthologs #6527. Best score 619 bits
Score difference with first non-orthologous sequence - *A. fumigatus*:619 *A. fischeri*:619
afum|EAL84684.1 100.00% afis|EAW20950.1 100.00%
Bootstrap support for afum|EAL84684.1 as seed ortholog is 100%.
Bootstrap support for afis|EAW20950.1 as seed ortholog is 100%.

---

Group of orthologs #6528. Best score 619 bits
Score difference with first non-orthologous sequence - *A. fumigatus*:619 *A. fischeri*:619
afum|EAL87064.1 100.00% afis|EAW19430.1 100.00%
Bootstrap support for afum|EAL87064.1 as seed ortholog is 100%.
Bootstrap support for afis|EAW19430.1 as seed ortholog is 100%.

---

Group of orthologs #6529. Best score 619 bits
Score difference with first non-orthologous sequence - *A. fumigatus*:619 *A. fischeri*:619
afum|EAL91667.1 100.00% afis|EAW17859.1 100.00%

Bootstrap support for afum|EAL91667.1 as seed ortholog is 100%.
Bootstrap support for afis|EAW17859.1 as seed ortholog is 100%.

---

Group of orthologs #6530. Best score 619 bits
Score difference with first non-orthologous sequence - *A. fumigatus*:619 *A. fischeri*:619
afum|EAL88461.1 100.00% afis|EAW23021.1 100.00%
Bootstrap support for afum|EAL88461.1 as seed ortholog is 100%.
Bootstrap support for afis|EAW23021.1 as seed ortholog is 100%.

---

Group of orthologs #6531. Best score 619 bits
Score difference with first non-orthologous sequence - *A. fumigatus*:619 *A. fischeri*:619
afum|EAL87997.1 100.00% afis|EAW23469.1 100.00%
Bootstrap support for afum|EAL87997.1 as seed ortholog is 100%.
Bootstrap support for afis|EAW23469.1 as seed ortholog is 100%.

---

Group of orthologs #6532. Best score 619 bits
Score difference with first non-orthologous sequence - *A. fumigatus*:619 *A. fischeri*:619
afum|EAL93864.1 100.00% afis|EAW19132.1 100.00%
Bootstrap support for afum|EAL93864.1 as seed ortholog is 100%.
Bootstrap support for afis|EAW19132.1 as seed ortholog is 100%.

---

Group of orthologs #6533. Best score 619 bits
Score difference with first non-orthologous sequence - *A. fumigatus*:619 *A. fischeri*:619
afum|EAL90620.2 100.00% afis|EAW22578.1 100.00%
Bootstrap support for afum|EAL90620.2 as seed ortholog is 100%.
Bootstrap support for afis|EAW22578.1 as seed ortholog is 100%.

---

Group of orthologs #6534. Best score 619 bits
Score difference with first non-orthologous sequence - *A. fumigatus*:619 *A. fischeri*:619
afum|EAL92333.1 100.00% afis|EAW21316.1 100.00%
Bootstrap support for afum|EAL92333.1 as seed ortholog is 100%.
Bootstrap support for afis|EAW21316.1 as seed ortholog is 100%.

---

Group of orthologs #6535. Best score 619 bits
Score difference with first non-orthologous sequence - *A. fumigatus*:619 *A. fischeri*:619
afum|EAL92817.1 100.00% afis|EAW21792.1 100.00%
Bootstrap support for afum|EAL92817.1 as seed ortholog is 100%.
Bootstrap support for afis|EAW21792.1 as seed ortholog is 100%.

---

Group of orthologs #6536. Best score 618 bits
Score difference with first non-orthologous sequence - *A. fumigatus*:618 *A. fischeri*:618
afum|EAL88747.1 100.00% afis|EAW16116.1 100.00%
Bootstrap support for afum|EAL88747.1 as seed ortholog is 100%.
Bootstrap support for afis|EAW16116.1 as seed ortholog is 100%.

---

Group of orthologs #6537. Best score 618 bits
Score difference with first non-orthologous sequence - *A. fumigatus*:618 *A. fischeri*:618
afum|EAL85290.1 100.00% afis|EAW20172.1 100.00%
Bootstrap support for afum|EAL85290.1 as seed ortholog is 100%.
Bootstrap support for afis|EAW20172.1 as seed ortholog is 100%.

---

Group of orthologs #6538. Best score 618 bits
Score difference with first non-orthologous sequence - *A. fumigatus*:618 *A. fischeri*:618
afum|EAL87721.1 100.00% afis|EAW18357.1 100.00%
Bootstrap support for afum|EAL87721.1 as seed ortholog is 100%.
Bootstrap support for afis|EAW18357.1 as seed ortholog is 100%.

Group of orthologs #6539. Best score 618 bits
Score difference with first non-orthologous sequence - *A. fumigatus*:454 *A. fischeri*:424
afum|EAL91234.2 100.00% afis|EAW17396.1 100.00%
Bootstrap support for afum|EAL91234.2 as seed ortholog is 100%.
Bootstrap support for afis|EAW17396.1 as seed ortholog is 100%.

---

Group of orthologs #6540. Best score 618 bits
Score difference with first non-orthologous sequence - *A. fumigatus*:618 *A. fischeri*:618
afum|EAL91327.1 100.00% afis|EAW17505.1 100.00%
Bootstrap support for afum|EAL91327.1 as seed ortholog is 100%.
Bootstrap support for afis|EAW17505.1 as seed ortholog is 100%.

---

Group of orthologs #6541. Best score 618 bits
Score difference with first non-orthologous sequence - *A. fumigatus*:618 *A. fischeri*:618
afum|EAL93154.1 100.00% afis|EAW18468.1 100.00%
Bootstrap support for afum|EAL93154.1 as seed ortholog is 100%.
Bootstrap support for afis|EAW18468.1 as seed ortholog is 100%.

---

Group of orthologs #6542. Best score 618 bits
Score difference with first non-orthologous sequence - *A. fumigatus*:618 *A. fischeri*:618
afum|EAL89806.2 100.00% afis|EAW25135.1 100.00%
Bootstrap support for afum|EAL89806.2 as seed ortholog is 100%.
Bootstrap support for afis|EAW25135.1 as seed ortholog is 100%.

---

Group of orthologs #6543. Best score 617 bits
Score difference with first non-orthologous sequence - *A. fumigatus*:617 *A. fischeri*:617
afum|EAL89029.2 100.00% afis|EAW16398.1 100.00%
Bootstrap support for afum|EAL89029.2 as seed ortholog is 100%.
Bootstrap support for afis|EAW16398.1 as seed ortholog is 100%.

---

Group of orthologs #6544. Best score 617 bits
Score difference with first non-orthologous sequence - *A. fumigatus*:617 *A. fischeri*:617
afum|EAL85229.1 100.00% afis|EAW20112.1 100.00%
Bootstrap support for afum|EAL85229.1 as seed ortholog is 100%.
Bootstrap support for afis|EAW20112.1 as seed ortholog is 100%.

---

Group of orthologs #6545. Best score 617 bits
Score difference with first non-orthologous sequence - *A. fumigatus*:617 *A. fischeri*:617
afum|EAL87277.1 100.00% afis|EAW23849.1 100.00%
Bootstrap support for afum|EAL87277.1 as seed ortholog is 100%.
Bootstrap support for afis|EAW23849.1 as seed ortholog is 100%.

---

Group of orthologs #6546. Best score 617 bits
Score difference with first non-orthologous sequence - *A. fumigatus*:543 *A. fischeri*:445
afum|EAL87391.2 100.00% afis|EAW23962.1 100.00%
Bootstrap support for afum|EAL87391.2 as seed ortholog is 100%.
Bootstrap support for afis|EAW23962.1 as seed ortholog is 100%.

---

Group of orthologs #6547. Best score 617 bits
Score difference with first non-orthologous sequence - *A. fumigatus*:617 *A. fischeri*:617
afum|EAL93909.1 100.00% afis|EAW19179.1 100.00%
Bootstrap support for afum|EAL93909.1 as seed ortholog is 100%.
Bootstrap support for afis|EAW19179.1 as seed ortholog is 100%.

---

Group of orthologs #6548. Best score 616 bits

Score difference with first non-orthologous sequence - *A. fumigatus*:616   *A. fischeri*:616
afum|EAL86577.1   100.00%   afis|EAW17115.1   100.00%
Bootstrap support for afum|EAL86577.1 as seed ortholog is 100%.
Bootstrap support for afis|EAW17115.1 as seed ortholog is 100%.

---

Group of orthologs #6549. Best score 616 bits
Score difference with first non-orthologous sequence - *A. fumigatus*:616   *A. fischeri*:616
afum|EAL88853.1   100.00%   afis|EAW16230.1   100.00%
Bootstrap support for afum|EAL88853.1 as seed ortholog is 100%.
Bootstrap support for afis|EAW16230.1 as seed ortholog is 100%.

---

Group of orthologs #6550. Best score 616 bits
Score difference with first non-orthologous sequence - *A. fumigatus*:616   *A. fischeri*:616
afum|EAL85120.1   100.00%   afis|EAW19775.1   100.00%
Bootstrap support for afum|EAL85120.1 as seed ortholog is 100%.
Bootstrap support for afis|EAW19775.1 as seed ortholog is 100%.

---

Group of orthologs #6551. Best score 616 bits
Score difference with first non-orthologous sequence - *A. fumigatus*:616   *A. fischeri*:616
afum|EAL91873.1   100.00%   afis|EAW18062.1   100.00%
Bootstrap support for afum|EAL91873.1 as seed ortholog is 100%.
Bootstrap support for afis|EAW18062.1 as seed ortholog is 100%.

---

Group of orthologs #6552. Best score 616 bits
Score difference with first non-orthologous sequence - *A. fumigatus*:616   *A. fischeri*:616
afum|EAL87345.1   100.00%   afis|EAW23919.1   100.00%
Bootstrap support for afum|EAL87345.1 as seed ortholog is 100%.
Bootstrap support for afis|EAW23919.1 as seed ortholog is 100%.

---

Group of orthologs #6553. Best score 616 bits
Score difference with first non-orthologous sequence - *A. fumigatus*:616   *A. fischeri*:616
afum|EAL93298.1   100.00%   afis|EAW18591.1   100.00%
Bootstrap support for afum|EAL93298.1 as seed ortholog is 100%.
Bootstrap support for afis|EAW18591.1 as seed ortholog is 100%.

---

Group of orthologs #6554. Best score 616 bits
Score difference with first non-orthologous sequence - *A. fumigatus*:616   *A. fischeri*:616
afum|EAL90835.1   100.00%   afis|EAW22356.1   100.00%
Bootstrap support for afum|EAL90835.1 as seed ortholog is 100%.
Bootstrap support for afis|EAW22356.1 as seed ortholog is 100%.

---

Group of orthologs #6555. Best score 616 bits
Score difference with first non-orthologous sequence - *A. fumigatus*:616   *A. fischeri*:616
afum|EAL91933.1   100.00%   afis|EAW24078.1   100.00%
Bootstrap support for afum|EAL91933.1 as seed ortholog is 100%.
Bootstrap support for afis|EAW24078.1 as seed ortholog is 100%.

---

Group of orthologs #6556. Best score 616 bits
Score difference with first non-orthologous sequence - *A. fumigatus*:616   *A. fischeri*:616
afum|EAL91977.1   100.00%   afis|EAW24121.1   100.00%
Bootstrap support for afum|EAL91977.1 as seed ortholog is 100%.
Bootstrap support for afis|EAW24121.1 as seed ortholog is 100%.

---

Group of orthologs #6557. Best score 615 bits
Score difference with first non-orthologous sequence - *A. fumigatus*:615   *A. fischeri*:615
afum|EAL85471.1   100.00%   afis|EAW20353.1   100.00%

Bootstrap support for afum|EAL85471.1 as seed ortholog is 100%.
Bootstrap support for afis|EAW20353.1 as seed ortholog is 100%.

---

Group of orthologs #6558. Best score 615 bits
Score difference with first non-orthologous sequence - *A. fumigatus*:615 *A. fischeri*:615
afum|EAL91720.1 100.00% afis|EAW17911.1 100.00%
Bootstrap support for afum|EAL91720.1 as seed ortholog is 100%.
Bootstrap support for afis|EAW17911.1 as seed ortholog is 100%.

---

Group of orthologs #6559. Best score 615 bits
Score difference with first non-orthologous sequence - *A. fumigatus*:615 *A. fischeri*:615
afum|EAL93816.1 100.00% afis|EAW19086.1 100.00%
Bootstrap support for afum|EAL93816.1 as seed ortholog is 100%.
Bootstrap support for afis|EAW19086.1 as seed ortholog is 100%.

---

Group of orthologs #6560. Best score 615 bits
Score difference with first non-orthologous sequence - *A. fumigatus*:615 *A. fischeri*:615
afum|EAL90640.1 100.00% afis|EAW22555.1 100.00%
Bootstrap support for afum|EAL90640.1 as seed ortholog is 100%.
Bootstrap support for afis|EAW22555.1 as seed ortholog is 100%.

---

Group of orthologs #6561. Best score 615 bits
Score difference with first non-orthologous sequence - *A. fumigatus*:615 *A. fischeri*:615
afum|EAL92158.2 100.00% afis|EAW21139.1 100.00%
Bootstrap support for afum|EAL92158.2 as seed ortholog is 100%.
Bootstrap support for afis|EAW21139.1 as seed ortholog is 100%.

---

Group of orthologs #6562. Best score 615 bits
Score difference with first non-orthologous sequence - *A. fumigatus*:615 *A. fischeri*:615
afum|EAL92475.1 100.00% afis|EAW21456.1 100.00%
Bootstrap support for afum|EAL92475.1 as seed ortholog is 100%.
Bootstrap support for afis|EAW21456.1 as seed ortholog is 100%.

---

Group of orthologs #6563. Best score 615 bits
Score difference with first non-orthologous sequence - *A. fumigatus*:266 *A. fischeri*:615
afum|EAL89569.1 100.00% afis|EAW24915.1 100.00%
Bootstrap support for afum|EAL89569.1 as seed ortholog is 100%.
Bootstrap support for afis|EAW24915.1 as seed ortholog is 100%.

---

Group of orthologs #6564. Best score 615 bits
Score difference with first non-orthologous sequence - *A. fumigatus*:615 *A. fischeri*:615
afum|EAL91959.1 100.00% afis|EAW24105.1 100.00%
Bootstrap support for afum|EAL91959.1 as seed ortholog is 100%.
Bootstrap support for afis|EAW24105.1 as seed ortholog is 100%.

---

Group of orthologs #6565. Best score 615 bits
Score difference with first non-orthologous sequence - *A. fumigatus*:615 *A. fischeri*:615
afum|EBA27346.1 100.00% afis|EAW23019.1 100.00%
Bootstrap support for afum|EBA27346.1 as seed ortholog is 100%.
Bootstrap support for afis|EAW23019.1 as seed ortholog is 100%.

---

Group of orthologs #6566. Best score 614 bits
Score difference with first non-orthologous sequence - *A. fumigatus*:614 *A. fischeri*:614
afum|EAL88749.1 100.00% afis|EAW16118.1 100.00%
Bootstrap support for afum|EAL88749.1 as seed ortholog is 100%.
Bootstrap support for afis|EAW16118.1 as seed ortholog is 100%.

Group of orthologs #6567. Best score 614 bits
Score difference with first non-orthologous sequence - *A. fumigatus*:564 *A. fischeri*:614
afum|EAL85322.1 100.00% afis|EAW20202.1 100.00%
Bootstrap support for afum|EAL85322.1 as seed ortholog is 100%.
Bootstrap support for afis|EAW20202.1 as seed ortholog is 100%.

---

Group of orthologs #6568. Best score 614 bits
Score difference with first non-orthologous sequence - *A. fumigatus*:614 *A. fischeri*:614
afum|EAL86072.1 100.00% afis|EAW24401.1 100.00%
Bootstrap support for afum|EAL86072.1 as seed ortholog is 100%.
Bootstrap support for afis|EAW24401.1 as seed ortholog is 100%.

---

Group of orthologs #6569. Best score 614 bits
Score difference with first non-orthologous sequence - *A. fumigatus*:614 *A. fischeri*:614
afum|EAL87259.1 100.00% afis|EAW23827.1 100.00%
Bootstrap support for afum|EAL87259.1 as seed ortholog is 100%.
Bootstrap support for afis|EAW23827.1 as seed ortholog is 100%.

---

Group of orthologs #6570. Best score 614 bits
Score difference with first non-orthologous sequence - *A. fumigatus*:512 *A. fischeri*:451
afum|EAL91031.1 100.00% afis|EAW22159.1 100.00%
Bootstrap support for afum|EAL91031.1 as seed ortholog is 100%.
Bootstrap support for afis|EAW22159.1 as seed ortholog is 100%.

---

Group of orthologs #6571. Best score 614 bits
Score difference with first non-orthologous sequence - *A. fumigatus*:614 *A. fischeri*:614
afum|EAL90989.1 100.00% afis|EAW22203.1 100.00%
Bootstrap support for afum|EAL90989.1 as seed ortholog is 100%.
Bootstrap support for afis|EAW22203.1 as seed ortholog is 100%.

---

Group of orthologs #6572. Best score 614 bits
Score difference with first non-orthologous sequence - *A. fumigatus*:614 *A. fischeri*:614
afum|EAL90598.1 100.00% afis|EAW22600.1 100.00%
Bootstrap support for afum|EAL90598.1 as seed ortholog is 100%.
Bootstrap support for afis|EAW22600.1 as seed ortholog is 100%.

---

Group of orthologs #6573. Best score 614 bits
Score difference with first non-orthologous sequence - *A. fumigatus*:614 *A. fischeri*:488
afum|EAL89761.2 100.00% afis|EAW25091.1 100.00%
Bootstrap support for afum|EAL89761.2 as seed ortholog is 100%.
Bootstrap support for afis|EAW25091.1 as seed ortholog is 100%.

---

Group of orthologs #6574. Best score 613 bits
Score difference with first non-orthologous sequence - *A. fumigatus*:613 *A. fischeri*:613
afum|EAL88973.1 100.00% afis|EAW16348.1 100.00%
Bootstrap support for afum|EAL88973.1 as seed ortholog is 100%.
Bootstrap support for afis|EAW16348.1 as seed ortholog is 100%.

---

Group of orthologs #6575. Best score 613 bits
Score difference with first non-orthologous sequence - *A. fumigatus*:560 *A. fischeri*:508
afum|EAL86811.1 100.00% afis|EAW19716.1 100.00%
Bootstrap support for afum|EAL86811.1 as seed ortholog is 100%.
Bootstrap support for afis|EAW19716.1 as seed ortholog is 100%.

---

Group of orthologs #6576. Best score 613 bits

Score difference with first non-orthologous sequence - *A. fumigatus*:613 *A. fischeri*:525
afum|EAL87389.1 100.00% afis|EAW23960.1 100.00%
Bootstrap support for afum|EAL87389.1 as seed ortholog is 100%.
Bootstrap support for afis|EAW23960.1 as seed ortholog is 100%.

---

Group of orthologs #6577. Best score 613 bits
Score difference with first non-orthologous sequence - *A. fumigatus*:613 *A. fischeri*:252
afum|EAL89343.1 100.00% afis|EAW24701.1 100.00%
Bootstrap support for afum|EAL89343.1 as seed ortholog is 100%.
Bootstrap support for afis|EAW24701.1 as seed ortholog is 100%.

---

Group of orthologs #6578. Best score 613 bits
Score difference with first non-orthologous sequence - *A. fumigatus*:613 *A. fischeri*:613
afum|EAL91978.1 100.00% afis|EAW24122.1 100.00%
Bootstrap support for afum|EAL91978.1 as seed ortholog is 100%.
Bootstrap support for afis|EAW24122.1 as seed ortholog is 100%.

---

Group of orthologs #6579. Best score 612 bits
Score difference with first non-orthologous sequence - *A. fumigatus*:612 *A. fischeri*:612
afum|EAL86966.1 100.00% afis|EAW19542.1 100.00%
Bootstrap support for afum|EAL86966.1 as seed ortholog is 100%.
Bootstrap support for afis|EAW19542.1 as seed ortholog is 100%.

---

Group of orthologs #6580. Best score 612 bits
Score difference with first non-orthologous sequence - *A. fumigatus*:612 *A. fischeri*:612
afum|EAL85874.2 100.00% afis|EAW24204.1 100.00%
Bootstrap support for afum|EAL85874.2 as seed ortholog is 100%.
Bootstrap support for afis|EAW24204.1 as seed ortholog is 100%.

---

Group of orthologs #6581. Best score 612 bits
Score difference with first non-orthologous sequence - *A. fumigatus*:612 *A. fischeri*:612
afum|EAL92466.1 100.00% afis|EAW21447.1 100.00%
Bootstrap support for afum|EAL92466.1 as seed ortholog is 100%.
Bootstrap support for afis|EAW21447.1 as seed ortholog is 100%.

---

Group of orthologs #6582. Best score 612 bits
Score difference with first non-orthologous sequence - *A. fumigatus*:612 *A. fischeri*:612
afum|EAL89540.1 100.00% afis|EAW24889.1 100.00%
Bootstrap support for afum|EAL89540.1 as seed ortholog is 100%.
Bootstrap support for afis|EAW24889.1 as seed ortholog is 100%.

---

Group of orthologs #6583. Best score 611 bits
Score difference with first non-orthologous sequence - *A. fumigatus*:611 *A. fischeri*:485
afum|EAL91516.1 100.00% afis|EAW17701.1 100.00%
Bootstrap support for afum|EAL91516.1 as seed ortholog is 100%.
Bootstrap support for afis|EAW17701.1 as seed ortholog is 100%.

---

Group of orthologs #6584. Best score 611 bits
Score difference with first non-orthologous sequence - *A. fumigatus*:611 *A. fischeri*:611
afum|EAL93879.1 100.00% afis|EAW19148.1 100.00%
Bootstrap support for afum|EAL93879.1 as seed ortholog is 100%.
Bootstrap support for afis|EAW19148.1 as seed ortholog is 100%.

---

Group of orthologs #6585. Best score 611 bits
Score difference with first non-orthologous sequence - *A. fumigatus*:503 *A. fischeri*:512
afum|EAL90770.1 100.00% afis|EAW22422.1 100.00%

Bootstrap support for afum|EAL90770.1 as seed ortholog is 100%.
Bootstrap support for afis|EAW22422.1 as seed ortholog is 100%.

---

Group of orthologs #6586. Best score 611 bits
Score difference with first non-orthologous sequence - *A. fumigatus*:526 *A. fischeri*:457
afum|EAL94078.1 100.00% afis|EAW19358.1 100.00%
Bootstrap support for afum|EAL94078.1 as seed ortholog is 100%.
Bootstrap support for afis|EAW19358.1 as seed ortholog is 100%.

---

Group of orthologs #6587. Best score 611 bits
Score difference with first non-orthologous sequence - *A. fumigatus*:611 *A. fischeri*:611
afum|EAL92783.1 100.00% afis|EAW21753.1 100.00%
Bootstrap support for afum|EAL92783.1 as seed ortholog is 100%.
Bootstrap support for afis|EAW21753.1 as seed ortholog is 100%.

---

Group of orthologs #6588. Best score 611 bits
Score difference with first non-orthologous sequence - *A. fumigatus*:611 *A. fischeri*:611
afum|EAL89959.2 100.00% afis|EAW25296.1 100.00%
Bootstrap support for afum|EAL89959.2 as seed ortholog is 100%.
Bootstrap support for afis|EAW25296.1 as seed ortholog is 100%.

---

Group of orthologs #6589. Best score 610 bits
Score difference with first non-orthologous sequence - *A. fumigatus*:443 *A. fischeri*:406
afum|EAL84503.1 100.00% afis|EAW20558.1 100.00%
Bootstrap support for afum|EAL84503.1 as seed ortholog is 100%.
Bootstrap support for afis|EAW20558.1 as seed ortholog is 100%.

---

Group of orthologs #6590. Best score 610 bits
Score difference with first non-orthologous sequence - *A. fumigatus*:610 *A. fischeri*:610
afum|EAL84809.1 100.00% afis|EAW20820.1 100.00%
Bootstrap support for afum|EAL84809.1 as seed ortholog is 100%.
Bootstrap support for afis|EAW20820.1 as seed ortholog is 100%.

---

Group of orthologs #6591. Best score 610 bits
Score difference with first non-orthologous sequence - *A. fumigatus*:610 *A. fischeri*:610
afum|EAL87603.1 100.00% afis|EAW18236.1 100.00%
Bootstrap support for afum|EAL87603.1 as seed ortholog is 100%.
Bootstrap support for afis|EAW18236.1 as seed ortholog is 100%.

---

Group of orthologs #6592. Best score 610 bits
Score difference with first non-orthologous sequence - *A. fumigatus*:610 *A. fischeri*:610
afum|EAL86890.1 100.00% afis|EAW19632.1 100.00%
Bootstrap support for afum|EAL86890.1 as seed ortholog is 100%.
Bootstrap support for afis|EAW19632.1 as seed ortholog is 100%.

---

Group of orthologs #6593. Best score 610 bits
Score difference with first non-orthologous sequence - *A. fumigatus*:610 *A. fischeri*:610
afum|EBA27237.1 100.00% afis|EAW15734.1 100.00%
Bootstrap support for afum|EBA27237.1 as seed ortholog is 100%.
Bootstrap support for afis|EAW15734.1 as seed ortholog is 100%.

---

Group of orthologs #6594. Best score 610 bits
Score difference with first non-orthologous sequence - *A. fumigatus*:610 *A. fischeri*:610
afum|EAL91866.1 100.00% afis|EAW18055.1 100.00%
Bootstrap support for afum|EAL91866.1 as seed ortholog is 100%.
Bootstrap support for afis|EAW18055.1 as seed ortholog is 100%.

Group of orthologs #6595. Best score 610 bits
Score difference with first non-orthologous sequence - *A. fumigatus*:610 *A. fischeri*:610
afum|EAL90063.1 100.00% afis|EAW25404.1 100.00%
Bootstrap support for afum|EAL90063.1 as seed ortholog is 100%.
Bootstrap support for afis|EAW25404.1 as seed ortholog is 100%.

---

Group of orthologs #6596. Best score 609 bits
Score difference with first non-orthologous sequence - *A. fumigatus*:609 *A. fischeri*:609
afum|EAL84746.1 100.00% afis|EAW20887.1 100.00%
Bootstrap support for afum|EAL84746.1 as seed ortholog is 100%.
Bootstrap support for afis|EAW20887.1 as seed ortholog is 100%.

---

Group of orthologs #6597. Best score 609 bits
Score difference with first non-orthologous sequence - *A. fumigatus*:609 *A. fischeri*:609
afum|EAL91310.1 100.00% afis|EAW17487.1 100.00%
Bootstrap support for afum|EAL91310.1 as seed ortholog is 100%.
Bootstrap support for afis|EAW17487.1 as seed ortholog is 100%.

---

Group of orthologs #6598. Best score 609 bits
Score difference with first non-orthologous sequence - *A. fumigatus*:609 *A. fischeri*:609
afum|EAL87916.1 100.00% afis|EAW23550.1 100.00%
Bootstrap support for afum|EAL87916.1 as seed ortholog is 100%.
Bootstrap support for afis|EAW23550.1 as seed ortholog is 100%.

---

Group of orthologs #6599. Best score 609 bits
Score difference with first non-orthologous sequence - *A. fumigatus*:609 *A. fischeri*:609
afum|EAL90301.1 100.00% afis|EAW22897.1 100.00%
Bootstrap support for afum|EAL90301.1 as seed ortholog is 100%.
Bootstrap support for afis|EAW22897.1 as seed ortholog is 100%.

---

Group of orthologs #6600. Best score 608 bits
Score difference with first non-orthologous sequence - *A. fumigatus*:265 *A. fischeri*:608
afum|EAL86525.1 100.00% afis|EAW17000.1 100.00%
Bootstrap support for afum|EAL86525.1 as seed ortholog is 100%.
Bootstrap support for afis|EAW17000.1 as seed ortholog is 100%.

---

Group of orthologs #6601. Best score 608 bits
Score difference with first non-orthologous sequence - *A. fumigatus*:608 *A. fischeri*:608
afum|EAL88857.1 100.00% afis|EAW16236.1 100.00%
Bootstrap support for afum|EAL88857.1 as seed ortholog is 100%.
Bootstrap support for afis|EAW16236.1 as seed ortholog is 100%.

---

Group of orthologs #6602. Best score 608 bits
Score difference with first non-orthologous sequence - *A. fumigatus*:608 *A. fischeri*:153
afum|EAL84959.1 100.00% afis|EAW19955.1 100.00%
Bootstrap support for afum|EAL84959.1 as seed ortholog is 100%.
Bootstrap support for afis|EAW19955.1 as seed ortholog is 99%.

---

Group of orthologs #6603. Best score 608 bits
Score difference with first non-orthologous sequence - *A. fumigatus*:608 *A. fischeri*:608
afum|EAL84442.1 100.00% afis|EAW20628.1 100.00%
Bootstrap support for afum|EAL84442.1 as seed ortholog is 100%.
Bootstrap support for afis|EAW20628.1 as seed ortholog is 100%.

---

Group of orthologs #6604. Best score 608 bits

Score difference with first non-orthologous sequence - *A. fumigatus*:608 *A. fischeri*:608
afum|EAL91470.1 100.00% afis|EAW17655.1 100.00%
Bootstrap support for afum|EAL91470.1 as seed ortholog is 100%.
Bootstrap support for afis|EAW17655.1 as seed ortholog is 100%.

---

Group of orthologs #6605. Best score 608 bits
Score difference with first non-orthologous sequence - *A. fumigatus*:608 *A. fischeri*:608
afum|EAL87390.1 100.00% afis|EAW23961.1 100.00%
Bootstrap support for afum|EAL87390.1 as seed ortholog is 100%.
Bootstrap support for afis|EAW23961.1 as seed ortholog is 100%.

---

Group of orthologs #6606. Best score 608 bits
Score difference with first non-orthologous sequence - *A. fumigatus*:608 *A. fischeri*:608
afum|EAL87998.1 100.00% afis|EAW23468.1 100.00%
Bootstrap support for afum|EAL87998.1 as seed ortholog is 100%.
Bootstrap support for afis|EAW23468.1 as seed ortholog is 100%.

---

Group of orthologs #6607. Best score 608 bits
Score difference with first non-orthologous sequence - *A. fumigatus*:546 *A. fischeri*:526
afum|EAL88037.1 100.00% afis|EAW24602.1 100.00%
Bootstrap support for afum|EAL88037.1 as seed ortholog is 100%.
Bootstrap support for afis|EAW24602.1 as seed ortholog is 100%.

---

Group of orthologs #6608. Best score 608 bits
Score difference with first non-orthologous sequence - *A. fumigatus*:608 *A. fischeri*:608
afum|EAL92451.1 100.00% afis|EAW21436.1 100.00%
Bootstrap support for afum|EAL92451.1 as seed ortholog is 100%.
Bootstrap support for afis|EAW21436.1 as seed ortholog is 100%.

---

Group of orthologs #6609. Best score 608 bits
Score difference with first non-orthologous sequence - *A. fumigatus*:608 *A. fischeri*:608
afum|EAL92720.1 100.00% afis|EAW21686.1 100.00%
Bootstrap support for afum|EAL92720.1 as seed ortholog is 100%.
Bootstrap support for afis|EAW21686.1 as seed ortholog is 100%.

---

Group of orthologs #6610. Best score 608 bits
Score difference with first non-orthologous sequence - *A. fumigatus*:608 *A. fischeri*:608
afum|EAL92749.1 100.00% afis|EAW21714.1 100.00%
Bootstrap support for afum|EAL92749.1 as seed ortholog is 100%.
Bootstrap support for afis|EAW21714.1 as seed ortholog is 100%.

---

Group of orthologs #6611. Best score 608 bits
Score difference with first non-orthologous sequence - *A. fumigatus*:608 *A. fischeri*:608
afum|EAL93022.1 100.00% afis|EAW22007.1 100.00%
Bootstrap support for afum|EAL93022.1 as seed ortholog is 100%.
Bootstrap support for afis|EAW22007.1 as seed ortholog is 100%.

---

Group of orthologs #6612. Best score 607 bits
Score difference with first non-orthologous sequence - *A. fumigatus*:607 *A. fischeri*:607
afum|EAL87972.1 100.00% afis|EAW23491.1 100.00%
Bootstrap support for afum|EAL87972.1 as seed ortholog is 100%.
Bootstrap support for afis|EAW23491.1 as seed ortholog is 100%.

---

Group of orthologs #6613. Best score 606 bits
Score difference with first non-orthologous sequence - *A. fumigatus*:606 *A. fischeri*:606
afum|EAL88547.2 100.00% afis|EAW15912.1 100.00%

Bootstrap support for afum|EAL88547.2 as seed ortholog is 100%.
Bootstrap support for afis|EAW15912.1 as seed ortholog is 100%.

---

Group of orthologs #6614. Best score 606 bits
Score difference with first non-orthologous sequence - *A. fumigatus*:606 *A. fischeri*:557
afum|EAL84984.2 100.00% afis|EAW19925.1 100.00%
Bootstrap support for afum|EAL84984.2 as seed ortholog is 100%.
Bootstrap support for afis|EAW19925.1 as seed ortholog is 100%.

---

Group of orthologs #6615. Best score 606 bits
Score difference with first non-orthologous sequence - *A. fumigatus*:606 *A. fischeri*:606
afum|EAL84895.1 100.00% afis|EAW20022.1 100.00%
Bootstrap support for afum|EAL84895.1 as seed ortholog is 100%.
Bootstrap support for afis|EAW20022.1 as seed ortholog is 100%.

---

Group of orthologs #6616. Best score 606 bits
Score difference with first non-orthologous sequence - *A. fumigatus*:606 *A. fischeri*:606
afum|EAL91373.1 100.00% afis|EAW17559.1 100.00%
Bootstrap support for afum|EAL91373.1 as seed ortholog is 100%.
Bootstrap support for afis|EAW17559.1 as seed ortholog is 100%.

---

Group of orthologs #6617. Best score 606 bits
Score difference with first non-orthologous sequence - *A. fumigatus*:606 *A. fischeri*:606
afum|EAL91778.1 100.00% afis|EAW17969.1 100.00%
Bootstrap support for afum|EAL91778.1 as seed ortholog is 100%.
Bootstrap support for afis|EAW17969.1 as seed ortholog is 100%.

---

Group of orthologs #6618. Best score 606 bits
Score difference with first non-orthologous sequence - *A. fumigatus*:606 *A. fischeri*:606
afum|EAL90289.1 100.00% afis|EAW22908.1 100.00%
Bootstrap support for afum|EAL90289.1 as seed ortholog is 100%.
Bootstrap support for afis|EAW22908.1 as seed ortholog is 100%.

---

Group of orthologs #6619. Best score 606 bits
Score difference with first non-orthologous sequence - *A. fumigatus*:606 *A. fischeri*:606
afum|EAL92507.1 100.00% afis|EAW21488.1 100.00%
Bootstrap support for afum|EAL92507.1 as seed ortholog is 100%.
Bootstrap support for afis|EAW21488.1 as seed ortholog is 100%.

---

Group of orthologs #6620. Best score 606 bits
Score difference with first non-orthologous sequence - *A. fumigatus*:606 *A. fischeri*:606
afum|EAL89967.2 100.00% afis|EAW25304.1 100.00%
Bootstrap support for afum|EAL89967.2 as seed ortholog is 100%.
Bootstrap support for afis|EAW25304.1 as seed ortholog is 100%.

---

Group of orthologs #6621. Best score 605 bits
Score difference with first non-orthologous sequence - *A. fumigatus*:605 *A. fischeri*:605
afum|EAL84719.1 100.00% afis|EAW20915.1 100.00%
Bootstrap support for afum|EAL84719.1 as seed ortholog is 100%.
Bootstrap support for afis|EAW20915.1 as seed ortholog is 100%.

---

Group of orthologs #6622. Best score 605 bits
Score difference with first non-orthologous sequence - *A. fumigatus*:605 *A. fischeri*:605
afum|EAL91510.1 100.00% afis|EAW17696.1 100.00%
Bootstrap support for afum|EAL91510.1 as seed ortholog is 100%.
Bootstrap support for afis|EAW17696.1 as seed ortholog is 100%.

---

Group of orthologs #6623. Best score 605 bits
Score difference with first non-orthologous sequence - *A. fumigatus*:605 *A. fischeri*:605

| | | | |
|---|---|---|---|
| afum\|EAL87394.1 | 100.00% | afis\|EAW23964.1 | 100.00% |

Bootstrap support for afum|EAL87394.1 as seed ortholog is 100%.
Bootstrap support for afis|EAW23964.1 as seed ortholog is 100%.

---

Group of orthologs #6624. Best score 605 bits
Score difference with first non-orthologous sequence - *A. fumigatus*:535 *A. fischeri*:530

| | | | |
|---|---|---|---|
| afum\|EAL93741.1 | 100.00% | afis\|EAW19007.1 | 100.00% |

Bootstrap support for afum|EAL93741.1 as seed ortholog is 100%.
Bootstrap support for afis|EAW19007.1 as seed ortholog is 100%.

---

Group of orthologs #6625. Best score 605 bits
Score difference with first non-orthologous sequence - *A. fumigatus*:389 *A. fischeri*:465

| | | | |
|---|---|---|---|
| afum\|EAL91098.2 | 100.00% | afis\|EAW22072.1 | 100.00% |

Bootstrap support for afum|EAL91098.2 as seed ortholog is 100%.
Bootstrap support for afis|EAW22072.1 as seed ortholog is 100%.

---

Group of orthologs #6626. Best score 605 bits
Score difference with first non-orthologous sequence - *A. fumigatus*:605 *A. fischeri*:605

| | | | |
|---|---|---|---|
| afum\|EAL90457.1 | 100.00% | afis\|EAW22738.1 | 100.00% |

Bootstrap support for afum|EAL90457.1 as seed ortholog is 100%.
Bootstrap support for afis|EAW22738.1 as seed ortholog is 100%.

---

Group of orthologs #6627. Best score 604 bits
Score difference with first non-orthologous sequence - *A. fumigatus*:604 *A. fischeri*:604

| | | | |
|---|---|---|---|
| afum\|EAL88660.1 | 100.00% | afis\|EAW16029.1 | 100.00% |

Bootstrap support for afum|EAL88660.1 as seed ortholog is 100%.
Bootstrap support for afis|EAW16029.1 as seed ortholog is 100%.

---

Group of orthologs #6628. Best score 604 bits
Score difference with first non-orthologous sequence - *A. fumigatus*:604 *A. fischeri*:604

| | | | |
|---|---|---|---|
| afum\|EAL88756.1 | 100.00% | afis\|EAW16126.1 | 100.00% |

Bootstrap support for afum|EAL88756.1 as seed ortholog is 100%.
Bootstrap support for afis|EAW16126.1 as seed ortholog is 100%.

---

Group of orthologs #6629. Best score 604 bits
Score difference with first non-orthologous sequence - *A. fumigatus*:604 *A. fischeri*:604

| | | | |
|---|---|---|---|
| afum\|EAL88943.1 | 100.00% | afis\|EAW16319.1 | 100.00% |

Bootstrap support for afum|EAL88943.1 as seed ortholog is 100%.
Bootstrap support for afis|EAW16319.1 as seed ortholog is 100%.

---

Group of orthologs #6630. Best score 604 bits
Score difference with first non-orthologous sequence - *A. fumigatus*:604 *A. fischeri*:604

| | | | |
|---|---|---|---|
| afum\|EAL87515.1 | 100.00% | afis\|EAW18156.1 | 100.00% |

Bootstrap support for afum|EAL87515.1 as seed ortholog is 100%.
Bootstrap support for afis|EAW18156.1 as seed ortholog is 100%.

---

Group of orthologs #6631. Best score 604 bits
Score difference with first non-orthologous sequence - *A. fumigatus*:496 *A. fischeri*:468

| | | | |
|---|---|---|---|
| afum\|EAL86018.1 | 100.00% | afis\|EAW24348.1 | 100.00% |

Bootstrap support for afum|EAL86018.1 as seed ortholog is 100%.
Bootstrap support for afis|EAW24348.1 as seed ortholog is 100%.

---

Group of orthologs #6632. Best score 604 bits

Score difference with first non-orthologous sequence - *A. fumigatus*:510 *A. fischeri*:304
afum|EAL94026.2 100.00% afis|EAW19309.1 100.00%
Bootstrap support for afum|EAL94026.2 as seed ortholog is 100%.
Bootstrap support for afis|EAW19309.1 as seed ortholog is 100%.

---

Group of orthologs #6633. Best score 604 bits
Score difference with first non-orthologous sequence - *A. fumigatus*:604 *A. fischeri*:604
afum|EAL92047.1 100.00% afis|EAW24187.1 100.00%
Bootstrap support for afum|EAL92047.1 as seed ortholog is 100%.
Bootstrap support for afis|EAW24187.1 as seed ortholog is 100%.

---

Group of orthologs #6634. Best score 603 bits
Score difference with first non-orthologous sequence - *A. fumigatus*:603 *A. fischeri*:603
afum|EAL85763.1 100.00% afis|EAW15570.1 100.00%
Bootstrap support for afum|EAL85763.1 as seed ortholog is 100%.
Bootstrap support for afis|EAW15570.1 as seed ortholog is 100%.

---

Group of orthologs #6635. Best score 603 bits
Score difference with first non-orthologous sequence - *A. fumigatus*:603 *A. fischeri*:603
afum|EAL86404.1 100.00% afis|EAW16865.1 100.00%
Bootstrap support for afum|EAL86404.1 as seed ortholog is 100%.
Bootstrap support for afis|EAW16865.1 as seed ortholog is 100%.

---

Group of orthologs #6636. Best score 603 bits
Score difference with first non-orthologous sequence - *A. fumigatus*:603 *A. fischeri*:603
afum|EAL85465.1 100.00% afis|EAW20345.1 100.00%
Bootstrap support for afum|EAL85465.1 as seed ortholog is 100%.
Bootstrap support for afis|EAW20345.1 as seed ortholog is 100%.

---

Group of orthologs #6637. Best score 603 bits
Score difference with first non-orthologous sequence - *A. fumigatus*:603 *A. fischeri*:438
afum|EAL87122.2 100.00% afis|EAW23679.1 100.00%
Bootstrap support for afum|EAL87122.2 as seed ortholog is 100%.
Bootstrap support for afis|EAW23679.1 as seed ortholog is 100%.

---

Group of orthologs #6638. Best score 603 bits
Score difference with first non-orthologous sequence - *A. fumigatus*:543 *A. fischeri*:603
afum|EAL92176.1 100.00% afis|EAW21158.1 100.00%
Bootstrap support for afum|EAL92176.1 as seed ortholog is 100%.
Bootstrap support for afis|EAW21158.1 as seed ortholog is 100%.

---

Group of orthologs #6639. Best score 603 bits
Score difference with first non-orthologous sequence - *A. fumigatus*:603 *A. fischeri*:543
afum|EAL92239.2 100.00% afis|EAW21227.1 100.00%
Bootstrap support for afum|EAL92239.2 as seed ortholog is 100%.
Bootstrap support for afis|EAW21227.1 as seed ortholog is 100%.

---

Group of orthologs #6640. Best score 603 bits
Score difference with first non-orthologous sequence - *A. fumigatus*:553 *A. fischeri*:532
afum|EAL89315.1 100.00% afis|EAW24655.1 100.00%
Bootstrap support for afum|EAL89315.1 as seed ortholog is 100%.
Bootstrap support for afis|EAW24655.1 as seed ortholog is 100%.

---

Group of orthologs #6641. Best score 603 bits
Score difference with first non-orthologous sequence - *A. fumigatus*:603 *A. fischeri*:603
afum|EAL89546.1 100.00% afis|EAW24893.1 100.00%

Bootstrap support for afum|EAL89546.1 as seed ortholog is 100%.
Bootstrap support for afis|EAW24893.1 as seed ortholog is 100%.

---

Group of orthologs #6642. Best score 603 bits
Score difference with first non-orthologous sequence - *A. fumigatus*:603 *A. fischeri*:282
afum|EAL90002.1 100.00% afis|EAW25339.1 100.00%
Bootstrap support for afum|EAL90002.1 as seed ortholog is 100%.
Bootstrap support for afis|EAW25339.1 as seed ortholog is 100%.

---

Group of orthologs #6643. Best score 603 bits
Score difference with first non-orthologous sequence - *A. fumigatus*:603 *A. fischeri*:603
afum|EAL90058.1 100.00% afis|EAW25399.1 100.00%
Bootstrap support for afum|EAL90058.1 as seed ortholog is 100%.
Bootstrap support for afis|EAW25399.1 as seed ortholog is 100%.

---

Group of orthologs #6644. Best score 602 bits
Score difference with first non-orthologous sequence - *A. fumigatus*:602 *A. fischeri*:602
afum|EAL85397.1 100.00% afis|EAW17757.1 100.00%
Bootstrap support for afum|EAL85397.1 as seed ortholog is 100%.
Bootstrap support for afis|EAW17757.1 as seed ortholog is 100%.

---

Group of orthologs #6645. Best score 602 bits
Score difference with first non-orthologous sequence - *A. fumigatus*:602 *A. fischeri*:602
afum|EAL84472.1 100.00% afis|EAW20589.1 100.00%
Bootstrap support for afum|EAL84472.1 as seed ortholog is 100%.
Bootstrap support for afis|EAW20589.1 as seed ortholog is 100%.

---

Group of orthologs #6646. Best score 602 bits
Score difference with first non-orthologous sequence - *A. fumigatus*:602 *A. fischeri*:602
afum|EAL93686.1 100.00% afis|EAW18951.1 100.00%
Bootstrap support for afum|EAL93686.1 as seed ortholog is 100%.
Bootstrap support for afis|EAW18951.1 as seed ortholog is 100%.

---

Group of orthologs #6647. Best score 602 bits
Score difference with first non-orthologous sequence - *A. fumigatus*:602 *A. fischeri*:602
afum|EAL92989.1 100.00% afis|EAW21974.1 100.00%
Bootstrap support for afum|EAL92989.1 as seed ortholog is 100%.
Bootstrap support for afis|EAW21974.1 as seed ortholog is 100%.

---

Group of orthologs #6648. Best score 601 bits
Score difference with first non-orthologous sequence - *A. fumigatus*:601 *A. fischeri*:601
afum|EAL84664.1 100.00% afis|EAW20970.1 100.00%
Bootstrap support for afum|EAL84664.1 as seed ortholog is 100%.
Bootstrap support for afis|EAW20970.1 as seed ortholog is 100%.

---

Group of orthologs #6649. Best score 601 bits
Score difference with first non-orthologous sequence - *A. fumigatus*:601 *A. fischeri*:452
afum|EAL86229.1 100.00% afis|EAW24566.1 100.00%
Bootstrap support for afum|EAL86229.1 as seed ortholog is 100%.
Bootstrap support for afis|EAW24566.1 as seed ortholog is 100%.

---

Group of orthologs #6650. Best score 601 bits
Score difference with first non-orthologous sequence - *A. fumigatus*:601 *A. fischeri*:601
afum|EAL93839.1 100.00% afis|EAW19108.1 100.00%
Bootstrap support for afum|EAL93839.1 as seed ortholog is 100%.
Bootstrap support for afis|EAW19108.1 as seed ortholog is 100%.

Group of orthologs #6651. Best score 601 bits
Score difference with first non-orthologous sequence - *A. fumigatus*:418 *A. fischeri*:340
afum|EAL94053.1 100.00% afis|EAW19336.1 100.00%
Bootstrap support for afum|EAL94053.1 as seed ortholog is 100%.
Bootstrap support for afis|EAW19336.1 as seed ortholog is 100%.

---

Group of orthologs #6652. Best score 600 bits
Score difference with first non-orthologous sequence - *A. fumigatus*:600 *A. fischeri*:600
afum|EAL86298.1 100.00% afis|EAW16724.1 100.00%
Bootstrap support for afum|EAL86298.1 as seed ortholog is 100%.
Bootstrap support for afis|EAW16724.1 as seed ortholog is 100%.

---

Group of orthologs #6653. Best score 600 bits
Score difference with first non-orthologous sequence - *A. fumigatus*:600 *A. fischeri*:496
afum|EAL92076.2 100.00% afis|EAW21037.1 100.00%
Bootstrap support for afum|EAL92076.2 as seed ortholog is 100%.
Bootstrap support for afis|EAW21037.1 as seed ortholog is 100%.

---

Group of orthologs #6654. Best score 600 bits
Score difference with first non-orthologous sequence - *A. fumigatus*:600 *A. fischeri*:600
afum|EAL90383.1 100.00% afis|EAW22818.1 100.00%
Bootstrap support for afum|EAL90383.1 as seed ortholog is 100%.
Bootstrap support for afis|EAW22818.1 as seed ortholog is 100%.

---

Group of orthologs #6655. Best score 600 bits
Score difference with first non-orthologous sequence - *A. fumigatus*:600 *A. fischeri*:551
afum|EAL94085.1 100.00% afis|EAW19366.1 100.00%
Bootstrap support for afum|EAL94085.1 as seed ortholog is 100%.
Bootstrap support for afis|EAW19366.1 as seed ortholog is 100%.

---

Group of orthologs #6656. Best score 600 bits
Score difference with first non-orthologous sequence - *A. fumigatus*:600 *A. fischeri*:600
afum|EAL92580.1 100.00% afis|EAW21561.1 100.00%
Bootstrap support for afum|EAL92580.1 as seed ortholog is 100%.
Bootstrap support for afis|EAW21561.1 as seed ortholog is 100%.

---

Group of orthologs #6657. Best score 600 bits
Score difference with first non-orthologous sequence - *A. fumigatus*:600 *A. fischeri*:600
afum|EAL89864.1 100.00% afis|EAW25201.1 100.00%
Bootstrap support for afum|EAL89864.1 as seed ortholog is 100%.
Bootstrap support for afis|EAW25201.1 as seed ortholog is 100%.

---

Group of orthologs #6658. Best score 599 bits
Score difference with first non-orthologous sequence - *A. fumigatus*:599 *A. fischeri*:599
afum|EAL88831.1 100.00% afis|EAW16203.1 100.00%
Bootstrap support for afum|EAL88831.1 as seed ortholog is 100%.
Bootstrap support for afis|EAW16203.1 as seed ortholog is 100%.

---

Group of orthologs #6659. Best score 599 bits
Score difference with first non-orthologous sequence - *A. fumigatus*:599 *A. fischeri*:599
afum|EAL85056.1 100.00% afis|EAW19836.1 100.00%
Bootstrap support for afum|EAL85056.1 as seed ortholog is 100%.
Bootstrap support for afis|EAW19836.1 as seed ortholog is 100%.

---

Group of orthologs #6660. Best score 599 bits

Score difference with first non-orthologous sequence - *A. fumigatus*:599 *A. fischeri*:599
afum|EAL84926.1 100.00% afis|EAW19989.1 100.00%
Bootstrap support for afum|EAL84926.1 as seed ortholog is 100%.
Bootstrap support for afis|EAW19989.1 as seed ortholog is 100%.

---

Group of orthologs #6661. Best score 599 bits
Score difference with first non-orthologous sequence - *A. fumigatus*:599 *A. fischeri*:599
afum|EAL91335.1 100.00% afis|EAW17517.1 100.00%
Bootstrap support for afum|EAL91335.1 as seed ortholog is 100%.
Bootstrap support for afis|EAW17517.1 as seed ortholog is 100%.

---

Group of orthologs #6662. Best score 599 bits
Score difference with first non-orthologous sequence - *A. fumigatus*:599 *A. fischeri*:599
afum|EAL91892.1 100.00% afis|EAW18080.1 100.00%
Bootstrap support for afum|EAL91892.1 as seed ortholog is 100%.
Bootstrap support for afis|EAW18080.1 as seed ortholog is 100%.

---

Group of orthologs #6663. Best score 599 bits
Score difference with first non-orthologous sequence - *A. fumigatus*:599 *A. fischeri*:599
afum|EAL92839.1 100.00% afis|EAW21823.1 100.00%
Bootstrap support for afum|EAL92839.1 as seed ortholog is 100%.
Bootstrap support for afis|EAW21823.1 as seed ortholog is 100%.

---

Group of orthologs #6664. Best score 598 bits
Score difference with first non-orthologous sequence - *A. fumigatus*:598 *A. fischeri*:598
afum|EAL84219.1 100.00% afis|EAW15425.1 100.00%
Bootstrap support for afum|EAL84219.1 as seed ortholog is 100%.
Bootstrap support for afis|EAW15425.1 as seed ortholog is 100%.

---

Group of orthologs #6665. Best score 598 bits
Score difference with first non-orthologous sequence - *A. fumigatus*:598 *A. fischeri*:598
afum|EAL84489.1 100.00% afis|EAW20576.1 100.00%
Bootstrap support for afum|EAL84489.1 as seed ortholog is 100%.
Bootstrap support for afis|EAW20576.1 as seed ortholog is 100%.

---

Group of orthologs #6666. Best score 598 bits
Score difference with first non-orthologous sequence - *A. fumigatus*:598 *A. fischeri*:598
afum|EAL86823.1 100.00% afis|EAW19703.1 100.00%
Bootstrap support for afum|EAL86823.1 as seed ortholog is 100%.
Bootstrap support for afis|EAW19703.1 as seed ortholog is 100%.

---

Group of orthologs #6667. Best score 598 bits
Score difference with first non-orthologous sequence - *A. fumigatus*:598 *A. fischeri*:598
afum|EAL91331.1 100.00% afis|EAW17512.1 100.00%
Bootstrap support for afum|EAL91331.1 as seed ortholog is 100%.
Bootstrap support for afis|EAW17512.1 as seed ortholog is 100%.

---

Group of orthologs #6668. Best score 598 bits
Score difference with first non-orthologous sequence - *A. fumigatus*:146 *A. fischeri*:412
afum|EAL89354.1 100.00% afis|EAW20704.1 100.00%
Bootstrap support for afum|EAL89354.1 as seed ortholog is 97%.
Bootstrap support for afis|EAW20704.1 as seed ortholog is 100%.

---

Group of orthologs #6669. Best score 598 bits
Score difference with first non-orthologous sequence - *A. fumigatus*:598 *A. fischeri*:598
afum|EAL86089.1 100.00% afis|EAW24420.1 100.00%

Bootstrap support for afum|EAL86089.1 as seed ortholog is 100%.
Bootstrap support for afis|EAW24420.1 as seed ortholog is 100%.

---

Group of orthologs #6670. Best score 598 bits
Score difference with first non-orthologous sequence - *A. fumigatus*:598 *A. fischeri*:598
afum|EAL93780.1 100.00% afis|EAW19051.1 100.00%
Bootstrap support for afum|EAL93780.1 as seed ortholog is 100%.
Bootstrap support for afis|EAW19051.1 as seed ortholog is 100%.

---

Group of orthologs #6671. Best score 598 bits
Score difference with first non-orthologous sequence - *A. fumigatus*:598 *A. fischeri*:365
afum|EAL90907.1 100.00% afis|EAW22274.1 100.00%
Bootstrap support for afum|EAL90907.1 as seed ortholog is 100%.
Bootstrap support for afis|EAW22274.1 as seed ortholog is 100%.

---

Group of orthologs #6672. Best score 598 bits
Score difference with first non-orthologous sequence - *A. fumigatus*:598 *A. fischeri*:598
afum|EAL90658.1 100.00% afis|EAW22536.1 100.00%
Bootstrap support for afum|EAL90658.1 as seed ortholog is 100%.
Bootstrap support for afis|EAW22536.1 as seed ortholog is 100%.

---

Group of orthologs #6673. Best score 598 bits
Score difference with first non-orthologous sequence - *A. fumigatus*:598 *A. fischeri*:598
afum|EAL92276.1 100.00% afis|EAW21261.1 100.00%
Bootstrap support for afum|EAL92276.1 as seed ortholog is 100%.
Bootstrap support for afis|EAW21261.1 as seed ortholog is 100%.

---

Group of orthologs #6674. Best score 597 bits
Score difference with first non-orthologous sequence - *A. fumigatus*:597 *A. fischeri*:597
afum|EAL84751.1 100.00% afis|EAW20881.1 100.00%
Bootstrap support for afum|EAL84751.1 as seed ortholog is 100%.
Bootstrap support for afis|EAW20881.1 as seed ortholog is 100%.

---

Group of orthologs #6675. Best score 597 bits
Score difference with first non-orthologous sequence - *A. fumigatus*:597 *A. fischeri*:505
afum|EAL86019.2 100.00% afis|EAW24349.1 100.00%
Bootstrap support for afum|EAL86019.2 as seed ortholog is 100%.
Bootstrap support for afis|EAW24349.1 as seed ortholog is 100%.

---

Group of orthologs #6676. Best score 597 bits
Score difference with first non-orthologous sequence - *A. fumigatus*:509 *A. fischeri*:490
afum|EAL87238.1 100.00% afis|EAW23804.1 100.00%
Bootstrap support for afum|EAL87238.1 as seed ortholog is 100%.
Bootstrap support for afis|EAW23804.1 as seed ortholog is 100%.

---

Group of orthologs #6677. Best score 597 bits
Score difference with first non-orthologous sequence - *A. fumigatus*:597 *A. fischeri*:597
afum|EAL93355.1 100.00% afis|EAW18641.1 100.00%
Bootstrap support for afum|EAL93355.1 as seed ortholog is 100%.
Bootstrap support for afis|EAW18641.1 as seed ortholog is 100%.

---

Group of orthologs #6678. Best score 597 bits
Score difference with first non-orthologous sequence - *A. fumigatus*:484 *A. fischeri*:489
afum|EBA27437.1 100.00% afis|EAW17673.1 100.00%
Bootstrap support for afum|EBA27437.1 as seed ortholog is 100%.
Bootstrap support for afis|EAW17673.1 as seed ortholog is 100%.

---

Group of orthologs #6679. Best score 597 bits
Score difference with first non-orthologous sequence - *A. fumigatus*:597 *A. fischeri*:597

| | | | |
|---|---|---|---|
| afum\|EAL93473.1 | 100.00% | afis\|EAW18747.1 | 100.00% |

Bootstrap support for afum|EAL93473.1 as seed ortholog is 100%.
Bootstrap support for afis|EAW18747.1 as seed ortholog is 100%.

---

Group of orthologs #6680. Best score 597 bits
Score difference with first non-orthologous sequence - *A. fumigatus*:597 *A. fischeri*:597

| | | | |
|---|---|---|---|
| afum\|EAL89510.1 | 100.00% | afis\|EAW24861.1 | 100.00% |

Bootstrap support for afum|EAL89510.1 as seed ortholog is 100%.
Bootstrap support for afis|EAW24861.1 as seed ortholog is 100%.

---

Group of orthologs #6681. Best score 597 bits
Score difference with first non-orthologous sequence - *A. fumigatus*:597 *A. fischeri*:597

| | | | |
|---|---|---|---|
| afum\|EAL92901.1 | 100.00% | afis\|EAW21887.1 | 100.00% |

Bootstrap support for afum|EAL92901.1 as seed ortholog is 100%.
Bootstrap support for afis|EAW21887.1 as seed ortholog is 100%.

---

Group of orthologs #6682. Best score 597 bits
Score difference with first non-orthologous sequence - *A. fumigatus*:597 *A. fischeri*:597

| | | | |
|---|---|---|---|
| afum\|EAL89945.1 | 100.00% | afis\|EAW25282.1 | 100.00% |

Bootstrap support for afum|EAL89945.1 as seed ortholog is 100%.
Bootstrap support for afis|EAW25282.1 as seed ortholog is 100%.

---

Group of orthologs #6683. Best score 597 bits
Score difference with first non-orthologous sequence - *A. fumigatus*:249 *A. fischeri*:597

| | | | |
|---|---|---|---|
| afum\|EAL90121.1 | 100.00% | afis\|EAW25460.1 | 100.00% |

Bootstrap support for afum|EAL90121.1 as seed ortholog is 100%.
Bootstrap support for afis|EAW25460.1 as seed ortholog is 100%.

---

Group of orthologs #6684. Best score 596 bits
Score difference with first non-orthologous sequence - *A. fumigatus*:596 *A. fischeri*:532

| | | | |
|---|---|---|---|
| afum\|EAL86430.2 | 100.00% | afis\|EAW16892.1 | 100.00% |

Bootstrap support for afum|EAL86430.2 as seed ortholog is 100%.
Bootstrap support for afis|EAW16892.1 as seed ortholog is 100%.

---

Group of orthologs #6685. Best score 596 bits
Score difference with first non-orthologous sequence - *A. fumigatus*:372 *A. fischeri*:433

| | | | |
|---|---|---|---|
| afum\|EAL84504.1 | 100.00% | afis\|EAW20557.1 | 100.00% |

Bootstrap support for afum|EAL84504.1 as seed ortholog is 100%.
Bootstrap support for afis|EAW20557.1 as seed ortholog is 100%.

---

Group of orthologs #6686. Best score 596 bits
Score difference with first non-orthologous sequence - *A. fumigatus*:184 *A. fischeri*:49

| | | | |
|---|---|---|---|
| afum\|EAL84279.2 | 100.00% | afis\|EAW25556.1 | 100.00% |

Bootstrap support for afum|EAL84279.2 as seed ortholog is 100%.
Bootstrap support for afis|EAW25556.1 as seed ortholog is 99%.

---

Group of orthologs #6687. Best score 596 bits
Score difference with first non-orthologous sequence - *A. fumigatus*:596 *A. fischeri*:596

| | | | |
|---|---|---|---|
| afum\|EAL87212.1 | 100.00% | afis\|EAW23778.1 | 100.00% |

Bootstrap support for afum|EAL87212.1 as seed ortholog is 100%.
Bootstrap support for afis|EAW23778.1 as seed ortholog is 100%.

---

Group of orthologs #6688. Best score 596 bits

Score difference with first non-orthologous sequence - *A. fumigatus*:596   *A. fischeri*:596
afum|EAL87395.1 100.00% afis|EAW23965.1 100.00%
Bootstrap support for afum|EAL87395.1 as seed ortholog is 100%.
Bootstrap support for afis|EAW23965.1 as seed ortholog is 100%.

---

Group of orthologs #6689. Best score 596 bits
Score difference with first non-orthologous sequence - *A. fumigatus*:596   *A. fischeri*:493
afum|EAL88325.1 100.00% afis|EAW23158.1 100.00%
Bootstrap support for afum|EAL88325.1 as seed ortholog is 100%.
Bootstrap support for afis|EAW23158.1 as seed ortholog is 100%.

---

Group of orthologs #6690. Best score 596 bits
Score difference with first non-orthologous sequence - *A. fumigatus*:596   *A. fischeri*:596
afum|EAL92654.1 100.00% afis|EAW21618.1 100.00%
Bootstrap support for afum|EAL92654.1 as seed ortholog is 100%.
Bootstrap support for afis|EAW21618.1 as seed ortholog is 100%.

---

Group of orthologs #6691. Best score 595 bits
Score difference with first non-orthologous sequence - *A. fumigatus*:595   *A. fischeri*:595
afum|EAL87097.1 100.00% afis|EAW19397.1 100.00%
Bootstrap support for afum|EAL87097.1 as seed ortholog is 100%.
Bootstrap support for afis|EAW19397.1 as seed ortholog is 100%.

---

Group of orthologs #6692. Best score 595 bits
Score difference with first non-orthologous sequence - *A. fumigatus*:595   *A. fischeri*:595
afum|EAL90555.1 100.00% afis|EAW22637.1 100.00%
Bootstrap support for afum|EAL90555.1 as seed ortholog is 100%.
Bootstrap support for afis|EAW22637.1 as seed ortholog is 100%.

---

Group of orthologs #6693. Best score 595 bits
Score difference with first non-orthologous sequence - *A. fumigatus*:595   *A. fischeri*:595
afum|EAL89604.1 100.00% afis|EAW24947.1 100.00%
Bootstrap support for afum|EAL89604.1 as seed ortholog is 100%.
Bootstrap support for afis|EAW24947.1 as seed ortholog is 100%.

---

Group of orthologs #6694. Best score 594 bits
Score difference with first non-orthologous sequence - *A. fumigatus*:594   *A. fischeri*:515
afum|EAL85008.1 100.00% afis|EAW18941.1 100.00%
Bootstrap support for afum|EAL85008.1 as seed ortholog is 100%.
Bootstrap support for afis|EAW18941.1 as seed ortholog is 100%.

---

Group of orthologs #6695. Best score 594 bits
Score difference with first non-orthologous sequence - *A. fumigatus*:495   *A. fischeri*:479
afum|EAL88764.1 100.00% afis|EAW16134.1 100.00%
Bootstrap support for afum|EAL88764.1 as seed ortholog is 100%.
Bootstrap support for afis|EAW16134.1 as seed ortholog is 100%.

---

Group of orthologs #6696. Best score 594 bits
Score difference with first non-orthologous sequence - *A. fumigatus*:594   *A. fischeri*:594
afum|EAL85084.1 100.00% afis|EAW19808.1 100.00%
Bootstrap support for afum|EAL85084.1 as seed ortholog is 100%.
Bootstrap support for afis|EAW19808.1 as seed ortholog is 100%.

---

Group of orthologs #6697. Best score 594 bits
Score difference with first non-orthologous sequence - *A. fumigatus*:594   *A. fischeri*:594
afum|EAL85017.1 100.00% afis|EAW19881.1 100.00%

Bootstrap support for afum|EAL85017.1 as seed ortholog is 100%.
Bootstrap support for afis|EAW19881.1 as seed ortholog is 100%.

---

Group of orthologs #6698. Best score 594 bits
Score difference with first non-orthologous sequence - *A. fumigatus*:594 *A. fischeri*:594
afum|EAL85355.1 100.00% afis|EAW20237.1 100.00%
Bootstrap support for afum|EAL85355.1 as seed ortholog is 100%.
Bootstrap support for afis|EAW20237.1 as seed ortholog is 100%.

---

Group of orthologs #6699. Best score 594 bits
Score difference with first non-orthologous sequence - *A. fumigatus*:594 *A. fischeri*:594
afum|EAL91730.1 100.00% afis|EAW17919.1 100.00%
Bootstrap support for afum|EAL91730.1 as seed ortholog is 100%.
Bootstrap support for afis|EAW17919.1 as seed ortholog is 100%.

---

Group of orthologs #6700. Best score 594 bits
Score difference with first non-orthologous sequence - *A. fumigatus*:594 *A. fischeri*:594
afum|EAL88152.1 100.00% afis|EAW23327.1 100.00%
Bootstrap support for afum|EAL88152.1 as seed ortholog is 100%.
Bootstrap support for afis|EAW23327.1 as seed ortholog is 100%.

---

Group of orthologs #6701. Best score 594 bits
Score difference with first non-orthologous sequence - *A. fumigatus*:594 *A. fischeri*:594
afum|EAL90169.2 100.00% afis|EAW25503.1 100.00%
Bootstrap support for afum|EAL90169.2 as seed ortholog is 100%.
Bootstrap support for afis|EAW25503.1 as seed ortholog is 100%.

---

Group of orthologs #6702. Best score 593 bits
Score difference with first non-orthologous sequence - *A. fumigatus*:518 *A. fischeri*:479
afum|EAL88848.1 100.00% afis|EAW16220.1 100.00%
Bootstrap support for afum|EAL88848.1 as seed ortholog is 100%.
Bootstrap support for afis|EAW16220.1 as seed ortholog is 100%.

---

Group of orthologs #6703. Best score 593 bits
Score difference with first non-orthologous sequence - *A. fumigatus*:593 *A. fischeri*:593
afum|EAL91356.1 100.00% afis|EAW17542.1 100.00%
Bootstrap support for afum|EAL91356.1 as seed ortholog is 100%.
Bootstrap support for afis|EAW17542.1 as seed ortholog is 100%.

---

Group of orthologs #6704. Best score 593 bits
Score difference with first non-orthologous sequence - *A. fumigatus*:593 *A. fischeri*:593
afum|EAL91518.1 100.00% afis|EAW17703.1 100.00%
Bootstrap support for afum|EAL91518.1 as seed ortholog is 100%.
Bootstrap support for afis|EAW17703.1 as seed ortholog is 100%.

---

Group of orthologs #6705. Best score 593 bits
Score difference with first non-orthologous sequence - *A. fumigatus*:593 *A. fischeri*:593
afum|EAL91535.2 100.00% afis|EAW17720.1 100.00%
Bootstrap support for afum|EAL91535.2 as seed ortholog is 100%.
Bootstrap support for afis|EAW17720.1 as seed ortholog is 100%.

---

Group of orthologs #6706. Best score 593 bits
Score difference with first non-orthologous sequence - *A. fumigatus*:593 *A. fischeri*:593
afum|EAL89232.1 100.00% afis|EAW20412.1 100.00%
Bootstrap support for afum|EAL89232.1 as seed ortholog is 100%.
Bootstrap support for afis|EAW20412.1 as seed ortholog is 100%.

Group of orthologs #6707. Best score 593 bits
Score difference with first non-orthologous sequence - *A. fumigatus*:593 *A. fischeri*:593
afum|EAL91027.1 100.00% afis|EAW22164.1 100.00%
Bootstrap support for afum|EAL91027.1 as seed ortholog is 100%.
Bootstrap support for afis|EAW22164.1 as seed ortholog is 100%.

Group of orthologs #6708. Best score 593 bits
Score difference with first non-orthologous sequence - *A. fumigatus*:593 *A. fischeri*:593
afum|EAL92000.1 100.00% afis|EAW24145.1 100.00%
Bootstrap support for afum|EAL92000.1 as seed ortholog is 100%.
Bootstrap support for afis|EAW24145.1 as seed ortholog is 100%.

Group of orthologs #6709. Best score 592 bits
Score difference with first non-orthologous sequence - *A. fumigatus*:592 *A. fischeri*:592
afum|EAL88712.1 100.00% afis|EAW16080.1 100.00%
Bootstrap support for afum|EAL88712.1 as seed ortholog is 100%.
Bootstrap support for afis|EAW16080.1 as seed ortholog is 100%.

Group of orthologs #6710. Best score 592 bits
Score difference with first non-orthologous sequence - *A. fumigatus*:592 *A. fischeri*:432
afum|EAL87472.1 100.00% afis|EAW18115.1 100.00%
Bootstrap support for afum|EAL87472.1 as seed ortholog is 100%.
Bootstrap support for afis|EAW18115.1 as seed ortholog is 100%.

Group of orthologs #6711. Best score 592 bits
Score difference with first non-orthologous sequence - *A. fumigatus*:592 *A. fischeri*:592
afum|EAL91634.1 100.00% afis|EAW17824.1 100.00%
Bootstrap support for afum|EAL91634.1 as seed ortholog is 100%.
Bootstrap support for afis|EAW17824.1 as seed ortholog is 100%.

Group of orthologs #6712. Best score 592 bits
Score difference with first non-orthologous sequence - *A. fumigatus*:592 *A. fischeri*:542
afum|EAL93072.1 100.00% afis|EAW18396.1 100.00%
Bootstrap support for afum|EAL93072.1 as seed ortholog is 100%.
Bootstrap support for afis|EAW18396.1 as seed ortholog is 100%.

Group of orthologs #6713. Best score 592 bits
Score difference with first non-orthologous sequence - *A. fumigatus*:592 *A. fischeri*:592
afum|EAL88286.1 100.00% afis|EAW23197.1 100.00%
Bootstrap support for afum|EAL88286.1 as seed ortholog is 100%.
Bootstrap support for afis|EAW23197.1 as seed ortholog is 100%.

Group of orthologs #6714. Best score 592 bits
Score difference with first non-orthologous sequence - *A. fumigatus*:592 *A. fischeri*:461
afum|EAL93955.1 100.00% afis|EAW19224.1 100.00%
Bootstrap support for afum|EAL93955.1 as seed ortholog is 100%.
Bootstrap support for afis|EAW19224.1 as seed ortholog is 100%.

Group of orthologs #6715. Best score 592 bits
Score difference with first non-orthologous sequence - *A. fumigatus*:592 *A. fischeri*:592
afum|EAL92541.1 100.00% afis|EAW21520.1 100.00%
Bootstrap support for afum|EAL92541.1 as seed ortholog is 100%.
Bootstrap support for afis|EAW21520.1 as seed ortholog is 100%.

Group of orthologs #6716. Best score 591 bits

Score difference with first non-orthologous sequence - *A. fumigatus*:591 *A. fischeri*:591
afum|EAL84649.2 100.00% afis|EAW20981.1 100.00%
Bootstrap support for afum|EAL84649.2 as seed ortholog is 100%.
Bootstrap support for afis|EAW20981.1 as seed ortholog is 100%.

---

Group of orthologs #6717. Best score 591 bits
Score difference with first non-orthologous sequence - *A. fumigatus*:591 *A. fischeri*:591
afum|EAL85447.1 100.00% afis|EAW20319.1 100.00%
Bootstrap support for afum|EAL85447.1 as seed ortholog is 100%.
Bootstrap support for afis|EAW20319.1 as seed ortholog is 100%.

---

Group of orthologs #6718. Best score 591 bits
Score difference with first non-orthologous sequence - *A. fumigatus*:591 *A. fischeri*:591
afum|EAL87826.1 100.00% afis|EAW23634.1 100.00%
Bootstrap support for afum|EAL87826.1 as seed ortholog is 100%.
Bootstrap support for afis|EAW23634.1 as seed ortholog is 100%.

---

Group of orthologs #6719. Best score 591 bits
Score difference with first non-orthologous sequence - *A. fumigatus*:591 *A. fischeri*:591
afum|EAL90467.1 100.00% afis|EAW22728.1 100.00%
Bootstrap support for afum|EAL90467.1 as seed ortholog is 100%.
Bootstrap support for afis|EAW22728.1 as seed ortholog is 100%.

---

Group of orthologs #6720. Best score 591 bits
Score difference with first non-orthologous sequence - *A. fumigatus*:591 *A. fischeri*:591
afum|EAL90414.1 100.00% afis|EAW22785.1 100.00%
Bootstrap support for afum|EAL90414.1 as seed ortholog is 100%.
Bootstrap support for afis|EAW22785.1 as seed ortholog is 100%.

---

Group of orthologs #6721. Best score 591 bits
Score difference with first non-orthologous sequence - *A. fumigatus*:591 *A. fischeri*:591
afum|EBA27183.1 100.00% afis|EAW20555.1 100.00%
Bootstrap support for afum|EBA27183.1 as seed ortholog is 100%.
Bootstrap support for afis|EAW20555.1 as seed ortholog is 100%.

---

Group of orthologs #6722. Best score 591 bits
Score difference with first non-orthologous sequence - *A. fumigatus*:591 *A. fischeri*:591
afum|EAL89744.1 100.00% afis|EAW25075.1 100.00%
Bootstrap support for afum|EAL89744.1 as seed ortholog is 100%.
Bootstrap support for afis|EAW25075.1 as seed ortholog is 100%.

---

Group of orthologs #6723. Best score 590 bits
Score difference with first non-orthologous sequence - *A. fumigatus*:282 *A. fischeri*:415
afum|EAL86534.1 100.00% afis|EAW16825.1 100.00%
Bootstrap support for afum|EAL86534.1 as seed ortholog is 100%.
Bootstrap support for afis|EAW16825.1 as seed ortholog is 100%.

---

Group of orthologs #6724. Best score 590 bits
Score difference with first non-orthologous sequence - *A. fumigatus*:590 *A. fischeri*:590
afum|EAL88557.2 100.00% afis|EAW15921.1 100.00%
Bootstrap support for afum|EAL88557.2 as seed ortholog is 100%.
Bootstrap support for afis|EAW15921.1 as seed ortholog is 100%.

---

Group of orthologs #6725. Best score 590 bits
Score difference with first non-orthologous sequence - *A. fumigatus*:590 *A. fischeri*:455
afum|EAL85127.1 100.00% afis|EAW19773.1 100.00%

Bootstrap support for afum|EAL85127.1 as seed ortholog is 100%.
Bootstrap support for afis|EAW19773.1 as seed ortholog is 100%.

---

Group of orthologs #6726. Best score 590 bits
Score difference with first non-orthologous sequence - *A. fumigatus*:590 *A. fischeri*:590
afum|EAL84536.1 100.00% afis|EAW20521.1 100.00%
Bootstrap support for afum|EAL84536.1 as seed ortholog is 100%.
Bootstrap support for afis|EAW20521.1 as seed ortholog is 100%.

---

Group of orthologs #6727. Best score 590 bits
Score difference with first non-orthologous sequence - *A. fumigatus*:590 *A. fischeri*:590
afum|EAL85244.1 100.00% afis|EAW20126.1 100.00%
Bootstrap support for afum|EAL85244.1 as seed ortholog is 100%.
Bootstrap support for afis|EAW20126.1 as seed ortholog is 100%.

---

Group of orthologs #6728. Best score 590 bits
Score difference with first non-orthologous sequence - *A. fumigatus*:590 *A. fischeri*:590
afum|EAL87155.1 100.00% afis|EAW23711.1 100.00%
Bootstrap support for afum|EAL87155.1 as seed ortholog is 100%.
Bootstrap support for afis|EAW23711.1 as seed ortholog is 100%.

---

Group of orthologs #6729. Best score 589 bits
Score difference with first non-orthologous sequence - *A. fumigatus*:589 *A. fischeri*:589
afum|EAL88492.1 100.00% afis|EAW15856.1 100.00%
Bootstrap support for afum|EAL88492.1 as seed ortholog is 100%.
Bootstrap support for afis|EAW15856.1 as seed ortholog is 100%.

---

Group of orthologs #6730. Best score 589 bits
Score difference with first non-orthologous sequence - *A. fumigatus*:589 *A. fischeri*:589
afum|EAL88645.1 100.00% afis|EAW16016.1 100.00%
Bootstrap support for afum|EAL88645.1 as seed ortholog is 100%.
Bootstrap support for afis|EAW16016.1 as seed ortholog is 100%.

---

Group of orthologs #6731. Best score 589 bits
Score difference with first non-orthologous sequence - *A. fumigatus*:589 *A. fischeri*:589
afum|EAL89202.2 100.00% afis|EAW16605.1 100.00%
Bootstrap support for afum|EAL89202.2 as seed ortholog is 100%.
Bootstrap support for afis|EAW16605.1 as seed ortholog is 100%.

---

Group of orthologs #6732. Best score 589 bits
Score difference with first non-orthologous sequence - *A. fumigatus*:589 *A. fischeri*:589
afum|EAL87660.1 100.00% afis|EAW18294.1 100.00%
Bootstrap support for afum|EAL87660.1 as seed ortholog is 100%.
Bootstrap support for afis|EAW18294.1 as seed ortholog is 100%.

---

Group of orthologs #6733. Best score 589 bits
Score difference with first non-orthologous sequence - *A. fumigatus*:589 *A. fischeri*:589
afum|EAL87074.1 100.00% afis|EAW19420.1 100.00%
Bootstrap support for afum|EAL87074.1 as seed ortholog is 100%.
Bootstrap support for afis|EAW19420.1 as seed ortholog is 100%.

---

Group of orthologs #6734. Best score 589 bits
Score difference with first non-orthologous sequence - *A. fumigatus*:589 *A. fischeri*:589
afum|EBA27379.1 100.00% afis|EAW24739.1 100.00%
Bootstrap support for afum|EBA27379.1 as seed ortholog is 100%.
Bootstrap support for afis|EAW24739.1 as seed ortholog is 100%.

---

Group of orthologs #6735. Best score 588 bits
Score difference with first non-orthologous sequence - *A. fumigatus*:309 *A. fischeri*:588
afum|EAL89114.1 100.00% afis|EAW16519.1 100.00%
Bootstrap support for afum|EAL89114.1 as seed ortholog is 100%.
Bootstrap support for afis|EAW16519.1 as seed ortholog is 100%.

---

Group of orthologs #6736. Best score 588 bits
Score difference with first non-orthologous sequence - *A. fumigatus*:588 *A. fischeri*:588
afum|EAL87650.1 100.00% afis|EAW18284.1 100.00%
Bootstrap support for afum|EAL87650.1 as seed ortholog is 100%.
Bootstrap support for afis|EAW18284.1 as seed ortholog is 100%.

---

Group of orthologs #6737. Best score 588 bits
Score difference with first non-orthologous sequence - *A. fumigatus*:452 *A. fischeri*:588
afum|EAL91150.1 100.00% afis|EAW17314.1 100.00%
Bootstrap support for afum|EAL91150.1 as seed ortholog is 100%.
Bootstrap support for afis|EAW17314.1 as seed ortholog is 100%.

---

Group of orthologs #6738. Best score 588 bits
Score difference with first non-orthologous sequence - *A. fumigatus*:588 *A. fischeri*:588
afum|EAL86167.1 100.00% afis|EAW24497.1 100.00%
Bootstrap support for afum|EAL86167.1 as seed ortholog is 100%.
Bootstrap support for afis|EAW24497.1 as seed ortholog is 100%.

---

Group of orthologs #6739. Best score 588 bits
Score difference with first non-orthologous sequence - *A. fumigatus*:588 *A. fischeri*:588
afum|EAL88183.1 100.00% afis|EAW23295.1 100.00%
Bootstrap support for afum|EAL88183.1 as seed ortholog is 100%.
Bootstrap support for afis|EAW23295.1 as seed ortholog is 100%.

---

Group of orthologs #6740. Best score 588 bits
Score difference with first non-orthologous sequence - *A. fumigatus*:588 *A. fischeri*:588
afum|EBA27233.1 100.00% afis|EAW20336.1 100.00%
Bootstrap support for afum|EBA27233.1 as seed ortholog is 100%.
Bootstrap support for afis|EAW20336.1 as seed ortholog is 100%.

---

Group of orthologs #6741. Best score 588 bits
Score difference with first non-orthologous sequence - *A. fumigatus*:588 *A. fischeri*:588
afum|EAL92784.1 100.00% afis|EAW21754.1 100.00%
Bootstrap support for afum|EAL92784.1 as seed ortholog is 100%.
Bootstrap support for afis|EAW21754.1 as seed ortholog is 100%.

---

Group of orthologs #6742. Best score 587 bits
Score difference with first non-orthologous sequence - *A. fumigatus*:587 *A. fischeri*:587
afum|EAL87563.1 100.00% afis|EAW18198.1 100.00%
Bootstrap support for afum|EAL87563.1 as seed ortholog is 100%.
Bootstrap support for afis|EAW18198.1 as seed ortholog is 100%.

---

Group of orthologs #6743. Best score 587 bits
Score difference with first non-orthologous sequence - *A. fumigatus*:587 *A. fischeri*:587
afum|EAL85983.1 100.00% afis|EAW24315.1 100.00%
Bootstrap support for afum|EAL85983.1 as seed ortholog is 100%.
Bootstrap support for afis|EAW24315.1 as seed ortholog is 100%.

---

Group of orthologs #6744. Best score 587 bits

Score difference with first non-orthologous sequence - *A. fumigatus*:587   *A. fischeri*:587
afum|EAL90272.1 100.00% afis|EAW22925.1 100.00%
Bootstrap support for afum|EAL90272.1 as seed ortholog is 100%.
Bootstrap support for afis|EAW22925.1 as seed ortholog is 100%.

---

Group of orthologs #6745. Best score 587 bits
Score difference with first non-orthologous sequence - *A. fumigatus*:587   *A. fischeri*:587
afum|EBA27200.1 100.00% afis|EAW20427.1 100.00%
Bootstrap support for afum|EBA27200.1 as seed ortholog is 100%.
Bootstrap support for afis|EAW20427.1 as seed ortholog is 100%.

---

Group of orthologs #6746. Best score 587 bits
Score difference with first non-orthologous sequence - *A. fumigatus*:587   *A. fischeri*:587
afum|EAL92996.1 100.00% afis|EAW21981.1 100.00%
Bootstrap support for afum|EAL92996.1 as seed ortholog is 100%.
Bootstrap support for afis|EAW21981.1 as seed ortholog is 100%.

---

Group of orthologs #6747. Best score 587 bits
Score difference with first non-orthologous sequence - *A. fumigatus*:587   *A. fischeri*:587
afum|EAL93024.1 100.00% afis|EAW22009.1 100.00%
Bootstrap support for afum|EAL93024.1 as seed ortholog is 100%.
Bootstrap support for afis|EAW22009.1 as seed ortholog is 100%.

---

Group of orthologs #6748. Best score 586 bits
Score difference with first non-orthologous sequence - *A. fumigatus*:586   *A. fischeri*:313
afum|EAL87214.1 100.00% afis|EAW23780.1 100.00%
Bootstrap support for afum|EAL87214.1 as seed ortholog is 100%.
Bootstrap support for afis|EAW23780.1 as seed ortholog is 100%.

---

Group of orthologs #6749. Best score 586 bits
Score difference with first non-orthologous sequence - *A. fumigatus*:586   *A. fischeri*:586
afum|EAL87286.1 100.00% afis|EAW23858.1 100.00%
Bootstrap support for afum|EAL87286.1 as seed ortholog is 100%.
Bootstrap support for afis|EAW23858.1 as seed ortholog is 100%.

---

Group of orthologs #6750. Best score 586 bits
Score difference with first non-orthologous sequence - *A. fumigatus*:6   *A. fischeri*:586
afum|EAL91117.1 100.00% afis|EAW20298.1 100.00%
Bootstrap support for afum|EAL91117.1 as seed ortholog is 57%. Alternative seed ortholog is afum|EAL85427.1 (6 bits away from this cluster)
Bootstrap support for afis|EAW20298.1 as seed ortholog is 100%.

---

Group of orthologs #6751. Best score 585 bits
Score difference with first non-orthologous sequence - *A. fumigatus*:585   *A. fischeri*:585
afum|EAL85623.1 100.00% afis|EAW15724.1 100.00%
Bootstrap support for afum|EAL85623.1 as seed ortholog is 100%.
Bootstrap support for afis|EAW15724.1 as seed ortholog is 100%.

---

Group of orthologs #6752. Best score 585 bits
Score difference with first non-orthologous sequence - *A. fumigatus*:585   *A. fischeri*:585
afum|EAL91269.1 100.00% afis|EAW17443.1 100.00%
Bootstrap support for afum|EAL91269.1 as seed ortholog is 100%.
Bootstrap support for afis|EAW17443.1 as seed ortholog is 100%.

---

Group of orthologs #6753. Best score 585 bits
Score difference with first non-orthologous sequence - *A. fumigatus*:585   *A. fischeri*:585

afum|EAL84351.1 100.00% afis|EAW25681.1 100.00%
Bootstrap support for afum|EAL84351.1 as seed ortholog is 100%.
Bootstrap support for afis|EAW25681.1 as seed ortholog is 100%.

---

Group of orthologs #6754. Best score 585 bits
Score difference with first non-orthologous sequence - *A. fumigatus*:585 *A. fischeri*:537
afum|EAL93074.1 100.00% afis|EAW18397.1 100.00%
Bootstrap support for afum|EAL93074.1 as seed ortholog is 100%.
Bootstrap support for afis|EAW18397.1 as seed ortholog is 100%.

---

Group of orthologs #6755. Best score 585 bits
Score difference with first non-orthologous sequence - *A. fumigatus*:585 *A. fischeri*:585
afum|EAL88465.1 100.00% afis|EAW23016.1 100.00%
Bootstrap support for afum|EAL88465.1 as seed ortholog is 100%.
Bootstrap support for afis|EAW23016.1 as seed ortholog is 100%.

---

Group of orthologs #6756. Best score 585 bits
Score difference with first non-orthologous sequence - *A. fumigatus*:585 *A. fischeri*:585
afum|EAL87968.1 100.00% afis|EAW23495.1 100.00%
Bootstrap support for afum|EAL87968.1 as seed ortholog is 100%.
Bootstrap support for afis|EAW23495.1 as seed ortholog is 100%.

---

Group of orthologs #6757. Best score 585 bits
Score difference with first non-orthologous sequence - *A. fumigatus*:585 *A. fischeri*:585
afum|EAL89794.1 100.00% afis|EAW25123.1 100.00%
Bootstrap support for afum|EAL89794.1 as seed ortholog is 100%.
Bootstrap support for afis|EAW25123.1 as seed ortholog is 100%.

---

Group of orthologs #6758. Best score 584 bits
Score difference with first non-orthologous sequence - *A. fumigatus*:584 *A. fischeri*:584
afum|EAL85570.1 100.00% afis|EAW15779.1 100.00%
Bootstrap support for afum|EAL85570.1 as seed ortholog is 100%.
Bootstrap support for afis|EAW15779.1 as seed ortholog is 100%.

---

Group of orthologs #6759. Best score 584 bits
Score difference with first non-orthologous sequence - *A. fumigatus*:584 *A. fischeri*:584
afum|EAL88512.1 100.00% afis|EAW15878.1 100.00%
Bootstrap support for afum|EAL88512.1 as seed ortholog is 100%.
Bootstrap support for afis|EAW15878.1 as seed ortholog is 100%.

---

Group of orthologs #6760. Best score 584 bits
Score difference with first non-orthologous sequence - *A. fumigatus*:584 *A. fischeri*:584
afum|EAL86930.1 100.00% afis|EAW19588.1 100.00%
Bootstrap support for afum|EAL86930.1 as seed ortholog is 100%.
Bootstrap support for afis|EAW19588.1 as seed ortholog is 100%.

---

Group of orthologs #6761. Best score 584 bits
Score difference with first non-orthologous sequence - *A. fumigatus*:584 *A. fischeri*:584
afum|EAL91528.1 100.00% afis|EAW17713.1 100.00%
Bootstrap support for afum|EAL91528.1 as seed ortholog is 100%.
Bootstrap support for afis|EAW17713.1 as seed ortholog is 100%.

---

Group of orthologs #6762. Best score 584 bits
Score difference with first non-orthologous sequence - *A. fumigatus*:584 *A. fischeri*:584
afum|EAL91683.1 100.00% afis|EAW17875.1 100.00%
Bootstrap support for afum|EAL91683.1 as seed ortholog is 100%.

Bootstrap support for afis|EAW17875.1 as seed ortholog is 100%.

---

Group of orthologs #6763. Best score 584 bits
Score difference with first non-orthologous sequence - *A. fumigatus*:584 *A. fischeri*:584
afum|EAL91693.1 100.00% afis|EAW17885.1 100.00%
Bootstrap support for afum|EAL91693.1 as seed ortholog is 100%.
Bootstrap support for afis|EAW17885.1 as seed ortholog is 100%.

---

Group of orthologs #6764. Best score 584 bits
Score difference with first non-orthologous sequence - *A. fumigatus*:531 *A. fischeri*:79
afum|EAL84374.1 100.00% afis|EAW25701.1 100.00%
Bootstrap support for afum|EAL84374.1 as seed ortholog is 100%.
Bootstrap support for afis|EAW25701.1 as seed ortholog is 99%.

---

Group of orthologs #6765. Best score 584 bits
Score difference with first non-orthologous sequence - *A. fumigatus*:584 *A. fischeri*:584
afum|EAL93185.1 100.00% afis|EAW18495.1 100.00%
Bootstrap support for afum|EAL93185.1 as seed ortholog is 100%.
Bootstrap support for afis|EAW18495.1 as seed ortholog is 100%.

---

Group of orthologs #6766. Best score 584 bits
Score difference with first non-orthologous sequence - *A. fumigatus*:584 *A. fischeri*:584
afum|EAL88054.1 100.00% afis|EAW23415.1 100.00%
Bootstrap support for afum|EAL88054.1 as seed ortholog is 100%.
Bootstrap support for afis|EAW23415.1 as seed ortholog is 100%.

---

Group of orthologs #6767. Best score 584 bits
Score difference with first non-orthologous sequence - *A. fumigatus*:584 *A. fischeri*:584
afum|EAL93265.2 100.00% afis|EAW18563.1 100.00%
Bootstrap support for afum|EAL93265.2 as seed ortholog is 100%.
Bootstrap support for afis|EAW18563.1 as seed ortholog is 100%.

---

Group of orthologs #6768. Best score 584 bits
Score difference with first non-orthologous sequence - *A. fumigatus*:584 *A. fischeri*:584
afum|EAL93734.1 100.00% afis|EAW18999.1 100.00%
Bootstrap support for afum|EAL93734.1 as seed ortholog is 100%.
Bootstrap support for afis|EAW18999.1 as seed ortholog is 100%.

---

Group of orthologs #6769. Best score 584 bits
Score difference with first non-orthologous sequence - *A. fumigatus*:584 *A. fischeri*:584
afum|EAL90979.1 100.00% afis|EAW22215.1 100.00%
Bootstrap support for afum|EAL90979.1 as seed ortholog is 100%.
Bootstrap support for afis|EAW22215.1 as seed ortholog is 100%.

---

Group of orthologs #6770. Best score 584 bits
Score difference with first non-orthologous sequence - *A. fumigatus*:584 *A. fischeri*:584
afum|EAL90346.1 100.00% afis|EAW22854.1 100.00%
Bootstrap support for afum|EAL90346.1 as seed ortholog is 100%.
Bootstrap support for afis|EAW22854.1 as seed ortholog is 100%.

---

Group of orthologs #6771. Best score 584 bits
Score difference with first non-orthologous sequence - *A. fumigatus*:584 *A. fischeri*:584
afum|EAL92485.1 100.00% afis|EAW21465.1 100.00%
Bootstrap support for afum|EAL92485.1 as seed ortholog is 100%.
Bootstrap support for afis|EAW21465.1 as seed ortholog is 100%.

---

Group of orthologs #6772. Best score 584 bits
Score difference with first non-orthologous sequence - *A. fumigatus*:584 *A. fischeri*:584
afum|EAL91993.1 100.00% afis|EAW24138.1 100.00%
Bootstrap support for afum|EAL91993.1 as seed ortholog is 100%.
Bootstrap support for afis|EAW24138.1 as seed ortholog is 100%.

---

Group of orthologs #6773. Best score 583 bits
Score difference with first non-orthologous sequence - *A. fumigatus*:583 *A. fischeri*:583
afum|EAL86356.1 100.00% afis|EAW16786.1 100.00%
Bootstrap support for afum|EAL86356.1 as seed ortholog is 100%.
Bootstrap support for afis|EAW16786.1 as seed ortholog is 100%.

---

Group of orthologs #6774. Best score 583 bits
Score difference with first non-orthologous sequence - *A. fumigatus*:583 *A. fischeri*:515
afum|EAL88820.2 100.00% afis|EAW16189.1 100.00%
Bootstrap support for afum|EAL88820.2 as seed ortholog is 100%.
Bootstrap support for afis|EAW16189.1 as seed ortholog is 100%.

---

Group of orthologs #6775. Best score 583 bits
Score difference with first non-orthologous sequence - *A. fumigatus*:583 *A. fischeri*:583
afum|EAL85130.1 100.00% afis|EAW19770.1 100.00%
Bootstrap support for afum|EAL85130.1 as seed ortholog is 100%.
Bootstrap support for afis|EAW19770.1 as seed ortholog is 100%.

---

Group of orthologs #6776. Best score 583 bits
Score difference with first non-orthologous sequence - *A. fumigatus*:501 *A. fischeri*:498
afum|EAL86151.2 100.00% afis|EAW24480.1 100.00%
Bootstrap support for afum|EAL86151.2 as seed ortholog is 100%.
Bootstrap support for afis|EAW24480.1 as seed ortholog is 100%.

---

Group of orthologs #6777. Best score 583 bits
Score difference with first non-orthologous sequence - *A. fumigatus*:583 *A. fischeri*:583
afum|EAL86245.1 100.00% afis|EAW24616.1 100.00%
Bootstrap support for afum|EAL86245.1 as seed ortholog is 100%.
Bootstrap support for afis|EAW24616.1 as seed ortholog is 100%.

---

Group of orthologs #6778. Best score 583 bits
Score difference with first non-orthologous sequence - *A. fumigatus*:583 *A. fischeri*:583
afum|EAL88290.1 100.00% afis|EAW23193.1 100.00%
Bootstrap support for afum|EAL88290.1 as seed ortholog is 100%.
Bootstrap support for afis|EAW23193.1 as seed ortholog is 100%.

---

Group of orthologs #6779. Best score 583 bits
Score difference with first non-orthologous sequence - *A. fumigatus*:583 *A. fischeri*:583
afum|EAL93504.1 100.00% afis|EAW18772.1 100.00%
Bootstrap support for afum|EAL93504.1 as seed ortholog is 100%.
Bootstrap support for afis|EAW18772.1 as seed ortholog is 100%.

---

Group of orthologs #6780. Best score 583 bits
Score difference with first non-orthologous sequence - *A. fumigatus*:583 *A. fischeri*:583
afum|EAL92757.1 100.00% afis|EAW21724.1 100.00%
Bootstrap support for afum|EAL92757.1 as seed ortholog is 100%.
Bootstrap support for afis|EAW21724.1 as seed ortholog is 100%.

---

Group of orthologs #6781. Best score 583 bits
Score difference with first non-orthologous sequence - *A. fumigatus*:583 *A. fischeri*:583

afum|EAL89786.1 100.00% afis|EAW25115.1 100.00%
Bootstrap support for afum|EAL89786.1 as seed ortholog is 100%.
Bootstrap support for afis|EAW25115.1 as seed ortholog is 100%.

---

Group of orthologs #6782. Best score 583 bits
Score difference with first non-orthologous sequence - *A. fumigatus*:517 *A. fischeri*:583
afum|EBA27445.1 100.00% afis|EAW24045.1 100.00%
Bootstrap support for afum|EBA27445.1 as seed ortholog is 100%.
Bootstrap support for afis|EAW24045.1 as seed ortholog is 100%.

---

Group of orthologs #6783. Best score 582 bits
Score difference with first non-orthologous sequence - *A. fumigatus*:536 *A. fischeri*:518
afum|EAL88881.1 100.00% afis|EAW16260.1 100.00%
Bootstrap support for afum|EAL88881.1 as seed ortholog is 100%.
Bootstrap support for afis|EAW16260.1 as seed ortholog is 100%.

---

Group of orthologs #6784. Best score 582 bits
Score difference with first non-orthologous sequence - *A. fumigatus*:582 *A. fischeri*:582
afum|EAL85075.1 100.00% afis|EAW19816.1 100.00%
Bootstrap support for afum|EAL85075.1 as seed ortholog is 100%.
Bootstrap support for afis|EAW19816.1 as seed ortholog is 100%.

---

Group of orthologs #6785. Best score 582 bits
Score difference with first non-orthologous sequence - *A. fumigatus*:582 *A. fischeri*:582
afum|EAL91395.1 100.00% afis|EAW17582.1 100.00%
Bootstrap support for afum|EAL91395.1 as seed ortholog is 100%.
Bootstrap support for afis|EAW17582.1 as seed ortholog is 100%.

---

Group of orthologs #6786. Best score 582 bits
Score difference with first non-orthologous sequence - *A. fumigatus*:582 *A. fischeri*:582
afum|EAL91545.1 100.00% afis|EAW17730.1 100.00%
Bootstrap support for afum|EAL91545.1 as seed ortholog is 100%.
Bootstrap support for afis|EAW17730.1 as seed ortholog is 100%.

---

Group of orthologs #6787. Best score 582 bits
Score difference with first non-orthologous sequence - *A. fumigatus*:582 *A. fischeri*:582
afum|EAL84278.1 100.00% afis|EAW25554.1 100.00%
Bootstrap support for afum|EAL84278.1 as seed ortholog is 100%.
Bootstrap support for afis|EAW25554.1 as seed ortholog is 100%.

---

Group of orthologs #6788. Best score 582 bits
Score difference with first non-orthologous sequence - *A. fumigatus*:582 *A. fischeri*:521
afum|EAL90745.1 100.00% afis|EAW22446.1 100.00%
Bootstrap support for afum|EAL90745.1 as seed ortholog is 100%.
Bootstrap support for afis|EAW22446.1 as seed ortholog is 100%.

---

Group of orthologs #6789. Best score 582 bits
Score difference with first non-orthologous sequence - *A. fumigatus*:582 *A. fischeri*:463
afum|EAL90138.2 100.00% afis|EAW25475.1 100.00%
Bootstrap support for afum|EAL90138.2 as seed ortholog is 100%.
Bootstrap support for afis|EAW25475.1 as seed ortholog is 100%.

---

Group of orthologs #6790. Best score 582 bits
Score difference with first non-orthologous sequence - *A. fumigatus*:582 *A. fischeri*:582
afum|EBA27462.1 100.00% afis|EAW21534.1 100.00%
Bootstrap support for afum|EBA27462.1 as seed ortholog is 100%.

Bootstrap support for afis|EAW21534.1 as seed ortholog is 100%.

---

Group of orthologs #6791. Best score 581 bits
Score difference with first non-orthologous sequence - *A. fumigatus*:581 *A. fischeri*:581
afum|EAL86594.1 100.00% afis|EAW17150.1 100.00%
Bootstrap support for afum|EAL86594.1 as seed ortholog is 100%.
Bootstrap support for afis|EAW17150.1 as seed ortholog is 100%.

---

Group of orthologs #6792. Best score 581 bits
Score difference with first non-orthologous sequence - *A. fumigatus*:336 *A. fischeri*:486
afum|EAL88548.2 100.00% afis|EAW15913.1 100.00%
Bootstrap support for afum|EAL88548.2 as seed ortholog is 100%.
Bootstrap support for afis|EAW15913.1 as seed ortholog is 100%.

---

Group of orthologs #6793. Best score 581 bits
Score difference with first non-orthologous sequence - *A. fumigatus*:581 *A. fischeri*:581
afum|EAL88770.2 100.00% afis|EAW16141.1 100.00%
Bootstrap support for afum|EAL88770.2 as seed ortholog is 100%.
Bootstrap support for afis|EAW16141.1 as seed ortholog is 100%.

---

Group of orthologs #6794. Best score 581 bits
Score difference with first non-orthologous sequence - *A. fumigatus*:581 *A. fischeri*:581
afum|EAL84589.1 100.00% afis|EAW20464.1 100.00%
Bootstrap support for afum|EAL84589.1 as seed ortholog is 100%.
Bootstrap support for afis|EAW20464.1 as seed ortholog is 100%.

---

Group of orthologs #6795. Best score 581 bits
Score difference with first non-orthologous sequence - *A. fumigatus*:581 *A. fischeri*:581
afum|EAL88989.1 100.00% afis|EAW16364.1 100.00%
Bootstrap support for afum|EAL88989.1 as seed ortholog is 100%.
Bootstrap support for afis|EAW16364.1 as seed ortholog is 100%.

---

Group of orthologs #6796. Best score 581 bits
Score difference with first non-orthologous sequence - *A. fumigatus*:492 *A. fischeri*:391
afum|EAL86272.1 100.00% afis|EAW20694.1 100.00%
Bootstrap support for afum|EAL86272.1 as seed ortholog is 100%.
Bootstrap support for afis|EAW20694.1 as seed ortholog is 100%.

---

Group of orthologs #6797. Best score 581 bits
Score difference with first non-orthologous sequence - *A. fumigatus*:581 *A. fischeri*:581
afum|EAL91500.1 100.00% afis|EAW17686.1 100.00%
Bootstrap support for afum|EAL91500.1 as seed ortholog is 100%.
Bootstrap support for afis|EAW17686.1 as seed ortholog is 100%.

---

Group of orthologs #6798. Best score 581 bits
Score difference with first non-orthologous sequence - *A. fumigatus*:492 *A. fischeri*:430
afum|EAL85922.1 100.00% afis|EAW24254.1 100.00%
Bootstrap support for afum|EAL85922.1 as seed ortholog is 100%.
Bootstrap support for afis|EAW24254.1 as seed ortholog is 100%.

---

Group of orthologs #6799. Best score 581 bits
Score difference with first non-orthologous sequence - *A. fumigatus*:581 *A. fischeri*:581
afum|EAL86015.1 100.00% afis|EAW24344.1 100.00%
Bootstrap support for afum|EAL86015.1 as seed ortholog is 100%.
Bootstrap support for afis|EAW24344.1 as seed ortholog is 100%.

---

Group of orthologs #6800. Best score 581 bits
Score difference with first non-orthologous sequence - *A. fumigatus*:581 *A. fischeri*:247
afum|EAL86249.1 100.00% afis|EAW24621.1 100.00%
Bootstrap support for afum|EAL86249.1 as seed ortholog is 100%.
Bootstrap support for afis|EAW24621.1 as seed ortholog is 100%.

---

Group of orthologs #6801. Best score 581 bits
Score difference with first non-orthologous sequence - *A. fumigatus*:581 *A. fischeri*:581
afum|EAL93056.1 100.00% afis|EAW18381.1 100.00%
Bootstrap support for afum|EAL93056.1 as seed ortholog is 100%.
Bootstrap support for afis|EAW18381.1 as seed ortholog is 100%.

---

Group of orthologs #6802. Best score 581 bits
Score difference with first non-orthologous sequence - *A. fumigatus*:581 *A. fischeri*:581
afum|EAL93507.1 100.00% afis|EAW18775.1 100.00%
Bootstrap support for afum|EAL93507.1 as seed ortholog is 100%.
Bootstrap support for afis|EAW18775.1 as seed ortholog is 100%.

---

Group of orthologs #6803. Best score 581 bits
Score difference with first non-orthologous sequence - *A. fumigatus*:581 *A. fischeri*:581
afum|EAL92075.1 100.00% afis|EAW21036.1 100.00%
Bootstrap support for afum|EAL92075.1 as seed ortholog is 100%.
Bootstrap support for afis|EAW21036.1 as seed ortholog is 100%.

---

Group of orthologs #6804. Best score 581 bits
Score difference with first non-orthologous sequence - *A. fumigatus*:581 *A. fischeri*:581
afum|EAL90894.1 100.00% afis|EAW22287.1 100.00%
Bootstrap support for afum|EAL90894.1 as seed ortholog is 100%.
Bootstrap support for afis|EAW22287.1 as seed ortholog is 100%.

---

Group of orthologs #6805. Best score 581 bits
Score difference with first non-orthologous sequence - *A. fumigatus*:360 *A. fischeri*:581
afum|EAL94073.1 100.00% afis|EAW19353.1 100.00%
Bootstrap support for afum|EAL94073.1 as seed ortholog is 100%.
Bootstrap support for afis|EAW19353.1 as seed ortholog is 100%.

---

Group of orthologs #6806. Best score 581 bits
Score difference with first non-orthologous sequence - *A. fumigatus*:581 *A. fischeri*:581
afum|EBA27294.1 100.00% afis|EAW19623.1 100.00%
Bootstrap support for afum|EBA27294.1 as seed ortholog is 100%.
Bootstrap support for afis|EAW19623.1 as seed ortholog is 100%.

---

Group of orthologs #6807. Best score 581 bits
Score difference with first non-orthologous sequence - *A. fumigatus*:420 *A. fischeri*:363
afum|EAL92665.1 100.00% afis|EAW21632.1 100.00%
Bootstrap support for afum|EAL92665.1 as seed ortholog is 100%.
Bootstrap support for afis|EAW21632.1 as seed ortholog is 100%.

---

Group of orthologs #6808. Best score 581 bits
Score difference with first non-orthologous sequence - *A. fumigatus*:581 *A. fischeri*:506
afum|EAL92896.1 100.00% afis|EAW21882.1 100.00%
Bootstrap support for afum|EAL92896.1 as seed ortholog is 100%.
Bootstrap support for afis|EAW21882.1 as seed ortholog is 100%.

---

Group of orthologs #6809. Best score 580 bits
Score difference with first non-orthologous sequence - *A. fumigatus*:580 *A. fischeri*:580

afum|EAL85469.1 100.00% afis|EAW20351.1 100.00%
Bootstrap support for afum|EAL85469.1 as seed ortholog is 100%.
Bootstrap support for afis|EAW20351.1 as seed ortholog is 100%.

---

Group of orthologs #6810. Best score 580 bits
Score difference with first non-orthologous sequence - *A. fumigatus*:580 *A. fischeri*:580
afum|EAL86074.1 100.00% afis|EAW24403.1 100.00%
Bootstrap support for afum|EAL86074.1 as seed ortholog is 100%.
Bootstrap support for afis|EAW24403.1 as seed ortholog is 100%.

---

Group of orthologs #6811. Best score 580 bits
Score difference with first non-orthologous sequence - *A. fumigatus*:580 *A. fischeri*:580
afum|EAL86201.1 100.00% afis|EAW24537.1 100.00%
Bootstrap support for afum|EAL86201.1 as seed ortholog is 100%.
Bootstrap support for afis|EAW24537.1 as seed ortholog is 100%.

---

Group of orthologs #6812. Best score 580 bits
Score difference with first non-orthologous sequence - *A. fumigatus*:580 *A. fischeri*:292
afum|EAL89344.2 100.00% afis|EAW24702.1 100.00%
Bootstrap support for afum|EAL89344.2 as seed ortholog is 100%.
Bootstrap support for afis|EAW24702.1 as seed ortholog is 100%.

---

Group of orthologs #6813. Best score 579 bits
Score difference with first non-orthologous sequence - *A. fumigatus*:579 *A. fischeri*:579
afum|EAL88497.1 100.00% afis|EAW15862.1 100.00%
Bootstrap support for afum|EAL88497.1 as seed ortholog is 100%.
Bootstrap support for afis|EAW15862.1 as seed ortholog is 100%.

---

Group of orthologs #6814. Best score 579 bits
Score difference with first non-orthologous sequence - *A. fumigatus*:579 *A. fischeri*:579
afum|EAL87011.1 100.00% afis|EAW19484.1 100.00%
Bootstrap support for afum|EAL87011.1 as seed ortholog is 100%.
Bootstrap support for afis|EAW19484.1 as seed ortholog is 100%.

---

Group of orthologs #6815. Best score 579 bits
Score difference with first non-orthologous sequence - *A. fumigatus*:319 *A. fischeri*:509
afum|EAL91219.1 100.00% afis|EAW17384.1 100.00%
Bootstrap support for afum|EAL91219.1 as seed ortholog is 100%.
Bootstrap support for afis|EAW17384.1 as seed ortholog is 100%.

---

Group of orthologs #6816. Best score 579 bits
Score difference with first non-orthologous sequence - *A. fumigatus*:579 *A. fischeri*:579
afum|EAL90794.1 100.00% afis|EAW22397.1 100.00%
Bootstrap support for afum|EAL90794.1 as seed ortholog is 100%.
Bootstrap support for afis|EAW22397.1 as seed ortholog is 100%.

---

Group of orthologs #6817. Best score 579 bits
Score difference with first non-orthologous sequence - *A. fumigatus*:579 *A. fischeri*:579
afum|EAL89765.2 100.00% afis|EAW25094.1 100.00%
Bootstrap support for afum|EAL89765.2 as seed ortholog is 100%.
Bootstrap support for afis|EAW25094.1 as seed ortholog is 100%.

---

Group of orthologs #6818. Best score 578 bits
Score difference with first non-orthologous sequence - *A. fumigatus*:578 *A. fischeri*:578
afum|EAL86345.1 100.00% afis|EAW16776.1 100.00%
Bootstrap support for afum|EAL86345.1 as seed ortholog is 100%.

Bootstrap support for afis|EAW16776.1 as seed ortholog is 100%.

---

Group of orthologs #6819. Best score 578 bits
Score difference with first non-orthologous sequence - *A. fumigatus*:578 *A. fischeri*:578
afum|EAL89951.1 100.00% afis|EAW25288.1 100.00%
Bootstrap support for afum|EAL89951.1 as seed ortholog is 100%.
Bootstrap support for afis|EAW25288.1 as seed ortholog is 100%.

---

Group of orthologs #6820. Best score 577 bits
Score difference with first non-orthologous sequence - *A. fumigatus*:577 *A. fischeri*:577
afum|EAL91719.2 100.00% afis|EAW17910.1 100.00%
Bootstrap support for afum|EAL91719.2 as seed ortholog is 100%.
Bootstrap support for afis|EAW17910.1 as seed ortholog is 100%.

---

Group of orthologs #6821. Best score 577 bits
Score difference with first non-orthologous sequence - *A. fumigatus*:577 *A. fischeri*:577
afum|EAL84313.2 100.00% afis|EAW25625.1 100.00%
Bootstrap support for afum|EAL84313.2 as seed ortholog is 100%.
Bootstrap support for afis|EAW25625.1 as seed ortholog is 100%.

---

Group of orthologs #6822. Best score 577 bits
Score difference with first non-orthologous sequence - *A. fumigatus*:577 *A. fischeri*:577
afum|EAL87899.1 100.00% afis|EAW23568.1 100.00%
Bootstrap support for afum|EAL87899.1 as seed ortholog is 100%.
Bootstrap support for afis|EAW23568.1 as seed ortholog is 100%.

---

Group of orthologs #6823. Best score 577 bits
Score difference with first non-orthologous sequence - *A. fumigatus*:577 *A. fischeri*:577
afum|EAL89521.1 100.00% afis|EAW24871.1 100.00%
Bootstrap support for afum|EAL89521.1 as seed ortholog is 100%.
Bootstrap support for afis|EAW24871.1 as seed ortholog is 100%.

---

Group of orthologs #6824. Best score 576 bits
Score difference with first non-orthologous sequence - *A. fumigatus*:576 *A. fischeri*:534
afum|EAL85761.1 100.00% afis|EAW15573.1 100.00%
Bootstrap support for afum|EAL85761.1 as seed ortholog is 100%.
Bootstrap support for afis|EAW15573.1 as seed ortholog is 100%.

---

Group of orthologs #6825. Best score 576 bits
Score difference with first non-orthologous sequence - *A. fumigatus*:576 *A. fischeri*:576
afum|EAL84359.1 100.00% afis|EAW25690.1 100.00%
Bootstrap support for afum|EAL84359.1 as seed ortholog is 100%.
Bootstrap support for afis|EAW25690.1 as seed ortholog is 100%.

---

Group of orthologs #6826. Best score 576 bits
Score difference with first non-orthologous sequence - *A. fumigatus*:576 *A. fischeri*:576
afum|EAL87246.1 100.00% afis|EAW23811.1 100.00%
Bootstrap support for afum|EAL87246.1 as seed ortholog is 100%.
Bootstrap support for afis|EAW23811.1 as seed ortholog is 100%.

---

Group of orthologs #6827. Best score 576 bits
Score difference with first non-orthologous sequence - *A. fumigatus*:515 *A. fischeri*:576
afum|EAL87902.1 100.00% afis|EAW23566.1 100.00%
Bootstrap support for afum|EAL87902.1 as seed ortholog is 100%.
Bootstrap support for afis|EAW23566.1 as seed ortholog is 100%.

---

Group of orthologs #6828. Best score 576 bits
Score difference with first non-orthologous sequence - *A. fumigatus*:477 *A. fischeri*:484
afum|EAL92052.1 100.00% afis|EAW21014.1 100.00%
Bootstrap support for afum|EAL92052.1 as seed ortholog is 100%.
Bootstrap support for afis|EAW21014.1 as seed ortholog is 100%.

---

Group of orthologs #6829. Best score 576 bits
Score difference with first non-orthologous sequence - *A. fumigatus*:576 *A. fischeri*:576
afum|EAL92715.1 100.00% afis|EAW21681.1 100.00%
Bootstrap support for afum|EAL92715.1 as seed ortholog is 100%.
Bootstrap support for afis|EAW21681.1 as seed ortholog is 100%.

---

Group of orthologs #6830. Best score 576 bits
Score difference with first non-orthologous sequence - *A. fumigatus*:471 *A. fischeri*:576
afum|EAL89666.1 100.00% afis|EAW25029.1 100.00%
Bootstrap support for afum|EAL89666.1 as seed ortholog is 100%.
Bootstrap support for afis|EAW25029.1 as seed ortholog is 100%.

---

Group of orthologs #6831. Best score 576 bits
Score difference with first non-orthologous sequence - *A. fumigatus*:576 *A. fischeri*:576
afum|EAL90185.1 100.00% afis|EAW25520.1 100.00%
Bootstrap support for afum|EAL90185.1 as seed ortholog is 100%.
Bootstrap support for afis|EAW25520.1 as seed ortholog is 100%.

---

Group of orthologs #6832. Best score 575 bits
Score difference with first non-orthologous sequence - *A. fumigatus*:575 *A. fischeri*:575
afum|EAL84192.1 100.00% afis|EAW15453.1 100.00%
Bootstrap support for afum|EAL84192.1 as seed ortholog is 100%.
Bootstrap support for afis|EAW15453.1 as seed ortholog is 100%.

---

Group of orthologs #6833. Best score 575 bits
Score difference with first non-orthologous sequence - *A. fumigatus*:575 *A. fischeri*:575
afum|EAL86705.1 100.00% afis|EAW17287.1 100.00%
Bootstrap support for afum|EAL86705.1 as seed ortholog is 100%.
Bootstrap support for afis|EAW17287.1 as seed ortholog is 100%.

---

Group of orthologs #6834. Best score 575 bits
Score difference with first non-orthologous sequence - *A. fumigatus*:575 *A. fischeri*:575
afum|EAL88575.1 100.00% afis|EAW15941.1 100.00%
Bootstrap support for afum|EAL88575.1 as seed ortholog is 100%.
Bootstrap support for afis|EAW15941.1 as seed ortholog is 100%.

---

Group of orthologs #6835. Best score 575 bits
Score difference with first non-orthologous sequence - *A. fumigatus*:575 *A. fischeri*:507
afum|EAL85047.1 100.00% afis|EAW19847.1 100.00%
Bootstrap support for afum|EAL85047.1 as seed ortholog is 100%.
Bootstrap support for afis|EAW19847.1 as seed ortholog is 100%.

---

Group of orthologs #6836. Best score 575 bits
Score difference with first non-orthologous sequence - *A. fumigatus*:575 *A. fischeri*:575
afum|EAL89037.1 100.00% afis|EAW16413.1 100.00%
Bootstrap support for afum|EAL89037.1 as seed ortholog is 100%.
Bootstrap support for afis|EAW16413.1 as seed ortholog is 100%.

---

Group of orthologs #6837. Best score 575 bits
Score difference with first non-orthologous sequence - *A. fumigatus*:575 *A. fischeri*:575

afum|EAL85470.1 100.00% afis|EAW20352.1 100.00%
Bootstrap support for afum|EAL85470.1 as seed ortholog is 100%.
Bootstrap support for afis|EAW20352.1 as seed ortholog is 100%.

---

Group of orthologs #6838. Best score 575 bits
Score difference with first non-orthologous sequence - *A. fumigatus*:575 *A. fischeri*:575
afum|EAL91602.1 100.00% afis|EAW17793.1 100.00%
Bootstrap support for afum|EAL91602.1 as seed ortholog is 100%.
Bootstrap support for afis|EAW17793.1 as seed ortholog is 100%.

---

Group of orthologs #6839. Best score 575 bits
Score difference with first non-orthologous sequence - *A. fumigatus*:575 *A. fischeri*:575
afum|EAL88372.1 100.00% afis|EAW23107.1 100.00%
Bootstrap support for afum|EAL88372.1 as seed ortholog is 100%.
Bootstrap support for afis|EAW23107.1 as seed ortholog is 100%.

---

Group of orthologs #6840. Best score 575 bits
Score difference with first non-orthologous sequence - *A. fumigatus*:575 *A. fischeri*:575
afum|EAL87778.1 100.00% afis|EAW24661.1 100.00%
Bootstrap support for afum|EAL87778.1 as seed ortholog is 100%.
Bootstrap support for afis|EAW24661.1 as seed ortholog is 100%.

---

Group of orthologs #6841. Best score 575 bits
Score difference with first non-orthologous sequence - *A. fumigatus*:575 *A. fischeri*:575
afum|EAL90960.1 100.00% afis|EAW22233.1 100.00%
Bootstrap support for afum|EAL90960.1 as seed ortholog is 100%.
Bootstrap support for afis|EAW22233.1 as seed ortholog is 100%.

---

Group of orthologs #6842. Best score 575 bits
Score difference with first non-orthologous sequence - *A. fumigatus*:575 *A. fischeri*:575
afum|EAL89556.1 100.00% afis|EAW24902.1 100.00%
Bootstrap support for afum|EAL89556.1 as seed ortholog is 100%.
Bootstrap support for afis|EAW24902.1 as seed ortholog is 100%.

---

Group of orthologs #6843. Best score 574 bits
Score difference with first non-orthologous sequence - *A. fumigatus*:574 *A. fischeri*:574
afum|EAL88701.2 100.00% afis|EAW16069.1 100.00%
Bootstrap support for afum|EAL88701.2 as seed ortholog is 100%.
Bootstrap support for afis|EAW16069.1 as seed ortholog is 100%.

---

Group of orthologs #6844. Best score 574 bits
Score difference with first non-orthologous sequence - *A. fumigatus*:574 *A. fischeri*:574
afum|EAL87824.1 100.00% afis|EAW23636.1 100.00%
Bootstrap support for afum|EAL87824.1 as seed ortholog is 100%.
Bootstrap support for afis|EAW23636.1 as seed ortholog is 100%.

---

Group of orthologs #6845. Best score 574 bits
Score difference with first non-orthologous sequence - *A. fumigatus*:574 *A. fischeri*:574
afum|EAL93758.1 100.00% afis|EAW19028.1 100.00%
Bootstrap support for afum|EAL93758.1 as seed ortholog is 100%.
Bootstrap support for afis|EAW19028.1 as seed ortholog is 100%.

---

Group of orthologs #6846. Best score 574 bits
Score difference with first non-orthologous sequence - *A. fumigatus*:574 *A. fischeri*:574
afum|EAL93993.1 100.00% afis|EAW19266.1 100.00%
Bootstrap support for afum|EAL93993.1 as seed ortholog is 100%.

Bootstrap support for afis|EAW19266.1 as seed ortholog is 100%.

---

Group of orthologs #6847. Best score 573 bits
Score difference with first non-orthologous sequence - *A. fumigatus*:573 *A. fischeri*:573
afum|EAL84502.1 100.00% afis|EAW20559.1 100.00%
Bootstrap support for afum|EAL84502.1 as seed ortholog is 100%.
Bootstrap support for afis|EAW20559.1 as seed ortholog is 100%.

---

Group of orthologs #6848. Best score 573 bits
Score difference with first non-orthologous sequence - *A. fumigatus*:573 *A. fischeri*:573
afum|EAL89150.1 100.00% afis|EAW16552.1 100.00%
Bootstrap support for afum|EAL89150.1 as seed ortholog is 100%.
Bootstrap support for afis|EAW16552.1 as seed ortholog is 100%.

---

Group of orthologs #6849. Best score 573 bits
Score difference with first non-orthologous sequence - *A. fumigatus*:573 *A. fischeri*:573
afum|EAL91263.1 100.00% afis|EAW17436.1 100.00%
Bootstrap support for afum|EAL91263.1 as seed ortholog is 100%.
Bootstrap support for afis|EAW17436.1 as seed ortholog is 100%.

---

Group of orthologs #6850. Best score 573 bits
Score difference with first non-orthologous sequence - *A. fumigatus*:573 *A. fischeri*:573
afum|EAL89464.1 100.00% afis|EAW24820.1 100.00%
Bootstrap support for afum|EAL89464.1 as seed ortholog is 100%.
Bootstrap support for afis|EAW24820.1 as seed ortholog is 100%.

---

Group of orthologs #6851. Best score 572 bits
Score difference with first non-orthologous sequence - *A. fumigatus*:572 *A. fischeri*:572
afum|EAL84240.1 100.00% afis|EAW15406.1 100.00%
Bootstrap support for afum|EAL84240.1 as seed ortholog is 100%.
Bootstrap support for afis|EAW15406.1 as seed ortholog is 100%.

---

Group of orthologs #6852. Best score 572 bits
Score difference with first non-orthologous sequence - *A. fumigatus*:485 *A. fischeri*:491
afum|EAL86358.1 100.00% afis|EAW16788.1 100.00%
Bootstrap support for afum|EAL86358.1 as seed ortholog is 100%.
Bootstrap support for afis|EAW16788.1 as seed ortholog is 100%.

---

Group of orthologs #6853. Best score 572 bits
Score difference with first non-orthologous sequence - *A. fumigatus*:572 *A. fischeri*:572
afum|EAL88763.1 100.00% afis|EAW16133.1 100.00%
Bootstrap support for afum|EAL88763.1 as seed ortholog is 100%.
Bootstrap support for afis|EAW16133.1 as seed ortholog is 100%.

---

Group of orthologs #6854. Best score 572 bits
Score difference with first non-orthologous sequence - *A. fumigatus*:572 *A. fischeri*:572
afum|EAL85040.1 100.00% afis|EAW19855.1 100.00%
Bootstrap support for afum|EAL85040.1 as seed ortholog is 100%.
Bootstrap support for afis|EAW19855.1 as seed ortholog is 100%.

---

Group of orthologs #6855. Best score 572 bits
Score difference with first non-orthologous sequence - *A. fumigatus*:572 *A. fischeri*:572
afum|EAL84962.1 100.00% afis|EAW19953.1 100.00%
Bootstrap support for afum|EAL84962.1 as seed ortholog is 100%.
Bootstrap support for afis|EAW19953.1 as seed ortholog is 100%.

---

Group of orthologs #6856. Best score 572 bits
Score difference with first non-orthologous sequence - *A. fumigatus*:572 *A. fischeri*:572
afum|EAL85342.1 100.00% afis|EAW20225.1 100.00%
Bootstrap support for afum|EAL85342.1 as seed ortholog is 100%.
Bootstrap support for afis|EAW20225.1 as seed ortholog is 100%.

---

Group of orthologs #6857. Best score 572 bits
Score difference with first non-orthologous sequence - *A. fumigatus*:572 *A. fischeri*:572
afum|EAL91771.1 100.00% afis|EAW17960.1 100.00%
Bootstrap support for afum|EAL91771.1 as seed ortholog is 100%.
Bootstrap support for afis|EAW17960.1 as seed ortholog is 100%.

---

Group of orthologs #6858. Best score 572 bits
Score difference with first non-orthologous sequence - *A. fumigatus*:572 *A. fischeri*:572
afum|EAL88139.1 100.00% afis|EAW23339.1 100.00%
Bootstrap support for afum|EAL88139.1 as seed ortholog is 100%.
Bootstrap support for afis|EAW23339.1 as seed ortholog is 100%.

---

Group of orthologs #6859. Best score 572 bits
Score difference with first non-orthologous sequence - *A. fumigatus*:572 *A. fischeri*:572
afum|EAL90543.1 100.00% afis|EAW22649.1 100.00%
Bootstrap support for afum|EAL90543.1 as seed ortholog is 100%.
Bootstrap support for afis|EAW22649.1 as seed ortholog is 100%.

---

Group of orthologs #6860. Best score 572 bits
Score difference with first non-orthologous sequence - *A. fumigatus*:572 *A. fischeri*:572
afum|EAL92486.1 100.00% afis|EAW21466.1 100.00%
Bootstrap support for afum|EAL92486.1 as seed ortholog is 100%.
Bootstrap support for afis|EAW21466.1 as seed ortholog is 100%.

---

Group of orthologs #6861. Best score 572 bits
Score difference with first non-orthologous sequence - *A. fumigatus*:572 *A. fischeri*:446
afum|EAL89943.1 100.00% afis|EAW25280.1 100.00%
Bootstrap support for afum|EAL89943.1 as seed ortholog is 100%.
Bootstrap support for afis|EAW25280.1 as seed ortholog is 100%.

---

Group of orthologs #6862. Best score 571 bits
Score difference with first non-orthologous sequence - *A. fumigatus*:318 *A. fischeri*:348
afum|EAL88500.1 100.00% afis|EAW15866.1 100.00%
Bootstrap support for afum|EAL88500.1 as seed ortholog is 100%.
Bootstrap support for afis|EAW15866.1 as seed ortholog is 100%.

---

Group of orthologs #6863. Best score 571 bits
Score difference with first non-orthologous sequence - *A. fumigatus*:571 *A. fischeri*:571
afum|EAL88577.1 100.00% afis|EAW15943.1 100.00%
Bootstrap support for afum|EAL88577.1 as seed ortholog is 100%.
Bootstrap support for afis|EAW15943.1 as seed ortholog is 100%.

---

Group of orthologs #6864. Best score 571 bits
Score difference with first non-orthologous sequence - *A. fumigatus*:571 *A. fischeri*:571
afum|EAL93277.1 100.00% afis|EAW18574.1 100.00%
Bootstrap support for afum|EAL93277.1 as seed ortholog is 100%.
Bootstrap support for afis|EAW18574.1 as seed ortholog is 100%.

---

Group of orthologs #6865. Best score 571 bits
Score difference with first non-orthologous sequence - *A. fumigatus*:571 *A. fischeri*:571

afum|EAL90926.1 100.00% afis|EAW22248.1 100.00%
Bootstrap support for afum|EAL90926.1 as seed ortholog is 100%.
Bootstrap support for afis|EAW22248.1 as seed ortholog is 100%.

---

Group of orthologs #6866. Best score 571 bits
Score difference with first non-orthologous sequence - *A. fumigatus*:571 *A. fischeri*:571
afum|EAL92281.1 100.00% afis|EAW21266.1 100.00%
Bootstrap support for afum|EAL92281.1 as seed ortholog is 100%.
Bootstrap support for afis|EAW21266.1 as seed ortholog is 100%.

---

Group of orthologs #6867. Best score 570 bits
Score difference with first non-orthologous sequence - *A. fumigatus*:570 *A. fischeri*:570
afum|EAL91159.1 100.00% afis|EAW17323.1 100.00%
Bootstrap support for afum|EAL91159.1 as seed ortholog is 100%.
Bootstrap support for afis|EAW17323.1 as seed ortholog is 100%.

---

Group of orthologs #6868. Best score 570 bits
Score difference with first non-orthologous sequence - *A. fumigatus*:570 *A. fischeri*:570
afum|EAL84291.1 100.00% afis|EAW25580.1 100.00%
Bootstrap support for afum|EAL84291.1 as seed ortholog is 100%.
Bootstrap support for afis|EAW25580.1 as seed ortholog is 100%.

---

Group of orthologs #6869. Best score 570 bits
Score difference with first non-orthologous sequence - *A. fumigatus*:570 *A. fischeri*:570
afum|EAL87231.2 100.00% afis|EAW23797.1 100.00%
Bootstrap support for afum|EAL87231.2 as seed ortholog is 100%.
Bootstrap support for afis|EAW23797.1 as seed ortholog is 100%.

---

Group of orthologs #6870. Best score 570 bits
Score difference with first non-orthologous sequence - *A. fumigatus*:570 *A. fischeri*:404
afum|EAL88184.1 100.00% afis|EAW23294.1 100.00%
Bootstrap support for afum|EAL88184.1 as seed ortholog is 100%.
Bootstrap support for afis|EAW23294.1 as seed ortholog is 100%.

---

Group of orthologs #6871. Best score 570 bits
Score difference with first non-orthologous sequence - *A. fumigatus*:570 *A. fischeri*:570
afum|EAL93759.1 100.00% afis|EAW19029.1 100.00%
Bootstrap support for afum|EAL93759.1 as seed ortholog is 100%.
Bootstrap support for afis|EAW19029.1 as seed ortholog is 100%.

---

Group of orthologs #6872. Best score 569 bits
Score difference with first non-orthologous sequence - *A. fumigatus*:569 *A. fischeri*:569
afum|EAL88530.1 100.00% afis|EAW15895.1 100.00%
Bootstrap support for afum|EAL88530.1 as seed ortholog is 100%.
Bootstrap support for afis|EAW15895.1 as seed ortholog is 100%.

---

Group of orthologs #6873. Best score 569 bits
Score difference with first non-orthologous sequence - *A. fumigatus*:569 *A. fischeri*:569
afum|EAL88353.1 100.00% afis|EAW23127.1 100.00%
Bootstrap support for afum|EAL88353.1 as seed ortholog is 100%.
Bootstrap support for afis|EAW23127.1 as seed ortholog is 100%.

---

Group of orthologs #6874. Best score 569 bits
Score difference with first non-orthologous sequence - *A. fumigatus*:569 *A. fischeri*:569
afum|EAL88192.1 100.00% afis|EAW23287.1 100.00%
Bootstrap support for afum|EAL88192.1 as seed ortholog is 100%.

Bootstrap support for afis|EAW23287.1 as seed ortholog is 100%.

---

Group of orthologs #6875. Best score 569 bits
Score difference with first non-orthologous sequence - *A. fumigatus*:569 *A. fischeri*:569
afum|EAL93438.1 100.00% afis|EAW18713.1 100.00%
Bootstrap support for afum|EAL93438.1 as seed ortholog is 100%.
Bootstrap support for afis|EAW18713.1 as seed ortholog is 100%.

---

Group of orthologs #6876. Best score 569 bits
Score difference with first non-orthologous sequence - *A. fumigatus*:569 *A. fischeri*:569
afum|EAL90519.1 100.00% afis|EAW22673.1 100.00%
Bootstrap support for afum|EAL90519.1 as seed ortholog is 100%.
Bootstrap support for afis|EAW22673.1 as seed ortholog is 100%.

---

Group of orthologs #6877. Best score 569 bits
Score difference with first non-orthologous sequence - *A. fumigatus*:569 *A. fischeri*:277
afum|EAL90117.1 100.00% afis|EAW25457.1 100.00%
Bootstrap support for afum|EAL90117.1 as seed ortholog is 100%.
Bootstrap support for afis|EAW25457.1 as seed ortholog is 100%.

---

Group of orthologs #6878. Best score 569 bits
Score difference with first non-orthologous sequence - *A. fumigatus*:569 *A. fischeri*:569
afum|EBA27419.1 100.00% afis|EAW22198.1 100.00%
Bootstrap support for afum|EBA27419.1 as seed ortholog is 100%.
Bootstrap support for afis|EAW22198.1 as seed ortholog is 100%.

---

Group of orthologs #6879. Best score 568 bits
Score difference with first non-orthologous sequence - *A. fumigatus*:568 *A. fischeri*:568
afum|EAL86213.1 100.00% afis|EAW24549.1 100.00%
Bootstrap support for afum|EAL86213.1 as seed ortholog is 100%.
Bootstrap support for afis|EAW24549.1 as seed ortholog is 100%.

---

Group of orthologs #6880. Best score 568 bits
Score difference with first non-orthologous sequence - *A. fumigatus*:568 *A. fischeri*:568
afum|EAL93424.1 100.00% afis|EAW18701.1 100.00%
Bootstrap support for afum|EAL93424.1 as seed ortholog is 100%.
Bootstrap support for afis|EAW18701.1 as seed ortholog is 100%.

---

Group of orthologs #6881. Best score 568 bits
Score difference with first non-orthologous sequence - *A. fumigatus*:568 *A. fischeri*:568
afum|EAL93598.1 100.00% afis|EAW18866.1 100.00%
Bootstrap support for afum|EAL93598.1 as seed ortholog is 100%.
Bootstrap support for afis|EAW18866.1 as seed ortholog is 100%.

---

Group of orthologs #6882. Best score 567 bits
Score difference with first non-orthologous sequence - *A. fumigatus*:567 *A. fischeri*:567
afum|EAL86385.1 100.00% afis|EAW16837.1 100.00%
Bootstrap support for afum|EAL86385.1 as seed ortholog is 100%.
Bootstrap support for afis|EAW16837.1 as seed ortholog is 100%.

---

Group of orthologs #6883. Best score 567 bits
Score difference with first non-orthologous sequence - *A. fumigatus*:567 *A. fischeri*:567
afum|EAL89709.1 100.00% afis|EAW25041.1 100.00%
Bootstrap support for afum|EAL89709.1 as seed ortholog is 100%.
Bootstrap support for afis|EAW25041.1 as seed ortholog is 100%.

---

Group of orthologs #6884. Best score 566 bits
Score difference with first non-orthologous sequence - *A. fumigatus*:566 *A. fischeri*:566
afum|EAL85728.1 100.00% afis|EAW15605.1 100.00%
Bootstrap support for afum|EAL85728.1 as seed ortholog is 100%.
Bootstrap support for afis|EAW15605.1 as seed ortholog is 100%.

---

Group of orthologs #6885. Best score 566 bits
Score difference with first non-orthologous sequence - *A. fumigatus*:566 *A. fischeri*:566
afum|EAL88714.1 100.00% afis|EAW16082.1 100.00%
Bootstrap support for afum|EAL88714.1 as seed ortholog is 100%.
Bootstrap support for afis|EAW16082.1 as seed ortholog is 100%.

---

Group of orthologs #6886. Best score 566 bits
Score difference with first non-orthologous sequence - *A. fumigatus*:566 *A. fischeri*:566
afum|EAL89118.1 100.00% afis|EAW16523.1 100.00%
Bootstrap support for afum|EAL89118.1 as seed ortholog is 100%.
Bootstrap support for afis|EAW16523.1 as seed ortholog is 100%.

---

Group of orthologs #6887. Best score 566 bits
Score difference with first non-orthologous sequence - *A. fumigatus*:566 *A. fischeri*:566
afum|EAL85361.1 100.00% afis|EAW20244.1 100.00%
Bootstrap support for afum|EAL85361.1 as seed ortholog is 100%.
Bootstrap support for afis|EAW20244.1 as seed ortholog is 100%.

---

Group of orthologs #6888. Best score 566 bits
Score difference with first non-orthologous sequence - *A. fumigatus*:566 *A. fischeri*:566
afum|EAL93063.1 100.00% afis|EAW18388.1 100.00%
Bootstrap support for afum|EAL93063.1 as seed ortholog is 100%.
Bootstrap support for afis|EAW18388.1 as seed ortholog is 100%.

---

Group of orthologs #6889. Best score 566 bits
Score difference with first non-orthologous sequence - *A. fumigatus*:566 *A. fischeri*:566
afum|EAL93830.1 100.00% afis|EAW19098.1 100.00%
Bootstrap support for afum|EAL93830.1 as seed ortholog is 100%.
Bootstrap support for afis|EAW19098.1 as seed ortholog is 100%.

---

Group of orthologs #6890. Best score 566 bits
Score difference with first non-orthologous sequence - *A. fumigatus*:566 *A. fischeri*:566
afum|EAL93937.1 100.00% afis|EAW19207.1 100.00%
Bootstrap support for afum|EAL93937.1 as seed ortholog is 100%.
Bootstrap support for afis|EAW19207.1 as seed ortholog is 100%.

---

Group of orthologs #6891. Best score 566 bits
Score difference with first non-orthologous sequence - *A. fumigatus*:566 *A. fischeri*:566
afum|EAL92106.1 100.00% afis|EAW21084.1 100.00%
Bootstrap support for afum|EAL92106.1 as seed ortholog is 100%.
Bootstrap support for afis|EAW21084.1 as seed ortholog is 100%.

---

Group of orthologs #6892. Best score 566 bits
Score difference with first non-orthologous sequence - *A. fumigatus*:566 *A. fischeri*:566
afum|EAL89777.1 100.00% afis|EAW25106.1 100.00%
Bootstrap support for afum|EAL89777.1 as seed ortholog is 100%.
Bootstrap support for afis|EAW25106.1 as seed ortholog is 100%.

---

Group of orthologs #6893. Best score 565 bits
Score difference with first non-orthologous sequence - *A. fumigatus*:509 *A. fischeri*:509

afum|EAL86520.1 100.00% afis|EAW16996.1 100.00%
Bootstrap support for afum|EAL86520.1 as seed ortholog is 100%.
Bootstrap support for afis|EAW16996.1 as seed ortholog is 100%.

---

Group of orthologs #6894. Best score 565 bits
Score difference with first non-orthologous sequence - *A. fumigatus*:565 *A. fischeri*:565
afum|EAL84519.2 100.00% afis|EAW20539.1 100.00%
Bootstrap support for afum|EAL84519.2 as seed ortholog is 100%.
Bootstrap support for afis|EAW20539.1 as seed ortholog is 100%.

---

Group of orthologs #6895. Best score 565 bits
Score difference with first non-orthologous sequence - *A. fumigatus*:285 *A. fischeri*:565
afum|EAL85426.1 100.00% afis|EAW20297.1 100.00%
Bootstrap support for afum|EAL85426.1 as seed ortholog is 100%.
Bootstrap support for afis|EAW20297.1 as seed ortholog is 100%.

---

Group of orthologs #6896. Best score 565 bits
Score difference with first non-orthologous sequence - *A. fumigatus*:565 *A. fischeri*:565
afum|EAL86906.2 100.00% afis|EAW19616.1 100.00%
Bootstrap support for afum|EAL86906.2 as seed ortholog is 100%.
Bootstrap support for afis|EAW19616.1 as seed ortholog is 100%.

---

Group of orthologs #6897. Best score 565 bits
Score difference with first non-orthologous sequence - *A. fumigatus*:565 *A. fischeri*:565
afum|EAL89583.1 100.00% afis|EAW24927.1 100.00%
Bootstrap support for afum|EAL89583.1 as seed ortholog is 100%.
Bootstrap support for afis|EAW24927.1 as seed ortholog is 100%.

---

Group of orthologs #6898. Best score 565 bits
Score difference with first non-orthologous sequence - *A. fumigatus*:565 *A. fischeri*:565
afum|EAL90197.1 100.00% afis|EAW25533.1 100.00%
Bootstrap support for afum|EAL90197.1 as seed ortholog is 100%.
Bootstrap support for afis|EAW25533.1 as seed ortholog is 100%.

---

Group of orthologs #6899. Best score 564 bits
Score difference with first non-orthologous sequence - *A. fumigatus*:564 *A. fischeri*:564
afum|EAL89139.1 100.00% afis|EAW16542.1 100.00%
Bootstrap support for afum|EAL89139.1 as seed ortholog is 100%.
Bootstrap support for afis|EAW16542.1 as seed ortholog is 100%.

---

Group of orthologs #6900. Best score 564 bits
Score difference with first non-orthologous sequence - *A. fumigatus*:564 *A. fischeri*:564
afum|EAL86903.1 100.00% afis|EAW19619.1 100.00%
Bootstrap support for afum|EAL86903.1 as seed ortholog is 100%.
Bootstrap support for afis|EAW19619.1 as seed ortholog is 100%.

---

Group of orthologs #6901. Best score 564 bits
Score difference with first non-orthologous sequence - *A. fumigatus*:564 *A. fischeri*:564
afum|EAL86883.1 100.00% afis|EAW19639.1 100.00%
Bootstrap support for afum|EAL86883.1 as seed ortholog is 100%.
Bootstrap support for afis|EAW19639.1 as seed ortholog is 100%.

---

Group of orthologs #6902. Best score 564 bits
Score difference with first non-orthologous sequence - *A. fumigatus*:564 *A. fischeri*:564
afum|EAL86832.1 100.00% afis|EAW19695.1 100.00%
Bootstrap support for afum|EAL86832.1 as seed ortholog is 100%.

Bootstrap support for afis|EAW19695.1 as seed ortholog is 100%.

---

Group of orthologs #6903. Best score 564 bits
Score difference with first non-orthologous sequence - *A. fumigatus*:564 *A. fischeri*:564
afum|EAL91243.1 100.00% afis|EAW17409.1 100.00%
Bootstrap support for afum|EAL91243.1 as seed ortholog is 100%.
Bootstrap support for afis|EAW17409.1 as seed ortholog is 100%.

---

Group of orthologs #6904. Best score 564 bits
Score difference with first non-orthologous sequence - *A. fumigatus*:564 *A. fischeri*:564
afum|EAL84300.1 100.00% afis|EAW25593.1 100.00%
Bootstrap support for afum|EAL84300.1 as seed ortholog is 100%.
Bootstrap support for afis|EAW25593.1 as seed ortholog is 100%.

---

Group of orthologs #6905. Best score 564 bits
Score difference with first non-orthologous sequence - *A. fumigatus*:564 *A. fischeri*:335
afum|EAL93407.1 100.00% afis|EAW18686.1 100.00%
Bootstrap support for afum|EAL93407.1 as seed ortholog is 100%.
Bootstrap support for afis|EAW18686.1 as seed ortholog is 100%.

---

Group of orthologs #6906. Best score 564 bits
Score difference with first non-orthologous sequence - *A. fumigatus*:564 *A. fischeri*:321
afum|EAL93702.1 100.00% afis|EAW18969.1 100.00%
Bootstrap support for afum|EAL93702.1 as seed ortholog is 100%.
Bootstrap support for afis|EAW18969.1 as seed ortholog is 100%.

---

Group of orthologs #6907. Best score 564 bits
Score difference with first non-orthologous sequence - *A. fumigatus*:564 *A. fischeri*:564
afum|EAL89634.1 100.00% afis|EAW24974.1 100.00%
Bootstrap support for afum|EAL89634.1 as seed ortholog is 100%.
Bootstrap support for afis|EAW24974.1 as seed ortholog is 100%.

---

Group of orthologs #6908. Best score 564 bits
Score difference with first non-orthologous sequence - *A. fumigatus*:564 *A. fischeri*:564
afum|EBA27331.1 100.00% afis|EAW23450.1 100.00%
Bootstrap support for afum|EBA27331.1 as seed ortholog is 100%.
Bootstrap support for afis|EAW23450.1 as seed ortholog is 100%.

---

Group of orthologs #6909. Best score 563 bits
Score difference with first non-orthologous sequence - *A. fumigatus*:563 *A. fischeri*:563
afum|EAL86161.1 100.00% afis|EAW24492.1 100.00%
Bootstrap support for afum|EAL86161.1 as seed ortholog is 100%.
Bootstrap support for afis|EAW24492.1 as seed ortholog is 100%.

---

Group of orthologs #6910. Best score 563 bits
Score difference with first non-orthologous sequence - *A. fumigatus*:563 *A. fischeri*:563
afum|EAL88257.1 100.00% afis|EAW23226.1 100.00%
Bootstrap support for afum|EAL88257.1 as seed ortholog is 100%.
Bootstrap support for afis|EAW23226.1 as seed ortholog is 100%.

---

Group of orthologs #6911. Best score 562 bits
Score difference with first non-orthologous sequence - *A. fumigatus*:562 *A. fischeri*:562
afum|EAL86297.2 100.00% afis|EAW16723.1 100.00%
Bootstrap support for afum|EAL86297.2 as seed ortholog is 100%.
Bootstrap support for afis|EAW16723.1 as seed ortholog is 100%.

---

Group of orthologs #6912. Best score 562 bits
Score difference with first non-orthologous sequence - *A. fumigatus*:371 *A. fischeri*:562
afum|EAL86408.2 100.00% afis|EAW16869.1 100.00%
Bootstrap support for afum|EAL86408.2 as seed ortholog is 100%.
Bootstrap support for afis|EAW16869.1 as seed ortholog is 100%.

---

Group of orthologs #6913. Best score 562 bits
Score difference with first non-orthologous sequence - *A. fumigatus*:562 *A. fischeri*:562
afum|EAL87071.1 100.00% afis|EAW19423.1 100.00%
Bootstrap support for afum|EAL87071.1 as seed ortholog is 100%.
Bootstrap support for afis|EAW19423.1 as seed ortholog is 100%.

---

Group of orthologs #6914. Best score 562 bits
Score difference with first non-orthologous sequence - *A. fumigatus*:562 *A. fischeri*:562
afum|EAL91305.1 100.00% afis|EAW17476.1 100.00%
Bootstrap support for afum|EAL91305.1 as seed ortholog is 100%.
Bootstrap support for afis|EAW17476.1 as seed ortholog is 100%.

---

Group of orthologs #6915. Best score 562 bits
Score difference with first non-orthologous sequence - *A. fumigatus*:562 *A. fischeri*:562
afum|EAL91501.1 100.00% afis|EAW17687.1 100.00%
Bootstrap support for afum|EAL91501.1 as seed ortholog is 100%.
Bootstrap support for afis|EAW17687.1 as seed ortholog is 100%.

---

Group of orthologs #6916. Best score 562 bits
Score difference with first non-orthologous sequence - *A. fumigatus*:562 *A. fischeri*:562
afum|EAL87258.1 100.00% afis|EAW23826.1 100.00%
Bootstrap support for afum|EAL87258.1 as seed ortholog is 100%.
Bootstrap support for afis|EAW23826.1 as seed ortholog is 100%.

---

Group of orthologs #6917. Best score 562 bits
Score difference with first non-orthologous sequence - *A. fumigatus*:562 *A. fischeri*:562
afum|EAL88312.2 100.00% afis|EAW23171.1 100.00%
Bootstrap support for afum|EAL88312.2 as seed ortholog is 100%.
Bootstrap support for afis|EAW23171.1 as seed ortholog is 100%.

---

Group of orthologs #6918. Best score 562 bits
Score difference with first non-orthologous sequence - *A. fumigatus*:562 *A. fischeri*:562
afum|EAL93482.1 100.00% afis|EAW18755.1 100.00%
Bootstrap support for afum|EAL93482.1 as seed ortholog is 100%.
Bootstrap support for afis|EAW18755.1 as seed ortholog is 100%.

---

Group of orthologs #6919. Best score 562 bits
Score difference with first non-orthologous sequence - *A. fumigatus*:562 *A. fischeri*:562
afum|EAL93611.1 100.00% afis|EAW18879.1 100.00%
Bootstrap support for afum|EAL93611.1 as seed ortholog is 100%.
Bootstrap support for afis|EAW18879.1 as seed ortholog is 100%.

---

Group of orthologs #6920. Best score 562 bits
Score difference with first non-orthologous sequence - *A. fumigatus*:562 *A. fischeri*:562
afum|EAL93642.1 100.00% afis|EAW18911.1 100.00%
Bootstrap support for afum|EAL93642.1 as seed ortholog is 100%.
Bootstrap support for afis|EAW18911.1 as seed ortholog is 100%.

---

Group of orthologs #6921. Best score 561 bits
Score difference with first non-orthologous sequence - *A. fumigatus*:561 *A. fischeri*:561

afum|EAL85224.1 100.00% afis|EAW20107.1 100.00%
Bootstrap support for afum|EAL85224.1 as seed ortholog is 100%.
Bootstrap support for afis|EAW20107.1 as seed ortholog is 100%.

---

Group of orthologs #6922. Best score 561 bits
Score difference with first non-orthologous sequence - *A. fumigatus*:561 *A. fischeri*:257
afum|EAL86857.1 100.00% afis|EAW19666.1 100.00%
Bootstrap support for afum|EAL86857.1 as seed ortholog is 100%.
Bootstrap support for afis|EAW19666.1 as seed ortholog is 100%.

---

Group of orthologs #6923. Best score 561 bits
Score difference with first non-orthologous sequence - *A. fumigatus*:561 *A. fischeri*:561
afum|EAL90755.1 100.00% afis|EAW22436.1 100.00%
Bootstrap support for afum|EAL90755.1 as seed ortholog is 100%.
Bootstrap support for afis|EAW22436.1 as seed ortholog is 100%.

---

Group of orthologs #6924. Best score 561 bits
Score difference with first non-orthologous sequence - *A. fumigatus*:561 *A. fischeri*:561
afum|EAL89696.1 100.00% afis|EAW25002.1 100.00%
Bootstrap support for afum|EAL89696.1 as seed ortholog is 100%.
Bootstrap support for afis|EAW25002.1 as seed ortholog is 100%.

---

Group of orthologs #6925. Best score 561 bits
Score difference with first non-orthologous sequence - *A. fumigatus*:561 *A. fischeri*:561
afum|EAL89992.1 100.00% afis|EAW25330.1 100.00%
Bootstrap support for afum|EAL89992.1 as seed ortholog is 100%.
Bootstrap support for afis|EAW25330.1 as seed ortholog is 100%.

---

Group of orthologs #6926. Best score 560 bits
Score difference with first non-orthologous sequence - *A. fumigatus*:560 *A. fischeri*:560
afum|EAL84209.1 100.00% afis|EAW15436.1 100.00%
Bootstrap support for afum|EAL84209.1 as seed ortholog is 100%.
Bootstrap support for afis|EAW15436.1 as seed ortholog is 100%.

---

Group of orthologs #6927. Best score 560 bits
Score difference with first non-orthologous sequence - *A. fumigatus*:464 *A. fischeri*:419
afum|EAL85077.1 100.00% afis|EAW19814.1 100.00%
Bootstrap support for afum|EAL85077.1 as seed ortholog is 100%.
Bootstrap support for afis|EAW19814.1 as seed ortholog is 100%.

---

Group of orthologs #6928. Best score 560 bits
Score difference with first non-orthologous sequence - *A. fumigatus*:560 *A. fischeri*:560
afum|EAL89708.1 100.00% afis|EAW25040.1 100.00%
Bootstrap support for afum|EAL89708.1 as seed ortholog is 100%.
Bootstrap support for afis|EAW25040.1 as seed ortholog is 100%.

---

Group of orthologs #6929. Best score 559 bits
Score difference with first non-orthologous sequence - *A. fumigatus*:559 *A. fischeri*:559
afum|EAL85779.1 100.00% afis|EAW15554.1 100.00%
Bootstrap support for afum|EAL85779.1 as seed ortholog is 100%.
Bootstrap support for afis|EAW15554.1 as seed ortholog is 100%.

---

Group of orthologs #6930. Best score 559 bits
Score difference with first non-orthologous sequence - *A. fumigatus*:559 *A. fischeri*:559
afum|EAL88075.1 100.00% afis|EAW23395.1 100.00%
Bootstrap support for afum|EAL88075.1 as seed ortholog is 100%.

Bootstrap support for afis|EAW23395.1 as seed ortholog is 100%.

---

Group of orthologs #6931. Best score 559 bits
Score difference with first non-orthologous sequence - *A. fumigatus*:559 *A. fischeri*:559
afum|EAL93675.1 100.00% afis|EAW18942.1 100.00%
Bootstrap support for afum|EAL93675.1 as seed ortholog is 100%.
Bootstrap support for afis|EAW18942.1 as seed ortholog is 100%.

---

Group of orthologs #6932. Best score 559 bits
Score difference with first non-orthologous sequence - *A. fumigatus*:559 *A. fischeri*:559
afum|EAL92137.1 100.00% afis|EAW21114.1 100.00%
Bootstrap support for afum|EAL92137.1 as seed ortholog is 100%.
Bootstrap support for afis|EAW21114.1 as seed ortholog is 100%.

---

Group of orthologs #6933. Best score 559 bits
Score difference with first non-orthologous sequence - *A. fumigatus*:559 *A. fischeri*:559
afum|EAL90385.1 100.00% afis|EAW22816.1 100.00%
Bootstrap support for afum|EAL90385.1 as seed ortholog is 100%.
Bootstrap support for afis|EAW22816.1 as seed ortholog is 100%.

---

Group of orthologs #6934. Best score 559 bits
Score difference with first non-orthologous sequence - *A. fumigatus*:559 *A. fischeri*:559
afum|EAL90371.2 100.00% afis|EAW22831.1 100.00%
Bootstrap support for afum|EAL90371.2 as seed ortholog is 100%.
Bootstrap support for afis|EAW22831.1 as seed ortholog is 100%.

---

Group of orthologs #6935. Best score 559 bits
Score difference with first non-orthologous sequence - *A. fumigatus*:559 *A. fischeri*:559
afum|EAL91960.1 100.00% afis|EAW24106.1 100.00%
Bootstrap support for afum|EAL91960.1 as seed ortholog is 100%.
Bootstrap support for afis|EAW24106.1 as seed ortholog is 100%.

---

Group of orthologs #6936. Best score 558 bits
Score difference with first non-orthologous sequence - *A. fumigatus*:558 *A. fischeri*:382
afum|EAL86316.1 100.00% afis|EAW16741.1 100.00%
Bootstrap support for afum|EAL86316.1 as seed ortholog is 100%.
Bootstrap support for afis|EAW16741.1 as seed ortholog is 100%.

---

Group of orthologs #6937. Best score 558 bits
Score difference with first non-orthologous sequence - *A. fumigatus*:558 *A. fischeri*:558
afum|EAL88627.1 100.00% afis|EAW15995.1 100.00%
Bootstrap support for afum|EAL88627.1 as seed ortholog is 100%.
Bootstrap support for afis|EAW15995.1 as seed ortholog is 100%.

---

Group of orthologs #6938. Best score 558 bits
Score difference with first non-orthologous sequence - *A. fumigatus*:558 *A. fischeri*:558
afum|EAL86241.1 100.00% afis|EAW24611.1 100.00%
Bootstrap support for afum|EAL86241.1 as seed ortholog is 100%.
Bootstrap support for afis|EAW24611.1 as seed ortholog is 100%.

---

Group of orthologs #6939. Best score 558 bits
Score difference with first non-orthologous sequence - *A. fumigatus*:558 *A. fischeri*:558
afum|EAL88445.1 100.00% afis|EAW23036.1 100.00%
Bootstrap support for afum|EAL88445.1 as seed ortholog is 100%.
Bootstrap support for afis|EAW23036.1 as seed ortholog is 100%.

---

Group of orthologs #6940. Best score 558 bits
Score difference with first non-orthologous sequence - *A. fumigatus*:558 *A. fischeri*:558
afum|EAL93517.2 100.00% afis|EAW18785.1 100.00%
Bootstrap support for afum|EAL93517.2 as seed ortholog is 100%.
Bootstrap support for afis|EAW18785.1 as seed ortholog is 100%.

---

Group of orthologs #6941. Best score 558 bits
Score difference with first non-orthologous sequence - *A. fumigatus*:558 *A. fischeri*:558
afum|EAL90659.1 100.00% afis|EAW22535.1 100.00%
Bootstrap support for afum|EAL90659.1 as seed ortholog is 100%.
Bootstrap support for afis|EAW22535.1 as seed ortholog is 100%.

---

Group of orthologs #6942. Best score 558 bits
Score difference with first non-orthologous sequence - *A. fumigatus*:558 *A. fischeri*:558
afum|EAL90190.2 100.00% afis|EAW25525.1 100.00%
Bootstrap support for afum|EAL90190.2 as seed ortholog is 100%.
Bootstrap support for afis|EAW25525.1 as seed ortholog is 100%.

---

Group of orthologs #6943. Best score 557 bits
Score difference with first non-orthologous sequence - *A. fumigatus*:557 *A. fischeri*:557
afum|EAL84389.2 100.00% afis|EAW25715.1 100.00%
Bootstrap support for afum|EAL84389.2 as seed ortholog is 100%.
Bootstrap support for afis|EAW25715.1 as seed ortholog is 100%.

---

Group of orthologs #6944. Best score 557 bits
Score difference with first non-orthologous sequence - *A. fumigatus*:557 *A. fischeri*:557
afum|EAL90321.1 100.00% afis|EAW22878.1 100.00%
Bootstrap support for afum|EAL90321.1 as seed ortholog is 100%.
Bootstrap support for afis|EAW22878.1 as seed ortholog is 100%.

---

Group of orthologs #6945. Best score 557 bits
Score difference with first non-orthologous sequence - *A. fumigatus*:468 *A. fischeri*:557
afum|EBA27230.1 100.00% afis|EAW20284.1 100.00%
Bootstrap support for afum|EBA27230.1 as seed ortholog is 100%.
Bootstrap support for afis|EAW20284.1 as seed ortholog is 100%.

---

Group of orthologs #6946. Best score 557 bits
Score difference with first non-orthologous sequence - *A. fumigatus*:557 *A. fischeri*:432
afum|EAL89825.1 100.00% afis|EAW25150.1 100.00%
Bootstrap support for afum|EAL89825.1 as seed ortholog is 100%.
Bootstrap support for afis|EAW25150.1 as seed ortholog is 100%.

---

Group of orthologs #6947. Best score 556 bits
Score difference with first non-orthologous sequence - *A. fumigatus*:556 *A. fischeri*:147
afum|EAL89282.1 100.00% afis|EAW16677.1 100.00%
Bootstrap support for afum|EAL89282.1 as seed ortholog is 100%.
Bootstrap support for afis|EAW16677.1 as seed ortholog is 100%.

---

Group of orthologs #6948. Best score 556 bits
Score difference with first non-orthologous sequence - *A. fumigatus*:556 *A. fischeri*:380
afum|EAL92161.1 100.00% afis|EAW21142.1 100.00%
Bootstrap support for afum|EAL92161.1 as seed ortholog is 100%.
Bootstrap support for afis|EAW21142.1 as seed ortholog is 100%.

---

Group of orthologs #6949. Best score 556 bits
Score difference with first non-orthologous sequence - *A. fumigatus*:556 *A. fischeri*:556

afum|EBA27186.1 100.00% afis|EAW20551.1 100.00%
Bootstrap support for afum|EBA27186.1 as seed ortholog is 100%.
Bootstrap support for afis|EAW20551.1 as seed ortholog is 100%.

---

Group of orthologs #6950. Best score 555 bits
Score difference with first non-orthologous sequence - *A. fumigatus*:555 *A. fischeri*:555
afum|EAL91418.1 100.00% afis|EAW17605.1 100.00%
Bootstrap support for afum|EAL91418.1 as seed ortholog is 100%.
Bootstrap support for afis|EAW17605.1 as seed ortholog is 100%.

---

Group of orthologs #6951. Best score 555 bits
Score difference with first non-orthologous sequence - *A. fumigatus*:481 *A. fischeri*:479
afum|EAL91067.1 100.00% afis|EAW22119.1 100.00%
Bootstrap support for afum|EAL91067.1 as seed ortholog is 100%.
Bootstrap support for afis|EAW22119.1 as seed ortholog is 100%.

---

Group of orthologs #6952. Best score 554 bits
Score difference with first non-orthologous sequence - *A. fumigatus*:554 *A. fischeri*:554
afum|EAL87003.1 100.00% afis|EAW19491.1 100.00%
Bootstrap support for afum|EAL87003.1 as seed ortholog is 100%.
Bootstrap support for afis|EAW19491.1 as seed ortholog is 100%.

---

Group of orthologs #6953. Best score 554 bits
Score difference with first non-orthologous sequence - *A. fumigatus*:554 *A. fischeri*:554
afum|EAL86912.1 100.00% afis|EAW19610.1 100.00%
Bootstrap support for afum|EAL86912.1 as seed ortholog is 100%.
Bootstrap support for afis|EAW19610.1 as seed ortholog is 100%.

---

Group of orthologs #6954. Best score 554 bits
Score difference with first non-orthologous sequence - *A. fumigatus*:554 *A. fischeri*:554
afum|EAL91271.1 100.00% afis|EAW17445.1 100.00%
Bootstrap support for afum|EAL91271.1 as seed ortholog is 100%.
Bootstrap support for afis|EAW17445.1 as seed ortholog is 100%.

---

Group of orthologs #6955. Best score 554 bits
Score difference with first non-orthologous sequence - *A. fumigatus*:554 *A. fischeri*:554
afum|EAL91746.1 100.00% afis|EAW17935.1 100.00%
Bootstrap support for afum|EAL91746.1 as seed ortholog is 100%.
Bootstrap support for afis|EAW17935.1 as seed ortholog is 100%.

---

Group of orthologs #6956. Best score 554 bits
Score difference with first non-orthologous sequence - *A. fumigatus*:554 *A. fischeri*:554
afum|EAL85864.1 100.00% afis|EAW24191.1 100.00%
Bootstrap support for afum|EAL85864.1 as seed ortholog is 100%.
Bootstrap support for afis|EAW24191.1 as seed ortholog is 100%.

---

Group of orthologs #6957. Best score 554 bits
Score difference with first non-orthologous sequence - *A. fumigatus*:554 *A. fischeri*:554
afum|EAL84345.1 100.00% afis|EAW25675.1 100.00%
Bootstrap support for afum|EAL84345.1 as seed ortholog is 100%.
Bootstrap support for afis|EAW25675.1 as seed ortholog is 100%.

---

Group of orthologs #6958. Best score 554 bits
Score difference with first non-orthologous sequence - *A. fumigatus*:554 *A. fischeri*:554
afum|EAL88249.1 100.00% afis|EAW23233.1 100.00%
Bootstrap support for afum|EAL88249.1 as seed ortholog is 100%.

Bootstrap support for afis|EAW23233.1 as seed ortholog is 100%.

---

Group of orthologs #6959. Best score 554 bits
Score difference with first non-orthologous sequence - *A. fumigatus*:554 *A. fischeri*:298
afum|EAL90343.1 100.00% afis|EAW22857.1 100.00%
Bootstrap support for afum|EAL90343.1 as seed ortholog is 100%.
Bootstrap support for afis|EAW22857.1 as seed ortholog is 100%.

---

Group of orthologs #6960. Best score 553 bits
Score difference with first non-orthologous sequence - *A. fumigatus*:553 *A. fischeri*:553
afum|EAL85308.1 100.00% afis|EAW20189.1 100.00%
Bootstrap support for afum|EAL85308.1 as seed ortholog is 100%.
Bootstrap support for afis|EAW20189.1 as seed ortholog is 100%.

---

Group of orthologs #6961. Best score 553 bits
Score difference with first non-orthologous sequence - *A. fumigatus*:493 *A. fischeri*:509
afum|EAL85445.1 100.00% afis|EAW20317.1 100.00%
Bootstrap support for afum|EAL85445.1 as seed ortholog is 100%.
Bootstrap support for afis|EAW20317.1 as seed ortholog is 100%.

---

Group of orthologs #6962. Best score 553 bits
Score difference with first non-orthologous sequence - *A. fumigatus*:553 *A. fischeri*:553
afum|EAL91899.1 100.00% afis|EAW18087.1 100.00%
Bootstrap support for afum|EAL91899.1 as seed ortholog is 100%.
Bootstrap support for afis|EAW18087.1 as seed ortholog is 100%.

---

Group of orthologs #6963. Best score 553 bits
Score difference with first non-orthologous sequence - *A. fumigatus*:553 *A. fischeri*:553
afum|EAL93086.1 100.00% afis|EAW18406.1 100.00%
Bootstrap support for afum|EAL93086.1 as seed ortholog is 100%.
Bootstrap support for afis|EAW18406.1 as seed ortholog is 100%.

---

Group of orthologs #6964. Best score 553 bits
Score difference with first non-orthologous sequence - *A. fumigatus*:553 *A. fischeri*:553
afum|EAL88433.1 100.00% afis|EAW23047.1 100.00%
Bootstrap support for afum|EAL88433.1 as seed ortholog is 100%.
Bootstrap support for afis|EAW23047.1 as seed ortholog is 100%.

---

Group of orthologs #6965. Best score 553 bits
Score difference with first non-orthologous sequence - *A. fumigatus*:457 *A. fischeri*:553
afum|EAL92430.1 100.00% afis|EAW21415.1 100.00%
Bootstrap support for afum|EAL92430.1 as seed ortholog is 100%.
Bootstrap support for afis|EAW21415.1 as seed ortholog is 100%.

---

Group of orthologs #6966. Best score 553 bits
Score difference with first non-orthologous sequence - *A. fumigatus*:553 *A. fischeri*:438
afum|EBA27286.1 100.00% afis|EAW23710.1 100.00%
Bootstrap support for afum|EBA27286.1 as seed ortholog is 100%.
Bootstrap support for afis|EAW23710.1 as seed ortholog is 100%.

---

Group of orthologs #6967. Best score 552 bits
Score difference with first non-orthologous sequence - *A. fumigatus*:552 *A. fischeri*:552
afum|EAL91495.1 100.00% afis|EAW17681.1 100.00%
Bootstrap support for afum|EAL91495.1 as seed ortholog is 100%.
Bootstrap support for afis|EAW17681.1 as seed ortholog is 100%.

---

Group of orthologs #6968. Best score 552 bits
Score difference with first non-orthologous sequence - *A. fumigatus*:552 *A. fischeri*:552
afum|EAL93052.1 100.00% afis|EAW18377.1 100.00%
Bootstrap support for afum|EAL93052.1 as seed ortholog is 100%.
Bootstrap support for afis|EAW18377.1 as seed ortholog is 100%.

---

Group of orthologs #6969. Best score 552 bits
Score difference with first non-orthologous sequence - *A. fumigatus*:552 *A. fischeri*:552
afum|EAL92693.1 100.00% afis|EAW21658.1 100.00%
Bootstrap support for afum|EAL92693.1 as seed ortholog is 100%.
Bootstrap support for afis|EAW21658.1 as seed ortholog is 100%.

---

Group of orthologs #6970. Best score 552 bits
Score difference with first non-orthologous sequence - *A. fumigatus*:205 *A. fischeri*:552
afum|EAL89588.1 100.00% afis|EAW24932.1 100.00%
Bootstrap support for afum|EAL89588.1 as seed ortholog is 100%.
Bootstrap support for afis|EAW24932.1 as seed ortholog is 100%.

---

Group of orthologs #6971. Best score 552 bits
Score difference with first non-orthologous sequence - *A. fumigatus*:552 *A. fischeri*:552
afum|EAL90049.1 100.00% afis|EAW25390.1 100.00%
Bootstrap support for afum|EAL90049.1 as seed ortholog is 100%.
Bootstrap support for afis|EAW25390.1 as seed ortholog is 100%.

---

Group of orthologs #6972. Best score 551 bits
Score difference with first non-orthologous sequence - *A. fumigatus*:551 *A. fischeri*:551
afum|EAL88864.1 100.00% afis|EAW16242.1 100.00%
Bootstrap support for afum|EAL88864.1 as seed ortholog is 100%.
Bootstrap support for afis|EAW16242.1 as seed ortholog is 100%.

---

Group of orthologs #6973. Best score 551 bits
Score difference with first non-orthologous sequence - *A. fumigatus*:551 *A. fischeri*:551
afum|EAL84699.1 100.00% afis|EAW20935.1 100.00%
Bootstrap support for afum|EAL84699.1 as seed ortholog is 100%.
Bootstrap support for afis|EAW20935.1 as seed ortholog is 100%.

---

Group of orthologs #6974. Best score 551 bits
Score difference with first non-orthologous sequence - *A. fumigatus*:551 *A. fischeri*:551
afum|EAL93217.1 100.00% afis|EAW18522.1 100.00%
Bootstrap support for afum|EAL93217.1 as seed ortholog is 100%.
Bootstrap support for afis|EAW18522.1 as seed ortholog is 100%.

---

Group of orthologs #6975. Best score 551 bits
Score difference with first non-orthologous sequence - *A. fumigatus*:551 *A. fischeri*:551
afum|EBA27249.1 100.00% afis|EAW24223.1 100.00%
Bootstrap support for afum|EBA27249.1 as seed ortholog is 100%.
Bootstrap support for afis|EAW24223.1 as seed ortholog is 100%.

---

Group of orthologs #6976. Best score 550 bits
Score difference with first non-orthologous sequence - *A. fumigatus*:550 *A. fischeri*:550
afum|EAL86670.1 100.00% afis|EAW17251.1 100.00%
Bootstrap support for afum|EAL86670.1 as seed ortholog is 100%.
Bootstrap support for afis|EAW17251.1 as seed ortholog is 100%.

---

Group of orthologs #6977. Best score 550 bits
Score difference with first non-orthologous sequence - *A. fumigatus*:550 *A. fischeri*:550

afum|EAL84929.1 100.00% afis|EAW19986.1 100.00%
Bootstrap support for afum|EAL84929.1 as seed ortholog is 100%.
Bootstrap support for afis|EAW19986.1 as seed ortholog is 100%.

---

Group of orthologs #6978. Best score 550 bits
Score difference with first non-orthologous sequence - *A. fumigatus*:550 *A. fischeri*:550
afum|EAL87151.1 100.00% afis|EAW23707.1 100.00%
Bootstrap support for afum|EAL87151.1 as seed ortholog is 100%.
Bootstrap support for afis|EAW23707.1 as seed ortholog is 100%.

---

Group of orthologs #6979. Best score 550 bits
Score difference with first non-orthologous sequence - *A. fumigatus*:550 *A. fischeri*:440
afum|EAL93462.1 100.00% afis|EAW18737.1 100.00%
Bootstrap support for afum|EAL93462.1 as seed ortholog is 100%.
Bootstrap support for afis|EAW18737.1 as seed ortholog is 100%.

---

Group of orthologs #6980. Best score 549 bits
Score difference with first non-orthologous sequence - *A. fumigatus*:303 *A. fischeri*:549
afum|EAL89281.1 100.00% afis|EAW16676.1 100.00%
Bootstrap support for afum|EAL89281.1 as seed ortholog is 100%.
Bootstrap support for afis|EAW16676.1 as seed ortholog is 100%.

---

Group of orthologs #6981. Best score 549 bits
Score difference with first non-orthologous sequence - *A. fumigatus*:549 *A. fischeri*:549
afum|EAL89283.1 100.00% afis|EAW16679.1 100.00%
Bootstrap support for afum|EAL89283.1 as seed ortholog is 100%.
Bootstrap support for afis|EAW16679.1 as seed ortholog is 100%.

---

Group of orthologs #6982. Best score 549 bits
Score difference with first non-orthologous sequence - *A. fumigatus*:549 *A. fischeri*:549
afum|EAL84321.1 100.00% afis|EAW25633.1 100.00%
Bootstrap support for afum|EAL84321.1 as seed ortholog is 100%.
Bootstrap support for afis|EAW25633.1 as seed ortholog is 100%.

---

Group of orthologs #6983. Best score 549 bits
Score difference with first non-orthologous sequence - *A. fumigatus*:549 *A. fischeri*:447
afum|EAL86037.1 100.00% afis|EAW24365.1 100.00%
Bootstrap support for afum|EAL86037.1 as seed ortholog is 100%.
Bootstrap support for afis|EAW24365.1 as seed ortholog is 100%.

---

Group of orthologs #6984. Best score 548 bits
Score difference with first non-orthologous sequence - *A. fumigatus*:548 *A. fischeri*:548
afum|EAL85828.1 100.00% afis|EAW15495.1 100.00%
Bootstrap support for afum|EAL85828.1 as seed ortholog is 100%.
Bootstrap support for afis|EAW15495.1 as seed ortholog is 100%.

---

Group of orthologs #6985. Best score 548 bits
Score difference with first non-orthologous sequence - *A. fumigatus*:548 *A. fischeri*:548
afum|EAL91420.1 100.00% afis|EAW17607.1 100.00%
Bootstrap support for afum|EAL91420.1 as seed ortholog is 100%.
Bootstrap support for afis|EAW17607.1 as seed ortholog is 100%.

---

Group of orthologs #6986. Best score 548 bits
Score difference with first non-orthologous sequence - *A. fumigatus*:548 *A. fischeri*:548
afum|EAL93183.1 100.00% afis|EAW18493.1 100.00%
Bootstrap support for afum|EAL93183.1 as seed ortholog is 100%.

Bootstrap support for afis|EAW18493.1 as seed ortholog is 100%.

---

Group of orthologs #6987. Best score 548 bits
Score difference with first non-orthologous sequence - *A. fumigatus*:548 *A. fischeri*:548
afum|EAL92355.2 100.00% afis|EAW21337.1 100.00%
Bootstrap support for afum|EAL92355.2 as seed ortholog is 100%.
Bootstrap support for afis|EAW21337.1 as seed ortholog is 100%.

---

Group of orthologs #6988. Best score 548 bits
Score difference with first non-orthologous sequence - *A. fumigatus*:548 *A. fischeri*:548
afum|EAL92698.1 100.00% afis|EAW21663.1 100.00%
Bootstrap support for afum|EAL92698.1 as seed ortholog is 100%.
Bootstrap support for afis|EAW21663.1 as seed ortholog is 100%.

---

Group of orthologs #6989. Best score 548 bits
Score difference with first non-orthologous sequence - *A. fumigatus*:548 *A. fischeri*:548
afum|EAL92046.1 100.00% afis|EAW24186.1 100.00%
Bootstrap support for afum|EAL92046.1 as seed ortholog is 100%.
Bootstrap support for afis|EAW24186.1 as seed ortholog is 100%.

---

Group of orthologs #6990. Best score 547 bits
Score difference with first non-orthologous sequence - *A. fumigatus*:547 *A. fischeri*:547
afum|EAL85560.1 100.00% afis|EAW15790.1 100.00%
Bootstrap support for afum|EAL85560.1 as seed ortholog is 100%.
Bootstrap support for afis|EAW15790.1 as seed ortholog is 100%.

---

Group of orthologs #6991. Best score 547 bits
Score difference with first non-orthologous sequence - *A. fumigatus*:547 *A. fischeri*:547
afum|EAL87467.1 100.00% afis|EAW18110.1 100.00%
Bootstrap support for afum|EAL87467.1 as seed ortholog is 100%.
Bootstrap support for afis|EAW18110.1 as seed ortholog is 100%.

---

Group of orthologs #6992. Best score 547 bits
Score difference with first non-orthologous sequence - *A. fumigatus*:547 *A. fischeri*:547
afum|EAL91581.1 100.00% afis|EAW17772.1 100.00%
Bootstrap support for afum|EAL91581.1 as seed ortholog is 100%.
Bootstrap support for afis|EAW17772.1 as seed ortholog is 100%.

---

Group of orthologs #6993. Best score 547 bits
Score difference with first non-orthologous sequence - *A. fumigatus*:547 *A. fischeri*:547
afum|EAL88335.1 100.00% afis|EAW23146.1 100.00%
Bootstrap support for afum|EAL88335.1 as seed ortholog is 100%.
Bootstrap support for afis|EAW23146.1 as seed ortholog is 100%.

---

Group of orthologs #6994. Best score 547 bits
Score difference with first non-orthologous sequence - *A. fumigatus*:547 *A. fischeri*:547
afum|EAL92243.1 100.00% afis|EAW21231.1 100.00%
Bootstrap support for afum|EAL92243.1 as seed ortholog is 100%.
Bootstrap support for afis|EAW21231.1 as seed ortholog is 100%.

---

Group of orthologs #6995. Best score 546 bits
Score difference with first non-orthologous sequence - *A. fumigatus*:546 *A. fischeri*:546
afum|EAL86523.1 100.00% afis|EAW16998.1 100.00%
Bootstrap support for afum|EAL86523.1 as seed ortholog is 100%.
Bootstrap support for afis|EAW16998.1 as seed ortholog is 100%.

---

Group of orthologs #6996. Best score 546 bits
Score difference with first non-orthologous sequence - *A. fumigatus*:546 *A. fischeri*:546
afum|EAL85163.1 100.00% afis|EAW20043.1 100.00%
Bootstrap support for afum|EAL85163.1 as seed ortholog is 100%.
Bootstrap support for afis|EAW20043.1 as seed ortholog is 100%.

---

Group of orthologs #6997. Best score 546 bits
Score difference with first non-orthologous sequence - *A. fumigatus*:546 *A. fischeri*:546
afum|EAL87636.1 100.00% afis|EAW18271.1 100.00%
Bootstrap support for afum|EAL87636.1 as seed ortholog is 100%.
Bootstrap support for afis|EAW18271.1 as seed ortholog is 100%.

---

Group of orthologs #6998. Best score 546 bits
Score difference with first non-orthologous sequence - *A. fumigatus*:546 *A. fischeri*:546
afum|EAL93732.1 100.00% afis|EAW18997.1 100.00%
Bootstrap support for afum|EAL93732.1 as seed ortholog is 100%.
Bootstrap support for afis|EAW18997.1 as seed ortholog is 100%.

---

Group of orthologs #6999. Best score 545 bits
Score difference with first non-orthologous sequence - *A. fumigatus*:545 *A. fischeri*:545
afum|EAL85508.1 100.00% afis|EAW15843.1 100.00%
Bootstrap support for afum|EAL85508.1 as seed ortholog is 100%.
Bootstrap support for afis|EAW15843.1 as seed ortholog is 100%.

---

Group of orthologs #7000. Best score 545 bits
Score difference with first non-orthologous sequence - *A. fumigatus*:545 *A. fischeri*:200
afum|EAL86741.1 100.00% afis|EAW20666.1 100.00%
Bootstrap support for afum|EAL86741.1 as seed ortholog is 100%.
Bootstrap support for afis|EAW20666.1 as seed ortholog is 100%.

---

Group of orthologs #7001. Best score 545 bits
Score difference with first non-orthologous sequence - *A. fumigatus*:545 *A. fischeri*:500
afum|EAL87845.1 100.00% afis|EAW23617.1 100.00%
Bootstrap support for afum|EAL87845.1 as seed ortholog is 100%.
Bootstrap support for afis|EAW23617.1 as seed ortholog is 100%.

---

Group of orthologs #7002. Best score 544 bits
Score difference with first non-orthologous sequence - *A. fumigatus*:544 *A. fischeri*:544
afum|EAL85759.1 100.00% afis|EAW15575.1 100.00%
Bootstrap support for afum|EAL85759.1 as seed ortholog is 100%.
Bootstrap support for afis|EAW15575.1 as seed ortholog is 100%.

---

Group of orthologs #7003. Best score 544 bits
Score difference with first non-orthologous sequence - *A. fumigatus*:544 *A. fischeri*:368
afum|EAL85109.1 100.00% afis|EAW19780.1 100.00%
Bootstrap support for afum|EAL85109.1 as seed ortholog is 100%.
Bootstrap support for afis|EAW19780.1 as seed ortholog is 100%.

---

Group of orthologs #7004. Best score 544 bits
Score difference with first non-orthologous sequence - *A. fumigatus*:544 *A. fischeri*:412
afum|EAL84275.1 100.00% afis|EAW25552.1 100.00%
Bootstrap support for afum|EAL84275.1 as seed ortholog is 100%.
Bootstrap support for afis|EAW25552.1 as seed ortholog is 100%.

---

Group of orthologs #7005. Best score 544 bits
Score difference with first non-orthologous sequence - *A. fumigatus*:544 *A. fischeri*:544

afum|EAL86198.1 100.00% afis|EAW24532.1 100.00%
Bootstrap support for afum|EAL86198.1 as seed ortholog is 100%.
Bootstrap support for afis|EAW24532.1 as seed ortholog is 100%.

---

Group of orthologs #7006. Best score 544 bits
Score difference with first non-orthologous sequence - *A. fumigatus*:431 *A. fischeri*:381
afum|EAL86199.1 100.00% afis|EAW24533.1 100.00%
Bootstrap support for afum|EAL86199.1 as seed ortholog is 100%.
Bootstrap support for afis|EAW24533.1 as seed ortholog is 100%.

---

Group of orthologs #7007. Best score 544 bits
Score difference with first non-orthologous sequence - *A. fumigatus*:544 *A. fischeri*:185
afum|EAL87328.1 100.00% afis|EAW23901.1 100.00%
Bootstrap support for afum|EAL87328.1 as seed ortholog is 100%.
Bootstrap support for afis|EAW23901.1 as seed ortholog is 100%.

---

Group of orthologs #7008. Best score 544 bits
Score difference with first non-orthologous sequence - *A. fumigatus*:544 *A. fischeri*:462
afum|EAL93650.2 100.00% afis|EAW18919.1 100.00%
Bootstrap support for afum|EAL93650.2 as seed ortholog is 100%.
Bootstrap support for afis|EAW18919.1 as seed ortholog is 100%.

---

Group of orthologs #7009. Best score 544 bits
Score difference with first non-orthologous sequence - *A. fumigatus*:544 *A. fischeri*:544
afum|EAL92018.1 100.00% afis|EAW24162.1 100.00%
Bootstrap support for afum|EAL92018.1 as seed ortholog is 100%.
Bootstrap support for afis|EAW24162.1 as seed ortholog is 100%.

---

Group of orthologs #7010. Best score 543 bits
Score difference with first non-orthologous sequence - *A. fumigatus*:543 *A. fischeri*:543
afum|EAL85351.1 100.00% afis|EAW20234.1 100.00%
Bootstrap support for afum|EAL85351.1 as seed ortholog is 100%.
Bootstrap support for afis|EAW20234.1 as seed ortholog is 100%.

---

Group of orthologs #7011. Best score 543 bits
Score difference with first non-orthologous sequence - *A. fumigatus*:543 *A. fischeri*:543
afum|EAL87028.1 100.00% afis|EAW19468.1 100.00%
Bootstrap support for afum|EAL87028.1 as seed ortholog is 100%.
Bootstrap support for afis|EAW19468.1 as seed ortholog is 100%.

---

Group of orthologs #7012. Best score 543 bits
Score difference with first non-orthologous sequence - *A. fumigatus*:543 *A. fischeri*:543
afum|EAL91558.1 100.00% afis|EAW17746.1 100.00%
Bootstrap support for afum|EAL91558.1 as seed ortholog is 100%.
Bootstrap support for afis|EAW17746.1 as seed ortholog is 100%.

---

Group of orthologs #7013. Best score 543 bits
Score difference with first non-orthologous sequence - *A. fumigatus*:543 *A. fischeri*:543
afum|EAL86096.1 100.00% afis|EAW24428.1 100.00%
Bootstrap support for afum|EAL86096.1 as seed ortholog is 100%.
Bootstrap support for afis|EAW24428.1 as seed ortholog is 100%.

---

Group of orthologs #7014. Best score 543 bits
Score difference with first non-orthologous sequence - *A. fumigatus*:543 *A. fischeri*:543
afum|EAL87352.1 100.00% afis|EAW23927.1 100.00%
Bootstrap support for afum|EAL87352.1 as seed ortholog is 100%.

Bootstrap support for afis|EAW23927.1 as seed ortholog is 100%.

---

Group of orthologs #7015. Best score 543 bits
Score difference with first non-orthologous sequence - *A. fumigatus*:543 *A. fischeri*:543
afum|EAL93409.1 100.00% afis|EAW18688.1 100.00%
Bootstrap support for afum|EAL93409.1 as seed ortholog is 100%.
Bootstrap support for afis|EAW18688.1 as seed ortholog is 100%.

---

Group of orthologs #7016. Best score 543 bits
Score difference with first non-orthologous sequence - *A. fumigatus*:543 *A. fischeri*:543
afum|EAL92421.1 100.00% afis|EAW21407.1 100.00%
Bootstrap support for afum|EAL92421.1 as seed ortholog is 100%.
Bootstrap support for afis|EAW21407.1 as seed ortholog is 100%.

---

Group of orthologs #7017. Best score 542 bits
Score difference with first non-orthologous sequence - *A. fumigatus*:542 *A. fischeri*:542
afum|EAL90378.2 100.00% afis|EAW22822.1 100.00%
Bootstrap support for afum|EAL90378.2 as seed ortholog is 100%.
Bootstrap support for afis|EAW22822.1 as seed ortholog is 100%.

---

Group of orthologs #7018. Best score 541 bits
Score difference with first non-orthologous sequence - *A. fumigatus*:541 *A. fischeri*:541
afum|EAL86350.1 100.00% afis|EAW16779.1 100.00%
Bootstrap support for afum|EAL86350.1 as seed ortholog is 100%.
Bootstrap support for afis|EAW16779.1 as seed ortholog is 100%.

---

Group of orthologs #7019. Best score 541 bits
Score difference with first non-orthologous sequence - *A. fumigatus*:541 *A. fischeri*:541
afum|EAL86677.1 100.00% afis|EAW17260.1 100.00%
Bootstrap support for afum|EAL86677.1 as seed ortholog is 100%.
Bootstrap support for afis|EAW17260.1 as seed ortholog is 100%.

---

Group of orthologs #7020. Best score 541 bits
Score difference with first non-orthologous sequence - *A. fumigatus*:541 *A. fischeri*:541
afum|EAL88619.1 100.00% afis|EAW15987.1 100.00%
Bootstrap support for afum|EAL88619.1 as seed ortholog is 100%.
Bootstrap support for afis|EAW15987.1 as seed ortholog is 100%.

---

Group of orthologs #7021. Best score 541 bits
Score difference with first non-orthologous sequence - *A. fumigatus*:541 *A. fischeri*:541
afum|EAL89220.1 100.00% afis|EAW16616.1 100.00%
Bootstrap support for afum|EAL89220.1 as seed ortholog is 100%.
Bootstrap support for afis|EAW16616.1 as seed ortholog is 100%.

---

Group of orthologs #7022. Best score 541 bits
Score difference with first non-orthologous sequence - *A. fumigatus*:541 *A. fischeri*:475
afum|EAL91609.1 100.00% afis|EAW17801.1 100.00%
Bootstrap support for afum|EAL91609.1 as seed ortholog is 100%.
Bootstrap support for afis|EAW17801.1 as seed ortholog is 100%.

---

Group of orthologs #7023. Best score 541 bits
Score difference with first non-orthologous sequence - *A. fumigatus*:541 *A. fischeri*:541
afum|EAL93054.1 100.00% afis|EAW18379.1 100.00%
Bootstrap support for afum|EAL93054.1 as seed ortholog is 100%.
Bootstrap support for afis|EAW18379.1 as seed ortholog is 100%.

---

Group of orthologs #7024. Best score 541 bits
Score difference with first non-orthologous sequence - *A. fumigatus*:541   *A. fischeri*:541
afum|EAL87337.1 100.00% afis|EAW23909.1 100.00%
Bootstrap support for afum|EAL87337.1 as seed ortholog is 100%.
Bootstrap support for afis|EAW23909.1 as seed ortholog is 100%.

---

Group of orthologs #7025. Best score 541 bits
Score difference with first non-orthologous sequence - *A. fumigatus*:541   *A. fischeri*:465
afum|EAL90958.1 100.00% afis|EAW22235.1 100.00%
Bootstrap support for afum|EAL90958.1 as seed ortholog is 100%.
Bootstrap support for afis|EAW22235.1 as seed ortholog is 100%.

---

Group of orthologs #7026. Best score 541 bits
Score difference with first non-orthologous sequence - *A. fumigatus*:541   *A. fischeri*:302
afum|EAL89342.1 100.00% afis|EAW24700.1 100.00%
Bootstrap support for afum|EAL89342.1 as seed ortholog is 100%.
Bootstrap support for afis|EAW24700.1 as seed ortholog is 100%.

---

Group of orthologs #7027. Best score 540 bits
Score difference with first non-orthologous sequence - *A. fumigatus*:540   *A. fischeri*:540
afum|EAL88974.1 100.00% afis|EAW16349.1 100.00%
Bootstrap support for afum|EAL88974.1 as seed ortholog is 100%.
Bootstrap support for afis|EAW16349.1 as seed ortholog is 100%.

---

Group of orthologs #7028. Best score 540 bits
Score difference with first non-orthologous sequence - *A. fumigatus*:540   *A. fischeri*:448
afum|EAL84630.1 100.00% afis|EAW20419.1 100.00%
Bootstrap support for afum|EAL84630.1 as seed ortholog is 100%.
Bootstrap support for afis|EAW20419.1 as seed ortholog is 100%.

---

Group of orthologs #7029. Best score 540 bits
Score difference with first non-orthologous sequence - *A. fumigatus*:540   *A. fischeri*:540
afum|EAL85300.1 100.00% afis|EAW20182.1 100.00%
Bootstrap support for afum|EAL85300.1 as seed ortholog is 100%.
Bootstrap support for afis|EAW20182.1 as seed ortholog is 100%.

---

Group of orthologs #7030. Best score 540 bits
Score difference with first non-orthologous sequence - *A. fumigatus*:540   *A. fischeri*:540
afum|EAL85441.1 100.00% afis|EAW20315.1 100.00%
Bootstrap support for afum|EAL85441.1 as seed ortholog is 100%.
Bootstrap support for afis|EAW20315.1 as seed ortholog is 100%.

---

Group of orthologs #7031. Best score 540 bits
Score difference with first non-orthologous sequence - *A. fumigatus*:540   *A. fischeri*:540
afum|EAL91266.1 100.00% afis|EAW17440.1 100.00%
Bootstrap support for afum|EAL91266.1 as seed ortholog is 100%.
Bootstrap support for afis|EAW17440.1 as seed ortholog is 100%.

---

Group of orthologs #7032. Best score 540 bits
Score difference with first non-orthologous sequence - *A. fumigatus*:452   *A. fischeri*:449
afum|EAL84315.1 100.00% afis|EAW25627.1 100.00%
Bootstrap support for afum|EAL84315.1 as seed ortholog is 100%.
Bootstrap support for afis|EAW25627.1 as seed ortholog is 100%.

---

Group of orthologs #7033. Best score 540 bits
Score difference with first non-orthologous sequence - *A. fumigatus*:540   *A. fischeri*:540

afum|EAL90847.1 100.00% afis|EAW22344.1 100.00%
Bootstrap support for afum|EAL90847.1 as seed ortholog is 100%.
Bootstrap support for afis|EAW22344.1 as seed ortholog is 100%.

---

Group of orthologs #7034. Best score 540 bits
Score difference with first non-orthologous sequence - *A. fumigatus*:540 *A. fischeri*:294
afum|EAL92674.1 100.00% afis|EAW21640.1 100.00%
Bootstrap support for afum|EAL92674.1 as seed ortholog is 100%.
Bootstrap support for afis|EAW21640.1 as seed ortholog is 100%.

---

Group of orthologs #7035. Best score 539 bits
Score difference with first non-orthologous sequence - *A. fumigatus*:539 *A. fischeri*:539
afum|EAL85594.1 100.00% afis|EAW15755.1 100.00%
Bootstrap support for afum|EAL85594.1 as seed ortholog is 100%.
Bootstrap support for afis|EAW15755.1 as seed ortholog is 100%.

---

Group of orthologs #7036. Best score 539 bits
Score difference with first non-orthologous sequence - *A. fumigatus*:539 *A. fischeri*:539
afum|EAL86364.1 100.00% afis|EAW16795.1 100.00%
Bootstrap support for afum|EAL86364.1 as seed ortholog is 100%.
Bootstrap support for afis|EAW16795.1 as seed ortholog is 100%.

---

Group of orthologs #7037. Best score 539 bits
Score difference with first non-orthologous sequence - *A. fumigatus*:539 *A. fischeri*:539
afum|EAL89083.1 100.00% afis|EAW16489.1 100.00%
Bootstrap support for afum|EAL89083.1 as seed ortholog is 100%.
Bootstrap support for afis|EAW16489.1 as seed ortholog is 100%.

---

Group of orthologs #7038. Best score 539 bits
Score difference with first non-orthologous sequence - *A. fumigatus*:539 *A. fischeri*:539
afum|EAL84807.1 100.00% afis|EAW20822.1 100.00%
Bootstrap support for afum|EAL84807.1 as seed ortholog is 100%.
Bootstrap support for afis|EAW20822.1 as seed ortholog is 100%.

---

Group of orthologs #7039. Best score 539 bits
Score difference with first non-orthologous sequence - *A. fumigatus*:539 *A. fischeri*:539
afum|EAL87641.1 100.00% afis|EAW18275.1 100.00%
Bootstrap support for afum|EAL87641.1 as seed ortholog is 100%.
Bootstrap support for afis|EAW18275.1 as seed ortholog is 100%.

---

Group of orthologs #7040. Best score 539 bits
Score difference with first non-orthologous sequence - *A. fumigatus*:539 *A. fischeri*:539
afum|EAL93868.1 100.00% afis|EAW19136.1 100.00%
Bootstrap support for afum|EAL93868.1 as seed ortholog is 100%.
Bootstrap support for afis|EAW19136.1 as seed ortholog is 100%.

---

Group of orthologs #7041. Best score 539 bits
Score difference with first non-orthologous sequence - *A. fumigatus*:539 *A. fischeri*:212
afum|EAL90200.1 100.00% afis|EAW25536.1 100.00%
Bootstrap support for afum|EAL90200.1 as seed ortholog is 100%.
Bootstrap support for afis|EAW25536.1 as seed ortholog is 100%.

---

Group of orthologs #7042. Best score 538 bits
Score difference with first non-orthologous sequence - *A. fumigatus*:538 *A. fischeri*:538
afum|EAL84919.1 100.00% afis|EAW20000.1 100.00%
Bootstrap support for afum|EAL84919.1 as seed ortholog is 100%.

Bootstrap support for afis|EAW20000.1 as seed ortholog is 100%.

---

Group of orthologs #7043. Best score 538 bits
Score difference with first non-orthologous sequence - *A. fumigatus*:538 *A. fischeri*:538
afum|EAL84794.2 100.00% afis|EAW20839.1 100.00%
Bootstrap support for afum|EAL84794.2 as seed ortholog is 100%.
Bootstrap support for afis|EAW20839.1 as seed ortholog is 100%.

---

Group of orthologs #7044. Best score 538 bits
Score difference with first non-orthologous sequence - *A. fumigatus*:538 *A. fischeri*:538
afum|EAL86788.1 100.00% afis|EAW19741.1 100.00%
Bootstrap support for afum|EAL86788.1 as seed ortholog is 100%.
Bootstrap support for afis|EAW19741.1 as seed ortholog is 100%.

---

Group of orthologs #7045. Best score 538 bits
Score difference with first non-orthologous sequence - *A. fumigatus*:538 *A. fischeri*:538
afum|EAL87157.1 100.00% afis|EAW23713.1 100.00%
Bootstrap support for afum|EAL87157.1 as seed ortholog is 100%.
Bootstrap support for afis|EAW23713.1 as seed ortholog is 100%.

---

Group of orthologs #7046. Best score 538 bits
Score difference with first non-orthologous sequence - *A. fumigatus*:494 *A. fischeri*:538
afum|EBA27282.1 100.00% afis|EAW17199.1 100.00%
Bootstrap support for afum|EBA27282.1 as seed ortholog is 100%.
Bootstrap support for afis|EAW17199.1 as seed ortholog is 100%.

---

Group of orthologs #7047. Best score 538 bits
Score difference with first non-orthologous sequence - *A. fumigatus*:538 *A. fischeri*:538
afum|EAL87407.1 100.00% afis|EAW23977.1 100.00%
Bootstrap support for afum|EAL87407.1 as seed ortholog is 100%.
Bootstrap support for afis|EAW23977.1 as seed ortholog is 100%.

---

Group of orthologs #7048. Best score 538 bits
Score difference with first non-orthologous sequence - *A. fumigatus*:538 *A. fischeri*:368
afum|EAL93726.1 100.00% afis|EAW18990.1 100.00%
Bootstrap support for afum|EAL93726.1 as seed ortholog is 100%.
Bootstrap support for afis|EAW18990.1 as seed ortholog is 100%.

---

Group of orthologs #7049. Best score 538 bits
Score difference with first non-orthologous sequence - *A. fumigatus*:538 *A. fischeri*:538
afum|EAL90675.1 100.00% afis|EAW22518.1 100.00%
Bootstrap support for afum|EAL90675.1 as seed ortholog is 100%.
Bootstrap support for afis|EAW22518.1 as seed ortholog is 100%.

---

Group of orthologs #7050. Best score 537 bits
Score difference with first non-orthologous sequence - *A. fumigatus*:537 *A. fischeri*:351
afum|EAL88744.1 100.00% afis|EAW16113.1 100.00%
Bootstrap support for afum|EAL88744.1 as seed ortholog is 100%.
Bootstrap support for afis|EAW16113.1 as seed ortholog is 100%.

---

Group of orthologs #7051. Best score 537 bits
Score difference with first non-orthologous sequence - *A. fumigatus*:537 *A. fischeri*:537
afum|EAL85015.1 100.00% afis|EAW19884.1 100.00%
Bootstrap support for afum|EAL85015.1 as seed ortholog is 100%.
Bootstrap support for afis|EAW19884.1 as seed ortholog is 100%.

---

Group of orthologs #7052. Best score 537 bits
Score difference with first non-orthologous sequence - *A. fumigatus*:537 *A. fischeri*:537
afum|EAL87518.1 100.00% afis|EAW18159.1 100.00%
Bootstrap support for afum|EAL87518.1 as seed ortholog is 100%.
Bootstrap support for afis|EAW18159.1 as seed ortholog is 100%.

---

Group of orthologs #7053. Best score 537 bits
Score difference with first non-orthologous sequence - *A. fumigatus*:537 *A. fischeri*:537
afum|EAL93918.1 100.00% afis|EAW19187.1 100.00%
Bootstrap support for afum|EAL93918.1 as seed ortholog is 100%.
Bootstrap support for afis|EAW19187.1 as seed ortholog is 100%.

---

Group of orthologs #7054. Best score 537 bits
Score difference with first non-orthologous sequence - *A. fumigatus*:537 *A. fischeri*:537
afum|EAL89833.1 100.00% afis|EAW25158.1 100.00%
Bootstrap support for afum|EAL89833.1 as seed ortholog is 100%.
Bootstrap support for afis|EAW25158.1 as seed ortholog is 100%.

---

Group of orthologs #7055. Best score 536 bits
Score difference with first non-orthologous sequence - *A. fumigatus*:536 *A. fischeri*:536
afum|EAL88549.1 100.00% afis|EAW15914.1 100.00%
Bootstrap support for afum|EAL88549.1 as seed ortholog is 100%.
Bootstrap support for afis|EAW15914.1 as seed ortholog is 100%.

---

Group of orthologs #7056. Best score 535 bits
Score difference with first non-orthologous sequence - *A. fumigatus*:318 *A. fischeri*:373
afum|EAL86974.1 100.00% afis|EAW19532.1 100.00%
Bootstrap support for afum|EAL86974.1 as seed ortholog is 100%.
Bootstrap support for afis|EAW19532.1 as seed ortholog is 100%.

---

Group of orthologs #7057. Best score 535 bits
Score difference with first non-orthologous sequence - *A. fumigatus*:479 *A. fischeri*:474
afum|EAL91207.2 100.00% afis|EAW17370.1 100.00%
Bootstrap support for afum|EAL91207.2 as seed ortholog is 100%.
Bootstrap support for afis|EAW17370.1 as seed ortholog is 100%.

---

Group of orthologs #7058. Best score 535 bits
Score difference with first non-orthologous sequence - *A. fumigatus*:535 *A. fischeri*:535
afum|EAL90362.2 100.00% afis|EAW22841.1 100.00%
Bootstrap support for afum|EAL90362.2 as seed ortholog is 100%.
Bootstrap support for afis|EAW22841.1 as seed ortholog is 100%.

---

Group of orthologs #7059. Best score 534 bits
Score difference with first non-orthologous sequence - *A. fumigatus*:534 *A. fischeri*:418
afum|EAL86425.2 100.00% afis|EAW16887.1 100.00%
Bootstrap support for afum|EAL86425.2 as seed ortholog is 100%.
Bootstrap support for afis|EAW16887.1 as seed ortholog is 100%.

---

Group of orthologs #7060. Best score 534 bits
Score difference with first non-orthologous sequence - *A. fumigatus*:534 *A. fischeri*:534
afum|EAL86647.1 100.00% afis|EAW17225.1 100.00%
Bootstrap support for afum|EAL86647.1 as seed ortholog is 100%.
Bootstrap support for afis|EAW17225.1 as seed ortholog is 100%.

---

Group of orthologs #7061. Best score 534 bits
Score difference with first non-orthologous sequence - *A. fumigatus*:534 *A. fischeri*:534

afum|EAL90929.1 100.00% afis|EAW16824.1 100.00%
Bootstrap support for afum|EAL90929.1 as seed ortholog is 100%.
Bootstrap support for afis|EAW16824.1 as seed ortholog is 100%.

---

Group of orthologs #7062. Best score 534 bits
Score difference with first non-orthologous sequence - *A. fumigatus*:534 *A. fischeri*:534
afum|EAL91228.1 100.00% afis|EAW17391.1 100.00%
Bootstrap support for afum|EAL91228.1 as seed ortholog is 100%.
Bootstrap support for afis|EAW17391.1 as seed ortholog is 100%.

---

Group of orthologs #7063. Best score 534 bits
Score difference with first non-orthologous sequence - *A. fumigatus*:534 *A. fischeri*:534
afum|EAL91275.2 100.00% afis|EAW17448.1 100.00%
Bootstrap support for afum|EAL91275.2 as seed ortholog is 100%.
Bootstrap support for afis|EAW17448.1 as seed ortholog is 100%.

---

Group of orthologs #7064. Best score 534 bits
Score difference with first non-orthologous sequence - *A. fumigatus*:534 *A. fischeri*:534
afum|EAL93366.1 100.00% afis|EAW18650.1 100.00%
Bootstrap support for afum|EAL93366.1 as seed ortholog is 100%.
Bootstrap support for afis|EAW18650.1 as seed ortholog is 100%.

---

Group of orthologs #7065. Best score 534 bits
Score difference with first non-orthologous sequence - *A. fumigatus*:534 *A. fischeri*:534
afum|EAL90297.2 100.00% afis|EAW22900.1 100.00%
Bootstrap support for afum|EAL90297.2 as seed ortholog is 100%.
Bootstrap support for afis|EAW22900.1 as seed ortholog is 100%.

---

Group of orthologs #7066. Best score 534 bits
Score difference with first non-orthologous sequence - *A. fumigatus*:534 *A. fischeri*:534
afum|EAL89541.2 100.00% afis|EAW24890.1 100.00%
Bootstrap support for afum|EAL89541.2 as seed ortholog is 100%.
Bootstrap support for afis|EAW24890.1 as seed ortholog is 100%.

---

Group of orthologs #7067. Best score 534 bits
Score difference with first non-orthologous sequence - *A. fumigatus*:534 *A. fischeri*:534
afum|EBA27418.1 100.00% afis|EAW22199.1 100.00%
Bootstrap support for afum|EBA27418.1 as seed ortholog is 100%.
Bootstrap support for afis|EAW22199.1 as seed ortholog is 100%.

---

Group of orthologs #7068. Best score 533 bits
Score difference with first non-orthologous sequence - *A. fumigatus*:533 *A. fischeri*:533
afum|EAL88858.1 100.00% afis|EAW16237.1 100.00%
Bootstrap support for afum|EAL88858.1 as seed ortholog is 100%.
Bootstrap support for afis|EAW16237.1 as seed ortholog is 100%.

---

Group of orthologs #7069. Best score 533 bits
Score difference with first non-orthologous sequence - *A. fumigatus*:533 *A. fischeri*:533
afum|EAL93596.1 100.00% afis|EAW18864.1 100.00%
Bootstrap support for afum|EAL93596.1 as seed ortholog is 100%.
Bootstrap support for afis|EAW18864.1 as seed ortholog is 100%.

---

Group of orthologs #7070. Best score 533 bits
Score difference with first non-orthologous sequence - *A. fumigatus*:533 *A. fischeri*:533
afum|EAL90446.2 100.00% afis|EAW22753.1 100.00%
Bootstrap support for afum|EAL90446.2 as seed ortholog is 100%.

Bootstrap support for afis|EAW22753.1 as seed ortholog is 100%.

---

Group of orthologs #7071. Best score 533 bits
Score difference with first non-orthologous sequence - *A. fumigatus*:533 *A. fischeri*:533
afum|EAL92958.1 100.00% afis|EAW21944.1 100.00%
Bootstrap support for afum|EAL92958.1 as seed ortholog is 100%.
Bootstrap support for afis|EAW21944.1 as seed ortholog is 100%.

---

Group of orthologs #7072. Best score 532 bits
Score difference with first non-orthologous sequence - *A. fumigatus*:532 *A. fischeri*:532
afum|EAL89153.1 100.00% afis|EAW16555.1 100.00%
Bootstrap support for afum|EAL89153.1 as seed ortholog is 100%.
Bootstrap support for afis|EAW16555.1 as seed ortholog is 100%.

---

Group of orthologs #7073. Best score 532 bits
Score difference with first non-orthologous sequence - *A. fumigatus*:532 *A. fischeri*:532
afum|EAL91152.1 100.00% afis|EAW17316.1 100.00%
Bootstrap support for afum|EAL91152.1 as seed ortholog is 100%.
Bootstrap support for afis|EAW17316.1 as seed ortholog is 100%.

---

Group of orthologs #7074. Best score 532 bits
Score difference with first non-orthologous sequence - *A. fumigatus*:532 *A. fischeri*:532
afum|EAL87936.1 100.00% afis|EAW23530.1 100.00%
Bootstrap support for afum|EAL87936.1 as seed ortholog is 100%.
Bootstrap support for afis|EAW23530.1 as seed ortholog is 100%.

---

Group of orthologs #7075. Best score 532 bits
Score difference with first non-orthologous sequence - *A. fumigatus*:532 *A. fischeri*:532
afum|EAL92952.2 100.00% afis|EAW22061.1 100.00%
Bootstrap support for afum|EAL92952.2 as seed ortholog is 100%.
Bootstrap support for afis|EAW22061.1 as seed ortholog is 100%.

---

Group of orthologs #7076. Best score 531 bits
Score difference with first non-orthologous sequence - *A. fumigatus*:531 *A. fischeri*:531
afum|EAL87034.1 100.00% afis|EAW19462.1 100.00%
Bootstrap support for afum|EAL87034.1 as seed ortholog is 100%.
Bootstrap support for afis|EAW19462.1 as seed ortholog is 100%.

---

Group of orthologs #7077. Best score 531 bits
Score difference with first non-orthologous sequence - *A. fumigatus*:531 *A. fischeri*:531
afum|EAL84334.1 100.00% afis|EAW25660.1 100.00%
Bootstrap support for afum|EAL84334.1 as seed ortholog is 100%.
Bootstrap support for afis|EAW25660.1 as seed ortholog is 100%.

---

Group of orthologs #7078. Best score 531 bits
Score difference with first non-orthologous sequence - *A. fumigatus*:432 *A. fischeri*:531
afum|EAL87153.1 100.00% afis|EAW23709.1 100.00%
Bootstrap support for afum|EAL87153.1 as seed ortholog is 100%.
Bootstrap support for afis|EAW23709.1 as seed ortholog is 100%.

---

Group of orthologs #7079. Best score 531 bits
Score difference with first non-orthologous sequence - *A. fumigatus*:460 *A. fischeri*:117
afum|EAL92066.1 100.00% afis|EAW21129.1 100.00%
Bootstrap support for afum|EAL92066.1 as seed ortholog is 100%.
Bootstrap support for afis|EAW21129.1 as seed ortholog is 96%.

---

Group of orthologs #7080. Best score 531 bits
Score difference with first non-orthologous sequence - *A. fumigatus*:531 *A. fischeri*:531
afum|EAL92930.1 100.00% afis|EAW21918.1 100.00%
Bootstrap support for afum|EAL92930.1 as seed ortholog is 100%.
Bootstrap support for afis|EAW21918.1 as seed ortholog is 100%.

---

Group of orthologs #7081. Best score 530 bits
Score difference with first non-orthologous sequence - *A. fumigatus*:530 *A. fischeri*:530
afum|EAL87479.1 100.00% afis|EAW18121.1 100.00%
Bootstrap support for afum|EAL87479.1 as seed ortholog is 100%.
Bootstrap support for afis|EAW18121.1 as seed ortholog is 100%.

---

Group of orthologs #7082. Best score 530 bits
Score difference with first non-orthologous sequence - *A. fumigatus*:530 *A. fischeri*:530
afum|EAL93783.1 100.00% afis|EAW19054.1 100.00%
Bootstrap support for afum|EAL93783.1 as seed ortholog is 100%.
Bootstrap support for afis|EAW19054.1 as seed ortholog is 100%.

---

Group of orthologs #7083. Best score 529 bits
Score difference with first non-orthologous sequence - *A. fumigatus*:529 *A. fischeri*:529
afum|EAL85608.1 100.00% afis|EAW15740.1 100.00%
Bootstrap support for afum|EAL85608.1 as seed ortholog is 100%.
Bootstrap support for afis|EAW15740.1 as seed ortholog is 100%.

---

Group of orthologs #7084. Best score 528 bits
Score difference with first non-orthologous sequence - *A. fumigatus*:528 *A. fischeri*:311
afum|EAL86410.2 100.00% afis|EAW16872.1 100.00%
Bootstrap support for afum|EAL86410.2 as seed ortholog is 100%.
Bootstrap support for afis|EAW16872.1 as seed ortholog is 100%.

---

Group of orthologs #7085. Best score 528 bits
Score difference with first non-orthologous sequence - *A. fumigatus*:528 *A. fischeri*:528
afum|EAL85411.1 100.00% afis|EAW20277.1 100.00%
Bootstrap support for afum|EAL85411.1 as seed ortholog is 100%.
Bootstrap support for afis|EAW20277.1 as seed ortholog is 100%.

---

Group of orthologs #7086. Best score 528 bits
Score difference with first non-orthologous sequence - *A. fumigatus*:216 *A. fischeri*:528
afum|EAL91151.1 100.00% afis|EAW17315.1 100.00%
Bootstrap support for afum|EAL91151.1 as seed ortholog is 100%.
Bootstrap support for afis|EAW17315.1 as seed ortholog is 100%.

---

Group of orthologs #7087. Best score 528 bits
Score difference with first non-orthologous sequence - *A. fumigatus*:528 *A. fischeri*:528
afum|EAL85877.1 100.00% afis|EAW24207.1 100.00%
Bootstrap support for afum|EAL85877.1 as seed ortholog is 100%.
Bootstrap support for afis|EAW24207.1 as seed ortholog is 100%.

---

Group of orthologs #7088. Best score 528 bits
Score difference with first non-orthologous sequence - *A. fumigatus*:528 *A. fischeri*:528
afum|EAL85960.1 100.00% afis|EAW24292.1 100.00%
Bootstrap support for afum|EAL85960.1 as seed ortholog is 100%.
Bootstrap support for afis|EAW24292.1 as seed ortholog is 100%.

---

Group of orthologs #7089. Best score 528 bits
Score difference with first non-orthologous sequence - *A. fumigatus*:528 *A. fischeri*:528

afum|EAL87891.1 100.00% afis|EAW23574.1 100.00%
Bootstrap support for afum|EAL87891.1 as seed ortholog is 100%.
Bootstrap support for afis|EAW23574.1 as seed ortholog is 100%.

---

Group of orthologs #7090. Best score 528 bits
Score difference with first non-orthologous sequence - *A. fumigatus*:528 *A. fischeri*:528
afum|EAL89352.1 100.00% afis|EAW24711.1 100.00%
Bootstrap support for afum|EAL89352.1 as seed ortholog is 100%.
Bootstrap support for afis|EAW24711.1 as seed ortholog is 100%.

---

Group of orthologs #7091. Best score 528 bits
Score difference with first non-orthologous sequence - *A. fumigatus*:528 *A. fischeri*:528
afum|EAL89849.1 100.00% afis|EAW25176.1 100.00%
Bootstrap support for afum|EAL89849.1 as seed ortholog is 100%.
Bootstrap support for afis|EAW25176.1 as seed ortholog is 100%.

---

Group of orthologs #7092. Best score 527 bits
Score difference with first non-orthologous sequence - *A. fumigatus*:527 *A. fischeri*:470
afum|EAL88476.2 100.00% afis|EAW23004.1 100.00%
Bootstrap support for afum|EAL88476.2 as seed ortholog is 100%.
Bootstrap support for afis|EAW23004.1 as seed ortholog is 100%.

---

Group of orthologs #7093. Best score 527 bits
Score difference with first non-orthologous sequence - *A. fumigatus*:527 *A. fischeri*:527
afum|EAL88108.1 100.00% afis|EAW23365.1 100.00%
Bootstrap support for afum|EAL88108.1 as seed ortholog is 100%.
Bootstrap support for afis|EAW23365.1 as seed ortholog is 100%.

---

Group of orthologs #7094. Best score 527 bits
Score difference with first non-orthologous sequence - *A. fumigatus*:527 *A. fischeri*:527
afum|EAL92679.1 100.00% afis|EAW21645.1 100.00%
Bootstrap support for afum|EAL92679.1 as seed ortholog is 100%.
Bootstrap support for afis|EAW21645.1 as seed ortholog is 100%.

---

Group of orthologs #7095. Best score 527 bits
Score difference with first non-orthologous sequence - *A. fumigatus*:527 *A. fischeri*:527
afum|EBA27377.1 100.00% afis|EAW24728.1 100.00%
Bootstrap support for afum|EBA27377.1 as seed ortholog is 100%.
Bootstrap support for afis|EAW24728.1 as seed ortholog is 100%.

---

Group of orthologs #7096. Best score 526 bits
Score difference with first non-orthologous sequence - *A. fumigatus*:526 *A. fischeri*:526
afum|EAL84816.1 100.00% afis|EAW20812.1 100.00%
Bootstrap support for afum|EAL84816.1 as seed ortholog is 100%.
Bootstrap support for afis|EAW20812.1 as seed ortholog is 100%.

---

Group of orthologs #7097. Best score 526 bits
Score difference with first non-orthologous sequence - *A. fumigatus*:526 *A. fischeri*:526
afum|EAL91828.1 100.00% afis|EAW18016.1 100.00%
Bootstrap support for afum|EAL91828.1 as seed ortholog is 100%.
Bootstrap support for afis|EAW18016.1 as seed ortholog is 100%.

---

Group of orthologs #7098. Best score 526 bits
Score difference with first non-orthologous sequence - *A. fumigatus*:526 *A. fischeri*:526
afum|EAL87283.1 100.00% afis|EAW23855.1 100.00%
Bootstrap support for afum|EAL87283.1 as seed ortholog is 100%.

Bootstrap support for afis|EAW23855.1 as seed ortholog is 100%.

---

Group of orthologs #7099. Best score 526 bits
Score difference with first non-orthologous sequence - *A. fumigatus*:526 *A. fischeri*:526
afum|EAL92271.1 100.00% afis|EAW21256.1 100.00%
Bootstrap support for afum|EAL92271.1 as seed ortholog is 100%.
Bootstrap support for afis|EAW21256.1 as seed ortholog is 100%.

---

Group of orthologs #7100. Best score 525 bits
Score difference with first non-orthologous sequence - *A. fumigatus*:525 *A. fischeri*:525
afum|EAL89023.1 100.00% afis|EAW16405.1 100.00%
Bootstrap support for afum|EAL89023.1 as seed ortholog is 100%.
Bootstrap support for afis|EAW16405.1 as seed ortholog is 100%.

---

Group of orthologs #7101. Best score 525 bits
Score difference with first non-orthologous sequence - *A. fumigatus*:525 *A. fischeri*:525
afum|EAL89245.1 100.00% afis|EAW16642.1 100.00%
Bootstrap support for afum|EAL89245.1 as seed ortholog is 100%.
Bootstrap support for afis|EAW16642.1 as seed ortholog is 100%.

---

Group of orthologs #7102. Best score 525 bits
Score difference with first non-orthologous sequence - *A. fumigatus*:525 *A. fischeri*:525
afum|EAL91762.1 100.00% afis|EAW17952.1 100.00%
Bootstrap support for afum|EAL91762.1 as seed ortholog is 100%.
Bootstrap support for afis|EAW17952.1 as seed ortholog is 100%.

---

Group of orthologs #7103. Best score 525 bits
Score difference with first non-orthologous sequence - *A. fumigatus*:525 *A. fischeri*:525
afum|EAL86041.1 100.00% afis|EAW24369.1 100.00%
Bootstrap support for afum|EAL86041.1 as seed ortholog is 100%.
Bootstrap support for afis|EAW24369.1 as seed ortholog is 100%.

---

Group of orthologs #7104. Best score 525 bits
Score difference with first non-orthologous sequence - *A. fumigatus*:525 *A. fischeri*:525
afum|EAL88268.1 100.00% afis|EAW23216.1 100.00%
Bootstrap support for afum|EAL88268.1 as seed ortholog is 100%.
Bootstrap support for afis|EAW23216.1 as seed ortholog is 100%.

---

Group of orthologs #7105. Best score 525 bits
Score difference with first non-orthologous sequence - *A. fumigatus*:525 *A. fischeri*:525
afum|EAL90738.1 100.00% afis|EAW22453.1 100.00%
Bootstrap support for afum|EAL90738.1 as seed ortholog is 100%.
Bootstrap support for afis|EAW22453.1 as seed ortholog is 100%.

---

Group of orthologs #7106. Best score 525 bits
Score difference with first non-orthologous sequence - *A. fumigatus*:525 *A. fischeri*:525
afum|EAL90786.1 100.00% afis|EAW22406.1 100.00%
Bootstrap support for afum|EAL90786.1 as seed ortholog is 100%.
Bootstrap support for afis|EAW22406.1 as seed ortholog is 100%.

---

Group of orthologs #7107. Best score 525 bits
Score difference with first non-orthologous sequence - *A. fumigatus*:525 *A. fischeri*:525
afum|EAL90299.1 100.00% afis|EAW22898.1 100.00%
Bootstrap support for afum|EAL90299.1 as seed ortholog is 100%.
Bootstrap support for afis|EAW22898.1 as seed ortholog is 100%.

---

Group of orthologs #7108. Best score 525 bits
Score difference with first non-orthologous sequence - *A. fumigatus*:525 *A. fischeri*:525
afum|EAL89395.2 100.00% afis|EAW24750.1 100.00%
Bootstrap support for afum|EAL89395.2 as seed ortholog is 100%.
Bootstrap support for afis|EAW24750.1 as seed ortholog is 100%.

---

Group of orthologs #7109. Best score 525 bits
Score difference with first non-orthologous sequence - *A. fumigatus*:525 *A. fischeri*:392
afum|EAL89506.2 100.00% afis|EAW24858.1 100.00%
Bootstrap support for afum|EAL89506.2 as seed ortholog is 100%.
Bootstrap support for afis|EAW24858.1 as seed ortholog is 100%.

---

Group of orthologs #7110. Best score 525 bits
Score difference with first non-orthologous sequence - *A. fumigatus*:525 *A. fischeri*:471
afum|EAL89984.1 100.00% afis|EAW25323.1 100.00%
Bootstrap support for afum|EAL89984.1 as seed ortholog is 100%.
Bootstrap support for afis|EAW25323.1 as seed ortholog is 100%.

---

Group of orthologs #7111. Best score 525 bits
Score difference with first non-orthologous sequence - *A. fumigatus*:525 *A. fischeri*:525
afum|EAL89999.1 100.00% afis|EAW25336.1 100.00%
Bootstrap support for afum|EAL89999.1 as seed ortholog is 100%.
Bootstrap support for afis|EAW25336.1 as seed ortholog is 100%.

---

Group of orthologs #7112. Best score 524 bits
Score difference with first non-orthologous sequence - *A. fumigatus*:524 *A. fischeri*:524
afum|EAL84596.1 100.00% afis|EAW20456.1 100.00%
Bootstrap support for afum|EAL84596.1 as seed ortholog is 100%.
Bootstrap support for afis|EAW20456.1 as seed ortholog is 100%.

---

Group of orthologs #7113. Best score 524 bits
Score difference with first non-orthologous sequence - *A. fumigatus*:524 *A. fischeri*:524
afum|EAL91645.1 100.00% afis|EAW17835.1 100.00%
Bootstrap support for afum|EAL91645.1 as seed ortholog is 100%.
Bootstrap support for afis|EAW17835.1 as seed ortholog is 100%.

---

Group of orthologs #7114. Best score 524 bits
Score difference with first non-orthologous sequence - *A. fumigatus*:524 *A. fischeri*:524
afum|EAL93039.1 100.00% afis|EAW18365.1 100.00%
Bootstrap support for afum|EAL93039.1 as seed ortholog is 100%.
Bootstrap support for afis|EAW18365.1 as seed ortholog is 100%.

---

Group of orthologs #7115. Best score 523 bits
Score difference with first non-orthologous sequence - *A. fumigatus*:523 *A. fischeri*:523
afum|EAL84663.1 100.00% afis|EAW20971.1 100.00%
Bootstrap support for afum|EAL84663.1 as seed ortholog is 100%.
Bootstrap support for afis|EAW20971.1 as seed ortholog is 100%.

---

Group of orthologs #7116. Best score 523 bits
Score difference with first non-orthologous sequence - *A. fumigatus*:363 *A. fischeri*:394
afum|EAL86773.1 100.00% afis|EAW20294.1 100.00%
Bootstrap support for afum|EAL86773.1 as seed ortholog is 100%.
Bootstrap support for afis|EAW20294.1 as seed ortholog is 100%.

---

Group of orthologs #7117. Best score 523 bits
Score difference with first non-orthologous sequence - *A. fumigatus*:523 *A. fischeri*:481

afum|EAL84378.1 100.00% afis|EAW25705.1 100.00%
Bootstrap support for afum|EAL84378.1 as seed ortholog is 100%.
Bootstrap support for afis|EAW25705.1 as seed ortholog is 100%.

---

Group of orthologs #7118. Best score 523 bits
Score difference with first non-orthologous sequence - *A. fumigatus*:523 *A. fischeri*:523
afum|EAL86127.1 100.00% afis|EAW24457.1 100.00%
Bootstrap support for afum|EAL86127.1 as seed ortholog is 100%.
Bootstrap support for afis|EAW24457.1 as seed ortholog is 100%.

---

Group of orthologs #7119. Best score 523 bits
Score difference with first non-orthologous sequence - *A. fumigatus*:523 *A. fischeri*:523
afum|EAL93777.1 100.00% afis|EAW19048.1 100.00%
Bootstrap support for afum|EAL93777.1 as seed ortholog is 100%.
Bootstrap support for afis|EAW19048.1 as seed ortholog is 100%.

---

Group of orthologs #7120. Best score 523 bits
Score difference with first non-orthologous sequence - *A. fumigatus*:523 *A. fischeri*:523
afum|EAL93833.1 100.00% afis|EAW19101.1 100.00%
Bootstrap support for afum|EAL93833.1 as seed ortholog is 100%.
Bootstrap support for afis|EAW19101.1 as seed ortholog is 100%.

---

Group of orthologs #7121. Best score 523 bits
Score difference with first non-orthologous sequence - *A. fumigatus*:523 *A. fischeri*:523
afum|EAL91036.1 100.00% afis|EAW22154.1 100.00%
Bootstrap support for afum|EAL91036.1 as seed ortholog is 100%.
Bootstrap support for afis|EAW22154.1 as seed ortholog is 100%.

---

Group of orthologs #7122. Best score 523 bits
Score difference with first non-orthologous sequence - *A. fumigatus*:523 *A. fischeri*:452
afum|EAL90397.1 100.00% afis|EAW22802.1 100.00%
Bootstrap support for afum|EAL90397.1 as seed ortholog is 100%.
Bootstrap support for afis|EAW22802.1 as seed ortholog is 100%.

---

Group of orthologs #7123. Best score 523 bits
Score difference with first non-orthologous sequence - *A. fumigatus*:523 *A. fischeri*:523
afum|EAL89931.1 100.00% afis|EAW25268.1 100.00%
Bootstrap support for afum|EAL89931.1 as seed ortholog is 100%.
Bootstrap support for afis|EAW25268.1 as seed ortholog is 100%.

---

Group of orthologs #7124. Best score 522 bits
Score difference with first non-orthologous sequence - *A. fumigatus*:522 *A. fischeri*:219
afum|EAL86921.1 100.00% afis|EAW19597.1 100.00%
Bootstrap support for afum|EAL86921.1 as seed ortholog is 100%.
Bootstrap support for afis|EAW19597.1 as seed ortholog is 100%.

---

Group of orthologs #7125. Best score 522 bits
Score difference with first non-orthologous sequence - *A. fumigatus*:522 *A. fischeri*:522
afum|EAL90213.1 100.00% afis|EAW22986.1 100.00%
Bootstrap support for afum|EAL90213.1 as seed ortholog is 100%.
Bootstrap support for afis|EAW22986.1 as seed ortholog is 100%.

---

Group of orthologs #7126. Best score 522 bits
Score difference with first non-orthologous sequence - *A. fumigatus*:522 *A. fischeri*:522
afum|EAL89458.2 100.00% afis|EAW24814.1 100.00%
Bootstrap support for afum|EAL89458.2 as seed ortholog is 100%.

Bootstrap support for afis|EAW24814.1 as seed ortholog is 100%.

---

Group of orthologs #7127. Best score 522 bits
Score difference with first non-orthologous sequence - *A. fumigatus*:522 *A. fischeri*:522
afum|EAL92862.1 100.00% afis|EAW21849.1 100.00%
Bootstrap support for afum|EAL92862.1 as seed ortholog is 100%.
Bootstrap support for afis|EAW21849.1 as seed ortholog is 100%.

---

Group of orthologs #7128. Best score 522 bits
Score difference with first non-orthologous sequence - *A. fumigatus*:522 *A. fischeri*:522
afum|EAL89737.2 100.00% afis|EAW25068.1 100.00%
Bootstrap support for afum|EAL89737.2 as seed ortholog is 100%.
Bootstrap support for afis|EAW25068.1 as seed ortholog is 100%.

---

Group of orthologs #7129. Best score 521 bits
Score difference with first non-orthologous sequence - *A. fumigatus*:521 *A. fischeri*:521
afum|EAL86444.1 100.00% afis|EAW16908.1 100.00%
Bootstrap support for afum|EAL86444.1 as seed ortholog is 100%.
Bootstrap support for afis|EAW16908.1 as seed ortholog is 100%.

---

Group of orthologs #7130. Best score 521 bits
Score difference with first non-orthologous sequence - *A. fumigatus*:386 *A. fischeri*:317
afum|EAL86597.1 100.00% afis|EAW17160.1 100.00%
Bootstrap support for afum|EAL86597.1 as seed ortholog is 100%.
Bootstrap support for afis|EAW17160.1 as seed ortholog is 100%.

---

Group of orthologs #7131. Best score 521 bits
Score difference with first non-orthologous sequence - *A. fumigatus*:521 *A. fischeri*:521
afum|EAL84594.2 100.00% afis|EAW20459.1 100.00%
Bootstrap support for afum|EAL84594.2 as seed ortholog is 100%.
Bootstrap support for afis|EAW20459.1 as seed ortholog is 100%.

---

Group of orthologs #7132. Best score 521 bits
Score difference with first non-orthologous sequence - *A. fumigatus*:521 *A. fischeri*:521
afum|EAL89189.1 100.00% afis|EAW16593.1 100.00%
Bootstrap support for afum|EAL89189.1 as seed ortholog is 100%.
Bootstrap support for afis|EAW16593.1 as seed ortholog is 100%.

---

Group of orthologs #7133. Best score 521 bits
Score difference with first non-orthologous sequence - *A. fumigatus*:521 *A. fischeri*:521
afum|EAL87586.1 100.00% afis|EAW18220.1 100.00%
Bootstrap support for afum|EAL87586.1 as seed ortholog is 100%.
Bootstrap support for afis|EAW18220.1 as seed ortholog is 100%.

---

Group of orthologs #7134. Best score 521 bits
Score difference with first non-orthologous sequence - *A. fumigatus*:521 *A. fischeri*:521
afum|EAL88357.1 100.00% afis|EAW23123.1 100.00%
Bootstrap support for afum|EAL88357.1 as seed ortholog is 100%.
Bootstrap support for afis|EAW23123.1 as seed ortholog is 100%.

---

Group of orthologs #7135. Best score 521 bits
Score difference with first non-orthologous sequence - *A. fumigatus*:521 *A. fischeri*:521
afum|EAL92995.1 100.00% afis|EAW21980.1 100.00%
Bootstrap support for afum|EAL92995.1 as seed ortholog is 100%.
Bootstrap support for afis|EAW21980.1 as seed ortholog is 100%.

---

Group of orthologs #7136. Best score 520 bits
Score difference with first non-orthologous sequence - *A. fumigatus*:520 *A. fischeri*:520
afum|EAL90351.1 100.00% afis|EAW22849.1 100.00%
Bootstrap support for afum|EAL90351.1 as seed ortholog is 100%.
Bootstrap support for afis|EAW22849.1 as seed ortholog is 100%.

---

Group of orthologs #7137. Best score 520 bits
Score difference with first non-orthologous sequence - *A. fumigatus*:473 *A. fischeri*:462
afum|EAL92710.1 100.00% afis|EAW21676.1 100.00%
Bootstrap support for afum|EAL92710.1 as seed ortholog is 100%.
Bootstrap support for afis|EAW21676.1 as seed ortholog is 100%.

---

Group of orthologs #7138. Best score 520 bits
Score difference with first non-orthologous sequence - *A. fumigatus*:520 *A. fischeri*:520
afum|EAL90013.1 100.00% afis|EAW25350.1 100.00%
Bootstrap support for afum|EAL90013.1 as seed ortholog is 100%.
Bootstrap support for afis|EAW25350.1 as seed ortholog is 100%.

---

Group of orthologs #7139. Best score 520 bits
Score difference with first non-orthologous sequence - *A. fumigatus*:520 *A. fischeri*:520
afum|EAL90132.1 100.00% afis|EAW25468.1 100.00%
Bootstrap support for afum|EAL90132.1 as seed ortholog is 100%.
Bootstrap support for afis|EAW25468.1 as seed ortholog is 100%.

---

Group of orthologs #7140. Best score 519 bits
Score difference with first non-orthologous sequence - *A. fumigatus*:519 *A. fischeri*:519
afum|EAL85506.2 100.00% afis|EAW15845.1 100.00%
Bootstrap support for afum|EAL85506.2 as seed ortholog is 100%.
Bootstrap support for afis|EAW15845.1 as seed ortholog is 100%.

---

Group of orthologs #7141. Best score 519 bits
Score difference with first non-orthologous sequence - *A. fumigatus*:519 *A. fischeri*:519
afum|EAL88590.1 100.00% afis|EAW15957.1 100.00%
Bootstrap support for afum|EAL88590.1 as seed ortholog is 100%.
Bootstrap support for afis|EAW15957.1 as seed ortholog is 100%.

---

Group of orthologs #7142. Best score 519 bits
Score difference with first non-orthologous sequence - *A. fumigatus*:519 *A. fischeri*:519
afum|EAL89149.1 100.00% afis|EAW16551.1 100.00%
Bootstrap support for afum|EAL89149.1 as seed ortholog is 100%.
Bootstrap support for afis|EAW16551.1 as seed ortholog is 100%.

---

Group of orthologs #7143. Best score 519 bits
Score difference with first non-orthologous sequence - *A. fumigatus*:519 *A. fischeri*:519
afum|EAL91488.1 100.00% afis|EAW17675.1 100.00%
Bootstrap support for afum|EAL91488.1 as seed ortholog is 100%.
Bootstrap support for afis|EAW17675.1 as seed ortholog is 100%.

---

Group of orthologs #7144. Best score 519 bits
Score difference with first non-orthologous sequence - *A. fumigatus*:519 *A. fischeri*:519
afum|EAL90544.1 100.00% afis|EAW22648.1 100.00%
Bootstrap support for afum|EAL90544.1 as seed ortholog is 100%.
Bootstrap support for afis|EAW22648.1 as seed ortholog is 100%.

---

Group of orthologs #7145. Best score 519 bits
Score difference with first non-orthologous sequence - *A. fumigatus*:519 *A. fischeri*:519

afum|EAL89715.1 100.00% afis|EAW25047.1 100.00%
Bootstrap support for afum|EAL89715.1 as seed ortholog is 100%.
Bootstrap support for afis|EAW25047.1 as seed ortholog is 100%.

---

Group of orthologs #7146. Best score 518 bits
Score difference with first non-orthologous sequence - *A. fumigatus*:518 *A. fischeri*:518
afum|EAL86847.2 100.00% afis|EAW19674.1 100.00%
Bootstrap support for afum|EAL86847.2 as seed ortholog is 100%.
Bootstrap support for afis|EAW19674.1 as seed ortholog is 100%.

---

Group of orthologs #7147. Best score 518 bits
Score difference with first non-orthologous sequence - *A. fumigatus*:518 *A. fischeri*:518
afum|EAL86795.1 100.00% afis|EAW19734.1 100.00%
Bootstrap support for afum|EAL86795.1 as seed ortholog is 100%.
Bootstrap support for afis|EAW19734.1 as seed ortholog is 100%.

---

Group of orthologs #7148. Best score 518 bits
Score difference with first non-orthologous sequence - *A. fumigatus*:518 *A. fischeri*:518
afum|EAL87237.1 100.00% afis|EAW23803.1 100.00%
Bootstrap support for afum|EAL87237.1 as seed ortholog is 100%.
Bootstrap support for afis|EAW23803.1 as seed ortholog is 100%.

---

Group of orthologs #7149. Best score 518 bits
Score difference with first non-orthologous sequence - *A. fumigatus*:518 *A. fischeri*:518
afum|EAL88096.1 100.00% afis|EAW23375.1 100.00%
Bootstrap support for afum|EAL88096.1 as seed ortholog is 100%.
Bootstrap support for afis|EAW23375.1 as seed ortholog is 100%.

---

Group of orthologs #7150. Best score 518 bits
Score difference with first non-orthologous sequence - *A. fumigatus*:518 *A. fischeri*:518
afum|EAL90546.1 100.00% afis|EAW22646.1 100.00%
Bootstrap support for afum|EAL90546.1 as seed ortholog is 100%.
Bootstrap support for afis|EAW22646.1 as seed ortholog is 100%.

---

Group of orthologs #7151. Best score 518 bits
Score difference with first non-orthologous sequence - *A. fumigatus*:518 *A. fischeri*:518
afum|EAL92262.1 100.00% afis|EAW21249.1 100.00%
Bootstrap support for afum|EAL92262.1 as seed ortholog is 100%.
Bootstrap support for afis|EAW21249.1 as seed ortholog is 100%.

---

Group of orthologs #7152. Best score 518 bits
Score difference with first non-orthologous sequence - *A. fumigatus*:518 *A. fischeri*:518
afum|EAL89948.1 100.00% afis|EAW25285.1 100.00%
Bootstrap support for afum|EAL89948.1 as seed ortholog is 100%.
Bootstrap support for afis|EAW25285.1 as seed ortholog is 100%.

---

Group of orthologs #7153. Best score 518 bits
Score difference with first non-orthologous sequence - *A. fumigatus*:518 *A. fischeri*:518
afum|EBA27411.1 100.00% afis|EAW22579.1 100.00%
Bootstrap support for afum|EBA27411.1 as seed ortholog is 100%.
Bootstrap support for afis|EAW22579.1 as seed ortholog is 100%.

---

Group of orthologs #7154. Best score 518 bits
Score difference with first non-orthologous sequence - *A. fumigatus*:518 *A. fischeri*:368
afum|EBA27382.1 100.00% afis|EAW24776.1 100.00%
Bootstrap support for afum|EBA27382.1 as seed ortholog is 100%.

Bootstrap support for afis|EAW24776.1 as seed ortholog is 100%.

---

Group of orthologs #7155. Best score 517 bits
Score difference with first non-orthologous sequence - *A. fumigatus*:517 *A. fischeri*:517
afum|EAL92526.1 100.00% afis|EAW21506.1 100.00%
Bootstrap support for afum|EAL92526.1 as seed ortholog is 100%.
Bootstrap support for afis|EAW21506.1 as seed ortholog is 100%.

---

Group of orthologs #7156. Best score 516 bits
Score difference with first non-orthologous sequence - *A. fumigatus*:516 *A. fischeri*:447
afum|EAL86301.1 100.00% afis|EAW16727.1 100.00%
Bootstrap support for afum|EAL86301.1 as seed ortholog is 100%.
Bootstrap support for afis|EAW16727.1 as seed ortholog is 100%.

---

Group of orthologs #7157. Best score 516 bits
Score difference with first non-orthologous sequence - *A. fumigatus*:516 *A. fischeri*:516
afum|EAL88612.1 100.00% afis|EAW15980.1 100.00%
Bootstrap support for afum|EAL88612.1 as seed ortholog is 100%.
Bootstrap support for afis|EAW15980.1 as seed ortholog is 100%.

---

Group of orthologs #7158. Best score 516 bits
Score difference with first non-orthologous sequence - *A. fumigatus*:516 *A. fischeri*:516
afum|EAL91900.1 100.00% afis|EAW18088.1 100.00%
Bootstrap support for afum|EAL91900.1 as seed ortholog is 100%.
Bootstrap support for afis|EAW18088.1 as seed ortholog is 100%.

---

Group of orthologs #7159. Best score 516 bits
Score difference with first non-orthologous sequence - *A. fumigatus*:516 *A. fischeri*:516
afum|EAL94004.1 100.00% afis|EAW19278.1 100.00%
Bootstrap support for afum|EAL94004.1 as seed ortholog is 100%.
Bootstrap support for afis|EAW19278.1 as seed ortholog is 100%.

---

Group of orthologs #7160. Best score 516 bits
Score difference with first non-orthologous sequence - *A. fumigatus*:516 *A. fischeri*:516
afum|EAL90330.2 100.00% afis|EAW22869.1 100.00%
Bootstrap support for afum|EAL90330.2 as seed ortholog is 100%.
Bootstrap support for afis|EAW22869.1 as seed ortholog is 100%.

---

Group of orthologs #7161. Best score 516 bits
Score difference with first non-orthologous sequence - *A. fumigatus*:516 *A. fischeri*:516
afum|EAL92768.1 100.00% afis|EAW21735.1 100.00%
Bootstrap support for afum|EAL92768.1 as seed ortholog is 100%.
Bootstrap support for afis|EAW21735.1 as seed ortholog is 100%.

---

Group of orthologs #7162. Best score 516 bits
Score difference with first non-orthologous sequence - *A. fumigatus*:516 *A. fischeri*:516
afum|EAL89690.1 100.00% afis|EAW25007.1 100.00%
Bootstrap support for afum|EAL89690.1 as seed ortholog is 100%.
Bootstrap support for afis|EAW25007.1 as seed ortholog is 100%.

---

Group of orthologs #7163. Best score 515 bits
Score difference with first non-orthologous sequence - *A. fumigatus*:515 *A. fischeri*:515
afum|EAL84640.2 100.00% afis|EAW20989.1 100.00%
Bootstrap support for afum|EAL84640.2 as seed ortholog is 100%.
Bootstrap support for afis|EAW20989.1 as seed ortholog is 100%.

---

Group of orthologs #7164. Best score 515 bits
Score difference with first non-orthologous sequence - *A. fumigatus*:515 *A. fischeri*:515
afum|EAL87007.1 100.00% afis|EAW19487.1 100.00%
Bootstrap support for afum|EAL87007.1 as seed ortholog is 100%.
Bootstrap support for afis|EAW19487.1 as seed ortholog is 100%.

---

Group of orthologs #7165. Best score 515 bits
Score difference with first non-orthologous sequence - *A. fumigatus*:515 *A. fischeri*:515
afum|EAL85979.1 100.00% afis|EAW24311.1 100.00%
Bootstrap support for afum|EAL85979.1 as seed ortholog is 100%.
Bootstrap support for afis|EAW24311.1 as seed ortholog is 100%.

---

Group of orthologs #7166. Best score 515 bits
Score difference with first non-orthologous sequence - *A. fumigatus*:515 *A. fischeri*:515
afum|EAL93614.1 100.00% afis|EAW18882.1 100.00%
Bootstrap support for afum|EAL93614.1 as seed ortholog is 100%.
Bootstrap support for afis|EAW18882.1 as seed ortholog is 100%.

---

Group of orthologs #7167. Best score 515 bits
Score difference with first non-orthologous sequence - *A. fumigatus*:515 *A. fischeri*:515
afum|EAL91038.1 100.00% afis|EAW22152.1 100.00%
Bootstrap support for afum|EAL91038.1 as seed ortholog is 100%.
Bootstrap support for afis|EAW22152.1 as seed ortholog is 100%.

---

Group of orthologs #7168. Best score 515 bits
Score difference with first non-orthologous sequence - *A. fumigatus*:418 *A. fischeri*:515
afum|EAL92222.1 100.00% afis|EAW21211.1 100.00%
Bootstrap support for afum|EAL92222.1 as seed ortholog is 100%.
Bootstrap support for afis|EAW21211.1 as seed ortholog is 100%.

---

Group of orthologs #7169. Best score 515 bits
Score difference with first non-orthologous sequence - *A. fumigatus*:515 *A. fischeri*:515
afum|EAL89474.1 100.00% afis|EAW24829.1 100.00%
Bootstrap support for afum|EAL89474.1 as seed ortholog is 100%.
Bootstrap support for afis|EAW24829.1 as seed ortholog is 100%.

---

Group of orthologs #7170. Best score 515 bits
Score difference with first non-orthologous sequence - *A. fumigatus*:515 *A. fischeri*:515
afum|EAL92815.1 100.00% afis|EAW21789.1 100.00%
Bootstrap support for afum|EAL92815.1 as seed ortholog is 100%.
Bootstrap support for afis|EAW21789.1 as seed ortholog is 100%.

---

Group of orthologs #7171. Best score 514 bits
Score difference with first non-orthologous sequence - *A. fumigatus*:514 *A. fischeri*:514
afum|EAL85856.1 100.00% afis|EAW15466.1 100.00%
Bootstrap support for afum|EAL85856.1 as seed ortholog is 100%.
Bootstrap support for afis|EAW15466.1 as seed ortholog is 100%.

---

Group of orthologs #7172. Best score 514 bits
Score difference with first non-orthologous sequence - *A. fumigatus*:514 *A. fischeri*:514
afum|EAL87213.1 100.00% afis|EAW23779.1 100.00%
Bootstrap support for afum|EAL87213.1 as seed ortholog is 100%.
Bootstrap support for afis|EAW23779.1 as seed ortholog is 100%.

---

Group of orthologs #7173. Best score 513 bits
Score difference with first non-orthologous sequence - *A. fumigatus*:513 *A. fischeri*:513

afum|EAL87048.1 100.00% afis|EAW19447.1 100.00%
Bootstrap support for afum|EAL87048.1 as seed ortholog is 100%.
Bootstrap support for afis|EAW19447.1 as seed ortholog is 100%.

---

Group of orthologs #7174. Best score 513 bits
Score difference with first non-orthologous sequence - *A. fumigatus*:513 *A. fischeri*:513
afum|EAL87217.2 100.00% afis|EAW23783.1 100.00%
Bootstrap support for afum|EAL87217.2 as seed ortholog is 100%.
Bootstrap support for afis|EAW23783.1 as seed ortholog is 100%.

---

Group of orthologs #7175. Best score 513 bits
Score difference with first non-orthologous sequence - *A. fumigatus*:513 *A. fischeri*:513
afum|EAL93756.1 100.00% afis|EAW19025.1 100.00%
Bootstrap support for afum|EAL93756.1 as seed ortholog is 100%.
Bootstrap support for afis|EAW19025.1 as seed ortholog is 100%.

---

Group of orthologs #7176. Best score 513 bits
Score difference with first non-orthologous sequence - *A. fumigatus*:513 *A. fischeri*:513
afum|EBA27490.1 100.00% afis|EAW18662.1 100.00%
Bootstrap support for afum|EBA27490.1 as seed ortholog is 100%.
Bootstrap support for afis|EAW18662.1 as seed ortholog is 100%.

---

Group of orthologs #7177. Best score 513 bits
Score difference with first non-orthologous sequence - *A. fumigatus*:513 *A. fischeri*:513
afum|EAL93908.1 100.00% afis|EAW19178.1 100.00%
Bootstrap support for afum|EAL93908.1 as seed ortholog is 100%.
Bootstrap support for afis|EAW19178.1 as seed ortholog is 100%.

---

Group of orthologs #7178. Best score 513 bits
Score difference with first non-orthologous sequence - *A. fumigatus*:513 *A. fischeri*:513
afum|EBA27391.1 100.00% afis|EAW25262.1 100.00%
Bootstrap support for afum|EBA27391.1 as seed ortholog is 100%.
Bootstrap support for afis|EAW25262.1 as seed ortholog is 100%.

---

Group of orthologs #7179. Best score 512 bits
Score difference with first non-orthologous sequence - *A. fumigatus*:512 *A. fischeri*:512
afum|EAL91241.1 100.00% afis|EAW17406.1 100.00%
Bootstrap support for afum|EAL91241.1 as seed ortholog is 100%.
Bootstrap support for afis|EAW17406.1 as seed ortholog is 100%.

---

Group of orthologs #7180. Best score 512 bits
Score difference with first non-orthologous sequence - *A. fumigatus*:512 *A. fischeri*:512
afum|EAL93299.2 100.00% afis|EAW18592.1 100.00%
Bootstrap support for afum|EAL93299.2 as seed ortholog is 100%.
Bootstrap support for afis|EAW18592.1 as seed ortholog is 100%.

---

Group of orthologs #7181. Best score 512 bits
Score difference with first non-orthologous sequence - *A. fumigatus*:512 *A. fischeri*:512
afum|EAL90689.1 100.00% afis|EAW22504.1 100.00%
Bootstrap support for afum|EAL90689.1 as seed ortholog is 100%.
Bootstrap support for afis|EAW22504.1 as seed ortholog is 100%.

---

Group of orthologs #7182. Best score 512 bits
Score difference with first non-orthologous sequence - *A. fumigatus*:512 *A. fischeri*:512
afum|EAL90000.1 100.00% afis|EAW25337.1 100.00%
Bootstrap support for afum|EAL90000.1 as seed ortholog is 100%.

Bootstrap support for afis|EAW25337.1 as seed ortholog is 100%.

---

Group of orthologs #7183. Best score 511 bits
Score difference with first non-orthologous sequence - *A. fumigatus*:511 *A. fischeri*:511
afum|EAL88694.1 100.00% afis|EAW16062.1 100.00%
Bootstrap support for afum|EAL88694.1 as seed ortholog is 100%.
Bootstrap support for afis|EAW16062.1 as seed ortholog is 100%.

---

Group of orthologs #7184. Best score 511 bits
Score difference with first non-orthologous sequence - *A. fumigatus*:511 *A. fischeri*:511
afum|EAL84538.1 100.00% afis|EAW20519.1 100.00%
Bootstrap support for afum|EAL84538.1 as seed ortholog is 100%.
Bootstrap support for afis|EAW20519.1 as seed ortholog is 100%.

---

Group of orthologs #7185. Best score 511 bits
Score difference with first non-orthologous sequence - *A. fumigatus*:511 *A. fischeri*:511
afum|EAL85305.1 100.00% afis|EAW20186.1 100.00%
Bootstrap support for afum|EAL85305.1 as seed ortholog is 100%.
Bootstrap support for afis|EAW20186.1 as seed ortholog is 100%.

---

Group of orthologs #7186. Best score 511 bits
Score difference with first non-orthologous sequence - *A. fumigatus*:511 *A. fischeri*:511
afum|EAL91251.1 100.00% afis|EAW17423.1 100.00%
Bootstrap support for afum|EAL91251.1 as seed ortholog is 100%.
Bootstrap support for afis|EAW17423.1 as seed ortholog is 100%.

---

Group of orthologs #7187. Best score 511 bits
Score difference with first non-orthologous sequence - *A. fumigatus*:511 *A. fischeri*:511
afum|EAL87944.1 100.00% afis|EAW23520.1 100.00%
Bootstrap support for afum|EAL87944.1 as seed ortholog is 100%.
Bootstrap support for afis|EAW23520.1 as seed ortholog is 100%.

---

Group of orthologs #7188. Best score 511 bits
Score difference with first non-orthologous sequence - *A. fumigatus*:511 *A. fischeri*:511
afum|EAL87848.1 100.00% afis|EAW23614.1 100.00%
Bootstrap support for afum|EAL87848.1 as seed ortholog is 100%.
Bootstrap support for afis|EAW23614.1 as seed ortholog is 100%.

---

Group of orthologs #7189. Best score 511 bits
Score difference with first non-orthologous sequence - *A. fumigatus*:511 *A. fischeri*:511
afum|EAL89578.2 100.00% afis|EAW24923.1 100.00%
Bootstrap support for afum|EAL89578.2 as seed ortholog is 100%.
Bootstrap support for afis|EAW24923.1 as seed ortholog is 100%.

---

Group of orthologs #7190. Best score 511 bits
Score difference with first non-orthologous sequence - *A. fumigatus*:511 *A. fischeri*:511
afum|EAL89857.1 100.00% afis|EAW25183.1 100.00%
Bootstrap support for afum|EAL89857.1 as seed ortholog is 100%.
Bootstrap support for afis|EAW25183.1 as seed ortholog is 100%.

---

Group of orthologs #7191. Best score 510 bits
Score difference with first non-orthologous sequence - *A. fumigatus*:510 *A. fischeri*:510
afum|EAL85565.1 100.00% afis|EAW15785.1 100.00%
Bootstrap support for afum|EAL85565.1 as seed ortholog is 100%.
Bootstrap support for afis|EAW15785.1 as seed ortholog is 100%.

---

Group of orthologs #7192. Best score 510 bits
Score difference with first non-orthologous sequence - *A. fumigatus*:430 *A. fischeri*:510
afum|EAL86620.1 100.00% afis|EAW17192.1 100.00%
Bootstrap support for afum|EAL86620.1 as seed ortholog is 100%.
Bootstrap support for afis|EAW17192.1 as seed ortholog is 100%.

---

Group of orthologs #7193. Best score 510 bits
Score difference with first non-orthologous sequence - *A. fumigatus*:510 *A. fischeri*:510
afum|EAL91145.1 100.00% afis|EAW17310.1 100.00%
Bootstrap support for afum|EAL91145.1 as seed ortholog is 100%.
Bootstrap support for afis|EAW17310.1 as seed ortholog is 100%.

---

Group of orthologs #7194. Best score 510 bits
Score difference with first non-orthologous sequence - *A. fumigatus*:510 *A. fischeri*:510
afum|EAL87937.1 100.00% afis|EAW23529.1 100.00%
Bootstrap support for afum|EAL87937.1 as seed ortholog is 100%.
Bootstrap support for afis|EAW23529.1 as seed ortholog is 100%.

---

Group of orthologs #7195. Best score 510 bits
Score difference with first non-orthologous sequence - *A. fumigatus*:510 *A. fischeri*:510
afum|EAL93025.1 100.00% afis|EAW22010.1 100.00%
Bootstrap support for afum|EAL93025.1 as seed ortholog is 100%.
Bootstrap support for afis|EAW22010.1 as seed ortholog is 100%.

---

Group of orthologs #7196. Best score 510 bits
Score difference with first non-orthologous sequence - *A. fumigatus*:350 *A. fischeri*:321
afum|EAL89818.1 100.00% afis|EAW25145.1 100.00%
Bootstrap support for afum|EAL89818.1 as seed ortholog is 100%.
Bootstrap support for afis|EAW25145.1 as seed ortholog is 100%.

---

Group of orthologs #7197. Best score 510 bits
Score difference with first non-orthologous sequence - *A. fumigatus*:510 *A. fischeri*:510
afum|EAL90175.2 100.00% afis|EAW25509.1 100.00%
Bootstrap support for afum|EAL90175.2 as seed ortholog is 100%.
Bootstrap support for afis|EAW25509.1 as seed ortholog is 100%.

---

Group of orthologs #7198. Best score 509 bits
Score difference with first non-orthologous sequence - *A. fumigatus*:509 *A. fischeri*:509
afum|EAL84208.2 100.00% afis|EAW15437.1 100.00%
Bootstrap support for afum|EAL84208.2 as seed ortholog is 100%.
Bootstrap support for afis|EAW15437.1 as seed ortholog is 100%.

---

Group of orthologs #7199. Best score 509 bits
Score difference with first non-orthologous sequence - *A. fumigatus*:509 *A. fischeri*:454
afum|EAL84601.1 100.00% afis|EAW20452.1 100.00%
Bootstrap support for afum|EAL84601.1 as seed ortholog is 100%.
Bootstrap support for afis|EAW20452.1 as seed ortholog is 100%.

---

Group of orthologs #7200. Best score 509 bits
Score difference with first non-orthologous sequence - *A. fumigatus*:509 *A. fischeri*:509
afum|EAL85357.1 100.00% afis|EAW20239.1 100.00%
Bootstrap support for afum|EAL85357.1 as seed ortholog is 100%.
Bootstrap support for afis|EAW20239.1 as seed ortholog is 100%.

---

Group of orthologs #7201. Best score 509 bits
Score difference with first non-orthologous sequence - *A. fumigatus*:509 *A. fischeri*:509

afum|EAL93534.1 100.00% afis|EAW18798.1 100.00%
Bootstrap support for afum|EAL93534.1 as seed ortholog is 100%.
Bootstrap support for afis|EAW18798.1 as seed ortholog is 100%.

---

Group of orthologs #7202. Best score 509 bits
Score difference with first non-orthologous sequence - *A. fumigatus*:509 *A. fischeri*:509
afum|EAL90714.1 100.00% afis|EAW22481.1 100.00%
Bootstrap support for afum|EAL90714.1 as seed ortholog is 100%.
Bootstrap support for afis|EAW22481.1 as seed ortholog is 100%.

---

Group of orthologs #7203. Best score 509 bits
Score difference with first non-orthologous sequence - *A. fumigatus*:509 *A. fischeri*:509
afum|EAL92602.1 100.00% afis|EAW21584.1 100.00%
Bootstrap support for afum|EAL92602.1 as seed ortholog is 100%.
Bootstrap support for afis|EAW21584.1 as seed ortholog is 100%.

---

Group of orthologs #7204. Best score 508 bits
Score difference with first non-orthologous sequence - *A. fumigatus*:508 *A. fischeri*:508
afum|EAL88994.1 100.00% afis|EAW16369.1 100.00%
Bootstrap support for afum|EAL88994.1 as seed ortholog is 100%.
Bootstrap support for afis|EAW16369.1 as seed ortholog is 100%.

---

Group of orthologs #7205. Best score 508 bits
Score difference with first non-orthologous sequence - *A. fumigatus*:508 *A. fischeri*:508
afum|EBA27426.1 100.00% afis|EAW17385.1 100.00%
Bootstrap support for afum|EBA27426.1 as seed ortholog is 100%.
Bootstrap support for afis|EAW17385.1 as seed ortholog is 100%.

---

Group of orthologs #7206. Best score 508 bits
Score difference with first non-orthologous sequence - *A. fumigatus*:508 *A. fischeri*:508
afum|EAL88201.1 100.00% afis|EAW23278.1 100.00%
Bootstrap support for afum|EAL88201.1 as seed ortholog is 100%.
Bootstrap support for afis|EAW23278.1 as seed ortholog is 100%.

---

Group of orthologs #7207. Best score 508 bits
Score difference with first non-orthologous sequence - *A. fumigatus*:508 *A. fischeri*:508
afum|EAL88254.1 100.00% afis|EAW23229.1 100.00%
Bootstrap support for afum|EAL88254.1 as seed ortholog is 100%.
Bootstrap support for afis|EAW23229.1 as seed ortholog is 100%.

---

Group of orthologs #7208. Best score 508 bits
Score difference with first non-orthologous sequence - *A. fumigatus*:508 *A. fischeri*:508
afum|EAL88141.1 100.00% afis|EAW23337.1 100.00%
Bootstrap support for afum|EAL88141.1 as seed ortholog is 100%.
Bootstrap support for afis|EAW23337.1 as seed ortholog is 100%.

---

Group of orthologs #7209. Best score 508 bits
Score difference with first non-orthologous sequence - *A. fumigatus*:508 *A. fischeri*:508
afum|EAL93362.1 100.00% afis|EAW18647.1 100.00%
Bootstrap support for afum|EAL93362.1 as seed ortholog is 100%.
Bootstrap support for afis|EAW18647.1 as seed ortholog is 100%.

---

Group of orthologs #7210. Best score 508 bits
Score difference with first non-orthologous sequence - *A. fumigatus*:508 *A. fischeri*:508
afum|EAL91026.1 100.00% afis|EAW22165.1 100.00%
Bootstrap support for afum|EAL91026.1 as seed ortholog is 100%.

Bootstrap support for afis|EAW22165.1 as seed ortholog is 100%.

---

Group of orthologs #7211. Best score 508 bits
Score difference with first non-orthologous sequence - *A. fumigatus*:508 *A. fischeri*:508
afum|EAL90869.1 100.00% afis|EAW22322.1 100.00%
Bootstrap support for afum|EAL90869.1 as seed ortholog is 100%.
Bootstrap support for afis|EAW22322.1 as seed ortholog is 100%.

---

Group of orthologs #7212. Best score 508 bits
Score difference with first non-orthologous sequence - *A. fumigatus*:508 *A. fischeri*:508
afum|EAL90833.1 100.00% afis|EAW22358.1 100.00%
Bootstrap support for afum|EAL90833.1 as seed ortholog is 100%.
Bootstrap support for afis|EAW22358.1 as seed ortholog is 100%.

---

Group of orthologs #7213. Best score 507 bits
Score difference with first non-orthologous sequence - *A. fumigatus*:507 *A. fischeri*:507
afum|EAL87673.1 100.00% afis|EAW18307.1 100.00%
Bootstrap support for afum|EAL87673.1 as seed ortholog is 100%.
Bootstrap support for afis|EAW18307.1 as seed ortholog is 100%.

---

Group of orthologs #7214. Best score 507 bits
Score difference with first non-orthologous sequence - *A. fumigatus*:507 *A. fischeri*:507
afum|EAL87278.2 100.00% afis|EAW23850.1 100.00%
Bootstrap support for afum|EAL87278.2 as seed ortholog is 100%.
Bootstrap support for afis|EAW23850.1 as seed ortholog is 100%.

---

Group of orthologs #7215. Best score 507 bits
Score difference with first non-orthologous sequence - *A. fumigatus*:507 *A. fischeri*:507
afum|EAL90499.1 100.00% afis|EAW22694.1 100.00%
Bootstrap support for afum|EAL90499.1 as seed ortholog is 100%.
Bootstrap support for afis|EAW22694.1 as seed ortholog is 100%.

---

Group of orthologs #7216. Best score 506 bits
Score difference with first non-orthologous sequence - *A. fumigatus*:506 *A. fischeri*:506
afum|EAL88519.1 100.00% afis|EAW15885.1 100.00%
Bootstrap support for afum|EAL88519.1 as seed ortholog is 100%.
Bootstrap support for afis|EAW15885.1 as seed ortholog is 100%.

---

Group of orthologs #7217. Best score 506 bits
Score difference with first non-orthologous sequence - *A. fumigatus*:506 *A. fischeri*:506
afum|EAL88428.2 100.00% afis|EAW23052.1 100.00%
Bootstrap support for afum|EAL88428.2 as seed ortholog is 100%.
Bootstrap support for afis|EAW23052.1 as seed ortholog is 100%.

---

Group of orthologs #7218. Best score 506 bits
Score difference with first non-orthologous sequence - *A. fumigatus*:506 *A. fischeri*:506
afum|EAL88369.1 100.00% afis|EAW23110.1 100.00%
Bootstrap support for afum|EAL88369.1 as seed ortholog is 100%.
Bootstrap support for afis|EAW23110.1 as seed ortholog is 100%.

---

Group of orthologs #7219. Best score 506 bits
Score difference with first non-orthologous sequence - *A. fumigatus*:506 *A. fischeri*:506
afum|EAL93349.1 100.00% afis|EAW18636.1 100.00%
Bootstrap support for afum|EAL93349.1 as seed ortholog is 100%.
Bootstrap support for afis|EAW18636.1 as seed ortholog is 100%.

---

Group of orthologs #7220. Best score 506 bits
Score difference with first non-orthologous sequence - *A. fumigatus*:352 *A. fischeri*:410

| | | | |
|---|---|---|---|
| afum\|EAL91093.2 | 100.00% | afis\|EAW22077.1 | 100.00% |

Bootstrap support for afum|EAL91093.2 as seed ortholog is 100%.
Bootstrap support for afis|EAW22077.1 as seed ortholog is 100%.

---

Group of orthologs #7221. Best score 506 bits
Score difference with first non-orthologous sequence - *A. fumigatus*:506 *A. fischeri*:506

| | | | |
|---|---|---|---|
| afum\|EAL90670.1 | 100.00% | afis\|EAW22524.1 | 100.00% |

Bootstrap support for afum|EAL90670.1 as seed ortholog is 100%.
Bootstrap support for afis|EAW22524.1 as seed ortholog is 100%.

---

Group of orthologs #7222. Best score 506 bits
Score difference with first non-orthologous sequence - *A. fumigatus*:506 *A. fischeri*:506

| | | | |
|---|---|---|---|
| afum\|EAL90409.1 | 100.00% | afis\|EAW22790.1 | 100.00% |

Bootstrap support for afum|EAL90409.1 as seed ortholog is 100%.
Bootstrap support for afis|EAW22790.1 as seed ortholog is 100%.

---

Group of orthologs #7223. Best score 506 bits
Score difference with first non-orthologous sequence - *A. fumigatus*:506 *A. fischeri*:463

| | | | |
|---|---|---|---|
| afum\|EAL89720.1 | 100.00% | afis\|EAW25052.1 | 100.00% |

Bootstrap support for afum|EAL89720.1 as seed ortholog is 100%.
Bootstrap support for afis|EAW25052.1 as seed ortholog is 100%.

---

Group of orthologs #7224. Best score 504 bits
Score difference with first non-orthologous sequence - *A. fumigatus*:504 *A. fischeri*:347

| | | | |
|---|---|---|---|
| afum\|EAL89062.1 | 100.00% | afis\|EAW16466.1 | 100.00% |

Bootstrap support for afum|EAL89062.1 as seed ortholog is 100%.
Bootstrap support for afis|EAW16466.1 as seed ortholog is 100%.

---

Group of orthologs #7225. Best score 504 bits
Score difference with first non-orthologous sequence - *A. fumigatus*:504 *A. fischeri*:504

| | | | |
|---|---|---|---|
| afum\|EAL86799.1 | 100.00% | afis\|EAW19728.1 | 100.00% |

Bootstrap support for afum|EAL86799.1 as seed ortholog is 100%.
Bootstrap support for afis|EAW19728.1 as seed ortholog is 100%.

---

Group of orthologs #7226. Best score 504 bits
Score difference with first non-orthologous sequence - *A. fumigatus*:504 *A. fischeri*:504

| | | | |
|---|---|---|---|
| afum\|EAL91712.1 | 100.00% | afis\|EAW17904.1 | 100.00% |

Bootstrap support for afum|EAL91712.1 as seed ortholog is 100%.
Bootstrap support for afis|EAW17904.1 as seed ortholog is 100%.

---

Group of orthologs #7227. Best score 504 bits
Score difference with first non-orthologous sequence - *A. fumigatus*:504 *A. fischeri*:353

| | | | |
|---|---|---|---|
| afum\|EAL92062.1 | 100.00% | afis\|EAW21023.1 | 100.00% |

Bootstrap support for afum|EAL92062.1 as seed ortholog is 100%.
Bootstrap support for afis|EAW21023.1 as seed ortholog is 100%.

---

Group of orthologs #7228. Best score 503 bits
Score difference with first non-orthologous sequence - *A. fumigatus*:503 *A. fischeri*:503

| | | | |
|---|---|---|---|
| afum\|EAL85571.1 | 100.00% | afis\|EAW15778.1 | 100.00% |

Bootstrap support for afum|EAL85571.1 as seed ortholog is 100%.
Bootstrap support for afis|EAW15778.1 as seed ortholog is 100%.

---

Group of orthologs #7229. Best score 503 bits
Score difference with first non-orthologous sequence - *A. fumigatus*:503 *A. fischeri*:503

afum|EAL88898.1 100.00% afis|EAW16277.1 100.00%
Bootstrap support for afum|EAL88898.1 as seed ortholog is 100%.
Bootstrap support for afis|EAW16277.1 as seed ortholog is 100%.

---

Group of orthologs #7230. Best score 503 bits
Score difference with first non-orthologous sequence - *A. fumigatus*:503 *A. fischeri*:503
afum|EAL84629.1 100.00% afis|EAW20421.1 100.00%
Bootstrap support for afum|EAL84629.1 as seed ortholog is 100%.
Bootstrap support for afis|EAW20421.1 as seed ortholog is 100%.

---

Group of orthologs #7231. Best score 503 bits
Score difference with first non-orthologous sequence - *A. fumigatus*:503 *A. fischeri*:503
afum|EAL91448.1 100.00% afis|EAW17633.1 100.00%
Bootstrap support for afum|EAL91448.1 as seed ortholog is 100%.
Bootstrap support for afis|EAW17633.1 as seed ortholog is 100%.

---

Group of orthologs #7232. Best score 503 bits
Score difference with first non-orthologous sequence - *A. fumigatus*:414 *A. fischeri*:503
afum|EAL90215.1 100.00% afis|EAW22983.1 100.00%
Bootstrap support for afum|EAL90215.1 as seed ortholog is 100%.
Bootstrap support for afis|EAW22983.1 as seed ortholog is 100%.

---

Group of orthologs #7233. Best score 503 bits
Score difference with first non-orthologous sequence - *A. fumigatus*:503 *A. fischeri*:503
afum|EAL90211.1 100.00% afis|EAW22988.1 100.00%
Bootstrap support for afum|EAL90211.1 as seed ortholog is 100%.
Bootstrap support for afis|EAW22988.1 as seed ortholog is 100%.

---

Group of orthologs #7234. Best score 503 bits
Score difference with first non-orthologous sequence - *A. fumigatus*:503 *A. fischeri*:503
afum|EAL90085.1 100.00% afis|EAW25426.1 100.00%
Bootstrap support for afum|EAL90085.1 as seed ortholog is 100%.
Bootstrap support for afis|EAW25426.1 as seed ortholog is 100%.

---

Group of orthologs #7235. Best score 502 bits
Score difference with first non-orthologous sequence - *A. fumigatus*:502 *A. fischeri*:502
afum|EAL85830.1 100.00% afis|EAW15493.1 100.00%
Bootstrap support for afum|EAL85830.1 as seed ortholog is 100%.
Bootstrap support for afis|EAW15493.1 as seed ortholog is 100%.

---

Group of orthologs #7236. Best score 502 bits
Score difference with first non-orthologous sequence - *A. fumigatus*:502 *A. fischeri*:502
afum|EAL85043.1 100.00% afis|EAW19850.1 100.00%
Bootstrap support for afum|EAL85043.1 as seed ortholog is 100%.
Bootstrap support for afis|EAW19850.1 as seed ortholog is 100%.

---

Group of orthologs #7237. Best score 502 bits
Score difference with first non-orthologous sequence - *A. fumigatus*:502 *A. fischeri*:502
afum|EAL86139.1 100.00% afis|EAW24467.1 100.00%
Bootstrap support for afum|EAL86139.1 as seed ortholog is 100%.
Bootstrap support for afis|EAW24467.1 as seed ortholog is 100%.

---

Group of orthologs #7238. Best score 501 bits
Score difference with first non-orthologous sequence - *A. fumigatus*:501 *A. fischeri*:450
afum|EAL87825.1 100.00% afis|EAW23635.1 100.00%
Bootstrap support for afum|EAL87825.1 as seed ortholog is 100%.

Bootstrap support for afis|EAW23635.1 as seed ortholog is 100%.

---

Group of orthologs #7239. Best score 501 bits
Score difference with first non-orthologous sequence - *A. fumigatus*:127 *A. fischeri*:501
afum|EAL92056.1 100.00% afis|EAW21018.1 100.00%
Bootstrap support for afum|EAL92056.1 as seed ortholog is 99%.
Bootstrap support for afis|EAW21018.1 as seed ortholog is 100%.

---

Group of orthologs #7240. Best score 500 bits
Score difference with first non-orthologous sequence - *A. fumigatus*:500 *A. fischeri*:500
afum|EAL88572.1 100.00% afis|EAW15936.1 100.00%
Bootstrap support for afum|EAL88572.1 as seed ortholog is 100%.
Bootstrap support for afis|EAW15936.1 as seed ortholog is 100%.

---

Group of orthologs #7241. Best score 500 bits
Score difference with first non-orthologous sequence - *A. fumigatus*:500 *A. fischeri*:500
afum|EAL84920.1 100.00% afis|EAW19996.1 100.00%
Bootstrap support for afum|EAL84920.1 as seed ortholog is 100%.
Bootstrap support for afis|EAW19996.1 as seed ortholog is 100%.

---

Group of orthologs #7242. Best score 500 bits
Score difference with first non-orthologous sequence - *A. fumigatus*:500 *A. fischeri*:500
afum|EAL91438.1 100.00% afis|EAW17624.1 100.00%
Bootstrap support for afum|EAL91438.1 as seed ortholog is 100%.
Bootstrap support for afis|EAW17624.1 as seed ortholog is 100%.

---

Group of orthologs #7243. Best score 500 bits
Score difference with first non-orthologous sequence - *A. fumigatus*:500 *A. fischeri*:500
afum|EAL84346.1 100.00% afis|EAW25676.1 100.00%
Bootstrap support for afum|EAL84346.1 as seed ortholog is 100%.
Bootstrap support for afis|EAW25676.1 as seed ortholog is 100%.

---

Group of orthologs #7244. Best score 500 bits
Score difference with first non-orthologous sequence - *A. fumigatus*:500 *A. fischeri*:500
afum|EAL90036.1 100.00% afis|EAW25377.1 100.00%
Bootstrap support for afum|EAL90036.1 as seed ortholog is 100%.
Bootstrap support for afis|EAW25377.1 as seed ortholog is 100%.

---

Group of orthologs #7245. Best score 499 bits
Score difference with first non-orthologous sequence - *A. fumigatus*:499 *A. fischeri*:499
afum|EAL89168.1 100.00% afis|EAW16571.1 100.00%
Bootstrap support for afum|EAL89168.1 as seed ortholog is 100%.
Bootstrap support for afis|EAW16571.1 as seed ortholog is 100%.

---

Group of orthologs #7246. Best score 499 bits
Score difference with first non-orthologous sequence - *A. fumigatus*:408 *A. fischeri*:400
afum|EAL91195.1 100.00% afis|EAW17353.1 100.00%
Bootstrap support for afum|EAL91195.1 as seed ortholog is 100%.
Bootstrap support for afis|EAW17353.1 as seed ortholog is 100%.

---

Group of orthologs #7247. Best score 499 bits
Score difference with first non-orthologous sequence - *A. fumigatus*:499 *A. fischeri*:499
afum|EAL91521.2 100.00% afis|EAW17707.1 100.00%
Bootstrap support for afum|EAL91521.2 as seed ortholog is 100%.
Bootstrap support for afis|EAW17707.1 as seed ortholog is 100%.

---

Group of orthologs #7248. Best score 499 bits
Score difference with first non-orthologous sequence - *A. fumigatus*:399 *A. fischeri*:231
afum|EAL91611.1 100.00% afis|EAW17803.1 100.00%
Bootstrap support for afum|EAL91611.1 as seed ortholog is 100%.
Bootstrap support for afis|EAW17803.1 as seed ortholog is 100%.

---

Group of orthologs #7249. Best score 499 bits
Score difference with first non-orthologous sequence - *A. fumigatus*:499 *A. fischeri*:215
afum|EAL92093.2 100.00% afis|EAW21058.1 100.00%
Bootstrap support for afum|EAL92093.2 as seed ortholog is 100%.
Bootstrap support for afis|EAW21058.1 as seed ortholog is 100%.

---

Group of orthologs #7250. Best score 499 bits
Score difference with first non-orthologous sequence - *A. fumigatus*:499 *A. fischeri*:499
afum|EAL90093.1 100.00% afis|EAW25435.1 100.00%
Bootstrap support for afum|EAL90093.1 as seed ortholog is 100%.
Bootstrap support for afis|EAW25435.1 as seed ortholog is 100%.

---

Group of orthologs #7251. Best score 498 bits
Score difference with first non-orthologous sequence - *A. fumigatus*:498 *A. fischeri*:498
afum|EAL85057.1 100.00% afis|EAW19835.1 100.00%
Bootstrap support for afum|EAL85057.1 as seed ortholog is 100%.
Bootstrap support for afis|EAW19835.1 as seed ortholog is 100%.

---

Group of orthologs #7252. Best score 498 bits
Score difference with first non-orthologous sequence - *A. fumigatus*:498 *A. fischeri*:498
afum|EAL93854.1 100.00% afis|EAW19122.1 100.00%
Bootstrap support for afum|EAL93854.1 as seed ortholog is 100%.
Bootstrap support for afis|EAW19122.1 as seed ortholog is 100%.

---

Group of orthologs #7253. Best score 498 bits
Score difference with first non-orthologous sequence - *A. fumigatus*:498 *A. fischeri*:498
afum|EAL90474.1 100.00% afis|EAW22721.1 100.00%
Bootstrap support for afum|EAL90474.1 as seed ortholog is 100%.
Bootstrap support for afis|EAW22721.1 as seed ortholog is 100%.

---

Group of orthologs #7254. Best score 498 bits
Score difference with first non-orthologous sequence - *A. fumigatus*:498 *A. fischeri*:498
afum|EAL90238.1 100.00% afis|EAW22959.1 100.00%
Bootstrap support for afum|EAL90238.1 as seed ortholog is 100%.
Bootstrap support for afis|EAW22959.1 as seed ortholog is 100%.

---

Group of orthologs #7255. Best score 498 bits
Score difference with first non-orthologous sequence - *A. fumigatus*:498 *A. fischeri*:498
afum|EBA27480.1 100.00% afis|EAW21905.1 100.00%
Bootstrap support for afum|EBA27480.1 as seed ortholog is 100%.
Bootstrap support for afis|EAW21905.1 as seed ortholog is 100%.

---

Group of orthologs #7256. Best score 497 bits
Score difference with first non-orthologous sequence - *A. fumigatus*:497 *A. fischeri*:497
afum|EAL86628.1 100.00% afis|EAW17201.1 100.00%
Bootstrap support for afum|EAL86628.1 as seed ortholog is 100%.
Bootstrap support for afis|EAW17201.1 as seed ortholog is 100%.

---

Group of orthologs #7257. Best score 497 bits
Score difference with first non-orthologous sequence - *A. fumigatus*:497 *A. fischeri*:497

afum|EAL84675.1 100.00% afis|EAW20958.1 100.00%
Bootstrap support for afum|EAL84675.1 as seed ortholog is 100%.
Bootstrap support for afis|EAW20958.1 as seed ortholog is 100%.

---

Group of orthologs #7258. Best score 497 bits
Score difference with first non-orthologous sequence - *A. fumigatus*:497 *A. fischeri*:497
afum|EAL87270.1 100.00% afis|EAW23839.1 100.00%
Bootstrap support for afum|EAL87270.1 as seed ortholog is 100%.
Bootstrap support for afis|EAW23839.1 as seed ortholog is 100%.

---

Group of orthologs #7259. Best score 497 bits
Score difference with first non-orthologous sequence - *A. fumigatus*:497 *A. fischeri*:497
afum|EAL90407.2 100.00% afis|EAW22792.1 100.00%
Bootstrap support for afum|EAL90407.2 as seed ortholog is 100%.
Bootstrap support for afis|EAW22792.1 as seed ortholog is 100%.

---

Group of orthologs #7260. Best score 497 bits
Score difference with first non-orthologous sequence - *A. fumigatus*:497 *A. fischeri*:497
afum|EAL89887.1 100.00% afis|EAW25226.1 100.00%
Bootstrap support for afum|EAL89887.1 as seed ortholog is 100%.
Bootstrap support for afis|EAW25226.1 as seed ortholog is 100%.

---

Group of orthologs #7261. Best score 496 bits
Score difference with first non-orthologous sequence - *A. fumigatus*:496 *A. fischeri*:158
afum|EAL85707.1 100.00% afis|EAW15627.1 100.00%
Bootstrap support for afum|EAL85707.1 as seed ortholog is 100%.
Bootstrap support for afis|EAW15627.1 as seed ortholog is 100%.

---

Group of orthologs #7262. Best score 496 bits
Score difference with first non-orthologous sequence - *A. fumigatus*:496 *A. fischeri*:496
afum|EAL87471.1 100.00% afis|EAW18114.1 100.00%
Bootstrap support for afum|EAL87471.1 as seed ortholog is 100%.
Bootstrap support for afis|EAW18114.1 as seed ortholog is 100%.

---

Group of orthologs #7263. Best score 496 bits
Score difference with first non-orthologous sequence - *A. fumigatus*:496 *A. fischeri*:496
afum|EAL89273.1 100.00% afis|EAW16669.1 100.00%
Bootstrap support for afum|EAL89273.1 as seed ortholog is 100%.
Bootstrap support for afis|EAW16669.1 as seed ortholog is 100%.

---

Group of orthologs #7264. Best score 496 bits
Score difference with first non-orthologous sequence - *A. fumigatus*:496 *A. fischeri*:496
afum|EAL86878.1 100.00% afis|EAW19644.1 100.00%
Bootstrap support for afum|EAL86878.1 as seed ortholog is 100%.
Bootstrap support for afis|EAW19644.1 as seed ortholog is 100%.

---

Group of orthologs #7265. Best score 496 bits
Score difference with first non-orthologous sequence - *A. fumigatus*:445 *A. fischeri*:279
afum|EAL84327.2 100.00% afis|EAW25640.1 100.00%
Bootstrap support for afum|EAL84327.2 as seed ortholog is 100%.
Bootstrap support for afis|EAW25640.1 as seed ortholog is 100%.

---

Group of orthologs #7266. Best score 496 bits
Score difference with first non-orthologous sequence - *A. fumigatus*:496 *A. fischeri*:496
afum|EAL93235.1 100.00% afis|EAW18537.1 100.00%
Bootstrap support for afum|EAL93235.1 as seed ortholog is 100%.

Bootstrap support for afis|EAW18537.1 as seed ortholog is 100%.

---

Group of orthologs #7267. Best score 496 bits
Score difference with first non-orthologous sequence - *A. fumigatus*:496 *A. fischeri*:360
afum|EAL93941.1 100.00% afis|EAW19211.1 100.00%
Bootstrap support for afum|EAL93941.1 as seed ortholog is 100%.
Bootstrap support for afis|EAW19211.1 as seed ortholog is 100%.

---

Group of orthologs #7268. Best score 496 bits
Score difference with first non-orthologous sequence - *A. fumigatus*:496 *A. fischeri*:496
afum|EAL92101.1 100.00% afis|EAW21079.1 100.00%
Bootstrap support for afum|EAL92101.1 as seed ortholog is 100%.
Bootstrap support for afis|EAW21079.1 as seed ortholog is 100%.

---

Group of orthologs #7269. Best score 496 bits
Score difference with first non-orthologous sequence - *A. fumigatus*:496 *A. fischeri*:400
afum|EAL91042.1 100.00% afis|EAW22148.1 100.00%
Bootstrap support for afum|EAL91042.1 as seed ortholog is 100%.
Bootstrap support for afis|EAW22148.1 as seed ortholog is 100%.

---

Group of orthologs #7270. Best score 496 bits
Score difference with first non-orthologous sequence - *A. fumigatus*:496 *A. fischeri*:496
afum|EAL92279.1 100.00% afis|EAW21264.1 100.00%
Bootstrap support for afum|EAL92279.1 as seed ortholog is 100%.
Bootstrap support for afis|EAW21264.1 as seed ortholog is 100%.

---

Group of orthologs #7271. Best score 496 bits
Score difference with first non-orthologous sequence - *A. fumigatus*:496 *A. fischeri*:496
afum|EAL92483.1 100.00% afis|EAW21463.1 100.00%
Bootstrap support for afum|EAL92483.1 as seed ortholog is 100%.
Bootstrap support for afis|EAW21463.1 as seed ortholog is 100%.

---

Group of orthologs #7272. Best score 496 bits
Score difference with first non-orthologous sequence - *A. fumigatus*:496 *A. fischeri*:496
afum|EAL92587.1 100.00% afis|EAW21568.1 100.00%
Bootstrap support for afum|EAL92587.1 as seed ortholog is 100%.
Bootstrap support for afis|EAW21568.1 as seed ortholog is 100%.

---

Group of orthologs #7273. Best score 495 bits
Score difference with first non-orthologous sequence - *A. fumigatus*:495 *A. fischeri*:495
afum|EAL88177.1 100.00% afis|EAW23302.1 100.00%
Bootstrap support for afum|EAL88177.1 as seed ortholog is 100%.
Bootstrap support for afis|EAW23302.1 as seed ortholog is 100%.

---

Group of orthologs #7274. Best score 495 bits
Score difference with first non-orthologous sequence - *A. fumigatus*:495 *A. fischeri*:495
afum|EAL90809.1 100.00% afis|EAW22382.1 100.00%
Bootstrap support for afum|EAL90809.1 as seed ortholog is 100%.
Bootstrap support for afis|EAW22382.1 as seed ortholog is 100%.

---

Group of orthologs #7275. Best score 494 bits
Score difference with first non-orthologous sequence - *A. fumigatus*:494 *A. fischeri*:494
afum|EAL91120.1 100.00% afis|EAW15511.1 100.00%
afum|EAL90936.1 20.13%
Bootstrap support for afum|EAL91120.1 as seed ortholog is 100%.
Bootstrap support for afis|EAW15511.1 as seed ortholog is 100%.

Group of orthologs #7276. Best score 494 bits
Score difference with first non-orthologous sequence - *A. fumigatus*:494 *A. fischeri*:494
afum|EAL85657.1 100.00% afis|EAW15691.1 100.00%
Bootstrap support for afum|EAL85657.1 as seed ortholog is 100%.
Bootstrap support for afis|EAW15691.1 as seed ortholog is 100%.

Group of orthologs #7277. Best score 494 bits
Score difference with first non-orthologous sequence - *A. fumigatus*:402 *A. fischeri*:400
afum|EAL92762.1 100.00% afis|EAW21729.1 100.00%
Bootstrap support for afum|EAL92762.1 as seed ortholog is 100%.
Bootstrap support for afis|EAW21729.1 as seed ortholog is 100%.

Group of orthologs #7278. Best score 494 bits
Score difference with first non-orthologous sequence - *A. fumigatus*:494 *A. fischeri*:494
afum|EAL89822.1 100.00% afis|EAW25147.1 100.00%
Bootstrap support for afum|EAL89822.1 as seed ortholog is 100%.
Bootstrap support for afis|EAW25147.1 as seed ortholog is 100%.

Group of orthologs #7279. Best score 493 bits
Score difference with first non-orthologous sequence - *A. fumigatus*:493 *A. fischeri*:493
afum|EAL86162.1 100.00% afis|EAW24493.1 100.00%
Bootstrap support for afum|EAL86162.1 as seed ortholog is 100%.
Bootstrap support for afis|EAW24493.1 as seed ortholog is 100%.

Group of orthologs #7280. Best score 492 bits
Score difference with first non-orthologous sequence - *A. fumigatus*:492 *A. fischeri*:492
afum|EAL89102.1 100.00% afis|EAW16507.1 100.00%
Bootstrap support for afum|EAL89102.1 as seed ortholog is 100%.
Bootstrap support for afis|EAW16507.1 as seed ortholog is 100%.

Group of orthologs #7281. Best score 492 bits
Score difference with first non-orthologous sequence - *A. fumigatus*:492 *A. fischeri*:492
afum|EBA27269.1 100.00% afis|EAW16790.1 100.00%
Bootstrap support for afum|EBA27269.1 as seed ortholog is 100%.
Bootstrap support for afis|EAW16790.1 as seed ortholog is 100%.

Group of orthologs #7282. Best score 492 bits
Score difference with first non-orthologous sequence - *A. fumigatus*:492 *A. fischeri*:492
afum|EAL89889.1 100.00% afis|EAW25228.1 100.00%
Bootstrap support for afum|EAL89889.1 as seed ortholog is 100%.
Bootstrap support for afis|EAW25228.1 as seed ortholog is 100%.

Group of orthologs #7283. Best score 491 bits
Score difference with first non-orthologous sequence - *A. fumigatus*:491 *A. fischeri*:491
afum|EAL91774.1 100.00% afis|EAW17964.1 100.00%
Bootstrap support for afum|EAL91774.1 as seed ortholog is 100%.
Bootstrap support for afis|EAW17964.1 as seed ortholog is 100%.

Group of orthologs #7284. Best score 491 bits
Score difference with first non-orthologous sequence - *A. fumigatus*:491 *A. fischeri*:491
afum|EBA27435.1 100.00% afis|EAW17661.1 100.00%
Bootstrap support for afum|EBA27435.1 as seed ortholog is 100%.
Bootstrap support for afis|EAW17661.1 as seed ortholog is 100%.

Group of orthologs #7285. Best score 491 bits

Score difference with first non-orthologous sequence - *A. fumigatus*:491   *A. fischeri*:491
afum|EAL93545.1 100.00% afis|EAW18809.1 100.00%
Bootstrap support for afum|EAL93545.1 as seed ortholog is 100%.
Bootstrap support for afis|EAW18809.1 as seed ortholog is 100%.

---

Group of orthologs #7286. Best score 491 bits
Score difference with first non-orthologous sequence - *A. fumigatus*:491   *A. fischeri*:491
afum|EAL93630.1 100.00% afis|EAW18898.1 100.00%
Bootstrap support for afum|EAL93630.1 as seed ortholog is 100%.
Bootstrap support for afis|EAW18898.1 as seed ortholog is 100%.

---

Group of orthologs #7287. Best score 491 bits
Score difference with first non-orthologous sequence - *A. fumigatus*:491   *A. fischeri*:491
afum|EAL92397.1 100.00% afis|EAW21382.1 100.00%
Bootstrap support for afum|EAL92397.1 as seed ortholog is 100%.
Bootstrap support for afis|EAW21382.1 as seed ortholog is 100%.

---

Group of orthologs #7288. Best score 491 bits
Score difference with first non-orthologous sequence - *A. fumigatus*:491   *A. fischeri*:341
afum|EAL92938.1 100.00% afis|EAW21925.1 100.00%
Bootstrap support for afum|EAL92938.1 as seed ortholog is 100%.
Bootstrap support for afis|EAW21925.1 as seed ortholog is 100%.

---

Group of orthologs #7289. Best score 491 bits
Score difference with first non-orthologous sequence - *A. fumigatus*:491   *A. fischeri*:491
afum|EAL93017.1 100.00% afis|EAW22002.1 100.00%
Bootstrap support for afum|EAL93017.1 as seed ortholog is 100%.
Bootstrap support for afis|EAW22002.1 as seed ortholog is 100%.

---

Group of orthologs #7290. Best score 490 bits
Score difference with first non-orthologous sequence - *A. fumigatus*:394   *A. fischeri*:368
afum|EAL91685.1 100.00% afis|EAW17877.1 100.00%
Bootstrap support for afum|EAL91685.1 as seed ortholog is 100%.
Bootstrap support for afis|EAW17877.1 as seed ortholog is 100%.

---

Group of orthologs #7291. Best score 490 bits
Score difference with first non-orthologous sequence - *A. fumigatus*:490   *A. fischeri*:490
afum|EAL91758.1 100.00% afis|EAW17948.1 100.00%
Bootstrap support for afum|EAL91758.1 as seed ortholog is 100%.
Bootstrap support for afis|EAW17948.1 as seed ortholog is 100%.

---

Group of orthologs #7292. Best score 490 bits
Score difference with first non-orthologous sequence - *A. fumigatus*:490   *A. fischeri*:490
afum|EAL88679.1 100.00% afis|EAW21810.1 100.00%
Bootstrap support for afum|EAL88679.1 as seed ortholog is 100%.
Bootstrap support for afis|EAW21810.1 as seed ortholog is 100%.

---

Group of orthologs #7293. Best score 490 bits
Score difference with first non-orthologous sequence - *A. fumigatus*:490   *A. fischeri*:418
afum|EAL87161.1 100.00% afis|EAW23721.1 100.00%
Bootstrap support for afum|EAL87161.1 as seed ortholog is 100%.
Bootstrap support for afis|EAW23721.1 as seed ortholog is 100%.

---

Group of orthologs #7294. Best score 489 bits
Score difference with first non-orthologous sequence - *A. fumigatus*:489   *A. fischeri*:489
afum|EAL88834.1 100.00% afis|EAW16206.1 100.00%

Bootstrap support for afum|EAL88834.1 as seed ortholog is 100%.
Bootstrap support for afis|EAW16206.1 as seed ortholog is 100%.

---

Group of orthologs #7295. Best score 489 bits
Score difference with first non-orthologous sequence - *A. fumigatus*:489 *A. fischeri*:489
afum|EAL85233.1 100.00% afis|EAW20116.1 100.00%
Bootstrap support for afum|EAL85233.1 as seed ortholog is 100%.
Bootstrap support for afis|EAW20116.1 as seed ortholog is 100%.

---

Group of orthologs #7296. Best score 489 bits
Score difference with first non-orthologous sequence - *A. fumigatus*:489 *A. fischeri*:489
afum|EAL92320.1 100.00% afis|EAW21303.1 100.00%
Bootstrap support for afum|EAL92320.1 as seed ortholog is 100%.
Bootstrap support for afis|EAW21303.1 as seed ortholog is 100%.

---

Group of orthologs #7297. Best score 489 bits
Score difference with first non-orthologous sequence - *A. fumigatus*:489 *A. fischeri*:489
afum|EAL90145.1 100.00% afis|EAW25480.1 100.00%
Bootstrap support for afum|EAL90145.1 as seed ortholog is 100%.
Bootstrap support for afis|EAW25480.1 as seed ortholog is 100%.

---

Group of orthologs #7298. Best score 488 bits
Score difference with first non-orthologous sequence - *A. fumigatus*:488 *A. fischeri*:488
afum|EAL85624.1 100.00% afis|EAW15723.1 100.00%
Bootstrap support for afum|EAL85624.1 as seed ortholog is 100%.
Bootstrap support for afis|EAW15723.1 as seed ortholog is 100%.

---

Group of orthologs #7299. Best score 488 bits
Score difference with first non-orthologous sequence - *A. fumigatus*:488 *A. fischeri*:488
afum|EAL89136.1 100.00% afis|EAW16539.1 100.00%
Bootstrap support for afum|EAL89136.1 as seed ortholog is 100%.
Bootstrap support for afis|EAW16539.1 as seed ortholog is 100%.

---

Group of orthologs #7300. Best score 488 bits
Score difference with first non-orthologous sequence - *A. fumigatus*:488 *A. fischeri*:488
afum|EAL85495.1 100.00% afis|EAW20382.1 100.00%
Bootstrap support for afum|EAL85495.1 as seed ortholog is 100%.
Bootstrap support for afis|EAW20382.1 as seed ortholog is 100%.

---

Group of orthologs #7301. Best score 488 bits
Score difference with first non-orthologous sequence - *A. fumigatus*:488 *A. fischeri*:488
afum|EAL91681.1 100.00% afis|EAW17873.1 100.00%
Bootstrap support for afum|EAL91681.1 as seed ortholog is 100%.
Bootstrap support for afis|EAW17873.1 as seed ortholog is 100%.

---

Group of orthologs #7302. Best score 488 bits
Score difference with first non-orthologous sequence - *A. fumigatus*:488 *A. fischeri*:488
afum|EAL87305.1 100.00% afis|EAW23877.1 100.00%
Bootstrap support for afum|EAL87305.1 as seed ortholog is 100%.
Bootstrap support for afis|EAW23877.1 as seed ortholog is 100%.

---

Group of orthologs #7303. Best score 488 bits
Score difference with first non-orthologous sequence - *A. fumigatus*:488 *A. fischeri*:488
afum|EAL90224.1 100.00% afis|EAW22973.1 100.00%
Bootstrap support for afum|EAL90224.1 as seed ortholog is 100%.
Bootstrap support for afis|EAW22973.1 as seed ortholog is 100%.

---
Group of orthologs #7304. Best score 488 bits
Score difference with first non-orthologous sequence - *A. fumigatus*:488 *A. fischeri*:488
afum|EAL92174.1 100.00% afis|EAW21156.1 100.00%
Bootstrap support for afum|EAL92174.1 as seed ortholog is 100%.
Bootstrap support for afis|EAW21156.1 as seed ortholog is 100%.

---
Group of orthologs #7305. Best score 487 bits
Score difference with first non-orthologous sequence - *A. fumigatus*:487 *A. fischeri*:487
afum|EAL91756.1 100.00% afis|EAW17945.1 100.00%
Bootstrap support for afum|EAL91756.1 as seed ortholog is 100%.
Bootstrap support for afis|EAW17945.1 as seed ortholog is 100%.

---
Group of orthologs #7306. Best score 487 bits
Score difference with first non-orthologous sequence - *A. fumigatus*:487 *A. fischeri*:487
afum|EAL87413.1 100.00% afis|EAW23984.1 100.00%
Bootstrap support for afum|EAL87413.1 as seed ortholog is 100%.
Bootstrap support for afis|EAW23984.1 as seed ortholog is 100%.

---
Group of orthologs #7307. Best score 487 bits
Score difference with first non-orthologous sequence - *A. fumigatus*:487 *A. fischeri*:487
afum|EAL87994.1 100.00% afis|EAW23472.1 100.00%
Bootstrap support for afum|EAL87994.1 as seed ortholog is 100%.
Bootstrap support for afis|EAW23472.1 as seed ortholog is 100%.

---
Group of orthologs #7308. Best score 487 bits
Score difference with first non-orthologous sequence - *A. fumigatus*:421 *A. fischeri*:345
afum|EBA27320.1 100.00% afis|EAW18238.1 100.00%
Bootstrap support for afum|EBA27320.1 as seed ortholog is 100%.
Bootstrap support for afis|EAW18238.1 as seed ortholog is 100%.

---
Group of orthologs #7309. Best score 486 bits
Score difference with first non-orthologous sequence - *A. fumigatus*:486 *A. fischeri*:486
afum|EAL85359.1 100.00% afis|EAW20242.1 100.00%
Bootstrap support for afum|EAL85359.1 as seed ortholog is 100%.
Bootstrap support for afis|EAW20242.1 as seed ortholog is 100%.

---
Group of orthologs #7310. Best score 486 bits
Score difference with first non-orthologous sequence - *A. fumigatus*:486 *A. fischeri*:486
afum|EAL86895.1 100.00% afis|EAW19626.1 100.00%
Bootstrap support for afum|EAL86895.1 as seed ortholog is 100%.
Bootstrap support for afis|EAW19626.1 as seed ortholog is 100%.

---
Group of orthologs #7311. Best score 486 bits
Score difference with first non-orthologous sequence - *A. fumigatus*:486 *A. fischeri*:486
afum|EAL91178.1 100.00% afis|EAW17335.1 100.00%
Bootstrap support for afum|EAL91178.1 as seed ortholog is 100%.
Bootstrap support for afis|EAW17335.1 as seed ortholog is 100%.

---
Group of orthologs #7312. Best score 486 bits
Score difference with first non-orthologous sequence - *A. fumigatus*:486 *A. fischeri*:486
afum|EAL91832.1 100.00% afis|EAW18020.1 100.00%
Bootstrap support for afum|EAL91832.1 as seed ortholog is 100%.
Bootstrap support for afis|EAW18020.1 as seed ortholog is 100%.

---
Group of orthologs #7313. Best score 486 bits

Score difference with first non-orthologous sequence - *A. fumigatus*:486 *A. fischeri*:486
afum|EAL88311.2 100.00% afis|EAW23172.1 100.00%
Bootstrap support for afum|EAL88311.2 as seed ortholog is 100%.
Bootstrap support for afis|EAW23172.1 as seed ortholog is 100%.

---

Group of orthologs #7314. Best score 486 bits
Score difference with first non-orthologous sequence - *A. fumigatus*:486 *A. fischeri*:486
afum|EAL91006.2 100.00% afis|EAW22183.1 100.00%
Bootstrap support for afum|EAL91006.2 as seed ortholog is 100%.
Bootstrap support for afis|EAW22183.1 as seed ortholog is 100%.

---

Group of orthologs #7315. Best score 486 bits
Score difference with first non-orthologous sequence - *A. fumigatus*:486 *A. fischeri*:486
afum|EAL90736.1 100.00% afis|EAW22455.1 100.00%
Bootstrap support for afum|EAL90736.1 as seed ortholog is 100%.
Bootstrap support for afis|EAW22455.1 as seed ortholog is 100%.

---

Group of orthologs #7316. Best score 486 bits
Score difference with first non-orthologous sequence - *A. fumigatus*:486 *A. fischeri*:486
afum|EAL90267.1 100.00% afis|EAW22930.1 100.00%
Bootstrap support for afum|EAL90267.1 as seed ortholog is 100%.
Bootstrap support for afis|EAW22930.1 as seed ortholog is 100%.

---

Group of orthologs #7317. Best score 486 bits
Score difference with first non-orthologous sequence - *A. fumigatus*:486 *A. fischeri*:486
afum|EAL90294.1 100.00% afis|EAW22903.1 100.00%
Bootstrap support for afum|EAL90294.1 as seed ortholog is 100%.
Bootstrap support for afis|EAW22903.1 as seed ortholog is 100%.

---

Group of orthologs #7318. Best score 486 bits
Score difference with first non-orthologous sequence - *A. fumigatus*:486 *A. fischeri*:486
afum|EAL89382.1 100.00% afis|EAW24735.1 100.00%
Bootstrap support for afum|EAL89382.1 as seed ortholog is 100%.
Bootstrap support for afis|EAW24735.1 as seed ortholog is 100%.

---

Group of orthologs #7319. Best score 486 bits
Score difference with first non-orthologous sequence - *A. fumigatus*:486 *A. fischeri*:486
afum|EAL89815.2 100.00% afis|EAW25143.1 100.00%
Bootstrap support for afum|EAL89815.2 as seed ortholog is 100%.
Bootstrap support for afis|EAW25143.1 as seed ortholog is 100%.

---

Group of orthologs #7320. Best score 485 bits
Score difference with first non-orthologous sequence - *A. fumigatus*:485 *A. fischeri*:485
afum|EAL86688.1 100.00% afis|EAW17271.1 100.00%
Bootstrap support for afum|EAL86688.1 as seed ortholog is 100%.
Bootstrap support for afis|EAW17271.1 as seed ortholog is 100%.

---

Group of orthologs #7321. Best score 485 bits
Score difference with first non-orthologous sequence - *A. fumigatus*:485 *A. fischeri*:485
afum|EAL85246.1 100.00% afis|EAW20130.1 100.00%
Bootstrap support for afum|EAL85246.1 as seed ortholog is 100%.
Bootstrap support for afis|EAW20130.1 as seed ortholog is 100%.

---

Group of orthologs #7322. Best score 485 bits
Score difference with first non-orthologous sequence - *A. fumigatus*:485 *A. fischeri*:485
afum|EAL85916.1 100.00% afis|EAW24248.1 100.00%

Bootstrap support for afum|EAL85916.1 as seed ortholog is 100%.
Bootstrap support for afis|EAW24248.1 as seed ortholog is 100%.

---

Group of orthologs #7323. Best score 485 bits
Score difference with first non-orthologous sequence - *A. fumigatus*:485 *A. fischeri*:485
afum|EAL88142.1 100.00% afis|EAW23336.1 100.00%
Bootstrap support for afum|EAL88142.1 as seed ortholog is 100%.
Bootstrap support for afis|EAW23336.1 as seed ortholog is 100%.

---

Group of orthologs #7324. Best score 485 bits
Score difference with first non-orthologous sequence - *A. fumigatus*:485 *A. fischeri*:485
afum|EAL89539.2 100.00% afis|EAW24888.1 100.00%
Bootstrap support for afum|EAL89539.2 as seed ortholog is 100%.
Bootstrap support for afis|EAW24888.1 as seed ortholog is 100%.

---

Group of orthologs #7325. Best score 484 bits
Score difference with first non-orthologous sequence - *A. fumigatus*:484 *A. fischeri*:484
afum|EAL84469.1 100.00% afis|EAW20592.1 100.00%
Bootstrap support for afum|EAL84469.1 as seed ortholog is 100%.
Bootstrap support for afis|EAW20592.1 as seed ortholog is 100%.

---

Group of orthologs #7326. Best score 484 bits
Score difference with first non-orthologous sequence - *A. fumigatus*:484 *A. fischeri*:484
afum|EAL85298.1 100.00% afis|EAW20180.1 100.00%
Bootstrap support for afum|EAL85298.1 as seed ortholog is 100%.
Bootstrap support for afis|EAW20180.1 as seed ortholog is 100%.

---

Group of orthologs #7327. Best score 484 bits
Score difference with first non-orthologous sequence - *A. fumigatus*:484 *A. fischeri*:484
afum|EAL84373.1 100.00% afis|EAW25698.1 100.00%
Bootstrap support for afum|EAL84373.1 as seed ortholog is 100%.
Bootstrap support for afis|EAW25698.1 as seed ortholog is 100%.

---

Group of orthologs #7328. Best score 484 bits
Score difference with first non-orthologous sequence - *A. fumigatus*:484 *A. fischeri*:484
afum|EAL93271.1 100.00% afis|EAW18569.1 100.00%
Bootstrap support for afum|EAL93271.1 as seed ortholog is 100%.
Bootstrap support for afis|EAW18569.1 as seed ortholog is 100%.

---

Group of orthologs #7329. Best score 484 bits
Score difference with first non-orthologous sequence - *A. fumigatus*:484 *A. fischeri*:484
afum|EAL89378.1 100.00% afis|EAW24731.1 100.00%
Bootstrap support for afum|EAL89378.1 as seed ortholog is 100%.
Bootstrap support for afis|EAW24731.1 as seed ortholog is 100%.

---

Group of orthologs #7330. Best score 483 bits
Score difference with first non-orthologous sequence - *A. fumigatus*:483 *A. fischeri*:483
afum|EAL88567.1 100.00% afis|EAW15931.1 100.00%
Bootstrap support for afum|EAL88567.1 as seed ortholog is 100%.
Bootstrap support for afis|EAW15931.1 as seed ortholog is 100%.

---

Group of orthologs #7331. Best score 483 bits
Score difference with first non-orthologous sequence - *A. fumigatus*:483 *A. fischeri*:483
afum|EAL91394.1 100.00% afis|EAW17581.1 100.00%
Bootstrap support for afum|EAL91394.1 as seed ortholog is 100%.
Bootstrap support for afis|EAW17581.1 as seed ortholog is 100%.

---

Group of orthologs #7332. Best score 482 bits
Score difference with first non-orthologous sequence - *A. fumigatus*:482 *A. fischeri*:482
afum|EAL88894.1 100.00% afis|EAW16273.1 100.00%
Bootstrap support for afum|EAL88894.1 as seed ortholog is 100%.
Bootstrap support for afis|EAW16273.1 as seed ortholog is 100%.

---

Group of orthologs #7333. Best score 482 bits
Score difference with first non-orthologous sequence - *A. fumigatus*:482 *A. fischeri*:376
afum|EAL89392.1 100.00% afis|EAW18104.1 100.00%
Bootstrap support for afum|EAL89392.1 as seed ortholog is 100%.
Bootstrap support for afis|EAW18104.1 as seed ortholog is 100%.

---

Group of orthologs #7334. Best score 482 bits
Score difference with first non-orthologous sequence - *A. fumigatus*:482 *A. fischeri*:482
afum|EAL88446.1 100.00% afis|EAW23035.1 100.00%
Bootstrap support for afum|EAL88446.1 as seed ortholog is 100%.
Bootstrap support for afis|EAW23035.1 as seed ortholog is 100%.

---

Group of orthologs #7335. Best score 482 bits
Score difference with first non-orthologous sequence - *A. fumigatus*:482 *A. fischeri*:482
afum|EAL93714.1 100.00% afis|EAW18978.1 100.00%
Bootstrap support for afum|EAL93714.1 as seed ortholog is 100%.
Bootstrap support for afis|EAW18978.1 as seed ortholog is 100%.

---

Group of orthologs #7336. Best score 482 bits
Score difference with first non-orthologous sequence - *A. fumigatus*:482 *A. fischeri*:482
afum|EAL92560.1 100.00% afis|EAW21540.1 100.00%
Bootstrap support for afum|EAL92560.1 as seed ortholog is 100%.
Bootstrap support for afis|EAW21540.1 as seed ortholog is 100%.

---

Group of orthologs #7337. Best score 482 bits
Score difference with first non-orthologous sequence - *A. fumigatus*:482 *A. fischeri*:482
afum|EAL91999.1 100.00% afis|EAW24144.1 100.00%
Bootstrap support for afum|EAL91999.1 as seed ortholog is 100%.
Bootstrap support for afis|EAW24144.1 as seed ortholog is 100%.

---

Group of orthologs #7338. Best score 482 bits
Score difference with first non-orthologous sequence - *A. fumigatus*:482 *A. fischeri*:482
afum|EBA27255.1 100.00% afis|EAW24367.1 100.00%
Bootstrap support for afum|EBA27255.1 as seed ortholog is 100%.
Bootstrap support for afis|EAW24367.1 as seed ortholog is 100%.

---

Group of orthologs #7339. Best score 481 bits
Score difference with first non-orthologous sequence - *A. fumigatus*:481 *A. fischeri*:481
afum|EAL86591.1 100.00% afis|EAW17131.1 100.00%
Bootstrap support for afum|EAL86591.1 as seed ortholog is 100%.
Bootstrap support for afis|EAW17131.1 as seed ortholog is 100%.

---

Group of orthologs #7340. Best score 481 bits
Score difference with first non-orthologous sequence - *A. fumigatus*:481 *A. fischeri*:481
afum|EAL89110.2 100.00% afis|EAW16515.1 100.00%
Bootstrap support for afum|EAL89110.2 as seed ortholog is 100%.
Bootstrap support for afis|EAW16515.1 as seed ortholog is 100%.

---

Group of orthologs #7341. Best score 481 bits

Score difference with first non-orthologous sequence - *A. fumigatus*:350   *A. fischeri*:481
afum|EAL93139.1   100.00%   afis|EAW18454.1   100.00%
Bootstrap support for afum|EAL93139.1 as seed ortholog is 100%.
Bootstrap support for afis|EAW18454.1 as seed ortholog is 100%.

---

Group of orthologs #7342. Best score 481 bits
Score difference with first non-orthologous sequence - *A. fumigatus*:481   *A. fischeri*:481
afum|EAL92807.1   100.00%   afis|EAW21778.1   100.00%
Bootstrap support for afum|EAL92807.1 as seed ortholog is 100%.
Bootstrap support for afis|EAW21778.1 as seed ortholog is 100%.

---

Group of orthologs #7343. Best score 481 bits
Score difference with first non-orthologous sequence - *A. fumigatus*:481   *A. fischeri*:421
afum|EAL89832.2   100.00%   afis|EAW25157.1   100.00%
Bootstrap support for afum|EAL89832.2 as seed ortholog is 100%.
Bootstrap support for afis|EAW25157.1 as seed ortholog is 100%.

---

Group of orthologs #7344. Best score 481 bits
Score difference with first non-orthologous sequence - *A. fumigatus*:481   *A. fischeri*:481
afum|EAL90127.1   100.00%   afis|EAW25464.1   100.00%
Bootstrap support for afum|EAL90127.1 as seed ortholog is 100%.
Bootstrap support for afis|EAW25464.1 as seed ortholog is 100%.

---

Group of orthologs #7345. Best score 480 bits
Score difference with first non-orthologous sequence - *A. fumigatus*:389   *A. fischeri*:380
afum|EAL91487.1   100.00%   afis|EAW17674.1   100.00%
Bootstrap support for afum|EAL91487.1 as seed ortholog is 100%.
Bootstrap support for afis|EAW17674.1 as seed ortholog is 100%.

---

Group of orthologs #7346. Best score 480 bits
Score difference with first non-orthologous sequence - *A. fumigatus*:480   *A. fischeri*:480
afum|EAL90841.1   100.00%   afis|EAW22351.1   100.00%
Bootstrap support for afum|EAL90841.1 as seed ortholog is 100%.
Bootstrap support for afis|EAW22351.1 as seed ortholog is 100%.

---

Group of orthologs #7347. Best score 480 bits
Score difference with first non-orthologous sequence - *A. fumigatus*:480   *A. fischeri*:480
afum|EAL92838.1   100.00%   afis|EAW21822.1   100.00%
Bootstrap support for afum|EAL92838.1 as seed ortholog is 100%.
Bootstrap support for afis|EAW21822.1 as seed ortholog is 100%.

---

Group of orthologs #7348. Best score 479 bits
Score difference with first non-orthologous sequence - *A. fumigatus*:479   *A. fischeri*:479
afum|EAL85626.1   100.00%   afis|EAW15721.1   100.00%
Bootstrap support for afum|EAL85626.1 as seed ortholog is 100%.
Bootstrap support for afis|EAW15721.1 as seed ortholog is 100%.

---

Group of orthologs #7349. Best score 479 bits
Score difference with first non-orthologous sequence - *A. fumigatus*:479   *A. fischeri*:479
afum|EAL89068.1   100.00%   afis|EAW16474.1   100.00%
Bootstrap support for afum|EAL89068.1 as seed ortholog is 100%.
Bootstrap support for afis|EAW16474.1 as seed ortholog is 100%.

---

Group of orthologs #7350. Best score 479 bits
Score difference with first non-orthologous sequence - *A. fumigatus*:479   *A. fischeri*:479
afum|EAL91473.1   100.00%   afis|EAW17658.1   100.00%

Bootstrap support for afum|EAL91473.1 as seed ortholog is 100%.
Bootstrap support for afis|EAW17658.1 as seed ortholog is 100%.

---

Group of orthologs #7351. Best score 479 bits
Score difference with first non-orthologous sequence - *A. fumigatus*:479 *A. fischeri*:479
afum|EAL91750.1 100.00% afis|EAW17939.1 100.00%
Bootstrap support for afum|EAL91750.1 as seed ortholog is 100%.
Bootstrap support for afis|EAW17939.1 as seed ortholog is 100%.

---

Group of orthologs #7352. Best score 479 bits
Score difference with first non-orthologous sequence - *A. fumigatus*:479 *A. fischeri*:479
afum|EAL92758.1 100.00% afis|EAW21725.1 100.00%
Bootstrap support for afum|EAL92758.1 as seed ortholog is 100%.
Bootstrap support for afis|EAW21725.1 as seed ortholog is 100%.

---

Group of orthologs #7353. Best score 479 bits
Score difference with first non-orthologous sequence - *A. fumigatus*:479 *A. fischeri*:479
afum|EAL89745.1 100.00% afis|EAW25076.1 100.00%
Bootstrap support for afum|EAL89745.1 as seed ortholog is 100%.
Bootstrap support for afis|EAW25076.1 as seed ortholog is 100%.

---

Group of orthologs #7354. Best score 479 bits
Score difference with first non-orthologous sequence - *A. fumigatus*:479 *A. fischeri*:337
afum|EBA27474.1 100.00% afis|EAW21783.1 100.00%
Bootstrap support for afum|EBA27474.1 as seed ortholog is 100%.
Bootstrap support for afis|EAW21783.1 as seed ortholog is 100%.

---

Group of orthologs #7355. Best score 478 bits
Score difference with first non-orthologous sequence - *A. fumigatus*:478 *A. fischeri*:478
afum|EAL88525.2 100.00% afis|EAW15890.1 100.00%
Bootstrap support for afum|EAL88525.2 as seed ortholog is 100%.
Bootstrap support for afis|EAW15890.1 as seed ortholog is 100%.

---

Group of orthologs #7356. Best score 478 bits
Score difference with first non-orthologous sequence - *A. fumigatus*:478 *A. fischeri*:478
afum|EAL89060.1 100.00% afis|EAW16464.1 100.00%
Bootstrap support for afum|EAL89060.1 as seed ortholog is 100%.
Bootstrap support for afis|EAW16464.1 as seed ortholog is 100%.

---

Group of orthologs #7357. Best score 478 bits
Score difference with first non-orthologous sequence - *A. fumigatus*:478 *A. fischeri*:478
afum|EAL88121.1 100.00% afis|EAW23355.1 100.00%
Bootstrap support for afum|EAL88121.1 as seed ortholog is 100%.
Bootstrap support for afis|EAW23355.1 as seed ortholog is 100%.

---

Group of orthologs #7358. Best score 478 bits
Score difference with first non-orthologous sequence - *A. fumigatus*:478 *A. fischeri*:478
afum|EAL93471.1 100.00% afis|EAW18745.1 100.00%
Bootstrap support for afum|EAL93471.1 as seed ortholog is 100%.
Bootstrap support for afis|EAW18745.1 as seed ortholog is 100%.

---

Group of orthologs #7359. Best score 478 bits
Score difference with first non-orthologous sequence - *A. fumigatus*:478 *A. fischeri*:478
afum|EAL92508.1 100.00% afis|EAW21489.1 100.00%
Bootstrap support for afum|EAL92508.1 as seed ortholog is 100%.
Bootstrap support for afis|EAW21489.1 as seed ortholog is 100%.

Group of orthologs #7360. Best score 477 bits
Score difference with first non-orthologous sequence - *A. fumigatus*:477 *A. fischeri*:477
afum|EAL85854.1 100.00% afis|EAW15468.1 100.00%
Bootstrap support for afum|EAL85854.1 as seed ortholog is 100%.
Bootstrap support for afis|EAW15468.1 as seed ortholog is 100%.

Group of orthologs #7361. Best score 477 bits
Score difference with first non-orthologous sequence - *A. fumigatus*:477 *A. fischeri*:477
afum|EAL86907.1 100.00% afis|EAW19615.1 100.00%
Bootstrap support for afum|EAL86907.1 as seed ortholog is 100%.
Bootstrap support for afis|EAW19615.1 as seed ortholog is 100%.

Group of orthologs #7362. Best score 477 bits
Score difference with first non-orthologous sequence - *A. fumigatus*:477 *A. fischeri*:477
afum|EBA27278.1 100.00% afis|EAW17056.1 100.00%
Bootstrap support for afum|EBA27278.1 as seed ortholog is 100%.
Bootstrap support for afis|EAW17056.1 as seed ortholog is 100%.

Group of orthologs #7363. Best score 477 bits
Score difference with first non-orthologous sequence - *A. fumigatus*:477 *A. fischeri*:477
afum|EAL93214.1 100.00% afis|EAW18519.1 100.00%
Bootstrap support for afum|EAL93214.1 as seed ortholog is 100%.
Bootstrap support for afis|EAW18519.1 as seed ortholog is 100%.

Group of orthologs #7364. Best score 477 bits
Score difference with first non-orthologous sequence - *A. fumigatus*:477 *A. fischeri*:477
afum|EAL93520.1 100.00% afis|EAW18787.1 100.00%
Bootstrap support for afum|EAL93520.1 as seed ortholog is 100%.
Bootstrap support for afis|EAW18787.1 as seed ortholog is 100%.

Group of orthologs #7365. Best score 476 bits
Score difference with first non-orthologous sequence - *A. fumigatus*:476 *A. fischeri*:476
afum|EAL85206.1 100.00% afis|EAW20087.1 100.00%
Bootstrap support for afum|EAL85206.1 as seed ortholog is 100%.
Bootstrap support for afis|EAW20087.1 as seed ortholog is 100%.

Group of orthologs #7366. Best score 476 bits
Score difference with first non-orthologous sequence - *A. fumigatus*:476 *A. fischeri*:476
afum|EAL88229.1 100.00% afis|EAW23252.1 100.00%
Bootstrap support for afum|EAL88229.1 as seed ortholog is 100%.
Bootstrap support for afis|EAW23252.1 as seed ortholog is 100%.

Group of orthologs #7367. Best score 476 bits
Score difference with first non-orthologous sequence - *A. fumigatus*:476 *A. fischeri*:476
afum|EAL94003.1 100.00% afis|EAW19277.1 100.00%
Bootstrap support for afum|EAL94003.1 as seed ortholog is 100%.
Bootstrap support for afis|EAW19277.1 as seed ortholog is 100%.

Group of orthologs #7368. Best score 476 bits
Score difference with first non-orthologous sequence - *A. fumigatus*:476 *A. fischeri*:476
afum|EAL92799.1 100.00% afis|EAW21770.1 100.00%
Bootstrap support for afum|EAL92799.1 as seed ortholog is 100%.
Bootstrap support for afis|EAW21770.1 as seed ortholog is 100%.

Group of orthologs #7369. Best score 476 bits

Score difference with first non-orthologous sequence - *A. fumigatus*:476 *A. fischeri*:476
afum|EAL92961.1 100.00% afis|EAW21947.1 100.00%
Bootstrap support for afum|EAL92961.1 as seed ortholog is 100%.
Bootstrap support for afis|EAW21947.1 as seed ortholog is 100%.

---

Group of orthologs #7370. Best score 475 bits
Score difference with first non-orthologous sequence - *A. fumigatus*:475 *A. fischeri*:475
afum|EAL87663.1 100.00% afis|EAW18297.1 100.00%
Bootstrap support for afum|EAL87663.1 as seed ortholog is 100%.
Bootstrap support for afis|EAW18297.1 as seed ortholog is 100%.

---

Group of orthologs #7371. Best score 475 bits
Score difference with first non-orthologous sequence - *A. fumigatus*:475 *A. fischeri*:475
afum|EAL90963.1 100.00% afis|EAW22230.1 100.00%
Bootstrap support for afum|EAL90963.1 as seed ortholog is 100%.
Bootstrap support for afis|EAW22230.1 as seed ortholog is 100%.

---

Group of orthologs #7372. Best score 475 bits
Score difference with first non-orthologous sequence - *A. fumigatus*:475 *A. fischeri*:475
afum|EAL90983.2 100.00% afis|EAW22211.1 100.00%
Bootstrap support for afum|EAL90983.2 as seed ortholog is 100%.
Bootstrap support for afis|EAW22211.1 as seed ortholog is 100%.

---

Group of orthologs #7373. Best score 475 bits
Score difference with first non-orthologous sequence - *A. fumigatus*:475 *A. fischeri*:475
afum|EAL90152.1 100.00% afis|EAW25487.1 100.00%
Bootstrap support for afum|EAL90152.1 as seed ortholog is 100%.
Bootstrap support for afis|EAW25487.1 as seed ortholog is 100%.

---

Group of orthologs #7374. Best score 474 bits
Score difference with first non-orthologous sequence - *A. fumigatus*:474 *A. fischeri*:387
afum|EAL86506.1 100.00% afis|EAW16979.1 100.00%
Bootstrap support for afum|EAL86506.1 as seed ortholog is 100%.
Bootstrap support for afis|EAW16979.1 as seed ortholog is 100%.

---

Group of orthologs #7375. Best score 474 bits
Score difference with first non-orthologous sequence - *A. fumigatus*:474 *A. fischeri*:474
afum|EAL87398.2 100.00% afis|EAW23968.1 100.00%
Bootstrap support for afum|EAL87398.2 as seed ortholog is 100%.
Bootstrap support for afis|EAW23968.1 as seed ortholog is 100%.

---

Group of orthologs #7376. Best score 474 bits
Score difference with first non-orthologous sequence - *A. fumigatus*:474 *A. fischeri*:474
afum|EAL88060.1 100.00% afis|EAW23410.1 100.00%
Bootstrap support for afum|EAL88060.1 as seed ortholog is 100%.
Bootstrap support for afis|EAW23410.1 as seed ortholog is 100%.

---

Group of orthologs #7377. Best score 474 bits
Score difference with first non-orthologous sequence - *A. fumigatus*:474 *A. fischeri*:474
afum|EAL87931.1 100.00% afis|EAW23535.1 100.00%
Bootstrap support for afum|EAL87931.1 as seed ortholog is 100%.
Bootstrap support for afis|EAW23535.1 as seed ortholog is 100%.

---

Group of orthologs #7378. Best score 474 bits
Score difference with first non-orthologous sequence - *A. fumigatus*:474 *A. fischeri*:474
afum|EAL90699.1 100.00% afis|EAW22494.1 100.00%

Bootstrap support for afum|EAL90699.1 as seed ortholog is 100%.
Bootstrap support for afis|EAW22494.1 as seed ortholog is 100%.

---

Group of orthologs #7379. Best score 474 bits
Score difference with first non-orthologous sequence - *A. fumigatus*:474 *A. fischeri*:474
afum|EAL92708.1 100.00% afis|EAW21674.1 100.00%
Bootstrap support for afum|EAL92708.1 as seed ortholog is 100%.
Bootstrap support for afis|EAW21674.1 as seed ortholog is 100%.

---

Group of orthologs #7380. Best score 473 bits
Score difference with first non-orthologous sequence - *A. fumigatus*:473 *A. fischeri*:473
afum|EAL89004.1 100.00% afis|EAW16379.1 100.00%
Bootstrap support for afum|EAL89004.1 as seed ortholog is 100%.
Bootstrap support for afis|EAW16379.1 as seed ortholog is 100%.

---

Group of orthologs #7381. Best score 473 bits
Score difference with first non-orthologous sequence - *A. fumigatus*:473 *A. fischeri*:473
afum|EAL90610.1 100.00% afis|EAW22589.1 100.00%
Bootstrap support for afum|EAL90610.1 as seed ortholog is 100%.
Bootstrap support for afis|EAW22589.1 as seed ortholog is 100%.

---

Group of orthologs #7382. Best score 473 bits
Score difference with first non-orthologous sequence - *A. fumigatus*:473 *A. fischeri*:473
afum|EAL90440.1 100.00% afis|EAW22759.1 100.00%
Bootstrap support for afum|EAL90440.1 as seed ortholog is 100%.
Bootstrap support for afis|EAW22759.1 as seed ortholog is 100%.

---

Group of orthologs #7383. Best score 473 bits
Score difference with first non-orthologous sequence - *A. fumigatus*:473 *A. fischeri*:473
afum|EAL90277.1 100.00% afis|EAW22920.1 100.00%
Bootstrap support for afum|EAL90277.1 as seed ortholog is 100%.
Bootstrap support for afis|EAW22920.1 as seed ortholog is 100%.

---

Group of orthologs #7384. Best score 473 bits
Score difference with first non-orthologous sequence - *A. fumigatus*:473 *A. fischeri*:473
afum|EAL89919.1 100.00% afis|EAW25258.1 100.00%
Bootstrap support for afum|EAL89919.1 as seed ortholog is 100%.
Bootstrap support for afis|EAW25258.1 as seed ortholog is 100%.

---

Group of orthologs #7385. Best score 472 bits
Score difference with first non-orthologous sequence - *A. fumigatus*:472 *A. fischeri*:326
afum|EAL87665.1 100.00% afis|EAW18299.1 100.00%
Bootstrap support for afum|EAL87665.1 as seed ortholog is 100%.
Bootstrap support for afis|EAW18299.1 as seed ortholog is 100%.

---

Group of orthologs #7386. Best score 472 bits
Score difference with first non-orthologous sequence - *A. fumigatus*:472 *A. fischeri*:472
afum|EAL89807.2 100.00% afis|EAW25136.1 100.00%
Bootstrap support for afum|EAL89807.2 as seed ortholog is 100%.
Bootstrap support for afis|EAW25136.1 as seed ortholog is 100%.

---

Group of orthologs #7387. Best score 471 bits
Score difference with first non-orthologous sequence - *A. fumigatus*:471 *A. fischeri*:471
afum|EAL86502.1 100.00% afis|EAW17527.1 100.00%
Bootstrap support for afum|EAL86502.1 as seed ortholog is 100%.
Bootstrap support for afis|EAW17527.1 as seed ortholog is 100%.

Group of orthologs #7388. Best score 471 bits
Score difference with first non-orthologous sequence - *A. fumigatus*:471 *A. fischeri*:471
afum|EAL85167.1 100.00% afis|EAW20046.1 100.00%
Bootstrap support for afum|EAL85167.1 as seed ortholog is 100%.
Bootstrap support for afis|EAW20046.1 as seed ortholog is 100%.

Group of orthologs #7389. Best score 471 bits
Score difference with first non-orthologous sequence - *A. fumigatus*:471 *A. fischeri*:471
afum|EAL91550.1 100.00% afis|EAW17735.1 100.00%
Bootstrap support for afum|EAL91550.1 as seed ortholog is 100%.
Bootstrap support for afis|EAW17735.1 as seed ortholog is 100%.

Group of orthologs #7390. Best score 471 bits
Score difference with first non-orthologous sequence - *A. fumigatus*:471 *A. fischeri*:471
afum|EAL91631.1 100.00% afis|EAW17821.1 100.00%
Bootstrap support for afum|EAL91631.1 as seed ortholog is 100%.
Bootstrap support for afis|EAW17821.1 as seed ortholog is 100%.

Group of orthologs #7391. Best score 471 bits
Score difference with first non-orthologous sequence - *A. fumigatus*:471 *A. fischeri*:414
afum|EBA27433.1 100.00% afis|EAW17597.1 100.00%
Bootstrap support for afum|EBA27433.1 as seed ortholog is 100%.
Bootstrap support for afis|EAW17597.1 as seed ortholog is 100%.

Group of orthologs #7392. Best score 471 bits
Score difference with first non-orthologous sequence - *A. fumigatus*:471 *A. fischeri*:471
afum|EAL93853.1 100.00% afis|EAW19121.1 100.00%
Bootstrap support for afum|EAL93853.1 as seed ortholog is 100%.
Bootstrap support for afis|EAW19121.1 as seed ortholog is 100%.

Group of orthologs #7393. Best score 471 bits
Score difference with first non-orthologous sequence - *A. fumigatus*:471 *A. fischeri*:471
afum|EAL90411.1 100.00% afis|EAW22788.1 100.00%
Bootstrap support for afum|EAL90411.1 as seed ortholog is 100%.
Bootstrap support for afis|EAW22788.1 as seed ortholog is 100%.

Group of orthologs #7394. Best score 471 bits
Score difference with first non-orthologous sequence - *A. fumigatus*:471 *A. fischeri*:471
afum|EAL92204.1 100.00% afis|EAW21183.1 100.00%
Bootstrap support for afum|EAL92204.1 as seed ortholog is 100%.
Bootstrap support for afis|EAW21183.1 as seed ortholog is 100%.

Group of orthologs #7395. Best score 471 bits
Score difference with first non-orthologous sequence - *A. fumigatus*:471 *A. fischeri*:471
afum|EAL89453.2 100.00% afis|EAW24810.1 100.00%
Bootstrap support for afum|EAL89453.2 as seed ortholog is 100%.
Bootstrap support for afis|EAW24810.1 as seed ortholog is 100%.

Group of orthologs #7396. Best score 470 bits
Score difference with first non-orthologous sequence - *A. fumigatus*:470 *A. fischeri*:470
afum|EAL87422.1 100.00% afis|EAW23992.1 100.00%
Bootstrap support for afum|EAL87422.1 as seed ortholog is 100%.
Bootstrap support for afis|EAW23992.1 as seed ortholog is 100%.

Group of orthologs #7397. Best score 470 bits

Score difference with first non-orthologous sequence - *A. fumigatus*:470 *A. fischeri*:470
afum|EAL93877.1 100.00% afis|EAW19146.1 100.00%
Bootstrap support for afum|EAL93877.1 as seed ortholog is 100%.
Bootstrap support for afis|EAW19146.1 as seed ortholog is 100%.

---

Group of orthologs #7398. Best score 470 bits
Score difference with first non-orthologous sequence - *A. fumigatus*:470 *A. fischeri*:470
afum|EAL90314.1 100.00% afis|EAW22884.1 100.00%
Bootstrap support for afum|EAL90314.1 as seed ortholog is 100%.
Bootstrap support for afis|EAW22884.1 as seed ortholog is 100%.

---

Group of orthologs #7399. Best score 469 bits
Score difference with first non-orthologous sequence - *A. fumigatus*:469 *A. fischeri*:469
afum|EAL88569.1 100.00% afis|EAW15933.1 100.00%
Bootstrap support for afum|EAL88569.1 as seed ortholog is 100%.
Bootstrap support for afis|EAW15933.1 as seed ortholog is 100%.

---

Group of orthologs #7400. Best score 469 bits
Score difference with first non-orthologous sequence - *A. fumigatus*:469 *A. fischeri*:469
afum|EAL89169.2 100.00% afis|EAW16572.1 100.00%
Bootstrap support for afum|EAL89169.2 as seed ortholog is 100%.
Bootstrap support for afis|EAW16572.1 as seed ortholog is 100%.

---

Group of orthologs #7401. Best score 469 bits
Score difference with first non-orthologous sequence - *A. fumigatus*:469 *A. fischeri*:469
afum|EAL93095.1 100.00% afis|EAW18414.1 100.00%
Bootstrap support for afum|EAL93095.1 as seed ortholog is 100%.
Bootstrap support for afis|EAW18414.1 as seed ortholog is 100%.

---

Group of orthologs #7402. Best score 469 bits
Score difference with first non-orthologous sequence - *A. fumigatus*:469 *A. fischeri*:469
afum|EAL88001.1 100.00% afis|EAW23465.1 100.00%
Bootstrap support for afum|EAL88001.1 as seed ortholog is 100%.
Bootstrap support for afis|EAW23465.1 as seed ortholog is 100%.

---

Group of orthologs #7403. Best score 469 bits
Score difference with first non-orthologous sequence - *A. fumigatus*:469 *A. fischeri*:469
afum|EAL92624.1 100.00% afis|EAW21605.1 100.00%
Bootstrap support for afum|EAL92624.1 as seed ortholog is 100%.
Bootstrap support for afis|EAW21605.1 as seed ortholog is 100%.

---

Group of orthologs #7404. Best score 468 bits
Score difference with first non-orthologous sequence - *A. fumigatus*:468 *A. fischeri*:468
afum|EAL84222.1 100.00% afis|EAW15422.1 100.00%
Bootstrap support for afum|EAL84222.1 as seed ortholog is 100%.
Bootstrap support for afis|EAW15422.1 as seed ortholog is 100%.

---

Group of orthologs #7405. Best score 468 bits
Score difference with first non-orthologous sequence - *A. fumigatus*:468 *A. fischeri*:376
afum|EAL85332.1 100.00% afis|EAW20217.1 100.00%
Bootstrap support for afum|EAL85332.1 as seed ortholog is 100%.
Bootstrap support for afis|EAW20217.1 as seed ortholog is 100%.

---

Group of orthologs #7406. Best score 468 bits
Score difference with first non-orthologous sequence - *A. fumigatus*:468 *A. fischeri*:468
afum|EAL87574.1 100.00% afis|EAW18208.1 100.00%

Bootstrap support for afum|EAL87574.1 as seed ortholog is 100%.
Bootstrap support for afis|EAW18208.1 as seed ortholog is 100%.

---

Group of orthologs #7407. Best score 468 bits
Score difference with first non-orthologous sequence - *A. fumigatus*:284 *A. fischeri*:256
afum|EAL84697.1 100.00% afis|EAW20937.1 100.00%
Bootstrap support for afum|EAL84697.1 as seed ortholog is 100%.
Bootstrap support for afis|EAW20937.1 as seed ortholog is 100%.

---

Group of orthologs #7408. Best score 468 bits
Score difference with first non-orthologous sequence - *A. fumigatus*:468 *A. fischeri*:468
afum|EAL84302.1 100.00% afis|EAW25596.1 100.00%
Bootstrap support for afum|EAL84302.1 as seed ortholog is 100%.
Bootstrap support for afis|EAW25596.1 as seed ortholog is 100%.

---

Group of orthologs #7409. Best score 468 bits
Score difference with first non-orthologous sequence - *A. fumigatus*:468 *A. fischeri*:468
afum|EAL85955.1 100.00% afis|EAW24286.1 100.00%
Bootstrap support for afum|EAL85955.1 as seed ortholog is 100%.
Bootstrap support for afis|EAW24286.1 as seed ortholog is 100%.

---

Group of orthologs #7410. Best score 468 bits
Score difference with first non-orthologous sequence - *A. fumigatus*:468 *A. fischeri*:468
afum|EAL90984.2 100.00% afis|EAW22210.1 100.00%
Bootstrap support for afum|EAL90984.2 as seed ortholog is 100%.
Bootstrap support for afis|EAW22210.1 as seed ortholog is 100%.

---

Group of orthologs #7411. Best score 468 bits
Score difference with first non-orthologous sequence - *A. fumigatus*:468 *A. fischeri*:468
afum|EAL92372.1 100.00% afis|EAW21357.1 100.00%
Bootstrap support for afum|EAL92372.1 as seed ortholog is 100%.
Bootstrap support for afis|EAW21357.1 as seed ortholog is 100%.

---

Group of orthologs #7412. Best score 468 bits
Score difference with first non-orthologous sequence - *A. fumigatus*:468 *A. fischeri*:468
afum|EAL89564.1 100.00% afis|EAW24910.1 100.00%
Bootstrap support for afum|EAL89564.1 as seed ortholog is 100%.
Bootstrap support for afis|EAW24910.1 as seed ortholog is 100%.

---

Group of orthologs #7413. Best score 468 bits
Score difference with first non-orthologous sequence - *A. fumigatus*:468 *A. fischeri*:468
afum|EAL89835.1 100.00% afis|EAW25160.1 100.00%
Bootstrap support for afum|EAL89835.1 as seed ortholog is 100%.
Bootstrap support for afis|EAW25160.1 as seed ortholog is 100%.

---

Group of orthologs #7414. Best score 467 bits
Score difference with first non-orthologous sequence - *A. fumigatus*:467 *A. fischeri*:467
afum|EAL86409.1 100.00% afis|EAW16871.1 100.00%
Bootstrap support for afum|EAL86409.1 as seed ortholog is 100%.
Bootstrap support for afis|EAW16871.1 as seed ortholog is 100%.

---

Group of orthologs #7415. Best score 467 bits
Score difference with first non-orthologous sequence - *A. fumigatus*:467 *A. fischeri*:241
afum|EAL88545.1 100.00% afis|EAW15910.1 100.00%
Bootstrap support for afum|EAL88545.1 as seed ortholog is 100%.
Bootstrap support for afis|EAW15910.1 as seed ortholog is 100%.

---

Group of orthologs #7416. Best score 467 bits
Score difference with first non-orthologous sequence - *A. fumigatus*:467 *A. fischeri*:467

| | | | |
|---|---|---|---|
| afum\|EAL88759.1 | 100.00% | afis\|EAW16129.1 | 100.00% |

Bootstrap support for afum|EAL88759.1 as seed ortholog is 100%.
Bootstrap support for afis|EAW16129.1 as seed ortholog is 100%.

---

Group of orthologs #7417. Best score 467 bits
Score difference with first non-orthologous sequence - *A. fumigatus*:467 *A. fischeri*:467

| | | | |
|---|---|---|---|
| afum\|EAL89718.1 | 100.00% | afis\|EAW25050.1 | 100.00% |

Bootstrap support for afum|EAL89718.1 as seed ortholog is 100%.
Bootstrap support for afis|EAW25050.1 as seed ortholog is 100%.

---

Group of orthologs #7418. Best score 467 bits
Score difference with first non-orthologous sequence - *A. fumigatus*:467 *A. fischeri*:467

| | | | |
|---|---|---|---|
| afum\|EAL90194.1 | 100.00% | afis\|EAW25529.1 | 100.00% |

Bootstrap support for afum|EAL90194.1 as seed ortholog is 100%.
Bootstrap support for afis|EAW25529.1 as seed ortholog is 100%.

---

Group of orthologs #7419. Best score 466 bits
Score difference with first non-orthologous sequence - *A. fumigatus*:466 *A. fischeri*:95

| | | | |
|---|---|---|---|
| afum\|EAL85371.1 | 100.00% | afis\|EAW18106.1 | 100.00% |

Bootstrap support for afum|EAL85371.1 as seed ortholog is 100%.
Bootstrap support for afis|EAW18106.1 as seed ortholog is 99%.

---

Group of orthologs #7420. Best score 466 bits
Score difference with first non-orthologous sequence - *A. fumigatus*:466 *A. fischeri*:466

| | | | |
|---|---|---|---|
| afum\|EAL89694.1 | 100.00% | afis\|EAW25004.1 | 100.00% |

Bootstrap support for afum|EAL89694.1 as seed ortholog is 100%.
Bootstrap support for afis|EAW25004.1 as seed ortholog is 100%.

---

Group of orthologs #7421. Best score 466 bits
Score difference with first non-orthologous sequence - *A. fumigatus*:466 *A. fischeri*:466

| | | | |
|---|---|---|---|
| afum\|EAL89958.1 | 100.00% | afis\|EAW25295.1 | 100.00% |

Bootstrap support for afum|EAL89958.1 as seed ortholog is 100%.
Bootstrap support for afis|EAW25295.1 as seed ortholog is 100%.

---

Group of orthologs #7422. Best score 465 bits
Score difference with first non-orthologous sequence - *A. fumigatus*:465 *A. fischeri*:465

| | | | |
|---|---|---|---|
| afum\|EAL85835.1 | 100.00% | afis\|EAW15488.1 | 100.00% |

Bootstrap support for afum|EAL85835.1 as seed ortholog is 100%.
Bootstrap support for afis|EAW15488.1 as seed ortholog is 100%.

---

Group of orthologs #7423. Best score 465 bits
Score difference with first non-orthologous sequence - *A. fumigatus*:465 *A. fischeri*:465

| | | | |
|---|---|---|---|
| afum\|EAL85704.1 | 100.00% | afis\|EAW15630.1 | 100.00% |

Bootstrap support for afum|EAL85704.1 as seed ortholog is 100%.
Bootstrap support for afis|EAW15630.1 as seed ortholog is 100%.

---

Group of orthologs #7424. Best score 465 bits
Score difference with first non-orthologous sequence - *A. fumigatus*:465 *A. fischeri*:465

| | | | |
|---|---|---|---|
| afum\|EAL86608.1 | 100.00% | afis\|EAW17178.1 | 100.00% |

Bootstrap support for afum|EAL86608.1 as seed ortholog is 100%.
Bootstrap support for afis|EAW17178.1 as seed ortholog is 100%.

---

Group of orthologs #7425. Best score 465 bits

Score difference with first non-orthologous sequence - *A. fumigatus*:465   *A. fischeri*:465
afum|EAL90773.1 100.00% afis|EAW22419.1 100.00%
Bootstrap support for afum|EAL90773.1 as seed ortholog is 100%.
Bootstrap support for afis|EAW22419.1 as seed ortholog is 100%.

---

Group of orthologs #7426. Best score 465 bits
Score difference with first non-orthologous sequence - *A. fumigatus*:465   *A. fischeri*:465
afum|EAL92548.1 100.00% afis|EAW21527.1 100.00%
Bootstrap support for afum|EAL92548.1 as seed ortholog is 100%.
Bootstrap support for afis|EAW21527.1 as seed ortholog is 100%.

---

Group of orthologs #7427. Best score 465 bits
Score difference with first non-orthologous sequence - *A. fumigatus*:465   *A. fischeri*:465
afum|EBA27397.1 100.00% afis|EAW25530.1 100.00%
Bootstrap support for afum|EBA27397.1 as seed ortholog is 100%.
Bootstrap support for afis|EAW25530.1 as seed ortholog is 100%.

---

Group of orthologs #7428. Best score 464 bits
Score difference with first non-orthologous sequence - *A. fumigatus*:464   *A. fischeri*:464
afum|EAL84834.1 100.00% afis|EAW20792.1 100.00%
Bootstrap support for afum|EAL84834.1 as seed ortholog is 100%.
Bootstrap support for afis|EAW20792.1 as seed ortholog is 100%.

---

Group of orthologs #7429. Best score 463 bits
Score difference with first non-orthologous sequence - *A. fumigatus*:463   *A. fischeri*:463
afum|EAL88501.1 100.00% afis|EAW15867.1 100.00%
Bootstrap support for afum|EAL88501.1 as seed ortholog is 100%.
Bootstrap support for afis|EAW15867.1 as seed ortholog is 100%.

---

Group of orthologs #7430. Best score 463 bits
Score difference with first non-orthologous sequence - *A. fumigatus*:359   *A. fischeri*:360
afum|EAL84808.1 100.00% afis|EAW20821.1 100.00%
Bootstrap support for afum|EAL84808.1 as seed ortholog is 100%.
Bootstrap support for afis|EAW20821.1 as seed ortholog is 100%.

---

Group of orthologs #7431. Best score 463 bits
Score difference with first non-orthologous sequence - *A. fumigatus*:463   *A. fischeri*:463
afum|EAL91257.1 100.00% afis|EAW17429.1 100.00%
Bootstrap support for afum|EAL91257.1 as seed ortholog is 100%.
Bootstrap support for afis|EAW17429.1 as seed ortholog is 100%.

---

Group of orthologs #7432. Best score 463 bits
Score difference with first non-orthologous sequence - *A. fumigatus*:463   *A. fischeri*:463
afum|EAL91533.1 100.00% afis|EAW17718.1 100.00%
Bootstrap support for afum|EAL91533.1 as seed ortholog is 100%.
Bootstrap support for afis|EAW17718.1 as seed ortholog is 100%.

---

Group of orthologs #7433. Best score 463 bits
Score difference with first non-orthologous sequence - *A. fumigatus*:463   *A. fischeri*:373
afum|EAL87114.1 100.00% afis|EAW23669.1 100.00%
Bootstrap support for afum|EAL87114.1 as seed ortholog is 100%.
Bootstrap support for afis|EAW23669.1 as seed ortholog is 100%.

---

Group of orthologs #7434. Best score 463 bits
Score difference with first non-orthologous sequence - *A. fumigatus*:463   *A. fischeri*:463
afum|EAL88205.1 100.00% afis|EAW23274.1 100.00%

Bootstrap support for afum|EAL88205.1 as seed ortholog is 100%.
Bootstrap support for afis|EAW23274.1 as seed ortholog is 100%.

---

Group of orthologs #7435. Best score 463 bits
Score difference with first non-orthologous sequence - *A. fumigatus*:463 *A. fischeri*:463
afum|EAL92220.1 100.00% afis|EAW21209.1 100.00%
Bootstrap support for afum|EAL92220.1 as seed ortholog is 100%.
Bootstrap support for afis|EAW21209.1 as seed ortholog is 100%.

---

Group of orthologs #7436. Best score 463 bits
Score difference with first non-orthologous sequence - *A. fumigatus*:463 *A. fischeri*:463
afum|EAL92511.1 100.00% afis|EAW21492.1 100.00%
Bootstrap support for afum|EAL92511.1 as seed ortholog is 100%.
Bootstrap support for afis|EAW21492.1 as seed ortholog is 100%.

---

Group of orthologs #7437. Best score 463 bits
Score difference with first non-orthologous sequence - *A. fumigatus*:463 *A. fischeri*:463
afum|EAL89989.1 100.00% afis|EAW25328.1 100.00%
Bootstrap support for afum|EAL89989.1 as seed ortholog is 100%.
Bootstrap support for afis|EAW25328.1 as seed ortholog is 100%.

---

Group of orthologs #7438. Best score 462 bits
Score difference with first non-orthologous sequence - *A. fumigatus*:462 *A. fischeri*:462
afum|EAL85637.1 100.00% afis|EAW15710.1 100.00%
Bootstrap support for afum|EAL85637.1 as seed ortholog is 100%.
Bootstrap support for afis|EAW15710.1 as seed ortholog is 100%.

---

Group of orthologs #7439. Best score 462 bits
Score difference with first non-orthologous sequence - *A. fumigatus*:462 *A. fischeri*:462
afum|EAL90492.1 100.00% afis|EAW22700.1 100.00%
Bootstrap support for afum|EAL90492.1 as seed ortholog is 100%.
Bootstrap support for afis|EAW22700.1 as seed ortholog is 100%.

---

Group of orthologs #7440. Best score 461 bits
Score difference with first non-orthologous sequence - *A. fumigatus*:461 *A. fischeri*:461
afum|EAL88953.1 100.00% afis|EAW16329.1 100.00%
Bootstrap support for afum|EAL88953.1 as seed ortholog is 100%.
Bootstrap support for afis|EAW16329.1 as seed ortholog is 100%.

---

Group of orthologs #7441. Best score 461 bits
Score difference with first non-orthologous sequence - *A. fumigatus*:461 *A. fischeri*:461
afum|EAL87303.1 100.00% afis|EAW23875.1 100.00%
Bootstrap support for afum|EAL87303.1 as seed ortholog is 100%.
Bootstrap support for afis|EAW23875.1 as seed ortholog is 100%.

---

Group of orthologs #7442. Best score 461 bits
Score difference with first non-orthologous sequence - *A. fumigatus*:461 *A. fischeri*:461
afum|EAL93388.1 100.00% afis|EAW18670.1 100.00%
Bootstrap support for afum|EAL93388.1 as seed ortholog is 100%.
Bootstrap support for afis|EAW18670.1 as seed ortholog is 100%.

---

Group of orthologs #7443. Best score 461 bits
Score difference with first non-orthologous sequence - *A. fumigatus*:461 *A. fischeri*:461
afum|EAL90046.2 100.00% afis|EAW25387.1 100.00%
Bootstrap support for afum|EAL90046.2 as seed ortholog is 100%.
Bootstrap support for afis|EAW25387.1 as seed ortholog is 100%.

Group of orthologs #7444. Best score 460 bits
Score difference with first non-orthologous sequence - *A. fumigatus*:460 *A. fischeri*:460
afum|EAL87943.1 100.00% afis|EAW23521.1 100.00%
Bootstrap support for afum|EAL87943.1 as seed ortholog is 100%.
Bootstrap support for afis|EAW23521.1 as seed ortholog is 100%.

Group of orthologs #7445. Best score 460 bits
Score difference with first non-orthologous sequence - *A. fumigatus*:460 *A. fischeri*:460
afum|EAL93954.1 100.00% afis|EAW19222.1 100.00%
Bootstrap support for afum|EAL93954.1 as seed ortholog is 100%.
Bootstrap support for afis|EAW19222.1 as seed ortholog is 100%.

Group of orthologs #7446. Best score 460 bits
Score difference with first non-orthologous sequence - *A. fumigatus*:460 *A. fischeri*:460
afum|EAL91113.1 100.00% afis|EAW22299.1 100.00%
Bootstrap support for afum|EAL91113.1 as seed ortholog is 100%.
Bootstrap support for afis|EAW22299.1 as seed ortholog is 100%.

Group of orthologs #7447. Best score 459 bits
Score difference with first non-orthologous sequence - *A. fumigatus*:459 *A. fischeri*:459
afum|EAL85586.1 100.00% afis|EAW15762.1 100.00%
Bootstrap support for afum|EAL85586.1 as seed ortholog is 100%.
Bootstrap support for afis|EAW15762.1 as seed ortholog is 100%.

Group of orthologs #7448. Best score 459 bits
Score difference with first non-orthologous sequence - *A. fumigatus*:459 *A. fischeri*:459
afum|EAL88906.1 100.00% afis|EAW16285.1 100.00%
Bootstrap support for afum|EAL88906.1 as seed ortholog is 100%.
Bootstrap support for afis|EAW16285.1 as seed ortholog is 100%.

Group of orthologs #7449. Best score 459 bits
Score difference with first non-orthologous sequence - *A. fumigatus*:459 *A. fischeri*:459
afum|EAL84667.1 100.00% afis|EAW20966.1 100.00%
Bootstrap support for afum|EAL84667.1 as seed ortholog is 100%.
Bootstrap support for afis|EAW20966.1 as seed ortholog is 100%.

Group of orthologs #7450. Best score 459 bits
Score difference with first non-orthologous sequence - *A. fumigatus*:459 *A. fischeri*:459
afum|EAL93330.1 100.00% afis|EAW18620.1 100.00%
Bootstrap support for afum|EAL93330.1 as seed ortholog is 100%.
Bootstrap support for afis|EAW18620.1 as seed ortholog is 100%.

Group of orthologs #7451. Best score 459 bits
Score difference with first non-orthologous sequence - *A. fumigatus*:459 *A. fischeri*:459
afum|EBA27408.1 100.00% afis|EAW22693.1 100.00%
Bootstrap support for afum|EBA27408.1 as seed ortholog is 100%.
Bootstrap support for afis|EAW22693.1 as seed ortholog is 100%.

Group of orthologs #7452. Best score 458 bits
Score difference with first non-orthologous sequence - *A. fumigatus*:458 *A. fischeri*:458
afum|EAL85683.1 100.00% afis|EAW15665.1 100.00%
Bootstrap support for afum|EAL85683.1 as seed ortholog is 100%.
Bootstrap support for afis|EAW15665.1 as seed ortholog is 100%.

Group of orthologs #7453. Best score 458 bits

Score difference with first non-orthologous sequence - *A. fumigatus*:458   *A. fischeri*:458
afum|EAL91226.1 100.00% afis|EAW17390.1 100.00%
Bootstrap support for afum|EAL91226.1 as seed ortholog is 100%.
Bootstrap support for afis|EAW17390.1 as seed ortholog is 100%.

---

Group of orthologs #7454. Best score 458 bits
Score difference with first non-orthologous sequence - *A. fumigatus*:458   *A. fischeri*:458
afum|EAL91606.1 100.00% afis|EAW17798.1 100.00%
Bootstrap support for afum|EAL91606.1 as seed ortholog is 100%.
Bootstrap support for afis|EAW17798.1 as seed ortholog is 100%.

---

Group of orthologs #7455. Best score 458 bits
Score difference with first non-orthologous sequence - *A. fumigatus*:458   *A. fischeri*:458
afum|EAL87196.1 100.00% afis|EAW23763.1 100.00%
Bootstrap support for afum|EAL87196.1 as seed ortholog is 100%.
Bootstrap support for afis|EAW23763.1 as seed ortholog is 100%.

---

Group of orthologs #7456. Best score 458 bits
Score difference with first non-orthologous sequence - *A. fumigatus*:458   *A. fischeri*:458
afum|EAL93168.1 100.00% afis|EAW18479.1 100.00%
Bootstrap support for afum|EAL93168.1 as seed ortholog is 100%.
Bootstrap support for afis|EAW18479.1 as seed ortholog is 100%.

---

Group of orthologs #7457. Best score 458 bits
Score difference with first non-orthologous sequence - *A. fumigatus*:458   *A. fischeri*:458
afum|EAL93405.1 100.00% afis|EAW18684.1 100.00%
Bootstrap support for afum|EAL93405.1 as seed ortholog is 100%.
Bootstrap support for afis|EAW18684.1 as seed ortholog is 100%.

---

Group of orthologs #7458. Best score 458 bits
Score difference with first non-orthologous sequence - *A. fumigatus*:458   *A. fischeri*:296
afum|EAL92547.1 100.00% afis|EAW21526.1 100.00%
Bootstrap support for afum|EAL92547.1 as seed ortholog is 100%.
Bootstrap support for afis|EAW21526.1 as seed ortholog is 100%.

---

Group of orthologs #7459. Best score 457 bits
Score difference with first non-orthologous sequence - *A. fumigatus*:457   *A. fischeri*:457
afum|EAL91534.2 100.00% afis|EAW17719.1 100.00%
Bootstrap support for afum|EAL91534.2 as seed ortholog is 100%.
Bootstrap support for afis|EAW17719.1 as seed ortholog is 100%.

---

Group of orthologs #7460. Best score 457 bits
Score difference with first non-orthologous sequence - *A. fumigatus*:457   *A. fischeri*:457
afum|EAL86022.1 100.00% afis|EAW24352.1 100.00%
Bootstrap support for afum|EAL86022.1 as seed ortholog is 100%.
Bootstrap support for afis|EAW24352.1 as seed ortholog is 100%.

---

Group of orthologs #7461. Best score 457 bits
Score difference with first non-orthologous sequence - *A. fumigatus*:457   *A. fischeri*:457
afum|EAL87865.1 100.00% afis|EAW23602.1 100.00%
Bootstrap support for afum|EAL87865.1 as seed ortholog is 100%.
Bootstrap support for afis|EAW23602.1 as seed ortholog is 100%.

---

Group of orthologs #7462. Best score 456 bits
Score difference with first non-orthologous sequence - *A. fumigatus*:456   *A. fischeri*:456
afum|EAL84618.1 100.00% afis|EAW20433.1 100.00%

Bootstrap support for afum|EAL84618.1 as seed ortholog is 100%.
Bootstrap support for afis|EAW20433.1 as seed ortholog is 100%.

---

Group of orthologs #7463. Best score 456 bits
Score difference with first non-orthologous sequence - *A. fumigatus*:456 *A. fischeri*:456
afum|EAL91591.2 100.00% afis|EAW17783.1 100.00%
Bootstrap support for afum|EAL91591.2 as seed ortholog is 100%.
Bootstrap support for afis|EAW17783.1 as seed ortholog is 100%.

---

Group of orthologs #7464. Best score 456 bits
Score difference with first non-orthologous sequence - *A. fumigatus*:456 *A. fischeri*:456
afum|EAL84390.2 100.00% afis|EAW25716.1 100.00%
Bootstrap support for afum|EAL84390.2 as seed ortholog is 100%.
Bootstrap support for afis|EAW25716.1 as seed ortholog is 100%.

---

Group of orthologs #7465. Best score 456 bits
Score difference with first non-orthologous sequence - *A. fumigatus*:456 *A. fischeri*:456
afum|EAL93128.1 100.00% afis|EAW18444.1 100.00%
Bootstrap support for afum|EAL93128.1 as seed ortholog is 100%.
Bootstrap support for afis|EAW18444.1 as seed ortholog is 100%.

---

Group of orthologs #7466. Best score 456 bits
Score difference with first non-orthologous sequence - *A. fumigatus*:456 *A. fischeri*:456
afum|EAL87431.1 100.00% afis|EAW24001.1 100.00%
Bootstrap support for afum|EAL87431.1 as seed ortholog is 100%.
Bootstrap support for afis|EAW24001.1 as seed ortholog is 100%.

---

Group of orthologs #7467. Best score 456 bits
Score difference with first non-orthologous sequence - *A. fumigatus*:456 *A. fischeri*:456
afum|EAL90239.1 100.00% afis|EAW22958.1 100.00%
Bootstrap support for afum|EAL90239.1 as seed ortholog is 100%.
Bootstrap support for afis|EAW22958.1 as seed ortholog is 100%.

---

Group of orthologs #7468. Best score 456 bits
Score difference with first non-orthologous sequence - *A. fumigatus*:456 *A. fischeri*:456
afum|EAL89550.1 100.00% afis|EAW24896.1 100.00%
Bootstrap support for afum|EAL89550.1 as seed ortholog is 100%.
Bootstrap support for afis|EAW24896.1 as seed ortholog is 100%.

---

Group of orthologs #7469. Best score 456 bits
Score difference with first non-orthologous sequence - *A. fumigatus*:456 *A. fischeri*:456
afum|EBA27172.1 100.00% afis|EAW25601.1 100.00%
Bootstrap support for afum|EBA27172.1 as seed ortholog is 100%.
Bootstrap support for afis|EAW25601.1 as seed ortholog is 100%.

---

Group of orthologs #7470. Best score 455 bits
Score difference with first non-orthologous sequence - *A. fumigatus*:455 *A. fischeri*:455
afum|EBA27240.1 100.00% afis|EAW15596.1 100.00%
Bootstrap support for afum|EBA27240.1 as seed ortholog is 100%.
Bootstrap support for afis|EAW15596.1 as seed ortholog is 100%.

---

Group of orthologs #7471. Best score 455 bits
Score difference with first non-orthologous sequence - *A. fumigatus*:455 *A. fischeri*:455
afum|EAL90229.1 100.00% afis|EAW22968.1 100.00%
Bootstrap support for afum|EAL90229.1 as seed ortholog is 100%.
Bootstrap support for afis|EAW22968.1 as seed ortholog is 100%.

---

Group of orthologs #7472. Best score 455 bits
Score difference with first non-orthologous sequence - *A. fumigatus*:455 *A. fischeri*:455
afum|EAL90278.1 100.00% afis|EAW22919.1 100.00%
Bootstrap support for afum|EAL90278.1 as seed ortholog is 100%.
Bootstrap support for afis|EAW22919.1 as seed ortholog is 100%.

---

Group of orthologs #7473. Best score 454 bits
Score difference with first non-orthologous sequence - *A. fumigatus*:454 *A. fischeri*:454
afum|EAL84896.2 100.00% afis|EAW20021.1 100.00%
Bootstrap support for afum|EAL84896.2 as seed ortholog is 100%.
Bootstrap support for afis|EAW20021.1 as seed ortholog is 100%.

---

Group of orthologs #7474. Best score 454 bits
Score difference with first non-orthologous sequence - *A. fumigatus*:454 *A. fischeri*:454
afum|EAL92145.1 100.00% afis|EAW21122.1 100.00%
Bootstrap support for afum|EAL92145.1 as seed ortholog is 100%.
Bootstrap support for afis|EAW21122.1 as seed ortholog is 100%.

---

Group of orthologs #7475. Best score 454 bits
Score difference with first non-orthologous sequence - *A. fumigatus*:454 *A. fischeri*:454
afum|EAL89377.1 100.00% afis|EAW24730.1 100.00%
Bootstrap support for afum|EAL89377.1 as seed ortholog is 100%.
Bootstrap support for afis|EAW24730.1 as seed ortholog is 100%.

---

Group of orthologs #7476. Best score 453 bits
Score difference with first non-orthologous sequence - *A. fumigatus*:453 *A. fischeri*:453
afum|EAL91790.1 100.00% afis|EAW17981.1 100.00%
Bootstrap support for afum|EAL91790.1 as seed ortholog is 100%.
Bootstrap support for afis|EAW17981.1 as seed ortholog is 100%.

---

Group of orthologs #7477. Best score 453 bits
Score difference with first non-orthologous sequence - *A. fumigatus*:453 *A. fischeri*:453
afum|EAL87368.1 100.00% afis|EAW23943.1 100.00%
Bootstrap support for afum|EAL87368.1 as seed ortholog is 100%.
Bootstrap support for afis|EAW23943.1 as seed ortholog is 100%.

---

Group of orthologs #7478. Best score 453 bits
Score difference with first non-orthologous sequence - *A. fumigatus*:453 *A. fischeri*:453
afum|EAL90956.1 100.00% afis|EAW22237.1 100.00%
Bootstrap support for afum|EAL90956.1 as seed ortholog is 100%.
Bootstrap support for afis|EAW22237.1 as seed ortholog is 100%.

---

Group of orthologs #7479. Best score 453 bits
Score difference with first non-orthologous sequence - *A. fumigatus*:453 *A. fischeri*:453
afum|EAL90270.1 100.00% afis|EAW22927.1 100.00%
Bootstrap support for afum|EAL90270.1 as seed ortholog is 100%.
Bootstrap support for afis|EAW22927.1 as seed ortholog is 100%.

---

Group of orthologs #7480. Best score 453 bits
Score difference with first non-orthologous sequence - *A. fumigatus*:453 *A. fischeri*:295
afum|EAL92299.1 100.00% afis|EAW21282.1 100.00%
Bootstrap support for afum|EAL92299.1 as seed ortholog is 100%.
Bootstrap support for afis|EAW21282.1 as seed ortholog is 100%.

---

Group of orthologs #7481. Best score 453 bits

Score difference with first non-orthologous sequence - *A. fumigatus*:453 *A. fischeri*:453
afum|EAL92356.1 100.00% afis|EAW21338.1 100.00%
Bootstrap support for afum|EAL92356.1 as seed ortholog is 100%.
Bootstrap support for afis|EAW21338.1 as seed ortholog is 100%.

---

Group of orthologs #7482. Best score 453 bits
Score difference with first non-orthologous sequence - *A. fumigatus*:453 *A. fischeri*:453
afum|EAL89489.1 100.00% afis|EAW24843.1 100.00%
Bootstrap support for afum|EAL89489.1 as seed ortholog is 100%.
Bootstrap support for afis|EAW24843.1 as seed ortholog is 100%.

---

Group of orthologs #7483. Best score 453 bits
Score difference with first non-orthologous sequence - *A. fumigatus*:453 *A. fischeri*:453
afum|EAL90187.1 100.00% afis|EAW25522.1 100.00%
Bootstrap support for afum|EAL90187.1 as seed ortholog is 100%.
Bootstrap support for afis|EAW25522.1 as seed ortholog is 100%.

---

Group of orthologs #7484. Best score 452 bits
Score difference with first non-orthologous sequence - *A. fumigatus*:452 *A. fischeri*:452
afum|EAL86398.1 100.00% afis|EAW16851.1 100.00%
Bootstrap support for afum|EAL86398.1 as seed ortholog is 100%.
Bootstrap support for afis|EAW16851.1 as seed ortholog is 100%.

---

Group of orthologs #7485. Best score 452 bits
Score difference with first non-orthologous sequence - *A. fumigatus*:452 *A. fischeri*:452
afum|EAL86112.1 100.00% afis|EAW24443.1 100.00%
Bootstrap support for afum|EAL86112.1 as seed ortholog is 100%.
Bootstrap support for afis|EAW24443.1 as seed ortholog is 100%.

---

Group of orthologs #7486. Best score 452 bits
Score difference with first non-orthologous sequence - *A. fumigatus*:452 *A. fischeri*:452
afum|EAL93191.2 100.00% afis|EAW18500.1 100.00%
Bootstrap support for afum|EAL93191.2 as seed ortholog is 100%.
Bootstrap support for afis|EAW18500.1 as seed ortholog is 100%.

---

Group of orthologs #7487. Best score 452 bits
Score difference with first non-orthologous sequence - *A. fumigatus*:452 *A. fischeri*:452
afum|EAL88180.1 100.00% afis|EAW23299.1 100.00%
Bootstrap support for afum|EAL88180.1 as seed ortholog is 100%.
Bootstrap support for afis|EAW23299.1 as seed ortholog is 100%.

---

Group of orthologs #7488. Best score 452 bits
Score difference with first non-orthologous sequence - *A. fumigatus*:452 *A. fischeri*:452
afum|EAL93781.1 100.00% afis|EAW19052.1 100.00%
Bootstrap support for afum|EAL93781.1 as seed ortholog is 100%.
Bootstrap support for afis|EAW19052.1 as seed ortholog is 100%.

---

Group of orthologs #7489. Best score 451 bits
Score difference with first non-orthologous sequence - *A. fumigatus*:451 *A. fischeri*:451
afum|EAL85793.1 100.00% afis|EAW15535.1 100.00%
Bootstrap support for afum|EAL85793.1 as seed ortholog is 100%.
Bootstrap support for afis|EAW15535.1 as seed ortholog is 100%.

---

Group of orthologs #7490. Best score 451 bits
Score difference with first non-orthologous sequence - *A. fumigatus*:451 *A. fischeri*:451
afum|EAL84451.1 100.00% afis|EAW20609.1 100.00%

Bootstrap support for afum|EAL84451.1 as seed ortholog is 100%.
Bootstrap support for afis|EAW20609.1 as seed ortholog is 100%.

---

Group of orthologs #7491. Best score 451 bits
Score difference with first non-orthologous sequence - *A. fumigatus*:451 *A. fischeri*:451
afum|EAL93464.1 100.00% afis|EAW18738.1 100.00%
Bootstrap support for afum|EAL93464.1 as seed ortholog is 100%.
Bootstrap support for afis|EAW18738.1 as seed ortholog is 100%.

---

Group of orthologs #7492. Best score 450 bits
Score difference with first non-orthologous sequence - *A. fumigatus*:450 *A. fischeri*:450
afum|EAL88674.1 100.00% afis|EAW16042.1 100.00%
Bootstrap support for afum|EAL88674.1 as seed ortholog is 100%.
Bootstrap support for afis|EAW16042.1 as seed ortholog is 100%.

---

Group of orthologs #7493. Best score 450 bits
Score difference with first non-orthologous sequence - *A. fumigatus*:450 *A. fischeri*:450
afum|EAL90370.1 100.00% afis|EAW22832.1 100.00%
Bootstrap support for afum|EAL90370.1 as seed ortholog is 100%.
Bootstrap support for afis|EAW22832.1 as seed ortholog is 100%.

---

Group of orthologs #7494. Best score 450 bits
Score difference with first non-orthologous sequence - *A. fumigatus*:450 *A. fischeri*:450
afum|EAL92931.1 100.00% afis|EAW21919.1 100.00%
Bootstrap support for afum|EAL92931.1 as seed ortholog is 100%.
Bootstrap support for afis|EAW21919.1 as seed ortholog is 100%.

---

Group of orthologs #7495. Best score 450 bits
Score difference with first non-orthologous sequence - *A. fumigatus*:450 *A. fischeri*:409
afum|EAL92044.1 100.00% afis|EAW24184.1 100.00%
Bootstrap support for afum|EAL92044.1 as seed ortholog is 100%.
Bootstrap support for afis|EAW24184.1 as seed ortholog is 100%.

---

Group of orthologs #7496. Best score 450 bits
Score difference with first non-orthologous sequence - *A. fumigatus*:450 *A. fischeri*:450
afum|EBA27247.1 100.00% afis|EAW24200.1 100.00%
Bootstrap support for afum|EBA27247.1 as seed ortholog is 100%.
Bootstrap support for afis|EAW24200.1 as seed ortholog is 100%.

---

Group of orthologs #7497. Best score 449 bits
Score difference with first non-orthologous sequence - *A. fumigatus*:449 *A. fischeri*:449
afum|EAL91544.1 100.00% afis|EAW17729.1 100.00%
Bootstrap support for afum|EAL91544.1 as seed ortholog is 100%.
Bootstrap support for afis|EAW17729.1 as seed ortholog is 100%.

---

Group of orthologs #7498. Best score 449 bits
Score difference with first non-orthologous sequence - *A. fumigatus*:449 *A. fischeri*:449
afum|EAL92315.1 100.00% afis|EAW21298.1 100.00%
Bootstrap support for afum|EAL92315.1 as seed ortholog is 100%.
Bootstrap support for afis|EAW21298.1 as seed ortholog is 100%.

---

Group of orthologs #7499. Best score 449 bits
Score difference with first non-orthologous sequence - *A. fumigatus*:449 *A. fischeri*:449
afum|EAL92401.1 100.00% afis|EAW21386.1 100.00%
Bootstrap support for afum|EAL92401.1 as seed ortholog is 100%.
Bootstrap support for afis|EAW21386.1 as seed ortholog is 100%.

---

Group of orthologs #7500. Best score 449 bits
Score difference with first non-orthologous sequence - *A. fumigatus*:449 *A. fischeri*:449
afum|EBA27251.1 100.00% afis|EAW24289.1 100.00%
Bootstrap support for afum|EBA27251.1 as seed ortholog is 100%.
Bootstrap support for afis|EAW24289.1 as seed ortholog is 100%.

---

Group of orthologs #7501. Best score 448 bits
Score difference with first non-orthologous sequence - *A. fumigatus*:448 *A. fischeri*:448
afum|EAL88917.1 100.00% afis|EAW16295.1 100.00%
Bootstrap support for afum|EAL88917.1 as seed ortholog is 100%.
Bootstrap support for afis|EAW16295.1 as seed ortholog is 100%.

---

Group of orthologs #7502. Best score 448 bits
Score difference with first non-orthologous sequence - *A. fumigatus*:448 *A. fischeri*:448
afum|EAL84773.1 100.00% afis|EAW20862.1 100.00%
Bootstrap support for afum|EAL84773.1 as seed ortholog is 100%.
Bootstrap support for afis|EAW20862.1 as seed ortholog is 100%.

---

Group of orthologs #7503. Best score 448 bits
Score difference with first non-orthologous sequence - *A. fumigatus*:448 *A. fischeri*:448
afum|EAL88140.1 100.00% afis|EAW23338.1 100.00%
Bootstrap support for afum|EAL88140.1 as seed ortholog is 100%.
Bootstrap support for afis|EAW23338.1 as seed ortholog is 100%.

---

Group of orthologs #7504. Best score 448 bits
Score difference with first non-orthologous sequence - *A. fumigatus*:448 *A. fischeri*:350
afum|EAL91035.1 100.00% afis|EAW22155.1 100.00%
Bootstrap support for afum|EAL91035.1 as seed ortholog is 100%.
Bootstrap support for afis|EAW22155.1 as seed ortholog is 100%.

---

Group of orthologs #7505. Best score 447 bits
Score difference with first non-orthologous sequence - *A. fumigatus*:447 *A. fischeri*:447
afum|EAL86371.1 100.00% afis|EAW16821.1 100.00%
Bootstrap support for afum|EAL86371.1 as seed ortholog is 100%.
Bootstrap support for afis|EAW16821.1 as seed ortholog is 100%.

---

Group of orthologs #7506. Best score 447 bits
Score difference with first non-orthologous sequence - *A. fumigatus*:294 *A. fischeri*:447
afum|EAL87841.1 100.00% afis|EAW17482.1 100.00%
Bootstrap support for afum|EAL87841.1 as seed ortholog is 100%.
Bootstrap support for afis|EAW17482.1 as seed ortholog is 100%.

---

Group of orthologs #7507. Best score 447 bits
Score difference with first non-orthologous sequence - *A. fumigatus*:447 *A. fischeri*:447
afum|EAL89723.2 100.00% afis|EAW25055.1 100.00%
Bootstrap support for afum|EAL89723.2 as seed ortholog is 100%.
Bootstrap support for afis|EAW25055.1 as seed ortholog is 100%.

---

Group of orthologs #7508. Best score 447 bits
Score difference with first non-orthologous sequence - *A. fumigatus*:447 *A. fischeri*:447
afum|EAL89877.1 100.00% afis|EAW25216.1 100.00%
Bootstrap support for afum|EAL89877.1 as seed ortholog is 100%.
Bootstrap support for afis|EAW25216.1 as seed ortholog is 100%.

---

Group of orthologs #7509. Best score 447 bits

Score difference with first non-orthologous sequence - *A. fumigatus*:447   *A. fischeri*:447
afum|EBA27460.1 100.00% afis|EAW21340.1 100.00%
Bootstrap support for afum|EBA27460.1 as seed ortholog is 100%.
Bootstrap support for afis|EAW21340.1 as seed ortholog is 100%.

---

Group of orthologs #7510. Best score 446 bits
Score difference with first non-orthologous sequence - *A. fumigatus*:446   *A. fischeri*:446
afum|EAL85813.1 100.00% afis|EAW15516.1 100.00%
Bootstrap support for afum|EAL85813.1 as seed ortholog is 100%.
Bootstrap support for afis|EAW15516.1 as seed ortholog is 100%.

---

Group of orthologs #7511. Best score 446 bits
Score difference with first non-orthologous sequence - *A. fumigatus*:446   *A. fischeri*:446
afum|EAL86697.1 100.00% afis|EAW17279.1 100.00%
Bootstrap support for afum|EAL86697.1 as seed ortholog is 100%.
Bootstrap support for afis|EAW17279.1 as seed ortholog is 100%.

---

Group of orthologs #7512. Best score 446 bits
Score difference with first non-orthologous sequence - *A. fumigatus*:446   *A. fischeri*:446
afum|EAL88945.1 100.00% afis|EAW16322.1 100.00%
Bootstrap support for afum|EAL88945.1 as seed ortholog is 100%.
Bootstrap support for afis|EAW16322.1 as seed ortholog is 100%.

---

Group of orthologs #7513. Best score 446 bits
Score difference with first non-orthologous sequence - *A. fumigatus*:446   *A. fischeri*:446
afum|EAL91253.1 100.00% afis|EAW17425.1 100.00%
Bootstrap support for afum|EAL91253.1 as seed ortholog is 100%.
Bootstrap support for afis|EAW17425.1 as seed ortholog is 100%.

---

Group of orthologs #7514. Best score 446 bits
Score difference with first non-orthologous sequence - *A. fumigatus*:446   *A. fischeri*:446
afum|EAL91296.1 100.00% afis|EAW17467.1 100.00%
Bootstrap support for afum|EAL91296.1 as seed ortholog is 100%.
Bootstrap support for afis|EAW17467.1 as seed ortholog is 100%.

---

Group of orthologs #7515. Best score 446 bits
Score difference with first non-orthologous sequence - *A. fumigatus*:446   *A. fischeri*:446
afum|EAL87247.1 100.00% afis|EAW23812.1 100.00%
Bootstrap support for afum|EAL87247.1 as seed ortholog is 100%.
Bootstrap support for afis|EAW23812.1 as seed ortholog is 100%.

---

Group of orthologs #7516. Best score 446 bits
Score difference with first non-orthologous sequence - *A. fumigatus*:446   *A. fischeri*:446
afum|EAL92834.1 100.00% afis|EAW21818.1 100.00%
Bootstrap support for afum|EAL92834.1 as seed ortholog is 100%.
Bootstrap support for afis|EAW21818.1 as seed ortholog is 100%.

---

Group of orthologs #7517. Best score 445 bits
Score difference with first non-orthologous sequence - *A. fumigatus*:139   *A. fischeri*:150
afum|EAL86843.1 100.00% afis|EAW23523.1 100.00%
afis|EAW16853.1 10.20%
afis|EAW15552.1 7.28%
Bootstrap support for afum|EAL86843.1 as seed ortholog is 90%.
Bootstrap support for afis|EAW23523.1 as seed ortholog is 97%.

---

Group of orthologs #7518. Best score 445 bits

Score difference with first non-orthologous sequence - *A. fumigatus*:445   *A. fischeri*:445
afum|EAL86590.1   100.00%   afis|EAW17130.1   100.00%
Bootstrap support for afum|EAL86590.1 as seed ortholog is 100%.
Bootstrap support for afis|EAW17130.1 as seed ortholog is 100%.

---

Group of orthologs #7519. Best score 445 bits
Score difference with first non-orthologous sequence - *A. fumigatus*:445   *A. fischeri*:445
afum|EAL85335.1   100.00%   afis|EAW20221.1   100.00%
Bootstrap support for afum|EAL85335.1 as seed ortholog is 100%.
Bootstrap support for afis|EAW20221.1 as seed ortholog is 100%.

---

Group of orthologs #7520. Best score 445 bits
Score difference with first non-orthologous sequence - *A. fumigatus*:445   *A. fischeri*:445
afum|EAL91333.1   100.00%   afis|EAW17515.1   100.00%
Bootstrap support for afum|EAL91333.1 as seed ortholog is 100%.
Bootstrap support for afis|EAW17515.1 as seed ortholog is 100%.

---

Group of orthologs #7521. Best score 445 bits
Score difference with first non-orthologous sequence - *A. fumigatus*:445   *A. fischeri*:445
afum|EAL90501.1   100.00%   afis|EAW22691.1   100.00%
Bootstrap support for afum|EAL90501.1 as seed ortholog is 100%.
Bootstrap support for afis|EAW22691.1 as seed ortholog is 100%.

---

Group of orthologs #7522. Best score 445 bits
Score difference with first non-orthologous sequence - *A. fumigatus*:223   *A. fischeri*:128
afum|EAL89025.2   100.00%   afis|EAW24555.1   100.00%
Bootstrap support for afum|EAL89025.2 as seed ortholog is 100%.
Bootstrap support for afis|EAW24555.1 as seed ortholog is 99%.

---

Group of orthologs #7523. Best score 445 bits
Score difference with first non-orthologous sequence - *A. fumigatus*:358   *A. fischeri*:238
afum|EAL92037.1   100.00%   afis|EAW24175.1   100.00%
Bootstrap support for afum|EAL92037.1 as seed ortholog is 100%.
Bootstrap support for afis|EAW24175.1 as seed ortholog is 100%.

---

Group of orthologs #7524. Best score 444 bits
Score difference with first non-orthologous sequence - *A. fumigatus*:444   *A. fischeri*:444
afum|EAL85984.1   100.00%   afis|EAW24316.1   100.00%
Bootstrap support for afum|EAL85984.1 as seed ortholog is 100%.
Bootstrap support for afis|EAW24316.1 as seed ortholog is 100%.

---

Group of orthologs #7525. Best score 444 bits
Score difference with first non-orthologous sequence - *A. fumigatus*:444   *A. fischeri*:444
afum|EAL86108.1   100.00%   afis|EAW24440.1   100.00%
Bootstrap support for afum|EAL86108.1 as seed ortholog is 100%.
Bootstrap support for afis|EAW24440.1 as seed ortholog is 100%.

---

Group of orthologs #7526. Best score 444 bits
Score difference with first non-orthologous sequence - *A. fumigatus*:444   *A. fischeri*:444
afum|EAL93529.1   100.00%   afis|EAW18793.1   100.00%
Bootstrap support for afum|EAL93529.1 as seed ortholog is 100%.
Bootstrap support for afis|EAW18793.1 as seed ortholog is 100%.

---

Group of orthologs #7527. Best score 443 bits
Score difference with first non-orthologous sequence - *A. fumigatus*:443   *A. fischeri*:443
afum|EAL85758.1   100.00%   afis|EAW15576.1   100.00%

Bootstrap support for afum|EAL85758.1 as seed ortholog is 100%.
Bootstrap support for afis|EAW15576.1 as seed ortholog is 100%.

---

Group of orthologs #7528. Best score 443 bits
Score difference with first non-orthologous sequence - *A. fumigatus*:443 *A. fischeri*:443
afum|EAL86335.1 100.00% afis|EAW16764.1 100.00%
Bootstrap support for afum|EAL86335.1 as seed ortholog is 100%.
Bootstrap support for afis|EAW16764.1 as seed ortholog is 100%.

---

Group of orthologs #7529. Best score 443 bits
Score difference with first non-orthologous sequence - *A. fumigatus*:443 *A. fischeri*:443
afum|EAL85254.1 100.00% afis|EAW20139.1 100.00%
Bootstrap support for afum|EAL85254.1 as seed ortholog is 100%.
Bootstrap support for afis|EAW20139.1 as seed ortholog is 100%.

---

Group of orthologs #7530. Best score 443 bits
Score difference with first non-orthologous sequence - *A. fumigatus*:443 *A. fischeri*:443
afum|EAL85476.1 100.00% afis|EAW20359.1 100.00%
Bootstrap support for afum|EAL85476.1 as seed ortholog is 100%.
Bootstrap support for afis|EAW20359.1 as seed ortholog is 100%.

---

Group of orthologs #7531. Best score 443 bits
Score difference with first non-orthologous sequence - *A. fumigatus*:443 *A. fischeri*:443
afum|EAL86812.1 100.00% afis|EAW19715.1 100.00%
Bootstrap support for afum|EAL86812.1 as seed ortholog is 100%.
Bootstrap support for afis|EAW19715.1 as seed ortholog is 100%.

---

Group of orthologs #7532. Best score 443 bits
Score difference with first non-orthologous sequence - *A. fumigatus*:443 *A. fischeri*:443
afum|EAL91427.1 100.00% afis|EAW17614.1 100.00%
Bootstrap support for afum|EAL91427.1 as seed ortholog is 100%.
Bootstrap support for afis|EAW17614.1 as seed ortholog is 100%.

---

Group of orthologs #7533. Best score 443 bits
Score difference with first non-orthologous sequence - *A. fumigatus*:443 *A. fischeri*:443
afum|EAL86071.1 100.00% afis|EAW24400.1 100.00%
Bootstrap support for afum|EAL86071.1 as seed ortholog is 100%.
Bootstrap support for afis|EAW24400.1 as seed ortholog is 100%.

---

Group of orthologs #7534. Best score 443 bits
Score difference with first non-orthologous sequence - *A. fumigatus*:443 *A. fischeri*:443
afum|EAL88398.1 100.00% afis|EAW23081.1 100.00%
Bootstrap support for afum|EAL88398.1 as seed ortholog is 100%.
Bootstrap support for afis|EAW23081.1 as seed ortholog is 100%.

---

Group of orthologs #7535. Best score 443 bits
Score difference with first non-orthologous sequence - *A. fumigatus*:443 *A. fischeri*:443
afum|EAL94034.1 100.00% afis|EAW19317.1 100.00%
Bootstrap support for afum|EAL94034.1 as seed ortholog is 100%.
Bootstrap support for afis|EAW19317.1 as seed ortholog is 100%.

---

Group of orthologs #7536. Best score 443 bits
Score difference with first non-orthologous sequence - *A. fumigatus*:443 *A. fischeri*:443
afum|EAL90673.1 100.00% afis|EAW22520.1 100.00%
Bootstrap support for afum|EAL90673.1 as seed ortholog is 100%.
Bootstrap support for afis|EAW22520.1 as seed ortholog is 100%.

Group of orthologs #7537. Best score 443 bits
Score difference with first non-orthologous sequence - *A. fumigatus*:443 *A. fischeri*:443
afum|EAL92540.1 100.00% afis|EAW21519.1 100.00%
Bootstrap support for afum|EAL92540.1 as seed ortholog is 100%.
Bootstrap support for afis|EAW21519.1 as seed ortholog is 100%.

---

Group of orthologs #7538. Best score 442 bits
Score difference with first non-orthologous sequence - *A. fumigatus*:442 *A. fischeri*:442
afum|EAL85517.1 100.00% afis|EAW15835.1 100.00%
Bootstrap support for afum|EAL85517.1 as seed ortholog is 100%.
Bootstrap support for afis|EAW15835.1 as seed ortholog is 100%.

---

Group of orthologs #7539. Best score 442 bits
Score difference with first non-orthologous sequence - *A. fumigatus*:442 *A. fischeri*:442
afum|EAL92250.1 100.00% afis|EAW21238.1 100.00%
Bootstrap support for afum|EAL92250.1 as seed ortholog is 100%.
Bootstrap support for afis|EAW21238.1 as seed ortholog is 100%.

---

Group of orthologs #7540. Best score 442 bits
Score difference with first non-orthologous sequence - *A. fumigatus*:442 *A. fischeri*:442
afum|EAL92375.1 100.00% afis|EAW21359.1 100.00%
Bootstrap support for afum|EAL92375.1 as seed ortholog is 100%.
Bootstrap support for afis|EAW21359.1 as seed ortholog is 100%.

---

Group of orthologs #7541. Best score 442 bits
Score difference with first non-orthologous sequence - *A. fumigatus*:442 *A. fischeri*:442
afum|EAL89975.1 100.00% afis|EAW25314.1 100.00%
Bootstrap support for afum|EAL89975.1 as seed ortholog is 100%.
Bootstrap support for afis|EAW25314.1 as seed ortholog is 100%.

---

Group of orthologs #7542. Best score 441 bits
Score difference with first non-orthologous sequence - *A. fumigatus*:441 *A. fischeri*:441
afum|EAL85597.1 100.00% afis|EAW15752.1 100.00%
Bootstrap support for afum|EAL85597.1 as seed ortholog is 100%.
Bootstrap support for afis|EAW15752.1 as seed ortholog is 100%.

---

Group of orthologs #7543. Best score 441 bits
Score difference with first non-orthologous sequence - *A. fumigatus*:441 *A. fischeri*:441
afum|EAL89141.1 100.00% afis|EAW16543.1 100.00%
Bootstrap support for afum|EAL89141.1 as seed ortholog is 100%.
Bootstrap support for afis|EAW16543.1 as seed ortholog is 100%.

---

Group of orthologs #7544. Best score 441 bits
Score difference with first non-orthologous sequence - *A. fumigatus*:441 *A. fischeri*:394
afum|EAL87136.1 100.00% afis|EAW23693.1 100.00%
Bootstrap support for afum|EAL87136.1 as seed ortholog is 100%.
Bootstrap support for afis|EAW23693.1 as seed ortholog is 100%.

---

Group of orthologs #7545. Best score 441 bits
Score difference with first non-orthologous sequence - *A. fumigatus*:441 *A. fischeri*:441
afum|EBA27319.1 100.00% afis|EAW18234.1 100.00%
Bootstrap support for afum|EBA27319.1 as seed ortholog is 100%.
Bootstrap support for afis|EAW18234.1 as seed ortholog is 100%.

---

Group of orthologs #7546. Best score 441 bits

Score difference with first non-orthologous sequence - *A. fumigatus*:441   *A. fischeri*:441
afum|EAL89425.1   100.00%   afis|EAW24786.1   100.00%
Bootstrap support for afum|EAL89425.1 as seed ortholog is 100%.
Bootstrap support for afis|EAW24786.1 as seed ortholog is 100%.

---

Group of orthologs #7547. Best score 440 bits
Score difference with first non-orthologous sequence - *A. fumigatus*:381   *A. fischeri*:386
afum|EAL84190.1   100.00%   afis|EAW15455.1   100.00%
Bootstrap support for afum|EAL84190.1 as seed ortholog is 100%.
Bootstrap support for afis|EAW15455.1 as seed ortholog is 100%.

---

Group of orthologs #7548. Best score 440 bits
Score difference with first non-orthologous sequence - *A. fumigatus*:440   *A. fischeri*:440
afum|EAL86333.1   100.00%   afis|EAW16762.1   100.00%
Bootstrap support for afum|EAL86333.1 as seed ortholog is 100%.
Bootstrap support for afis|EAW16762.1 as seed ortholog is 100%.

---

Group of orthologs #7549. Best score 440 bits
Score difference with first non-orthologous sequence - *A. fumigatus*:440   *A. fischeri*:440
afum|EAL85295.1   100.00%   afis|EAW20177.1   100.00%
Bootstrap support for afum|EAL85295.1 as seed ortholog is 100%.
Bootstrap support for afis|EAW20177.1 as seed ortholog is 100%.

---

Group of orthologs #7550. Best score 440 bits
Score difference with first non-orthologous sequence - *A. fumigatus*:440   *A. fischeri*:440
afum|EAL93727.1   100.00%   afis|EAW18991.1   100.00%
Bootstrap support for afum|EAL93727.1 as seed ortholog is 100%.
Bootstrap support for afis|EAW18991.1 as seed ortholog is 100%.

---

Group of orthologs #7551. Best score 440 bits
Score difference with first non-orthologous sequence - *A. fumigatus*:440   *A. fischeri*:440
afum|EAL94002.1   100.00%   afis|EAW19276.1   100.00%
Bootstrap support for afum|EAL94002.1 as seed ortholog is 100%.
Bootstrap support for afis|EAW19276.1 as seed ortholog is 100%.

---

Group of orthologs #7552. Best score 440 bits
Score difference with first non-orthologous sequence - *A. fumigatus*:440   *A. fischeri*:440
afum|EAL92664.1   100.00%   afis|EAW21630.1   100.00%
Bootstrap support for afum|EAL92664.1 as seed ortholog is 100%.
Bootstrap support for afis|EAW21630.1 as seed ortholog is 100%.

---

Group of orthologs #7553. Best score 439 bits
Score difference with first non-orthologous sequence - *A. fumigatus*:439   *A. fischeri*:439
afum|EAL85577.1   100.00%   afis|EAW15771.1   100.00%
Bootstrap support for afum|EAL85577.1 as seed ortholog is 100%.
Bootstrap support for afis|EAW15771.1 as seed ortholog is 100%.

---

Group of orthologs #7554. Best score 439 bits
Score difference with first non-orthologous sequence - *A. fumigatus*:439   *A. fischeri*:439
afum|EAL93153.1   100.00%   afis|EAW18467.1   100.00%
Bootstrap support for afum|EAL93153.1 as seed ortholog is 100%.
Bootstrap support for afis|EAW18467.1 as seed ortholog is 100%.

---

Group of orthologs #7555. Best score 438 bits
Score difference with first non-orthologous sequence - *A. fumigatus*:438   *A. fischeri*:438
afum|EAL85574.1   100.00%   afis|EAW15775.1   100.00%

Bootstrap support for afum|EAL85574.1 as seed ortholog is 100%.
Bootstrap support for afis|EAW15775.1 as seed ortholog is 100%.

---

Group of orthologs #7556. Best score 438 bits
Score difference with first non-orthologous sequence - *A. fumigatus*:438 *A. fischeri*:438
afum|EAL90251.1 100.00% afis|EAW22946.1 100.00%
Bootstrap support for afum|EAL90251.1 as seed ortholog is 100%.
Bootstrap support for afis|EAW22946.1 as seed ortholog is 100%.

---

Group of orthologs #7557. Best score 438 bits
Score difference with first non-orthologous sequence - *A. fumigatus*:173 *A. fischeri*:273
afum|EBA27291.1 100.00% afis|EAW19707.1 100.00%
Bootstrap support for afum|EBA27291.1 as seed ortholog is 100%.
Bootstrap support for afis|EAW19707.1 as seed ortholog is 100%.

---

Group of orthologs #7558. Best score 438 bits
Score difference with first non-orthologous sequence - *A. fumigatus*:438 *A. fischeri*:397
afum|EAL89384.1 100.00% afis|EAW24737.1 100.00%
Bootstrap support for afum|EAL89384.1 as seed ortholog is 100%.
Bootstrap support for afis|EAW24737.1 as seed ortholog is 100%.

---

Group of orthologs #7559. Best score 437 bits
Score difference with first non-orthologous sequence - *A. fumigatus*:437 *A. fischeri*:437
afum|EAL85705.1 100.00% afis|EAW15629.1 100.00%
Bootstrap support for afum|EAL85705.1 as seed ortholog is 100%.
Bootstrap support for afis|EAW15629.1 as seed ortholog is 100%.

---

Group of orthologs #7560. Best score 437 bits
Score difference with first non-orthologous sequence - *A. fumigatus*:437 *A. fischeri*:437
afum|EAL84923.1 100.00% afis|EAW19994.1 100.00%
Bootstrap support for afum|EAL84923.1 as seed ortholog is 100%.
Bootstrap support for afis|EAW19994.1 as seed ortholog is 100%.

---

Group of orthologs #7561. Best score 437 bits
Score difference with first non-orthologous sequence - *A. fumigatus*:437 *A. fischeri*:437
afum|EAL87555.1 100.00% afis|EAW18196.1 100.00%
Bootstrap support for afum|EAL87555.1 as seed ortholog is 100%.
Bootstrap support for afis|EAW18196.1 as seed ortholog is 100%.

---

Group of orthologs #7562. Best score 437 bits
Score difference with first non-orthologous sequence - *A. fumigatus*:437 *A. fischeri*:437
afum|EAL85494.1 100.00% afis|EAW20381.1 100.00%
Bootstrap support for afum|EAL85494.1 as seed ortholog is 100%.
Bootstrap support for afis|EAW20381.1 as seed ortholog is 100%.

---

Group of orthologs #7563. Best score 437 bits
Score difference with first non-orthologous sequence - *A. fumigatus*:437 *A. fischeri*:285
afum|EAL91633.1 100.00% afis|EAW17823.1 100.00%
Bootstrap support for afum|EAL91633.1 as seed ortholog is 100%.
Bootstrap support for afis|EAW17823.1 as seed ortholog is 100%.

---

Group of orthologs #7564. Best score 437 bits
Score difference with first non-orthologous sequence - *A. fumigatus*:437 *A. fischeri*:437
afum|EAL86084.1 100.00% afis|EAW24415.1 100.00%
Bootstrap support for afum|EAL86084.1 as seed ortholog is 100%.
Bootstrap support for afis|EAW24415.1 as seed ortholog is 100%.

---

Group of orthologs #7565. Best score 437 bits
Score difference with first non-orthologous sequence - *A. fumigatus*:437 *A. fischeri*:437
afum|EAL87234.2 100.00% afis|EAW23799.1 100.00%
Bootstrap support for afum|EAL87234.2 as seed ortholog is 100%.
Bootstrap support for afis|EAW23799.1 as seed ortholog is 100%.

---

Group of orthologs #7566. Best score 437 bits
Score difference with first non-orthologous sequence - *A. fumigatus*:437 *A. fischeri*:379
afum|EAL87904.1 100.00% afis|EAW23564.1 100.00%
Bootstrap support for afum|EAL87904.1 as seed ortholog is 100%.
Bootstrap support for afis|EAW23564.1 as seed ortholog is 100%.

---

Group of orthologs #7567. Best score 437 bits
Score difference with first non-orthologous sequence - *A. fumigatus*:437 *A. fischeri*:437
afum|EAL94049.1 100.00% afis|EAW19332.1 100.00%
Bootstrap support for afum|EAL94049.1 as seed ortholog is 100%.
Bootstrap support for afis|EAW19332.1 as seed ortholog is 100%.

---

Group of orthologs #7568. Best score 437 bits
Score difference with first non-orthologous sequence - *A. fumigatus*:437 *A. fischeri*:437
afum|EAL90552.2 100.00% afis|EAW22640.1 100.00%
Bootstrap support for afum|EAL90552.2 as seed ortholog is 100%.
Bootstrap support for afis|EAW22640.1 as seed ortholog is 100%.

---

Group of orthologs #7569. Best score 437 bits
Score difference with first non-orthologous sequence - *A. fumigatus*:437 *A. fischeri*:437
afum|EAL89424.1 100.00% afis|EAW24785.1 100.00%
Bootstrap support for afum|EAL89424.1 as seed ortholog is 100%.
Bootstrap support for afis|EAW24785.1 as seed ortholog is 100%.

---

Group of orthologs #7570. Best score 437 bits
Score difference with first non-orthologous sequence - *A. fumigatus*:437 *A. fischeri*:437
afum|EBA27398.1 100.00% afis|EAW25531.1 100.00%
Bootstrap support for afum|EBA27398.1 as seed ortholog is 100%.
Bootstrap support for afis|EAW25531.1 as seed ortholog is 100%.

---

Group of orthologs #7571. Best score 436 bits
Score difference with first non-orthologous sequence - *A. fumigatus*:436 *A. fischeri*:436
afum|EAL86475.1 100.00% afis|EAW16948.1 100.00%
Bootstrap support for afum|EAL86475.1 as seed ortholog is 100%.
Bootstrap support for afis|EAW16948.1 as seed ortholog is 100%.

---

Group of orthologs #7572. Best score 436 bits
Score difference with first non-orthologous sequence - *A. fumigatus*:436 *A. fischeri*:436
afum|EAL89156.1 100.00% afis|EAW16558.1 100.00%
Bootstrap support for afum|EAL89156.1 as seed ortholog is 100%.
Bootstrap support for afis|EAW16558.1 as seed ortholog is 100%.

---

Group of orthologs #7573. Best score 436 bits
Score difference with first non-orthologous sequence - *A. fumigatus*:436 *A. fischeri*:436
afum|EAL93036.1 100.00% afis|EAW18363.1 100.00%
Bootstrap support for afum|EAL93036.1 as seed ortholog is 100%.
Bootstrap support for afis|EAW18363.1 as seed ortholog is 100%.

---

Group of orthologs #7574. Best score 436 bits

Score difference with first non-orthologous sequence - *A. fumigatus*:436 *A. fischeri*:436
afum|EAL93561.1 100.00% afis|EAW18827.1 100.00%
Bootstrap support for afum|EAL93561.1 as seed ortholog is 100%.
Bootstrap support for afis|EAW18827.1 as seed ortholog is 100%.

---

Group of orthologs #7575. Best score 436 bits
Score difference with first non-orthologous sequence - *A. fumigatus*:436 *A. fischeri*:436
afum|EAL92447.1 100.00% afis|EAW21432.1 100.00%
Bootstrap support for afum|EAL92447.1 as seed ortholog is 100%.
Bootstrap support for afis|EAW21432.1 as seed ortholog is 100%.

---

Group of orthologs #7576. Best score 435 bits
Score difference with first non-orthologous sequence - *A. fumigatus*:435 *A. fischeri*:435
afum|EAL85711.1 100.00% afis|EAW15623.1 100.00%
Bootstrap support for afum|EAL85711.1 as seed ortholog is 100%.
Bootstrap support for afis|EAW15623.1 as seed ortholog is 100%.

---

Group of orthologs #7577. Best score 435 bits
Score difference with first non-orthologous sequence - *A. fumigatus*:435 *A. fischeri*:435
afum|EAL86338.1 100.00% afis|EAW16770.1 100.00%
Bootstrap support for afum|EAL86338.1 as seed ortholog is 100%.
Bootstrap support for afis|EAW16770.1 as seed ortholog is 100%.

---

Group of orthologs #7578. Best score 435 bits
Score difference with first non-orthologous sequence - *A. fumigatus*:435 *A. fischeri*:435
afum|EAL88859.1 100.00% afis|EAW16238.1 100.00%
Bootstrap support for afum|EAL88859.1 as seed ortholog is 100%.
Bootstrap support for afis|EAW16238.1 as seed ortholog is 100%.

---

Group of orthologs #7579. Best score 435 bits
Score difference with first non-orthologous sequence - *A. fumigatus*:435 *A. fischeri*:435
afum|EAL84737.2 100.00% afis|EAW20896.1 100.00%
Bootstrap support for afum|EAL84737.2 as seed ortholog is 100%.
Bootstrap support for afis|EAW20896.1 as seed ortholog is 100%.

---

Group of orthologs #7580. Best score 435 bits
Score difference with first non-orthologous sequence - *A. fumigatus*:435 *A. fischeri*:435
afum|EAL90647.1 100.00% afis|EAW22548.1 100.00%
Bootstrap support for afum|EAL90647.1 as seed ortholog is 100%.
Bootstrap support for afis|EAW22548.1 as seed ortholog is 100%.

---

Group of orthologs #7581. Best score 434 bits
Score difference with first non-orthologous sequence - *A. fumigatus*:434 *A. fischeri*:434
afum|EAL85610.1 100.00% afis|EAW15738.1 100.00%
Bootstrap support for afum|EAL85610.1 as seed ortholog is 100%.
Bootstrap support for afis|EAW15738.1 as seed ortholog is 100%.

---

Group of orthologs #7582. Best score 434 bits
Score difference with first non-orthologous sequence - *A. fumigatus*:434 *A. fischeri*:434
afum|EAL85509.1 100.00% afis|EAW15842.1 100.00%
Bootstrap support for afum|EAL85509.1 as seed ortholog is 100%.
Bootstrap support for afis|EAW15842.1 as seed ortholog is 100%.

---

Group of orthologs #7583. Best score 434 bits
Score difference with first non-orthologous sequence - *A. fumigatus*:434 *A. fischeri*:434
afum|EAL88503.1 100.00% afis|EAW15869.1 100.00%

Bootstrap support for afum|EAL88503.1 as seed ortholog is 100%.
Bootstrap support for afis|EAW15869.1 as seed ortholog is 100%.

---

Group of orthologs #7584. Best score 434 bits
Score difference with first non-orthologous sequence - *A. fumigatus*:434 *A. fischeri*:434
afum|EAL88908.1 100.00% afis|EAW16286.1 100.00%
Bootstrap support for afum|EAL88908.1 as seed ortholog is 100%.
Bootstrap support for afis|EAW16286.1 as seed ortholog is 100%.

---

Group of orthologs #7585. Best score 434 bits
Score difference with first non-orthologous sequence - *A. fumigatus*:434 *A. fischeri*:434
afum|EAL84795.1 100.00% afis|EAW20838.1 100.00%
Bootstrap support for afum|EAL84795.1 as seed ortholog is 100%.
Bootstrap support for afis|EAW20838.1 as seed ortholog is 100%.

---

Group of orthologs #7586. Best score 434 bits
Score difference with first non-orthologous sequence - *A. fumigatus*:434 *A. fischeri*:434
afum|EAL92277.1 100.00% afis|EAW21262.1 100.00%
Bootstrap support for afum|EAL92277.1 as seed ortholog is 100%.
Bootstrap support for afis|EAW21262.1 as seed ortholog is 100%.

---

Group of orthologs #7587. Best score 434 bits
Score difference with first non-orthologous sequence - *A. fumigatus*:434 *A. fischeri*:306
afum|EAL92899.1 100.00% afis|EAW21885.1 100.00%
Bootstrap support for afum|EAL92899.1 as seed ortholog is 100%.
Bootstrap support for afis|EAW21885.1 as seed ortholog is 100%.

---

Group of orthologs #7588. Best score 433 bits
Score difference with first non-orthologous sequence - *A. fumigatus*:433 *A. fischeri*:433
afum|EAL90700.1 100.00% afis|EAW22493.1 100.00%
Bootstrap support for afum|EAL90700.1 as seed ortholog is 100%.
Bootstrap support for afis|EAW22493.1 as seed ortholog is 100%.

---

Group of orthologs #7589. Best score 433 bits
Score difference with first non-orthologous sequence - *A. fumigatus*:433 *A. fischeri*:433
afum|EAL92247.1 100.00% afis|EAW21235.1 100.00%
Bootstrap support for afum|EAL92247.1 as seed ortholog is 100%.
Bootstrap support for afis|EAW21235.1 as seed ortholog is 100%.

---

Group of orthologs #7590. Best score 432 bits
Score difference with first non-orthologous sequence - *A. fumigatus*:432 *A. fischeri*:432
afum|EAL85035.1 100.00% afis|EAW19860.1 100.00%
Bootstrap support for afum|EAL85035.1 as seed ortholog is 100%.
Bootstrap support for afis|EAW19860.1 as seed ortholog is 100%.

---

Group of orthologs #7591. Best score 432 bits
Score difference with first non-orthologous sequence - *A. fumigatus*:432 *A. fischeri*:432
afum|EAL84419.1 100.00% afis|EAW24043.1 100.00%
Bootstrap support for afum|EAL84419.1 as seed ortholog is 100%.
Bootstrap support for afis|EAW24043.1 as seed ortholog is 100%.

---

Group of orthologs #7592. Best score 432 bits
Score difference with first non-orthologous sequence - *A. fumigatus*:432 *A. fischeri*:432
afum|EAL88464.1 100.00% afis|EAW23017.1 100.00%
Bootstrap support for afum|EAL88464.1 as seed ortholog is 100%.
Bootstrap support for afis|EAW23017.1 as seed ortholog is 100%.

Group of orthologs #7593. Best score 431 bits
Score difference with first non-orthologous sequence - *A. fumigatus*:431 *A. fischeri*:431
afum|EAL84854.1 100.00% afis|EAW20765.1 100.00%
Bootstrap support for afum|EAL84854.1 as seed ortholog is 100%.
Bootstrap support for afis|EAW20765.1 as seed ortholog is 100%.

---

Group of orthologs #7594. Best score 431 bits
Score difference with first non-orthologous sequence - *A. fumigatus*:221 *A. fischeri*:139
afum|EAL86960.1 100.00% afis|EAW19549.1 100.00%
Bootstrap support for afum|EAL86960.1 as seed ortholog is 100%.
Bootstrap support for afis|EAW19549.1 as seed ortholog is 99%.

---

Group of orthologs #7595. Best score 431 bits
Score difference with first non-orthologous sequence - *A. fumigatus*:431 *A. fischeri*:431
afum|EAL88346.1 100.00% afis|EAW23135.1 100.00%
Bootstrap support for afum|EAL88346.1 as seed ortholog is 100%.
Bootstrap support for afis|EAW23135.1 as seed ortholog is 100%.

---

Group of orthologs #7596. Best score 431 bits
Score difference with first non-orthologous sequence - *A. fumigatus*:431 *A. fischeri*:431
afum|EBA27431.1 100.00% afis|EAW17568.1 100.00%
Bootstrap support for afum|EBA27431.1 as seed ortholog is 100%.
Bootstrap support for afis|EAW17568.1 as seed ortholog is 100%.

---

Group of orthologs #7597. Best score 431 bits
Score difference with first non-orthologous sequence - *A. fumigatus*:431 *A. fischeri*:431
afum|EAL91043.1 100.00% afis|EAW22147.1 100.00%
Bootstrap support for afum|EAL91043.1 as seed ortholog is 100%.
Bootstrap support for afis|EAW22147.1 as seed ortholog is 100%.

---

Group of orthologs #7598. Best score 431 bits
Score difference with first non-orthologous sequence - *A. fumigatus*:431 *A. fischeri*:431
afum|EAL92036.2 100.00% afis|EAW24174.1 100.00%
Bootstrap support for afum|EAL92036.2 as seed ortholog is 100%.
Bootstrap support for afis|EAW24174.1 as seed ortholog is 100%.

---

Group of orthologs #7599. Best score 430 bits
Score difference with first non-orthologous sequence - *A. fumigatus*:430 *A. fischeri*:390
afum|EAL85741.1 100.00% afis|EAW15591.1 100.00%
Bootstrap support for afum|EAL85741.1 as seed ortholog is 100%.
Bootstrap support for afis|EAW15591.1 as seed ortholog is 100%.

---

Group of orthologs #7600. Best score 430 bits
Score difference with first non-orthologous sequence - *A. fumigatus*:430 *A. fischeri*:430
afum|EAL86584.1 100.00% afis|EAW17123.1 100.00%
Bootstrap support for afum|EAL86584.1 as seed ortholog is 100%.
Bootstrap support for afis|EAW17123.1 as seed ortholog is 100%.

---

Group of orthologs #7601. Best score 430 bits
Score difference with first non-orthologous sequence - *A. fumigatus*:430 *A. fischeri*:430
afum|EAL84465.1 100.00% afis|EAW20596.1 100.00%
Bootstrap support for afum|EAL84465.1 as seed ortholog is 100%.
Bootstrap support for afis|EAW20596.1 as seed ortholog is 100%.

---

Group of orthologs #7602. Best score 430 bits

Score difference with first non-orthologous sequence - *A. fumigatus*:430 *A. fischeri*:430
afum|EAL84332.1 100.00% afis|EAW25646.1 100.00%
Bootstrap support for afum|EAL84332.1 as seed ortholog is 100%.
Bootstrap support for afis|EAW25646.1 as seed ortholog is 100%.

---

Group of orthologs #7603. Best score 430 bits
Score difference with first non-orthologous sequence - *A. fumigatus*:430 *A. fischeri*:430
afum|EAL88125.2 100.00% afis|EAW23351.1 100.00%
Bootstrap support for afum|EAL88125.2 as seed ortholog is 100%.
Bootstrap support for afis|EAW23351.1 as seed ortholog is 100%.

---

Group of orthologs #7604. Best score 430 bits
Score difference with first non-orthologous sequence - *A. fumigatus*:430 *A. fischeri*:430
afum|EAL88081.1 100.00% afis|EAW23390.1 100.00%
Bootstrap support for afum|EAL88081.1 as seed ortholog is 100%.
Bootstrap support for afis|EAW23390.1 as seed ortholog is 100%.

---

Group of orthologs #7605. Best score 429 bits
Score difference with first non-orthologous sequence - *A. fumigatus*:429 *A. fischeri*:429
afum|EAL89249.1 100.00% afis|EAW16645.1 100.00%
Bootstrap support for afum|EAL89249.1 as seed ortholog is 100%.
Bootstrap support for afis|EAW16645.1 as seed ortholog is 100%.

---

Group of orthologs #7606. Best score 429 bits
Score difference with first non-orthologous sequence - *A. fumigatus*:429 *A. fischeri*:429
afum|EAL86558.1 100.00% afis|EAW20707.1 100.00%
Bootstrap support for afum|EAL86558.1 as seed ortholog is 100%.
Bootstrap support for afis|EAW20707.1 as seed ortholog is 100%.

---

Group of orthologs #7607. Best score 429 bits
Score difference with first non-orthologous sequence - *A. fumigatus*:429 *A. fischeri*:429
afum|EAL91664.1 100.00% afis|EAW17856.1 100.00%
Bootstrap support for afum|EAL91664.1 as seed ortholog is 100%.
Bootstrap support for afis|EAW17856.1 as seed ortholog is 100%.

---

Group of orthologs #7608. Best score 429 bits
Score difference with first non-orthologous sequence - *A. fumigatus*:429 *A. fischeri*:429
afum|EAL87336.1 100.00% afis|EAW23908.1 100.00%
Bootstrap support for afum|EAL87336.1 as seed ortholog is 100%.
Bootstrap support for afis|EAW23908.1 as seed ortholog is 100%.

---

Group of orthologs #7609. Best score 429 bits
Score difference with first non-orthologous sequence - *A. fumigatus*:429 *A. fischeri*:429
afum|EAL89824.1 100.00% afis|EAW25149.1 100.00%
Bootstrap support for afum|EAL89824.1 as seed ortholog is 100%.
Bootstrap support for afis|EAW25149.1 as seed ortholog is 100%.

---

Group of orthologs #7610. Best score 429 bits
Score difference with first non-orthologous sequence - *A. fumigatus*:429 *A. fischeri*:429
afum|EAL90014.1 100.00% afis|EAW25351.1 100.00%
Bootstrap support for afum|EAL90014.1 as seed ortholog is 100%.
Bootstrap support for afis|EAW25351.1 as seed ortholog is 100%.

---

Group of orthologs #7611. Best score 428 bits
Score difference with first non-orthologous sequence - *A. fumigatus*:428 *A. fischeri*:428
afum|EAL85512.1 100.00% afis|EAW15839.1 100.00%

Bootstrap support for afum|EAL85512.1 as seed ortholog is 100%.
Bootstrap support for afis|EAW15839.1 as seed ortholog is 100%.

---

Group of orthologs #7612. Best score 428 bits
Score difference with first non-orthologous sequence - *A. fumigatus*:428 *A. fischeri*:428
afum|EAL86378.1 100.00% afis|EAW16830.1 100.00%
Bootstrap support for afum|EAL86378.1 as seed ortholog is 100%.
Bootstrap support for afis|EAW16830.1 as seed ortholog is 100%.

---

Group of orthologs #7613. Best score 428 bits
Score difference with first non-orthologous sequence - *A. fumigatus*:428 *A. fischeri*:428
afum|EAL85900.1 100.00% afis|EAW24231.1 100.00%
Bootstrap support for afum|EAL85900.1 as seed ortholog is 100%.
Bootstrap support for afis|EAW24231.1 as seed ortholog is 100%.

---

Group of orthologs #7614. Best score 428 bits
Score difference with first non-orthologous sequence - *A. fumigatus*:428 *A. fischeri*:428
afum|EAL91003.1 100.00% afis|EAW22187.1 100.00%
Bootstrap support for afum|EAL91003.1 as seed ortholog is 100%.
Bootstrap support for afis|EAW22187.1 as seed ortholog is 100%.

---

Group of orthologs #7615. Best score 428 bits
Score difference with first non-orthologous sequence - *A. fumigatus*:428 *A. fischeri*:428
afum|EAL90164.1 100.00% afis|EAW25498.1 100.00%
Bootstrap support for afum|EAL90164.1 as seed ortholog is 100%.
Bootstrap support for afis|EAW25498.1 as seed ortholog is 100%.

---

Group of orthologs #7616. Best score 427 bits
Score difference with first non-orthologous sequence - *A. fumigatus*:427 *A. fischeri*:427
afum|EAL88513.1 100.00% afis|EAW15879.1 100.00%
Bootstrap support for afum|EAL88513.1 as seed ortholog is 100%.
Bootstrap support for afis|EAW15879.1 as seed ortholog is 100%.

---

Group of orthologs #7617. Best score 427 bits
Score difference with first non-orthologous sequence - *A. fumigatus*:427 *A. fischeri*:427
afum|EAL91648.1 100.00% afis|EAW17838.1 100.00%
Bootstrap support for afum|EAL91648.1 as seed ortholog is 100%.
Bootstrap support for afis|EAW17838.1 as seed ortholog is 100%.

---

Group of orthologs #7618. Best score 427 bits
Score difference with first non-orthologous sequence - *A. fumigatus*:427 *A. fischeri*:427
afum|EAL93576.2 100.00% afis|EAW17531.1 100.00%
Bootstrap support for afum|EAL93576.2 as seed ortholog is 100%.
Bootstrap support for afis|EAW17531.1 as seed ortholog is 100%.

---

Group of orthologs #7619. Best score 427 bits
Score difference with first non-orthologous sequence - *A. fumigatus*:427 *A. fischeri*:427
afum|EAL87910.1 100.00% afis|EAW23556.1 100.00%
Bootstrap support for afum|EAL87910.1 as seed ortholog is 100%.
Bootstrap support for afis|EAW23556.1 as seed ortholog is 100%.

---

Group of orthologs #7620. Best score 427 bits
Score difference with first non-orthologous sequence - *A. fumigatus*:427 *A. fischeri*:427
afum|EAL87847.2 100.00% afis|EAW23615.1 100.00%
Bootstrap support for afum|EAL87847.2 as seed ortholog is 100%.
Bootstrap support for afis|EAW23615.1 as seed ortholog is 100%.

Group of orthologs #7621. Best score 427 bits
Score difference with first non-orthologous sequence - *A. fumigatus*:427 *A. fischeri*:427
afum|EAL93474.1 100.00% afis|EAW18748.1 100.00%
Bootstrap support for afum|EAL93474.1 as seed ortholog is 100%.
Bootstrap support for afis|EAW18748.1 as seed ortholog is 100%.

Group of orthologs #7622. Best score 427 bits
Score difference with first non-orthologous sequence - *A. fumigatus*:427 *A. fischeri*:427
afum|EAL92255.1 100.00% afis|EAW21242.1 100.00%
Bootstrap support for afum|EAL92255.1 as seed ortholog is 100%.
Bootstrap support for afis|EAW21242.1 as seed ortholog is 100%.

Group of orthologs #7623. Best score 426 bits
Score difference with first non-orthologous sequence - *A. fumigatus*:426 *A. fischeri*:426
afum|EAL85703.1 100.00% afis|EAW15634.1 100.00%
Bootstrap support for afum|EAL85703.1 as seed ortholog is 100%.
Bootstrap support for afis|EAW15634.1 as seed ortholog is 100%.

Group of orthologs #7624. Best score 426 bits
Score difference with first non-orthologous sequence - *A. fumigatus*:426 *A. fischeri*:426
afum|EAL88766.2 100.00% afis|EAW16136.1 100.00%
Bootstrap support for afum|EAL88766.2 as seed ortholog is 100%.
Bootstrap support for afis|EAW16136.1 as seed ortholog is 100%.

Group of orthologs #7625. Best score 426 bits
Score difference with first non-orthologous sequence - *A. fumigatus*:426 *A. fischeri*:426
afum|EAL84485.2 100.00% afis|EAW20579.1 100.00%
Bootstrap support for afum|EAL84485.2 as seed ortholog is 100%.
Bootstrap support for afis|EAW20579.1 as seed ortholog is 100%.

Group of orthologs #7626. Best score 426 bits
Score difference with first non-orthologous sequence - *A. fumigatus*:426 *A. fischeri*:353
afum|EAL84428.1 100.00% afis|EAW20645.1 100.00%
Bootstrap support for afum|EAL84428.1 as seed ortholog is 100%.
Bootstrap support for afis|EAW20645.1 as seed ortholog is 100%.

Group of orthologs #7627. Best score 426 bits
Score difference with first non-orthologous sequence - *A. fumigatus*:426 *A. fischeri*:426
afum|EAL93159.2 100.00% afis|EAW18472.1 100.00%
Bootstrap support for afum|EAL93159.2 as seed ortholog is 100%.
Bootstrap support for afis|EAW18472.1 as seed ortholog is 100%.

Group of orthologs #7628. Best score 426 bits
Score difference with first non-orthologous sequence - *A. fumigatus*:426 *A. fischeri*:426
afum|EAL92298.1 100.00% afis|EAW21281.1 100.00%
Bootstrap support for afum|EAL92298.1 as seed ortholog is 100%.
Bootstrap support for afis|EAW21281.1 as seed ortholog is 100%.

Group of orthologs #7629. Best score 426 bits
Score difference with first non-orthologous sequence - *A. fumigatus*:426 *A. fischeri*:426
afum|EAL89976.1 100.00% afis|EAW25315.1 100.00%
Bootstrap support for afum|EAL89976.1 as seed ortholog is 100%.
Bootstrap support for afis|EAW25315.1 as seed ortholog is 100%.

Group of orthologs #7630. Best score 426 bits

Score difference with first non-orthologous sequence - *A. fumigatus*:426 *A. fischeri*:426
afum|EAL91964.1 100.00% afis|EAW24110.1 100.00%
Bootstrap support for afum|EAL91964.1 as seed ortholog is 100%.
Bootstrap support for afis|EAW24110.1 as seed ortholog is 100%.

---

Group of orthologs #7631. Best score 425 bits
Score difference with first non-orthologous sequence - *A. fumigatus*:425 *A. fischeri*:425
afum|EAL93151.2 100.00% afis|EAW18464.1 100.00%
Bootstrap support for afum|EAL93151.2 as seed ortholog is 100%.
Bootstrap support for afis|EAW18464.1 as seed ortholog is 100%.

---

Group of orthologs #7632. Best score 425 bits
Score difference with first non-orthologous sequence - *A. fumigatus*:425 *A. fischeri*:425
afum|EAL94038.1 100.00% afis|EAW19320.1 100.00%
Bootstrap support for afum|EAL94038.1 as seed ortholog is 100%.
Bootstrap support for afis|EAW19320.1 as seed ortholog is 100%.

---

Group of orthologs #7633. Best score 425 bits
Score difference with first non-orthologous sequence - *A. fumigatus*:425 *A. fischeri*:425
afum|EAL90743.1 100.00% afis|EAW22448.1 100.00%
Bootstrap support for afum|EAL90743.1 as seed ortholog is 100%.
Bootstrap support for afis|EAW22448.1 as seed ortholog is 100%.

---

Group of orthologs #7634. Best score 425 bits
Score difference with first non-orthologous sequence - *A. fumigatus*:425 *A. fischeri*:425
afum|EAL90252.1 100.00% afis|EAW22945.1 100.00%
Bootstrap support for afum|EAL90252.1 as seed ortholog is 100%.
Bootstrap support for afis|EAW22945.1 as seed ortholog is 100%.

---

Group of orthologs #7635. Best score 425 bits
Score difference with first non-orthologous sequence - *A. fumigatus*:425 *A. fischeri*:425
afum|EAL92734.1 100.00% afis|EAW21700.1 100.00%
Bootstrap support for afum|EAL92734.1 as seed ortholog is 100%.
Bootstrap support for afis|EAW21700.1 as seed ortholog is 100%.

---

Group of orthologs #7636. Best score 425 bits
Score difference with first non-orthologous sequence - *A. fumigatus*:425 *A. fischeri*:425
afum|EAL92743.1 100.00% afis|EAW21708.1 100.00%
Bootstrap support for afum|EAL92743.1 as seed ortholog is 100%.
Bootstrap support for afis|EAW21708.1 as seed ortholog is 100%.

---

Group of orthologs #7637. Best score 424 bits
Score difference with first non-orthologous sequence - *A. fumigatus*:424 *A. fischeri*:424
afum|EAL88523.1 100.00% afis|EAW15889.1 100.00%
Bootstrap support for afum|EAL88523.1 as seed ortholog is 100%.
Bootstrap support for afis|EAW15889.1 as seed ortholog is 100%.

---

Group of orthologs #7638. Best score 424 bits
Score difference with first non-orthologous sequence - *A. fumigatus*:424 *A. fischeri*:424
afum|EAL84784.1 100.00% afis|EAW20849.1 100.00%
Bootstrap support for afum|EAL84784.1 as seed ortholog is 100%.
Bootstrap support for afis|EAW20849.1 as seed ortholog is 100%.

---

Group of orthologs #7639. Best score 424 bits
Score difference with first non-orthologous sequence - *A. fumigatus*:424 *A. fischeri*:424
afum|EAL88370.1 100.00% afis|EAW23109.1 100.00%

Bootstrap support for afum|EAL88370.1 as seed ortholog is 100%.
Bootstrap support for afis|EAW23109.1 as seed ortholog is 100%.

---

Group of orthologs #7640. Best score 424 bits
Score difference with first non-orthologous sequence - *A. fumigatus*:424 *A. fischeri*:367
afum|EAL93434.1 100.00% afis|EAW18711.1 100.00%
Bootstrap support for afum|EAL93434.1 as seed ortholog is 100%.
Bootstrap support for afis|EAW18711.1 as seed ortholog is 100%.

---

Group of orthologs #7641. Best score 424 bits
Score difference with first non-orthologous sequence - *A. fumigatus*:424 *A. fischeri*:424
afum|EAL92452.2 100.00% afis|EAW21437.1 100.00%
Bootstrap support for afum|EAL92452.2 as seed ortholog is 100%.
Bootstrap support for afis|EAW21437.1 as seed ortholog is 100%.

---

Group of orthologs #7642. Best score 424 bits
Score difference with first non-orthologous sequence - *A. fumigatus*:424 *A. fischeri*:424
afum|EAL89383.2 100.00% afis|EAW24736.1 100.00%
Bootstrap support for afum|EAL89383.2 as seed ortholog is 100%.
Bootstrap support for afis|EAW24736.1 as seed ortholog is 100%.

---

Group of orthologs #7643. Best score 424 bits
Score difference with first non-orthologous sequence - *A. fumigatus*:424 *A. fischeri*:424
afum|EAL92823.2 100.00% afis|EAW21798.1 100.00%
Bootstrap support for afum|EAL92823.2 as seed ortholog is 100%.
Bootstrap support for afis|EAW21798.1 as seed ortholog is 100%.

---

Group of orthologs #7644. Best score 424 bits
Score difference with first non-orthologous sequence - *A. fumigatus*:424 *A. fischeri*:424
afum|EAL89630.2 100.00% afis|EAW24971.1 100.00%
Bootstrap support for afum|EAL89630.2 as seed ortholog is 100%.
Bootstrap support for afis|EAW24971.1 as seed ortholog is 100%.

---

Group of orthologs #7645. Best score 423 bits
Score difference with first non-orthologous sequence - *A. fumigatus*:423 *A. fischeri*:423
afum|EAL88826.2 100.00% afis|EAW16195.1 100.00%
Bootstrap support for afum|EAL88826.2 as seed ortholog is 100%.
Bootstrap support for afis|EAW16195.1 as seed ortholog is 100%.

---

Group of orthologs #7646. Best score 423 bits
Score difference with first non-orthologous sequence - *A. fumigatus*:423 *A. fischeri*:423
afum|EAL84841.1 100.00% afis|EAW20782.1 100.00%
Bootstrap support for afum|EAL84841.1 as seed ortholog is 100%.
Bootstrap support for afis|EAW20782.1 as seed ortholog is 100%.

---

Group of orthologs #7647. Best score 423 bits
Score difference with first non-orthologous sequence - *A. fumigatus*:362 *A. fischeri*:423
afum|EAL86231.1 100.00% afis|EAW24568.1 100.00%
Bootstrap support for afum|EAL86231.1 as seed ortholog is 100%.
Bootstrap support for afis|EAW24568.1 as seed ortholog is 100%.

---

Group of orthologs #7648. Best score 423 bits
Score difference with first non-orthologous sequence - *A. fumigatus*:423 *A. fischeri*:423
afum|EAL87228.2 100.00% afis|EAW23794.1 100.00%
Bootstrap support for afum|EAL87228.2 as seed ortholog is 100%.
Bootstrap support for afis|EAW23794.1 as seed ortholog is 100%.

Group of orthologs #7649. Best score 423 bits
Score difference with first non-orthologous sequence - *A. fumigatus*:423 *A. fischeri*:423
afum|EAL88005.1 100.00% afis|EAW23461.1 100.00%
Bootstrap support for afum|EAL88005.1 as seed ortholog is 100%.
Bootstrap support for afis|EAW23461.1 as seed ortholog is 100%.

Group of orthologs #7650. Best score 423 bits
Score difference with first non-orthologous sequence - *A. fumigatus*:423 *A. fischeri*:423
afum|EAL93393.2 100.00% afis|EAW18673.1 100.00%
Bootstrap support for afum|EAL93393.2 as seed ortholog is 100%.
Bootstrap support for afis|EAW18673.1 as seed ortholog is 100%.

Group of orthologs #7651. Best score 423 bits
Score difference with first non-orthologous sequence - *A. fumigatus*:423 *A. fischeri*:423
afum|EAL92162.2 100.00% afis|EAW21143.1 100.00%
Bootstrap support for afum|EAL92162.2 as seed ortholog is 100%.
Bootstrap support for afis|EAW21143.1 as seed ortholog is 100%.

Group of orthologs #7652. Best score 423 bits
Score difference with first non-orthologous sequence - *A. fumigatus*:423 *A. fischeri*:423
afum|EAL89813.1 100.00% afis|EAW25141.1 100.00%
Bootstrap support for afum|EAL89813.1 as seed ortholog is 100%.
Bootstrap support for afis|EAW25141.1 as seed ortholog is 100%.

Group of orthologs #7653. Best score 422 bits
Score difference with first non-orthologous sequence - *A. fumigatus*:422 *A. fischeri*:422
afum|EAL89090.1 100.00% afis|EAW16496.1 100.00%
Bootstrap support for afum|EAL89090.1 as seed ortholog is 100%.
Bootstrap support for afis|EAW16496.1 as seed ortholog is 100%.

Group of orthologs #7654. Best score 422 bits
Score difference with first non-orthologous sequence - *A. fumigatus*:422 *A. fischeri*:422
afum|EAL88348.1 100.00% afis|EAW23133.1 100.00%
Bootstrap support for afum|EAL88348.1 as seed ortholog is 100%.
Bootstrap support for afis|EAW23133.1 as seed ortholog is 100%.

Group of orthologs #7655. Best score 422 bits
Score difference with first non-orthologous sequence - *A. fumigatus*:381 *A. fischeri*:422
afum|EAL92612.1 100.00% afis|EAW21594.1 100.00%
Bootstrap support for afum|EAL92612.1 as seed ortholog is 100%.
Bootstrap support for afis|EAW21594.1 as seed ortholog is 100%.

Group of orthologs #7656. Best score 422 bits
Score difference with first non-orthologous sequence - *A. fumigatus*:422 *A. fischeri*:422
afum|EAL90027.1 100.00% afis|EAW25367.1 100.00%
Bootstrap support for afum|EAL90027.1 as seed ortholog is 100%.
Bootstrap support for afis|EAW25367.1 as seed ortholog is 100%.

Group of orthologs #7657. Best score 421 bits
Score difference with first non-orthologous sequence - *A. fumigatus*:421 *A. fischeri*:421
afum|EAL91573.1 100.00% afis|EAW17763.1 100.00%
Bootstrap support for afum|EAL91573.1 as seed ortholog is 100%.
Bootstrap support for afis|EAW17763.1 as seed ortholog is 100%.

Group of orthologs #7658. Best score 421 bits

Score difference with first non-orthologous sequence - *A. fumigatus*:421 *A. fischeri*:421
afum|EAL86118.1 100.00% afis|EAW24449.1 100.00%
Bootstrap support for afum|EAL86118.1 as seed ortholog is 100%.
Bootstrap support for afis|EAW24449.1 as seed ortholog is 100%.

---

Group of orthologs #7659. Best score 420 bits
Score difference with first non-orthologous sequence - *A. fumigatus*:420 *A. fischeri*:420
afum|EAL85648.1 100.00% afis|EAW15700.1 100.00%
Bootstrap support for afum|EAL85648.1 as seed ortholog is 100%.
Bootstrap support for afis|EAW15700.1 as seed ortholog is 100%.

---

Group of orthologs #7660. Best score 420 bits
Score difference with first non-orthologous sequence - *A. fumigatus*:420 *A. fischeri*:420
afum|EAL85651.2 100.00% afis|EAW15697.1 100.00%
Bootstrap support for afum|EAL85651.2 as seed ortholog is 100%.
Bootstrap support for afis|EAW15697.1 as seed ortholog is 100%.

---

Group of orthologs #7661. Best score 420 bits
Score difference with first non-orthologous sequence - *A. fumigatus*:420 *A. fischeri*:420
afum|EAL87256.1 100.00% afis|EAW23821.1 100.00%
Bootstrap support for afum|EAL87256.1 as seed ortholog is 100%.
Bootstrap support for afis|EAW23821.1 as seed ortholog is 100%.

---

Group of orthologs #7662. Best score 420 bits
Score difference with first non-orthologous sequence - *A. fumigatus*:420 *A. fischeri*:420
afum|EAL93803.1 100.00% afis|EAW19075.1 100.00%
Bootstrap support for afum|EAL93803.1 as seed ortholog is 100%.
Bootstrap support for afis|EAW19075.1 as seed ortholog is 100%.

---

Group of orthologs #7663. Best score 420 bits
Score difference with first non-orthologous sequence - *A. fumigatus*:420 *A. fischeri*:420
afum|EBA27332.1 100.00% afis|EAW23449.1 100.00%
Bootstrap support for afum|EBA27332.1 as seed ortholog is 100%.
Bootstrap support for afis|EAW23449.1 as seed ortholog is 100%.

---

Group of orthologs #7664. Best score 419 bits
Score difference with first non-orthologous sequence - *A. fumigatus*:419 *A. fischeri*:419
afum|EAL91112.2 100.00% afis|EAW17303.1 100.00%
Bootstrap support for afum|EAL91112.2 as seed ortholog is 100%.
Bootstrap support for afis|EAW17303.1 as seed ortholog is 100%.

---

Group of orthologs #7665. Best score 419 bits
Score difference with first non-orthologous sequence - *A. fumigatus*:419 *A. fischeri*:419
afum|EAL93932.1 100.00% afis|EAW19202.1 100.00%
Bootstrap support for afum|EAL93932.1 as seed ortholog is 100%.
Bootstrap support for afis|EAW19202.1 as seed ortholog is 100%.

---

Group of orthologs #7666. Best score 418 bits
Score difference with first non-orthologous sequence - *A. fumigatus*:418 *A. fischeri*:418
afum|EAL88488.1 100.00% afis|EAW15852.1 100.00%
Bootstrap support for afum|EAL88488.1 as seed ortholog is 100%.
Bootstrap support for afis|EAW15852.1 as seed ortholog is 100%.

---

Group of orthologs #7667. Best score 418 bits
Score difference with first non-orthologous sequence - *A. fumigatus*:418 *A. fischeri*:418
afum|EAL84437.1 100.00% afis|EAW20630.1 100.00%

Bootstrap support for afum|EAL84437.1 as seed ortholog is 100%.
Bootstrap support for afis|EAW20630.1 as seed ortholog is 100%.

---

Group of orthologs #7668. Best score 418 bits
Score difference with first non-orthologous sequence - *A. fumigatus*:418 *A. fischeri*:418
afum|EAL85328.1 100.00% afis|EAW20211.1 100.00%
Bootstrap support for afum|EAL85328.1 as seed ortholog is 100%.
Bootstrap support for afis|EAW20211.1 as seed ortholog is 100%.

---

Group of orthologs #7669. Best score 417 bits
Score difference with first non-orthologous sequence - *A. fumigatus*:417 *A. fischeri*:417
afum|EAL84603.1 100.00% afis|EAW20450.1 100.00%
Bootstrap support for afum|EAL84603.1 as seed ortholog is 100%.
Bootstrap support for afis|EAW20450.1 as seed ortholog is 100%.

---

Group of orthologs #7670. Best score 416 bits
Score difference with first non-orthologous sequence - *A. fumigatus*:314 *A. fischeri*:138
afum|EAL85780.1 100.00% afis|EAW16713.1 100.00%
Bootstrap support for afum|EAL85780.1 as seed ortholog is 100%.
Bootstrap support for afis|EAW16713.1 as seed ortholog is 99%.

---

Group of orthologs #7671. Best score 416 bits
Score difference with first non-orthologous sequence - *A. fumigatus*:416 *A. fischeri*:369
afum|EAL91704.1 100.00% afis|EAW17896.1 100.00%
Bootstrap support for afum|EAL91704.1 as seed ortholog is 100%.
Bootstrap support for afis|EAW17896.1 as seed ortholog is 100%.

---

Group of orthologs #7672. Best score 416 bits
Score difference with first non-orthologous sequence - *A. fumigatus*:416 *A. fischeri*:416
afum|EAL91014.1 100.00% afis|EAW22175.1 100.00%
Bootstrap support for afum|EAL91014.1 as seed ortholog is 100%.
Bootstrap support for afis|EAW22175.1 as seed ortholog is 100%.

---

Group of orthologs #7673. Best score 415 bits
Score difference with first non-orthologous sequence - *A. fumigatus*:415 *A. fischeri*:415
afum|EBA27182.1 100.00% afis|EAW20571.1 100.00%
afis|EAW15461.1 53.82%
Bootstrap support for afum|EBA27182.1 as seed ortholog is 100%.
Bootstrap support for afis|EAW20571.1 as seed ortholog is 100%.

---

Group of orthologs #7674. Best score 415 bits
Score difference with first non-orthologous sequence - *A. fumigatus*:415 *A. fischeri*:415
afum|EAL88591.1 100.00% afis|EAW15958.1 100.00%
Bootstrap support for afum|EAL88591.1 as seed ortholog is 100%.
Bootstrap support for afis|EAW15958.1 as seed ortholog is 100%.

---

Group of orthologs #7675. Best score 415 bits
Score difference with first non-orthologous sequence - *A. fumigatus*:415 *A. fischeri*:305
afum|EAL84831.1 100.00% afis|EAW20798.1 100.00%
Bootstrap support for afum|EAL84831.1 as seed ortholog is 100%.
Bootstrap support for afis|EAW20798.1 as seed ortholog is 100%.

---

Group of orthologs #7676. Best score 415 bits
Score difference with first non-orthologous sequence - *A. fumigatus*:44 *A. fischeri*:415
afum|EAL90927.1 100.00% afis|EAW17066.1 100.00%
Bootstrap support for afum|EAL90927.1 as seed ortholog is 99%.

Bootstrap support for afis|EAW17066.1 as seed ortholog is 100%.

---

Group of orthologs #7677. Best score 415 bits
Score difference with first non-orthologous sequence - *A. fumigatus*:415 *A. fischeri*:415
afum|EAL91576.1 100.00% afis|EAW17767.1 100.00%
Bootstrap support for afum|EAL91576.1 as seed ortholog is 100%.
Bootstrap support for afis|EAW17767.1 as seed ortholog is 100%.

---

Group of orthologs #7678. Best score 415 bits
Score difference with first non-orthologous sequence - *A. fumigatus*:415 *A. fischeri*:415
afum|EAL91652.1 100.00% afis|EAW17843.1 100.00%
Bootstrap support for afum|EAL91652.1 as seed ortholog is 100%.
Bootstrap support for afis|EAW17843.1 as seed ortholog is 100%.

---

Group of orthologs #7679. Best score 415 bits
Score difference with first non-orthologous sequence - *A. fumigatus*:415 *A. fischeri*:415
afum|EAL92676.1 100.00% afis|EAW21642.1 100.00%
Bootstrap support for afum|EAL92676.1 as seed ortholog is 100%.
Bootstrap support for afis|EAW21642.1 as seed ortholog is 100%.

---

Group of orthologs #7680. Best score 415 bits
Score difference with first non-orthologous sequence - *A. fumigatus*:415 *A. fischeri*:415
afum|EAL89466.1 100.00% afis|EAW24822.1 100.00%
Bootstrap support for afum|EAL89466.1 as seed ortholog is 100%.
Bootstrap support for afis|EAW24822.1 as seed ortholog is 100%.

---

Group of orthologs #7681. Best score 415 bits
Score difference with first non-orthologous sequence - *A. fumigatus*:415 *A. fischeri*:415
afum|EAL92810.1 100.00% afis|EAW21782.1 100.00%
Bootstrap support for afum|EAL92810.1 as seed ortholog is 100%.
Bootstrap support for afis|EAW21782.1 as seed ortholog is 100%.

---

Group of orthologs #7682. Best score 414 bits
Score difference with first non-orthologous sequence - *A. fumigatus*:414 *A. fischeri*:414
afum|EAL87928.1 100.00% afis|EAW23538.1 100.00%
Bootstrap support for afum|EAL87928.1 as seed ortholog is 100%.
Bootstrap support for afis|EAW23538.1 as seed ortholog is 100%.

---

Group of orthologs #7683. Best score 414 bits
Score difference with first non-orthologous sequence - *A. fumigatus*:414 *A. fischeri*:414
afum|EAL93346.1 100.00% afis|EAW18633.1 100.00%
Bootstrap support for afum|EAL93346.1 as seed ortholog is 100%.
Bootstrap support for afis|EAW18633.1 as seed ortholog is 100%.

---

Group of orthologs #7684. Best score 413 bits
Score difference with first non-orthologous sequence - *A. fumigatus*:413 *A. fischeri*:413
afum|EBA27270.1 100.00% afis|EAW16819.1 100.00%
Bootstrap support for afum|EBA27270.1 as seed ortholog is 100%.
Bootstrap support for afis|EAW16819.1 as seed ortholog is 100%.

---

Group of orthologs #7685. Best score 413 bits
Score difference with first non-orthologous sequence - *A. fumigatus*:413 *A. fischeri*:413
afum|EAL87236.1 100.00% afis|EAW23802.1 100.00%
Bootstrap support for afum|EAL87236.1 as seed ortholog is 100%.
Bootstrap support for afis|EAW23802.1 as seed ortholog is 100%.

---

Group of orthologs #7686. Best score 413 bits
Score difference with first non-orthologous sequence - *A. fumigatus*:413 *A. fischeri*:413
afum|EAL89758.2 100.00% afis|EAW25089.1 100.00%
Bootstrap support for afum|EAL89758.2 as seed ortholog is 100%.
Bootstrap support for afis|EAW25089.1 as seed ortholog is 100%.

---

Group of orthologs #7687. Best score 412 bits
Score difference with first non-orthologous sequence - *A. fumigatus*:412 *A. fischeri*:412
afum|EAL86681.1 100.00% afis|EAW17264.1 100.00%
Bootstrap support for afum|EAL86681.1 as seed ortholog is 100%.
Bootstrap support for afis|EAW17264.1 as seed ortholog is 100%.

---

Group of orthologs #7688. Best score 412 bits
Score difference with first non-orthologous sequence - *A. fumigatus*:412 *A. fischeri*:412
afum|EAL84617.1 100.00% afis|EAW20435.1 100.00%
Bootstrap support for afum|EAL84617.1 as seed ortholog is 100%.
Bootstrap support for afis|EAW20435.1 as seed ortholog is 100%.

---

Group of orthologs #7689. Best score 412 bits
Score difference with first non-orthologous sequence - *A. fumigatus*:412 *A. fischeri*:412
afum|EAL85201.1 100.00% afis|EAW20082.1 100.00%
Bootstrap support for afum|EAL85201.1 as seed ortholog is 100%.
Bootstrap support for afis|EAW20082.1 as seed ortholog is 100%.

---

Group of orthologs #7690. Best score 412 bits
Score difference with first non-orthologous sequence - *A. fumigatus*:412 *A. fischeri*:412
afum|EAL84747.1 100.00% afis|EAW20886.1 100.00%
Bootstrap support for afum|EAL84747.1 as seed ortholog is 100%.
Bootstrap support for afis|EAW20886.1 as seed ortholog is 100%.

---

Group of orthologs #7691. Best score 412 bits
Score difference with first non-orthologous sequence - *A. fumigatus*:412 *A. fischeri*:412
afum|EAL91735.2 100.00% afis|EAW17924.1 100.00%
Bootstrap support for afum|EAL91735.2 as seed ortholog is 100%.
Bootstrap support for afis|EAW17924.1 as seed ortholog is 100%.

---

Group of orthologs #7692. Best score 412 bits
Score difference with first non-orthologous sequence - *A. fumigatus*:412 *A. fischeri*:412
afum|EAL92166.1 100.00% afis|EAW21147.1 100.00%
Bootstrap support for afum|EAL92166.1 as seed ortholog is 100%.
Bootstrap support for afis|EAW21147.1 as seed ortholog is 100%.

---

Group of orthologs #7693. Best score 411 bits
Score difference with first non-orthologous sequence - *A. fumigatus*:266 *A. fischeri*:411
afum|EAL84430.1 100.00% afis|EAW20639.1 100.00%
Bootstrap support for afum|EAL84430.1 as seed ortholog is 100%.
Bootstrap support for afis|EAW20639.1 as seed ortholog is 100%.

---

Group of orthologs #7694. Best score 411 bits
Score difference with first non-orthologous sequence - *A. fumigatus*:411 *A. fischeri*:411
afum|EAL90234.1 100.00% afis|EAW22963.1 100.00%
Bootstrap support for afum|EAL90234.1 as seed ortholog is 100%.
Bootstrap support for afis|EAW22963.1 as seed ortholog is 100%.

---

Group of orthologs #7695. Best score 411 bits
Score difference with first non-orthologous sequence - *A. fumigatus*:411 *A. fischeri*:411

afum|EAL89322.1 100.00% afis|EAW24664.1 100.00%
Bootstrap support for afum|EAL89322.1 as seed ortholog is 100%.
Bootstrap support for afis|EAW24664.1 as seed ortholog is 100%.

---

Group of orthologs #7696. Best score 410 bits
Score difference with first non-orthologous sequence - *A. fumigatus*:410 *A. fischeri*:410
afum|EAL85257.1 100.00% afis|EAW20142.1 100.00%
Bootstrap support for afum|EAL85257.1 as seed ortholog is 100%.
Bootstrap support for afis|EAW20142.1 as seed ortholog is 100%.

---

Group of orthologs #7697. Best score 410 bits
Score difference with first non-orthologous sequence - *A. fumigatus*:410 *A. fischeri*:410
afum|EAL86236.1 100.00% afis|EAW20690.1 100.00%
Bootstrap support for afum|EAL86236.1 as seed ortholog is 100%.
Bootstrap support for afis|EAW20690.1 as seed ortholog is 100%.

---

Group of orthologs #7698. Best score 410 bits
Score difference with first non-orthologous sequence - *A. fumigatus*:410 *A. fischeri*:410
afum|EBA27463.1 100.00% afis|EAW21535.1 100.00%
Bootstrap support for afum|EBA27463.1 as seed ortholog is 100%.
Bootstrap support for afis|EAW21535.1 as seed ortholog is 100%.

---

Group of orthologs #7699. Best score 409 bits
Score difference with first non-orthologous sequence - *A. fumigatus*:409 *A. fischeri*:409
afum|EAL91547.1 100.00% afis|EAW17732.1 100.00%
Bootstrap support for afum|EAL91547.1 as seed ortholog is 100%.
Bootstrap support for afis|EAW17732.1 as seed ortholog is 100%.

---

Group of orthologs #7700. Best score 409 bits
Score difference with first non-orthologous sequence - *A. fumigatus*:409 *A. fischeri*:409
afum|EAL91788.1 100.00% afis|EAW17979.1 100.00%
Bootstrap support for afum|EAL91788.1 as seed ortholog is 100%.
Bootstrap support for afis|EAW17979.1 as seed ortholog is 100%.

---

Group of orthologs #7701. Best score 409 bits
Score difference with first non-orthologous sequence - *A. fumigatus*:409 *A. fischeri*:409
afum|EAL87211.1 100.00% afis|EAW23777.1 100.00%
Bootstrap support for afum|EAL87211.1 as seed ortholog is 100%.
Bootstrap support for afis|EAW23777.1 as seed ortholog is 100%.

---

Group of orthologs #7702. Best score 409 bits
Score difference with first non-orthologous sequence - *A. fumigatus*:409 *A. fischeri*:409
afum|EAL87411.1 100.00% afis|EAW23981.1 100.00%
Bootstrap support for afum|EAL87411.1 as seed ortholog is 100%.
Bootstrap support for afis|EAW23981.1 as seed ortholog is 100%.

---

Group of orthologs #7703. Best score 409 bits
Score difference with first non-orthologous sequence - *A. fumigatus*:409 *A. fischeri*:409
afum|EAL88450.1 100.00% afis|EAW23032.1 100.00%
Bootstrap support for afum|EAL88450.1 as seed ortholog is 100%.
Bootstrap support for afis|EAW23032.1 as seed ortholog is 100%.

---

Group of orthologs #7704. Best score 409 bits
Score difference with first non-orthologous sequence - *A. fumigatus*:409 *A. fischeri*:409
afum|EAL90537.1 100.00% afis|EAW22655.1 100.00%
Bootstrap support for afum|EAL90537.1 as seed ortholog is 100%.

Bootstrap support for afis|EAW22655.1 as seed ortholog is 100%.

---

Group of orthologs #7705. Best score 408 bits
Score difference with first non-orthologous sequence - *A. fumigatus*:408 *A. fischeri*:408
afum|EAL85811.1 100.00% afis|EAW15518.1 100.00%
Bootstrap support for afum|EAL85811.1 as seed ortholog is 100%.
Bootstrap support for afis|EAW15518.1 as seed ortholog is 100%.

---

Group of orthologs #7706. Best score 408 bits
Score difference with first non-orthologous sequence - *A. fumigatus*:408 *A. fischeri*:408
afum|EAL88516.1 100.00% afis|EAW15882.1 100.00%
Bootstrap support for afum|EAL88516.1 as seed ortholog is 100%.
Bootstrap support for afis|EAW15882.1 as seed ortholog is 100%.

---

Group of orthologs #7707. Best score 408 bits
Score difference with first non-orthologous sequence - *A. fumigatus*:408 *A. fischeri*:408
afum|EAL86093.1 100.00% afis|EAW24425.1 100.00%
Bootstrap support for afum|EAL86093.1 as seed ortholog is 100%.
Bootstrap support for afis|EAW24425.1 as seed ortholog is 100%.

---

Group of orthologs #7708. Best score 408 bits
Score difference with first non-orthologous sequence - *A. fumigatus*:408 *A. fischeri*:408
afum|EAL93429.1 100.00% afis|EAW18706.1 100.00%
Bootstrap support for afum|EAL93429.1 as seed ortholog is 100%.
Bootstrap support for afis|EAW18706.1 as seed ortholog is 100%.

---

Group of orthologs #7709. Best score 408 bits
Score difference with first non-orthologous sequence - *A. fumigatus*:408 *A. fischeri*:408
afum|EAL93029.1 100.00% afis|EAW22014.1 100.00%
Bootstrap support for afum|EAL93029.1 as seed ortholog is 100%.
Bootstrap support for afis|EAW22014.1 as seed ortholog is 100%.

---

Group of orthologs #7710. Best score 407 bits
Score difference with first non-orthologous sequence - *A. fumigatus*:407 *A. fischeri*:407
afum|EAL84446.1 100.00% afis|EAW20615.1 100.00%
Bootstrap support for afum|EAL84446.1 as seed ortholog is 100%.
Bootstrap support for afis|EAW20615.1 as seed ortholog is 100%.

---

Group of orthologs #7711. Best score 407 bits
Score difference with first non-orthologous sequence - *A. fumigatus*:407 *A. fischeri*:407
afum|EAL93411.2 100.00% afis|EAW18690.1 100.00%
Bootstrap support for afum|EAL93411.2 as seed ortholog is 100%.
Bootstrap support for afis|EAW18690.1 as seed ortholog is 100%.

---

Group of orthologs #7712. Best score 407 bits
Score difference with first non-orthologous sequence - *A. fumigatus*:407 *A. fischeri*:407
afum|EAL92598.1 100.00% afis|EAW21580.1 100.00%
Bootstrap support for afum|EAL92598.1 as seed ortholog is 100%.
Bootstrap support for afis|EAW21580.1 as seed ortholog is 100%.

---

Group of orthologs #7713. Best score 407 bits
Score difference with first non-orthologous sequence - *A. fumigatus*:407 *A. fischeri*:407
afum|EAL92605.1 100.00% afis|EAW21587.1 100.00%
Bootstrap support for afum|EAL92605.1 as seed ortholog is 100%.
Bootstrap support for afis|EAW21587.1 as seed ortholog is 100%.

---

Group of orthologs #7714. Best score 406 bits
Score difference with first non-orthologous sequence - *A. fumigatus*:406 *A. fischeri*:406
afum|EAL86332.1 100.00% afis|EAW16761.1 100.00%
Bootstrap support for afum|EAL86332.1 as seed ortholog is 100%.
Bootstrap support for afis|EAW16761.1 as seed ortholog is 100%.

---

Group of orthologs #7715. Best score 406 bits
Score difference with first non-orthologous sequence - *A. fumigatus*:406 *A. fischeri*:406
afum|EAL86619.2 100.00% afis|EAW17191.1 100.00%
Bootstrap support for afum|EAL86619.2 as seed ortholog is 100%.
Bootstrap support for afis|EAW17191.1 as seed ortholog is 100%.

---

Group of orthologs #7716. Best score 406 bits
Score difference with first non-orthologous sequence - *A. fumigatus*:406 *A. fischeri*:406
afum|EAL90022.1 100.00% afis|EAW25360.1 100.00%
Bootstrap support for afum|EAL90022.1 as seed ortholog is 100%.
Bootstrap support for afis|EAW25360.1 as seed ortholog is 100%.

---

Group of orthologs #7717. Best score 405 bits
Score difference with first non-orthologous sequence - *A. fumigatus*:405 *A. fischeri*:405
afum|EAL88716.1 100.00% afis|EAW16084.1 100.00%
Bootstrap support for afum|EAL88716.1 as seed ortholog is 100%.
Bootstrap support for afis|EAW16084.1 as seed ortholog is 100%.

---

Group of orthologs #7718. Best score 405 bits
Score difference with first non-orthologous sequence - *A. fumigatus*:405 *A. fischeri*:405
afum|EAL84847.1 100.00% afis|EAW20772.1 100.00%
Bootstrap support for afum|EAL84847.1 as seed ortholog is 100%.
Bootstrap support for afis|EAW20772.1 as seed ortholog is 100%.

---

Group of orthologs #7719. Best score 405 bits
Score difference with first non-orthologous sequence - *A. fumigatus*:405 *A. fischeri*:405
afum|EAL86813.1 100.00% afis|EAW19714.1 100.00%
Bootstrap support for afum|EAL86813.1 as seed ortholog is 100%.
Bootstrap support for afis|EAW19714.1 as seed ortholog is 100%.

---

Group of orthologs #7720. Best score 405 bits
Score difference with first non-orthologous sequence - *A. fumigatus*:405 *A. fischeri*:405
afum|EAL92134.1 100.00% afis|EAW21111.1 100.00%
Bootstrap support for afum|EAL92134.1 as seed ortholog is 100%.
Bootstrap support for afis|EAW21111.1 as seed ortholog is 100%.

---

Group of orthologs #7721. Best score 405 bits
Score difference with first non-orthologous sequence - *A. fumigatus*:405 *A. fischeri*:405
afum|EBA27327.1 100.00% afis|EAW23528.1 100.00%
Bootstrap support for afum|EBA27327.1 as seed ortholog is 100%.
Bootstrap support for afis|EAW23528.1 as seed ortholog is 100%.

---

Group of orthologs #7722. Best score 404 bits
Score difference with first non-orthologous sequence - *A. fumigatus*:404 *A. fischeri*:404
afum|EAL89513.1 100.00% afis|EAW24863.1 100.00%
Bootstrap support for afum|EAL89513.1 as seed ortholog is 100%.
Bootstrap support for afis|EAW24863.1 as seed ortholog is 100%.

---

Group of orthologs #7723. Best score 404 bits
Score difference with first non-orthologous sequence - *A. fumigatus*:404 *A. fischeri*:404

afum|EAL89600.1 100.00% afis|EAW24943.1 100.00%
Bootstrap support for afum|EAL89600.1 as seed ortholog is 100%.
Bootstrap support for afis|EAW24943.1 as seed ortholog is 100%.

---

Group of orthologs #7724. Best score 404 bits
Score difference with first non-orthologous sequence - *A. fumigatus*:404 *A. fischeri*:404
afum|EAL92871.1 100.00% afis|EAW21858.1 100.00%
Bootstrap support for afum|EAL92871.1 as seed ortholog is 100%.
Bootstrap support for afis|EAW21858.1 as seed ortholog is 100%.

---

Group of orthologs #7725. Best score 403 bits
Score difference with first non-orthologous sequence - *A. fumigatus*:403 *A. fischeri*:403
afum|EAL88983.1 100.00% afis|EAW16358.1 100.00%
Bootstrap support for afum|EAL88983.1 as seed ortholog is 100%.
Bootstrap support for afis|EAW16358.1 as seed ortholog is 100%.

---

Group of orthologs #7726. Best score 403 bits
Score difference with first non-orthologous sequence - *A. fumigatus*:403 *A. fischeri*:403
afum|EAL85360.1 100.00% afis|EAW20243.1 100.00%
Bootstrap support for afum|EAL85360.1 as seed ortholog is 100%.
Bootstrap support for afis|EAW20243.1 as seed ortholog is 100%.

---

Group of orthologs #7727. Best score 403 bits
Score difference with first non-orthologous sequence - *A. fumigatus*:403 *A. fischeri*:403
afum|EAL86871.1 100.00% afis|EAW19651.1 100.00%
Bootstrap support for afum|EAL86871.1 as seed ortholog is 100%.
Bootstrap support for afis|EAW19651.1 as seed ortholog is 100%.

---

Group of orthologs #7728. Best score 403 bits
Score difference with first non-orthologous sequence - *A. fumigatus*:403 *A. fischeri*:403
afum|EBA27476.1 100.00% afis|EAW21839.1 100.00%
Bootstrap support for afum|EBA27476.1 as seed ortholog is 100%.
Bootstrap support for afis|EAW21839.1 as seed ortholog is 100%.

---

Group of orthologs #7729. Best score 403 bits
Score difference with first non-orthologous sequence - *A. fumigatus*:403 *A. fischeri*:403
afum|EBA27406.1 100.00% afis|EAW22808.1 100.00%
Bootstrap support for afum|EBA27406.1 as seed ortholog is 100%.
Bootstrap support for afis|EAW22808.1 as seed ortholog is 100%.

---

Group of orthologs #7730. Best score 402 bits
Score difference with first non-orthologous sequence - *A. fumigatus*:402 *A. fischeri*:402
afum|EAL91209.1 100.00% afis|EAW17371.1 100.00%
Bootstrap support for afum|EAL91209.1 as seed ortholog is 100%.
Bootstrap support for afis|EAW17371.1 as seed ortholog is 100%.

---

Group of orthologs #7731. Best score 402 bits
Score difference with first non-orthologous sequence - *A. fumigatus*:402 *A. fischeri*:402
afum|EAL93081.1 100.00% afis|EAW18403.1 100.00%
Bootstrap support for afum|EAL93081.1 as seed ortholog is 100%.
Bootstrap support for afis|EAW18403.1 as seed ortholog is 100%.

---

Group of orthologs #7732. Best score 402 bits
Score difference with first non-orthologous sequence - *A. fumigatus*:402 *A. fischeri*:402
afum|EAL88048.2 100.00% afis|EAW23421.1 100.00%
Bootstrap support for afum|EAL88048.2 as seed ortholog is 100%.

Bootstrap support for afis|EAW23421.1 as seed ortholog is 100%.

---

Group of orthologs #7733. Best score 402 bits
Score difference with first non-orthologous sequence - *A. fumigatus*:402 *A. fischeri*:402
afum|EAL93609.1 100.00% afis|EAW18877.1 100.00%
Bootstrap support for afum|EAL93609.1 as seed ortholog is 100%.
Bootstrap support for afis|EAW18877.1 as seed ortholog is 100%.

---

Group of orthologs #7734. Best score 402 bits
Score difference with first non-orthologous sequence - *A. fumigatus*:402 *A. fischeri*:402
afum|EAL92544.1 100.00% afis|EAW21523.1 100.00%
Bootstrap support for afum|EAL92544.1 as seed ortholog is 100%.
Bootstrap support for afis|EAW21523.1 as seed ortholog is 100%.

---

Group of orthologs #7735. Best score 402 bits
Score difference with first non-orthologous sequence - *A. fumigatus*:402 *A. fischeri*:402
afum|EAL89760.2 100.00% afis|EAW25090.1 100.00%
Bootstrap support for afum|EAL89760.2 as seed ortholog is 100%.
Bootstrap support for afis|EAW25090.1 as seed ortholog is 100%.

---

Group of orthologs #7736. Best score 402 bits
Score difference with first non-orthologous sequence - *A. fumigatus*:402 *A. fischeri*:402
afum|EAL91995.1 100.00% afis|EAW24140.1 100.00%
Bootstrap support for afum|EAL91995.1 as seed ortholog is 100%.
Bootstrap support for afis|EAW24140.1 as seed ortholog is 100%.

---

Group of orthologs #7737. Best score 401 bits
Score difference with first non-orthologous sequence - *A. fumigatus*:401 *A. fischeri*:401
afum|EAL85802.1 100.00% afis|EAW15526.1 100.00%
Bootstrap support for afum|EAL85802.1 as seed ortholog is 100%.
Bootstrap support for afis|EAW15526.1 as seed ortholog is 100%.

---

Group of orthologs #7738. Best score 401 bits
Score difference with first non-orthologous sequence - *A. fumigatus*:401 *A. fischeri*:401
afum|EAL86798.1 100.00% afis|EAW19729.1 100.00%
Bootstrap support for afum|EAL86798.1 as seed ortholog is 100%.
Bootstrap support for afis|EAW19729.1 as seed ortholog is 100%.

---

Group of orthologs #7739. Best score 401 bits
Score difference with first non-orthologous sequence - *A. fumigatus*:23 *A. fischeri*:77
afum|EAL91132.1 100.00% afis|EAW16922.1 100.00%
Bootstrap support for afum|EAL91132.1 as seed ortholog is 67%. Alternative seed ortholog is afum|EAL91922.1 (23 bits away from this cluster)
Bootstrap support for afis|EAW16922.1 as seed ortholog is 94%.

---

Group of orthologs #7740. Best score 401 bits
Score difference with first non-orthologous sequence - *A. fumigatus*:401 *A. fischeri*:401
afum|EAL88227.1 100.00% afis|EAW23254.1 100.00%
Bootstrap support for afum|EAL88227.1 as seed ortholog is 100%.
Bootstrap support for afis|EAW23254.1 as seed ortholog is 100%.

---

Group of orthologs #7741. Best score 401 bits
Score difference with first non-orthologous sequence - *A. fumigatus*:401 *A. fischeri*:401
afum|EAL92462.2 100.00% afis|EAW21444.1 100.00%
Bootstrap support for afum|EAL92462.2 as seed ortholog is 100%.
Bootstrap support for afis|EAW21444.1 as seed ortholog is 100%.

---

Group of orthologs #7742. Best score 401 bits
Score difference with first non-orthologous sequence - *A. fumigatus*:292 *A. fischeri*:262
afum|EBA27363.1 100.00% afis|EAW21811.1 100.00%
Bootstrap support for afum|EBA27363.1 as seed ortholog is 99%.
Bootstrap support for afis|EAW21811.1 as seed ortholog is 99%.

---

Group of orthologs #7743. Best score 400 bits
Score difference with first non-orthologous sequence - *A. fumigatus*:400 *A. fischeri*:400
afum|EAL86160.1 100.00% afis|EAW24491.1 100.00%
Bootstrap support for afum|EAL86160.1 as seed ortholog is 100%.
Bootstrap support for afis|EAW24491.1 as seed ortholog is 100%.

---

Group of orthologs #7744. Best score 400 bits
Score difference with first non-orthologous sequence - *A. fumigatus*:400 *A. fischeri*:400
afum|EAL93480.1 100.00% afis|EAW18753.1 100.00%
Bootstrap support for afum|EAL93480.1 as seed ortholog is 100%.
Bootstrap support for afis|EAW18753.1 as seed ortholog is 100%.

---

Group of orthologs #7745. Best score 400 bits
Score difference with first non-orthologous sequence - *A. fumigatus*:400 *A. fischeri*:400
afum|EAL93689.2 100.00% afis|EAW18953.1 100.00%
Bootstrap support for afum|EAL93689.2 as seed ortholog is 100%.
Bootstrap support for afis|EAW18953.1 as seed ortholog is 100%.

---

Group of orthologs #7746. Best score 399 bits
Score difference with first non-orthologous sequence - *A. fumigatus*:399 *A. fischeri*:399
afum|EAL85774.1 100.00% afis|EAW15558.1 100.00%
Bootstrap support for afum|EAL85774.1 as seed ortholog is 100%.
Bootstrap support for afis|EAW15558.1 as seed ortholog is 100%.

---

Group of orthologs #7747. Best score 399 bits
Score difference with first non-orthologous sequence - *A. fumigatus*:399 *A. fischeri*:399
afum|EAL85240.1 100.00% afis|EAW20122.1 100.00%
Bootstrap support for afum|EAL85240.1 as seed ortholog is 100%.
Bootstrap support for afis|EAW20122.1 as seed ortholog is 100%.

---

Group of orthologs #7748. Best score 399 bits
Score difference with first non-orthologous sequence - *A. fumigatus*:399 *A. fischeri*:399
afum|EAL89106.1 100.00% afis|EAW16511.1 100.00%
Bootstrap support for afum|EAL89106.1 as seed ortholog is 100%.
Bootstrap support for afis|EAW16511.1 as seed ortholog is 100%.

---

Group of orthologs #7749. Best score 399 bits
Score difference with first non-orthologous sequence - *A. fumigatus*:399 *A. fischeri*:399
afum|EAL85277.1 100.00% afis|EAW20160.1 100.00%
Bootstrap support for afum|EAL85277.1 as seed ortholog is 100%.
Bootstrap support for afis|EAW20160.1 as seed ortholog is 100%.

---

Group of orthologs #7750. Best score 399 bits
Score difference with first non-orthologous sequence - *A. fumigatus*:399 *A. fischeri*:399
afum|EAL86824.1 100.00% afis|EAW19702.1 100.00%
Bootstrap support for afum|EAL86824.1 as seed ortholog is 100%.
Bootstrap support for afis|EAW19702.1 as seed ortholog is 100%.

---

Group of orthologs #7751. Best score 399 bits

Score difference with first non-orthologous sequence - *A. fumigatus*:399   *A. fischeri*:399
afum|EAL84331.1 100.00% afis|EAW25645.1 100.00%
Bootstrap support for afum|EAL84331.1 as seed ortholog is 100%.
Bootstrap support for afis|EAW25645.1 as seed ortholog is 100%.

---

Group of orthologs #7752. Best score 399 bits
Score difference with first non-orthologous sequence - *A. fumigatus*:399   *A. fischeri*:399
afum|EAL85977.2 100.00% afis|EAW24309.1 100.00%
Bootstrap support for afum|EAL85977.2 as seed ortholog is 100%.
Bootstrap support for afis|EAW24309.1 as seed ortholog is 100%.

---

Group of orthologs #7753. Best score 399 bits
Score difference with first non-orthologous sequence - *A. fumigatus*:399   *A. fischeri*:399
afum|EAL93720.1 100.00% afis|EAW18984.1 100.00%
Bootstrap support for afum|EAL93720.1 as seed ortholog is 100%.
Bootstrap support for afis|EAW18984.1 as seed ortholog is 100%.

---

Group of orthologs #7754. Best score 399 bits
Score difference with first non-orthologous sequence - *A. fumigatus*:399   *A. fischeri*:399
afum|EAL93848.2 100.00% afis|EAW19116.1 100.00%
Bootstrap support for afum|EAL93848.2 as seed ortholog is 100%.
Bootstrap support for afis|EAW19116.1 as seed ortholog is 100%.

---

Group of orthologs #7755. Best score 399 bits
Score difference with first non-orthologous sequence - *A. fumigatus*:399   *A. fischeri*:399
afum|EAL92439.1 100.00% afis|EAW21424.1 100.00%
Bootstrap support for afum|EAL92439.1 as seed ortholog is 100%.
Bootstrap support for afis|EAW21424.1 as seed ortholog is 100%.

---

Group of orthologs #7756. Best score 399 bits
Score difference with first non-orthologous sequence - *A. fumigatus*:399   *A. fischeri*:399
afum|EAL92727.1 100.00% afis|EAW21693.1 100.00%
Bootstrap support for afum|EAL92727.1 as seed ortholog is 100%.
Bootstrap support for afis|EAW21693.1 as seed ortholog is 100%.

---

Group of orthologs #7757. Best score 399 bits
Score difference with first non-orthologous sequence - *A. fumigatus*:399   *A. fischeri*:399
afum|EAL90150.1 100.00% afis|EAW25485.1 100.00%
Bootstrap support for afum|EAL90150.1 as seed ortholog is 100%.
Bootstrap support for afis|EAW25485.1 as seed ortholog is 100%.

---

Group of orthologs #7758. Best score 398 bits
Score difference with first non-orthologous sequence - *A. fumigatus*:398   *A. fischeri*:398
afum|EAL87612.1 100.00% afis|EAW18247.1 100.00%
Bootstrap support for afum|EAL87612.1 as seed ortholog is 100%.
Bootstrap support for afis|EAW18247.1 as seed ortholog is 100%.

---

Group of orthologs #7759. Best score 398 bits
Score difference with first non-orthologous sequence - *A. fumigatus*:398   *A. fischeri*:195
afum|EAL94089.1 100.00% afis|EAW19370.1 100.00%
Bootstrap support for afum|EAL94089.1 as seed ortholog is 100%.
Bootstrap support for afis|EAW19370.1 as seed ortholog is 100%.

---

Group of orthologs #7760. Best score 398 bits
Score difference with first non-orthologous sequence - *A. fumigatus*:398   *A. fischeri*:398
afum|EAL89724.1 100.00% afis|EAW25056.1 100.00%

Bootstrap support for afum|EAL89724.1 as seed ortholog is 100%.
Bootstrap support for afis|EAW25056.1 as seed ortholog is 100%.

---

Group of orthologs #7761. Best score 398 bits
Score difference with first non-orthologous sequence - *A. fumigatus*:398 *A. fischeri*:398
afum|EAL90088.1 100.00% afis|EAW25429.1 100.00%
Bootstrap support for afum|EAL90088.1 as seed ortholog is 100%.
Bootstrap support for afis|EAW25429.1 as seed ortholog is 100%.

---

Group of orthologs #7762. Best score 397 bits
Score difference with first non-orthologous sequence - *A. fumigatus*:397 *A. fischeri*:397
afum|EAL88801.1 100.00% afis|EAW16166.1 100.00%
Bootstrap support for afum|EAL88801.1 as seed ortholog is 100%.
Bootstrap support for afis|EAW16166.1 as seed ortholog is 100%.

---

Group of orthologs #7763. Best score 397 bits
Score difference with first non-orthologous sequence - *A. fumigatus*:397 *A. fischeri*:229
afum|EAL94055.1 100.00% afis|EAW19337.1 100.00%
Bootstrap support for afum|EAL94055.1 as seed ortholog is 100%.
Bootstrap support for afis|EAW19337.1 as seed ortholog is 100%.

---

Group of orthologs #7764. Best score 397 bits
Score difference with first non-orthologous sequence - *A. fumigatus*:397 *A. fischeri*:397
afum|EAL90437.1 100.00% afis|EAW22762.1 100.00%
Bootstrap support for afum|EAL90437.1 as seed ortholog is 100%.
Bootstrap support for afis|EAW22762.1 as seed ortholog is 100%.

---

Group of orthologs #7765. Best score 397 bits
Score difference with first non-orthologous sequence - *A. fumigatus*:397 *A. fischeri*:397
afum|EAL91957.1 100.00% afis|EAW24103.1 100.00%
Bootstrap support for afum|EAL91957.1 as seed ortholog is 100%.
Bootstrap support for afis|EAW24103.1 as seed ortholog is 100%.

---

Group of orthologs #7766. Best score 397 bits
Score difference with first non-orthologous sequence - *A. fumigatus*:397 *A. fischeri*:397
afum|EBA27312.1 100.00% afis|EAW23921.1 100.00%
Bootstrap support for afum|EBA27312.1 as seed ortholog is 100%.
Bootstrap support for afis|EAW23921.1 as seed ortholog is 100%.

---

Group of orthologs #7767. Best score 396 bits
Score difference with first non-orthologous sequence - *A. fumigatus*:396 *A. fischeri*:396
afum|EAL85825.1 100.00% afis|EAW15498.1 100.00%
Bootstrap support for afum|EAL85825.1 as seed ortholog is 100%.
Bootstrap support for afis|EAW15498.1 as seed ortholog is 100%.

---

Group of orthologs #7768. Best score 396 bits
Score difference with first non-orthologous sequence - *A. fumigatus*:396 *A. fischeri*:396
afum|EAL89063.1 100.00% afis|EAW16469.1 100.00%
Bootstrap support for afum|EAL89063.1 as seed ortholog is 100%.
Bootstrap support for afis|EAW16469.1 as seed ortholog is 100%.

---

Group of orthologs #7769. Best score 396 bits
Score difference with first non-orthologous sequence - *A. fumigatus*:396 *A. fischeri*:396
afum|EAL87466.1 100.00% afis|EAW18109.1 100.00%
Bootstrap support for afum|EAL87466.1 as seed ortholog is 100%.
Bootstrap support for afis|EAW18109.1 as seed ortholog is 100%.

Group of orthologs #7770. Best score 396 bits
Score difference with first non-orthologous sequence - *A. fumigatus*:396 *A. fischeri*:396
afum|EAL85479.1 100.00% afis|EAW20363.1 100.00%
Bootstrap support for afum|EAL85479.1 as seed ortholog is 100%.
Bootstrap support for afis|EAW20363.1 as seed ortholog is 100%.

Group of orthologs #7771. Best score 396 bits
Score difference with first non-orthologous sequence - *A. fumigatus*:396 *A. fischeri*:396
afum|EAL87084.2 100.00% afis|EAW19410.1 100.00%
Bootstrap support for afum|EAL87084.2 as seed ortholog is 100%.
Bootstrap support for afis|EAW19410.1 as seed ortholog is 100%.

Group of orthologs #7772. Best score 396 bits
Score difference with first non-orthologous sequence - *A. fumigatus*:396 *A. fischeri*:396
afum|EAL87325.1 100.00% afis|EAW23898.1 100.00%
Bootstrap support for afum|EAL87325.1 as seed ortholog is 100%.
Bootstrap support for afis|EAW23898.1 as seed ortholog is 100%.

Group of orthologs #7773. Best score 396 bits
Score difference with first non-orthologous sequence - *A. fumigatus*:396 *A. fischeri*:305
afum|EBA27305.1 100.00% afis|EAW23761.1 100.00%
Bootstrap support for afum|EBA27305.1 as seed ortholog is 100%.
Bootstrap support for afis|EAW23761.1 as seed ortholog is 100%.

Group of orthologs #7774. Best score 395 bits
Score difference with first non-orthologous sequence - *A. fumigatus*:395 *A. fischeri*:395
afum|EAL87159.1 100.00% afis|EAW23714.1 100.00%
Bootstrap support for afum|EAL87159.1 as seed ortholog is 100%.
Bootstrap support for afis|EAW23714.1 as seed ortholog is 100%.

Group of orthologs #7775. Best score 395 bits
Score difference with first non-orthologous sequence - *A. fumigatus*:395 *A. fischeri*:395
afum|EBA27296.1 100.00% afis|EAW19535.1 100.00%
Bootstrap support for afum|EBA27296.1 as seed ortholog is 100%.
Bootstrap support for afis|EAW19535.1 as seed ortholog is 100%.

Group of orthologs #7776. Best score 394 bits
Score difference with first non-orthologous sequence - *A. fumigatus*:394 *A. fischeri*:394
afum|EAL88765.1 100.00% afis|EAW16135.1 100.00%
Bootstrap support for afum|EAL88765.1 as seed ortholog is 100%.
Bootstrap support for afis|EAW16135.1 as seed ortholog is 100%.

Group of orthologs #7777. Best score 394 bits
Score difference with first non-orthologous sequence - *A. fumigatus*:394 *A. fischeri*:394
afum|EAL86976.2 100.00% afis|EAW19528.1 100.00%
Bootstrap support for afum|EAL86976.2 as seed ortholog is 100%.
Bootstrap support for afis|EAW19528.1 as seed ortholog is 100%.

Group of orthologs #7778. Best score 394 bits
Score difference with first non-orthologous sequence - *A. fumigatus*:394 *A. fischeri*:394
afum|EAL90737.1 100.00% afis|EAW22454.1 100.00%
Bootstrap support for afum|EAL90737.1 as seed ortholog is 100%.
Bootstrap support for afis|EAW22454.1 as seed ortholog is 100%.

Group of orthologs #7779. Best score 394 bits

Score difference with first non-orthologous sequence - *A. fumigatus*:394   *A. fischeri*:394
afum|EAL90044.1 100.00% afis|EAW25385.1 100.00%
Bootstrap support for afum|EAL90044.1 as seed ortholog is 100%.
Bootstrap support for afis|EAW25385.1 as seed ortholog is 100%.

---

Group of orthologs #7780. Best score 393 bits
Score difference with first non-orthologous sequence - *A. fumigatus*:393   *A. fischeri*:393
afum|EAL85764.1 100.00% afis|EAW15569.1 100.00%
Bootstrap support for afum|EAL85764.1 as seed ortholog is 100%.
Bootstrap support for afis|EAW15569.1 as seed ortholog is 100%.

---

Group of orthologs #7781. Best score 393 bits
Score difference with first non-orthologous sequence - *A. fumigatus*:393   *A. fischeri*:393
afum|EAL91232.1 100.00% afis|EAW17394.1 100.00%
Bootstrap support for afum|EAL91232.1 as seed ortholog is 100%.
Bootstrap support for afis|EAW17394.1 as seed ortholog is 100%.

---

Group of orthologs #7782. Best score 393 bits
Score difference with first non-orthologous sequence - *A. fumigatus*:393   *A. fischeri*:393
afum|EAL88471.1 100.00% afis|EAW23009.1 100.00%
Bootstrap support for afum|EAL88471.1 as seed ortholog is 100%.
Bootstrap support for afis|EAW23009.1 as seed ortholog is 100%.

---

Group of orthologs #7783. Best score 393 bits
Score difference with first non-orthologous sequence - *A. fumigatus*:393   *A. fischeri*:393
afum|EAL87863.2 100.00% afis|EAW23603.1 100.00%
Bootstrap support for afum|EAL87863.2 as seed ortholog is 100%.
Bootstrap support for afis|EAW23603.1 as seed ortholog is 100%.

---

Group of orthologs #7784. Best score 393 bits
Score difference with first non-orthologous sequence - *A. fumigatus*:393   *A. fischeri*:393
afum|EAL93232.1 100.00% afis|EAW18534.1 100.00%
Bootstrap support for afum|EAL93232.1 as seed ortholog is 100%.
Bootstrap support for afis|EAW18534.1 as seed ortholog is 100%.

---

Group of orthologs #7785. Best score 393 bits
Score difference with first non-orthologous sequence - *A. fumigatus*:393   *A. fischeri*:393
afum|EAL90337.1 100.00% afis|EAW22863.1 100.00%
Bootstrap support for afum|EAL90337.1 as seed ortholog is 100%.
Bootstrap support for afis|EAW22863.1 as seed ortholog is 100%.

---

Group of orthologs #7786. Best score 393 bits
Score difference with first non-orthologous sequence - *A. fumigatus*:393   *A. fischeri*:393
afum|EAL92791.1 100.00% afis|EAW21761.1 100.00%
Bootstrap support for afum|EAL92791.1 as seed ortholog is 100%.
Bootstrap support for afis|EAW21761.1 as seed ortholog is 100%.

---

Group of orthologs #7787. Best score 392 bits
Score difference with first non-orthologous sequence - *A. fumigatus*:392   *A. fischeri*:392
afum|EAL90888.1 100.00% afis|EAW22293.1 100.00%
Bootstrap support for afum|EAL90888.1 as seed ortholog is 100%.
Bootstrap support for afis|EAW22293.1 as seed ortholog is 100%.

---

Group of orthologs #7788. Best score 392 bits
Score difference with first non-orthologous sequence - *A. fumigatus*:392   *A. fischeri*:392
afum|EAL89399.1 100.00% afis|EAW24757.1 100.00%

Bootstrap support for afum|EAL89399.1 as seed ortholog is 100%.
Bootstrap support for afis|EAW24757.1 as seed ortholog is 100%.

---

Group of orthologs #7789. Best score 392 bits
Score difference with first non-orthologous sequence - *A. fumigatus*:392 *A. fischeri*:392
afum|EAL89479.1 100.00% afis|EAW24834.1 100.00%
Bootstrap support for afum|EAL89479.1 as seed ortholog is 100%.
Bootstrap support for afis|EAW24834.1 as seed ortholog is 100%.

---

Group of orthologs #7790. Best score 392 bits
Score difference with first non-orthologous sequence - *A. fumigatus*:392 *A. fischeri*:156
afum|EAL89644.1 100.00% afis|EAW24983.1 100.00%
Bootstrap support for afum|EAL89644.1 as seed ortholog is 100%.
Bootstrap support for afis|EAW24983.1 as seed ortholog is 100%.

---

Group of orthologs #7791. Best score 391 bits
Score difference with first non-orthologous sequence - *A. fumigatus*:391 *A. fischeri*:391
afum|EAL85306.1 100.00% afis|EAW20187.1 100.00%
Bootstrap support for afum|EAL85306.1 as seed ortholog is 100%.
Bootstrap support for afis|EAW20187.1 as seed ortholog is 100%.

---

Group of orthologs #7792. Best score 391 bits
Score difference with first non-orthologous sequence - *A. fumigatus*:391 *A. fischeri*:391
afum|EAL91400.1 100.00% afis|EAW17587.1 100.00%
Bootstrap support for afum|EAL91400.1 as seed ortholog is 100%.
Bootstrap support for afis|EAW17587.1 as seed ortholog is 100%.

---

Group of orthologs #7793. Best score 391 bits
Score difference with first non-orthologous sequence - *A. fumigatus*:144 *A. fischeri*:267
afum|EAL86541.1 100.00% afis|EAW23604.1 100.00%
Bootstrap support for afum|EAL86541.1 as seed ortholog is 99%.
Bootstrap support for afis|EAW23604.1 as seed ortholog is 100%.

---

Group of orthologs #7794. Best score 391 bits
Score difference with first non-orthologous sequence - *A. fumigatus*:391 *A. fischeri*:391
afum|EAL87320.1 100.00% afis|EAW23892.1 100.00%
Bootstrap support for afum|EAL87320.1 as seed ortholog is 100%.
Bootstrap support for afis|EAW23892.1 as seed ortholog is 100%.

---

Group of orthologs #7795. Best score 391 bits
Score difference with first non-orthologous sequence - *A. fumigatus*:391 *A. fischeri*:391
afum|EAL91088.1 100.00% afis|EAW22102.1 100.00%
Bootstrap support for afum|EAL91088.1 as seed ortholog is 100%.
Bootstrap support for afis|EAW22102.1 as seed ortholog is 100%.

---

Group of orthologs #7796. Best score 391 bits
Score difference with first non-orthologous sequence - *A. fumigatus*:391 *A. fischeri*:391
afum|EAL89732.1 100.00% afis|EAW25063.1 100.00%
Bootstrap support for afum|EAL89732.1 as seed ortholog is 100%.
Bootstrap support for afis|EAW25063.1 as seed ortholog is 100%.

---

Group of orthologs #7797. Best score 390 bits
Score difference with first non-orthologous sequence - *A. fumigatus*:390 *A. fischeri*:390
afum|EAL85280.1 100.00% afis|EAW20162.1 100.00%
Bootstrap support for afum|EAL85280.1 as seed ortholog is 100%.
Bootstrap support for afis|EAW20162.1 as seed ortholog is 100%.

---

Group of orthologs #7798. Best score 390 bits
Score difference with first non-orthologous sequence - *A. fumigatus*:390 *A. fischeri*:390

| | | | |
|---|---|---|---|
| afum\|EAL89178.1 | 100.00% | afis\|EAW16583.1 | 100.00% |

Bootstrap support for afum|EAL89178.1 as seed ortholog is 100%.
Bootstrap support for afis|EAW16583.1 as seed ortholog is 100%.

---

Group of orthologs #7799. Best score 390 bits
Score difference with first non-orthologous sequence - *A. fumigatus*:390 *A. fischeri*:390

| | | | |
|---|---|---|---|
| afum\|EAL93044.1 | 100.00% | afis\|EAW18369.1 | 100.00% |

Bootstrap support for afum|EAL93044.1 as seed ortholog is 100%.
Bootstrap support for afis|EAW18369.1 as seed ortholog is 100%.

---

Group of orthologs #7800. Best score 390 bits
Score difference with first non-orthologous sequence - *A. fumigatus*:390 *A. fischeri*:390

| | | | |
|---|---|---|---|
| afum\|EAL88373.1 | 100.00% | afis\|EAW23106.1 | 100.00% |

Bootstrap support for afum|EAL88373.1 as seed ortholog is 100%.
Bootstrap support for afis|EAW23106.1 as seed ortholog is 100%.

---

Group of orthologs #7801. Best score 390 bits
Score difference with first non-orthologous sequence - *A. fumigatus*:390 *A. fischeri*:302

| | | | |
|---|---|---|---|
| afum\|EAL90225.1 | 100.00% | afis\|EAW22972.1 | 100.00% |

Bootstrap support for afum|EAL90225.1 as seed ortholog is 100%.
Bootstrap support for afis|EAW22972.1 as seed ortholog is 100%.

---

Group of orthologs #7802. Best score 390 bits
Score difference with first non-orthologous sequence - *A. fumigatus*:390 *A. fischeri*:390

| | | | |
|---|---|---|---|
| afum\|EAL89469.1 | 100.00% | afis\|EAW24825.1 | 100.00% |

Bootstrap support for afum|EAL89469.1 as seed ortholog is 100%.
Bootstrap support for afis|EAW24825.1 as seed ortholog is 100%.

---

Group of orthologs #7803. Best score 389 bits
Score difference with first non-orthologous sequence - *A. fumigatus*:389 *A. fischeri*:303

| | | | |
|---|---|---|---|
| afum\|EAL86488.2 | 100.00% | afis\|EAW16960.1 | 100.00% |

Bootstrap support for afum|EAL86488.2 as seed ortholog is 100%.
Bootstrap support for afis|EAW16960.1 as seed ortholog is 100%.

---

Group of orthologs #7804. Best score 389 bits
Score difference with first non-orthologous sequence - *A. fumigatus*:389 *A. fischeri*:389

| | | | |
|---|---|---|---|
| afum\|EAL91577.1 | 100.00% | afis\|EAW17769.1 | 100.00% |

Bootstrap support for afum|EAL91577.1 as seed ortholog is 100%.
Bootstrap support for afis|EAW17769.1 as seed ortholog is 100%.

---

Group of orthologs #7805. Best score 389 bits
Score difference with first non-orthologous sequence - *A. fumigatus*:389 *A. fischeri*:389

| | | | |
|---|---|---|---|
| afum\|EAL91747.1 | 100.00% | afis\|EAW17936.1 | 100.00% |

Bootstrap support for afum|EAL91747.1 as seed ortholog is 100%.
Bootstrap support for afis|EAW17936.1 as seed ortholog is 100%.

---

Group of orthologs #7806. Best score 389 bits
Score difference with first non-orthologous sequence - *A. fumigatus*:389 *A. fischeri*:389

| | | | |
|---|---|---|---|
| afum\|EAL93605.1 | 100.00% | afis\|EAW18873.1 | 100.00% |

Bootstrap support for afum|EAL93605.1 as seed ortholog is 100%.
Bootstrap support for afis|EAW18873.1 as seed ortholog is 100%.

---

Group of orthologs #7807. Best score 389 bits

Score difference with first non-orthologous sequence - *A. fumigatus*:389 *A. fischeri*:389
afum|EAL91988.1 100.00% afis|EAW24133.1 100.00%
Bootstrap support for afum|EAL91988.1 as seed ortholog is 100%.
Bootstrap support for afis|EAW24133.1 as seed ortholog is 100%.

---

Group of orthologs #7808. Best score 388 bits
Score difference with first non-orthologous sequence - *A. fumigatus*:388 *A. fischeri*:388
afum|EAL92055.1 100.00% afis|EAW21017.1 100.00%
Bootstrap support for afum|EAL92055.1 as seed ortholog is 100%.
Bootstrap support for afis|EAW21017.1 as seed ortholog is 100%.

---

Group of orthologs #7809. Best score 387 bits
Score difference with first non-orthologous sequence - *A. fumigatus*:387 *A. fischeri*:387
afum|EAL93729.1 100.00% afis|EAW18994.1 100.00%
Bootstrap support for afum|EAL93729.1 as seed ortholog is 100%.
Bootstrap support for afis|EAW18994.1 as seed ortholog is 100%.

---

Group of orthologs #7810. Best score 387 bits
Score difference with first non-orthologous sequence - *A. fumigatus*:387 *A. fischeri*:387
afum|EAL90105.1 100.00% afis|EAW25447.1 100.00%
Bootstrap support for afum|EAL90105.1 as seed ortholog is 100%.
Bootstrap support for afis|EAW25447.1 as seed ortholog is 100%.

---

Group of orthologs #7811. Best score 387 bits
Score difference with first non-orthologous sequence - *A. fumigatus*:387 *A. fischeri*:387
afum|EAL91963.1 100.00% afis|EAW24109.1 100.00%
Bootstrap support for afum|EAL91963.1 as seed ortholog is 100%.
Bootstrap support for afis|EAW24109.1 as seed ortholog is 100%.

---

Group of orthologs #7812. Best score 386 bits
Score difference with first non-orthologous sequence - *A. fumigatus*:386 *A. fischeri*:386
afum|EAL87138.1 100.00% afis|EAW16171.1 100.00%
Bootstrap support for afum|EAL87138.1 as seed ortholog is 100%.
Bootstrap support for afis|EAW16171.1 as seed ortholog is 100%.

---

Group of orthologs #7813. Best score 386 bits
Score difference with first non-orthologous sequence - *A. fumigatus*:386 *A. fischeri*:386
afum|EAL86496.1 100.00% afis|EAW16968.1 100.00%
Bootstrap support for afum|EAL86496.1 as seed ortholog is 100%.
Bootstrap support for afis|EAW16968.1 as seed ortholog is 100%.

---

Group of orthologs #7814. Best score 386 bits
Score difference with first non-orthologous sequence - *A. fumigatus*:386 *A. fischeri*:386
afum|EAL86166.1 100.00% afis|EAW24496.1 100.00%
Bootstrap support for afum|EAL86166.1 as seed ortholog is 100%.
Bootstrap support for afis|EAW24496.1 as seed ortholog is 100%.

---

Group of orthologs #7815. Best score 386 bits
Score difference with first non-orthologous sequence - *A. fumigatus*:386 *A. fischeri*:386
afum|EAL93637.1 100.00% afis|EAW18905.1 100.00%
Bootstrap support for afum|EAL93637.1 as seed ortholog is 100%.
Bootstrap support for afis|EAW18905.1 as seed ortholog is 100%.

---

Group of orthologs #7816. Best score 386 bits
Score difference with first non-orthologous sequence - *A. fumigatus*:386 *A. fischeri*:386
afum|EAL92079.1 100.00% afis|EAW21040.1 100.00%

Bootstrap support for afum|EAL92079.1 as seed ortholog is 100%.
Bootstrap support for afis|EAW21040.1 as seed ortholog is 100%.

---

Group of orthologs #7817. Best score 386 bits
Score difference with first non-orthologous sequence - *A. fumigatus*:386 *A. fischeri*:386
afum|EAL90629.1 100.00% afis|EAW22569.1 100.00%
Bootstrap support for afum|EAL90629.1 as seed ortholog is 100%.
Bootstrap support for afis|EAW22569.1 as seed ortholog is 100%.

---

Group of orthologs #7818. Best score 386 bits
Score difference with first non-orthologous sequence - *A. fumigatus*:386 *A. fischeri*:386
afum|EAL92025.1 100.00% afis|EAW24167.1 100.00%
Bootstrap support for afum|EAL92025.1 as seed ortholog is 100%.
Bootstrap support for afis|EAW24167.1 as seed ortholog is 100%.

---

Group of orthologs #7819. Best score 386 bits
Score difference with first non-orthologous sequence - *A. fumigatus*:386 *A. fischeri*:386
afum|EBA27392.1 100.00% afis|EAW25341.1 100.00%
Bootstrap support for afum|EBA27392.1 as seed ortholog is 100%.
Bootstrap support for afis|EAW25341.1 as seed ortholog is 100%.

---

Group of orthologs #7820. Best score 385 bits
Score difference with first non-orthologous sequence - *A. fumigatus*:385 *A. fischeri*:385
afum|EBA27235.1 100.00% afis|EAW15780.1 100.00%
Bootstrap support for afum|EBA27235.1 as seed ortholog is 100%.
Bootstrap support for afis|EAW15780.1 as seed ortholog is 100%.

---

Group of orthologs #7821. Best score 385 bits
Score difference with first non-orthologous sequence - *A. fumigatus*:385 *A. fischeri*:385
afum|EAL87423.1 100.00% afis|EAW23993.1 100.00%
Bootstrap support for afum|EAL87423.1 as seed ortholog is 100%.
Bootstrap support for afis|EAW23993.1 as seed ortholog is 100%.

---

Group of orthologs #7822. Best score 384 bits
Score difference with first non-orthologous sequence - *A. fumigatus*:384 *A. fischeri*:384
afum|EAL85795.1 100.00% afis|EAW15533.1 100.00%
Bootstrap support for afum|EAL85795.1 as seed ortholog is 100%.
Bootstrap support for afis|EAW15533.1 as seed ortholog is 100%.

---

Group of orthologs #7823. Best score 384 bits
Score difference with first non-orthologous sequence - *A. fumigatus*:331 *A. fischeri*:384
afum|EAL84997.1 100.00% afis|EAW19909.1 100.00%
Bootstrap support for afum|EAL84997.1 as seed ortholog is 100%.
Bootstrap support for afis|EAW19909.1 as seed ortholog is 100%.

---

Group of orthologs #7824. Best score 384 bits
Score difference with first non-orthologous sequence - *A. fumigatus*:384 *A. fischeri*:384
afum|EBA27271.1 100.00% afis|EAW16820.1 100.00%
Bootstrap support for afum|EBA27271.1 as seed ortholog is 100%.
Bootstrap support for afis|EAW16820.1 as seed ortholog is 100%.

---

Group of orthologs #7825. Best score 384 bits
Score difference with first non-orthologous sequence - *A. fumigatus*:384 *A. fischeri*:384
afum|EAL93688.1 100.00% afis|EAW18952.1 100.00%
Bootstrap support for afum|EAL93688.1 as seed ortholog is 100%.
Bootstrap support for afis|EAW18952.1 as seed ortholog is 100%.

Group of orthologs #7826. Best score 384 bits
Score difference with first non-orthologous sequence - *A. fumigatus*:384 *A. fischeri*:384
afum|EAL92872.1 100.00% afis|EAW21859.1 100.00%
Bootstrap support for afum|EAL92872.1 as seed ortholog is 100%.
Bootstrap support for afis|EAW21859.1 as seed ortholog is 100%.

Group of orthologs #7827. Best score 383 bits
Score difference with first non-orthologous sequence - *A. fumigatus*:383 *A. fischeri*:383
afum|EAL85039.1 100.00% afis|EAW19856.1 100.00%
Bootstrap support for afum|EAL85039.1 as seed ortholog is 100%.
Bootstrap support for afis|EAW19856.1 as seed ortholog is 100%.

Group of orthologs #7828. Best score 383 bits
Score difference with first non-orthologous sequence - *A. fumigatus*:383 *A. fischeri*:383
afum|EAL91848.1 100.00% afis|EAW18037.1 100.00%
Bootstrap support for afum|EAL91848.1 as seed ortholog is 100%.
Bootstrap support for afis|EAW18037.1 as seed ortholog is 100%.

Group of orthologs #7829. Best score 383 bits
Score difference with first non-orthologous sequence - *A. fumigatus*:383 *A. fischeri*:383
afum|EAL93733.1 100.00% afis|EAW18998.1 100.00%
Bootstrap support for afum|EAL93733.1 as seed ortholog is 100%.
Bootstrap support for afis|EAW18998.1 as seed ortholog is 100%.

Group of orthologs #7830. Best score 383 bits
Score difference with first non-orthologous sequence - *A. fumigatus*:383 *A. fischeri*:383
afum|EAL93933.1 100.00% afis|EAW19203.1 100.00%
Bootstrap support for afum|EAL93933.1 as seed ortholog is 100%.
Bootstrap support for afis|EAW19203.1 as seed ortholog is 100%.

Group of orthologs #7831. Best score 383 bits
Score difference with first non-orthologous sequence - *A. fumigatus*:383 *A. fischeri*:383
afum|EAL89829.2 100.00% afis|EAW25154.1 100.00%
Bootstrap support for afum|EAL89829.2 as seed ortholog is 100%.
Bootstrap support for afis|EAW25154.1 as seed ortholog is 100%.

Group of orthologs #7832. Best score 382 bits
Score difference with first non-orthologous sequence - *A. fumigatus*:382 *A. fischeri*:382
afum|EAL88895.1 100.00% afis|EAW16274.1 100.00%
Bootstrap support for afum|EAL88895.1 as seed ortholog is 100%.
Bootstrap support for afis|EAW16274.1 as seed ortholog is 100%.

Group of orthologs #7833. Best score 382 bits
Score difference with first non-orthologous sequence - *A. fumigatus*:382 *A. fischeri*:382
afum|EBA27229.1 100.00% afis|EAW20269.1 100.00%
Bootstrap support for afum|EBA27229.1 as seed ortholog is 100%.
Bootstrap support for afis|EAW20269.1 as seed ortholog is 100%.

Group of orthologs #7834. Best score 381 bits
Score difference with first non-orthologous sequence - *A. fumigatus*:381 *A. fischeri*:381
afum|EAL84658.1 100.00% afis|EAW20975.1 100.00%
Bootstrap support for afum|EAL84658.1 as seed ortholog is 100%.
Bootstrap support for afis|EAW20975.1 as seed ortholog is 100%.

Group of orthologs #7835. Best score 381 bits

Score difference with first non-orthologous sequence - *A. fumigatus*:381 *A. fischeri*:381
afum|EAL87073.1 100.00% afis|EAW19421.1 100.00%
Bootstrap support for afum|EAL87073.1 as seed ortholog is 100%.
Bootstrap support for afis|EAW19421.1 as seed ortholog is 100%.

---

Group of orthologs #7836. Best score 381 bits
Score difference with first non-orthologous sequence - *A. fumigatus*:381 *A. fischeri*:381
afum|EAL86868.1 100.00% afis|EAW19654.1 100.00%
Bootstrap support for afum|EAL86868.1 as seed ortholog is 100%.
Bootstrap support for afis|EAW19654.1 as seed ortholog is 100%.

---

Group of orthologs #7837. Best score 381 bits
Score difference with first non-orthologous sequence - *A. fumigatus*:381 *A. fischeri*:381
afum|EAL93664.1 100.00% afis|EAW18932.1 100.00%
Bootstrap support for afum|EAL93664.1 as seed ortholog is 100%.
Bootstrap support for afis|EAW18932.1 as seed ortholog is 100%.

---

Group of orthologs #7838. Best score 381 bits
Score difference with first non-orthologous sequence - *A. fumigatus*:301 *A. fischeri*:381
afum|EAL92364.1 100.00% afis|EAW21347.1 100.00%
Bootstrap support for afum|EAL92364.1 as seed ortholog is 100%.
Bootstrap support for afis|EAW21347.1 as seed ortholog is 100%.

---

Group of orthologs #7839. Best score 381 bits
Score difference with first non-orthologous sequence - *A. fumigatus*:381 *A. fischeri*:381
afum|EAL89652.1 100.00% afis|EAW24992.1 100.00%
Bootstrap support for afum|EAL89652.1 as seed ortholog is 100%.
Bootstrap support for afis|EAW24992.1 as seed ortholog is 100%.

---

Group of orthologs #7840. Best score 380 bits
Score difference with first non-orthologous sequence - *A. fumigatus*:380 *A. fischeri*:380
afum|EAL89257.1 100.00% afis|EAW16653.1 100.00%
Bootstrap support for afum|EAL89257.1 as seed ortholog is 100%.
Bootstrap support for afis|EAW16653.1 as seed ortholog is 100%.

---

Group of orthologs #7841. Best score 380 bits
Score difference with first non-orthologous sequence - *A. fumigatus*:380 *A. fischeri*:380
afum|EAL88291.1 100.00% afis|EAW23192.1 100.00%
Bootstrap support for afum|EAL88291.1 as seed ortholog is 100%.
Bootstrap support for afis|EAW23192.1 as seed ortholog is 100%.

---

Group of orthologs #7842. Best score 380 bits
Score difference with first non-orthologous sequence - *A. fumigatus*:380 *A. fischeri*:380
afum|EAL92197.1 100.00% afis|EAW21177.1 100.00%
Bootstrap support for afum|EAL92197.1 as seed ortholog is 100%.
Bootstrap support for afis|EAW21177.1 as seed ortholog is 100%.

---

Group of orthologs #7843. Best score 380 bits
Score difference with first non-orthologous sequence - *A. fumigatus*:380 *A. fischeri*:380
afum|EAL92346.2 100.00% afis|EAW21328.1 100.00%
Bootstrap support for afum|EAL92346.2 as seed ortholog is 100%.
Bootstrap support for afis|EAW21328.1 as seed ortholog is 100%.

---

Group of orthologs #7844. Best score 380 bits
Score difference with first non-orthologous sequence - *A. fumigatus*:380 *A. fischeri*:380
afum|EAL89455.1 100.00% afis|EAW24812.1 100.00%

Bootstrap support for afum|EAL89455.1 as seed ortholog is 100%.
Bootstrap support for afis|EAW24812.1 as seed ortholog is 100%.

---

Group of orthologs #7845. Best score 380 bits
Score difference with first non-orthologous sequence - *A. fumigatus*:380 *A. fischeri*:380
afum|EAL92882.1 100.00% afis|EAW21868.1 100.00%
Bootstrap support for afum|EAL92882.1 as seed ortholog is 100%.
Bootstrap support for afis|EAW21868.1 as seed ortholog is 100%.

---

Group of orthologs #7846. Best score 379 bits
Score difference with first non-orthologous sequence - *A. fumigatus*:234 *A. fischeri*:379
afum|EAL85433.1 100.00% afis|EAW20305.1 100.00%
Bootstrap support for afum|EAL85433.1 as seed ortholog is 100%.
Bootstrap support for afis|EAW20305.1 as seed ortholog is 100%.

---

Group of orthologs #7847. Best score 379 bits
Score difference with first non-orthologous sequence - *A. fumigatus*:379 *A. fischeri*:379
afum|EAL91255.1 100.00% afis|EAW17427.1 100.00%
Bootstrap support for afum|EAL91255.1 as seed ortholog is 100%.
Bootstrap support for afis|EAW17427.1 as seed ortholog is 100%.

---

Group of orthologs #7848. Best score 379 bits
Score difference with first non-orthologous sequence - *A. fumigatus*:379 *A. fischeri*:379
afum|EBA27279.1 100.00% afis|EAW17081.1 100.00%
Bootstrap support for afum|EBA27279.1 as seed ortholog is 100%.
Bootstrap support for afis|EAW17081.1 as seed ortholog is 100%.

---

Group of orthologs #7849. Best score 378 bits
Score difference with first non-orthologous sequence - *A. fumigatus*:378 *A. fischeri*:378
afum|EAL84656.1 100.00% afis|EAW20977.1 100.00%
Bootstrap support for afum|EAL84656.1 as seed ortholog is 100%.
Bootstrap support for afis|EAW20977.1 as seed ortholog is 100%.

---

Group of orthologs #7850. Best score 378 bits
Score difference with first non-orthologous sequence - *A. fumigatus*:378 *A. fischeri*:315
afum|EAL85873.2 100.00% afis|EAW24203.1 100.00%
Bootstrap support for afum|EAL85873.2 as seed ortholog is 100%.
Bootstrap support for afis|EAW24203.1 as seed ortholog is 100%.

---

Group of orthologs #7851. Best score 378 bits
Score difference with first non-orthologous sequence - *A. fumigatus*:378 *A. fischeri*:378
afum|EAL88022.1 100.00% afis|EAW23440.1 100.00%
Bootstrap support for afum|EAL88022.1 as seed ortholog is 100%.
Bootstrap support for afis|EAW23440.1 as seed ortholog is 100%.

---

Group of orthologs #7852. Best score 378 bits
Score difference with first non-orthologous sequence - *A. fumigatus*:378 *A. fischeri*:378
afum|EAL90892.1 100.00% afis|EAW22289.1 100.00%
Bootstrap support for afum|EAL90892.1 as seed ortholog is 100%.
Bootstrap support for afis|EAW22289.1 as seed ortholog is 100%.

---

Group of orthologs #7853. Best score 378 bits
Score difference with first non-orthologous sequence - *A. fumigatus*:378 *A. fischeri*:378
afum|EAL92965.1 100.00% afis|EAW21951.1 100.00%
Bootstrap support for afum|EAL92965.1 as seed ortholog is 100%.
Bootstrap support for afis|EAW21951.1 as seed ortholog is 100%.

Group of orthologs #7854. Best score 377 bits
Score difference with first non-orthologous sequence - *A. fumigatus*:377 *A. fischeri*:377
afum|EAL85658.1 100.00% afis|EAW15690.1 100.00%
Bootstrap support for afum|EAL85658.1 as seed ortholog is 100%.
Bootstrap support for afis|EAW15690.1 as seed ortholog is 100%.

Group of orthologs #7855. Best score 377 bits
Score difference with first non-orthologous sequence - *A. fumigatus*:377 *A. fischeri*:197
afum|EAL91643.1 100.00% afis|EAW17833.1 100.00%
Bootstrap support for afum|EAL91643.1 as seed ortholog is 100%.
Bootstrap support for afis|EAW17833.1 as seed ortholog is 100%.

Group of orthologs #7856. Best score 377 bits
Score difference with first non-orthologous sequence - *A. fumigatus*:377 *A. fischeri*:377
afum|EAL92532.1 100.00% afis|EAW21512.1 100.00%
Bootstrap support for afum|EAL92532.1 as seed ortholog is 100%.
Bootstrap support for afis|EAW21512.1 as seed ortholog is 100%.

Group of orthologs #7857. Best score 377 bits
Score difference with first non-orthologous sequence - *A. fumigatus*:377 *A. fischeri*:377
afum|EAL92650.1 100.00% afis|EAW21616.1 100.00%
Bootstrap support for afum|EAL92650.1 as seed ortholog is 100%.
Bootstrap support for afis|EAW21616.1 as seed ortholog is 100%.

Group of orthologs #7858. Best score 377 bits
Score difference with first non-orthologous sequence - *A. fumigatus*:377 *A. fischeri*:377
afum|EAL92828.1 100.00% afis|EAW21802.1 100.00%
Bootstrap support for afum|EAL92828.1 as seed ortholog is 100%.
Bootstrap support for afis|EAW21802.1 as seed ortholog is 100%.

Group of orthologs #7859. Best score 377 bits
Score difference with first non-orthologous sequence - *A. fumigatus*:377 *A. fischeri*:377
afum|EAL89725.1 100.00% afis|EAW25057.1 100.00%
Bootstrap support for afum|EAL89725.1 as seed ortholog is 100%.
Bootstrap support for afis|EAW25057.1 as seed ortholog is 100%.

Group of orthologs #7860. Best score 376 bits
Score difference with first non-orthologous sequence - *A. fumigatus*:376 *A. fischeri*:376
afum|EAL88233.1 100.00% afis|EAW23248.1 100.00%
Bootstrap support for afum|EAL88233.1 as seed ortholog is 100%.
Bootstrap support for afis|EAW23248.1 as seed ortholog is 100%.

Group of orthologs #7861. Best score 376 bits
Score difference with first non-orthologous sequence - *A. fumigatus*:376 *A. fischeri*:376
afum|EAL90853.1 100.00% afis|EAW22339.1 100.00%
Bootstrap support for afum|EAL90853.1 as seed ortholog is 100%.
Bootstrap support for afis|EAW22339.1 as seed ortholog is 100%.

Group of orthologs #7862. Best score 375 bits
Score difference with first non-orthologous sequence - *A. fumigatus*:375 *A. fischeri*:375
afum|EAL85797.1 100.00% afis|EAW15531.1 100.00%
Bootstrap support for afum|EAL85797.1 as seed ortholog is 100%.
Bootstrap support for afis|EAW15531.1 as seed ortholog is 100%.

Group of orthologs #7863. Best score 375 bits

Score difference with first non-orthologous sequence - *A. fumigatus*:375 *A. fischeri*:375
afum|EAL85519.1 100.00% afis|EAW15833.1 100.00%
Bootstrap support for afum|EAL85519.1 as seed ortholog is 100%.
Bootstrap support for afis|EAW15833.1 as seed ortholog is 100%.

---

Group of orthologs #7864. Best score 375 bits
Score difference with first non-orthologous sequence - *A. fumigatus*:375 *A. fischeri*:375
afum|EAL85957.1 100.00% afis|EAW24288.1 100.00%
Bootstrap support for afum|EAL85957.1 as seed ortholog is 100%.
Bootstrap support for afis|EAW24288.1 as seed ortholog is 100%.

---

Group of orthologs #7865. Best score 375 bits
Score difference with first non-orthologous sequence - *A. fumigatus*:375 *A. fischeri*:375
afum|EAL86029.1 100.00% afis|EAW24359.1 100.00%
Bootstrap support for afum|EAL86029.1 as seed ortholog is 100%.
Bootstrap support for afis|EAW24359.1 as seed ortholog is 100%.

---

Group of orthologs #7866. Best score 375 bits
Score difference with first non-orthologous sequence - *A. fumigatus*:375 *A. fischeri*:375
afum|EAL90255.1 100.00% afis|EAW22941.1 100.00%
Bootstrap support for afum|EAL90255.1 as seed ortholog is 100%.
Bootstrap support for afis|EAW22941.1 as seed ortholog is 100%.

---

Group of orthologs #7867. Best score 375 bits
Score difference with first non-orthologous sequence - *A. fumigatus*:375 *A. fischeri*:375
afum|EAL92455.1 100.00% afis|EAW21439.1 100.00%
Bootstrap support for afum|EAL92455.1 as seed ortholog is 100%.
Bootstrap support for afis|EAW21439.1 as seed ortholog is 100%.

---

Group of orthologs #7868. Best score 374 bits
Score difference with first non-orthologous sequence - *A. fumigatus*:374 *A. fischeri*:374
afum|EAL86379.1 100.00% afis|EAW16831.1 100.00%
Bootstrap support for afum|EAL86379.1 as seed ortholog is 100%.
Bootstrap support for afis|EAW16831.1 as seed ortholog is 100%.

---

Group of orthologs #7869. Best score 374 bits
Score difference with first non-orthologous sequence - *A. fumigatus*:374 *A. fischeri*:374
afum|EAL91401.1 100.00% afis|EAW17588.1 100.00%
Bootstrap support for afum|EAL91401.1 as seed ortholog is 100%.
Bootstrap support for afis|EAW17588.1 as seed ortholog is 100%.

---

Group of orthologs #7870. Best score 374 bits
Score difference with first non-orthologous sequence - *A. fumigatus*:374 *A. fischeri*:374
afum|EAL84301.1 100.00% afis|EAW25595.1 100.00%
Bootstrap support for afum|EAL84301.1 as seed ortholog is 100%.
Bootstrap support for afis|EAW25595.1 as seed ortholog is 100%.

---

Group of orthologs #7871. Best score 374 bits
Score difference with first non-orthologous sequence - *A. fumigatus*:374 *A. fischeri*:374
afum|EAL94060.1 100.00% afis|EAW19341.1 100.00%
Bootstrap support for afum|EAL94060.1 as seed ortholog is 100%.
Bootstrap support for afis|EAW19341.1 as seed ortholog is 100%.

---

Group of orthologs #7872. Best score 374 bits
Score difference with first non-orthologous sequence - *A. fumigatus*:374 *A. fischeri*:374
afum|EAL92182.1 100.00% afis|EAW21163.1 100.00%

Bootstrap support for afum|EAL92182.1 as seed ortholog is 100%.
Bootstrap support for afis|EAW21163.1 as seed ortholog is 100%.

---

Group of orthologs #7873. Best score 373 bits
Score difference with first non-orthologous sequence - *A. fumigatus*:373 *A. fischeri*:373
afum|EAL86396.1 100.00% afis|EAW16850.1 100.00%
Bootstrap support for afum|EAL86396.1 as seed ortholog is 100%.
Bootstrap support for afis|EAW16850.1 as seed ortholog is 100%.

---

Group of orthologs #7874. Best score 373 bits
Score difference with first non-orthologous sequence - *A. fumigatus*:373 *A. fischeri*:373
afum|EAL88306.1 100.00% afis|EAW23177.1 100.00%
Bootstrap support for afum|EAL88306.1 as seed ortholog is 100%.
Bootstrap support for afis|EAW23177.1 as seed ortholog is 100%.

---

Group of orthologs #7875. Best score 373 bits
Score difference with first non-orthologous sequence - *A. fumigatus*:373 *A. fischeri*:373
afum|EAL88118.1 100.00% afis|EAW23358.1 100.00%
Bootstrap support for afum|EAL88118.1 as seed ortholog is 100%.
Bootstrap support for afis|EAW23358.1 as seed ortholog is 100%.

---

Group of orthologs #7876. Best score 373 bits
Score difference with first non-orthologous sequence - *A. fumigatus*:373 *A. fischeri*:373
afum|EAL93876.1 100.00% afis|EAW19145.1 100.00%
Bootstrap support for afum|EAL93876.1 as seed ortholog is 100%.
Bootstrap support for afis|EAW19145.1 as seed ortholog is 100%.

---

Group of orthologs #7877. Best score 373 bits
Score difference with first non-orthologous sequence - *A. fumigatus*:373 *A. fischeri*:373
afum|EAL90081.1 100.00% afis|EAW25422.1 100.00%
Bootstrap support for afum|EAL90081.1 as seed ortholog is 100%.
Bootstrap support for afis|EAW25422.1 as seed ortholog is 100%.

---

Group of orthologs #7878. Best score 373 bits
Score difference with first non-orthologous sequence - *A. fumigatus*:373 *A. fischeri*:373
afum|EAL90191.2 100.00% afis|EAW25526.1 100.00%
Bootstrap support for afum|EAL90191.2 as seed ortholog is 100%.
Bootstrap support for afis|EAW25526.1 as seed ortholog is 100%.

---

Group of orthologs #7879. Best score 373 bits
Score difference with first non-orthologous sequence - *A. fumigatus*:373 *A. fischeri*:313
afum|EBA27176.1 100.00% afis|EAW25722.1 100.00%
Bootstrap support for afum|EBA27176.1 as seed ortholog is 100%.
Bootstrap support for afis|EAW25722.1 as seed ortholog is 100%.

---

Group of orthologs #7880. Best score 371 bits
Score difference with first non-orthologous sequence - *A. fumigatus*:314 *A. fischeri*:289
afum|EAL86417.1 100.00% afis|EAW16880.1 100.00%
Bootstrap support for afum|EAL86417.1 as seed ortholog is 100%.
Bootstrap support for afis|EAW16880.1 as seed ortholog is 100%.

---

Group of orthologs #7881. Best score 371 bits
Score difference with first non-orthologous sequence - *A. fumigatus*:371 *A. fischeri*:313
afum|EAL85485.1 100.00% afis|EAW20371.1 100.00%
Bootstrap support for afum|EAL85485.1 as seed ortholog is 100%.
Bootstrap support for afis|EAW20371.1 as seed ortholog is 100%.

---

Group of orthologs #7882. Best score 371 bits
Score difference with first non-orthologous sequence - *A. fumigatus*:371 *A. fischeri*:371
afum|EAL91230.1 100.00% afis|EAW17392.1 100.00%
Bootstrap support for afum|EAL91230.1 as seed ortholog is 100%.
Bootstrap support for afis|EAW17392.1 as seed ortholog is 100%.

---

Group of orthologs #7883. Best score 371 bits
Score difference with first non-orthologous sequence - *A. fumigatus*:371 *A. fischeri*:371
afum|EAL93628.1 100.00% afis|EAW18896.1 100.00%
Bootstrap support for afum|EAL93628.1 as seed ortholog is 100%.
Bootstrap support for afis|EAW18896.1 as seed ortholog is 100%.

---

Group of orthologs #7884. Best score 370 bits
Score difference with first non-orthologous sequence - *A. fumigatus*:370 *A. fischeri*:370
afum|EAL85158.1 100.00% afis|EAW20037.1 100.00%
Bootstrap support for afum|EAL85158.1 as seed ortholog is 100%.
Bootstrap support for afis|EAW20037.1 as seed ortholog is 100%.

---

Group of orthologs #7885. Best score 370 bits
Score difference with first non-orthologous sequence - *A. fumigatus*:370 *A. fischeri*:370
afum|EAL91880.1 100.00% afis|EAW18068.1 100.00%
Bootstrap support for afum|EAL91880.1 as seed ortholog is 100%.
Bootstrap support for afis|EAW18068.1 as seed ortholog is 100%.

---

Group of orthologs #7886. Best score 370 bits
Score difference with first non-orthologous sequence - *A. fumigatus*:370 *A. fischeri*:370
afum|EAL93592.1 100.00% afis|EAW18860.1 100.00%
Bootstrap support for afum|EAL93592.1 as seed ortholog is 100%.
Bootstrap support for afis|EAW18860.1 as seed ortholog is 100%.

---

Group of orthologs #7887. Best score 370 bits
Score difference with first non-orthologous sequence - *A. fumigatus*:370 *A. fischeri*:370
afum|EAL93952.1 100.00% afis|EAW19219.1 100.00%
Bootstrap support for afum|EAL93952.1 as seed ortholog is 100%.
Bootstrap support for afis|EAW19219.1 as seed ortholog is 100%.

---

Group of orthologs #7888. Best score 370 bits
Score difference with first non-orthologous sequence - *A. fumigatus*:241 *A. fischeri*:370
afum|EAL90751.1 100.00% afis|EAW22440.1 100.00%
Bootstrap support for afum|EAL90751.1 as seed ortholog is 100%.
Bootstrap support for afis|EAW22440.1 as seed ortholog is 100%.

---

Group of orthologs #7889. Best score 370 bits
Score difference with first non-orthologous sequence - *A. fumigatus*:370 *A. fischeri*:370
afum|EAL89852.1 100.00% afis|EAW25178.1 100.00%
Bootstrap support for afum|EAL89852.1 as seed ortholog is 100%.
Bootstrap support for afis|EAW25178.1 as seed ortholog is 100%.

---

Group of orthologs #7890. Best score 369 bits
Score difference with first non-orthologous sequence - *A. fumigatus*:369 *A. fischeri*:369
afum|EAL91332.1 100.00% afis|EAW17514.1 100.00%
Bootstrap support for afum|EAL91332.1 as seed ortholog is 100%.
Bootstrap support for afis|EAW17514.1 as seed ortholog is 100%.

---

Group of orthologs #7891. Best score 369 bits

Score difference with first non-orthologous sequence - *A. fumigatus*:369   *A. fischeri*:369
afum|EAL91549.1 100.00% afis|EAW17734.1 100.00%
Bootstrap support for afum|EAL91549.1 as seed ortholog is 100%.
Bootstrap support for afis|EAW17734.1 as seed ortholog is 100%.

---

Group of orthologs #7892. Best score 369 bits
Score difference with first non-orthologous sequence - *A. fumigatus*:369   *A. fischeri*:369
afum|EAL90423.1 100.00% afis|EAW22775.1 100.00%
Bootstrap support for afum|EAL90423.1 as seed ortholog is 100%.
Bootstrap support for afis|EAW22775.1 as seed ortholog is 100%.

---

Group of orthologs #7893. Best score 368 bits
Score difference with first non-orthologous sequence - *A. fumigatus*:368   *A. fischeri*:368
afum|EBA27243.1 100.00% afis|EAW15542.1 100.00%
Bootstrap support for afum|EBA27243.1 as seed ortholog is 100%.
Bootstrap support for afis|EAW15542.1 as seed ortholog is 100%.

---

Group of orthologs #7894. Best score 368 bits
Score difference with first non-orthologous sequence - *A. fumigatus*:368   *A. fischeri*:368
afum|EAL88419.1 100.00% afis|EAW23060.1 100.00%
Bootstrap support for afum|EAL88419.1 as seed ortholog is 100%.
Bootstrap support for afis|EAW23060.1 as seed ortholog is 100%.

---

Group of orthologs #7895. Best score 368 bits
Score difference with first non-orthologous sequence - *A. fumigatus*:368   *A. fischeri*:368
afum|EAL93772.1 100.00% afis|EAW19043.1 100.00%
Bootstrap support for afum|EAL93772.1 as seed ortholog is 100%.
Bootstrap support for afis|EAW19043.1 as seed ortholog is 100%.

---

Group of orthologs #7896. Best score 368 bits
Score difference with first non-orthologous sequence - *A. fumigatus*:368   *A. fischeri*:368
afum|EAL93824.1 100.00% afis|EAW19093.1 100.00%
Bootstrap support for afum|EAL93824.1 as seed ortholog is 100%.
Bootstrap support for afis|EAW19093.1 as seed ortholog is 100%.

---

Group of orthologs #7897. Best score 368 bits
Score difference with first non-orthologous sequence - *A. fumigatus*:368   *A. fischeri*:368
afum|EAL93003.1 100.00% afis|EAW21987.1 100.00%
Bootstrap support for afum|EAL93003.1 as seed ortholog is 100%.
Bootstrap support for afis|EAW21987.1 as seed ortholog is 100%.

---

Group of orthologs #7898. Best score 368 bits
Score difference with first non-orthologous sequence - *A. fumigatus*:368   *A. fischeri*:368
afum|EAL90118.2 100.00% afis|EAW25458.1 100.00%
Bootstrap support for afum|EAL90118.2 as seed ortholog is 100%.
Bootstrap support for afis|EAW25458.1 as seed ortholog is 100%.

---

Group of orthologs #7899. Best score 368 bits
Score difference with first non-orthologous sequence - *A. fumigatus*:368   *A. fischeri*:368
afum|EAL91930.2 100.00% afis|EAW24074.1 100.00%
Bootstrap support for afum|EAL91930.2 as seed ortholog is 100%.
Bootstrap support for afis|EAW24074.1 as seed ortholog is 100%.

---

Group of orthologs #7900. Best score 368 bits
Score difference with first non-orthologous sequence - *A. fumigatus*:368   *A. fischeri*:368
afum|EBA27301.1 100.00% afis|EAW23727.1 100.00%

Bootstrap support for afum|EBA27301.1 as seed ortholog is 100%.
Bootstrap support for afis|EAW23727.1 as seed ortholog is 100%.

---

Group of orthologs #7901. Best score 367 bits
Score difference with first non-orthologous sequence - *A. fumigatus*:279 *A. fischeri*:367
afum|EAL86537.1 100.00% afis|EAW17053.1 100.00%
Bootstrap support for afum|EAL86537.1 as seed ortholog is 100%.
Bootstrap support for afis|EAW17053.1 as seed ortholog is 100%.

---

Group of orthologs #7902. Best score 367 bits
Score difference with first non-orthologous sequence - *A. fumigatus*:367 *A. fischeri*:367
afum|EAL88608.1 100.00% afis|EAW15976.1 100.00%
Bootstrap support for afum|EAL88608.1 as seed ortholog is 100%.
Bootstrap support for afis|EAW15976.1 as seed ortholog is 100%.

---

Group of orthologs #7903. Best score 367 bits
Score difference with first non-orthologous sequence - *A. fumigatus*:367 *A. fischeri*:367
afum|EAL88197.2 100.00% afis|EAW23282.1 100.00%
Bootstrap support for afum|EAL88197.2 as seed ortholog is 100%.
Bootstrap support for afis|EAW23282.1 as seed ortholog is 100%.

---

Group of orthologs #7904. Best score 367 bits
Score difference with first non-orthologous sequence - *A. fumigatus*:367 *A. fischeri*:367
afum|EAL87461.1 100.00% afis|EAW24030.1 100.00%
Bootstrap support for afum|EAL87461.1 as seed ortholog is 100%.
Bootstrap support for afis|EAW24030.1 as seed ortholog is 100%.

---

Group of orthologs #7905. Best score 367 bits
Score difference with first non-orthologous sequence - *A. fumigatus*:367 *A. fischeri*:367
afum|EAL92827.1 100.00% afis|EAW21801.1 100.00%
Bootstrap support for afum|EAL92827.1 as seed ortholog is 100%.
Bootstrap support for afis|EAW21801.1 as seed ortholog is 100%.

---

Group of orthologs #7906. Best score 367 bits
Score difference with first non-orthologous sequence - *A. fumigatus*:367 *A. fischeri*:367
afum|EAL89844.1 100.00% afis|EAW25171.1 100.00%
Bootstrap support for afum|EAL89844.1 as seed ortholog is 100%.
Bootstrap support for afis|EAW25171.1 as seed ortholog is 100%.

---

Group of orthologs #7907. Best score 367 bits
Score difference with first non-orthologous sequence - *A. fumigatus*:367 *A. fischeri*:302
afum|EBA27389.1 100.00% afis|EAW25199.1 100.00%
Bootstrap support for afum|EBA27389.1 as seed ortholog is 100%.
Bootstrap support for afis|EAW25199.1 as seed ortholog is 100%.

---

Group of orthologs #7908. Best score 366 bits
Score difference with first non-orthologous sequence - *A. fumigatus*:366 *A. fischeri*:366
afum|EAL87049.1 100.00% afis|EAW19446.1 100.00%
Bootstrap support for afum|EAL87049.1 as seed ortholog is 100%.
Bootstrap support for afis|EAW19446.1 as seed ortholog is 100%.

---

Group of orthologs #7909. Best score 366 bits
Score difference with first non-orthologous sequence - *A. fumigatus*:196 *A. fischeri*:11
afum|EAL87770.2 100.00% afis|EAW19678.1 100.00%
Bootstrap support for afum|EAL87770.2 as seed ortholog is 100%.

Bootstrap support for afis|EAW19678.1 as seed ortholog is 60%. Alternative seed ortholog is afis|EAW24039.1 (11 bits away from this cluster)

---

Group of orthologs #7910. Best score 366 bits
Score difference with first non-orthologous sequence - *A. fumigatus*:366 *A. fischeri*:366
afum|EAL94022.1 100.00% afis|EAW19304.1 100.00%
Bootstrap support for afum|EAL94022.1 as seed ortholog is 100%.
Bootstrap support for afis|EAW19304.1 as seed ortholog is 100%.

---

Group of orthologs #7911. Best score 366 bits
Score difference with first non-orthologous sequence - *A. fumigatus*:366 *A. fischeri*:366
afum|EAL92331.1 100.00% afis|EAW21314.1 100.00%
Bootstrap support for afum|EAL92331.1 as seed ortholog is 100%.
Bootstrap support for afis|EAW21314.1 as seed ortholog is 100%.

---

Group of orthologs #7912. Best score 365 bits
Score difference with first non-orthologous sequence - *A. fumigatus*:365 *A. fischeri*:365
afum|EAL91677.1 100.00% afis|EAW17869.1 100.00%
Bootstrap support for afum|EAL91677.1 as seed ortholog is 100%.
Bootstrap support for afis|EAW17869.1 as seed ortholog is 100%.

---

Group of orthologs #7913. Best score 365 bits
Score difference with first non-orthologous sequence - *A. fumigatus*:365 *A. fischeri*:365
afum|EBA27326.1 100.00% afis|EAW17485.1 100.00%
Bootstrap support for afum|EBA27326.1 as seed ortholog is 100%.
Bootstrap support for afis|EAW17485.1 as seed ortholog is 100%.

---

Group of orthologs #7914. Best score 365 bits
Score difference with first non-orthologous sequence - *A. fumigatus*:365 *A. fischeri*:365
afum|EAL93304.2 100.00% afis|EAW18596.1 100.00%
Bootstrap support for afum|EAL93304.2 as seed ortholog is 100%.
Bootstrap support for afis|EAW18596.1 as seed ortholog is 100%.

---

Group of orthologs #7915. Best score 365 bits
Score difference with first non-orthologous sequence - *A. fumigatus*:365 *A. fischeri*:365
afum|EAL90280.1 100.00% afis|EAW22917.1 100.00%
Bootstrap support for afum|EAL90280.1 as seed ortholog is 100%.
Bootstrap support for afis|EAW22917.1 as seed ortholog is 100%.

---

Group of orthologs #7916. Best score 364 bits
Score difference with first non-orthologous sequence - *A. fumigatus*:69 *A. fischeri*:364
afum|EAL86288.2 100.00% afis|EAW16389.1 100.00%
afum|EAL89015.1 100.00%
Bootstrap support for afum|EAL86288.2 as seed ortholog is 99%.
Bootstrap support for afum|EAL89015.1 as seed ortholog is 99%.
Bootstrap support for afis|EAW16389.1 as seed ortholog is 100%.

---

Group of orthologs #7917. Best score 364 bits
Score difference with first non-orthologous sequence - *A. fumigatus*:364 *A. fischeri*:364
afum|EAL84991.1 100.00% afis|EAW19916.1 100.00%
Bootstrap support for afum|EAL84991.1 as seed ortholog is 100%.
Bootstrap support for afis|EAW19916.1 as seed ortholog is 100%.

---

Group of orthologs #7918. Best score 364 bits
Score difference with first non-orthologous sequence - *A. fumigatus*:364 *A. fischeri*:364
afum|EAL84992.1 100.00% afis|EAW19915.1 100.00%

Bootstrap support for afum|EAL84992.1 as seed ortholog is 100%.
Bootstrap support for afis|EAW19915.1 as seed ortholog is 100%.

---

Group of orthologs #7919. Best score 364 bits
Score difference with first non-orthologous sequence - *A. fumigatus*:364 *A. fischeri*:364
afum|EAL91313.1 100.00% afis|EAW17491.1 100.00%
Bootstrap support for afum|EAL91313.1 as seed ortholog is 100%.
Bootstrap support for afis|EAW17491.1 as seed ortholog is 100%.

---

Group of orthologs #7920. Best score 364 bits
Score difference with first non-orthologous sequence - *A. fumigatus*:194 *A. fischeri*:364
afum|EAL86105.1 100.00% afis|EAW24437.1 100.00%
Bootstrap support for afum|EAL86105.1 as seed ortholog is 100%.
Bootstrap support for afis|EAW24437.1 as seed ortholog is 100%.

---

Group of orthologs #7921. Best score 364 bits
Score difference with first non-orthologous sequence - *A. fumigatus*:364 *A. fischeri*:364
afum|EAL93133.2 100.00% afis|EAW18448.1 100.00%
Bootstrap support for afum|EAL93133.2 as seed ortholog is 100%.
Bootstrap support for afis|EAW18448.1 as seed ortholog is 100%.

---

Group of orthologs #7922. Best score 364 bits
Score difference with first non-orthologous sequence - *A. fumigatus*:364 *A. fischeri*:364
afum|EAL90559.1 100.00% afis|EAW22634.1 100.00%
Bootstrap support for afum|EAL90559.1 as seed ortholog is 100%.
Bootstrap support for afis|EAW22634.1 as seed ortholog is 100%.

---

Group of orthologs #7923. Best score 364 bits
Score difference with first non-orthologous sequence - *A. fumigatus*:364 *A. fischeri*:314
afum|EBA27512.1 100.00% afis|EAW19326.1 100.00%
Bootstrap support for afum|EBA27512.1 as seed ortholog is 100%.
Bootstrap support for afis|EAW19326.1 as seed ortholog is 100%.

---

Group of orthologs #7924. Best score 364 bits
Score difference with first non-orthologous sequence - *A. fumigatus*:364 *A. fischeri*:364
afum|EAL89867.1 100.00% afis|EAW25206.1 100.00%
Bootstrap support for afum|EAL89867.1 as seed ortholog is 100%.
Bootstrap support for afis|EAW25206.1 as seed ortholog is 100%.

---

Group of orthologs #7925. Best score 363 bits
Score difference with first non-orthologous sequence - *A. fumigatus*:363 *A. fischeri*:363
afum|EAL88514.1 100.00% afis|EAW15880.1 100.00%
Bootstrap support for afum|EAL88514.1 as seed ortholog is 100%.
Bootstrap support for afis|EAW15880.1 as seed ortholog is 100%.

---

Group of orthologs #7926. Best score 363 bits
Score difference with first non-orthologous sequence - *A. fumigatus*:363 *A. fischeri*:363
afum|EAL89056.2 100.00% afis|EAW16460.1 100.00%
Bootstrap support for afum|EAL89056.2 as seed ortholog is 100%.
Bootstrap support for afis|EAW16460.1 as seed ortholog is 100%.

---

Group of orthologs #7927. Best score 363 bits
Score difference with first non-orthologous sequence - *A. fumigatus*:363 *A. fischeri*:363
afum|EBA27246.1 100.00% afis|EAW15513.1 100.00%
Bootstrap support for afum|EBA27246.1 as seed ortholog is 100%.
Bootstrap support for afis|EAW15513.1 as seed ortholog is 100%.

Group of orthologs #7928. Best score 363 bits
Score difference with first non-orthologous sequence - *A. fumigatus*:363 *A. fischeri*:363
afum|EAL93339.1 100.00% afis|EAW18627.1 100.00%
Bootstrap support for afum|EAL93339.1 as seed ortholog is 100%.
Bootstrap support for afis|EAW18627.1 as seed ortholog is 100%.

Group of orthologs #7929. Best score 363 bits
Score difference with first non-orthologous sequence - *A. fumigatus*:363 *A. fischeri*:363
afum|EAL92070.1 100.00% afis|EAW21031.1 100.00%
Bootstrap support for afum|EAL92070.1 as seed ortholog is 100%.
Bootstrap support for afis|EAW21031.1 as seed ortholog is 100%.

Group of orthologs #7930. Best score 362 bits
Score difference with first non-orthologous sequence - *A. fumigatus*:362 *A. fischeri*:362
afum|EAL85564.1 100.00% afis|EAW15786.1 100.00%
Bootstrap support for afum|EAL85564.1 as seed ortholog is 100%.
Bootstrap support for afis|EAW15786.1 as seed ortholog is 100%.

Group of orthologs #7931. Best score 362 bits
Score difference with first non-orthologous sequence - *A. fumigatus*:362 *A. fischeri*:362
afum|EAL88762.1 100.00% afis|EAW16132.1 100.00%
Bootstrap support for afum|EAL88762.1 as seed ortholog is 100%.
Bootstrap support for afis|EAW16132.1 as seed ortholog is 100%.

Group of orthologs #7932. Best score 362 bits
Score difference with first non-orthologous sequence - *A. fumigatus*:362 *A. fischeri*:362
afum|EAL93540.1 100.00% afis|EAW18804.1 100.00%
Bootstrap support for afum|EAL93540.1 as seed ortholog is 100%.
Bootstrap support for afis|EAW18804.1 as seed ortholog is 100%.

Group of orthologs #7933. Best score 362 bits
Score difference with first non-orthologous sequence - *A. fumigatus*:362 *A. fischeri*:311
afum|EAL93557.1 100.00% afis|EAW18820.1 100.00%
Bootstrap support for afum|EAL93557.1 as seed ortholog is 100%.
Bootstrap support for afis|EAW18820.1 as seed ortholog is 100%.

Group of orthologs #7934. Best score 362 bits
Score difference with first non-orthologous sequence - *A. fumigatus*:362 *A. fischeri*:362
afum|EAL90196.2 100.00% afis|EAW25532.1 100.00%
Bootstrap support for afum|EAL90196.2 as seed ortholog is 100%.
Bootstrap support for afis|EAW25532.1 as seed ortholog is 100%.

Group of orthologs #7935. Best score 361 bits
Score difference with first non-orthologous sequence - *A. fumigatus*:361 *A. fischeri*:361
afum|EAL91430.1 100.00% afis|EAW17617.1 100.00%
Bootstrap support for afum|EAL91430.1 as seed ortholog is 100%.
Bootstrap support for afis|EAW17617.1 as seed ortholog is 100%.

Group of orthologs #7936. Best score 361 bits
Score difference with first non-orthologous sequence - *A. fumigatus*:361 *A. fischeri*:361
afum|EAL88216.1 100.00% afis|EAW23264.1 100.00%
Bootstrap support for afum|EAL88216.1 as seed ortholog is 100%.
Bootstrap support for afis|EAW23264.1 as seed ortholog is 100%.

Group of orthologs #7937. Best score 361 bits

Score difference with first non-orthologous sequence - *A. fumigatus*:361 *A. fischeri*:361
afum|EAL92922.2 100.00% afis|EAW21910.1 100.00%
Bootstrap support for afum|EAL92922.2 as seed ortholog is 100%.
Bootstrap support for afis|EAW21910.1 as seed ortholog is 100%.

---

Group of orthologs #7938. Best score 360 bits
Score difference with first non-orthologous sequence - *A. fumigatus*:360 *A. fischeri*:360
afum|EAL88163.1 100.00% afis|EAW23314.1 100.00%
Bootstrap support for afum|EAL88163.1 as seed ortholog is 100%.
Bootstrap support for afis|EAW23314.1 as seed ortholog is 100%.

---

Group of orthologs #7939. Best score 360 bits
Score difference with first non-orthologous sequence - *A. fumigatus*:360 *A. fischeri*:360
afum|EAL92714.1 100.00% afis|EAW21680.1 100.00%
Bootstrap support for afum|EAL92714.1 as seed ortholog is 100%.
Bootstrap support for afis|EAW21680.1 as seed ortholog is 100%.

---

Group of orthologs #7940. Best score 359 bits
Score difference with first non-orthologous sequence - *A. fumigatus*:359 *A. fischeri*:359
afum|EAL84197.1 100.00% afis|EAW15449.1 100.00%
Bootstrap support for afum|EAL84197.1 as seed ortholog is 100%.
Bootstrap support for afis|EAW15449.1 as seed ortholog is 100%.

---

Group of orthologs #7941. Best score 359 bits
Score difference with first non-orthologous sequence - *A. fumigatus*:359 *A. fischeri*:359
afum|EBA27353.1 100.00% afis|EAW16008.1 100.00%
Bootstrap support for afum|EBA27353.1 as seed ortholog is 100%.
Bootstrap support for afis|EAW16008.1 as seed ortholog is 100%.

---

Group of orthologs #7942. Best score 359 bits
Score difference with first non-orthologous sequence - *A. fumigatus*:359 *A. fischeri*:359
afum|EAL90790.1 100.00% afis|EAW22401.1 100.00%
Bootstrap support for afum|EAL90790.1 as seed ortholog is 100%.
Bootstrap support for afis|EAW22401.1 as seed ortholog is 100%.

---

Group of orthologs #7943. Best score 359 bits
Score difference with first non-orthologous sequence - *A. fumigatus*:359 *A. fischeri*:359
afum|EAL92880.2 100.00% afis|EAW21866.1 100.00%
Bootstrap support for afum|EAL92880.2 as seed ortholog is 100%.
Bootstrap support for afis|EAW21866.1 as seed ortholog is 100%.

---

Group of orthologs #7944. Best score 359 bits
Score difference with first non-orthologous sequence - *A. fumigatus*:359 *A. fischeri*:359
afum|EAL89783.2 100.00% afis|EAW25112.1 100.00%
Bootstrap support for afum|EAL89783.2 as seed ortholog is 100%.
Bootstrap support for afis|EAW25112.1 as seed ortholog is 100%.

---

Group of orthologs #7945. Best score 358 bits
Score difference with first non-orthologous sequence - *A. fumigatus*:358 *A. fischeri*:358
afum|EAL86359.1 100.00% afis|EAW16789.1 100.00%
Bootstrap support for afum|EAL86359.1 as seed ortholog is 100%.
Bootstrap support for afis|EAW16789.1 as seed ortholog is 100%.

---

Group of orthologs #7946. Best score 358 bits
Score difference with first non-orthologous sequence - *A. fumigatus*:358 *A. fischeri*:358
afum|EAL88637.2 100.00% afis|EAW16005.1 100.00%

Bootstrap support for afum|EAL88637.2 as seed ortholog is 100%.
Bootstrap support for afis|EAW16005.1 as seed ortholog is 100%.

---

Group of orthologs #7947. Best score 358 bits
Score difference with first non-orthologous sequence - *A. fumigatus*:358 *A. fischeri*:358
afum|EAL85213.1 100.00% afis|EAW20095.1 100.00%
Bootstrap support for afum|EAL85213.1 as seed ortholog is 100%.
Bootstrap support for afis|EAW20095.1 as seed ortholog is 100%.

---

Group of orthologs #7948. Best score 358 bits
Score difference with first non-orthologous sequence - *A. fumigatus*:358 *A. fischeri*:358
afum|EAL87725.1 100.00% afis|EAW23649.1 100.00%
Bootstrap support for afum|EAL87725.1 as seed ortholog is 100%.
Bootstrap support for afis|EAW23649.1 as seed ortholog is 100%.

---

Group of orthologs #7949. Best score 358 bits
Score difference with first non-orthologous sequence - *A. fumigatus*:291 *A. fischeri*:279
afum|EAL92030.1 100.00% afis|EAW23114.1 100.00%
Bootstrap support for afum|EAL92030.1 as seed ortholog is 100%.
Bootstrap support for afis|EAW23114.1 as seed ortholog is 100%.

---

Group of orthologs #7950. Best score 358 bits
Score difference with first non-orthologous sequence - *A. fumigatus*:358 *A. fischeri*:358
afum|EBA27313.1 100.00% afis|EAW23983.1 100.00%
Bootstrap support for afum|EBA27313.1 as seed ortholog is 100%.
Bootstrap support for afis|EAW23983.1 as seed ortholog is 100%.

---

Group of orthologs #7951. Best score 357 bits
Score difference with first non-orthologous sequence - *A. fumigatus*:357 *A. fischeri*:357
afum|EAL87712.2 100.00% afis|EAW18348.1 100.00%
Bootstrap support for afum|EAL87712.2 as seed ortholog is 100%.
Bootstrap support for afis|EAW18348.1 as seed ortholog is 100%.

---

Group of orthologs #7952. Best score 357 bits
Score difference with first non-orthologous sequence - *A. fumigatus*:357 *A. fischeri*:357
afum|EAL87935.1 100.00% afis|EAW23531.1 100.00%
Bootstrap support for afum|EAL87935.1 as seed ortholog is 100%.
Bootstrap support for afis|EAW23531.1 as seed ortholog is 100%.

---

Group of orthologs #7953. Best score 357 bits
Score difference with first non-orthologous sequence - *A. fumigatus*:357 *A. fischeri*:357
afum|EAL90685.1 100.00% afis|EAW22508.1 100.00%
Bootstrap support for afum|EAL90685.1 as seed ortholog is 100%.
Bootstrap support for afis|EAW22508.1 as seed ortholog is 100%.

---

Group of orthologs #7954. Best score 356 bits
Score difference with first non-orthologous sequence - *A. fumigatus*:356 *A. fischeri*:356
afum|EBA27383.1 100.00% afis|EAW24777.1 100.00%
Bootstrap support for afum|EBA27383.1 as seed ortholog is 100%.
Bootstrap support for afis|EAW24777.1 as seed ortholog is 100%.

---

Group of orthologs #7955. Best score 355 bits
Score difference with first non-orthologous sequence - *A. fumigatus*:355 *A. fischeri*:355
afum|EAL90489.1 100.00% afis|EAW22703.1 100.00%
Bootstrap support for afum|EAL90489.1 as seed ortholog is 100%.
Bootstrap support for afis|EAW22703.1 as seed ortholog is 100%.

Group of orthologs #7956. Best score 355 bits
Score difference with first non-orthologous sequence - *A. fumigatus*:355 *A. fischeri*:355
afum|EAL92444.1 100.00% afis|EAW21429.1 100.00%
Bootstrap support for afum|EAL92444.1 as seed ortholog is 100%.
Bootstrap support for afis|EAW21429.1 as seed ortholog is 100%.

Group of orthologs #7957. Best score 355 bits
Score difference with first non-orthologous sequence - *A. fumigatus*:355 *A. fischeri*:355
afum|EAL89611.1 100.00% afis|EAW24954.1 100.00%
Bootstrap support for afum|EAL89611.1 as seed ortholog is 100%.
Bootstrap support for afis|EAW24954.1 as seed ortholog is 100%.

Group of orthologs #7958. Best score 354 bits
Score difference with first non-orthologous sequence - *A. fumigatus*:354 *A. fischeri*:354
afum|EAL87521.1 100.00% afis|EAW18161.1 100.00%
Bootstrap support for afum|EAL87521.1 as seed ortholog is 100%.
Bootstrap support for afis|EAW18161.1 as seed ortholog is 100%.

Group of orthologs #7959. Best score 354 bits
Score difference with first non-orthologous sequence - *A. fumigatus*:354 *A. fischeri*:354
afum|EAL86095.1 100.00% afis|EAW24427.1 100.00%
Bootstrap support for afum|EAL86095.1 as seed ortholog is 100%.
Bootstrap support for afis|EAW24427.1 as seed ortholog is 100%.

Group of orthologs #7960. Best score 354 bits
Score difference with first non-orthologous sequence - *A. fumigatus*:354 *A. fischeri*:354
afum|EAL93621.1 100.00% afis|EAW18888.1 100.00%
Bootstrap support for afum|EAL93621.1 as seed ortholog is 100%.
Bootstrap support for afis|EAW18888.1 as seed ortholog is 100%.

Group of orthologs #7961. Best score 354 bits
Score difference with first non-orthologous sequence - *A. fumigatus*:354 *A. fischeri*:354
afum|EAL94020.1 100.00% afis|EAW19301.1 100.00%
Bootstrap support for afum|EAL94020.1 as seed ortholog is 100%.
Bootstrap support for afis|EAW19301.1 as seed ortholog is 100%.

Group of orthologs #7962. Best score 353 bits
Score difference with first non-orthologous sequence - *A. fumigatus*:353 *A. fischeri*:353
afum|EAL84623.1 100.00% afis|EAW20426.1 100.00%
Bootstrap support for afum|EAL84623.1 as seed ortholog is 100%.
Bootstrap support for afis|EAW20426.1 as seed ortholog is 100%.

Group of orthologs #7963. Best score 353 bits
Score difference with first non-orthologous sequence - *A. fumigatus*:353 *A. fischeri*:353
afum|EAL86942.1 100.00% afis|EAW19575.1 100.00%
Bootstrap support for afum|EAL86942.1 as seed ortholog is 100%.
Bootstrap support for afis|EAW19575.1 as seed ortholog is 100%.

Group of orthologs #7964. Best score 353 bits
Score difference with first non-orthologous sequence - *A. fumigatus*:353 *A. fischeri*:353
afum|EAL87254.1 100.00% afis|EAW23819.1 100.00%
Bootstrap support for afum|EAL87254.1 as seed ortholog is 100%.
Bootstrap support for afis|EAW23819.1 as seed ortholog is 100%.

Group of orthologs #7965. Best score 353 bits

Score difference with first non-orthologous sequence - *A. fumigatus*:353   *A. fischeri*:353
afum|EAL88051.2 100.00% afis|EAW23418.1 100.00%
Bootstrap support for afum|EAL88051.2 as seed ortholog is 100%.
Bootstrap support for afis|EAW23418.1 as seed ortholog is 100%.

---

Group of orthologs #7966. Best score 353 bits
Score difference with first non-orthologous sequence - *A. fumigatus*:353   *A. fischeri*:180
afum|EAL89348.2 100.00% afis|EAW24706.1 100.00%
Bootstrap support for afum|EAL89348.2 as seed ortholog is 100%.
Bootstrap support for afis|EAW24706.1 as seed ortholog is 100%.

---

Group of orthologs #7967. Best score 353 bits
Score difference with first non-orthologous sequence - *A. fumigatus*:353   *A. fischeri*:353
afum|EAL89391.1 100.00% afis|EAW24745.1 100.00%
Bootstrap support for afum|EAL89391.1 as seed ortholog is 100%.
Bootstrap support for afis|EAW24745.1 as seed ortholog is 100%.

---

Group of orthologs #7968. Best score 353 bits
Score difference with first non-orthologous sequence - *A. fumigatus*:353   *A. fischeri*:353
afum|EAL92772.1 100.00% afis|EAW21738.1 100.00%
Bootstrap support for afum|EAL92772.1 as seed ortholog is 100%.
Bootstrap support for afis|EAW21738.1 as seed ortholog is 100%.

---

Group of orthologs #7969. Best score 353 bits
Score difference with first non-orthologous sequence - *A. fumigatus*:353   *A. fischeri*:353
afum|EBA27378.1 100.00% afis|EAW24729.1 100.00%
Bootstrap support for afum|EBA27378.1 as seed ortholog is 100%.
Bootstrap support for afis|EAW24729.1 as seed ortholog is 100%.

---

Group of orthologs #7970. Best score 352 bits
Score difference with first non-orthologous sequence - *A. fumigatus*:352   *A. fischeri*:352
afum|EAL85788.1 100.00% afis|EAW15543.1 100.00%
Bootstrap support for afum|EAL85788.1 as seed ortholog is 100%.
Bootstrap support for afis|EAW15543.1 as seed ortholog is 100%.

---

Group of orthologs #7971. Best score 352 bits
Score difference with first non-orthologous sequence - *A. fumigatus*:352   *A. fischeri*:352
afum|EAL88261.1 100.00% afis|EAW23223.1 100.00%
Bootstrap support for afum|EAL88261.1 as seed ortholog is 100%.
Bootstrap support for afis|EAW23223.1 as seed ortholog is 100%.

---

Group of orthologs #7972. Best score 351 bits
Score difference with first non-orthologous sequence - *A. fumigatus*:351   *A. fischeri*:351
afum|EAL86877.1 100.00% afis|EAW19645.1 100.00%
Bootstrap support for afum|EAL86877.1 as seed ortholog is 100%.
Bootstrap support for afis|EAW19645.1 as seed ortholog is 100%.

---

Group of orthologs #7973. Best score 351 bits
Score difference with first non-orthologous sequence - *A. fumigatus*:351   *A. fischeri*:351
afum|EBA27275.1 100.00% afis|EAW16984.1 100.00%
Bootstrap support for afum|EBA27275.1 as seed ortholog is 100%.
Bootstrap support for afis|EAW16984.1 as seed ortholog is 100%.

---

Group of orthologs #7974. Best score 351 bits
Score difference with first non-orthologous sequence - *A. fumigatus*:351   *A. fischeri*:351
afum|EAL93602.1 100.00% afis|EAW18870.1 100.00%

Bootstrap support for afum|EAL93602.1 as seed ortholog is 100%.
Bootstrap support for afis|EAW18870.1 as seed ortholog is 100%.

---

Group of orthologs #7975. Best score 351 bits
Score difference with first non-orthologous sequence - *A. fumigatus*:351 *A. fischeri*:351
afum|EAL92102.1 100.00% afis|EAW21081.1 100.00%
Bootstrap support for afum|EAL92102.1 as seed ortholog is 100%.
Bootstrap support for afis|EAW21081.1 as seed ortholog is 100%.

---

Group of orthologs #7976. Best score 350 bits
Score difference with first non-orthologous sequence - *A. fumigatus*:350 *A. fischeri*:350
afum|EAL94042.1 100.00% afis|EAW19324.1 100.00%
Bootstrap support for afum|EAL94042.1 as seed ortholog is 100%.
Bootstrap support for afis|EAW19324.1 as seed ortholog is 100%.

---

Group of orthologs #7977. Best score 350 bits
Score difference with first non-orthologous sequence - *A. fumigatus*:350 *A. fischeri*:350
afum|EAL90334.1 100.00% afis|EAW22865.1 100.00%
Bootstrap support for afum|EAL90334.1 as seed ortholog is 100%.
Bootstrap support for afis|EAW22865.1 as seed ortholog is 100%.

---

Group of orthologs #7978. Best score 350 bits
Score difference with first non-orthologous sequence - *A. fumigatus*:350 *A. fischeri*:350
afum|EAL90111.1 100.00% afis|EAW25452.1 100.00%
Bootstrap support for afum|EAL90111.1 as seed ortholog is 100%.
Bootstrap support for afis|EAW25452.1 as seed ortholog is 100%.

---

Group of orthologs #7979. Best score 349 bits
Score difference with first non-orthologous sequence - *A. fumigatus*:349 *A. fischeri*:349
afum|EAL86323.1 100.00% afis|EAW16750.1 100.00%
Bootstrap support for afum|EAL86323.1 as seed ortholog is 100%.
Bootstrap support for afis|EAW16750.1 as seed ortholog is 100%.

---

Group of orthologs #7980. Best score 349 bits
Score difference with first non-orthologous sequence - *A. fumigatus*:349 *A. fischeri*:349
afum|EAL90733.1 100.00% afis|EAW22459.1 100.00%
Bootstrap support for afum|EAL90733.1 as seed ortholog is 100%.
Bootstrap support for afis|EAW22459.1 as seed ortholog is 100%.

---

Group of orthologs #7981. Best score 349 bits
Score difference with first non-orthologous sequence - *A. fumigatus*:349 *A. fischeri*:349
afum|EAL90273.1 100.00% afis|EAW22924.1 100.00%
Bootstrap support for afum|EAL90273.1 as seed ortholog is 100%.
Bootstrap support for afis|EAW22924.1 as seed ortholog is 100%.

---

Group of orthologs #7982. Best score 348 bits
Score difference with first non-orthologous sequence - *A. fumigatus*:348 *A. fischeri*:348
afum|EAL85547.1 100.00% afis|EAW15803.1 100.00%
Bootstrap support for afum|EAL85547.1 as seed ortholog is 100%.
Bootstrap support for afis|EAW15803.1 as seed ortholog is 100%.

---

Group of orthologs #7983. Best score 348 bits
Score difference with first non-orthologous sequence - *A. fumigatus*:222 *A. fischeri*:173
afum|EAL89047.1 100.00% afis|EAW16447.1 100.00%
Bootstrap support for afum|EAL89047.1 as seed ortholog is 100%.
Bootstrap support for afis|EAW16447.1 as seed ortholog is 99%.

---

Group of orthologs #7984. Best score 348 bits
Score difference with first non-orthologous sequence - *A. fumigatus*:348 *A. fischeri*:348
afum|EAL87306.1 100.00% afis|EAW23878.1 100.00%
Bootstrap support for afum|EAL87306.1 as seed ortholog is 100%.
Bootstrap support for afis|EAW23878.1 as seed ortholog is 100%.

---

Group of orthologs #7985. Best score 348 bits
Score difference with first non-orthologous sequence - *A. fumigatus*:348 *A. fischeri*:348
afum|EAL92949.2 100.00% afis|EAW21936.1 100.00%
Bootstrap support for afum|EAL92949.2 as seed ortholog is 100%.
Bootstrap support for afis|EAW21936.1 as seed ortholog is 100%.

---

Group of orthologs #7986. Best score 347 bits
Score difference with first non-orthologous sequence - *A. fumigatus*:347 *A. fischeri*:347
afum|EAL87507.1 100.00% afis|EAW18149.1 100.00%
Bootstrap support for afum|EAL87507.1 as seed ortholog is 100%.
Bootstrap support for afis|EAW18149.1 as seed ortholog is 100%.

---

Group of orthologs #7987. Best score 347 bits
Score difference with first non-orthologous sequence - *A. fumigatus*:347 *A. fischeri*:347
afum|EAL93076.2 100.00% afis|EAW18399.1 100.00%
Bootstrap support for afum|EAL93076.2 as seed ortholog is 100%.
Bootstrap support for afis|EAW18399.1 as seed ortholog is 100%.

---

Group of orthologs #7988. Best score 346 bits
Score difference with first non-orthologous sequence - *A. fumigatus*:294 *A. fischeri*:346
afum|EAL85414.1 100.00% afis|EAW20279.1 100.00%
Bootstrap support for afum|EAL85414.1 as seed ortholog is 100%.
Bootstrap support for afis|EAW20279.1 as seed ortholog is 100%.

---

Group of orthologs #7989. Best score 346 bits
Score difference with first non-orthologous sequence - *A. fumigatus*:346 *A. fischeri*:346
afum|EAL86953.1 100.00% afis|EAW19558.1 100.00%
Bootstrap support for afum|EAL86953.1 as seed ortholog is 100%.
Bootstrap support for afis|EAW19558.1 as seed ortholog is 100%.

---

Group of orthologs #7990. Best score 346 bits
Score difference with first non-orthologous sequence - *A. fumigatus*:346 *A. fischeri*:346
afum|EAL86899.2 100.00% afis|EAW19622.1 100.00%
Bootstrap support for afum|EAL86899.2 as seed ortholog is 100%.
Bootstrap support for afis|EAW19622.1 as seed ortholog is 100%.

---

Group of orthologs #7991. Best score 346 bits
Score difference with first non-orthologous sequence - *A. fumigatus*:346 *A. fischeri*:346
afum|EBA27369.1 100.00% afis|EAW16576.1 100.00%
Bootstrap support for afum|EBA27369.1 as seed ortholog is 100%.
Bootstrap support for afis|EAW16576.1 as seed ortholog is 100%.

---

Group of orthologs #7992. Best score 346 bits
Score difference with first non-orthologous sequence - *A. fumigatus*:264 *A. fischeri*:346
afum|EAL87360.1 100.00% afis|EAW23935.1 100.00%
Bootstrap support for afum|EAL87360.1 as seed ortholog is 100%.
Bootstrap support for afis|EAW23935.1 as seed ortholog is 100%.

---

Group of orthologs #7993. Best score 345 bits

Score difference with first non-orthologous sequence - *A. fumigatus*:345   *A. fischeri*:345
afum|EAL88836.1 100.00% afis|EAW16209.1 100.00%
Bootstrap support for afum|EAL88836.1 as seed ortholog is 100%.
Bootstrap support for afis|EAW16209.1 as seed ortholog is 100%.

---

Group of orthologs #7994. Best score 344 bits
Score difference with first non-orthologous sequence - *A. fumigatus*:344   *A. fischeri*:344
afum|EAL84760.1 100.00% afis|EAW20874.1 100.00%
Bootstrap support for afum|EAL84760.1 as seed ortholog is 100%.
Bootstrap support for afis|EAW20874.1 as seed ortholog is 100%.

---

Group of orthologs #7995. Best score 344 bits
Score difference with first non-orthologous sequence - *A. fumigatus*:344   *A. fischeri*:344
afum|EAL90245.1 100.00% afis|EAW22952.1 100.00%
Bootstrap support for afum|EAL90245.1 as seed ortholog is 100%.
Bootstrap support for afis|EAW22952.1 as seed ortholog is 100%.

---

Group of orthologs #7996. Best score 343 bits
Score difference with first non-orthologous sequence - *A. fumigatus*:85   *A. fischeri*:192
afum|EAL86719.2 100.00% afis|EAW20668.1 100.00%
afum|EAL92228.1 24.18%
Bootstrap support for afum|EAL86719.2 as seed ortholog is 39%. Alternative seed ortholog is afum|EAL84364.1 (85 bits away from this cluster)
Bootstrap support for afis|EAW20668.1 as seed ortholog is 90%.

---

Group of orthologs #7997. Best score 343 bits
Score difference with first non-orthologous sequence - *A. fumigatus*:343   *A. fischeri*:343
afum|EAL86519.2 100.00% afis|EAW16995.1 100.00%
Bootstrap support for afum|EAL86519.2 as seed ortholog is 100%.
Bootstrap support for afis|EAW16995.1 as seed ortholog is 100%.

---

Group of orthologs #7998. Best score 343 bits
Score difference with first non-orthologous sequence - *A. fumigatus*:343   *A. fischeri*:343
afum|EAL93345.1 100.00% afis|EAW18632.1 100.00%
Bootstrap support for afum|EAL93345.1 as seed ortholog is 100%.
Bootstrap support for afis|EAW18632.1 as seed ortholog is 100%.

---

Group of orthologs #7999. Best score 343 bits
Score difference with first non-orthologous sequence - *A. fumigatus*:343   *A. fischeri*:343
afum|EAL93401.1 100.00% afis|EAW18680.1 100.00%
Bootstrap support for afum|EAL93401.1 as seed ortholog is 100%.
Bootstrap support for afis|EAW18680.1 as seed ortholog is 100%.

---

Group of orthologs #8000. Best score 343 bits
Score difference with first non-orthologous sequence - *A. fumigatus*:343   *A. fischeri*:343
afum|EAL89859.1 100.00% afis|EAW25184.1 100.00%
Bootstrap support for afum|EAL89859.1 as seed ortholog is 100%.
Bootstrap support for afis|EAW25184.1 as seed ortholog is 100%.

---

Group of orthologs #8001. Best score 343 bits
Score difference with first non-orthologous sequence - *A. fumigatus*:343   *A. fischeri*:343
afum|EBA27417.1 100.00% afis|EAW22208.1 100.00%
Bootstrap support for afum|EBA27417.1 as seed ortholog is 100%.
Bootstrap support for afis|EAW22208.1 as seed ortholog is 100%.

---

Group of orthologs #8002. Best score 342 bits

Score difference with first non-orthologous sequence - *A. fumigatus*:342 *A. fischeri*:342
afum|EAL91267.1 100.00% afis|EAW17441.1 100.00%
Bootstrap support for afum|EAL91267.1 as seed ortholog is 100%.
Bootstrap support for afis|EAW17441.1 as seed ortholog is 100%.

---

Group of orthologs #8003. Best score 342 bits
Score difference with first non-orthologous sequence - *A. fumigatus*:342 *A. fischeri*:342
afum|EAL91498.1 100.00% afis|EAW17684.1 100.00%
Bootstrap support for afum|EAL91498.1 as seed ortholog is 100%.
Bootstrap support for afis|EAW17684.1 as seed ortholog is 100%.

---

Group of orthologs #8004. Best score 342 bits
Score difference with first non-orthologous sequence - *A. fumigatus*:342 *A. fischeri*:342
afum|EAL93949.1 100.00% afis|EAW19216.1 100.00%
Bootstrap support for afum|EAL93949.1 as seed ortholog is 100%.
Bootstrap support for afis|EAW19216.1 as seed ortholog is 100%.

---

Group of orthologs #8005. Best score 341 bits
Score difference with first non-orthologous sequence - *A. fumigatus*:285 *A. fischeri*:83
afum|EAL89347.1 100.00% afis|EAW24705.1 100.00%
Bootstrap support for afum|EAL89347.1 as seed ortholog is 100%.
Bootstrap support for afis|EAW24705.1 as seed ortholog is 99%.

---

Group of orthologs #8006. Best score 341 bits
Score difference with first non-orthologous sequence - *A. fumigatus*:341 *A. fischeri*:341
afum|EBA27484.1 100.00% afis|EAW21990.1 100.00%
Bootstrap support for afum|EBA27484.1 as seed ortholog is 100%.
Bootstrap support for afis|EAW21990.1 as seed ortholog is 100%.

---

Group of orthologs #8007. Best score 340 bits
Score difference with first non-orthologous sequence - *A. fumigatus*:340 *A. fischeri*:340
afum|EAL85368.1 100.00% afis|EAW20250.1 100.00%
Bootstrap support for afum|EAL85368.1 as seed ortholog is 100%.
Bootstrap support for afis|EAW20250.1 as seed ortholog is 100%.

---

Group of orthologs #8008. Best score 340 bits
Score difference with first non-orthologous sequence - *A. fumigatus*:340 *A. fischeri*:340
afum|EAL91352.1 100.00% afis|EAW17538.1 100.00%
Bootstrap support for afum|EAL91352.1 as seed ortholog is 100%.
Bootstrap support for afis|EAW17538.1 as seed ortholog is 100%.

---

Group of orthologs #8009. Best score 340 bits
Score difference with first non-orthologous sequence - *A. fumigatus*:272 *A. fischeri*:263
afum|EAL93826.1 100.00% afis|EAW19095.1 100.00%
Bootstrap support for afum|EAL93826.1 as seed ortholog is 100%.
Bootstrap support for afis|EAW19095.1 as seed ortholog is 100%.

---

Group of orthologs #8010. Best score 339 bits
Score difference with first non-orthologous sequence - *A. fumigatus*:339 *A. fischeri*:339
afum|EAL86380.1 100.00% afis|EAW16832.1 100.00%
Bootstrap support for afum|EAL86380.1 as seed ortholog is 100%.
Bootstrap support for afis|EAW16832.1 as seed ortholog is 100%.

---

Group of orthologs #8011. Best score 339 bits
Score difference with first non-orthologous sequence - *A. fumigatus*:339 *A. fischeri*:339
afum|EBA27503.1 100.00% afis|EAW19017.1 100.00%

Bootstrap support for afum|EBA27503.1 as seed ortholog is 100%.
Bootstrap support for afis|EAW19017.1 as seed ortholog is 100%.

---

Group of orthologs #8012. Best score 338 bits
Score difference with first non-orthologous sequence - *A. fumigatus*:338 *A. fischeri*:254
afum|EAL91668.1 100.00% afis|EAW17860.1 100.00%
Bootstrap support for afum|EAL91668.1 as seed ortholog is 100%.
Bootstrap support for afis|EAW17860.1 as seed ortholog is 100%.

---

Group of orthologs #8013. Best score 338 bits
Score difference with first non-orthologous sequence - *A. fumigatus*:338 *A. fischeri*:338
afum|EAL91743.1 100.00% afis|EAW17932.1 100.00%
Bootstrap support for afum|EAL91743.1 as seed ortholog is 100%.
Bootstrap support for afis|EAW17932.1 as seed ortholog is 100%.

---

Group of orthologs #8014. Best score 338 bits
Score difference with first non-orthologous sequence - *A. fumigatus*:338 *A. fischeri*:338
afum|EAL92354.1 100.00% afis|EAW21336.1 100.00%
Bootstrap support for afum|EAL92354.1 as seed ortholog is 100%.
Bootstrap support for afis|EAW21336.1 as seed ortholog is 100%.

---

Group of orthologs #8015. Best score 337 bits
Score difference with first non-orthologous sequence - *A. fumigatus*:337 *A. fischeri*:271
afum|EAL88705.1 100.00% afis|EAW16073.1 100.00%
Bootstrap support for afum|EAL88705.1 as seed ortholog is 100%.
Bootstrap support for afis|EAW16073.1 as seed ortholog is 100%.

---

Group of orthologs #8016. Best score 337 bits
Score difference with first non-orthologous sequence - *A. fumigatus*:280 *A. fischeri*:337
afum|EBA27178.1 100.00% afis|EAW20642.1 100.00%
Bootstrap support for afum|EBA27178.1 as seed ortholog is 100%.
Bootstrap support for afis|EAW20642.1 as seed ortholog is 100%.

---

Group of orthologs #8017. Best score 336 bits
Score difference with first non-orthologous sequence - *A. fumigatus*:336 *A. fischeri*:336
afum|EAL84206.1 100.00% afis|EAW15440.1 100.00%
Bootstrap support for afum|EAL84206.1 as seed ortholog is 100%.
Bootstrap support for afis|EAW15440.1 as seed ortholog is 100%.

---

Group of orthologs #8018. Best score 336 bits
Score difference with first non-orthologous sequence - *A. fumigatus*:336 *A. fischeri*:336
afum|EAL86426.1 100.00% afis|EAW16888.1 100.00%
Bootstrap support for afum|EAL86426.1 as seed ortholog is 100%.
Bootstrap support for afis|EAW16888.1 as seed ortholog is 100%.

---

Group of orthologs #8019. Best score 336 bits
Score difference with first non-orthologous sequence - *A. fumigatus*:336 *A. fischeri*:336
afum|EAL86629.2 100.00% afis|EAW17203.1 100.00%
Bootstrap support for afum|EAL86629.2 as seed ortholog is 100%.
Bootstrap support for afis|EAW17203.1 as seed ortholog is 100%.

---

Group of orthologs #8020. Best score 336 bits
Score difference with first non-orthologous sequence - *A. fumigatus*:336 *A. fischeri*:336
afum|EAL84892.1 100.00% afis|EAW20025.1 100.00%
Bootstrap support for afum|EAL84892.1 as seed ortholog is 100%.
Bootstrap support for afis|EAW20025.1 as seed ortholog is 100%.

---
Group of orthologs #8021. Best score 336 bits
Score difference with first non-orthologous sequence - *A. fumigatus*:336 *A. fischeri*:336
afum|EAL86097.1 100.00% afis|EAW24429.1 100.00%
Bootstrap support for afum|EAL86097.1 as seed ortholog is 100%.
Bootstrap support for afis|EAW24429.1 as seed ortholog is 100%.

---
Group of orthologs #8022. Best score 336 bits
Score difference with first non-orthologous sequence - *A. fumigatus*:336 *A. fischeri*:336
afum|EAL89827.1 100.00% afis|EAW25152.1 100.00%
Bootstrap support for afum|EAL89827.1 as seed ortholog is 100%.
Bootstrap support for afis|EAW25152.1 as seed ortholog is 100%.

---
Group of orthologs #8023. Best score 335 bits
Score difference with first non-orthologous sequence - *A. fumigatus*:335 *A. fischeri*:335
afum|EAL85579.1 100.00% afis|EAW15769.1 100.00%
Bootstrap support for afum|EAL85579.1 as seed ortholog is 100%.
Bootstrap support for afis|EAW15769.1 as seed ortholog is 100%.

---
Group of orthologs #8024. Best score 335 bits
Score difference with first non-orthologous sequence - *A. fumigatus*:335 *A. fischeri*:335
afum|EAL84946.1 100.00% afis|EAW19965.1 100.00%
Bootstrap support for afum|EAL84946.1 as seed ortholog is 100%.
Bootstrap support for afis|EAW19965.1 as seed ortholog is 100%.

---
Group of orthologs #8025. Best score 334 bits
Score difference with first non-orthologous sequence - *A. fumigatus*:334 *A. fischeri*:334
afum|EAL86642.1 100.00% afis|EAW17216.1 100.00%
Bootstrap support for afum|EAL86642.1 as seed ortholog is 100%.
Bootstrap support for afis|EAW17216.1 as seed ortholog is 100%.

---
Group of orthologs #8026. Best score 334 bits
Score difference with first non-orthologous sequence - *A. fumigatus*:334 *A. fischeri*:334
afum|EAL85088.2 100.00% afis|EAW19805.1 100.00%
Bootstrap support for afum|EAL85088.2 as seed ortholog is 100%.
Bootstrap support for afis|EAW19805.1 as seed ortholog is 100%.

---
Group of orthologs #8027. Best score 333 bits
Score difference with first non-orthologous sequence - *A. fumigatus*:333 *A. fischeri*:333
afum|EAL84853.1 100.00% afis|EAW20766.1 100.00%
Bootstrap support for afum|EAL84853.1 as seed ortholog is 100%.
Bootstrap support for afis|EAW20766.1 as seed ortholog is 100%.

---
Group of orthologs #8028. Best score 333 bits
Score difference with first non-orthologous sequence - *A. fumigatus*:333 *A. fischeri*:333
afum|EBA27434.1 100.00% afis|EAW17601.1 100.00%
Bootstrap support for afum|EBA27434.1 as seed ortholog is 100%.
Bootstrap support for afis|EAW17601.1 as seed ortholog is 100%.

---
Group of orthologs #8029. Best score 333 bits
Score difference with first non-orthologous sequence - *A. fumigatus*:333 *A. fischeri*:333
afum|EAL93494.1 100.00% afis|EAW18762.1 100.00%
Bootstrap support for afum|EAL93494.1 as seed ortholog is 100%.
Bootstrap support for afis|EAW18762.1 as seed ortholog is 100%.

---
Group of orthologs #8030. Best score 333 bits

Score difference with first non-orthologous sequence - *A. fumigatus*:333   *A. fischeri*:333
afum|EAL90232.1 100.00% afis|EAW22965.1 100.00%
Bootstrap support for afum|EAL90232.1 as seed ortholog is 100%.
Bootstrap support for afis|EAW22965.1 as seed ortholog is 100%.

---

Group of orthologs #8031. Best score 333 bits
Score difference with first non-orthologous sequence - *A. fumigatus*:333   *A. fischeri*:333
afum|EAL90354.1 100.00% afis|EAW22848.1 100.00%
Bootstrap support for afum|EAL90354.1 as seed ortholog is 100%.
Bootstrap support for afis|EAW22848.1 as seed ortholog is 100%.

---

Group of orthologs #8032. Best score 332 bits
Score difference with first non-orthologous sequence - *A. fumigatus*:332   *A. fischeri*:332
afum|EAL84204.1 100.00% afis|EAW15443.1 100.00%
Bootstrap support for afum|EAL84204.1 as seed ortholog is 100%.
Bootstrap support for afis|EAW15443.1 as seed ortholog is 100%.

---

Group of orthologs #8033. Best score 332 bits
Score difference with first non-orthologous sequence - *A. fumigatus*:332   *A. fischeri*:332
afum|EAL88584.1 100.00% afis|EAW15951.1 100.00%
Bootstrap support for afum|EAL88584.1 as seed ortholog is 100%.
Bootstrap support for afis|EAW15951.1 as seed ortholog is 100%.

---

Group of orthologs #8034. Best score 332 bits
Score difference with first non-orthologous sequence - *A. fumigatus*:332   *A. fischeri*:332
afum|EAL85036.1 100.00% afis|EAW19859.1 100.00%
Bootstrap support for afum|EAL85036.1 as seed ortholog is 100%.
Bootstrap support for afis|EAW19859.1 as seed ortholog is 100%.

---

Group of orthologs #8035. Best score 332 bits
Score difference with first non-orthologous sequence - *A. fumigatus*:332   *A. fischeri*:332
afum|EAL85258.1 100.00% afis|EAW20143.1 100.00%
Bootstrap support for afum|EAL85258.1 as seed ortholog is 100%.
Bootstrap support for afis|EAW20143.1 as seed ortholog is 100%.

---

Group of orthologs #8036. Best score 332 bits
Score difference with first non-orthologous sequence - *A. fumigatus*:332   *A. fischeri*:332
afum|EAL93028.1 100.00% afis|EAW22013.1 100.00%
Bootstrap support for afum|EAL93028.1 as seed ortholog is 100%.
Bootstrap support for afis|EAW22013.1 as seed ortholog is 100%.

---

Group of orthologs #8037. Best score 331 bits
Score difference with first non-orthologous sequence - *A. fumigatus*:331   *A. fischeri*:331
afum|EAL93627.1 100.00% afis|EAW18895.1 100.00%
Bootstrap support for afum|EAL93627.1 as seed ortholog is 100%.
Bootstrap support for afis|EAW18895.1 as seed ortholog is 100%.

---

Group of orthologs #8038. Best score 331 bits
Score difference with first non-orthologous sequence - *A. fumigatus*:331   *A. fischeri*:331
afum|EAL90578.1 100.00% afis|EAW22618.1 100.00%
Bootstrap support for afum|EAL90578.1 as seed ortholog is 100%.
Bootstrap support for afis|EAW22618.1 as seed ortholog is 100%.

---

Group of orthologs #8039. Best score 331 bits
Score difference with first non-orthologous sequence - *A. fumigatus*:331   *A. fischeri*:331
afum|EBA27401.1 100.00% afis|EAW22894.1 100.00%

Bootstrap support for afum|EBA27401.1 as seed ortholog is 100%.
Bootstrap support for afis|EAW22894.1 as seed ortholog is 100%.

---

Group of orthologs #8040. Best score 330 bits
Score difference with first non-orthologous sequence - *A. fumigatus*:330 *A. fischeri*:330
afum|EAL86580.1 100.00% afis|EAW17118.1 100.00%
Bootstrap support for afum|EAL86580.1 as seed ortholog is 100%.
Bootstrap support for afis|EAW17118.1 as seed ortholog is 100%.

---

Group of orthologs #8041. Best score 329 bits
Score difference with first non-orthologous sequence - *A. fumigatus*:329 *A. fischeri*:284
afum|EAL84491.1 100.00% afis|EAW20574.1 100.00%
Bootstrap support for afum|EAL84491.1 as seed ortholog is 100%.
Bootstrap support for afis|EAW20574.1 as seed ortholog is 100%.

---

Group of orthologs #8042. Best score 329 bits
Score difference with first non-orthologous sequence - *A. fumigatus*:329 *A. fischeri*:329
afum|EAL88277.1 100.00% afis|EAW23206.1 100.00%
Bootstrap support for afum|EAL88277.1 as seed ortholog is 100%.
Bootstrap support for afis|EAW23206.1 as seed ortholog is 100%.

---

Group of orthologs #8043. Best score 329 bits
Score difference with first non-orthologous sequence - *A. fumigatus*:329 *A. fischeri*:329
afum|EAL88211.1 100.00% afis|EAW23268.1 100.00%
Bootstrap support for afum|EAL88211.1 as seed ortholog is 100%.
Bootstrap support for afis|EAW23268.1 as seed ortholog is 100%.

---

Group of orthologs #8044. Best score 329 bits
Score difference with first non-orthologous sequence - *A. fumigatus*:329 *A. fischeri*:329
afum|EAL93896.1 100.00% afis|EAW19166.1 100.00%
Bootstrap support for afum|EAL93896.1 as seed ortholog is 100%.
Bootstrap support for afis|EAW19166.1 as seed ortholog is 100%.

---

Group of orthologs #8045. Best score 329 bits
Score difference with first non-orthologous sequence - *A. fumigatus*:329 *A. fischeri*:329
afum|EAL92411.1 100.00% afis|EAW21397.1 100.00%
Bootstrap support for afum|EAL92411.1 as seed ortholog is 100%.
Bootstrap support for afis|EAW21397.1 as seed ortholog is 100%.

---

Group of orthologs #8046. Best score 328 bits
Score difference with first non-orthologous sequence - *A. fumigatus*:328 *A. fischeri*:328
afum|EAL88680.1 100.00% afis|EAW16047.1 100.00%
Bootstrap support for afum|EAL88680.1 as seed ortholog is 100%.
Bootstrap support for afis|EAW16047.1 as seed ortholog is 100%.

---

Group of orthologs #8047. Best score 328 bits
Score difference with first non-orthologous sequence - *A. fumigatus*:328 *A. fischeri*:328
afum|EAL84463.1 100.00% afis|EAW20598.1 100.00%
Bootstrap support for afum|EAL84463.1 as seed ortholog is 100%.
Bootstrap support for afis|EAW20598.1 as seed ortholog is 100%.

---

Group of orthologs #8048. Best score 328 bits
Score difference with first non-orthologous sequence - *A. fumigatus*:328 *A. fischeri*:171
afum|EAL85466.1 100.00% afis|EAW20346.1 100.00%
Bootstrap support for afum|EAL85466.1 as seed ortholog is 100%.
Bootstrap support for afis|EAW20346.1 as seed ortholog is 100%.

Group of orthologs #8049. Best score 328 bits
Score difference with first non-orthologous sequence - *A. fumigatus*:328 *A. fischeri*:328
afum|EAL93449.1 100.00% afis|EAW18724.1 100.00%
Bootstrap support for afum|EAL93449.1 as seed ortholog is 100%.
Bootstrap support for afis|EAW18724.1 as seed ortholog is 100%.

Group of orthologs #8050. Best score 328 bits
Score difference with first non-orthologous sequence - *A. fumigatus*:328 *A. fischeri*:328
afum|EAL92322.1 100.00% afis|EAW21306.1 100.00%
Bootstrap support for afum|EAL92322.1 as seed ortholog is 100%.
Bootstrap support for afis|EAW21306.1 as seed ortholog is 100%.

Group of orthologs #8051. Best score 327 bits
Score difference with first non-orthologous sequence - *A. fumigatus*:327 *A. fischeri*:327
afum|EAL84553.1 100.00% afis|EAW20503.1 100.00%
Bootstrap support for afum|EAL84553.1 as seed ortholog is 100%.
Bootstrap support for afis|EAW20503.1 as seed ortholog is 100%.

Group of orthologs #8052. Best score 327 bits
Score difference with first non-orthologous sequence - *A. fumigatus*:327 *A. fischeri*:327
afum|EAL88245.1 100.00% afis|EAW23237.1 100.00%
Bootstrap support for afum|EAL88245.1 as seed ortholog is 100%.
Bootstrap support for afis|EAW23237.1 as seed ortholog is 100%.

Group of orthologs #8053. Best score 327 bits
Score difference with first non-orthologous sequence - *A. fumigatus*:327 *A. fischeri*:327
afum|EAL87833.1 100.00% afis|EAW23627.1 100.00%
Bootstrap support for afum|EAL87833.1 as seed ortholog is 100%.
Bootstrap support for afis|EAW23627.1 as seed ortholog is 100%.

Group of orthologs #8054. Best score 327 bits
Score difference with first non-orthologous sequence - *A. fumigatus*:327 *A. fischeri*:327
afum|EAL90155.1 100.00% afis|EAW25490.1 100.00%
Bootstrap support for afum|EAL90155.1 as seed ortholog is 100%.
Bootstrap support for afis|EAW25490.1 as seed ortholog is 100%.

Group of orthologs #8055. Best score 326 bits
Score difference with first non-orthologous sequence - *A. fumigatus*:249 *A. fischeri*:253
afum|EAL88998.1 100.00% afis|EAW16374.1 100.00%
Bootstrap support for afum|EAL88998.1 as seed ortholog is 100%.
Bootstrap support for afis|EAW16374.1 as seed ortholog is 100%.

Group of orthologs #8056. Best score 326 bits
Score difference with first non-orthologous sequence - *A. fumigatus*:326 *A. fischeri*:326
afum|EAL89166.1 100.00% afis|EAW16569.1 100.00%
Bootstrap support for afum|EAL89166.1 as seed ortholog is 100%.
Bootstrap support for afis|EAW16569.1 as seed ortholog is 100%.

Group of orthologs #8057. Best score 326 bits
Score difference with first non-orthologous sequence - *A. fumigatus*:326 *A. fischeri*:326
afum|EAL89196.1 100.00% afis|EAW16600.1 100.00%
Bootstrap support for afum|EAL89196.1 as seed ortholog is 100%.
Bootstrap support for afis|EAW16600.1 as seed ortholog is 100%.

Group of orthologs #8058. Best score 326 bits

Score difference with first non-orthologous sequence - *A. fumigatus*:326 *A. fischeri*:326
afum|EAL86749.1 100.00% afis|EAW24648.1 100.00%
Bootstrap support for afum|EAL86749.1 as seed ortholog is 100%.
Bootstrap support for afis|EAW24648.1 as seed ortholog is 100%.

---

Group of orthologs #8059. Best score 326 bits
Score difference with first non-orthologous sequence - *A. fumigatus*:326 *A. fischeri*:326
afum|EAL87458.1 100.00% afis|EAW24027.1 100.00%
Bootstrap support for afum|EAL87458.1 as seed ortholog is 100%.
Bootstrap support for afis|EAW24027.1 as seed ortholog is 100%.

---

Group of orthologs #8060. Best score 326 bits
Score difference with first non-orthologous sequence - *A. fumigatus*:326 *A. fischeri*:326
afum|EAL90747.1 100.00% afis|EAW22444.1 100.00%
Bootstrap support for afum|EAL90747.1 as seed ortholog is 100%.
Bootstrap support for afis|EAW22444.1 as seed ortholog is 100%.

---

Group of orthologs #8061. Best score 326 bits
Score difference with first non-orthologous sequence - *A. fumigatus*:326 *A. fischeri*:326
afum|EAL89905.1 100.00% afis|EAW25243.1 100.00%
Bootstrap support for afum|EAL89905.1 as seed ortholog is 100%.
Bootstrap support for afis|EAW25243.1 as seed ortholog is 100%.

---

Group of orthologs #8062. Best score 326 bits
Score difference with first non-orthologous sequence - *A. fumigatus*:326 *A. fischeri*:326
afum|EAL90001.1 100.00% afis|EAW25338.1 100.00%
Bootstrap support for afum|EAL90001.1 as seed ortholog is 100%.
Bootstrap support for afis|EAW25338.1 as seed ortholog is 100%.

---

Group of orthologs #8063. Best score 326 bits
Score difference with first non-orthologous sequence - *A. fumigatus*:326 *A. fischeri*:326
afum|EAL92031.1 100.00% afis|EAW24169.1 100.00%
Bootstrap support for afum|EAL92031.1 as seed ortholog is 100%.
Bootstrap support for afis|EAW24169.1 as seed ortholog is 100%.

---

Group of orthologs #8064. Best score 325 bits
Score difference with first non-orthologous sequence - *A. fumigatus*:325 *A. fischeri*:325
afum|EAL91809.2 100.00% afis|EAW17998.1 100.00%
Bootstrap support for afum|EAL91809.2 as seed ortholog is 100%.
Bootstrap support for afis|EAW17998.1 as seed ortholog is 100%.

---

Group of orthologs #8065. Best score 324 bits
Score difference with first non-orthologous sequence - *A. fumigatus*:324 *A. fischeri*:324
afum|EAL87055.1 100.00% afis|EAW19441.1 100.00%
Bootstrap support for afum|EAL87055.1 as seed ortholog is 100%.
Bootstrap support for afis|EAW19441.1 as seed ortholog is 100%.

---

Group of orthologs #8066. Best score 324 bits
Score difference with first non-orthologous sequence - *A. fumigatus*:324 *A. fischeri*:324
afum|EAL85943.1 100.00% afis|EAW24275.1 100.00%
Bootstrap support for afum|EAL85943.1 as seed ortholog is 100%.
Bootstrap support for afis|EAW24275.1 as seed ortholog is 100%.

---

Group of orthologs #8067. Best score 324 bits
Score difference with first non-orthologous sequence - *A. fumigatus*:324 *A. fischeri*:173
afum|EAL86109.1 100.00% afis|EAW24441.1 100.00%

Bootstrap support for afum|EAL86109.1 as seed ortholog is 100%.
Bootstrap support for afis|EAW24441.1 as seed ortholog is 100%.

---

Group of orthologs #8068. Best score 324 bits
Score difference with first non-orthologous sequence - *A. fumigatus*:324 *A. fischeri*:237
afum|EAL91055.1 100.00% afis|EAW22135.1 100.00%
Bootstrap support for afum|EAL91055.1 as seed ortholog is 100%.
Bootstrap support for afis|EAW22135.1 as seed ortholog is 100%.

---

Group of orthologs #8069. Best score 324 bits
Score difference with first non-orthologous sequence - *A. fumigatus*:324 *A. fischeri*:324
afum|EAL90711.1 100.00% afis|EAW22483.1 100.00%
Bootstrap support for afum|EAL90711.1 as seed ortholog is 100%.
Bootstrap support for afis|EAW22483.1 as seed ortholog is 100%.

---

Group of orthologs #8070. Best score 324 bits
Score difference with first non-orthologous sequence - *A. fumigatus*:324 *A. fischeri*:324
afum|EAL90310.1 100.00% afis|EAW22888.1 100.00%
Bootstrap support for afum|EAL90310.1 as seed ortholog is 100%.
Bootstrap support for afis|EAW22888.1 as seed ortholog is 100%.

---

Group of orthologs #8071. Best score 323 bits
Score difference with first non-orthologous sequence - *A. fumigatus*:323 *A. fischeri*:323
afum|EAL88132.1 100.00% afis|EAW23345.1 100.00%
Bootstrap support for afum|EAL88132.1 as seed ortholog is 100%.
Bootstrap support for afis|EAW23345.1 as seed ortholog is 100%.

---

Group of orthologs #8072. Best score 323 bits
Score difference with first non-orthologous sequence - *A. fumigatus*:323 *A. fischeri*:323
afum|EAL93939.2 100.00% afis|EAW19209.1 100.00%
Bootstrap support for afum|EAL93939.2 as seed ortholog is 100%.
Bootstrap support for afis|EAW19209.1 as seed ortholog is 100%.

---

Group of orthologs #8073. Best score 322 bits
Score difference with first non-orthologous sequence - *A. fumigatus*:272 *A. fischeri*:233
afum|EAL86589.1 100.00% afis|EAW17129.1 100.00%
Bootstrap support for afum|EAL86589.1 as seed ortholog is 100%.
Bootstrap support for afis|EAW17129.1 as seed ortholog is 100%.

---

Group of orthologs #8074. Best score 322 bits
Score difference with first non-orthologous sequence - *A. fumigatus*:322 *A. fischeri*:322
afum|EAL86603.1 100.00% afis|EAW17172.1 100.00%
Bootstrap support for afum|EAL86603.1 as seed ortholog is 100%.
Bootstrap support for afis|EAW17172.1 as seed ortholog is 100%.

---

Group of orthologs #8075. Best score 322 bits
Score difference with first non-orthologous sequence - *A. fumigatus*:322 *A. fischeri*:322
afum|EAL90687.1 100.00% afis|EAW22506.1 100.00%
Bootstrap support for afum|EAL90687.1 as seed ortholog is 100%.
Bootstrap support for afis|EAW22506.1 as seed ortholog is 100%.

---

Group of orthologs #8076. Best score 321 bits
Score difference with first non-orthologous sequence - *A. fumigatus*:321 *A. fischeri*:321
afum|EAL87729.1 100.00% afis|EAW16109.1 100.00%
afum|EAL88789.1 86.13%
Bootstrap support for afum|EAL87729.1 as seed ortholog is 100%.

Bootstrap support for afis|EAW16109.1 as seed ortholog is 100%.

---

Group of orthologs #8077. Best score 321 bits
Score difference with first non-orthologous sequence - *A. fumigatus*:321 *A. fischeri*:321
afum|EAL92832.1 100.00% afis|EAW21816.1 100.00%
Bootstrap support for afum|EAL92832.1 as seed ortholog is 100%.
Bootstrap support for afis|EAW21816.1 as seed ortholog is 100%.

---

Group of orthologs #8078. Best score 320 bits
Score difference with first non-orthologous sequence - *A. fumigatus*:320 *A. fischeri*:320
afum|EAL84532.1 100.00% afis|EAW20525.1 100.00%
Bootstrap support for afum|EAL84532.1 as seed ortholog is 100%.
Bootstrap support for afis|EAW20525.1 as seed ortholog is 100%.

---

Group of orthologs #8079. Best score 320 bits
Score difference with first non-orthologous sequence - *A. fumigatus*:320 *A. fischeri*:320
afum|EAL87907.1 100.00% afis|EAW23559.1 100.00%
Bootstrap support for afum|EAL87907.1 as seed ortholog is 100%.
Bootstrap support for afis|EAW23559.1 as seed ortholog is 100%.

---

Group of orthologs #8080. Best score 320 bits
Score difference with first non-orthologous sequence - *A. fumigatus*:320 *A. fischeri*:320
afum|EAL93244.1 100.00% afis|EAW18544.1 100.00%
Bootstrap support for afum|EAL93244.1 as seed ortholog is 100%.
Bootstrap support for afis|EAW18544.1 as seed ortholog is 100%.

---

Group of orthologs #8081. Best score 320 bits
Score difference with first non-orthologous sequence - *A. fumigatus*:320 *A. fischeri*:320
afum|EAL93935.1 100.00% afis|EAW19205.1 100.00%
Bootstrap support for afum|EAL93935.1 as seed ortholog is 100%.
Bootstrap support for afis|EAW19205.1 as seed ortholog is 100%.

---

Group of orthologs #8082. Best score 319 bits
Score difference with first non-orthologous sequence - *A. fumigatus*:125 *A. fischeri*:319
afum|EAL86973.1 100.00% afis|EAW19533.1 100.00%
Bootstrap support for afum|EAL86973.1 as seed ortholog is 99%.
Bootstrap support for afis|EAW19533.1 as seed ortholog is 100%.

---

Group of orthologs #8083. Best score 319 bits
Score difference with first non-orthologous sequence - *A. fumigatus*:319 *A. fischeri*:319
afum|EAL87178.1 100.00% afis|EAW23741.1 100.00%
Bootstrap support for afum|EAL87178.1 as seed ortholog is 100%.
Bootstrap support for afis|EAW23741.1 as seed ortholog is 100%.

---

Group of orthologs #8084. Best score 319 bits
Score difference with first non-orthologous sequence - *A. fumigatus*:319 *A. fischeri*:319
afum|EAL93325.1 100.00% afis|EAW18615.1 100.00%
Bootstrap support for afum|EAL93325.1 as seed ortholog is 100%.
Bootstrap support for afis|EAW18615.1 as seed ortholog is 100%.

---

Group of orthologs #8085. Best score 319 bits
Score difference with first non-orthologous sequence - *A. fumigatus*:319 *A. fischeri*:319
afum|EAL90514.1 100.00% afis|EAW22678.1 100.00%
Bootstrap support for afum|EAL90514.1 as seed ortholog is 100%.
Bootstrap support for afis|EAW22678.1 as seed ortholog is 100%.

---

Group of orthologs #8086. Best score 319 bits
Score difference with first non-orthologous sequence - *A. fumigatus*:319 *A. fischeri*:319
afum|EAL89617.2 100.00% afis|EAW24959.1 100.00%
Bootstrap support for afum|EAL89617.2 as seed ortholog is 100%.
Bootstrap support for afis|EAW24959.1 as seed ortholog is 100%.

---

Group of orthologs #8087. Best score 319 bits
Score difference with first non-orthologous sequence - *A. fumigatus*:319 *A. fischeri*:319
afum|EAL90056.1 100.00% afis|EAW25397.1 100.00%
Bootstrap support for afum|EAL90056.1 as seed ortholog is 100%.
Bootstrap support for afis|EAW25397.1 as seed ortholog is 100%.

---

Group of orthologs #8088. Best score 318 bits
Score difference with first non-orthologous sequence - *A. fumigatus*:318 *A. fischeri*:318
afum|EAL84680.2 100.00% afis|EAW20954.1 100.00%
Bootstrap support for afum|EAL84680.2 as seed ortholog is 100%.
Bootstrap support for afis|EAW20954.1 as seed ortholog is 100%.

---

Group of orthologs #8089. Best score 318 bits
Score difference with first non-orthologous sequence - *A. fumigatus*:318 *A. fischeri*:318
afum|EAL91484.1 100.00% afis|EAW17670.1 100.00%
Bootstrap support for afum|EAL91484.1 as seed ortholog is 100%.
Bootstrap support for afis|EAW17670.1 as seed ortholog is 100%.

---

Group of orthologs #8090. Best score 318 bits
Score difference with first non-orthologous sequence - *A. fumigatus*:318 *A. fischeri*:318
afum|EAL84298.2 100.00% afis|EAW25590.1 100.00%
Bootstrap support for afum|EAL84298.2 as seed ortholog is 100%.
Bootstrap support for afis|EAW25590.1 as seed ortholog is 100%.

---

Group of orthologs #8091. Best score 318 bits
Score difference with first non-orthologous sequence - *A. fumigatus*:318 *A. fischeri*:318
afum|EAL91997.1 100.00% afis|EAW24142.1 100.00%
Bootstrap support for afum|EAL91997.1 as seed ortholog is 100%.
Bootstrap support for afis|EAW24142.1 as seed ortholog is 100%.

---

Group of orthologs #8092. Best score 317 bits
Score difference with first non-orthologous sequence - *A. fumigatus*:317 *A. fischeri*:317
afum|EAL88972.1 100.00% afis|EAW16347.1 100.00%
Bootstrap support for afum|EAL88972.1 as seed ortholog is 100%.
Bootstrap support for afis|EAW16347.1 as seed ortholog is 100%.

---

Group of orthologs #8093. Best score 317 bits
Score difference with first non-orthologous sequence - *A. fumigatus*:317 *A. fischeri*:317
afum|EAL88345.2 100.00% afis|EAW23136.1 100.00%
Bootstrap support for afum|EAL88345.2 as seed ortholog is 100%.
Bootstrap support for afis|EAW23136.1 as seed ortholog is 100%.

---

Group of orthologs #8094. Best score 316 bits
Score difference with first non-orthologous sequence - *A. fumigatus*:316 *A. fischeri*:316
afum|EAL87481.1 100.00% afis|EAW18123.1 100.00%
Bootstrap support for afum|EAL87481.1 as seed ortholog is 100%.
Bootstrap support for afis|EAW18123.1 as seed ortholog is 100%.

---

Group of orthologs #8095. Best score 315 bits
Score difference with first non-orthologous sequence - *A. fumigatus*:315 *A. fischeri*:315

afum|EAL89108.1 100.00% afis|EAW16513.1 100.00%
Bootstrap support for afum|EAL89108.1 as seed ortholog is 100%.
Bootstrap support for afis|EAW16513.1 as seed ortholog is 100%.

---

Group of orthologs #8096. Best score 315 bits
Score difference with first non-orthologous sequence - *A. fumigatus*:315 *A. fischeri*:315
afum|EAL91306.1 100.00% afis|EAW17477.1 100.00%
Bootstrap support for afum|EAL91306.1 as seed ortholog is 100%.
Bootstrap support for afis|EAW17477.1 as seed ortholog is 100%.

---

Group of orthologs #8097. Best score 315 bits
Score difference with first non-orthologous sequence - *A. fumigatus*:315 *A. fischeri*:315
afum|EAL88256.1 100.00% afis|EAW23227.1 100.00%
Bootstrap support for afum|EAL88256.1 as seed ortholog is 100%.
Bootstrap support for afis|EAW23227.1 as seed ortholog is 100%.

---

Group of orthologs #8098. Best score 315 bits
Score difference with first non-orthologous sequence - *A. fumigatus*:315 *A. fischeri*:315
afum|EAL90374.2 100.00% afis|EAW22826.1 100.00%
Bootstrap support for afum|EAL90374.2 as seed ortholog is 100%.
Bootstrap support for afis|EAW22826.1 as seed ortholog is 100%.

---

Group of orthologs #8099. Best score 315 bits
Score difference with first non-orthologous sequence - *A. fumigatus*:315 *A. fischeri*:315
afum|EBA27510.1 100.00% afis|EAW19270.1 100.00%
Bootstrap support for afum|EBA27510.1 as seed ortholog is 100%.
Bootstrap support for afis|EAW19270.1 as seed ortholog is 100%.

---

Group of orthologs #8100. Best score 314 bits
Score difference with first non-orthologous sequence - *A. fumigatus*:314 *A. fischeri*:314
afum|EAL85656.1 100.00% afis|EAW15692.1 100.00%
Bootstrap support for afum|EAL85656.1 as seed ortholog is 100%.
Bootstrap support for afis|EAW15692.1 as seed ortholog is 100%.

---

Group of orthologs #8101. Best score 314 bits
Score difference with first non-orthologous sequence - *A. fumigatus*:314 *A. fischeri*:314
afum|EAL93148.2 100.00% afis|EAW18462.1 100.00%
Bootstrap support for afum|EAL93148.2 as seed ortholog is 100%.
Bootstrap support for afis|EAW18462.1 as seed ortholog is 100%.

---

Group of orthologs #8102. Best score 314 bits
Score difference with first non-orthologous sequence - *A. fumigatus*:314 *A. fischeri*:314
afum|EAL92054.1 100.00% afis|EAW21016.1 100.00%
Bootstrap support for afum|EAL92054.1 as seed ortholog is 100%.
Bootstrap support for afis|EAW21016.1 as seed ortholog is 100%.

---

Group of orthologs #8103. Best score 314 bits
Score difference with first non-orthologous sequence - *A. fumigatus*:314 *A. fischeri*:314
afum|EAL90901.1 100.00% afis|EAW22280.1 100.00%
Bootstrap support for afum|EAL90901.1 as seed ortholog is 100%.
Bootstrap support for afis|EAW22280.1 as seed ortholog is 100%.

---

Group of orthologs #8104. Best score 313 bits
Score difference with first non-orthologous sequence - *A. fumigatus*:247 *A. fischeri*:313
afum|EAL86579.1 100.00% afis|EAW17117.1 100.00%
Bootstrap support for afum|EAL86579.1 as seed ortholog is 100%.

Bootstrap support for afis|EAW17117.1 as seed ortholog is 100%.

---

Group of orthologs #8105. Best score 313 bits
Score difference with first non-orthologous sequence - *A. fumigatus*:313 *A. fischeri*:313
afum|EAL88431.1 100.00% afis|EAW23049.1 100.00%
Bootstrap support for afum|EAL88431.1 as seed ortholog is 100%.
Bootstrap support for afis|EAW23049.1 as seed ortholog is 100%.

---

Group of orthologs #8106. Best score 313 bits
Score difference with first non-orthologous sequence - *A. fumigatus*:313 *A. fischeri*:313
afum|EAL92304.1 100.00% afis|EAW21287.1 100.00%
Bootstrap support for afum|EAL92304.1 as seed ortholog is 100%.
Bootstrap support for afis|EAW21287.1 as seed ortholog is 100%.

---

Group of orthologs #8107. Best score 312 bits
Score difference with first non-orthologous sequence - *A. fumigatus*:312 *A. fischeri*:312
afum|EAL85616.1 100.00% afis|EAW15731.1 100.00%
Bootstrap support for afum|EAL85616.1 as seed ortholog is 100%.
Bootstrap support for afis|EAW15731.1 as seed ortholog is 100%.

---

Group of orthologs #8108. Best score 312 bits
Score difference with first non-orthologous sequence - *A. fumigatus*:312 *A. fischeri*:312
afum|EAL85002.1 100.00% afis|EAW19905.1 100.00%
Bootstrap support for afum|EAL85002.1 as seed ortholog is 100%.
Bootstrap support for afis|EAW19905.1 as seed ortholog is 100%.

---

Group of orthologs #8109. Best score 312 bits
Score difference with first non-orthologous sequence - *A. fumigatus*:312 *A. fischeri*:260
afum|EAL84471.1 100.00% afis|EAW20590.1 100.00%
Bootstrap support for afum|EAL84471.1 as seed ortholog is 100%.
Bootstrap support for afis|EAW20590.1 as seed ortholog is 100%.

---

Group of orthologs #8110. Best score 312 bits
Score difference with first non-orthologous sequence - *A. fumigatus*:312 *A. fischeri*:312
afum|EAL92426.1 100.00% afis|EAW21412.1 100.00%
Bootstrap support for afum|EAL92426.1 as seed ortholog is 100%.
Bootstrap support for afis|EAW21412.1 as seed ortholog is 100%.

---

Group of orthologs #8111. Best score 312 bits
Score difference with first non-orthologous sequence - *A. fumigatus*:312 *A. fischeri*:312
afum|EAL92918.2 100.00% afis|EAW21906.1 100.00%
Bootstrap support for afum|EAL92918.2 as seed ortholog is 100%.
Bootstrap support for afis|EAW21906.1 as seed ortholog is 100%.

---

Group of orthologs #8112. Best score 312 bits
Score difference with first non-orthologous sequence - *A. fumigatus*:312 *A. fischeri*:312
afum|EAL89803.1 100.00% afis|EAW25132.1 100.00%
Bootstrap support for afum|EAL89803.1 as seed ortholog is 100%.
Bootstrap support for afis|EAW25132.1 as seed ortholog is 100%.

---

Group of orthologs #8113. Best score 312 bits
Score difference with first non-orthologous sequence - *A. fumigatus*:312 *A. fischeri*:312
afum|EBA27253.1 100.00% afis|EAW24355.1 100.00%
Bootstrap support for afum|EBA27253.1 as seed ortholog is 100%.
Bootstrap support for afis|EAW24355.1 as seed ortholog is 100%.

---

Group of orthologs #8114. Best score 311 bits
Score difference with first non-orthologous sequence - *A. fumigatus*:311 *A. fischeri*:311
afum|EAL86407.1 100.00% afis|EAW16868.1 100.00%
Bootstrap support for afum|EAL86407.1 as seed ortholog is 100%.
Bootstrap support for afis|EAW16868.1 as seed ortholog is 100%.

---

Group of orthologs #8115. Best score 311 bits
Score difference with first non-orthologous sequence - *A. fumigatus*:311 *A. fischeri*:311
afum|EAL87565.1 100.00% afis|EAW18200.1 100.00%
Bootstrap support for afum|EAL87565.1 as seed ortholog is 100%.
Bootstrap support for afis|EAW18200.1 as seed ortholog is 100%.

---

Group of orthologs #8116. Best score 311 bits
Score difference with first non-orthologous sequence - *A. fumigatus*:311 *A. fischeri*:311
afum|EAL85902.1 100.00% afis|EAW24233.1 100.00%
Bootstrap support for afum|EAL85902.1 as seed ortholog is 100%.
Bootstrap support for afis|EAW24233.1 as seed ortholog is 100%.

---

Group of orthologs #8117. Best score 311 bits
Score difference with first non-orthologous sequence - *A. fumigatus*:311 *A. fischeri*:311
afum|EAL91068.1 100.00% afis|EAW22118.1 100.00%
Bootstrap support for afum|EAL91068.1 as seed ortholog is 100%.
Bootstrap support for afis|EAW22118.1 as seed ortholog is 100%.

---

Group of orthologs #8118. Best score 310 bits
Score difference with first non-orthologous sequence - *A. fumigatus*:310 *A. fischeri*:310
afum|EAL85031.1 100.00% afis|EAW19865.1 100.00%
Bootstrap support for afum|EAL85031.1 as seed ortholog is 100%.
Bootstrap support for afis|EAW19865.1 as seed ortholog is 100%.

---

Group of orthologs #8119. Best score 310 bits
Score difference with first non-orthologous sequence - *A. fumigatus*:310 *A. fischeri*:149
afum|EAL85928.1 100.00% afis|EAW24260.1 100.00%
Bootstrap support for afum|EAL85928.1 as seed ortholog is 100%.
Bootstrap support for afis|EAW24260.1 as seed ortholog is 100%.

---

Group of orthologs #8120. Best score 310 bits
Score difference with first non-orthologous sequence - *A. fumigatus*:310 *A. fischeri*:310
afum|EAL93459.1 100.00% afis|EAW18734.1 100.00%
Bootstrap support for afum|EAL93459.1 as seed ortholog is 100%.
Bootstrap support for afis|EAW18734.1 as seed ortholog is 100%.

---

Group of orthologs #8121. Best score 310 bits
Score difference with first non-orthologous sequence - *A. fumigatus*:310 *A. fischeri*:310
afum|EAL90855.1 100.00% afis|EAW22337.1 100.00%
Bootstrap support for afum|EAL90855.1 as seed ortholog is 100%.
Bootstrap support for afis|EAW22337.1 as seed ortholog is 100%.

---

Group of orthologs #8122. Best score 309 bits
Score difference with first non-orthologous sequence - *A. fumigatus*:309 *A. fischeri*:309
afum|EAL88093.1 100.00% afis|EAW23378.1 100.00%
Bootstrap support for afum|EAL88093.1 as seed ortholog is 100%.
Bootstrap support for afis|EAW23378.1 as seed ortholog is 100%.

---

Group of orthologs #8123. Best score 309 bits
Score difference with first non-orthologous sequence - *A. fumigatus*:309 *A. fischeri*:309

afum|EAL93270.1 100.00% afis|EAW18568.1 100.00%
Bootstrap support for afum|EAL93270.1 as seed ortholog is 100%.
Bootstrap support for afis|EAW18568.1 as seed ortholog is 100%.

---

Group of orthologs #8124. Best score 309 bits
Score difference with first non-orthologous sequence - *A. fumigatus*:309 *A. fischeri*:207
afum|EBA27185.1 100.00% afis|EAW20552.1 100.00%
Bootstrap support for afum|EBA27185.1 as seed ortholog is 100%.
Bootstrap support for afis|EAW20552.1 as seed ortholog is 100%.

---

Group of orthologs #8125. Best score 308 bits
Score difference with first non-orthologous sequence - *A. fumigatus*:308 *A. fischeri*:308
afum|EAL88200.1 100.00% afis|EAW23279.1 100.00%
Bootstrap support for afum|EAL88200.1 as seed ortholog is 100%.
Bootstrap support for afis|EAW23279.1 as seed ortholog is 100%.

---

Group of orthologs #8126. Best score 308 bits
Score difference with first non-orthologous sequence - *A. fumigatus*:308 *A. fischeri*:308
afum|EBA27335.1 100.00% afis|EAW23360.1 100.00%
Bootstrap support for afum|EBA27335.1 as seed ortholog is 100%.
Bootstrap support for afis|EAW23360.1 as seed ortholog is 100%.

---

Group of orthologs #8127. Best score 308 bits
Score difference with first non-orthologous sequence - *A. fumigatus*:308 *A. fischeri*:308
afum|EBA27328.1 100.00% afis|EAW23527.1 100.00%
Bootstrap support for afum|EBA27328.1 as seed ortholog is 100%.
Bootstrap support for afis|EAW23527.1 as seed ortholog is 100%.

---

Group of orthologs #8128. Best score 307 bits
Score difference with first non-orthologous sequence - *A. fumigatus*:307 *A. fischeri*:307
afum|EAL84949.1 100.00% afis|EAW19967.1 100.00%
Bootstrap support for afum|EAL84949.1 as seed ortholog is 100%.
Bootstrap support for afis|EAW19967.1 as seed ortholog is 100%.

---

Group of orthologs #8129. Best score 307 bits
Score difference with first non-orthologous sequence - *A. fumigatus*:229 *A. fischeri*:307
afum|EBA27194.1 100.00% afis|EAW20470.1 100.00%
Bootstrap support for afum|EBA27194.1 as seed ortholog is 100%.
Bootstrap support for afis|EAW20470.1 as seed ortholog is 100%.

---

Group of orthologs #8130. Best score 307 bits
Score difference with first non-orthologous sequence - *A. fumigatus*:307 *A. fischeri*:142
afum|EAL90012.1 100.00% afis|EAW25349.1 100.00%
Bootstrap support for afum|EAL90012.1 as seed ortholog is 100%.
Bootstrap support for afis|EAW25349.1 as seed ortholog is 100%.

---

Group of orthologs #8131. Best score 307 bits
Score difference with first non-orthologous sequence - *A. fumigatus*:307 *A. fischeri*:307
afum|EAL90162.1 100.00% afis|EAW25496.1 100.00%
Bootstrap support for afum|EAL90162.1 as seed ortholog is 100%.
Bootstrap support for afis|EAW25496.1 as seed ortholog is 100%.

---

Group of orthologs #8132. Best score 307 bits
Score difference with first non-orthologous sequence - *A. fumigatus*:240 *A. fischeri*:307
afum|EAL92015.1 100.00% afis|EAW24159.1 100.00%
Bootstrap support for afum|EAL92015.1 as seed ortholog is 100%.

Bootstrap support for afis|EAW24159.1 as seed ortholog is 100%.

---

Group of orthologs #8133. Best score 306 bits
Score difference with first non-orthologous sequence - *A. fumigatus*:306 *A. fischeri*:306
afum|EAL89193.1 100.00% afis|EAW16597.1 100.00%
Bootstrap support for afum|EAL89193.1 as seed ortholog is 100%.
Bootstrap support for afis|EAW16597.1 as seed ortholog is 100%.

---

Group of orthologs #8134. Best score 306 bits
Score difference with first non-orthologous sequence - *A. fumigatus*:306 *A. fischeri*:306
afum|EAL87296.1 100.00% afis|EAW23868.1 100.00%
Bootstrap support for afum|EAL87296.1 as seed ortholog is 100%.
Bootstrap support for afis|EAW23868.1 as seed ortholog is 100%.

---

Group of orthologs #8135. Best score 305 bits
Score difference with first non-orthologous sequence - *A. fumigatus*:305 *A. fischeri*:305
afum|EAL85520.1 100.00% afis|EAW15832.1 100.00%
Bootstrap support for afum|EAL85520.1 as seed ortholog is 100%.
Bootstrap support for afis|EAW15832.1 as seed ortholog is 100%.

---

Group of orthologs #8136. Best score 305 bits
Score difference with first non-orthologous sequence - *A. fumigatus*:207 *A. fischeri*:222
afum|EAL85068.1 100.00% afis|EAW19824.1 100.00%
Bootstrap support for afum|EAL85068.1 as seed ortholog is 99%.
Bootstrap support for afis|EAW19824.1 as seed ortholog is 100%.

---

Group of orthologs #8137. Best score 305 bits
Score difference with first non-orthologous sequence - *A. fumigatus*:305 *A. fischeri*:305
afum|EAL84648.2 100.00% afis|EAW20982.1 100.00%
Bootstrap support for afum|EAL84648.2 as seed ortholog is 100%.
Bootstrap support for afis|EAW20982.1 as seed ortholog is 100%.

---

Group of orthologs #8138. Best score 304 bits
Score difference with first non-orthologous sequence - *A. fumigatus*:304 *A. fischeri*:304
afum|EAL84999.1 100.00% afis|EAW19907.1 100.00%
Bootstrap support for afum|EAL84999.1 as seed ortholog is 100%.
Bootstrap support for afis|EAW19907.1 as seed ortholog is 100%.

---

Group of orthologs #8139. Best score 304 bits
Score difference with first non-orthologous sequence - *A. fumigatus*:304 *A. fischeri*:304
afum|EAL93593.2 100.00% afis|EAW18861.1 100.00%
Bootstrap support for afum|EAL93593.2 as seed ortholog is 100%.
Bootstrap support for afis|EAW18861.1 as seed ortholog is 100%.

---

Group of orthologs #8140. Best score 304 bits
Score difference with first non-orthologous sequence - *A. fumigatus*:304 *A. fischeri*:304
afum|EAL90449.1 100.00% afis|EAW22750.1 100.00%
Bootstrap support for afum|EAL90449.1 as seed ortholog is 100%.
Bootstrap support for afis|EAW22750.1 as seed ortholog is 100%.

---

Group of orthologs #8141. Best score 303 bits
Score difference with first non-orthologous sequence - *A. fumigatus*:303 *A. fischeri*:303
afum|EAL86337.1 100.00% afis|EAW16769.1 100.00%
Bootstrap support for afum|EAL86337.1 as seed ortholog is 100%.
Bootstrap support for afis|EAW16769.1 as seed ortholog is 100%.

---

Group of orthologs #8142. Best score 303 bits
Score difference with first non-orthologous sequence - *A. fumigatus*:303 *A. fischeri*:303
afum|EAL88413.1 100.00% afis|EAW23066.1 100.00%
Bootstrap support for afum|EAL88413.1 as seed ortholog is 100%.
Bootstrap support for afis|EAW23066.1 as seed ortholog is 100%.

---

Group of orthologs #8143. Best score 303 bits
Score difference with first non-orthologous sequence - *A. fumigatus*:303 *A. fischeri*:303
afum|EAL93958.1 100.00% afis|EAW19228.1 100.00%
Bootstrap support for afum|EAL93958.1 as seed ortholog is 100%.
Bootstrap support for afis|EAW19228.1 as seed ortholog is 100%.

---

Group of orthologs #8144. Best score 303 bits
Score difference with first non-orthologous sequence - *A. fumigatus*:303 *A. fischeri*:303
afum|EAL92518.1 100.00% afis|EAW21499.1 100.00%
Bootstrap support for afum|EAL92518.1 as seed ortholog is 100%.
Bootstrap support for afis|EAW21499.1 as seed ortholog is 100%.

---

Group of orthologs #8145. Best score 302 bits
Score difference with first non-orthologous sequence - *A. fumigatus*:302 *A. fischeri*:302
afum|EAL85562.1 100.00% afis|EAW15788.1 100.00%
Bootstrap support for afum|EAL85562.1 as seed ortholog is 100%.
Bootstrap support for afis|EAW15788.1 as seed ortholog is 100%.

---

Group of orthologs #8146. Best score 302 bits
Score difference with first non-orthologous sequence - *A. fumigatus*:302 *A. fischeri*:302
afum|EAL88835.1 100.00% afis|EAW16208.1 100.00%
Bootstrap support for afum|EAL88835.1 as seed ortholog is 100%.
Bootstrap support for afis|EAW16208.1 as seed ortholog is 100%.

---

Group of orthologs #8147. Best score 302 bits
Score difference with first non-orthologous sequence - *A. fumigatus*:302 *A. fischeri*:189
afum|EAL84660.1 100.00% afis|EAW20973.1 100.00%
Bootstrap support for afum|EAL84660.1 as seed ortholog is 100%.
Bootstrap support for afis|EAW20973.1 as seed ortholog is 100%.

---

Group of orthologs #8148. Best score 302 bits
Score difference with first non-orthologous sequence - *A. fumigatus*:302 *A. fischeri*:302
afum|EAL86873.1 100.00% afis|EAW19649.1 100.00%
Bootstrap support for afum|EAL86873.1 as seed ortholog is 100%.
Bootstrap support for afis|EAW19649.1 as seed ortholog is 100%.

---

Group of orthologs #8149. Best score 302 bits
Score difference with first non-orthologous sequence - *A. fumigatus*:302 *A. fischeri*:302
afum|EAL85923.1 100.00% afis|EAW24255.1 100.00%
Bootstrap support for afum|EAL85923.1 as seed ortholog is 100%.
Bootstrap support for afis|EAW24255.1 as seed ortholog is 100%.

---

Group of orthologs #8150. Best score 302 bits
Score difference with first non-orthologous sequence - *A. fumigatus*:302 *A. fischeri*:302
afum|EAL92535.1 100.00% afis|EAW21515.1 100.00%
Bootstrap support for afum|EAL92535.1 as seed ortholog is 100%.
Bootstrap support for afis|EAW21515.1 as seed ortholog is 100%.

---

Group of orthologs #8151. Best score 302 bits
Score difference with first non-orthologous sequence - *A. fumigatus*:302 *A. fischeri*:302

afum|EAL89756.1 100.00% afis|EAW25087.1 100.00%
Bootstrap support for afum|EAL89756.1 as seed ortholog is 100%.
Bootstrap support for afis|EAW25087.1 as seed ortholog is 100%.

---

Group of orthologs #8152. Best score 302 bits
Score difference with first non-orthologous sequence - *A. fumigatus*:302 *A. fischeri*:302
afum|EBA27324.1 100.00% afis|EAW23631.1 100.00%
Bootstrap support for afum|EBA27324.1 as seed ortholog is 100%.
Bootstrap support for afis|EAW23631.1 as seed ortholog is 100%.

---

Group of orthologs #8153. Best score 301 bits
Score difference with first non-orthologous sequence - *A. fumigatus*:301 *A. fischeri*:301
afum|EAL92718.2 100.00% afis|EAW21684.1 100.00%
Bootstrap support for afum|EAL92718.2 as seed ortholog is 100%.
Bootstrap support for afis|EAW21684.1 as seed ortholog is 100%.

---

Group of orthologs #8154. Best score 300 bits
Score difference with first non-orthologous sequence - *A. fumigatus*:300 *A. fischeri*:300
afum|EAL91143.1 100.00% afis|EAW17308.1 100.00%
Bootstrap support for afum|EAL91143.1 as seed ortholog is 100%.
Bootstrap support for afis|EAW17308.1 as seed ortholog is 100%.

---

Group of orthologs #8155. Best score 300 bits
Score difference with first non-orthologous sequence - *A. fumigatus*:300 *A. fischeri*:300
afum|EAL88388.1 100.00% afis|EAW23091.1 100.00%
Bootstrap support for afum|EAL88388.1 as seed ortholog is 100%.
Bootstrap support for afis|EAW23091.1 as seed ortholog is 100%.

---

Group of orthologs #8156. Best score 300 bits
Score difference with first non-orthologous sequence - *A. fumigatus*:300 *A. fischeri*:300
afum|EBA27376.1 100.00% afis|EAW24725.1 100.00%
Bootstrap support for afum|EBA27376.1 as seed ortholog is 100%.
Bootstrap support for afis|EAW24725.1 as seed ortholog is 100%.

---

Group of orthologs #8157. Best score 299 bits
Score difference with first non-orthologous sequence - *A. fumigatus*:299 *A. fischeri*:299
afum|EAL87686.1 100.00% afis|EAW18319.1 100.00%
Bootstrap support for afum|EAL87686.1 as seed ortholog is 100%.
Bootstrap support for afis|EAW18319.1 as seed ortholog is 100%.

---

Group of orthologs #8158. Best score 299 bits
Score difference with first non-orthologous sequence - *A. fumigatus*:299 *A. fischeri*:299
afum|EAL93496.2 100.00% afis|EAW18764.1 100.00%
Bootstrap support for afum|EAL93496.2 as seed ortholog is 100%.
Bootstrap support for afis|EAW18764.1 as seed ortholog is 100%.

---

Group of orthologs #8159. Best score 298 bits
Score difference with first non-orthologous sequence - *A. fumigatus*:89 *A. fischeri*:298
afum|EAL84921.1 100.00% afis|EAW15395.1 100.00%
Bootstrap support for afum|EAL84921.1 as seed ortholog is 96%.
Bootstrap support for afis|EAW15395.1 as seed ortholog is 100%.

---

Group of orthologs #8160. Best score 298 bits
Score difference with first non-orthologous sequence - *A. fumigatus*:298 *A. fischeri*:298
afum|EAL85746.1 100.00% afis|EAW15586.1 100.00%
Bootstrap support for afum|EAL85746.1 as seed ortholog is 100%.

Bootstrap support for afis|EAW15586.1 as seed ortholog is 100%.

---

Group of orthologs #8161. Best score 298 bits
Score difference with first non-orthologous sequence - *A. fumigatus*:298 *A. fischeri*:298
afum|EAL87648.1 100.00% afis|EAW18282.1 100.00%
Bootstrap support for afum|EAL87648.1 as seed ortholog is 100%.
Bootstrap support for afis|EAW18282.1 as seed ortholog is 100%.

---

Group of orthologs #8162. Best score 298 bits
Score difference with first non-orthologous sequence - *A. fumigatus*:298 *A. fischeri*:298
afum|EAL91082.1 100.00% afis|EAW16966.1 100.00%
Bootstrap support for afum|EAL91082.1 as seed ortholog is 100%.
Bootstrap support for afis|EAW16966.1 as seed ortholog is 100%.

---

Group of orthologs #8163. Best score 298 bits
Score difference with first non-orthologous sequence - *A. fumigatus*:298 *A. fischeri*:298
afum|EAL85974.1 100.00% afis|EAW24306.1 100.00%
Bootstrap support for afum|EAL85974.1 as seed ortholog is 100%.
Bootstrap support for afis|EAW24306.1 as seed ortholog is 100%.

---

Group of orthologs #8164. Best score 298 bits
Score difference with first non-orthologous sequence - *A. fumigatus*:298 *A. fischeri*:298
afum|EAL88246.1 100.00% afis|EAW23236.1 100.00%
Bootstrap support for afum|EAL88246.1 as seed ortholog is 100%.
Bootstrap support for afis|EAW23236.1 as seed ortholog is 100%.

---

Group of orthologs #8165. Best score 298 bits
Score difference with first non-orthologous sequence - *A. fumigatus*:298 *A. fischeri*:298
afum|EBA27430.1 100.00% afis|EAW17503.1 100.00%
Bootstrap support for afum|EBA27430.1 as seed ortholog is 100%.
Bootstrap support for afis|EAW17503.1 as seed ortholog is 100%.

---

Group of orthologs #8166. Best score 298 bits
Score difference with first non-orthologous sequence - *A. fumigatus*:298 *A. fischeri*:298
afum|EAL93428.1 100.00% afis|EAW18705.1 100.00%
Bootstrap support for afum|EAL93428.1 as seed ortholog is 100%.
Bootstrap support for afis|EAW18705.1 as seed ortholog is 100%.

---

Group of orthologs #8167. Best score 298 bits
Score difference with first non-orthologous sequence - *A. fumigatus*:298 *A. fischeri*:298
afum|EAL90828.1 100.00% afis|EAW22363.1 100.00%
Bootstrap support for afum|EAL90828.1 as seed ortholog is 100%.
Bootstrap support for afis|EAW22363.1 as seed ortholog is 100%.

---

Group of orthologs #8168. Best score 298 bits
Score difference with first non-orthologous sequence - *A. fumigatus*:298 *A. fischeri*:298
afum|EAL90824.1 100.00% afis|EAW22368.1 100.00%
Bootstrap support for afum|EAL90824.1 as seed ortholog is 100%.
Bootstrap support for afis|EAW22368.1 as seed ortholog is 100%.

---

Group of orthologs #8169. Best score 297 bits
Score difference with first non-orthologous sequence - *A. fumigatus*:221 *A. fischeri*:176
afum|EAL91185.1 100.00% afis|EAW17340.1 100.00%
Bootstrap support for afum|EAL91185.1 as seed ortholog is 100%.
Bootstrap support for afis|EAW17340.1 as seed ortholog is 100%.

---

Group of orthologs #8170. Best score 297 bits
Score difference with first non-orthologous sequence - *A. fumigatus*:297 *A. fischeri*:297
afum|EAL87898.2 100.00% afis|EAW23569.1 100.00%
Bootstrap support for afum|EAL87898.2 as seed ortholog is 100%.
Bootstrap support for afis|EAW23569.1 as seed ortholog is 100%.

---

Group of orthologs #8171. Best score 297 bits
Score difference with first non-orthologous sequence - *A. fumigatus*:297 *A. fischeri*:297
afum|EAL93308.1 100.00% afis|EAW18600.1 100.00%
Bootstrap support for afum|EAL93308.1 as seed ortholog is 100%.
Bootstrap support for afis|EAW18600.1 as seed ortholog is 100%.

---

Group of orthologs #8172. Best score 296 bits
Score difference with first non-orthologous sequence - *A. fumigatus*:296 *A. fischeri*:296
afum|EAL89263.1 100.00% afis|EAW16659.1 100.00%
Bootstrap support for afum|EAL89263.1 as seed ortholog is 100%.
Bootstrap support for afis|EAW16659.1 as seed ortholog is 100%.

---

Group of orthologs #8173. Best score 296 bits
Score difference with first non-orthologous sequence - *A. fumigatus*:296 *A. fischeri*:296
afum|EBA27169.1 100.00% afis|EAW15435.1 100.00%
Bootstrap support for afum|EBA27169.1 as seed ortholog is 100%.
Bootstrap support for afis|EAW15435.1 as seed ortholog is 100%.

---

Group of orthologs #8174. Best score 296 bits
Score difference with first non-orthologous sequence - *A. fumigatus*:296 *A. fischeri*:296
afum|EAL88070.1 100.00% afis|EAW23400.1 100.00%
Bootstrap support for afum|EAL88070.1 as seed ortholog is 100%.
Bootstrap support for afis|EAW23400.1 as seed ortholog is 100%.

---

Group of orthologs #8175. Best score 296 bits
Score difference with first non-orthologous sequence - *A. fumigatus*:296 *A. fischeri*:296
afum|EBA27438.1 100.00% afis|EAW17894.1 100.00%
Bootstrap support for afum|EBA27438.1 as seed ortholog is 100%.
Bootstrap support for afis|EAW17894.1 as seed ortholog is 100%.

---

Group of orthologs #8176. Best score 296 bits
Score difference with first non-orthologous sequence - *A. fumigatus*:296 *A. fischeri*:231
afum|EAL90475.1 100.00% afis|EAW22719.1 100.00%
Bootstrap support for afum|EAL90475.1 as seed ortholog is 100%.
Bootstrap support for afis|EAW22719.1 as seed ortholog is 100%.

---

Group of orthologs #8177. Best score 296 bits
Score difference with first non-orthologous sequence - *A. fumigatus*:296 *A. fischeri*:296
afum|EAL92167.1 100.00% afis|EAW21148.1 100.00%
Bootstrap support for afum|EAL92167.1 as seed ortholog is 100%.
Bootstrap support for afis|EAW21148.1 as seed ortholog is 100%.

---

Group of orthologs #8178. Best score 296 bits
Score difference with first non-orthologous sequence - *A. fumigatus*:296 *A. fischeri*:296
afum|EAL90073.1 100.00% afis|EAW25414.1 100.00%
Bootstrap support for afum|EAL90073.1 as seed ortholog is 100%.
Bootstrap support for afis|EAW25414.1 as seed ortholog is 100%.

---

Group of orthologs #8179. Best score 295 bits
Score difference with first non-orthologous sequence - *A. fumigatus*:295 *A. fischeri*:295

afum|EAL91119.1 100.00% afis|EAW20300.1 100.00%
afum|EAL90937.1 25.00%
Bootstrap support for afum|EAL91119.1 as seed ortholog is 100%.
Bootstrap support for afis|EAW20300.1 as seed ortholog is 100%.

---

Group of orthologs #8180. Best score 295 bits
Score difference with first non-orthologous sequence - *A. fumigatus*:295 *A. fischeri*:295
afum|EAL88349.1 100.00% afis|EAW23132.1 100.00%
Bootstrap support for afum|EAL88349.1 as seed ortholog is 100%.
Bootstrap support for afis|EAW23132.1 as seed ortholog is 100%.

---

Group of orthologs #8181. Best score 295 bits
Score difference with first non-orthologous sequence - *A. fumigatus*:295 *A. fischeri*:295
afum|EBA27215.1 100.00% afis|EAW19923.1 100.00%
Bootstrap support for afum|EBA27215.1 as seed ortholog is 100%.
Bootstrap support for afis|EAW19923.1 as seed ortholog is 100%.

---

Group of orthologs #8182. Best score 295 bits
Score difference with first non-orthologous sequence - *A. fumigatus*:295 *A. fischeri*:295
afum|EAL92719.1 100.00% afis|EAW21685.1 100.00%
Bootstrap support for afum|EAL92719.1 as seed ortholog is 100%.
Bootstrap support for afis|EAW21685.1 as seed ortholog is 100%.

---

Group of orthologs #8183. Best score 294 bits
Score difference with first non-orthologous sequence - *A. fumigatus*:294 *A. fischeri*:294
afum|EAL91247.1 100.00% afis|EAW17422.1 100.00%
Bootstrap support for afum|EAL91247.1 as seed ortholog is 100%.
Bootstrap support for afis|EAW17422.1 as seed ortholog is 100%.

---

Group of orthologs #8184. Best score 294 bits
Score difference with first non-orthologous sequence - *A. fumigatus*:294 *A. fischeri*:294
afum|EAL86224.1 100.00% afis|EAW24563.1 100.00%
Bootstrap support for afum|EAL86224.1 as seed ortholog is 100%.
Bootstrap support for afis|EAW24563.1 as seed ortholog is 100%.

---

Group of orthologs #8185. Best score 294 bits
Score difference with first non-orthologous sequence - *A. fumigatus*:294 *A. fischeri*:294
afum|EAL93835.1 100.00% afis|EAW19103.1 100.00%
Bootstrap support for afum|EAL93835.1 as seed ortholog is 100%.
Bootstrap support for afis|EAW19103.1 as seed ortholog is 100%.

---

Group of orthologs #8186. Best score 293 bits
Score difference with first non-orthologous sequence - *A. fumigatus*:293 *A. fischeri*:293
afum|EAL88539.1 100.00% afis|EAW15904.1 100.00%
Bootstrap support for afum|EAL88539.1 as seed ortholog is 100%.
Bootstrap support for afis|EAW15904.1 as seed ortholog is 100%.

---

Group of orthologs #8187. Best score 293 bits
Score difference with first non-orthologous sequence - *A. fumigatus*:293 *A. fischeri*:293
afum|EAL89163.1 100.00% afis|EAW16566.1 100.00%
Bootstrap support for afum|EAL89163.1 as seed ortholog is 100%.
Bootstrap support for afis|EAW16566.1 as seed ortholog is 100%.

---

Group of orthologs #8188. Best score 293 bits
Score difference with first non-orthologous sequence - *A. fumigatus*:293 *A. fischeri*:293
afum|EAL93728.1 100.00% afis|EAW18993.1 100.00%

Bootstrap support for afum|EAL93728.1 as seed ortholog is 100%.
Bootstrap support for afis|EAW18993.1 as seed ortholog is 100%.

---

Group of orthologs #8189. Best score 292 bits
Score difference with first non-orthologous sequence - *A. fumigatus*:292 *A. fischeri*:292
afum|EAL85535.1 100.00% afis|EAW15816.1 100.00%
Bootstrap support for afum|EAL85535.1 as seed ortholog is 100%.
Bootstrap support for afis|EAW15816.1 as seed ortholog is 100%.

---

Group of orthologs #8190. Best score 292 bits
Score difference with first non-orthologous sequence - *A. fumigatus*:292 *A. fischeri*:292
afum|EAL89222.2 100.00% afis|EAW16614.1 100.00%
Bootstrap support for afum|EAL89222.2 as seed ortholog is 100%.
Bootstrap support for afis|EAW16614.1 as seed ortholog is 100%.

---

Group of orthologs #8191. Best score 292 bits
Score difference with first non-orthologous sequence - *A. fumigatus*:292 *A. fischeri*:292
afum|EAL89911.1 100.00% afis|EAW25249.1 100.00%
Bootstrap support for afum|EAL89911.1 as seed ortholog is 100%.
Bootstrap support for afis|EAW25249.1 as seed ortholog is 100%.

---

Group of orthologs #8192. Best score 292 bits
Score difference with first non-orthologous sequence - *A. fumigatus*:292 *A. fischeri*:292
afum|EBA27393.1 100.00% afis|EAW25366.1 100.00%
Bootstrap support for afum|EBA27393.1 as seed ortholog is 100%.
Bootstrap support for afis|EAW25366.1 as seed ortholog is 100%.

---

Group of orthologs #8193. Best score 291 bits
Score difference with first non-orthologous sequence - *A. fumigatus*:291 *A. fischeri*:291
afum|EAL84551.1 100.00% afis|EAW20505.1 100.00%
Bootstrap support for afum|EAL84551.1 as seed ortholog is 100%.
Bootstrap support for afis|EAW20505.1 as seed ortholog is 100%.

---

Group of orthologs #8194. Best score 291 bits
Score difference with first non-orthologous sequence - *A. fumigatus*:291 *A. fischeri*:291
afum|EAL91834.1 100.00% afis|EAW18022.1 100.00%
Bootstrap support for afum|EAL91834.1 as seed ortholog is 100%.
Bootstrap support for afis|EAW18022.1 as seed ortholog is 100%.

---

Group of orthologs #8195. Best score 291 bits
Score difference with first non-orthologous sequence - *A. fumigatus*:291 *A. fischeri*:173
afum|EAL87734.1 100.00% afis|EAW22033.1 100.00%
Bootstrap support for afum|EAL87734.1 as seed ortholog is 100%.
Bootstrap support for afis|EAW22033.1 as seed ortholog is 99%.

---

Group of orthologs #8196. Best score 291 bits
Score difference with first non-orthologous sequence - *A. fumigatus*:291 *A. fischeri*:291
afum|EAL90389.1 100.00% afis|EAW22812.1 100.00%
Bootstrap support for afum|EAL90389.1 as seed ortholog is 100%.
Bootstrap support for afis|EAW22812.1 as seed ortholog is 100%.

---

Group of orthologs #8197. Best score 291 bits
Score difference with first non-orthologous sequence - *A. fumigatus*:291 *A. fischeri*:291
afum|EAL90379.2 100.00% afis|EAW22821.1 100.00%
Bootstrap support for afum|EAL90379.2 as seed ortholog is 100%.
Bootstrap support for afis|EAW22821.1 as seed ortholog is 100%.

Group of orthologs #8198. Best score 291 bits
Score difference with first non-orthologous sequence - *A. fumigatus*:291 *A. fischeri*:291
afum|EAL90283.1 100.00% afis|EAW22914.1 100.00%
Bootstrap support for afum|EAL90283.1 as seed ortholog is 100%.
Bootstrap support for afis|EAW22914.1 as seed ortholog is 100%.

Group of orthologs #8199. Best score 290 bits
Score difference with first non-orthologous sequence - *A. fumigatus*:290 *A. fischeri*:290
afum|EAL88455.1 100.00% afis|EAW23027.1 100.00%
Bootstrap support for afum|EAL88455.1 as seed ortholog is 100%.
Bootstrap support for afis|EAW23027.1 as seed ortholog is 100%.

Group of orthologs #8200. Best score 290 bits
Score difference with first non-orthologous sequence - *A. fumigatus*:290 *A. fischeri*:290
afum|EAL92732.1 100.00% afis|EAW21698.1 100.00%
Bootstrap support for afum|EAL92732.1 as seed ortholog is 100%.
Bootstrap support for afis|EAW21698.1 as seed ortholog is 100%.

Group of orthologs #8201. Best score 289 bits
Score difference with first non-orthologous sequence - *A. fumigatus*:289 *A. fischeri*:289
afum|EAL84976.2 100.00% afis|EAW19933.1 100.00%
Bootstrap support for afum|EAL84976.2 as seed ortholog is 100%.
Bootstrap support for afis|EAW19933.1 as seed ortholog is 100%.

Group of orthologs #8202. Best score 289 bits
Score difference with first non-orthologous sequence - *A. fumigatus*:289 *A. fischeri*:289
afum|EAL87604.2 100.00% afis|EAW18237.1 100.00%
Bootstrap support for afum|EAL87604.2 as seed ortholog is 100%.
Bootstrap support for afis|EAW18237.1 as seed ortholog is 100%.

Group of orthologs #8203. Best score 289 bits
Score difference with first non-orthologous sequence - *A. fumigatus*:289 *A. fischeri*:289
afum|EAL92603.2 100.00% afis|EAW21585.1 100.00%
Bootstrap support for afum|EAL92603.2 as seed ortholog is 100%.
Bootstrap support for afis|EAW21585.1 as seed ortholog is 100%.

Group of orthologs #8204. Best score 289 bits
Score difference with first non-orthologous sequence - *A. fumigatus*:289 *A. fischeri*:289
afum|EBA27193.1 100.00% afis|EAW20474.1 100.00%
Bootstrap support for afum|EBA27193.1 as seed ortholog is 100%.
Bootstrap support for afis|EAW20474.1 as seed ortholog is 100%.

Group of orthologs #8205. Best score 289 bits
Score difference with first non-orthologous sequence - *A. fumigatus*:289 *A. fischeri*:289
afum|EAL89571.1 100.00% afis|EAW24916.1 100.00%
Bootstrap support for afum|EAL89571.1 as seed ortholog is 100%.
Bootstrap support for afis|EAW24916.1 as seed ortholog is 100%.

Group of orthologs #8206. Best score 289 bits
Score difference with first non-orthologous sequence - *A. fumigatus*:59 *A. fischeri*:90
afum|EBA27227.1 100.00% afis|EAW24094.1 100.00%
Bootstrap support for afum|EBA27227.1 as seed ortholog is 85%.
Bootstrap support for afis|EAW24094.1 as seed ortholog is 93%.

Group of orthologs #8207. Best score 288 bits

Score difference with first non-orthologous sequence - *A. fumigatus*:288 *A. fischeri*:96
afum|EAL85154.2 100.00% afis|EAW23663.1 100.00%
Bootstrap support for afum|EAL85154.2 as seed ortholog is 100%.
Bootstrap support for afis|EAW23663.1 as seed ortholog is 97%.

---

Group of orthologs #8208. Best score 288 bits
Score difference with first non-orthologous sequence - *A. fumigatus*:288 *A. fischeri*:288
afum|EAL91726.1 100.00% afis|EAW17916.1 100.00%
Bootstrap support for afum|EAL91726.1 as seed ortholog is 100%.
Bootstrap support for afis|EAW17916.1 as seed ortholog is 100%.

---

Group of orthologs #8209. Best score 288 bits
Score difference with first non-orthologous sequence - *A. fumigatus*:288 *A. fischeri*:288
afum|EAL86187.1 100.00% afis|EAW24522.1 100.00%
Bootstrap support for afum|EAL86187.1 as seed ortholog is 100%.
Bootstrap support for afis|EAW24522.1 as seed ortholog is 100%.

---

Group of orthologs #8210. Best score 288 bits
Score difference with first non-orthologous sequence - *A. fumigatus*:288 *A. fischeri*:288
afum|EBA27446.1 100.00% afis|EAW24075.1 100.00%
Bootstrap support for afum|EBA27446.1 as seed ortholog is 100%.
Bootstrap support for afis|EAW24075.1 as seed ortholog is 100%.

---

Group of orthologs #8211. Best score 287 bits
Score difference with first non-orthologous sequence - *A. fumigatus*:287 *A. fischeri*:287
afum|EAL88746.1 100.00% afis|EAW16115.1 100.00%
Bootstrap support for afum|EAL88746.1 as seed ortholog is 100%.
Bootstrap support for afis|EAW16115.1 as seed ortholog is 100%.

---

Group of orthologs #8212. Best score 287 bits
Score difference with first non-orthologous sequence - *A. fumigatus*:287 *A. fischeri*:287
afum|EAL88451.2 100.00% afis|EAW23031.1 100.00%
Bootstrap support for afum|EAL88451.2 as seed ortholog is 100%.
Bootstrap support for afis|EAW23031.1 as seed ortholog is 100%.

---

Group of orthologs #8213. Best score 287 bits
Score difference with first non-orthologous sequence - *A. fumigatus*:287 *A. fischeri*:26
afum|EAL92224.2 100.00% afis|EAW21213.1 100.00%
Bootstrap support for afum|EAL92224.2 as seed ortholog is 100%.
Bootstrap support for afis|EAW21213.1 as seed ortholog is 88%.

---

Group of orthologs #8214. Best score 286 bits
Score difference with first non-orthologous sequence - *A. fumigatus*:286 *A. fischeri*:286
afum|EAL89237.1 100.00% afis|EAW16635.1 100.00%
Bootstrap support for afum|EAL89237.1 as seed ortholog is 100%.
Bootstrap support for afis|EAW16635.1 as seed ortholog is 100%.

---

Group of orthologs #8215. Best score 286 bits
Score difference with first non-orthologous sequence - *A. fumigatus*:286 *A. fischeri*:286
afum|EAL84635.1 100.00% afis|EAW20993.1 100.00%
Bootstrap support for afum|EAL84635.1 as seed ortholog is 100%.
Bootstrap support for afis|EAW20993.1 as seed ortholog is 100%.

---

Group of orthologs #8216. Best score 286 bits
Score difference with first non-orthologous sequence - *A. fumigatus*:286 *A. fischeri*:286
afum|EAL90536.1 100.00% afis|EAW22656.1 100.00%

Bootstrap support for afum|EAL90536.1 as seed ortholog is 100%.
Bootstrap support for afis|EAW22656.1 as seed ortholog is 100%.

---

Group of orthologs #8217. Best score 285 bits
Score difference with first non-orthologous sequence - *A. fumigatus*:285 *A. fischeri*:285
afum|EAL87435.1 100.00% afis|EAW24005.1 100.00%
Bootstrap support for afum|EAL87435.1 as seed ortholog is 100%.
Bootstrap support for afis|EAW24005.1 as seed ortholog is 100%.

---

Group of orthologs #8218. Best score 285 bits
Score difference with first non-orthologous sequence - *A. fumigatus*:285 *A. fischeri*:285
afum|EAL92344.1 100.00% afis|EAW21326.1 100.00%
Bootstrap support for afum|EAL92344.1 as seed ortholog is 100%.
Bootstrap support for afis|EAW21326.1 as seed ortholog is 100%.

---

Group of orthologs #8219. Best score 284 bits
Score difference with first non-orthologous sequence - *A. fumigatus*:284 *A. fischeri*:284
afum|EAL87425.1 100.00% afis|EAW23995.1 100.00%
Bootstrap support for afum|EAL87425.1 as seed ortholog is 100%.
Bootstrap support for afis|EAW23995.1 as seed ortholog is 100%.

---

Group of orthologs #8220. Best score 284 bits
Score difference with first non-orthologous sequence - *A. fumigatus*:284 *A. fischeri*:284
afum|EAL90618.1 100.00% afis|EAW22581.1 100.00%
Bootstrap support for afum|EAL90618.1 as seed ortholog is 100%.
Bootstrap support for afis|EAW22581.1 as seed ortholog is 100%.

---

Group of orthologs #8221. Best score 284 bits
Score difference with first non-orthologous sequence - *A. fumigatus*:284 *A. fischeri*:284
afum|EAL89853.2 100.00% afis|EAW25179.1 100.00%
Bootstrap support for afum|EAL89853.2 as seed ortholog is 100%.
Bootstrap support for afis|EAW25179.1 as seed ortholog is 100%.

---

Group of orthologs #8222. Best score 283 bits
Score difference with first non-orthologous sequence - *A. fumigatus*:283 *A. fischeri*:283
afum|EAL85104.1 100.00% afis|EAW19837.1 100.00%
Bootstrap support for afum|EAL85104.1 as seed ortholog is 100%.
Bootstrap support for afis|EAW19837.1 as seed ortholog is 100%.

---

Group of orthologs #8223. Best score 282 bits
Score difference with first non-orthologous sequence - *A. fumigatus*:282 *A. fischeri*:282
afum|EAL88095.1 100.00% afis|EAW23376.1 100.00%
Bootstrap support for afum|EAL88095.1 as seed ortholog is 100%.
Bootstrap support for afis|EAW23376.1 as seed ortholog is 100%.

---

Group of orthologs #8224. Best score 281 bits
Score difference with first non-orthologous sequence - *A. fumigatus*:281 *A. fischeri*:281
afum|EAL87950.1 100.00% afis|EAW23513.1 100.00%
Bootstrap support for afum|EAL87950.1 as seed ortholog is 100%.
Bootstrap support for afis|EAW23513.1 as seed ortholog is 100%.

---

Group of orthologs #8225. Best score 281 bits
Score difference with first non-orthologous sequence - *A. fumigatus*:281 *A. fischeri*:281
afum|EAL92905.1 100.00% afis|EAW21891.1 100.00%
Bootstrap support for afum|EAL92905.1 as seed ortholog is 100%.
Bootstrap support for afis|EAW21891.1 as seed ortholog is 100%.

Group of orthologs #8226. Best score 280 bits
Score difference with first non-orthologous sequence - *A. fumigatus*:280 *A. fischeri*:280
afum|EAL84894.1 100.00% afis|EAW20023.1 100.00%
Bootstrap support for afum|EAL84894.1 as seed ortholog is 100%.
Bootstrap support for afis|EAW20023.1 as seed ortholog is 100%.

---

Group of orthologs #8227. Best score 280 bits
Score difference with first non-orthologous sequence - *A. fumigatus*:280 *A. fischeri*:280
afum|EAL89224.2 100.00% afis|EAW16629.1 100.00%
Bootstrap support for afum|EAL89224.2 as seed ortholog is 100%.
Bootstrap support for afis|EAW16629.1 as seed ortholog is 100%.

---

Group of orthologs #8228. Best score 280 bits
Score difference with first non-orthologous sequence - *A. fumigatus*:280 *A. fischeri*:280
afum|EAL87312.1 100.00% afis|EAW23884.1 100.00%
Bootstrap support for afum|EAL87312.1 as seed ortholog is 100%.
Bootstrap support for afis|EAW23884.1 as seed ortholog is 100%.

---

Group of orthologs #8229. Best score 280 bits
Score difference with first non-orthologous sequence - *A. fumigatus*:280 *A. fischeri*:280
afum|EAL93725.1 100.00% afis|EAW18989.1 100.00%
Bootstrap support for afum|EAL93725.1 as seed ortholog is 100%.
Bootstrap support for afis|EAW18989.1 as seed ortholog is 100%.

---

Group of orthologs #8230. Best score 280 bits
Score difference with first non-orthologous sequence - *A. fumigatus*:280 *A. fischeri*:280
afum|EAL92154.1 100.00% afis|EAW21135.1 100.00%
Bootstrap support for afum|EAL92154.1 as seed ortholog is 100%.
Bootstrap support for afis|EAW21135.1 as seed ortholog is 100%.

---

Group of orthologs #8231. Best score 279 bits
Score difference with first non-orthologous sequence - *A. fumigatus*:279 *A. fischeri*:279
afum|EAL93760.2 100.00% afis|EAW19031.1 100.00%
Bootstrap support for afum|EAL93760.2 as seed ortholog is 100%.
Bootstrap support for afis|EAW19031.1 as seed ortholog is 100%.

---

Group of orthologs #8232. Best score 279 bits
Score difference with first non-orthologous sequence - *A. fumigatus*:279 *A. fischeri*:279
afum|EAL90284.1 100.00% afis|EAW22913.1 100.00%
Bootstrap support for afum|EAL90284.1 as seed ortholog is 100%.
Bootstrap support for afis|EAW22913.1 as seed ortholog is 100%.

---

Group of orthologs #8233. Best score 279 bits
Score difference with first non-orthologous sequence - *A. fumigatus*:279 *A. fischeri*:279
afum|EAL89798.1 100.00% afis|EAW25127.1 100.00%
Bootstrap support for afum|EAL89798.1 as seed ortholog is 100%.
Bootstrap support for afis|EAW25127.1 as seed ortholog is 100%.

---

Group of orthologs #8234. Best score 278 bits
Score difference with first non-orthologous sequence - *A. fumigatus*:278 *A. fischeri*:278
afum|EAL87324.1 100.00% afis|EAW23896.1 100.00%
Bootstrap support for afum|EAL87324.1 as seed ortholog is 100%.
Bootstrap support for afis|EAW23896.1 as seed ortholog is 100%.

---

Group of orthologs #8235. Best score 278 bits

Score difference with first non-orthologous sequence - *A. fumigatus*:278 *A. fischeri*:278
afum|EAL90271.1 100.00% afis|EAW22926.1 100.00%
Bootstrap support for afum|EAL90271.1 as seed ortholog is 100%.
Bootstrap support for afis|EAW22926.1 as seed ortholog is 100%.

---

Group of orthologs #8236. Best score 278 bits
Score difference with first non-orthologous sequence - *A. fumigatus*:278 *A. fischeri*:278
afum|EBA27501.1 100.00% afis|EAW19014.1 100.00%
Bootstrap support for afum|EBA27501.1 as seed ortholog is 100%.
Bootstrap support for afis|EAW19014.1 as seed ortholog is 100%.

---

Group of orthologs #8237. Best score 277 bits
Score difference with first non-orthologous sequence - *A. fumigatus*:277 *A. fischeri*:277
afum|EAL93513.1 100.00% afis|EAW18781.1 100.00%
Bootstrap support for afum|EAL93513.1 as seed ortholog is 100%.
Bootstrap support for afis|EAW18781.1 as seed ortholog is 100%.

---

Group of orthologs #8238. Best score 277 bits
Score difference with first non-orthologous sequence - *A. fumigatus*:277 *A. fischeri*:277
afum|EAL89764.1 100.00% afis|EAW25093.1 100.00%
Bootstrap support for afum|EAL89764.1 as seed ortholog is 100%.
Bootstrap support for afis|EAW25093.1 as seed ortholog is 100%.

---

Group of orthologs #8239. Best score 276 bits
Score difference with first non-orthologous sequence - *A. fumigatus*:276 *A. fischeri*:276
afum|EAL84789.1 100.00% afis|EAW20844.1 100.00%
Bootstrap support for afum|EAL84789.1 as seed ortholog is 100%.
Bootstrap support for afis|EAW20844.1 as seed ortholog is 100%.

---

Group of orthologs #8240. Best score 276 bits
Score difference with first non-orthologous sequence - *A. fumigatus*:276 *A. fischeri*:276
afum|EAL87602.2 100.00% afis|EAW18235.1 100.00%
Bootstrap support for afum|EAL87602.2 as seed ortholog is 100%.
Bootstrap support for afis|EAW18235.1 as seed ortholog is 100%.

---

Group of orthologs #8241. Best score 276 bits
Score difference with first non-orthologous sequence - *A. fumigatus*:276 *A. fischeri*:276
afum|EBA27478.1 100.00% afis|EAW21895.1 100.00%
Bootstrap support for afum|EBA27478.1 as seed ortholog is 100%.
Bootstrap support for afis|EAW21895.1 as seed ortholog is 100%.

---

Group of orthologs #8242. Best score 275 bits
Score difference with first non-orthologous sequence - *A. fumigatus*:275 *A. fischeri*:275
afum|EAL89058.1 100.00% afis|EAW16462.1 100.00%
Bootstrap support for afum|EAL89058.1 as seed ortholog is 100%.
Bootstrap support for afis|EAW16462.1 as seed ortholog is 100%.

---

Group of orthologs #8243. Best score 275 bits
Score difference with first non-orthologous sequence - *A. fumigatus*:275 *A. fischeri*:275
afum|EAL87581.1 100.00% afis|EAW18215.1 100.00%
Bootstrap support for afum|EAL87581.1 as seed ortholog is 100%.
Bootstrap support for afis|EAW18215.1 as seed ortholog is 100%.

---

Group of orthologs #8244. Best score 275 bits
Score difference with first non-orthologous sequence - *A. fumigatus*:275 *A. fischeri*:275
afum|EAL86980.2 100.00% afis|EAW19517.1 100.00%

Bootstrap support for afum|EAL86980.2 as seed ortholog is 100%.
Bootstrap support for afis|EAW19517.1 as seed ortholog is 100%.

---

Group of orthologs #8245. Best score 275 bits
Score difference with first non-orthologous sequence - *A. fumigatus*:275 *A. fischeri*:275
afum|EBA27356.1 100.00% afis|EAW16137.1 100.00%
Bootstrap support for afum|EBA27356.1 as seed ortholog is 100%.
Bootstrap support for afis|EAW16137.1 as seed ortholog is 100%.

---

Group of orthologs #8246. Best score 275 bits
Score difference with first non-orthologous sequence - *A. fumigatus*:275 *A. fischeri*:275
afum|EBA27284.1 100.00% afis|EAW17244.1 100.00%
Bootstrap support for afum|EBA27284.1 as seed ortholog is 100%.
Bootstrap support for afis|EAW17244.1 as seed ortholog is 100%.

---

Group of orthologs #8247. Best score 275 bits
Score difference with first non-orthologous sequence - *A. fumigatus*:275 *A. fischeri*:275
afum|EAL93116.1 100.00% afis|EAW18435.1 100.00%
Bootstrap support for afum|EAL93116.1 as seed ortholog is 100%.
Bootstrap support for afis|EAW18435.1 as seed ortholog is 100%.

---

Group of orthologs #8248. Best score 274 bits
Score difference with first non-orthologous sequence - *A. fumigatus*:274 *A. fischeri*:274
afum|EAL85294.1 100.00% afis|EAW20176.1 100.00%
Bootstrap support for afum|EAL85294.1 as seed ortholog is 100%.
Bootstrap support for afis|EAW20176.1 as seed ortholog is 100%.

---

Group of orthologs #8249. Best score 274 bits
Score difference with first non-orthologous sequence - *A. fumigatus*:274 *A. fischeri*:274
afum|EAL87340.1 100.00% afis|EAW23911.1 100.00%
Bootstrap support for afum|EAL87340.1 as seed ortholog is 100%.
Bootstrap support for afis|EAW23911.1 as seed ortholog is 100%.

---

Group of orthologs #8250. Best score 274 bits
Score difference with first non-orthologous sequence - *A. fumigatus*:274 *A. fischeri*:274
afum|EAL87379.1 100.00% afis|EAW23951.1 100.00%
Bootstrap support for afum|EAL87379.1 as seed ortholog is 100%.
Bootstrap support for afis|EAW23951.1 as seed ortholog is 100%.

---

Group of orthologs #8251. Best score 274 bits
Score difference with first non-orthologous sequence - *A. fumigatus*:274 *A. fischeri*:274
afum|EAL92656.1 100.00% afis|EAW21620.1 100.00%
Bootstrap support for afum|EAL92656.1 as seed ortholog is 100%.
Bootstrap support for afis|EAW21620.1 as seed ortholog is 100%.

---

Group of orthologs #8252. Best score 274 bits
Score difference with first non-orthologous sequence - *A. fumigatus*:274 *A. fischeri*:274
afum|EBA27188.1 100.00% afis|EAW20545.1 100.00%
Bootstrap support for afum|EBA27188.1 as seed ortholog is 100%.
Bootstrap support for afis|EAW20545.1 as seed ortholog is 100%.

---

Group of orthologs #8253. Best score 274 bits
Score difference with first non-orthologous sequence - *A. fumigatus*:274 *A. fischeri*:274
afum|EAL92942.1 100.00% afis|EAW21930.1 100.00%
Bootstrap support for afum|EAL92942.1 as seed ortholog is 100%.
Bootstrap support for afis|EAW21930.1 as seed ortholog is 100%.

Group of orthologs #8254. Best score 273 bits
Score difference with first non-orthologous sequence - *A. fumigatus*:192 *A. fischeri*:273
afum|EAL86212.1 100.00% afis|EAW24548.1 100.00%
Bootstrap support for afum|EAL86212.1 as seed ortholog is 100%.
Bootstrap support for afis|EAW24548.1 as seed ortholog is 100%.

Group of orthologs #8255. Best score 273 bits
Score difference with first non-orthologous sequence - *A. fumigatus*:273 *A. fischeri*:273
afum|EAL93210.1 100.00% afis|EAW18515.1 100.00%
Bootstrap support for afum|EAL93210.1 as seed ortholog is 100%.
Bootstrap support for afis|EAW18515.1 as seed ortholog is 100%.

Group of orthologs #8256. Best score 273 bits
Score difference with first non-orthologous sequence - *A. fumigatus*:273 *A. fischeri*:273
afum|EAL90401.1 100.00% afis|EAW22798.1 100.00%
Bootstrap support for afum|EAL90401.1 as seed ortholog is 100%.
Bootstrap support for afis|EAW22798.1 as seed ortholog is 100%.

Group of orthologs #8257. Best score 273 bits
Score difference with first non-orthologous sequence - *A. fumigatus*:273 *A. fischeri*:273
afum|EAL92175.1 100.00% afis|EAW21157.1 100.00%
Bootstrap support for afum|EAL92175.1 as seed ortholog is 100%.
Bootstrap support for afis|EAW21157.1 as seed ortholog is 100%.

Group of orthologs #8258. Best score 272 bits
Score difference with first non-orthologous sequence - *A. fumigatus*:272 *A. fischeri*:272
afum|EAL85754.1 100.00% afis|EAW15580.1 100.00%
Bootstrap support for afum|EAL85754.1 as seed ortholog is 100%.
Bootstrap support for afis|EAW15580.1 as seed ortholog is 100%.

Group of orthologs #8259. Best score 272 bits
Score difference with first non-orthologous sequence - *A. fumigatus*:272 *A. fischeri*:272
afum|EAL85756.1 100.00% afis|EAW15578.1 100.00%
Bootstrap support for afum|EAL85756.1 as seed ortholog is 100%.
Bootstrap support for afis|EAW15578.1 as seed ortholog is 100%.

Group of orthologs #8260. Best score 272 bits
Score difference with first non-orthologous sequence - *A. fumigatus*:272 *A. fischeri*:272
afum|EAL91946.2 100.00% afis|EAW24091.1 100.00%
Bootstrap support for afum|EAL91946.2 as seed ortholog is 100%.
Bootstrap support for afis|EAW24091.1 as seed ortholog is 100%.

Group of orthologs #8261. Best score 271 bits
Score difference with first non-orthologous sequence - *A. fumigatus*:271 *A. fischeri*:271
afum|EAL91926.1 100.00% afis|EAW24070.1 100.00%
Bootstrap support for afum|EAL91926.1 as seed ortholog is 100%.
Bootstrap support for afis|EAW24070.1 as seed ortholog is 100%.

Group of orthologs #8262. Best score 270 bits
Score difference with first non-orthologous sequence - *A. fumigatus*:270 *A. fischeri*:270
afum|EAL89171.2 100.00% afis|EAW16574.1 100.00%
Bootstrap support for afum|EAL89171.2 as seed ortholog is 100%.
Bootstrap support for afis|EAW16574.1 as seed ortholog is 100%.

Group of orthologs #8263. Best score 270 bits

Score difference with first non-orthologous sequence - *A. fumigatus*:270 *A. fischeri*:270
afum|EAL89260.1 100.00% afis|EAW16656.1 100.00%
Bootstrap support for afum|EAL89260.1 as seed ortholog is 100%.
Bootstrap support for afis|EAW16656.1 as seed ortholog is 100%.

---

Group of orthologs #8264. Best score 269 bits
Score difference with first non-orthologous sequence - *A. fumigatus*:269 *A. fischeri*:269
afum|EAL84804.1 100.00% afis|EAW20825.1 100.00%
Bootstrap support for afum|EAL84804.1 as seed ortholog is 100%.
Bootstrap support for afis|EAW20825.1 as seed ortholog is 100%.

---

Group of orthologs #8265. Best score 269 bits
Score difference with first non-orthologous sequence - *A. fumigatus*:269 *A. fischeri*:269
afum|EAL91324.1 100.00% afis|EAW17501.1 100.00%
Bootstrap support for afum|EAL91324.1 as seed ortholog is 100%.
Bootstrap support for afis|EAW17501.1 as seed ortholog is 100%.

---

Group of orthologs #8266. Best score 269 bits
Score difference with first non-orthologous sequence - *A. fumigatus*:269 *A. fischeri*:269
afum|EAL93536.1 100.00% afis|EAW18800.1 100.00%
Bootstrap support for afum|EAL93536.1 as seed ortholog is 100%.
Bootstrap support for afis|EAW18800.1 as seed ortholog is 100%.

---

Group of orthologs #8267. Best score 269 bits
Score difference with first non-orthologous sequence - *A. fumigatus*:269 *A. fischeri*:223
afum|EBA27306.1 100.00% afis|EAW23805.1 100.00%
Bootstrap support for afum|EBA27306.1 as seed ortholog is 100%.
Bootstrap support for afis|EAW23805.1 as seed ortholog is 100%.

---

Group of orthologs #8268. Best score 268 bits
Score difference with first non-orthologous sequence - *A. fumigatus*:268 *A. fischeri*:268
afum|EAL85849.1 100.00% afis|EAW15472.1 100.00%
Bootstrap support for afum|EAL85849.1 as seed ortholog is 100%.
Bootstrap support for afis|EAW15472.1 as seed ortholog is 100%.

---

Group of orthologs #8269. Best score 268 bits
Score difference with first non-orthologous sequence - *A. fumigatus*:218 *A. fischeri*:20
afum|EAL86481.2 100.00% afis|EAW16954.1 100.00%
Bootstrap support for afum|EAL86481.2 as seed ortholog is 100%.
Bootstrap support for afis|EAW16954.1 as seed ortholog is 77%.

---

Group of orthologs #8270. Best score 267 bits
Score difference with first non-orthologous sequence - *A. fumigatus*:267 *A. fischeri*:267
afum|EAL84453.1 100.00% afis|EAW20607.1 100.00%
Bootstrap support for afum|EAL84453.1 as seed ortholog is 100%.
Bootstrap support for afis|EAW20607.1 as seed ortholog is 100%.

---

Group of orthologs #8271. Best score 267 bits
Score difference with first non-orthologous sequence - *A. fumigatus*:267 *A. fischeri*:267
afum|EAL89557.1 100.00% afis|EAW24904.1 100.00%
Bootstrap support for afum|EAL89557.1 as seed ortholog is 100%.
Bootstrap support for afis|EAW24904.1 as seed ortholog is 100%.

---

Group of orthologs #8272. Best score 266 bits
Score difference with first non-orthologous sequence - *A. fumigatus*:266 *A. fischeri*:266
afum|EAL88084.1 100.00% afis|EAW23387.1 100.00%

Bootstrap support for afum|EAL88084.1 as seed ortholog is 100%.
Bootstrap support for afis|EAW23387.1 as seed ortholog is 100%.

---

Group of orthologs #8273. Best score 265 bits
Score difference with first non-orthologous sequence - *A. fumigatus*:265 *A. fischeri*:265
afum|EAL84742.1 100.00% afis|EAW20891.1 100.00%
Bootstrap support for afum|EAL84742.1 as seed ortholog is 100%.
Bootstrap support for afis|EAW20891.1 as seed ortholog is 100%.

---

Group of orthologs #8274. Best score 265 bits
Score difference with first non-orthologous sequence - *A. fumigatus*:265 *A. fischeri*:265
afum|EAL93785.1 100.00% afis|EAW19056.1 100.00%
Bootstrap support for afum|EAL93785.1 as seed ortholog is 100%.
Bootstrap support for afis|EAW19056.1 as seed ortholog is 100%.

---

Group of orthologs #8275. Best score 264 bits
Score difference with first non-orthologous sequence - *A. fumigatus*:264 *A. fischeri*:264
afum|EBA27473.1 100.00% afis|EAW21751.1 100.00%
Bootstrap support for afum|EBA27473.1 as seed ortholog is 100%.
Bootstrap support for afis|EAW21751.1 as seed ortholog is 100%.

---

Group of orthologs #8276. Best score 263 bits
Score difference with first non-orthologous sequence - *A. fumigatus*:263 *A. fischeri*:263
afum|EAL93865.1 100.00% afis|EAW19133.1 100.00%
Bootstrap support for afum|EAL93865.1 as seed ortholog is 100%.
Bootstrap support for afis|EAW19133.1 as seed ortholog is 100%.

---

Group of orthologs #8277. Best score 263 bits
Score difference with first non-orthologous sequence - *A. fumigatus*:263 *A. fischeri*:263
afum|EAL90149.1 100.00% afis|EAW25484.1 100.00%
Bootstrap support for afum|EAL90149.1 as seed ortholog is 100%.
Bootstrap support for afis|EAW25484.1 as seed ortholog is 100%.

---

Group of orthologs #8278. Best score 263 bits
Score difference with first non-orthologous sequence - *A. fumigatus*:263 *A. fischeri*:263
afum|EAL91986.1 100.00% afis|EAW24130.1 100.00%
Bootstrap support for afum|EAL91986.1 as seed ortholog is 100%.
Bootstrap support for afis|EAW24130.1 as seed ortholog is 100%.

---

Group of orthologs #8279. Best score 262 bits
Score difference with first non-orthologous sequence - *A. fumigatus*:131 *A. fischeri*:69
afum|EAL86286.2 100.00% afis|EAW17202.1 100.00%
Bootstrap support for afum|EAL86286.2 as seed ortholog is 99%.
Bootstrap support for afis|EAW17202.1 as seed ortholog is 95%.

---

Group of orthologs #8280. Best score 262 bits
Score difference with first non-orthologous sequence - *A. fumigatus*:262 *A. fischeri*:262
afum|EAL84499.1 100.00% afis|EAW20562.1 100.00%
Bootstrap support for afum|EAL84499.1 as seed ortholog is 100%.
Bootstrap support for afis|EAW20562.1 as seed ortholog is 100%.

---

Group of orthologs #8281. Best score 262 bits
Score difference with first non-orthologous sequence - *A. fumigatus*:262 *A. fischeri*:262
afum|EAL85483.1 100.00% afis|EAW20367.1 100.00%
Bootstrap support for afum|EAL85483.1 as seed ortholog is 100%.
Bootstrap support for afis|EAW20367.1 as seed ortholog is 100%.

---

Group of orthologs #8282. Best score 261 bits
Score difference with first non-orthologous sequence - *A. fumigatus*:261 *A. fischeri*:261

| | | | |
|---|---|---|---|
| afum\|EAL85514.1 | 100.00% | afis\|EAW15837.1 | 100.00% |

Bootstrap support for afum|EAL85514.1 as seed ortholog is 100%.
Bootstrap support for afis|EAW15837.1 as seed ortholog is 100%.

---

Group of orthologs #8283. Best score 261 bits
Score difference with first non-orthologous sequence - *A. fumigatus*:261 *A. fischeri*:261

| | | | |
|---|---|---|---|
| afum\|EAL84943.1 | 100.00% | afis\|EAW19962.1 | 100.00% |

Bootstrap support for afum|EAL84943.1 as seed ortholog is 100%.
Bootstrap support for afis|EAW19962.1 as seed ortholog is 100%.

---

Group of orthologs #8284. Best score 261 bits
Score difference with first non-orthologous sequence - *A. fumigatus*:261 *A. fischeri*:261

| | | | |
|---|---|---|---|
| afum\|EAL88294.2 | 100.00% | afis\|EAW23189.1 | 100.00% |

Bootstrap support for afum|EAL88294.2 as seed ortholog is 100%.
Bootstrap support for afis|EAW23189.1 as seed ortholog is 100%.

---

Group of orthologs #8285. Best score 261 bits
Score difference with first non-orthologous sequence - *A. fumigatus*:261 *A. fischeri*:261

| | | | |
|---|---|---|---|
| afum\|EAL93550.2 | 100.00% | afis\|EAW18814.1 | 100.00% |

Bootstrap support for afum|EAL93550.2 as seed ortholog is 100%.
Bootstrap support for afis|EAW18814.1 as seed ortholog is 100%.

---

Group of orthologs #8286. Best score 261 bits
Score difference with first non-orthologous sequence - *A. fumigatus*:261 *A. fischeri*:261

| | | | |
|---|---|---|---|
| afum\|EAL92143.1 | 100.00% | afis\|EAW21119.1 | 100.00% |

Bootstrap support for afum|EAL92143.1 as seed ortholog is 100%.
Bootstrap support for afis|EAW21119.1 as seed ortholog is 100%.

---

Group of orthologs #8287. Best score 260 bits
Score difference with first non-orthologous sequence - *A. fumigatus*:260 *A. fischeri*:260

| | | | |
|---|---|---|---|
| afum\|EAL89356.1 | 100.00% | afis\|EAW19608.1 | 100.00% |

Bootstrap support for afum|EAL89356.1 as seed ortholog is 100%.
Bootstrap support for afis|EAW19608.1 as seed ortholog is 100%.

---

Group of orthologs #8288. Best score 260 bits
Score difference with first non-orthologous sequence - *A. fumigatus*:260 *A. fischeri*:260

| | | | |
|---|---|---|---|
| afum\|EAL91772.1 | 100.00% | afis\|EAW17962.1 | 100.00% |

Bootstrap support for afum|EAL91772.1 as seed ortholog is 100%.
Bootstrap support for afis|EAW17962.1 as seed ortholog is 100%.

---

Group of orthologs #8289. Best score 260 bits
Score difference with first non-orthologous sequence - *A. fumigatus*:260 *A. fischeri*:260

| | | | |
|---|---|---|---|
| afum\|EAL87341.1 | 100.00% | afis\|EAW23912.1 | 100.00% |

Bootstrap support for afum|EAL87341.1 as seed ortholog is 100%.
Bootstrap support for afis|EAW23912.1 as seed ortholog is 100%.

---

Group of orthologs #8290. Best score 259 bits
Score difference with first non-orthologous sequence - *A. fumigatus*:259 *A. fischeri*:259

| | | | |
|---|---|---|---|
| afum\|EAL91479.1 | 100.00% | afis\|EAW17665.1 | 100.00% |

Bootstrap support for afum|EAL91479.1 as seed ortholog is 100%.
Bootstrap support for afis|EAW17665.1 as seed ortholog is 100%.

---

Group of orthologs #8291. Best score 259 bits

Score difference with first non-orthologous sequence - *A. fumigatus*:259 *A. fischeri*:259
afum|EAL93184.2 100.00% afis|EAW18494.1 100.00%
Bootstrap support for afum|EAL93184.2 as seed ortholog is 100%.
Bootstrap support for afis|EAW18494.1 as seed ortholog is 100%.

---

Group of orthologs #8292. Best score 259 bits
Score difference with first non-orthologous sequence - *A. fumigatus*:259 *A. fischeri*:259
afum|EAL88354.1 100.00% afis|EAW23126.1 100.00%
Bootstrap support for afum|EAL88354.1 as seed ortholog is 100%.
Bootstrap support for afis|EAW23126.1 as seed ortholog is 100%.

---

Group of orthologs #8293. Best score 259 bits
Score difference with first non-orthologous sequence - *A. fumigatus*:259 *A. fischeri*:259
afum|EAL88150.1 100.00% afis|EAW23329.1 100.00%
Bootstrap support for afum|EAL88150.1 as seed ortholog is 100%.
Bootstrap support for afis|EAW23329.1 as seed ortholog is 100%.

---

Group of orthologs #8294. Best score 258 bits
Score difference with first non-orthologous sequence - *A. fumigatus*:258 *A. fischeri*:258
afum|EAL93110.2 100.00% afis|EAW18430.1 100.00%
Bootstrap support for afum|EAL93110.2 as seed ortholog is 100%.
Bootstrap support for afis|EAW18430.1 as seed ortholog is 100%.

---

Group of orthologs #8295. Best score 258 bits
Score difference with first non-orthologous sequence - *A. fumigatus*:258 *A. fischeri*:258
afum|EAL87401.1 100.00% afis|EAW23971.1 100.00%
Bootstrap support for afum|EAL87401.1 as seed ortholog is 100%.
Bootstrap support for afis|EAW23971.1 as seed ortholog is 100%.

---

Group of orthologs #8296. Best score 258 bits
Score difference with first non-orthologous sequence - *A. fumigatus*:258 *A. fischeri*:258
afum|EAL93878.1 100.00% afis|EAW19147.1 100.00%
Bootstrap support for afum|EAL93878.1 as seed ortholog is 100%.
Bootstrap support for afis|EAW19147.1 as seed ortholog is 100%.

---

Group of orthologs #8297. Best score 258 bits
Score difference with first non-orthologous sequence - *A. fumigatus*:258 *A. fischeri*:258
afum|EAL92326.1 100.00% afis|EAW21310.1 100.00%
Bootstrap support for afum|EAL92326.1 as seed ortholog is 100%.
Bootstrap support for afis|EAW21310.1 as seed ortholog is 100%.

---

Group of orthologs #8298. Best score 258 bits
Score difference with first non-orthologous sequence - *A. fumigatus*:258 *A. fischeri*:258
afum|EAL92353.1 100.00% afis|EAW21335.1 100.00%
Bootstrap support for afum|EAL92353.1 as seed ortholog is 100%.
Bootstrap support for afis|EAW21335.1 as seed ortholog is 100%.

---

Group of orthologs #8299. Best score 257 bits
Score difference with first non-orthologous sequence - *A. fumigatus*:257 *A. fischeri*:257
afum|EAL93305.1 100.00% afis|EAW18597.1 100.00%
Bootstrap support for afum|EAL93305.1 as seed ortholog is 100%.
Bootstrap support for afis|EAW18597.1 as seed ortholog is 100%.

---

Group of orthologs #8300. Best score 257 bits
Score difference with first non-orthologous sequence - *A. fumigatus*:257 *A. fischeri*:257
afum|EAL92089.1 100.00% afis|EAW21053.1 100.00%

Bootstrap support for afum|EAL92089.1 as seed ortholog is 100%.
Bootstrap support for afis|EAW21053.1 as seed ortholog is 100%.

---

Group of orthologs #8301. Best score 256 bits
Score difference with first non-orthologous sequence - *A. fumigatus*:256 *A. fischeri*:256
afum|EAL86319.2 100.00% afis|EAW16744.1 100.00%
Bootstrap support for afum|EAL86319.2 as seed ortholog is 100%.
Bootstrap support for afis|EAW16744.1 as seed ortholog is 100%.

---

Group of orthologs #8302. Best score 256 bits
Score difference with first non-orthologous sequence - *A. fumigatus*:256 *A. fischeri*:256
afum|EAL93562.1 100.00% afis|EAW18828.1 100.00%
Bootstrap support for afum|EAL93562.1 as seed ortholog is 100%.
Bootstrap support for afis|EAW18828.1 as seed ortholog is 100%.

---

Group of orthologs #8303. Best score 256 bits
Score difference with first non-orthologous sequence - *A. fumigatus*:256 *A. fischeri*:256
afum|EAL92420.1 100.00% afis|EAW21406.1 100.00%
Bootstrap support for afum|EAL92420.1 as seed ortholog is 100%.
Bootstrap support for afis|EAW21406.1 as seed ortholog is 100%.

---

Group of orthologs #8304. Best score 255 bits
Score difference with first non-orthologous sequence - *A. fumigatus*:255 *A. fischeri*:255
afum|EAL92701.1 100.00% afis|EAW21666.1 100.00%
Bootstrap support for afum|EAL92701.1 as seed ortholog is 100%.
Bootstrap support for afis|EAW21666.1 as seed ortholog is 100%.

---

Group of orthologs #8305. Best score 255 bits
Score difference with first non-orthologous sequence - *A. fumigatus*:255 *A. fischeri*:255
afum|EAL92928.1 100.00% afis|EAW24536.1 100.00%
Bootstrap support for afum|EAL92928.1 as seed ortholog is 100%.
Bootstrap support for afis|EAW24536.1 as seed ortholog is 100%.

---

Group of orthologs #8306. Best score 254 bits
Score difference with first non-orthologous sequence - *A. fumigatus*:254 *A. fischeri*:254
afum|EAL86328.1 100.00% afis|EAW16756.1 100.00%
Bootstrap support for afum|EAL86328.1 as seed ortholog is 100%.
Bootstrap support for afis|EAW16756.1 as seed ortholog is 100%.

---

Group of orthologs #8307. Best score 254 bits
Score difference with first non-orthologous sequence - *A. fumigatus*:254 *A. fischeri*:254
afum|EAL88394.1 100.00% afis|EAW23085.1 100.00%
Bootstrap support for afum|EAL88394.1 as seed ortholog is 100%.
Bootstrap support for afis|EAW23085.1 as seed ortholog is 100%.

---

Group of orthologs #8308. Best score 254 bits
Score difference with first non-orthologous sequence - *A. fumigatus*:254 *A. fischeri*:254
afum|EAL92739.1 100.00% afis|EAW21705.1 100.00%
Bootstrap support for afum|EAL92739.1 as seed ortholog is 100%.
Bootstrap support for afis|EAW21705.1 as seed ortholog is 100%.

---

Group of orthologs #8309. Best score 253 bits
Score difference with first non-orthologous sequence - *A. fumigatus*:253 *A. fischeri*:253
afum|EAL85745.1 100.00% afis|EAW15587.1 100.00%
Bootstrap support for afum|EAL85745.1 as seed ortholog is 100%.
Bootstrap support for afis|EAW15587.1 as seed ortholog is 100%.

Group of orthologs #8310. Best score 253 bits
Score difference with first non-orthologous sequence - *A. fumigatus*:253 *A. fischeri*:253
afum|EAL84956.1 100.00% afis|EAW19959.1 100.00%
Bootstrap support for afum|EAL84956.1 as seed ortholog is 100%.
Bootstrap support for afis|EAW19959.1 as seed ortholog is 100%.

---

Group of orthologs #8311. Best score 253 bits
Score difference with first non-orthologous sequence - *A. fumigatus*:253 *A. fischeri*:253
afum|EAL88903.1 100.00% afis|EAW16282.1 100.00%
Bootstrap support for afum|EAL88903.1 as seed ortholog is 100%.
Bootstrap support for afis|EAW16282.1 as seed ortholog is 100%.

---

Group of orthologs #8312. Best score 253 bits
Score difference with first non-orthologous sequence - *A. fumigatus*:253 *A. fischeri*:253
afum|EAL89585.1 100.00% afis|EAW24929.1 100.00%
Bootstrap support for afum|EAL89585.1 as seed ortholog is 100%.
Bootstrap support for afis|EAW24929.1 as seed ortholog is 100%.

---

Group of orthologs #8313. Best score 251 bits
Score difference with first non-orthologous sequence - *A. fumigatus*:251 *A. fischeri*:251
afum|EAL87819.1 100.00% afis|EAW16225.1 100.00%
Bootstrap support for afum|EAL87819.1 as seed ortholog is 100%.
Bootstrap support for afis|EAW16225.1 as seed ortholog is 100%.

---

Group of orthologs #8314. Best score 251 bits
Score difference with first non-orthologous sequence - *A. fumigatus*:251 *A. fischeri*:251
afum|EAL88880.1 100.00% afis|EAW16259.1 100.00%
Bootstrap support for afum|EAL88880.1 as seed ortholog is 100%.
Bootstrap support for afis|EAW16259.1 as seed ortholog is 100%.

---

Group of orthologs #8315. Best score 251 bits
Score difference with first non-orthologous sequence - *A. fumigatus*:251 *A. fischeri*:251
afum|EAL91763.1 100.00% afis|EAW17953.1 100.00%
Bootstrap support for afum|EAL91763.1 as seed ortholog is 100%.
Bootstrap support for afis|EAW17953.1 as seed ortholog is 100%.

---

Group of orthologs #8316. Best score 251 bits
Score difference with first non-orthologous sequence - *A. fumigatus*:251 *A. fischeri*:251
afum|EAL93216.1 100.00% afis|EAW18521.1 100.00%
Bootstrap support for afum|EAL93216.1 as seed ortholog is 100%.
Bootstrap support for afis|EAW18521.1 as seed ortholog is 100%.

---

Group of orthologs #8317. Best score 251 bits
Score difference with first non-orthologous sequence - *A. fumigatus*:251 *A. fischeri*:251
afum|EAL92208.1 100.00% afis|EAW21186.1 100.00%
Bootstrap support for afum|EAL92208.1 as seed ortholog is 100%.
Bootstrap support for afis|EAW21186.1 as seed ortholog is 100%.

---

Group of orthologs #8318. Best score 250 bits
Score difference with first non-orthologous sequence - *A. fumigatus*:250 *A. fischeri*:250
afum|EAL87367.1 100.00% afis|EAW23942.1 100.00%
Bootstrap support for afum|EAL87367.1 as seed ortholog is 100%.
Bootstrap support for afis|EAW23942.1 as seed ortholog is 100%.

---

Group of orthologs #8319. Best score 250 bits

Score difference with first non-orthologous sequence - *A. fumigatus*:250 *A. fischeri*:250
afum|EAL90531.1 100.00% afis|EAW22661.1 100.00%
Bootstrap support for afum|EAL90531.1 as seed ortholog is 100%.
Bootstrap support for afis|EAW22661.1 as seed ortholog is 100%.

---

Group of orthologs #8320. Best score 250 bits
Score difference with first non-orthologous sequence - *A. fumigatus*:250 *A. fischeri*:250
afum|EAL90447.1 100.00% afis|EAW22752.1 100.00%
Bootstrap support for afum|EAL90447.1 as seed ortholog is 100%.
Bootstrap support for afis|EAW22752.1 as seed ortholog is 100%.

---

Group of orthologs #8321. Best score 249 bits
Score difference with first non-orthologous sequence - *A. fumigatus*:249 *A. fischeri*:249
afum|EAL86433.1 100.00% afis|EAW16896.1 100.00%
Bootstrap support for afum|EAL86433.1 as seed ortholog is 100%.
Bootstrap support for afis|EAW16896.1 as seed ortholog is 100%.

---

Group of orthologs #8322. Best score 248 bits
Score difference with first non-orthologous sequence - *A. fumigatus*:92 *A. fischeri*:248
afum|EAL86767.1 100.00% afis|EAW20712.1 100.00%
afum|EAL86746.1 52.43%
afum|EAL90935.1 9.24%
afum|EAL91121.1 8.93%
Bootstrap support for afum|EAL86767.1 as seed ortholog is 97%.
Bootstrap support for afis|EAW20712.1 as seed ortholog is 100%.

---

Group of orthologs #8323. Best score 248 bits
Score difference with first non-orthologous sequence - *A. fumigatus*:248 *A. fischeri*:248
afum|EAL86845.1 100.00% afis|EAW19677.1 100.00%
Bootstrap support for afum|EAL86845.1 as seed ortholog is 100%.
Bootstrap support for afis|EAW19677.1 as seed ortholog is 100%.

---

Group of orthologs #8324. Best score 248 bits
Score difference with first non-orthologous sequence - *A. fumigatus*:248 *A. fischeri*:248
afum|EAL91595.1 100.00% afis|EAW19747.1 100.00%
Bootstrap support for afum|EAL91595.1 as seed ortholog is 100%.
Bootstrap support for afis|EAW19747.1 as seed ortholog is 100%.

---

Group of orthologs #8325. Best score 248 bits
Score difference with first non-orthologous sequence - *A. fumigatus*:248 *A. fischeri*:248
afum|EAL89499.1 100.00% afis|EAW24852.1 100.00%
Bootstrap support for afum|EAL89499.1 as seed ortholog is 100%.
Bootstrap support for afis|EAW24852.1 as seed ortholog is 100%.

---

Group of orthologs #8326. Best score 248 bits
Score difference with first non-orthologous sequence - *A. fumigatus*:248 *A. fischeri*:206
afum|EAL89727.1 100.00% afis|EAW25059.1 100.00%
Bootstrap support for afum|EAL89727.1 as seed ortholog is 100%.
Bootstrap support for afis|EAW25059.1 as seed ortholog is 100%.

---

Group of orthologs #8327. Best score 248 bits
Score difference with first non-orthologous sequence - *A. fumigatus*:248 *A. fischeri*:248
afum|EAL89956.1 100.00% afis|EAW25293.1 100.00%
Bootstrap support for afum|EAL89956.1 as seed ortholog is 100%.
Bootstrap support for afis|EAW25293.1 as seed ortholog is 100%.

---

Group of orthologs #8328. Best score 248 bits
Score difference with first non-orthologous sequence - *A. fumigatus*:248 *A. fischeri*:248
afum|EBA27177.1 100.00% afis|EAW25727.1 100.00%
Bootstrap support for afum|EBA27177.1 as seed ortholog is 100%.
Bootstrap support for afis|EAW25727.1 as seed ortholog is 100%.

---

Group of orthologs #8329. Best score 247 bits
Score difference with first non-orthologous sequence - *A. fumigatus*:247 *A. fischeri*:247
afum|EAL85394.1 100.00% afis|EAW20659.1 100.00%
Bootstrap support for afum|EAL85394.1 as seed ortholog is 100%.
Bootstrap support for afis|EAW20659.1 as seed ortholog is 100%.

---

Group of orthologs #8330. Best score 247 bits
Score difference with first non-orthologous sequence - *A. fumigatus*:247 *A. fischeri*:247
afum|EAL93136.1 100.00% afis|EAW18451.1 100.00%
Bootstrap support for afum|EAL93136.1 as seed ortholog is 100%.
Bootstrap support for afis|EAW18451.1 as seed ortholog is 100%.

---

Group of orthologs #8331. Best score 246 bits
Score difference with first non-orthologous sequence - *A. fumigatus*:246 *A. fischeri*:246
afum|EAL86856.1 100.00% afis|EAW19667.1 100.00%
Bootstrap support for afum|EAL86856.1 as seed ortholog is 100%.
Bootstrap support for afis|EAW19667.1 as seed ortholog is 100%.

---

Group of orthologs #8332. Best score 246 bits
Score difference with first non-orthologous sequence - *A. fumigatus*:14 *A. fischeri*:4
afum|EAL90710.1 100.00% afis|EAW22484.1 100.00%
Bootstrap support for afum|EAL90710.1 as seed ortholog is 100%.
Bootstrap support for afis|EAW22484.1 as seed ortholog is 75%.

---

Group of orthologs #8333. Best score 246 bits
Score difference with first non-orthologous sequence - *A. fumigatus*:246 *A. fischeri*:161
afum|EBA27187.1 100.00% afis|EAW20549.1 100.00%
Bootstrap support for afum|EBA27187.1 as seed ortholog is 100%.
Bootstrap support for afis|EAW20549.1 as seed ortholog is 100%.

---

Group of orthologs #8334. Best score 245 bits
Score difference with first non-orthologous sequence - *A. fumigatus*:245 *A. fischeri*:245
afum|EAL86209.1 100.00% afis|EAW24545.1 100.00%
Bootstrap support for afum|EAL86209.1 as seed ortholog is 100%.
Bootstrap support for afis|EAW24545.1 as seed ortholog is 100%.

---

Group of orthologs #8335. Best score 245 bits
Score difference with first non-orthologous sequence - *A. fumigatus*:245 *A. fischeri*:245
afum|EAL93569.1 100.00% afis|EAW18833.1 100.00%
Bootstrap support for afum|EAL93569.1 as seed ortholog is 100%.
Bootstrap support for afis|EAW18833.1 as seed ortholog is 100%.

---

Group of orthologs #8336. Best score 245 bits
Score difference with first non-orthologous sequence - *A. fumigatus*:245 *A. fischeri*:245
afum|EAL93594.1 100.00% afis|EAW18862.1 100.00%
Bootstrap support for afum|EAL93594.1 as seed ortholog is 100%.
Bootstrap support for afis|EAW18862.1 as seed ortholog is 100%.

---

Group of orthologs #8337. Best score 245 bits
Score difference with first non-orthologous sequence - *A. fumigatus*:245 *A. fischeri*:245

afum|EAL92564.1 100.00% afis|EAW21544.1 100.00%
Bootstrap support for afum|EAL92564.1 as seed ortholog is 100%.
Bootstrap support for afis|EAW21544.1 as seed ortholog is 100%.

---

Group of orthologs #8338. Best score 244 bits
Score difference with first non-orthologous sequence - *A. fumigatus*:244 *A. fischeri*:244
afum|EAL88490.1 100.00% afis|EAW15854.1 100.00%
Bootstrap support for afum|EAL88490.1 as seed ortholog is 100%.
Bootstrap support for afis|EAW15854.1 as seed ortholog is 100%.

---

Group of orthologs #8339. Best score 244 bits
Score difference with first non-orthologous sequence - *A. fumigatus*:244 *A. fischeri*:180
afum|EAL86190.1 100.00% afis|EAW24526.1 100.00%
Bootstrap support for afum|EAL86190.1 as seed ortholog is 100%.
Bootstrap support for afis|EAW24526.1 as seed ortholog is 100%.

---

Group of orthologs #8340. Best score 244 bits
Score difference with first non-orthologous sequence - *A. fumigatus*:244 *A. fischeri*:244
afum|EAL90481.1 100.00% afis|EAW22713.1 100.00%
Bootstrap support for afum|EAL90481.1 as seed ortholog is 100%.
Bootstrap support for afis|EAW22713.1 as seed ortholog is 100%.

---

Group of orthologs #8341. Best score 244 bits
Score difference with first non-orthologous sequence - *A. fumigatus*:244 *A. fischeri*:244
afum|EAL92521.1 100.00% afis|EAW21502.1 100.00%
Bootstrap support for afum|EAL92521.1 as seed ortholog is 100%.
Bootstrap support for afis|EAW21502.1 as seed ortholog is 100%.

---

Group of orthologs #8342. Best score 242 bits
Score difference with first non-orthologous sequence - *A. fumigatus*:242 *A. fischeri*:242
afum|EAL92121.1 100.00% afis|EAW21098.1 100.00%
Bootstrap support for afum|EAL92121.1 as seed ortholog is 100%.
Bootstrap support for afis|EAW21098.1 as seed ortholog is 100%.

---

Group of orthologs #8343. Best score 242 bits
Score difference with first non-orthologous sequence - *A. fumigatus*:242 *A. fischeri*:242
afum|EAL92014.2 100.00% afis|EAW24158.1 100.00%
Bootstrap support for afum|EAL92014.2 as seed ortholog is 100%.
Bootstrap support for afis|EAW24158.1 as seed ortholog is 100%.

---

Group of orthologs #8344. Best score 241 bits
Score difference with first non-orthologous sequence - *A. fumigatus*:241 *A. fischeri*:241
afum|EAL89132.1 100.00% afis|EAW16535.1 100.00%
Bootstrap support for afum|EAL89132.1 as seed ortholog is 100%.
Bootstrap support for afis|EAW16535.1 as seed ortholog is 100%.

---

Group of orthologs #8345. Best score 241 bits
Score difference with first non-orthologous sequence - *A. fumigatus*:241 *A. fischeri*:241
afum|EAL90761.1 100.00% afis|EAW22430.1 100.00%
Bootstrap support for afum|EAL90761.1 as seed ortholog is 100%.
Bootstrap support for afis|EAW22430.1 as seed ortholog is 100%.

---

Group of orthologs #8346. Best score 240 bits
Score difference with first non-orthologous sequence - *A. fumigatus*:240 *A. fischeri*:240
afum|EAL85684.1 100.00% afis|EAW15664.1 100.00%
Bootstrap support for afum|EAL85684.1 as seed ortholog is 100%.

Bootstrap support for afis|EAW15664.1 as seed ortholog is 100%.

---

Group of orthologs #8347. Best score 240 bits
Score difference with first non-orthologous sequence - *A. fumigatus*:240 *A. fischeri*:240
afum|EAL85698.2 100.00% afis|EAW15652.1 100.00%
Bootstrap support for afum|EAL85698.2 as seed ortholog is 100%.
Bootstrap support for afis|EAW15652.1 as seed ortholog is 100%.

---

Group of orthologs #8348. Best score 240 bits
Score difference with first non-orthologous sequence - *A. fumigatus*:240 *A. fischeri*:240
afum|EAL89061.1 100.00% afis|EAW16465.1 100.00%
Bootstrap support for afum|EAL89061.1 as seed ortholog is 100%.
Bootstrap support for afis|EAW16465.1 as seed ortholog is 100%.

---

Group of orthologs #8349. Best score 240 bits
Score difference with first non-orthologous sequence - *A. fumigatus*:240 *A. fischeri*:240
afum|EAL91421.1 100.00% afis|EAW17608.1 100.00%
Bootstrap support for afum|EAL91421.1 as seed ortholog is 100%.
Bootstrap support for afis|EAW17608.1 as seed ortholog is 100%.

---

Group of orthologs #8350. Best score 239 bits
Score difference with first non-orthologous sequence - *A. fumigatus*:239 *A. fischeri*:239
afum|EAL85401.1 100.00% afis|EAW21807.1 100.00%
afum|EAL86716.1 14.44% afis|EAW20671.1 14.29%
afis|EAW18099.1 11.56%
Bootstrap support for afum|EAL85401.1 as seed ortholog is 100%.
Bootstrap support for afis|EAW21807.1 as seed ortholog is 100%.

---

Group of orthologs #8351. Best score 239 bits
Score difference with first non-orthologous sequence - *A. fumigatus*:175 *A. fischeri*:239
afum|EAL87692.1 100.00% afis|EAW18326.1 100.00%
Bootstrap support for afum|EAL87692.1 as seed ortholog is 100%.
Bootstrap support for afis|EAW18326.1 as seed ortholog is 100%.

---

Group of orthologs #8352. Best score 239 bits
Score difference with first non-orthologous sequence - *A. fumigatus*:239 *A. fischeri*:239
afum|EAL87396.1 100.00% afis|EAW23966.1 100.00%
Bootstrap support for afum|EAL87396.1 as seed ortholog is 100%.
Bootstrap support for afis|EAW23966.1 as seed ortholog is 100%.

---

Group of orthologs #8353. Best score 239 bits
Score difference with first non-orthologous sequence - *A. fumigatus*:239 *A. fischeri*:239
afum|EBA27459.1 100.00% afis|EAW21339.1 100.00%
Bootstrap support for afum|EBA27459.1 as seed ortholog is 100%.
Bootstrap support for afis|EAW21339.1 as seed ortholog is 100%.

---

Group of orthologs #8354. Best score 239 bits
Score difference with first non-orthologous sequence - *A. fumigatus*:239 *A. fischeri*:239
afum|EAL91941.1 100.00% afis|EAW24086.1 100.00%
Bootstrap support for afum|EAL91941.1 as seed ortholog is 100%.
Bootstrap support for afis|EAW24086.1 as seed ortholog is 100%.

---

Group of orthologs #8355. Best score 238 bits
Score difference with first non-orthologous sequence - *A. fumigatus*:238 *A. fischeri*:238
afum|EAL87642.1 100.00% afis|EAW18276.1 100.00%
Bootstrap support for afum|EAL87642.1 as seed ortholog is 100%.

Bootstrap support for afis|EAW18276.1 as seed ortholog is 100%.

---

Group of orthologs #8356. Best score 238 bits
Score difference with first non-orthologous sequence - *A. fumigatus*:238 *A. fischeri*:238
afum|EAL92583.1 100.00% afis|EAW21564.1 100.00%
Bootstrap support for afum|EAL92583.1 as seed ortholog is 100%.
Bootstrap support for afis|EAW21564.1 as seed ortholog is 100%.

---

Group of orthologs #8357. Best score 238 bits
Score difference with first non-orthologous sequence - *A. fumigatus*:238 *A. fischeri*:123
afum|EBA27260.1 100.00% afis|EAW24512.1 100.00%
Bootstrap support for afum|EBA27260.1 as seed ortholog is 100%.
Bootstrap support for afis|EAW24512.1 as seed ortholog is 100%.

---

Group of orthologs #8358. Best score 237 bits
Score difference with first non-orthologous sequence - *A. fumigatus*:237 *A. fischeri*:237
afum|EAL84380.1 100.00% afis|EAW25707.1 100.00%
Bootstrap support for afum|EAL84380.1 as seed ortholog is 100%.
Bootstrap support for afis|EAW25707.1 as seed ortholog is 100%.

---

Group of orthologs #8359. Best score 237 bits
Score difference with first non-orthologous sequence - *A. fumigatus*:237 *A. fischeri*:237
afum|EAL88321.1 100.00% afis|EAW23162.1 100.00%
Bootstrap support for afum|EAL88321.1 as seed ortholog is 100%.
Bootstrap support for afis|EAW23162.1 as seed ortholog is 100%.

---

Group of orthologs #8360. Best score 236 bits
Score difference with first non-orthologous sequence - *A. fumigatus*:236 *A. fischeri*:236
afum|EAL88196.2 100.00% afis|EAW23283.1 100.00%
Bootstrap support for afum|EAL88196.2 as seed ortholog is 100%.
Bootstrap support for afis|EAW23283.1 as seed ortholog is 100%.

---

Group of orthologs #8361. Best score 235 bits
Score difference with first non-orthologous sequence - *A. fumigatus*:235 *A. fischeri*:235
afum|EAL85096.2 100.00% afis|EAW19794.1 100.00%
Bootstrap support for afum|EAL85096.2 as seed ortholog is 100%.
Bootstrap support for afis|EAW19794.1 as seed ortholog is 100%.

---

Group of orthologs #8362. Best score 235 bits
Score difference with first non-orthologous sequence - *A. fumigatus*:235 *A. fischeri*:235
afum|EAL84449.1 100.00% afis|EAW20612.1 100.00%
Bootstrap support for afum|EAL84449.1 as seed ortholog is 100%.
Bootstrap support for afis|EAW20612.1 as seed ortholog is 100%.

---

Group of orthologs #8363. Best score 235 bits
Score difference with first non-orthologous sequence - *A. fumigatus*:235 *A. fischeri*:235
afum|EAL92735.1 100.00% afis|EAW21701.1 100.00%
Bootstrap support for afum|EAL92735.1 as seed ortholog is 100%.
Bootstrap support for afis|EAW21701.1 as seed ortholog is 100%.

---

Group of orthologs #8364. Best score 234 bits
Score difference with first non-orthologous sequence - *A. fumigatus*:234 *A. fischeri*:234
afum|EAL86414.1 100.00% afis|EAW16877.1 100.00%
Bootstrap support for afum|EAL86414.1 as seed ortholog is 100%.
Bootstrap support for afis|EAW16877.1 as seed ortholog is 100%.

---

Group of orthologs #8365. Best score 234 bits
Score difference with first non-orthologous sequence - *A. fumigatus*:234 *A. fischeri*:234
afum|EAL92892.1 100.00% afis|EAW21878.1 100.00%
Bootstrap support for afum|EAL92892.1 as seed ortholog is 100%.
Bootstrap support for afis|EAW21878.1 as seed ortholog is 100%.

---

Group of orthologs #8366. Best score 233 bits
Score difference with first non-orthologous sequence - *A. fumigatus*:233 *A. fischeri*:233
afum|EBA27364.1 100.00% afis|EAW16321.1 100.00%
Bootstrap support for afum|EBA27364.1 as seed ortholog is 100%.
Bootstrap support for afis|EAW16321.1 as seed ortholog is 100%.

---

Group of orthologs #8367. Best score 233 bits
Score difference with first non-orthologous sequence - *A. fumigatus*:233 *A. fischeri*:233
afum|EAL86222.1 100.00% afis|EAW24561.1 100.00%
Bootstrap support for afum|EAL86222.1 as seed ortholog is 100%.
Bootstrap support for afis|EAW24561.1 as seed ortholog is 100%.

---

Group of orthologs #8368. Best score 233 bits
Score difference with first non-orthologous sequence - *A. fumigatus*:233 *A. fischeri*:233
afum|EAL90495.1 100.00% afis|EAW22697.1 100.00%
Bootstrap support for afum|EAL90495.1 as seed ortholog is 100%.
Bootstrap support for afis|EAW22697.1 as seed ortholog is 100%.

---

Group of orthologs #8369. Best score 233 bits
Score difference with first non-orthologous sequence - *A. fumigatus*:233 *A. fischeri*:233
afum|EAL92195.1 100.00% afis|EAW21175.1 100.00%
Bootstrap support for afum|EAL92195.1 as seed ortholog is 100%.
Bootstrap support for afis|EAW21175.1 as seed ortholog is 100%.

---

Group of orthologs #8370. Best score 232 bits
Score difference with first non-orthologous sequence - *A. fumigatus*:232 *A. fischeri*:232
afum|EAL84452.1 100.00% afis|EAW20608.1 100.00%
Bootstrap support for afum|EAL84452.1 as seed ortholog is 100%.
Bootstrap support for afis|EAW20608.1 as seed ortholog is 100%.

---

Group of orthologs #8371. Best score 232 bits
Score difference with first non-orthologous sequence - *A. fumigatus*:232 *A. fischeri*:232
afum|EAL88157.1 100.00% afis|EAW23319.1 100.00%
Bootstrap support for afum|EAL88157.1 as seed ortholog is 100%.
Bootstrap support for afis|EAW23319.1 as seed ortholog is 100%.

---

Group of orthologs #8372. Best score 232 bits
Score difference with first non-orthologous sequence - *A. fumigatus*:232 *A. fischeri*:232
afum|EBA27262.1 100.00% afis|EAW24541.1 100.00%
Bootstrap support for afum|EBA27262.1 as seed ortholog is 100%.
Bootstrap support for afis|EAW24541.1 as seed ortholog is 100%.

---

Group of orthologs #8373. Best score 231 bits
Score difference with first non-orthologous sequence - *A. fumigatus*:231 *A. fischeri*:231
afum|EBA27292.1 100.00% afis|EAW19628.1 100.00%
Bootstrap support for afum|EBA27292.1 as seed ortholog is 100%.
Bootstrap support for afis|EAW19628.1 as seed ortholog is 100%.

---

Group of orthologs #8374. Best score 230 bits
Score difference with first non-orthologous sequence - *A. fumigatus*:230 *A. fischeri*:230

afum|EAL88920.1 100.00% afis|EAW16298.1 100.00%
Bootstrap support for afum|EAL88920.1 as seed ortholog is 100%.
Bootstrap support for afis|EAW16298.1 as seed ortholog is 100%.

---

Group of orthologs #8375. Best score 228 bits
Score difference with first non-orthologous sequence - *A. fumigatus*:228 *A. fischeri*:228
afum|EAL85742.1 100.00% afis|EAW15590.1 100.00%
Bootstrap support for afum|EAL85742.1 as seed ortholog is 100%.
Bootstrap support for afis|EAW15590.1 as seed ortholog is 100%.

---

Group of orthologs #8376. Best score 228 bits
Score difference with first non-orthologous sequence - *A. fumigatus*:228 *A. fischeri*:228
afum|EBA27482.1 100.00% afis|EAW21914.1 100.00%
Bootstrap support for afum|EBA27482.1 as seed ortholog is 100%.
Bootstrap support for afis|EAW21914.1 as seed ortholog is 100%.

---

Group of orthologs #8377. Best score 227 bits
Score difference with first non-orthologous sequence - *A. fumigatus*:227 *A. fischeri*:227
afum|EAL93090.1 100.00% afis|EAW18410.1 100.00%
Bootstrap support for afum|EAL93090.1 as seed ortholog is 100%.
Bootstrap support for afis|EAW18410.1 as seed ortholog is 100%.

---

Group of orthologs #8378. Best score 226 bits
Score difference with first non-orthologous sequence - *A. fumigatus*:226 *A. fischeri*:226
afum|EAL86874.1 100.00% afis|EAW19648.1 100.00%
Bootstrap support for afum|EAL86874.1 as seed ortholog is 100%.
Bootstrap support for afis|EAW19648.1 as seed ortholog is 100%.

---

Group of orthologs #8379. Best score 225 bits
Score difference with first non-orthologous sequence - *A. fumigatus*:225 *A. fischeri*:225
afum|EAL88232.2 100.00% afis|EAW23249.1 100.00%
Bootstrap support for afum|EAL88232.2 as seed ortholog is 100%.
Bootstrap support for afis|EAW23249.1 as seed ortholog is 100%.

---

Group of orthologs #8380. Best score 225 bits
Score difference with first non-orthologous sequence - *A. fumigatus*:225 *A. fischeri*:225
afum|EAL92763.1 100.00% afis|EAW21730.1 100.00%
Bootstrap support for afum|EAL92763.1 as seed ortholog is 100%.
Bootstrap support for afis|EAW21730.1 as seed ortholog is 100%.

---

Group of orthologs #8381. Best score 224 bits
Score difference with first non-orthologous sequence - *A. fumigatus*:160 *A. fischeri*:224
afum|EAL91154.1 100.00% afis|EAW17318.1 100.00%
Bootstrap support for afum|EAL91154.1 as seed ortholog is 100%.
Bootstrap support for afis|EAW17318.1 as seed ortholog is 100%.

---

Group of orthologs #8382. Best score 223 bits
Score difference with first non-orthologous sequence - *A. fumigatus*:223 *A. fischeri*:223
afum|EAL85890.1 100.00% afis|EAW24221.1 100.00%
Bootstrap support for afum|EAL85890.1 as seed ortholog is 100%.
Bootstrap support for afis|EAW24221.1 as seed ortholog is 100%.

---

Group of orthologs #8383. Best score 223 bits
Score difference with first non-orthologous sequence - *A. fumigatus*:223 *A. fischeri*:223
afum|EAL93917.1 100.00% afis|EAW19186.1 100.00%
Bootstrap support for afum|EAL93917.1 as seed ortholog is 100%.

Bootstrap support for afis|EAW19186.1 as seed ortholog is 100%.

---

Group of orthologs #8384. Best score 223 bits
Score difference with first non-orthologous sequence - *A. fumigatus*:10 *A. fischeri*:223
afum|EAL90453.1 100.00% afis|EAW22742.1 100.00%
Bootstrap support for afum|EAL90453.1 as seed ortholog is 76%.
Bootstrap support for afis|EAW22742.1 as seed ortholog is 100%.

---

Group of orthologs #8385. Best score 223 bits
Score difference with first non-orthologous sequence - *A. fumigatus*:223 *A. fischeri*:223
afum|EBA27412.1 100.00% afis|EAW22543.1 100.00%
Bootstrap support for afum|EBA27412.1 as seed ortholog is 100%.
Bootstrap support for afis|EAW22543.1 as seed ortholog is 100%.

---

Group of orthologs #8386. Best score 222 bits
Score difference with first non-orthologous sequence - *A. fumigatus*:222 *A. fischeri*:222
afum|EAL86507.1 100.00% afis|EAW16980.1 100.00%
Bootstrap support for afum|EAL86507.1 as seed ortholog is 100%.
Bootstrap support for afis|EAW16980.1 as seed ortholog is 100%.

---

Group of orthologs #8387. Best score 222 bits
Score difference with first non-orthologous sequence - *A. fumigatus*:222 *A. fischeri*:222
afum|EAL90541.1 100.00% afis|EAW22651.1 100.00%
Bootstrap support for afum|EAL90541.1 as seed ortholog is 100%.
Bootstrap support for afis|EAW22651.1 as seed ortholog is 100%.

---

Group of orthologs #8388. Best score 220 bits
Score difference with first non-orthologous sequence - *A. fumigatus*:220 *A. fischeri*:220
afum|EAL85059.1 100.00% afis|EAW19833.1 100.00%
Bootstrap support for afum|EAL85059.1 as seed ortholog is 100%.
Bootstrap support for afis|EAW19833.1 as seed ortholog is 100%.

---

Group of orthologs #8389. Best score 220 bits
Score difference with first non-orthologous sequence - *A. fumigatus*:220 *A. fischeri*:220
afum|EAL85353.1 100.00% afis|EAW20236.1 100.00%
Bootstrap support for afum|EAL85353.1 as seed ortholog is 100%.
Bootstrap support for afis|EAW20236.1 as seed ortholog is 100%.

---

Group of orthologs #8390. Best score 220 bits
Score difference with first non-orthologous sequence - *A. fumigatus*:220 *A. fischeri*:220
afum|EAL87203.1 100.00% afis|EAW23769.1 100.00%
Bootstrap support for afum|EAL87203.1 as seed ortholog is 100%.
Bootstrap support for afis|EAW23769.1 as seed ortholog is 100%.

---

Group of orthologs #8391. Best score 220 bits
Score difference with first non-orthologous sequence - *A. fumigatus*:220 *A. fischeri*:174
afum|EAL87941.1 100.00% afis|EAW23524.1 100.00%
Bootstrap support for afum|EAL87941.1 as seed ortholog is 100%.
Bootstrap support for afis|EAW23524.1 as seed ortholog is 100%.

---

Group of orthologs #8392. Best score 220 bits
Score difference with first non-orthologous sequence - *A. fumigatus*:220 *A. fischeri*:220
afum|EAL90412.1 100.00% afis|EAW22787.1 100.00%
Bootstrap support for afum|EAL90412.1 as seed ortholog is 100%.
Bootstrap support for afis|EAW22787.1 as seed ortholog is 100%.

---

Group of orthologs #8393. Best score 220 bits
Score difference with first non-orthologous sequence - *A. fumigatus*:220   *A. fischeri*:220
afum|EBA27261.1 100.00% afis|EAW24513.1 100.00%
Bootstrap support for afum|EBA27261.1 as seed ortholog is 100%.
Bootstrap support for afis|EAW24513.1 as seed ortholog is 100%.

---

Group of orthologs #8394. Best score 219 bits
Score difference with first non-orthologous sequence - *A. fumigatus*:219   *A. fischeri*:219
afum|EAL89089.1 100.00% afis|EAW16495.1 100.00%
Bootstrap support for afum|EAL89089.1 as seed ortholog is 100%.
Bootstrap support for afis|EAW16495.1 as seed ortholog is 100%.

---

Group of orthologs #8395. Best score 219 bits
Score difference with first non-orthologous sequence - *A. fumigatus*:219   *A. fischeri*:9
afum|EAL88871.1 100.00% afis|EAW21127.1 100.00%
Bootstrap support for afum|EAL88871.1 as seed ortholog is 100%.
Bootstrap support for afis|EAW21127.1 as seed ortholog is 56%. Alternative seed ortholog is afis|EAW21125.1 (9 bits away from this cluster)

---

Group of orthologs #8396. Best score 218 bits
Score difference with first non-orthologous sequence - *A. fumigatus*:218   *A. fischeri*:218
afum|EAL91858.1 100.00% afis|EAW18046.1 100.00%
Bootstrap support for afum|EAL91858.1 as seed ortholog is 100%.
Bootstrap support for afis|EAW18046.1 as seed ortholog is 100%.

---

Group of orthologs #8397. Best score 218 bits
Score difference with first non-orthologous sequence - *A. fumigatus*:218   *A. fischeri*:218
afum|EAL89740.1 100.00% afis|EAW25071.1 100.00%
Bootstrap support for afum|EAL89740.1 as seed ortholog is 100%.
Bootstrap support for afis|EAW25071.1 as seed ortholog is 100%.

---

Group of orthologs #8398. Best score 217 bits
Score difference with first non-orthologous sequence - *A. fumigatus*:217   *A. fischeri*:217
afum|EAL88484.1 100.00% afis|EAW15848.1 100.00%
Bootstrap support for afum|EAL88484.1 as seed ortholog is 100%.
Bootstrap support for afis|EAW15848.1 as seed ortholog is 100%.

---

Group of orthologs #8399. Best score 217 bits
Score difference with first non-orthologous sequence - *A. fumigatus*:217   *A. fischeri*:217
afum|EAL87655.1 100.00% afis|EAW18290.1 100.00%
Bootstrap support for afum|EAL87655.1 as seed ortholog is 100%.
Bootstrap support for afis|EAW18290.1 as seed ortholog is 100%.

---

Group of orthologs #8400. Best score 217 bits
Score difference with first non-orthologous sequence - *A. fumigatus*:217   *A. fischeri*:217
afum|EBA27241.1 100.00% afis|EAW15564.1 100.00%
Bootstrap support for afum|EBA27241.1 as seed ortholog is 100%.
Bootstrap support for afis|EAW15564.1 as seed ortholog is 100%.

---

Group of orthologs #8401. Best score 217 bits
Score difference with first non-orthologous sequence - *A. fumigatus*:217   *A. fischeri*:217
afum|EAL93149.1 100.00% afis|EAW18463.1 100.00%
Bootstrap support for afum|EAL93149.1 as seed ortholog is 100%.
Bootstrap support for afis|EAW18463.1 as seed ortholog is 100%.

---

Group of orthologs #8402. Best score 217 bits

Score difference with first non-orthologous sequence - *A. fumigatus*:217   *A. fischeri*:217
afum|EAL87454.1 100.00% afis|EAW24023.1 100.00%
Bootstrap support for afum|EAL87454.1 as seed ortholog is 100%.
Bootstrap support for afis|EAW24023.1 as seed ortholog is 100%.

---

Group of orthologs #8403. Best score 215 bits
Score difference with first non-orthologous sequence - *A. fumigatus*:215   *A. fischeri*:91
afum|EAL85363.2 100.00% afis|EAW20245.1 100.00%
Bootstrap support for afum|EAL85363.2 as seed ortholog is 100%.
Bootstrap support for afis|EAW20245.1 as seed ortholog is 100%.

---

Group of orthologs #8404. Best score 214 bits
Score difference with first non-orthologous sequence - *A. fumigatus*:214   *A. fischeri*:77
afum|EAL84825.1 100.00% afis|EAW20803.1 100.00%
Bootstrap support for afum|EAL84825.1 as seed ortholog is 100%.
Bootstrap support for afis|EAW20803.1 as seed ortholog is 99%.

---

Group of orthologs #8405. Best score 213 bits
Score difference with first non-orthologous sequence - *A. fumigatus*:213   *A. fischeri*:213
afum|EAL86740.1 100.00% afis|EAW16818.1 100.00%
Bootstrap support for afum|EAL86740.1 as seed ortholog is 100%.
Bootstrap support for afis|EAW16818.1 as seed ortholog is 100%.

---

Group of orthologs #8406. Best score 213 bits
Score difference with first non-orthologous sequence - *A. fumigatus*:213   *A. fischeri*:213
afum|EAL88076.1 100.00% afis|EAW23394.1 100.00%
Bootstrap support for afum|EAL88076.1 as seed ortholog is 100%.
Bootstrap support for afis|EAW23394.1 as seed ortholog is 100%.

---

Group of orthologs #8407. Best score 213 bits
Score difference with first non-orthologous sequence - *A. fumigatus*:213   *A. fischeri*:213
afum|EAL93818.1 100.00% afis|EAW19088.1 100.00%
Bootstrap support for afum|EAL93818.1 as seed ortholog is 100%.
Bootstrap support for afis|EAW19088.1 as seed ortholog is 100%.

---

Group of orthologs #8408. Best score 213 bits
Score difference with first non-orthologous sequence - *A. fumigatus*:213   *A. fischeri*:213
afum|EAL90980.2 100.00% afis|EAW22214.1 100.00%
Bootstrap support for afum|EAL90980.2 as seed ortholog is 100%.
Bootstrap support for afis|EAW22214.1 as seed ortholog is 100%.

---

Group of orthologs #8409. Best score 213 bits
Score difference with first non-orthologous sequence - *A. fumigatus*:213   *A. fischeri*:213
afum|EAL90023.1 100.00% afis|EAW25361.1 100.00%
Bootstrap support for afum|EAL90023.1 as seed ortholog is 100%.
Bootstrap support for afis|EAW25361.1 as seed ortholog is 100%.

---

Group of orthologs #8410. Best score 212 bits
Score difference with first non-orthologous sequence - *A. fumigatus*:212   *A. fischeri*:212
afum|EAL91193.1 100.00% afis|EAW17350.1 100.00%
Bootstrap support for afum|EAL91193.1 as seed ortholog is 100%.
Bootstrap support for afis|EAW17350.1 as seed ortholog is 100%.

---

Group of orthologs #8411. Best score 212 bits
Score difference with first non-orthologous sequence - *A. fumigatus*:212   *A. fischeri*:118
afum|EAL91905.2 100.00% afis|EAW18097.1 100.00%

Bootstrap support for afum|EAL91905.2 as seed ortholog is 100%.
Bootstrap support for afis|EAW18097.1 as seed ortholog is 100%.

---

Group of orthologs #8412. Best score 212 bits
Score difference with first non-orthologous sequence - *A. fumigatus*:212 *A. fischeri*:212
afum|EBA27322.1 100.00% afis|EAW18289.1 100.00%
Bootstrap support for afum|EBA27322.1 as seed ortholog is 100%.
Bootstrap support for afis|EAW18289.1 as seed ortholog is 100%.

---

Group of orthologs #8413. Best score 210 bits
Score difference with first non-orthologous sequence - *A. fumigatus*:210 *A. fischeri*:210
afum|EAL93987.1 100.00% afis|EAW19260.1 100.00%
Bootstrap support for afum|EAL93987.1 as seed ortholog is 100%.
Bootstrap support for afis|EAW19260.1 as seed ortholog is 100%.

---

Group of orthologs #8414. Best score 210 bits
Score difference with first non-orthologous sequence - *A. fumigatus*:210 *A. fischeri*:210
afum|EAL92646.1 100.00% afis|EAW21612.1 100.00%
Bootstrap support for afum|EAL92646.1 as seed ortholog is 100%.
Bootstrap support for afis|EAW21612.1 as seed ortholog is 100%.

---

Group of orthologs #8415. Best score 209 bits
Score difference with first non-orthologous sequence - *A. fumigatus*:209 *A. fischeri*:209
afum|EAL86583.1 100.00% afis|EAW17122.1 100.00%
Bootstrap support for afum|EAL86583.1 as seed ortholog is 100%.
Bootstrap support for afis|EAW17122.1 as seed ortholog is 100%.

---

Group of orthologs #8416. Best score 209 bits
Score difference with first non-orthologous sequence - *A. fumigatus*:209 *A. fischeri*:209
afum|EBA27404.1 100.00% afis|EAW22839.1 100.00%
Bootstrap support for afum|EBA27404.1 as seed ortholog is 100%.
Bootstrap support for afis|EAW22839.1 as seed ortholog is 100%.

---

Group of orthologs #8417. Best score 208 bits
Score difference with first non-orthologous sequence - *A. fumigatus*:208 *A. fischeri*:208
afum|EAL86944.1 100.00% afis|EAW19573.1 100.00%
Bootstrap support for afum|EAL86944.1 as seed ortholog is 100%.
Bootstrap support for afis|EAW19573.1 as seed ortholog is 100%.

---

Group of orthologs #8418. Best score 208 bits
Score difference with first non-orthologous sequence - *A. fumigatus*:208 *A. fischeri*:208
afum|EAL84404.1 100.00% afis|EAW25744.1 100.00%
Bootstrap support for afum|EAL84404.1 as seed ortholog is 100%.
Bootstrap support for afis|EAW25744.1 as seed ortholog is 100%.

---

Group of orthologs #8419. Best score 206 bits
Score difference with first non-orthologous sequence - *A. fumigatus*:206 *A. fischeri*:206
afum|EAL86054.1 100.00% afis|EAW24382.1 100.00%
Bootstrap support for afum|EAL86054.1 as seed ortholog is 100%.
Bootstrap support for afis|EAW24382.1 as seed ortholog is 100%.

---

Group of orthologs #8420. Best score 206 bits
Score difference with first non-orthologous sequence - *A. fumigatus*:206 *A. fischeri*:206
afum|EAL87974.1 100.00% afis|EAW23489.1 100.00%
Bootstrap support for afum|EAL87974.1 as seed ortholog is 100%.
Bootstrap support for afis|EAW23489.1 as seed ortholog is 100%.

Group of orthologs #8421. Best score 206 bits
Score difference with first non-orthologous sequence - *A. fumigatus*:206 *A. fischeri*:206
afum|EAL90109.2 100.00% afis|EAW25450.1 100.00%
Bootstrap support for afum|EAL90109.2 as seed ortholog is 100%.
Bootstrap support for afis|EAW25450.1 as seed ortholog is 100%.

---

Group of orthologs #8422. Best score 205 bits
Score difference with first non-orthologous sequence - *A. fumigatus*:205 *A. fischeri*:205
afum|EAL85578.1 100.00% afis|EAW15770.1 100.00%
Bootstrap support for afum|EAL85578.1 as seed ortholog is 100%.
Bootstrap support for afis|EAW15770.1 as seed ortholog is 100%.

---

Group of orthologs #8423. Best score 204 bits
Score difference with first non-orthologous sequence - *A. fumigatus*:204 *A. fischeri*:204
afum|EAL88281.1 100.00% afis|EAW23202.1 100.00%
Bootstrap support for afum|EAL88281.1 as seed ortholog is 100%.
Bootstrap support for afis|EAW23202.1 as seed ortholog is 100%.

---

Group of orthologs #8424. Best score 204 bits
Score difference with first non-orthologous sequence - *A. fumigatus*:204 *A. fischeri*:204
afum|EBA27311.1 100.00% afis|EAW23920.1 100.00%
Bootstrap support for afum|EBA27311.1 as seed ortholog is 100%.
Bootstrap support for afis|EAW23920.1 as seed ortholog is 100%.

---

Group of orthologs #8425. Best score 203 bits
Score difference with first non-orthologous sequence - *A. fumigatus*:203 *A. fischeri*:203
afum|EAL85636.1 100.00% afis|EAW15712.1 100.00%
Bootstrap support for afum|EAL85636.1 as seed ortholog is 100%.
Bootstrap support for afis|EAW15712.1 as seed ortholog is 100%.

---

Group of orthologs #8426. Best score 203 bits
Score difference with first non-orthologous sequence - *A. fumigatus*:203 *A. fischeri*:203
afum|EAL84316.1 100.00% afis|EAW25628.1 100.00%
Bootstrap support for afum|EAL84316.1 as seed ortholog is 100%.
Bootstrap support for afis|EAW25628.1 as seed ortholog is 100%.

---

Group of orthologs #8427. Best score 203 bits
Score difference with first non-orthologous sequence - *A. fumigatus*:203 *A. fischeri*:203
afum|EAL92904.1 100.00% afis|EAW21890.1 100.00%
Bootstrap support for afum|EAL92904.1 as seed ortholog is 100%.
Bootstrap support for afis|EAW21890.1 as seed ortholog is 100%.

---

Group of orthologs #8428. Best score 203 bits
Score difference with first non-orthologous sequence - *A. fumigatus*:203 *A. fischeri*:203
afum|EAL90129.2 100.00% afis|EAW25466.1 100.00%
Bootstrap support for afum|EAL90129.2 as seed ortholog is 100%.
Bootstrap support for afis|EAW25466.1 as seed ortholog is 100%.

---

Group of orthologs #8429. Best score 202 bits
Score difference with first non-orthologous sequence - *A. fumigatus*:202 *A. fischeri*:202
afum|EAL93069.1 100.00% afis|EAW18393.1 100.00%
Bootstrap support for afum|EAL93069.1 as seed ortholog is 100%.
Bootstrap support for afis|EAW18393.1 as seed ortholog is 100%.

---

Group of orthologs #8430. Best score 201 bits

Score difference with first non-orthologous sequence - *A. fumigatus*:73 *A. fischeri*:59
afum|EAL85018.1 100.00% afis|EAW19880.1 100.00%
Bootstrap support for afum|EAL85018.1 as seed ortholog is 51%. Alternative seed ortholog is afum|EAL87816.2 (73 bits away from this cluster)
Bootstrap support for afis|EAW19880.1 as seed ortholog is 76%.

---

Group of orthologs #8431. Best score 201 bits
Score difference with first non-orthologous sequence - *A. fumigatus*:201 *A. fischeri*:201
afum|EAL91701.1 100.00% afis|EAW17892.1 100.00%
Bootstrap support for afum|EAL91701.1 as seed ortholog is 100%.
Bootstrap support for afis|EAW17892.1 as seed ortholog is 100%.

---

Group of orthologs #8432. Best score 200 bits
Score difference with first non-orthologous sequence - *A. fumigatus*:200 *A. fischeri*:200
afum|EAL89085.1 100.00% afis|EAW16491.1 100.00%
Bootstrap support for afum|EAL89085.1 as seed ortholog is 100%.
Bootstrap support for afis|EAW16491.1 as seed ortholog is 100%.

---

Group of orthologs #8433. Best score 198 bits
Score difference with first non-orthologous sequence - *A. fumigatus*:198 *A. fischeri*:198
afum|EAL84692.1 100.00% afis|EAW20942.1 100.00%
Bootstrap support for afum|EAL84692.1 as seed ortholog is 100%.
Bootstrap support for afis|EAW20942.1 as seed ortholog is 100%.

---

Group of orthologs #8434. Best score 198 bits
Score difference with first non-orthologous sequence - *A. fumigatus*:198 *A. fischeri*:198
afum|EAL89736.1 100.00% afis|EAW25067.1 100.00%
Bootstrap support for afum|EAL89736.1 as seed ortholog is 100%.
Bootstrap support for afis|EAW25067.1 as seed ortholog is 100%.

---

Group of orthologs #8435. Best score 198 bits
Score difference with first non-orthologous sequence - *A. fumigatus*:198 *A. fischeri*:198
afum|EAL89842.1 100.00% afis|EAW25169.1 100.00%
Bootstrap support for afum|EAL89842.1 as seed ortholog is 100%.
Bootstrap support for afis|EAW25169.1 as seed ortholog is 100%.

---

Group of orthologs #8436. Best score 196 bits
Score difference with first non-orthologous sequence - *A. fumigatus*:196 *A. fischeri*:196
afum|EAL89128.1 100.00% afis|EAW16531.1 100.00%
Bootstrap support for afum|EAL89128.1 as seed ortholog is 100%.
Bootstrap support for afis|EAW16531.1 as seed ortholog is 100%.

---

Group of orthologs #8437. Best score 196 bits
Score difference with first non-orthologous sequence - *A. fumigatus*:196 *A. fischeri*:196
afum|EAL93113.1 100.00% afis|EAW18432.1 100.00%
Bootstrap support for afum|EAL93113.1 as seed ortholog is 100%.
Bootstrap support for afis|EAW18432.1 as seed ortholog is 100%.

---

Group of orthologs #8438. Best score 196 bits
Score difference with first non-orthologous sequence - *A. fumigatus*:196 *A. fischeri*:196
afum|EAL92730.1 100.00% afis|EAW21696.1 100.00%
Bootstrap support for afum|EAL92730.1 as seed ortholog is 100%.
Bootstrap support for afis|EAW21696.1 as seed ortholog is 100%.

---

Group of orthologs #8439. Best score 194 bits
Score difference with first non-orthologous sequence - *A. fumigatus*:194 *A. fischeri*:194

afum|EAL92092.2 100.00% afis|EAW21057.1 100.00%
Bootstrap support for afum|EAL92092.2 as seed ortholog is 100%.
Bootstrap support for afis|EAW21057.1 as seed ortholog is 100%.

---

Group of orthologs #8440. Best score 193 bits
Score difference with first non-orthologous sequence - *A. fumigatus*:193 *A. fischeri*:76
afum|EAL84242.1 100.00% afis|EAW17742.1 100.00%
Bootstrap support for afum|EAL84242.1 as seed ortholog is 100%.
Bootstrap support for afis|EAW17742.1 as seed ortholog is 96%.

---

Group of orthologs #8441. Best score 193 bits
Score difference with first non-orthologous sequence - *A. fumigatus*:193 *A. fischeri*:193
afum|EAL85323.1 100.00% afis|EAW20203.1 100.00%
Bootstrap support for afum|EAL85323.1 as seed ortholog is 100%.
Bootstrap support for afis|EAW20203.1 as seed ortholog is 100%.

---

Group of orthologs #8442. Best score 193 bits
Score difference with first non-orthologous sequence - *A. fumigatus*:193 *A. fischeri*:193
afum|EAL90811.1 100.00% afis|EAW22380.1 100.00%
Bootstrap support for afum|EAL90811.1 as seed ortholog is 100%.
Bootstrap support for afis|EAW22380.1 as seed ortholog is 100%.

---

Group of orthologs #8443. Best score 193 bits
Score difference with first non-orthologous sequence - *A. fumigatus*:193 *A. fischeri*:193
afum|EAL90318.1 100.00% afis|EAW22881.1 100.00%
Bootstrap support for afum|EAL90318.1 as seed ortholog is 100%.
Bootstrap support for afis|EAW22881.1 as seed ortholog is 100%.

---

Group of orthologs #8444. Best score 192 bits
Score difference with first non-orthologous sequence - *A. fumigatus*:192 *A. fischeri*:192
afum|EAL90680.1 100.00% afis|EAW22513.1 100.00%
Bootstrap support for afum|EAL90680.1 as seed ortholog is 100%.
Bootstrap support for afis|EAW22513.1 as seed ortholog is 100%.

---

Group of orthologs #8445. Best score 191 bits
Score difference with first non-orthologous sequence - *A. fumigatus*:191 *A. fischeri*:191
afum|EAL85862.1 100.00% afis|EAW24189.1 100.00%
Bootstrap support for afum|EAL85862.1 as seed ortholog is 100%.
Bootstrap support for afis|EAW24189.1 as seed ortholog is 100%.

---

Group of orthologs #8446. Best score 191 bits
Score difference with first non-orthologous sequence - *A. fumigatus*:191 *A. fischeri*:191
afum|EAL90003.1 100.00% afis|EAW25340.1 100.00%
Bootstrap support for afum|EAL90003.1 as seed ortholog is 100%.
Bootstrap support for afis|EAW25340.1 as seed ortholog is 100%.

---

Group of orthologs #8447. Best score 190 bits
Score difference with first non-orthologous sequence - *A. fumigatus*:190 *A. fischeri*:190
afum|EAL85718.1 100.00% afis|EAW15617.1 100.00%
Bootstrap support for afum|EAL85718.1 as seed ortholog is 100%.
Bootstrap support for afis|EAW15617.1 as seed ortholog is 100%.

---

Group of orthologs #8448. Best score 190 bits
Score difference with first non-orthologous sequence - *A. fumigatus*:190 *A. fischeri*:190
afum|EAL91490.1 100.00% afis|EAW17677.1 100.00%
Bootstrap support for afum|EAL91490.1 as seed ortholog is 100%.

Bootstrap support for afis|EAW17677.1 as seed ortholog is 100%.

---

Group of orthologs #8449. Best score 189 bits
Score difference with first non-orthologous sequence - *A. fumigatus*:189 *A. fischeri*:189
afum|EBA27447.1 100.00% afis|EAW21044.1 100.00%
afis|EAW20263.1 12.64%
Bootstrap support for afum|EBA27447.1 as seed ortholog is 100%.
Bootstrap support for afis|EAW21044.1 as seed ortholog is 100%.

---

Group of orthologs #8450. Best score 188 bits
Score difference with first non-orthologous sequence - *A. fumigatus*:188 *A. fischeri*:188
afum|EAL85263.1 100.00% afis|EAW20148.1 100.00%
Bootstrap support for afum|EAL85263.1 as seed ortholog is 100%.
Bootstrap support for afis|EAW20148.1 as seed ortholog is 100%.

---

Group of orthologs #8451. Best score 188 bits
Score difference with first non-orthologous sequence - *A. fumigatus*:188 *A. fischeri*:188
afum|EAL88014.1 100.00% afis|EAW23452.1 100.00%
Bootstrap support for afum|EAL88014.1 as seed ortholog is 100%.
Bootstrap support for afis|EAW23452.1 as seed ortholog is 100%.

---

Group of orthologs #8452. Best score 188 bits
Score difference with first non-orthologous sequence - *A. fumigatus*:188 *A. fischeri*:188
afum|EAL90217.1 100.00% afis|EAW22980.1 100.00%
Bootstrap support for afum|EAL90217.1 as seed ortholog is 100%.
Bootstrap support for afis|EAW22980.1 as seed ortholog is 100%.

---

Group of orthologs #8453. Best score 188 bits
Score difference with first non-orthologous sequence - *A. fumigatus*:188 *A. fischeri*:188
afum|EAL92804.1 100.00% afis|EAW21775.1 100.00%
Bootstrap support for afum|EAL92804.1 as seed ortholog is 100%.
Bootstrap support for afis|EAW21775.1 as seed ortholog is 100%.

---

Group of orthologs #8454. Best score 188 bits
Score difference with first non-orthologous sequence - *A. fumigatus*:188 *A. fischeri*:188
afum|EBA27472.1 100.00% afis|EAW21748.1 100.00%
Bootstrap support for afum|EBA27472.1 as seed ortholog is 100%.
Bootstrap support for afis|EAW21748.1 as seed ortholog is 100%.

---

Group of orthologs #8455. Best score 187 bits
Score difference with first non-orthologous sequence - *A. fumigatus*:187 *A. fischeri*:187
afum|EAL87490.1 100.00% afis|EAW18131.1 100.00%
Bootstrap support for afum|EAL87490.1 as seed ortholog is 100%.
Bootstrap support for afis|EAW18131.1 as seed ortholog is 100%.

---

Group of orthologs #8456. Best score 186 bits
Score difference with first non-orthologous sequence - *A. fumigatus*:186 *A. fischeri*:186
afum|EBA27232.1 100.00% afis|EAW20335.1 100.00%
Bootstrap support for afum|EBA27232.1 as seed ortholog is 100%.
Bootstrap support for afis|EAW20335.1 as seed ortholog is 100%.

---

Group of orthologs #8457. Best score 185 bits
Score difference with first non-orthologous sequence - *A. fumigatus*:185 *A. fischeri*:185
afum|EAL93242.1 100.00% afis|EAW18542.1 100.00%
Bootstrap support for afum|EAL93242.1 as seed ortholog is 100%.
Bootstrap support for afis|EAW18542.1 as seed ortholog is 100%.

---
Group of orthologs #8458. Best score 184 bits
Score difference with first non-orthologous sequence - *A. fumigatus*:184 *A. fischeri*:184
afum|EBA27252.1 100.00% afis|EAW24310.1 100.00%
Bootstrap support for afum|EBA27252.1 as seed ortholog is 100%.
Bootstrap support for afis|EAW24310.1 as seed ortholog is 100%.

---
Group of orthologs #8459. Best score 182 bits
Score difference with first non-orthologous sequence - *A. fumigatus*:182 *A. fischeri*:182
afum|EAL86839.1 100.00% afis|EAW19685.1 100.00%
Bootstrap support for afum|EAL86839.1 as seed ortholog is 100%.
Bootstrap support for afis|EAW19685.1 as seed ortholog is 100%.

---
Group of orthologs #8460. Best score 182 bits
Score difference with first non-orthologous sequence - *A. fumigatus*:182 *A. fischeri*:182
afum|EAL89801.1 100.00% afis|EAW25130.1 100.00%
Bootstrap support for afum|EAL89801.1 as seed ortholog is 100%.
Bootstrap support for afis|EAW25130.1 as seed ortholog is 100%.

---
Group of orthologs #8461. Best score 181 bits
Score difference with first non-orthologous sequence - *A. fumigatus*:181 *A. fischeri*:181
afum|EBA27168.1 100.00% afis|EAW15439.1 100.00%
Bootstrap support for afum|EBA27168.1 as seed ortholog is 100%.
Bootstrap support for afis|EAW15439.1 as seed ortholog is 100%.

---
Group of orthologs #8462. Best score 181 bits
Score difference with first non-orthologous sequence - *A. fumigatus*:181 *A. fischeri*:181
afum|EAL90606.1 100.00% afis|EAW22593.1 100.00%
Bootstrap support for afum|EAL90606.1 as seed ortholog is 100%.
Bootstrap support for afis|EAW22593.1 as seed ortholog is 100%.

---
Group of orthologs #8463. Best score 181 bits
Score difference with first non-orthologous sequence - *A. fumigatus*:181 *A. fischeri*:181
afum|EAL92800.1 100.00% afis|EAW21771.1 100.00%
Bootstrap support for afum|EAL92800.1 as seed ortholog is 100%.
Bootstrap support for afis|EAW21771.1 as seed ortholog is 100%.

---
Group of orthologs #8464. Best score 180 bits
Score difference with first non-orthologous sequence - *A. fumigatus*:180 *A. fischeri*:180
afum|EAL88437.1 100.00% afis|EAW23044.1 100.00%
Bootstrap support for afum|EAL88437.1 as seed ortholog is 100%.
Bootstrap support for afis|EAW23044.1 as seed ortholog is 100%.

---
Group of orthologs #8465. Best score 180 bits
Score difference with first non-orthologous sequence - *A. fumigatus*:180 *A. fischeri*:180
afum|EAL93847.2 100.00% afis|EAW19115.1 100.00%
Bootstrap support for afum|EAL93847.2 as seed ortholog is 100%.
Bootstrap support for afis|EAW19115.1 as seed ortholog is 100%.

---
Group of orthologs #8466. Best score 180 bits
Score difference with first non-orthologous sequence - *A. fumigatus*:180 *A. fischeri*:180
afum|EAL90715.1 100.00% afis|EAW22477.1 100.00%
Bootstrap support for afum|EAL90715.1 as seed ortholog is 100%.
Bootstrap support for afis|EAW22477.1 as seed ortholog is 100%.

---
Group of orthologs #8467. Best score 179 bits

Score difference with first non-orthologous sequence - *A. fumigatus*:179   *A. fischeri*:179
afum|EBA27352.1 100.00% afis|EAW15924.1 100.00%
Bootstrap support for afum|EBA27352.1 as seed ortholog is 100%.
Bootstrap support for afis|EAW15924.1 as seed ortholog is 100%.

---

Group of orthologs #8468. Best score 178 bits
Score difference with first non-orthologous sequence - *A. fumigatus*:178   *A. fischeri*:178
afum|EAL87683.1 100.00% afis|EAW18316.1 100.00%
Bootstrap support for afum|EAL87683.1 as seed ortholog is 100%.
Bootstrap support for afis|EAW18316.1 as seed ortholog is 100%.

---

Group of orthologs #8469. Best score 178 bits
Score difference with first non-orthologous sequence - *A. fumigatus*:178   *A. fischeri*:178
afum|EAL89303.1 100.00% afis|EAW22066.1 100.00%
Bootstrap support for afum|EAL89303.1 as seed ortholog is 100%.
Bootstrap support for afis|EAW22066.1 as seed ortholog is 100%.

---

Group of orthologs #8470. Best score 177 bits
Score difference with first non-orthologous sequence - *A. fumigatus*:177   *A. fischeri*:177
afum|EAL89041.1 100.00% afis|EAW16438.1 100.00%
Bootstrap support for afum|EAL89041.1 as seed ortholog is 100%.
Bootstrap support for afis|EAW16438.1 as seed ortholog is 100%.

---

Group of orthologs #8471. Best score 176 bits
Score difference with first non-orthologous sequence - *A. fumigatus*:176   *A. fischeri*:176
afum|EAL86954.1 100.00% afis|EAW19557.1 100.00%
Bootstrap support for afum|EAL86954.1 as seed ortholog is 100%.
Bootstrap support for afis|EAW19557.1 as seed ortholog is 100%.

---

Group of orthologs #8472. Best score 176 bits
Score difference with first non-orthologous sequence - *A. fumigatus*:176   *A. fischeri*:176
afum|EAL91358.1 100.00% afis|EAW17544.1 100.00%
Bootstrap support for afum|EAL91358.1 as seed ortholog is 100%.
Bootstrap support for afis|EAW17544.1 as seed ortholog is 100%.

---

Group of orthologs #8473. Best score 176 bits
Score difference with first non-orthologous sequence - *A. fumigatus*:176   *A. fischeri*:176
afum|EAL87188.1 100.00% afis|EAW23751.1 100.00%
Bootstrap support for afum|EAL87188.1 as seed ortholog is 100%.
Bootstrap support for afis|EAW23751.1 as seed ortholog is 100%.

---

Group of orthologs #8474. Best score 176 bits
Score difference with first non-orthologous sequence - *A. fumigatus*:176   *A. fischeri*:176
afum|EAL93297.1 100.00% afis|EAW18590.1 100.00%
Bootstrap support for afum|EAL93297.1 as seed ortholog is 100%.
Bootstrap support for afis|EAW18590.1 as seed ortholog is 100%.

---

Group of orthologs #8475. Best score 176 bits
Score difference with first non-orthologous sequence - *A. fumigatus*:176   *A. fischeri*:176
afum|EAL91066.1 100.00% afis|EAW22120.1 100.00%
Bootstrap support for afum|EAL91066.1 as seed ortholog is 100%.
Bootstrap support for afis|EAW22120.1 as seed ortholog is 100%.

---

Group of orthologs #8476. Best score 174 bits
Score difference with first non-orthologous sequence - *A. fumigatus*:11   *A. fischeri*:174
afum|EAL86216.1 100.00% afis|EAW24551.1 100.00%

Bootstrap support for afum|EAL86216.1 as seed ortholog is 68%. Alternative seed ortholog is afum|EAL91136.1 (11 bits away from this cluster)
Bootstrap support for afis|EAW24551.1 as seed ortholog is 100%.

---

Group of orthologs #8477. Best score 174 bits
Score difference with first non-orthologous sequence - *A. fumigatus*:174 *A. fischeri*:174
afum|EAL93866.1 100.00% afis|EAW19134.1 100.00%
Bootstrap support for afum|EAL93866.1 as seed ortholog is 100%.
Bootstrap support for afis|EAW19134.1 as seed ortholog is 100%.

---

Group of orthologs #8478. Best score 173 bits
Score difference with first non-orthologous sequence - *A. fumigatus*:173 *A. fischeri*:173
afum|EAL93227.1 100.00% afis|EAW18530.1 100.00%
Bootstrap support for afum|EAL93227.1 as seed ortholog is 100%.
Bootstrap support for afis|EAW18530.1 as seed ortholog is 100%.

---

Group of orthologs #8479. Best score 172 bits
Score difference with first non-orthologous sequence - *A. fumigatus*:172 *A. fischeri*:172
afum|EBA27242.1 100.00% afis|EAW15545.1 100.00%
Bootstrap support for afum|EBA27242.1 as seed ortholog is 100%.
Bootstrap support for afis|EAW15545.1 as seed ortholog is 100%.

---

Group of orthologs #8480. Best score 170 bits
Score difference with first non-orthologous sequence - *A. fumigatus*:170 *A. fischeri*:170
afum|EAL86978.1 100.00% afis|EAW19519.1 100.00%
Bootstrap support for afum|EAL86978.1 as seed ortholog is 100%.
Bootstrap support for afis|EAW19519.1 as seed ortholog is 100%.

---

Group of orthologs #8481. Best score 170 bits
Score difference with first non-orthologous sequence - *A. fumigatus*:170 *A. fischeri*:170
afum|EAL87791.1 100.00% afis|EAW20710.1 100.00%
Bootstrap support for afum|EAL87791.1 as seed ortholog is 100%.
Bootstrap support for afis|EAW20710.1 as seed ortholog is 100%.

---

Group of orthologs #8482. Best score 170 bits
Score difference with first non-orthologous sequence - *A. fumigatus*:170 *A. fischeri*:170
afum|EAL92429.1 100.00% afis|EAW22413.1 100.00%
Bootstrap support for afum|EAL92429.1 as seed ortholog is 100%.
Bootstrap support for afis|EAW22413.1 as seed ortholog is 100%.

---

Group of orthologs #8483. Best score 169 bits
Score difference with first non-orthologous sequence - *A. fumigatus*:169 *A. fischeri*:169
afum|EAL88970.1 100.00% afis|EAW16345.1 100.00%
Bootstrap support for afum|EAL88970.1 as seed ortholog is 100%.
Bootstrap support for afis|EAW16345.1 as seed ortholog is 100%.

---

Group of orthologs #8484. Best score 169 bits
Score difference with first non-orthologous sequence - *A. fumigatus*:169 *A. fischeri*:169
afum|EAL89596.1 100.00% afis|EAW24939.1 100.00%
Bootstrap support for afum|EAL89596.1 as seed ortholog is 100%.
Bootstrap support for afis|EAW24939.1 as seed ortholog is 100%.

---

Group of orthologs #8485. Best score 169 bits
Score difference with first non-orthologous sequence - *A. fumigatus*:169 *A. fischeri*:169
afum|EBA27175.1 100.00% afis|EAW25720.1 100.00%
Bootstrap support for afum|EBA27175.1 as seed ortholog is 100%.

Bootstrap support for afis|EAW25720.1 as seed ortholog is 100%.

---

Group of orthologs #8486. Best score 168 bits
Score difference with first non-orthologous sequence - *A. fumigatus*:168 *A. fischeri*:168
afum|EAL86478.1 100.00% afis|EAW16951.1 100.00%
Bootstrap support for afum|EAL86478.1 as seed ortholog is 100%.
Bootstrap support for afis|EAW16951.1 as seed ortholog is 100%.

---

Group of orthologs #8487. Best score 166 bits
Score difference with first non-orthologous sequence - *A. fumigatus*:166 *A. fischeri*:166
afum|EAL85352.1 100.00% afis|EAW20235.1 100.00%
Bootstrap support for afum|EAL85352.1 as seed ortholog is 100%.
Bootstrap support for afis|EAW20235.1 as seed ortholog is 100%.

---

Group of orthologs #8488. Best score 166 bits
Score difference with first non-orthologous sequence - *A. fumigatus*:166 *A. fischeri*:166
afum|EAL88023.1 100.00% afis|EAW23439.1 100.00%
Bootstrap support for afum|EAL88023.1 as seed ortholog is 100%.
Bootstrap support for afis|EAW23439.1 as seed ortholog is 100%.

---

Group of orthologs #8489. Best score 164 bits
Score difference with first non-orthologous sequence - *A. fumigatus*:164 *A. fischeri*:164
afum|EAL84205.1 100.00% afis|EAW15442.1 100.00%
Bootstrap support for afum|EAL84205.1 as seed ortholog is 100%.
Bootstrap support for afis|EAW15442.1 as seed ortholog is 100%.

---

Group of orthologs #8490. Best score 164 bits
Score difference with first non-orthologous sequence - *A. fumigatus*:164 *A. fischeri*:164
afum|EAL85660.1 100.00% afis|EAW15688.1 100.00%
Bootstrap support for afum|EAL85660.1 as seed ortholog is 100%.
Bootstrap support for afis|EAW15688.1 as seed ortholog is 100%.

---

Group of orthologs #8491. Best score 164 bits
Score difference with first non-orthologous sequence - *A. fumigatus*:164 *A. fischeri*:164
afum|EAL90944.1 100.00% afis|EAW24645.1 100.00%
Bootstrap support for afum|EAL90944.1 as seed ortholog is 100%.
Bootstrap support for afis|EAW24645.1 as seed ortholog is 100%.

---

Group of orthologs #8492. Best score 163 bits
Score difference with first non-orthologous sequence - *A. fumigatus*:163 *A. fischeri*:163
afum|EAL91198.1 100.00% afis|EAW17361.1 100.00%
Bootstrap support for afum|EAL91198.1 as seed ortholog is 100%.
Bootstrap support for afis|EAW17361.1 as seed ortholog is 100%.

---

Group of orthologs #8493. Best score 163 bits
Score difference with first non-orthologous sequence - *A. fumigatus*:163 *A. fischeri*:163
afum|EAL89551.1 100.00% afis|EAW24897.1 100.00%
Bootstrap support for afum|EAL89551.1 as seed ortholog is 100%.
Bootstrap support for afis|EAW24897.1 as seed ortholog is 100%.

---

Group of orthologs #8494. Best score 161 bits
Score difference with first non-orthologous sequence - *A. fumigatus*:161 *A. fischeri*:161
afum|EBA27204.1 100.00% afis|EAW20851.1 100.00%
Bootstrap support for afum|EBA27204.1 as seed ortholog is 100%.
Bootstrap support for afis|EAW20851.1 as seed ortholog is 100%.

---

Group of orthologs #8495. Best score 159 bits
Score difference with first non-orthologous sequence - *A. fumigatus*:159 *A. fischeri*:159
afum|EAL92627.1 100.00% afis|EAW19670.1 100.00%
Bootstrap support for afum|EAL92627.1 as seed ortholog is 100%.
Bootstrap support for afis|EAW19670.1 as seed ortholog is 100%.

---

Group of orthologs #8496. Best score 159 bits
Score difference with first non-orthologous sequence - *A. fumigatus*:159 *A. fischeri*:159
afum|EAL92144.1 100.00% afis|EAW21121.1 100.00%
Bootstrap support for afum|EAL92144.1 as seed ortholog is 100%.
Bootstrap support for afis|EAW21121.1 as seed ortholog is 100%.

---

Group of orthologs #8497. Best score 158 bits
Score difference with first non-orthologous sequence - *A. fumigatus*:158 *A. fischeri*:158
afum|EAL85982.1 100.00% afis|EAW24314.1 100.00%
Bootstrap support for afum|EAL85982.1 as seed ortholog is 100%.
Bootstrap support for afis|EAW24314.1 as seed ortholog is 100%.

---

Group of orthologs #8498. Best score 158 bits
Score difference with first non-orthologous sequence - *A. fumigatus*:158 *A. fischeri*:158
afum|EBA27206.1 100.00% afis|EAW20817.1 100.00%
Bootstrap support for afum|EBA27206.1 as seed ortholog is 100%.
Bootstrap support for afis|EAW20817.1 as seed ortholog is 100%.

---

Group of orthologs #8499. Best score 157 bits
Score difference with first non-orthologous sequence - *A. fumigatus*:157 *A. fischeri*:157
afum|EAL84824.1 100.00% afis|EAW20804.1 100.00%
Bootstrap support for afum|EAL84824.1 as seed ortholog is 100%.
Bootstrap support for afis|EAW20804.1 as seed ortholog is 100%.

---

Group of orthologs #8500. Best score 157 bits
Score difference with first non-orthologous sequence - *A. fumigatus*:157 *A. fischeri*:157
afum|EAL90590.1 100.00% afis|EAW22608.1 100.00%
Bootstrap support for afum|EAL90590.1 as seed ortholog is 100%.
Bootstrap support for afis|EAW22608.1 as seed ortholog is 100%.

---

Group of orthologs #8501. Best score 157 bits
Score difference with first non-orthologous sequence - *A. fumigatus*:157 *A. fischeri*:157
afum|EAL89517.2 100.00% afis|EAW24867.1 100.00%
Bootstrap support for afum|EAL89517.2 as seed ortholog is 100%.
Bootstrap support for afis|EAW24867.1 as seed ortholog is 100%.

---

Group of orthologs #8502. Best score 155 bits
Score difference with first non-orthologous sequence - *A. fumigatus*:155 *A. fischeri*:155
afum|EAL89795.1 100.00% afis|EAW25124.1 100.00%
Bootstrap support for afum|EAL89795.1 as seed ortholog is 100%.
Bootstrap support for afis|EAW25124.1 as seed ortholog is 100%.

---

Group of orthologs #8503. Best score 154 bits
Score difference with first non-orthologous sequence - *A. fumigatus*:154 *A. fischeri*:154
afum|EAL94109.1 100.00% afis|EAW25759.1 100.00%
afum|EAL92048.1 69.55% afis|EAW24638.1 44.99%
afum|EAL87107.1 38.53%
Bootstrap support for afum|EAL94109.1 as seed ortholog is 100%.
Bootstrap support for afis|EAW25759.1 as seed ortholog is 100%.

---

Group of orthologs #8504. Best score 154 bits
Score difference with first non-orthologous sequence - *A. fumigatus*:154 *A. fischeri*:154
afum|EAL91083.1 100.00% afis|EAW19744.1 100.00%
Bootstrap support for afum|EAL91083.1 as seed ortholog is 100%.
Bootstrap support for afis|EAW19744.1 as seed ortholog is 100%.

---

Group of orthologs #8505. Best score 152 bits
Score difference with first non-orthologous sequence - *A. fumigatus*:152 *A. fischeri*:152
afum|EAL87156.2 100.00% afis|EAW23712.1 100.00%
Bootstrap support for afum|EAL87156.2 as seed ortholog is 100%.
Bootstrap support for afis|EAW23712.1 as seed ortholog is 100%.

---

Group of orthologs #8506. Best score 152 bits
Score difference with first non-orthologous sequence - *A. fumigatus*:152 *A. fischeri*:152
afum|EAL87416.1 100.00% afis|EAW23987.1 100.00%
Bootstrap support for afum|EAL87416.1 as seed ortholog is 100%.
Bootstrap support for afis|EAW23987.1 as seed ortholog is 100%.

---

Group of orthologs #8507. Best score 151 bits
Score difference with first non-orthologous sequence - *A. fumigatus*:151 *A. fischeri*:151
afum|EAL87942.1 100.00% afis|EAW23522.1 100.00%
Bootstrap support for afum|EAL87942.1 as seed ortholog is 100%.
Bootstrap support for afis|EAW23522.1 as seed ortholog is 100%.

---

Group of orthologs #8508. Best score 151 bits
Score difference with first non-orthologous sequence - *A. fumigatus*:151 *A. fischeri*:151
afum|EAL92122.1 100.00% afis|EAW21101.1 100.00%
Bootstrap support for afum|EAL92122.1 as seed ortholog is 100%.
Bootstrap support for afis|EAW21101.1 as seed ortholog is 100%.

---

Group of orthologs #8509. Best score 150 bits
Score difference with first non-orthologous sequence - *A. fumigatus*:150 *A. fischeri*:150
afum|EAL87043.1 100.00% afis|EAW19453.1 100.00%
Bootstrap support for afum|EAL87043.1 as seed ortholog is 100%.
Bootstrap support for afis|EAW19453.1 as seed ortholog is 100%.

---

Group of orthologs #8510. Best score 149 bits
Score difference with first non-orthologous sequence - *A. fumigatus*:149 *A. fischeri*:149
afum|EAL92878.2 100.00% afis|EAW21864.1 100.00%
Bootstrap support for afum|EAL92878.2 as seed ortholog is 100%.
Bootstrap support for afis|EAW21864.1 as seed ortholog is 100%.

---

Group of orthologs #8511. Best score 148 bits
Score difference with first non-orthologous sequence - *A. fumigatus*:148 *A. fischeri*:148
afum|EAL89112.1 100.00% afis|EAW16517.1 100.00%
Bootstrap support for afum|EAL89112.1 as seed ortholog is 100%.
Bootstrap support for afis|EAW16517.1 as seed ortholog is 100%.

---

Group of orthologs #8512. Best score 148 bits
Score difference with first non-orthologous sequence - *A. fumigatus*:148 *A. fischeri*:148
afum|EAL85975.1 100.00% afis|EAW24307.1 100.00%
Bootstrap support for afum|EAL85975.1 as seed ortholog is 100%.
Bootstrap support for afis|EAW24307.1 as seed ortholog is 100%.

---

Group of orthologs #8513. Best score 148 bits
Score difference with first non-orthologous sequence - *A. fumigatus*:148 *A. fischeri*:148

afum|EAL92530.1 100.00% afis|EAW21510.1 100.00%
Bootstrap support for afum|EAL92530.1 as seed ortholog is 100%.
Bootstrap support for afis|EAW21510.1 as seed ortholog is 100%.

---

Group of orthologs #8514. Best score 147 bits
Score difference with first non-orthologous sequence - *A. fumigatus*:147 *A. fischeri*:147
afum|EAL87185.1 100.00% afis|EAW23689.1 100.00%
Bootstrap support for afum|EAL87185.1 as seed ortholog is 100%.
Bootstrap support for afis|EAW23689.1 as seed ortholog is 100%.

---

Group of orthologs #8515. Best score 146 bits
Score difference with first non-orthologous sequence - *A. fumigatus*:146 *A. fischeri*:146
afum|EAL92348.1 100.00% afis|EAW21330.1 100.00%
Bootstrap support for afum|EAL92348.1 as seed ortholog is 100%.
Bootstrap support for afis|EAW21330.1 as seed ortholog is 100%.

---

Group of orthologs #8516. Best score 144 bits
Score difference with first non-orthologous sequence - *A. fumigatus*:144 *A. fischeri*:144
afum|EAL84259.1 100.00% afis|EAW17040.1 100.00%
Bootstrap support for afum|EAL84259.1 as seed ortholog is 100%.
Bootstrap support for afis|EAW17040.1 as seed ortholog is 100%.

---

Group of orthologs #8517. Best score 144 bits
Score difference with first non-orthologous sequence - *A. fumigatus*:144 *A. fischeri*:144
afum|EAL84704.1 100.00% afis|EAW20930.1 100.00%
Bootstrap support for afum|EAL84704.1 as seed ortholog is 100%.
Bootstrap support for afis|EAW20930.1 as seed ortholog is 100%.

---

Group of orthologs #8518. Best score 144 bits
Score difference with first non-orthologous sequence - *A. fumigatus*:144 *A. fischeri*:144
afum|EAL90154.1 100.00% afis|EAW25489.1 100.00%
Bootstrap support for afum|EAL90154.1 as seed ortholog is 100%.
Bootstrap support for afis|EAW25489.1 as seed ortholog is 100%.

---

Group of orthologs #8519. Best score 141 bits
Score difference with first non-orthologous sequence - *A. fumigatus*:141 *A. fischeri*:141
afum|EAL86588.1 100.00% afis|EAW17128.1 100.00%
Bootstrap support for afum|EAL86588.1 as seed ortholog is 100%.
Bootstrap support for afis|EAW17128.1 as seed ortholog is 100%.

---

Group of orthologs #8520. Best score 141 bits
Score difference with first non-orthologous sequence - *A. fumigatus*:42 *A. fischeri*:47
afum|EAL92223.1 100.00% afis|EAW16436.1 100.00%
Bootstrap support for afum|EAL92223.1 as seed ortholog is 83%.
Bootstrap support for afis|EAW16436.1 as seed ortholog is 90%.

---

Group of orthologs #8521. Best score 141 bits
Score difference with first non-orthologous sequence - *A. fumigatus*:141 *A. fischeri*:141
afum|EAL93804.1 100.00% afis|EAW19076.1 100.00%
Bootstrap support for afum|EAL93804.1 as seed ortholog is 100%.
Bootstrap support for afis|EAW19076.1 as seed ortholog is 100%.

---

Group of orthologs #8522. Best score 141 bits
Score difference with first non-orthologous sequence - *A. fumigatus*:141 *A. fischeri*:141
afum|EBA27310.1 100.00% afis|EAW23897.1 100.00%
Bootstrap support for afum|EBA27310.1 as seed ortholog is 100%.

Bootstrap support for afis|EAW23897.1 as seed ortholog is 100%.

---

Group of orthologs #8523. Best score 140 bits
Score difference with first non-orthologous sequence - *A. fumigatus*:140 *A. fischeri*:140
afum|EAL84744.1 100.00% afis|EAW20889.1 100.00%
Bootstrap support for afum|EAL84744.1 as seed ortholog is 100%.
Bootstrap support for afis|EAW20889.1 as seed ortholog is 100%.

---

Group of orthologs #8524. Best score 138 bits
Score difference with first non-orthologous sequence - *A. fumigatus*:138 *A. fischeri*:138
afum|EAL90973.1 100.00% afis|EAW22221.1 100.00%
Bootstrap support for afum|EAL90973.1 as seed ortholog is 100%.
Bootstrap support for afis|EAW22221.1 as seed ortholog is 100%.

---

Group of orthologs #8525. Best score 137 bits
Score difference with first non-orthologous sequence - *A. fumigatus*:137 *A. fischeri*:137
afum|EAL85161.2 100.00% afis|EAW20041.1 100.00%
Bootstrap support for afum|EAL85161.2 as seed ortholog is 100%.
Bootstrap support for afis|EAW20041.1 as seed ortholog is 100%.

---

Group of orthologs #8526. Best score 137 bits
Score difference with first non-orthologous sequence - *A. fumigatus*:137 *A. fischeri*:137
afum|EAL87799.1 100.00% afis|EAW19105.1 100.00%
Bootstrap support for afum|EAL87799.1 as seed ortholog is 100%.
Bootstrap support for afis|EAW19105.1 as seed ortholog is 100%.

---

Group of orthologs #8527. Best score 137 bits
Score difference with first non-orthologous sequence - *A. fumigatus*:137 *A. fischeri*:137
afum|EBA27489.1 100.00% afis|EAW18639.1 100.00%
Bootstrap support for afum|EBA27489.1 as seed ortholog is 100%.
Bootstrap support for afis|EAW18639.1 as seed ortholog is 100%.

---

Group of orthologs #8528. Best score 136 bits
Score difference with first non-orthologous sequence - *A. fumigatus*:136 *A. fischeri*:136
afum|EAL85794.1 100.00% afis|EAW15534.1 100.00%
Bootstrap support for afum|EAL85794.1 as seed ortholog is 100%.
Bootstrap support for afis|EAW15534.1 as seed ortholog is 100%.

---

Group of orthologs #8529. Best score 134 bits
Score difference with first non-orthologous sequence - *A. fumigatus*:134 *A. fischeri*:134
afum|EAL87142.1 100.00% afis|EAW23701.1 100.00%
Bootstrap support for afum|EAL87142.1 as seed ortholog is 100%.
Bootstrap support for afis|EAW23701.1 as seed ortholog is 100%.

---

Group of orthologs #8530. Best score 132 bits
Score difference with first non-orthologous sequence - *A. fumigatus*:132 *A. fischeri*:57
afum|EAL87067.1 100.00% afis|EAW19427.1 100.00%
Bootstrap support for afum|EAL87067.1 as seed ortholog is 100%.
Bootstrap support for afis|EAW19427.1 as seed ortholog is 99%.

---

Group of orthologs #8531. Best score 131 bits
Score difference with first non-orthologous sequence - *A. fumigatus*:131 *A. fischeri*:131
afum|EBA27234.1 100.00% afis|EAW15830.1 100.00%
Bootstrap support for afum|EBA27234.1 as seed ortholog is 100%.
Bootstrap support for afis|EAW15830.1 as seed ortholog is 100%.

---

Group of orthologs #8532. Best score 131 bits
Score difference with first non-orthologous sequence - *A. fumigatus*:131 *A. fischeri*:131
afum|EBA27390.1 100.00% afis|EAW25261.1 100.00%
Bootstrap support for afum|EBA27390.1 as seed ortholog is 100%.
Bootstrap support for afis|EAW25261.1 as seed ortholog is 100%.

---

Group of orthologs #8533. Best score 129 bits
Score difference with first non-orthologous sequence - *A. fumigatus*:129 *A. fischeri*:129
afum|EAL90102.1 100.00% afis|EAW25444.1 100.00%
Bootstrap support for afum|EAL90102.1 as seed ortholog is 100%.
Bootstrap support for afis|EAW25444.1 as seed ortholog is 100%.

---

Group of orthologs #8534. Best score 127 bits
Score difference with first non-orthologous sequence - *A. fumigatus*:127 *A. fischeri*:127
afum|EAL94065.1 100.00% afis|EAW19345.1 100.00%
Bootstrap support for afum|EAL94065.1 as seed ortholog is 100%.
Bootstrap support for afis|EAW19345.1 as seed ortholog is 100%.

---

Group of orthologs #8535. Best score 126 bits
Score difference with first non-orthologous sequence - *A. fumigatus*:74 *A. fischeri*:2
afum|EAL86781.1 100.00% afis|EAW25743.1 100.00%
afis|EAW19901.1 19.14%
Bootstrap support for afum|EAL86781.1 as seed ortholog is 97%.
Bootstrap support for afis|EAW25743.1 as seed ortholog is 54%. Alternative seed ortholog is afis|EAW19890.1 (2 bits away from this cluster)

---

Group of orthologs #8536. Best score 125 bits
Score difference with first non-orthologous sequence - *A. fumigatus*:125 *A. fischeri*:125
afum|EAL89698.1 100.00% afis|EAW25000.1 100.00%
Bootstrap support for afum|EAL89698.1 as seed ortholog is 100%.
Bootstrap support for afis|EAW25000.1 as seed ortholog is 100%.

---

Group of orthologs #8537. Best score 123 bits
Score difference with first non-orthologous sequence - *A. fumigatus*:123 *A. fischeri*:123
afum|EAL90672.1 100.00% afis|EAW22522.1 100.00%
Bootstrap support for afum|EAL90672.1 as seed ortholog is 100%.
Bootstrap support for afis|EAW22522.1 as seed ortholog is 100%.

---

Group of orthologs #8538. Best score 120 bits
Score difference with first non-orthologous sequence - *A. fumigatus*:120 *A. fischeri*:120
afum|EBA27272.1 100.00% afis|EAW16863.1 100.00%
Bootstrap support for afum|EBA27272.1 as seed ortholog is 100%.
Bootstrap support for afis|EAW16863.1 as seed ortholog is 100%.

---

Group of orthologs #8539. Best score 118 bits
Score difference with first non-orthologous sequence - *A. fumigatus*:118 *A. fischeri*:118
afum|EAL86402.1 100.00% afis|EAW16406.1 100.00%
afum|EBA27374.1 100.00%
Bootstrap support for afum|EAL86402.1 as seed ortholog is 100%.
Bootstrap support for afum|EBA27374.1 as seed ortholog is 100%.
Bootstrap support for afis|EAW16406.1 as seed ortholog is 100%.

---

Group of orthologs #8540. Best score 118 bits
Score difference with first non-orthologous sequence - *A. fumigatus*:25 *A. fischeri*:118
afum|EAL85379.1 100.00% afis|EAW19387.1 100.00%
afum|EAL87817.1 73.48%

Bootstrap support for afum|EAL85379.1 as seed ortholog is 75%.
Bootstrap support for afis|EAW19387.1 as seed ortholog is 100%.

---

Group of orthologs #8541. Best score 118 bits
Score difference with first non-orthologous sequence - *A. fumigatus*:118 *A. fischeri*:118
afum|EAL88458.1 100.00% afis|EAW23024.1 100.00%
Bootstrap support for afum|EAL88458.1 as seed ortholog is 100%.
Bootstrap support for afis|EAW23024.1 as seed ortholog is 100%.

---

Group of orthologs #8542. Best score 115 bits
Score difference with first non-orthologous sequence - *A. fumigatus*:115 *A. fischeri*:115
afum|EAL87023.1 100.00% afis|EAW19473.1 100.00%
Bootstrap support for afum|EAL87023.1 as seed ortholog is 100%.
Bootstrap support for afis|EAW19473.1 as seed ortholog is 100%.

---

Group of orthologs #8543. Best score 113 bits
Score difference with first non-orthologous sequence - *A. fumigatus*:113 *A. fischeri*:113
afum|EAL90934.1 100.00% afis|EAW20714.1 100.00%
afum|EAL91122.1 50.31%
Bootstrap support for afum|EAL90934.1 as seed ortholog is 100%.
Bootstrap support for afis|EAW20714.1 as seed ortholog is 100%.

---

Group of orthologs #8544. Best score 106 bits
Score difference with first non-orthologous sequence - *A. fumigatus*:106 *A. fischeri*:106
afum|EAL91212.1 100.00% afis|EAW17377.1 100.00%
Bootstrap support for afum|EAL91212.1 as seed ortholog is 100%.
Bootstrap support for afis|EAW17377.1 as seed ortholog is 100%.

---

Group of orthologs #8545. Best score 101 bits
Score difference with first non-orthologous sequence - *A. fumigatus*:101 *A. fischeri*:101
afum|EAL87512.1 100.00% afis|EAW18153.1 100.00%
Bootstrap support for afum|EAL87512.1 as seed ortholog is 100%.
Bootstrap support for afis|EAW18153.1 as seed ortholog is 100%.

---

Group of orthologs #8546. Best score 100 bits
Score difference with first non-orthologous sequence - *A. fumigatus*:100 *A. fischeri*:100
afum|EAL92635.1 100.00% afis|EAW16912.1 100.00%
Bootstrap support for afum|EAL92635.1 as seed ortholog is 100%.
Bootstrap support for afis|EAW16912.1 as seed ortholog is 100%.

---

Group of orthologs #8547. Best score 97 bits
Score difference with first non-orthologous sequence - *A. fumigatus*:97 *A. fischeri*:97
afum|EBA27210.1 100.00% afis|EAW19982.1 100.00%
Bootstrap support for afum|EBA27210.1 as seed ortholog is 100%.
Bootstrap support for afis|EAW19982.1 as seed ortholog is 100%.

---

Group of orthologs #8548. Best score 93 bits
Score difference with first non-orthologous sequence - *A. fumigatus*:93 *A. fischeri*:93
afum|EAL86553.1 100.00% afis|EAW17057.1 100.00%
Bootstrap support for afum|EAL86553.1 as seed ortholog is 100%.
Bootstrap support for afis|EAW17057.1 as seed ortholog is 100%.

---

Group of orthologs #8549. Best score 88 bits
Score difference with first non-orthologous sequence - *A. fumigatus*:88 *A. fischeri*:88
afum|EAL84371.1 100.00% afis|EAW19872.1 100.00%
Bootstrap support for afum|EAL84371.1 as seed ortholog is 100%.

Bootstrap support for afis|EAW19872.1 as seed ortholog is 100%.

---

Group of orthologs #8550. Best score 86 bits
Score difference with first non-orthologous sequence - *A. fumigatus*:86 *A. fischeri*:86
afum|EBA27450.1 100.00% afis|EAW17120.1 100.00%
afis|EAW17354.1 9.70%
Bootstrap support for afum|EBA27450.1 as seed ortholog is 100%.
Bootstrap support for afis|EAW17120.1 as seed ortholog is 100%.

---

Group of orthologs #8551. Best score 85 bits
Score difference with first non-orthologous sequence - *A. fumigatus*:85 *A. fischeri*:85
afum|EAL92339.1 100.00% afis|EAW21322.1 100.00%
Bootstrap support for afum|EAL92339.1 as seed ortholog is 100%.
Bootstrap support for afis|EAW21322.1 as seed ortholog is 100%.

---

Group of orthologs #8552. Best score 83 bits
Score difference with first non-orthologous sequence - *A. fumigatus*:83 *A. fischeri*:83
afum|EAL87807.1 100.00% afis|EAW22023.1 100.00%
afum|EAL86764.1 30.82%
Bootstrap support for afum|EAL87807.1 as seed ortholog is 100%.
Bootstrap support for afis|EAW22023.1 as seed ortholog is 100%.

---

Group of orthologs #8553. Best score 83 bits
Score difference with first non-orthologous sequence - *A. fumigatus*:83 *A. fischeri*:37
afum|EAL93575.1 100.00% afis|EAW24573.1 100.00%
Bootstrap support for afum|EAL93575.1 as seed ortholog is 100%.
Bootstrap support for afis|EAW24573.1 as seed ortholog is 77%.

---

Group of orthologs #8554. Best score 82 bits
Score difference with first non-orthologous sequence - *A. fumigatus*:82 *A. fischeri*:82
afum|EAL84497.1 100.00% afis|EAW20564.1 100.00%
Bootstrap support for afum|EAL84497.1 as seed ortholog is 100%.
Bootstrap support for afis|EAW20564.1 as seed ortholog is 100%.

---

Group of orthologs #8555. Best score 68 bits
Score difference with first non-orthologous sequence - *A. fumigatus*:68 *A. fischeri*:68
afum|EAL89871.1 100.00% afis|EAW22091.1 100.00%
Bootstrap support for afum|EAL89871.1 as seed ortholog is 100%.
Bootstrap support for afis|EAW22091.1 as seed ortholog is 100%.

---

Group of orthologs #8556. Best score 63 bits
Score difference with first non-orthologous sequence - *A. fumigatus*:63 *A. fischeri*:63
afum|EAL84803.1 100.00% afis|EAW17752.1 100.00%
Bootstrap support for afum|EAL84803.1 as seed ortholog is 100%.
Bootstrap support for afis|EAW17752.1 as seed ortholog is 100%.

---

Group of orthologs #8557. Best score 63 bits
Score difference with first non-orthologous sequence - *A. fumigatus*:63 *A. fischeri*:63
afum|EAL93071.1 100.00% afis|EAW18395.1 100.00%
Bootstrap support for afum|EAL93071.1 as seed ortholog is 100%.
Bootstrap support for afis|EAW18395.1 as seed ortholog is 100%.

---

Group of orthologs #8558. Best score 60 bits
Score difference with first non-orthologous sequence - *A. fumigatus*:60 *A. fischeri*:60
afum|EAL91589.1 100.00% afis|EAW18844.1 100.00%
Bootstrap support for afum|EAL91589.1 as seed ortholog is 100%.

Bootstrap support for afis|EAW18844.1 as seed ortholog is 100%.

---

Group of orthologs #8559. Best score 60 bits
Score difference with first non-orthologous sequence - *A. fumigatus*:60 *A. fischeri*:60
afum|EBA27361.1 100.00% afis|EAW20663.1 100.00%
Bootstrap support for afum|EBA27361.1 as seed ortholog is 100%.
Bootstrap support for afis|EAW20663.1 as seed ortholog is 100%.

---

Group of orthologs #8560. Best score 59 bits
Score difference with first non-orthologous sequence - *A. fumigatus*:59 *A. fischeri*:59
afum|EAL85009.1 100.00% afis|EAW22095.1 100.00%
Bootstrap support for afum|EAL85009.1 as seed ortholog is 100%.
Bootstrap support for afis|EAW22095.1 as seed ortholog is 100%.

---

Group of orthologs #8561. Best score 53 bits
Score difference with first non-orthologous sequence - *A. fumigatus*:53 *A. fischeri*:53
afum|EAL90928.1 100.00% afis|EAW16894.1 100.00%
Bootstrap support for afum|EAL90928.1 as seed ortholog is 100%.
Bootstrap support for afis|EAW16894.1 as seed ortholog is 100%.

---

Group of orthologs #8562. Best score 53 bits
Score difference with first non-orthologous sequence - *A. fumigatus*:53 *A. fischeri*:53
afum|EBA27449.1 100.00% afis|EAW21152.1 100.00%
Bootstrap support for afum|EBA27449.1 as seed ortholog is 100%.
Bootstrap support for afis|EAW21152.1 as seed ortholog is 100%.

--------------------------------------------------------------------------------

Abbreviations: afum - *A. fumigatus*; afis - *A. fischeri*
