## Additional file 3 for "*Aspergillus fumigatus* versus Genus *Aspergillus*: Conservation, adaptive evolution and specific virulence genes"

### This PDF file includes:

**1. Supplemental methods**

- **(i) Test for recombination**
- **(ii) Tests of functional category enrichment**
- **(iii) Inference of positive selection in *A. fumigatus* virulence genes**

**References**

### Other Additional file for this manuscript includes the following:

**Additional file 1**

**Additional file 2**

* Corresponding author

### Supplementary methods

#### Test for recombination

Recombination can inflate the rate of false positives in studies aimed at detecting positive selection based on dN/dS ratios, because the common methods used for detecting positive selection assume that all sites share the same underlying phylogeny [1]. Therefore, we eliminated any genes in the analysis that showed evidence for recombination signals in aligned columns, using a statistical test implemented in GARD program [2] under HyPhy package (Hypothesis testing using Phylogenies) [3]. GARD implements a genetic algorithm for detecting the evidence for potential recombination events from MSA of homologous sequences. Briefly, the method compares a likelihood model assuming a single recombination breakpoint with two different topologies on each side of the breakpoint, with a model that assumes no recombination. If the model with recombination was supported by the Akaike Information Criterion [4], a likelihood ratio test based on the parametric bootstrapping [5] is used to determine whether the model with no recombination can be rejected in favor of a model with recombination. Based on the outcome of the analysis, a level of support is assigned and expressed as a breakpoint placement score [2]. Identified breakpoints by GARD were then assessed for statistical significance by using SH test (Shimodaira-Hasegawa test) [6]. Evidence of significant recombination events were found in 879 alignments (table S15, for details, see Additional file 1 Supplemental Note "Recombination breakpoints"). These clusters where we found statistical evidence of significant breakpoints in alignment, were discarded from further analysis.

#### Tests of functional category enrichment

GO annotations of *A. fumigatus* single copy ortholog proteins were performed with Blast2GO package [7]. To identify significantly over-represented functional categories, present in the data sets used in this study, Fisher's Exact Test [8] was used. The reference set was constituted of all *A. fumigatus* protein coding genes with GO annotations. To acquire a systematic overview of the PSGs functionalities, we performed the sequence similarity search with BlastP [9] against the KOG database [10]. We used Integrating Network Objects with Hierarchies (INOH) database signal transduction pathway database [11] at InnateDB platform [12] to perform the pathway over-representation analysis of PSGs. DAVID tool [13] was used to calculate whether a certain GO category occurred more often in the PSGs then to be expected looking at all *A.*

*fumigatus* protein-coding gene annotations at the background, as we did in our recent work [14]. We used level 3 GO terms and a very conservative EASE Score (a modified Fisher Exact p-Value; [15]) threshold 0.05 to identify the significantly over-represented pathways in *A. fumigatus* PSGs. Fisher's exact test was applied to calculate the statistical significance of the over-represented GO categories. The p-value was calculated for Blast2GO, InnateDB and DAVID analysis using hypergeometric tests, and Benjamini-Hochberg adjustment was used for multiple test correction to estimate the proportion of enriched gene sets that would occur by chance given the number of tested gene sets.

#### Inference of positive selection in *A. fumigatus* virulence genes

A total of 1931 virulence-related genes were collected from literature [16, 17]. A total of 1498 genes in non-SCOs were analysed for positive selection. For reasons of accuracy and computational power, a size reduction approach was followed. In orthogroups with high number of sequences and lower sequence similarity, inference of positive selection is less powerful and less accurate [18] and removal of divergent sequences from the orthogroup for better accuracy can eliminate the high false positive rate caused by highly divergent sequences. Therefore, removal of divergent sequences was performed until the size of 18 sequences was reached. Resulting orthogroups are then aligned first at amino acid level, which are then converted into codon alignments and filtered to remove gap-rich regions. Resulting alignments are then used to infer the phylogenetic gene trees and positive selection with PAML branch-site models.
